# Downregulation of Neurodevelopmental Gene Expression in iPSC-Derived Cerebral Organoids Upon Infection by Human Cytomegalovirus

**DOI:** 10.1101/2021.08.01.454651

**Authors:** Benjamin S. O’Brien, Rebekah L. Mokry, Megan L. Schumacher, Kirthi Pulakanti, Sridhar Rao, Scott S. Terhune, Allison D. Ebert

## Abstract

Human cytomegalovirus (HCMV) is a beta herpesvirus that, upon congenital infection, can cause severe birth defects including vision and hearing loss, microcephaly, and seizures. Currently, no approved treatment options exist for in utero infections. We previously demonstrated that HCMV infection decreases calcium signaling responses and alters neuronal differentiation in induced pluripotent stem cell (iPSC) derived neural progenitor cells (NPCs). Here we aimed to determine the impact of infection on the transcriptome in developing human neurons using iPSC-derived 3-dimensional cerebral organoids. We infected iPSC-derived cerebral organoids with HCMV encoding eGFP and sorted cell populations based on GFP signal strength. Significant transcriptional downregulation was observed including in key neurodevelopmental gene pathways in both the GFP (+) and intermediate groups. Interestingly, the GFP (-) group also showed downregulation of the same targets indicating a mismatch between GFP expression and viral infection. Using a modified HCMV virus destabilizing IE 1 and 2 proteins, we still observed significant downregulation of neurodevelopmental gene expression in infected neural progenitor cells. Together, these data indicate that IE viral proteins are not the main drivers of neurodevelopmental gene dysregulation in HCMV infected neural tissues suggesting therapeutically targeting IE gene expression is insufficient to restore neural differentiation and function.

## Introduction

Human cytomegalovirus (HCMV) is a beta-herpesvirus with a 235-kbp double-stranded DNA (dsDNA) genome and the potential to express more than 700 proteins (Mocarski and Kemble 1996; Mocarski 2007). The virus infects a majority of the world’s population with seroprevalence of 40-100% depending on age, socio-economic status and geographic location (Sinzger et al. 2008; Griffiths et al. 2015). Infection by HCMV is lifelong and can result in a range of conditions. Vertical transmission can also occur causing congenital CMV (cCMV) infection (Mocarski and Kemble 1996; Griffiths et al. 2015). A portion of babies born with cCMV infection will have long-term health problems including vision and hearing loss, microcephaly, and seizures. Neurological symptoms are likely the result of infection of neural progenitor cells (NPCs) (Luo et al. 2008; Hindley et al. 2012; Sun et al. 2020). In vitro, NPCs derived from both induced pluripotent stem cells (iPSCs) and fetal stem cells are fully permissive for HCMV infection, though susceptibility depends upon degree of NPC terminal differentiation (Odeberg et al. 2006; Luo et al. 2010; Pan et al. 2013; Gonzalez-Sanchez et al. 2015). In the fetal brain, NPCs are located in the bilateral subventricular zone aligned with the developing ventricle, and these cells express a number of key transcription factors that are required for maintenance of the progenitor cell pool and subsequent differentiation. NPCs can differentiate into the multiple neuronal and glial lineages found in the central nervous system (CNS). HCMV infection of NPCs has been shown to alter differentiation and function (Luo et al. 2010; Liu et al. 2017a; Wu et al. 2018; Brown et al. 2019). However, the mechanisms controlling this remain to be fully established.

Upon infection, the HCMV genes are expressed in three sequential steps known as immediate early (IE), early (E), and late (L). Viral IE gene expression can be detected within hours of infection and total viral replication occurs within approximately 96 hours post infection (hpi) (Mocarski 2007; Marcinowski et al. 2012; Griffiths et al. 2015). In general, IE proteins are responsible for inhibiting intrinsic and innate host cell responses and initiating the transcription of viral early genes. Viral early genes then regulate host cell function to allow for viral genome replication and packaging (Griffiths et al. 2015; Xu et al. 2018). Late viral genes are expressed after viral DNA replication has begun and encode mostly structural proteins required for egress (Mocarski 2007; Perng et al. 2011; D’Aiuto et al. 2012; Tirosh et al. 2015). Following primary infection, the virus may remain lytic, replicating and activating the immune system, or enter a latent stage lacking replication and immune system activation (Griffiths et al. 2015). HCMV can replicate in differentiated cell types to varying degrees (Marcinowski et al. 2012; Pan et al. 2013; Gonzalez-Sanchez et al. 2015). In symptomatic children and adults, HCMV infection can be managed using (val)ganciclovir, cidofovir, and/or foscarnet, all of which inhibit viral DNA synthesis, or letermovir, which targets viral DNA packaging into nucleocapsids (Britt and Prichard 2018; Pass and Arav-Boger 2018; Altman et al. 2019; Steingruber and Marschall 2020). There is no FDA-approved therapy for treating expecting mothers, making vaccine development a major public health priority. Clinical trials are being conducted for use of valganciclovir in infants and children with confirmed symptomatic and asymptomatic cCMV infections.

Initial human microarray experiments using RNA isolated from HCMV-infected NPCs identified several downregulated neurodevelopmental genes such as NES, SOX2/4, and OLIG1 along with disruptions to differentiation (Luo et al. 2010; D’Aiuto et al. 2012). A subsequent mechanistic study in NPCs demonstrated the ability for viral protein IE1 to cause the loss of SOX2 expression through trapping of unphosphorylated STAT3 in the nucleus (Wu et al. 2018). Meanwhile, recent RNAseq studies conducted in HCMV-infected cerebral organoids support these findings identifying SOX2, OLIG1, ENO2, ALDOC, and FEZF2 among other downregulated neurodevelopmental genes (Brown et al. 2019; Sun et al. 2020). These studies also found upregulation in ontology terms interleukin, inflammatory, and immune response with downregulation in ontology terms axonogenesis, calcium regulation, and glycolytic processes (Sun et al. 2020). Further, this work identifies the potential interaction of the PDGFRa receptor and viral pentameric complex in promoting virus entry into NPCs (Rak et al. 2018; Sun et al. 2020). It also found that administration of HCMV neutralizing antibody (NAbs) was sufficient to prevent infection and disruptions to structural development within the organoid (Sun et al. 2020). Additional work from us and others using the cerebral organoid model have found consistent disruptions to neuronal differentiation, cortical layering, calcium signaling, and electrical activity after HCMV infection (Brown et al. 2019; Sison et al. 2019; Sun et al. 2020).

In the current study we aim to identify neurodevelopmental transcriptional networks that are altered in response to HCMV infection and whether specific viral proteins contribute to gene misregulation. We found transcriptional downregulation among many genes which was not limited to cells expressing high levels of GFP, a surrogate marker of viral replication and was observed throughout the infected tissue. Previous studies have suggested that HCMV IE1 viral protein expression can regulate host gene expression (Marcinowski et al. 2012; Khan et al. 2014; Pignoloni et al. 2016; Liu et al. 2017b; Adamson and Nevels 2020). We found this to be the case for some gene targets, while regulation of others appeared to be independent of IE1/2 protein expression suggesting that other HCMV-related mechanisms are involved. Together, this work reveals that limiting IE gene expression offers minimal benefit on neural differentiation and function suggesting that therapeutic development against cCMV infection will require a more comprehensive approach.

## Results

### Cells from infected neural organoids exhibit reduced transcriptional profiles despite varying levels of viral gene expression

Human iPSC-derived cerebral organoids were generated from two independent healthy control iPSC lines (Ebert et al. 2013; Sison et al. 2019) and infected after 30 days of development with a recombinant HCMV strain TB40/E expressing EGFP. Because tissues varied in size and likely cell numbers, we infected organoids using 500 infectious units per μg of tissue (IU/µg) and allowed infection to progress for two weeks. We detected GFP fluorescence by 4 days post infection (dpi), increased at 8 dpi, and continued to be present at 12 dpi indicating viral spread (Fig. 1A). At 14 dpi, HCMV-infected and mock-treated uninfected organoids (Fig. 1B) were dissociated into single cell suspensions, and cells were sorted based on levels of GFP fluorescence as an indirect measurement of infection (Fig. 1C). The gating parameters for live cells were set based on cells isolated from uninfected mock-treated organoids (Fig. S1). This approach is routinely used to investigate latency in HCMV-infected hematopoietic cells (Rak et al. 2018). The EGFP gene is present within the viral genome and expressed using an SV40 promoter, and fluorescence is postulated to be proportional to transcriptionally competent HCMV genomes (Rak et al. 2018; Collins-McMillen et al. 2019). We sorted cells into groups exhibiting high, intermediate, and undetectable levels of fluorescence (Fig. 1C). We defined these populations as GFP positive (+), GFP intermediate (Inter), and GFP negative (-). For example, the total number of live cells from a representative experiment was approximately 3.5 x10^6^ which consisted of 0.5 x10^6^ GFP (-) cells, 2 x10^6^ GFP (Inter) cells, and 1x10^6^ GFP (+) cells (Fig. 1C). Infection did not cause large scale cell death as we observed that the number of live cells within each infected organoid group were comparable to the uninfected mock group (Fig. 1D). High expression of GFP likely represents a population of cells exhibiting high viral genome copies and viral gene expression. In contrast, cells isolated based on the absence of fluorescence are anticipated to be uninfected, or perhaps newly infected with fluorescence below the threshold of detection. To address these differences, we quantified levels of HCMV gene classes, immediate early (IE; UL123, UL122), early-late (E-L; UL44) and late (L; UL99) with late expression dependent on viral genome synthesis (Chambers et al. 1999). Cell populations of GFP (+) and GFP (Inter) exhibited high levels of viral gene expression in all classes with differences consistent with their levels of fluorescence (Fig. 1E). In the GFP (-) group, we did observe low but statistically significant levels of viral IE RNAs (Fig.1E) likely representing a mixture of uninfected and newly infected cells within the tissues.

**Figure 1.**
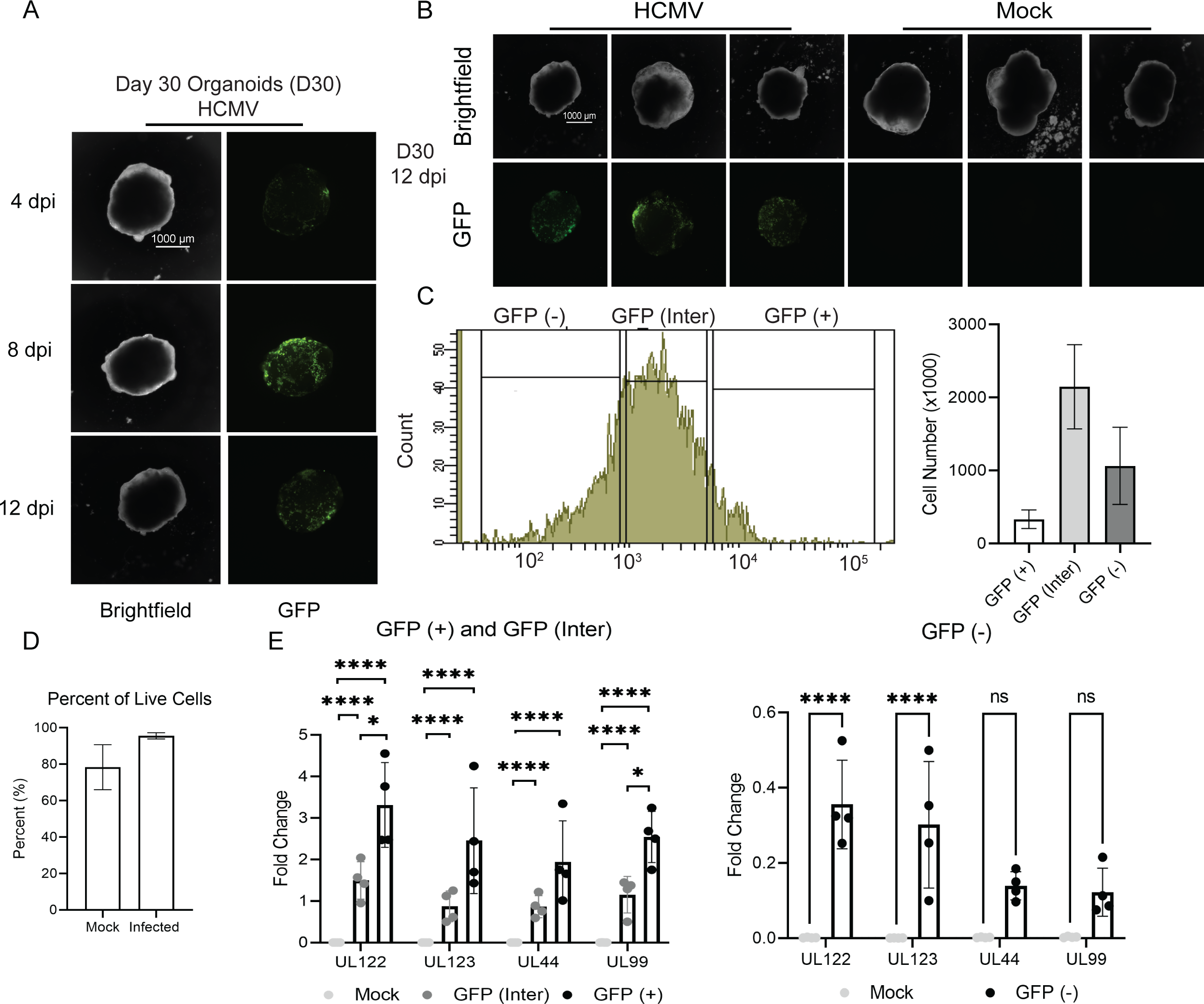
Three-dimensional cerebral organoid infection with HCMV. (A) On Day 30 of differentiation, organoids were weighed and infected at 500 IU/ug with HCMV-TB40EGFP. Bright-field and fluorescent images of one representative infected organoid taken at 4, 8, and 12 dpi are shown. (B) Images at 12 dpi of infected and uninfected organoids are displayed, one representative organoid from each sorted group is shown. (C) Representative GFP intensity plot from FACs analysis of an infected organoid and percentage of living cells in each organoid group as determined by FACs each infected organoid group was made up of three pooled organoids. (D) Percentage of live cells within whole infected and uninfected organoids across samples. (E) qPCR analysis of viral gene expression within mock compared to GFP (+) and GFP intermediate subpopulations. (F) qPCR analysis of viral gene expression within mock compared to the GFP (-) subpopulation.

We quantified infection-induced changes to host cell gene expression using RNA sequencing on sorted cell populations from infected organoids and sorted cells isolated from uninfected Mock-treated tissues. These studies were completed using three biological replicate experiments that were sequenced separately with multiple organoids combined for each condition. Mock and infected samples were mixed for each biological replicate resulting in the pooled library. Samples had at least 2.5 million aligned reads and the External RNA Controls Consortium (ERCC) spike-in was used to normalize variation in RNA expression across samples. After trimming, 80% of the bases had a quality score of 30 indicating the probability of an incorrect base call was 1 in 1000. One of the GFP (+) replicates was removed from our subsequent analysis due to a large drop off in total number of mapped reads likely related to technical error in the sequencing reaction. Initially, we performed principal component analysis (PCA) on the data sets which revealed two groups (Fig. 2A). One group contained the Mock-treated samples, and the other contained samples isolated from infected organoids regardless of GFP signal strength (Fig. 2A). This was initially surprising as we hypothesized that the subpopulations from infected organoids would cluster separately based on varying states of infection. However, as noted above, we did detect low levels of viral transcripts in the GFP (-) group indicating this population contains some infected cells. These data also suggest a possible indirect effect of HCMV on uninfected cells in the tissues.

**Figure 2.**
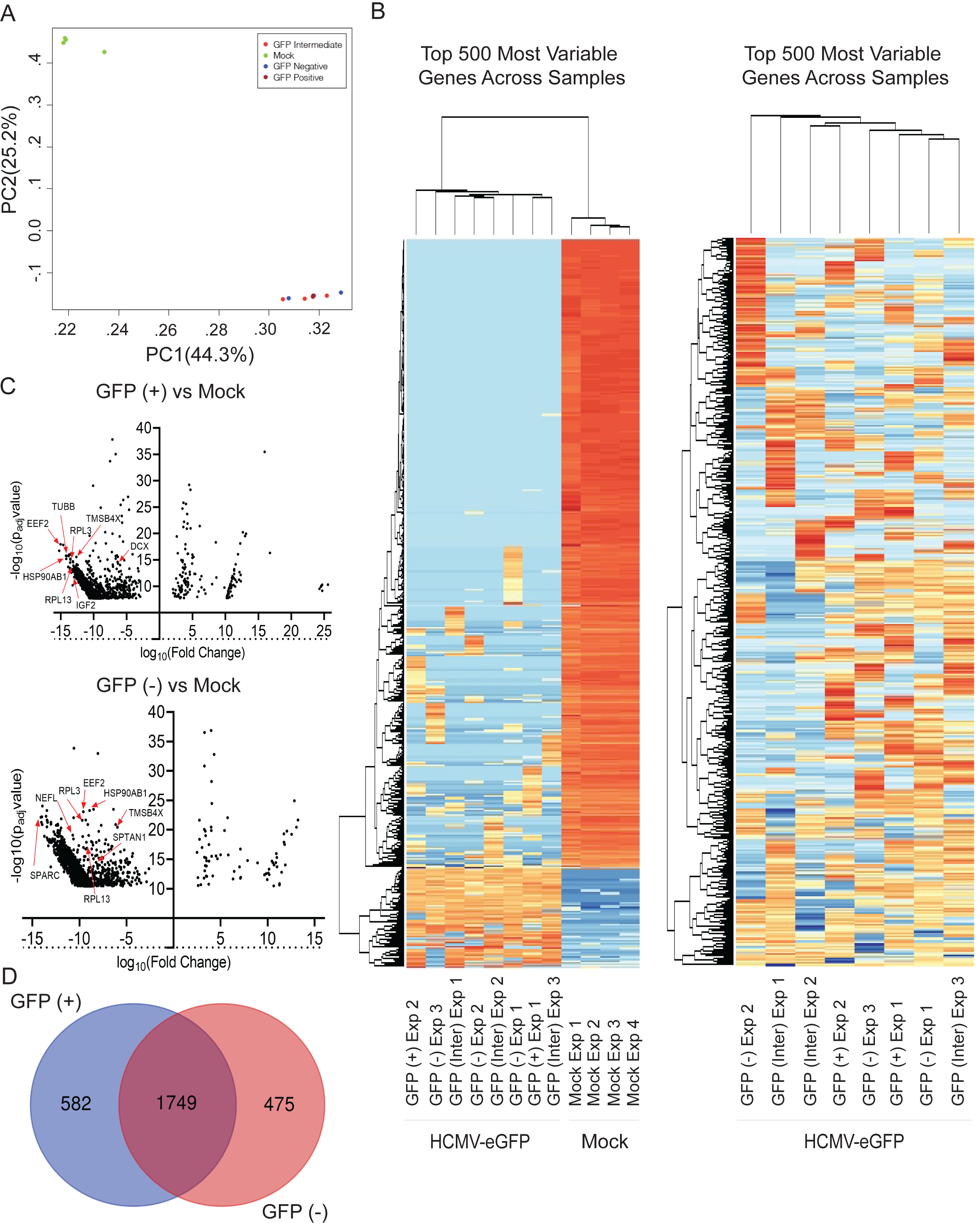
All infected organoid subpopulations cluster distinctly from mock samples with many genes being downregulated. (A) PCA with components being all mapped genes across samples split by axis determined by adj p-value and compared between each sample. (B) Hierarchical cluster analysis of all sequenced samples for the top 500 most variable genes according to read counts assigned. Hierarchical cluster analysis of only infected samples for the top 500 most variable genes according to read counts assigned. (C) Volcano plot of the top 2,500 differentially expressed genes comparing GFP (-) vs. Mock as determined by adj p-value < .01. Volcano plot of the top 2,500 differentially expressed genes comparing GFP (+) vs Mock as determined by adj p-value. Arrows denote representative genes downregulated in both groups. (D) Venn diagram comparing the 2,500 most differentially expressed genes in GFP (-) vs. Mock and GFP (+) vs. Mock.

We identified 1,222 cellular transcripts upregulated, 17,696 downregulated, and 2,882 not differentially expressed in GFP (+) cells from HCMV-infected tissues compared to cells from Mock tissues, using an adjusted p-value < 0.05 and 3-fold cutoff. In the GFP (-) population, we observed 911 transcripts upregulated and 18,379 downregulated compared to Mock. Table S1 and S2 contain full gene lists from DESEQ2 analysis along with a gene list of those not differentially expressed for GFP (+) vs Mock and GFP (-) vs Mock, respectively. Completion of hierarchical cluster analysis on the top 500 variable genes across samples showed distinct clustering between infected samples and mock samples (Fig. 2B, left), and we did not observe any additional clustering based on GFP designation following the removal of the mock data sets (Fig. 2B, right). These differences are shown in volcano plots for GFP (+) versus Mock and GFP (-) versus Mock (Fig. 2C) illustrating the disproportional number of genes being downregulated. For both GFP (+) and GFP (-) groups, over 90% of significantly differentially expressed genes were downregulated compared to cells isolated from uninfected Mock tissues. These observations demonstrate that HCMV infection profoundly impacts transcription of infected cells in neural tissues and supports a hypothesis that changes are occurring at very early times of infection (i.e., GFP (-) populations) and likely also involves indirect effects on uninfected cells from the neighboring GFP (+) infected cells.

### Cell populations from HCMV-infected organoids display downregulation in critical neurodevelopmental and signaling genes

To determine which biological processes are disproportionately altered in response to infection, we began by analyzing the complete data sets using Gene Set Enrichment Analysis (GSEA) that defines significant differences between two biological states in hallmark gene sets related to specific biological processes. Full gene lists for each analyzed gene set and rank metrics can be found in Supplemental Table 3. Compared to HCMV-infected GFP (+) cells, GSEA identified enrichment in genes up-regulated in response to alpha interferon proteins (Fig. 3A). We have included a representative heatmap of 40 genes out of the 97 hallmark genes in this category (Fig. 3A). This interferon response is known to occur during HCMV infection (Boyle et al. 1999; Paulus et al. 2006). Although GFP (+) and GFP (-) populations clustered together, this hallmark gene set was not significantly enriched in HCMV-infected GFP (-) compared to Mock (Fig. 3A). In contrast, HCMV-infected GFP (+) cells exhibited negative correlation and dysregulation in several sets (Fig.3B). This included genes up-regulated during the unfolded protein response, a subgroup of genes regulated by MYC, genes involved in the G2/M checkpoint, genes up-regulated in response to TGFß1, and genes up-regulated by activation of the PI3K/AKT/mTOR pathway. Disruption to genes involved with MYC, TGFß1, and AKT signaling in the GFP (+) population is likely related to infection causing an arrest in cell proliferation. We have included heatmaps of 40 genes within each hallmark set to further demonstrate that most genes were downregulated upon infection (Fig 3B). Our lab and others have demonstrated HCMV-mediated dysregulating of several of these processes in 2D cultures of primary human fibroblasts with many of the activities dysregulated by HCMV immediate early genes (Michelson et al. 1994; Jault et al. 1995; Kudchodkar et al. 2006; Moorman et al. 2008).

**Figure 3.**
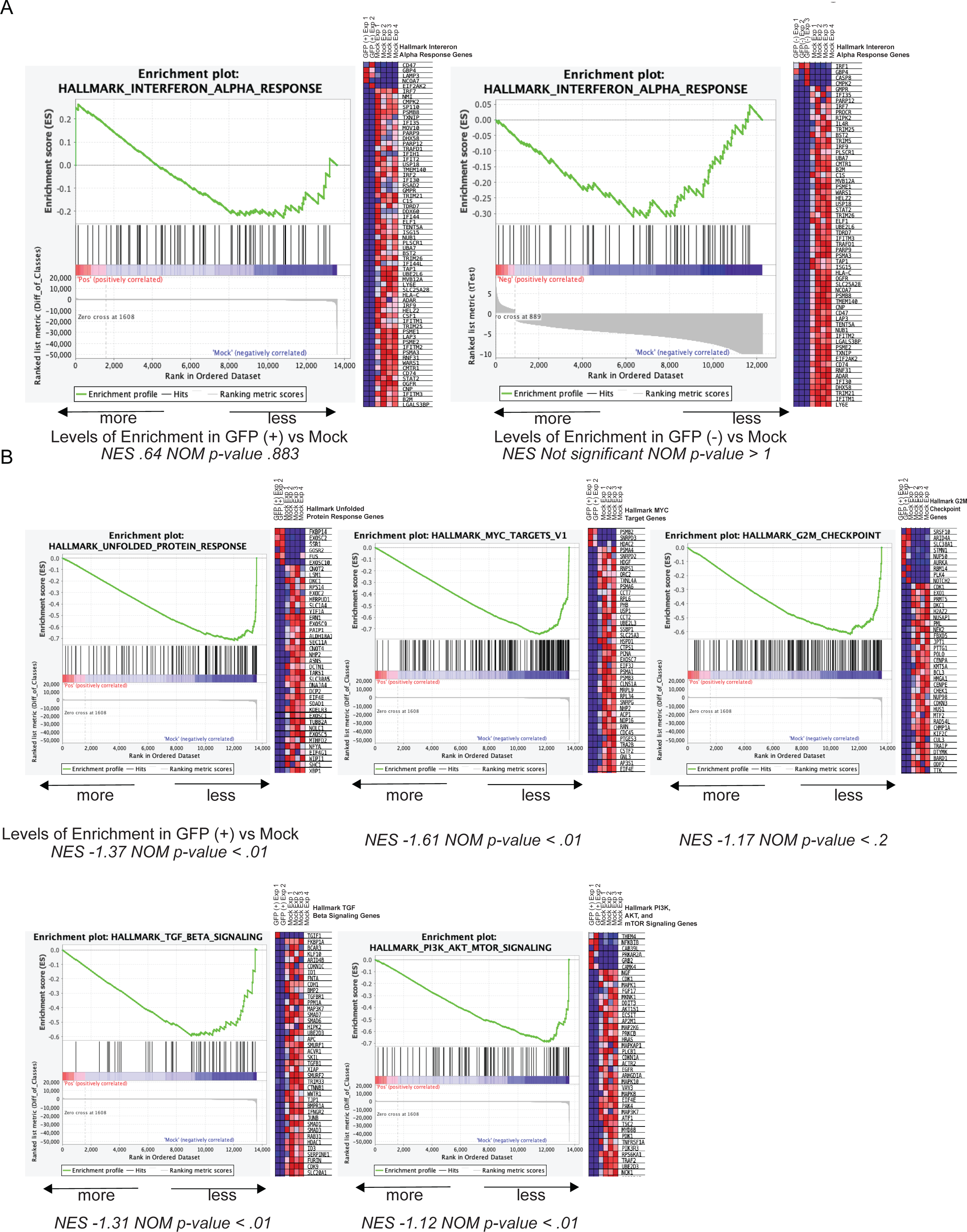
Gene set enrichment analysis (GSEA) plots comparing infected organoid populations to mock. GSEA using the hallmark gene set on a list of all differentially expressed genes with fold change ± 3 and adj p-value < .05. (A) Hallmark interferon alpha response enrichment plots and corresponding heat maps from GFP (+) vs Mock and GFP (-) vs Mock respectively. (B) Additional enrichment plots and heat maps (40 representative genes each) from GSEA comparing GFP (+) vs Mock revealing significant de-enrichment in GFP (+) groups. A normalized enrichment score (NES) and normalized p-value (NOM p-value) for each plot are shown below, a cutoff of NOM p-value < .05 was used to establish significance.

Next, we used gene ontology enrichment analysis to more precisely map specific processes that are impacted during infection. We analyzed the top 3,000 differentially expressed genes, both increased and decreased, between cells isolated from HCMV-infected GFP (+) and Mock-treated organoids. As noted earlier, most expression was downregulated in infected populations. Using a p-value cutoff of < 0.01, gene ontology (GO) analysis identified significant changes in 804 terms related to biological processes, 95 molecular functions, and 201 cellular components. The top five statistically significant terms are shown in Figure 5A with full lists in Table S5 and additional enriched terms in Figure S3 (additional ontology analysis performed using DAVID is found in Figure S3). The GO terms Ion Signaling, Cell-Cell Communication, and Junction Signaling were all significantly affected in HCMV GFP (+) cells (Fig. 4B). For example, we observed a 10.9-fold reduction in expression of GJA1, 7.8-fold reduction in CACNA1C, 5.7-fold reduction in CACNA1G, 5.4-fold for CAMKV, and 5.8-fold for KCNF1. We observed similar changes in the GFP (-) group compared to Mock, with GJA1 reduced by 11.2-fold, CACNA1C by 4.7-fold, CACNA1G by 5.6-fold and CAMKV by 7.1-fold (data not shown). KCNF1 was also downregulated but not significantly in the GFP (-) group. These data are consistent with previous studies demonstrating reduced expression of neuron-specific ion channel subunits and gap junctions (Luo et al. 2010; Khan et al. 2014) as well as our previous studies demonstrating reduced Ca^2+^ signaling in 2D NPC cultures and cells derived from HCMV-infected organoids (Sison et al. 2019).

**Figure 4.**
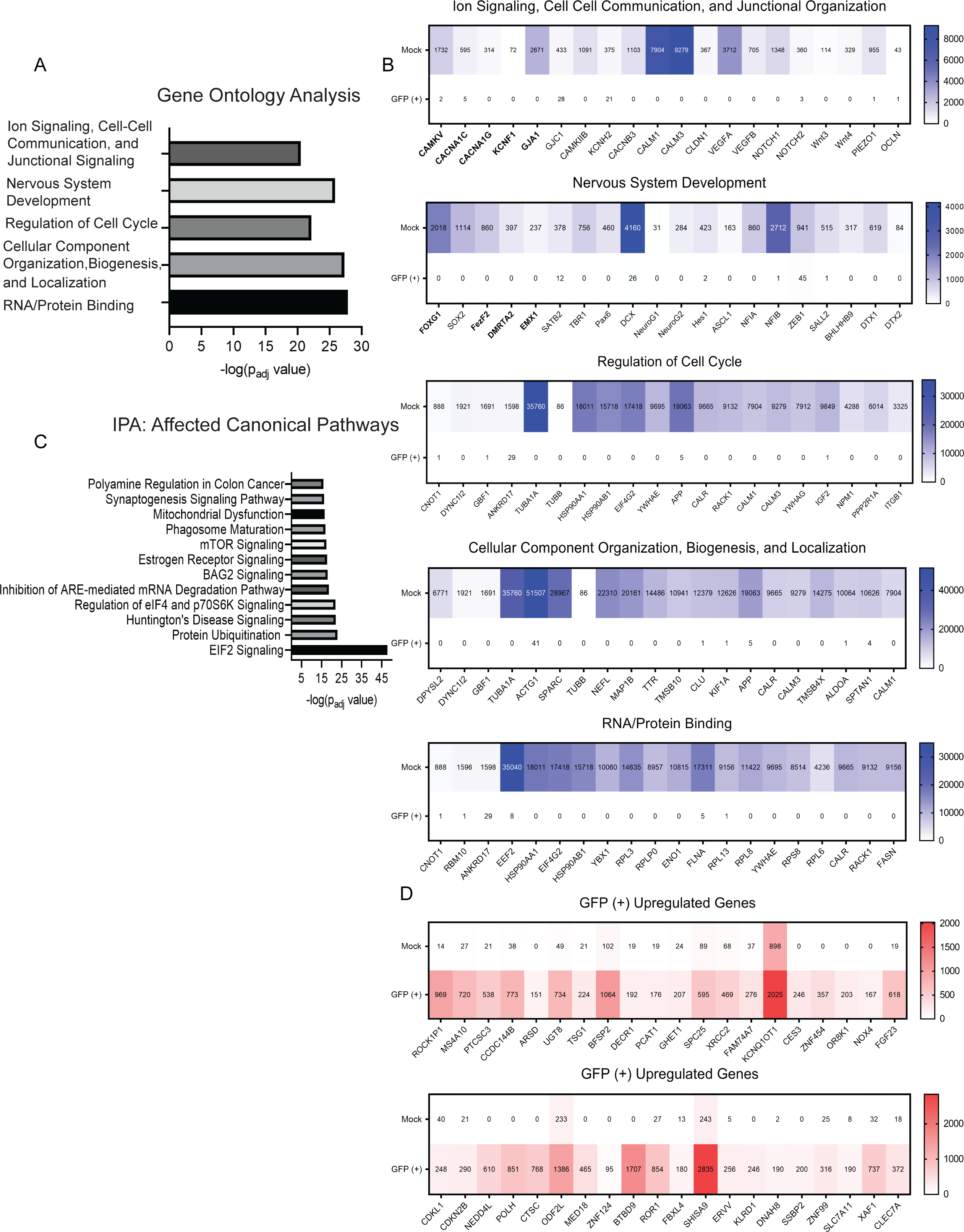
Pathway and ontology analysis conducted on GFP (+) vs Mock and heat maps displaying up and downregulated genes. (A) Gene Ontology analysis of the top 3,000 differentially expressed genes when comparing GFP (+) vs. Mock using fold change ± 3 and adj p-value < .01. Terms listed were the five most significant across categories molecular function, biological process, and cellular components. (B) Heat maps showing 20 representative genes within the ontology classifications from (A), the values in the heat map are the average raw read count number across GFP (+) or Mock samples. (C) Most significantly impacted canonical pathways as determined by ingenuity pathway analysis using the same gene list as in (A). (D) Node diagram generated from IPA core analysis using the same gene list as in (A) organized based on most significantly affected canonical pathway regulators, how they interconnect and effect downstream functions (Solid lines indicate direct interaction and dashed lines indicate indirect interaction). (E) Heat maps containing 20 of the most significantly upregulated genes in GFP (+) and GFP (-) groups vs Mock determined by adj p-value and fold change. For ontology and IPA significance is displayed as -log(adj p-value) (Bolded genes within the heat maps identify those used in subsequent shield-1 experiments).

**Figure 5.**
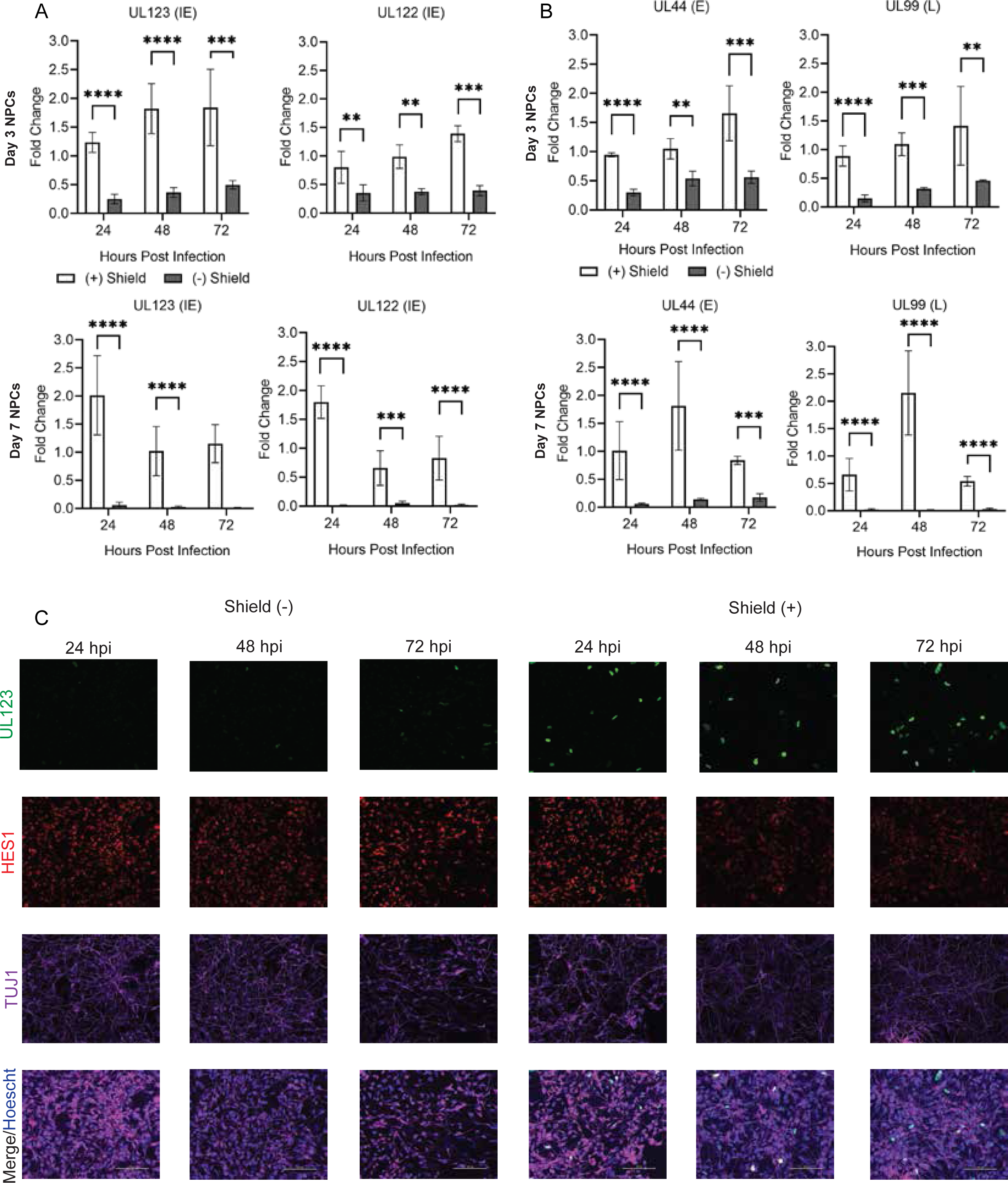
Infection of NPCs with TB40r mGFP-IE-FKBP virus results in limited expression of viral proteins UL122/UL123 unless shield-1 is administered. NPCs were infected at 3- or 7-days post plate down with **TB40r mGFP-IE-FKBP** at an MOI of 1. One group was then supplemented with 1 uM shield-1 daily (shield +) while the other was given vehicle (shield -). (A) Immunofluorescent images at 24, 48, and 72 hpi from NPCs infected 3 days post plate down with HCMV-IE1/IE2-ddFKBP and probed for HCMV IE gene IE1 (green), neuronal specific beta-tubulin Tuj1 (purple), neurodevelopmental transcription factor Hes1 (red), Hoescht (blue), and merge (All images taken at 20X, Using Nikon TL-1). (B-C) Expression of HCMV viral genes (UL122 (IE), UL123 (IE), UL44 (E), and UL99 (L)) within NPCs infected 3- or 7-days post plate down at 24, 48, and 72 hours post infection (hpi) as determined by qPCR. Stars were assigned based on level of significance as determined by T-test: * = p ≤ .05, ** = p ≤ .01, *** = p ≤ .001, and **** = p ≤ .00001.

We observed significant changes in the GO category Nervous System Development (Fig. 4B). In the GFP (+) cells compared to Mock, we quantified a 9.7-fold reduction in SOX2, 8.3-fold in FOXG1, 8.4-fold in DMRTA2, 7.8-fold in EMX1, and 9.5-fold in FEZF2. In GFP (-) cells, SOX2 decreased by 9.1-fold, FOXG1 by 7.5-fold, and FEZF2 by 8.5-fold while no significant changes occurred in DMRTA and EMX. Numerous additional developmental genes that were significantly reduced in both GFP (+) and GFP (-) include HES1, PAX6, and NEUROG1 and G2 (Fig. 4B). The decrease in several of these transcription factors has been observations made in HCMV-infected NPCs (Luo et al. 2008; Luo et al. 2010; Li et al. 2015), and we have now expanded the observations to changes occurring within the context of a more complex 3D tissue model system.

The remaining most significantly enriched terms included cell cycle, cellular component organization, and RNA/protein binding (Fig. 5A) with several genes spanning multiple categories. Examples of genes related to cell cycle included NPM1 and PP2R1A, to cellular organization include NEFL and DPYSL2, and to RNA-protein binding include EEF2 and RPL3 (Fig. 4B). A complete list of GO terms enriched using the top 5,000 differentially expressed genes for GFP (+) and GFP (-) vs Mock as determined by adjusted p value of < 0.025 and fold change of > 5 are presented in Figure S3. We observed conservation of enriched terms between GFP (+) and GFP (-) populations including processes regulating protein expression, neuron differentiation, cellular component organization, and cell cycle.

Finally, we sought to define specific pathways using the Ingenuity Pathway Analysis tool. We analyzed the top 3,000 differentially expressed genes which defined 12 pathways as significantly altered during infection (Fig. 4C). Most pathway components are downregulated and consistent with results from the GO analysis. Several of these pathways involve regulating protein translation and degradation including protein ubiquitination, signaling relating to BAG2, EIF2 signaling, regulation of EIF4 and p70S6K, and mTOR signaling (Fig. S4A). Others related to neurodevelopmental processes such as Synaptogenesis Signaling Pathway and Huntington’s Disease signaling (Fig. S4A). Meanwhile, RHOGDI signaling was one of a few pathways containing several upregulated components and many of the unchanged genes between GFP (+) and Mock related to GPCR signaling (Fig. S4B). Finally, IPA’s core analysis tool generated a node diagram connecting regulators from the major affected canonical pathways (Fig. S5). Several of these regulators fall into identified ontology classifications (TGFB1, RICTOR, NGF, and SP1) and have impacts on functional pathways that match our previous analyses. Complete analyses from IPA comparing GFP (+) vs Mock can be found in Supplemental Table S4.

### Impact of HCMV replication on neurodevelopment gene expression

The GFP (-) cell populations isolated from infected neural organoids exhibit profound changes in host transcriptomes. These GFP (-) cells exhibit low levels of HCMV IE gene expression, and (Wu et al. 2018) have demonstrated that expression of HCMV IE1 disrupts SOX2 expression in a STAT3-dependent mechanism (Reitsma et al. 2013). Therefore, we hypothesized that expression of viral proteins IE1 and possibly IE2 might be necessary to downregulate other key neurodevelopmental genes identified by our RNA seq experiments. To test their contributions, we infected cultures of NPCs using TB40r mGFP-IE-FKBP virus that expresses IE1 and IE2 with a destabilization domain (DD) tag in a shared exon (Glass et al. 2009). Addition of Shield1 ligand to the culture media prevents DD-dependent IE1 and IE2 degradation and increases its steady-state levels (Glass et al. 2009). We cultured neurosphere-derived NPCs for 3 and 7 days prior to infection to mimic developmental states. Cells were infected using TB40r mGFP-IE-FKBP at an MOI of 1.0 IU/cell, based on titers from fibroblasts, and we added 1 μM Shield1 ligand (Shield (+)) or vehicle (Shield (-)) starting at 2 hpi. We quantified changes in HCMV gene expression and observed significant reductions in IE RNAs UL123 (IE1) and UL122 (IE2) at all time points in Shield (-) compared to Shield (+) (Fig. 5A). Reductions in RNA levels for UL44 and UL99 also occurred (Fig. 5B), and these differences were observed for infections done using Day 7 NPCs (Fig. 5A, B). We observed IE1-positive nuclei using immunofluorescence in Shield (+) but not Shield (-), albeit lower than the anticipated number of infected cells (Fig. 5C). Shield (+) IE1-positive cultures also exhibited reductions in TUJ1 and HES1, which is consistent with published studies (Liu et al. 2017b; Brown et al. 2019; Sun et al. 2020). Our data demonstrate that NPCs infected in the absence of Shield1 exhibit reduced HCMV IE1 protein and gene expression by 24 hpi as well as disruptions to additional viral gene classes.

We next asked whether destabilizing IE1 and IE2 in Shield (-) conditions could limit the downregulation of key developmental and transcription factors which we observed in GFP (+) and GFP (-) organoid populations. We infected NPCs as described above and measured differences in cellular gene expression of the nervous system developmental genes FEZF2, FOXG1, DMRTA1, and EMX1 in the Shield (+) and Shield (-) conditions (Fig. 6). We observed significant reductions in FEZF2 and FOXG1 RNA levels by 24 hpi in Day 3 NPCs, and by 48 hpi in Day 7 NPCs regardless of Shield1 (Fig. 6A). Changes in DMRTA2 and EMX1 occurred late during infection of Day 3 NPCs but was detectible by 24 hpi in Day 7 NPCs (Fig. 6B). There were some differences between Shield (+) and (-) conditions, but our data show that limiting IE1/IE2 protein expression was not sufficient to prevent the downregulation of any of the key neurodevelopmental targets explored. We next investigated a set of ion signaling, cell-cell communication and junctional organizational genes (Fig. 4B), specifically KCNF1, CACNA1C, CAMKV, CACNA1G, and GJA1 (Fig. 7). We consistently observed small decreases in calcium channel genes, KCNF1 and CACNA1C at late time points independent of Shield addition (Fig. 7A). Likewise, we observed decreases in CAMKV, CACNA1G, and GJA1. However, a Shield-dependent effect was noted particularly in the case of GJA1 and CAMKV (Fig. 7B,C). Together with our previous results (Sison et al. 2019), our data confirm that HCMV mediated a disruption of several key neurodevelopmental and functional genes (Fig. 7D), but this effect has limited dependence on expression of IE1 and IE2.

**Figure 6.**
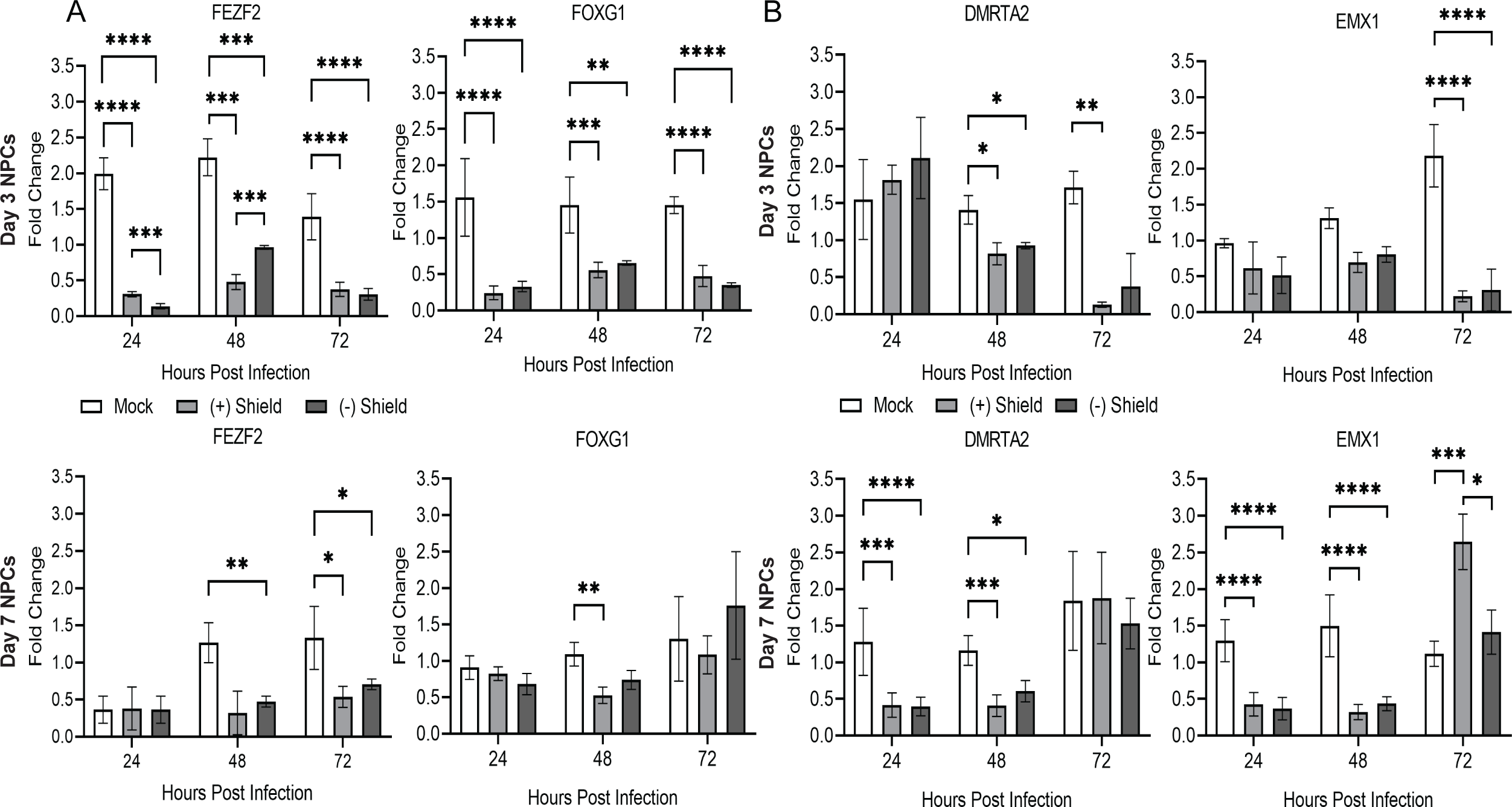
Neurodevelopmental gene targets are downregulated within NPCs infected with TB40r mGFP-IE-FKBP independent of shield-1 administration. (A-B) qPCRs were performed for several key neurodevelopmental transcription factors (FezF2, FOXG1, DMRTA2, and EMX1) within the HCMV-IE1/IE2-ddFKBP infected (shield +) and (shield –) groups plus uninfected (mock) NPCs at both 3- and 7-days post plate down infected NPCs. No effect of shield administration was observed for these targets. Stars were assigned based on level of significance as determined by ANOVA: * = p ≤ .05, ** = p ≤ .01, *** = p ≤ .001, and **** = p ≤ .00001.

**Figure 7.**
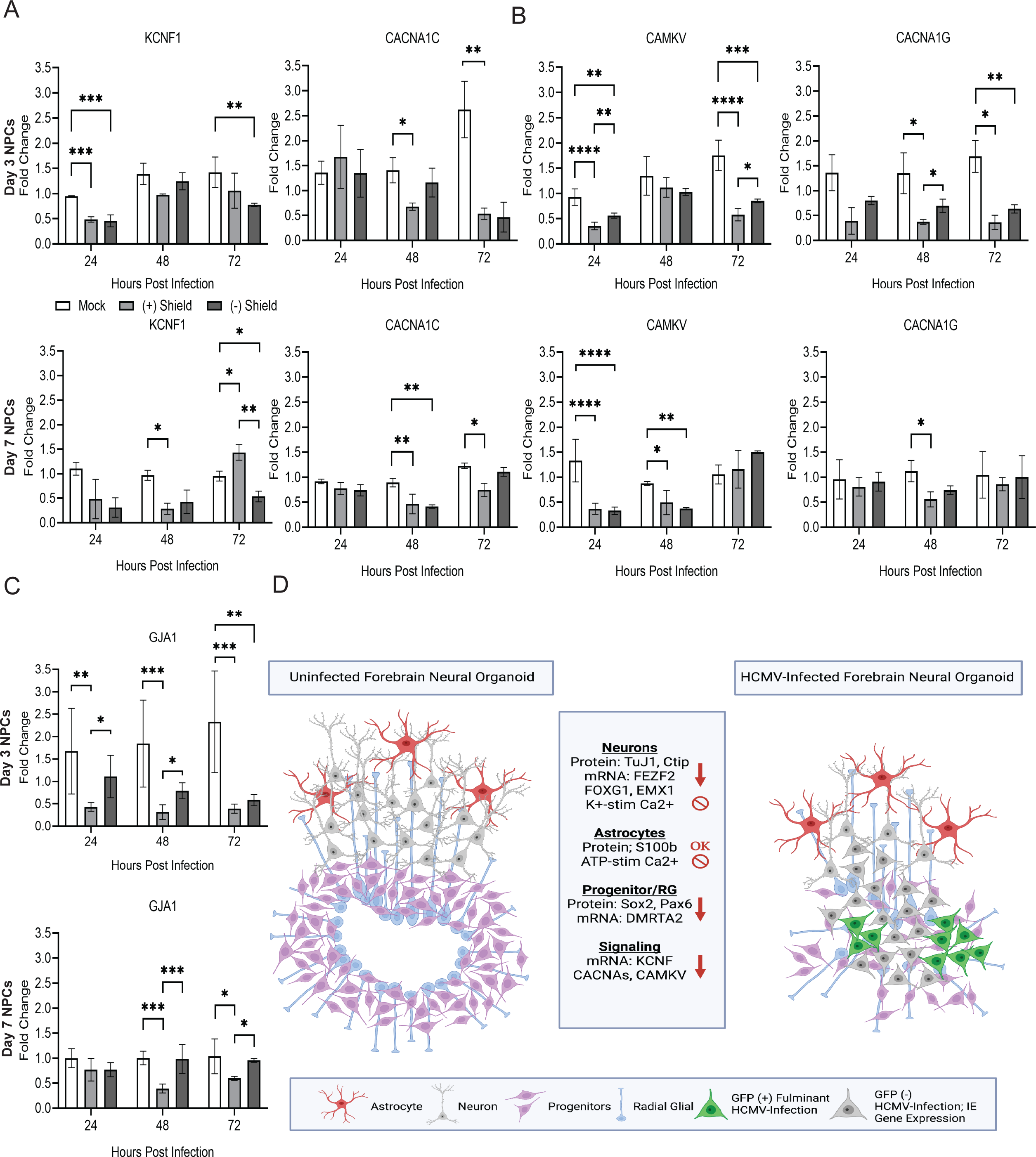
Signaling and cell-cell communication gene targets are downregulated within NPCs infected with TB40r mGFP-IE-FKBP depending on shield-1 administration. Following the same parameters as Figure 6, qPCRs were conducted for expression of signaling, cell-cell communication, and junctional genes (CACNA1C, CACNA1G, CAMKV, KCNF1, and GJA1). (A) For these targets shield administration had no impact on gene expression levels. (B,C) A shield dependent effect was observed when comparing shield (-) and shield (+) infected groups for these targets. Stars were assigned based on level of significance as determined by ANOVA: * = p ≤ .05, ** = p ≤ .01, *** = p ≤ .001, and **** = p ≤ .00001. (D) Graphical summary integrating current literature with this study’s findings regarding HCMVs impact on development within NPCs and cerebral organoids.

## Discussion

Stem cell-derived organoids provide a unique model system in which to study the developing human brain. For example, their 3D structure more closely represents the developing brain compared to 2D monolayer culture system and allows for the generation of a layered cortical structure expressing a wide variety of the markers associated with the human brain (Lancaster et al. 2013; Xu et al. 2018; Sison et al. 2019). We and others have previously demonstrated that cerebral organoids can be infected by HCMV with infection substantially altering organoid structure and function (Brown et al. 2019; Sison et al. 2019; Sun et al. 2020) (Figure 7D). However, a more thorough examination of how the transcriptional landscape is altered upon HCMV infection and what role viral proteins play in this process is needed to better understand the overall impact of HCMV infection on neural development.

Neurodevelopment is a highly complex process dependent on both spatial and temporal cues that activate large transcription factor networks within the NPC population. Ultimately, these cues cause NPCs to develop into a variety of terminally differentiated CNS cell types. RNA seq analysis of Day 30 organoids sorted for GFP signal strength as a proxy for the level of infection revealed downregulation of developmental transcription factors DMRTA2, FEZF2, EMX1, and FOXG1 in all populations regardless of GFP signal strength. These transcription factors are involved with the development of the telencephalon and regulation of early neuronal cell fate decision making. FEZF2 is typically expressed in the early cortical progenitor cell population and is critical for fate specification of subcerebral projection neurons through activation of downstream transcription factor networks (Chen et al. 2008; Wang et al. 2011; Guo et al. 2013). FOXG1 knock-out mice experience severe microcephaly, a common phenotype of HCMV infection (Martynoga et al. 2005; Kumamoto and Hanashima 2017). In humans, FOXG1 syndrome, caused by mutations or deletions in the long (q) arm of chromosome 14, is characterized by impaired development and structural brain abnormalities (Tohyama et al. 2011; Chiola et al. 2019). DMRTA2 along with EMX1 are highly expressed in the developing dorsal telencephalon where research in mice reports reduced cortex size upon mutation (Shinozaki et al. 2004; Konno et al. 2012; Young et al. 2017). In addition to these roles, several of these transcription factors are involved in key signaling pathways with wide-ranging impacts. For example, DMRTA2 is cross regulated by the Wnt/β-catenin signaling pathway, FOXG1 is associated with the SHH pathway, and FEZF2 is an upstream regulator of cTIP2 and SATB2 regulating fate decisions (Chen et al. 2008; Kuwahara et al. 2010; Lui et al. 2011). Interestingly, despite no overt indication of active viral replication, the GFP (-) population exhibit similar transcriptional effects as the GFP (+) and intermediate populations. This suggests that viral infection has more widespread consequences that likely go beyond active viral replication. Although further studies are needed to elucidate the mechanisms underlying the global effect on the transcriptional profile, disruption of cell-cell communication processes, spread of viral proteins, or distribution of tegument proteins that disrupt cellular differentiation and function could be potential mechanisms.

Several members of the basic helix loop helix (bHLH) transcription factor family, including HES1, ASCL1, NEUROG1/2, OLIG 2, and N-MYC were also significantly downregulated across infected organoid groups. This family overlaps with DMRTA2 in protecting NPC maintenance and has additional roles in controlling the timing behind NPC proliferation and differentiation (Young et al. 2017). Previous research also indicates cross regulation of bHLH factors by FEZF2 and DMRTA2 (Seo et al. 2007; Yang et al. 2012; Genin et al. 2014; Liu et al. 2017a; Dennis et al. 2019). Further, bHLH factors like HES1 have previously been identified as downregulated in NPCs following HCMV infection (Liu et al. 2017a; Liu et al. 2017b; Dennis et al. 2019). N-MYC regulates fating and division of intermediate neural progenitors, has influence on the expression of other family members through downstream cascades, and is cross regulated by SHH and Wnt (Kuwahara et al. 2010). The exact mechanism by which HCMV impacts bHLH factors is not fully elucidated, but studies indicate that HCMV proteins can act directly on genes to affect transcription or translation, and viral kinases can also function as pseudo kinases to impart post-translational modifications on host proteins and transcription factors (Mocarski 2007; Marcinowski et al. 2012; Steingruber and Marschall 2020). Specifically, the viral kinase UL97 can act as CDK1, which post-translationally modifies many of the bHLH factors altering their functionality (Gill et al. 2012). The wide-ranging functions within the bHLH family and their involvement within many common signaling cascades highlights the complexity of the potential avenues by which HCMV infection can affect neurodevelopmental pathways (Li et al. 2015).

It is well established that NPCs exhibit impaired signaling, differentiation, and even apoptosis following HCMV infection (Luo et al. 2010; Brown et al. 2019; Sison et al. 2019; Sun et al. 2020) (Figure 7C). NPCs are the cell type most permissive to infection (Luo et al. 2008) and found abundantly within cultured cerebral organoids. NPCs form early developmental structures and differentiate into a wide range of cell types in the human brain. Previous work has shown that IE1 traps unphosphorylated STAT3 in the nucleus and contributes to the reduction in SOX2 expression HCMV infected NPCs (Wu et al. 2018). However, little else is known about how HCMV downregulates key neurodevelopmental targets. Data from fibroblast cultures indicate the ability of HCMV infection to transcriptionally regulate host genes as well as degrade their protein products (Cohen and Stern-Ginossar 2014; Khan et al. 2014). Further, overexpression of IE1 or 2 alone in healthy fibroblasts infected with mutated virus unable to transcribe either IE gene demonstrated rescue in gene targets previously downregulated by infection (Luo 2010, Khan 2013). Therefore, we used a conditional approach to assess the role of IE1/2 on additional neurodevelopmental gene expression. Initially, infections were conducted at 7 days post plate down; however, we observed a lack of viral protein replication and low expression of viral proteins (Figure S2). These results were not entirely unexpected as other groups have reported that NPC differentiation is inversely correlated to infectability (Odeberg et al. 2006; Gonzalez-Sanchez et al. 2015). While our experiments using the TB40r mGFP-IE-FKBP virus support this idea, our data also show that infection at 7 days post plate down still results in transcript level downregulation despite incomplete viral replication (Figure 4, S2). Although this downregulation was not as significant or sustained as what was observed in the 72-hour post plate down infection, these data indicate that viral material and/or partially replicating virus is capable of inducing acute transcriptional changes in more differentiated NPCs. The neurodevelopmental targets assessed following infection with the TB40r mGFP-IE-FKBP virus to control IE1/2 expression were clearly downregulated. However, the results indicate that transcriptional downregulation was largely independent of IE1/2 expression, which further highlights the ability of other viral components to impact transcription and translation in infected NPCs.

Previous studies from us and others have shown that HCMV infection disrupts calcium and potassium signaling in infected NPCs and organoids (Brown et al. 2019; Sison et al. 2019; Sun et al. 2020). We therefore hypothesized that these functional deficits could be due to downregulation of channel encoding genes responsible for ion transport and maintaining membrane potential.

Several such targets showed downregulation within the GFP (+) and intermediate groups (Fig. 3G). CACNA1C and CACNA1G are genes that encode unique voltage gated calcium channel subtypes. CACNA1G expression gives rise to T-type calcium channels that belong to the low voltage activated subgroup. Recently, mutations in CACNA1G have been shown to be associated with early onset cerebellar ataxia and epilepsy (Calhoun et al. 2016; Barresi et al. 2020). The gene CACNA1C encodes a critical subunit leading to the formation of L-type calcium channels. L-type channels have a prominent role in controlling gene expression through coupling membrane depolarization with cAMP response element-binding protein (CREB) phosphorylation via local Ca2+/calmodulin-dependent protein kinase II (CAMKII) signaling; interestingly, both CAMKIIB and Calmodulin 1/3 (CALM1/3) are downregulated following infection (Zhang et al. 2005; Liang et al. 2016) thus limiting the ability of CREB to act as the transcriptional regulator of nearly 4,000 downstream gene targets (Wheeler et al. 2012; Chiola et al. 2019). In mice it has been demonstrated that reduced expression of CACNA1C during development leads to a reduction in neurogenesis (Lee et al. 2016; Moon et al. 2018). This demonstrates that alterations to the expression of either CACNA1G or CACNA1C can cause phenotypes similar to those associated with congenital HCMV infection. CAMKV, while not involved in the activation of CREB, is a Ca^2+^/calmodulin pseudo kinase that is required for the activity dependent maintenance of dendritic spines (Saneyoshi et al. 2010; Liang et al. 2016). It is unsurprising then that HCMV infected organoids and NPCs that display decreased neuronal firing and ability to transport calcium have CAMKV deficits as many of these processes are cross regulated (Odeberg et al. 2006; Brown et al. 2019; Sison et al. 2019; Sun et al. 2020). Meanwhile, several potassium channel encoding genes were downregulated in the GFP (+) and intermediate groups. These genes are important for potassium ion transport leading to neurotransmitter release and neural excitation. KCNF1, which encodes a member of the voltage-gated potassium channel subfamily F, was particularly interesting because previous studies had shown it to be downregulated upon HCMV infection in NPCs (Luo et al. 2010; Young et al. 2011; Brown et al. 2019), and mutations in it and other family members have been linked to seizures and epilepsy, phenotypes of congenital HCMV infection (Kohling and Wolfart 2016).

Finally, GJA1, which encodes protein connexin-43, was downregulated in both GFP (+) and intermediate organoid groups compared to mock (Fig. 3,4). In addition to ion channels and calcium sensors, NPCs rely on gap and tight junctions to communicate with each other through intercellular ion transport (Wei et al. 2004; Goodenough and Paul 2009; Zhou and Jiang 2014). This type of communication can inform neural cell fating by cuing NPCs to continue proliferating or to begin differentiating (Lemcke and Kuznetsov 2013; Zhou and Jiang 2014). There is some evidence to suggest that HCMV hijacks the existing junctional machinery to spread viral material (Silva et al. 2005; Cohen and Stern-Ginossar 2014; Khan et al. 2014). In HCMV infected fibroblasts, there is clear evidence that expression of IE genes specifically leads to decreased expression of GJA1 and GJC1 (Luo et al. 2008; Khan et al. 2014). Intriguingly, our data suggest that this is also the case in NPCs (Fig. 6D). As such, we postulate that remodeling of the cell-cell communication network could be a way by which the virus impacts both signaling and development in the host and promotes cell-to-cell viral spread early in infection.

Despite the differences in viral strain, viral titre, and 2D vs 3D culture models, the shield (-) NPCs and isolated GFP (-) organoid cells have some interesting similarities. Previous studies show that the TB40r mGFP-IE-FKBP construct maintains a low level of viral transcript expression (Pan et al. 2016; Rak et al. 2018); however, our data suggest that this level of expression is likely not sufficient to cause full replication (Fig. 4, S2). Similarly, IE viral transcripts were still significantly increased in the isolated GFP (-) cells compared to mock conditions despite a lack of robust GFP expression (Fig. 1D, 4). It is possible that if the GFP (-) cells were given more time in culture that GFP expression and E-L and L viral transcripts would be more evident. Nevertheless, significant downregulation of several gene targets was still observed in the absence of robust viral replication (Fig. 2,3,5,6) suggesting that the presence of some viral transcripts and/or viral material is sufficient to induce significant gene expression changes in neural tissues.

Taken together, we demonstrate that neurodevelopmental gene networks and critical neural signaling pathways are not dependent on IE1/2 protein expression indicating that other HCMV-related mechanisms are involved. Therefore, these data suggest that therapeutics designed to solely limit viral gene and protein expression may be insufficient to impact the widespread neural differentiation and functional deficits induced by congenital HCMV infection.

## Materials and Methods

### Cell culture and viruses

MRC-5 fibroblasts were cultured in Dulbecco’s modified Eagle medium (DMEM) (ThermoFisher Scientific) containing 7% fetal bovine serum (FBS) (Atlanta Biologicals) and 1% penicillin-streptomycin (ThermoFisher Scientific). Cells were plated at 1.0×10^4^ cells per well of a 24-well plate onto Matrigel coated coverslips. Matrigel was diluted in DMEM, placed on coverslips for approximately 12 hr, and aspirated off prior to plating cells. Virus stocks were prepared by infecting MRC-5 fibroblasts (ATCC) with HCMV strain TB40/E encoding EGFP (Umashankar et al. 2011). Cell culture medium was collected and pelleted through a sorbitol cushion (20% sorbitol, 50 mM Tris-HCl, pH 7.2, 1 mM MgCl2) at 55,000 × g for 1 h in a Sorvall WX-90 ultracentrifuge and SureSpin 630 rotor (ThermoFisher Scientific). The TB40r mGFP-IE-FKBP BAC was kindly provided by E. Borst (unpublished). The virus was cultured in the present of 1 µM Shield-1 (AOBIUS #AOB1848) and replaced every 24 hr. Titers of viral stocks were determined by a limited dilution assay with the 50% tissue culture infectious dose (TCID50) in MRC-5 cells in a 96-well dish. At 2 weeks post infection, HCMV IE1-positive cells were counted to determine viral titers, reported as the number of infectious units (IU) per milliliter. IE1-positive cells were determined using a mouse anti-HCMV IE1 antibody (clone 1B12; generously provided by Tom Shenk, Princeton University, Princeton, NJ).

Two independent iPSC lines were used (4.2 and 21.5/8) (Ebert et al. 2013). The iPSCs were grown and maintained in Essential 8 Media (ThermoFisher Scientific) and were cultured in feeder-free conditions on Matrigel (Corning). Neural progenitor cells (NPCs) were differentiated and maintained as neurospheres (EZ spheres) in Stemline (Millipore Sigma) supplemented with 0.5% N-2 supplement (ThermoFisher Scientific), 100 ng/ml EGF (Miltenyi Biotech), 100 ng/ml fibroblast growth factor (FGF; Stem Cell Technologies), and 5 ug/ml heparin (Millipore Sigma) as described previously (Ebert et al., 2013). Plated NPCs were grown in Neurobasal medium (ThermoFisher Scientific) supplemented with 2% B-27 (ThermoFisher Scientific) and 1% antibiotic-antimycotic (ThermoFisher Scientific). Cells were plated at 1.0×10^4^ cells per well of a 24-well plate onto Matrigel coated coverslips. Matrigel was diluted in DMEM, placed on coverslips for approximately 12 hr, and aspirated off prior to plating cells.

### Cerebral Organoids

Cerebral organoid cultures were differentiated from iPSCs according to the specification of the cerebral organoid kit from StemCell Technologies (#08570) that relies on an established protocol (Lancaster et al. 2013). These organoids were made from either of the two independent WT stem cell lines (4.2 and 21.5/8), Infected 1 was derived from the 21.5/8 line, Infected 2 and 3 were derived from the 4.2 line, Mock 1 and 2 were derived from the 21.5/8 line, and Mock 3 and 4 were derived from the 4.2 line. (Ebert et al. 2013). Briefly, iPSCs were seeded at 9 × 10^3^ cells per well onto 96-well ultralow attachment plates for embryoid body (EB) formation and grown in EB formation media (StemCell Technologies). At day 5, the induction of neural epithelium was initiated by moving the EBs into an ultralow attachment 24-well plate where they were then fed with induction media (StemCell Technologies). On day 7, neural tissues were embedded in Matrigel droplets and moved to ultralow attachment 6-well plates and fed expansion media (Stemcell Technologies). Then at day 10, the plate of developing organoids was transferred to a rocker which elicits the circulation of nutrients and prevents organoids from sticking to the dish. From this point on the organoids were fed maturation media (StemCell Technologies) every 3 days. At day 30, organoids were weighed and infected with 200-500 infectious units/ug HCMV strain TB40/E encoding GFP (Collins-McMillen et al. 2018; Rak et al. 2018; Collins-McMillen et al. 2019). Media was changed every 3-4 days. At 14 days post infection, organoids were either fixed for cryosectioning, prepped for FACs sorting, or dissociated in accutase enzyme and lysed for protein or RNA isolation. Three organoids containing approximately 1-2 million cells each were combined prior to sorting to ensure sufficient cell numbers in each population for whole transcriptome analysis.

### FACs Sorting

Organoids were dissociated using the enzyme accutase at 37°C for x min, and then washed in PBS. Organoid sorting buffer (1% FBS and 2 mM EDTA in DPBS) was then added to the tube and gently blown at the organoid until it completely dissociated. This cell suspension was then filtered to remove any remaining cell clumps. Finally, the filtered suspension was placed on ice until ready for sorting. Fluorescence Activated Cell Sorting (FACs) was performed on BD FACS Diva 8.0.1. Live cell gating was established using the forward and side scatter plots from an uninfected organoid. Infected organoids were then further sorted based on GFP fluorescence signal into GFP (+), GFP intermediate, or GFP (-) that were consistent across replicates. Gating analysis and plots were generated using Flowjo.

### RNA Sequencing and qRT-PCR

Total RNA was isolated from organoid samples following FACs sorting for live cells and GFP expression. To insure a high enough RNA concentration, 3 infected organoids were combined for each N. After isolation RNA was analyzed on the tape station using screen tape from Agilent (#5067-5576) for quantity and quality. External RNA Controls Consortium (ERCC) spike-in control from Invitrogen (#4456740) was added to each sample to minimize external sources of variability. cDNA libraries were then generated for each sample using the NEBNext Poly(A) mRNA Magnetic Isolation Module (NEB #E7490). Libraries were then analyzed by qPCR generating CT values using NEBNEXT Library Quant Kit (#E7630) and by tape station with D1000 screen tape from Agilent (#5067-5582) to calculate fragment base pair size. These measures were then analyzed by the NEB bio-calculator tool to get an estimated library concentration. Libraries were then diluted to 4 nM and pooled. Another qPCR was run with the pooled library to confirm concentration and a new dilution was performed if necessary. The diluted pooled library (25 pM) was then combined with diluted NEXTSeq PhiX control (#FC-110-3002). The PhiX control and library were then combined and loaded into the cartridge. Sequencing was performed using the NextSeq 500/550 High Output Kit v2.5 (#20024906) from Illumina. Preliminary analysis was then performed using the online platform Basepair Technologies.

Total RNA was isolated, and reverse transcribed into cDNA using the Promega RT Kit (#A3500). q-RT-PCR was performed using specific primer sequences as outlined in (Table S5). The resulting CT values were normalized to GAPDH and the Delta Delta CT method was employed for analysis.

### Immunofluorescence and Western Blot Analysis

Neural progenitor cells were fixed on coverslips with 4% paraformaldehyde (PFA) for 20 minutes at 4°C, washed with phosphate-buffered saline (PBS), and placed in PBS. For immunofluorescence, coverslips were blocked with 5% normal donkey serum (S30; Sigma) and 0.1% Triton in PBS for 30 min, incubated in primary antibodies overnight at 4°C, and incubated in secondary antibodies for 1 h at room temperature. The nuclear stain Hoechst was used to label nuclei. Primary antibodies used were HES1 (rabbit, PA5-28802; Thermo Fisher), UL123 (mouse;), and Tuj1 (chicken, GTX85469; GeneTex). Species-appropriate fluorescent secondary antibodies were used. An upright TS100 Nikon fluorescence microscope and NIS Elements were used for imaging and analysis.

Protein was isolated from NPCs or MRC5 fibroblasts by lifting the cells from the plate with trypsin (Invitrogen #15400-054) and lysing the cells in nuclear lysis buffer (10 mM NaCl, 50 mM Tris-HCl, pH 7.5, 1 mM EDTA, 1% SDS). Samples were then sonicated to ensure lyses and quantified using a Pierce BCA assay (ThermoFisher #23225). Equal amounts of protein (20 μg) were then loaded onto a 10% SDS-PAGE made with 2,2,2-trichloroethanol and run at 130 volts for 1 hour. Total protein was visualized via stain free detection on the Biorad ChemiDoc MP. Gels were transferred to .45 μm Amershan Protran nitrocellulose membranes (Cytiva #10600002) using a BioRad Trans-Blot Turbo semi dry transfer system. Membranes were probed for IE1 (1:1000, clone 1B12, Shenk Lab), IE2 (1:1000, clone 3A9, Shenk Lab), UL44(1:10000, Virusys #CA006-100), and pp28 (1:1000 clone 10B4-29, Shenk Lab), using a goat α mouse HRP secondary (Jackson Immunoresearch Laboratories #115-035-003) detected by Novex ECL (Invitrogen #WP20005).

### Data analysis

All statistical analysis was performed using Graph Pad Version 9. Data were collected from 3 independent organoid differentiations and infections and analyzed by one-way ANOVA or Student’s T-test as appropriate with Tukey post hoc test for qPCR and western blot. p<0.05 was considered significant.

### Bioinformatics

Fastq files were exported from Illumina sequence hub following sequencing then optical duplicates were removed and reference genome created using the raw fasta sequences for hg19. The raw RNA Seq reads are mapped and quantified to combined reference genome using Salmon v1.4.0. The read count matrix for all transcripts were utilized in R to make PCA plots and to run differential expression analysis. DESeq2 in R was used to perform the comparisons against mock samples to get significant up regulated and down regulated genes at adjusted P value < 0.05 and fold change of 3. Basepair Tech software and expression count analysis of the trimmed reads was conducted using STAR. Differentially expressed genes were identified by DESeq2 analysis using cutoffs of adjusted p-value <.05 and log2 fold change of ± 3. Gene ontology analysis was subsequently performed using G-profiler and DAVID. Additional RNA seq analysis was performed using ingenuity pathway analysis from Qiagen and Gene Set Enrichment Analysis tool from UC San Diego and the Broad Institute. For both G-profiler and IPA analysis a list of the 3,000 most differentially expressed genes across samples analyzed were used. All these genes met the threshold of adj p-value < .01 and log2 fold change ± 9 as determined by DESeq2 analysis. For GSEA analysis the entire list of differentially expressed genes identified by DESeq2 meeting the adj p-value < .05 and log2 fold change of ± 3 cutoffs were used.

### GSEA

Gene set enrichment analysis was performed using GSEA version 3.0 software with default parameters, classic enrichment, and phenotype permutations at 1000 times. GSEA was performed on differentially expressed genes determined from DESeq2 analysis of GFP (+) vs Mock and GFP (-) vs Mock samples using genes that met the cutoff of adj p-value of < .05 and log2 fold change ± 3. The enrichment plots displayed in Figure 3 were generated from analysis of GFP (+) vs Mock or GFP (-) vs Mock using the software’s hallmark gene set

## Acknowledgements

The authors would like to thank Tom Shenk for providing antibodies against HCMV proteins and Eva Borst for providing the TB40r mGFP-IE-FKBP virus. We thank Benedetta Bonacci in the Versiti-Blood Research Institute Flow Cytometry Core, and Emma Thomas for the artwork included in this work. Creation of this art is supported by the VI4-ArtLab Artist-in-Residence program. Finally, we thank members of the Terhune, Hudson, Rao, and Ebert laboratories for helpful discussions and input on the project.

Research reported in this publication was supported by the National Institute of Allergy and Infectious Diseases division of the National Institutes of Health under award number R01AI132414 (S.S.T. and A.D.E.) and R01CA2042031 (S.R.). The content is solely the responsibility of the authors and does not necessarily represent the official views of the National Institutes of Health.

All authors declare no conflicts of interests related to the manuscript. B.S.O, R.L.M, S.S.T. and A.D.E. were responsible for study design. B.S.O., R.L.M., M.L.S., K.P., S.R. S.S.T., and A.D.E. conducted experiments, analyzed data, and/or interpreted data. S.R., S.S.T., and A.D.E. provided resources. B.S.O. R.L.M, and K.P. contributed to writing the manuscript. B.S.O, S.S.T., and A.D.E. wrote and edited the final manuscript.

## Supplementary

**Figure S1. Representative FACs traces from an HCMV-TB40EGFP infected organoid.** (A) Set of three scatter analysis plots used to determine the live cell population within the organoid, live cell gates were set based on the uninfected condition. (B) Table breaking down relative cell counts within each gated population. (C) Plot of GFP signal intensity against total number of sorted cells, GFP gates were set based on previous infected organoid sorting. (D) Relative number of cells within each infected subpopulation, broken down by replicate.

**Figure S2. Representative FACs traces from an uninfected organoid.** (A) Set of three scatter analysis plots used to determine the live cell population within the organoid, live cell gates were set based on the uninfected condition. (B-C) Side scatter versus GFP plot in an uninfected organoid showing lack of GFP signal. (D) Percentage of live cells across all organoids sorted for sequencing.

**Figure S3. Additional gene ontology analysis for GFP (+) vs. Mock and GFP (-) vs. Mock.** (A) Ontology conducted using the same 3,000 gene list as in Figure 3A with the platform DAVID instead of G-profiler. Results shown are the top 20 most significant terms.

**Figure S4. Network diagrams generated from IPA, summarize three key affected canonical pathways that were differentially affected by infection.** (A) Synaptogenesis signaling pathway-log(adj p-value) = 16.367 majority of pathway components were downregulated. (B) mTOR signaling -log(adj p-value) = 17.479 majority of pathway components were downregulated. (C) RHOGDI signaling -log(adj p-value) = 11.123 mixture of upregulated and downregulated pathway components.

**Figure S5. Ingenuity pathway core analysis diagram for GFP (+) vs. Mock.** (A) Node diagram generated using IPA to connect major affected pathways and functional outcomes within the GFP (+) samples to regulators, done using the same gene list as IPA from Figure 3.

