## Supplemental Table 1 for "Downregulation of Neurodevelopmental Gene Expression in iPSC-Derived Cerebral Organoids Upon Infection by Human Cytomegalovirus"

| | Transcript ID | Gene | baseMean | log2FoldChai | lfcSE |
|---|---|---|---|---|---|
| 1 | NR_125368.1 | LOC100129434 | 3249.78269 | 4.90846741 | 0.22614162 |
| 2 | NR_047679.1 | UGDH-AS1 | 4240.14496 | 4.40373721 | 0.25243388 |
| 3 | NR_126435.1 | LINC00504 | 7301.41713 | 4.21173606 | 0.25645715 |
| 4 | NR_125837.1 | LOC101926898 | 1190.42614 | 4.66547485 | 0.31653415 |
| 5 | NR_046088.1 | LINC01776 | 193.231486 | 5.40785391 | 0.36787743 |
| 6 | NM_001402.5 | #N/A | 33648.0805 | -7.1670099 | 0.52604771 |
| 7 | NM_001190881.2 | ORC4 | 2019.7174 | 15.9594456 | 1.2076091 |
| 8 | NM_000599.3 | #N/A | 3888.47051 | -6.671084 | 0.50811128 |
| 9 | NM_002568.3 | #N/A | 2710.01066 | -7.5054552 | 0.58263873 |
| 10 | NM_001164436.1 | #N/A | 785.868599 | 4.49517537 | 0.37289743 |
| 11 | NM_000088.3 | COL1A1 | 14089.3145 | -10.086921 | 0.83971668 |
| 12 | NR_049735.2 | PTCSC3 | 192.995625 | 4.69027508 | 0.3955378 |
| 13 | NR_027451.1 | NORAD | 2119.24645 | -4.7925399 | 0.4134876 |
| 14 | NM_002332.2 | #N/A | 2045.52274 | -5.5179658 | 0.48120944 |
| 15 | NR_034022.1 | LOC100131257 | 1815.7193 | 3.61660474 | 0.31848146 |
| 16 | NR_110133.1 | LOC101927374 | 1757.80909 | 4.00879143 | 0.35462112 |
| 17 | NM_006597.5 | #N/A | 5029.85763 | -5.9081719 | 0.52453822 |
| 18 | NM_002046.5 | #N/A | 7293.23457 | -8.9254718 | 0.8006075 |
| 19 | NR_002728.3 | KCNQ1OT1 | 3940.58133 | 3.48174464 | 0.31340367 |
| 20 | NM_005934.3 | #N/A | 744.801673 | -4.6223299 | 0.41796171 |
| 21 | NR_130914.1 | LINC01949 | 411.983906 | 4.18518337 | 0.38443753 |
| 22 | NM_004371.3 | #N/A | 1014.85616 | -5.7398269 | 0.53126293 |
| 23 | NR_047532.1 | CDKN2B-AS1 | 111.087569 | 3.73131146 | 0.35312379 |
| 24 | NM_002136.3 | #N/A | 4300.51106 | -5.6812199 | 0.53972714 |
| 25 | NM_009589.3 | #N/A | 100.218756 | 4.24653919 | 0.40499185 |
| 26 | NR_033770.1 | ROCK1P1 | 332.430227 | 6.09886541 | 0.58781299 |
| 27 | NR_130931.1 | LINC01873 | 1410.93141 | 3.60293507 | 0.34874809 |
| 28 | NM_206893.3 | #N/A | 258.48343 | 4.70886875 | 0.45830557 |
| 29 | NR_047469.1 | LINC00506 | 4080.27165 | 4.1388784 | 0.40357376 |
| 30 | NM_004134.6 | #N/A | 1141.94576 | -8.2519311 | 0.81660028 |
| 31 | NM_001243960.1 | #N/A | 204.998823 | 12.658191 | 1.25345347 |
| 32 | NM_001291969.1 | #N/A | 283.658487 | 13.1300565 | 1.30877276 |
| 33 | NM_001940.3 | #N/A | 1370.96149 | -7.1532752 | 0.71323692 |
| 34 | NM_001101.3 | #N/A | 16063.6579 | -10.620527 | 1.06819986 |
| 35 | NM_001114173.2 | #N/A | 256.046554 | 12.9824694 | 1.3084077 |
| 36 | NM_006600.3 | #N/A | 569.182755 | -6.068081 | 0.61286083 |
| 37 | NM_020729.2 | #N/A | 613.711445 | 2.60807594 | 0.26336265 |
| 38 | NM_178564.3 | #N/A | 422.937371 | -5.5430364 | 0.57313469 |
| 39 | NR_130727.1 | LOC401357 | 84.6012645 | 8.53778287 | 0.88607258 |
| 40 | NM_015276.1 | #N/A | 2443.86277 | -9.7192624 | 1.01251574 |
| 41 | NM_001127350.1 | #N/A | 155.082034 | 12.2554699 | 1.27992699 |
| 42 | NM_001190470.1 | #N/A | 15365.6855 | -8.1486001 | 0.85099192 |

| | | | | | |
|---|---|---|---|---|---|
| 44 | NR_029405.1 | LOC643406 | 729.965393 | 3.92236936 | 0.41142494 |
| 43 | NM_001614.3 | #N/A | 14499.7517 | -15.034364 | 1.57726774 |
| 45 | NM_001961.3 | #N/A | 10923.4795 | -14.625773 | 1.5393484 |
| 46 | NR_027053.2 | LOC646214 | 1040.84735 | 3.89584042 | 0.41839686 |
| 47 | NM_001387.2 | #N/A | 4404.97825 | -10.098988 | 1.09347982 |
| 48 | NM_006009.3 | #N/A | 16687.4587 | -15.237095 | 1.65172338 |
| 49 | NM_005507.2 | #N/A | 3057.90291 | -7.2294738 | 0.78890607 |
| 50 | NM_152733.2 | BTBD9 | 3518.43182 | 16.7596061 | 1.83747253 |
| 51 | NM_005431.1 | #N/A | 201.596476 | 2.77267951 | 0.30461625 |
| 52 | NR_036647.1 | CCDC144B | 283.224401 | 4.32577591 | 0.47640803 |
| 53 | NR_109904.1 | LINC02108 | 215.287097 | 4.25859063 | 0.46943527 |
| 55 | NM_002111.8 | HTT | 412.185331 | -4.1057943 | 0.45367999 |
| 54 | NM_001206491.1 | MAVS | 121.314793 | 11.9033825 | 1.31544716 |
| 56 | NM_007355.3 | #N/A | 5256.99967 | -13.57066 | 1.50223744 |
| 57 | NM_001297567.1 | ZNF124 | 132.617435 | 4.98207335 | 0.55212402 |
| 58 | NM_005348.3 | #N/A | 6017.22032 | -13.765509 | 1.52634084 |
| 59 | NM_005722.3 | ACTR2 | 1149.86459 | -6.4609915 | 0.71946479 |
| 60 | NM_003380.3 | #N/A | 7480.09621 | -10.56751 | 1.17856171 |
| 61 | NM_003118.3 | #N/A | 8188.12875 | -14.209967 | 1.58599223 |
| 62 | NM_178014.3 | #N/A | 7596.19423 | -14.10169 | 1.57455309 |
| 63 | NM_002032.2 | #N/A | 6191.95282 | -13.806822 | 1.54316214 |
| 64 | NM_021009.6 | #N/A | 5598.05674 | -6.6673278 | 0.74533639 |
| 65 | NR_135520.1 | LINC01684 | 643.104614 | 5.15429523 | 0.57903026 |
| 66 | NM_020738.2 | #N/A | 1840.17338 | -6.3335283 | 0.71167219 |
| 67 | NM_021109.3 | #N/A | 7404.06422 | -12.643244 | 1.42146032 |
| 68 | NM_001286561.1 | NUMA1 | 1141.7868 | -9.2726687 | 1.04472028 |
| 69 | NR_110665.1 | ROR1-AS1 | 302.832286 | 4.99247665 | 0.56383927 |
| 70 | NM_178153.2 | #N/A | 1503.20433 | -6.6660235 | 0.75660354 |
| 72 | NR_144568.1 | MALAT1 | 7908.31863 | -14.159749 | 1.61017258 |
| 71 | NM_001418.3 | #N/A | 5216.33659 | -13.559445 | 1.54202131 |
| 73 | NM_000967.3 | #N/A | 5108.41754 | -13.529276 | 1.54412604 |
| 74 | NM_001010.2 | #N/A | 2596.13284 | -7.9002237 | 0.90238154 |
| 75 | NR_110130.1 | LINC02444 | 100.38705 | 3.29559861 | 0.37736098 |
| 76 | NM_001278559.1 | #N/A | 573.153822 | -5.3586729 | 0.61465613 |
| 77 | NR_040244.1 | LINC00547 | 136.197432 | 3.52081769 | 0.40423142 |
| 78 | NM_004395.3 | DBN1 | 2124.51549 | -9.0248944 | 1.03972409 |
| 79 | NM_000968.3 | #N/A | 4613.79925 | -10.735437 | 1.23813607 |
| 80 | NM_013318.3 | PRRC2B | 2468.56281 | -11.14944 | 1.28710914 |
| 81 | NM_001428.3 | #N/A | 3912.28145 | -13.144368 | 1.51791041 |
| 82 | NM_006082.2 | #N/A | 5036.773 | -10.835059 | 1.25324766 |
| 83 | NM_005027.3 | #N/A | 853.228382 | -6.3879822 | 0.74048484 |
| 84 | NR_126447.1 | WARS2-IT1 | 81.1090933 | 3.49033907 | 0.40494931 |
| 85 | NM_001110556.1 | #N/A | 3900.81897 | -13.140246 | 1.52558963 |

| | | | | | |
|---|---|---|---|---|---|
| 86 | NM_005909.4 | #N/A | 5250.48998 | -13.568841 | 1.5793652 |
| 87 | NM_005381.2 | #N/A | 2170.40205 | -10.967213 | 1.28190307 |
| 89 | NR_038326.1 | LINC01541 | 51.1861717 | 4.93678613 | 0.57815189 |
| 88 | NM_000972.2 | #N/A | 3030.83825 | -6.311691 | 0.73920534 |
| 90 | NM_032591.2 | #N/A | 95.6592116 | 11.5561012 | 1.35988032 |
| 92 | NR_037845.1 | LOC100506023 | 143.525825 | 3.82324461 | 0.4503498 |
| 91 | NM_184041.2 | #N/A | 2661.36791 | -12.588586 | 1.48296349 |
| 93 | NM_001190702.1 | #N/A | 17862.1708 | -7.1051 | 0.83895686 |
| 94 | NM_002654.5 | #N/A | 2375.94298 | -12.425029 | 1.46873312 |
| 95 | NM_001329222.1 | GANAB | 397.295719 | -6.0008283 | 0.70998513 |
| 96 | NM_001848.2 | #N/A | 1778.8707 | -10.691004 | 1.26609826 |
| 97 | NM_000146.3 | #N/A | 3778.63633 | -13.0943 | 1.55450641 |
| 98 | NM_003299.2 | #N/A | 2852.79953 | -12.688849 | 1.5108691 |
| 99 | NM_013402.4 | #N/A | 818.473716 | -7.0796051 | 0.84653271 |
| 100 | NM_006158.4 | NEFL | 6526.59065 | -13.882749 | 1.66180261 |
| 101 | NM_005382.2 | NEFM | 6280.85769 | -13.827383 | 1.65634194 |
| 102 | NM_006445.3 | #N/A | 1468.35607 | -10.424664 | 1.25019923 |
| 103 | NM_003360.4 | UGT8 | 277.330411 | 3.9183672 | 0.47019036 |
| 104 | NM_000973.4 | #N/A | 3493.29043 | -12.980981 | 1.55800677 |
| 105 | NR_125813.1 | LOC101929415 | 204.306239 | 3.5317896 | 0.42486632 |
| 106 | NR_103837.1 | #N/A | 68.6904413 | 3.84154864 | 0.46262088 |
| 107 | NM_003334.3 | #N/A | 2387.89667 | -12.432242 | 1.50297135 |
| 108 | NM_004559.4 | #N/A | 5202.73414 | -13.555661 | 1.63901588 |
| 109 | NR_144567.1 | MALAT1 | 2361.87083 | -12.416409 | 1.5015925 |
| 110 | NM_001684.4 | #N/A | 341.759224 | -3.2842193 | 0.39726126 |
| 111 | NM_000597.2 | #N/A | 2904.7148 | -12.71473 | 1.53902994 |
| 112 | NM_004343.3 | CALR | 2887.54801 | -12.70632 | 1.53875326 |
| 113 | NM_007085.4 | #N/A | 3694.6722 | -13.061904 | 1.58363732 |
| 114 | NM_007114.2 | #N/A | 246.957936 | -4.5768814 | 0.55514638 |
| 115 | NM_152924.4 | #N/A | 469.533376 | -6.5040975 | 0.7899259 |
| 116 | NM_005346.4 | #N/A | 729.053492 | -7.8500331 | 0.95444186 |
| 117 | NM_000093.4 | #N/A | 2072.72379 | -10.872626 | 1.32338802 |
| 118 | NM_001080409.2 | #N/A | 121.425336 | 3.76305774 | 0.45806953 |
| 119 | NM_001845.5 | #N/A | 2139.07056 | -8.299677 | 1.01018392 |
| 120 | NM_178012.4 | #N/A | 5387.05878 | -13.605893 | 1.65573119 |
| 121 | NM_001185176.1 | CES3 | 82.0139301 | 11.3352785 | 1.38097039 |
| 122 | NM_002778.3 | #N/A | 2082.40872 | -12.234915 | 1.4923877 |
| 123 | NM_000291.3 | #N/A | 1648.38276 | -10.564698 | 1.28952968 |
| 124 | NM_001002.3 | #N/A | 4272.05462 | -13.271319 | 1.62033058 |
| 125 | NM_005324.4 | #N/A | 2192.52588 | -12.308908 | 1.50314889 |
| 126 | NM_001746.3 | #N/A | 2452.33764 | -12.470581 | 1.52493967 |
| 127 | NM_001831.3 | #N/A | 3445.43616 | -12.961136 | 1.5863361 |
| 128 | NM_002823.4 | PTMA | 3300.8678 | -7.7857849 | 0.9532177 |

| | | | | | |
|---|---|---|---|---|---|
| 129 | NM_004396.4 | #N/A | 1997.71781 | -10.818168 | 1.32492872 |
| 130 | NM_001975.2 | #N/A | 2238.93685 | -12.339158 | 1.51564544 |
| 131 | NM_003107.2 | #N/A | 3219.53077 | -10.339107 | 1.27096646 |
| 132 | NR_046479.1 | #N/A | 81.9419246 | 11.3389175 | 1.394569 |
| 133 | NM_012423.3 | #N/A | 3265.82582 | -12.883828 | 1.58623373 |
| 134 | NR_106781.1 | #N/A | 5008.06306 | -13.5007 | 1.66340584 |
| 135 | NM_006986.3 | #N/A | 1859.25486 | -12.071253 | 1.48866764 |
| 136 | NM_003262.3 | #N/A | 664.39255 | -7.5786964 | 0.93514603 |
| 137 | NM_000970.4 | #N/A | 2973.56486 | -12.748573 | 1.5739011 |
| 138 | NM_006766.4 | #N/A | 357.862973 | -5.8839851 | 0.72700135 |
| 139 | NM_004423.3 | #N/A | 781.59792 | -3.1879089 | 0.39403767 |
| 140 | NM_006761.4 | #N/A | 3121.78477 | -12.818765 | 1.58603877 |
| 141 | NM_004321.7 | KIF1A | 3313.61223 | -12.904811 | 1.59813482 |
| 142 | NM_000977.3 | RPL13 | 3664.59612 | -13.050038 | 1.6177646 |
| 143 | NM_006931.2 | #N/A | 353.918585 | -6.1623922 | 0.76466141 |
| 144 | NM_000516.5 | #N/A | 3802.32282 | -13.10328 | 1.62615636 |
| 145 | NM_006386.4 | #N/A | 1582.96989 | -11.839231 | 1.46985059 |
| 146 | NM_000175.4 | #N/A | 1681.28609 | -11.92592 | 1.48134864 |
| 147 | NM_006098.4 | #N/A | 2885.66417 | -12.705305 | 1.5790755 |
| 148 | NM_153692.3 | #N/A | 74.2505796 | 4.57654368 | 0.56874276 |
| 149 | NM_198589.2 | #N/A | 1792.1874 | -12.018183 | 1.49634145 |
| 150 | NM_004104.4 | #N/A | 2816.4825 | -12.670308 | 1.57837822 |
| 151 | NM_001278593.1 | #N/A | 142.655537 | 12.1298531 | 1.51157997 |
| 152 | NM_003128.2 | #N/A | 1714.869 | -10.600028 | 1.32176715 |
| 153 | NM_001686.3 | #N/A | 2375.37044 | -12.424532 | 1.54956706 |
| 154 | NM_005184.3 | #N/A | 2859.04279 | -12.691933 | 1.58413321 |
| 155 | NM_002074.4 | #N/A | 2413.00182 | -12.447202 | 1.55402031 |
| 156 | NM_001145204.2 | #N/A | 1107.25441 | 3.54621919 | 0.44279958 |
| 157 | NM_007029.3 | #N/A | 4725.12091 | -13.416757 | 1.67903168 |
| 158 | NM_001012.1 | #N/A | 3062.03487 | -12.790868 | 1.60238343 |
| 159 | NM_002473.5 | MYH9 | 1901.73063 | -12.103915 | 1.51888137 |
| 160 | NM_000971.3 | #N/A | 2592.26038 | -12.55059 | 1.57595916 |
| 161 | NM_004404.4 | #N/A | 1112.19553 | -10.052843 | 1.26365655 |
| 162 | NM_000918.3 | #N/A | 1955.50285 | -12.144036 | 1.52681459 |
| 163 | NM_201413.2 | #N/A | 1485.33385 | -11.747537 | 1.4789066 |
| 164 | NR_109833.1 | PRNCR1 | 151.042039 | 3.97360229 | 0.50050674 |
| 165 | NM_000202.7 | #N/A | 1662.47681 | -11.90971 | 1.50152986 |
| 166 | NR_125956.1 | LOC101928565 | 50.3334401 | 5.28556942 | 0.66699676 |
| 167 | NM_152473.2 | ERVV-1 | 84.0366885 | 5.41931198 | 0.6841182 |
| 168 | NM_000969.3 | #N/A | 2596.0335 | -12.552698 | 1.58543524 |
| 172 | NM_007334.2 | KLRD1 | 82.0859357 | 11.3324149 | 1.43185797 |
| 169 | NM_001493.2 | #N/A | 2523.2785 | -12.511697 | 1.58091883 |
| 170 | NM_003088.3 | #N/A | 2107.90405 | -12.252208 | 1.54825006 |

| | | | | | |
|---|---|---|---|---|---|
| 171 | NM_003217.2 | #N/A | 1936.03577 | -12.12951 | 1.53285479 |
| 173 | NM_002824.5 | #N/A | 3193.40617 | -12.851474 | 1.62636388 |
| 174 | NM_001130438.2 | #N/A | 2653.56723 | -12.584298 | 1.59314734 |
| 175 | NM_000852.3 | #N/A | 1295.42368 | -10.274673 | 1.30090212 |
| 176 | NM_002356.6 | #N/A | 2517.6091 | -12.508397 | 1.58422524 |
| 177 | NM_015092.4 | SMG1 | 445.443845 | -5.7593448 | 0.72987639 |
| 178 | NM_006888.4 | #N/A | 2582.55777 | -12.545162 | 1.59013137 |
| 179 | NR_038380.1 | SLC7A11-AS1 | 68.7960754 | 4.59390874 | 0.58276844 |
| 181 | NM_002520.6 | NPM1 | 1997.58198 | -12.174658 | 1.54592886 |
| 180 | NM_001330681.1 | #N/A | 2078.45139 | -10.960816 | 1.39187282 |
| 182 | NM_012401.3 | #N/A | 1422.39366 | -11.684969 | 1.48394891 |
| 183 | NM_001291311.1 | PLD3 | 2090.94737 | -12.240671 | 1.55557831 |
| 184 | NM_002156.4 | #N/A | 1418.35122 | -11.68076 | 1.48470816 |
| 185 | NM_004651.3 | #N/A | 1562.93698 | -11.820766 | 1.50368054 |
| 186 | NM_130445.3 | #N/A | 1723.71321 | -11.962205 | 1.52320976 |
| 188 | NR_109994.1 | LOC101926889 | 311.546339 | 3.11124251 | 0.3963775 |
| 187 | NM_014014.4 | #N/A | 1315.76829 | -11.572432 | 1.47431321 |
| 189 | NM_007158.5 | #N/A | 1741.12489 | -11.976395 | 1.52598202 |
| 190 | NM_001317897.1 | #N/A | 162.03285 | 12.3243234 | 1.57056873 |
| 191 | NM_000612.5 | #N/A | 2319.73021 | -12.390481 | 1.58024567 |
| 192 | NM_003752.4 | EIF3C | 1778.70038 | -12.007239 | 1.53454384 |
| 193 | NM_004046.5 | #N/A | 1756.3735 | -11.989033 | 1.53453997 |
| 194 | NM_001042465.2 | #N/A | 1971.76117 | -12.155942 | 1.55727625 |
| 195 | NM_001417.6 | #N/A | 1581.11173 | -11.83724 | 1.51730909 |
| 196 | NM_007126.3 | #N/A | 1552.6264 | -11.811195 | 1.51443048 |
| 197 | NM_023009.6 | #N/A | 3040.87723 | -12.780868 | 1.6404508 |
| 198 | NM_014423.3 | #N/A | 593.613514 | -8.268029 | 1.06140713 |
| 199 | NM_018955.3 | #N/A | 1827.9797 | -12.046638 | 1.5464996 |
| 200 | NM_005762.2 | #N/A | 1854.11459 | -12.067095 | 1.55129239 |
| 201 | NM_018834.5 | #N/A | 2008.4663 | -12.182433 | 1.5666892 |
| 202 | NM_001823.4 | #N/A | 2405.59174 | -12.442713 | 1.60040277 |
| 203 | NM_012479.3 | #N/A | 2513.82712 | -12.50628 | 1.6088295 |
| 204 | NM_001743.5 | #N/A | 2527.54292 | -12.514134 | 1.6101357 |
| 205 | NM_000998.4 | #N/A | 2304.85212 | -12.38106 | 1.59344562 |
| 206 | NM_001846.3 | #N/A | 2683.04807 | -12.600329 | 1.62168504 |
| 207 | NM_001321678.1 | #N/A | 67.5098918 | 11.05619 | 1.42348275 |
| 208 | NR_104668.1 | LOC101928505 | 36.1185875 | 6.36642452 | 0.81992233 |
| 209 | NM_016841.4 | #N/A | 2317.43344 | -10.100514 | 1.30094855 |
| 210 | NM_001309840.1 | #N/A | 2229.70612 | -12.33333 | 1.58933022 |
| 211 | NM_212482.2 | #N/A | 1486.81553 | -11.748861 | 1.51542585 |
| 212 | NR_015362.1 | #N/A | 88.4911222 | 3.43262266 | 0.44278881 |
| 213 | NM_015001.2 | #N/A | 449.150694 | -7.2944824 | 0.94117305 |
| 214 | NM_001306129.1 | FN1 | 1416.73495 | -11.679348 | 1.50724815 |

| | | | | | |
|---|---|---|---|---|---|
| 215 | NM_001923.4 | #N/A | 1284.35829 | -11.537603 | 1.48962561 |
| 216 | NM_007363.4 | #N/A | 1568.71615 | -11.825945 | 1.52688938 |
| 217 | NM_201414.2 | #N/A | 3020.05896 | -12.770983 | 1.65071915 |
| 218 | NM_001270427.1 | #N/A | 71.6491388 | 11.1370981 | 1.43986524 |
| 220 | NR_135276.1 | BFSP2-AS1 | 422.735737 | 3.37452779 | 0.43665706 |
| 219 | NM_006826.3 | #N/A | 2368.74218 | -12.420503 | 1.60717886 |
| 221 | NM_016395.2 | #N/A | 878.974883 | -8.924649 | 1.15512434 |
| 222 | NM_005801.3 | #N/A | 1913.87267 | -12.112875 | 1.56792769 |
| 223 | NM_003737.3 | #N/A | 1056.58533 | -9.038195 | 1.17107012 |
| 224 | NM_003011.3 | #N/A | 1889.53235 | -12.094385 | 1.56776371 |
| 226 | NM_022051.2 | EGLN1 | 315.652191 | -5.9873437 | 0.77643247 |
| 225 | NM_001039580.1 | #N/A | 297.384851 | -4.3413663 | 0.56301715 |
| 227 | NM_000365.5 | #N/A | 1722.93725 | -11.961134 | 1.55199808 |
| 228 | NM_014918.4 | #N/A | 193.768193 | -4.7104127 | 0.6112116 |
| 229 | NM_021103.3 | #N/A | 3472.59132 | -12.972407 | 1.68360919 |
| 230 | NM_032727.3 | #N/A | 2947.84012 | -12.736076 | 1.65567774 |
| 231 | NM_001416.3 | #N/A | 1898.36544 | -12.101161 | 1.57345796 |
| 232 | NM_006039.4 | #N/A | 1584.33473 | -10.568531 | 1.37508342 |
| 233 | NM_000981.3 | #N/A | 1908.5017 | -10.70574 | 1.39335788 |
| 234 | NM_001253383.1 | #N/A | 1735.3818 | -11.971608 | 1.5588471 |
| 235 | NM_014225.5 | #N/A | 1889.68036 | -12.094539 | 1.57485853 |
| 236 | NM_012204.3 | #N/A | 258.078187 | -5.0535233 | 0.65819369 |
| 237 | NM_001315537.1 | LDHB | 1999.01599 | -12.175675 | 1.58607045 |
| 238 | NM_001206927.1 | #N/A | 63.6374671 | 8.61931971 | 1.12305212 |
| 239 | NM_001009944.2 | #N/A | 1344.25268 | -11.603285 | 1.51277096 |
| 240 | NM_002117.5 | #N/A | 562.628652 | -6.8143023 | 0.88923887 |
| 241 | NM_012433.3 | SF3B1 | 1116.75516 | -11.335834 | 1.4801249 |
| 242 | NM_005313.4 | #N/A | 1550.38982 | -11.809139 | 1.54217018 |
| 243 | NM_020755.3 | #N/A | 1941.74457 | -10.72549 | 1.40101233 |
| 244 | NM_001256736.2 | SSBP2 | 64.5279498 | 10.9918962 | 1.43669379 |
| 245 | NR_135293.1 | LINC02280 | 148.0749 | 4.2610469 | 0.55791021 |
| 246 | NM_001256651.1 | #N/A | 78.3751821 | 11.2781347 | 1.477056 |
| 247 | NM_001256802.1 | #N/A | 2006.46348 | -12.181042 | 1.59622424 |
| 248 | NM_001005271.2 | #N/A | 1861.97368 | -12.073121 | 1.5839777 |
| 249 | NM_006317.4 | #N/A | 2771.18257 | -11.193359 | 1.46853481 |
| 250 | NM_194434.2 | #N/A | 522.035779 | -6.8059981 | 0.89287469 |
| 251 | NM_002128.5 | #N/A | 1729.58259 | -11.966744 | 1.57143999 |
| 252 | NM_000937.4 | #N/A | 1070.16318 | -11.274216 | 1.48069572 |
| 253 | NM_001282236.1 | #N/A | 110.965947 | 2.52581592 | 0.33180333 |
| 254 | NM_004068.3 | #N/A | 1538.74404 | -11.798164 | 1.55032059 |
| 255 | NM_003295.3 | #N/A | 3908.75024 | -9.5481029 | 1.25504937 |
| 256 | NM_013336.3 | #N/A | 1414.01779 | -11.67629 | 1.5354516 |
| 257 | NM_001007.4 | #N/A | 1860.14071 | -12.071809 | 1.58804674 |

| | | | | | |
|---|---|---|---|---|---|
| 258 | NM_001006.4 | #N/A | 3555.41551 | -7.9848463 | 1.05067573 |
| 259 | NM_001002907.1 | OR8K1 | 67.5221877 | 11.0513998 | 1.45453079 |
| 260 | NM_000979.3 | #N/A | 1879.69894 | -12.086878 | 1.59149371 |
| 261 | NM_005563.3 | #N/A | 2700.54138 | -12.609637 | 1.6606997 |
| 262 | NM_006088.5 | #N/A | 994.403754 | -11.168753 | 1.47087631 |
| 263 | NM_002211.3 | #N/A | 1597.36651 | -11.852202 | 1.56105369 |
| 264 | NM_000975.3 | #N/A | 1749.76206 | -11.983525 | 1.57991242 |
| 265 | NM_004181.4 | #N/A | 2487.12912 | -12.490881 | 1.64670053 |
| 266 | NM_001207.4 | BTF3 | 818.003752 | -6.0747842 | 0.8010493 |
| 267 | NM_006791.3 | #N/A | 1347.16706 | -11.606389 | 1.53040986 |
| 268 | NM_000615.6 | #N/A | 2561.08357 | -11.083888 | 1.46210099 |
| 269 | NM_004039.2 | #N/A | 1570.50564 | -11.827748 | 1.56077483 |
| 270 | NM_198576.3 | #N/A | 1254.14497 | -11.50331 | 1.5187156 |
| 271 | NM_001003.2 | #N/A | 2320.01545 | -12.390518 | 1.63612786 |
| 272 | NM_020338.3 | #N/A | 1556.05903 | -11.81431 | 1.56042188 |
| 273 | NM_003255.4 | #N/A | 1558.79917 | -10.424544 | 1.37716725 |
| 274 | NR_046398.1 | #N/A | 167.792443 | 3.83874284 | 0.5071278 |
| 277 | NM_177401.4 | MIDN | 762.954451 | -6.3765837 | 0.84297455 |
| 275 | NM_001854.3 | #N/A | 1210.20681 | -11.451904 | 1.51377592 |
| 276 | NM_016127.5 | #N/A | 1528.62594 | -11.788651 | 1.55847533 |
| 278 | NM_001303624.1 | #N/A | 1814.16033 | -12.035733 | 1.59133452 |
| 279 | NM_005885.3 | #N/A | 1142.68161 | -11.368762 | 1.50473396 |
| 280 | NM_015626.8 | #N/A | 1115.68496 | -11.33414 | 1.50045173 |
| 281 | NM_001021.4 | #N/A | 2105.04039 | -12.250223 | 1.62196067 |
| 282 | NM_003373.3 | #N/A | 688.621232 | -8.0409637 | 1.06479157 |
| 283 | NM_001261826.1 | #N/A | 1100.97156 | -11.315265 | 1.49899297 |
| 284 | NM_002972.3 | #N/A | 1405.4483 | -11.667341 | 1.54579896 |
| 285 | NM_001539.3 | #N/A | 1381.18157 | -11.642297 | 1.54324826 |
| 286 | NM_001658.3 | #N/A | 1342.73041 | -11.601604 | 1.5378247 |
| 288 | NR_110765.1 | L3MBTL4-AS1 | 141.832111 | 6.16666738 | 0.81764065 |
| 287 | NM_000398.6 | #N/A | 1134.93482 | -11.359097 | 1.50603876 |
| 289 | NM_000430.3 | #N/A | 1359.6737 | -11.619634 | 1.54097735 |
| 290 | NM_002375.4 | #N/A | 1062.23603 | -11.263984 | 1.49371898 |
| 291 | NM_005100.3 | #N/A | 1116.95779 | -11.336177 | 1.50338466 |
| 292 | NM_006842.2 | #N/A | 1002.55466 | -11.180164 | 1.48286268 |
| 293 | NM_006035.3 | #N/A | 914.730121 | -9.7149044 | 1.28884426 |
| 294 | NM_021999.4 | #N/A | 1042.36435 | -11.236468 | 1.49118199 |
| 295 | NM_014106.3 | #N/A | 361.337824 | -4.5076794 | 0.59836047 |
| 296 | NM_001030006.1 | #N/A | 1334.03599 | -11.59218 | 1.54022942 |
| 297 | NM_005911.5 | #N/A | 950.437809 | -11.10306 | 1.47549168 |
| 298 | NM_001304998.1 | TMEM170A | 135.55894 | 12.0560815 | 1.60224995 |
| 299 | NM_002096.2 | #N/A | 489.826189 | -7.524687 | 1.00029234 |
| 300 | NM_001258401.2 | SLC4A8 | 90.9105724 | 11.4927397 | 1.52807725 |

| | | | | | |
|---|---|---|---|---|---|
| 301 | NR_027409.1 | #N/A | 988.727411 | -11.159686 | 1.4872626 |
| 302 | NM_012293.2 | #N/A | 1157.20263 | -11.387245 | 1.51844993 |
| 303 | NM_031844.2 | #N/A | 1041.87059 | -11.235325 | 1.49834493 |
| 304 | NM_001206799.1 | #N/A | 1232.75426 | -10.105109 | 1.34853591 |
| 305 | NM_012215.3 | #N/A | 996.067005 | -11.17056 | 1.4909776 |
| 306 | NM_002629.3 | #N/A | 1439.70312 | -11.702098 | 1.56223383 |
| 307 | NM_001330262.1 | #N/A | 1171.7831 | -11.405257 | 1.52331516 |
| 308 | NM_003601.3 | #N/A | 779.756764 | -3.1287632 | 0.41811436 |
| 309 | NM_015315.4 | #N/A | 1095.547 | -11.308096 | 1.51127229 |
| 310 | NM_006805.3 | #N/A | 1470.00434 | -5.810066 | 0.77664436 |
| 311 | NM_001903.4 | #N/A | 886.816053 | -11.003579 | 1.471174 |
| 312 | NM_004265.3 | #N/A | 1912.05528 | -12.111526 | 1.62008014 |
| 313 | NM_000978.3 | #N/A | 1581.34802 | -11.837529 | 1.58380666 |
| 314 | NM_001020.5 | #N/A | 1741.70357 | -11.976851 | 1.60247066 |
| 315 | NM_005566.3 | #N/A | 1303.1608 | -11.558171 | 1.54658555 |
| 316 | NM_001873.3 | #N/A | 1578.33921 | -11.834772 | 1.58394852 |
| 317 | NM_001839.4 | #N/A | 1073.32584 | -11.278712 | 1.51026111 |
| 318 | NM_000992.2 | #N/A | 1739.14619 | -11.974746 | 1.6037435 |
| 319 | NM_002295.5 | #N/A | 2365.60471 | -6.6403088 | 0.88973004 |
| 320 | NM_003290.2 | #N/A | 1285.74163 | -11.539143 | 1.54633651 |
| 321 | NM_022748.11 | #N/A | 499.049294 | -4.5498506 | 0.60987531 |
| 322 | NM_001909.4 | #N/A | 982.736305 | -11.151623 | 1.49510712 |
| 323 | NM_031992.1 | #N/A | 1090.00475 | -11.300724 | 1.51533714 |
| 324 | NM_005506.3 | #N/A | 1099.24435 | -11.312911 | 1.51752801 |
| 325 | NM_001113202.1 | #N/A | 1524.81983 | -11.784971 | 1.58136527 |
| 326 | NM_005724.5 | #N/A | 1175.29997 | -11.409451 | 1.53105628 |
| 327 | NR_110867.1 | LOC101928936 | 375.467039 | 3.69936043 | 0.49653196 |
| 328 | NM_024830.4 | #N/A | 569.043661 | -5.1739168 | 0.69483123 |
| 329 | NM_153828.2 | #N/A | 1388.51648 | -11.649975 | 1.56453353 |
| 330 | NM_005560.4 | #N/A | 985.628969 | -9.8040222 | 1.31675218 |
| 331 | NM_033022.3 | #N/A | 2025.91859 | -12.194931 | 1.63824379 |
| 332 | NM_016609.4 | SLC22A17 | 876.777444 | -10.986954 | 1.47690998 |
| 333 | NM_007371.3 | #N/A | 547.992504 | -5.3072393 | 0.71353895 |
| 334 | NM_001013.3 | #N/A | 1414.62039 | -11.676755 | 1.57094963 |
| 335 | NM_021141.3 | #N/A | 1163.59654 | -10.021908 | 1.34825136 |
| 336 | NM_033426.2 | #N/A | 213.573328 | -3.8016094 | 0.51153936 |
| 337 | NM_014819.4 | #N/A | 1359.90521 | -11.619831 | 1.56380135 |
| 338 | NM_002444.2 | #N/A | 933.455021 | -9.8028183 | 1.32049536 |
| 339 | NM_000986.3 | #N/A | 1726.84847 | -11.964499 | 1.61214621 |
| 340 | NM_004731.4 | #N/A | 162.819602 | 12.3310927 | 1.66172968 |
| 341 | NM_001297721.1 | C1orf43 | 60.8540841 | 10.9102671 | 1.47075297 |
| 342 | NM_002265.5 | #N/A | 911.520508 | -11.043045 | 1.48867937 |
| 343 | NM_001035006.2 | #N/A | 1652.72627 | -10.484353 | 1.41348106 |

| | | | | | |
|---|---|---|---|---|---|
| 344 | NM_001252007.1 | PAPOLA | 73.1195922 | 3.80041051 | 0.51250452 |
| 345 | NM_000089.3 | #N/A | 8150.26652 | -7.0796539 | 0.95490951 |
| 346 | NM_000598.4 | #N/A | 1967.62964 | -10.884931 | 1.46825237 |
| 347 | NM_004140.3 | #N/A | 590.557769 | -3.7249004 | 0.50247657 |
| 348 | NM_002045.3 | #N/A | 2223.7002 | -12.329373 | 1.66358704 |
| 349 | NM_003973.4 | #N/A | 1976.38735 | -12.159238 | 1.64107093 |
| 350 | NM_000188.2 | HK1 | 1065.5761 | -11.268004 | 1.52089604 |
| 351 | NM_014286.3 | #N/A | 1125.60662 | -7.8941761 | 1.06625424 |
| 352 | NR_120621.1 | LINC02681 | 77.4226865 | 4.03035918 | 0.54465472 |
| 353 | NM_000983.3 | #N/A | 1479.95369 | -11.741888 | 1.58726117 |
| 354 | NM_001664.3 | #N/A | 946.000027 | -11.096378 | 1.50043192 |
| 355 | NM_017952.5 | #N/A | 251.134908 | -5.7460651 | 0.77745227 |
| 356 | NM_012117.2 | #N/A | 923.753879 | -11.061651 | 1.49701078 |
| 357 | NM_001195605.1 | ZNF865 | 229.93974 | -4.0877856 | 0.55387481 |
| 361 | NM_014944.4 | CLSTN1 | 1266.50056 | -11.517324 | 1.56104697 |
| 358 | NM_000661.4 | RPL9 | 2203.81193 | -12.316372 | 1.66937338 |
| 359 | NM_002808.4 | #N/A | 1018.31654 | -11.202632 | 1.51819495 |
| 360 | NM_005654.5 | #N/A | 898.067163 | -11.021107 | 1.49371061 |
| 362 | NM_182810.2 | ATF4 | 1179.20055 | -11.414246 | 1.54728125 |
| 363 | NM_016577.3 | #N/A | 831.730963 | -8.1335877 | 1.10271889 |
| 365 | NR_120387.1 | LOC339059 | 109.04488 | 4.05412966 | 0.54969973 |
| 364 | NM_004521.2 | #N/A | 882.488232 | -9.6496176 | 1.30833284 |
| 366 | NM_003486.6 | #N/A | 1258.26838 | -11.507893 | 1.5606516 |
| 367 | NM_003633.3 | ENC1 | 1693.45052 | -11.93633 | 1.61898653 |
| 368 | NM_020795.3 | #N/A | 1050.35349 | -11.247305 | 1.52638011 |
| 369 | NM_004501.3 | HNRNPU | 1032.9634 | -11.223177 | 1.52372207 |
| 370 | NM_152436.2 | GLIPR1L2 | 102.991316 | 4.27702516 | 0.58097235 |
| 371 | NM_005617.3 | #N/A | 1749.91972 | -11.983657 | 1.62803925 |
| 372 | NM_001273.3 | #N/A | 821.60378 | -10.893197 | 1.48011652 |
| 373 | NM_001207065.1 | #N/A | 106.82482 | 11.7112944 | 1.59135373 |
| 374 | NM_000701.7 | #N/A | 963.838126 | -11.123346 | 1.51254636 |
| 375 | NM_001286968.1 | #N/A | 1053.77893 | -4.0646951 | 0.55269867 |
| 376 | NM_001033026.1 | #N/A | 997.314271 | -11.172389 | 1.5196023 |
| 377 | NM_001023.3 | RPS20 | 1481.79971 | -11.743729 | 1.59761969 |
| 378 | NM_001969.4 | #N/A | 924.260129 | -11.062739 | 1.5055378 |
| 379 | NM_004356.3 | CD81 | 1170.28726 | -11.403316 | 1.55207517 |
| 380 | NM_000393.4 | #N/A | 1360.48297 | -11.620667 | 1.58230702 |
| 381 | NM_052925.3 | #N/A | 1004.52992 | -11.183302 | 1.52283813 |
| 382 | NM_017491.3 | #N/A | 924.225864 | -9.7072739 | 1.32209185 |
| 383 | NM_001136262.1 | #N/A | 1075.69883 | -11.281638 | 1.53689065 |
| 384 | NM_005517.3 | #N/A | 1333.44914 | -11.59146 | 1.57956645 |
| 385 | NM_170665.3 | #N/A | 1104.97671 | -11.320444 | 1.54300844 |
| 386 | NM_000987.3 | #N/A | 1679.46954 | -11.924382 | 1.62565111 |

| | | | | | |
|---|---|---|---|---|---|
| 387 | NM_001014809.2 | #N/A | 2149.31582 | -12.280304 | 1.67454365 |
| 388 | NM_006148.3 | #N/A | 982.657319 | -11.151213 | 1.5205719 |
| 389 | NM_206917.2 | #N/A | 1113.16241 | -11.331023 | 1.54578363 |
| 390 | NM_001313.4 | #N/A | 2083.3604 | -12.235284 | 1.67003909 |
| 391 | NM_006429.3 | #N/A | 1122.46548 | -11.342988 | 1.54851708 |
| 392 | NM_006373.3 | #N/A | 1307.58614 | -11.563394 | 1.57911864 |
| 393 | NM_004152.3 | OAZ1 | 1361.93124 | -11.622006 | 1.5873328 |
| 394 | NM_000484.3 | #N/A | 1108.04181 | -11.324759 | 1.54683041 |
| 395 | NM_005626.4 | #N/A | 417.2598 | -5.2387207 | 0.71571495 |
| 396 | NR_002940.2 | LRRC37A4P | 169.802 | -3.546183 | 0.48457461 |
| 397 | NM_005610.2 | #N/A | 960.385946 | -11.117913 | 1.51983913 |
| 398 | NM_005022.3 | #N/A | 1102.76957 | -11.317623 | 1.54772878 |
| 399 | NM_006054.3 | #N/A | 1655.84121 | -11.903943 | 1.62785306 |
| 400 | NM_001184735.1 | ASTN2 | 929.716472 | 3.82009426 | 0.5224576 |
| 401 | NM_021960.4 | #N/A | 829.10934 | -10.90623 | 1.49175021 |
| 402 | NM_144570.2 | #N/A | 367.251686 | -5.3630599 | 0.73356775 |
| 403 | NM_001130.5 | #N/A | 1412.05815 | -11.674116 | 1.59735199 |
| 404 | NM_002687.3 | #N/A | 749.265918 | -10.760418 | 1.47262156 |
| 405 | NM_197949.2 | #N/A | 136.649567 | 4.34947155 | 0.59530749 |
| 406 | NM_020675.3 | #N/A | 261.778946 | 2.78051499 | 0.38072748 |
| 407 | NM_006617.1 | #N/A | 1463.71863 | -10.30976 | 1.41250349 |
| 408 | NM_007184.3 | #N/A | 1001.20482 | -11.178239 | 1.53219785 |
| 409 | NM_000993.4 | #N/A | 1454.64571 | -11.717026 | 1.60613567 |
| 410 | NM_002743.3 | PRKCSH | 832.595806 | -9.5637806 | 1.31134067 |
| 411 | NM_004500.3 | HNRNPC | 1094.9108 | -11.306972 | 1.55067215 |
| 412 | NM_002948.3 | #N/A | 1591.07574 | -11.84636 | 1.62519495 |
| 413 | NM_006835.2 | #N/A | 1270.05757 | -11.521175 | 1.58085472 |
| 414 | NM_152293.2 | #N/A | 231.431289 | -5.065117 | 0.69532235 |
| 415 | NM_133259.3 | #N/A | 689.41967 | -5.2498989 | 0.72075503 |
| 416 | NM_004736.3 | #N/A | 1634.91778 | -11.885594 | 1.63238955 |
| 417 | NM_006559.2 | #N/A | 1111.05016 | -11.328191 | 1.55610795 |
| 418 | NM_018847.3 | #N/A | 360.766091 | -5.3622224 | 0.73702851 |
| 419 | NM_003400.3 | #N/A | 920.228711 | -11.056323 | 1.51985773 |
| 420 | NR_026712.1 | RPL13AP5 | 1480.86347 | -11.742796 | 1.61453195 |
| 421 | NM_006946.2 | #N/A | 896.068657 | -8.9714767 | 1.23363286 |
| 422 | NM_001013251.2 | #N/A | 921.195995 | -11.058163 | 1.52089315 |
| 423 | NM_002825.6 | #N/A | 773.9562 | -10.806777 | 1.48718414 |
| 424 | NM_152221.2 | #N/A | 1131.57501 | -11.35463 | 1.56268055 |
| 425 | NM_002626.5 | #N/A | 741.225276 | -10.744333 | 1.47926809 |
| 426 | NM_014766.4 | #N/A | 1149.21589 | -11.376916 | 1.56660695 |
| 427 | NM_001320965.1 | #N/A | 761.434155 | -10.782829 | 1.4851667 |
| 428 | NM_001028.2 | #N/A | 1242.05458 | -11.489105 | 1.58321613 |
| 429 | NM_001064.3 | #N/A | 1070.93691 | -11.275321 | 1.55423075 |

| | | | | | |
|---|---|---|---|---|---|
| 430 | NM_001145667.1 | #N/A | 906.096028 | -11.034144 | 1.52131971 |
| 431 | NM_004932.3 | #N/A | 433.703892 | -3.5790898 | 0.49346324 |
| 432 | NM_005066.2 | #N/A | 959.733809 | -11.116966 | 1.53302812 |
| 433 | NM_001018006.1 | #N/A | 1275.74137 | -11.527935 | 1.59008653 |
| 434 | NM_001970.4 | #N/A | 1156.77628 | -11.386464 | 1.57098547 |
| 435 | NM_002567.3 | #N/A | 1369.95865 | -11.630496 | 1.60479921 |
| 436 | NM_001759.3 | #N/A | 1173.54463 | -11.407114 | 1.57421025 |
| 437 | NM_001297654.1 | DDR1 | 741.475797 | -10.745383 | 1.48332512 |
| 438 | NM_001792.4 | #N/A | 1065.64036 | -11.268274 | 1.55554101 |
| 439 | NM_004484.3 | #N/A | 954.063019 | -11.108952 | 1.53347664 |
| 440 | NM_005742.3 | #N/A | 960.849701 | -11.118898 | 1.53476978 |
| 441 | NM_006029.4 | #N/A | 1035.33802 | -11.226465 | 1.54973167 |
| 442 | NM_001128204.1 | #N/A | 1007.1267 | -11.187105 | 1.54465985 |
| 443 | NM_032242.3 | PLXNA1 | 1326.58029 | -11.584121 | 1.59967985 |
| 444 | NM_003559.4 | PIP4K2B | 1222.64543 | -11.466398 | 1.5835427 |
| 445 | NM_001271938.1 | #N/A | 914.219914 | -11.046999 | 1.52575714 |
| 446 | NM_001205251.1 | ACSL6 | 55.9403818 | 10.7812288 | 1.48971884 |
| 448 | NM_006943.3 | SOX12 | 1001.40053 | -11.178335 | 1.54497707 |
| 447 | NM_004637.5 | #N/A | 911.897649 | -11.043426 | 1.52628283 |
| 449 | NM_001901.2 | #N/A | 1599.05404 | -11.853711 | 1.63889121 |
| 450 | NM_004924.5 | #N/A | 877.472441 | -10.988218 | 1.51965346 |
| 451 | NM_004995.3 | #N/A | 861.681385 | -10.962159 | 1.51656621 |
| 452 | NM_005809.5 | #N/A | 1096.41917 | -9.919688 | 1.37233904 |
| 453 | NM_001193416.2 | #N/A | 778.185186 | -10.814451 | 1.49642252 |
| 454 | NM_001018.4 | RPS15 | 1417.63005 | -11.67981 | 1.61626243 |
| 455 | NM_000976.3 | #N/A | 1314.00317 | -11.570359 | 1.60143889 |
| 456 | NM_030926.5 | #N/A | 797.516968 | -10.850139 | 1.50235646 |
| 459 | NM_005737.3 | ARL4C | 800.85539 | -10.856379 | 1.50354826 |
| 457 | NM_000989.3 | #N/A | 1270.49726 | -11.521711 | 1.5956717 |
| 458 | NM_004859.3 | #N/A | 1963.01041 | -6.3741963 | 0.88282246 |
| 460 | NM_016091.3 | #N/A | 1151.77796 | -11.380229 | 1.57632391 |
| 461 | NM_001297698.1 | #N/A | 54.7830768 | 10.7572061 | 1.49041618 |
| 462 | NM_033296.2 | MRFAP1 | 991.765643 | -11.164429 | 1.54746474 |
| 463 | NM_002901.2 | #N/A | 802.169923 | -10.858599 | 1.50576344 |
| 465 | NM_005318.3 | H1-0 | 1107.86415 | -9.9311619 | 1.3772394 |
| 464 | NM_005273.3 | #N/A | 1074.58727 | -11.280149 | 1.56438472 |
| 466 | NM_177433.2 | #N/A | 1150.86155 | -11.379238 | 1.57816308 |
| 467 | NR_003594.1 | REXO1L2P | 57.0951827 | 7.98160508 | 1.10705835 |
| 468 | NM_001009.3 | #N/A | 1721.8903 | -10.513251 | 1.45840289 |
| 469 | NM_005166.4 | APLP1 | 1531.44847 | -11.791317 | 1.63592577 |
| 470 | NM_001011.3 | #N/A | 1368.73199 | -11.629183 | 1.61367429 |
| 471 | NM_006083.3 | #N/A | 480.848042 | -7.8016437 | 1.08376913 |
| 472 | NM_001321483.1 | #N/A | 1403.18529 | -11.66505 | 1.62073394 |

| | | | | | |
|---|---|---|---|---|---|
| 473 | NM_007209.3 | #N/A | 1381.62487 | -11.642684 | 1.61764122 |
| 474 | NM_002415.1 | #N/A | 1285.90534 | -11.539087 | 1.60333192 |
| 475 | NM_002130.7 | #N/A | 1393.94194 | -11.655618 | 1.61983271 |
| 476 | NM_178819.3 | #N/A | 501.38395 | -7.577842 | 1.05312415 |
| 477 | NM_002734.4 | PRKAR1A | 694.099168 | -10.649785 | 1.48056084 |
| 478 | NM_006289.3 | #N/A | 762.415426 | -10.785682 | 1.49986691 |
| 479 | NR_002315.1 | H3P6 | 1194.02495 | -11.4322 | 1.5906732 |
| 480 | NM_024536.5 | #N/A | 712.019875 | -10.686835 | 1.48756589 |
| 481 | NM_000165.4 | #N/A | 825.236635 | -10.899858 | 1.51754022 |
| 482 | NM_002228.3 | #N/A | 729.32986 | -9.3763359 | 1.30544216 |
| 483 | NM_001142495.1 | #N/A | 912.969753 | -6.0472001 | 0.8419947 |
| 484 | NM_001904.3 | #N/A | 681.467876 | -10.622977 | 1.47924286 |
| 485 | NM_001849.3 | #N/A | 901.929513 | -11.02798 | 1.53594005 |
| 487 | NM_001780.5 | CD63 | 888.840727 | -11.006482 | 1.53363174 |
| 486 | NM_001155.4 | #N/A | 727.249256 | -10.717637 | 1.49340835 |
| 488 | NM_012090.5 | MACF1 | 924.103626 | -11.062544 | 1.54160254 |
| 489 | NR_028052.1 | NGRN | 1137.90939 | -11.36272 | 1.5835032 |
| 490 | NR_104643.1 | FLJ32255 | 45.4437525 | 3.99397595 | 0.55673629 |
| 491 | NM_016548.3 | #N/A | 917.971626 | -11.052922 | 1.54092953 |
| 492 | NM_001014.4 | #N/A | 1608.678 | -11.862234 | 1.65402824 |
| 493 | NM_144665.3 | #N/A | 842.156895 | -10.928672 | 1.52390185 |
| 494 | NM_178537.4 | #N/A | 1137.38806 | -4.2435629 | 0.59172766 |
| 495 | NM_138383.2 | #N/A | 814.42381 | -10.880085 | 1.51733591 |
| 496 | NM_001037333.2 | #N/A | 958.411259 | -11.115091 | 1.55031998 |
| 497 | NM_000996.2 | #N/A | 1079.62857 | -11.286843 | 1.57436751 |
| 498 | NR_034017.1 | LINC01502 | 116.857715 | 25.5737188 | 3.56754933 |
| 499 | NM_025219.2 | #N/A | 735.777065 | -9.3862937 | 1.30947766 |
| 500 | NM_001142277.1 | #N/A | 1117.46261 | -11.336925 | 1.58215108 |
| 501 | NM_031372.3 | HNRNPDL | 960.572883 | -11.118201 | 1.55176535 |
| 502 | NM_007375.3 | TARDBP | 588.287533 | -9.1342305 | 1.27494174 |
| 503 | NM_002635.3 | #N/A | 1046.02187 | -11.241299 | 1.56919755 |
| 504 | NM_001004.3 | #N/A | 1506.72198 | -11.767764 | 1.6428114 |
| 505 | NM_006015.4 | #N/A | 717.284369 | -10.696938 | 1.49346244 |
| 506 | NM_015161.2 | #N/A | 1073.0093 | -11.277952 | 1.57460167 |
| 507 | NM_030571.3 | #N/A | 1403.11787 | -10.237 | 1.42970162 |
| 509 | NM_005216.4 | DDOST | 862.733878 | -10.963524 | 1.53236605 |
| 508 | NM_001346792.1 | TSR2 | 62.1167425 | 24.6939215 | 3.45150424 |
| 510 | NM_006927.3 | #N/A | 286.059684 | -6.2388665 | 0.87200844 |
| 511 | NM_006454.2 | #N/A | 490.124913 | -7.96106 | 1.11287844 |
| 512 | NM_003186.3 | #N/A | 1803.4567 | -12.027221 | 1.68150811 |
| 513 | NM_021019.4 | #N/A | 1179.40236 | -11.414487 | 1.59611283 |
| 514 | NM_000451.3 | SHOX | 129.756602 | 4.35734643 | 0.60934206 |
| 515 | NM_173595.3 | #N/A | 491.156265 | -7.9639921 | 1.11377636 |

| | | | | | |
|---|---|---|---|---|---|
| 516 | NM_024911.6 | #N/A | 818.217431 | -10.887446 | 1.52288831 |
| 517 | NM_031263.3 | #N/A | 649.882588 | -10.554136 | 1.47662009 |
| 518 | NM_006026.3 | #N/A | 1558.15878 | -10.373496 | 1.45161484 |
| 519 | NM_001280.2 | CIRBP | 832.9225 | -10.91248 | 1.52793982 |
| 520 | NM_004539.3 | #N/A | 662.437319 | -9.3067406 | 1.30312993 |
| 521 | NM_005964.3 | #N/A | 1278.39371 | -10.112036 | 1.41577543 |
| 522 | NM_033643.2 | #N/A | 1047.18106 | -9.2093538 | 1.28938133 |
| 523 | NR_138054.1 | #N/A | 59.3877049 | 10.8642065 | 1.52117923 |
| 524 | NM_001293798.2 | DUX4 | 75.5003635 | 11.2255682 | 1.5722506 |
| 525 | NM_001008938.3 | #N/A | 847.869589 | -10.938323 | 1.5327461 |
| 526 | NM_006601.6 | #N/A | 1057.77943 | -11.2573 | 1.57735593 |
| 527 | NM_018243.3 | #N/A | 1179.2663 | -5.7355826 | 0.80370241 |
| 528 | NM_139323.3 | #N/A | 1292.88048 | -11.54696 | 1.61807234 |
| 529 | NM_145296.1 | #N/A | 835.185043 | -8.8721025 | 1.24333274 |
| 530 | NM_014390.3 | #N/A | 829.379123 | -10.906632 | 1.52970759 |
| 531 | NM_004040.3 | #N/A | 829.277909 | -5.8126291 | 0.81549986 |
| 532 | NM_005561.3 | #N/A | 1012.42514 | -11.194344 | 1.57063829 |
| 533 | NM_001257293.1 | #N/A | 763.416852 | -10.786577 | 1.51362846 |
| 534 | NM_005567.3 | #N/A | 746.541773 | -10.755253 | 1.50942517 |
| 535 | NM_003344.3 | #N/A | 783.835286 | -10.824949 | 1.51948978 |
| 536 | NM_000702.3 | #N/A | 298.677398 | -6.3000505 | 0.88436522 |
| 537 | NM_006825.3 | #N/A | 924.773028 | -11.063708 | 1.55334978 |
| 538 | NM_001016.3 | #N/A | 1439.56981 | -11.702022 | 1.64309095 |
| 539 | NM_005514.7 | #N/A | 822.279842 | -10.894345 | 1.52983445 |
| 540 | NM_002374.3 | MAP2 | 1435.60443 | -11.698025 | 1.64289739 |
| 541 | NM_004161.4 | #N/A | 825.692088 | -10.900063 | 1.5309184 |
| 542 | NM_000371.3 | TTR | 4050.86997 | -13.194674 | 1.85359451 |
| 543 | NM_001219.4 | #N/A | 861.391732 | -10.961385 | 1.54011387 |
| 544 | NM_001270366.1 | PLPPR3 | 1266.92241 | -11.517708 | 1.61883308 |
| 545 | NM_001655.4 | #N/A | 666.932766 | -10.592232 | 1.48870833 |
| 546 | NM_019106.5 | SEPTIN3 | 1641.05548 | -11.890984 | 1.67152955 |
| 547 | NM_001323310.1 | #N/A | 59.540472 | 24.6332469 | 3.46321213 |
| 548 | NM_001030.4 | #N/A | 1208.00179 | -11.448948 | 1.60971343 |
| 549 | NM_021149.3 | #N/A | 964.499332 | -11.124333 | 1.56416858 |
| 550 | NM_001009931.2 | HRNR | 52.6721875 | 10.6949923 | 1.50409244 |
| 551 | NM_001109662.3 | #N/A | 654.975602 | -10.565724 | 1.48615878 |
| 552 | NM_001762.3 | #N/A | 913.100207 | -11.045149 | 1.55378374 |
| 553 | NM_030801.3 | MAGED4B | 935.264097 | -11.079689 | 1.55874406 |
| 554 | NM_001012614.1 | #N/A | 753.496882 | -10.768013 | 1.51509756 |
| 555 | NM_000859.2 | #N/A | 1313.60624 | -11.569949 | 1.62844797 |
| 556 | NM_006908.4 | #N/A | 1020.01347 | -11.204964 | 1.57748733 |
| 557 | NM_004309.5 | #N/A | 1048.5217 | -11.244806 | 1.58342087 |
| 558 | NM_005898.4 | #N/A | 830.864166 | -10.90899 | 1.53620008 |

| | | | | | |
|---|---|---|---|---|---|
| 559 | NM_006819.2 | #N/A | 794.901665 | -10.845335 | 1.52764097 |
| 560 | NM_002081.2 | #N/A | 1111.40123 | -11.328752 | 1.59596169 |
| 561 | NM_006201.4 | #N/A | 782.073094 | -10.821907 | 1.52452351 |
| 563 | NM_001330700.1 | TOP2B | 812.42158 | -10.876505 | 1.53281853 |
| 562 | NM_001110781.2 | #N/A | 739.248619 | -9.3841076 | 1.32248862 |
| 564 | NM_012096.2 | #N/A | 243.222944 | -4.6376154 | 0.6536335 |
| 565 | NM_031157.3 | #N/A | 871.341125 | -10.977617 | 1.54721391 |
| 566 | NR_102755.1 | FLJ31104 | 94.1392862 | 3.38099147 | 0.47669552 |
| 567 | NM_006013.4 | RPL10 | 1213.52595 | -11.455461 | 1.615289 |
| 568 | NM_015295.2 | SMCHD1 | 342.092096 | -5.2643945 | 0.74230953 |
| 569 | NM_001527.3 | #N/A | 1035.02136 | -11.225966 | 1.58306863 |
| 570 | NM_001291861.2 | #N/A | 753.473699 | -9.409129 | 1.32710056 |
| 571 | NM_002271.4 | #N/A | 812.502101 | -10.876737 | 1.53447692 |
| 572 | NM_002160.3 | #N/A | 992.022685 | -5.510719 | 0.77762691 |
| 573 | NM_001694.3 | #N/A | 1106.66634 | -11.322651 | 1.59812011 |
| 574 | NM_018247.3 | #N/A | 807.634694 | -10.868265 | 1.53407835 |
| 575 | NM_004530.5 | #N/A | 752.858599 | -10.767373 | 1.51997573 |
| 576 | NM_170679.2 | #N/A | 1021.70541 | -11.207347 | 1.58269318 |
| 577 | NM_139072.3 | #N/A | 720.306701 | -8.1359424 | 1.14907067 |
| 578 | NM_014739.2 | #N/A | 651.291819 | -10.557648 | 1.49142653 |
| 579 | NM_001278116.1 | #N/A | 1571.19524 | -11.828257 | 1.6715117 |
| 580 | NM_018407.4 | #N/A | 950.209438 | -9.7127169 | 1.37283658 |
| 582 | NR_046687.1 | KLHL6-AS1 | 76.1844706 | 4.89999332 | 0.6926578 |
| 581 | NM_021643.3 | TRIB2 | 672.712279 | -10.604686 | 1.49906136 |
| 583 | NM_000942.4 | PPIB | 902.642273 | -11.028784 | 1.55922232 |
| 584 | NM_000984.5 | #N/A | 1055.31493 | -11.254111 | 1.59128518 |
| 585 | NM_001033044.3 | #N/A | 571.3223 | -10.368738 | 1.46609835 |
| 586 | NM_003768.4 | #N/A | 1470.41251 | -11.732602 | 1.65950841 |
| 587 | NM_001537.3 | #N/A | 794.594946 | -10.844697 | 1.53428846 |
| 588 | NM_016424.4 | #N/A | 618.135318 | -10.482363 | 1.48319081 |
| 589 | NM_181689.2 | #N/A | 909.567474 | -11.039485 | 1.56204263 |
| 590 | NM_004339.3 | #N/A | 976.993079 | -11.142877 | 1.57763913 |
| 591 | NM_003129.3 | #N/A | 1020.21217 | -11.205278 | 1.58678972 |
| 592 | NM_001920.4 | #N/A | 431.892926 | -5.7631881 | 0.81634318 |
| 593 | NM_002373.5 | #N/A | 880.70318 | -10.99323 | 1.55765264 |
| 594 | NM_000389.4 | #N/A | 768.872303 | -10.797382 | 1.53016241 |
| 595 | NM_207318.3 | #N/A | 356.121203 | -5.7582479 | 0.81635275 |
| 596 | NM_001195532.1 | #N/A | 702.100831 | -10.66681 | 1.51250735 |
| 597 | NM_001099294.1 | #N/A | 222.078772 | -5.5651228 | 0.78921594 |
| 598 | NM_004747.3 | #N/A | 697.53686 | -10.656833 | 1.51132481 |
| 599 | NM_001018108.3 | #N/A | 704.650386 | -10.671447 | 1.51377007 |
| 600 | NM_001101417.3 | #N/A | 55.8191925 | 24.5460604 | 3.48207149 |
| 601 | NM_004807.2 | #N/A | 853.373615 | -10.947579 | 1.55322175 |

| | | | | | | |
|---|---|---|---|---|---|---|
| 602 | NR_126166.1 | FAM74A7 | | 117.047997 | 2.90279422 | 0.41187557 |
| 603 | NR_033320.2 | MIAT | | 826.233435 | -10.900915 | 1.54711162 |
| 604 | NM_000362.4 | TIMP3 | | 700.013847 | -10.662472 | 1.5137479 |
| 605 | NM_005730.3 | | #N/A | 601.496476 | -10.443316 | 1.48314578 |
| 606 | NM_178031.2 | | #N/A | 879.657595 | -10.991406 | 1.56102263 |
| 607 | NM_001322271.1 | | #N/A | 55.1764524 | 10.7711844 | 1.52994827 |
| 608 | NM_004640.6 | | #N/A | 590.142649 | -10.414947 | 1.48014184 |
| 609 | NM_006667.4 | | #N/A | 916.915758 | -11.051227 | 1.57064481 |
| 610 | NM_004368.3 | | #N/A | 748.984317 | -10.759572 | 1.52948823 |
| 611 | NM_014335.2 | | #N/A | 915.38489 | -11.04878 | 1.5712979 |
| 612 | NM_002516.3 | | #N/A | 518.081952 | -6.1980524 | 0.8814977 |
| 614 | NM_003133.5 | SRP9 | | 983.447113 | -11.152244 | 1.58703169 |
| 613 | NM_001320214.1 | | #N/A | 547.951876 | -10.308236 | 1.46687164 |
| 615 | NM_006184.5 | | #N/A | 703.154573 | -10.66863 | 1.51816347 |
| 616 | NM_001908.4 | | #N/A | 835.790065 | -10.91779 | 1.55382545 |
| 617 | NM_152409.3 | C5orf24 | | 90.0395196 | 11.4792035 | 1.63403707 |
| 618 | NM_001797.3 | | #N/A | 889.734823 | -11.008122 | 1.56714092 |
| 619 | NM_005918.3 | | #N/A | 673.45254 | -10.605981 | 1.50985478 |
| 620 | NM_020123.3 | | #N/A | 720.779968 | -10.704015 | 1.52388724 |
| 621 | NM_032221.4 | | #N/A | 290.921738 | -6.5484743 | 0.9322999 |
| 622 | NM_017590.5 | | #N/A | 835.176057 | -10.916562 | 1.55429136 |
| 623 | NM_014765.2 | | #N/A | 978.860258 | -11.145454 | 1.58707764 |
| 624 | NM_183236.2 | | #N/A | 54.3879312 | 24.5072227 | 3.49001822 |
| 625 | NM_020532.4 | | #N/A | 1411.23277 | -11.673332 | 1.6624668 |
| 626 | NM_001010858.2 | | #N/A | 1067.66813 | -11.270793 | 1.60526641 |
| 627 | NM_001012478.1 | U2AF2 | | 712.678975 | -10.687667 | 1.52245008 |
| 628 | NM_001958.3 | | #N/A | 1540.59169 | -11.799844 | 1.68109601 |
| 629 | NM_002951.4 | | #N/A | 748.227266 | -10.75817 | 1.53297333 |
| 630 | NM_004462.4 | | #N/A | 1286.08504 | -11.539366 | 1.64447788 |
| 631 | NM_005993.4 | | #N/A | 599.108598 | -10.437284 | 1.48742136 |
| 632 | NM_006431.2 | | #N/A | 982.750772 | -11.151216 | 1.58901568 |
| 633 | NM_014938.5 | | #N/A | 447.656881 | -3.4217131 | 0.48762809 |
| 634 | NM_001017392.4 | | #N/A | 568.022147 | -10.360726 | 1.47698539 |
| 637 | NM_206876.1 | PPP1CB | | 465.960184 | -6.4671129 | 0.9221533 |
| 635 | NM_001154.3 | | #N/A | 739.692258 | -10.741744 | 1.53168467 |
| 636 | NM_013291.2 | | #N/A | 699.766279 | -10.661377 | 1.52026774 |
| 638 | NM_003900.4 | | #N/A | 735.899659 | -10.734355 | 1.53084082 |
| 639 | NM_014232.2 | | #N/A | 918.355434 | -9.0191507 | 1.28619889 |
| 640 | NR_073534.1 | | #N/A | 53.8154266 | 24.4949544 | 3.49331524 |
| 641 | NM_006496.3 | | #N/A | 479.162584 | -8.0463765 | 1.1476747 |
| 643 | NM_017644.3 | KLHL24 | | 221.671439 | -5.8724061 | 0.83765139 |
| 642 | NM_002631.3 | | #N/A | 917.526951 | -11.052244 | 1.57648518 |
| 644 | NM_002402.3 | | #N/A | 1035.17196 | -11.226571 | 1.60171629 |

| | | | | | |
|---|---|---|---|---|---|
| 645 | NM_003644.2 | #N/A | 61.2843474 | 10.9083637 | 1.55630085 |
| 646 | NR_040039.1 | LOC157273 | 114.040563 | 4.24886206 | 0.60639203 |
| 647 | NM_004643.3 | PABPN1 | 603.058994 | -10.446994 | 1.49128228 |
| 648 | NM_015710.4 | #N/A | 812.910136 | -10.877457 | 1.55284876 |
| 649 | NM_001533.2 | HNRNPL | 830.194927 | -10.907934 | 1.55739979 |
| 650 | NM_001618.3 | #N/A | 762.683508 | -10.785385 | 1.5400236 |
| 651 | NM_001494.3 | #N/A | 906.359327 | -11.034535 | 1.57569153 |
| 652 | NM_021648.4 | #N/A | 714.621564 | -9.4181549 | 1.34543602 |
| 653 | NM_015059.2 | TLN2 | 269.621976 | -5.0217312 | 0.71747517 |
| 654 | NM_001164097.1 | VCAN | 1119.49616 | -11.339289 | 1.62018618 |
| 655 | NM_004800.2 | #N/A | 642.447585 | -10.538057 | 1.50578357 |
| 656 | NM_001195553.1 | #N/A | 1505.59848 | -11.766728 | 1.68158846 |
| 657 | NM_001318188.1 | HNRNPK | 621.603305 | -10.489855 | 1.49952321 |
| 658 | NM_018128.4 | #N/A | 236.615228 | -5.2045512 | 0.74400568 |
| 659 | NM_001142423.1 | MORF4L2 | 675.415599 | -10.610251 | 1.51783059 |
| 660 | NM_001300995.1 | NOX4 | 55.5470061 | 10.7675922 | 1.54043371 |
| 661 | NM_000358.2 | #N/A | 1232.72293 | -8.7937504 | 1.25830195 |
| 662 | NM_139215.2 | #N/A | 656.727766 | -10.56984 | 1.51256378 |
| 663 | NM_006644.3 | #N/A | 621.328436 | -10.489627 | 1.50149641 |
| 664 | NM_003146.2 | #N/A | 733.795418 | -10.729759 | 1.53599553 |
| 665 | NR_033966.1 | LINC01761 | 71.4561194 | 3.33968186 | 0.47819611 |
| 666 | NM_001301047.2 | #N/A | 57.9792655 | 10.8441066 | 1.55302093 |
| 667 | NM_032737.3 | #N/A | 683.984764 | -10.628363 | 1.5221515 |
| 668 | NM_001089.2 | #N/A | 792.574876 | -9.5687176 | 1.37055822 |
| 669 | NM_017758.3 | #N/A | 455.239271 | -8.7632078 | 1.25519852 |
| 670 | NM_000988.3 | #N/A | 1381.87018 | -11.642976 | 1.66799818 |
| 671 | NM_003097.4 | #N/A | 939.5143 | -11.086331 | 1.5883524 |
| 673 | NM_194247.3 | HNRNPA3 | 758.927786 | -10.778404 | 1.54462093 |
| 672 | NM_080840.2 | PTPRA | 594.225702 | -10.425239 | 1.49405144 |
| 674 | NM_012094.4 | #N/A | 818.601177 | -10.887555 | 1.56086714 |
| 675 | NM_001001701.3 | #N/A | 343.308867 | -7.5479593 | 1.08215749 |
| 676 | NM_005721.4 | #N/A | 722.256903 | -10.706939 | 1.5351017 |
| 677 | NM_001204077.1 | #N/A | 274.844839 | -4.6266686 | 0.66344403 |
| 678 | NM_003681.4 | #N/A | 558.483367 | -10.336148 | 1.48216238 |
| 679 | NM_206852.2 | RTN1 | 1453.99339 | -11.7164 | 1.68019671 |
| 680 | NM_002741.3 | #N/A | 732.169544 | -9.3568734 | 1.34204119 |
| 681 | NM_018023.4 | #N/A | 427.21789 | -7.7618824 | 1.11324304 |
| 682 | NM_019020.3 | #N/A | 752.226917 | -10.765664 | 1.54425433 |
| 683 | NM_001098.2 | #N/A | 601.504776 | -10.443224 | 1.49821557 |
| 684 | NM_002154.3 | #N/A | 675.723463 | -10.610977 | 1.52251601 |
| 685 | NM_000302.3 | #N/A | 639.950524 | -10.53283 | 1.51181157 |
| 686 | NM_006621.5 | #N/A | 538.989051 | -10.284966 | 1.47639096 |
| 687 | NM_015447.3 | #N/A | 645.167658 | -9.2698447 | 1.3306593 |

| | | | | | | |
|---|---|---|---|---|---|---|
| 688 | NM_018638.4 | | #N/A | 390.718131 | -6.3850739 | 0.91654343 |
| 689 | NM_030752.2 | | #N/A | 953.035185 | -11.106937 | 1.59454734 |
| 690 | NM_005552.4 | KLC1 | | 1386.79956 | -11.648129 | 1.67286636 |
| 691 | NM_005998.4 | | #N/A | 877.422656 | -10.987633 | 1.5780088 |
| 692 | NM_001009184.1 | | #N/A | 707.172384 | -10.676592 | 1.53359448 |
| 693 | NM_013436.4 | | #N/A | 604.711034 | -10.450084 | 1.50117399 |
| 694 | NM_018206.5 | | #N/A | 651.211505 | -10.557708 | 1.51692624 |
| 695 | NM_001844.4 | | #N/A | 936.645309 | -11.082172 | 1.59249617 |
| 696 | NM_014730.3 | | #N/A | 688.33314 | -10.637674 | 1.52876396 |
| 697 | NM_001305105.1 | | #N/A | 895.476011 | -11.017069 | 1.58347512 |
| 698 | NM_014718.3 | | #N/A | 902.177755 | -9.6299594 | 1.38425897 |
| 699 | NM_030649.2 | | #N/A | 953.820906 | -11.108118 | 1.59687778 |
| 700 | NM_033161.3 | | #N/A | 718.85928 | -10.700289 | 1.53833508 |
| 701 | NM_002391.4 | | #N/A | 769.375381 | -10.798297 | 1.55249882 |
| 702 | NM_018675.2 | | #N/A | 3569.31064 | 6.38138053 | 0.91758195 |
| 703 | NM_002139.3 | | #N/A | 670.276321 | -10.599047 | 1.5245518 |
| 704 | NM_017994.4 | | #N/A | 338.883769 | -5.3883118 | 0.7754705 |
| 705 | NM_203499.2 | | #N/A | 465.017621 | -8.794414 | 1.26579047 |
| 706 | NM_002293.3 | | #N/A | 617.82827 | -10.482237 | 1.50896722 |
| 707 | NM_002874.4 | | #N/A | 747.276373 | -10.756008 | 1.54849901 |
| 708 | NM_001127228.1 | | #N/A | 756.924461 | -10.774553 | 1.55132596 |
| 709 | NM_002868.3 | | #N/A | 631.847731 | -10.513924 | 1.51377301 |
| 710 | NM_016185.3 | | #N/A | 1401.08327 | -11.662889 | 1.67919382 |
| 711 | NM_001271.3 | | #N/A | 549.805027 | -10.31312 | 1.48496184 |
| 712 | NM_002821.4 | | #N/A | 632.019557 | -10.514482 | 1.51400707 |
| 713 | NM_006430.3 | | #N/A | 787.934751 | -10.832413 | 1.56072896 |
| 714 | NM_001126340.2 | | #N/A | 529.256218 | -10.258931 | 1.47824883 |
| 715 | NM_006836.1 | | #N/A | 588.790102 | -10.412165 | 1.50040408 |
| 716 | NM_001098272.2 | | #N/A | 1093.27352 | -11.304909 | 1.62916402 |
| 717 | NM_002270.3 | TNPO1 | | 723.215516 | -10.708671 | 1.54374398 |
| 718 | NM_003036.3 | | #N/A | 454.436901 | -5.3059528 | 0.76488666 |
| 719 | NM_018926.2 | PCDHGB6 | | 670.064996 | -10.598924 | 1.52806277 |
| 720 | NM_001199780.1 | PSMB2 | | 108.828017 | 9.83504097 | 1.41801275 |
| 721 | NM_003089.5 | | #N/A | 616.611981 | -10.478377 | 1.51083072 |
| 722 | NM_015338.5 | | #N/A | 561.448054 | -10.343406 | 1.49137345 |
| 723 | NM_002107.4 | | #N/A | 1016.32199 | -11.199566 | 1.61512012 |
| 724 | NM_001080424.1 | | #N/A | 648.849759 | -9.1929288 | 1.32623436 |
| 725 | NM_021029.5 | | #N/A | 1101.85092 | -11.316238 | 1.63261559 |
| 726 | NM_021228.2 | | #N/A | 681.195936 | -5.5553056 | 0.80160602 |
| 727 | NM_014141.5 | | #N/A | 702.247818 | -7.0306589 | 1.01478363 |
| 728 | NM_004218.3 | | #N/A | 719.234591 | -10.700912 | 1.54524859 |
| 729 | NM_004494.2 | | #N/A | 594.143009 | -10.424912 | 1.50543487 |
| 730 | NM_001606.4 | | #N/A | 575.311031 | -10.378858 | 1.49908264 |

| | | | | | |
|---|---|---|---|---|---|
| 731 | NM_003134.5 | #N/A | 764.642365 | -10.789242 | 1.55843195 |
| 732 | NM_001278742.1 | TSPAN6 | 48.1395649 | 10.5682572 | 1.5265684 |
| 733 | NM_001101662.1 | NRDC | 603.987162 | -10.448971 | 1.50953929 |
| 734 | NM_002862.3 | #N/A | 792.385454 | -9.4549385 | 1.36591818 |
| 735 | NM_012469.3 | #N/A | 660.625798 | -10.578182 | 1.52849362 |
| 736 | NM_001183.5 | #N/A | 864.153577 | -10.965823 | 1.58474902 |
| 737 | NM_001386.5 | #N/A | 1035.12216 | -11.226163 | 1.62300837 |
| 738 | NM_006815.3 | #N/A | 663.050874 | -10.583533 | 1.53030314 |
| 739 | NM_018975.3 | #N/A | 881.606657 | -10.994504 | 1.58975336 |
| 740 | NM_003286.3 | #N/A | 563.748961 | -10.349447 | 1.49658355 |
| 741 | NM_005642.2 | #N/A | 618.707571 | -10.48352 | 1.5163851 |
| 742 | NM_001030009.1 | #N/A | 1254.75322 | -11.503753 | 1.66425809 |
| 743 | NM_000999.3 | #N/A | 1005.02043 | -11.183551 | 1.61860155 |
| 744 | NM_024408.3 | #N/A | 509.208821 | -10.203624 | 1.47675054 |
| 745 | NM_014868.4 | #N/A | 678.057661 | -10.615799 | 1.53706203 |
| 746 | NM_001025368.2 | #N/A | 797.290763 | -10.849545 | 1.57103067 |
| 747 | NM_004990.3 | #N/A | 811.029222 | -10.874182 | 1.57479145 |
| 748 | NM_206926.1 | #N/A | 600.138856 | -10.440197 | 1.51208058 |
| 749 | NM_001304275.1 | RAPGEF1 | 616.667919 | -10.478882 | 1.51776547 |
| 750 | NM_001069.2 | #N/A | 2992.20587 | -7.4672752 | 1.08175905 |
| 751 | NM_021980.4 | OPTN | 338.742052 | -6.8679398 | 0.99514829 |
| 752 | NM_000997.4 | #N/A | 1053.48701 | -11.251539 | 1.63054826 |
| 753 | NM_001659.2 | #N/A | 836.390504 | -10.918572 | 1.58229004 |
| 754 | NM_014607.3 | #N/A | 511.115367 | -10.20825 | 1.47930845 |
| 755 | NM_000551.3 | VHL | 637.993425 | -10.527618 | 1.52574475 |
| 756 | NM_033446.2 | #N/A | 445.7453 | -5.9633373 | 0.86429599 |
| 757 | NM_002715.2 | #N/A | 792.084667 | -10.840052 | 1.57134037 |
| 758 | NM_053056.2 | #N/A | 631.945672 | -10.514894 | 1.52443884 |
| 761 | NM_203504.2 | G3BP2 | 681.367996 | -10.622733 | 1.54038141 |
| 759 | NM_002745.4 | MAPK1 | 602.329429 | -10.444936 | 1.51458847 |
| 760 | NM_003072.3 | #N/A | 542.229248 | -10.293376 | 1.49257883 |
| 762 | NM_018036.6 | #N/A | 163.873696 | -4.5874428 | 0.66526139 |
| 763 | NR_134873.1 | LOC100419170 | 198.865806 | 3.10458841 | 0.4503469 |
| 764 | NM_006854.3 | #N/A | 653.105599 | -10.561878 | 1.53214162 |
| 765 | NM_014713.4 | #N/A | 647.859066 | -10.550331 | 1.53054331 |
| 766 | NM_148174.3 | #N/A | 824.142657 | -10.897271 | 1.58083797 |
| 767 | NM_012325.2 | #N/A | 720.437263 | -10.703249 | 1.55278627 |
| 768 | NM_003769.2 | #N/A | 590.253807 | -10.415608 | 1.51130352 |
| 769 | NR_135154.1 | #N/A | 527.513935 | -10.253661 | 1.48797467 |
| 770 | NM_152862.2 | #N/A | 793.282587 | -10.842267 | 1.57394504 |
| 771 | NM_013974.2 | #N/A | 801.307955 | -10.856713 | 1.57613707 |
| 772 | NM_002586.4 | #N/A | 472.397642 | -8.7705094 | 1.27382975 |
| 773 | NM_006870.3 | #N/A | 961.336776 | -11.119462 | 1.61499475 |

| | | | | | |
|---|---|---|---|---|---|
| 774 | NM_000445.4 | #N/A | 646.56851 | -10.547398 | 1.53202954 |
| 775 | NM_017797.3 | #N/A | 636.681435 | -10.524886 | 1.5287302 |
| 776 | NM_002603.3 | #N/A | 57.7299009 | 10.8219888 | 1.57211506 |
| 777 | NM_002809.3 | #N/A | 677.10366 | -10.613836 | 1.54229276 |
| 778 | NM_198475.2 | #N/A | 795.477043 | -10.846121 | 1.57602637 |
| 779 | NM_181697.2 | #N/A | 712.455476 | -10.687267 | 1.55306966 |
| 780 | NM_024718.4 | #N/A | 676.829781 | -10.613043 | 1.54271157 |
| 781 | NM_001321743.1 | FAM168B | 194.218638 | -3.8707352 | 0.5627849 |
| 782 | NM_002539.2 | #N/A | 1163.56611 | -11.394785 | 1.65670259 |
| 783 | NM_006141.2 | #N/A | 656.51096 | -10.569301 | 1.5371043 |
| 784 | NM_006391.2 | #N/A | 678.981105 | -10.617704 | 1.54412932 |
| 785 | NM_006924.4 | #N/A | 718.424018 | -10.699186 | 1.55593192 |
| 786 | NM_018011.3 | #N/A | 509.223692 | -10.202573 | 1.48403984 |
| 787 | NM_001129.4 | #N/A | 561.309323 | -10.344178 | 1.50476583 |
| 788 | NM_000018.3 | #N/A | 588.600788 | -10.41223 | 1.51495288 |
| 789 | NM_025081.2 | #N/A | 489.614664 | -10.146959 | 1.4766654 |
| 790 | NM_002899.3 | #N/A | 740.958605 | -10.744137 | 1.56398285 |
| 791 | NM_001688.4 | #N/A | 456.096647 | -6.0406088 | 0.87937121 |
| 792 | NM_014972.2 | #N/A | 582.822439 | -10.397648 | 1.51364901 |
| 793 | NM_003816.2 | #N/A | 642.598353 | -10.538706 | 1.53439097 |
| 794 | NM_002819.4 | #N/A | 560.811474 | -10.342162 | 1.50613258 |
| 795 | NM_001568.2 | #N/A | 761.910857 | -10.783961 | 1.57065354 |
| 796 | NM_080677.2 | #N/A | 768.655262 | -10.796863 | 1.57306537 |
| 797 | NM_001080432.2 | #N/A | 357.207743 | -6.5491485 | 0.9542793 |
| 798 | NM_012218.3 | #N/A | 652.098843 | -10.559582 | 1.53884945 |
| 799 | NM_001128619.2 | #N/A | 782.643618 | -10.822755 | 1.57735838 |
| 800 | NM_002086.4 | #N/A | 782.01528 | -10.821598 | 1.57767176 |
| 801 | NM_007065.3 | #N/A | 633.548643 | -10.517932 | 1.53343123 |
| 802 | NM_013271.4 | #N/A | 981.302205 | -11.149175 | 1.62595391 |
| 803 | NM_001320547.1 | WDR6 | 527.781996 | -10.254939 | 1.49568257 |
| 804 | NM_004930.4 | #N/A | 720.402472 | -10.703267 | 1.5612276 |
| 805 | NM_001015.4 | #N/A | 2923.8325 | -8.5753161 | 1.25088064 |
| 806 | NR_002206.3 | GTF2IP1 | 841.615129 | -10.927519 | 1.59465705 |
| 807 | NM_006904.6 | #N/A | 568.82545 | -10.362349 | 1.5124036 |
| 808 | NM_001111020.2 | #N/A | 497.607857 | -10.169272 | 1.48480812 |
| 809 | NM_007278.1 | #N/A | 804.781785 | -10.86313 | 1.58616336 |
| 810 | NM_001305275.1 | #N/A | 845.980026 | -10.934947 | 1.59691111 |
| 811 | NM_007040.5 | #N/A | 510.898624 | -10.207202 | 1.49064683 |
| 812 | NM_001199085.1 | #N/A | 46.0866154 | 24.2799353 | 3.54592068 |
| 813 | NM_017811.3 | #N/A | 577.732789 | -10.384722 | 1.51675227 |
| 814 | NM_004719.2 | #N/A | 267.772593 | -7.1795664 | 1.04885227 |
| 815 | NM_013236.3 | #N/A | 755.602114 | -10.772017 | 1.57392927 |
| 816 | NM_018031.4 | #N/A | 521.384403 | -10.237524 | 1.49589015 |

| | | | | | |
|---|---|---|---|---|---|
| 822 | NR_103797.1 | SON | 495.040377 | -10.162001 | 1.48500178 |
| 817 | NM_001102.3 | #N/A | 922.567358 | -11.060286 | 1.6164377 |
| 818 | NM_001540.3 | #N/A | 595.386198 | -10.428565 | 1.52412099 |
| 819 | NM_006310.3 | #N/A | 577.480582 | -10.384132 | 1.51757572 |
| 820 | NM_015013.3 | #N/A | 578.17046 | -9.1115353 | 1.33151504 |
| 821 | NM_018448.4 | #N/A | 729.081634 | -10.720414 | 1.5666813 |
| 823 | NM_198390.2 | CMIP | 685.004864 | -10.630448 | 1.55421082 |
| 824 | NM_004331.2 | #N/A | 732.228157 | -10.726388 | 1.56851283 |
| 825 | NM_001286366.1 | #N/A | 904.836274 | -11.032185 | 1.61329437 |
| 826 | NR_003249.2 | HNRNPDL | 551.647588 | -10.317787 | 1.50889316 |
| 827 | NM_032632.4 | PAPOLA | 549.058604 | -10.311184 | 1.50814179 |
| 828 | NM_007359.4 | #N/A | 566.333746 | -10.355935 | 1.51487491 |
| 829 | NM_015980.4 | #N/A | 1484.88752 | -11.746702 | 1.71844721 |
| 830 | NM_015526.2 | #N/A | 878.000815 | -10.98862 | 1.6080345 |
| 831 | NM_012181.4 | #N/A | 702.023492 | -10.665988 | 1.5610304 |
| 832 | NM_002660.2 | #N/A | 667.830417 | -10.594015 | 1.55065594 |
| 833 | NM_006801.2 | #N/A | 564.546301 | -10.351857 | 1.51529694 |
| 834 | NM_030662.3 | MAP2K2 | 630.510174 | -10.511021 | 1.53893413 |
| 835 | NM_005051.2 | #N/A | 493.784179 | -10.158766 | 1.48786951 |
| 836 | NM_001320967.1 | #N/A | 528.296651 | -10.256745 | 1.50243763 |
| 837 | NM_001136135.1 | #N/A | 763.428981 | -10.786944 | 1.58024187 |
| 838 | NR_033902.1 | #N/A | 603.070806 | -10.446738 | 1.53047458 |
| 839 | NM_001102420.2 | #N/A | 616.340766 | -10.477987 | 1.53526519 |
| 840 | NM_001193370.1 | #N/A | 46.3623417 | 10.5144362 | 1.54058371 |
| 841 | NM_001660.3 | #N/A | 780.401944 | -10.818641 | 1.58520802 |
| 842 | NR_023312.2 | #N/A | 507.1473 | -10.19601 | 1.49418361 |
| 843 | NR_027001.1 | GOLGA2P7 | 496.590971 | -10.166284 | 1.49027392 |
| 844 | NR_037946.1 | HNRNPUL2-BSCL2 | 509.642459 | -10.204172 | 1.4961678 |
| 845 | NM_001394.6 | #N/A | 987.379423 | -11.158143 | 1.63617377 |
| 846 | NM_001164319.1 | #N/A | 583.388827 | -10.399559 | 1.52509542 |
| 847 | NM_183422.3 | #N/A | 475.045127 | -10.103321 | 1.48186378 |
| 848 | NM_006224.3 | #N/A | 554.183586 | -10.32498 | 1.51443722 |
| 849 | NM_003836.6 | DLK1 | 1104.59015 | -11.320142 | 1.6606063 |
| 850 | NM_153649.3 | #N/A | 544.847544 | -10.299953 | 1.51102561 |
| 851 | NM_000938.2 | #N/A | 593.324411 | -10.423188 | 1.52919084 |
| 852 | NM_080426.3 | #N/A | 580.745521 | -10.392594 | 1.52501853 |
| 853 | NM_006227.3 | #N/A | 553.544867 | -10.32403 | 1.5150132 |
| 854 | NM_053064.4 | #N/A | 1017.96805 | -11.201943 | 1.64391971 |
| 856 | NM_014325.3 | CORO1C | 883.269804 | -10.997214 | 1.6139612 |
| 855 | NM_003079.4 | #N/A | 563.996919 | -7.8704497 | 1.15505789 |
| 857 | NM_001012241.1 | #N/A | 586.673343 | -10.406682 | 1.52733607 |
| 858 | NM_016176.3 | #N/A | 590.849097 | -10.417376 | 1.52930035 |
| 859 | NM_022098.3 | #N/A | 579.936632 | 2.05828531 | 0.30219236 |

| | | | | | | |
|---|---|---|---|---|---|---|
| 860 | NM_002254.6 | | #N/A | 1155.9955 | -11.385485 | 1.67217665 |
| 861 | NM_003074.3 | | #N/A | 577.25889 | -10.383544 | 1.52492994 |
| 862 | NM_003170.4 | | #N/A | 478.114258 | -10.112538 | 1.48518915 |
| 863 | NM_138615.2 | | #N/A | 465.960801 | -7.8639577 | 1.15495037 |
| 864 | NM_001031.4 | | #N/A | 896.006809 | -11.017945 | 1.61844241 |
| 865 | NM_001693.3 | | #N/A | 725.335529 | -10.713091 | 1.57393528 |
| 866 | NM_017620.2 | ILF3 | | 490.365432 | -10.14824 | 1.49118113 |
| 867 | NM_001005332.1 | | #N/A | 640.239942 | -5.3998734 | 0.79357873 |
| 868 | NM_003142.4 | | #N/A | 525.966076 | -10.249344 | 1.50629252 |
| 869 | NM_012268.3 | | #N/A | 615.932264 | -10.477085 | 1.53968834 |
| 870 | NM_022574.4 | | #N/A | 479.64019 | -10.116456 | 1.48675215 |
| 871 | NM_002037.5 | FYN | | 698.63756 | -10.658919 | 1.56657831 |
| 872 | NM_005968.4 | | #N/A | 550.300532 | -10.314767 | 1.51602694 |
| 873 | NM_003017.4 | | #N/A | 705.561001 | -10.673043 | 1.56887524 |
| 874 | NM_004427.3 | | #N/A | 486.57183 | -10.136984 | 1.4901174 |
| 875 | NM_004168.3 | | #N/A | 435.415994 | -9.9767713 | 1.46679072 |
| 876 | NM_020117.10 | | #N/A | 528.561011 | -10.256122 | 1.50802563 |
| 877 | NM_005385.3 | | #N/A | 451.400956 | -10.028524 | 1.47465808 |
| 878 | NM_080391.3 | | #N/A | 557.912686 | -10.334339 | 1.51977725 |
| 879 | NM_003405.3 | | #N/A | 982.791348 | -11.15132 | 1.63999059 |
| 880 | NM_001997.4 | | #N/A | 982.048361 | -11.150207 | 1.64050247 |
| 881 | NM_003062.3 | | #N/A | 505.160312 | -5.4830136 | 0.80675105 |
| 882 | NM_004515.3 | | #N/A | 729.381409 | -10.721093 | 1.57806176 |
| 883 | NM_006585.3 | | #N/A | 676.37158 | -10.612254 | 1.56229751 |
| 884 | NM_015894.3 | | #N/A | 957.90059 | -11.114346 | 1.63627663 |
| 885 | NM_000289.5 | | #N/A | 708.795233 | -10.67972 | 1.57263972 |
| 886 | NM_002340.5 | | #N/A | 824.858295 | -10.898592 | 1.60509284 |
| 887 | NM_001190708.1 | | #N/A | 1261.93559 | -11.512196 | 1.69563581 |
| 888 | NM_001996.3 | | #N/A | 589.085264 | -10.413503 | 1.53394768 |
| 889 | NM_178862.2 | | #N/A | 665.861842 | -10.589569 | 1.56015173 |
| 890 | NM_004048.2 | | #N/A | 580.369225 | -10.391845 | 1.53108271 |
| 891 | NM_003754.2 | | #N/A | 725.9202 | -10.714216 | 1.57862358 |
| 892 | NM_025235.3 | | #N/A | 404.116989 | -4.8815116 | 0.71937347 |
| 893 | NM_004910.2 | | #N/A | 441.973191 | -9.9993199 | 1.47361468 |
| 894 | NM_001032221.3 | | #N/A | 953.033157 | -11.106922 | 1.63729199 |
| 895 | NM_016368.4 | | #N/A | 449.090493 | -10.021969 | 1.47755955 |
| 896 | NR_024074.1 | GOLGA8IP | | 84.801481 | 4.59105429 | 0.67688593 |
| 897 | NR_039980.1 | LOC286437 | | 477.2254 | 3.42258917 | 0.50475852 |
| 898 | NM_004793.3 | | #N/A | 481.352356 | -10.121111 | 1.49281867 |
| 899 | NM_152713.4 | | #N/A | 481.718747 | -10.122935 | 1.49307556 |
| 901 | NR_110493.1 | TUG1 | | 572.746497 | -10.372343 | 1.53015428 |
| 900 | NM_013438.4 | | #N/A | 666.103251 | -10.589826 | 1.56222267 |
| 902 | NM_001893.4 | | #N/A | 529.467483 | -10.258797 | 1.51357773 |

| | | | | | |
|---|---|---|---|---|---|
| 903 | NM_017514.4 | #N/A | 658.366465 | -10.57342 | 1.55997101 |
| 904 | NM_001162371.2 | #N/A | 745.3148 | -10.752332 | 1.58651086 |
| 905 | NM_005877.5 | #N/A | 483.769486 | -10.128838 | 1.49452189 |
| 906 | NM_007245.3 | ATXN2L | 496.455483 | -10.166801 | 1.50022248 |
| 907 | NM_002764.3 | #N/A | 428.233421 | -9.9528756 | 1.46883539 |
| 908 | NM_015154.2 | #N/A | 211.463187 | -3.9620195 | 0.58473958 |
| 909 | NM_017510.5 | #N/A | 580.376044 | -10.391691 | 1.53364424 |
| 910 | NM_003470.2 | #N/A | 536.448453 | -10.277872 | 1.51721371 |
| 911 | NM_005896.3 | IDH1 | 709.281411 | -10.680845 | 1.57701934 |
| 912 | NM_006389.4 | #N/A | 539.837548 | -10.286897 | 1.51908298 |
| 913 | NM_001546.3 | #N/A | 774.319336 | -10.807358 | 1.59624511 |
| 914 | NM_001037675.3 | NBPF9 | 440.598898 | -9.9942934 | 1.47619725 |
| 915 | NM_001197293.2 | #N/A | 957.147304 | -11.113185 | 1.64151359 |
| 916 | NM_020960.4 | #N/A | 538.169034 | -10.282678 | 1.51899734 |
| 917 | NR_033976.1 | LINC02232 | 37.585689 | 6.78374433 | 1.00247377 |
| 918 | NM_021943.2 | #N/A | 519.310887 | -10.231078 | 1.51207173 |
| 919 | NM_001134231.1 | #N/A | 798.823182 | -10.852379 | 1.60406496 |
| 920 | NM_007178.3 | #N/A | 884.028562 | -10.998493 | 1.62613634 |
| 921 | NR_130107.1 | GHET1 | 85.2882493 | 3.05133656 | 0.45116539 |
| 922 | NM_015382.3 | #N/A | 488.830195 | -10.144167 | 1.4999703 |
| 923 | NM_006513.3 | #N/A | 641.099165 | -10.53506 | 1.55796447 |
| 924 | NM_015134.3 | #N/A | 619.054921 | -10.483808 | 1.55042473 |
| 925 | NM_021034.2 | #N/A | 526.445682 | -10.251703 | 1.51652835 |
| 926 | NM_003366.3 | #N/A | 621.839589 | -10.490833 | 1.55209004 |
| 927 | NM_003463.4 | PTP4A1 | 746.849453 | -10.755107 | 1.59128075 |
| 928 | NM_032888.3 | #N/A | 229.223709 | -6.4082726 | 0.94812257 |
| 929 | NM_001272053.1 | MRFAP1 | 449.800174 | -10.023279 | 1.48306369 |
| 930 | NM_130853.2 | #N/A | 964.049706 | -9.6918325 | 1.43430729 |
| 931 | NM_020335.2 | #N/A | 617.464566 | -10.480725 | 1.55143207 |
| 932 | NM_004712.4 | #N/A | 513.493412 | -10.214822 | 1.5121668 |
| 933 | NM_031989.4 | #N/A | 648.453354 | -10.551579 | 1.56213715 |
| 934 | NM_005456.3 | #N/A | 687.685001 | -10.636168 | 1.57501076 |
| 935 | NM_014394.2 | #N/A | 592.024857 | -9.1468833 | 1.35455752 |
| 936 | NM_000454.4 | SOD1 | 550.315617 | -10.314753 | 1.52780785 |
| 937 | NM_152879.2 | #N/A | 456.121561 | -10.043834 | 1.48769981 |
| 938 | NM_001098534.1 | #N/A | 618.880537 | -10.484348 | 1.55337674 |
| 939 | NM_030665.3 | #N/A | 509.82756 | -10.204574 | 1.51193449 |
| 940 | NM_004599.3 | #N/A | 2250.85851 | -5.737474 | 0.85011222 |
| 941 | NM_002628.4 | #N/A | 515.180413 | -10.21969 | 1.51469024 |
| 942 | NM_001206654.1 | #N/A | 525.892134 | -10.249313 | 1.51932142 |
| 943 | NM_018215.3 | #N/A | 498.263026 | -10.171249 | 1.50772786 |
| 944 | NM_031370.2 | #N/A | 440.847921 | -9.9953592 | 1.481769 |
| 945 | NM_003378.3 | #N/A | 1120.20808 | -11.340205 | 1.68141833 |

| | | | | | |
|---|---|---|---|---|---|
| 946 | NM_001142641.1 | #N/A | 474.990982 | -10.10201 | 1.49787901 |
| 947 | NM_001093725.1 | #N/A | 739.63441 | -10.741187 | 1.59295347 |
| 948 | NM_004663.4 | #N/A | 617.607534 | -10.481161 | 1.55449382 |
| 949 | NM_014262.4 | #N/A | 322.115462 | -7.4627949 | 1.10687491 |
| 950 | NM_001000.3 | #N/A | 1039.84331 | -11.232684 | 1.66610722 |
| 951 | NM_000786.3 | #N/A | 710.032018 | -10.682317 | 1.58466701 |
| 952 | NR_026581.1 | #N/A | 645.980662 | -10.545808 | 1.56440292 |
| 953 | NM_001458.4 | FLNC | 491.779955 | -10.15295 | 1.50661523 |
| 954 | NM_001145543.1 | ZSCAN18 | 430.399029 | -9.9593817 | 1.47799004 |
| 955 | NM_014931.3 | #N/A | 458.045933 | -10.050342 | 1.49148133 |
| 956 | NM_006364.3 | #N/A | 437.275337 | -9.9840224 | 1.48176579 |
| 957 | NM_198334.2 | #N/A | 812.663315 | -10.877205 | 1.6146899 |
| 958 | NM_001319682.1 | #N/A | 276.481666 | 2.4502755 | 0.36376684 |
| 959 | NM_014862.3 | #N/A | 588.913323 | -10.412594 | 1.545947 |
| 960 | NM_001018069.1 | #N/A | 503.218724 | -10.185024 | 1.51260546 |
| 961 | NM_001078171.1 | #N/A | 696.723926 | -9.2624228 | 1.37567296 |
| 962 | NM_016604.3 | #N/A | 427.871792 | -9.9519331 | 1.47805666 |
| 963 | NM_001017.2 | #N/A | 1087.13847 | -11.296886 | 1.67829669 |
| 964 | NM_006420.2 | #N/A | 188.491223 | -4.4668233 | 0.66370778 |
| 965 | NM_001338.4 | #N/A | 571.849764 | -10.369837 | 1.54095512 |
| 966 | NM_012478.3 | #N/A | 529.911632 | -10.260321 | 1.52478026 |
| 967 | NM_024595.2 | #N/A | 480.368911 | -10.118508 | 1.50384427 |
| 968 | NM_001553.2 | #N/A | 642.683702 | -10.539135 | 1.56645971 |
| 969 | NM_017892.3 | #N/A | 436.447896 | -8.6473873 | 1.2853373 |
| 970 | NM_001204.6 | #N/A | 562.402283 | -5.8478262 | 0.86932585 |
| 971 | NM_015015.2 | #N/A | 625.084081 | -9.1188263 | 1.35572712 |
| 972 | NM_003746.2 | #N/A | 585.943662 | -9.1319861 | 1.35778838 |
| 973 | NM_004595.4 | #N/A | 673.314018 | -10.605647 | 1.57691609 |
| 974 | NM_005969.3 | #N/A | 650.125186 | -10.555086 | 1.5695067 |
| 975 | NM_006472.5 | #N/A | 432.552673 | -9.9673303 | 1.48260162 |
| 976 | NM_001319236.1 | RPL34 | 975.335101 | -11.140299 | 1.65718899 |
| 977 | NM_001141980.1 | #N/A | 469.314627 | -10.085333 | 1.50101244 |
| 978 | NM_016201.3 | #N/A | 492.487377 | -10.154741 | 1.51228854 |
| 979 | NM_019589.2 | #N/A | 481.081238 | -10.120683 | 1.50729221 |
| 980 | NM_031462.3 | #N/A | 608.680625 | -10.460377 | 1.55786562 |
| 981 | NM_206833.3 | #N/A | 704.573342 | -10.671183 | 1.58921593 |
| 983 | NR_027166.1 | SNHG29 | 840.306838 | -10.925295 | 1.62764107 |
| 982 | NM_012073.4 | #N/A | 680.970956 | -10.621833 | 1.58245838 |
| 985 | NR_027420.1 | LOC389834 | 102.483449 | 3.92072959 | 0.58430516 |
| 984 | NM_016297.3 | #N/A | 457.316073 | -5.2440491 | 0.7815198 |
| 986 | NM_001791.3 | #N/A | 637.514084 | -10.526884 | 1.56901047 |
| 987 | NM_019058.3 | #N/A | 422.01985 | -9.9317324 | 1.48033952 |
| 988 | NM_032595.4 | #N/A | 640.073939 | -10.532599 | 1.57009817 |

| | | | | | | |
|---|---|---|---|---|---|---|
| 989 | NM_014263.3 | | #N/A | 461.784774 | -10.061061 | 1.49985874 |
| 990 | NM_001193495.1 | | #N/A | 376.157354 | -8.4498531 | 1.25977807 |
| 991 | NM_003179.2 | SYP | | 959.262042 | -11.116394 | 1.65756051 |
| 992 | NM_013286.4 | | #N/A | 501.206156 | -8.9051166 | 1.32796877 |
| 993 | NM_001085460.1 | | #N/A | 499.101043 | -10.174317 | 1.51757904 |
| 994 | NM_001166110.1 | | #N/A | 610.651896 | -9.0854149 | 1.35586389 |
| 995 | NM_001358.2 | | #N/A | 469.213243 | -10.084544 | 1.50527349 |
| 996 | NM_053024.3 | | #N/A | 847.942587 | -10.938394 | 1.63277521 |
| 997 | NM_001080425.3 | | #N/A | 611.29319 | -10.466035 | 1.56232985 |
| 998 | NM_020137.4 | | #N/A | 457.537738 | -10.04875 | 1.50007348 |
| 999 | NM_002695.3 | | #N/A | 526.234409 | -10.250108 | 1.53037438 |
| 1000 | NM_002792.3 | | #N/A | 684.193 | -10.62882 | 1.58703271 |
| 1001 | NM_024299.2 | | #N/A | 649.323719 | -10.553429 | 1.57575639 |
| 1002 | NM_184043.2 | ALDOA | | 530.019669 | -10.260053 | 1.53214271 |
| 1003 | NM_001068.3 | TOP2B | | 492.209246 | -10.153011 | 1.51622242 |
| 1004 | NM_006305.3 | | #N/A | 727.450221 | -10.717086 | 1.60046447 |
| 1005 | NM_015125.4 | CIC | | 565.477437 | -10.353749 | 1.54631404 |
| 1006 | NR_036625.1 | | #N/A | 537.58364 | -10.280576 | 1.53565161 |
| 1007 | NM_004295.3 | | #N/A | 428.681992 | -8.6201811 | 1.28776714 |
| 1008 | NM_002893.3 | | #N/A | 501.735652 | -10.181505 | 1.52109827 |
| 1009 | NM_003144.4 | | #N/A | 502.001702 | -10.182261 | 1.52146991 |
| 1010 | NM_032271.2 | | #N/A | 484.356355 | -10.130499 | 1.5137442 |
| 1011 | NM_001271148.1 | ST3GAL6 | | 44.0723235 | 10.4396499 | 1.56002939 |
| 1012 | NM_001190836.1 | DCTN1 | | 469.97283 | -10.087614 | 1.5084928 |
| 1013 | NM_004984.2 | | #N/A | 881.966798 | -10.9952 | 1.64449579 |
| 1014 | NM_006745.4 | | #N/A | 636.433184 | -10.524411 | 1.57408847 |
| 1015 | NM_018837.3 | SULF2 | | 561.647783 | -10.344475 | 1.54723952 |
| 1016 | NM_004402.3 | | #N/A | 62.6453732 | 24.7020497 | 3.69486372 |
| 1017 | NM_139276.2 | STAT3 | | 485.824279 | -10.135619 | 1.5166091 |
| 1018 | NM_001080398.1 | | #N/A | 448.425611 | -10.019369 | 1.49926951 |
| 1019 | NM_005744.3 | | #N/A | 360.687377 | -8.4271711 | 1.26107507 |
| 1020 | NM_004518.5 | | #N/A | 880.895365 | -10.993359 | 1.64519757 |
| 1021 | NM_014413.3 | | #N/A | 457.371187 | -10.047603 | 1.50369455 |
| 1022 | NM_014329.4 | | #N/A | 477.111632 | -10.108732 | 1.51297223 |
| 1023 | NM_175852.3 | | #N/A | 362.09939 | -8.3977056 | 1.25689882 |
| 1024 | NM_182557.2 | | #N/A | 430.88147 | -9.9615391 | 1.49104022 |
| 1025 | NM_019613.3 | | #N/A | 462.454779 | -10.063706 | 1.50642067 |
| 1026 | NM_198318.4 | PRMT1 | | 612.889301 | -10.470058 | 1.56734479 |
| 1027 | NM_004327.3 | | #N/A | 587.075971 | -10.408036 | 1.55815062 |
| 1028 | NM_003021.3 | | #N/A | 600.204323 | -10.439834 | 1.56356672 |
| 1029 | NM_007235.4 | | #N/A | 496.740189 | -10.166722 | 1.52284637 |
| 1030 | NM_006812.3 | | #N/A | 545.323849 | -10.301318 | 1.54312558 |
| 1031 | NM_001748.4 | | #N/A | 454.119855 | -10.038451 | 1.50397911 |

| | | | | | |
|---|---|---|---|---|---|
| 1032 | NM_001307924.2 | #N/A | 531.715311 | -10.264788 | 1.53794051 |
| 1033 | NM_001098210.1 | #N/A | 471.989871 | -10.093346 | 1.51237066 |
| 1034 | NM_001304.4 | #N/A | 465.029202 | -10.072331 | 1.50925576 |
| 1035 | NM_004689.3 | #N/A | 482.804178 | -10.125656 | 1.51730669 |
| 1036 | NM_030810.4 | #N/A | 530.044545 | -10.260803 | 1.53755136 |
| 1037 | NM_004187.3 | #N/A | 469.840234 | -10.085994 | 1.51152601 |
| 1038 | NM_018240.6 | #N/A | 471.1937 | -10.091242 | 1.51231049 |
| 1039 | NM_005736.3 | #N/A | 847.542641 | -9.5117511 | 1.42549823 |
| 1040 | NM_007118.3 | #N/A | 561.392201 | -10.343342 | 1.55026161 |
| 1041 | NM_001429.3 | #N/A | 390.594032 | -9.820136 | 1.47188586 |
| 1042 | NM_001022.3 | #N/A | 988.824637 | -11.160149 | 1.67280944 |
| 1043 | NM_001440.3 | #N/A | 447.972016 | -10.017828 | 1.50162355 |
| 1044 | NM_030915.3 | #N/A | 595.239213 | -10.427948 | 1.56337033 |
| 1045 | NM_001042681.1 | #N/A | 460.097661 | -10.056086 | 1.50778524 |
| 1046 | NM_004157.3 | #N/A | 433.621473 | -9.9709187 | 1.49508325 |
| 1048 | NM_020151.3 | STARD7 | 557.254972 | -10.332659 | 1.54940743 |
| 1047 | NM_005137.2 | #N/A | 572.919745 | -10.37271 | 1.55539191 |
| 1049 | NM_007103.3 | #N/A | 543.755722 | -10.297376 | 1.54423644 |
| 1050 | NM_014599.5 | #N/A | 593.185222 | -10.422502 | 1.56316344 |
| 1051 | NM_006618.4 | #N/A | 609.032637 | -10.460901 | 1.56907958 |
| 1052 | NM_001105079.2 | #N/A | 403.7376 | -9.8690003 | 1.48033835 |
| 1053 | NM_002807.3 | #N/A | 565.9492 | -10.355175 | 1.55346771 |
| 1054 | NM_004630.3 | #N/A | 438.014913 | -9.9846159 | 1.49787328 |
| 1055 | NM_024293.5 | #N/A | 535.400931 | -10.275156 | 1.54170742 |
| 1056 | NR_028272.1 | NEAT1 | 580.259043 | -10.39172 | 1.55959004 |
| 1057 | NM_003632.2 | #N/A | 476.348914 | -10.107042 | 1.51699526 |
| 1058 | NM_014891.6 | #N/A | 620.946048 | -5.4267692 | 0.81452904 |
| 1059 | NM_002585.3 | #N/A | 881.285428 | -10.993992 | 1.65018147 |
| 1060 | NM_000382.2 | #N/A | 287.023382 | -4.5905425 | 0.68905858 |
| 1061 | NM_005338.6 | #N/A | 434.300139 | -9.973335 | 1.49723733 |
| 1062 | NM_006295.2 | #N/A | 558.042922 | -9.0618605 | 1.36041088 |
| 1063 | NM_144997.5 | #N/A | 245.286688 | -4.5341037 | 0.68067885 |
| 1064 | NM_005973.4 | #N/A | 329.310977 | -7.4989152 | 1.12581321 |
| 1065 | NM_080616.4 | #N/A | 357.207675 | -7.0919184 | 1.06481383 |
| 1066 | NM_130787.2 | #N/A | 402.143589 | -9.8628132 | 1.48082466 |
| 1067 | NM_032408.3 | #N/A | 473.66035 | -10.098231 | 1.51649883 |
| 1068 | NM_001080457.1 | #N/A | 636.212928 | -10.523731 | 1.58066252 |
| 1069 | NM_002106.3 | #N/A | 670.47815 | -10.59948 | 1.59209549 |
| 1070 | NM_004286.4 | #N/A | 410.353673 | -8.6148567 | 1.29436308 |
| 1071 | NM_013444.3 | UBQLN2 | 457.048882 | -8.6993438 | 1.30725871 |
| 1072 | NM_001895.3 | #N/A | 453.287486 | -10.034719 | 1.50811992 |
| 1073 | NM_005803.3 | #N/A | 648.036826 | -10.550549 | 1.58606804 |
| 1074 | NR_045262.1 | #N/A | 71.4736919 | 3.19232792 | 0.4798973 |

| | | | | | |
|---|---|---|---|---|---|
| 1075 | NM_002428.3 | #N/A | 816.94359 | -10.884756 | 1.6363452 |
| 1076 | NM_130854.2 | #N/A | 389.126034 | -9.8153329 | 1.47646231 |
| 1077 | NM_003753.3 | #N/A | 482.816018 | -10.125846 | 1.52333884 |
| 1078 | NM_002092.3 | #N/A | 431.322794 | -9.9632308 | 1.49893601 |
| 1079 | NM_012319.3 | #N/A | 701.344396 | -10.664462 | 1.60464653 |
| 1080 | NM_001252102.1 | #N/A | 878.016527 | -10.988535 | 1.65359255 |
| 1081 | NM_002414.4 | #N/A | 638.819261 | -10.530076 | 1.58467241 |
| 1082 | NM_014866.1 | #N/A | 386.05802 | -9.8033476 | 1.4753786 |
| 1083 | NM_002266.3 | #N/A | 609.957653 | -10.463217 | 1.57516672 |
| 1084 | NM_002085.4 | #N/A | 510.498694 | -10.206433 | 1.53656554 |
| 1085 | NM_000099.3 | #N/A | 451.105545 | -10.028192 | 1.50982635 |
| 1086 | NM_001144831.1 | #N/A | 586.129042 | -10.40556 | 1.56665925 |
| 1087 | NM_002094.3 | #N/A | 476.226589 | -10.106129 | 1.52159345 |
| 1088 | NM_020531.2 | #N/A | 556.369262 | -10.330461 | 1.55533912 |
| 1089 | NM_058243.2 | BRD4 | 373.88174 | -9.7571333 | 1.46935259 |
| 1090 | NM_001244701.1 | ZFP36L1 | 398.415026 | -9.8498826 | 1.4833614 |
| 1091 | NM_003333.4 | #N/A | 649.72884 | -10.554236 | 1.58946483 |
| 1092 | NM_014275.4 | #N/A | 459.583239 | -10.055099 | 1.51446617 |
| 1093 | NM_016463.8 | #N/A | 467.488265 | -10.080038 | 1.51829403 |
| 1094 | NM_006979.2 | #N/A | 423.516074 | -9.9374989 | 1.49693552 |
| 1095 | NM_177983.2 | #N/A | 556.18188 | -9.0572255 | 1.36447764 |
| 1096 | NM_001199456.1 | BRD2 | 464.251702 | -10.069093 | 1.51718987 |
| 1099 | NM_207577.1 | MAP6 | 605.611265 | -10.452885 | 1.57528057 |
| 1097 | NM_001009552.1 | #N/A | 528.355535 | -10.255773 | 1.54560923 |
| 1098 | NM_020782.1 | #N/A | 308.549428 | -5.4329364 | 0.81878493 |
| 1100 | NM_001330682.1 | #N/A | 568.425164 | -10.361366 | 1.5616509 |
| 1101 | NM_002673.5 | PLXNB1 | 437.590764 | -9.9829269 | 1.50504329 |
| 1102 | NM_001330575.1 | DECR1 | 76.2000061 | 3.36588936 | 0.50752443 |
| 1103 | NM_025181.4 | #N/A | 367.728092 | -9.73329 | 1.46771152 |
| 1104 | NM_002147.3 | HOXB5 | 721.84679 | -10.706275 | 1.61468476 |
| 1105 | NM_173602.2 | #N/A | 336.358051 | -7.5318589 | 1.13606037 |
| 1106 | NM_001319.6 | #N/A | 448.135018 | -10.018448 | 1.51115655 |
| 1107 | NM_000489.4 | #N/A | 450.719317 | -10.026743 | 1.51256302 |
| 1108 | NM_139207.4 | #N/A | 471.436413 | -10.09069 | 1.52224126 |
| 1109 | NM_001282963.1 | HJURP | 55.5101184 | 10.7827573 | 1.62668598 |
| 1110 | NM_002047.3 | #N/A | 497.446041 | -10.168973 | 1.53419698 |
| 1111 | NR_038419.1 | SLC39A12-AS1 | 65.1705355 | 5.92098228 | 0.89333195 |
| 1112 | NM_002865.2 | #N/A | 616.427964 | -10.478272 | 1.58114545 |
| 1113 | NM_001098722.1 | #N/A | 664.923955 | -10.587496 | 1.59783032 |
| 1114 | NM_015409.4 | #N/A | 431.82055 | -9.9644761 | 1.5038868 |
| 1115 | NM_022463.4 | #N/A | 441.622189 | -9.9983143 | 1.5090662 |
| 1116 | NM_005766.3 | #N/A | 606.900936 | -10.455913 | 1.5782282 |
| 1117 | NR_034009.1 | #N/A | 423.371489 | -9.9363149 | 1.5002504 |

| | | | | | | |
|---|---|---|---|---|---|---|
| 1118 | NM_006016.5 | | #N/A | 425.556581 | -9.9447674 | 1.50169467 |
| 1119 | NM_017519.2 | ARID1B | | 247.073016 | -7.0000563 | 1.05706625 |
| 1120 | NM_003012.4 | | #N/A | 660.11705 | -10.576847 | 1.59726245 |
| 1121 | NM_005443.4 | | #N/A | 496.483281 | -10.166104 | 1.53543185 |
| 1122 | NM_014847.3 | | #N/A | 430.978391 | -9.9623776 | 1.50470107 |
| 1123 | NM_005271.4 | | #N/A | 468.897502 | -10.083678 | 1.52338852 |
| 1124 | NM_001040715.1 | | #N/A | 587.17337 | -10.408154 | 1.57251739 |
| 1125 | NM_002911.3 | | #N/A | 420.497039 | -9.9262134 | 1.49971958 |
| 1126 | NM_001332.3 | | #N/A | 762.204998 | -10.784384 | 1.62973237 |
| 1127 | NM_016337.2 | | #N/A | 549.511107 | -10.31275 | 1.55860385 |
| 1128 | NM_007033.4 | | #N/A | 462.925478 | -10.065116 | 1.52150659 |
| 1129 | NM_058004.3 | | #N/A | 500.034254 | -10.176323 | 1.53872471 |
| 1130 | NM_003343.5 | | #N/A | 406.811961 | -9.8783713 | 1.49392181 |
| 1131 | NM_012414.3 | | #N/A | 365.505993 | -9.7245132 | 1.47064612 |
| 1132 | NM_015270.4 | | #N/A | 433.047427 | -8.6937845 | 1.31485001 |
| 1133 | NM_019116.2 | | #N/A | 492.869136 | -10.155765 | 1.53629722 |
| 1134 | NM_003564.2 | | #N/A | 399.618044 | -9.8535886 | 1.49062601 |
| 1135 | NM_006371.4 | | #N/A | 393.891768 | -8.4991684 | 1.28589901 |
| 1136 | NM_015904.3 | | #N/A | 564.2943 | -10.350724 | 1.56615098 |
| 1137 | NM_001681.3 | | #N/A | 551.719918 | -10.318348 | 1.56189661 |
| 1138 | NM_004593.2 | | #N/A | 545.888816 | -10.302811 | 1.5597128 |
| 1139 | NM_002161.5 | IARS1 | | 413.801912 | -9.9034849 | 1.49941255 |
| 1140 | NM_001190274.1 | FBXO11 | | 462.923145 | -10.064977 | 1.52402571 |
| 1141 | NM_003365.2 | | #N/A | 545.91184 | -10.303145 | 1.56040646 |
| 1142 | NM_001304717.2 | | #N/A | 448.575363 | -8.744729 | 1.32451924 |
| 1143 | NM_006044.3 | | #N/A | 433.439965 | -9.9708094 | 1.51027067 |
| 1144 | NR_135469.1 | | #N/A | 43.2732764 | 10.4108212 | 1.57698717 |
| 1145 | NM_001867.2 | | #N/A | 625.18856 | -10.498698 | 1.59033272 |
| 1146 | NM_003791.3 | | #N/A | 446.294789 | -10.012756 | 1.51699108 |
| 1147 | NM_138444.3 | | #N/A | 417.909132 | -9.9168144 | 1.50244946 |
| 1148 | NM_005386.3 | | #N/A | 447.761223 | -10.016138 | 1.51760441 |
| 1149 | NM_005445.3 | SMC3 | | 418.091336 | -9.9180316 | 1.50333987 |
| 1150 | NM_006555.3 | | #N/A | 633.352712 | -9.1199409 | 1.38251801 |
| 1151 | NM_016107.4 | | #N/A | 477.200879 | -10.108897 | 1.53243588 |
| 1152 | NM_001012761.2 | | #N/A | 584.335427 | -10.401312 | 1.57717616 |
| 1153 | NM_020713.2 | | #N/A | 483.888543 | -10.129161 | 1.53602177 |
| 1154 | NM_014287.3 | | #N/A | 538.815691 | -10.284357 | 1.55959251 |
| 1155 | NM_014849.4 | | #N/A | 664.383921 | -10.586621 | 1.60573061 |
| 1156 | NM_004538.5 | | #N/A | 671.311796 | -10.6014 | 1.60804801 |
| 1157 | NM_006411.3 | | #N/A | 581.151459 | -10.393298 | 1.57658531 |
| 1158 | NM_013390.2 | | #N/A | 454.942803 | -10.040542 | 1.52313786 |
| 1159 | NM_004826.3 | | #N/A | 410.374792 | -9.8926205 | 1.50097529 |
| 1160 | NM_003009.2 | | #N/A | 526.067002 | -10.249592 | 1.55526949 |

| | | | | | |
|---|---|---|---|---|---|
| 1161 | NM_015322.4 | #N/A | 471.934419 | -10.09281 | 1.53146371 |
| 1162 | NM_014292.4 | #N/A | 398.376057 | -9.8482435 | 1.49445131 |
| 1163 | NM_015043.3 | #N/A | 410.47402 | -9.8917449 | 1.50118485 |
| 1164 | NM_002710.3 | #N/A | 439.830317 | -9.9910968 | 1.51652863 |
| 1165 | NM_031243.2 | #N/A | 440.995061 | -9.9953181 | 1.51720877 |
| 1166 | NM_052905.3 | #N/A | 407.873074 | -9.8817696 | 1.50002748 |
| 1167 | NM_004638.3 | #N/A | 454.949995 | -10.039502 | 1.52417089 |
| 1168 | NM_003590.4 | #N/A | 423.18675 | -9.9357482 | 1.50848886 |
| 1169 | NM_130440.3 | PTPRF | 436.141885 | -9.9803675 | 1.51604264 |
| 1170 | NM_000671.4 | ADH5 | 540.045874 | -10.287497 | 1.56277239 |
| 1171 | NM_001145263.1 | NCOA4 | 498.608468 | -10.172487 | 1.54533007 |
| 1172 | NM_003565.2 | #N/A | 483.154409 | -10.126834 | 1.53855281 |
| 1173 | NM_001543.4 | #N/A | 543.310648 | -10.296171 | 1.56437934 |
| 1174 | NM_002140.4 | HNRNPK | 383.695687 | -9.7955833 | 1.48835767 |
| 1175 | NM_004434.2 | #N/A | 217.478959 | -6.8284886 | 1.03756451 |
| 1176 | NM_001076785.2 | #N/A | 341.867566 | -5.1768488 | 0.7866768 |
| 1177 | NM_006401.2 | #N/A | 446.040727 | -10.011647 | 1.52173329 |
| 1178 | NM_015392.3 | #N/A | 631.934926 | -10.514162 | 1.59811713 |
| 1179 | NM_020297.3 | ABCC9 | 752.354761 | 3.2007748 | 0.48655111 |
| 1180 | NM_002184.3 | #N/A | 548.878772 | -10.311233 | 1.56749896 |
| 1181 | NM_002537.3 | OAZ2 | 468.049476 | -10.081079 | 1.53260546 |
| 1182 | NM_004537.6 | #N/A | 569.59716 | -10.364246 | 1.57563508 |
| 1183 | NM_003906.4 | #N/A | 404.492698 | -9.8706742 | 1.50070805 |
| 1184 | NM_021102.3 | #N/A | 385.954318 | -9.8023874 | 1.49034138 |
| 1185 | NM_006876.2 | #N/A | 471.952769 | -10.093164 | 1.5349088 |
| 1186 | NM_014268.3 | #N/A | 726.405312 | -10.715187 | 1.62948012 |
| 1187 | NM_001171613.1 | PREPL | 598.895744 | -10.43664 | 1.58734524 |
| 1188 | NM_001145638.2 | #N/A | 588.526215 | -10.411402 | 1.58389181 |
| 1189 | NR_110825.1 | LINC01482 | 174.276131 | 4.29076635 | 0.65279489 |
| 1190 | NM_032430.1 | #N/A | 772.098333 | -10.803146 | 1.64364017 |
| 1191 | NM_001099415.2 | #N/A | 392.556125 | -9.8277789 | 1.49543907 |
| 1192 | NM_002213.4 | #N/A | 425.671939 | -9.9451444 | 1.51329703 |
| 1193 | NM_002952.3 | #N/A | 5593.16572 | -7.7982227 | 1.18670082 |
| 1194 | NM_198486.3 | #N/A | 538.10586 | -10.282071 | 1.56483424 |
| 1197 | NR_026811.3 | GOLGA2P10 | 390.839699 | -9.8199776 | 1.49465564 |
| 1195 | NM_002954.5 | RPS27A | 823.594177 | -10.896298 | 1.6584966 |
| 1196 | NM_005570.3 | #N/A | 387.219455 | -9.8088889 | 1.49299065 |
| 1198 | NM_001001563.3 | #N/A | 406.469876 | -9.8773615 | 1.50352397 |
| 1199 | NM_004342.6 | #N/A | 942.816299 | -11.091572 | 1.68836277 |
| 1200 | NM_174909.4 | #N/A | 574.651564 | -10.377104 | 1.57954685 |
| 1201 | NM_001077488.3 | #N/A | 780.123647 | -10.81822 | 1.6469324 |
| 1202 | NM_005095.2 | #N/A | 365.87367 | -5.1812094 | 0.78876153 |
| 1203 | NM_001788.5 | SEPTIN7 | 583.490165 | -10.399205 | 1.58327246 |

| | | | | | |
|---|---|---|---|---|---|
| 1204 | NM_057161.3 | #N/A | 542.652863 | -10.294316 | 1.56734631 |
| 1205 | NM_003747.2 | #N/A | 452.665618 | -10.03269 | 1.5276922 |
| 1206 | NM_002812.4 | PSMD8 | 551.819012 | -9.0464876 | 1.37754879 |
| 1207 | NM_001641.3 | #N/A | 482.57138 | -10.124959 | 1.54199469 |
| 1208 | NM_017921.3 | #N/A | 394.810971 | -9.8356865 | 1.4980064 |
| 1209 | NM_001172437.2 | PEG10 | 628.624512 | -10.506628 | 1.60036853 |
| 1211 | NM_014330.3 | PPP1R15A | 375.041895 | -9.7621504 | 1.48723295 |
| 1210 | NM_001530.3 | #N/A | 470.457724 | -10.088444 | 1.53693098 |
| 1212 | NM_006387.5 | #N/A | 420.726431 | -9.926793 | 1.51245984 |
| 1213 | NM_013253.4 | DKK3 | 696.308343 | -10.654152 | 1.62333985 |
| 1214 | NM_031266.2 | #N/A | 541.298767 | -10.290708 | 1.56800405 |
| 1215 | NM_022087.3 | #N/A | 395.905702 | -9.8401866 | 1.49955451 |
| 1216 | NM_015720.3 | #N/A | 906.1124 | -11.034075 | 1.6815301 |
| 1217 | NM_014615.3 | #N/A | 504.453612 | -10.189432 | 1.55297997 |
| 1218 | NM_000801.4 | #N/A | 541.404471 | -10.291069 | 1.56875133 |
| 1219 | NM_005231.3 | #N/A | 447.964799 | -10.018516 | 1.52733014 |
| 1220 | NM_001152.4 | #N/A | 543.743252 | -10.2974 | 1.57005935 |
| 1221 | NM_004819.2 | #N/A | 431.774519 | -9.9646338 | 1.51931991 |
| 1222 | NM_005765.2 | #N/A | 536.619974 | -10.278331 | 1.56717176 |
| 1223 | NM_004703.5 | #N/A | 462.253724 | -10.062909 | 1.53449205 |
| 1224 | NM_002317.6 | #N/A | 468.27295 | -10.082341 | 1.53785118 |
| 1225 | NM_004787.3 | #N/A | 412.393344 | -9.8996349 | 1.5099857 |
| 1226 | NM_014045.4 | #N/A | 428.23807 | -9.9537568 | 1.5183845 |
| 1227 | NM_001448.2 | #N/A | 707.848261 | -10.678 | 1.62908318 |
| 1228 | NM_020746.4 | #N/A | 353.90117 | -9.677565 | 1.47647394 |
| 1229 | NM_022818.4 | #N/A | 612.545973 | -10.469303 | 1.59728503 |
| 1230 | NM_000014.5 | A2M | 797.079415 | -10.84932 | 1.65538233 |
| 1231 | NM_004394.2 | #N/A | 370.992563 | -9.7465159 | 1.48750872 |
| 1232 | NM_001193417.2 | #N/A | 367.969792 | -9.7344227 | 1.48571844 |
| 1236 | NR_003580.2 | GTF2IP4 | 629.512648 | -10.508641 | 1.60406279 |
| 1233 | NM_002461.2 | #N/A | 788.920359 | -10.834369 | 1.65380728 |
| 1234 | NM_018509.3 | #N/A | 474.290343 | -10.100202 | 1.54178541 |
| 1235 | NM_032800.2 | #N/A | 546.907959 | -10.305943 | 1.57316348 |
| 1237 | NM_006516.2 | #N/A | 870.066412 | -10.975829 | 1.67554128 |
| 1238 | NM_015002.2 | #N/A | 360.854287 | -9.7053059 | 1.48172989 |
| 1239 | NM_018462.4 | #N/A | 508.482681 | -10.200626 | 1.55754823 |
| 1240 | NM_033008.2 | #N/A | 486.926847 | -10.138234 | 1.54806732 |
| 1241 | NM_014649.2 | #N/A | 375.059731 | -9.7613503 | 1.49057503 |
| 1242 | NM_001271641.1 | MTCH1 | 644.844029 | -10.543307 | 1.61003387 |
| 1243 | NM_006708.2 | #N/A | 452.962296 | -10.033794 | 1.53255838 |
| 1244 | NM_014445.3 | #N/A | 474.971495 | -10.102057 | 1.54322617 |
| 1245 | NM_015253.1 | WSCD1 | 436.675794 | -9.9810647 | 1.52479186 |
| 1246 | NM_004905.2 | #N/A | 441.303005 | -9.9962392 | 1.52730845 |

| | | | | | | |
|---|---|---|---|---|---|---|
| 1247 | NM_002487.2 | | #N/A | 648.099366 | -10.550495 | 1.61204333 |
| 1248 | NM_005751.4 | AKAP9 | | 417.331853 | -9.9151453 | 1.51506946 |
| 1249 | NM_001040456.2 | | #N/A | 432.271051 | -9.9667582 | 1.52346623 |
| 1250 | NM_014548.3 | | #N/A | 803.476949 | -10.860678 | 1.66025298 |
| 1251 | NM_152437.2 | | #N/A | 444.708277 | -10.007825 | 1.52988572 |
| 1252 | NM_201599.2 | | #N/A | 369.65084 | -9.7418036 | 1.48929276 |
| 1253 | NM_001619.3 | | #N/A | 469.185526 | -10.084304 | 1.54186148 |
| 1254 | NM_004656.3 | | #N/A | 512.663909 | -10.212343 | 1.56162314 |
| 1255 | NM_014971.1 | | #N/A | 650.485541 | -10.556019 | 1.61429295 |
| 1256 | NM_052875.4 | | #N/A | 589.616195 | -10.414095 | 1.59262927 |
| 1257 | NM_080836.3 | | #N/A | 188.491677 | -6.1192902 | 0.93585522 |
| 1258 | NM_004960.3 | FUS | | 825.032957 | -10.898771 | 1.6668666 |
| 1259 | NM_002087.3 | GRN | | 559.108982 | -10.337901 | 1.58112516 |
| 1260 | NM_002714.3 | | #N/A | 365.739791 | -9.7248114 | 1.48742141 |
| 1261 | NM_003119.3 | | #N/A | 395.868028 | -9.8395 | 1.50548223 |
| 1262 | NM_001096.2 | | #N/A | 606.802402 | -10.455528 | 1.59990996 |
| 1263 | NM_024038.3 | | #N/A | 564.251155 | -10.350658 | 1.58388821 |
| 1264 | NM_018639.4 | WSB2 | | 766.167584 | -10.792002 | 1.65154578 |
| 1266 | NR_002196.2 | H19 | | 780.417547 | -10.818848 | 1.65582872 |
| 1265 | NM_001300905.1 | | #N/A | 367.718911 | -9.7319472 | 1.48947968 |
| 1267 | NM_004450.2 | | #N/A | 475.426784 | -8.0560983 | 1.23322753 |
| 1268 | NM_001288976.1 | | #N/A | 547.55625 | -10.307121 | 1.57784989 |
| 1269 | NM_001170634.1 | FUS | | 563.625659 | -10.34944 | 1.58455936 |
| 1270 | NM_002688.5 | | #N/A | 627.108607 | -10.502942 | 1.60806099 |
| 1271 | NM_014680.3 | | #N/A | 524.481063 | -10.24606 | 1.56875031 |
| 1272 | NM_031203.3 | | #N/A | 540.625132 | -10.288784 | 1.57527432 |
| 1273 | NM_001287.5 | | #N/A | 439.503374 | -9.9904156 | 1.52996446 |
| 1274 | NM_002796.2 | | #N/A | 504.035828 | -10.187918 | 1.56021225 |
| 1275 | NM_014812.2 | | #N/A | 554.708276 | -10.326033 | 1.58166799 |
| 1276 | NM_014389.2 | | #N/A | 460.591329 | -10.057862 | 1.54077194 |
| 1278 | NM_020791.2 | TAOK1 | | 480.736667 | -10.11926 | 1.5502471 |
| 1277 | NM_003061.2 | | #N/A | 686.084494 | -9.2155239 | 1.41184535 |
| 1279 | NM_024757.4 | | #N/A | 397.680209 | -9.8457982 | 1.50839169 |
| 1280 | NM_003908.4 | | #N/A | 428.685073 | -9.9542579 | 1.52506395 |
| 1281 | NM_004738.4 | | #N/A | 387.380681 | -9.8085319 | 1.50282392 |
| 1282 | NM_018060.3 | | #N/A | 352.435904 | -9.6715395 | 1.48188026 |
| 1283 | NM_001040134.1 | | #N/A | 799.798169 | -9.4162013 | 1.44288159 |
| 1284 | NM_001294.3 | | #N/A | 555.163291 | -10.327463 | 1.58264982 |
| 1285 | NM_001031711.2 | | #N/A | 523.794656 | -10.243625 | 1.56983922 |
| 1286 | NM_053275.3 | RPLP0 | | 695.20279 | -10.651772 | 1.63246425 |
| 1287 | NM_032287.2 | | #N/A | 459.29507 | -10.053663 | 1.54091805 |
| 1288 | NM_000581.3 | | #N/A | 479.32004 | -10.115325 | 1.5506364 |
| 1289 | NM_001554.4 | | #N/A | 623.238244 | -10.494478 | 1.60881483 |

| | | | | | |
|---|---|---|---|---|---|
| 1290 | NM_003498.5 | #N/A | 624.224918 | -9.0913251 | 1.39368775 |
| 1291 | NM_001142276.1 | #N/A | 499.901391 | -10.176108 | 1.5603759 |
| 1292 | NM_014719.2 | #N/A | 552.381856 | -10.320241 | 1.58248725 |
| 1294 | NM_017457.5 | CYTH2 | 607.493327 | -10.4575 | 1.60369438 |
| 1293 | NM_005520.2 | #N/A | 794.835628 | -5.4639833 | 0.83790623 |
| 1295 | NM_001145056.1 | #N/A | 499.875734 | -10.176256 | 1.56067406 |
| 1296 | NM_003680.3 | YARS1 | 527.144302 | -10.252669 | 1.57256426 |
| 1297 | NM_005529.6 | #N/A | 410.099103 | -9.8915112 | 1.51716142 |
| 1298 | NM_012435.2 | SHC2 | 548.725379 | -10.310546 | 1.58165142 |
| 1299 | NM_002014.3 | #N/A | 688.557994 | -10.637912 | 1.63228108 |
| 1300 | NM_003756.2 | #N/A | 622.315477 | -10.491983 | 1.61018437 |
| 1301 | NM_004175.4 | #N/A | 448.67357 | -10.01999 | 1.53779455 |
| 1302 | NM_145716.3 | #N/A | 604.361487 | -10.449449 | 1.60371671 |
| 1303 | NM_199141.1 | #N/A | 454.566851 | -10.03897 | 1.54101323 |
| 1304 | NM_032208.2 | ANTXR1 | 386.913217 | -9.8070078 | 1.50549695 |
| 1305 | NM_001282611.1 | #N/A | 846.048931 | -10.935209 | 1.67878216 |
| 1306 | NM_001024808.2 | #N/A | 363.681192 | -6.5676801 | 1.0084101 |
| 1307 | NM_005759.5 | #N/A | 366.046382 | -9.7257274 | 1.49331513 |
| 1308 | NM_023015.4 | #N/A | 376.329747 | -9.7666703 | 1.49956046 |
| 1309 | NM_000391.3 | #N/A | 354.639256 | -8.3504183 | 1.28221336 |
| 1310 | NM_013403.2 | #N/A | 449.996288 | -10.023721 | 1.53915191 |
| 1311 | NM_032808.6 | #N/A | 517.291447 | -10.225293 | 1.57009252 |
| 1314 | NR_003491.3 | MIAT | 451.45815 | -10.028088 | 1.54022167 |
| 1312 | NM_015902.5 | #N/A | 410.520288 | -9.8916479 | 1.51923822 |
| 1313 | NM_030981.2 | #N/A | 441.210276 | -9.9961072 | 1.53529526 |
| 1315 | NM_003106.3 | #N/A | 370.294102 | -9.7423231 | 1.49641094 |
| 1317 | NR_002594.1 | SLC7A5P2 | 139.859532 | 2.08006517 | 0.31956309 |
| 1316 | NM_003969.3 | #N/A | 518.877818 | -10.229801 | 1.57160068 |
| 1318 | NM_014396.3 | #N/A | 354.428465 | -9.6798437 | 1.48715657 |
| 1319 | NM_001143998.1 | #N/A | 395.21136 | -9.837055 | 1.51159697 |
| 1320 | NM_016732.2 | #N/A | 479.652783 | -10.116406 | 1.55458785 |
| 1321 | NM_030939.4 | #N/A | 459.839509 | -10.055269 | 1.54557026 |
| 1322 | NM_005629.3 | #N/A | 595.809973 | -10.429568 | 1.60323301 |
| 1323 | NM_138370.2 | #N/A | 436.116128 | -9.9797548 | 1.53412117 |
| 1324 | NM_014294.5 | #N/A | 362.394354 | -9.7131064 | 1.49320117 |
| 1325 | NM_001308373.1 | #N/A | 457.559183 | -10.048241 | 1.54487006 |
| 1326 | NM_152742.2 | #N/A | 735.019537 | -10.732218 | 1.65049741 |
| 1327 | NM_001042544.1 | LTBP4 | 382.785224 | -9.7917478 | 1.50594995 |
| 1328 | NM_014786.3 | #N/A | 389.369672 | -9.8163759 | 1.50990479 |
| 1329 | NM_001419.2 | #N/A | 441.931848 | -9.998066 | 1.53865224 |
| 1330 | NM_000884.2 | #N/A | 485.423856 | -10.133607 | 1.55973151 |
| 1331 | NM_001033930.2 | #N/A | 509.98545 | -10.204843 | 1.5707296 |
| 1332 | NM_003339.2 | #N/A | 524.515826 | -10.245373 | 1.57699035 |

| | | | | | |
|---|---|---|---|---|---|
| 1333 | NM_013245.2 | #N/A | 513.455708 | -10.21457 | 1.57240987 |
| 1334 | NM_015327.2 | #N/A | 404.58404 | -9.8705658 | 1.51949979 |
| 1335 | NM_015332.3 | #N/A | 514.904563 | -10.218764 | 1.57337513 |
| 1336 | NM_031449.3 | #N/A | 383.686152 | -9.7944259 | 1.5081183 |
| 1337 | NM_001009993.3 | #N/A | 590.956045 | -10.417221 | 1.6041699 |
| 1338 | NM_031412.2 | #N/A | 499.004804 | -10.173491 | 1.56671274 |
| 1339 | NM_001079864.2 | TAX1BP1 | 378.738381 | -9.7759076 | 1.50556032 |
| 1340 | NM_005077.4 | #N/A | 392.149624 | -9.8257112 | 1.51335527 |
| 1341 | NM_032039.3 | #N/A | 390.646406 | -9.82085 | 1.51280223 |
| 1342 | NM_005465.4 | #N/A | 716.064516 | -10.694312 | 1.64738594 |
| 1343 | NM_021259.2 | #N/A | 339.598366 | -8.2909789 | 1.27723159 |
| 1344 | NM_006947.3 | #N/A | 438.704616 | -9.9875304 | 1.53871997 |
| 1345 | NM_016322.3 | #N/A | 494.164387 | -10.159389 | 1.56518587 |
| 1346 | NM_005340.6 | #N/A | 578.834322 | -10.387567 | 1.60039051 |
| 1347 | NM_007107.4 | SSR3 | 446.554494 | -10.013276 | 1.54289616 |
| 1348 | NM_000527.4 | #N/A | 555.302576 | -10.328031 | 1.59146857 |
| 1349 | NM_005275.3 | GNL1 | 437.972394 | -9.9851467 | 1.53866037 |
| 1350 | NM_198968.3 | #N/A | 356.664013 | -9.689369 | 1.49313001 |
| 1351 | NM_007353.2 | #N/A | 456.497177 | -10.044992 | 1.54802039 |
| 1352 | NM_002906.3 | RDX | 381.349577 | -9.7852722 | 1.50803919 |
| 1353 | NM_031845.2 | MAP2 | 974.042805 | -11.138365 | 1.71678357 |
| 1354 | NM_014742.3 | #N/A | 408.668955 | -9.8853998 | 1.52392361 |
| 1355 | NM_002383.3 | #N/A | 855.86871 | -10.95171 | 1.68863831 |
| 1356 | NM_003025.3 | #N/A | 420.446814 | -9.9262841 | 1.53054364 |
| 1357 | NM_001001520.2 | #N/A | 408.044367 | -9.882771 | 1.52398877 |
| 1358 | NM_001330071.1 | #N/A | 685.723289 | -10.631973 | 1.63969356 |
| 1359 | NM_004136.3 | #N/A | 364.314454 | -9.7192247 | 1.49902869 |
| 1360 | NM_014935.4 | #N/A | 377.32175 | -8.4955387 | 1.31038913 |
| 1361 | NM_015578.2 | #N/A | 340.526885 | -9.6213392 | 1.48412631 |
| 1362 | NM_015177.1 | #N/A | 395.689477 | -9.8386808 | 1.51791632 |
| 1363 | NM_004516.3 | #N/A | 511.782639 | -10.209686 | 1.57557068 |
| 1364 | NM_002794.4 | #N/A | 483.147759 | -10.12684 | 1.5628704 |
| 1365 | NM_182931.2 | #N/A | 369.911917 | -9.7412634 | 1.50345265 |
| 1366 | NM_001100590.1 | KIAA0232 | 379.673089 | -9.7792737 | 1.5095079 |
| 1367 | NM_004247.3 | #N/A | 432.378552 | -9.9668206 | 1.53846693 |
| 1368 | NM_033000.3 | #N/A | 399.68677 | -9.8520782 | 1.52071356 |
| 1369 | NM_182498.3 | #N/A | 634.377695 | -10.519707 | 1.62408185 |
| 1370 | NM_015130.2 | #N/A | 356.914381 | -9.691185 | 1.49628124 |
| 1371 | NM_014970.3 | #N/A | 542.038883 | -10.292857 | 1.58922009 |
| 1372 | NM_174911.4 | #N/A | 378.238851 | -9.774333 | 1.50917506 |
| 1373 | NM_001322410.1 | PNISR | 366.301895 | -9.7268187 | 1.50198218 |
| 1374 | NM_014639.3 | TTC37 | 371.866786 | -9.749377 | 1.50560459 |
| 1375 | NM_033133.4 | #N/A | 421.424892 | -9.9299245 | 1.53349381 |

| | | | | | |
|---|---|---|---|---|---|
| 1376 | NM_006107.3 | LUC7L3 | 431.245558 | -9.9625797 | 1.53866101 |
| 1377 | NM_001032296.3 | STK24 | 327.226261 | -9.5657047 | 1.47740761 |
| 1378 | NM_001195427.1 | #N/A | 454.964617 | -10.039949 | 1.55085485 |
| 1379 | NM_001198802.1 | #N/A | 401.095376 | -9.8589448 | 1.52302838 |
| 1380 | NM_002709.2 | #N/A | 488.366427 | -10.142416 | 1.56686574 |
| 1381 | NM_020719.2 | #N/A | 371.024072 | -9.7457493 | 1.50562983 |
| 1382 | NM_016292.2 | #N/A | 412.26995 | -9.8979825 | 1.52929902 |
| 1383 | NM_018426.2 | #N/A | 376.990287 | -9.7690715 | 1.5094206 |
| 1384 | NM_004475.2 | #N/A | 406.392625 | -9.8775804 | 1.52635053 |
| 1385 | NM_006135.2 | #N/A | 399.071932 | -9.8511624 | 1.52227106 |
| 1387 | NM_020474.3 | GALNT1 | 378.230779 | -8.4980994 | 1.31341746 |
| 1386 | NM_018649.2 | #N/A | 514.407884 | -10.217279 | 1.57913704 |
| 1388 | NM_001039550.1 | #N/A | 349.209355 | -9.6581837 | 1.49294672 |
| 1389 | NM_020461.3 | #N/A | 359.575877 | -5.0602154 | 0.78223818 |
| 1390 | NM_001037663.1 | EEF1B2 | 605.078857 | -10.451432 | 1.61599557 |
| 1391 | NM_014631.2 | SH3PXD2A | 341.945363 | -9.6283566 | 1.48876956 |
| 1392 | NM_001100623.1 | PLEKHB2 | 401.661342 | -9.8593191 | 1.52457036 |
| 1393 | NM_001286134.1 | RIC8A | 444.486672 | -10.006687 | 1.54742021 |
| 1394 | NM_006698.3 | #N/A | 499.594599 | -10.175278 | 1.57377899 |
| 1395 | NM_001127173.2 | #N/A | 768.833798 | -10.797096 | 1.67000813 |
| 1396 | NM_022731.4 | #N/A | 1557.103 | -6.0167904 | 0.93076326 |
| 1397 | NM_015331.2 | #N/A | 390.510503 | -9.8201509 | 1.51916728 |
| 1398 | NM_001033853.1 | #N/A | 533.081759 | -10.269018 | 1.58865509 |
| 1399 | NM_022754.6 | #N/A | 430.420514 | -9.9598224 | 1.54107094 |
| 1400 | NM_001164782.1 | #N/A | 46.148095 | 10.4987906 | 1.62449409 |
| 1401 | NM_003640.4 | #N/A | 337.046094 | -9.6061268 | 1.48673826 |
| 1402 | NM_017871.5 | #N/A | 364.421121 | -9.7197282 | 1.50439959 |
| 1403 | NM_001954.4 | DDR1 | 451.829962 | -10.030802 | 1.55257979 |
| 1404 | NM_144703.2 | #N/A | 439.352156 | -9.9898179 | 1.54627135 |
| 1405 | NM_003040.3 | #N/A | 398.539933 | -9.850064 | 1.52477804 |
| 1406 | NM_021079.4 | #N/A | 408.992823 | -9.886812 | 1.53056656 |
| 1407 | NM_001197294.1 | #N/A | 498.092042 | -10.171049 | 1.5746759 |
| 1408 | NM_003913.4 | #N/A | 328.635358 | -9.5708501 | 1.48173595 |
| 1409 | NM_006379.3 | #N/A | 614.534743 | -10.474061 | 1.62163481 |
| 1410 | NM_017737.4 | #N/A | 696.772025 | -9.2266759 | 1.4284744 |
| 1411 | NM_198896.1 | #N/A | 646.637098 | -10.547341 | 1.63295726 |
| 1412 | NM_000636.3 | #N/A | 346.391647 | -9.6466825 | 1.49361287 |
| 1413 | NM_012111.2 | #N/A | 393.519415 | -9.8309548 | 1.52217745 |
| 1414 | NM_019844.3 | #N/A | 25.6219267 | 6.96540325 | 1.07863845 |
| 1415 | NM_004583.3 | #N/A | 472.459672 | -10.094473 | 1.56325305 |
| 1416 | NM_014298.4 | #N/A | 405.203001 | -9.8736177 | 1.52907931 |
| 1417 | NM_001251971.1 | #N/A | 405.756144 | -5.3131669 | 0.82284919 |
| 1418 | NM_001098504.1 | #N/A | 413.041772 | -9.9002215 | 1.53338876 |

| | | | | | |
|---|---|---|---|---|---|
| 1419 | NM_003045.4 | #N/A | 347.334626 | -9.6514975 | 1.49487965 |
| 1420 | NM_015164.2 | #N/A | 394.352819 | -9.8339491 | 1.5234269 |
| 1421 | NM_005735.3 | #N/A | 370.796647 | -9.7446748 | 1.50963852 |
| 1422 | NM_024103.2 | #N/A | 427.109748 | -9.9492521 | 1.5416902 |
| 1423 | NM_022151.4 | #N/A | 448.431293 | -7.4530992 | 1.1549274 |
| 1424 | NM_018026.3 | #N/A | 463.767609 | -10.06796 | 1.56030055 |
| 1425 | NM_002273.3 | #N/A | 464.171046 | -10.069628 | 1.56069382 |
| 1426 | NM_177438.2 | DICER1 | 350.349894 | -9.6643129 | 1.49793533 |
| 1427 | NM_015097.2 | #N/A | 462.886042 | -10.065025 | 1.56015074 |
| 1428 | NM_002613.4 | #N/A | 356.214037 | -9.6866869 | 1.50163286 |
| 1429 | NM_003200.3 | #N/A | 425.735865 | -9.9439069 | 1.54154253 |
| 1430 | NM_006990.4 | #N/A | 342.021866 | -9.6299203 | 1.49293133 |
| 1431 | NM_004687.4 | #N/A | 484.474085 | -10.130695 | 1.57062175 |
| 1432 | NM_006047.5 | #N/A | 370.229877 | -9.7420486 | 1.51053446 |
| 1433 | NM_005778.3 | #N/A | 349.829195 | -9.6598656 | 1.49788482 |
| 1434 | NM_032223.3 | PCNX3 | 365.411206 | -9.7240643 | 1.50821275 |
| 1435 | NM_013446.3 | #N/A | 422.435296 | -9.9330594 | 1.54065619 |
| 1436 | NM_001014283.1 | #N/A | 149.851574 | -5.075134 | 0.78731241 |
| 1437 | NM_001025091.1 | #N/A | 398.501758 | -9.8491169 | 1.5279866 |
| 1438 | NM_002442.3 | #N/A | 350.4044 | -7.2499221 | 1.12472676 |
| 1439 | NM_014764.3 | #N/A | 375.581837 | -9.7627907 | 1.51460846 |
| 1440 | NM_015966.2 | #N/A | 369.062349 | -9.7393891 | 1.51103768 |
| 1441 | NM_001130144.2 | LTBP3 | 318.500989 | -9.5252136 | 1.47809823 |
| 1442 | NM_001678.4 | #N/A | 328.113284 | -9.5677963 | 1.48503107 |
| 1446 | NR_046252.1 | FGD5-AS1 | 429.754415 | -9.957723 | 1.54569474 |
| 1443 | NM_006022.3 | #N/A | 626.151945 | -10.500856 | 1.62999374 |
| 1444 | NM_006767.3 | #N/A | 415.552509 | -9.9094593 | 1.53821465 |
| 1445 | NM_016312.2 | #N/A | 369.831884 | -9.7409394 | 1.51201125 |
| 1447 | NM_006623.3 | #N/A | 557.230937 | -7.7481104 | 1.2029499 |
| 1448 | NM_015245.2 | #N/A | 416.799444 | -9.9140051 | 1.5392261 |
| 1449 | NM_001199383.1 | #N/A | 411.433169 | -9.8951877 | 1.53651025 |
| 1450 | NM_004514.3 | #N/A | 475.984115 | -10.105195 | 1.56924125 |
| 1451 | NM_001271803.1 | #N/A | 527.053431 | -10.25231 | 1.59227844 |
| 1452 | NM_015388.3 | #N/A | 435.678248 | -9.9776178 | 1.54959658 |
| 1453 | NM_001110.3 | #N/A | 346.212119 | -9.6459993 | 1.49833388 |
| 1454 | NM_002210.4 | #N/A | 432.783857 | -9.967932 | 1.54833461 |
| 1455 | NM_002810.2 | #N/A | 492.192035 | -10.153712 | 1.57709246 |
| 1456 | NM_004768.3 | #N/A | 369.287296 | -9.739296 | 1.51278 |
| 1457 | NM_015356.4 | #N/A | 362.773312 | -9.7140455 | 1.50892297 |
| 1458 | NM_194285.2 | #N/A | 127.131584 | -4.496392 | 0.69842431 |
| 1459 | NM_001105570.1 | NUDT19 | 238.633151 | 2.36752459 | 0.36778515 |
| 1460 | NM_001136200.1 | #N/A | 134.720727 | -3.6759158 | 0.57103995 |
| 1461 | NM_000414.3 | #N/A | 319.788394 | -9.5318074 | 1.48081442 |

| | | | | | |
|---|---|---|---|---|---|
| 1462 | NM_003792.3 | #N/A | 517.391528 | -10.225573 | 1.58861283 |
| 1463 | NM_006367.3 | CAP1 | 489.22339 | -10.144712 | 1.57624332 |
| 1464 | NM_000179.2 | MSH6 | 382.632532 | -9.790066 | 1.52123112 |
| 1467 | NR_046228.1 | ANKRD20A12P | 51.77685 | 4.0710501 | 0.63278535 |
| 1465 | NM_001098209.1 | #N/A | 340.442299 | -9.6231199 | 1.49580791 |
| 1466 | NM_006546.3 | #N/A | 409.84824 | -9.8899095 | 1.53725722 |
| 1468 | NM_152280.4 | #N/A | 1413.01731 | -4.7832017 | 0.74352345 |
| 1469 | NM_006369.4 | LRRC41 | 358.341565 | -9.6961407 | 1.50730275 |
| 1470 | NM_001042424.2 | #N/A | 434.191881 | -9.9725438 | 1.55038317 |
| 1471 | NM_001626.5 | #N/A | 308.840749 | -9.4812524 | 1.4740737 |
| 1472 | NM_181672.2 | #N/A | 378.718075 | -9.7751268 | 1.51981425 |
| 1473 | NM_001270975.1 | #N/A | 378.914344 | -9.7762574 | 1.52005389 |
| 1474 | NM_002333.3 | #N/A | 613.885663 | -10.472321 | 1.62837743 |
| 1475 | NM_004691.4 | #N/A | 458.089035 | -10.05028 | 1.5629199 |
| 1478 | NM_015874.4 | RBPJ | 445.946127 | -10.010625 | 1.5568133 |
| 1476 | NM_014502.4 | #N/A | 443.048551 | -10.001886 | 1.5554743 |
| 1477 | NM_015306.2 | #N/A | 339.543804 | -9.6177501 | 1.4957468 |
| 1479 | NM_001360.2 | DHCR7 | 617.364751 | -10.480671 | 1.62998501 |
| 1480 | NM_005121.2 | #N/A | 255.953392 | -7.1293935 | 1.10887191 |
| 1481 | NM_002225.3 | #N/A | 306.633388 | -9.4719545 | 1.47330681 |
| 1486 | NR_026656.1 | LINC02381 | 704.745653 | -10.671709 | 1.66006809 |
| 1482 | NM_001316.3 | #N/A | 463.231126 | -10.065938 | 1.56576449 |
| 1483 | NM_006795.3 | #N/A | 401.007349 | -9.8583429 | 1.53348365 |
| 1484 | NM_007144.2 | #N/A | 346.452036 | -8.3710909 | 1.30217774 |
| 1485 | NM_007270.4 | #N/A | 356.556794 | -9.6889407 | 1.50717381 |
| 1487 | NM_001288732.1 | #N/A | 323.064812 | -9.547595 | 1.48529352 |
| 1488 | NM_016057.2 | #N/A | 443.914671 | -10.004508 | 1.55646084 |
| 1489 | NM_000687.3 | #N/A | 430.099671 | -9.9594525 | 1.54947861 |
| 1490 | NM_022003.3 | #N/A | 850.83018 | -10.943245 | 1.70256147 |
| 1491 | NM_014521.2 | #N/A | 373.858269 | -9.7574216 | 1.51822739 |
| 1492 | NM_015658.3 | #N/A | 450.319288 | -10.025329 | 1.5602458 |
| 1493 | NM_001131016.1 | #N/A | 356.433044 | -9.687624 | 1.50787693 |
| 1494 | NM_003755.4 | #N/A | 454.425092 | -10.038423 | 1.56249252 |
| 1495 | NM_004373.3 | #N/A | 694.970669 | -10.651351 | 1.65795324 |
| 1496 | NM_014663.2 | #N/A | 383.110563 | -9.7921205 | 1.52431376 |
| 1497 | NM_018998.3 | #N/A | 417.20931 | -9.9150378 | 1.54344574 |
| 1498 | NM_014856.2 | #N/A | 371.746374 | -9.7491442 | 1.51767437 |
| 1499 | NM_007165.4 | #N/A | 433.940148 | -9.9718522 | 1.55242938 |
| 1500 | NM_019884.2 | #N/A | 461.526887 | -10.060833 | 1.56626705 |
| 1501 | NM_001281454.1 | #N/A | 481.198188 | -10.120914 | 1.5757157 |
| 1502 | NM_006628.5 | #N/A | 518.167996 | -10.227638 | 1.59232675 |
| 1507 | NM_006329.3 | FBLN5 | 634.34534 | -10.519931 | 1.63820245 |
| 1504 | NM_002804.4 | PSMC3 | 445.350596 | -10.009385 | 1.55867054 |

| | | | | | |
|---|---|---|---|---|---|
| 1503 | NM_001243127.2 | #N/A | 211.545322 | -4.818847 | 0.75045233 |
| 1505 | NM_004958.3 | #N/A | 355.202321 | -9.6830019 | 1.50789292 |
| 1506 | NM_005146.4 | #N/A | 358.038156 | -9.6945207 | 1.50967438 |
| 1508 | NM_014003.3 | #N/A | 350.19316 | -9.6623822 | 1.50474539 |
| 1509 | NM_014909.4 | #N/A | 534.959163 | -10.273569 | 1.59981866 |
| 1510 | NM_020440.3 | #N/A | 481.410107 | -10.121727 | 1.57629546 |
| 1511 | NM_176812.4 | #N/A | 403.336466 | -9.8663014 | 1.53653365 |
| 1512 | NM_130839.2 | #N/A | 401.592437 | -9.8601256 | 1.53589509 |
| 1513 | NM_001689.4 | #N/A | 485.261018 | -10.133216 | 1.57864662 |
| 1514 | NM_002482.3 | #N/A | 454.270942 | -10.037913 | 1.56433961 |
| 1515 | NM_002730.3 | #N/A | 505.476397 | -10.192244 | 1.58838101 |
| 1516 | NM_002080.3 | #N/A | 427.603969 | -9.9507741 | 1.5508442 |
| 1517 | NM_005777.2 | #N/A | 324.243384 | -9.5509549 | 1.48851079 |
| 1518 | NM_006464.3 | #N/A | 408.51607 | -9.8848044 | 1.54055796 |
| 1519 | NM_012305.3 | #N/A | 409.98822 | -9.8898207 | 1.54140626 |
| 1520 | NM_003403.4 | #N/A | 345.80494 | -9.6439772 | 1.50314987 |
| 1521 | NM_032493.3 | #N/A | 415.197357 | -9.9082782 | 1.54440916 |
| 1522 | NM_031262.3 | HNRNPK | 368.427574 | -9.7355793 | 1.51751903 |
| 1523 | NM_003076.4 | #N/A | 432.058096 | -9.9652731 | 1.55349163 |
| 1524 | NM_001007553.2 | #N/A | 342.883672 | -9.6318102 | 1.50156238 |
| 1525 | NM_003145.3 | #N/A | 491.249046 | -10.150792 | 1.58276911 |
| 1527 | NM_001135586.1 | C5orf24 | 357.227811 | -9.690859 | 1.51114781 |
| 1526 | NM_001130991.2 | #N/A | 560.872588 | -10.342465 | 1.6127744 |
| 1528 | NM_003692.4 | #N/A | 645.267496 | -10.544298 | 1.64421987 |
| 1529 | NM_021807.3 | #N/A | 348.580816 | -9.6554396 | 1.5056893 |
| 1530 | NM_004380.2 | #N/A | 319.648089 | -9.532089 | 1.48648853 |
| 1531 | NM_005870.4 | SAP18 | 412.12248 | -9.8974865 | 1.54358674 |
| 1532 | NM_015965.6 | #N/A | 504.456067 | -10.189141 | 1.5892368 |
| 1533 | NM_001164.4 | #N/A | 582.500295 | -10.396526 | 1.62161657 |
| 1534 | NM_001098833.1 | ATXN7L3 | 453.008039 | -10.034072 | 1.56525877 |
| 1535 | NM_004052.3 | BNIP3 | 546.811081 | -10.304984 | 1.6077824 |
| 1536 | NR_123722.1 | PKD1P6-NPIPP1 | 305.113904 | -8.1395937 | 1.26998553 |
| 1537 | NM_003345.4 | #N/A | 453.207182 | -10.03439 | 1.56573091 |
| 1538 | NM_006312.5 | #N/A | 422.953565 | -9.9347872 | 1.55035451 |
| 1539 | NM_022744.3 | #N/A | 367.989675 | -9.7340488 | 1.51923122 |
| 1540 | NM_001042539.2 | #N/A | 491.59831 | -10.151634 | 1.5844828 |
| 1541 | NM_001281720.1 | UBB | 648.721994 | -10.552148 | 1.64758237 |
| 1542 | NM_014997.3 | #N/A | 311.947625 | -8.2185726 | 1.28329401 |
| 1543 | NM_002430.2 | #N/A | 612.18024 | -8.4403329 | 1.31800426 |
| 1544 | NM_017934.6 | #N/A | 384.297711 | -8.5214015 | 1.33081286 |
| 1545 | NM_004888.3 | #N/A | 491.671212 | -10.152166 | 1.58552739 |
| 1546 | NM_181891.2 | #N/A | 457.628716 | -10.048255 | 1.56947314 |
| 1547 | NM_001145418.1 | TTC28 | 469.835011 | -3.9886227 | 0.62307847 |

| | | | | | |
|---|---|---|---|---|---|
| 1548 | NM_002802.2 | #N/A | 427.197614 | -9.9493228 | 1.55433246 |
| 1549 | NM_025185.3 | #N/A | 389.803584 | -9.8169542 | 1.53369158 |
| 1550 | NM_014820.4 | #N/A | 389.69697 | -9.8164653 | 1.53402156 |
| 1551 | NM_015171.3 | #N/A | 534.539938 | -5.3514871 | 0.83637366 |
| 1552 | NM_015208.4 | #N/A | 551.088415 | -10.316476 | 1.61240191 |
| 1553 | NM_014823.2 | #N/A | 349.304705 | -9.6592792 | 1.51028298 |
| 1554 | NM_005176.5 | #N/A | 521.45388 | -10.236876 | 1.60077301 |
| 1555 | NM_022075.4 | #N/A | 368.043205 | -9.7348357 | 1.5229275 |
| 1556 | NM_001302883.1 | CLSTN1 | 406.586492 | -9.8782912 | 1.54555381 |
| 1557 | NM_030912.2 | TRIM8 | 316.632075 | -9.5170619 | 1.48931161 |
| 1558 | NM_139062.2 | #N/A | 329.187076 | -9.5737644 | 1.49820946 |
| 1559 | NM_001009998.3 | #N/A | 328.807797 | -9.5715743 | 1.49790728 |
| 1560 | NM_012286.2 | #N/A | 390.199814 | -9.8183251 | 1.53672845 |
| 1561 | NM_015039.3 | #N/A | 536.947874 | -10.27913 | 1.60885149 |
| 1562 | NM_020198.2 | #N/A | 333.037125 | -9.5901499 | 1.50109478 |
| 1563 | NM_001015048.2 | BAG5 | 158.472168 | -4.820803 | 0.75461106 |
| 1564 | NM_025241.2 | #N/A | 424.938929 | -9.9414396 | 1.55627318 |
| 1565 | NM_001348.2 | #N/A | 371.052139 | -9.746041 | 1.52583964 |
| 1566 | NM_005415.4 | #N/A | 316.75485 | -9.5189054 | 1.49030539 |
| 1567 | NM_002959.6 | #N/A | 582.11647 | -10.395808 | 1.62767669 |
| 1568 | NM_006572.5 | #N/A | 362.752569 | -9.7130008 | 1.52083624 |
| 1570 | NM_005128.3 | DOP1B | 41.1975049 | 10.3492574 | 1.62049165 |
| 1569 | NM_005119.3 | #N/A | 337.394541 | -9.6082685 | 1.50448694 |
| 1571 | NM_005500.2 | #N/A | 492.140049 | -10.153418 | 1.5899502 |
| 1572 | NM_004355.3 | CD74 | 780.704994 | -10.819383 | 1.69426208 |
| 1573 | NM_022765.3 | #N/A | 503.154762 | -10.18544 | 1.59523921 |
| 1574 | NM_001267782.1 | #N/A | 133.258809 | -4.0252925 | 0.6305619 |
| 1575 | NM_006426.2 | #N/A | 664.86679 | -9.15508 | 1.43420603 |
| 1576 | NM_006793.4 | #N/A | 353.965431 | -9.6780939 | 1.51617159 |
| 1577 | NM_001039960.2 | #N/A | 458.949967 | -10.05246 | 1.57496718 |
| 1578 | NM_018976.4 | #N/A | 637.709618 | -10.527245 | 1.64940806 |
| 1579 | NM_013325.4 | #N/A | 351.960447 | -9.669415 | 1.51503905 |
| 1580 | NM_014344.3 | #N/A | 603.743636 | -10.448332 | 1.63742098 |
| 1581 | NR_046456.1 | LINC00907 | 69.2417055 | 3.98678758 | 0.62482849 |
| 1582 | NM_019609.4 | #N/A | 333.003158 | -9.5913963 | 1.50340652 |
| 1583 | NM_003472.3 | #N/A | 394.322304 | -9.833068 | 1.54132629 |
| 1584 | NM_022912.2 | #N/A | 661.5757 | -10.580219 | 1.65870786 |
| 1585 | NM_004943.1 | #N/A | 337.07584 | -9.6078752 | 1.50632192 |
| 1586 | NM_005631.4 | #N/A | 312.042765 | -9.4974549 | 1.48907084 |
| 1587 | NR_110292.1 | SNHG31 | 84.270979 | 2.88886798 | 0.45295692 |
| 1588 | NM_005063.4 | #N/A | 3880.31315 | -6.8662647 | 1.07665255 |
| 1589 | NM_030579.2 | #N/A | 343.370579 | -7.0494177 | 1.10537571 |
| 1590 | NM_174921.2 | #N/A | 318.031276 | -9.523282 | 1.49333403 |

| | | | | | |
|---|---|---|---|---|---|
| 1591 | NM_015850.3 | #N/A | 290.525447 | -9.3934927 | 1.47307226 |
| 1592 | NM_021961.5 | #N/A | 297.783931 | -8.1053195 | 1.27107712 |
| 1593 | NM_031463.4 | #N/A | 466.524126 | -10.076163 | 1.58014109 |
| 1594 | NM_024042.3 | #N/A | 319.274833 | -9.5296224 | 1.49457528 |
| 1595 | NM_001039705.2 | #N/A | 430.993537 | -9.9624326 | 1.56250284 |
| 1596 | NM_033546.3 | MYL12B | 522.802742 | -10.240761 | 1.60625111 |
| 1597 | NM_012207.2 | HNRNPH3 | 377.845059 | -9.7718941 | 1.53309016 |
| 1598 | NM_014675.4 | #N/A | 299.747242 | -9.4388108 | 1.48090872 |
| 1599 | NM_007260.2 | #N/A | 307.064831 | -9.474188 | 1.48651034 |
| 1600 | NM_005255.3 | #N/A | 312.057489 | -9.4962456 | 1.49000013 |
| 1601 | NM_014392.4 | NSG1 | 651.921845 | -10.559191 | 1.65684042 |
| 1602 | NM_006336.3 | #N/A | 328.766727 | -4.6578259 | 0.73089321 |
| 1603 | NM_005339.4 | #N/A | 448.894962 | -10.020649 | 1.57244232 |
| 1604 | NM_001145108.1 | NELL2 | 756.919301 | -10.77436 | 1.6907747 |
| 1605 | NM_001289123.1 | TUBB4A | 594.75926 | -10.426775 | 1.63624993 |
| 1606 | NM_015466.3 | #N/A | 332.169577 | -9.5863027 | 1.50434914 |
| 1607 | NM_014740.3 | #N/A | 348.614497 | -9.6558943 | 1.51531411 |
| 1608 | NM_001024807.2 | APLP1 | 518.090152 | -10.227665 | 1.60528233 |
| 1609 | NM_021814.4 | #N/A | 420.175284 | -9.9254367 | 1.55786863 |
| 1610 | NM_001306131.1 | FN1 | 461.849 | -10.062361 | 1.57945083 |
| 1611 | NM_003715.3 | USO1 | 299.154226 | -9.4363041 | 1.48122099 |
| 1612 | NM_002902.2 | #N/A | 482.238472 | -10.124125 | 1.58959334 |
| 1613 | NM_001037984.2 | #N/A | 376.943879 | -9.7693281 | 1.53391486 |
| 1614 | NM_001278244.1 | ZFAND5 | 44.000318 | 10.4466291 | 1.64043516 |
| 1615 | NM_001810.5 | #N/A | 384.93453 | -9.7990057 | 1.53893768 |
| 1616 | NM_024612.4 | #N/A | 288.036019 | -9.3796883 | 1.47325028 |
| 1617 | NM_012477.3 | #N/A | 351.852954 | -9.6693211 | 1.51881733 |
| 1618 | NM_012197.3 | #N/A | 369.859787 | -8.4664046 | 1.33006244 |
| 1619 | NM_024516.3 | #N/A | 311.748865 | -9.4958505 | 1.49184947 |
| 1620 | NM_001080533.2 | #N/A | 286.899253 | -9.3758084 | 1.47317857 |
| 1621 | NM_020410.2 | #N/A | 356.578733 | -9.6891895 | 1.52269723 |
| 1622 | NM_001099784.2 | FBXL19 | 483.725145 | -10.128698 | 1.59179327 |
| 1623 | NM_003139.3 | #N/A | 341.924186 | -9.6278367 | 1.5131858 |
| 1624 | NM_006476.4 | #N/A | 470.040976 | -10.08714 | 1.58548033 |
| 1625 | NM_001662.3 | #N/A | 497.863938 | -10.170115 | 1.59857212 |
| 1626 | NM_015456.4 | #N/A | 411.074504 | -9.8936339 | 1.55532129 |
| 1627 | NM_001017995.2 | #N/A | 358.605494 | -9.6967627 | 1.52467193 |
| 1628 | NM_004907.2 | #N/A | 297.520376 | -9.427584 | 1.48248605 |
| 1629 | NM_003481.2 | #N/A | 389.516575 | -9.8159202 | 1.54365838 |
| 1630 | NM_012235.3 | #N/A | 360.663732 | -9.7050261 | 1.52622528 |
| 1631 | NM_014700.3 | #N/A | 345.578794 | -9.6431979 | 1.51645844 |
| 1632 | NM_198241.2 | #N/A | 465.223912 | -10.073144 | 1.58437313 |
| 1633 | NM_020898.2 | #N/A | 307.052896 | -9.472888 | 1.49002846 |

| | | | | | |
|---|---|---|---|---|---|
| 1634 | NM_001029.3 | #N/A | 700.344565 | -10.662447 | 1.67742981 |
| 1635 | NM_001101372.1 | #N/A | 519.854979 | -8.9617088 | 1.40987862 |
| 1636 | NM_001469.4 | #N/A | 392.979675 | -9.8289325 | 1.54627526 |
| 1637 | NM_003418.4 | #N/A | 389.355229 | -9.8162586 | 1.54431247 |
| 1638 | NM_001007097.2 | #N/A | 295.276558 | -8.1406846 | 1.28093019 |
| 1639 | NM_145290.3 | #N/A | 285.940733 | -9.3707868 | 1.47446561 |
| 1640 | NM_001005.4 | #N/A | 488.891948 | -10.143962 | 1.59633509 |
| 1641 | NM_002883.3 | #N/A | 463.272467 | -10.066215 | 1.5841682 |
| 1644 | NR_026682.1 | LOC100268168 | 75.4864402 | 3.73921146 | 0.5884895 |
| 1642 | NM_003100.3 | #N/A | 251.850526 | -3.1539183 | 0.49637457 |
| 1643 | NM_005780.2 | #N/A | 233.640962 | -6.9977048 | 1.101305 |
| 1645 | NM_001006935.2 | TCEAL4 | 443.954851 | -10.004628 | 1.57484016 |
| 1646 | NM_001148.4 | #N/A | 640.27356 | -10.533053 | 1.65802581 |
| 1647 | NM_004341.4 | #N/A | 292.710843 | -9.4046699 | 1.48075162 |
| 1648 | NM_014747.2 | #N/A | 531.740536 | -10.265311 | 1.61633639 |
| 1649 | NM_015213.3 | #N/A | 395.569792 | -9.838192 | 1.54923721 |
| 1650 | NM_006345.3 | #N/A | 393.71926 | -9.8313852 | 1.54822335 |
| 1651 | NM_138423.3 | #N/A | 356.185817 | -9.6872179 | 1.52553328 |
| 1652 | NM_001420.3 | #N/A | 852.306626 | -10.945722 | 1.72379096 |
| 1653 | NM_002572.3 | #N/A | 502.280628 | -10.182674 | 1.60361693 |
| 1654 | NM_015151.3 | #N/A | 278.054081 | -9.3298742 | 1.46934723 |
| 1655 | NM_001144.5 | #N/A | 327.123994 | -9.5644757 | 1.50667618 |
| 1656 | NM_005032.6 | #N/A | 283.452362 | -9.3586017 | 1.47433803 |
| 1657 | NM_005016.5 | #N/A | 429.026499 | -9.9545004 | 1.56832922 |
| 1658 | NM_001552.2 | #N/A | 321.304003 | -9.5399072 | 1.50319089 |
| 1659 | NM_017905.4 | #N/A | 298.828103 | -9.4352075 | 1.48669499 |
| 1660 | NM_015584.4 | #N/A | 393.552316 | -9.8309361 | 1.54908499 |
| 1661 | NM_012262.3 | #N/A | 208.95487 | -6.7536732 | 1.06422159 |
| 1662 | NM_005891.2 | #N/A | 620.746475 | -10.488472 | 1.65276963 |
| 1663 | NM_025029.4 | #N/A | 490.00733 | -10.147072 | 1.59905177 |
| 1664 | NM_177972.2 | TUB | 478.720768 | -10.113361 | 1.59404024 |
| 1665 | NM_001284214.1 | #N/A | 323.206105 | -9.5482904 | 1.50501719 |
| 1666 | NM_001316374.1 | #N/A | 429.403642 | -9.9566583 | 1.56944271 |
| 1667 | NM_001711.5 | #N/A | 372.664872 | -9.7533138 | 1.53737818 |
| 1668 | NM_138501.5 | #N/A | 359.740757 | -9.7016536 | 1.52936318 |
| 1669 | NM_001282659.1 | #N/A | 347.667908 | -9.6524558 | 1.52194739 |
| 1670 | NM_006769.3 | #N/A | 448.717485 | -10.020242 | 1.57998706 |
| 1671 | NM_001104647.1 | #N/A | 360.174625 | -9.7026747 | 1.53012064 |
| 1672 | NM_012255.4 | #N/A | 323.064308 | -9.5461805 | 1.50556726 |
| 1673 | NM_003347.3 | #N/A | 396.679625 | -9.8422277 | 1.55229073 |
| 1674 | NM_002834.4 | PTPN11 | 538.765348 | -3.1489709 | 0.49665935 |
| 1675 | NM_003190.4 | TAPBP | 329.945354 | -9.5778311 | 1.51073103 |
| 1676 | NM_004359.1 | #N/A | 382.597473 | -9.7900637 | 1.54429286 |

| | | | | | |
|---|---|---|---|---|---|
| 1677 | NM_058164.3 | #N/A | 558.22945 | -10.335117 | 1.63029359 |
| 1678 | NM_014246.1 | #N/A | 371.059963 | -9.7457678 | 1.53738908 |
| 1679 | NM_058179.3 | #N/A | 457.385502 | -10.047775 | 1.58514984 |
| 1680 | NM_014372.4 | #N/A | 484.825339 | -10.1318 | 1.59846615 |
| 1681 | NM_080821.2 | #N/A | 256.850955 | -5.7259888 | 0.9034298 |
| 1682 | NM_003795.5 | #N/A | 415.758714 | -9.910162 | 1.56365658 |
| 1683 | NM_001276252.1 | #N/A | 304.314989 | -9.4599883 | 1.49270984 |
| 1684 | NM_145899.2 | #N/A | 513.724987 | -10.215242 | 1.61192134 |
| 1685 | NM_007346.3 | #N/A | 285.876967 | -9.3694102 | 1.47871698 |
| 1686 | NM_012264.4 | #N/A | 309.964861 | -9.4866347 | 1.49724369 |
| 1687 | NM_014914.4 | #N/A | 484.142893 | -10.12948 | 1.59888727 |
| 1688 | NM_003367.3 | #N/A | 427.439642 | -9.9499304 | 1.57070715 |
| 1689 | NM_005157.5 | #N/A | 365.26016 | -9.7242086 | 1.535114 |
| 1690 | NM_138967.3 | #N/A | 560.157042 | -10.340039 | 1.63231372 |
| 1691 | NM_014500.4 | #N/A | 386.822692 | -9.8056899 | 1.54822813 |
| 1692 | NM_016578.3 | RSF1 | 312.825863 | -9.4994437 | 1.50032637 |
| 1693 | NM_001020658.1 | #N/A | 310.486544 | -9.4879622 | 1.49894631 |
| 1694 | NM_152988.2 | #N/A | 294.751231 | -9.415449 | 1.48750463 |
| 1695 | NM_001856.3 | #N/A | 271.215822 | -8.0169974 | 1.26662506 |
| 1696 | NM_017870.3 | #N/A | 594.786427 | -10.426744 | 1.64739457 |
| 1697 | NM_001130965.2 | #N/A | 296.850051 | -9.4237086 | 1.48898862 |
| 1698 | NM_004860.3 | #N/A | 369.215078 | -9.7388362 | 1.53909478 |
| 1699 | NM_006816.2 | #N/A | 359.764887 | -9.7015788 | 1.53323649 |
| 1700 | NM_005625.3 | #N/A | 315.553857 | -9.5126676 | 1.50347781 |
| 1701 | NM_004385.4 | #N/A | 373.886042 | -9.7570778 | 1.54237584 |
| 1702 | NM_002305.3 | #N/A | 485.976657 | -10.135702 | 1.60232662 |
| 1703 | NM_002768.4 | #N/A | 383.86032 | -9.7947465 | 1.54853969 |
| 1704 | NM_032409.2 | #N/A | 421.842331 | -9.931175 | 1.57021889 |
| 1705 | NM_030790.4 | #N/A | 437.831981 | -9.9847156 | 1.5787825 |
| 1706 | NM_001098509.1 | #N/A | 436.705523 | -9.9811662 | 1.57841473 |
| 1707 | NM_032108.3 | #N/A | 464.748668 | -7.0064784 | 1.10799175 |
| 1708 | NM_001833.3 | #N/A | 384.039467 | -9.7957079 | 1.54911822 |
| 1711 | NM_015541.2 | LRIG1 | 301.683394 | -8.1150108 | 1.28344718 |
| 1709 | NM_004739.3 | #N/A | 432.123725 | -9.9656399 | 1.57618983 |
| 1710 | NM_005153.2 | #N/A | 345.305695 | -9.6421039 | 1.52500882 |
| 1712 | NM_015570.3 | #N/A | 683.277682 | -10.626755 | 1.68075546 |
| 1714 | NM_003663.3 | CGGBP1 | 392.631113 | -9.8272683 | 1.55444381 |
| 1713 | NM_001146688.1 | KDM3A | 338.467055 | -9.6127194 | 1.52052284 |
| 1715 | NM_001048183.1 | #N/A | 299.692001 | -9.4372082 | 1.4929591 |
| 1716 | NM_015113.3 | #N/A | 292.768164 | -9.4041084 | 1.48839025 |
| 1717 | NM_004986.3 | #N/A | 323.383656 | -9.5484921 | 1.51136929 |
| 1718 | NM_006409.3 | #N/A | 407.183568 | -9.8801247 | 1.56400304 |
| 1719 | NM_002410.4 | #N/A | 460.601784 | -10.058014 | 1.59221657 |

| | | | | | |
|---|---|---|---|---|---|
| 1720 | NM_015423.2 | #N/A | 499.437115 | -10.174714 | 1.61068755 |
| 1722 | NM_015292.2 | ESYT1 | 317.737755 | -9.5226521 | 1.50760025 |
| 1721 | NM_014738.5 | #N/A | 281.229545 | -9.3449335 | 1.47947977 |
| 1723 | NM_001136025.4 | #N/A | 394.414658 | -9.8351952 | 1.55716003 |
| 1724 | NM_001320153.1 | #N/A | 362.933398 | -9.7140357 | 1.53834716 |
| 1725 | NM_004370.5 | #N/A | 383.31839 | -9.7935266 | 1.55122362 |
| 1726 | NM_004866.5 | #N/A | 381.03145 | -9.7839949 | 1.54970281 |
| 1727 | NM_001010924.1 | #N/A | 529.345199 | -8.8477422 | 1.40147953 |
| 1728 | NM_000941.2 | #N/A | 377.522089 | -8.4063076 | 1.33159897 |
| 1729 | NM_016073.3 | #N/A | 540.129263 | -10.287554 | 1.62958774 |
| 1730 | NM_006247.3 | #N/A | 376.825778 | -9.7683017 | 1.54744839 |
| 1731 | NM_002793.3 | #N/A | 384.220081 | -9.7963451 | 1.55191656 |
| 1732 | NM_004783.3 | #N/A | 354.481008 | -9.6803125 | 1.53396362 |
| 1733 | NM_032421.2 | #N/A | 332.2846 | -9.5881192 | 1.51935388 |
| 1734 | NM_015482.1 | SLC22A23 | 462.280256 | -10.062981 | 1.59472693 |
| 1735 | NM_017911.2 | #N/A | 120.634715 | -3.851601 | 0.61042303 |
| 1736 | NM_001042646.2 | #N/A | 347.673334 | -9.652062 | 1.52975382 |
| 1737 | NM_006502.2 | #N/A | 287.058303 | -9.3761047 | 1.48602993 |
| 1738 | NM_016019.4 | #N/A | 323.927914 | -9.549627 | 1.51377776 |
| 1739 | NM_006925.4 | #N/A | 270.594044 | -9.2915332 | 1.47292457 |
| 1740 | NM_015138.4 | RTF1 | 671.833204 | -4.7054633 | 0.74596356 |
| 1741 | NM_014828.3 | #N/A | 334.777899 | -9.5974391 | 1.52160583 |
| 1742 | NM_001170755.2 | NBPF15 | 313.308731 | -9.5019228 | 1.5065474 |
| 1743 | NM_004393.5 | #N/A | 367.720484 | -9.7336621 | 1.54322849 |
| 1744 | NM_005999.2 | #N/A | 166.519465 | -4.5093001 | 0.71495347 |
| 1745 | NM_022117.3 | #N/A | 295.915613 | -9.4197104 | 1.49349579 |
| 1746 | NM_016480.4 | #N/A | 377.026646 | -9.7690473 | 1.54906622 |
| 1747 | NM_006392.3 | #N/A | 354.067886 | -9.6785029 | 1.53473335 |
| 1748 | NM_001286242.1 | #N/A | 269.973095 | -9.2882209 | 1.47293533 |
| 1749 | NM_138734.2 | #N/A | 483.873004 | -10.128926 | 1.60634006 |
| 1750 | NM_022833.3 | #N/A | 305.243831 | -9.4639126 | 1.50106609 |
| 1751 | NM_145754.3 | #N/A | 509.118868 | -10.202293 | 1.61831745 |
| 1752 | NM_003959.2 | #N/A | 521.228626 | -10.236138 | 1.6239208 |
| 1753 | NM_004285.3 | #N/A | 270.147003 | -9.289325 | 1.47377765 |
| 1754 | NM_003439.3 | #N/A | 345.7672 | -9.644486 | 1.53029576 |
| 1755 | NM_017747.2 | #N/A | 294.59134 | -9.4139848 | 1.49376344 |
| 1756 | NM_024524.3 | ATP13A3 | 305.186823 | -9.4638543 | 1.50200612 |
| 1757 | NM_001040158.1 | MACROH2A1 | 415.223796 | -9.9076728 | 1.57253453 |
| 1758 | NM_152758.5 | #N/A | 294.148153 | -9.4102351 | 1.49364387 |
| 1759 | NM_015995.3 | #N/A | 344.489735 | -9.6382724 | 1.53006339 |
| 1760 | NM_002896.3 | RBM4 | 412.851947 | -9.9000553 | 1.57190323 |
| 1761 | NM_004761.4 | #N/A | 301.3894 | -9.4476707 | 1.50032312 |
| 1762 | NM_005054.2 | #N/A | 288.812276 | -9.3836542 | 1.49014475 |

| | | | | | |
|---|---|---|---|---|---|
| 1763 | NM_030920.4 | #N/A | 288.234771 | -9.3801832 | 1.48966626 |
| 1764 | NM_001288.4 | CLIC1 | 317.495889 | -9.5217795 | 1.51228088 |
| 1765 | NM_015360.4 | #N/A | 296.285262 | -9.4215748 | 1.49638738 |
| 1766 | NM_005345.5 | #N/A | 387.78395 | -9.8096478 | 1.55815042 |
| 1767 | NM_025179.3 | #N/A | 583.543265 | -10.399055 | 1.65176399 |
| 1768 | NM_199054.2 | MKNK2 | 281.669568 | -9.3494281 | 1.48512442 |
| 1769 | NM_006666.2 | #N/A | 317.080181 | -9.5193163 | 1.51217156 |
| 1770 | NM_001007237.2 | #N/A | 556.179064 | -10.329666 | 1.64105234 |
| 1771 | NM_004302.4 | #N/A | 456.770096 | -10.045749 | 1.5959661 |
| 1772 | NM_001134422.1 | #N/A | 337.63513 | -9.6097464 | 1.52675809 |
| 1773 | NM_006918.4 | #N/A | 545.949885 | -8.8854601 | 1.4118 |
| 1774 | NM_001278912.1 | #N/A | 400.916042 | -9.8576703 | 1.56652757 |
| 1775 | NM_002088.4 | GRIK5 | 419.129869 | -9.9214788 | 1.57676441 |
| 1776 | NM_199511.2 | CCDC80 | 442.118368 | -9.9992798 | 1.58931211 |
| 1777 | NM_024835.4 | #N/A | 287.900043 | -9.3804967 | 1.49111534 |
| 1778 | NM_182729.2 | #N/A | 354.805709 | -9.6810596 | 1.53887736 |
| 1779 | NM_030755.4 | #N/A | 162.639994 | -5.0040049 | 0.79544801 |
| 1780 | NM_018667.3 | #N/A | 245.764099 | -7.0765319 | 1.12494086 |
| 1781 | NM_015277.5 | #N/A | 603.023929 | -10.446449 | 1.66082142 |
| 1782 | NM_001173431.1 | #N/A | 506.045694 | -4.9984907 | 0.79470428 |
| 1783 | NM_022173.2 | #N/A | 406.436251 | -8.6048481 | 1.36808212 |
| 1784 | NM_001297595.1 | #N/A | 364.429337 | -9.7200119 | 1.54556971 |
| 1785 | NM_021239.2 | #N/A | 646.049631 | -4.4874979 | 0.71360385 |
| 1786 | NM_006823.3 | #N/A | 636.049919 | -10.52344 | 1.67346665 |
| 1788 | NM_020721.1 | KIAA1210 | 72.0258245 | 2.80741079 | 0.44647192 |
| 1787 | NM_017617.4 | #N/A | 484.420054 | -10.130151 | 1.6110464 |
| 1791 | NM_020722.1 | CRACD | 440.858618 | -9.99465 | 1.58956152 |
| 1793 | NM_173614.2 | NOMO2 | 462.881728 | -10.065365 | 1.60085862 |
| 1789 | NM_006756.3 | #N/A | 403.320065 | -9.866101 | 1.5691666 |
| 1790 | NM_014874.3 | #N/A | 384.785962 | -9.7977096 | 1.55825695 |
| 1792 | NM_022089.3 | #N/A | 454.04501 | -10.037026 | 1.59627048 |
| 1794 | NM_031991.3 | #N/A | 272.521362 | -9.3006885 | 1.47928354 |
| 1795 | NM_000284.3 | #N/A | 331.08457 | -9.5815399 | 1.52397629 |
| 1796 | NM_002627.4 | #N/A | 444.359777 | -10.006514 | 1.59164064 |
| 1797 | NM_018071.4 | #N/A | 332.409714 | -9.5880869 | 1.52534009 |
| 1798 | NM_005669.4 | #N/A | 417.509181 | -9.916194 | 1.57756422 |
| 1799 | NM_001324320.1 | MAST2 | 453.630276 | -10.035676 | 1.59664051 |
| 1800 | NM_001613.2 | #N/A | 620.346806 | -10.487713 | 1.66858545 |
| 1801 | NM_183376.2 | #N/A | 277.375599 | -9.3270765 | 1.48417627 |
| 1802 | NM_000875.4 | #N/A | 345.203234 | -9.6407337 | 1.53414352 |
| 1803 | NM_002392.5 | MDM2 | 275.386315 | -9.3160154 | 1.48256772 |
| 1804 | NM_015705.5 | #N/A | 324.255712 | -9.5512373 | 1.52015927 |
| 1805 | NM_016263.3 | #N/A | 370.828093 | -9.745019 | 1.55102873 |

| | | | | | | |
|---|---|---|---|---|---|---|
| 1806 | NM_006048.4 | | #N/A | 285.248108 | -9.3670737 | 1.49104279 |
| 1807 | NM_001315507.1 | | #N/A | 312.424666 | -9.497626 | 1.51201047 |
| 1808 | NM_017455.3 | | #N/A | 383.378719 | -9.7932799 | 1.55917499 |
| 1809 | NM_001636.3 | | #N/A | 1474.74285 | -6.1000891 | 0.97128867 |
| 1810 | NM_002409.4 | | #N/A | 412.407906 | -9.8983887 | 1.57610266 |
| 1811 | NM_012324.4 | | #N/A | 579.26229 | -10.38859 | 1.65408529 |
| 1812 | NM_020444.4 | | #N/A | 331.452297 | -9.5827066 | 1.52578989 |
| 1813 | NM_021137.4 | | #N/A | 314.349718 | -9.5072068 | 1.51381607 |
| 1814 | NM_001278389.1 | | #N/A | 654.373264 | -10.564455 | 1.68221676 |
| 1816 | NM_001303275.1 | ACLY | | 325.311381 | -9.5574544 | 1.5220207 |
| 1815 | NM_001012426.1 | FOXP4 | | 290.292915 | -9.3907323 | 1.49546337 |
| 1817 | NM_015271.4 | | #N/A | 473.54906 | -8.0619756 | 1.28386907 |
| 1818 | NM_013233.2 | | #N/A | 267.342249 | -9.2745214 | 1.4769969 |
| 1819 | NM_004992.3 | MECP2 | | 345.90398 | -7.1858454 | 1.14448733 |
| 1820 | NM_003592.2 | | #N/A | 325.125553 | -9.5551598 | 1.52207238 |
| 1821 | NM_006145.2 | | #N/A | 363.246958 | -9.7157084 | 1.54784649 |
| 1822 | NM_006544.3 | | #N/A | 381.420979 | -9.7854601 | 1.55900571 |
| 1823 | NM_001520.3 | | #N/A | 312.991442 | -9.5008512 | 1.513721 |
| 1824 | NM_004093.3 | | #N/A | 709.457998 | -10.680957 | 1.70186033 |
| 1825 | NM_019094.5 | | #N/A | 312.512379 | -9.4976054 | 1.51334993 |
| 1826 | NM_002489.3 | | #N/A | 555.48045 | -10.328119 | 1.64574954 |
| 1827 | NM_144498.2 | | #N/A | 268.042677 | -9.276812 | 1.47821438 |
| 1828 | NM_006178.3 | | #N/A | 479.785002 | -10.116764 | 1.61213693 |
| 1829 | NM_006802.3 | | #N/A | 333.780059 | -9.5934217 | 1.52873398 |
| 1830 | NM_016404.2 | | #N/A | 346.58935 | -9.6473557 | 1.53758325 |
| 1831 | NM_002817.3 | | #N/A | 411.551409 | -9.8953379 | 1.57715495 |
| 1832 | NM_152486.2 | | #N/A | 488.713475 | -10.143742 | 1.61676142 |
| 1833 | NM_015229.3 | | #N/A | 319.995668 | -9.5326026 | 1.51941905 |
| 1834 | NM_007263.3 | | #N/A | 395.383114 | -9.8375484 | 1.56811232 |
| 1835 | NM_138689.2 | | #N/A | 556.777282 | -10.331384 | 1.64681701 |
| 1836 | NM_015933.5 | | #N/A | 386.658798 | -9.8053349 | 1.56314521 |
| 1837 | NM_004699.3 | | #N/A | 361.210455 | -9.7072738 | 1.54758708 |
| 1838 | NM_016308.2 | | #N/A | 382.490958 | -9.7894207 | 1.56068244 |
| 1839 | NM_003861.2 | | #N/A | 328.496162 | -9.5703811 | 1.52580927 |
| 1840 | NM_015047.2 | | #N/A | 300.757804 | -9.4420715 | 1.50537897 |
| 1841 | NM_015656.1 | KIF26A | | 563.281181 | -10.348053 | 1.65001881 |
| 1842 | NM_001115113.2 | | #N/A | 349.340328 | -9.6586042 | 1.5401154 |
| 1843 | NM_001312.3 | | #N/A | 416.317323 | -9.9118755 | 1.58079071 |
| 1844 | NM_032801.4 | | #N/A | 320.12736 | -9.5333092 | 1.52050587 |
| 1845 | NM_006110.2 | | #N/A | 356.899293 | -9.6899543 | 1.54551643 |
| 1846 | NM_001087.4 | | #N/A | 364.191338 | -8.3536037 | 1.33240456 |
| 1847 | NM_003352.4 | | #N/A | 384.994601 | -9.7990706 | 1.56302324 |
| 1848 | NM_001024.3 | | #N/A | 666.865998 | -10.591762 | 1.68957695 |

| | | | | | |
|---|---|---|---|---|---|
| 1849 | NM_016289.3 | #N/A | 381.699406 | -9.7865921 | 1.5612268 |
| 1850 | NM_001287054.2 | #N/A | 65.8889756 | 11.0117117 | 1.75676145 |
| 1851 | NM_005087.3 | #N/A | 282.247754 | -9.3511939 | 1.49196178 |
| 1852 | NM_001032396.2 | #N/A | 340.843824 | -9.6234331 | 1.5355389 |
| 1853 | NM_003496.3 | TRRAP | 275.176505 | -9.3160804 | 1.48662571 |
| 1854 | NM_005151.3 | #N/A | 318.409502 | -9.5250584 | 1.51999268 |
| 1855 | NM_007262.4 | #N/A | 411.621764 | -9.8956836 | 1.57907792 |
| 1856 | NM_181442.3 | #N/A | 380.786382 | -9.7826981 | 1.56110474 |
| 1857 | NM_001322434.1 | HNRNPH3 | 326.332083 | -9.5603463 | 1.52581286 |
| 1858 | NM_002168.3 | IDH2 | 372.028808 | -9.7499921 | 1.55602838 |
| 1859 | NM_079834.3 | #N/A | 269.929544 | -9.2880594 | 1.48232656 |
| 1860 | NM_080645.2 | #N/A | 776.284003 | -10.811134 | 1.72542688 |
| 1861 | NM_003801.3 | #N/A | 385.281888 | -9.8004532 | 1.5643054 |
| 1862 | NM_178026.2 | #N/A | 386.649674 | -9.8053295 | 1.56552154 |
| 1863 | NR_002578.2 | #N/A | 533.567511 | -10.270022 | 1.63976996 |
| 1864 | NM_004549.5 | #N/A | 443.501142 | -10.003186 | 1.59737348 |
| 1865 | NM_014706.3 | #N/A | 306.116572 | -9.4679988 | 1.51205832 |
| 1866 | NM_001294336.1 | #N/A | 505.536949 | -10.191891 | 1.62775381 |
| 1867 | NM_001640.3 | #N/A | 349.925438 | -9.6618134 | 1.54307553 |
| 1868 | NM_014608.4 | #N/A | 260.164626 | -9.23438 | 1.4748231 |
| 1869 | NM_002646.3 | PIK3C2B | 268.896281 | -6.1328881 | 0.97960559 |
| 1870 | NM_005243.3 | #N/A | 368.157723 | -9.7346051 | 1.5549259 |
| 1871 | NM_001122955.3 | BSCL2 | 488.948519 | -10.144124 | 1.62036273 |
| 1872 | NM_006713.3 | #N/A | 526.916803 | -10.25181 | 1.63782838 |
| 1873 | NM_001159673.1 | SYNCRIP | 326.185303 | -9.5595732 | 1.52739457 |
| 1874 | NM_002860.3 | #N/A | 314.066911 | -9.5048536 | 1.51902393 |
| 1875 | NM_001135653.1 | #N/A | 271.389209 | -9.2955215 | 1.48574444 |
| 1876 | NM_020959.2 | #N/A | 394.40296 | -9.8340328 | 1.5718854 |
| 1877 | NM_003860.3 | #N/A | 402.190566 | -9.8621706 | 1.57650706 |
| 1878 | NM_001001391.1 | #N/A | 673.797021 | -10.606892 | 1.69579359 |
| 1879 | NM_006407.3 | #N/A | 408.572207 | -9.8849455 | 1.58038588 |
| 1880 | NM_020992.3 | #N/A | 443.536229 | -10.003928 | 1.59951054 |
| 1881 | NM_001077489.3 | #N/A | 344.037028 | -9.6364607 | 1.54099201 |
| 1882 | NM_012287.5 | #N/A | 176.252642 | -6.5745289 | 1.05134481 |
| 1883 | NM_005103.4 | #N/A | 717.560462 | -9.2364738 | 1.47706488 |
| 1884 | NM_001242559.1 | #N/A | 466.827394 | -10.07692 | 1.61149293 |
| 1885 | NM_001679.3 | #N/A | 390.134635 | -9.8182433 | 1.5701693 |
| 1886 | NM_014172.5 | #N/A | 363.027477 | -9.7144515 | 1.55359904 |
| 1887 | NM_021078.2 | #N/A | 289.385172 | -9.3871779 | 1.50146771 |
| 1888 | NM_006486.2 | #N/A | 279.912277 | -9.3410364 | 1.49414477 |
| 1889 | NM_018235.2 | #N/A | 338.909575 | -9.6154964 | 1.53810046 |
| 1890 | NM_052916.2 | #N/A | 459.199522 | -10.053341 | 1.60873309 |
| 1891 | NM_001317940.1 | #N/A | 38.0013162 | 10.22245 | 1.63595713 |

| | | | | | |
|---|---|---|---|---|---|
| 1892 | NM_017761.3 | #N/A | 282.885835 | -8.0781662 | 1.29279887 |
| 1893 | NM_001042368.2 | #N/A | 444.656993 | -10.007122 | 1.60158853 |
| 1894 | NM_198321.3 | #N/A | 304.487925 | -9.4623009 | 1.51444958 |
| 1895 | NM_001193646.1 | ATF5 | 358.423246 | -9.6959488 | 1.55202563 |
| 1896 | NM_006817.3 | #N/A | 367.286488 | -9.7313233 | 1.55786712 |
| 1897 | NM_145012.5 | #N/A | 310.781445 | -9.4901338 | 1.51923671 |
| 1898 | NM_016080.3 | GLOD4 | 319.506199 | -9.5302509 | 1.5257092 |
| 1899 | NM_006709.4 | #N/A | 391.457792 | -9.8230819 | 1.57277476 |
| 1900 | NM_006989.5 | RASA4 | 381.650456 | -9.7859464 | 1.56692323 |
| 1901 | NM_001242758.1 | HLA-A | 318.639665 | -9.5277494 | 1.52563556 |
| 1902 | NM_002827.3 | #N/A | 323.298858 | -9.5476386 | 1.52909287 |
| 1903 | NM_004822.2 | #N/A | 601.406214 | -9.1745535 | 1.46942669 |
| 1904 | NM_005342.3 | #N/A | 391.064707 | -9.8217412 | 1.5731057 |
| 1905 | NM_002811.4 | #N/A | 326.205498 | -9.5601168 | 1.53133625 |
| 1906 | NM_014613.2 | #N/A | 295.084082 | -9.4156358 | 1.50820363 |
| 1907 | NM_173562.4 | #N/A | 318.441783 | -9.5252314 | 1.52572758 |
| 1908 | NM_014188.2 | #N/A | 351.01224 | -9.6659607 | 1.54832929 |
| 1909 | NM_006838.3 | #N/A | 328.521479 | -9.5704144 | 1.53312057 |
| 1910 | NM_033200.2 | #N/A | 330.262221 | -9.5781088 | 1.53460791 |
| 1911 | NM_001017405.2 | #N/A | 297.811144 | -9.4291548 | 1.51077611 |
| 1912 | NR_036592.1 | EBLN3P | 282.782824 | -9.3536105 | 1.49870462 |
| 1913 | NM_001037806.3 | #N/A | 293.66379 | -9.4092923 | 1.50766329 |
| 1914 | NM_001005335.1 | #N/A | 288.90965 | -6.9827388 | 1.11895776 |
| 1915 | NM_002775.4 | #N/A | 415.495771 | -9.9097672 | 1.58812906 |
| 1916 | NM_002314.3 | #N/A | 439.399532 | -9.990079 | 1.60166219 |
| 1917 | NM_004074.2 | #N/A | 387.124241 | -9.8072672 | 1.57233289 |
| 1918 | NM_152793.2 | #N/A | 475.363314 | -10.10312 | 1.61983639 |
| 1919 | NM_001398.2 | #N/A | 286.138776 | -9.3718426 | 1.50288772 |
| 1921 | NM_139156.3 | AMPD2 | 249.900705 | -7.8585612 | 1.26034475 |
| 1920 | NM_032281.2 | #N/A | 567.722531 | -10.359443 | 1.66142739 |
| 1922 | NM_001272082.1 | #N/A | 439.627124 | -9.9906156 | 1.60232936 |
| 1923 | NM_006325.4 | #N/A | 1123.12882 | -7.7561254 | 1.24419569 |
| 1924 | NM_021224.5 | #N/A | 304.345152 | -9.4597439 | 1.51767869 |
| 1925 | NM_014889.3 | #N/A | 270.381492 | -9.2894557 | 1.49051982 |
| 1926 | NM_001330063.1 | #N/A | 298.408199 | -9.4327089 | 1.51364092 |
| 1927 | NM_015527.3 | #N/A | 352.067492 | -9.6701616 | 1.55174214 |
| 1928 | NM_153273.3 | #N/A | 304.651357 | -9.4618609 | 1.51828119 |
| 1929 | NM_004429.4 | EFNB1 | 327.177593 | -9.5637191 | 1.53483525 |
| 1933 | NR_037612.1 | SEPT5-GP1BB | 496.335668 | -10.165611 | 1.63152891 |
| 1930 | NM_004776.3 | #N/A | 385.318042 | -9.800561 | 1.57292667 |
| 1931 | NM_006315.5 | #N/A | 382.682416 | -8.4132257 | 1.35024646 |
| 1932 | NM_030927.2 | #N/A | 421.509609 | -9.9298009 | 1.59362057 |
| 1934 | NM_001318734.1 | #N/A | 418.271134 | -9.9184999 | 1.59193612 |

| | | | | | |
|---|---|---|---|---|---|
| 1935 | NM_005407.2 | #N/A | 303.829909 | -9.4584459 | 1.51813357 |
| 1936 | NM_015374.2 | #N/A | 526.934344 | -10.251875 | 1.64545729 |
| 1937 | NM_016605.2 | #N/A | 307.346568 | -9.4740543 | 1.52062437 |
| 1938 | NM_003099.4 | #N/A | 284.317479 | -9.361991 | 1.50273925 |
| 1939 | NM_006324.2 | #N/A | 352.684692 | -9.6726262 | 1.5526064 |
| 1940 | NM_022130.3 | #N/A | 314.101209 | -9.5055207 | 1.52582588 |
| 1941 | NM_001113239.2 | #N/A | 331.51195 | -9.5832886 | 1.53842963 |
| 1942 | NM_001518.4 | #N/A | 321.41059 | -9.5396583 | 1.53142303 |
| 1943 | NM_002018.3 | #N/A | 307.116276 | -9.4724752 | 1.52060798 |
| 1944 | NM_001008895.2 | #N/A | 248.651263 | -9.1692692 | 1.47200943 |
| 1945 | NM_000637.3 | #N/A | 265.906359 | -9.2654313 | 1.487539 |
| 1946 | NM_006262.3 | #N/A | 495.198628 | -10.162709 | 1.63169229 |
| 1947 | NM_004730.3 | #N/A | 319.080997 | -9.5280761 | 1.52993407 |
| 1948 | NM_031912.4 | #N/A | 164.677665 | 2.32510497 | 0.37334828 |
| 1950 | NM_014183.3 | DYNLRB1 | 358.620032 | -9.6969547 | 1.55712256 |
| 1949 | NM_000690.3 | #N/A | 328.657303 | -9.5709698 | 1.53690113 |
| 1951 | NM_001112736.1 | #N/A | 249.534242 | -9.1728679 | 1.47308489 |
| 1952 | NM_003971.5 | #N/A | 295.241806 | -9.4156122 | 1.51228114 |
| 1953 | NM_015909.3 | #N/A | 270.120827 | -9.2881623 | 1.4920172 |
| 1954 | NM_001330520.1 | #N/A | 283.127489 | -9.3558363 | 1.5029084 |
| 1955 | NM_006833.4 | #N/A | 395.303472 | -9.8372368 | 1.58039157 |
| 1956 | NM_015057.4 | #N/A | 462.875978 | -10.065067 | 1.61712704 |
| 1957 | NM_006442.3 | #N/A | 432.038234 | -9.965551 | 1.60131093 |
| 1958 | NM_022479.2 | #N/A | 419.174448 | -5.5296142 | 0.88857074 |
| 1959 | NM_005614.3 | #N/A | 393.057097 | -9.828961 | 1.57960905 |
| 1960 | NM_002708.3 | #N/A | 336.482748 | -9.6050158 | 1.54365189 |
| 1961 | NM_018179.4 | ATF7IP | 374.765626 | -9.7593948 | 1.568489 |
| 1962 | NM_016021.2 | #N/A | 343.647655 | -9.635519 | 1.54886424 |
| 1963 | NM_001284280.1 | PPP4R3A | 276.906842 | -9.3232122 | 1.49874102 |
| 1964 | NM_001030055.1 | #N/A | 289.937606 | -9.3917431 | 1.50978775 |
| 1965 | NM_001344.3 | #N/A | 349.899073 | -9.6615465 | 1.55335097 |
| 1966 | NM_001134382.2 | #N/A | 280.492166 | -9.3430197 | 1.5021789 |
| 1967 | NM_002609.3 | #N/A | 308.428536 | -9.4807414 | 1.52446198 |
| 1968 | NM_018084.4 | #N/A | 349.952852 | -9.6616731 | 1.55393725 |
| 1969 | NM_004344.1 | #N/A | 258.654181 | -9.2269767 | 1.48408738 |
| 1970 | NM_001330221.1 | #N/A | 561.489436 | -10.343592 | 1.66382582 |
| 1971 | NM_001673.4 | #N/A | 418.236912 | -9.9186038 | 1.59550195 |
| 1972 | NM_013265.3 | #N/A | 393.978984 | -9.8321696 | 1.58165718 |
| 1973 | NM_016147.2 | #N/A | 318.061479 | -9.5237154 | 1.5320851 |
| 1974 | NM_006302.2 | #N/A | 276.175968 | -9.320675 | 1.49954397 |
| 1975 | NM_032389.4 | #N/A | 298.802592 | -9.4338442 | 1.51773365 |
| 1976 | NM_001102397.2 | #N/A | 310.319395 | -9.4879567 | 1.52655165 |
| 1977 | NM_001330395.1 | #N/A | 375.958453 | -9.7647993 | 1.57112108 |

| | | | | | |
|---|---|---|---|---|---|
| 1978 | NM_015544.2 | #N/A | 264.453014 | -9.2582504 | 1.4897749 |
| 1979 | NM_001320.6 | CSNK2B | 344.24392 | -9.6380664 | 1.55094107 |
| 1980 | NM_054016.3 | #N/A | 291.432759 | -9.3969398 | 1.51215133 |
| 1981 | NM_005186.3 | #N/A | 362.416253 | -9.7116464 | 1.5629796 |
| 1982 | NM_002855.4 | #N/A | 298.3011 | -9.4310036 | 1.5178649 |
| 1983 | NM_021033.6 | #N/A | 480.03736 | -8.706077 | 1.40119543 |
| 1984 | NM_020135.2 | #N/A | 279.45378 | -9.3367196 | 1.50283149 |
| 1985 | NM_207122.1 | EXT2 | 341.129867 | -9.6249864 | 1.54925506 |
| 1986 | NM_033089.6 | #N/A | 255.717458 | -9.2087511 | 1.48278947 |
| 1987 | NM_014306.4 | #N/A | 323.76622 | -9.549425 | 1.53783749 |
| 1988 | NM_004092.3 | #N/A | 408.596612 | -9.8853286 | 1.59196193 |
| 1989 | NM_003273.3 | #N/A | 318.443939 | -9.5255613 | 1.53443185 |
| 1990 | NM_000990.4 | #N/A | 2009.66883 | -4.1533503 | 0.66911706 |
| 1991 | NM_018013.3 | #N/A | 316.513561 | -7.4609262 | 1.20198195 |
| 1992 | NM_022149.4 | #N/A | 328.28223 | -9.569274 | 1.54162963 |
| 1993 | NR_039988.1 | #N/A | 78.3798392 | 2.61057305 | 0.42056039 |
| 1994 | NM_004718.3 | #N/A | 403.565584 | -9.8670423 | 1.58968354 |
| 1995 | NM_001039479.1 | #N/A | 345.912822 | -9.644849 | 1.55396473 |
| 1996 | NM_001113363.1 | #N/A | 437.199763 | -9.9822293 | 1.60834121 |
| 1997 | NM_005785.3 | #N/A | 312.239574 | -9.4961517 | 1.52998979 |
| 1998 | NM_016255.2 | #N/A | 289.139822 | -9.385941 | 1.51231714 |
| 1999 | NM_000075.3 | #N/A | 391.303708 | -9.8226645 | 1.58270717 |
| 2000 | NM_001060.5 | #N/A | 143.901419 | 2.87377933 | 0.46305984 |
| 2001 | NM_004397.5 | #N/A | 372.326664 | -9.7501237 | 1.57118366 |
| 2002 | NM_032333.4 | #N/A | 277.756581 | -9.3274669 | 1.50306207 |
| 2003 | NM_014371.3 | #N/A | 284.152547 | -9.3623677 | 1.50878176 |
| 2004 | NM_031286.3 | #N/A | 308.731888 | -9.4809907 | 1.52790901 |
| 2005 | NM_005124.3 | #N/A | 278.924793 | -9.334686 | 1.50449401 |
| 2006 | NM_006510.4 | #N/A | 317.234978 | -9.5198517 | 1.53432066 |
| 2007 | NM_016142.2 | #N/A | 288.443055 | -9.3824919 | 1.5127358 |
| 2008 | NM_001349.3 | #N/A | 277.72732 | -9.3269515 | 1.5038273 |
| 2009 | NM_003581.4 | #N/A | 323.233362 | -9.5467019 | 1.53929318 |
| 2010 | NM_021158.4 | TRIB3 | 170.323456 | -5.5929511 | 0.90186282 |
| 2011 | NM_007002.3 | #N/A | 334.458011 | -9.5959447 | 1.54759836 |
| 2012 | NM_020182.4 | #N/A | 386.657515 | -9.8056977 | 1.58148064 |
| 2013 | NM_001206938.1 | #N/A | 281.903319 | -9.3485551 | 1.50785867 |
| 2014 | NM_001286050.1 | #N/A | 519.141126 | -10.230286 | 1.65033963 |
| 2015 | NM_005984.4 | #N/A | 327.200184 | -9.5641357 | 1.54286997 |
| 2016 | NM_002512.3 | #N/A | 494.206083 | -10.159446 | 1.63907425 |
| 2017 | NM_003634.3 | #N/A | 384.004289 | -9.7954749 | 1.58036842 |
| 2018 | NM_138565.2 | CTTN | 229.91989 | -7.7435174 | 1.24941794 |
| 2019 | NM_002105.2 | H2AX | 379.989628 | -9.7800451 | 1.57806273 |
| 2020 | NM_177924.4 | #N/A | 398.64707 | -9.849769 | 1.58930561 |

| | | | | | | |
|---|---|---|---|---|---|---|
| 2021 | NM_001212.3 | | #N/A | 358.804645 | -9.6975234 | 1.56482066 |
| 2022 | NM_021116.2 | | #N/A | 432.592194 | -5.4079517 | 0.87273269 |
| 2023 | NM_015241.2 | | #N/A | 380.6583 | -8.4015902 | 1.35589984 |
| 2024 | NM_021070.4 | LTBP3 | | 313.76728 | -9.5054451 | 1.53409582 |
| 2025 | NM_020524.3 | | #N/A | 311.037394 | -9.4928699 | 1.53211454 |
| 2026 | NM_014077.3 | | #N/A | 361.767464 | -9.7094543 | 1.56710534 |
| 2027 | NM_001182.4 | | #N/A | 278.279856 | -9.3313291 | 1.50615144 |
| 2028 | NM_012321.4 | | #N/A | 402.426012 | -9.8629315 | 1.5920527 |
| 2029 | NM_003331.4 | | #N/A | 256.687526 | -9.2152134 | 1.48756231 |
| 2030 | NM_004060.3 | | #N/A | 317.021087 | -9.5178593 | 1.53638848 |
| 2031 | NM_032960.3 | | #N/A | 279.434525 | -9.3370735 | 1.50722383 |
| 2032 | NM_002227.3 | | #N/A | 265.88427 | -9.264626 | 1.49563215 |
| 2033 | NM_006517.4 | | #N/A | 276.981431 | -7.9867668 | 1.28942673 |
| 2034 | NM_012316.4 | | #N/A | 475.066484 | -2.9263328 | 0.47245229 |
| 2035 | NM_021252.4 | | #N/A | 365.826447 | -9.7254097 | 1.57013527 |
| 2036 | NM_002404.2 | | #N/A | 425.194994 | -9.9428687 | 1.60529641 |
| 2037 | NM_001136157.1 | | #N/A | 287.390708 | -9.378914 | 1.51435883 |
| 2038 | NM_006230.3 | POLD2 | | 314.786198 | -9.5087926 | 1.53560376 |
| 2039 | NM_017896.2 | | #N/A | 377.222889 | -7.2504753 | 1.17098714 |
| 2040 | NM_006772.2 | | #N/A | 257.412216 | -9.2176281 | 1.48883256 |
| 2041 | NM_015348.1 | | #N/A | 304.106377 | -9.4587119 | 1.52778551 |
| 2042 | NM_001113407.2 | | #N/A | 435.251561 | -9.976268 | 1.61159007 |
| 2043 | NM_016026.3 | | #N/A | 449.591156 | -10.023082 | 1.61913941 |
| 2044 | NM_001282539.1 | GNB1 | | 640.816877 | -10.534323 | 1.70178459 |
| 2045 | NM_001242560.1 | | #N/A | 547.526106 | -10.30721 | 1.66520409 |
| 2046 | NM_001130072.1 | | #N/A | 329.747842 | -9.5756356 | 1.54712599 |
| 2047 | NM_001261.3 | | #N/A | 260.723088 | -9.2363239 | 1.49242273 |
| 2048 | NM_001029858.3 | | #N/A | 419.14422 | -6.8567964 | 1.10799907 |
| 2049 | NM_032853.3 | | #N/A | 293.509719 | -9.4079838 | 1.52025496 |
| 2050 | NM_007185.6 | | #N/A | 499.532606 | -10.174841 | 1.6442244 |
| 2051 | NM_006380.3 | | #N/A | 268.06831 | -9.2776189 | 1.49943336 |
| 2052 | NM_001130138.1 | ISLR2 | | 800.070329 | -7.4530507 | 1.2046926 |
| 2053 | NM_006245.3 | | #N/A | 340.009433 | -9.6198521 | 1.5549968 |
| 2054 | NM_005789.3 | | #N/A | 341.612459 | -9.6269003 | 1.55629914 |
| 2055 | NM_002116.7 | | #N/A | 334.500616 | -9.5969371 | 1.55152051 |
| 2056 | NM_003736.2 | | #N/A | 366.273425 | -9.7275925 | 1.57274912 |
| 2057 | NM_004047.4 | | #N/A | 477.103447 | -10.108618 | 1.63443598 |
| 2058 | NM_020865.2 | | #N/A | 323.315348 | -9.5469147 | 1.54363271 |
| 2059 | NM_004326.3 | | #N/A | 288.36063 | -9.381946 | 1.51697882 |
| 2060 | NM_017582.6 | | #N/A | 309.442338 | -9.4842325 | 1.53353909 |
| 2061 | NM_001082968.1 | | #N/A | 297.852563 | -9.428583 | 1.52493223 |
| 2062 | NM_001278556.1 | | #N/A | 435.631049 | -9.9773856 | 1.61383055 |
| 2063 | NM_016407.4 | RTF2 | | 338.494571 | -9.6133099 | 1.55497289 |

| | | | | | |
|---|---|---|---|---|---|
| 2064 | NM_005455.4 | #N/A | 304.762406 | -9.4615271 | 1.53051587 |
| 2067 | NM_030789.3 | HM13 | 346.035772 | -9.645527 | 1.56047379 |
| 2065 | NM_003751.3 | #N/A | 280.489378 | -9.3426502 | 1.5114812 |
| 2066 | NM_005861.3 | #N/A | 333.813634 | -9.5934878 | 1.55205458 |
| 2068 | NM_021008.3 | DEAF1 | 372.985429 | -9.7534266 | 1.57799273 |
| 2069 | NM_022826.3 | #N/A | 267.761116 | -9.2740267 | 1.50046043 |
| 2070 | NM_007046.3 | #N/A | 311.717961 | -9.4958563 | 1.53642601 |
| 2071 | NM_005207.3 | #N/A | 402.126134 | -4.8415521 | 0.78340539 |
| 2072 | NM_015219.4 | #N/A | 342.731821 | -9.6326228 | 1.55859959 |
| 2073 | NM_024656.3 | #N/A | 395.829346 | -4.1998392 | 0.67957141 |
| 2074 | NM_032772.5 | #N/A | 266.561545 | -9.2706872 | 1.50008427 |
| 2075 | NM_001270520.1 | #N/A | 325.178799 | -9.5551507 | 1.5464074 |
| 2076 | NM_001082579.1 | #N/A | 510.86091 | -10.207909 | 1.65221918 |
| 2077 | NM_024671.3 | #N/A | 287.235763 | -9.3766191 | 1.51766666 |
| 2078 | NM_015571.3 | #N/A | 270.097245 | -9.2880112 | 1.50346008 |
| 2079 | NM_004615.3 | TSPAN7 | 394.301061 | -9.8337711 | 1.59184191 |
| 2081 | NM_001128217.2 | PSIP1 | 262.938896 | -9.2475078 | 1.49702461 |
| 2080 | NM_001031684.2 | #N/A | 334.694879 | -9.5972092 | 1.55361791 |
| 2082 | NM_002318.2 | #N/A | 412.016481 | -9.8976129 | 1.60230619 |
| 2083 | NM_016320.4 | #N/A | 246.7016 | -9.1576774 | 1.48272539 |
| 2084 | NM_006423.2 | #N/A | 412.029766 | -9.8970931 | 1.60265581 |
| 2085 | NM_001318727.1 | #N/A | 373.673297 | -9.756227 | 1.58014316 |
| 2086 | NM_002805.5 | #N/A | 334.263982 | -9.5953514 | 1.55426051 |
| 2087 | NM_015144.2 | #N/A | 286.850321 | -9.3742674 | 1.51851163 |
| 2088 | NR_027410.1 | GOLGA8B | 374.622519 | -4.670929 | 0.75670553 |
| 2089 | NM_001697.2 | #N/A | 426.811821 | -9.9480447 | 1.61169837 |
| 2090 | NM_001190476.1 | #N/A | 404.792293 | -7.0785324 | 1.14692463 |
| 2091 | NM_001271741.1 | #N/A | 277.844173 | -9.3296498 | 1.51165649 |
| 2092 | NM_006660.4 | #N/A | 242.833176 | -9.1342749 | 1.48000542 |
| 2093 | NM_030782.4 | #N/A | 334.149214 | -9.5950417 | 1.55465112 |
| 2094 | NM_020944.2 | #N/A | 279.270124 | -9.3359856 | 1.51282501 |
| 2095 | NM_018117.11 | #N/A | 239.204916 | -9.1138862 | 1.47693436 |
| 2096 | NM_032311.4 | #N/A | 336.804179 | -9.6057453 | 1.55664757 |
| 2097 | NM_012326.3 | #N/A | 396.010799 | -9.8399165 | 1.59469376 |
| 2098 | NM_002803.3 | #N/A | 297.83525 | -9.4291498 | 1.5282838 |
| 2099 | NM_006096.3 | #N/A | 398.977425 | -9.8506063 | 1.59659949 |
| 2100 | NM_181449.2 | #N/A | 45.0671505 | 5.72047685 | 0.92721539 |
| 2102 | NM_001002811.2 | PDE4DIP | 243.328131 | -9.1380004 | 1.48122228 |
| 2101 | NM_001001998.2 | #N/A | 257.917794 | -9.2215701 | 1.49475991 |
| 2103 | NM_014685.3 | #N/A | 286.094843 | -9.3707327 | 1.51892833 |
| 2104 | NM_001242836.1 | NDRG4 | 568.006648 | -9.0920196 | 1.47378779 |
| 2105 | NM_003842.4 | #N/A | 263.255349 | -9.250133 | 1.49943863 |
| 2106 | NM_003676.3 | #N/A | 285.99585 | -9.3709337 | 1.5190477 |

| | | | | | |
|---|---|---|---|---|---|
| 2107 | NM_004273.4 | #N/A | 260.157221 | -9.2355112 | 1.49711872 |
| 2108 | NM_001146334.1 | #N/A | 445.414191 | -10.009609 | 1.62279018 |
| 2109 | NM_133443.3 | #N/A | 328.878321 | -9.5713998 | 1.55180307 |
| 2110 | NM_007362.3 | #N/A | 340.290363 | -9.6208244 | 1.55985239 |
| 2111 | NM_005109.2 | #N/A | 278.621897 | -9.3324234 | 1.51314264 |
| 2113 | NM_172349.2 | NSD1 | 247.864454 | -9.1651895 | 1.48613416 |
| 2112 | NM_007106.3 | #N/A | 301.161874 | -9.4449399 | 1.53149389 |
| 2114 | NM_004598.3 | #N/A | 573.712057 | -8.932474 | 1.44846785 |
| 2115 | NM_138457.2 | #N/A | 295.862601 | -5.3722771 | 0.87115098 |
| 2116 | NM_001304743.1 | #N/A | 232.487936 | -9.0719523 | 1.47111951 |
| 2117 | NM_152346.2 | #N/A | 259.268857 | -9.2296461 | 1.49692311 |
| 2118 | NM_000108.4 | #N/A | 275.051628 | -9.3142696 | 1.51068976 |
| 2119 | NM_022749.5 | #N/A | 263.406914 | -9.2517963 | 1.50057589 |
| 2120 | NM_001252100.1 | #N/A | 580.798448 | -10.39241 | 1.68601483 |
| 2121 | NM_005226.3 | #N/A | 200.348086 | -5.0941587 | 0.82649269 |
| 2122 | NM_001323619.1 | UNC45A | 261.66683 | -9.2421603 | 1.49954802 |
| 2123 | NM_002593.3 | #N/A | 334.057117 | -8.3215808 | 1.35022146 |
| 2124 | NM_206538.3 | #N/A | 493.676559 | -10.157894 | 1.64851189 |
| 2125 | NM_001128833.1 | #N/A | 248.197802 | -9.1658685 | 1.48775219 |
| 2126 | NM_002078.4 | #N/A | 252.889357 | -9.1928318 | 1.49212505 |
| 2127 | NM_002869.4 | #N/A | 254.714257 | -9.202687 | 1.49373545 |
| 2128 | NM_003799.2 | #N/A | 262.750073 | -9.2474146 | 1.50107273 |
| 2129 | NM_005493.2 | #N/A | 307.456737 | -9.4747108 | 1.53801645 |
| 2130 | NM_001204467.1 | #N/A | 283.880545 | -9.3598599 | 1.5194268 |
| 2132 | NM_016456.4 | TMEM9 | 325.010274 | -9.5546944 | 1.55136975 |
| 2131 | NM_016272.3 | #N/A | 249.340155 | -6.8019652 | 1.1044274 |
| 2133 | NM_006503.3 | #N/A | 362.480166 | -9.7124647 | 1.57712311 |
| 2134 | NM_017789.4 | #N/A | 386.667578 | -9.8054877 | 1.59223781 |
| 2135 | NM_001193483.2 | #N/A | 258.969843 | -9.2279836 | 1.49855989 |
| 2136 | NM_002398.2 | #N/A | 377.979439 | -5.9157236 | 0.96072669 |
| 2137 | NM_003469.4 | #N/A | 697.397445 | -9.1855934 | 1.49184707 |
| 2138 | NM_005419.3 | #N/A | 271.394313 | -9.2943356 | 1.50950917 |
| 2139 | NM_006622.3 | #N/A | 425.375799 | -9.9430493 | 1.61483673 |
| 2140 | NM_003131.3 | #N/A | 281.785828 | -9.3489089 | 1.51869108 |
| 2141 | NM_016628.4 | #N/A | 266.15076 | -9.2675767 | 1.50548158 |
| 2142 | NM_006814.4 | #N/A | 321.3227 | -9.5382738 | 1.54973504 |
| 2143 | NM_022356.3 | #N/A | 307.108812 | -9.4737767 | 1.53930953 |
| 2144 | NM_001135699.1 | YWHAZ | 400.899622 | -9.8577425 | 1.60176652 |
| 2145 | NM_014983.2 | #N/A | 249.337782 | -9.1729701 | 1.49051606 |
| 2146 | NM_012398.2 | #N/A | 386.04386 | -9.802895 | 1.59291107 |
| 2147 | NM_012137.3 | #N/A | 289.554533 | -9.3883349 | 1.52574599 |
| 2148 | NM_020928.1 | #N/A | 230.252651 | -6.8647197 | 1.11569109 |
| 2149 | NM_001297776.1 | #N/A | 281.128185 | -9.3447362 | 1.51882577 |

| | | | | | |
|---|---|---|---|---|---|
| 2150 | NM_001318728.1 | #N/A | 296.112002 | -9.4195672 | 1.53101933 |
| 2151 | NM_005526.3 | #N/A | 334.491464 | -9.5963135 | 1.55989736 |
| 2152 | NM_001323674.1 | #N/A | 39.360572 | 10.2856044 | 1.67232948 |
| 2153 | NM_025222.3 | #N/A | 1133.14547 | -6.4907887 | 1.05543389 |
| 2154 | NM_006548.5 | #N/A | 268.223784 | -9.2773657 | 1.50872692 |
| 2155 | NM_012247.4 | #N/A | 290.239103 | -9.3914519 | 1.52729692 |
| 2156 | NM_001146156.1 | #N/A | 390.015031 | -9.8176834 | 1.59673574 |
| 2157 | NM_020830.4 | #N/A | 249.946835 | -9.1755309 | 1.49232624 |
| 2158 | NR_110936.1 | #N/A | 51.1460987 | 10.6451674 | 1.73146504 |
| 2159 | NM_002396.4 | #N/A | 259.969083 | -9.2322425 | 1.50179746 |
| 2160 | NM_014671.2 | #N/A | 281.582203 | -9.348031 | 1.52068379 |
| 2161 | NM_018476.3 | #N/A | 607.841292 | -10.458065 | 1.70124024 |
| 2162 | NR_138470.1 | #N/A | 381.314252 | -9.7850637 | 1.59176994 |
| 2163 | NM_005004.3 | #N/A | 361.413741 | -9.7081002 | 1.57928817 |
| 2164 | NM_004711.4 | #N/A | 335.147914 | -9.5989189 | 1.56155197 |
| 2165 | NM_032476.3 | #N/A | 237.532388 | -9.1040458 | 1.48110307 |
| 2166 | NM_005104.3 | #N/A | 294.324019 | -9.412682 | 1.53161599 |
| 2167 | NM_016626.4 | #N/A | 299.868874 | -9.4383988 | 1.53586863 |
| 2168 | NM_001492.5 | #N/A | 374.752303 | -9.7603388 | 1.58828219 |
| 2169 | NM_001243332.1 | SEZ6L2 | 338.397336 | -9.6126042 | 1.56426553 |
| 2170 | NM_145686.3 | MAP4K4 | 368.489921 | -9.7358957 | 1.58439155 |
| 2171 | NM_022735.3 | #N/A | 238.997786 | -9.1116519 | 1.48285686 |
| 2172 | NM_000274.3 | #N/A | 270.264067 | -9.2889915 | 1.51177509 |
| 2173 | NM_005412.5 | #N/A | 468.293899 | -10.081458 | 1.64079527 |
| 2174 | NM_024294.3 | #N/A | 276.034376 | -9.3198366 | 1.51684587 |
| 2175 | NM_001170937.1 | FUS | 401.30785 | -9.859618 | 1.60479199 |
| 2176 | NM_015339.4 | #N/A | 243.855975 | -9.1417422 | 1.48794679 |
| 2177 | NM_004378.2 | #N/A | 379.429985 | -9.7786266 | 1.5918532 |
| 2178 | NM_014280.2 | #N/A | 355.075498 | -9.6825575 | 1.57639284 |
| 2179 | NM_001291694.1 | #N/A | 276.121932 | -8.0426508 | 1.30948326 |
| 2180 | NM_018489.2 | #N/A | 255.712957 | -7.8749409 | 1.28217147 |
| 2181 | NM_025069.2 | #N/A | 380.166969 | -9.7811341 | 1.59255741 |
| 2182 | NM_022737.2 | #N/A | 455.397023 | -10.041557 | 1.63510451 |
| 2183 | NM_138459.3 | #N/A | 308.523196 | -9.4795674 | 1.54365717 |
| 2184 | NM_001830.3 | #N/A | 421.548434 | -9.9298337 | 1.61703856 |
| 2185 | NM_016222.3 | #N/A | 268.873633 | -9.2813856 | 1.51144067 |
| 2186 | NM_014061.4 | #N/A | 314.312253 | -9.5063414 | 1.54824979 |
| 2187 | NM_001130111.1 | #N/A | 355.61652 | -9.6845432 | 1.57734064 |
| 2188 | NM_032236.6 | #N/A | 269.693866 | -9.2855453 | 1.51234282 |
| 2189 | NM_018490.3 | #N/A | 342.091581 | -9.6292402 | 1.56843478 |
| 2190 | NM_001136528.1 | #N/A | 378.992725 | -9.7767423 | 1.59291518 |
| 2191 | NM_015321.2 | #N/A | 278.302272 | -9.330631 | 1.52026227 |
| 2192 | NM_001144995.1 | #N/A | 241.40994 | -9.1263521 | 1.48707202 |

| | | | | | |
|---|---|---|---|---|---|
| 2193 | NM_001318509.1 | #N/A | 436.266995 | -9.9796759 | 1.62619178 |
| 2194 | NM_001031835.2 | #N/A | 279.923388 | -9.3397912 | 1.52195384 |
| 2195 | NR_040046.1 | LINC01531 | 148.750152 | 3.21114406 | 0.52330072 |
| 2196 | NM_024334.2 | TMEM43 | 340.212022 | -9.6209477 | 1.56825107 |
| 2197 | NM_014976.1 | #N/A | 286.692533 | -9.3740749 | 1.52807828 |
| 2198 | NM_006191.2 | #N/A | 547.745485 | -7.6417861 | 1.24586318 |
| 2199 | NM_001627.3 | #N/A | 250.461977 | -9.1793365 | 1.49657067 |
| 2200 | NM_080598.5 | #N/A | 450.103073 | -10.024548 | 1.63437262 |
| 2201 | NM_001032288.2 | #N/A | 395.040256 | -9.8364923 | 1.60377689 |
| 2202 | NM_001271875.1 | #N/A | 260.132118 | -9.234146 | 1.5055935 |
| 2203 | NM_021098.2 | #N/A | 372.726793 | -9.7520668 | 1.5901058 |
| 2204 | NM_032704.4 | #N/A | 302.624655 | -9.4521927 | 1.54140163 |
| 2205 | NM_139279.5 | #N/A | 273.273804 | -9.3062807 | 1.51767721 |
| 2206 | NM_003461.4 | #N/A | 324.963888 | -9.5550175 | 1.55830687 |
| 2207 | NM_001407.2 | #N/A | 563.445807 | -8.9037198 | 1.45218967 |
| 2208 | NM_004597.6 | SNRPD2 | 404.188701 | -9.8693298 | 1.60980438 |
| 2209 | NM_001261420.1 | OS9 | 313.936252 | -9.505724 | 1.55052682 |
| 2210 | NM_002866.4 | #N/A | 657.206671 | -10.570749 | 1.72451211 |
| 2211 | NM_173587.3 | #N/A | 463.422624 | -10.06628 | 1.64232352 |
| 2212 | NM_020337.2 | #N/A | 269.039391 | -9.2815655 | 1.51448058 |
| 2213 | NM_001199161.1 | USP19 | 233.951362 | -9.0797435 | 1.48156922 |
| 2214 | NM_018715.3 | #N/A | 255.878635 | -9.2092929 | 1.50291264 |
| 2215 | NM_020132.4 | #N/A | 279.899262 | -9.3396617 | 1.5242657 |
| 2216 | NM_001142481.1 | NREP | 330.15466 | -9.5775872 | 1.56312765 |
| 2217 | NM_032812.8 | #N/A | 232.284036 | -9.0711975 | 1.48054614 |
| 2218 | NM_006320.4 | #N/A | 349.481985 | -9.659682 | 1.57662126 |
| 2219 | NM_020131.4 | #N/A | 268.999517 | -9.2814248 | 1.51495767 |
| 2220 | NM_000346.3 | #N/A | 239.554555 | -9.1144535 | 1.48777529 |
| 2221 | NM_017745.5 | #N/A | 259.099704 | -9.2279216 | 1.50629881 |
| 2222 | NM_207116.2 | #N/A | 208.498502 | -4.1151838 | 0.67174262 |
| 2223 | NM_015046.5 | #N/A | 245.502572 | -9.1505748 | 1.49384251 |
| 2224 | NM_139321.2 | #N/A | 374.082215 | -9.7577595 | 1.59300195 |
| 2225 | NM_018174.5 | #N/A | 291.102716 | -9.3960267 | 1.53417939 |
| 2226 | NM_001330249.1 | HNRNPA3 | 249.963278 | -9.1780329 | 1.49876714 |
| 2227 | NM_019845.2 | #N/A | 523.326462 | -10.242315 | 1.67258958 |
| 2228 | NR_001591.1 | TPTEP1 | 67.7755491 | 3.09828056 | 0.50596512 |
| 2229 | NM_015170.2 | SULF1 | 302.505706 | -9.4526346 | 1.54369835 |
| 2230 | NM_004879.4 | #N/A | 333.335391 | -9.5912532 | 1.56641842 |
| 2231 | NM_006109.4 | #N/A | 281.922572 | -9.3496187 | 1.52718232 |
| 2232 | NM_001128148.2 | #N/A | 304.378508 | -9.4605229 | 1.54536428 |
| 2233 | NM_015640.3 | #N/A | 359.936321 | -9.7030543 | 1.58536484 |
| 2234 | NM_015185.2 | #N/A | 278.985836 | -9.3340851 | 1.52511426 |
| 2235 | NM_024551.2 | #N/A | 303.046573 | -9.4536197 | 1.5446763 |

| | | | | | |
|---|---|---|---|---|---|
| 2236 | NM_001257291.1 | #N/A | 248.128307 | -9.1661478 | 1.49777737 |
| 2238 | NM_030973.3 | MED25 | 230.191853 | -9.0587694 | 1.48025465 |
| 2237 | NM_014838.2 | #N/A | 240.166013 | -6.1655202 | 1.00749063 |
| 2239 | NM_053023.4 | #N/A | 314.948916 | -9.5096205 | 1.55422217 |
| 2240 | NM_000696.3 | ALDH9A1 | 225.300177 | -9.0275241 | 1.47563202 |
| 2241 | NM_002887.3 | #N/A | 260.718245 | -9.2373186 | 1.50996471 |
| 2242 | NM_145173.3 | #N/A | 446.923911 | -10.014275 | 1.63705831 |
| 2243 | NM_003714.2 | STC2 | 247.564701 | -9.1622575 | 1.49780424 |
| 2244 | NM_006360.5 | #N/A | 343.938272 | -9.6363732 | 1.57541035 |
| 2245 | NM_001436.3 | #N/A | 375.92423 | -9.7645239 | 1.59644453 |
| 2246 | NM_173701.1 | #N/A | 311.231233 | -9.4923664 | 1.55198087 |
| 2247 | NM_178502.3 | #N/A | 294.669806 | -9.4134277 | 1.53912026 |
| 2248 | NM_001300792.1 | #N/A | 464.633055 | -10.070281 | 1.64658374 |
| 2249 | NM_014737.2 | #N/A | 279.930304 | -9.339313 | 1.52711646 |
| 2250 | NM_015056.2 | #N/A | 284.05521 | -9.3600943 | 1.53056577 |
| 2251 | NM_001077516.1 | #N/A | 276.712043 | -9.3230981 | 1.5246366 |
| 2252 | NM_001032.4 | RPS29 | 572.3001 | -10.371179 | 1.6963196 |
| 2253 | NM_001286.3 | #N/A | 327.154008 | -9.5645239 | 1.56436614 |
| 2254 | NR_136702.1 | SFPQ | 373.902818 | -9.7568791 | 1.59589846 |
| 2255 | NM_022080.2 | NAPB | 429.827836 | -9.958335 | 1.6289178 |
| 2256 | NM_003329.3 | #N/A | 384.552904 | -9.7975931 | 1.6026655 |
| 2257 | NM_133452.2 | #N/A | 225.580782 | -9.0281776 | 1.47699484 |
| 2258 | NM_033140.3 | #N/A | 259.053894 | -9.2264986 | 1.50950138 |
| 2259 | NM_001080517.2 | #N/A | 349.503881 | -9.6608576 | 1.58065875 |
| 2260 | NM_024298.4 | #N/A | 264.899639 | -9.2597128 | 1.51512484 |
| 2261 | NM_001878.3 | #N/A | 284.055614 | -9.3621714 | 1.53195237 |
| 2262 | NM_019102.3 | #N/A | 448.904938 | -10.021014 | 1.6397961 |
| 2263 | NM_004607.2 | TBCA | 350.726805 | -9.6646393 | 1.58152168 |
| 2264 | NM_182972.2 | IRF2BP2 | 297.272563 | -9.4254367 | 1.54246898 |
| 2265 | NM_014633.4 | #N/A | 250.958885 | -9.1819849 | 1.50266023 |
| 2266 | NM_005006.6 | #N/A | 253.206837 | -9.1943154 | 1.50491206 |
| 2267 | NM_206915.2 | BEX3 | 360.64185 | -9.7046527 | 1.58847843 |
| 2268 | NM_001308176.1 | #N/A | 284.42587 | -8.0866681 | 1.32374941 |
| 2269 | NM_001321075.1 | #N/A | 332.821383 | -9.5884423 | 1.56962335 |
| 2270 | NM_006788.3 | #N/A | 262.717786 | -7.9070845 | 1.29447367 |
| 2271 | NM_007146.2 | #N/A | 260.899843 | -9.2369526 | 1.51225556 |
| 2272 | NM_004064.4 | CDKN1B | 396.531271 | -9.8413498 | 1.61127061 |
| 2273 | NM_002797.4 | #N/A | 353.891806 | -9.6775315 | 1.58456283 |
| 2274 | NM_000528.3 | #N/A | 251.490072 | -9.1856745 | 1.50404651 |
| 2275 | NM_006598.2 | #N/A | 250.916801 | -6.79312 | 1.11239566 |
| 2276 | NM_138795.3 | #N/A | 453.801191 | -10.036395 | 1.64350263 |
| 2277 | NM_001283054.1 | #N/A | 552.743896 | -10.320975 | 1.69015004 |
| 2279 | NR_026943.1 | LINC00205 | 324.042511 | -9.550374 | 1.56401594 |

| | | | | | |
|---|---|---|---|---|---|
| 2278 | NM_001142551.1 | WDR47 | 416.492903 | -9.9125347 | 1.62330675 |
| 2280 | NM_006646.5 | #N/A | 350.789201 | -9.6646899 | 1.58283392 |
| 2281 | NM_014630.2 | #N/A | 248.648468 | -9.1689931 | 1.50164099 |
| 2282 | NM_001009984.2 | C20orf194 | 313.51033 | -9.5025725 | 1.55639318 |
| 2283 | NM_003091.3 | #N/A | 324.651389 | -9.5529499 | 1.56461692 |
| 2284 | NM_020764.3 | #N/A | 351.564202 | -9.6682691 | 1.58352848 |
| 2285 | NM_002157.2 | #N/A | 303.353559 | -9.4551849 | 1.54866064 |
| 2286 | NM_004844.4 | #N/A | 331.812941 | -9.5847198 | 1.56989597 |
| 2287 | NM_015251.2 | #N/A | 345.794106 | -9.6442157 | 1.57970952 |
| 2288 | NM_001034.3 | #N/A | 140.550903 | -4.5323222 | 0.74240591 |
| 2290 | NM_017670.2 | OTUB1 | 307.121558 | -9.4731874 | 1.55211514 |
| 2289 | NM_001277.2 | #N/A | 239.077505 | -9.1117053 | 1.49288007 |
| 2291 | NM_007315.3 | STAT1 | 273.657102 | -9.3080831 | 1.52508433 |
| 2292 | NM_005548.2 | KARS1 | 342.730895 | -9.631395 | 1.57814328 |
| 2293 | NM_019044.4 | #N/A | 256.808696 | -9.2146185 | 1.50985285 |
| 2294 | NM_020777.2 | #N/A | 241.436159 | -9.1279804 | 1.49572748 |
| 2295 | NM_006804.3 | #N/A | 243.476393 | -9.1383311 | 1.49765558 |
| 2296 | NM_001863.4 | #N/A | 410.094607 | -8.4857893 | 1.39075587 |
| 2297 | NM_017542.4 | #N/A | 301.983259 | -9.4486956 | 1.54857279 |
| 2298 | NM_001001973.2 | #N/A | 337.741972 | -9.6099548 | 1.57509527 |
| 2299 | NM_004090.3 | #N/A | 293.309258 | -9.4068776 | 1.54180311 |
| 2300 | NM_012308.2 | #N/A | 218.060992 | -8.9787616 | 1.47165308 |
| 2301 | NM_015537.4 | NSMF | 240.533949 | -9.1219019 | 1.49519154 |
| 2302 | NM_001099285.1 | PTMA | 411.604477 | -9.8954738 | 1.62212999 |
| 2303 | NM_014752.2 | #N/A | 326.921937 | -9.5634272 | 1.567772 |
| 2304 | NM_004420.2 | #N/A | 494.560603 | -10.160449 | 1.66583931 |
| 2305 | NM_019592.6 | #N/A | 269.662349 | -8.0098326 | 1.31322863 |
| 2306 | NM_001004431.2 | #N/A | 236.280553 | -9.0963357 | 1.4914215 |
| 2307 | NM_001164803.1 | #N/A | 300.83442 | -9.4429596 | 1.54826037 |
| 2308 | NM_001170629.1 | #N/A | 263.756217 | -9.2525802 | 1.51702855 |
| 2309 | NM_001137673.1 | #N/A | 241.060457 | -9.1235787 | 1.49594115 |
| 2310 | NM_005544.2 | IRS1 | 241.979481 | -7.7989067 | 1.27878651 |
| 2311 | NM_001320596.1 | #N/A | 383.523071 | -9.793957 | 1.60598418 |
| 2312 | NM_019121.1 | #N/A | 269.689766 | -9.2856273 | 1.52296717 |
| 2313 | NM_080732.3 | #N/A | 333.454502 | -9.5918645 | 1.57318779 |
| 2314 | NM_004461.2 | #N/A | 320.584737 | -9.535045 | 1.56411605 |
| 2315 | NM_001010972.1 | #N/A | 317.045503 | -9.5195317 | 1.56162012 |
| 2316 | NM_001042603.2 | #N/A | 240.020106 | -9.1164844 | 1.49563751 |
| 2317 | NM_016025.4 | #N/A | 301.679646 | -9.4475276 | 1.55011918 |
| 2318 | NM_003051.3 | SLC16A1 | 386.384275 | -7.2461348 | 1.18895513 |
| 2319 | NM_001202502.1 | NIPSNAP1 | 327.17054 | -9.5647979 | 1.56943596 |
| 2320 | NM_013333.3 | EPN1 | 436.713798 | -9.9810835 | 1.63776498 |
| 2321 | NM_001321208.1 | #N/A | 313.017624 | -9.5006036 | 1.55941644 |

| | | | | | |
|---|---|---|---|---|---|
| 2322 | NM_001806.3 | #N/A | 337.898805 | -9.6105061 | 1.57745117 |
| 2323 | NM_013379.2 | #N/A | 233.816656 | -9.0794855 | 1.49030983 |
| 2324 | NM_002736.2 | #N/A | 516.04667 | -10.221746 | 1.67793242 |
| 2325 | NM_017595.5 | #N/A | 283.539303 | -9.358488 | 1.53622886 |
| 2326 | NM_052876.3 | #N/A | 309.492416 | -9.4844448 | 1.55699784 |
| 2327 | NM_015457.2 | #N/A | 255.958496 | -9.2111999 | 1.51216061 |
| 2328 | NM_001163285.1 | #N/A | 230.803819 | -9.0607389 | 1.48749181 |
| 2330 | NM_145230.3 | ATP6V0E2 | 471.819491 | -10.09254 | 1.65697961 |
| 2329 | NM_006328.3 | #N/A | 276.553073 | -9.3220439 | 1.53046468 |
| 2331 | NM_020851.2 | #N/A | 728.765456 | -10.71978 | 1.76001103 |
| 2332 | NM_006471.3 | MYL12A | 267.727614 | -9.2758377 | 1.52301886 |
| 2333 | NM_001284215.1 | #N/A | 276.532448 | -9.3209334 | 1.53052771 |
| 2334 | NM_001011553.3 | SEPTIN7 | 386.178569 | -9.803225 | 1.6100093 |
| 2335 | NM_001032283.2 | TMPO | 255.338608 | -9.2053954 | 1.51188509 |
| 2336 | NM_014726.2 | TBKBP1 | 322.872899 | -9.5453079 | 1.56784286 |
| 2337 | NM_015135.2 | #N/A | 251.804751 | -9.1862999 | 1.50894099 |
| 2338 | NM_022462.4 | HIF3A | 41.8771328 | 10.3767561 | 1.70453621 |
| 2339 | NM_133642.3 | #N/A | 330.180385 | -8.2018051 | 1.34734692 |
| 2340 | NM_003136.3 | #N/A | 250.834537 | -9.1806507 | 1.50820434 |
| 2341 | NM_005436.4 | #N/A | 239.804013 | -9.1163771 | 1.4977163 |
| 2342 | NM_001304815.1 | CIC | 232.35206 | -9.0712926 | 1.49039057 |
| 2343 | NM_004448.3 | ERBB2 | 270.004628 | -9.2891338 | 1.52623505 |
| 2344 | NM_021095.2 | #N/A | 254.208384 | -9.2021492 | 1.51195454 |
| 2345 | NM_170707.3 | #N/A | 237.366266 | -9.1029891 | 1.4957526 |
| 2346 | NM_001199698.1 | BAG6 | 298.248736 | -9.4318982 | 1.54982856 |
| 2347 | NM_138346.2 | #N/A | 253.246235 | -9.1953032 | 1.51096636 |
| 2348 | NM_015178.2 | RHOBTB2 | 226.143579 | -9.0311486 | 1.48417753 |
| 2349 | NM_015160.2 | #N/A | 246.904682 | -9.1584133 | 1.50520299 |
| 2350 | NM_012121.4 | #N/A | 239.681884 | -9.1149964 | 1.49810829 |
| 2352 | NM_001128846.1 | SMARCA4 | 283.819016 | -9.3581262 | 1.5381373 |
| 2351 | NM_001080509.2 | TSPAN11 | 229.570337 | -9.0550892 | 1.48835235 |
| 2353 | NM_004628.4 | XPC | 218.665301 | -8.9846006 | 1.47675577 |
| 2354 | NM_020239.3 | #N/A | 226.100405 | -6.8337735 | 1.12322015 |
| 2355 | NM_003375.4 | VDAC2 | 395.303956 | -9.8372359 | 1.61742903 |
| 2356 | NM_018230.2 | #N/A | 253.653866 | -9.1971022 | 1.51220772 |
| 2357 | NM_006003.2 | #N/A | 303.529185 | -9.4562212 | 1.55484473 |
| 2358 | NM_001321473.1 | #N/A | 253.811509 | -9.1992296 | 1.51263758 |
| 2359 | NM_001081550.1 | #N/A | 262.558802 | -7.9015617 | 1.29929644 |
| 2360 | NM_004587.2 | RRBP1 | 407.274034 | -4.6674112 | 0.76749692 |
| 2361 | NM_001286077.1 | #N/A | 344.139222 | -9.6372672 | 1.58500758 |
| 2362 | NM_002786.3 | PSMA1 | 318.205196 | -9.5243568 | 1.56654184 |
| 2363 | NM_001080421.2 | UNC13A | 417.285433 | -7.8823498 | 1.29659681 |
| 2364 | NM_002616.2 | #N/A | 233.893391 | -9.0816275 | 1.49385576 |

| | | | | | |
|---|---|---|---|---|---|
| 2370 | NR_130934.1 | LINC01602 | 75.8521092 | 3.71429122 | 0.61106995 |
| 2368 | NM_012446.4 | SSBP2 | 303.638036 | -9.4559362 | 1.55571688 |
| 2365 | NM_001077198.2 | #N/A | 279.861006 | -9.3393447 | 1.53649987 |
| 2366 | NM_001843.3 | #N/A | 449.772828 | -10.023297 | 1.64907733 |
| 2367 | NM_004627.4 | #N/A | 266.629062 | -9.268816 | 1.52492624 |
| 2369 | NM_030809.2 | #N/A | 313.862681 | -9.5043402 | 1.56365644 |
| 2371 | NM_001142478.1 | #N/A | 458.768641 | -10.052076 | 1.65386735 |
| 2372 | NM_003827.3 | #N/A | 302.555603 | -9.4515529 | 1.55505948 |
| 2373 | NM_014412.2 | #N/A | 297.56095 | -9.4274001 | 1.55118794 |
| 2374 | NM_021005.3 | #N/A | 307.365922 | -9.4751999 | 1.55904396 |
| 2375 | NM_012104.4 | BACE1 | 280.316045 | -9.3420392 | 1.53717731 |
| 2376 | NM_006886.3 | #N/A | 412.972118 | -9.9004879 | 1.62923438 |
| 2377 | NM_000094.3 | COL7A1 | 273.262962 | -9.3052819 | 1.53132689 |
| 2378 | NM_178126.3 | #N/A | 238.667008 | -9.1096868 | 1.49921821 |
| 2379 | NM_005120.2 | #N/A | 224.104522 | -9.0204366 | 1.48461825 |
| 2380 | NM_002957.5 | #N/A | 237.767424 | -7.7726371 | 1.27929337 |
| 2381 | NM_014272.4 | #N/A | 322.830361 | -9.545605 | 1.5712785 |
| 2382 | NM_198976.2 | #N/A | 278.998821 | -9.3346306 | 1.53662451 |
| 2383 | NM_004381.4 | #N/A | 298.13986 | -9.4304197 | 1.5524896 |
| 2384 | NM_130809.4 | #N/A | 222.85377 | -9.0099916 | 1.48335523 |
| 2385 | NM_016462.3 | #N/A | 267.177519 | -9.2722754 | 1.52668615 |
| 2386 | NM_003247.3 | THBS2 | 245.51323 | -9.152055 | 1.50703125 |
| 2387 | NM_004756.4 | #N/A | 281.656691 | -9.3476232 | 1.53920276 |
| 2388 | NR_109905.1 | #N/A | 37.0705539 | 10.1981078 | 1.67926356 |
| 2389 | NM_021815.4 | #N/A | 459.400832 | -10.054324 | 1.65574848 |
| 2390 | NM_032268.4 | #N/A | 340.925221 | -7.3771521 | 1.2148792 |
| 2391 | NM_003325.3 | #N/A | 282.466673 | -9.3524158 | 1.54027412 |
| 2392 | NM_002319.4 | #N/A | 223.47334 | -9.0162738 | 1.48501897 |
| 2393 | NM_014864.3 | #N/A | 279.315739 | -9.3352184 | 1.53756854 |
| 2394 | NM_005826.4 | #N/A | 315.313253 | -9.5107307 | 1.56656159 |
| 2395 | NM_005494.2 | #N/A | 437.954695 | -8.566896 | 1.41113742 |
| 2396 | NM_015379.4 | #N/A | 325.152528 | -9.5556695 | 1.57408209 |
| 2397 | NM_022720.6 | #N/A | 259.285724 | -9.2290437 | 1.52028773 |
| 2398 | NM_005234.3 | NR2F6 | 252.668978 | -9.192821 | 1.51434268 |
| 2399 | NM_031490.4 | #N/A | 212.585531 | -8.9416875 | 1.47306389 |
| 2400 | NM_000046.3 | #N/A | 185.757879 | -5.6031021 | 0.92311982 |
| 2401 | NM_006929.4 | #N/A | 256.695086 | -7.871261 | 1.29680765 |
| 2402 | NM_012290.4 | #N/A | 212.659881 | -8.9434373 | 1.47350625 |
| 2403 | NM_002634.3 | #N/A | 327.941752 | -9.5677087 | 1.57639238 |
| 2404 | NM_001031714.3 | #N/A | 243.204735 | -9.1366869 | 1.50541792 |
| 2405 | NM_001406.3 | #N/A | 252.21871 | -9.1890451 | 1.51407926 |
| 2406 | NM_014216.5 | #N/A | 319.048743 | -9.5277449 | 1.56992577 |
| 2407 | NM_000216.3 | #N/A | 374.873519 | -9.761148 | 1.60845509 |

| | | | | | |
|---|---|---|---|---|---|
| 2408 | NM_002938.4 | #N/A | 260.716875 | -9.2365439 | 1.52202561 |
| 2409 | NM_001819.2 | #N/A | 459.931114 | -10.055927 | 1.65708611 |
| 2410 | NM_014399.3 | #N/A | 562.741463 | -10.346832 | 1.70512349 |
| 2411 | NM_001128912.1 | #N/A | 301.141249 | -9.4436537 | 1.55630302 |
| 2412 | NM_004300.3 | #N/A | 277.969909 | -9.3289874 | 1.53746885 |
| 2413 | NM_005633.3 | SOS1 | 262.487611 | -9.245865 | 1.52381876 |
| 2414 | NM_001105203.1 | #N/A | 380.81383 | -9.7833519 | 1.61245874 |
| 2415 | NM_004962.4 | #N/A | 372.347026 | -9.751404 | 1.60719854 |
| 2416 | NM_015214.2 | #N/A | 235.136047 | -9.0875274 | 1.49778332 |
| 2417 | NM_032905.4 | #N/A | 248.977718 | -9.1704345 | 1.51147689 |
| 2418 | NM_001318187.1 | HNRNPK | 250.018687 | -9.1776347 | 1.51293147 |
| 2419 | NM_001172623.1 | NEO1 | 377.4435 | -9.7708586 | 1.6109563 |
| 2420 | NM_006079.4 | #N/A | 285.211432 | -9.3658503 | 1.54414953 |
| 2421 | NM_006565.3 | #N/A | 246.671216 | -9.1565223 | 1.50961866 |
| 2422 | NM_012203.1 | #N/A | 296.182705 | -9.42098 | 1.55323686 |
| 2423 | NM_206962.3 | #N/A | 267.693112 | -9.2744156 | 1.52909304 |
| 2424 | NM_001083913.1 | #N/A | 246.387315 | -9.1554628 | 1.50953597 |
| 2425 | NM_177535.2 | MAGED4B | 370.118797 | -9.7433768 | 1.60657854 |
| 2426 | NM_006197.3 | #N/A | 269.817526 | -9.2864138 | 1.53131512 |
| 2427 | NM_001135999.1 | RUSC2 | 386.674304 | -9.8054845 | 1.61700836 |
| 2428 | NM_002480.2 | #N/A | 229.55138 | -9.0533587 | 1.49307115 |
| 2429 | NM_002541.3 | #N/A | 231.957028 | -9.07007 | 1.49582961 |
| 2430 | NM_033201.2 | #N/A | 418.858003 | -9.920757 | 1.63622901 |
| 2431 | NM_001861.4 | #N/A | 495.275104 | -10.162549 | 1.67616918 |
| 2432 | NM_207368.4 | #N/A | 385.418317 | -9.8008328 | 1.61699501 |
| 2434 | NM_018969.5 | GPR173 | 273.954021 | -5.5954169 | 0.9232118 |
| 2433 | NM_001128847.1 | #N/A | 295.403876 | -9.4171023 | 1.55376698 |
| 2435 | NM_178152.2 | #N/A | 692.856864 | -10.64683 | 1.75666378 |
| 2436 | NM_001318100.1 | #N/A | 268.968556 | -9.2814516 | 1.53154732 |
| 2437 | NM_001291411.1 | SON | 260.767516 | -9.2379789 | 1.52443875 |
| 2438 | NM_002073.2 | #N/A | 427.924794 | -8.5351635 | 1.40849405 |
| 2439 | NM_005076.4 | #N/A | 249.806518 | -9.1735179 | 1.51384812 |
| 2440 | NM_001039538.1 | MAP2 | 384.709984 | -9.797773 | 1.61694222 |
| 2441 | NM_003872.2 | #N/A | 351.177711 | -9.6668005 | 1.59538329 |
| 2442 | NM_152493.2 | #N/A | 347.336456 | -9.6506947 | 1.59271876 |
| 2443 | NM_001009999.2 | #N/A | 318.399501 | -9.5250951 | 1.57204666 |
| 2444 | NM_015908.5 | #N/A | 240.148697 | -9.1174079 | 1.50479051 |
| 2445 | NM_013341.4 | #N/A | 405.618085 | -9.8743753 | 1.62981975 |
| 2446 | NM_003901.3 | #N/A | 392.671346 | -3.5645821 | 0.58841042 |
| 2447 | NM_001130021.1 | #N/A | 535.639685 | -10.275627 | 1.6962673 |
| 2448 | NM_001136195.1 | #N/A | 308.371964 | -9.4785778 | 1.56474585 |
| 2449 | NM_005717.3 | #N/A | 484.404092 | -8.6963361 | 1.43562363 |
| 2450 | NM_001172630.1 | #N/A | 401.197542 | -9.858592 | 1.62752196 |

| | | | | | |
|---|---|---|---|---|---|
| 2451 | NM_001142289.2 | MGRN1 | 274.360322 | -9.3102848 | 1.53702239 |
| 2452 | NM_007313.2 | ABL1 | 225.373719 | -9.0254589 | 1.49007318 |
| 2453 | NM_001080434.1 | LMTK3 | 443.072893 | -10.001881 | 1.65130137 |
| 2454 | NM_001164831.2 | #N/A | 481.33945 | -10.121413 | 1.67108225 |
| 2455 | NM_031484.3 | #N/A | 231.013704 | -9.0641189 | 1.49665562 |
| 2456 | NM_001242318.2 | #N/A | 224.967853 | -9.0233955 | 1.48999386 |
| 2457 | NM_052899.2 | #N/A | 476.714595 | -10.107499 | 1.66899496 |
| 2458 | NM_001202515.1 | CFLAR | 738.73038 | 3.12497843 | 0.51602186 |
| 2459 | NM_015375.2 | #N/A | 219.740452 | -8.9888802 | 1.48448323 |
| 2460 | NM_001177413.1 | RPS27A | 399.784427 | -8.5828599 | 1.41760788 |
| 2461 | NM_002814.3 | #N/A | 269.363132 | -9.2829829 | 1.53339535 |
| 2462 | NM_183001.4 | #N/A | 276.290965 | -9.3205501 | 1.53973732 |
| 2463 | NM_003150.3 | #N/A | 258.365501 | -9.2230696 | 1.52372411 |
| 2464 | NM_001512.3 | #N/A | 351.189947 | -9.666417 | 1.59707561 |
| 2465 | NM_032233.2 | #N/A | 214.782507 | -8.9580286 | 1.48011444 |
| 2466 | NM_023018.4 | #N/A | 261.371926 | -9.2407796 | 1.52691246 |
| 2467 | NM_173517.5 | #N/A | 278.363486 | -9.3312777 | 1.54197577 |
| 2468 | NM_016306.5 | DNAJB11 | 299.320502 | -9.436423 | 1.5595216 |
| 2469 | NM_016148.3 | #N/A | 456.119413 | -10.043826 | 1.6599314 |
| 2470 | NM_004850.4 | #N/A | 222.424343 | -9.0070521 | 1.48861307 |
| 2471 | NM_001291964.1 | AFDN | 227.87296 | -9.0418141 | 1.49445735 |
| 2472 | NM_002267.3 | #N/A | 330.80822 | -9.5802445 | 1.58348153 |
| 2473 | NM_012336.3 | #N/A | 297.943123 | -9.4284894 | 1.55845584 |
| 2474 | NM_003342.4 | UBE2G1 | 288.413788 | -9.3822434 | 1.55088975 |
| 2475 | NM_018660.2 | #N/A | 195.643379 | -6.6391134 | 1.09750156 |
| 2476 | NM_001136196.1 | #N/A | 262.792588 | -7.9715956 | 1.31783885 |
| 2477 | NM_018584.5 | #N/A | 643.953157 | -10.541295 | 1.7428136 |
| 2478 | NM_016032.3 | #N/A | 297.60809 | -9.4282155 | 1.55880581 |
| 2479 | NM_007348.3 | #N/A | 269.715256 | -9.2857408 | 1.53543553 |
| 2480 | NM_015999.5 | #N/A | 222.203838 | -9.0073976 | 1.48941219 |
| 2481 | NM_001159390.1 | GUK1 | 376.65416 | -9.7675213 | 1.61515388 |
| 2482 | NM_007145.2 | ZNF146 | 277.863614 | -9.3274143 | 1.5424452 |
| 2483 | NM_001079514.2 | #N/A | 222.2846 | -9.007884 | 1.48967637 |
| 2484 | NM_207013.2 | #N/A | 376.626357 | -9.7673326 | 1.61529209 |
| 2485 | NM_018979.3 | #N/A | 255.709373 | -9.2090154 | 1.52298288 |
| 2486 | NM_004592.3 | #N/A | 267.399516 | -9.2729821 | 1.53367551 |
| 2487 | NM_032442.2 | #N/A | 294.017303 | -9.4099335 | 1.55636174 |
| 2488 | NM_022370.3 | #N/A | 237.829134 | -7.7697167 | 1.28512805 |
| 2489 | NM_001190987.1 | #N/A | 294.501705 | -9.4121 | 1.55701913 |
| 2490 | NM_003010.3 | #N/A | 295.966993 | -9.4197478 | 1.558336 |
| 2491 | NM_021135.5 | #N/A | 349.495271 | -9.6597741 | 1.59814 |
| 2492 | NM_001329.3 | #N/A | 258.326547 | -9.2240668 | 1.52607224 |
| 2493 | NM_014641.2 | #N/A | 232.021257 | -9.0688125 | 1.50046184 |

| | | | | | |
|---|---|---|---|---|---|
| 2494 | NM_022841.5 | #N/A | 216.611431 | -6.3628622 | 1.05288841 |
| 2495 | NM_001301059.1 | #N/A | 368.440964 | -9.7357577 | 1.6111641 |
| 2496 | NM_004255.3 | #N/A | 389.166618 | -9.8146862 | 1.62424688 |
| 2497 | NM_022760.4 | #N/A | 221.069475 | -8.9998828 | 1.48940778 |
| 2498 | NM_015629.3 | #N/A | 262.418117 | -9.2464662 | 1.53033173 |
| 2501 | NM_032361.3 | THOC3 | 221.065885 | -8.9988705 | 1.48944037 |
| 2499 | NM_015197.3 | #N/A | 593.33313 | -7.7068408 | 1.27558636 |
| 2500 | NM_016410.5 | #N/A | 285.723033 | -9.368775 | 1.5506366 |
| 2502 | NM_017873.3 | #N/A | 208.964951 | -8.9174596 | 1.47609824 |
| 2503 | NM_004162.4 | #N/A | 254.158249 | -9.1999441 | 1.52290636 |
| 2504 | NM_006755.1 | #N/A | 340.57074 | -9.6223492 | 1.59297401 |
| 2505 | NM_030796.4 | #N/A | 441.481849 | -9.9964172 | 1.65493721 |
| 2506 | NM_144599.4 | #N/A | 291.270269 | -9.396678 | 1.55596448 |
| 2507 | NM_017588.2 | #N/A | 253.810541 | -9.1976229 | 1.52328614 |
| 2508 | NM_002856.2 | #N/A | 315.545828 | -9.5125884 | 1.57549535 |
| 2509 | NM_198935.2 | #N/A | 313.480937 | -9.5021902 | 1.57394826 |
| 2510 | NM_001865.3 | #N/A | 451.351268 | -10.028463 | 1.66115127 |
| 2511 | NM_004481.4 | #N/A | 846.503679 | -6.8681033 | 1.13777383 |
| 2512 | NM_022491.2 | #N/A | 250.55051 | -9.1793475 | 1.52103133 |
| 2513 | NM_015670.5 | #N/A | 291.739532 | -9.3991348 | 1.55747515 |
| 2514 | NM_007285.6 | #N/A | 392.184106 | -9.8258489 | 1.6282871 |
| 2515 | NM_000700.2 | #N/A | 296.092247 | -9.421298 | 1.56144104 |
| 2516 | NM_014811.3 | #N/A | 226.46453 | -9.034004 | 1.49725325 |
| 2517 | NM_020443.4 | NAV1 | 336.143126 | -9.6034434 | 1.59170257 |
| 2518 | NM_001321092.1 | #N/A | 309.558983 | -9.4844422 | 1.57208789 |
| 2519 | NM_020211.2 | #N/A | 547.799367 | -10.308044 | 1.70860071 |
| 2520 | NM_002746.2 | #N/A | 269.149189 | -9.2827744 | 1.53874019 |
| 2521 | NM_004612.3 | #N/A | 361.006977 | -9.7069512 | 1.60913706 |
| 2522 | NM_007257.5 | #N/A | 510.046002 | -3.742069 | 0.62038833 |
| 2523 | NM_032272.4 | #N/A | 302.588711 | -9.4514171 | 1.56709313 |
| 2524 | NM_001130691.1 | #N/A | 328.759883 | -9.5709328 | 1.58698836 |
| 2525 | NM_012156.2 | EPB41L1 | 428.578123 | -9.9539465 | 1.65062408 |
| 2526 | NM_001039355.2 | #N/A | 266.373828 | -9.2672592 | 1.53680545 |
| 2527 | NM_024731.3 | #N/A | 218.114737 | -8.9798554 | 1.48914879 |
| 2528 | NM_016039.2 | #N/A | 305.911081 | -9.4674007 | 1.57006208 |
| 2529 | NM_006136.2 | #N/A | 319.230527 | -9.5286569 | 1.58029262 |
| 2530 | NM_015037.3 | ZSWIM8 | 286.802876 | -9.3758404 | 1.55498854 |
| 2531 | NM_014313.3 | #N/A | 296.693735 | -9.4230259 | 1.56311931 |
| 2532 | NM_020062.3 | #N/A | 237.656729 | -9.1032561 | 1.51017553 |
| 2533 | NM_001190467.1 | #N/A | 345.823769 | -9.6440914 | 1.60016089 |
| 2534 | NM_133373.4 | #N/A | 175.916483 | -4.3046591 | 0.71424438 |
| 2535 | NM_004172.4 | #N/A | 228.038343 | -9.0421841 | 1.50047623 |
| 2536 | NM_003250.5 | #N/A | 418.847975 | -9.9203475 | 1.6462628 |

| | | | | | |
|---|---|---|---|---|---|
| 2537 | NM_018650.4 | #N/A | 218.514046 | -8.9804789 | 1.49027865 |
| 2538 | NM_020733.1 | #N/A | 352.563104 | -8.2789327 | 1.37385311 |
| 2539 | NR_001568.1 | BCYRN1 | 1057.90292 | -7.0937568 | 1.17726146 |
| 2540 | NM_001143775.1 | #N/A | 324.241789 | -9.5512761 | 1.5852158 |
| 2541 | NM_018936.3 | #N/A | 292.857891 | -9.4048535 | 1.56097643 |
| 2542 | NM_002791.2 | #N/A | 372.830406 | -9.7528525 | 1.6188505 |
| 2543 | NM_012106.3 | #N/A | 271.986618 | -9.2979524 | 1.54337343 |
| 2544 | NM_181042.4 | #N/A | 306.649366 | -9.4712704 | 1.5722703 |
| 2545 | NM_002508.2 | #N/A | 241.047055 | -9.1255234 | 1.51490743 |
| 2546 | NM_021942.5 | #N/A | 219.030332 | -8.9845237 | 1.4915656 |
| 2547 | NM_138387.3 | G6PC3 | 265.159525 | -9.2619464 | 1.53766135 |
| 2548 | NM_018083.4 | #N/A | 331.389067 | -9.582838 | 1.59097537 |
| 2549 | NM_033088.3 | #N/A | 328.848341 | -9.5718142 | 1.58913552 |
| 2550 | NM_001004333.4 | RNASEK | 313.27787 | -9.5019129 | 1.57764013 |
| 2551 | NM_005439.2 | MLF2 | 292.997512 | -9.4054611 | 1.56169569 |
| 2552 | NM_001345840.1 | #N/A | 393.330995 | -9.8301683 | 1.63228158 |
| 2553 | NM_005481.2 | #N/A | 313.360285 | -7.000324 | 1.16251462 |
| 2554 | NM_001137560.1 | #N/A | 526.858174 | -10.251679 | 1.70249473 |
| 2555 | NM_181741.3 | #N/A | 325.098667 | -9.5542757 | 1.58666202 |
| 2556 | NM_031954.4 | #N/A | 303.772261 | -9.4573781 | 1.57066024 |
| 2557 | NM_014341.2 | MTCH1 | 252.241362 | -9.1906995 | 1.52645688 |
| 2558 | NM_178167.3 | #N/A | 253.119816 | -9.194145 | 1.52704016 |
| 2559 | NR_024077.2 | WASH2P | 217.250032 | -8.9754744 | 1.49095642 |
| 2560 | NM_003780.4 | B4GALT2 | 396.22113 | -9.8404803 | 1.63468575 |
| 2561 | NM_152265.4 | #N/A | 348.712553 | -9.6561655 | 1.60420222 |
| 2562 | NR_134878.1 | POTEC | 64.3587685 | 5.73989952 | 0.95362662 |
| 2563 | NM_145725.2 | #N/A | 222.12837 | -9.0074837 | 1.49655335 |
| 2564 | NM_006221.3 | PIN1 | 317.705292 | -9.5221085 | 1.58225209 |
| 2565 | NM_001113493.1 | #N/A | 268.624372 | -9.2798684 | 1.54206951 |
| 2566 | NM_014516.3 | #N/A | 229.87329 | -9.0554629 | 1.50477968 |
| 2567 | NM_057166.4 | #N/A | 315.943438 | -9.5151159 | 1.5812264 |
| 2568 | NM_001119.4 | #N/A | 307.241158 | -8.0993743 | 1.34598995 |
| 2569 | NM_001198832.2 | PDE4DIP | 301.981065 | -9.4488913 | 1.57029795 |
| 2570 | NM_005628.2 | #N/A | 283.817966 | -9.3592837 | 1.55547809 |
| 2571 | NM_001170687.1 | #N/A | 214.978842 | -8.9588491 | 1.48904336 |
| 2572 | NM_198880.2 | #N/A | 238.986516 | -9.1121294 | 1.51465595 |
| 2573 | NM_001080417.2 | #N/A | 224.945484 | -9.0235371 | 1.5000642 |
| 2574 | NM_006451.4 | #N/A | 361.802622 | -9.7093573 | 1.61408888 |
| 2575 | NM_005080.3 | XBP1 | 265.195141 | -9.2623439 | 1.53983321 |
| 2576 | NM_182649.1 | #N/A | 260.208473 | -9.233202 | 1.53508828 |
| 2577 | NM_014780.4 | CUL7 | 224.376796 | -9.0220949 | 1.50000607 |
| 2578 | NM_001100423.1 | #N/A | 214.324322 | -8.9557925 | 1.48936686 |
| 2579 | NM_015318.3 | #N/A | 243.437649 | -9.1373966 | 1.51979677 |

| | | | | | |
|---|---|---|---|---|---|
| 2580 | NR_034144.1 | LINC00670 | 104.861503 | 4.17542106 | 0.69449814 |
| 2581 | NM_000127.2 | EXT1 | 242.940983 | -9.1358014 | 1.51966275 |
| 2582 | NM_001291986.1 | #N/A | 230.068907 | -9.0581292 | 1.50676634 |
| 2583 | NM_015050.2 | #N/A | 244.870417 | -9.146725 | 1.52152545 |
| 2584 | NM_016219.4 | #N/A | 301.515904 | -9.4465795 | 1.57149375 |
| 2585 | NM_006004.3 | #N/A | 360.940773 | -9.7060921 | 1.61469911 |
| 2586 | NM_001001.4 | #N/A | 270.830686 | -9.2919283 | 1.54584054 |
| 2587 | NM_001287491.1 | #N/A | 281.923183 | -9.3491822 | 1.55553679 |
| 2588 | NM_001322257.1 | RAD54L2 | 40.3036303 | 10.3018287 | 1.71418294 |
| 2589 | NM_003028.2 | #N/A | 252.340649 | -9.1893965 | 1.52908661 |
| 2590 | NM_005072.4 | #N/A | 212.041163 | -8.9405284 | 1.48777622 |
| 2591 | NM_007111.4 | #N/A | 225.537527 | -9.0271182 | 1.50224445 |
| 2592 | NM_006595.3 | #N/A | 210.21875 | -8.9250115 | 1.48529168 |
| 2593 | NM_002740.5 | #N/A | 238.430479 | -9.1072458 | 1.51567721 |
| 2594 | NM_003651.4 | #N/A | 237.506092 | -9.1040223 | 1.51523516 |
| 2595 | NM_018471.2 | #N/A | 305.151532 | -9.4637433 | 1.57516905 |
| 2596 | NM_032288.6 | #N/A | 347.335214 | -9.650525 | 1.60635196 |
| 2597 | NM_019015.2 | CHPF2 | 239.799122 | -9.1169432 | 1.517617 |
| 2598 | NM_032876.5 | #N/A | 230.445058 | -9.0607226 | 1.50824719 |
| 2599 | NM_006861.6 | #N/A | 290.757292 | -9.3938176 | 1.56386702 |
| 2600 | NM_001829.3 | #N/A | 250.878373 | -9.1807661 | 1.52847539 |
| 2601 | NM_003371.3 | #N/A | 212.590564 | -8.9407832 | 1.48850285 |
| 2602 | NM_138473.2 | #N/A | 198.975454 | -8.8475709 | 1.47300632 |
| 2603 | NM_003038.4 | #N/A | 333.633781 | -9.5926825 | 1.59707968 |
| 2604 | NM_001634.5 | AMD1 | 228.790497 | -9.0470008 | 1.50630358 |
| 2605 | NM_015419.3 | #N/A | 354.458388 | -7.1265151 | 1.18662877 |
| 2606 | NM_020873.6 | #N/A | 222.368888 | -7.7310101 | 1.28740504 |
| 2607 | NM_006717.2 | #N/A | 741.478961 | -5.3341838 | 0.88837544 |
| 2608 | NM_152787.4 | #N/A | 202.55319 | -8.8720232 | 1.47762292 |
| 2609 | NM_006519.3 | #N/A | 301.410466 | -9.445947 | 1.57326568 |
| 2610 | NM_006460.2 | #N/A | 233.343658 | -6.6845247 | 1.11337259 |
| 2611 | NM_025238.3 | #N/A | 316.315132 | -9.5152423 | 1.58485977 |
| 2612 | NM_006244.3 | #N/A | 357.947056 | -9.6939314 | 1.61508283 |
| 2613 | NM_020196.2 | #N/A | 275.01846 | -9.3135336 | 1.55173373 |
| 2614 | NM_016617.3 | #N/A | 252.457702 | -9.1890971 | 1.53102905 |
| 2615 | NM_033198.3 | #N/A | 274.052186 | -7.9478483 | 1.32433796 |
| 2616 | NM_004872.4 | #N/A | 252.565836 | -9.1925467 | 1.53178362 |
| 2617 | NM_020925.2 | #N/A | 215.704574 | -8.9649282 | 1.4938913 |
| 2618 | NM_005875.2 | #N/A | 299.296317 | -9.4358383 | 1.57238284 |
| 2619 | NM_001199281.1 | CABIN1 | 208.020713 | -8.9119137 | 1.48512352 |
| 2620 | NM_001308038.1 | #N/A | 66.2246664 | 3.74739983 | 0.62448844 |
| 2621 | NM_021210.4 | #N/A | 278.240732 | -9.3306598 | 1.55518248 |
| 2623 | NM_152295.4 | TARS1 | 297.902762 | -9.429686 | 1.5717577 |

| | | | | | |
|---|---|---|---|---|---|
| 2622 | NM_020133.2 | #N/A | 297.904783 | -9.4288902 | 1.57163892 |
| 2624 | NM_014366.4 | #N/A | 241.866421 | -9.1285667 | 1.52168091 |
| 2625 | NM_025137.3 | #N/A | 205.205188 | -8.8924309 | 1.48259449 |
| 2626 | NM_001170718.1 | #N/A | 253.006549 | -9.1932489 | 1.53291284 |
| 2627 | NM_014643.3 | #N/A | 170.874731 | -3.3786504 | 0.56338162 |
| 2628 | NM_004124.2 | #N/A | 336.763922 | -9.605786 | 1.60181305 |
| 2629 | NM_020342.2 | #N/A | 270.497943 | -9.2897408 | 1.54911031 |
| 2630 | NM_012470.3 | TNPO3 | 268.03276 | -9.2765976 | 1.54699592 |
| 2631 | NM_015153.3 | #N/A | 204.830224 | -8.8888412 | 1.48234472 |
| 2632 | NM_002129.3 | #N/A | 299.85056 | -9.438516 | 1.5740839 |
| 2633 | NM_018045.7 | #N/A | 296.366779 | -9.4215745 | 1.57128407 |
| 2634 | NM_030913.4 | #N/A | 307.759808 | -9.4759822 | 1.58035934 |
| 2635 | NM_021975.3 | #N/A | 240.780169 | -9.1221721 | 1.52156059 |
| 2636 | NR_130745.1 | WASH8P | 207.049071 | -8.9043161 | 1.48531873 |
| 2637 | NM_017739.3 | POMGNT1 | 279.405449 | -9.3371252 | 1.55753568 |
| 2638 | NM_020699.3 | #N/A | 284.24688 | -9.3608263 | 1.56162086 |
| 2639 | NM_001079872.1 | CUL4B | 256.279273 | -9.2128512 | 1.53724231 |
| 2640 | NM_015695.2 | #N/A | 221.113744 | -7.6676835 | 1.27940704 |
| 2641 | NM_007190.3 | #N/A | 204.977878 | -8.8908941 | 1.48360698 |
| 2642 | NM_014748.3 | #N/A | 269.334787 | -9.2839561 | 1.54933513 |
| 2643 | NM_020223.3 | #N/A | 341.651935 | -9.6269964 | 1.60661487 |
| 2644 | NM_002940.2 | #N/A | 279.59709 | -9.3372855 | 1.55833151 |
| 2645 | NM_033386.3 | #N/A | 301.770804 | -9.4480048 | 1.57684745 |
| 2646 | NM_001085454.1 | #N/A | 516.351246 | -10.222618 | 1.70624749 |
| 2647 | NM_000143.3 | FH | 258.313339 | -9.2235156 | 1.5395503 |
| 2648 | NM_015336.2 | #N/A | 298.280098 | -9.4305782 | 1.57438248 |
| 2649 | NM_002103.4 | #N/A | 213.305493 | -8.9483878 | 1.49395366 |
| 2650 | NM_005916.4 | #N/A | 340.249066 | -9.6205256 | 1.60632242 |
| 2651 | NM_001284389.1 | #N/A | 252.881095 | -9.1911377 | 1.53472816 |
| 2652 | NM_015560.2 | OPA1 | 259.646786 | -9.2302885 | 1.54129527 |
| 2653 | NM_001008701.2 | #N/A | 573.199428 | -10.373321 | 1.73220207 |
| 2654 | NM_052844.3 | WDR34 | 208.576267 | -8.9160816 | 1.48896255 |
| 2655 | NM_002816.4 | #N/A | 284.445945 | -9.3625151 | 1.56353404 |
| 2656 | NM_001317824.1 | #N/A | 45.4684671 | 10.4750001 | 1.7493636 |
| 2657 | NM_000251.2 | #N/A | 224.575244 | -9.0210776 | 1.50669181 |
| 2658 | NM_033222.4 | #N/A | 303.410576 | -9.4551087 | 1.57919481 |
| 2659 | NM_138962.3 | #N/A | 256.544735 | -9.214558 | 1.53902666 |
| 2660 | NM_001012329.1 | #N/A | 270.384899 | -7.061241 | 1.17940415 |
| 2661 | NM_052897.3 | #N/A | 205.411227 | -8.8929681 | 1.48556391 |
| 2662 | NM_017449.4 | EPHB2 | 378.131219 | -9.7731329 | 1.63290617 |
| 2663 | NM_001184823.1 | VDAC2 | 282.300975 | -9.3516501 | 1.56260349 |
| 2664 | NM_015159.2 | #N/A | 348.054147 | -9.6543703 | 1.61322489 |
| 2665 | NM_000994.3 | #N/A | 1611.53593 | -6.2902925 | 1.05113137 |

| | | | | | |
|---|---|---|---|---|---|
| 2666 | NM_000041.3 | #N/A | 284.316794 | -9.3632241 | 1.56466789 |
| 2667 | NM_006068.4 | TLR6 | 39.8794635 | 3.13950025 | 0.52469834 |
| 2668 | NM_004631.4 | #N/A | 217.304923 | -8.9760774 | 1.5002477 |
| 2669 | NM_001144062.1 | COPB1 | 242.580061 | -9.1344036 | 1.5267645 |
| 2670 | NM_022575.3 | #N/A | 268.540532 | -9.2795175 | 1.55107583 |
| 2671 | NM_006034.4 | #N/A | 370.312851 | -9.7428205 | 1.62865082 |
| 2672 | NM_001009881.2 | #N/A | 209.165584 | -8.9205695 | 1.4912955 |
| 2673 | NM_001244604.1 | #N/A | 176.472487 | -22.483533 | 3.7587701 |
| 2674 | NM_058163.2 | #N/A | 213.002778 | -8.9450254 | 1.4954269 |
| 2676 | NM_005336.5 | HDLBP | 249.595667 | -9.1754719 | 1.53403794 |
| 2675 | NM_003238.4 | #N/A | 343.579799 | -9.6353498 | 1.61093205 |
| 2677 | NM_199368.1 | TRPC4AP | 217.564415 | -7.6454238 | 1.27828437 |
| 2678 | NM_014603.2 | #N/A | 205.661945 | -8.8963848 | 1.4875167 |
| 2679 | NM_001077180.2 | METTL9 | 317.255568 | -9.5195769 | 1.59188926 |
| 2680 | NM_014071.4 | #N/A | 302.508196 | -9.4508189 | 1.58044456 |
| 2681 | NM_012234.6 | RYBP | 268.950599 | -9.2812279 | 1.55221699 |
| 2682 | NM_201274.3 | #N/A | 397.270346 | -9.8452384 | 1.64654456 |
| 2683 | NM_152421.3 | #N/A | 282.837596 | -9.354254 | 1.56447265 |
| 2684 | NM_020920.3 | #N/A | 239.595255 | -9.1162948 | 1.52472191 |
| 2685 | NM_001289823.1 | FURIN | 229.236996 | -9.0519677 | 1.51397996 |
| 2686 | NM_016441.2 | #N/A | 346.320968 | -9.64671 | 1.61364734 |
| 2687 | NM_182917.4 | EIF4G1 | 252.318698 | -9.1908373 | 1.53743337 |
| 2688 | NM_002835.3 | #N/A | 259.980922 | -9.232356 | 1.54444893 |
| 2689 | NM_023080.2 | #N/A | 279.435616 | -9.3366092 | 1.56201009 |
| 2690 | NM_004114.3 | #N/A | 411.721164 | -9.8960852 | 1.65587792 |
| 2691 | NM_002789.5 | #N/A | 313.48276 | -9.502535 | 1.59009503 |
| 2692 | NM_004622.2 | #N/A | 295.57097 | -9.4175973 | 1.57587734 |
| 2693 | NM_012112.4 | #N/A | 236.309344 | -9.0945411 | 1.5218345 |
| 2694 | NM_016303.2 | #N/A | 217.480634 | -8.976126 | 1.50206002 |
| 2695 | NM_003655.2 | #N/A | 263.590652 | -9.2526109 | 1.54850027 |
| 2696 | NM_014712.2 | #N/A | 202.201074 | -8.8701331 | 1.48455609 |
| 2697 | NM_007172.3 | #N/A | 230.580061 | -9.0591183 | 1.51629442 |
| 2698 | NM_138352.2 | #N/A | 375.287617 | -9.762047 | 1.63398471 |
| 2699 | NM_020315.4 | #N/A | 386.507604 | -9.8046153 | 1.64128566 |
| 2700 | NM_001159702.2 | #N/A | 369.301479 | -9.7393949 | 1.63046101 |
| 2701 | NM_001663.3 | #N/A | 282.913681 | -7.9846187 | 1.33670369 |
| 2702 | NM_152429.4 | #N/A | 436.558077 | -9.980652 | 1.6710219 |
| 2703 | NM_006339.2 | #N/A | 219.192399 | -8.9867512 | 1.50474195 |
| 2704 | NM_001005388.2 | NFASC | 457.771228 | -10.049022 | 1.6826696 |
| 2706 | NM_016211.4 | SEC31A | 194.601881 | -8.8142386 | 1.4758813 |
| 2705 | NM_002858.3 | #N/A | 248.860019 | -9.1692601 | 1.53534162 |
| 2707 | NM_015284.3 | #N/A | 209.167269 | -8.9187059 | 1.49350417 |
| 2708 | NM_005639.2 | #N/A | 484.453415 | -10.130501 | 1.69681198 |

| | | | | | |
|---|---|---|---|---|---|
| 2709 | NM_001320882.1 | #N/A | 124.070898 | -22.00739 | 3.68624711 |
| 2710 | NM_030630.2 | #N/A | 242.946733 | -9.1351089 | 1.53016048 |
| 2711 | NM_001164829.1 | #N/A | 90.6901189 | 3.48779423 | 0.58425446 |
| 2712 | NM_020307.3 | CCNL1 | 202.436299 | -8.871694 | 1.48630927 |
| 2713 | NM_000791.3 | #N/A | 251.763645 | -9.1857516 | 1.53903214 |
| 2714 | NM_015937.5 | #N/A | 290.681495 | -9.3939193 | 1.57419755 |
| 2715 | NM_016016.3 | SLC25A39 | 243.491407 | -9.1376384 | 1.53131854 |
| 2716 | NM_014153.3 | #N/A | 211.187555 | -8.9329034 | 1.4970848 |
| 2717 | NM_005899.4 | #N/A | 244.346479 | -9.1415313 | 1.53208383 |
| 2718 | NM_183079.2 | #N/A | 207.28761 | -8.9053449 | 1.49253034 |
| 2719 | NM_003902.4 | FUBP1 | 241.021379 | -9.124279 | 1.52928436 |
| 2720 | NM_005928.3 | #N/A | 337.019178 | -9.6078091 | 1.61034676 |
| 2721 | NM_012179.3 | #N/A | 219.761803 | -8.990197 | 1.50683634 |
| 2722 | NM_003557.2 | #N/A | 222.093532 | -9.003799 | 1.50940071 |
| 2723 | NM_025248.2 | #N/A | 317.220608 | -9.5198668 | 1.59606841 |
| 2724 | NM_004261.4 | #N/A | 255.239349 | -9.2058505 | 1.54346036 |
| 2725 | NM_001082959.1 | #N/A | 201.226515 | -8.8625181 | 1.48594161 |
| 2726 | NM_006710.4 | #N/A | 277.2667 | -9.3253835 | 1.563574 |
| 2727 | NM_000310.3 | PPT1 | 288.897909 | -9.38496 | 1.57364896 |
| 2728 | NM_001025370.2 | #N/A | 233.650418 | -9.0794271 | 1.52244258 |
| 2729 | NM_003341.4 | #N/A | 276.551887 | -9.3220083 | 1.56312372 |
| 2730 | NR_003561.2 | DPY19L2P2 | 242.160593 | 2.85645793 | 0.47906342 |
| 2731 | NM_000183.2 | #N/A | 202.464361 | -8.8737631 | 1.48832861 |
| 2732 | NM_152277.2 | #N/A | 273.755279 | -8.0330877 | 1.34737022 |
| 2733 | NM_080594.3 | #N/A | 325.464659 | -9.5566872 | 1.60296291 |
| 2734 | NM_001271223.2 | OBSCN | 199.383213 | -8.8495168 | 1.48437785 |
| 2735 | NM_001164816.1 | #N/A | 291.121802 | -9.3967668 | 1.57620643 |
| 2736 | NM_015386.2 | #N/A | 256.246714 | -9.2118031 | 1.54517454 |
| 2737 | NM_001099402.1 | #N/A | 221.382007 | -9.0018807 | 1.50998675 |
| 2738 | NM_014757.4 | #N/A | 198.337542 | -7.563817 | 1.26884416 |
| 2739 | NM_032590.4 | #N/A | 279.804504 | -9.3379187 | 1.56644125 |
| 2740 | NM_006055.2 | #N/A | 266.095875 | -9.2665764 | 1.55457463 |
| 2741 | NM_015027.3 | #N/A | 226.879622 | -9.035917 | 1.51594475 |
| 2742 | NM_001153484.1 | SCOC | 328.619251 | -9.5705591 | 1.60568604 |
| 2743 | NM_001199417.1 | #N/A | 271.841237 | -9.296392 | 1.55982413 |
| 2744 | NM_003075.4 | #N/A | 212.11467 | -8.9383823 | 1.49980532 |
| 2745 | NM_004438.4 | #N/A | 320.54907 | -9.5344431 | 1.59984409 |
| 2746 | NM_005088.2 | #N/A | 217.812724 | -8.9783054 | 1.5065115 |
| 2747 | NM_016343.3 | #N/A | 240.13027 | -7.7693927 | 1.30366047 |
| 2748 | NM_001289135.2 | #N/A | 256.355133 | -9.2135006 | 1.546012 |
| 2749 | NM_031459.4 | #N/A | 225.914799 | -9.0302406 | 1.51529714 |
| 2750 | NM_015905.2 | #N/A | 224.144835 | -9.0195761 | 1.51352822 |
| 2752 | NM_019026.4 | TMCO1 | 242.545216 | -9.132516 | 1.53254544 |

| | | | | | |
|---|---|---|---|---|---|
| 2751 | NM_012295.3 | #N/A | 225.21021 | -9.0273905 | 1.51490446 |
| 2753 | NM_005497.3 | #N/A | 216.488922 | -8.9679068 | 1.50515959 |
| 2754 | NM_015678.4 | NBEA | 298.997583 | -9.4336932 | 1.58360484 |
| 2755 | NM_015100.3 | #N/A | 202.955567 | -8.8749289 | 1.4898846 |
| 2756 | NM_014888.2 | #N/A | 290.010138 | -9.3892009 | 1.57625855 |
| 2757 | NM_016531.5 | #N/A | 209.957139 | -8.9254682 | 1.49852415 |
| 2758 | NM_152743.3 | #N/A | 233.719256 | -9.0792596 | 1.52437103 |
| 2759 | NM_000090.3 | COL3A1 | 8392.80323 | -8.1372951 | 1.36630716 |
| 2760 | NM_001288738.1 | #N/A | 309.404575 | -8.0990752 | 1.35989422 |
| 2761 | NM_004094.4 | #N/A | 259.704765 | -7.9561856 | 1.33592825 |
| 2762 | NM_003607.3 | #N/A | 265.376062 | -9.2620058 | 1.5552756 |
| 2763 | NM_023008.3 | #N/A | 151.39493 | -3.2213407 | 0.54097689 |
| 2764 | NM_002167.4 | #N/A | 220.405703 | -8.9963863 | 1.51093268 |
| 2765 | NM_014303.3 | #N/A | 249.036426 | -9.1710144 | 1.54030364 |
| 2766 | NM_001318852.1 | MAPK8IP3 | 495.695712 | -10.163955 | 1.70711531 |
| 2767 | NM_019088.3 | #N/A | 247.461774 | -9.1619952 | 1.53886384 |
| 2768 | NM_002806.3 | #N/A | 253.139488 | -9.1940494 | 1.54431954 |
| 2769 | NM_006703.3 | #N/A | 307.60247 | -9.4752081 | 1.59153709 |
| 2770 | NM_003812.3 | #N/A | 234.77479 | -9.0851961 | 1.52612538 |
| 2771 | NM_182978.3 | #N/A | 344.412492 | -9.6386125 | 1.61919444 |
| 2772 | NM_001001555.2 | #N/A | 191.163739 | -8.7891235 | 1.47662842 |
| 2773 | NM_002717.3 | #N/A | 238.294282 | -9.1068382 | 1.52998595 |
| 2774 | NM_006913.3 | #N/A | 323.725694 | -9.5490211 | 1.60429523 |
| 2775 | NM_003195.4 | #N/A | 255.802623 | -9.2094151 | 1.54727646 |
| 2776 | NM_003713.4 | #N/A | 191.915119 | -8.7950259 | 1.47769266 |
| 2777 | NM_013237.3 | #N/A | 362.877095 | -8.3053443 | 1.39571774 |
| 2778 | NM_152345.4 | #N/A | 414.487354 | -9.9055452 | 1.66466789 |
| 2779 | NM_002109.5 | #N/A | 291.666321 | -9.3985944 | 1.57962147 |
| 2780 | NM_003660.3 | #N/A | 401.006411 | -9.85792 | 1.65681511 |
| 2781 | NM_023019.3 | DCTN1 | 482.09265 | -10.12371 | 1.70163182 |
| 2782 | NM_007208.3 | #N/A | 272.359026 | -9.2995224 | 1.56323857 |
| 2783 | NM_018064.3 | #N/A | 294.120721 | -9.4104023 | 1.58190199 |
| 2784 | NM_018332.4 | #N/A | 227.675293 | -9.0417345 | 1.51993183 |
| 2785 | NM_020783.3 | #N/A | 811.61408 | -8.0478661 | 1.35295132 |
| 2786 | NM_001010942.2 | #N/A | 282.157216 | -9.3504153 | 1.57205139 |
| 2787 | NR_024019.3 | #N/A | 199.691696 | -8.8526121 | 1.48854812 |
| 2788 | NM_001379.3 | #N/A | 232.161861 | -9.0691845 | 1.52504802 |
| 2789 | NM_021003.4 | #N/A | 279.011798 | -9.3341928 | 1.56960498 |
| 2790 | NM_021254.2 | #N/A | 200.250614 | -8.8581759 | 1.48967268 |
| 2791 | NM_020988.2 | #N/A | 533.62529 | -10.270178 | 1.72719406 |
| 2792 | NM_001282925.1 | #N/A | 430.517717 | -9.9602492 | 1.67511006 |
| 2793 | NM_001270362.1 | RAD23A | 209.629856 | -8.9225923 | 1.50067416 |
| 2794 | NM_003073.4 | #N/A | 334.717242 | -9.5970257 | 1.61414956 |

| | | | | | |
|---|---|---|---|---|---|
| 2795 | NM_138373.4 | #N/A | 222.677402 | -9.0103607 | 1.51550695 |
| 2796 | NM_014089.3 | #N/A | 252.353652 | -9.188846 | 1.54556686 |
| 2797 | NM_014892.4 | SCAF8 | 244.099116 | -7.7887367 | 1.3101174 |
| 2798 | NM_001318797.1 | #N/A | 223.962381 | -9.0186027 | 1.51712047 |
| 2799 | NM_004231.3 | #N/A | 345.431043 | -9.6427543 | 1.62217056 |
| 2800 | NM_001286665.1 | #N/A | 366.708995 | -9.7294655 | 1.63684435 |
| 2801 | NM_002206.2 | #N/A | 242.88556 | -9.1340084 | 1.53665379 |
| 2802 | NM_030811.3 | #N/A | 202.154497 | -8.8696608 | 1.49217996 |
| 2803 | NM_015103.2 | PLXND1 | 437.490174 | -6.5406026 | 1.10038959 |
| 2804 | NM_019082.3 | DDX56 | 244.883074 | -9.1464484 | 1.53885142 |
| 2805 | NM_025082.3 | #N/A | 250.73621 | -9.1803866 | 1.5446371 |
| 2806 | NM_016132.4 | #N/A | 203.242187 | -8.8791692 | 1.49397197 |
| 2807 | NM_003819.3 | #N/A | 233.556136 | -9.0783109 | 1.52753664 |
| 2808 | NM_018999.3 | #N/A | 155.335855 | -22.308911 | 3.75381209 |
| 2809 | NM_016208.3 | #N/A | 289.052311 | -9.3857411 | 1.57943638 |
| 2810 | NM_014647.3 | #N/A | 213.397769 | -8.9466667 | 1.505591 |
| 2811 | NM_018127.6 | #N/A | 247.631946 | -9.1623155 | 1.54191922 |
| 2812 | NM_006156.2 | #N/A | 312.756685 | -9.4995886 | 1.59870557 |
| 2813 | NM_017526.4 | #N/A | 197.127055 | -8.835135 | 1.48689284 |
| 2814 | NM_005760.2 | #N/A | 219.078673 | -8.9852972 | 1.51222219 |
| 2815 | NM_144723.2 | #N/A | 297.841235 | -9.4289291 | 1.58689558 |
| 2816 | NM_080703.2 | #N/A | 238.163614 | -9.1078793 | 1.53305238 |
| 2817 | NM_007005.4 | TLE4 | 312.775335 | -9.5 | 1.59929951 |
| 2818 | NM_006097.4 | #N/A | 287.755961 | -9.3801277 | 1.57936011 |
| 2819 | NM_001284307.1 | #N/A | 204.743475 | -8.8890091 | 1.49669656 |
| 2820 | NM_003055.2 | #N/A | 389.772288 | -9.817423 | 1.65329734 |
| 2821 | NM_003507.1 | FZD7 | 222.299498 | -9.0065116 | 1.51685013 |
| 2822 | NM_181833.2 | NF2 | 40.9586664 | 10.3454418 | 1.74240951 |
| 2823 | NM_005572.3 | LMNA | 242.432741 | -9.1338693 | 1.53840264 |
| 2824 | NM_000925.3 | #N/A | 220.015743 | -8.9912516 | 1.51452102 |
| 2825 | NM_014949.3 | #N/A | 200.492207 | -8.8576409 | 1.49214537 |
| 2826 | NM_133171.4 | #N/A | 268.256564 | -9.2780798 | 1.56306791 |
| 2827 | NM_001193489.1 | #N/A | 234.059683 | -9.0798776 | 1.52972748 |
| 2829 | NM_031866.2 | FZD8 | 201.150305 | -8.8608103 | 1.49287162 |
| 2828 | NM_015048.1 | #N/A | 204.201554 | -8.883594 | 1.49671823 |
| 2830 | NM_016235.2 | #N/A | 222.38533 | -9.008321 | 1.51784344 |
| 2831 | NM_006236.2 | #N/A | 589.04857 | -10.412527 | 1.75451241 |
| 2832 | NM_001082576.1 | #N/A | 357.24729 | -9.6905361 | 1.6329924 |
| 2833 | NM_001278216.1 | CDK5RAP3 | 197.044417 | -8.8341992 | 1.48880491 |
| 2834 | NM_153377.4 | LRIG3 | 260.561006 | -9.2365761 | 1.55665326 |
| 2838 | NM_031314.2 | HNRNPC | 317.750326 | -9.5218821 | 1.60487338 |
| 2835 | NM_001142397.1 | #N/A | 283.27157 | -9.3562995 | 1.57698952 |
| 2836 | NM_001142864.3 | #N/A | 272.675774 | -7.9323317 | 1.33697121 |

| | | | | | |
|---|---|---|---|---|---|
| 2837 | NM_002336.2 | #N/A | 229.939343 | -9.0550055 | 1.52618298 |
| 2839 | NM_001127231.2 | #N/A | 327.834471 | -9.5664167 | 1.61257328 |
| 2840 | NM_001731.2 | #N/A | 249.07236 | -9.1702797 | 1.54582042 |
| 2841 | NM_004463.2 | #N/A | 243.198905 | -9.1363861 | 1.54020012 |
| 2842 | NM_005596.3 | NFIB | 735.242707 | -10.732415 | 1.80928601 |
| 2843 | NM_006319.4 | #N/A | 250.305988 | -9.1778343 | 1.54746537 |
| 2844 | NM_080861.3 | #N/A | 275.658434 | -9.3166854 | 1.5708895 |
| 2845 | NM_001031710.2 | #N/A | 262.708004 | -9.2477258 | 1.55933714 |
| 2846 | NM_015246.3 | #N/A | 204.865737 | -8.8892902 | 1.49889317 |
| 2847 | NM_001278390.1 | SLC38A1 | 50.8459726 | 3.06696471 | 0.5171584 |
| 2848 | NM_006406.1 | #N/A | 298.323615 | -9.4311005 | 1.59029851 |
| 2849 | NM_005040.3 | #N/A | 219.663 | -8.9904272 | 1.5162891 |
| 2850 | NM_001098507.1 | ZNF207 | 233.081901 | -9.0732027 | 1.53028392 |
| 2851 | NM_025176.5 | #N/A | 229.165123 | -9.051282 | 1.52664348 |
| 2852 | NM_152727.5 | #N/A | 187.506503 | -8.7624903 | 1.47806295 |
| 2853 | NM_002166.4 | #N/A | 455.641653 | -10.042163 | 1.69406615 |
| 2854 | NM_032246.4 | #N/A | 333.881853 | -9.5931842 | 1.61835887 |
| 2855 | NM_015084.2 | #N/A | 220.103104 | -8.9923222 | 1.51705295 |
| 2856 | NM_001168370.1 | CUL7 | 184.900065 | -8.741161 | 1.47492654 |
| 2857 | NM_173689.6 | #N/A | 209.754611 | -8.9252607 | 1.50606459 |
| 2858 | NM_003483.4 | HMGA2 | 395.044803 | -8.4123236 | 1.41963224 |
| 2859 | NM_032102.3 | SRSF8 | 242.397848 | -9.1316625 | 1.54104034 |
| 2860 | NM_020225.2 | #N/A | 206.697543 | -8.9007139 | 1.50220443 |
| 2861 | NM_017727.4 | #N/A | 212.530936 | -8.9429015 | 1.50945822 |
| 2862 | NM_015949.2 | #N/A | 258.134849 | -9.2221592 | 1.55669881 |
| 2863 | NM_004713.5 | #N/A | 240.736788 | -9.1214524 | 1.5397289 |
| 2864 | NM_002403.3 | #N/A | 219.86012 | -8.9917779 | 1.51793303 |
| 2865 | NM_001304809.1 | BRD1 | 224.255263 | -9.0192402 | 1.52263101 |
| 2866 | NM_031892.2 | #N/A | 247.034707 | -9.1596224 | 1.54636584 |
| 2867 | NM_080670.3 | SLC35A4 | 238.009816 | -9.1055667 | 1.53729318 |
| 2868 | NM_001142565.1 | #N/A | 255.278934 | -9.2077945 | 1.55456735 |
| 2869 | NM_018383.4 | #N/A | 201.6582 | -8.8663046 | 1.49694451 |
| 2870 | NM_005655.3 | #N/A | 182.880116 | -8.7253207 | 1.4732498 |
| 2871 | NM_001013649.3 | #N/A | 131.374539 | -5.5118643 | 0.9307685 |
| 2872 | NM_182977.2 | #N/A | 227.484548 | -9.0401362 | 1.52662147 |
| 2873 | NM_052852.3 | #N/A | 168.603245 | 2.12268808 | 0.35848374 |
| 2874 | NM_018257.2 | #N/A | 232.23428 | -9.0692647 | 1.53169354 |
| 2875 | NM_020362.4 | #N/A | 307.141761 | -9.4729494 | 1.59990846 |
| 2876 | NM_005732.3 | #N/A | 217.046765 | -8.9712713 | 1.51520254 |
| 2877 | NM_001256095.1 | MYH10 | 353.04631 | -9.6748233 | 1.63416534 |
| 2878 | NM_002221.3 | ITPKB | 264.314097 | -9.2578616 | 1.56380644 |
| 2879 | NM_015638.2 | #N/A | 301.892582 | -9.4484266 | 1.59605934 |
| 2880 | NM_182976.3 | #N/A | 198.095307 | -8.840632 | 1.49345268 |

| | | | | | |
|---|---|---|---|---|---|
| 2881 | NM_003724.3 | #N/A | 163.506996 | -5.9368526 | 1.00294151 |
| 2882 | NM_006428.4 | #N/A | 236.083756 | -9.0933191 | 1.53617868 |
| 2883 | NM_016529.5 | #N/A | 333.144405 | -9.5905775 | 1.62019389 |
| 2884 | NM_000268.3 | NF2 | 200.422603 | -8.8591007 | 1.4966613 |
| 2885 | NM_001008529.2 | #N/A | 410.204559 | -9.8909888 | 1.67104684 |
| 2886 | NM_001282850.1 | #N/A | 241.838159 | -9.1265743 | 1.54188645 |
| 2887 | NM_004566.3 | #N/A | 282.692015 | -9.3536361 | 1.58019317 |
| 2888 | NM_012317.3 | #N/A | 296.596332 | -9.4229528 | 1.59197583 |
| 2889 | NM_014300.3 | #N/A | 241.117603 | -9.1242811 | 1.5415055 |
| 2890 | NM_152734.3 | #N/A | 194.950669 | -8.815977 | 1.48946822 |
| 2891 | NM_001286572.2 | #N/A | 515.851445 | -6.7499324 | 1.14044942 |
| 2893 | NM_031896.4 | CACNG7 | 453.653548 | -10.035888 | 1.69572177 |
| 2892 | NM_001271958.1 | #N/A | 222.741809 | -9.0104129 | 1.52244927 |
| 2894 | NM_002826.4 | #N/A | 280.037686 | -9.3406582 | 1.57834908 |
| 2895 | NM_017707.3 | #N/A | 221.144732 | -9.0001696 | 1.52080753 |
| 2896 | NM_006736.5 | #N/A | 323.964766 | -9.55008 | 1.61386982 |
| 2897 | NM_007173.5 | #N/A | 365.877734 | -9.726016 | 1.6436454 |
| 2898 | NM_198252.2 | #N/A | 196.87663 | -8.8331007 | 1.49274376 |
| 2899 | NM_003629.3 | #N/A | 317.681595 | -9.5222924 | 1.60932669 |
| 2900 | NR_073460.1 | BMS1P17 | 71.7447555 | 5.5672464 | 0.94098993 |
| 2901 | NM_148415.2 | ATXN2L | 224.669407 | -9.0211306 | 1.52491771 |
| 2902 | NM_000100.3 | CSTB | 198.659473 | -8.845366 | 1.49530448 |
| 2903 | NM_020801.3 | #N/A | 278.067537 | -9.3296865 | 1.57719169 |
| 2904 | NM_022156.4 | #N/A | 202.868713 | -8.8753568 | 1.50047236 |
| 2905 | NM_198268.2 | #N/A | 257.984515 | -9.2216908 | 1.55902283 |
| 2906 | NM_014233.3 | #N/A | 234.877712 | -9.0854235 | 1.53610013 |
| 2907 | NM_001128916.1 | TOMM40 | 242.232673 | -9.1301918 | 1.54384927 |
| 2908 | NM_021962.4 | #N/A | 321.655683 | -9.5393929 | 1.61320332 |
| 2909 | NM_207012.3 | #N/A | 208.533476 | -8.9155452 | 1.50772715 |
| 2910 | NM_019023.2 | #N/A | 183.408548 | -8.7296393 | 1.4763683 |
| 2911 | NM_005461.4 | #N/A | 355.7271 | -9.6854302 | 1.63827234 |
| 2912 | NM_004082.4 | DCTN1 | 347.537275 | -9.6516492 | 1.63291525 |
| 2913 | NM_004227.3 | #N/A | 215.417516 | -8.9617475 | 1.51621486 |
| 2914 | NM_022566.2 | #N/A | 222.610536 | -9.0090365 | 1.5242841 |
| 2915 | NM_015934.4 | #N/A | 267.733183 | -9.2747262 | 1.56928737 |
| 2916 | NR_027406.1 | LOC100129034 | 299.921503 | -8.0525479 | 1.36253526 |
| 2917 | NM_181336.3 | #N/A | 248.488802 | -9.1675272 | 1.55122021 |
| 2918 | NM_013366.3 | #N/A | 203.321926 | -8.8783514 | 1.50257585 |
| 2919 | NM_005000.4 | #N/A | 322.884023 | -9.5450321 | 1.61545182 |
| 2920 | NM_004499.3 | #N/A | 289.884781 | -9.3896864 | 1.58922942 |
| 2921 | NM_006268.4 | #N/A | 236.785947 | -9.0982303 | 1.53991792 |
| 2922 | NM_004431.4 | #N/A | 190.996691 | -8.7897229 | 1.48779685 |
| 2923 | NM_005278.4 | GPM6B | 342.407685 | -9.6296387 | 1.63003373 |

| | | | | | | |
|---|---|---|---|---|---|---|
| 2924 | NM_000919.3 | PAM | | 206.883296 | -8.9036865 | 1.50725458 |
| 2925 | NM_006830.3 | | #N/A | 336.997684 | -9.6070374 | 1.62633868 |
| 2926 | NM_000404.3 | | #N/A | 205.658744 | -8.8966203 | 1.50617256 |
| 2927 | NM_005782.3 | | #N/A | 296.545295 | -9.422287 | 1.59523071 |
| 2928 | NM_015994.3 | | #N/A | 258.048693 | -9.2218905 | 1.56127975 |
| 2929 | NR_109842.1 | | #N/A | 143.244292 | 3.48253251 | 0.58960451 |
| 2930 | NR_037600.1 | LINC01578 | | 301.415334 | -9.445756 | 1.59922967 |
| 2931 | NM_015045.3 | | #N/A | 202.598738 | -8.8722343 | 1.50216095 |
| 2932 | NM_005414.4 | | #N/A | 210.381016 | -8.9274717 | 1.51153558 |
| 2933 | NM_177424.2 | | #N/A | 357.342252 | -9.6916262 | 1.64094308 |
| 2934 | NM_002355.3 | | #N/A | 274.712575 | -9.3124898 | 1.57677364 |
| 2935 | NM_014772.2 | | #N/A | 356.221265 | -9.68728 | 1.64032124 |
| 2936 | NM_014570.4 | | #N/A | 204.561202 | -8.8859718 | 1.5046788 |
| 2937 | NM_024667.2 | | #N/A | 258.350527 | -9.2228818 | 1.56179509 |
| 2938 | NM_006910.4 | | #N/A | 242.163322 | -9.1288529 | 1.54603604 |
| 2939 | NM_001081563.2 | DMPK | | 207.493168 | -8.9091317 | 1.5088885 |
| 2940 | NM_001321291.1 | USP36 | | 190.819319 | -8.7862006 | 1.48823569 |
| 2941 | NM_002669.3 | | #N/A | 237.273331 | -9.1004013 | 1.54145587 |
| 2942 | NM_016002.2 | | #N/A | 234.034143 | -9.0816979 | 1.53824775 |
| 2943 | NR_135167.1 | | #N/A | 255.106744 | -9.206553 | 1.5594338 |
| 2944 | NM_001145358.1 | | #N/A | 217.970857 | -8.9776403 | 1.5207497 |
| 2945 | NM_014781.4 | | #N/A | 248.744591 | -9.1679702 | 1.55299555 |
| 2946 | NM_001329443.1 | | #N/A | 231.362628 | -9.0637778 | 1.53536838 |
| 2947 | NM_001300.5 | | #N/A | 251.483021 | -9.185088 | 1.55604748 |
| 2948 | NM_001012339.2 | | #N/A | 132.395314 | -5.2217282 | 0.88466894 |
| 2949 | NM_018645.5 | | #N/A | 489.746998 | -10.146017 | 1.7190972 |
| 2950 | NM_033063.1 | MAP6 | | 202.192767 | -8.8715122 | 1.50331672 |
| 2951 | NM_203304.3 | MEX3D | | 250.556427 | -9.1797549 | 1.55571511 |
| 2952 | NM_001330330.1 | | #N/A | 242.046643 | -9.1290037 | 1.54716982 |
| 2953 | NM_015168.1 | | #N/A | 222.561881 | -9.0082492 | 1.52675518 |
| 2954 | NM_003688.3 | CASK | | 262.231876 | -9.2451369 | 1.56699495 |
| 2955 | NM_033081.2 | DIDO1 | | 186.725926 | -8.7536793 | 1.48370771 |
| 2957 | NM_176801.2 | ADD1 | | 270.459477 | -9.2884504 | 1.57448352 |
| 2956 | NM_001862.2 | | #N/A | 342.055853 | -9.6285058 | 1.63212232 |
| 2958 | NM_184234.2 | | #N/A | 218.317486 | -8.9828372 | 1.52276439 |
| 2959 | NM_001204961.1 | PBX1 | | 306.032394 | -9.4687665 | 1.60528875 |
| 2960 | NM_001282582.1 | MXRA8 | | 254.46754 | -9.2028401 | 1.56022783 |
| 2961 | NM_005796.2 | | #N/A | 370.850986 | -4.8503837 | 0.82234202 |
| 2962 | NM_177949.3 | | #N/A | 251.541668 | -9.1853749 | 1.55736064 |
| 2963 | NM_182811.1 | | #N/A | 192.133501 | -8.7973839 | 1.49160917 |
| 2964 | NM_001265612.1 | | #N/A | 139.201464 | -22.163392 | 3.75811581 |
| 2965 | NM_003758.3 | | #N/A | 318.285838 | -9.5243846 | 1.61512976 |
| 2966 | NM_015525.3 | | #N/A | 201.203685 | -8.8629002 | 1.50295922 |

| | | | | | |
|---|---|---|---|---|---|
| 2967 | NM_000308.3 | CTSA | 242.463547 | -9.132902 | 1.54891795 |
| 2968 | NM_016040.4 | #N/A | 185.258431 | -8.7434213 | 1.48295873 |
| 2969 | NM_002069.5 | #N/A | 344.504601 | -9.6386838 | 1.63483498 |
| 2970 | NM_001161726.1 | #N/A | 276.748601 | -9.3224715 | 1.58131178 |
| 2971 | NM_005746.2 | #N/A | 222.377492 | -9.0072355 | 1.52799597 |
| 2972 | NM_017721.4 | #N/A | 210.738681 | -8.9298697 | 1.51489474 |
| 2973 | NM_013986.3 | #N/A | 320.499987 | -9.5348169 | 1.61758594 |
| 2974 | NM_014520.3 | #N/A | 223.882118 | -9.0169785 | 1.52972961 |
| 2975 | NM_002475.4 | #N/A | 365.069454 | -9.7224964 | 1.64960721 |
| 2976 | NM_015476.3 | #N/A | 307.680028 | -9.4752982 | 1.60768443 |
| 2977 | NR_037611.1 | SEPT5-GP1BB | 198.283156 | -8.8423971 | 1.50031672 |
| 2978 | NM_004544.3 | #N/A | 266.76555 | -9.2695162 | 1.57289675 |
| 2979 | NM_003198.2 | ELOA | 219.816979 | -8.9901584 | 1.52556912 |
| 2980 | NM_015509.3 | #N/A | 427.054314 | -9.9486509 | 1.68823956 |
| 2982 | NR_145444.1 | FAM30C | 46.6556135 | 3.68836315 | 0.62591409 |
| 2981 | NM_021136.2 | #N/A | 225.71984 | -9.0292683 | 1.53227615 |
| 2983 | NM_001042466.2 | #N/A | 282.163213 | -9.35097 | 1.58698926 |
| 2984 | NM_000487.5 | #N/A | 186.84495 | -8.75505 | 1.48608914 |
| 2985 | NM_004229.3 | #N/A | 186.6741 | -8.756677 | 1.48636057 |
| 2986 | NM_201575.3 | #N/A | 382.463418 | -9.7898443 | 1.66171164 |
| 2987 | NM_001281.2 | #N/A | 330.377251 | -9.5783754 | 1.62599383 |
| 2988 | NM_016641.3 | #N/A | 257.553367 | -9.2189416 | 1.56504677 |
| 2990 | NR_033321.2 | MIAT | 382.932447 | -8.5221946 | 1.44687715 |
| 2989 | NM_006356.2 | #N/A | 284.242546 | -9.3614281 | 1.58935486 |
| 2991 | NM_019112.3 | #N/A | 231.678486 | -9.0667203 | 1.53941949 |
| 2992 | NM_001003796.1 | #N/A | 285.334557 | -9.3670193 | 1.59045727 |
| 2993 | NM_080820.5 | #N/A | 327.343366 | -9.5650877 | 1.62415833 |
| 2994 | NM_002013.3 | #N/A | 322.445121 | -9.5433473 | 1.62050346 |
| 2995 | NM_001136203.1 | #N/A | 297.672349 | -9.4279073 | 1.60093999 |
| 2996 | NM_019009.3 | #N/A | 282.036706 | -9.3505151 | 1.58778348 |
| 2997 | NM_006690.3 | #N/A | 320.729776 | -9.5356181 | 1.61925057 |
| 2998 | NM_000117.2 | #N/A | 268.930254 | -9.281332 | 1.57624363 |
| 2999 | NM_006377.3 | #N/A | 201.932282 | -8.8669744 | 1.50607266 |
| 3000 | NM_017825.2 | #N/A | 185.370273 | -8.7446597 | 1.48542547 |
| 3001 | NM_020825.3 | #N/A | 223.572414 | -9.0145972 | 1.53126947 |
| 3002 | NM_005650.3 | TCF20 | 281.863934 | -9.3489343 | 1.58812677 |
| 3003 | NM_017969.2 | #N/A | 250.021251 | -7.8125148 | 1.32716009 |
| 3004 | NM_005165.2 | #N/A | 253.805593 | -9.1975037 | 1.56248831 |
| 3005 | NM_018124.3 | #N/A | 129.054251 | -5.4840898 | 0.93175977 |
| 3006 | NM_032228.5 | #N/A | 273.068205 | -9.3033259 | 1.58066296 |
| 3007 | NM_006011.3 | #N/A | 372.465708 | -9.7515184 | 1.65686358 |
| 3009 | NM_014593.3 | CXXC1 | 224.361565 | -9.0198961 | 1.53260833 |
| 3008 | NM_005589.3 | #N/A | 195.552107 | -8.8226886 | 1.4991038 |

| | | | | | | |
|---|---|---|---|---|---|---|
| 3010 | NM_020695.3 | | #N/A | 272.766959 | -7.9251818 | 1.34669551 |
| 3011 | NM_001127.3 | AP1B1 | | 277.352547 | -9.3265283 | 1.5849013 |
| 3012 | NM_001261413.1 | DCTN2 | | 272.018785 | -9.2973444 | 1.57994667 |
| 3013 | NM_030938.4 | | #N/A | 284.147121 | -9.3610274 | 1.59086889 |
| 3014 | NM_052920.1 | | #N/A | 216.669032 | -8.9694308 | 1.52435546 |
| 3015 | NM_001318119.1 | SEC31A | | 197.187146 | -8.8322875 | 1.50106203 |
| 3016 | NM_015032.3 | | #N/A | 239.442671 | -7.7556276 | 1.31810251 |
| 3017 | NM_013234.3 | | #N/A | 301.345067 | -9.445481 | 1.60547377 |
| 3018 | NM_021639.4 | | #N/A | 188.42028 | -8.769031 | 1.49052308 |
| 3019 | NM_170746.3 | | #N/A | 251.043125 | -9.1819266 | 1.56074498 |
| 3020 | NM_001170535.1 | | #N/A | 220.573127 | -8.9952403 | 1.52905999 |
| 3021 | NM_001330247.1 | HNRNPA3 | | 191.047756 | -8.7890779 | 1.49405411 |
| 3022 | NM_032932.5 | | #N/A | 344.216881 | -9.6375161 | 1.63836915 |
| 3023 | NM_145239.2 | | #N/A | 358.265639 | -6.2848901 | 1.06844153 |
| 3024 | NM_003156.3 | STIM1 | | 212.981717 | -8.9461111 | 1.52090847 |
| 3025 | NM_001040061.2 | | #N/A | 140.088915 | 4.01757377 | 0.68302739 |
| 3026 | NM_005883.2 | APC2 | | 1827.98038 | -6.4010465 | 1.08828769 |
| 3027 | NM_000287.3 | | #N/A | 187.908643 | -8.763861 | 1.49002891 |
| 3028 | NM_007213.2 | | #N/A | 284.980757 | -9.3655354 | 1.59236575 |
| 3029 | NM_020310.2 | | #N/A | 206.911544 | -8.903398 | 1.5138401 |
| 3030 | NM_139030.3 | CD151 | | 283.042102 | -9.3557853 | 1.5908664 |
| 3031 | NM_019858.1 | | #N/A | 215.052649 | -8.958058 | 1.52339986 |
| 3033 | NM_178150.2 | FBH1 | | 199.082826 | -8.8479588 | 1.50471364 |
| 3032 | NM_005754.2 | | #N/A | 224.478199 | -9.0211777 | 1.53416454 |
| 3034 | NM_018061.2 | | #N/A | 218.921055 | -8.9842387 | 1.5279126 |
| 3035 | NM_001347721.1 | | #N/A | 219.198126 | -8.9856655 | 1.52824375 |
| 3036 | NM_024036.4 | | #N/A | 375.878077 | -9.7645221 | 1.6606973 |
| 3037 | NM_145733.2 | | #N/A | 498.108389 | -10.170784 | 1.72980437 |
| 3038 | NM_014245.4 | | #N/A | 220.21972 | -8.99353 | 1.52968298 |
| 3039 | NM_002787.4 | | #N/A | 291.398662 | -8.1245437 | 1.3821473 |
| 3040 | NM_015150.1 | | #N/A | 208.749516 | -8.9180651 | 1.51715743 |
| 3041 | NM_001040260.3 | | #N/A | 300.313839 | -9.4404424 | 1.60605548 |
| 3042 | NM_002446.3 | | #N/A | 353.728966 | -9.6769108 | 1.64630143 |
| 3043 | NM_198551.3 | | #N/A | 190.582767 | -8.7832994 | 1.49428519 |
| 3044 | NM_001346581.1 | | #N/A | 247.937874 | -9.1636746 | 1.55901601 |
| 3045 | NM_032840.2 | | #N/A | 217.531996 | -8.9758944 | 1.52714705 |
| 3046 | NM_032864.3 | | #N/A | 180.725274 | -8.7077612 | 1.4815555 |
| 3047 | NM_001282875.1 | | #N/A | 240.310465 | -9.1209821 | 1.5520035 |
| 3048 | NM_001242898.1 | | #N/A | 242.418642 | -9.1314404 | 1.55387601 |
| 3049 | NM_013243.3 | | #N/A | 458.326224 | -10.05069 | 1.71031536 |
| 3050 | NM_001198665.1 | | #N/A | 254.790913 | -7.8367856 | 1.33360477 |
| 3051 | NM_033505.3 | | #N/A | 290.065763 | -9.3905803 | 1.59806428 |
| 3052 | NM_005676.4 | | #N/A | 125.969094 | -22.029284 | 3.74896317 |

| | | | | | | |
|---|---|---|---|---|---|---|
| 3053 | NM_001048241.2 | | #N/A | 301.62269 | -9.4468762 | 1.60772729 |
| 3054 | NM_004436.2 | | #N/A | 191.313178 | -8.7909601 | 1.49615849 |
| 3055 | NM_001777.3 | CD47 | | 372.908043 | -9.7529828 | 1.65995247 |
| 3056 | NM_005451.4 | | #N/A | 210.540954 | -8.9298258 | 1.51989335 |
| 3057 | NM_014316.3 | | #N/A | 329.753396 | -9.5754807 | 1.62978993 |
| 3058 | NM_016141.3 | | #N/A | 288.769285 | -9.3841565 | 1.59736522 |
| 3059 | NM_182553.2 | | #N/A | 341.281337 | -9.6253153 | 1.63860305 |
| 3060 | NM_006449.4 | | #N/A | 219.31336 | -8.9869891 | 1.52996704 |
| 3061 | NM_018209.3 | | #N/A | 192.377522 | -8.7986572 | 1.49804823 |
| 3062 | NM_020820.3 | | #N/A | 253.767274 | -9.1964947 | 1.56580291 |
| 3063 | NM_018972.2 | | #N/A | 307.616412 | -9.4754203 | 1.61336792 |
| 3064 | NM_153831.3 | | #N/A | 202.084001 | -8.8685642 | 1.51010988 |
| 3065 | NM_015179.3 | | #N/A | 174.675707 | -8.6600198 | 1.47462203 |
| 3066 | NM_017607.3 | | #N/A | 191.967427 | -8.7948901 | 1.49763342 |
| 3067 | NM_001103146.1 | | #N/A | 216.428211 | -8.9693662 | 1.52736755 |
| 3068 | NM_024906.2 | SCD5 | | 101.093584 | 2.68610989 | 0.45744417 |
| 3069 | NM_019597.4 | | #N/A | 275.477634 | -9.316132 | 1.58668216 |
| 3070 | NM_170782.2 | | #N/A | 33.0156084 | 10.0186366 | 1.70639551 |
| 3071 | NM_001330123.1 | | #N/A | 243.803524 | -9.1405931 | 1.55686479 |
| 3072 | NM_032575.2 | GLIS2 | | 181.549026 | -8.7136774 | 1.48444629 |
| 3073 | NM_001984.1 | | #N/A | 249.912042 | -9.175537 | 1.56318057 |
| 3074 | NM_001007257.2 | | #N/A | 208.129496 | -8.9122884 | 1.51840568 |
| 3075 | NM_003941.3 | | #N/A | 197.535174 | -8.836426 | 1.50547746 |
| 3076 | NM_001033.4 | | #N/A | 269.302332 | -9.2831434 | 1.58170134 |
| 3077 | NM_002296.3 | | #N/A | 249.719226 | -9.1746119 | 1.56321597 |
| 3078 | NM_058167.2 | | #N/A | 215.910427 | -8.964411 | 1.52752952 |
| 3079 | NM_020199.2 | | #N/A | 200.366723 | -8.8585572 | 1.50973658 |
| 3080 | NM_020919.3 | | #N/A | 190.848121 | -8.787027 | 1.49759803 |
| 3081 | NM_153718.3 | | #N/A | 234.345351 | -9.0817365 | 1.54781922 |
| 3082 | NM_015461.2 | | #N/A | 193.173122 | -8.806474 | 1.50094292 |
| 3083 | NM_015094.2 | | #N/A | 226.759377 | -9.0362479 | 1.5402978 |
| 3084 | NM_004964.2 | | #N/A | 198.035636 | -8.8421351 | 1.50731915 |
| 3085 | NM_001285448.1 | PDXDC1 | | 131.447616 | -3.6907788 | 0.62921955 |
| 3086 | NM_001184970.1 | | #N/A | 186.412683 | -8.7544482 | 1.49264265 |
| 3087 | NM_020761.2 | | #N/A | 201.579773 | -8.8657963 | 1.51192823 |
| 3088 | NM_003022.2 | | #N/A | 233.135507 | -9.0757632 | 1.54777487 |
| 3089 | NM_001018003.2 | | #N/A | 216.861025 | -8.9718055 | 1.53012035 |
| 3090 | NM_001271977.1 | | #N/A | 277.233148 | -9.3253493 | 1.59048509 |
| 3091 | NM_005967.3 | | #N/A | 279.310702 | -9.3357782 | 1.59226321 |
| 3092 | NM_152299.3 | | #N/A | 268.71047 | -9.2803623 | 1.5828409 |
| 3093 | NM_015175.2 | | #N/A | 177.060864 | -6.0642598 | 1.03432212 |
| 3094 | NM_012458.3 | | #N/A | 240.501873 | -9.120414 | 1.55560932 |
| 3095 | NM_198335.3 | | #N/A | 190.421968 | -8.7842119 | 1.4983741 |

| | | | | | |
|---|---|---|---|---|---|
| 3096 | NM_018683.3 | #N/A | 210.244443 | -8.9259095 | 1.52283337 |
| 3097 | NM_005811.4 | #N/A | 199.545096 | -8.8508961 | 1.51005391 |
| 3098 | NM_001136139.2 | #N/A | 193.179276 | -8.805452 | 1.50232796 |
| 3099 | NM_016061.2 | #N/A | 274.22784 | -9.3095404 | 1.58834035 |
| 3100 | NM_002890.2 | #N/A | 202.084498 | -8.8689776 | 1.5133355 |
| 3101 | NM_016038.2 | #N/A | 247.896413 | -9.1639508 | 1.56380696 |
| 3102 | NM_000123.3 | ERCC5 | 195.905293 | -8.8237049 | 1.50588623 |
| 3103 | NM_015140.3 | #N/A | 233.640112 | -9.0780021 | 1.54927321 |
| 3104 | NM_001145161.2 | #N/A | 361.719703 | -9.70912 | 1.65713222 |
| 3105 | NM_001302633.1 | #N/A | 266.099355 | -9.2670107 | 1.58169299 |
| 3106 | NR_125977.1 | WARS2-AS1 | 352.680083 | 3.98160784 | 0.67959389 |
| 3107 | NM_001135822.1 | #N/A | 286.743567 | -9.3743971 | 1.60017072 |
| 3108 | NM_024522.2 | #N/A | 403.790589 | -9.8678498 | 1.68447817 |
| 3109 | NM_001293696.1 | #N/A | 281.404672 | -9.3466998 | 1.59557776 |
| 3110 | NM_007214.4 | #N/A | 532.343166 | -3.9739558 | 0.67839492 |
| 3111 | NM_002226.4 | #N/A | 229.150916 | -9.0503704 | 1.54506056 |
| 3112 | NM_015320.3 | #N/A | 187.824384 | -8.7660102 | 1.4965918 |
| 3113 | NM_138773.3 | #N/A | 275.266141 | -9.3147204 | 1.59032699 |
| 3114 | NM_174913.2 | #N/A | 235.231678 | -9.0882498 | 1.55164699 |
| 3115 | NM_001283.3 | #N/A | 356.260155 | -9.687182 | 1.65400017 |
| 3116 | NM_014041.3 | #N/A | 226.623265 | -9.0348756 | 1.5426573 |
| 3118 | NM_018447.3 | EMC3 | 172.142568 | -4.6222865 | 0.78929438 |
| 3117 | NM_001098635.1 | #N/A | 450.419313 | -10.025596 | 1.71196003 |
| 3119 | NM_022081.5 | HPS4 | 178.622278 | -8.6919614 | 1.48430739 |
| 3120 | NM_016274.5 | #N/A | 299.484307 | -9.4367215 | 1.61152112 |
| 3121 | NM_019107.3 | #N/A | 278.215092 | -9.3308917 | 1.59351824 |
| 3122 | NM_001291345.1 | #N/A | 234.228562 | -9.0814119 | 1.55106072 |
| 3123 | NM_001042682.1 | #N/A | 238.509753 | -9.1100741 | 1.5559879 |
| 3124 | NM_001924.3 | #N/A | 187.518643 | -8.761006 | 1.49638656 |
| 3125 | NM_024569.4 | #N/A | 316.346705 | -9.5157767 | 1.62541053 |
| 3126 | NM_138774.3 | #N/A | 287.845373 | -8.1067638 | 1.38484076 |
| 3127 | NM_014758.2 | SNX19 | 191.750297 | -8.7926444 | 1.5020403 |
| 3128 | NM_006547.2 | #N/A | 201.129587 | -8.8631763 | 1.51410506 |
| 3129 | NM_000611.5 | CD59 | 239.892251 | -9.1165438 | 1.55756489 |
| 3130 | NM_152588.2 | #N/A | 260.937313 | -9.2382864 | 1.57835907 |
| 3131 | NM_001126055.2 | CASK | 192.666819 | -8.800898 | 1.50378204 |
| 3132 | NM_001134647.1 | AFAP1 | 378.018283 | -9.7724615 | 1.66986046 |
| 3133 | NM_152419.2 | #N/A | 258.46908 | -9.2249013 | 1.57632753 |
| 3134 | NM_152699.4 | #N/A | 223.376156 | -9.0134897 | 1.54032224 |
| 3135 | NM_001470.3 | #N/A | 309.835431 | -9.4863763 | 1.62153649 |
| 3136 | NM_014283.4 | #N/A | 251.032139 | -9.182115 | 1.56951689 |
| 3137 | NM_014975.2 | #N/A | 583.347524 | -5.8661781 | 1.00272479 |
| 3138 | NM_020796.4 | #N/A | 268.177204 | -9.2775043 | 1.58597196 |

| | | | | | |
|---|---|---|---|---|---|
| 3139 | NM_000122.1 | #N/A | 215.756542 | -7.6158607 | 1.30193867 |
| 3140 | NM_015404.3 | #N/A | 206.148391 | -8.8995941 | 1.52156406 |
| 3141 | NM_177947.2 | #N/A | 288.00322 | -9.3814438 | 1.60394512 |
| 3142 | NM_001195278.1 | #N/A | 310.97684 | -9.4910341 | 1.62300298 |
| 3143 | NM_016030.5 | #N/A | 195.617273 | -8.8210729 | 1.50846271 |
| 3144 | NM_005347.4 | #N/A | 2576.25339 | -6.0799865 | 1.03974206 |
| 3145 | NM_003086.3 | SNAPC4 | 170.808534 | -8.6265128 | 1.47540689 |
| 3146 | NM_001014794.2 | #N/A | 201.832125 | -8.8680072 | 1.51680814 |
| 3147 | NM_001696.3 | #N/A | 265.899553 | -9.2649511 | 1.58475786 |
| 3148 | NM_000311.3 | #N/A | 247.615148 | -9.1638829 | 1.56750497 |
| 3149 | NM_001964.2 | #N/A | 363.179008 | -9.7156066 | 1.66196834 |
| 3150 | NM_003254.2 | #N/A | 271.610626 | -9.2967127 | 1.59040288 |
| 3151 | NM_003907.2 | #N/A | 206.402556 | -8.8996647 | 1.52249141 |
| 3152 | NM_006280.2 | #N/A | 300.880069 | -9.4434887 | 1.615512 |
| 3153 | NM_006482.2 | #N/A | 209.148218 | -8.9181481 | 1.52563522 |
| 3154 | NM_001287190.1 | #N/A | 252.731958 | -9.1934453 | 1.5727979 |
| 3155 | NM_016485.4 | #N/A | 332.834125 | -9.5887171 | 1.64043172 |
| 3156 | NM_015662.2 | #N/A | 212.473335 | -8.9428164 | 1.52997293 |
| 3157 | NM_004279.2 | #N/A | 198.398911 | -8.8429879 | 1.51306444 |
| 3158 | NM_014774.2 | #N/A | 170.942049 | -8.6292526 | 1.47656243 |
| 3159 | NM_052924.2 | #N/A | 171.780516 | -8.6363193 | 1.47783341 |
| 3160 | NM_006575.4 | #N/A | 199.485001 | -8.8502685 | 1.51449075 |
| 3161 | NM_018049.2 | #N/A | 257.214828 | -9.2171789 | 1.57734873 |
| 3162 | NM_005829.4 | #N/A | 190.652338 | -8.7845391 | 1.50349716 |
| 3163 | NM_006773.3 | #N/A | 282.055104 | -5.1780469 | 0.88625729 |
| 3164 | NM_013417.3 | IARS1 | 235.764689 | -9.0912399 | 1.55604923 |
| 3165 | NM_005534.3 | IFNGR2 | 168.442247 | -5.8495271 | 1.00132306 |
| 3166 | NM_014236.3 | #N/A | 230.239171 | -9.0572085 | 1.55041417 |
| 3167 | NM_005611.3 | #N/A | 198.248359 | -8.8412707 | 1.51351209 |
| 3168 | NM_000153.3 | #N/A | 174.82553 | -8.6614831 | 1.4827553 |
| 3169 | NM_001165036.1 | #N/A | 170.271835 | -8.6231727 | 1.47622426 |
| 3170 | NM_015299.2 | #N/A | 191.66612 | -8.7941523 | 1.50562298 |
| 3171 | NM_006521.5 | #N/A | 210.84783 | -8.9315231 | 1.52943396 |
| 3172 | NR_028325.1 | LOC100132062 | 176.462275 | -8.675302 | 1.48557216 |
| 3173 | NM_002388.4 | #N/A | 214.915289 | -8.9574596 | 1.53391788 |
| 3174 | NM_130465.4 | #N/A | 287.153177 | -9.376236 | 1.60566412 |
| 3175 | NM_001317122.1 | AGO1 | 214.280282 | -8.9527632 | 1.53317529 |
| 3176 | NM_003184.3 | #N/A | 134.369801 | -5.2459539 | 0.89838104 |
| 3177 | NM_002839.3 | #N/A | 186.705781 | -8.7567202 | 1.49990868 |
| 3178 | NM_031442.3 | #N/A | 215.672268 | -8.9636801 | 1.5353535 |
| 3179 | NM_017575.4 | #N/A | 186.113064 | -8.7515335 | 1.49903854 |
| 3180 | NM_000214.2 | #N/A | 194.43857 | -8.8147529 | 1.51006333 |
| 3181 | NM_001033723.2 | #N/A | 231.051171 | -6.8373328 | 1.17134128 |

| | | | | | |
|---|---|---|---|---|---|
| 3182 | NM_001040445.2 | #N/A | 199.026872 | -8.8467232 | 1.51558569 |
| 3183 | NM_001291813.1 | #N/A | 175.728308 | -8.6671946 | 1.48481087 |
| 3184 | NM_001321.2 | #N/A | 431.528273 | -9.9634404 | 1.7069295 |
| 3185 | NM_002138.3 | HNRNPD | 435.872792 | -9.9780819 | 1.70971809 |
| 3186 | NM_024735.4 | #N/A | 215.607804 | -8.9628457 | 1.53575808 |
| 3187 | NM_181521.2 | #N/A | 178.453689 | -8.6894608 | 1.48889274 |
| 3188 | NM_015577.2 | #N/A | 245.850338 | -9.1510732 | 1.56810565 |
| 3189 | NM_022832.3 | #N/A | 169.665872 | -8.617423 | 1.47681199 |
| 3190 | NM_147187.2 | #N/A | 223.831645 | -9.0183129 | 1.54554512 |
| 3191 | NM_015294.4 | #N/A | 240.556744 | -9.1198688 | 1.56305562 |
| 3192 | NM_006753.5 | #N/A | 201.552782 | -8.8651579 | 1.51942243 |
| 3193 | NM_001172673.1 | #N/A | 392.704825 | -7.8033079 | 1.33746874 |
| 3194 | NM_003016.4 | SRSF2 | 182.931051 | -8.7268621 | 1.49584527 |
| 3195 | NM_013433.4 | #N/A | 333.127712 | -9.590488 | 1.64390437 |
| 3196 | NM_001136018.3 | EPHX1 | 213.232984 | -8.947777 | 1.53376543 |
| 3197 | NM_138471.2 | #N/A | 203.173253 | -8.8773607 | 1.52168365 |
| 3198 | NM_021074.4 | #N/A | 247.495244 | -9.1619022 | 1.57070009 |
| 3199 | NM_017999.4 | #N/A | 190.084396 | -8.7811389 | 1.50544252 |
| 3200 | NM_022917.4 | #N/A | 223.650094 | -9.0157083 | 1.54570434 |
| 3201 | NM_144628.3 | #N/A | 262.510518 | -3.8852542 | 0.66611396 |
| 3202 | NM_015089.3 | #N/A | 174.905061 | -8.6608092 | 1.48505175 |
| 3203 | NM_002850.3 | #N/A | 282.207254 | -9.3510054 | 1.60353391 |
| 3204 | NM_004432.3 | #N/A | 346.003703 | -9.6449645 | 1.65397192 |
| 3205 | NM_004716.3 | #N/A | 287.780189 | -9.3792982 | 1.60853816 |
| 3206 | NM_000895.2 | #N/A | 194.762144 | -8.8164725 | 1.5120392 |
| 3207 | NM_182548.3 | #N/A | 48.3617806 | 4.51666538 | 0.77461683 |
| 3208 | NM_006561.3 | CELF2 | 209.80976 | -8.9234354 | 1.53043437 |
| 3209 | NM_018907.3 | #N/A | 290.925083 | -9.3954137 | 1.61138887 |
| 3210 | NM_003270.3 | #N/A | 169.010378 | -8.6126067 | 1.47720517 |
| 3211 | NM_002913.4 | #N/A | 188.785528 | -8.7696699 | 1.504222 |
| 3212 | NM_013382.5 | POMT2 | 197.592822 | -8.8376875 | 1.51592162 |
| 3213 | NM_001304840.1 | #N/A | 31.918013 | 9.97860981 | 1.71170558 |
| 3214 | NM_001256378.1 | MCMBP | 175.434092 | -8.6649817 | 1.4863844 |
| 3215 | NM_001095.3 | #N/A | 305.948129 | -9.4674967 | 1.62425082 |
| 3216 | NM_002841.3 | #N/A | 264.817138 | -6.7614009 | 1.16000263 |
| 3217 | NM_014913.3 | #N/A | 200.442802 | -8.8574575 | 1.51961582 |
| 3218 | NM_003689.3 | AKR7A2 | 168.934708 | -8.609985 | 1.47728127 |
| 3219 | NM_001321688.1 | #N/A | 200.585958 | -8.8597251 | 1.52022155 |
| 3220 | NM_024818.4 | UBA5 | 181.282446 | -8.7109476 | 1.49471198 |
| 3221 | NM_017824.4 | #N/A | 237.404309 | -9.101476 | 1.56176685 |
| 3222 | NM_003137.4 | #N/A | 245.08341 | -9.1473038 | 1.56989471 |
| 3223 | NM_003562.4 | #N/A | 254.616203 | -9.2025082 | 1.57949398 |
| 3224 | NM_015496.4 | #N/A | 189.545781 | -8.7772651 | 1.50654969 |

| | | | | | |
|---|---|---|---|---|---|
| 3225 | NM_031946.6 | #N/A | 251.810749 | -9.1861329 | 1.57674473 |
| 3226 | NM_006844.4 | #N/A | 252.293832 | -9.1894843 | 1.5775537 |
| 3227 | NM_044472.2 | #N/A | 403.687392 | -9.8676058 | 1.69423918 |
| 3228 | NM_001330564.1 | #N/A | 221.629684 | -9.0038334 | 1.54602311 |
| 3229 | NM_016237.4 | #N/A | 285.518722 | -9.3675866 | 1.60851387 |
| 3230 | NM_033274.4 | #N/A | 478.676517 | -8.6464287 | 1.4846825 |
| 3231 | NM_001122659.2 | #N/A | 224.346284 | -9.0206332 | 1.54927496 |
| 3232 | NM_007166.3 | #N/A | 344.088481 | -9.6369616 | 1.65512991 |
| 3233 | NM_014452.4 | #N/A | 234.615057 | -6.0850299 | 1.04510783 |
| 3234 | NM_175614.4 | #N/A | 326.238115 | -9.5601736 | 1.64196063 |
| 3235 | NM_001024455.3 | #N/A | 229.600704 | -9.0536501 | 1.55504983 |
| 3236 | NM_002720.2 | #N/A | 239.595355 | -9.1145062 | 1.56593438 |
| 3239 | NM_002970.3 | SAT1 | 178.862574 | -8.6928042 | 1.49361208 |
| 3238 | NM_001303524.1 | TUBB6 | 247.916205 | -9.1644475 | 1.57464573 |
| 3237 | NM_001164665.1 | #N/A | 188.531137 | -8.7691755 | 1.50671328 |
| 3240 | NM_015317.2 | #N/A | 242.465524 | -9.1318462 | 1.56898534 |
| 3241 | NM_016630.6 | #N/A | 209.043555 | -8.9174915 | 1.53219083 |
| 3242 | NM_203364.2 | #N/A | 500.04383 | -7.4514081 | 1.28031318 |
| 3243 | NM_002798.2 | #N/A | 292.484951 | -9.4028228 | 1.61566902 |
| 3244 | NM_138369.2 | #N/A | 249.253813 | -9.1719682 | 1.57603353 |
| 3245 | NM_024662.2 | #N/A | 187.722276 | -8.7618422 | 1.50567606 |
| 3246 | NM_006372.4 | #N/A | 193.484957 | -8.8061198 | 1.51329992 |
| 3247 | NM_182762.3 | #N/A | 98.0022555 | 2.81877832 | 0.48441508 |
| 3248 | NM_005397.3 | #N/A | 326.728381 | -9.5630545 | 1.6434835 |
| 3249 | NM_001037131.2 | #N/A | 215.193927 | -8.9602144 | 1.53991412 |
| 3250 | NM_032940.2 | #N/A | 222.640652 | -9.0090902 | 1.5483151 |
| 3251 | NM_015005.2 | #N/A | 259.636188 | -9.2313065 | 1.58656967 |
| 3252 | NM_015599.2 | #N/A | 170.604706 | -8.6256777 | 1.48252607 |
| 3253 | NM_007224.3 | #N/A | 295.783538 | -8.0208589 | 1.37863615 |
| 3254 | NR_110744.1 | ZIM2-AS1 | 92.2019189 | 3.96257584 | 0.68111032 |
| 3255 | NM_003575.3 | #N/A | 216.665395 | -8.9695128 | 1.54183044 |
| 3256 | NR_136326.1 | #N/A | 176.489324 | -8.6760534 | 1.49167627 |
| 3257 | NM_001320883.1 | #N/A | 328.184557 | -9.5682864 | 1.645128 |
| 3258 | NM_022766.5 | #N/A | 597.031 | -5.5276736 | 0.95045543 |
| 3259 | NM_001077397.1 | IRF2BP2 | 195.263232 | -8.8211234 | 1.51679151 |
| 3260 | NM_001519.3 | #N/A | 211.845339 | -8.9361429 | 1.53656189 |
| 3261 | NM_001130067.1 | #N/A | 328.377314 | -9.5697835 | 1.64560366 |
| 3262 | NM_002499.3 | #N/A | 165.572474 | -22.394547 | 3.85095972 |
| 3263 | NM_003388.4 | #N/A | 410.567565 | -9.89181 | 1.70099462 |
| 3264 | NM_032875.2 | #N/A | 180.654759 | -8.7058511 | 1.49705627 |
| 3265 | NM_016530.2 | #N/A | 215.336722 | -8.9598271 | 1.54075311 |
| 3266 | NM_003875.2 | #N/A | 197.555735 | -8.8357687 | 1.51948787 |
| 3267 | NM_006758.2 | U2AF1 | 241.175044 | -9.124441 | 1.56926734 |

| | | | | | |
|---|---|---|---|---|---|
| 3268 | NM_183419.3 | #N/A | 182.585153 | -8.7242184 | 1.50048509 |
| 3269 | NM_014815.3 | #N/A | 201.434473 | -8.8662022 | 1.52495506 |
| 3270 | NM_001300860.1 | #N/A | 256.460288 | -9.2123479 | 1.58460819 |
| 3271 | NM_012341.2 | #N/A | 207.669173 | -8.9085081 | 1.53235359 |
| 3272 | NM_005503.3 | APBA2 | 412.629334 | -9.8991782 | 1.70287839 |
| 3273 | NM_207311.2 | BICDL1 | 239.292472 | -9.1125642 | 1.56756348 |
| 3274 | NM_013442.2 | #N/A | 224.22883 | -9.0182753 | 1.55138172 |
| 3275 | NM_001303256.2 | #N/A | 234.432056 | -9.0832592 | 1.56263692 |
| 3276 | NM_018981.2 | #N/A | 281.822907 | -9.3494605 | 1.608425 |
| 3277 | NM_001320219.1 | #N/A | 351.810271 | -9.6688905 | 1.66344276 |
| 3278 | NM_004444.4 | #N/A | 198.972528 | -8.8485885 | 1.52232726 |
| 3279 | NM_002971.5 | #N/A | 185.26072 | -8.7416253 | 1.50405725 |
| 3280 | NM_000168.5 | #N/A | 201.398675 | -8.8633102 | 1.52504583 |
| 3281 | NM_145693.2 | #N/A | 212.975084 | -8.945488 | 1.53917994 |
| 3282 | NM_001024401.2 | #N/A | 1491.31256 | -4.7617659 | 0.81938306 |
| 3283 | NM_025146.3 | #N/A | 239.127861 | -9.1118217 | 1.56795738 |
| 3284 | NM_001449.4 | #N/A | 299.063028 | -9.4347126 | 1.62356214 |
| 3285 | NM_001308068.1 | FLYWCH1 | 229.694292 | -9.0550514 | 1.55859603 |
| 3286 | NM_004528.3 | #N/A | 303.325507 | -8.1835946 | 1.40861145 |
| 3287 | NM_003001.3 | SDHC | 220.239484 | -8.9935502 | 1.54813292 |
| 3288 | NM_014117.2 | #N/A | 206.768728 | -8.9025413 | 1.53250922 |
| 3289 | NM_017437.2 | #N/A | 183.139798 | -8.7259325 | 1.50209933 |
| 3290 | NM_001931.4 | #N/A | 186.393242 | -8.7531128 | 1.50679947 |
| 3291 | NM_138609.2 | MACROH2A1 | 221.00905 | -8.9985455 | 1.54918383 |
| 3292 | NM_006435.2 | IFITM2 | 182.632355 | -8.7257384 | 1.50231432 |
| 3293 | NM_030645.1 | #N/A | 218.553239 | -8.9825688 | 1.54657215 |
| 3294 | NM_006765.3 | #N/A | 205.991258 | -8.8980012 | 1.53205786 |
| 3295 | NM_001293.2 | #N/A | 214.714029 | -8.9564992 | 1.5421848 |
| 3296 | NM_001330351.1 | #N/A | 222.988349 | -9.0118303 | 1.5518402 |
| 3297 | NM_016139.3 | #N/A | 206.727003 | -8.9037196 | 1.53321086 |
| 3298 | NM_005685.3 | #N/A | 182.959292 | -8.7237719 | 1.50230659 |
| 3299 | NM_000386.3 | #N/A | 226.225882 | -9.03252 | 1.55564849 |
| 3300 | NM_014060.2 | #N/A | 196.193308 | -8.8262754 | 1.52010767 |
| 3301 | NM_144586.6 | #N/A | 243.783339 | -9.1396563 | 1.57409327 |
| 3302 | NM_006620.3 | #N/A | 202.353414 | -8.8712115 | 1.52796895 |
| 3303 | NM_002204.3 | #N/A | 282.420723 | -9.352367 | 1.61104209 |
| 3304 | NM_014282.3 | #N/A | 246.846625 | -9.158057 | 1.57770517 |
| 3305 | NM_006700.2 | #N/A | 230.092292 | -9.0562671 | 1.56018626 |
| 3306 | NM_001079803.2 | #N/A | 233.379009 | -9.0762584 | 1.56377518 |
| 3307 | NM_005781.4 | TNK2 | 249.869545 | -9.1748719 | 1.58093989 |
| 3308 | NM_001128219.2 | #N/A | 196.37861 | -8.828517 | 1.52142682 |
| 3309 | NM_001137675.3 | #N/A | 170.747229 | -8.6273362 | 1.48675329 |
| 3310 | NM_013285.2 | #N/A | 206.699032 | -8.9022309 | 1.53422594 |

| | | | | | |
|---|---|---|---|---|---|
| 3311 | NM_022064.4 | #N/A | 216.290514 | -8.9669557 | 1.54538693 |
| 3312 | NM_020378.3 | #N/A | 207.93173 | -8.9108315 | 1.5358349 |
| 3313 | NM_130847.2 | #N/A | 199.497201 | -8.8522876 | 1.52589581 |
| 3314 | NM_002573.3 | #N/A | 407.976996 | -9.8827519 | 1.70362365 |
| 3315 | NM_001031732.3 | #N/A | 181.260015 | -8.7130473 | 1.50201355 |
| 3316 | NM_024649.4 | #N/A | 179.793685 | -8.7029766 | 1.50036048 |
| 3317 | NM_017592.2 | #N/A | 207.381874 | -8.9081861 | 1.53592444 |
| 3318 | NM_016374.5 | #N/A | 194.983375 | -8.8173459 | 1.52029936 |
| 3319 | NM_022166.3 | #N/A | 218.468266 | -7.7067674 | 1.32881808 |
| 3320 | NM_100264.2 | #N/A | 238.697751 | -9.1089309 | 1.5706303 |
| 3321 | NM_002496.3 | #N/A | 251.49364 | -9.1848103 | 1.58375981 |
| 3322 | NM_007277.4 | #N/A | 194.299864 | -8.8133045 | 1.51975843 |
| 3323 | NM_018170.4 | #N/A | 264.122338 | -9.2552333 | 1.59600027 |
| 3324 | NR_126027.1 | EXTL3-AS1 | 133.338408 | 3.19696302 | 0.55132095 |
| 3325 | NM_201269.2 | #N/A | 186.561063 | -8.7534532 | 1.50970844 |
| 3326 | NM_001031689.2 | #N/A | 202.580591 | -8.8719259 | 1.53017681 |
| 3327 | NM_001242792.1 | SNAP91 | 366.975829 | -9.7298911 | 1.67823914 |
| 3328 | NM_018233.3 | #N/A | 188.358703 | -8.7688856 | 1.51253214 |
| 3329 | NM_005864.3 | #N/A | 195.359438 | -8.819095 | 1.52125321 |
| 3330 | NM_000402.4 | G6PD | 231.157213 | -9.0628924 | 1.56332319 |
| 3331 | NM_001256488.1 | GOLGB1 | 186.483968 | -8.7553713 | 1.51032659 |
| 3332 | NM_182547.3 | #N/A | 182.593977 | -8.7234622 | 1.50483438 |
| 3333 | NM_001033561.1 | #N/A | 190.656756 | -8.7858301 | 1.51565849 |
| 3334 | NM_015649.2 | #N/A | 221.923894 | -9.0042503 | 1.5533384 |
| 3335 | NM_001145415.1 | #N/A | 179.649642 | -8.6989885 | 1.50069804 |
| 3336 | NM_004504.4 | AGFG1 | 174.303202 | -8.6559884 | 1.49334593 |
| 3337 | NM_001257181.1 | SLC20A2 | 191.18905 | -8.7916904 | 1.516833 |
| 3338 | NM_005197.3 | #N/A | 283.019325 | -9.3546211 | 1.61394298 |
| 3339 | NM_018303.5 | #N/A | 213.534045 | -8.9485325 | 1.54387325 |
| 3340 | NR_037142.1 | LOC100289473 | 65.4641888 | 6.46850082 | 1.11617829 |
| 3341 | NM_152902.4 | #N/A | 302.883096 | -9.4527039 | 1.63120896 |
| 3342 | NM_017740.2 | #N/A | 184.540002 | -8.7386579 | 1.50811335 |
| 3343 | NM_178000.2 | #N/A | 236.885139 | -9.0998781 | 1.57054929 |
| 3344 | NM_001201339.1 | #N/A | 341.225676 | -9.6241248 | 1.6613487 |
| 3345 | NM_198076.5 | #N/A | 284.741354 | -9.3636162 | 1.61651121 |
| 3346 | NM_006832.2 | #N/A | 208.085548 | -8.9115718 | 1.53868356 |
| 3347 | NM_003249.4 | #N/A | 214.729008 | -8.9567706 | 1.54654607 |
| 3348 | NM_020839.3 | #N/A | 206.54268 | -8.9003916 | 1.53687399 |
| 3349 | NM_015631.5 | #N/A | 173.993875 | -8.6544244 | 1.49465985 |
| 3350 | NM_024836.2 | #N/A | 192.549496 | -8.7998926 | 1.51977138 |
| 3351 | NM_014754.2 | #N/A | 269.877271 | -9.2865133 | 1.60394561 |
| 3352 | NM_001030007.1 | #N/A | 307.779706 | -9.4766056 | 1.63689898 |
| 3353 | NM_032514.3 | #N/A | 246.198644 | -9.1539346 | 1.58116691 |

| | | | | | |
|---|---|---|---|---|---|
| 3354 | NM_001127266.1 | #N/A | 197.365507 | -8.834972 | 1.52620742 |
| 3355 | NM_002833.3 | #N/A | 280.677434 | -4.9266811 | 0.85108676 |
| 3356 | NM_001025107.2 | #N/A | 354.401888 | -9.6802488 | 1.67232094 |
| 3357 | NM_020831.4 | #N/A | 221.825505 | -9.0034637 | 1.55543593 |
| 3358 | NM_148921.3 | EPN2 | 207.068025 | -8.9038799 | 1.53828512 |
| 3359 | NM_006659.3 | TUBGCP2 | 307.091895 | -9.4728957 | 1.63662877 |
| 3360 | NM_138300.3 | #N/A | 198.968619 | -8.8470717 | 1.52856204 |
| 3361 | NM_173546.2 | #N/A | 221.703578 | -9.0045321 | 1.55577662 |
| 3362 | NM_001081755.2 | #N/A | 204.121095 | -8.8829439 | 1.53480255 |
| 3363 | NM_001301044.1 | #N/A | 270.480342 | -9.2884706 | 1.60488384 |
| 3364 | NM_001031690.2 | #N/A | 404.683339 | -9.8712229 | 1.70560955 |
| 3365 | NM_001163.3 | #N/A | 368.729915 | -9.7368963 | 1.68244291 |
| 3366 | NM_005723.3 | #N/A | 296.276138 | -9.4209372 | 1.62787093 |
| 3367 | NM_019892.5 | #N/A | 184.944935 | -8.7416749 | 1.51055371 |
| 3368 | NM_018425.3 | PI4K2A | 204.747444 | -8.8886526 | 1.53600155 |
| 3369 | NM_014501.2 | #N/A | 585.29055 | -5.5350946 | 0.95652829 |
| 3370 | NM_020148.2 | #N/A | 316.332277 | -9.5151215 | 1.64461792 |
| 3371 | NM_001286365.1 | #N/A | 201.946748 | -8.8685932 | 1.53291709 |
| 3372 | NM_002851.2 | #N/A | 429.737758 | -9.957439 | 1.72113701 |
| 3373 | NM_001120.4 | MFSD10 | 178.819512 | -8.6932169 | 1.50275043 |
| 3374 | NM_002588.3 | #N/A | 181.309281 | -8.7149687 | 1.50655894 |
| 3375 | NM_003359.3 | #N/A | 167.470293 | -8.6005725 | 1.48677809 |
| 3376 | NM_001040000.2 | #N/A | 386.820625 | -9.8056753 | 1.69516901 |
| 3377 | NM_015104.2 | #N/A | 175.8087 | -8.6682613 | 1.49855015 |
| 3378 | NM_001317231.1 | MTCH2 | 272.008779 | -9.2977099 | 1.60743896 |
| 3379 | NM_002726.4 | #N/A | 221.918552 | -9.0044459 | 1.55674307 |
| 3380 | NM_025161.5 | #N/A | 201.094208 | -8.8622868 | 1.53220593 |
| 3381 | NM_133635.4 | #N/A | 197.227623 | -8.8363328 | 1.52772654 |
| 3382 | NM_007030.2 | #N/A | 193.262519 | -8.8049239 | 1.52232801 |
| 3383 | NM_014669.4 | #N/A | 213.167001 | -8.9462653 | 1.54677755 |
| 3384 | NM_015906.3 | #N/A | 185.060722 | -8.7417763 | 1.51139881 |
| 3385 | NM_004186.4 | #N/A | 159.970952 | -8.532041 | 1.4752104 |
| 3386 | NM_020314.5 | #N/A | 194.648918 | -8.8165884 | 1.5244114 |
| 3387 | NM_001794.4 | #N/A | 247.171197 | -9.1596189 | 1.58378665 |
| 3388 | NM_000254.2 | MTR | 185.488859 | -8.7466747 | 1.51245055 |
| 3389 | NM_001242488.1 | #N/A | 192.896948 | -8.8022741 | 1.52207906 |
| 3390 | NM_003274.4 | #N/A | 198.393281 | -8.8422338 | 1.52901551 |
| 3391 | NM_006005.3 | WFS1 | 195.030747 | -8.817027 | 1.52468049 |
| 3392 | NM_003930.4 | #N/A | 289.435567 | -9.3877918 | 1.62349758 |
| 3393 | NM_018375.4 | #N/A | 210.122596 | -8.9241932 | 1.54339904 |
| 3394 | NM_006010.5 | #N/A | 208.804959 | -8.9168843 | 1.54227518 |
| 3395 | NM_001001410.2 | #N/A | 243.237361 | -9.1365574 | 1.58043547 |
| 3396 | NM_001071.2 | #N/A | 188.685607 | -8.7693699 | 1.51693663 |

| | | | | | | |
|---|---|---|---|---|---|---|
| 3397 | NM_001322119.1 | PDPR | | 225.314097 | -3.1528068 | 0.54538241 |
| 3398 | NM_017780.3 | | #N/A | 329.985372 | -3.190917 | 0.55199887 |
| 3399 | NM_018227.5 | | #N/A | 236.662603 | -9.0966092 | 1.57368617 |
| 3400 | NM_019066.4 | | #N/A | 167.592443 | -8.5974667 | 1.48733005 |
| 3401 | NM_017838.3 | NHP2 | | 268.265085 | -9.2779663 | 1.60536075 |
| 3402 | NM_018184.2 | | #N/A | 322.559507 | -8.1277132 | 1.40633108 |
| 3403 | NM_023011.3 | | #N/A | 227.229142 | -9.0376742 | 1.5639316 |
| 3404 | NM_001040438.2 | | #N/A | 215.72597 | -8.9636326 | 1.55125416 |
| 3405 | NM_013373.3 | | #N/A | 205.062127 | -8.890473 | 1.53868684 |
| 3406 | NM_004947.4 | | #N/A | 249.075636 | -7.8969575 | 1.3667987 |
| 3407 | NM_001301045.1 | | #N/A | 326.463304 | -9.5610793 | 1.65502327 |
| 3408 | NR_130152.1 | | #N/A | 170.59956 | -8.6242702 | 1.49286622 |
| 3409 | NM_004548.2 | | #N/A | 238.850684 | -9.110318 | 1.57702311 |
| 3410 | NM_001032363.3 | | #N/A | 212.341062 | -8.9409799 | 1.54774813 |
| 3411 | NM_018403.6 | | #N/A | 173.360971 | -8.6463838 | 1.49703225 |
| 3412 | NM_053043.2 | | #N/A | 326.645955 | -4.4670654 | 0.77346607 |
| 3413 | NM_002822.4 | | #N/A | 213.41176 | -8.9467312 | 1.54918973 |
| 3414 | NM_015026.2 | | #N/A | 195.913256 | -8.8232539 | 1.52787513 |
| 3415 | NM_018010.3 | | #N/A | 175.087519 | -8.6637651 | 1.50040109 |
| 3416 | NM_019096.4 | | #N/A | 190.748532 | -8.784969 | 1.52141739 |
| 3417 | NM_001113348.1 | | #N/A | 198.70224 | -8.8455005 | 1.53198389 |
| 3418 | NM_014547.4 | | #N/A | 170.724482 | -8.6256627 | 1.49392804 |
| 3419 | NM_020680.3 | | #N/A | 182.576669 | -8.7232787 | 1.51082498 |
| 3420 | NM_018677.3 | | #N/A | 243.039757 | -9.1356407 | 1.58227826 |
| 3421 | NM_001092.4 | | #N/A | 234.506975 | -9.0856705 | 1.57365071 |
| 3422 | NM_005219.4 | | #N/A | 216.773687 | -8.9705966 | 1.55376631 |
| 3423 | NM_001134673.3 | | #N/A | 240.607957 | -9.1200434 | 1.57969914 |
| 3424 | NM_203372.1 | | #N/A | 303.482718 | -9.4560636 | 1.6380823 |
| 3425 | NM_018222.4 | | #N/A | 258.34437 | -4.487384 | 0.77737173 |
| 3426 | NM_016121.4 | | #N/A | 169.196879 | -8.6130464 | 1.49214468 |
| 3427 | NM_033480.2 | | #N/A | 198.179871 | -8.8414858 | 1.53181265 |
| 3428 | NM_152753.3 | | #N/A | 171.778466 | -8.6367077 | 1.49632239 |
| 3429 | NM_001134232.1 | | #N/A | 221.379918 | -9.000598 | 1.55945942 |
| 3430 | NM_001127671.1 | | #N/A | 234.839022 | -9.085347 | 1.57419056 |
| 3431 | NM_003406.3 | | #N/A | 248.50967 | -9.1677511 | 1.58847981 |
| 3432 | NM_002070.3 | | #N/A | 808.300952 | -5.4816137 | 0.94988794 |
| 3436 | NM_153033.4 | KCTD7 | | 254.887621 | -9.2040752 | 1.59509071 |
| 3433 | NM_001243998.1 | SPIB | | 149.294949 | 2.66644854 | 0.4620996 |
| 3434 | NM_003569.2 | | #N/A | 330.797923 | -9.580164 | 1.66027183 |
| 3435 | NM_004192.3 | | #N/A | 190.40175 | -7.4426568 | 1.28981917 |
| 3437 | NM_014315.2 | | #N/A | 240.565991 | -9.1204648 | 1.58068653 |
| 3438 | NM_182895.4 | | #N/A | 181.138723 | -8.7118176 | 1.50985962 |
| 3439 | NM_004204.3 | | #N/A | 195.948597 | -8.8257597 | 1.52975284 |

| | | | | | |
|---|---|---|---|---|---|
| 3442 | NM_015203.4 | RPRD2 | 206.924038 | -8.9051696 | 1.54360282 |
| 3440 | NM_001085365.1 | #N/A | 250.574554 | -9.1794838 | 1.59114944 |
| 3441 | NM_007062.2 | #N/A | 239.853561 | -9.1162724 | 1.5801966 |
| 3443 | NM_025138.4 | #N/A | 198.439131 | -8.8423863 | 1.53270121 |
| 3444 | NM_014612.4 | #N/A | 210.892963 | -8.9328707 | 1.54846877 |
| 3445 | NM_016952.4 | CDON | 220.173056 | -6.5492301 | 1.13536302 |
| 3446 | NM_006609.4 | #N/A | 143.956886 | -6.2924789 | 1.0908936 |
| 3447 | NM_001039141.2 | #N/A | 193.123517 | -8.8036403 | 1.52635335 |
| 3448 | NM_001070.4 | #N/A | 184.081649 | -8.7349769 | 1.51447298 |
| 3449 | NM_005688.3 | #N/A | 167.859816 | -8.6028651 | 1.49158537 |
| 3450 | NM_015680.5 | #N/A | 234.29825 | -9.0826468 | 1.57478913 |
| 3451 | NM_014884.4 | #N/A | 269.400986 | -9.2841997 | 1.60976 |
| 3452 | NM_018368.3 | #N/A | 170.697535 | -8.6261454 | 1.49568235 |
| 3453 | NM_016275.3 | #N/A | 312.621245 | -9.4988294 | 1.64704152 |
| 3454 | NM_004417.3 | #N/A | 184.925447 | -8.7412045 | 1.51571395 |
| 3455 | NM_005009.2 | #N/A | 209.764506 | -8.9234149 | 1.54737883 |
| 3456 | NM_152395.2 | #N/A | 168.779804 | -8.6100624 | 1.49308733 |
| 3457 | NM_025160.6 | WDR26 | 171.787009 | -8.6370869 | 1.49783685 |
| 3458 | NM_005805.5 | #N/A | 246.86556 | -9.1580405 | 1.58824438 |
| 3459 | NM_001135865.1 | NPIPB5 | 278.943043 | -9.3354134 | 1.61903079 |
| 3460 | NM_005850.4 | #N/A | 212.246725 | -8.9401574 | 1.55049648 |
| 3461 | NM_012161.3 | #N/A | 230.491213 | -9.0587301 | 1.57111216 |
| 3462 | NM_003910.3 | #N/A | 239.806651 | -9.1160086 | 1.5810765 |
| 3463 | NM_052874.4 | #N/A | 338.523062 | -5.1265161 | 0.88915868 |
| 3464 | NM_001277325.1 | #N/A | 179.446999 | -8.6978149 | 1.50862774 |
| 3465 | NM_001080516.1 | GRXCR2 | 26.8811932 | 9.00077679 | 1.56127733 |
| 3466 | NM_004319.2 | #N/A | 270.479747 | -9.2897303 | 1.61140136 |
| 3467 | NM_018841.5 | #N/A | 213.274249 | -8.9477712 | 1.55207493 |
| 3468 | NM_016282.3 | #N/A | 193.406801 | -8.8056965 | 1.52747464 |
| 3469 | NM_001204298.1 | #N/A | 271.179895 | -7.8994474 | 1.37055993 |
| 3470 | NM_001098616.2 | #N/A | 290.341911 | -9.3916989 | 1.62949223 |
| 3471 | NM_001081491.1 | #N/A | 172.662106 | -8.6445393 | 1.49995117 |
| 3472 | NM_001303448.1 | #N/A | 211.508257 | -8.9330945 | 1.54999529 |
| 3473 | NM_001308208.1 | #N/A | 175.729281 | -8.6681122 | 1.5040427 |
| 3474 | NM_001321096.2 | #N/A | 212.390944 | -8.9402328 | 1.55126215 |
| 3475 | NM_014810.4 | #N/A | 178.963729 | -8.6931888 | 1.50837967 |
| 3476 | NM_004077.2 | #N/A | 728.139289 | -5.320365 | 0.9231924 |
| 3477 | NM_004078.2 | #N/A | 289.796268 | -9.3897876 | 1.62931034 |
| 3478 | NM_004705.3 | #N/A | 205.71474 | -8.8941908 | 1.54335123 |
| 3479 | NM_000849.4 | #N/A | 231.621875 | -9.0655133 | 1.57309303 |
| 3480 | NM_018994.2 | #N/A | 205.146789 | -8.8907073 | 1.54281077 |
| 3481 | NM_030581.3 | #N/A | 176.378901 | -8.6733134 | 1.50510376 |
| 3482 | NM_001009925.1 | TMEM230 | 203.584327 | -8.8815423 | 1.5413988 |

| | | | | | |
|---|---|---|---|---|---|
| 3483 | NM_145802.3 | #N/A | 304.885606 | -9.4627746 | 1.64235514 |
| 3484 | NM_147164.2 | #N/A | 286.378146 | -9.3720218 | 1.62662801 |
| 3485 | NM_152716.2 | #N/A | 167.998372 | -8.6027088 | 1.49313336 |
| 3486 | NM_182620.3 | #N/A | 237.159364 | -9.1000107 | 1.57948946 |
| 3487 | NM_024100.3 | #N/A | 210.621256 | -8.9288148 | 1.5498662 |
| 3488 | NM_006370.2 | #N/A | 229.22045 | -9.0512556 | 1.57121912 |
| 3489 | NM_005573.3 | #N/A | 259.836981 | -9.2316641 | 1.60255643 |
| 3490 | NM_130811.3 | SNAP25 | 398.904026 | -9.8506162 | 1.71002623 |
| 3492 | NR_003929.2 | PPIEL | 144.753236 | 1.82140915 | 0.31619615 |
| 3491 | NM_007234.4 | #N/A | 276.18649 | -9.3198205 | 1.61792153 |
| 3493 | NM_000814.5 | #N/A | 340.429788 | -9.6216092 | 1.67040683 |
| 3494 | NM_001098416.3 | #N/A | 183.180646 | -8.7277259 | 1.51521028 |
| 3495 | NM_015344.2 | #N/A | 226.263423 | -9.0320083 | 1.56811862 |
| 3496 | NM_006808.2 | #N/A | 247.72058 | -9.1630073 | 1.5908788 |
| 3497 | NM_001289076.1 | #N/A | 39.5045831 | 10.2717033 | 1.78340488 |
| 3498 | NM_181892.3 | UBE2D3 | 175.02498 | -8.6638908 | 1.50441029 |
| 3499 | NM_014679.4 | #N/A | 178.489526 | -8.6885396 | 1.50870701 |
| 3500 | NM_003707.2 | #N/A | 215.219813 | -8.9594649 | 1.55580005 |
| 3501 | NM_001286771.2 | #N/A | 217.241342 | -8.9735155 | 1.55826365 |
| 3502 | NM_014939.3 | #N/A | 175.177695 | -8.6621489 | 1.50420176 |
| 3504 | NM_001102653.1 | OTUD4 | 162.54339 | -8.555828 | 1.48585678 |
| 3503 | NM_001017969.2 | #N/A | 214.087614 | -8.9517665 | 1.5546156 |
| 3505 | NM_001483.2 | #N/A | 169.025744 | -8.6125689 | 1.49569664 |
| 3506 | NM_018089.2 | #N/A | 177.167948 | -8.6776365 | 1.50698873 |
| 3507 | NM_001001323.1 | #N/A | 369.930741 | -9.7417116 | 1.69194178 |
| 3508 | NM_018202.5 | #N/A | 250.636402 | -9.1796052 | 1.59430821 |
| 3509 | NM_020655.3 | #N/A | 364.59958 | -9.7207103 | 1.68841248 |
| 3510 | NM_001098787.1 | BET1L | 178.061107 | -8.6888679 | 1.50920219 |
| 3511 | NM_015959.3 | #N/A | 229.23548 | -9.0510193 | 1.5721479 |
| 3512 | NM_024658.3 | #N/A | 181.815959 | -8.7193337 | 1.51453911 |
| 3513 | NM_016075.3 | #N/A | 210.303492 | -8.9265036 | 1.55058122 |
| 3514 | NM_022733.2 | #N/A | 267.629005 | -9.2743376 | 1.61133987 |
| 3515 | NM_006091.4 | #N/A | 425.243454 | -9.9423725 | 1.72748353 |
| 3517 | NM_017895.7 | DDX27 | 170.540253 | -8.6247076 | 1.498671 |
| 3516 | NM_015209.2 | KAZN | 165.811163 | -8.5858891 | 1.49193081 |
| 3518 | NM_198963.2 | #N/A | 195.401284 | -8.8201128 | 1.53268331 |
| 3519 | NM_017755.5 | #N/A | 183.085809 | -8.7262816 | 1.51644003 |
| 3520 | NM_024006.5 | #N/A | 184.271088 | -8.7364793 | 1.51834242 |
| 3522 | NR_046251.1 | FGD5-AS1 | 212.118913 | -8.9379959 | 1.55342698 |
| 3521 | NM_001276275.1 | #N/A | 221.144203 | -9.0007787 | 1.56434145 |
| 3523 | NM_004786.2 | #N/A | 262.0754 | -9.244225 | 1.60686887 |
| 3524 | NM_001145856.1 | #N/A | 284.193463 | -9.3606839 | 1.62720768 |
| 3525 | NM_001281300.1 | #N/A | 217.999397 | -8.9779766 | 1.56069553 |

| | | | | | |
|---|---|---|---|---|---|
| 3526 | NM_004112.3 | #N/A | 185.560772 | -8.7460145 | 1.52041523 |
| 3527 | NM_024682.2 | #N/A | 203.268674 | -8.8774003 | 1.54324878 |
| 3528 | NM_001950.3 | #N/A | 193.809741 | -8.8085108 | 1.5313614 |
| 3529 | NM_020896.3 | #N/A | 221.669557 | -9.0024374 | 1.56509136 |
| 3530 | NM_032019.5 | #N/A | 198.14197 | -8.841484 | 1.53713613 |
| 3531 | NM_015483.2 | #N/A | 221.202526 | -7.6340856 | 1.32725805 |
| 3532 | NM_020343.3 | #N/A | 188.011202 | -8.7638131 | 1.52366159 |
| 3533 | NM_003653.3 | #N/A | 244.302505 | -9.1423619 | 1.58954013 |
| 3534 | NM_000917.3 | P4HA1 | 237.978129 | -9.1061474 | 1.58328457 |
| 3535 | NM_006887.4 | #N/A | 254.518625 | -9.2031333 | 1.60017795 |
| 3537 | NM_021242.5 | MID1IP1 | 180.174417 | -8.7062656 | 1.51396342 |
| 3536 | NM_002790.3 | #N/A | 243.296688 | -9.1368505 | 1.58883545 |
| 3538 | NM_183006.3 | #N/A | 190.134887 | -8.7813085 | 1.52705892 |
| 3539 | NM_018644.3 | B3GAT1 | 622.408617 | -10.492136 | 1.82463583 |
| 3540 | NM_024407.4 | #N/A | 254.087577 | -9.1994678 | 1.59985795 |
| 3541 | NM_003315.3 | #N/A | 163.15824 | -8.5594386 | 1.48860936 |
| 3542 | NM_001300828.1 | #N/A | 206.134077 | -8.8977472 | 1.54747631 |
| 3543 | NM_004728.3 | #N/A | 325.934487 | -6.9562175 | 1.20983231 |
| 3544 | NM_014824.2 | #N/A | 202.872105 | -8.874299 | 1.54356562 |
| 3545 | NM_001670.2 | #N/A | 452.347726 | -3.3561847 | 0.58387003 |
| 3546 | NM_001933.4 | #N/A | 234.59031 | -9.0850097 | 1.58055926 |
| 3547 | NM_032016.3 | #N/A | 302.819116 | -9.452889 | 1.64459719 |
| 3548 | NM_001134693.1 | #N/A | 234.034656 | -9.0813464 | 1.58006134 |
| 3549 | NM_012399.4 | #N/A | 227.484888 | -9.0390868 | 1.57278083 |
| 3550 | NM_020962.2 | #N/A | 186.382636 | -8.7544751 | 1.52358308 |
| 3551 | NM_004252.4 | #N/A | 168.193227 | -8.6069797 | 1.49794323 |
| 3552 | NM_032025.4 | #N/A | 219.777908 | -8.9897367 | 1.56479848 |
| 3554 | NM_130439.3 | MXI1 | 200.232881 | -8.8547982 | 1.54150335 |
| 3553 | NM_030648.3 | SETD7 | 221.082097 | -8.9984711 | 1.56652201 |
| 3555 | NM_021129.3 | #N/A | 366.273349 | -9.7271119 | 1.69342839 |
| 3556 | NM_152600.2 | #N/A | 198.227754 | -8.8415048 | 1.53928984 |
| 3557 | NM_003883.3 | #N/A | 205.803107 | -8.8953512 | 1.54888109 |
| 3558 | NM_002038.3 | #N/A | 259.508202 | -7.8414521 | 1.36542803 |
| 3559 | NM_002501.3 | #N/A | 284.411554 | -9.3615446 | 1.6303129 |
| 3560 | NM_004720.5 | #N/A | 254.92996 | -9.204281 | 1.60312532 |
| 3561 | NM_016608.1 | #N/A | 235.807377 | -9.0918518 | 1.58361133 |
| 3562 | NM_013400.3 | REPIN1 | 182.245734 | -8.7209607 | 1.51909245 |
| 3563 | NM_001122630.1 | #N/A | 208.875972 | -8.9187901 | 1.5535682 |
| 3564 | NM_001297569.1 | ZNF124 | 31.8705993 | 9.96643408 | 1.73625091 |
| 3569 | NM_016022.3 | APH1A | 184.264964 | -8.7369732 | 1.52213066 |
| 3565 | NM_001034996.2 | #N/A | 298.636782 | -9.4324803 | 1.64330286 |
| 3566 | NM_001199973.1 | #N/A | 158.732126 | -8.5203394 | 1.48434623 |
| 3567 | NM_002079.2 | #N/A | 252.634224 | -7.8055888 | 1.35985491 |

| | | | | | |
|---|---|---|---|---|---|
| 3568 | NM_015540.3 | #N/A | 174.909317 | -8.6611804 | 1.50892247 |
| 3570 | NM_001079673.1 | #N/A | 166.047864 | -8.586363 | 1.49590766 |
| 3571 | NM_001242866.2 | #N/A | 176.612556 | -8.673312 | 1.51108825 |
| 3572 | NM_001004304.3 | #N/A | 180.537774 | -8.7057058 | 1.51678305 |
| 3573 | NM_000381.3 | #N/A | 174.726152 | -8.6569456 | 1.50831259 |
| 3574 | NM_144982.4 | #N/A | 170.621601 | -8.626486 | 1.50305427 |
| 3575 | NM_001126.3 | #N/A | 186.511151 | -8.7532181 | 1.52538849 |
| 3576 | NM_014660.3 | #N/A | 292.527168 | -9.4025944 | 1.63864265 |
| 3577 | NM_001040450.2 | #N/A | 203.231324 | -8.8764883 | 1.54698732 |
| 3578 | NM_004674.4 | #N/A | 177.336517 | -8.6812597 | 1.51295584 |
| 3579 | NM_182692.2 | #N/A | 252.388318 | -9.1900317 | 1.60166181 |
| 3580 | NM_001195518.1 | #N/A | 223.439743 | -9.013879 | 1.57099703 |
| 3581 | NM_145056.2 | #N/A | 278.623534 | -9.3326442 | 1.62667873 |
| 3582 | NM_005632.2 | #N/A | 183.78503 | -8.7322863 | 1.52207329 |
| 3583 | NM_001199179.1 | #N/A | 235.280193 | -9.0881895 | 1.58420563 |
| 3584 | NM_001960.5 | #N/A | 296.354872 | -9.4215865 | 1.6423191 |
| 3585 | NM_002731.3 | PRKACB | 268.903161 | -9.281302 | 1.61793826 |
| 3586 | NM_020376.3 | #N/A | 169.311236 | -8.6159097 | 1.50203259 |
| 3587 | NM_005997.2 | #N/A | 275.289355 | -9.315129 | 1.62396577 |
| 3588 | NM_024295.5 | #N/A | 225.469054 | -9.0267642 | 1.57380538 |
| 3589 | NM_015255.2 | UBR2 | 164.102132 | -8.5691044 | 1.49405106 |
| 3590 | NM_001143667.1 | #N/A | 182.235554 | -8.7201513 | 1.52046886 |
| 3591 | NM_001193389.1 | #N/A | 35.2582128 | 10.1086114 | 1.76284015 |
| 3593 | NM_015878.5 | AZIN1 | 223.042097 | -9.0107748 | 1.57141195 |
| 3592 | NM_004374.3 | #N/A | 283.539453 | -9.3579457 | 1.63196697 |
| 3594 | NM_033549.4 | #N/A | 191.104142 | -8.7882216 | 1.53262918 |
| 3595 | NR_134469.1 | #N/A | 230.980574 | -9.0621815 | 1.58042187 |
| 3596 | NM_006474.4 | PDPN | 240.497786 | -9.1207061 | 1.59074187 |
| 3597 | NM_001080419.2 | #N/A | 167.469883 | -8.5985805 | 1.49976311 |
| 3598 | NM_003953.5 | #N/A | 220.06241 | -8.9931348 | 1.56863728 |
| 3599 | NM_001199378.1 | #N/A | 110.48391 | -21.838248 | 3.80935203 |
| 3600 | NM_001080429.2 | #N/A | 279.193475 | -9.3357433 | 1.62849312 |
| 3601 | NM_001015053.1 | #N/A | 257.79174 | -9.2215395 | 1.6086147 |
| 3602 | NM_016076.4 | #N/A | 159.030467 | -8.5239408 | 1.48705073 |
| 3603 | NM_001446.4 | #N/A | 589.433827 | -10.413628 | 1.8167511 |
| 3604 | NM_001197026.1 | #N/A | 221.270958 | -9.0005669 | 1.57061475 |
| 3605 | NM_017836.3 | SLC41A3 | 215.024641 | -8.9585262 | 1.56338175 |
| 3606 | NM_004069.4 | #N/A | 252.052746 | -9.1879827 | 1.60344864 |
| 3607 | NM_025114.3 | #N/A | 200.868121 | -8.8593238 | 1.54630683 |
| 3608 | NM_015542.3 | #N/A | 168.338309 | -8.6060966 | 1.50218043 |
| 3609 | NM_014038.2 | #N/A | 246.34708 | -9.154908 | 1.59807081 |
| 3610 | NM_013354.6 | #N/A | 214.574175 | -8.9552149 | 1.5633706 |
| 3611 | NM_001303512.1 | PDZD4 | 271.743175 | -9.2964133 | 1.62304273 |

| | | | | | |
|---|---|---|---|---|---|
| 3612 | NR_027260.1 | RN7SL2 | 565.645696 | -7.1790462 | 1.25338674 |
| 3613 | NM_020745.3 | #N/A | 177.149216 | -4.4406329 | 0.77529979 |
| 3614 | NR_036610.1 | SRSF3 | 217.34472 | -8.9739866 | 1.56686463 |
| 3615 | NM_015201.4 | #N/A | 296.175601 | -9.4207749 | 1.644952 |
| 3616 | NM_002197.2 | #N/A | 177.653498 | -8.6834554 | 1.51634933 |
| 3617 | NM_004618.4 | #N/A | 160.943011 | -8.5416503 | 1.49169935 |
| 3618 | NR_110303.1 | LOC101929295 | 83.0862893 | 3.59091298 | 0.6271331 |
| 3619 | NM_018142.2 | #N/A | 167.557992 | -8.5989181 | 1.50183897 |
| 3620 | NM_004684.5 | SPARCL1 | 166.712407 | -8.5939847 | 1.50101981 |
| 3621 | NM_000501.3 | #N/A | 96.1745739 | -21.662862 | 3.78379126 |
| 3622 | NM_001172640.1 | CELF1 | 247.072596 | -9.1578311 | 1.59966582 |
| 3623 | NM_003350.2 | #N/A | 256.688477 | -9.2140951 | 1.60950866 |
| 3624 | NM_007027.3 | #N/A | 217.796798 | -8.9770187 | 1.56815855 |
| 3625 | NM_213636.2 | PDLIM7 | 203.215046 | -8.8784202 | 1.55107332 |
| 3627 | NM_199166.2 | ALAS1 | 154.208438 | -8.4807905 | 1.481639 |
| 3626 | NM_005858.3 | #N/A | 152.249826 | -8.4615274 | 1.47826737 |
| 3628 | NM_001270377.1 | #N/A | 268.793671 | -9.2806637 | 1.62140945 |
| 3629 | NM_003372.6 | #N/A | 272.309566 | -9.2993332 | 1.62473497 |
| 3630 | NM_176096.2 | #N/A | 219.465487 | -8.987729 | 1.57039458 |
| 3632 | NM_012402.3 | ARFIP2 | 172.902462 | -8.6445523 | 1.51045921 |
| 3631 | NM_002721.4 | #N/A | 200.670167 | -8.85915 | 1.54795536 |
| 3633 | NR_027692.1 | #N/A | 228.242237 | -9.043444 | 1.58027327 |
| 3634 | NM_006494.3 | #N/A | 160.59138 | -8.5398827 | 1.49229739 |
| 3635 | NM_018992.3 | #N/A | 209.044468 | -8.9178952 | 1.55840715 |
| 3636 | NM_001141973.2 | #N/A | 275.865973 | -9.3187096 | 1.62850423 |
| 3637 | NM_024665.5 | #N/A | 232.061262 | -9.0691311 | 1.58501099 |
| 3638 | NM_000665.4 | #N/A | 266.432738 | -9.2683259 | 1.62006296 |
| 3639 | NM_000546.5 | TP53 | 203.576752 | -8.8800055 | 1.55234709 |
| 3640 | NM_016848.5 | #N/A | 316.20984 | -9.5151848 | 1.66339896 |
| 3641 | NM_001099625.1 | #N/A | 196.073799 | -8.8257383 | 1.54298973 |
| 3642 | NM_001024660.4 | KALRN | 181.831914 | -6.6485252 | 1.16240695 |
| 3643 | NM_181787.2 | #N/A | 156.782869 | -3.8725692 | 0.67706415 |
| 3644 | NM_001144774.2 | #N/A | 324.773861 | -9.5535707 | 1.67039924 |
| 3645 | NM_022458.3 | #N/A | 204.780487 | -8.8885799 | 1.55414796 |
| 3646 | NM_015080.3 | #N/A | 423.055182 | -9.9350907 | 1.73718251 |
| 3647 | NM_001206652.1 | #N/A | 297.794559 | -9.4282756 | 1.64860259 |
| 3648 | NM_001204063.1 | CHURC1 | 157.534502 | -8.5114011 | 1.48835844 |
| 3649 | NM_015691.3 | WWC3 | 160.482237 | -8.5368401 | 1.49280606 |
| 3650 | NM_022913.3 | #N/A | 199.369251 | -8.8475556 | 1.54714856 |
| 3651 | NM_020946.1 | DENND1A | 162.455554 | -8.5554813 | 1.49616786 |
| 3652 | NM_024034.4 | #N/A | 345.495169 | -9.6429906 | 1.68634183 |
| 3653 | NM_005399.4 | #N/A | 199.097035 | -8.8475264 | 1.54727854 |
| 3654 | NM_020747.2 | ZNF608 | 208.501372 | -7.5532365 | 1.32095043 |

| | | | | | | |
|---|---|---|---|---|---|---|
| 3655 | NM_012087.3 | | #N/A | 220.386755 | -8.9946668 | 1.57305466 |
| 3656 | NM_001172774.1 | | #N/A | 196.57973 | -8.8289573 | 1.54415058 |
| 3657 | NM_032991.2 | CASP3 | | 168.951423 | -8.6098373 | 1.50605948 |
| 3658 | NM_019059.3 | | #N/A | 279.874211 | -9.3388271 | 1.63362481 |
| 3659 | NM_002657.3 | PLAGL2 | | 157.435752 | -7.2277853 | 1.26439556 |
| 3660 | NM_182970.3 | | #N/A | 287.710937 | -6.4439717 | 1.12727348 |
| 3662 | NR_003700.1 | PI4KAP2 | | 219.234322 | -4.3209303 | 0.75591706 |
| 3661 | NM_003560.2 | | #N/A | 162.474323 | -8.5571354 | 1.49701893 |
| 3663 | NM_000400.3 | ERCC2 | | 198.939539 | -8.8471173 | 1.54790341 |
| 3664 | NM_005474.4 | | #N/A | 331.134441 | -9.5811083 | 1.67634638 |
| 3665 | NM_013260.7 | | #N/A | 213.747214 | -8.9501921 | 1.5660156 |
| 3666 | NM_015173.3 | | #N/A | 151.645479 | -8.4559533 | 1.47954626 |
| 3667 | NM_006241.7 | | #N/A | 199.631126 | -8.851861 | 1.54884444 |
| 3668 | NM_030650.2 | | #N/A | 232.245387 | -9.0695016 | 1.58701514 |
| 3669 | NM_006023.2 | | #N/A | 231.264354 | -9.0639844 | 1.58624719 |
| 3670 | NM_004927.3 | | #N/A | 171.989135 | -8.6367094 | 1.51155463 |
| 3671 | NM_004424.4 | | #N/A | 198.987682 | -8.8465036 | 1.54843705 |
| 3672 | NM_016172.2 | | #N/A | 204.485301 | -8.8861162 | 1.55536291 |
| 3673 | NM_018047.2 | | #N/A | 201.544902 | -8.8654073 | 1.5518121 |
| 3674 | NM_001015877.1 | | #N/A | 158.853789 | -8.5223785 | 1.49190554 |
| 3675 | NM_005741.4 | | #N/A | 182.158836 | -8.7185992 | 1.52656026 |
| 3676 | NM_199436.1 | SPAST | | 215.020395 | -7.6843002 | 1.345678 |
| 3677 | NM_001253384.1 | | #N/A | 250.750132 | 2.51095012 | 0.43972642 |
| 3678 | NM_001013635.3 | | #N/A | 321.753646 | -9.5404625 | 1.6707977 |
| 3679 | NM_004690.3 | | #N/A | 167.210702 | -8.5943418 | 1.50529338 |
| 3680 | NM_153029.3 | | #N/A | 201.194339 | -6.4363036 | 1.12758207 |
| 3681 | NM_001282858.1 | | #N/A | 181.864448 | -8.7175707 | 1.52739824 |
| 3682 | NM_004075.4 | | #N/A | 163.528724 | -8.5633637 | 1.50044698 |
| 3683 | NM_022902.4 | | #N/A | 155.057931 | -8.4883862 | 1.48736353 |
| 3684 | NM_001127211.2 | | #N/A | 162.963292 | -8.5582649 | 1.49966981 |
| 3685 | NM_015475.4 | | #N/A | 174.956785 | -8.6611166 | 1.51779233 |
| 3686 | NM_015199.3 | | #N/A | 201.259949 | -8.8624818 | 1.55318768 |
| 3687 | NM_001174116.1 | | #N/A | 209.591082 | -8.9200939 | 1.56332378 |
| 3688 | NM_003458.3 | | #N/A | 285.402678 | -9.3672789 | 1.64184798 |
| 3689 | NM_024016.3 | | #N/A | 267.452361 | -9.2744679 | 1.62588942 |
| 3690 | NM_001270.2 | CHD1 | | 172.994058 | -8.6439759 | 1.51544404 |
| 3691 | NM_152925.2 | | #N/A | 157.695555 | -8.5108589 | 1.49216304 |
| 3693 | NM_004383.2 | CSK | | 241.06043 | -9.1234197 | 1.59963419 |
| 3692 | NM_001076788.1 | | #N/A | 223.704148 | -9.0154115 | 1.58070626 |
| 3694 | NM_001033057.1 | | #N/A | 156.764245 | -8.5010282 | 1.49077642 |
| 3695 | NM_003916.4 | | #N/A | 181.90927 | -8.7192154 | 1.52905264 |
| 3696 | NM_001319957.1 | | #N/A | 218.402169 | -8.9811124 | 1.57504153 |
| 3697 | NM_001323342.1 | | #N/A | 172.417242 | -8.6426814 | 1.51571117 |

| | | | | | |
|---|---|---|---|---|---|
| 3698 | NM_080667.6 | CFAP36 | 178.489468 | -8.6895994 | 1.52395179 |
| 3699 | NM_000232.4 | SGCB | 402.173029 | -3.1539756 | 0.55314495 |
| 3700 | NM_020428.3 | #N/A | 203.156717 | -8.8771725 | 1.55687798 |
| 3701 | NM_014203.2 | #N/A | 374.173469 | -9.7580915 | 1.71139373 |
| 3702 | NM_014345.2 | #N/A | 198.499197 | -7.4874918 | 1.31319052 |
| 3703 | NM_016406.3 | #N/A | 184.631004 | -8.7393597 | 1.5327581 |
| 3704 | NM_023924.4 | #N/A | 186.750198 | -8.755191 | 1.53555438 |
| 3705 | NM_005576.3 | #N/A | 177.65613 | -8.6858561 | 1.52350881 |
| 3706 | NM_006195.5 | #N/A | 347.782354 | -9.6527926 | 1.69312229 |
| 3707 | NM_013242.2 | #N/A | 202.617779 | -8.872716 | 1.5563861 |
| 3708 | NM_000368.4 | #N/A | 184.746047 | -8.7403412 | 1.53328373 |
| 3709 | NM_024535.4 | #N/A | 238.58286 | -9.1081551 | 1.5979604 |
| 3710 | NM_005132.2 | #N/A | 155.873644 | -8.4964378 | 1.49086618 |
| 3711 | NM_001195555.1 | #N/A | 169.83897 | -8.6160289 | 1.51189371 |
| 3712 | NM_005013.2 | #N/A | 164.846222 | -8.5753063 | 1.50478897 |
| 3713 | NM_014351.3 | #N/A | 309.795516 | -9.4854467 | 1.66449573 |
| 3714 | NM_139315.2 | #N/A | 191.386991 | -8.7899649 | 1.54246204 |
| 3715 | NM_024953.3 | #N/A | 160.616009 | -8.5365608 | 1.49805808 |
| 3716 | NM_012306.3 | #N/A | 368.110949 | -9.7343667 | 1.70839959 |
| 3717 | NM_001199802.1 | RPL11 | 225.868729 | -9.030022 | 1.58482549 |
| 3718 | NR_016021.1 | #N/A | 210.360277 | -3.5297415 | 0.61951779 |
| 3719 | NM_001170794.1 | BACH2 | 392.608571 | -8.5596975 | 1.50238317 |
| 3720 | NM_181453.3 | #N/A | 183.361195 | -8.7302695 | 1.53235375 |
| 3721 | NM_004207.3 | SLC16A3 | 209.378155 | -8.9192251 | 1.56562706 |
| 3722 | NM_019852.4 | #N/A | 163.028556 | -8.5610394 | 1.50280014 |
| 3723 | NM_001321358.1 | #N/A | 221.176737 | -8.9989675 | 1.57974487 |
| 3724 | NM_001414.3 | #N/A | 217.58666 | -8.9753136 | 1.57562613 |
| 3725 | NM_007024.4 | #N/A | 185.542993 | -8.7464795 | 1.53552735 |
| 3726 | NM_001550.3 | IFRD1 | 259.630554 | -9.2306165 | 1.6205446 |
| 3727 | NM_003709.3 | #N/A | 293.076853 | -9.4053889 | 1.65134469 |
| 3728 | NM_144973.3 | #N/A | 246.772732 | -9.1569649 | 1.60774496 |
| 3729 | NR_003563.1 | PI4KAP1 | 188.530058 | -8.768315 | 1.53970578 |
| 3730 | NM_018028.3 | SAMD4B | 161.35976 | -8.5467512 | 1.50086752 |
| 3731 | NM_025128.4 | #N/A | 217.308567 | -8.9740821 | 1.57594239 |
| 3732 | NM_003478.3 | #N/A | 172.800802 | -8.6434975 | 1.51790592 |
| 3733 | NR_110257.1 | LINC01814 | 509.373125 | 3.89947979 | 0.68487208 |
| 3734 | NM_003379.4 | #N/A | 241.493805 | -9.1264696 | 1.60292846 |
| 3735 | NM_015387.4 | #N/A | 251.757973 | -9.1857474 | 1.61334064 |
| 3736 | NM_001035267.1 | #N/A | 244.307912 | -9.1428057 | 1.60586052 |
| 3737 | NM_005698.3 | #N/A | 190.319624 | -8.7829771 | 1.54273427 |
| 3738 | NM_004896.4 | #N/A | 192.268017 | -8.7979616 | 1.54543057 |
| 3739 | NM_007006.2 | #N/A | 595.78808 | -5.2333557 | 0.91930291 |
| 3740 | NM_152390.2 | #N/A | 245.343538 | -9.1496895 | 1.6073013 |

| | | | | | |
|---|---|---|---|---|---|
| 3741 | NM_005953.3 | #N/A | 257.321707 | -9.2176697 | 1.61928727 |
| 3742 | NM_015124.4 | #N/A | 216.648057 | -8.9692222 | 1.57570718 |
| 3743 | NM_182641.3 | #N/A | 178.507845 | -8.6917865 | 1.52702242 |
| 3744 | NM_018091.5 | #N/A | 158.765519 | -8.5203147 | 1.49694471 |
| 3745 | NM_002870.3 | #N/A | 179.255654 | -8.6971025 | 1.52803842 |
| 3746 | NM_152531.4 | #N/A | 203.758327 | -8.8812168 | 1.56042112 |
| 3747 | NM_016068.2 | #N/A | 227.124266 | -9.0377013 | 1.58817289 |
| 3748 | NM_001135998.2 | NDUFB11 | 258.880781 | -9.2266248 | 1.62160266 |
| 3749 | NM_016417.2 | #N/A | 188.109085 | -8.7658104 | 1.54062326 |
| 3750 | NM_080491.2 | #N/A | 229.971783 | -9.0556472 | 1.59166121 |
| 3751 | NM_001080410.2 | #N/A | 304.313514 | -9.4597012 | 1.66272754 |
| 3752 | NM_033254.3 | #N/A | 151.505111 | -8.4517944 | 1.48575362 |
| 3753 | NM_001191061.1 | #N/A | 221.306314 | -8.9998363 | 1.58220748 |
| 3754 | NM_138440.2 | #N/A | 236.969812 | -5.6045516 | 0.98532413 |
| 3755 | NM_032015.4 | #N/A | 183.633278 | -8.7320603 | 1.53536445 |
| 3756 | NM_024541.2 | #N/A | 133.400887 | -5.5207662 | 0.97073484 |
| 3757 | NM_006688.4 | #N/A | 249.954891 | -9.1767313 | 1.6136395 |
| 3758 | NM_022892.1 | NAIP | 173.312774 | -7.3683931 | 1.29582118 |
| 3759 | NM_001040455.1 | #N/A | 169.205125 | -7.2792858 | 1.28019629 |
| 3760 | NM_005663.4 | #N/A | 200.924948 | -7.5007003 | 1.31924943 |
| 3761 | NM_022763.3 | #N/A | 181.868386 | -8.7167192 | 1.53317128 |
| 3762 | NM_005113.3 | #N/A | 170.064197 | -8.6209027 | 1.51636402 |
| 3763 | NM_005817.4 | #N/A | 188.572083 | -8.770139 | 1.54270648 |
| 3765 | NM_024589.2 | ROGDI | 245.449266 | -9.1494351 | 1.60945782 |
| 3764 | NM_015062.4 | #N/A | 160.739349 | -8.5381907 | 1.50192592 |
| 3766 | NM_006111.2 | #N/A | 152.003097 | -8.4608067 | 1.48839978 |
| 3767 | NM_000071.2 | CBS | 280.225566 | -9.3404932 | 1.64333592 |
| 3768 | NM_033251.2 | RPL13 | 167.041544 | -8.5938194 | 1.5122317 |
| 3769 | NM_001628.3 | #N/A | 272.91992 | -7.8969211 | 1.3896188 |
| 3770 | NM_006285.2 | #N/A | 208.258776 | -8.9127786 | 1.56840989 |
| 3771 | NM_145185.3 | #N/A | 164.244595 | -8.569056 | 1.50801565 |
| 3772 | NM_033001.3 | #N/A | 255.354098 | -9.2066855 | 1.62028518 |
| 3773 | NM_000269.2 | NME1 | 381.717884 | -8.3310006 | 1.46637138 |
| 3774 | NM_018948.3 | #N/A | 152.598847 | -8.4653416 | 1.49001139 |
| 3775 | NM_015226.2 | #N/A | 205.013783 | -8.8900419 | 1.5648265 |
| 3776 | NM_018360.2 | #N/A | 176.138006 | -8.6696207 | 1.52609994 |
| 3777 | NM_006297.2 | XRCC1 | 181.784144 | -8.7168618 | 1.53443192 |
| 3778 | NM_001110303.3 | #N/A | 186.83951 | -8.7553875 | 1.54124034 |
| 3779 | NM_018008.3 | #N/A | 313.168791 | -9.5007189 | 1.67270496 |
| 3780 | NM_002882.3 | #N/A | 243.785291 | -9.1394658 | 1.60927617 |
| 3781 | NM_019885.3 | CYP26B1 | 300.114309 | -9.4402265 | 1.66251561 |
| 3782 | NM_001282116.1 | RFX3 | 155.232138 | -8.4902021 | 1.4952864 |
| 3783 | NM_005233.5 | #N/A | 191.323726 | -8.7896182 | 1.54812704 |

| | | | | | |
|---|---|---|---|---|---|
| 3784 | NM_016291.3 | #N/A | 214.603625 | -8.9550234 | 1.57731328 |
| 3785 | NM_000285.3 | #N/A | 198.7763 | -8.8457569 | 1.55824881 |
| 3786 | NM_003087.2 | #N/A | 313.315278 | -9.5023933 | 1.67399328 |
| 3787 | NM_004698.2 | #N/A | 171.367003 | -7.2924215 | 1.28466478 |
| 3788 | NM_006337.4 | #N/A | 222.344568 | -9.0065193 | 1.58663157 |
| 3789 | NM_014160.4 | #N/A | 160.922903 | -8.5410366 | 1.50461797 |
| 3790 | NM_021128.4 | #N/A | 194.666893 | -6.2251256 | 1.09664719 |
| 3791 | NM_017623.4 | #N/A | 157.59638 | -8.509255 | 1.49908624 |
| 3792 | NM_001667.3 | #N/A | 186.207465 | -8.7520102 | 1.54202092 |
| 3793 | NM_001535.4 | PRMT2 | 210.52518 | -8.928907 | 1.57325685 |
| 3794 | NM_000434.3 | #N/A | 206.362018 | -7.5343067 | 1.32769485 |
| 3795 | NM_014255.5 | #N/A | 185.474539 | -8.7457663 | 1.54123691 |
| 3796 | NM_018903.3 | #N/A | 223.134512 | -9.012817 | 1.58835648 |
| 3797 | NM_207354.2 | #N/A | 177.666925 | -8.6820897 | 1.53020148 |
| 3798 | NM_001042472.2 | #N/A | 226.647895 | -9.0349491 | 1.59253216 |
| 3799 | NM_005704.4 | #N/A | 360.333103 | -9.7036674 | 1.71056805 |
| 3800 | NM_001321967.1 | ATAD1 | 216.097006 | -8.9658146 | 1.58053739 |
| 3801 | NM_000956.3 | #N/A | 33.0040793 | 4.7466389 | 0.83678552 |
| 3803 | NM_017452.3 | STAU1 | 202.269612 | -8.8703905 | 1.56383695 |
| 3802 | NM_001031804.2 | #N/A | 196.182695 | -8.8284783 | 1.55644516 |
| 3805 | NM_002857.3 | PEX19 | 167.747549 | -8.6008154 | 1.51634683 |
| 3804 | NM_001112726.2 | #N/A | 171.576027 | -8.632746 | 1.52196514 |
| 3806 | NM_006349.2 | #N/A | 226.126889 | -9.0312402 | 1.59225884 |
| 3807 | NM_018046.4 | #N/A | 167.315458 | -8.5955759 | 1.51551373 |
| 3808 | NM_153350.3 | #N/A | 262.170174 | -9.244808 | 1.63002886 |
| 3810 | NM_033644.2 | FBXW11 | 185.689163 | -8.7447826 | 1.54219005 |
| 3809 | NM_002212.3 | #N/A | 213.553569 | -8.9488614 | 1.57817749 |
| 3811 | NM_006351.3 | #N/A | 198.954922 | -8.8467748 | 1.56030706 |
| 3812 | NM_005716.3 | #N/A | 170.258971 | -8.6223286 | 1.5207341 |
| 3813 | NM_013263.4 | #N/A | 145.900314 | -8.4009962 | 1.48175852 |
| 3814 | NM_007364.3 | #N/A | 230.941997 | -9.0617702 | 1.5983932 |
| 3815 | NM_015907.2 | LAP3 | 152.00054 | -8.4607151 | 1.49242879 |
| 3816 | NM_033656.3 | #N/A | 164.774295 | -8.57335 | 1.512317 |
| 3817 | NM_001040006.1 | #N/A | 249.386992 | -7.7781945 | 1.37208673 |
| 3818 | NM_194291.2 | TMEM65 | 175.895254 | -8.6691282 | 1.52934749 |
| 3819 | NM_020638.2 | #N/A | 218.973524 | 4.99281925 | 0.88081094 |
| 3820 | NM_006405.6 | #N/A | 155.834396 | -8.4964951 | 1.49900685 |
| 3821 | NM_001042616.2 | PIGY | 224.554763 | -9.0211321 | 1.59162661 |
| 3822 | NM_014497.4 | #N/A | 305.999222 | -9.4667654 | 1.67026696 |
| 3823 | NM_006304.1 | SEM1 | 176.778021 | -8.6758341 | 1.5308066 |
| 3824 | NM_016265.3 | #N/A | 146.00377 | -8.4015016 | 1.48252672 |
| 3825 | NM_033121.1 | #N/A | 145.644964 | -5.4866007 | 0.96821017 |
| 3826 | NM_003897.3 | #N/A | 165.826735 | -8.5864422 | 1.5152534 |

| | | | | | |
|---|---|---|---|---|---|
| 3827 | NM_020313.3 | #N/A | 207.917895 | -8.9103869 | 1.57253865 |
| 3828 | NM_014573.2 | #N/A | 295.847467 | -8.1489727 | 1.43832931 |
| 3829 | NM_004901.4 | #N/A | 217.315252 | -8.9744917 | 1.58408327 |
| 3830 | NM_001077484.1 | #N/A | 257.996642 | -9.2211136 | 1.6276287 |
| 3831 | NM_153225.3 | #N/A | 77.326193 | 2.69480136 | 0.47568132 |
| 3832 | NM_032486.3 | #N/A | 331.324591 | -9.5832669 | 1.69174194 |
| 3833 | NM_080817.4 | #N/A | 835.338451 | 4.91098263 | 0.86702817 |
| 3834 | NM_001145522.1 | #N/A | 313.595707 | -9.5039365 | 1.67794435 |
| 3835 | NM_017926.3 | #N/A | 178.36583 | -8.6867745 | 1.53375856 |
| 3836 | NM_004553.4 | #N/A | 226.332456 | -9.032701 | 1.59491503 |
| 3837 | NM_001136472.1 | LITAF | 168.129555 | -7.3288775 | 1.29410987 |
| 3838 | NM_001892.5 | #N/A | 288.648545 | -9.3834899 | 1.65693554 |
| 3839 | NM_002408.3 | #N/A | 160.808695 | -8.5407409 | 1.50823888 |
| 3840 | NM_001304504.1 | #N/A | 188.544797 | -8.7688681 | 1.54862521 |
| 3841 | NM_001031698.2 | PRPF40B | 176.867528 | -8.676495 | 1.53246426 |
| 3842 | NM_003412.3 | #N/A | 280.803819 | -9.3448441 | 1.65110223 |
| 3843 | NM_003873.5 | #N/A | 156.523496 | -8.5032857 | 1.50243861 |
| 3844 | NM_001408.2 | #N/A | 786.782663 | -5.4982341 | 0.9714995 |
| 3845 | NM_017946.3 | #N/A | 154.978532 | -8.4886751 | 1.49988993 |
| 3846 | NM_006099.3 | PIAS3 | 163.064056 | -8.5625144 | 1.51302255 |
| 3848 | NM_016479.4 | SHISA5 | 236.77055 | -9.099335 | 1.60786197 |
| 3847 | NM_015347.4 | #N/A | 168.996855 | -8.6120492 | 1.52174741 |
| 3849 | NM_022132.4 | #N/A | 144.815538 | -7.1092199 | 1.25622038 |
| 3850 | NM_001244008.1 | KIF1A | 230.526458 | -9.0598813 | 1.6009772 |
| 3851 | NM_001014840.1 | #N/A | 185.265812 | -8.7439959 | 1.54517741 |
| 3852 | NM_181716.2 | #N/A | 302.519767 | -9.4509817 | 1.6701693 |
| 3853 | NM_016200.4 | #N/A | 192.778946 | -7.5243405 | 1.3297337 |
| 3854 | NM_006712.4 | #N/A | 194.861311 | -8.8160044 | 1.55805081 |
| 3855 | NM_024996.5 | GFM1 | 187.132239 | -8.7571941 | 1.54780144 |
| 3856 | NM_022805.3 | #N/A | 263.766215 | -9.2536405 | 1.63574134 |
| 3857 | NM_001330701.1 | AGTPBP1 | 198.581167 | -8.8424165 | 1.56317878 |
| 3858 | NM_001617.3 | #N/A | 218.561826 | -8.9817729 | 1.58787635 |
| 3859 | NM_001098536.1 | #N/A | 335.831269 | -9.6022443 | 1.69761909 |
| 3860 | NM_020357.2 | #N/A | 242.482213 | -9.1319335 | 1.61448092 |
| 3861 | NM_020659.3 | #N/A | 276.698488 | -9.3219818 | 1.64820099 |
| 3862 | NR_002599.1 | #N/A | 273.037333 | -9.3030346 | 1.64492908 |
| 3863 | NM_024329.5 | #N/A | 176.770943 | -8.6772896 | 1.53439038 |
| 3864 | NM_004818.2 | #N/A | 412.102559 | -4.7162219 | 0.83400819 |
| 3865 | NM_018201.4 | #N/A | 195.989654 | -8.8258524 | 1.56075311 |
| 3866 | NM_001130040.1 | #N/A | 245.090839 | -9.1491536 | 1.61797783 |
| 3867 | NM_004317.3 | #N/A | 232.523662 | -9.0717734 | 1.60431068 |
| 3868 | NM_015634.3 | #N/A | 187.857767 | -8.7639396 | 1.54987444 |
| 3869 | NM_001167738.1 | #N/A | 350.241842 | -4.6596045 | 0.8240443 |

| | | | | | |
|---|---|---|---|---|---|
| 3870 | NM_015311.2 | #N/A | 217.0812 | -8.9734689 | 1.58695796 |
| 3871 | NM_015532.4 | #N/A | 185.453066 | -8.7453873 | 1.5466447 |
| 3872 | NM_001134870.1 | #N/A | 221.581042 | -9.002477 | 1.59234076 |
| 3873 | NM_001282547.1 | #N/A | 319.946668 | -6.9063266 | 1.22161 |
| 3874 | NM_016062.3 | #N/A | 237.804844 | -9.1041087 | 1.6103534 |
| 3875 | NM_012473.3 | #N/A | 210.379522 | -8.927332 | 1.57911876 |
| 3876 | NM_001001794.3 | #N/A | 149.385937 | -8.4339359 | 1.49192123 |
| 3877 | NM_007274.3 | ACOT7 | 316.859873 | -9.5181983 | 1.68378452 |
| 3878 | NM_006415.3 | #N/A | 181.991664 | -8.7186467 | 1.54235479 |
| 3879 | NM_006742.2 | #N/A | 158.885507 | -7.1929158 | 1.27251372 |
| 3880 | NM_014967.4 | #N/A | 165.991883 | -8.5858395 | 1.51908305 |
| 3881 | NM_015574.1 | #N/A | 110.182271 | -21.20221 | 3.75126359 |
| 3882 | NM_175907.5 | #N/A | 148.724254 | -8.4277433 | 1.4911246 |
| 3883 | NM_015665.5 | #N/A | 165.992209 | -8.5852954 | 1.51909184 |
| 3884 | NM_213595.3 | ISCU | 177.795716 | -8.6850575 | 1.53683496 |
| 3886 | NM_001289405.1 | MMS19 | 200.469088 | -8.8578726 | 1.56744088 |
| 3885 | NM_001201428.1 | #N/A | 194.864335 | -8.8174111 | 1.56028741 |
| 3887 | NM_017693.3 | #N/A | 176.316412 | -8.6725191 | 1.53487771 |
| 3888 | NM_014982.2 | #N/A | 226.323525 | -9.0324848 | 1.59860196 |
| 3890 | NM_145730.2 | AP1B1 | 154.746877 | -8.4831765 | 1.50144771 |
| 3889 | NM_001291978.1 | #N/A | 153.572701 | -7.1946993 | 1.2733888 |
| 3891 | NM_014734.3 | #N/A | 146.076353 | -8.4019563 | 1.4873907 |
| 3892 | NM_001303513.1 | #N/A | 255.985435 | -9.2098634 | 1.63045303 |
| 3893 | NM_003132.2 | #N/A | 312.95374 | -8.0688387 | 1.42856069 |
| 3894 | NM_002696.2 | #N/A | 229.759972 | -9.0541557 | 1.60309309 |
| 3895 | NM_032404.2 | TMPRSS3 | 34.5799346 | 3.59499648 | 0.6365489 |
| 3896 | NM_016497.3 | #N/A | 219.036625 | -8.9853516 | 1.59115412 |
| 3897 | NM_001001487.1 | ATP2C1 | 146.740079 | -8.4101496 | 1.48941528 |
| 3898 | NM_031229.3 | #N/A | 180.679952 | -8.7090577 | 1.54234082 |
| 3899 | NM_017801.2 | #N/A | 158.13617 | -8.5162089 | 1.50835594 |
| 3900 | NM_017830.3 | OCIAD1 | 172.371786 | -8.6398121 | 1.53034764 |
| 3901 | NM_006403.3 | #N/A | 174.018717 | -8.6531107 | 1.53273288 |
| 3902 | NM_016099.2 | GOLGA7 | 216.181549 | -8.9664065 | 1.58833032 |
| 3903 | NM_001170543.1 | #N/A | 168.672696 | -8.6074831 | 1.52479473 |
| 3904 | NM_138425.3 | #N/A | 355.613777 | -9.6843992 | 1.71572529 |
| 3905 | NM_007176.3 | #N/A | 213.976206 | -8.9521082 | 1.58604681 |
| 3906 | NM_021117.3 | #N/A | 172.076617 | -8.6381451 | 1.53056325 |
| 3907 | NM_016243.2 | #N/A | 155.19407 | -8.4897926 | 1.50430055 |
| 3908 | NM_015701.4 | #N/A | 174.062963 | -8.6551263 | 1.53361258 |
| 3909 | NM_001261406.1 | MPDZ | 179.713657 | -8.6977934 | 1.54121942 |
| 3910 | NM_003169.3 | SUPT5H | 157.114427 | -8.5077481 | 1.50753519 |
| 3911 | NM_017560.1 | #N/A | 204.265834 | -7.6097896 | 1.34842021 |
| 3912 | NM_016561.2 | #N/A | 175.77828 | -8.6685035 | 1.53616485 |

| | | | | | |
|---|---|---|---|---|---|
| 3913 | NM_021183.4 | #N/A | 207.308053 | -5.0963324 | 0.90316136 |
| 3914 | NM_003936.4 | #N/A | 288.384307 | -9.382439 | 1.66281108 |
| 3915 | NM_001291303.1 | FAT4 | 260.468254 | -9.2346931 | 1.63664006 |
| 3916 | NM_001136035.2 | #N/A | 183.888288 | -8.7330745 | 1.54778474 |
| 3917 | NM_175610.3 | #N/A | 138.761209 | -6.2468685 | 1.10721042 |
| 3918 | NR_102436.2 | #N/A | 83.1686391 | 2.02231971 | 0.35843867 |
| 3919 | NM_004522.2 | #N/A | 2629.04241 | -7.6745945 | 1.36028377 |
| 3920 | NM_001135575.1 | #N/A | 290.271913 | -9.3914938 | 1.66474881 |
| 3921 | NM_006368.4 | #N/A | 214.157325 | -8.9527737 | 1.58702817 |
| 3922 | NM_001288997.1 | LIN54 | 52.8021328 | 10.711873 | 1.89892311 |
| 3923 | NM_021991.2 | #N/A | 201.978829 | -8.8689746 | 1.57242013 |
| 3924 | NM_003670.2 | #N/A | 156.84387 | -8.5063963 | 1.5082282 |
| 3925 | NM_005134.3 | #N/A | 198.769055 | -8.84418 | 1.56815643 |
| 3926 | NM_002633.2 | #N/A | 144.603454 | -8.3849596 | 1.4868701 |
| 3927 | NM_005148.3 | #N/A | 146.375465 | -8.4038763 | 1.49019841 |
| 3928 | NM_015702.2 | #N/A | 220.868363 | -8.9975192 | 1.59550015 |
| 3929 | NM_172231.3 | #N/A | 187.125854 | -8.7577306 | 1.55297478 |
| 3930 | NM_015380.4 | #N/A | 201.245215 | -8.8632766 | 1.57186168 |
| 3931 | NM_003368.4 | #N/A | 166.319797 | -8.5866523 | 1.52292242 |
| 3932 | NR_111971.1 | #N/A | 210.650641 | -8.927918 | 1.58356232 |
| 3934 | NM_002735.2 | PRKAR1B | 258.463392 | -9.2241987 | 1.63619077 |
| 3935 | NM_017893.3 | SEMA4G | 166.873132 | -8.5931976 | 1.5242438 |
| 3933 | NM_001431.3 | #N/A | 190.741407 | -8.7873505 | 1.5586958 |
| 3936 | NM_031902.4 | #N/A | 170.524877 | -8.6239974 | 1.52982001 |
| 3937 | NM_001048200.2 | #N/A | 148.328546 | -8.4252224 | 1.49457908 |
| 3938 | NM_031886.2 | KCNA7 | 85.3551261 | 3.0287055 | 0.53729582 |
| 3939 | NM_001018050.3 | #N/A | 201.669003 | -8.8661615 | 1.57301364 |
| 3940 | NM_001301226.1 | #N/A | 213.307118 | -8.948583 | 1.58780416 |
| 3941 | NM_005834.4 | #N/A | 188.196549 | -8.7670553 | 1.55559243 |
| 3942 | NM_001346813.1 | ARID1B | 204.644285 | -8.8878776 | 1.57710505 |
| 3943 | NM_022831.2 | #N/A | 249.124766 | -7.7717845 | 1.37907132 |
| 3944 | NM_014363.5 | #N/A | 220.096249 | -8.9925779 | 1.5958053 |
| 3945 | NM_003796.3 | URI1 | 161.239544 | -8.5421042 | 1.51589402 |
| 3946 | NM_021627.2 | #N/A | 155.584844 | -8.4927942 | 1.50720527 |
| 3947 | NM_033070.2 | #N/A | 233.596647 | -9.0776577 | 1.61100871 |
| 3948 | NM_024755.3 | #N/A | 186.628504 | -8.7542285 | 1.55366214 |
| 3949 | NM_000723.4 | #N/A | 250.915952 | -9.1814233 | 1.62961757 |
| 3950 | NM_001162383.1 | #N/A | 272.367298 | -9.2997163 | 1.65061162 |
| 3951 | NM_001076683.1 | #N/A | 223.148873 | -7.6301882 | 1.35437396 |
| 3952 | NM_032195.2 | SON | 175.746176 | -8.6660124 | 1.53849609 |
| 3953 | NM_001171944.1 | #N/A | 143.147328 | -8.3708693 | 1.48612811 |
| 3954 | NM_006837.2 | #N/A | 215.201259 | -8.9599554 | 1.59071837 |
| 3955 | NM_152558.4 | #N/A | 200.938856 | -8.8594657 | 1.57291592 |

| | | | | | |
|---|---|---|---|---|---|
| 3956 | NM_001080543.1 | #N/A | 165.830995 | -8.5836651 | 1.5240114 |
| 3957 | NM_018268.3 | #N/A | 214.828281 | -8.9578771 | 1.59044895 |
| 3958 | NM_015565.2 | #N/A | 167.348909 | -8.596518 | 1.5265533 |
| 3959 | NM_005646.3 | #N/A | 153.180848 | -7.1899061 | 1.27694948 |
| 3960 | NM_005866.3 | #N/A | 198.710189 | -8.8458341 | 1.57110661 |
| 3961 | NM_004578.3 | #N/A | 238.149231 | -9.1062799 | 1.61749146 |
| 3962 | NM_000126.3 | #N/A | 169.756096 | -8.6179048 | 1.53081108 |
| 3963 | NR_036569.1 | STAG3L5P-PVRIG2P-PI | 181.967724 | -8.7183673 | 1.54870882 |
| 3964 | NM_017548.4 | #N/A | 176.331152 | -8.6716998 | 1.5405039 |
| 3965 | NM_007367.3 | #N/A | 180.356053 | -8.7056721 | 1.54666008 |
| 3966 | NM_194071.3 | #N/A | 173.058659 | -8.6481216 | 1.53659282 |
| 3967 | NM_032478.3 | #N/A | 184.961449 | -5.4636541 | 0.97085773 |
| 3968 | NM_004890.2 | #N/A | 191.511355 | -8.7923322 | 1.56241129 |
| 3969 | NM_003646.3 | #N/A | 209.760022 | -8.9228833 | 1.58570201 |
| 3970 | NM_032436.3 | #N/A | 160.082362 | -8.5346062 | 1.51677377 |
| 3971 | NM_032558.2 | #N/A | 155.472981 | -8.4920452 | 1.50937845 |
| 3972 | NM_015913.3 | #N/A | 189.798679 | -8.7785869 | 1.56039757 |
| 3973 | NM_001199867.1 | #N/A | 164.200704 | -8.5723459 | 1.5237686 |
| 3974 | NM_032621.3 | #N/A | 279.587481 | -9.3374164 | 1.65979805 |
| 3975 | NM_001134493.1 | TOMM6 | 224.004988 | -9.0176965 | 1.6030234 |
| 3976 | NM_025135.4 | #N/A | 222.773318 | -9.009059 | 1.60152522 |
| 3977 | NM_018191.3 | #N/A | 181.109171 | -8.7110274 | 1.54858648 |
| 3978 | NM_152726.2 | #N/A | 167.37697 | -8.5969455 | 1.52830414 |
| 3979 | NM_001801.2 | #N/A | 176.184664 | -8.6726605 | 1.54179987 |
| 3981 | NM_022745.4 | ATPAF1 | 159.108836 | -8.5232683 | 1.51532776 |
| 3980 | NM_004854.4 | #N/A | 233.937089 | -9.0799107 | 1.61428492 |
| 3982 | NM_138638.4 | #N/A | 200.869646 | -8.8607728 | 1.57543494 |
| 3983 | NM_001142640.1 | TNRC6C | 170.804516 | -8.6261911 | 1.53375667 |
| 3984 | NM_001320595.1 | #N/A | 174.701405 | -8.6589845 | 1.5396061 |
| 3985 | NM_003435.4 | #N/A | 153.291053 | -8.4705566 | 1.50619816 |
| 3986 | NM_019048.2 | #N/A | 189.495685 | -8.7762009 | 1.56062157 |
| 3987 | NM_001033117.2 | #N/A | 358.469356 | -9.6962892 | 1.7243076 |
| 3988 | NM_015859.3 | #N/A | 161.052502 | -8.5413188 | 1.51892492 |
| 3989 | NM_153816.5 | #N/A | 169.876798 | -8.618519 | 1.53273459 |
| 3990 | NM_005819.5 | #N/A | 188.727812 | -8.770225 | 1.55978025 |
| 3993 | NM_197968.3 | ZMYM2 | 201.204471 | -8.8613707 | 1.57616234 |
| 3991 | NM_016199.2 | #N/A | 242.650086 | -9.1329495 | 1.62444971 |
| 3992 | NM_139245.3 | #N/A | 263.699697 | -9.2527493 | 1.64577686 |
| 3994 | NM_020651.3 | #N/A | 199.419887 | -8.8498762 | 1.5742247 |
| 3995 | NM_001080485.3 | ZNF275 | 143.700322 | -8.3800546 | 1.49082877 |
| 3996 | NM_001256447.1 | #N/A | 177.391073 | -8.6821665 | 1.54471351 |
| 3997 | NM_031469.3 | #N/A | 144.808925 | -8.3906704 | 1.49294613 |
| 3998 | NM_001163547.2 | #N/A | 30.6535847 | 9.92393325 | 1.76577374 |

| | | | | | |
|---|---|---|---|---|---|
| 3999 | NM_004267.4 | #N/A | 188.673883 | -8.7703119 | 1.56061895 |
| 4000 | NM_006012.2 | #N/A | 214.707999 | -8.9563937 | 1.59372272 |
| 4001 | NM_174916.2 | #N/A | 160.181866 | -8.5343902 | 1.51863024 |
| 4002 | NM_015444.2 | #N/A | 152.57208 | -8.4618439 | 1.50574124 |
| 4003 | NM_001173464.1 | #N/A | 368.592469 | -9.7363955 | 1.73259073 |
| 4004 | NM_016491.3 | MRPL37 | 194.626885 | -8.8151432 | 1.56867285 |
| 4005 | NM_004852.2 | #N/A | 408.72984 | -8.4098901 | 1.49661728 |
| 4006 | NM_014665.3 | #N/A | 167.781148 | -8.5990796 | 1.53028385 |
| 4007 | NM_023012.5 | #N/A | 203.949336 | -8.8823593 | 1.58080895 |
| 4008 | NM_004443.3 | #N/A | 152.683784 | -8.4674344 | 1.50710088 |
| 4009 | NM_018686.5 | #N/A | 194.592998 | -8.8140648 | 1.56879141 |
| 4010 | NM_152892.2 | #N/A | 170.240478 | -8.6213913 | 1.53448728 |
| 4011 | NM_001282697.1 | #N/A | 190.198699 | -8.7812538 | 1.56313873 |
| 4012 | NM_020466.4 | #N/A | 156.601146 | -8.5006444 | 1.51323191 |
| 4013 | NM_133266.4 | #N/A | 165.052265 | -8.5750157 | 1.52649858 |
| 4014 | NM_012194.2 | #N/A | 244.179518 | -6.5793792 | 1.17128537 |
| 4015 | NM_004550.4 | NDUFS2 | 312.207541 | -6.8683901 | 1.22283752 |
| 4016 | NR_033711.1 | TP73-AS1 | 139.027581 | -8.3308761 | 1.48321991 |
| 4017 | NM_014691.2 | #N/A | 148.552973 | -8.424839 | 1.4999699 |
| 4018 | NM_018255.3 | #N/A | 167.591779 | -8.5966843 | 1.53067595 |
| 4019 | NM_002615.6 | #N/A | 390.461209 | -9.8201384 | 1.74853146 |
| 4020 | NM_004547.5 | #N/A | 230.356763 | -9.0579321 | 1.61283678 |
| 4021 | NM_020441.2 | #N/A | 162.873153 | -8.5596312 | 1.52412189 |
| 4022 | NM_052948.3 | #N/A | 228.326837 | -9.0450834 | 1.61056677 |
| 4023 | NM_003850.2 | SUCLA2 | 194.586189 | -8.81459 | 1.56959283 |
| 4024 | NM_016166.2 | PIAS1 | 188.36846 | -7.4071258 | 1.31899358 |
| 4025 | NM_001190706.1 | #N/A | 528.354777 | -10.25622 | 1.82637603 |
| 4026 | NM_031287.2 | #N/A | 221.383014 | -9.0010073 | 1.60308257 |
| 4027 | NM_032806.5 | #N/A | 158.983405 | -8.523597 | 1.51807351 |
| 4028 | NM_003430.3 | #N/A | 209.74894 | -8.9210076 | 1.58887273 |
| 4029 | NM_004318.3 | #N/A | 160.867663 | -8.542285 | 1.52143005 |
| 4030 | NM_021244.4 | #N/A | 171.560246 | -8.632524 | 1.53757349 |
| 4031 | NM_001690.3 | #N/A | 711.157331 | -2.9710672 | 0.52920923 |
| 4032 | NM_012421.3 | #N/A | 188.163412 | -8.7655486 | 1.56144818 |
| 4033 | NM_001018004.1 | #N/A | 191.165547 | -8.789218 | 1.56568062 |
| 4034 | NM_020689.3 | #N/A | 270.40377 | -9.2899834 | 1.65489365 |
| 4035 | NM_024297.2 | #N/A | 171.259242 | -8.6307331 | 1.53752658 |
| 4036 | NM_198530.3 | #N/A | 155.501716 | -8.4929629 | 1.51297592 |
| 4037 | NM_001003713.2 | #N/A | 214.259097 | -8.9537507 | 1.5951048 |
| 4038 | NM_001039877.1 | #N/A | 233.21147 | -9.0766673 | 1.61703781 |
| 4039 | NM_001278563.2 | #N/A | 213.689745 | -8.9511314 | 1.59476043 |
| 4040 | NM_194460.2 | #N/A | 211.205234 | -8.9323917 | 1.59145815 |
| 4041 | NM_015049.2 | #N/A | 164.066085 | -8.5685824 | 1.52669646 |

| | | | | | |
|---|---|---|---|---|---|
| 4042 | NM_001930.3 | #N/A | 225.035013 | -9.024066 | 1.60788925 |
| 4043 | NM_018708.2 | #N/A | 171.621066 | -8.6334261 | 1.5382751 |
| 4044 | NM_001017962.2 | P4HA1 | 158.948955 | -8.5220233 | 1.51847353 |
| 4045 | NM_147223.2 | #N/A | 175.858195 | -8.6661962 | 1.54431326 |
| 4046 | NM_004552.2 | #N/A | 239.506958 | -9.1144326 | 1.6242907 |
| 4047 | NM_032434.3 | #N/A | 186.583835 | -8.7537828 | 1.56001684 |
| 4048 | NM_006402.2 | LAMTOR5 | 212.654148 | -8.9425278 | 1.59370181 |
| 4049 | NM_173564.3 | #N/A | 343.069964 | -8.1818399 | 1.45815875 |
| 4050 | NM_001130026.2 | TCAF2 | 120.820546 | 2.55808899 | 0.45594001 |
| 4051 | NM_001042405.1 | CEP170 | 310.375494 | -9.4877823 | 1.69107072 |
| 4052 | NM_025240.2 | CD276 | 236.77505 | -9.0984554 | 1.62169933 |
| 4053 | NM_021204.4 | #N/A | 236.506193 | -9.0952286 | 1.62121085 |
| 4054 | NM_001330653.1 | #N/A | 229.335724 | -9.0521873 | 1.61364312 |
| 4055 | NM_002097.2 | #N/A | 280.023379 | -9.3398156 | 1.6651193 |
| 4056 | NM_021090.3 | #N/A | 167.040702 | -8.5971894 | 1.53272468 |
| 4057 | NM_005542.5 | #N/A | 307.446707 | -9.4746456 | 1.6892136 |
| 4058 | NM_001042537.1 | SLC9A6 | 300.166703 | -9.440271 | 1.6831222 |
| 4059 | NM_001098511.2 | KIF2A | 275.939535 | -9.3182917 | 1.66155127 |
| 4060 | NM_020442.5 | VARS2 | 155.145109 | -8.4904749 | 1.51398683 |
| 4061 | NM_173834.3 | #N/A | 166.036566 | -8.5850499 | 1.53089169 |
| 4062 | NR_134537.1 | C7orf50 | 226.449626 | -9.0332936 | 1.61085992 |
| 4063 | NM_004938.3 | #N/A | 168.896974 | -8.6109308 | 1.53569611 |
| 4064 | NM_145728.2 | #N/A | 181.075404 | -8.7114714 | 1.55365284 |
| 4065 | NM_005729.3 | #N/A | 146.46645 | -8.4064801 | 1.49928012 |
| 4066 | NM_001282790.1 | #N/A | 183.133921 | -8.7255718 | 1.55623165 |
| 4067 | NM_015267.3 | #N/A | 225.968795 | -9.0302737 | 1.6106202 |
| 4068 | NM_001300971.1 | #N/A | 241.00702 | -9.123258 | 1.62727933 |
| 4069 | NM_001346768.1 | #N/A | 234.015286 | -9.0807085 | 1.61991068 |
| 4070 | NM_003211.4 | #N/A | 227.294143 | -7.6487147 | 1.3645378 |
| 4071 | NM_024865.3 | #N/A | 177.190179 | -8.6817428 | 1.54885299 |
| 4072 | NM_020895.4 | #N/A | 355.087957 | -9.6824005 | 1.7275242 |
| 4073 | NM_003794.3 | #N/A | 184.11525 | -8.7343078 | 1.55849735 |
| 4074 | NM_021205.5 | #N/A | 139.693976 | -8.3389412 | 1.4879776 |
| 4075 | NM_002795.3 | #N/A | 212.813724 | -8.9438658 | 1.59593384 |
| 4076 | NM_001002800.2 | #N/A | 151.668679 | -8.4555457 | 1.50902666 |
| 4077 | NM_001080414.3 | #N/A | 149.087046 | -8.4292678 | 1.50434499 |
| 4078 | NM_015235.2 | #N/A | 174.718966 | -8.6590695 | 1.54537357 |
| 4079 | NM_002788.3 | #N/A | 187.65601 | -8.7618413 | 1.56381991 |
| 4081 | NR_110051.1 | LOC101927770 | 175.568898 | 3.79757689 | 0.67783879 |
| 4080 | NM_003336.3 | #N/A | 216.08054 | -8.9653829 | 1.60026416 |
| 4082 | NM_025083.3 | #N/A | 181.247028 | -8.7121197 | 1.55509763 |
| 4083 | NM_002904.5 | #N/A | 203.007358 | -8.8755971 | 1.58440286 |
| 4084 | NM_030798.4 | #N/A | 158.049267 | -8.5160681 | 1.52026195 |

| | | | | | |
|---|---|---|---|---|---|
| 4088 | NR_038967.1 | LOC100630923 | 152.775299 | -8.4651819 | 1.5112392 |
| 4085 | NM_007358.3 | #N/A | 180.471203 | -8.7050412 | 1.55405257 |
| 4086 | NM_022777.3 | #N/A | 145.009719 | -8.3903822 | 1.49788715 |
| 4087 | NM_144711.5 | #N/A | 215.563844 | -8.9625949 | 1.60004746 |
| 4089 | NM_000821.6 | #N/A | 144.476282 | -8.3826698 | 1.49662434 |
| 4090 | NM_005653.4 | #N/A | 183.097882 | -8.7275355 | 1.55840077 |
| 4091 | NM_153613.2 | #N/A | 210.230607 | -8.9268125 | 1.59417939 |
| 4092 | NM_023083.3 | #N/A | 184.729349 | -8.7391914 | 1.56085015 |
| 4093 | NM_001409.3 | #N/A | 206.149765 | -6.4256048 | 1.14768022 |
| 4094 | NM_001128827.2 | #N/A | 211.834516 | -8.9366995 | 1.59632726 |
| 4095 | NM_013388.5 | #N/A | 178.126886 | -8.6868189 | 1.55170746 |
| 4096 | NM_033257.3 | #N/A | 188.918232 | -8.7717751 | 1.56691641 |
| 4097 | NM_203434.2 | #N/A | 210.906104 | -8.9311257 | 1.59539259 |
| 4098 | NM_014678.4 | #N/A | 162.556168 | -8.5546362 | 1.52827418 |
| 4099 | NM_173354.4 | #N/A | 141.039719 | -8.348912 | 1.49155358 |
| 4100 | NM_018997.3 | MRPS21 | 180.42513 | -8.7062379 | 1.55541768 |
| 4101 | NM_005180.8 | #N/A | 213.197488 | -8.945797 | 1.59834782 |
| 4102 | NM_198402.4 | #N/A | 171.027665 | -8.6275863 | 1.5415871 |
| 4103 | NM_001135643.1 | DCTN4 | 293.360758 | -9.4065518 | 1.68089147 |
| 4104 | NM_003849.3 | #N/A | 204.882147 | -8.8890569 | 1.58847036 |
| 4105 | NM_003905.3 | #N/A | 192.756145 | -8.8002677 | 1.57269863 |
| 4106 | NM_001272060.1 | RPS6KB1 | 161.726632 | -8.5474793 | 1.52756376 |
| 4107 | NM_001265605.1 | MORF4L1 | 307.957683 | -9.4768047 | 1.69368564 |
| 4108 | NM_001174063.1 | #N/A | 154.650409 | -8.4809287 | 1.51577598 |
| 4109 | NM_001318729.1 | GPM6B | 247.620417 | -9.1618137 | 1.63754272 |
| 4110 | NM_015326.4 | SRGAP2 | 263.191973 | -9.2509989 | 1.65354224 |
| 4111 | NM_139067.3 | #N/A | 569.251293 | -5.43125 | 0.97080394 |
| 4112 | NM_001114183.1 | GRIA1 | 504.678952 | -10.189622 | 1.82139179 |
| 4113 | NM_021227.3 | #N/A | 217.475615 | -8.9754674 | 1.60436618 |
| 4114 | NM_001330177.1 | #N/A | 138.671075 | -8.3268975 | 1.48854031 |
| 4115 | NM_033630.2 | SCAND1 | 212.726457 | -8.9432589 | 1.59898014 |
| 4116 | NM_001915.3 | #N/A | 143.119329 | -8.3741047 | 1.49726346 |
| 4117 | NM_033116.5 | #N/A | 174.274099 | -8.6537974 | 1.54730433 |
| 4118 | NM_001308023.1 | TNS1 | 142.483087 | -8.3644425 | 1.49558276 |
| 4119 | NM_014947.4 | #N/A | 144.112703 | -8.3833812 | 1.49897684 |
| 4120 | NM_006284.3 | #N/A | 204.826544 | -8.8885122 | 1.58931241 |
| 4121 | NM_001193533.1 | #N/A | 28.9957807 | 9.83983904 | 1.75944882 |
| 4122 | NM_013346.3 | #N/A | 207.095444 | -8.9047858 | 1.59230637 |
| 4123 | NM_033129.3 | #N/A | 231.655755 | -9.0657884 | 1.62116473 |
| 4124 | NM_002912.4 | REV3L | 225.846553 | -7.637998 | 1.36586013 |
| 4125 | NM_001271776.1 | #N/A | 175.415792 | -8.6652258 | 1.54960494 |
| 4126 | NM_032041.2 | #N/A | 304.312536 | -9.460273 | 1.69188584 |
| 4127 | NM_021219.3 | #N/A | 162.473128 | -8.5539925 | 1.52982611 |

| | | | | | |
|---|---|---|---|---|---|
| 4128 | NM_001040446.2 | #N/A | 137.584453 | -8.3123642 | 1.48666883 |
| 4129 | NM_014814.2 | PSMD6 | 189.422567 | -8.7750617 | 1.56943493 |
| 4130 | NM_078485.3 | #N/A | 149.602277 | -8.4375871 | 1.50928911 |
| 4131 | NM_014047.2 | #N/A | 197.599047 | -8.8367793 | 1.58084545 |
| 4132 | NM_003920.4 | #N/A | 158.639203 | -8.5188673 | 1.52409548 |
| 4133 | NR_026550.2 | #N/A | 200.72042 | -8.8594145 | 1.5850181 |
| 4134 | NM_001346445.1 | #N/A | 415.316609 | -7.1763752 | 1.28411054 |
| 4135 | NM_030940.3 | #N/A | 194.210052 | -8.8111844 | 1.57668394 |
| 4136 | NM_001042545.1 | #N/A | 150.21492 | -8.4439633 | 1.51125564 |
| 4137 | NM_002847.4 | #N/A | 137.586872 | -8.3158351 | 1.48838888 |
| 4138 | NM_207113.2 | #N/A | 209.248595 | -8.9191958 | 1.5964038 |
| 4139 | NM_003340.6 | UBE2D3 | 151.482065 | -8.4525198 | 1.51288976 |
| 4141 | NM_001144061.1 | COPB1 | 201.581419 | -8.8660062 | 1.58705633 |
| 4140 | NM_001008397.3 | #N/A | 156.753516 | -7.2276261 | 1.29377101 |
| 4142 | NM_024927.4 | #N/A | 145.76064 | -8.3956401 | 1.50291332 |
| 4143 | NM_004781.3 | #N/A | 152.700691 | -8.466942 | 1.51569012 |
| 4144 | NM_203350.2 | #N/A | 161.495789 | -8.5462642 | 1.53004531 |
| 4145 | NM_000038.5 | #N/A | 390.124463 | -9.8182431 | 1.75781427 |
| 4146 | NM_004421.2 | #N/A | 174.784377 | -8.6612597 | 1.55068801 |
| 4147 | NM_001284239.1 | SARAF | 172.742921 | -8.6425879 | 1.54738242 |
| 4148 | NM_001277333.1 | #N/A | 95.6169947 | 3.28284582 | 0.58784843 |
| 4149 | NM_052906.4 | #N/A | 213.399501 | -8.9468099 | 1.60214658 |
| 4152 | NR_027873.1 | PAXBP1 | 141.992149 | -8.3612893 | 1.49748181 |
| 4150 | NM_015470.2 | #N/A | 147.907994 | -6.2228227 | 1.11448512 |
| 4151 | NM_198526.3 | #N/A | 165.244481 | -8.5788823 | 1.53644811 |
| 4153 | NM_014870.3 | #N/A | 150.53375 | -8.445011 | 1.51270985 |
| 4154 | NM_001198774.1 | #N/A | 187.234433 | -8.7592375 | 1.56901993 |
| 4155 | NM_032569.3 | #N/A | 185.007833 | -8.7439561 | 1.56639332 |
| 4156 | NM_001802.1 | #N/A | 139.755501 | -8.3394906 | 1.49410052 |
| 4157 | NM_005253.3 | #N/A | 185.255825 | -8.7450666 | 1.56677332 |
| 4158 | NM_173809.4 | #N/A | 186.162974 | -8.7511463 | 1.56794406 |
| 4159 | NM_005005.2 | #N/A | 290.054098 | -9.3906686 | 1.6826627 |
| 4160 | NM_005085.3 | #N/A | 163.66578 | -8.5637608 | 1.53448527 |
| 4161 | NM_024602.5 | #N/A | 186.94267 | -8.7569049 | 1.56921637 |
| 4162 | NM_002338.4 | #N/A | 302.177649 | -4.0353175 | 0.72316836 |
| 4163 | NM_001166007.1 | #N/A | 140.802313 | -8.3475345 | 1.49600868 |
| 4164 | NM_001242362.1 | MAGED4B | 148.430753 | -8.4239036 | 1.50979026 |
| 4165 | NM_014887.2 | #N/A | 144.86646 | -8.3887126 | 1.50384465 |
| 4166 | NM_138391.4 | #N/A | 175.314967 | -8.6641339 | 1.55327265 |
| 4167 | NM_007317.2 | #N/A | 171.865991 | -8.6349395 | 1.5480681 |
| 4168 | NM_007100.3 | #N/A | 188.84593 | -8.7717206 | 1.57266249 |
| 4169 | NM_207330.2 | #N/A | 57.1574381 | 2.94303874 | 0.52764841 |
| 4170 | NM_001135597.1 | #N/A | 283.442825 | -9.3566577 | 1.67756497 |

| | | | | | |
|---|---|---|---|---|---|
| 4171 | NM_015726.3 | #N/A | 171.692269 | -7.2812857 | 1.30564141 |
| 4172 | NM_002214.2 | #N/A | 221.107857 | -3.0607653 | 0.5488482 |
| 4173 | NM_001959.3 | #N/A | 205.331259 | -8.8926654 | 1.594765 |
| 4174 | NM_015189.2 | #N/A | 175.046829 | -8.6613741 | 1.55338949 |
| 4175 | NM_003764.3 | #N/A | 75.7300129 | 2.77950329 | 0.4984997 |
| 4176 | NM_004546.2 | #N/A | 239.330419 | -9.113356 | 1.63449809 |
| 4177 | NM_000297.3 | #N/A | 173.444528 | -8.6497372 | 1.5513704 |
| 4178 | NM_178832.3 | #N/A | 170.192653 | -8.621433 | 1.54646076 |
| 4179 | NM_001004439.1 | #N/A | 337.588412 | -9.6099576 | 1.72393412 |
| 4180 | NM_001008273.1 | TAGLN3 | 324.302997 | -8.1034487 | 1.45370525 |
| 4181 | NM_004882.3 | #N/A | 138.027148 | -8.31862 | 1.49229594 |
| 4182 | NM_005476.5 | #N/A | 168.267569 | -8.6065321 | 1.54403468 |
| 4183 | NM_006231.3 | #N/A | 234.577954 | -5.049331 | 0.90589752 |
| 4184 | NM_001319239.1 | ZC3H11A | 155.129467 | -8.488232 | 1.52292383 |
| 4185 | NM_018428.2 | #N/A | 157.31944 | -8.5071537 | 1.52637336 |
| 4186 | NM_138394.3 | #N/A | 161.958817 | -8.5489786 | 1.53388501 |
| 4187 | NM_001042678.1 | RHOC | 210.392742 | -8.9286363 | 1.60207322 |
| 4188 | NM_198236.2 | #N/A | 193.983456 | -8.8113961 | 1.58103902 |
| 4189 | NM_001077628.2 | #N/A | 169.999064 | -8.619828 | 1.54668805 |
| 4190 | NM_152903.4 | #N/A | 290.701689 | -3.3710009 | 0.60487521 |
| 4192 | NM_001110514.1 | EBF4 | 168.74056 | -8.6114904 | 1.54528514 |
| 4191 | NM_001001852.3 | #N/A | 197.206027 | -8.8338918 | 1.58519806 |
| 4193 | NM_016146.5 | #N/A | 192.0306 | -8.7956218 | 1.57835052 |
| 4194 | NM_031283.2 | #N/A | 135.47075 | -8.2946781 | 1.48848553 |
| 4195 | NM_005370.4 | #N/A | 159.71108 | -8.5300577 | 1.53074442 |
| 4196 | NM_014248.3 | #N/A | 230.213181 | -9.0571882 | 1.62535534 |
| 4197 | NM_006813.2 | #N/A | 178.653319 | -8.6908971 | 1.55969247 |
| 4198 | NM_012394.3 | #N/A | 261.998606 | -9.2435487 | 1.65891453 |
| 4199 | NM_201568.2 | #N/A | 155.949805 | -8.4948012 | 1.52463414 |
| 4200 | NM_001035005.3 | #N/A | 241.299461 | -9.1252545 | 1.63782644 |
| 4201 | NM_016079.3 | #N/A | 187.519131 | -8.7602479 | 1.5723133 |
| 4202 | NM_024333.2 | #N/A | 262.906265 | -9.2487507 | 1.66000751 |
| 4203 | NM_001985.2 | #N/A | 161.607511 | -8.546659 | 1.53411394 |
| 4204 | NM_007218.3 | RNF139 | 157.398917 | -8.5083291 | 1.52730599 |
| 4205 | NM_022140.4 | #N/A | 142.825433 | -7.0897834 | 1.27270913 |
| 4206 | NM_020470.2 | #N/A | 188.459712 | -8.7685741 | 1.57414205 |
| 4207 | NM_006824.2 | EBNA1BP2 | 204.643137 | -8.8872028 | 1.59558178 |
| 4208 | NM_015368.3 | #N/A | 191.971046 | -8.7949257 | 1.57902998 |
| 4209 | NM_006636.3 | #N/A | 211.956106 | -8.9379943 | 1.6049086 |
| 4210 | NM_031431.3 | #N/A | 162.684126 | -8.5560549 | 1.53633763 |
| 4211 | NM_014937.3 | #N/A | 201.200853 | -8.8628966 | 1.59149968 |
| 4212 | NM_031417.3 | #N/A | 248.792922 | -9.1688656 | 1.64646019 |
| 4213 | NM_020856.3 | #N/A | 232.318765 | -9.0704784 | 1.62884455 |

| | | | | | |
|---|---|---|---|---|---|
| 4214 | NM_032846.3 | #N/A | 180.904561 | -8.7112445 | 1.56435687 |
| 4215 | NM_182947.3 | #N/A | 256.515672 | -9.2129449 | 1.65453804 |
| 4216 | NM_002966.2 | #N/A | 241.955179 | -9.1294871 | 1.63962771 |
| 4217 | NM_002892.3 | #N/A | 193.616544 | -8.8061809 | 1.58159182 |
| 4218 | NM_020447.4 | #N/A | 151.81549 | -8.4579901 | 1.51910125 |
| 4219 | NM_015132.4 | #N/A | 144.086548 | -8.380552 | 1.50521767 |
| 4220 | NM_001195001.1 | #N/A | 156.447795 | -8.5022847 | 1.52711923 |
| 4221 | NM_021202.2 | #N/A | 167.831453 | -8.6034077 | 1.54535932 |
| 4222 | NM_022062.2 | #N/A | 140.462521 | -7.0676692 | 1.26966259 |
| 4223 | NM_001007246.2 | BRWD1 | 243.197128 | -9.1361246 | 1.64137283 |
| 4224 | NM_000725.3 | #N/A | 215.398097 | -8.9612896 | 1.60997994 |
| 4225 | NM_021633.3 | #N/A | 175.492815 | -8.6656167 | 1.55688752 |
| 4226 | NM_003093.2 | #N/A | 212.351886 | -8.9408347 | 1.60636041 |
| 4227 | NM_017432.4 | #N/A | 144.167427 | -8.3792253 | 1.50550234 |
| 4228 | NM_001834.3 | #N/A | 166.321307 | -8.5896796 | 1.54334642 |
| 4229 | NM_004171.3 | #N/A | 200.073318 | -8.8535212 | 1.59079533 |
| 4230 | NM_001567.3 | #N/A | 505.01787 | -4.5671729 | 0.82076272 |
| 4231 | NM_017859.3 | #N/A | 164.033539 | -8.5678464 | 1.53982822 |
| 4232 | NM_182776.2 | MCM7 | 161.551568 | -8.5458443 | 1.53590539 |
| 4233 | NM_030624.2 | #N/A | 122.268974 | -6.0522136 | 1.08777482 |
| 4234 | NM_000709.3 | #N/A | 163.007113 | -8.5579072 | 1.53819284 |
| 4235 | NM_032999.3 | #N/A | 293.419431 | -9.4076816 | 1.69107515 |
| 4236 | NM_014155.4 | ZBTB44 | 141.87236 | -8.3573863 | 1.50231422 |
| 4237 | NM_001085411.2 | #N/A | 171.545679 | -8.6320645 | 1.55174398 |
| 4238 | NM_013365.4 | #N/A | 161.671798 | -8.5464228 | 1.53636395 |
| 4239 | NM_002926.3 | RGS12 | 193.779595 | -8.8081877 | 1.58343794 |
| 4240 | NM_199168.3 | #N/A | 259.202053 | -9.2289111 | 1.65929824 |
| 4241 | NM_005697.4 | #N/A | 146.189465 | -8.4040046 | 1.51100729 |
| 4242 | NM_015411.2 | #N/A | 203.231187 | -8.8777547 | 1.59638592 |
| 4243 | NM_012329.2 | #N/A | 260.6369 | -9.2361308 | 1.66085134 |
| 4244 | NM_004147.3 | #N/A | 190.958757 | -8.7875461 | 1.58020411 |
| 4245 | NM_002622.4 | #N/A | 231.255312 | -9.0639013 | 1.62992727 |
| 4246 | NM_012088.2 | #N/A | 187.216199 | -8.7589809 | 1.57517546 |
| 4247 | NM_145294.4 | #N/A | 153.456918 | -8.4737779 | 1.52388483 |
| 4248 | NM_001123377.1 | PARK7 | 229.188243 | -9.0513182 | 1.62785318 |
| 4249 | NM_001853.3 | #N/A | 209.364313 | -8.9213245 | 1.60448161 |
| 4250 | NM_022101.3 | #N/A | 141.771176 | -8.3580569 | 1.50318707 |
| 4251 | NM_001300801.1 | PITPNM2 | 186.290109 | -8.7518717 | 1.57413799 |
| 4252 | NM_001329140.1 | PTPRF | 209.364408 | -8.9190733 | 1.60421404 |
| 4253 | NM_018051.4 | #N/A | 176.902466 | -7.3186828 | 1.31646985 |
| 4254 | NM_004596.4 | #N/A | 194.916391 | -7.542606 | 1.35679495 |
| 4255 | NM_001042663.1 | #N/A | 159.474877 | -8.527002 | 1.53404375 |
| 4256 | NM_001992.4 | #N/A | 403.548489 | -7.135112 | 1.28371142 |

| | | | | | |
|---|---|---|---|---|---|
| 4257 | NM_032775.3 | #N/A | 182.659496 | -8.7234156 | 1.56951166 |
| 4258 | NM_003893.4 | #N/A | 143.362783 | -8.3747771 | 1.50696326 |
| 4259 | NM_001300821.2 | #N/A | 165.720399 | -8.5834163 | 1.5445401 |
| 4260 | NM_015190.4 | #N/A | 192.536267 | -8.7992143 | 1.58336057 |
| 4261 | NM_001048218.1 | #N/A | 162.576503 | -8.556958 | 1.53986519 |
| 4262 | NM_015516.3 | #N/A | 135.319012 | -8.2922992 | 1.49232781 |
| 4264 | NM_004239.4 | TRIP11 | 141.936534 | -8.3604513 | 1.50484071 |
| 4263 | NM_001330597.1 | #N/A | 249.206585 | -9.1718669 | 1.65090089 |
| 4265 | NM_000255.3 | #N/A | 158.175142 | -8.5156077 | 1.53279041 |
| 4266 | NM_005594.5 | #N/A | 190.380164 | -8.7829539 | 1.58100155 |
| 4267 | NM_005052.2 | #N/A | 225.828213 | -9.0293121 | 1.62538418 |
| 4268 | NM_006390.3 | #N/A | 142.698915 | -8.3681851 | 1.50637843 |
| 4269 | NM_001318975.1 | BCKDHB | 45.3595738 | 10.4935829 | 1.88927105 |
| 4270 | NR_119374.1 | LOC101927560 | 52.8275678 | 5.08401184 | 0.91532413 |
| 4271 | NM_000642.2 | #N/A | 127.975156 | -8.2105152 | 1.47827052 |
| 4272 | NM_019593.3 | #N/A | 206.735002 | -4.0511533 | 0.72943481 |
| 4273 | NM_031461.5 | #N/A | 182.664799 | -8.7241637 | 1.5709519 |
| 4274 | NM_032813.3 | #N/A | 159.72427 | -8.5287093 | 1.53584315 |
| 4275 | NM_001128172.2 | PAK3 | 38.0733218 | 10.2177803 | 1.84031554 |
| 4276 | NM_001291459.1 | #N/A | 159.521518 | -8.5281876 | 1.53605956 |
| 4277 | NM_001363.4 | #N/A | 149.52567 | -8.4332151 | 1.51910456 |
| 4278 | NM_001014839.1 | NCDN | 130.685165 | -8.2384948 | 1.48423557 |
| 4279 | NM_004649.7 | #N/A | 187.449887 | -8.7599689 | 1.57829582 |
| 4280 | NM_004214.4 | #N/A | 236.637436 | -9.0969351 | 1.63903694 |
| 4281 | NM_024527.4 | #N/A | 168.32012 | -8.6051249 | 1.55058155 |
| 4282 | NM_175748.3 | #N/A | 157.137842 | -8.5056876 | 1.53267507 |
| 4285 | NM_213619.2 | ATP6V1H | 211.64749 | -8.935982 | 1.61025283 |
| 4283 | NM_015325.2 | #N/A | 229.434268 | -6.5191017 | 1.17473873 |
| 4284 | NM_017546.4 | #N/A | 135.039419 | -8.2873346 | 1.49335435 |
| 4286 | NM_019606.5 | #N/A | 137.928253 | -8.3181082 | 1.49894648 |
| 4288 | NR_135515.1 | CYYR1-AS1 | 35.3934944 | 3.30582045 | 0.59576562 |
| 4287 | NM_004894.2 | #N/A | 220.936757 | -8.9977891 | 1.62156448 |
| 4289 | NM_004208.3 | #N/A | 128.683997 | -8.2200014 | 1.48147231 |
| 4290 | NM_173798.3 | #N/A | 207.84332 | -8.9101039 | 1.60600647 |
| 4291 | NM_032299.3 | #N/A | 198.506293 | -8.8433474 | 1.59406899 |
| 4292 | NM_152833.2 | #N/A | 159.529097 | -8.5279137 | 1.53729006 |
| 4293 | NM_001242818.1 | #N/A | 140.330715 | -8.341307 | 1.50373026 |
| 4294 | NM_021215.3 | #N/A | 134.478923 | -8.2822144 | 1.49308039 |
| 4295 | NM_006362.4 | #N/A | 162.631267 | -8.5556629 | 1.54267541 |
| 4296 | NM_007293.2 | #N/A | 302.277464 | -9.4508379 | 1.70408488 |
| 4297 | NM_006999.4 | #N/A | 146.149264 | -8.4005143 | 1.5147282 |
| 4298 | NM_003092.4 | #N/A | 193.693522 | -8.8076261 | 1.58823272 |
| 4299 | NM_017864.3 | #N/A | 144.176299 | -8.3807962 | 1.51125939 |

| | | | | | |
|---|---|---|---|---|---|
| 4300 | NM_016118.4 | NUB1 | 141.876116 | -7.0804107 | 1.2767825 |
| 4301 | NM_198836.2 | ACACA | 372.279873 | -9.750859 | 1.75836607 |
| 4302 | NM_001033519.1 | #N/A | 154.548285 | -8.4816475 | 1.5296056 |
| 4303 | NM_002072.4 | #N/A | 586.096622 | -4.0436765 | 0.72935937 |
| 4304 | NM_021976.4 | #N/A | 165.624558 | -8.5839544 | 1.54831566 |
| 4305 | NM_001042590.2 | #N/A | 160.675246 | -8.5385567 | 1.54018712 |
| 4306 | NM_033082.3 | #N/A | 195.091026 | -8.8178847 | 1.59056521 |
| 4307 | NM_001164273.1 | MGA | 144.886269 | -8.3915858 | 1.51375086 |
| 4308 | NM_006465.3 | #N/A | 154.998867 | -8.4874002 | 1.53104553 |
| 4309 | NR_003098.1 | #N/A | 178.88043 | -8.69285 | 1.56812858 |
| 4310 | NM_001610.3 | #N/A | 175.610446 | -8.6670464 | 1.56349795 |
| 4311 | NM_030632.2 | #N/A | 145.662868 | -8.395041 | 1.51446057 |
| 4312 | NM_213674.1 | TPM2 | 256.831587 | -5.985963 | 1.07987506 |
| 4314 | NM_007275.2 | TUSC2 | 174.99243 | -8.6619709 | 1.56283774 |
| 4313 | NM_004529.3 | #N/A | 212.177148 | -8.9396868 | 1.61294357 |
| 4315 | NR_046331.1 | #N/A | 200.348322 | -8.8564162 | 1.59793236 |
| 4316 | NM_001913.4 | #N/A | 135.655238 | -8.2968237 | 1.4971051 |
| 4317 | NM_001166347.1 | #N/A | 134.547366 | -8.2836035 | 1.49481327 |
| 4318 | NM_003130.3 | SRI | 177.65642 | -8.6830862 | 1.56695197 |
| 4319 | NM_005763.3 | #N/A | 156.233921 | -7.2197377 | 1.30288149 |
| 4320 | NM_144732.4 | #N/A | 159.856341 | -7.1854496 | 1.2966763 |
| 4321 | NM_025073.2 | #N/A | 165.327383 | -8.5803673 | 1.54844783 |
| 4322 | NM_004389.3 | CTNNA2 | 180.598398 | -8.7082534 | 1.57157056 |
| 4323 | NM_032635.3 | #N/A | 189.308704 | -8.7754939 | 1.58394538 |
| 4324 | NM_145166.3 | #N/A | 143.785019 | -8.3782324 | 1.51223541 |
| 4325 | NM_016155.5 | #N/A | 216.04837 | -8.9655698 | 1.61828482 |
| 4326 | NM_004708.3 | #N/A | 193.815898 | -8.8086337 | 1.5899874 |
| 4327 | NM_012328.2 | #N/A | 146.824428 | -8.4078994 | 1.51771327 |
| 4328 | NM_022106.2 | #N/A | 135.114237 | -8.2858639 | 1.49570867 |
| 4329 | NM_002229.2 | #N/A | 164.462826 | -8.5737357 | 1.54769046 |
| 4330 | NM_000436.3 | #N/A | 219.743816 | -8.9899042 | 1.62298049 |
| 4331 | NM_006848.2 | #N/A | 205.409789 | -8.8921586 | 1.60535586 |
| 4332 | NM_004489.4 | #N/A | 183.477511 | -8.7299043 | 1.57613212 |
| 4333 | NM_001995.3 | #N/A | 148.392598 | -8.4237795 | 1.52094442 |
| 4334 | NR_024345.1 | HNF1A-AS1 | 146.202056 | 3.41839918 | 0.6172199 |
| 4335 | NM_014795.3 | #N/A | 310.138247 | -9.4866752 | 1.71292594 |
| 4336 | NM_001343.3 | #N/A | 173.339737 | -8.6502414 | 1.56194969 |
| 4337 | NM_001320845.1 | SAT2 | 148.347859 | -8.4221071 | 1.52092056 |
| 4338 | NM_012086.4 | #N/A | 147.398372 | -8.4134939 | 1.51936029 |
| 4339 | NM_001035254.2 | #N/A | 146.278359 | -7.0689575 | 1.27659862 |
| 4340 | NM_017489.2 | #N/A | 149.280796 | -8.4323016 | 1.52286335 |
| 4341 | NM_020857.2 | #N/A | 194.015996 | -7.4345874 | 1.34269874 |
| 4342 | NM_020750.2 | #N/A | 316.531448 | -6.8587687 | 1.23880086 |

| | | | | | |
|---|---|---|---|---|---|
| 4343 | NM_003587.4 | #N/A | 155.806853 | -8.4959496 | 1.53452163 |
| 4344 | NM_032975.3 | DTNA | 130.313231 | -8.2369868 | 1.48794658 |
| 4345 | NM_019594.3 | LRRC8A | 211.154454 | -8.9335229 | 1.61382532 |
| 4346 | NM_014618.2 | #N/A | 244.903576 | -9.1467836 | 1.65242655 |
| 4347 | NM_020145.3 | SH3GLB2 | 150.749018 | -8.449064 | 1.52642009 |
| 4348 | NM_001278196.1 | ZBTB18 | 386.477255 | -9.804396 | 1.77130171 |
| 4349 | NM_023936.1 | #N/A | 215.95624 | -8.9649069 | 1.61971985 |
| 4350 | NM_001276380.1 | ESF1 | 140.855215 | -8.3469041 | 1.50810162 |
| 4351 | NM_001039697.1 | SNAPC3 | 226.732074 | -3.0416148 | 0.54957137 |
| 4352 | NM_003188.3 | #N/A | 217.820792 | -8.9772303 | 1.62208278 |
| 4353 | NM_001317812.1 | #N/A | 133.89626 | -8.2745724 | 1.49517312 |
| 4354 | NM_001654.4 | #N/A | 132.699513 | -8.264579 | 1.49339504 |
| 4355 | NM_001199172.1 | #N/A | 225.907187 | -9.0299342 | 1.63172386 |
| 4356 | NM_001256405.1 | #N/A | 149.763871 | -8.4391293 | 1.52495668 |
| 4357 | NM_207044.1 | ENSA | 229.555001 | -9.0538605 | 1.63626252 |
| 4358 | NM_001444.2 | #N/A | 263.198131 | -9.2503891 | 1.67184886 |
| 4359 | NM_000754.3 | #N/A | 184.84043 | -8.7410769 | 1.57983498 |
| 4360 | NM_001128913.1 | PCBP2 | 212.334303 | -8.941078 | 1.61619842 |
| 4361 | NM_004089.3 | #N/A | 145.381942 | -8.3960116 | 1.51773009 |
| 4362 | NM_002881.2 | #N/A | 154.677829 | -8.483412 | 1.53359598 |
| 4363 | NM_020933.4 | #N/A | 142.691626 | -8.3649919 | 1.51220252 |
| 4364 | NM_021160.2 | #N/A | 202.459067 | -8.8713769 | 1.60373121 |
| 4365 | NM_024067.3 | #N/A | 173.622969 | -8.6500284 | 1.56369603 |
| 4366 | NM_181784.2 | #N/A | 157.379816 | -8.5094452 | 1.5383156 |
| 4367 | NM_014899.3 | #N/A | 348.06447 | -3.666666 | 0.66288466 |
| 4368 | NM_175866.4 | #N/A | 153.655285 | -8.4766449 | 1.53259607 |
| 4369 | NM_015156.3 | #N/A | 150.595155 | -8.4443027 | 1.52680728 |
| 4370 | NM_181575.4 | #N/A | 202.738814 | -8.8746244 | 1.60471952 |
| 4371 | NM_053045.1 | #N/A | 164.168699 | -8.5691294 | 1.54965756 |
| 4372 | NM_004508.3 | IDI1 | 251.690585 | -9.1860389 | 1.66132971 |
| 4373 | NM_024721.4 | #N/A | 140.525855 | -8.3457788 | 1.50955722 |
| 4374 | NM_001244950.1 | #N/A | 204.296445 | -8.8866773 | 1.60746421 |
| 4375 | NM_016029.3 | #N/A | 171.194415 | -8.6304135 | 1.5611097 |
| 4376 | NM_017586.3 | #N/A | 205.015461 | -8.8896333 | 1.60799318 |
| 4377 | NM_018971.2 | #N/A | 214.146188 | -8.9522168 | 1.61945802 |
| 4378 | NM_020240.2 | CDC42SE2 | 135.328774 | -8.2879197 | 1.49932193 |
| 4379 | NM_005920.3 | #N/A | 174.92393 | -8.6600773 | 1.56668228 |
| 4380 | NM_017528.4 | #N/A | 155.886594 | -8.4959893 | 1.53703173 |
| 4381 | NM_018085.4 | #N/A | 783.950354 | -2.5051863 | 0.45322598 |
| 4382 | NM_001256126.1 | #N/A | 395.663244 | -9.8384695 | 1.77997957 |
| 4383 | NM_020319.2 | #N/A | 150.972994 | -8.4483642 | 1.52866596 |
| 4384 | NM_001136108.2 | #N/A | 197.912092 | -8.8385967 | 1.59931723 |
| 4385 | NM_014302.3 | #N/A | 199.791949 | -8.8527548 | 1.60198912 |

| | | | | | |
|---|---|---|---|---|---|
| 4386 | NM_001569.3 | #N/A | 183.71314 | -8.7314322 | 1.58015978 |
| 4387 | NM_006527.3 | #N/A | 180.011906 | -8.7018565 | 1.57483198 |
| 4388 | NM_052903.4 | #N/A | 96.9980911 | -4.7539672 | 0.86051299 |
| 4389 | NM_199002.1 | #N/A | 140.419309 | -8.3443188 | 1.51051133 |
| 4390 | NM_006899.4 | #N/A | 153.475176 | -8.4730934 | 1.53389159 |
| 4391 | NM_002818.2 | #N/A | 155.354147 | -8.4903526 | 1.53703445 |
| 4392 | NM_001135690.2 | #N/A | 131.572397 | -8.2518157 | 1.4938863 |
| 4393 | NM_000848.3 | #N/A | 157.411474 | -8.5101311 | 1.54081393 |
| 4394 | NM_001244926.1 | PRPF4 | 149.803103 | -8.4370986 | 1.52761925 |
| 4395 | NM_001040151.1 | SCN3B | 464.78253 | -4.5989023 | 0.83269985 |
| 4396 | NM_003165.3 | #N/A | 264.670533 | -9.2585035 | 1.67659389 |
| 4397 | NM_015559.2 | #N/A | 261.190202 | -7.815369 | 1.41530679 |
| 4398 | NM_020827.2 | #N/A | 158.136266 | -8.5146354 | 1.54197482 |
| 4399 | NM_002146.4 | HOXB3 | 216.583708 | -8.9700634 | 1.6245358 |
| 4400 | NM_014773.4 | #N/A | 151.166475 | -8.4516992 | 1.53067543 |
| 4401 | NM_001291412.1 | #N/A | 159.779675 | -8.5297828 | 1.54492764 |
| 4402 | NM_005983.3 | #N/A | 129.258515 | -8.2253605 | 1.48988535 |
| 4403 | NM_001178139.1 | #N/A | 137.298441 | -8.3088283 | 1.50502326 |
| 4404 | NM_001319113.1 | RBM42 | 185.280363 | -8.7444171 | 1.58402022 |
| 4405 | NM_020372.3 | SLC22A17 | 130.171487 | -8.2359808 | 1.49190204 |
| 4406 | NR_145450.1 | TNK2-AS1 | 92.7455864 | 2.56070297 | 0.4638635 |
| 4407 | NM_004644.4 | #N/A | 199.05393 | -8.8475004 | 1.60286656 |
| 4408 | NM_005927.4 | #N/A | 170.318012 | -8.622472 | 1.5621756 |
| 4409 | NM_000718.3 | #N/A | 204.99307 | -8.8903045 | 1.61105321 |
| 4410 | NM_016355.3 | #N/A | 170.490128 | -8.6232429 | 1.56266444 |
| 4411 | NM_002419.3 | #N/A | 132.480278 | -8.2614403 | 1.49718815 |
| 4412 | NM_020382.4 | #N/A | 151.692525 | -8.4557891 | 1.53247858 |
| 4413 | NM_022457.6 | #N/A | 148.262827 | -8.4219828 | 1.5265285 |
| 4414 | NM_016224.4 | #N/A | 139.291877 | -8.3337164 | 1.51059284 |
| 4415 | NM_002766.2 | #N/A | 147.340755 | -8.4123053 | 1.52491348 |
| 4416 | NM_006246.3 | #N/A | 191.242741 | -8.7883925 | 1.59337113 |
| 4417 | NM_000755.4 | #N/A | 150.330933 | -8.4442053 | 1.53107206 |
| 4418 | NM_198317.2 | #N/A | 167.1527 | -8.5957307 | 1.55853615 |
| 4419 | NM_014110.4 | #N/A | 185.462747 | -8.7452236 | 1.58567991 |
| 4420 | NM_014956.4 | #N/A | 136.328919 | -8.3012719 | 1.50531759 |
| 4421 | NM_020940.3 | #N/A | 154.560043 | -8.4822818 | 1.53825058 |
| 4422 | NM_194356.2 | #N/A | 135.007878 | -8.2875692 | 1.50293763 |
| 4425 | NM_016607.3 | ARMCX3 | 197.861957 | -8.8376585 | 1.60275113 |
| 4423 | NM_001257399.1 | #N/A | 145.356964 | -8.3949535 | 1.52243307 |
| 4424 | NM_016151.3 | #N/A | 140.749064 | -8.3479899 | 1.51393196 |
| 4426 | NM_018159.3 | #N/A | 214.61585 | -8.9557468 | 1.62416797 |
| 4427 | NM_001078172.1 | RTL8A | 196.925816 | -8.8315411 | 1.60169788 |
| 4428 | NM_004414.6 | #N/A | 263.453556 | -9.2514758 | 1.67796674 |

| | | | | | | |
|---|---|---|---|---|---|---|
| 4429 | NM_001194955.1 | MATR3 | | 165.487888 | -8.5799471 | 1.55621254 |
| 4430 | NM_001284.3 | | #N/A | 169.051055 | -8.6111347 | 1.56185976 |
| 4431 | NM_007198.3 | | #N/A | 135.738651 | -8.2951847 | 1.50460716 |
| 4432 | NM_024548.3 | | #N/A | 143.206924 | -8.3700142 | 1.51822978 |
| 4433 | NM_174908.3 | | #N/A | 375.081958 | -7.0396484 | 1.2769298 |
| 4434 | NM_016145.3 | | #N/A | 208.801582 | -8.9164676 | 1.61742026 |
| 4435 | NM_012453.2 | | #N/A | 147.010328 | -8.4103999 | 1.52565344 |
| 4436 | NM_080927.3 | | #N/A | 288.112517 | -6.7489542 | 1.2242981 |
| 4437 | NM_016100.4 | | #N/A | 135.327158 | -8.2923624 | 1.50443822 |
| 4438 | NM_005094.3 | | #N/A | 217.980561 | -8.9787631 | 1.6289966 |
| 4439 | NM_018216.2 | | #N/A | 155.034282 | -8.4866617 | 1.53975999 |
| 4440 | NM_015076.4 | | #N/A | 167.227445 | -8.5949683 | 1.55963819 |
| 4441 | NM_032611.2 | | #N/A | 126.181392 | -8.1900808 | 1.48623177 |
| 4442 | NM_020154.2 | | #N/A | 185.351003 | -8.7445766 | 1.58697331 |
| 4443 | NM_004223.4 | | #N/A | 177.23236 | -8.6803275 | 1.57539132 |
| 4444 | NM_007373.3 | SHOC2 | | 189.982904 | -8.7801816 | 1.59353192 |
| 4445 | NM_080546.4 | | #N/A | 124.89768 | -8.176053 | 1.48389825 |
| 4446 | NM_015630.3 | | #N/A | 173.790814 | -8.6512453 | 1.57028515 |
| 4447 | NM_181443.3 | BTBD3 | | 128.618128 | -8.2152568 | 1.49116922 |
| 4448 | NM_001983.3 | | #N/A | 131.950891 | -8.2532287 | 1.4981637 |
| 4449 | NM_025015.2 | | #N/A | 299.501476 | -9.4372085 | 1.71307757 |
| 4450 | NM_007121.5 | | #N/A | 132.764734 | -8.264442 | 1.50023706 |
| 4451 | NM_005453.4 | | #N/A | 157.98947 | -8.5133827 | 1.54549375 |
| 4452 | NM_006031.5 | | #N/A | 160.329122 | -8.5349313 | 1.54956195 |
| 4453 | NM_024843.3 | | #N/A | 155.98199 | -8.4976211 | 1.54278326 |
| 4454 | NM_003377.4 | | #N/A | 148.341898 | -8.4238748 | 1.52945226 |
| 4455 | NM_001079520.1 | | #N/A | 309.561406 | -9.4840488 | 1.72196858 |
| 4456 | NM_016129.2 | | #N/A | 172.768807 | -8.6430448 | 1.56931062 |
| 4457 | NM_001131024.1 | PEX5 | | 172.353167 | -8.6387811 | 1.56859229 |
| 4458 | NM_003626.3 | | #N/A | 155.093574 | -8.4887229 | 1.54135324 |
| 4459 | NM_004027.2 | | #N/A | 228.79937 | -9.0480264 | 1.64298066 |
| 4460 | NM_014278.3 | | #N/A | 126.834495 | -8.1959835 | 1.48827557 |
| 4461 | NM_174983.4 | | #N/A | 168.08463 | -8.6033588 | 1.56224506 |
| 4462 | NM_153042.3 | | #N/A | 129.903319 | -8.2297891 | 1.49447509 |
| 4463 | NM_003123.4 | | #N/A | 267.520119 | 2.67552819 | 0.48587649 |
| 4464 | NM_138392.3 | | #N/A | 142.428261 | -6.2906913 | 1.1423926 |
| 4465 | NM_012157.3 | | #N/A | 145.005093 | -8.3913837 | 1.52394166 |
| 4466 | NM_152609.2 | | #N/A | 152.748751 | -8.4659117 | 1.5374791 |
| 4467 | NM_001001486.1 | ATP2C1 | | 126.541227 | -8.1941916 | 1.48817139 |
| 4468 | NM_001166286.1 | | #N/A | 184.187163 | -8.7357627 | 1.58662723 |
| 4469 | NM_022787.3 | | #N/A | 319.27881 | 2.91009109 | 0.52853992 |
| 4470 | NM_173611.3 | | #N/A | 181.589674 | -8.7138383 | 1.58271215 |
| 4471 | NM_006196.3 | | #N/A | 1129.17883 | -4.8089612 | 0.87347815 |

| | | | | | | |
|---|---|---|---|---|---|---|
| 4472 | NM_005768.5 | | #N/A | 135.891438 | -4.3382209 | 0.78798198 |
| 4473 | NM_001290474.1 | | #N/A | 162.691183 | -8.5556026 | 1.55410022 |
| 4476 | NM_198097.3 | CCZ1B | | 163.877204 | -8.5654161 | 1.55599536 |
| 4474 | NM_014916.3 | | #N/A | 218.742903 | -3.8841466 | 0.70559513 |
| 4475 | NM_017922.3 | | #N/A | 146.654102 | -8.4059069 | 1.5270226 |
| 4477 | NM_001105568.2 | | #N/A | 126.066196 | -8.1910381 | 1.48801195 |
| 4478 | NM_175075.4 | TDRP | | 129.932273 | -8.2337141 | 1.4957767 |
| 4479 | NM_007320.2 | | #N/A | 145.430792 | -8.3933604 | 1.52480205 |
| 4480 | NM_004834.4 | | #N/A | 151.328212 | -8.4537981 | 1.53584919 |
| 4481 | NM_024678.5 | | #N/A | 138.211599 | -8.3197904 | 1.51165363 |
| 4482 | NM_001329431.1 | | #N/A | 132.478188 | -8.2577584 | 1.50045338 |
| 4483 | NM_018052.3 | | #N/A | 175.295768 | -8.6643976 | 1.57440966 |
| 4484 | NM_006421.4 | | #N/A | 278.736426 | -4.0650644 | 0.73873301 |
| 4485 | NM_003313.3 | | #N/A | 172.303895 | -8.6391608 | 1.57007621 |
| 4486 | NM_152641.3 | | #N/A | 159.377891 | -8.5255376 | 1.54947879 |
| 4487 | NM_019843.3 | | #N/A | 135.68395 | -8.2932518 | 1.50730956 |
| 4488 | NM_012138.3 | | #N/A | 146.888908 | -8.4087441 | 1.52836104 |
| 4489 | NM_021970.3 | | #N/A | 194.307052 | -8.8119891 | 1.60165954 |
| 4490 | NM_001321381.1 | | #N/A | 161.792636 | -8.5469349 | 1.55365543 |
| 4491 | NM_033028.4 | | #N/A | 131.191296 | -8.2457022 | 1.49896075 |
| 4492 | NM_001102421.2 | | #N/A | 179.212924 | -8.6954093 | 1.58080047 |
| 4493 | NM_004330.3 | | #N/A | 138.372625 | -8.322915 | 1.51315776 |
| 4494 | NM_017643.2 | | #N/A | 155.753808 | -8.4923153 | 1.54395356 |
| 4495 | NM_032168.2 | | #N/A | 149.250916 | -8.4317919 | 1.53306752 |
| 4496 | NR_040722.1 | | #N/A | 192.208518 | -8.7950982 | 1.59914887 |
| 4497 | NM_015971.3 | | #N/A | 160.276162 | -8.5345601 | 1.55182569 |
| 4498 | NM_001145437.1 | LSS | | 222.617429 | -9.009236 | 1.63815686 |
| 4499 | NM_001184979.1 | NDUFS5 | | 249.774154 | -9.1748245 | 1.66828213 |
| 4500 | NM_014845.5 | FIG4 | | 162.979932 | -8.5591078 | 1.55635325 |
| 4501 | NM_006988.4 | | #N/A | 256.858292 | -9.2163213 | 1.67589333 |
| 4502 | NM_001330420.1 | LUC7L | | 153.82517 | -8.4755463 | 1.54124919 |
| 4503 | NM_006366.2 | | #N/A | 207.680464 | -8.9083076 | 1.61997699 |
| 4504 | NM_001039618.2 | | #N/A | 130.892906 | -8.2440881 | 1.49920621 |
| 4505 | NR_136519.1 | | #N/A | 77.6038 | 4.12217055 | 0.74963159 |
| 4506 | NM_006560.3 | | #N/A | 188.561159 | -8.7703503 | 1.59494929 |
| 4507 | NM_153487.3 | | #N/A | 138.524169 | -8.3266859 | 1.51427779 |
| 4508 | NM_032013.3 | | #N/A | 186.860694 | -8.7554226 | 1.59231666 |
| 4509 | NR_024281.1 | LINC00599 | | 213.522951 | -8.9484531 | 1.62744424 |
| 4511 | NM_001166.4 | BIRC2 | | 159.083109 | -8.5232594 | 1.55015281 |
| 4510 | NM_000405.4 | | #N/A | 138.492002 | -8.3246126 | 1.51402981 |
| 4512 | NM_001275.3 | | #N/A | 287.20261 | -9.3764297 | 1.70531561 |
| 4513 | NM_020909.3 | | #N/A | 137.826095 | -8.3180189 | 1.51286594 |
| 4514 | NM_001330782.1 | | #N/A | 250.648681 | -9.1807816 | 1.669867 |

| | | | | | |
|---|---|---|---|---|---|
| 4515 | NM_152624.5 | #N/A | 129.468184 | -8.2286027 | 1.49668975 |
| 4516 | NM_001164814.1 | ACIN1 | 138.308776 | -8.3212798 | 1.51357793 |
| 4517 | NM_021821.3 | #N/A | 171.24018 | -8.6299617 | 1.56972905 |
| 4518 | NM_032204.4 | #N/A | 150.50873 | -8.4449218 | 1.53610174 |
| 4519 | NM_005189.2 | #N/A | 173.05037 | -8.644799 | 1.57249367 |
| 4520 | NM_018161.4 | #N/A | 138.417034 | -8.3243172 | 1.51436296 |
| 4521 | NM_006784.2 | #N/A | 151.612111 | -8.4545103 | 1.53850113 |
| 4522 | NM_020312.3 | #N/A | 134.811792 | -8.286249 | 1.50788524 |
| 4523 | NM_016396.2 | #N/A | 174.614021 | -8.6573953 | 1.5754366 |
| 4524 | NM_015889.4 | #N/A | 181.819902 | -8.715967 | 1.58614696 |
| 4525 | NM_014149.3 | #N/A | 123.282997 | -5.1389291 | 0.93526147 |
| 4526 | NM_001177701.2 | #N/A | 141.521598 | -8.3559912 | 1.52091217 |
| 4527 | NM_003869.5 | #N/A | 143.137758 | -8.3742134 | 1.52425446 |
| 4528 | NM_001284230.1 | MAP3K9 | 176.53794 | -8.6740036 | 1.57884173 |
| 4529 | NM_001282974.1 | MTSS1 | 316.901995 | -9.5180554 | 1.73254394 |
| 4530 | NM_001281534.1 | #N/A | 233.398843 | -9.0763419 | 1.65245441 |
| 4531 | NM_016454.3 | #N/A | 167.389028 | -8.5977744 | 1.56532868 |
| 4532 | NM_002645.3 | #N/A | 146.451448 | -8.4069833 | 1.53071657 |
| 4533 | NM_017653.3 | #N/A | 155.581668 | -8.4919675 | 1.54617998 |
| 4534 | NM_000476.2 | AK1 | 240.092589 | -9.1182654 | 1.66025496 |
| 4535 | NM_015069.4 | ZNF423 | 201.93185 | -8.8676172 | 1.61468606 |
| 4536 | NM_016400.3 | #N/A | 172.22832 | -8.63853 | 1.57297168 |
| 4537 | NM_030627.3 | #N/A | 165.556029 | -8.5806649 | 1.56241712 |
| 4538 | NM_015131.2 | #N/A | 147.414544 | -8.4137374 | 1.53206088 |
| 4539 | NM_001206736.1 | NSFL1C | 255.141793 | -9.2055143 | 1.67650521 |
| 4540 | NM_003164.4 | #N/A | 148.113774 | -8.4213482 | 1.53382439 |
| 4541 | NM_006353.2 | #N/A | 182.810604 | -8.7243039 | 1.58905747 |
| 4542 | NM_001164462.1 | MUC12 | 31.8036895 | 4.24995096 | 0.77411538 |
| 4543 | NM_001142930.1 | #N/A | 180.186015 | -8.703817 | 1.58546548 |
| 4544 | NM_015462.4 | #N/A | 168.089397 | -8.6026482 | 1.56703163 |
| 4545 | NM_001289972.1 | ZNF84 | 131.66746 | -8.2483692 | 1.50255308 |
| 4546 | NM_001029863.2 | C6orf120 | 150.160623 | -8.4402009 | 1.53760028 |
| 4547 | NM_201624.2 | USP33 | 229.686779 | -9.0535186 | 1.64933919 |
| 4548 | NM_015107.2 | #N/A | 125.046444 | -8.174957 | 1.48931548 |
| 4549 | NM_016823.3 | #N/A | 166.465925 | -8.5896709 | 1.56487056 |
| 4550 | NM_001098794.1 | #N/A | 121.801327 | -8.1379543 | 1.4826548 |
| 4551 | NM_002843.3 | #N/A | 135.56433 | -8.2937691 | 1.51108032 |
| 4552 | NM_032303.4 | #N/A | 121.861107 | -8.1410964 | 1.48326741 |
| 4553 | NM_001190452.1 | MTRNR2L1 | 5235.29027 | -7.2553375 | 1.32197591 |
| 4554 | NM_006357.3 | #N/A | 253.740258 | -7.7731337 | 1.41645914 |
| 4555 | NM_205842.2 | #N/A | 288.026757 | -9.3807063 | 1.70950207 |
| 4556 | NM_014771.3 | #N/A | 127.016289 | -8.2004499 | 1.49447353 |
| 4557 | NM_058246.3 | #N/A | 154.013122 | -8.4773693 | 1.54493333 |

| | | | | | |
|---|---|---|---|---|---|
| 4558 | NR_104307.2 | #N/A | 146.844575 | -8.4070891 | 1.53215825 |
| 4559 | NM_004352.3 | #N/A | 121.588193 | -8.1371335 | 1.48300079 |
| 4560 | NM_001184772.2 | BCORL1 | 122.3358 | -8.1470646 | 1.48490073 |
| 4561 | NM_139246.4 | #N/A | 159.542789 | -8.5276256 | 1.55437004 |
| 4562 | NM_002767.3 | #N/A | 193.19302 | -8.803729 | 1.60496067 |
| 4563 | NM_001167582.1 | #N/A | 149.608841 | -8.4369502 | 1.53812084 |
| 4564 | NM_006293.3 | #N/A | 139.931741 | -8.3395062 | 1.52035392 |
| 4565 | NM_000466.2 | #N/A | 130.093737 | -8.2327737 | 1.50092408 |
| 4566 | NM_004725.3 | #N/A | 212.468077 | -8.9414429 | 1.63011215 |
| 4567 | NM_001077446.3 | #N/A | 129.129499 | -8.2232026 | 1.49930004 |
| 4569 | NR_046661.1 | ATP11A-AS1 | 40.3600696 | 4.26185637 | 0.7771715 |
| 4568 | NM_019030.3 | #N/A | 147.27318 | -8.4107089 | 1.53372914 |
| 4570 | NM_024552.2 | #N/A | 165.617225 | -8.5817793 | 1.56497507 |
| 4571 | NM_175885.3 | #N/A | 134.504692 | -8.2791719 | 1.50980089 |
| 4572 | NM_012079.5 | #N/A | 194.168132 | -8.8113728 | 1.60688259 |
| 4573 | NM_001145774.2 | #N/A | 208.520675 | -8.9127011 | 1.6255956 |
| 4574 | NR_001564.2 | XIST | 253.553309 | -9.1976919 | 1.67759922 |
| 4575 | NM_030808.4 | #N/A | 159.129411 | -8.5242563 | 1.55487368 |
| 4576 | NM_003968.3 | #N/A | 160.370236 | -8.5354737 | 1.55698024 |
| 4577 | NM_003078.3 | #N/A | 201.816703 | -8.8675094 | 1.61760414 |
| 4578 | NM_016205.2 | #N/A | 187.678451 | -8.7621531 | 1.59841046 |
| 4579 | NM_001100112.1 | MYH2 | 20.5998752 | 5.1897054 | 0.94677154 |
| 4580 | NM_017534.5 | #N/A | 20.5998752 | 5.1897054 | 0.94677154 |
| 4581 | NM_018946.3 | #N/A | 143.399483 | -8.3749007 | 1.52791925 |
| 4582 | NM_004184.3 | #N/A | 210.48253 | -8.9284119 | 1.62899197 |
| 4583 | NM_001316342.1 | #N/A | 164.380316 | -8.5713237 | 1.56388342 |
| 4584 | NM_005749.3 | #N/A | 126.265726 | -8.1928999 | 1.49485193 |
| 4585 | NM_025205.4 | #N/A | 169.115601 | -7.3365008 | 1.33877834 |
| 4586 | NM_001199753.1 | CPT1C | 172.955169 | -8.6439606 | 1.57741344 |
| 4587 | NM_153333.2 | #N/A | 163.92587 | -8.5669944 | 1.5634333 |
| 4588 | NM_177968.3 | #N/A | 120.05128 | -6.0345322 | 1.10130468 |
| 4589 | NM_015274.2 | #N/A | 168.072636 | -8.6034898 | 1.57017566 |
| 4590 | NM_138348.5 | #N/A | 90.6106902 | -3.9960202 | 0.72931414 |
| 4591 | NM_001144072.1 | #N/A | 155.291154 | -8.489044 | 1.54939915 |
| 4592 | NM_182706.4 | #N/A | 238.987287 | -9.1106361 | 1.66288659 |
| 4593 | NM_001135821.1 | #N/A | 189.316813 | -8.7753437 | 1.60172282 |
| 4594 | NM_032989.2 | #N/A | 187.731315 | -8.7625284 | 1.59947978 |
| 4595 | NM_001185076.1 | FMR1 | 157.604036 | -8.5104883 | 1.55352978 |
| 4596 | NM_003201.2 | #N/A | 158.643466 | -8.5187566 | 1.55507432 |
| 4597 | NM_014760.3 | #N/A | 303.519831 | -6.7948682 | 1.24037519 |
| 4598 | NM_001134398.1 | #N/A | 209.214776 | -8.9195954 | 1.62836104 |
| 4599 | NM_002165.3 | #N/A | 128.094602 | -8.2138492 | 1.49959839 |
| 4600 | NM_000305.2 | #N/A | 166.138508 | -8.5875956 | 1.5681624 |

| | | | | | |
|---|---|---|---|---|---|
| 4601 | NM_015694.2 | #N/A | 141.338924 | -8.3531277 | 1.52534752 |
| 4603 | NM_017550.2 | MIER2 | 149.328255 | -8.4325914 | 1.53995349 |
| 4602 | NM_002345.3 | #N/A | 242.893597 | -7.7154308 | 1.40898133 |
| 4604 | NM_177402.4 | #N/A | 233.960957 | -7.8100935 | 1.42634662 |
| 4605 | NM_001278628.1 | CRNKL1 | 123.879352 | -8.1625548 | 1.49074919 |
| 4606 | NM_003157.4 | #N/A | 126.745327 | -8.1955571 | 1.49675826 |
| 4607 | NM_015353.2 | #N/A | 206.064014 | -8.8970389 | 1.62488339 |
| 4608 | NM_181050.2 | #N/A | 133.610926 | -8.271182 | 1.51063997 |
| 4609 | NM_005956.3 | #N/A | 210.209314 | -7.5283157 | 1.37502428 |
| 4610 | NM_015323.4 | #N/A | 122.759413 | -8.1509684 | 1.48881573 |
| 4611 | NM_014456.4 | #N/A | 212.461031 | -8.9431507 | 1.63357877 |
| 4612 | NM_015886.4 | #N/A | 237.899096 | -9.1050947 | 1.66316227 |
| 4613 | NM_003945.3 | #N/A | 158.349446 | -8.518058 | 1.55597699 |
| 4614 | NM_007108.3 | #N/A | 252.420026 | -9.1901944 | 1.67883213 |
| 4615 | NM_001031723.3 | #N/A | 144.852356 | -8.3880841 | 1.53233743 |
| 4616 | NM_006701.4 | #N/A | 158.518038 | -8.518618 | 1.55621624 |
| 4617 | NM_001321439.1 | #N/A | 152.01833 | -8.4575888 | 1.54511969 |
| 4618 | NM_022157.3 | #N/A | 161.240056 | -8.5439878 | 1.56092368 |
| 4619 | NM_001037954.3 | #N/A | 148.258234 | -8.4210122 | 1.5384945 |
| 4620 | NM_006589.2 | #N/A | 161.278023 | -8.5453802 | 1.56125961 |
| 4621 | NM_001128852.1 | #N/A | 192.983783 | -8.8041762 | 1.60868052 |
| 4622 | NM_001330574.1 | #N/A | 188.081152 | -8.7650252 | 1.6015284 |
| 4623 | NM_020189.5 | #N/A | 170.197118 | -8.6209519 | 1.57534236 |
| 4624 | NM_018434.5 | #N/A | 140.399046 | -8.3445059 | 1.52488977 |
| 4625 | NM_032141.3 | #N/A | 127.04227 | -8.1994874 | 1.49842155 |
| 4626 | NM_004098.3 | #N/A | 171.907706 | -8.6348517 | 1.57808486 |
| 4627 | NM_004952.4 | #N/A | 234.819046 | -9.0857905 | 1.66059989 |
| 4628 | NM_015443.3 | KANSL1 | 169.676781 | -7.2500476 | 1.32510181 |
| 4629 | NM_007186.5 | #N/A | 136.700117 | -8.3027968 | 1.5175814 |
| 4630 | NR_108066.1 | #N/A | 97.3772742 | 3.6133681 | 0.660521 |
| 4631 | NM_138820.3 | #N/A | 141.02745 | -8.3506001 | 1.52654297 |
| 4632 | NM_001128636.2 | #N/A | 145.408069 | -8.3949632 | 1.53468523 |
| 4633 | NM_018840.4 | #N/A | 172.867998 | -8.6434394 | 1.58037838 |
| 4634 | NM_015258.1 | #N/A | 131.203267 | -8.24702 | 1.50795282 |
| 4636 | NM_015120.4 | ALMS1 | 138.290275 | -8.3204125 | 1.5213991 |
| 4635 | NM_001243765.1 | #N/A | 157.490673 | -8.5096413 | 1.55599534 |
| 4637 | NM_018226.4 | #N/A | 149.967734 | -8.4394148 | 1.54319273 |
| 4638 | NM_173854.5 | #N/A | 195.327829 | -7.4336077 | 1.3593918 |
| 4639 | NM_001252006.1 | PAPOLA | 144.967907 | -8.3886514 | 1.5342436 |
| 4640 | NM_015187.4 | #N/A | 243.348488 | -9.1370149 | 1.67115184 |
| 4641 | NM_153702.3 | #N/A | 145.723939 | -8.398138 | 1.53601837 |
| 4642 | NM_015123.2 | #N/A | 134.786621 | -8.2820382 | 1.51482749 |
| 4643 | NM_001243186.1 | #N/A | 132.109273 | -8.2569069 | 1.51030928 |

| | | | | | |
|---|---|---|---|---|---|
| 4644 | NM_001080855.2 | #N/A | 166.008144 | -8.5856824 | 1.5704922 |
| 4645 | NM_001142299.1 | #N/A | 126.973808 | -8.1998392 | 1.49991844 |
| 4646 | NM_144572.1 | TBC1D2B | 117.786605 | -8.0902209 | 1.47988975 |
| 4647 | NM_002006.4 | #N/A | 122.91793 | -8.1548672 | 1.49185868 |
| 4648 | NM_133369.2 | #N/A | 223.344522 | -9.0127655 | 1.64882101 |
| 4649 | NM_022756.5 | #N/A | 283.756346 | -9.358575 | 1.71212411 |
| 4650 | NM_001040653.3 | ZXDC | 135.053621 | -8.2870729 | 1.51619807 |
| 4651 | NM_013328.3 | #N/A | 159.523556 | -8.5280775 | 1.56042506 |
| 4652 | NM_005862.2 | #N/A | 166.875319 | -7.2292835 | 1.32283245 |
| 4653 | NR_026752.1 | CROCCP2 | 153.291243 | -8.4704716 | 1.55006873 |
| 4654 | NM_020170.3 | #N/A | 154.176648 | -8.4806576 | 1.55202096 |
| 4656 | NR_109783.1 | LINC02078 | 141.186602 | 2.87262942 | 0.52572054 |
| 4655 | NM_139045.3 | #N/A | 171.527182 | -8.631052 | 1.57957049 |
| 4657 | NM_018952.4 | #N/A | 226.422946 | -9.0339358 | 1.65350198 |
| 4658 | NM_004357.4 | #N/A | 166.81693 | -8.5930559 | 1.57295019 |
| 4659 | NM_003302.2 | #N/A | 139.712765 | -8.3388682 | 1.52646626 |
| 4660 | NM_199069.1 | #N/A | 142.364016 | -8.3640562 | 1.53109836 |
| 4661 | NM_007051.2 | #N/A | 172.37098 | -8.6397908 | 1.58159661 |
| 4662 | NM_001100170.1 | IMMT | 227.551282 | -9.040073 | 1.65489413 |
| 4663 | NM_003324.4 | #N/A | 125.74653 | -8.1823634 | 1.49791784 |
| 4664 | NM_015975.4 | #N/A | 133.824258 | -8.2746053 | 1.51483323 |
| 4665 | NM_001024921.3 | RPL9 | 215.789085 | -8.9635349 | 1.64100336 |
| 4666 | NM_032926.2 | #N/A | 171.697894 | -8.6338527 | 1.58067807 |
| 4667 | NM_003952.2 | #N/A | 143.216939 | -8.3735221 | 1.53303291 |
| 4668 | NM_001288730.1 | #N/A | 156.302621 | -8.498639 | 1.55605192 |
| 4669 | NM_002706.5 | #N/A | 125.576639 | -8.1832214 | 1.49831667 |
| 4670 | NM_001193628.1 | #N/A | 142.120363 | 2.83926628 | 0.51990448 |
| 4671 | NM_015932.5 | #N/A | 237.670031 | -7.6845712 | 1.40720759 |
| 4674 | NM_019008.5 | MIEF1 | 178.37354 | -8.6870777 | 1.59088173 |
| 4672 | NM_001142573.1 | #N/A | 144.789326 | -8.3890114 | 1.53629951 |
| 4673 | NM_014498.4 | #N/A | 174.958885 | -8.6631947 | 1.58651331 |
| 4675 | NM_015497.4 | #N/A | 155.319316 | -8.488871 | 1.55460346 |
| 4676 | NM_005843.5 | #N/A | 132.066008 | -6.1781659 | 1.13150448 |
| 4677 | NM_004661.3 | #N/A | 133.671659 | -8.272994 | 1.51522025 |
| 4678 | NM_001317784.1 | SCYL2 | 192.164114 | -6.5516888 | 1.20004971 |
| 4679 | NM_005274.2 | #N/A | 151.195002 | -8.4510766 | 1.54795073 |
| 4680 | NM_006453.2 | #N/A | 174.38957 | -8.6566043 | 1.58583781 |
| 4681 | NM_015088.2 | #N/A | 139.781829 | -8.3355064 | 1.52709133 |
| 4682 | NM_006868.3 | #N/A | 194.541493 | -8.8148678 | 1.615004 |
| 4683 | NM_145214.2 | #N/A | 127.078865 | -8.2001711 | 1.50245796 |
| 4684 | NR_034031.1 | LOC389906 | 176.383009 | -8.6721042 | 1.58896288 |
| 4685 | NM_018110.3 | #N/A | 204.110887 | -8.8838758 | 1.62780435 |
| 4686 | NM_001127358.1 | #N/A | 157.720539 | -8.5113536 | 1.55956008 |

| | | | | | | |
|---|---|---|---|---|---|---|
| 4687 | NR_073503.1 | | #N/A | 170.560136 | -8.623724 | 1.58016107 |
| 4688 | NM_006586.4 | | #N/A | 137.916075 | -8.318863 | 1.52435467 |
| 4689 | NM_018449.3 | UBAP2 | | 152.605799 | -8.4641527 | 1.55099968 |
| 4690 | NM_030806.3 | | #N/A | 250.580565 | -5.3069742 | 0.9724969 |
| 4691 | NM_001001521.1 | UGP2 | | 219.90085 | -7.5836449 | 1.38970935 |
| 4692 | NM_004753.6 | | #N/A | 150.870635 | -8.4496932 | 1.54842051 |
| 4693 | NM_139202.2 | MLC1 | | 284.475408 | -9.3629426 | 1.71582831 |
| 4694 | NM_052902.3 | | #N/A | 119.067546 | -8.1083914 | 1.48615098 |
| 4696 | NM_003995.3 | NPR2 | | 127.171666 | -8.2040897 | 1.5037833 |
| 4695 | NM_000903.2 | | #N/A | 143.06823 | -8.3739764 | 1.53492212 |
| 4697 | NM_015569.4 | | #N/A | 228.839162 | -9.0485328 | 1.65855106 |
| 4698 | NM_024725.3 | | #N/A | 146.750548 | -8.4064535 | 1.5408587 |
| 4699 | NM_004055.4 | | #N/A | 178.05278 | -8.6865294 | 1.59229001 |
| 4700 | NM_173551.4 | | #N/A | 118.845454 | -8.1032008 | 1.48537502 |
| 4701 | NM_138812.3 | DYDC1 | | 68.2483137 | 3.81305926 | 0.69903375 |
| 4702 | NR_033680.1 | CARD8 | | 39.0620238 | 10.278566 | 1.88446225 |
| 4703 | NM_001113482.1 | MANEAL | | 220.439448 | -8.9945693 | 1.6490883 |
| 4704 | NM_014703.2 | | #N/A | 121.994769 | -8.1408446 | 1.49257811 |
| 4705 | NM_030914.3 | | #N/A | 187.770274 | -8.7628943 | 1.60668014 |
| 4706 | NM_001277196.1 | | #N/A | 138.05106 | -8.3201147 | 1.52552751 |
| 4707 | NM_021959.2 | | #N/A | 212.055677 | -7.6667816 | 1.40586866 |
| 4708 | NM_001317955.1 | | #N/A | 137.395381 | -8.3112477 | 1.52412683 |
| 4709 | NM_000681.3 | | #N/A | 140.977362 | -8.347086 | 1.53119184 |
| 4710 | NM_016169.3 | SUFU | | 124.266515 | -8.1669319 | 1.49816628 |
| 4711 | NM_004838.3 | | #N/A | 145.345986 | -8.3945069 | 1.53994677 |
| 4712 | NM_001048249.3 | | #N/A | 153.09239 | -8.4686438 | 1.55357346 |
| 4713 | NM_000858.5 | | #N/A | 163.229002 | -8.5603681 | 1.57048827 |
| 4714 | NM_007241.3 | | #N/A | 189.976716 | -8.7796859 | 1.611058 |
| 4715 | NM_001248005.1 | | #N/A | 68.140336 | 11.0775401 | 2.03274414 |
| 4716 | NM_015881.5 | DKK3 | | 170.812956 | -8.6279228 | 1.5833704 |
| 4718 | NM_005010.4 | NRCAM | | 217.489476 | -8.9752297 | 1.64715917 |
| 4717 | NM_001303614.1 | SAMD4B | | 126.271734 | -8.1913618 | 1.50331037 |
| 4719 | NM_002691.3 | | #N/A | 141.264362 | -8.3514821 | 1.53285352 |
| 4720 | NM_001348129.1 | | #N/A | 164.016218 | -8.5679365 | 1.57267177 |
| 4721 | NM_000985.4 | | #N/A | 127.011801 | -8.2008796 | 1.50533286 |
| 4722 | NM_004939.2 | | #N/A | 540.924461 | -4.776364 | 0.87676376 |
| 4723 | NM_194332.2 | RNF38 | | 141.15866 | -8.3501119 | 1.53278865 |
| 4724 | NM_014487.5 | | #N/A | 125.407371 | -8.1804529 | 1.50179922 |
| 4725 | NM_024598.3 | | #N/A | 208.907483 | -8.9169639 | 1.6370757 |
| 4726 | NM_001130674.2 | | #N/A | 132.248443 | -8.2595148 | 1.51673312 |
| 4727 | NM_001029882.3 | AHDC1 | | 337.951921 | -2.0838275 | 0.38267329 |
| 4728 | NM_007188.4 | | #N/A | 161.232315 | -8.5430846 | 1.56900964 |
| 4729 | NM_001037283.1 | | #N/A | 333.488018 | -6.1348802 | 1.12673757 |

| | | | | | |
|---|---|---|---|---|---|
| 4730 | NM_201412.2 | LUC7L | 198.094796 | -8.8400704 | 1.62359162 |
| 4731 | NM_016240.2 | #N/A | 155.135476 | -8.4880528 | 1.55899263 |
| 4732 | NM_031966.3 | #N/A | 149.423609 | -8.4342867 | 1.54931706 |
| 4733 | NM_005872.2 | #N/A | 172.944337 | -8.6442483 | 1.58790045 |
| 4734 | NM_001321691.1 | #N/A | 217.564974 | -7.5686429 | 1.39047011 |
| 4735 | NM_153812.2 | #N/A | 124.074618 | -8.1676366 | 1.50051793 |
| 4736 | NM_001005336.2 | #N/A | 181.327032 | -8.7128947 | 1.60091856 |
| 4737 | NM_019063.4 | #N/A | 124.54196 | -8.1730866 | 1.50177188 |
| 4738 | NM_016423.2 | #N/A | 174.937208 | -8.6609244 | 1.59142239 |
| 4739 | NM_012133.5 | #N/A | 130.405797 | -8.2371025 | 1.51357177 |
| 4740 | NM_030816.4 | #N/A | 178.302175 | -7.4135942 | 1.36231687 |
| 4745 | NM_138342.3 | GLB1L2 | 141.944355 | -7.0153701 | 1.28918849 |
| 4741 | NM_000264.3 | #N/A | 218.558899 | -8.983136 | 1.65077415 |
| 4742 | NM_001006634.2 | #N/A | 127.664714 | -8.206848 | 1.50814338 |
| 4743 | NM_001271618.1 | #N/A | 191.768063 | -8.7933748 | 1.61588794 |
| 4744 | NM_004749.3 | #N/A | 141.097944 | -8.3512031 | 1.53465534 |
| 4746 | NM_020216.3 | #N/A | 133.341435 | -8.2699895 | 1.51988682 |
| 4747 | NM_004493.2 | #N/A | 171.071141 | -8.6288677 | 1.58590129 |
| 4748 | NM_004913.2 | #N/A | 176.642837 | -8.6751799 | 1.59445807 |
| 4749 | NM_024537.3 | #N/A | 157.060095 | -8.5054923 | 1.56330256 |
| 4750 | NM_052850.3 | #N/A | 190.451142 | -8.7836814 | 1.61443256 |
| 4751 | NM_013387.3 | #N/A | 178.583429 | -8.6910824 | 1.59744867 |
| 4752 | NM_005902.3 | #N/A | 179.70165 | -8.7016452 | 1.59952602 |
| 4753 | NM_003330.3 | #N/A | 159.377932 | -8.526403 | 1.56737894 |
| 4754 | NM_014453.3 | #N/A | 155.182125 | -8.4879646 | 1.56033498 |
| 4756 | NR_034113.1 | DNAJC27-AS1 | 109.551432 | 2.03813252 | 0.37471628 |
| 4755 | NM_004567.3 | #N/A | 148.529919 | -8.4237767 | 1.54872923 |
| 4758 | NM_017778.2 | NSD3 | 148.981565 | -8.4283749 | 1.54963997 |
| 4757 | NM_014319.4 | #N/A | 172.848331 | -8.6433113 | 1.5891507 |
| 4759 | NM_020780.1 | #N/A | 240.663688 | -9.1207242 | 1.6770397 |
| 4760 | NM_001142806.1 | SLC6A8 | 204.40299 | -8.887222 | 1.63412375 |
| 4761 | NM_001291310.1 | #N/A | 166.201658 | -8.586786 | 1.57892697 |
| 4762 | NM_022461.4 | #N/A | 144.070129 | -8.3802989 | 1.54098834 |
| 4763 | NM_004870.3 | #N/A | 172.238932 | -8.6389815 | 1.58861908 |
| 4764 | NM_014674.2 | #N/A | 137.182421 | -8.3094367 | 1.52830046 |
| 4765 | NM_016083.4 | #N/A | 291.236074 | -9.3962938 | 1.72822884 |
| 4766 | NM_001139441.1 | BCAP31 | 192.307297 | -8.7980998 | 1.61825593 |
| 4767 | NM_001146069.1 | #N/A | 186.187591 | -8.7518823 | 1.60975742 |
| 4768 | NM_015622.5 | #N/A | 178.235263 | -8.6885514 | 1.59809559 |
| 4770 | NM_006185.3 | NUMA1 | 158.87496 | -8.5242099 | 1.56793917 |
| 4769 | NM_001303264.1 | #N/A | 168.06569 | -7.3290764 | 1.34811334 |
| 4771 | NM_014806.4 | #N/A | 185.712453 | -8.7476402 | 1.60903726 |
| 4772 | NM_001099695.1 | #N/A | 188.888619 | -7.3841805 | 1.358282 |

| | | | | | |
|---|---|---|---|---|---|
| 4773 | NM_005776.2 | #N/A | 197.340247 | -8.8349342 | 1.62521247 |
| 4774 | NM_005581.4 | #N/A | 146.993863 | -8.4127477 | 1.54762128 |
| 4775 | NM_032804.5 | #N/A | 168.976732 | -8.6111236 | 1.58417569 |
| 4776 | NM_004830.3 | #N/A | 136.360741 | -8.3000905 | 1.52705794 |
| 4777 | NM_005539.4 | #N/A | 157.389437 | -8.5082263 | 1.56535843 |
| 4778 | NM_012342.2 | #N/A | 142.95258 | -8.3725092 | 1.54039466 |
| 4779 | NM_019022.3 | #N/A | 180.367543 | -7.3245108 | 1.34757524 |
| 4780 | NM_006348.3 | COG5 | 152.950312 | -8.4666709 | 1.55776118 |
| 4781 | NM_178314.4 | #N/A | 140.501522 | -8.3444462 | 1.53526818 |
| 4782 | NM_005488.2 | #N/A | 136.860081 | -8.3070952 | 1.52847087 |
| 4783 | NM_032023.3 | #N/A | 137.050829 | -8.308795 | 1.5287904 |
| 4784 | NM_015111.1 | #N/A | 120.856235 | -8.127781 | 1.49551046 |
| 4785 | NM_024112.3 | #N/A | 209.355754 | -8.9200536 | 1.64131976 |
| 4786 | NM_005711.4 | #N/A | 286.845302 | -9.3752536 | 1.72524294 |
| 4787 | NM_001198999.1 | #N/A | 242.926502 | -9.1348992 | 1.68102434 |
| 4788 | NM_001199174.1 | #N/A | 155.253284 | -8.4889749 | 1.56224336 |
| 4789 | NM_022095.3 | #N/A | 139.48481 | -6.9926807 | 1.28688937 |
| 4790 | NM_000178.3 | #N/A | 153.62008 | -8.4736082 | 1.55946948 |
| 4791 | NM_000532.4 | #N/A | 128.640396 | -8.2173629 | 1.51232905 |
| 4792 | NM_001080412.2 | #N/A | 121.845144 | -8.1383344 | 1.4979116 |
| 4793 | NM_138360.3 | #N/A | 280.091432 | -9.3399371 | 1.71910667 |
| 4794 | NM_199040.3 | #N/A | 184.124656 | -8.7337774 | 1.6075532 |
| 4795 | NM_173829.3 | #N/A | 149.913031 | -5.1469872 | 0.94746505 |
| 4796 | NM_006624.5 | ZMYND11 | 165.179091 | -8.5774391 | 1.57898992 |
| 4797 | NM_032259.3 | #N/A | 130.500597 | -8.2383786 | 1.51657644 |
| 4798 | NM_138367.1 | #N/A | 131.103178 | -8.2445416 | 1.51777445 |
| 4799 | NM_024117.3 | MAPKAP1 | 154.381394 | -8.4814988 | 1.56147692 |
| 4800 | NM_001024948.2 | FNBP1L | 120.203072 | -8.1171994 | 1.49442588 |
| 4801 | NM_001516.4 | #N/A | 124.110784 | -8.1674182 | 1.50368364 |
| 4802 | NM_000345.3 | #N/A | 226.034922 | -9.0306524 | 1.66264172 |
| 4803 | NM_001256494.1 | MAN2C1 | 141.594588 | -8.356273 | 1.5385126 |
| 4804 | NM_002492.3 | #N/A | 145.66144 | -8.3954192 | 1.54573332 |
| 4805 | NM_001346143.1 | #N/A | 136.279709 | -8.3030053 | 1.52883541 |
| 4806 | NM_018133.3 | #N/A | 135.596125 | -8.2924786 | 1.5269141 |
| 4807 | NM_032520.4 | #N/A | 156.247798 | -8.4982008 | 1.56493891 |
| 4808 | NM_001135655.1 | #N/A | 285.094016 | -9.3655823 | 1.72474374 |
| 4809 | NM_033064.4 | #N/A | 1049.64088 | -3.3724676 | 0.62107183 |
| 4810 | NM_001172773.1 | #N/A | 123.891092 | -8.1608025 | 1.50293526 |
| 4811 | NM_001668.3 | #N/A | 119.943399 | -8.1191996 | 1.49526715 |
| 4812 | NM_020967.2 | #N/A | 134.213945 | -8.2760399 | 1.52437039 |
| 4813 | NM_182767.5 | #N/A | 169.869765 | -8.6184516 | 1.5876204 |
| 4814 | NM_006342.2 | #N/A | 143.954885 | -8.3799145 | 1.5438723 |
| 4815 | NM_182552.4 | #N/A | 91.5714585 | -3.7949445 | 0.69915647 |

| | | | | | |
|---|---|---|---|---|---|
| 4816 | NM_001321856.1 | JAK1 | 129.945526 | -8.2345079 | 1.51713987 |
| 4817 | NM_002235.3 | KCNA6 | 194.259159 | -8.8115695 | 1.62347864 |
| 4818 | NM_001318052.1 | DENND2A | 127.111485 | -8.2002862 | 1.51093613 |
| 4819 | NM_002973.3 | #N/A | 125.853969 | -8.1851377 | 1.50814694 |
| 4820 | NM_080686.2 | #N/A | 1549.56016 | -6.6939066 | 1.23338521 |
| 4821 | NM_001015881.1 | #N/A | 173.438343 | -8.648414 | 1.59356603 |
| 4822 | NM_004329.2 | BMPR1A | 151.660933 | -8.4559407 | 1.55823026 |
| 4823 | NM_033554.3 | HLA-DPA1 | 266.321602 | -9.2679217 | 1.70789292 |
| 4824 | NM_005839.3 | #N/A | 160.278755 | -8.5372027 | 1.57330718 |
| 4825 | NM_001123329.1 | ZBTB1 | 115.175778 | -8.057483 | 1.48493881 |
| 4826 | NM_032020.4 | #N/A | 128.562848 | -8.2175886 | 1.51455598 |
| 4827 | NM_001143835.1 | #N/A | 118.277434 | -8.0940186 | 1.49186447 |
| 4828 | NM_001271184.1 | #N/A | 181.751514 | -8.7157301 | 1.60644871 |
| 4829 | NM_003690.4 | #N/A | 173.497452 | -8.6495606 | 1.59425559 |
| 4830 | NM_021115.4 | #N/A | 229.734762 | -9.0541105 | 1.66910436 |
| 4831 | NM_003473.3 | #N/A | 135.457962 | -8.2912417 | 1.52854567 |
| 4832 | NM_003034.3 | ST8SIA1 | 165.471239 | -8.5807484 | 1.5820549 |
| 4833 | NM_018067.4 | #N/A | 178.179062 | -8.6876323 | 1.60175486 |
| 4834 | NM_020829.3 | #N/A | 123.263211 | -8.1544512 | 1.50345421 |
| 4835 | NM_014016.4 | #N/A | 129.834354 | -8.2284918 | 1.51717112 |
| 4836 | NM_001130969.1 | NSMF | 573.268852 | -10.373504 | 1.91286019 |
| 4837 | NM_002149.3 | #N/A | 212.370649 | -8.9409609 | 1.64869364 |
| 4838 | NM_001330231.1 | #N/A | 163.743679 | -8.5649524 | 1.5794742 |
| 4839 | NM_004209.5 | #N/A | 220.68491 | -8.9962051 | 1.65900187 |
| 4840 | NM_022048.4 | #N/A | 131.222269 | -8.2457038 | 1.52067989 |
| 4841 | NM_005786.5 | #N/A | 122.133482 | -8.1446316 | 1.50206725 |
| 4842 | NM_003363.3 | #N/A | 143.951375 | -8.3800931 | 1.54563492 |
| 4843 | NM_022767.3 | #N/A | 165.477418 | -8.5805275 | 1.58262182 |
| 4844 | NM_006263.3 | #N/A | 140.062718 | -8.3405338 | 1.53843513 |
| 4845 | NM_001156.4 | #N/A | 133.625392 | -8.2745195 | 1.52632664 |
| 4846 | NM_016403.3 | #N/A | 163.882215 | -8.5672193 | 1.58032035 |
| 4847 | NM_001330161.1 | #N/A | 86.8312259 | -3.5272841 | 0.65066415 |
| 4848 | NM_138389.3 | #N/A | 119.163811 | -8.10877 | 1.4958231 |
| 4849 | NM_001126131.1 | #N/A | 136.37669 | -6.9607744 | 1.28409076 |
| 4850 | NM_001128431.2 | #N/A | 138.976875 | -8.3310132 | 1.53686498 |
| 4851 | NM_001146213.1 | #N/A | 163.06077 | -8.5582442 | 1.57877845 |
| 4852 | NM_001031702.3 | #N/A | 174.697929 | -8.6590004 | 1.59743224 |
| 4853 | NM_020412.4 | #N/A | 258.151741 | -4.18496 | 0.7720497 |
| 4854 | NM_030969.4 | #N/A | 150.291082 | -8.4413838 | 1.55743324 |
| 4855 | NM_001256798.1 | #N/A | 142.93911 | -8.3683877 | 1.54397981 |
| 4856 | NM_018927.3 | #N/A | 223.65595 | -9.0153247 | 1.66340877 |
| 4857 | NM_005662.6 | #N/A | 145.45045 | -8.3976818 | 1.54946114 |
| 4858 | NM_022834.4 | #N/A | 140.475598 | -8.3446282 | 1.53969149 |

| | | | | | |
|---|---|---|---|---|---|
| 4859 | NM_001130831.1 | #N/A | 194.668559 | -8.814031 | 1.6263403 |
| 4860 | NR_028103.1 | #N/A | 142.679399 | -8.3699571 | 1.54475891 |
| 4861 | NM_031309.5 | #N/A | 221.847226 | -9.0036717 | 1.66176081 |
| 4862 | NM_020142.3 | NDUFA4L2 | 168.77921 | -8.6080005 | 1.58877879 |
| 4863 | NM_001083956.1 | ZNF655 | 124.807363 | -8.1771155 | 1.50926654 |
| 4864 | NM_001308321.1 | PARP15 | 816.409826 | 3.67160814 | 0.67770158 |
| 4865 | NM_016258.2 | #N/A | 162.427637 | -8.551986 | 1.57880703 |
| 4866 | NM_003168.2 | #N/A | 162.052025 | -8.5510646 | 1.57875677 |
| 4867 | NM_015922.2 | #N/A | 165.080416 | -8.5781154 | 1.58376907 |
| 4868 | NM_021637.2 | #N/A | 379.578056 | -8.281124 | 1.52902499 |
| 4869 | NM_001282142.1 | #N/A | 132.089009 | -8.2573498 | 1.52466216 |
| 4870 | NM_016231.4 | #N/A | 172.740524 | -6.5882248 | 1.21646626 |
| 4871 | NM_181866.2 | #N/A | 230.304674 | -9.0582553 | 1.67261075 |
| 4872 | NM_001046.2 | #N/A | 147.87098 | -8.4196762 | 1.55473856 |
| 4873 | NM_138408.3 | #N/A | 140.40255 | -8.3439362 | 1.54074762 |
| 4874 | NM_001170418.1 | #N/A | 75.4904364 | 3.98459435 | 0.73579709 |
| 4875 | NM_000521.3 | #N/A | 151.886653 | -7.1016855 | 1.31150448 |
| 4876 | NM_138730.2 | #N/A | 118.184239 | -8.0943158 | 1.49483979 |
| 4877 | NM_001302632.1 | #N/A | 129.128268 | -8.2262652 | 1.51933836 |
| 4878 | NM_014397.5 | NEK6 | 145.595939 | -8.3969919 | 1.55089992 |
| 4879 | NM_023037.2 | #N/A | 226.506183 | -9.0340591 | 1.6685534 |
| 4880 | NM_001821.3 | CHML | 229.598775 | -5.9836507 | 1.10520279 |
| 4881 | NR_125921.1 | LINC02261 | 136.031004 | 3.74752301 | 0.6922046 |
| 4882 | NM_001198934.1 | #N/A | 142.826799 | -8.3695727 | 1.54600467 |
| 4883 | NM_018158.2 | SLC4A1AP | 136.872882 | -8.3076085 | 1.5345955 |
| 4884 | NM_001164748.1 | #N/A | 141.421522 | -8.3566131 | 1.54375654 |
| 4885 | NM_024324.4 | #N/A | 150.073496 | -8.4393685 | 1.55903797 |
| 4886 | NM_001135694.2 | VDAC3 | 338.951188 | -9.6153287 | 1.77644111 |
| 4887 | NM_015014.3 | #N/A | 140.660034 | -8.3458692 | 1.54199904 |
| 4888 | NM_001033581.1 | #N/A | 188.759851 | -8.7700239 | 1.62047426 |
| 4889 | NM_182789.3 | PAIP1 | 180.407026 | -8.7048594 | 1.60850101 |
| 4890 | NM_002751.6 | #N/A | 229.353348 | -9.0516627 | 1.67260395 |
| 4891 | NM_024666.4 | #N/A | 133.229914 | -8.2711591 | 1.52861885 |
| 4892 | NM_182565.3 | #N/A | 152.062232 | -8.4582578 | 1.56322704 |
| 4893 | NM_021729.5 | VPS11 | 133.715118 | -8.2734309 | 1.52914481 |
| 4894 | NM_017736.4 | #N/A | 131.192419 | -8.2455144 | 1.5239999 |
| 4895 | NM_022484.5 | #N/A | 114.545862 | -8.0483601 | 1.48768297 |
| 4896 | NM_002485.4 | NBN | 121.013 | -4.1564837 | 0.76831722 |
| 4897 | NM_014755.2 | #N/A | 124.023188 | -8.1647452 | 1.50926351 |
| 4898 | NM_005452.5 | #N/A | 165.823481 | -8.5842358 | 1.58683656 |
| 4899 | NM_015191.2 | #N/A | 124.977256 | -8.1792865 | 1.51201279 |
| 4900 | NM_002752.4 | #N/A | 149.719106 | -8.4361615 | 1.5596084 |
| 4901 | NM_001282918.1 | #N/A | 161.042072 | -8.5415239 | 1.57913827 |

| | | | | | |
|---|---|---|---|---|---|
| 4902 | NM_005831.4 | #N/A | 116.776166 | -8.0806212 | 1.49395708 |
| 4903 | NR_038170.1 | #N/A | 171.945687 | -8.6363595 | 1.59670225 |
| 4904 | NM_001079862.2 | #N/A | 181.695328 | -8.7159319 | 1.61149826 |
| 4905 | NM_006991.4 | #N/A | 119.17027 | -8.1087561 | 1.49932524 |
| 4906 | NM_013392.3 | #N/A | 130.328688 | -8.2350877 | 1.52272015 |
| 4907 | NM_006363.4 | #N/A | 112.747668 | -8.0287873 | 1.48463329 |
| 4908 | NM_003101.5 | #N/A | 122.370698 | -8.1454619 | 1.50628475 |
| 4909 | NM_004044.6 | #N/A | 164.641387 | -8.5736782 | 1.58557805 |
| 4910 | NM_001271874.1 | #N/A | 130.843291 | -8.2425534 | 1.52443558 |
| 4911 | NM_015268.3 | #N/A | 119.930692 | -8.1155745 | 1.5010364 |
| 4912 | NM_015963.5 | #N/A | 152.944882 | -8.4667735 | 1.56600992 |
| 4913 | NM_001311162.1 | #N/A | 240.123863 | -9.1182986 | 1.68659532 |
| 4914 | NM_018120.5 | #N/A | 180.304819 | -8.7044668 | 1.61009203 |
| 4915 | NM_001040441.2 | #N/A | 148.292165 | -8.4222726 | 1.55793106 |
| 4916 | NM_001310.3 | #N/A | 125.835699 | -5.9887941 | 1.10780687 |
| 4917 | NM_020457.2 | #N/A | 145.443193 | -8.3941202 | 1.55277493 |
| 4918 | NM_001040202.1 | #N/A | 192.363698 | -8.7978364 | 1.62759792 |
| 4919 | NM_022900.4 | #N/A | 173.400482 | -8.6487286 | 1.60002212 |
| 4920 | NM_013232.3 | #N/A | 136.263586 | -8.2979703 | 1.53514279 |
| 4921 | NM_198563.2 | #N/A | 141.733945 | -8.3579103 | 1.54625305 |
| 4922 | NM_015340.3 | #N/A | 131.460972 | -8.2505003 | 1.52642864 |
| 4923 | NM_020385.3 | #N/A | 148.675749 | -8.42567 | 1.5595394 |
| 4924 | NM_001164161.1 | #N/A | 157.147454 | -8.5046816 | 1.57419313 |
| 4925 | NM_032477.2 | #N/A | 178.841251 | -8.6927602 | 1.60901749 |
| 4926 | NM_016238.2 | ANAPC7 | 176.23685 | -8.6712402 | 1.60506071 |
| 4927 | NM_001328682.1 | #N/A | 302.39343 | -9.4505508 | 1.74951426 |
| 4928 | NR_034076.1 | THAP9-AS1 | 126.568181 | -8.1940688 | 1.5169504 |
| 4929 | NM_016072.4 | #N/A | 169.577364 | -8.6158488 | 1.59504999 |
| 4930 | NM_001749.3 | #N/A | 172.436696 | -8.6400326 | 1.59954285 |
| 4931 | NM_001193511.1 | #N/A | 143.92826 | -8.3818347 | 1.55184175 |
| 4932 | NM_002246.2 | #N/A | 224.193772 | -6.4650839 | 1.19697095 |
| 4933 | NM_016938.4 | #N/A | 154.679526 | -8.4830798 | 1.57057609 |
| 4934 | NM_145886.3 | #N/A | 116.739206 | -8.0776292 | 1.49556981 |
| 4935 | NM_001134364.1 | #N/A | 162.024327 | -8.5491239 | 1.58293925 |
| 4936 | NM_001173523.1 | #N/A | 191.812999 | -8.7948065 | 1.62845577 |
| 4937 | NM_004664.3 | LIN7A | 168.202103 | -8.6027307 | 1.59290772 |
| 4938 | NM_002956.2 | CLIP1 | 217.941579 | -8.9773014 | 1.66229796 |
| 4939 | NM_022131.2 | #N/A | 303.091423 | -4.4829166 | 0.83010655 |
| 4940 | NM_001174096.1 | ZEB1 | 119.313725 | -8.1110468 | 1.50197757 |
| 4941 | NM_001291879.1 | #N/A | 111.362912 | -5.8304634 | 1.07973169 |
| 4942 | NM_006650.3 | #N/A | 272.529209 | -9.3008661 | 1.72239953 |
| 4943 | NM_015713.4 | #N/A | 163.699526 | -8.5648581 | 1.58620442 |
| 4944 | NM_213611.2 | #N/A | 156.922146 | -8.504881 | 1.57512247 |

| | | | | | |
|---|---|---|---|---|---|
| 4945 | NM_024069.3 | #N/A | 170.751579 | -8.6260925 | 1.59763567 |
| 4946 | NM_005620.1 | #N/A | 175.026794 | -8.6628915 | 1.60449232 |
| 4947 | NM_182488.3 | #N/A | 164.609134 | -8.5728573 | 1.58790352 |
| 4948 | NM_000026.3 | #N/A | 153.143832 | -8.4696765 | 1.56880343 |
| 4949 | NM_001025248.1 | #N/A | 144.673865 | -8.3859865 | 1.55345053 |
| 4950 | NM_015310.3 | #N/A | 173.134995 | -8.6449365 | 1.60149469 |
| 4951 | NM_032353.3 | #N/A | 156.292637 | -8.4987521 | 1.57447516 |
| 4952 | NM_000474.3 | #N/A | 194.049475 | -8.8117569 | 1.63256064 |
| 4953 | NM_001195626.1 | #N/A | 122.369513 | -8.1440957 | 1.50892791 |
| 4954 | NM_001242352.1 | ODF2 | 150.586212 | -8.4448029 | 1.56478619 |
| 4955 | NM_001303274.1 | ACLY | 286.694645 | -9.3734092 | 1.736935 |
| 4956 | NM_012384.4 | #N/A | 113.274943 | -8.0336112 | 1.48870636 |
| 4957 | NM_018427.4 | RRN3 | 188.055424 | -8.76528 | 1.62438436 |
| 4958 | NM_006590.3 | #N/A | 163.591457 | -8.5634069 | 1.58705297 |
| 4962 | NR_104644.1 | LINC01204 | 98.6582216 | 3.15707019 | 0.585126 |
| 4961 | NM_020379.3 | MAN1C1 | 115.750613 | -8.068279 | 1.49536216 |
| 4959 | NM_001080394.3 | #N/A | 132.352379 | -8.2586558 | 1.53063621 |
| 4960 | NM_017482.3 | #N/A | 240.373767 | -9.1192629 | 1.69012135 |
| 4963 | NM_014159.6 | SETD2 | 371.158641 | -4.33446 | 0.80345717 |
| 4964 | NM_138396.5 | #N/A | 195.185375 | -8.819058 | 1.63477505 |
| 4965 | NM_078471.3 | #N/A | 137.259032 | -8.3129276 | 1.54098964 |
| 4966 | NM_001316772.1 | GARS1 | 219.328865 | -8.9877793 | 1.66634725 |
| 4967 | NM_022776.4 | #N/A | 122.218666 | -8.1446551 | 1.51011895 |
| 4968 | NM_003716.3 | #N/A | 236.145601 | -9.0940956 | 1.68620499 |
| 4969 | NM_145049.4 | #N/A | 147.984841 | -5.4324676 | 1.00727707 |
| 4970 | NM_001304782.1 | #N/A | 66.7539026 | 2.79883728 | 0.51896512 |
| 4971 | NM_003374.2 | #N/A | 154.061389 | -8.4782651 | 1.57204687 |
| 4972 | NM_138797.2 | #N/A | 118.85247 | -8.1016008 | 1.5022497 |
| 4973 | NM_017865.3 | #N/A | 154.89529 | -8.4845237 | 1.57328986 |
| 4974 | NM_020368.2 | #N/A | 128.310858 | -8.2150368 | 1.52331269 |
| 4975 | NM_001142674.1 | #N/A | 126.35272 | -8.1912312 | 1.51894795 |
| 4976 | NM_012448.3 | #N/A | 205.231069 | -6.3504313 | 1.17769003 |
| 4977 | NM_206853.2 | #N/A | 429.732476 | -4.4703749 | 0.82915256 |
| 4978 | NM_003977.3 | #N/A | 177.472727 | -8.6818622 | 1.61033025 |
| 4979 | NM_001135944.1 | #N/A | 177.370951 | -8.6809512 | 1.61022777 |
| 4980 | NM_032538.2 | #N/A | 213.013322 | -8.9450351 | 1.65920454 |
| 4981 | NM_001145337.2 | #N/A | 141.894047 | -8.3561308 | 1.55008067 |
| 4982 | NM_001271592.1 | #N/A | 182.587356 | -8.7222689 | 1.61799408 |
| 4983 | NM_006579.2 | #N/A | 191.204245 | -8.7894053 | 1.63050441 |
| 4984 | NM_001018068.1 | #N/A | 226.826746 | -9.0346701 | 1.67601783 |
| 4985 | NM_022060.2 | ABHD4 | 158.375971 | -7.1513689 | 1.32666671 |
| 4986 | NM_017907.2 | #N/A | 203.046692 | -8.8761671 | 1.64670836 |
| 4987 | NM_024041.3 | #N/A | 129.429203 | -8.2253324 | 1.52598381 |

| | | | | | |
|---|---|---|---|---|---|
| 4988 | NM_001282412.1 | PPP1R7 | 155.593593 | -8.4918321 | 1.575531 |
| 4989 | NM_001134479.1 | #N/A | 159.524308 | -8.5278014 | 1.58225743 |
| 4990 | NM_000543.4 | #N/A | 189.405603 | -8.7757648 | 1.62828782 |
| 4991 | NM_006094.4 | #N/A | 131.591647 | -8.2497644 | 1.53073255 |
| 4992 | NM_201442.3 | #N/A | 128.848652 | -8.2226582 | 1.52578145 |
| 4993 | NR_024461.1 | RAMP2-AS1 | 146.446303 | 2.03865508 | 0.37829758 |
| 4994 | NM_020751.2 | #N/A | 112.981364 | -8.0322047 | 1.49070799 |
| 4995 | NM_173050.3 | #N/A | 160.501235 | -8.5361882 | 1.58423662 |
| 4998 | NM_144985.3 | CDH24 | 168.1213 | -8.6033397 | 1.59673675 |
| 4996 | NM_001321882.1 | NAA35 | 127.682731 | -8.2048266 | 1.52277803 |
| 4997 | NM_001328.2 | #N/A | 209.235038 | -7.5087298 | 1.39359118 |
| 4999 | NM_014923.4 | #N/A | 135.309316 | -8.2920067 | 1.53906517 |
| 5000 | NM_001079839.2 | OCIAD1 | 152.247989 | -8.4614115 | 1.57052506 |
| 5001 | NM_016010.2 | #N/A | 182.155231 | -8.7191902 | 1.61841489 |
| 5002 | NM_203388.2 | #N/A | 152.603087 | -8.4642192 | 1.57122373 |
| 5003 | NM_005252.3 | #N/A | 167.850087 | -8.6029993 | 1.59705359 |
| 5004 | NM_198398.1 | #N/A | 257.23704 | -9.217067 | 1.71106489 |
| 5005 | NR_036443.1 | HERC2P9 | 139.728603 | -8.3373727 | 1.54782401 |
| 5006 | NM_001025159.2 | CD74 | 263.499068 | -9.2525683 | 1.71777019 |
| 5007 | NM_001170689.1 | #N/A | 123.153512 | -8.1554099 | 1.51407537 |
| 5008 | NM_014905.4 | #N/A | 204.543034 | -8.8869411 | 1.64991636 |
| 5009 | NM_002945.3 | #N/A | 327.925817 | -6.1015798 | 1.13280576 |
| 5011 | NM_003955.4 | SOCS3 | 134.189477 | -8.2813599 | 1.53756971 |
| 5010 | NM_001037442.3 | #N/A | 114.416358 | -8.0514071 | 1.49486246 |
| 5012 | NM_005458.7 | #N/A | 331.106022 | -5.6529816 | 1.04956662 |
| 5013 | NM_001105530.1 | CAP1 | 233.797963 | -7.6517485 | 1.42076169 |
| 5014 | NM_170674.4 | #N/A | 239.165001 | -9.1130352 | 1.69211444 |
| 5015 | NM_024663.3 | #N/A | 109.400592 | -7.9849299 | 1.48267293 |
| 5017 | NM_030775.2 | WNT5B | 112.827713 | -8.0270164 | 1.49055533 |
| 5016 | NM_003337.3 | #N/A | 181.444104 | -8.7132871 | 1.6179821 |
| 5018 | NM_006939.3 | #N/A | 138.234527 | -8.3214636 | 1.54525619 |
| 5019 | NM_024015.4 | #N/A | 182.188461 | -8.7204093 | 1.61938333 |
| 5020 | NM_032286.2 | #N/A | 161.512211 | -8.5456332 | 1.58691725 |
| 5021 | NM_030819.3 | #N/A | 156.963584 | -8.504654 | 1.57943456 |
| 5022 | NM_001206426.1 | #N/A | 213.929902 | -8.9512492 | 1.66241601 |
| 5023 | NM_016589.3 | #N/A | 167.836285 | -7.3266453 | 1.36069814 |
| 5024 | NM_003904.4 | #N/A | 171.521814 | -8.6324982 | 1.6032502 |
| 5025 | NM_004483.4 | #N/A | 182.947408 | -8.7255648 | 1.62060773 |
| 5026 | NM_001321178.1 | FAM156B | 118.228465 | -6.8184466 | 1.26648107 |
| 5027 | NM_016045.2 | #N/A | 142.193247 | -8.3615377 | 1.55309095 |
| 5028 | NM_021727.4 | #N/A | 150.149595 | -8.4411826 | 1.56788465 |
| 5029 | NM_024523.5 | #N/A | 117.919444 | -8.0931089 | 1.50325373 |
| 5030 | NM_022830.2 | #N/A | 127.138862 | -8.20017 | 1.5232081 |

| | | | | | |
|---|---|---|---|---|---|
| 5031 | NM_006985.2 | #N/A | 142.89456 | -8.3691694 | 1.55474798 |
| 5032 | NM_022726.3 | #N/A | 193.469951 | -8.8062582 | 1.6359945 |
| 5033 | NM_018394.3 | #N/A | 137.954591 | -8.3175468 | 1.5452216 |
| 5034 | NM_014374.3 | #N/A | 138.854549 | -8.3298023 | 1.54756213 |
| 5035 | NR_024344.1 | MIR4697HG | 184.185492 | -8.7350488 | 1.62289436 |
| 5036 | NM_001145468.3 | SPECC1L | 239.60037 | -9.1154387 | 1.69359872 |
| 5038 | NM_002488.4 | NDUFA2 | 166.636933 | -8.5907111 | 1.59613264 |
| 5037 | NM_001079518.1 | #N/A | 120.428639 | -8.1228013 | 1.50920404 |
| 5039 | NM_005530.2 | #N/A | 135.153843 | -8.2881872 | 1.53998317 |
| 5040 | NM_018717.4 | #N/A | 147.131325 | -8.411912 | 1.56300878 |
| 5041 | NM_016207.3 | #N/A | 143.475633 | -8.3737806 | 1.55605089 |
| 5043 | NM_052834.2 | WDR7 | 143.80232 | -8.3792867 | 1.55712399 |
| 5042 | NM_018445.5 | #N/A | 144.368176 | -8.3842747 | 1.55805187 |
| 5044 | NM_199359.2 | #N/A | 139.164221 | -8.3311597 | 1.54817876 |
| 5045 | NM_014851.3 | #N/A | 122.021422 | -6.8135453 | 1.26622968 |
| 5046 | NM_001677.3 | #N/A | 584.921506 | -5.8177019 | 1.08120336 |
| 5047 | NM_001981.2 | #N/A | 118.031528 | -8.096207 | 1.50465028 |
| 5048 | NM_014804.2 | #N/A | 109.417901 | -7.9843053 | 1.48388837 |
| 5049 | NM_001375.2 | #N/A | 101.140463 | -3.60791 | 0.67057846 |
| 5050 | NM_004603.3 | #N/A | 208.721091 | -8.9158393 | 1.65719192 |
| 5051 | NM_020897.2 | #N/A | 201.853213 | -8.8676899 | 1.64824806 |
| 5052 | NM_003047.4 | #N/A | 128.462146 | -8.2164581 | 1.52723122 |
| 5053 | NM_004814.2 | #N/A | 145.019273 | -8.3901878 | 1.55952398 |
| 5054 | NM_001321694.1 | #N/A | 190.095289 | -8.7803743 | 1.63206479 |
| 5055 | NM_005667.3 | #N/A | 137.585036 | -8.3143008 | 1.54551571 |
| 5056 | NM_033414.2 | #N/A | 155.054738 | -8.4873463 | 1.57770234 |
| 5057 | NM_024077.4 | #N/A | 145.08802 | -8.3894646 | 1.55966654 |
| 5058 | NM_001005463.2 | #N/A | 126.881424 | -8.2001079 | 1.5245066 |
| 5059 | NM_007122.4 | #N/A | 220.054088 | -8.9917293 | 1.67167767 |
| 5061 | NM_053041.2 | COMMD7 | 162.959239 | -8.5590954 | 1.59127852 |
| 5060 | NM_016468.6 | #N/A | 164.606758 | -8.5728906 | 1.59384091 |
| 5062 | NM_030628.1 | #N/A | 123.504821 | -8.1594732 | 1.51716783 |
| 5063 | NM_030621.4 | DICER1 | 122.511141 | -8.1443814 | 1.51455543 |
| 5064 | NM_000609.6 | #N/A | 204.713598 | -8.8887645 | 1.65305033 |
| 5065 | NM_014367.3 | #N/A | 142.840721 | -8.3672148 | 1.55611095 |
| 5066 | NM_198560.2 | #N/A | 221.923257 | -9.0046491 | 1.6749014 |
| 5067 | NM_020840.2 | #N/A | 168.367076 | -8.6058013 | 1.60087006 |
| 5068 | NM_013437.4 | LRP12 | 128.799533 | -8.2200109 | 1.5294211 |
| 5069 | NM_001261434.1 | #N/A | 161.006713 | -8.5413462 | 1.58923484 |
| 5070 | NM_004642.3 | #N/A | 253.134326 | -9.194318 | 1.71083959 |
| 5071 | NM_175875.4 | #N/A | 121.384196 | -8.1364013 | 1.5140054 |
| 5073 | NM_173470.2 | MMGT1 | 171.29892 | -8.6307291 | 1.60607789 |
| 5072 | NM_002292.3 | #N/A | 878.010483 | -6.4583511 | 1.20182063 |

| | | | | | |
|---|---|---|---|---|---|
| 5074 | NM_033271.2 | #N/A | 203.695325 | -7.4718486 | 1.39046873 |
| 5076 | NM_001285878.1 | CEBPB | 118.414788 | -8.0957414 | 1.50663709 |
| 5075 | NM_001042371.2 | #N/A | 161.985791 | -7.2756706 | 1.35401786 |
| 5077 | NM_014408.4 | #N/A | 159.212268 | -8.5252004 | 1.586641 |
| 5078 | NM_001199839.1 | BCL2L2 | 132.550433 | -8.2609191 | 1.53749524 |
| 5079 | NM_021903.2 | GABBR1 | 202.876915 | -8.8741142 | 1.65164551 |
| 5080 | NM_152926.2 | #N/A | 181.816788 | -8.7186967 | 1.62275106 |
| 5081 | NM_014222.2 | #N/A | 163.827568 | -8.566492 | 1.59455596 |
| 5082 | NM_016494.3 | #N/A | 156.430915 | -8.4997492 | 1.58231856 |
| 5083 | NM_024923.3 | #N/A | 152.719328 | -7.1014498 | 1.32200444 |
| 5084 | NM_153232.3 | #N/A | 158.479035 | -8.5181712 | 1.58574485 |
| 5085 | NM_020850.2 | #N/A | 108.860207 | -7.9776101 | 1.48513666 |
| 5086 | NM_001308096.1 | GLYR1 | 150.147412 | -8.439031 | 1.57111163 |
| 5087 | NM_005163.2 | AKT1 | 138.558735 | -8.3228647 | 1.54950447 |
| 5088 | NM_030792.6 | #N/A | 189.56066 | -8.7772349 | 1.63411378 |
| 5089 | NM_032306.3 | #N/A | 173.967263 | -8.6526664 | 1.61097152 |
| 5090 | NM_001751.5 | #N/A | 166.704193 | -8.5918875 | 1.59983625 |
| 5091 | NM_001129994.1 | #N/A | 143.545528 | -8.3770963 | 1.55985726 |
| 5092 | NM_002747.3 | #N/A | 120.926074 | -8.1307838 | 1.51401509 |
| 5093 | NM_003291.3 | #N/A | 166.876092 | -7.2139041 | 1.34341466 |
| 5094 | NM_018457.3 | #N/A | 136.55676 | -8.3029001 | 1.54620489 |
| 5095 | NM_016143.4 | NSFL1C | 149.012191 | -8.4317324 | 1.57021721 |
| 5096 | NR_037608.1 | SLX1A-SULT1A3 | 162.901307 | -8.5576964 | 1.59375198 |
| 5097 | NM_006768.4 | #N/A | 130.922736 | -8.2420387 | 1.53498745 |
| 5098 | NM_020529.2 | NFKBIA | 109.319104 | -7.9842931 | 1.48713311 |
| 5099 | NM_001142416.1 | #N/A | 128.478548 | -8.2138619 | 1.52997909 |
| 5100 | NM_152278.3 | #N/A | 238.185014 | -9.106248 | 1.69620446 |
| 5102 | NR_033465.1 | NT5C3B | 177.426779 | -8.6813198 | 1.61713842 |
| 5101 | NM_007284.3 | #N/A | 152.840018 | -8.466029 | 1.57703506 |
| 5103 | NM_001002029.3 | C4B | 160.332205 | -8.5377192 | 1.59042377 |
| 5105 | NM_203331.2 | CD59 | 189.068943 | -8.7741911 | 1.63447795 |
| 5104 | NM_002999.3 | #N/A | 100.560798 | -4.593074 | 0.85560773 |
| 5106 | NM_005003.2 | #N/A | 199.460661 | -8.8500713 | 1.64884885 |
| 5107 | NM_000271.4 | #N/A | 138.808173 | -8.3275715 | 1.55157112 |
| 5109 | NM_001330618.1 | CYHR1 | 148.317329 | -8.4216499 | 1.56921951 |
| 5108 | NM_001163474.1 | ZNF746 | 122.66282 | -8.1461896 | 1.51791485 |
| 5110 | NM_014333.3 | #N/A | 275.482393 | -9.317274 | 1.73612054 |
| 5111 | NM_201521.2 | #N/A | 164.412234 | -8.5711666 | 1.59710898 |
| 5112 | NM_013396.4 | #N/A | 113.424797 | -8.0371008 | 1.49771752 |
| 5114 | NR_027982.1 | RASAL2-AS1 | 94.636337 | 4.23681437 | 0.78957048 |
| 5113 | NM_030576.3 | #N/A | 195.29364 | -8.8196047 | 1.64362579 |
| 5115 | NM_012425.3 | #N/A | 198.876886 | -8.845976 | 1.64861814 |
| 5116 | NM_001142685.1 | ARHGAP32 | 164.726485 | -8.575638 | 1.59825062 |

| | | | | | |
|---|---|---|---|---|---|
| 5117 | NM_005659.6 | #N/A | 151.743214 | -8.4562622 | 1.57609965 |
| 5118 | NM_015833.3 | #N/A | 137.635243 | -8.3174929 | 1.55026807 |
| 5119 | NM_001114382.2 | TSC2 | 267.118601 | -9.2715884 | 1.72824905 |
| 5120 | NM_194326.3 | #N/A | 160.407406 | -8.5362554 | 1.59117379 |
| 5121 | NM_005316.3 | #N/A | 125.776813 | -8.1858363 | 1.52596124 |
| 5122 | NM_025010.4 | #N/A | 142.313204 | -8.3639484 | 1.55919328 |
| 5123 | NM_003358.2 | #N/A | 201.256327 | -7.455474 | 1.38985194 |
| 5124 | NM_018171.3 | #N/A | 133.093933 | -8.2641779 | 1.54065957 |
| 5125 | NM_007368.3 | #N/A | 129.693206 | -8.2292557 | 1.53419565 |
| 5126 | NM_021640.3 | #N/A | 153.752688 | -8.4747108 | 1.57995105 |
| 5127 | NM_033540.2 | #N/A | 132.41358 | -8.2596202 | 1.53985786 |
| 5128 | NM_023072.2 | #N/A | 123.856189 | -8.1625624 | 1.52184264 |
| 5129 | NM_015534.5 | #N/A | 131.262739 | -8.2464548 | 1.53749707 |
| 5130 | NM_020243.4 | #N/A | 182.128397 | -8.7190484 | 1.62574758 |
| 5131 | NM_001282431.1 | #N/A | 129.763013 | -8.230723 | 1.53473386 |
| 5132 | NM_015533.3 | #N/A | 114.371584 | -8.0465318 | 1.50042457 |
| 5133 | NM_018144.3 | #N/A | 178.402477 | -8.6893172 | 1.62038772 |
| 5134 | NM_020546.2 | #N/A | 181.486497 | -6.1410964 | 1.14521057 |
| 5135 | NM_012208.3 | #N/A | 127.793197 | -8.2086949 | 1.53081495 |
| 5136 | NM_001967.3 | #N/A | 1840.7067 | -6.9259699 | 1.29167926 |
| 5137 | NM_018439.3 | #N/A | 118.252519 | -6.7688942 | 1.2624846 |
| 5138 | NM_005047.3 | #N/A | 117.093101 | -8.0825334 | 1.50760145 |
| 5139 | NM_020824.3 | #N/A | 789.396483 | -3.692964 | 0.68884098 |
| 5140 | NM_002982.3 | #N/A | 175.635906 | -8.6677874 | 1.61692182 |
| 5141 | NM_002027.2 | #N/A | 139.707584 | -8.3356726 | 1.55501048 |
| 5142 | NM_176869.2 | #N/A | 122.2607 | -8.1435389 | 1.51924161 |
| 5143 | NM_003462.3 | #N/A | 207.343395 | -8.9075205 | 1.66184713 |
| 5144 | NM_001130056.3 | #N/A | 150.077606 | -8.4401595 | 1.57469016 |
| 5145 | NM_001142334.1 | RBFOX1 | 235.057655 | -9.0878699 | 1.69563741 |
| 5146 | NM_006809.4 | #N/A | 115.19081 | -8.0604374 | 1.50393965 |
| 5147 | NM_015202.3 | #N/A | 128.624506 | -8.2191597 | 1.53357737 |
| 5148 | NM_021189.4 | #N/A | 260.500187 | -9.2362181 | 1.72337932 |
| 5149 | NR_038135.2 | MGAT4EP | 37.7416793 | 4.835678 | 0.90229843 |
| 5150 | NM_001304438.1 | #N/A | 227.688249 | -9.0414361 | 1.68707271 |
| 5151 | NM_016603.3 | FAM13B | 144.920422 | -8.3897116 | 1.56557915 |
| 5152 | NM_001145076.2 | #N/A | 170.104948 | -8.6208784 | 1.60880627 |
| 5153 | NM_004871.2 | #N/A | 165.275024 | -8.5780603 | 1.600806 |
| 5154 | NM_032116.4 | #N/A | 140.626584 | -8.3480747 | 1.5580288 |
| 5155 | NM_012097.3 | #N/A | 131.920986 | -8.2523736 | 1.54021221 |
| 5156 | NM_001919.3 | #N/A | 126.654615 | -8.1948047 | 1.5295497 |
| 5157 | NM_006233.4 | #N/A | 145.948276 | -8.3999811 | 1.56784717 |
| 5158 | NM_001755.2 | #N/A | 124.789895 | -8.1737352 | 1.52576111 |
| 5159 | NM_012105.4 | #N/A | 105.960773 | -7.937229 | 1.48162465 |

| | | | | | | |
|---|---|---|---|---|---|---|
| 5160 | NM_016613.6 | | #N/A | 186.217219 | -8.7519263 | 1.63371842 |
| 5161 | NM_080862.2 | | #N/A | 188.963929 | -8.7728142 | 1.63781678 |
| 5162 | NM_001195478.1 | TFG | | 135.556368 | -8.294661 | 1.54863237 |
| 5163 | NM_006898.4 | | #N/A | 212.976368 | -7.674453 | 1.43285578 |
| 5164 | NM_022340.3 | | #N/A | 117.582799 | -8.0873913 | 1.51000053 |
| 5166 | NM_007195.2 | POLI | | 125.447417 | -6.9017553 | 1.28870577 |
| 5165 | NM_001287135.1 | | #N/A | 118.845667 | -8.1037751 | 1.51314495 |
| 5167 | NM_001278203.1 | CLCC1 | | 43.009846 | 10.4171449 | 1.9451674 |
| 5168 | NM_014028.3 | | #N/A | 193.102257 | -6.7603291 | 1.26234159 |
| 5169 | NM_002949.3 | | #N/A | 160.840118 | -8.539642 | 1.59468673 |
| 5170 | NM_001271978.1 | | #N/A | 145.849386 | -8.3999218 | 1.56873011 |
| 5171 | NM_006330.3 | | #N/A | 127.568916 | -8.2057517 | 1.53249639 |
| 5172 | NM_001306144.1 | | #N/A | 130.251086 | -8.2351553 | 1.53806109 |
| 5173 | NM_007266.3 | | #N/A | 150.483905 | -8.4439611 | 1.57714194 |
| 5174 | NM_022495.5 | PCNX4 | | 135.235626 | -6.9415672 | 1.29660872 |
| 5175 | NM_024876.3 | | #N/A | 110.938219 | -8.0071998 | 1.49566126 |
| 5176 | NM_015420.6 | | #N/A | 153.304699 | -8.4701436 | 1.58219841 |
| 5177 | NM_022553.5 | | #N/A | 121.300468 | -8.1313337 | 1.51901106 |
| 5178 | NM_001322243.1 | | #N/A | 130.13094 | -8.2363868 | 1.53867613 |
| 5179 | NM_145804.2 | | #N/A | 125.260385 | -8.1783979 | 1.52792101 |
| 5180 | NM_173510.3 | | #N/A | 117.646349 | -8.0888688 | 1.51132306 |
| 5181 | NM_013321.3 | | #N/A | 125.966725 | -8.1883206 | 1.53003491 |
| 5182 | NM_001166412.1 | | #N/A | 135.838445 | -8.2988337 | 1.55074042 |
| 5184 | NM_001281501.1 | FADS2 | | 194.86473 | -8.8176933 | 1.64772755 |
| 5183 | NM_001024924.1 | | #N/A | 153.408252 | -8.4716181 | 1.58306784 |
| 5185 | NM_004281.3 | BAG3 | | 145.025929 | -7.031922 | 1.3140582 |
| 5186 | NM_018557.2 | | #N/A | 113.05361 | -8.0346516 | 1.50143045 |
| 5187 | NM_012474.4 | | #N/A | 183.320004 | -8.7281172 | 1.63112382 |
| 5188 | NM_021260.3 | | #N/A | 133.280664 | -8.2691361 | 1.54538966 |
| 5189 | NM_020418.3 | | #N/A | 162.017867 | -8.5508537 | 1.59811054 |
| 5190 | NM_000921.4 | PDE3A | | 189.239482 | -8.7750209 | 1.64014216 |
| 5191 | NM_002205.4 | | #N/A | 172.894255 | -8.6452853 | 1.6160751 |
| 5192 | NM_153757.3 | | #N/A | 193.857239 | -8.8091094 | 1.64676139 |
| 5193 | NM_006963.4 | | #N/A | 144.522414 | -8.3841677 | 1.56742171 |
| 5194 | NM_001320044.1 | | #N/A | 172.666446 | -8.6418728 | 1.61564517 |
| 5196 | NM_032382.4 | COG8 | | 117.885713 | -8.0931872 | 1.51308658 |
| 5195 | NM_014281.4 | | #N/A | 188.505415 | -8.7670417 | 1.63907531 |
| 5197 | NM_012158.2 | | #N/A | 124.330577 | -8.1685636 | 1.52725829 |
| 5198 | NM_000238.3 | | #N/A | 182.076915 | -8.7190157 | 1.6301971 |
| 5199 | NM_000633.2 | BCL2 | | 122.743922 | -8.1503754 | 1.52389126 |
| 5200 | NM_005102.2 | | #N/A | 119.215984 | -8.109286 | 1.51638061 |
| 5201 | NM_001048210.2 | CLCC1 | | 118.617288 | -8.0990139 | 1.51448528 |
| 5202 | NM_032342.2 | | #N/A | 185.554457 | -8.7461661 | 1.63561526 |

| | | | | | |
|---|---|---|---|---|---|
| 5203 | NM_138799.3 | #N/A | 159.138786 | -8.5241062 | 1.59413432 |
| 5204 | NM_016622.3 | #N/A | 145.401332 | -8.3940556 | 1.56987709 |
| 5205 | NM_203390.2 | #N/A | 118.757819 | -8.1008216 | 1.51508181 |
| 5206 | NM_003731.2 | #N/A | 166.728125 | -8.591764 | 1.60699575 |
| 5207 | NM_001326.2 | #N/A | 135.659176 | -8.2942091 | 1.55137586 |
| 5208 | NM_006218.3 | #N/A | 141.146333 | -8.3506354 | 1.56192942 |
| 5209 | NM_004041.4 | #N/A | 180.501001 | -8.7051907 | 1.62828484 |
| 5210 | NM_000421.3 | #N/A | 140.14535 | -8.3415509 | 1.56035899 |
| 5211 | NM_024047.4 | #N/A | 119.207086 | -8.1081672 | 1.51669897 |
| 5212 | NM_201379.2 | #N/A | 272.521047 | -9.3019036 | 1.74007212 |
| 5213 | NM_194278.3 | ELMSAN1 | 114.610432 | -8.0516016 | 1.50620693 |
| 5214 | NM_173216.2 | ST6GAL1 | 190.063681 | -8.7810143 | 1.64270324 |
| 5215 | NM_001321305.1 | #N/A | 117.619081 | -8.0919101 | 1.51386703 |
| 5216 | NM_152267.3 | #N/A | 121.558211 | -8.1364468 | 1.52228293 |
| 5217 | NM_213569.2 | NEBL | 111.03722 | -8.0066511 | 1.49801096 |
| 5218 | NM_001145135.1 | #N/A | 111.953733 | -8.0164106 | 1.49985871 |
| 5219 | NM_001098785.1 | #N/A | 254.197301 | -9.1999657 | 1.72132612 |
| 5220 | NM_003090.3 | #N/A | 135.838527 | -8.2955669 | 1.55220693 |
| 5221 | NM_024909.3 | #N/A | 266.93716 | -9.270588 | 1.73470046 |
| 5222 | NM_001002909.3 | #N/A | 181.50822 | -8.7140526 | 1.63059475 |
| 5224 | NM_212471.2 | PRKAR1A | 155.972404 | -8.4974805 | 1.59020298 |
| 5223 | NM_001320851.1 | #N/A | 109.548421 | -7.9849572 | 1.4942941 |
| 5225 | NM_032431.2 | #N/A | 124.549681 | -8.1715494 | 1.52932807 |
| 5226 | NM_018244.4 | #N/A | 126.891499 | -8.1979924 | 1.53433306 |
| 5227 | NM_000263.3 | #N/A | 131.019814 | -8.2451503 | 1.54328279 |
| 5228 | NR_024384.1 | #N/A | 167.764721 | -8.5999626 | 1.60970417 |
| 5229 | NM_005900.2 | #N/A | 168.144833 | -8.6036945 | 1.61055696 |
| 5230 | NM_001077415.2 | #N/A | 156.345396 | -8.4987469 | 1.5909508 |
| 5231 | NM_001035507.2 | #N/A | 123.710139 | -8.1633556 | 1.52820271 |
| 5232 | NM_006378.3 | #N/A | 169.326875 | -8.6137495 | 1.6125746 |
| 5233 | NM_020416.3 | #N/A | 199.615996 | -8.8510762 | 1.65701988 |
| 5234 | NM_031209.2 | #N/A | 152.145468 | -8.4594122 | 1.5838165 |
| 5235 | NM_001866.2 | #N/A | 188.966289 | -8.7727358 | 1.64256155 |
| 5236 | NM_001003801.1 | #N/A | 157.313173 | -8.5064263 | 1.59272958 |
| 5237 | NM_022092.2 | #N/A | 151.761551 | -7.0872983 | 1.32704596 |
| 5238 | NM_005002.4 | #N/A | 167.116752 | -8.5954864 | 1.60957368 |
| 5239 | NM_002513.2 | NME3 | 128.764354 | -8.2177906 | 1.53891574 |
| 5240 | NM_153832.2 | GPR161 | 135.986739 | -8.2958874 | 1.55355256 |
| 5241 | NM_012304.4 | #N/A | 104.125525 | -7.9128959 | 1.48194336 |
| 5242 | NM_022730.3 | #N/A | 155.936514 | -8.4949619 | 1.59096567 |
| 5243 | NM_001042498.2 | #N/A | 108.381784 | -7.9718767 | 1.4930825 |
| 5244 | NM_015484.4 | #N/A | 126.222663 | -8.1895534 | 1.53389409 |
| 5245 | NR_028389.1 | ACVR2B-AS1 | 69.79264 | 3.41937899 | 0.64046393 |

| | | | | | | |
|---|---|---|---|---|---|---|
| 5246 | NM_130434.4 | | #N/A | 123.726432 | -8.1624135 | 1.52893592 |
| 5247 | NM_004541.3 | | #N/A | 155.573404 | -8.492053 | 1.59071198 |
| 5248 | NM_016059.4 | | #N/A | 175.139348 | -8.6627243 | 1.6226729 |
| 5249 | NM_001003712.1 | | #N/A | 143.329374 | -8.3750343 | 1.56881534 |
| 5250 | NM_130475.2 | | #N/A | 203.09569 | -8.8764212 | 1.66279907 |
| 5251 | NM_007286.5 | | #N/A | 119.842038 | -5.9211162 | 1.10922546 |
| 5252 | NM_001142353.2 | PPP3CB | | 221.85576 | -9.0034238 | 1.68679616 |
| 5253 | NM_002547.2 | | #N/A | 789.259215 | 2.19341442 | 0.41094483 |
| 5254 | NM_001080395.2 | | #N/A | 128.060628 | -8.2102583 | 1.53825035 |
| 5255 | NM_001130921.1 | | #N/A | 142.77429 | -8.3696675 | 1.56811055 |
| 5256 | NM_001124758.2 | | #N/A | 151.135387 | -8.4497036 | 1.58322585 |
| 5257 | NM_012143.3 | | #N/A | 144.377376 | -8.3826138 | 1.57065029 |
| 5258 | NM_014648.3 | | #N/A | 289.717908 | -2.5610308 | 0.47985842 |
| 5259 | NM_025074.6 | | #N/A | 144.521857 | -8.3870786 | 1.57151993 |
| 5260 | NM_001286577.1 | C2CD3 | | 110.392907 | -7.9988515 | 1.49878937 |
| 5261 | NM_148977.2 | PANK1 | | 108.099834 | -7.9638405 | 1.49223792 |
| 5262 | NM_003491.3 | | #N/A | 170.672289 | -8.6258592 | 1.61635735 |
| 5263 | NM_153367.3 | | #N/A | 130.358117 | -6.9621535 | 1.30464751 |
| 5264 | NM_001035235.3 | SRA1 | | 126.069852 | -8.1881071 | 1.53442388 |
| 5265 | NM_003760.4 | EIF4G3 | | 177.385215 | -8.6801221 | 1.62668532 |
| 5266 | NM_138462.2 | | #N/A | 170.387283 | -8.6227199 | 1.61595566 |
| 5267 | NM_181491.2 | MED22 | | 119.100306 | -8.1076272 | 1.51945923 |
| 5268 | NM_152527.4 | | #N/A | 169.819864 | -8.6175757 | 1.61508173 |
| 5269 | NM_005327.4 | | #N/A | 132.794589 | -8.2635668 | 1.54876095 |
| 5270 | NM_018094.4 | | #N/A | 114.697958 | -8.051257 | 1.50899513 |
| 5271 | NR_038863.2 | MROCKI | | 63.0521557 | 5.36758979 | 1.00610619 |
| 5272 | NM_001142421.1 | MORF4L2 | | 133.062381 | -8.2681966 | 1.54988492 |
| 5273 | NM_001168241.1 | | #N/A | 218.506868 | -8.9820223 | 1.68368317 |
| 5274 | NM_015852.3 | | #N/A | 83.1609646 | -4.5278606 | 0.84875917 |
| 5275 | NM_032227.3 | | #N/A | 121.732262 | -8.1390259 | 1.52568879 |
| 5276 | NM_139126.3 | | #N/A | 114.683809 | -8.0517679 | 1.50935673 |
| 5277 | NM_001300959.1 | ZMIZ2 | | 130.764058 | -8.2399291 | 1.54470756 |
| 5278 | NM_153633.2 | | #N/A | 192.135841 | -8.7968829 | 1.6492408 |
| 5279 | NM_030593.2 | | #N/A | 150.719929 | -8.4467596 | 1.58361596 |
| 5280 | NM_001347.3 | | #N/A | 186.037518 | -7.3491042 | 1.37786406 |
| 5281 | NM_182487.3 | | #N/A | 118.930076 | -8.1071566 | 1.51999192 |
| 5282 | NM_024544.2 | | #N/A | 138.037452 | -8.320507 | 1.56038921 |
| 5283 | NM_015099.3 | | #N/A | 155.980589 | -8.4957378 | 1.59335392 |
| 5284 | NM_001615.3 | | #N/A | 219.051083 | -8.9859849 | 1.68531439 |
| 5285 | NM_020841.4 | | #N/A | 238.536643 | -9.1078058 | 1.70820418 |
| 5286 | NM_001085466.1 | | #N/A | 130.884944 | -8.2428727 | 1.54600205 |
| 5287 | NM_152795.3 | | #N/A | 125.677131 | -8.1835434 | 1.53515353 |
| 5288 | NM_002196.2 | | #N/A | 146.293746 | -8.4007359 | 1.57598333 |

| | | | | | |
|---|---|---|---|---|---|
| 5289 | NM_174938.5 | #N/A | 177.783501 | -8.6845293 | 1.62928178 |
| 5290 | NM_005884.3 | #N/A | 107.085355 | -7.9523504 | 1.49198258 |
| 5291 | NM_001321431.1 | FBN3 | 163.211575 | -8.5604288 | 1.60620946 |
| 5292 | NM_007254.3 | #N/A | 120.726156 | -8.1266253 | 1.52481507 |
| 5293 | NM_006033.3 | #N/A | 132.985487 | -8.2653833 | 1.55087533 |
| 5294 | NM_015884.3 | #N/A | 149.910955 | -8.4386884 | 1.58346397 |
| 5296 | NM_001320714.1 | DOP1B | 220.476107 | -8.9941997 | 1.68783607 |
| 5295 | NM_001282507.1 | NPIPA7 | 161.985372 | -8.5486389 | 1.60422002 |
| 5297 | NM_001037163.1 | #N/A | 161.609864 | -8.5464019 | 1.6039354 |
| 5298 | NM_052937.3 | #N/A | 109.318384 | -7.9835058 | 1.49835679 |
| 5299 | NM_006234.4 | #N/A | 143.963089 | -8.3794822 | 1.57269581 |
| 5300 | NM_033018.3 | #N/A | 122.524233 | -8.1479048 | 1.52943466 |
| 5301 | NM_006540.3 | #N/A | 124.713612 | -8.1709502 | 1.53378137 |
| 5302 | NM_032508.3 | #N/A | 135.943933 | -8.2958469 | 1.55722433 |
| 5303 | NM_030974.3 | #N/A | 138.695123 | -8.3257948 | 1.56290599 |
| 5304 | NM_000199.3 | #N/A | 122.160092 | -8.144452 | 1.52888037 |
| 5305 | NM_014940.3 | #N/A | 120.716901 | -8.1266051 | 1.52580865 |
| 5306 | NR_046593.1 | LINC01840 | 203.798599 | 3.14370817 | 0.59025015 |
| 5307 | NM_005586.3 | #N/A | 150.036491 | -8.4392187 | 1.5845946 |
| 5308 | NM_014572.2 | #N/A | 119.981372 | -8.1196689 | 1.52463625 |
| 5309 | NM_182715.2 | #N/A | 117.920663 | -8.0935056 | 1.51972731 |
| 5310 | NM_017412.3 | #N/A | 243.783705 | -9.1390806 | 1.71608734 |
| 5311 | NM_001100427.1 | #N/A | 141.47753 | -8.353292 | 1.56860102 |
| 5312 | NM_025224.3 | #N/A | 128.8923 | -8.2219309 | 1.54405131 |
| 5313 | NM_002230.2 | #N/A | 161.767111 | -8.5486135 | 1.60543099 |
| 5314 | NM_025233.6 | #N/A | 130.457696 | -8.2366388 | 1.54711133 |
| 5315 | NM_015143.2 | #N/A | 154.197706 | -8.4785095 | 1.59279312 |
| 5316 | NM_001031623.2 | #N/A | 108.147846 | -7.9677568 | 1.49714328 |
| 5317 | NM_001304808.1 | #N/A | 111.182241 | -8.0066253 | 1.50458588 |
| 5318 | NM_001240.3 | #N/A | 105.167278 | -7.9250811 | 1.48928177 |
| 5319 | NM_001199302.1 | CNOT2 | 185.852537 | -6.7031881 | 1.25971945 |
| 5320 | NM_006432.3 | #N/A | 265.455568 | -4.4799147 | 0.84198858 |
| 5321 | NM_003896.3 | #N/A | 157.880513 | -8.5138236 | 1.60018933 |
| 5322 | NM_022752.5 | #N/A | 128.901387 | -8.2208141 | 1.54516225 |
| 5323 | NM_001042414.2 | #N/A | 158.255276 | -8.5157378 | 1.60063461 |
| 5324 | NM_004111.5 | #N/A | 127.186165 | -8.2008205 | 1.54145994 |
| 5325 | NR_103804.1 | #N/A | 125.312297 | -8.1831112 | 1.53831336 |
| 5326 | NM_032515.4 | #N/A | 132.25856 | -8.2583524 | 1.5524993 |
| 5327 | NR_033227.1 | #N/A | 143.616986 | -8.3755451 | 1.57466503 |
| 5328 | NM_006675.4 | #N/A | 187.623421 | -8.7638312 | 1.64767683 |
| 5329 | NM_000031.5 | #N/A | 127.528907 | -8.2033281 | 1.5425307 |
| 5330 | NM_153827.4 | #N/A | 112.371087 | -8.0260812 | 1.50921989 |
| 5331 | NM_000819.4 | #N/A | 160.685355 | -8.5390432 | 1.60575422 |

| | | | | | |
|---|---|---|---|---|---|
| 5332 | NM_152341.4 | #N/A | 129.34228 | -8.2247627 | 1.54666812 |
| 5333 | NM_017945.3 | #N/A | 155.308456 | -8.4893205 | 1.59648441 |
| 5334 | NM_001171617.1 | PREPL | 119.574536 | -8.1153712 | 1.52626979 |
| 5335 | NM_014963.2 | #N/A | 114.167258 | -8.0442697 | 1.51296594 |
| 5336 | NM_001271044.2 | #N/A | 263.463782 | -9.2520039 | 1.74013132 |
| 5337 | NM_024792.2 | #N/A | 109.869478 | -7.9896865 | 1.50273855 |
| 5338 | NM_001025105.2 | #N/A | 113.87829 | -8.0388425 | 1.51214357 |
| 5339 | NM_001289022.1 | #N/A | 156.389194 | -8.4994552 | 1.59880369 |
| 5340 | NM_004492.2 | #N/A | 156.938217 | -8.5041254 | 1.59977483 |
| 5341 | NM_018357.3 | #N/A | 132.51612 | -8.2606585 | 1.55399255 |
| 5342 | NM_001321166.1 | #N/A | 103.301075 | -7.9014761 | 1.48649008 |
| 5343 | NM_024102.3 | #N/A | 150.564447 | -8.4450163 | 1.58874161 |
| 5344 | NM_014846.3 | #N/A | 140.930597 | -8.3492895 | 1.5707485 |
| 5345 | NM_138499.3 | #N/A | 104.333808 | -7.9172317 | 1.48951577 |
| 5346 | NM_001130414.1 | APBA2 | 138.854831 | -8.3292913 | 1.56707183 |
| 5347 | NM_001011713.2 | #N/A | 136.201369 | -8.2991104 | 1.5614643 |
| 5348 | NM_013386.4 | #N/A | 120.765478 | -8.1263147 | 1.52905755 |
| 5349 | NM_022488.4 | #N/A | 149.942398 | -8.4386317 | 1.58796674 |
| 5350 | NM_001114617.1 | MGAT1 | 134.844841 | -8.2871779 | 1.55950182 |
| 5351 | NM_001277062.1 | #N/A | 149.448829 | -8.4325784 | 1.58686634 |
| 5352 | NM_015316.2 | #N/A | 149.945226 | -8.4390484 | 1.58812417 |
| 5353 | NM_183415.2 | #N/A | 109.821244 | -7.9870149 | 1.50313043 |
| 5354 | NM_014434.3 | NDOR1 | 136.210221 | -8.3026386 | 1.56300566 |
| 5355 | NM_020245.4 | #N/A | 157.284467 | -8.5081668 | 1.60170916 |
| 5356 | NM_001017371.4 | #N/A | 140.20088 | -8.3409919 | 1.57024929 |
| 5357 | NM_001040431.2 | #N/A | 162.058644 | -8.5507593 | 1.61000789 |
| 5358 | NM_018433.5 | #N/A | 114.325262 | -8.0445433 | 1.51476162 |
| 5359 | NM_003683.5 | #N/A | 133.098404 | -8.2669985 | 1.55677278 |
| 5360 | NM_001121.3 | #N/A | 116.216935 | -8.0745895 | 1.52055219 |
| 5361 | NM_024812.2 | #N/A | 160.972816 | -8.5408204 | 1.60839509 |
| 5362 | NM_031215.2 | CABLES2 | 128.462907 | -8.2148084 | 1.54722057 |
| 5363 | NM_183357.2 | ADCY5 | 197.602894 | -8.8360546 | 1.66425031 |
| 5364 | NM_014886.4 | #N/A | 185.477025 | -8.7453877 | 1.64720553 |
| 5365 | NM_001318942.1 | MEA1 | 142.17302 | -8.3619719 | 1.57501678 |
| 5366 | NM_001346113.1 | #N/A | 32.3956901 | 9.98435083 | 1.88074373 |
| 5367 | NM_003111.4 | SP3 | 121.115504 | -8.1281906 | 1.53111409 |
| 5368 | NM_006118.3 | #N/A | 157.70694 | -8.5109902 | 1.60324889 |
| 5369 | NM_138761.3 | #N/A | 143.315641 | -8.3737115 | 1.57751153 |
| 5370 | NM_001321365.1 | #N/A | 149.492599 | -8.4330187 | 1.5887362 |
| 5371 | NM_016097.4 | #N/A | 149.257281 | -8.4313346 | 1.58840971 |
| 5372 | NM_014142.3 | #N/A | 121.087112 | -8.1293692 | 1.53156164 |
| 5373 | NM_031454.1 | #N/A | 139.32166 | -8.3320404 | 1.56982151 |
| 5374 | NM_001009813.2 | #N/A | 208.193179 | -8.9119608 | 1.67912914 |

| | | | | | | |
|---|---|---|---|---|---|---|
| 5375 | NM_173542.3 | | #N/A | 171.540179 | -8.6331036 | 1.62659372 |
| 5376 | NM_001077183.2 | TSC2 | | 155.659912 | -8.4915561 | 1.60002307 |
| 5377 | NM_172353.2 | | #N/A | 115.71488 | -8.0636625 | 1.51946698 |
| 5378 | NM_018695.3 | | #N/A | 126.53655 | -8.1913782 | 1.54357792 |
| 5379 | NM_001258437.1 | TARS1 | | 132.729733 | -8.2608737 | 1.55673171 |
| 5380 | NM_004582.3 | | #N/A | 142.108454 | -8.3614686 | 1.57597389 |
| 5381 | NM_000757.5 | | #N/A | 110.836692 | -8.0058411 | 1.50902413 |
| 5382 | NM_001453.2 | | #N/A | 142.529362 | -8.3679551 | 1.57732699 |
| 5383 | NM_005860.2 | | #N/A | 135.097556 | -8.2892498 | 1.56250998 |
| 5384 | NM_017971.3 | | #N/A | 158.847801 | -8.5217283 | 1.60636312 |
| 5385 | NM_006938.3 | | #N/A | 187.600142 | -8.7614134 | 1.65162513 |
| 5386 | NM_002056.3 | | #N/A | 604.456729 | -2.0548629 | 0.38737473 |
| 5387 | NM_012227.3 | GTPBP6 | | 160.367236 | -8.5352137 | 1.60904434 |
| 5388 | NM_001282149.1 | | #N/A | 273.958079 | -9.3074242 | 1.75487343 |
| 5389 | NM_001040402.2 | | #N/A | 140.606882 | -8.343719 | 1.5732856 |
| 5390 | NM_004457.3 | | #N/A | 153.792648 | -7.2001555 | 1.35776971 |
| 5391 | NM_001323964.1 | TIAL1 | | 100.430251 | -7.8601779 | 1.48247843 |
| 5392 | NM_014415.3 | | #N/A | 115.061257 | -8.0562695 | 1.51950751 |
| 5393 | NM_000374.4 | | #N/A | 147.433912 | -8.4145276 | 1.58715387 |
| 5394 | NM_002137.3 | | #N/A | 2633.83921 | -5.7999331 | 1.09398799 |
| 5395 | NM_177441.2 | | #N/A | 127.245642 | -8.2022155 | 1.54720308 |
| 5396 | NM_005903.6 | | #N/A | 149.791492 | -8.4378968 | 1.5916801 |
| 5397 | NM_001347217.1 | | #N/A | 167.149971 | -8.5951272 | 1.62141924 |
| 5398 | NM_017660.3 | | #N/A | 132.383141 | -8.256922 | 1.55767216 |
| 5399 | NM_138793.3 | | #N/A | 133.491863 | -8.2701459 | 1.56028486 |
| 5400 | NM_001144382.1 | | #N/A | 123.767097 | -8.1599834 | 1.53953925 |
| 5401 | NM_052947.3 | | #N/A | 216.965045 | -8.9722904 | 1.69281132 |
| 5402 | NM_001003703.1 | ATP5PF | | 135.828196 | -8.2959629 | 1.56537127 |
| 5403 | NM_001040661.1 | | #N/A | 126.006445 | -8.1874279 | 1.54490063 |
| 5404 | NM_001164817.1 | | #N/A | 156.878954 | -8.5018246 | 1.60434541 |
| 5405 | NM_001256864.1 | | #N/A | 187.604324 | -8.7635756 | 1.65373648 |
| 5406 | NM_006384.3 | | #N/A | 110.160729 | -7.9963974 | 1.5089608 |
| 5407 | NM_003314.2 | | #N/A | 139.446891 | -8.3345504 | 1.57303761 |
| 5408 | NM_007056.2 | | #N/A | 103.965356 | -7.9077543 | 1.49281511 |
| 5409 | NM_001301073.1 | | #N/A | 133.391732 | -8.2700971 | 1.56148686 |
| 5410 | NM_018266.2 | | #N/A | 102.361111 | -7.8906208 | 1.48998875 |
| 5411 | NM_001099272.1 | | #N/A | 106.817934 | -7.9482625 | 1.50099596 |
| 5412 | NR_003038.2 | SNHG5 | | 186.09566 | -8.7503484 | 1.65252088 |
| 5413 | NR_138033.1 | EMC2 | | 32.1919694 | 9.99960499 | 1.88849227 |
| 5414 | NM_024113.4 | | #N/A | 139.519646 | -8.3347839 | 1.57414068 |
| 5415 | NM_002063.3 | | #N/A | 217.990137 | -8.9787294 | 1.69586879 |
| 5416 | NM_015603.2 | | #N/A | 102.537479 | -7.8929655 | 1.49085688 |
| 5417 | NM_001098204.1 | HNRNPF | | 159.989193 | -8.5319788 | 1.61164783 |

| | | | | | |
|---|---|---|---|---|---|
| 5418 | NM_012200.3 | #N/A | 153.631703 | -8.4732124 | 1.60054812 |
| 5419 | NM_152857.2 | WTAP | 132.845254 | -8.2637929 | 1.56103076 |
| 5420 | NR_027700.2 | #N/A | 118.236077 | -8.0975491 | 1.52965935 |
| 5421 | NM_206900.2 | #N/A | 223.795794 | -9.0162704 | 1.70324696 |
| 5422 | NM_001142726.1 | C1orf122 | 155.950739 | -8.4951796 | 1.60484739 |
| 5423 | NR_125376.1 | LOC100419583 | 140.070526 | -8.3409622 | 1.57571088 |
| 5424 | NM_001042462.1 | #N/A | 117.243051 | -8.0830244 | 1.5270417 |
| 5425 | NM_033315.3 | #N/A | 147.046187 | -8.4109568 | 1.58908089 |
| 5426 | NM_016505.3 | ZCCHC17 | 148.342081 | -8.4227961 | 1.59133542 |
| 5427 | NR_109948.1 | LOC101929645 | 42.8597994 | 3.25405124 | 0.61481742 |
| 5428 | NM_005239.5 | #N/A | 150.674779 | -7.1726469 | 1.35529745 |
| 5429 | NM_016733.2 | #N/A | 173.710492 | -8.6519239 | 1.63481747 |
| 5431 | NM_020718.3 | USP31 | 126.805721 | -8.1957859 | 1.54886356 |
| 5430 | NM_001127229.1 | #N/A | 123.138691 | -8.1543122 | 1.54102795 |
| 5432 | NM_020141.3 | #N/A | 125.688849 | -8.18276 | 1.54641699 |
| 5433 | NM_032360.3 | #N/A | 205.546792 | -7.4708716 | 1.4118927 |
| 5434 | NR_038241.1 | VEZT | 118.970732 | -8.101604 | 1.53119475 |
| 5436 | NR_024417.1 | LINC01634 | 46.4296012 | 2.84677061 | 0.53804717 |
| 5435 | NM_018238.3 | #N/A | 128.468505 | -8.2154689 | 1.55274775 |
| 5437 | NM_005190.3 | #N/A | 132.603898 | -8.2602359 | 1.56128754 |
| 5438 | NM_023934.3 | #N/A | 175.776944 | -8.6676808 | 1.63830717 |
| 5439 | NM_001130158.2 | #N/A | 159.262399 | -8.5253742 | 1.61153491 |
| 5440 | NM_052848.2 | #N/A | 122.023043 | -8.1419269 | 1.5391539 |
| 5441 | NM_016573.3 | #N/A | 135.651393 | -8.2939727 | 1.56799112 |
| 5442 | NM_017875.2 | #N/A | 126.874411 | -8.197262 | 1.54972183 |
| 5443 | NM_138428.5 | #N/A | 117.999402 | -6.0164243 | 1.13743232 |
| 5444 | NM_022456.4 | #N/A | 111.144342 | -8.0058705 | 1.51359269 |
| 5445 | NM_017594.4 | #N/A | 178.850203 | -8.6931379 | 1.64374545 |
| 5446 | NM_001517.4 | #N/A | 132.255423 | -8.257879 | 1.56148957 |
| 5447 | NM_001244014.1 | #N/A | 153.142479 | -8.4704927 | 1.60175262 |
| 5448 | NM_001039703.5 | #N/A | 145.817954 | -8.396813 | 1.58784778 |
| 5449 | NM_014173.3 | #N/A | 154.951705 | -8.4870559 | 1.60517614 |
| 5450 | NM_014206.3 | #N/A | 162.307099 | -8.5532752 | 1.61772133 |
| 5451 | NM_001079521.1 | #N/A | 110.627014 | -7.9971781 | 1.51259003 |
| 5452 | NM_031903.2 | #N/A | 129.169528 | -8.2229917 | 1.55537451 |
| 5453 | NM_001168243.1 | C4orf48 | 244.483804 | -9.1438705 | 1.72966485 |
| 5456 | NR_134301.1 | IDI1 | 194.230565 | -8.8119975 | 1.66689299 |
| 5454 | NM_019070.4 | #N/A | 167.554737 | -8.5992775 | 1.62664322 |
| 5455 | NM_139280.3 | #N/A | 123.180558 | -8.1555021 | 1.54269502 |
| 5457 | NM_080607.2 | #N/A | 417.904221 | -7.0992648 | 1.34301683 |
| 5458 | NM_001880.3 | #N/A | 188.306767 | -8.7664562 | 1.65844345 |
| 5459 | NM_014251.2 | #N/A | 108.725563 | -7.9782416 | 1.50941546 |
| 5460 | NM_134442.4 | #N/A | 132.616067 | -8.2596692 | 1.56265771 |

| | | | | | |
|---|---|---|---|---|---|
| 5461 | NM_006129.4 | #N/A | 156.821277 | -8.5040784 | 1.60894228 |
| 5462 | NM_033138.3 | #N/A | 129.946302 | -8.2344388 | 1.55791886 |
| 5463 | NM_001277335.1 | #N/A | 131.933478 | -8.2520922 | 1.56134672 |
| 5464 | NM_004906.4 | #N/A | 162.75874 | -8.555901 | 1.61890114 |
| 5465 | NM_003355.2 | UCP2 | 111.053084 | -8.0053954 | 1.51475454 |
| 5466 | NM_001319943.1 | PAM | 114.116325 | -8.0453343 | 1.52235282 |
| 5467 | NM_006400.4 | DCTN2 | 210.581884 | -8.9283561 | 1.68946168 |
| 5468 | NM_012461.2 | #N/A | 130.201837 | -8.2351969 | 1.55831321 |
| 5469 | NM_005504.6 | #N/A | 190.674521 | -8.7848382 | 1.66257342 |
| 5470 | NM_016096.3 | #N/A | 153.883473 | -8.4757108 | 1.60412715 |
| 5471 | NM_001278579.1 | #N/A | 170.029315 | -8.6206322 | 1.63182798 |
| 5472 | NM_022493.2 | #N/A | 132.098743 | -8.2556746 | 1.56284675 |
| 5473 | NR_036605.1 | #N/A | 126.949034 | -8.198907 | 1.55210337 |
| 5474 | NM_023928.4 | #N/A | 107.487507 | -7.9568529 | 1.50638959 |
| 5475 | NM_175876.4 | #N/A | 103.238932 | -7.9002426 | 1.49566618 |
| 5476 | NM_033278.3 | #N/A | 158.022676 | -8.5140859 | 1.6119131 |
| 5477 | NM_015528.2 | #N/A | 122.600978 | -8.1459146 | 1.54224736 |
| 5478 | NM_025092.4 | #N/A | 119.151187 | -8.1105841 | 1.53561981 |
| 5479 | NM_001199140.1 | AMMECR1L | 130.249923 | -8.2374022 | 1.55966733 |
| 5481 | NM_198389.2 | PDPN | 186.372046 | -8.7534717 | 1.65737805 |
| 5480 | NM_016033.2 | #N/A | 115.712695 | -8.0646672 | 1.52696515 |
| 5482 | NM_015117.2 | #N/A | 127.884379 | -8.2086807 | 1.55426649 |
| 5483 | NM_013293.4 | #N/A | 154.234038 | -8.478696 | 1.60543598 |
| 5484 | NM_032457.3 | #N/A | 125.051388 | -8.17744 | 1.54838838 |
| 5485 | NM_001173463.1 | #N/A | 183.121873 | -8.72554 | 1.65222858 |
| 5486 | NM_001039690.3 | #N/A | 123.044316 | -8.1517832 | 1.54369264 |
| 5487 | NM_000416.2 | IFNGR1 | 121.520502 | -8.1365819 | 1.5412006 |
| 5488 | NM_002647.3 | #N/A | 105.605484 | -7.9325967 | 1.50292351 |
| 5489 | NM_015000.3 | #N/A | 134.188951 | -6.9196346 | 1.31103591 |
| 5490 | NM_001164094.1 | #N/A | 131.272013 | -8.2466409 | 1.56262863 |
| 5491 | NM_014961.4 | #N/A | 268.46483 | -9.2787416 | 1.75819558 |
| 5492 | NM_000016.5 | ACADM | 117.465165 | -8.0893206 | 1.53303667 |
| 5493 | NM_022762.4 | #N/A | 132.137694 | -8.2562374 | 1.56468995 |
| 5494 | NM_207299.1 | #N/A | 226.426616 | -9.0331869 | 1.71192616 |
| 5495 | NM_001278208.1 | DMC1 | 80.7987425 | 3.92435414 | 0.74375528 |
| 5496 | NM_005702.3 | #N/A | 156.551776 | -8.5007793 | 1.61109095 |
| 5497 | NM_004401.2 | #N/A | 108.096926 | -7.968725 | 1.51028118 |
| 5498 | NM_198525.2 | #N/A | 118.229566 | -8.0956929 | 1.53435549 |
| 5499 | NM_014377.1 | #N/A | 118.428477 | -8.0980344 | 1.53484469 |
| 5500 | NM_021178.4 | #N/A | 153.125421 | -8.4686794 | 1.6051591 |
| 5501 | NM_153321.2 | #N/A | 198.268978 | -8.8421627 | 1.67597516 |
| 5502 | NM_001145845.1 | ROBO1 | 116.206654 | -8.0678496 | 1.52949731 |
| 5503 | NM_024831.7 | #N/A | 110.817803 | -8.0021914 | 1.51708185 |

| | | | | | |
|---|---|---|---|---|---|
| 5504 | NM_020932.2 | #N/A | 222.048484 | -7.5676283 | 1.43479099 |
| 5505 | NM_001320546.1 | #N/A | 122.122912 | -8.1416666 | 1.54368203 |
| 5506 | NM_001127649.2 | #N/A | 152.456276 | -8.4635838 | 1.60490669 |
| 5507 | NM_016286.3 | #N/A | 160.578917 | -8.5373498 | 1.61891173 |
| 5508 | NM_153326.2 | #N/A | 118.492791 | -8.1016887 | 1.53640379 |
| 5509 | NM_001321324.1 | MOV10 | 123.524366 | -8.1620312 | 1.54788935 |
| 5510 | NM_032014.2 | #N/A | 160.722054 | -8.5386455 | 1.61938677 |
| 5511 | NM_183394.2 | #N/A | 143.145619 | -8.371884 | 1.58777189 |
| 5512 | NM_001301191.1 | #N/A | 201.318939 | -8.8632472 | 1.68097772 |
| 5513 | NM_003214.3 | #N/A | 122.43106 | -8.1481979 | 1.54553577 |
| 5514 | NM_002294.2 | #N/A | 117.851812 | -8.0938445 | 1.53524894 |
| 5515 | NM_001217.4 | #N/A | 140.497515 | -8.3439346 | 1.58281681 |
| 5516 | NM_018364.4 | RSBN1 | 151.108845 | -8.4490162 | 1.60283423 |
| 5517 | NM_170692.2 | #N/A | 112.972814 | -8.0328121 | 1.52387177 |
| 5518 | NM_021132.3 | #N/A | 194.563729 | -8.8144449 | 1.67219066 |
| 5519 | NM_005937.3 | #N/A | 459.123516 | -3.7359085 | 0.70877445 |
| 5520 | NM_022121.4 | #N/A | 112.218362 | -8.0239547 | 1.52241657 |
| 5521 | NM_001635.3 | #N/A | 149.668468 | -8.4358055 | 1.60071176 |
| 5522 | NM_001031685.2 | #N/A | 150.205471 | -8.4396537 | 1.60153352 |
| 5523 | NM_201998.2 | #N/A | 105.209052 | -7.9283499 | 1.50450322 |
| 5524 | NM_018092.4 | #N/A | 231.061209 | -9.0625166 | 1.71974398 |
| 5525 | NM_020921.3 | NIN | 146.48009 | -8.4043947 | 1.59491341 |
| 5526 | NM_001330211.1 | #N/A | 128.900581 | -8.2200788 | 1.55998775 |
| 5527 | NM_020383.3 | #N/A | 182.664504 | -8.7227786 | 1.65548228 |
| 5528 | NM_001282730.1 | SUPT7L | 111.802636 | -8.0146916 | 1.52113127 |
| 5529 | NR_015353.1 | #N/A | 104.79476 | -7.9220141 | 1.50364839 |
| 5530 | NM_016047.3 | #N/A | 159.396952 | -8.5265246 | 1.61844212 |
| 5531 | NM_001294338.1 | #N/A | 110.693681 | -8.0007815 | 1.51867013 |
| 5532 | NR_024359.2 | #N/A | 130.194409 | -6.8796763 | 1.30589081 |
| 5533 | NM_017953.3 | #N/A | 115.351431 | -8.0608499 | 1.53011727 |
| 5535 | NM_001271287.1 | ZNF512 | 129.268454 | -8.2234047 | 1.56099161 |
| 5534 | NM_001199097.1 | #N/A | 252.105794 | -9.1891597 | 1.74432508 |
| 5536 | NM_016551.2 | #N/A | 266.558813 | -3.4448597 | 0.65393992 |
| 5537 | NM_005618.3 | #N/A | 137.882825 | -8.3167259 | 1.57881016 |
| 5538 | NM_017645.4 | #N/A | 123.738789 | -8.1612464 | 1.54929861 |
| 5539 | NM_004193.2 | #N/A | 125.993056 | -8.1909144 | 1.55499558 |
| 5540 | NM_021203.3 | #N/A | 205.009882 | -7.4638552 | 1.41696275 |
| 5541 | NM_001199954.1 | #N/A | 190.70232 | -8.7854826 | 1.66790088 |
| 5542 | NM_001286060.1 | #N/A | 140.758462 | -8.3457276 | 1.58453375 |
| 5543 | NM_005049.2 | #N/A | 116.943338 | -8.0803552 | 1.53415943 |
| 5544 | NM_022061.3 | #N/A | 116.012549 | -8.0677708 | 1.53176501 |
| 5545 | NM_014753.3 | #N/A | 277.088234 | -5.8799729 | 1.11643515 |
| 5546 | NM_018405.3 | #N/A | 117.298521 | -8.0852596 | 1.53520742 |

| | | | | | |
|---|---|---|---|---|---|
| 5547 | NM_001012732.1 | DCTD | 112.015552 | -8.0206286 | 1.52297242 |
| 5548 | NM_018400.3 | SCN3B | 195.174996 | -8.8188551 | 1.67453278 |
| 5549 | NM_001134381.1 | TBC1D5 | 154.305955 | -8.4802489 | 1.6102836 |
| 5550 | NM_022464.4 | #N/A | 108.3145 | -7.970966 | 1.51360429 |
| 5551 | NM_003591.3 | #N/A | 141.411165 | -8.3544098 | 1.58650143 |
| 5552 | NM_001127511.2 | #N/A | 124.896376 | -8.1739867 | 1.55227089 |
| 5553 | NM_005053.3 | #N/A | 182.512695 | -8.7217412 | 1.65646661 |
| 5554 | NM_032283.2 | #N/A | 146.815543 | -8.4075879 | 1.5968464 |
| 5555 | NM_170601.4 | #N/A | 121.76422 | -6.7945678 | 1.29054439 |
| 5556 | NM_022739.3 | #N/A | 118.424021 | -8.0969062 | 1.53798117 |
| 5557 | NM_001097615.2 | POLR2J3 | 137.419609 | -8.3139977 | 1.57926375 |
| 5558 | NM_001003722.1 | GLE1 | 118.407621 | -8.0970767 | 1.53820225 |
| 5559 | NM_015252.3 | #N/A | 158.763073 | -8.5205357 | 1.61869225 |
| 5560 | NM_001025603.1 | #N/A | 140.025969 | -8.3398078 | 1.58448777 |
| 5561 | NM_012266.5 | #N/A | 216.764483 | -8.9700685 | 1.70424936 |
| 5562 | NM_015018.3 | DOP1A | 117.360005 | -8.0827343 | 1.53569906 |
| 5563 | NM_001294335.1 | #N/A | 252.527229 | -9.190211 | 1.74614162 |
| 5564 | NM_000744.6 | CHRNA4 | 225.606755 | -9.0278575 | 1.71532691 |
| 5565 | NM_001005920.2 | #N/A | 122.149034 | -8.140436 | 1.54670738 |
| 5566 | NM_152371.3 | #N/A | 121.425438 | -8.1358355 | 1.5458427 |
| 5567 | NM_001281482.1 | #N/A | 138.383066 | -8.3244486 | 1.58188525 |
| 5568 | NM_002084.4 | #N/A | 166.109023 | -8.5871891 | 1.63180407 |
| 5569 | NM_017802.3 | #N/A | 122.369972 | -8.1461402 | 1.54801931 |
| 5570 | NM_033318.4 | #N/A | 169.115397 | -8.6121517 | 1.63670576 |
| 5571 | NM_001272004.1 | #N/A | 100.961752 | -7.8685073 | 1.49541762 |
| 5572 | NM_001130440.1 | SRP9 | 159.071048 | -8.5228073 | 1.62015623 |
| 5573 | NM_001271789.1 | #N/A | 182.736166 | -8.7252606 | 1.65868624 |
| 5574 | NM_016063.2 | #N/A | 112.580168 | -8.0263145 | 1.52593783 |
| 5575 | NM_001322393.1 | ENTPD6 | 120.497648 | -8.1257457 | 1.54486011 |
| 5576 | NM_001007099.2 | #N/A | 130.187391 | -8.2345439 | 1.5655755 |
| 5577 | NM_005082.4 | #N/A | 124.260671 | -8.1699586 | 1.55330253 |
| 5578 | NM_016930.3 | #N/A | 108.605419 | -7.9757132 | 1.5163806 |
| 5579 | NM_003917.4 | #N/A | 101.965777 | -7.881161 | 1.4984341 |
| 5580 | NM_014384.2 | ACAD8 | 118.372852 | -8.0971549 | 1.53958771 |
| 5581 | NM_016034.4 | #N/A | 140.608091 | -7.0695318 | 1.34422488 |
| 5582 | NM_001080826.2 | #N/A | 173.163536 | -7.3733075 | 1.40204473 |
| 5583 | NM_003174.3 | SVIL | 107.129489 | -7.9557135 | 1.51281603 |
| 5584 | NM_001159969.1 | #N/A | 220.475588 | -8.9947283 | 1.71042357 |
| 5585 | NM_152415.2 | #N/A | 123.179324 | -8.1534077 | 1.55046207 |
| 5586 | NM_015346.3 | #N/A | 114.87322 | -8.0548322 | 1.5318047 |
| 5587 | NM_014904.2 | #N/A | 110.357015 | -7.9955082 | 1.52059599 |
| 5588 | NM_001286611.1 | REPS1 | 170.486802 | -8.624382 | 1.64026963 |
| 5589 | NM_001042601.2 | #N/A | 99.5015289 | -7.8498894 | 1.49312754 |

| | | | | | |
|---|---|---|---|---|---|
| 5590 | NM_014393.2 | STAU2 | 157.994842 | -8.5132039 | 1.61933227 |
| 5591 | NM_001040181.1 | #N/A | 123.882518 | -8.1631725 | 1.55277191 |
| 5592 | NM_001002251.2 | ARL6IP4 | 117.621894 | -8.0874756 | 1.53852309 |
| 5593 | NM_003031.3 | #N/A | 158.738177 | -8.5202051 | 1.62094239 |
| 5594 | NM_033624.2 | #N/A | 113.547411 | -8.0392582 | 1.52945072 |
| 5595 | NM_001270965.1 | PSD | 191.089307 | -8.7868854 | 1.67169895 |
| 5596 | NM_001661.3 | #N/A | 165.110455 | -8.5767992 | 1.63180969 |
| 5597 | NR_028038.1 | #N/A | 159.243368 | -8.5250921 | 1.62204999 |
| 5598 | NM_001080453.2 | #N/A | 799.517027 | -6.3216617 | 1.20281636 |
| 5599 | NM_006045.2 | #N/A | 1146.29389 | -6.800988 | 1.29403925 |
| 5600 | NM_144568.3 | PIP4P1 | 137.903825 | -8.3180285 | 1.58274409 |
| 5601 | NR_104615.1 | LOC100506688 | 156.276061 | 3.68872431 | 0.70189496 |
| 5602 | NM_007318.2 | #N/A | 139.104195 | -6.9643313 | 1.32525869 |
| 5603 | NM_001278640.1 | #N/A | 170.703866 | -8.6255831 | 1.64151198 |
| 5604 | NM_018346.2 | #N/A | 135.837812 | -8.2952635 | 1.57864971 |
| 5605 | NM_024958.2 | #N/A | 139.291213 | -8.3306362 | 1.58556446 |
| 5606 | NM_001297707.1 | #N/A | 257.327621 | -9.2179177 | 1.7545232 |
| 5607 | NM_005343.3 | #N/A | 177.080798 | -8.6784623 | 1.65189238 |
| 5608 | NM_003887.2 | #N/A | 165.301364 | -8.5811275 | 1.63342052 |
| 5609 | NM_018137.2 | #N/A | 113.426051 | -6.7582665 | 1.28649055 |
| 5610 | NM_023071.3 | SPATS2 | 188.96754 | -8.77172 | 1.66983413 |
| 5611 | NM_016639.2 | #N/A | 180.673739 | -8.7084917 | 1.65786454 |
| 5612 | NM_006818.3 | #N/A | 1507.76093 | -5.9458822 | 1.13213294 |
| 5613 | NM_025134.4 | #N/A | 154.696768 | -7.0970858 | 1.35134239 |
| 5614 | NM_015330.4 | #N/A | 133.867954 | -8.2722778 | 1.57529714 |
| 5615 | NM_002703.4 | #N/A | 129.815535 | -8.2302264 | 1.56732509 |
| 5616 | NM_001023567.4 | #N/A | 237.197021 | -9.1002353 | 1.73304881 |
| 5617 | NM_016621.3 | #N/A | 204.947786 | -7.4602628 | 1.42073617 |
| 5618 | NM_002437.4 | #N/A | 116.415551 | -8.0730154 | 1.5374572 |
| 5619 | NM_001329201.1 | IAPP | 179.467855 | 4.66855982 | 0.88911917 |
| 5620 | NM_005647.3 | #N/A | 147.555509 | -8.4161925 | 1.60291561 |
| 5621 | NM_015722.3 | #N/A | 107.464174 | -7.9604106 | 1.51616048 |
| 5622 | NM_145295.3 | #N/A | 133.686499 | -8.2726415 | 1.5756724 |
| 5623 | NM_017750.3 | #N/A | 108.400651 | -7.9710421 | 1.51842974 |
| 5624 | NM_001814.5 | #N/A | 110.299926 | -7.999406 | 1.52386017 |
| 5625 | NM_001308142.1 | #N/A | 103.311493 | -7.8992857 | 1.50485509 |
| 5626 | NM_000788.2 | #N/A | 161.837966 | -8.5482491 | 1.62854806 |
| 5627 | NM_000495.4 | #N/A | 175.29201 | -8.6648 | 1.65077455 |
| 5628 | NM_013301.2 | CCDC106 | 128.77543 | -8.2182047 | 1.56574109 |
| 5629 | NM_003000.2 | #N/A | 145.042928 | -8.3901533 | 1.59853847 |
| 5630 | NM_015989.4 | #N/A | 125.970881 | -8.1848974 | 1.55950076 |
| 5632 | NM_138821.2 | PAM | 112.334236 | -8.0255259 | 1.52917361 |
| 5631 | NM_015261.2 | #N/A | 112.995404 | -8.029309 | 1.52989499 |

| | | | | | | |
|---|---|---|---|---|---|---|
| 5633 | NM_001178030.1 | SF1 | | 108.411393 | -7.9725275 | 1.51911831 |
| 5634 | NM_138792.3 | | #N/A | 118.803063 | -8.1028604 | 1.54397272 |
| 5635 | NM_144772.2 | | #N/A | 153.216827 | -8.4701696 | 1.61404401 |
| 5636 | NM_003942.2 | | #N/A | 98.5519514 | -7.8316846 | 1.492423 |
| 5637 | NM_004789.3 | | #N/A | 332.051985 | -9.58533 | 1.82660337 |
| 5638 | NM_033547.3 | | #N/A | 119.627224 | -8.1115983 | 1.54581866 |
| 5639 | NM_001198916.1 | | #N/A | 111.183649 | -8.0105163 | 1.52657102 |
| 5640 | NM_015355.3 | | #N/A | 152.299069 | -8.4608068 | 1.61247995 |
| 5641 | NR_134559.1 | LINC00649 | | 176.987257 | 3.20711592 | 0.61123593 |
| 5642 | NM_001037172.2 | | #N/A | 166.495956 | -8.5895732 | 1.63709157 |
| 5643 | NM_005643.3 | | #N/A | 136.918726 | -8.3077881 | 1.5834243 |
| 5644 | NR_027418.1 | LINC00674 | | 407.938892 | -1.8765895 | 0.35769179 |
| 5645 | NM_032110.2 | | #N/A | 142.115829 | -8.3598213 | 1.59356567 |
| 5646 | NM_012166.2 | | #N/A | 135.095374 | -8.288653 | 1.58008462 |
| 5647 | NM_001131025.1 | PEX5 | | 110.260051 | -7.9919544 | 1.52361076 |
| 5648 | NM_019095.5 | | #N/A | 123.630535 | -8.160402 | 1.55571725 |
| 5649 | NM_014043.3 | | #N/A | 122.205578 | -8.1427426 | 1.55241775 |
| 5650 | NM_001171135.1 | ZBED1 | | 119.018354 | -8.1043092 | 1.54515247 |
| 5651 | NM_004301.4 | | #N/A | 121.458526 | -8.132498 | 1.55054672 |
| 5652 | NM_014830.2 | | #N/A | 115.558702 | -8.0622641 | 1.53720835 |
| 5653 | NR_024542.1 | SNHG7 | | 140.185541 | -8.341331 | 1.59054327 |
| 5654 | NM_080759.5 | | #N/A | 126.014289 | -8.1857462 | 1.56089727 |
| 5655 | NM_001324331.1 | | #N/A | 154.596869 | -8.4821042 | 1.6174672 |
| 5656 | NM_133471.3 | | #N/A | 135.37093 | -8.2905368 | 1.58093977 |
| 5657 | NM_012412.4 | H2AZ2 | | 133.015041 | -8.264707 | 1.57617195 |
| 5658 | NM_001243028.2 | | #N/A | 107.598552 | -7.9600085 | 1.51816287 |
| 5659 | NM_014655.3 | | #N/A | 159.48116 | -8.5278736 | 1.62648764 |
| 5660 | NM_018231.2 | | #N/A | 133.549735 | -8.2724399 | 1.57776285 |
| 5661 | NM_015530.4 | | #N/A | 158.586244 | -7.1290752 | 1.35975342 |
| 5662 | NM_003878.2 | | #N/A | 148.773628 | -8.4270986 | 1.60739301 |
| 5663 | NM_024604.2 | | #N/A | 116.431626 | -8.0714707 | 1.53970947 |
| 5664 | NM_001176.3 | | #N/A | 141.184342 | -8.3520552 | 1.59333915 |
| 5665 | NM_003013.2 | | #N/A | 135.264323 | -6.9263942 | 1.32144702 |
| 5666 | NM_001142479.1 | NREP | | 197.612709 | -8.8366122 | 1.68606687 |
| 5667 | NM_018077.2 | | #N/A | 103.720386 | -7.9069318 | 1.50867923 |
| 5668 | NM_001077418.2 | | #N/A | 127.24577 | -8.2037216 | 1.56532252 |
| 5669 | NM_033380.2 | | #N/A | 348.301957 | -9.6550701 | 1.84243022 |
| 5670 | NM_001172665.1 | RAB40C | | 122.938316 | -8.1516422 | 1.55555938 |
| 5671 | NM_001320852.1 | ARHGEF7 | | 127.449357 | -8.2028831 | 1.56546687 |
| 5672 | NR_033436.1 | | #N/A | 116.777096 | -8.077318 | 1.54152209 |
| 5673 | NM_001040437.2 | | #N/A | 145.598789 | -8.3945227 | 1.60207793 |
| 5674 | NM_001289971.1 | | #N/A | 192.038328 | -8.7945515 | 1.678498 |
| 5675 | NM_203418.2 | | #N/A | 195.944137 | -8.8256134 | 1.68446855 |

| | | | | | |
|---|---|---|---|---|---|
| 5676 | NM_020690.5 | ANKHD1-EIF4EBP3 | 113.183806 | -8.0321559 | 1.53305919 |
| 5677 | NM_017567.4 | #N/A | 135.897244 | -8.2958283 | 1.58346268 |
| 5678 | NM_032377.3 | ELOF1 | 134.988296 | -7.0111006 | 1.33826734 |
| 5679 | NM_012142.4 | #N/A | 116.315049 | -8.0753311 | 1.54148035 |
| 5680 | NM_172351.2 | #N/A | 120.027436 | -8.1153074 | 1.549148 |
| 5681 | NM_018180.2 | #N/A | 100.118368 | -7.8594974 | 1.50041817 |
| 5682 | NM_080797.3 | #N/A | 118.12246 | -8.0961476 | 1.54563138 |
| 5683 | NM_014953.4 | #N/A | 149.168231 | -8.431416 | 1.6097002 |
| 5684 | NM_052859.3 | #N/A | 107.854544 | -3.3463421 | 0.63887278 |
| 5685 | NM_023010.3 | UPF3B | 136.486645 | -8.3021741 | 1.58543072 |
| 5686 | NM_001111322.1 | #N/A | 128.969044 | -8.2207868 | 1.57002134 |
| 5687 | NM_001172173.1 | #N/A | 141.928913 | -8.3591359 | 1.59650109 |
| 5688 | NM_198859.3 | #N/A | 126.860697 | -8.1975619 | 1.56563273 |
| 5689 | NM_003664.4 | AP3B1 | 126.040167 | -8.1913348 | 1.56450833 |
| 5690 | NM_006706.3 | #N/A | 140.418585 | -8.3419337 | 1.59342108 |
| 5691 | NM_024948.3 | #N/A | 126.834969 | -8.1966748 | 1.56582128 |
| 5692 | NM_153223.3 | CEP120 | 106.407299 | -7.9415428 | 1.51710211 |
| 5693 | NM_017814.2 | #N/A | 112.414982 | -8.0239754 | 1.53291612 |
| 5694 | NM_175061.3 | #N/A | 124.917914 | -8.1758152 | 1.56201529 |
| 5695 | NM_006122.3 | #N/A | 136.785104 | -8.3067591 | 1.58707967 |
| 5696 | NM_006554.4 | #N/A | 158.127426 | -8.5149292 | 1.62686847 |
| 5697 | NM_025136.3 | #N/A | 484.08899 | 1.33871363 | 0.25577583 |
| 5699 | NM_181675.3 | PPP2R2B | 184.359864 | -8.7374168 | 1.66944674 |
| 5698 | NM_012141.2 | #N/A | 106.2391 | -7.9397531 | 1.51703862 |
| 5700 | NM_182540.4 | #N/A | 114.975097 | -8.0554293 | 1.53917921 |
| 5701 | NM_018149.6 | SMG8 | 110.448162 | -7.9977937 | 1.5282317 |
| 5702 | NM_183377.1 | ASIC2 | 193.459361 | -8.8065444 | 1.68294122 |
| 5703 | NM_001297779.1 | #N/A | 150.33752 | 2.77370772 | 0.53013028 |
| 5704 | NM_015946.4 | #N/A | 119.333344 | -8.1108504 | 1.55019353 |
| 5705 | NM_020464.1 | #N/A | 127.121764 | -8.198234 | 1.56696654 |
| 5706 | NM_025099.5 | #N/A | 118.521519 | -8.0979521 | 1.54782413 |
| 5707 | NM_012129.4 | #N/A | 117.965345 | -8.0897313 | 1.54629959 |
| 5708 | NM_198426.2 | #N/A | 126.121685 | -8.1899095 | 1.56552267 |
| 5709 | NM_001287060.1 | #N/A | 114.095173 | -8.0455744 | 1.53800292 |
| 5710 | NM_024753.4 | #N/A | 99.0352069 | -7.8409106 | 1.49894538 |
| 5711 | NM_005827.2 | #N/A | 122.075371 | -8.1421884 | 1.55664629 |
| 5712 | NM_001170635.1 | #N/A | 143.947205 | -8.3804346 | 1.60230476 |
| 5713 | NM_001190965.2 | #N/A | 213.600546 | -8.9491668 | 1.71104005 |
| 5714 | NM_001136204.2 | #N/A | 249.874694 | -7.7121808 | 1.47458997 |
| 5715 | NM_033280.3 | #N/A | 200.063254 | -8.8544582 | 1.69316472 |
| 5716 | NM_002466.3 | #N/A | 137.782883 | -8.3164559 | 1.59037969 |
| 5717 | NM_006782.3 | #N/A | 122.059251 | -8.1416448 | 1.55711469 |
| 5718 | NM_017822.3 | #N/A | 138.612168 | -8.3243404 | 1.59206838 |

| | | | | | |
|---|---|---|---|---|---|
| 5719 | NM_004645.2 | #N/A | 114.739613 | -8.050933 | 1.53983049 |
| 5720 | NM_198868.2 | #N/A | 200.469987 | -8.8577915 | 1.69415372 |
| 5721 | NM_175859.2 | #N/A | 117.516981 | -8.0890229 | 1.54723276 |
| 5722 | NM_198242.2 | #N/A | 198.069894 | -8.8395173 | 1.69082281 |
| 5723 | NM_003390.3 | #N/A | 122.984306 | -8.1520493 | 1.55938183 |
| 5724 | NM_001722.2 | #N/A | 114.675665 | -8.052859 | 1.5404264 |
| 5725 | NM_004526.3 | #N/A | 245.479082 | -6.4854497 | 1.24062648 |
| 5726 | NM_006693.3 | #N/A | 126.288884 | -8.1895189 | 1.56660806 |
| 5727 | NM_014426.3 | #N/A | 101.515284 | -7.8761823 | 1.50669619 |
| 5728 | NM_006651.3 | #N/A | 179.511245 | -8.6984521 | 1.66404805 |
| 5729 | NM_001018020.1 | #N/A | 107.655256 | -7.9605238 | 1.5229103 |
| 5730 | NM_001308319.1 | CHD9 | 148.41832 | -8.4252723 | 1.61185028 |
| 5731 | NM_022070.4 | #N/A | 104.104929 | -7.9118785 | 1.51367098 |
| 5732 | NM_133372.2 | #N/A | 121.09192 | -8.1292271 | 1.55549894 |
| 5733 | NM_001254952.2 | #N/A | 189.666547 | -8.7779699 | 1.67973723 |
| 5734 | NM_006074.4 | #N/A | 107.095108 | -7.9567892 | 1.5226089 |
| 5735 | NM_000390.3 | #N/A | 100.133843 | -7.8580768 | 1.503755 |
| 5736 | NM_018090.4 | #N/A | 105.002445 | -7.9260932 | 1.51680585 |
| 5737 | NM_020320.4 | #N/A | 98.0682145 | -7.8231411 | 1.49711621 |
| 5738 | NM_001076677.2 | #N/A | 198.240264 | -8.8417642 | 1.69208582 |
| 5739 | NM_001005373.3 | LRSAM1 | 130.148273 | -8.2332482 | 1.57573489 |
| 5740 | NM_006909.2 | #N/A | 140.633481 | -6.2884788 | 1.20360268 |
| 5741 | NM_014853.2 | #N/A | 105.437565 | -7.9296016 | 1.51772159 |
| 5742 | NM_014175.3 | #N/A | 137.49198 | -8.3138587 | 1.5913753 |
| 5743 | NM_138785.3 | #N/A | 106.364596 | -7.9446653 | 1.52079391 |
| 5744 | NM_007202.3 | #N/A | 96.5410405 | -7.8040437 | 1.49390818 |
| 5745 | NM_015472.4 | #N/A | 133.142473 | -8.2690985 | 1.58293929 |
| 5746 | NM_018229.3 | #N/A | 117.130145 | -8.081893 | 1.54710137 |
| 5747 | NM_025078.4 | #N/A | 114.484593 | -8.048866 | 1.54079468 |
| 5748 | NM_005125.1 | #N/A | 124.716089 | -8.172839 | 1.56453853 |
| 5749 | NM_004073.3 | #N/A | 115.441946 | -8.0624335 | 1.54346756 |
| 5750 | NM_006856.2 | #N/A | 115.052348 | -8.0565746 | 1.54235319 |
| 5751 | NM_012080.4 | #N/A | 104.145933 | -7.9138591 | 1.51505987 |
| 5752 | NM_013399.2 | #N/A | 170.286475 | -8.6214709 | 1.6505287 |
| 5753 | NM_000479.4 | #N/A | 135.768131 | -8.2958771 | 1.58822886 |
| 5755 | NR_003615.2 | LOC728554 | 98.6397709 | -7.8371566 | 1.50044598 |
| 5754 | NM_199129.2 | #N/A | 118.121382 | -8.0954487 | 1.54989096 |
| 5756 | NM_001042576.1 | RRBP1 | 129.654007 | -8.2317801 | 1.57603402 |
| 5757 | NR_125375.1 | SLC9A3-AS1 | 140.980216 | -6.9765974 | 1.33578905 |
| 5760 | NM_183373.3 | PXDC1 | 125.434665 | -8.182471 | 1.56685832 |
| 5758 | NM_018838.4 | #N/A | 149.701288 | -8.435944 | 1.61539663 |
| 5759 | NM_020246.3 | #N/A | 153.630076 | -8.4738339 | 1.62264747 |
| 5761 | NM_000318.2 | #N/A | 95.85144 | -7.791594 | 1.49203078 |

| | | | | | |
|---|---|---|---|---|---|
| 5762 | NM_004095.3 | #N/A | 183.350817 | -8.7287338 | 1.67151562 |
| 5763 | NM_015885.3 | #N/A | 114.222302 | -8.044027 | 1.54040051 |
| 5764 | NM_007372.3 | DDX42 | 99.2935665 | -7.8428141 | 1.50188408 |
| 5765 | NM_001077195.1 | #N/A | 115.482262 | -8.0633953 | 1.54419372 |
| 5766 | NM_015935.4 | #N/A | 120.744412 | -8.1258134 | 1.55626752 |
| 5767 | NM_133265.2 | #N/A | 104.252786 | -7.9156263 | 1.51600674 |
| 5768 | NM_000930.4 | #N/A | 132.466755 | -8.2604263 | 1.58206694 |
| 5770 | NM_003185.3 | TAF4 | 113.55026 | -8.036688 | 1.53925808 |
| 5769 | NM_001255.2 | #N/A | 113.689393 | -8.0381669 | 1.53953234 |
| 5771 | NM_001242860.2 | PRKACB | 189.151612 | -8.7738009 | 1.68045119 |
| 5772 | NM_178863.4 | #N/A | 144.787835 | -8.3883287 | 1.60663643 |
| 5773 | NM_001206665.2 | IGSF8 | 168.22799 | -8.604113 | 1.64816888 |
| 5774 | NM_022450.3 | #N/A | 108.771882 | -7.9784868 | 1.52836065 |
| 5775 | NM_203284.2 | RBPJ | 110.238804 | -7.9944941 | 1.53175901 |
| 5777 | NM_020862.1 | LRFN1 | 177.451787 | -8.6816575 | 1.6634984 |
| 5776 | NM_016041.4 | #N/A | 113.822303 | -8.039785 | 1.54050176 |
| 5778 | NR_038888.1 | LOC284950 | 98.0751118 | 3.46919614 | 0.66474043 |
| 5779 | NM_001757.3 | #N/A | 109.422017 | -7.9836585 | 1.5299028 |
| 5780 | NM_000098.2 | #N/A | 100.126391 | -7.8596906 | 1.50615693 |
| 5781 | NM_006461.3 | #N/A | 116.801266 | -8.0777729 | 1.54796654 |
| 5782 | NM_178511.5 | #N/A | 101.316475 | -7.8737312 | 1.50886692 |
| 5783 | NM_001286117.1 | CCDC90B | 139.634065 | -8.3354713 | 1.59747636 |
| 5784 | NM_001018139.2 | NME2 | 134.850023 | -8.2865482 | 1.58818708 |
| 5786 | NM_018113.3 | LMBR1L | 146.606904 | -8.4059439 | 1.61129695 |
| 5785 | NM_001160226.1 | #N/A | 188.445934 | -8.7684619 | 1.68078504 |
| 5787 | NM_001170880.1 | #N/A | 149.950424 | -8.4381457 | 1.61753432 |
| 5788 | NM_001142782.1 | #N/A | 119.408368 | -8.1092652 | 1.55451211 |
| 5789 | NM_003642.3 | #N/A | 109.148512 | -7.9785719 | 1.52949928 |
| 5790 | NM_002505.4 | #N/A | 180.75768 | -8.707935 | 1.66937579 |
| 5791 | NM_003239.4 | TGFB3 | 110.849833 | -8.0064735 | 1.53511233 |
| 5792 | NM_078469.2 | #N/A | 127.304252 | -8.2023362 | 1.57271437 |
| 5793 | NM_018138.4 | #N/A | 92.5088298 | -7.7421364 | 1.48459758 |
| 5794 | NM_024092.2 | #N/A | 146.603353 | -7.0254698 | 1.34720577 |
| 5795 | NM_013364.5 | #N/A | 98.6730363 | -7.8379956 | 1.50306962 |
| 5796 | NM_003014.3 | #N/A | 154.96052 | -8.4872579 | 1.62762667 |
| 5797 | NM_012250.5 | #N/A | 118.639991 | -8.100022 | 1.55336695 |
| 5798 | NM_024624.5 | SMC6 | 115.374684 | -8.0581107 | 1.54534346 |
| 5799 | NM_001042522.2 | SPRED3 | 135.654269 | -8.2946897 | 1.59073526 |
| 5800 | NM_007039.3 | #N/A | 113.486289 | -8.0368355 | 1.54129682 |
| 5802 | NM_177996.2 | EPB41L1 | 163.652346 | -8.5659263 | 1.64285492 |
| 5801 | NM_014247.2 | #N/A | 276.655271 | -5.864835 | 1.12481387 |
| 5803 | NM_001080.3 | ALDH5A1 | 173.190674 | -4.4507349 | 0.85363468 |
| 5804 | NM_000808.3 | #N/A | 221.372082 | -6.1908333 | 1.18740404 |

| | | | | | | |
|---|---|---|---|---|---|---|
| 5805 | NM_014681.5 | | #N/A | 112.989476 | -8.0302691 | 1.54032148 |
| 5806 | NM_004260.3 | RECQL4 | | 140.941343 | -8.3488209 | 1.60143417 |
| 5807 | NM_001001485.2 | ATP2C1 | | 99.9302963 | -7.8521554 | 1.50618359 |
| 5808 | NM_001278270.1 | TADA3 | | 131.750027 | -6.9773676 | 1.3384111 |
| 5809 | NM_012238.4 | | #N/A | 120.626045 | -8.1244516 | 1.55860517 |
| 5810 | NM_001905.3 | | #N/A | 161.613199 | -8.5469584 | 1.63968567 |
| 5811 | NM_001305043.1 | | #N/A | 168.364675 | -8.6043732 | 1.65077509 |
| 5812 | NM_139048.2 | | #N/A | 109.026726 | -6.6428193 | 1.27445299 |
| 5813 | NM_014802.2 | C2CD5 | | 101.815137 | -7.878695 | 1.51159104 |
| 5815 | NM_006095.2 | ATP8A1 | | 169.818924 | -8.6179658 | 1.65352454 |
| 5814 | NM_001080123.1 | | #N/A | 143.565321 | -8.3768895 | 1.60728975 |
| 5816 | NM_007013.3 | | #N/A | 118.381918 | -6.8203539 | 1.30863146 |
| 5817 | NM_014458.3 | | #N/A | 144.274262 | -8.3830974 | 1.60847267 |
| 5818 | NM_000142.4 | FGFR3 | | 300.745821 | -9.4423408 | 1.81173386 |
| 5820 | NM_178151.2 | DCX | | 145.372514 | -8.3939222 | 1.6106382 |
| 5819 | NM_018145.2 | | #N/A | 116.313257 | -8.0727474 | 1.54900181 |
| 5821 | NM_001211.5 | BUB1B | | 97.2042103 | -7.8147601 | 1.49952645 |
| 5823 | NR_040093.1 | STARD4-AS1 | | 117.912316 | -6.742081 | 1.29378975 |
| 5822 | NM_020431.3 | | #N/A | 154.040388 | -8.4773763 | 1.62679864 |
| 5824 | NM_173852.3 | KRTCAP2 | | 132.193346 | -8.2575576 | 1.58478709 |
| 5825 | NM_000270.3 | PNP | | 133.041504 | -8.2668585 | 1.58661848 |
| 5826 | NM_001009996.2 | | #N/A | 103.844174 | -7.9108513 | 1.5183474 |
| 5827 | NM_001498.3 | | #N/A | 149.841252 | -8.4388959 | 1.61970225 |
| 5828 | NM_024509.1 | | #N/A | 129.864272 | -8.2312143 | 1.57990187 |
| 5829 | NM_014858.3 | | #N/A | 187.951463 | -8.7642991 | 1.68234119 |
| 5830 | NM_181715.2 | | #N/A | 98.9727563 | -7.8407799 | 1.50516856 |
| 5831 | NM_001243374.1 | CLCN3 | | 231.104567 | -5.3906775 | 1.03485487 |
| 5832 | NM_012232.5 | | #N/A | 652.439763 | -3.354543 | 0.64403219 |
| 5833 | NM_001242575.2 | | #N/A | 95.9344656 | 3.26427904 | 0.62670772 |
| 5834 | NM_001113490.1 | AMOT | | 108.213377 | -7.9705039 | 1.53031515 |
| 5835 | NM_003187.4 | | #N/A | 162.744258 | -8.5565981 | 1.64283942 |
| 5836 | NM_002360.3 | | #N/A | 124.996244 | -6.8189579 | 1.30924301 |
| 5837 | NM_206855.2 | | #N/A | 115.604341 | -8.0631344 | 1.54819021 |
| 5839 | NM_018724.3 | IL20 | | 18.8912083 | 8.47028821 | 1.62644902 |
| 5838 | NM_014402.4 | | #N/A | 286.852988 | -5.1282761 | 0.98472092 |
| 5840 | NR_036659.1 | | #N/A | 164.762602 | -8.5742248 | 1.64647057 |
| 5841 | NM_004263.4 | | #N/A | 176.029709 | -8.6703074 | 1.66500413 |
| 5842 | NM_001323685.1 | | #N/A | 130.684536 | -8.2407035 | 1.58253808 |
| 5843 | NM_001190.3 | | #N/A | 107.028969 | -7.9542648 | 1.52762999 |
| 5844 | NM_022757.4 | | #N/A | 120.393709 | -6.7671607 | 1.29972997 |
| 5845 | NM_001142467.1 | | #N/A | 194.717097 | -8.8153841 | 1.6933164 |
| 5846 | NM_015607.3 | | #N/A | 123.954408 | -8.1665187 | 1.56876046 |
| 5847 | NM_001029885.1 | | #N/A | 125.43261 | -8.1813013 | 1.57162945 |

| | | | | | |
|---|---|---|---|---|---|
| 5848 | NM_033112.3 | #N/A | 134.293636 | -8.2799506 | 1.59059367 |
| 5849 | NM_001142424.1 | MORF4L2 | 123.327568 | -8.1567238 | 1.56696909 |
| 5850 | NM_130463.3 | #N/A | 219.281146 | -8.9868062 | 1.72646156 |
| 5851 | NM_017966.4 | #N/A | 109.949157 | -7.9911947 | 1.535232 |
| 5852 | NM_001245005.1 | #N/A | 130.257636 | -8.2338259 | 1.58194309 |
| 5853 | NM_152359.2 | CPT1C | 101.527221 | -7.8795095 | 1.51391916 |
| 5854 | NM_001001790.2 | #N/A | 165.050817 | -8.5769527 | 1.64799343 |
| 5855 | NM_016302.3 | CRBN | 129.658707 | -8.2301819 | 1.58138998 |
| 5856 | NM_031432.3 | #N/A | 136.241588 | -8.2992903 | 1.59475037 |
| 5857 | NM_005627.3 | #N/A | 135.278362 | -8.2913209 | 1.59339204 |
| 5858 | NR_109911.1 | CTD-3080P12 | 34.9930575 | 4.4543997 | 0.85603979 |
| 5859 | NR_027856.1 | #N/A | 102.261559 | -7.8873186 | 1.51600831 |
| 5860 | NM_033415.3 | #N/A | 145.47878 | -8.3952802 | 1.6137939 |
| 5861 | NM_018285.3 | IMP3 | 140.91124 | -8.3484229 | 1.60480281 |
| 5862 | NM_018119.3 | #N/A | 126.297535 | -8.1910651 | 1.57467555 |
| 5863 | NM_007357.2 | #N/A | 95.0933266 | -7.7818762 | 1.49625 |
| 5865 | NM_004287.4 | GOSR2 | 125.667766 | -8.1852734 | 1.57384913 |
| 5864 | NM_002813.6 | #N/A | 127.686262 | -8.2075904 | 1.57814089 |
| 5866 | NM_014783.5 | #N/A | 102.946081 | -7.8954907 | 1.51811772 |
| 5867 | NM_000170.2 | GLDC | 123.207228 | -8.1545663 | 1.56806745 |
| 5868 | NM_032439.3 | #N/A | 181.732018 | -8.7161727 | 1.67613175 |
| 5869 | NM_001320704.1 | #N/A | 113.124389 | -8.0344722 | 1.54521516 |
| 5870 | NM_001080495.2 | TNRC18 | 996.567349 | -6.3183202 | 1.21519275 |
| 5871 | NM_006062.2 | #N/A | 132.479278 | -8.2597466 | 1.58865421 |
| 5872 | NM_003443.2 | ZBTB17 | 140.061289 | -8.3392942 | 1.60397144 |
| 5873 | NM_001242809.1 | #N/A | 105.963188 | -7.9395103 | 1.52710244 |
| 5874 | NM_016309.2 | #N/A | 154.288232 | -8.4800157 | 1.63136612 |
| 5875 | NM_001139490.1 | TRAF3IP1 | 123.090357 | -6.7983 | 1.30785433 |
| 5876 | NM_001007225.2 | #N/A | 117.928988 | -8.0949307 | 1.5573103 |
| 5877 | NM_014015.3 | #N/A | 112.717702 | -8.0272881 | 1.54434376 |
| 5878 | NM_015721.2 | #N/A | 109.786185 | -7.9891786 | 1.53704485 |
| 5879 | NM_001242659.1 | #N/A | 119.189396 | -8.108118 | 1.55996649 |
| 5880 | NM_080725.2 | #N/A | 126.535619 | -6.8333499 | 1.31474313 |
| 5881 | NM_001008272.1 | #N/A | 165.377876 | -8.5799607 | 1.65082239 |
| 5882 | NM_001142426.1 | MORF4L2 | 222.444683 | -9.0079415 | 1.73325565 |
| 5883 | NM_015921.2 | #N/A | 123.005786 | -8.1539705 | 1.56902537 |
| 5884 | NM_001331037.1 | GPBP1 | 114.521008 | -8.0514056 | 1.54941526 |
| 5885 | NM_014673.4 | #N/A | 111.426921 | -8.0126421 | 1.54198725 |
| 5886 | NM_021934.4 | ATG101 | 127.046983 | -8.1992767 | 1.57804807 |
| 5887 | NM_174902.3 | #N/A | 137.935989 | -8.3184989 | 1.60099601 |
| 5888 | NM_001042479.1 | #N/A | 101.751611 | -7.8764395 | 1.51607552 |
| 5889 | NM_001048.3 | #N/A | 165.349676 | -8.5806613 | 1.65177819 |
| 5890 | NM_001123385.1 | #N/A | 114.711027 | -8.0518533 | 1.54997422 |

| | | | | | |
|---|---|---|---|---|---|
| 5891 | NM_016614.2 | #N/A | 113.245807 | -8.0320787 | 1.5461958 |
| 5892 | NR_110386.1 | LOC101927476 | 89.4430451 | 3.12192387 | 0.60102535 |
| 5893 | NM_001321149.1 | #N/A | 123.406423 | -8.1607023 | 1.57109689 |
| 5894 | NM_178860.4 | #N/A | 256.8204 | -9.2147034 | 1.77409511 |
| 5895 | NM_198098.3 | AQP1 | 732.365498 | -10.727193 | 2.06537634 |
| 5896 | NM_030667.2 | #N/A | 171.130403 | -8.6286546 | 1.66137977 |
| 5897 | NM_001136265.1 | #N/A | 148.61601 | -7.1514559 | 1.37700828 |
| 5898 | NM_138704.3 | #N/A | 118.281052 | -8.0964363 | 1.55898676 |
| 5899 | NM_015040.3 | #N/A | 242.327891 | -4.122424 | 0.79378877 |
| 5900 | NM_001010000.2 | #N/A | 128.567582 | -8.216968 | 1.5822643 |
| 5901 | NM_014417.4 | #N/A | 136.976869 | -8.3077265 | 1.59976362 |
| 5902 | NM_001322246.1 | #N/A | 182.4978 | -8.7220765 | 1.67959859 |
| 5903 | NM_001135110.2 | #N/A | 124.913063 | -8.1734722 | 1.57404793 |
| 5904 | NM_001300759.1 | #N/A | 131.829831 | -8.2520885 | 1.58922873 |
| 5905 | NM_001135190.1 | ARAP1 | 101.586227 | -7.8792217 | 1.51758036 |
| 5906 | NM_002532.5 | #N/A | 123.742588 | -8.160609 | 1.5717691 |
| 5907 | NM_001128226.2 | #N/A | 108.955969 | -7.976918 | 1.536442 |
| 5908 | NM_006505.4 | #N/A | 141.757234 | -8.3584297 | 1.60992952 |
| 5909 | NM_001099270.1 | #N/A | 106.563522 | -7.9449832 | 1.530323 |
| 5910 | NM_001301268.1 | #N/A | 102.065183 | -7.8828913 | 1.51878257 |
| 5911 | NM_001251974.1 | RCAN2 | 154.257738 | -8.4805472 | 1.63398013 |
| 5912 | NM_006473.3 | #N/A | 117.717687 | -8.0884581 | 1.55854136 |
| 5913 | NM_001330658.1 | #N/A | 110.867898 | -8.003744 | 1.54225434 |
| 5914 | NM_031862.3 | #N/A | 144.677245 | -8.3895467 | 1.61677487 |
| 5915 | NM_000320.2 | #N/A | 169.437142 | -8.6147891 | 1.6602153 |
| 5916 | NM_006571.3 | #N/A | 132.756214 | -8.263306 | 1.59248527 |
| 5917 | NM_001198972.1 | #N/A | 173.412479 | -8.6496085 | 1.66699021 |
| 5918 | NM_025195.3 | #N/A | 94.9647257 | -7.7831899 | 1.50002316 |
| 5919 | NM_030780.4 | #N/A | 106.53181 | -7.9459281 | 1.53146899 |
| 5920 | NM_014051.3 | #N/A | 148.241659 | -8.4222175 | 1.62333621 |
| 5921 | NM_001287520.1 | CDC25B | 99.4492466 | -7.8437502 | 1.51191312 |
| 5922 | NM_012420.2 | #N/A | 101.623926 | -7.8782433 | 1.51865401 |
| 5923 | NM_001300846.1 | #N/A | 118.73137 | -8.1022542 | 1.56194009 |
| 5924 | NM_001136263.1 | #N/A | 117.126493 | -8.0823119 | 1.55824874 |
| 5925 | NM_022338.3 | #N/A | 122.79098 | -8.1515825 | 1.57163674 |
| 5926 | NM_001852.3 | #N/A | 120.411415 | -8.123268 | 1.56627991 |
| 5927 | NM_170744.4 | #N/A | 152.86628 | -8.4682924 | 1.63289311 |
| 5928 | NM_001174100.1 | #N/A | 149.345503 | -8.4325967 | 1.62607324 |
| 5929 | NM_001199058.1 | #N/A | 129.073208 | -8.2224917 | 1.58562448 |
| 5930 | NM_001242824.1 | #N/A | 157.805082 | -8.5128339 | 1.64165873 |
| 5931 | NM_001079818.2 | #N/A | 120.969541 | -8.1320736 | 1.56825574 |
| 5932 | NM_024632.5 | #N/A | 109.132028 | -7.9785185 | 1.53872582 |
| 5933 | NM_021076.3 | #N/A | 147.630697 | -8.4168604 | 1.62333119 |

| | | | | | |
|---|---|---|---|---|---|
| 5934 | NM_033158.4 | HYAL2 | 105.43658 | -7.9284901 | 1.52915505 |
| 5935 | NM_030793.4 | #N/A | 120.404981 | -8.1223327 | 1.56655977 |
| 5936 | NM_018192.3 | #N/A | 120.633536 | -8.1272817 | 1.56754451 |
| 5937 | NM_033087.3 | #N/A | 95.3120645 | -7.7850947 | 1.50169884 |
| 5938 | NM_001161841.1 | SULF2 | 114.94053 | -8.0585604 | 1.55448547 |
| 5939 | NR_036570.1 | STAG3L5P-PVRIG2P-PI | 106.973332 | -7.9533485 | 1.53437058 |
| 5940 | NM_004232.3 | #N/A | 110.012456 | -7.9925404 | 1.54206619 |
| 5941 | NM_001142749.2 | #N/A | 95.914212 | -7.7945589 | 1.50410012 |
| 5942 | NM_004225.2 | #N/A | 141.534972 | -8.3544803 | 1.61237328 |
| 5943 | NM_024096.1 | #N/A | 157.613514 | -8.5103835 | 1.64258546 |
| 5944 | NM_001101802.1 | #N/A | 93.2885772 | -7.7562234 | 1.49728563 |
| 5945 | NM_006323.4 | #N/A | 115.780113 | -8.0625524 | 1.5564444 |
| 5946 | NM_001098784.1 | #N/A | 134.947646 | -8.2865474 | 1.59975075 |
| 5947 | NM_021216.4 | ZNF71 | 119.74668 | -8.1134668 | 1.56648948 |
| 5948 | NM_021826.4 | #N/A | 105.889121 | -7.9360513 | 1.53230123 |
| 5949 | NR_073080.1 | PHLDA3 | 154.90516 | -8.4861068 | 1.63857793 |
| 5950 | NM_006739.3 | #N/A | 130.018222 | -8.2324162 | 1.58963909 |
| 5951 | NR_130935.1 | C11orf72 | 386.530617 | 1.75941405 | 0.33975017 |
| 5952 | NM_005550.3 | #N/A | 213.566892 | -8.948766 | 1.72809054 |
| 5953 | NM_000431.3 | #N/A | 139.020803 | -8.329599 | 1.60864045 |
| 5954 | NM_001007468.2 | #N/A | 135.149787 | -8.290898 | 1.60124225 |
| 5955 | NM_001065.3 | #N/A | 141.278201 | -8.3545253 | 1.61356154 |
| 5956 | NM_001002027.1 | #N/A | 160.526038 | -8.5370513 | 1.64888831 |
| 5957 | NM_001174061.1 | #N/A | 150.621595 | -8.4432141 | 1.63081842 |
| 5958 | NM_018151.4 | #N/A | 124.427066 | -8.1667241 | 1.57742136 |
| 5959 | NM_003081.4 | SNAP25 | 277.141879 | -9.3245988 | 1.80112261 |
| 5963 | NM_145021.5 | MARCHF8 | 97.3355875 | -7.8130071 | 1.50920792 |
| 5960 | NM_015391.3 | #N/A | 143.159241 | -8.3720128 | 1.61717666 |
| 5961 | NM_018262.3 | #N/A | 128.321806 | -8.2158802 | 1.58702932 |
| 5962 | NM_022719.2 | #N/A | 130.054479 | -8.23302 | 1.59030253 |
| 5964 | NR_045058.1 | #N/A | 120.444716 | -8.1207314 | 1.56864249 |
| 5965 | NM_001184985.1 | WNK1 | 93.9504017 | -7.7648689 | 1.49994367 |
| 5966 | NM_032320.6 | BTBD10 | 134.747051 | -8.2840173 | 1.60030747 |
| 5967 | NM_024095.4 | #N/A | 117.384187 | -8.0845368 | 1.56192227 |
| 5968 | NM_001033025.2 | #N/A | 124.449415 | -8.1704453 | 1.57861284 |
| 5969 | NM_138619.3 | #N/A | 122.202164 | -8.1424044 | 1.57324285 |
| 5970 | NM_001318849.1 | #N/A | 170.024433 | -8.6186383 | 1.66536928 |
| 5971 | NM_001136193.1 | #N/A | 135.247981 | -8.2873751 | 1.60185325 |
| 5973 | NR_002715.1 | RN7SL1 | 308.383339 | -7.9724501 | 1.54100343 |
| 5972 | NM_001282799.1 | #N/A | 114.80936 | -8.0515516 | 1.55629095 |
| 5974 | NM_007279.2 | U2AF2 | 101.129791 | -7.8693683 | 1.52108961 |
| 5975 | NM_001146054.1 | #N/A | 151.295468 | -8.4523088 | 1.63390812 |
| 5976 | NM_022875.2 | #N/A | 97.6585562 | -7.8223317 | 1.51212328 |

| | | | | | | |
|---|---|---|---|---|---|---|
| 5977 | NM_014181.2 | | #N/A | 124.26458 | -8.1668248 | 1.57881644 |
| 5978 | NM_012151.3 | F8A1 | | 96.7187979 | -7.8057941 | 1.5090466 |
| 5979 | NM_024031.3 | | #N/A | 141.212343 | -8.3505864 | 1.61436954 |
| 5980 | NM_018718.2 | | #N/A | 103.988925 | -7.9106855 | 1.52938236 |
| 5981 | NM_005460.3 | | #N/A | 105.403968 | -7.9287578 | 1.53290865 |
| 5982 | NM_006702.4 | | #N/A | 141.139831 | -8.3513076 | 1.61461056 |
| 5983 | NM_032038.2 | | #N/A | 153.659995 | -8.4727341 | 1.63811574 |
| 5984 | NM_001143981.1 | | #N/A | 188.640658 | -8.7697478 | 1.69562554 |
| 5985 | NM_003153.4 | | #N/A | 121.826855 | -8.1411507 | 1.57415947 |
| 5986 | NM_001004342.3 | | #N/A | 200.657987 | -8.8586192 | 1.71299273 |
| 5987 | NR_073148.1 | | #N/A | 103.170265 | -7.8997626 | 1.52758458 |
| 5988 | NM_001317724.1 | NOL7 | | 129.647816 | -8.2279769 | 1.59107225 |
| 5989 | NM_001100588.1 | | #N/A | 111.445279 | -8.0103559 | 1.54908406 |
| 5990 | NM_002523.2 | | #N/A | 137.485341 | -8.314475 | 1.60794788 |
| 5991 | NM_001317089.1 | | #N/A | 24.7617063 | 9.60339726 | 1.85730042 |
| 5992 | NM_022494.2 | | #N/A | 110.128017 | -7.9934468 | 1.54595922 |
| 5993 | NM_153280.2 | | #N/A | 131.617882 | -8.2504885 | 1.59579485 |
| 5995 | NM_020195.2 | SDR39U1 | | 113.689429 | -8.0388394 | 1.55509547 |
| 5994 | NM_003095.2 | | #N/A | 156.845894 | -8.5033148 | 1.64494544 |
| 5996 | NM_145034.4 | | #N/A | 189.251029 | 2.536897 | 0.49078413 |
| 5997 | NM_198291.2 | SRC | | 386.271756 | -4.649176 | 0.89949766 |
| 5998 | NM_005131.2 | | #N/A | 100.639189 | -7.8628454 | 1.5212944 |
| 5999 | NM_003404.4 | | #N/A | 123.122093 | -8.1527249 | 1.57742039 |
| 6000 | NM_004099.5 | | #N/A | 90.5222177 | -7.7100809 | 1.49180317 |
| 6001 | NM_147782.3 | | #N/A | 137.690297 | -8.3129344 | 1.60849158 |
| 6002 | NM_006133.2 | | #N/A | 141.345675 | -8.3533354 | 1.61639103 |
| 6003 | NM_020943.2 | | #N/A | 89.2361175 | -7.6897319 | 1.48797795 |
| 6004 | NM_004805.3 | | #N/A | 125.961106 | -6.9100026 | 1.3371128 |
| 6006 | NM_016451.4 | COPB1 | | 144.041966 | -8.3795905 | 1.62151736 |
| 6005 | NM_001166266.1 | LTBP1 | | 125.478858 | -8.1826956 | 1.58342419 |
| 6007 | NM_003924.3 | | #N/A | 183.646461 | -8.7319473 | 1.68981092 |
| 6008 | NM_005648.3 | | #N/A | 154.739852 | -8.4836033 | 1.64176007 |
| 6009 | NM_004441.4 | | #N/A | 196.725892 | -8.8299164 | 1.70882931 |
| 6010 | NM_001330542.1 | | #N/A | 111.623582 | -8.0113286 | 1.55042826 |
| 6011 | NM_003619.3 | | #N/A | 90.391443 | -7.7112706 | 1.49239221 |
| 6012 | NM_000837.1 | GRINA | | 117.403728 | -8.0860658 | 1.56499253 |
| 6013 | NM_033107.3 | | #N/A | 102.526185 | -7.8883566 | 1.52672689 |
| 6014 | NM_014906.4 | | #N/A | 133.823158 | -8.2760798 | 1.60179664 |
| 6015 | NM_013375.3 | | #N/A | 107.023002 | -7.9507539 | 1.53885806 |
| 6016 | NM_001204450.1 | CCPG1 | | 112.086597 | -8.0214224 | 1.55271632 |
| 6017 | NM_014600.2 | | #N/A | 126.405401 | -8.1929039 | 1.58598114 |
| 6018 | NM_015434.3 | | #N/A | 109.510455 | -7.9833546 | 1.54556975 |
| 6019 | NM_002610.4 | | #N/A | 130.817964 | -8.2395754 | 1.59542059 |

| | | | | | |
|---|---|---|---|---|---|
| 6020 | NM_020759.2 | #N/A | 101.75053 | -6.6004146 | 1.27803149 |
| 6021 | NR_045600.1 | #N/A | 131.985158 | -8.2552763 | 1.59855545 |
| 6022 | NM_001001975.1 | #N/A | 170.004381 | -8.6193047 | 1.66923677 |
| 6023 | NM_001946.3 | #N/A | 110.88527 | -8.0044955 | 1.55042353 |
| 6024 | NM_001080428.2 | #N/A | 105.571172 | -7.9322313 | 1.53650861 |
| 6025 | NM_020139.3 | #N/A | 94.8602079 | -7.7810251 | 1.50723067 |
| 6026 | NM_015449.3 | #N/A | 140.094357 | -8.3426811 | 1.61604157 |
| 6027 | NM_006457.4 | #N/A | 208.496799 | -8.9155454 | 1.72708823 |
| 6029 | NM_004799.3 | ZFYVE9 | 120.40653 | -8.1239634 | 1.57382056 |
| 6028 | NM_001007253.3 | #N/A | 110.826202 | -8.0029172 | 1.55037131 |
| 6030 | NM_001043351.1 | #N/A | 145.943777 | -8.4012383 | 1.6277717 |
| 6031 | NM_005871.3 | #N/A | 119.751455 | -8.1131947 | 1.57201755 |
| 6032 | NM_005067.5 | #N/A | 143.612996 | -6.3227198 | 1.22515764 |
| 6033 | NM_003992.4 | CLK3 | 99.0625128 | -7.8412221 | 1.51944304 |
| 6034 | NR_026922.1 | LOC150776 | 122.618418 | -8.1472496 | 1.57875248 |
| 6035 | NM_001170464.2 | #N/A | 153.873645 | -8.4750312 | 1.64235404 |
| 6036 | NM_015036.2 | #N/A | 140.085786 | -8.3409047 | 1.6165018 |
| 6037 | NM_138333.4 | #N/A | 87.5966361 | -7.6636689 | 1.48531346 |
| 6038 | NM_001190946.2 | #N/A | 128.386897 | -8.214368 | 1.59206984 |
| 6039 | NM_022480.3 | #N/A | 153.900272 | -8.4752115 | 1.64269332 |
| 6040 | NM_030583.3 | #N/A | 141.506975 | -8.3560554 | 1.61961244 |
| 6041 | NM_001047.3 | #N/A | 160.639167 | -8.5380699 | 1.65491624 |
| 6042 | NM_181671.2 | #N/A | 121.776186 | -8.1380917 | 1.57738869 |
| 6043 | NM_001037637.1 | BTF3 | 132.494488 | -8.2601158 | 1.60105587 |
| 6044 | NM_024639.4 | #N/A | 140.498293 | -8.3461308 | 1.61775212 |
| 6045 | NM_032880.4 | #N/A | 130.142452 | -8.2345103 | 1.59613154 |
| 6046 | NM_005944.6 | #N/A | 190.211922 | -8.7821036 | 1.70240971 |
| 6047 | NM_005483.2 | #N/A | 105.765832 | -7.935096 | 1.53828177 |
| 6048 | NM_133476.4 | #N/A | 110.856365 | -8.000662 | 1.55104423 |
| 6049 | NM_177537.2 | MAGED4B | 110.130957 | -7.9925361 | 1.54951836 |
| 6050 | NM_001252499.2 | #N/A | 112.256995 | -8.0193302 | 1.55481007 |
| 6051 | NM_001284254.1 | #N/A | 98.3051455 | -7.8332445 | 1.51882321 |
| 6052 | NM_032822.2 | #N/A | 103.170509 | -7.8992396 | 1.53164397 |
| 6053 | NM_001145306.1 | CDK6 | 107.51981 | -7.9572193 | 1.54292585 |
| 6054 | NM_002676.2 | #N/A | 131.146952 | -8.2453248 | 1.59878843 |
| 6055 | NM_014592.3 | #N/A | 120.172131 | -8.1196901 | 1.57457878 |
| 6056 | NM_080748.2 | #N/A | 144.479922 | -8.385016 | 1.62609006 |
| 6057 | NM_144631.5 | #N/A | 98.9449517 | -7.8388861 | 1.52030767 |
| 6058 | NM_133499.2 | SYN1 | 198.656934 | -8.8449457 | 1.71546489 |
| 6060 | NR_028327.1 | LOC100133331 | 90.8231714 | -7.7156267 | 1.49646064 |
| 6059 | NM_080430.3 | #N/A | 113.146013 | -8.0329374 | 1.55799779 |
| 6061 | NM_004554.4 | #N/A | 98.4738213 | -7.8341665 | 1.51949062 |
| 6062 | NM_024758.4 | #N/A | 172.993298 | 2.39279139 | 0.46410791 |

| | | | | | |
|---|---|---|---|---|---|
| 6063 | NM_001146227.1 | #N/A | 108.048722 | -5.34708 | 1.03716844 |
| 6064 | NM_015915.4 | ATL1 | 170.230441 | -8.6214525 | 1.67233947 |
| 6065 | NM_004284.4 | #N/A | 99.5460899 | -7.8504245 | 1.52283054 |
| 6066 | NM_001130053.3 | #N/A | 194.869151 | -8.8165941 | 1.7103206 |
| 6067 | NM_001321304.1 | #N/A | 112.466215 | -8.0262788 | 1.55706433 |
| 6068 | NM_004542.3 | #N/A | 140.438012 | -8.3442955 | 1.61883044 |
| 6069 | NM_001675.4 | ATF4 | 98.1933016 | -7.8243744 | 1.51819162 |
| 6070 | NM_018288.3 | #N/A | 91.5749781 | -7.7303814 | 1.50024738 |
| 6071 | NM_152334.2 | #N/A | 103.237845 | -7.8983835 | 1.53295867 |
| 6072 | NM_015453.2 | #N/A | 132.879747 | -8.2635181 | 1.60386078 |
| 6073 | NM_005384.2 | #N/A | 155.01888 | -8.4857434 | 1.64700241 |
| 6074 | NM_015063.2 | #N/A | 178.143465 | -8.6870418 | 1.68612661 |
| 6075 | NM_005828.4 | #N/A | 1065.85944 | -5.723301 | 1.11097896 |
| 6076 | NM_138439.2 | #N/A | 115.266911 | -8.0602195 | 1.56475689 |
| 6077 | NM_000663.4 | ABAT | 141.099762 | -8.3498329 | 1.62104717 |
| 6078 | NM_020652.2 | #N/A | 121.152301 | -8.1305249 | 1.57846503 |
| 6079 | NM_001896.2 | #N/A | 104.95011 | -7.9233056 | 1.53843133 |
| 6081 | NM_031206.4 | LAS1L | 115.356607 | -8.0600687 | 1.56507911 |
| 6080 | NM_001293233.1 | #N/A | 163.163336 | -8.5595716 | 1.66207084 |
| 6082 | NM_014860.2 | #N/A | 117.252118 | -8.0828765 | 1.56958866 |
| 6083 | NM_005700.4 | #N/A | 132.154504 | -8.2569137 | 1.60345776 |
| 6084 | NM_020923.2 | #N/A | 144.775531 | -8.3901087 | 1.62950579 |
| 6085 | NM_003104.5 | #N/A | 121.579034 | -8.1360675 | 1.58028528 |
| 6086 | NM_006327.3 | #N/A | 131.723185 | -8.2509428 | 1.60259361 |
| 6087 | NM_133173.2 | APBB3 | 174.290224 | -8.6552947 | 1.68123762 |
| 6088 | NM_006080.2 | #N/A | 183.912301 | -8.7333048 | 1.69655734 |
| 6089 | NM_001078645.2 | #N/A | 151.978304 | -8.4570093 | 1.64295652 |
| 6090 | NM_001256763.1 | #N/A | 164.101478 | -8.5685873 | 1.66462874 |
| 6091 | NM_002145.3 | HOXB2 | 133.999939 | -8.2778391 | 1.6081767 |
| 6092 | NM_015471.3 | #N/A | 110.04907 | -7.9902165 | 1.55238113 |
| 6093 | NM_002249.5 | #N/A | 93.268202 | -7.7511244 | 1.50598281 |
| 6094 | NM_001098205.1 | HNRNPF | 179.251202 | -8.6973694 | 1.68989753 |
| 6095 | NM_020961.3 | #N/A | 97.6492781 | -7.8184529 | 1.5191485 |
| 6098 | NR_110954.1 | LOC101927040 | 30.7959162 | 3.20351669 | 0.62248605 |
| 6096 | NM_007043.6 | #N/A | 111.230904 | -8.0065525 | 1.55576669 |
| 6097 | NM_012429.4 | #N/A | 95.0135989 | -7.784253 | 1.51258809 |
| 6099 | NM_001242868.1 | #N/A | 152.696566 | -8.4639115 | 1.64474288 |
| 6100 | NM_001160372.3 | #N/A | 138.571414 | -8.3258042 | 1.61795914 |
| 6101 | NM_022465.3 | #N/A | 108.705064 | -7.9746475 | 1.54981597 |
| 6102 | NM_001320803.1 | ORMDL3 | 92.0796852 | -7.7341491 | 1.50311086 |
| 6103 | NR_036684.1 | #N/A | 107.256036 | -7.9531028 | 1.54565962 |
| 6104 | NM_024083.3 | #N/A | 114.866904 | -8.0530164 | 1.56509462 |
| 6105 | NM_014239.3 | #N/A | 119.082747 | -8.1061889 | 1.57561552 |

| | | | | | |
|---|---|---|---|---|---|
| 6106 | NM_001136053.2 | #N/A | 113.45215 | -8.0366566 | 1.56214846 |
| 6107 | NM_004998.3 | #N/A | 97.7499252 | -7.8239169 | 1.52084222 |
| 6108 | NM_012124.2 | #N/A | 101.108488 | -7.8701169 | 1.52982395 |
| 6109 | NM_001134225.1 | #N/A | 97.4444548 | -7.8186041 | 1.51989072 |
| 6110 | NM_014166.3 | #N/A | 118.022439 | -8.0930016 | 1.57322522 |
| 6111 | NM_014049.4 | #N/A | 99.2163785 | -7.8439492 | 1.52497787 |
| 6112 | NM_182640.2 | #N/A | 98.7568142 | -7.8373541 | 1.52369252 |
| 6113 | NM_033395.1 | #N/A | 100.809547 | -7.8655577 | 1.52930347 |
| 6114 | NM_005908.3 | #N/A | 109.686099 | -6.7100471 | 1.30476651 |
| 6115 | NM_001318785.1 | RARS2 | 123.596928 | -8.1593334 | 1.58664806 |
| 6116 | NM_014190.3 | #N/A | 433.27391 | -9.9697415 | 1.93870132 |
| 6117 | NM_014688.3 | #N/A | 102.578632 | -7.8906041 | 1.53439507 |
| 6118 | NM_014640.4 | #N/A | 222.226803 | -9.0073198 | 1.75174597 |
| 6119 | NM_013283.4 | #N/A | 95.566285 | -7.7869188 | 1.51457909 |
| 6120 | NM_015061.3 | #N/A | 89.2521591 | -7.6891756 | 1.49558772 |
| 6121 | NM_001040440.2 | #N/A | 185.429513 | -8.7446157 | 1.7008958 |
| 6122 | NM_016579.3 | #N/A | 146.620641 | -8.4063399 | 1.63513545 |
| 6123 | NM_001039708.1 | #N/A | 104.25277 | -7.9124914 | 1.53914551 |
| 6124 | NM_001270454.1 | #N/A | 99.0433405 | -7.8417373 | 1.52539009 |
| 6125 | NM_002946.4 | #N/A | 129.967389 | -8.2315807 | 1.60123171 |
| 6126 | NR_130156.1 | #N/A | 93.2807967 | -7.7532569 | 1.50829598 |
| 6127 | NM_032167.4 | #N/A | 109.243796 | -7.9820261 | 1.55293672 |
| 6128 | NM_013448.2 | #N/A | 101.687084 | -7.8789183 | 1.53302418 |
| 6129 | NM_002918.4 | #N/A | 98.6526873 | -7.8346724 | 1.52448683 |
| 6130 | NM_022823.2 | #N/A | 161.065613 | -7.2691925 | 1.41469261 |
| 6131 | NM_002077.3 | #N/A | 127.097094 | -6.8288891 | 1.32907857 |
| 6132 | NM_207111.3 | #N/A | 104.510401 | -7.9143948 | 1.54044818 |
| 6133 | NM_001135050.1 | #N/A | 155.199842 | -8.4879528 | 1.65216402 |
| 6134 | NM_005089.3 | ZRSR2 | 107.866211 | -7.9620023 | 1.54982613 |
| 6135 | NM_033394.2 | #N/A | 144.585945 | -8.3878422 | 1.6328011 |
| 6136 | NM_152313.3 | #N/A | 121.037173 | -8.1291234 | 1.58243667 |
| 6137 | NM_018208.3 | #N/A | 108.723629 | -7.974968 | 1.55252515 |
| 6138 | NM_016035.4 | #N/A | 118.964353 | -6.743888 | 1.3129199 |
| 6139 | NM_173552.3 | #N/A | 91.2400393 | -7.7231381 | 1.50359665 |
| 6140 | NM_032882.5 | PNMA6A | 107.325456 | -7.9563236 | 1.54906538 |
| 6141 | NM_001019.4 | #N/A | 144.554115 | -8.385491 | 1.63273838 |
| 6142 | NM_001130142.1 | #N/A | 93.5709076 | -7.7582311 | 1.51073704 |
| 6143 | NM_001916.4 | #N/A | 358.431313 | -6.1483641 | 1.19727315 |
| 6144 | NM_001134375.1 | #N/A | 101.780788 | -7.877752 | 1.53407351 |
| 6145 | NR_135599.1 | #N/A | 115.709752 | -8.0636284 | 1.57038636 |
| 6146 | NM_152735.3 | #N/A | 100.762868 | -7.867216 | 1.53228304 |
| 6147 | NM_032603.4 | #N/A | 93.4656856 | -7.758935 | 1.51121968 |
| 6148 | NM_000021.3 | #N/A | 148.252859 | -8.4220671 | 1.64052544 |

| | | | | | |
|---|---|---|---|---|---|
| 6149 | NM_018335.4 | #N/A | 91.9140788 | -7.7301004 | 1.50576461 |
| 6150 | NM_144607.5 | #N/A | 98.705382 | -7.837641 | 1.526757 |
| 6151 | NM_001008215.2 | #N/A | 105.57575 | -7.9319063 | 1.54526427 |
| 6152 | NM_031288.3 | #N/A | 88.6830435 | -7.6829454 | 1.49683443 |
| 6153 | NM_001258429.1 | #N/A | 142.214339 | -8.3620704 | 1.6291939 |
| 6154 | NM_015675.3 | #N/A | 123.169368 | -8.1563583 | 1.58912249 |
| 6155 | NM_001135651.2 | #N/A | 94.4789856 | -7.7728535 | 1.51445856 |
| 6156 | NM_032301.2 | #N/A | 89.0785377 | -7.6893891 | 1.4982082 |
| 6157 | NM_000367.4 | TPMT | 89.5419645 | -7.6943011 | 1.49917504 |
| 6158 | NM_002455.4 | MTX1 | 103.519385 | -7.9055925 | 1.54040309 |
| 6159 | NM_001551.2 | #N/A | 178.77305 | -7.2655562 | 1.41572521 |
| 6160 | NM_032380.4 | #N/A | 115.705187 | -8.0654131 | 1.57161612 |
| 6161 | NM_018648.3 | NOP10 | 146.539795 | -8.4061657 | 1.63804272 |
| 6162 | NM_170784.2 | MKKS | 105.240299 | -7.9279206 | 1.54488809 |
| 6163 | NM_016001.2 | #N/A | 122.519364 | -8.1459501 | 1.58756729 |
| 6164 | NM_025158.4 | #N/A | 104.686522 | -7.9174339 | 1.54307116 |
| 6165 | NM_182905.4 | #N/A | 100.505353 | -7.8611833 | 1.53217563 |
| 6166 | NM_015448.2 | #N/A | 85.7958913 | -7.6323533 | 1.48760329 |
| 6167 | NM_005981.4 | #N/A | 97.3768477 | -7.8134852 | 1.5230653 |
| 6168 | NM_006849.2 | #N/A | 109.937218 | -7.9923584 | 1.55793187 |
| 6169 | NM_032169.4 | #N/A | 112.469221 | -8.0248136 | 1.56425032 |
| 6170 | NM_146387.1 | MRPL4 | 128.267496 | -8.2134191 | 1.60109524 |
| 6171 | NM_003370.3 | #N/A | 106.537123 | -7.9461602 | 1.54906088 |
| 6172 | NM_175921.5 | #N/A | 135.62726 | -7.0186934 | 1.36827825 |
| 6173 | NM_001278.4 | #N/A | 98.5357413 | -7.8331633 | 1.52711084 |
| 6174 | NM_001244808.1 | #N/A | 118.173561 | -8.0960745 | 1.57846819 |
| 6175 | NR_045209.1 | #N/A | 117.269732 | -8.0821073 | 1.57586993 |
| 6176 | NM_058229.3 | #N/A | 153.348695 | -8.471532 | 1.65181371 |
| 6177 | NM_001307943.1 | TTC17 | 91.155506 | -7.7197192 | 1.5053397 |
| 6178 | NM_001005909.2 | IP6K2 | 128.104578 | -8.213281 | 1.60160483 |
| 6180 | NM_032445.2 | MEGF11 | 115.817185 | -8.0673768 | 1.5731881 |
| 6179 | NM_001320941.1 | #N/A | 25.0953723 | 9.63599895 | 1.87908649 |
| 6181 | NM_152721.5 | #N/A | 249.183313 | -7.6906556 | 1.49976492 |
| 6182 | NM_003825.3 | #N/A | 92.0501521 | -7.7380886 | 1.50907918 |
| 6183 | NM_130468.3 | CHST14 | 99.5157255 | -7.8505294 | 1.53105693 |
| 6184 | NM_004775.3 | #N/A | 87.3062897 | -7.6605197 | 1.49400978 |
| 6185 | NM_198244.2 | #N/A | 118.807769 | -8.1043338 | 1.58064934 |
| 6186 | NM_032145.4 | #N/A | 96.3433115 | -7.7997364 | 1.52128279 |
| 6187 | NM_017812.3 | #N/A | 92.191323 | -7.7348712 | 1.50868694 |
| 6188 | NM_001134338.2 | #N/A | 127.53755 | -8.2058494 | 1.60070155 |
| 6189 | NM_003369.3 | #N/A | 138.198106 | -6.9336461 | 1.35258852 |
| 6190 | NM_183393.2 | #N/A | 167.86663 | -8.6018468 | 1.6781252 |
| 6191 | NM_001128228.2 | #N/A | 102.474993 | -7.8897368 | 1.53923843 |

| | | | | | | |
|---|---|---|---|---|---|---|
| 6192 | NM_152470.2 | | #N/A | 110.099432 | -7.9917079 | 1.55924195 |
| 6193 | NM_032313.3 | | #N/A | 99.7523655 | -7.8502674 | 1.53167658 |
| 6194 | NM_001163735.1 | MYO19 | | 101.501281 | -7.8747494 | 1.53648263 |
| 6195 | NM_001172624.1 | NEO1 | | 278.04393 | -9.3305153 | 1.82059827 |
| 6196 | NR_028104.1 | | #N/A | 105.356943 | -7.9271626 | 1.54682913 |
| 6197 | NM_001017408.2 | | #N/A | 115.403011 | -8.0577062 | 1.57232885 |
| 6198 | NM_144949.2 | | #N/A | 95.0972048 | -7.7814505 | 1.51848836 |
| 6199 | NM_001957.3 | | #N/A | 104.682605 | -7.9227119 | 1.5460782 |
| 6200 | NM_005198.4 | CHKB | | 111.197855 | -6.654812 | 1.2988563 |
| 6201 | NM_016156.5 | MTMR2 | | 141.922785 | -8.3598838 | 1.63166147 |
| 6202 | NM_003705.4 | | #N/A | 132.919177 | -8.264407 | 1.61304511 |
| 6203 | NM_001077199.2 | | #N/A | 169.094811 | -8.6117817 | 1.68087655 |
| 6204 | NM_020662.3 | | #N/A | 98.1804627 | -7.8267862 | 1.52781195 |
| 6205 | NM_014062.2 | | #N/A | 109.990034 | -6.639248 | 1.29601938 |
| 6206 | NM_199287.2 | | #N/A | 102.196031 | -7.885405 | 1.53928103 |
| 6207 | NM_002830.3 | | #N/A | 121.209627 | -8.1291855 | 1.58687912 |
| 6208 | NM_014604.3 | | #N/A | 137.853956 | -8.3172732 | 1.62375376 |
| 6209 | NM_001013843.2 | | #N/A | 113.692836 | -8.0386787 | 1.56938581 |
| 6210 | NM_001042610.2 | | #N/A | 164.389918 | -8.5716353 | 1.67349626 |
| 6211 | NM_001199260.1 | | #N/A | 92.0434755 | -7.7355488 | 1.51032031 |
| 6212 | NM_138384.2 | | #N/A | 116.60702 | -8.0763893 | 1.57690365 |
| 6213 | NM_003477.2 | | #N/A | 111.105649 | -8.0051837 | 1.56304508 |
| 6214 | NM_145245.4 | | #N/A | 275.966969 | -6.5808693 | 1.28494246 |
| 6215 | NM_032124.4 | | #N/A | 118.76584 | -8.1013267 | 1.5818349 |
| 6216 | NM_004416.2 | DTX1 | | 196.247282 | -8.8267791 | 1.72355013 |
| 6217 | NM_014629.3 | | #N/A | 88.3945088 | -7.6763473 | 1.49891387 |
| 6218 | NM_013306.4 | | #N/A | 127.374517 | -8.2031651 | 1.60185121 |
| 6219 | NM_001330103.1 | | #N/A | 100.091169 | -4.939545 | 0.96463293 |
| 6220 | NM_032656.3 | | #N/A | 110.164125 | -7.9940538 | 1.56120116 |
| 6221 | NM_001798.4 | | #N/A | 98.7648255 | -7.8339765 | 1.53010844 |
| 6222 | NM_198467.2 | | #N/A | 115.983903 | -8.0672587 | 1.57568849 |
| 6223 | NM_206831.2 | | #N/A | 136.852874 | -8.3065995 | 1.62244675 |
| 6224 | NM_001321209.1 | | #N/A | 107.804804 | -7.9621056 | 1.55517879 |
| 6225 | NM_001002914.2 | KCTD11 | | 91.2873771 | -7.7264751 | 1.509252 |
| 6226 | NM_014034.2 | | #N/A | 129.670013 | -8.2284884 | 1.6073854 |
| 6227 | NM_003617.3 | | #N/A | 159.662475 | -8.5301209 | 1.66632856 |
| 6228 | NM_005779.2 | | #N/A | 107.232402 | -7.958637 | 1.55478345 |
| 6229 | NM_005110.3 | | #N/A | 92.5430833 | -7.7457465 | 1.51328563 |
| 6230 | NM_005718.4 | ARPC4 | | 139.439782 | -8.3343072 | 1.62836651 |
| 6231 | NM_001252078.1 | | #N/A | 122.009726 | -8.1415955 | 1.59075119 |
| 6232 | NM_019012.5 | | #N/A | 95.5025536 | -7.7850336 | 1.52113706 |
| 6233 | NM_001201338.1 | | #N/A | 179.031798 | -8.695689 | 1.69926687 |
| 6234 | NM_022343.3 | | #N/A | 159.00876 | -8.5240054 | 1.66575547 |

| | | | | | |
|---|---|---|---|---|---|
| 6235 | NM_001039465.1 | #N/A | 96.0136141 | -7.7953501 | 1.52343448 |
| 6236 | NM_016174.4 | #N/A | 118.649012 | -8.1014904 | 1.58337717 |
| 6237 | NR_110492.1 | TUG1 | 152.510412 | -8.4637164 | 1.65422657 |
| 6238 | NM_020784.2 | #N/A | 128.485265 | -8.2154091 | 1.60578291 |
| 6239 | NM_003798.3 | #N/A | 89.1304233 | -7.6914716 | 1.5034402 |
| 6240 | NM_002495.3 | #N/A | 120.831331 | -8.1275743 | 1.5887677 |
| 6241 | NM_006803.3 | #N/A | 147.852561 | -7.1443568 | 1.39666439 |
| 6242 | NM_020456.3 | #N/A | 95.8239117 | -7.790106 | 1.52292877 |
| 6243 | NM_016056.3 | TMBIM4 | 94.0994321 | -7.7669968 | 1.51843215 |
| 6244 | NR_028024.1 | #N/A | 109.251936 | -7.9799236 | 1.56011481 |
| 6245 | NM_022753.3 | #N/A | 101.40312 | -7.8763632 | 1.53994373 |
| 6246 | NM_005802.4 | #N/A | 90.0449369 | -7.7017798 | 1.50593033 |
| 6247 | NM_002035.2 | #N/A | 209.482546 | -3.2793273 | 0.64122955 |
| 6248 | NM_001206612.1 | CHTOP | 172.059273 | -8.6361356 | 1.68870243 |
| 6249 | NM_015698.5 | #N/A | 100.854014 | -7.8675394 | 1.53841866 |
| 6250 | NM_001130089.1 | KARS1 | 126.518906 | -8.1933898 | 1.60217942 |
| 6251 | NM_031420.3 | #N/A | 117.281998 | -8.0840159 | 1.58082117 |
| 6252 | NM_005606.6 | #N/A | 151.033229 | -8.4491245 | 1.65232103 |
| 6253 | NM_015503.2 | SH2B1 | 98.6995885 | -7.8363429 | 1.53256011 |
| 6254 | NM_014784.3 | #N/A | 160.845722 | -8.5392906 | 1.670093 |
| 6255 | NM_001305626.1 | CALM2 | 91.5425239 | -7.7305304 | 1.51193497 |
| 6256 | NM_012192.3 | #N/A | 122.89977 | -6.7809314 | 1.3263684 |
| 6257 | NM_002897.4 | #N/A | 124.52574 | -8.1729608 | 1.59876368 |
| 6258 | NM_025140.2 | #N/A | 112.845235 | -8.0279632 | 1.57085822 |
| 6259 | NM_004922.3 | #N/A | 98.4924331 | -7.8332854 | 1.53277481 |
| 6260 | NM_005400.2 | #N/A | 172.724341 | -8.6424561 | 1.69116121 |
| 6261 | NM_181349.2 | #N/A | 96.0073106 | -7.7927118 | 1.5249674 |
| 6262 | NM_017813.4 | #N/A | 435.001594 | -4.555135 | 0.89149546 |
| 6263 | NR_024565.1 | ZNF271P | 113.180748 | -8.0325207 | 1.57210103 |
| 6264 | NM_001004356.2 | #N/A | 113.396112 | -8.0363642 | 1.57297095 |
| 6265 | NM_001160103.1 | #N/A | 87.2529053 | -7.6606547 | 1.49942677 |
| 6266 | NM_004259.6 | #N/A | 91.259621 | -7.7241113 | 1.51187898 |
| 6267 | NM_032383.4 | #N/A | 92.1762787 | -7.7385063 | 1.51472364 |
| 6268 | NM_024066.2 | #N/A | 112.818239 | -8.0263717 | 1.57125202 |
| 6269 | NM_001168618.1 | IFT46 | 85.3725985 | -7.6290261 | 1.493534 |
| 6271 | NM_001281533.1 | EPB41L3 | 172.182158 | -8.6375055 | 1.69107699 |
| 6270 | NM_000836.2 | GRIN2D | 116.026105 | -8.0688786 | 1.57975757 |
| 6272 | NM_001938.2 | #N/A | 279.01101 | -3.9494665 | 0.77325324 |
| 6273 | NM_001079841.2 | OCIAD1 | 163.080903 | -8.5599086 | 1.67603227 |
| 6274 | NM_181783.3 | #N/A | 105.684697 | -7.9338752 | 1.55349167 |
| 6275 | NM_139055.3 | ADAMTS15 | 104.536419 | -7.9206823 | 1.55092147 |
| 6276 | NM_001171506.2 | MCFD2 | 94.6846686 | -7.7757082 | 1.52257983 |
| 6277 | NM_005881.3 | #N/A | 98.0109802 | -7.8283106 | 1.53291543 |

| | | | | | |
|---|---|---|---|---|---|
| 6278 | NM_001015052.2 | #N/A | 118.769372 | -8.1022062 | 1.58670579 |
| 6279 | NM_002928.3 | #N/A | 105.5238 | -7.9292349 | 1.5529631 |
| 6280 | NM_006219.2 | #N/A | 97.9265938 | -7.8234033 | 1.53223893 |
| 6281 | NM_005380.7 | #N/A | 100.869694 | -7.8699769 | 1.54146776 |
| 6282 | NM_024686.4 | #N/A | 84.3262479 | -7.6071778 | 1.4900315 |
| 6283 | NM_018093.3 | WDR74 | 104.645781 | -7.9191738 | 1.55133517 |
| 6284 | NM_001382.3 | #N/A | 102.180033 | -7.8878216 | 1.5452056 |
| 6285 | NM_001278480.1 | SUPT20H | 103.584651 | -7.9036208 | 1.54833755 |
| 6286 | NM_014223.4 | #N/A | 137.073155 | -8.3088457 | 1.62773846 |
| 6287 | NM_003171.4 | #N/A | 91.5912333 | -7.726468 | 1.51368855 |
| 6288 | NM_000868.3 | HTR2C | 161.286917 | -8.5453055 | 1.6743532 |
| 6289 | NM_004517.3 | #N/A | 109.571807 | -7.9860002 | 1.5648659 |
| 6290 | NM_004228.6 | #N/A | 134.58275 | -8.283234 | 1.62322982 |
| 6291 | NR_135468.1 | #N/A | 108.078279 | -7.9654527 | 1.56096991 |
| 6292 | NM_001271836.1 | #N/A | 88.4196157 | -7.6748962 | 1.50408149 |
| 6293 | NM_001282689.1 | #N/A | 91.9962914 | -7.7362998 | 1.51622544 |
| 6294 | NM_012280.3 | #N/A | 133.606546 | -8.2719994 | 1.62121817 |
| 6295 | NM_003765.2 | #N/A | 99.9180045 | -7.8538125 | 1.53931441 |
| 6296 | NM_003685.2 | #N/A | 955.775138 | -6.2533041 | 1.22565334 |
| 6297 | NM_004426.2 | #N/A | 656.692171 | -6.0453953 | 1.18490126 |
| 6298 | NM_205768.2 | ZBTB18 | 301.975257 | -9.4483755 | 1.85197528 |
| 6299 | NM_173680.3 | #N/A | 104.425197 | -7.9153767 | 1.55150811 |
| 6300 | NM_007067.4 | #N/A | 113.782965 | -8.0433468 | 1.57661526 |
| 6301 | NM_199341.3 | #N/A | 97.319278 | -7.8150777 | 1.53187338 |
| 6302 | NM_001127496.1 | #N/A | 119.105307 | -8.1068281 | 1.58908304 |
| 6303 | NM_003704.3 | FAM193A | 122.39265 | -8.1446041 | 1.59649863 |
| 6304 | NM_001288973.1 | #N/A | 218.840385 | -6.3215056 | 1.23916977 |
| 6305 | NM_017638.2 | #N/A | 85.3169735 | -7.6263892 | 1.49500586 |
| 6306 | NM_001455.3 | #N/A | 112.728466 | -8.0252798 | 1.57328516 |
| 6307 | NM_001040284.2 | #N/A | 84.9021817 | -7.6190304 | 1.49367368 |
| 6308 | NM_006852.3 | #N/A | 115.490575 | -8.0598737 | 1.5801229 |
| 6309 | NR_033302.1 | #N/A | 88.5839739 | -7.679069 | 1.50548293 |
| 6310 | NM_005869.3 | #N/A | 105.013394 | -7.9236008 | 1.55344134 |
| 6311 | NM_001007559.2 | #N/A | 96.5486419 | -7.8070587 | 1.53071361 |
| 6312 | NM_001098638.1 | #N/A | 92.1257947 | -7.7353363 | 1.51689037 |
| 6313 | NM_033301.2 | #N/A | 102.505987 | -7.8906095 | 1.54749743 |
| 6314 | NM_017667.3 | #N/A | 115.599447 | -8.0627672 | 1.5812884 |
| 6315 | NM_001242783.1 | #N/A | 106.545528 | -7.9455809 | 1.55841384 |
| 6316 | NM_018431.4 | #N/A | 147.489454 | -8.4138961 | 1.65040052 |
| 6317 | NM_152868.2 | #N/A | 111.341922 | -8.0076484 | 1.57074499 |
| 6318 | NM_003321.4 | #N/A | 681.249399 | -4.9344759 | 0.9679369 |
| 6319 | NM_001077352.1 | #N/A | 86.8392367 | -7.6507775 | 1.50079007 |
| 6320 | NM_002611.4 | #N/A | 106.445922 | -7.9459921 | 1.55870058 |

| | | | | | |
|---|---|---|---|---|---|
| 6321 | NM_025132.3 | #N/A | 120.436252 | -8.1239231 | 1.59363634 |
| 6322 | NM_002202.2 | #N/A | 139.475058 | -8.3349795 | 1.63519847 |
| 6323 | NM_032289.2 | #N/A | 189.258819 | -7.3327965 | 1.43862148 |
| 6324 | NM_181842.2 | #N/A | 121.472599 | -8.1349511 | 1.59605667 |
| 6325 | NM_024814.3 | #N/A | 111.942692 | -8.0162289 | 1.57277339 |
| 6326 | NM_001206528.2 | CRTAC1 | 173.58668 | -8.650486 | 1.69738601 |
| 6327 | NM_020871.3 | #N/A | 113.765746 | -8.03895 | 1.57741525 |
| 6328 | NM_003748.3 | #N/A | 160.14228 | -8.5345228 | 1.67485803 |
| 6329 | NM_001034841.3 | ITPRIPL2 | 100.276782 | -7.8617058 | 1.54283558 |
| 6330 | NM_003110.5 | #N/A | 85.9356471 | -7.6375644 | 1.49888432 |
| 6331 | NM_018116.3 | MSTO1 | 107.711606 | -7.9620993 | 1.56260566 |
| 6333 | NM_001168303.1 | KLHL13 | 106.601122 | -7.9479487 | 1.55985561 |
| 6332 | NM_001044305.2 | #N/A | 263.124535 | -9.2496558 | 1.81532058 |
| 6334 | NM_017849.3 | TMEM127 | 149.739684 | -8.4348519 | 1.6554371 |
| 6335 | NR_024113.1 | #N/A | 134.042048 | -8.2755596 | 1.62418245 |
| 6336 | NM_032982.3 | #N/A | 109.475514 | -7.9873234 | 1.56785866 |
| 6337 | NM_033114.3 | #N/A | 214.709302 | -7.4938675 | 1.47100324 |
| 6338 | NM_006593.3 | #N/A | 410.959148 | -9.8930952 | 1.94212855 |
| 6339 | NM_001400.4 | #N/A | 87.7392864 | -7.6651772 | 1.50508735 |
| 6340 | NM_032193.3 | #N/A | 106.297929 | -7.9430461 | 1.5596857 |
| 6341 | NM_002093.3 | #N/A | 120.18259 | -8.1204699 | 1.5946561 |
| 6342 | NR_102706.1 | #N/A | 123.23953 | -8.1564705 | 1.60178282 |
| 6344 | NM_001242882.1 | NAXD | 118.509602 | -8.097848 | 1.59039207 |
| 6343 | NM_001136159.1 | #N/A | 90.0186641 | -7.7024237 | 1.51271867 |
| 6345 | NM_024779.4 | #N/A | 101.393436 | -7.8737214 | 1.54637411 |
| 6346 | NM_001242833.1 | NDRG4 | 159.295161 | -8.5266187 | 1.67465064 |
| 6347 | NM_018325.4 | #N/A | 127.458303 | -8.2030606 | 1.61109777 |
| 6348 | NM_001013836.1 | #N/A | 120.860296 | -8.1272309 | 1.59623205 |
| 6349 | NM_020187.2 | #N/A | 98.8257255 | -7.834995 | 1.53889897 |
| 6350 | NM_024619.3 | #N/A | 122.355176 | -6.8685196 | 1.34923228 |
| 6351 | NM_001278380.1 | ARL6IP4 | 95.2320335 | -7.7837907 | 1.52905053 |
| 6352 | NM_006858.3 | #N/A | 97.1513175 | -7.8145691 | 1.53518134 |
| 6353 | NM_001134367.3 | SLC6A6 | 87.5008805 | -7.659513 | 1.50478399 |
| 6354 | NM_001609.3 | #N/A | 95.8322872 | -7.7912893 | 1.53077621 |
| 6355 | NM_014805.3 | #N/A | 422.625811 | -2.5339953 | 0.49786407 |
| 6356 | NM_031450.3 | #N/A | 104.361287 | -7.9160141 | 1.55532913 |
| 6357 | NM_003635.3 | #N/A | 90.8291179 | -7.7171064 | 1.51633588 |
| 6358 | NM_022073.3 | #N/A | 102.154208 | -7.8832498 | 1.54910802 |
| 6359 | NM_016581.4 | #N/A | 135.486016 | -8.2915385 | 1.62944644 |
| 6360 | NM_012279.3 | #N/A | 91.7470653 | -7.7281226 | 1.51881233 |
| 6361 | NM_181659.2 | #N/A | 91.6995185 | -7.7317854 | 1.51955826 |
| 6362 | NR_108090.1 | LINC01447 | 77.1327068 | 3.63077115 | 0.71359653 |
| 6363 | NM_015529.3 | #N/A | 125.444086 | -8.1820092 | 1.60813994 |

| | | | | | |
|---|---|---|---|---|---|
| 6364 | NM_025196.3 | #N/A | 106.304481 | -7.9424014 | 1.5611519 |
| 6365 | NM_175884.5 | #N/A | 114.991653 | -8.0546953 | 1.58330265 |
| 6366 | NM_017867.2 | #N/A | 120.133677 | -8.1178755 | 1.59579632 |
| 6367 | NM_032349.3 | NUDT16L1 | 129.130324 | -8.2227563 | 1.61644251 |
| 6369 | NR_003089.1 | OTUB1 | 89.9846912 | -7.7027505 | 1.51435334 |
| 6368 | NM_003598.1 | #N/A | 101.483717 | -7.8742952 | 1.54808785 |
| 6370 | NM_007344.3 | #N/A | 87.6082049 | -7.6616125 | 1.50632385 |
| 6371 | NM_001032382.1 | PQBP1 | 113.988204 | -8.0413413 | 1.58100273 |
| 6372 | NM_025234.2 | #N/A | 119.066235 | -8.1058348 | 1.59368584 |
| 6373 | NM_147175.3 | #N/A | 137.954841 | -8.3175977 | 1.63533149 |
| 6374 | NM_015454.2 | LARP7 | 98.2199565 | -7.8258185 | 1.5386592 |
| 6375 | NM_018387.4 | #N/A | 144.974212 | -4.1988126 | 0.82554401 |
| 6376 | NM_004937.2 | #N/A | 89.6043564 | -7.6963554 | 1.51333188 |
| 6377 | NM_182924.3 | #N/A | 104.82803 | -7.9239055 | 1.55807169 |
| 6378 | NM_018330.6 | #N/A | 197.264192 | -8.8340044 | 1.73707581 |
| 6379 | NM_001135734.2 | ZNF384 | 105.308629 | -7.9284854 | 1.55913981 |
| 6380 | NR_144474.1 | DPH1 | 135.221031 | -8.2916726 | 1.63064129 |
| 6381 | NM_014817.3 | #N/A | 103.18854 | -7.9029297 | 1.55423059 |
| 6382 | NM_004836.6 | #N/A | 89.6709548 | -7.6946207 | 1.51327607 |
| 6383 | NM_020774.3 | MIB1 | 554.520719 | -3.972129 | 0.78119514 |
| 6384 | NM_014602.2 | #N/A | 91.7428045 | -7.7291792 | 1.52020017 |
| 6385 | NM_001099409.1 | #N/A | 86.869992 | -7.6509318 | 1.50482001 |
| 6386 | NM_001270448.1 | ACADVL | 97.2578402 | -7.8180788 | 1.53778033 |
| 6387 | NM_052854.3 | #N/A | 95.1705969 | -7.7852575 | 1.53138822 |
| 6388 | NM_001114118.2 | #N/A | 288.248305 | -3.1577775 | 0.62118437 |
| 6389 | NM_001193582.1 | NRCAM | 273.800981 | -9.3071353 | 1.83102739 |
| 6390 | NM_033542.3 | #N/A | 88.5435329 | -7.6804895 | 1.51107976 |
| 6391 | NM_152704.3 | AMER2 | 137.816632 | -8.3161488 | 1.63619819 |
| 6392 | NM_001077207.3 | #N/A | 151.24144 | -8.4513463 | 1.66300201 |
| 6393 | NM_001331012.1 | #N/A | 129.553111 | -8.2287896 | 1.61926499 |
| 6394 | NM_001171510.2 | #N/A | 204.410236 | 7.74121429 | 1.5233374 |
| 6395 | NM_152287.3 | #N/A | 87.2203615 | -4.1913552 | 0.82481137 |
| 6396 | NM_002941.3 | #N/A | 150.318701 | -8.4418085 | 1.66127822 |
| 6398 | NR_024448.2 | GUSBP11 | 96.0661267 | -7.7953048 | 1.53407196 |
| 6397 | NM_012323.3 | #N/A | 87.3724499 | -7.6599606 | 1.50743841 |
| 6399 | NM_014822.3 | #N/A | 131.383596 | -8.2483936 | 1.62331176 |
| 6400 | NM_001256574.1 | ENC1 | 104.444791 | -7.9207313 | 1.55898539 |
| 6401 | NM_001206998.1 | #N/A | 94.1713625 | -7.76724 | 1.52884434 |
| 6403 | NM_001171093.1 | FAM131A | 109.961444 | -7.9909905 | 1.57290159 |
| 6402 | NM_001145670.1 | SORBS2 | 94.3592686 | -7.7708241 | 1.52956864 |
| 6404 | NM_001286646.1 | SLC45A4 | 102.037761 | -6.5395344 | 1.2872461 |
| 6405 | NM_001197297.1 | #N/A | 95.2886684 | -7.783689 | 1.53216949 |
| 6406 | NM_130799.2 | MEN1 | 104.962705 | -7.9231908 | 1.55970999 |

| | | | | | |
|---|---|---|---|---|---|
| 6407 | NM_001253389.1 | ISYNA1 | 84.9523503 | -7.6222455 | 1.50051592 |
| 6408 | NM_018275.4 | #N/A | 128.825275 | -8.2199114 | 1.61839045 |
| 6409 | NM_006084.4 | #N/A | 103.717648 | -7.9101241 | 1.55745539 |
| 6410 | NM_000660.6 | #N/A | 106.919434 | -6.6756673 | 1.31451513 |
| 6411 | NM_020975.4 | #N/A | 163.226628 | -8.561671 | 1.68589342 |
| 6412 | NM_001003940.1 | #N/A | 109.050832 | -7.9821084 | 1.57181936 |
| 6413 | NM_003687.3 | #N/A | 89.6319038 | -7.6980788 | 1.51588567 |
| 6414 | NM_005034.3 | #N/A | 142.647925 | -8.3663195 | 1.64763583 |
| 6415 | NM_001204386.1 | DLG1 | 94.8451353 | -7.7738322 | 1.53098226 |
| 6416 | NM_003071.3 | #N/A | 94.8629534 | -7.7777217 | 1.53177133 |
| 6417 | NM_001199344.1 | #N/A | 111.333684 | -8.0112382 | 1.57778716 |
| 6418 | NM_001288655.1 | #N/A | 86.1628532 | -7.6403726 | 1.50475875 |
| 6419 | NM_016094.3 | #N/A | 257.53903 | -3.0283899 | 0.59646792 |
| 6420 | NM_001142730.2 | #N/A | 89.3533355 | -7.6928028 | 1.51518536 |
| 6421 | NM_001286379.1 | FAM120B | 95.037165 | -7.7797573 | 1.53235859 |
| 6422 | NM_206854.2 | #N/A | 128.59903 | -8.2178132 | 1.61868972 |
| 6423 | NM_002718.4 | #N/A | 133.063798 | -8.2655763 | 1.62814229 |
| 6424 | NM_005986.2 | #N/A | 107.159836 | -7.9527427 | 1.56661055 |
| 6425 | NM_016216.3 | #N/A | 84.8419596 | -7.6175255 | 1.50060262 |
| 6426 | NM_016070.3 | #N/A | 120.850241 | -8.1273714 | 1.60109149 |
| 6427 | NM_022482.4 | #N/A | 98.694778 | -7.8341343 | 1.54333343 |
| 6428 | NM_033027.3 | #N/A | 83.1069534 | -7.586999 | 1.49464714 |
| 6429 | NM_004961.3 | #N/A | 131.650285 | -8.2524283 | 1.62587323 |
| 6430 | NM_024907.6 | #N/A | 88.2698682 | -7.674229 | 1.51195469 |
| 6431 | NM_006869.3 | #N/A | 158.474098 | -8.5186512 | 1.67848741 |
| 6432 | NM_001139500.1 | #N/A | 130.712358 | -8.2412847 | 1.62386525 |
| 6433 | NR_073024.1 | RPL13A | 113.579451 | -8.0385531 | 1.58400055 |
| 6434 | NM_016525.4 | #N/A | 113.185098 | -8.0322005 | 1.58276077 |
| 6435 | NM_003162.3 | #N/A | 125.299952 | -6.7996893 | 1.33991965 |
| 6436 | NM_005569.3 | #N/A | 85.3108221 | -7.621523 | 1.50187414 |
| 6437 | NM_032151.4 | #N/A | 215.011208 | 2.27536172 | 0.44839539 |
| 6438 | NM_206920.2 | #N/A | 98.8090457 | -7.8368207 | 1.54444341 |
| 6439 | NM_020779.3 | #N/A | 96.8229113 | -7.8096421 | 1.53910334 |
| 6440 | NM_003851.2 | #N/A | 126.834268 | -6.9213693 | 1.36412957 |
| 6441 | NM_004096.4 | #N/A | 702.789261 | -3.6597045 | 0.7213076 |
| 6442 | NM_080605.3 | #N/A | 291.016899 | -4.6480694 | 0.91617561 |
| 6443 | NM_021933.3 | #N/A | 151.028358 | -8.4487357 | 1.66537822 |
| 6444 | NM_001040439.1 | #N/A | 257.868531 | -9.2218252 | 1.81782433 |
| 6445 | NM_003143.2 | #N/A | 120.740816 | -8.1256447 | 1.60183019 |
| 6446 | NM_181501.1 | #N/A | 148.07323 | -8.4213731 | 1.66016653 |
| 6447 | NR_037162.1 | #N/A | 87.4574658 | -7.66411 | 1.51128993 |
| 6448 | NM_013449.3 | #N/A | 123.323172 | -8.1583467 | 1.60882191 |
| 6449 | NM_053051.3 | #N/A | 100.86763 | -7.8646935 | 1.55093653 |

| | | | | | | |
|---|---|---|---|---|---|---|
| 6450 | NM_001242911.1 | ZFAND6 | | 98.8174123 | -7.8371202 | 1.54564592 |
| 6451 | NM_001039844.2 | | #N/A | 249.097132 | 1.92319664 | 0.37933791 |
| 6452 | NM_021237.4 | | #N/A | 129.160856 | -8.2227383 | 1.6219414 |
| 6453 | NM_001198869.1 | CAPN1 | | 135.710284 | -8.2970574 | 1.63665393 |
| 6454 | NM_012239.5 | | #N/A | 88.4337701 | -7.6779605 | 1.51489544 |
| 6455 | NM_004447.5 | | #N/A | 100.880262 | -7.8666884 | 1.55219468 |
| 6456 | NR_073599.1 | | #N/A | 112.005793 | -8.0203673 | 1.58259631 |
| 6457 | NM_001025203.1 | U2AF1 | | 108.582838 | -7.9713908 | 1.57297421 |
| 6458 | NM_015373.3 | CBY1 | | 85.9746852 | -7.6388938 | 1.50744554 |
| 6459 | NM_152243.2 | | #N/A | 126.12017 | -8.1902133 | 1.61625229 |
| 6460 | NM_014299.2 | BRD4 | | 108.02394 | -7.9651111 | 1.57187006 |
| 6461 | NM_133502.2 | | #N/A | 104.161473 | -7.9116053 | 1.56139315 |
| 6462 | NM_013235.4 | | #N/A | 109.640581 | -7.9865924 | 1.57649601 |
| 6463 | NM_018125.3 | | #N/A | 126.928822 | -8.1989919 | 1.61844822 |
| 6464 | NM_001039469.2 | MARK2 | | 88.7744934 | -7.6817211 | 1.51635997 |
| 6465 | NM_001135245.1 | | #N/A | 118.768946 | -8.1001319 | 1.59904331 |
| 6466 | NM_001128920.2 | MARK3 | | 115.840648 | -8.0693765 | 1.59302208 |
| 6467 | NM_020121.3 | | #N/A | 96.2706568 | -4.7811102 | 0.94395276 |
| 6468 | NM_032830.2 | | #N/A | 121.08535 | -8.1290025 | 1.60505167 |
| 6469 | NM_016458.3 | | #N/A | 118.000339 | -8.09315 | 1.5980324 |
| 6470 | NM_033117.3 | | #N/A | 134.410382 | -8.2806819 | 1.63505734 |
| 6471 | NM_000748.2 | | #N/A | 166.716347 | -8.5915406 | 1.69645952 |
| 6473 | NM_001760.4 | CCND3 | | 117.966967 | -8.092282 | 1.59798826 |
| 6472 | NM_001031812.3 | CSNK1G3 | | 93.96235 | -7.7645536 | 1.5332794 |
| 6474 | NM_001014433.2 | CUTA | | 115.33923 | -8.0605076 | 1.59195615 |
| 6475 | NM_030768.2 | | #N/A | 114.120548 | -8.0438937 | 1.58867621 |
| 6476 | NM_001001872.3 | | #N/A | 96.725999 | -7.8052695 | 1.54157932 |
| 6477 | NM_017943.3 | | #N/A | 105.305922 | -7.9301018 | 1.56625259 |
| 6478 | NM_013393.1 | | #N/A | 100.224807 | -7.857811 | 1.55226501 |
| 6479 | NM_018321.3 | | #N/A | 98.1081117 | -7.8246322 | 1.54575257 |
| 6480 | NM_015239.2 | AGTPBP1 | | 107.624645 | -7.9612878 | 1.57277397 |
| 6481 | NM_030885.3 | | #N/A | 396.250581 | -4.4202872 | 0.87325041 |
| 6482 | NM_032836.2 | | #N/A | 107.187747 | -7.9535527 | 1.57129238 |
| 6483 | NM_005262.2 | | #N/A | 142.420359 | -6.962424 | 1.37557431 |
| 6484 | NM_002515.2 | | #N/A | 133.328905 | -8.2688317 | 1.63378807 |
| 6485 | NM_001136223.1 | | #N/A | 131.171681 | -8.2453269 | 1.62921582 |
| 6486 | NM_000156.5 | | #N/A | 130.940915 | -8.24334 | 1.62888034 |
| 6487 | NM_178865.4 | | #N/A | 143.122508 | -8.3708813 | 1.65422119 |
| 6488 | NM_152298.3 | | #N/A | 91.0105609 | -7.7142439 | 1.52458186 |
| 6489 | NM_001687.4 | | #N/A | 119.89774 | -8.1162151 | 1.60408169 |
| 6490 | NM_138709.2 | DAB2IP | | 92.0728976 | -7.7357457 | 1.52894105 |
| 6491 | NM_004565.2 | | #N/A | 106.037968 | -7.9390381 | 1.56924312 |
| 6492 | NM_017728.3 | | #N/A | 109.766956 | -7.9894471 | 1.57924627 |

| | | | | | |
|---|---|---|---|---|---|
| 6493 | NM_021170.3 | #N/A | 121.812833 | -8.1374346 | 1.60860219 |
| 6494 | NM_014169.3 | #N/A | 100.758478 | -7.8651329 | 1.55479642 |
| 6495 | NM_021130.4 | #N/A | 2512.01966 | -5.531984 | 1.09369162 |
| 6496 | NR_111973.1 | GS1-124K5 | 109.133713 | 2.57914755 | 0.50991487 |
| 6497 | NM_182705.2 | RFLNB | 118.17395 | -6.7241026 | 1.32945645 |
| 6498 | NM_001256442.1 | #N/A | 149.230108 | -8.4334375 | 1.66755124 |
| 6499 | NM_170774.1 | RASSF2 | 27.0043107 | 9.72188648 | 1.9223461 |
| 6500 | NM_001786.4 | #N/A | 103.886943 | -7.9080664 | 1.56377988 |
| 6501 | NM_006556.3 | #N/A | 131.530877 | -8.2491469 | 1.63150207 |
| 6502 | NM_001322040.1 | #N/A | 97.4854114 | -7.8184224 | 1.54635829 |
| 6503 | NM_001167734.1 | VARS2 | 144.88401 | -8.3880417 | 1.65904744 |
| 6504 | NM_000366.5 | #N/A | 103.146686 | -7.8976293 | 1.56209914 |
| 6505 | NM_004176.4 | SREBF1 | 100.698647 | -7.8639556 | 1.55547711 |
| 6506 | NM_183050.3 | #N/A | 80.9332812 | -7.5505417 | 1.49368108 |
| 6507 | NM_175931.2 | #N/A | 119.086681 | -8.1071473 | 1.60383575 |
| 6508 | NM_003115.4 | UAP1 | 101.692666 | -7.8782964 | 1.55859088 |
| 6509 | NM_001656.3 | #N/A | 103.652189 | -7.904853 | 1.5639064 |
| 6510 | NM_001136044.1 | #N/A | 120.940359 | -8.1279869 | 1.60811133 |
| 6511 | NM_014871.4 | #N/A | 152.028573 | -3.9754253 | 0.78654926 |
| 6512 | NM_001257359.1 | #N/A | 305.925467 | -6.676473 | 1.3209874 |
| 6513 | NM_006341.3 | #N/A | 103.51707 | -7.9037427 | 1.56380667 |
| 6514 | NR_109983.1 | LOC729218 | 86.6754752 | -7.6510995 | 1.51384108 |
| 6515 | NR_136715.1 | #N/A | 87.4212713 | -7.6612492 | 1.51586227 |
| 6516 | NM_001282597.2 | CTNNA2 | 236.526366 | -9.0961104 | 1.79985591 |
| 6517 | NM_021214.1 | #N/A | 138.288081 | -8.3217957 | 1.64665549 |
| 6518 | NM_032118.3 | #N/A | 98.0427125 | -7.8258164 | 1.54857214 |
| 6519 | NM_024046.4 | #N/A | 374.562854 | -9.759315 | 1.93129931 |
| 6520 | NR_047519.1 | LINC01128 | 111.343428 | -8.0091156 | 1.58495856 |
| 6521 | NR_003662.3 | #N/A | 113.017199 | -8.0341034 | 1.58991551 |
| 6522 | NM_002595.4 | #N/A | 92.1820248 | -7.7392069 | 1.53164035 |
| 6523 | NM_002749.3 | #N/A | 109.412003 | -7.9825823 | 1.57983682 |
| 6524 | NM_018246.2 | #N/A | 102.287729 | -7.8836016 | 1.56031781 |
| 6525 | NM_002121.5 | #N/A | 165.417065 | -8.580948 | 1.6984959 |
| 6526 | NM_019108.3 | #N/A | 184.170398 | -2.8096313 | 0.55614068 |
| 6527 | NM_013247.4 | HTRA2 | 101.83354 | -7.8782576 | 1.55945562 |
| 6528 | NM_012098.2 | #N/A | 103.533035 | -7.9057367 | 1.56497156 |
| 6529 | NM_017495.5 | #N/A | 87.7567681 | -7.6686163 | 1.51836622 |
| 6530 | NM_018297.3 | #N/A | 86.3966932 | -7.6468249 | 1.51404616 |
| 6531 | NM_001032295.1 | #N/A | 89.5459927 | -7.6985734 | 1.52441064 |
| 6532 | NM_001012991.2 | #N/A | 101.582973 | -7.8766116 | 1.55972527 |
| 6533 | NM_172316.2 | MEIS2 | 186.055372 | -8.7501687 | 1.73273582 |
| 6534 | NM_020692.2 | #N/A | 114.593188 | -8.051586 | 1.59444076 |
| 6535 | NM_007247.5 | #N/A | 105.684671 | -7.936423 | 1.57172347 |

| | | | | | |
|---|---|---|---|---|---|
| 6536 | NM_001377.2 | #N/A | 102.877339 | -7.898024 | 1.56416209 |
| 6537 | NM_002372.3 | MAN2A1 | 120.091982 | -6.744013 | 1.33581726 |
| 6538 | NM_001270529.1 | #N/A | 107.306752 | -7.9529085 | 1.57534355 |
| 6539 | NM_181481.4 | LDLRAD4 | 117.939289 | -8.0949406 | 1.60351513 |
| 6540 | NM_001607.3 | #N/A | 114.269867 | -8.0468359 | 1.59402345 |
| 6541 | NM_003020.4 | SCG5 | 166.785808 | -8.5916809 | 1.70202201 |
| 6542 | NM_020351.3 | #N/A | 124.585667 | -8.1714782 | 1.6189114 |
| 6543 | NM_006058.4 | #N/A | 156.111004 | -8.4953753 | 1.68322964 |
| 6544 | NM_001008493.2 | #N/A | 140.53091 | -8.3434182 | 1.65323002 |
| 6545 | NM_004313.3 | #N/A | 152.942598 | -8.4672835 | 1.67794558 |
| 6546 | NM_001130445.2 | #N/A | 91.5789119 | -7.7230856 | 1.53060118 |
| 6547 | NM_003383.4 | #N/A | 96.9809377 | -7.81049 | 1.54793819 |
| 6548 | NM_025072.6 | #N/A | 114.893721 | -8.0542788 | 1.59645453 |
| 6549 | NM_001278379.1 | ARL6IP4 | 135.452663 | -8.2928412 | 1.64376666 |
| 6550 | NM_018115.3 | #N/A | 98.7337403 | -7.8343361 | 1.55288715 |
| 6551 | NM_001322898.1 | #N/A | 129.966294 | -8.2327709 | 1.63241298 |
| 6552 | NM_022453.2 | #N/A | 103.320733 | -7.900961 | 1.56679446 |
| 6553 | NM_023926.4 | ZSCAN18 | 83.1340906 | -7.584497 | 1.50409118 |
| 6554 | NM_182943.2 | #N/A | 144.124104 | -8.3827041 | 1.66248128 |
| 6555 | NM_152326.2 | #N/A | 81.7973553 | -7.5671585 | 1.50076451 |
| 6556 | NM_001142302.1 | CCDC113 | 39.1709171 | 10.2584439 | 2.03457042 |
| 6557 | NM_004435.2 | ENDOG | 117.857557 | -8.0911026 | 1.60478918 |
| 6558 | NM_001114172.1 | PIK3R3 | 142.484257 | -8.3656347 | 1.65925548 |
| 6559 | NM_024419.4 | #N/A | 88.6908312 | -7.6784281 | 1.52299894 |
| 6560 | NM_015137.5 | #N/A | 87.5138057 | -7.665505 | 1.52056127 |
| 6561 | NM_003002.3 | #N/A | 94.4167271 | -7.770616 | 1.54146732 |
| 6562 | NR_026556.1 | ANKRD26P1 | 34.2968484 | 8.76926661 | 1.73961171 |
| 6563 | NM_199342.3 | #N/A | 133.912148 | -8.2749202 | 1.64163438 |
| 6564 | NM_016436.4 | #N/A | 265.130953 | -4.5424495 | 0.90122083 |
| 6565 | NM_001163280.1 | HTATSF1 | 103.503027 | -7.9043969 | 1.56825255 |
| 6566 | NM_001324188.1 | #N/A | 89.5291029 | -7.6951821 | 1.52683436 |
| 6567 | NM_033416.2 | #N/A | 112.061169 | -8.0197663 | 1.5914981 |
| 6568 | NM_001427.3 | #N/A | 174.703473 | -8.6596339 | 1.71882907 |
| 6569 | NM_002263.3 | #N/A | 99.1708172 | -7.8411143 | 1.55639373 |
| 6570 | NM_005842.3 | #N/A | 108.71776 | -7.9746461 | 1.58301177 |
| 6571 | NM_012265.2 | #N/A | 115.367255 | -8.0603622 | 1.60004995 |
| 6572 | NR_144402.1 | LOC108783654 | 88.5003885 | -7.6783482 | 1.52428676 |
| 6573 | NM_198273.2 | LYSMD3 | 81.721832 | -7.5668387 | 1.50223207 |
| 6574 | NM_015205.2 | ATP11A | 105.262744 | -7.9265142 | 1.57374474 |
| 6575 | NM_145858.2 | #N/A | 88.9222916 | -7.6848902 | 1.52585931 |
| 6576 | NM_001042412.1 | #N/A | 105.693112 | -7.9330861 | 1.57518325 |
| 6577 | NM_005270.4 | #N/A | 96.4624006 | -7.8022403 | 1.54938601 |
| 6578 | NM_130443.3 | #N/A | 106.043013 | -7.9389397 | 1.57653307 |

| | | | | | |
|---|---|---|---|---|---|
| 6579 | NM_001198536.1 | MEFV | 225.811666 | 3.82187765 | 0.75898483 |
| 6580 | NM_001163424.1 | #N/A | 156.68366 | -8.5022314 | 1.68846996 |
| 6581 | NM_003597.4 | #N/A | 90.1250821 | -7.7018821 | 1.52956263 |
| 6582 | NM_002028.3 | #N/A | 92.7215012 | -7.7436599 | 1.53793615 |
| 6583 | NM_001278710.1 | NRN1 | 122.687812 | -8.1479851 | 1.61824932 |
| 6584 | NM_004911.4 | #N/A | 758.630836 | -4.7727486 | 0.94791492 |
| 6585 | NM_153689.5 | #N/A | 104.234037 | -7.9127402 | 1.57155875 |
| 6588 | NM_007124.2 | UTRN | 142.719464 | -5.8536643 | 1.16271601 |
| 6586 | NM_001031727.3 | #N/A | 82.2688037 | -7.5705443 | 1.50372405 |
| 6587 | NM_003180.2 | #N/A | 129.95823 | -8.2318591 | 1.63508951 |
| 6589 | NM_019080.2 | NDFIP2 | 147.131482 | -8.4108493 | 1.67087153 |
| 6590 | NM_032511.3 | #N/A | 129.971274 | -8.2334932 | 1.63565385 |
| 6591 | NM_015474.3 | SAMHD1 | 101.846008 | -7.8832071 | 1.56607702 |
| 6592 | NM_022835.2 | #N/A | 579.99389 | -2.4639359 | 0.48954756 |
| 6593 | NR_033230.2 | #N/A | 111.85065 | -5.9491665 | 1.18214094 |
| 6594 | NM_015463.2 | #N/A | 148.184367 | -8.421361 | 1.6734496 |
| 6595 | NM_152316.2 | #N/A | 107.829202 | -7.9625369 | 1.58230627 |
| 6596 | NM_078470.5 | #N/A | 108.645923 | -7.9728067 | 1.58438713 |
| 6597 | NM_005536.3 | #N/A | 93.0996605 | -7.7514679 | 1.54043004 |
| 6598 | NM_017769.4 | #N/A | 108.215651 | -7.966342 | 1.58321538 |
| 6599 | NM_019045.4 | #N/A | 90.997268 | -7.7195758 | 1.53416739 |
| 6600 | NM_139169.4 | #N/A | 114.951665 | -8.0546453 | 1.60085281 |
| 6601 | NM_005851.4 | #N/A | 96.6302356 | -7.8048818 | 1.55126094 |
| 6602 | NR_125816.1 | LINC02099 | 74.1667321 | 3.82918349 | 0.76111618 |
| 6603 | NM_022143.4 | #N/A | 98.4913519 | -7.8327688 | 1.55691778 |
| 6604 | NM_001024649.1 | #N/A | 234.266158 | -9.0836176 | 1.80579522 |
| 6605 | NM_017553.2 | #N/A | 106.526595 | -7.9442745 | 1.57938343 |
| 6606 | NM_017746.3 | #N/A | 106.395305 | -7.9432976 | 1.57933704 |
| 6607 | NM_032314.3 | #N/A | 112.128963 | -8.0185721 | 1.59430999 |
| 6608 | NM_020798.2 | #N/A | 103.646671 | -7.9062805 | 1.5719962 |
| 6609 | NM_001267040.1 | JARID2 | 117.799453 | -8.0930419 | 1.60919927 |
| 6610 | NM_138349.3 | #N/A | 92.8240851 | -7.7480405 | 1.54076601 |
| 6611 | NM_001278304.1 | #N/A | 90.2815822 | -5.5116458 | 1.09604709 |
| 6612 | NM_001316990.1 | #N/A | 87.2187943 | -7.6542651 | 1.52216731 |
| 6613 | NM_019109.4 | #N/A | 110.844356 | -8.0023701 | 1.59141232 |
| 6614 | NM_024296.4 | CCDC28B | 125.65272 | -8.1835342 | 1.62746564 |
| 6615 | NM_000920.3 | #N/A | 99.5540907 | -7.8462584 | 1.5604678 |
| 6616 | NM_005433.3 | #N/A | 254.262335 | -1.7782114 | 0.35366411 |
| 6617 | NM_178454.4 | #N/A | 79.6561383 | -7.5239196 | 1.49648266 |
| 6618 | NM_015950.4 | #N/A | 95.6859132 | -7.7891545 | 1.54924607 |
| 6619 | NM_004714.2 | #N/A | 95.1501962 | -7.7842961 | 1.54829313 |
| 6620 | NM_001305424.1 | #N/A | 123.866626 | -8.1616409 | 1.62351628 |
| 6621 | NM_001330418.1 | #N/A | 114.239473 | -8.0458507 | 1.60053472 |

| | | | | | |
|---|---|---|---|---|---|
| 6622 | NM_031212.3 | #N/A | 95.0635103 | -7.7796263 | 1.54764715 |
| 6623 | NM_001998.2 | #N/A | 130.001263 | -6.841882 | 1.36110978 |
| 6624 | NM_016098.3 | #N/A | 122.107324 | -8.142218 | 1.6198412 |
| 6625 | NM_016271.4 | #N/A | 107.894072 | -7.9628482 | 1.58424422 |
| 6626 | NM_032339.4 | #N/A | 132.940934 | -8.2649845 | 1.6443473 |
| 6627 | NM_033025.5 | #N/A | 104.685059 | -7.921925 | 1.57612716 |
| 6628 | NM_012175.3 | #N/A | 111.914652 | -8.01677 | 1.59501604 |
| 6629 | NM_201628.2 | #N/A | 203.584306 | -8.8793802 | 1.76667565 |
| 6630 | NM_024105.3 | #N/A | 91.1085808 | -7.7215179 | 1.53644224 |
| 6631 | NR_073110.1 | #N/A | 157.664299 | -8.5108147 | 1.6935055 |
| 6632 | NM_001991.4 | #N/A | 107.982373 | -7.9674614 | 1.58542579 |
| 6633 | NM_001174072.2 | #N/A | 274.650076 | -4.5117352 | 0.89783425 |
| 6634 | NM_004707.3 | #N/A | 125.828925 | -8.1863523 | 1.62916277 |
| 6635 | NM_022552.4 | DNMT3A | 90.198818 | -7.7013279 | 1.53273093 |
| 6636 | NM_006157.4 | #N/A | 126.708948 | -8.1955142 | 1.63109449 |
| 6637 | NM_175732.2 | #N/A | 82.9399512 | -7.5867003 | 1.5100851 |
| 6638 | NM_153329.3 | #N/A | 98.1052041 | -7.8264944 | 1.55791488 |
| 6639 | NM_024571.3 | #N/A | 118.436066 | -8.09841 | 1.61208452 |
| 6640 | NM_007157.3 | ZXDB | 84.511423 | -7.6102058 | 1.51508069 |
| 6641 | NM_006785.3 | #N/A | 93.1234616 | -7.7527132 | 1.54346993 |
| 6642 | NM_016221.3 | DCTN4 | 159.205636 | -6.4892321 | 1.29208709 |
| 6643 | NM_002853.3 | #N/A | 93.8566564 | -7.7614901 | 1.54549961 |
| 6644 | NM_005740.2 | #N/A | 88.4063321 | -7.6765197 | 1.5285854 |
| 6645 | NM_005733.2 | #N/A | 95.9286841 | -7.7957895 | 1.55240636 |
| 6647 | NM_001040025.2 | ARL16 | 107.559751 | -7.9588411 | 1.58492982 |
| 6646 | NM_000157.3 | #N/A | 113.53782 | -8.0373335 | 1.60055646 |
| 6648 | NM_006912.5 | #N/A | 102.362675 | -7.887817 | 1.57077928 |
| 6649 | NM_018359.3 | #N/A | 87.4335799 | -7.6626277 | 1.52597562 |
| 6650 | NM_022077.3 | #N/A | 98.5623756 | -7.8328545 | 1.55989759 |
| 6651 | NM_000154.1 | #N/A | 82.1753806 | -7.5699992 | 1.50760692 |
| 6652 | NM_021035.2 | #N/A | 160.901159 | -2.5052783 | 0.49894266 |
| 6653 | NM_178191.2 | ATP5IF1 | 90.7241237 | -7.7137639 | 1.5363587 |
| 6654 | NM_001308210.1 | TSHZ1 | 88.4449963 | -7.679596 | 1.52956579 |
| 6655 | NM_172170.4 | #N/A | 102.864198 | -7.8920041 | 1.57199134 |
| 6656 | NM_005245.3 | #N/A | 827.408795 | -3.9083314 | 0.77865228 |
| 6657 | NM_001282874.1 | #N/A | 180.599399 | -8.7073178 | 1.73476846 |
| 6658 | NM_152783.4 | #N/A | 88.2846969 | -7.6788346 | 1.52987696 |
| 6659 | NM_006038.3 | #N/A | 83.059726 | -7.5903102 | 1.51227384 |
| 6660 | NM_001141969.1 | #N/A | 103.279036 | -7.8990324 | 1.57383789 |
| 6661 | NM_001284364.2 | #N/A | 125.893987 | -8.1864243 | 1.63109452 |
| 6662 | NM_024809.4 | #N/A | 96.9925334 | -7.8096587 | 1.55603698 |
| 6663 | NM_015523.3 | #N/A | 97.8258331 | -7.8222479 | 1.55857492 |
| 6664 | NM_004765.3 | #N/A | 113.775712 | -8.0406674 | 1.60219222 |

| | | | | | |
|---|---|---|---|---|---|
| 6665 | NM_005236.2 | #N/A | 80.7328405 | -3.8308772 | 0.76335673 |
| 6666 | NM_003718.4 | #N/A | 94.7227897 | -7.7753088 | 1.54944332 |
| 6667 | NM_001032381.1 | PQBP1 | 104.516632 | -7.9169611 | 1.57769519 |
| 6668 | NM_006537.3 | #N/A | 107.701123 | -7.961287 | 1.58656749 |
| 6669 | NM_013254.3 | #N/A | 81.780672 | -7.5636973 | 1.50748928 |
| 6670 | NM_030625.2 | #N/A | 84.1298817 | -7.6036296 | 1.51546885 |
| 6671 | NM_000249.3 | MLH1 | 103.808913 | -7.9068589 | 1.57601875 |
| 6672 | NM_018030.4 | OSBPL1A | 116.075604 | -8.0710828 | 1.60875089 |
| 6673 | NM_001291107.1 | #N/A | 208.725932 | -8.9153523 | 1.7770524 |
| 6674 | NM_001282281.1 | #N/A | 124.858789 | -8.1740771 | 1.62962635 |
| 6675 | NM_001319944.1 | #N/A | 97.6071822 | -7.818194 | 1.55867621 |
| 6676 | NM_033428.2 | #N/A | 125.759793 | -8.1847349 | 1.63174321 |
| 6677 | NM_018389.4 | #N/A | 106.82689 | -7.9506331 | 1.58517053 |
| 6678 | NM_001079537.1 | #N/A | 115.921538 | -8.0662216 | 1.60825005 |
| 6679 | NM_032451.1 | #N/A | 174.279083 | -3.7704209 | 0.75174578 |
| 6680 | NM_001042618.1 | PARP2 | 111.642342 | -8.0137509 | 1.59787442 |
| 6681 | NM_025106.3 | #N/A | 116.987088 | -8.0817274 | 1.61143567 |
| 6682 | NM_003824.3 | FADD | 98.757358 | -7.8363746 | 1.56262145 |
| 6683 | NM_020812.3 | #N/A | 79.4373425 | -7.5251148 | 1.50059207 |
| 6684 | NM_033100.3 | #N/A | 123.225963 | -6.7711175 | 1.35028076 |
| 6685 | NM_001278356.1 | #N/A | 92.4364952 | -7.7391734 | 1.54334392 |
| 6686 | NM_001745.3 | #N/A | 334.826832 | -3.5570769 | 0.70935444 |
| 6687 | NM_005675.4 | #N/A | 103.748265 | -7.9100841 | 1.57747823 |
| 6688 | NM_153631.2 | HOXA3 | 132.879693 | -8.2655828 | 1.64845053 |
| 6689 | NM_001007228.1 | #N/A | 143.101118 | -8.3710642 | 1.66958704 |
| 6690 | NM_178148.3 | #N/A | 103.219258 | -7.8999172 | 1.5756158 |
| 6691 | NM_001308317.1 | #N/A | 133.951645 | -8.2752637 | 1.65055861 |
| 6692 | NM_181701.3 | #N/A | 209.6896 | -3.7721716 | 0.75243956 |
| 6693 | NM_001205294.1 | #N/A | 195.09302 | -8.818017 | 1.75895906 |
| 6694 | NM_020912.1 | FLYWCH1 | 82.8882885 | -7.5843465 | 1.51307279 |
| 6695 | NM_001252512.1 | SOAT1 | 23.3778587 | 9.53211107 | 1.90166551 |
| 6696 | NM_005202.3 | #N/A | 88.7012496 | -7.6849151 | 1.53322448 |
| 6697 | NM_014463.2 | #N/A | 132.85921 | -8.2639462 | 1.64884118 |
| 6698 | NM_001122772.2 | AGAP2 | 312.665619 | -7.9613976 | 1.58849383 |
| 6699 | NM_175839.2 | #N/A | 88.2635785 | -7.6736886 | 1.53114554 |
| 6700 | NM_006764.4 | IFRD2 | 97.9420598 | -7.8239001 | 1.56113035 |
| 6701 | NM_144611.3 | #N/A | 118.113227 | -4.3411529 | 0.86621064 |
| 6702 | NM_022893.3 | #N/A | 347.477246 | -9.6510163 | 1.92572834 |
| 6703 | NM_020971.2 | #N/A | 153.271974 | -8.4696517 | 1.69009324 |
| 6704 | NM_001166114.1 | #N/A | 100.497296 | -7.8606376 | 1.56859716 |
| 6705 | NM_001300896.1 | #N/A | 196.336666 | 2.13643024 | 0.42632856 |
| 6706 | NM_078473.2 | #N/A | 121.967342 | -6.8668822 | 1.37029459 |
| 6707 | NM_001134337.2 | #N/A | 107.968345 | -7.9632704 | 1.58911002 |

| | | | | | |
|---|---|---|---|---|---|
| 6708 | NM_030937.4 | #N/A | 109.171003 | -7.9805324 | 1.59265344 |
| 6709 | NM_152482.2 | #N/A | 115.736209 | -8.0644719 | 1.60942168 |
| 6710 | NM_001199873.1 | B4GALT3 | 115.529438 | -8.0613422 | 1.6091833 |
| 6711 | NM_001301056.1 | #N/A | 133.34056 | -8.2699879 | 1.65086512 |
| 6712 | NM_001315.2 | MAPK14 | 94.607091 | -7.7714309 | 1.5513669 |
| 6713 | NM_153207.4 | #N/A | 91.0524287 | -7.7187979 | 1.54090935 |
| 6714 | NM_020162.3 | #N/A | 84.4606203 | -7.6118094 | 1.51957648 |
| 6715 | NM_001277074.1 | CCDC12 | 104.779904 | -7.9210603 | 1.58135302 |
| 6716 | NM_004398.3 | #N/A | 106.045188 | -7.9373613 | 1.58460359 |
| 6717 | NM_006070.5 | #N/A | 112.78697 | -8.0256319 | 1.60228784 |
| 6718 | NM_015399.3 | #N/A | 110.747124 | -8.000943 | 1.59737584 |
| 6719 | NM_020536.4 | KAT14 | 89.0095352 | -7.6866335 | 1.53470261 |
| 6720 | NM_130464.2 | #N/A | 368.872151 | -4.8668412 | 0.97174195 |
| 6721 | NM_001319956.1 | RNF220 | 128.140591 | -8.2134895 | 1.64005983 |
| 6722 | NM_198285.2 | #N/A | 85.8164773 | -7.6347128 | 1.52455753 |
| 6723 | NM_016475.4 | #N/A | 80.5460413 | -7.5441952 | 1.50660455 |
| 6724 | NM_001244262.1 | HBP1 | 78.0107253 | -7.4935651 | 1.49656968 |
| 6725 | NM_001286626.1 | #N/A | 88.040557 | -7.667823 | 1.53142892 |
| 6726 | NM_004741.4 | #N/A | 219.952847 | -8.9922092 | 1.79595609 |
| 6727 | NM_001003681.2 | #N/A | 111.862343 | -8.0138347 | 1.60063381 |
| 6728 | NM_002406.3 | #N/A | 107.142975 | -7.9567451 | 1.58928026 |
| 6729 | NR_136158.1 | #N/A | 75.7933209 | -4.8034896 | 0.95944775 |
| 6730 | NM_005957.4 | #N/A | 104.279502 | -7.9167609 | 1.5813892 |
| 6731 | NM_001007793.2 | BUB3 | 96.4161481 | -7.8022023 | 1.55855493 |
| 6733 | NM_001324304.1 | RPN2 | 151.240607 | -8.4507926 | 1.6881082 |
| 6732 | NM_001319661.1 | TMEM263 | 99.376035 | -7.8478897 | 1.56767029 |
| 6734 | NM_004220.2 | #N/A | 108.235881 | -7.9702341 | 1.59217718 |
| 6735 | NM_002276.4 | #N/A | 88.0332889 | -7.6727128 | 1.53280764 |
| 6736 | NM_020640.3 | #N/A | 119.762674 | -5.8782976 | 1.17435232 |
| 6737 | NM_139273.3 | #N/A | 86.5016792 | -7.6464115 | 1.5276634 |
| 6738 | NM_032390.4 | #N/A | 124.609411 | -6.8960117 | 1.37779019 |
| 6739 | NM_001130709.1 | RUFY3 | 95.1463283 | -7.7825451 | 1.55496493 |
| 6740 | NM_145251.3 | #N/A | 91.4372754 | -7.7243044 | 1.54345839 |
| 6741 | NM_012071.3 | #N/A | 124.287004 | -8.1671925 | 1.63208653 |
| 6742 | NM_017924.3 | #N/A | 131.363293 | -4.3288987 | 0.86510366 |
| 6743 | NM_024747.5 | #N/A | 96.3537991 | -7.8012773 | 1.55910114 |
| 6744 | NM_006334.3 | #N/A | 180.787714 | -8.7085534 | 1.74049024 |
| 6745 | NM_001330740.1 | #N/A | 155.847618 | -8.4943346 | 1.69771205 |
| 6748 | NM_015441.2 | OLFML2B | 82.9179807 | -7.5883745 | 1.51667287 |
| 6746 | NM_001002292.3 | WLS | 122.358347 | -8.1464693 | 1.62821701 |
| 6747 | NM_001172.3 | #N/A | 151.955607 | -8.4577859 | 1.69043103 |
| 6749 | NM_015964.3 | #N/A | 182.663782 | -8.7246523 | 1.74388957 |
| 6750 | NM_001282307.1 | STK25 | 138.325043 | -8.3212751 | 1.66332572 |

| | | | | | |
|---|---|---|---|---|---|
| 6751 | NM_005725.5 | #N/A | 85.0908571 | -7.6180291 | 1.52282075 |
| 6752 | NM_001317919.1 | #N/A | 109.388197 | -7.9865996 | 1.5966222 |
| 6753 | NM_001319193.1 | FBF1 | 76.2821749 | -7.4618186 | 1.49176589 |
| 6754 | NM_148957.3 | #N/A | 146.297837 | -8.4031936 | 1.68014353 |
| 6755 | NM_173500.3 | #N/A | 78.0343411 | -4.847373 | 0.96921846 |
| 6756 | NM_002435.2 | #N/A | 78.9847263 | -7.5145109 | 1.50252504 |
| 6757 | NM_005800.4 | #N/A | 94.0358517 | -7.7653677 | 1.55270441 |
| 6758 | NR_015404.1 | #N/A | 101.4815 | -7.873927 | 1.57442055 |
| 6759 | NM_182923.3 | KLC1 | 105.18644 | -7.9255081 | 1.58482859 |
| 6760 | NM_080592.3 | #N/A | 120.381995 | -8.1214566 | 1.62402913 |
| 6761 | NM_015361.3 | #N/A | 132.61293 | -8.26093 | 1.65196302 |
| 6762 | NM_024116.3 | #N/A | 80.6384995 | -7.5444188 | 1.50878579 |
| 6763 | NM_030818.3 | #N/A | 78.169232 | -7.4970745 | 1.49940671 |
| 6764 | NM_003467.2 | #N/A | 81.2662763 | -7.5563238 | 1.51127444 |
| 6765 | NR_003260.1 | DNM1P46 | 55.6720473 | 2.2847685 | 0.45697491 |
| 6766 | NM_015188.1 | #N/A | 107.796081 | -7.9615884 | 1.59241965 |
| 6767 | NM_001127183.2 | #N/A | 95.1618107 | -7.7855106 | 1.55727202 |
| 6768 | NM_001321179.1 | FAM156B | 81.1613415 | -7.5565955 | 1.51153431 |
| 6769 | NM_002490.4 | #N/A | 255.713557 | -6.4588273 | 1.29199628 |
| 6770 | NM_001164315.1 | #N/A | 124.190506 | -8.165663 | 1.63345546 |
| 6771 | NM_007192.3 | #N/A | 733.724334 | -2.626245 | 0.52541978 |
| 6772 | NM_007240.2 | #N/A | 104.159757 | -7.9122497 | 1.58301999 |
| 6773 | NM_014157.3 | #N/A | 116.778208 | -8.0794721 | 1.61662507 |
| 6774 | NM_025191.3 | #N/A | 110.77609 | -8.0052785 | 1.60201162 |
| 6775 | NM_000147.4 | FUCA1 | 92.847742 | -6.4703327 | 1.29487456 |
| 6776 | NM_006067.4 | #N/A | 114.352299 | -8.0464722 | 1.61029904 |
| 6777 | NM_024591.4 | #N/A | 101.4082 | -7.875101 | 1.57603019 |
| 6778 | NM_001300908.1 | #N/A | 143.769729 | -8.3773398 | 1.676552 |
| 6779 | NM_145698.4 | #N/A | 105.239704 | -7.9272967 | 1.58661669 |
| 6780 | NM_001080452.1 | GPR108 | 88.0482078 | -7.6710562 | 1.53541375 |
| 6781 | NM_001320663.1 | #N/A | 123.477377 | -8.1572605 | 1.63280448 |
| 6782 | NM_003423.3 | #N/A | 90.4761004 | -7.7060009 | 1.54251121 |
| 6783 | NM_004128.2 | #N/A | 125.004601 | -6.9012252 | 1.38144667 |
| 6784 | NM_018135.3 | MRPS18A | 75.5428077 | -7.4507958 | 1.49158257 |
| 6785 | NM_001278530.1 | #N/A | 99.2734277 | -7.8450874 | 1.57061472 |
| 6786 | NM_032832.5 | #N/A | 227.088833 | -5.5856209 | 1.11833267 |
| 6790 | NR_026773.1 | LINC01558 | 77.8767061 | 4.47807581 | 0.89669089 |
| 6787 | NM_016218.3 | POLK | 83.2260207 | -7.5872755 | 1.51925302 |
| 6789 | NM_181485.2 | ZGPAT | 97.1380876 | -7.8117759 | 1.56421738 |
| 6788 | NM_031480.2 | #N/A | 118.941077 | -8.1030247 | 1.62254316 |
| 6791 | NM_007246.3 | #N/A | 104.664044 | -7.9187371 | 1.58567021 |
| 6792 | NM_018590.4 | #N/A | 120.291635 | -8.1211361 | 1.62624209 |
| 6793 | NM_016188.4 | #N/A | 191.150259 | -8.7888525 | 1.759974 |

| | | | | | |
|---|---|---|---|---|---|
| 6794 | NM_080390.3 | #N/A | 180.046056 | -8.702333 | 1.74274864 |
| 6795 | NM_032663.4 | #N/A | 106.702627 | -7.945517 | 1.59120042 |
| 6796 | NM_007066.4 | #N/A | 87.9765068 | -7.66926 | 1.53593518 |
| 6797 | NM_001288811.1 | ARL17A | 84.7749629 | 2.27489373 | 0.45561251 |
| 6798 | NR_028025.1 | #N/A | 89.7101464 | -7.6981952 | 1.54184456 |
| 6799 | NM_032351.5 | #N/A | 106.373565 | -7.9439489 | 1.59123172 |
| 6800 | NM_002061.3 | #N/A | 122.982012 | -8.1537306 | 1.63332377 |
| 6801 | NM_001254738.1 | RND3 | 109.721286 | -7.9844621 | 1.59958466 |
| 6802 | NM_052951.2 | #N/A | 101.956607 | -7.8827593 | 1.57921232 |
| 6803 | NM_001146070.1 | #N/A | 77.8577882 | -7.4947143 | 1.50151597 |
| 6804 | NM_001346494.1 | #N/A | 117.765272 | -8.0898764 | 1.62075877 |
| 6805 | NM_003958.3 | #N/A | 93.7472004 | -7.7606789 | 1.55483805 |
| 6806 | NM_000107.2 | DDB2 | 82.9414032 | -7.5809667 | 1.51888181 |
| 6807 | NM_001005407.1 | CACNA1H | 265.759277 | -3.3264165 | 0.66654707 |
| 6808 | NM_001101663.4 | NBPF11 | 105.969091 | -5.8668233 | 1.17566463 |
| 6809 | NM_015442.2 | #N/A | 95.6971048 | -7.7889348 | 1.56084096 |
| 6810 | NM_003276.2 | TMPO | 113.107293 | -8.0311873 | 1.60944557 |
| 6811 | NM_001330289.1 | #N/A | 245.385628 | -9.1486837 | 1.83344019 |
| 6812 | NM_002493.4 | #N/A | 128.131966 | -8.210959 | 1.64552424 |
| 6813 | NM_001114618.1 | MGAT1 | 99.9572183 | -7.8499832 | 1.5732013 |
| 6814 | NM_006208.2 | #N/A | 104.753004 | 2.27085778 | 0.45513028 |
| 6815 | NM_020644.2 | #N/A | 107.98532 | -7.9650514 | 1.5964814 |
| 6816 | NM_025154.5 | #N/A | 96.9021327 | -5.7355644 | 1.14971089 |
| 6817 | NM_004107.4 | #N/A | 93.0881703 | -7.7539443 | 1.55431042 |
| 6818 | NM_030803.6 | #N/A | 106.329315 | -7.9421035 | 1.59219507 |
| 6819 | NM_005070.3 | #N/A | 130.940542 | -8.2436002 | 1.65273889 |
| 6820 | NM_024319.3 | #N/A | 102.451159 | -7.8883066 | 1.58151585 |
| 6821 | NM_015436.3 | RCHY1 | 110.269883 | -7.9957362 | 1.60311585 |
| 6822 | NM_032112.2 | #N/A | 123.177794 | -8.154449 | 1.63493552 |
| 6823 | NM_024583.4 | #N/A | 95.0069699 | -7.7797302 | 1.55983714 |
| 6824 | NM_006922.3 | #N/A | 174.072066 | -8.6540115 | 1.73535724 |
| 6825 | NM_024701.3 | #N/A | 104.261179 | -7.9165906 | 1.58758899 |
| 6826 | NM_144664.4 | #N/A | 87.9486117 | -7.6698045 | 1.53816018 |
| 6827 | NM_016539.3 | #N/A | 94.0151041 | -7.7629598 | 1.55694863 |
| 6828 | NM_173694.4 | #N/A | 267.300311 | -9.2734363 | 1.85992999 |
| 6829 | NM_000033.3 | #N/A | 107.920502 | -7.9652015 | 1.59768862 |
| 6830 | NM_017549.4 | #N/A | 99.6107692 | -7.8516249 | 1.57509855 |
| 6831 | NM_001281775.2 | #N/A | 94.0328261 | -7.7691472 | 1.55870518 |
| 6832 | NM_007097.3 | #N/A | 211.148848 | -8.9322214 | 1.79204215 |
| 6833 | NM_025194.2 | #N/A | 91.2110781 | -7.7235271 | 1.54960253 |
| 6834 | NM_001346231.1 | #N/A | 109.794832 | -7.9891509 | 1.60292242 |
| 6835 | NM_032848.2 | #N/A | 82.6852184 | -7.5811477 | 1.52106818 |
| 6836 | NM_001143887.1 | #N/A | 122.317355 | -8.1440897 | 1.63405492 |

| | | | | | |
|---|---|---|---|---|---|
| 6837 | NM_145201.5 | #N/A | 90.7413259 | -7.7177962 | 1.54856928 |
| 6838 | NM_000303.2 | #N/A | 86.6634734 | -7.6488174 | 1.53476621 |
| 6839 | NM_002095.4 | #N/A | 87.4020103 | -7.659559 | 1.53695248 |
| 6840 | NR_027293.2 | BMS1P20 | 82.0485551 | -7.5686866 | 1.51883281 |
| 6841 | NM_172081.2 | #N/A | 129.860813 | -8.2288976 | 1.65135155 |
| 6842 | NM_001312673.1 | #N/A | 78.9044222 | -7.5153939 | 1.50822223 |
| 6843 | NM_006950.3 | SYN1 | 159.337395 | -8.5257373 | 1.71110072 |
| 6844 | NM_030955.3 | #N/A | 74.1882167 | -7.4248391 | 1.49027121 |
| 6845 | NM_183075.2 | #N/A | 79.0588646 | -7.5165462 | 1.50875218 |
| 6846 | NM_001042559.2 | EIF4G2 | 164.077208 | -8.5706306 | 1.72034909 |
| 6847 | NM_004272.4 | #N/A | 112.560197 | -8.0238647 | 1.61063875 |
| 6848 | NM_024692.5 | #N/A | 75.6430849 | -7.4525315 | 1.49597967 |
| 6849 | NR_037676.1 | FRMD6-AS1 | 277.178739 | 4.21557228 | 0.84623656 |
| 6850 | NR_110730.1 | PTOV1-AS2 | 84.2690155 | -7.6091349 | 1.52753887 |
| 6851 | NM_007187.4 | #N/A | 104.179344 | -7.9115814 | 1.58841796 |
| 6852 | NR_024463.1 | #N/A | 37.1461648 | 3.76820795 | 0.75655306 |
| 6853 | NM_021992.2 | #N/A | 178.31641 | -8.6884131 | 1.74443252 |
| 6854 | NM_024326.3 | FBXL15 | 118.149714 | -8.0945295 | 1.62522256 |
| 6855 | NM_015568.3 | #N/A | 148.52829 | -8.425017 | 1.69169954 |
| 6857 | NM_006711.4 | RNPS1 | 115.260873 | -8.0586017 | 1.61817519 |
| 6856 | NM_001198801.1 | #N/A | 103.412189 | -7.9049961 | 1.5873453 |
| 6858 | NM_017807.3 | #N/A | 94.7706576 | -7.7768536 | 1.56160507 |
| 6859 | NM_017799.3 | #N/A | 79.4191724 | -7.5224723 | 1.51062161 |
| 6860 | NM_001079524.1 | #N/A | 637.461585 | -2.5634763 | 0.51478894 |
| 6861 | NM_001146699.1 | #N/A | 78.2164241 | -7.4963835 | 1.50554097 |
| 6862 | NM_001128430.1 | #N/A | 117.887257 | -8.0907046 | 1.62502395 |
| 6863 | NM_005604.3 | #N/A | 110.076255 | -7.9911283 | 1.60511146 |
| 6864 | NM_001320977.1 | #N/A | 109.192111 | -7.9807154 | 1.60307301 |
| 6865 | NM_024090.2 | #N/A | 110.642457 | -8.001732 | 1.60732067 |
| 6866 | NM_001282112.1 | #N/A | 91.6797431 | -7.7298347 | 1.55282504 |
| 6867 | NM_003192.2 | #N/A | 115.954575 | -5.0821846 | 1.02098651 |
| 6868 | NM_032810.3 | ATAD1 | 134.81233 | -8.2848644 | 1.66448295 |
| 6869 | NM_201542.4 | #N/A | 106.490919 | -7.9451281 | 1.59628675 |
| 6870 | NM_001001329.2 | PRKCSH | 84.9238683 | -7.6209308 | 1.53117581 |
| 6871 | NM_205861.2 | #N/A | 90.1834257 | -7.7044262 | 1.54794886 |
| 6872 | NM_005050.3 | #N/A | 88.1536266 | -7.6704053 | 1.54126763 |
| 6873 | NR_130929.1 | LOC644285 | 87.0795609 | 3.24237894 | 0.65153107 |
| 6874 | NM_001142420.1 | MORF4L2 | 117.896609 | -8.0918147 | 1.62610083 |
| 6875 | NM_024417.4 | #N/A | 79.3369261 | -7.5193002 | 1.51109371 |
| 6876 | NR_104318.1 | #N/A | 133.156815 | -8.2669679 | 1.66140981 |
| 6877 | NM_001033859.2 | ACADVL | 93.5687145 | -7.762301 | 1.56002551 |
| 6878 | NM_001322118.1 | PDPR | 123.680333 | -8.1613685 | 1.6402712 |
| 6879 | NM_001290261.1 | ZNF337 | 129.815185 | -8.2283745 | 1.65375827 |

| | | | | | |
|---|---|---|---|---|---|
| 6880 | NM_000034.3 | #N/A | 79.1139472 | -7.5131933 | 1.51008382 |
| 6881 | NM_001290058.1 | #N/A | 23.3304449 | 9.51520212 | 1.91252889 |
| 6882 | NM_003596.3 | #N/A | 99.2205596 | -7.8439592 | 1.5766408 |
| 6883 | NM_004897.4 | #N/A | 100.898243 | -7.8679524 | 1.58152094 |
| 6884 | NM_001195446.1 | SRSF7 | 93.6054819 | -7.7560463 | 1.55908412 |
| 6886 | NM_138357.2 | MCU | 89.513082 | -7.6950679 | 1.54692092 |
| 6885 | NM_001114394.1 | #N/A | 79.933976 | -7.5284428 | 1.51342201 |
| 6887 | NM_001297600.1 | #N/A | 89.2437977 | -7.6936541 | 1.54665546 |
| 6888 | NM_012417.3 | #N/A | 173.960533 | -8.6532723 | 1.73996316 |
| 6889 | NM_023039.4 | #N/A | 85.3559811 | -7.6237139 | 1.53297933 |
| 6890 | NM_016437.2 | #N/A | 112.064587 | -6.6408513 | 1.3355172 |
| 6891 | NM_001082486.1 | #N/A | 103.479828 | -7.9027338 | 1.58932106 |
| 6892 | NM_001136049.2 | LMLN | 73.9869252 | -7.4207392 | 1.49245069 |
| 6893 | NM_006645.2 | STARD10 | 81.8217804 | -7.5649587 | 1.52156801 |
| 6894 | NM_001301085.1 | #N/A | 92.5616986 | -7.745365 | 1.55791978 |
| 6895 | NM_001001433.2 | #N/A | 90.3866806 | -7.7093193 | 1.55068957 |
| 6896 | NM_002370.3 | #N/A | 87.3009779 | -7.6568973 | 1.54015015 |
| 6897 | NM_130466.3 | #N/A | 89.8666974 | -7.7016476 | 1.54918191 |
| 6898 | NM_001260.2 | #N/A | 93.0661851 | -7.7480143 | 1.55853212 |
| 6899 | NR_110248.1 | LOC101929319 | 32.7425141 | 3.23110358 | 0.65000868 |
| 6900 | NM_014285.6 | #N/A | 86.5086166 | -7.6426427 | 1.53766095 |
| 6901 | NM_019052.3 | #N/A | 78.2238711 | -7.5000872 | 1.50897922 |
| 6902 | NM_001318005.1 | ZNF514 | 85.9907288 | -7.6362078 | 1.53643818 |
| 6903 | NM_005154.4 | #N/A | 74.2230953 | -7.4224912 | 1.49360655 |
| 6904 | NM_018453.3 | #N/A | 93.8634717 | -7.7631851 | 1.56224341 |
| 6905 | NM_018207.2 | #N/A | 95.8116195 | -7.7933843 | 1.56848242 |
| 6906 | NM_005509.5 | #N/A | 97.2374301 | -7.8130838 | 1.57260015 |
| 6907 | NM_021184.3 | #N/A | 82.6662249 | -7.5806417 | 1.52583019 |
| 6908 | NM_001293697.1 | #N/A | 147.671314 | -8.4156741 | 1.69396339 |
| 6909 | NM_004053.3 | #N/A | 99.8434383 | -7.8523101 | 1.58063143 |
| 6910 | NM_152374.1 | #N/A | 227.879128 | -7.5481412 | 1.51953014 |
| 6911 | NM_001142419.1 | MORF4L2 | 90.5766015 | -7.710808 | 1.55230691 |
| 6912 | NM_033637.3 | #N/A | 126.500938 | -8.1941398 | 1.64965342 |
| 6913 | NM_001113434.4 | #N/A | 86.9306601 | -6.3689658 | 1.28223071 |
| 6914 | NM_006468.7 | #N/A | 100.777916 | -7.8660433 | 1.58371482 |
| 6915 | NM_001128206.1 | #N/A | 110.976163 | -8.0065743 | 1.6120897 |
| 6916 | NM_014460.3 | #N/A | 157.654108 | -8.511051 | 1.7136882 |
| 6917 | NM_001388.4 | #N/A | 100.1892 | -7.85729 | 1.58211676 |
| 6918 | NM_021120.3 | #N/A | 151.287604 | -8.4510263 | 1.70168278 |
| 6919 | NR_123740.1 | #N/A | 105.036166 | -7.923749 | 1.59552317 |
| 6920 | NM_201433.1 | #N/A | 182.970132 | -8.7269581 | 1.7573542 |
| 6921 | NM_145246.4 | #N/A | 91.1985636 | -7.7202276 | 1.55465784 |
| 6922 | NM_016353.4 | #N/A | 102.175707 | -6.6058067 | 1.33034783 |

| | | | | | |
|---|---|---|---|---|---|
| 6923 | NM_153485.2 | #N/A | 84.5972498 | -7.6110616 | 1.53281112 |
| 6924 | NM_002719.3 | #N/A | 97.1215753 | -7.8135031 | 1.57360841 |
| 6925 | NM_018674.5 | ASIC4 | 100.864544 | -7.8673482 | 1.58450811 |
| 6926 | NM_019887.5 | #N/A | 109.293569 | -7.9822336 | 1.60764182 |
| 6927 | NM_001011649.2 | #N/A | 100.921583 | -7.8657665 | 1.5842333 |
| 6928 | NM_032182.3 | #N/A | 90.9544936 | -7.7162515 | 1.55414026 |
| 6929 | NM_025065.6 | #N/A | 90.0241125 | -7.7018763 | 1.55132315 |
| 6930 | NR_109843.1 | #N/A | 79.578049 | -7.5244702 | 1.51558786 |
| 6931 | NM_001199135.1 | #N/A | 89.7790044 | -7.6979488 | 1.55054568 |
| 6932 | NM_014946.3 | SPAST | 169.403653 | -8.6137321 | 1.73504751 |
| 6933 | NM_021218.2 | #N/A | 98.7301657 | -7.834051 | 1.57809284 |
| 6934 | NM_001014286.2 | #N/A | 98.2364134 | -7.8285573 | 1.57702125 |
| 6935 | NM_033182.5 | #N/A | 89.2113641 | -7.6853323 | 1.54817971 |
| 6936 | NM_001301064.1 | RRN3 | 101.254565 | -7.8691061 | 1.58543006 |
| 6937 | NM_006043.1 | #N/A | 141.801758 | -8.3585614 | 1.68408892 |
| 6938 | NR_104013.1 | NSUN5P1 | 78.0220567 | -7.4998565 | 1.51119465 |
| 6939 | NR_135223.1 | PCNP | 88.2208432 | -7.6736392 | 1.54627092 |
| 6940 | NM_152227.2 | #N/A | 86.7938075 | -7.6499667 | 1.54164709 |
| 6941 | NM_018337.3 | #N/A | 94.3525311 | -7.7697461 | 1.56580279 |
| 6942 | NM_004865.3 | #N/A | 104.433668 | -7.9163296 | 1.59535219 |
| 6943 | NM_001186.3 | #N/A | 129.210457 | -8.2234596 | 1.65731681 |
| 6944 | NM_007255.2 | #N/A | 107.199988 | -7.9548398 | 1.60323125 |
| 6945 | NM_001130860.3 | #N/A | 106.896709 | -7.9525453 | 1.6027933 |
| 6946 | NM_002346.2 | #N/A | 123.598855 | -8.1627113 | 1.645223 |
| 6947 | NM_015342.3 | #N/A | 81.3838343 | -7.554984 | 1.52282405 |
| 6948 | NM_015972.3 | #N/A | 110.984246 | -8.0041584 | 1.61341328 |
| 6950 | NR_037701.1 | LINC01963 | 176.088877 | -7.2172713 | 1.45507987 |
| 6949 | NM_004928.2 | #N/A | 78.3058762 | -7.498446 | 1.51177439 |
| 6951 | NM_022063.2 | #N/A | 93.8004367 | -7.759428 | 1.5644102 |
| 6952 | NM_030954.3 | #N/A | 82.3549837 | -7.577489 | 1.52773093 |
| 6953 | NM_018356.2 | #N/A | 98.9686177 | -7.8417004 | 1.5811659 |
| 6954 | NM_025164.4 | #N/A | 89.1034204 | -7.6877831 | 1.55018784 |
| 6955 | NM_018685.4 | #N/A | 92.8955877 | -7.7477382 | 1.56235153 |
| 6956 | NM_014186.3 | #N/A | 116.21164 | -8.0702446 | 1.62741972 |
| 6957 | NM_001330729.1 | #N/A | 123.493715 | -8.1589049 | 1.64532168 |
| 6958 | NM_153358.2 | #N/A | 69.2373478 | -4.5725286 | 0.92209221 |
| 6959 | NR_038886.1 | LINC01426 | 83.2234775 | 5.26501787 | 1.06176608 |
| 6960 | NM_001206916.1 | #N/A | 136.583211 | -8.3029664 | 1.67459447 |
| 6961 | NM_001105539.2 | #N/A | 124.693637 | -8.1703662 | 1.64800902 |
| 6962 | NM_001482.2 | #N/A | 88.8369421 | -7.6805693 | 1.54948721 |
| 6963 | NR_015370.2 | EPB41L4A-AS1 | 107.419451 | -7.9565119 | 1.60519585 |
| 6964 | NM_153365.2 | #N/A | 97.8757159 | -7.8238239 | 1.57849759 |
| 6965 | NM_001166278.1 | DLG3 | 96.2077815 | -7.7981535 | 1.57341358 |

| | | | | | |
|---|---|---|---|---|---|
| 6966 | NM_016133.3 | #N/A | 77.2775408 | -7.4865649 | 1.51056883 |
| 6967 | NM_001034023.1 | #N/A | 91.8983959 | -7.7344216 | 1.56081358 |
| 6968 | NM_014966.3 | DHX30 | 108.722234 | -7.9718475 | 1.60891224 |
| 6969 | NM_001161346.1 | #N/A | 108.37032 | -7.9686891 | 1.60832739 |
| 6970 | NM_021963.3 | #N/A | 107.840297 | -7.9632729 | 1.60732796 |
| 6971 | NM_021134.3 | #N/A | 117.212402 | -8.0831209 | 1.63152914 |
| 6972 | NM_013337.3 | #N/A | 102.416914 | -7.8892025 | 1.59240471 |
| 6973 | NR_125900.1 | LOC101928551 | 36.3818858 | 2.539522 | 0.51264286 |
| 6974 | NM_003427.4 | #N/A | 78.5984447 | -7.5068732 | 1.5154731 |
| 6975 | NM_014920.3 | #N/A | 121.250431 | -5.5432005 | 1.11905627 |
| 6976 | NR_024408.1 | #N/A | 84.4939061 | -6.3330636 | 1.27855032 |
| 6977 | NM_001206999.1 | #N/A | 82.947307 | -7.5814827 | 1.53077575 |
| 6978 | NM_001025436.2 | SPAG16 | 88.2726762 | -6.3925332 | 1.29077191 |
| 6980 | NM_001309242.1 | MYO15B | 104.775019 | -7.9230267 | 1.59982247 |
| 6979 | NM_001300865.1 | #N/A | 110.162899 | -7.9945953 | 1.61427885 |
| 6981 | NM_032194.2 | #N/A | 86.1695736 | -7.6373056 | 1.54215481 |
| 6982 | NM_006159.2 | NELL2 | 173.191288 | -8.6448745 | 1.74566025 |
| 6983 | NM_016570.2 | #N/A | 105.437522 | -7.9296045 | 1.60128177 |
| 6984 | NM_001278554.1 | UBE2E3 | 113.974247 | -8.0420281 | 1.62403552 |
| 6985 | NM_144683.3 | #N/A | 107.296017 | -7.9552227 | 1.60650057 |
| 6986 | NM_032305.2 | #N/A | 100.397779 | -7.8581973 | 1.58696806 |
| 6987 | NR_073396.1 | ZNF818P | 258.555292 | 3.02888403 | 0.61168979 |
| 6988 | NM_020724.1 | #N/A | 84.2569514 | -7.6061841 | 1.53610753 |
| 6989 | NM_001313998.1 | #N/A | 101.913644 | -7.8796843 | 1.59135145 |
| 6990 | NM_002559.3 | #N/A | 131.133443 | -8.2459651 | 1.66543497 |
| 6991 | NM_016564.3 | #N/A | 140.771587 | -8.347897 | 1.68640573 |
| 6992 | NM_198545.3 | #N/A | 122.372779 | -8.1442628 | 1.64527682 |
| 6993 | NM_006225.3 | #N/A | 93.100048 | -7.7524458 | 1.56614541 |
| 6994 | NM_001286625.1 | RNPS1 | 114.5276 | -8.0500539 | 1.62632054 |
| 6995 | NM_005105.4 | RBM8A | 721.077268 | -3.1081027 | 0.62795421 |
| 6996 | NM_016504.2 | #N/A | 100.59962 | -7.8624674 | 1.58852318 |
| 6997 | NM_017415.2 | #N/A | 72.4207818 | -7.3880961 | 1.49268434 |
| 6998 | NM_001256695.1 | PRDM11 | 75.0458733 | -7.4435091 | 1.50399384 |
| 6999 | NM_020813.3 | #N/A | 92.6947048 | -7.7430546 | 1.56457374 |
| 7000 | NM_183238.3 | #N/A | 93.2610281 | -7.7542991 | 1.56687193 |
| 7001 | NM_001198905.1 | YY1AP1 | 94.4937777 | -7.7705849 | 1.57030278 |
| 7002 | NM_003098.2 | #N/A | 95.0492455 | -6.4278185 | 1.29897067 |
| 7003 | NM_003338.4 | #N/A | 121.406681 | -8.1338806 | 1.64389152 |
| 7004 | NM_001282146.1 | NLGN4X | 89.8427029 | -7.6979347 | 1.55581133 |
| 7005 | NM_001199054.1 | #N/A | 109.596127 | -7.9889931 | 1.61468845 |
| 7006 | NM_152441.2 | #N/A | 90.9999499 | -7.7168898 | 1.55968671 |
| 7007 | NM_174893.3 | #N/A | 91.7644792 | -7.7278863 | 1.56195786 |
| 7008 | NM_017411.3 | #N/A | 81.1771628 | -7.5568644 | 1.52742715 |

| | | | | | |
|---|---|---|---|---|---|
| 7009 | NM_001145199.1 | #N/A | 85.2170994 | -7.6237047 | 1.54114529 |
| 7010 | NM_201589.3 | #N/A | 152.874488 | -8.4673284 | 1.71169942 |
| 7011 | NM_172226.2 | CAMKK2 | 291.482239 | -9.3976673 | 1.90007855 |
| 7012 | NR_037901.1 | ARHGEF26-AS1 | 156.058632 | 3.04617727 | 0.61591755 |
| 7013 | NM_173348.1 | #N/A | 81.1220343 | -7.5529395 | 1.52734069 |
| 7014 | NM_014733.4 | #N/A | 159.843808 | -8.5316804 | 1.72531138 |
| 7015 | NM_138578.2 | #N/A | 109.083906 | -7.9780626 | 1.61338457 |
| 7016 | NM_001003398.1 | #N/A | 79.0881707 | -7.514038 | 1.51956125 |
| 7017 | NM_017931.2 | #N/A | 76.6409955 | -7.4732909 | 1.51136308 |
| 7018 | NR_110439.1 | OTUD6B-AS1 | 111.112942 | -8.0050328 | 1.61891651 |
| 7019 | NM_001962.2 | #N/A | 139.811882 | -6.9198359 | 1.39952637 |
| 7020 | NM_015305.3 | #N/A | 83.5329878 | -7.5960072 | 1.53631063 |
| 7021 | NR_104473.1 | DUSP22 | 104.322224 | -7.9147217 | 1.60083193 |
| 7023 | NR_034041.1 | SRSF1 | 132.743134 | -5.234382 | 1.05878889 |
| 7022 | NM_153813.2 | #N/A | 92.0567684 | -7.7348547 | 1.56457787 |
| 7024 | NM_145253.2 | #N/A | 89.7150363 | -7.698031 | 1.55717993 |
| 7025 | NM_001166005.1 | EPB41 | 144.454061 | -8.3844216 | 1.69611459 |
| 7026 | NM_016946.4 | #N/A | 72.3907319 | -7.3895454 | 1.49488247 |
| 7027 | NM_020247.4 | #N/A | 73.801632 | -7.4187531 | 1.50092076 |
| 7028 | NM_024690.2 | MUC16 | 25.6128775 | 5.71435924 | 1.15612333 |
| 7029 | NR_026785.1 | RHPN1-AS1 | 53.2896831 | 3.64025218 | 0.73651209 |
| 7031 | NM_015141.3 | GPD1L | 124.897194 | -8.1738953 | 1.65381482 |
| 7030 | NM_007265.2 | #N/A | 87.3645808 | -7.6586116 | 1.54956592 |
| 7032 | NM_001083614.1 | #N/A | 81.1304545 | -7.5521025 | 1.52809457 |
| 7033 | NM_001018056.2 | #N/A | 124.447985 | -8.1696932 | 1.65308374 |
| 7034 | NR_103529.1 | #N/A | 127.353163 | -8.203292 | 1.6599211 |
| 7035 | NM_022346.4 | #N/A | 111.801133 | -8.0132279 | 1.62181621 |
| 7036 | NM_001304997.1 | #N/A | 77.3610313 | -7.4826912 | 1.51469215 |
| 7037 | NM_001257361.1 | AMPD2 | 79.480577 | -7.5236958 | 1.52300204 |
| 7038 | NR_003286.2 | #N/A | 10369.8449 | -7.0692883 | 1.43108158 |
| 7039 | NM_175744.4 | RHOC | 120.18409 | -8.1190782 | 1.6438447 |
| 7040 | NM_001197109.1 | SCHIP1 | 139.961508 | -8.3387703 | 1.68840422 |
| 7041 | NM_015955.3 | #N/A | 88.4200351 | -7.6762568 | 1.55428311 |
| 7042 | NM_139056.3 | #N/A | 162.614727 | -8.5568356 | 1.73266861 |
| 7043 | NM_001136499.1 | ZNF841 | 85.7532331 | -7.6290638 | 1.54489091 |
| 7044 | NM_030949.2 | #N/A | 87.0334062 | -7.6532961 | 1.54981645 |
| 7045 | NM_001113378.1 | FANCI | 90.0067462 | -7.6994928 | 1.55921824 |
| 7046 | NM_032357.3 | #N/A | 116.158809 | -8.0692406 | 1.63411817 |
| 7047 | NM_001099857.2 | #N/A | 89.0363246 | -7.689111 | 1.55720021 |
| 7048 | NM_033449.2 | #N/A | 84.8170418 | -7.6179213 | 1.54279363 |
| 7049 | NM_001024662.2 | #N/A | 105.569487 | -7.9343583 | 1.60690865 |
| 7050 | NM_017819.3 | #N/A | 104.594487 | -7.9184893 | 1.60371121 |
| 7051 | NM_001006617.1 | #N/A | 101.543014 | -7.8732784 | 1.59458231 |

| | | | | | |
|---|---|---|---|---|---|
| 7052 | NM_170713.2 | #N/A | 88.7671376 | -7.6827616 | 1.55600377 |
| 7053 | NM_031452.3 | #N/A | 126.180594 | -6.7871273 | 1.37469092 |
| 7054 | NM_003863.3 | DPM2 | 95.7358083 | -7.7913735 | 1.57818559 |
| 7055 | NM_001039848.3 | #N/A | 93.7526647 | -7.7619044 | 1.57224808 |
| 7056 | NM_145306.2 | #N/A | 123.444662 | -8.1578905 | 1.65251593 |
| 7057 | NM_001348098.1 | #N/A | 106.314572 | -7.9417595 | 1.60877562 |
| 7058 | NM_030797.3 | #N/A | 319.610432 | -9.5303058 | 1.93058726 |
| 7059 | NM_003362.3 | #N/A | 107.466023 | -7.9574274 | 1.61200108 |
| 7060 | NM_002969.5 | #N/A | 133.078863 | -8.2669992 | 1.67477823 |
| 7061 | NM_001102564.1 | IFT43 | 80.7865909 | -7.5449639 | 1.52852659 |
| 7062 | NM_032403.2 | #N/A | 159.462024 | -8.5284004 | 1.72782648 |
| 7063 | NM_198925.2 | #N/A | 83.1787649 | -7.5884276 | 1.53755262 |
| 7064 | NM_032375.5 | AKT1S1 | 94.2428186 | -7.7690844 | 1.57420044 |
| 7065 | NM_024713.2 | #N/A | 97.9549081 | -7.8240072 | 1.58533975 |
| 7066 | NM_001287000.1 | ST6GALNAC6 | 89.5395265 | -7.6924325 | 1.55870505 |
| 7067 | NM_001144990.1 | #N/A | 144.796796 | -8.3889481 | 1.69985856 |
| 7068 | NM_001135812.1 | #N/A | 212.610213 | -2.6547087 | 0.53802603 |
| 7069 | NM_001199763.1 | #N/A | 129.743068 | -8.2312536 | 1.66824573 |
| 7070 | NM_001330645.1 | #N/A | 168.883936 | -8.6115259 | 1.74541006 |
| 7071 | NM_000194.2 | #N/A | 83.7148286 | -7.5954766 | 1.53959833 |
| 7072 | NM_001130720.2 | #N/A | 148.915316 | -8.4284774 | 1.70861053 |
| 7073 | NM_001145524.1 | #N/A | 98.3784535 | -7.8296143 | 1.58758439 |
| 7074 | NM_004604.4 | #N/A | 82.6510324 | -7.58002 | 1.53722391 |
| 7075 | NM_020159.4 | #N/A | 132.029762 | -8.2556049 | 1.67432173 |
| 7076 | NM_017615.2 | #N/A | 87.1462878 | -7.6560106 | 1.55283692 |
| 7077 | NM_004635.4 | MAPKAPK3 | 76.2673998 | -7.4654735 | 1.51427413 |
| 7078 | NM_021974.4 | #N/A | 109.846532 | -7.9890133 | 1.62046467 |
| 7079 | NM_015646.5 | #N/A | 104.206623 | -7.9144031 | 1.6053844 |
| 7080 | NM_006108.3 | #N/A | 89.7524672 | -7.7017378 | 1.56241484 |
| 7081 | NM_001329857.1 | #N/A | 106.318606 | -7.9404264 | 1.61085108 |
| 7082 | NM_001017440.2 | #N/A | 72.3647341 | -7.3875335 | 1.49870645 |
| 7083 | NM_030634.2 | ZNF436 | 93.4743269 | -7.7576099 | 1.57388742 |
| 7084 | NR_109988.1 | #N/A | 90.0027936 | -7.7001344 | 1.56236479 |
| 7085 | NM_002231.3 | #N/A | 106.55986 | -7.9484445 | 1.61285291 |
| 7086 | NM_023007.2 | JMJD4 | 85.7003831 | -7.6317968 | 1.54879761 |
| 7087 | NM_052950.3 | #N/A | 71.4219176 | -7.3676567 | 1.49519738 |
| 7088 | NM_013330.4 | #N/A | 73.1331533 | -7.4008642 | 1.50196261 |
| 7089 | NM_001135733.1 | #N/A | 126.65529 | -8.1932178 | 1.66291536 |
| 7090 | NM_001100913.2 | #N/A | 89.158868 | -7.6866611 | 1.56027545 |
| 7091 | NM_003426.3 | #N/A | 138.282348 | -8.3202031 | 1.68889589 |
| 7092 | NM_000155.3 | #N/A | 77.484122 | -7.4874603 | 1.52005814 |
| 7093 | NM_001193315.1 | VIPAS39 | 93.2682485 | -7.7537379 | 1.57414159 |
| 7094 | NM_152272.4 | #N/A | 104.114019 | -7.9119741 | 1.60626542 |

| | | | | | |
|---|---|---|---|---|---|
| 7095 | NM_001198557.1 | #N/A | 99.8190749 | -7.8488956 | 1.5935988 |
| 7096 | NM_198055.1 | #N/A | 74.5949141 | -7.4342164 | 1.50944066 |
| 7097 | NM_001008220.1 | #N/A | 199.780039 | -8.8520088 | 1.79736042 |
| 7098 | NM_001134773.2 | #N/A | 87.3476901 | -7.6583637 | 1.55500504 |
| 7099 | NM_139058.2 | #N/A | 95.0821313 | -7.7776718 | 1.57926739 |
| 7100 | NM_003096.3 | #N/A | 119.859762 | -8.1144655 | 1.64773948 |
| 7101 | NM_006238.4 | #N/A | 95.0190239 | -7.7800583 | 1.57987291 |
| 7102 | NM_001114093.1 | #N/A | 123.746263 | -8.1620609 | 1.65756421 |
| 7103 | NM_003069.4 | #N/A | 124.106121 | -8.1682955 | 1.65891965 |
| 7104 | NM_016524.3 | #N/A | 78.4761591 | -7.5073148 | 1.5247745 |
| 7105 | NM_001242810.2 | BRDT | 71.6099949 | 4.5452397 | 0.92319854 |
| 7106 | NM_031267.3 | CDK13 | 129.072032 | -5.7114987 | 1.16012322 |
| 7107 | NM_020409.2 | #N/A | 104.171086 | -7.9131572 | 1.60735864 |
| 7108 | NM_145798.2 | #N/A | 84.9375814 | -7.619525 | 1.54776826 |
| 7109 | NR_046258.1 | FAR2P2 | 74.6643451 | -7.4314951 | 1.50967931 |
| 7110 | NM_138376.2 | #N/A | 84.7408836 | -7.6167483 | 1.54747108 |
| 7111 | NM_001163435.2 | #N/A | 93.2971857 | -7.7523577 | 1.57504268 |
| 7112 | NR_033936.3 | GOLGA2P10 | 85.1996838 | -7.6234949 | 1.5489095 |
| 7113 | NM_014601.3 | #N/A | 183.222903 | -3.9526559 | 0.80311051 |
| 7114 | NM_022167.3 | #N/A | 98.1391659 | -7.8304148 | 1.5910207 |
| 7115 | NM_000281.3 | #N/A | 95.0441447 | -7.7821449 | 1.58132081 |
| 7116 | NM_012310.4 | #N/A | 77.822543 | -7.4920512 | 1.5223784 |
| 7117 | NM_052853.3 | #N/A | 109.781277 | -7.987782 | 1.62323532 |
| 7118 | NM_139265.3 | #N/A | 79.4864829 | -7.5262385 | 1.5295463 |
| 7119 | NM_001323572.1 | #N/A | 133.32004 | -8.2685722 | 1.68043657 |
| 7120 | NM_001173408.1 | #N/A | 265.972038 | -4.5172255 | 0.9180945 |
| 7121 | NM_001042454.2 | #N/A | 100.03252 | -7.8560372 | 1.59673687 |
| 7122 | NM_001099694.1 | #N/A | 102.1995 | 1.7457769 | 0.35482805 |
| 7123 | NM_153252.4 | #N/A | 137.277405 | -5.7713861 | 1.17307293 |
| 7124 | NM_033453.3 | #N/A | 85.7689739 | -7.6335373 | 1.55164572 |
| 7125 | NM_152367.2 | MAB21L3 | 594.159545 | 3.50504253 | 0.71247621 |
| 7126 | NM_001040627.1 | NCALD | 84.6799523 | -7.6161123 | 1.54820463 |
| 7127 | NM_001134670.1 | HOGA1 | 79.7821452 | 1.97071204 | 0.40064048 |
| 7128 | NM_006633.3 | #N/A | 80.456504 | -7.5415511 | 1.53321509 |
| 7129 | NM_022039.3 | #N/A | 72.8991657 | -7.3964322 | 1.50379587 |
| 7130 | NM_031216.3 | SEH1L | 111.111257 | -8.0053133 | 1.62768347 |
| 7131 | NM_006443.2 | #N/A | 74.8659792 | -7.4376409 | 1.5123167 |
| 7132 | NM_001082967.2 | #N/A | 139.367106 | -8.3325992 | 1.69436179 |
| 7133 | NM_001024809.3 | #N/A | 92.3885168 | -7.7426105 | 1.57448602 |
| 7134 | NM_024844.4 | #N/A | 99.9701659 | -7.8520591 | 1.59683606 |
| 7135 | NM_014962.3 | #N/A | 88.2710894 | -7.6763604 | 1.56124059 |
| 7136 | NM_001171941.2 | #N/A | 79.6717123 | -7.5303059 | 1.53156244 |
| 7137 | NM_004846.3 | #N/A | 91.428744 | -7.7236122 | 1.5709402 |

| | | | | | |
|---|---|---|---|---|---|
| 7138 | NM_016154.4 | #N/A | 110.387291 | -7.9963907 | 1.62647168 |
| 7139 | NM_005388.4 | #N/A | 111.153639 | -6.62054 | 1.34668432 |
| 7140 | NM_003033.3 | #N/A | 86.1756067 | -7.6421438 | 1.55461224 |
| 7141 | NM_001330167.1 | RAB11FIP2 | 94.8286214 | -7.779376 | 1.5826447 |
| 7142 | NM_138431.2 | #N/A | 90.137924 | -7.7022643 | 1.5670113 |
| 7143 | NM_015939.4 | #N/A | 85.8597492 | -7.6324331 | 1.55285652 |
| 7144 | NM_001330125.1 | VAMP2 | 166.691768 | -8.5902855 | 1.74786995 |
| 7145 | NM_032350.5 | C7orf50 | 103.485812 | -7.9035289 | 1.60820573 |
| 7146 | NM_014836.4 | #N/A | 82.3344037 | -7.5780534 | 1.54204892 |
| 7147 | NM_181507.1 | #N/A | 70.3266828 | -7.3463853 | 1.49491902 |
| 7148 | NM_007223.2 | #N/A | 94.6796414 | -7.7763141 | 1.58242251 |
| 7149 | NM_018953.3 | HOXC5 | 120.327259 | -8.122309 | 1.65285262 |
| 7150 | NM_001020820.2 | #N/A | 82.3585126 | -7.5764462 | 1.54179286 |
| 7151 | NM_001130967.2 | #N/A | 71.4147353 | -7.3677405 | 1.4993818 |
| 7152 | NM_183353.2 | #N/A | 98.4146046 | -7.8273112 | 1.59294224 |
| 7153 | NM_021925.3 | #N/A | 86.5894939 | -7.6461767 | 1.55612499 |
| 7154 | NM_001999.3 | #N/A | 652.745913 | -6.0153545 | 1.22425961 |
| 7155 | NM_001079559.2 | #N/A | 560.610563 | -4.5222222 | 0.92038641 |
| 7156 | NM_152913.2 | #N/A | 146.291819 | -8.4036338 | 1.71054903 |
| 7157 | NM_005308.2 | #N/A | 113.757578 | -8.0402961 | 1.63660686 |
| 7158 | NM_178833.5 | #N/A | 77.7528738 | -7.4912278 | 1.52498907 |
| 7159 | NM_018034.3 | #N/A | 99.0313967 | -7.8402183 | 1.59604485 |
| 7160 | NR_003285.2 | #N/A | 174.023209 | -8.6527394 | 1.76179744 |
| 7161 | NM_003193.4 | #N/A | 103.277567 | -7.8996645 | 1.60850882 |
| 7162 | NM_001205293.1 | #N/A | 172.366311 | -8.6393195 | 1.75914902 |
| 7163 | NM_001011552.3 | #N/A | 231.184872 | 3.89399936 | 0.79291668 |
| 7164 | NM_014908.3 | #N/A | 96.1510334 | -7.7995743 | 1.58819338 |
| 7165 | NM_005195.3 | #N/A | 82.1511741 | -7.5750686 | 1.5425028 |
| 7166 | NM_015412.3 | #N/A | 76.4833912 | -7.4681284 | 1.52076881 |
| 7167 | NM_138610.2 | MACROH2A1 | 106.532106 | -7.9481383 | 1.6185526 |
| 7168 | NM_001571.5 | #N/A | 74.566795 | -7.4281253 | 1.51276737 |
| 7169 | NM_007271.3 | #N/A | 106.062685 | -7.9355009 | 1.6162685 |
| 7170 | NM_025250.2 | #N/A | 2371.25257 | -5.5993957 | 1.14045884 |
| 7171 | NR_033339.1 | CTBP1-DT | 84.9187905 | -4.1509504 | 0.84547597 |
| 7172 | NM_203495.3 | COMMD6 | 140.758254 | -8.3473208 | 1.70022757 |
| 7173 | NM_001286635.1 | #N/A | 154.858667 | -8.4858631 | 1.72852615 |
| 7174 | NM_001321926.1 | CKMT1A | 143.488234 | -8.3751635 | 1.70604875 |
| 7175 | NM_018930.3 | #N/A | 90.4667948 | -7.7100707 | 1.57056994 |
| 7176 | NM_001077443.1 | HNRNPC | 621.101473 | -5.9492977 | 1.21196902 |
| 7177 | NM_005326.4 | #N/A | 108.752207 | -7.9750219 | 1.6247345 |
| 7178 | NM_018452.5 | #N/A | 112.500416 | -8.0240063 | 1.63481386 |
| 7179 | NR_036624.1 | #N/A | 102.016464 | -7.8863859 | 1.60690792 |
| 7180 | NM_206836.2 | ECI2 | 72.808476 | -7.3993833 | 1.5077667 |

| | | | | | |
|---|---|---|---|---|---|
| 7181 | NM_032368.4 | #N/A | 100.620811 | -7.8664241 | 1.60303906 |
| 7182 | NM_016101.4 | #N/A | 71.3704362 | -7.3657711 | 1.501244 |
| 7183 | NM_001308263.1 | #N/A | 93.7198446 | -7.7634224 | 1.58239958 |
| 7184 | NM_001188.3 | #N/A | 76.0891214 | -7.4621219 | 1.52107048 |
| 7185 | NR_103449.1 | LINC01544 | 30.3551847 | 4.21173418 | 0.85852811 |
| 7186 | NM_015176.3 | #N/A | 95.1693661 | -7.7817141 | 1.58626616 |
| 7187 | NM_004710.4 | #N/A | 78.7464539 | -7.5091648 | 1.53073095 |
| 7188 | NM_004864.3 | #N/A | 86.0870692 | -7.6424684 | 1.55790074 |
| 7189 | NM_021241.2 | #N/A | 100.443914 | -7.8617986 | 1.60282492 |
| 7190 | NM_023937.3 | #N/A | 95.9068963 | -7.7946835 | 1.58915738 |
| 7191 | NM_001098790.1 | #N/A | 81.9294121 | -7.5648361 | 1.54232396 |
| 7192 | NM_000743.4 | #N/A | 116.335675 | -8.0734403 | 1.64607036 |
| 7193 | NM_001100418.1 | REX1BD | 98.9574093 | -7.8380054 | 1.59813363 |
| 7194 | NR_131917.1 | LOC105747689 | 78.9708986 | 2.84927456 | 0.5809935 |
| 7195 | NM_001198838.1 | #N/A | 81.54932 | -7.5641561 | 1.54254216 |
| 7196 | NM_021809.6 | #N/A | 78.1758853 | -7.495039 | 1.52845817 |
| 7197 | NM_017832.3 | #N/A | 69.9532832 | -7.3387018 | 1.49663728 |
| 7198 | NM_001323289.1 | #N/A | 93.7938768 | -7.759979 | 1.58257226 |
| 7199 | NM_001330074.1 | #N/A | 133.56624 | -8.2741308 | 1.68754839 |
| 7200 | NM_001042683.2 | #N/A | 82.4202683 | -7.572773 | 1.54483836 |
| 7201 | NM_006923.3 | #N/A | 93.9287951 | -7.7644017 | 1.5840204 |
| 7202 | NM_003859.2 | DPM1 | 83.9011109 | -7.5991274 | 1.55031578 |
| 7203 | NM_015055.3 | #N/A | 87.0747301 | -7.6536535 | 1.56161008 |
| 7204 | NM_001316973.1 | #N/A | 158.391733 | -8.5163042 | 1.73765148 |
| 7205 | NM_006014.4 | LAGE3 | 102.682711 | -7.8917247 | 1.61035641 |
| 7206 | NM_078483.3 | #N/A | 107.596498 | -7.960475 | 1.62451506 |
| 7207 | NM_153712.4 | #N/A | 367.888758 | -6.1299132 | 1.25098237 |
| 7208 | NM_022769.4 | #N/A | 106.098915 | -7.9385193 | 1.62012111 |
| 7209 | NM_005028.4 | #N/A | 100.986135 | -7.8679256 | 1.60580945 |
| 7210 | NM_002124.3 | HLA-DRB1 | 147.758906 | -8.418333 | 1.71822068 |
| 7211 | NM_194255.2 | #N/A | 81.0616705 | -7.5512082 | 1.54126612 |
| 7212 | NM_016103.3 | #N/A | 113.289823 | -8.034619 | 1.64013767 |
| 7213 | NM_004704.4 | #N/A | 103.845741 | -7.908023 | 1.61465998 |
| 7214 | NM_001193630.1 | ZNF705B | 21.7797643 | 9.42028566 | 1.92345055 |
| 7215 | NM_001270941.1 | #N/A | 205.393795 | -8.8923966 | 1.81572827 |
| 7216 | NM_004045.3 | #N/A | 117.759756 | -8.090597 | 1.65206846 |
| 7217 | NM_022443.4 | #N/A | 83.465307 | -7.5926163 | 1.55057048 |
| 7218 | NM_001193360.1 | EXD2 | 102.543141 | -7.887598 | 1.61086001 |
| 7219 | NR_076392.1 | #N/A | 132.713238 | -8.2637312 | 1.68777975 |
| 7220 | NM_012225.3 | #N/A | 123.282846 | -8.156026 | 1.66589189 |
| 7221 | NM_017854.1 | #N/A | 120.187306 | -8.1188 | 1.65830714 |
| 7222 | NM_002536.2 | #N/A | 104.167735 | -7.9128277 | 1.61632946 |
| 7223 | NM_001282772.1 | #N/A | 76.0829528 | -7.4598919 | 1.52388105 |

| | | | | | |
|---|---|---|---|---|---|
| 7224 | NM_017637.5 | #N/A | 77.2286716 | -7.4837864 | 1.52887948 |
| 7225 | NM_170663.4 | #N/A | 218.379737 | -8.980657 | 1.83488295 |
| 7226 | NM_133489.2 | SLC26A10 | 113.46222 | -8.0355943 | 1.64188014 |
| 7227 | NM_004519.3 | #N/A | 196.775064 | -8.8301329 | 1.80425925 |
| 7228 | NM_001146068.1 | CAPN2 | 73.3641382 | -7.4044923 | 1.51305723 |
| 7229 | NM_032795.2 | #N/A | 94.6370029 | -7.7736907 | 1.58851897 |
| 7230 | NM_001318040.1 | ZNF618 | 161.589222 | -8.5462057 | 1.74644557 |
| 7231 | NM_001242837.1 | AP2A2 | 118.480377 | -8.099069 | 1.65509356 |
| 7232 | NR_135125.1 | #N/A | 103.297738 | -7.9022551 | 1.61487998 |
| 7233 | NM_134323.1 | #N/A | 86.7640567 | -7.6490051 | 1.56319554 |
| 7234 | NR_037178.1 | LOC100294145 | 158.226543 | -7.0678853 | 1.44444592 |
| 7235 | NM_015367.3 | #N/A | 86.5326225 | -7.6451844 | 1.56244982 |
| 7237 | NM_016338.4 | IPO11 | 90.0300075 | -6.4229473 | 1.31270021 |
| 7236 | NM_001330988.1 | SLC25A25 | 116.18067 | -8.0697171 | 1.64925865 |
| 7239 | NM_183381.2 | RNF13 | 118.269768 | -6.6966774 | 1.36867062 |
| 7238 | NM_001174101.1 | #N/A | 89.9951115 | -7.70122 | 1.57397229 |
| 7241 | NM_025147.4 | COQ10B | 76.9324532 | -7.4739312 | 1.52759419 |
| 7240 | NM_004686.4 | #N/A | 112.471182 | -8.0235482 | 1.63993551 |
| 7242 | NM_001256265.1 | #N/A | 108.244997 | -7.9696337 | 1.62894835 |
| 7243 | NM_017572.3 | #N/A | 76.3246442 | -7.4652977 | 1.525859 |
| 7244 | NR_003265.3 | SDHAP2 | 68.6881833 | -7.3129979 | 1.49476375 |
| 7245 | NM_001042535.3 | #N/A | 134.808526 | -8.2846839 | 1.69343203 |
| 7246 | NM_001284498.1 | #N/A | 86.4886826 | -7.6475268 | 1.56321939 |
| 7247 | NM_001199818.1 | ARMCX5-GPRASP2 | 103.195674 | -7.8988913 | 1.61461908 |
| 7248 | NM_001842.4 | #N/A | 107.447961 | -7.9571762 | 1.62654084 |
| 7249 | NM_003730.4 | #N/A | 71.997233 | -7.3813627 | 1.50889457 |
| 7250 | NM_015469.2 | #N/A | 115.567847 | -8.0626747 | 1.64825227 |
| 7251 | NM_021145.3 | DMTF1 | 88.7213219 | -7.677114 | 1.56946825 |
| 7252 | NM_020902.1 | #N/A | 117.03274 | -8.078686 | 1.65157424 |
| 7253 | NM_001800.3 | #N/A | 142.279794 | -8.3625011 | 1.70968659 |
| 7254 | NM_175736.4 | #N/A | 105.392582 | -7.9304127 | 1.62140242 |
| 7255 | NM_003840.4 | #N/A | 92.7282847 | -6.4661623 | 1.32214009 |
| 7256 | NM_178559.5 | #N/A | 42.30469 | 2.85942321 | 0.58473634 |
| 7257 | NM_001172688.1 | #N/A | 69.9653411 | -7.3405185 | 1.5011328 |
| 7258 | NM_017841.2 | SDHAF2 | 105.064517 | -7.9250859 | 1.6206967 |
| 7259 | NM_001135629.2 | #N/A | 94.4382884 | -7.7705879 | 1.58924914 |
| 7260 | NM_004366.5 | #N/A | 73.8813406 | -7.4160836 | 1.51675734 |
| 7261 | NM_032860.4 | #N/A | 78.6054894 | -7.507029 | 1.53538908 |
| 7262 | NM_020191.2 | #N/A | 82.0876924 | -7.5694784 | 1.54823408 |
| 7263 | NM_153687.3 | #N/A | 72.7962254 | -7.3961008 | 1.51277382 |
| 7264 | NM_002131.3 | #N/A | 118.848271 | -8.1042498 | 1.65765321 |
| 7265 | NM_022037.2 | #N/A | 147.379699 | -8.4118977 | 1.72073177 |
| 7266 | NM_013300.2 | #N/A | 98.4254855 | -7.8311739 | 1.60200408 |

| | | | | | |
|---|---|---|---|---|---|
| 7267 | NM_033550.3 | #N/A | 125.177625 | -4.8051839 | 0.98303263 |
| 7269 | NM_001303103.1 | TLE1 | 120.231698 | -8.1175049 | 1.66071839 |
| 7268 | NM_001161662.1 | #N/A | 86.3804622 | -7.6469768 | 1.5644589 |
| 7270 | NM_001042635.1 | #N/A | 95.0279962 | -7.7802934 | 1.59177997 |
| 7271 | NM_016048.2 | ISOC1 | 96.740483 | -7.8062932 | 1.59718379 |
| 7272 | NM_001297705.1 | #N/A | 84.6016853 | -7.6151908 | 1.55815692 |
| 7273 | NM_001826.2 | #N/A | 85.1782122 | -7.6212605 | 1.55956325 |
| 7274 | NM_031426.3 | #N/A | 89.2047328 | -7.6875717 | 1.57314399 |
| 7275 | NM_021126.5 | #N/A | 113.664049 | -8.0383648 | 1.6449568 |
| 7276 | NR_027383.1 | #N/A | 93.2186761 | -7.7505042 | 1.58609294 |
| 7277 | NM_139157.2 | DENND2B | 112.935074 | -6.6367511 | 1.35829663 |
| 7278 | NR_003264.2 | SDHAP1 | 69.1651966 | -7.3215759 | 1.49860492 |
| 7279 | NR_029435.1 | PSMA3-AS1 | 85.0496087 | -7.6166867 | 1.55911618 |
| 7280 | NM_001136156.1 | #N/A | 88.2747651 | -7.6733581 | 1.57078123 |
| 7281 | NM_001037335.2 | HELZ2 | 107.507704 | -7.9604578 | 1.62962701 |
| 7282 | NM_001587.3 | #N/A | 212.655934 | -6.9351849 | 1.41981224 |
| 7283 | NM_024640.3 | #N/A | 102.133494 | -7.884016 | 1.61406474 |
| 7285 | NR_026771.1 | LINC02249 | 70.6823773 | 3.31950863 | 0.67959996 |
| 7284 | NM_001007156.2 | NTRK3 | 114.11883 | -8.0447503 | 1.64699525 |
| 7286 | NM_001317934.1 | #N/A | 76.9305586 | -7.4782362 | 1.53118062 |
| 7290 | NM_032450.2 | MROH1 | 99.5499126 | -7.8503587 | 1.60743774 |
| 7287 | NM_003259.3 | #N/A | 76.9185513 | -7.4740317 | 1.53035791 |
| 7288 | NM_014606.2 | #N/A | 102.023286 | -7.8806442 | 1.61362365 |
| 7289 | NM_017866.5 | #N/A | 87.9429422 | -7.6682734 | 1.57014751 |
| 7291 | NR_103464.1 | #N/A | 93.2207969 | -7.7531457 | 1.58756676 |
| 7292 | NM_181837.2 | ORC3 | 74.6119981 | -7.431233 | 1.52170995 |
| 7293 | NM_001024845.2 | #N/A | 106.357062 | -7.9422072 | 1.62639738 |
| 7294 | NM_005001.4 | #N/A | 89.803665 | -7.6989803 | 1.57661468 |
| 7295 | NM_001214909.1 | #N/A | 114.011508 | -8.0415817 | 1.64679664 |
| 7296 | NM_002075.3 | #N/A | 74.6137975 | -7.4354983 | 1.52270235 |
| 7297 | NM_020803.4 | #N/A | 93.0548122 | -7.7520519 | 1.58753068 |
| 7298 | NM_181514.1 | #N/A | 94.4442332 | -7.7717942 | 1.59171658 |
| 7299 | NM_001111067.2 | #N/A | 101.229063 | -7.8724809 | 1.61252909 |
| 7300 | NM_004584.2 | #N/A | 73.8304623 | -7.4198567 | 1.51981422 |
| 7301 | NM_018697.3 | LANCL2 | 228.309809 | -6.2971491 | 1.28990473 |
| 7302 | NM_030918.5 | #N/A | 95.8789059 | -7.7927283 | 1.59625055 |
| 7303 | NM_203458.4 | NOTCH2NLA | 82.0938193 | -7.571512 | 1.55099097 |
| 7304 | NM_001321191.1 | RUVBL2 | 92.0819444 | -7.7346752 | 1.58445975 |
| 7305 | NM_153000.4 | #N/A | 80.2298432 | -7.5374334 | 1.54417545 |
| 7306 | NM_006358.3 | #N/A | 94.2380704 | -7.7683263 | 1.59156017 |
| 7307 | NM_015566.2 | #N/A | 81.9303173 | -7.5652892 | 1.54998958 |
| 7308 | NM_020184.3 | #N/A | 85.2566883 | -7.623547 | 1.56198298 |
| 7309 | NM_001330437.1 | PTPN11 | 75.6605965 | -7.4504728 | 1.52661556 |

| | | | | | |
|---|---|---|---|---|---|
| 7310 | NM_021045.2 | ZNF248 | 129.821054 | -8.2297745 | 1.68632607 |
| 7311 | NM_001005209.2 | #N/A | 116.668686 | -8.0755991 | 1.65479106 |
| 7312 | NM_018367.6 | #N/A | 129.508893 | -8.2282716 | 1.68629756 |
| 7313 | NM_001084.4 | #N/A | 251.078625 | -4.2627833 | 0.87364952 |
| 7314 | NM_001330218.1 | CCNYL1 | 76.7588162 | -7.4693453 | 1.53085765 |
| 7315 | NM_019007.3 | #N/A | 90.7016855 | -7.7146455 | 1.58127125 |
| 7316 | NM_032998.2 | #N/A | 108.610045 | -7.9726699 | 1.6342152 |
| 7317 | NM_001042679.1 | RHOC | 86.4024573 | -7.6442058 | 1.56692154 |
| 7318 | NM_001282670.1 | FAAP20 | 129.670363 | 2.70734624 | 0.55496259 |
| 7319 | NM_016474.4 | #N/A | 79.849295 | -7.5283618 | 1.54332526 |
| 7320 | NM_033215.4 | #N/A | 92.8344814 | -7.7463912 | 1.58802422 |
| 7321 | NM_001174128.1 | #N/A | 74.7086103 | -7.4359194 | 1.52447413 |
| 7322 | NM_020175.2 | #N/A | 89.2083832 | -7.6887986 | 1.57675876 |
| 7323 | NM_001301228.1 | HMGB3 | 102.904661 | -7.895194 | 1.61910517 |
| 7324 | NM_006086.3 | #N/A | 4145.96823 | -7.1076814 | 1.45760416 |
| 7325 | NM_024617.3 | #N/A | 81.6402768 | -7.5594036 | 1.55028762 |
| 7326 | NM_018316.2 | #N/A | 78.9438444 | -7.5124864 | 1.54068515 |
| 7327 | NM_001144757.2 | SCG5 | 105.970401 | -7.9388798 | 1.62815345 |
| 7328 | NM_001084392.1 | DDT | 115.479337 | -8.061687 | 1.65337184 |
| 7330 | NM_001584.2 | MPPED2 | 149.64677 | -8.4356254 | 1.7301503 |
| 7329 | NM_000713.2 | #N/A | 83.7093866 | -7.5973579 | 1.55821954 |
| 7331 | NM_006541.4 | #N/A | 132.018497 | -8.2546456 | 1.69302873 |
| 7332 | NM_017900.2 | #N/A | 103.564778 | -7.9038316 | 1.62106965 |
| 7333 | NM_000375.2 | #N/A | 102.914633 | -7.8956832 | 1.61960185 |
| 7334 | NM_003316.3 | #N/A | 101.293009 | -7.8724133 | 1.6149637 |
| 7335 | NM_003585.4 | #N/A | 73.0967429 | -7.4054445 | 1.51920527 |
| 7336 | NR_015433.1 | #N/A | 90.7325504 | -7.7125642 | 1.5822854 |
| 7337 | NM_001025780.2 | ABHD17B | 133.53259 | -2.8061641 | 0.57571228 |
| 7338 | NR_038841.1 | AFF1-AS1 | 143.379087 | 3.02512014 | 0.6206613 |
| 7339 | NR_028408.1 | LINC00938 | 99.1212326 | -7.8409421 | 1.60872849 |
| 7340 | NM_016226.4 | #N/A | 112.305914 | -8.0217112 | 1.64592691 |
| 7341 | NM_007010.4 | #N/A | 112.046197 | -8.019344 | 1.64548379 |
| 7342 | NM_022497.4 | #N/A | 82.5805708 | -7.5797506 | 1.55537052 |
| 7343 | NM_012258.3 | #N/A | 133.202452 | -8.266457 | 1.69629948 |
| 7344 | NM_152769.2 | #N/A | 149.205802 | -6.9900671 | 1.43444366 |
| 7345 | NM_002180.2 | IGHMBP2 | 99.0146264 | -7.8398053 | 1.60889605 |
| 7346 | NM_001167.3 | XIAP | 113.112454 | -8.0301851 | 1.64801338 |
| 7347 | NM_052879.4 | #N/A | 88.1208781 | -7.6692055 | 1.5739667 |
| 7349 | NR_144445.1 | ERVK3-1 | 140.970715 | -6.9178108 | 1.41984078 |
| 7348 | NR_002809.2 | #N/A | 78.810803 | -7.5102511 | 1.54143129 |
| 7351 | NM_001163809.1 | WDR81 | 71.1172348 | -7.3606151 | 1.51077035 |
| 7350 | NM_001029991.1 | #N/A | 73.6413038 | -7.4110182 | 1.5211207 |
| 7352 | NM_012230.3 | POMZP3 | 109.50318 | -7.9847086 | 1.63898895 |

| | | | | | |
|---|---|---|---|---|---|
| 7353 | NM_031937.2 | #N/A | 75.5339193 | -7.4497184 | 1.52925575 |
| 7354 | NM_015898.3 | #N/A | 101.882718 | -6.4994787 | 1.33422176 |
| 7355 | NM_001323934.1 | SNAP47 | 91.2038618 | -7.7227092 | 1.58545029 |
| 7356 | NM_001281749.1 | #N/A | 72.5869666 | -7.3926356 | 1.51789984 |
| 7357 | NM_020773.2 | #N/A | 68.2144513 | -7.3032748 | 1.49956687 |
| 7358 | NM_003609.4 | #N/A | 97.9392882 | -7.8225177 | 1.60619504 |
| 7359 | NM_014320.2 | #N/A | 94.1436743 | -7.7671048 | 1.5949873 |
| 7361 | NR_036750.2 | NPEPPSP1 | 71.5042214 | -7.3742479 | 1.51437435 |
| 7360 | NM_001171624.1 | VEGFA | 137.957198 | -8.3187013 | 1.70832487 |
| 7362 | NM_032340.3 | #N/A | 123.096213 | -8.1536627 | 1.67444982 |
| 7363 | NM_018264.3 | #N/A | 80.3837935 | -7.5384115 | 1.54819474 |
| 7364 | NM_001008395.3 | #N/A | 96.8436288 | -7.8092747 | 1.60400815 |
| 7365 | NM_153611.4 | #N/A | 85.0512141 | -7.6237458 | 1.56595079 |
| 7366 | NM_001347423.1 | #N/A | 2746.64851 | -5.4756687 | 1.12474359 |
| 7367 | NM_001257.4 | #N/A | 139.785868 | -8.3374682 | 1.71271692 |
| 7368 | NM_001270401.1 | #N/A | 97.3683143 | -7.8121514 | 1.60480347 |
| 7369 | NM_138732.2 | #N/A | 252.572365 | -6.4051535 | 1.31581351 |
| 7370 | NM_152407.3 | #N/A | 96.0480873 | -7.7952513 | 1.6016555 |
| 7371 | NM_001005210.2 | #N/A | 139.436403 | -8.3332947 | 1.71225436 |
| 7372 | NM_020754.3 | #N/A | 75.0400608 | -7.4434491 | 1.52942008 |
| 7373 | NM_001048205.1 | #N/A | 78.3244988 | -7.5048698 | 1.54210866 |
| 7374 | NM_014621.2 | #N/A | 123.974978 | -8.1650425 | 1.67787031 |
| 7375 | NM_001237.4 | #N/A | 93.8815403 | -7.7627823 | 1.59527735 |
| 7376 | NM_203352.2 | #N/A | 90.5431072 | -7.7131023 | 1.58516294 |
| 7377 | NM_152271.4 | #N/A | 143.669526 | -8.3759493 | 1.7214092 |
| 7378 | NM_001136191.2 | #N/A | 85.9314544 | -7.639578 | 1.57022003 |
| 7379 | NM_020230.6 | #N/A | 73.5914925 | -7.4086245 | 1.52284391 |
| 7380 | NM_002861.4 | #N/A | 140.899639 | -8.3486578 | 1.71613464 |
| 7383 | NR_073527.1 | PPP2R2B | 110.420985 | -7.9948725 | 1.6435157 |
| 7381 | NM_014379.3 | #N/A | 142.340285 | 3.16885049 | 0.65142304 |
| 7382 | NM_016128.3 | #N/A | 747.926244 | -5.9016519 | 1.21321369 |
| 7384 | NM_018107.4 | #N/A | 94.1191 | -7.7696967 | 1.59727002 |
| 7385 | NM_001330290.1 | #N/A | 76.2292718 | -7.4605844 | 1.53374993 |
| 7386 | NM_001194946.1 | #N/A | 385.899952 | -9.8025253 | 2.01523843 |
| 7387 | NM_000158.3 | #N/A | 75.7186088 | -7.4564454 | 1.53294142 |
| 7388 | NM_006715.3 | #N/A | 88.499457 | -7.6771854 | 1.57834759 |
| 7390 | NM_005174.3 | ATP5F1C | 97.198823 | -7.8156162 | 1.60686651 |
| 7389 | NM_001142295.1 | SPART | 67.9808454 | -7.3004193 | 1.50094853 |
| 7391 | NM_004487.4 | #N/A | 164.945521 | -8.5756507 | 1.76337505 |
| 7392 | NM_001404.4 | #N/A | 6334.07037 | -7.1500075 | 1.47026064 |
| 7393 | NM_001193655.1 | CYBC1 | 77.8362069 | -7.4914995 | 1.54064616 |
| 7394 | NM_014017.3 | #N/A | 92.8091926 | -7.746166 | 1.59324807 |
| 7395 | NM_025150.4 | #N/A | 89.9831372 | -7.6978809 | 1.58338246 |

| | | | | | |
|---|---|---|---|---|---|
| 7396 | NM_032783.4 | #N/A | 84.2672049 | -7.6049708 | 1.56445763 |
| 7397 | NM_001330660.1 | #N/A | 89.3371521 | -7.694954 | 1.58298085 |
| 7398 | NM_006252.3 | #N/A | 74.4316226 | -4.663842 | 0.95944764 |
| 7399 | NM_194314.2 | ZBTB41 | 129.053382 | -6.8025885 | 1.39944059 |
| 7400 | NM_030569.6 | #N/A | 93.8339739 | -7.7655571 | 1.59758353 |
| 7401 | NM_001146345.1 | #N/A | 74.3527403 | -7.4285896 | 1.52833506 |
| 7402 | NM_005822.3 | RCAN2 | 142.880614 | -8.3690744 | 1.72187427 |
| 7403 | NM_001077351.1 | #N/A | 80.9129115 | -7.5462089 | 1.55268922 |
| 7404 | NM_001099220.2 | #N/A | 127.711059 | -5.6769884 | 1.16808703 |
| 7405 | NM_006893.2 | #N/A | 75.3807595 | -7.4486505 | 1.53267281 |
| 7406 | NM_015028.3 | #N/A | 214.837515 | -8.957511 | 1.84324908 |
| 7407 | NM_001912.4 | #N/A | 75.1901314 | -7.4424459 | 1.53156315 |
| 7408 | NM_006085.5 | #N/A | 98.9868801 | -7.8374537 | 1.61284397 |
| 7409 | NM_018310.3 | #N/A | 85.4455038 | -7.6268743 | 1.5695356 |
| 7410 | NM_001256575.1 | ENC1 | 135.775243 | -8.2949678 | 1.70703488 |
| 7411 | NM_033288.3 | ZNF160 | 91.5182767 | -7.7243162 | 1.58970629 |
| 7412 | NM_052907.3 | TMEM132B | 102.729134 | -7.8915563 | 1.62417146 |
| 7413 | NM_145269.4 | #N/A | 90.2417132 | -7.7049111 | 1.58582119 |
| 7414 | NM_001013839.3 | #N/A | 132.295405 | -8.2557272 | 1.69938834 |
| 7415 | NM_001145445.1 | SCAF4 | 76.2267606 | -7.4664595 | 1.53697736 |
| 7416 | NM_003750.2 | #N/A | 1055.39692 | -4.3164371 | 0.88858467 |
| 7417 | NR_024031.2 | DANCR | 102.167781 | -7.884512 | 1.62321498 |
| 7418 | NM_032291.3 | #N/A | 139.722212 | -8.3365042 | 1.71653564 |
| 7419 | NM_003310.3 | #N/A | 101.136578 | -7.8695896 | 1.62041264 |
| 7420 | NM_006996.2 | #N/A | 78.5725958 | -7.5053501 | 1.54546644 |
| 7421 | NM_015917.2 | #N/A | 74.2708004 | -7.4273456 | 1.52944715 |
| 7422 | NM_001105549.1 | ZNF83 | 74.2585732 | -7.429242 | 1.52995668 |
| 7423 | NM_152773.4 | #N/A | 89.5276389 | -7.6927549 | 1.58432366 |
| 7424 | NM_001002926.1 | #N/A | 79.8216028 | -7.5295122 | 1.55075118 |
| 7425 | NM_032837.2 | FAM104A | 87.0484282 | -7.6567259 | 1.57701162 |
| 7426 | NM_001463.3 | #N/A | 86.7737443 | -7.6513783 | 1.57594343 |
| 7427 | NM_002237.3 | #N/A | 145.981511 | -8.3993584 | 1.73011901 |
| 7428 | NR_027928.2 | CHKB-CPT1B | 70.0683663 | -7.3376225 | 1.51150154 |
| 7429 | NM_021248.2 | #N/A | 89.2896049 | -7.6917682 | 1.58453264 |
| 7430 | NM_139012.2 | MAPK14 | 79.5996707 | -7.52835 | 1.55090824 |
| 7431 | NM_021104.1 | #N/A | 2920.3928 | -6.7293452 | 1.38633374 |
| 7432 | NM_018211.3 | #N/A | 78.8788583 | -7.5102689 | 1.54730841 |
| 7433 | NM_152551.3 | #N/A | 80.0109552 | -7.5320607 | 1.55182696 |
| 7434 | NM_001038707.1 | CDC42SE1 | 77.7773787 | -7.4928911 | 1.54379212 |
| 7435 | NM_001077665.2 | AGAP6 | 108.526999 | -4.0280597 | 0.82994268 |
| 7436 | NM_207115.1 | #N/A | 114.497827 | -8.0496683 | 1.65857236 |
| 7437 | NM_031925.2 | #N/A | 109.949874 | -7.9907688 | 1.64647629 |
| 7438 | NM_152283.4 | ZFP62 | 122.524907 | -8.1470442 | 1.67872018 |

| | | | | | |
|---|---|---|---|---|---|
| 7439 | NM_001261395.1 | #N/A | 35.73412 | 10.1507814 | 2.09168656 |
| 7440 | NM_194280.3 | #N/A | 87.4337162 | -7.6604098 | 1.57876109 |
| 7441 | NR_027768.1 | DPY19L2P2 | 80.2180237 | -7.5381434 | 1.55359266 |
| 7442 | NM_024309.3 | #N/A | 77.3426648 | -7.4815186 | 1.54194125 |
| 7443 | NM_004240.3 | #N/A | 76.7572703 | -7.4758716 | 1.54081737 |
| 7444 | NM_153747.1 | #N/A | 88.6774697 | -7.6822694 | 1.58336092 |
| 7445 | NM_001310137.1 | #N/A | 147.992948 | -5.178038 | 1.06735961 |
| 7446 | NM_033208.3 | #N/A | 73.6297567 | -7.4108902 | 1.52768779 |
| 7447 | NM_024165.2 | #N/A | 95.8313639 | -7.796328 | 1.6071914 |
| 7448 | NM_001193388.3 | #N/A | 66.1487392 | -7.2566066 | 1.49596048 |
| 7449 | NM_148414.2 | ATXN2L | 95.026582 | -7.7819071 | 1.60435109 |
| 7450 | NM_017423.2 | #N/A | 95.1863806 | -7.7834749 | 1.60471623 |
| 7451 | NM_014177.2 | #N/A | 93.1288903 | -7.7509408 | 1.59802917 |
| 7452 | NM_001319837.1 | ARHGEF10L | 84.7904337 | -7.6170132 | 1.57043993 |
| 7453 | NM_017906.2 | #N/A | 84.6180835 | -7.612428 | 1.56956837 |
| 7454 | NM_014989.5 | RIMS1 | 128.846607 | -8.2203552 | 1.6949913 |
| 7455 | NM_005682.6 | #N/A | 177.577088 | -8.6827173 | 1.79037103 |
| 7456 | NM_058169.5 | #N/A | 71.8997812 | -7.3770135 | 1.52120742 |
| 7457 | NM_033211.3 | #N/A | 111.164035 | -8.006975 | 1.65130879 |
| 7458 | NM_004760.2 | #N/A | 97.8283043 | -7.8213183 | 1.61307647 |
| 7459 | NM_001018072.1 | #N/A | 73.1840777 | -7.4077542 | 1.52789129 |
| 7460 | NM_001323523.1 | #N/A | 144.268349 | -8.3832882 | 1.72911185 |
| 7461 | NR_110912.1 | #N/A | 83.2281355 | -7.5893272 | 1.56537342 |
| 7462 | NM_032873.4 | #N/A | 86.0580299 | -7.6368986 | 1.57523868 |
| 7463 | NM_016446.3 | TMEM8B | 81.0382401 | -7.5506564 | 1.55747206 |
| 7464 | NM_005649.2 | #N/A | 81.8168804 | -7.5629716 | 1.56015034 |
| 7465 | NM_024528.3 | #N/A | 76.1742315 | -7.4620999 | 1.53932484 |
| 7466 | NM_148956.3 | #N/A | 97.3558766 | -7.8152164 | 1.61217979 |
| 7467 | NR_111980.1 | #N/A | 556.5668 | 3.19126746 | 0.65833971 |
| 7468 | NM_022365.3 | #N/A | 121.052802 | -6.7186675 | 1.38608116 |
| 7469 | NR_033688.1 | #N/A | 78.6047712 | -7.5100556 | 1.54936099 |
| 7470 | NM_022827.3 | #N/A | 86.7780589 | -7.6516562 | 1.57864186 |
| 7471 | NM_014723.3 | #N/A | 113.870519 | -8.0419822 | 1.65923402 |
| 7472 | NM_170686.2 | #N/A | 77.6276384 | -7.4863982 | 1.54475762 |
| 7473 | NM_016651.5 | #N/A | 150.924529 | -8.4475889 | 1.74321659 |
| 7474 | NM_030759.4 | #N/A | 81.0565605 | -7.5511071 | 1.55828696 |
| 7475 | NR_126511.1 | #N/A | 88.8393806 | -7.6818282 | 1.58527849 |
| 7476 | NM_001248008.1 | SKIL | 102.564169 | -7.8897865 | 1.62822586 |
| 7477 | NM_018136.4 | #N/A | 84.8369443 | -7.6157477 | 1.57168407 |
| 7478 | NM_001040101.1 | NSG1 | 171.87822 | -8.6356216 | 1.78220502 |
| 7479 | NM_032881.2 | #N/A | 103.920885 | -7.9099259 | 1.63254787 |
| 7480 | NM_014667.3 | #N/A | 90.0673099 | -7.7063766 | 1.59055478 |
| 7481 | NM_002617.3 | PEX10 | 84.4934913 | -7.6108989 | 1.57088556 |

| | | | | | |
|---|---|---|---|---|---|
| 7482 | NM_001008495.3 | #N/A | 72.2266672 | -7.3883785 | 1.52497986 |
| 7483 | NM_015004.3 | #N/A | 77.9787985 | -7.4954348 | 1.54718606 |
| 7484 | NM_001300909.1 | #N/A | 78.3681383 | -7.5061329 | 1.54941183 |
| 7485 | NM_144679.2 | #N/A | 84.2989627 | -7.6073289 | 1.57030534 |
| 7486 | NM_152522.5 | #N/A | 85.2621806 | -7.6226063 | 1.57348615 |
| 7487 | NM_000373.3 | #N/A | 90.7440006 | -7.7132064 | 1.59219706 |
| 7488 | NM_018197.2 | #N/A | 79.0124598 | -7.5119928 | 1.55067411 |
| 7489 | NM_001321077.1 | #N/A | 107.380686 | -7.956383 | 1.64243883 |
| 7490 | NM_025126.3 | RNF34 | 91.3138886 | -7.7217926 | 1.59411695 |
| 7491 | NM_001082575.2 | RBFOX3 | 263.370779 | -6.4452755 | 1.33062588 |
| 7492 | NM_014362.3 | #N/A | 67.0183713 | -7.2779108 | 1.50259376 |
| 7493 | NM_017762.2 | #N/A | 68.5080773 | -7.3103017 | 1.50929977 |
| 7494 | NM_022373.4 | #N/A | 118.888042 | -3.6048743 | 0.74428501 |
| 7495 | NM_001300815.1 | #N/A | 76.3184477 | -7.4627961 | 1.54108592 |
| 7496 | NM_001002762.2 | #N/A | 75.8877783 | -7.4571001 | 1.53993584 |
| 7497 | NM_032420.3 | #N/A | 114.392694 | -8.0481433 | 1.66205543 |
| 7498 | NM_024098.3 | #N/A | 83.4197418 | -7.5931378 | 1.56810952 |
| 7499 | NM_001002262.3 | #N/A | 89.2144075 | -7.6899345 | 1.58822602 |
| 7500 | NM_001134937.1 | FIP1L1 | 79.1063283 | -7.5120583 | 1.55155693 |
| 7501 | NM_001321646.1 | #N/A | 103.093042 | -7.8967096 | 1.63119829 |
| 7502 | NM_006346.2 | #N/A | 70.4512293 | -7.3478188 | 1.51784575 |
| 7503 | NM_032862.4 | #N/A | 83.9601287 | -7.6007989 | 1.57013302 |
| 7504 | NM_001007069.1 | SDCBP | 87.8593225 | -7.6638246 | 1.583198 |
| 7505 | NM_016484.4 | #N/A | 104.39363 | -7.9167715 | 1.63552319 |
| 7506 | NM_006696.3 | #N/A | 91.4708431 | -7.7275025 | 1.59643327 |
| 7507 | NM_019857.4 | #N/A | 86.3631572 | -7.6392758 | 1.57826931 |
| 7508 | NM_002522.3 | #N/A | 199.747855 | -8.8519421 | 1.82889985 |
| 7509 | NM_145109.2 | #N/A | 70.5325192 | -7.3502499 | 1.51863976 |
| 7511 | NR_027754.2 | CEACAM22P | 95.8444962 | 3.34495045 | 0.69117949 |
| 7510 | NM_145177.2 | #N/A | 69.9118655 | -7.340055 | 1.5166991 |
| 7512 | NM_032816.4 | #N/A | 65.5919882 | -7.2444449 | 1.49698198 |
| 7513 | NM_032859.2 | #N/A | 83.5899337 | -7.593993 | 1.56921842 |
| 7514 | NM_006182.2 | #N/A | 70.9381277 | -7.3636487 | 1.52169342 |
| 7515 | NM_032773.3 | LRCH3 | 68.5523208 | -7.3100868 | 1.51063802 |
| 7516 | NM_001276318.1 | #N/A | 80.7765052 | -7.5462978 | 1.55946152 |
| 7517 | NM_080655.2 | MSANTD3 | 97.0182362 | -7.8099391 | 1.61396115 |
| 7518 | NM_198243.2 | #N/A | 74.6585663 | -7.4301356 | 1.5354936 |
| 7520 | NM_138766.2 | PAM | 105.118583 | -7.9269009 | 1.63850618 |
| 7519 | NM_025075.3 | #N/A | 97.6862299 | -7.8200278 | 1.61641991 |
| 7521 | NM_199044.3 | NSUN4 | 74.5726573 | -7.4287986 | 1.53557102 |
| 7522 | NM_001080522.2 | CC2D2A | 86.7874784 | -7.6509913 | 1.58158345 |
| 7523 | NM_030935.4 | #N/A | 87.4585411 | -7.6635052 | 1.58423945 |
| 7524 | NM_004824.3 | #N/A | 76.0014699 | -7.460928 | 1.54244856 |

| | | | | | |
|---|---|---|---|---|---|
| 7525 | NM_001286245.1 | #N/A | 93.3114831 | -7.7527106 | 1.60282695 |
| 7526 | NM_032051.1 | #N/A | 67.2338066 | -7.2827883 | 1.50575941 |
| 7527 | NM_025212.3 | #N/A | 95.7648915 | -7.794282 | 1.61166474 |
| 7528 | NM_020244.2 | #N/A | 87.5913401 | -7.6628452 | 1.58461574 |
| 7529 | NM_145868.1 | #N/A | 68.4907623 | -7.3090405 | 1.51149007 |
| 7530 | NM_001256155.2 | ARMCX4 | 76.4412072 | -7.4673842 | 1.54441923 |
| 7531 | NM_007112.4 | #N/A | 110.959127 | -8.0069425 | 1.65613641 |
| 7532 | NM_018710.2 | #N/A | 109.525869 | -7.985414 | 1.65175004 |
| 7533 | NM_003611.2 | #N/A | 84.9547196 | -7.6154073 | 1.57523869 |
| 7534 | NM_012256.3 | #N/A | 67.5633897 | -7.289947 | 1.50791 |
| 7535 | NM_144736.4 | #N/A | 67.776169 | -7.2947519 | 1.50893339 |
| 7536 | NM_001010917.2 | #N/A | 161.110427 | -8.5421177 | 1.76708896 |
| 7537 | NM_006775.2 | #N/A | 68.0224358 | -7.2986438 | 1.50987821 |
| 7538 | NM_001300826.1 | #N/A | 96.0085542 | -7.7931435 | 1.61222151 |
| 7539 | NM_002335.3 | #N/A | 297.069263 | -2.9577193 | 0.6118831 |
| 7540 | NM_004973.3 | #N/A | 122.797026 | -6.7322284 | 1.39275215 |
| 7541 | NM_016277.4 | #N/A | 96.350662 | -7.7997813 | 1.61368178 |
| 7542 | NM_033390.1 | #N/A | 76.4881646 | -7.4639848 | 1.54424045 |
| 7543 | NM_001126111.2 | #N/A | 76.7083159 | -7.471241 | 1.54576743 |
| 7544 | NM_000714.5 | #N/A | 125.977519 | -8.1866928 | 1.69380301 |
| 7545 | NM_152240.2 | #N/A | 70.5375697 | -7.3476346 | 1.52036669 |
| 7546 | NM_198216.1 | #N/A | 92.3548008 | -7.7397416 | 1.6016247 |
| 7547 | NM_003392.4 | WNT5A | 97.6124841 | -7.8220983 | 1.61871288 |
| 7548 | NR_033323.3 | NSUN5P2 | 70.8578875 | -7.3608921 | 1.52335176 |
| 7549 | NM_007349.3 | #N/A | 78.4262239 | -7.5030387 | 1.55288029 |
| 7550 | NM_003612.4 | #N/A | 123.288716 | -8.1555946 | 1.68796391 |
| 7551 | NM_004694.4 | #N/A | 72.6818099 | -7.3968414 | 1.5309523 |
| 7552 | NM_001321645.1 | #N/A | 66.2532511 | -7.2575299 | 1.50216409 |
| 7553 | NM_032043.2 | #N/A | 289.624602 | 2.46631493 | 0.51048915 |
| 7554 | NM_005952.3 | #N/A | 128.343371 | -8.2141993 | 1.70029069 |
| 7555 | NM_173484.3 | #N/A | 38.1451852 | 2.94390283 | 0.60939004 |
| 7556 | NM_021873.3 | CDC25B | 70.7813932 | -7.3516588 | 1.52192227 |
| 7557 | NM_152625.2 | #N/A | 51.7950546 | 3.79708134 | 0.78608015 |
| 7558 | NM_001288767.1 | #N/A | 85.4333867 | -7.6249923 | 1.57873187 |
| 7559 | NM_171982.4 | #N/A | 85.8269695 | -7.6315065 | 1.58013239 |
| 7561 | NM_001322384.1 | ENTPD6 | 88.3056493 | -7.6745853 | 1.58947639 |
| 7560 | NM_001303503.1 | #N/A | 77.6118415 | -7.4871444 | 1.55065498 |
| 7562 | NM_032627.4 | #N/A | 71.9620922 | -7.3761313 | 1.52774059 |
| 7563 | NM_001204142.1 | #N/A | 77.098696 | -7.4785263 | 1.54910407 |
| 7564 | NM_145030.3 | #N/A | 66.9837551 | -7.2761288 | 1.50720821 |
| 7565 | NM_001242374.1 | ANAPC13 | 84.6250382 | -7.6126469 | 1.5769707 |
| 7566 | NM_003271.4 | #N/A | 78.9808149 | -7.5098991 | 1.55577765 |
| 7567 | NM_002668.2 | #N/A | 67.0122 | -7.2795291 | 1.50811833 |

| | | | | | |
|---|---|---|---|---|---|
| 7568 | NM_001185077.2 | #N/A | 78.0989646 | -7.5004448 | 1.55397132 |
| 7569 | NR_038366.1 | HOTAIRM1 | 122.487323 | -8.1478349 | 1.68817151 |
| 7570 | NM_017456.3 | #N/A | 134.79068 | -8.2850971 | 1.71677026 |
| 7571 | NM_003957.3 | #N/A | 123.580398 | -8.1592112 | 1.69091417 |
| 7572 | NM_144695.3 | #N/A | 73.690492 | -7.4132919 | 1.53633332 |
| 7573 | NM_021163.3 | #N/A | 91.9362638 | -7.7359616 | 1.60324445 |
| 7574 | NM_001204478.1 | #N/A | 64.6974292 | -7.2247205 | 1.49738379 |
| 7575 | NM_017636.3 | #N/A | 84.9717037 | -7.6192304 | 1.57924163 |
| 7577 | NM_001256828.1 | SMG6 | 74.9859648 | -7.4422255 | 1.54257767 |
| 7576 | NM_001160259.1 | #N/A | 120.018106 | -8.1181018 | 1.68266879 |
| 7578 | NM_001012979.2 | #N/A | 148.918118 | -8.4282736 | 1.74701254 |
| 7579 | NM_015192.3 | #N/A | 84.0461535 | -7.6023515 | 1.57581148 |
| 7580 | NM_001128631.2 | #N/A | 74.717776 | -7.4344276 | 1.54106972 |
| 7581 | NM_054025.2 | #N/A | 159.580887 | -8.5276489 | 1.76777086 |
| 7582 | NM_025080.3 | #N/A | 118.496969 | -8.0982993 | 1.67884165 |
| 7583 | NM_001145028.1 | #N/A | 115.568424 | -8.0634202 | 1.67182633 |
| 7584 | NM_178507.2 | #N/A | 82.9970258 | -7.5895549 | 1.57357782 |
| 7585 | NM_007222.4 | #N/A | 119.318801 | -8.108718 | 1.68123175 |
| 7586 | NM_001270764.1 | #N/A | 97.7375636 | -7.8222604 | 1.62212198 |
| 7587 | NM_001083600.2 | #N/A | 95.5748446 | -7.7882825 | 1.61516749 |
| 7588 | NM_018688.4 | #N/A | 74.0584425 | -7.4206521 | 1.5389325 |
| 7589 | NM_014008.4 | #N/A | 77.1936628 | -7.4819394 | 1.55165135 |
| 7590 | NM_001079670.2 | #N/A | 87.5255699 | -7.6620912 | 1.5890274 |
| 7591 | NM_015238.2 | #N/A | 142.379289 | -4.3119454 | 0.89426755 |
| 7592 | NM_015567.1 | SLITRK5 | 66.8216378 | -7.2724216 | 1.50825845 |
| 7593 | NM_001322477.1 | HPS1 | 79.4277746 | -7.5234807 | 1.56050124 |
| 7594 | NM_005686.2 | #N/A | 73.030549 | -7.405637 | 1.53618025 |
| 7595 | NM_014473.2 | #N/A | 93.4468394 | -7.7548028 | 1.60863229 |
| 7596 | NM_001198777.1 | #N/A | 83.5936078 | -7.5941284 | 1.57538772 |
| 7597 | NM_014902.5 | #N/A | 82.0371046 | -7.5687729 | 1.57013217 |
| 7598 | NM_024816.2 | #N/A | 73.509225 | -7.4131087 | 1.53783206 |
| 7599 | NM_012162.3 | #N/A | 88.6747222 | -7.6810436 | 1.59351204 |
| 7600 | NM_024637.4 | #N/A | 124.261977 | -4.9025919 | 1.01710446 |
| 7601 | NM_022170.1 | #N/A | 84.9651613 | -7.6230567 | 1.58159442 |
| 7602 | NM_152864.3 | #N/A | 73.4305119 | -7.4087203 | 1.53712762 |
| 7603 | NM_198516.2 | #N/A | 85.3269336 | -7.6279924 | 1.58279336 |
| 7604 | NM_005837.2 | #N/A | 130.171932 | -8.2344332 | 1.7087666 |
| 7605 | NM_001145073.2 | #N/A | 69.4917809 | -7.3320447 | 1.52153193 |
| 7606 | NM_014594.2 | #N/A | 92.4056226 | -7.7376083 | 1.60572797 |
| 7607 | NM_001278198.1 | #N/A | 81.0193583 | -7.548586 | 1.56661303 |
| 7608 | NM_002582.3 | #N/A | 82.6122667 | -7.5766193 | 1.57245015 |
| 7609 | NM_001111.4 | #N/A | 140.119504 | -8.3389755 | 1.73081464 |
| 7610 | NM_205767.2 | MICOS13 | 100.486864 | -7.8600616 | 1.63148289 |

| | | | | | |
|---|---|---|---|---|---|
| 7611 | NR_135260.1 | C1orf61 | 148.592597 | -8.4247904 | 1.74872292 |
| 7612 | NR_024271.1 | SEPTIN7P2 | 71.2345745 | -7.3625376 | 1.52824638 |
| 7613 | NM_006937.3 | #N/A | 886.484562 | -4.5097611 | 0.93610284 |
| 7614 | NM_001258277.1 | TMEM200A | 95.2591532 | -7.7854427 | 1.61613478 |
| 7615 | NM_053050.4 | #N/A | 90.0304734 | -7.7032639 | 1.59907458 |
| 7616 | NM_006857.2 | #N/A | 106.71005 | -7.9471189 | 1.64971537 |
| 7617 | NM_018328.4 | MBD5 | 77.0887145 | -7.4779512 | 1.55242356 |
| 7618 | NM_001168401.1 | #N/A | 131.688529 | -8.2508391 | 1.71295416 |
| 7619 | NM_001282129.1 | #N/A | 91.2992827 | -7.7187245 | 1.60250468 |
| 7620 | NM_000876.3 | IGF2R | 748.077567 | -2.6691345 | 0.55423136 |
| 7621 | NM_002624.3 | #N/A | 547.8661 | -6.1347668 | 1.27384448 |
| 7622 | NM_001199198.2 | #N/A | 112.572732 | -8.0246027 | 1.66631604 |
| 7623 | NM_018318.4 | #N/A | 79.5686279 | -7.5238658 | 1.56250252 |
| 7624 | NM_031438.3 | #N/A | 64.813437 | -7.2270891 | 1.50096393 |
| 7625 | NM_134269.2 | #N/A | 91.2213574 | -7.7239803 | 1.60421853 |
| 7626 | NM_004929.3 | #N/A | 88.5475353 | -7.6819128 | 1.59565737 |
| 7627 | NM_015617.2 | #N/A | 70.4089008 | -7.3507234 | 1.52688771 |
| 7628 | NM_152705.2 | #N/A | 90.0127548 | -7.7024862 | 1.59997825 |
| 7629 | NM_001161576.2 | SAMD4A | 69.8810267 | -7.340255 | 1.52485169 |
| 7630 | NM_001318934.1 | HLTF | 74.5691647 | -7.4258544 | 1.54268299 |
| 7631 | NM_001025077.2 | #N/A | 266.317979 | -9.2670331 | 1.92523763 |
| 7633 | NM_203505.2 | G3BP2 | 105.250432 | -7.9291439 | 1.64733086 |
| 7632 | NM_004242.3 | #N/A | 66.3335687 | -7.2596828 | 1.50824486 |
| 7634 | NM_019056.6 | NDUFB11 | 94.8001225 | -7.7767918 | 1.61574409 |
| 7635 | NM_080621.4 | #N/A | 98.0833086 | -7.8269825 | 1.62623265 |
| 7636 | NM_004545.3 | #N/A | 107.844625 | -7.9631334 | 1.65466383 |
| 7637 | NM_006649.3 | #N/A | 69.3019005 | -7.3255268 | 1.52220088 |
| 7638 | NM_024954.4 | #N/A | 77.5665271 | -7.4885967 | 1.55614861 |
| 7639 | NM_001122679.1 | #N/A | 112.637956 | -8.0281537 | 1.66831923 |
| 7640 | NM_001135702.1 | #N/A | 80.8778083 | -7.5504527 | 1.56904394 |
| 7641 | NM_007041.3 | #N/A | 81.7181076 | -7.5590561 | 1.57084209 |
| 7642 | NM_001297597.1 | SIN3B | 69.807114 | -7.3342875 | 1.52422801 |
| 7643 | NM_017840.3 | #N/A | 80.6894353 | -7.5418379 | 1.56739515 |
| 7644 | NM_015204.2 | #N/A | 180.963891 | -7.2253288 | 1.50164657 |
| 7645 | NM_001008239.2 | C18orf25 | 71.1183954 | -7.3642568 | 1.53060689 |
| 7646 | NM_147188.2 | #N/A | 83.5971131 | -7.5932471 | 1.57847015 |
| 7647 | NM_000593.5 | TAP1 | 77.3811391 | -7.4831591 | 1.55562868 |
| 7648 | NM_001761.2 | #N/A | 91.2843797 | -7.7252057 | 1.60599989 |
| 7649 | NM_032251.5 | #N/A | 68.2705803 | -7.30288 | 1.51819545 |
| 7650 | NM_001014841.1 | NCDN | 194.999617 | -8.8179454 | 1.83327143 |
| 7652 | NM_005389.2 | PCMT1 | 161.241642 | -8.5436784 | 1.77658195 |
| 7651 | NM_001330698.1 | #N/A | 114.334294 | -8.047226 | 1.67335278 |
| 7653 | NM_018256.3 | #N/A | 92.138302 | -7.7359838 | 1.60862641 |

| | | | | | | |
|---|---|---|---|---|---|---|
| 7654 | NM_001319050.1 | BRPF1 | | 72.9379015 | -7.4000418 | 1.53879053 |
| 7655 | NM_000182.4 | | #N/A | 551.9796 | -5.7824862 | 1.20245337 |
| 7656 | NM_007084.3 | | #N/A | 111.131599 | -8.0047237 | 1.6645723 |
| 7657 | NM_005496.3 | SMC4 | | 113.413523 | -8.0343131 | 1.67080601 |
| 7658 | NM_182691.2 | SRPK2 | | 93.6498126 | -7.7567457 | 1.61307966 |
| 7659 | NM_015313.2 | | #N/A | 88.7264963 | -7.6786361 | 1.59688633 |
| 7660 | NM_021626.2 | | #N/A | 78.7860944 | -7.5129539 | 1.56247339 |
| 7661 | NM_001130107.1 | KLC1 | | 87.9150487 | -7.6710019 | 1.59549027 |
| 7662 | NM_001098577.2 | RPL31 | | 65.92316 | -7.256665 | 1.50934962 |
| 7663 | NM_002828.3 | | #N/A | 63.6809238 | -7.2015013 | 1.49798759 |
| 7664 | NM_017426.3 | | #N/A | 71.1438788 | -7.3668383 | 1.53245621 |
| 7665 | NM_001142568.2 | | #N/A | 116.739309 | -8.0799097 | 1.68086116 |
| 7666 | NM_178439.4 | | #N/A | 94.5988124 | -4.0105367 | 0.83433769 |
| 7667 | NM_033400.2 | | #N/A | 66.9767127 | -7.2798926 | 1.51455074 |
| 7668 | NM_021904.3 | GABBR1 | | 146.610107 | -8.4057637 | 1.74897743 |
| 7669 | NM_004470.3 | | #N/A | 78.9865383 | -7.514523 | 1.56356138 |
| 7670 | NR_135195.1 | LINC02177 | | 60.3718069 | 3.04781177 | 0.63417446 |
| 7671 | NM_006243.3 | | #N/A | 63.1046145 | -7.1902561 | 1.49614187 |
| 7672 | NM_001286086.1 | C11orf98 | | 90.5918292 | -7.7123611 | 1.60484786 |
| 7673 | NR_024057.2 | ZNF542P | | 96.3328702 | -7.8012529 | 1.62336654 |
| 7674 | NM_014380.2 | | #N/A | 64.7243189 | -7.226995 | 1.50395292 |
| 7675 | NM_202468.2 | | #N/A | 127.486863 | -8.2040858 | 1.70735578 |
| 7676 | NM_001347701.1 | SYNE1 | | 65.1040391 | -7.2372665 | 1.50621441 |
| 7677 | NM_024827.3 | | #N/A | 72.5546754 | -7.3931943 | 1.53874836 |
| 7678 | NM_001135156.2 | DPF1 | | 223.526819 | -9.0145294 | 1.87627634 |
| 7679 | NR_047665.1 | EPS15L1 | | 85.6323547 | -7.6275221 | 1.58765895 |
| 7680 | NM_003888.3 | | #N/A | 111.677195 | -8.0146868 | 1.66839525 |
| 7681 | NR_003256.2 | | #N/A | 85.3310963 | -7.6271679 | 1.58782737 |
| 7682 | NM_021947.2 | | #N/A | 93.9616769 | -7.7631728 | 1.61622587 |
| 7683 | NM_005598.3 | | #N/A | 115.952561 | -8.0657112 | 1.67925581 |
| 7684 | NM_012120.2 | | #N/A | 72.7234145 | -7.3972821 | 1.54010087 |
| 7685 | NM_015608.2 | EDRF1 | | 82.6464491 | -7.5785856 | 1.57787755 |
| 7686 | NM_001308120.1 | | #N/A | 66.6298656 | -7.2709911 | 1.51390646 |
| 7687 | NM_025203.2 | | #N/A | 80.2626853 | -7.5357507 | 1.56904316 |
| 7688 | NM_014808.3 | | #N/A | 70.4378775 | -7.3467399 | 1.52982629 |
| 7689 | NM_024573.2 | | #N/A | 79.5734283 | -7.5250458 | 1.56702886 |
| 7690 | NM_001177.5 | | #N/A | 229.821636 | -3.7145909 | 0.77356266 |
| 7691 | NM_004935.3 | | #N/A | 122.095308 | -8.1418728 | 1.69558635 |
| 7692 | NM_022100.2 | | #N/A | 84.073573 | -7.6025722 | 1.58337119 |
| 7693 | NM_001281956.1 | CSMD2 | | 109.548148 | -7.9861578 | 1.66337708 |
| 7694 | NM_178565.4 | | #N/A | 74.7785337 | -7.4332535 | 1.5482215 |
| 7695 | NM_001184727.1 | | #N/A | 94.6605061 | -7.7715706 | 1.61878546 |
| 7696 | NM_030771.1 | | #N/A | 72.3036975 | -7.3853792 | 1.53838873 |

| | | | | | |
|---|---|---|---|---|---|
| 7697 | NM_015974.2 | #N/A | 66.1577582 | -7.2603288 | 1.51247382 |
| 7698 | NM_000313.3 | #N/A | 103.070011 | -6.5072069 | 1.35572063 |
| 7699 | NM_001304550.1 | #N/A | 68.4818012 | -7.3054687 | 1.52206783 |
| 7700 | NM_012062.4 | #N/A | 194.314748 | -8.8117064 | 1.83601284 |
| 7701 | NM_006397.2 | RNASEH2A | 85.1402364 | -7.6205954 | 1.58789082 |
| 7702 | NM_024078.2 | #N/A | 90.5801241 | -7.7105619 | 1.60663367 |
| 7703 | NM_018684.3 | #N/A | 101.445592 | -7.8746565 | 1.64084689 |
| 7704 | NM_004589.3 | #N/A | 92.2179919 | -6.3627367 | 1.32586162 |
| 7705 | NM_012263.4 | #N/A | 82.5170111 | -7.5755159 | 1.57862172 |
| 7706 | NM_024685.3 | #N/A | 73.6013051 | -7.4114203 | 1.54449046 |
| 7707 | NM_003194.4 | #N/A | 75.3220032 | -7.4477845 | 1.55208503 |
| 7708 | NM_001478.4 | #N/A | 98.6380595 | -7.8344542 | 1.6327091 |
| 7709 | NM_030777.3 | #N/A | 95.5331595 | -5.1873829 | 1.08109311 |
| 7710 | NM_032492.3 | #N/A | 90.3222034 | -7.7073971 | 1.60637133 |
| 7712 | NM_201592.2 | GPM6A | 128.662823 | -8.2162625 | 1.71248443 |
| 7711 | NM_001322885.1 | #N/A | 91.3750085 | -7.7249345 | 1.6100724 |
| 7713 | NM_020706.2 | SCAF4 | 97.5682915 | -7.8161322 | 1.6291227 |
| 7714 | NM_148170.4 | #N/A | 65.1401757 | -7.2360735 | 1.50822787 |
| 7715 | NM_173469.3 | #N/A | 95.6402528 | -7.7899396 | 1.62384793 |
| 7716 | NM_033023.4 | PDGFA | 509.654139 | 4.11582356 | 0.85799818 |
| 7717 | NM_001018138.1 | NME2 | 107.946432 | -7.9642906 | 1.66031351 |
| 7718 | NM_001285987.1 | POU5F1 | 114.433653 | 2.81919693 | 0.58773389 |
| 7719 | NR_024356.2 | #N/A | 132.055031 | -8.2554008 | 1.72104741 |
| 7720 | NM_004362.2 | #N/A | 89.1453698 | -6.3182335 | 1.31727496 |
| 7721 | NM_023948.4 | #N/A | 84.0710068 | -7.604178 | 1.585825 |
| 7722 | NR_110241.1 | LINC01920 | 27.4099823 | 9.74144307 | 2.03160906 |
| 7723 | NM_001172681.1 | #N/A | 98.7180394 | -7.8346709 | 1.63403481 |
| 7724 | NM_001025088.1 | #N/A | 148.487194 | -8.4226695 | 1.75676658 |
| 7725 | NM_001145301.2 | #N/A | 71.543999 | -7.3686165 | 1.53694275 |
| 7726 | NM_001130106.1 | HSPBP1 | 98.2367925 | -7.8269389 | 1.63256123 |
| 7727 | NM_002393.4 | #N/A | 95.8935052 | -7.7910795 | 1.62513001 |
| 7728 | NM_002141.4 | #N/A | 103.114609 | -7.899106 | 1.64768073 |
| 7729 | NM_015492.4 | #N/A | 65.2594667 | -7.2392314 | 1.51006237 |
| 7730 | NM_001031703.2 | #N/A | 85.4729929 | -7.6276973 | 1.59116311 |
| 7731 | NM_018214.4 | #N/A | 65.3323501 | -7.2412781 | 1.51060537 |
| 7733 | NM_201997.2 | SF1 | 86.3558325 | -7.6386552 | 1.59368528 |
| 7732 | NM_018444.3 | #N/A | 96.855256 | -7.8090703 | 1.62924116 |
| 7734 | NM_020709.2 | #N/A | 87.9634905 | -7.6685876 | 1.59997119 |
| 7735 | NM_024644.4 | #N/A | 79.7174374 | -7.5275637 | 1.57059028 |
| 7736 | NR_037646.1 | TMX2-CTNND1 | 92.0214123 | -7.7307842 | 1.61301096 |
| 7737 | NM_001199022.2 | #N/A | 69.2626121 | -7.3254371 | 1.52864176 |
| 7738 | NM_198219.2 | #N/A | 85.9517077 | -7.6322515 | 1.59267838 |
| 7739 | NM_001004067.3 | #N/A | 254.223435 | -4.2904854 | 0.89533546 |

| | | | | | | |
|---|---|---|---|---|---|---|
| 7740 | NM_147150.2 | | #N/A | 88.3452634 | -7.6757417 | 1.60180152 |
| 7741 | NM_004623.4 | | #N/A | 87.096937 | -7.6541779 | 1.59731247 |
| 7742 | NM_001033910.2 | | #N/A | 62.389096 | -7.1755288 | 1.49747483 |
| 7743 | NM_001130848.1 | | #N/A | 101.834572 | -7.8836863 | 1.64530874 |
| 7744 | NM_018464.4 | | #N/A | 95.7900099 | -7.7912797 | 1.62601499 |
| 7745 | NM_001242854.1 | ARFIP2 | | 63.463961 | -7.1959756 | 1.50185241 |
| 7746 | NM_012219.4 | MRAS | | 76.031456 | -7.454223 | 1.5558463 |
| 7747 | NM_032138.5 | | #N/A | 73.8767851 | -7.4151083 | 1.5477878 |
| 7748 | NM_001142333.1 | | #N/A | 106.965004 | -7.9501265 | 1.65948834 |
| 7749 | NM_015635.3 | | #N/A | 62.2119177 | -7.1702592 | 1.4966969 |
| 7750 | NM_020347.3 | | #N/A | 78.8999815 | -7.5126153 | 1.56827015 |
| 7751 | NM_032889.4 | | #N/A | 86.2724445 | -7.6405451 | 1.59501696 |
| 7752 | NM_017887.2 | | #N/A | 88.1096405 | -7.6714573 | 1.60163059 |
| 7753 | NM_032111.3 | | #N/A | 85.6811508 | -7.6305625 | 1.59308435 |
| 7754 | NM_001319139.1 | | #N/A | 92.2713028 | -7.7345796 | 1.61483712 |
| 7755 | NM_001256891.1 | SRI | | 68.2794983 | -7.3070225 | 1.52558417 |
| 7756 | NM_032731.3 | | #N/A | 101.371993 | -7.8736815 | 1.64440818 |
| 7757 | NM_152383.4 | DIS3L2 | | 70.5253564 | -7.3507769 | 1.53525732 |
| 7758 | NM_153603.3 | | #N/A | 108.235619 | -3.3817933 | 0.70632939 |
| 7759 | NM_001321706.1 | | #N/A | 132.809257 | -8.2639813 | 1.72617911 |
| 7760 | NM_015194.2 | | #N/A | 82.9389358 | -7.5870101 | 1.58480814 |
| 7761 | NR_102326.1 | HCG18 | | 66.6654826 | -7.2701432 | 1.51865565 |
| 7762 | NM_004629.1 | FANCG | | 84.6413893 | -7.6131645 | 1.59032975 |
| 7763 | NM_014187.3 | | #N/A | 82.0077188 | -7.5690745 | 1.58113549 |
| 7765 | NM_033632.3 | FBXW7 | | 79.283395 | -7.5182603 | 1.57069971 |
| 7764 | NM_006395.2 | | #N/A | 69.120874 | -7.322706 | 1.52983882 |
| 7766 | NM_004257.5 | | #N/A | 114.665182 | -8.0523294 | 1.6823334 |
| 7767 | NM_001199764.1 | | #N/A | 153.024092 | -8.4693487 | 1.76962453 |
| 7768 | NM_018270.5 | | #N/A | 82.9776592 | -7.5824236 | 1.58431407 |
| 7769 | NM_138401.3 | | #N/A | 87.2943423 | -7.6575518 | 1.60002113 |
| 7770 | NM_001114600.2 | | #N/A | 98.390225 | -7.8310605 | 1.63637432 |
| 7771 | NM_001164463.1 | RGPD8 | | 76.2135968 | -7.4669991 | 1.56031499 |
| 7772 | NM_031280.3 | | #N/A | 121.331879 | -6.7108738 | 1.40241402 |
| 7773 | NM_001001132.1 | | #N/A | 154.172688 | -8.4779011 | 1.77176171 |
| 7774 | NM_018224.3 | COA1 | | 97.3611467 | -7.8145953 | 1.63324511 |
| 7775 | NM_016024.2 | | #N/A | 81.9325807 | -7.5666297 | 1.58146088 |
| 7776 | NM_001308248.1 | EVI5 | | 78.7155941 | -7.5089227 | 1.56943832 |
| 7777 | NM_014293.3 | | #N/A | 471.152128 | -3.7521066 | 0.78422797 |
| 7778 | NM_052917.3 | | #N/A | 120.469158 | -8.122381 | 1.69765663 |
| 7779 | NM_001330178.1 | | #N/A | 66.8001724 | -7.2673737 | 1.5190893 |
| 7780 | NM_001308036.1 | NCBP2 | | 84.4375248 | -7.6099431 | 1.59088009 |
| 7781 | NM_031284.4 | | #N/A | 92.971957 | -7.7503601 | 1.62024619 |
| 7782 | NM_001278642.1 | EDIL3 | | 185.624903 | -7.2522355 | 1.51616552 |

| | | | | | |
|---|---|---|---|---|---|
| 7783 | NM_001329984.1 | #N/A | 242.50965 | -9.1314983 | 1.90905669 |
| 7784 | NM_139070.2 | #N/A | 103.608946 | -7.9047342 | 1.6525864 |
| 7785 | NM_020533.2 | #N/A | 98.8244364 | -7.8376555 | 1.63858054 |
| 7786 | NM_019091.3 | #N/A | 79.7322354 | -7.5271705 | 1.57370886 |
| 7787 | NM_015381.6 | #N/A | 87.9228882 | -7.6691776 | 1.60351428 |
| 7788 | NM_004813.2 | #N/A | 92.6781983 | -7.7438841 | 1.61919973 |
| 7789 | NR_135475.1 | #N/A | 62.0918198 | -7.1688402 | 1.49897221 |
| 7790 | NM_004898.3 | #N/A | 87.4377417 | -6.2923654 | 1.31577879 |
| 7791 | NM_024122.4 | #N/A | 89.1210505 | -7.6860656 | 1.60724296 |
| 7792 | NM_015087.4 | #N/A | 103.016222 | -7.9002813 | 1.65204847 |
| 7793 | NM_144658.3 | DOCK11 | 83.7968491 | -7.6004446 | 1.58949582 |
| 7794 | NM_173824.3 | #N/A | 77.3215049 | -7.4818323 | 1.56473706 |
| 7795 | NM_000235.3 | #N/A | 78.4478692 | -7.4996551 | 1.56850744 |
| 7796 | NM_139283.1 | #N/A | 76.7178621 | -7.4703452 | 1.56250437 |
| 7797 | NM_017853.2 | #N/A | 70.8256713 | -7.3533336 | 1.53812829 |
| 7798 | NM_017544.3 | #N/A | 81.4940468 | -7.5576531 | 1.58088242 |
| 7799 | NM_017659.3 | #N/A | 80.9948485 | -7.5495369 | 1.57925912 |
| 7800 | NM_032985.4 | #N/A | 92.6464386 | -6.4622855 | 1.35197097 |
| 7801 | NM_001006607.2 | LRRC37A2 | 65.0441375 | -7.2365793 | 1.51407501 |
| 7805 | NR_036521.1 | ZNF667-AS1 | 89.3663461 | -7.6912927 | 1.60921602 |
| 7802 | NM_001012968.2 | #N/A | 69.1728637 | -7.3228535 | 1.53210793 |
| 7803 | NM_001284351.2 | #N/A | 68.2711463 | -7.3065258 | 1.52870388 |
| 7804 | NM_152307.2 | #N/A | 79.6709698 | -7.5264925 | 1.57473417 |
| 7806 | NM_019005.3 | #N/A | 97.1653828 | -6.4260166 | 1.34453768 |
| 7807 | NM_080476.4 | PIGU | 104.46186 | -6.5189489 | 1.36408473 |
| 7808 | NR_033183.1 | #N/A | 95.2795547 | -7.7847965 | 1.62897496 |
| 7809 | NM_198316.1 | #N/A | 70.733364 | -7.3545856 | 1.53896341 |
| 7811 | NR_027701.1 | LINC00346 | 62.0724064 | 3.19101998 | 0.66775681 |
| 7810 | NM_000097.5 | #N/A | 77.5970901 | -7.4890519 | 1.567169 |
| 7812 | NM_181705.3 | #N/A | 109.985393 | -7.9931062 | 1.67280785 |
| 7813 | NM_001321701.1 | #N/A | 79.8127494 | -7.5329101 | 1.57651169 |
| 7814 | NM_138447.2 | #N/A | 84.4444626 | -7.6096873 | 1.59274278 |
| 7815 | NM_032338.3 | #N/A | 89.862604 | -6.3250093 | 1.32387492 |
| 7816 | NM_022460.3 | #N/A | 75.7496452 | -7.4538694 | 1.56018239 |
| 7817 | NM_001010866.3 | #N/A | 86.6534565 | -7.6483619 | 1.60091829 |
| 7818 | NM_016399.2 | #N/A | 100.739238 | -7.8636491 | 1.64599009 |
| 7819 | NM_138701.3 | #N/A | 84.0756399 | -7.6033858 | 1.5915053 |
| 7820 | NM_152737.3 | #N/A | 70.7771273 | -7.3551081 | 1.53967245 |
| 7821 | NM_144609.2 | #N/A | 85.2927115 | -7.6239956 | 1.59597667 |
| 7822 | NM_006416.4 | #N/A | 92.8133242 | -7.7462173 | 1.62160349 |
| 7823 | NM_014050.3 | #N/A | 108.579214 | -6.6971049 | 1.40215796 |
| 7824 | NM_020701.3 | #N/A | 87.673704 | -7.6679804 | 1.60543658 |
| 7825 | NM_001112732.2 | #N/A | 363.121216 | -6.0886405 | 1.27479959 |

| | | | | | |
|---|---|---|---|---|---|
| 7826 | NM_001674.3 | #N/A | 76.7933256 | -7.4752781 | 1.56524156 |
| 7827 | NM_006286.4 | #N/A | 105.485376 | -7.9331137 | 1.66121138 |
| 7828 | NM_003954.4 | #N/A | 65.0140568 | -7.2377255 | 1.5156602 |
| 7829 | NM_016162.3 | #N/A | 83.3252541 | -7.5913565 | 1.58984466 |
| 7830 | NM_033113.2 | #N/A | 66.2458921 | -7.2579609 | 1.52003104 |
| 7831 | NM_001145365.1 | #N/A | 106.422519 | -7.9417827 | 1.66328332 |
| 7832 | NM_173547.3 | #N/A | 98.8090452 | -7.8353993 | 1.64101041 |
| 7833 | NM_052840.4 | #N/A | 104.296799 | -7.9128259 | 1.65725127 |
| 7834 | NM_001100165.1 | #N/A | 77.5513126 | -7.4905914 | 1.5688719 |
| 7835 | NM_000455.4 | #N/A | 359.028554 | -5.5905822 | 1.17099235 |
| 7837 | NM_001286045.1 | RBM7 | 61.3890981 | -7.1491432 | 1.4977513 |
| 7836 | NM_001110199.2 | #N/A | 125.006019 | -8.1755414 | 1.7127853 |
| 7838 | NM_001316314.1 | #N/A | 73.4110618 | -7.4077048 | 1.55193108 |
| 7839 | NM_001300862.1 | MPND | 81.8394611 | -7.5638287 | 1.58467774 |
| 7840 | NM_003927.4 | #N/A | 69.8093852 | -7.3370632 | 1.53716535 |
| 7841 | NM_001204375.1 | #N/A | 120.204636 | -8.1177818 | 1.7008887 |
| 7842 | NM_001099658.1 | #N/A | 144.158218 | -8.3816721 | 1.75626903 |
| 7843 | NM_006653.4 | #N/A | 89.5684135 | -7.6946896 | 1.61231609 |
| 7844 | NM_139025.4 | ADAMTS13 | 68.3611009 | -7.3060472 | 1.53108063 |
| 7845 | NM_001099439.1 | #N/A | 82.0562687 | -7.5690959 | 1.58621646 |
| 7846 | NM_001077497.2 | #N/A | 88.8304189 | -7.6822809 | 1.60995809 |
| 7847 | NM_032781.3 | #N/A | 82.8136209 | -7.5846905 | 1.58971327 |
| 7848 | NM_000181.3 | #N/A | 82.3913746 | -7.5754005 | 1.58780455 |
| 7849 | NM_003932.4 | #N/A | 1253.9325 | -6.1532782 | 1.28976514 |
| 7850 | NM_000485.2 | #N/A | 77.3186766 | -7.4835857 | 1.56862974 |
| 7851 | NM_001258023.1 | GRIA1 | 167.947098 | -8.6016419 | 1.80301287 |
| 7852 | NM_001347219.1 | #N/A | 73.3285909 | -7.4108551 | 1.55345077 |
| 7853 | NM_020465.3 | NDRG4 | 135.512981 | -8.2924328 | 1.73827445 |
| 7854 | NM_024529.4 | CDC73 | 123.975272 | -5.613879 | 1.17679775 |
| 7855 | NM_001195305.1 | #N/A | 84.6122022 | -7.6124384 | 1.5958556 |
| 7856 | NM_003641.3 | #N/A | 120.903798 | -8.1300015 | 1.70436903 |
| 7857 | NM_145345.2 | UBXN11 | 71.4671456 | -7.3717061 | 1.54544281 |
| 7858 | NM_001143826.2 | MAPRE2 | 132.567354 | -8.26178 | 1.73206563 |
| 7859 | NM_052868.5 | #N/A | 89.8936629 | -7.7030712 | 1.61492881 |
| 7860 | NM_018307.4 | RHOT1 | 82.3670414 | -7.5747958 | 1.58805156 |
| 7861 | NM_001263.3 | #N/A | 63.8915466 | -7.2098566 | 1.51194395 |
| 7862 | NM_032036.2 | #N/A | 86.1506569 | -7.6388928 | 1.60206304 |
| 7863 | NM_001144826.1 | RUNDC3A | 155.465586 | -8.4902249 | 1.78065931 |
| 7864 | NR_111952.1 | LINC00869 | 77.8098509 | -7.4921574 | 1.57143577 |
| 7865 | NM_015282.2 | #N/A | 111.538556 | -8.0114102 | 1.68053508 |
| 7866 | NM_001198690.1 | #N/A | 83.1887999 | -7.5899084 | 1.5922229 |
| 7867 | NM_001130475.2 | #N/A | 68.759399 | -7.3149826 | 1.53456992 |
| 7868 | NM_013241.2 | #N/A | 65.6707636 | -7.2493562 | 1.52083198 |

| | | | | | |
|---|---|---|---|---|---|
| 7869 | NM_014587.4 | #N/A | 80.3704827 | -7.5354471 | 1.58084437 |
| 7870 | NM_033055.2 | #N/A | 223.011967 | -5.5068718 | 1.1552689 |
| 7871 | NM_001195479.1 | TFG | 96.0459314 | -7.7973468 | 1.63615382 |
| 7872 | NR_108087.1 | LINC01087 | 74.3090202 | 3.60189717 | 0.75586018 |
| 7873 | NM_173575.3 | #N/A | 113.207063 | -8.0325503 | 1.68564461 |
| 7874 | NR_135109.1 | LINC02547 | 90.4855967 | 3.01855585 | 0.63345524 |
| 7875 | NM_001128610.2 | #N/A | 62.8045605 | -7.1873051 | 1.50838326 |
| 7876 | NM_144710.4 | #N/A | 66.2675554 | -7.2566985 | 1.52296449 |
| 7877 | NM_152344.3 | #N/A | 203.76177 | -4.1896227 | 0.87928309 |
| 7878 | NM_020360.3 | #N/A | 100.626032 | -7.8639504 | 1.65051206 |
| 7879 | NM_033641.3 | #N/A | 220.145761 | -8.9933312 | 1.88756573 |
| 7880 | NM_004283.3 | #N/A | 142.004674 | -4.7224935 | 0.99119774 |
| 7881 | NM_018079.4 | #N/A | 67.4823717 | -4.5233912 | 0.94949977 |
| 7882 | NM_172080.2 | #N/A | 190.852245 | -8.7866629 | 1.84445464 |
| 7883 | NM_001163484.1 | DCAF11 | 92.5775986 | -7.7462909 | 1.62607823 |
| 7884 | NM_001167740.1 | #N/A | 94.8444829 | -7.7777837 | 1.63287987 |
| 7885 | NM_001514.5 | #N/A | 84.5901061 | -7.6124388 | 1.59817545 |
| 7886 | NM_017908.3 | #N/A | 67.161081 | -7.2793352 | 1.52825034 |
| 7887 | NR_034035.2 | #N/A | 72.3970059 | -7.3887785 | 1.55132739 |
| 7888 | NM_022742.4 | #N/A | 89.454291 | -7.6942228 | 1.61574435 |
| 7889 | NM_003085.4 | #N/A | 150.231758 | -8.4408462 | 1.77256451 |
| 7890 | NM_001297704.1 | #N/A | 123.49888 | -8.1569825 | 1.71297532 |
| 7891 | NM_152740.3 | #N/A | 65.7434161 | -7.2540812 | 1.52361383 |
| 7892 | NR_001434.4 | HLA-H | 73.7881322 | -7.4186156 | 1.55831145 |
| 7893 | NM_012424.4 | #N/A | 83.9739312 | -7.6003108 | 1.5965158 |
| 7894 | NM_001077269.1 | WIPF1 | 72.4229122 | -7.3937491 | 1.55324319 |
| 7895 | NM_016192.3 | #N/A | 111.96983 | -8.017625 | 1.68444747 |
| 7896 | NM_001276293.1 | #N/A | 66.3920275 | -7.2650681 | 1.52636921 |
| 7897 | NM_207009.3 | #N/A | 80.4611344 | -7.5402555 | 1.58418955 |
| 7898 | NM_006626.5 | #N/A | 85.3877583 | -7.6247157 | 1.60195145 |
| 7899 | NM_001144758.2 | #N/A | 138.07431 | -8.3196601 | 1.7480183 |
| 7900 | NM_032646.5 | #N/A | 62.7354444 | -7.1805996 | 1.5087291 |
| 7901 | NM_003425.3 | #N/A | 63.4535515 | -7.1994015 | 1.51269925 |
| 7902 | NM_012320.3 | #N/A | 113.895464 | -6.7685148 | 1.4221664 |
| 7903 | NM_020238.2 | #N/A | 112.286229 | -6.6072531 | 1.38831267 |
| 7904 | NM_014462.2 | #N/A | 87.1479574 | -7.6553148 | 1.60898583 |
| 7905 | NM_004815.3 | #N/A | 74.3540779 | -7.429211 | 1.561613 |
| 7906 | NM_018934.3 | PCDHB14 | 83.2907816 | -7.589391 | 1.59533952 |
| 7908 | NR_033464.1 | NT5C3B | 105.69089 | -7.9351868 | 1.66810748 |
| 7907 | NM_016004.4 | #N/A | 93.5890032 | -7.7556012 | 1.63035578 |
| 7909 | NM_001313959.1 | AFF1 | 110.330716 | -7.9969096 | 1.68122718 |
| 7910 | NM_001023563.3 | #N/A | 73.7443478 | -7.4134734 | 1.55861319 |
| 7911 | NM_003155.2 | #N/A | 62.8987848 | -7.1862872 | 1.51084821 |

| | | | | | |
|---|---|---|---|---|---|
| 7912 | NM_181503.2 | #N/A | 86.4442922 | -7.642685 | 1.60680489 |
| 7913 | NM_002357.3 | #N/A | 81.4817854 | -7.5599582 | 1.58947851 |
| 7914 | NM_001080482.2 | #N/A | 83.3765856 | -7.5922026 | 1.5964294 |
| 7915 | NM_004264.4 | #N/A | 87.7556785 | -7.6644235 | 1.61168902 |
| 7916 | NM_207435.2 | C12orf76 | 88.7957664 | -4.9011798 | 1.03070429 |
| 7917 | NM_001487.3 | BLOC1S1 | 105.997907 | -7.9379631 | 1.66935272 |
| 7918 | NM_001006613.1 | #N/A | 74.8601177 | -7.4327715 | 1.56319459 |
| 7919 | NM_001348022.1 | #N/A | 68.0213885 | -7.2936182 | 1.53393385 |
| 7920 | NM_048368.3 | #N/A | 71.4988371 | -7.3672803 | 1.5494777 |
| 7921 | NM_001300947.1 | #N/A | 65.2951601 | -7.2380884 | 1.52233764 |
| 7922 | NM_006253.4 | #N/A | 176.971661 | -5.9781842 | 1.25745168 |
| 7923 | NR_027039.1 | LINC00410 | 40.0689112 | 5.40017169 | 1.13593707 |
| 7924 | NM_001322956.1 | TMEM183A | 121.204273 | -8.1311519 | 1.71046186 |
| 7925 | NM_002504.5 | #N/A | 77.1095131 | -7.479117 | 1.57331446 |
| 7926 | NM_182533.2 | #N/A | 79.5540265 | -7.5243174 | 1.58282379 |
| 7927 | NM_032900.5 | #N/A | 69.0174497 | -7.316994 | 1.53930354 |
| 7928 | NM_183398.2 | #N/A | 72.7872578 | -7.3936547 | 1.55549206 |
| 7929 | NM_018566.3 | YOD1 | 80.0101443 | -6.1729621 | 1.2986886 |
| 7930 | NM_001166293.1 | #N/A | 73.4222309 | -7.4101577 | 1.55903704 |
| 7931 | NM_033513.2 | #N/A | 93.0123322 | -7.7491389 | 1.63041437 |
| 7932 | NM_033259.2 | #N/A | 121.756684 | -8.138293 | 1.71252609 |
| 7933 | NM_005726.5 | #N/A | 76.047507 | -7.4599497 | 1.56983491 |
| 7934 | NM_014161.4 | #N/A | 99.5114782 | -7.8480148 | 1.65154192 |
| 7935 | NM_017872.4 | #N/A | 63.9577962 | -7.209656 | 1.51733575 |
| 7936 | NM_031443.3 | CCM2 | 94.0862642 | -7.7657244 | 1.63443947 |
| 7937 | NR_024058.1 | YWHAE | 84.8366102 | -7.6159468 | 1.60300174 |
| 7938 | NM_054035.2 | UNC119 | 61.3988012 | -7.1530665 | 1.50579165 |
| 7939 | NM_001163546.2 | #N/A | 94.2154344 | -7.7677725 | 1.63526278 |
| 7940 | NM_001143993.1 | #N/A | 66.3616103 | -7.2593219 | 1.52837159 |
| 7941 | NM_178841.3 | #N/A | 88.7748676 | -7.6803728 | 1.61705274 |
| 7942 | NM_145685.2 | BRF1 | 71.8445315 | -7.3761767 | 1.55306035 |
| 7943 | NM_001253792.1 | #N/A | 63.8224649 | -7.2063129 | 1.51731373 |
| 7944 | NM_001301072.1 | MAP3K4 | 61.6643286 | -7.1580867 | 1.50718116 |
| 7945 | NM_024805.2 | #N/A | 90.7072527 | -7.7128133 | 1.62398152 |
| 7946 | NM_007242.5 | #N/A | 84.1017666 | -7.6031431 | 1.60095899 |
| 7947 | NM_001099289.2 | #N/A | 107.230426 | -7.9540598 | 1.67501431 |
| 7948 | NM_014284.2 | #N/A | 83.6015028 | -7.5958026 | 1.59978443 |
| 7949 | NR_110555.1 | LOC102724814 | 59.8378421 | -7.1148383 | 1.49849936 |
| 7950 | NM_001145191.1 | #N/A | 122.840603 | -6.1105048 | 1.28701952 |
| 7951 | NR_122111.1 | DUXAP9 | 73.7755657 | 4.16534375 | 0.87732913 |
| 7952 | NM_001303514.1 | PDZD4 | 245.347848 | -7.8858436 | 1.66104116 |
| 7953 | NM_198573.2 | #N/A | 155.549333 | -8.491211 | 1.78854515 |
| 7954 | NM_020218.1 | ATXN7L3 | 96.952072 | -7.8089964 | 1.64488595 |

| | | | | | |
|---|---|---|---|---|---|
| 7955 | NM_003203.4 | #N/A | 66.2969821 | -7.2606276 | 1.52948131 |
| 7956 | NM_001290039.1 | #N/A | 107.785494 | -7.9598972 | 1.67682288 |
| 7957 | NM_017777.3 | #N/A | 70.8746412 | -7.3581359 | 1.5500749 |
| 7958 | NM_016500.3 | #N/A | 73.8525865 | -7.417799 | 1.56267961 |
| 7959 | NM_001160329.1 | #N/A | 113.423874 | -8.0354438 | 1.69281911 |
| 7960 | NM_003502.3 | #N/A | 62.2636185 | -7.1745774 | 1.51146205 |
| 7961 | NM_017820.4 | #N/A | 61.1558279 | -7.1430843 | 1.50484794 |
| 7962 | NM_001042529.2 | #N/A | 62.3308812 | -7.1757517 | 1.51179808 |
| 7963 | NM_017742.5 | #N/A | 61.0082804 | -7.1402952 | 1.50435277 |
| 7964 | NM_018074.5 | #N/A | 76.269222 | -7.4619335 | 1.57227731 |
| 7965 | NR_049730.1 | ASB16-AS1 | 40.3527404 | 2.50906231 | 0.52871363 |
| 7966 | NM_018668.4 | VPS33B | 89.682046 | -7.699908 | 1.62266044 |
| 7967 | NM_021928.3 | #N/A | 368.591702 | -3.8246103 | 0.80605979 |
| 7968 | NM_001286657.1 | #N/A | 63.5308439 | -7.2018576 | 1.51802604 |
| 7969 | NM_004957.5 | #N/A | 111.531372 | -8.0106091 | 1.68853976 |
| 7970 | NM_007342.2 | #N/A | 76.4551924 | -7.4667207 | 1.57400722 |
| 7971 | NM_012168.5 | #N/A | 63.9984396 | -7.2132767 | 1.52058516 |
| 7972 | NM_001287147.1 | #N/A | 32.2189614 | 4.27938933 | 0.90212799 |
| 7973 | NM_001330431.1 | #N/A | 84.3170434 | -7.6074752 | 1.60378607 |
| 7974 | NR_024493.2 | #N/A | 76.0487857 | -7.459772 | 1.57265377 |
| 7975 | NR_133680.1 | #N/A | 70.0554988 | 5.11485602 | 1.07832685 |
| 7976 | NM_000191.2 | #N/A | 63.7229842 | -7.2089185 | 1.51985032 |
| 7977 | NM_006845.3 | #N/A | 64.7176102 | -7.2286672 | 1.52403708 |
| 7978 | NM_022828.4 | #N/A | 70.5996776 | -7.3512124 | 1.54992956 |
| 7979 | NM_006574.3 | #N/A | 96.153437 | -7.7980828 | 1.64416037 |
| 7980 | NM_004831.3 | #N/A | 74.7600076 | -6.0842108 | 1.28285937 |
| 7981 | NM_018164.2 | #N/A | 87.0956603 | -7.6544663 | 1.61405825 |
| 7982 | NM_032352.3 | #N/A | 81.321682 | -7.5540305 | 1.59308422 |
| 7983 | NM_001264573.1 | #N/A | 69.8214861 | -7.3364868 | 1.54730762 |
| 7984 | NM_001173462.1 | #N/A | 88.5360776 | -7.6779821 | 1.61936209 |
| 7985 | NM_058174.2 | #N/A | 66.3784867 | -7.268002 | 1.53294337 |
| 7986 | NM_001292034.2 | #N/A | 260.268853 | -3.2599077 | 0.68757593 |
| 7987 | NM_024572.3 | #N/A | 94.6233958 | -7.7757169 | 1.6401111 |
| 7988 | NM_032042.5 | #N/A | 81.8323535 | -7.5661688 | 1.59591053 |
| 7989 | NM_153340.4 | #N/A | 91.7564904 | -7.7296496 | 1.63042039 |
| 7990 | NM_001005367.2 | #N/A | 95.4191673 | -7.7867209 | 1.64247811 |
| 7991 | NM_006810.3 | #N/A | 82.1219429 | -7.5710526 | 1.59698465 |
| 7992 | NM_004109.4 | #N/A | 64.7494389 | -7.2287149 | 1.52485283 |
| 7993 | NM_001078173.1 | #N/A | 86.6478247 | -7.6469516 | 1.6131498 |
| 7994 | NM_012297.4 | G3BP2 | 95.2906928 | -7.7850261 | 1.64232562 |
| 7995 | NM_030934.4 | #N/A | 69.2177422 | -7.3217798 | 1.54462085 |
| 7996 | NM_002533.3 | #N/A | 65.9545012 | -7.2488669 | 1.52929591 |
| 7997 | NM_001127217.2 | SMAD9 | 77.1457854 | -7.4837852 | 1.57887146 |

| | | | | | |
|---|---|---|---|---|---|
| 7998 | NM_017955.3 | #N/A | 68.7673736 | -7.3146196 | 1.54328329 |
| 7999 | NM_012244.3 | #N/A | 242.352183 | -9.1321014 | 1.92691508 |
| 8000 | NM_003773.4 | #N/A | 99.4376872 | -7.8479144 | 1.65597289 |
| 8001 | NM_001040458.1 | #N/A | 63.5518597 | -7.2051586 | 1.52056638 |
| 8003 | NM_001282313.1 | EXOC7 | 143.178986 | -8.3720026 | 1.76690117 |
| 8004 | NM_018928.2 | PCDHGC4 | 96.596444 | -7.8049016 | 1.64720354 |
| 8002 | NM_001010922.2 | #N/A | 124.012484 | 2.51954237 | 0.53174577 |
| 8005 | NM_001304963.1 | #N/A | 100.088627 | -6.5791136 | 1.38854985 |
| 8006 | NM_016167.4 | #N/A | 70.8260182 | -7.3571045 | 1.55275609 |
| 8007 | NM_001171155.1 | #N/A | 96.3126089 | -7.8001878 | 1.64628863 |
| 8008 | NM_001303142.1 | #N/A | 60.3957517 | -7.1280145 | 1.50442921 |
| 8009 | NM_001002030.1 | #N/A | 86.2722298 | -7.6412551 | 1.61292151 |
| 8010 | NM_015679.2 | #N/A | 88.9898216 | -7.6850729 | 1.62216332 |
| 8011 | NM_015655.3 | #N/A | 76.2271339 | -7.4664231 | 1.57611413 |
| 8013 | NR_134241.1 | LOC101930370 | 32.8208979 | 2.93339051 | 0.61923713 |
| 8012 | NM_032995.2 | #N/A | 66.1391871 | -7.2583904 | 1.53223663 |
| 8014 | NM_019037.2 | #N/A | 83.0271152 | -7.5846347 | 1.60118132 |
| 8015 | NM_001271822.1 | #N/A | 63.8226178 | -7.2101938 | 1.52216408 |
| 8016 | NM_003998.3 | #N/A | 62.8274171 | -7.1859606 | 1.51710368 |
| 8017 | NM_005482.2 | #N/A | 97.9493796 | -7.8238015 | 1.65183562 |
| 8018 | NM_023004.5 | #N/A | 91.1405723 | -7.719311 | 1.62980712 |
| 8019 | NM_001198843.1 | RBM4 | 77.297483 | -7.4823352 | 1.57992604 |
| 8020 | NM_018410.4 | #N/A | 71.6511113 | -7.3715707 | 1.55653223 |
| 8021 | NM_005720.3 | #N/A | 91.6939786 | -6.3478142 | 1.34049082 |
| 8022 | NM_001198852.1 | FOXJ3 | 121.403286 | -6.091922 | 1.286474 |
| 8023 | NR_028502.1 | MIR22HG | 67.9934457 | -7.3025398 | 1.54217361 |
| 8024 | NM_006591.2 | #N/A | 62.4345454 | -7.176846 | 1.51563828 |
| 8025 | NM_001271999.1 | DOCK7 | 82.1096068 | -7.5663248 | 1.59789936 |
| 8026 | NR_104586.1 | #N/A | 67.9699016 | -7.2996738 | 1.54160654 |
| 8027 | NM_001014763.1 | ETFB | 101.485571 | -7.8752521 | 1.66326584 |
| 8028 | NM_018178.5 | #N/A | 60.9356037 | -7.1438563 | 1.5090799 |
| 8029 | NM_052857.3 | #N/A | 74.1409647 | -7.4215655 | 1.56773863 |
| 8030 | NM_001137667.1 | #N/A | 91.76699 | -7.7276091 | 1.63244746 |
| 8031 | NM_030965.2 | #N/A | 82.960163 | -7.5847502 | 1.60229721 |
| 8032 | NM_001008892.2 | NIPA2 | 88.8971465 | -7.6831761 | 1.62321215 |
| 8033 | NM_031297.4 | RNF208 | 98.3385895 | -7.8299994 | 1.65432764 |
| 8034 | NM_001320485.1 | #N/A | 88.7747962 | -7.6807263 | 1.62281934 |
| 8035 | NM_032376.3 | #N/A | 68.7819893 | -7.3148663 | 1.54554134 |
| 8036 | NM_030805.3 | LMAN2L | 69.4215921 | -7.3271622 | 1.5481795 |
| 8037 | NM_007221.3 | PMF1 | 72.1074117 | -7.3813404 | 1.55966245 |
| 8038 | NM_015335.4 | MED13L | 440.52704 | -5.8419314 | 1.2345257 |
| 8039 | NR_024110.1 | SBDSP1 | 68.8241198 | -7.3109299 | 1.54497786 |
| 8040 | NM_173537.3 | #N/A | 65.6861987 | -4.1941939 | 0.88639598 |

| | | | | | |
|---|---|---|---|---|---|
| 8041 | NM_001130912.2 | #N/A | 69.0305056 | -7.3192873 | 1.54691535 |
| 8042 | NM_032315.2 | #N/A | 75.1299523 | -7.4406982 | 1.57273504 |
| 8043 | NM_001008392.1 | #N/A | 64.9916583 | -7.2292426 | 1.52810327 |
| 8044 | NM_002693.2 | POLG | 91.9131017 | -7.733001 | 1.63459759 |
| 8045 | NM_015500.1 | #N/A | 67.1097955 | -7.2837129 | 1.53966064 |
| 8046 | NM_177538.2 | #N/A | 768.449968 | 1.63711299 | 0.34609603 |
| 8047 | NM_001199620.1 | #N/A | 66.1672333 | -7.259601 | 1.53486814 |
| 8048 | NM_001243159.1 | #N/A | 66.2929981 | -7.2629526 | 1.53559732 |
| 8049 | NM_139179.3 | #N/A | 93.0212931 | -7.7510062 | 1.63892844 |
| 8050 | NM_005605.4 | PPP3CC | 77.6288991 | -7.4877312 | 1.58328227 |
| 8051 | NM_022839.4 | #N/A | 80.7003185 | -7.5451059 | 1.59551735 |
| 8052 | NM_007110.4 | #N/A | 73.8241467 | -7.4136242 | 1.56774711 |
| 8053 | NM_203298.2 | #N/A | 84.2614092 | -7.6063369 | 1.60850537 |
| 8054 | NM_005830.3 | #N/A | 74.3928039 | -7.4253615 | 1.57026877 |
| 8055 | NM_139162.3 | #N/A | 87.7260122 | -7.6631422 | 1.62055857 |
| 8056 | NR_051963.1 | MRPS31P5 | 63.4293487 | -7.1961815 | 1.52183685 |
| 8057 | NR_130944.1 | WDR73 | 69.2767529 | -7.3213374 | 1.54837285 |
| 8058 | NM_018100.3 | #N/A | 89.5217253 | -7.6954859 | 1.62788409 |
| 8059 | NM_020979.4 | #N/A | 97.7025253 | -7.8201352 | 1.65439237 |
| 8060 | NM_014431.2 | #N/A | 67.4961972 | -7.2852193 | 1.54127277 |
| 8061 | NM_001114632.1 | #N/A | 60.5532504 | -7.1323773 | 1.50899621 |
| 8062 | NM_032828.3 | #N/A | 143.318512 | -8.372423 | 1.77141165 |
| 8063 | NM_001082969.1 | #N/A | 90.7837686 | -7.7128308 | 1.63189171 |
| 8064 | NM_014996.2 | #N/A | 75.8947974 | -7.4554674 | 1.57743813 |
| 8065 | NR_103814.2 | LINC00276 | 64.0868318 | 5.67296138 | 1.20030418 |
| 8066 | NM_014055.3 | #N/A | 60.8379122 | -7.1410863 | 1.51098261 |
| 8067 | NM_020232.4 | #N/A | 80.3708736 | -7.5388099 | 1.59518565 |
| 8068 | NM_032831.3 | #N/A | 69.9378814 | -7.3423792 | 1.55366596 |
| 8069 | NM_004459.6 | #N/A | 325.87286 | -5.4615261 | 1.15573325 |
| 8070 | NM_001077399.2 | #N/A | 75.2704969 | -7.4450677 | 1.57549373 |
| 8071 | NM_152261.3 | #N/A | 110.395399 | -7.9964587 | 1.69228627 |
| 8072 | NM_172215.2 | #N/A | 30.5815791 | 9.92714844 | 2.10096579 |
| 8073 | NM_006427.3 | #N/A | 85.41787 | -7.6267965 | 1.6142727 |
| 8074 | NM_001304762.1 | #N/A | 61.5918416 | -7.1579541 | 1.51507046 |
| 8075 | NM_024576.4 | #N/A | 69.7742928 | -7.3396629 | 1.55354421 |
| 8076 | NM_001130690.2 | PDE10A | 140.275475 | -8.3423199 | 1.76588438 |
| 8077 | NM_016440.3 | #N/A | 64.246395 | -7.2126646 | 1.5267856 |
| 8078 | NM_001300989.1 | #N/A | 124.89931 | -8.1743647 | 1.730474 |
| 8079 | NM_001304420.2 | SLMAP | 70.0974449 | -7.3371793 | 1.55327097 |
| 8080 | NM_148893.2 | #N/A | 73.039653 | -7.3992318 | 1.56641926 |
| 8081 | NM_017975.4 | #N/A | 91.8911269 | -7.7309544 | 1.63670578 |
| 8082 | NM_001130970.1 | NSMF | 340.494203 | -9.6218533 | 2.0372654 |
| 8083 | NM_020844.2 | #N/A | 70.814188 | -7.3537162 | 1.55705527 |

| | | | | | |
|---|---|---|---|---|---|
| 8084 | NM_024075.4 | #N/A | 85.2750835 | -7.623001 | 1.61427003 |
| 8085 | NM_001142418.1 | MORF4L2 | 79.0980359 | -7.5136056 | 1.59114598 |
| 8086 | NM_016836.3 | #N/A | 108.758984 | -7.9763851 | 1.68914294 |
| 8087 | NM_004336.4 | #N/A | 75.9780698 | -7.4567813 | 1.57915756 |
| 8088 | NM_015508.4 | #N/A | 62.1294191 | -7.1717442 | 1.51879107 |
| 8089 | NM_007331.1 | NSD2 | 73.3875972 | -7.4068908 | 1.56867812 |
| 8090 | NM_172230.2 | #N/A | 100.154106 | -7.857738 | 1.66433552 |
| 8091 | NR_103741.1 | IPO5P1 | 93.2191274 | -7.7507009 | 1.64170539 |
| 8092 | NM_015307.1 | FAM189A1 | 80.6297427 | -7.5436243 | 1.59792873 |
| 8093 | NM_198256.3 | #N/A | 60.0411364 | -7.1184181 | 1.5078828 |
| 8094 | NM_019886.3 | #N/A | 86.0339984 | -7.6386331 | 1.61814761 |
| 8095 | NM_001318730.1 | #N/A | 100.643528 | -7.8614486 | 1.66539757 |
| 8096 | NM_016115.4 | #N/A | 75.7038024 | -7.4516527 | 1.5787684 |
| 8097 | NM_001114.4 | #N/A | 86.1752905 | -7.6400522 | 1.61877637 |
| 8098 | NM_001271314.1 | ZNF266 | 81.676467 | -7.5610719 | 1.6021415 |
| 8099 | NM_001024226.1 | ARF1 | 108.443305 | -7.9694065 | 1.68868242 |
| 8100 | NM_016044.2 | #N/A | 72.025673 | -7.3801052 | 1.5638341 |
| 8101 | NM_020823.1 | TMEM181 | 290.752052 | -3.4848693 | 0.73844507 |
| 8102 | NM_005229.4 | ELK1 | 80.8361949 | -7.546689 | 1.59938317 |
| 8103 | NM_007026.3 | #N/A | 71.9218084 | -7.3782756 | 1.56372362 |
| 8104 | NM_001271634.1 | #N/A | 71.8023416 | -7.3799335 | 1.56410161 |
| 8105 | NM_001144766.2 | DNAJC7 | 115.292706 | -8.0595546 | 1.70820314 |
| 8106 | NM_015058.1 | #N/A | 75.4071238 | -4.8223085 | 1.02214356 |
| 8107 | NM_001286566.1 | KIAA0513 | 176.439077 | -8.6730973 | 1.83837484 |
| 8108 | NM_002601.3 | #N/A | 93.859072 | -7.7624344 | 1.64542385 |
| 8109 | NM_014478.4 | #N/A | 127.370761 | -8.2021956 | 1.738667 |
| 8110 | NM_018141.3 | #N/A | 140.753387 | -3.0475253 | 0.64604747 |
| 8111 | NM_021177.4 | #N/A | 104.028356 | -7.9108883 | 1.67706298 |
| 8112 | NM_018019.2 | #N/A | 82.2903814 | -6.292182 | 1.33406891 |
| 8113 | NM_014878.4 | #N/A | 76.4171519 | -7.4667485 | 1.58328955 |
| 8114 | NM_002923.3 | #N/A | 83.6715716 | -7.5981846 | 1.61119953 |
| 8115 | NM_016542.3 | #N/A | 60.3270062 | -7.1284304 | 1.51164511 |
| 8116 | NM_152652.2 | #N/A | 82.9129084 | -7.5838938 | 1.6082442 |
| 8117 | NM_007212.3 | #N/A | 134.07109 | -6.8254135 | 1.44747508 |
| 8118 | NM_001130924.2 | #N/A | 61.7504852 | -7.1587852 | 1.51834689 |
| 8119 | NM_201559.2 | #N/A | 211.910837 | -4.0814954 | 0.86568966 |
| 8120 | NM_001079855.1 | #N/A | 114.559656 | -8.050042 | 1.70745976 |
| 8121 | NM_016370.2 | #N/A | 104.950481 | -7.9232023 | 1.68056251 |
| 8122 | NM_021194.2 | #N/A | 70.4391488 | -7.3516376 | 1.55941274 |
| 8123 | NM_004975.3 | #N/A | 91.735539 | -5.4938564 | 1.165453 |
| 8124 | NM_001194956.1 | MATR3 | 162.508575 | -5.4300664 | 1.15199502 |
| 8125 | NM_016467.4 | #N/A | 61.1305934 | -7.1450702 | 1.51611326 |
| 8126 | NM_145648.3 | #N/A | 82.9986457 | -7.5836557 | 1.60920186 |

| | | | | | |
|---|---|---|---|---|---|
| 8127 | NM_005904.3 | SMAD7 | 59.1094976 | -7.0941886 | 1.50546769 |
| 8128 | NM_013943.2 | #N/A | 599.880848 | -4.1966984 | 0.89059098 |
| 8129 | NM_024519.3 | #N/A | 363.494739 | -4.0086852 | 0.85072947 |
| 8130 | NM_144693.2 | #N/A | 81.1317196 | -7.5517512 | 1.60264902 |
| 8131 | NM_001286831.1 | RCBTB2 | 69.8716946 | -7.3385869 | 1.55744164 |
| 8133 | NM_133631.3 | ROBO1 | 104.295407 | -7.9155565 | 1.67998594 |
| 8132 | NM_001287387.1 | #N/A | 102.563383 | -7.891469 | 1.67487002 |
| 8134 | NM_001017528.2 | PRR5 | 59.323252 | -7.0999469 | 1.50689276 |
| 8135 | NM_001300952.1 | SRGAP2 | 70.5607047 | -7.3525658 | 1.56053 |
| 8136 | NM_001099922.2 | #N/A | 72.8442733 | -7.3937668 | 1.56942893 |
| 8137 | NM_001010898.3 | #N/A | 126.873999 | -8.1975918 | 1.74016939 |
| 8138 | NM_001203258.1 | #N/A | 89.3246213 | -7.6926794 | 1.63319957 |
| 8139 | NM_031921.5 | #N/A | 86.460616 | -7.6424319 | 1.6225241 |
| 8140 | NM_003595.3 | #N/A | 62.1461042 | -7.1727734 | 1.52290121 |
| 8141 | NM_145273.3 | #N/A | 45.5676042 | 4.18770612 | 0.8892553 |
| 8142 | NM_021937.4 | #N/A | 92.7883759 | -6.4699351 | 1.37398823 |
| 8143 | NM_001099403.1 | #N/A | 211.15147 | -8.9321205 | 1.89697513 |
| 8144 | NM_001005862.2 | ERBB2 | 81.8328766 | -7.5657808 | 1.6069439 |
| 8145 | NM_004477.2 | #N/A | 85.5900939 | -7.6296834 | 1.62055482 |
| 8146 | NM_018122.4 | #N/A | 68.8774189 | -7.3169832 | 1.55415666 |
| 8147 | NM_001256627.1 | #N/A | 91.2191856 | -7.7211952 | 1.64006978 |
| 8148 | NM_001258329.1 | EPB41L1 | 109.583762 | -7.9875188 | 1.69666006 |
| 8149 | NM_004579.4 | #N/A | 64.4793772 | -7.2216445 | 1.53399131 |
| 8150 | NM_024653.3 | #N/A | 95.6159735 | -5.7327542 | 1.21781371 |
| 8151 | NM_020693.3 | #N/A | 66.9894032 | -7.2726843 | 1.54501417 |
| 8152 | NM_014517.4 | #N/A | 97.3862149 | -7.8139505 | 1.66011766 |
| 8153 | NM_020905.3 | #N/A | 66.2252805 | -7.258946 | 1.5423382 |
| 8154 | NM_001114090.1 | #N/A | 75.6096948 | -7.4484139 | 1.5826805 |
| 8155 | NM_004386.2 | #N/A | 808.256569 | -5.9138008 | 1.2566029 |
| 8156 | NM_014867.2 | #N/A | 395.856363 | -4.0586342 | 0.86242439 |
| 8157 | NM_004832.2 | #N/A | 75.6904371 | -7.4526778 | 1.58367408 |
| 8158 | NM_182961.3 | #N/A | 89.8694389 | -7.697778 | 1.63574944 |
| 8159 | NM_017818.3 | #N/A | 75.065629 | -7.4382867 | 1.58063471 |
| 8160 | NM_001165966.1 | #N/A | 20.348503 | 9.31952786 | 1.98051375 |
| 8161 | NR_110000.1 | SAMMSON | 83.8982119 | 2.91583091 | 0.61966274 |
| 8162 | NM_001827.2 | #N/A | 98.8594569 | -7.836694 | 1.66548709 |
| 8163 | NM_182765.4 | #N/A | 78.9965095 | -7.513315 | 1.59677499 |
| 8164 | NM_003449.4 | #N/A | 63.2840611 | -7.1968188 | 1.52959638 |
| 8165 | NM_001127691.2 | #N/A | 68.1705544 | 2.33606194 | 0.4965228 |
| 8166 | NM_017970.3 | #N/A | 59.8932848 | -7.1154176 | 1.51237492 |
| 8167 | NM_017719.4 | #N/A | 88.0505481 | -7.6706457 | 1.63048241 |
| 8168 | NM_001145928.1 | SAP130 | 65.0820218 | -7.237643 | 1.53846946 |
| 8169 | NM_001128214.1 | #N/A | 60.0786899 | -7.1167178 | 1.51295982 |

| | | | | | |
|---|---|---|---|---|---|
| 8170 | NM_021255.2 | #N/A | 101.086742 | -7.8675809 | 1.67258096 |
| 8171 | NM_032164.3 | #N/A | 58.464755 | -7.0783312 | 1.50484362 |
| 8172 | NM_052860.2 | ZNF300 | 127.232236 | -8.2005773 | 1.74345459 |
| 8173 | NM_006283.2 | #N/A | 69.3861173 | -7.3318196 | 1.55885232 |
| 8174 | NM_198082.2 | #N/A | 59.8258486 | -7.1108689 | 1.51197518 |
| 8175 | NM_001198633.1 | RUNX1T1 | 107.218668 | -7.9533734 | 1.6912075 |
| 8176 | NM_018446.3 | GLT8D1 | 65.4506113 | -7.2444884 | 1.54048358 |
| 8177 | NM_018176.3 | #N/A | 89.143224 | -7.6894815 | 1.63511783 |
| 8178 | NM_032631.3 | #N/A | 105.206783 | -7.9282017 | 1.68588813 |
| 8179 | NM_022065.4 | THADA | 69.6765311 | -7.3332165 | 1.55954783 |
| 8180 | NM_005941.4 | #N/A | 65.073082 | -7.2348032 | 1.53868336 |
| 8181 | NM_001330124.1 | #N/A | 70.7856332 | -7.3561756 | 1.56453607 |
| 8182 | NM_006577.5 | #N/A | 64.4191092 | -7.2165643 | 1.53493924 |
| 8183 | NM_030767.5 | AKNA | 103.468541 | -7.9027574 | 1.68090148 |
| 8184 | NM_017640.5 | #N/A | 65.9336435 | -7.2483876 | 1.54187348 |
| 8185 | NM_007074.3 | CORO1A | 122.576399 | -8.1473372 | 1.73330314 |
| 8186 | NM_001289103.1 | PRKCSH | 83.6389384 | -7.597794 | 1.61648378 |
| 8187 | NR_109840.1 | #N/A | 29.7228223 | 9.88621559 | 2.10336234 |
| 8188 | NR_029407.1 | LOC642361 | 79.804518 | -7.5289648 | 1.60186995 |
| 8189 | NM_001243965.1 | HLA-DRB1 | 121.331095 | -8.1339109 | 1.73078646 |
| 8190 | NM_006489.2 | #N/A | 159.164253 | -5.4026829 | 1.1496324 |
| 8191 | NM_001318798.1 | #N/A | 82.4072399 | -6.2037003 | 1.3201412 |
| 8192 | NM_001105251.2 | #N/A | 64.7738674 | -7.2231695 | 1.53714118 |
| 8193 | NM_016003.3 | #N/A | 114.745103 | -8.052245 | 1.71368312 |
| 8194 | NM_001321770.1 | #N/A | 79.3689396 | -7.5238236 | 1.60131986 |
| 8195 | NM_003778.3 | #N/A | 64.3984515 | -7.2234351 | 1.53742217 |
| 8196 | NM_018269.3 | #N/A | 103.162631 | -7.8986857 | 1.68114577 |
| 8197 | NM_001329545.1 | #N/A | 59.5777101 | -7.106885 | 1.51267087 |
| 8198 | NM_001330518.1 | #N/A | 85.8432132 | -7.6345502 | 1.62498724 |
| 8199 | NM_005679.3 | #N/A | 110.953336 | -8.0067048 | 1.70418175 |
| 8200 | NM_206933.2 | #N/A | 26.1028474 | 6.61568625 | 1.40811629 |
| 8201 | NM_001318875.1 | FLOT1 | 77.8653808 | -7.4956381 | 1.59547498 |
| 8202 | NM_032842.3 | #N/A | 81.9736862 | -7.5710005 | 1.61166513 |
| 8203 | NM_001329451.1 | KANK2 | 65.7385678 | -7.2497447 | 1.54335054 |
| 8204 | NM_138980.3 | #N/A | 101.182038 | -7.8699541 | 1.67545236 |
| 8205 | NM_001198953.1 | ZFYVE21 | 71.6844998 | -7.3720851 | 1.56947209 |
| 8206 | NM_017650.2 | PPP1R9A | 80.8472323 | -7.5473751 | 1.60684404 |
| 8207 | NM_001317920.1 | #N/A | 110.263882 | -7.9938165 | 1.70194787 |
| 8208 | NM_138453.3 | #N/A | 534.522958 | -4.5689771 | 0.97278129 |
| 8209 | NM_005690.4 | DNM1L | 91.1867756 | -7.7192196 | 1.64353608 |
| 8210 | NM_001324276.1 | #N/A | 70.9453903 | -7.3590526 | 1.56689483 |
| 8211 | NM_001135005.2 | #N/A | 104.015387 | -7.9114134 | 1.6845674 |
| 8212 | NM_198446.2 | #N/A | 96.1651602 | -7.7965579 | 1.6601458 |

| | | | | | |
|---|---|---|---|---|---|
| 8213 | NM_013299.3 | #N/A | 114.249626 | -6.7732389 | 1.44229828 |
| 8214 | NM_001932.4 | #N/A | 105.764952 | -7.9356656 | 1.68988075 |
| 8215 | NM_016538.2 | #N/A | 76.1115729 | -6.1004121 | 1.29911588 |
| 8216 | NM_001136050.2 | #N/A | 64.7039817 | -7.2291121 | 1.5396349 |
| 8217 | NM_153265.2 | #N/A | 73.1487177 | -7.4068944 | 1.57753355 |
| 8218 | NM_016640.3 | #N/A | 69.9413026 | -7.3378447 | 1.56292577 |
| 8219 | NM_012212.3 | PTGR1 | 65.7324024 | -7.2520271 | 1.54467585 |
| 8220 | NM_002491.2 | #N/A | 99.3588884 | -7.8450332 | 1.6710378 |
| 8221 | NR_028482.1 | #N/A | 67.2959409 | -7.2823522 | 1.55126742 |
| 8223 | NM_015242.4 | ARAP1 | 148.607412 | -8.4264799 | 1.79502035 |
| 8222 | NM_001305066.1 | TMEM59 | 75.5505259 | -7.4487829 | 1.58674658 |
| 8224 | NM_023947.3 | #N/A | 107.976732 | -7.9652643 | 1.69677789 |
| 8225 | NM_001080396.2 | #N/A | 102.590037 | -7.8909546 | 1.68100586 |
| 8226 | NM_032466.3 | #N/A | 76.5930183 | -7.4663179 | 1.59068152 |
| 8227 | NM_006024.6 | TAX1BP1 | 80.0887322 | -7.533191 | 1.60503728 |
| 8228 | NM_001300867.1 | MIA3 | 71.4656376 | -7.3681632 | 1.56996319 |
| 8229 | NM_014067.3 | #N/A | 77.9652522 | -7.4934083 | 1.59669237 |
| 8230 | NM_001330091.1 | NRXN1 | 107.376878 | -7.9561568 | 1.69531526 |
| 8231 | NM_032869.3 | #N/A | 72.9864859 | -7.4002266 | 1.57688791 |
| 8232 | NM_001282471.1 | PRORY | 57.8062861 | 5.67774969 | 1.20987389 |
| 8233 | NM_004182.3 | UXT | 83.1403062 | -7.5877373 | 1.61687409 |
| 8234 | NM_020860.3 | #N/A | 101.235679 | -7.8713486 | 1.67730158 |
| 8235 | NM_001242928.1 | #N/A | 60.6039757 | -7.1289159 | 1.51912827 |
| 8236 | NR_015432.1 | #N/A | 68.9689983 | -7.3176462 | 1.5593573 |
| 8237 | NM_138278.3 | #N/A | 72.7368517 | 3.29416308 | 0.70197582 |
| 8238 | NM_138782.2 | #N/A | 62.5961416 | -7.1781819 | 1.5297348 |
| 8239 | NM_016339.4 | #N/A | 64.281223 | -7.2125583 | 1.5370696 |
| 8240 | NM_002953.3 | #N/A | 60.2873159 | -7.128213 | 1.51910483 |
| 8241 | NM_015669.4 | #N/A | 109.727654 | -7.9870862 | 1.70219045 |
| 8242 | NM_001135244.1 | #N/A | 80.0679368 | -7.5332465 | 1.60548666 |
| 8243 | NM_181351.4 | #N/A | 211.546779 | -8.9355278 | 1.90459084 |
| 8244 | NM_018293.2 | #N/A | 64.9552237 | -7.231088 | 1.54131223 |
| 8245 | NM_001099269.2 | #N/A | 85.6969666 | -7.6292265 | 1.62627768 |
| 8246 | NM_144684.3 | #N/A | 67.188888 | -7.2766719 | 1.55115027 |
| 8247 | NR_015434.1 | MRPL20-AS1 | 68.4714048 | -7.3102691 | 1.55833808 |
| 8248 | NR_036608.1 | #N/A | 71.9242641 | -7.3794617 | 1.57315296 |
| 8249 | NM_001099652.1 | #N/A | 85.0681371 | -7.6207969 | 1.62488912 |
| 8250 | NM_001131017.1 | #N/A | 85.6364228 | -7.6292202 | 1.62669857 |
| 8251 | NM_176819.3 | #N/A | 67.158475 | 3.03531185 | 0.64720811 |
| 8252 | NR_026903.1 | AMZ2P1 | 72.5426778 | -7.3901689 | 1.5760298 |
| 8253 | NM_201267.1 | #N/A | 74.7006822 | -7.4367402 | 1.58616657 |
| 8254 | NM_001037738.2 | #N/A | 145.283965 | -2.9743395 | 0.63451629 |
| 8255 | NM_152617.3 | #N/A | 184.238247 | -3.6936369 | 0.78797848 |

| | | | | | | |
|---|---|---|---|---|---|---|
| 8256 | NM_016478.4 | | #N/A | 74.7762198 | -7.4352642 | 1.58621876 |
| 8257 | NM_080743.4 | | #N/A | 100.497563 | -5.6087073 | 1.19671847 |
| 8258 | NM_212492.2 | | #N/A | 77.1821283 | -7.4789935 | 1.59582172 |
| 8259 | NM_032304.3 | | #N/A | 70.1465497 | -7.342683 | 1.56694101 |
| 8260 | NM_004170.5 | | #N/A | 87.5875135 | -7.6627305 | 1.6353679 |
| 8261 | NM_000355.3 | | #N/A | 58.0725835 | -7.071518 | 1.50921999 |
| 8262 | NM_002386.3 | MC1R | | 64.1128028 | -7.214281 | 1.53974423 |
| 8263 | NM_175575.5 | | #N/A | 285.89518 | -9.3703809 | 1.99997751 |
| 8264 | NM_178128.5 | | #N/A | 58.7520003 | 2.50394768 | 0.53443111 |
| 8265 | NM_021927.2 | | #N/A | 98.2399531 | -7.8292278 | 1.67113474 |
| 8266 | NM_080668.3 | | #N/A | 71.7537544 | -7.377243 | 1.57470769 |
| 8267 | NM_001142298.1 | | #N/A | 75.8310018 | -7.4518742 | 1.59066234 |
| 8268 | NM_001130864.1 | | #N/A | 58.3227031 | -7.0726754 | 1.50976406 |
| 8269 | NM_015384.4 | | #N/A | 65.1810625 | -7.2417838 | 1.54596774 |
| 8270 | NM_001013642.2 | | #N/A | 110.88953 | -8.0030438 | 1.70857169 |
| 8271 | NM_016333.3 | | #N/A | 2366.63161 | -6.3064261 | 1.34639433 |
| 8272 | NM_005761.2 | | #N/A | 71.9804464 | -7.3818861 | 1.57611859 |
| 8273 | NM_001113547.1 | | #N/A | 89.3426689 | -7.6918265 | 1.64256125 |
| 8274 | NM_001769.3 | | #N/A | 60.4584384 | -7.1315299 | 1.52291966 |
| 8275 | NM_001317898.1 | | #N/A | 102.23167 | -7.8854124 | 1.6839353 |
| 8276 | NM_017556.2 | | #N/A | 122.8666 | -8.1515422 | 1.74077202 |
| 8277 | NM_001040092.2 | | #N/A | 80.2989366 | -7.5407423 | 1.61037166 |
| 8278 | NM_022728.3 | | #N/A | 638.328477 | -10.52862 | 2.24848923 |
| 8279 | NM_017585.3 | | #N/A | 93.9565903 | -7.7656199 | 1.65846712 |
| 8280 | NM_005469.3 | | #N/A | 75.8887698 | -7.4546431 | 1.59210712 |
| 8281 | NM_016304.2 | | #N/A | 379.428555 | -6.1242706 | 1.30813308 |
| 8282 | NM_178570.2 | | #N/A | 99.677515 | -7.8485241 | 1.67652126 |
| 8283 | NM_001362.3 | DIO3 | | 68.4573511 | -7.3094606 | 1.56140676 |
| 8284 | NM_017446.3 | | #N/A | 68.3566267 | -7.3037578 | 1.56020667 |
| 8285 | NM_001145297.3 | | #N/A | 67.9532133 | -7.3011582 | 1.55967212 |
| 8286 | NM_182543.3 | | #N/A | 57.6985161 | -7.05879 | 1.50799938 |
| 8287 | NM_001083620.1 | GRIA2 | | 130.212771 | -8.2348412 | 1.75928304 |
| 8288 | NM_002765.4 | PRPS2 | | 62.9182782 | -7.1856151 | 1.53512746 |
| 8289 | NM_004472.2 | | #N/A | 141.926796 | -4.1485483 | 0.88631676 |
| 8290 | NM_001243963.1 | LRCH2 | | 117.563614 | -8.0865148 | 1.72772213 |
| 8291 | NM_001010983.2 | GLT8D1 | | 62.3488754 | -7.1772132 | 1.53349634 |
| 8292 | NM_006885.3 | | #N/A | 72.0177877 | -7.3841054 | 1.57776078 |
| 8293 | NM_001100420.1 | | #N/A | 58.8195871 | -7.0877652 | 1.51451155 |
| 8294 | NM_032312.3 | | #N/A | 100.47787 | -6.4520001 | 1.37875938 |
| 8295 | NM_001145436.1 | | #N/A | 124.679261 | -8.1744164 | 1.74684539 |
| 8296 | NM_006383.3 | | #N/A | 90.6274175 | -7.7112112 | 1.647883 |
| 8297 | NM_023075.5 | | #N/A | 63.7581767 | -7.2016473 | 1.53902035 |
| 8298 | NM_001306130.1 | FN1 | | 105.395181 | -7.9326162 | 1.69525412 |

| | | | | | | |
|---|---|---|---|---|---|---|
| 8299 | NM_198507.2 | | #N/A | 61.2815859 | -7.1479803 | 1.52757586 |
| 8300 | NM_001143944.1 | LEMD2 | | 80.8694118 | -7.5471989 | 1.61294493 |
| 8301 | NR_134455.1 | | #N/A | 65.4348103 | -7.2424148 | 1.54791187 |
| 8302 | NM_000954.5 | | #N/A | 90.9652704 | -7.7187898 | 1.64978684 |
| 8303 | NM_024111.4 | | #N/A | 78.8456357 | -7.5105953 | 1.6053822 |
| 8304 | NM_182607.4 | | #N/A | 127.7479 | 2.85293134 | 0.60984656 |
| 8305 | NM_194318.3 | | #N/A | 61.3635776 | -7.1541211 | 1.52941516 |
| 8306 | NM_001242537.1 | | #N/A | 53.3623414 | 10.7251163 | 2.29299822 |
| 8307 | NM_001314042.1 | KDM5B | | 209.705014 | -4.7521752 | 1.01609447 |
| 8308 | NM_173832.5 | | #N/A | 75.6423666 | -7.4507423 | 1.59325098 |
| 8309 | NM_001321387.2 | | #N/A | 75.2038742 | -7.4454269 | 1.59218418 |
| 8310 | NM_001122.3 | | #N/A | 57.3029223 | -7.0504539 | 1.50775428 |
| 8311 | NR_040091.1 | LOC100506990 | | 95.1158893 | 1.78707913 | 0.38218376 |
| 8312 | NM_001034173.3 | | #N/A | 122.935047 | -8.1518934 | 1.74338723 |
| 8313 | NM_198204.1 | | #N/A | 82.9690228 | -7.5840859 | 1.62222531 |
| 8314 | NM_030930.3 | | #N/A | 63.9081318 | -7.2131907 | 1.54291027 |
| 8315 | NM_001145966.1 | | #N/A | 163.024013 | -5.8631518 | 1.25414353 |
| 8316 | NM_001300883.2 | | #N/A | 66.2449969 | -7.2590826 | 1.55289389 |
| 8317 | NM_022371.3 | | #N/A | 95.9486689 | -6.392628 | 1.36753767 |
| 8318 | NM_001100429.1 | | #N/A | 129.694208 | -8.2288845 | 1.76049332 |
| 8319 | NM_022821.3 | | #N/A | 80.577586 | -7.5446609 | 1.6142316 |
| 8320 | NM_003135.2 | | #N/A | 84.0560422 | -7.6036119 | 1.62698013 |
| 8321 | NM_001695.4 | | #N/A | 316.735792 | -2.7352705 | 0.58530397 |
| 8322 | NM_006662.2 | | #N/A | 459.776766 | -2.8319084 | 0.60603556 |
| 8323 | NM_001173476.1 | | #N/A | 100.256809 | -7.8574226 | 1.68155253 |
| 8324 | NM_015958.2 | | #N/A | 75.8156655 | -7.4544717 | 1.59532539 |
| 8325 | NM_003766.4 | BECN1 | | 64.8686942 | -7.2320505 | 1.54779772 |
| 8326 | NM_176867.3 | PPA2 | | 66.8679383 | 8.27323702 | 1.77066912 |
| 8327 | NM_198537.3 | | #N/A | 64.6642024 | -7.2216651 | 1.54561051 |
| 8328 | NM_001086521.1 | | #N/A | 94.5362046 | -7.772861 | 1.66371899 |
| 8329 | NM_016228.3 | | #N/A | 78.1141934 | -7.498164 | 1.60509705 |
| 8330 | NM_002510.2 | | #N/A | 90.2533952 | -7.7086641 | 1.650204 |
| 8331 | NR_033327.1 | | #N/A | 140.337015 | -8.3444635 | 1.78633581 |
| 8332 | NM_001195044.1 | | #N/A | 80.4002537 | -7.5351713 | 1.61311748 |
| 8333 | NM_152996.2 | | #N/A | 82.9989138 | -7.5851028 | 1.62391896 |
| 8334 | NM_001330327.1 | | #N/A | 58.9780033 | -7.0970576 | 1.51944099 |
| 8335 | NM_152616.4 | | #N/A | 19.7162888 | 9.27687222 | 1.98630044 |
| 8336 | NM_170709.2 | | #N/A | 52.8882042 | 10.6954361 | 2.29008731 |
| 8337 | NM_016138.4 | | #N/A | 60.115101 | -7.1184133 | 1.52423469 |
| 8338 | NM_178526.4 | | #N/A | 66.8742822 | -7.2719242 | 1.55713386 |
| 8339 | NM_001254757.1 | | #N/A | 58.0184538 | -7.067012 | 1.51334744 |
| 8340 | NM_032428.1 | | #N/A | 105.496181 | -5.4155211 | 1.1597075 |
| 8342 | NR_027436.2 | LOC283788 | | 63.661244 | -7.2080276 | 1.54360897 |

| | | | | | |
|---|---|---|---|---|---|
| 8341 | NM_000317.2 | #N/A | 84.0057472 | -7.6037478 | 1.62834477 |
| 8343 | NM_022804.2 | #N/A | 79.6488652 | -7.5253475 | 1.61183921 |
| 8344 | NM_019556.2 | #N/A | 56.3671556 | -7.0285317 | 1.50546932 |
| 8345 | NM_001532.2 | #N/A | 71.2097973 | -7.3633452 | 1.57720705 |
| 8346 | NM_001321350.1 | #N/A | 81.7424406 | -7.5611439 | 1.61976804 |
| 8347 | NM_001320815.1 | #N/A | 99.4462707 | -7.844942 | 1.68064085 |
| 8348 | NM_006821.5 | ACOT2 | 56.4143048 | -7.0258414 | 1.50520428 |
| 8349 | NM_032412.3 | #N/A | 82.9392568 | -6.208243 | 1.3300386 |
| 8350 | NM_001006684.1 | TCEAL8 | 74.8879316 | -7.4370248 | 1.59332551 |
| 8351 | NM_001968.4 | EIF4E | 113.074292 | -8.0328078 | 1.72101037 |
| 8352 | NM_016656.3 | #N/A | 78.1132437 | -7.4960739 | 1.60601956 |
| 8353 | NM_012388.3 | BLOC1S6 | 92.5285192 | -7.7390656 | 1.65816408 |
| 8354 | NM_015879.2 | #N/A | 113.588871 | -6.7665375 | 1.44984141 |
| 8355 | NR_003604.2 | #N/A | 89.6204747 | -7.6956634 | 1.64903474 |
| 8356 | NM_001301169.1 | #N/A | 126.561743 | -8.1934169 | 1.75583847 |
| 8357 | NM_177556.2 | #N/A | 91.0452845 | -7.7194065 | 1.65437384 |
| 8358 | NM_001293215.1 | #N/A | 79.8824764 | -7.5310504 | 1.61408677 |
| 8359 | NM_138358.2 | #N/A | 66.4188303 | -7.2635561 | 1.5567537 |
| 8360 | NM_017657.4 | #N/A | 80.5622652 | -7.540618 | 1.61614586 |
| 8361 | NM_016360.3 | #N/A | 66.1101854 | -7.2575776 | 1.55552017 |
| 8363 | NM_032833.4 | PPP1R15B | 312.421162 | -2.5556906 | 0.54780331 |
| 8362 | NM_006064.4 | #N/A | 73.9060398 | -7.4203607 | 1.590526 |
| 8364 | NM_024839.2 | #N/A | 84.0213744 | -7.6023029 | 1.62954173 |
| 8365 | NM_014154.4 | ARMC8 | 76.6475429 | -7.4688088 | 1.60094189 |
| 8366 | NM_022755.5 | #N/A | 64.7120863 | -7.2266792 | 1.54915439 |
| 8367 | NM_014409.3 | TAF5L | 83.0395848 | -7.5851726 | 1.62601733 |
| 8368 | NM_001034850.2 | RETREG1 | 110.83904 | -4.7367981 | 1.01544224 |
| 8369 | NM_024089.2 | #N/A | 67.9827777 | -7.2996822 | 1.56484961 |
| 8370 | NM_001330556.1 | #N/A | 122.329154 | -8.1442249 | 1.74593097 |
| 8371 | NM_001282459.1 | #N/A | 50.6125223 | 3.41414785 | 0.73194048 |
| 8372 | NM_001271052.1 | ADGRE2 | 20.0622507 | 9.29851776 | 1.99356166 |
| 8373 | NM_001328675.1 | #N/A | 111.486157 | -8.0103326 | 1.71742523 |
| 8374 | NM_145057.3 | #N/A | 62.3295532 | -7.1773547 | 1.53891222 |
| 8375 | NM_000932.2 | #N/A | 83.9061948 | -7.6004274 | 1.6298012 |
| 8376 | NM_020935.2 | #N/A | 94.860053 | -5.1846736 | 1.11182699 |
| 8377 | NR_135550.1 | DENND6A-DT | 94.0534653 | 4.15354534 | 0.89074394 |
| 8378 | NM_001451.2 | #N/A | 100.922725 | -7.8684698 | 1.68743634 |
| 8379 | NM_004845.4 | #N/A | 107.660069 | -7.9597947 | 1.70704729 |
| 8380 | NM_183413.2 | FBXO44 | 122.799551 | -8.1511642 | 1.74824761 |
| 8381 | NR_027164.1 | SNHG29 | 116.505302 | -8.075238 | 1.73203494 |
| 8382 | NM_024028.3 | #N/A | 109.760635 | -7.9888866 | 1.71369562 |
| 8383 | NM_017647.3 | #N/A | 209.45984 | -3.7667338 | 0.80819475 |
| 8384 | NM_003457.3 | ZNF207 | 118.456692 | -8.1001506 | 1.73804495 |

| | | | | | |
|---|---|---|---|---|---|
| 8385 | NM_001014447.2 | #N/A | 68.0216063 | -7.3003044 | 1.56650281 |
| 8386 | NM_080656.2 | #N/A | 116.839028 | -4.0137376 | 0.86127968 |
| 8387 | NM_001145416.1 | #N/A | 62.2804302 | -7.1724576 | 1.53912782 |
| 8388 | NM_006594.4 | #N/A | 80.5473092 | -7.5384912 | 1.61776188 |
| 8389 | NM_181482.4 | #N/A | 77.6976002 | -7.4872979 | 1.60678684 |
| 8390 | NM_002894.2 | #N/A | 58.1004653 | -7.0733277 | 1.51811245 |
| 8391 | NM_004965.6 | #N/A | 949.69504 | -5.968045 | 1.28090829 |
| 8392 | NM_005126.4 | #N/A | 72.9588688 | -7.3969524 | 1.58758751 |
| 8393 | NR_037713.1 | #N/A | 92.3408774 | -7.7383735 | 1.66092162 |
| 8394 | NM_001318819.1 | #N/A | 70.1919514 | -7.3390049 | 1.57535546 |
| 8395 | NM_021190.3 | PTBP2 | 124.451565 | -8.1693339 | 1.75369329 |
| 8396 | NM_001305077.1 | #N/A | 80.2865115 | -7.5398954 | 1.61863877 |
| 8397 | NM_005859.4 | #N/A | 76.6170584 | -6.1026794 | 1.31017033 |
| 8398 | NM_201398.2 | #N/A | 88.2295431 | -7.6728311 | 1.64726373 |
| 8399 | NM_032359.3 | #N/A | 99.456838 | -7.8460021 | 1.68452213 |
| 8400 | NM_016535.3 | #N/A | 71.3404761 | -7.3676482 | 1.58183375 |
| 8401 | NM_001190810.1 | AGAP9 | 56.8056072 | -7.0370525 | 1.51089355 |
| 8402 | NR_036573.1 | LOC155060 | 65.3414567 | -3.2934519 | 0.7071383 |
| 8403 | NM_001039613.2 | #N/A | 68.7812606 | -7.3136929 | 1.5703387 |
| 8404 | NM_017503.4 | #N/A | 73.4814379 | -7.4089683 | 1.59079992 |
| 8405 | NR_003697.1 | #N/A | 79.5905811 | -7.5244296 | 1.61561762 |
| 8406 | NM_001319228.1 | #N/A | 77.0943603 | -7.4740514 | 1.60481112 |
| 8407 | NM_001130963.1 | #N/A | 70.1102783 | -7.3373663 | 1.57547338 |
| 8408 | NM_017765.2 | #N/A | 60.9691132 | -7.1432662 | 1.53386899 |
| 8409 | NM_133496.4 | #N/A | 65.9741354 | -7.2539123 | 1.5577166 |
| 8410 | NM_002248.4 | KCNN1 | 99.4263723 | -7.8442873 | 1.68453265 |
| 8411 | NM_015038.1 | #N/A | 65.1095588 | -3.8748605 | 0.83215988 |
| 8412 | NM_182502.3 | TMPRSS11B | 35.3720389 | 3.00791278 | 0.64600558 |
| 8413 | NM_021075.3 | #N/A | 58.0432818 | -7.0713194 | 1.51878269 |
| 8414 | NM_032458.2 | #N/A | 65.4168308 | -7.2421548 | 1.5556201 |
| 8415 | NM_153756.2 | #N/A | 61.0398697 | -7.1427668 | 1.53436012 |
| 8416 | NM_001083601.2 | #N/A | 69.1861872 | -7.3196181 | 1.57252608 |
| 8417 | NM_182527.2 | #N/A | 96.7805643 | -7.8041464 | 1.67676854 |
| 8418 | NM_145918.2 | #N/A | 79.8072676 | -7.533026 | 1.61861455 |
| 8419 | NM_032324.2 | #N/A | 72.1168349 | -7.3813483 | 1.58606217 |
| 8420 | NM_002380.4 | #N/A | 87.263637 | -7.6599498 | 1.64612834 |
| 8421 | NM_021644.3 | HNRNPH3 | 82.9634609 | -7.5818014 | 1.62935789 |
| 8422 | NR_003584.3 | SNHG8 | 84.0463954 | -7.6021528 | 1.63374532 |
| 8423 | NM_021121.3 | EEF1B2 | 84.1958705 | -7.6085647 | 1.63520139 |
| 8424 | NM_182913.2 | #N/A | 94.2968246 | -7.7667342 | 1.66931419 |
| 8425 | NM_016527.3 | #N/A | 21.5741153 | 8.63190567 | 1.85537268 |
| 8426 | NM_025129.4 | #N/A | 68.2820555 | -7.3021073 | 1.5695683 |
| 8427 | NM_004058.4 | #N/A | 97.6469394 | -7.8216592 | 1.68133078 |

| | | | | | |
|---|---|---|---|---|---|
| 8428 | NM_001318197.1 | #N/A | 76.0940419 | 2.93947358 | 0.63195641 |
| 8429 | NM_014903.5 | #N/A | 175.275762 | -6.3319877 | 1.36133612 |
| 8430 | NM_002015.3 | #N/A | 77.6727984 | -7.4928991 | 1.61110894 |
| 8431 | NM_001322402.1 | #N/A | 90.8148568 | -7.7155044 | 1.65902092 |
| 8432 | NM_001322897.1 | #N/A | 23.484982 | 9.54598395 | 2.05274325 |
| 8433 | NM_201632.4 | #N/A | 36.2469149 | 10.1708856 | 2.187128 |
| 8434 | NM_001137550.1 | #N/A | 79.5151826 | -7.524764 | 1.618343 |
| 8435 | NM_001286760.1 | CERCAM | 94.6477498 | -7.7761728 | 1.67246458 |
| 8436 | NM_004874.3 | #N/A | 75.3648817 | -7.442227 | 1.60075691 |
| 8437 | NM_001248004.1 | #N/A | 60.664901 | -7.1294725 | 1.53353528 |
| 8438 | NM_001033504.1 | TMEM98 | 66.6556755 | -7.2698106 | 1.56373827 |
| 8439 | NM_001142964.1 | #N/A | 69.1206994 | -7.3224414 | 1.57511442 |
| 8440 | NM_031231.3 | #N/A | 62.4291125 | -7.178374 | 1.54418292 |
| 8441 | NM_207336.2 | #N/A | 83.3451947 | -7.5887244 | 1.63250097 |
| 8442 | NM_001033045.3 | #N/A | 65.3284285 | -7.2390395 | 1.55768553 |
| 8443 | NM_001144888.1 | #N/A | 69.7070606 | -7.337791 | 1.57895319 |
| 8444 | NM_001145044.1 | SLCO3A1 | 61.9614222 | -7.163929 | 1.54167733 |
| 8445 | NM_001314063.1 | #N/A | 180.482151 | -8.7055836 | 1.87353679 |
| 8446 | NM_001143841.1 | TMEM106C | 65.116351 | -7.2318621 | 1.55639404 |
| 8447 | NM_015926.5 | #N/A | 65.8802507 | -7.2543255 | 1.56134866 |
| 8448 | NM_016246.2 | #N/A | 64.1860035 | -7.2181508 | 1.55366042 |
| 8449 | NM_153451.2 | #N/A | 74.3204878 | -2.9048728 | 0.62525298 |
| 8450 | NM_152715.3 | #N/A | 59.1380489 | -7.0969205 | 1.52759253 |
| 8451 | NM_024339.4 | #N/A | 88.4056478 | -7.6744627 | 1.6521213 |
| 8452 | NM_018941.3 | #N/A | 171.854962 | -2.9525992 | 0.63569211 |
| 8453 | NM_018622.6 | #N/A | 73.4601056 | -7.4085749 | 1.59521556 |
| 8454 | NM_153690.4 | #N/A | 81.4475243 | -7.5603676 | 1.62790105 |
| 8455 | NM_001004106.2 | #N/A | 100.713826 | -7.8659691 | 1.69381837 |
| 8456 | NR_003272.2 | #N/A | 77.322527 | -7.4841913 | 1.61169076 |
| 8457 | NM_006627.2 | #N/A | 73.467302 | -7.4094705 | 1.59565539 |
| 8458 | NM_001113361.1 | TBC1D14 | 57.6256824 | -7.0564579 | 1.51964486 |
| 8459 | NM_001039803.2 | CDK20 | 75.0799049 | -7.4413462 | 1.60254667 |
| 8461 | NM_001282684.1 | KCTD17 | 73.7285108 | -7.4107543 | 1.59598804 |
| 8460 | NM_001247.4 | #N/A | 59.214895 | -7.1035036 | 1.52982485 |
| 8462 | NM_001130996.1 | HMGCR | 132.616614 | -8.2616121 | 1.77931163 |
| 8463 | NM_001304562.1 | #N/A | 100.119145 | -7.856119 | 1.6920877 |
| 8464 | NM_023078.4 | #N/A | 58.9394195 | -7.0919014 | 1.52762085 |
| 8465 | NM_153348.2 | #N/A | 57.4474556 | -7.0549052 | 1.51965832 |
| 8466 | NR_122035.1 | ARRDC1-AS1 | 80.258578 | -7.5353401 | 1.62316981 |
| 8469 | NM_032968.4 | PCDH11X | 142.581928 | -8.3659743 | 1.80214932 |
| 8467 | NM_001042497.1 | #N/A | 77.4094388 | -7.4803414 | 1.61138474 |
| 8468 | NM_014943.3 | #N/A | 54.4343851 | -6.9787807 | 1.50334117 |
| 8470 | NM_005637.3 | SS18 | 178.359521 | -2.8594833 | 0.61598437 |

| | | | | | |
|---|---|---|---|---|---|
| 8471 | NM_018087.4 | #N/A | 76.4518086 | -7.4664956 | 1.60844081 |
| 8472 | NM_000965.4 | RARB | 55.5298699 | -7.0093057 | 1.50997866 |
| 8473 | NM_139015.4 | #N/A | 410.296357 | -3.5362092 | 0.76184365 |
| 8474 | NM_001199163.1 | PSMC5 | 63.0909282 | -7.1899917 | 1.54911261 |
| 8475 | NM_001018064.2 | #N/A | 134.100743 | -6.8122643 | 1.46802011 |
| 8476 | NM_201280.2 | #N/A | 61.8926756 | -7.1588037 | 1.54271557 |
| 8477 | NR_038861.1 | TUNAR | 64.268726 | -7.2203626 | 1.55622442 |
| 8478 | NM_001122606.1 | LAMP2 | 112.191321 | -8.0205609 | 1.72876832 |
| 8479 | NM_030674.3 | #N/A | 109.370751 | -7.9836268 | 1.72083899 |
| 8480 | NM_004724.3 | #N/A | 63.9775873 | -7.2109345 | 1.5543028 |
| 8481 | NM_024635.3 | #N/A | 66.1596262 | -7.26076 | 1.5650561 |
| 8482 | NM_178587.2 | #N/A | 70.9019592 | -7.3593574 | 1.58638681 |
| 8483 | NM_005249.4 | #N/A | 787.698635 | -10.832002 | 2.33521095 |
| 8484 | NM_001031740.2 | MANEAL | 74.7712665 | -7.4369477 | 1.60330491 |
| 8485 | NR_033322.3 | NSUN5P1 | 55.4226837 | -6.9998582 | 1.50909049 |
| 8486 | NM_001289950.2 | #N/A | 77.3966769 | -7.4810465 | 1.61296107 |
| 8487 | NM_020648.5 | #N/A | 234.840716 | -2.8965103 | 0.62452213 |
| 8488 | NM_001321018.1 | #N/A | 79.416556 | -7.5203512 | 1.62152929 |
| 8489 | NR_027435.1 | ARRDC3-AS1 | 69.9228154 | 2.52853094 | 0.54520718 |
| 8490 | NM_016422.3 | #N/A | 217.462899 | -2.187063 | 0.47159062 |
| 8491 | NM_007048.5 | #N/A | 53.9096989 | -6.9630305 | 1.50143783 |
| 8492 | NM_078468.2 | #N/A | 60.0163124 | -7.1158495 | 1.53443921 |
| 8493 | NM_001135740.1 | PNCK | 68.026725 | -7.2960507 | 1.5733813 |
| 8494 | NM_001257363.2 | #N/A | 67.3929463 | -7.2824681 | 1.57044813 |
| 8495 | NM_144567.4 | #N/A | 66.1280773 | -7.2581404 | 1.56524558 |
| 8496 | NM_001291867.1 | #N/A | 72.5513652 | -7.3942311 | 1.59485954 |
| 8497 | NM_001166373.1 | MARCHF1 | 104.220592 | -7.9136459 | 1.70709469 |
| 8498 | NM_177938.2 | P4HTM | 63.1238976 | -7.1889641 | 1.55078915 |
| 8499 | NM_001202413.1 | AKR1A1 | 135.500575 | -8.292222 | 1.78898275 |
| 8500 | NM_001029896.1 | #N/A | 80.7420948 | -7.5462331 | 1.62806229 |
| 8501 | NM_001081562.2 | #N/A | 65.3991749 | -7.2386559 | 1.56170064 |
| 8502 | NM_020676.6 | ABHD6 | 94.3065132 | -7.7679448 | 1.67605127 |
| 8503 | NR_038269.1 | GIHCG | 59.8685317 | -7.1112835 | 1.53456477 |
| 8504 | NM_001276270.1 | #N/A | 64.5072174 | -7.2185362 | 1.55775962 |
| 8505 | NM_024051.3 | #N/A | 63.6873169 | -7.200569 | 1.55398724 |
| 8506 | NM_002623.3 | #N/A | 103.287938 | -7.8998784 | 1.70494821 |
| 8507 | NM_006463.4 | #N/A | 70.0095584 | -7.3406116 | 1.58428724 |
| 8508 | NM_022078.2 | #N/A | 65.2217235 | -7.2367841 | 1.5619191 |
| 8509 | NR_046084.1 | SH3BP5-AS1 | 55.6090505 | -7.0072468 | 1.51238955 |
| 8510 | NM_001128150.1 | #N/A | 74.6090967 | -7.4311207 | 1.60389344 |
| 8511 | NM_014798.2 | #N/A | 62.5872039 | -7.1788497 | 1.54953011 |
| 8512 | NM_032459.2 | #N/A | 71.3524433 | -7.3657339 | 1.5898628 |
| 8513 | NM_004365.3 | CETN3 | 60.5141693 | -7.1294226 | 1.53888138 |

| | | | | | |
|---|---|---|---|---|---|
| 8514 | NM_203371.1 | #N/A | 109.260294 | -7.9831477 | 1.72320839 |
| 8515 | NM_033286.3 | KNSTRN | 76.6626858 | -7.4703769 | 1.61254951 |
| 8516 | NM_001135993.1 | #N/A | 63.4490783 | -7.1951527 | 1.55315886 |
| 8517 | NM_032175.3 | #N/A | 61.71183 | -7.1530755 | 1.54412852 |
| 8518 | NM_021705.3 | #N/A | 66.6343499 | -7.2676448 | 1.5689679 |
| 8519 | NM_152269.4 | #N/A | 80.4706592 | -7.542964 | 1.62845683 |
| 8520 | NM_001203247.1 | #N/A | 79.1286647 | -7.5126596 | 1.62207766 |
| 8521 | NM_001199847.1 | #N/A | 76.7073746 | -6.0988141 | 1.31685715 |
| 8522 | NM_024336.2 | #N/A | 88.0724394 | -7.6718545 | 1.65661987 |
| 8523 | NM_005033.2 | #N/A | 64.0875712 | -7.2108801 | 1.55718996 |
| 8524 | NM_001270393.1 | PKIB | 103.568361 | -6.6320382 | 1.43222534 |
| 8525 | NM_001184971.1 | PACSIN2 | 72.6372505 | -7.3940031 | 1.5968634 |
| 8526 | NM_004269.3 | #N/A | 83.1442911 | -7.5867824 | 1.63858083 |
| 8527 | NM_013326.4 | #N/A | 72.2912474 | -7.3878072 | 1.59570129 |
| 8528 | NM_001178085.1 | ZFX | 67.663435 | -7.2935379 | 1.57538473 |
| 8529 | NM_025201.4 | #N/A | 58.0484356 | -7.0714665 | 1.52741019 |
| 8530 | NR_037601.1 | LINC01578 | 76.5743058 | -7.466869 | 1.6129015 |
| 8531 | NR_024550.1 | DMTF1 | 100.474451 | -7.8578165 | 1.69739333 |
| 8532 | NM_001130136.1 | ISLR2 | 153.481204 | -8.4721997 | 1.83022286 |
| 8533 | NM_003772.3 | #N/A | 59.0439582 | -7.096322 | 1.53311225 |
| 8534 | NM_002996.4 | #N/A | 76.1679382 | -7.4655282 | 1.61289895 |
| 8535 | NM_001042548.1 | #N/A | 95.6377537 | -7.788909 | 1.6828519 |
| 8536 | NM_001261843.1 | #N/A | 72.2909095 | -7.3887348 | 1.59641181 |
| 8537 | NM_001011725.1 | #N/A | 64.746498 | -7.2246016 | 1.56097313 |
| 8538 | NM_001001396.2 | ATP2B4 | 93.5227336 | -7.7571861 | 1.67625404 |
| 8539 | NM_003659.3 | #N/A | 267.2953 | -5.6860823 | 1.22870219 |
| 8540 | NR_003610.1 | PDXDC2P-NPIPB14P | 193.893492 | -3.5086412 | 0.75823663 |
| 8541 | NM_006747.3 | #N/A | 74.2657659 | -6.0562802 | 1.3089292 |
| 8542 | NM_001112734.3 | #N/A | 63.4181707 | -7.1941592 | 1.55487023 |
| 8543 | NM_013320.2 | #N/A | 84.5685643 | -7.6112255 | 1.64505961 |
| 8544 | NM_020821.2 | #N/A | 103.558876 | -7.9034137 | 1.70821458 |
| 8545 | NM_001321076.1 | #N/A | 143.369263 | -8.3733965 | 1.80990749 |
| 8546 | NM_001011880.2 | CLEC18B | 102.576919 | -6.6186323 | 1.43074998 |
| 8548 | NM_001330637.1 | MPDZ | 96.0142937 | -7.7981006 | 1.68578505 |
| 8547 | NM_001127190.1 | #N/A | 72.6653321 | -7.3957995 | 1.59881538 |
| 8549 | NM_006917.4 | #N/A | 87.1694499 | -7.6572082 | 1.65534953 |
| 8550 | NM_002417.4 | #N/A | 80.3892787 | -7.543344 | 1.63078253 |
| 8551 | NM_001164381.1 | STAU2 | 89.779244 | -7.6982331 | 1.66439062 |
| 8552 | NM_003884.4 | #N/A | 58.2481768 | -7.0734103 | 1.52935578 |
| 8553 | NR_002211.1 | MEIS3P1 | 64.3934509 | -7.2241707 | 1.56211435 |
| 8554 | NM_001172646.1 | PLCB4 | 59.413582 | -7.1060296 | 1.53660223 |
| 8555 | NR_027155.2 | SMG1P3 | 61.2557517 | -7.1467901 | 1.54543633 |
| 8556 | NM_014869.6 | #N/A | 56.3959773 | -7.0275464 | 1.51974963 |

| | | | | | |
|---|---|---|---|---|---|
| 8557 | NM_000144.4 | #N/A | 207.115502 | 1.77249296 | 0.38334077 |
| 8558 | NM_024093.2 | #N/A | 76.5103315 | -7.4649868 | 1.61447192 |
| 8559 | NM_001282757.1 | TRA2A | 97.71726 | -3.5587306 | 0.76967751 |
| 8560 | NM_001297551.1 | #N/A | 65.7862098 | -7.2494095 | 1.56792378 |
| 8561 | NM_003277.3 | #N/A | 116.162624 | -8.0719382 | 1.745863 |
| 8562 | NR_132749.1 | #N/A | 85.3617065 | -7.6240346 | 1.64903626 |
| 8563 | NM_033375.4 | #N/A | 123.369198 | -8.1588253 | 1.76490085 |
| 8564 | NM_031890.3 | #N/A | 103.278709 | -7.9008444 | 1.70913978 |
| 8565 | NM_022727.5 | #N/A | 73.2443434 | -7.4068018 | 1.60247446 |
| 8566 | NM_001190438.1 | NCOR1 | 79.9448486 | -7.5292503 | 1.6291833 |
| 8567 | NM_000321.2 | RB1 | 205.188249 | -5.3745692 | 1.16297204 |
| 8568 | NM_025182.3 | #N/A | 70.8193453 | -7.3568594 | 1.59212553 |
| 8569 | NM_031210.5 | #N/A | 91.7692676 | -7.7301737 | 1.67297063 |
| 8570 | NM_004976.4 | KCNC1 | 95.7678263 | -7.7915929 | 1.68638519 |
| 8571 | NM_152479.5 | TTC9B | 146.935035 | -8.4091996 | 1.82016076 |
| 8572 | NM_002979.4 | #N/A | 56.3136179 | -7.0298172 | 1.52170047 |
| 8573 | NM_018075.4 | #N/A | 92.0622314 | -7.735529 | 1.67459015 |
| 8574 | NR_110834.1 | #N/A | 85.6695408 | -7.6335662 | 1.65251045 |
| 8575 | NM_001130971.1 | NSMF | 63.3807782 | -7.1917208 | 1.55694396 |
| 8576 | NM_004563.3 | #N/A | 80.3451498 | -7.5386034 | 1.63206622 |
| 8577 | NM_000745.3 | CHRNA5 | 72.8261417 | -7.3965379 | 1.60159846 |
| 8578 | NM_024580.5 | #N/A | 56.5063667 | -7.0275815 | 1.52176402 |
| 8579 | NM_000349.2 | #N/A | 103.126923 | 2.44902429 | 0.5303449 |
| 8580 | NM_032427.3 | #N/A | 62.7424804 | -7.1870852 | 1.55645858 |
| 8582 | NM_020408.5 | LYRM4 | 86.9415598 | -7.6521579 | 1.65741998 |
| 8581 | NM_001047160.2 | #N/A | 60.8974963 | -7.1352271 | 1.54544803 |
| 8583 | NM_201430.2 | #N/A | 95.0126578 | -7.7791117 | 1.68505914 |
| 8584 | NM_145080.3 | #N/A | 97.069751 | -7.8104821 | 1.6920205 |
| 8585 | NM_004763.4 | #N/A | 72.9989747 | -7.399142 | 1.60294815 |
| 8586 | NR_130915.1 | LCAL1 | 31.7069159 | 6.25493506 | 1.35512996 |
| 8587 | NM_003921.4 | #N/A | 63.5156263 | -7.1966012 | 1.55920393 |
| 8588 | NM_001184720.1 | GYG1 | 74.6331889 | -7.4333965 | 1.61055326 |
| 8589 | NM_001199155.1 | #N/A | 70.108878 | -7.3370256 | 1.58970709 |
| 8590 | NM_001297696.1 | CCDC125 | 100.248858 | 2.57682216 | 0.55833701 |
| 8591 | NM_006270.4 | #N/A | 61.4188916 | -7.1538079 | 1.55017546 |
| 8592 | NR_038335.1 | CLU | 109.308992 | -7.9843495 | 1.73031619 |
| 8593 | NM_002507.3 | #N/A | 889.525516 | -5.1044601 | 1.10627201 |
| 8594 | NM_001799.3 | #N/A | 54.6679811 | -6.9794401 | 1.51274507 |
| 8595 | NM_004849.3 | #N/A | 76.1795192 | -7.4595549 | 1.61680633 |
| 8596 | NM_016507.3 | #N/A | 124.477952 | -8.1695617 | 1.77075209 |
| 8597 | NM_001014432.1 | AKT1 | 85.1492736 | -7.6180235 | 1.65128701 |
| 8598 | NM_001010862.2 | #N/A | 100.290046 | -5.8096459 | 1.2594675 |
| 8599 | NM_001145408.1 | #N/A | 71.8602128 | -7.3751972 | 1.59888004 |

| | | | | | |
|---|---|---|---|---|---|
| 8600 | NM_021805.2 | #N/A | 57.1433874 | -7.0445962 | 1.52736011 |
| 8601 | NM_002958.3 | RYK | 94.4209543 | -7.7686143 | 1.68453946 |
| 8602 | NM_001017919.1 | RCCD1 | 64.1714218 | -7.2117524 | 1.5639244 |
| 8603 | NR_033677.1 | #N/A | 78.8435262 | -7.509827 | 1.62866425 |
| 8604 | NM_005484.3 | PARP2 | 63.2021181 | -7.1875707 | 1.55878674 |
| 8605 | NM_001145206.1 | KIAA1671 | 91.7183589 | -4.939989 | 1.07137625 |
| 8606 | NM_020650.2 | #N/A | 67.2862503 | -7.2864932 | 1.58032563 |
| 8607 | NM_006807.4 | #N/A | 71.0191996 | -7.3602865 | 1.59642167 |
| 8608 | NM_001320360.1 | #N/A | 90.9003152 | -7.7168837 | 1.67395526 |
| 8609 | NM_001321612.1 | #N/A | 315.082313 | -5.3993206 | 1.1712492 |
| 8610 | NM_005940.4 | #N/A | 88.3596862 | -7.6764262 | 1.66528861 |
| 8611 | NM_014057.4 | #N/A | 75.4257228 | -7.4507142 | 1.6163183 |
| 8612 | NM_133458.3 | #N/A | 103.858791 | -7.9066218 | 1.71535889 |
| 8613 | NM_020190.3 | #N/A | 78.6915498 | -7.5116986 | 1.62972772 |
| 8614 | NM_014585.5 | SLC40A1 | 70.8992872 | -7.3601836 | 1.59686551 |
| 8615 | NR_104023.1 | #N/A | 175.395542 | -5.1938156 | 1.12696408 |
| 8616 | NM_032582.3 | #N/A | 315.758931 | -3.6093963 | 0.78328557 |
| 8617 | NM_001145077.1 | #N/A | 80.6154471 | -7.5429606 | 1.63696929 |
| 8618 | NM_013349.4 | #N/A | 66.5962394 | -7.2669102 | 1.57715957 |
| 8619 | NM_001040167.1 | #N/A | 70.0625869 | -7.3405584 | 1.59336234 |
| 8620 | NM_021971.2 | #N/A | 55.649284 | -7.0105308 | 1.52176944 |
| 8621 | NM_018413.5 | #N/A | 108.994296 | -7.9773917 | 1.73172162 |
| 8622 | NM_053002.5 | MED12L | 64.3923776 | -7.2202578 | 1.56756489 |
| 8623 | NM_173793.4 | C22orf39 | 71.6101996 | -7.3714511 | 1.60041629 |
| 8624 | NM_015227.4 | #N/A | 66.8985429 | -7.2783685 | 1.58022172 |
| 8625 | NM_001171887.1 | #N/A | 77.1981354 | -7.4792542 | 1.62394094 |
| 8626 | NM_002863.4 | #N/A | 63.2285565 | -7.1982912 | 1.56318922 |
| 8627 | NM_005502.3 | #N/A | 72.6624767 | -6.1191517 | 1.32884201 |
| 8628 | NM_000203.4 | #N/A | 67.3239506 | -7.2885077 | 1.58284091 |
| 8629 | NM_032140.2 | #N/A | 52.4353053 | -6.9241325 | 1.5037274 |
| 8630 | NM_020649.2 | #N/A | 59.72216 | -7.1081504 | 1.54378978 |
| 8631 | NM_172109.2 | #N/A | 90.1921397 | -7.7033757 | 1.67310109 |
| 8632 | NM_002481.3 | #N/A | 98.7861456 | -7.8365657 | 1.70212416 |
| 8633 | NM_015987.4 | #N/A | 61.8347243 | -7.1608463 | 1.55535809 |
| 8634 | NM_018203.2 | #N/A | 64.2118282 | -7.2138332 | 1.5668965 |
| 8635 | NM_020236.3 | #N/A | 90.3676358 | -7.7074884 | 1.67420923 |
| 8636 | NM_032765.3 | #N/A | 60.4607999 | -7.1266161 | 1.5480936 |
| 8637 | NM_031298.3 | #N/A | 76.7503286 | -7.4728624 | 1.62339384 |
| 8638 | NM_000190.3 | #N/A | 71.8243589 | -7.376899 | 1.60262494 |
| 8639 | NM_001304996.1 | #N/A | 62.465976 | -7.1701526 | 1.55773702 |
| 8640 | NR_144318.1 | #N/A | 64.3560723 | -7.2155262 | 1.56796326 |
| 8641 | NM_001161009.2 | #N/A | 78.6478716 | -7.506314 | 1.63120566 |
| 8642 | NM_001625.3 | #N/A | 68.2608894 | -4.2737873 | 0.92874292 |

| | | | | | |
|---|---|---|---|---|---|
| 8643 | NM_001127325.1 | MAD2L2 | 94.7060791 | -7.7749252 | 1.68960962 |
| 8644 | NM_001164722.2 | #N/A | 102.998538 | 2.97154251 | 0.64577573 |
| 8645 | NM_144969.2 | #N/A | 81.4810455 | -7.5584329 | 1.64269039 |
| 8646 | NM_001300799.1 | #N/A | 119.883354 | -8.1147714 | 1.76385185 |
| 8647 | NM_001127896.1 | #N/A | 104.753601 | -7.92155 | 1.72209523 |
| 8648 | NM_033267.4 | #N/A | 62.3775071 | -7.1754351 | 1.55990063 |
| 8649 | NM_002436.3 | #N/A | 62.2074496 | -7.1681567 | 1.55841721 |
| 8650 | NM_002716.4 | #N/A | 68.6106403 | -7.3089235 | 1.58907256 |
| 8651 | NM_138446.1 | #N/A | 64.3141266 | -7.2169523 | 1.56911456 |
| 8652 | NM_197964.4 | #N/A | 56.5274819 | -7.0308538 | 1.52864968 |
| 8653 | NR_038460.1 | #N/A | 61.862137 | 2.25317118 | 0.48988146 |
| 8655 | NM_201574.2 | SLC4A3 | 68.5321554 | -7.3093983 | 1.58927375 |
| 8654 | NM_001345882.1 | #N/A | 66.9454325 | -7.2712557 | 1.58097985 |
| 8656 | NM_001289175.1 | VPS52 | 70.9913794 | -7.3603909 | 1.60039078 |
| 8657 | NM_001330272.1 | #N/A | 59.7485269 | -7.1139098 | 1.5470261 |
| 8658 | NM_001168368.1 | #N/A | 68.8432657 | -7.3190605 | 1.59168318 |
| 8659 | NM_002599.4 | #N/A | 123.957953 | -8.1625712 | 1.77512974 |
| 8660 | NM_033512.2 | #N/A | 76.067832 | -6.0833344 | 1.32299059 |
| 8661 | NM_152484.2 | #N/A | 57.5270979 | -7.0529324 | 1.5340273 |
| 8662 | NM_199478.2 | #N/A | 74.9315528 | -7.4370453 | 1.61763526 |
| 8663 | NM_001199803.2 | #N/A | 56.0827565 | -7.0190468 | 1.52697 |
| 8664 | NM_014890.3 | FILIP1L | 84.0491864 | -7.6049233 | 1.65449853 |
| 8665 | NM_145914.2 | #N/A | 58.7006206 | -7.0839595 | 1.54129507 |
| 8666 | NM_021100.4 | #N/A | 62.6825204 | -7.1803411 | 1.56230613 |
| 8667 | NM_181471.2 | #N/A | 79.1222335 | -7.5132845 | 1.63477152 |
| 8668 | NM_201378.3 | #N/A | 140.764919 | -8.3454285 | 1.8160036 |
| 8669 | NM_013323.2 | SNX11 | 70.4277526 | -7.3500783 | 1.59942351 |
| 8670 | NM_006741.3 | #N/A | 93.9831349 | -7.7643922 | 1.68962477 |
| 8671 | NM_145112.2 | #N/A | 72.768566 | -7.3955973 | 1.60937089 |
| 8672 | NM_001320752.1 | STS | 77.4609225 | -7.4844942 | 1.62885333 |
| 8673 | NM_007361.3 | #N/A | 169.149979 | -8.61363 | 1.8747639 |
| 8674 | NM_198552.2 | #N/A | 62.0785257 | -7.1703721 | 1.56063176 |
| 8675 | NM_033238.2 | #N/A | 61.2706058 | -7.1497973 | 1.55624312 |
| 8677 | NM_020657.3 | ZNF304 | 65.7790378 | -7.2497903 | 1.5780314 |
| 8676 | NM_001122821.1 | #N/A | 60.0166858 | -7.1228437 | 1.55040716 |
| 8678 | NM_202001.2 | #N/A | 67.3308352 | -7.28543 | 1.58578047 |
| 8679 | NM_181718.3 | #N/A | 95.497959 | -7.7873889 | 1.6952168 |
| 8680 | NM_032711.3 | #N/A | 61.5810688 | -7.1538252 | 1.5573184 |
| 8681 | NM_148172.2 | #N/A | 73.3357694 | -7.4093667 | 1.61297057 |
| 8682 | NM_007019.3 | #N/A | 68.663682 | -7.3105408 | 1.59160386 |
| 8683 | NM_203464.2 | AK4 | 73.093817 | -7.4036224 | 1.61189207 |
| 8684 | NM_001002857.1 | #N/A | 80.2965186 | -7.5412936 | 1.64217421 |
| 8685 | NM_001080430.3 | #N/A | 60.9302739 | -7.1348082 | 1.55367697 |

| | | | | | |
|---|---|---|---|---|---|
| 8686 | NM_001005861.2 | RYK | 87.3094217 | -7.6603969 | 1.66816641 |
| 8688 | NM_153264.6 | COL6A5 | 24.2506924 | 6.4325424 | 1.4008315 |
| 8687 | NM_001003787.2 | #N/A | 64.2843045 | -7.2169081 | 1.57164658 |
| 8689 | NM_001323631.1 | #N/A | 58.1036197 | -7.0723002 | 1.54016585 |
| 8690 | NM_022773.2 | #N/A | 60.9790552 | -7.1438462 | 1.5558665 |
| 8691 | NM_001040457.2 | RHBDD2 | 55.1685251 | -6.9929694 | 1.5230266 |
| 8692 | NM_198513.1 | PHF20L1 | 132.774899 | -6.9929253 | 1.52303837 |
| 8693 | NM_017847.5 | #N/A | 67.2200897 | -7.2839404 | 1.58657106 |
| 8694 | NM_138783.3 | #N/A | 60.6846784 | -7.1323573 | 1.55361605 |
| 8695 | NM_001014765.1 | FBXO44 | 52.006792 | -6.9099582 | 1.50527192 |
| 8696 | NM_001318477.1 | TSKU | 109.892149 | 1.68970715 | 0.36808903 |
| 8697 | NM_000980.3 | #N/A | 2510.85455 | -6.5066176 | 1.41742447 |
| 8698 | NM_013231.5 | #N/A | 69.9245277 | -7.3340023 | 1.59784722 |
| 8699 | NM_016195.3 | KIF20B | 55.9220462 | -7.0112324 | 1.52758965 |
| 8700 | NM_198183.2 | UBE2L6 | 84.6480949 | -7.6127867 | 1.65871615 |
| 8701 | NM_006978.2 | #N/A | 67.9109208 | -7.2939797 | 1.58936669 |
| 8702 | NM_001229.4 | #N/A | 56.0655043 | -7.0148384 | 1.52858313 |
| 8703 | NM_001243198.2 | #N/A | 20.1096644 | 9.31811475 | 2.03058299 |
| 8704 | NM_001303523.1 | #N/A | 20.1096644 | 9.31811475 | 2.03058299 |
| 8705 | NR_024388.1 | #N/A | 112.938666 | -5.4677858 | 1.19161069 |
| 8706 | NM_145255.3 | #N/A | 78.1096488 | -7.4973324 | 1.63400122 |
| 8707 | NM_001317078.1 | MED19 | 82.1208779 | -7.5694113 | 1.6497397 |
| 8708 | NM_001321155.1 | #N/A | 66.3622697 | -7.2629305 | 1.58295028 |
| 8709 | NM_001001994.2 | #N/A | 158.428253 | -7.2514582 | 1.58053473 |
| 8710 | NR_033997.1 | RNF144A-AS1 | 76.9060643 | 2.88950347 | 0.62984481 |
| 8711 | NM_001290187.1 | KRBA1 | 63.5436063 | -7.1990026 | 1.5692398 |
| 8712 | NM_001348121.1 | TRIM16 | 26.2877951 | 9.70959008 | 2.11654819 |
| 8713 | NM_001243953.1 | KIF2A | 74.1105042 | -7.4228683 | 1.61808912 |
| 8714 | NM_001261439.1 | #N/A | 41.6877401 | 3.98467188 | 0.86870199 |
| 8715 | NM_003437.4 | #N/A | 56.6738706 | -7.0354339 | 1.53380925 |
| 8716 | NM_003580.3 | #N/A | 57.8421517 | -7.066748 | 1.54070488 |
| 8717 | NM_032634.3 | #N/A | 56.3728433 | -7.0282707 | 1.53233258 |
| 8718 | NM_020832.2 | #N/A | 65.8930472 | -7.2570336 | 1.58236253 |
| 8719 | NR_028326.1 | #N/A | 56.3585244 | -7.0321792 | 1.53335636 |
| 8720 | NM_016546.3 | #N/A | 60.7057611 | -7.1382155 | 1.5566069 |
| 8721 | NM_001128126.2 | #N/A | 80.7237086 | -7.5451118 | 1.64544493 |
| 8722 | NM_001243042.1 | HLA-C | 101.367823 | -4.350289 | 0.94875119 |
| 8723 | NM_012291.4 | #N/A | 60.8909932 | -7.1393968 | 1.55704519 |
| 8724 | NM_032961.2 | #N/A | 158.761525 | -5.3769813 | 1.17271222 |
| 8725 | NM_001122764.1 | #N/A | 66.2774556 | -7.2601216 | 1.58351182 |
| 8727 | NM_018057.6 | SLC6A15 | 79.8182034 | -7.5268936 | 1.64171695 |
| 8726 | NM_001329429.1 | #N/A | 82.6082028 | -7.5780741 | 1.65287337 |
| 8728 | NM_021647.7 | #N/A | 63.6118402 | -7.19935 | 1.57034436 |

| | | | | | |
|---|---|---|---|---|---|
| 8729 | NM_005585.4 | #N/A | 59.7553506 | -7.1168708 | 1.55237052 |
| 8730 | NM_014957.2 | #N/A | 68.6094147 | -7.3102069 | 1.59455531 |
| 8731 | NM_001193465.1 | KANSL1 | 75.0872671 | -7.4433707 | 1.62362083 |
| 8732 | NM_001024678.3 | #N/A | 74.8884789 | -7.4364877 | 1.62236159 |
| 8733 | NM_145261.3 | #N/A | 57.2743402 | -7.0497532 | 1.53798724 |
| 8734 | NM_012437.5 | #N/A | 71.7791283 | -7.375753 | 1.60914472 |
| 8735 | NM_003216.3 | #N/A | 59.0373126 | -7.0906229 | 1.5471879 |
| 8736 | NM_002871.4 | #N/A | 82.3184601 | -7.5728936 | 1.65245967 |
| 8737 | NM_001076778.2 | #N/A | 97.2994739 | -7.8124511 | 1.7047503 |
| 8738 | NM_003876.2 | TMEM11 | 65.5384106 | -7.2465386 | 1.5813091 |
| 8739 | NM_001318740.1 | #N/A | 58.1610037 | 5.82761929 | 1.27181707 |
| 8740 | NM_001297659.1 | #N/A | 72.0191706 | -7.3803031 | 1.61069453 |
| 8741 | NM_138316.3 | #N/A | 96.2398121 | 3.1813602 | 0.69435166 |
| 8742 | NM_014243.2 | #N/A | 75.2020364 | -7.4413364 | 1.62463212 |
| 8743 | NM_001945.2 | #N/A | 84.8082454 | -7.6171016 | 1.66303605 |
| 8744 | NM_001142784.2 | IL11RA | 65.4542688 | -7.2434671 | 1.58157212 |
| 8745 | NM_001164326.1 | BRD8 | 59.9103641 | -7.1175834 | 1.55410943 |
| 8747 | NM_147184.3 | TP53I3 | 55.449738 | -7.006649 | 1.52990312 |
| 8746 | NM_053279.2 | #N/A | 56.571389 | -7.037822 | 1.53671 |
| 8748 | NM_057169.4 | #N/A | 57.2436875 | -7.0499389 | 1.53943994 |
| 8749 | NR_033787.2 | DNM1P41 | 133.700258 | -2.0055772 | 0.43794505 |
| 8750 | NM_014790.4 | #N/A | 75.8093631 | -7.4583819 | 1.62868424 |
| 8751 | NM_018102.4 | #N/A | 54.3084849 | -6.9690551 | 1.52182773 |
| 8752 | NM_001164213.1 | #N/A | 82.3275882 | -7.5720391 | 1.65366504 |
| 8753 | NM_001330665.1 | #N/A | 65.2787101 | -7.2413454 | 1.5814391 |
| 8754 | NM_020414.3 | #N/A | 812.557237 | -5.7484537 | 1.25542078 |
| 8755 | NM_017827.3 | #N/A | 59.755874 | -7.1099664 | 1.55282282 |
| 8756 | NM_001330170.1 | #N/A | 67.4850246 | -7.2871852 | 1.59162415 |
| 8757 | NM_003584.2 | DUSP11 | 54.0959267 | -6.9683112 | 1.52202736 |
| 8758 | NM_203389.2 | #N/A | 85.33567 | -7.6249213 | 1.66545442 |
| 8759 | NM_001286219.1 | TMEM132B | 33.0981399 | 10.0405007 | 2.1932148 |
| 8760 | NM_003656.4 | #N/A | 90.4588674 | -7.7093764 | 1.68406704 |
| 8761 | NM_018412.3 | #N/A | 82.9739148 | -7.5845719 | 1.65681637 |
| 8762 | NM_005921.1 | #N/A | 65.750796 | -5.972085 | 1.30462259 |
| 8763 | NM_203459.2 | #N/A | 151.22819 | -8.4501109 | 1.84604548 |
| 8764 | NM_001135564.1 | HSF2 | 119.772659 | -5.5072374 | 1.20322867 |
| 8765 | NM_015169.3 | #N/A | 66.4845628 | -7.2651543 | 1.58742978 |
| 8766 | NM_001199933.1 | SESN1 | 52.6964402 | -6.9339771 | 1.5151461 |
| 8767 | NM_012475.4 | #N/A | 77.7850225 | -7.4915094 | 1.63710885 |
| 8768 | NR_073007.1 | #N/A | 83.388478 | -7.5913782 | 1.65898533 |
| 8769 | NM_000240.3 | MAOA | 58.1750828 | -7.0754394 | 1.54626367 |
| 8770 | NR_038397.2 | DNM3OS | 80.1969497 | -7.5375362 | 1.64737115 |
| 8771 | NM_001134389.1 | #N/A | 61.6065132 | -7.1535722 | 1.5635273 |

| | | | | | |
|---|---|---|---|---|---|
| 8772 | NM_017882.2 | #N/A | 73.1211622 | -7.404112 | 1.6183025 |
| 8773 | NM_005147.5 | #N/A | 74.1392719 | -7.4240907 | 1.62269214 |
| 8774 | NM_020888.2 | #N/A | 121.999229 | -8.140981 | 1.77946096 |
| 8775 | NM_052886.2 | #N/A | 85.9261806 | -7.6334918 | 1.66855826 |
| 8776 | NM_025141.3 | TM2D3 | 90.5145344 | -7.7110712 | 1.68558874 |
| 8777 | NM_012202.4 | #N/A | 250.244505 | -6.3085895 | 1.37905679 |
| 8778 | NM_015480.2 | #N/A | 68.9712273 | -7.3183178 | 1.59982985 |
| 8779 | NM_016357.4 | #N/A | 55.1669725 | -6.9989997 | 1.53003373 |
| 8780 | NM_017668.2 | #N/A | 53.0088215 | -6.9379793 | 1.51681696 |
| 8781 | NM_021949.3 | ATP2B3 | 150.182681 | -8.4417606 | 1.84582219 |
| 8782 | NM_007104.4 | #N/A | 1447.86094 | -6.1630565 | 1.34758352 |
| 8783 | NM_152339.3 | #N/A | 79.1594468 | -6.1288787 | 1.34011877 |
| 8784 | NM_017661.3 | ZNF280D | 54.5370833 | -6.9837646 | 1.52726072 |
| 8785 | NM_001243733.1 | #N/A | 64.8315254 | -7.2318706 | 1.58170163 |
| 8786 | NM_000152.4 | #N/A | 60.0822088 | -7.117051 | 1.55661112 |
| 8787 | NM_004889.3 | #N/A | 72.5703485 | -7.3911126 | 1.61657593 |
| 8788 | NM_138570.3 | #N/A | 57.6918879 | -7.0616727 | 1.54461757 |
| 8789 | NM_001039614.2 | INSYN1 | 90.563628 | -7.7103407 | 1.68651296 |
| 8790 | NM_000553.4 | #N/A | 62.9584862 | -5.9052341 | 1.29184407 |
| 8791 | NM_182587.3 | #N/A | 56.4728331 | -7.0332621 | 1.53898998 |
| 8792 | NM_078480.2 | #N/A | 267.054049 | -5.671974 | 1.24128301 |
| 8793 | NM_021032.4 | FGF12 | 80.421657 | -7.5395479 | 1.65003356 |
| 8794 | NM_015074.3 | KIF1B | 1452.98839 | -3.7563442 | 0.82208895 |
| 8795 | NM_174889.4 | #N/A | 78.628957 | -7.5057446 | 1.64278307 |
| 8796 | NM_004083.5 | DDIT3 | 88.2232078 | -7.6725374 | 1.67982728 |
| 8797 | NM_032294.2 | #N/A | 72.7072557 | -7.391343 | 1.6182549 |
| 8798 | NM_016627.4 | #N/A | 90.3585429 | -7.706964 | 1.6874845 |
| 8799 | NM_001282602.1 | #N/A | 100.088799 | -7.8533807 | 1.71975698 |
| 8800 | NM_001300981.1 | #N/A | 55.7388136 | -7.0144172 | 1.53621767 |
| 8801 | NM_014180.3 | #N/A | 58.8816911 | -7.0905259 | 1.55288631 |
| 8802 | NM_018082.5 | #N/A | 69.35056 | -7.3255322 | 1.60435051 |
| 8803 | NM_018569.5 | #N/A | 94.4876516 | -7.7694021 | 1.70168566 |
| 8804 | NM_078474.2 | #N/A | 85.9473572 | -7.633372 | 1.67199127 |
| 8805 | NM_152932.2 | GLT8D1 | 63.6643935 | -7.2072883 | 1.57867599 |
| 8806 | NM_199337.2 | #N/A | 60.7378778 | -7.1371024 | 1.56330646 |
| 8807 | NM_001098811.1 | #N/A | 107.709455 | -7.9608621 | 1.74377689 |
| 8808 | NM_001278463.1 | DNM1L | 111.864607 | -8.0179453 | 1.75651531 |
| 8809 | NM_022551.2 | #N/A | 3227.55528 | -6.743068 | 1.47724747 |
| 8810 | NM_003864.3 | #N/A | 74.2887811 | -7.4237894 | 1.62641594 |
| 8811 | NM_006288.4 | #N/A | 107.916416 | -7.9639458 | 1.74474626 |
| 8812 | NM_017734.4 | #N/A | 78.7551154 | -7.5109182 | 1.6459757 |
| 8813 | NM_139076.2 | #N/A | 55.310814 | -6.9999607 | 1.5340323 |
| 8814 | NM_001204214.1 | NOS1 | 104.035849 | 2.1552012 | 0.47234101 |

| | | | | | |
|---|---|---|---|---|---|
| 8815 | NM_018066.3 | #N/A | 61.0961994 | -7.144154 | 1.56579451 |
| 8816 | NM_024928.4 | #N/A | 64.0311673 | -7.2144238 | 1.581302 |
| 8817 | NM_024942.3 | #N/A | 64.6215169 | -7.2231372 | 1.58331846 |
| 8818 | NM_198207.2 | #N/A | 64.2119123 | -7.2171944 | 1.58201373 |
| 8819 | NM_012099.1 | #N/A | 216.01445 | 2.41186645 | 0.52873702 |
| 8820 | NM_025232.3 | #N/A | 55.4866066 | -7.0072272 | 1.53622301 |
| 8821 | NM_018958.2 | #N/A | 42.1615216 | 2.75998764 | 0.60509072 |
| 8822 | NM_130843.3 | #N/A | 97.1150729 | -7.813385 | 1.71310263 |
| 8823 | NM_025189.3 | #N/A | 59.096647 | -7.0995345 | 1.55659928 |
| 8824 | NM_001130825.1 | SUPT5H | 97.1543305 | -7.8129399 | 1.71315533 |
| 8825 | NM_024664.3 | #N/A | 72.3518462 | -7.3842857 | 1.61917516 |
| 8826 | NR_135198.1 | LOC729652 | 147.264774 | 2.38507838 | 0.52309709 |
| 8827 | NM_020365.4 | #N/A | 85.3116126 | -7.6242148 | 1.67223634 |
| 8828 | NM_001822.5 | #N/A | 73.55137 | -7.4127079 | 1.62590456 |
| 8829 | NM_022478.3 | CDH24 | 71.5542254 | -7.3694354 | 1.61649914 |
| 8830 | NM_033419.4 | #N/A | 63.5542079 | -7.2021129 | 1.57980271 |
| 8831 | NM_001283044.2 | #N/A | 72.0487163 | -7.37686 | 1.61821965 |
| 8832 | NM_015478.6 | #N/A | 50.2269537 | -6.8627061 | 1.50544667 |
| 8833 | NM_004440.3 | #N/A | 79.9112028 | -7.533409 | 1.65272452 |
| 8834 | NM_001083617.1 | RB1CC1 | 164.858847 | -7.0592432 | 1.5487128 |
| 8835 | NM_024787.3 | RNF122 | 87.3874288 | -6.3829479 | 1.40040626 |
| 8836 | NM_002836.3 | PTPRA | 71.5615896 | -7.3706029 | 1.61712152 |
| 8837 | NM_203344.2 | #N/A | 57.3214383 | -7.0569066 | 1.54837652 |
| 8838 | NM_018320.4 | #N/A | 75.3997378 | -7.4471006 | 1.63406747 |
| 8839 | NM_001031801.1 | #N/A | 64.7762984 | -7.2274018 | 1.5858899 |
| 8840 | NM_001130058.1 | #N/A | 83.0995008 | -7.5865828 | 1.66493696 |
| 8841 | NM_203290.3 | #N/A | 60.3860186 | -7.1251873 | 1.56371403 |
| 8842 | NM_001321312.1 | #N/A | 66.4448943 | -7.2602404 | 1.5934073 |
| 8843 | NM_030647.1 | #N/A | 98.8277826 | -6.4069418 | 1.40618067 |
| 8844 | NM_030791.3 | #N/A | 71.2789419 | -7.3651254 | 1.61691004 |
| 8845 | NM_052941.4 | #N/A | 147.876237 | 3.44443949 | 0.75619135 |
| 8846 | NM_206925.2 | CA12 | 82.1058137 | -7.5685809 | 1.66164375 |
| 8847 | NM_023940.2 | #N/A | 80.4915083 | -7.5394776 | 1.65537407 |
| 8848 | NM_016116.2 | #N/A | 43.8737361 | 5.25292457 | 1.15338952 |
| 8849 | NM_017522.4 | #N/A | 87.140535 | -6.2458621 | 1.37145599 |
| 8850 | NM_001271675.1 | LOC441155 | 30.6294579 | 6.19681609 | 1.3607069 |
| 8851 | NM_001033505.1 | CSTF3 | 56.1263677 | -7.0204449 | 1.54160589 |
| 8852 | NM_014167.4 | #N/A | 64.2198253 | -7.2122951 | 1.58379494 |
| 8853 | NM_022114.3 | #N/A | 145.36123 | -8.3947819 | 1.84348166 |
| 8854 | NM_014018.2 | #N/A | 59.6185836 | -7.1072888 | 1.56078335 |
| 8855 | NR_134562.1 | #N/A | 53.5358845 | -6.9512511 | 1.52652991 |
| 8856 | NM_022006.1 | #N/A | 87.6768373 | -7.6654206 | 1.68343555 |
| 8857 | NM_001161780.1 | #N/A | 106.781658 | -7.9487449 | 1.74572188 |

| | | | | | |
|---|---|---|---|---|---|
| 8858 | NM_012392.3 | #N/A | 76.1433168 | -7.4594661 | 1.6383695 |
| 8859 | NM_207363.2 | #N/A | 52.2718447 | -6.9175763 | 1.51948806 |
| 8860 | NM_194279.3 | #N/A | 58.9859057 | -7.0920963 | 1.55807046 |
| 8861 | NM_003793.3 | #N/A | 90.7636368 | -3.9576494 | 0.86958234 |
| 8862 | NM_014595.2 | #N/A | 59.9595861 | -7.1166916 | 1.56368708 |
| 8863 | NM_018263.4 | #N/A | 53.9909432 | -6.9700777 | 1.53152848 |
| 8864 | NM_018899.5 | #N/A | 89.3339266 | -7.6903444 | 1.68979855 |
| 8865 | NM_001319674.1 | #N/A | 54.6801874 | -6.9776302 | 1.53326615 |
| 8866 | NM_172315.2 | #N/A | 210.957807 | -5.1560981 | 1.13301348 |
| 8867 | NM_001303095.1 | ISG20L2 | 250.965659 | -1.4439927 | 0.31731636 |
| 8868 | NM_001321463.1 | #N/A | 145.322618 | -8.3930586 | 1.84458147 |
| 8869 | NM_178448.3 | #N/A | 77.4501109 | -7.4829936 | 1.64475964 |
| 8870 | NM_139078.2 | #N/A | 88.864252 | -7.683814 | 1.68893589 |
| 8871 | NR_026874.2 | LINC02593 | 78.6098257 | -7.5083995 | 1.65077876 |
| 8872 | NM_207320.2 | #N/A | 28.7176902 | 3.47318059 | 0.76361547 |
| 8873 | NM_006153.5 | #N/A | 68.5763594 | -7.3106641 | 1.60739201 |
| 8874 | NM_002067.4 | #N/A | 470.140085 | -4.9380643 | 1.08576434 |
| 8875 | NM_001330323.1 | HOXB3 | 74.637191 | -7.434475 | 1.63469499 |
| 8876 | NM_001130137.1 | ISLR2 | 49.9329957 | -6.8525817 | 1.50679216 |
| 8877 | NM_016283.4 | #N/A | 61.5532174 | -7.1547928 | 1.57339399 |
| 8878 | NM_017634.3 | #N/A | 56.4127934 | -7.0294486 | 1.54586438 |
| 8879 | NM_019896.2 | #N/A | 84.0424579 | -7.6028194 | 1.67210844 |
| 8880 | NR_144509.1 | #N/A | 75.6576829 | -7.4531552 | 1.63920754 |
| 8881 | NM_145691.3 | #N/A | 57.8681441 | -7.0635289 | 1.55355317 |
| 8882 | NM_002192.3 | #N/A | 142.937045 | -6.0516869 | 1.33113503 |
| 8883 | NM_147128.3 | #N/A | 54.1256385 | -6.9655696 | 1.53215666 |
| 8884 | NM_001004105.2 | #N/A | 63.4054028 | -7.195897 | 1.58284339 |
| 8885 | NM_001744.5 | #N/A | 139.689433 | -8.3346329 | 1.83336795 |
| 8886 | NM_021222.2 | #N/A | 64.3450954 | -7.2217751 | 1.58860564 |
| 8887 | NM_024955.5 | #N/A | 112.895245 | -8.0313719 | 1.76672917 |
| 8888 | NR_132421.1 | COX20 | 18.7381127 | 9.21056453 | 2.02614048 |
| 8889 | NM_031220.3 | #N/A | 62.4600072 | -7.1772505 | 1.57890621 |
| 8890 | NM_138328.2 | #N/A | 67.6133096 | -7.2901618 | 1.60383311 |
| 8891 | NM_003440.3 | ZNF140 | 60.5840486 | -7.1275367 | 1.56812773 |
| 8892 | NM_013401.3 | #N/A | 55.8664039 | -7.0198163 | 1.54447213 |
| 8893 | NM_005206.4 | #N/A | 215.514155 | -3.1249668 | 0.68756726 |
| 8894 | NM_024065.4 | #N/A | 64.76909 | -7.2272182 | 1.59017337 |
| 8895 | NM_014339.6 | #N/A | 64.5634888 | -7.2214034 | 1.58894993 |
| 8896 | NM_001128224.2 | TMCC1 | 81.0091202 | -7.5502551 | 1.66133262 |
| 8897 | NM_152601.3 | #N/A | 66.4335148 | -5.8995969 | 1.29818473 |
| 8898 | NM_016565.2 | #N/A | 69.4755324 | -7.3291788 | 1.61282807 |
| 8899 | NM_032378.5 | #N/A | 68.6185079 | -7.3119675 | 1.60913977 |
| 8900 | NM_014978.2 | #N/A | 81.5489602 | -7.5594118 | 1.6636169 |

| | | | | | |
|---|---|---|---|---|---|
| 8901 | NM_002916.3 | #N/A | 74.2815434 | -7.4241612 | 1.63389517 |
| 8902 | NM_003384.2 | #N/A | 94.7177567 | -7.7748917 | 1.71120437 |
| 8903 | NM_174891.3 | #N/A | 50.8872009 | -6.8836831 | 1.51506752 |
| 8904 | NM_203374.1 | #N/A | 52.2050307 | -6.9145697 | 1.52191414 |
| 8906 | NM_133475.1 | ANKRD24 | 69.0532806 | -7.3184964 | 1.61084573 |
| 8905 | NM_016408.3 | #N/A | 69.1782443 | -7.3187296 | 1.61089854 |
| 8907 | NM_000237.2 | #N/A | 119.464145 | -8.1095002 | 1.78496249 |
| 8908 | NM_033120.3 | #N/A | 52.0443538 | -6.9146364 | 1.5220351 |
| 8909 | NM_017910.3 | #N/A | 73.1694053 | -7.4034985 | 1.62966774 |
| 8910 | NM_018130.2 | #N/A | 51.1906088 | -6.8904216 | 1.51681663 |
| 8911 | NR_037599.1 | PDCD6IPP2 | 59.6857772 | 2.77877253 | 0.61171309 |
| 8912 | NM_024562.1 | #N/A | 64.1742453 | -7.2130108 | 1.58788107 |
| 8913 | NM_014628.2 | #N/A | 68.2631714 | -7.30183 | 1.60745459 |
| 8914 | NM_001321690.1 | BZW1 | 99.6773196 | -7.8483624 | 1.72779119 |
| 8915 | NR_132355.1 | BLOC1S6 | 67.3127629 | -7.2842808 | 1.60366619 |
| 8916 | NM_005932.3 | #N/A | 54.3297285 | -6.9773854 | 1.53611474 |
| 8917 | NM_001256545.1 | #N/A | 72.8112525 | -7.3992491 | 1.62902183 |
| 8918 | NM_001321146.1 | #N/A | 54.4102446 | -6.9726314 | 1.53521043 |
| 8919 | NM_001947.3 | #N/A | 98.5301441 | -6.4016512 | 1.40949142 |
| 8921 | NM_052910.2 | SLITRK1 | 90.2835106 | -7.7055568 | 1.69663287 |
| 8920 | NM_015250.3 | #N/A | 73.7680487 | -7.4181567 | 1.63336017 |
| 8922 | NM_003870.3 | #N/A | 355.833944 | -1.5977625 | 0.35182373 |
| 8923 | NR_028040.1 | CASTOR3 | 81.9778003 | -7.5649759 | 1.66583229 |
| 8924 | NM_177533.4 | #N/A | 76.7726705 | -7.4711733 | 1.64545462 |
| 8925 | NM_001207051.1 | #N/A | 74.1074863 | -7.4243341 | 1.63518362 |
| 8926 | NM_001004318.2 | #N/A | 55.6623477 | 2.00190115 | 0.44095183 |
| 8927 | NM_001168300.1 | KLHL13 | 89.9721659 | -7.7026806 | 1.69685951 |
| 8928 | NM_005007.3 | #N/A | 67.2073589 | -7.2790982 | 1.60365999 |
| 8929 | NM_017626.4 | #N/A | 64.1505492 | -7.2135314 | 1.58927824 |
| 8930 | NM_018463.3 | #N/A | 62.6687908 | -7.1790684 | 1.581718 |
| 8931 | NM_174905.3 | FAM98C | 71.364835 | -7.3656975 | 1.6229054 |
| 8932 | NM_001085375.1 | #N/A | 51.9979823 | -6.9139616 | 1.52339339 |
| 8933 | NM_001206559.1 | #N/A | 64.3456806 | -7.2144504 | 1.58975185 |
| 8934 | NM_001320464.1 | #N/A | 71.2887044 | -7.3674578 | 1.6235641 |
| 8935 | NM_145265.2 | #N/A | 60.7353573 | -7.1347474 | 1.57227409 |
| 8936 | NM_001079843.2 | #N/A | 67.1516493 | -5.914479 | 1.30347122 |
| 8937 | NM_019065.2 | NECAB2 | 63.7308462 | -7.2054332 | 1.5880113 |
| 8938 | NM_152653.3 | #N/A | 90.6185445 | -6.2953224 | 1.38756815 |
| 8939 | NM_002269.2 | #N/A | 58.7953466 | -7.0865527 | 1.56202979 |
| 8940 | NM_001330111.1 | #N/A | 64.6654961 | -7.2276234 | 1.59323015 |
| 8941 | NM_203318.1 | #N/A | 68.0060786 | -7.2983455 | 1.6089571 |
| 8942 | NM_014666.3 | #N/A | 88.1637619 | -7.6757772 | 1.69220873 |
| 8943 | NM_001201483.1 | #N/A | 59.2290148 | -7.0997097 | 1.56529582 |

| | | | | | |
|---|---|---|---|---|---|
| 8944 | NM_015302.1 | #N/A | 162.949799 | -1.9310178 | 0.42580029 |
| 8945 | NM_013312.2 | #N/A | 78.6479519 | -7.5053396 | 1.65506038 |
| 8946 | NM_006388.3 | #N/A | 57.9442118 | -7.0671298 | 1.55849987 |
| 8947 | NM_053005.5 | MOB2 | 66.8795111 | -7.2722205 | 1.60374215 |
| 8948 | NM_020188.3 | #N/A | 86.8865706 | -7.6503768 | 1.6871506 |
| 8949 | NM_178496.3 | #N/A | 76.7906562 | -7.472846 | 1.64801219 |
| 8950 | NM_054013.3 | MGAT4B | 53.4485904 | -6.9488529 | 1.53246472 |
| 8951 | NM_206909.2 | #N/A | 94.0025162 | -7.7656598 | 1.71263813 |
| 8952 | NM_014717.2 | #N/A | 49.1597347 | -6.8283656 | 1.50602365 |
| 8953 | NM_001316311.1 | #N/A | 71.3095366 | -7.3673479 | 1.62497018 |
| 8954 | NM_007200.4 | #N/A | 72.6234805 | -7.3949959 | 1.63128302 |
| 8955 | NM_001321022.1 | #N/A | 21.3863887 | 9.38385584 | 2.07022768 |
| 8956 | NM_001079539.1 | XBP1 | 73.3876851 | -7.4059096 | 1.63396146 |
| 8957 | NM_004794.2 | #N/A | 89.280042 | -7.6903369 | 1.69679853 |
| 8958 | NM_181861.1 | #N/A | 80.5416773 | -7.5387252 | 1.66335477 |
| 8959 | NR_040582.2 | STAG3L3 | 52.313002 | -6.9170413 | 1.52624255 |
| 8960 | NM_001278376.2 | #N/A | 95.199817 | -6.51258 | 1.43700715 |
| 8961 | NM_018474.4 | #N/A | 49.4115573 | -6.8351403 | 1.50819745 |
| 8962 | NM_018317.3 | #N/A | 57.6716312 | -7.057844 | 1.55735885 |
| 8963 | NM_001252065.1 | SYT7 | 88.3042901 | -7.6753414 | 1.69381502 |
| 8964 | NM_001253380.1 | #N/A | 86.9025338 | -7.6546821 | 1.68948571 |
| 8965 | NM_025152.2 | #N/A | 52.6877841 | -6.9351571 | 1.5306812 |
| 8966 | NM_015945.11 | #N/A | 82.9049366 | -7.5839625 | 1.67395347 |
| 8967 | NM_001079669.2 | #N/A | 101.207404 | -7.8722878 | 1.73773482 |
| 8968 | NM_001039673.2 | #N/A | 118.503367 | -8.0995109 | 1.78790931 |
| 8969 | NM_152789.3 | #N/A | 80.878066 | -7.5445455 | 1.66549349 |
| 8970 | NM_001242916.1 | ZFAND6 | 81.6864093 | -7.5602265 | 1.66919823 |
| 8971 | NM_153636.2 | #N/A | 92.0214511 | -7.7347229 | 1.70771958 |
| 8972 | NM_133328.3 | #N/A | 57.7421933 | -7.0610128 | 1.55907796 |
| 8973 | NM_017599.3 | #N/A | 181.071987 | -7.1722989 | 1.58381589 |
| 8974 | NM_005641.3 | #N/A | 70.4419352 | -7.3519191 | 1.62355323 |
| 8975 | NM_138493.2 | #N/A | 76.7819106 | -7.4719552 | 1.6500773 |
| 8976 | NM_145119.3 | #N/A | 64.5213507 | -7.223475 | 1.59524619 |
| 8977 | NM_172014.3 | TNFSF14 | 86.5984161 | 2.9665687 | 0.65522479 |
| 8978 | NM_173467.4 | #N/A | 56.4198479 | -7.0274909 | 1.55220153 |
| 8979 | NM_001042413.1 | #N/A | 52.018837 | -6.9107198 | 1.52663955 |
| 8980 | NM_053277.2 | #N/A | 280.791318 | -9.3443458 | 2.06425346 |
| 8981 | NM_001278728.1 | RFXANK | 58.7880125 | -7.0880308 | 1.5658623 |
| 8982 | NM_001082537.2 | #N/A | 57.6877043 | -7.0583659 | 1.55934223 |
| 8983 | NM_173540.2 | #N/A | 54.0852851 | -6.9672267 | 1.53942515 |
| 8984 | NM_001080488.1 | #N/A | 95.4726201 | -7.7881284 | 1.72095235 |
| 8985 | NM_002412.4 | #N/A | 54.0941278 | -6.9686953 | 1.53992888 |
| 8986 | NM_002931.3 | #N/A | 295.701737 | -5.3053955 | 1.17243608 |

| | | | | | |
|---|---|---|---|---|---|
| 8987 | NM_006558.2 | #N/A | 203.02504 | -7.3170556 | 1.61701489 |
| 8988 | NM_015102.4 | #N/A | 61.4057094 | -7.1520363 | 1.58057577 |
| 8989 | NM_001256523.1 | #N/A | 79.2556472 | -7.5206972 | 1.66213176 |
| 8990 | NM_025155.2 | #N/A | 62.6302361 | -7.1772377 | 1.58621882 |
| 8991 | NM_001284513.1 | #N/A | 62.4522047 | -7.1728667 | 1.58527942 |
| 8992 | NR_073502.1 | #N/A | 83.8101189 | -7.601787 | 1.68007777 |
| 8993 | NM_016175.3 | #N/A | 49.9803559 | -6.852887 | 1.51459791 |
| 8994 | NM_006375.3 | ENOX2 | 58.2624537 | -7.0740241 | 1.56351984 |
| 8995 | NM_006087.3 | #N/A | 113.808385 | -8.0407809 | 1.77722443 |
| 8996 | NM_001308325.1 | AKTIP | 70.0999219 | -7.3409404 | 1.62260798 |
| 8997 | NM_001301699.1 | ERI3 | 72.1861226 | -7.3851725 | 1.63268091 |
| 8998 | NM_001348015.1 | ZNF83 | 107.791804 | -5.4014679 | 1.1942724 |
| 9000 | NM_006676.7 | USP20 | 69.6964369 | -7.3346599 | 1.62173345 |
| 8999 | NM_001006605.4 | #N/A | 69.7676143 | -7.3344986 | 1.62169519 |
| 9001 | NM_001003700.1 | #N/A | 19.2509076 | 9.25429773 | 2.04648664 |
| 9002 | NM_173555.3 | #N/A | 50.8541148 | -6.8741971 | 1.52038842 |
| 9003 | NM_001102575.1 | #N/A | 136.075335 | -3.3604835 | 0.74326997 |
| 9004 | NM_001297741.1 | BTBD10 | 84.6808838 | -7.6153021 | 1.68437391 |
| 9005 | NM_017983.6 | #N/A | 68.3261026 | -7.308799 | 1.61661842 |
| 9006 | NM_001014446.2 | #N/A | 88.1526414 | -7.6731819 | 1.69726326 |
| 9008 | NM_017757.2 | ZNF407 | 53.2212678 | -6.9467318 | 1.53663386 |
| 9007 | NM_001080209.1 | #N/A | 55.5042231 | -7.0039254 | 1.5492799 |
| 9009 | NM_021101.4 | #N/A | 106.210038 | -7.9430336 | 1.75722441 |
| 9010 | NM_001319121.1 | DNPEP | 66.5277548 | -7.2677395 | 1.60786868 |
| 9011 | NM_001277324.1 | NPIPA2 | 61.9335867 | -7.1595145 | 1.58395139 |
| 9012 | NM_001083914.2 | #N/A | 77.394895 | -7.483723 | 1.65577058 |
| 9014 | NM_025251.2 | ARHGAP39 | 65.9267 | -7.256573 | 1.60566816 |
| 9013 | NM_002690.2 | #N/A | 113.727842 | -6.7688454 | 1.49774071 |
| 9015 | NM_001278442.1 | ILK | 56.04898 | -7.0202376 | 1.55354308 |
| 9016 | NM_001271507.1 | #N/A | 54.3729119 | -6.9694368 | 1.54243667 |
| 9017 | NM_001077525.2 | #N/A | 60.0338514 | -7.116155 | 1.57494019 |
| 9018 | NM_015155.2 | #N/A | 268.314686 | -3.6717241 | 0.81264392 |
| 9019 | NR_103715.1 | LINC00269 | 58.8787577 | 2.99742659 | 0.66342966 |
| 9020 | NM_153208.2 | IQCK | 79.8190225 | -7.5266033 | 1.66598166 |
| 9021 | NM_001111125.2 | #N/A | 72.3545141 | -7.3852958 | 1.6347279 |
| 9022 | NM_145690.2 | #N/A | 2544.97309 | -5.5782147 | 1.2347394 |
| 9023 | NM_001145426.1 | #N/A | 57.5492856 | -7.0617654 | 1.56318333 |
| 9024 | NM_014205.3 | #N/A | 56.6926152 | -7.0358615 | 1.55745543 |
| 9026 | NM_057167.3 | COL6A3 | 100.702268 | -7.8663855 | 1.74145457 |
| 9025 | NM_001142273.1 | #N/A | 154.808059 | -8.4854035 | 1.87849368 |
| 9027 | NM_001278719.1 | SV2A | 73.5846338 | -7.4135735 | 1.64126135 |
| 9028 | NM_201222.2 | #N/A | 71.9018598 | -7.3828432 | 1.63446947 |
| 9029 | NM_001039590.2 | #N/A | 60.0289881 | -7.1172819 | 1.57576414 |

| | | | | | |
|---|---|---|---|---|---|
| 9030 | NM_001145023.1 | #N/A | 53.1998753 | -6.9480184 | 1.53839728 |
| 9031 | NM_005012.3 | #N/A | 56.1634587 | -7.0254707 | 1.55560275 |
| 9032 | NM_020205.3 | #N/A | 68.3971566 | -5.936201 | 1.31442394 |
| 9033 | NM_003721.3 | #N/A | 64.8636157 | -7.2284943 | 1.60058003 |
| 9034 | NM_014346.3 | #N/A | 59.1998129 | -7.0976326 | 1.57161481 |
| 9035 | NM_004071.3 | #N/A | 80.6769114 | -7.5439236 | 1.67067978 |
| 9036 | NM_006759.3 | UGP2 | 88.5568494 | -7.6800351 | 1.70093825 |
| 9037 | NM_025103.2 | #N/A | 56.0327482 | -7.0197883 | 1.55480173 |
| 9038 | NM_001260492.1 | RDX | 77.1088597 | -7.4791945 | 1.65662121 |
| 9039 | NM_015693.3 | #N/A | 52.1709822 | -6.9172083 | 1.53214419 |
| 9040 | NM_001018136.2 | #N/A | 82.5940274 | -7.5783432 | 1.6786942 |
| 9041 | NM_019053.5 | #N/A | 59.7814899 | -7.1114642 | 1.5752845 |
| 9042 | NM_057159.2 | #N/A | 54.1055611 | -6.9689372 | 1.54379556 |
| 9043 | NR_002909.2 | SNHG3 | 63.1508879 | -7.1938769 | 1.59369335 |
| 9044 | NM_004280.4 | #N/A | 70.2125823 | -7.3439089 | 1.6269715 |
| 9045 | NM_015166.3 | MLC1 | 98.6797261 | -7.8345588 | 1.73569241 |
| 9046 | NM_001308154.1 | #N/A | 162.479915 | -8.5546802 | 1.89525953 |
| 9047 | NM_032689.4 | #N/A | 52.9472356 | -6.936201 | 1.53672858 |
| 9048 | NM_006806.4 | #N/A | 109.728055 | -5.3944916 | 1.19521612 |
| 9049 | NM_001495.4 | #N/A | 63.839694 | -7.2057674 | 1.59655661 |
| 9050 | NM_001291285.1 | FAT4 | 66.0092555 | -7.2562847 | 1.60798852 |
| 9051 | NM_001031677.3 | #N/A | 66.9068055 | -7.2758519 | 1.61238585 |
| 9052 | NM_001035258.1 | #N/A | 88.9620624 | -7.6858978 | 1.70327336 |
| 9053 | NM_032802.3 | #N/A | 55.2879246 | -7.0022068 | 1.55175742 |
| 9054 | NM_001135570.1 | #N/A | 84.6909347 | -7.6136715 | 1.68728535 |
| 9055 | NM_016050.4 | #N/A | 60.2665427 | -7.1218147 | 1.57828871 |
| 9056 | NM_001321116.1 | #N/A | 13.2302209 | -18.987232 | 4.20792815 |
| 9057 | NR_045072.1 | #N/A | 103.390026 | -7.901846 | 1.75154683 |
| 9058 | NM_017941.5 | C17orf80 | 72.8100267 | -7.3925199 | 1.63889909 |
| 9059 | NM_032630.2 | #N/A | 60.7735777 | -7.1368406 | 1.58250916 |
| 9060 | NM_001122636.1 | #N/A | 86.6528141 | -7.647475 | 1.69583076 |
| 9061 | NM_023933.2 | #N/A | 69.9918443 | -7.3381182 | 1.62725076 |
| 9062 | NM_004289.6 | #N/A | 83.8114553 | -6.1930135 | 1.37334837 |
| 9063 | NR_027948.2 | #N/A | 117.546656 | -8.0848493 | 1.7928753 |
| 9064 | NR_109997.1 | PRKAR2A-AS1 | 415.314919 | 2.55555606 | 0.56673124 |
| 9065 | NM_014044.6 | UNC50 | 71.82867 | -7.3765125 | 1.63587536 |
| 9066 | NM_001139498.1 | #N/A | 101.904814 | -7.8806726 | 1.7477047 |
| 9067 | NM_020904.2 | #N/A | 72.9890802 | -7.402786 | 1.64211347 |
| 9071 | NR_110745.1 | LINC01858 | 52.8864981 | 2.92382 | 0.64859782 |
| 9070 | NM_007150.3 | ZNF185 | 13.0384786 | -18.968904 | 4.20795564 |
| 9068 | NM_001005747.3 | #N/A | 13.0384786 | -18.968904 | 4.20795564 |
| 9069 | NM_002840.4 | #N/A | 100.895556 | -7.8644788 | 1.74460251 |
| 9072 | NM_024337.3 | #N/A | 69.8404333 | -7.3387172 | 1.62818841 |

| | | | | | |
|---|---|---|---|---|---|
| 9073 | NM_013262.3 | #N/A | 54.6273513 | -6.984064 | 1.54957085 |
| 9074 | NM_144563.2 | #N/A | 58.8023459 | -7.0886977 | 1.57283264 |
| 9075 | NM_001099696.2 | REPIN1 | 85.9259839 | -7.6382776 | 1.69480188 |
| 9076 | NM_003983.5 | #N/A | 104.70808 | -7.9224635 | 1.7580383 |
| 9077 | NM_194298.2 | SLC16A9 | 105.434185 | -7.932797 | 1.76047501 |
| 9078 | NM_001127320.2 | UBAP2L | 235.55812 | -4.0431848 | 0.89728585 |
| 9079 | NM_003272.3 | #N/A | 59.7883335 | -7.111528 | 1.57823947 |
| 9080 | NM_032758.3 | #N/A | 95.7475785 | -6.359052 | 1.41131212 |
| 9081 | NR_026865.1 | #N/A | 63.6146377 | -7.2004772 | 1.59806468 |
| 9082 | NM_001135036.1 | TVP23C | 187.34644 | 1.96793439 | 0.43678654 |
| 9083 | NM_001185082.1 | #N/A | 49.8226172 | -6.8459956 | 1.51981617 |
| 9085 | NM_001243657.1 | NUDT7 | 55.1350756 | 2.68567092 | 0.59624273 |
| 9084 | NM_000052.6 | #N/A | 61.0487457 | -7.147065 | 1.58670469 |
| 9086 | NM_001500.3 | #N/A | 71.2303402 | -7.3647386 | 1.63525046 |
| 9087 | NM_001166426.2 | #N/A | 49.7758606 | -6.8508633 | 1.52116579 |
| 9088 | NM_152499.2 | #N/A | 47.8338261 | -6.7907235 | 1.50782607 |
| 9089 | NM_000824.4 | #N/A | 68.985015 | -7.3170582 | 1.6250224 |
| 9090 | NM_001283053.1 | #N/A | 72.9775905 | -7.3987304 | 1.64328091 |
| 9091 | NM_152296.4 | #N/A | 1923.22287 | -6.3995687 | 1.42138967 |
| 9092 | NM_001146175.1 | #N/A | 77.2116435 | -7.4805881 | 1.66151783 |
| 9093 | NM_032178.2 | #N/A | 54.7153732 | -6.9832961 | 1.55115971 |
| 9094 | NM_001252053.1 | PCMT1 | 66.1312824 | -7.2609153 | 1.61289561 |
| 9095 | NM_006232.4 | POLR2H | 59.0062984 | -7.0954747 | 1.57616429 |
| 9096 | NM_002039.3 | #N/A | 71.2299759 | -5.9830849 | 1.32912271 |
| 9097 | NR_037941.1 | #N/A | 77.5396559 | -7.4907402 | 1.6641056 |
| 9098 | NM_181723.2 | #N/A | 74.6263915 | -7.4319771 | 1.65107045 |
| 9099 | NM_138356.2 | #N/A | 82.2800689 | -7.5730358 | 1.6824314 |
| 9100 | NM_001282736.1 | #N/A | 18.2253178 | 9.16491504 | 2.0361004 |
| 9101 | NM_145887.3 | #N/A | 62.0417786 | -7.1646925 | 1.59179308 |
| 9102 | NM_002915.3 | #N/A | 59.4436817 | -7.1009813 | 1.57766441 |
| 9103 | NM_001242840.1 | #N/A | 78.8225574 | -7.5114595 | 1.66920404 |
| 9104 | NM_001331078.1 | #N/A | 48.759875 | -6.8168178 | 1.51483695 |
| 9105 | NM_001001331.2 | #N/A | 158.170817 | -8.5167552 | 1.89292994 |
| 9106 | NM_001199340.1 | #N/A | 54.5865128 | -6.976639 | 1.55073682 |
| 9107 | NM_171998.3 | #N/A | 96.5896865 | -5.7598297 | 1.28026895 |
| 9108 | NM_001322215.1 | #N/A | 57.8564047 | -7.0666546 | 1.57083631 |
| 9109 | NM_016498.4 | #N/A | 51.2287469 | -6.8865442 | 1.53091638 |
| 9110 | NM_001009921.2 | VPS8 | 131.357701 | -8.2466476 | 1.83332895 |
| 9111 | NM_182642.2 | #N/A | 68.5066463 | -7.311692 | 1.62566569 |
| 9112 | NR_040104.1 | NAXD | 59.4128475 | -7.1012675 | 1.57896343 |
| 9113 | NM_001258392.2 | #N/A | 64.845835 | -7.2278262 | 1.60720511 |
| 9114 | NM_005360.4 | #N/A | 61.5721821 | -7.1590272 | 1.59201461 |
| 9115 | NM_001145939.1 | PAFAH1B3 | 103.630194 | -7.9045115 | 1.75790665 |

| | | | | | |
|---|---|---|---|---|---|
| 9116 | NM_001789.2 | #N/A | 61.0036335 | -7.1394113 | 1.58779259 |
| 9117 | NM_012154.3 | #N/A | 47.7386421 | -6.7855038 | 1.50923573 |
| 9118 | NM_006612.5 | #N/A | 419.695954 | -1.7730784 | 0.39438109 |
| 9119 | NM_020806.4 | GPHN | 55.4738725 | -7.0031505 | 1.55777531 |
| 9120 | NM_001135556.1 | DYNC1I1 | 110.264341 | -7.9952468 | 1.77860513 |
| 9121 | NM_052963.2 | #N/A | 66.9588751 | -7.2747367 | 1.61834636 |
| 9123 | NM_001142471.1 | INTS12 | 62.2083423 | -7.1679081 | 1.59463988 |
| 9122 | NM_001122842.2 | #N/A | 75.4221453 | -7.4457705 | 1.65645001 |
| 9124 | NM_001303457.1 | #N/A | 109.103136 | -5.382024 | 1.19740858 |
| 9125 | NM_001318860.1 | #N/A | 75.5434383 | -7.4489807 | 1.65732885 |
| 9126 | NM_017454.3 | STAU1 | 60.3036869 | -7.126963 | 1.58578622 |
| 9127 | NM_001605.2 | AARS1 | 754.2161 | -5.6398913 | 1.25505002 |
| 9128 | NM_014078.5 | #N/A | 70.1447904 | -7.3412604 | 1.6336622 |
| 9129 | NM_014421.2 | #N/A | 56.5006567 | -7.0361191 | 1.56577118 |
| 9130 | NM_018663.2 | #N/A | 65.5144001 | -7.2426844 | 1.61173089 |
| 9131 | NM_183065.2 | #N/A | 49.2710484 | -6.8301395 | 1.51993375 |
| 9132 | NR_036434.1 | #N/A | 60.6834839 | -7.1315548 | 1.58707578 |
| 9133 | NM_017998.2 | #N/A | 73.5659418 | -7.4106809 | 1.64930795 |
| 9134 | NM_012330.3 | #N/A | 61.1842365 | -7.1415807 | 1.58951941 |
| 9135 | NM_001040427.1 | #N/A | 68.0339407 | -7.2986078 | 1.62452282 |
| 9136 | NM_033026.5 | #N/A | 91.4040436 | -7.7236326 | 1.71915598 |
| 9137 | NM_013332.3 | #N/A | 105.089884 | -3.3157535 | 0.73805691 |
| 9138 | NM_138465.3 | #N/A | 61.5253523 | -7.1516441 | 1.59190611 |
| 9139 | NM_032052.1 | #N/A | 61.2576236 | -7.1435824 | 1.59028713 |
| 9140 | NM_001017926.2 | ZHX1 | 63.4527211 | -7.1973012 | 1.60239632 |
| 9141 | NM_005966.3 | #N/A | 79.9729062 | -7.528531 | 1.67612916 |
| 9142 | NM_014650.3 | #N/A | 69.2084388 | -7.3261275 | 1.63105779 |
| 9143 | NM_022736.2 | #N/A | 67.3157817 | -7.2805621 | 1.62092584 |
| 9144 | NM_001080496.2 | #N/A | 48.028291 | -6.7997404 | 1.5139554 |
| 9145 | NM_001029875.2 | #N/A | 53.3419722 | -6.9467099 | 1.54683988 |
| 9146 | NM_018366.2 | #N/A | 71.86885 | -6.0988304 | 1.35805125 |
| 9147 | NM_001135746.1 | #N/A | 59.8065154 | -7.1098967 | 1.58321798 |
| 9148 | NM_201446.2 | #N/A | 51.9678785 | -6.9083555 | 1.53845083 |
| 9149 | NM_001122769.2 | #N/A | 55.2934325 | -7.0006733 | 1.55919783 |
| 9150 | NM_004556.2 | #N/A | 57.6068307 | -7.0580365 | 1.57198834 |
| 9151 | NM_153701.2 | #N/A | 102.810757 | 1.72871069 | 0.38503033 |
| 9152 | NM_014035.2 | #N/A | 59.6594905 | -5.8272033 | 1.29797487 |
| 9153 | NM_013338.4 | #N/A | 71.1045645 | -7.3620274 | 1.63987795 |
| 9154 | NM_024114.3 | #N/A | 23.5212459 | 7.28834178 | 1.62347738 |
| 9155 | NM_032829.2 | #N/A | 61.9210294 | -7.1664819 | 1.59639306 |
| 9156 | NM_001276405.1 | DALRD3 | 55.6528515 | -7.0054249 | 1.5605338 |
| 9157 | NM_001077366.1 | #N/A | 80.1593574 | -7.5369288 | 1.67927296 |
| 9158 | NM_001253382.1 | #N/A | 110.015153 | -7.9918704 | 1.78064773 |

| | | | | | |
|---|---|---|---|---|---|
| 9159 | NM_004853.2 | #N/A | 66.6564388 | -7.269363 | 1.61978075 |
| 9160 | NM_152405.4 | #N/A | 141.636018 | -2.4600599 | 0.54816822 |
| 9161 | NM_005035.3 | #N/A | 327.943077 | -4.0486555 | 0.90225122 |
| 9162 | NM_001080418.2 | #N/A | 91.9496517 | -7.7329204 | 1.72332319 |
| 9164 | NM_032507.3 | PGBD1 | 72.4906126 | -7.3895835 | 1.64690552 |
| 9163 | NM_001145432.1 | SMIM20 | 58.2557446 | -7.0732389 | 1.57640699 |
| 9165 | NM_006570.4 | #N/A | 356.870026 | -5.5350771 | 1.23370774 |
| 9166 | NM_001001664.2 | #N/A | 67.2782712 | -5.9053691 | 1.31631014 |
| 9167 | NM_181839.2 | #N/A | 83.5449065 | -7.5957222 | 1.6931152 |
| 9168 | NM_002040.3 | #N/A | 50.0008915 | -6.8498569 | 1.52686872 |
| 9169 | NM_001080423.3 | #N/A | 91.1503323 | -7.7202199 | 1.72090309 |
| 9170 | NM_018204.4 | #N/A | 111.018728 | -8.0067207 | 1.78494693 |
| 9171 | NM_017948.5 | #N/A | 88.5423188 | -7.6780898 | 1.71186687 |
| 9172 | NM_018050.2 | #N/A | 55.6685109 | -7.0084769 | 1.56255874 |
| 9173 | NM_021198.2 | #N/A | 54.3243507 | -6.9708025 | 1.55416658 |
| 9174 | NM_145806.3 | #N/A | 55.7150202 | -7.0098615 | 1.56286992 |
| 9175 | NM_002579.2 | #N/A | 68.9675768 | -7.3226233 | 1.63263565 |
| 9176 | NM_001324094.1 | #N/A | 90.9770847 | -7.7154686 | 1.72039343 |
| 9177 | NM_001010875.3 | #N/A | 56.4900322 | -7.0304773 | 1.56799199 |
| 9178 | NR_047657.1 | #N/A | 76.6309958 | -7.4668082 | 1.66531951 |
| 9179 | NM_020311.2 | #N/A | 72.910579 | -7.3987106 | 1.65024443 |
| 9180 | NM_052918.4 | #N/A | 130.400169 | -8.2360631 | 1.83714219 |
| 9181 | NM_001239.3 | #N/A | 65.0532905 | -7.2327531 | 1.61341069 |
| 9182 | NM_006089.2 | #N/A | 54.0486811 | -6.9621802 | 1.5531223 |
| 9183 | NM_001204160.2 | #N/A | 76.4289796 | -7.4654465 | 1.6655958 |
| 9184 | NM_182916.2 | #N/A | 62.3093469 | -7.1709659 | 1.59991258 |
| 9185 | NM_006299.4 | #N/A | 51.0425808 | -6.8855305 | 1.53638255 |
| 9186 | NM_020437.4 | #N/A | 72.3057936 | -7.386313 | 1.64815569 |
| 9187 | NM_018217.2 | #N/A | 55.8669118 | -7.0185505 | 1.56611417 |
| 9188 | NM_001321368.1 | #N/A | 60.7811964 | -7.1376753 | 1.59275289 |
| 9189 | NM_020197.2 | #N/A | 97.8309251 | -5.2684296 | 1.175675 |
| 9190 | NM_032343.2 | #N/A | 129.595977 | -8.2280277 | 1.83615335 |
| 9191 | NM_001098208.1 | HNRNPF | 93.1551379 | -7.7514862 | 1.72986125 |
| 9192 | NM_001100814.2 | PIP4P1 | 64.4347615 | -7.2182207 | 1.61087097 |
| 9193 | NM_001114614.2 | #N/A | 63.2002118 | -7.1933764 | 1.60532468 |
| 9194 | NM_007276.4 | #N/A | 57.2435515 | -7.0479599 | 1.57299547 |
| 9195 | NM_032479.3 | #N/A | 67.645143 | -7.2900056 | 1.62703225 |
| 9196 | NM_001144950.1 | #N/A | 63.1496908 | -7.1950033 | 1.60611481 |
| 9197 | NM_015696.4 | #N/A | 51.490487 | -6.9007208 | 1.54042598 |
| 9198 | NM_001083602.1 | #N/A | 80.813257 | -7.5474722 | 1.68482993 |
| 9199 | NM_001173984.2 | #N/A | 87.7965661 | -7.665885 | 1.71127965 |
| 9200 | NM_001174118.1 | #N/A | 69.9966174 | -7.3355502 | 1.63754875 |
| 9201 | NM_001128220.2 | #N/A | 53.2206345 | -6.9452014 | 1.55046266 |

| | | | | | |
|---|---|---|---|---|---|
| 9202 | NM_207645.3 | #N/A | 103.755514 | -7.9065126 | 1.76518244 |
| 9203 | NM_006123.4 | #N/A | 53.6650505 | -6.9558785 | 1.55309594 |
| 9204 | NM_001142272.1 | #N/A | 74.3303097 | -7.4267903 | 1.65826274 |
| 9205 | NM_016341.3 | #N/A | 55.8895732 | -7.0110746 | 1.56555216 |
| 9206 | NM_002399.3 | MEIS2 | 78.9186756 | -7.5119906 | 1.67744313 |
| 9207 | NM_014759.3 | #N/A | 143.229554 | -8.3716453 | 1.86940337 |
| 9208 | NM_003797.4 | #N/A | 67.9979151 | -7.2953791 | 1.62913737 |
| 9209 | NM_001323036.2 | #N/A | 29.3768604 | 9.86934031 | 2.20398159 |
| 9210 | NM_004732.3 | KCNAB3 | 60.3037901 | -7.1189402 | 1.58997038 |
| 9211 | NM_002846.3 | #N/A | 112.828501 | -8.0259512 | 1.79256064 |
| 9212 | NR_144937.1 | #N/A | 48.1316147 | 4.85011435 | 1.08325811 |
| 9213 | NM_003720.3 | #N/A | 57.5060411 | -7.0565732 | 1.57616862 |
| 9214 | NM_005845.4 | ABCC4 | 55.0945496 | -6.993457 | 1.56214855 |
| 9215 | NM_005559.3 | #N/A | 177.813076 | -4.0680852 | 0.90871581 |
| 9216 | NM_019051.2 | #N/A | 63.3774548 | -7.1951874 | 1.60725149 |
| 9217 | NM_201591.2 | GPM6A | 1319.98085 | -4.4504177 | 0.99431086 |
| 9218 | NR_036564.1 | CCT3 | 60.8221507 | -7.1362435 | 1.59439239 |
| 9219 | NM_001320846.1 | SAT2 | 81.4436247 | -7.5585573 | 1.68884833 |
| 9220 | NM_001145797.1 | SH2B1 | 73.6329414 | -7.4113301 | 1.65596298 |
| 9221 | NM_015041.2 | #N/A | 74.4469125 | -7.4322194 | 1.6606571 |
| 9222 | NM_012267.4 | #N/A | 58.2961319 | -7.0770439 | 1.58133586 |
| 9223 | NM_006851.2 | #N/A | 74.038839 | -7.4218187 | 1.65846754 |
| 9224 | NM_024838.4 | #N/A | 53.9332935 | -6.9604782 | 1.55549871 |
| 9225 | NM_001356.4 | DDX3X | 47.9792911 | -6.7945579 | 1.51842754 |
| 9226 | NM_001008393.3 | #N/A | 57.482114 | -7.0533228 | 1.5762642 |
| 9227 | NM_203486.2 | #N/A | 91.6968621 | -7.7275939 | 1.72709041 |
| 9228 | NM_014793.4 | #N/A | 63.9390353 | -7.2068265 | 1.61078412 |
| 9229 | NM_207351.4 | #N/A | 83.2194614 | -2.9311942 | 0.65514899 |
| 9230 | NM_015957.3 | #N/A | 59.441302 | -7.1005221 | 1.58705116 |
| 9231 | NM_001080466.1 | #N/A | 47.3187129 | -6.7741589 | 1.51420954 |
| 9232 | NM_004569.3 | #N/A | 66.3007798 | -7.2605917 | 1.62294712 |
| 9233 | NM_015704.2 | #N/A | 307.716067 | -2.4075506 | 0.53824605 |
| 9234 | NM_000712.3 | #N/A | 89.4326272 | -7.6929679 | 1.71989952 |
| 9235 | NM_024525.4 | #N/A | 67.2027699 | -7.2786285 | 1.62729639 |
| 9236 | NM_001276414.1 | #N/A | 51.6737486 | -6.9067235 | 1.54419314 |
| 9237 | NM_152305.2 | #N/A | 64.5722406 | -7.2241793 | 1.61517078 |
| 9238 | NM_003422.2 | MZF1 | 51.6152598 | -6.9046796 | 1.54383278 |
| 9239 | NM_032122.4 | DTNBP1 | 64.7292758 | -7.2241196 | 1.61530041 |
| 9240 | NM_003060.3 | #N/A | 53.0446422 | -6.9433607 | 1.55277451 |
| 9242 | NM_152754.2 | SEMA3D | 82.615029 | -7.5795011 | 1.69511586 |
| 9241 | NM_001242487.1 | #N/A | 96.2629949 | -7.7957691 | 1.74347924 |
| 9243 | NM_001011655.2 | #N/A | 58.682009 | -7.0847195 | 1.58465439 |
| 9244 | NM_017566.3 | #N/A | 61.1439596 | -7.1402623 | 1.59712955 |

| | | | | | |
|---|---|---|---|---|---|
| 9245 | NM_017914.3 | #N/A | 68.6984437 | -7.3130756 | 1.63579892 |
| 9246 | NM_001320558.1 | ZSCAN16 | 19.7085924 | 4.14237057 | 0.92661366 |
| 9247 | NM_173601.1 | #N/A | 68.4748381 | -7.3084252 | 1.63497716 |
| 9248 | NM_001160258.1 | #N/A | 91.7774155 | -7.7310619 | 1.72955676 |
| 9249 | NM_017846.4 | #N/A | 59.9905961 | -7.1158648 | 1.59192261 |
| 9250 | NM_021930.5 | #N/A | 72.9992712 | -7.4032446 | 1.65623148 |
| 9251 | NM_003711.3 | #N/A | 60.5503623 | -7.1304897 | 1.59527318 |
| 9252 | NM_002358.3 | #N/A | 67.2734845 | -7.2796262 | 1.6287739 |
| 9253 | NM_018315.4 | #N/A | 62.3810269 | -7.1702254 | 1.60442362 |
| 9254 | NM_023943.3 | #N/A | 89.6269526 | -7.6932293 | 1.7215262 |
| 9255 | NR_023384.1 | RNF216P1 | 64.0823257 | -7.2096328 | 1.61334777 |
| 9256 | NM_207332.2 | #N/A | 50.8222969 | -6.8812859 | 1.54009581 |
| 9257 | NM_001129995.1 | #N/A | 201.718551 | -5.3159716 | 1.18977749 |
| 9258 | NR_130945.1 | WDR73 | 53.2250568 | -6.9392101 | 1.55312496 |
| 9259 | NR_073365.1 | #N/A | 61.8961285 | -7.1650411 | 1.60377705 |
| 9260 | NM_000935.2 | PLOD2 | 104.988574 | -6.4676941 | 1.44793742 |
| 9261 | NR_037718.1 | #N/A | 80.5737653 | -7.544989 | 1.6891338 |
| 9262 | NM_001256443.1 | #N/A | 51.1344082 | -6.8924172 | 1.54321231 |
| 9263 | NM_001258330.1 | EPB41L1 | 50.6178173 | -6.877717 | 1.54008712 |
| 9264 | NM_032366.4 | #N/A | 58.4047664 | -7.0758226 | 1.58444793 |
| 9265 | NM_001001567.1 | PDE9A | 67.2846553 | -7.2816549 | 1.63082067 |
| 9266 | NM_018691.3 | #N/A | 51.4925005 | -6.8959014 | 1.54451289 |
| 9267 | NM_015351.1 | #N/A | 163.359297 | -5.7954982 | 1.29815508 |
| 9268 | NM_001130020.1 | #N/A | 88.97593 | -7.686444 | 1.72174337 |
| 9269 | NM_173659.3 | #N/A | 68.2673606 | -7.3028861 | 1.63592664 |
| 9270 | NM_001345868.1 | GUF1 | 62.5856636 | 10.9404198 | 2.45133879 |
| 9271 | NM_130786.3 | #N/A | 62.9641033 | -7.1852153 | 1.60996135 |
| 9272 | NM_001243076.2 | E2F3 | 71.5757209 | -5.3044465 | 1.18856014 |
| 9273 | NM_012241.4 | #N/A | 52.1802917 | -6.9160073 | 1.54982632 |
| 9274 | NM_134268.4 | #N/A | 46.9484971 | -6.7656691 | 1.51613213 |
| 9275 | NM_001163788.2 | #N/A | 97.1409416 | -7.8115125 | 1.75062003 |
| 9276 | NM_001308293.1 | SH2B1 | 51.576316 | -6.9007329 | 1.5467373 |
| 9277 | NM_032199.2 | #N/A | 55.2134255 | -7.0013602 | 1.56931948 |
| 9278 | NM_006166.3 | #N/A | 148.968734 | -5.694282 | 1.276392 |
| 9279 | NM_001010867.3 | #N/A | 69.6447669 | -7.3284051 | 1.64272458 |
| 9280 | NM_001017964.1 | #N/A | 66.8693207 | -7.2729112 | 1.6303091 |
| 9281 | NM_001164160.1 | #N/A | 82.1585934 | -7.574006 | 1.69785391 |
| 9282 | NM_020346.2 | #N/A | 63.7394394 | -7.2029227 | 1.61473954 |
| 9283 | NM_001114633.1 | PLA2G4B | 122.172845 | -5.4842886 | 1.22962491 |
| 9284 | NM_025125.3 | #N/A | 64.1847672 | -7.2119627 | 1.6169942 |
| 9285 | NM_001100818.1 | #N/A | 78.3614489 | -7.502408 | 1.68212953 |
| 9287 | NR_002930.2 | PTPRVP | 23.6993657 | 3.98039221 | 0.89247882 |
| 9286 | NM_001184902.1 | #N/A | 61.7424581 | -7.1531318 | 1.60386779 |

| | | | | | |
|---|---|---|---|---|---|
| 9288 | NM_014936.4 | #N/A | 56.8787688 | -7.040415 | 1.57874739 |
| 9289 | NM_001897.4 | #N/A | 60.0372472 | -7.1230641 | 1.59729851 |
| 9290 | NM_145893.2 | #N/A | 82.2503715 | -7.5702679 | 1.69758366 |
| 9291 | NM_001162536.2 | #N/A | 75.5234251 | -7.4458366 | 1.66974194 |
| 9292 | NM_015163.5 | TRIM9 | 91.8406454 | -7.7300179 | 1.73357896 |
| 9293 | NR_027946.2 | #N/A | 110.266035 | -7.9969958 | 1.79344696 |
| 9294 | NM_021983.4 | HLA-DRB4 | 98.8050838 | -7.8378028 | 1.75795583 |
| 9295 | NM_018372.3 | #N/A | 50.8570495 | -6.8773928 | 1.54256291 |
| 9296 | NM_212551.4 | #N/A | 66.356087 | -7.2613757 | 1.62870707 |
| 9297 | NM_001243740.2 | ZNF124 | 68.5321028 | 1.93200245 | 0.43336917 |
| 9298 | NM_018491.3 | CBWD1 | 67.4984331 | -7.2889967 | 1.63508321 |
| 9299 | NM_001201372.1 | #N/A | 79.0086104 | -7.5128598 | 1.68534246 |
| 9300 | NM_002521.2 | #N/A | 90.1511586 | -7.7056706 | 1.7288267 |
| 9301 | NM_152342.3 | #N/A | 57.201506 | -7.0460433 | 1.58097398 |
| 9302 | NM_018977.3 | #N/A | 62.420488 | -7.1767071 | 1.61032424 |
| 9303 | NM_001329631.1 | #N/A | 64.3865929 | -7.2226863 | 1.62066962 |
| 9304 | NM_022451.10 | #N/A | 54.1704062 | -6.9673283 | 1.56340776 |
| 9305 | NM_004996.3 | #N/A | 259.380577 | -4.6505416 | 1.04363937 |
| 9306 | NM_001024218.1 | GPHN | 121.040207 | -8.129558 | 1.82462969 |
| 9307 | NM_020931.3 | #N/A | 64.0751858 | -7.2089564 | 1.61803029 |
| 9308 | NR_110396.1 | LOC101928336 | 176.723475 | 3.16805562 | 0.71108768 |
| 9309 | NM_007238.4 | #N/A | 65.5008508 | -7.2433029 | 1.6258123 |
| 9310 | NM_001079823.1 | #N/A | 88.3726632 | -6.4046444 | 1.43759224 |
| 9311 | NM_022044.2 | #N/A | 58.0220999 | -7.0708166 | 1.58727898 |
| 9312 | NM_001012978.2 | #N/A | 92.7392111 | -7.7455315 | 1.738777 |
| 9313 | NM_001278651.1 | #N/A | 60.4397826 | -7.1266373 | 1.59995301 |
| 9314 | NM_032437.3 | #N/A | 54.9439374 | -6.9868234 | 1.56859326 |
| 9315 | NM_001002755.2 | #N/A | 68.4393041 | -7.3069815 | 1.64053601 |
| 9316 | NM_138443.3 | #N/A | 64.6634568 | -7.2240655 | 1.62195062 |
| 9317 | NM_001142533.1 | SH2D3C | 68.203548 | -7.2992937 | 1.63885728 |
| 9318 | NM_001083613.1 | #N/A | 61.6072202 | -7.1552595 | 1.60664187 |
| 9319 | NM_032047.4 | #N/A | 77.9807782 | -6.0886812 | 1.36730301 |
| 9320 | NM_001288629.1 | #N/A | 112.659038 | -8.0240684 | 1.80199456 |
| 9321 | NM_001203245.2 | #N/A | 63.5303765 | -7.2026018 | 1.61757421 |
| 9322 | NM_004468.4 | #N/A | 56.6867025 | -7.0363034 | 1.58022036 |
| 9323 | NM_018486.2 | #N/A | 53.9896605 | -6.9655181 | 1.56434975 |
| 9324 | NM_020210.3 | #N/A | 46.7806044 | -6.7548523 | 1.51731725 |
| 9325 | NM_014064.3 | NTMT1 | 70.5311936 | -7.3490192 | 1.65103697 |
| 9326 | NR_027122.1 | ESRG | 221.903193 | 2.37427209 | 0.53343087 |
| 9327 | NM_005694.1 | COX17 | 62.5781839 | -7.1783643 | 1.61288055 |
| 9328 | NM_004503.3 | #N/A | 83.9785499 | -7.6039908 | 1.70882105 |
| 9329 | NM_032620.3 | #N/A | 75.7908318 | -7.4520597 | 1.67481702 |
| 9330 | NM_001008910.3 | #N/A | 50.1559057 | -6.8549626 | 1.54063234 |

| | | | | | |
|---|---|---|---|---|---|
| 9331 | NM_002553.3 | #N/A | 65.7047228 | -7.2462521 | 1.62865738 |
| 9332 | NM_005238.3 | #N/A | 53.281056 | -6.9503079 | 1.56216913 |
| 9333 | NM_001166037.1 | #N/A | 73.6626963 | -7.4125585 | 1.66613645 |
| 9334 | NM_001127195.1 | #N/A | 106.733706 | -7.9482996 | 1.78673382 |
| 9335 | NM_018440.3 | #N/A | 227.957995 | -3.1718383 | 0.71313264 |
| 9336 | NM_001324116.1 | UAP1 | 50.3745555 | -6.8624644 | 1.5430945 |
| 9337 | NM_001330474.1 | #N/A | 70.9454646 | -7.3582937 | 1.65480932 |
| 9338 | NM_002382.4 | #N/A | 70.2479309 | -7.3463545 | 1.65229594 |
| 9339 | NM_173527.2 | #N/A | 62.0755576 | -7.1663024 | 1.6118761 |
| 9340 | NM_030653.3 | #N/A | 58.113693 | -7.0731782 | 1.59110135 |
| 9341 | NM_001003891.2 | #N/A | 59.7810858 | -7.1170018 | 1.60107277 |
| 9342 | NM_005692.4 | LOC114483834 | 59.6618716 | -7.1082888 | 1.59924412 |
| 9343 | NM_024418.2 | #N/A | 132.831498 | -6.9987623 | 1.574615 |
| 9344 | NM_000138.4 | FBN1 | 668.891739 | -3.57991 | 0.80544421 |
| 9345 | NR_037668.2 | SCNN1D | 50.7175466 | -6.8706072 | 1.54584327 |
| 9346 | NM_016586.2 | #N/A | 57.3525828 | -7.0507035 | 1.58639053 |
| 9347 | NM_182709.2 | #N/A | 61.5829069 | -7.1500924 | 1.60882001 |
| 9348 | NM_001145125.1 | MKRN1 | 87.4852925 | -7.6608642 | 1.72384711 |
| 9349 | NM_203288.1 | #N/A | 69.018292 | -7.318174 | 1.6467611 |
| 9350 | NM_003501.2 | #N/A | 51.9184831 | -6.9049367 | 1.55390524 |
| 9351 | NM_017573.4 | PCSK4 | 50.1244212 | -6.8619985 | 1.54432629 |
| 9352 | NM_012407.3 | #N/A | 70.5515924 | -7.3522336 | 1.65472814 |
| 9353 | NM_001102368.2 | DGLUCY | 59.9195735 | -7.1171825 | 1.60190376 |
| 9354 | NM_015692.2 | #N/A | 57.8657602 | -4.2961442 | 0.96698868 |
| 9355 | NM_001288766.1 | #N/A | 51.4696849 | -6.9018166 | 1.55352697 |
| 9356 | NM_024512.4 | #N/A | 123.810721 | 2.15305274 | 0.48463011 |
| 9357 | NM_020166.4 | #N/A | 57.9994049 | -7.0657601 | 1.59047075 |
| 9358 | NR_002824.3 | HERC2P2 | 688.314977 | -4.3799297 | 0.98591713 |
| 9359 | NM_001286835.1 | CBWD5 | 54.8592139 | -6.9872463 | 1.57296033 |
| 9360 | NM_002484.3 | #N/A | 51.2322973 | -6.8900748 | 1.55112231 |
| 9361 | NM_001323600.1 | SVIL | 49.3195254 | -6.8357819 | 1.53909031 |
| 9362 | NR_026580.1 | TFDP1 | 61.3635772 | -7.1509455 | 1.61004759 |
| 9363 | NM_004210.4 | #N/A | 80.1722516 | -7.5344354 | 1.69641393 |
| 9364 | NM_001277842.1 | #N/A | 73.3127067 | -7.4049792 | 1.66746655 |
| 9365 | NM_000683.3 | #N/A | 54.5601594 | -6.9827943 | 1.57253453 |
| 9366 | NM_031300.3 | #N/A | 55.7722874 | -7.0086629 | 1.57852561 |
| 9367 | NM_203406.1 | #N/A | 58.5456619 | -7.0804674 | 1.59470826 |
| 9368 | NM_006214.3 | #N/A | 55.5701844 | -7.0072787 | 1.57823357 |
| 9369 | NM_001085049.2 | #N/A | 140.991002 | -8.3501907 | 1.88072007 |
| 9370 | NM_206949.2 | #N/A | 47.6885444 | -6.789268 | 1.52915511 |
| 9371 | NM_148920.2 | #N/A | 72.4906164 | -7.3899988 | 1.66463083 |
| 9372 | NM_032959.5 | #N/A | 69.8324989 | -7.3318612 | 1.65174993 |
| 9373 | NM_007014.4 | #N/A | 62.1788899 | -7.1695919 | 1.61521171 |

| | | | | | |
|---|---|---|---|---|---|
| 9374 | NM_002859.3 | #N/A | 139.305503 | -3.4112142 | 0.76851798 |
| 9375 | NM_001300958.1 | #N/A | 56.5883378 | -7.0356182 | 1.58533893 |
| 9376 | NM_006466.3 | #N/A | 49.5512512 | -6.8416391 | 1.54163861 |
| 9377 | NM_004886.3 | #N/A | 54.9300136 | -6.9880616 | 1.57479262 |
| 9378 | NM_001320836.1 | #N/A | 50.0959949 | -6.8558477 | 1.54503642 |
| 9379 | NM_014580.4 | #N/A | 67.8880764 | -7.2967366 | 1.64441699 |
| 9380 | NM_019002.3 | #N/A | 64.1385642 | -5.8360962 | 1.31526908 |
| 9381 | NM_001304347.1 | #N/A | 82.5152479 | -7.5782719 | 1.70794624 |
| 9382 | NM_198897.1 | #N/A | 50.7695013 | -6.8747696 | 1.54952885 |
| 9383 | NM_000067.2 | #N/A | 154.912386 | -8.4867799 | 1.91289329 |
| 9384 | NM_001258374.1 | #N/A | 65.6556979 | -7.2497911 | 1.63413017 |
| 9385 | NM_173548.2 | #N/A | 49.9605948 | -6.8524058 | 1.54462237 |
| 9386 | NM_021243.2 | #N/A | 105.192807 | -7.9262051 | 1.78670267 |
| 9387 | NM_001308115.1 | #N/A | 48.1139799 | -6.7997605 | 1.53282997 |
| 9388 | NM_001079910.1 | #N/A | 70.9427575 | -7.3611293 | 1.65965729 |
| 9389 | NM_001290117.1 | TIMM23B | 47.9818178 | -6.7995677 | 1.53305563 |
| 9390 | NM_001109903.1 | RNFT2 | 85.4924586 | -7.6270518 | 1.71970621 |
| 9391 | NM_024900.4 | #N/A | 53.5272504 | -6.95347 | 1.56787698 |
| 9392 | NM_001256012.1 | #N/A | 121.479417 | -8.1364304 | 1.83464562 |
| 9393 | NM_016023.3 | #N/A | 62.2934823 | -7.1716439 | 1.61714164 |
| 9394 | NM_001320193.1 | #N/A | 75.6914912 | -7.4526444 | 1.68057202 |
| 9395 | NM_017956.3 | #N/A | 52.7107026 | -6.9286386 | 1.56241975 |
| 9396 | NM_001083893.1 | #N/A | 60.7197175 | -7.1336374 | 1.60868328 |
| 9397 | NM_013335.3 | #N/A | 51.2533462 | -6.8891101 | 1.55358072 |
| 9398 | NM_000899.4 | #N/A | 86.2753566 | -7.6415194 | 1.72330219 |
| 9399 | NM_001005333.1 | #N/A | 46.524415 | -6.7466177 | 1.52153655 |
| 9400 | NM_001330566.1 | #N/A | 83.5819911 | -7.5937092 | 1.71257121 |
| 9401 | NM_001017963.2 | #N/A | 87.2076044 | -6.3822698 | 1.43937731 |
| 9402 | NM_001305145.1 | TMEFF2 | 13.1976755 | 5.7719374 | 1.30177514 |
| 9403 | NM_001207062.1 | C22orf23 | 79.2891365 | -7.519261 | 1.69589181 |
| 9405 | NM_003244.3 | TGIF1 | 46.318622 | -6.7442992 | 1.52111728 |
| 9404 | NM_002997.4 | #N/A | 76.2704992 | -6.1877586 | 1.39560171 |
| 9406 | NM_022486.4 | #N/A | 61.9771887 | -7.1618728 | 1.61531114 |
| 9407 | NM_005788.3 | #N/A | 51.7236566 | -6.9000043 | 1.55633177 |
| 9408 | NM_001190242.1 | IFT80 | 74.4264366 | -6.1471561 | 1.38654777 |
| 9409 | NM_001425.2 | #N/A | 73.9759467 | -7.4211921 | 1.67397108 |
| 9410 | NM_018059.4 | #N/A | 51.2854953 | -6.89071 | 1.55430944 |
| 9411 | NM_006223.3 | #N/A | 60.5107528 | -7.1278315 | 1.60792954 |
| 9412 | NM_005480.3 | #N/A | 57.5928493 | -7.0572954 | 1.59209466 |
| 9413 | NM_019848.4 | #N/A | 49.5168328 | -6.8460016 | 1.54460944 |
| 9414 | NM_001127213.1 | LY6E | 87.4793746 | -7.6581223 | 1.7279387 |
| 9415 | NR_038353.1 | #N/A | 170.615453 | 3.34981207 | 0.75586801 |
| 9416 | NM_015167.2 | JMJD6 | 49.502054 | -6.8391124 | 1.54321994 |

| | | | | | |
|---|---|---|---|---|---|
| 9417 | NM_001112802.1 | #N/A | 54.6493149 | -6.9774842 | 1.57466788 |
| 9418 | NR_034181.1 | SUV39H2 | 57.8729378 | -7.0638996 | 1.59432893 |
| 9419 | NM_001039577.3 | #N/A | 56.9311249 | -7.0465794 | 1.59043251 |
| 9420 | NR_073000.1 | #N/A | 58.7047372 | -7.0886019 | 1.59997784 |
| 9421 | NM_006875.3 | #N/A | 59.1693754 | -7.099588 | 1.60301364 |
| 9422 | NM_017883.5 | #N/A | 69.3513362 | -7.3294558 | 1.65496974 |
| 9423 | NM_014590.3 | ERVW-1 | 53.1218529 | 5.69593504 | 1.28613682 |
| 9424 | NM_001283020.1 | NAPB | 63.0773653 | -7.1848679 | 1.6223734 |
| 9425 | NM_001317956.1 | #N/A | 50.6803968 | -6.8728006 | 1.55192051 |
| 9426 | NM_153607.2 | #N/A | 50.4039173 | -6.8609025 | 1.5493193 |
| 9427 | NM_015416.4 | #N/A | 67.2249761 | -7.278297 | 1.64360613 |
| 9428 | NM_001093767.2 | #N/A | 98.9483142 | -7.8380482 | 1.77004193 |
| 9429 | NM_182914.2 | SYNE2 | 162.975269 | -8.5578945 | 1.93293131 |
| 9430 | NM_001207067.1 | #N/A | 94.1507487 | -7.7658055 | 1.7543215 |
| 9433 | NM_172171.2 | CAMK2G | 105.030791 | -7.9247322 | 1.79043911 |
| 9431 | NM_001256007.2 | #N/A | 54.9982686 | -6.9944284 | 1.58025474 |
| 9432 | NM_004219.3 | #N/A | 56.9640437 | -7.0424646 | 1.59109236 |
| 9434 | NM_057092.2 | FKBP2 | 49.3974735 | -6.834788 | 1.54421846 |
| 9435 | NM_001265609.1 | TRAK1 | 47.0972336 | -6.7630266 | 1.5280402 |
| 9436 | NM_001346413.1 | #N/A | 55.6342676 | -7.0078922 | 1.58349007 |
| 9437 | NM_130836.2 | #N/A | 64.2037109 | -7.2112048 | 1.62944744 |
| 9438 | NM_033427.2 | #N/A | 145.425817 | -6.8731036 | 1.55308323 |
| 9439 | NM_001178040.1 | #N/A | 91.1710476 | -6.2808419 | 1.41928195 |
| 9440 | NM_025202.3 | #N/A | 47.0412925 | -6.7696185 | 1.52975918 |
| 9441 | NM_003981.3 | #N/A | 53.9166842 | -6.9593137 | 1.57275992 |
| 9442 | NM_001001436.2 | #N/A | 71.762414 | -7.3735016 | 1.66640963 |
| 9443 | NM_001163213.1 | FGFR3 | 60.09943 | -7.117245 | 1.60869863 |
| 9444 | NM_014751.5 | MTSS1 | 61.8387859 | -7.1643969 | 1.61937493 |
| 9445 | NM_001166392.1 | TPBG | 107.688404 | -7.9628953 | 1.80005915 |
| 9446 | NM_001291941.1 | #N/A | 88.9874549 | -7.685009 | 1.73746278 |
| 9447 | NM_001283009.1 | #N/A | 77.7368004 | -7.4904948 | 1.69350745 |
| 9448 | NM_001282749.1 | TLE4 | 103.651193 | -7.9048227 | 1.78719201 |
| 9449 | NM_152729.2 | #N/A | 71.6478529 | -7.3756962 | 1.66780668 |
| 9450 | NM_203437.3 | AFTPH | 68.1443727 | -7.299595 | 1.65064804 |
| 9451 | NM_004307.1 | #N/A | 69.5835635 | -7.3343791 | 1.65879378 |
| 9452 | NM_001199861.1 | #N/A | 45.5319813 | -6.7210731 | 1.52014569 |
| 9453 | NM_000037.3 | #N/A | 90.9609193 | -7.7179712 | 1.74573412 |
| 9454 | NM_001193268.1 | #N/A | 71.2055808 | -7.3629858 | 1.66561477 |
| 9455 | NM_006480.4 | #N/A | 45.4982911 | -6.7186831 | 1.51990657 |
| 9456 | NM_001282680.1 | #N/A | 65.7561347 | -7.2541386 | 1.64105156 |
| 9457 | NM_001330112.1 | #N/A | 57.5675635 | -7.058812 | 1.5972561 |
| 9458 | NM_001289912.1 | #N/A | 46.6522744 | -6.7559084 | 1.52877309 |
| 9459 | NM_001319990.1 | #N/A | 58.2820365 | -7.0803114 | 1.60222233 |

| | | | | | |
|---|---|---|---|---|---|
| 9460 | NM_145689.2 | #N/A | 69.0587836 | -7.3240891 | 1.65744732 |
| 9461 | NR_003579.1 | #N/A | 71.0630835 | -5.9656912 | 1.35008251 |
| 9462 | NM_001278677.1 | #N/A | 59.9064921 | -7.1167769 | 1.61069262 |
| 9463 | NM_005907.3 | #N/A | 65.3215693 | -4.6231471 | 1.04635361 |
| 9464 | NM_001039503.2 | PRSS53 | 48.4585997 | -6.8064275 | 1.54072773 |
| 9465 | NM_005171.4 | #N/A | 44.9479246 | -6.7014783 | 1.51700862 |
| 9466 | NM_000019.3 | #N/A | 185.328285 | -3.9765575 | 0.90022117 |
| 9467 | NM_032536.2 | #N/A | 80.8286459 | -7.5458143 | 1.70832928 |
| 9468 | NM_001243738.1 | RGL2 | 62.5362381 | -7.1717776 | 1.62366326 |
| 9469 | NM_001189.3 | #N/A | 81.2138062 | -7.5555836 | 1.71073977 |
| 9470 | NM_024675.3 | #N/A | 48.4533776 | -6.8077896 | 1.54146272 |
| 9471 | NM_052990.2 | #N/A | 55.7194996 | -7.0109151 | 1.58750461 |
| 9472 | NM_015260.3 | #N/A | 62.553359 | -7.1807836 | 1.62598716 |
| 9473 | NM_021030.2 | #N/A | 46.5530169 | -6.74849 | 1.52814699 |
| 9474 | NM_006352.4 | ZBTB18 | 105.115952 | -7.9240176 | 1.79437339 |
| 9475 | NM_001042551.1 | #N/A | 91.3675791 | -7.722437 | 1.74886088 |
| 9476 | NM_001177387.1 | ATXN7 | 50.1730338 | -6.8530543 | 1.55206365 |
| 9477 | NM_170750.2 | PSMD10 | 67.6588366 | -7.2904557 | 1.65115074 |
| 9478 | NM_014945.4 | ABLIM3 | 78.8917155 | -7.5115271 | 1.70128569 |
| 9479 | NM_001206856.2 | CAPRIN2 | 51.2391504 | -6.884035 | 1.55924359 |
| 9480 | NM_001006639.1 | #N/A | 61.3096523 | -7.1449949 | 1.61836243 |
| 9481 | NM_153688.3 | #N/A | 63.0627326 | -7.1913419 | 1.62890941 |
| 9482 | NM_012309.4 | SHANK2 | 54.6739348 | -6.9807899 | 1.58123616 |
| 9483 | NM_013248.2 | #N/A | 61.2989166 | -7.1466444 | 1.61899373 |
| 9484 | NM_176824.2 | #N/A | 47.0626544 | -6.7669742 | 1.53298104 |
| 9485 | NM_005804.3 | #N/A | 245.201028 | -5.5346513 | 1.25383686 |
| 9486 | NM_001198800.2 | #N/A | 59.1854954 | -7.0965685 | 1.60769574 |
| 9487 | NM_001127660.1 | MFN2 | 142.137827 | -8.3629335 | 1.89462073 |
| 9488 | NM_004116.4 | #N/A | 88.5400945 | -7.6789751 | 1.73974931 |
| 9489 | NM_020438.4 | #N/A | 70.8229553 | -7.3551773 | 1.66640967 |
| 9490 | NM_001010906.1 | #N/A | 88.8268215 | 1.89492331 | 0.42933021 |
| 9491 | NM_018341.2 | #N/A | 50.2852158 | -6.8593312 | 1.55416585 |
| 9492 | NM_178140.3 | #N/A | 50.2693038 | -6.8591059 | 1.55419693 |
| 9493 | NM_001100594.1 | SNRK | 64.3142031 | -7.2133552 | 1.6344775 |
| 9494 | NM_001320633.1 | #N/A | 55.527409 | -7.0009308 | 1.58637803 |
| 9495 | NM_176814.4 | #N/A | 44.4654632 | -6.6856568 | 1.51493913 |
| 9496 | NM_003275.3 | #N/A | 76.2334536 | -7.4629313 | 1.69131927 |
| 9497 | NM_032928.3 | #N/A | 63.0972888 | -7.1902871 | 1.62960647 |
| 9498 | NM_207644.2 | #N/A | 56.164633 | -7.024276 | 1.59198647 |
| 9499 | NR_027237.1 | LOC728743 | 73.230063 | -7.4026021 | 1.67778692 |
| 9500 | NM_001184777.1 | #N/A | 78.4421226 | -7.5014975 | 1.70021955 |
| 9501 | NM_005038.2 | #N/A | 221.442286 | -2.2989892 | 0.5210697 |
| 9502 | NM_032687.3 | #N/A | 52.9612914 | -6.937449 | 1.57244465 |

| | | | | | |
|---|---|---|---|---|---|
| 9503 | NM_001130012.2 | #N/A | 64.8995107 | -7.2315649 | 1.6392334 |
| 9504 | NM_001166159.1 | NDUFS2 | 47.7329868 | -6.7890058 | 1.53904953 |
| 9505 | NM_001124759.1 | #N/A | 32.2572901 | 5.65620865 | 1.28225995 |
| 9506 | NM_199363.2 | #N/A | 65.2297972 | -7.2379302 | 1.64110776 |
| 9507 | NM_001430.4 | #N/A | 151.807077 | -8.4575274 | 1.9176956 |
| 9508 | NM_001127453.1 | #N/A | 71.8002968 | -7.3753258 | 1.67237682 |
| 9509 | NM_005792.2 | MPHOSPH6 | 63.3902429 | -7.1960042 | 1.63177879 |
| 9510 | NM_001167671.2 | LPP | 96.5210903 | -7.8011564 | 1.76914955 |
| 9511 | NM_080669.5 | #N/A | 49.9310131 | -6.8531631 | 1.55418038 |
| 9512 | NM_001083.3 | #N/A | 78.209766 | -7.5033091 | 1.70165336 |
| 9513 | NM_001303.3 | #N/A | 53.9728366 | -6.966067 | 1.57987608 |
| 9514 | NM_001352.4 | #N/A | 50.5949459 | -6.8692154 | 1.55796172 |
| 9515 | NM_001003897.1 | #N/A | 55.9073578 | -7.0141022 | 1.5909898 |
| 9516 | NM_001105573.1 | #N/A | 213.286865 | -2.5949243 | 0.5886078 |
| 9517 | NM_004979.5 | #N/A | 46.3262525 | -6.7435069 | 1.52963919 |
| 9518 | NM_001164387.1 | #N/A | 62.9451116 | -7.1847741 | 1.62975061 |
| 9519 | NM_001018005.1 | #N/A | 86.3731278 | -7.6452735 | 1.73451142 |
| 9520 | NR_120685.1 | LINC01503 | 62.326637 | -7.1721896 | 1.62728854 |
| 9521 | NM_001289746.1 | #N/A | 101.074415 | -6.4083757 | 1.45427155 |
| 9522 | NM_015986.3 | #N/A | 48.9448601 | -6.8203566 | 1.54785636 |
| 9523 | NM_004316.3 | #N/A | 57.4013988 | -7.0487688 | 1.59976309 |
| 9524 | NM_020374.3 | #N/A | 76.5309055 | -7.4655846 | 1.6945669 |
| 9525 | NM_199413.2 | #N/A | 80.373966 | -7.5376243 | 1.71091303 |
| 9526 | NM_005706.3 | #N/A | 67.2310975 | -7.2788003 | 1.6522339 |
| 9527 | NM_005288.3 | #N/A | 159.042283 | -7.2586734 | 1.64789571 |
| 9528 | NM_012198.4 | #N/A | 79.3880745 | -7.5196136 | 1.70712795 |
| 9529 | NM_080872.3 | #N/A | 49.2317361 | -6.8356066 | 1.55185112 |
| 9530 | NM_002134.3 | #N/A | 87.7531388 | -7.6656871 | 1.74041392 |
| 9531 | NM_024040.2 | #N/A | 302.881572 | -4.1981558 | 0.95314359 |
| 9532 | NM_144489.3 | RGS3 | 43.8366973 | -6.6669599 | 1.51369599 |
| 9533 | NM_001034852.2 | #N/A | 61.4274988 | -7.1494105 | 1.62327261 |
| 9534 | NM_001142444.1 | #N/A | 51.3821936 | -6.8959302 | 1.56608622 |
| 9535 | NM_001007527.1 | #N/A | 60.3668706 | -7.1244745 | 1.618143 |
| 9536 | NM_007059.3 | #N/A | 57.811798 | -7.0618134 | 1.60401317 |
| 9537 | NM_022898.2 | #N/A | 139.801501 | -8.3372634 | 1.89412311 |
| 9538 | NM_014425.4 | #N/A | 48.747519 | -6.8181118 | 1.54904751 |
| 9539 | NM_176806.3 | #N/A | 98.7215735 | -6.3791166 | 1.44938224 |
| 9540 | NM_001300750.1 | LMBR1L | 51.4056926 | -6.8991201 | 1.56774197 |
| 9541 | NM_001348078.1 | #N/A | 45.6612681 | -6.7287412 | 1.5290503 |
| 9542 | NM_014637.3 | #N/A | 48.9683946 | -6.8239831 | 1.55069204 |
| 9543 | NM_017842.2 | #N/A | 143.826497 | -2.9625594 | 0.67321936 |
| 9544 | NM_014782.6 | #N/A | 60.1444372 | -7.1230483 | 1.6187226 |
| 9545 | NM_001100122.1 | #N/A | 81.6865275 | -7.5632988 | 1.71881666 |

| | | | | | |
|---|---|---|---|---|---|
| 9546 | NM_012426.4 | #N/A | 705.086089 | -2.1550496 | 0.48974963 |
| 9547 | NM_207370.2 | #N/A | 274.57704 | -3.8203083 | 0.86829778 |
| 9548 | NM_001301007.1 | #N/A | 82.3195358 | -7.5710778 | 1.72081433 |
| 9549 | NM_006347.3 | #N/A | 61.6342137 | -7.1557464 | 1.62644519 |
| 9550 | NM_016464.4 | #N/A | 46.0770066 | -6.7368283 | 1.53122862 |
| 9551 | NM_007327.3 | #N/A | 116.087363 | -8.0682563 | 1.83388401 |
| 9552 | NM_175609.2 | #N/A | 94.0328824 | -7.7642462 | 1.7648276 |
| 9553 | NM_001005745.1 | NUMB | 104.939898 | -6.4548451 | 1.46728167 |
| 9554 | NM_032509.3 | #N/A | 66.0073066 | -5.9752377 | 1.35831285 |
| 9555 | NM_213720.2 | #N/A | 46.3151153 | -6.7399539 | 1.53216227 |
| 9556 | NR_036485.1 | SBF2-AS1 | 80.4917191 | 2.95036087 | 0.67072513 |
| 9557 | NM_001042469.1 | #N/A | 49.4907191 | -6.8431816 | 1.55573228 |
| 9558 | NM_001702.2 | #N/A | 215.209376 | -4.0552218 | 0.92195583 |
| 9559 | NM_001042388.2 | #N/A | 73.8812506 | -7.4205683 | 1.68711435 |
| 9560 | NM_005680.2 | #N/A | 43.4191487 | -6.653386 | 1.51272615 |
| 9561 | NM_001323543.1 | #N/A | 49.4452894 | -6.8405943 | 1.55540238 |
| 9562 | NM_001304792.1 | SENP6 | 57.7526774 | -5.7815034 | 1.31467155 |
| 9563 | NM_001256550.1 | #N/A | 54.5048376 | -6.9792584 | 1.58705868 |
| 9564 | NM_024833.2 | #N/A | 48.8050658 | -6.8154236 | 1.54980314 |
| 9565 | NM_005994.3 | #N/A | 185.797251 | -5.9166067 | 1.34545477 |
| 9566 | NM_152264.4 | SLC39A13 | 70.5537153 | -7.3507237 | 1.67164008 |
| 9567 | NM_198722.2 | #N/A | 47.7972237 | -6.7867136 | 1.54338549 |
| 9568 | NM_002395.5 | #N/A | 64.2946596 | -7.2177979 | 1.64142947 |
| 9569 | NM_003582.2 | #N/A | 45.5560494 | -6.7229468 | 1.52892951 |
| 9570 | NM_001009185.2 | #N/A | 60.2592657 | -7.1204086 | 1.61944403 |
| 9571 | NM_015896.3 | #N/A | 111.073404 | -8.0072814 | 1.82131713 |
| 9572 | NM_198080.3 | #N/A | 72.9012925 | -7.3971628 | 1.68268313 |
| 9573 | NM_001201461.1 | #N/A | 72.9601822 | -7.3962836 | 1.682511 |
| 9574 | NM_001142446.1 | #N/A | 77.4195107 | -7.4857943 | 1.70328121 |
| 9575 | NM_001279354.1 | #N/A | 90.0205653 | -7.7036735 | 1.75285354 |
| 9576 | NM_006855.3 | #N/A | 44.7027981 | -6.6960532 | 1.52374892 |
| 9577 | NM_080678.2 | UBE2F | 66.9350658 | -7.2735155 | 1.65520837 |
| 9578 | NM_013251.3 | #N/A | 89.5025276 | -7.6946866 | 1.75109828 |
| 9579 | NM_015690.4 | #N/A | 118.91727 | -8.1061948 | 1.84482964 |
| 9580 | NM_001564.3 | #N/A | 55.9056788 | -7.0117938 | 1.59580371 |
| 9581 | NM_012063.3 | DNM1L | 58.3944874 | -7.082232 | 1.61189761 |
| 9582 | NM_001160302.1 | SYNJ1 | 73.7121192 | -7.4132592 | 1.68725451 |
| 9583 | NR_038322.1 | #N/A | 99.6135902 | -3.7667552 | 0.85747318 |
| 9584 | NM_181504.3 | #N/A | 111.499082 | -8.0110312 | 1.82380801 |
| 9585 | NM_014974.2 | #N/A | 378.368105 | -4.0560108 | 0.92345393 |
| 9586 | NM_006606.2 | #N/A | 120.92436 | -3.5652103 | 0.81181316 |
| 9587 | NM_001199563.1 | #N/A | 45.5958225 | -6.7181335 | 1.52987976 |
| 9588 | NM_001127196.1 | #N/A | 399.9193 | -2.7706683 | 0.63096301 |

| | | | | | |
|---|---|---|---|---|---|
| 9589 | NM_198458.2 | #N/A | 52.3228461 | -6.9190291 | 1.57579233 |
| 9590 | NM_001376.4 | #N/A | 2321.97464 | -6.159534 | 1.4029187 |
| 9591 | NM_017547.3 | #N/A | 71.1270993 | -7.3611601 | 1.67661919 |
| 9592 | NR_045106.2 | #N/A | 43.8658582 | -6.661712 | 1.51731328 |
| 9594 | NR_037709.1 | TEN1-CDK3 | 43.8460918 | -6.6659842 | 1.51833032 |
| 9593 | NM_001304749.1 | #N/A | 45.3329464 | -6.7117682 | 1.52876153 |
| 9595 | NM_006993.2 | NPM3 | 69.3488381 | -7.3258315 | 1.66871409 |
| 9596 | NM_080416.3 | SEPTIN4 | 56.1057466 | -7.0194921 | 1.59895023 |
| 9597 | NM_001329628.1 | #N/A | 48.911579 | -6.8240022 | 1.5545099 |
| 9598 | NM_004954.4 | #N/A | 74.1731996 | -7.4209259 | 1.69062942 |
| 9599 | NM_017828.4 | #N/A | 43.3694014 | -6.6463118 | 1.51425865 |
| 9600 | NM_001136201.1 | #N/A | 59.2077893 | -7.0995623 | 1.61768759 |
| 9601 | NR_024403.1 | LOC730101 | 47.8542198 | -6.7886289 | 1.5468974 |
| 9602 | NM_003762.4 | #N/A | 57.4706437 | -7.0535841 | 1.60740402 |
| 9603 | NM_006907.3 | #N/A | 62.6100534 | -7.1792882 | 1.63613355 |
| 9604 | NR_038971.1 | LINC00641 | 52.9587819 | -6.9344667 | 1.58048229 |
| 9606 | NR_102759.1 | TRIM52-AS1 | 49.8425222 | -6.8486087 | 1.56094978 |
| 9605 | NM_001545.2 | #N/A | 59.0017652 | -7.0927603 | 1.61660096 |
| 9607 | NM_001198569.1 | ATP6V0C | 56.358514 | -7.0313141 | 1.60308291 |
| 9608 | NM_004804.2 | #N/A | 532.066682 | -2.1244423 | 0.48436961 |
| 9609 | NM_002910.5 | #N/A | 48.3409548 | -6.8080193 | 1.55226958 |
| 9610 | NR_038849.1 | LINC01133 | 306.281275 | 4.01128554 | 0.91473715 |
| 9611 | NM_138740.3 | #N/A | 63.0430395 | -7.1873456 | 1.63901913 |
| 9612 | NM_017816.2 | #N/A | 52.8622609 | -6.9333419 | 1.58114956 |
| 9613 | NM_021221.2 | LY6G5B | 43.758041 | -6.6632831 | 1.51972291 |
| 9614 | NM_030761.4 | #N/A | 96.8254901 | -6.3517545 | 1.44867933 |
| 9615 | NR_033855.1 | LINC00485 | 77.6884045 | 4.07058351 | 0.92843928 |
| 9616 | NM_001320817.1 | #N/A | 43.9268047 | -6.6701688 | 1.52137499 |
| 9617 | NM_001172509.1 | #N/A | 94.8752283 | -7.7769855 | 1.77387756 |
| 9618 | NM_001345873.1 | #N/A | 73.8510449 | -7.4157169 | 1.69158717 |
| 9619 | NR_110837.1 | LINC02080 | 321.73276 | 2.93822746 | 0.67024534 |
| 9620 | NM_002053.2 | #N/A | 49.7354191 | -6.8521479 | 1.56322212 |
| 9621 | NM_145232.3 | #N/A | 67.5024912 | -7.286082 | 1.66237535 |
| 9622 | NM_001330669.1 | SARS1 | 81.4761015 | -7.557159 | 1.72431019 |
| 9623 | NM_005644.3 | #N/A | 55.2762143 | -6.9978967 | 1.59681025 |
| 9624 | NM_024330.1 | #N/A | 57.4395892 | -7.0526153 | 1.60944947 |
| 9625 | NM_001130528.2 | #N/A | 73.2537394 | -7.4090476 | 1.69087684 |
| 9626 | NR_135091.1 | LINC02746 | 33.9906305 | 2.76937975 | 0.6321731 |
| 9627 | NM_001031827.2 | BOLA2 | 68.5410052 | -7.3101199 | 1.66871928 |
| 9628 | NM_001318525.1 | #N/A | 73.1235771 | -7.4030958 | 1.68993623 |
| 9629 | NM_006834.4 | #N/A | 58.9277006 | -7.0912589 | 1.61881832 |
| 9630 | NM_001286836.1 | CBWD5 | 54.1330511 | -6.9689697 | 1.59094564 |
| 9631 | NM_001256335.1 | PTGES2 | 57.0256427 | -7.0442082 | 1.60833255 |

| | | | | | |
|---|---|---|---|---|---|
| 9632 | NM_006275.5 | #N/A | 4189.0934 | 3.89952214 | 0.89034219 |
| 9633 | NM_033212.3 | #N/A | 45.9017029 | -6.7366889 | 1.53813845 |
| 9634 | NM_052811.3 | #N/A | 77.0224114 | -7.4769219 | 1.707505 |
| 9635 | NM_001166242.1 | C22orf39 | 71.0499062 | -6.0811487 | 1.38885723 |
| 9636 | NM_014930.2 | ZNF510 | 65.8802719 | -7.2482789 | 1.6554731 |
| 9637 | NM_006412.3 | #N/A | 43.6764242 | -6.6629529 | 1.52202948 |
| 9638 | NM_021138.3 | #N/A | 67.4797826 | -6.0082533 | 1.3724931 |
| 9639 | NM_018555.5 | #N/A | 43.1522694 | -6.6418688 | 1.51737708 |
| 9640 | NM_130442.3 | ELMO1 | 79.2336766 | -7.5184232 | 1.71767543 |
| 9641 | NM_001321983.1 | #N/A | 26.2280854 | 9.70655974 | 2.21762571 |
| 9642 | NM_001017979.2 | #N/A | 56.9585131 | -7.0411224 | 1.60871442 |
| 9643 | NM_001286177.1 | C2CD5 | 74.5539901 | -7.4291027 | 1.69747636 |
| 9644 | NR_045121.1 | DGCR5 | 46.9206952 | -6.7686939 | 1.54667838 |
| 9645 | NM_001163417.1 | FAM172A | 81.3629798 | -7.5534971 | 1.72617306 |
| 9646 | NM_001031725.5 | #N/A | 45.5723397 | -6.7237068 | 1.53671363 |
| 9647 | NM_001142282.1 | RPS24 | 68.0244485 | -7.3021307 | 1.6689647 |
| 9648 | NM_017622.2 | #N/A | 52.9558421 | -6.9347214 | 1.5849995 |
| 9649 | NM_016301.3 | #N/A | 55.760335 | -7.010146 | 1.60228301 |
| 9650 | NM_030957.3 | #N/A | 291.024325 | -4.1970955 | 0.9593248 |
| 9651 | NM_001011658.3 | #N/A | 54.334342 | -6.9726223 | 1.59380649 |
| 9652 | NM_014000.2 | VCL | 47.8120431 | -6.7958344 | 1.55346543 |
| 9653 | NM_001987.4 | #N/A | 95.7817705 | -6.3351055 | 1.44815574 |
| 9654 | NM_006396.2 | #N/A | 62.2923941 | -7.169611 | 1.63898569 |
| 9655 | NM_005868.5 | #N/A | 53.6750101 | -6.9573873 | 1.59080504 |
| 9656 | NM_138284.1 | IL17D | 57.3188052 | -7.0501359 | 1.61232169 |
| 9657 | NM_203453.3 | #N/A | 64.5698589 | -7.2220546 | 1.65175605 |
| 9658 | NM_012243.2 | #N/A | 60.8016598 | -7.1415408 | 1.63342917 |
| 9659 | NM_015997.3 | #N/A | 45.0638412 | -6.7037043 | 1.53342684 |
| 9660 | NM_001006946.1 | SDC1 | 64.8125108 | -7.2296981 | 1.65379235 |
| 9661 | NM_001008703.2 | #N/A | 69.2616866 | -7.3245611 | 1.67579967 |
| 9662 | NM_001297573.1 | TRERF1 | 61.6394224 | -7.1604362 | 1.63836163 |
| 9663 | NM_001316979.1 | #N/A | 60.9590721 | -7.1402918 | 1.63388377 |
| 9664 | NM_001318069.1 | #N/A | 59.6544441 | -7.1110136 | 1.62724096 |
| 9665 | NM_182491.2 | #N/A | 63.5483771 | -7.2003029 | 1.64769406 |
| 9666 | NM_014231.4 | #N/A | 74.6193778 | -7.4317693 | 1.70068684 |
| 9667 | NM_006670.4 | TPBG | 113.354381 | -8.0364983 | 1.83936684 |
| 9668 | NM_001286631.1 | RBM26 | 73.9316274 | -7.4174282 | 1.69772437 |
| 9669 | NM_001083908.1 | DNAAF2 | 46.8683855 | -6.7550401 | 1.54613703 |
| 9670 | NM_005023.3 | #N/A | 43.8829436 | -6.6640731 | 1.52557266 |
| 9671 | NM_001047434.2 | #N/A | 54.3791205 | -6.9789989 | 1.59777846 |
| 9672 | NM_178229.4 | #N/A | 71.3879213 | -5.3072512 | 1.21505247 |
| 9673 | NR_026984.1 | #N/A | 70.0349656 | -3.6188585 | 0.82851618 |
| 9674 | NM_001178074.1 | #N/A | 85.630634 | -7.628896 | 1.74670576 |

| | | | | | |
|---|---|---|---|---|---|
| 9675 | NM_016588.2 | #N/A | 119.500412 | -6.6132627 | 1.51418781 |
| 9676 | NM_006499.4 | LGALS8 | 53.7539896 | -6.953502 | 1.59209855 |
| 9677 | NM_001267046.1 | #N/A | 52.7612709 | -6.9289518 | 1.5865013 |
| 9678 | NM_016008.3 | #N/A | 46.1884221 | -6.7457946 | 1.54456505 |
| 9679 | NM_001010938.1 | #N/A | 68.7840565 | -7.3141001 | 1.67471888 |
| 9680 | NM_016173.4 | #N/A | 65.7566665 | -7.2456061 | 1.65902798 |
| 9681 | NM_001032291.2 | PSRC1 | 77.4099377 | -7.4850215 | 1.71396202 |
| 9682 | NM_024629.3 | #N/A | 61.5730744 | -7.1519245 | 1.63774605 |
| 9683 | NM_152581.3 | #N/A | 51.576121 | -6.8987669 | 1.57978942 |
| 9684 | NM_024581.5 | #N/A | 42.8039716 | -6.6331477 | 1.5192506 |
| 9685 | NM_021211.3 | #N/A | 64.9015405 | -7.2288956 | 1.65579753 |
| 9686 | NM_000252.2 | MTM1 | 42.9289733 | -6.6307418 | 1.51895306 |
| 9687 | NM_001330731.1 | SLC44A1 | 48.1836123 | -6.8004271 | 1.55791953 |
| 9688 | NM_001330580.1 | #N/A | 52.5533729 | -6.9255307 | 1.58658987 |
| 9689 | NM_016069.9 | #N/A | 66.6523165 | -7.268167 | 1.6651652 |
| 9690 | NM_016648.3 | #N/A | 60.4727367 | -7.1292808 | 1.63347411 |
| 9691 | NM_001286706.1 | #N/A | 57.560872 | -5.6878254 | 1.3032526 |
| 9692 | NM_001271662.1 | SEMA4F | 60.3083985 | -7.1257646 | 1.63281148 |
| 9693 | NM_022758.5 | #N/A | 53.5535047 | -6.9546697 | 1.59362362 |
| 9694 | NR_028138.1 | ATP2B1-AS1 | 63.6713594 | -7.2031295 | 1.6507549 |
| 9695 | NM_001297774.1 | #N/A | 61.3279472 | -7.1471525 | 1.63812114 |
| 9696 | NM_001330373.1 | FGD4 | 78.6918392 | -7.5053743 | 1.72026978 |
| 9697 | NM_175617.3 | #N/A | 80.7909766 | -7.5467652 | 1.72976586 |
| 9699 | NR_026659.1 | LOC727896 | 86.5020209 | 1.946678 | 0.44619727 |
| 9698 | NM_031940.3 | #N/A | 57.570246 | -7.0534658 | 1.61671948 |
| 9700 | NM_001153.4 | #N/A | 101.908189 | -7.8832739 | 1.80696037 |
| 9701 | NM_148923.3 | #N/A | 61.4295344 | -7.1510705 | 1.63925259 |
| 9702 | NM_001288953.1 | #N/A | 45.1317371 | -6.7007333 | 1.53605246 |
| 9703 | NM_000136.2 | #N/A | 59.9349263 | -7.1170696 | 1.63151228 |
| 9704 | NM_020739.3 | #N/A | 81.0736831 | -6.1203063 | 1.40301024 |
| 9705 | NM_006531.4 | #N/A | 47.1887276 | -6.7698738 | 1.55201315 |
| 9706 | NM_007148.4 | #N/A | 126.15158 | -8.1885315 | 1.87726448 |
| 9707 | NM_001010911.2 | #N/A | 54.5441894 | -6.9792827 | 1.60012096 |
| 9708 | NM_032326.3 | #N/A | 54.1556806 | -6.9684098 | 1.59766687 |
| 9709 | NM_052965.3 | #N/A | 68.7382896 | -7.3122647 | 1.67676736 |
| 9710 | NM_003289.3 | TPM2 | 63.4693814 | -7.2017009 | 1.65147628 |
| 9711 | NM_016352.3 | CPA4 | 52.3787588 | -6.923595 | 1.58771223 |
| 9712 | NR_003263.2 | SDHAP3 | 41.7354167 | -6.5951766 | 1.51255095 |
| 9713 | NM_014883.3 | #N/A | 55.1121721 | -6.9895338 | 1.60317693 |
| 9714 | NM_006439.4 | #N/A | 55.1454149 | -6.9980095 | 1.6051441 |
| 9715 | NM_001099642.1 | #N/A | 51.0188327 | -6.8822904 | 1.5786709 |
| 9716 | NM_172016.2 | TRIM39 | 45.9348195 | -6.7340398 | 1.54493149 |
| 9717 | NM_015009.2 | #N/A | 71.9196674 | -7.3822225 | 1.69367128 |

| | | | | | |
|---|---|---|---|---|---|
| 9718 | NM_001099638.1 | ZNF146 | 75.4408763 | -7.4512711 | 1.7097075 |
| 9719 | NR_132761.1 | STOX2 | 52.5879742 | -6.9251327 | 1.58908387 |
| 9720 | NR_037861.1 | PPT2-EGFL8 | 46.6566797 | -6.7582229 | 1.55090394 |
| 9721 | NM_001286659.2 | TBC1D4 | 62.2690204 | -7.1706058 | 1.64561851 |
| 9722 | NM_001303471.2 | #N/A | 61.5517561 | -7.1523335 | 1.64150723 |
| 9724 | NR_026758.2 | FAR2P1 | 83.1461393 | -7.5902655 | 1.74219568 |
| 9723 | NM_016373.3 | #N/A | 47.3406605 | -6.7733751 | 1.55469349 |
| 9725 | NM_004206.3 | #N/A | 50.0361664 | -6.8537032 | 1.57320697 |
| 9726 | NM_001002252.2 | ARL6IP4 | 54.1299792 | -6.9645947 | 1.598751 |
| 9727 | NM_001206747.1 | #N/A | 47.4092486 | -6.7781399 | 1.55596278 |
| 9728 | NM_005783.3 | #N/A | 70.3296279 | -7.3467094 | 1.68650511 |
| 9729 | NM_003763.5 | #N/A | 91.9209474 | -7.7354284 | 1.77580008 |
| 9730 | NM_004733.3 | #N/A | 50.8272818 | -6.8744337 | 1.57813794 |
| 9731 | NM_001165414.1 | LDHA | 73.2002584 | -7.4042458 | 1.69985054 |
| 9732 | NR_027303.1 | #N/A | 115.467977 | -6.5673286 | 1.50781972 |
| 9733 | NM_001198941.1 | #N/A | 58.6014797 | -7.0878433 | 1.62734988 |
| 9734 | NM_139286.3 | #N/A | 53.9618539 | -6.9623712 | 1.59857671 |
| 9735 | NM_001301168.1 | #N/A | 54.6282303 | -6.9833214 | 1.60344647 |
| 9736 | NM_152285.3 | #N/A | 48.2367967 | -6.8067679 | 1.5630013 |
| 9738 | NM_032941.2 | RECQL | 47.2732991 | -6.771257 | 1.55500056 |
| 9737 | NM_022163.3 | #N/A | 55.4247291 | -7.0029632 | 1.60821682 |
| 9739 | NM_001282198.1 | SLC25A14 | 43.3917219 | -6.6469632 | 1.5264902 |
| 9740 | NM_152414.4 | #N/A | 168.067666 | -8.6028943 | 1.97566696 |
| 9741 | NM_024913.4 | #N/A | 53.4389088 | -6.9540587 | 1.59712721 |
| 9742 | NM_002519.2 | #N/A | 50.1577941 | -6.8598607 | 1.57550168 |
| 9743 | NM_001289969.1 | PPP3CB | 65.7938445 | -7.247012 | 1.6644756 |
| 9745 | NR_040072.1 | NDRG4 | 44.6464641 | -6.6940567 | 1.53758916 |
| 9744 | NM_017950.3 | #N/A | 93.0350423 | -7.7519269 | 1.78057222 |
| 9746 | NM_013280.4 | FLRT1 | 44.7285562 | -6.6914277 | 1.53702132 |
| 9747 | NR_125362.1 | LOC100506679 | 57.7562674 | 2.17669674 | 0.49998779 |
| 9748 | NM_001256141.1 | #N/A | 48.9589828 | -6.8196192 | 1.56660093 |
| 9749 | NM_000750.3 | #N/A | 65.1571567 | -7.236887 | 1.66250606 |
| 9750 | NM_130802.2 | MEN1 | 54.5080577 | -6.9758187 | 1.60259746 |
| 9751 | NM_002101.4 | #N/A | 58.577955 | -7.0841839 | 1.62750369 |
| 9752 | NM_032907.4 | #N/A | 59.6850283 | -7.1099708 | 1.63344055 |
| 9753 | NM_198850.3 | PHLDB3 | 46.9234527 | -6.7612462 | 1.55355379 |
| 9754 | NM_003939.4 | #N/A | 87.781677 | -6.2186187 | 1.42888712 |
| 9755 | NM_001164440.1 | #N/A | 69.5347559 | -5.9225015 | 1.36097949 |
| 9756 | NM_006506.3 | #N/A | 65.4119687 | -7.2403479 | 1.66388087 |
| 9757 | NM_001127510.2 | #N/A | 68.3891356 | -7.309746 | 1.68005321 |
| 9758 | NM_024585.3 | #N/A | 50.7044435 | -6.8750396 | 1.58019688 |
| 9759 | NM_001017998.3 | #N/A | 73.370376 | -7.4082893 | 1.70282427 |
| 9760 | NM_014736.5 | #N/A | 59.3942318 | -7.0995712 | 1.63195286 |

| | | | | | |
|---|---|---|---|---|---|
| 9761 | NR_126432.1 | #N/A | 63.7263773 | -7.2061721 | 1.65646629 |
| 9762 | NM_004632.3 | #N/A | 57.6919598 | -7.0573638 | 1.62232046 |
| 9763 | NM_018467.3 | #N/A | 61.8417589 | -7.1605466 | 1.64606512 |
| 9764 | NM_001167675.1 | #N/A | 95.1185947 | -7.780759 | 1.78870908 |
| 9765 | NM_001018109.2 | #N/A | 44.6662307 | -6.6966533 | 1.53952681 |
| 9766 | NM_080717.2 | #N/A | 36.0699347 | 8.06949986 | 1.85517742 |
| 9767 | NM_001128205.1 | #N/A | 59.3847475 | -7.1037947 | 1.63325269 |
| 9768 | NM_017551.2 | #N/A | 150.317112 | -5.262914 | 1.21001354 |
| 9769 | NM_177477.3 | #N/A | 56.6902398 | -7.0376135 | 1.61813204 |
| 9770 | NM_016329.3 | #N/A | 46.4373719 | -6.7456483 | 1.55108967 |
| 9771 | NR_144366.1 | #N/A | 52.4555116 | -6.9185022 | 1.59091373 |
| 9772 | NM_003450.2 | #N/A | 44.0997933 | -6.6702293 | 1.53386098 |
| 9773 | NR_037882.1 | RTEL1-TNFRSF6B | 46.9642315 | -6.7668801 | 1.55623698 |
| 9774 | NM_012301.3 | #N/A | 95.5648956 | -7.7886471 | 1.79135725 |
| 9775 | NM_018319.3 | #N/A | 43.3704295 | -6.6515879 | 1.52983809 |
| 9776 | NM_003668.3 | #N/A | 48.3845491 | -6.8069005 | 1.56559976 |
| 9777 | NM_015329.3 | #N/A | 318.042679 | -2.7903623 | 0.64179664 |
| 9778 | NM_014007.3 | #N/A | 60.9356717 | -7.1356957 | 1.64126228 |
| 9779 | NM_032793.4 | #N/A | 78.2294617 | -7.4991062 | 1.72496648 |
| 9780 | NR_052003.1 | #N/A | 52.6352884 | -6.9318593 | 1.59460609 |
| 9781 | NM_022492.4 | #N/A | 45.0933837 | -6.7028977 | 1.54199596 |
| 9782 | NM_032593.2 | #N/A | 61.5318037 | -7.1529752 | 1.64569077 |
| 9783 | NM_012213.2 | #N/A | 49.1426398 | -6.8325892 | 1.57200644 |
| 9784 | NM_024828.3 | #N/A | 47.5062419 | -6.7793842 | 1.55986862 |
| 9785 | NM_004415.3 | #N/A | 88.6834155 | -7.6832383 | 1.76805659 |
| 9786 | NM_018381.3 | #N/A | 61.3273883 | -7.1522483 | 1.64589957 |
| 9787 | NR_135763.1 | #N/A | 55.7761209 | -7.0097736 | 1.61326341 |
| 9788 | NM_003599.3 | #N/A | 45.0018868 | -6.7045048 | 1.54304173 |
| 9789 | NM_001002002.2 | #N/A | 51.8234176 | -6.9041083 | 1.58903138 |
| 9790 | NM_001040152.1 | #N/A | 2137.17791 | -5.2288426 | 1.20348214 |
| 9791 | NR_110505.1 | LOC100508046 | 17.0803087 | 9.06873424 | 2.08729507 |
| 9792 | NM_001294340.1 | #N/A | 42.7080412 | -6.623356 | 1.52447101 |
| 9793 | NR_026827.1 | LINC00839 | 23.2407313 | 3.06048064 | 0.70449643 |
| 9794 | NM_002852.3 | #N/A | 118.77136 | -6.6017203 | 1.51984434 |
| 9795 | NM_018294.5 | #N/A | 60.0729329 | -7.1170928 | 1.63851731 |
| 9796 | NM_153260.2 | #N/A | 282.162187 | 1.74830623 | 0.40251516 |
| 9797 | NM_001306195.1 | ARPP19 | 178.515111 | -4.3519938 | 1.00204178 |
| 9798 | NM_198043.2 | ZDHHC16 | 61.4973614 | -7.1537877 | 1.64729109 |
| 9799 | NM_018719.4 | #N/A | 58.2629276 | -7.07131 | 1.6283642 |
| 9800 | NM_021830.4 | #N/A | 52.4456979 | -6.9250444 | 1.5949224 |
| 9801 | NM_001008860.2 | NIPA2 | 62.6236884 | -7.1793049 | 1.65350698 |
| 9802 | NM_000538.3 | RFXAP | 48.0974791 | -6.7967924 | 1.56552378 |
| 9803 | NM_173485.5 | #N/A | 64.546427 | -7.2264353 | 1.66451012 |

| | | | | | |
|---|---|---|---|---|---|
| 9804 | NM_001042463.2 | TMEM80 | 46.6250687 | -6.7502796 | 1.55488181 |
| 9805 | NM_022126.3 | #N/A | 47.3327336 | -6.7721281 | 1.56004601 |
| 9806 | NM_213622.2 | #N/A | 55.27379 | -6.9958431 | 1.61157736 |
| 9807 | NM_001145099.1 | #N/A | 63.2257459 | -7.1919112 | 1.65684079 |
| 9808 | NM_145043.2 | #N/A | 45.8824198 | -6.7276553 | 1.5500163 |
| 9809 | NM_033196.2 | #N/A | 64.6448825 | -7.2226579 | 1.6640938 |
| 9810 | NM_001130519.2 | #N/A | 94.1296552 | -7.7646454 | 1.7893068 |
| 9811 | NM_001040118.2 | ARAP1 | 45.9613086 | -6.7387664 | 1.5530342 |
| 9812 | NR_024117.2 | MSTO2P | 41.2509859 | -6.578454 | 1.51633172 |
| 9813 | NM_016067.3 | #N/A | 53.5353767 | -6.9508879 | 1.60222713 |
| 9814 | NM_001145450.1 | MORN2 | 46.7124837 | -6.7622852 | 1.55885776 |
| 9815 | NM_001299.5 | #N/A | 71.8326321 | -7.37889 | 1.70099236 |
| 9816 | NM_001004343.2 | #N/A | 64.7754317 | 2.72191553 | 0.62749214 |
| 9817 | NR_132422.1 | PCCA-DT | 68.4090099 | 2.26029279 | 0.52113477 |
| 9818 | NM_022474.3 | #N/A | 64.7422268 | -7.2289358 | 1.66695762 |
| 9819 | NM_022872.2 | #N/A | 53.9428132 | -6.9670539 | 1.60662429 |
| 9821 | NM_020459.1 | PAIP2B | 85.5499459 | -4.6574061 | 1.07425969 |
| 9820 | NM_001145443.2 | #N/A | 80.8694289 | -7.5462913 | 1.74060395 |
| 9822 | NM_000229.1 | #N/A | 43.5146321 | -6.6586243 | 1.53593283 |
| 9824 | NR_133003.1 | MIR9-3HG | 78.5131844 | -7.5015252 | 1.73040922 |
| 9823 | NM_001170766.1 | #N/A | 65.8068749 | -7.2495929 | 1.6723025 |
| 9825 | NM_001706.4 | #N/A | 44.335164 | -6.6775715 | 1.5403877 |
| 9826 | NM_032211.6 | #N/A | 48.100128 | -6.8046807 | 1.56972103 |
| 9827 | NM_203500.1 | #N/A | 49.2755778 | -6.8349995 | 1.57675859 |
| 9828 | NR_103777.1 | LRIG2-DT | 204.173208 | 2.67349449 | 0.61675134 |
| 9831 | NR_145429.1 | LINC01181 | 27.4437049 | 6.14380932 | 1.41771311 |
| 9829 | NM_001270998.1 | ZFAND2B | 58.156305 | -7.0705752 | 1.6315718 |
| 9830 | NM_012113.2 | #N/A | 50.041088 | -6.8606824 | 1.5831245 |
| 9832 | NM_001243313.1 | #N/A | 78.6534758 | -7.505518 | 1.7319899 |
| 9833 | NM_001044370.1 | #N/A | 222.137117 | -9.0055754 | 2.0784173 |
| 9834 | NM_004650.2 | #N/A | 45.8125514 | -6.7268021 | 1.55262628 |
| 9835 | NM_000169.2 | GLA | 52.2246685 | -6.9175656 | 1.59681531 |
| 9836 | NM_015127.4 | CLCC1 | 49.4274651 | -6.8407751 | 1.57921955 |
| 9837 | NM_207335.2 | #N/A | 41.1580269 | 2.40605228 | 0.55550613 |
| 9838 | NM_001024956.2 | SC5D | 44.1185885 | -6.6692162 | 1.53983237 |
| 9839 | NM_001323408.1 | #N/A | 49.834102 | -6.853774 | 1.5825269 |
| 9840 | NM_001099271.1 | #N/A | 48.0607569 | -6.7963888 | 1.56929843 |
| 9841 | NM_022555.3 | HLA-DRB3 | 83.6023357 | -7.5971868 | 1.75430481 |
| 9842 | NM_018210.3 | #N/A | 47.1541805 | -6.7740362 | 1.56441114 |
| 9843 | NM_145911.2 | #N/A | 46.044677 | -6.7329463 | 1.55493603 |
| 9844 | NM_001282791.1 | #N/A | 66.7411844 | -7.2714786 | 1.67933147 |
| 9845 | NM_017724.2 | LRRFIP2 | 59.9688396 | -7.1208564 | 1.64466199 |
| 9846 | NM_003083.3 | #N/A | 63.6529287 | -7.2019662 | 1.6634253 |

| | | | | | |
|---|---|---|---|---|---|
| 9847 | NM_001002248.2 | ANAPC11 | 65.8646719 | -7.2502943 | 1.67482368 |
| 9848 | NM_016028.4 | KMT5B | 81.5871444 | -7.5584431 | 1.74603745 |
| 9849 | NM_032125.2 | TMEM222 | 143.695915 | -5.6108695 | 1.29615326 |
| 9850 | NM_013410.3 | #N/A | 70.1480559 | -7.3423484 | 1.69615352 |
| 9851 | NR_132426.1 | ACAD11 | 60.0589602 | -7.1218131 | 1.645414 |
| 9852 | NM_004127.5 | #N/A | 71.1751022 | -7.3650962 | 1.70168096 |
| 9853 | NM_022822.2 | #N/A | 94.8613509 | -7.7803379 | 1.7977072 |
| 9854 | NM_001122670.1 | PHKA1 | 47.7580938 | -6.7882064 | 1.56852065 |
| 9855 | NM_025045.5 | #N/A | 41.9940423 | -6.6076506 | 1.52683795 |
| 9856 | NM_001252049.1 | PCMT1 | 79.2168896 | -7.5140388 | 1.73635023 |
| 9857 | NM_001178094.1 | #N/A | 84.534769 | -7.6083817 | 1.75824633 |
| 9858 | NM_022717.3 | #N/A | 51.0960531 | -6.8847911 | 1.5910428 |
| 9859 | NM_005116.5 | #N/A | 529.753378 | -3.9645504 | 0.91625429 |
| 9860 | NM_001010935.2 | RAP1A | 46.0339697 | -6.7414306 | 1.55808095 |
| 9861 | NM_003149.2 | STAC | 71.5085263 | -7.3705285 | 1.70350251 |
| 9862 | NM_152766.3 | #N/A | 59.2633966 | -7.0999275 | 1.64101741 |
| 9863 | NM_001004128.2 | QSOX1 | 66.7194009 | -7.2703174 | 1.68074175 |
| 9864 | NM_173664.5 | #N/A | 59.8021086 | -7.1115421 | 1.6440822 |
| 9865 | NM_014850.3 | #N/A | 143.654821 | -3.9747464 | 0.91896582 |
| 9867 | NM_001287045.1 | SH3GLB2 | 96.7129978 | -7.8051786 | 1.80465499 |
| 9866 | NM_001267547.2 | #N/A | 57.5572195 | -7.0551139 | 1.63122733 |
| 9868 | NM_006212.2 | PFKFB2 | 56.7250665 | -3.1869028 | 0.73688137 |
| 9869 | NM_001284325.1 | #N/A | 49.201733 | -6.837171 | 1.5809693 |
| 9870 | NM_001098816.2 | #N/A | 292.85908 | -4.7878743 | 1.1071572 |
| 9871 | NM_183057.2 | VPS28 | 56.4943691 | -7.027958 | 1.62525177 |
| 9872 | NM_032026.3 | #N/A | 51.0365755 | -6.8839171 | 1.59197644 |
| 9873 | NM_152791.4 | #N/A | 55.9014073 | -7.0103901 | 1.62126326 |
| 9874 | NM_153266.3 | #N/A | 73.3307338 | -7.4058621 | 1.71272176 |
| 9875 | NM_001195144.1 | #N/A | 79.2572877 | -7.5171888 | 1.73864975 |
| 9876 | NM_001129993.2 | #N/A | 60.4233156 | -7.1302372 | 1.64917163 |
| 9877 | NM_014066.3 | #N/A | 55.6021649 | -7.0064501 | 1.62062306 |
| 9878 | NM_014653.3 | #N/A | 53.2340976 | -6.9437227 | 1.60610622 |
| 9879 | NM_007147.3 | #N/A | 42.3315688 | -6.6168055 | 1.53051371 |
| 9880 | NM_020143.3 | #N/A | 61.8224284 | -7.160361 | 1.65625474 |
| 9881 | NM_001042724.1 | #N/A | 181.356715 | -5.1570281 | 1.19288264 |
| 9882 | NM_016223.4 | #N/A | 47.763997 | -6.7862195 | 1.56977877 |
| 9883 | NM_032501.3 | #N/A | 52.3907549 | -6.9188701 | 1.60068074 |
| 9884 | NM_003516.2 | H2AC18 | 41.6029309 | -6.5904626 | 1.52494416 |
| 9885 | NM_207291.2 | #N/A | 70.6338731 | -7.3546732 | 1.70181655 |
| 9886 | NM_032018.6 | #N/A | 41.5896244 | -6.5900807 | 1.52507365 |
| 9887 | NR_033243.2 | #N/A | 50.5245809 | -6.8657592 | 1.58890896 |
| 9888 | NM_198686.2 | #N/A | 64.2126621 | -7.2138696 | 1.66954333 |
| 9889 | NM_001286722.1 | #N/A | 72.8756596 | -7.3936422 | 1.71141058 |

| | | | | | |
|---|---|---|---|---|---|
| 9890 | NM_012174.1 | FBXW8 | 73.0732931 | -5.9828532 | 1.38496785 |
| 9891 | NM_018004.2 | #N/A | 48.9280811 | -6.827608 | 1.58056917 |
| 9892 | NM_019099.4 | #N/A | 64.0748788 | -7.2103423 | 1.66916315 |
| 9893 | NM_001328650.1 | PIK3R3 | 76.2149398 | -7.4590808 | 1.72681813 |
| 9894 | NM_001199580.1 | #N/A | 64.1403664 | -7.2149068 | 1.67041017 |
| 9895 | NR_135199.1 | FAM78B | 69.559838 | -7.3303455 | 1.69719342 |
| 9896 | NM_145062.2 | #N/A | 61.3726632 | -7.1494853 | 1.65532844 |
| 9897 | NM_001318784.1 | #N/A | 99.5379718 | -7.8465778 | 1.81673673 |
| 9898 | NM_001346218.1 | #N/A | 73.9334055 | -7.416445 | 1.7172482 |
| 9899 | NM_018944.2 | #N/A | 61.8445647 | -5.8796589 | 1.36156721 |
| 9900 | NM_213590.2 | #N/A | 49.2102447 | -6.8255775 | 1.58061151 |
| 9901 | NM_182836.2 | RABGGTA | 50.6441064 | -6.8715356 | 1.5913235 |
| 9902 | NM_013342.3 | TFPT | 57.6355158 | -7.0570846 | 1.63436347 |
| 9903 | NM_001330232.1 | #N/A | 91.2941281 | -7.7241771 | 1.7890638 |
| 9904 | NM_004774.3 | #N/A | 269.9369 | -5.1545322 | 1.19389982 |
| 9905 | NM_199245.2 | #N/A | 42.0465544 | -6.6066754 | 1.5303229 |
| 9906 | NM_013443.4 | ST6GALNAC6 | 52.2322896 | -6.9185495 | 1.60270503 |
| 9907 | NM_019104.2 | #N/A | 60.1361602 | -7.1207744 | 1.6496311 |
| 9908 | NM_000201.2 | #N/A | 64.2408629 | -5.9430343 | 1.37682807 |
| 9909 | NM_024646.2 | #N/A | 352.942469 | -3.113736 | 0.72138503 |
| 9910 | NM_001204197.1 | NBEA | 119.364484 | -6.6018571 | 1.52965212 |
| 9911 | NM_001235.3 | #N/A | 1546.65823 | -5.0125745 | 1.16141876 |
| 9912 | NM_032174.5 | #N/A | 68.7889466 | -7.3150194 | 1.6951315 |
| 9913 | NM_001007466.2 | #N/A | 57.7350071 | -7.0587508 | 1.63577826 |
| 9914 | NM_001005415.1 | MARCHF2 | 63.3212347 | -7.1942113 | 1.66731103 |
| 9915 | NM_001271581.1 | #N/A | 57.6297935 | -7.0543988 | 1.63498796 |
| 9916 | NM_032332.3 | #N/A | 47.1948585 | -6.76587 | 1.56823198 |
| 9917 | NM_213618.1 | #N/A | 81.4281734 | -7.5609056 | 1.75250767 |
| 9918 | NM_001242858.2 | PRKACB | 81.6706201 | -7.5614797 | 1.75270204 |
| 9919 | NM_001105244.1 | PTPRM | 91.2820912 | -7.7201062 | 1.78951404 |
| 9920 | NM_020809.3 | ARHGAP20 | 44.7427399 | -6.6963458 | 1.55245385 |
| 9921 | NM_001159676.1 | SYNCRIP | 51.973322 | -6.9082753 | 1.60170578 |
| 9922 | NM_005645.3 | #N/A | 50.0150204 | -6.8560247 | 1.58959849 |
| 9923 | NM_144778.3 | #N/A | 56.4739177 | -7.0281019 | 1.62951837 |
| 9924 | NM_001009894.2 | #N/A | 62.2544755 | -7.1683313 | 1.6621127 |
| 9925 | NM_006198.2 | #N/A | 52.3602754 | -6.9233603 | 1.60543064 |
| 9926 | NM_001039548.2 | KLHL35 | 62.4434159 | -7.1725751 | 1.66329926 |
| 9927 | NM_145160.2 | #N/A | 52.3611499 | -6.9235703 | 1.60556155 |
| 9928 | NM_001289930.1 | ZNF718 | 48.8094464 | -6.8156457 | 1.58058122 |
| 9929 | NM_015101.3 | #N/A | 62.1299183 | -7.1705712 | 1.66297852 |
| 9930 | NM_004117.3 | FKBP5 | 45.6618202 | -6.729762 | 1.56081879 |
| 9931 | NM_001166271.2 | #N/A | 61.8508467 | -7.157502 | 1.66016356 |
| 9932 | NM_004560.3 | #N/A | 61.7518758 | -7.1623443 | 1.66131647 |

| | | | | | |
|---|---|---|---|---|---|
| 9933 | NR_046462.1 | FAM169A | 65.2999247 | -7.2384759 | 1.67899563 |
| 9935 | NM_198441.2 | PRR20A | 26.0384306 | 9.66645459 | 2.24225897 |
| 9934 | NM_001168.2 | #N/A | 64.970234 | -7.2299379 | 1.67707567 |
| 9936 | NM_181889.2 | UBE2D3 | 67.4365628 | -7.2853884 | 1.68995606 |
| 9937 | NM_021105.2 | #N/A | 49.9282141 | -6.857574 | 1.59072756 |
| 9938 | NM_153271.1 | #N/A | 41.6969891 | -6.5967185 | 1.53029948 |
| 9939 | NM_006051.3 | APBB3 | 54.5376429 | -6.9822898 | 1.61978313 |
| 9940 | NM_138335.2 | #N/A | 45.5182553 | -6.7149029 | 1.55780823 |
| 9941 | NM_015659.2 | #N/A | 572.55336 | -4.8119209 | 1.11634714 |
| 9942 | NM_007057.3 | #N/A | 60.7297019 | -7.133984 | 1.65524036 |
| 9943 | NM_001012759.2 | #N/A | 63.9531255 | -7.2086254 | 1.67269513 |
| 9944 | NM_015960.2 | #N/A | 43.9764168 | -6.6684059 | 1.54736716 |
| 9945 | NM_002581.4 | #N/A | 97.8722954 | -6.3505202 | 1.4738774 |
| 9946 | NM_005876.4 | #N/A | 233.640322 | -3.1632716 | 0.73420991 |
| 9947 | NM_001142616.1 | #N/A | 77.6033248 | -7.4866853 | 1.73785094 |
| 9948 | NM_182760.3 | #N/A | 55.0124333 | -6.9915562 | 1.62296037 |
| 9949 | NR_024006.1 | #N/A | 48.3376195 | -6.8036304 | 1.57936019 |
| 9950 | NM_015052.4 | #N/A | 74.3641317 | -7.426361 | 1.72403481 |
| 9951 | NM_018477.2 | #N/A | 367.739648 | -3.6414126 | 0.8453638 |
| 9952 | NM_012343.3 | #N/A | 57.3134983 | -7.0484127 | 1.63632701 |
| 9953 | NM_173686.2 | #N/A | 43.7799949 | -6.6676906 | 1.54797916 |
| 9954 | NM_016287.3 | #N/A | 863.132351 | -2.7305116 | 0.63394163 |
| 9955 | NM_001014834.2 | #N/A | 41.7710713 | -6.5985836 | 1.5320486 |
| 9956 | NM_001309443.1 | #N/A | 149.112866 | -6.8828881 | 1.59811757 |
| 9957 | NM_001330785.1 | #N/A | 90.611803 | -7.7102512 | 1.79034448 |
| 9958 | NM_022165.2 | #N/A | 74.4191245 | -7.4280735 | 1.72493976 |
| 9959 | NM_001098633.3 | #N/A | 56.7153669 | -7.0334578 | 1.63382203 |
| 9960 | NM_001099679.1 | #N/A | 68.0079562 | -7.3006851 | 1.69597402 |
| 9961 | NR_049729.1 | ASB16-AS1 | 43.4599055 | -6.6540512 | 1.54579418 |
| 9962 | NM_001321228.1 | #N/A | 44.6531397 | -6.6872648 | 1.55351808 |
| 9963 | NM_181643.5 | #N/A | 97.5095095 | -7.8196557 | 1.81667626 |
| 9964 | NM_001322206.1 | #N/A | 67.5536027 | -7.2863248 | 1.6929702 |
| 9965 | NM_032777.9 | #N/A | 551.55737 | -5.4377185 | 1.26357745 |
| 9966 | NM_015490.3 | #N/A | 127.986956 | -2.9363415 | 0.68239291 |
| 9967 | NM_003783.3 | B3GALT2 | 83.6471632 | -7.5959961 | 1.76548576 |
| 9968 | NM_016950.2 | SPOCK3 | 69.3625651 | -7.3296345 | 1.70407223 |
| 9969 | NM_001002009.2 | NT5C3A | 57.786092 | -7.0593177 | 1.64146882 |
| 9970 | NM_024681.3 | KCTD17 | 50.5930439 | -6.8765413 | 1.59905203 |
| 9971 | NM_003800.4 | #N/A | 77.3083038 | -7.4854362 | 1.74080186 |
| 9972 | NM_004065.2 | CDR1 | 67.8288399 | -7.2907089 | 1.69558753 |
| 9973 | NM_001040108.1 | MLH3 | 51.8336325 | -6.8998798 | 1.60479849 |
| 9974 | NM_001321693.1 | #N/A | 69.2065147 | -6.0482283 | 1.40676756 |
| 9975 | NM_001161345.1 | CHFR | 72.0458678 | -7.3825705 | 1.71763682 |

| | | | | | |
|---|---|---|---|---|---|
| 9976 | NR_026717.1 | STK19 | 59.9560567 | -7.1125548 | 1.65485573 |
| 9977 | NM_181652.2 | #N/A | 64.9095701 | -7.2313386 | 1.68255244 |
| 9978 | NM_018913.2 | PCDHGA10 | 63.6038965 | -7.2011363 | 1.67570263 |
| 9979 | NM_005441.2 | #N/A | 41.6948251 | -6.5898412 | 1.53358209 |
| 9980 | NM_016213.4 | #N/A | 53.0488366 | -6.9381289 | 1.61479002 |
| 9981 | NM_000086.2 | CLN3 | 58.4559038 | -7.0787284 | 1.64761301 |
| 9982 | NM_001329139.1 | PTPRF | 126.743313 | -8.1964855 | 1.90778281 |
| 9983 | NM_001080469.1 | #N/A | 49.9405722 | -6.852217 | 1.59490556 |
| 9984 | NM_014184.3 | #N/A | 52.0726049 | -6.9137824 | 1.60927354 |
| 9985 | NM_001184819.1 | #N/A | 65.0556383 | -7.2383569 | 1.68510747 |
| 9986 | NM_020666.2 | #N/A | 59.9006486 | -5.7242632 | 1.33271893 |
| 9987 | NM_177966.6 | #N/A | 103.568791 | -3.5246382 | 0.82063706 |
| 9988 | NM_001282391.1 | #N/A | 67.4459316 | -7.2863642 | 1.69673642 |
| 9989 | NR_131336.1 | #N/A | 60.2471883 | -7.1255013 | 1.6592724 |
| 9990 | NM_032847.2 | #N/A | 49.3720878 | -6.8363335 | 1.59202216 |
| 9991 | NM_005772.4 | #N/A | 42.8654738 | -6.6323221 | 1.54452172 |
| 9992 | NM_024638.3 | #N/A | 43.6243745 | -6.6541169 | 1.54962135 |
| 9993 | NM_006165.3 | NFRKB | 51.0324478 | -6.8882325 | 1.6042141 |
| 9994 | NM_024603.3 | #N/A | 62.5071132 | -7.1742077 | 1.67085844 |
| 9995 | NM_182633.2 | #N/A | 272.838984 | 1.42234948 | 0.33128503 |
| 9996 | NM_018025.2 | #N/A | 55.4502786 | -7.000244 | 1.63052662 |
| 9997 | NM_030919.2 | #N/A | 49.7488299 | -6.8467974 | 1.59482403 |
| 9998 | NM_001321328.1 | #N/A | 42.8000676 | -6.6261913 | 1.54345683 |
| 9999 | NM_001160246.1 | RPAIN | 71.0358117 | -7.359825 | 1.71440934 |
| 10000 | NM_001127448.1 | #N/A | 163.894786 | -8.5680544 | 1.99586793 |
| 10001 | NM_018983.3 | #N/A | 57.0102991 | -7.0420664 | 1.64055825 |
| 10002 | NM_001039887.2 | #N/A | 42.0839595 | -6.6096821 | 1.53986132 |
| 10003 | NM_005853.5 | #N/A | 58.4601811 | -7.0810343 | 1.64966494 |
| 10005 | NR_045196.2 | SNHG18 | 40.5892007 | -6.5569478 | 1.52761515 |
| 10004 | NM_001005476.2 | #N/A | 72.8270553 | -7.3993477 | 1.72387831 |
| 10006 | NM_023106.2 | #N/A | 82.9984473 | -7.587022 | 1.76770637 |
| 10007 | NM_004715.4 | CTDP1 | 59.1260776 | -7.0933227 | 1.65269116 |
| 10008 | NR_145420.1 | LINC00355 | 56.8366467 | 3.93101486 | 0.91592374 |
| 10009 | NM_002514.3 | #N/A | 56.7468028 | -5.6578286 | 1.31833261 |
| 10010 | NM_021047.2 | #N/A | 44.6933608 | -6.6879383 | 1.55838399 |
| 10011 | NM_001415.3 | #N/A | 994.808444 | -2.4708838 | 0.57577461 |
| 10012 | NM_145169.2 | #N/A | 43.3999371 | -6.6510904 | 1.5498935 |
| 10013 | NM_001288634.1 | #N/A | 64.281872 | -7.2206193 | 1.68262838 |
| 10014 | NM_153617.1 | #N/A | 54.3515363 | -6.9761442 | 1.62570645 |
| 10015 | NM_001256617.1 | TUBGCP2 | 90.303475 | -6.2460775 | 1.45562824 |
| 10016 | NM_198149.2 | SHISA4 | 57.3277787 | -7.0516007 | 1.64339377 |
| 10017 | NM_006190.4 | #N/A | 50.0702303 | -6.8551742 | 1.59762595 |
| 10018 | NM_001077440.1 | BCLAF1 | 61.6796856 | -7.1561005 | 1.66783433 |

| | | | | | |
|---|---|---|---|---|---|
| 10019 | NM_201277.2 | #N/A | 90.7681702 | 2.69453735 | 0.62804366 |
| 10020 | NM_001278604.1 | #N/A | 53.585706 | -6.9509764 | 1.62022661 |
| 10021 | NM_018334.4 | #N/A | 54.2328764 | -6.9738982 | 1.62570802 |
| 10022 | NM_001301726.1 | #N/A | 83.4416161 | -3.7659347 | 0.8780672 |
| 10023 | NM_145176.2 | #N/A | 87.7067244 | -6.2084896 | 1.44790901 |
| 10024 | NM_152222.1 | #N/A | 48.4518167 | -6.8086195 | 1.58787555 |
| 10025 | NM_003944.3 | #N/A | 47.9876911 | -6.8013315 | 1.58619528 |
| 10026 | NM_005600.2 | #N/A | 53.7717361 | -6.9620532 | 1.62373134 |
| 10027 | NM_001330495.1 | #N/A | 67.136171 | -7.277332 | 1.69731262 |
| 10028 | NM_007131.4 | #N/A | 39.9321765 | -6.5340255 | 1.52395148 |
| 10029 | NM_001294348.1 | #N/A | 81.3925375 | -7.5542604 | 1.7619442 |
| 10030 | NM_015941.3 | #N/A | 57.1261299 | -7.0470099 | 1.64362798 |
| 10031 | NM_024884.2 | #N/A | 688.133955 | 2.23330046 | 0.52092202 |
| 10032 | NM_001316943.1 | #N/A | 44.9341394 | -6.6957246 | 1.56180544 |
| 10033 | NM_024613.3 | #N/A | 42.2751761 | -6.6068207 | 1.54108393 |
| 10034 | NM_001128161.1 | UBP1 | 58.7375759 | -7.0892766 | 1.65378325 |
| 10035 | NM_001029884.2 | #N/A | 85.3632655 | -7.6276629 | 1.77949891 |
| 10036 | NR_024458.1 | TPT1-AS1 | 88.5155982 | -2.7225734 | 0.6351731 |
| 10037 | NM_001348149.1 | #N/A | 54.3524444 | -6.9747757 | 1.62725795 |
| 10038 | NM_001331036.1 | #N/A | 51.5739839 | -6.8982974 | 1.60943863 |
| 10039 | NR_108047.1 | RNF139-AS1 | 99.0109762 | 2.86350228 | 0.66809711 |
| 10040 | NM_001303007.1 | #N/A | 41.8727807 | -6.6010023 | 1.54036557 |
| 10041 | NM_201543.2 | #N/A | 45.8652728 | -6.7330021 | 1.57150447 |
| 10042 | NM_001174084.1 | #N/A | 43.2589122 | -6.6446466 | 1.55096754 |
| 10043 | NM_004037.7 | #N/A | 49.8476566 | -6.8541416 | 1.59992054 |
| 10044 | NM_020235.6 | #N/A | 70.193754 | -7.3388089 | 1.71311236 |
| 10045 | NM_213594.2 | RFX4 | 64.98841 | -7.2308875 | 1.68796088 |
| 10046 | NM_001319066.1 | #N/A | 55.9963365 | -7.0156046 | 1.63805984 |
| 10047 | NM_173632.3 | #N/A | 104.632853 | -5.2592384 | 1.22803727 |
| 10048 | NM_138361.5 | LRSAM1 | 53.3189202 | -6.9433462 | 1.62142188 |
| 10049 | NM_016457.4 | PRKD2 | 56.1428579 | -7.0256383 | 1.64073749 |
| 10050 | NM_001206941.1 | #N/A | 182.777243 | -5.8650763 | 1.36971247 |
| 10051 | NM_005540.2 | #N/A | 58.2114458 | -7.0774139 | 1.65287928 |
| 10052 | NM_001199562.1 | #N/A | 44.6710556 | -6.6981766 | 1.56435935 |
| 10053 | NM_001135654.1 | #N/A | 77.7586946 | -7.4903025 | 1.74972972 |
| 10054 | NM_014729.2 | #N/A | 56.0145468 | -7.0170834 | 1.63928287 |
| 10055 | NM_001039211.2 | #N/A | 318.983863 | 3.06019333 | 0.71495604 |
| 10056 | NM_001114100.2 | #N/A | 127.4719 | -8.2042538 | 1.916788 |
| 10057 | NM_018095.5 | #N/A | 43.2355966 | -6.6463868 | 1.55299984 |
| 10058 | NM_002431.3 | #N/A | 46.4958447 | -6.7450507 | 1.57608115 |
| 10059 | NM_020200.6 | #N/A | 54.9947587 | -6.9878358 | 1.63299912 |
| 10060 | NM_004568.5 | SERPINB6 | 53.0737907 | -6.941479 | 1.62229004 |
| 10061 | NM_005836.2 | #N/A | 41.3578178 | -6.5755296 | 1.53675909 |

| | | | | | | |
|---|---|---|---|---|---|---|
| 10062 | NM_001948.3 | | #N/A | 50.0909593 | -6.8591767 | 1.60313127 |
| 10063 | NM_002907.3 | | #N/A | 59.1691675 | -7.0956536 | 1.65846955 |
| 10064 | NM_001278231.1 | | #N/A | 56.9221329 | -7.0419019 | 1.64614826 |
| 10065 | NM_018479.3 | ECHDC1 | | 72.2377147 | -7.3866185 | 1.72677906 |
| 10066 | NM_000280.4 | PAX6 | | 65.8957628 | -7.2478376 | 1.69451455 |
| 10067 | NM_005708.4 | | #N/A | 263.860646 | -5.1191109 | 1.19683354 |
| 10068 | NM_199437.1 | | #N/A | 53.0025978 | -6.9376601 | 1.62231797 |
| 10069 | NM_012145.3 | | #N/A | 54.430322 | -6.9759432 | 1.63139719 |
| 10070 | NM_021080.3 | | #N/A | 85.8631635 | -7.633135 | 1.78514034 |
| 10071 | NM_033281.5 | | #N/A | 58.5987974 | -7.0831958 | 1.65674245 |
| 10072 | NM_020192.4 | | #N/A | 46.0961169 | -6.7358897 | 1.57556662 |
| 10073 | NM_000387.5 | | #N/A | 51.397589 | -6.894066 | 1.61272883 |
| 10074 | NM_003815.4 | | #N/A | 51.3965343 | -6.8957497 | 1.61331604 |
| 10075 | NM_001038618.2 | | #N/A | 39.0883962 | -6.5012463 | 1.52105292 |
| 10076 | NR_033669.1 | | #N/A | 50.2997782 | -6.8621025 | 1.60556585 |
| 10077 | NM_207446.2 | | #N/A | 52.1679522 | -6.9183221 | 1.61877418 |
| 10078 | NM_014895.3 | | #N/A | 40.6977135 | -6.5545652 | 1.53376472 |
| 10079 | NM_006056.4 | | #N/A | 82.8313017 | 3.0718391 | 0.71884147 |
| 10080 | NM_001265596.1 | PFDN6 | | 56.5842297 | -7.0346192 | 1.64619979 |
| 10081 | NM_001286834.1 | | #N/A | 87.885142 | -3.7395139 | 0.87513846 |
| 10082 | NR_024122.1 | | #N/A | 63.7725362 | -7.2058041 | 1.68634418 |
| 10083 | NM_016401.3 | | #N/A | 74.2418108 | -7.422814 | 1.73730821 |
| 10084 | NM_000408.4 | | #N/A | 60.4312668 | -7.125671 | 1.66827629 |
| 10085 | NM_080881.2 | DBN1 | | 50.4080134 | -6.8684038 | 1.60806996 |
| 10086 | NM_006483.2 | | #N/A | 56.4923867 | -7.0282415 | 1.64555265 |
| 10087 | NM_001201366.1 | | #N/A | 83.0456754 | -7.5873371 | 1.77646728 |
| 10088 | NM_001161358.1 | | #N/A | 60.9916614 | -7.1388781 | 1.67162444 |
| 10089 | NM_172056.2 | KCNH2 | | 54.3638027 | -6.9741029 | 1.63316979 |
| 10090 | NM_001538.3 | | #N/A | 53.1173176 | -6.9355337 | 1.62416433 |
| 10091 | NM_001289413.1 | EHMT2 | | 77.2593428 | -7.4815611 | 1.75207671 |
| 10092 | NM_021127.2 | | #N/A | 40.0987152 | -6.5407959 | 1.53177256 |
| 10093 | NM_001271441.1 | | #N/A | 39.3990402 | -6.5127875 | 1.52525072 |
| 10094 | NM_001127218.2 | POLD2 | | 52.8747496 | -6.9318133 | 1.62344201 |
| 10095 | NM_004701.3 | | #N/A | 69.2398119 | -6.0471101 | 1.41642614 |
| 10096 | NM_016299.3 | | #N/A | 45.5733445 | -6.7201169 | 1.57417936 |
| 10097 | NM_001113525.1 | | #N/A | 44.4352896 | -6.683418 | 1.56579559 |
| 10098 | NM_016332.3 | MSRB1 | | 50.8595628 | -6.8787193 | 1.61171615 |
| 10099 | NM_005613.5 | RGS4 | | 48.9638209 | -6.8259112 | 1.59938011 |
| 10100 | NM_015428.3 | | #N/A | 50.6084907 | -6.8664289 | 1.60904441 |
| 10101 | NM_004581.5 | RABGGTA | | 50.746786 | -6.8745757 | 1.61100338 |
| 10102 | NR_046186.1 | | #N/A | 64.4619547 | 2.98864541 | 0.70040034 |
| 10103 | NM_001085354.2 | | #N/A | 62.9853157 | -7.1863553 | 1.68422138 |
| 10104 | NR_024491.1 | | #N/A | 17.6577763 | 4.54784666 | 1.0659431 |

| | | | | | |
|---|---|---|---|---|---|
| 10105 | NM_001257096.1 | PAX1 | 74.0502134 | -7.4218877 | 1.73962322 |
| 10106 | NM_198264.1 | #N/A | 49.7346408 | -6.8451943 | 1.6044675 |
| 10107 | NM_001111077.1 | #N/A | 452.96699 | -4.079049 | 0.95611123 |
| 10108 | NM_006530.3 | #N/A | 59.9053368 | -7.1125863 | 1.66723459 |
| 10111 | NM_001322412.1 | PNISR | 66.9920617 | -7.2764357 | 1.70576883 |
| 10109 | NM_001184700.1 | #N/A | 19.4177406 | 9.2726139 | 2.17374829 |
| 10110 | NM_001297736.1 | #N/A | 19.4177406 | 9.2726139 | 2.17374829 |
| 10112 | NM_032207.3 | #N/A | 42.0268035 | -6.6025448 | 1.54781639 |
| 10113 | NM_001316951.1 | GPALPP1 | 41.0294232 | -6.5666171 | 1.53942529 |
| 10114 | NM_181332.2 | #N/A | 116.205277 | -8.0709836 | 1.89209994 |
| 10115 | NM_177925.3 | H2AJ | 47.4917101 | -6.7802737 | 1.58954897 |
| 10116 | NR_024440.1 | LOC729609 | 18.1857002 | 3.37029851 | 0.79014874 |
| 10117 | NM_001277285.1 | #N/A | 47.7475537 | -6.7869793 | 1.59118119 |
| 10118 | NR_027405.1 | MTHFD2 | 61.9117033 | -7.1651623 | 1.67988326 |
| 10119 | NM_001330760.1 | #N/A | 77.2899351 | -7.4801276 | 1.75374314 |
| 10120 | NM_001278431.1 | #N/A | 64.0430859 | -7.2149472 | 1.69178014 |
| 10121 | NM_001077427.3 | #N/A | 68.5946304 | -7.3115671 | 1.71445865 |
| 10122 | NM_001164277.1 | SLC37A4 | 38.8630804 | -6.4876142 | 1.52145437 |
| 10123 | NM_001281789.1 | CNEP1R1 | 62.0897358 | -7.1641871 | 1.68025265 |
| 10124 | NM_002439.4 | #N/A | 92.1871577 | -3.2333007 | 0.75842007 |
| 10125 | NM_001004354.2 | #N/A | 51.5057389 | -6.8919732 | 1.61665748 |
| 10126 | NM_024574.3 | #N/A | 56.4451327 | -7.0327134 | 1.6496723 |
| 10128 | NM_001243246.1 | P3H1 | 42.1556831 | -6.6124699 | 1.55114692 |
| 10127 | NM_001220473.2 | #N/A | 67.2925177 | -7.2783957 | 1.70736291 |
| 10131 | NR_133948.1 | TOR1B | 74.1828909 | -7.4206706 | 1.74092875 |
| 10129 | NM_001297553.1 | #N/A | 567.464822 | -4.2663027 | 1.00089908 |
| 10130 | NM_018451.4 | #N/A | 55.0888255 | -6.9917965 | 1.64030846 |
| 10132 | NM_001196.3 | BID | 54.4035536 | -6.9754545 | 1.63665431 |
| 10133 | NR_137176.1 | MIR100HG | 96.8774726 | -7.8085292 | 1.83219149 |
| 10134 | NM_001013625.3 | #N/A | 63.8207118 | -7.2095141 | 1.69168186 |
| 10135 | NM_145739.2 | #N/A | 48.1500909 | 10.5764434 | 2.48190981 |
| 10136 | NM_152760.2 | #N/A | 67.0566279 | -7.2763349 | 1.70750332 |
| 10137 | NM_017775.3 | #N/A | 246.849377 | -3.8520242 | 0.90396368 |
| 10138 | NM_004275.4 | #N/A | 49.2412649 | -6.8333529 | 1.60370958 |
| 10139 | NM_001145860.1 | #N/A | 53.0804635 | -6.9392656 | 1.62871284 |
| 10140 | NM_172251.2 | #N/A | 59.2296308 | -7.0983466 | 1.6660721 |
| 10141 | NM_001146343.1 | #N/A | 48.3238363 | -6.8039383 | 1.59700125 |
| 10142 | NR_132442.1 | LUADT1 | 135.907412 | 5.07148674 | 1.19040998 |
| 10143 | NM_138421.2 | #N/A | 53.0789144 | -6.9386837 | 1.62875009 |
| 10144 | NM_001142529.1 | BHLHB9 | 41.7811445 | -6.5942587 | 1.54791805 |
| 10145 | NM_015564.2 | #N/A | 76.2646938 | -6.1866288 | 1.45239291 |
| 10146 | NM_005133.2 | #N/A | 54.8419352 | -6.9856532 | 1.64008203 |
| 10147 | NM_001286440.1 | NME4 | 51.7507574 | -6.9071551 | 1.62183683 |

| | | | | | |
|---|---|---|---|---|---|
| 10148 | NM_001008662.2 | KYAT3 | 61.690402 | -5.7556881 | 1.35149308 |
| 10149 | NM_032027.2 | #N/A | 62.4308381 | -7.1718845 | 1.68413995 |
| 10150 | NM_025215.5 | #N/A | 53.4582016 | -6.9474726 | 1.6314858 |
| 10151 | NM_181506.4 | #N/A | 18.5191074 | 4.34197056 | 1.01967713 |
| 10152 | NM_025112.4 | #N/A | 40.9995419 | -6.5677153 | 1.54239871 |
| 10153 | NM_017444.5 | #N/A | 42.5232988 | -6.6143652 | 1.55337222 |
| 10154 | NM_181776.2 | #N/A | 128.997871 | 3.81328239 | 0.89553816 |
| 10155 | NM_014698.2 | #N/A | 44.0199701 | -6.6763599 | 1.56808694 |
| 10157 | NM_003848.3 | SUCLG2 | 54.8092546 | -5.7099886 | 1.3411571 |
| 10156 | NM_001029859.2 | #N/A | 62.9977418 | -4.1765307 | 0.98098255 |
| 10158 | NR_036442.1 | #N/A | 76.4068814 | -6.0263111 | 1.41546919 |
| 10160 | NM_001698.2 | AUH | 47.2360331 | -6.7785232 | 1.59221488 |
| 10159 | NM_001243349.1 | #N/A | 54.0599327 | -6.968236 | 1.6367737 |
| 10161 | NM_021948.4 | #N/A | 69.9779838 | -7.338981 | 1.72398758 |
| 10162 | NM_032790.3 | ORAI1 | 38.9928704 | -6.4975145 | 1.52646717 |
| 10163 | NM_001318082.1 | #N/A | 52.8176884 | -6.9293023 | 1.62799257 |
| 10164 | NM_030758.3 | #N/A | 47.9133421 | -6.7895948 | 1.59527348 |
| 10166 | NM_174976.2 | ZDHHC22 | 65.9781448 | -7.2551174 | 1.70471869 |
| 10165 | NM_004748.4 | #N/A | 47.5164487 | -6.783945 | 1.5940027 |
| 10167 | NR_144477.1 | #N/A | 54.673163 | -6.9870421 | 1.64178951 |
| 10168 | NM_001253387.1 | CHL1 | 89.3318483 | -7.6907766 | 1.80742421 |
| 10169 | NM_005531.2 | #N/A | 45.322207 | -6.7143518 | 1.57794653 |
| 10170 | NR_130704.1 | #N/A | 48.851608 | -6.8215261 | 1.60316889 |
| 10171 | NM_001197113.1 | IQCJ-SCHIP1 | 58.4243463 | -7.082051 | 1.66443943 |
| 10172 | NM_032355.3 | MON1A | 63.0054927 | -7.1875216 | 1.68927053 |
| 10173 | NM_016567.3 | #N/A | 55.330977 | -6.9976271 | 1.64465115 |
| 10174 | NM_022481.5 | #N/A | 50.7049535 | -6.8677957 | 1.61415167 |
| 10175 | NR_027671.2 | #N/A | 56.7841033 | -7.0400073 | 1.65462331 |
| 10176 | NM_018640.4 | #N/A | 138.692075 | -8.3255545 | 1.95681861 |
| 10177 | NR_024209.1 | #N/A | 51.4752131 | -6.8916733 | 1.6199664 |
| 10178 | NM_001330611.1 | #N/A | 44.3942257 | -6.6849052 | 1.57137839 |
| 10179 | NM_001288580.1 | BROX | 55.6836427 | -7.0048312 | 1.64667566 |
| 10180 | NM_001077664.2 | URGCP | 119.737546 | -8.1140724 | 1.90744882 |
| 10181 | NM_001023560.3 | ZSCAN26 | 55.3410093 | -6.9975013 | 1.64497826 |
| 10182 | NM_012257.3 | #N/A | 55.460823 | -7.0066396 | 1.64720471 |
| 10183 | NR_036477.1 | TMEM198B | 43.9528202 | -6.6663561 | 1.56726856 |
| 10184 | NM_001024858.2 | #N/A | 54.3374278 | -6.9735104 | 1.63951997 |
| 10185 | NM_152275.3 | #N/A | 43.0872804 | -6.6439319 | 1.5620419 |
| 10186 | NM_001135774.1 | #N/A | 57.4688836 | -7.0578185 | 1.65950012 |
| 10187 | NM_001201362.1 | #N/A | 54.6482831 | -6.9851876 | 1.64242571 |
| 10188 | NM_001346363.1 | #N/A | 44.2361379 | -6.6827631 | 1.5714107 |
| 10189 | NM_001007075.2 | KLHL5 | 57.9357319 | -7.0684433 | 1.66211954 |
| 10190 | NM_032346.1 | #N/A | 58.1909921 | -7.0718072 | 1.66309268 |

| | | | | | |
|---|---|---|---|---|---|
| 10191 | NM_005101.3 | #N/A | 46.9098578 | -6.7672735 | 1.59154415 |
| 10192 | NM_032824.2 | #N/A | 37.8573254 | -6.4552947 | 1.5182927 |
| 10193 | NM_003678.4 | #N/A | 62.933501 | -7.1856513 | 1.6901588 |
| 10194 | NM_001301861.1 | #N/A | 50.9809006 | -6.8880986 | 1.6201952 |
| 10195 | NM_003431.3 | ZNF124 | 38.2143527 | -6.4660735 | 1.52096182 |
| 10196 | NM_001025235.1 | TSPAN4 | 50.2981081 | -6.8669526 | 1.61526776 |
| 10197 | NM_016011.3 | #N/A | 59.3813053 | -5.8256974 | 1.37037257 |
| 10198 | NM_014624.3 | #N/A | 48.950307 | -6.8253937 | 1.60578446 |
| 10199 | NR_036640.1 | #N/A | 56.1159133 | -7.0191283 | 1.65135884 |
| 10200 | NM_016108.3 | #N/A | 75.8977213 | -6.017608 | 1.41577824 |
| 10201 | NM_001042475.2 | #N/A | 84.7213421 | -7.6141278 | 1.79164292 |
| 10202 | NM_017735.4 | #N/A | 53.4512067 | -6.9503545 | 1.63557417 |
| 10203 | NM_138691.2 | TMC1 | 38.1232394 | 5.04597743 | 1.18750641 |
| 10204 | NM_024308.3 | #N/A | 39.9504841 | -6.5335676 | 1.53768166 |
| 10205 | NM_080649.2 | #N/A | 56.1384239 | -7.0175291 | 1.65158532 |
| 10206 | NM_001284388.1 | NOLC1 | 56.7113902 | -7.0305574 | 1.65467518 |
| 10207 | NM_001256577.2 | RPL10 | 88.7760356 | -7.683375 | 1.80842745 |
| 10208 | NM_032319.2 | #N/A | 52.2152666 | -6.916397 | 1.62802346 |
| 10209 | NM_024819.6 | #N/A | 50.4127687 | -6.8643393 | 1.61579491 |
| 10210 | NR_046330.1 | TUBGCP2 | 43.5551194 | -6.660323 | 1.56788552 |
| 10211 | NM_001200003.1 | #N/A | 55.429527 | -7.0007123 | 1.6482889 |
| 10212 | NM_000688.5 | #N/A | 55.5695562 | -7.0081097 | 1.65013387 |
| 10213 | NM_001199111.1 | MDH1 | 91.4552441 | -7.7251728 | 1.81902342 |
| 10214 | NM_012237.3 | #N/A | 71.3761919 | -7.3668313 | 1.73467809 |
| 10215 | NM_001002010.2 | #N/A | 53.2611006 | -6.9464037 | 1.63572721 |
| 10216 | NM_138420.2 | #N/A | 267.406989 | -3.8417407 | 0.90471178 |
| 10217 | NM_005522.4 | HOXA1 | 73.1686923 | -7.4045345 | 1.74393251 |
| 10218 | NM_001206602.1 | #N/A | 54.2397459 | -6.9738864 | 1.64270508 |
| 10219 | NM_014254.2 | #N/A | 48.6875378 | -6.8153126 | 1.60538731 |
| 10220 | NM_134446.3 | #N/A | 62.0768565 | -7.1648381 | 1.68783703 |
| 10221 | NM_144720.3 | #N/A | 86.3384415 | -7.6409848 | 1.80023541 |
| 10222 | NM_022079.2 | #N/A | 69.4684006 | -7.3252213 | 1.72594259 |
| 10223 | NM_052954.3 | #N/A | 40.1183478 | -6.5393098 | 1.54076952 |
| 10224 | NM_001319842.1 | CAPRIN2 | 45.5448104 | -6.7172765 | 1.58274705 |
| 10225 | NM_001322815.1 | #N/A | 22.7930582 | 9.50484882 | 2.2396641 |
| 10226 | NM_001308230.1 | ATXN2L | 47.2478009 | -6.7673609 | 1.59467324 |
| 10227 | NM_001145678.1 | #N/A | 299.243576 | 3.9604163 | 0.93329182 |
| 10228 | NM_181523.2 | #N/A | 232.275804 | -4.9546476 | 1.16761114 |
| 10230 | NM_001198812.1 | MSANTD3-TMEFF1 | 80.4267095 | -7.5402063 | 1.77695008 |
| 10229 | NM_001037495.1 | #N/A | 57.2829849 | -7.0487945 | 1.66113634 |
| 10231 | NM_172359.2 | #N/A | 57.180082 | -7.0499625 | 1.66149727 |
| 10232 | NM_133277.3 | #N/A | 185.344352 | 3.41065289 | 0.80382964 |
| 10233 | NM_001330488.1 | PHC2 | 62.6139291 | -7.1778794 | 1.69205513 |

| | | | | | |
|---|---|---|---|---|---|
| 10234 | NM_005071.2 | SLC1A6 | 64.3766558 | -7.2185007 | 1.70166723 |
| 10235 | NR_044998.1 | #N/A | 46.6099772 | -6.7496652 | 1.59117015 |
| 10236 | NM_145729.2 | #N/A | 53.5912437 | -6.9546537 | 1.63957723 |
| 10237 | NM_033489.2 | #N/A | 41.3161413 | -6.5810305 | 1.551507 |
| 10238 | NM_001243108.1 | #N/A | 44.0135741 | -6.6682866 | 1.57209011 |
| 10239 | NM_001163287.1 | #N/A | 44.5210315 | -6.6928151 | 1.57804642 |
| 10240 | NM_012406.3 | #N/A | 265.752291 | -3.1082514 | 0.73286689 |
| 10241 | NM_001008695.1 | THAP7 | 43.6508198 | -6.6542184 | 1.56897842 |
| 10242 | NM_001278074.1 | COL5A1 | 85.7665351 | -6.1711368 | 1.45510907 |
| 10243 | NM_003630.2 | #N/A | 50.136032 | -6.855601 | 1.61650154 |
| 10244 | NM_001017958.2 | OS9 | 82.6546264 | -7.5822519 | 1.78788102 |
| 10245 | NM_006120.3 | #N/A | 71.4396612 | -7.3702093 | 1.73791269 |
| 10246 | NM_001013663.1 | PTRHD1 | 56.5749304 | -7.0314984 | 1.65806106 |
| 10247 | NM_001081677.1 | #N/A | 124.595872 | -8.1715609 | 1.92692205 |
| 10249 | NM_001130679.2 | EIF4E | 39.7359145 | -6.5172611 | 1.53694587 |
| 10250 | NM_007171.3 | POMT1 | 69.4196985 | -7.3230961 | 1.72698538 |
| 10248 | NM_000982.3 | RPL21 | 1432.28657 | -5.972974 | 1.40859465 |
| 10252 | NR_003239.1 | SNHG11 | 46.8055551 | -6.7603137 | 1.59433933 |
| 10251 | NM_001322006.1 | #N/A | 70.2181878 | -7.3418285 | 1.73147552 |
| 10253 | NM_001467.5 | #N/A | 53.0573539 | -6.9365503 | 1.63595736 |
| 10254 | NM_174858.2 | #N/A | 60.9250381 | -7.1397582 | 1.68391557 |
| 10255 | NM_001733.4 | #N/A | 137.499817 | -3.4997311 | 0.82542167 |
| 10256 | NM_019069.3 | #N/A | 58.4345017 | -3.7344903 | 0.88079655 |
| 10257 | NM_001167604.1 | XPNPEP1 | 49.8584381 | -6.8515002 | 1.61604342 |
| 10258 | NM_001303426.1 | #N/A | 52.9285302 | -6.9343904 | 1.63567304 |
| 10259 | NM_001321512.1 | #N/A | 43.4279751 | -6.6524124 | 1.56917822 |
| 10260 | NM_001135610.1 | #N/A | 49.804157 | -6.8507002 | 1.6160191 |
| 10261 | NM_001317990.1 | #N/A | 65.5857073 | -7.2494858 | 1.71013527 |
| 10262 | NR_132419.1 | COX20 | 62.2165487 | -7.1708368 | 1.69162584 |
| 10263 | NM_001317343.1 | #N/A | 66.0013643 | -5.8348979 | 1.37651353 |
| 10264 | NM_001329991.1 | #N/A | 50.4591549 | -6.866954 | 1.62018641 |
| 10265 | NM_012450.3 | #N/A | 96.4740362 | -7.8042955 | 1.84135478 |
| 10267 | NM_001136541.1 | APOL1 | 179.636893 | 3.35076167 | 0.79059384 |
| 10266 | NM_001005910.2 | #N/A | 46.069356 | -6.7291565 | 1.58770191 |
| 10268 | NM_001042492.2 | #N/A | 97.7450276 | -7.8207429 | 1.84538791 |
| 10269 | NM_194249.2 | #N/A | 44.0277307 | -6.6696784 | 1.57379092 |
| 10270 | NM_016472.4 | #N/A | 40.1835393 | -6.5451721 | 1.54455701 |
| 10271 | NM_199351.2 | #N/A | 98.7191649 | -5.1842791 | 1.22344679 |
| 10272 | NR_027158.1 | SNHG29 | 77.6792044 | -7.4911531 | 1.76798913 |
| 10273 | NM_001330762.1 | #N/A | 51.311646 | -6.8878904 | 1.62562362 |
| 10274 | NM_015129.5 | #N/A | 45.5362604 | -6.7253363 | 1.58733766 |
| 10275 | NM_012430.4 | #N/A | 69.2313748 | -5.8975385 | 1.39196906 |
| 10276 | NR_046413.1 | POLD4 | 40.8115701 | 2.34065995 | 0.55248587 |

| | | | | | |
|---|---|---|---|---|---|
| 10277 | NM_152943.2 | #N/A | 58.1458395 | -7.0689444 | 1.66856487 |
| 10278 | NM_001145033.1 | #N/A | 77.2117659 | -7.4789307 | 1.76536945 |
| 10279 | NM_001002245.2 | ANAPC11 | 53.2556932 | -6.946174 | 1.63962604 |
| 10280 | NM_018139.2 | DNAAF2 | 47.4994663 | -6.7860839 | 1.60184773 |
| 10281 | NM_001270399.1 | TUBA1A | 57.8936677 | -7.0622047 | 1.66707736 |
| 10283 | NM_020978.4 | AMY2B | 43.7766078 | -6.6606363 | 1.57236977 |
| 10282 | NM_002023.4 | #N/A | 48.4222633 | -6.8122339 | 1.60815813 |
| 10284 | NM_001256661.1 | #N/A | 38.0767957 | -6.4626455 | 1.52566316 |
| 10285 | NM_203413.2 | ELP5 | 47.1504575 | -6.7712104 | 1.59858008 |
| 10286 | NR_036462.1 | #N/A | 37.9427865 | -6.4570182 | 1.5244018 |
| 10287 | NM_018112.2 | #N/A | 49.1170675 | -6.8260579 | 1.61177037 |
| 10288 | NM_014799.3 | #N/A | 78.455151 | -7.5044793 | 1.7719997 |
| 10289 | NM_139242.3 | #N/A | 283.76344 | 1.8934543 | 0.44710998 |
| 10290 | NM_177452.3 | #N/A | 66.7678028 | -7.2716463 | 1.71710276 |
| 10291 | NM_006716.3 | #N/A | 49.4614067 | -6.835695 | 1.61429202 |
| 10292 | NM_001303269.1 | POLR2B | 45.6326148 | -6.715307 | 1.5859638 |
| 10293 | NR_039969.1 | MIR5047 | 40.0162294 | -6.5370535 | 1.54396572 |
| 10294 | NM_001083591.1 | CELF2 | 131.899396 | -8.252737 | 1.94923879 |
| 10295 | NR_028027.1 | #N/A | 45.6747908 | -6.7276543 | 1.58906262 |
| 10296 | NM_173573.2 | #N/A | 82.3407895 | -7.5754082 | 1.78934587 |
| 10297 | NM_001320942.1 | #N/A | 40.1071373 | -6.5311994 | 1.54277549 |
| 10298 | NM_001224.4 | CASP2 | 54.9805032 | -6.9878677 | 1.6507734 |
| 10299 | NM_002390.5 | #N/A | 72.0837435 | -7.3810553 | 1.74375652 |
| 10300 | NM_021195.4 | #N/A | 67.893797 | -7.2963168 | 1.7237856 |
| 10301 | NM_024769.3 | #N/A | 67.7861548 | -5.2328135 | 1.2362808 |
| 10302 | NM_003312.5 | #N/A | 37.8807153 | -6.4528239 | 1.52455248 |
| 10303 | NM_032192.3 | #N/A | 44.3266657 | -6.6747747 | 1.57731722 |
| 10304 | NM_001323393.1 | #N/A | 76.5395761 | -7.4678638 | 1.76485118 |
| 10305 | NM_001287816.1 | UBAP2L | 65.7869342 | -7.2520943 | 1.71387073 |
| 10306 | NM_001256875.1 | #N/A | 49.5434867 | -6.840202 | 1.6165439 |
| 10307 | NM_000945.3 | #N/A | 504.759505 | -4.03437 | 0.95344745 |
| 10308 | NR_033426.1 | ACAD9 | 41.9045771 | -6.6047098 | 1.56111471 |
| 10309 | NM_001286808.1 | #N/A | 47.605139 | -6.7839276 | 1.60359491 |
| 10310 | NM_019087.2 | #N/A | 63.8991922 | -7.2069599 | 1.70362868 |
| 10311 | NM_001002257.2 | #N/A | 47.4190195 | -6.7732951 | 1.60123621 |
| 10312 | NM_001325.2 | CSTF2 | 43.5771056 | -6.6606238 | 1.57461145 |
| 10313 | NM_001740.4 | #N/A | 60.1637899 | -7.1213092 | 1.68372801 |
| 10314 | NM_016618.2 | #N/A | 47.5108089 | -6.7818779 | 1.6035276 |
| 10315 | NM_015575.3 | #N/A | 68.1776878 | -7.2972014 | 1.72566101 |
| 10316 | NM_030763.2 | #N/A | 44.6011784 | -6.6831562 | 1.580537 |
| 10317 | NM_001174120.1 | ZFYVE27 | 57.1413341 | -7.0417844 | 1.66536241 |
| 10318 | NM_018086.3 | #N/A | 42.6965648 | -6.6332343 | 1.56875037 |
| 10319 | NM_001199282.2 | LRBA | 62.3938092 | -7.1731215 | 1.69647074 |

| | | | | | |
|---|---|---|---|---|---|
| 10322 | NR_134485.1 | LMNB1-DT | 20.439469 | 6.72088596 | 1.58954426 |
| 10320 | NM_138555.3 | #N/A | 43.6304011 | -6.6534139 | 1.57358298 |
| 10321 | NM_175063.5 | #N/A | 51.2253095 | -6.8905086 | 1.62966066 |
| 10323 | NM_001323627.1 | #N/A | 45.1836232 | -6.7129607 | 1.58771241 |
| 10324 | NM_058180.4 | #N/A | 39.8076089 | -6.5296182 | 1.54439137 |
| 10325 | NM_024630.2 | #N/A | 54.9632819 | -6.9921386 | 1.65416138 |
| 10326 | NM_025077.3 | #N/A | 49.8158318 | -6.8497574 | 1.62057064 |
| 10327 | NM_001002246.2 | ANAPC11 | 50.2327726 | -6.8592614 | 1.62299627 |
| 10328 | NM_175085.2 | #N/A | 82.0525201 | -6.1093719 | 1.44580592 |
| 10329 | NR_026808.1 | ANP32A-IT1 | 75.7958853 | 2.35256436 | 0.55677498 |
| 10330 | NR_132737.1 | #N/A | 100.653791 | -6.5944321 | 1.56068463 |
| 10331 | NM_022734.2 | #N/A | 55.2887502 | -7.0016269 | 1.65710223 |
| 10332 | NM_001114748.1 | #N/A | 46.9472934 | -6.7602671 | 1.60002506 |
| 10333 | NM_001319998.1 | #N/A | 46.8516627 | -6.7632407 | 1.60079842 |
| 10334 | NM_001293186.1 | METTL5 | 47.6297928 | -6.7833777 | 1.60562588 |
| 10335 | NM_001171162.1 | ZMYM3 | 40.4257112 | -6.5499431 | 1.55055008 |
| 10336 | NM_003112.4 | #N/A | 58.1834414 | -7.0685189 | 1.6734767 |
| 10337 | NM_005713.2 | #N/A | 86.7935389 | -7.6484248 | 1.81080121 |
| 10338 | NM_017633.2 | #N/A | 42.6350465 | -6.6305804 | 1.56983989 |
| 10339 | NM_016046.4 | #N/A | 47.2152019 | -6.7734655 | 1.60373164 |
| 10340 | NM_001142798.2 | #N/A | 64.0159988 | -7.2103952 | 1.7072925 |
| 10341 | NM_013281.3 | FLRT3 | 57.6338108 | -5.6663451 | 1.34176273 |
| 10342 | NM_001143779.1 | #N/A | 53.2641969 | -6.9394903 | 1.64329553 |
| 10343 | NM_014727.2 | KMT2B | 393.049059 | -3.8573127 | 0.91349273 |
| 10344 | NM_015279.1 | #N/A | 67.6149706 | -7.2881777 | 1.72600751 |
| 10345 | NM_001191057.3 | #N/A | 52.2856973 | -6.9193645 | 1.63883087 |
| 10346 | NM_001130415.1 | #N/A | 43.6372738 | -6.6566896 | 1.57670029 |
| 10347 | NM_004335.3 | #N/A | 59.4444016 | -7.1067189 | 1.68328561 |
| 10348 | NM_001099733.1 | #N/A | 64.8588076 | -7.2310907 | 1.71290871 |
| 10350 | NR_024173.1 | LINC00923 | 76.9394136 | 2.21338516 | 0.52431819 |
| 10351 | NR_110395.1 | LOC101928335 | 66.4994826 | 3.72248738 | 0.88180842 |
| 10349 | NM_001319245.1 | MAST2 | 70.3776084 | -6.0743382 | 1.43893138 |
| 10352 | NM_173812.4 | #N/A | 51.9565182 | -6.9056361 | 1.63621494 |
| 10353 | NM_001031734.3 | #N/A | 57.0216561 | -7.0444392 | 1.6691739 |
| 10354 | NM_001011663.1 | #N/A | 41.3547416 | -6.5801183 | 1.55922654 |
| 10355 | NM_015589.5 | #N/A | 43.725478 | -6.6669849 | 1.57988855 |
| 10356 | NM_001303257.2 | MORC2 | 45.5347911 | -6.7227462 | 1.59312059 |
| 10357 | NM_014324.5 | #N/A | 48.4998391 | -6.8129817 | 1.61452011 |
| 10358 | NM_024079.4 | #N/A | 56.200647 | -7.0245041 | 1.66470684 |
| 10359 | NM_012245.2 | #N/A | 228.684272 | -4.9307563 | 1.1686095 |
| 10360 | NM_001287526.1 | IFT22 | 96.8214689 | 4.79214271 | 1.13581713 |
| 10361 | NM_173814.5 | #N/A | 81.5025896 | -6.2895547 | 1.49075898 |
| 10362 | NM_152565.1 | ATP6V0D2 | 29.542475 | 3.02395489 | 0.71676898 |

| | | | | | |
|---|---|---|---|---|---|
| 10363 | NM_016085.4 | #N/A | 44.8880176 | -6.6984414 | 1.58786005 |
| 10364 | NR_073499.1 | #N/A | 59.6326267 | -7.1081947 | 1.68507365 |
| 10365 | NM_015976.4 | #N/A | 47.752311 | -6.7851667 | 1.60853627 |
| 10366 | NM_001171509.2 | #N/A | 91.9075905 | -7.7319652 | 1.83301305 |
| 10367 | NM_001201556.1 | ZNF821 | 50.2805774 | -6.8614472 | 1.62665208 |
| 10368 | NM_014932.3 | #N/A | 53.3236989 | -6.9434982 | 1.64616435 |
| 10369 | NR_004858.1 | #N/A | 41.3202291 | -6.5783537 | 1.55959738 |
| 10370 | NM_001142725.1 | #N/A | 38.4254589 | -6.4706244 | 1.53434064 |
| 10371 | NM_001326295.1 | #N/A | 42.5856854 | -6.6228476 | 1.57047427 |
| 10372 | NM_001243571.1 | #N/A | 55.1819922 | -7.0004549 | 1.66012541 |
| 10373 | NR_027107.2 | TENM3-AS1 | 38.7151141 | -6.4877891 | 1.53863208 |
| 10374 | NM_001319143.1 | AK2 | 49.2296472 | -6.8309191 | 1.62024057 |
| 10375 | NR_027512.1 | BEND3P3 | 54.3508553 | -5.6960714 | 1.35118457 |
| 10376 | NM_178326.2 | #N/A | 54.1697834 | -6.9743501 | 1.65443971 |
| 10377 | NM_001170649.1 | LAS1L | 38.6302057 | -6.4787156 | 1.53693622 |
| 10378 | NM_001330333.1 | #N/A | 66.5610224 | -7.2642241 | 1.72329265 |
| 10379 | NM_001288786.1 | #N/A | 64.8461671 | -7.2323026 | 1.71579418 |
| 10380 | NM_012260.3 | #N/A | 39.6703667 | -6.5262167 | 1.54837186 |
| 10381 | NM_007080.2 | #N/A | 48.0521156 | -6.7984516 | 1.61300646 |
| 10382 | NM_000512.4 | #N/A | 53.7806193 | -6.9591958 | 1.65124655 |
| 10383 | NM_032261.4 | #N/A | 46.9727957 | -6.7653481 | 1.6053079 |
| 10384 | NM_144635.4 | #N/A | 65.5672585 | -7.2439734 | 1.71892852 |
| 10385 | NM_001270771.1 | #N/A | 40.9477955 | -6.5665626 | 1.55819676 |
| 10386 | NM_001290047.1 | #N/A | 105.949914 | -4.2424258 | 1.00671424 |
| 10387 | NM_001142351.1 | #N/A | 74.1633374 | -7.4245394 | 1.76188717 |
| 10388 | NM_001256348.1 | SEMA5B | 48.4459988 | -6.8047718 | 1.61493942 |
| 10389 | NM_001013703.3 | #N/A | 277.811797 | -3.1166829 | 0.73970275 |
| 10390 | NM_001282474.1 | #N/A | 46.6695248 | -6.7578389 | 1.60408085 |
| 10391 | NM_020760.2 | #N/A | 47.3011119 | -6.7742671 | 1.60804547 |
| 10392 | NM_005302.3 | #N/A | 63.5336226 | -7.199475 | 1.70909981 |
| 10393 | NM_001113496.1 | #N/A | 55.8228986 | -7.0154984 | 1.66554223 |
| 10394 | NM_004978.4 | #N/A | 61.1191599 | -7.1434501 | 1.69591379 |
| 10395 | NM_015090.3 | NFASC | 88.4244723 | -7.6762014 | 1.82276968 |
| 10396 | NM_001319115.1 | #N/A | 48.9038788 | -6.8223997 | 1.62023663 |
| 10397 | NM_014369.3 | #N/A | 65.3060409 | -7.2376512 | 1.71898031 |
| 10398 | NM_006303.3 | #N/A | 57.213308 | -7.0463593 | 1.67367482 |
| 10399 | NM_001031737.2 | #N/A | 44.3994806 | -6.688337 | 1.58866643 |
| 10400 | NM_000784.3 | #N/A | 132.459951 | -8.2609865 | 1.9622608 |
| 10401 | NM_012249.3 | #N/A | 327.768867 | -5.0246122 | 1.19367082 |
| 10402 | NM_001321012.1 | #N/A | 59.8860509 | -7.1130372 | 1.68986234 |
| 10403 | NM_001048230.1 | #N/A | 65.0167281 | -7.232675 | 1.71861364 |
| 10404 | NM_000795.3 | #N/A | 60.2353343 | -7.1229163 | 1.6926075 |
| 10405 | NR_038301.1 | LINC00888 | 45.6815583 | -6.7234196 | 1.59770234 |

| | | | | | |
|---|---|---|---|---|---|
| 10406 | NM_018035.2 | #N/A | 60.0646543 | -7.1168258 | 1.6911994 |
| 10407 | NM_144579.2 | #N/A | 37.4234731 | -6.4316851 | 1.52852742 |
| 10408 | NM_001145165.1 | #N/A | 62.3015445 | -7.168417 | 1.70369133 |
| 10409 | NM_199191.2 | BABAM2 | 38.0714398 | -6.4651046 | 1.53664546 |
| 10410 | NM_201526.1 | #N/A | 67.9387814 | -7.2985563 | 1.73473615 |
| 10411 | NM_006928.4 | PMEL | 53.4032913 | -6.9525165 | 1.65258093 |
| 10412 | NR_104457.1 | FAM214A | 38.9406228 | -6.489005 | 1.54242713 |
| 10413 | NM_001206840.1 | #N/A | 43.5895062 | -6.6624981 | 1.58380699 |
| 10414 | NM_001297442.1 | #N/A | 79.0116983 | -7.5159799 | 1.78669661 |
| 10415 | NM_013409.2 | #N/A | 100.262254 | -4.9295761 | 1.17185744 |
| 10416 | NM_153354.4 | TMEM161B | 53.5110489 | -6.9497873 | 1.6523079 |
| 10417 | NM_198971.2 | #N/A | 66.0980015 | -4.9974224 | 1.18815558 |
| 10418 | NM_001280787.1 | SURF1 | 55.0270521 | -6.9890081 | 1.66180307 |
| 10419 | NM_003846.2 | #N/A | 69.4729716 | -7.32903 | 1.74289994 |
| 10420 | NM_013322.2 | SNX10 | 79.9237353 | -7.5299197 | 1.7907901 |
| 10422 | NM_183380.3 | DST | 97.726324 | -7.8223138 | 1.86045955 |
| 10421 | NM_032839.2 | #N/A | 50.4939326 | -6.8665862 | 1.6331537 |
| 10423 | NM_001206646.1 | #N/A | 48.47885 | -6.8146248 | 1.6210794 |
| 10424 | NM_016371.3 | HSD17B7 | 72.396752 | -4.0146762 | 0.95502893 |
| 10425 | NM_001348136.1 | CCNI | 98.0212817 | -7.8258735 | 1.86167371 |
| 10426 | NM_001004125.2 | TUSC1 | 111.027934 | -5.291242 | 1.2588151 |
| 10427 | NM_018480.4 | #N/A | 52.2934157 | -6.918909 | 1.64605811 |
| 10428 | NR_027456.1 | LINC00894 | 76.9632498 | -7.4784712 | 1.77924597 |
| 10429 | NM_005434.4 | #N/A | 65.5335001 | 1.9639476 | 0.46727942 |
| 10430 | NM_001122823.1 | #N/A | 38.6948252 | -6.4835199 | 1.54264976 |
| 10431 | NM_005938.3 | #N/A | 36.76735 | -6.4138045 | 1.52609524 |
| 10432 | NM_019625.3 | #N/A | 59.6665314 | -7.1060711 | 1.6908728 |
| 10433 | NM_033529.3 | #N/A | 52.7625839 | -6.9360222 | 1.65045044 |
| 10434 | NM_001288800.1 | #N/A | 44.0659065 | -6.6703939 | 1.58734899 |
| 10435 | NM_001166112.1 | PNPLA6 | 70.3829982 | -7.3508801 | 1.74932628 |
| 10436 | NM_001039840.2 | #N/A | 45.3408209 | -6.7137177 | 1.59776398 |
| 10437 | NM_002560.2 | #N/A | 38.1514281 | -6.4668821 | 1.53909186 |
| 10438 | NM_139033.2 | MAPK7 | 43.2960773 | -6.6413604 | 1.58069034 |
| 10439 | NM_145810.2 | #N/A | 50.6681483 | -6.8736682 | 1.63613556 |
| 10440 | NM_001199863.1 | #N/A | 49.8749971 | -6.8477555 | 1.63020313 |
| 10441 | NM_001256166.1 | #N/A | 50.8165308 | -6.8799668 | 1.6379258 |
| 10442 | NM_138632.2 | TRIOBP | 40.3809614 | -6.547662 | 1.5589899 |
| 10443 | NM_001317961.1 | UIMC1 | 47.6671499 | -6.7825439 | 1.61504522 |
| 10445 | NR_038367.1 | HOTAIRM1 | 61.1276064 | -7.1442966 | 1.70130734 |
| 10444 | NM_153047.3 | #N/A | 42.2783291 | -6.6103643 | 1.57415459 |
| 10446 | NM_001167934.1 | CCM2 | 51.8188716 | -6.9051581 | 1.64446076 |
| 10447 | NR_130898.1 | THAP12 | 47.9837812 | -6.7946872 | 1.61814857 |
| 10448 | NM_001199267.1 | #N/A | 53.3035128 | -6.9485171 | 1.65483501 |

| | | | | | |
|---|---|---|---|---|---|
| 10449 | NM_001077188.1 | #N/A | 87.3949721 | -7.6602212 | 1.82463209 |
| 10450 | NM_052936.4 | #N/A | 44.3784816 | -6.6773397 | 1.5907094 |
| 10451 | NM_015011.1 | #N/A | 60.9833745 | -5.7307027 | 1.3652385 |
| 10452 | NM_001348096.1 | GFRA1 | 58.7167534 | -7.0900406 | 1.68911657 |
| 10453 | NM_001329943.1 | #N/A | 48.0033735 | -6.7919178 | 1.61817372 |
| 10454 | NR_026982.1 | RPL23AP82 | 58.158411 | -7.0720249 | 1.68493945 |
| 10455 | NM_012139.3 | #N/A | 41.0842832 | -6.5656818 | 1.56435447 |
| 10456 | NM_021021.3 | #N/A | 60.6961389 | -7.1371436 | 1.70063585 |
| 10457 | NM_003094.3 | #N/A | 176.588687 | -3.7503116 | 0.89364037 |
| 10458 | NM_024969.3 | CSRNP3 | 84.2829525 | -7.6063065 | 1.81263496 |
| 10459 | NM_012300.2 | FBXW11 | 60.7731876 | -7.1340785 | 1.70013038 |
| 10460 | NM_015158.3 | #N/A | 98.5369535 | -6.3370184 | 1.5101764 |
| 10461 | NM_005333.4 | #N/A | 53.8897927 | -6.9660954 | 1.66014947 |
| 10462 | NM_017990.4 | #N/A | 55.3065133 | -7.0049054 | 1.66960814 |
| 10463 | NM_004237.3 | #N/A | 41.81939 | -6.5952803 | 1.57209859 |
| 10464 | NM_007069.3 | PLAAT3 | 45.1968434 | -6.7116496 | 1.5999122 |
| 10465 | NM_001003697.1 | ATP5PF | 50.5820096 | -6.8714286 | 1.63810751 |
| 10466 | NM_014264.4 | #N/A | 40.314316 | -6.5428972 | 1.55988919 |
| 10467 | NM_001039583.1 | PICK1 | 61.511545 | -7.14847 | 1.70433035 |
| 10468 | NM_001198793.1 | ARPC4-TTLL3 | 55.3666122 | -6.9989019 | 1.66871381 |
| 10469 | NM_001317395.1 | #N/A | 55.6958378 | -7.0082996 | 1.6710773 |
| 10470 | NM_018353.4 | #N/A | 43.1020511 | -6.6357617 | 1.58225928 |
| 10471 | NM_001083965.1 | #N/A | 54.5560918 | -6.976962 | 1.6638635 |
| 10472 | NM_003442.5 | #N/A | 39.5222501 | -6.5092363 | 1.55246177 |
| 10473 | NM_001281453.1 | #N/A | 57.535374 | -7.0535527 | 1.68239714 |
| 10474 | NM_001330164.1 | #N/A | 58.1617414 | -7.075731 | 1.6876791 |
| 10475 | NM_001310159.1 | PAX6 | 37.3477397 | -6.4336362 | 1.53465871 |
| 10476 | NM_001014443.2 | #N/A | 43.0742523 | -6.640766 | 1.58423586 |
| 10477 | NM_006078.3 | #N/A | 81.2682384 | -6.2829236 | 1.49895533 |
| 10478 | NM_024019.3 | #N/A | 108.154238 | -7.9662313 | 1.90059259 |
| 10479 | NM_001033088.1 | #N/A | 104.956743 | -6.6572803 | 1.58856101 |
| 10480 | NM_001278141.1 | NBPF12 | 218.885505 | -4.8714055 | 1.16243407 |
| 10481 | NM_138364.3 | #N/A | 36.6248195 | -6.4040235 | 1.52815746 |
| 10482 | NM_017440.4 | #N/A | 42.8013417 | -6.6234806 | 1.58079828 |
| 10483 | NM_001077238.1 | #N/A | 48.2624734 | -6.8018171 | 1.62345131 |
| 10484 | NM_001300731.1 | #N/A | 64.4068358 | -7.2203787 | 1.7234494 |
| 10485 | NM_001206836.1 | #N/A | 16.5675138 | 9.01869966 | 2.15283699 |
| 10486 | NM_001184897.1 | PHF8 | 36.4792337 | -6.4030818 | 1.52849673 |
| 10487 | NM_001243351.1 | NUB1 | 49.2805105 | -6.8353672 | 1.63183442 |
| 10488 | NR_002217.1 | PMS2CL | 43.5571565 | -6.6559837 | 1.58902526 |
| 10489 | NM_000017.3 | #N/A | 38.318935 | -6.4657066 | 1.54375015 |
| 10490 | NM_001289789.1 | GPI | 77.2314272 | -7.4809731 | 1.78638709 |
| 10491 | NM_001301098.1 | #N/A | 41.0844963 | -6.569292 | 1.56928501 |

| | | | | | |
|---|---|---|---|---|---|
| 10492 | NM_003242.5 | #N/A | 42.2821647 | -6.6168132 | 1.58070032 |
| 10493 | NM_003622.3 | #N/A | 47.7545906 | -6.785341 | 1.62096236 |
| 10494 | NM_020341.3 | #N/A | 64.6379308 | -7.2227276 | 1.72545814 |
| 10495 | NM_001098206.1 | HNRNPF | 45.0601229 | -6.704521 | 1.6018691 |
| 10496 | NM_031434.3 | #N/A | 38.4953174 | -6.476433 | 1.54736847 |
| 10497 | NM_000204.4 | CFI | 62.6120777 | -7.1832211 | 1.71625783 |
| 10498 | NM_152640.4 | #N/A | 40.6084149 | -6.558214 | 1.56695754 |
| 10499 | NM_203436.2 | ASCL4 | 12.1994123 | 7.85624612 | 1.87712422 |
| 10500 | NM_001171195.1 | #N/A | 78.6583107 | -7.5080133 | 1.79416154 |
| 10501 | NM_031891.3 | #N/A | 43.9086189 | -6.6662254 | 1.59300785 |
| 10502 | NR_104164.1 | LINC01443 | 78.9664943 | 3.4908977 | 0.83427528 |
| 10503 | NM_001042384.1 | #N/A | 42.4689797 | -6.614903 | 1.58099191 |
| 10504 | NM_006242.3 | #N/A | 47.782617 | -6.790637 | 1.62298966 |
| 10505 | NM_024698.5 | #N/A | 54.7922587 | -6.9902908 | 1.67087925 |
| 10506 | NM_031904.4 | #N/A | 51.9819916 | -6.9158441 | 1.65326041 |
| 10507 | NM_002268.4 | #N/A | 504.779173 | -2.1307132 | 0.50937361 |
| 10508 | NM_153335.5 | #N/A | 44.2032833 | -6.6764453 | 1.59613353 |
| 10509 | NM_004251.4 | #N/A | 52.3196911 | -6.9227484 | 1.65516853 |
| 10510 | NM_001199346.1 | #N/A | 81.0145942 | -7.5484189 | 1.80478046 |
| 10511 | NM_017443.4 | #N/A | 103.974517 | -7.9115186 | 1.8917306 |
| 10512 | NM_014026.4 | #N/A | 36.7371622 | -6.4081604 | 1.5322759 |
| 10513 | NM_052839.3 | #N/A | 38.2122963 | -6.4645282 | 1.54576053 |
| 10514 | NM_033411.4 | #N/A | 42.0910453 | -6.6059371 | 1.57960132 |
| 10515 | NR_110049.2 | TMED2-DT | 110.732331 | 2.01905427 | 0.48288519 |
| 10516 | NM_001288765.1 | DMPK | 65.3491909 | -7.2384067 | 1.73118101 |
| 10517 | NR_038383.1 | LINC02453 | 32.6421909 | 4.34506646 | 1.03923973 |
| 10519 | NM_080593.2 | H2BC12 | 37.1346552 | -6.431038 | 1.53834585 |
| 10518 | NM_002486.4 | #N/A | 277.991066 | -2.5461321 | 0.60905238 |
| 10520 | NM_152526.5 | #N/A | 37.330556 | -6.4336193 | 1.53901045 |
| 10521 | NM_177939.2 | #N/A | 166.12066 | -3.358162 | 0.80336329 |
| 10522 | NM_005634.2 | SOX3 | 68.651211 | -7.3085011 | 1.74852345 |
| 10523 | NM_199442.1 | #N/A | 59.0766134 | -7.0941991 | 1.69768747 |
| 10524 | NM_003567.3 | #N/A | 42.7279708 | -6.6332891 | 1.58746667 |
| 10525 | NM_001216.2 | #N/A | 106.690256 | -7.9464678 | 1.90174726 |
| 10526 | NM_001122633.2 | #N/A | 39.3018832 | -6.5109472 | 1.55828397 |
| 10527 | NM_206866.2 | #N/A | 52.8443781 | -6.9378426 | 1.66066573 |
| 10528 | NM_138330.2 | #N/A | 53.5054738 | -6.9492699 | 1.66342824 |
| 10529 | NM_144580.2 | #N/A | 51.0837666 | -6.8880076 | 1.64887425 |
| 10530 | NM_000439.4 | #N/A | 58.9750574 | -7.0928938 | 1.69806829 |
| 10531 | NM_006704.4 | #N/A | 60.2061196 | -7.1227361 | 1.7052745 |
| 10532 | NR_136734.1 | RNF217 | 74.71652 | -2.3566086 | 0.56422502 |
| 10533 | NM_000437.3 | #N/A | 45.7219341 | -3.5423104 | 0.84812852 |
| 10534 | NM_001001976.2 | ATE1 | 48.0420466 | -6.799014 | 1.62794275 |

| | | | | | |
|---|---|---|---|---|---|
| 10535 | NM_001270966.1 | PSD | 67.5345632 | -7.2885391 | 1.74534698 |
| 10536 | NM_144638.2 | #N/A | 41.5050154 | -6.5835048 | 1.57657433 |
| 10537 | NM_018352.2 | #N/A | 47.0972857 | -6.7637057 | 1.61978164 |
| 10538 | NM_001242793.1 | SNAP91 | 53.4018819 | -6.9507201 | 1.66478816 |
| 10539 | NM_032730.5 | RTN4IP1 | 38.0248104 | -6.4550298 | 1.54608056 |
| 10540 | NM_001308195.1 | #N/A | 50.4423008 | -6.8692331 | 1.64531747 |
| 10541 | NM_022366.2 | #N/A | 40.7169792 | -6.5537141 | 1.56976521 |
| 10542 | NM_001033719.2 | #N/A | 41.1055907 | -6.5693509 | 1.57353396 |
| 10543 | NR_003613.1 | PMS2P1 | 43.9681255 | -6.670356 | 1.59778542 |
| 10544 | NR_109792.1 | MIR124-2HG | 174.472394 | -4.5634941 | 1.09313376 |
| 10545 | NM_001008390.1 | #N/A | 49.8427623 | -6.8494248 | 1.64082977 |
| 10546 | NM_173685.2 | #N/A | 43.0623876 | -6.6387461 | 1.59039148 |
| 10547 | NM_001134369.2 | LRRFIP2 | 54.9209299 | -6.9851403 | 1.67345324 |
| 10549 | NR_045026.1 | SNHG29 | 39.3403247 | -6.5071505 | 1.55893179 |
| 10548 | NM_006478.4 | #N/A | 62.3545939 | -7.1736826 | 1.71862614 |
| 10550 | NM_001195738.1 | #N/A | 71.0579392 | -7.3589155 | 1.76301435 |
| 10551 | NR_033399.1 | DDX12P | 43.403258 | -6.6445737 | 1.5919205 |
| 10552 | NM_007370.6 | #N/A | 44.9567759 | -6.6995869 | 1.60512346 |
| 10553 | NM_001324.2 | #N/A | 39.2397402 | -6.5063153 | 1.55887725 |
| 10554 | NM_000124.3 | #N/A | 50.0065232 | -6.8541878 | 1.64225526 |
| 10555 | NM_001256469.1 | KAT6B | 171.54581 | -3.3653703 | 0.80634715 |
| 10556 | NM_198514.3 | #N/A | 47.4309717 | -3.5984614 | 0.86220678 |
| 10557 | NM_001122957.2 | #N/A | 54.2615749 | -6.9703285 | 1.6701329 |
| 10558 | NM_001345869.1 | GUF1 | 53.1257468 | -6.937715 | 1.66234483 |
| 10559 | NM_177951.2 | #N/A | 42.0360575 | -6.6040928 | 1.58273144 |
| 10560 | NM_003818.3 | #N/A | 490.105649 | -3.1763328 | 0.76126893 |
| 10561 | NM_032317.2 | #N/A | 55.5354794 | -4.9319115 | 1.18202667 |
| 10562 | NM_030917.3 | #N/A | 43.7569231 | -6.6652188 | 1.59761663 |
| 10563 | NM_032490.4 | #N/A | 35.4840118 | -6.3567028 | 1.52369839 |
| 10564 | NM_144721.5 | #N/A | 38.6644581 | -6.487647 | 1.55510147 |
| 10565 | NR_134476.1 | LOC101929165 | 135.956364 | 2.47623771 | 0.59367018 |
| 10566 | NM_001127245.1 | #N/A | 57.2860477 | -7.0483761 | 1.68986898 |
| 10567 | NM_017632.3 | #N/A | 55.4815123 | -7.0040289 | 1.67924504 |
| 10568 | NM_022836.3 | #N/A | 37.3144754 | -6.4348383 | 1.54279079 |
| 10570 | NM_001199693.1 | SLC4A2 | 67.9848307 | -7.3007973 | 1.75043309 |
| 10569 | NM_001001991.2 | #N/A | 39.6748763 | -6.5244865 | 1.56430499 |
| 10571 | NM_001008216.1 | #N/A | 62.2988351 | -7.1705835 | 1.71947955 |
| 10572 | NM_001172705.1 | EIF4G2 | 44.159827 | -6.6752359 | 1.60071743 |
| 10573 | NM_001304999.1 | #N/A | 42.2993747 | -6.6130252 | 1.58584182 |
| 10574 | NM_013238.2 | #N/A | 66.9067747 | -4.4701277 | 1.07209233 |
| 10575 | NM_007055.3 | #N/A | 322.071408 | -2.084981 | 0.50009575 |
| 10577 | NM_001136004.3 | MICAL3 | 79.7767599 | -6.063892 | 1.45453992 |
| 10576 | NM_001013694.2 | #N/A | 45.7967276 | -6.7260991 | 1.61337527 |

| | | | | | |
|---|---|---|---|---|---|
| 10578 | NR_026956.1 | LINC00663 | 178.111856 | 1.78328448 | 0.42777565 |
| 10579 | NM_015510.4 | DHRS7B | 44.7046245 | -6.6929421 | 1.60555058 |
| 10580 | NM_016520.2 | #N/A | 301.04764 | -2.1805557 | 0.52313167 |
| 10581 | NM_004936.3 | #N/A | 55.2144014 | -7.0012938 | 1.6796977 |
| 10582 | NM_001330332.1 | #N/A | 203.62476 | 3.13419371 | 0.75199219 |
| 10583 | NM_003172.3 | #N/A | 58.7670443 | -7.0870155 | 1.70045036 |
| 10584 | NM_003618.3 | #N/A | 103.798568 | -7.906657 | 1.89716091 |
| 10585 | NM_006200.5 | #N/A | 68.3908001 | -7.307812 | 1.75349933 |
| 10586 | NM_001004127.2 | #N/A | 39.1938667 | -6.5053598 | 1.56098089 |
| 10587 | NM_005748.5 | YAF2 | 60.3198013 | -7.1230588 | 1.70929959 |
| 10588 | NM_020399.3 | #N/A | 133.957011 | -8.2764486 | 1.9861078 |
| 10589 | NM_001039672.2 | #N/A | 49.9534235 | -6.8482038 | 1.64343898 |
| 10590 | NM_012134.2 | #N/A | 39.8998491 | -6.5343111 | 1.56830689 |
| 10591 | NR_034105.3 | CRNDE | 45.7468669 | -6.728951 | 1.6151201 |
| 10592 | NM_032424.2 | #N/A | 53.0265703 | -6.9343802 | 1.66447127 |
| 10593 | NM_198799.3 | BCAS4 | 37.9289272 | -6.4548681 | 1.54939915 |
| 10594 | NM_004277.4 | #N/A | 39.4044043 | -6.5175041 | 1.56453394 |
| 10595 | NM_006117.2 | ECI2 | 51.5917563 | -6.8985559 | 1.6562505 |
| 10596 | NM_033657.2 | DAP3 | 67.0363071 | -7.2774169 | 1.7474019 |
| 10597 | NM_001127322.1 | CBX5 | 44.2768972 | -6.6789589 | 1.60374647 |
| 10598 | NM_001067.3 | #N/A | 445.857608 | -5.1373738 | 1.23360319 |
| 10599 | NR_125838.1 | LOC101926915 | 33.3922599 | 2.69886697 | 0.64812839 |
| 10600 | NM_001014838.1 | #N/A | 47.6919764 | -6.7875952 | 1.630077 |
| 10601 | NM_001272051.1 | #N/A | 48.7248931 | -6.8142208 | 1.63650033 |
| 10602 | NM_021111.2 | #N/A | 72.82963 | -5.9488042 | 1.42891401 |
| 10603 | NM_001143912.1 | #N/A | 41.9544853 | -6.6008488 | 1.58564701 |
| 10604 | NM_015082.1 | #N/A | 46.2277356 | -6.7428558 | 1.61986808 |
| 10605 | NM_001286414.2 | #N/A | 65.0315262 | -7.2357049 | 1.73841203 |
| 10606 | NM_001011551.2 | #N/A | 40.9011468 | -6.5634923 | 1.57693169 |
| 10607 | NM_152396.3 | METTL6 | 45.206659 | -6.7088617 | 1.6120059 |
| 10608 | NM_079423.3 | #N/A | 744.053827 | -5.5956357 | 1.34457276 |
| 10609 | NM_001024934.1 | WASF1 | 57.0090377 | -7.0442546 | 1.69294471 |
| 10610 | NM_005475.2 | #N/A | 46.4775675 | -6.7505072 | 1.62256354 |
| 10611 | NM_022838.3 | #N/A | 53.5965118 | -6.952514 | 1.67119426 |
| 10612 | NM_001127716.1 | ETFA | 58.2961853 | -7.077577 | 1.70132468 |
| 10613 | NM_001649.3 | #N/A | 69.9739383 | -7.3417398 | 1.76482338 |
| 10615 | NM_001316920.1 | TVP23B | 43.2505816 | -6.6424905 | 1.59680058 |
| 10614 | NM_001290326.1 | USP38 | 48.6131417 | -6.8136029 | 1.63792684 |
| 10616 | NM_001077181.1 | #N/A | 37.3538489 | -6.4329507 | 1.54660341 |
| 10617 | NM_014481.3 | #N/A | 37.4674251 | -6.4354755 | 1.54724351 |
| 10618 | NM_178826.3 | #N/A | 37.9211006 | -6.4617777 | 1.55361094 |
| 10619 | NM_138355.3 | #N/A | 43.3601146 | -6.6434309 | 1.5974885 |
| 10620 | NM_022359.5 | #N/A | 60.7479117 | -7.1330814 | 1.71540612 |

| | | | | | |
|---|---|---|---|---|---|
| 10621 | NM_178336.2 | #N/A | 49.2938943 | -6.8317701 | 1.64302758 |
| 10622 | NM_006831.2 | #N/A | 46.1376685 | -6.7349719 | 1.61989861 |
| 10623 | NM_001164317.1 | #N/A | 53.6344109 | -6.9574726 | 1.67354958 |
| 10624 | NR_074083.1 | #N/A | 41.823472 | -6.5926506 | 1.58583088 |
| 10625 | NM_001330360.1 | #N/A | 45.3115336 | -6.7149762 | 1.6152736 |
| 10626 | NM_024641.3 | #N/A | 49.4374864 | -4.7553199 | 1.14394253 |
| 10627 | NM_001682.2 | #N/A | 328.443532 | -3.270272 | 0.78676392 |
| 10628 | NM_001128843.2 | TNIP3 | 37.9979057 | 4.4509203 | 1.07088089 |
| 10629 | NM_018398.2 | #N/A | 40.8339099 | -6.5624872 | 1.5789253 |
| 10630 | NM_001271781.1 | PSMD6 | 49.5576695 | -6.8435433 | 1.64662568 |
| 10631 | NM_001270392.1 | RASSF6 | 62.8976462 | 6.45597762 | 1.55358651 |
| 10632 | NM_006677.2 | USP19 | 75.0806923 | -7.4429401 | 1.79112306 |
| 10634 | NR_027906.1 | AFDN-DT | 43.3861388 | -6.6461443 | 1.59947415 |
| 10633 | NM_001008212.1 | #N/A | 58.0583484 | -7.0736879 | 1.7023711 |
| 10635 | NM_024706.4 | #N/A | 45.9186948 | -6.7280975 | 1.61928507 |
| 10636 | NM_001006115.2 | IP6K1 | 69.0860441 | -7.320213 | 1.76181447 |
| 10637 | NM_002528.6 | #N/A | 52.3931624 | -6.9191843 | 1.66543187 |
| 10639 | NR_109900.1 | LINC02225 | 69.8015467 | 4.92164991 | 1.18467479 |
| 10638 | NM_001271562.1 | #N/A | 42.2892228 | -6.6114182 | 1.59141234 |
| 10640 | NM_018465.3 | #N/A | 50.011462 | -6.8535228 | 1.64973788 |
| 10641 | NM_001323960.1 | RBCK1 | 39.7303564 | -6.5183122 | 1.56911961 |
| 10642 | NM_001172225.2 | #N/A | 44.0145244 | -6.6701564 | 1.60569137 |
| 10643 | NR_134528.1 | LAMTOR4 | 57.2313595 | -7.0456755 | 1.69611497 |
| 10644 | NM_207520.1 | #N/A | 68.103512 | -7.3017028 | 1.75780015 |
| 10645 | NM_001330502.1 | CAP1 | 95.1605834 | -7.7849284 | 1.87435151 |
| 10646 | NM_022768.4 | RBM15 | 37.035569 | -6.4182299 | 1.54533005 |
| 10648 | NM_019032.5 | ADAMTSL4 | 58.4291866 | -7.0826387 | 1.7053318 |
| 10647 | NM_015954.3 | #N/A | 41.8045207 | -6.5950786 | 1.587947 |
| 10649 | NM_001297418.1 | #N/A | 58.2626427 | -7.0759316 | 1.70393147 |
| 10650 | NR_037188.1 | #N/A | 39.06831 | -6.5043704 | 1.56642418 |
| 10651 | NM_001128917.1 | #N/A | 51.3788625 | -6.8951258 | 1.66085606 |
| 10652 | NM_017576.2 | #N/A | 63.5483398 | -7.2013202 | 1.73466367 |
| 10653 | NR_104025.1 | #N/A | 48.5949515 | -6.8101068 | 1.6404784 |
| 10654 | NM_001290118.1 | TIMM23B | 20.7892923 | 9.37247853 | 2.25775477 |
| 10655 | NM_001002258.4 | #N/A | 38.161049 | -6.4671346 | 1.55795297 |
| 10656 | NM_000853.3 | GSTT1 | 44.6300608 | -6.6843264 | 1.61037688 |
| 10657 | NM_032309.3 | #N/A | 51.5200764 | -6.8941595 | 1.66092924 |
| 10658 | NM_130797.3 | #N/A | 76.7713676 | -7.4710616 | 1.79991304 |
| 10659 | NM_001303062.1 | #N/A | 39.0045733 | -6.5020919 | 1.56656379 |
| 10660 | NM_001317026.1 | #N/A | 57.6882448 | -7.0598946 | 1.70100262 |
| 10662 | NR_131900.1 | AGAP1-IT1 | 17.6209241 | 5.7234009 | 1.37917548 |
| 10661 | NR_109807.1 | #N/A | 36.5886606 | -6.4067541 | 1.5438392 |
| 10663 | NM_017988.5 | #N/A | 49.4263596 | -6.8362161 | 1.64741252 |

| | | | | | |
|---|---|---|---|---|---|
| 10664 | NM_001288645.1 | #N/A | 95.9177614 | -7.793769 | 1.87821766 |
| 10665 | NM_001025253.2 | #N/A | 80.5875156 | -7.541425 | 1.81769954 |
| 10666 | NM_001330213.1 | #N/A | 44.5474823 | -6.6909737 | 1.61272165 |
| 10667 | NM_182511.3 | #N/A | 53.0059076 | -6.9392661 | 1.6726195 |
| 10668 | NM_018000.2 | #N/A | 144.353645 | 1.71976512 | 0.41460168 |
| 10669 | NM_001267536.2 | #N/A | 52.6439773 | -5.6440803 | 1.36079219 |
| 10670 | NM_001267065.1 | PLEKHB2 | 41.7161552 | -6.5866719 | 1.5882681 |
| 10671 | NM_001122752.1 | SERPINI1 | 72.081612 | -7.3804291 | 1.7798103 |
| 10672 | NR_024241.1 | FAM86DP | 36.7768318 | -6.4136081 | 1.54680044 |
| 10673 | NM_078629.3 | #N/A | 54.1931846 | -6.970285 | 1.6811109 |
| 10674 | NM_153334.6 | #N/A | 61.7437626 | -7.1628515 | 1.72761455 |
| 10675 | NM_004614.4 | #N/A | 48.6480787 | -6.8097774 | 1.64255136 |
| 10676 | NM_002975.2 | #N/A | 35.099219 | -6.347724 | 1.53113569 |
| 10677 | NM_024741.2 | #N/A | 49.691721 | -6.8490464 | 1.65212446 |
| 10678 | NM_001174108.2 | ZBED6 | 36.900036 | -6.4220103 | 1.54914659 |
| 10679 | NM_003993.3 | #N/A | 48.3540298 | -6.8077719 | 1.64237246 |
| 10680 | NM_001098484.2 | #N/A | 37.6155504 | -6.4373945 | 1.55312342 |
| 10681 | NM_138414.2 | #N/A | 49.3771073 | -6.8355287 | 1.6491867 |
| 10682 | NM_032012.3 | #N/A | 502.696874 | -3.520344 | 0.84937177 |
| 10683 | NM_080629.2 | COL11A1 | 48.6802768 | -6.8156593 | 1.64460108 |
| 10684 | NM_001171137.1 | #N/A | 104.232772 | -4.2249604 | 1.01957525 |
| 10685 | NM_001304833.1 | OGFOD2 | 38.0861209 | -6.4667868 | 1.56087071 |
| 10686 | NM_001278158.1 | ZNF205 | 37.0303341 | -6.4192772 | 1.54941634 |
| 10687 | NM_052923.1 | #N/A | 41.001068 | -6.5672926 | 1.58524202 |
| 10688 | NM_001143688.2 | #N/A | 42.951355 | -6.6408065 | 1.60313835 |
| 10689 | NM_032901.3 | #N/A | 43.8114745 | -6.6614804 | 1.6081628 |
| 10690 | NM_001319206.1 | #N/A | 66.6901295 | -7.2677775 | 1.75454654 |
| 10691 | NM_014583.3 | #N/A | 47.2340714 | -6.7732616 | 1.63520935 |
| 10692 | NM_014337.3 | #N/A | 60.6245225 | -7.1360461 | 1.72282368 |
| 10693 | NM_002312.3 | LIG4 | 60.4195121 | -7.1308043 | 1.72163224 |
| 10694 | NM_078476.3 | #N/A | 58.7510244 | -7.0842868 | 1.71054829 |
| 10695 | NM_012322.2 | #N/A | 65.1995751 | -7.2366705 | 1.7475259 |
| 10696 | NM_001025071.1 | #N/A | 45.0945878 | -6.7048055 | 1.61921225 |
| 10697 | NM_001317968.1 | ARRDC1 | 36.4491358 | -6.3971154 | 1.54508008 |
| 10698 | NM_020165.3 | #N/A | 46.1623863 | -6.7367042 | 1.6272004 |
| 10699 | NR_102705.1 | #N/A | 44.4473813 | -6.686731 | 1.61517686 |
| 10700 | NM_001159291.1 | MICAL1 | 54.3135462 | -6.9718528 | 1.68421381 |
| 10701 | NM_001329639.1 | #N/A | 39.7876818 | -6.5293776 | 1.57732629 |
| 10702 | NM_032936.3 | #N/A | 43.5516928 | -6.658525 | 1.60860331 |
| 10704 | NR_102432.2 | C13orf42 | 19.1134594 | 3.80791677 | 0.9199601 |
| 10703 | NM_001080529.2 | WIPF3 | 80.2337507 | -6.2627515 | 1.51301984 |
| 10705 | NM_145231.3 | #N/A | 39.9520084 | -6.5365717 | 1.57919589 |
| 10706 | NM_001014436.2 | #N/A | 248.810303 | -3.9645923 | 0.95782663 |

| | | | | | |
|---|---|---|---|---|---|
| 10707 | NM_005251.2 | #N/A | 47.6781201 | -6.7885519 | 1.64020978 |
| 10708 | NM_030636.2 | #N/A | 47.2270599 | -6.7699625 | 1.63574363 |
| 10709 | NM_001098497.2 | #N/A | 42.4274153 | -6.6167145 | 1.59878655 |
| 10710 | NM_001319846.1 | #N/A | 64.8184006 | -7.226105 | 1.74615834 |
| 10713 | NR_029408.1 | LINC00863 | 52.2282026 | -6.9156197 | 1.67126395 |
| 10711 | NM_001143787.1 | #N/A | 55.1039197 | -6.9911815 | 1.68952677 |
| 10712 | NM_003258.4 | #N/A | 54.7796627 | -6.986297 | 1.68835092 |
| 10714 | NM_005044.4 | #N/A | 388.675148 | -1.4206288 | 0.34333217 |
| 10715 | NM_001007563.2 | #N/A | 1062.0099 | -3.3892904 | 0.81918672 |
| 10716 | NM_199444.1 | #N/A | 49.6196129 | -6.8467922 | 1.65485232 |
| 10717 | NM_012405.3 | #N/A | 370.521662 | -1.8843127 | 0.45544983 |
| 10718 | NM_016125.3 | #N/A | 36.148589 | -6.3880364 | 1.54408101 |
| 10719 | NM_014466.2 | #N/A | 83.1346337 | -7.5894702 | 1.83451784 |
| 10720 | NM_000164.3 | #N/A | 57.7445396 | -7.0640988 | 1.70761523 |
| 10721 | NM_014750.4 | #N/A | 35.9950892 | -6.3841839 | 1.54325425 |
| 10722 | NM_017425.3 | SPA17 | 43.0982718 | -6.643659 | 1.60605941 |
| 10723 | NM_001142395.1 | #N/A | 44.8380289 | -6.6963559 | 1.61887975 |
| 10724 | NM_001199.3 | #N/A | 50.9842688 | -6.885597 | 1.66473636 |
| 10725 | NM_001089584.2 | #N/A | 55.6945343 | -7.0128985 | 1.69556999 |
| 10726 | NR_045286.1 | #N/A | 73.3361993 | -7.4056475 | 1.79066968 |
| 10730 | NR_033904.1 | LINC00921 | 56.7763295 | 2.33947355 | 0.56576505 |
| 10727 | NM_001144037.1 | TMEM25 | 41.6786089 | -6.5919064 | 1.5941518 |
| 10728 | NM_001319999.1 | #N/A | 84.3455651 | -7.6070562 | 1.83964276 |
| 10729 | NM_080764.3 | #N/A | 39.3172319 | -6.5028015 | 1.57259944 |
| 10731 | NM_001320537.1 | #N/A | 47.6227691 | -6.7848884 | 1.64083401 |
| 10732 | NM_001142599.1 | #N/A | 36.7414271 | -6.4141662 | 1.55122016 |
| 10733 | NM_013367.2 | #N/A | 60.3503607 | -7.1238064 | 1.72303862 |
| 10734 | NM_144617.2 | #N/A | 38.3926834 | -6.4743513 | 1.56609417 |
| 10735 | NM_001077700.2 | #N/A | 43.7915216 | -6.6643681 | 1.61216822 |
| 10736 | NM_001184749.2 | #N/A | 38.4930517 | -6.4741761 | 1.56618969 |
| 10737 | NM_001269.4 | #N/A | 46.2900871 | -6.741433 | 1.63083725 |
| 10738 | NM_024615.3 | #N/A | 42.1265975 | -6.6061689 | 1.59829592 |
| 10739 | NM_019013.2 | #N/A | 60.6233455 | -7.1307435 | 1.72535883 |
| 10740 | NM_130906.2 | #N/A | 55.2106145 | -6.9956087 | 1.6926715 |
| 10741 | NM_001310315.1 | #N/A | 53.0692878 | -6.93966 | 1.67917722 |
| 10742 | NM_006949.3 | #N/A | 42.3176988 | -6.6177746 | 1.60140387 |
| 10743 | NM_001042486.3 | #N/A | 420.319007 | -5.3409697 | 1.29250174 |
| 10744 | NM_145271.3 | ZNF688 | 37.9485625 | -6.4527807 | 1.56161259 |
| 10745 | NM_000964.3 | #N/A | 36.0252869 | -6.383966 | 1.5449797 |
| 10746 | NM_012460.3 | #N/A | 51.5040135 | -6.8941942 | 1.66852222 |
| 10747 | NR_003246.4 | GOLGA6L5P | 38.855669 | -6.4951798 | 1.57198602 |
| 10748 | NM_054027.4 | #N/A | 201.852316 | -3.8440045 | 0.93035224 |
| 10749 | NM_003779.3 | #N/A | 48.3432823 | -6.8100636 | 1.64830066 |

| | | | | | |
|---|---|---|---|---|---|
| 10750 | NM_006113.4 | #N/A | 38.784108 | -6.4900677 | 1.57099552 |
| 10751 | NM_203475.2 | #N/A | 76.0956643 | -7.4590069 | 1.8055404 |
| 10752 | NM_001033714.2 | #N/A | 40.4306768 | -6.5443475 | 1.58424696 |
| 10753 | NM_021269.2 | #N/A | 42.1162719 | -6.6030221 | 1.59849432 |
| 10754 | NM_001017524.2 | SIRT3 | 35.9378695 | -6.3823909 | 1.54509685 |
| 10755 | NM_001242819.1 | DEF8 | 59.1241149 | -7.0982329 | 1.71858387 |
| 10756 | NM_001164473.2 | FNBP1L | 115.643583 | -8.0627713 | 1.95216006 |
| 10757 | NM_030799.8 | #N/A | 267.833368 | -2.7429421 | 0.66423726 |
| 10758 | NM_018053.3 | #N/A | 34.7847423 | -6.3286306 | 1.53268141 |
| 10759 | NM_001278182.1 | #N/A | 112.655893 | -8.024426 | 1.94349304 |
| 10760 | NM_001322324.1 | #N/A | 93.8838521 | -6.2611897 | 1.51659968 |
| 10761 | NM_000635.3 | #N/A | 55.7504576 | -7.014451 | 1.69916255 |
| 10762 | NM_019891.3 | #N/A | 37.0857095 | -6.4175868 | 1.55459449 |
| 10763 | NR_028324.1 | LINC01002 | 40.7079287 | -6.5583528 | 1.58870628 |
| 10764 | NM_003885.2 | #N/A | 884.359697 | -5.8321642 | 1.41292098 |
| 10765 | NM_182679.2 | GPATCH4 | 63.7594078 | -7.2051565 | 1.74561796 |
| 10766 | NR_037148.1 | DNAJC25 | 44.0410201 | -6.6657518 | 1.61498723 |
| 10767 | NM_152309.2 | #N/A | 48.4085913 | -6.8089646 | 1.64973813 |
| 10768 | NR_015439.1 | UBA6-AS1 | 48.0968665 | -6.7984414 | 1.64729385 |
| 10769 | NM_001286485.1 | #N/A | 44.1831745 | -6.6798514 | 1.61860693 |
| 10770 | NM_017680.4 | #N/A | 36.9083435 | -6.4231784 | 1.55647774 |
| 10771 | NM_152540.3 | #N/A | 71.5959034 | -5.9181423 | 1.4341644 |
| 10772 | NM_019083.2 | TRMT13 | 38.9321717 | -6.4945968 | 1.57391565 |
| 10773 | NR_033416.1 | CCT6P3 | 43.2601705 | -6.6425193 | 1.60977525 |
| 10774 | NR_027387.1 | LINC01003 | 54.6443024 | -6.9813618 | 1.69203703 |
| 10775 | NM_003503.3 | #N/A | 60.7796693 | -7.131751 | 1.72851905 |
| 10776 | NM_001144994.1 | #N/A | 55.7017154 | -7.0071506 | 1.69845782 |
| 10777 | NM_001170670.1 | ZNHIT6 | 38.4912893 | 10.2356134 | 2.48109382 |
| 10778 | NM_001145727.2 | #N/A | 57.0649516 | 2.77859392 | 0.67365273 |
| 10779 | NM_001199856.1 | AK3 | 58.1302185 | -7.0722531 | 1.71485829 |
| 10780 | NM_001001556.2 | #N/A | 38.0056042 | -6.4637445 | 1.56733517 |
| 10781 | NM_139178.3 | #N/A | 38.8091459 | -6.4890614 | 1.57366974 |
| 10782 | NM_001031696.3 | PLD3 | 44.6762874 | -6.697857 | 1.62431716 |
| 10783 | NM_001242462.1 | GHR | 33.9586667 | 10.0533544 | 2.43823433 |
| 10784 | NR_045861.1 | #N/A | 35.0826875 | -6.3416268 | 1.53820603 |
| 10785 | NM_015918.3 | #N/A | 48.0341042 | -6.7939183 | 1.64798446 |
| 10786 | NM_001014445.2 | NLE1 | 68.828749 | -7.3145804 | 1.77440525 |
| 10787 | NM_001142287.1 | #N/A | 49.3405924 | -6.8328226 | 1.65770818 |
| 10788 | NM_001042588.1 | SNUPN | 40.6783584 | -6.557407 | 1.59091162 |
| 10789 | NM_133455.3 | #N/A | 60.809428 | -7.1335135 | 1.73071288 |
| 10790 | NM_001009932.2 | DNASE1L1 | 35.8105552 | -6.3772372 | 1.54732839 |
| 10791 | NM_016257.3 | #N/A | 74.2852734 | -7.4247339 | 1.80154263 |
| 10793 | NR_135262.1 | C1orf61 | 96.3157301 | -7.7989593 | 1.89283141 |

| | | | | | |
|---|---|---|---|---|---|
| 10792 | NM_178003.2 | #N/A | 89.2023182 | -7.6877312 | 1.86583149 |
| 10794 | NM_001281297.1 | #N/A | 49.9054024 | -6.8535532 | 1.66352559 |
| 10795 | NM_000282.3 | #N/A | 57.2199664 | -7.0508964 | 1.71201156 |
| 10796 | NM_001145847.1 | #N/A | 51.6327669 | -6.8944726 | 1.67409434 |
| 10797 | NM_206824.2 | VKORC1 | 46.4939149 | -6.7499227 | 1.63904074 |
| 10798 | NM_198150.2 | #N/A | 46.4823857 | -5.4671276 | 1.32758739 |
| 10799 | NM_004233.3 | #N/A | 42.1225118 | -6.6040106 | 1.60379253 |
| 10800 | NM_001318122.1 | #N/A | 43.2664133 | -6.6489531 | 1.61477289 |
| 10801 | NM_152288.2 | #N/A | 56.5478743 | -3.6999721 | 0.89867722 |
| 10802 | NM_198887.2 | #N/A | 182.023465 | -2.0587774 | 0.50004976 |
| 10803 | NM_182563.3 | BRICD5 | 38.0567755 | -6.4594379 | 1.56903005 |
| 10804 | NM_001163297.1 | #N/A | 41.5932073 | -6.5934377 | 1.60163251 |
| 10805 | NM_152516.3 | COMMD1 | 48.4793949 | -6.8110212 | 1.65466845 |
| 10806 | NM_018196.3 | #N/A | 37.7805896 | -6.4468026 | 1.5662181 |
| 10807 | NM_014182.4 | #N/A | 47.4374767 | -6.7793821 | 1.64707009 |
| 10808 | NM_002705.4 | #N/A | 53.8989287 | -6.9663846 | 1.6925351 |
| 10810 | NR_024333.1 | ILF3-DT | 183.655639 | -2.2081161 | 0.53650532 |
| 10809 | NM_178834.4 | LAYN | 55.8472612 | -7.0173092 | 1.70500222 |
| 10811 | NM_133467.2 | #N/A | 36.6708579 | -6.4110006 | 1.55776751 |
| 10812 | NM_001291994.1 | #N/A | 84.879204 | -7.6171203 | 1.85088115 |
| 10813 | NM_015871.4 | #N/A | 44.880751 | -6.6969787 | 1.62769431 |
| 10814 | NM_001098844.2 | #N/A | 77.3245364 | 2.72980675 | 0.66355899 |
| 10815 | NM_001160037.1 | RHOBTB2 | 57.0104408 | -7.0460616 | 1.71276651 |
| 10816 | NM_001039885.2 | #N/A | 47.8742162 | -6.7945251 | 1.65163468 |
| 10817 | NM_001166105.2 | #N/A | 46.0448266 | -6.7362021 | 1.63761262 |
| 10818 | NM_004419.3 | #N/A | 57.3302848 | -7.0495168 | 1.71379819 |
| 10819 | NM_181836.5 | #N/A | 317.426658 | -3.309573 | 0.80463734 |
| 10820 | NR_110291.1 | KANSL1L-AS1 | 24.8062461 | 3.69728922 | 0.89899841 |
| 10821 | NM_001330308.1 | #N/A | 41.1505261 | -6.572041 | 1.5980203 |
| 10822 | NM_024052.4 | #N/A | 57.9837485 | -4.8156399 | 1.17094223 |
| 10823 | NM_001195563.1 | #N/A | 36.6131798 | -6.3995085 | 1.55621546 |
| 10824 | NM_024944.2 | #N/A | 49.9084272 | -6.851548 | 1.66642885 |
| 10825 | NM_001277145.1 | #N/A | 36.6426901 | -6.4118714 | 1.55951356 |
| 10826 | NM_001284201.1 | JKAMP | 42.2741842 | -6.6130075 | 1.60845737 |
| 10827 | NM_174923.2 | #N/A | 66.7076824 | -7.2699362 | 1.76823162 |
| 10828 | NM_001322437.1 | HNRNPH3 | 36.0800509 | -6.3805692 | 1.55205125 |
| 10829 | NM_133334.2 | NSD2 | 275.128969 | -3.2031307 | 0.77932358 |
| 10830 | NM_001244705.1 | #N/A | 40.4940223 | -6.5483626 | 1.59361817 |
| 10831 | NM_001272049.1 | #N/A | 66.0792568 | -7.2547539 | 1.76565612 |
| 10832 | NM_007324.3 | #N/A | 73.7112306 | -5.9497898 | 1.44810633 |
| 10833 | NM_001002247.2 | ANAPC11 | 65.6308714 | -7.2452759 | 1.76343612 |
| 10834 | NM_001317902.1 | #N/A | 34.3569108 | 7.48038001 | 1.82065874 |
| 10835 | NM_174890.3 | #N/A | 33.9940648 | -6.2980733 | 1.53296716 |

| | | | | | |
|---|---|---|---|---|---|
| 10836 | NM_016937.3 | #N/A | 49.0280752 | -6.8235987 | 1.66104148 |
| 10839 | NR_024024.1 | ACAA1 | 38.705073 | -6.4881415 | 1.57948659 |
| 10837 | NM_001199388.2 | EPB41L2 | 37.8370698 | -6.4553102 | 1.57149731 |
| 10838 | NM_001202546.2 | #N/A | 49.2100853 | -6.8303024 | 1.66277583 |
| 10840 | NM_001198719.1 | #N/A | 55.2285425 | -6.9971206 | 1.7034088 |
| 10841 | NM_001127202.3 | #N/A | 49.4761547 | -6.8402576 | 1.66529082 |
| 10842 | NM_001143780.2 | #N/A | 43.5298654 | -6.6558739 | 1.62040851 |
| 10843 | NM_001294339.1 | #N/A | 39.7911 | -6.5238595 | 1.58843874 |
| 10844 | NR_028301.1 | DUBR | 51.3407733 | -6.8912269 | 1.67790256 |
| 10845 | NM_017891.4 | #N/A | 47.080717 | -6.768128 | 1.64796727 |
| 10846 | NM_001085426.2 | #N/A | 49.2841341 | -6.839027 | 1.66540951 |
| 10847 | NR_027783.2 | SAT1 | 44.2785624 | -6.6746426 | 1.62563835 |
| 10848 | NM_005892.3 | #N/A | 49.1457398 | -6.8272931 | 1.66289603 |
| 10849 | NM_003455.3 | #N/A | 50.3557828 | -6.8588327 | 1.67065418 |
| 10850 | NM_194282.3 | #N/A | 43.2099867 | -6.64522 | 1.61877283 |
| 10851 | NM_019555.2 | #N/A | 54.6271969 | -6.9812683 | 1.70068083 |
| 10852 | NM_198155.4 | #N/A | 66.039049 | -7.2572866 | 1.76791688 |
| 10853 | NM_138417.2 | #N/A | 42.7909073 | -6.6291706 | 1.61510686 |
| 10854 | NM_001193523.1 | #N/A | 37.0123551 | -6.4225901 | 1.56483626 |
| 10855 | NM_152914.2 | #N/A | 121.593683 | -2.6057861 | 0.63489899 |
| 10856 | NM_016334.3 | #N/A | 42.4161061 | -6.6168499 | 1.61221576 |
| 10857 | NM_194330.2 | RNF38 | 68.5287211 | -7.305694 | 1.7801549 |
| 10858 | NM_001018161.1 | PON2 | 44.1733277 | -6.6725735 | 1.62603358 |
| 10859 | NM_012214.2 | #N/A | 133.111507 | -3.9460693 | 0.96161262 |
| 10860 | NM_213649.1 | #N/A | 48.3493668 | -6.8033816 | 1.65791971 |
| 10861 | NM_002125.3 | #N/A | 70.5546385 | -7.3521333 | 1.79178495 |
| 10862 | NM_000137.2 | #N/A | 40.9541915 | -6.5701072 | 1.60125649 |
| 10863 | NR_027321.1 | LINC00964 | 38.63842 | 3.41507817 | 0.83237424 |
| 10864 | NM_001317238.1 | #N/A | 76.2193675 | -5.1709459 | 1.26040313 |
| 10865 | NM_152260.2 | #N/A | 59.2496701 | -5.6744203 | 1.38313297 |
| 10866 | NR_037947.1 | #N/A | 43.5732466 | -6.6502511 | 1.62099889 |
| 10867 | NM_001042723.1 | #N/A | 39.840917 | -6.51874 | 1.58905548 |
| 10868 | NM_001134405.1 | #N/A | 56.6759635 | -7.0333699 | 1.71459205 |
| 10869 | NM_001198664.1 | #N/A | 55.1712628 | -6.9916252 | 1.70441148 |
| 10870 | NM_153685.3 | #N/A | 70.3473447 | -7.3435659 | 1.79029022 |
| 10871 | NM_001671.4 | #N/A | 50.0141073 | -6.8529351 | 1.67079089 |
| 10872 | NM_002583.2 | #N/A | 38.3424262 | -6.4761866 | 1.57911504 |
| 10873 | NM_002309.4 | #N/A | 62.8790663 | -7.1852137 | 1.75242327 |
| 10874 | NM_001113498.2 | MDGA2 | 46.7151145 | -6.7557449 | 1.64775379 |
| 10875 | NM_032892.4 | #N/A | 63.1770127 | -7.1938469 | 1.75470149 |
| 10876 | NM_002801.3 | #N/A | 49.258742 | -6.8333767 | 1.66706402 |
| 10877 | NM_144660.2 | #N/A | 49.3620343 | -6.8308823 | 1.66660027 |
| 10878 | NM_001172574.1 | MCPH1 | 47.4623747 | -6.7794562 | 1.65406429 |

| | | | | | |
|---|---|---|---|---|---|
| 10879 | NM_001177711.1 | #N/A | 49.5703305 | -6.8423696 | 1.66957834 |
| 10880 | NM_014472.4 | #N/A | 34.0531175 | -6.3017444 | 1.53774309 |
| 10881 | NM_153719.3 | #N/A | 51.021064 | -6.8870304 | 1.68064279 |
| 10882 | NM_001164407.1 | #N/A | 419.386934 | 2.05348088 | 0.50117323 |
| 10883 | NM_004773.3 | #N/A | 43.4620457 | -6.6453824 | 1.62193914 |
| 10884 | NR_033897.1 | EPHA1-AS1 | 59.9888823 | 3.56362963 | 0.86981124 |
| 10885 | NM_001039366.1 | ATP6V1E1 | 60.7744159 | -7.1381098 | 1.74238561 |
| 10886 | NM_001143853.1 | PYM1 | 52.6863671 | -6.9293948 | 1.69143327 |
| 10887 | NM_015358.2 | #N/A | 68.0439268 | -7.3008217 | 1.78211713 |
| 10888 | NM_021168.4 | #N/A | 69.2621094 | -7.3252299 | 1.78818464 |
| 10889 | NM_002291.2 | #N/A | 955.899537 | -4.4475266 | 1.08578652 |
| 10890 | NM_016434.3 | #N/A | 41.5560535 | -6.5896103 | 1.6087918 |
| 10891 | NR_036636.1 | #N/A | 43.5267565 | -6.6534689 | 1.62443715 |
| 10892 | NM_001127715.2 | STXBP5 | 58.4751566 | -7.0749607 | 1.72739854 |
| 10893 | NM_194324.2 | #N/A | 58.1547783 | -7.0707778 | 1.72638597 |
| 10894 | NR_015453.2 | NOP14-AS1 | 46.2858354 | -6.7421581 | 1.64619434 |
| 10895 | NM_016647.2 | #N/A | 52.5728281 | -6.925089 | 1.69096029 |
| 10896 | NM_001330104.1 | #N/A | 38.188124 | -6.4606466 | 1.57756906 |
| 10897 | NM_174922.4 | #N/A | 55.5674289 | -7.0051476 | 1.7105883 |
| 10898 | NM_001001974.2 | #N/A | 45.4034241 | -6.7142276 | 1.63958715 |
| 10899 | NR_033710.1 | TP73-AS1 | 40.0872395 | -6.5352683 | 1.59607228 |
| 10900 | NR_126169.1 | PRR29-AS1 | 66.4014091 | 3.20265715 | 0.78229557 |
| 10901 | NM_001166279.1 | #N/A | 57.4324388 | -7.0517164 | 1.72254135 |
| 10902 | NM_014672.3 | #N/A | 50.3912468 | -6.8628494 | 1.67660233 |
| 10903 | NM_130901.2 | #N/A | 57.3988252 | -5.7735052 | 1.41067524 |
| 10904 | NM_017974.3 | #N/A | 43.4797564 | -6.6558991 | 1.62636819 |
| 10905 | NM_001198938.1 | #N/A | 33.7856752 | -6.2939471 | 1.53799984 |
| 10906 | NM_207340.1 | #N/A | 38.5213991 | -6.4786419 | 1.5831322 |
| 10907 | NM_177937.2 | GOLM1 | 37.9571029 | -6.4500796 | 1.57640944 |
| 10908 | NM_018154.2 | #N/A | 38.5139638 | -6.4769849 | 1.58310089 |
| 10909 | NR_003578.1 | ZNF702P | 39.5662574 | -6.5113774 | 1.59163731 |
| 10910 | NM_001098525.2 | CKAP2 | 58.8809188 | -7.0853093 | 1.73209464 |
| 10911 | NM_001309842.1 | #N/A | 65.3854 | -7.2390121 | 1.76967466 |
| 10912 | NM_024852.3 | #N/A | 37.3158151 | -6.4280041 | 1.57141879 |
| 10913 | NM_001288582.1 | #N/A | 38.3450335 | -6.4711958 | 1.58198977 |
| 10914 | NR_136240.1 | ATXN1-AS1 | 68.3151727 | 2.11797099 | 0.51777574 |
| 10915 | NM_152544.2 | #N/A | 39.8983917 | -6.5279328 | 1.59604663 |
| 10916 | NM_001172679.1 | #N/A | 56.6251855 | -5.6151026 | 1.37296544 |
| 10917 | NM_001135635.1 | #N/A | 43.8275018 | -6.6665577 | 1.63025693 |
| 10918 | NM_001318192.1 | #N/A | 159.44621 | -8.5281868 | 2.08559742 |
| 10919 | NM_001318234.1 | #N/A | 84.4152517 | -7.6088427 | 1.86093069 |
| 10920 | NM_001317915.1 | #N/A | 57.5922964 | -7.0592814 | 1.72656341 |
| 10921 | NM_178858.4 | #N/A | 102.099595 | 2.40022162 | 0.58706552 |

| | | | | | |
|---|---|---|---|---|---|
| 10922 | NM_014636.2 | #N/A | 74.8672976 | -7.4333813 | 1.81813935 |
| 10923 | NM_001001522.1 | #N/A | 41.1223496 | -6.5769866 | 1.60871764 |
| 10924 | NM_001037281.1 | #N/A | 54.2940836 | -6.9691358 | 1.70463482 |
| 10925 | NM_001331007.1 | #N/A | 68.9716939 | -7.3212092 | 1.79076256 |
| 10926 | NM_016013.3 | #N/A | 46.6268536 | -6.753463 | 1.65189872 |
| 10927 | NM_201522.2 | #N/A | 40.0732285 | -6.5363595 | 1.59883455 |
| 10928 | NM_014550.3 | #N/A | 36.9985103 | -6.4253162 | 1.57168424 |
| 10929 | NM_001114123.2 | ELK1 | 56.362272 | -7.0261665 | 1.71884057 |
| 10930 | NM_001287439.1 | #N/A | 54.6005318 | -6.9763541 | 1.70665807 |
| 10931 | NM_024063.2 | #N/A | 43.6161941 | -6.6541526 | 1.62785563 |
| 10932 | NM_030751.5 | #N/A | 66.9281353 | -7.2713073 | 1.77883434 |
| 10933 | NM_006630.2 | #N/A | 45.8128937 | -6.7250673 | 1.64523059 |
| 10934 | NM_006602.3 | #N/A | 40.3408782 | -6.545884 | 1.60146389 |
| 10935 | NM_032827.6 | #N/A | 34.8599398 | -6.3384934 | 1.55086344 |
| 10936 | NM_000027.3 | #N/A | 35.3746452 | -6.3483668 | 1.55334685 |
| 10937 | NM_001271783.1 | #N/A | 65.9093854 | -7.2524664 | 1.77458738 |
| 10938 | NM_001300984.1 | #N/A | 33.5290784 | -6.2756101 | 1.53560929 |
| 10939 | NM_001100119.1 | #N/A | 45.0208926 | -6.6960321 | 1.63850167 |
| 10940 | NM_024908.3 | #N/A | 51.5326719 | -6.8957396 | 1.68744617 |
| 10941 | NM_001198904.1 | YY1AP1 | 38.1953327 | -6.4659207 | 1.58243951 |
| 10942 | NM_002744.4 | #N/A | 38.6276103 | -6.4876651 | 1.58776953 |
| 10943 | NM_005011.4 | #N/A | 44.3353336 | -6.6783772 | 1.63448407 |
| 10944 | NR_038446.1 | LINC00861 | 32.2609608 | 3.87423055 | 0.94821473 |
| 10945 | NM_001276290.1 | PRKAR1A | 40.489778 | -6.556492 | 1.60478534 |
| 10946 | NM_001031738.2 | #N/A | 44.7893775 | -6.7011352 | 1.64023019 |
| 10947 | NM_001113402.1 | #N/A | 54.1682673 | -6.9665156 | 1.70538887 |
| 10948 | NM_030795.3 | #N/A | 66.3686833 | -7.2631679 | 1.77810087 |
| 10949 | NM_001178034.1 | DPP10 | 57.5616374 | -7.0559802 | 1.72769029 |
| 10950 | NM_173804.4 | #N/A | 33.7621643 | -6.2905663 | 1.54060023 |
| 10951 | NM_001114107.4 | PDLIM3 | 42.2197339 | -6.6135927 | 1.6197343 |
| 10952 | NM_022152.5 | #N/A | 36.7441438 | -6.4139147 | 1.57084236 |
| 10953 | NM_016310.4 | #N/A | 46.2846955 | -6.7409622 | 1.65101489 |
| 10954 | NM_001951.3 | #N/A | 51.8367841 | -6.906997 | 1.69176994 |
| 10955 | NM_007045.3 | #N/A | 65.282474 | -7.2419298 | 1.77418592 |
| 10957 | NM_194455.1 | KRIT1 | 37.6580963 | -6.4433291 | 1.57860481 |
| 10956 | NM_001321953.1 | #N/A | 36.6633625 | -6.4116443 | 1.5708369 |
| 10958 | NM_001135179.1 | #N/A | 236.866772 | -2.7597714 | 0.67615369 |
| 10959 | NM_024597.3 | #N/A | 36.5283128 | -6.4068945 | 1.56998773 |
| 10960 | NM_001297599.1 | MIER3 | 79.1235348 | -7.5146941 | 1.8415548 |
| 10961 | NM_203456.2 | PPIE | 42.6554596 | -6.6248899 | 1.62373533 |
| 10962 | NM_001270551.1 | #N/A | 53.8140102 | -6.9603174 | 1.70613899 |
| 10963 | NM_001318508.1 | #N/A | 38.4226557 | -6.4805289 | 1.58854118 |
| 10964 | NM_015110.3 | #N/A | 224.607925 | -2.8731207 | 0.70430981 |

| | | | | | |
|---|---|---|---|---|---|
| 10965 | NR_120513.1 | LOC101928861 | 30.7132796 | 3.95273559 | 0.968983 |
| 10966 | NM_001163636.2 | #N/A | 69.7419831 | -7.3358859 | 1.79836251 |
| 10967 | NM_001146051.1 | HSDL1 | 37.5587569 | 10.2198757 | 2.50544188 |
| 10968 | NM_003224.5 | #N/A | 41.0431451 | -6.5730091 | 1.61144544 |
| 10969 | NM_001320036.1 | #N/A | 35.3327994 | -6.3586383 | 1.55900969 |
| 10970 | NM_202469.2 | #N/A | 51.3070839 | -6.8937014 | 1.69025773 |
| 10971 | NM_001322929.1 | STAU1 | 54.2033146 | -6.9656859 | 1.70798424 |
| 10972 | NM_001267783.1 | #N/A | 52.1808246 | -6.9171728 | 1.6961741 |
| 10973 | NM_001282759.1 | TRA2A | 36.7402235 | -6.4121527 | 1.57234703 |
| 10974 | NR_037802.1 | ZNF625-ZNF20 | 37.0951744 | -6.4163457 | 1.57348769 |
| 10975 | NM_001199461.1 | PDCD2 | 48.4191711 | -6.8052402 | 1.66897482 |
| 10976 | NM_031446.4 | #N/A | 38.5804962 | -6.4798713 | 1.58917931 |
| 10977 | NM_001105520.1 | #N/A | 36.3034119 | -6.3892743 | 1.56697288 |
| 10978 | NM_015148.3 | #N/A | 52.2206072 | -6.9132705 | 1.69550821 |
| 10979 | NM_001244580.1 | TRRAP | 65.9617468 | -7.2523573 | 1.77870566 |
| 10980 | NM_152515.4 | #N/A | 38.149007 | -6.4702748 | 1.58709677 |
| 10981 | NM_014291.3 | GCAT | 49.6652091 | -6.8452736 | 1.67912109 |
| 10982 | NM_001130845.1 | #N/A | 41.5176796 | -6.5886152 | 1.61643466 |
| 10983 | NM_001142651.2 | #N/A | 77.9102361 | -7.4934475 | 1.83864806 |
| 10984 | NM_005612.4 | #N/A | 63.862626 | -7.2098199 | 1.76908791 |
| 10985 | NM_001128918.2 | #N/A | 51.2400096 | -6.890592 | 1.69077117 |
| 10986 | NM_001100117.2 | #N/A | 54.8529702 | -6.9887115 | 1.7149891 |
| 10987 | NR_038361.1 | APTR | 33.7892827 | -6.2888067 | 1.54328058 |
| 10988 | NM_015545.3 | #N/A | 154.686244 | -2.1653693 | 0.53139819 |
| 10989 | NR_034034.1 | LOC100288748 | 46.964891 | 2.47460252 | 0.60729668 |
| 10990 | NM_138317.2 | #N/A | 39.3437208 | -6.5038688 | 1.59620116 |
| 10991 | NM_002594.4 | #N/A | 44.384744 | -6.678932 | 1.63935662 |
| 10992 | NM_001039111.2 | #N/A | 43.0552222 | -6.639894 | 1.62980271 |
| 10993 | NM_004059.4 | #N/A | 44.920092 | -6.694312 | 1.64322444 |
| 10994 | NM_007109.2 | #N/A | 40.335221 | -6.5366498 | 1.60462907 |
| 10995 | NM_001304331.1 | #N/A | 99.0608572 | -7.8401973 | 1.92479268 |
| 10996 | NM_000380.3 | XPA | 64.0889923 | -5.7664355 | 1.41576598 |
| 10997 | NM_020703.2 | #N/A | 39.6697252 | -6.517327 | 1.60030965 |
| 10998 | NM_004966.3 | HNRNPF | 49.5970119 | -6.8423985 | 1.68024275 |
| 10999 | NM_005880.3 | #N/A | 351.40267 | -3.3552942 | 0.82397782 |
| 11000 | NM_001346890.1 | WHRN | 39.1741085 | -6.5066235 | 1.59788556 |
| 11001 | NM_002207.2 | #N/A | 38.3625603 | -6.4720376 | 1.58942966 |
| 11002 | NM_016592.3 | #N/A | 56.2168557 | -7.0248891 | 1.72526086 |
| 11003 | NM_001171136.1 | #N/A | 74.9091854 | -7.4340259 | 1.82592281 |
| 11004 | NM_001136232.2 | SEC13 | 38.8197958 | -6.4962554 | 1.59572484 |
| 11005 | NM_001205228.1 | SORT1 | 46.7221823 | -6.7568687 | 1.65974321 |
| 11006 | NM_001142305.1 | #N/A | 52.273694 | -6.9205061 | 1.70003916 |
| 11007 | NM_001291878.1 | #N/A | 133.02472 | -8.2657966 | 2.03054795 |

| | | | | | |
|---|---|---|---|---|---|
| 11008 | NM_145859.1 | PDCD10 | 40.4235846 | -6.5486413 | 1.60878392 |
| 11009 | NM_001257389.1 | CD63 | 55.0649455 | -6.9927204 | 1.71804725 |
| 11010 | NM_007281.3 | #N/A | 65.2987508 | -7.2406919 | 1.77908141 |
| 11011 | NM_002010.2 | #N/A | 52.9319279 | -6.936 | 1.70431895 |
| 11012 | NM_003335.2 | #N/A | 54.7036544 | -5.5716918 | 1.3690992 |
| 11013 | NM_001318327.1 | DNMBP | 37.2003471 | -6.4281155 | 1.57987349 |
| 11014 | NM_003068.4 | #N/A | 42.2140298 | -6.6140299 | 1.62557464 |
| 11016 | NR_045014.1 | LMO3 | 151.371717 | -8.452021 | 2.07748374 |
| 11015 | NM_080685.2 | PTPN13 | 41.4884769 | -6.5775561 | 1.61674341 |
| 11017 | NM_005410.2 | #N/A | 39.2437613 | -6.5081926 | 1.59979524 |
| 11018 | NR_021489.1 | #N/A | 52.6654296 | -6.9310826 | 1.7038425 |
| 11019 | NM_001145648.1 | #N/A | 63.3892676 | -7.1949573 | 1.76872551 |
| 11020 | NM_001174051.2 | #N/A | 74.6077012 | -7.4303214 | 1.82661285 |
| 11021 | NR_047594.1 | #N/A | 39.7073799 | -6.5199603 | 1.60293398 |
| 11022 | NM_173515.2 | #N/A | 64.1069607 | -7.2150877 | 1.77397476 |
| 11023 | NM_018129.3 | #N/A | 293.5313 | 2.3053019 | 0.56684488 |
| 11024 | NM_004454.2 | #N/A | 109.281431 | -5.2255522 | 1.28491264 |
| 11025 | NM_001282764.1 | CDV3 | 46.5195617 | -6.7477224 | 1.65922699 |
| 11026 | NR_037193.1 | #N/A | 45.7860256 | -6.7242823 | 1.65348278 |
| 11027 | NM_022055.1 | #N/A | 62.7894242 | -7.1816653 | 1.76607115 |
| 11028 | NM_001290293.2 | SDR39U1 | 34.9676487 | -6.3341458 | 1.55785982 |
| 11029 | NM_173514.3 | #N/A | 44.2044394 | -6.6798387 | 1.64290809 |
| 11030 | NM_001256495.1 | MAN2C1 | 43.1603972 | -6.6394097 | 1.63298877 |
| 11031 | NM_020158.3 | #N/A | 56.9889751 | -7.041277 | 1.73185849 |
| 11032 | NM_144574.3 | #N/A | 35.3815097 | -6.3508084 | 1.56221962 |
| 11033 | NM_024855.3 | #N/A | 51.5204967 | -5.6158985 | 1.38147205 |
| 11034 | NM_001134938.1 | FIP1L1 | 46.5761897 | -6.751278 | 1.6608533 |
| 11035 | NM_002445.3 | #N/A | 40.4736706 | 4.3635811 | 1.07346322 |
| 11036 | NM_018916.3 | #N/A | 76.1929896 | -7.460249 | 1.83535339 |
| 11037 | NM_001098207.1 | HNRNPF | 51.8325653 | -6.9050908 | 1.69899708 |
| 11039 | NR_015454.1 | MAFG-DT | 36.8373976 | -6.4151814 | 1.57844469 |
| 11038 | NM_001323956.1 | RBCK1 | 71.629028 | -7.3724445 | 1.81398971 |
| 11040 | NM_181655.3 | #N/A | 41.2183258 | -6.5714783 | 1.61693298 |
| 11041 | NM_006101.2 | #N/A | 38.6044349 | -6.4784042 | 1.59408035 |
| 11042 | NM_001008897.1 | TCP1 | 60.2179365 | -7.1190616 | 1.75175112 |
| 11043 | NM_001321099.1 | #N/A | 89.9411431 | -7.6982847 | 1.89437743 |
| 11044 | NM_001134450.1 | #N/A | 41.0155229 | -6.5613186 | 1.61462506 |
| 11045 | NM_018429.2 | #N/A | 228.223891 | -2.8636405 | 0.7047126 |
| 11046 | NM_052880.4 | #N/A | 50.8694818 | -6.8828312 | 1.69394374 |
| 11047 | NM_021922.2 | #N/A | 44.1239085 | -6.6724875 | 1.64220322 |
| 11048 | NR_002186.1 | DKFZP586I1420 | 36.9602397 | -6.4145182 | 1.57885762 |
| 11049 | NM_001102426.2 | TBC1D8 | 37.488527 | -6.4453901 | 1.58647995 |
| 11050 | NM_006090.4 | #N/A | 39.5898435 | -6.5122504 | 1.60326356 |

| | | | | | |
|---|---|---|---|---|---|
| 11051 | NM_019591.3 | #N/A | 41.6262102 | -6.5838784 | 1.62097035 |
| 11052 | NM_001080849.2 | #N/A | 50.7334416 | -6.87353 | 1.69238683 |
| 11053 | NM_001456.3 | FLNA | 1205.68678 | -5.448453 | 1.34155963 |
| 11054 | NR_046357.1 | RNF165 | 71.166079 | -7.3622628 | 1.81286063 |
| 11055 | NM_001218.4 | #N/A | 96.007182 | -7.796396 | 1.91987861 |
| 11056 | NM_005524.3 | #N/A | 135.229804 | -4.751079 | 1.1699916 |
| 11058 | NR_034146.1 | SIRT2 | 44.5609131 | -5.4118575 | 1.33274456 |
| 11057 | NM_147686.3 | #N/A | 41.4837675 | -6.5878974 | 1.6223585 |
| 11060 | NM_145063.3 | OARD1 | 35.8068818 | -6.3665643 | 1.56795023 |
| 11059 | NM_001330322.1 | #N/A | 36.9184283 | -6.4092674 | 1.57846722 |
| 11061 | NM_003741.3 | CHRD | 36.7985318 | -6.4124414 | 1.57940034 |
| 11062 | NM_001271006.1 | #N/A | 44.7637111 | -6.6911015 | 1.64866311 |
| 11063 | NM_001130182.1 | DNAJA4 | 35.8140179 | -6.3670475 | 1.56883991 |
| 11064 | NR_136703.1 | SFPQ | 37.5606608 | -6.4371627 | 1.58614737 |
| 11065 | NM_006169.2 | #N/A | 63.3493316 | -7.1962081 | 1.77386565 |
| 11066 | NR_037672.1 | SERF2 | 38.4472182 | -6.4811559 | 1.59762135 |
| 11067 | NM_006154.3 | #N/A | 41.0997376 | -6.5738022 | 1.62054209 |
| 11068 | NM_001278225.1 | RUFY2 | 39.7151838 | -6.5232683 | 1.60810007 |
| 11069 | NM_014494.2 | #N/A | 92.4987408 | -7.7419207 | 1.90854337 |
| 11070 | NM_080822.2 | #N/A | 49.6921411 | -6.8438439 | 1.68718245 |
| 11071 | NM_001317951.1 | #N/A | 39.4081592 | -6.5122624 | 1.60574958 |
| 11072 | NM_030802.3 | #N/A | 40.6920529 | -6.562482 | 1.61821556 |
| 11073 | NM_001080397.2 | SLC45A1 | 55.4080481 | -7.0025643 | 1.72674394 |
| 11074 | NM_001142448.1 | #N/A | 50.8916601 | -6.8847343 | 1.69777749 |
| 11075 | NM_015421.3 | #N/A | 37.7670463 | -6.4490375 | 1.5906621 |
| 11076 | NR_027241.2 | #N/A | 44.4148783 | -6.6800209 | 1.64771373 |
| 11077 | NM_001184774.1 | #N/A | 43.2383289 | -6.6419734 | 1.6383939 |
| 11078 | NM_001256277.1 | #N/A | 47.8132163 | -6.791649 | 1.67535181 |
| 11079 | NR_037630.1 | LOC645513 | 35.9550448 | -6.3758024 | 1.5728462 |
| 11080 | NM_022490.2 | #N/A | 94.5513881 | -3.1498726 | 0.77708278 |
| 11081 | NM_138717.2 | #N/A | 33.4127082 | -6.2778397 | 1.54881156 |
| 11082 | NM_201541.1 | NDRG2 | 37.9362462 | -6.459126 | 1.59355616 |
| 11083 | NM_001323607.1 | SYNDIG1 | 49.6060368 | -6.8424016 | 1.68815067 |
| 11084 | NM_006658.4 | #N/A | 34.0342258 | -6.2952523 | 1.5532405 |
| 11085 | NM_148965.1 | TNFRSF25 | 42.2184319 | -6.6061169 | 1.63002518 |
| 11086 | NM_001145639.1 | GPSM1 | 49.5715973 | -6.8431017 | 1.68859469 |
| 11087 | NM_001329615.1 | #N/A | 37.2908335 | -6.4247193 | 1.58557449 |
| 11088 | NM_018163.2 | #N/A | 39.1143834 | -6.4983551 | 1.60374208 |
| 11089 | NR_125936.1 | LINC02506 | 19.6136136 | 3.87665729 | 0.95681479 |
| 11090 | NM_001143819.2 | #N/A | 56.5080195 | -7.0311664 | 1.73553729 |
| 11091 | NR_037688.1 | #N/A | 1578.18916 | -4.9783943 | 1.22888473 |
| 11092 | NM_014762.3 | #N/A | 1897.92861 | -5.8621936 | 1.44705609 |
| 11093 | NR_040252.1 | #N/A | 39.117562 | -6.4967812 | 1.60371232 |

| | | | | | |
|---|---|---|---|---|---|
| 11094 | NM_001324266.1 | #N/A | 53.6622871 | -6.9518619 | 1.71616337 |
| 11095 | NM_001160147.1 | #N/A | 58.9196079 | -7.0896114 | 1.75039966 |
| 11096 | NM_024813.2 | #N/A | 44.2936969 | -6.6759619 | 1.6483492 |
| 11097 | NM_001001660.2 | #N/A | 49.5120331 | -6.839006 | 1.68866726 |
| 11098 | NM_006321.3 | #N/A | 87.6960697 | -7.6639572 | 1.89241225 |
| 11099 | NM_001100878.1 | #N/A | 31.7679955 | -6.2031939 | 1.53173323 |
| 11100 | NM_130783.4 | #N/A | 218.86309 | -3.5298929 | 0.87172136 |
| 11101 | NM_001317770.1 | #N/A | 62.9812762 | -4.5910948 | 1.13379669 |
| 11102 | NM_006440.4 | #N/A | 36.1495485 | -6.3880703 | 1.57759315 |
| 11103 | NM_005392.3 | #N/A | 414.133681 | -3.4297212 | 0.84712382 |
| 11104 | NM_016955.3 | #N/A | 32.4436798 | -6.2290048 | 1.53864864 |
| 11105 | NM_001949.4 | #N/A | 52.2820774 | -6.916054 | 1.70841047 |
| 11106 | NM_080571.1 | C20orf96 | 73.7649182 | -7.4141478 | 1.83156928 |
| 11107 | NM_001190487.2 | MTRNR2L6 | 1410.07098 | -4.9756318 | 1.22917787 |
| 11108 | NM_001008707.1 | #N/A | 98.420024 | -7.8299581 | 1.93443334 |
| 11109 | NM_017917.3 | PPP2R3C | 43.9412087 | -6.6664966 | 1.64713169 |
| 11110 | NM_024940.7 | #N/A | 34.0028678 | -6.3028197 | 1.55734378 |
| 11111 | NM_182501.3 | #N/A | 38.0403845 | -6.4544356 | 1.59507099 |
| 11112 | NM_005393.2 | #N/A | 49.5677628 | -6.84148 | 1.69102494 |
| 11113 | NR_034170.1 | EIF3J-DT | 36.1911022 | -6.3920397 | 1.58002147 |
| 11114 | NM_007099.3 | ACP1 | 41.9675809 | -6.6070616 | 1.63320778 |
| 11115 | NM_001031720.3 | GSTCD | 35.6580176 | -6.3710789 | 1.57489796 |
| 11116 | NM_004839.3 | #N/A | 37.1192953 | -6.4316562 | 1.58988306 |
| 11117 | NM_080611.4 | DUSP15 | 38.7606081 | -6.4873172 | 1.60369708 |
| 11118 | NM_012180.2 | #N/A | 38.4945898 | -6.4764969 | 1.60109252 |
| 11119 | NM_001171894.2 | #N/A | 48.972348 | -6.8279125 | 1.68800114 |
| 11120 | NM_001037340.2 | PDE4B | 52.7709838 | -6.9321306 | 1.7138145 |
| 11121 | NM_153741.1 | #N/A | 45.6561456 | -6.7226127 | 1.66209029 |
| 11122 | NM_016093.3 | RPL26L1 | 64.2349156 | -7.2138084 | 1.78372425 |
| 11123 | NM_198220.2 | #N/A | 44.6604967 | -6.6897807 | 1.65430285 |
| 11124 | NM_001080477.3 | #N/A | 459.383451 | -4.0126534 | 0.99234205 |
| 11125 | NM_002194.3 | #N/A | 42.4439616 | -6.6164988 | 1.63631658 |
| 11126 | NM_016202.2 | ZNF580 | 32.9905558 | -6.2526212 | 1.54645937 |
| 11127 | NM_004451.4 | #N/A | 54.7247105 | -6.9828136 | 1.72709059 |
| 11128 | NM_173473.3 | #N/A | 66.3081461 | -7.2595252 | 1.7955873 |
| 11130 | NR_026899.1 | ARHGAP27P1-BPTFP1- | 37.6618595 | -6.4456715 | 1.59433555 |
| 11129 | NM_017918.4 | #N/A | 35.6036756 | -6.3581341 | 1.57267787 |
| 11131 | NM_001884.3 | #N/A | 57.5161891 | -7.0569532 | 1.74571828 |
| 11132 | NM_025188.3 | #N/A | 39.1951642 | -6.5006017 | 1.60809975 |
| 11133 | NM_006002.4 | UCHL3 | 47.1363797 | -6.7678369 | 1.67429343 |
| 11134 | NM_015149.4 | #N/A | 37.52615 | -6.438614 | 1.59308253 |
| 11135 | NM_144576.3 | #N/A | 42.2612231 | -6.6046691 | 1.63431153 |
| 11136 | NM_001077350.2 | #N/A | 67.9139076 | -7.2935879 | 1.80506976 |

| | | | | | |
|---|---|---|---|---|---|
| 11137 | NM_001144775.2 | #N/A | 73.2476109 | -7.4058597 | 1.83287753 |
| 11138 | NR_028039.1 | CASTOR3 | 48.8685235 | -6.8257574 | 1.68942611 |
| 11139 | NM_138775.2 | #N/A | 39.1559223 | -6.5004132 | 1.60891771 |
| 11140 | NM_001278222.1 | C11orf49 | 38.6997517 | -6.4854593 | 1.60532271 |
| 11141 | NM_001320973.1 | #N/A | 42.9568514 | -6.6319315 | 1.64175002 |
| 11142 | NM_003972.2 | #N/A | 318.33634 | -3.0832309 | 0.76336082 |
| 11143 | NM_001303143.1 | #N/A | 49.2925962 | -6.8278502 | 1.6907887 |
| 11144 | NM_017679.3 | #N/A | 37.6295816 | -6.4415592 | 1.59516308 |
| 11145 | NM_001198951.1 | RNF103 | 36.3942273 | -6.392726 | 1.58325261 |
| 11146 | NM_001145205.1 | #N/A | 42.0803308 | -6.6059515 | 1.63622047 |
| 11147 | NM_017793.2 | #N/A | 48.4111268 | -6.8045683 | 1.6854212 |
| 11148 | NR_036435.1 | #N/A | 38.9461438 | -6.4863031 | 1.60657521 |
| 11149 | NM_145052.3 | #N/A | 38.7100978 | -6.4827448 | 1.60576454 |
| 11150 | NM_213566.1 | #N/A | 333.770659 | 1.43389057 | 0.3551981 |
| 11151 | NM_001258218.1 | MIS12 | 37.7270972 | -6.4449276 | 1.5965463 |
| 11152 | NM_001098398.1 | #N/A | 88.5067423 | -7.6799484 | 1.90265059 |
| 11153 | NM_003173.3 | #N/A | 37.9704608 | -6.4551295 | 1.59946551 |
| 11154 | NM_005505.4 | #N/A | 47.8005072 | -6.7936242 | 1.68336854 |
| 11155 | NM_024578.2 | #N/A | 32.2339422 | -6.2224692 | 1.54188392 |
| 11156 | NM_016104.3 | #N/A | 49.3644497 | -6.8336331 | 1.69337672 |
| 11157 | NM_001185011.1 | #N/A | 44.7699043 | -6.6898188 | 1.65791183 |
| 11158 | NM_001023587.2 | #N/A | 42.5825103 | -6.6209881 | 1.64093003 |
| 11159 | NM_003420.3 | #N/A | 41.8653975 | -6.5963728 | 1.63485776 |
| 11160 | NM_032899.6 | FAM83A | 20.3803049 | 3.84601241 | 0.95327142 |
| 11161 | NM_001243203.1 | #N/A | 56.8481901 | -7.0377158 | 1.74439013 |
| 11162 | NM_006933.6 | #N/A | 391.803553 | -4.6439806 | 1.15118657 |
| 11163 | NM_032565.4 | #N/A | 44.45335 | -6.6855058 | 1.65736964 |
| 11164 | NM_006331.7 | #N/A | 46.5078259 | -6.7488183 | 1.67315263 |
| 11165 | NM_001986.2 | #N/A | 45.8695992 | -6.7307775 | 1.66873343 |
| 11166 | NR_126059.1 | LINC01560 | 32.6564314 | -6.2363321 | 1.54619805 |
| 11167 | NR_145128.1 | #N/A | 42.6607685 | -6.621991 | 1.64186789 |
| 11168 | NR_134984.1 | #N/A | 40.5644818 | -5.2675871 | 1.30610453 |
| 11169 | NM_001307936.1 | #N/A | 46.275126 | -6.7360185 | 1.67023098 |
| 11170 | NM_172373.3 | #N/A | 42.0540395 | -6.6105741 | 1.63914211 |
| 11171 | NM_001452.1 | #N/A | 82.2986795 | -6.0755223 | 1.50658295 |
| 11172 | NM_201286.3 | USP51 | 37.2773285 | -6.4336731 | 1.59544535 |
| 11173 | NR_033268.1 | #N/A | 34.4504152 | -6.3194109 | 1.56731233 |
| 11174 | NM_001348124.1 | TRIM16 | 52.9994001 | -6.9347637 | 1.72003167 |
| 11176 | NM_001288656.1 | UBN1 | 51.0937205 | -6.8873398 | 1.70834418 |
| 11175 | NM_001033050.2 | #N/A | 44.4975739 | -6.6821095 | 1.65744641 |
| 11177 | NM_024057.3 | #N/A | 35.7502516 | -6.3686716 | 1.57997169 |
| 11178 | NR_024386.2 | PLEKHM1P1 | 35.237766 | -6.3444225 | 1.57402219 |
| 11179 | NM_003182.2 | #N/A | 156.663949 | -5.6095241 | 1.39203243 |

| | | | | | |
|---|---|---|---|---|---|
| 11180 | NM_001267774.1 | #N/A | 46.5136555 | -6.7486496 | 1.67488024 |
| 11181 | NM_021831.5 | #N/A | 40.3397016 | -6.5452724 | 1.6244025 |
| 11182 | NM_016245.4 | #N/A | 42.4101109 | -6.6167553 | 1.64227255 |
| 11183 | NR_038994.1 | VSTM2A-OT1 | 43.3758565 | 4.39938097 | 1.09193102 |
| 11184 | NM_021825.4 | CCDC90B | 43.2233552 | -6.6448386 | 1.64942593 |
| 11185 | NM_001168647.2 | #N/A | 44.7870751 | -6.6935874 | 1.66154184 |
| 11186 | NM_005079.3 | #N/A | 46.8131057 | -6.7637386 | 1.6791583 |
| 11187 | NM_001707.3 | #N/A | 46.3785105 | -6.7462722 | 1.67483512 |
| 11188 | NM_030922.6 | NIPA2 | 55.7249111 | -7.0089729 | 1.74029063 |
| 11189 | NM_001282529.1 | #N/A | 44.454212 | -6.6847504 | 1.65983063 |
| 11190 | NM_021630.5 | #N/A | 47.1139782 | -6.7698118 | 1.68118817 |
| 11191 | NR_103498.1 | UBE2F | 39.9824677 | -6.5329908 | 1.62245374 |
| 11192 | NM_016042.3 | #N/A | 37.1480555 | -6.4251187 | 1.59571171 |
| 11193 | NM_014449.1 | #N/A | 59.1019723 | -7.0936649 | 1.76178358 |
| 11194 | NR_073008.1 | CDK2AP1 | 36.8582769 | -6.4109093 | 1.59224725 |
| 11195 | NM_004785.5 | #N/A | 33.5443173 | -6.2763573 | 1.55884865 |
| 11196 | NM_001145312.1 | #N/A | 32.476163 | -6.233289 | 1.54836929 |
| 11197 | NM_001130029.1 | RELL2 | 68.3474536 | -7.305072 | 1.81462675 |
| 11198 | NM_004667.5 | HERC2 | 521.495363 | -3.2374554 | 0.80421047 |
| 11199 | NM_012339.3 | #N/A | 34.1757448 | -6.3099443 | 1.56748106 |
| 11200 | NR_045720.1 | #N/A | 44.3817911 | -6.6852282 | 1.66076076 |
| 11201 | NM_017982.3 | #N/A | 46.3361919 | -6.7436844 | 1.67531704 |
| 11202 | NM_001297723.1 | C1orf43 | 32.5524499 | -6.2374311 | 1.54961815 |
| 11203 | NM_145111.3 | #N/A | 49.4622547 | -4.7606437 | 1.18274518 |
| 11204 | NM_207334.2 | #N/A | 37.8846663 | -6.4506578 | 1.60262743 |
| 11205 | NM_001286439.1 | NME4 | 34.5216744 | -6.3191977 | 1.57013142 |
| 11206 | NM_001006625.1 | #N/A | 14.8974139 | 8.88289707 | 2.20716427 |
| 11207 | NM_001291379.1 | AGAP4 | 51.084637 | -6.8817396 | 1.71002315 |
| 11208 | NM_001301771.1 | #N/A | 40.4473301 | -6.5496201 | 1.62755939 |
| 11209 | NM_000456.2 | #N/A | 33.1756387 | -6.2589819 | 1.55549944 |
| 11210 | NM_001226.3 | #N/A | 39.7726787 | -6.5227975 | 1.6210657 |
| 11211 | NM_178539.4 | TAFA2 | 75.4226727 | -5.9624556 | 1.48191139 |
| 11212 | NM_000885.5 | #N/A | 37.3018859 | -6.4384221 | 1.60022258 |
| 11213 | NM_022090.4 | #N/A | 37.2474383 | -6.4250257 | 1.59712886 |
| 11214 | NM_001166660.1 | NLGN3 | 81.4739685 | -7.5587582 | 1.87916508 |
| 11215 | NM_198309.3 | TTC8 | 33.9923147 | -6.303264 | 1.56712664 |
| 11216 | NM_016510.5 | #N/A | 36.411045 | -6.3914545 | 1.58912443 |
| 11217 | NM_001271969.1 | HSP90AB1 | 61.0014259 | -7.1428795 | 1.77603557 |
| 11218 | NM_001099338.1 | #N/A | 41.2331708 | -6.5768903 | 1.63564018 |
| 11219 | NM_053035.2 | #N/A | 41.6107563 | -6.5896271 | 1.63886141 |
| 11220 | NR_038292.1 | LINC01012 | 139.816631 | 2.78394918 | 0.6923889 |
| 11221 | NM_006042.2 | #N/A | 42.8373318 | -6.6333933 | 1.64983101 |
| 11222 | NM_004215.4 | #N/A | 31.9396602 | -6.2094479 | 1.54448852 |

| | | | | | |
|---|---|---|---|---|---|
| 11223 | NM_153339.2 | #N/A | 42.4806994 | -6.6168284 | 1.64608258 |
| 11224 | NM_001303525.1 | TUBB6 | 38.0108138 | -6.4650875 | 1.60835979 |
| 11225 | NM_001190440.1 | NCOR1 | 33.3017756 | -6.2621462 | 1.55790111 |
| 11226 | NM_001001438.2 | LSS | 53.829156 | -6.9584048 | 1.73114838 |
| 11227 | NM_017563.4 | #N/A | 438.435158 | -3.778317 | 0.94014026 |
| 11228 | NM_052935.4 | #N/A | 41.8112704 | -6.5944322 | 1.64114427 |
| 11229 | NM_001281460.1 | #N/A | 35.1230545 | -6.3376957 | 1.57729366 |
| 11230 | NM_001256513.1 | SSBP1 | 47.204578 | -6.7720837 | 1.68545246 |
| 11231 | NM_148908.3 | #N/A | 57.1244775 | -7.0443188 | 1.75322177 |
| 11232 | NM_004453.3 | #N/A | 36.1655451 | -6.3882142 | 1.59007031 |
| 11233 | NR_036576.1 | #N/A | 39.0941937 | -6.4989492 | 1.61764743 |
| 11234 | NM_001278228.1 | RUSC1 | 46.9337087 | -6.7618321 | 1.6831511 |
| 11235 | NM_080916.2 | #N/A | 35.6875442 | -6.364891 | 1.58439475 |
| 11236 | NM_001286069.1 | #N/A | 45.7916755 | -6.7245639 | 1.67397846 |
| 11237 | NM_031905.4 | #N/A | 42.2810105 | -6.6146459 | 1.64667552 |
| 11238 | NM_177948.2 | ARMCX3 | 45.7507953 | -6.7301786 | 1.67549459 |
| 11239 | NM_031407.6 | #N/A | 956.655437 | -4.3525348 | 1.08358378 |
| 11240 | NM_001330523.1 | PYCR1 | 40.9856897 | -6.5701895 | 1.63572024 |
| 11241 | NM_001322030.1 | #N/A | 36.57432 | -6.4008065 | 1.59366535 |
| 11242 | NM_024496.3 | #N/A | 914.373378 | -3.7600772 | 0.93633573 |
| 11243 | NM_001322263.2 | #N/A | 43.5542131 | -6.6545775 | 1.65714742 |
| 11244 | NM_145309.5 | LRTOMT | 45.8653231 | -6.7300914 | 1.6761921 |
| 11245 | NM_016540.3 | #N/A | 32.8866401 | -6.2444485 | 1.55526492 |
| 11246 | NM_001098576.1 | TMBIM6 | 47.9494896 | -6.7915897 | 1.69168164 |
| 11247 | NM_001206641.2 | #N/A | 41.2138651 | -6.5751128 | 1.63785641 |
| 11248 | NM_001291106.1 | #N/A | 70.2664492 | -7.3472405 | 1.83021134 |
| 11249 | NM_198205.1 | MLX | 45.3808367 | -6.7154135 | 1.67285339 |
| 11250 | NM_001302378.1 | #N/A | 191.004466 | -2.5655216 | 0.6391506 |
| 11251 | NM_003468.3 | #N/A | 38.4194115 | -6.4769392 | 1.61366149 |
| 11252 | NM_138736.2 | #N/A | 321.004316 | -5.3250663 | 1.32671528 |
| 11253 | NR_036614.1 | #N/A | 63.5662368 | -7.2031431 | 1.79486634 |
| 11254 | NM_133177.3 | #N/A | 68.744346 | -7.3138879 | 1.82252409 |
| 11255 | NM_004390.4 | #N/A | 35.3356965 | -6.3607154 | 1.58502492 |
| 11256 | NM_138559.1 | #N/A | 89.365637 | -7.6913215 | 1.91678902 |
| 11257 | NM_001243962.1 | HLA-DQB1 | 63.1784033 | -7.1925215 | 1.79251539 |
| 11258 | NM_152385.2 | #N/A | 42.1339957 | -6.6034783 | 1.64571955 |
| 11259 | NR_024127.1 | #N/A | 40.3876818 | -6.5446625 | 1.63112211 |
| 11260 | NM_032667.6 | BSCL2 | 99.8029717 | -3.4003182 | 0.84748703 |
| 11261 | NM_015186.3 | VPS13A | 49.4965361 | -6.8349207 | 1.70388435 |
| 11262 | NM_020458.3 | #N/A | 45.3514927 | -6.7193991 | 1.67522111 |
| 11263 | NM_016144.3 | #N/A | 40.6629417 | -6.5547226 | 1.63422474 |
| 11264 | NM_022485.4 | #N/A | 33.6820772 | -6.289623 | 1.56822664 |
| 11265 | NM_173642.3 | #N/A | 174.655896 | -3.8763101 | 0.96650481 |

| | | | | | |
|---|---|---|---|---|---|
| 11266 | NM_001297419.1 | RAD52 | 36.7437844 | -6.4047866 | 1.5969542 |
| 11267 | NR_049762.1 | #N/A | 56.1811565 | -7.0173556 | 1.74977332 |
| 11268 | NM_001042399.1 | CEP83 | 38.6671747 | -6.4856176 | 1.61725059 |
| 11269 | NM_002779.4 | #N/A | 87.9936286 | -7.6705896 | 1.9128269 |
| 11270 | NM_001271043.2 | NFIX | 84.2007088 | -7.6057898 | 1.89675686 |
| 11271 | NR_029411.1 | LOC100133091 | 32.6969723 | -6.2458203 | 1.55765792 |
| 11272 | NM_014238.1 | #N/A | 151.836522 | -3.1559416 | 0.78709126 |
| 11273 | NM_001333.3 | #N/A | 34.5554545 | -6.3258253 | 1.57769475 |
| 11274 | NM_001297658.1 | #N/A | 35.7506868 | -6.3678017 | 1.58872037 |
| 11275 | NM_001127230.1 | #N/A | 67.1913378 | -7.2808814 | 1.81656224 |
| 11276 | NM_001305115.1 | DUSP26 | 66.8904673 | -7.271652 | 1.81428588 |
| 11277 | NM_152282.4 | PXYLP1 | 34.8563973 | -6.3364583 | 1.5810622 |
| 11278 | NM_001330612.1 | #N/A | 46.6331268 | -6.7483953 | 1.68390346 |
| 11279 | NM_032673.2 | #N/A | 37.4290161 | -6.4346777 | 1.60570343 |
| 11280 | NR_136642.1 | CT75 | 25.8899507 | 3.32844032 | 0.83067031 |
| 11281 | NM_001303253.1 | ACAT2 | 58.7228975 | -7.0860263 | 1.76846482 |
| 11282 | NM_001017390.2 | SULT1A4 | 33.6731421 | -6.2813 | 1.56771147 |
| 11283 | NM_013284.3 | #N/A | 33.2498364 | -6.2724755 | 1.56551258 |
| 11284 | NR_120613.1 | NUTM2B-AS1 | 44.102369 | -6.6743919 | 1.66600881 |
| 11285 | NM_004772.2 | #N/A | 1822.99456 | -5.0616096 | 1.26349514 |
| 11286 | NM_004299.4 | #N/A | 35.4006457 | -6.360298 | 1.58770946 |
| 11287 | NM_012385.2 | #N/A | 42.9682743 | -6.6383117 | 1.6573246 |
| 11288 | NM_018043.5 | #N/A | 46.0916177 | -6.7401201 | 1.68278626 |
| 11289 | NM_018475.4 | #N/A | 161.104286 | -3.6186118 | 0.90357883 |
| 11291 | NM_001144000.1 | AGAP5 | 31.5656335 | -6.1902474 | 1.54588744 |
| 11290 | NM_000425.4 | #N/A | 57.2379472 | -7.0527337 | 1.76127387 |
| 11292 | NM_004292.2 | #N/A | 36.8937226 | -6.4179231 | 1.6027857 |
| 11293 | NM_001003652.3 | #N/A | 78.9604395 | -7.5125415 | 1.8767585 |
| 11294 | NM_001300907.1 | KMT5B | 41.5263865 | -6.5821352 | 1.64440335 |
| 11295 | NM_006020.2 | #N/A | 36.6702691 | -6.4066389 | 1.60058312 |
| 11296 | NR_073526.1 | #N/A | 71.1781779 | -7.3610044 | 1.83924712 |
| 11297 | NM_001145374.1 | #N/A | 48.5504671 | -6.8131315 | 1.70242964 |
| 11298 | NR_103819.1 | SNRPD3 | 42.2418079 | 6.67001563 | 1.66673365 |
| 11299 | NM_018492.3 | #N/A | 42.7112758 | -6.6233206 | 1.65514286 |
| 11300 | NM_001083909.2 | ADGRA1 | 34.9141982 | -6.3378237 | 1.58389501 |
| 11301 | NM_152511.4 | #N/A | 34.7080024 | -6.3195702 | 1.5793971 |
| 11302 | NM_000309.3 | #N/A | 33.5501027 | -6.2854193 | 1.57089587 |
| 11303 | NM_001317786.1 | #N/A | 32.4115259 | -6.2269358 | 1.55654835 |
| 11304 | NM_000904.4 | #N/A | 45.5153633 | -6.7176555 | 1.67951665 |
| 11305 | NM_001142677.1 | #N/A | 47.0324913 | -6.7671345 | 1.69192141 |
| 11306 | NM_001320616.1 | #N/A | 52.446722 | -6.922244 | 1.730823 |
| 11307 | NM_001042734.3 | SEC24B | 36.9282615 | -6.4232105 | 1.60615925 |
| 11308 | NM_001174164.1 | PRRT4 | 50.7265687 | -6.8742752 | 1.71900811 |

| | | | | | | |
|---|---|---|---|---|---|---|
| 11309 | NR_073520.1 | | #N/A | 31.5229598 | -6.1903427 | 1.54807534 |
| 11310 | NM_015719.3 | | #N/A | 71.7703381 | -7.3778122 | 1.84506145 |
| 11311 | NM_024540.3 | | #N/A | 42.8288824 | -6.6292274 | 1.6581076 |
| 11312 | NM_005833.3 | | #N/A | 38.5478861 | -6.4778929 | 1.62027979 |
| 11313 | NM_016429.3 | | #N/A | 32.9516424 | -6.2586126 | 1.56544444 |
| 11314 | NM_001142296.1 | | #N/A | 57.2712297 | -7.0457842 | 1.76242255 |
| 11315 | NM_018048.4 | | #N/A | 50.8045218 | -6.8745734 | 1.71965724 |
| 11316 | NR_038854.1 | NRAV | | 30.6038376 | -6.149156 | 1.53840698 |
| 11317 | NM_001172418.1 | BTBD9 | | 39.0540492 | -6.5003716 | 1.62629744 |
| 11318 | NM_001278230.1 | | #N/A | 35.0076423 | -6.3380185 | 1.5857207 |
| 11319 | NM_005745.7 | BCAP31 | | 37.1026144 | -6.4190376 | 1.60611072 |
| 11320 | NM_002452.3 | NUDT1 | | 54.8096211 | -6.9835194 | 1.74746758 |
| 11321 | NM_138972.3 | BACE1 | | 87.1084151 | -7.6541952 | 1.91542393 |
| 11322 | NM_022351.4 | | #N/A | 46.0949459 | -6.7365348 | 1.68582041 |
| 11323 | NM_024055.4 | | #N/A | 31.3269052 | -6.1806324 | 1.54683301 |
| 11324 | NM_006281.3 | | #N/A | 31.8315546 | -6.2090359 | 1.55406768 |
| 11325 | NM_014901.4 | | #N/A | 467.290721 | -4.009455 | 1.00357767 |
| 11326 | NM_001012974.2 | LRRC73 | | 39.1408067 | -6.5058967 | 1.62857853 |
| 11327 | NM_001308028.1 | | #N/A | 58.7839329 | -7.0855656 | 1.77368616 |
| 11328 | NM_012333.4 | | #N/A | 30.823944 | -6.1584264 | 1.5416188 |
| 11329 | NM_001258420.1 | | #N/A | 40.5489973 | -6.5501886 | 1.63974633 |
| 11330 | NM_020116.4 | | #N/A | 43.8221104 | -6.6679996 | 1.66924222 |
| 11331 | NM_198471.2 | | #N/A | 31.4742386 | -6.1873602 | 1.54913968 |
| 11332 | NM_001318134.1 | | #N/A | 43.0116447 | -6.6402725 | 1.66255423 |
| 11333 | NM_001134388.1 | ZDHHC4 | | 36.6254179 | -6.4006214 | 1.60268114 |
| 11334 | NM_015365.2 | | #N/A | 39.2788508 | -6.5038233 | 1.62855623 |
| 11335 | NM_003948.4 | | #N/A | 39.1782844 | -6.4953551 | 1.62644641 |
| 11336 | NM_019110.4 | | #N/A | 47.5564064 | -5.3754273 | 1.34609942 |
| 11337 | NM_144566.2 | | #N/A | 35.9751253 | -6.375606 | 1.59658833 |
| 11338 | NM_001144058.1 | NTM | | 42.3056936 | -6.6194101 | 1.65788189 |
| 11339 | NM_001324150.1 | ZNF41 | | 35.7975505 | -6.3693281 | 1.5952612 |
| 11340 | NM_007034.4 | | #N/A | 31.8269002 | -6.2044721 | 1.55397957 |
| 11341 | NM_020350.4 | | #N/A | 42.1335905 | -6.6125479 | 1.65621285 |
| 11342 | NM_173630.3 | | #N/A | 46.5735142 | -6.7465039 | 1.68977824 |
| 11343 | NM_001278789.1 | | #N/A | 37.7507568 | -6.4527367 | 1.616388 |
| 11344 | NM_016305.3 | SS18L2 | | 52.9878859 | -6.9362594 | 1.73756083 |
| 11345 | NM_002208.4 | | #N/A | 46.0683711 | -6.7365534 | 1.6876468 |
| 11346 | NM_015557.2 | | #N/A | 205.213923 | -5.2250209 | 1.3092252 |
| 11347 | NM_032947.4 | SMIM3 | | 31.1314751 | -6.1752016 | 1.5473318 |
| 11348 | NM_017703.2 | | #N/A | 47.1421691 | -6.7666116 | 1.69564054 |
| 11349 | NM_001012754.3 | | #N/A | 41.0644341 | -6.5752285 | 1.6477312 |
| 11350 | NM_001282533.1 | | #N/A | 317.972141 | -4.0879803 | 1.024661 |
| 11351 | NM_001012643.3 | | #N/A | 42.011925 | -6.600457 | 1.65446504 |

| | | | | | |
|---|---|---|---|---|---|
| 11352 | NM_005787.5 | #N/A | 46.5886097 | -6.7553549 | 1.69330408 |
| 11353 | NR_027318.2 | LINC00906 | 14.5574975 | 6.02762571 | 1.51095849 |
| 11354 | NM_152850.3 | PIGO | 33.392551 | -6.2713929 | 1.57209283 |
| 11355 | NM_001282671.1 | FAAP20 | 36.9247409 | -6.4159683 | 1.60848773 |
| 11356 | NM_024834.3 | MCMBP | 82.998213 | -7.5847056 | 1.90148454 |
| 11357 | NM_173828.4 | #N/A | 49.0584188 | -6.8252053 | 1.71119697 |
| 11358 | NM_018157.3 | #N/A | 67.9259937 | -5.8253918 | 1.46063928 |
| 11359 | NM_001198868.1 | #N/A | 56.9538904 | -7.0452605 | 1.76652503 |
| 11361 | NM_017711.3 | GDPD2 | 39.4830564 | -6.5146909 | 1.63377112 |
| 11360 | NM_001033678.3 | #N/A | 32.625961 | -6.2338183 | 1.56333618 |
| 11362 | NM_138477.2 | #N/A | 128.912256 | -3.1664929 | 0.79414887 |
| 11363 | NM_152998.2 | #N/A | 41.7218263 | -6.5906766 | 1.65294094 |
| 11364 | NM_017518.7 | HAUS7 | 41.2261577 | -6.5751752 | 1.64927373 |
| 11365 | NR_037183.2 | SPANXA2-OT1 | 24.8923604 | 4.25396601 | 1.06705101 |
| 11366 | NM_001276312.1 | #N/A | 45.7924567 | -6.7300598 | 1.68821426 |
| 11367 | NM_133462.3 | #N/A | 41.9609818 | -6.600733 | 1.65582918 |
| 11368 | NM_001013436.2 | #N/A | 46.5236249 | -6.7486075 | 1.69298179 |
| 11369 | NM_004388.2 | #N/A | 37.824927 | -6.4565145 | 1.61987133 |
| 11370 | NM_001033554.2 | #N/A | 97.6540914 | 1.61240237 | 0.40453881 |
| 11371 | NM_021967.2 | SERF1A | 48.1016726 | -6.8010165 | 1.70638339 |
| 11372 | NM_001207055.1 | #N/A | 32.8940912 | -6.2450075 | 1.56713814 |
| 11373 | NM_016466.5 | #N/A | 42.1958832 | -6.6081445 | 1.65834603 |
| 11374 | NM_178040.3 | #N/A | 74.7431221 | -7.433498 | 1.86562748 |
| 11375 | NM_001099858.1 | C7orf25 | 34.7590549 | -6.3332837 | 1.5895886 |
| 11376 | NM_001289745.1 | #N/A | 30.4357239 | -6.1340605 | 1.53961042 |
| 11377 | NM_001265589.1 | #N/A | 38.2292161 | -6.4600112 | 1.62175233 |
| 11378 | NM_001161376.1 | #N/A | 31.1752972 | -6.171253 | 1.54934991 |
| 11380 | NR_027504.1 | MST1P2 | 32.8905047 | -6.2514331 | 1.56955683 |
| 11379 | NM_024313.2 | #N/A | 89.5281781 | -2.7521377 | 0.69098122 |
| 11381 | NM_001902.5 | #N/A | 29.6802314 | -6.101478 | 1.53193931 |
| 11382 | NR_073562.1 | RABGGTB | 30.4007344 | -6.1386645 | 1.54134295 |
| 11383 | NM_001300962.1 | #N/A | 70.2205985 | -7.3465622 | 1.84465044 |
| 11384 | NM_003141.3 | #N/A | 31.4747034 | -6.1910943 | 1.55469558 |
| 11385 | NM_001128167.2 | PAK3 | 95.045944 | -7.7783032 | 1.95343045 |
| 11387 | NM_201429.2 | RTN3 | 91.5564425 | -7.7255915 | 1.94049052 |
| 11386 | NM_004766.2 | #N/A | 589.020268 | -5.055481 | 1.26982198 |
| 11388 | NM_175039.3 | #N/A | 36.5677451 | -6.4016665 | 1.60821296 |
| 11389 | NM_004332.3 | #N/A | 43.8306183 | -6.6641809 | 1.67440291 |
| 11390 | NR_133665.1 | #N/A | 8.7409393 | 5.60092422 | 1.40726925 |
| 11391 | NM_001301267.1 | #N/A | 52.3558661 | -6.9216666 | 1.73914757 |
| 11392 | NM_052858.5 | #N/A | 59.0214827 | 2.81638285 | 0.70771098 |
| 11394 | NR_034075.1 | THAP9-AS1 | 39.2762787 | -6.4980652 | 1.63291489 |
| 11393 | NM_001286734.1 | #N/A | 59.3142053 | -5.650864 | 1.42002372 |

| | | | | | |
|---|---|---|---|---|---|
| 11395 | NM_000044.3 | #N/A | 42.8508733 | -6.6319043 | 1.66662863 |
| 11396 | NR_110864.1 | #N/A | 42.5996612 | -6.6284233 | 1.66583123 |
| 11397 | NM_001032387.1 | #N/A | 39.6836867 | -6.5243066 | 1.63967467 |
| 11398 | NR_052010.1 | IL11RA | 39.7299237 | -6.5272348 | 1.6404794 |
| 11399 | NM_007086.3 | #N/A | 46.4362146 | -6.7454468 | 1.6953891 |
| 11400 | NM_001042432.1 | CLN3 | 35.6102593 | -6.3656329 | 1.60015309 |
| 11401 | NM_138285.4 | #N/A | 36.0390957 | -6.379715 | 1.60377909 |
| 11402 | NM_001135058.1 | COCH | 48.5117655 | -6.8074456 | 1.71138342 |
| 11403 | NM_005638.5 | #N/A | 52.0129449 | -6.9138885 | 1.73819356 |
| 11404 | NM_013264.4 | #N/A | 45.8910322 | -6.7290492 | 1.69185216 |
| 11405 | NM_001287059.1 | #N/A | 74.9474879 | -6.166636 | 1.55047859 |
| 11406 | NM_021723.4 | #N/A | 42.1440214 | -6.6016239 | 1.65987127 |
| 11407 | NM_001267062.1 | PLEKHB2 | 54.5554574 | -6.9799849 | 1.75524648 |
| 11408 | NM_019590.4 | #N/A | 61.2356819 | -7.1488051 | 1.79772016 |
| 11409 | NM_001003945.2 | ALAD | 42.21858 | -6.6142408 | 1.66352529 |
| 11410 | NM_001100912.1 | BEND7 | 37.1994537 | -6.4287614 | 1.61694615 |
| 11411 | NM_014767.2 | SPOCK2 | 449.210064 | -3.5883776 | 0.9026035 |
| 11412 | NM_001304463.1 | #N/A | 49.6904828 | -6.8436635 | 1.72145363 |
| 11413 | NM_058230.2 | #N/A | 35.1505634 | -6.3415218 | 1.59520751 |
| 11414 | NM_001042552.2 | #N/A | 50.6029553 | -6.8666855 | 1.72754393 |
| 11415 | NM_001282750.1 | #N/A | 46.8660432 | -6.7559663 | 1.69971659 |
| 11416 | NM_001310121.1 | ATXN2 | 31.9742571 | -6.2025702 | 1.56055876 |
| 11419 | NR_024374.1 | LOC642846 | 36.5496952 | -6.4024376 | 1.61088285 |
| 11417 | NM_001040058.1 | #N/A | 46.0732866 | -6.7347445 | 1.69449754 |
| 11418 | NM_014297.4 | #N/A | 44.1713214 | -6.6734563 | 1.67906915 |
| 11420 | NM_004217.3 | #N/A | 39.9217815 | -6.5277952 | 1.64251264 |
| 11421 | NM_024740.2 | ALG9 | 52.3597499 | -6.9155969 | 1.74016039 |
| 11422 | NM_145330.2 | MRPL33 | 44.4462556 | -6.6849143 | 1.6822097 |
| 11423 | NM_021089.2 | #N/A | 37.3777872 | -6.429085 | 1.61790155 |
| 11424 | NM_007137.3 | ZNF81 | 44.4699009 | -6.68544 | 1.68255506 |
| 11425 | NM_001002913.2 | #N/A | 31.8749647 | -6.2104605 | 1.56319918 |
| 11426 | NM_002387.2 | #N/A | 70.1809318 | -2.9393033 | 0.73984206 |
| 11427 | NM_144966.5 | FREM1 | 30.6101083 | -6.1507702 | 1.54821686 |
| 11428 | NM_031218.3 | #N/A | 38.4976749 | -6.4746801 | 1.62983031 |
| 11429 | NM_014174.2 | #N/A | 38.0664543 | -6.4594103 | 1.62606175 |
| 11430 | NM_001258022.1 | GRIA1 | 76.0832801 | -7.4600127 | 1.87805097 |
| 11431 | NM_001018000.3 | KAZN | 41.9824554 | -6.5996553 | 1.66172811 |
| 11432 | NM_003675.3 | #N/A | 63.6511587 | -5.7370239 | 1.44463946 |
| 11433 | NM_001113528.1 | #N/A | 51.0019768 | -6.8871864 | 1.73429378 |
| 11434 | NM_001099693.1 | RPL31 | 35.6662433 | -6.3625611 | 1.6021992 |
| 11435 | NM_001318089.1 | SWI5 | 39.8673713 | -6.5249164 | 1.64309527 |
| 11436 | NM_001006109.1 | HMCES | 41.5065429 | -6.5886412 | 1.65923411 |
| 11437 | NM_001322033.1 | #N/A | 271.710011 | -4.7474066 | 1.19557916 |

| | | | | | |
|---|---|---|---|---|---|
| 11438 | NM_133468.4 | #N/A | 30.8243496 | -6.1625932 | 1.55201311 |
| 11439 | NM_005269.2 | #N/A | 33.2160875 | -6.2694454 | 1.57901899 |
| 11440 | NM_012083.2 | #N/A | 42.9981249 | -6.6354729 | 1.6713646 |
| 11442 | NR_034125.1 | C10orf143 | 74.0064289 | 3.00072354 | 0.75585883 |
| 11441 | NR_034053.2 | TNPO3 | 44.0819223 | -6.6704348 | 1.68023025 |
| 11443 | NM_002643.3 | #N/A | 39.6400614 | -6.5165371 | 1.64168933 |
| 11444 | NM_001270875.1 | #N/A | 81.1495377 | -7.5551909 | 1.90340587 |
| 11445 | NM_002889.3 | #N/A | 86.496496 | -7.6466652 | 1.92664944 |
| 11446 | NM_001017922.1 | #N/A | 32.743525 | -6.2488888 | 1.57449443 |
| 11447 | NR_134867.1 | #N/A | 50.9891181 | -6.8847328 | 1.73471919 |
| 11448 | NM_020702.4 | #N/A | 271.603485 | 1.41839118 | 0.35741369 |
| 11449 | NM_145283.2 | #N/A | 48.9322474 | 3.57671048 | 0.90146979 |
| 11450 | NM_001136494.1 | #N/A | 67.400418 | -7.2870566 | 1.83665321 |
| 11451 | NM_001193350.1 | #N/A | 46.6745684 | -6.7566154 | 1.70304381 |
| 11452 | NM_134324.2 | #N/A | 36.2671705 | -6.3899903 | 1.61068033 |
| 11453 | NM_024316.2 | #N/A | 40.1740601 | -6.5385737 | 1.64821548 |
| 11454 | NM_001320542.1 | #N/A | 39.4904638 | -6.5187331 | 1.64324692 |
| 11455 | NM_207366.2 | #N/A | 68.2667679 | 2.28095914 | 0.57498351 |
| 11456 | NM_014279.4 | #N/A | 62.1171087 | -7.1636265 | 1.80632533 |
| 11457 | NM_001282551.1 | BTBD3 | 46.050234 | -6.7368331 | 1.69881869 |
| 11458 | NM_001348013.1 | ZNF136 | 31.262977 | 9.93368104 | 2.50500008 |
| 11459 | NM_015237.3 | #N/A | 41.1921668 | -6.5725037 | 1.65740238 |
| 11460 | NR_145460.1 | #N/A | 14.034287 | 4.10223993 | 1.03454491 |
| 11461 | NM_012327.5 | #N/A | 45.3153176 | -6.7139901 | 1.69327837 |
| 11462 | NM_001993.4 | #N/A | 86.0368887 | -7.6392104 | 1.92668575 |
| 11463 | NM_015910.5 | #N/A | 29.6606187 | -6.1045944 | 1.5397873 |
| 11464 | NM_001144759.2 | #N/A | 409.424511 | -5.0017732 | 1.26166724 |
| 11465 | NM_012338.3 | #N/A | 40.044865 | -6.5352901 | 1.64882769 |
| 11466 | NM_005491.4 | MAMLD1 | 32.897329 | -6.2423766 | 1.57510074 |
| 11467 | NM_001271895.1 | #N/A | 39.2672345 | -6.5096675 | 1.64258973 |
| 11468 | NM_001085400.1 | #N/A | 69.616176 | -2.5917864 | 0.65400692 |
| 11469 | NM_005114.2 | #N/A | 37.0324371 | -6.4197216 | 1.62008051 |
| 11470 | NM_001304536.1 | EPHA4 | 66.5618574 | -7.2695142 | 1.83459254 |
| 11471 | NM_001146314.1 | ABHD14B | 30.481547 | -6.137095 | 1.54912743 |
| 11472 | NM_207327.4 | #N/A | 35.2545827 | -6.3464883 | 1.60216231 |
| 11473 | NM_001085347.2 | #N/A | 34.1745619 | -6.3055149 | 1.59184666 |
| 11474 | NM_183235.2 | #N/A | 40.7989369 | -6.5628011 | 1.656807 |
| 11476 | NM_001135599.3 | TGFB2 | 49.4666273 | -6.8416769 | 1.72725079 |
| 11475 | NM_001006622.2 | WDR33 | 39.2898658 | -6.5063782 | 1.64260629 |
| 11477 | NM_001172896.1 | #N/A | 50.9065486 | -6.8821917 | 1.73748198 |
| 11478 | NM_147173.2 | #N/A | 43.6864973 | -6.6600944 | 1.68141736 |
| 11479 | NM_005840.2 | SPRY3 | 37.1481129 | -6.4240132 | 1.62195772 |
| 11480 | NM_001160243.1 | RPAIN | 34.0398924 | -6.3057991 | 1.59243138 |

| | | | | | |
|---|---|---|---|---|---|
| 11481 | NM_001142475.1 | #N/A | 64.166757 | -7.2155636 | 1.82221548 |
| 11482 | NM_138415.4 | #N/A | 65.4325967 | -7.2396201 | 1.82835041 |
| 11483 | NM_001303099.1 | #N/A | 39.5859358 | -6.5170352 | 1.64589848 |
| 11484 | NR_135227.1 | PCNP | 30.5934655 | -6.14291 | 1.5514269 |
| 11485 | NM_001164811.1 | #N/A | 37.7174836 | -6.4519206 | 1.62952252 |
| 11486 | NM_031448.4 | #N/A | 45.1920486 | -6.7024039 | 1.69312356 |
| 11487 | NM_001322224.1 | #N/A | 40.0970033 | -6.5319155 | 1.65008476 |
| 11488 | NR_030737.1 | TTC3P1 | 36.4169804 | -6.3955476 | 1.61570191 |
| 11489 | NM_005854.2 | #N/A | 31.2709786 | -6.17394 | 1.55982387 |
| 11490 | NM_021229.3 | #N/A | 37.9544529 | -6.4593771 | 1.63196164 |
| 11492 | NM_080796.3 | DIDO1 | 77.3796637 | -4.8288024 | 1.22034842 |
| 11491 | NM_001270805.1 | SYT6 | 49.5735684 | -6.8415576 | 1.72902499 |
| 11493 | NM_018401.2 | #N/A | 41.1486691 | -6.5744548 | 1.66153725 |
| 11494 | NM_001195226.1 | #N/A | 74.3086983 | -7.4277178 | 1.87734362 |
| 11495 | NR_002570.3 | #N/A | 37.9968366 | -6.454576 | 1.63145795 |
| 11496 | NM_001282438.1 | BLOC1S2 | 71.9631736 | 5.28279779 | 1.3353116 |
| 11497 | NM_001012662.2 | #N/A | 37.9671435 | -6.4593426 | 1.63292956 |
| 11498 | NM_006176.2 | #N/A | 42.4940766 | -6.6195287 | 1.67343455 |
| 11499 | NM_054033.3 | #N/A | 50.6685486 | -6.8736332 | 1.73778367 |
| 11500 | NM_001001890.2 | #N/A | 41.2200331 | -6.5787591 | 1.66346366 |
| 11501 | NM_178580.2 | HM13 | 50.1373781 | -6.855046 | 1.73336094 |
| 11502 | NR_132964.1 | SNORA109 | 892.95526 | -4.85529 | 1.22775464 |
| 11503 | NM_001195215.1 | #N/A | 41.1340451 | -6.5760727 | 1.66298702 |
| 11504 | NM_018056.3 | #N/A | 44.971493 | -6.7021079 | 1.69500374 |
| 11505 | NM_022341.1 | #N/A | 33.5684784 | -6.2757918 | 1.58720502 |
| 11506 | NM_001114121.2 | CHEK1 | 31.0237184 | -6.167152 | 1.55973822 |
| 11507 | NM_015221.3 | #N/A | 35.7296829 | -6.3761887 | 1.61263735 |
| 11508 | NM_022845.2 | #N/A | 74.1054928 | -5.1095765 | 1.29249859 |
| 11509 | NM_006822.2 | #N/A | 67.6199597 | -3.159768 | 0.79932454 |
| 11510 | NM_016126.3 | HSPB11 | 31.4106916 | -6.1852903 | 1.56487646 |
| 11511 | NM_001177651.1 | #N/A | 40.2331814 | -6.5361741 | 1.65375596 |
| 11512 | NM_001243756.1 | PXN | 34.7080744 | -6.3339466 | 1.60267818 |
| 11513 | NM_005962.4 | #N/A | 35.6751301 | -6.373878 | 1.61285227 |
| 11514 | NM_001278170.1 | #N/A | 21.5883395 | 9.42691206 | 2.38565604 |
| 11515 | NM_001039591.2 | #N/A | 579.210893 | -3.7642079 | 0.95264681 |
| 11516 | NM_001099673.2 | #N/A | 39.6672691 | -6.521325 | 1.65066151 |
| 11517 | NM_005086.4 | SSPN | 30.4207178 | -6.1417804 | 1.55481517 |
| 11518 | NM_003408.2 | #N/A | 40.44862 | -6.5416133 | 1.65614827 |
| 11519 | NM_201999.2 | #N/A | 50.5511751 | -6.8680219 | 1.73878445 |
| 11520 | NM_130808.2 | CPNE4 | 52.0258943 | -6.9124666 | 1.75016835 |
| 11521 | NM_014353.4 | #N/A | 38.9087953 | -6.4858207 | 1.6421969 |
| 11522 | NR_026846.1 | LOC285074 | 58.1484338 | -5.6202718 | 1.42317696 |
| 11523 | NM_001142315.1 | #N/A | 33.5019003 | -6.2716511 | 1.58823648 |

| | | | | | |
|---|---|---|---|---|---|
| 11524 | NM_016053.3 | #N/A | 52.2538775 | -6.9158371 | 1.75152859 |
| 11525 | NM_019114.4 | EPB41L4B | 35.1296129 | -6.3488932 | 1.6079535 |
| 11526 | NM_020445.5 | #N/A | 86.2452632 | -7.6392193 | 1.93477454 |
| 11527 | NM_022124.5 | #N/A | 74.8230429 | -7.4375843 | 1.88374276 |
| 11528 | NR_038190.1 | #N/A | 32.710578 | -6.2363941 | 1.57955915 |
| 11529 | NM_014214.2 | #N/A | 30.129415 | -6.1264418 | 1.55178531 |
| 11530 | NM_001252092.1 | #N/A | 39.2308715 | -6.5076257 | 1.64835947 |
| 11531 | NM_001491.2 | GCNT2 | 43.8942052 | -6.6652113 | 1.6883653 |
| 11532 | NM_001184876.2 | #N/A | 73.475114 | -7.4067115 | 1.87622153 |
| 11533 | NM_003826.2 | #N/A | 167.255851 | -4.9682808 | 1.25860441 |
| 11534 | NM_013397.5 | #N/A | 33.4723769 | -6.2751103 | 1.58973595 |
| 11535 | NM_201551.2 | NELL1 | 69.4804078 | -7.3301986 | 1.85712425 |
| 11536 | NM_001320627.1 | BOLA2-SMG1P6 | 42.4064381 | -6.6206966 | 1.67748827 |
| 11537 | NM_001242859.2 | PRKACB | 113.350277 | -8.0354496 | 2.0359871 |
| 11538 | NM_001143975.1 | UBTFL1 | 20.3574181 | 7.81453418 | 1.98003691 |
| 11539 | NR_135104.1 | #N/A | 36.168504 | -6.379009 | 1.61638598 |
| 11540 | NM_001284281.1 | PPP4R3A | 45.6834551 | -5.440103 | 1.37857772 |
| 11541 | NM_012383.4 | #N/A | 42.1854463 | -6.6079811 | 1.67452406 |
| 11542 | NM_001291931.1 | #N/A | 65.1516127 | -4.1749715 | 1.05801446 |
| 11543 | NM_145870.2 | #N/A | 36.5646609 | -6.4019917 | 1.62242881 |
| 11544 | NM_001136575.1 | LANCL1 | 121.186627 | -6.5400847 | 1.65754835 |
| 11545 | NM_172177.3 | #N/A | 30.3465401 | -6.1364163 | 1.55540304 |
| 11546 | NM_001320848.1 | FAHD2B | 33.2411818 | -6.2690427 | 1.58924922 |
| 11547 | NM_020423.6 | #N/A | 35.2201333 | -6.3403044 | 1.60737341 |
| 11548 | NM_006811.3 | #N/A | 462.479515 | -2.1930552 | 0.55598648 |
| 11549 | NM_182746.2 | #N/A | 318.100122 | -2.5628548 | 0.64978135 |
| 11550 | NM_001083112.2 | #N/A | 48.042848 | -6.7922429 | 1.72224886 |
| 11551 | NM_006951.4 | #N/A | 40.1182711 | -6.535145 | 1.65707442 |
| 11552 | NM_020744.3 | MTA3 | 32.4071879 | -6.2333338 | 1.5805648 |
| 11553 | NM_138635.3 | H2AZ2 | 452.048534 | -5.1331569 | 1.30173619 |
| 11554 | NM_173666.3 | #N/A | 39.2095963 | -6.4977457 | 1.64778121 |
| 11555 | NR_134855.1 | #N/A | 34.4486034 | -6.3173586 | 1.60207432 |
| 11556 | NM_012140.4 | #N/A | 38.8040193 | -6.4948562 | 1.64709739 |
| 11557 | NM_180982.2 | #N/A | 31.1701722 | -6.1695208 | 1.56462675 |
| 11558 | NM_017879.1 | #N/A | 32.4067902 | -6.2260893 | 1.57928247 |
| 11559 | NM_015077.3 | #N/A | 346.50213 | -3.9477574 | 1.00139476 |
| 11560 | NM_032949.2 | #N/A | 41.7317827 | -6.5905417 | 1.67182559 |
| 11561 | NM_014562.3 | #N/A | 32.4341285 | -6.2275667 | 1.57989466 |
| 11562 | NM_020993.4 | #N/A | 79.721447 | -7.5240478 | 1.90887116 |
| 11563 | NM_001330983.1 | LIMCH1 | 108.0025 | -7.9661801 | 2.02117731 |
| 11564 | NM_001171740.2 | #N/A | 34.0507683 | -6.2930782 | 1.5967682 |
| 11565 | NM_001319664.1 | TMEM263 | 32.621057 | -6.2310277 | 1.58121313 |
| 11566 | NM_002498.2 | #N/A | 34.6137669 | -6.3223611 | 1.60440179 |

| | | | | | |
|---|---|---|---|---|---|
| 11567 | NM_007032.5 | TRIOBP | 188.540171 | -1.8364482 | 0.466042 |
| 11569 | NM_182546.3 | VSTM2A | 46.5880106 | -6.7556328 | 1.71441318 |
| 11568 | NM_153322.2 | #N/A | 51.3714637 | -6.8964553 | 1.75014929 |
| 11570 | NM_001515.3 | GTF2H2 | 42.5553768 | -6.6178861 | 1.67955748 |
| 11571 | NM_001331030.1 | #N/A | 24.6071692 | 9.58514888 | 2.43273872 |
| 11572 | NR_033797.2 | MIR4453HG | 39.7226892 | -6.5200863 | 1.65485458 |
| 11573 | NM_152750.4 | #N/A | 41.8900668 | -6.5913883 | 1.67297943 |
| 11574 | NR_134456.1 | LINC02671 | 40.2150079 | 2.7769245 | 0.70487796 |
| 11575 | NM_020160.2 | #N/A | 46.6627618 | -6.7531705 | 1.71420614 |
| 11576 | NM_201265.1 | UBL7 | 47.8147983 | -6.7894578 | 1.72352833 |
| 11577 | NM_001321072.1 | CBS | 64.4780484 | -5.7501656 | 1.45977575 |
| 11578 | NM_000933.3 | PLCB4 | 38.6649123 | -6.4898071 | 1.64757999 |
| 11579 | NM_001130035.1 | #N/A | 30.8752047 | -6.1607165 | 1.56406425 |
| 11580 | NM_032811.2 | #N/A | 45.8341334 | -6.7283159 | 1.70816181 |
| 11581 | NM_183043.2 | RNF6 | 43.9136403 | -6.6603234 | 1.69094341 |
| 11582 | NM_020186.2 | #N/A | 35.4393856 | -6.3597321 | 1.61470128 |
| 11583 | NM_033148.3 | DTNB | 34.6749874 | -6.3265575 | 1.60648677 |
| 11584 | NM_032844.4 | #N/A | 36.7510449 | -6.4100571 | 1.62769934 |
| 11585 | NM_017450.2 | #N/A | 206.61943 | -2.9718355 | 0.75464798 |
| 11586 | NM_001330724.1 | #N/A | 34.9055843 | -6.3381711 | 1.60993129 |
| 11587 | NM_001012393.2 | #N/A | 126.77969 | -8.1955178 | 2.08173548 |
| 11588 | NM_016941.3 | #N/A | 46.1785953 | -6.7360194 | 1.71102061 |
| 11589 | NR_038270.1 | #N/A | 31.1627919 | -6.1752092 | 1.5685925 |
| 11590 | NM_033452.2 | #N/A | 29.4330087 | -6.0953634 | 1.54832075 |
| 11591 | NR_138095.1 | #N/A | 34.6792965 | -6.3236231 | 1.6064126 |
| 11592 | NM_001001550.2 | #N/A | 45.8784105 | -6.7321574 | 1.71027232 |
| 11593 | NM_002477.1 | MYL5 | 33.1984577 | -6.2605842 | 1.59052774 |
| 11594 | NM_153026.2 | #N/A | 66.8326796 | -7.2741076 | 1.84823961 |
| 11595 | NM_001134707.1 | #N/A | 30.1774911 | -6.1313322 | 1.5579351 |
| 11596 | NR_110533.1 | DGCR5 | 31.3492098 | -6.1735206 | 1.5686882 |
| 11597 | NM_001005914.2 | SEMA3B | 60.6031436 | -7.1352172 | 1.81326187 |
| 11598 | NM_001171161.1 | #N/A | 118.144807 | 2.5245661 | 0.64158379 |
| 11599 | NM_178558.4 | #N/A | 34.632803 | -6.3204198 | 1.60636108 |
| 11600 | NM_032231.5 | #N/A | 37.6275911 | -6.4471332 | 1.63866031 |
| 11601 | NM_001076678.2 | #N/A | 62.6279503 | -7.1761961 | 1.8240543 |
| 11602 | NM_173576.2 | #N/A | 46.6694271 | -6.7566119 | 1.71742659 |
| 11603 | NR_136649.1 | #N/A | 54.7590439 | -6.9877978 | 1.77618286 |
| 11604 | NM_001190849.1 | #N/A | 34.7505937 | -6.3279326 | 1.6085245 |
| 11605 | NM_001321708.1 | DGKI | 42.8890802 | -6.6295504 | 1.68529909 |
| 11606 | NM_033309.2 | #N/A | 38.8855153 | -6.4961076 | 1.65155717 |
| 11607 | NM_001202412.1 | ZNF559 | 38.9742025 | -6.4880627 | 1.64969493 |
| 11608 | NM_001284499.1 | SLC43A2 | 83.0026441 | 7.18354473 | 1.8265665 |
| 11609 | NM_152658.2 | #N/A | 32.9667951 | -6.2531202 | 1.59009179 |

| | | | | | |
|---|---|---|---|---|---|
| 11610 | NM_001307941.1 | #N/A | 46.1663962 | -6.7411137 | 1.71423527 |
| 11611 | NM_001105541.1 | #N/A | 50.2916004 | -6.8601113 | 1.74450763 |
| 11612 | NM_001033549.2 | BABAM1 | 46.0516289 | -6.728835 | 1.7111798 |
| 11613 | NR_038279.1 | LINC00665 | 38.6459063 | -6.4808197 | 1.648161 |
| 11614 | NR_036542.1 | #N/A | 38.9347856 | -6.4864366 | 1.64968462 |
| 11615 | NM_001145347.1 | #N/A | 33.4954445 | -6.2771302 | 1.59649577 |
| 11616 | NM_000817.2 | #N/A | 40.0758442 | -6.5395186 | 1.66325387 |
| 11617 | NM_003456.2 | #N/A | 35.2226556 | -6.3408789 | 1.61284384 |
| 11618 | NM_182665.3 | #N/A | 34.8595531 | -6.3374015 | 1.61196374 |
| 11619 | NM_001561.5 | #N/A | 107.327992 | 1.54150792 | 0.39212917 |
| 11620 | NM_001270468.1 | #N/A | 46.0111458 | -6.7321502 | 1.71256489 |
| 11621 | NM_020421.3 | #N/A | 32.9954226 | -6.2496208 | 1.58985258 |
| 11622 | NR_045680.1 | HEIH | 51.2063749 | -4.8243569 | 1.22731524 |
| 11623 | NM_016230.3 | #N/A | 42.0905139 | -6.6036031 | 1.67998234 |
| 11624 | NR_048569.1 | #N/A | 63.3168573 | -7.1928556 | 1.829881 |
| 11625 | NM_001107.4 | #N/A | 35.6773565 | -6.3664697 | 1.61970044 |
| 11626 | NM_003838.4 | FPGT | 29.8746398 | -6.1143191 | 1.55576379 |
| 11627 | NM_003594.3 | #N/A | 43.8273215 | -4.586512 | 1.1672561 |
| 11628 | NM_017787.4 | #N/A | 37.3545963 | -6.4391291 | 1.63882744 |
| 11629 | NM_019071.2 | #N/A | 37.899746 | -6.4506688 | 1.64182696 |
| 11630 | NM_207303.4 | ATRNL1 | 131.634478 | -5.3806476 | 1.36952464 |
| 11631 | NM_001318828.1 | CD200 | 48.6469884 | -6.8114486 | 1.73377767 |
| 11632 | NM_003999.2 | #N/A | 55.2975356 | -7.0008251 | 1.7820281 |
| 11633 | NM_005715.2 | #N/A | 91.3926083 | -4.9871463 | 1.26949391 |
| 11634 | NR_024330.1 | SPACA6 | 36.7901692 | -6.4171567 | 1.63359942 |
| 11635 | NM_001272012.1 | ITPRIP | 43.5785679 | -6.6505363 | 1.69318825 |
| 11636 | NM_018942.2 | #N/A | 57.9197975 | -7.0658964 | 1.79902487 |
| 11637 | NM_004891.3 | #N/A | 34.8069306 | -6.3282711 | 1.61136811 |
| 11638 | NR_132353.1 | BLOC1S6 | 36.5718811 | -6.4000979 | 1.62989672 |
| 11639 | NM_000943.4 | #N/A | 37.8505605 | -6.4589257 | 1.64490991 |
| 11640 | NM_004361.3 | #N/A | 54.3091721 | -6.9715476 | 1.77545698 |
| 11641 | NM_020371.2 | #N/A | 38.3535829 | -6.4736177 | 1.64865617 |
| 11642 | NM_130384.2 | #N/A | 34.5246976 | -6.3178723 | 1.60917299 |
| 11643 | NM_001323384.1 | #N/A | 49.4331866 | -6.8348705 | 1.74092479 |
| 11644 | NR_045587.1 | SRA1 | 52.6860593 | -6.9306614 | 1.76535883 |
| 11645 | NM_003950.3 | #N/A | 53.5823491 | 1.94989819 | 0.49673913 |
| 11646 | NM_015086.1 | #N/A | 100.400811 | -7.8588262 | 2.00215061 |
| 11647 | NM_017880.1 | #N/A | 32.9160212 | -6.2522211 | 1.59287469 |
| 11648 | NM_001170416.1 | #N/A | 29.6029251 | -6.100101 | 1.55418352 |
| 11649 | NM_133640.4 | #N/A | 57.3315891 | -7.0515615 | 1.7966309 |
| 11650 | NR_073090.1 | #N/A | 39.3999604 | -6.5044291 | 1.65722316 |
| 11651 | NM_203384.1 | #N/A | 54.5704744 | -6.9782763 | 1.77798259 |
| 11652 | NM_032449.2 | CC2D1B | 50.3897545 | -6.8655343 | 1.74933497 |

| | | | | | |
|---|---|---|---|---|---|
| 11653 | NR_125382.1 | LOC103611081 | 36.0347496 | -6.3825447 | 1.62642452 |
| 11654 | NM_005591.3 | MRE11 | 30.1904456 | -6.1324332 | 1.56280793 |
| 11655 | NM_001190991.1 | #N/A | 31.8840437 | -6.2043993 | 1.58131364 |
| 11656 | NM_006981.3 | #N/A | 70.2767673 | -7.3445837 | 1.87218979 |
| 11657 | NM_022658.3 | #N/A | 58.0406979 | -7.0704339 | 1.80233137 |
| 11658 | NM_153256.3 | #N/A | 29.3947901 | -6.0901549 | 1.55245889 |
| 11659 | NM_001122964.2 | #N/A | 59.6493731 | -7.1117975 | 1.81320263 |
| 11660 | NM_021824.3 | NIF3L1 | 32.5584457 | -6.2372371 | 1.59024354 |
| 11661 | NM_173618.2 | #N/A | 46.338827 | -6.7428003 | 1.71919523 |
| 11662 | NM_152683.3 | #N/A | 28.9053203 | -6.0588207 | 1.54490404 |
| 11664 | NR_003226.1 | AFG3L1P | 35.1614938 | -6.3534431 | 1.62020001 |
| 11663 | NM_021724.4 | #N/A | 37.9409506 | -6.4528849 | 1.64556342 |
| 11665 | NM_001256449.1 | #N/A | 37.4549153 | -6.4391848 | 1.64218993 |
| 11666 | NM_203411.1 | #N/A | 30.2705401 | -6.1376332 | 1.56529659 |
| 11667 | NM_001317924.1 | #N/A | 47.0532768 | -6.7643675 | 1.72532883 |
| 11668 | NM_001097641.1 | #N/A | 30.6887025 | 9.93009845 | 2.53285956 |
| 11669 | NM_080648.2 | #N/A | 60.3204474 | -7.1277165 | 1.81816211 |
| 11670 | NM_001136501.2 | #N/A | 30.8209616 | -6.1533718 | 1.56969402 |
| 11671 | NM_001083588.1 | #N/A | 29.3751264 | -6.0848905 | 1.55258831 |
| 11672 | NM_001323302.1 | #N/A | 53.4416316 | -6.9458239 | 1.77231449 |
| 11673 | NM_001204890.1 | ISY1-RAB43 | 60.6280777 | -7.1303913 | 1.81949354 |
| 11674 | NM_001307976.1 | #N/A | 31.9973442 | -6.213692 | 1.58559918 |
| 11675 | NM_207395.2 | #N/A | 34.3904979 | -6.313914 | 1.61130967 |
| 11676 | NM_001168399.1 | BEX2 | 41.2132575 | -6.5720382 | 1.67727545 |
| 11677 | NM_001348081.1 | #N/A | 43.7454224 | -6.6630658 | 1.70059024 |
| 11678 | NM_015627.2 | #N/A | 40.6220659 | -6.5599955 | 1.6744229 |
| 11679 | NM_015439.2 | CCDC28A | 31.4960835 | -6.1914786 | 1.58046951 |
| 11680 | NM_138353.3 | DCAF15 | 221.771957 | -3.5646083 | 0.90992901 |
| 11681 | NM_001317813.1 | #N/A | 175.452249 | 4.67348827 | 1.19312301 |
| 11682 | NM_001165972.1 | NRG3 | 53.1862464 | -6.9406463 | 1.77193644 |
| 11683 | NM_015398.2 | #N/A | 37.4573296 | -6.431375 | 1.64195691 |
| 11684 | NM_001321958.1 | #N/A | 38.6595549 | -6.4785667 | 1.65409087 |
| 11685 | NM_001481.2 | #N/A | 36.8931666 | -6.4129887 | 1.63758944 |
| 11686 | NM_001040454.1 | SLC26A6 | 33.8501336 | -6.2892122 | 1.606008 |
| 11687 | NM_003776.3 | MRPL40 | 47.5009729 | -6.7797587 | 1.73149863 |
| 11688 | NM_021939.3 | #N/A | 991.783271 | -3.8709095 | 0.98869687 |
| 11689 | NM_019018.2 | #N/A | 59.8046572 | -4.1281947 | 1.05445271 |
| 11690 | NM_001282778.1 | #N/A | 49.4977067 | -6.8363557 | 1.74627917 |
| 11691 | NM_016576.4 | #N/A | 30.6617009 | -6.1555242 | 1.57262673 |
| 11692 | NM_173872.3 | #N/A | 39.0840947 | -6.504031 | 1.66168644 |
| 11693 | NM_001271629.1 | #N/A | 29.8265814 | -6.1085983 | 1.5606768 |
| 11694 | NM_001168682.1 | #N/A | 28.1946057 | -6.0304242 | 1.5407814 |
| 11695 | NM_198705.2 | #N/A | 38.9188548 | -6.489166 | 1.65815303 |

| | | | | | |
|---|---|---|---|---|---|
| 11696 | NM_003728.3 | #N/A | 102.419722 | -5.1015038 | 1.30393245 |
| 11697 | NM_018105.2 | #N/A | 36.9333789 | -6.4168507 | 1.64015285 |
| 11698 | NM_015651.2 | #N/A | 42.0541722 | -6.6026473 | 1.68780013 |
| 11699 | NM_001282787.1 | NFIB | 336.433141 | -6.2206792 | 1.59022093 |
| 11700 | NM_000290.3 | #N/A | 29.4290014 | -6.0889596 | 1.5566617 |
| 11701 | NR_104223.1 | SH3YL1 | 33.5102731 | -6.2755019 | 1.60436623 |
| 11702 | NM_001256732.2 | SSBP2 | 82.7488369 | -7.579864 | 1.93795847 |
| 11703 | NM_020990.4 | CKMT1B | 47.4377816 | -6.7797747 | 1.73388068 |
| 11704 | NM_001040143.1 | #N/A | 73.9873779 | -7.4182133 | 1.89718993 |
| 11705 | NM_001078.3 | #N/A | 33.876295 | -6.2938951 | 1.60979369 |
| 11706 | NM_001093756.1 | TRAPPC13 | 37.6128835 | -6.4414393 | 1.64765199 |
| 11707 | NM_001002877.1 | #N/A | 38.467011 | -6.4744544 | 1.65616152 |
| 11709 | NR_047521.1 | LINC01128 | 51.3761399 | -6.8957943 | 1.76412613 |
| 11708 | NM_025268.3 | #N/A | 30.7698201 | -6.1538409 | 1.57431275 |
| 11710 | NM_014382.3 | #N/A | 62.1975972 | -7.1702423 | 1.83441674 |
| 11711 | NM_033046.2 | RTKN | 32.2319699 | -6.2165972 | 1.5905865 |
| 11712 | NM_001300751.1 | LMBR1L | 68.6091442 | -7.3097598 | 1.8703823 |
| 11714 | NR_033397.4 | PRMT1 | 35.0989619 | -6.3449805 | 1.62355129 |
| 11713 | NM_001321266.1 | #N/A | 44.372963 | -6.6788265 | 1.70897406 |
| 11715 | NM_001267828.1 | ZNF839 | 33.3861621 | 10.0296052 | 2.56672809 |
| 11716 | NM_001135213.2 | #N/A | 49.9601656 | -6.847538 | 1.75262208 |
| 11717 | NM_001204859.1 | #N/A | 44.6166959 | -6.6885281 | 1.71195599 |
| 11718 | NM_001190992.1 | SLC33A1 | 30.6634628 | -6.1546329 | 1.57554834 |
| 11719 | NM_001297423.1 | #N/A | 39.0683545 | -6.4944657 | 1.66271537 |
| 11720 | NR_132104.1 | PPP3CB-AS1 | 97.6147784 | 3.40122847 | 0.87080094 |
| 11721 | NM_001291814.1 | #N/A | 44.2617882 | -6.6732513 | 1.70861773 |
| 11722 | NM_001115.2 | #N/A | 30.4077651 | -6.1421584 | 1.57265145 |
| 11723 | NM_001040429.2 | #N/A | 166.164915 | -3.9428682 | 1.00969713 |
| 11724 | NM_024590.3 | #N/A | 46.3778905 | -6.7473897 | 1.72791407 |
| 11725 | NM_001025930.3 | #N/A | 35.447945 | -6.3631795 | 1.62968807 |
| 11726 | NM_001323295.1 | #N/A | 50.7085078 | -6.8691497 | 1.75934235 |
| 11727 | NM_001006945.1 | LRIF1 | 37.0421112 | -6.419859 | 1.64429 |
| 11728 | NM_001011718.1 | #N/A | 63.0634329 | -5.148743 | 1.31872747 |
| 11729 | NM_001322820.1 | #N/A | 57.6583311 | -7.0626601 | 1.8091232 |
| 11730 | NM_003454.3 | #N/A | 31.2071233 | -6.1730983 | 1.58151827 |
| 11731 | NM_001311178.1 | SLMAP | 30.6983378 | -6.1553007 | 1.5771452 |
| 11732 | NM_001243794.1 | #N/A | 42.3251879 | -6.6137955 | 1.69483719 |
| 11733 | NM_024570.3 | #N/A | 51.4769832 | -6.8944257 | 1.7667687 |
| 11734 | NM_001024939.3 | #N/A | 27.9957532 | -6.0213317 | 1.54312458 |
| 11735 | NM_001330617.1 | ZNF17 | 32.231382 | -6.2170076 | 1.59330367 |
| 11736 | NM_001102657.1 | #N/A | 31.7725078 | -6.1992744 | 1.5888289 |
| 11737 | NR_003654.2 | SCAND2P | 53.1884342 | -2.8002309 | 0.71772125 |
| 11738 | NM_144998.3 | #N/A | 43.7171363 | -6.6610742 | 1.70756339 |

| | | | | | |
|---|---|---|---|---|---|
| 11739 | NM_001139442.1 | TTLL11 | 37.6229014 | -6.4449469 | 1.65226885 |
| 11740 | NM_001253816.1 | SLC52A2 | 57.2004457 | -7.0478996 | 1.80686777 |
| 11741 | NM_021994.2 | #N/A | 51.6385167 | -4.0917684 | 1.04902328 |
| 11742 | NM_015942.4 | #N/A | 43.9609672 | -6.6677742 | 1.70953871 |
| 11743 | NM_001136498.1 | #N/A | 37.1569277 | -6.4246096 | 1.64724115 |
| 11744 | NM_001292018.1 | CYTH1 | 48.7438265 | -6.8154993 | 1.74752321 |
| 11745 | NM_005739.3 | #N/A | 43.2288564 | -6.6398301 | 1.70248671 |
| 11746 | NM_153182.3 | #N/A | 34.0880086 | -6.2938143 | 1.61375595 |
| 11747 | NM_001316984.1 | FAM207A | 33.8795702 | -6.2841477 | 1.61131304 |
| 11748 | NR_015361.1 | #N/A | 34.8220359 | 2.31110009 | 0.59261114 |
| 11749 | NM_001044385.2 | #N/A | 58.5555791 | -7.086038 | 1.81715238 |
| 11750 | NM_016654.4 | #N/A | 43.5619511 | -6.654721 | 1.70654447 |
| 11751 | NM_199424.2 | #N/A | 52.9097529 | -6.9350019 | 1.77843956 |
| 11752 | NM_198536.2 | #N/A | 38.2116607 | -6.465674 | 1.65822585 |
| 11753 | NM_001300778.1 | ZNF140 | 36.3362545 | -6.3915089 | 1.63929034 |
| 11754 | NM_000221.2 | #N/A | 45.5975438 | -6.7201165 | 1.72373184 |
| 11755 | NM_018988.3 | #N/A | 40.5079773 | -6.5504783 | 1.68042521 |
| 11756 | NM_148175.2 | #N/A | 34.8791927 | -6.3344845 | 1.62509753 |
| 11757 | NM_001172432.1 | PHKG2 | 41.496153 | -6.586769 | 1.68983112 |
| 11758 | NM_002026.3 | FN1 | 841.920548 | -10.928108 | 2.80385054 |
| 11759 | NR_045020.1 | #N/A | 34.1936119 | -6.3014261 | 1.61680036 |
| 11760 | NM_001123040.1 | #N/A | 77.2381558 | 1.57966183 | 0.40531204 |
| 11761 | NM_000797.3 | #N/A | 28.014029 | -6.0138533 | 1.54308086 |
| 11765 | NR_104010.1 | ZNHIT3 | 51.8116942 | -6.9050817 | 1.77187582 |
| 11762 | NM_001173982.1 | #N/A | 66.2576189 | -5.7749934 | 1.48188117 |
| 11763 | NM_003108.3 | #N/A | 2373.60595 | -4.0344347 | 1.03525939 |
| 11764 | NM_182697.2 | #N/A | 64.9032229 | -7.2326077 | 1.85591477 |
| 11766 | NM_006052.1 | #N/A | 176.666905 | -1.6799446 | 0.43110942 |
| 11767 | NM_033036.2 | #N/A | 75.9300025 | -5.9431027 | 1.52517236 |
| 11768 | NM_138340.4 | #N/A | 28.6549903 | -6.04877 | 1.55250364 |
| 11769 | NM_001330404.1 | #N/A | 36.5374023 | -6.3983672 | 1.64230719 |
| 11770 | NM_015888.4 | #N/A | 71.6094034 | -5.8711342 | 1.50699825 |
| 11771 | NR_132739.1 | #N/A | 138.767656 | -4.7355351 | 1.21554255 |
| 11772 | NM_001202414.1 | #N/A | 34.9635503 | -6.3304565 | 1.62497753 |
| 11773 | NM_001666.4 | #N/A | 29.2453385 | -6.0736432 | 1.55908489 |
| 11775 | NM_182835.2 | SCFD1 | 48.2858202 | -6.8037468 | 1.74659942 |
| 11776 | NR_023392.1 | ZNF252P | 70.686251 | -2.6955553 | 0.69197647 |
| 11774 | NM_004491.4 | #N/A | 829.947854 | -5.1884639 | 1.33192731 |
| 11777 | NM_025163.3 | #N/A | 33.8555701 | -6.2937737 | 1.61569316 |
| 11778 | NM_144589.2 | #N/A | 45.4537876 | -6.7162013 | 1.72429475 |
| 11779 | NM_001300965.1 | #N/A | 39.4795406 | -6.5132242 | 1.67227986 |
| 11780 | NM_133436.3 | ASNS | 40.5166622 | -6.5554676 | 1.68316797 |
| 11781 | NM_001194986.1 | #N/A | 39.7138295 | -6.5280561 | 1.67615966 |

| | | | | | |
|---|---|---|---|---|---|
| 11782 | NR_033403.1 | #N/A | 28.6103794 | -6.051788 | 1.55388244 |
| 11783 | NM_001317232.1 | MTCH2 | 48.7884663 | -6.82249 | 1.7520284 |
| 11784 | NR_130947.1 | WDR73 | 15.7684667 | 8.94163659 | 2.29623451 |
| 11785 | NM_001276271.1 | MBD4 | 35.8711876 | -6.3747994 | 1.63713954 |
| 11786 | NM_002658.4 | #N/A | 34.0209651 | -6.2969906 | 1.6174444 |
| 11787 | NM_000857.3 | #N/A | 67.6807694 | -7.2896332 | 1.87244466 |
| 11788 | NM_020737.2 | #N/A | 49.7793099 | -6.8477638 | 1.75905488 |
| 11789 | NM_003616.2 | GEMIN2 | 37.5491833 | -6.435814 | 1.65330254 |
| 11790 | NM_001308027.1 | #N/A | 35.9086645 | -6.3748245 | 1.63773925 |
| 11791 | NM_024556.3 | #N/A | 44.1925875 | -6.6790774 | 1.71602732 |
| 11792 | NM_001085.4 | #N/A | 64.0331132 | -7.2134359 | 1.85343608 |
| 11793 | NM_022909.3 | #N/A | 55.3167499 | -5.7208934 | 1.46996837 |
| 11794 | NM_001282306.1 | #N/A | 41.1206644 | -6.5758386 | 1.6899123 |
| 11795 | NM_138386.2 | #N/A | 28.0345754 | -6.0239718 | 1.54817239 |
| 11796 | NM_001276373.1 | #N/A | 25.5256357 | 9.63873376 | 2.47721063 |
| 11797 | NM_001200016.1 | #N/A | 29.0035973 | -6.0687485 | 1.55977624 |
| 11798 | NM_001280794.2 | EPHB6 | 49.5613597 | -6.8393309 | 1.75805496 |
| 11799 | NM_001321744.1 | FAM168B | 69.550554 | -7.3277243 | 1.8837888 |
| 11800 | NM_001308440.1 | ZFP28 | 42.4314163 | -6.6157781 | 1.7008332 |
| 11801 | NR_125755.1 | #N/A | 29.7520611 | -6.1105503 | 1.57108956 |
| 11802 | NM_001127392.2 | #N/A | 30.3519185 | -6.1289021 | 1.57582037 |
| 11803 | NM_018418.4 | #N/A | 32.9159816 | -6.2442978 | 1.60566662 |
| 11804 | NM_001199723.1 | CRABP2 | 40.3948508 | -6.5518811 | 1.68479571 |
| 11805 | NR_130726.1 | LINC00243 | 37.4160714 | 3.01268926 | 0.77471808 |
| 11806 | NM_148887.2 | #N/A | 41.8103537 | -5.3152973 | 1.36684685 |
| 11807 | NR_015449.1 | RNF216P1 | 38.4267996 | -6.4790547 | 1.66628645 |
| 11808 | NM_001204468.1 | RBM10 | 67.3226568 | -7.2854816 | 1.87371387 |
| 11809 | NM_001256067.1 | #N/A | 28.553435 | -6.0515125 | 1.5563513 |
| 11810 | NM_005341.3 | #N/A | 38.6861467 | -6.4793167 | 1.66639448 |
| 11811 | NM_198794.2 | #N/A | 51.5275325 | -6.896707 | 1.77394232 |
| 11812 | NM_002036.3 | #N/A | 45.4351589 | -6.7156462 | 1.72751144 |
| 11813 | NM_001303413.1 | RPRD1A | 34.8997248 | -6.3309674 | 1.62869648 |
| 11814 | NM_000286.2 | #N/A | 30.5814427 | -6.1497573 | 1.58209914 |
| 11815 | NR_135249.1 | #N/A | 36.7729688 | -6.417868 | 1.65128701 |
| 11816 | NM_000916.3 | OXTR | 28.603508 | -6.0484277 | 1.5562597 |
| 11817 | NM_001202519.1 | #N/A | 42.4847043 | -6.6202657 | 1.70342078 |
| 11818 | NM_001145434.1 | #N/A | 36.0775123 | -6.3802564 | 1.64185554 |
| 11819 | NR_003287.2 | #N/A | 32352.3662 | -7.0118943 | 1.8044317 |
| 11820 | NM_001258039.1 | SPRY1 | 32.4177786 | -6.2224689 | 1.60147533 |
| 11821 | NM_001291281.2 | #N/A | 228.24359 | -6.0128769 | 1.54760619 |
| 11822 | NM_018696.2 | #N/A | 32.9612934 | -6.2545497 | 1.60995968 |
| 11823 | NM_013355.4 | #N/A | 32.2684059 | -6.2235397 | 1.60211796 |
| 11824 | NR_026649.2 | BPHL | 30.3221641 | -6.1296941 | 1.57798698 |

| | | | | | |
|---|---|---|---|---|---|
| 11825 | NM_079421.2 | #N/A | 162.480763 | -5.6073386 | 1.44353371 |
| 11826 | NM_001003694.1 | #N/A | 36.4986168 | -6.4003859 | 1.64780009 |
| 11827 | NM_020765.2 | #N/A | 685.040349 | -5.0787717 | 1.30769601 |
| 11828 | NR_103803.1 | EBPL | 43.6522624 | -6.6544359 | 1.71355349 |
| 11829 | NM_001278689.1 | #N/A | 31.6249755 | -6.192186 | 1.59456788 |
| 11830 | NR_135786.1 | #N/A | 35.3428961 | -6.3497258 | 1.63543964 |
| 11831 | NM_001025242.1 | #N/A | 60.0192286 | -7.1209572 | 1.83419113 |
| 11832 | NM_016487.4 | #N/A | 33.9823133 | -6.295412 | 1.62163149 |
| 11833 | NM_022901.2 | #N/A | 51.3684124 | 2.98819937 | 0.76976396 |
| 11834 | NR_121570.1 | LOC101928381 | 44.178308 | 4.55829995 | 1.17433679 |
| 11835 | NM_001300977.1 | #N/A | 35.8557498 | -6.3745704 | 1.64227632 |
| 11836 | NM_001330449.1 | #N/A | 35.024074 | -6.3397735 | 1.63341947 |
| 11837 | NM_001014342.2 | #N/A | 58.2865063 | 3.12424703 | 0.80495736 |
| 11838 | NM_001039396.1 | #N/A | 16.0162054 | 5.2946385 | 1.36416056 |
| 11839 | NM_139075.3 | #N/A | 51.1709732 | -5.4448291 | 1.40286786 |
| 11840 | NM_001134445.1 | #N/A | 53.4464813 | -6.9505196 | 1.79100826 |
| 11841 | NR_038911.1 | MIF-AS1 | 32.882072 | -6.2469114 | 1.60980244 |
| 11842 | NM_001006943.2 | #N/A | 27.2487227 | -5.9784589 | 1.54063305 |
| 11843 | NM_181303.1 | #N/A | 30.0155777 | -6.1209989 | 1.5774392 |
| 11844 | NM_001424.5 | #N/A | 363.773616 | 1.50547335 | 0.38800905 |
| 11846 | NR_126480.1 | SALRNA1 | 16.6272809 | 4.75060894 | 1.22439845 |
| 11845 | NM_032345.2 | #N/A | 28.4049619 | -6.0368976 | 1.5559271 |
| 11847 | NM_152274.4 | #N/A | 41.8274652 | -6.5950806 | 1.69997583 |
| 11848 | NM_001310123.1 | ATXN2 | 106.688311 | -3.530851 | 0.91018917 |
| 11849 | NM_001085384.2 | #N/A | 32.0494304 | -6.2196064 | 1.60351648 |
| 11850 | NM_017704.2 | #N/A | 35.5233094 | -6.3588104 | 1.63955037 |
| 11851 | NM_001330305.1 | #N/A | 44.676621 | -6.6883628 | 1.72477148 |
| 11852 | NM_001011671.2 | #N/A | 31.1409471 | -6.1680107 | 1.59071019 |
| 11853 | NM_001034025.1 | ERP29 | 50.7110606 | -6.8747023 | 1.77301047 |
| 11854 | NR_045553.1 | #N/A | 35.8453182 | -6.3761337 | 1.64443439 |
| 11855 | NM_001321702.1 | #N/A | 121.606271 | -8.1362265 | 2.09871647 |
| 11856 | NR_132114.1 | SNHG19 | 34.1992038 | -6.3054195 | 1.62661765 |
| 11857 | NM_006779.3 | #N/A | 36.8209692 | -6.4155975 | 1.65510695 |
| 11858 | NR_002813.1 | KCNIP4-IT1 | 25.9261429 | 5.64655267 | 1.45671924 |
| 11859 | NM_004570.5 | PIK3C2G | 24.4940357 | 8.24378692 | 2.12680746 |
| 11860 | NM_016307.3 | #N/A | 35.1876851 | -6.3497031 | 1.63817373 |
| 11861 | NR_138575.1 | #N/A | 35.5462679 | -6.3675404 | 1.64291446 |
| 11862 | NM_001258026.1 | TPI1 | 35.4295447 | -6.3564986 | 1.64017951 |
| 11863 | NM_001270977.1 | #N/A | 51.547149 | -6.901955 | 1.78094245 |
| 11864 | NM_001807.4 | #N/A | 27.550674 | -5.9948087 | 1.54689275 |
| 11865 | NM_014735.4 | #N/A | 36.7483282 | -6.4141509 | 1.65510944 |
| 11866 | NM_001320717.1 | #N/A | 36.0844112 | -6.378788 | 1.64602279 |
| 11867 | NR_110479.1 | MIRLET7BHG | 80.5812211 | 2.10610037 | 0.54355194 |

| | | | | | |
|---|---|---|---|---|---|
| 11868 | NM_000328.2 | #N/A | 71.9040243 | -7.380017 | 1.90481618 |
| 11869 | NM_022916.4 | #N/A | 157.987522 | -3.1547589 | 0.81430452 |
| 11870 | NM_001271953.1 | #N/A | 35.2634144 | -6.3493874 | 1.63902343 |
| 11871 | NM_001033002.3 | RPAIN | 45.2834655 | -6.7106137 | 1.73241842 |
| 11872 | NR_135077.1 | CSGALNACT2 | 38.2539724 | -6.4646818 | 1.66900866 |
| 11873 | NM_001317177.1 | CHCHD3 | 66.4134724 | -7.2643188 | 1.87562163 |
| 11874 | NM_194281.3 | #N/A | 28.9536642 | -6.0708357 | 1.56751754 |
| 11875 | NM_001033602.2 | #N/A | 27.8634775 | -6.0123665 | 1.55272767 |
| 11876 | NM_001003665.3 | #N/A | 60.4371259 | -5.8525351 | 1.51148273 |
| 11877 | NM_024657.4 | #N/A | 59.8897507 | -7.1180504 | 1.83869371 |
| 11878 | NM_004315.5 | #N/A | 40.2189694 | -6.533675 | 1.6878193 |
| 11879 | NM_198455.2 | #N/A | 27.2052532 | -5.9785169 | 1.54442251 |
| 11880 | NR_135778.1 | RSRP1 | 36.7643142 | -6.4112246 | 1.65623189 |
| 11881 | NM_001320126.1 | #N/A | 30.7474688 | -6.1444869 | 1.58736206 |
| 11882 | NM_006578.3 | #N/A | 282.163612 | -3.5205708 | 0.90957525 |
| 11883 | NM_152379.3 | #N/A | 28.7339707 | -6.051505 | 1.56361276 |
| 11884 | NM_001301837.1 | C12orf57 | 40.2127011 | -6.54292 | 1.69083629 |
| 11885 | NM_001100169.1 | IMMT | 53.8116351 | -6.9623063 | 1.79925475 |
| 11886 | NM_012345.2 | #N/A | 56.1340021 | -5.5585658 | 1.43654654 |
| 11887 | NM_001002018.2 | HCFC1R1 | 38.1770613 | -6.4604113 | 1.6696723 |
| 11888 | NR_046392.1 | LINC00507 | 82.7141628 | 4.06899899 | 1.05161992 |
| 11889 | NM_001012279.2 | #N/A | 1010.57326 | -4.6234193 | 1.19501289 |
| 11890 | NM_001135662.1 | #N/A | 42.1191874 | -6.60192 | 1.70643368 |
| 11891 | NM_001289153.1 | #N/A | 30.726967 | -6.1436868 | 1.58799283 |
| 11892 | NM_005319.3 | H1-2 | 34.5958439 | -6.3229328 | 1.63435488 |
| 11893 | NM_033364.3 | #N/A | 52.3432919 | -6.9225511 | 1.78935878 |
| 11894 | NM_001042400.1 | #N/A | 44.7120597 | -5.415735 | 1.40008257 |
| 11895 | NM_001315530.1 | #N/A | 36.3881994 | -6.3986461 | 1.65423191 |
| 11896 | NM_001321035.1 | SEPTIN2 | 46.7957395 | -6.7537235 | 1.74610098 |
| 11897 | NM_014338.3 | PISD | 54.6418792 | -6.9791953 | 1.80446318 |
| 11898 | NR_144431.1 | PEG13 | 45.157441 | -6.7082768 | 1.73444107 |
| 11899 | NM_024109.3 | #N/A | 30.0061367 | -6.1233515 | 1.58323091 |
| 11900 | NM_016606.3 | #N/A | 66.3424412 | -7.2652014 | 1.87857418 |
| 11901 | NM_001122898.2 | #N/A | 43.7008284 | -6.660148 | 1.72220906 |
| 11902 | NR_104481.1 | #N/A | 14.2651998 | 8.82455661 | 2.28197108 |
| 11903 | NM_019854.4 | #N/A | 46.3985456 | -6.7457299 | 1.74446915 |
| 11904 | NM_001130677.1 | #N/A | 45.8739265 | -6.7274651 | 1.74007149 |
| 11905 | NR_036623.1 | #N/A | 32.9871878 | -6.2541164 | 1.61772441 |
| 11906 | NM_001286420.1 | #N/A | 42.411321 | -6.6151277 | 1.71122094 |
| 11907 | NM_020972.2 | #N/A | 33.5501892 | -6.2847479 | 1.62590411 |
| 11908 | NM_001018024.2 | #N/A | 33.0234469 | -6.2539482 | 1.6182019 |
| 11909 | NM_030578.3 | #N/A | 28.0785801 | -6.0255612 | 1.55911031 |
| 11910 | NR_036508.1 | LOC100128398 | 58.9111653 | -2.4476391 | 0.63341833 |

| | | | | | |
|---|---|---|---|---|---|
| 11911 | NM_001006623.2 | #N/A | 34.1681695 | -6.3031484 | 1.63120881 |
| 11912 | NM_001204051.1 | #N/A | 30.1987934 | -6.1193182 | 1.58365718 |
| 11913 | NM_001305840.1 | TCEAL4 | 37.1982554 | -6.4252767 | 1.6629395 |
| 11914 | NM_001290263.1 | #N/A | 34.0598976 | -6.3073638 | 1.63247192 |
| 11915 | NM_001284295.1 | #N/A | 41.8495908 | -5.31364 | 1.37534962 |
| 11916 | NM_001200049.2 | #N/A | 83.0110278 | -7.5872285 | 1.96397787 |
| 11917 | NM_198042.3 | #N/A | 36.1791219 | -6.3912003 | 1.65444472 |
| 11918 | NM_172115.2 | CAMK2D | 29.1216499 | -6.0799819 | 1.57389112 |
| 11921 | NR_037846.1 | MSH5-SAPCD1 | 33.5927788 | -6.2717707 | 1.62358455 |
| 11920 | NM_057180.2 | VPS29 | 41.0550022 | -6.5670091 | 1.70002467 |
| 11919 | NM_001024736.1 | #N/A | 339.586312 | -3.6125337 | 0.93519008 |
| 11922 | NM_001256799.2 | #N/A | 37.6012644 | -6.4438241 | 1.66849789 |
| 11923 | NM_001010872.2 | #N/A | 81.9225389 | 3.22280593 | 0.83457488 |
| 11924 | NM_001921.2 | DCTD | 45.5422926 | -6.7169765 | 1.7394907 |
| 11925 | NM_003915.5 | CPNE1 | 36.6363654 | -6.4040933 | 1.65861317 |
| 11926 | NM_004655.3 | #N/A | 132.451924 | -3.4790366 | 0.90109221 |
| 11927 | NM_001164093.1 | #N/A | 40.0159381 | -6.5265913 | 1.69052517 |
| 11928 | NM_025108.2 | #N/A | 44.4084758 | -6.6815735 | 1.73079086 |
| 11929 | NM_001282930.1 | #N/A | 64.7979541 | -7.2272092 | 1.87220843 |
| 11930 | NM_001286616.1 | ANO4 | 31.5872634 | -6.1905331 | 1.60370214 |
| 11932 | NM_198141.2 | GANC | 27.7769916 | -6.0092685 | 1.55688144 |
| 11931 | NM_020300.4 | #N/A | 33.3809096 | -6.27258 | 1.62509672 |
| 11934 | NR_028330.1 | ST20-AS1 | 34.0138105 | -6.2985272 | 1.63184948 |
| 11933 | NM_016315.3 | #N/A | 39.1422201 | -6.5020491 | 1.68457872 |
| 11935 | NM_001276469.1 | B4GALNT1 | 57.408154 | -7.0514736 | 1.82709608 |
| 11937 | NR_034108.1 | TRAF3IP2-AS1 | 29.8687212 | -6.1115154 | 1.58353805 |
| 11936 | NM_032273.3 | #N/A | 44.0569817 | -6.6694283 | 1.72808939 |
| 11938 | NM_015341.4 | #N/A | 37.5835451 | -6.4445602 | 1.66991727 |
| 11939 | NM_133445.2 | #N/A | 42.6825693 | -6.627858 | 1.7175949 |
| 11940 | NM_001313905.1 | EMP3 | 27.5982593 | -5.9949786 | 1.55371747 |
| 11941 | NM_020422.5 | #N/A | 29.8265388 | -6.1035437 | 1.58185134 |
| 11942 | NM_001136114.1 | POMT1 | 40.3745845 | -6.5424512 | 1.69565331 |
| 11943 | NM_194292.2 | #N/A | 32.453092 | -6.2288576 | 1.61447851 |
| 11944 | NM_033515.2 | #N/A | 39.5459134 | -6.5203538 | 1.69024012 |
| 11945 | NR_027782.1 | #N/A | 33.9873807 | -6.2929869 | 1.63138092 |
| 11946 | NM_001270482.1 | #N/A | 39.1399155 | -6.5050928 | 1.68644425 |
| 11947 | NR_105056.2 | LOC101926905 | 26.3008534 | 7.5330515 | 1.95302817 |
| 11948 | NM_152263.3 | #N/A | 190.57059 | 1.77219201 | 0.45951272 |
| 11949 | NM_001124.2 | #N/A | 28.7435163 | -6.0595813 | 1.57141381 |
| 11950 | NM_003026.3 | #N/A | 196.69908 | -5.2787369 | 1.36894836 |
| 11951 | NM_001143942.1 | #N/A | 27.2945521 | -5.98151 | 1.55123944 |
| 11952 | NM_001184792.1 | PARD3 | 29.8790918 | -6.1156459 | 1.58607238 |
| 11953 | NM_001714.2 | #N/A | 32.4584363 | -6.2272383 | 1.61509684 |

| | | | | | |
|---|---|---|---|---|---|
| 11954 | NM_001128922.1 | #N/A | 38.1463381 | -6.4683706 | 1.67768046 |
| 11955 | NM_001164796.1 | SLC25A26 | 34.3102924 | -6.308796 | 1.63645133 |
| 11956 | NR_038834.1 | #N/A | 129.751288 | 3.15712789 | 0.81903541 |
| 11957 | NM_004101.3 | #N/A | 28.2795306 | -6.0359568 | 1.56588279 |
| 11958 | NM_016653.2 | #N/A | 29.7044993 | -6.1032247 | 1.58348969 |
| 11959 | NR_002784.2 | #N/A | 17.0169008 | 8.33503935 | 2.16278228 |
| 11960 | NM_001170721.1 | #N/A | 27.9134459 | -6.0197061 | 1.56203024 |
| 11961 | NM_198195.1 | #N/A | 46.9030455 | -6.7597408 | 1.75405887 |
| 11962 | NM_177995.2 | #N/A | 33.2596223 | -6.262829 | 1.6251666 |
| 11963 | NM_016209.4 | #N/A | 37.8156069 | -6.4452237 | 1.67282263 |
| 11964 | NM_001348050.1 | #N/A | 81.8784755 | -4.843693 | 1.25716754 |
| 11965 | NM_020890.2 | #N/A | 31.8384235 | -6.2040657 | 1.61029326 |
| 11966 | NR_038334.1 | LOC100505715 | 28.8976927 | -6.0579204 | 1.57253177 |
| 11967 | NM_172218.2 | SPAG1 | 49.1719381 | -6.8332514 | 1.77381137 |
| 11968 | NM_001185118.1 | #N/A | 43.2263008 | -6.6373824 | 1.72298531 |
| 11969 | NR_024048.2 | #N/A | 28.3768284 | -6.0314946 | 1.56573187 |
| 11970 | NR_046480.1 | ROGDI | 33.746986 | -6.290836 | 1.63321689 |
| 11971 | NM_017803.4 | #N/A | 31.93219 | -6.207741 | 1.6116833 |
| 11972 | NM_000212.2 | ITGB3 | 39.7878036 | -6.5285517 | 1.69509226 |
| 11973 | NM_001346590.1 | INSIG1 | 32.5788047 | -6.2399389 | 1.62050269 |
| 11974 | NM_001330454.1 | #N/A | 52.394817 | -6.9192745 | 1.79694135 |
| 11975 | NM_022840.4 | #N/A | 33.8030171 | -6.2858085 | 1.6324775 |
| 11976 | NM_001039374.4 | #N/A | 32.1264483 | -6.2202102 | 1.61567461 |
| 11977 | NR_135107.1 | S100PBP | 28.112432 | 9.80436345 | 2.54737333 |
| 11978 | NM_001322949.1 | #N/A | 38.6284896 | -6.4789709 | 1.68338452 |
| 11979 | NM_001013659.2 | #N/A | 639.350452 | 1.52989142 | 0.39751982 |
| 11980 | NM_001200.3 | #N/A | 38.3794104 | -6.4777057 | 1.68327948 |
| 11981 | NM_032387.4 | #N/A | 30.2131339 | -6.1297915 | 1.59293611 |
| 11982 | NM_021165.3 | #N/A | 33.9405643 | -6.2940168 | 1.6356689 |
| 11983 | NM_001205.2 | #N/A | 35.8954035 | -6.3755507 | 1.65690424 |
| 11984 | NM_001171797.1 | #N/A | 58.3263967 | -7.0750642 | 1.83885074 |
| 11985 | NR_036577.1 | APTX | 30.5391011 | -6.1486355 | 1.59831184 |
| 11986 | NM_015291.3 | #N/A | 50.0884987 | -6.8576773 | 1.78264698 |
| 11987 | NM_001321112.1 | #N/A | 29.2002268 | -6.0784121 | 1.58009468 |
| 11988 | NM_022169.4 | #N/A | 33.9336463 | -6.2884023 | 1.63472788 |
| 11989 | NM_006743.4 | #N/A | 830.30503 | -2.3963417 | 0.62297642 |
| 11990 | NM_001292004.1 | HEXB | 28.7331265 | -6.0544276 | 1.57399867 |
| 11991 | NM_001320194.1 | #N/A | 32.8272774 | -6.2416522 | 1.62269039 |
| 11992 | NM_001253823.1 | BLVRA | 27.3467826 | -5.9774054 | 1.55408777 |
| 11993 | NM_001303112.1 | #N/A | 39.5272686 | -6.5096771 | 1.69250981 |
| 11994 | NM_014948.3 | #N/A | 36.7636311 | -6.4176095 | 1.66863332 |
| 11995 | NM_006568.2 | #N/A | 35.8610294 | -6.3732705 | 1.65730382 |
| 11996 | NM_001114396.1 | #N/A | 27.9644734 | -6.0197923 | 1.56550341 |

| | | | | | |
|---|---|---|---|---|---|
| 11997 | NM_001752.3 | #N/A | 69.2617373 | -7.3235368 | 1.90478193 |
| 11998 | NM_001282673.1 | FAAP20 | 55.0423809 | -6.9923538 | 1.8188556 |
| 11999 | NM_006491.2 | NOVA1 | 30.5467899 | -6.1423414 | 1.59774134 |
| 12000 | NM_007156.4 | ZXDA | 30.7236201 | -6.147283 | 1.59903059 |
| 12001 | NM_003743.4 | #N/A | 64.8895641 | -7.2312171 | 1.88118105 |
| 12002 | NM_001322427.1 | #N/A | 33.6945266 | -6.2853671 | 1.63519878 |
| 12003 | NM_006668.1 | #N/A | 35.4410209 | -6.3588175 | 1.65458703 |
| 12004 | NM_001171132.2 | #N/A | 35.39832 | -6.3587144 | 1.65458667 |
| 12005 | NM_000641.3 | #N/A | 45.3079352 | -6.712447 | 1.74673883 |
| 12006 | NR_045557.1 | URI1 | 40.3129159 | -6.548407 | 1.70418653 |
| 12007 | NM_015215.3 | #N/A | 382.268969 | -4.027076 | 1.04806978 |
| 12008 | NM_001136258.1 | SGMS2 | 45.2015221 | -6.7118467 | 1.74687483 |
| 12009 | NM_001002031.2 | #N/A | 28.2680401 | -6.0368379 | 1.57120124 |
| 12010 | NR_038832.1 | HOXA-AS3 | 79.1700576 | -5.9847062 | 1.55770492 |
| 12011 | NM_014290.2 | #N/A | 33.5543702 | -6.2736155 | 1.63292503 |
| 12012 | NM_001329990.1 | #N/A | 56.6540702 | -7.0331397 | 1.83071073 |
| 12013 | NR_045574.1 | #N/A | 30.7974058 | -6.1612251 | 1.60377286 |
| 12014 | NM_001257191.2 | DDX6 | 300.250967 | -3.6841659 | 0.95909047 |
| 12015 | NM_024091.3 | #N/A | 31.2217666 | -6.1690437 | 1.60601286 |
| 12016 | NM_001004722.3 | NCK2 | 38.6850564 | -6.4831405 | 1.68779463 |
| 12017 | NM_003223.2 | #N/A | 56.628633 | -5.5639464 | 1.44869849 |
| 12018 | NM_014881.4 | #N/A | 30.0388388 | -6.1265846 | 1.59527114 |
| 12020 | NR_135812.1 | CBLB | 51.2627475 | -3.8760782 | 1.00930193 |
| 12019 | NM_001268.3 | #N/A | 36.5505573 | -6.4049778 | 1.66780916 |
| 12021 | NM_001287215.1 | AMD1 | 32.65969 | -6.2392637 | 1.62467024 |
| 12022 | NM_002663.4 | #N/A | 39.8348859 | -6.5319102 | 1.7010641 |
| 12023 | NM_203451.2 | #N/A | 29.6326569 | -6.1031223 | 1.58941129 |
| 12025 | NR_026967.1 | LOC143666 | 36.8586344 | -6.410505 | 1.66949539 |
| 12024 | NM_001003684.1 | UQCR10 | 32.4796764 | -6.2307696 | 1.62269261 |
| 12026 | NR_110281.1 | LOC101927884 | 76.3317811 | 4.853615 | 1.26406958 |
| 12027 | NM_020717.3 | SHROOM4 | 28.01917 | -6.0261521 | 1.56945298 |
| 12028 | NR_015358.2 | LINC00643 | 52.0368045 | -6.9093942 | 1.79956272 |
| 12029 | NM_033119.4 | #N/A | 66.7027226 | -4.4451363 | 1.15775114 |
| 12030 | NM_001013438.2 | #N/A | 238.164554 | -2.4560333 | 0.63977447 |
| 12031 | NM_001485.3 | #N/A | 41.171074 | -6.5759595 | 1.71303912 |
| 12032 | NM_021800.2 | #N/A | 45.6372642 | -6.7215417 | 1.75101608 |
| 12033 | NM_001114176.1 | #N/A | 32.1724084 | -6.2173913 | 1.61973879 |
| 12034 | NM_001284382.1 | #N/A | 54.4798089 | -6.9762052 | 1.81758248 |
| 12035 | NM_018322.2 | #N/A | 35.6163436 | -6.3626288 | 1.65778858 |
| 12036 | NM_018078.3 | #N/A | 32.1510795 | -6.2146723 | 1.61958782 |
| 12037 | NM_153376.2 | #N/A | 31.2526586 | -6.1779652 | 1.61012085 |
| 12038 | NM_001128615.1 | #N/A | 35.6117834 | -6.3664036 | 1.65924957 |
| 12039 | NM_001282686.1 | KCTD17 | 38.015203 | -6.4633999 | 1.68454529 |

| | | | | | |
|---|---|---|---|---|---|
| 12040 | NM_001308301.1 | MYLK3 | 133.619092 | 1.8347191 | 0.47832996 |
| 12041 | NM_001130858.2 | PPIP5K1 | 36.4889368 | -6.3969536 | 1.66779047 |
| 12042 | NM_197941.3 | #N/A | 36.6524694 | -5.1278138 | 1.33701309 |
| 12043 | NM_021931.3 | #N/A | 52.9974232 | -6.9375577 | 1.80894018 |
| 12044 | NM_000841.3 | #N/A | 43.8772407 | -6.6662632 | 1.73823513 |
| 12045 | NM_001039707.1 | #N/A | 31.9512993 | -6.2007283 | 1.61699138 |
| 12046 | NM_001018078.2 | FPGS | 33.6038972 | -6.2798926 | 1.63764859 |
| 12047 | NM_004782.3 | #N/A | 176.700775 | -2.715524 | 0.70815901 |
| 12048 | NM_001161352.1 | #N/A | 38.5363994 | -6.4793436 | 1.68974033 |
| 12049 | NM_144976.3 | #N/A | 29.3147629 | -6.088115 | 1.58772034 |
| 12050 | NM_153605.3 | #N/A | 34.4815248 | -6.3227248 | 1.64890335 |
| 12051 | NM_032423.2 | #N/A | 65.2135179 | -5.7381614 | 1.49650494 |
| 12052 | NM_001164380.1 | #N/A | 37.7564399 | -6.4473524 | 1.681597 |
| 12053 | NM_030786.2 | #N/A | 40.9139798 | -6.5664475 | 1.71277883 |
| 12054 | NM_001037494.1 | #N/A | 40.9694591 | -6.5660013 | 1.71269513 |
| 12055 | NM_001272067.1 | SHISA5 | 35.1870978 | -6.3509922 | 1.65679547 |
| 12056 | NR_110290.1 | SLC22A17 | 36.1526708 | -6.3934102 | 1.66790529 |
| 12057 | NM_002518.3 | #N/A | 40.5466386 | -6.55497 | 1.71063501 |
| 12058 | NM_003774.4 | #N/A | 33.0989719 | -6.2668445 | 1.63554361 |
| 12059 | NM_001285829.1 | CEBPA | 29.0164408 | -6.068019 | 1.5836912 |
| 12060 | NM_012081.5 | #N/A | 43.541984 | -4.5824496 | 1.19598534 |
| 12061 | NM_033169.2 | B3GALNT1 | 31.0393961 | -6.1617183 | 1.60831496 |
| 12062 | NM_001256829.1 | #N/A | 29.9925148 | -6.1176682 | 1.59694706 |
| 12063 | NM_152911.3 | #N/A | 32.1223973 | -6.2179521 | 1.62321584 |
| 12064 | NR_073415.2 | HSP90AB4P | 58.5311473 | 3.59224494 | 0.93785435 |
| 12065 | NM_001329846.1 | #N/A | 46.3128032 | -6.7394679 | 1.75959077 |
| 12066 | NM_025180.3 | #N/A | 33.7192038 | -6.27819 | 1.63920964 |
| 12067 | NM_022369.3 | #N/A | 44.545758 | -6.6840232 | 1.74534437 |
| 12068 | NM_001242921.1 | PLTP | 30.90707 | -6.1634867 | 1.60961205 |
| 12069 | NM_000573.3 | #N/A | 62.7778442 | 2.96503481 | 0.77436428 |
| 12070 | NM_001319655.1 | #N/A | 36.5268011 | -6.4035595 | 1.67250189 |
| 12071 | NM_002632.5 | PGF | 36.189118 | -6.3937424 | 1.66997736 |
| 12072 | NM_152698.2 | #N/A | 51.9247806 | -6.9039732 | 1.80363725 |
| 12073 | NM_001190201.1 | #N/A | 28.4101926 | -6.0371015 | 1.57750669 |
| 12074 | NM_015414.3 | #N/A | 38.0187955 | -6.4613994 | 1.68871564 |
| 12075 | NR_144529.1 | LOC100507599 | 29.8773047 | -6.1120777 | 1.59759662 |
| 12076 | NM_001008394.2 | #N/A | 19.0331018 | 3.15864971 | 0.82563327 |
| 12077 | NM_152447.4 | LRFN5 | 41.0028799 | -6.5648808 | 1.71603699 |
| 12078 | NM_198227.1 | #N/A | 73.9463405 | -7.4190842 | 1.93931567 |
| 12079 | NM_000186.3 | #N/A | 28.5180766 | -6.0508198 | 1.58178078 |
| 12080 | NR_134862.1 | MRPL18 | 36.5785385 | -6.401134 | 1.6733678 |
| 12081 | NM_199250.2 | C19orf48 | 44.2009001 | -6.6708565 | 1.74399446 |
| 12082 | NM_001282338.1 | #N/A | 62.1433556 | -7.1673313 | 1.87400177 |

| | | | | | |
|---|---|---|---|---|---|
| 12083 | NM_001288764.1 | #N/A | 43.2841777 | -6.6516453 | 1.73930253 |
| 12084 | NM_004102.4 | #N/A | 53.8552296 | -6.960809 | 1.82025855 |
| 12085 | NM_024676.4 | #N/A | 40.3997747 | -6.547739 | 1.71235867 |
| 12086 | NM_017784.4 | OSBPL10 | 35.9402521 | -6.3756312 | 1.66740046 |
| 12087 | NR_024084.1 | #N/A | 33.3541362 | -6.2672509 | 1.63920211 |
| 12088 | NM_014454.2 | #N/A | 57.3141284 | -5.5748147 | 1.45842408 |
| 12089 | NM_001136469.2 | TMEM108 | 44.1574277 | -6.6790543 | 1.74746582 |
| 12090 | NM_001321532.1 | #N/A | 53.1803001 | -6.9409709 | 1.81615828 |
| 12091 | NM_001177549.2 | #N/A | 53.6029603 | 2.57126386 | 0.67280957 |
| 12092 | NM_194259.2 | #N/A | 39.7308298 | -6.5194975 | 1.70622578 |
| 12093 | NM_002183.3 | #N/A | 28.0841491 | -6.0248592 | 1.57691335 |
| 12094 | NM_001015049.2 | #N/A | 44.5936649 | -6.6905118 | 1.75127289 |
| 12095 | NM_001145783.1 | #N/A | 49.9781881 | -6.8523028 | 1.79369597 |
| 12096 | NM_198156.2 | #N/A | 237.849606 | 1.66225896 | 0.43513207 |
| 12097 | NM_001201459.1 | ZNF513 | 37.7161454 | -6.4446415 | 1.68720556 |
| 12098 | NM_020134.3 | #N/A | 1865.31476 | -5.6325072 | 1.47459836 |
| 12099 | NM_001166600.2 | FAM122B | 27.1264821 | -5.9790245 | 1.56544349 |
| 12100 | NR_045839.1 | #N/A | 31.8088097 | -6.2019688 | 1.62384034 |
| 12101 | NM_030651.3 | #N/A | 45.0735026 | -6.7027602 | 1.75512338 |
| 12103 | NM_004200.3 | SYT7 | 98.486071 | -4.6349567 | 1.21382237 |
| 12102 | NM_001346810.1 | #N/A | 47.3316602 | -6.7746149 | 1.77416668 |
| 12104 | NM_001123378.2 | ADSL | 35.7182512 | -6.3720939 | 1.66892856 |
| 12105 | NM_001316766.1 | #N/A | 30.7943519 | -6.155304 | 1.61213398 |
| 12106 | NM_015892.4 | #N/A | 55.1655738 | -5.5231737 | 1.446584 |
| 12107 | NM_001252393.1 | #N/A | 33.2216746 | -6.2717977 | 1.64290571 |
| 12108 | NM_001190981.1 | #N/A | 47.5945305 | -4.1634932 | 1.09076078 |
| 12109 | NM_031301.3 | #N/A | 59.2738972 | -7.102845 | 1.86116967 |
| 12110 | NM_005065.5 | #N/A | 491.388883 | -3.4776478 | 0.91135314 |
| 12111 | NM_001330521.1 | #N/A | 30.8443301 | -6.1650278 | 1.61572024 |
| 12112 | NM_001164715.1 | #N/A | 34.4280139 | -6.3079448 | 1.65322347 |
| 12113 | NM_015278.4 | #N/A | 39.5191033 | -6.5196124 | 1.70889629 |
| 12114 | NM_003428.5 | ZNF84 | 15.3504992 | 8.93492871 | 2.3420889 |
| 12115 | NM_031307.3 | #N/A | 31.0708873 | -6.1647556 | 1.61601676 |
| 12116 | NM_001199708.1 | #N/A | 32.5497238 | -6.2262769 | 1.63222681 |
| 12117 | NM_001113324.2 | #N/A | 29.1196686 | -6.0745556 | 1.59252501 |
| 12118 | NM_001818.3 | #N/A | 26.967423 | 9.74474335 | 2.55481626 |
| 12119 | NM_001412.3 | #N/A | 639.307805 | -2.0601512 | 0.54015168 |
| 12121 | NM_206825.1 | GNL3 | 36.9791942 | -6.4131384 | 1.68155744 |
| 12120 | NM_001290325.1 | USP38 | 42.3371012 | -5.1963934 | 1.36251783 |
| 12122 | NR_134612.1 | #N/A | 43.6090399 | -6.6562207 | 1.74541602 |
| 12123 | NM_001123363.3 | RGPD6 | 43.1098779 | -6.6412877 | 1.74161186 |
| 12124 | NM_001080514.2 | #N/A | 31.4004446 | -6.186201 | 1.6223885 |
| 12125 | NM_001145862.1 | #N/A | 28.5443181 | -6.0507007 | 1.58690971 |

| | | | | | |
|---|---|---|---|---|---|
| 12127 | NR_028344.1 | LINC00632 | 45.5500819 | -4.6480731 | 1.21918017 |
| 12126 | NM_174934.3 | SCN4B | 33.3631356 | -6.2754044 | 1.64602135 |
| 12128 | NR_033709.1 | TP73-AS1 | 34.3370162 | -6.3066136 | 1.65429002 |
| 12129 | NR_024492.1 | FLJ42627 | 119.71169 | -1.8758151 | 0.49205869 |
| 12130 | NM_001008537.2 | #N/A | 124.993778 | -5.2843326 | 1.38619859 |
| 12131 | NM_024709.4 | #N/A | 57.9247034 | -5.5856002 | 1.46533122 |
| 12132 | NM_001313893.1 | HMGB1 | 49.8283468 | -6.848471 | 1.79697257 |
| 12133 | NM_024832.4 | #N/A | 28.8221042 | -6.0671697 | 1.59207828 |
| 12134 | NR_120648.1 | LOC101928994 | 29.1149931 | 5.81010518 | 1.52466762 |
| 12135 | NM_033402.4 | #N/A | 61.0449019 | -5.6499067 | 1.48265654 |
| 12136 | NM_001007464.2 | #N/A | 34.0996542 | -6.3066481 | 1.65517792 |
| 12137 | NR_024397.1 | NUTM2A-AS1 | 34.2743019 | -6.3093913 | 1.65597407 |
| 12138 | NM_001323599.1 | #N/A | 30.5934153 | -6.1504821 | 1.61435738 |
| 12139 | NM_152835.4 | #N/A | 54.5836607 | -6.9768007 | 1.83133262 |
| 12140 | NM_001282285.1 | DYRK4 | 33.3772079 | -6.2756593 | 1.64738178 |
| 12141 | NM_012434.4 | #N/A | 121.319742 | -3.1601502 | 0.82958226 |
| 12142 | NM_016049.3 | #N/A | 30.7476019 | -6.1552888 | 1.61587014 |
| 12143 | NR_134608.1 | #N/A | 44.642619 | -6.6889178 | 1.75595454 |
| 12144 | NM_020939.1 | #N/A | 39.34199 | -6.5026054 | 1.70707023 |
| 12145 | NM_144999.3 | #N/A | 135.836952 | -3.4366328 | 0.90221441 |
| 12146 | NM_030753.4 | #N/A | 34.6307507 | -6.3261507 | 1.66083929 |
| 12147 | NM_002122.3 | #N/A | 49.5215165 | -6.8428175 | 1.79653821 |
| 12148 | NM_001199397.1 | #N/A | 49.6125098 | -6.845093 | 1.797249 |
| 12149 | NM_001130926.2 | MEF2A | 34.6516424 | -6.3203029 | 1.65964217 |
| 12150 | NM_153369.2 | #N/A | 43.9205482 | -3.3062221 | 0.86823176 |
| 12151 | NR_045019.1 | #N/A | 30.9903608 | -6.1654599 | 1.61925075 |
| 12152 | NR_102748.1 | LOC727751 | 27.0923693 | -5.9694925 | 1.56780782 |
| 12153 | NM_001320309.1 | #N/A | 48.7876145 | -6.8170252 | 1.79046394 |
| 12154 | NM_001008.3 | #N/A | 43.3113222 | -6.6482757 | 1.74620235 |
| 12155 | NM_016545.4 | #N/A | 258.037297 | -2.8571287 | 0.75060577 |
| 12156 | NM_182984.4 | #N/A | 30.3652454 | -6.1424875 | 1.61391035 |
| 12157 | NM_024654.4 | #N/A | 496.892755 | 0.91728015 | 0.24102057 |
| 12158 | NM_001345880.1 | #N/A | 40.8680178 | -6.5603375 | 1.72435413 |
| 12159 | NM_001291324.1 | #N/A | 341.785717 | -3.7355637 | 0.98188781 |
| 12160 | NM_016331.2 | ZNF639 | 78.9368638 | -4.7875198 | 1.2584601 |
| 12161 | NM_018694.3 | ARL6IP4 | 31.7946884 | -6.2003469 | 1.62998793 |
| 12162 | NM_033157.3 | #N/A | 40.084187 | -6.5351896 | 1.71813237 |
| 12163 | NM_001145642.4 | RUBCN | 78.9439871 | -7.5135235 | 1.97560513 |
| 12164 | NR_109758.1 | LINC02315 | 36.4619303 | 2.93864384 | 0.77278926 |
| 12165 | NR_135312.1 | SMG1P6 | 75.2798614 | 2.93349132 | 0.7714522 |
| 12166 | NM_001204864.1 | ELOC | 19.8784979 | 8.06552461 | 2.12123168 |
| 12167 | NM_001322019.1 | NDUFA10 | 30.500617 | -6.1479966 | 1.61698085 |
| 12168 | NM_022469.3 | #N/A | 42.4474433 | -6.6173863 | 1.74055055 |

| | | | | | |
|---|---|---|---|---|---|
| 12169 | NM_006656.5 | #N/A | 28.6424045 | -6.052047 | 1.59186293 |
| 12170 | NM_006155.2 | SEPTIN2 | 30.6117738 | -6.1422085 | 1.61558902 |
| 12171 | NM_001123.3 | ADK | 30.4833807 | -6.1390633 | 1.61478731 |
| 12172 | NM_152328.4 | ADSS1 | 35.6904974 | -6.3725653 | 1.67627588 |
| 12173 | NM_001257413.1 | #N/A | 298.9892 | 2.54934658 | 0.67067974 |
| 12174 | NM_001322414.1 | #N/A | 37.2970399 | -6.4254281 | 1.69039083 |
| 12175 | NM_133444.2 | #N/A | 28.7981357 | -6.0663451 | 1.59618946 |
| 12176 | NM_001267727.1 | ARSG | 27.8038207 | -6.0096638 | 1.58136637 |
| 12177 | NR_038285.1 | MELTF-AS1 | 30.4253003 | -6.1370663 | 1.61499682 |
| 12178 | NM_017738.3 | #N/A | 34.7476811 | -5.0366298 | 1.32589987 |
| 12179 | NM_024871.2 | #N/A | 30.117146 | -6.1207539 | 1.61138889 |
| 12180 | NM_152478.2 | #N/A | 42.4934884 | -6.6177027 | 1.74222087 |
| 12181 | NM_001272013.1 | #N/A | 51.9857286 | -6.9141985 | 1.82030921 |
| 12182 | NR_015384.2 | LOC100126784 | 48.1187509 | -6.7983169 | 1.78981311 |
| 12183 | NM_001127400.1 | #N/A | 26.6537134 | -5.9460203 | 1.5655379 |
| 12186 | NR_110787.1 | DDX59-AS1 | 219.077367 | 3.25953986 | 0.85825669 |
| 12184 | NM_001122824.1 | #N/A | 51.2785607 | -3.5632041 | 0.93821421 |
| 12185 | NM_138794.4 | #N/A | 41.607727 | -6.5845304 | 1.73374472 |
| 12187 | NM_001204872.1 | #N/A | 32.2581926 | -6.2286237 | 1.64021509 |
| 12188 | NR_134302.1 | LOC101927752 | 29.5700254 | -6.093259 | 1.60477175 |
| 12189 | NM_001144823.2 | DENND4A | 55.1798648 | -6.9920765 | 1.84167307 |
| 12190 | NM_206839.2 | MORF4L1 | 48.9687866 | -6.8227069 | 1.79709983 |
| 12191 | NM_001166345.1 | MDFIC | 93.3092723 | -2.2275111 | 0.58674516 |
| 12194 | NM_019895.2 | CLDND1 | 51.1126959 | -6.8880244 | 1.8144142 |
| 12193 | NM_001324020.1 | PDXDC1 | 44.9979982 | -6.6971857 | 1.76413329 |
| 12192 | NM_001167607.1 | #N/A | 26.3949184 | 9.71397891 | 2.55880377 |
| 12195 | NM_152446.3 | #N/A | 25.3834944 | -5.8766489 | 1.54800906 |
| 12196 | NM_001289808.1 | #N/A | 39.4179175 | -6.5101335 | 1.71510635 |
| 12197 | NM_001162429.2 | PDCD6IP | 54.6336335 | -6.981284 | 1.83956696 |
| 12198 | NM_012471.2 | TRPC5 | 40.740407 | -6.560846 | 1.72900577 |
| 12199 | NM_001113567.2 | #N/A | 41.2472604 | -6.5713638 | 1.73183584 |
| 12200 | NM_021615.4 | #N/A | 303.020832 | 2.08870432 | 0.55048424 |
| 12201 | NM_001135055.2 | TKT | 39.901014 | -6.5265277 | 1.72012218 |
| 12202 | NM_020848.2 | #N/A | 65.6990894 | -4.4201062 | 1.16498845 |
| 12203 | NM_032373.4 | #N/A | 56.8832155 | -7.0385132 | 1.85511144 |
| 12204 | NM_001258038.1 | #N/A | 35.4026829 | -6.361385 | 1.67673181 |
| 12205 | NM_014984.3 | #N/A | 46.0550606 | -6.740691 | 1.77670028 |
| 12206 | NM_032857.4 | #N/A | 27.3047969 | -5.9812148 | 1.57655142 |
| 12207 | NR_024170.1 | LAMTOR3 | 46.9800393 | -6.7681535 | 1.78409439 |
| 12208 | NM_001320220.1 | #N/A | 67.3576171 | -7.2814549 | 1.91959049 |
| 12209 | NM_178006.3 | #N/A | 28.9904893 | -6.0701824 | 1.6003258 |
| 12210 | NM_016354.3 | #N/A | 28.1389208 | -6.0275935 | 1.58917353 |
| 12211 | NM_001330260.1 | #N/A | 50.2376367 | -6.8624702 | 1.80942915 |

| | | | | | |
|---|---|---|---|---|---|
| 12212 | NM_001243247.1 | NPRL3 | 41.1770587 | -6.5800451 | 1.73506555 |
| 12213 | NM_001308295.1 | #N/A | 43.3854171 | -6.6488008 | 1.75335276 |
| 12214 | NM_001130913.1 | ZNF720 | 31.6073657 | -6.1976043 | 1.63458046 |
| 12215 | NM_001270523.1 | #N/A | 71.5735572 | -7.3734903 | 1.94513286 |
| 12216 | NM_001286752.1 | #N/A | 26.1565093 | -5.9194996 | 1.56172723 |
| 12217 | NM_001286129.1 | #N/A | 26.7726438 | -5.9517502 | 1.57075095 |
| 12218 | NM_024514.4 | CYP2R1 | 31.7936309 | -6.1988973 | 1.63598622 |
| 12219 | NR_046438.2 | MEIS1-AS3 | 77.0216562 | -5.9373396 | 1.5671079 |
| 12220 | NM_001082480.2 | ZNF623 | 27.834581 | -6.0075773 | 1.58567902 |
| 12221 | NM_001103167.1 | ZGLP1 | 29.9963463 | -6.1179817 | 1.61492864 |
| 12222 | NM_018845.3 | #N/A | 29.6007334 | -6.0898176 | 1.60751679 |
| 12223 | NM_001030010.2 | #N/A | 27.2614177 | -5.9750182 | 1.57743871 |
| 12224 | NM_021082.3 | #N/A | 33.7945027 | -6.2937668 | 1.66167409 |
| 12225 | NM_018015.5 | #N/A | 34.0822616 | -6.3075504 | 1.66548178 |
| 12226 | NM_001163079.1 | #N/A | 37.1379445 | -6.4246643 | 1.69647622 |
| 12227 | NM_001943.4 | #N/A | 165.469918 | -1.5446266 | 0.40789359 |
| 12228 | NM_001203.2 | #N/A | 34.909635 | -6.3314798 | 1.67205501 |
| 12229 | NM_016196.3 | #N/A | 41.1886465 | -6.5720717 | 1.73582167 |
| 12230 | NM_001039888.3 | #N/A | 43.1909829 | -6.6413882 | 1.75425937 |
| 12231 | NM_022137.5 | #N/A | 63.7809555 | -7.2061903 | 1.90403028 |
| 12232 | NM_007117.4 | #N/A | 27.6189784 | -6.0016282 | 1.58579485 |
| 12233 | NM_181553.2 | #N/A | 106.334225 | -3.310522 | 0.8747638 |
| 12234 | NM_001330766.1 | ZNF415 | 31.8637833 | -6.2117833 | 1.64145407 |
| 12235 | NR_135823.1 | ZNF616 | 54.9938239 | -3.9388565 | 1.04085053 |
| 12236 | NM_001317991.1 | #N/A | 36.9331422 | -6.4182535 | 1.69616331 |
| 12237 | NM_001085447.1 | #N/A | 40.1872909 | -6.5419437 | 1.72894344 |
| 12238 | NM_152457.2 | #N/A | 36.8761361 | -6.4101152 | 1.69415834 |
| 12239 | NM_001289399.1 | SMARCA2 | 32.6531255 | -6.2465543 | 1.65103824 |
| 12240 | NM_001321187.1 | #N/A | 28.8237144 | -6.0614474 | 1.60213742 |
| 12241 | NM_003285.2 | #N/A | 42.981029 | -6.6366319 | 1.75422298 |
| 12242 | NM_145165.3 | CHURC1 | 33.3049111 | -6.2628293 | 1.65568632 |
| 12243 | NM_138966.3 | #N/A | 43.9555305 | -6.6684628 | 1.76295931 |
| 12244 | NR_027840.1 | #N/A | 32.4702144 | -6.236544 | 1.64890711 |
| 12245 | NR_131187.1 | LOC105376360 | 32.0633335 | 2.85293184 | 0.75431002 |
| 12246 | NM_001042473.3 | #N/A | 31.2845958 | -6.1797488 | 1.63403356 |
| 12247 | NM_031232.3 | #N/A | 30.5558267 | -6.1385374 | 1.6231778 |
| 12248 | NM_032789.4 | #N/A | 50.0786977 | -6.8589435 | 1.81393513 |
| 12249 | NM_003080.2 | #N/A | 25.3793546 | -5.8773737 | 1.55437521 |
| 12250 | NM_001319238.1 | ZC3H11A | 119.209818 | -5.2221849 | 1.3811807 |
| 12251 | NM_001008661.2 | #N/A | 31.6734508 | -6.1992132 | 1.63960458 |
| 12252 | NM_001348137.1 | CCNI | 69.1243067 | -7.3229899 | 1.93698351 |
| 12253 | NM_018380.3 | #N/A | 47.8165888 | -5.3396558 | 1.41242356 |
| 12254 | NM_001122838.1 | #N/A | 33.1557342 | -6.2535907 | 1.65446373 |

| | | | | | |
|---|---|---|---|---|---|
| 12255 | NR_033861.1 | LINC00514 | 80.3048682 | 2.09351 | 0.55389349 |
| 12256 | NM_025139.5 | #N/A | 38.2761256 | -6.4728552 | 1.71258762 |
| 12257 | NM_003070.4 | #N/A | 38.219224 | -6.4681723 | 1.71136408 |
| 12258 | NM_004562.2 | #N/A | 32.3751184 | -6.2224035 | 1.64694128 |
| 12259 | NM_001080449.2 | #N/A | 32.6591965 | -6.2381117 | 1.65110826 |
| 12260 | NM_014999.3 | #N/A | 335.046 | -2.8514239 | 0.75481556 |
| 12262 | NM_001163147.1 | ETV1 | 39.5883498 | -6.5191297 | 1.72574421 |
| 12261 | NM_001135095.1 | #N/A | 53.5822633 | -5.6853218 | 1.50501172 |
| 12263 | NM_000816.3 | GABRG2 | 57.8783187 | -7.0645849 | 1.87040855 |
| 12264 | NM_001243784.1 | PDE2A | 32.4281615 | -6.227383 | 1.64880026 |
| 12265 | NM_005159.4 | #N/A | 31.5709806 | -6.1954729 | 1.64045406 |
| 12266 | NM_017649.4 | #N/A | 42.3442124 | -6.6164837 | 1.75199797 |
| 12267 | NR_046106.1 | NOL8 | 27.1995275 | -5.9745809 | 1.58235252 |
| 12268 | NM_153015.2 | #N/A | 37.9697043 | -6.4535318 | 1.70923962 |
| 12269 | NM_001297652.1 | DDR1 | 31.0052258 | -6.1701727 | 1.63426811 |
| 12270 | NR_039979.1 | LINC00967 | 25.1003141 | 4.71663183 | 1.24930946 |
| 12271 | NM_052864.2 | #N/A | 33.0969225 | -6.2544367 | 1.65692717 |
| 12272 | NM_198282.3 | #N/A | 50.5199492 | -6.8721717 | 1.82060106 |
| 12273 | NM_004670.3 | #N/A | 31.2249774 | -6.181354 | 1.63765644 |
| 12274 | NM_144671.4 | #N/A | 62.168155 | -7.1680017 | 1.89935283 |
| 12275 | NM_001345876.1 | #N/A | 34.6778234 | -6.3293308 | 1.67743192 |
| 12276 | NM_176818.2 | #N/A | 249.046538 | -3.7514205 | 0.99427252 |
| 12277 | NM_015450.2 | #N/A | 34.1827553 | -6.307559 | 1.67182224 |
| 12278 | NM_001171134.2 | FAM3A | 25.9801301 | -5.9114033 | 1.56707594 |
| 12279 | NM_033083.6 | #N/A | 213.210058 | -2.4254688 | 0.64299292 |
| 12280 | NM_001037317.1 | #N/A | 46.7226433 | -6.7557097 | 1.79095376 |
| 12281 | NM_024672.5 | #N/A | 28.1963844 | -6.0222137 | 1.59681035 |
| 12282 | NM_001278214.1 | LRRC20 | 60.9246701 | -7.1364482 | 1.8922803 |
| 12283 | NM_015193.4 | #N/A | 56.6786614 | -7.0332017 | 1.86494231 |
| 12284 | NM_022746.3 | #N/A | 30.153969 | -6.1206624 | 1.62313161 |
| 12285 | NM_001318239.1 | ZNF821 | 30.1437851 | -6.1298261 | 1.62571954 |
| 12286 | NR_134983.1 | #N/A | 30.6581173 | -6.1561919 | 1.63274607 |
| 12287 | NM_001130683.3 | #N/A | 29.0947812 | -6.0769421 | 1.61178407 |
| 12288 | NM_001321547.1 | #N/A | 30.9050986 | -6.1558518 | 1.63272773 |
| 12289 | NM_001276282.3 | #N/A | 33.3275554 | -6.2630768 | 1.66133426 |
| 12290 | NM_001835.3 | #N/A | 30.0755351 | -6.1247817 | 1.62482257 |
| 12291 | NM_001142541.2 | SLC35B3 | 29.1444737 | -6.0676486 | 1.60968838 |
| 12292 | NM_001195263.1 | #N/A | 34.7732393 | -6.3302216 | 1.6795879 |
| 12293 | NM_016077.4 | PTRH2 | 35.5861394 | -6.3656216 | 1.68922411 |
| 12294 | NM_001278698.1 | #N/A | 28.6636097 | -6.0454101 | 1.60434061 |
| 12295 | NM_001308392.1 | #N/A | 47.9080432 | -6.790279 | 1.8021563 |
| 12296 | NM_005657.2 | #N/A | 64.4041853 | -7.2188435 | 1.91589997 |
| 12297 | NM_001321053.1 | #N/A | 22.5382442 | 2.61393484 | 0.69381951 |

| | | | | | |
|---|---|---|---|---|---|
| 12298 | NM_001286373.1 | TOMM40L | 31.4226478 | -6.1777382 | 1.63986808 |
| 12299 | NM_052845.3 | #N/A | 158.1247 | -2.7803857 | 0.73808958 |
| 12300 | NM_181711.3 | #N/A | 28.0871717 | -6.0174208 | 1.59778903 |
| 12301 | NM_001195131.1 | TSTD3 | 87.2799123 | 1.75978676 | 0.46730003 |
| 12302 | NM_001144936.1 | #N/A | 852.950761 | -4.3060925 | 1.14349027 |
| 12303 | NM_133503.3 | DCN | 48.5481178 | -6.8153245 | 1.80986963 |
| 12304 | NM_173039.2 | #N/A | 29.9710694 | -6.1142664 | 1.62391945 |
| 12305 | NM_001289061.1 | #N/A | 31.7035488 | -6.2014152 | 1.64716938 |
| 12306 | NM_145715.2 | TIGD2 | 25.3501946 | -5.8805965 | 1.56213607 |
| 12307 | NM_003745.1 | SOCS1 | 29.5818274 | -6.0971516 | 1.61981782 |
| 12308 | NM_017877.3 | #N/A | 283.274267 | -1.6134918 | 0.42867115 |
| 12309 | NM_001267042.2 | #N/A | 34.3667818 | -6.3065817 | 1.67555226 |
| 12310 | NM_002577.4 | PAK2 | 579.701066 | -3.4995019 | 0.92976687 |
| 12311 | NM_147168.1 | #N/A | 44.3233326 | -6.6830679 | 1.77578778 |
| 12312 | NM_032709.2 | #N/A | 25.8543929 | -5.9071411 | 1.56982052 |
| 12313 | NM_001282450.1 | ESRRA | 29.3445525 | -6.0851967 | 1.61733146 |
| 12314 | NM_033407.3 | DOCK7 | 26.950052 | -5.9628611 | 1.58484453 |
| 12315 | NM_016084.4 | #N/A | 42.3390322 | -6.6123142 | 1.75756877 |
| 12316 | NR_023314.4 | THTPA | 40.4068632 | -6.5439842 | 1.73966272 |
| 12317 | NR_046176.2 | #N/A | 29.9519057 | -6.1191018 | 1.62676046 |
| 12318 | NM_001166357.1 | SHMT2 | 48.8279768 | -6.8191537 | 1.81310781 |
| 12319 | NM_198315.2 | VWA5A | 26.4157793 | -5.9409389 | 1.57961443 |
| 12320 | NM_181726.3 | #N/A | 25.6309719 | -5.8911485 | 1.56649391 |
| 12321 | NM_001184766.1 | ODF2L | 25.9855782 | -5.9161101 | 1.57316741 |
| 12322 | NM_001253875.1 | UXS1 | 60.2737353 | -7.1191252 | 1.89307776 |
| 12323 | NM_004294.2 | #N/A | 26.477673 | -5.9381341 | 1.57907856 |
| 12324 | NM_080676.5 | #N/A | 34.7453152 | -6.3282559 | 1.68291886 |
| 12325 | NM_177559.2 | #N/A | 25.392336 | -5.8737373 | 1.56217999 |
| 12326 | NM_183372.5 | #N/A | 28.2374973 | -6.0240511 | 1.60233258 |
| 12327 | NM_178169.3 | #N/A | 39.9407303 | -6.5342775 | 1.73810576 |
| 12328 | NM_017613.3 | #N/A | 117.474086 | -3.1091801 | 0.82712441 |
| 12329 | NR_002781.1 | TSPY26P | 27.336269 | -5.9813112 | 1.59125474 |
| 12330 | NM_001289045.1 | #N/A | 26.6306339 | -5.9488952 | 1.58275423 |
| 12331 | NM_012109.2 | #N/A | 702.568739 | -4.5578312 | 1.21264814 |
| 12332 | NR_027487.2 | ARHGAP27P1 | 29.8956798 | -6.1128912 | 1.62659641 |
| 12334 | NR_015380.2 | A1BG-AS1 | 25.7782358 | -5.8967877 | 1.56933932 |
| 12333 | NM_020800.2 | #N/A | 50.5313139 | -6.8735132 | 1.82927509 |
| 12335 | NM_012261.3 | #N/A | 26.5433331 | -5.9434409 | 1.58183597 |
| 12336 | NM_000082.3 | ERCC8 | 34.3597186 | -6.3136111 | 1.68038728 |
| 12337 | NM_001048265.1 | #N/A | 28.7054513 | -6.0608045 | 1.61311422 |
| 12338 | NM_003260.4 | #N/A | 45.0611446 | -6.698402 | 1.78293057 |
| 12339 | NM_153759.3 | DNMT3A | 129.080375 | -3.0669171 | 0.8165163 |
| 12340 | NM_001282755.1 | MTA3 | 26.0953993 | -5.9174139 | 1.575512 |

| | | | | | |
|---|---|---|---|---|---|
| 12341 | NM_006509.3 | #N/A | 30.2052984 | -6.1345728 | 1.63332252 |
| 12342 | NM_001013437.1 | #N/A | 32.7146207 | -6.2424114 | 1.66219725 |
| 12343 | NM_001318054.1 | #N/A | 24.6265122 | -5.8313237 | 1.55272789 |
| 12344 | NM_001171025.1 | PTPRD | 42.1656305 | -6.6131817 | 1.76100482 |
| 12345 | NR_034107.2 | LOC257396 | 52.660279 | 2.49667257 | 0.66486554 |
| 12346 | NM_001276336.1 | EIF4E2 | 54.3841197 | -6.9704236 | 1.85636992 |
| 12347 | NR_027149.1 | #N/A | 28.9943919 | -6.066258 | 1.61571532 |
| 12348 | NM_022097.3 | #N/A | 27.6378995 | 3.71728074 | 0.99010067 |
| 12349 | NM_014619.4 | GRIK4 | 29.5969083 | -6.1047533 | 1.62606761 |
| 12351 | NM_032823.5 | AOPEP | 29.1955946 | -6.0808936 | 1.61987911 |
| 12350 | NM_018348.5 | #N/A | 26.3763355 | -5.9298182 | 1.57963722 |
| 12352 | NM_001105207.2 | #N/A | 42.4482118 | -6.6187935 | 1.76318613 |
| 12353 | NM_001262.2 | CDKN2C | 31.5866692 | -6.1911139 | 1.6496891 |
| 12354 | NR_034132.1 | LINC00939 | 26.0559366 | 3.6358152 | 0.96882133 |
| 12355 | NM_001331010.1 | #N/A | 37.6177757 | -6.4448807 | 1.71735373 |
| 12356 | NM_005026.3 | #N/A | 130.962534 | -2.2776663 | 0.60695532 |
| 12357 | NM_032440.3 | #N/A | 64.816827 | -7.230187 | 1.92676705 |
| 12358 | NR_103729.1 | #N/A | 47.7242364 | -6.7827778 | 1.80766361 |
| 12359 | NM_001098510.1 | PELI3 | 28.8310207 | -6.0546902 | 1.61368809 |
| 12360 | NM_001281504.1 | #N/A | 30.907952 | -6.1522156 | 1.63982102 |
| 12361 | NM_015633.2 | #N/A | 34.1530108 | -6.2995707 | 1.67927872 |
| 12362 | NM_001301871.1 | #N/A | 41.1113457 | -6.5738387 | 1.75253269 |
| 12363 | NR_136405.1 | #N/A | 31.8477642 | -6.2115212 | 1.65597501 |
| 12364 | NM_006859.3 | #N/A | 27.0183259 | -5.9687487 | 1.59127154 |
| 12365 | NM_152402.2 | #N/A | 31.452008 | -6.180071 | 1.64771253 |
| 12366 | NM_001077490.2 | GNAS | 31.4895502 | -6.1920767 | 1.65101644 |
| 12367 | NM_033331.2 | #N/A | 40.3308193 | -6.5449632 | 1.74522773 |
| 12368 | NM_001159279.1 | ZNF716 | 162.792768 | 3.16699248 | 0.84450073 |
| 12369 | NM_000159.3 | #N/A | 36.4419584 | -6.3970553 | 1.70592273 |
| 12370 | NR_126510.1 | #N/A | 35.2290294 | -6.3477835 | 1.6927988 |
| 12371 | NM_052969.1 | #N/A | 36.801242 | -6.4095868 | 1.7093621 |
| 12372 | NR_136651.1 | #N/A | 29.5524095 | -6.0976124 | 1.62626209 |
| 12373 | NR_135601.1 | #N/A | 13.6452815 | 8.73516695 | 2.32981445 |
| 12374 | NM_001003788.2 | #N/A | 24.7673519 | -5.8392899 | 1.55771707 |
| 12375 | NR_134535.1 | PHF21B | 34.5583476 | -6.3248839 | 1.68743113 |
| 12376 | NM_001267606.1 | ZNF248 | 26.2173964 | -5.9216657 | 1.57991694 |
| 12377 | NM_004983.2 | #N/A | 51.7023372 | -6.8999779 | 1.84093888 |
| 12378 | NM_152912.4 | MTIF3 | 29.7555579 | -6.1106812 | 1.63043541 |
| 12379 | NM_032532.2 | #N/A | 77.258839 | -3.7805489 | 1.00878317 |
| 12380 | NM_152730.5 | #N/A | 107.13947 | 1.84635121 | 0.49268433 |
| 12381 | NM_002143.2 | #N/A | 75.7311746 | -7.4520118 | 1.98858965 |
| 12382 | NM_001287429.1 | #N/A | 43.4949129 | -6.6552337 | 1.7760395 |
| 12383 | NR_026551.1 | #N/A | 34.1777176 | -6.3021394 | 1.68206032 |

| | | | | | |
|---|---|---|---|---|---|
| 12384 | NR_046418.2 | #N/A | 25.495049 | -5.8765946 | 1.56848587 |
| 12385 | NM_001141945.2 | ACTA2 | 34.4975696 | -6.3246683 | 1.68815725 |
| 12386 | NM_024321.4 | #N/A | 107.073228 | -3.1264892 | 0.83455555 |
| 12387 | NM_004654.3 | #N/A | 40.8576476 | -6.5649151 | 1.75247203 |
| 12388 | NM_015969.2 | #N/A | 36.0060693 | -6.3808409 | 1.70335667 |
| 12389 | NM_007008.2 | #N/A | 28.7650121 | -6.0609087 | 1.61797413 |
| 12390 | NM_015604.3 | #N/A | 36.1283724 | -6.3811841 | 1.70347781 |
| 12391 | NM_001040667.2 | #N/A | 41.8157829 | -6.6005149 | 1.762168 |
| 12392 | NM_198353.2 | #N/A | 27.7540847 | -6.0090412 | 1.6043826 |
| 12393 | NM_001040616.2 | #N/A | 30.0177361 | -6.1172284 | 1.63342638 |
| 12394 | NR_104624.1 | LINC01647 | 48.4437942 | 4.91039425 | 1.31127557 |
| 12395 | NM_005138.2 | SCO2 | 25.8375235 | -5.9073188 | 1.57767177 |
| 12396 | NM_145065.2 | #N/A | 30.1241554 | -6.1281343 | 1.63686017 |
| 12397 | NM_020714.2 | #N/A | 245.25649 | 1.20492499 | 0.32184838 |
| 12398 | NM_022113.5 | #N/A | 26.349697 | -5.9374535 | 1.58597322 |
| 12399 | NM_007098.3 | #N/A | 30.2901653 | -6.1310199 | 1.63771446 |
| 12400 | NR_046329.1 | #N/A | 35.6151366 | -6.3677331 | 1.70122786 |
| 12401 | NM_020528.2 | PCBP3 | 38.1829111 | -6.4665871 | 1.72765752 |
| 12402 | NR_033519.1 | #N/A | 39.692711 | -6.5163056 | 1.7410042 |
| 12404 | NR_003262.1 | FDPSP2 | 153.664329 | 2.5257526 | 0.67483644 |
| 12403 | NM_001039476.2 | NPRL3 | 48.9487008 | -6.8233755 | 1.82308121 |
| 12405 | NM_182524.3 | #N/A | 33.7976927 | -6.2818461 | 1.67850245 |
| 12406 | NM_001142425.1 | MORF4L2 | 47.0140571 | -6.7692686 | 1.80893621 |
| 12407 | NM_012331.4 | #N/A | 28.0299331 | -6.0223654 | 1.60940184 |
| 12408 | NM_001290295.1 | #N/A | 35.8273108 | -6.3726612 | 1.70318392 |
| 12409 | NM_001320042.1 | ISCU | 26.1405417 | -5.9122726 | 1.58020097 |
| 12410 | NM_000361.2 | #N/A | 25.2731736 | -5.8722493 | 1.56970801 |
| 12411 | NM_033132.3 | #N/A | 56.2466542 | -7.0263989 | 1.87828244 |
| 12412 | NM_005773.2 | #N/A | 27.6938315 | -5.9990727 | 1.60379899 |
| 12413 | NM_138393.2 | #N/A | 28.0636374 | -6.0286288 | 1.6117394 |
| 12414 | NM_031485.3 | #N/A | 131.894957 | -3.3109796 | 0.88520397 |
| 12415 | NR_109846.1 | KCNA3 | 31.521808 | -6.1839268 | 1.65337527 |
| 12416 | NM_001177465.2 | #N/A | 29.7858313 | -6.0995431 | 1.63109317 |
| 12417 | NM_001007248.2 | #N/A | 25.0476646 | -5.8589553 | 1.56677025 |
| 12418 | NM_001284431.1 | UTP15 | 24.6774048 | 9.61752186 | 2.57197593 |
| 12419 | NM_020317.4 | #N/A | 28.3739068 | -6.0400249 | 1.61544547 |
| 12420 | NR_049753.2 | #N/A | 28.6041117 | -6.0553997 | 1.61976933 |
| 12421 | NM_006936.2 | #N/A | 427.616683 | -3.8529017 | 1.03064663 |
| 12422 | NM_170725.2 | #N/A | 24.4764328 | -5.828826 | 1.55937794 |
| 12423 | NM_001308237.1 | ZZZ3 | 25.8078041 | -5.8968634 | 1.5776547 |
| 12424 | NM_001243749.1 | CYREN | 26.3003186 | -5.930927 | 1.58693078 |
| 12425 | NR_002165.1 | HMGB3P1 | 64.6549963 | 2.23511001 | 0.59807129 |
| 12426 | NM_001145850.1 | #N/A | 24.4374292 | -5.8259013 | 1.55892156 |

| | | | | | |
|---|---|---|---|---|---|
| 12427 | NM_002699.3 | #N/A | 39.2419757 | -6.5012031 | 1.73966596 |
| 12428 | NM_006980.4 | #N/A | 26.5028369 | -5.9397434 | 1.58942689 |
| 12429 | NM_001256455.1 | ZBTB7B | 25.684759 | -5.8975933 | 1.57829164 |
| 12432 | NR_120332.1 | EHD4-AS1 | 28.0670267 | 3.94914003 | 1.05699813 |
| 12431 | NR_033908.1 | LOC100288069 | 39.4969728 | -5.2388407 | 1.40218613 |
| 12430 | NM_182572.3 | #N/A | 24.0646749 | -5.7986444 | 1.55202039 |
| 12433 | NR_120346.1 | THSD4-AS2 | 27.8966496 | 3.79534444 | 1.01588909 |
| 12434 | NM_001100389.1 | #N/A | 138.169068 | -2.0528615 | 0.54951898 |
| 12435 | NM_001077692.1 | #N/A | 26.5051651 | -5.9350832 | 1.58893456 |
| 12436 | NM_152245.2 | #N/A | 35.8283947 | -6.3764077 | 1.70722569 |
| 12437 | NM_178123.4 | #N/A | 441.973616 | -3.8735479 | 1.03715593 |
| 12438 | NM_145008.2 | #N/A | 29.9234332 | -6.1088329 | 1.63569627 |
| 12439 | NM_004856.6 | #N/A | 32.87804 | -6.2552713 | 1.67494656 |
| 12440 | NM_001318794.1 | #N/A | 30.0741167 | -6.1246225 | 1.63997988 |
| 12441 | NM_199298.1 | #N/A | 37.2054555 | -6.4258836 | 1.72084414 |
| 12442 | NM_013441.3 | #N/A | 26.5862317 | -5.9448211 | 1.59208358 |
| 12443 | NM_022572.4 | PNKD | 32.2438585 | -6.2270682 | 1.66774046 |
| 12444 | NR_135527.1 | DLG4 | 34.1273733 | -5.0156167 | 1.34344383 |
| 12445 | NM_001257971.1 | #N/A | 30.3768208 | -6.1374508 | 1.64399172 |
| 12446 | NM_001318915.1 | #N/A | 39.8658656 | -6.525215 | 1.74791574 |
| 12447 | NM_001536.5 | PRMT1 | 34.9953422 | -6.3359094 | 1.6974001 |
| 12448 | NM_001319106.1 | OPCML | 44.3872102 | -6.6825681 | 1.7904731 |
| 12449 | NM_005541.4 | #N/A | 51.101665 | -6.8885375 | 1.84568223 |
| 12450 | NM_145291.3 | #N/A | 27.0420722 | -5.9662534 | 1.59860326 |
| 12451 | NM_001010852.3 | #N/A | 74.035461 | -4.8914803 | 1.31069974 |
| 12452 | NM_133460.2 | #N/A | 27.3831576 | -5.9785487 | 1.60198055 |
| 12453 | NM_198597.2 | #N/A | 493.556558 | -4.7921726 | 1.28408265 |
| 12454 | NM_001271816.1 | #N/A | 32.9962945 | -6.2468641 | 1.67407864 |
| 12455 | NM_000245.3 | #N/A | 36.4117337 | -6.3979698 | 1.71465345 |
| 12456 | NM_001172303.2 | #N/A | 36.3912594 | -6.3951429 | 1.71415434 |
| 12457 | NM_198892.1 | BMP2K | 27.2841125 | -5.975389 | 1.60170125 |
| 12458 | NM_000520.5 | #N/A | 262.391459 | -4.6673336 | 1.25108906 |
| 12459 | NR_033328.1 | #N/A | 40.9500183 | -6.5647792 | 1.75971266 |
| 12460 | NR_038118.1 | #N/A | 24.6303846 | -5.8369595 | 1.56463534 |
| 12461 | NM_001171168.1 | #N/A | 34.3920309 | -6.3083482 | 1.69113503 |
| 12462 | NM_016060.2 | #N/A | 30.7047546 | -6.1498872 | 1.64877962 |
| 12463 | NM_001008223.1 | #N/A | 36.6141039 | -6.4071532 | 1.71780979 |
| 12464 | NM_001130997.2 | #N/A | 31.0875348 | -6.1691797 | 1.65421274 |
| 12465 | NM_033224.4 | #N/A | 311.2342 | -2.5337397 | 0.67943758 |
| 12466 | NR_015382.2 | CD27-AS1 | 81.455355 | -4.7061141 | 1.2619896 |
| 12467 | NM_006497.3 | #N/A | 37.47712 | -6.4436511 | 1.72793893 |
| 12468 | NM_201266.1 | #N/A | 46.0033831 | -6.7358999 | 1.80635147 |
| 12469 | NM_001142702.1 | DLG2 | 72.4468466 | -7.3871171 | 1.98116424 |

| | | | | | |
|---|---|---|---|---|---|
| 12470 | NM_173690.4 | #N/A | 90.0846384 | -7.7021554 | 2.06606858 |
| 12471 | NM_002555.5 | #N/A | 31.9451329 | -6.2105035 | 1.66600518 |
| 12472 | NM_006019.3 | #N/A | 31.862044 | -6.2090794 | 1.66567864 |
| 12473 | NM_001033555.2 | #N/A | 63.8334363 | -5.9368884 | 1.59282382 |
| 12474 | NM_000944.4 | #N/A | 86.8913852 | -7.650693 | 2.05267317 |
| 12475 | NM_001278438.1 | FNDC3A | 35.3162037 | -6.3475589 | 1.70327985 |
| 12476 | NM_001318339.1 | #N/A | 89.2419256 | -7.6876148 | 2.06291586 |
| 12477 | NM_014229.2 | #N/A | 31.5704043 | -6.1876709 | 1.66060576 |
| 12478 | NR_024159.1 | DGCR9 | 27.2809312 | -5.9804632 | 1.60505932 |
| 12479 | NM_022483.4 | #N/A | 38.6684339 | -5.2018225 | 1.39619755 |
| 12480 | NM_031207.5 | #N/A | 34.353321 | -6.3089014 | 1.69341265 |
| 12481 | NM_000966.5 | #N/A | 29.1542948 | -6.0826645 | 1.63310146 |
| 12482 | NM_018393.3 | #N/A | 41.9823435 | -6.5988862 | 1.77182044 |
| 12483 | NM_001330234.1 | #N/A | 27.1814501 | -5.9693042 | 1.60280018 |
| 12484 | NM_001024457.3 | RGPD1 | 73.1597174 | -7.4017079 | 1.98742862 |
| 12485 | NM_207343.3 | #N/A | 50.8197646 | -6.8744795 | 1.84586856 |
| 12486 | NM_052943.3 | #N/A | 26.4456073 | -5.9436968 | 1.59595831 |
| 12487 | NM_176880.5 | #N/A | 36.1783643 | -6.3859661 | 1.71473589 |
| 12488 | NM_001164283.2 | #N/A | 278.065232 | 2.42398044 | 0.65090756 |
| 12489 | NM_001318083.1 | #N/A | 24.7838278 | -5.8396264 | 1.56831444 |
| 12490 | NR_033998.1 | LOC729970 | 27.7685169 | -6.0114228 | 1.61466812 |
| 12491 | NM_020770.2 | #N/A | 26.3628766 | -5.9383693 | 1.59525769 |
| 12492 | NM_001075098.3 | #N/A | 25.6456027 | -5.8933902 | 1.58327584 |
| 12493 | NM_016027.2 | #N/A | 25.4294335 | -5.8799103 | 1.57965364 |
| 12494 | NM_001195608.1 | PLEKHA1 | 34.06361 | -6.2948862 | 1.69149025 |
| 12495 | NM_024817.2 | THSD4 | 26.1139676 | -5.9124497 | 1.58900833 |
| 12496 | NR_024247.1 | #N/A | 36.3629418 | -6.3935777 | 1.71855678 |
| 12497 | NM_004769.3 | #N/A | 28.9415714 | -6.0590385 | 1.62865828 |
| 12498 | NM_001308117.1 | RBFOX1 | 30.051163 | -6.1218679 | 1.64591914 |
| 12499 | NM_173810.3 | #N/A | 29.9862101 | -6.1145828 | 1.64404323 |
| 12500 | NM_001025247.1 | #N/A | 25.7834748 | -5.8982277 | 1.58606065 |
| 12501 | NM_001031746.4 | #N/A | 72.8934898 | -3.7144915 | 0.99884308 |
| 12502 | NM_138771.3 | #N/A | 34.6674042 | -6.3246004 | 1.70074091 |
| 12503 | NM_001198805.1 | MSANTD3 | 28.0763403 | -6.0217456 | 1.61933844 |
| 12504 | NM_001204159.2 | #N/A | 45.971091 | -6.7362704 | 1.81156732 |
| 12505 | NM_019100.4 | #N/A | 29.2518493 | -6.0860076 | 1.63672248 |
| 12506 | NR_015381.1 | TP53TG1 | 27.6299188 | -5.9998144 | 1.6135832 |
| 12507 | NM_025211.3 | #N/A | 34.4654751 | -6.3103679 | 1.69712894 |
| 12508 | NR_108072.1 | #N/A | 27.560675 | -5.9892964 | 1.61089869 |
| 12509 | NM_138720.2 | H2BC5 | 26.5107221 | -5.9437367 | 1.59884035 |
| 12510 | NM_001172713.1 | #N/A | 119.92754 | -2.9921239 | 0.80489033 |
| 12511 | NM_001201480.1 | #N/A | 32.7996604 | -6.2493236 | 1.6811767 |
| 12512 | NM_001242483.1 | #N/A | 35.898254 | -6.3807419 | 1.71681791 |

| | | | | | |
|---|---|---|---|---|---|
| 12513 | NM_014857.4 | RABGAP1L | 24.7652121 | -5.83921 | 1.57120973 |
| 12514 | NM_001164278.1 | #N/A | 31.8800775 | -6.2083581 | 1.6706154 |
| 12515 | NM_001083619.1 | GRIA2 | 132.061753 | -5.3162067 | 1.43067673 |
| 12516 | NM_001109938.2 | #N/A | 30.0170315 | -6.1193221 | 1.646885 |
| 12517 | NM_001320147.1 | #N/A | 251.599372 | -5.4245732 | 1.4600108 |
| 12518 | NM_003706.2 | #N/A | 28.8848403 | -6.0614537 | 1.63151737 |
| 12519 | NM_199076.2 | #N/A | 43.4469797 | -5.206393 | 1.40143168 |
| 12521 | NM_032534.3 | KRBA1 | 29.835554 | -6.1155198 | 1.64626676 |
| 12520 | NM_001128914.1 | PCBP2 | 463.61131 | -4.0129935 | 1.08027579 |
| 12522 | NM_152686.3 | #N/A | 264.837559 | -2.9687024 | 0.79919105 |
| 12523 | NM_001297429.1 | #N/A | 16.7466427 | 9.02628296 | 2.43001746 |
| 12524 | NR_003255.2 | TSIX | 445.75297 | 1.75984583 | 0.47378892 |
| 12525 | NM_001301243.1 | #N/A | 32.517087 | -6.2267647 | 1.67644493 |
| 12526 | NM_001146082.2 | #N/A | 25.4727325 | -5.8795906 | 1.58310873 |
| 12527 | NM_018502.4 | TMCO6 | 30.6948377 | -6.1441824 | 1.65438741 |
| 12528 | NM_000465.3 | #N/A | 28.2515785 | -6.0217959 | 1.62151422 |
| 12529 | NR_073536.1 | SHANK2-AS3 | 63.0709636 | 2.58747741 | 0.69676854 |
| 12530 | NM_004425.3 | #N/A | 28.4835705 | -6.0423083 | 1.62740704 |
| 12532 | NR_033248.1 | GCSHP3 | 29.8638872 | -6.110043 | 1.64568246 |
| 12531 | NM_007215.3 | #N/A | 35.494246 | -4.9492103 | 1.333021 |
| 12533 | NM_031922.4 | REPS1 | 57.0889851 | -7.0405865 | 1.89634774 |
| 12534 | NM_001170640.1 | CDK20 | 24.7790093 | -5.8474722 | 1.575098 |
| 12535 | NM_001321351.1 | #N/A | 45.2938768 | -6.708448 | 1.8070088 |
| 12536 | NM_001242795.1 | #N/A | 70.4143532 | -3.6288479 | 0.97752247 |
| 12537 | NM_000478.5 | #N/A | 25.4685167 | -5.8889971 | 1.58654702 |
| 12538 | NM_021956.4 | GRIK2 | 44.1913659 | -6.6788803 | 1.7993791 |
| 12539 | NM_152432.2 | #N/A | 31.9466218 | -4.9245101 | 1.32677575 |
| 12540 | NM_004915.3 | ABCG1 | 28.9790431 | -6.0668025 | 1.63471552 |
| 12541 | NM_014711.5 | #N/A | 30.1914961 | -6.1189826 | 1.64884905 |
| 12542 | NM_001305051.1 | TMEM59 | 61.3639189 | -7.1487996 | 1.92662503 |
| 12543 | NM_007038.4 | #N/A | 37.0212255 | -4.3413035 | 1.1700535 |
| 12544 | NM_018672.4 | ABCA5 | 31.7146979 | -6.1922934 | 1.66903907 |
| 12545 | NM_013377.3 | #N/A | 25.6559286 | -5.8908552 | 1.5878136 |
| 12546 | NM_013259.2 | TAGLN3 | 40.947624 | -6.5656079 | 1.77028362 |
| 12547 | NM_001348092.1 | #N/A | 23.818648 | 9.56674669 | 2.579519 |
| 12548 | NM_019111.4 | #N/A | 624.761479 | -5.3208143 | 1.43473452 |
| 12549 | NM_001184991.1 | NDUFC1 | 37.3930855 | -6.4315032 | 1.73443987 |
| 12550 | NM_201554.1 | DGKA | 32.1610081 | -6.2132359 | 1.67558895 |
| 12552 | NR_039985.1 | FLJ22447 | 8.57924599 | 6.36662406 | 1.7169807 |
| 12551 | NM_033363.1 | MRPS12 | 31.5300361 | -6.1931029 | 1.67017707 |
| 12553 | NM_024039.2 | MIS12 | 29.521471 | -6.0954359 | 1.64391624 |
| 12554 | NM_000329.2 | #N/A | 41.6465242 | -6.5943996 | 1.77852715 |
| 12555 | NM_138467.2 | #N/A | 26.3555532 | -5.9330519 | 1.60023829 |

| | | | | | |
|---|---|---|---|---|---|
| 12556 | NM_001037536.2 | #N/A | 39.5534854 | -6.5180927 | 1.75821914 |
| 12557 | NM_001099334.2 | #N/A | 44.1861383 | -6.674763 | 1.80047429 |
| 12558 | NM_012135.2 | #N/A | 27.281931 | -5.9764382 | 1.61209801 |
| 12559 | NM_001173524.1 | #N/A | 23.6767096 | -5.7756771 | 1.55807128 |
| 12560 | NM_053013.3 | #N/A | 24.2191265 | -5.8109436 | 1.56786651 |
| 12561 | NM_001297712.1 | #N/A | 22.829946 | 9.47699097 | 2.55717498 |
| 12562 | NM_001168320.1 | TSPAN9 | 93.9330667 | -5.3754093 | 1.45050905 |
| 12563 | NM_001307981.1 | DNAJC4 | 24.1496517 | -5.8097904 | 1.56786989 |
| 12564 | NM_052865.3 | #N/A | 36.6055134 | -6.4037414 | 1.72817876 |
| 12565 | NM_001288666.1 | HCFC1R1 | 44.2435527 | -6.6797729 | 1.80268358 |
| 12566 | NR_037945.1 | STX16-NPEPL1 | 30.397052 | -6.1305406 | 1.65448714 |
| 12567 | NM_018295.4 | #N/A | 25.8209173 | -5.9037369 | 1.59330806 |
| 12568 | NM_001098671.1 | #N/A | 30.8377617 | -6.1531081 | 1.66079416 |
| 12569 | NM_003202.4 | #N/A | 37.036686 | -6.4249988 | 1.73419283 |
| 12570 | NM_002123.4 | #N/A | 38.5037617 | -6.4791492 | 1.748893 |
| 12571 | NM_005575.2 | #N/A | 26.814467 | -5.9550039 | 1.60759215 |
| 12572 | NM_022895.2 | #N/A | 53.9232505 | -6.9599978 | 1.87896271 |
| 12573 | NM_032792.3 | #N/A | 27.8748739 | -6.0125517 | 1.62330801 |
| 12574 | NM_001300847.1 | #N/A | 131.980413 | -4.6262509 | 1.24921481 |
| 12575 | NM_014714.3 | #N/A | 321.499304 | -4.6483532 | 1.25517784 |
| 12576 | NM_001301097.1 | KIRREL3 | 58.8688788 | -7.0862303 | 1.91363652 |
| 12577 | NM_001242823.2 | C4B_2 | 94.4458145 | -7.7732208 | 2.09922579 |
| 12579 | NR_037868.1 | SLFNL1-AS1 | 167.790506 | 1.96404415 | 0.53042351 |
| 12578 | NM_006343.2 | #N/A | 35.4414984 | -6.3622678 | 1.71822934 |
| 12580 | NR_040586.1 | STAG3L4 | 37.7800802 | -6.4523134 | 1.74259324 |
| 12581 | NM_207310.2 | #N/A | 32.0594165 | -6.2126006 | 1.67791497 |
| 12582 | NM_001008225.2 | CNOT4 | 24.8594157 | -5.8446284 | 1.57892238 |
| 12583 | NM_001003698.3 | #N/A | 30.5791298 | -6.1475259 | 1.66085458 |
| 12584 | NM_005258.2 | #N/A | 25.4845249 | -5.8884388 | 1.59096948 |
| 12585 | NM_194283.3 | #N/A | 29.7638827 | -6.0973003 | 1.64746247 |
| 12586 | NM_001405.3 | #N/A | 671.373919 | 3.43403124 | 0.92787221 |
| 12587 | NR_033379.1 | LINC01623 | 21.5504685 | 4.80666322 | 1.29876846 |
| 12588 | NM_004460.4 | #N/A | 28.9658487 | -6.0726826 | 1.64087128 |
| 12589 | NR_109790.1 | RAB24 | 45.480808 | -6.713589 | 1.81409005 |
| 12590 | NM_152832.2 | #N/A | 43.139159 | -6.6412339 | 1.79465069 |
| 12591 | NM_001257281.1 | #N/A | 29.0644735 | -6.071625 | 1.64079833 |
| 12592 | NR_036641.1 | #N/A | 35.4614647 | -6.3646466 | 1.71998629 |
| 12593 | NM_001018070.2 | CORO1B | 55.906182 | -7.0135207 | 1.8956377 |
| 12594 | NR_125916.1 | BHLHE40-AS1 | 68.2282529 | 2.99395124 | 0.8092509 |
| 12595 | NM_001289978.1 | DPF1 | 44.5669062 | -6.6870017 | 1.80783589 |
| 12596 | NM_058216.2 | #N/A | 31.0057514 | -6.1628297 | 1.66612889 |
| 12597 | NM_001320925.2 | NAA38 | 33.6517181 | -6.2779672 | 1.69726583 |
| 12598 | NM_001011724.1 | #N/A | 31.8256632 | -6.1983556 | 1.67587281 |

| | | | | | |
|---|---|---|---|---|---|
| 12599 | NR_103535.1 | NEXN-AS1 | 109.491619 | 3.71000904 | 1.00310088 |
| 12600 | NM_006520.2 | #N/A | 74.1275926 | -4.6821446 | 1.26596375 |
| 12601 | NM_006258.3 | #N/A | 24.6680195 | -5.8414125 | 1.57943673 |
| 12602 | NM_001017370.2 | NHLRC3 | 26.652081 | -5.9445166 | 1.60735429 |
| 12603 | NM_001198534.1 | OXR1 | 44.8077909 | -6.69377 | 1.81012458 |
| 12604 | NM_001134335.1 | EIF2AK1 | 76.19896 | -4.7112349 | 1.27401887 |
| 12605 | NM_000411.6 | #N/A | 33.724853 | -6.2903702 | 1.70110133 |
| 12606 | NM_001168407.1 | #N/A | 28.7439165 | -6.0580997 | 1.63835647 |
| 12607 | NR_123713.1 | RAD52 | 28.2753114 | -6.0300157 | 1.63081174 |
| 12608 | NR_026680.3 | LINC02210 | 33.0462174 | -6.2557004 | 1.69197816 |
| 12609 | NM_014234.4 | #N/A | 23.7586069 | -5.7868087 | 1.56518204 |
| 12610 | NM_001318736.1 | CCDC82 | 30.3438884 | -6.1417392 | 1.66124577 |
| 12611 | NM_003781.3 | #N/A | 34.5437568 | -6.3163936 | 1.708555 |
| 12612 | NM_001001992.1 | USP16 | 75.1450892 | -4.6950457 | 1.27004678 |
| 12613 | NM_001276294.1 | #N/A | 34.7963791 | -6.3381222 | 1.71477073 |
| 12614 | NM_007210.3 | #N/A | 106.020377 | 3.13958799 | 0.84943131 |
| 12615 | NM_000689.4 | #N/A | 42.5003921 | -6.6229244 | 1.79188357 |
| 12616 | NM_001199013.1 | #N/A | 33.4940109 | -6.2787335 | 1.69890184 |
| 12617 | NR_003227.1 | AFG3L1P | 40.7429397 | -6.5567074 | 1.77417369 |
| 12618 | NM_001134423.2 | CDV3 | 41.9312452 | -6.6043746 | 1.78713903 |
| 12619 | NM_198535.2 | #N/A | 232.616613 | 1.78953521 | 0.48427305 |
| 12620 | NM_004203.4 | #N/A | 36.1172206 | -6.3816126 | 1.72714082 |
| 12621 | NM_016281.3 | #N/A | 30.2236984 | -6.1209927 | 1.65708726 |
| 12622 | NM_001011546.1 | #N/A | 37.7819022 | -6.4501734 | 1.74651637 |
| 12623 | NM_201555.1 | #N/A | 40.7803781 | -3.928979 | 1.06389252 |
| 12624 | NM_001118885.1 | GLRA2 | 50.2354496 | -6.8617324 | 1.85813084 |
| 12625 | NM_020805.2 | #N/A | 46.5010985 | -6.7462853 | 1.82686391 |
| 12626 | NM_214462.4 | #N/A | 28.8030049 | -6.0661811 | 1.6428073 |
| 12627 | NM_001101672.1 | #N/A | 28.9004588 | -6.0643633 | 1.64236279 |
| 12628 | NM_001270433.1 | #N/A | 27.8302028 | -6.0169227 | 1.62958879 |
| 12629 | NM_020734.3 | #N/A | 43.523171 | -6.6547395 | 1.8023431 |
| 12630 | NR_029377.1 | #N/A | 38.0283318 | -6.4550567 | 1.74826319 |
| 12631 | NM_212476.2 | FN1 | 501.327738 | -10.180181 | 2.75741231 |
| 12632 | NM_001007794.2 | CEPT1 | 27.3875755 | -5.9945035 | 1.62376051 |
| 12633 | NR_015394.1 | LINC00636 | 35.9400446 | 8.47120394 | 2.29462792 |
| 12634 | NM_001204082.1 | MICOS10 | 70.6354958 | 5.13136475 | 1.39001925 |
| 12635 | NM_001304961.1 | #N/A | 49.7822954 | -6.8457578 | 1.85449833 |
| 12636 | NM_001143978.2 | #N/A | 50.7708925 | -5.394432 | 1.46142086 |
| 12637 | NM_001037666.2 | CASTOR1 | 23.9153298 | -5.7967718 | 1.57064612 |
| 12638 | NM_015697.7 | #N/A | 32.7712686 | -6.2422229 | 1.69136522 |
| 12639 | NM_001278280.1 | GLT8D1 | 40.9929188 | -6.5643159 | 1.77896359 |
| 12640 | NM_001278913.1 | #N/A | 41.6352088 | -6.5936648 | 1.7869575 |
| 12641 | NM_003946.6 | NOL3 | 31.5513923 | -6.1820731 | 1.67558548 |

| | | | | | |
|---|---|---|---|---|---|
| 12642 | NM_052841.3 | #N/A | 25.1126747 | -5.8688689 | 1.59069317 |
| 12643 | NM_144974.3 | #N/A | 68.5906261 | -2.4356708 | 0.66017277 |
| 12644 | NM_001300797.1 | RAN | 30.7200159 | -6.1540043 | 1.66841194 |
| 12645 | NM_178011.4 | #N/A | 40.1887104 | -6.5398315 | 1.77319708 |
| 12646 | NM_005440.4 | #N/A | 347.392936 | -2.3033197 | 0.6245299 |
| 12647 | NR_135321.1 | LINC01043 | 9.62891179 | 6.22581239 | 1.68828402 |
| 12648 | NR_027460.1 | RRN3P3 | 26.6276385 | -5.9433662 | 1.61187013 |
| 12649 | NM_017570.4 | #N/A | 45.8620313 | -3.3924457 | 0.92014454 |
| 12650 | NM_153704.5 | #N/A | 26.6651676 | -5.9504054 | 1.61397499 |
| 12651 | NM_001177984.2 | SCN8A | 109.89011 | -3.8636169 | 1.04810165 |
| 12652 | NM_032341.4 | #N/A | 25.404243 | -5.8808448 | 1.5954191 |
| 12653 | NM_015703.4 | #N/A | 446.642977 | -1.2445724 | 0.33764954 |
| 12654 | NM_024936.2 | #N/A | 41.276606 | -3.9554833 | 1.07326758 |
| 12655 | NM_001297754.1 | #N/A | 26.891491 | -5.9587545 | 1.61685092 |
| 12656 | NM_001278495.1 | NUTM2B | 33.4951391 | -6.2759884 | 1.70301511 |
| 12657 | NR_131192.1 | LINC01671 | 33.8613884 | 6.1165912 | 1.6598129 |
| 12658 | NM_000189.4 | #N/A | 502.897249 | -2.6602933 | 0.72194822 |
| 12659 | NM_001317938.1 | CCDC192 | 33.7541588 | 6.83517955 | 1.85512482 |
| 12660 | NM_173615.4 | #N/A | 95.2142719 | -7.7848057 | 2.11286963 |
| 12661 | NM_032017.2 | #N/A | 29.0510264 | -6.0732483 | 1.64847345 |
| 12662 | NM_004869.3 | #N/A | 177.24345 | -2.5728627 | 0.69837696 |
| 12663 | NM_016134.3 | #N/A | 25.7395775 | -5.9048472 | 1.6028624 |
| 12664 | NM_000101.3 | #N/A | 28.2723375 | -6.0327121 | 1.63770346 |
| 12665 | NM_052901.4 | SLC25A25 | 35.8407977 | -5.0924175 | 1.38246271 |
| 12666 | NM_014301.4 | ISCU | 28.9513248 | -6.0564434 | 1.64426248 |
| 12667 | NM_001005242.2 | PKP2 | 34.5372266 | -6.3191647 | 1.71561386 |
| 12668 | NR_135534.1 | FLNB-AS1 | 25.4736048 | -5.8788482 | 1.5961333 |
| 12669 | NR_024593.1 | POM121L10P | 124.15151 | 2.44808513 | 0.66467205 |
| 12670 | NM_001288984.1 | SDAD1 | 43.1541344 | -6.6421675 | 1.80342859 |
| 12671 | NM_001204080.1 | #N/A | 44.3763681 | -6.6790895 | 1.81377702 |
| 12672 | NM_001271442.1 | CFAP410 | 22.9548995 | -5.7294534 | 1.55594866 |
| 12673 | NM_014683.3 | #N/A | 281.231391 | -1.7485605 | 0.47489834 |
| 12674 | NM_144633.2 | #N/A | 53.6985934 | -5.4621583 | 1.48371378 |
| 12675 | NM_016558.3 | SCAND1 | 26.3290622 | -5.9348897 | 1.61249206 |
| 12676 | NM_001083607.1 | #N/A | 29.916106 | -6.1042406 | 1.65860096 |
| 12677 | NM_005664.3 | #N/A | 28.3732314 | -6.0321126 | 1.63903742 |
| 12678 | NM_002576.4 | PAK1 | 333.215537 | -3.7805547 | 1.02746336 |
| 12679 | NM_001040199.1 | CLDND1 | 25.9190201 | -5.908693 | 1.60599481 |
| 12680 | NM_138436.3 | #N/A | 32.7896612 | -6.2434508 | 1.69698338 |
| 12681 | NM_032131.5 | #N/A | 27.9266741 | -6.0154837 | 1.63523146 |
| 12682 | NM_001308287.1 | PTPN2 | 60.041952 | 7.41422031 | 2.01549845 |
| 12683 | NM_001321109.1 | #N/A | 24.6451788 | -5.8406862 | 1.58775372 |
| 12684 | NM_021083.3 | #N/A | 24.4342932 | -5.8227346 | 1.58302011 |

| | | | | | |
|---|---|---|---|---|---|
| 12685 | NM_005371.5 | #N/A | 29.0679393 | -6.074227 | 1.65140543 |
| 12686 | NR_033255.1 | #N/A | 27.8760672 | -6.0040173 | 1.63233584 |
| 12687 | NR_038450.1 | LOC100130849 | 25.474769 | 4.09113117 | 1.11240658 |
| 12688 | NM_020768.3 | #N/A | 27.3203739 | -5.9844127 | 1.62728786 |
| 12689 | NM_201650.2 | #N/A | 34.5896316 | -6.3252019 | 1.72004844 |
| 12690 | NM_002606.2 | #N/A | 32.8444588 | -6.2408066 | 1.69714474 |
| 12691 | NM_003834.2 | #N/A | 30.9997341 | -6.1569814 | 1.67444058 |
| 12692 | NM_018259.5 | #N/A | 267.10531 | -3.385375 | 0.92069396 |
| 12693 | NM_181468.2 | EIF6 | 28.9066951 | -6.0696049 | 1.65072934 |
| 12694 | NM_001709.4 | #N/A | 32.2112491 | -6.2190483 | 1.69146722 |
| 12695 | NM_001286208.1 | GAS8 | 26.6345744 | -5.9396448 | 1.61548429 |
| 12696 | NM_001242485.1 | #N/A | 31.5472412 | -6.1850616 | 1.68246184 |
| 12697 | NR_135043.1 | #N/A | 36.1844948 | -6.3872961 | 1.73748072 |
| 12698 | NM_001293212.1 | #N/A | 32.2836087 | -6.2172253 | 1.69138851 |
| 12699 | NM_194286.3 | #N/A | 322.741019 | -4.2415078 | 1.15392037 |
| 12700 | NM_001170881.1 | #N/A | 36.6030955 | -6.4085576 | 1.74351295 |
| 12701 | NM_001025593.2 | ARFIP1 | 31.9571457 | -6.1992466 | 1.68664684 |
| 12702 | NR_034006.2 | #N/A | 25.1640182 | -5.8636298 | 1.59536369 |
| 12703 | NM_014825.2 | #N/A | 267.703992 | -1.6703748 | 0.45448213 |
| 12704 | NM_030762.2 | #N/A | 24.1637102 | -5.8102436 | 1.58099899 |
| 12705 | NM_002065.6 | GLUL | 33.2011967 | -6.2682753 | 1.70567395 |
| 12706 | NM_001198846.1 | RBM14-RBM4 | 23.2988048 | -5.7572448 | 1.56681588 |
| 12707 | NM_001284206.1 | DCAF5 | 40.970144 | -6.5634147 | 1.78628982 |
| 12708 | NM_001631.4 | #N/A | 11.957746 | 7.06935947 | 1.92417385 |
| 12709 | NM_001258373.1 | SPATA20 | 35.5219893 | -6.3651613 | 1.73269937 |
| 12711 | NR_104359.1 | GPR1-AS | 51.3839087 | 2.84092542 | 0.77340054 |
| 12710 | NM_001320283.1 | #N/A | 31.5052935 | -6.1875881 | 1.6844737 |
| 12712 | NM_018901.3 | #N/A | 51.3611949 | -6.8922457 | 1.87633363 |
| 12713 | NM_001130085.1 | #N/A | 31.6018084 | -6.1947964 | 1.68648653 |
| 12714 | NM_001289861.1 | PER3 | 35.0038054 | -6.3375617 | 1.72543254 |
| 12717 | NR_104096.1 | KIR3DX1 | 50.2214243 | 2.38578614 | 0.64961386 |
| 12715 | NM_016544.2 | #N/A | 27.0750747 | -5.9740248 | 1.62665308 |
| 12716 | NM_145266.5 | #N/A | 84.317062 | -2.5190886 | 0.68591299 |
| 12718 | NM_001319190.1 | SCIMP | 152.144619 | 2.04515198 | 0.55701178 |
| 12719 | NM_001013672.4 | #N/A | 27.0904985 | -5.9758455 | 1.62764425 |
| 12720 | NM_001097612.1 | #N/A | 41.0745955 | -6.5721029 | 1.79019802 |
| 12721 | NM_001286582.1 | #N/A | 69.437512 | -7.3245827 | 1.99526554 |
| 12722 | NM_003385.4 | #N/A | 60.7668709 | -7.1339783 | 1.94335596 |
| 12723 | NM_001136103.2 | TMEM132C | 43.2801211 | -6.6456856 | 1.81049532 |
| 12724 | NM_004270.4 | #N/A | 31.365851 | -6.1805414 | 1.68377009 |
| 12725 | NM_152259.3 | #N/A | 33.8307425 | -6.2846261 | 1.7121141 |
| 12726 | NR_134875.1 | TMEM241 | 32.2979057 | -6.2268106 | 1.69639097 |
| 12727 | NM_021908.2 | #N/A | 33.8529258 | -6.2923612 | 1.71426259 |

| | | | | | |
|---|---|---|---|---|---|
| 12728 | NM_015676.1 | #N/A | 30.3293097 | -6.1306203 | 1.67021399 |
| 12729 | NR_024052.2 | HCG18 | 43.8959227 | -6.6643914 | 1.81574137 |
| 12730 | NR_033334.1 | TMEM70 | 22.8835515 | -5.7286 | 1.56077746 |
| 12731 | NM_005057.3 | #N/A | 54.6965448 | -6.9841716 | 1.90362398 |
| 12732 | NM_145305.2 | #N/A | 22.5164992 | -5.7035793 | 1.55461632 |
| 12733 | NM_001077441.1 | #N/A | 29.7872022 | -6.1048999 | 1.6642639 |
| 12734 | NM_023112.3 | #N/A | 35.5875201 | -4.9444052 | 1.34839786 |
| 12735 | NM_001145544.1 | #N/A | 40.0429089 | -6.5390516 | 1.78345081 |
| 12736 | NM_018365.2 | #N/A | 46.2328298 | -6.7443108 | 1.83944847 |
| 12737 | NM_001293228.1 | #N/A | 40.7851831 | -6.5578724 | 1.78862412 |
| 12738 | NM_001142284.1 | RPS24 | 65.9567129 | -7.2532888 | 1.97844873 |
| 12739 | NM_001304947.1 | #N/A | 26.2255817 | -5.9238993 | 1.61603849 |
| 12740 | NM_000846.4 | #N/A | 24.4257284 | 8.87519529 | 2.42119866 |
| 12741 | NM_006721.3 | #N/A | 38.8338425 | -6.4899541 | 1.77076544 |
| 12742 | NM_001129996.1 | #N/A | 27.3259922 | -5.9803142 | 1.63174212 |
| 12743 | NM_001127258.2 | HHIPL1 | 23.5885108 | -5.76778 | 1.57381978 |
| 12744 | NM_032664.3 | FUT10 | 23.6133637 | -5.7739987 | 1.57554318 |
| 12745 | NM_052837.2 | #N/A | 33.2353173 | -6.2683065 | 1.71057256 |
| 12746 | NM_022786.2 | #N/A | 39.5986824 | -6.5181176 | 1.77879046 |
| 12747 | NM_001242784.1 | #N/A | 51.7439716 | -3.1090346 | 0.8485331 |
| 12748 | NR_036556.1 | #N/A | 30.8130454 | -6.1637261 | 1.68225382 |
| 12749 | NM_001330663.1 | #N/A | 26.683144 | -5.946922 | 1.62318843 |
| 12750 | NM_001258306.1 | #N/A | 25.4943922 | -5.8817777 | 1.60545808 |
| 12751 | NM_002885.2 | #N/A | 53.5220661 | -6.9540636 | 1.89818437 |
| 12752 | NM_144964.3 | #N/A | 35.6613678 | -6.3671841 | 1.73803118 |
| 12753 | NR_136669.1 | #N/A | 29.2341755 | -6.0718994 | 1.65748793 |
| 12754 | NR_138046.1 | #N/A | 34.010738 | -6.2934911 | 1.71808549 |
| 12755 | NM_001256585.1 | MGLL | 29.5226329 | -6.0925227 | 1.66333289 |
| 12756 | NM_021004.3 | #N/A | 23.5803255 | -5.7649094 | 1.57412686 |
| 12758 | NR_002786.1 | CIDECP1 | 31.9936567 | -6.2093193 | 1.69554995 |
| 12757 | NM_016448.3 | #N/A | 37.7727143 | -6.4494073 | 1.76111466 |
| 12759 | NM_145860.1 | PDCD10 | 29.6390839 | -6.1001888 | 1.66581451 |
| 12760 | NM_001330763.1 | #N/A | 28.73415 | -6.0450473 | 1.65088756 |
| 12761 | NM_001243227.1 | #N/A | 68.4565608 | -7.3036768 | 1.99463732 |
| 12762 | NR_073387.1 | ALG1L9P | 94.4557798 | 2.7640774 | 0.75514017 |
| 12763 | NM_014676.2 | #N/A | 175.104317 | -3.6394178 | 0.99432023 |
| 12764 | NR_132272.1 | #N/A | 38.8195625 | -6.4926689 | 1.7739733 |
| 12765 | NM_016256.3 | #N/A | 37.4466923 | -6.4359456 | 1.75851924 |
| 12766 | NM_001127898.3 | #N/A | 34.0917782 | -6.2979536 | 1.72094186 |
| 12767 | NM_001098797.1 | #N/A | 41.0671421 | -6.5678865 | 1.79493776 |
| 12768 | NM_001271886.1 | AAGAB | 32.7065215 | -6.2398349 | 1.70532185 |
| 12769 | NM_000055.3 | #N/A | 29.8069437 | -6.1062329 | 1.66908956 |
| 12770 | NM_194429.2 | #N/A | 47.5383578 | -6.7783876 | 1.85282204 |

| | | | | | |
|---|---|---|---|---|---|
| 12771 | NM_003943.4 | STBD1 | 23.3951007 | -5.7555107 | 1.57328468 |
| 12772 | NM_198568.2 | #N/A | 33.2781981 | 4.57450738 | 1.25045535 |
| 12773 | NM_001204267.1 | #N/A | 31.0650678 | 4.45993398 | 1.21915266 |
| 12774 | NM_138690.2 | #N/A | 40.5606034 | -6.5562746 | 1.79229509 |
| 12775 | NM_001304466.1 | #N/A | 26.7193064 | -5.9557799 | 1.62825787 |
| 12776 | NM_001286564.1 | #N/A | 44.1116759 | -6.6672257 | 1.82300972 |
| 12777 | NM_001194998.1 | #N/A | 24.0306684 | -5.7922391 | 1.58378852 |
| 12778 | NR_110761.1 | LOC102723769 | 45.3178492 | 3.15836884 | 0.86360894 |
| 12779 | NR_137177.1 | MIR100HG | 49.305246 | -6.8314779 | 1.86817366 |
| 12780 | NR_027455.5 | LINC00893 | 45.8691414 | -6.7327771 | 1.84128661 |
| 12781 | NM_033629.4 | #N/A | 22.2897375 | -5.6916767 | 1.55657955 |
| 12782 | NM_177959.2 | #N/A | 42.3928545 | -6.6150078 | 1.80926086 |
| 12783 | NM_001350.4 | #N/A | 34.3200476 | -6.3052915 | 1.72482709 |
| 12784 | NM_017760.6 | #N/A | 88.3310386 | -3.1152474 | 0.85218458 |
| 12785 | NR_120415.1 | LINC00368 | 62.7113737 | 3.63719757 | 0.99503737 |
| 12786 | NM_018347.2 | #N/A | 28.3548938 | -6.0440546 | 1.65387009 |
| 12787 | NM_175040.3 | ST6GALNAC4 | 25.0757301 | -5.8538495 | 1.60189248 |
| 12788 | NM_001195755.1 | #N/A | 65.3765437 | 2.22423953 | 0.60866825 |
| 12789 | NM_018846.4 | #N/A | 32.8163139 | -6.2437852 | 1.7090064 |
| 12790 | NM_006567.4 | #N/A | 25.7783757 | -5.8968772 | 1.61408673 |
| 12791 | NM_005008.3 | #N/A | 108.341079 | -5.0717285 | 1.38831229 |
| 12792 | NM_001320893.1 | DNMT3A | 37.5173433 | -6.4456584 | 1.76443071 |
| 12793 | NM_018702.3 | #N/A | 42.6800378 | -5.1667514 | 1.41437275 |
| 12794 | NR_027282.1 | LOC399815 | 16.3399218 | 4.24590829 | 1.1623162 |
| 12796 | NR_003570.2 | DMBT1P1 | 49.7273025 | 2.86554581 | 0.78448235 |
| 12795 | NM_001146698.1 | #N/A | 33.2810526 | -6.2709403 | 1.7167621 |
| 12797 | NM_012089.2 | #N/A | 95.9203148 | -3.1477089 | 0.86175405 |
| 12798 | NM_014652.3 | #N/A | 244.348755 | -2.2004379 | 0.6024324 |
| 12799 | NM_005568.4 | #N/A | 24.4789809 | -5.8304374 | 1.5963944 |
| 12800 | NM_001301016.2 | #N/A | 27.0593067 | -5.9747772 | 1.63599931 |
| 12801 | NR_103834.1 | #N/A | 35.9070036 | -6.3712895 | 1.74470588 |
| 12802 | NM_007365.2 | #N/A | 41.3421786 | -6.5818834 | 1.80249212 |
| 12803 | NM_001276723.1 | C11orf74 | 29.9380517 | -6.1092512 | 1.67306895 |
| 12804 | NM_139284.2 | #N/A | 57.2410785 | -7.0515999 | 1.93120365 |
| 12805 | NM_198040.2 | PHC2 | 39.5903604 | -6.518889 | 1.7854729 |
| 12806 | NM_001144035.1 | TMEM25 | 29.1692484 | -6.0784971 | 1.66505097 |
| 12807 | NM_032626.5 | RBBP6 | 22.7585229 | -5.722276 | 1.56749601 |
| 12808 | NM_001286512.1 | SLC35B2 | 43.3957249 | -6.6470296 | 1.82086805 |
| 12809 | NM_001303037.1 | #N/A | 103.53342 | -2.9046121 | 0.79571047 |
| 12810 | NM_032564.4 | #N/A | 28.7933562 | -6.056198 | 1.65914556 |
| 12811 | NM_001117.4 | #N/A | 29.4704905 | -6.0912677 | 1.66878323 |
| 12813 | NM_001497.3 | B4GALT1 | 177.59377 | -3.1738794 | 0.86955603 |
| 12812 | NM_001201397.1 | EDNRB | 28.9108793 | -6.0673545 | 1.66228804 |

| | | | | | |
|---|---|---|---|---|---|
| 12814 | NM_001324337.1 | SHOC2 | 37.1200847 | -6.4190602 | 1.75867766 |
| 12815 | NM_005139.2 | #N/A | 24.8037947 | -5.8504652 | 1.60289239 |
| 12816 | NM_138811.3 | #N/A | 23.1527985 | -5.7452273 | 1.5742462 |
| 12817 | NM_001242915.1 | ZFAND6 | 49.1356152 | -6.8309904 | 1.87182144 |
| 12818 | NR_026678.1 | TMEM79 | 26.657534 | -5.9474205 | 1.63021759 |
| 12819 | NM_003757.3 | EIF3I | 535.35753 | -3.4553675 | 0.94715527 |
| 12820 | NR_103446.1 | TRAM2-AS1 | 25.1405452 | -5.8667257 | 1.60818363 |
| 12821 | NM_022036.2 | #N/A | 25.0104221 | -5.8595666 | 1.60634425 |
| 12822 | NR_003494.1 | #N/A | 24.178461 | -5.8125236 | 1.59385584 |
| 12823 | NM_001130028.1 | CLK3 | 33.820794 | -6.2873406 | 1.72417887 |
| 12824 | NM_019024.2 | #N/A | 192.389796 | -2.8762204 | 0.78877258 |
| 12825 | NR_033925.1 | FENDRR | 34.0282499 | -6.3021461 | 1.72833209 |
| 12826 | NR_027052.1 | THAP7-AS1 | 23.3291798 | -5.7559259 | 1.57859355 |
| 12827 | NM_001142928.1 | #N/A | 36.9460657 | -6.4135471 | 1.75897127 |
| 12828 | NM_001142650.1 | HNRNPLL | 36.9458383 | -6.4161705 | 1.75973857 |
| 12829 | NM_001244606.1 | #N/A | 26.8224227 | -5.9564918 | 1.63391019 |
| 12830 | NM_021151.3 | #N/A | 44.4966736 | -4.4061676 | 1.20867191 |
| 12831 | NM_001145065.1 | CCSER1 | 29.3379978 | -6.0789853 | 1.66770894 |
| 12832 | NM_015020.3 | PHLPP2 | 39.2371312 | -6.5003521 | 1.78330512 |
| 12833 | NR_004053.3 | SOX2-OT | 23.0111086 | -5.7354722 | 1.57347995 |
| 12834 | NM_001258276.1 | TMEM200A | 30.6476603 | -6.1560911 | 1.68895502 |
| 12835 | NM_032878.3 | ALKBH6 | 27.6045804 | -5.9879087 | 1.64294581 |
| 12836 | NM_021938.3 | #N/A | 37.0567604 | -6.4205715 | 1.76175869 |
| 12837 | NM_001136039.2 | NIF3L1 | 28.0782458 | -6.0167227 | 1.65117893 |
| 12838 | NM_153200.2 | EDF1 | 28.1003414 | -6.0240416 | 1.65328281 |
| 12839 | NM_002531.2 | #N/A | 46.1617315 | -6.7363696 | 1.84881294 |
| 12840 | NM_052883.1 | #N/A | 22.8679913 | -5.729128 | 1.57243298 |
| 12841 | NM_005307.2 | #N/A | 29.358346 | -6.0817616 | 1.66923292 |
| 12842 | NM_138476.3 | #N/A | 28.5203816 | -6.0456202 | 1.65941322 |
| 12843 | NM_006417.4 | #N/A | 36.7710462 | -6.4134056 | 1.76050576 |
| 12844 | NM_006994.4 | #N/A | 27.109247 | -5.9790838 | 1.64129622 |
| 12845 | NM_001257180.1 | #N/A | 56.9261843 | -7.043338 | 1.93352575 |
| 12846 | NM_145267.2 | #N/A | 28.2784254 | -6.0326295 | 1.65606508 |
| 12847 | NM_001135004.2 | DNAJB5 | 48.2005524 | -6.8000457 | 1.8669009 |
| 12848 | NR_026850.1 | MTMR9LP | 24.7706959 | -5.8389274 | 1.60305891 |
| 12849 | NM_172229.2 | #N/A | 27.4502868 | -5.9823101 | 1.64266854 |
| 12850 | NM_001159287.1 | TPI1 | 56.2064155 | -7.0206271 | 1.92798069 |
| 12851 | NM_005374.4 | #N/A | 48.8080472 | -6.8201847 | 1.87297482 |
| 12852 | NM_145061.5 | #N/A | 25.5967602 | -5.8852429 | 1.61677912 |
| 12853 | NM_001008727.3 | #N/A | 23.4089193 | -5.7528178 | 1.58048969 |
| 12854 | NM_001308445.1 | #N/A | 64.1589909 | -3.7728829 | 1.03663051 |
| 12855 | NM_005107.3 | #N/A | 35.8157658 | -6.3725754 | 1.75092525 |
| 12856 | NM_153018.2 | #N/A | 30.2300116 | -6.1257264 | 1.68309339 |

| | | | | | |
|---|---|---|---|---|---|
| 12857 | NM_001199570.1 | #N/A | 79.2804262 | -5.9394806 | 1.63197784 |
| 12858 | NM_025040.3 | #N/A | 70.0309318 | -4.5998797 | 1.26389693 |
| 12859 | NM_001042406.1 | HMGCLL1 | 35.9153879 | -6.3760502 | 1.75212819 |
| 12860 | NM_001301131.1 | #N/A | 34.5592184 | -6.3209784 | 1.73707778 |
| 12861 | NM_001198670.1 | #N/A | 29.9486993 | -6.1056481 | 1.67807166 |
| 12862 | NM_001197122.1 | #N/A | 30.5910728 | -6.1520887 | 1.69085668 |
| 12863 | NM_001319042.1 | FABP7 | 58.4565235 | -7.0773208 | 1.94536539 |
| 12864 | NR_131782.1 | CCEPR | 32.1032917 | -6.2131719 | 1.70785322 |
| 12865 | NM_001301076.1 | #N/A | 29.1070384 | -6.0776058 | 1.6708871 |
| 12866 | NM_005879.2 | #N/A | 27.6734131 | -5.9973055 | 1.64884224 |
| 12867 | NM_014365.2 | HSPB8 | 37.0069701 | -6.4215192 | 1.76550028 |
| 12868 | NM_001243795.1 | #N/A | 27.212823 | -5.9762996 | 1.64319919 |
| 12869 | NM_015550.3 | #N/A | 25.1261446 | -5.8587734 | 1.61102378 |
| 12870 | NM_001304476.1 | STXBP6 | 28.1952992 | -6.0201597 | 1.65551554 |
| 12871 | NR_110204.1 | CHROMR | 25.082223 | -5.859214 | 1.61172193 |
| 12872 | NM_001271869.1 | #N/A | 50.2710227 | -6.863814 | 1.88824195 |
| 12873 | NM_003603.6 | SORBS2 | 32.8928212 | -6.2440474 | 1.71779903 |
| 12874 | NM_001130914.1 | #N/A | 25.7019761 | -5.8948568 | 1.62176975 |
| 12875 | NR_003661.2 | UBE2Q2P1 | 164.028913 | 2.15986939 | 0.59435612 |
| 12876 | NM_002055.4 | #N/A | 672.678368 | -5.2160558 | 1.4354553 |
| 12877 | NM_004278.3 | #N/A | 23.6659265 | -5.7828417 | 1.59151482 |
| 12878 | NM_001127394.3 | #N/A | 35.7582744 | -6.3719198 | 1.75374261 |
| 12879 | NM_001318230.1 | #N/A | 41.4330227 | -6.579092 | 1.81077386 |
| 12880 | NR_046724.1 | TM4SF19-AS1 | 88.7270947 | 3.6687347 | 1.0097678 |
| 12881 | NM_001146218.1 | #N/A | 29.295562 | -6.0851642 | 1.67496568 |
| 12882 | NM_031488.4 | #N/A | 167.051995 | -4.4279461 | 1.21881003 |
| 12883 | NM_080422.2 | PTPN2 | 27.5674984 | -5.9933253 | 1.64976769 |
| 12884 | NM_001031694.2 | SCMH1 | 25.1435563 | -5.8702717 | 1.61594263 |
| 12885 | NM_007243.2 | NRM | 22.8175286 | -5.722409 | 1.57533395 |
| 12886 | NM_001136020.2 | #N/A | 36.8659564 | -6.4094092 | 1.76450033 |
| 12887 | NM_020799.3 | #N/A | 30.5538781 | -6.1437745 | 1.69136702 |
| 12888 | NM_014638.3 | #N/A | 45.2737412 | -6.7094885 | 1.84714445 |
| 12889 | NM_001278346.1 | PHYKPL | 26.0194074 | -5.9097318 | 1.62697878 |
| 12890 | NM_022342.4 | #N/A | 24.1154361 | -5.8110316 | 1.59995815 |
| 12891 | NM_005187.5 | #N/A | 27.156933 | -5.971993 | 1.64443321 |
| 12892 | NM_032310.4 | #N/A | 42.3331064 | -6.6128534 | 1.82094144 |
| 12893 | NM_020201.3 | #N/A | 44.2754648 | -5.4017744 | 1.48754604 |
| 12894 | NM_013275.5 | #N/A | 357.176992 | -9.6912233 | 2.66886311 |
| 12895 | NM_152695.5 | #N/A | 38.1302633 | -5.0238054 | 1.38355651 |
| 12896 | NM_005418.3 | DENND2B | 34.3081643 | -6.3047565 | 1.7364766 |
| 12897 | NM_002033.3 | FUT4 | 37.0499073 | -3.7755327 | 1.03987968 |
| 12898 | NR_125926.1 | CXXC4-AS1 | 51.4955313 | 2.48887592 | 0.68591565 |
| 12899 | NR_138039.1 | #N/A | 62.3088934 | 2.04558947 | 0.56380684 |

| | | | | | |
|---|---|---|---|---|---|
| 12900 | NM_001267868.1 | #N/A | 31.4174679 | -4.8975613 | 1.34990242 |
| 12901 | NM_001066.2 | #N/A | 32.0863353 | 2.15453774 | 0.59385569 |
| 12902 | NM_032551.4 | #N/A | 40.6494659 | -6.558369 | 1.80783887 |
| 12903 | NM_016453.3 | #N/A | 49.9842788 | -6.849972 | 1.88823612 |
| 12904 | NM_001318750.1 | MSANTD4 | 85.0402174 | -4.7979499 | 1.32259404 |
| 12905 | NM_001301700.1 | ERI3 | 30.1678158 | -6.1275395 | 1.68923065 |
| 12906 | NM_025246.2 | #N/A | 31.5187563 | -6.1860618 | 1.70537085 |
| 12907 | NM_007018.4 | #N/A | 43.332899 | -5.1838053 | 1.42907768 |
| 12908 | NM_003643.3 | #N/A | 52.9727574 | 2.55338123 | 0.70400021 |
| 12909 | NM_001277066.1 | #N/A | 40.2366795 | -6.5395319 | 1.80304677 |
| 12910 | NM_016391.7 | #N/A | 30.1007482 | -6.1233284 | 1.68835085 |
| 12911 | NR_038244.1 | LOC100507557 | 21.9508768 | -5.6636477 | 1.56190987 |
| 12912 | NM_001164638.1 | #N/A | 25.578865 | -5.8891754 | 1.62415583 |
| 12913 | NM_203346.4 | #N/A | 343.189917 | -9.633599 | 2.65695291 |
| 12914 | NM_001322248.1 | #N/A | 33.0848093 | -6.2510846 | 1.72415065 |
| 12915 | NM_022145.4 | #N/A | 26.5204675 | -5.9459028 | 1.64005169 |
| 12916 | NM_025221.5 | #N/A | 37.4855722 | -6.4393157 | 1.7762589 |
| 12917 | NM_001321829.1 | CCDC186 | 49.0194079 | -6.8244073 | 1.88254533 |
| 12918 | NR_130139.1 | NPLOC4 | 49.6079584 | -6.8426994 | 1.88771697 |
| 12919 | NR_110064.1 | LINC01441 | 99.6211604 | 3.10797676 | 0.85750543 |
| 12920 | NM_144994.7 | #N/A | 25.5683574 | -5.882022 | 1.62292787 |
| 12921 | NM_004498.2 | #N/A | 33.0455167 | -6.2583059 | 1.72678015 |
| 12922 | NM_016486.3 | #N/A | 69.3862995 | -4.5843992 | 1.26520007 |
| 12923 | NR_110743.1 | LINC01929 | 19.4307989 | 7.0993287 | 1.95928432 |
| 12924 | NM_017837.3 | PIGV | 21.9414713 | -5.667398 | 1.56440475 |
| 12925 | NM_001033583.2 | #N/A | 30.015949 | -6.1154124 | 1.68820682 |
| 12926 | NR_029434.1 | PSMA3-AS1 | 50.5361461 | -3.3792445 | 0.93291205 |
| 12927 | NM_001031717.3 | #N/A | 28.4577344 | -6.0450983 | 1.66891228 |
| 12928 | NM_001146071.1 | TDRD3 | 23.9490704 | -5.7841674 | 1.59691392 |
| 12929 | NM_007022.4 | #N/A | 33.5168654 | -6.2724827 | 1.73204728 |
| 12930 | NM_001135674.1 | #N/A | 28.5517923 | -6.0397391 | 1.6678496 |
| 12931 | NM_004346.3 | #N/A | 31.0771672 | -6.1750212 | 1.70520101 |
| 12932 | NM_001143774.1 | PHYHIPL | 21.5286298 | 9.42185555 | 2.60205914 |
| 12934 | NM_006168.2 | NKX6-1 | 33.442068 | -6.2742381 | 1.73287386 |
| 12933 | NM_001170820.3 | #N/A | 58.7024181 | -5.5613101 | 1.53597035 |
| 12935 | NR_049755.1 | #N/A | 28.7634179 | -6.0572122 | 1.67297762 |
| 12936 | NM_002245.3 | #N/A | 38.0291751 | -6.4622618 | 1.78499621 |
| 12937 | NR_026641.1 | #N/A | 27.6148519 | -5.9945467 | 1.65581017 |
| 12938 | NR_103456.1 | ZSCAN16-AS1 | 24.3650989 | -5.8163192 | 1.60667532 |
| 12939 | NM_014398.3 | #N/A | 76.4254801 | 1.82458658 | 0.50401896 |
| 12940 | NM_003621.3 | #N/A | 32.8320467 | -6.251741 | 1.72705766 |
| 12941 | NM_006983.1 | #N/A | 22.5148382 | -5.7028911 | 1.57545532 |
| 12942 | NM_001113561.2 | RNF180 | 25.3981873 | -5.8752499 | 1.62310996 |

| | | | | | |
|---|---|---|---|---|---|
| 12943 | NM_005227.2 | #N/A | 25.8338442 | -5.9007911 | 1.63031614 |
| 12944 | NM_018205.3 | #N/A | 61.9447421 | -7.1623511 | 1.97888929 |
| 12945 | NM_014720.3 | #N/A | 24.3271845 | -5.8072245 | 1.60454885 |
| 12946 | NM_001347716.1 | #N/A | 44.1570678 | -6.6776457 | 1.84506853 |
| 12947 | NR_136214.1 | KCNIP1-AS1 | 13.3266952 | 5.63068177 | 1.55585436 |
| 12948 | NM_001282982.1 | #N/A | 36.2273858 | -6.3837485 | 1.76417547 |
| 12949 | NM_001271951.1 | TPGS2 | 29.013525 | -6.0622827 | 1.6756026 |
| 12950 | NM_001193464.1 | #N/A | 31.5837888 | -6.1854047 | 1.70974511 |
| 12951 | NM_001329609.1 | ARL16 | 27.2521388 | -5.9805764 | 1.65339935 |
| 12952 | NM_001079811.2 | GLB1 | 30.9503647 | -6.167358 | 1.70541269 |
| 12953 | NM_001320672.1 | RUNX3 | 12.0240435 | 3.46784275 | 0.95894573 |
| 12954 | NM_001099627.1 | MTFR1L | 41.4889773 | -6.5882438 | 1.82198815 |
| 12955 | NM_006841.5 | #N/A | 21.6298699 | -5.6473092 | 1.56176842 |
| 12956 | NM_017458.3 | MVP | 48.5650958 | -6.8079539 | 1.8827687 |
| 12957 | NM_001143960.2 | C2orf74 | 25.2176097 | -5.8640405 | 1.62192298 |
| 12958 | NM_001174153.1 | RABEPK | 36.2341637 | -6.3843639 | 1.76597218 |
| 12959 | NM_152586.3 | USP54 | 32.8670415 | -6.2449967 | 1.72741571 |
| 12960 | NM_001292032.1 | #N/A | 31.7706389 | -6.2044395 | 1.7162306 |
| 12961 | NM_006143.2 | #N/A | 36.599914 | -6.4042717 | 1.7715485 |
| 12962 | NM_001170686.1 | #N/A | 36.2211867 | -6.3839575 | 1.76600742 |
| 12963 | NM_032482.2 | #N/A | 581.708838 | -3.7658618 | 1.04176449 |
| 12964 | NM_001093726.1 | #N/A | 25.0868128 | -5.859882 | 1.62110043 |
| 12965 | NM_001271948.1 | PPP2R2B | 23.6399915 | -5.778446 | 1.59858665 |
| 12966 | NM_001318810.1 | #N/A | 32.3582334 | -6.2290059 | 1.72345168 |
| 12967 | NM_001306206.1 | CSTF2 | 36.5348105 | -6.396351 | 1.76987602 |
| 12968 | NM_001281521.1 | MCM8 | 27.4473298 | -5.9968185 | 1.65939046 |
| 12969 | NR_109899.1 | KCNIP1-OT1 | 37.8711225 | 5.17668509 | 1.43251901 |
| 12970 | NM_001030059.2 | PLPP4 | 23.6722238 | -5.781517 | 1.60012592 |
| 12971 | NM_020787.3 | #N/A | 26.3735889 | -5.9293533 | 1.64107535 |
| 12972 | NM_001261430.1 | PTGES3L | 23.7679547 | -5.7743317 | 1.59826431 |
| 12973 | NM_003677.4 | #N/A | 418.992976 | -2.5343953 | 0.70152094 |
| 12974 | NM_001270876.1 | #N/A | 96.4086638 | 7.80566896 | 2.16079438 |
| 12975 | NM_003215.2 | #N/A | 28.1034629 | 4.33439835 | 1.19989348 |
| 12976 | NM_004999.3 | #N/A | 61.4450748 | -7.1537832 | 1.98046549 |
| 12977 | NM_001177519.2 | #N/A | 31.3641292 | -6.1880869 | 1.71314993 |
| 12978 | NM_001267580.1 | #N/A | 46.1304983 | -6.7395772 | 1.8658331 |
| 12979 | NR_046347.1 | #N/A | 27.0013614 | -5.9721901 | 1.65338018 |
| 12980 | NM_001323529.1 | #N/A | 38.0129997 | -6.4614102 | 1.78889183 |
| 12981 | NM_001258040.2 | HARS1 | 26.5921824 | -5.9493959 | 1.6472065 |
| 12982 | NM_001347919.1 | SNX19 | 38.0777006 | -6.459334 | 1.78847603 |
| 12983 | NM_001080501.2 | #N/A | 38.3944092 | -5.0270602 | 1.39207963 |
| 12984 | NM_020730.2 | #N/A | 24.5853345 | -5.8356637 | 1.61598798 |
| 12985 | NM_001130102.2 | #N/A | 24.3154507 | -5.8143766 | 1.61020218 |

| | | | | | |
|---|---|---|---|---|---|
| 12986 | NR_024080.1 | ACP1 | 29.6798388 | -6.0941788 | 1.68775167 |
| 12987 | NM_001277200.1 | #N/A | 26.0195052 | -5.914306 | 1.6380082 |
| 12988 | NM_001003407.1 | #N/A | 33.6579837 | -6.2821569 | 1.73992489 |
| 12989 | NM_001003676.2 | C11orf49 | 25.2383217 | -5.8636377 | 1.62404925 |
| 12990 | NM_001278740.1 | #N/A | 28.0092916 | -6.0201068 | 1.66756027 |
| 12991 | NM_144564.4 | #N/A | 134.034901 | -3.4378005 | 0.95230687 |
| 12992 | NM_032521.2 | #N/A | 245.36244 | 2.2373605 | 0.61985281 |
| 12993 | NM_001322042.1 | #N/A | 36.5794364 | -6.3989439 | 1.77297511 |
| 12994 | NM_001242761.1 | DLGAP1 | 35.4853482 | -6.3554202 | 1.76113298 |
| 12995 | NM_003213.3 | #N/A | 26.9310594 | -5.9582117 | 1.65109813 |
| 12996 | NM_001098503.1 | #N/A | 92.1969831 | 2.78491534 | 0.77194178 |
| 12997 | NM_005602.5 | #N/A | 53.9749274 | -6.9670492 | 1.93131927 |
| 12998 | NM_001329917.1 | #N/A | 556.387351 | 2.52545228 | 0.70011467 |
| 12999 | NM_020872.2 | CNTN3 | 34.1728173 | -6.3078212 | 1.74869487 |
| 13000 | NM_001330512.1 | RPS6KA6 | 25.3894184 | -5.8828231 | 1.6309796 |
| 13001 | NM_152354.5 | #N/A | 30.397625 | -6.1301218 | 1.69953912 |
| 13002 | NR_033342.1 | #N/A | 34.4159452 | -6.3132223 | 1.75039889 |
| 13003 | NR_038456.1 | #N/A | 35.3470499 | -6.3569939 | 1.76282649 |
| 13004 | NM_001127204.1 | HMOX2 | 33.3987089 | -6.278251 | 1.74107933 |
| 13006 | NM_001331022.1 | VPS26C | 29.4663733 | -6.0831872 | 1.68698358 |
| 13005 | NM_001271661.1 | #N/A | 190.717568 | 3.347345 | 0.92828614 |
| 13008 | NR_109845.1 | KCNA3 | 15.2907896 | 8.93137126 | 2.4770057 |
| 13007 | NM_014551.4 | NCAPH2 | 23.9445956 | -5.7891311 | 1.60553892 |
| 13009 | NM_138770.2 | #N/A | 28.1449581 | -6.0301211 | 1.67247195 |
| 13010 | NM_032222.2 | #N/A | 25.4623709 | -5.8839301 | 1.63210693 |
| 13012 | NM_024742.2 | ARMC5 | 31.6706846 | -4.9135394 | 1.36322165 |
| 13011 | NM_013975.3 | #N/A | 152.671607 | -3.0256809 | 0.83945413 |
| 13013 | NM_001271420.1 | #N/A | 22.1780489 | -5.6819209 | 1.57642904 |
| 13014 | NM_001008844.2 | #N/A | 24.6926508 | -5.8268038 | 1.61668412 |
| 13015 | NM_003489.3 | #N/A | 227.136336 | -2.4558849 | 0.68153337 |
| 13016 | NM_213601.2 | #N/A | 192.460288 | -2.768156 | 0.76822436 |
| 13017 | NM_018024.2 | #N/A | 27.1316039 | -5.9754388 | 1.65832622 |
| 13018 | NM_023927.2 | #N/A | 41.9516372 | -6.605459 | 1.83318333 |
| 13019 | NM_032622.2 | LNX1 | 32.8170986 | -6.2397948 | 1.73173055 |
| 13020 | NM_054026.3 | CNOT7 | 26.1399271 | -5.916707 | 1.64238796 |
| 13021 | NR_046360.1 | #N/A | 35.3486625 | -6.3512805 | 1.76318032 |
| 13022 | NM_000946.2 | #N/A | 32.5326076 | -6.2292359 | 1.72955904 |
| 13023 | NM_001318716.1 | KBTBD4 | 24.4036246 | -5.8147077 | 1.61453086 |
| 13024 | NR_026921.1 | LOC202181 | 23.5454748 | -5.7763016 | 1.60387319 |
| 13025 | NM_001244584.2 | FMC1-LUC7L2 | 49.686234 | -6.8442745 | 1.90068581 |
| 13026 | NR_015363.2 | GLIDR | 40.2190105 | -5.0814719 | 1.41116366 |
| 13027 | NM_030907.3 | #N/A | 23.0328015 | -5.7393643 | 1.59388728 |
| 13028 | NM_032367.3 | #N/A | 21.7888992 | -5.6537265 | 1.57028603 |

| | | | | | |
|---|---|---|---|---|---|
| 13029 | NM_012082.3 | #N/A | 91.1710189 | -6.1017137 | 1.69474135 |
| 13030 | NM_001164496.1 | #N/A | 75.4357897 | -7.4492534 | 2.06919908 |
| 13031 | NM_199190.2 | #N/A | 40.2512647 | -6.5360956 | 1.81561404 |
| 13032 | NM_024899.3 | #N/A | 24.9840214 | -5.8561632 | 1.62680637 |
| 13033 | NM_001317818.1 | #N/A | 32.4390291 | 5.74262825 | 1.59538875 |
| 13034 | NM_001199878.1 | #N/A | 23.5564294 | -5.7771269 | 1.60512715 |
| 13035 | NM_001122835.2 | #N/A | 43.4097787 | -6.6514475 | 1.8484278 |
| 13036 | NR_033935.1 | DCTN1 | 27.3598664 | -5.9863766 | 1.66362319 |
| 13037 | NM_001348090.1 | C20orf204 | 28.8158164 | -6.057612 | 1.68348939 |
| 13038 | NM_018131.4 | #N/A | 26.7212236 | -5.948364 | 1.65314238 |
| 13039 | NM_001044264.2 | #N/A | 25.0065656 | -5.8526643 | 1.62657832 |
| 13040 | NM_152618.2 | #N/A | 21.1112047 | -5.61002 | 1.55940656 |
| 13041 | NR_026658.1 | LOC100240735 | 31.9920089 | -6.2129785 | 1.72713543 |
| 13042 | NR_134482.1 | #N/A | 38.2013773 | -5.012767 | 1.39350347 |
| 13043 | NR_034134.1 | LINC01249 | 10.7082011 | 5.85602038 | 1.62800651 |
| 13044 | NM_001135146.1 | #N/A | 27.6549315 | -6.0077605 | 1.67047662 |
| 13045 | NM_004881.4 | #N/A | 27.9618075 | -6.0237725 | 1.67508627 |
| 13046 | NM_014839.4 | PLPPR4 | 73.3815939 | -7.4071102 | 2.05979419 |
| 13048 | NM_003504.4 | CDC45 | 30.9521478 | -6.1599723 | 1.71327605 |
| 13047 | NM_001202234.1 | NR4A1 | 99.414924 | -1.8371242 | 0.51095972 |
| 13049 | NR_003370.2 | RRN3P1 | 24.2436092 | -5.8096867 | 1.61586718 |
| 13050 | NR_132352.1 | BLOC1S6 | 31.5956789 | -6.1899006 | 1.72175802 |
| 13051 | NM_002306.3 | #N/A | 21.7215531 | -5.6544399 | 1.57283427 |
| 13052 | NM_001085457.1 | CBWD6 | 32.2175352 | -6.2199213 | 1.73021622 |
| 13053 | NM_015122.2 | #N/A | 31.8520993 | -6.2005707 | 1.72520822 |
| 13054 | NR_028383.1 | #N/A | 27.5730616 | -5.9989442 | 1.66921539 |
| 13055 | NR_110835.1 | #N/A | 39.8642206 | -6.5299119 | 1.81694802 |
| 13056 | NM_006569.5 | #N/A | 41.5323009 | -6.5851529 | 1.83243759 |
| 13057 | NM_001122731.2 | MICAL3 | 29.7340498 | 5.23358442 | 1.45634902 |
| 13058 | NM_001164095.2 | #N/A | 24.7542714 | -5.8348134 | 1.62383025 |
| 13059 | NM_001665.3 | #N/A | 42.8774575 | -5.1635197 | 1.43711026 |
| 13060 | NM_001144942.1 | #N/A | 38.1555227 | -6.4691793 | 1.80069887 |
| 13061 | NM_020153.3 | IFT46 | 22.8704236 | -5.7299844 | 1.5949889 |
| 13062 | NR_135607.1 | #N/A | 29.0246359 | -6.0766496 | 1.69165916 |
| 13063 | NM_001322131.1 | ZNF160 | 31.2787586 | -6.1687694 | 1.71731544 |
| 13064 | NM_001238.3 | #N/A | 23.8425484 | -5.7791463 | 1.60885792 |
| 13065 | NM_033510.2 | #N/A | 176.754045 | -3.5230425 | 0.98079044 |
| 13066 | NM_001330354.1 | UNC50 | 24.0256592 | -5.7908902 | 1.61222823 |
| 13067 | NR_109820.1 | NR2F1-AS1 | 37.5215716 | -6.4368561 | 1.79218578 |
| 13068 | NM_005949.3 | #N/A | 25.9931422 | -5.9089269 | 1.64531926 |
| 13069 | NM_000062.2 | #N/A | 28.6205077 | -6.0562626 | 1.68639508 |
| 13070 | NR_103551.2 | #N/A | 38.7846244 | -6.4842576 | 1.80647955 |
| 13071 | NM_080622.3 | ABHD16B | 26.4565971 | -5.9341613 | 1.6532468 |

| | | | | | |
|---|---|---|---|---|---|
| 13072 | NM_001270639.1 | JOSD2 | 25.3197113 | -5.8646434 | 1.63397691 |
| 13073 | NM_001199635.1 | ADCYAP1R1 | 39.7186019 | -6.5234667 | 1.81759499 |
| 13074 | NM_001135934.1 | #N/A | 40.9814329 | -6.5670679 | 1.82986874 |
| 13075 | NM_001130688.1 | HMGB2 | 46.5846396 | -6.7479352 | 1.88036173 |
| 13076 | NR_047465.1 | LINC00501 | 45.9915085 | 2.40788844 | 0.67097247 |
| 13077 | NM_006820.3 | #N/A | 79.7877222 | -5.934581 | 1.65407473 |
| 13078 | NM_130800.2 | MEN1 | 34.8661502 | -6.3325296 | 1.76505519 |
| 13079 | NM_015070.5 | #N/A | 29.4920473 | -6.0982684 | 1.69981804 |
| 13080 | NR_028051.1 | MTERF4 | 24.7335467 | -5.8445632 | 1.62911385 |
| 13081 | NM_198391.2 | #N/A | 25.0923762 | -5.8643887 | 1.63471965 |
| 13082 | NM_002842.4 | #N/A | 27.0641522 | -5.9751179 | 1.66564332 |
| 13083 | NM_199051.2 | #N/A | 34.0006942 | -6.2984423 | 1.75579777 |
| 13084 | NM_001005368.2 | ZNF32 | 22.1231034 | -5.671904 | 1.58125473 |
| 13085 | NM_001199165.2 | #N/A | 28.6054419 | -6.0550107 | 1.68805408 |
| 13086 | NM_001304509.1 | #N/A | 27.3401093 | -5.9745573 | 1.66576377 |
| 13087 | NM_018273.3 | #N/A | 24.7549436 | -5.8390449 | 1.62798464 |
| 13088 | NM_014079.3 | #N/A | 157.246338 | 3.98631867 | 1.11144653 |
| 13089 | NM_001003795.2 | #N/A | 145.170279 | -4.2454155 | 1.18370811 |
| 13090 | NM_001257234.1 | ALG13 | 34.1712414 | -6.2996328 | 1.75676518 |
| 13091 | NM_032435.2 | #N/A | 21.3935441 | -5.6275779 | 1.56936508 |
| 13092 | NM_001205254.1 | #N/A | 44.5817079 | -6.6919079 | 1.8662635 |
| 13093 | NM_001130442.2 | HRAS | 29.0432145 | -6.073441 | 1.69415521 |
| 13094 | NM_173558.3 | #N/A | 18.8226743 | 3.26468288 | 0.91083348 |
| 13095 | NM_020130.4 | #N/A | 34.6435531 | -6.3220732 | 1.76388754 |
| 13096 | NM_032229.2 | #N/A | 39.5732376 | -5.0624135 | 1.41246546 |
| 13097 | NM_014145.4 | TMEM230 | 27.7110152 | -5.9933831 | 1.67228374 |
| 13098 | NM_014323.2 | #N/A | 22.1706988 | -5.6779166 | 1.58434435 |
| 13099 | NM_016055.5 | #N/A | 37.8852956 | -6.4533641 | 1.80105582 |
| 13100 | NM_024518.2 | #N/A | 200.869013 | 1.46980555 | 0.41021717 |
| 13101 | NM_001161476.1 | #N/A | 38.9308179 | -5.0387126 | 1.40641139 |
| 13102 | NM_001242929.1 | ANKS3 | 35.3302101 | -6.3550163 | 1.77386191 |
| 13103 | NM_002350.3 | #N/A | 24.1051129 | -5.8101208 | 1.62179269 |
| 13104 | NM_001098634.1 | #N/A | 107.110115 | -7.9543419 | 2.22055986 |
| 13105 | NM_018153.3 | ANTXR1 | 28.8996962 | -6.0688767 | 1.69425859 |
| 13106 | NM_007232.2 | #N/A | 36.4066209 | -6.3978707 | 1.78624283 |
| 13107 | NM_000991.4 | #N/A | 771.222893 | -3.5678786 | 0.99614794 |
| 13108 | NM_001077690.1 | ALG9 | 119.37002 | -2.6827472 | 0.74905303 |
| 13109 | NR_138249.1 | #N/A | 17.3819313 | 7.22050886 | 2.01605755 |
| 13110 | NM_001134831.1 | #N/A | 43.217159 | -6.6485785 | 1.85639872 |
| 13111 | NM_001324366.1 | #N/A | 34.1840026 | -6.3020601 | 1.75975054 |
| 13112 | NR_135677.1 | #N/A | 50.8311935 | -6.8747925 | 1.91985659 |
| 13113 | NM_001098502.1 | #N/A | 26.4237308 | -5.9405189 | 1.65901878 |
| 13114 | NM_006610.3 | #N/A | 22.792897 | -5.7266716 | 1.59985372 |

| | | | | | |
|---|---|---|---|---|---|
| 13115 | NM_001206427.1 | #N/A | 30.1782639 | -6.1267207 | 1.71179531 |
| 13116 | NM_001199263.1 | #N/A | 29.2596379 | -6.0774076 | 1.69809756 |
| 13117 | NM_021944.3 | #N/A | 23.5216 | -5.7658639 | 1.61120307 |
| 13118 | NM_153347.2 | #N/A | 22.6582001 | -5.7122702 | 1.59623964 |
| 13119 | NR_122073.1 | LINC01879 | 13.4186112 | 3.1315157 | 0.87509514 |
| 13120 | NM_016071.3 | #N/A | 26.4479092 | -5.9331432 | 1.65812899 |
| 13121 | NM_001193451.1 | #N/A | 92.4150926 | -3.9546951 | 1.10525257 |
| 13122 | NM_001303507.1 | #N/A | 39.1878199 | -6.5070905 | 1.8187281 |
| 13123 | NM_001267616.1 | #N/A | 59.833659 | -7.1114963 | 1.98796633 |
| 13124 | NM_001318093.1 | #N/A | 38.2571854 | -6.4645314 | 1.80710943 |
| 13125 | NR_002188.3 | GBAP1 | 26.8942781 | -5.9577648 | 1.66550281 |
| 13126 | NM_152836.2 | #N/A | 22.6986257 | -5.7061952 | 1.5953439 |
| 13127 | NM_001199379.1 | #N/A | 279.938393 | -9.3397445 | 2.61171448 |
| 13128 | NM_001201427.1 | #N/A | 69.6429022 | -3.2643854 | 0.91296335 |
| 13129 | NM_020772.2 | #N/A | 440.173843 | -2.1435457 | 0.59949569 |
| 13130 | NM_014470.3 | #N/A | 35.2905346 | -6.3479975 | 1.77540195 |
| 13131 | NM_153836.3 | #N/A | 33.3173367 | -6.2691064 | 1.75344677 |
| 13132 | NM_016284.4 | #N/A | 274.222309 | -9.3099807 | 2.60417685 |
| 13133 | NM_001330227.1 | #N/A | 29.7268542 | -6.1061598 | 1.70805666 |
| 13134 | NR_144640.1 | #N/A | 26.081762 | -5.9094842 | 1.65307822 |
| 13135 | NM_005599.3 | NHLH2 | 35.0088387 | -5.0622995 | 1.4162024 |
| 13136 | NM_001142799.2 | #N/A | 29.2662558 | -6.0839309 | 1.70241255 |
| 13137 | NM_001284350.1 | #N/A | 10.4099506 | 4.08975343 | 1.14449246 |
| 13138 | NM_001290307.2 | CTNNA1 | 23.5045693 | -5.7743097 | 1.61603098 |
| 13139 | NM_000352.4 | #N/A | 35.5181043 | -6.3610965 | 1.78050241 |
| 13140 | NM_001007227.1 | SPOP | 26.0441895 | -5.9191621 | 1.65687422 |
| 13141 | NM_001134321.1 | #N/A | 28.3993272 | -6.0387239 | 1.69045258 |
| 13142 | NM_001164465.3 | GOLGA6L10 | 20.7359867 | -5.5826363 | 1.56323257 |
| 13143 | NM_024763.4 | #N/A | 32.5055337 | -6.2375312 | 1.74661092 |
| 13144 | NM_001303506.1 | #N/A | 22.1497867 | -5.6718707 | 1.5883252 |
| 13145 | NM_004455.2 | #N/A | 20.8230931 | -5.5898783 | 1.56543178 |
| 13146 | NR_003605.1 | #N/A | 37.6321589 | -6.4440679 | 1.80483617 |
| 13147 | NM_001195200.1 | CCDC107 | 29.3541313 | -6.0833898 | 1.7038477 |
| 13148 | NM_002873.1 | RAD17 | 27.9948976 | -6.0097094 | 1.6833467 |
| 13149 | NM_001204284.1 | #N/A | 33.2483581 | -6.2669566 | 1.75543122 |
| 13150 | NM_001297624.1 | ZNF480 | 25.8715976 | 9.66012418 | 2.70593766 |
| 13151 | NM_001242935.2 | SNCAIP | 31.7799628 | -6.2053812 | 1.73831039 |
| 13152 | NM_001007169.3 | #N/A | 152.494752 | 2.82279811 | 0.79075955 |
| 13153 | NM_153361.3 | #N/A | 22.7549378 | -5.7117382 | 1.60015765 |
| 13154 | NM_001242846.1 | RNF146 | 29.5722107 | -6.1012164 | 1.70933456 |
| 13155 | NM_014176.3 | #N/A | 62.7298943 | -5.6280884 | 1.57697859 |
| 13156 | NM_001289095.1 | PXK | 20.9710994 | -5.6021865 | 1.56974881 |
| 13157 | NM_001271849.1 | #N/A | 21.7385178 | -5.6596506 | 1.58601019 |

| | | | | | |
|---|---|---|---|---|---|
| 13158 | NM_001294333.1 | #N/A | 23.8884495 | -5.7966413 | 1.62439332 |
| 13159 | NM_001322152.1 | #N/A | 63.2003272 | -7.1887254 | 2.01451818 |
| 13160 | NM_006800.3 | #N/A | 21.8667398 | -5.6650015 | 1.58756032 |
| 13161 | NM_001302818.1 | #N/A | 26.1874443 | 8.31707373 | 2.33095695 |
| 13162 | NM_001287747.1 | FDFT1 | 26.9142198 | -5.9611203 | 1.67077608 |
| 13163 | NM_001130420.2 | #N/A | 39.9997477 | -6.5302804 | 1.83052501 |
| 13164 | NM_203386.2 | #N/A | 36.0322526 | -6.3750567 | 1.78710849 |
| 13165 | NM_001037305.1 | #N/A | 25.6292333 | -5.8869892 | 1.65030536 |
| 13166 | NR_103486.1 | DNM3OS | 31.3296596 | -6.1800138 | 1.73257022 |
| 13167 | NM_001321091.1 | #N/A | 25.5417334 | -5.882039 | 1.64913691 |
| 13168 | NM_053004.2 | #N/A | 32.4965792 | -6.2309887 | 1.74706464 |
| 13169 | NR_111915.1 | LOC101929057 | 60.9262854 | 2.54595133 | 0.71384641 |
| 13170 | NM_001329877.1 | FUT1 | 81.03462 | 2.29386448 | 0.64318278 |
| 13171 | NR_002166.1 | #N/A | 26.5392684 | -5.9404953 | 1.66569422 |
| 13172 | NM_019004.1 | #N/A | 292.860059 | -4.0056053 | 1.1234485 |
| 13173 | NM_020251.3 | #N/A | 46.4435395 | -6.7515529 | 1.89363488 |
| 13174 | NM_004328.4 | #N/A | 28.3036429 | -6.0305611 | 1.69145992 |
| 13175 | NM_020924.3 | #N/A | 23.7757505 | -5.7806345 | 1.62140277 |
| 13176 | NM_194272.2 | #N/A | 21.8181623 | -5.6654841 | 1.58952367 |
| 13177 | NM_020700.1 | #N/A | 120.687255 | -3.0234248 | 0.84845316 |
| 13178 | NM_138779.4 | #N/A | 24.585455 | -5.825195 | 1.63476054 |
| 13179 | NM_001001683.3 | #N/A | 23.6298405 | -5.7690992 | 1.61906666 |
| 13180 | NM_021973.2 | #N/A | 36.123998 | -6.3858982 | 1.79253558 |
| 13181 | NM_015832.4 | #N/A | 29.0197271 | -6.0665432 | 1.7032419 |
| 13182 | NR_026991.1 | H1FX-AS1 | 29.0507796 | -4.7891034 | 1.3447925 |
| 13183 | NM_001290050.1 | UHRF1 | 144.545434 | -2.4773879 | 0.69567326 |
| 13184 | NM_016095.2 | #N/A | 51.6179393 | -5.3832181 | 1.51167697 |
| 13185 | NM_007001.2 | #N/A | 27.711631 | -6.0097272 | 1.68772385 |
| 13186 | NM_001306188.1 | #N/A | 22.0902617 | -5.6814231 | 1.59553577 |
| 13187 | NM_001286119.1 | CCDC90B | 23.223951 | -5.7400973 | 1.61221491 |
| 13188 | NM_001198798.2 | ASCC1 | 29.8787474 | -6.1123517 | 1.71692842 |
| 13189 | NM_152355.2 | #N/A | 69.1005703 | -4.5547377 | 1.27950728 |
| 13190 | NM_025027.4 | ZNF606 | 27.3179119 | -5.9868431 | 1.68182996 |
| 13191 | NM_013381.2 | TRHDE | 27.0686326 | -5.9692213 | 1.67707675 |
| 13192 | NM_001270378.1 | #N/A | 21.2324603 | -5.6211423 | 1.57930855 |
| 13193 | NM_001271933.1 | CEP164 | 26.8865879 | -5.9604889 | 1.67466094 |
| 13194 | NM_001290144.1 | #N/A | 43.6100117 | -6.6542862 | 1.86980369 |
| 13195 | NM_005486.2 | #N/A | 35.7865059 | -6.3749044 | 1.79137427 |
| 13196 | NM_024683.3 | #N/A | 22.2512987 | -5.6817279 | 1.59659551 |
| 13197 | NM_023070.2 | #N/A | 29.5615385 | -6.0940642 | 1.7125438 |
| 13198 | NM_004841.3 | #N/A | 59.3269777 | -5.5565971 | 1.56158577 |
| 13199 | NM_004177.4 | #N/A | 35.590596 | -6.3702097 | 1.79056847 |
| 13200 | NM_030671.2 | #N/A | 43.2216086 | -6.6416133 | 1.86691523 |

| | | | | | |
|---|---|---|---|---|---|
| 13201 | NM_001031849.2 | MASP1 | 45.4971025 | -6.7148425 | 1.88772805 |
| 13202 | NM_005334.2 | #N/A | 508.369606 | -4.6409813 | 1.30472184 |
| 13203 | NM_002758.3 | #N/A | 23.799533 | -5.7824683 | 1.62587092 |
| 13204 | NR_126040.1 | LOC101927188 | 14.4215884 | 4.49114053 | 1.26302672 |
| 13205 | NM_002334.3 | #N/A | 150.193549 | -4.2804311 | 1.20384531 |
| 13206 | NM_182743.2 | #N/A | 37.5355253 | -6.4463481 | 1.81330456 |
| 13207 | NM_145897.2 | PFDN5 | 54.6092029 | -6.9834029 | 1.96446468 |
| 13208 | NM_212552.2 | #N/A | 23.8160932 | -5.7877955 | 1.62825232 |
| 13209 | NR_034101.1 | NCAM1-AS1 | 32.904354 | -6.2514975 | 1.75878051 |
| 13210 | NM_001139457.2 | BCAP31 | 24.4281692 | -5.8205782 | 1.63760033 |
| 13211 | NR_073449.1 | #N/A | 33.0269957 | -6.2595726 | 1.76112208 |
| 13212 | NM_000505.3 | F12 | 36.1937419 | -6.3855362 | 1.7967696 |
| 13213 | NM_001185093.1 | NIT1 | 22.3868837 | -5.6915597 | 1.60149854 |
| 13214 | NM_007023.3 | #N/A | 24.4898561 | -5.8235792 | 1.63867336 |
| 13215 | NM_022372.4 | #N/A | 33.012844 | -6.2510074 | 1.75923466 |
| 13216 | NM_001319670.1 | #N/A | 23.8732589 | -5.7964338 | 1.63135958 |
| 13217 | NM_001330419.1 | #N/A | 32.7031615 | -6.2413442 | 1.7567898 |
| 13218 | NM_001320470.1 | #N/A | 23.1083985 | -5.7402427 | 1.61578912 |
| 13219 | NM_001031806.1 | #N/A | 26.9517736 | -5.9656532 | 1.67937546 |
| 13220 | NM_001128848.1 | SMARCA4 | 373.333912 | -9.7549147 | 2.7465204 |
| 13221 | NM_000121.3 | #N/A | 23.5045217 | -5.7699967 | 1.62464054 |
| 13222 | NM_212539.1 | #N/A | 27.814453 | -6.0142063 | 1.6934913 |
| 13223 | NM_001101667.1 | ACOX3 | 25.3861746 | -5.8839535 | 1.6569815 |
| 13224 | NR_125434.1 | LOC100505915 | 114.0774 | 1.4680676 | 0.41342197 |
| 13225 | NM_016335.4 | #N/A | 56.263781 | -7.0270207 | 1.979061 |
| 13226 | NM_001142310.1 | TMEM169 | 41.7293567 | -6.5923308 | 1.85669598 |
| 13227 | NM_198123.1 | #N/A | 33.8632742 | -6.293877 | 1.77265313 |
| 13228 | NM_001323682.1 | #N/A | 28.9707821 | -6.0713082 | 1.71051015 |
| 13229 | NM_001002880.1 | CBY1 | 21.9451979 | -5.6663994 | 1.59645137 |
| 13230 | NM_001199355.1 | RPL17-C18orf32 | 37.5472745 | -6.4391601 | 1.81422338 |
| 13231 | NM_001346167.1 | ZNF287 | 32.1926925 | -6.2100961 | 1.74970777 |
| 13232 | NR_033349.1 | RAI2 | 28.0566398 | -6.0107345 | 1.693653 |
| 13233 | NM_145243.4 | #N/A | 21.2379064 | -5.625541 | 1.58531582 |
| 13234 | NM_001286606.1 | #N/A | 25.337034 | -5.8779842 | 1.65646765 |
| 13235 | NM_001685.4 | ATP5PF | 45.3332171 | -6.7109697 | 1.89139079 |
| 13236 | NM_001195007.1 | PPIE | 22.9325828 | -5.7202 | 1.61222807 |
| 13237 | NM_001206540.2 | #N/A | 23.8609246 | -5.7867527 | 1.63113801 |
| 13238 | NM_001329138.1 | PTPRF | 55.6331762 | -7.0047519 | 1.97451735 |
| 13239 | NM_001334.2 | #N/A | 31.2431354 | -6.1809428 | 1.7423037 |
| 13240 | NM_003551.2 | #N/A | 26.033811 | -5.9190908 | 1.66853416 |
| 13241 | NM_016526.4 | BET1L | 23.6606478 | -5.7707778 | 1.62680603 |
| 13242 | NM_003811.3 | #N/A | 20.7294683 | -5.5888539 | 1.5756065 |
| 13243 | NM_005619.4 | #N/A | 35.5165945 | -6.35749 | 1.7925143 |

| | | | | | |
|---|---|---|---|---|---|
| 13244 | NR_038885.1 | LINC01426 | 50.5382048 | 4.59110187 | 1.29453369 |
| 13245 | NM_016090.3 | RBM7 | 31.7500712 | -6.2002565 | 1.74866694 |
| 13246 | NM_004360.4 | #N/A | 23.1122926 | -5.7441713 | 1.62024165 |
| 13247 | NM_001321367.1 | #N/A | 121.81711 | 2.37071679 | 0.66874388 |
| 13248 | NM_001289404.1 | MMS19 | 27.6718662 | -6.0027064 | 1.69329655 |
| 13249 | NM_001305624.1 | CALM2 | 35.5143042 | -6.360109 | 1.7944187 |
| 13250 | NM_001317777.1 | RIPPLY3 | 55.9000787 | 2.40932627 | 0.67975709 |
| 13251 | NM_016020.3 | #N/A | 30.7144861 | -6.1491962 | 1.73502683 |
| 13252 | NR_027855.1 | #N/A | 39.7057727 | -6.5203203 | 1.83995096 |
| 13253 | NM_052898.1 | #N/A | 33.3425728 | -6.2702952 | 1.76952344 |
| 13254 | NR_131765.1 | #N/A | 25.7697464 | -5.8957402 | 1.66388542 |
| 13255 | NM_020845.2 | PITPNM2 | 45.396935 | -6.7175487 | 1.89684055 |
| 13256 | NM_024071.3 | ZFYVE21 | 38.10244 | -6.4659667 | 1.82605001 |
| 13257 | NM_021253.3 | #N/A | 30.9260188 | -4.8780674 | 1.37764464 |
| 13258 | NM_020156.4 | #N/A | 133.907189 | -1.8677304 | 0.52750803 |
| 13259 | NM_024610.5 | #N/A | 23.8191061 | -5.7827135 | 1.63322122 |
| 13260 | NM_177454.3 | #N/A | 264.063375 | -4.6493811 | 1.31313404 |
| 13261 | NM_004126.3 | #N/A | 24.7832538 | -5.8374041 | 1.64878896 |
| 13262 | NR_136746.1 | #N/A | 40.8243682 | -6.5601867 | 1.85309531 |
| 13263 | NM_001320990.1 | #N/A | 21.970212 | -5.6621275 | 1.59949243 |
| 13264 | NM_201434.2 | #N/A | 30.5361415 | -4.8576911 | 1.37225995 |
| 13265 | NM_032726.3 | #N/A | 20.4648685 | -5.5661572 | 1.57270694 |
| 13266 | NM_001128302.2 | #N/A | 32.4140232 | -6.2312203 | 1.76067619 |
| 13267 | NM_001330241.1 | #N/A | 26.8011782 | -5.9542422 | 1.68253678 |
| 13268 | NM_020468.5 | SNX14 | 31.5201791 | -6.1946342 | 1.75052387 |
| 13269 | NM_003416.3 | #N/A | 32.5649214 | -6.2330557 | 1.76146027 |
| 13270 | NM_001330596.1 | #N/A | 26.1724846 | -5.9215212 | 1.67349379 |
| 13271 | NR_104273.1 | FBXW7-AS1 | 29.0699901 | 8.16702096 | 2.30829977 |
| 13272 | NR_036541.1 | GPR89A | 22.4956622 | -5.6940274 | 1.60965887 |
| 13273 | NR_073074.1 | #N/A | 29.3840214 | -6.0894033 | 1.72143227 |
| 13274 | NM_018339.5 | #N/A | 250.735613 | -3.2036349 | 0.90582279 |
| 13275 | NR_047674.1 | LRRC4C | 26.085766 | -5.9116583 | 1.67207422 |
| 13277 | NM_202494.2 | GIPC1 | 25.9735238 | -5.9138611 | 1.67297131 |
| 13276 | NM_022103.3 | ZNF667 | 26.0228709 | -5.9053355 | 1.67056248 |
| 13278 | NR_051991.1 | SENP1 | 24.3490429 | -5.8144552 | 1.64487117 |
| 13279 | NM_145697.2 | #N/A | 29.3574788 | -6.0871338 | 1.72250411 |
| 13280 | NM_004087.2 | DLG1 | 33.9827342 | -6.3035564 | 1.78377425 |
| 13281 | NM_030573.2 | #N/A | 22.9988378 | -5.7379323 | 1.62405737 |
| 13282 | NM_001282179.1 | #N/A | 44.9861649 | -6.7006331 | 1.89681445 |
| 13283 | NR_038879.1 | PAXBP1-AS1 | 20.8529013 | -5.5942137 | 1.5836326 |
| 13284 | NM_032505.2 | #N/A | 21.2971471 | -5.615396 | 1.58975217 |
| 13285 | NM_001322342.1 | DDX31 | 33.0110634 | -4.969432 | 1.40688429 |
| 13286 | NM_001347902.1 | #N/A | 29.2469705 | -6.0809129 | 1.72167372 |

| | | | | | |
|---|---|---|---|---|---|
| 13287 | NM_001271512.3 | ACBD5 | 26.4414332 | -5.932614 | 1.6799361 |
| 13288 | NM_018134.2 | #N/A | 86.353633 | 2.6375367 | 0.74694014 |
| 13289 | NM_005512.2 | LRRC32 | 27.3661952 | -5.9851664 | 1.69511523 |
| 13290 | NM_017940.4 | #N/A | 412.315603 | -2.0888289 | 0.59160382 |
| 13291 | NR_028028.1 | #N/A | 13.4187389 | 8.70630301 | 2.46583202 |
| 13292 | NM_001336.3 | #N/A | 38.718852 | -6.4890359 | 1.83792808 |
| 13293 | NM_000161.2 | #N/A | 30.9598647 | -6.1621506 | 1.74542912 |
| 13294 | NM_017845.4 | #N/A | 31.9681973 | -6.2053747 | 1.75777163 |
| 13295 | NR_104080.1 | RNU6-9 | 29.1008317 | -6.0703224 | 1.71960291 |
| 13296 | NM_058237.1 | #N/A | 19.975297 | -5.5312943 | 1.56694779 |
| 13297 | NM_001267776.1 | #N/A | 23.1503366 | -5.749931 | 1.62893533 |
| 13298 | NM_001243160.1 | #N/A | 20.773484 | -5.5897725 | 1.58365879 |
| 13299 | NR_138463.1 | #N/A | 23.8831369 | -5.7798789 | 1.63751999 |
| 13300 | NM_152411.3 | #N/A | 74.2477925 | -3.0790932 | 0.87253538 |
| 13301 | NM_002394.5 | #N/A | 26.1208995 | -5.9190634 | 1.67735918 |
| 13302 | NM_004085.3 | #N/A | 26.4100325 | -5.9339776 | 1.68158755 |
| 13303 | NM_005671.3 | #N/A | 22.2972479 | -5.6904563 | 1.6125954 |
| 13304 | NM_001319119.1 | #N/A | 26.5172391 | -5.9301781 | 1.68070018 |
| 13305 | NM_198531.4 | #N/A | 50.8991002 | -6.8772388 | 1.94932071 |
| 13306 | NM_001145784.1 | #N/A | 20.4968472 | -5.5704928 | 1.5792046 |
| 13307 | NM_145206.3 | VTI1A | 27.1592423 | -5.9755428 | 1.69418267 |
| 13308 | NM_144651.4 | #N/A | 60.1997764 | -5.5688705 | 1.57889384 |
| 13309 | NM_001242534.1 | #N/A | 25.7815759 | -5.9050028 | 1.67421174 |
| 13310 | NR_144476.1 | DPH1 | 27.36497 | -5.9821872 | 1.69616201 |
| 13311 | NM_005261.3 | #N/A | 35.0096801 | -6.34237 | 1.79845305 |
| 13312 | NM_001284528.1 | ZSCAN32 | 29.1571444 | -6.0723526 | 1.7219734 |
| 13313 | NM_015426.4 | #N/A | 21.6289817 | -5.6357972 | 1.59830218 |
| 13314 | NR_029406.1 | RPL23AP87 | 66.6710851 | 2.0009911 | 0.56754877 |
| 13315 | NM_001271573.1 | #N/A | 21.1492408 | -5.6085036 | 1.59081809 |
| 13316 | NM_001278647.1 | #N/A | 22.9102627 | -5.7301415 | 1.62566183 |
| 13317 | NM_001145313.2 | #N/A | 25.4716642 | -5.8872772 | 1.67038479 |
| 13318 | NM_015060.2 | #N/A | 347.398904 | -1.7991271 | 0.51054019 |
| 13319 | NR_027749.1 | #N/A | 24.5236414 | -5.8316911 | 1.65503166 |
| 13320 | NR_132313.1 | PCA3 | 36.4593916 | 3.25195733 | 0.92302112 |
| 13321 | NM_001136113.1 | POMT1 | 23.5781082 | -5.7731878 | 1.63865207 |
| 13322 | NM_002381.4 | #N/A | 22.3990792 | -5.6971892 | 1.61739707 |
| 13323 | NR_048551.1 | SLC4A3 | 44.6948518 | -6.691191 | 1.89969671 |
| 13324 | NR_036492.1 | PEX19 | 29.9946485 | -6.1215903 | 1.73801293 |
| 13325 | NR_015445.1 | BREA2 | 63.5969358 | 3.89269172 | 1.1052382 |
| 13326 | NM_001256167.1 | ZNF620 | 38.9979463 | -6.4919257 | 1.84332316 |
| 13327 | NM_001160328.1 | SYT3 | 29.3250157 | -6.082429 | 1.72739333 |
| 13328 | NM_001009181.1 | #N/A | 19.2386117 | 9.26066354 | 2.63028493 |
| 13329 | NM_001077243.2 | GRIA4 | 29.5377862 | -6.098909 | 1.73265872 |

| | | | | | |
|---|---|---|---|---|---|
| 13331 | NM_033313.2 | CDC14A | 18.9266017 | 6.75660707 | 1.91962765 |
| 13330 | NM_001320608.1 | #N/A | 22.3249867 | -5.6906456 | 1.61677765 |
| 13332 | NM_004507.3 | #N/A | 22.3845931 | -5.6931789 | 1.61753722 |
| 13333 | NM_199122.2 | TBRG4 | 27.5770059 | -5.9953178 | 1.70353504 |
| 13334 | NM_053052.3 | SNAP47 | 35.3089994 | -6.3493243 | 1.80418454 |
| 13335 | NR_003501.1 | EARS2 | 23.1878356 | -5.7453387 | 1.63292359 |
| 13336 | NM_080651.3 | #N/A | 23.4155001 | -5.7582141 | 1.63701367 |
| 13337 | NR_037653.1 | #N/A | 21.2013538 | -5.6156917 | 1.59651861 |
| 13338 | NM_000674.2 | #N/A | 39.5904308 | -6.5182745 | 1.85318919 |
| 13339 | NM_001164637.2 | #N/A | 28.0845637 | -6.0256675 | 1.71314169 |
| 13340 | NM_025057.2 | #N/A | 45.5641591 | -5.2222152 | 1.48484762 |
| 13341 | NM_001285439.1 | #N/A | 35.1100568 | -6.3390195 | 1.8024152 |
| 13342 | NM_015319.2 | TNS2 | 23.8107055 | -5.7746354 | 1.64198807 |
| 13343 | NM_001302348.1 | #N/A | 22.8961878 | -5.7252255 | 1.62796599 |
| 13344 | NM_004867.4 | #N/A | 27.2361436 | -5.9801905 | 1.70049739 |
| 13345 | NM_003394.3 | #N/A | 24.2765691 | -5.8028684 | 1.65016489 |
| 13346 | NM_006511.2 | RSC1A1 | 21.9277585 | -5.6650211 | 1.61106095 |
| 13347 | NM_016594.2 | #N/A | 21.5747155 | -5.6474865 | 1.60630868 |
| 13348 | NM_006902.4 | #N/A | 34.2984211 | -6.3148295 | 1.79617341 |
| 13349 | NM_031439.3 | #N/A | 47.2571246 | 2.44046261 | 0.69419682 |
| 13351 | NR_004847.3 | UBE2Q2P2 | 22.6580118 | -5.7136451 | 1.62536447 |
| 13350 | NM_017414.3 | #N/A | 23.675904 | -5.7816148 | 1.64469928 |
| 13352 | NM_152376.4 | #N/A | 54.7969343 | -6.9885083 | 1.98805201 |
| 13353 | NM_153213.4 | #N/A | 21.8433845 | -5.657038 | 1.60942464 |
| 13354 | NM_001322194.1 | JAK2 | 30.9140167 | -6.1657101 | 1.75418772 |
| 13355 | NM_018530.2 | GSDMB | 21.3171572 | -5.6168483 | 1.59806303 |
| 13356 | NM_020647.3 | #N/A | 27.4180699 | -5.9834974 | 1.70259198 |
| 13357 | NM_006748.3 | #N/A | 92.5892629 | -7.7425644 | 2.20322228 |
| 13358 | NM_017857.3 | #N/A | 29.8277872 | -6.1134877 | 1.73981307 |
| 13359 | NM_001128845.1 | SMARCA4 | 331.964405 | -9.5854275 | 2.72802611 |
| 13360 | NM_001256946.1 | FAAP20 | 24.4025672 | -5.8133855 | 1.65451297 |
| 13361 | NM_145017.2 | #N/A | 50.4022273 | -6.86789 | 1.95468629 |
| 13362 | NM_001085377.1 | #N/A | 20.2542993 | -5.545253 | 1.57827168 |
| 13363 | NM_001320310.1 | #N/A | 50.478095 | -6.8649492 | 1.95413816 |
| 13364 | NM_023074.3 | #N/A | 64.0647265 | -4.4579238 | 1.26909416 |
| 13365 | NM_003309.3 | TSPYL1 | 823.621897 | -2.9405086 | 0.8371858 |
| 13366 | NM_001310160.1 | PAX6 | 26.7461977 | -5.9534089 | 1.69501234 |
| 13367 | NM_017815.2 | #N/A | 27.5084448 | -5.9925273 | 1.70616858 |
| 13369 | NM_001172702.1 | SLC38A6 | 21.4493746 | -5.6385974 | 1.6054421 |
| 13368 | NM_001043352.1 | #N/A | 27.311376 | -5.9875004 | 1.7047907 |
| 13370 | NM_001320586.1 | #N/A | 23.3856744 | -5.7655134 | 1.64156198 |
| 13371 | NR_134893.1 | #N/A | 26.350736 | -5.926043 | 1.68728808 |
| 13372 | NM_018190.3 | BBS7 | 65.7518973 | -3.3417137 | 0.95163251 |

| | | | | | |
|---|---|---|---|---|---|
| 13373 | NM_003789.3 | #N/A | 20.6325638 | -5.5782663 | 1.58865647 |
| 13374 | NM_001300733.1 | #N/A | 21.4534498 | -5.6320531 | 1.60399437 |
| 13375 | NM_001142648.1 | #N/A | 38.4816273 | -6.4795568 | 1.84538718 |
| 13376 | NM_001243101.1 | NTRK3 | 27.7340224 | -6.0018554 | 1.70954118 |
| 13377 | NM_001199942.1 | DOP1A | 42.1588316 | -6.6107081 | 1.88359727 |
| 13378 | NR_104225.1 | SH3YL1 | 25.7721654 | -5.8918751 | 1.67880861 |
| 13379 | NM_001330393.1 | #N/A | 24.5939815 | -5.8340956 | 1.66239267 |
| 13380 | NM_014569.3 | #N/A | 27.5576245 | -6.0018471 | 1.71022428 |
| 13381 | NM_001174124.1 | GOLGA7 | 37.4001116 | -6.4286345 | 1.832086 |
| 13382 | NM_178009.4 | #N/A | 29.1495519 | -6.0692395 | 1.72971544 |
| 13383 | NM_001134875.1 | TEDC1 | 22.4564881 | -5.6907308 | 1.62185573 |
| 13384 | NM_001080433.1 | #N/A | 35.3614811 | -6.3530343 | 1.81062263 |
| 13385 | NM_016377.3 | #N/A | 23.9690855 | -5.7956982 | 1.65197771 |
| 13386 | NM_014668.3 | #N/A | 254.471381 | -2.0750389 | 0.59146639 |
| 13387 | NM_022761.2 | #N/A | 27.5033478 | -5.9921686 | 1.70806051 |
| 13388 | NM_001291420.1 | GOLGA6L9 | 111.21766 | -4.3737871 | 1.24684258 |
| 13389 | NM_001197123.1 | #N/A | 20.6945193 | -5.5836112 | 1.59176044 |
| 13390 | NM_005815.4 | #N/A | 21.1861632 | -5.6161565 | 1.6011191 |
| 13391 | NM_001320640.1 | #N/A | 45.7040973 | -6.7258644 | 1.9175492 |
| 13392 | NM_001164757.1 | #N/A | 40.0844053 | -6.5347316 | 1.86307494 |
| 13393 | NM_001002814.2 | #N/A | 41.6168052 | -6.5923697 | 1.87968198 |
| 13394 | NM_017665.3 | #N/A | 26.2097111 | -5.9161602 | 1.68693588 |
| 13395 | NM_031951.4 | #N/A | 27.9821385 | 6.69114771 | 1.90810875 |
| 13396 | NM_006040.2 | #N/A | 77.1299356 | -5.8707644 | 1.67419738 |
| 13397 | NM_001293187.1 | METTL5 | 32.1990334 | -6.2189677 | 1.77357788 |
| 13398 | NM_020843.2 | #N/A | 162.467776 | -3.2008762 | 0.91292372 |
| 13399 | NM_001331039.1 | #N/A | 26.834023 | -5.9474364 | 1.69642285 |
| 13400 | NM_001321697.1 | #N/A | 23.2908402 | -5.7558499 | 1.64233379 |
| 13401 | NM_001321221.1 | #N/A | 188.213186 | 2.32148105 | 0.6624003 |
| 13402 | NM_001102594.1 | DTX2 | 20.376676 | -5.5568258 | 1.58578922 |
| 13403 | NM_001039712.1 | DEDD | 25.6880972 | -5.9011119 | 1.68415855 |
| 13404 | NM_032786.2 | #N/A | 26.4697501 | -5.9371186 | 1.69459653 |
| 13405 | NM_176793.1 | #N/A | 22.1394894 | -5.6742615 | 1.61960978 |
| 13406 | NR_134519.1 | #N/A | 31.5558015 | -6.1941095 | 1.76798383 |
| 13407 | NM_001143785.1 | #N/A | 34.5378621 | 5.04225599 | 1.43923275 |
| 13408 | NM_001347869.1 | #N/A | 19.4268801 | -5.4895418 | 1.56700021 |
| 13409 | NM_145313.3 | #N/A | 33.6824019 | -4.8368693 | 1.38072762 |
| 13410 | NM_175066.3 | #N/A | 329.437858 | -1.5076585 | 0.43041321 |
| 13411 | NM_001220778.1 | CDKN1A | 28.1993185 | -6.0342506 | 1.72284722 |
| 13412 | NM_178013.3 | #N/A | 19.8904463 | -5.5244279 | 1.5772804 |
| 13413 | NM_024946.2 | #N/A | 340.979797 | -3.1735048 | 0.90627753 |
| 13414 | NM_001203263.1 | #N/A | 25.2710434 | -5.859001 | 1.67336149 |
| 13415 | NM_203446.2 | #N/A | 36.8348924 | -6.41521 | 1.83224415 |

| | | | | | |
|---|---|---|---|---|---|
| 13416 | NM_001320247.1 | #N/A | 50.1679495 | -6.8569484 | 1.95859577 |
| 13417 | NM_145331.2 | #N/A | 44.1498147 | -6.6780028 | 1.90752929 |
| 13418 | NR_024249.1 | FAM86C2P | 20.8958929 | -5.5877557 | 1.59628284 |
| 13420 | NM_001286265.1 | MRS2 | 27.4557665 | -5.9917947 | 1.71178436 |
| 13419 | NM_001042665.1 | PLEKHG5 | 25.2604785 | -5.8733027 | 1.67793536 |
| 13421 | NM_014620.5 | #N/A | 37.2228332 | -6.4321428 | 1.83767457 |
| 13423 | NM_001288754.1 | RHOT1 | 47.0242784 | -6.7626909 | 1.93215695 |
| 13422 | NM_001282866.1 | #N/A | 25.2409054 | -5.8728858 | 1.67793936 |
| 13424 | NM_004502.3 | #N/A | 28.6519557 | -6.0527234 | 1.72934616 |
| 13425 | NM_000177.4 | #N/A | 51.3213707 | -6.8949279 | 1.97005569 |
| 13426 | NR_037642.1 | LHX4-AS1 | 23.5374971 | -5.7637541 | 1.64688424 |
| 13427 | NM_001302082.1 | BEND5 | 30.288535 | -6.1308547 | 1.75180507 |
| 13428 | NM_001164760.1 | #N/A | 29.9611571 | -6.1181099 | 1.74821341 |
| 13429 | NM_001127584.1 | #N/A | 43.687703 | -6.653299 | 1.90120127 |
| 13430 | NM_001753.4 | #N/A | 32.1700376 | -6.2214459 | 1.77791712 |
| 13431 | NR_003682.1 | MGC70870 | 35.239267 | 3.078317 | 0.87972591 |
| 13432 | NM_001318129.1 | ZMAT5 | 33.430351 | -6.2699426 | 1.79189861 |
| 13433 | NM_019042.4 | #N/A | 29.3935943 | -6.0937283 | 1.74153811 |
| 13434 | NM_005954.2 | #N/A | 34.3271071 | -6.307741 | 1.80275232 |
| 13435 | NM_001319235.1 | RPL34 | 40.6470385 | -6.5553551 | 1.87359221 |
| 13436 | NM_006393.2 | NEBL | 22.0582353 | -5.6768882 | 1.62279317 |
| 13437 | NM_000196.3 | #N/A | 27.2676001 | -5.9817682 | 1.71000469 |
| 13438 | NR_002718.2 | TDGF1P3 | 52.186921 | 4.38525069 | 1.25379143 |
| 13439 | NR_102281.1 | KCTD21-AS1 | 47.3190202 | 5.44550823 | 1.55701849 |
| 13440 | NM_000746.5 | #N/A | 37.9439181 | -6.456628 | 1.84628544 |
| 13441 | NM_001197222.1 | #N/A | 45.5224838 | -6.7151552 | 1.92034784 |
| 13442 | NM_001144872.1 | #N/A | 38.4854699 | -6.4803273 | 1.85326058 |
| 13443 | NM_001242850.1 | RNF146 | 38.6327076 | -6.4767464 | 1.85225639 |
| 13444 | NM_001323981.1 | SLC16A9 | 36.1675704 | -6.381243 | 1.82511955 |
| 13445 | NM_018140.3 | #N/A | 23.4735268 | -5.7573345 | 1.64718624 |
| 13446 | NM_001330416.1 | ST8SIA2 | 111.665306 | -3.5373399 | 1.01212131 |
| 13447 | NM_001318909.1 | GTF3C2 | 24.9466593 | -5.8602368 | 1.67692577 |
| 13448 | NM_213603.2 | #N/A | 22.3609505 | -5.6908609 | 1.62867756 |
| 13449 | NM_020526.4 | #N/A | 23.84341 | -5.7865693 | 1.65608125 |
| 13450 | NM_181353.2 | #N/A | 21.3829454 | -5.6309418 | 1.61155709 |
| 13451 | NR_002773.1 | AOC4P | 42.830428 | 3.6712226 | 1.05088798 |
| 13452 | NM_001323312.1 | #N/A | 26.3542566 | -5.9283215 | 1.69704017 |
| 13453 | NM_001319220.1 | #N/A | 59.1773972 | 3.30452189 | 0.94596533 |
| 13454 | NM_152905.3 | #N/A | 43.1621166 | -6.6451875 | 1.90233369 |
| 13455 | NM_170783.3 | #N/A | 26.6872152 | -5.9434677 | 1.70161313 |
| 13456 | NR_131907.1 | ABALON | 23.3392353 | -5.7498905 | 1.64645466 |
| 13457 | NM_001127398.2 | ERLEC1 | 30.4374101 | -6.1318684 | 1.7558584 |
| 13458 | NM_032530.1 | #N/A | 55.4699125 | -3.8738925 | 1.10929958 |

| | | | | | | |
|---|---|---|---|---|---|---|
| 13459 | NM_001258.2 | CDK3 | | 24.5179515 | -5.8240884 | 1.66781963 |
| 13460 | NM_001267574.2 | EIF3C | | 30.3646118 | -6.1352936 | 1.75761724 |
| 13461 | NM_001330185.1 | | #N/A | 38.9799468 | -6.4994804 | 1.86203575 |
| 13462 | NM_014170.2 | | #N/A | 25.2269598 | -5.8663547 | 1.68072879 |
| 13463 | NM_018725.3 | | #N/A | 23.2829531 | -5.7492991 | 1.64722009 |
| 13464 | NM_014682.2 | ST18 | | 61.4245069 | -5.5822207 | 1.59945794 |
| 13465 | NM_001271610.1 | | #N/A | 21.6260978 | -5.6441048 | 1.61725253 |
| 13466 | NM_152774.3 | TMEM196 | | 31.8118441 | -6.2014623 | 1.77698254 |
| 13467 | NM_001304421.2 | SLMAP | | 29.0549844 | -6.0737719 | 1.740645 |
| 13468 | NM_001195150.2 | | #N/A | 27.7846874 | -6.0086993 | 1.72214764 |
| 13469 | NM_000239.2 | | #N/A | 219.057408 | 2.93590623 | 0.84149089 |
| 13470 | NM_023931.3 | ZNF747 | | 27.8795497 | -6.0113278 | 1.72299655 |
| 13471 | NM_001164382.1 | STAU2 | | 23.0439275 | -5.7356896 | 1.64402028 |
| 13472 | NM_148912.3 | ABHD11 | | 31.4313337 | -6.1908365 | 1.77454216 |
| 13473 | NM_032127.3 | | #N/A | 28.4016385 | -6.0416865 | 1.73185451 |
| 13474 | NM_003490.3 | | #N/A | 53.3829547 | -6.9487683 | 1.99200515 |
| 13475 | NM_001322207.1 | | #N/A | 33.064191 | -6.2595132 | 1.79448768 |
| 13476 | NR_033823.2 | | #N/A | 21.2549196 | -5.6284505 | 1.61374233 |
| 13477 | NM_001195328.1 | RAB9A | | 23.8243002 | -5.7789658 | 1.65725522 |
| 13478 | NR_027780.1 | | #N/A | 36.5374289 | -6.4065354 | 1.83750986 |
| 13479 | NM_006992.3 | | #N/A | 22.1992467 | -5.6887992 | 1.63172118 |
| 13480 | NM_212543.1 | | #N/A | 20.6832118 | -5.5726926 | 1.59855596 |
| 13481 | NM_032712.2 | | #N/A | 33.8500984 | -6.2939302 | 1.80549795 |
| 13482 | NM_003292.2 | | #N/A | 542.672237 | -3.1824714 | 0.91313939 |
| 13483 | NM_002155.4 | | #N/A | 31.7136398 | -6.1984506 | 1.77867205 |
| 13485 | NM_205845.2 | AKR1C2 | | 100.510039 | -4.9305167 | 1.41487757 |
| 13484 | NM_018947.5 | | #N/A | 904.561474 | -2.083069 | 0.59776726 |
| 13486 | NM_001006635.2 | | #N/A | 32.0902167 | -3.5655489 | 1.02322081 |
| 13487 | NM_144657.4 | | #N/A | 27.8561905 | -6.0061036 | 1.723772 |
| 13488 | NM_006333.3 | | #N/A | 21.193002 | -5.617048 | 1.61220993 |
| 13489 | NM_001106.3 | | #N/A | 324.308498 | -3.495677 | 1.00336905 |
| 13490 | NM_152308.2 | | #N/A | 24.1877214 | -5.8047584 | 1.6663379 |
| 13491 | NM_001267708.1 | ZNF706 | | 21.2204077 | -5.6252043 | 1.61494789 |
| 13492 | NM_001320755.1 | | #N/A | 32.2612583 | -6.2248047 | 1.78710937 |
| 13493 | NM_001300756.1 | | #N/A | 27.5022062 | -5.9934518 | 1.72077298 |
| 13494 | NM_031311.4 | | #N/A | 26.1356765 | -5.9220605 | 1.70081657 |
| 13495 | NM_001286488.1 | TMEM14B | | 22.6833351 | -5.7049466 | 1.63866998 |
| 13496 | NM_001184882.1 | CD84 | | 71.4202891 | 2.47197758 | 0.71009574 |
| 13497 | NR_027040.1 | ATP6V0E2-AS1 | | 31.2144493 | -6.1714056 | 1.77295021 |
| 13498 | NM_001306177.1 | | #N/A | 69.4755249 | -4.530249 | 1.30156857 |
| 13499 | NR_002451.2 | ABCA11P | | 20.4905409 | -5.5746175 | 1.60177769 |
| 13500 | NR_104321.1 | SCHLAP1 | | 61.9485272 | 4.39084299 | 1.26163954 |
| 13501 | NM_001033578.2 | | #N/A | 25.5829726 | -5.8912921 | 1.69282106 |

| | | | | | |
|---|---|---|---|---|---|
| 13502 | NM_130470.2 | #N/A | 30.0356427 | -6.1112969 | 1.75615994 |
| 13503 | NM_001287763.1 | NSG1 | 68.4295669 | -7.3053195 | 2.09938393 |
| 13504 | NM_006714.4 | #N/A | 24.216934 | -5.8020054 | 1.6673734 |
| 13505 | NM_001166109.1 | #N/A | 212.292157 | -3.706277 | 1.06511336 |
| 13506 | NM_032344.3 | #N/A | 27.0668273 | -5.9625404 | 1.71363814 |
| 13507 | NM_001303451.1 | #N/A | 18.3798549 | 9.1952144 | 2.64273238 |
| 13508 | NR_135192.1 | LOC101928595 | 29.4077879 | -6.0871588 | 1.749494 |
| 13509 | NM_001318873.1 | #N/A | 25.5183556 | -5.8791577 | 1.68975625 |
| 13512 | NM_206997.1 | GPR152 | 22.7273384 | 8.18104252 | 2.35137206 |
| 13510 | NM_001317836.1 | #N/A | 33.5772061 | -6.2760836 | 1.80387382 |
| 13511 | NM_178232.3 | #N/A | 29.0630692 | -6.0741701 | 1.74583505 |
| 13513 | NM_001304434.1 | #N/A | 37.6718106 | -6.4400254 | 1.85113425 |
| 13514 | NM_001317077.1 | CHTOP | 33.9750436 | -6.2995136 | 1.81091529 |
| 13515 | NM_033343.3 | #N/A | 29.4863613 | -6.0957796 | 1.75238201 |
| 13516 | NM_001304835.1 | #N/A | 19.8501731 | -5.5167 | 1.58597487 |
| 13517 | NR_135229.1 | PCNP | 43.7059777 | -6.6556174 | 1.91362566 |
| 13518 | NM_001330466.1 | #N/A | 21.8272198 | -5.6654404 | 1.62899487 |
| 13519 | NM_001323377.1 | #N/A | 13.5732759 | 8.75978176 | 2.51895446 |
| 13520 | NM_001014972.2 | ZNF638 | 223.591349 | -9.0155744 | 2.59260467 |
| 13521 | NR_073201.3 | #N/A | 41.831467 | -6.5914697 | 1.89550921 |
| 13522 | NM_001242397.1 | #N/A | 49.4355673 | -6.8382468 | 1.96661935 |
| 13523 | NM_006735.3 | #N/A | 27.9110057 | -6.0137113 | 1.72989146 |
| 13524 | NM_001142476.1 | NREP | 34.0843753 | -6.2994673 | 1.8122473 |
| 13525 | NM_001005386.2 | ACTR2 | 28.5654864 | -6.0511549 | 1.7408451 |
| 13526 | NM_001271907.1 | SPATA33 | 22.8717258 | -5.73333 | 1.64946633 |
| 13527 | NM_001270408.1 | #N/A | 42.4959038 | -2.519075 | 0.72474443 |
| 13528 | NM_001719.2 | #N/A | 473.183487 | -4.5352385 | 1.30484533 |
| 13529 | NM_144606.5 | #N/A | 20.0423811 | -5.5289589 | 1.59114405 |
| 13530 | NM_001322978.1 | #N/A | 42.2496281 | -6.6091467 | 1.90211215 |
| 13531 | NM_001184744.1 | PTH1R | 29.1128128 | -6.081014 | 1.75026622 |
| 13532 | NM_001346516.1 | #N/A | 21.3863648 | -5.6385069 | 1.62296256 |
| 13533 | NM_015035.3 | #N/A | 176.620609 | -2.1059977 | 0.60628646 |
| 13534 | NM_138418.3 | #N/A | 26.5081161 | -5.9303616 | 1.7073384 |
| 13535 | NM_001301210.1 | #N/A | 225.838933 | -3.2368997 | 0.93193062 |
| 13536 | NM_001297561.1 | ZNF708 | 159.04069 | 2.24859412 | 0.64739539 |
| 13537 | NM_001329717.1 | #N/A | 30.1166595 | -6.1247304 | 1.76339921 |
| 13538 | NR_103844.1 | NPTN-IT1 | 23.6588359 | -5.7721683 | 1.66191709 |
| 13539 | NM_001317771.1 | #N/A | 22.9239201 | -5.7354136 | 1.65167669 |
| 13540 | NM_003492.2 | #N/A | 22.6645975 | -5.7148084 | 1.64574392 |
| 13541 | NM_001320210.1 | UBA5 | 27.6115351 | -5.9913471 | 1.72557365 |
| 13542 | NM_012229.4 | NT5C2 | 153.921811 | -3.1761093 | 0.91479953 |
| 13543 | NM_001303412.1 | #N/A | 27.1026567 | -5.9732825 | 1.72055944 |
| 13544 | NM_003805.4 | #N/A | 22.5664234 | -5.7042156 | 1.6431723 |

| | | | | | |
|---|---|---|---|---|---|
| 13545 | NM_001080825.2 | TMEM120B | 313.624673 | -1.2451825 | 0.35870654 |
| 13546 | NR_033704.1 | #N/A | 23.6137879 | -5.772546 | 1.66304688 |
| 13547 | NM_001136005.1 | GART | 19.5103828 | -5.4922963 | 1.58232099 |
| 13548 | NM_019605.3 | #N/A | 25.8718371 | -5.9068856 | 1.70177535 |
| 13549 | NM_001318391.1 | UTP4 | 26.866396 | -5.9663284 | 1.71912025 |
| 13550 | NM_207406.3 | #N/A | 43.1931696 | -6.6404377 | 1.91355622 |
| 13551 | NM_058190.3 | #N/A | 23.3014992 | -5.7483202 | 1.65655427 |
| 13552 | NM_173474.3 | #N/A | 26.6458837 | -5.9410262 | 1.7121683 |
| 13553 | NM_033362.3 | MRPS12 | 37.3557156 | -6.432235 | 1.85391851 |
| 13554 | NM_014661.3 | #N/A | 281.426407 | -4.4421826 | 1.28077203 |
| 13555 | NM_002753.4 | #N/A | 33.7856049 | -6.2941177 | 1.81473746 |
| 13556 | NM_001297653.1 | DDR1 | 299.587627 | -9.4373311 | 2.72105085 |
| 13557 | NM_001004317.3 | #N/A | 56.4067495 | -5.4729731 | 1.57810557 |
| 13558 | NM_001199697.1 | BAG6 | 37.4891414 | -6.4432383 | 1.85793943 |
| 13559 | NM_203397.2 | #N/A | 22.0666685 | -5.6719981 | 1.6355503 |
| 13560 | NM_203349.3 | #N/A | 48.2890006 | -5.2802017 | 1.52258896 |
| 13561 | NM_181869.1 | APAF1 | 128.680312 | 1.96537657 | 0.56680343 |
| 13562 | NR_136234.1 | #N/A | 27.937888 | -6.0141401 | 1.73444593 |
| 13563 | NM_001242415.1 | #N/A | 23.3381232 | -5.7484921 | 1.65800961 |
| 13564 | NM_002799.3 | #N/A | 427.031648 | -3.8346993 | 1.10602295 |
| 13565 | NM_001242821.1 | DEF8 | 33.3983853 | -6.268505 | 1.80818015 |
| 13566 | NM_001145145.1 | #N/A | 23.4497742 | -5.7597669 | 1.66158003 |
| 13567 | NM_002238.3 | #N/A | 27.6696004 | -6.0005198 | 1.73105663 |
| 13568 | NM_001282864.1 | SLC2A11 | 22.2191794 | -5.6833255 | 1.63975568 |
| 13569 | NM_145238.3 | #N/A | 19.6449994 | -5.5063056 | 1.58890418 |
| 13570 | NM_024605.3 | #N/A | 20.1326643 | -5.5454602 | 1.60022371 |
| 13571 | NR_045634.1 | BOLA3-AS1 | 107.476888 | -4.3227263 | 1.24748046 |
| 13573 | NR_144555.1 | RN7SL832P | 104.083127 | 2.92692654 | 0.84471026 |
| 13572 | NM_001308229.1 | #N/A | 33.0097423 | -3.6046922 | 1.04031168 |
| 13574 | NM_000073.2 | CD3G | 28.158847 | 2.25339309 | 0.65035942 |
| 13575 | NM_001321133.1 | #N/A | 24.3316323 | -5.8124801 | 1.67757934 |
| 13576 | NR_138092.1 | #N/A | 221.615709 | 1.11004354 | 0.32038868 |
| 13577 | NM_130393.3 | PTPRD | 62.8425141 | -5.6057263 | 1.61799781 |
| 13578 | NM_001136019.2 | #N/A | 21.7685967 | -5.6642052 | 1.63490425 |
| 13579 | NM_001033081.2 | #N/A | 50.0979806 | -6.854355 | 1.97846246 |
| 13580 | NM_001040214.2 | #N/A | 76.107373 | -7.4599893 | 2.15362376 |
| 13581 | NM_199185.3 | NPM1 | 43.6168986 | -6.660123 | 1.92294031 |
| 13582 | NM_201647.2 | #N/A | 27.6061489 | -6.0043324 | 1.73367959 |
| 13583 | NM_015435.4 | RNF19A | 35.1965095 | -6.351705 | 1.83402804 |
| 13584 | NM_001166168.1 | NEK6 | 22.4783401 | -5.7082326 | 1.64836699 |
| 13585 | NM_025193.3 | #N/A | 21.6985606 | -5.6462392 | 1.63051596 |
| 13586 | NM_031455.3 | #N/A | 31.2507651 | -6.1814863 | 1.78509796 |
| 13587 | NM_001127692.2 | PCCA | 26.0751328 | -5.9158783 | 1.70848349 |

| | | | | | |
|---|---|---|---|---|---|
| 13588 | NM_001319218.1 | ALDH4A1 | 25.8509773 | -5.8964048 | 1.70290554 |
| 13589 | NM_001314025.1 | #N/A | 24.0902231 | -5.7939165 | 1.67350428 |
| 13590 | NM_004234.4 | ZNF235 | 20.9820282 | -5.6002618 | 1.61766547 |
| 13591 | NM_178839.4 | #N/A | 27.867343 | -6.0147715 | 1.73755972 |
| 13592 | NM_001284418.1 | #N/A | 29.846231 | -6.1098914 | 1.76518886 |
| 13593 | NM_003847.2 | #N/A | 20.5212025 | -5.5693667 | 1.60904589 |
| 13594 | NM_153341.3 | #N/A | 26.7702545 | -5.9597309 | 1.72186288 |
| 13595 | NM_198993.3 | #N/A | 33.6484719 | -4.830746 | 1.39576419 |
| 13596 | NR_027007.3 | THUMPD3-AS1 | 28.2533584 | -6.0381294 | 1.744812 |
| 13597 | NR_125372.1 | #N/A | 23.7480202 | -5.7894008 | 1.67329065 |
| 13598 | NM_001242906.1 | #N/A | 31.7760232 | -6.2026737 | 1.79280445 |
| 13599 | NM_001080506.1 | #N/A | 25.7291816 | -5.897392 | 1.70481186 |
| 13600 | NR_024181.1 | #N/A | 18.9589362 | -5.4517116 | 1.57597821 |
| 13601 | NM_020425.5 | SMIM8 | 23.5402446 | -5.7741576 | 1.66925706 |
| 13602 | NM_001128309.1 | #N/A | 29.1995014 | 9.83703423 | 2.84421126 |
| 13603 | NM_024097.3 | #N/A | 26.3816019 | -5.9295836 | 1.71460229 |
| 13604 | NM_152420.2 | #N/A | 32.3080836 | -6.2180218 | 1.79801494 |
| 13606 | NR_135681.1 | LOC105370792 | 27.6446102 | -4.7204597 | 1.36509396 |
| 13605 | NM_001277064.1 | #N/A | 35.2482178 | -6.3518466 | 1.83686424 |
| 13608 | NM_198793.2 | CD47 | 1658.33038 | 3.10256067 | 0.89724067 |
| 13607 | NM_015257.2 | #N/A | 30.1665669 | -6.1288739 | 1.77243302 |
| 13609 | NM_001283049.1 | #N/A | 38.521764 | -4.99866 | 1.44576672 |
| 13610 | NM_001015509.2 | PTRH2 | 23.3719315 | -5.7504977 | 1.66342677 |
| 13611 | NM_024081.5 | #N/A | 110.484105 | 2.3938482 | 0.69248068 |
| 13612 | NR_109841.1 | LINC01205 | 18.5041589 | 2.4395668 | 0.70572441 |
| 13613 | NR_026884.1 | LINC00652 | 36.1154859 | 3.18159282 | 0.92046273 |
| 13614 | NR_120378.1 | LOC102724034 | 20.1907543 | -5.5451491 | 1.60426656 |
| 13615 | NM_014001.4 | #N/A | 35.7397878 | -6.374697 | 1.8444785 |
| 13616 | NM_001303058.1 | TMED4 | 320.966155 | 1.42417952 | 0.41214266 |
| 13617 | NM_001040023.1 | #N/A | 32.9495739 | -6.2497647 | 1.80862926 |
| 13618 | NM_001345944.1 | PFKP | 30.3336353 | -6.1337679 | 1.77507498 |
| 13619 | NM_001143679.1 | GSTK1 | 23.2971795 | -5.7496689 | 1.66407354 |
| 13620 | NM_001303012.1 | #N/A | 28.5880405 | -6.0424129 | 1.74888622 |
| 13621 | NM_001025236.1 | TSPAN4 | 31.2403088 | -6.1759872 | 1.78770164 |
| 13622 | NR_033714.1 | #N/A | 33.4515573 | -6.2701947 | 1.8150653 |
| 13623 | NM_021912.4 | #N/A | 26.4041465 | -5.9251753 | 1.71532297 |
| 13624 | NM_006294.4 | UQCRB | 498.605831 | -3.6590024 | 1.05931321 |
| 13625 | NM_001032284.2 | #N/A | 185.140913 | -2.9884492 | 0.86524693 |
| 13626 | NM_001128591.1 | #N/A | 25.4142588 | -5.8773845 | 1.70179305 |
| 13627 | NM_001204404.1 | #N/A | 61.7072827 | -5.5790559 | 1.61543363 |
| 13628 | NM_018032.4 | #N/A | 25.8136684 | -5.8991862 | 1.70832375 |
| 13629 | NM_016210.4 | #N/A | 21.3666139 | -5.6303689 | 1.63064212 |
| 13630 | NM_175872.4 | #N/A | 18.991901 | -5.452682 | 1.57936091 |

| | | | | | |
|---|---|---|---|---|---|
| 13631 | NR_030717.1 | #N/A | 21.1831583 | -5.611072 | 1.62523855 |
| 13632 | NM_001319184.1 | #N/A | 26.3220018 | -5.9268695 | 1.71672559 |
| 13633 | NM_032494.2 | #N/A | 40.359851 | -4.2417489 | 1.22864415 |
| 13634 | NM_144641.3 | #N/A | 26.5940286 | -5.9499185 | 1.7234937 |
| 13635 | NR_001445.2 | RN7SK | 35.9397985 | -6.3739097 | 1.84648122 |
| 13636 | NR_036467.1 | #N/A | 24.2459351 | -5.8093144 | 1.68332466 |
| 13637 | NM_000462.3 | #N/A | 44.2223068 | -6.6733696 | 1.93387753 |
| 13638 | NM_194317.3 | #N/A | 26.0704662 | -5.9122111 | 1.71333494 |
| 13639 | NM_173620.2 | #N/A | 24.5670831 | -5.8301176 | 1.68964022 |
| 13640 | NM_001199251.2 | #N/A | 20.7737692 | -5.5824677 | 1.61789562 |
| 13641 | NM_025049.3 | #N/A | 27.6584922 | -6.0013955 | 1.7395999 |
| 13642 | NR_028351.1 | LINC02145 | 47.8902694 | 3.16710158 | 0.91806521 |
| 13643 | NR_015360.1 | #N/A | 20.0955108 | -5.5355932 | 1.60476901 |
| 13644 | NR_109819.1 | NR2F1-AS1 | 32.0437934 | -4.926421 | 1.42832564 |
| 13645 | NM_004675.3 | #N/A | 22.5030015 | -5.7093448 | 1.65539125 |
| 13646 | NM_001127203.3 | #N/A | 34.6189193 | -6.3175293 | 1.83177946 |
| 13647 | NM_001330993.1 | #N/A | 20.1903042 | -5.5433879 | 1.60746052 |
| 13648 | NR_003922.1 | PMCHL2 | 27.4424683 | 2.65119745 | 0.76883889 |
| 13649 | NM_012123.3 | #N/A | 39.7655766 | -6.5235274 | 1.89182056 |
| 13650 | NR_130154.1 | #N/A | 36.2268225 | -6.3874171 | 1.85241507 |
| 13651 | NM_001314057.1 | #N/A | 29.7993283 | -4.8135241 | 1.39637445 |
| 13652 | NM_006037.3 | HDAC4 | 144.400076 | -1.8718115 | 0.543017 |
| 13653 | NM_001014812.2 | CIAO2A | 29.3709729 | -6.0831391 | 1.76475946 |
| 13654 | NM_003579.3 | #N/A | 23.731505 | -5.7705539 | 1.67413305 |
| 13655 | NM_002193.3 | #N/A | 28.7261049 | -6.0480726 | 1.75468402 |
| 13656 | NM_001317163.1 | #N/A | 20.1296979 | -5.5501682 | 1.61036361 |
| 13657 | NM_001112800.1 | #N/A | 56.0599906 | -7.0202321 | 2.03694548 |
| 13658 | NM_001242313.1 | TMEM191B | 23.2418364 | -5.7442396 | 1.66675441 |
| 13659 | NM_152517.2 | #N/A | 25.4853368 | -5.8888572 | 1.70874028 |
| 13660 | NM_001130082.2 | #N/A | 217.677851 | -8.9769656 | 2.60500791 |
| 13661 | NR_027064.2 | #N/A | 78.1589574 | 2.19202443 | 0.63616065 |
| 13662 | NM_001008737.1 | #N/A | 63.769892 | 2.37328542 | 0.68886395 |
| 13663 | NM_178238.3 | PILRB | 26.2268732 | -5.9176122 | 1.71781253 |
| 13664 | NM_001136158.1 | OTUD5 | 28.0323675 | -6.0269757 | 1.74959175 |
| 13665 | NM_203414.2 | ELP5 | 75.8130352 | -4.6060304 | 1.33710857 |
| 13666 | NM_001143761.1 | EIF5A | 28.0047065 | -6.0188969 | 1.74727149 |
| 13667 | NM_001207014.1 | #N/A | 21.4018839 | -5.6212874 | 1.63198058 |
| 13668 | NM_001199875.1 | SERF2 | 19.821375 | -5.5277925 | 1.60485015 |
| 13669 | NM_017421.3 | #N/A | 31.1309345 | -6.1667825 | 1.79055187 |
| 13670 | NM_001309474.1 | #N/A | 21.4760645 | -5.6383482 | 1.63719328 |
| 13671 | NM_001308420.1 | #N/A | 21.4255 | -5.6391039 | 1.63751088 |
| 13672 | NM_001199662.1 | PMF1-BGLAP | 24.5595996 | -5.8273632 | 1.69240283 |
| 13674 | NR_038956.1 | NUP50-DT | 27.1160179 | -5.9733838 | 1.73483235 |

| | | | | | |
|---|---|---|---|---|---|
| 13673 | NR_037669.1 | #N/A | 24.4973813 | -5.8257978 | 1.69196288 |
| 13675 | NR_045018.1 | #N/A | 24.4780766 | -5.8272567 | 1.69253499 |
| 13676 | NR_036511.1 | LOC100129917 | 21.9871113 | -5.6679784 | 1.64647096 |
| 13677 | NR_024534.1 | ALG3 | 27.1587813 | -5.9751902 | 1.73597534 |
| 13678 | NM_016598.2 | #N/A | 28.0247338 | -6.0143431 | 1.7475363 |
| 13679 | NM_015836.3 | WARS2 | 28.3416636 | -6.0417595 | 1.75555525 |
| 13680 | NM_001319652.1 | #N/A | 33.5589127 | -6.2706234 | 1.82209063 |
| 13681 | NM_001316952.1 | #N/A | 23.2443984 | -5.7427941 | 1.66895423 |
| 13682 | NM_014690.4 | #N/A | 34.3831661 | -6.3104373 | 1.83391491 |
| 13683 | NM_006827.5 | #N/A | 889.000374 | -3.7722506 | 1.09646023 |
| 13684 | NM_004884.3 | #N/A | 241.779739 | -3.4518969 | 1.00350154 |
| 13685 | NR_015366.4 | #N/A | 19.6921003 | -5.5161413 | 1.6036686 |
| 13686 | NM_018044.4 | NSUN5 | 29.2509396 | -6.0767508 | 1.76669509 |
| 13687 | NM_144689.4 | #N/A | 34.5880505 | -6.3165296 | 1.83642109 |
| 13688 | NR_135305.1 | SMG1P2 | 27.3993243 | -5.9796949 | 1.7386984 |
| 13689 | NM_001018057.1 | DKK3 | 573.201829 | -4.0627847 | 1.1816766 |
| 13690 | NM_148178.2 | #N/A | 23.4317789 | -5.7606284 | 1.67583159 |
| 13691 | NR_038192.1 | DHPS | 45.6404961 | -6.7205754 | 1.95512493 |
| 13692 | NM_001242739.1 | #N/A | 19.0399334 | -5.4617712 | 1.58898732 |
| 13693 | NM_024328.5 | #N/A | 21.5979811 | -5.6426223 | 1.64161345 |
| 13694 | NM_001256876.1 | #N/A | 31.586646 | -4.912348 | 1.42932385 |
| 13695 | NM_033388.1 | #N/A | 26.9772105 | -5.9649367 | 1.73576941 |
| 13696 | NM_002608.3 | #N/A | 31.4171225 | -6.1841404 | 1.79978978 |
| 13697 | NM_003771.4 | KRT36 | 17.5210981 | 9.1266245 | 2.65632912 |
| 13698 | NR_134539.1 | #N/A | 24.5343281 | -5.8205261 | 1.69415362 |
| 13699 | NM_020233.4 | #N/A | 24.0172706 | -5.7925039 | 1.68613255 |
| 13700 | NM_020804.4 | #N/A | 21.9134331 | -5.6527793 | 1.64587227 |
| 13701 | NM_003414.5 | #N/A | 21.0324768 | -5.5991287 | 1.63038265 |
| 13702 | NR_027624.1 | #N/A | 28.7901774 | -6.0542564 | 1.76294558 |
| 13703 | NM_018354.2 | TMEM74B | 21.9783035 | -5.662502 | 1.64926235 |
| 13704 | NM_001256470.1 | #N/A | 44.4560876 | -6.6824987 | 1.94649059 |
| 13705 | NM_004146.5 | #N/A | 255.345097 | -3.5407685 | 1.03150711 |
| 13706 | NM_018323.3 | #N/A | 30.4080542 | -6.1347466 | 1.78728001 |
| 13707 | NM_001242336.1 | #N/A | 26.8153937 | -5.9492902 | 1.73331428 |
| 13708 | NM_001287533.1 | #N/A | 33.8853543 | -4.2117893 | 1.22718354 |
| 13709 | NM_170606.2 | #N/A | 328.927245 | -3.4554829 | 1.00685757 |
| 13710 | NM_001160305.2 | #N/A | 29.8119936 | -6.1048921 | 1.77954417 |
| 13711 | NM_014646.2 | LPIN2 | 311.746132 | -4.3390483 | 1.26488596 |
| 13712 | NM_005985.3 | #N/A | 45.3120973 | -5.1939915 | 1.51425644 |
| 13713 | NM_001198621.2 | #N/A | 17.1639863 | 6.83406047 | 1.99258338 |
| 13714 | NM_001100419.1 | REX1BD | 35.4784814 | -6.3601572 | 1.85451544 |
| 13715 | NM_174899.4 | #N/A | 26.7739415 | -4.6694159 | 1.36167181 |
| 13716 | NM_001243879.1 | #N/A | 33.6355048 | -6.2820393 | 1.83196681 |

| | | | | | |
|---|---|---|---|---|---|
| 13717 | NM_153695.3 | #N/A | 26.2191205 | -5.918328 | 1.72590757 |
| 13718 | NM_183425.2 | #N/A | 39.0211098 | -6.4959262 | 1.89432552 |
| 13719 | NM_001256630.1 | BRSK2 | 25.8820717 | -5.897013 | 1.71979259 |
| 13720 | NM_001005290.3 | PSRC1 | 28.5085135 | -6.0488514 | 1.76412336 |
| 13721 | NM_133491.4 | #N/A | 30.6068082 | -6.1510983 | 1.79395312 |
| 13722 | NM_018242.2 | #N/A | 60.1754371 | 2.13840416 | 0.62381978 |
| 13723 | NM_178221.2 | ATG4C | 27.1960921 | -5.9683667 | 1.74139052 |
| 13724 | NM_001271696.1 | #N/A | 22.680576 | -5.7094336 | 1.66622464 |
| 13725 | NM_001204088.1 | #N/A | 21.6380475 | -5.6408586 | 1.64641469 |
| 13726 | NM_001006640.1 | #N/A | 27.0659863 | -5.9676574 | 1.74207931 |
| 13727 | NR_046292.1 | MSTO1 | 22.9189776 | -5.7375499 | 1.67498031 |
| 13728 | NR_033967.1 | SLC2A1-AS1 | 23.0937147 | 2.50250413 | 0.73061285 |
| 13729 | NM_001303534.1 | #N/A | 21.9112956 | -5.664861 | 1.65395564 |
| 13730 | NM_024642.4 | #N/A | 38.104125 | -6.46589 | 1.88807298 |
| 13731 | NM_020685.4 | #N/A | 39.5750645 | -6.5149527 | 1.90255202 |
| 13732 | NM_152268.3 | #N/A | 19.8638761 | -5.5299386 | 1.61499484 |
| 13733 | NM_018234.2 | #N/A | 30.3140273 | -4.855572 | 1.41820826 |
| 13734 | NM_001167676.1 | FAM229A | 18.4299675 | -5.4111935 | 1.58065963 |
| 13735 | NM_152709.4 | #N/A | 27.0388853 | -5.9710935 | 1.74422741 |
| 13736 | NM_023079.4 | #N/A | 383.294445 | -3.5177906 | 1.02761845 |
| 13737 | NM_018221.4 | #N/A | 194.454329 | -8.8141967 | 2.57484397 |
| 13738 | NM_001142560.1 | #N/A | 24.0163013 | -5.7969112 | 1.6935697 |
| 13739 | NR_135106.1 | S100PBP | 21.6107187 | -5.6327901 | 1.64573398 |
| 13740 | NR_024583.1 | POM121L8P | 90.9849916 | 4.84244067 | 1.41501834 |
| 13741 | NM_001286696.1 | ZNF79 | 25.6351085 | -5.879886 | 1.71875013 |
| 13742 | NM_002236.4 | #N/A | 24.504869 | -5.819756 | 1.7013164 |
| 13743 | NM_152679.3 | #N/A | 25.2775024 | -5.868319 | 1.71565259 |
| 13744 | NM_004843.3 | #N/A | 43.0095544 | -3.3179181 | 0.97027436 |
| 13745 | NM_000048.3 | #N/A | 26.3198987 | -5.9351416 | 1.73566362 |
| 13746 | NM_006332.4 | #N/A | 26.2962274 | -5.9331164 | 1.73520731 |
| 13748 | NR_108099.1 | LOC101928401 | 53.4183917 | 2.03553663 | 0.59535309 |
| 13747 | NM_001271918.1 | SEC11A | 24.0852986 | -5.8037672 | 1.69747441 |
| 13749 | NM_001305163.1 | #N/A | 36.6762402 | -6.4007354 | 1.87245939 |
| 13750 | NM_001318733.1 | RHBDL1 | 19.9048519 | -5.5276249 | 1.6170832 |
| 13751 | NM_024989.3 | #N/A | 196.061036 | -8.8260285 | 2.5822437 |
| 13752 | NM_001145095.1 | HHLA1 | 35.6830632 | 5.75871046 | 1.68521161 |
| 13753 | NR_109913.1 | SLC35B3 | 19.7025642 | -5.509718 | 1.61250043 |
| 13754 | NM_001303031.1 | #N/A | 25.741202 | -5.9000756 | 1.72681975 |
| 13755 | NM_198333.2 | P2RY10 | 81.2428938 | 4.51250673 | 1.3207303 |
| 13756 | NM_015417.4 | #N/A | 37.4267675 | -6.4399682 | 1.8851031 |
| 13757 | NM_001300914.1 | DNAJB1 | 25.3530896 | -5.881431 | 1.72170975 |
| 13758 | NM_006515.3 | SETMAR | 21.5123526 | -5.6341371 | 1.64947268 |
| 13759 | NM_001167575.1 | CAAP1 | 21.2375647 | -5.627198 | 1.64749531 |

| | | | | | |
|---|---|---|---|---|---|
| 13760 | NR_104246.1 | #N/A | 36.0004741 | -6.3808687 | 1.86850794 |
| 13761 | NM_139343.2 | #N/A | 268.086324 | -9.2770709 | 2.71685493 |
| 13762 | NM_001572.3 | #N/A | 22.2200499 | -5.6911656 | 1.6667397 |
| 13763 | NM_000850.4 | #N/A | 21.8141801 | -5.6625534 | 1.65839859 |
| 13764 | NM_001302545.1 | #N/A | 32.6134232 | -6.2435219 | 1.82861945 |
| 13765 | NM_015286.5 | #N/A | 62.5542796 | -5.5891497 | 1.63697765 |
| 13766 | NR_026694.2 | #N/A | 23.6396117 | -5.7661175 | 1.6891516 |
| 13767 | NM_001001740.3 | COP1 | 28.0595481 | -6.0205213 | 1.76373357 |
| 13768 | NR_144421.1 | #N/A | 21.8241594 | -5.6658383 | 1.6598228 |
| 13769 | NM_015033.2 | #N/A | 340.222567 | -4.4569099 | 1.30577108 |
| 13770 | NM_001201536.1 | TAF1A | 23.5222202 | -5.7595636 | 1.68750363 |
| 13771 | NM_015397.3 | #N/A | 211.714422 | -2.6866939 | 0.78718475 |
| 13772 | NM_001306147.1 | #N/A | 34.4916203 | -6.3174302 | 1.85118513 |
| 13773 | NM_001033521.1 | CSTF1 | 25.5263025 | -5.8800804 | 1.72315431 |
| 13774 | NM_001267609.1 | GPR161 | 31.136898 | -6.1730212 | 1.80910198 |
| 13775 | NM_024717.5 | MCTP1 | 20.4913147 | -5.5578826 | 1.62916429 |
| 13776 | NR_110071.1 | LOC101928075 | 15.1809928 | 5.75035769 | 1.68565798 |
| 13777 | NM_001330237.1 | #N/A | 25.9600137 | -5.9078672 | 1.7319791 |
| 13778 | NR_037719.1 | TMEM256-PLSCR3 | 22.2463163 | -5.6786796 | 1.66480419 |
| 13779 | NM_001099646.1 | #N/A | 24.3915335 | -5.8236693 | 1.70733754 |
| 13780 | NM_052958.2 | #N/A | 22.6813669 | -5.7118295 | 1.67455712 |
| 13781 | NM_001103161.1 | #N/A | 26.6915136 | -5.9471041 | 1.74375299 |
| 13782 | NM_001079512.3 | #N/A | 36.4245023 | -3.5715938 | 1.04733431 |
| 13783 | NM_173627.4 | #N/A | 24.9097098 | -5.8510625 | 1.71579799 |
| 13784 | NM_001098728.1 | #N/A | 57.9869495 | -3.8292182 | 1.12307701 |
| 13785 | NM_001127895.1 | #N/A | 22.5012996 | -5.7053155 | 1.67338788 |
| 13786 | NM_001271285.1 | #N/A | 25.4610385 | -5.8826592 | 1.72539782 |
| 13787 | NM_001293104.1 | #N/A | 23.5404984 | -5.7723866 | 1.69307117 |
| 13788 | NM_001320884.1 | #N/A | 56.9523014 | -7.0428121 | 2.06578212 |
| 13789 | NM_001284352.1 | #N/A | 31.6084264 | -6.1963052 | 1.81750283 |
| 13790 | NM_001172477.1 | RRM2B | 28.5758798 | -6.0505287 | 1.7747952 |
| 13791 | NM_001130057.3 | EEF1D | 31.2919597 | -6.1806141 | 1.81310551 |
| 13792 | NM_000728.3 | #N/A | 29.3268938 | -6.0862147 | 1.78554942 |
| 13793 | NM_001113207.1 | #N/A | 24.7253381 | -5.8322392 | 1.71131777 |
| 13794 | NM_001167608.1 | #N/A | 27.0693916 | -5.9732261 | 1.75279436 |
| 13795 | NM_001322089.1 | CNOT7 | 27.7305734 | -6.0020656 | 1.76132427 |
| 13796 | NM_022344.3 | #N/A | 609.10694 | 0.99007408 | 0.29064279 |
| 13797 | NM_001317998.1 | ZNF677 | 19.9209332 | -5.5203622 | 1.62068651 |
| 13798 | NR_038253.1 | LOC100288637 | 20.9249144 | -5.5930525 | 1.64223085 |
| 13799 | NR_144434.1 | RSPH3 | 199.760882 | 1.32150417 | 0.38803878 |
| 13800 | NM_001289401.1 | ZNF135 | 30.5879368 | -6.1396128 | 1.80282776 |
| 13801 | NR_073405.1 | PRKXP1 | 111.135049 | 1.72329015 | 0.50607238 |
| 13802 | NR_105019.1 | TMEM161B-AS1 | 21.5096359 | -5.6335517 | 1.65494514 |

| | | | | | |
|---|---|---|---|---|---|
| 13803 | NM_005701.3 | #N/A | 19.8811825 | -5.5302818 | 1.62473385 |
| 13804 | NM_001312920.1 | MSANTD2 | 18.269316 | -5.4076447 | 1.58881025 |
| 13805 | NR_047673.1 | LRRC4C | 27.7069338 | -6.0039817 | 1.76403673 |
| 13806 | NM_002665.4 | PLGLB2 | 18.0987643 | -5.3909554 | 1.58396113 |
| 13807 | NM_006260.4 | #N/A | 236.923017 | -1.7610962 | 0.51744562 |
| 13808 | NM_013363.3 | #N/A | 18.2782149 | -5.3970806 | 1.58579152 |
| 13809 | NR_138077.1 | #N/A | 42.4987463 | -6.6194453 | 1.94498211 |
| 13810 | NM_001324309.1 | #N/A | 24.0799809 | -5.8093871 | 1.70712016 |
| 13811 | NR_130125.1 | #N/A | 25.9137023 | 7.51008292 | 2.20689673 |
| 13812 | NR_003698.1 | FAHD2CP | 20.1386146 | -5.5420116 | 1.62860577 |
| 13813 | NM_183323.2 | PAIP1 | 23.3931419 | -5.7604762 | 1.69294476 |
| 13814 | NM_001320179.1 | #N/A | 25.8683411 | -5.8963782 | 1.7331058 |
| 13815 | NM_001318475.1 | ZFP1 | 25.3471901 | -5.8691194 | 1.72533295 |
| 13816 | NM_001278622.1 | RAD17 | 25.7225408 | -5.9039474 | 1.73563786 |
| 13817 | NM_000662.7 | #N/A | 22.7617264 | 5.07777934 | 1.49282896 |
| 13819 | NR_109890.1 | LINC02202 | 77.6312866 | 2.77149031 | 0.81498934 |
| 13818 | NM_001258315.1 | #N/A | 20.3215967 | -5.5533131 | 1.63301997 |
| 13820 | NM_001346393.1 | #N/A | 27.046753 | -5.958117 | 1.75250819 |
| 13821 | NM_001286124.1 | MARK1 | 36.6484561 | -6.4083082 | 1.88494214 |
| 13822 | NM_001048223.2 | DBNDD2 | 27.385671 | -5.9834636 | 1.76001044 |
| 13823 | NM_003222.3 | #N/A | 36.8012438 | -6.4092082 | 1.88525877 |
| 13824 | NM_001014795.2 | #N/A | 22.2122229 | -5.6802542 | 1.67101611 |
| 13825 | NR_135579.1 | #N/A | 26.2916107 | -5.9296956 | 1.7445722 |
| 13826 | NM_198212.2 | #N/A | 43.322213 | 7.04535688 | 2.07289954 |
| 13827 | NM_001346141.1 | PPAN | 24.8803433 | -5.8414228 | 1.7187001 |
| 13828 | NR_033805.1 | WAC-AS1 | 237.982217 | -3.0394505 | 0.8943815 |
| 13829 | NM_001286272.1 | TPT1 | 21.5621164 | -5.6382052 | 1.65912662 |
| 13830 | NR_045033.1 | APH1A | 27.7394883 | -6.0039253 | 1.76680622 |
| 13831 | NR_036478.1 | TMEM198B | 20.0052469 | -5.5379454 | 1.62977824 |
| 13832 | NM_020398.3 | #N/A | 26.7843485 | 3.51287372 | 1.03389722 |
| 13833 | NM_001014440.3 | ODF3B | 56.8404194 | -7.0412805 | 2.07263162 |
| 13834 | NM_002126.4 | #N/A | 19.8395028 | -5.5160653 | 1.62415425 |
| 13835 | NR_132117.1 | DNAJC3-DT | 23.4295409 | -5.7599695 | 1.69604761 |
| 13836 | NM_001098173.1 | PRDM7 | 56.5923264 | 5.62085775 | 1.65509897 |
| 13837 | NM_001174168.2 | #N/A | 38.5353485 | 4.64193288 | 1.36696165 |
| 13838 | NM_001304274.1 | #N/A | 21.8942892 | -5.6620186 | 1.66755494 |
| 13839 | NM_001323072.1 | PFKP | 21.292238 | -5.6189861 | 1.6549371 |
| 13840 | NM_001145453.2 | GFRA1 | 212.358099 | -8.9412329 | 2.63368542 |
| 13841 | NM_001308330.1 | #N/A | 24.0085838 | -5.7956527 | 1.70725455 |
| 13842 | NM_024848.2 | #N/A | 25.1614943 | -5.8665914 | 1.7285624 |
| 13843 | NM_001323511.1 | #N/A | 124.797043 | -3.3548141 | 0.9885049 |
| 13844 | NR_110729.1 | LOC101928295 | 34.2396133 | 2.09926922 | 0.6186256 |
| 13845 | NM_024501.2 | #N/A | 26.8179617 | -5.9560191 | 1.75519173 |

| | | | | | |
|---|---|---|---|---|---|
| 13846 | NM_016532.3 | #N/A | 107.375323 | -4.3024015 | 1.26798186 |
| 13847 | NM_025142.1 | TAOK1 | 13.7647008 | 8.74164836 | 2.57696775 |
| 13848 | NM_002188.2 | #N/A | 23.9174493 | 7.88760755 | 2.32523182 |
| 13849 | NM_033015.3 | FASTK | 34.149937 | -6.3005704 | 1.85757685 |
| 13854 | NR_103523.1 | AMN1 | 33.6879136 | -6.2816027 | 1.85200438 |
| 13853 | NR_027295.2 | LINC02210 | 20.0883025 | -5.5352676 | 1.63198864 |
| 13850 | NM_001127362.1 | #N/A | 18.971879 | -5.4617328 | 1.61028914 |
| 13851 | NM_001185181.2 | #N/A | 25.5245997 | -5.8888312 | 1.73622798 |
| 13852 | NM_174936.3 | #N/A | 74.2605362 | -2.9693125 | 0.87545989 |
| 13855 | NM_002939.3 | #N/A | 22.7004098 | -5.7061019 | 1.6823927 |
| 13856 | NM_002689.3 | #N/A | 83.1020058 | -2.3087129 | 0.68073556 |
| 13857 | NM_001146206.1 | RASGRP4 | 41.7513533 | 2.68266509 | 0.79104823 |
| 13858 | NM_001270381.1 | #N/A | 22.6940142 | -5.7189246 | 1.68635751 |
| 13859 | NM_001193524.1 | #N/A | 26.1193572 | -5.9233372 | 1.74668866 |
| 13860 | NM_174869.2 | #N/A | 18.7513848 | -5.4379978 | 1.60359675 |
| 13861 | NM_001330189.1 | #N/A | 19.5209558 | -5.4867588 | 1.61799617 |
| 13862 | NM_001145440.2 | #N/A | 19.4639321 | -5.4933469 | 1.61998922 |
| 13863 | NM_001271099.1 | #N/A | 35.4502624 | -6.356324 | 1.87476016 |
| 13864 | NM_001026.4 | RPS24 | 27.6898821 | -6.0054739 | 1.77139956 |
| 13865 | NR_046104.1 | LINC02297 | 25.9462938 | 7.90071858 | 2.33046055 |
| 13866 | NR_134303.1 | LOC101927752 | 16.6225706 | 7.38846953 | 2.17939231 |
| 13867 | NM_005929.5 | #N/A | 21.4841074 | -5.6413445 | 1.66414025 |
| 13868 | NM_001130487.1 | NDRG4 | 26.2031646 | -5.929672 | 1.74920891 |
| 13869 | NM_001324339.1 | #N/A | 16.6623413 | 9.05459284 | 2.67145342 |
| 13870 | NM_001301838.1 | C12orf57 | 24.6840945 | -5.8285499 | 1.71966716 |
| 13871 | NM_003804.4 | #N/A | 84.2445535 | -2.9605104 | 0.87357059 |
| 13872 | NR_024433.1 | LINC00926 | 46.1208619 | 2.34320975 | 0.69143677 |
| 13873 | NM_001253852.2 | #N/A | 31.0027604 | -6.1689457 | 1.82036201 |
| 13874 | NR_135781.1 | RSRP1 | 23.7776145 | -5.7893714 | 1.70838469 |
| 13875 | NM_018014.3 | #N/A | 33.7814046 | -6.2825203 | 1.85400887 |
| 13876 | NM_001287492.2 | #N/A | 26.5609687 | -5.9453038 | 1.75465497 |
| 13877 | NR_134623.1 | LOC105373021 | 48.1419097 | 3.06083313 | 0.90337475 |
| 13878 | NM_001204199.1 | #N/A | 24.7335642 | -5.8361464 | 1.72271731 |
| 13879 | NM_002748.3 | #N/A | 673.288577 | -3.837192 | 1.13266723 |
| 13880 | NM_001146256.1 | ZDHHC15 | 14.6831672 | 8.83528914 | 2.60818801 |
| 13881 | NM_001271471.2 | LRTOMT | 21.7697806 | -5.6445751 | 1.66673067 |
| 13882 | NM_001010879.3 | #N/A | 22.434232 | -5.6990475 | 1.68301297 |
| 13883 | NM_001014842.2 | #N/A | 21.5988832 | -5.6461637 | 1.66749782 |
| 13884 | NM_205850.2 | #N/A | 19.5432792 | -5.5000998 | 1.62441791 |
| 13885 | NM_017715.3 | #N/A | 22.4306375 | -5.6937446 | 1.68162901 |
| 13886 | NR_144335.1 | #N/A | 22.1393264 | -5.6722132 | 1.67553694 |
| 13887 | NM_001134651.1 | #N/A | 56.7856193 | -5.4577507 | 1.61237134 |
| 13888 | NM_001242481.1 | #N/A | 23.3737441 | -5.7484849 | 1.69826464 |

| | | | | | |
|---|---|---|---|---|---|
| 13889 | NM_001012271.1 | #N/A | 43.0897038 | -3.5592608 | 1.05154506 |
| 13890 | NM_001033054.2 | #N/A | 53.7988618 | 3.02205544 | 0.89283701 |
| 13891 | NM_001267723.1 | #N/A | 19.1082428 | -5.4689746 | 1.61580776 |
| 13892 | NR_036444.1 | FENDRR | 29.1138757 | -6.0764217 | 1.79562344 |
| 13893 | NM_080867.2 | #N/A | 134.748218 | -2.663867 | 0.78724352 |
| 13894 | NM_020309.3 | #N/A | 456.4896 | -5.1708519 | 1.5282392 |
| 13895 | NM_032347.2 | #N/A | 22.1963293 | -5.6765283 | 1.67778313 |
| 13896 | NM_001134340.1 | #N/A | 28.3174201 | -6.0354027 | 1.7838708 |
| 13897 | NM_006000.2 | #N/A | 25.2008906 | -5.8677065 | 1.73448356 |
| 13898 | NM_201995.2 | SF1 | 27.9268244 | -6.0177038 | 1.77885828 |
| 13899 | NM_001321847.1 | #N/A | 35.3343857 | -6.3513421 | 1.87769549 |
| 13900 | NM_003729.3 | RTCA | 215.985553 | -2.0403741 | 0.60323591 |
| 13901 | NM_174937.3 | #N/A | 60.7615328 | -7.1340412 | 2.10937467 |
| 13902 | NR_028099.1 | DCAF11 | 25.6515843 | -5.8999332 | 1.74463399 |
| 13904 | NM_201538.1 | NDRG2 | 33.7908933 | -6.2823043 | 1.85836985 |
| 13903 | NM_001008712.2 | RBPMS | 24.1943143 | -5.8106155 | 1.718846 |
| 13905 | NM_001308382.1 | POLL | 19.9068538 | -5.5298921 | 1.63625542 |
| 13906 | NR_110858.1 | EDRF1 | 21.6581422 | -5.6571062 | 1.67424455 |
| 13907 | NM_001291858.1 | #N/A | 177.788915 | -8.684974 | 2.57039186 |
| 13908 | NM_001280792.1 | SURF4 | 25.4872909 | -5.888985 | 1.74293704 |
| 13909 | NM_032050.1 | #N/A | 209.715344 | -4.3223522 | 1.27934867 |
| 13910 | NR_015419.2 | LOC145783 | 25.0136251 | -5.8554909 | 1.73332469 |
| 13911 | NM_001323413.1 | #N/A | 23.5190813 | -5.7610199 | 1.70544604 |
| 13912 | NM_000141.4 | FGFR2 | 201.515371 | -8.8656181 | 2.6247356 |
| 13913 | NM_018237.3 | #N/A | 177.305193 | -8.6810358 | 2.5701148 |
| 13914 | NM_138381.3 | #N/A | 18.4157186 | -5.414704 | 1.60307403 |
| 13915 | NM_018659.2 | #N/A | 29.413465 | -6.0901022 | 1.80305747 |
| 13916 | NM_198310.3 | TTC8 | 22.9723111 | -5.7399291 | 1.69942425 |
| 13917 | NM_002526.3 | #N/A | 24.9100847 | -5.8546801 | 1.73360834 |
| 13918 | NR_029193.1 | SH3RF3-AS1 | 26.3075778 | -5.9233819 | 1.75403316 |
| 13919 | NM_025115.3 | #N/A | 26.0841828 | -5.9197198 | 1.75297616 |
| 13921 | NM_181640.2 | CKLF | 21.2298439 | -5.6214639 | 1.6648791 |
| 13920 | NM_001025356.2 | #N/A | 181.334288 | -8.7134604 | 2.58061193 |
| 13922 | NM_022475.2 | #N/A | 50.06577 | -5.3029153 | 1.57080357 |
| 13923 | NM_006036.4 | PREPL | 24.1653772 | -5.8112395 | 1.72159354 |
| 13924 | NR_133646.1 | #N/A | 22.9963906 | -5.7338043 | 1.69874899 |
| 13925 | NM_014578.3 | #N/A | 19.2536019 | -5.4838732 | 1.624805 |
| 13926 | NM_001291428.1 | #N/A | 20.8761959 | -5.5868094 | 1.65532397 |
| 13927 | NR_003606.2 | #N/A | 30.5370405 | -6.1449366 | 1.82096436 |
| 13928 | NR_132751.1 | #N/A | 19.8244511 | -5.5222818 | 1.63659388 |
| 13929 | NM_001005362.2 | DNM2 | 179.788908 | -8.7011334 | 2.57892472 |
| 13930 | NM_152732.4 | #N/A | 23.3280245 | -5.757762 | 1.70659795 |
| 13931 | NM_031293.2 | #N/A | 28.9810062 | -6.0678153 | 1.79870363 |

| | | | | | | |
|---|---|---|---|---|---|---|
| 13932 | NR_015364.2 | | #N/A | 17.8069629 | -5.3709441 | 1.59226745 |
| 13933 | NM_004617.3 | | #N/A | 27.8897624 | -6.0141385 | 1.78304578 |
| 13934 | NM_001008222.2 | | #N/A | 21.7209596 | -5.6477308 | 1.67457263 |
| 13935 | NM_001281425.1 | DCTN3 | | 43.2199104 | -6.6444924 | 1.97065062 |
| 13936 | NM_182734.2 | | #N/A | 26.102276 | -5.9149459 | 1.75455276 |
| 13937 | NM_001172309.1 | NEXN | | 23.0553324 | -5.745267 | 1.70435726 |
| 13938 | NM_001013653.2 | LRRC26 | | 22.7010252 | -5.7127308 | 1.69473269 |
| 13939 | NM_014260.3 | PFDN6 | | 33.9695775 | -6.2905241 | 1.86641778 |
| 13940 | NM_005479.3 | | #N/A | 23.721274 | -5.7792766 | 1.71483579 |
| 13941 | NM_016556.3 | | #N/A | 23.2541121 | -5.741778 | 1.70401702 |
| 13942 | NM_006576.3 | AVIL | | 18.9350596 | -5.4530091 | 1.61841428 |
| 13943 | NM_001013257.2 | BCAM | | 23.7449577 | -5.7879825 | 1.71806347 |
| 13944 | NM_145274.3 | | #N/A | 22.0256687 | -5.6734215 | 1.68413312 |
| 13945 | NM_018950.2 | HLA-F | | 18.7579608 | -5.4451713 | 1.61656542 |
| 13946 | NM_004189.3 | | #N/A | 27.078393 | -5.9717972 | 1.77306924 |
| 13947 | NM_017806.3 | | #N/A | 35.5480686 | -6.3548848 | 1.88694539 |
| 13948 | NM_212478.2 | FN1 | | 254.398525 | -9.2015993 | 2.73269766 |
| 13949 | NM_004023.2 | | #N/A | 32.5041875 | -6.2261052 | 1.84905792 |
| 13950 | NM_001160148.1 | | #N/A | 42.5339671 | -6.616536 | 1.96528021 |
| 13951 | NM_206937.1 | | #N/A | 43.1880897 | -6.6393242 | 1.97209166 |
| 13952 | NM_018937.4 | PCDHB3 | | 21.744255 | -5.6488771 | 1.67798715 |
| 13953 | NM_198179.2 | | #N/A | 39.2318304 | 2.70610011 | 0.80386794 |
| 13954 | NM_001127244.1 | LRRC8A | | 20.0207189 | -5.5261343 | 1.64166431 |
| 13955 | NM_001011554.2 | | #N/A | 25.9662176 | -5.8994124 | 1.75258577 |
| 13956 | NM_001346492.1 | | #N/A | 22.1882602 | -5.6837459 | 1.68874484 |
| 13957 | NM_001018053.1 | | #N/A | 18.8540893 | -5.4562771 | 1.6215429 |
| 13959 | NR_038103.1 | TECR | | 22.9211692 | -5.723485 | 1.7010089 |
| 13958 | NM_018653.3 | | #N/A | 22.3634317 | -5.6973405 | 1.69324638 |
| 13960 | NM_182700.5 | | #N/A | 27.5770306 | -5.9892737 | 1.78005244 |
| 13962 | NM_001198546.1 | NOL4 | | 59.5836975 | -7.1070671 | 2.11245756 |
| 13961 | NM_001099400.1 | | #N/A | 34.8583105 | -6.3325278 | 1.88224233 |
| 13963 | NR_072977.1 | | #N/A | 24.6345163 | -5.8329865 | 1.73374323 |
| 13964 | NM_001270550.1 | MDK | | 18.9947771 | -5.4573593 | 1.62219324 |
| 13965 | NM_001080461.1 | | #N/A | 20.0540526 | -5.5462895 | 1.64883388 |
| 13966 | NM_146388.1 | | #N/A | 58.8941712 | -4.3109221 | 1.28156909 |
| 13967 | NM_032854.3 | CORO6 | | 19.6354048 | -5.5149886 | 1.63998991 |
| 13968 | NM_175850.2 | DNMT3B | | 18.2927336 | -5.4080685 | 1.60830498 |
| 13969 | NM_001243288.1 | | #N/A | 18.2319586 | -5.3987067 | 1.60571465 |
| 13970 | NM_001171868.1 | TMEM200B | | 19.7926924 | -5.5071522 | 1.63806421 |
| 13971 | NR_073169.1 | ATG9B | | 22.5245452 | -5.7100449 | 1.69855367 |
| 13972 | NR_136664.1 | | #N/A | 21.828856 | -5.6584824 | 1.68334991 |
| 13973 | NM_001291641.1 | | #N/A | 25.6351355 | -5.8921161 | 1.75287738 |
| 13974 | NM_001286526.1 | | #N/A | 19.2558407 | -5.4782516 | 1.63004769 |

| | | | | | |
|---|---|---|---|---|---|
| 13975 | NM_004625.3 | #N/A | 34.3638748 | -6.3102491 | 1.87761274 |
| 13976 | NM_006492.2 | #N/A | 21.3903001 | -5.6323443 | 1.67590848 |
| 13979 | NM_012415.3 | RAD54B | 20.6734989 | -5.5730156 | 1.65836555 |
| 13977 | NM_001156474.1 | #N/A | 23.1383823 | -5.7480432 | 1.71045558 |
| 13978 | NM_006958.2 | #N/A | 35.3940659 | -4.8657352 | 1.4478964 |
| 13980 | NM_000051.3 | ATM | 290.725652 | -1.0386185 | 0.30908922 |
| 13981 | NM_001260508.1 | KCNJ3 | 140.898333 | 3.15599662 | 0.93935995 |
| 13982 | NM_005112.4 | #N/A | 25.1623302 | -5.8592478 | 1.74415872 |
| 13983 | NR_026934.1 | LINC02085 | 13.5696685 | 3.62782545 | 1.07998453 |
| 13984 | NM_001136230.2 | #N/A | 84.6626003 | 2.97447496 | 0.8855834 |
| 13985 | NM_024531.4 | #N/A | 23.7307088 | -5.7807852 | 1.72110218 |
| 13986 | NR_037166.1 | SOD2-OT1 | 24.2793139 | 2.07234771 | 0.61710855 |
| 13987 | NR_045513.1 | #N/A | 19.4642126 | -5.4874928 | 1.63427082 |
| 13988 | NM_001130714.2 | LEF1 | 50.2864643 | -6.8592265 | 2.04292712 |
| 13989 | NM_031217.3 | #N/A | 28.6807618 | -4.7701611 | 1.42084781 |
| 13990 | NM_001005619.1 | ITGB4 | 89.2000546 | -4.753709 | 1.41607513 |
| 13991 | NM_001007073.1 | RPL32 | 29.1993504 | -6.0791527 | 1.81139868 |
| 13992 | NM_001322800.1 | LOC100996842 | 28.8256319 | -6.0555554 | 1.80445241 |
| 13993 | NR_144376.1 | AKR1B1 | 18.4641445 | -5.4218985 | 1.61567163 |
| 13994 | NM_001256725.1 | #N/A | 20.3313569 | -5.5616038 | 1.65738031 |
| 13995 | NM_001005339.1 | #N/A | 27.3653945 | -5.9866974 | 1.78411878 |
| 13996 | NR_027274.1 | SRRM2-AS1 | 40.1001543 | 2.59606818 | 0.77370186 |
| 13997 | NM_017898.4 | #N/A | 24.662743 | -5.8372326 | 1.7397535 |
| 13998 | NR_028050.1 | MTERF4 | 28.8023205 | -6.0515586 | 1.80374712 |
| 13999 | NM_001276718.1 | #N/A | 28.052059 | -6.0178215 | 1.79379865 |
| 14000 | NM_032850.4 | #N/A | 19.6199792 | -5.5085822 | 1.6423035 |
| 14001 | NM_001323982.1 | #N/A | 17.4363327 | -5.3374969 | 1.59139376 |
| 14002 | NM_005590.3 | #N/A | 20.4872704 | -5.5759382 | 1.66274443 |
| 14003 | NM_001288727.1 | PPRC1 | 22.9867192 | -5.7325959 | 1.70955317 |
| 14004 | NM_080680.2 | #N/A | 24.1319347 | -5.8047906 | 1.73140422 |
| 14005 | NM_005295.2 | #N/A | 60.3893715 | -7.125419 | 2.12551318 |
| 14006 | NM_001145520.1 | RAI14 | 24.448515 | -5.8179178 | 1.73554541 |
| 14007 | NM_001173989.2 | #N/A | 18.9809753 | -5.4664419 | 1.63071825 |
| 14008 | NM_001170423.1 | #N/A | 24.5841262 | -5.8258138 | 1.73796721 |
| 14009 | NM_001048201.2 | #N/A | 37.2329771 | -6.4286678 | 1.91784902 |
| 14010 | NR_144632.1 | #N/A | 21.2076671 | -5.6237263 | 1.67779572 |
| 14011 | NR_131235.1 | PANTR1 | 59.5090416 | -7.1036066 | 2.11957215 |
| 14012 | NM_147202.1 | #N/A | 172.069295 | -8.637759 | 2.5778142 |
| 14013 | NM_001318170.1 | #N/A | 89.2206491 | 1.77003825 | 0.52827625 |
| 14014 | NM_052927.2 | #N/A | 24.9544928 | -5.8591864 | 1.74938637 |
| 14015 | NM_203291.1 | RBBP8 | 24.1746416 | -5.8069269 | 1.73383929 |
| 14016 | NM_001013619.3 | #N/A | 95.3886258 | 1.73865341 | 0.51917483 |
| 14017 | NM_145058.2 | RILPL2 | 19.0773257 | -5.4667953 | 1.63250075 |

| | | | | | |
|---|---|---|---|---|---|
| 14018 | NR_120405.1 | LOC400867 | 17.7463474 | 6.68245671 | 1.99572943 |
| 14019 | NM_001256600.1 | #N/A | 31.189187 | -6.1679279 | 1.84212966 |
| 14020 | NM_014235.4 | #N/A | 175.26668 | -3.2857566 | 0.98134691 |
| 14021 | NR_038331.1 | LINC01095 | 33.8228696 | 9.37411804 | 2.79976112 |
| 14022 | NR_036538.1 | LOC646903 | 19.7174633 | -5.5161469 | 1.64754758 |
| 14023 | NM_152731.2 | #N/A | 24.0192301 | -5.7970471 | 1.7314993 |
| 14024 | NM_203403.1 | #N/A | 21.8785884 | -5.6651772 | 1.69225575 |
| 14025 | NM_058175.2 | COL6A2 | 25.4673552 | -5.8864601 | 1.75843689 |
| 14026 | NR_024543.1 | SNHG7 | 27.4571083 | -5.9922446 | 1.79029235 |
| 14027 | NR_046437.1 | SLC30A10 | 18.4920618 | -5.4270098 | 1.62153383 |
| 14028 | NM_005019.4 | #N/A | 42.8332431 | -6.6284229 | 1.98054478 |
| 14029 | NM_001301820.1 | ARMC5 | 29.4202261 | -6.0931554 | 1.82085416 |
| 14030 | NM_001163315.2 | #N/A | 124.706057 | -3.4420578 | 1.02863348 |
| 14031 | NR_002934.3 | #N/A | 19.1013905 | -5.4667796 | 1.63397977 |
| 14032 | NM_057749.2 | CCNE2 | 21.4731537 | -5.6357559 | 1.6845791 |
| 14033 | NR_103783.1 | BLACAT1 | 20.0450463 | -5.5410549 | 1.65628578 |
| 14034 | NM_182703.5 | #N/A | 41.6895246 | -3.5975133 | 1.07537823 |
| 14035 | NM_024874.4 | #N/A | 401.798164 | -2.2551803 | 0.67414694 |
| 14036 | NM_000294.2 | #N/A | 145.407257 | -1.2217086 | 0.36526459 |
| 14037 | NM_031244.3 | SIRT5 | 21.659881 | -5.6417216 | 1.68683311 |
| 14039 | NR_109973.1 | DLEU1 | 18.6299082 | -5.4301346 | 1.62384555 |
| 14038 | NR_039997.1 | LOC100506083 | 18.5487672 | -5.4150909 | 1.61935087 |
| 14040 | NM_001433.3 | #N/A | 17.0050147 | -5.2956111 | 1.58363848 |
| 14041 | NM_013255.4 | #N/A | 189.737249 | -2.1009218 | 0.62831865 |
| 14042 | NM_001003679.3 | LEPR | 24.9764854 | -5.8582772 | 1.75223424 |
| 14043 | NM_024587.3 | #N/A | 24.4027326 | -5.8099732 | 1.73794293 |
| 14044 | NM_001282997.1 | #N/A | 22.8626736 | -5.7189298 | 1.7108111 |
| 14045 | NM_021159.4 | #N/A | 34.6425987 | -6.3287142 | 1.89328119 |
| 14046 | NM_001143937.1 | PSMA1 | 18.2971122 | -5.4016804 | 1.61602991 |
| 14047 | NM_001080503.2 | #N/A | 18.6917885 | -5.4412289 | 1.62791088 |
| 14048 | NM_000333.3 | ATXN7 | 26.433504 | -5.9416767 | 1.77766936 |
| 14049 | NR_015401.2 | LINC00957 | 41.4687422 | -5.0573789 | 1.51321743 |
| 14050 | NR_024322.1 | MSL3P1 | 19.2327193 | -5.4686558 | 1.63627073 |
| 14051 | NM_152233.3 | #N/A | 280.78747 | -3.4918122 | 1.04479616 |
| 14052 | NM_173794.3 | #N/A | 62.9365812 | -4.3709599 | 1.30788793 |
| 14053 | NM_001139467.1 | TBL1X | 37.5973029 | -6.4454855 | 1.92875089 |
| 14054 | NM_020147.3 | #N/A | 20.5996149 | -5.5732277 | 1.66779192 |
| 14055 | NM_199421.1 | #N/A | 25.3616536 | -5.8801813 | 1.7596819 |
| 14056 | NM_000839.3 | #N/A | 28.1733244 | -6.0223581 | 1.80226758 |
| 14057 | NM_001330616.1 | #N/A | 30.4841627 | -6.1364162 | 1.83648483 |
| 14058 | NM_001079802.1 | FKTN | 28.7371866 | -6.0590347 | 1.81426613 |
| 14059 | NM_001258213.2 | #N/A | 25.6881212 | -5.8891348 | 1.76353207 |
| 14060 | NM_001331028.1 | #N/A | 24.2946855 | -5.8171978 | 1.74220506 |

| | | | | | |
|---|---|---|---|---|---|
| 14062 | NR_024480.1 | LINC00887 | 39.5839733 | 2.45811278 | 0.73627891 |
| 14061 | NM_001324255.1 | #N/A | 1374.79641 | -7.937931 | 2.37765478 |
| 14063 | NM_001137551.1 | LRRFIP1 | 33.6450463 | -6.2811321 | 1.88144462 |
| 14064 | NM_177439.2 | #N/A | 22.2359121 | -5.6895954 | 1.70428903 |
| 14065 | NM_001166113.1 | PNPLA6 | 31.4031668 | -6.185497 | 1.85307091 |
| 14066 | NM_001271685.1 | #N/A | 32.636722 | -6.2310264 | 1.86672772 |
| 14067 | NM_020318.2 | #N/A | 28.7443444 | -6.0545361 | 1.81392274 |
| 14068 | NM_017752.2 | #N/A | 28.0518468 | -6.0245247 | 1.80510985 |
| 14069 | NM_173487.2 | #N/A | 27.493531 | -5.9885168 | 1.79439501 |
| 14070 | NM_001136137.1 | RPL28 | 22.9970386 | -5.730046 | 1.71701706 |
| 14071 | NM_001167947.2 | TIMM17B | 20.2821866 | -5.5506577 | 1.66337843 |
| 14072 | NM_001297566.1 | #N/A | 25.3887285 | -5.8793734 | 1.7619374 |
| 14073 | NR_046341.1 | #N/A | 84.9682603 | -2.548907 | 0.76390961 |
| 14074 | NM_001287487.1 | #N/A | 21.4114304 | -5.6266322 | 1.68655037 |
| 14075 | NR_073091.1 | #N/A | 23.1665996 | -5.74136 | 1.72100733 |
| 14076 | NM_145315.4 | #N/A | 20.277468 | -5.5477689 | 1.66300352 |
| 14077 | NM_001330602.1 | MRPL37 | 19.9421694 | -5.5236914 | 1.65600783 |
| 14078 | NM_001330692.1 | #N/A | 24.2752511 | -5.8130109 | 1.74286282 |
| 14079 | NM_001134433.1 | #N/A | 22.4685061 | -5.7004864 | 1.70919623 |
| 14080 | NM_030639.2 | #N/A | 40.0723939 | -6.5313825 | 1.95861865 |
| 14081 | NR_073083.1 | #N/A | 39.720982 | -6.526049 | 1.95708161 |
| 14082 | NM_001012507.3 | CENPW | 21.1249402 | -5.6094711 | 1.68224075 |
| 14084 | NM_145731.3 | SYNGR1 | 17.5815808 | -5.3501004 | 1.60447719 |
| 14083 | NM_001193478.1 | #N/A | 19.4695516 | -5.4936408 | 1.6475315 |
| 14085 | NM_001166348.1 | #N/A | 24.382591 | -5.8246573 | 1.74690755 |
| 14086 | NM_004165.2 | #N/A | 35.211175 | -6.3518745 | 1.90510861 |
| 14087 | NM_001278055.1 | #N/A | 146.484658 | -4.2039516 | 1.26089698 |
| 14088 | NM_001322310.1 | PHACTR1 | 27.2572957 | -5.983014 | 1.79456064 |
| 14089 | NM_002003.4 | #N/A | 113.169944 | 1.64581985 | 0.49375359 |
| 14090 | NR_027702.1 | #N/A | 59.9054442 | -5.5128596 | 1.65390559 |
| 14091 | NM_001018052.3 | POLR3H | 22.397772 | -5.6976173 | 1.70935283 |
| 14092 | NM_001206572.1 | #N/A | 26.8051148 | -5.9619446 | 1.78868538 |
| 14093 | NM_014709.3 | #N/A | 528.090393 | -3.5427572 | 1.06294885 |
| 14094 | NM_014696.3 | #N/A | 25.0959564 | -5.8584545 | 1.75780282 |
| 14095 | NM_001271838.1 | #N/A | 26.8196068 | -5.9468477 | 1.78440627 |
| 14096 | NM_001270731.1 | BCL2L13 | 31.2700205 | -6.1743476 | 1.85285297 |
| 14097 | NR_037717.1 | RNASEK-C17orf49 | 25.104348 | -5.8570542 | 1.75768113 |
| 14098 | NM_173662.2 | #N/A | 28.9389005 | -6.0626447 | 1.81948736 |
| 14099 | NR_104034.1 | GNRHR2 | 128.718063 | 3.03565233 | 0.91113015 |
| 14100 | NM_005192.3 | #N/A | 20.97448 | -5.593904 | 1.67911793 |
| 14101 | NM_001042510.1 | ZNF706 | 28.9433401 | -6.0624729 | 1.81981121 |
| 14103 | NM_001145805.1 | IRGM | 37.5300259 | 3.9532323 | 1.18671212 |
| 14102 | NM_001143990.1 | WRAP53 | 25.7443917 | -5.8861211 | 1.76694686 |

| | | | | | |
|---|---|---|---|---|---|
| 14104 | NM_001171888.1 | FGFR1OP2 | 17.124185 | -5.310711 | 1.59429484 |
| 14105 | NM_001270491.1 | RPL13A | 201.609762 | -8.866398 | 2.66184716 |
| 14106 | NR_003189.2 | SYS1-DBNDD2 | 26.895479 | -5.9568023 | 1.78875687 |
| 14107 | NM_001029840.2 | TCAIM | 27.8345362 | -6.0074197 | 1.80406174 |
| 14108 | NM_009590.3 | #N/A | 21.5636813 | -5.6395481 | 1.6936163 |
| 14109 | NM_207123.2 | #N/A | 24.5161107 | -5.8267708 | 1.7500145 |
| 14110 | NM_001184824.1 | #N/A | 23.921791 | -5.7992232 | 1.74184624 |
| 14111 | NM_001199752.2 | CPT1C | 20.6117665 | -5.5670645 | 1.67214835 |
| 14112 | NM_002199.3 | #N/A | 29.7226269 | -4.6475395 | 1.39621646 |
| 14113 | NM_194271.2 | RNF34 | 21.4265998 | -5.6254671 | 1.690095 |
| 14114 | NM_172057.2 | KCNH2 | 24.8933422 | -5.8534632 | 1.75867902 |
| 14115 | NR_134489.1 | #N/A | 20.7637323 | -5.5845773 | 1.67809648 |
| 14116 | NM_001135032.1 | #N/A | 22.0340874 | -5.6759839 | 1.70575912 |
| 14117 | NM_001018008.1 | #N/A | 38.7409519 | -4.9738139 | 1.49491559 |
| 14118 | NM_001321163.1 | #N/A | 21.3976172 | -5.624552 | 1.69056917 |
| 14119 | NM_199420.3 | #N/A | 17.5949595 | -5.3508922 | 1.60835742 |
| 14120 | NM_198833.1 | SERPINB8 | 25.0202394 | -5.8483052 | 1.75797956 |
| 14121 | NM_001199341.1 | #N/A | 27.8192144 | -6.010771 | 1.80686003 |
| 14122 | NM_001104544.1 | TMEM255A | 29.904015 | -6.1110833 | 1.83707007 |
| 14123 | NM_001242699.1 | #N/A | 33.0861844 | -6.262083 | 1.88270701 |
| 14124 | NR_024525.2 | #N/A | 17.5314521 | -5.3432873 | 1.60649007 |
| 14125 | NR_033297.1 | #N/A | 19.1865482 | -5.4693316 | 1.64455057 |
| 14126 | NM_018482.3 | #N/A | 296.118428 | -3.8240521 | 1.14991094 |
| 14127 | NM_001302688.1 | #N/A | 20.2113334 | -5.5401445 | 1.66602092 |
| 14128 | NM_014305.3 | #N/A | 17.8523043 | -5.3595782 | 1.61198012 |
| 14129 | NM_001301030.1 | GRIK5 | 19.7138567 | -5.516844 | 1.65942274 |
| 14130 | NM_001272083.1 | #N/A | 18.6299718 | -5.4256957 | 1.63206697 |
| 14131 | NM_020696.3 | #N/A | 288.55793 | -3.4238217 | 1.0299543 |
| 14132 | NM_025209.3 | #N/A | 23.4227116 | -5.7638727 | 1.73388251 |
| 14133 | NR_104064.1 | SMIM2-AS1 | 21.1568111 | 5.24537909 | 1.57815269 |
| 14134 | NM_001204212.1 | #N/A | 25.9895225 | 7.6065396 | 2.28860007 |
| 14135 | NM_080832.2 | #N/A | 31.9794529 | -4.7365111 | 1.42509711 |
| 14136 | NM_001145636.1 | #N/A | 19.1381593 | -5.4785044 | 1.64844677 |
| 14137 | NM_014518.3 | #N/A | 24.8182764 | -5.8441714 | 1.7587436 |
| 14138 | NR_024547.1 | TMEM11 | 20.3663456 | -5.5500408 | 1.67027643 |
| 14139 | NM_001242822.1 | #N/A | 22.8609592 | -5.7224735 | 1.72221568 |
| 14140 | NM_052932.2 | #N/A | 341.318439 | -4.248527 | 1.278761 |
| 14141 | NM_024086.3 | #N/A | 184.086914 | -1.8347888 | 0.55227073 |
| 14142 | NM_022171.2 | TCTA | 104.089807 | -2.4624578 | 0.7412253 |
| 14143 | NR_033194.1 | ZFAND1 | 36.3263201 | -4.8877034 | 1.47124935 |
| 14144 | NM_001032386.1 | #N/A | 30.1005575 | -6.1229881 | 1.84334456 |
| 14145 | NM_001318841.1 | #N/A | 26.6607639 | -5.9489521 | 1.79103313 |
| 14146 | NM_001042728.2 | #N/A | 21.4760657 | -5.6425898 | 1.69895166 |

| | | | | | |
|---|---|---|---|---|---|
| 14147 | NM_054022.3 | #N/A | 24.3806266 | 9.57526603 | 2.88350824 |
| 14148 | NM_001521.3 | GTF3C2 | 23.15625 | -5.7498549 | 1.73171048 |
| 14149 | NM_001037144.5 | #N/A | 23.0707083 | -5.7471421 | 1.7310531 |
| 14151 | NM_001286095.1 | TMEM9B | 37.3833597 | -6.4357755 | 1.93869351 |
| 14150 | NM_001258304.1 | #N/A | 20.9258521 | -5.6040694 | 1.68816215 |
| 14152 | NM_001297739.1 | #N/A | 22.2517833 | -5.6873784 | 1.71328421 |
| 14153 | NM_020911.1 | PLXNA4 | 362.200945 | -4.0169154 | 1.21014675 |
| 14154 | NR_024466.1 | #N/A | 16.9565491 | -5.2922962 | 1.59437926 |
| 14156 | NR_120466.1 | LINC01489 | 27.3418577 | 5.07349739 | 1.5285211 |
| 14155 | NM_199336.2 | #N/A | 22.0254151 | -5.6733535 | 1.70923722 |
| 14157 | NM_181463.2 | #N/A | 19.8588223 | -5.5173201 | 1.66229424 |
| 14158 | NM_001286581.1 | #N/A | 167.614323 | -8.5999547 | 2.59134211 |
| 14159 | NR_102371.1 | MFF-DT | 39.660583 | 2.81132178 | 0.8471532 |
| 14160 | NM_001006933.1 | TCEAL3 | 23.542226 | -5.7680535 | 1.73814073 |
| 14161 | NM_001151.3 | #N/A | 403.939336 | -2.8525165 | 0.85964474 |
| 14162 | NR_015341.2 | SMURF2P1-LRRC37BP1 | 180.795519 | -1.8230085 | 0.54939641 |
| 14163 | NM_001303241.1 | #N/A | 210.578272 | -3.1475056 | 0.94859201 |
| 14164 | NR_024469.1 | LOC100130987 | 19.6378796 | -5.5101399 | 1.66080376 |
| 14165 | NR_122038.1 | ZNF561-AS1 | 22.1658731 | -5.6785381 | 1.7116314 |
| 14166 | NM_017848.5 | #N/A | 20.4234547 | -5.5583328 | 1.67559485 |
| 14167 | NM_001290040.1 | #N/A | 34.0059551 | -6.2918927 | 1.89691068 |
| 14168 | NM_001278159.1 | AKT1S1 | 18.9493009 | -5.4570805 | 1.64524473 |
| 14169 | NR_024494.1 | BCRP3 | 18.9604188 | -5.4490328 | 1.64300125 |
| 14170 | NM_001076680.1 | #N/A | 36.1866648 | -4.8820713 | 1.47217232 |
| 14171 | NR_023318.1 | #N/A | 25.425558 | -5.8740964 | 1.77130961 |
| 14172 | NM_006796.2 | #N/A | 291.03593 | -4.2335517 | 1.27675387 |
| 14173 | NM_001288567.1 | TMEM9 | 25.7238595 | -5.9020976 | 1.78022349 |
| 14174 | NM_002674.3 | #N/A | 51.4422157 | -6.8961918 | 2.08007948 |
| 14175 | NM_001323635.1 | #N/A | 21.4181979 | -5.6337976 | 1.69933913 |
| 14176 | NR_126523.1 | C5orf64 | 31.5705952 | 2.93789389 | 0.88625305 |
| 14177 | NM_001330200.1 | RPL19 | 19.1031029 | -5.4616795 | 1.647708 |
| 14178 | NM_145055.3 | #N/A | 22.1643557 | -5.6696883 | 1.71053817 |
| 14180 | NM_020922.4 | WNK3 | 23.288771 | -5.7594251 | 1.73787921 |
| 14179 | NM_001318821.1 | #N/A | 156.805625 | -8.5038547 | 2.56599839 |
| 14181 | NM_002542.5 | OGG1 | 21.1915318 | -5.6145658 | 1.694298 |
| 14182 | NM_015688.1 | #N/A | 50.4425355 | -3.6029407 | 1.08732143 |
| 14183 | NR_111943.1 | #N/A | 27.2260304 | -5.9742019 | 1.80304095 |
| 14184 | NM_003940.2 | #N/A | 147.048109 | -1.1947835 | 0.36062973 |
| 14185 | NR_134858.1 | LRTOMT | 136.337893 | 2.72444699 | 0.82252546 |
| 14186 | NM_001135775.2 | #N/A | 30.4416 | -4.0629162 | 1.22690473 |
| 14187 | NM_001082578.1 | #N/A | 261.016337 | -9.2382706 | 2.79016638 |
| 14188 | NM_001184748.1 | PAFAH1B2 | 34.0718246 | 2.14394297 | 0.64754077 |
| 14189 | NM_030577.2 | #N/A | 22.844078 | -5.7260476 | 1.72948345 |

| | | | | | |
|---|---|---|---|---|---|
| 14190 | NM_001020819.2 | MYADM | 29.9298923 | -6.1184426 | 1.84809471 |
| 14191 | NR_033184.1 | #N/A | 32.5507938 | -6.230369 | 1.88205644 |
| 14192 | NM_001973.3 | #N/A | 119.586201 | -2.0754606 | 0.62707055 |
| 14193 | NM_001257999.2 | ANKFY1 | 20.7176686 | -5.5927052 | 1.68980823 |
| 14195 | NR_110005.1 | LRPAP1 | 145.931328 | 8.00665754 | 2.41918777 |
| 14194 | NM_001278784.1 | #N/A | 54.3037819 | -5.3852248 | 1.62712885 |
| 14196 | NM_001321705.1 | #N/A | 51.8875946 | -6.9097299 | 2.08784021 |
| 14197 | NM_014189.3 | ADD1 | 217.386602 | -8.9747137 | 2.7118703 |
| 14198 | NM_001324192.1 | PANK2 | 19.3795592 | -5.492016 | 1.65965686 |
| 14199 | NM_152900.2 | #N/A | 26.3324877 | -5.9359861 | 1.79423341 |
| 14200 | NM_001282989.1 | #N/A | 18.569724 | -5.4333276 | 1.64291114 |
| 14201 | NM_152654.2 | DAND5 | 178.235856 | 2.08514672 | 0.63053906 |
| 14202 | NM_001127711.2 | TMEM14B | 25.9989605 | -5.9050577 | 1.78571256 |
| 14203 | NM_001098512.2 | #N/A | 19.046792 | -5.4635839 | 1.65242955 |
| 14204 | NR_110003.1 | LINC01733 | 58.6002906 | 2.64369899 | 0.79969228 |
| 14206 | NR_027087.2 | LINC02800 | 29.6141759 | 5.16000126 | 1.56109526 |
| 14205 | NM_001289155.1 | #N/A | 41.0075058 | -3.9837298 | 1.20522959 |
| 14207 | NM_005422.2 | TECTA | 20.7120269 | -5.5841768 | 1.68947861 |
| 14208 | NM_080742.2 | #N/A | 26.7685403 | -5.9506071 | 1.80035423 |
| 14209 | NM_019607.2 | #N/A | 28.2349881 | -4.5722227 | 1.3833626 |
| 14210 | NR_103707.1 | TSPEAR-AS1 | 18.9187263 | -5.4491606 | 1.64870611 |
| 14211 | NM_001270732.1 | BCL2L13 | 20.4643806 | -5.5576789 | 1.68189618 |
| 14212 | NM_001330676.1 | PSMA4 | 17.844011 | -5.367222 | 1.62435654 |
| 14213 | NM_033406.2 | #N/A | 18.015195 | -5.3774964 | 1.62747347 |
| 14214 | NM_001286451.1 | #N/A | 21.2816937 | -5.6218534 | 1.70147879 |
| 14215 | NM_002347.4 | #N/A | 25.6295885 | -5.896383 | 1.78468063 |
| 14216 | NR_024611.2 | HINT1 | 26.8372357 | -5.9556243 | 1.80274624 |
| 14217 | NM_001290194.1 | #N/A | 13.4784485 | 8.71100606 | 2.63695583 |
| 14218 | NM_001040182.1 | CLDND1 | 26.6544466 | -5.9398212 | 1.79809012 |
| 14219 | NM_001172415.1 | #N/A | 224.705402 | -3.3438721 | 1.01227026 |
| 14220 | NM_148918.2 | SHMT1 | 21.6618932 | -5.6457235 | 1.70922153 |
| 14221 | NM_007207.5 | #N/A | 19.8559881 | -5.5255237 | 1.67301392 |
| 14222 | NM_001346100.1 | #N/A | 20.1507717 | -5.5517393 | 1.680983 |
| 14224 | NM_001282728.1 | PXYLP1 | 24.0412116 | -5.7985799 | 1.75581008 |
| 14223 | NM_001098533.2 | #N/A | 22.7763841 | -5.7236905 | 1.73313553 |
| 14225 | NM_020778.4 | ALPK3 | 28.0821624 | -6.0257339 | 1.82465305 |
| 14226 | NM_013451.3 | #N/A | 242.472368 | -9.1323995 | 2.76572775 |
| 14227 | NM_001136043.2 | #N/A | 30.2797051 | -6.127036 | 1.85627576 |
| 14228 | NM_021804.2 | #N/A | 29.4241463 | 6.10349818 | 1.84916936 |
| 14229 | NM_007191.4 | #N/A | 27.0912327 | -5.9739185 | 1.81011753 |
| 14230 | NM_001014764.2 | EMC6 | 20.8701692 | -5.59102 | 1.6941164 |
| 14231 | NM_001319993.1 | PARPBP | 18.4446674 | -5.4044983 | 1.63776688 |
| 14232 | NM_001141936.2 | #N/A | 27.7254506 | -6.0054711 | 1.82002604 |

| | | | | | |
|---|---|---|---|---|---|
| 14233 | NM_001330128.1 | MMS19 | 33.9415836 | -6.2923073 | 1.90704337 |
| 14234 | NM_130834.2 | #N/A | 26.5551077 | -5.9464938 | 1.80277108 |
| 14235 | NR_109929.1 | #N/A | 34.2854988 | -6.314685 | 1.91453641 |
| 14238 | NR_130172.1 | H2BW3P | 33.0839461 | -6.2567388 | 1.89701801 |
| 14236 | NM_001159699.1 | #N/A | 16.4961367 | -5.2546898 | 1.59318519 |
| 14237 | NM_003364.3 | #N/A | 27.9630847 | -6.0199767 | 1.82522547 |
| 14239 | NM_198128.2 | RNF138 | 23.1661948 | -5.7381427 | 1.73988932 |
| 14240 | NM_052836.3 | #N/A | 22.6891928 | -5.7141172 | 1.73295321 |
| 14241 | NM_000640.2 | #N/A | 21.3905813 | -5.6258034 | 1.70650755 |
| 14242 | NM_001193452.1 | NUDT16L1 | 17.5968585 | -5.3542421 | 1.62417951 |
| 14243 | NM_001306212.1 | THBS4 | 19.7120968 | -5.518834 | 1.67412305 |
| 14244 | NR_034055.1 | LINC00692 | 21.6231248 | 6.64674446 | 2.01633443 |
| 14245 | NM_001013838.1 | #N/A | 26.8293413 | -5.955107 | 1.80660814 |
| 14246 | NM_001007022.2 | #N/A | 31.4519855 | -6.1908953 | 1.87841642 |
| 14247 | NM_025207.4 | #N/A | 17.4530624 | -5.3345652 | 1.61867628 |
| 14248 | NM_001159397.1 | #N/A | 29.9747545 | 6.17894987 | 1.87495698 |
| 14249 | NM_001300929.1 | #N/A | 21.6910162 | -5.6449665 | 1.713026 |
| 14250 | NM_005912.2 | #N/A | 25.1916694 | 6.36539017 | 1.93174501 |
| 14251 | NM_001204143.1 | #N/A | 23.7925681 | -5.7916663 | 1.7576646 |
| 14252 | NM_017861.3 | #N/A | 246.424864 | -1.0461036 | 0.31749419 |
| 14253 | NM_001004196.3 | CD200 | 32.7332497 | -6.2380141 | 1.89336332 |
| 14254 | NM_001197223.1 | #N/A | 44.0357773 | -6.6675981 | 2.02378566 |
| 14255 | NM_018072.5 | #N/A | 177.642304 | -2.0734038 | 0.62934863 |
| 14256 | NR_110438.1 | OTUD6B-AS1 | 17.2966516 | -5.3258205 | 1.61677171 |
| 14257 | NM_002589.2 | #N/A | 25.1051198 | -5.8673558 | 1.78122131 |
| 14258 | NM_001195639.1 | #N/A | 21.9167651 | -5.6624904 | 1.71923985 |
| 14259 | NM_002448.3 | MSX1 | 1397.82913 | 4.38406898 | 1.33115984 |
| 14260 | NR_036447.1 | PKD1P1 | 1734.5612 | -4.159119 | 1.26294612 |
| 14261 | NM_033160.6 | ZNF658 | 19.5531329 | -5.509688 | 1.67316737 |
| 14262 | NM_201532.2 | #N/A | 24.0251818 | -5.7952724 | 1.75990126 |
| 14263 | NM_001099339.1 | COMMD7 | 18.3451038 | -5.4044801 | 1.64138 |
| 14264 | NM_001918.3 | #N/A | 276.949926 | 1.00947782 | 0.30665145 |
| 14265 | NM_006103.3 | #N/A | 16.9989443 | -5.3029483 | 1.61090711 |
| 14266 | NM_016361.4 | #N/A | 16.6794309 | -5.2739087 | 1.60209715 |
| 14267 | NM_001616.4 | #N/A | 127.780731 | -3.1600142 | 0.95996571 |
| 14268 | NR_024529.1 | RPL23AP7 | 20.4887521 | -5.5766352 | 1.69435717 |
| 14269 | NM_016078.5 | #N/A | 19.9823268 | -5.5283685 | 1.67983501 |
| 14270 | NM_016442.3 | #N/A | 18.9745558 | -5.4506215 | 1.65621806 |
| 14271 | NM_001291484.2 | #N/A | 68.4080489 | 2.3750435 | 0.72168379 |
| 14272 | NM_024307.2 | #N/A | 17.0624546 | -5.3038473 | 1.61168502 |
| 14273 | NM_003935.4 | #N/A | 18.2202705 | -5.3890103 | 1.63782223 |
| 14274 | NM_001135770.3 | PVR | 23.9232257 | -5.7913304 | 1.76054821 |
| 14275 | NM_004855.4 | #N/A | 17.6548306 | -5.3492177 | 1.62623996 |

| | | | | | |
|---|---|---|---|---|---|
| 14276 | NM_152449.3 | #N/A | 27.1926835 | -5.967767 | 1.81447719 |
| 14277 | NM_181712.4 | #N/A | 19.6265274 | -5.5093721 | 1.67521648 |
| 14278 | NM_001008530.2 | #N/A | 130.673078 | -4.0529018 | 1.23244102 |
| 14279 | NM_001077268.1 | #N/A | 19.5102155 | -5.5014024 | 1.67291301 |
| 14281 | NR_027345.1 | LINC00173 | 17.1487969 | -5.3018454 | 1.6123653 |
| 14280 | NM_005297.3 | MCHR1 | 22.763054 | -5.721334 | 1.73993447 |
| 14282 | NM_015030.1 | #N/A | 174.202257 | -2.5020208 | 0.76105771 |
| 14283 | NM_001232.3 | CASQ2 | 13.6595278 | 5.90931343 | 1.79755479 |
| 14284 | NM_003613.3 | #N/A | 47.2247458 | 4.17494863 | 1.27018968 |
| 14285 | NM_001037808.2 | EIF3C | 28.3046905 | -6.03848 | 1.83746912 |
| 14286 | NM_001032279.1 | #N/A | 18.2142605 | -5.4026875 | 1.64471746 |
| 14287 | NM_139164.2 | #N/A | 146.872192 | -8.4094536 | 2.5600748 |
| 14288 | NR_045147.1 | ITGB3BP | 25.4439824 | -5.8729012 | 1.78795934 |
| 14289 | NM_001322362.1 | #N/A | 42.5415478 | -6.6206222 | 2.01578259 |
| 14290 | NM_153046.2 | #N/A | 18.819818 | 5.14610001 | 1.56705865 |
| 14291 | NM_004130.3 | GYG1 | 20.7266297 | -5.5846617 | 1.70063945 |
| 14292 | NM_024094.2 | #N/A | 24.2107047 | -5.8050456 | 1.76782785 |
| 14293 | NM_001201543.1 | #N/A | 31.0702676 | -6.1732155 | 1.88004496 |
| 14294 | NM_145059.2 | #N/A | 79.6640045 | -3.3053183 | 1.00664347 |
| 14295 | NM_173497.3 | HECTD2 | 23.9913336 | -5.7901231 | 1.7634111 |
| 14296 | NM_001324047.1 | #N/A | 28.4116405 | -6.0412733 | 1.83997158 |
| 14297 | NR_002323.2 | TUG1 | 185.592282 | -8.7465366 | 2.66402699 |
| 14298 | NM_001290214.2 | #N/A | 36.8322633 | -6.4107591 | 1.95268499 |
| 14299 | NM_175062.3 | #N/A | 21.8350344 | -5.651493 | 1.72147523 |
| 14300 | NM_001164756.1 | ASPH | 27.7379113 | -6.001962 | 1.82824318 |
| 14301 | NM_017723.2 | #N/A | 26.731813 | -4.5012722 | 1.37129488 |
| 14302 | NM_030971.3 | SFXN3 | 375.07016 | -2.8849376 | 0.87893244 |
| 14303 | NR_040111.1 | LOC100505782 | 10.0252855 | 4.34446158 | 1.32364175 |
| 14304 | NM_183008.2 | UBXN11 | 37.5801309 | -6.4452345 | 1.96370531 |
| 14305 | NM_001330651.1 | ARHGAP36 | 24.3493039 | -5.8192432 | 1.77304021 |
| 14306 | NR_121580.1 | SMC2-AS1 | 33.5549522 | 3.43274703 | 1.04592635 |
| 14307 | NM_001102560.1 | #N/A | 16.6918652 | -5.2765847 | 1.60798054 |
| 14308 | NM_001164751.1 | #N/A | 19.0268657 | -5.4570124 | 1.66302372 |
| 14309 | NM_144988.3 | #N/A | 19.8129059 | -5.5235034 | 1.68328447 |
| 14310 | NM_017729.3 | EPS8L1 | 21.5139047 | -5.6439242 | 1.72014388 |
| 14311 | NM_173582.4 | #N/A | 843.020349 | -4.0198699 | 1.22520193 |
| 14312 | NR_027086.1 | #N/A | 17.9482579 | 3.59227503 | 1.09494381 |
| 14313 | NM_018106.3 | #N/A | 30.1484884 | -6.1279738 | 1.86801641 |
| 14314 | NM_001025237.1 | TSPAN4 | 18.6542415 | -5.4321728 | 1.65594152 |
| 14315 | NM_001134745.2 | LRRTM4 | 18.2982558 | -5.3996569 | 1.64610231 |
| 14316 | NM_001324167.1 | ZNF32 | 18.055843 | -5.3922391 | 1.64387722 |
| 14317 | NM_001100873.1 | C16orf46 | 17.7418072 | -5.3644513 | 1.6354329 |
| 14318 | NM_001318966.1 | EIF2B4 | 24.7756552 | -5.8423834 | 1.7811412 |

| | | | | | |
|---|---|---|---|---|---|
| 14319 | NM_001317094.1 | #N/A | 21.3790789 | -5.6308865 | 1.71678347 |
| 14320 | NM_001280559.1 | #N/A | 22.3136078 | -5.682199 | 1.73244412 |
| 14321 | NM_001287750.1 | FDFT1 | 20.0528831 | -5.5455398 | 1.69083459 |
| 14322 | NM_001008660.2 | #N/A | 145.90164 | -8.3999086 | 2.56191155 |
| 14323 | NR_036484.1 | FEZF1-AS1 | 41.3679402 | -6.5826797 | 2.00777074 |
| 14324 | NM_018362.3 | #N/A | 261.499215 | -2.8645969 | 0.8737419 |
| 14325 | NM_001320678.1 | SETMAR | 24.5439273 | -5.8232447 | 1.776352 |
| 14326 | NM_000958.2 | #N/A | 17.993623 | -5.3879903 | 1.64359069 |
| 14327 | NM_001165937.1 | STARD3 | 38.3614192 | -6.4718496 | 1.97430354 |
| 14328 | NM_001282931.1 | #N/A | 34.0510516 | -6.3036923 | 1.92327477 |
| 14329 | NR_036520.1 | LOC100287042 | 16.3598759 | -5.2464598 | 1.60073988 |
| 14330 | NM_001286821.1 | LETM2 | 22.1172978 | -4.3880539 | 1.33885169 |
| 14331 | NM_014746.4 | #N/A | 263.720862 | -3.283475 | 1.00189942 |
| 14332 | NM_017897.2 | #N/A | 16.0983336 | -5.2221762 | 1.59353405 |
| 14333 | NM_001142465.2 | #N/A | 22.7435215 | -5.716219 | 1.74431014 |
| 14334 | NM_001136225.1 | #N/A | 16.4438961 | -5.2541743 | 1.60334628 |
| 14335 | NM_001024936.1 | WASF1 | 366.74312 | -3.737742 | 1.14061654 |
| 14336 | NM_001270469.1 | #N/A | 21.9997554 | -5.6607794 | 1.727521 |
| 14337 | NM_018424.3 | EPB41L4B | 35.0369889 | -6.3439125 | 1.9362774 |
| 14338 | NM_001137553.1 | #N/A | 129.768146 | -2.7821731 | 0.84918376 |
| 14339 | NM_001311199.1 | CLNS1A | 19.8373771 | -5.5235646 | 1.68603809 |
| 14340 | NM_001276468.1 | #N/A | 35.8814656 | -6.3735571 | 1.94551753 |
| 14341 | NM_001013644.2 | #N/A | 18.2375097 | -5.4045194 | 1.64981649 |
| 14342 | NM_014230.3 | #N/A | 275.733437 | -2.662448 | 0.81275112 |
| 14343 | NM_133448.2 | #N/A | 22.6464901 | -5.7091657 | 1.7429526 |
| 14344 | NM_001320847.1 | #N/A | 22.5652003 | -5.7021881 | 1.74096597 |
| 14345 | NR_004859.1 | SCAND2P | 17.1181176 | -5.3102428 | 1.62142379 |
| 14346 | NM_001278251.2 | SLC25A45 | 74.8123483 | 2.36963206 | 0.72354567 |
| 14347 | NM_001286249.1 | #N/A | 17.5040106 | -5.3430761 | 1.63147726 |
| 14348 | NM_173194.2 | #N/A | 25.4623983 | -5.8756198 | 1.79424226 |
| 14349 | NM_001323951.1 | MBD1 | 18.7685614 | -5.4288838 | 1.65815704 |
| 14350 | NM_032552.3 | DAB2IP | 440.807306 | -4.42172 | 1.35059617 |
| 14351 | NM_001258324.1 | TP53I11 | 24.344111 | -5.8129291 | 1.77564055 |
| 14352 | NM_001102566.1 | #N/A | 17.4423753 | -5.325156 | 1.62686693 |
| 14353 | NM_001319054.1 | GEMIN7 | 20.7425547 | -5.5854706 | 1.7064773 |
| 14354 | NM_001178093.1 | #N/A | 144.973192 | -8.3906791 | 2.56393424 |
| 14355 | NR_027471.1 | LOC440173 | 26.1119308 | -4.6304659 | 1.41505438 |
| 14356 | NM_006734.3 | #N/A | 251.299331 | -3.9640782 | 1.2116275 |
| 14357 | NM_001305.4 | CLDN4 | 26.50076 | -5.9427588 | 1.81646906 |
| 14358 | NM_001136116.1 | #N/A | 22.7383269 | -5.713336 | 1.74639777 |
| 14359 | NM_004980.4 | KCND3 | 20.5598086 | -5.5730345 | 1.70372041 |
| 14360 | NR_026849.1 | LINC00526 | 19.4025387 | -5.4851697 | 1.67686115 |
| 14361 | NM_001321194.1 | #N/A | 35.8849578 | -6.3723849 | 1.94821202 |

| | | | | | | |
|---|---|---|---|---|---|---|
| 14362 | NM_001251989.1 | | #N/A | 19.8253808 | -5.5178369 | 1.68714965 |
| 14363 | NM_014241.3 | | #N/A | 23.9008071 | -5.789274 | 1.77016232 |
| 14364 | NM_198853.2 | | #N/A | 17.7981193 | -5.3708861 | 1.64231932 |
| 14365 | NM_004080.2 | | #N/A | 53.8285641 | -4.1814575 | 1.27865257 |
| 14366 | NM_001136202.1 | | #N/A | 18.5936333 | -5.4339514 | 1.66179408 |
| 14367 | NM_001099455.1 | | #N/A | 74.8111579 | 7.22344767 | 2.20934245 |
| 14368 | NM_001008213.1 | OPTN | | 23.6736213 | -5.7681036 | 1.76432391 |
| 14369 | NM_001324087.1 | VDAC2 | | 17.8347065 | -5.3682287 | 1.64222412 |
| 14370 | NM_001303529.1 | | #N/A | 17.3476495 | -5.3365518 | 1.63254804 |
| 14371 | NM_001329862.1 | TRUB2 | | 17.6557008 | -5.3458391 | 1.6354753 |
| 14372 | NM_032601.3 | | #N/A | 18.4153955 | -5.4116785 | 1.65576692 |
| 14373 | NM_001329494.1 | EMC2 | | 18.5238968 | -5.4252937 | 1.65997014 |
| 14374 | NM_001330235.1 | | #N/A | 29.9142183 | -4.6345393 | 1.41813409 |
| 14375 | NM_002234.3 | | #N/A | 22.015558 | -5.6774905 | 1.73748725 |
| 14376 | NM_001256192.1 | PDCD6IP | | 17.0643556 | -5.309946 | 1.6250242 |
| 14377 | NM_015488.4 | | #N/A | 27.1090465 | -5.970474 | 1.82723396 |
| 14378 | NM_032484.4 | | #N/A | 39.9440306 | -4.9939345 | 1.52837207 |
| 14379 | NR_024206.1 | | #N/A | 27.9910345 | -6.0188157 | 1.84206366 |
| 14380 | NM_006786.3 | | #N/A | 24.2010734 | -5.80756 | 1.77747594 |
| 14381 | NR_023915.1 | IPW | | 176.889533 | -4.0826028 | 1.24955602 |
| 14382 | NM_001032730.1 | | #N/A | 17.5117133 | -5.3406264 | 1.63462457 |
| 14383 | NM_145812.2 | AIFM1 | | 23.5798724 | -5.7713296 | 1.76654439 |
| 14384 | NM_138412.3 | | #N/A | 20.5195231 | -5.5644639 | 1.70343714 |
| 14385 | NM_005515.3 | | #N/A | 26.3743394 | -5.9335891 | 1.81655616 |
| 14386 | NM_001306080.1 | | #N/A | 27.8173025 | -6.0135727 | 1.84115016 |
| 14387 | NM_014037.2 | | #N/A | 15.9515226 | -5.2086763 | 1.59475635 |
| 14389 | NR_121644.1 | DBET | | 9.45952662 | 4.81842913 | 1.47536549 |
| 14388 | NR_110602.1 | | #N/A | 25.1322393 | -5.8541713 | 1.79250715 |
| 14390 | NM_001278374.1 | | #N/A | 30.229848 | -6.1269671 | 1.87661463 |
| 14391 | NM_001321345.1 | | #N/A | 21.5925063 | -5.6416012 | 1.72801792 |
| 14392 | NM_001271819.1 | | #N/A | 237.00739 | 6.15586974 | 1.88627878 |
| 14393 | NM_052872.3 | | #N/A | 25.4502494 | 8.09212422 | 2.48012356 |
| 14394 | NM_001278542.1 | DIXDC1 | | 21.3646008 | -5.6203768 | 1.7226557 |
| 14395 | NM_145719.2 | | #N/A | 21.8274326 | -5.6515843 | 1.73233732 |
| 14396 | NM_001318926.1 | RPE | | 17.329109 | -5.3298582 | 1.63377906 |
| 14397 | NM_000288.3 | | #N/A | 17.4206505 | -5.3350032 | 1.6354908 |
| 14398 | NM_003803.3 | MYOM1 | | 17.1672424 | -5.3212145 | 1.631622 |
| 14399 | NM_015024.4 | | #N/A | 405.408885 | -3.4790205 | 1.0668903 |
| 14400 | NM_001267612.1 | GPR161 | | 33.8366978 | -6.2951403 | 1.93084929 |
| 14401 | NR_036471.1 | | #N/A | 19.3716712 | -5.4889007 | 1.68362617 |
| 14402 | NM_175573.2 | ADRM1 | | 26.105276 | -5.9233123 | 1.81697949 |
| 14403 | NM_001031679.2 | | #N/A | 39.6625386 | -6.5226619 | 2.00118191 |
| 14404 | NM_182559.2 | | #N/A | 30.9835524 | 9.17247111 | 2.81430566 |

| | | | | | |
|---|---|---|---|---|---|
| 14405 | NM_003568.2 | #N/A | 17.3729235 | -5.3306716 | 1.63557305 |
| 14409 | NR_135263.1 | C1orf61 | 52.5551372 | -6.92475 | 2.12478864 |
| 14406 | NM_001142451.1 | #N/A | 20.7683379 | -5.5840473 | 1.71343179 |
| 14407 | NM_024108.2 | #N/A | 17.3238106 | -5.3307503 | 1.63570988 |
| 14408 | NM_177963.3 | #N/A | 16.9263752 | -5.2949132 | 1.62471459 |
| 14410 | NR_045214.1 | LOC100507373 | 16.3240248 | -5.2315732 | 1.60545167 |
| 14411 | NM_018487.2 | #N/A | 17.3022137 | -5.3307785 | 1.63592352 |
| 14412 | NM_001293562.1 | #N/A | 16.5054747 | -5.2480497 | 1.61060073 |
| 14413 | NR_003083.2 | SLC6A10P | 17.1223923 | -5.3124796 | 1.63040131 |
| 14414 | NM_032679.2 | ZNF577 | 19.2263962 | -5.4685038 | 1.67845237 |
| 14415 | NM_014584.2 | #N/A | 317.736877 | -1.9227004 | 0.59024372 |
| 14416 | NM_001008739.1 | #N/A | 23.3621227 | -5.7577846 | 1.76766332 |
| 14417 | NM_001164464.1 | TRIM43B | 27.8470182 | 8.37309804 | 2.57149295 |
| 14418 | NR_135065.1 | LINC02745 | 29.2432838 | 3.33758395 | 1.0250413 |
| 14419 | NM_001171109.1 | #N/A | 21.5676924 | -5.6381138 | 1.73179501 |
| 14420 | NM_001199760.1 | ST20-MTHFS | 36.5353799 | 6.68142462 | 2.05229845 |
| 14421 | NR_133652.1 | ATG9B | 18.4382422 | -5.4142576 | 1.66323459 |
| 14422 | NM_001040125.1 | #N/A | 21.9665589 | -5.6583338 | 1.73834551 |
| 14423 | NM_001166686.1 | #N/A | 17.849345 | -5.3590107 | 1.64662832 |
| 14424 | NM_001040451.2 | #N/A | 22.0811781 | -5.6796306 | 1.74515987 |
| 14425 | NM_001130698.1 | #N/A | 19.8810437 | -5.5228779 | 1.69705394 |
| 14426 | NM_001321111.1 | #N/A | 35.6569192 | -6.3673682 | 1.95702392 |
| 14427 | NM_001127695.2 | CTSA | 28.348453 | -6.0421289 | 1.85755215 |
| 14428 | NM_001098621.1 | #N/A | 18.3118576 | -5.3990455 | 1.65986843 |
| 14429 | NM_024766.4 | #N/A | 17.0685241 | -5.306226 | 1.63132255 |
| 14430 | NM_001547.4 | #N/A | 19.9995454 | -5.5425853 | 1.70431638 |
| 14431 | NM_178498.3 | #N/A | 92.6967185 | 3.76982771 | 1.15920845 |
| 14432 | NR_026698.1 | #N/A | 23.3784209 | -5.7642521 | 1.77253893 |
| 14433 | NM_032432.4 | #N/A | 20.9565162 | -5.5927774 | 1.71990078 |
| 14434 | NM_004097.2 | #N/A | 92.8493169 | -7.7466887 | 2.382401 |
| 14435 | NM_080869.1 | #N/A | 19.9099175 | 7.62523932 | 2.34529545 |
| 14436 | NM_001286522.1 | #N/A | 21.7834546 | -5.6470681 | 1.73702416 |
| 14437 | NM_003558.3 | #N/A | 33.2325025 | -6.2670744 | 1.92775468 |
| 14438 | NM_003287.3 | #N/A | 24.2075861 | -5.8078148 | 1.7865551 |
| 14439 | NM_181651.2 | #N/A | 41.0277677 | -6.5718971 | 2.02162239 |
| 14441 | NM_001165037.1 | GABRA5 | 27.0999666 | -5.9672577 | 1.83583613 |
| 14440 | NM_001122631.1 | #N/A | 20.17897 | -5.5511284 | 1.70780424 |
| 14442 | NM_001301083.1 | CNOT8 | 22.6845372 | -5.7094676 | 1.75660612 |
| 14443 | NR_144404.1 | #N/A | 31.3818217 | -4.6871625 | 1.44214943 |
| 14444 | NR_130740.1 | FAM86EP | 17.1209065 | -5.3112201 | 1.63427996 |
| 14445 | NM_001171743.2 | #N/A | 26.9159125 | -5.9595497 | 1.83386107 |
| 14446 | NM_020177.2 | #N/A | 154.752431 | -2.2915815 | 0.70522531 |
| 14447 | NM_004902.3 | #N/A | 188.57565 | -8.7695521 | 2.69894046 |

| | | | | | |
|---|---|---|---|---|---|
| 14448 | NM_016602.2 | #N/A | 16.1508251 | -5.2269692 | 1.60879781 |
| 14449 | NM_001243532.1 | PDIK1L | 43.7714639 | -6.6639694 | 2.05138953 |
| 14450 | NM_014941.3 | MORC2 | 17.3677576 | -5.3207724 | 1.63809382 |
| 14451 | NM_017987.4 | RUFY2 | 27.1327205 | -5.9708975 | 1.83833247 |
| 14452 | NM_001324058.1 | #N/A | 17.404478 | -5.3365991 | 1.6431892 |
| 14453 | NM_001242498.1 | MIF4GD | 17.2839191 | -5.3310049 | 1.64155546 |
| 14454 | NM_007075.3 | WDR45 | 56.8223322 | -2.4804645 | 0.76379966 |
| 14455 | NR_122047.1 | TBC1D22A-AS1 | 52.3125574 | 2.60233286 | 0.80133459 |
| 14456 | NM_005795.5 | #N/A | 17.3209275 | -5.3324387 | 1.6421316 |
| 14457 | NR_037884.1 | LOC100507053 | 17.304059 | -5.3252939 | 1.64031643 |
| 14458 | NM_001100388.1 | C11orf88 | 33.9166582 | -6.2977722 | 1.93996226 |
| 14460 | NR_003138.3 | SNHG10 | 18.6336495 | -5.4178279 | 1.66951693 |
| 14459 | NM_133325.2 | #N/A | 18.9728304 | -5.4553019 | 1.68107359 |
| 14461 | NM_080388.2 | #N/A | 21.528676 | -5.6427082 | 1.73898447 |
| 14462 | NM_001143830.1 | #N/A | 23.4675743 | -5.7621581 | 1.77582613 |
| 14463 | NM_001620.2 | #N/A | 148.991626 | -8.4299721 | 2.59836887 |
| 14464 | NM_002449.4 | #N/A | 30.3873085 | -6.1399918 | 1.8925682 |
| 14465 | NM_001143936.1 | #N/A | 24.9102141 | -5.8476828 | 1.80285199 |
| 14466 | NR_026847.1 | LOC286359 | 31.6689485 | 4.80534009 | 1.48161551 |
| 14467 | NM_005164.3 | #N/A | 24.5130096 | -5.8192179 | 1.79428324 |
| 14468 | NM_145020.4 | #N/A | 23.0598896 | -5.7416098 | 1.77063047 |
| 14470 | NM_001252204.1 | C4A | 10.9970055 | 8.45592165 | 2.60827156 |
| 14469 | NM_001012511.2 | GOSR2 | 21.1533847 | -5.6124111 | 1.73118392 |
| 14471 | NM_177531.4 | #N/A | 17.7695172 | 5.64914349 | 1.74290138 |
| 14472 | NM_018369.2 | #N/A | 25.1000386 | -5.8552682 | 1.80651242 |
| 14473 | NM_001166304.1 | #N/A | 18.9990161 | -5.4660815 | 1.68694356 |
| 14474 | NM_001303228.1 | #N/A | 25.7654363 | 2.20623406 | 0.68091323 |
| 14475 | NR_125752.1 | LINC00383 | 18.9626121 | 5.3339677 | 1.64626482 |
| 14476 | NM_000813.2 | #N/A | 56.7489093 | -5.7699006 | 1.78091942 |
| 14477 | NM_080627.3 | #N/A | 692.045403 | -2.0116957 | 0.62092645 |
| 14478 | NM_001024732.2 | #N/A | 21.3711322 | -5.6185386 | 1.73424077 |
| 14479 | NM_001330422.1 | #N/A | 29.379937 | -6.0889155 | 1.87941853 |
| 14481 | NM_024762.3 | ZNF552 | 26.5078431 | -4.4793336 | 1.38266038 |
| 14480 | NM_001172435.1 | #N/A | 46.1152413 | -6.7390291 | 2.08015611 |
| 14482 | NR_104041.1 | #N/A | 27.6375267 | -6.0005738 | 1.8522259 |
| 14484 | NR_037702.1 | RNF7 | 21.6427136 | -5.6449396 | 1.74253062 |
| 14483 | NM_001197181.1 | #N/A | 25.2515773 | -5.8635728 | 1.81001417 |
| 14485 | NM_001110503.2 | TMEM87A | 38.6546317 | 2.4367635 | 0.75230176 |
| 14486 | NM_002118.4 | #N/A | 27.0504163 | -5.96984 | 1.84307533 |
| 14487 | NM_001347828.1 | #N/A | 36.182064 | -6.3819267 | 1.97059032 |
| 14488 | NM_032107.4 | #N/A | 22.8825227 | -4.4383844 | 1.37049708 |
| 14489 | NM_006292.3 | #N/A | 257.757495 | -4.2969556 | 1.32689778 |
| 14492 | NM_001297697.1 | CCDC125 | 25.4820926 | -5.877713 | 1.81508834 |

| | | | | | |
|---|---|---|---|---|---|
| 14490 | NM_001080437.1 | #N/A | 27.4858132 | -5.9945926 | 1.85116289 |
| 14491 | NM_001257995.1 | #N/A | 30.1247956 | 9.13197606 | 2.82003462 |
| 14493 | NR_136526.1 | LINC01872 | 24.9022348 | 6.33094648 | 1.95507962 |
| 14494 | NM_015114.2 | #N/A | 319.902989 | -3.2693215 | 1.0096273 |
| 14495 | NM_006942.1 | #N/A | 16.6563651 | -5.2731831 | 1.62864592 |
| 14496 | NR_109949.1 | HNF4A-AS1 | 23.2098598 | 7.01332338 | 2.1669734 |
| 14497 | NM_001018066.2 | #N/A | 18.290667 | -5.4004426 | 1.66884018 |
| 14498 | NM_001271954.1 | TPGS2 | 20.3718108 | -5.568153 | 1.72073797 |
| 14499 | NM_006961.3 | #N/A | 16.0086484 | -5.2075092 | 1.60938683 |
| 14500 | NM_001320623.1 | BOLA2-SMG1P6 | 38.6833704 | -6.4829765 | 2.00375807 |
| 14501 | NM_182623.2 | #N/A | 16.8163308 | -5.2757642 | 1.63066148 |
| 14502 | NM_152387.3 | #N/A | 16.3228109 | -5.2410486 | 1.62002342 |
| 14503 | NM_004452.3 | ESRRB | 21.9265652 | -5.6651891 | 1.75134233 |
| 14504 | NM_175623.3 | #N/A | 25.9890485 | -5.9050155 | 1.82562396 |
| 14505 | NR_037576.1 | #N/A | 19.3697415 | -5.4867784 | 1.69661587 |
| 14506 | NM_145901.2 | HMGA1 | 21.3408076 | -5.6274741 | 1.74033672 |
| 14507 | NM_001199298.1 | #N/A | 34.1296106 | -6.3022275 | 1.94906391 |
| 14508 | NM_001346145.1 | FLRT2 | 37.9858894 | -6.4550305 | 1.99638914 |
| 14509 | NM_017858.2 | #N/A | 21.6759199 | -5.6429711 | 1.74526715 |
| 14510 | NM_001286458.1 | #N/A | 20.2586734 | -5.5551382 | 1.71823614 |
| 14511 | NM_000995.4 | RPL34 | 22.9837907 | -5.7317184 | 1.7729917 |
| 14512 | NR_111893.2 | CNTNAP3P2 | 66.8956546 | 1.6396466 | 0.50720413 |
| 14513 | NM_001131036.1 | ZCCHC9 | 23.2568533 | -5.7506764 | 1.7789453 |
| 14514 | NM_032452.2 | #N/A | 46.1282569 | -6.7369459 | 2.08416891 |
| 14515 | NM_001888.4 | #N/A | 32.2144108 | -3.6009897 | 1.11403836 |
| 14516 | NM_001101338.1 | ZNF883 | 21.0776291 | -5.6048346 | 1.73413884 |
| 14517 | NR_037807.1 | #N/A | 15.6053397 | -5.1704404 | 1.59985118 |
| 14518 | NR_023313.2 | #N/A | 150.33021 | -8.4430558 | 2.61275068 |
| 14519 | NM_001256421.1 | GPAT3 | 35.936664 | 4.43442135 | 1.37227697 |
| 14520 | NM_001284285.1 | #N/A | 26.4333212 | -5.9304615 | 1.83532902 |
| 14521 | NR_073574.1 | #N/A | 21.8587941 | -5.6665968 | 1.75385397 |
| 14522 | NM_001173454.1 | PDHA1 | 16.5359331 | -5.2663004 | 1.63008433 |
| 14523 | NM_001330336.1 | CBWD2 | 24.5108594 | -5.8292992 | 1.80440346 |
| 14524 | NM_001324195.1 | DDX3Y | 20.1066497 | -5.5347339 | 1.71327031 |
| 14525 | NM_001270530.1 | RNF111 | 29.6690201 | -6.1001591 | 1.88838215 |
| 14526 | NM_017774.3 | CDKAL1 | 98.6422528 | -2.2674902 | 0.70199947 |
| 14527 | NM_022782.3 | #N/A | 51.0614104 | -4.1112853 | 1.27292421 |
| 14528 | NM_003508.2 | #N/A | 19.994862 | -5.5290761 | 1.71232532 |
| 14529 | NM_145235.3 | #N/A | 32.8671077 | -6.2521944 | 1.93653555 |
| 14530 | NM_173628.3 | #N/A | 18.3097073 | -5.4092076 | 1.67544581 |
| 14531 | NM_024645.2 | #N/A | 19.1120595 | -5.4656622 | 1.69299203 |
| 14532 | NM_001127323.1 | GRM8 | 33.901869 | -6.2955916 | 1.95015901 |
| 14533 | NM_014317.4 | #N/A | 22.0244574 | -5.666467 | 1.75554816 |

| | | | | | |
|---|---|---|---|---|---|
| 14534 | NM_001123391.3 | TBC1D3 | 21.4992039 | -5.6397484 | 1.74746501 |
| 14535 | NM_001346818.1 | TPMT | 135.629486 | 5.89982865 | 1.82827287 |
| 14536 | NM_198281.2 | #N/A | 43.9802829 | -5.1024059 | 1.58138795 |
| 14537 | NM_001286436.1 | #N/A | 17.3846163 | -5.3388561 | 1.6548074 |
| 14538 | NM_152788.4 | ANKS1B | 23.0265811 | -5.7306696 | 1.77634112 |
| 14539 | NM_001203261.1 | #N/A | 19.030077 | -5.454315 | 1.69083867 |
| 14540 | NM_018249.5 | #N/A | 135.479493 | -8.2929448 | 2.57087293 |
| 14541 | NM_001330429.1 | FSD1 | 29.0430611 | -6.0727051 | 1.88358993 |
| 14542 | NM_003512.3 | H2AC6 | 16.2656307 | -5.2401786 | 1.62591848 |
| 14543 | NM_001261382.1 | #N/A | 21.5657149 | -5.6352251 | 1.74851192 |
| 14544 | NM_015312.3 | KIAA1109 | 420.367454 | -4.3515198 | 1.35023829 |
| 14545 | NM_017712.3 | #N/A | 132.584629 | -8.2618051 | 2.56366234 |
| 14546 | NM_001302084.1 | #N/A | 27.300625 | -5.9809397 | 1.85605529 |
| 14547 | NM_002815.3 | #N/A | 322.47734 | -3.3129078 | 1.02810277 |
| 14548 | NM_024025.2 | #N/A | 30.319379 | -6.136241 | 1.90431653 |
| 14549 | NM_001261467.1 | NSRP1 | 16.8601234 | -5.2801188 | 1.6388915 |
| 14550 | NM_004985.4 | #N/A | 304.103034 | -2.9278917 | 0.90882407 |
| 14551 | NM_032788.2 | #N/A | 73.1918544 | -2.7966804 | 0.86816601 |
| 14552 | NM_001324105.1 | KLF8 | 20.5703159 | -5.5712378 | 1.72951963 |
| 14553 | NM_001206943.1 | #N/A | 27.4478505 | -5.9856536 | 1.85822275 |
| 14554 | NM_001204856.1 | #N/A | 33.4478788 | -6.2777229 | 1.94890869 |
| 14555 | NM_001008237.2 | #N/A | 15.9541023 | -5.2029081 | 1.61547578 |
| 14556 | NR_109978.1 | #N/A | 24.3040461 | -5.8074927 | 1.80327329 |
| 14557 | NM_004143.3 | #N/A | 20.4709457 | -5.5636255 | 1.72768203 |
| 14558 | NR_002453.5 | SMG1P5 | 16.6602404 | -5.2740354 | 1.63786103 |
| 14559 | NR_138072.1 | #N/A | 31.5133148 | -6.1809773 | 1.91950412 |
| 14560 | NM_017676.2 | GIN1 | 19.3169756 | -5.4760174 | 1.70068236 |
| 14561 | NM_001421.3 | ELF4 | 18.9914582 | -5.461757 | 1.69627779 |
| 14563 | NR_033440.1 | PAM | 21.6421139 | -5.6491524 | 1.75452768 |
| 14562 | NM_006150.4 | #N/A | 23.1304219 | -5.7461254 | 1.7846431 |
| 14564 | NM_025165.2 | #N/A | 19.1922988 | -5.4807842 | 1.70238216 |
| 14565 | NM_001330306.1 | #N/A | 20.2839597 | -5.5483527 | 1.72349412 |
| 14566 | NM_019040.4 | #N/A | 22.259023 | -5.6857047 | 1.76626861 |
| 14567 | NR_073484.1 | #N/A | 25.3087527 | -5.8729547 | 1.82484308 |
| 14568 | NM_001321703.1 | #N/A | 31.1215378 | -6.1709549 | 1.91748828 |
| 14569 | NM_052929.1 | #N/A | 35.5519465 | -6.3651908 | 1.97784477 |
| 14570 | NM_001013439.2 | #N/A | 20.0631765 | -5.5298229 | 1.71856813 |
| 14571 | NM_001270574.1 | #N/A | 22.4967501 | -5.7006228 | 1.77174798 |
| 14572 | NR_135777.1 | RSRP1 | 19.0675117 | -5.4540776 | 1.69515215 |
| 14573 | NM_001256139.1 | CAPG | 29.3396378 | -6.0817721 | 1.89027918 |
| 14574 | NR_046308.1 | #N/A | 18.584188 | -5.425771 | 1.68661668 |
| 14575 | NM_031477.4 | #N/A | 17.7478361 | -5.3587928 | 1.66593166 |
| 14576 | NM_005527.3 | #N/A | 18.5439018 | -5.4189675 | 1.68471147 |

| | | | | | |
|---|---|---|---|---|---|
| 14577 | NM_001252641.1 | #N/A | 16.3360943 | -5.2335413 | 1.62717345 |
| 14578 | NM_016018.4 | #N/A | 33.5104979 | -6.2728231 | 1.95035671 |
| 14579 | NM_001322386.1 | ENTPD6 | 29.209388 | -6.0840491 | 1.89174784 |
| 14580 | NM_001324066.1 | #N/A | 35.1810727 | -6.3447853 | 1.97290159 |
| 14581 | NR_104228.1 | LINC00632 | 16.0097652 | -5.2100421 | 1.62019801 |
| 14582 | NM_001282394.1 | WNK2 | 185.280268 | -4.1350252 | 1.28609165 |
| 14583 | NM_022110.3 | #N/A | 20.5036143 | -5.5702011 | 1.73265837 |
| 14584 | NR_040016.1 | TMEM50B | 114.07944 | -3.8688378 | 1.20344562 |
| 14585 | NR_135150.1 | #N/A | 16.3424037 | -5.2419638 | 1.63078747 |
| 14586 | NM_001130929.1 | #N/A | 316.655961 | -2.4890721 | 0.77437622 |
| 14587 | NM_001161546.1 | #N/A | 15.4058659 | -5.1614135 | 1.60586652 |
| 14588 | NM_001080493.3 | #N/A | 17.1055019 | -5.3078946 | 1.65149156 |
| 14589 | NM_015308.3 | #N/A | 166.161888 | -7.337655 | 2.2831703 |
| 14590 | NM_014309.2 | #N/A | 320.495528 | -4.1577356 | 1.29385509 |
| 14591 | NM_199296.2 | #N/A | 33.9943135 | -6.3009776 | 1.96102211 |
| 14592 | NM_001321484.1 | #N/A | 22.643953 | -5.7073342 | 1.77628476 |
| 14593 | NM_001347722.1 | #N/A | 17.7400964 | -5.361051 | 1.66888774 |
| 14594 | NR_126522.1 | EXOC3-AS1 | 20.0036352 | -5.5309198 | 1.72197397 |
| 14595 | NR_029393.1 | KRT16P3 | 13.7998186 | 8.78435057 | 2.7349505 |
| 14596 | NR_033240.1 | SLC25A21-AS1 | 15.6014972 | -5.1718512 | 1.61035946 |
| 14597 | NM_173494.1 | #N/A | 18.9316077 | 2.39811681 | 0.7468779 |
| 14598 | NM_001161500.1 | ZNF611 | 25.3239101 | -5.8644944 | 1.82654927 |
| 14599 | NM_003205.3 | #N/A | 128.082606 | -8.2120326 | 2.55808742 |
| 14600 | NM_001039802.1 | #N/A | 117.486929 | -3.9042344 | 1.21636448 |
| 14601 | NR_038991.1 | RBM26-AS1 | 15.7116775 | -5.1873434 | 1.6162134 |
| 14602 | NM_019054.2 | #N/A | 18.1233158 | -5.3931948 | 1.68043113 |
| 14603 | NM_001243280.1 | #N/A | 40.5365869 | -5.0014341 | 1.5584615 |
| 14604 | NM_001321746.1 | #N/A | 153.860727 | -8.4764714 | 2.64170957 |
| 14605 | NM_001243609.1 | LMO3 | 210.323624 | -8.9266627 | 2.78218623 |
| 14606 | NM_015106.3 | #N/A | 178.61936 | -8.691169 | 2.70895036 |
| 14607 | NM_001032410.1 | USP16 | 38.8452563 | -6.4887766 | 2.02266162 |
| 14608 | NM_001031743.2 | #N/A | 22.0320688 | -5.6766159 | 1.76953109 |
| 14609 | NM_018386.4 | #N/A | 18.4107364 | -5.4210749 | 1.69004549 |
| 14610 | NM_020390.5 | #N/A | 41.6181992 | -2.2598659 | 0.70458465 |
| 14611 | NM_001002296.1 | #N/A | 25.7946884 | -5.9062769 | 1.84180889 |
| 14612 | NM_182626.3 | #N/A | 39.4626843 | 4.02331729 | 1.25496083 |
| 14613 | NM_003441.2 | #N/A | 16.4060292 | -5.2454673 | 1.63625355 |
| 14614 | NM_001267556.1 | #N/A | 30.4214816 | -6.1408499 | 1.91605614 |
| 14615 | NR_045631.1 | #N/A | 21.4245323 | -5.6267304 | 1.75615609 |
| 14616 | NM_017916.2 | #N/A | 172.729568 | -4.0414768 | 1.2614012 |
| 14617 | NM_001193538.2 | #N/A | 30.7503145 | -4.6528715 | 1.45229947 |
| 14618 | NM_001193535.1 | FBXL5 | 18.1505102 | -5.3806886 | 1.67950527 |
| 14619 | NM_004531.4 | #N/A | 32.58764 | -6.2332274 | 1.945646 |

| | | | | | |
|---|---|---|---|---|---|
| 14620 | NM_015093.5 | TAB2 | 17.6950592 | -5.346472 | 1.66886376 |
| 14621 | NM_153247.2 | #N/A | 131.940244 | -8.2547639 | 2.57814781 |
| 14622 | NR_046079.1 | SEMA5B | 20.7644856 | -5.5775916 | 1.74203715 |
| 14623 | NR_026562.3 | #N/A | 15.4493919 | -5.1653163 | 1.61340459 |
| 14624 | NR_135548.1 | #N/A | 18.1304495 | -5.3836404 | 1.68165326 |
| 14625 | NM_007076.2 | #N/A | 17.3085664 | -5.3224852 | 1.66257882 |
| 14626 | NR_134950.1 | #N/A | 16.0360114 | -5.207666 | 1.62671284 |
| 14627 | NM_213725.1 | #N/A | 164.812647 | -3.8250674 | 1.1948965 |
| 14628 | NM_001018067.1 | #N/A | 130.581129 | -8.2398658 | 2.57427401 |
| 14629 | NM_020381.3 | #N/A | 52.2146747 | -2.4975742 | 0.7802905 |
| 14630 | NM_001346700.1 | #N/A | 19.5223693 | -5.5061699 | 1.72030564 |
| 14631 | NM_033102.2 | #N/A | 16.7275308 | -5.2833687 | 1.65105434 |
| 14632 | NM_001308196.1 | SIMC1 | 46.6541069 | -2.7835144 | 0.86985725 |
| 14633 | NM_001129979.1 | #N/A | 16.9659436 | -5.2909862 | 1.65349989 |
| 14634 | NM_153336.2 | #N/A | 17.4507153 | -5.3338535 | 1.66703091 |
| 14635 | NM_170710.4 | #N/A | 18.4301799 | -5.4244684 | 1.69572993 |
| 14636 | NM_001357.4 | #N/A | 864.704136 | -3.8061997 | 1.18990735 |
| 14637 | NM_000419.4 | #N/A | 20.4150266 | -5.5675207 | 1.74059299 |
| 14638 | NM_001321301.1 | #N/A | 23.5551702 | -5.7620459 | 1.80180943 |
| 14639 | NM_001037161.1 | #N/A | 16.9047832 | -5.2868494 | 1.6533119 |
| 14640 | NM_001049.2 | #N/A | 23.1026618 | -5.7390542 | 1.79477332 |
| 14641 | NM_001256644.1 | LTA4H | 25.4706481 | -5.8744338 | 1.83720193 |
| 14642 | NM_001330385.1 | #N/A | 26.1589431 | -5.9186398 | 1.85138722 |
| 14643 | NM_001171816.1 | #N/A | 15.3619494 | -5.1501167 | 1.61123726 |
| 14644 | NM_005097.3 | #N/A | 30.4306777 | -6.1409985 | 1.92125721 |
| 14645 | NM_001134233.1 | #N/A | 15.7505293 | -5.1784548 | 1.62018342 |
| 14646 | NM_001301132.1 | #N/A | 17.9046163 | -5.3703758 | 1.68025891 |
| 14647 | NM_014840.2 | NUAK1 | 331.908953 | -3.5853873 | 1.12183496 |
| 14648 | NM_001347995.1 | FAM189A2 | 21.5559945 | -5.6463955 | 1.7668002 |
| 14649 | NM_145233.3 | #N/A | 15.9906709 | -5.2099785 | 1.63033103 |
| 14650 | NM_006724.3 | MAP3K4 | 44.2012662 | -5.1041149 | 1.59722665 |
| 14651 | NM_203376.1 | #N/A | 19.1780593 | -5.4678813 | 1.71106863 |
| 14652 | NR_120632.1 | LOC101927964 | 26.3669205 | 3.03434029 | 0.94960331 |
| 14653 | NM_001278648.1 | #N/A | 24.4859675 | -5.8258938 | 1.82331008 |
| 14654 | NM_001322084.1 | #N/A | 18.845969 | -5.4432089 | 1.70362748 |
| 14655 | NM_183374.2 | #N/A | 24.3780144 | 6.20379441 | 1.9416928 |
| 14656 | NR_046262.1 | NIPBL-DT | 206.348717 | -1.891343 | 0.59213555 |
| 14657 | NM_017751.4 | SMPD4 | 126.93689 | -8.1989998 | 2.56734565 |
| 14658 | NM_181673.2 | #N/A | 178.17028 | -8.6878158 | 2.72042749 |
| 14659 | NM_016492.4 | #N/A | 31.3801775 | -4.6747304 | 1.46384937 |
| 14660 | NM_032807.4 | FBH1 | 17.0528917 | -5.3062866 | 1.66166218 |
| 14661 | NR_104230.1 | EXOSC2 | 17.392459 | -5.3356908 | 1.67091849 |
| 14662 | NR_105020.1 | TMEM161B-AS1 | 55.0922123 | -6.9926726 | 2.1903759 |

| | | | | | |
|---|---|---|---|---|---|
| 14663 | NM_138424.1 | KIF12 | 36.1802171 | -6.3907941 | 2.00243383 |
| 14664 | NM_001265582.1 | URM1 | 18.3801817 | -5.4103022 | 1.69585022 |
| 14665 | NR_038988.1 | #N/A | 15.0781303 | -5.124089 | 1.60635296 |
| 14666 | NM_001282658.1 | #N/A | 18.3438522 | -5.4142108 | 1.69750001 |
| 14667 | NM_003407.3 | #N/A | 43.2215084 | -5.0744478 | 1.59099829 |
| 14668 | NM_000908.3 | #N/A | 166.163081 | -8.5869721 | 2.69277662 |
| 14669 | NM_001002000.2 | #N/A | 20.531932 | -5.5610052 | 1.74397185 |
| 14670 | NM_001012994.1 | #N/A | 239.063389 | -2.8429521 | 0.89160788 |
| 14671 | NM_198532.2 | #N/A | 39.7854538 | 2.09880511 | 0.65835794 |
| 14672 | NM_001098783.2 | RPP14 | 17.3241913 | -5.3265755 | 1.67097257 |
| 14673 | NM_001130156.2 | #N/A | 27.2069859 | -5.9817839 | 1.8766852 |
| 14674 | NM_000259.3 | MYO5A | 23.7841842 | -5.7838591 | 1.81471423 |
| 14675 | NM_001173466.1 | #N/A | 27.480986 | -5.9906842 | 1.87973015 |
| 14676 | NM_001134222.1 | #N/A | 27.0976747 | -5.9735547 | 1.87444344 |
| 14677 | NM_001304468.1 | #N/A | 19.347832 | -5.4743826 | 1.71792745 |
| 14678 | NM_014623.3 | #N/A | 23.0491528 | -5.7385185 | 1.80103543 |
| 14679 | NM_001314059.1 | #N/A | 23.8984627 | -5.7974329 | 1.81992537 |
| 14680 | NM_032855.3 | HSH2D | 50.9652063 | 2.05722338 | 0.64598453 |
| 14681 | NM_001319958.1 | #N/A | 16.6339632 | -5.2573712 | 1.65096448 |
| 14682 | NM_001145511.1 | #N/A | 45.3175132 | -6.7083774 | 2.1066514 |
| 14683 | NM_175900.3 | #N/A | 29.9324371 | 2.36164994 | 0.74177478 |
| 14684 | NM_144987.3 | #N/A | 17.0486714 | -5.3050615 | 1.66635158 |
| 14685 | NM_001161520.1 | COG5 | 15.0536265 | -5.1196967 | 1.60826121 |
| 14686 | NR_037926.1 | SENP3-EIF4A1 | 15.5254703 | -5.1593764 | 1.62080612 |
| 14687 | NM_001199653.1 | PMF1 | 18.5973833 | -5.4256453 | 1.70453601 |
| 14688 | NM_003105.5 | #N/A | 195.538329 | -3.5998581 | 1.1309365 |
| 14689 | NM_001300970.1 | #N/A | 22.2792019 | -5.6764895 | 1.78353455 |
| 14690 | NM_013434.4 | #N/A | 26.1931992 | -5.9167874 | 1.8590681 |
| 14692 | NR_137175.1 | MIR100HG | 40.5301102 | -6.5545781 | 2.05959462 |
| 14691 | NM_138927.2 | #N/A | 123.346512 | -8.1576152 | 2.56329906 |
| 14693 | NM_001025100.1 | #N/A | 16.5473071 | -5.2689805 | 1.65564807 |
| 14694 | NM_001100166.1 | #N/A | 16.2338478 | 8.98274241 | 2.82291511 |
| 14695 | NM_002546.3 | #N/A | 15.0435609 | -5.1265134 | 1.61116845 |
| 14696 | NM_152548.2 | #N/A | 31.223734 | -6.178764 | 1.94194083 |
| 14697 | NM_001173539.1 | #N/A | 22.2320564 | -5.6823377 | 1.7859949 |
| 14699 | NM_018925.2 | PCDHGB5 | 15.8987645 | -5.2028362 | 1.63531624 |
| 14698 | NM_016074.4 | #N/A | 17.9137453 | -5.3764183 | 1.68986756 |
| 14700 | NM_005537.5 | ING1 | 16.2802592 | -5.2434754 | 1.64830658 |
| 14701 | NM_005584.4 | #N/A | 27.8452298 | -4.5266756 | 1.42311977 |
| 14702 | NM_006265.2 | #N/A | 855.880374 | -4.4953739 | 1.41330714 |
| 14703 | NM_194315.1 | #N/A | 18.9576292 | -5.4654025 | 1.71833439 |
| 14704 | NM_003124.4 | #N/A | 17.1491527 | -5.3085268 | 1.66917427 |
| 14705 | NM_001321196.1 | SLC27A5 | 17.0143826 | -5.2990862 | 1.66622265 |

| | | | | | |
|---|---|---|---|---|---|
| 14706 | NM_001330240.1 | ARL17B | 16.6098608 | -5.2637285 | 1.65513533 |
| 14707 | NM_032834.3 | #N/A | 19.0703946 | -5.457543 | 1.71638984 |
| 14708 | NR_029383.1 | PAN3-AS1 | 14.8900204 | -5.1100535 | 1.60719651 |
| 14709 | NM_012233.2 | #N/A | 231.48084 | -3.0039867 | 0.94489308 |
| 14710 | NM_080672.4 | #N/A | 24.6310945 | -5.8403506 | 1.83775106 |
| 14711 | NM_001204827.1 | XPNPEP3 | 16.6659043 | -5.2685111 | 1.65785383 |
| 14712 | NM_003500.3 | #N/A | 21.5139734 | -5.6448519 | 1.7763364 |
| 14713 | NM_001099670.2 | RBIS | 25.4190111 | -5.8714726 | 1.8476984 |
| 14714 | NM_198920.2 | #N/A | 17.1537166 | -5.3086433 | 1.67099107 |
| 14715 | NM_004312.2 | #N/A | 18.5827273 | 6.16179057 | 1.93973645 |
| 14716 | NM_004792.2 | #N/A | 297.289676 | -3.2463871 | 1.02217438 |
| 14717 | NR_110751.1 | LINC02071 | 92.4433303 | 3.9879472 | 1.25595459 |
| 14718 | NM_018040.3 | #N/A | 50.0366709 | -1.9240544 | 0.60598705 |
| 14719 | NM_001320610.1 | #N/A | 22.0378756 | -5.6794973 | 1.78888281 |
| 14720 | NM_003899.3 | #N/A | 30.4197128 | -6.1420961 | 1.93464925 |
| 14721 | NM_015272.4 | #N/A | 29.274216 | -6.0808884 | 1.91561685 |
| 14722 | NM_001010926.3 | #N/A | 27.9352105 | -6.0075817 | 1.8927331 |
| 14723 | NR_026916.1 | ANKRD30BP2 | 49.0103805 | 2.87698012 | 0.90650186 |
| 14724 | NM_001037132.2 | #N/A | 42.4651523 | -6.619359 | 2.08587915 |
| 14725 | NM_019117.4 | #N/A | 22.2518131 | -5.6896798 | 1.79296051 |
| 14726 | NR_002838.1 | #N/A | 17.01657 | -5.2997898 | 1.67056862 |
| 14727 | NR_027130.1 | #N/A | 259.375102 | -1.7381261 | 0.54795765 |
| 14728 | NM_001128849.1 | #N/A | 25.7194407 | -5.9025007 | 1.86119212 |
| 14729 | NM_001317911.1 | #N/A | 13.227314 | 8.72358233 | 2.75079308 |
| 14730 | NM_000667.3 | #N/A | 12.613999 | 6.65719132 | 2.09922082 |
| 14731 | NM_001320038.1 | #N/A | 20.212794 | -5.5414691 | 1.74751527 |
| 14732 | NM_001319240.1 | LOC100287896 | 30.7622571 | -4.6454338 | 1.46500039 |
| 14733 | NM_003204.2 | #N/A | 146.189059 | -8.4022364 | 2.65000039 |
| 14734 | NM_001284501.1 | #N/A | 17.2625817 | -5.3139613 | 1.67604836 |
| 14735 | NR_134951.1 | #N/A | 16.8564605 | -5.2956618 | 1.67027357 |
| 14736 | NM_032429.3 | #N/A | 26.295743 | -5.9330635 | 1.87135786 |
| 14737 | NM_001278237.1 | RXYLT1 | 17.3895032 | -5.324129 | 1.67933904 |
| 14738 | NM_024697.2 | #N/A | 19.4880405 | -5.4940674 | 1.73305349 |
| 14739 | NM_017902.2 | #N/A | 227.908086 | -3.0517393 | 0.96266024 |
| 14740 | NM_003914.3 | #N/A | 19.5475677 | -5.4873868 | 1.7311684 |
| 14741 | NM_001258282.1 | #N/A | 21.7429224 | -4.3714863 | 1.37930962 |
| 14742 | NM_021249.4 | #N/A | 27.3194342 | -5.9867596 | 1.88896838 |
| 14743 | NM_001098500.2 | #N/A | 26.6482464 | -5.947921 | 1.87683564 |
| 14744 | NR_040070.1 | LOXL1-AS1 | 17.6055444 | -5.352867 | 1.68908945 |
| 14745 | NM_138796.3 | #N/A | 26.2763325 | -5.930543 | 1.87150629 |
| 14746 | NM_001301196.1 | TRIB3 | 26.5421446 | -5.9452535 | 1.87633125 |
| 14747 | NM_001282175.1 | CLPTM1 | 52.8685088 | -5.6692382 | 1.78981342 |
| 14748 | NM_001063.3 | #N/A | 15.4175558 | -5.1535017 | 1.62717047 |

| | | | | | |
|---|---|---|---|---|---|
| 14749 | NM_001145463.1 | #N/A | 29.4553155 | -4.5914932 | 1.44983994 |
| 14750 | NM_001288980.1 | #N/A | 24.4218926 | -5.8192507 | 1.83765095 |
| 14751 | NM_003786.3 | #N/A | 19.7771699 | -5.5197819 | 1.74309924 |
| 14752 | NR_046219.1 | EGFLAM-AS4 | 23.3285499 | 5.19853443 | 1.64229866 |
| 14753 | NM_007096.3 | #N/A | 32.0214108 | -6.2067533 | 1.96082938 |
| 14754 | NM_173826.3 | #N/A | 22.7871334 | -5.7241217 | 1.80837671 |
| 14755 | NR_038840.1 | LINC00693 | 20.8764378 | -5.5923572 | 1.76717671 |
| 14756 | NM_001301170.1 | #N/A | 27.4262716 | -5.9914352 | 1.89332227 |
| 14757 | NM_013309.5 | #N/A | 170.761567 | 1.21212697 | 0.38311668 |
| 14758 | NR_138604.1 | #N/A | 21.5776094 | -5.6456377 | 1.78442163 |
| 14759 | NM_001242501.1 | MIF4GD | 21.630608 | -5.6415924 | 1.78326264 |
| 14760 | NM_001242532.1 | #N/A | 24.6018486 | -5.8271334 | 1.84195075 |
| 14761 | NM_001174085.1 | POLL | 20.1710293 | -5.5403419 | 1.75134517 |
| 14762 | NR_134654.1 | LOC401261 | 14.902523 | -5.1034014 | 1.61327289 |
| 14763 | NM_001286137.1 | #N/A | 46.9506469 | 6.06734336 | 1.9180823 |
| 14764 | NR_144326.1 | PIGG | 15.6226979 | -5.1776777 | 1.63702575 |
| 14765 | NM_001301873.1 | #N/A | 37.2115845 | 3.97649265 | 1.25740251 |
| 14766 | NM_001301075.1 | #N/A | 21.3344654 | -5.6158078 | 1.77578111 |
| 14767 | NM_014994.2 | #N/A | 124.41051 | -8.1700528 | 2.58354789 |
| 14768 | NM_017860.4 | #N/A | 15.5005502 | -5.1711227 | 1.63523933 |
| 14769 | NR_051974.1 | #N/A | 20.693401 | -5.5812591 | 1.76506561 |
| 14770 | NR_038377.1 | LINC01667 | 27.545759 | 3.72210309 | 1.17711835 |
| 14771 | NR_104476.1 | #N/A | 36.6444853 | -6.4099645 | 2.02721521 |
| 14772 | NM_001099432.1 | #N/A | 34.4927374 | -6.3195337 | 1.99868682 |
| 14773 | NM_015600.4 | #N/A | 18.9120108 | -5.4550931 | 1.72535978 |
| 14774 | NM_001199196.1 | #N/A | 28.7209548 | -6.0541991 | 1.9151115 |
| 14775 | NM_133368.2 | #N/A | 120.686968 | -8.126225 | 2.5705458 |
| 14776 | NM_001284194.1 | FUT11 | 16.3288008 | -5.2327408 | 1.65531667 |
| 14777 | NM_004236.3 | #N/A | 145.859829 | -2.4008269 | 0.75961167 |
| 14778 | NM_022041.3 | GAN | 18.0587237 | -5.3797832 | 1.70257078 |
| 14779 | NM_182688.2 | #N/A | 29.0024548 | -6.0713794 | 1.92185316 |
| 14780 | NM_001166693.2 | AFF1 | 27.4801234 | -5.9958225 | 1.89796064 |
| 14781 | NM_001304723.1 | PLEKHO1 | 17.619829 | -5.3478849 | 1.69289775 |
| 14782 | NM_021140.3 | KDM6A | 25.7375506 | -5.9018562 | 1.86840292 |
| 14783 | NM_001346992.1 | #N/A | 16.0325752 | -5.2111582 | 1.64981799 |
| 14784 | NM_001284236.1 | #N/A | 147.474131 | -8.4152384 | 2.66443026 |
| 14785 | NM_001205028.1 | #N/A | 23.6134464 | -5.777666 | 1.8293475 |
| 14786 | NM_031219.3 | #N/A | 18.0478397 | -5.3880659 | 1.70605571 |
| 14787 | NM_001136205.2 | KCTD1 | 15.4540252 | -5.1691536 | 1.63676451 |
| 14788 | NM_001288668.1 | HCFC1R1 | 16.5790496 | -5.255421 | 1.66419539 |
| 14790 | NR_126481.1 | SALRNA3 | 15.2285277 | 6.37119188 | 2.01749587 |
| 14789 | NR_110191.1 | #N/A | 18.5923259 | -5.4351934 | 1.72111976 |
| 14791 | NM_001184940.1 | #N/A | 156.987185 | -8.5048654 | 2.69367464 |

| | | | | | |
|---|---|---|---|---|---|
| 14792 | NR_102752.1 | LOC100506639 | 15.1314135 | -5.124436 | 1.62313146 |
| 14793 | NM_012419.4 | #N/A | 22.5343624 | -5.7006125 | 1.80569771 |
| 14794 | NM_001306140.1 | QDPR | 24.2999096 | -5.8155491 | 1.84217179 |
| 14795 | NM_001317227.1 | CDH12 | 19.7574262 | -5.5096841 | 1.74544292 |
| 14796 | NM_181742.3 | ORC4 | 37.0882394 | -6.426483 | 2.03645397 |
| 14797 | NM_020226.3 | PRDM8 | 29.2444745 | -6.0747462 | 1.92510772 |
| 14798 | NM_182919.3 | TICAM1 | 15.220518 | -5.1296766 | 1.62568754 |
| 14799 | NM_001002033.2 | #N/A | 15.9947681 | -5.2195805 | 1.65437979 |
| 14800 | NM_001318316.1 | #N/A | 16.2120498 | -5.2193815 | 1.65452733 |
| 14801 | NM_001002836.3 | #N/A | 168.095473 | -2.9703063 | 0.94162644 |
| 14802 | NM_006603.4 | #N/A | 113.938212 | -8.0432381 | 2.54993107 |
| 14803 | NM_001162501.1 | #N/A | 31.2948155 | -6.1821946 | 1.96001312 |
| 14804 | NM_207352.3 | #N/A | 230.294555 | 2.33346905 | 0.73985131 |
| 14805 | NM_001457.3 | #N/A | 179.886139 | -8.7016964 | 2.75900955 |
| 14806 | NM_133644.3 | #N/A | 23.4825186 | -5.7716397 | 1.83013546 |
| 14807 | NM_001032393.2 | HNRNPH2 | 25.9154826 | -5.9099046 | 1.87407021 |
| 14808 | NM_018982.4 | #N/A | 31.7816162 | -4.6828648 | 1.48503281 |
| 14809 | NM_001002017.2 | HCFC1R1 | 23.1178579 | -5.7376213 | 1.81957674 |
| 14810 | NM_001324122.1 | #N/A | 27.9265367 | -6.0087751 | 1.90563071 |
| 14811 | NM_014951.2 | #N/A | 18.5957737 | -5.4167732 | 1.71798193 |
| 14812 | NM_001198780.1 | #N/A | 148.667047 | -8.4264664 | 2.67260958 |
| 14813 | NM_001253699.1 | #N/A | 17.9025055 | -5.3810398 | 1.70678406 |
| 14814 | NM_031948.4 | #N/A | 19.6701256 | -5.5116156 | 1.74832492 |
| 14815 | NM_019118.4 | #N/A | 21.4481465 | -5.6359252 | 1.78788076 |
| 14816 | NM_147134.3 | #N/A | 234.714266 | 1.12119302 | 0.35568825 |
| 14817 | NM_001025205.1 | AP2M1 | 22.9690189 | -5.7389417 | 1.821251 |
| 14818 | NM_020436.4 | #N/A | 18.2782042 | -5.4121348 | 1.7176169 |
| 14819 | NM_001297565.1 | #N/A | 57.5115104 | -4.19953 | 1.3329657 |
| 14820 | NM_001001557.3 | #N/A | 76.0117164 | -4.5029008 | 1.42936577 |
| 14821 | NM_133646.2 | #N/A | 160.347674 | -2.0311285 | 0.64475516 |
| 14823 | NM_014507.3 | MCAT | 26.4597015 | -4.4533484 | 1.41368655 |
| 14822 | NM_001142679.1 | #N/A | 22.6281355 | -5.7098213 | 1.8125485 |
| 14824 | NM_014770.3 | #N/A | 26.5749936 | -5.9424123 | 1.88636795 |
| 14825 | NM_006438.4 | #N/A | 38.6368847 | 5.9893639 | 1.90141211 |
| 14826 | NM_018291.3 | #N/A | 17.4186803 | -5.3412105 | 1.69588129 |
| 14827 | NM_001145268.1 | #N/A | 16.2950314 | -5.232155 | 1.66145323 |
| 14828 | NM_016381.5 | #N/A | 16.349961 | -5.2494476 | 1.66717524 |
| 14829 | NM_001134435.2 | IQCG | 48.8644208 | -6.8231348 | 2.16714788 |
| 14830 | NR_144485.1 | #N/A | 15.706163 | -5.1885739 | 1.64815753 |
| 14831 | NM_001144825.1 | #N/A | 215.48573 | -4.3260433 | 1.37439962 |
| 14832 | NM_016009.4 | #N/A | 116.588125 | -8.0764741 | 2.56602095 |
| 14833 | NM_014030.3 | #N/A | 105.610354 | -3.0932281 | 0.98291693 |
| 14834 | NR_024237.1 | #N/A | 16.2283095 | 5.4511616 | 1.73220133 |

| | | | | | |
|---|---|---|---|---|---|
| 14835 | NM_014002.3 | #N/A | 15.766261 | -5.1918999 | 1.65005621 |
| 14836 | NM_001185.3 | #N/A | 15.5895274 | 4.91234807 | 1.5612897 |
| 14837 | NM_001274.5 | CHEK1 | 20.0170356 | -5.5220793 | 1.75512817 |
| 14838 | NR_004845.1 | #N/A | 20.8490373 | 6.21155806 | 1.97435296 |
| 14839 | NM_001130478.1 | #N/A | 22.1056762 | -5.6772838 | 1.80460902 |
| 14840 | NM_032148.3 | #N/A | 41.0128517 | -2.8141057 | 0.89457859 |
| 14841 | NM_001282418.1 | #N/A | 54.4473911 | -6.9738509 | 2.21698608 |
| 14842 | NM_001303001.1 | #N/A | 25.6944481 | 7.04486141 | 2.23963913 |
| 14843 | NM_019855.4 | #N/A | 9.98402913 | 7.54346116 | 2.39830293 |
| 14844 | NM_001164211.1 | #N/A | 26.3586648 | -5.9264141 | 1.88428085 |
| 14845 | NM_001267594.1 | #N/A | 32.6214289 | -4.7150465 | 1.49918025 |
| 14846 | NM_006441.3 | MTHFS | 15.4601234 | -5.1686351 | 1.64372557 |
| 14847 | NM_001174138.2 | CCK | 26.9297132 | 2.99627606 | 0.95293085 |
| 14848 | NM_173811.3 | #N/A | 17.2432444 | -5.3196107 | 1.69188487 |
| 14849 | NM_033418.3 | #N/A | 15.5721787 | -5.1643832 | 1.64270899 |
| 14850 | NM_000262.2 | #N/A | 117.563304 | -3.8906991 | 1.23778933 |
| 14851 | NM_001145812.1 | SH2B1 | 30.3034175 | -6.1314202 | 1.95068151 |
| 14852 | NM_001288704.1 | #N/A | 85.7905656 | 3.78513147 | 1.20422775 |
| 14853 | NM_001454.3 | #N/A | 285.406143 | -3.6973319 | 1.17636296 |
| 14854 | NM_153268.3 | PLCXD2 | 22.4344569 | -5.6930554 | 1.81134669 |
| 14855 | NM_001300773.1 | SYT7 | 34.596758 | -6.3177629 | 2.01029035 |
| 14856 | NM_005677.3 | #N/A | 33.4473455 | -4.7444259 | 1.50970371 |
| 14857 | NM_153810.4 | #N/A | 244.205149 | -1.0752833 | 0.34219558 |
| 14858 | NM_001283021.1 | #N/A | 15.9571598 | -5.207287 | 1.65718194 |
| 14860 | NR_024284.1 | ZEB1-AS1 | 28.6164507 | -4.5499011 | 1.44821433 |
| 14859 | NM_017884.5 | #N/A | 28.9962529 | -6.0637153 | 1.93004767 |
| 14861 | NR_024566.1 | ZNF271P | 45.0217514 | -2.7650081 | 0.88013875 |
| 14862 | NM_001195737.1 | #N/A | 26.9089689 | -5.9585715 | 1.89680678 |
| 14863 | NM_002950.3 | #N/A | 865.304952 | -4.5097546 | 1.4359944 |
| 14864 | NM_139201.2 | #N/A | 15.3584587 | -5.1523746 | 1.64076388 |
| 14865 | NM_133450.3 | #N/A | 19.44567 | -5.484676 | 1.74673949 |
| 14866 | NM_173674.2 | DCBLD1 | 17.4606006 | -5.3388543 | 1.7005608 |
| 14867 | NM_001104631.1 | #N/A | 29.6108567 | -6.1002471 | 1.94329163 |
| 14868 | NM_001319945.1 | #N/A | 22.6819228 | -5.7175601 | 1.82158721 |
| 14869 | NM_001142292.1 | LMAN2L | 24.3858754 | -5.8248134 | 1.85577969 |
| 14870 | NM_138930.3 | DIABLO | 31.4282847 | -6.1853956 | 1.97076004 |
| 14871 | NM_016724.2 | #N/A | 23.258182 | -5.7546455 | 1.83375084 |
| 14872 | NM_001032282.3 | KLF10 | 16.4901425 | -5.2635706 | 1.67735451 |
| 14873 | NR_135268.1 | #N/A | 32.4500955 | -6.2262511 | 1.98428536 |
| 14874 | NM_080626.5 | #N/A | 17.096671 | -5.294474 | 1.68760944 |
| 14875 | NM_001243804.1 | #N/A | 21.9859182 | -5.6744053 | 1.8088013 |
| 14876 | NM_007175.6 | #N/A | 262.911498 | -2.4512894 | 0.78141462 |
| 14877 | NM_003744.5 | NUMB | 21.070238 | -5.6152867 | 1.7901615 |

| | | | | | |
|---|---|---|---|---|---|
| 14878 | NM_031483.6 | #N/A | 116.002354 | -8.0691363 | 2.57272011 |
| 14879 | NM_032188.2 | #N/A | 152.860895 | -2.9661928 | 0.94585443 |
| 14880 | NM_001291991.1 | KIAA1324L | 31.8447418 | -6.1998621 | 1.97708132 |
| 14881 | NM_020178.4 | #N/A | 28.006335 | -6.0150604 | 1.91835599 |
| 14882 | NR_073135.1 | HPS4 | 30.7716121 | -6.1521302 | 1.96221779 |
| 14883 | NM_024712.3 | #N/A | 15.8769813 | -5.2069726 | 1.66084209 |
| 14884 | NM_001130687.2 | GUCY1A1 | 16.4260116 | -5.2418403 | 1.67206286 |
| 14885 | NM_001001395.2 | LMO3 | 154.091056 | -8.4779003 | 2.70442863 |
| 14886 | NM_001330261.1 | #N/A | 116.769941 | -8.0785334 | 2.57720869 |
| 14887 | NM_024773.2 | #N/A | 17.4432803 | -5.3432421 | 1.70510875 |
| 14888 | NR_046412.1 | POLD4 | 19.5832699 | -5.5074087 | 1.75767309 |
| 14889 | NM_001321970.1 | #N/A | 27.5995596 | -5.9939809 | 1.91299593 |
| 14890 | NR_073424.1 | #N/A | 27.220144 | -5.978967 | 1.90824642 |
| 14891 | NR_130112.1 | #N/A | 18.1997994 | -5.3900432 | 1.72037922 |
| 14892 | NM_001128844.1 | #N/A | 22.6049402 | -5.7173442 | 1.82486319 |
| 14893 | NM_015393.3 | #N/A | 172.783502 | -4.0892781 | 1.30528975 |
| 14894 | NM_001278511.1 | AP3B2 | 77.8237715 | -4.5218821 | 1.44341783 |
| 14895 | NR_040724.1 | #N/A | 112.319025 | -8.0226145 | 2.56117458 |
| 14896 | NR_130778.1 | LINC02198 | 19.1456427 | 3.18347825 | 1.01638164 |
| 14897 | NM_145003.4 | #N/A | 14.4694449 | -5.0637015 | 1.61676368 |
| 14898 | NM_052953.3 | LRRC3B | 28.472086 | -6.0376132 | 1.92788948 |
| 14899 | NR_045585.1 | SDF2 | 18.9798116 | -5.4573891 | 1.7430227 |
| 14900 | NM_001002879.1 | THOC5 | 15.9993012 | -5.1983895 | 1.66042134 |
| 14901 | NR_136513.1 | #N/A | 19.5643083 | 6.51749359 | 2.08180326 |
| 14902 | NM_001328674.1 | #N/A | 150.669986 | -8.4456769 | 2.69777085 |
| 14903 | NM_001145316.1 | DSN1 | 17.6814621 | -5.3474815 | 1.70823053 |
| 14904 | NM_001282957.1 | #N/A | 34.5089916 | -6.3226528 | 2.01979915 |
| 14905 | NM_001287421.1 | ZSCAN26 | 16.8757687 | -5.2815626 | 1.68733805 |
| 14906 | NM_139034.2 | #N/A | 29.4016716 | -6.0915352 | 1.94630166 |
| 14907 | NM_001252124.1 | #N/A | 17.8363686 | -5.3752167 | 1.71750041 |
| 14908 | NM_007309.3 | DIAPH2 | 16.4497375 | -5.2417374 | 1.67493201 |
| 14909 | NR_046293.1 | MSTO1 | 17.8137516 | -5.3550874 | 1.71122588 |
| 14910 | NM_001244678.1 | THNSL2 | 18.4150598 | -5.4168647 | 1.73097824 |
| 14911 | NR_003086.1 | HSD17B7P2 | 15.2985828 | -5.1558627 | 1.64773273 |
| 14912 | NM_030923.4 | #N/A | 69.5794375 | -3.1663147 | 1.01208063 |
| 14913 | NM_001293167.1 | PANO1 | 34.5862709 | -2.63424 | 0.84205545 |
| 14914 | NR_037776.2 | #N/A | 18.8145374 | -5.4361187 | 1.7378088 |
| 14915 | NM_001286632.1 | RBM26 | 110.468635 | -7.9986349 | 2.55706644 |
| 14916 | NR_104599.1 | #N/A | 14.8009446 | -5.1009879 | 1.63080276 |
| 14917 | NM_001199917.1 | #N/A | 20.0899229 | -5.5334993 | 1.7691086 |
| 14918 | NM_022072.3 | #N/A | 16.4461121 | -5.2569644 | 1.6807671 |
| 14919 | NM_006475.2 | #N/A | 136.963395 | -8.3083483 | 2.65647231 |
| 14920 | NM_001135953.1 | GKAP1 | 21.2647422 | -5.6256383 | 1.79884663 |

| | | | | | |
|---|---|---|---|---|---|
| 14921 | NM_001323890.1 | #N/A | 20.5372204 | -5.5759849 | 1.78296745 |
| 14922 | NR_027775.2 | PMS2P5 | 18.0828242 | -5.3736827 | 1.71837061 |
| 14923 | NR_036480.1 | VPS9D1-AS1 | 15.5361178 | -5.175344 | 1.65497498 |
| 14924 | NM_033425.4 | DIXDC1 | 54.4090434 | -4.1342987 | 1.32208027 |
| 14925 | NR_033684.1 | #N/A | 22.3668852 | -5.6839076 | 1.81772815 |
| 14926 | NM_052978.4 | TRIM9 | 19.9098844 | -5.5237766 | 1.7665764 |
| 14927 | NM_016547.2 | SDF4 | 21.7771666 | -5.6619552 | 1.81086206 |
| 14928 | NR_120317.1 | #N/A | 43.6003003 | -4.2852901 | 1.3705557 |
| 14929 | NM_031968.2 | NARF | 14.5430347 | -5.0771368 | 1.62414051 |
| 14930 | NM_005387.6 | #N/A | 20.3011596 | -5.5550586 | 1.77710268 |
| 14931 | NM_001014979.2 | #N/A | 19.8309412 | -5.5253629 | 1.76774494 |
| 14932 | NM_001130413.3 | #N/A | 18.8777966 | -5.4578395 | 1.74614065 |
| 14933 | NM_001134848.1 | #N/A | 28.8169651 | -4.5518951 | 1.45638523 |
| 14935 | NR_036533.1 | LOC100499489 | 20.1274924 | -5.5381808 | 1.77207583 |
| 14934 | NM_001271286.1 | ZNF512 | 20.5304587 | -5.5795854 | 1.78531915 |
| 14936 | NM_006041.2 | #N/A | 92.6560868 | -1.6728774 | 0.53546847 |
| 14937 | NM_202002.2 | #N/A | 34.0551504 | -6.2984003 | 2.01618495 |
| 14938 | NM_001130009.2 | #N/A | 27.8973959 | -6.0084951 | 1.92362404 |
| 14940 | NM_001244973.1 | NRP1 | 152.270753 | -8.4611846 | 2.70892206 |
| 14939 | NM_001202431.1 | PRDX1 | 18.8389273 | -5.4399219 | 1.74163372 |
| 14941 | NM_002413.4 | #N/A | 14.7097836 | -5.0931276 | 1.63072648 |
| 14942 | NR_110840.1 | LOC101927136 | 32.040642 | 3.6557104 | 1.17053254 |
| 14943 | NM_001300732.1 | TMC7 | 56.2043831 | 3.40338657 | 1.08996232 |
| 14944 | NR_037185.1 | #N/A | 15.5462139 | -5.1726306 | 1.65664374 |
| 14945 | NR_138150.1 | #N/A | 14.1255565 | -5.028922 | 1.61084378 |
| 14946 | NR_024402.2 | SNAI3-AS1 | 18.1731884 | -5.3896445 | 1.72641693 |
| 14947 | NM_001321047.1 | #N/A | 24.2158612 | -5.8054983 | 1.85967957 |
| 14948 | NM_001206845.1 | #N/A | 144.459408 | 6.28064449 | 2.01194715 |
| 14949 | NM_001285450.1 | #N/A | 14.7383808 | -5.10078 | 1.63401221 |
| 14950 | NM_001168254.1 | OCIAD1 | 25.4389159 | -5.8746067 | 1.88196323 |
| 14951 | NM_021813.3 | #N/A | 32.0288852 | -6.2144739 | 1.9910802 |
| 14952 | NM_153769.2 | #N/A | 17.1527878 | -5.3077237 | 1.70055667 |
| 14953 | NM_001003692.1 | #N/A | 17.3758035 | -5.3377259 | 1.71034741 |
| 14954 | NM_152445.2 | #N/A | 99.5579942 | -2.9048759 | 0.93095579 |
| 14955 | NM_178422.5 | PAQR7 | 100.787244 | -1.5507603 | 0.49703225 |
| 14956 | NR_130789.1 | MIEF1 | 113.991506 | -8.0439114 | 2.57826211 |
| 14957 | NM_001319838.1 | ARHGEF10L | 18.551363 | -5.4236868 | 1.7385483 |
| 14958 | NM_021817.2 | #N/A | 22.6036142 | -5.7049176 | 1.82879395 |
| 14959 | NM_001109.4 | #N/A | 18.8788081 | -5.4516185 | 1.74766556 |
| 14960 | NM_001040165.2 | #N/A | 15.5638315 | -5.1650081 | 1.65589559 |
| 14961 | NM_001270643.1 | LUC7L2 | 26.4643698 | -5.9414973 | 1.9048947 |
| 14962 | NM_001167674.1 | #N/A | 23.0580349 | -5.7344549 | 1.83855248 |
| 14963 | NR_138032.1 | LOC107985033 | 61.4521023 | 1.92648647 | 0.61767173 |

| | | | | | |
|---|---|---|---|---|---|
| 14964 | NM_001255980.1 | #N/A | 14.2704369 | -5.0520524 | 1.6198346 |
| 14965 | NM_001256534.1 | #N/A | 21.9444762 | -5.6623408 | 1.81570142 |
| 14966 | NR_104345.1 | #N/A | 25.8716868 | -5.901107 | 1.89227225 |
| 14967 | NM_006354.3 | TADA3 | 144.22585 | -8.3825736 | 2.68811816 |
| 14968 | NM_001143782.1 | FKBP11 | 14.6059347 | -5.0825985 | 1.62993908 |
| 14969 | NR_024116.2 | RASA4CP | 20.5934936 | -5.5709821 | 1.78657912 |
| 14970 | NR_026981.1 | RPL23AP82 | 14.0129492 | -5.0217998 | 1.61047842 |
| 14971 | NM_001190316.1 | IP6K2 | 21.3761253 | -5.616512 | 1.80142598 |
| 14972 | NR_024034.1 | CLUHP3 | 16.3815787 | -5.2397237 | 1.6807593 |
| 14973 | NM_001282568.1 | ZCCHC17 | 17.7689631 | -5.3616986 | 1.72034688 |
| 14974 | NM_001292008.1 | #N/A | 35.7412634 | 6.76710943 | 2.17153332 |
| 14975 | NM_001256409.1 | #N/A | 32.5192513 | -6.231149 | 1.99968577 |
| 14976 | NR_003290.2 | EP400P1 | 24.7291452 | -3.7473607 | 1.20280402 |
| 14977 | NM_012347.4 | FBXO9 | 15.0291291 | 8.87058189 | 2.84750391 |
| 14978 | NM_001131023.1 | PEX5 | 25.131898 | -5.8602366 | 1.88118405 |
| 14979 | NM_001198763.1 | CD302 | 9.85199644 | 8.29669467 | 2.66350659 |
| 14980 | NM_024099.3 | LBHD1 | 14.7462956 | -5.0963274 | 1.63615799 |
| 14981 | NM_003852.3 | #N/A | 143.807648 | -8.3783478 | 2.69031477 |
| 14982 | NM_004333.4 | #N/A | 44.9460182 | -3.9346605 | 1.26344973 |
| 14983 | NM_018932.3 | #N/A | 22.97717 | -5.726189 | 1.83873116 |
| 14984 | NM_033204.3 | #N/A | 26.9894629 | -5.9680457 | 1.91650673 |
| 14985 | NR_037771.1 | #N/A | 39.3952184 | 3.86965802 | 1.24266155 |
| 14986 | NR_024018.1 | #N/A | 21.3867819 | -5.6359415 | 1.81014676 |
| 14987 | NM_172084.2 | #N/A | 11.9874775 | 8.54076176 | 2.74324287 |
| 14988 | NM_016553.4 | #N/A | 19.3973367 | -5.4848124 | 1.76178158 |
| 14989 | NM_001330683.1 | #N/A | 145.960784 | -8.3996208 | 2.69814078 |
| 14990 | NM_001346048.1 | #N/A | 21.1368202 | -5.6081882 | 1.80146934 |
| 14991 | NM_001280801.1 | #N/A | 24.3148496 | -5.8104067 | 1.86644378 |
| 14992 | NM_001304493.1 | #N/A | 32.6117832 | -4.9516408 | 1.59068281 |
| 14993 | NM_198120.2 | EBAG9 | 22.3059071 | -5.6853974 | 1.82695725 |
| 14994 | NM_001204103.1 | #N/A | 31.0041305 | -6.1588386 | 1.97938506 |
| 14995 | NM_001010897.2 | #N/A | 29.1566132 | -6.0744781 | 1.95230161 |
| 14996 | NM_024768.2 | EFCC1 | 16.5962155 | -5.262594 | 1.69140786 |
| 14997 | NM_199367.2 | #N/A | 20.7023877 | -4.2996205 | 1.38200587 |
| 14998 | NM_001202464.1 | ZMYND11 | 213.511551 | -4.0269776 | 1.29451448 |
| 14999 | NM_001198837.1 | #N/A | 25.7657525 | 7.78114141 | 2.50134644 |
| 15000 | NM_001570.3 | #N/A | 71.9766684 | 1.88941316 | 0.60741912 |
| 15002 | NR_024548.1 | CRCP | 90.1765671 | -2.7337657 | 0.87889528 |
| 15001 | NM_182612.3 | #N/A | 146.708484 | -8.4070651 | 2.70285546 |
| 15003 | NM_172020.4 | POM121 | 383.352729 | -3.272558 | 1.05226509 |
| 15004 | NM_001323032.2 | #N/A | 25.7851327 | -5.9039795 | 1.8984136 |
| 15006 | NM_001135163.1 | SIGLEC11 | 23.6853437 | 2.50822131 | 0.80654655 |
| 15005 | NM_001003800.1 | #N/A | 220.223265 | -3.0758908 | 0.98908084 |

| | | | | | |
|---|---|---|---|---|---|
| 15007 | NM_004034.3 | #N/A | 31.3186847 | -6.1734308 | 1.98514885 |
| 15008 | NR_036634.1 | #N/A | 25.5081223 | -5.8871648 | 1.89362444 |
| 15009 | NM_018833.2 | TAP2 | 18.9557144 | -5.464956 | 1.75811751 |
| 15010 | NM_030821.4 | #N/A | 172.063584 | -2.0680827 | 0.66531572 |
| 15011 | NR_028407.2 | #N/A | 14.2898109 | -5.0530652 | 1.62574216 |
| 15012 | NM_001288992.1 | #N/A | 19.7359962 | -5.5213916 | 1.77669517 |
| 15013 | NR_037860.1 | #N/A | 18.6168043 | -5.4304467 | 1.74756473 |
| 15014 | NM_024887.3 | DHDDS | 30.3837293 | -6.1343305 | 1.97419953 |
| 15015 | NM_001270616.1 | #N/A | 27.1079396 | -4.4714608 | 1.43913039 |
| 15016 | NM_022091.4 | ASCC3 | 14.4519441 | -5.0626875 | 1.6295018 |
| 15017 | NM_025130.3 | #N/A | 15.3667383 | -5.1503687 | 1.65774751 |
| 15018 | NM_001005417.2 | B4GALT2 | 22.7139319 | -5.7245619 | 1.84258776 |
| 15019 | NM_005188.3 | #N/A | 257.632349 | -2.3192785 | 0.74667319 |
| 15020 | NM_018212.5 | #N/A | 1133.1619 | -1.7221467 | 0.55442995 |
| 15021 | NM_005025.4 | SERPINI1 | 21.2250159 | -5.6123963 | 1.80698457 |
| 15022 | NM_001346372.1 | #N/A | 28.5308768 | -6.0428208 | 1.94631243 |
| 15023 | NM_018670.3 | MESP1 | 22.7459166 | -5.7123355 | 1.84015979 |
| 15024 | NM_001813.2 | #N/A | 19.9450444 | -5.522857 | 1.77922778 |
| 15025 | NR_136586.1 | #N/A | 36.9812927 | -6.4158911 | 2.06701834 |
| 15026 | NM_001282195.1 | #N/A | 24.1914451 | -5.8125525 | 1.87278719 |
| 15027 | NM_001308226.1 | RPS15 | 22.9170742 | -5.7284773 | 1.84606815 |
| 15028 | NM_001077526.2 | #N/A | 18.7002611 | -5.4373684 | 1.75236648 |
| 15029 | NM_001317334.1 | #N/A | 23.2162268 | -5.7500882 | 1.85326553 |
| 15030 | NR_120457.1 | #N/A | 19.2114466 | -5.47749 | 1.7654208 |
| 15031 | NM_018081.2 | WRAP53 | 17.1987403 | -5.3064546 | 1.7103083 |
| 15032 | NM_017672.5 | #N/A | 106.38706 | -7.9442879 | 2.56068698 |
| 15034 | NR_026544.1 | PCED1B-AS1 | 65.6003378 | 2.02286617 | 0.65205714 |
| 15033 | NM_001013628.2 | #N/A | 15.0068675 | -5.1227846 | 1.65130493 |
| 15035 | NM_024700.3 | #N/A | 122.443583 | -2.0165 | 0.65004722 |
| 15036 | NM_001100399.1 | #N/A | 399.650375 | -3.2912703 | 1.06103852 |
| 15037 | NM_001979.5 | #N/A | 23.1775209 | -5.7495699 | 1.853559 |
| 15038 | NM_024863.5 | TCEAL4 | 27.6052959 | -5.996341 | 1.93333698 |
| 15039 | NM_206907.3 | PRKAA1 | 32.9546243 | -6.2486224 | 2.0146942 |
| 15040 | NM_004486.4 | #N/A | 606.893254 | -2.111168 | 0.6806958 |
| 15041 | NM_012118.3 | #N/A | 61.0355101 | -2.5641263 | 0.82675915 |
| 15042 | NM_002024.5 | FMR1 | 19.4284162 | -5.4824874 | 1.76780666 |
| 15043 | NM_001166059.1 | HMGCL | 17.5677171 | -5.3479397 | 1.72452195 |
| 15044 | NM_004951.4 | #N/A | 21.798346 | 8.08841366 | 2.60830883 |
| 15045 | NM_207325.2 | #N/A | 26.3347727 | -5.9355042 | 1.91419496 |
| 15046 | NR_126327.1 | #N/A | 16.5140546 | -5.2613691 | 1.6968953 |
| 15048 | NR_038934.1 | CAMTA1 | 15.6374908 | -5.1802744 | 1.6708795 |
| 15047 | NM_001282397.1 | HPCAL4 | 41.1297337 | -4.9812759 | 1.60669789 |
| 15049 | NM_001261412.1 | DCTN2 | 141.58102 | -8.3560775 | 2.69526813 |

| | | | | | |
|---|---|---|---|---|---|
| 15050 | NM_203370.1 | #N/A | 21.1437402 | -5.6089877 | 1.80921117 |
| 15051 | NM_001206729.1 | AKT3 | 15.0613017 | -5.1324139 | 1.65563118 |
| 15052 | NR_130136.1 | #N/A | 26.7656176 | -5.9555218 | 1.9221389 |
| 15053 | NM_181785.3 | #N/A | 18.0206212 | -5.385408 | 1.73818123 |
| 15054 | NM_001166621.1 | TRAPPC1 | 21.0165185 | -5.6112226 | 1.81131017 |
| 15055 | NM_001128853.1 | #N/A | 110.674381 | -8.0013151 | 2.58299798 |
| 15056 | NM_005794.3 | #N/A | 15.9002007 | -5.2012692 | 1.67909793 |
| 15057 | NM_001003931.3 | PARP3 | 17.8386084 | -5.3657571 | 1.7322797 |
| 15058 | NM_030958.2 | #N/A | 25.0792686 | -5.8578077 | 1.89124697 |
| 15059 | NM_001301069.1 | #N/A | 535.947129 | -4.2527227 | 1.37307596 |
| 15060 | NM_153219.3 | #N/A | 16.6487117 | -5.2661603 | 1.70027795 |
| 15061 | NM_182832.2 | #N/A | 34.7460199 | 6.13173431 | 1.97986208 |
| 15062 | NM_017593.3 | #N/A | 14.6023875 | -5.080438 | 1.64053172 |
| 15063 | NM_145905.2 | HMGA1 | 19.1025556 | -5.4577962 | 1.76258791 |
| 15064 | NM_017577.4 | #N/A | 13.7397487 | -4.9883201 | 1.61114218 |
| 15066 | NR_103849.2 | CASC9 | 26.3195212 | 1.8832555 | 0.60832592 |
| 15065 | NM_025258.2 | #N/A | 19.6451832 | -5.5125503 | 1.78065716 |
| 15067 | NM_001278716.1 | #N/A | 16.4978024 | -5.2541661 | 1.69733082 |
| 15068 | NM_013959.3 | NRG1 | 21.098931 | -5.606321 | 1.81114029 |
| 15069 | NM_001324215.1 | #N/A | 34.3957926 | -6.3174591 | 2.04107038 |
| 15070 | NR_028057.1 | #N/A | 19.7603186 | -5.5236699 | 1.78504788 |
| 15071 | NR_036682.1 | #N/A | 19.6827659 | -5.5005504 | 1.77770643 |
| 15072 | NM_004535.2 | #N/A | 203.616279 | -3.200436 | 1.03438852 |
| 15073 | NM_001316994.1 | #N/A | 15.3989789 | -5.1576381 | 1.66703552 |
| 15074 | NR_126407.1 | LINC01392 | 23.4634094 | 7.28912643 | 2.35631352 |
| 15075 | NM_001284240.1 | #N/A | 118.255184 | -8.0966744 | 2.61751426 |
| 15076 | NM_014716.3 | #N/A | 17.4533923 | -5.3377583 | 1.72578176 |
| 15077 | NM_001040146.3 | #N/A | 31.7570035 | -4.6691431 | 1.50965992 |
| 15078 | NM_001146197.1 | #N/A | 19.3825148 | 7.28637059 | 2.35604013 |
| 15079 | NM_024959.3 | SLC8B1 | 16.9543419 | -5.2917029 | 1.7112788 |
| 15080 | NM_004723.3 | ARHGEF2 | 192.911348 | -3.4123251 | 1.10359617 |
| 15081 | NM_174945.2 | ZNF575 | 17.8239361 | -5.3628954 | 1.73447014 |
| 15082 | NM_001319666.1 | TMEM263 | 14.7632018 | -5.0975532 | 1.64871637 |
| 15083 | NM_001855.4 | #N/A | 47.4316228 | -5.1615922 | 1.66945665 |
| 15084 | NM_012416.3 | #N/A | 194.796572 | -3.9012695 | 1.26186803 |
| 15085 | NM_001164766.1 | #N/A | 93.593625 | -2.637781 | 0.85332514 |
| 15086 | NM_001346509.1 | SASH1 | 23.2875921 | -5.7493842 | 1.85995208 |
| 15087 | NM_024310.4 | #N/A | 15.361452 | -5.1606703 | 1.67004576 |
| 15088 | NM_001195733.1 | #N/A | 109.53452 | -7.9863332 | 2.58479126 |
| 15089 | NM_178033.1 | #N/A | 23.3231597 | -5.7519065 | 1.86165527 |
| 15091 | NR_033862.1 | DKFZp451B082 | 16.1176794 | 4.6803083 | 1.51491388 |
| 15090 | NM_001145304.1 | #N/A | 13.6659071 | -4.9817224 | 1.61247016 |
| 15092 | NM_001178119.1 | ABI1 | 26.8302731 | -5.9503694 | 1.92608875 |

| | | | | | |
|---|---|---|---|---|---|
| 15093 | NR_033773.1 | #N/A | 13.906117 | -5.0084195 | 1.62129164 |
| 15094 | NM_001128931.1 | #N/A | 14.2884321 | -5.0475463 | 1.63408798 |
| 15095 | NM_001044723.2 | CSNK1G3 | 26.7016718 | -5.9468718 | 1.92533816 |
| 15097 | NM_001287804.1 | ZDHHC16 | 15.386267 | -5.1563335 | 1.66950013 |
| 15096 | NM_001199877.1 | #N/A | 14.959858 | -5.1107042 | 1.65472956 |
| 15098 | NM_024815.3 | #N/A | 14.9324629 | -5.1064458 | 1.65334211 |
| 15099 | NM_003227.3 | #N/A | 14.2549915 | -5.0381039 | 1.63127969 |
| 15100 | NR_033761.1 | #N/A | 20.5093396 | -5.5583417 | 1.79972598 |
| 15101 | NM_020660.2 | #N/A | 19.8089791 | -5.5185854 | 1.78716921 |
| 15102 | NM_001320371.1 | #N/A | 18.1988745 | -5.3935258 | 1.74681804 |
| 15103 | NM_002755.3 | MAP2K1 | 268.885853 | -3.0586347 | 0.99088462 |
| 15104 | NM_001282696.1 | #N/A | 17.6153108 | -5.3468955 | 1.73221679 |
| 15105 | NM_001291398.1 | WASHC2A | 109.087415 | -7.9804712 | 2.58552809 |
| 15106 | NM_001184888.1 | NIPA2 | 17.0334752 | -5.311237 | 1.72102901 |
| 15107 | NM_001004051.3 | #N/A | 105.175064 | -7.9278475 | 2.56903713 |
| 15108 | NM_001271635.1 | CNOT9 | 16.6121257 | -5.2757008 | 1.70968669 |
| 15109 | NM_001136537.1 | BTBD19 | 25.0340901 | -5.8623643 | 1.89993795 |
| 15110 | NM_138345.2 | #N/A | 22.1256703 | -5.6739944 | 1.83888874 |
| 15111 | NM_001145673.1 | #N/A | 18.3490338 | -5.4168551 | 1.7557194 |
| 15112 | NM_001034194.1 | #N/A | 20.6096456 | -5.5737355 | 1.8065901 |
| 15113 | NM_016481.4 | #N/A | 16.2179151 | -5.2332086 | 1.69639945 |
| 15114 | NR_126046.1 | GLIDR | 22.1032607 | -4.378495 | 1.41935078 |
| 15115 | NM_031942.4 | #N/A | 31.6957152 | -6.1931848 | 2.0076259 |
| 15116 | NM_001010853.2 | #N/A | 140.116966 | -1.4830161 | 0.48075939 |
| 15117 | NR_049780.1 | #N/A | 16.8360833 | -5.2762369 | 1.71049871 |
| 15118 | NM_001165931.1 | RRM2 | 32.4646217 | -6.2286372 | 2.01931048 |
| 15119 | NR_104662.1 | LOC102467081 | 194.836649 | 2.77935333 | 0.90115221 |
| 15120 | NR_105005.1 | LOC101929528 | 12.082305 | 8.59369208 | 2.78670863 |
| 15121 | NM_000621.4 | HTR2A | 16.1458902 | -5.2246753 | 1.69425152 |
| 15122 | NM_001329953.1 | LTV1 | 15.1253174 | -5.1280747 | 1.6630026 |
| 15123 | NM_003413.3 | #N/A | 35.1008185 | -6.3469503 | 2.0582923 |
| 15124 | NM_001282404.1 | #N/A | 30.4933252 | -6.1440403 | 1.99262135 |
| 15125 | NM_001143906.1 | TRAFD1 | 17.2809624 | -5.309875 | 1.72231838 |
| 15126 | NM_001201407.1 | ZNF778 | 27.8015329 | -6.0016746 | 1.94673784 |
| 15127 | NM_198838.1 | ACACA | 20.1313492 | -5.5482349 | 1.79979117 |
| 15128 | NM_032237.4 | #N/A | 16.6522013 | -5.2568345 | 1.70585609 |
| 15129 | NM_012216.3 | #N/A | 33.0006967 | -6.2563912 | 2.03043795 |
| 15130 | NM_001024683.1 | ZNF688 | 16.6716785 | -5.2723279 | 1.7111154 |
| 15131 | NM_001318845.1 | CNPY3 | 31.6083005 | -6.1892585 | 2.00886673 |
| 15132 | NM_001172650.2 | #N/A | 15.2167847 | -5.1317097 | 1.66581327 |
| 15133 | NM_170677.4 | #N/A | 134.389802 | -8.2807503 | 2.68820114 |
| 15134 | NR_104131.1 | #N/A | 85.2916264 | 2.79381176 | 0.90698111 |
| 15135 | NM_000267.3 | NF1 | 283.579491 | -2.281614 | 0.74071106 |

| | | | | | |
|---|---|---|---|---|---|
| 15136 | NM_001306153.1 | CCT5 | 118.86614 | -8.104286 | 2.63110117 |
| 15137 | NM_004956.4 | #N/A | 25.9735582 | -5.9047061 | 1.91753412 |
| 15138 | NR_122105.1 | MIR570HG | 13.9761428 | -5.0088713 | 1.62676088 |
| 15139 | NM_001101339.2 | #N/A | 28.8796938 | 3.4107794 | 1.10776507 |
| 15140 | NM_206943.2 | #N/A | 330.767066 | -4.1841102 | 1.35902832 |
| 15141 | NM_017631.5 | #N/A | 34.237037 | -6.3105926 | 2.04983092 |
| 15142 | NM_001317171.1 | SNRPG | 23.1261806 | -5.7445514 | 1.86621604 |
| 15143 | NR_135629.1 | #N/A | 18.3708455 | -5.4040381 | 1.75633001 |
| 15144 | NR_110861.1 | RARA-AS1 | 20.4230283 | -5.5621387 | 1.80837988 |
| 15145 | NM_001304350.1 | ZNF558 | 18.7169876 | -5.4304631 | 1.76571579 |
| 15146 | NR_026595.2 | FAM226A | 17.5913697 | -5.3416924 | 1.73701905 |
| 15147 | NM_001282506.1 | #N/A | 14.2141813 | -5.0468913 | 1.64118529 |
| 15148 | NR_135622.1 | #N/A | 16.9147627 | -5.290171 | 1.72029929 |
| 15149 | NM_001330130.1 | #N/A | 15.7175129 | -5.1923148 | 1.68863216 |
| 15150 | NM_152400.2 | #N/A | 26.9628704 | -5.9584743 | 1.93795675 |
| 15151 | NR_138074.1 | #N/A | 18.9615416 | -5.4435136 | 1.77057817 |
| 15152 | NR_121647.1 | LINC01410 | 25.6598985 | 3.43409197 | 1.11710652 |
| 15153 | NM_001244889.1 | #N/A | 101.025756 | -4.8057483 | 1.56343896 |
| 15154 | NR_104295.1 | #N/A | 15.0666218 | -5.1300073 | 1.66908762 |
| 15155 | NM_001301240.1 | #N/A | 33.9338056 | -6.295651 | 2.04835594 |
| 15156 | NM_152398.3 | OCIAD2 | 32.2074284 | -6.2138571 | 2.02176146 |
| 15157 | NM_001286174.1 | #N/A | 24.5613437 | -5.8335301 | 1.8981143 |
| 15158 | NM_032217.4 | #N/A | 131.202804 | -8.246487 | 2.68328987 |
| 15159 | NM_006467.2 | #N/A | 15.2861672 | -5.1425807 | 1.67336198 |
| 15160 | NM_001017975.4 | #N/A | 15.6637657 | -5.1763069 | 1.68457345 |
| 15161 | NM_018385.2 | #N/A | 186.70289 | -2.2392066 | 0.72875552 |
| 15162 | NM_012289.3 | #N/A | 233.885515 | -3.1020292 | 1.00957026 |
| 15163 | NM_032512.3 | #N/A | 490.696441 | -4.4007272 | 1.43232589 |
| 15164 | NM_002942.4 | #N/A | 133.22913 | -8.2683268 | 2.69118452 |
| 15165 | NM_001145971.1 | #N/A | 16.158628 | -5.2267839 | 1.70124846 |
| 15166 | NM_001204193.1 | SRP19 | 18.0991188 | -5.3761002 | 1.74997067 |
| 15167 | NM_001319041.1 | #N/A | 24.6193385 | -5.8289524 | 1.89750417 |
| 15168 | NM_203394.2 | #N/A | 60.2739926 | -1.8547517 | 0.60380582 |
| 15169 | NM_001288633.1 | #N/A | 19.538325 | -5.5035274 | 1.79171439 |
| 15170 | NR_046325.1 | PCAT6 | 16.5805461 | -5.2630974 | 1.71371285 |
| 15171 | NM_001286585.1 | #N/A | 22.9814385 | -5.7274221 | 1.86499948 |
| 15172 | NM_145260.2 | #N/A | 23.6483628 | -5.7787268 | 1.88175347 |
| 15173 | NM_031918.3 | #N/A | 104.922839 | -3.7336682 | 1.21581964 |
| 15174 | NR_136241.1 | #N/A | 14.996146 | -5.1257496 | 1.66937454 |
| 15175 | NM_001172697.1 | TSFM | 20.5943647 | -5.5670168 | 1.8131431 |
| 15176 | NM_015716.4 | #N/A | 103.393825 | -7.903279 | 2.57451559 |
| 15177 | NM_182958.2 | #N/A | 16.3992797 | -5.2486079 | 1.71006549 |
| 15178 | NM_001315529.1 | #N/A | 65.0475142 | -2.8648446 | 0.93350719 |

| | | | | | |
|---|---|---|---|---|---|
| 15179 | NM_002524.4 | #N/A | 377.912708 | -1.45814 | 0.47517082 |
| 15180 | NM_018039.2 | #N/A | 13.8843346 | -5.0029027 | 1.6304389 |
| 15183 | NR_110650.1 | LINC02133 | 28.210797 | 4.84308893 | 1.57854107 |
| 15181 | NM_001330292.1 | #N/A | 16.8839783 | -5.2974421 | 1.72663471 |
| 15182 | NM_031421.3 | #N/A | 27.8214607 | -6.012163 | 1.95958269 |
| 15184 | NM_001308090.1 | #N/A | 100.827658 | -7.8669917 | 2.56426569 |
| 15185 | NM_024707.2 | #N/A | 14.7786203 | -5.0921434 | 1.66044784 |
| 15186 | NM_001017923.1 | #N/A | 17.499832 | -5.3399062 | 1.74135494 |
| 15187 | NM_001227.4 | #N/A | 14.1452435 | -5.0257359 | 1.63892248 |
| 15188 | NM_182609.3 | ZNF677 | 59.6033214 | 1.70471018 | 0.55599098 |
| 15189 | NR_109849.1 | LOC102606465 | 15.6844093 | -5.1815109 | 1.69005749 |
| 15190 | NM_001320793.1 | #N/A | 16.2356689 | -5.2386385 | 1.70883724 |
| 15191 | NM_004282.3 | #N/A | 27.5140513 | -4.4827369 | 1.46231424 |
| 15192 | NM_014168.3 | #N/A | 19.347724 | -5.4813393 | 1.78831402 |
| 15193 | NM_004555.3 | #N/A | 18.9622558 | -5.4650004 | 1.78309359 |
| 15194 | NM_152329.3 | #N/A | 24.424259 | -5.8167726 | 1.89796356 |
| 15195 | NM_005895.3 | #N/A | 727.911991 | -1.9575241 | 0.63876546 |
| 15196 | NM_001316307.1 | #N/A | 36.3244921 | -6.3924761 | 2.08596813 |
| 15197 | NM_001127208.2 | TET2 | 115.702364 | -2.7308034 | 0.89116193 |
| 15198 | NM_001809.3 | #N/A | 17.7094985 | -5.3602173 | 1.74957389 |
| 15199 | NM_182796.2 | MAT2B | 18.8656267 | -5.4574242 | 1.78150332 |
| 15200 | NM_001302694.1 | TRIM2 | 22.5271552 | -5.7033278 | 1.86185536 |
| 15201 | NM_153374.2 | #N/A | 18.6652953 | -5.4296659 | 1.77255955 |
| 15202 | NM_012428.3 | #N/A | 20.2637541 | -5.5442333 | 1.81062388 |
| 15203 | NM_001316309.1 | #N/A | 20.0720779 | -5.5421571 | 1.80999465 |
| 15204 | NM_176787.4 | #N/A | 17.5881211 | -5.343151 | 1.74509266 |
| 15205 | NM_005670.3 | #N/A | 34.796438 | -3.9811647 | 1.30036688 |
| 15206 | NM_001113756.2 | #N/A | 73.0488838 | -7.4022999 | 2.41830142 |
| 15207 | NM_001244015.1 | #N/A | 25.2998232 | -5.8773775 | 1.92015187 |
| 15208 | NR_125405.1 | LOC102723582 | 32.5214526 | 5.67109167 | 1.85284803 |
| 15209 | NR_038282.1 | PTPRG-AS1 | 16.6507831 | -5.2714709 | 1.72230537 |
| 15210 | NM_001127718.2 | #N/A | 11.7960527 | 8.55928516 | 2.79670359 |
| 15211 | NM_003246.3 | #N/A | 969.066207 | -3.4542497 | 1.12871698 |
| 15212 | NM_014992.2 | DAAM1 | 198.358562 | -2.7154654 | 0.88734073 |
| 15213 | NM_198240.1 | #N/A | 100.343629 | -7.8599429 | 2.56863468 |
| 15214 | NM_000771.3 | #N/A | 20.5936273 | 8.00517879 | 2.61630237 |
| 15216 | NM_001305654.1 | LIME1 | 20.6163893 | -5.5783652 | 1.8234093 |
| 15215 | NM_001252507.2 | #N/A | 1012.90544 | -3.8075665 | 1.24457969 |
| 15217 | NM_001319844.1 | #N/A | 31.1162759 | -4.8931269 | 1.59962969 |
| 15218 | NR_026950.1 | LOC283922 | 13.6282405 | -4.9854065 | 1.62990339 |
| 15219 | NR_104429.1 | PPIL6 | 237.503679 | 2.5711002 | 0.84057746 |
| 15220 | NM_001303048.1 | MYL12A | 30.5089999 | -6.143735 | 2.00861726 |
| 15221 | NM_014267.5 | #N/A | 823.251115 | -1.836013 | 0.60029096 |

| | | | | | |
|---|---|---|---|---|---|
| 15222 | NM_001033575.1 | DUSP28 | 17.2099721 | -5.3134022 | 1.73729859 |
| 15223 | NR_045022.1 | SNHG29 | 18.2346484 | -5.3920562 | 1.76309368 |
| 15224 | NM_001185090.1 | #N/A | 23.9958451 | -5.7861307 | 1.89229891 |
| 15225 | NM_004798.3 | #N/A | 624.536817 | -3.0940424 | 1.01207037 |
| 15226 | NM_001243942.1 | PRPSAP2 | 22.7984405 | -5.7179524 | 1.87070698 |
| 15227 | NM_033261.2 | #N/A | 14.8516094 | -5.1018248 | 1.66915296 |
| 15228 | NM_007325.4 | #N/A | 128.679723 | -8.2179395 | 2.68882019 |
| 15229 | NM_138410.3 | #N/A | 14.6330283 | -5.0889169 | 1.66511298 |
| 15230 | NM_180991.4 | #N/A | 29.7457248 | 3.25470871 | 1.06506129 |
| 15231 | NM_022716.3 | #N/A | 18.3003662 | -5.4059392 | 1.76903752 |
| 15232 | NM_006300.3 | #N/A | 25.9839587 | -4.4073648 | 1.44260831 |
| 15233 | NM_024738.2 | #N/A | 501.478176 | -2.9450174 | 0.96407073 |
| 15234 | NR_028514.1 | PITPNA-AS1 | 17.1546186 | -5.3132823 | 1.73939129 |
| 15235 | NR_104414.1 | C11orf98 | 14.7004485 | -5.0915381 | 1.66692579 |
| 15236 | NM_001256761.2 | HTR2C | 158.168686 | -8.5161956 | 2.78823194 |
| 15238 | NR_024489.1 | LOC100129534 | 14.6558335 | -5.09165 | 1.66701717 |
| 15237 | NM_002470.3 | #N/A | 14.9909564 | -5.1131111 | 1.67406085 |
| 15239 | NM_001195076.1 | #N/A | 16.8684187 | -5.2788462 | 1.72856505 |
| 15240 | NM_001122961.1 | #N/A | 20.3671904 | -5.5629708 | 1.8216341 |
| 15241 | NM_002009.3 | #N/A | 37.9086191 | 3.5924365 | 1.17644394 |
| 15242 | NM_001267804.1 | #N/A | 16.6477949 | -5.2578674 | 1.72187199 |
| 15243 | NM_001198696.1 | AIFM2 | 32.6495831 | -6.240737 | 2.0441127 |
| 15244 | NM_024693.4 | #N/A | 25.878232 | -5.9083796 | 1.93529109 |
| 15245 | NM_001304960.1 | #N/A | 16.2777992 | -5.2337827 | 1.71441831 |
| 15246 | NM_005906.5 | MAK | 20.7304528 | -5.5880366 | 1.83055934 |
| 15247 | NR_132415.1 | TRIT1 | 17.7370952 | -5.3495044 | 1.7524255 |
| 15248 | NM_018198.3 | #N/A | 264.18973 | -3.1283475 | 1.02489412 |
| 15249 | NM_001105199.1 | #N/A | 16.6234218 | -5.2586873 | 1.72293891 |
| 15250 | NM_001318802.1 | #N/A | 15.7555185 | -5.1894298 | 1.70034399 |
| 15251 | NR_136290.1 | CNIH3 | 18.5028963 | -5.415032 | 1.77478637 |
| 15252 | NM_001114734.1 | #N/A | 17.706247 | -5.3647579 | 1.75857333 |
| 15253 | NM_170733.3 | BDNF | 18.6826463 | -5.4369695 | 1.78235222 |
| 15254 | NR_024380.1 | LOC441666 | 229.932683 | 2.01570991 | 0.66081336 |
| 15255 | NM_144495.2 | PQBP1 | 17.9491176 | -5.3836165 | 1.7650153 |
| 15256 | NR_109909.1 | CTC-338M12 | 13.7575468 | -4.9896746 | 1.63608979 |
| 15257 | NM_001015055.1 | #N/A | 22.1621501 | -5.6851021 | 1.86417167 |
| 15258 | NM_001288770.1 | #N/A | 13.8479605 | -4.9951705 | 1.63813445 |
| 15259 | NM_024893.2 | #N/A | 30.5210756 | -6.142122 | 2.01425873 |
| 15260 | NM_139021.2 | #N/A | 201.244079 | -4.5679291 | 1.49837285 |
| 15262 | NM_025230.4 | DCAF11 | 127.001016 | -8.1989814 | 2.6895292 |
| 15263 | NM_145755.2 | TTC21A | 23.1327666 | -5.7372093 | 1.88200596 |
| 15261 | NM_001292030.1 | #N/A | 197.313382 | 5.86756881 | 1.9247568 |
| 15264 | NR_003521.1 | WHAMMP3 | 72.0852038 | 1.84596286 | 0.60555511 |

| | | | | | |
|---|---|---|---|---|---|
| 15265 | NR_047658.1 | #N/A | 18.9126631 | -5.458667 | 1.79104517 |
| 15266 | NM_172103.3 | EYA4 | 20.0468445 | -5.5387162 | 1.81735327 |
| 15267 | NM_001305005.2 | #N/A | 43.7732494 | -6.664148 | 2.18698294 |
| 15268 | NM_015982.3 | #N/A | 17.1823445 | -5.3137176 | 1.74448539 |
| 15269 | NM_001300946.1 | #N/A | 305.473763 | -3.1784069 | 1.04351617 |
| 15270 | NM_001320953.1 | ZNF232 | 16.3472129 | -5.2515892 | 1.72429469 |
| 15271 | NM_152927.2 | #N/A | 15.8608879 | -5.2053593 | 1.70937645 |
| 15272 | NM_000790.3 | #N/A | 17.2279815 | -5.3192572 | 1.74681296 |
| 15273 | NM_001286094.1 | TMEM9B | 19.2005576 | -5.4749111 | 1.7984106 |
| 15274 | NM_001003683.2 | PDE1A | 42.6211806 | -6.6220337 | 2.1752528 |
| 15275 | NM_001287244.1 | RALYL | 19.843929 | -5.5190175 | 1.81306812 |
| 15276 | NM_001318360.1 | #N/A | 93.0360321 | -7.7509987 | 2.54629929 |
| 15277 | NM_001303117.1 | #N/A | 17.0490682 | -5.3074874 | 1.74366072 |
| 15278 | NM_001319138.1 | #N/A | 16.4115606 | -5.2503143 | 1.72489145 |
| 15279 | NM_015840.3 | #N/A | 101.780212 | -7.8805643 | 2.58902467 |
| 15280 | NM_001330069.1 | #N/A | 24.4650909 | -5.8252622 | 1.91384131 |
| 15281 | NM_001321924.1 | #N/A | 13.3299048 | -4.9436132 | 1.62419352 |
| 15282 | NM_173480.2 | #N/A | 15.1535089 | -5.126776 | 1.68451851 |
| 15283 | NM_001006681.1 | SPIN2B | 24.0794853 | -5.7995735 | 1.90571803 |
| 15284 | NM_001135862.2 | #N/A | 124.381933 | -8.1691207 | 2.68453457 |
| 15285 | NM_001723.5 | #N/A | 15.0006583 | -5.118015 | 1.68189494 |
| 15286 | NM_206940.1 | #N/A | 49.1458894 | 2.73698734 | 0.89944771 |
| 15287 | NM_001566.2 | INPP4A | 17.1569419 | -5.3106264 | 1.74528912 |
| 15288 | NR_125408.1 | #N/A | 14.7010889 | -5.0892111 | 1.67291208 |
| 15289 | NM_000246.3 | CIITA | 22.5729615 | -5.7047081 | 1.87529562 |
| 15290 | NR_136596.1 | #N/A | 19.3315039 | -5.4891187 | 1.80454206 |
| 15291 | NM_133343.1 | RAD17 | 15.9107894 | -5.1925162 | 1.70704804 |
| 15292 | NM_001077701.2 | #N/A | 41.8156563 | -4.9864237 | 1.63943392 |
| 15293 | NM_001128921.2 | MARK3 | 19.4644167 | -5.4967805 | 1.80728133 |
| 15294 | NM_004585.4 | #N/A | 15.3360283 | -5.15725 | 1.69567897 |
| 15295 | NM_001199876.1 | SERF2 | 16.4404953 | -5.2528651 | 1.72736027 |
| 15296 | NM_152236.2 | #N/A | 31.2742362 | -6.1769362 | 2.03123162 |
| 15297 | NM_052909.3 | #N/A | 160.946342 | -2.3145554 | 0.76129015 |
| 15298 | NM_001256552.1 | NEIL1 | 14.107587 | -5.0374043 | 1.65697974 |
| 15299 | NM_001040020.1 | FAM3C | 108.945421 | -7.9785782 | 2.62482488 |
| 15300 | NM_024751.3 | GSTCD | 83.3146731 | 1.98615273 | 0.6534651 |
| 15301 | NM_001102654.1 | #N/A | 18.6144363 | -5.4359483 | 1.78852097 |
| 15302 | NM_001317843.1 | #N/A | 17.0628094 | -5.3002896 | 1.74397832 |
| 15303 | NM_001143757.1 | #N/A | 31.3643721 | -6.174101 | 2.03175256 |
| 15304 | NM_017831.3 | #N/A | 182.071908 | 1.78998641 | 0.58904445 |
| 15305 | NM_003548.2 | H4C14 | 15.3835948 | -5.1496915 | 1.69471095 |
| 15306 | NR_037675.1 | #N/A | 15.3346052 | -5.15898 | 1.69789038 |
| 15307 | NM_020204.2 | #N/A | 13.541259 | -4.9688856 | 1.63545692 |

| | | | | | |
|---|---|---|---|---|---|
| 15308 | NM_001001936.2 | #N/A | 19.165642 | -5.4678177 | 1.79987455 |
| 15309 | NM_020056.4 | #N/A | 25.5402758 | -5.8874979 | 1.93802625 |
| 15310 | NM_020981.3 | B3GALT1 | 13.9355338 | -5.0190577 | 1.65216893 |
| 15311 | NR_023353.1 | EXOSC7 | 18.2280208 | -5.3953745 | 1.77619885 |
| 15312 | NM_020439.2 | #N/A | 17.4817516 | -5.3404035 | 1.75813277 |
| 15313 | NM_022806.3 | #N/A | 27.0187484 | -5.9640393 | 1.96374888 |
| 15314 | NM_033495.3 | KLHL13 | 40.1199429 | -4.1631007 | 1.37080418 |
| 15315 | NM_007211.4 | #N/A | 27.1656862 | -4.4591604 | 1.46842472 |
| 15316 | NM_002397.4 | #N/A | 128.641645 | -8.2173998 | 2.70617306 |
| 15317 | NM_001313964.1 | DNAJB1 | 15.655852 | -5.1881748 | 1.70873218 |
| 15318 | NM_007225.2 | #N/A | 33.1158523 | -6.2574078 | 2.0608938 |
| 15319 | NM_001316362.1 | #N/A | 20.2552526 | -5.5394419 | 1.82464257 |
| 15320 | NM_001197260.1 | DOK1 | 17.7194419 | -5.3578987 | 1.76496294 |
| 15321 | NM_006053.3 | TCIRG1 | 15.9595585 | -5.2138475 | 1.71757969 |
| 15322 | NM_001008710.2 | #N/A | 18.8827887 | -5.4578969 | 1.79805242 |
| 15323 | NM_033050.5 | #N/A | 19.9150106 | -5.5295759 | 1.82175208 |
| 15324 | NM_001243611.1 | LMO3 | 18.0925607 | -5.3821013 | 1.77339981 |
| 15325 | NM_012464.4 | #N/A | 15.4475555 | -5.1652014 | 1.70196516 |
| 15326 | NM_024806.3 | #N/A | 19.4461186 | -5.4983635 | 1.811804 |
| 15327 | NM_145018.3 | #N/A | 20.9854177 | -4.3019217 | 1.4176281 |
| 15328 | NM_017823.4 | #N/A | 17.1210502 | -5.313764 | 1.75108122 |
| 15329 | NM_001730.4 | #N/A | 21.6929794 | -5.6549917 | 1.86358652 |
| 15330 | NM_001288667.1 | HCFC1R1 | 18.9158134 | -5.4478001 | 1.79547602 |
| 15331 | NM_178136.2 | #N/A | 95.9507605 | -7.7954095 | 2.56921511 |
| 15332 | NM_001318737.1 | #N/A | 19.2985111 | -5.4772416 | 1.80547719 |
| 15333 | NM_001319100.1 | #N/A | 92.4924266 | -7.7425262 | 2.55257927 |
| 15334 | NR_104006.1 | #N/A | 17.6249211 | -5.3389935 | 1.76036062 |
| 15335 | NM_001322240.1 | #N/A | 25.9680509 | -5.9012202 | 1.94595071 |
| 15336 | NM_173042.2 | IL18BP | 13.9851677 | -5.0068175 | 1.65104707 |
| 15337 | NM_001242597.1 | #N/A | 16.5658304 | -5.2579464 | 1.73388913 |
| 15338 | NR_135317.1 | SMG1P2 | 20.8172434 | -5.5868067 | 1.84273333 |
| 15339 | NM_001172428.1 | KLHL7 | 18.343193 | -5.408918 | 1.78419907 |
| 15340 | NR_138423.1 | #N/A | 15.4840485 | -5.1656826 | 1.7044149 |
| 15341 | NM_001193343.1 | MRPS18A | 19.6793932 | -5.5111523 | 1.81844897 |
| 15342 | NM_001199181.1 | #N/A | 31.9782599 | -6.2125063 | 2.04988082 |
| 15343 | NR_024455.1 | LINC01134 | 39.2275856 | 4.53083614 | 1.49500709 |
| 15344 | NM_152485.3 | #N/A | 14.7795845 | -5.0921118 | 1.68032162 |
| 15345 | NM_006648.3 | WNK2 | 270.439868 | -2.9815301 | 0.98402132 |
| 15346 | NM_001278262.1 | CARHSP1 | 14.6277337 | -5.0734934 | 1.67480286 |
| 15347 | NM_001080480.2 | #N/A | 28.4107324 | -6.0375294 | 1.99307493 |
| 15348 | NR_024014.1 | #N/A | 18.8220339 | -5.4411026 | 1.79625024 |
| 15349 | NM_152989.4 | #N/A | 39.1128404 | -6.4945294 | 2.14407624 |
| 15350 | NM_001198695.1 | #N/A | 25.5269238 | -5.8894914 | 1.94494915 |

| | | | | | |
|---|---|---|---|---|---|
| 15351 | NR_028137.1 | SLC26A4-AS1 | 44.9125295 | -6.697905 | 2.21196217 |
| 15352 | NM_020762.3 | #N/A | 16.2349067 | -5.2328698 | 1.72815157 |
| 15353 | NM_002937.4 | #N/A | 14.804688 | -5.1069248 | 1.68682752 |
| 15354 | NM_001286774.1 | OCIAD2 | 26.3822298 | -5.9306413 | 1.95904116 |
| 15356 | NR_132320.1 | FRG1FP | 28.0648236 | -4.4940247 | 1.4845978 |
| 15355 | NM_001170717.1 | #N/A | 27.2628182 | -5.9764398 | 1.97431705 |
| 15357 | NM_014657.2 | #N/A | 17.5836513 | -5.3567477 | 1.76991373 |
| 15358 | NM_001321226.1 | #N/A | 27.2341435 | -5.9840304 | 1.9773845 |
| 15359 | NM_001166269.1 | HAUS4 | 19.1515846 | -5.4726486 | 1.80841998 |
| 15360 | NM_024565.6 | #N/A | 16.0445566 | -5.2003333 | 1.71864687 |
| 15361 | NM_001300921.1 | #N/A | 18.7410536 | 3.66018755 | 1.20981388 |
| 15362 | NM_015548.4 | #N/A | 436.355972 | -3.3878364 | 1.11993262 |
| 15363 | NM_005045.3 | #N/A | 125.291019 | -8.1799898 | 2.704203 |
| 15364 | NM_031371.3 | #N/A | 19.5328056 | -5.5064619 | 1.8206281 |
| 15365 | NM_001282768.1 | KIAA1217 | 16.5129999 | -5.2547272 | 1.73742979 |
| 15366 | NM_001101340.1 | ADM5 | 13.4684035 | -4.9566234 | 1.63923596 |
| 15367 | NM_001163678.1 | SHOX2 | 14.4111526 | -5.0586368 | 1.67297062 |
| 15368 | NM_001303433.1 | FUBP1 | 70.7282382 | -3.2086367 | 1.06124199 |
| 15369 | NM_021916.3 | #N/A | 52.4575398 | -2.3933575 | 0.79162121 |
| 15370 | NM_001320825.1 | #N/A | 14.5025237 | -5.0635279 | 1.67485279 |
| 15371 | NM_152266.4 | #N/A | 15.3542177 | -5.1378089 | 1.69944237 |
| 15372 | NM_001080532.1 | #N/A | 19.2797261 | -5.4799165 | 1.81268651 |
| 15373 | NR_135105.1 | #N/A | 47.3004468 | -3.2793304 | 1.0848117 |
| 15374 | NM_198503.3 | #N/A | 31.3575199 | -6.1760597 | 2.04311911 |
| 15375 | NM_001288570.1 | TMEM9 | 17.98846 | -5.37618 | 1.77855135 |
| 15376 | NR_103718.1 | LINC00528 | 16.415802 | 4.63422624 | 1.53325176 |
| 15377 | NM_001282162.1 | #N/A | 14.6868754 | -5.0899398 | 1.68404842 |
| 15378 | NM_031478.5 | #N/A | 437.608738 | -3.805295 | 1.25924615 |
| 15379 | NR_045863.1 | #N/A | 14.1355528 | -5.0422483 | 1.66861477 |
| 15380 | NM_013304.2 | #N/A | 14.5797104 | -5.0801892 | 1.68144325 |
| 15381 | NM_012455.2 | #N/A | 14.8993942 | -5.1141198 | 1.69288733 |
| 15382 | NM_001199173.1 | #N/A | 16.0536033 | -5.2183786 | 1.72753262 |
| 15383 | NM_001170637.3 | SRGAP2 | 138.485183 | -8.3238186 | 2.75579653 |
| 15384 | NR_110900.1 | ZNF213-AS1 | 22.4379243 | -4.218376 | 1.39669001 |
| 15385 | NM_001206855.1 | #N/A | 18.7137487 | -5.4294934 | 1.79778198 |
| 15386 | NM_001172651.1 | ZNF177 | 17.3318174 | -5.3277004 | 1.76430796 |
| 15387 | NR_027157.1 | TMPO-AS1 | 16.4791204 | -5.2592451 | 1.74167684 |
| 15388 | NM_001320252.1 | #N/A | 15.4807529 | -5.1621387 | 1.70954278 |
| 15389 | NR_038393.1 | LOC100506178 | 25.066886 | 5.5962791 | 1.85341419 |
| 15390 | NM_018442.3 | DCAF6 | 19.941429 | -5.5320875 | 1.83233071 |
| 15391 | NM_001161586.1 | ME3 | 19.8368009 | -5.5122606 | 1.82601893 |
| 15392 | NM_001098832.1 | FAM104A | 24.3473082 | -5.8056612 | 1.92324999 |
| 15393 | NM_005710.2 | PQBP1 | 15.4503325 | -5.1561599 | 1.70830914 |

| | | | | | |
|---|---|---|---|---|---|
| 15394 | NM_153210.4 | #N/A | 17.5844165 | -5.35372 | 1.77378597 |
| 15395 | NR_138091.1 | ANKRD20A9P | 17.2283331 | 4.47250143 | 1.48224631 |
| 15396 | NM_001195831.2 | #N/A | 31.6279674 | -6.1962712 | 2.05370234 |
| 15397 | NR_133920.1 | #N/A | 27.7867641 | -2.7730615 | 0.91932129 |
| 15398 | NM_001039999.2 | #N/A | 13.791033 | -5.0054347 | 1.65949528 |
| 15399 | NM_001330187.1 | TBC1D30 | 15.9620876 | -5.2147543 | 1.72902813 |
| 15400 | NM_001206985.2 | METTL23 | 18.6554348 | -5.4224088 | 1.79795541 |
| 15401 | NM_001077654.2 | #N/A | 15.2527577 | -5.1466451 | 1.70668958 |
| 15402 | NM_001320048.1 | C1orf112 | 17.363157 | -5.3227118 | 1.76519788 |
| 15403 | NM_001288566.1 | TMEM9 | 17.4155562 | -5.3317029 | 1.76828458 |
| 15404 | NM_001330081.1 | #N/A | 27.6061105 | -5.994541 | 1.98818042 |
| 15405 | NM_001318808.1 | MAJIN | 14.7573766 | 7.19729689 | 2.38749468 |
| 15406 | NM_015146.1 | #N/A | 97.1530874 | -7.8132683 | 2.59200773 |
| 15407 | NM_001304458.1 | PRMT9 | 19.6440587 | -5.5016412 | 1.8253489 |
| 15408 | NR_033872.1 | PCBP1-AS1 | 30.694731 | -3.2923476 | 1.09237793 |
| 15409 | NM_001184889.1 | NIPA2 | 28.1740548 | -4.5021771 | 1.49382303 |
| 15410 | NM_001330374.1 | FGD4 | 9.23207818 | 8.16548688 | 2.7093487 |
| 15411 | NM_006532.3 | #N/A | 64.5030925 | -2.4236168 | 0.80419133 |
| 15412 | NR_131157.1 | NKILA | 18.1853434 | -5.4008226 | 1.79227775 |
| 15414 | NM_024490.3 | ATP10A | 17.2869111 | -5.3298578 | 1.76899021 |
| 15413 | NM_018373.2 | #N/A | 230.0097 | -1.0861987 | 0.36051044 |
| 15415 | NM_080798.3 | #N/A | 21.1668197 | -5.6090594 | 1.861936 |
| 15416 | NM_001281710.1 | #N/A | 19.6996907 | -5.4994614 | 1.82559267 |
| 15418 | NM_198044.2 | ZDHHC16 | 16.4442685 | -5.2479807 | 1.74213952 |
| 15417 | NM_153214.2 | #N/A | 13.8765615 | -5.0125438 | 1.66397967 |
| 15419 | NR_126012.1 | LINC01695 | 35.9655675 | 2.28200055 | 0.75756681 |
| 15420 | NM_001330381.1 | #N/A | 14.6953158 | -5.0950543 | 1.69173541 |
| 15421 | NM_000396.3 | #N/A | 28.4998547 | -3.9954931 | 1.32667471 |
| 15422 | NM_001642.2 | #N/A | 169.325268 | -8.614212 | 2.86046981 |
| 15423 | NM_024997.3 | #N/A | 15.8917187 | -5.1936385 | 1.72463687 |
| 15424 | NR_023390.1 | LINC00476 | 161.492197 | 1.11755831 | 0.37112804 |
| 15425 | NM_006862.3 | #N/A | 16.7866524 | -5.2830837 | 1.75471689 |
| 15426 | NM_001256486.1 | #N/A | 91.6590015 | -7.7294619 | 2.56747425 |
| 15427 | NM_033256.2 | #N/A | 12.9940839 | -4.9037554 | 1.62887954 |
| 15430 | NR_104317.1 | FAM234A | 13.2669039 | -4.9418719 | 1.64179654 |
| 15428 | NM_001256008.2 | #N/A | 118.436906 | -8.098213 | 2.69039252 |
| 15429 | NM_005444.2 | #N/A | 116.172311 | -8.0705075 | 2.68121308 |
| 15431 | NM_001288977.1 | #N/A | 94.226393 | -7.7692986 | 2.58195943 |
| 15432 | NM_001313990.1 | #N/A | 23.826004 | -5.7915921 | 1.92474749 |
| 15433 | NM_002739.4 | #N/A | 19.3571413 | -5.4766698 | 1.82029066 |
| 15434 | NM_001290222.1 | NQO2 | 13.3533502 | -4.9541256 | 1.64667065 |
| 15435 | NM_003396.2 | #N/A | 26.7239561 | 5.02227648 | 1.66953164 |
| 15436 | NR_136638.1 | CT75 | 12.9080766 | -4.8971283 | 1.62808573 |

| | | | | | |
|---|---|---|---|---|---|
| 15437 | NR_002330.1 | ST7-AS1 | 13.6425489 | -4.9764682 | 1.65451543 |
| 15438 | NM_001318943.1 | MEA1 | 13.082119 | -4.9182742 | 1.63522771 |
| 15439 | NM_021167.4 | #N/A | 469.774832 | 1.12698773 | 0.3747025 |
| 15440 | NM_005738.4 | #N/A | 19.2312131 | -5.4711465 | 1.81915428 |
| 15441 | NM_001204824.1 | KCNQ3 | 8.99323963 | 8.16429804 | 2.71464473 |
| 15442 | NM_001098531.2 | #N/A | 17.4634469 | -5.3362753 | 1.77456553 |
| 15443 | NM_001286056.1 | CNN3 | 15.4229771 | -5.1488579 | 1.71248998 |
| 15444 | NM_006877.3 | #N/A | 29.5897194 | -6.1011246 | 2.02921837 |
| 15445 | NM_002517.3 | #N/A | 15.1548193 | -5.1368177 | 1.70853057 |
| 15446 | NR_109947.1 | #N/A | 17.9783231 | -5.384475 | 1.7910424 |
| 15447 | NM_198830.1 | ACLY | 675.65995 | -4.3486188 | 1.44655264 |
| 15448 | NR_110141.1 | LOC101927870 | 8.34404188 | 3.43925773 | 1.14407236 |
| 15449 | NM_001267608.1 | #N/A | 20.7348754 | -5.5737 | 1.85430826 |
| 15450 | NM_025004.2 | #N/A | 12.871748 | -4.8966847 | 1.62909842 |
| 15452 | NR_003503.1 | GGT8P | 39.9214683 | 3.59787569 | 1.19704719 |
| 15451 | NM_145040.2 | #N/A | 14.8377094 | -5.1113294 | 1.70058369 |
| 15454 | NR_034137.1 | ZBED5-AS1 | 12.9062313 | -4.9070571 | 1.63266396 |
| 15453 | NR_015427.2 | #N/A | 281.661583 | -2.5762295 | 0.85715541 |
| 15455 | NR_040013.1 | LOC644554 | 15.3613734 | -5.1427631 | 1.71124179 |
| 15456 | NR_038344.1 | MAGI2-AS3 | 13.6196091 | -4.9858977 | 1.65909154 |
| 15457 | NM_024704.4 | #N/A | 16.6149303 | -5.2729413 | 1.75477029 |
| 15458 | NM_002592.2 | PCNA | 17.7247363 | -5.3553285 | 1.78230179 |
| 15459 | NM_005250.2 | #N/A | 37.5089982 | -4.8476225 | 1.61339091 |
| 15460 | NM_001243121.1 | PDE4A | 17.0485954 | -5.3125103 | 1.76815688 |
| 15461 | NM_176792.2 | #N/A | 16.7733362 | -5.2854942 | 1.75930338 |
| 15462 | NR_144490.1 | #N/A | 15.6557834 | -5.1780037 | 1.72351526 |
| 15463 | NM_001345907.1 | WDR53 | 12.9698357 | -4.9018542 | 1.63186293 |
| 15464 | NM_001136040.2 | #N/A | 122.793657 | -8.1502742 | 2.71337227 |
| 15465 | NM_001316981.1 | #N/A | 15.6476268 | -5.1725944 | 1.72211166 |
| 15466 | NM_001164384.1 | STAU2 | 17.7925343 | -5.3600895 | 1.78470407 |
| 15468 | NM_016620.3 | ZNF644 | 19.5178337 | -5.4883367 | 1.82739222 |
| 15467 | NM_007231.4 | #N/A | 44.6680298 | 4.46953466 | 1.4881741 |
| 15469 | NM_138622.3 | BCL2L11 | 15.1178655 | -5.1335723 | 1.70932697 |
| 15470 | NM_001142316.1 | #N/A | 17.5726031 | -5.3426661 | 1.77925489 |
| 15471 | NR_037144.1 | PLEKHA8P1 | 13.459551 | -4.9649352 | 1.65365849 |
| 15472 | NR_135179.1 | LOC102723692 | 18.3158389 | 7.31738808 | 2.43733185 |
| 15473 | NR_015395.2 | MIR4435-2HG | 17.8367206 | -5.3726984 | 1.78964166 |
| 15474 | NM_001319225.1 | #N/A | 16.7410692 | -5.2816218 | 1.75946843 |
| 15475 | NR_073113.1 | NNT-AS1 | 89.9947896 | -2.0575657 | 0.68558241 |
| 15476 | NM_017619.3 | #N/A | 63.5162467 | -1.7468132 | 0.58207121 |
| 15477 | NM_001271779.1 | PSMD6 | 13.4954991 | -4.9682966 | 1.65604055 |
| 15479 | NM_194252.2 | TTLL11 | 14.0417762 | -5.026465 | 1.67549242 |
| 15478 | NM_001330208.1 | #N/A | 25.4387295 | -5.872647 | 1.95755271 |

| | | | | | |
|---|---|---|---|---|---|
| 15480 | NM_001164883.1 | CTNNA2 | 19.2335539 | -5.474323 | 1.82485997 |
| 15481 | NM_001166006.1 | EPB41 | 14.9145816 | -5.1076858 | 1.70263982 |
| 15482 | NM_006952.3 | #N/A | 28.3054532 | 3.92715143 | 1.3091469 |
| 15483 | NM_001029989.2 | #N/A | 167.267727 | 4.44718876 | 1.48257778 |
| 15484 | NM_001316935.1 | HSPB11 | 16.6832078 | -5.2600608 | 1.75366097 |
| 15485 | NM_001319183.1 | #N/A | 12.7372731 | -4.8814377 | 1.62763056 |
| 15486 | NM_001142300.1 | CCNYL1 | 20.6993399 | -5.5798697 | 1.86052668 |
| 15487 | NM_000863.2 | #N/A | 19.5457195 | -5.5043838 | 1.83555573 |
| 15488 | NR_045999.1 | #N/A | 8.599864 | 8.0678452 | 2.69075527 |
| 15489 | NM_031369.2 | HNRNPD | 119.808915 | -8.1147678 | 2.70702041 |
| 15490 | NM_001170797.1 | ZFAND1 | 14.3774711 | -5.0543672 | 1.68630157 |
| 15491 | NM_001321180.1 | FAM156B | 13.1052671 | -4.9290144 | 1.64452766 |
| 15492 | NM_005061.2 | #N/A | 15.3938822 | 5.92197122 | 1.97581329 |
| 15493 | NM_145010.3 | #N/A | 17.5271192 | -5.3451668 | 1.78363714 |
| 15494 | NM_001253912.1 | #N/A | 16.7074311 | -5.2680812 | 1.75802346 |
| 15495 | NM_001184943.1 | #N/A | 128.404409 | -8.2148107 | 2.74157325 |
| 15496 | NR_136612.1 | LOC100130698 | 37.2332629 | 4.73681177 | 1.5808535 |
| 15498 | NR_138084.1 | HCG24 | 9.91134508 | 2.74989992 | 0.9177721 |
| 15497 | NR_125974.1 | WARS2-AS1 | 41.9149615 | 2.78518904 | 0.92955176 |
| 15499 | NR_034150.1 | LINC01136 | 58.6387951 | 3.76949624 | 1.25818945 |
| 15500 | NM_001317994.1 | #N/A | 17.9463859 | -5.3706268 | 1.79287138 |
| 15501 | NR_046366.2 | DDX39A | 23.7091131 | -5.7781967 | 1.92914913 |
| 15502 | NR_033819.1 | #N/A | 17.0250914 | -5.2919538 | 1.76686122 |
| 15503 | NM_001143854.1 | #N/A | 94.2116686 | -7.769193 | 2.59445456 |
| 15504 | NM_007049.4 | #N/A | 88.1792588 | -7.673737 | 2.56280136 |
| 15505 | NM_001082533.1 | CA10 | 18.3919113 | -5.4064622 | 1.80601677 |
| 15506 | NR_103478.1 | #N/A | 20.8532059 | -5.5981329 | 1.87007957 |
| 15507 | NM_014654.3 | #N/A | 1303.58055 | -2.8778119 | 0.96139226 |
| 15508 | NM_004759.4 | #N/A | 15.5297003 | -5.1760634 | 1.729252 |
| 15509 | NM_014616.2 | #N/A | 136.305389 | -3.6983368 | 1.23569449 |
| 15510 | NM_001323424.1 | #N/A | 15.8497966 | -5.1926071 | 1.73542897 |
| 15511 | NM_015275.2 | #N/A | 190.941994 | -3.3612905 | 1.12337535 |
| 15512 | NM_003673.3 | #N/A | 16.2597502 | -5.2304602 | 1.74810634 |
| 15513 | NM_001310321.1 | #N/A | 13.4437619 | -4.9566034 | 1.6567481 |
| 15514 | NR_033886.1 | RAD21-AS1 | 38.3255289 | 4.47319737 | 1.49547663 |
| 15515 | NM_012307.3 | #N/A | 16.1186588 | -5.2216175 | 1.74578564 |
| 15516 | NM_001144997.1 | ITGA7 | 20.3073537 | -5.5576557 | 1.85827988 |
| 15517 | NM_001303501.1 | #N/A | 20.1516033 | -5.5516056 | 1.85633484 |
| 15518 | NM_001279359.1 | #N/A | 30.403434 | -6.1312627 | 2.05043837 |
| 15519 | NR_024054.2 | GUSBP14 | 15.6699474 | -5.1902357 | 1.73588114 |
| 15520 | NM_001319086.1 | RUVBL1 | 15.0842998 | -5.1318751 | 1.71637621 |
| 15521 | NR_125383.1 | LINC02551 | 65.630464 | 1.88085404 | 0.62906772 |
| 15522 | NM_001306084.1 | #N/A | 24.4962244 | -5.8283319 | 1.94955079 |

| | | | | | |
|---|---|---|---|---|---|
| 15523 | NM_006309.3 | #N/A | 20.4808162 | -5.5652218 | 1.86155112 |
| 15524 | NM_004791.2 | #N/A | 21.6748686 | -5.6507976 | 1.89025551 |
| 15525 | NM_001271168.1 | #N/A | 14.8292871 | -5.0967145 | 1.70496308 |
| 15526 | NM_006469.4 | #N/A | 662.862992 | -3.5759826 | 1.19624108 |
| 15527 | NM_001320599.1 | #N/A | 18.7282624 | -5.4273837 | 1.81576444 |
| 15528 | NM_001322796.1 | IDE | 14.61834 | -5.0674918 | 1.69545108 |
| 15529 | NM_001321964.1 | #N/A | 17.3473558 | -5.3260523 | 1.78197608 |
| 15530 | NM_001130867.1 | ERCC2 | 15.0523231 | -5.1288511 | 1.71613717 |
| 15531 | NM_001164758.1 | PRKAR1B | 20.7247458 | -5.5902941 | 1.87077497 |
| 15532 | NM_054110.4 | #N/A | 18.1534629 | -5.3976176 | 1.80632137 |
| 15533 | NM_194457.1 | #N/A | 23.4342613 | -5.7581678 | 1.92701573 |
| 15534 | NM_007129.3 | #N/A | 224.306782 | -3.3031671 | 1.10571322 |
| 15536 | NM_001145364.2 | ABHD11 | 16.48371 | -5.2476242 | 1.75664536 |
| 15535 | NM_000508.4 | #N/A | 30.1630697 | 9.19299058 | 3.07736054 |
| 15537 | NR_135237.1 | LOC730100 | 24.5700458 | 3.36635057 | 1.1269382 |
| 15538 | NM_021913.4 | #N/A | 112.133692 | -8.0196444 | 2.68479902 |
| 15539 | NM_020336.3 | #N/A | 106.64572 | -7.9470804 | 2.66115947 |
| 15540 | NM_001134830.1 | AHI1 | 63.7825325 | -3.3682366 | 1.12803974 |
| 15541 | NM_001102398.2 | #N/A | 115.356536 | -8.060052 | 2.69947044 |
| 15542 | NM_001165884.2 | #N/A | 9.86429235 | 8.25854109 | 2.76618831 |
| 15544 | NM_024952.7 | DGLUCY | 15.8715479 | -5.2031943 | 1.74297769 |
| 15543 | NM_005888.3 | SLC25A3 | 15.7808647 | -5.1854067 | 1.73701737 |
| 15545 | NM_020677.4 | #N/A | 17.01141 | -5.2861813 | 1.77084012 |
| 15546 | NM_198947.3 | #N/A | 17.9898498 | -5.3874948 | 1.80540673 |
| 15547 | NM_001145409.1 | #N/A | 85.2742113 | -7.6254198 | 2.55576197 |
| 15548 | NR_110852.1 | LOC101928855 | 40.5674285 | 2.21903861 | 0.74379606 |
| 15549 | NM_001001894.2 | TTC3 | 90.1121045 | -7.7047668 | 2.58297173 |
| 15550 | NR_001282.2 | PCDHB19P | 26.4459726 | -3.8642719 | 1.29554947 |
| 15551 | NM_001303460.1 | #N/A | 14.768297 | -5.0931257 | 1.70766777 |
| 15552 | NM_001169574.1 | CXorf58 | 14.5494576 | 7.45993906 | 2.50127932 |
| 15553 | NM_001040142.1 | #N/A | 33.9379111 | -6.2950727 | 2.11077193 |
| 15554 | NR_002931.2 | CLEC4GP1 | 69.1991045 | 2.30353072 | 0.77242444 |
| 15555 | NR_132379.1 | LINC01022 | 49.1000107 | 2.83215328 | 0.94977473 |
| 15556 | NM_001199201.1 | ZBBX | 19.453074 | -5.4965458 | 1.84381661 |
| 15557 | NM_001321217.1 | #N/A | 13.5229664 | -4.9727813 | 1.66885773 |
| 15558 | NR_138050.1 | #N/A | 30.9435952 | 4.63308297 | 1.55500964 |
| 15559 | NM_001348097.1 | #N/A | 135.254406 | -8.289726 | 2.78287896 |
| 15560 | NR_134281.1 | LOC101927357 | 65.0678625 | 2.63098178 | 0.88323333 |
| 15561 | NM_017525.2 | #N/A | 31.9057448 | -6.2089634 | 2.08455863 |
| 15562 | NM_025076.4 | UXS1 | 87.0156465 | -7.6545392 | 2.57013708 |
| 15563 | NM_001278205.1 | HSPA14 | 23.2847332 | -4.2491194 | 1.42695556 |
| 15564 | NM_001083953.1 | THADA | 17.8390568 | -5.3720271 | 1.80420099 |
| 15565 | NM_001038628.1 | B3GALNT1 | 19.0174492 | -5.4518807 | 1.83106763 |

| | | | | | |
|---|---|---|---|---|---|
| 15566 | NM_001322881.1 | #N/A | 26.4114018 | 5.47721888 | 1.83963615 |
| 15567 | NM_014364.4 | #N/A | 14.4304135 | -5.053407 | 1.69747956 |
| 15568 | NM_004298.3 | #N/A | 19.1860856 | -5.4760567 | 1.83975627 |
| 15569 | NR_135502.1 | #N/A | 82.5890923 | -7.5791707 | 2.54649928 |
| 15570 | NM_006006.4 | #N/A | 13.0093458 | -4.9162725 | 1.65205196 |
| 15572 | NR_132978.1 | LOC105376331 | 15.2288735 | 4.58918148 | 1.54231893 |
| 15571 | NM_001300789.1 | PEX5 | 17.3475157 | -5.3359835 | 1.79330158 |
| 15573 | NM_001145109.1 | NELL2 | 88.4310457 | -7.6779399 | 2.58041135 |
| 15574 | NM_001136022.2 | NFATC4 | 89.3583088 | -7.6927156 | 2.5856522 |
| 15576 | NR_120449.1 | LOC101927267 | 13.2568105 | -4.9400405 | 1.6604865 |
| 15575 | NM_001124767.1 | #N/A | 15.0364399 | -5.1259175 | 1.72297518 |
| 15577 | NM_000384.2 | #N/A | 15.0102081 | 4.95961032 | 1.66739968 |
| 15578 | NR_052022.1 | #N/A | 18.7820453 | -5.443715 | 1.83021907 |
| 15579 | NM_024811.3 | CPSF7 | 16.7824982 | -5.2873367 | 1.77777208 |
| 15580 | NM_001101312.1 | #N/A | 15.0550268 | -5.1291684 | 1.72469679 |
| 15581 | NM_001100159.2 | #N/A | 15.8636502 | -5.2039951 | 1.75025965 |
| 15582 | NM_005950.2 | #N/A | 24.7445863 | -5.8348852 | 1.96246113 |
| 15583 | NM_001257197.1 | CUL3 | 27.6446396 | -5.9939962 | 2.01606908 |
| 15584 | NM_001324302.1 | RPN2 | 21.880057 | -5.6528241 | 1.90138049 |
| 15585 | NM_002692.3 | #N/A | 13.8583689 | -4.9966252 | 1.68066199 |
| 15586 | NM_001286084.1 | #N/A | 16.5602425 | -5.2656227 | 1.77122785 |
| 15587 | NM_001258462.1 | #N/A | 17.7023237 | -5.342921 | 1.79727905 |
| 15588 | NM_199242.2 | UNC13D | 13.450311 | -4.9476992 | 1.66442228 |
| 15589 | NM_001322886.1 | #N/A | 118.043606 | -8.0940083 | 2.7231142 |
| 15590 | NM_001130145.2 | #N/A | 85.479833 | -7.6288628 | 2.56675301 |
| 15591 | NM_001330443.1 | #N/A | 110.622127 | -8.0000786 | 2.69171969 |
| 15592 | NM_024684.3 | AAMDC | 15.0953257 | -5.1218728 | 1.72336498 |
| 15593 | NM_001164098.1 | VCAN | 115.334949 | -8.0604097 | 2.71214097 |
| 15594 | NM_005115.4 | #N/A | 84.7852366 | -7.6170219 | 2.5631388 |
| 15595 | NM_001199707.1 | #N/A | 13.6822868 | -4.9837986 | 1.67708402 |
| 15596 | NM_001282622.1 | #N/A | 88.6276193 | -7.6811476 | 2.58489419 |
| 15597 | NM_031916.4 | #N/A | 20.1795989 | -5.5485682 | 1.86722638 |
| 15598 | NM_001271788.1 | #N/A | 105.985606 | -7.9378339 | 2.67242 |
| 15599 | NM_198969.1 | TLE5 | 24.5552457 | -5.8321855 | 1.96358909 |
| 15600 | NM_175605.4 | #N/A | 16.0365975 | -5.2210127 | 1.75791499 |
| 15601 | NM_001137554.1 | #N/A | 298.024669 | 1.77855686 | 0.59886436 |
| 15602 | NM_001025778.1 | VRK3 | 14.9707741 | -5.1255023 | 1.72586545 |
| 15603 | NM_012177.4 | #N/A | 15.9649202 | -5.2150375 | 1.75620964 |
| 15604 | NM_024106.2 | #N/A | 107.380437 | -7.9570289 | 2.67960424 |
| 15605 | NM_180699.3 | SNRNP35 | 13.3389002 | -4.9564249 | 1.66922713 |
| 15606 | NM_001163942.1 | #N/A | 29.7811227 | 2.43627832 | 0.82049981 |
| 15607 | NR_104194.1 | #N/A | 18.7412369 | -5.4375563 | 1.83156071 |
| 15608 | NM_199440.1 | HSPD1 | 17.2223535 | -5.3269858 | 1.79435426 |

| | | | | | |
|---|---|---|---|---|---|
| 15609 | NR_039987.1 | LINC01005 | 14.272036 | 4.27703824 | 1.44070584 |
| 15610 | NM_001242607.1 | #N/A | 32.9175931 | -6.2479757 | 2.10483079 |
| 15611 | NM_001293274.1 | CACHD1 | 13.7251847 | -4.9938268 | 1.68247315 |
| 15612 | NM_001286589.1 | AIG1 | 13.0903857 | -4.9146545 | 1.65600741 |
| 15613 | NM_177400.2 | #N/A | 19.6070138 | -5.5056896 | 1.855693 |
| 15614 | NM_176875.3 | #N/A | 15.2522034 | -5.141489 | 1.73309174 |
| 15615 | NM_006954.1 | #N/A | 106.731955 | -7.9483124 | 2.67998491 |
| 15616 | NM_007050.5 | #N/A | 46.0626398 | -3.5064462 | 1.18229696 |
| 15617 | NR_002822.3 | #N/A | 13.846008 | -5.006259 | 1.68813171 |
| 15618 | NM_033413.3 | #N/A | 25.4535576 | -5.8848384 | 1.98463232 |
| 15619 | NM_001012301.3 | #N/A | 13.0923254 | -4.9294424 | 1.66294649 |
| 15620 | NM_153257.4 | #N/A | 24.2021154 | -5.8013477 | 1.9571171 |
| 15621 | NM_001168331.1 | #N/A | 12.9441822 | -4.9020365 | 1.65386488 |
| 15622 | NM_015459.4 | #N/A | 80.3581231 | -7.5397077 | 2.54376747 |
| 15623 | NM_130386.2 | #N/A | 379.152249 | -4.0285893 | 1.35916899 |
| 15624 | NR_026789.1 | FAM66A | 13.9116235 | -5.0012027 | 1.68734402 |
| 15625 | NM_015944.3 | #N/A | 16.9443169 | -5.2993301 | 1.78798861 |
| 15626 | NM_032276.3 | #N/A | 30.967471 | -4.5996034 | 1.55189312 |
| 15627 | NM_006207.2 | PDGFRL | 14.0205004 | -5.027378 | 1.69657833 |
| 15628 | NM_001005212.3 | #N/A | 9.96910379 | 7.56000594 | 2.55146392 |
| 15629 | NM_001207011.1 | CNTFR | 18.6755927 | -5.4375833 | 1.83523941 |
| 15630 | NR_136300.1 | #N/A | 56.2251512 | 2.61425276 | 0.88236462 |
| 15631 | NR_033753.2 | LRRC37A11P | 21.1190089 | 4.62027357 | 1.55955701 |
| 15632 | NM_133638.4 | #N/A | 43.3996262 | 3.78756826 | 1.27871437 |
| 15633 | NM_001284242.1 | #N/A | 12.6196917 | 8.61636255 | 2.9095047 |
| 15634 | NM_005108.3 | #N/A | 13.0840209 | -4.9147336 | 1.65956478 |
| 15635 | NM_001207020.1 | #N/A | 14.4964057 | -5.063133 | 1.70983042 |
| 15636 | NM_001198547.1 | NOL4 | 23.2993376 | -5.7515826 | 1.94240683 |
| 15637 | NM_015303.3 | VPS8 | 87.704482 | -7.6656629 | 2.58889463 |
| 15638 | NM_001113512.2 | ARHGEF7 | 32.578777 | -6.2335146 | 2.10539703 |
| 15639 | NM_001007067.1 | SDCBP | 18.5844012 | -5.43001 | 1.83443594 |
| 15640 | NM_148177.2 | FBXO32 | 32.7806046 | 6.77236578 | 2.28836501 |
| 15641 | NM_001030018.1 | #N/A | 31.2050665 | -6.1744779 | 2.08671866 |
| 15642 | NM_006272.2 | #N/A | 214.536473 | -2.2958396 | 0.77599634 |
| 15643 | NM_001079522.1 | #N/A | 109.052008 | -7.9791515 | 2.697099 |
| 15644 | NM_001163692.1 | UBAP1L | 12.9824821 | -4.9050602 | 1.65815225 |
| 15645 | NM_031923.3 | #N/A | 80.4247776 | -3.8414904 | 1.29884556 |
| 15646 | NR_040733.1 | #N/A | 18.0396473 | -5.3885794 | 1.82191955 |
| 15647 | NM_001145293.1 | NGLY1 | 20.1792397 | -5.5450317 | 1.87500052 |
| 15648 | NR_047510.2 | N6AMT1 | 14.1844196 | -5.0360245 | 1.70311449 |
| 15649 | NM_017559.2 | #N/A | 16.7071358 | -5.2697827 | 1.78220527 |
| 15650 | NM_001308358.1 | CIAPIN1 | 23.772761 | -5.7812593 | 1.95528609 |
| 15651 | NM_001004719.2 | OR4M2 | 20.9898017 | 8.60958212 | 2.91231733 |

| | | | | | |
|---|---|---|---|---|---|
| 15652 | NM_018923.2 | PCDHGB2 | 16.9789717 | -5.2989581 | 1.79249496 |
| 15653 | NM_001242618.1 | #N/A | 16.0236929 | -5.2101152 | 1.76255646 |
| 15654 | NR_024187.2 | SUZ12P1 | 18.2695743 | -5.3941185 | 1.82515547 |
| 15655 | NM_001318872.1 | FARS2 | 14.2248167 | -5.0499542 | 1.70874216 |
| 15656 | NM_015920.3 | #N/A | 196.446075 | -3.894892 | 1.31802858 |
| 15657 | NM_001320676.1 | SETMAR | 18.4338772 | -5.4177221 | 1.83341776 |
| 15658 | NM_198484.4 | #N/A | 104.436748 | -7.9168415 | 2.67952645 |
| 15659 | NR_033667.1 | GYG2P1 | 20.3410143 | 2.35554647 | 0.79730827 |
| 15660 | NM_152318.2 | #N/A | 19.7247308 | -5.5073747 | 1.86487818 |
| 15661 | NM_001171930.1 | #N/A | 20.5547613 | -5.5719447 | 1.88694237 |
| 15662 | NM_000224.2 | #N/A | 194.511069 | -3.8815077 | 1.31459034 |
| 15663 | NR_024071.1 | #N/A | 12.9461535 | -4.9041272 | 1.66092164 |
| 15664 | NR_132417.1 | MHENCR | 14.5326174 | -5.0788445 | 1.72026604 |
| 15665 | NM_000670.4 | ADH4 | 34.6075187 | 4.24505286 | 1.43803992 |
| 15667 | NR_120596.1 | LINC01315 | 14.5859847 | -5.0714375 | 1.71801016 |
| 15666 | NM_001104546.1 | #N/A | 35.8627509 | -3.403603 | 1.15300634 |
| 15668 | NR_003099.1 | ZNF273 | 20.7227075 | -5.5847329 | 1.89192992 |
| 15669 | NM_001304834.1 | #N/A | 13.7808932 | -5.0066169 | 1.69622865 |
| 15670 | NM_003102.2 | #N/A | 21.1212062 | -5.6165405 | 1.90295607 |
| 15671 | NM_001164446.2 | #N/A | 26.2618127 | 2.2809844 | 0.77287253 |
| 15672 | NM_181291.2 | #N/A | 17.6481248 | -5.3401768 | 1.80945041 |
| 15673 | NM_199037.4 | #N/A | 91.8475046 | 1.80897048 | 0.61297031 |
| 15674 | NM_016824.4 | #N/A | 84.7227955 | -7.6160234 | 2.5807884 |
| 15675 | NM_001330259.1 | #N/A | 305.288783 | 0.94929113 | 0.32168898 |
| 15676 | NM_018299.4 | #N/A | 85.5608678 | -7.6301409 | 2.58563789 |
| 15677 | NM_014722.4 | #N/A | 33.3848208 | -6.2668525 | 2.12388849 |
| 15678 | NM_001001709.2 | #N/A | 20.7170623 | 7.9327783 | 2.68883712 |
| 15679 | NM_001042599.1 | ERBB4 | 19.626287 | -5.5126241 | 1.86853529 |
| 15680 | NR_037631.1 | CRIM1-DT | 18.8096295 | -5.4431464 | 1.84502558 |
| 15681 | NM_006750.3 | SNTB2 | 127.916852 | -1.1384674 | 0.3859157 |
| 15682 | NM_001206847.1 | #N/A | 20.0702537 | -5.5344259 | 1.87612143 |
| 15683 | NM_001142482.1 | NREP | 88.775523 | -7.6832628 | 2.60477991 |
| 15684 | NM_130391.3 | PTPRD | 117.148448 | -8.0823071 | 2.74006832 |
| 15685 | NR_145426.1 | RUSC1-AS1 | 15.5357707 | -5.1712218 | 1.75315672 |
| 15686 | NR_027288.1 | LOC341056 | 20.6784707 | 8.5898619 | 2.91231521 |
| 15687 | NM_152682.3 | #N/A | 80.7790504 | -7.5472733 | 2.55914693 |
| 15688 | NM_001178079.1 | #N/A | 19.6247298 | -5.5022619 | 1.86601429 |
| 15689 | NM_001191033.1 | GAREM2 | 42.1681178 | -2.3218561 | 0.78752011 |
| 15690 | NM_024070.3 | PVRIG | 13.1872505 | -4.9368167 | 1.67452981 |
| 15691 | NM_001320572.1 | #N/A | 107.546863 | -7.9598627 | 2.7001107 |
| 15692 | NM_001136136.1 | RPL28 | 17.7628815 | -5.3562575 | 1.81724782 |
| 15693 | NM_001305182.1 | RSPRY1 | 13.1639311 | -4.9369548 | 1.67508042 |
| 15694 | NM_005577.2 | #N/A | 15.9815475 | 6.45705519 | 2.1908812 |

| | | | | | |
|---|---|---|---|---|---|
| 15695 | NM_172209.2 | #N/A | 265.360379 | 1.92732457 | 0.65397632 |
| 15696 | NM_001314014.1 | #N/A | 17.8066839 | -5.3542876 | 1.81689528 |
| 15697 | NM_001323954.1 | MBD1 | 19.1034432 | -5.4647595 | 1.85445905 |
| 15698 | NM_001308014.1 | #N/A | 25.381942 | 8.87492706 | 3.0119972 |
| 15699 | NM_001204837.1 | #N/A | 24.8567783 | 6.75963885 | 2.29433865 |
| 15700 | NR_138426.1 | #N/A | 21.1515339 | -5.6053759 | 1.90261513 |
| 15701 | NM_032918.2 | #N/A | 13.468113 | -4.9585348 | 1.68318723 |
| 15702 | NR_134869.1 | #N/A | 14.1335853 | -5.0241364 | 1.70557328 |
| 15703 | NR_046381.1 | #N/A | 16.5374175 | -5.2624373 | 1.78656928 |
| 15704 | NM_182542.2 | #N/A | 11.7935005 | 6.00263582 | 2.03793736 |
| 15705 | NM_139355.2 | #N/A | 14.0852159 | -5.0185351 | 1.70394689 |
| 15706 | NM_015343.4 | #N/A | 184.497979 | -2.3293032 | 0.79090455 |
| 15707 | NM_022908.2 | #N/A | 15.6329019 | -5.1725041 | 1.7564069 |
| 15708 | NM_003116.2 | #N/A | 18.0330936 | -5.3888273 | 1.8299275 |
| 15709 | NM_020937.3 | #N/A | 20.1663559 | -5.536674 | 1.88015104 |
| 15710 | NM_001243194.1 | INPP5F | 508.498522 | -3.5424464 | 1.20297798 |
| 15712 | NM_015044.4 | GGA2 | 418.427374 | -1.8743183 | 0.63651046 |
| 15711 | NM_001330383.1 | #N/A | 13.3164734 | -4.9479098 | 1.68029232 |
| 15713 | NM_001243228.1 | #N/A | 23.1459542 | -5.7425201 | 1.95024328 |
| 15714 | NM_001308393.1 | #N/A | 16.6799079 | -5.2811087 | 1.79390353 |
| 15715 | NM_175932.2 | #N/A | 17.6436709 | -5.3592994 | 1.82052813 |
| 15716 | NM_001142683.2 | CCDC121 | 14.3600082 | -5.0652574 | 1.7207157 |
| 15717 | NM_001297426.1 | TMEM175 | 17.7356148 | -5.3534061 | 1.81877159 |
| 15718 | NR_135060.1 | DCP1B | 63.4863826 | 2.28380774 | 0.77608585 |
| 15719 | NR_033385.1 | #N/A | 14.8633827 | -5.1122583 | 1.73733091 |
| 15720 | NM_181435.5 | #N/A | 21.5541912 | -5.6339499 | 1.91496204 |
| 15721 | NM_003482.3 | KMT2D | 396.420778 | -1.5653941 | 0.53210443 |
| 15723 | NR_102708.1 | LINC00678 | 20.081702 | 3.48174419 | 1.18350075 |
| 15722 | NM_015289.3 | #N/A | 460.648173 | -4.0281639 | 1.36923736 |
| 15724 | NM_001204423.1 | #N/A | 19.0495945 | -5.4535345 | 1.85384022 |
| 15725 | NM_080823.3 | SRMS | 13.0711366 | -4.9084409 | 1.66860006 |
| 15726 | NM_138468.5 | #N/A | 24.4986162 | -5.8173095 | 1.97781618 |
| 15727 | NM_005933.3 | #N/A | 363.503051 | -3.9670346 | 1.3489834 |
| 15728 | NM_031361.2 | #N/A | 78.8182739 | -7.5119169 | 2.55452499 |
| 15729 | NM_001304959.1 | #N/A | 35.1560907 | -4.7501103 | 1.61535679 |
| 15730 | NM_015133.4 | #N/A | 125.027892 | -8.175948 | 2.78102735 |
| 15731 | NM_001135212.1 | #N/A | 32.2224512 | -4.1803012 | 1.42202673 |
| 15732 | NM_001278173.1 | ZNF33A | 14.2048572 | -5.0423769 | 1.71540436 |
| 15734 | NM_183237.2 | RNF7 | 16.5702666 | -5.2627864 | 1.79067103 |
| 15733 | NM_001271427.1 | #N/A | 111.242066 | -8.0083217 | 2.7248323 |
| 15735 | NM_001168379.1 | ZIC4 | 29.5082858 | -6.09385 | 2.07384276 |
| 15736 | NM_006732.2 | #N/A | 23.4240758 | -5.7636491 | 1.96173763 |
| 15737 | NR_038281.1 | PTPRG-AS1 | 14.2332775 | -5.0285549 | 1.7115541 |

| | | | | | |
|---|---|---|---|---|---|
| 15738 | NM_001287349.1 | #N/A | 12.5105244 | -4.8610991 | 1.65465776 |
| 15739 | NM_001130846.3 | AIFM1 | 21.5998636 | -4.3673279 | 1.48661212 |
| 15740 | NM_172159.3 | KCNAB1 | 16.206886 | -5.2179003 | 1.77625879 |
| 15741 | NM_033290.3 | #N/A | 76.7822452 | -7.4740848 | 2.54437396 |
| 15742 | NR_038357.1 | ZSWIM8-AS1 | 12.2910264 | -4.8286312 | 1.64393179 |
| 15743 | NM_004485.3 | #N/A | 85.7305272 | 5.5047282 | 1.87418205 |
| 15744 | NM_001144989.1 | #N/A | 212.656509 | 1.03088803 | 0.35100116 |
| 15745 | NM_015465.4 | #N/A | 76.9361798 | -7.4770013 | 2.5458475 |
| 15746 | NM_134261.2 | #N/A | 21.8056709 | -5.6560693 | 1.9258677 |
| 15747 | NR_073411.1 | #N/A | 14.3526144 | -5.0393111 | 1.71594146 |
| 15748 | NM_001172677.1 | ZNF607 | 15.6029044 | -5.1600768 | 1.75740304 |
| 15749 | NM_138378.2 | #N/A | 20.8609301 | -5.5979669 | 1.90672843 |
| 15750 | NM_152688.3 | #N/A | 22.0088715 | -5.6670615 | 1.93032212 |
| 15751 | NR_003528.3 | ZNF658B | 12.9816951 | -4.9083912 | 1.67198522 |
| 15752 | NM_006264.2 | #N/A | 83.4001201 | -7.5932478 | 2.5866308 |
| 15753 | NM_001320034.1 | #N/A | 82.9423903 | -7.5852992 | 2.58433486 |
| 15754 | NM_001271950.1 | TPGS2 | 18.2546693 | -5.3954271 | 1.8382516 |
| 15755 | NM_001324086.1 | #N/A | 23.6577584 | -5.7652284 | 1.96463614 |
| 15756 | NM_006749.4 | #N/A | 18.2652255 | -5.4001979 | 1.84023602 |
| 15757 | NM_139354.2 | #N/A | 15.307317 | -5.1373441 | 1.75066341 |
| 15758 | NR_133914.1 | PCF11-AS1 | 11.9751889 | -4.796222 | 1.63457499 |
| 15759 | NR_034126.1 | LINC01160 | 85.7811896 | 1.5177368 | 0.51730587 |
| 15760 | NR_027027.1 | GUSBP1 | 15.8031426 | -5.1940667 | 1.77042592 |
| 15761 | NM_001002021.2 | #N/A | 14.0231408 | -5.0235907 | 1.7124119 |
| 15762 | NM_033177.3 | #N/A | 19.682746 | -5.5114513 | 1.87871599 |
| 15763 | NM_001271754.1 | #N/A | 21.1318026 | -5.613161 | 1.91398071 |
| 15764 | NM_152349.2 | #N/A | 20.2952274 | -5.5500961 | 1.8925903 |
| 15765 | NR_015415.1 | DICER1-AS1 | 21.2681934 | -4.1268292 | 1.4072994 |
| 15766 | NR_024199.2 | CBLL1 | 17.1398363 | -5.3146258 | 1.81247901 |
| 15767 | NM_000234.2 | #N/A | 77.3819943 | -7.4853061 | 2.55297481 |
| 15768 | NM_152373.3 | #N/A | 13.2471373 | -4.9305676 | 1.68163611 |
| 15769 | NM_001256165.1 | TMCO1 | 22.0189776 | -5.663636 | 1.93201192 |
| 15770 | NM_001144932.2 | PSMB5 | 15.2093775 | -5.1478002 | 1.75613603 |
| 15771 | NM_001291821.1 | CBWD3 | 18.0227401 | -5.3703998 | 1.83216123 |
| 15772 | NM_000602.4 | #N/A | 273.712578 | -2.9904551 | 1.02023108 |
| 15773 | NM_013327.4 | PARVB | 14.728912 | -5.0973504 | 1.73935804 |
| 15774 | NM_001142699.1 | DLG2 | 16.6802828 | -5.2650935 | 1.79681079 |
| 15775 | NM_001079685.2 | KIAA1191 | 99.4991236 | -7.8470145 | 2.67818923 |
| 15776 | NM_001297671.1 | #N/A | 77.0084503 | -7.4783321 | 2.55252094 |
| 15777 | NM_001321433.1 | TMBIM1 | 23.8388886 | -3.7160454 | 1.26847269 |
| 15778 | NM_080702.2 | #N/A | 135.073065 | -8.28754 | 2.8291667 |
| 15779 | NM_001280797.1 | GLS2 | 15.843591 | -5.1818482 | 1.76908661 |
| 15780 | NM_001145054.1 | C2orf81 | 12.2780737 | -4.8327679 | 1.65012791 |

| | | | | | |
|---|---|---|---|---|---|
| 15781 | NM_001286072.1 | MOV10 | 14.0368992 | -5.0314193 | 1.71819734 |
| 15782 | NR_024629.1 | IDI2-AS1 | 12.004014 | -4.7991949 | 1.63907088 |
| 15784 | NM_001330245.1 | SLC39A13 | 92.014935 | -7.7344181 | 2.64186333 |
| 15783 | NM_001306158.1 | #N/A | 84.0612144 | -7.6046857 | 2.59753591 |
| 15785 | NM_001039664.1 | TNFRSF25 | 12.5178159 | -4.8600401 | 1.66010454 |
| 15786 | NR_024355.1 | LINC00634 | 20.3980059 | -5.5560449 | 1.8980839 |
| 15787 | NM_001178087.1 | SERF1B | 81.2220513 | -7.5551572 | 2.58115076 |
| 15789 | NR_038219.1 | LINC01010 | 22.4499603 | 4.8980282 | 1.67339713 |
| 15788 | NM_001145710.1 | #N/A | 12.3107829 | -4.8309714 | 1.650487 |
| 15790 | NM_182513.3 | SPC24 | 186.487281 | 1.394184 | 0.47633865 |
| 15791 | NM_001318939.1 | #N/A | 13.1460356 | -4.9228659 | 1.681991 |
| 15792 | NM_001374.2 | #N/A | 17.3826369 | -5.3321596 | 1.8220332 |
| 15793 | NR_133644.1 | #N/A | 27.6526271 | -5.9940115 | 2.0482076 |
| 15794 | NM_001001669.2 | #N/A | 14.6077872 | -5.0876229 | 1.73854545 |
| 15795 | NM_001319658.1 | #N/A | 15.3977307 | -5.1441523 | 1.75787773 |
| 15796 | NM_206886.4 | CCDC18 | 14.079402 | -5.0182402 | 1.71489039 |
| 15797 | NR_125963.1 | LOC101928718 | 12.2335646 | -4.8229907 | 1.64832081 |
| 15798 | NM_001195156.1 | TMEM35B | 14.0865123 | -5.0372245 | 1.72162884 |
| 15799 | NM_001202560.2 | #N/A | 16.2597573 | -5.2256741 | 1.7862319 |
| 15800 | NM_174894.2 | #N/A | 19.1954819 | -5.4771294 | 1.87235316 |
| 15801 | NM_144722.3 | #N/A | 18.907896 | -5.4560974 | 1.86550872 |
| 15802 | NM_144653.4 | #N/A | 286.103274 | -3.1102989 | 1.06351826 |
| 15803 | NM_207514.2 | #N/A | 28.8301609 | -6.0641932 | 2.0737035 |
| 15804 | NR_027788.1 | ZNF767P | 64.6189457 | -2.5542186 | 0.87344668 |
| 15805 | NM_005296.2 | #N/A | 13.5504037 | -4.9750279 | 1.70132971 |
| 15806 | NM_001282144.1 | #N/A | 22.1423135 | -5.6860668 | 1.94454629 |
| 15807 | NM_020386.4 | #N/A | 12.1030869 | -4.8135029 | 1.64626827 |
| 15808 | NR_024252.1 | FAM86HP | 14.1753511 | -5.0353089 | 1.72243736 |
| 15809 | NM_001288623.1 | ICA1L | 105.952295 | -1.4404715 | 0.49274875 |
| 15810 | NM_002667.4 | #N/A | 32.2796689 | 6.92051416 | 2.36747194 |
| 15811 | NM_175733.3 | #N/A | 20.8737909 | -3.5029957 | 1.19852694 |
| 15812 | NM_004446.2 | #N/A | 670.116966 | -4.1330914 | 1.4141318 |
| 15814 | NR_144395.1 | SUZ12P1 | 329.795611 | 1.35557526 | 0.46388503 |
| 15813 | NM_001123384.1 | #N/A | 17.4326769 | -5.3396414 | 1.82724966 |
| 15816 | NR_036506.1 | PPIAP30 | 11.6374849 | 5.40493151 | 1.84962682 |
| 15815 | NM_001146106.1 | #N/A | 16.4118283 | -5.2557854 | 1.79858634 |
| 15818 | NR_136514.1 | LOC105371998 | 13.6466432 | -4.9706585 | 1.70118011 |
| 15817 | NR_033884.1 | #N/A | 153.418166 | 1.15511868 | 0.39533178 |
| 15819 | NM_001256197.1 | ARAF | 13.0142052 | -4.9094017 | 1.68023247 |
| 15820 | NM_001330446.1 | FAM118B | 28.9752183 | -6.0635952 | 2.07554025 |
| 15821 | NR_029401.1 | LINC01347 | 93.9534952 | -3.1075268 | 1.06387998 |
| 15823 | NR_110128.1 | ZNF276 | 18.0082298 | -5.3864156 | 1.84412766 |
| 15822 | NM_001330058.1 | #N/A | 97.8517629 | -7.8229457 | 2.67831451 |

| | | | | | |
|---|---|---|---|---|---|
| 15824 | NR_002216.1 | NXF4 | 14.761776 | 6.80195771 | 2.32895209 |
| 15826 | NM_001143964.3 | TBC1D7 | 21.2914707 | -5.6220954 | 1.92504577 |
| 15825 | NM_000266.3 | #N/A | 22.47442 | -5.6975869 | 1.95088424 |
| 15827 | NR_037902.1 | #N/A | 35.8551099 | 2.37112909 | 0.81194435 |
| 15828 | NM_022571.5 | #N/A | 16.6305335 | -5.2730922 | 1.80583175 |
| 15829 | NM_001198954.1 | RNF103-CHMP3 | 14.6524608 | -5.092982 | 1.74418102 |
| 15830 | NR_033179.1 | #N/A | 17.4686114 | -5.3274945 | 1.82449064 |
| 15831 | NM_015073.2 | #N/A | 171.995384 | -2.6785691 | 0.91762045 |
| 15832 | NM_020453.3 | #N/A | 74.3036483 | -2.3208918 | 0.79510152 |
| 15833 | NM_001204424.1 | #N/A | 18.5836272 | -5.4307771 | 1.86052162 |
| 15834 | NM_001323329.1 | #N/A | 101.711253 | -7.8784052 | 2.69952344 |
| 15835 | NM_001303111.1 | #N/A | 15.1243848 | -5.1218899 | 1.7550376 |
| 15836 | NM_001269054.1 | CTCFL | 25.4727231 | 3.26405314 | 1.11884148 |
| 15837 | NM_001145373.2 | OTUD1 | 57.7634299 | -2.727833 | 0.93510798 |
| 15838 | NM_080657.4 | #N/A | 27.3259437 | -5.9860887 | 2.05215705 |
| 15839 | NM_001271100.1 | #N/A | 25.4429491 | -5.8816661 | 2.01641703 |
| 15840 | NR_037862.1 | #N/A | 19.2270234 | -5.4805314 | 1.87897419 |
| 15841 | NR_024130.2 | #N/A | 25.2998024 | -3.5397196 | 1.21374204 |
| 15842 | NM_198480.3 | ZNF615 | 17.6113767 | -5.3388636 | 1.83087591 |
| 15843 | NM_001318203.1 | VTI1A | 29.6812008 | -6.1013617 | 2.09248888 |
| 15844 | NM_001080383.1 | GJC1 | 80.582369 | 6.71972162 | 2.30472492 |
| 15845 | NM_001284333.1 | #N/A | 20.1958609 | -5.5551715 | 1.90535211 |
| 15846 | NM_012242.2 | #N/A | 20.4317461 | -5.565499 | 1.90916277 |
| 15847 | NM_001270425.1 | #N/A | 72.8958602 | -7.3992127 | 2.53860133 |
| 15848 | NM_001129840.1 | #N/A | 97.5933658 | -7.8192927 | 2.68304316 |
| 15849 | NM_001282951.2 | COPS7B | 15.3120645 | -5.153054 | 1.76818593 |
| 15850 | NM_001135196.1 | #N/A | 13.039863 | 7.02064494 | 2.40917466 |
| 15851 | NM_001271082.1 | NKD2 | 14.3567941 | -5.0531925 | 1.73417464 |
| 15852 | NM_001935.3 | #N/A | 13.0677441 | -4.913088 | 1.68613627 |
| 15853 | NM_001282286.1 | DYRK4 | 22.3389587 | -5.6856133 | 1.95131432 |
| 15854 | NM_054014.3 | FKBP1A | 12.7068593 | -4.8773291 | 1.67395103 |
| 15855 | NM_001258379.1 | #N/A | 91.8485916 | -7.7318691 | 2.65370646 |
| 15856 | NM_005178.4 | #N/A | 22.5972758 | -4.2021496 | 1.44229488 |
| 15857 | NM_001321210.1 | #N/A | 13.0037461 | -4.9169045 | 1.68769095 |
| 15858 | NM_001146339.1 | VSTM2B | 38.183371 | -4.8406438 | 1.66158928 |
| 15859 | NM_001320966.1 | #N/A | 237.287733 | -7.1303876 | 2.44756524 |
| 15860 | NM_001321905.1 | #N/A | 42.9378404 | 10.4072836 | 3.57240671 |
| 15861 | NM_001203251.1 | #N/A | 23.8065339 | -5.7798126 | 1.98399482 |
| 15862 | NM_001012758.2 | #N/A | 12.9669794 | -4.8993247 | 1.68185327 |
| 15863 | NM_032307.4 | #N/A | 12.566767 | -4.8649034 | 1.67011552 |
| 15864 | NM_015198.4 | #N/A | 14.4397311 | -5.0731664 | 1.74170552 |
| 15865 | NM_176805.3 | #N/A | 50.1932593 | 4.3898015 | 1.50711677 |
| 15866 | NM_001130065.1 | #N/A | 106.653688 | -7.948025 | 2.72917501 |

| | | | | | |
|---|---|---|---|---|---|
| 15867 | NR_135218.1 | #N/A | 17.6378911 | -5.3608667 | 1.84105621 |
| 15868 | NM_183421.1 | #N/A | 23.2017418 | -5.7496763 | 1.97470356 |
| 15869 | NM_201566.2 | #N/A | 14.4395276 | -5.0596846 | 1.73786451 |
| 15870 | NM_023923.3 | #N/A | 108.88267 | 3.79370692 | 1.30307756 |
| 15871 | NM_001012455.1 | #N/A | 13.0311598 | -4.9093041 | 1.68636146 |
| 15872 | NM_145799.3 | #N/A | 15.7878351 | -5.1990334 | 1.78611085 |
| 15873 | NM_002552.4 | #N/A | 74.2854724 | -7.4264582 | 2.55147022 |
| 15874 | NM_033326.3 | SOX6 | 21.4727037 | -4.3368078 | 1.4901359 |
| 15875 | NM_001284301.2 | #N/A | 16.9677052 | -5.3044037 | 1.82272341 |
| 15876 | NM_004496.3 | #N/A | 17.5871348 | -5.3485808 | 1.8380261 |
| 15877 | NM_001330614.1 | DOCK7 | 98.319069 | -7.8300649 | 2.69085822 |
| 15878 | NM_001134363.2 | #N/A | 53.6584349 | -2.7495639 | 0.94493567 |
| 15879 | NM_033518.3 | #N/A | 15.1668273 | -5.1215786 | 1.7601358 |
| 15880 | NM_016445.2 | #N/A | 14.3341063 | 5.47255944 | 1.88103581 |
| 15881 | NR_052852.1 | #N/A | 143.309485 | -8.3738793 | 2.87844938 |
| 15882 | NM_001012967.2 | DDX60L | 12.0023222 | -4.798269 | 1.64940274 |
| 15883 | NM_003806.2 | #N/A | 15.5996559 | -5.1796513 | 1.78050662 |
| 15884 | NR_130972.1 | PPP2R3C | 17.5027608 | -5.34015 | 1.83575967 |
| 15885 | NR_002776.4 | MCM3AP-AS1 | 13.5120591 | -4.9567086 | 1.70396564 |
| 15886 | NM_002641.3 | PIGA | 17.6251254 | -5.3568434 | 1.84168694 |
| 15887 | NM_001024960.2 | ARPC4 | 111.780344 | -8.0151777 | 2.7557174 |
| 15888 | NM_001326580.1 | #N/A | 16.4995031 | -5.2403678 | 1.80173158 |
| 15889 | NM_001207044.1 | #N/A | 611.185232 | -3.7253451 | 1.28129451 |
| 15890 | NM_001195056.1 | DDIT3 | 15.6683428 | -5.1902158 | 1.78545369 |
| 15891 | NR_027175.1 | SNHG29 | 12.4975706 | -4.8478484 | 1.66775559 |
| 15892 | NM_001319297.1 | #N/A | 126.359611 | 2.67696931 | 0.92094 |
| 15893 | NM_030773.3 | #N/A | 16.1766419 | -5.219805 | 1.79579054 |
| 15894 | NM_001003811.1 | #N/A | 20.9371487 | 4.38563169 | 1.50888194 |
| 15895 | NM_144682.5 | #N/A | 47.3485179 | 2.48476032 | 0.85488672 |
| 15896 | NM_001128933.2 | #N/A | 30.3989209 | -3.7826017 | 1.3014272 |
| 15897 | NM_003488.3 | #N/A | 192.120124 | -3.2656367 | 1.12375487 |
| 15898 | NM_001253723.1 | DPYSL5 | 199.765325 | -3.7876553 | 1.30350111 |
| 15899 | NM_001184742.1 | #N/A | 14.3723878 | -5.0492052 | 1.73768427 |
| 15900 | NM_001167672.2 | #N/A | 498.751502 | 1.86565026 | 0.64219527 |
| 15901 | NM_001330640.1 | #N/A | 59.0325152 | -3.2518177 | 1.11949746 |
| 15902 | NM_205835.3 | LSR | 17.1393468 | -5.3123017 | 1.82899183 |
| 15903 | NM_014567.3 | #N/A | 74.4356247 | -7.4292653 | 2.55826811 |
| 15904 | NM_006866.3 | #N/A | 9.17897179 | 6.19655568 | 2.13389187 |
| 15905 | NR_040515.1 | #N/A | 151.351527 | 1.05025388 | 0.36169176 |
| 15906 | NM_138769.2 | #N/A | 440.930581 | -3.1973048 | 1.10113387 |
| 15907 | NM_006638.3 | #N/A | 16.5693655 | -5.254656 | 1.80984819 |
| 15908 | NM_201613.2 | #N/A | 13.1570032 | -4.9255584 | 1.69665248 |
| 15909 | NM_018156.3 | #N/A | 104.00256 | -7.9112007 | 2.72560492 |

| | | | | | |
|---|---|---|---|---|---|
| 15910 | NM_001034955.1 | SORBS1 | 15.00648 | -5.1073625 | 1.75992434 |
| 15911 | NM_001302621.1 | #N/A | 19.0757646 | -5.4691728 | 1.88463718 |
| 15912 | NM_004993.5 | #N/A | 73.58111 | -7.4126183 | 2.55455139 |
| 15913 | NR_136279.1 | LOC105375800 | 14.092932 | 3.66134831 | 1.26183037 |
| 15914 | NM_002557.3 | #N/A | 14.9543731 | -5.1041184 | 1.75909768 |
| 15915 | NM_001001342.2 | BLOC1S2 | 21.6440483 | -5.6485407 | 1.94690264 |
| 15916 | NM_006322.5 | #N/A | 94.1673963 | -7.7675607 | 2.67765094 |
| 15917 | NM_014873.2 | #N/A | 301.607081 | -2.1317178 | 0.73490573 |
| 15918 | NM_032298.2 | SYT3 | 25.9421246 | -5.9123296 | 2.0384057 |
| 15919 | NR_135480.1 | LOC102724532 | 15.0438751 | -5.129532 | 1.76864104 |
| 15920 | NR_046223.1 | #N/A | 15.3400013 | -5.1455133 | 1.77418378 |
| 15922 | NM_018396.2 | METTL2B | 143.634577 | -1.6509274 | 0.56927321 |
| 15921 | NM_001202.5 | #N/A | 21.8281965 | -5.6655973 | 1.95360594 |
| 15923 | NR_109848.1 | #N/A | 15.7500573 | -5.1838532 | 1.78765463 |
| 15924 | NM_024593.3 | #N/A | 25.4564823 | -5.8840474 | 2.02916227 |
| 15925 | NM_201400.3 | #N/A | 20.0535818 | -5.5438278 | 1.91189216 |
| 15926 | NM_015681.4 | #N/A | 22.6989482 | -4.2028979 | 1.44945962 |
| 15927 | NM_001320724.1 | RWDD2B | 17.8971974 | -5.3819274 | 1.85614787 |
| 15928 | NM_001039490.1 | CD151 | 27.6120695 | -5.999119 | 2.06922305 |
| 15929 | NR_038903.1 | RAB30-DT | 12.9679933 | -4.9026438 | 1.69154584 |
| 15930 | NM_152284.3 | #N/A | 48.0563341 | 2.69414401 | 0.92958194 |
| 15931 | NM_001346576.1 | DPH1 | 20.5654256 | -5.5623485 | 1.9194118 |
| 15932 | NM_001281766.2 | EPHA5 | 95.1884341 | -7.7829934 | 2.68592226 |
| 15933 | NM_197962.2 | GLRX2 | 14.4735054 | -5.07091 | 1.75058823 |
| 15934 | NM_033504.3 | #N/A | 14.9780324 | -5.1134294 | 1.76559396 |
| 15935 | NR_039992.2 | LINC00645 | 16.9114671 | -5.2902379 | 1.82672755 |
| 15936 | NM_001161779.1 | PDP1 | 26.156109 | -5.9142113 | 2.04257723 |
| 15937 | NR_103867.1 | #N/A | 77.2252597 | -7.4822956 | 2.5842768 |
| 15938 | NM_001145848.1 | #N/A | 71.0137605 | -7.3615503 | 2.54287575 |
| 15939 | NM_001300806.1 | #N/A | 14.3573785 | -5.0492107 | 1.74415318 |
| 15940 | NM_004490.2 | #N/A | 14.96864 | -5.1163322 | 1.76736851 |
| 15941 | NM_001278562.1 | #N/A | 13.1435997 | -4.9356719 | 1.70500948 |
| 15942 | NM_006296.6 | VRK2 | 14.5157322 | -5.0807116 | 1.75521125 |
| 15943 | NM_030613.3 | #N/A | 12.2948857 | -4.839081 | 1.67172488 |
| 15944 | NM_006940.5 | #N/A | 108.911898 | -7.977271 | 2.75597712 |
| 15945 | NM_003433.3 | #N/A | 12.0947637 | -4.8119807 | 1.66254318 |
| 15946 | NM_213613.3 | #N/A | 12.1151109 | -4.8063789 | 1.66083074 |
| 15947 | NM_031949.4 | #N/A | 24.1190345 | 5.19981665 | 1.79680951 |
| 15949 | NR_033932.1 | RGMB-AS1 | 13.4251373 | -4.9604142 | 1.71421096 |
| 15948 | NM_001098519.1 | #N/A | 25.0026358 | -5.8585407 | 2.02458759 |
| 15950 | NM_001318219.1 | CEP250 | 127.870327 | -3.5980181 | 1.24371163 |
| 15951 | NM_018042.4 | #N/A | 26.3062616 | 2.62942946 | 0.90904537 |
| 15952 | NR_029410.1 | LOC389765 | 17.2238774 | -4.0229404 | 1.39092453 |

| | | | | | |
|---|---|---|---|---|---|
| 15953 | NR_037858.1 | NIFK-AS1 | 13.5120732 | -4.969486 | 1.71817989 |
| 15954 | NM_001284261.1 | #N/A | 14.8193235 | -5.1068729 | 1.76575751 |
| 15955 | NM_022058.3 | #N/A | 74.249367 | -7.4255546 | 2.56771744 |
| 15956 | NM_001135241.2 | AKR1C2 | 17.1487207 | -5.3168034 | 1.83855207 |
| 15957 | NM_173471.3 | SLC25A26 | 14.4414702 | -5.0631724 | 1.7508863 |
| 15958 | NM_001039763.3 | #N/A | 13.0351411 | -4.9211074 | 1.70186678 |
| 15959 | NM_001282601.1 | #N/A | 32.5125786 | -6.2300268 | 2.15508302 |
| 15960 | NR_037715.1 | RNASEK | 12.386782 | -4.8376467 | 1.67356783 |
| 15961 | NM_032021.2 | #N/A | 11.9530826 | -4.7980489 | 1.66000797 |
| 15962 | NM_001284303.1 | TBC1D22A | 14.3454936 | -5.0588008 | 1.75055913 |
| 15963 | NM_001040712.2 | PTPRD | 78.4572482 | -7.505117 | 2.59719095 |
| 15964 | NM_001077206.3 | SEC31A | 94.5093704 | -7.7726661 | 2.6898123 |
| 15965 | NM_005490.2 | #N/A | 14.7021369 | 2.48161576 | 0.85891497 |
| 15966 | NM_004758.3 | #N/A | 103.436499 | -7.9024709 | 2.73516179 |
| 15967 | NM_001317946.1 | #N/A | 22.484167 | -5.6968527 | 1.9717793 |
| 15968 | NM_001331020.1 | #N/A | 29.9521988 | -4.5334891 | 1.56914206 |
| 15969 | NM_170724.2 | #N/A | 18.9393364 | 6.96443705 | 2.41078752 |
| 15970 | NM_139211.4 | HOPX | 16.7816096 | -5.2728394 | 1.82524483 |
| 15971 | NM_001301253.1 | #N/A | 41.5065791 | 10.3584792 | 3.58571323 |
| 15972 | NR_024233.1 | LINC00167 | 31.4844874 | 1.64297649 | 0.56879029 |
| 15973 | NM_001348099.1 | GFRA1 | 8.13448282 | 8.01823371 | 2.77610031 |
| 15974 | NM_030945.3 | #N/A | 19.4526714 | -5.4994131 | 1.90407049 |
| 15975 | NM_001002269.2 | EXOSC3 | 13.1686783 | -4.9381351 | 1.70986998 |
| 15976 | NM_002354.2 | #N/A | 19.5717859 | -5.5001018 | 1.90481804 |
| 15977 | NM_001291018.1 | PDE6D | 15.627778 | -5.1770769 | 1.79306839 |
| 15978 | NR_046337.1 | MELK | 34.9612568 | -3.0539911 | 1.05786464 |
| 15979 | NM_012476.2 | #N/A | 17.4305812 | -5.3311203 | 1.8468353 |
| 15980 | NM_183399.2 | RNF14 | 39.3382394 | 6.46435425 | 2.23949588 |
| 15981 | NM_001080515.2 | #N/A | 25.1997089 | -5.8627392 | 2.03129338 |
| 15982 | NM_005281.3 | #N/A | 13.5226501 | -4.9687874 | 1.72164838 |
| 15983 | NM_001329922.1 | CALM3 | 23.8029607 | -5.7896523 | 2.00611024 |
| 15984 | NM_025213.2 | #N/A | 13.6501962 | -4.982674 | 1.72656583 |
| 15985 | NM_174907.3 | #N/A | 275.102447 | -2.9902971 | 1.03639257 |
| 15986 | NR_137424.1 | A2M-AS1 | 11.6946097 | -4.7527606 | 1.64753573 |
| 15987 | NM_173092.2 | #N/A | 20.8742353 | 2.64115977 | 0.91557597 |
| 15988 | NM_145045.4 | #N/A | 15.7303189 | -5.1892989 | 1.79900409 |
| 15989 | NM_001955.4 | #N/A | 14.2267482 | -5.0369131 | 1.74621018 |
| 15990 | NM_001267698.1 | CD63 | 11.6342163 | -4.7537155 | 1.6482505 |
| 15991 | NR_135221.1 | LOC100130111 | 34.0012883 | 4.33612041 | 1.50353373 |
| 15992 | NR_135203.1 | FAM201B | 14.6917485 | -5.081351 | 1.76206061 |
| 15993 | NM_020424.4 | #N/A | 13.9655994 | -5.0079138 | 1.73678361 |
| 15994 | NM_001040629.1 | NCALD | 21.7365998 | -5.6426592 | 1.95703462 |
| 15995 | NM_207307.2 | #N/A | 24.1335647 | -5.8072561 | 2.01417848 |

| | | | | | |
|---|---|---|---|---|---|
| 15996 | NM_001284367.1 | NDUFB8 | 13.2250431 | -4.9426545 | 1.71431547 |
| 15997 | NM_001286565.1 | KIAA0513 | 238.659731 | -1.6187413 | 0.56153628 |
| 15998 | NM_024409.3 | #N/A | 12.1236452 | -4.8199452 | 1.67233472 |
| 15999 | NR_130910.1 | #N/A | 13.2398162 | -4.9445238 | 1.71564041 |
| 16000 | NR_136271.1 | LOC105375304 | 12.610179 | -4.8724245 | 1.69088368 |
| 16001 | NM_004969.3 | #N/A | 68.6576066 | -7.3128553 | 2.53833823 |
| 16002 | NR_104155.1 | LINC02764 | 18.9609571 | 8.46490013 | 2.93829567 |
| 16003 | NR_024254.1 | FAM86FP | 11.4932062 | -4.7346552 | 1.64352283 |
| 16005 | NM_201598.3 | EEF2KMT | 25.5672946 | -5.8837647 | 2.04250348 |
| 16004 | NM_014563.5 | TRAPPC2 | 243.881054 | 1.09249727 | 0.37925119 |
| 16006 | NR_046422.1 | #N/A | 48.2597862 | 1.64582102 | 0.57133067 |
| 16007 | NM_178557.3 | #N/A | 490.270828 | -4.1140541 | 1.42840408 |
| 16008 | NR_104239.1 | TLE4 | 15.5291595 | -5.1788151 | 1.79816612 |
| 16009 | NM_001127617.2 | ALDOA | 13.0320013 | -4.9081954 | 1.70439862 |
| 16010 | NM_173676.2 | PNPLA1 | 61.8276482 | 4.384091 | 1.52244688 |
| 16011 | NM_020807.2 | ZNF319 | 104.307882 | -2.5729838 | 0.89361359 |
| 16012 | NM_001282979.1 | #N/A | 105.960707 | -7.9382706 | 2.75708002 |
| 16013 | NM_001256172.1 | #N/A | 14.0364133 | -5.0190753 | 1.74326427 |
| 16014 | NR_126419.1 | P3H2-AS1 | 62.5638494 | 2.45032641 | 0.85108493 |
| 16015 | NM_001170714.1 | #N/A | 13.4715907 | -4.9566464 | 1.72206176 |
| 16016 | NM_001040274.2 | #N/A | 16.8515502 | -5.276578 | 1.83323938 |
| 16017 | NR_136632.1 | CFAP20 | 13.9686533 | -5.0227302 | 1.74507176 |
| 16018 | NR_003190.2 | USP32P1 | 13.5984332 | -4.9689027 | 1.72658914 |
| 16019 | NR_026904.1 | PABPC1P2 | 62.2795653 | 2.12372749 | 0.7379673 |
| 16020 | NR_027023.1 | #N/A | 13.4831104 | -4.9696751 | 1.72698295 |
| 16021 | NR_125338.1 | PIFO | 9.7922868 | 8.29195212 | 2.88155718 |
| 16022 | NM_001145064.2 | #N/A | 352.442528 | -2.059995 | 0.71589584 |
| 16023 | NM_201520.2 | #N/A | 13.2096335 | -4.9431822 | 1.71786397 |
| 16024 | NM_012413.3 | #N/A | 13.1145368 | -4.9291937 | 1.71303767 |
| 16025 | NM_001287821.1 | #N/A | 20.0746669 | -5.5409507 | 1.92588313 |
| 16026 | NM_021133.3 | #N/A | 13.2631345 | -4.9326745 | 1.71446354 |
| 16027 | NM_176797.2 | P2RY6 | 16.9900945 | 4.70253155 | 1.63451207 |
| 16028 | NM_001242895.1 | #N/A | 215.626016 | 2.41090513 | 0.83799556 |
| 16029 | NM_001135935.1 | POSTN | 86.1092676 | -7.6390096 | 2.65539236 |
| 16030 | NM_001243234.1 | #N/A | 69.5091852 | -7.3305312 | 2.54846024 |
| 16031 | NM_001194947.1 | EIF4G1 | 74.0634281 | -7.4222367 | 2.58049108 |
| 16032 | NM_015075.1 | #N/A | 17.5359897 | -5.3501957 | 1.8601785 |
| 16033 | NM_006634.2 | #N/A | 23.5848063 | -3.7052546 | 1.28840009 |
| 16034 | NM_022750.3 | #N/A | 17.0969392 | -5.3100646 | 1.84645305 |
| 16035 | NM_001001996.2 | GPM6B | 74.6563923 | -7.4337004 | 2.58495524 |
| 16036 | NM_152388.3 | #N/A | 111.380632 | -2.8946966 | 1.0066449 |
| 16037 | NM_152416.3 | #N/A | 16.2887072 | -5.2268996 | 1.81779741 |
| 16038 | NR_110138.1 | PCAT5 | 37.3952102 | 2.02298708 | 0.70357598 |

| | | | | | |
|---|---|---|---|---|---|
| 16039 | NM_203401.1 | STMN1 | 19.1740354 | -5.4698648 | 1.90238119 |
| 16040 | NM_144989.2 | #N/A | 15.1222764 | -5.1176069 | 1.78003526 |
| 16041 | NM_022770.3 | #N/A | 16.6437423 | -5.2625525 | 1.83069242 |
| 16042 | NM_001025.4 | #N/A | 2179.82215 | -4.5224258 | 1.57358042 |
| 16043 | NM_001160042.1 | #N/A | 23.5847391 | -5.7748823 | 2.00943976 |
| 16044 | NM_032280.2 | ZCCHC9 | 12.6140047 | -4.8611097 | 1.69167157 |
| 16045 | NM_144967.3 | #N/A | 18.2716992 | -5.393395 | 1.87716349 |
| 16046 | NR_046090.1 | LOC283214 | 10.0456053 | 4.42609562 | 1.54053066 |
| 16047 | NR_104128.1 | #N/A | 13.3163527 | -4.949086 | 1.72273374 |
| 16048 | NM_001166702.1 | FAM104B | 13.1941216 | -4.9202435 | 1.71285228 |
| 16049 | NM_001256187.1 | SNX12 | 17.9868685 | -5.3780103 | 1.87233258 |
| 16050 | NM_001159643.1 | MCTP2 | 11.414973 | 8.47036361 | 2.94916743 |
| 16051 | NR_109781.1 | NCMAP-DT | 12.6766697 | -4.8841409 | 1.70053845 |
| 16052 | NR_037665.1 | #N/A | 519.605118 | -2.6749972 | 0.93153536 |
| 16053 | NM_144668.5 | #N/A | 21.6481135 | -5.6510279 | 1.96827561 |
| 16054 | NM_000359.2 | #N/A | 13.5926888 | -4.9723151 | 1.73230703 |
| 16055 | NM_032120.3 | #N/A | 163.404038 | 1.53408629 | 0.53446998 |
| 16056 | NM_005510.3 | #N/A | 84.3664672 | -2.4817006 | 0.86474997 |
| 16057 | NM_001184743.1 | PGBD1 | 14.5521976 | -5.071536 | 1.76727592 |
| 16058 | NM_001323987.1 | CTNNA1 | 29.0120644 | -6.0675392 | 2.11442837 |
| 16059 | NM_001163811.1 | #N/A | 15.9167764 | -5.1946021 | 1.81028887 |
| 16060 | NR_110410.1 | RAP2C-AS1 | 20.8120087 | -3.4970792 | 1.21873425 |
| 16061 | NM_001198842.1 | #N/A | 13.8680143 | -4.999522 | 1.7426786 |
| 16062 | NM_033229.2 | #N/A | 18.6747048 | 8.4429762 | 2.94303557 |
| 16063 | NM_001050.2 | #N/A | 165.05412 | -1.9359383 | 0.67483349 |
| 16064 | NM_001195528.1 | #N/A | 17.2386 | -5.3179712 | 1.85396957 |
| 16065 | NM_001193493.1 | #N/A | 15.0334974 | -5.1282899 | 1.78799939 |
| 16066 | NR_024037.1 | #N/A | 11.9826074 | -4.7925007 | 1.67096892 |
| 16067 | NR_145328.1 | #N/A | 14.365033 | -5.0465842 | 1.75958649 |
| 16068 | NM_001195654.1 | ZGPAT | 22.9273207 | -5.7213065 | 1.99503274 |
| 16069 | NM_001134316.1 | #N/A | 11.8420253 | -4.7830993 | 1.66790618 |
| 16070 | NM_212535.2 | #N/A | 27.5412886 | -4.4311256 | 1.54524747 |
| 16071 | NM_178571.4 | TBC1D26 | 12.6603483 | -4.8643139 | 1.69633386 |
| 16072 | NM_001319960.1 | #N/A | 92.2852241 | -7.7380948 | 2.69872126 |
| 16073 | NM_001345993.1 | #N/A | 12.1207877 | -4.798478 | 1.67357815 |
| 16074 | NR_110106.1 | CRAT37 | 45.3138693 | 1.8211851 | 0.63520898 |
| 16075 | NM_001330342.1 | CBWD2 | 14.1686423 | -5.0214583 | 1.75147972 |
| 16076 | NM_001003690.1 | MAD2L1BP | 11.8650584 | -4.7799733 | 1.66727872 |
| 16077 | NM_001146264.2 | #N/A | 13.9818643 | -5.0193228 | 1.75115701 |
| 16078 | NM_001010860.1 | #N/A | 15.9914343 | -5.2161802 | 1.82000546 |
| 16079 | NM_001329629.1 | #N/A | 88.8215563 | -7.6833145 | 2.68100909 |
| 16080 | NR_037566.1 | BCRP2 | 15.3592921 | -5.1482828 | 1.79645067 |
| 16081 | NM_001128303.1 | #N/A | 11.8825196 | -4.7883496 | 1.67090038 |

| | | | | | |
|---|---|---|---|---|---|
| 16083 | NM_001270767.1 | NTAN1 | 15.4261079 | -5.1603835 | 1.80112056 |
| 16082 | NM_000924.3 | #N/A | 22.8921422 | -5.7308936 | 2.00026044 |
| 16084 | NM_001305102.1 | #N/A | 74.288667 | -7.4265668 | 2.59209589 |
| 16085 | NM_005358.5 | LMO7 | 14.0266551 | -5.0287797 | 1.75570181 |
| 16086 | NM_001244949.1 | #N/A | 84.4298802 | -7.6100787 | 2.65710781 |
| 16087 | NM_001286740.1 | UBL7 | 21.7281641 | -5.6458338 | 1.97131214 |
| 16088 | NR_110997.1 | MIR3936HG | 40.3945387 | 2.25527577 | 0.78761768 |
| 16089 | NR_121565.1 | LOC101927954 | 38.2630013 | 2.45087968 | 0.8559419 |
| 16090 | NM_001291163.1 | EPHX1 | 17.3246362 | -5.3314036 | 1.86217347 |
| 16091 | NM_020682.3 | #N/A | 255.501919 | 1.10611859 | 0.38636171 |
| 16092 | NM_024864.4 | #N/A | 14.3100647 | -5.0443905 | 1.76235724 |
| 16093 | NR_130911.1 | RSAD1 | 13.7499905 | -4.9952026 | 1.74543088 |
| 16094 | NM_018558.3 | #N/A | 16.255357 | -5.2415746 | 1.83165955 |
| 16095 | NM_145005.6 | C9orf72 | 11.4264073 | -4.7228979 | 1.65045802 |
| 16096 | NM_001128608.1 | #N/A | 87.9586261 | -7.6692578 | 2.68013851 |
| 16097 | NM_213604.2 | #N/A | 11.5878728 | -4.7508677 | 1.66027736 |
| 16098 | NM_001287188.1 | #N/A | 98.9444351 | -7.8390113 | 2.73953601 |
| 16099 | NR_033314.1 | BDNF-AS | 11.7398375 | -4.7713998 | 1.66774175 |
| 16100 | NM_001319120.1 | #N/A | 68.6079083 | -7.3116487 | 2.55585146 |
| 16101 | NM_024821.3 | #N/A | 12.1366731 | -4.8065396 | 1.68039577 |
| 16102 | NM_001014431.1 | #N/A | 557.977754 | -4.0669603 | 1.42185728 |
| 16103 | NM_001127181.2 | CENPL | 17.9794363 | -5.3705789 | 1.87768002 |
| 16104 | NM_001323324.1 | #N/A | 68.1237652 | -7.3015279 | 2.55287539 |
| 16105 | NM_001278730.1 | #N/A | 13.3560907 | -4.9555222 | 1.73281216 |
| 16106 | NM_133625.4 | #N/A | 13.9376753 | -5.0014874 | 1.74897185 |
| 16107 | NM_001184801.1 | #N/A | 20.2332428 | -4.0459179 | 1.41495343 |
| 16108 | NR_135131.1 | #N/A | 11.4269425 | -4.7282926 | 1.65362708 |
| 16110 | NR_027121.1 | SHROOM4 | 95.4544854 | 2.67743344 | 0.93640685 |
| 16109 | NM_001103176.1 | #N/A | 18.2129979 | -5.3975571 | 1.88773356 |
| 16111 | NM_001306.3 | #N/A | 21.9724049 | -5.6728058 | 1.9841064 |
| 16112 | NR_109990.1 | NRSN2-AS1 | 28.2244466 | -2.6641223 | 0.93180406 |
| 16113 | NM_007007.2 | #N/A | 463.654809 | -3.2392148 | 1.13311179 |
| 16114 | NM_005923.3 | #N/A | 11.6919051 | -4.7671846 | 1.66788582 |
| 16115 | NM_001078651.2 | #N/A | 19.4618556 | -5.4942952 | 1.92247551 |
| 16116 | NM_198566.3 | #N/A | 13.4871926 | -4.9543602 | 1.7336098 |
| 16117 | NM_001303421.1 | #N/A | 48.2789551 | -3.8938949 | 1.36256393 |
| 16118 | NM_001329914.1 | #N/A | 24.4624906 | -5.820959 | 2.03697276 |
| 16119 | NM_001286648.1 | SLC45A4 | 14.8013164 | -5.0965086 | 1.78387931 |
| 16120 | NM_030916.2 | #N/A | 27.2633962 | 2.82763027 | 0.98977422 |
| 16121 | NM_001256865.1 | DNAJC6 | 87.2379908 | -7.6571158 | 2.68042275 |
| 16122 | NM_001321086.1 | #N/A | 11.8578243 | -4.7721333 | 1.67061802 |
| 16123 | NM_148919.3 | #N/A | 11.2376482 | -4.7029915 | 1.64645978 |
| 16124 | NM_006867.3 | #N/A | 11.9120047 | -4.7899489 | 1.67742555 |

| | | | | | |
|---|---|---|---|---|---|
| 16125 | NR_003611.2 | BMS1P1 | 192.21206 | 1.17388322 | 0.41118322 |
| 16126 | NM_001204173.1 | #N/A | 12.0764721 | -4.8030562 | 1.68242207 |
| 16127 | NM_153236.3 | #N/A | 17.7927425 | 8.45231787 | 2.96088184 |
| 16128 | NR_144444.1 | LOC105372480 | 15.0479871 | -5.124512 | 1.79543575 |
| 16129 | NR_110917.1 | LINC02141 | 28.9378754 | 2.30280865 | 0.8068419 |
| 16130 | NM_005282.2 | #N/A | 19.3664092 | 3.36270567 | 1.17824444 |
| 16131 | NR_109773.1 | HERC2P4 | 38.3702875 | 2.93384544 | 1.02807019 |
| 16132 | NM_014005.4 | PCDHA9 | 92.5792553 | 1.86094788 | 0.65212952 |
| 16133 | NM_001290292.1 | SDR39U1 | 23.1094985 | -5.7446707 | 2.01326235 |
| 16134 | NR_046827.1 | LNX1-AS2 | 15.9988113 | 4.1425699 | 1.45201316 |
| 16135 | NM_000344.3 | SMN1 | 87.7744942 | -7.6658419 | 2.68699434 |
| 16136 | NM_001099678.1 | #N/A | 352.655151 | -1.1390324 | 0.39924587 |
| 16137 | NM_001288592.1 | #N/A | 12.3859219 | -4.8524635 | 1.70093602 |
| 16138 | NM_001198978.1 | SMAP2 | 15.8040655 | -5.2018999 | 1.82344698 |
| 16139 | NM_001305040.1 | #N/A | 15.5522349 | -5.1604617 | 1.80908141 |
| 16140 | NM_000029.3 | #N/A | 12.9831958 | -4.9158865 | 1.72336441 |
| 16141 | NM_001199622.1 | #N/A | 25.337201 | 7.07182067 | 2.47973101 |
| 16142 | NM_020866.2 | #N/A | 21.6465931 | -5.6422818 | 1.97878402 |
| 16143 | NM_019006.3 | #N/A | 15.7646863 | -5.1964974 | 1.82265496 |
| 16144 | NM_001031617.2 | #N/A | 154.6007 | -1.1140715 | 0.39077564 |
| 16145 | NM_001136026.2 | #N/A | 15.0645282 | -5.1119402 | 1.79310307 |
| 16146 | NR_046361.1 | #N/A | 13.7821914 | -5.0058444 | 1.75599184 |
| 16147 | NM_001282494.1 | GOLGA8N | 22.0771032 | -4.1519434 | 1.45650584 |
| 16148 | NM_173641.2 | EPHA10 | 17.9314952 | -4.0921316 | 1.43554978 |
| 16149 | NM_001135219.1 | #N/A | 15.989408 | -5.2078508 | 1.82698488 |
| 16150 | NM_000949.6 | #N/A | 13.3720722 | -4.9597996 | 1.74002335 |
| 16151 | NM_173483.3 | #N/A | 17.2194977 | 6.76930945 | 2.37484573 |
| 16152 | NM_001114120.2 | #N/A | 24.4517555 | -4.2709019 | 1.49847997 |
| 16153 | NM_001098612.1 | SIGLEC14 | 39.0598754 | 4.74572672 | 1.66560442 |
| 16154 | NM_018565.3 | SERINC2 | 17.8841969 | -5.3633871 | 1.88252406 |
| 16155 | NM_001199629.1 | #N/A | 20.770874 | -5.5839612 | 1.96014186 |
| 16156 | NM_012482.4 | #N/A | 76.2978609 | -7.4647431 | 2.62041917 |
| 16157 | NM_138809.3 | #N/A | 162.435733 | -2.6284415 | 0.92279488 |
| 16158 | NM_001282958.1 | EIF4E2 | 98.0909199 | -7.8269991 | 2.74846832 |
| 16159 | NM_001018038.2 | VPS13A | 19.800124 | -5.5247431 | 1.94031812 |
| 16160 | NM_001164386.1 | #N/A | 72.7178566 | -7.3956195 | 2.59832679 |
| 16161 | NM_198439.2 | #N/A | 11.8743567 | -4.7779314 | 1.67903911 |
| 16162 | NR_135008.1 | #N/A | 13.6012026 | -4.9875222 | 1.75281147 |
| 16163 | NM_001329942.1 | #N/A | 7.84823055 | 7.96599028 | 2.79960478 |
| 16164 | NM_006106.4 | #N/A | 65.6478217 | -7.2481255 | 2.54748011 |
| 16165 | NR_046342.1 | #N/A | 24.2122739 | -5.8023935 | 2.03984719 |
| 16166 | NM_001008800.2 | CCT3 | 90.6964564 | -7.7138318 | 2.71192567 |
| 16167 | NM_152335.2 | #N/A | 13.5950961 | -4.9648541 | 1.74550518 |

| | | | | | |
|---|---|---|---|---|---|
| 16168 | NM_145247.4 | SFR1 | 15.9153417 | -5.1965609 | 1.82740775 |
| 16169 | NM_001012333.2 | MDK | 16.2918431 | -5.2438506 | 1.84407436 |
| 16170 | NM_014406.4 | #N/A | 15.7998311 | 7.5405336 | 2.65183084 |
| 16171 | NM_001318760.1 | #N/A | 15.0491311 | -5.1307878 | 1.80450673 |
| 16172 | NM_006282.3 | #N/A | 195.455338 | -1.3641949 | 0.47980416 |
| 16173 | NM_021144.3 | PSIP1 | 85.0456778 | -7.6203878 | 2.68041704 |
| 16174 | NM_017881.2 | #N/A | 13.7108967 | -4.972981 | 1.74931633 |
| 16175 | NR_034002.1 | MCF2L-AS1 | 13.9766522 | -5.0109685 | 1.762706 |
| 16177 | NM_001290216.2 | RARB | 17.3971112 | -5.3389935 | 1.87840087 |
| 16176 | NM_000347.5 | #N/A | 27.3864848 | -4.7069496 | 1.65603176 |
| 16178 | NM_001392.4 | #N/A | 20.7345495 | -5.5880319 | 1.96611826 |
| 16179 | NM_001005753.2 | #N/A | 20.6117396 | -5.5837856 | 1.96483743 |
| 16180 | NR_046226.1 | CPB2-AS1 | 12.6518307 | -4.8633854 | 1.71136427 |
| 16181 | NM_013246.2 | #N/A | 12.7039872 | -4.877995 | 1.71653892 |
| 16182 | NR_110062.1 | LOC101927620 | 17.5064902 | 8.42906236 | 2.96618919 |
| 16183 | NR_136736.1 | MLLT10 | 12.7246021 | -4.8876924 | 1.72007136 |
| 16184 | NM_014217.3 | #N/A | 15.6968971 | -5.1765576 | 1.82179884 |
| 16185 | NM_021131.4 | #N/A | 117.037204 | -8.080917 | 2.84413091 |
| 16186 | NM_001144952.1 | #N/A | 226.015595 | -2.7914316 | 0.98249923 |
| 16187 | NR_038429.1 | RNASEH1-AS1 | 14.3732737 | -5.0444612 | 1.77551224 |
| 16188 | NR_102735.1 | DBH-AS1 | 12.0068833 | -4.8036081 | 1.69107889 |
| 16189 | NM_001190906.1 | #N/A | 43.614876 | 2.70263448 | 0.95148515 |
| 16190 | NM_001080379.1 | PACRG | 13.8422242 | -5.004805 | 1.7621083 |
| 16191 | NM_031965.2 | HASPIN | 12.4519356 | -4.8426358 | 1.70505781 |
| 16192 | NR_103782.1 | NUP54 | 15.7592229 | -5.1795891 | 1.82378463 |
| 16193 | NM_022164.2 | #N/A | 19.1021377 | -5.469025 | 1.92574296 |
| 16194 | NM_001039582.3 | PNCK | 112.727271 | -8.0276914 | 2.82685138 |
| 16195 | NM_000960.3 | #N/A | 27.3812339 | 4.54542368 | 1.60087219 |
| 16196 | NM_003578.3 | #N/A | 11.894854 | 6.88943741 | 2.42651365 |
| 16197 | NM_001012727.1 | #N/A | 11.3960064 | -4.7194534 | 1.66233559 |
| 16198 | NM_001278916.1 | #N/A | 69.6333728 | -7.3333499 | 2.58355744 |
| 16199 | NM_001080401.1 | PPM1N | 20.8414777 | 2.208626 | 0.77818955 |
| 16201 | NM_001287251.1 | NPIPB9 | 13.6913106 | -4.9722852 | 1.75221143 |
| 16200 | NM_001271711.1 | SLC2A8 | 18.3320242 | -5.3996299 | 1.90279816 |
| 16202 | NM_183009.2 | VIRMA | 13.7013013 | -4.9919384 | 1.75914248 |
| 16203 | NM_014865.3 | #N/A | 310.582115 | -3.0004431 | 1.05739663 |
| 16204 | NM_181887.2 | UBE2D3 | 85.1651897 | -7.622422 | 2.686335 |
| 16205 | NM_001346870.1 | NAIP | 12.331254 | -4.833762 | 1.70366949 |
| 16206 | NM_001202425.1 | ZNF559-ZNF177 | 15.6259249 | -5.1722178 | 1.82314929 |
| 16207 | NM_032868.5 | MPND | 21.3671354 | -5.6258095 | 1.9830845 |
| 16208 | NM_138571.4 | #N/A | 302.263741 | -3.0893368 | 1.08901187 |
| 16209 | NM_170676.4 | #N/A | 90.2964662 | -7.7066024 | 2.71699134 |
| 16210 | NM_018835.2 | #N/A | 11.2449704 | -4.7009312 | 1.6574795 |

| | | | | | |
|---|---|---|---|---|---|
| 16211 | NM_183352.2 | #N/A | 308.851598 | -3.8893286 | 1.37132708 |
| 16212 | NM_006314.2 | #N/A | 15.6506305 | -5.1724538 | 1.82381457 |
| 16213 | NM_001271957.1 | #N/A | 13.7862115 | -4.9986634 | 1.76276443 |
| 16214 | NM_001308429.1 | #N/A | 15.5674805 | -5.1621387 | 1.82067805 |
| 16215 | NM_001166419.1 | #N/A | 12.6622481 | -4.8588612 | 1.71379109 |
| 16216 | NM_001281293.1 | #N/A | 83.5515652 | -7.5945539 | 2.67881714 |
| 16217 | NM_138625.3 | #N/A | 42.8705664 | -2.3409137 | 0.82571002 |
| 16218 | NR_120660.1 | LOC101929431 | 61.2150742 | 3.17849035 | 1.12148699 |
| 16219 | NM_201595.2 | #N/A | 85.5904351 | -7.6297942 | 2.69220082 |
| 16220 | NM_001331009.1 | #N/A | 78.6994193 | -7.5087869 | 2.64965792 |
| 16221 | NR_110644.1 | ERICH2 | 12.5625639 | -4.8607448 | 1.71547621 |
| 16222 | NM_001277291.1 | ZNF891 | 13.770306 | -4.9811823 | 1.75809646 |
| 16223 | NM_001112718.3 | #N/A | 104.816137 | -7.9215903 | 2.79615302 |
| 16224 | NM_001168359.1 | TDRD6 | 20.6674814 | 5.39300038 | 1.90373627 |
| 16225 | NM_003288.3 | #N/A | 76.6217101 | -7.4702195 | 2.63781998 |
| 16226 | NM_001322884.1 | #N/A | 85.130527 | -7.6221006 | 2.6915778 |
| 16227 | NM_199282.2 | #N/A | 11.4001046 | -4.7213198 | 1.66788397 |
| 16228 | NM_173827.3 | #N/A | 63.56193 | -7.2017175 | 2.54431226 |
| 16229 | NR_136637.1 | LINC02604 | 12.1239296 | -4.8178346 | 1.70219734 |
| 16230 | NM_015653.4 | #N/A | 24.7389647 | -5.8426725 | 2.06441118 |
| 16231 | NM_001131035.1 | #N/A | 14.715114 | -5.0932782 | 1.79996704 |
| 16232 | NM_015984.3 | #N/A | 83.4357188 | -7.5928333 | 2.68354123 |
| 16233 | NM_020397.3 | #N/A | 77.1217435 | 1.2895786 | 0.45578726 |
| 16234 | NM_024754.4 | #N/A | 14.7481726 | -5.0995875 | 1.80247905 |
| 16235 | NM_001882.3 | #N/A | 18.610453 | -5.4300017 | 1.91932951 |
| 16236 | NM_177991.2 | #N/A | 19.5709713 | -5.5044061 | 1.94566787 |
| 16237 | NM_001318154.1 | #N/A | 82.5342042 | -7.5774078 | 2.6785879 |
| 16238 | NM_032561.4 | #N/A | 15.1034153 | -5.1344322 | 1.81504643 |
| 16239 | NM_006948.4 | #N/A | 444.139731 | -3.3148712 | 1.17186099 |
| 16240 | NM_003550.2 | #N/A | 14.1196834 | -5.0296515 | 1.77813225 |
| 16242 | NR_038839.1 | MIR3976HG | 17.220238 | 8.40542247 | 2.97165044 |
| 16241 | NM_002722.4 | #N/A | 17.220238 | 8.40542247 | 2.97165044 |
| 16243 | NR_126047.1 | LOC728673 | 11.487661 | -4.7379606 | 1.67526932 |
| 16244 | NM_005167.5 | PPM1J | 14.4556925 | -5.0650908 | 1.79098722 |
| 16245 | NM_001318833.1 | EHMT2 | 40.7284754 | -3.7066922 | 1.31077196 |
| 16246 | NM_012164.3 | #N/A | 325.190431 | -2.6168136 | 0.9254444 |
| 16247 | NM_033223.4 | #N/A | 14.975103 | -5.1171967 | 1.80970317 |
| 16248 | NM_001185075.1 | FMR1 | 50.598186 | 10.6368901 | 3.76217101 |
| 16249 | NM_145278.4 | #N/A | 24.1142233 | 4.75977328 | 1.68355902 |
| 16250 | NM_020775.4 | #N/A | 83.2064867 | -7.589141 | 2.68444021 |
| 16251 | NM_015513.4 | #N/A | 69.6703557 | -2.3038044 | 0.81497024 |
| 16252 | NM_001045556.2 | #N/A | 89.2603551 | -7.6898296 | 2.7206437 |
| 16253 | NM_001281778.2 | #N/A | 82.8566408 | -7.5829808 | 2.68319803 |

| | | | | | |
|---|---|---|---|---|---|
| 16254 | NM_001099666.1 | #N/A | 196.33195 | -3.6412758 | 1.28846663 |
| 16255 | NR_033896.1 | FLJ46906 | 12.3143766 | -4.8285002 | 1.70858432 |
| 16256 | NM_000447.2 | #N/A | 69.8111635 | -7.3370311 | 2.59631986 |
| 16257 | NR_134505.1 | CERNA2 | 16.0169948 | 2.61569104 | 0.9257359 |
| 16258 | NM_173808.2 | #N/A | 254.324117 | -4.2340742 | 1.49852771 |
| 16259 | NM_001754.4 | RUNX1 | 50.6077319 | 6.55220502 | 2.31913745 |
| 16260 | NM_001317157.1 | PLAGL1 | 11.4894776 | -4.7324543 | 1.67505353 |
| 16261 | NM_001135751.1 | #N/A | 10.8560654 | -4.6511991 | 1.64637516 |
| 16262 | NM_025231.2 | #N/A | 13.5009363 | -4.9756818 | 1.7617414 |
| 16263 | NM_001301011.1 | #N/A | 13.2579603 | -4.9368552 | 1.74822162 |
| 16264 | NM_001085481.1 | #N/A | 11.8837622 | -4.7721021 | 1.68994507 |
| 16265 | NM_001710.5 | #N/A | 34.804012 | -4.7048648 | 1.6661279 |
| 16266 | NR_027278.1 | FAM230C | 25.0993043 | 6.06605031 | 2.14827987 |
| 16267 | NR_073178.1 | LINC01993 | 28.6829407 | -3.055766 | 1.08243398 |
| 16268 | NR_133635.1 | TMEM234 | 12.2761263 | -4.8287613 | 1.71050536 |
| 16269 | NR_136399.1 | IDE | 17.7114451 | -5.3526621 | 1.89620242 |
| 16270 | NM_001083926.1 | #N/A | 15.8407322 | -5.206836 | 1.84470102 |
| 16271 | NM_199001.3 | #N/A | 12.4460765 | -4.8600336 | 1.72182976 |
| 16272 | NM_001242713.2 | #N/A | 14.9131696 | -5.1206759 | 1.81463484 |
| 16273 | NM_001167856.2 | #N/A | 112.504878 | -8.0244826 | 2.84469211 |
| 16274 | NM_001145207.1 | #N/A | 11.9245141 | -4.7839526 | 1.69614209 |
| 16275 | NM_001300742.2 | #N/A | 64.9625193 | 4.19073544 | 1.48586081 |
| 16276 | NM_001321471.1 | #N/A | 14.806671 | -5.085074 | 1.80297357 |
| 16277 | NM_032331.3 | EEF1AKMT4 | 14.9236104 | -5.1117449 | 1.81250438 |
| 16278 | NM_001312911.1 | #N/A | 13.8206486 | -4.9926758 | 1.7703251 |
| 16279 | NM_003199.2 | #N/A | 91.858936 | -7.7314303 | 2.74164039 |
| 16280 | NR_103768.1 | CH17-340M24 | 10.8766525 | -4.6546244 | 1.65086478 |
| 16281 | NM_001282424.2 | A2ML1 | 26.7380271 | 4.33298881 | 1.53686688 |
| 16282 | NM_207040.1 | #N/A | 80.8849391 | -7.5481122 | 2.67829182 |
| 16283 | NM_001322383.1 | ENTPD6 | 13.7627095 | -4.9889241 | 1.77059326 |
| 16284 | NM_005471.4 | #N/A | 188.805686 | -2.8068797 | 0.99653507 |
| 16285 | NM_020654.4 | #N/A | 65.7238877 | -7.2496807 | 2.57392896 |
| 16286 | NM_002880.3 | RAF1 | 555.466325 | -3.3247272 | 1.18043072 |
| 16287 | NM_001300949.1 | ZNF101 | 240.880452 | 2.42977933 | 0.86274382 |
| 16288 | NM_018287.6 | #N/A | 62.7974055 | -7.1840742 | 2.55092967 |
| 16289 | NM_004899.4 | BABAM2 | 14.1429212 | -5.0201185 | 1.782574 |
| 16290 | NM_015873.3 | VILL | 21.4541923 | -4.1034012 | 1.45709388 |
| 16291 | NM_013249.2 | #N/A | 13.6264626 | -4.9891121 | 1.77173988 |
| 16292 | NR_104432.1 | #N/A | 19.3740734 | -5.4885134 | 1.94910244 |
| 16293 | NR_130747.1 | #N/A | 14.6404653 | -5.0785247 | 1.80349447 |
| 16295 | NR_126343.1 | LOC171391 | 11.2981963 | -4.7107206 | 1.67304811 |
| 16294 | NR_073154.1 | OBSCN-AS1 | 13.0004014 | -4.9192197 | 1.74710193 |
| 16296 | NM_001080508.2 | #N/A | 10.9894689 | -4.6749915 | 1.66042416 |

| | | | | | |
|---|---|---|---|---|---|
| 16297 | NM_001083537.1 | #N/A | 16.328649 | -5.2311775 | 1.85805842 |
| 16298 | NM_022440.2 | MAL | 16.9339857 | 8.38138642 | 2.97727235 |
| 16299 | NM_000363.4 | #N/A | 21.9294543 | -4.128684 | 1.46680265 |
| 16300 | NM_003540.3 | H4C6 | 17.5547744 | 8.351687 | 2.9673095 |
| 16301 | NM_207317.2 | #N/A | 21.6859783 | -5.6536411 | 2.00883221 |
| 16302 | NR_026873.1 | LINC00174 | 22.4777573 | -3.0574163 | 1.08637373 |
| 16304 | NM_001098622.2 | NANOS3 | 13.6434666 | -4.9809005 | 1.76989955 |
| 16303 | NM_001035512.1 | #N/A | 18.1047943 | -5.3907769 | 1.9155376 |
| 16305 | NM_003628.4 | #N/A | 81.450958 | -7.5580129 | 2.6857286 |
| 16306 | NR_033738.1 | LOC440300 | 342.786495 | 1.13099234 | 0.40194856 |
| 16307 | NM_004311.3 | #N/A | 397.844333 | -1.8477609 | 0.65682193 |
| 16308 | NM_001033925.1 | TIAL1 | 65.5575173 | -7.2459928 | 2.57604382 |
| 16309 | NR_131170.1 | LGR4-AS1 | 6.42900985 | 3.74513806 | 1.33159696 |
| 16310 | NM_001289181.1 | ZNF302 | 14.5738528 | -5.062695 | 1.80017127 |
| 16311 | NM_139319.2 | #N/A | 62.6348605 | -7.1803986 | 2.55320369 |
| 16312 | NM_003196.2 | #N/A | 13.7100328 | -4.9961899 | 1.77656475 |
| 16313 | NM_213589.2 | #N/A | 252.038072 | -2.6959345 | 0.95864183 |
| 16314 | NM_001271983.1 | #N/A | 22.6562369 | 3.99169372 | 1.41944301 |
| 16315 | NM_022740.4 | #N/A | 65.9772889 | -7.2551739 | 2.57999808 |
| 16316 | NR_134895.1 | #N/A | 14.3906102 | -5.0663393 | 1.80180008 |
| 16317 | NM_001322031.1 | GPATCH2L | 23.5628897 | 8.38728244 | 2.98315322 |
| 16318 | NM_001163545.2 | #N/A | 62.4379125 | -7.1759182 | 2.55253751 |
| 16319 | NM_001169125.1 | #N/A | 90.0934111 | -7.7033709 | 2.74016395 |
| 16320 | NM_024010.2 | #N/A | 18.9886513 | -5.4550844 | 1.94045851 |
| 16321 | NM_153051.2 | #N/A | 99.6262901 | -7.8482214 | 2.79214552 |
| 16322 | NM_001316306.1 | #N/A | 16.0295191 | -5.2144641 | 1.85523218 |
| 16323 | NM_001009955.3 | #N/A | 342.676678 | -3.8746407 | 1.37877951 |
| 16324 | NM_004246.2 | #N/A | 16.4934941 | 5.18593068 | 1.84550188 |
| 16325 | NR_027340.1 | FKBP9P1 | 53.5994498 | 2.65784181 | 0.9459481 |
| 16326 | NM_144968.3 | #N/A | 27.5422841 | 5.1279446 | 1.82515374 |
| 16327 | NR_002944.2 | HNRNPA1P10 | 10.8880441 | -4.6599422 | 1.65863936 |
| 16328 | NM_001267066.1 | PLEKHB2 | 65.1752079 | -7.237963 | 2.57679509 |
| 16329 | NM_018374.3 | #N/A | 22.6568081 | -5.704649 | 2.03110843 |
| 16330 | NR_023319.1 | #N/A | 16.8432459 | -5.2903146 | 1.88380391 |
| 16331 | NR_038420.1 | ZRANB2-AS1 | 12.3588461 | -4.8425805 | 1.72447947 |
| 16332 | NM_001085430.2 | #N/A | 12.6047874 | -4.8666077 | 1.73355908 |
| 16333 | NM_001322272.1 | #N/A | 16.3101448 | -5.2399754 | 1.86659773 |
| 16334 | NM_016252.3 | BIRC6 | 320.776989 | -3.7804497 | 1.34671799 |
| 16335 | NM_080650.3 | #N/A | 14.6343135 | -5.086769 | 1.81255995 |
| 16336 | NR_027356.2 | LSP1P5 | 15.0863712 | -5.1211901 | 1.82509885 |
| 16337 | NM_001308123.1 | #N/A | 11.6148423 | -4.7516331 | 1.69343709 |
| 16338 | NM_001318739.1 | #N/A | 14.9965927 | -5.1196606 | 1.82490226 |
| 16339 | NM_001166262.1 | MTIF3 | 17.4865669 | -5.3308166 | 1.90020644 |

| | | | | | |
|---|---|---|---|---|---|
| 16340 | NM_001272000.1 | DOCK7 | 78.6367489 | -7.5075482 | 2.67660964 |
| 16341 | NM_018922.2 | PCDHGB1 | 12.00479 | -4.7975606 | 1.71053254 |
| 16342 | NR_027284.1 | LINC00602 | 17.0245729 | 7.36932436 | 2.62762583 |
| 16343 | NM_001271007.1 | #N/A | 17.8640696 | -5.3686825 | 1.9143024 |
| 16344 | NR_002305.1 | PDIA3P1 | 14.3929629 | -5.0600641 | 1.80438102 |
| 16345 | NR_110267.1 | LINC01923 | 19.7646228 | 2.17704552 | 0.77633215 |
| 16346 | NM_001142777.1 | #N/A | 13.071681 | -4.9295398 | 1.75824822 |
| 16347 | NR_003719.2 | NBPF22P | 36.9265428 | 10.1901764 | 3.63477288 |
| 16348 | NM_012411.5 | PTPN22 | 17.9760574 | 5.12584761 | 1.82846137 |
| 16349 | NR_045564.1 | ASCC1 | 15.8194729 | -5.1890329 | 1.85107727 |
| 16350 | NM_032483.3 | PLPP5 | 11.6333681 | -4.7541273 | 1.69597804 |
| 16353 | NM_001284335.1 | EIF3J | 18.2293879 | -5.4072335 | 1.9291624 |
| 16351 | NM_000413.3 | #N/A | 12.71179 | -4.8818247 | 1.74170289 |
| 16352 | NM_001118886.1 | #N/A | 11.4246601 | -4.7348741 | 1.68925866 |
| 16354 | NM_153028.3 | ZNF75A | 11.2813159 | -4.7150638 | 1.68238455 |
| 16355 | NM_001102596.1 | #N/A | 13.2460837 | -4.9344046 | 1.76066382 |
| 16356 | NM_001170460.1 | CDK16 | 20.3954102 | -5.5677249 | 1.98668772 |
| 16357 | NM_001101669.2 | INPP4B | 27.6475063 | -3.4222474 | 1.2212083 |
| 16358 | NM_004734.4 | #N/A | 18.2226895 | -5.3982052 | 1.92645745 |
| 16359 | NM_057162.2 | #N/A | 11.7422467 | -4.7619727 | 1.69939538 |
| 16360 | NM_015517.4 | #N/A | 24.6761527 | -5.8379347 | 2.08342256 |
| 16361 | NM_181581.2 | #N/A | 14.5735187 | -5.0615484 | 1.80642488 |
| 16362 | NM_001077365.1 | #N/A | 61.9997939 | -7.1658177 | 2.55750338 |
| 16363 | NM_001101401.2 | #N/A | 17.5402993 | 4.17095446 | 1.48878481 |
| 16364 | NM_001320636.1 | AQP9 | 17.2685222 | 8.32795627 | 2.97275305 |
| 16365 | NM_001297550.1 | #N/A | 24.5471289 | 4.15542004 | 1.48360801 |
| 16366 | NM_015196.3 | TMEM131L | 63.6811106 | -7.2044096 | 2.57224541 |
| 16367 | NM_001321590.1 | #N/A | 93.1987083 | -7.7520521 | 2.76812207 |
| 16368 | NM_013450.3 | #N/A | 59.6727943 | -7.1106565 | 2.5390785 |
| 16369 | NM_001323526.1 | #N/A | 84.640291 | -7.6131682 | 2.71885377 |
| 16370 | NR_136740.1 | #N/A | 74.8873451 | -7.4367016 | 2.65616205 |
| 16371 | NM_021952.4 | #N/A | 22.7674882 | -5.7176566 | 2.04261968 |
| 16372 | NR_135639.1 | LINC01970 | 73.0759547 | 2.11722929 | 0.7564089 |
| 16373 | NM_003353.3 | #N/A | 11.4212268 | -4.7290691 | 1.68975381 |
| 16374 | NR_027898.1 | COL6A4P2 | 80.3955522 | 3.0028466 | 1.07308115 |
| 16375 | NM_001159508.1 | #N/A | 200.266286 | 4.31202467 | 1.54096112 |
| 16376 | NM_001199319.1 | PEX26 | 75.5658529 | -2.0191339 | 0.72158854 |
| 16377 | NM_005426.2 | TP53BP2 | 59.5273206 | -7.1071263 | 2.54026675 |
| 16378 | NM_001297708.1 | #N/A | 80.1876496 | -7.536519 | 2.69395875 |
| 16379 | NR_026833.1 | LOC400940 | 17.1353009 | -5.3047512 | 1.89621831 |
| 16380 | NM_001385.2 | #N/A | 12.1982174 | 5.66511434 | 2.02511082 |
| 16381 | NR_145327.1 | #N/A | 15.9677172 | -5.2160065 | 1.86457064 |
| 16382 | NM_001242852.1 | RNF146 | 13.4699278 | -4.9718126 | 1.7774603 |

| | | | | | |
|---|---|---|---|---|---|
| 16383 | NM_001254748.1 | RGS5 | 14.4293768 | -5.0626514 | 1.8099821 |
| 16384 | NM_002272.3 | #N/A | 20.9612837 | 4.25382916 | 1.52089232 |
| 16385 | NM_032119.3 | #N/A | 60.0741576 | -7.1202603 | 2.54599795 |
| 16386 | NM_001143760.1 | EIF5A | 21.9824193 | -5.6720548 | 2.02829505 |
| 16387 | NR_126453.1 | IRAIN | 21.0912826 | 3.10490038 | 1.11029642 |
| 16388 | NM_178171.4 | #N/A | 65.5629094 | 2.5631591 | 0.91663036 |
| 16389 | NM_058166.4 | #N/A | 11.9679925 | -4.7792919 | 1.70938734 |
| 16390 | NM_001039846.1 | #N/A | 12.3651337 | -4.8474265 | 1.73387133 |
| 16391 | NM_033276.3 | #N/A | 13.4413049 | -4.9622584 | 1.77506563 |
| 16392 | NM_001100595.1 | #N/A | 74.2544011 | -1.8604659 | 0.66570109 |
| 16393 | NM_001278342.1 | DIABLO | 23.476865 | -5.7608382 | 2.06149195 |
| 16394 | NR_037918.2 | PRH1-PRR4 | 14.0314856 | -5.0282237 | 1.79936104 |
| 16395 | NR_027032.1 | AGAP2-AS1 | 15.230295 | -5.1340416 | 1.83731082 |
| 16396 | NM_004912.3 | #N/A | 22.5137052 | -5.704406 | 2.04162395 |
| 16397 | NM_005225.2 | #N/A | 100.692178 | -2.8067205 | 1.00463513 |
| 16398 | NM_001287749.1 | FDFT1 | 15.4194321 | -5.1627077 | 1.84796466 |
| 16399 | NM_001324216.1 | ELAVL4 | 17.4266129 | -5.336165 | 1.91023097 |
| 16400 | NM_006839.2 | IMMT | 63.2259359 | -7.1940141 | 2.57530757 |
| 16401 | NM_017782.4 | TASOR2 | 79.2466785 | -7.5182186 | 2.69161041 |
| 16402 | NM_001130081.2 | #N/A | 19.9930074 | -5.5364255 | 1.98224302 |
| 16403 | NR_040038.1 | LINC01907 | 68.5026571 | 2.20454239 | 0.78936706 |
| 16404 | NR_104049.1 | #N/A | 15.2691419 | -5.1344007 | 1.83844347 |
| 16405 | NM_001289172.1 | ZNF227 | 11.5455894 | -4.7281681 | 1.69301503 |
| 16406 | NM_005437.3 | NCOA4 | 28.6371475 | -6.0452943 | 2.16469753 |
| 16407 | NM_006731.2 | FKTN | 76.7827596 | -7.4731199 | 2.67674269 |
| 16408 | NM_015378.3 | #N/A | 74.5156538 | -7.4296696 | 2.661493 |
| 16409 | NM_001297549.1 | #N/A | 58.8788439 | -7.0914226 | 2.54043764 |
| 16410 | NM_001163391.1 | #N/A | 11.9696289 | -4.7906263 | 1.71639233 |
| 16411 | NM_133474.3 | #N/A | 152.917031 | -2.5717293 | 0.92150776 |
| 16412 | NM_015395.2 | #N/A | 376.429308 | -2.3915157 | 0.85695335 |
| 16413 | NM_001300927.1 | #N/A | 315.589941 | 1.90085161 | 0.6811616 |
| 16414 | NR_109952.1 | LOC101927051 | 7.14085054 | 4.70724126 | 1.68722825 |
| 16415 | NM_173074.2 | #N/A | 11.5305943 | -4.746726 | 1.70169025 |
| 16416 | NR_144394.1 | SUZ12P1 | 18.634603 | -5.4313087 | 1.94722949 |
| 16417 | NM_001346030.1 | #N/A | 11.5297162 | -4.7350479 | 1.69769799 |
| 16418 | NM_001146016.1 | TJAP1 | 24.7054799 | -5.8344588 | 2.09199586 |
| 16419 | NM_001256512.1 | SSBP1 | 15.432671 | -5.1643117 | 1.85188218 |
| 16420 | NM_002908.3 | #N/A | 48.3064712 | -2.2840618 | 0.81927855 |
| 16421 | NR_109961.1 | LOC101928107 | 16.3614812 | 8.33207284 | 2.98902739 |
| 16422 | NM_138341.2 | TMEM116 | 15.48642 | -5.1510428 | 1.847917 |
| 16423 | NM_001286371.1 | VTA1 | 13.211444 | -4.9388642 | 1.77188377 |
| 16424 | NR_040583.2 | STAG3L1 | 48.0176248 | -2.6272031 | 0.94255997 |
| 16425 | NM_001301175.1 | USP30 | 32.8460008 | -3.8578028 | 1.38473654 |

| | | | | | |
|---|---|---|---|---|---|
| 16426 | NM_025218.3 | #N/A | 14.1862244 | -5.0413827 | 1.80961495 |
| 16427 | NR_038924.1 | PXN-AS1 | 11.7679459 | -4.7699282 | 1.7122269 |
| 16428 | NM_152756.4 | #N/A | 57.9702775 | -7.0689682 | 2.53765065 |
| 16429 | NM_203383.1 | #N/A | 61.0852589 | -7.1443036 | 2.56468187 |
| 16430 | NM_024332.3 | BRCC3 | 12.5554596 | -4.8708977 | 1.7486405 |
| 16431 | NM_001270896.1 | #N/A | 16.0282948 | -5.2053424 | 1.86884758 |
| 16432 | NM_001143957.2 | #N/A | 26.3158123 | -3.3223363 | 1.19285633 |
| 16433 | NM_015511.4 | AAR2 | 24.5828155 | -5.8281973 | 2.09282763 |
| 16434 | NM_138403.4 | #N/A | 17.5050201 | 2.61557204 | 0.93926815 |
| 16435 | NR_073468.1 | #N/A | 76.6053074 | -7.4698988 | 2.68255047 |
| 16436 | NM_003922.3 | #N/A | 475.268029 | -3.9439082 | 1.41636006 |
| 16437 | NR_133638.1 | MIR194-2HG | 50.1770309 | 4.11426106 | 1.47760641 |
| 16438 | NR_110666.1 | ROR1-AS1 | 34.8395844 | 2.13757587 | 0.76779135 |
| 16439 | NM_005175.2 | #N/A | 12.9575185 | -4.9123081 | 1.76447589 |
| 16441 | NR_015389.1 | LINC00667 | 236.679217 | -2.0761075 | 0.74575137 |
| 16440 | NM_014059.2 | #N/A | 12.0969739 | -4.805347 | 1.72612279 |
| 16442 | NM_003625.4 | #N/A | 75.6970042 | -7.4524637 | 2.67712692 |
| 16443 | NR_038133.1 | #N/A | 21.5165308 | -5.6391717 | 2.02586856 |
| 16444 | NM_001323387.1 | #N/A | 25.0434055 | -5.8539652 | 2.10310129 |
| 16445 | NR_036595.1 | RBM15-AS1 | 42.0296008 | 3.10949407 | 1.11718373 |
| 16446 | NM_080744.1 | #N/A | 10.7016656 | -4.6326768 | 1.66465761 |
| 16447 | NM_001277348.1 | #N/A | 11.2061065 | -4.6973626 | 1.6880654 |
| 16448 | NM_206923.3 | #N/A | 12.7838853 | -4.8876497 | 1.75668797 |
| 16449 | NM_001287061.1 | #N/A | 15.7187273 | -5.1917151 | 1.86607841 |
| 16450 | NM_001330229.1 | #N/A | 14.7612206 | -5.0891827 | 1.82923721 |
| 16451 | NM_005244.4 | #N/A | 23.9097011 | -5.7938749 | 2.08252538 |
| 16452 | NM_001793.5 | #N/A | 61.6660753 | -7.158061 | 2.57314263 |
| 16453 | NM_004784.2 | #N/A | 22.0854645 | -3.6034253 | 1.29535612 |
| 16457 | NR_026868.1 | ANKRD19P | 13.9235681 | -5.0077337 | 1.80029006 |
| 16454 | NM_004235.5 | #N/A | 15.6303968 | -5.1835454 | 1.86349301 |
| 16455 | NM_014515.5 | #N/A | 58.3538338 | -7.0783234 | 2.54467125 |
| 16456 | NM_014809.3 | #N/A | 16.7585625 | -5.2778884 | 1.89737243 |
| 16458 | NM_144597.2 | #N/A | 44.5557806 | -1.6223309 | 0.58326897 |
| 16459 | NM_001145299.3 | #N/A | 75.6611994 | -7.4517215 | 2.68011544 |
| 16460 | NM_001253866.1 | YME1L1 | 148.836697 | -6.0976648 | 2.19328891 |
| 16461 | NM_001303494.1 | #N/A | 81.5500882 | -6.3083351 | 2.26922794 |
| 16463 | NR_002806.2 | FAM13A-AS1 | 13.8942197 | -4.9945837 | 1.79667969 |
| 16462 | NM_001177802.1 | #N/A | 20.2916042 | -5.5486492 | 1.99598325 |
| 16465 | NR_027085.1 | LINC01226 | 71.7199879 | 4.2109362 | 1.51488181 |
| 16464 | NM_001282206.1 | SIGMAR1 | 18.3087032 | -5.4035455 | 1.9439223 |
| 16466 | NM_001033569.1 | AMZ2 | 25.5151014 | -5.8827083 | 2.11663703 |
| 16467 | NM_001168236.1 | #N/A | 70.5461634 | -7.3511968 | 2.64575703 |
| 16468 | NM_001322980.1 | #N/A | 149.649801 | 4.19008377 | 1.5080898 |

| | | | | | |
|---|---|---|---|---|---|
| 16469 | NM_032986.3 | #N/A | 59.6590235 | -7.1102281 | 2.55927444 |
| 16470 | NR_137178.1 | MIR100HG | 13.2516875 | -4.9379627 | 1.77792892 |
| 16471 | NM_001100.3 | #N/A | 14.6568284 | -5.0878799 | 1.83193028 |
| 16472 | NM_018058.6 | #N/A | 25.6847966 | -4.623615 | 1.66488282 |
| 16473 | NR_026788.1 | FAM66C | 23.6721387 | -4.2156313 | 1.51809916 |
| 16474 | NM_001286198.1 | #N/A | 76.7439418 | 1.33013367 | 0.47901926 |
| 16475 | NM_001282907.1 | CCT8 | 12.9267461 | -4.9069053 | 1.76726035 |
| 16477 | NR_046213.1 | LINC00260 | 34.3919492 | 2.73701165 | 0.98593691 |
| 16476 | NM_152720.2 | NEK3 | 14.0937514 | -5.0353062 | 1.81382616 |
| 16478 | NM_001324091.1 | GLCE | 35.2477937 | 5.50087598 | 1.98164002 |
| 16479 | NM_001291157.1 | #N/A | 67.1225987 | -7.280475 | 2.62284719 |
| 16480 | NM_001330282.1 | #N/A | 17.5620876 | -5.3440553 | 1.92544243 |
| 16481 | NR_036510.1 | #N/A | 15.6576261 | -5.1884744 | 1.86950028 |
| 16482 | NM_001323916.1 | #N/A | 13.4261424 | -4.9593233 | 1.78716803 |
| 16483 | NR_026893.2 | #N/A | 77.4602437 | -7.4854555 | 2.69804907 |
| 16484 | NM_001017989.2 | #N/A | 10.94972 | -4.6682209 | 1.68280296 |
| 16486 | NM_199285.2 | PRR19 | 15.7183473 | -5.181886 | 1.8681557 |
| 16485 | NM_018195.3 | #N/A | 58.6947852 | -7.0869054 | 2.55494854 |
| 16487 | NM_001304435.1 | LDB2 | 12.8265935 | -4.8952292 | 1.76487155 |
| 16488 | NM_001142501.1 | MON1A | 16.0960864 | -5.2123061 | 1.87930732 |
| 16489 | NM_024963.5 | #N/A | 552.951042 | 1.41404238 | 0.50985169 |
| 16490 | NM_001115016.2 | #N/A | 74.0808327 | -7.4224134 | 2.67654619 |
| 16491 | NM_203462.2 | #N/A | 416.596946 | -3.1745716 | 1.14478785 |
| 16492 | NM_001031721.3 | #N/A | 13.1716506 | -4.9391157 | 1.781188 |
| 16493 | NM_000227.4 | #N/A | 11.0279095 | -4.6651559 | 1.6824635 |
| 16494 | NM_002898.3 | #N/A | 345.701938 | 1.3760919 | 0.49631311 |
| 16495 | NM_001242768.1 | MTHFD1L | 58.9779857 | -7.0939465 | 2.55867658 |
| 16496 | NM_031303.2 | #N/A | 14.5511778 | -5.0785942 | 1.83177287 |
| 16497 | NM_001305142.1 | #N/A | 19.8494335 | -5.511064 | 1.98797008 |
| 16498 | NM_002990.4 | #N/A | 58.0966779 | 2.15267093 | 0.77654275 |
| 16499 | NM_014829.3 | #N/A | 62.9040019 | -7.1863351 | 2.5923693 |
| 16500 | NM_152765.3 | #N/A | 14.6554615 | -5.0789529 | 1.83228594 |
| 16501 | NM_019600.3 | FAM214A | 66.3568778 | -2.3172293 | 0.83608933 |
| 16502 | NM_020448.4 | #N/A | 56.6524646 | -7.0357248 | 2.53875527 |
| 16503 | NM_017569.3 | SUPT20H | 11.7086873 | -4.7604897 | 1.71783861 |
| 16504 | NM_001282585.1 | MXRA8 | 14.0216928 | -5.0309493 | 1.81545549 |
| 16505 | NM_016932.4 | #N/A | 14.6513519 | -5.0923469 | 1.83764751 |
| 16506 | NR_045639.1 | UQCRB | 12.0919242 | -4.8164016 | 1.7381271 |
| 16507 | NM_001130055.3 | EEF1D | 12.8230505 | -4.8913088 | 1.76529742 |
| 16508 | NM_001304264.1 | LDLRAD3 | 25.1039036 | -4.2869919 | 1.54719494 |
| 16509 | NM_032728.3 | #N/A | 13.1469534 | -4.9272334 | 1.77831474 |
| 16510 | NM_001012456.1 | SEC61G | 14.2436752 | -5.047677 | 1.82188898 |
| 16511 | NM_001258217.1 | #N/A | 18.6879804 | -5.439734 | 1.96340612 |

| | | | | | |
|---|---|---|---|---|---|
| 16512 | NM_001321518.1 | COX11 | 16.1898671 | -5.225246 | 1.88616825 |
| 16513 | NM_001105521.3 | #N/A | 17.6789858 | -5.357885 | 1.93437908 |
| 16514 | NR_134283.1 | LOC100996419 | 11.0308421 | -4.6783057 | 1.6893626 |
| 16515 | NR_001459.2 | SNHG10 | 13.0816979 | -4.932252 | 1.78111541 |
| 16516 | NR_110875.1 | LINC01497 | 13.6021591 | 6.34952155 | 2.29340017 |
| 16517 | NR_136700.1 | GLYR1 | 14.8552537 | -5.1108392 | 1.84614166 |
| 16518 | NR_110706.1 | ZNF529-AS1 | 11.5244405 | -4.7289963 | 1.70824361 |
| 16519 | NM_001286432.1 | REV3L | 57.2935454 | -7.051956 | 2.54763412 |
| 16520 | NM_003549.3 | #N/A | 11.6531988 | -4.7593898 | 1.7194114 |
| 16521 | NM_001330239.1 | #N/A | 64.8922962 | -7.2312175 | 2.6124469 |
| 16522 | NM_032741.4 | #N/A | 59.4954748 | -7.1062518 | 2.56755944 |
| 16523 | NM_152744.3 | #N/A | 187.750132 | -3.5734823 | 1.29113748 |
| 16524 | NR_036461.1 | H2BC20P | 10.535326 | -4.6008851 | 1.66236537 |
| 16525 | NM_001083335.1 | #N/A | 22.7747072 | -5.7229391 | 2.06781193 |
| 16526 | NM_001130110.1 | #N/A | 12.9813155 | -4.9019668 | 1.77123423 |
| 16527 | NM_006116.2 | #N/A | 167.910093 | -2.7348101 | 0.98821774 |
| 16528 | NM_000828.4 | GRIA3 | 16.5047602 | -5.2498619 | 1.8971766 |
| 16529 | NR_024248.1 | TSSC2 | 25.1412764 | -4.283576 | 1.54799417 |
| 16530 | NM_001288985.1 | #N/A | 10.6099407 | -4.6215684 | 1.67015081 |
| 16531 | NM_133367.4 | #N/A | 119.860637 | -2.1966171 | 0.79386897 |
| 16532 | NM_001317941.2 | ENTPD6 | 75.2771456 | -7.4441612 | 2.69039777 |
| 16533 | NM_198524.2 | TEX9 | 16.5264426 | -5.2594373 | 1.90083454 |
| 16534 | NM_001190880.2 | HYI | 12.8531362 | -4.9022211 | 1.77199814 |
| 16535 | NM_001243750.1 | #N/A | 12.8837332 | -4.8888667 | 1.76721986 |
| 16536 | NM_001135937.2 | SMAD2 | 76.9192683 | -7.4758377 | 2.70285576 |
| 16537 | NM_001100819.2 | #N/A | 149.881868 | 1.97841643 | 0.71532802 |
| 16538 | NM_182925.4 | #N/A | 12.3350488 | -4.8325805 | 1.74728475 |
| 16539 | NR_104387.1 | #N/A | 11.253038 | -4.6991747 | 1.69908618 |
| 16540 | NM_004418.3 | #N/A | 15.0153472 | -5.1140858 | 1.84948587 |
| 16541 | NM_000020.2 | #N/A | 11.1003558 | -4.6877886 | 1.69533188 |
| 16542 | NM_031427.3 | #N/A | 319.739838 | 1.59823211 | 0.57807257 |
| 16543 | NR_040094.1 | LINC01193 | 15.7076463 | 6.78966479 | 2.45610986 |
| 16544 | NM_001184773.1 | #N/A | 67.9722905 | -7.298543 | 2.64040397 |
| 16545 | NM_001035534.2 | #N/A | 60.3345493 | -7.1266674 | 2.57832582 |
| 16546 | NR_024371.1 | PRDM16-DT | 21.3061092 | -5.6273216 | 2.03592442 |
| 16547 | NM_001290051.1 | #N/A | 16.4136515 | -5.2349046 | 1.89397171 |
| 16548 | NM_030641.3 | #N/A | 181.670091 | 2.5965939 | 0.93953249 |
| 16549 | NM_000076.2 | CDKN1C | 19.6295367 | -5.5129489 | 1.9948556 |
| 16550 | NM_144659.5 | #N/A | 12.2714719 | -4.8177291 | 1.74359338 |
| 16551 | NM_198181.3 | GOLGA6L9 | 57.0313007 | -7.0452956 | 2.55004808 |
| 16552 | NM_001328685.1 | #N/A | 74.7462334 | -7.4339749 | 2.69079153 |
| 16553 | NM_181352.1 | #N/A | 26.3251975 | -5.9328944 | 2.14750207 |
| 16554 | NM_003475.3 | #N/A | 18.0625656 | -5.3822946 | 1.94861259 |

| | | | | | |
|---|---|---|---|---|---|
| 16555 | NM_183240.2 | #N/A | 11.6384514 | -4.7621319 | 1.72437657 |
| 16556 | NM_001042717.2 | #N/A | 82.861952 | 5.6496863 | 2.04613449 |
| 16557 | NM_001243767.1 | BTF3L4 | 16.7927228 | -5.2798352 | 1.91238716 |
| 16558 | NM_001319134.1 | STAU1 | 75.6560439 | -7.451285 | 2.69897273 |
| 16559 | NM_001317960.1 | XRN2 | 17.2768599 | -5.3163949 | 1.92592638 |
| 16561 | NM_198546.1 | SPATA21 | 16.9418202 | 5.70356698 | 2.06626689 |
| 16560 | NM_001276277.2 | #N/A | 73.9346797 | -7.418847 | 2.68767864 |
| 16562 | NM_144680.3 | ZNF18 | 14.5043889 | -5.0584241 | 1.83275427 |
| 16563 | NM_001146155.1 | #N/A | 16.631264 | -5.2692793 | 1.90919901 |
| 16564 | NM_001289068.1 | #N/A | 17.1114879 | -5.3019273 | 1.92110401 |
| 16565 | NM_181456.2 | MRPL55 | 17.9602179 | -5.3690922 | 1.9455771 |
| 16566 | NR_038382.1 | LINC00865 | 17.8826395 | -5.3620366 | 1.94321987 |
| 16567 | NR_104452.1 | #N/A | 12.3787521 | -4.8323061 | 1.75187164 |
| 16568 | NM_153251.3 | ZDHHC20 | 88.9811636 | -7.6854527 | 2.78627073 |
| 16569 | NM_080879.2 | #N/A | 10.7902804 | -4.6434637 | 1.68345299 |
| 16570 | NM_001164755.1 | #N/A | 16.3753244 | -5.2542187 | 1.90501657 |
| 16571 | NR_110836.1 | LOC101928782 | 9.98904098 | 4.52006325 | 1.63898804 |
| 16572 | NM_001281496.1 | PHB | 19.1602789 | -5.4743915 | 1.98541675 |
| 16573 | NM_001163813.1 | #N/A | 11.9098815 | -4.7730434 | 1.73115436 |
| 16574 | NR_024360.1 | ZFHX4-AS1 | 10.8038822 | -4.6404074 | 1.68328655 |
| 16575 | NM_001159531.1 | BEGAIN | 56.9846946 | -7.0441804 | 2.55528047 |
| 16576 | NM_001286042.1 | ACADM | 21.7473224 | -5.6462889 | 2.04839719 |
| 16577 | NM_205847.2 | #N/A | 18.5511707 | -5.4306517 | 1.97028835 |
| 16578 | NR_120593.1 | #N/A | 49.2277606 | 2.0776113 | 0.75383313 |
| 16579 | NM_021902.3 | #N/A | 19.1603669 | -5.47593 | 1.98694848 |
| 16580 | NM_001177358.1 | GPR137 | 14.463112 | -5.0708845 | 1.84006947 |
| 16581 | NM_014844.4 | #N/A | 189.356548 | -3.5836497 | 1.30040825 |
| 16582 | NR_110331.1 | HOXB-AS3 | 14.2716372 | -5.0470507 | 1.83171949 |
| 16583 | NM_001099645.1 | #N/A | 11.9943807 | -4.8073612 | 1.74490654 |
| 16584 | NM_207362.2 | #N/A | 11.2130938 | -4.7063783 | 1.70837814 |
| 16585 | NM_144573.3 | NEXN | 11.9595465 | -4.7868854 | 1.73762669 |
| 16586 | NM_001284260.1 | #N/A | 13.5157175 | -4.9624299 | 1.80144225 |
| 16587 | NM_005609.3 | #N/A | 22.6581483 | -5.7170222 | 2.07555101 |
| 16588 | NM_001307929.1 | #N/A | 14.0586782 | -5.0279949 | 1.82552957 |
| 16589 | NM_005309.2 | #N/A | 10.9304996 | -4.6551971 | 1.69024727 |
| 16590 | NM_001282383.1 | PTTG1 | 16.1541528 | -5.2116242 | 1.89237258 |
| 16591 | NM_001243333.1 | SEZ6L2 | 70.2373424 | -7.3454264 | 2.66742472 |
| 16592 | NR_135160.1 | TM2D1 | 14.7225354 | -5.1011849 | 1.85248096 |
| 16593 | NM_032531.3 | #N/A | 65.4811716 | -7.2444328 | 2.6313964 |
| 16594 | NM_018933.3 | #N/A | 12.8517426 | -4.8933041 | 1.77749549 |
| 16595 | NR_144629.1 | LOC389602 | 29.0231467 | 4.70243727 | 1.70832619 |
| 16596 | NM_016203.3 | #N/A | 13.3027595 | -4.9349594 | 1.79282585 |
| 16597 | NR_029451.2 | LINC00477 | 24.631927 | 3.97508462 | 1.4441783 |

| | | | | | |
|---|---|---|---|---|---|
| 16598 | NM_080604.2 | #N/A | 17.6656276 | -5.3456268 | 1.94221293 |
| 16599 | NR_002821.2 | TPTEP2 | 10.8215821 | -4.6393261 | 1.6856669 |
| 16600 | NM_001318066.1 | #N/A | 69.6562717 | -7.3334151 | 2.66524571 |
| 16601 | NM_001271520.1 | #N/A | 11.9538743 | -4.7910791 | 1.74142016 |
| 16602 | NM_024960.5 | #N/A | 55.4165585 | -7.0040458 | 2.54578386 |
| 16603 | NM_001321949.1 | FGF14 | 13.0032118 | -4.9048083 | 1.78282834 |
| 16604 | NM_001171163.1 | ZMYM3 | 13.3442865 | -4.9500494 | 1.79938713 |
| 16605 | NM_001134999.1 | FERMT2 | 73.9265021 | -7.4187559 | 2.69700488 |
| 16606 | NM_022459.4 | XPO4 | 149.686369 | -2.1068745 | 0.76595936 |
| 16607 | NR_024148.1 | RNF121 | 13.5116701 | -4.9741607 | 1.80853715 |
| 16608 | NM_001195104.1 | #N/A | 34.3502723 | 10.0860846 | 3.66765204 |
| 16609 | NM_001097620.1 | #N/A | 10.7970775 | -4.6492215 | 1.69063198 |
| 16610 | NM_007280.1 | #N/A | 13.6723481 | -4.9862774 | 1.8134371 |
| 16611 | NM_001320628.1 | BOLA2-SMG1P6 | 18.6208125 | -5.419894 | 1.97146499 |
| 16612 | NR_125949.1 | LOC101928118 | 114.783501 | 4.11105625 | 1.49562669 |
| 16613 | NR_031675.1 | #N/A | 83.7512578 | 2.2134635 | 0.80529054 |
| 16614 | NM_153618.1 | #N/A | 13.0248785 | -4.9263496 | 1.79242899 |
| 16615 | NM_024053.4 | #N/A | 15.9726655 | -5.2069618 | 1.89478103 |
| 16616 | NR_023921.2 | DHRS4-AS1 | 15.1098681 | -5.1186388 | 1.86269402 |
| 16617 | NM_001079535.1 | #N/A | 14.0979345 | -5.0230992 | 1.82807789 |
| 16618 | NM_001320329.1 | NDRG2 | 12.5167325 | -4.8523759 | 1.76597385 |
| 16619 | NR_029385.1 | #N/A | 128.563375 | 2.78433767 | 1.01339666 |
| 16620 | NM_182894.2 | #N/A | 95.2016987 | -3.0602145 | 1.11387049 |
| 16621 | NM_001127364.2 | #N/A | 15.6017557 | -5.1712142 | 1.88327174 |
| 16622 | NM_001271959.1 | SLC39A1 | 15.5667653 | -5.1770513 | 1.88544619 |
| 16623 | NM_016270.3 | #N/A | 21.3213935 | -4.0774529 | 1.4849765 |
| 16624 | NM_152657.3 | #N/A | 11.3867149 | -4.7223945 | 1.72001552 |
| 16625 | NM_001347822.1 | OGDHL | 55.7343946 | -7.0122669 | 2.55440442 |
| 16626 | NM_001191009.2 | #N/A | 44.5745925 | 10.4536939 | 3.80830268 |
| 16627 | NR_103737.1 | SOX9-AS1 | 63.7990412 | 3.82167356 | 1.3922959 |
| 16628 | NM_001076786.2 | #N/A | 221.468701 | -3.594238 | 1.30985542 |
| 16629 | NM_001243719.1 | RAB6A | 34.0640201 | 10.0740407 | 3.67158486 |
| 16630 | NM_002285.2 | #N/A | 62.3109819 | -7.1732603 | 2.61437796 |
| 16631 | NM_005357.3 | #N/A | 49.4651042 | -3.8741805 | 1.41198763 |
| 16632 | NR_000011.1 | SNORA70 | 12.4583016 | -4.8554469 | 1.77012864 |
| 16633 | NR_045682.1 | #N/A | 67.1135946 | 2.23880974 | 0.81630344 |
| 16634 | NR_002163.3 | OR7E37P | 11.0932546 | 5.34749982 | 1.94983913 |
| 16635 | NM_002556.2 | #N/A | 337.025014 | -3.7015033 | 1.34981739 |
| 16637 | NR_027378.1 | NKAIN3-IT1 | 92.1723422 | -3.1013027 | 1.13097363 |
| 16636 | NM_001039842.2 | #N/A | 10.6191935 | -4.6091193 | 1.68083315 |
| 16638 | NM_015507.3 | #N/A | 15.5465869 | -5.1725373 | 1.8865134 |
| 16639 | NM_014448.3 | #N/A | 15.8223049 | -5.2013447 | 1.89725725 |
| 16640 | NR_125384.1 | LOC100507437 | 10.1860867 | -4.5582204 | 1.66274542 |

| | | | | | |
|---|---|---|---|---|---|
| 16641 | NM_001302371.1 | #N/A | 107.499759 | -6.05062 | 2.20725847 |
| 16642 | NR_036530.1 | LOC100289230 | 12.3571839 | -4.8292204 | 1.7618134 |
| 16643 | NR_073571.1 | #N/A | 37.1520517 | 8.11013021 | 2.95887374 |
| 16644 | NM_032369.3 | #N/A | 15.6090884 | -5.1806027 | 1.89034011 |
| 16645 | NM_206808.3 | CLYBL | 13.5017294 | -4.9743974 | 1.8151156 |
| 16646 | NM_001099335.1 | PHYHIP | 12.3168563 | -4.8417348 | 1.76675787 |
| 16647 | NM_033240.2 | #N/A | 10.5800186 | -4.6166168 | 1.68468685 |
| 16648 | NM_006059.3 | #N/A | 16.4044884 | -5.2539028 | 1.91728985 |
| 16650 | NR_038926.1 | PRKAG2-AS1 | 16.1009162 | -5.2157416 | 1.90359675 |
| 16649 | NM_005204.3 | #N/A | 16.97097 | -5.3025247 | 1.93527159 |
| 16651 | NM_013352.3 | #N/A | 54.3356081 | -6.9756555 | 2.54594602 |
| 16652 | NM_002048.2 | #N/A | 145.612894 | -2.5360372 | 0.92561519 |
| 16653 | NM_001102562.1 | #N/A | 16.1232506 | -5.2231947 | 1.90660382 |
| 16654 | NM_002503.4 | #N/A | 15.1933471 | -5.1360371 | 1.87481661 |
| 16655 | NR_034011.1 | SNHG8 | 13.670025 | -4.9844626 | 1.81965369 |
| 16656 | NM_001142958.1 | #N/A | 11.4208562 | -4.7127506 | 1.72050996 |
| 16657 | NM_001271096.1 | #N/A | 60.278546 | -7.125425 | 2.60190363 |
| 16658 | NM_001204090.1 | #N/A | 43.0743011 | 2.62913791 | 0.9600929 |
| 16659 | NM_001317930.1 | #N/A | 15.5906856 | -5.1779551 | 1.89104622 |
| 16660 | NR_026857.1 | BAIAP2-DT | 442.868949 | -2.6638012 | 0.9728826 |
| 16661 | NM_001320035.1 | #N/A | 86.8491914 | -3.1076129 | 1.13501674 |
| 16664 | NR_023382.1 | ZNF815P | 32.4618293 | -2.4880417 | 0.9087707 |
| 16662 | NM_001329686.1 | #N/A | 13.8019932 | -4.9954819 | 1.824614 |
| 16663 | NM_024761.4 | #N/A | 83.1775838 | -3.3698132 | 1.23084148 |
| 16665 | NM_019090.2 | MAP10 | 23.0182176 | -3.0980059 | 1.13187755 |
| 16666 | NM_001207012.1 | PGF | 13.9244481 | -4.9961747 | 1.8255866 |
| 16667 | NR_004382.1 | MESTIT1 | 39.7862093 | 2.99654578 | 1.09499586 |
| 16668 | NM_001127598.2 | #N/A | 16.830148 | -5.2822222 | 1.93027275 |
| 16669 | NM_001142552.1 | #N/A | 16.7009831 | -5.27013 | 1.92590154 |
| 16670 | NM_001286815.1 | #N/A | 11.1843242 | -4.6911376 | 1.71434598 |
| 16671 | NM_002983.2 | #N/A | 17.0477784 | 7.24727893 | 2.64877842 |
| 16672 | NM_001134877.1 | TEDC1 | 14.1763532 | -5.0436628 | 1.84388885 |
| 16673 | NR_001281.2 | PCDHB18P | 33.8896781 | -5.0221967 | 1.83624166 |
| 16674 | NM_001278711.1 | NRN1 | 13.3104595 | -4.9365543 | 1.80507386 |
| 16675 | NM_004510.3 | #N/A | 10.5624305 | -4.6094249 | 1.68546891 |
| 16680 | NR_144442.1 | AMZ2 | 12.6325334 | -4.8801644 | 1.78451849 |
| 16678 | NM_018166.2 | EVA1B | 13.8836927 | -5.0132469 | 1.83319194 |
| 16679 | NM_182592.2 | YIPF7 | 22.240676 | 6.27152326 | 2.29331623 |
| 16676 | NM_001272103.1 | #N/A | 14.6243421 | -5.0734647 | 1.85517829 |
| 16677 | NM_014506.2 | #N/A | 56.5273316 | -7.0325552 | 2.57159149 |
| 16681 | NR_038280.1 | ELOA-AS1 | 10.3518216 | -4.5763728 | 1.6735823 |
| 16682 | NM_001288965.1 | #N/A | 17.4455063 | -5.3358646 | 1.95191951 |
| 16683 | NM_030817.2 | #N/A | 73.513159 | 4.10036437 | 1.5002437 |

| | | | | | |
|---|---|---|---|---|---|
| 16684 | NM_001013845.1 | #N/A | 121.295567 | -3.50056 | 1.28089446 |
| 16685 | NM_000332.3 | ATXN1 | 71.4606904 | -7.3690998 | 2.69674782 |
| 16686 | NR_038937.1 | RPARP-AS1 | 11.038528 | -4.6609816 | 1.70581781 |
| 16687 | NM_001318927.1 | RPE | 18.5066816 | -5.4229762 | 1.9847969 |
| 16688 | NR_046757.1 | SNRK-AS1 | 13.1944884 | -4.9266299 | 1.803319 |
| 16689 | NM_001142929.1 | CHAT | 19.4512888 | -5.4900717 | 2.00959959 |
| 16690 | NM_001300943.1 | #N/A | 70.1240737 | -7.3418564 | 2.68755908 |
| 16691 | NM_021738.2 | #N/A | 10.742811 | -4.6436599 | 1.69992708 |
| 16694 | NM_003903.4 | CDC16 | 58.9852716 | -7.0942082 | 2.59750567 |
| 16692 | NM_001304724.1 | PLEKHO1 | 14.0027269 | -5.0030816 | 1.831852 |
| 16698 | NR_038458.1 | TOB1-AS1 | 15.3726827 | -5.1449893 | 1.88378409 |
| 16693 | NM_001322197.1 | #N/A | 33.4915155 | 10.0496464 | 3.67963326 |
| 16695 | NM_031960.2 | #N/A | 33.4915155 | 10.0496464 | 3.67963326 |
| 16696 | NM_032592.3 | #N/A | 11.572494 | -4.7496497 | 1.73901426 |
| 16697 | NR_027006.1 | #N/A | 17.7716039 | -5.3648528 | 1.96427332 |
| 16699 | NM_001111018.1 | NAV2 | 69.4234572 | -7.32743 | 2.68302603 |
| 16700 | NM_006914.3 | #N/A | 13.0112495 | -4.8968464 | 1.79315379 |
| 16701 | NM_138422.3 | #N/A | 11.5415934 | -4.7396167 | 1.73560112 |
| 16702 | NM_001322288.1 | ZNF391 | 14.930172 | -5.1199808 | 1.87529371 |
| 16703 | NM_152545.2 | #N/A | 72.9436668 | -7.3985526 | 2.70987089 |
| 16704 | NR_024373.2 | MIR4435-2HG | 13.9283692 | -5.01435 | 1.83669159 |
| 16705 | NM_024719.3 | #N/A | 10.9164875 | -4.6469542 | 1.70225349 |
| 16706 | NM_001258308.1 | #N/A | 64.7445617 | -7.2281385 | 2.64798322 |
| 16707 | NM_001177306.1 | PAM | 72.6724004 | -7.3941905 | 2.70917487 |
| 16708 | NR_024522.1 | #N/A | 28.6504312 | -4.4377778 | 1.62616141 |
| 16709 | NR_033839.1 | GSEC | 10.1280719 | -4.5474524 | 1.66641316 |
| 16710 | NM_145044.3 | ZNF501 | 15.3168505 | -5.1547344 | 1.88906077 |
| 16712 | NM_032927.3 | TMEM128 | 15.7948914 | -5.1840008 | 1.90019839 |
| 16711 | NM_001301189.1 | #N/A | 18.2757328 | -5.3998869 | 1.97932561 |
| 16713 | NM_001317782.1 | #N/A | 14.6117988 | -5.0831555 | 1.86329775 |
| 16714 | NM_001466.3 | #N/A | 340.211439 | -2.492559 | 0.91368113 |
| 16715 | NM_052813.4 | #N/A | 13.2256236 | -4.9312039 | 1.80759782 |
| 16716 | NM_014827.5 | ZC3H11A | 296.083032 | -2.9555788 | 1.08342717 |
| 16717 | NM_001144951.1 | GLYCTK | 10.7315898 | -4.6410069 | 1.70140949 |
| 16718 | NM_001137601.2 | #N/A | 9.99696282 | -4.532412 | 1.66163546 |
| 16719 | NM_004796.5 | #N/A | 11.7345716 | -4.7479354 | 1.7407939 |
| 16720 | NM_001248007.1 | TRIM3 | 11.595971 | -4.7474455 | 1.74068508 |
| 16721 | NM_183243.2 | #N/A | 17.940405 | -5.3815349 | 1.97330338 |
| 16722 | NM_015485.4 | #N/A | 17.3287715 | -5.3154548 | 1.94913523 |
| 16723 | NM_001256944.1 | CLCN4 | 43.3698738 | 10.4140902 | 3.81903913 |
| 16724 | NR_004862.1 | #N/A | 52.3282685 | -6.9213788 | 2.5382477 |
| 16725 | NM_001167890.1 | #N/A | 18.1691073 | -5.3981424 | 1.98003902 |
| 16726 | NM_001297763.1 | #N/A | 145.904402 | 2.30277844 | 0.84467928 |

| | | | | | |
|---|---|---|---|---|---|
| 16727 | NM_181838.1 | UBE2D2 | 11.7571309 | -4.7699015 | 1.74978049 |
| 16728 | NM_181466.2 | #N/A | 12.2708526 | -4.8265675 | 1.77097702 |
| 16729 | NM_184231.2 | #N/A | 52.148439 | -6.9163374 | 2.53780465 |
| 16730 | NM_001005739.1 | #N/A | 63.4003604 | -7.1967341 | 2.64109518 |
| 16731 | NM_001080556.1 | #N/A | 22.1406228 | -5.6818629 | 2.08515833 |
| 16733 | NM_004677.2 | XKRY | 33.2052633 | 10.0372926 | 3.6837516 |
| 16732 | NM_001144996.1 | #N/A | 53.9947145 | -6.9667032 | 2.55683227 |
| 16734 | NM_001001665.3 | CYP27C1 | 55.8804269 | -1.9935015 | 0.73169529 |
| 16735 | NM_001288662.1 | CRMP1 | 15.5790579 | -5.1768142 | 1.90015207 |
| 16736 | NM_001317100.1 | DNAJB4 | 13.2876742 | -4.9471922 | 1.81596476 |
| 16737 | NM_012231.4 | PRDM2 | 68.6929842 | -7.3122056 | 2.6843549 |
| 16738 | NM_001303458.2 | #N/A | 57.2183113 | -7.0499799 | 2.58813183 |
| 16739 | NM_016642.3 | #N/A | 12.8392243 | -4.8997501 | 1.79893104 |
| 16740 | NR_135301.1 | SEMA3F-AS1 | 46.1641567 | 2.39153017 | 0.87818258 |
| 16741 | NM_138813.3 | #N/A | 10.3116049 | -4.567249 | 1.67713435 |
| 16742 | NM_198465.3 | #N/A | 18.2608838 | -4.1193762 | 1.51273487 |
| 16743 | NM_001134836.1 | #N/A | 30.6395081 | 2.87375627 | 1.05543982 |
| 16744 | NM_001318336.1 | #N/A | 68.0018993 | -7.2989153 | 2.68064755 |
| 16745 | NM_001012973.1 | #N/A | 10.1464905 | -4.5564095 | 1.67382508 |
| 16746 | NM_001779.2 | #N/A | 12.5177949 | -4.853492 | 1.78295299 |
| 16747 | NR_045494.1 | CLU | 71.4623582 | -4.2863398 | 1.57473628 |
| 16748 | NM_014508.2 | #N/A | 12.6822744 | -4.8770606 | 1.79202114 |
| 16749 | NM_001324227.1 | STOML1 | 15.4836005 | -5.1646053 | 1.89780689 |
| 16750 | NM_019619.3 | #N/A | 51.8110115 | -6.9069663 | 2.5382198 |
| 16751 | NM_001198858.1 | #N/A | 8.36102545 | 8.06472243 | 2.96421694 |
| 16752 | NM_001321039.1 | #N/A | 53.6255074 | -1.5608313 | 0.57370381 |
| 16753 | NM_001017415.1 | USP1 | 13.5780334 | -4.9750461 | 1.82898207 |
| 16754 | NM_178310.3 | #N/A | 10.4503614 | -4.5925176 | 1.68858732 |
| 16755 | NM_001135243.1 | TCOF1 | 56.4436154 | -7.0305925 | 2.58505372 |
| 16756 | NM_152605.3 | #N/A | 24.6847659 | -4.246097 | 1.56169784 |
| 16757 | NR_144687.1 | #N/A | 11.3822044 | -4.7082994 | 1.73172563 |
| 16758 | NM_001318746.1 | RTN4IP1 | 32.919011 | 10.0248283 | 3.68793463 |
| 16759 | NM_006256.3 | #N/A | 62.5623485 | -7.1776445 | 2.64063144 |
| 16760 | NM_001174087.1 | #N/A | 72.4057899 | -7.3880447 | 2.71844223 |
| 16761 | NM_003782.3 | #N/A | 10.3218421 | -4.5825211 | 1.68618939 |
| 16762 | NR_135120.1 | MAGOHB | 12.4338839 | -4.8429552 | 1.78221393 |
| 16763 | NM_001320264.1 | #N/A | 18.4752344 | -5.4154514 | 1.99322329 |
| 16764 | NM_015266.2 | #N/A | 51.8963941 | -6.9092892 | 2.54339067 |
| 16765 | NM_001145319.1 | #N/A | 10.5510675 | -4.5992575 | 1.69306043 |
| 16766 | NM_016140.3 | #N/A | 10.4223095 | -4.5999544 | 1.6936848 |
| 16767 | NM_021007.2 | #N/A | 89.2091192 | -2.2988983 | 0.84650711 |
| 16768 | NM_001321276.1 | #N/A | 16.867046 | -5.2737926 | 1.94205954 |
| 16769 | NM_001293810.1 | GSTT1 | 14.8968836 | -5.1088107 | 1.88138659 |

| | | | | | |
|---|---|---|---|---|---|
| 16770 | NM_013356.2 | SLC16A8 | 9.95007307 | -4.526913 | 1.66726887 |
| 16771 | NM_001015051.3 | RUNX2 | 13.287714 | -4.9416655 | 1.82016373 |
| 16772 | NM_001190265.1 | #N/A | 11.2463826 | -4.7005056 | 1.73152145 |
| 16773 | NM_006161.2 | #N/A | 10.7675516 | -4.6355551 | 1.70764725 |
| 16774 | NM_015853.4 | #N/A | 13.9868474 | -5.0057197 | 1.84401576 |
| 16775 | NR_003595.1 | #N/A | 19.2278391 | 3.53580299 | 1.30256655 |
| 16776 | NM_153826.3 | #N/A | 57.5886608 | -7.0591804 | 2.60060246 |
| 16777 | NM_017951.4 | SMPD4 | 72.7196477 | -7.3941495 | 2.7241568 |
| 16778 | NM_001324362.1 | OGFOD1 | 23.4763578 | 3.5583672 | 1.31110106 |
| 16779 | NM_001450.3 | #N/A | 18.042366 | -5.3880685 | 1.98538682 |
| 16780 | NM_173495.2 | #N/A | 27.6728127 | -3.0052674 | 1.10750383 |
| 16781 | NM_001167959.1 | TMEM178A | 15.3142163 | -5.1452723 | 1.89615889 |
| 16782 | NM_004558.4 | NRTN | 11.2865351 | -4.7145951 | 1.73752448 |
| 16783 | NM_032180.2 | #N/A | 51.467243 | -6.8974404 | 2.54203111 |
| 16784 | NM_018027.4 | #N/A | 168.492634 | -6.6343207 | 2.445097 |
| 16785 | NM_001312909.1 | #N/A | 17.5849367 | -5.3510584 | 1.972279 |
| 16786 | NM_022918.3 | #N/A | 50.9827053 | -6.8837686 | 2.5374974 |
| 16787 | NM_003280.2 | #N/A | 11.2828938 | -4.7168431 | 1.73877589 |
| 16788 | NM_007314.3 | #N/A | 58.0601416 | -7.0709003 | 2.60665333 |
| 16789 | NR_028504.1 | MIR22HG | 12.4249761 | -4.8490908 | 1.78768125 |
| 16790 | NR_103866.1 | #N/A | 63.9049892 | -7.2078109 | 2.65736371 |
| 16791 | NM_001308232.1 | TMIGD2 | 27.7965853 | 4.13905887 | 1.52601228 |
| 16792 | NM_001243770.1 | #N/A | 10.6905583 | -4.6261224 | 1.70565028 |
| 16793 | NM_001321560.1 | #N/A | 93.2014094 | -7.7517549 | 2.85808452 |
| 16794 | NM_001172685.1 | CELF6 | 16.0638831 | -5.2162907 | 1.92354895 |
| 16795 | NM_170607.2 | MLX | 10.7045971 | -4.6260771 | 1.70600733 |
| 16796 | NM_001278494.1 | ANAPC15 | 11.5751032 | -4.728892 | 1.74404076 |
| 16797 | NR_027058.1 | SNHG20 | 198.638747 | 1.39201921 | 0.51338283 |
| 16798 | NM_004056.5 | #N/A | 12.5019524 | -4.8565855 | 1.79119008 |
| 16799 | NM_000047.2 | #N/A | 15.3526897 | -5.1565927 | 1.90186962 |
| 16800 | NM_001297578.1 | #N/A | 79.5947516 | -7.5255647 | 2.77585334 |
| 16801 | NM_018376.3 | #N/A | 11.8781297 | -4.7730901 | 1.76064448 |
| 16802 | NM_005096.3 | ZMYM3 | 63.660283 | -7.2025098 | 2.65682877 |
| 16803 | NM_020435.3 | #N/A | 12.8922111 | -4.8949463 | 1.80564682 |
| 16804 | NM_002252.4 | #N/A | 10.8084999 | -4.6526071 | 1.71670572 |
| 16805 | NR_133007.1 | #N/A | 25.9026914 | 4.84824817 | 1.78889534 |
| 16806 | NM_001303098.1 | TMEM198 | 15.8407213 | -5.2056779 | 1.92220369 |
| 16807 | NM_207510.3 | #N/A | 16.7521617 | -5.2778781 | 1.94887556 |
| 16808 | NR_103765.1 | CAND1 | 26.1333501 | 4.3374531 | 1.60165152 |
| 16809 | NM_001278175.1 | #N/A | 10.959967 | -4.6595825 | 1.72063409 |
| 16810 | NM_152572.2 | #N/A | 14.9928865 | -5.1232852 | 1.89192835 |
| 16811 | NM_001162426.1 | #N/A | 70.416369 | -7.348419 | 2.71379906 |
| 16812 | NM_001244766.1 | #N/A | 17.9167082 | -5.3721541 | 1.98394541 |

| | | | | | |
|---|---|---|---|---|---|
| 16813 | NM_180989.5 | #N/A | 68.4870321 | -1.9071778 | 0.70436992 |
| 16814 | NM_001282167.1 | #N/A | 11.6960913 | -4.7590626 | 1.75795537 |
| 16815 | NM_001320824.1 | #N/A | 17.2296483 | -5.3131996 | 1.96285298 |
| 16816 | NM_080866.2 | #N/A | 26.1960205 | 2.3345972 | 0.86251876 |
| 16817 | NM_002376.6 | MARK3 | 78.7618505 | -7.5091742 | 2.77429674 |
| 16818 | NM_006500.2 | #N/A | 203.721863 | -2.7964688 | 1.0332234 |
| 16819 | NR_033876.1 | LINC00511 | 10.1977952 | 5.93854254 | 2.19416685 |
| 16820 | NM_001005498.3 | RHBDF2 | 12.7595156 | -4.8939149 | 1.80846625 |
| 16821 | NM_006726.4 | LRBA | 47.8500531 | -3.2325141 | 1.19455819 |
| 16823 | NR_135591.1 | HDAC6 | 51.5984824 | -6.9009099 | 2.5502362 |
| 16822 | NR_030770.1 | #N/A | 12.9868063 | -4.9188256 | 1.81776563 |
| 16824 | NM_001277961.1 | STIM1 | 13.6062617 | -4.97655 | 1.83920724 |
| 16825 | NM_000833.4 | #N/A | 66.1627142 | -7.258435 | 2.6828214 |
| 16826 | NM_001318827.1 | TSC2 | 51.3447007 | -6.8939165 | 2.54833008 |
| 16827 | NM_001145317.1 | #N/A | 32.3465064 | 9.99958153 | 3.69650062 |
| 16828 | NM_001329126.1 | #N/A | 66.523051 | -7.2663692 | 2.68613567 |
| 16829 | NM_001146266.1 | GPR85 | 14.5806229 | -5.0755719 | 1.87629795 |
| 16830 | NM_001204458.1 | #N/A | 56.376587 | -7.0284161 | 2.59829778 |
| 16831 | NM_001146341.1 | #N/A | 14.5626372 | 6.68059696 | 2.46974291 |
| 16832 | NM_001320444.1 | TMEM102 | 11.8262847 | -4.7719343 | 1.76428306 |
| 16833 | NR_073409.1 | #N/A | 11.3054214 | -4.7219695 | 1.74582225 |
| 16834 | NR_110375.1 | THRIL | 40.8307406 | -2.273754 | 0.84074352 |
| 16835 | NM_001033046.3 | #N/A | 63.000473 | -7.187588 | 2.65775072 |
| 16836 | NM_004428.2 | #N/A | 15.5534996 | -5.1784436 | 1.91483274 |
| 16838 | NM_001289910.1 | IDH2 | 62.7219196 | -7.1811683 | 2.65606951 |
| 16837 | NM_001082973.2 | #N/A | 15.9666092 | -5.2123911 | 1.92787775 |
| 16839 | NR_036519.1 | #N/A | 10.9189257 | -4.6654291 | 1.72555943 |
| 16840 | NM_001101387.1 | #N/A | 11.5211897 | -4.7390413 | 1.75293948 |
| 16841 | NM_001130721.1 | #N/A | 66.6466451 | -7.2685333 | 2.68905001 |
| 16842 | NM_001317102.1 | DNAJB4 | 17.9794619 | -5.3755122 | 1.98883314 |
| 16844 | NR_047687.1 | NALCN-AS1 | 17.2122705 | 6.66550118 | 2.46621956 |
| 16843 | NM_153681.2 | PIGP | 10.7206073 | -4.6285337 | 1.71255715 |
| 16845 | NM_001320418.1 | MITD1 | 14.8444378 | -5.0906676 | 1.88385427 |
| 16846 | NM_016227.3 | SUCO | 11.453044 | -4.7282981 | 1.7498524 |
| 16847 | NM_001184987.1 | NDUFC1 | 65.366091 | -7.2408665 | 2.67980143 |
| 16848 | NM_001330131.1 | #N/A | 10.2828255 | -4.5740029 | 1.69322308 |
| 16849 | NM_001319300.1 | #N/A | 16.6835941 | -5.2673968 | 1.95008312 |
| 16850 | NM_001329539.1 | #N/A | 13.474245 | -4.9498754 | 1.83255501 |
| 16851 | NM_001300735.1 | #N/A | 51.9304328 | -6.910418 | 2.55846781 |
| 16852 | NM_173631.3 | ZNF547 | 19.4986504 | -3.4173606 | 1.26523775 |
| 16853 | NM_018283.3 | #N/A | 96.5983024 | -1.4037739 | 0.51976276 |
| 16854 | NM_198580.2 | #N/A | 185.890319 | -1.5415816 | 0.57091533 |
| 16855 | NM_003183.5 | #N/A | 214.666009 | -3.5464725 | 1.31354949 |

| | | | | | |
|---|---|---|---|---|---|
| 16856 | NM_001105194.1 | AMER3 | 67.5697465 | -7.2891342 | 2.70004751 |
| 16857 | NR_024264.1 | LOC145845 | 13.7320657 | -4.9921067 | 1.84920536 |
| 16858 | NM_001346102.1 | #N/A | 11.3232232 | -4.706528 | 1.74356773 |
| 16859 | NM_001292043.1 | #N/A | 344.172638 | -3.0745697 | 1.13918582 |
| 16860 | NM_001204832.2 | ANO10 | 12.3876087 | -4.8323695 | 1.7905106 |
| 16861 | NR_024372.1 | FLJ25758 | 32.0602542 | 9.98679023 | 3.70067196 |
| 16862 | NR_026900.1 | #N/A | 28.3440271 | -4.4107018 | 1.63466011 |
| 16863 | NM_133465.3 | #N/A | 52.1234216 | -6.9157693 | 2.56337203 |
| 16864 | NM_001177383.1 | #N/A | 65.6359187 | -7.2464815 | 2.68613169 |
| 16865 | NR_027417.1 | LOC644669 | 33.7653253 | 2.06067271 | 0.76389828 |
| 16866 | NM_054012.3 | #N/A | 275.658929 | -1.7741722 | 0.65774212 |
| 16867 | NM_001297669.1 | #N/A | 13.265874 | -4.9264605 | 1.82650179 |
| 16868 | NM_001017980.3 | VMA21 | 279.607834 | -1.5558891 | 0.57688924 |
| 16869 | NM_005977.3 | #N/A | 49.6882281 | -6.8466484 | 2.53936124 |
| 16870 | NM_001305564.1 | #N/A | 10.2021821 | -4.5620753 | 1.69217196 |
| 16871 | NM_001330635.1 | #N/A | 62.3890437 | -4.1159586 | 1.52672866 |
| 16872 | NM_021020.3 | #N/A | 352.157605 | -3.9054842 | 1.44879093 |
| 16873 | NM_153698.1 | #N/A | 12.3281082 | -4.8424849 | 1.79637537 |
| 16874 | NM_015842.2 | LMO7 | 67.7939187 | -7.2928527 | 2.70587453 |
| 16875 | NM_001018054.2 | #N/A | 69.5514811 | -7.3312997 | 2.72042419 |
| 16876 | NM_020936.2 | #N/A | 64.8288637 | -7.2288406 | 2.68256229 |
| 16877 | NM_203497.3 | COMMD6 | 31.0991139 | -6.1662149 | 2.28826192 |
| 16878 | NR_037888.1 | STX18-AS1 | 10.1280154 | -4.5407286 | 1.68519286 |
| 16879 | NM_006251.5 | PRKAA1 | 126.872424 | -1.4429234 | 0.53553494 |
| 16880 | NM_001128.5 | #N/A | 156.802889 | -2.4477953 | 0.90849956 |
| 16881 | NM_020794.2 | #N/A | 23.3283448 | -5.7525675 | 2.13518508 |
| 16882 | NM_001257392.1 | CD63 | 54.4663651 | -6.9793073 | 2.59076873 |
| 16883 | NM_001204857.1 | ELOC | 11.9992687 | -4.7980923 | 1.78112638 |
| 16884 | NM_001330205.1 | #N/A | 11.523731 | -4.7439379 | 1.76118372 |
| 16885 | NM_001127321.1 | CBX5 | 316.345351 | -1.2881357 | 0.47822379 |
| 16886 | NR_104103.1 | #N/A | 32.3210826 | 4.76067743 | 1.76773936 |
| 16887 | NM_014505.5 | #N/A | 180.705309 | -2.223677 | 0.82579089 |
| 16888 | NM_032088.1 | PCDHGA8 | 12.9780766 | -4.8999898 | 1.82019431 |
| 16889 | NM_001330539.1 | #N/A | 11.8710625 | -4.7832883 | 1.77685733 |
| 16892 | NR_038915.1 | PRR7-AS1 | 11.8806603 | -4.777004 | 1.77471313 |
| 16890 | NM_021069.4 | #N/A | 64.8542597 | -7.2297595 | 2.68597226 |
| 16891 | NM_022915.3 | #N/A | 142.493389 | -2.6416176 | 0.98140444 |
| 16893 | NM_145113.2 | #N/A | 15.6989576 | -5.1821226 | 1.92532336 |
| 16894 | NM_001321279.1 | TSEN2 | 11.4469676 | -4.7328336 | 1.75884346 |
| 16895 | NM_014023.3 | #N/A | 245.435291 | -3.5533159 | 1.32050313 |
| 16896 | NM_021188.2 | #N/A | 64.1638152 | -7.2138422 | 2.68143388 |
| 16897 | NR_121564.1 | LOC101927827 | 25.9808856 | 6.29047846 | 2.3382695 |
| 16898 | NM_005660.2 | #N/A | 11.3763371 | -4.72123 | 1.75531582 |

| | | | | | |
|---|---|---|---|---|---|
| 16899 | NM_006828.3 | #N/A | 192.365873 | -2.2481747 | 0.83588623 |
| 16900 | NM_001042461.2 | #N/A | 18.4903013 | -5.4199756 | 2.01542687 |
| 16901 | NM_001043229.1 | #N/A | 11.1719336 | -4.6880507 | 1.74329573 |
| 16902 | NM_001190972.1 | #N/A | 10.624476 | -4.6184568 | 1.717416 |
| 16903 | NM_018225.2 | #N/A | 399.71015 | -1.3269574 | 0.49343508 |
| 16904 | NM_001330572.1 | #N/A | 11.7522715 | -4.7491536 | 1.76605853 |
| 16905 | NM_207305.4 | FOXD4 | 12.6295142 | -4.8685258 | 1.81052806 |
| 16906 | NM_001165893.1 | #N/A | 49.9343793 | -6.8537539 | 2.54882284 |
| 16907 | NM_001005786.2 | DAZ2 | 40.9604363 | 10.3314678 | 3.84221759 |
| 16908 | NM_001346093.1 | #N/A | 40.9604363 | 10.3314678 | 3.84221759 |
| 16909 | NM_144650.2 | #N/A | 12.658187 | -4.8807416 | 1.81530303 |
| 16910 | NM_031910.3 | #N/A | 171.798981 | -0.9599979 | 0.35705878 |
| 16911 | NR_023389.1 | LINC00476 | 14.7664069 | -5.0933182 | 1.89453873 |
| 16912 | NM_001145335.1 | UBE2Q2 | 9.82572232 | -4.5088353 | 1.6772489 |
| 16913 | NR_144550.1 | SDAD1P1 | 10.5671511 | -4.6211784 | 1.71905717 |
| 16914 | NM_001033886.2 | CXCL12 | 14.9415316 | -5.1084361 | 1.90072199 |
| 16915 | NR_136244.1 | TSBP1-AS1 | 14.5038263 | 4.70151296 | 1.74939257 |
| 16916 | NM_001190317.1 | IP6K2 | 49.7496325 | -6.8485434 | 2.54838706 |
| 16917 | NM_001167741.1 | #N/A | 50.6004597 | -6.8727145 | 2.55753587 |
| 16918 | NM_018416.2 | #N/A | 153.887601 | -2.6811437 | 0.99794035 |
| 16919 | NM_015506.2 | #N/A | 37.1492049 | -1.9433546 | 0.72336445 |
| 16920 | NM_018929.2 | #N/A | 12.6891575 | -4.8730588 | 1.81403393 |
| 16921 | NM_004197.1 | STK19 | 50.061843 | -6.8573579 | 2.55282948 |
| 16922 | NM_003452.3 | #N/A | 17.1266961 | -5.3161456 | 1.9790951 |
| 16923 | NM_139241.3 | FGD4 | 63.412435 | -7.1969955 | 2.67969664 |
| 16924 | NM_001015050.2 | ALG1L | 11.1287207 | 8.43504886 | 3.1408166 |
| 16925 | NM_018397.4 | #N/A | 23.3937095 | -3.3968093 | 1.26483843 |
| 16926 | NM_012240.2 | SIRT4 | 13.3163059 | -4.9413075 | 1.84000101 |
| 16927 | NM_001265604.1 | MORF4L1 | 12.1569065 | -4.8272708 | 1.79758206 |
| 16928 | NM_017629.3 | #N/A | 226.528236 | -2.0485087 | 0.76300562 |
| 16929 | NR_104189.1 | SRGAP2-AS1 | 20.595082 | 4.83555921 | 1.80159847 |
| 16930 | NR_038875.2 | LINC01289 | 22.3390106 | 1.81475455 | 0.67616333 |
| 16931 | NM_001135217.1 | #N/A | 17.3988656 | -5.3373363 | 1.98871116 |
| 16932 | NM_001127605.2 | #N/A | 52.8732329 | -6.936472 | 2.58480058 |
| 16933 | NM_001142588.1 | #N/A | 13.102796 | -4.9210192 | 1.83389577 |
| 16934 | NM_058172.5 | #N/A | 70.4561853 | -7.3489877 | 2.73878227 |
| 16935 | NM_004205.4 | #N/A | 21.5498074 | -5.6442307 | 2.10363595 |
| 16936 | NM_004746.3 | #N/A | 64.3818534 | -7.2185351 | 2.69059332 |
| 16937 | NM_020836.3 | #N/A | 69.4130105 | -7.3266841 | 2.73115812 |
| 16938 | NR_037928.1 | P2RX5-TAX1BP3 | 233.435313 | 1.41771359 | 0.52856625 |
| 16939 | NM_001037165.1 | #N/A | 486.848695 | -0.8094809 | 0.30180898 |
| 16940 | NM_014872.2 | #N/A | 203.038923 | -2.6900412 | 1.00298058 |
| 16941 | NM_004523.3 | #N/A | 154.132443 | -1.4149077 | 0.52759643 |

| | | | | | |
|---|---|---|---|---|---|
| 16942 | NR_038338.2 | OSER1-DT | 14.8811279 | -5.1026861 | 1.90287305 |
| 16943 | NM_001318521.1 | CENPI | 12.6747578 | -4.8772349 | 1.81892912 |
| 16944 | NM_019596.5 | #N/A | 68.7671162 | -7.3144478 | 2.72806317 |
| 16945 | NM_001308422.1 | #N/A | 13.9252253 | -5.0181803 | 1.87176013 |
| 16947 | NR_137192.1 | MIR100HG | 226.135744 | -1.4210848 | 0.530136 |
| 16946 | NM_001320641.1 | #N/A | 70.4953277 | -7.3497382 | 2.74180696 |
| 16948 | NM_001322286.1 | #N/A | 14.9379211 | -3.8076788 | 1.42051182 |
| 16949 | NR_145419.1 | DCTN2 | 50.2414775 | -6.8626835 | 2.56041754 |
| 16950 | NM_207315.3 | CMPK2 | 13.2207675 | -4.9398324 | 1.84304457 |
| 16951 | NM_001190411.1 | RPH3AL | 14.9799502 | -5.1209273 | 1.9106783 |
| 16952 | NM_001037872.2 | #N/A | 62.450543 | -7.1743775 | 2.67740705 |
| 16953 | NM_001198995.1 | NADK | 13.2116375 | -4.9445032 | 1.84532663 |
| 16954 | NM_001184768.1 | ARMCX6 | 13.4493583 | -4.9531305 | 1.84874866 |
| 16955 | NR_046294.1 | MSTO1 | 18.1322433 | -5.39656 | 2.01434036 |
| 16956 | NM_001184750.2 | SLITRK4 | 269.006455 | 5.75173822 | 2.14698739 |
| 16957 | NM_152279.3 | #N/A | 164.893975 | 0.88187037 | 0.32918826 |
| 16958 | NM_001252079.1 | USP15 | 11.6390719 | -4.7642879 | 1.77853162 |
| 16959 | NM_005855.3 | #N/A | 11.1144143 | -4.688951 | 1.75048403 |
| 16960 | NR_046095.1 | DACT1 | 21.6981839 | -5.646989 | 2.10827142 |
| 16961 | NM_014921.4 | #N/A | 622.475997 | -3.8399824 | 1.43367513 |
| 16962 | NM_001321363.1 | #N/A | 54.4203279 | -6.9776423 | 2.60534753 |
| 16963 | NM_001323525.1 | #N/A | 65.1006665 | -7.2347013 | 2.70172654 |
| 16964 | NM_014310.3 | #N/A | 11.4554314 | -4.7253248 | 1.76470732 |
| 16965 | NR_073609.1 | #N/A | 58.7675581 | 1.48419763 | 0.55434486 |
| 16966 | NM_199248.2 | #N/A | 13.4580998 | -4.9461777 | 1.84754343 |
| 16967 | NM_001320584.1 | QRICH1 | 66.5980878 | -7.2674637 | 2.71472392 |
| 16968 | NR_111929.1 | #N/A | 11.9424105 | -4.7874579 | 1.78844864 |
| 16969 | NM_213599.2 | ANO5 | 58.5458971 | -7.0818673 | 2.64572381 |
| 16970 | NM_015601.3 | #N/A | 48.1871296 | -6.802574 | 2.54142565 |
| 16971 | NM_001621.4 | #N/A | 104.840057 | -2.6103846 | 0.97527811 |
| 16972 | NM_001321712.1 | #N/A | 60.5846737 | -7.1308485 | 2.66425049 |
| 16973 | NM_004887.4 | #N/A | 1043.91024 | 2.68999488 | 1.00507814 |
| 16974 | NM_152304.1 | #N/A | 30.9872504 | 5.09791254 | 1.90476514 |
| 16975 | NM_001102668.2 | PSMA4 | 13.0167434 | -4.9239843 | 1.84058813 |
| 16976 | NR_002612.1 | #N/A | 76.8558134 | 1.9867774 | 0.74286123 |
| 16977 | NM_032861.3 | #N/A | 94.6598627 | -2.6410531 | 0.98754613 |
| 16978 | NR_109796.1 | KIAA1191 | 63.2915307 | -7.1938901 | 2.69028418 |
| 16979 | NM_001289984.1 | #N/A | 14.3537333 | -5.060179 | 1.89249173 |
| 16980 | NM_015147.2 | #N/A | 70.1070349 | -7.3411427 | 2.74562465 |
| 16981 | NM_006973.2 | #N/A | 62.0943282 | -7.166625 | 2.68061435 |
| 16982 | NM_006239.2 | #N/A | 36.9562353 | 2.86008318 | 1.06981396 |
| 16983 | NM_024784.3 | #N/A | 103.057444 | 0.89720808 | 0.33560387 |
| 16984 | NM_032370.2 | #N/A | 11.4290435 | -4.722318 | 1.76666494 |

| | | | | | |
|---|---|---|---|---|---|
| 16985 | NM_144598.4 | #N/A | 67.2997053 | -7.2825361 | 2.72456614 |
| 16986 | NM_001318104.1 | #N/A | 10.7948691 | -4.6280315 | 1.73158371 |
| 16987 | NM_001287507.1 | PPCS | 10.324194 | -4.5813419 | 1.71416908 |
| 16988 | NM_001272095.1 | STX4 | 11.1098346 | -4.6964366 | 1.75731771 |
| 16989 | NM_002405.3 | #N/A | 14.3273532 | -5.0365134 | 1.8845721 |
| 16990 | NM_172312.1 | #N/A | 20.3269374 | -5.5599692 | 2.08078418 |
| 16991 | NM_033020.2 | #N/A | 167.523109 | -2.2021819 | 0.82424227 |
| 16992 | NM_145341.3 | PDCD4 | 10.3689854 | -4.5868562 | 1.71691663 |
| 16994 | NR_135825.1 | LOC646548 | 18.062357 | 3.8275818 | 1.43285063 |
| 16993 | NM_001190329.1 | #N/A | 11.2400021 | -4.7046858 | 1.76119365 |
| 16995 | NM_001039615.3 | ZNF705D | 30.9152451 | 9.93445335 | 3.7189905 |
| 16996 | NM_001282437.1 | BLOC1S2 | 18.6956198 | -5.4268876 | 2.03175471 |
| 16998 | NR_135516.1 | CYYR1-AS1 | 19.3206012 | 8.54606338 | 3.19962355 |
| 16997 | NM_005588.2 | #N/A | 19.3206012 | 8.54606338 | 3.19962355 |
| 16999 | NM_001031712.2 | #N/A | 62.1672391 | -3.4544705 | 1.29341067 |
| 17000 | NM_001282652.1 | STIP1 | 14.1511325 | -5.0266248 | 1.88209642 |
| 17001 | NM_001321230.1 | #N/A | 11.1913531 | -4.6966497 | 1.7589655 |
| 17002 | NM_001319307.1 | #N/A | 62.5347846 | -7.1770496 | 2.68798183 |
| 17003 | NR_120479.1 | LINC02367 | 33.5490355 | 4.83206223 | 1.80986219 |
| 17004 | NM_014692.1 | #N/A | 9.80837418 | -4.5121065 | 1.69008759 |
| 17006 | NM_207304.2 | MBNL2 | 47.6184257 | -6.7853802 | 2.54162638 |
| 17005 | NM_001042625.1 | #N/A | 19.2584955 | -5.4832842 | 2.05390214 |
| 17007 | NM_001003810.1 | HNRNPD | 123.266829 | -6.2228182 | 2.33095518 |
| 17008 | NM_001320698.1 | #N/A | 10.6884177 | -4.6391541 | 1.73776235 |
| 17009 | NM_001025434.1 | #N/A | 11.5478294 | -4.7358335 | 1.77402119 |
| 17010 | NM_001330199.1 | #N/A | 11.2301336 | -4.7058472 | 1.76290314 |
| 17011 | NR_110616.1 | LINC01355 | 12.0097325 | -4.7906208 | 1.79468066 |
| 17012 | NM_001346446.1 | #N/A | 48.4060174 | -6.8091035 | 2.55103702 |
| 17013 | NR_026757.1 | #N/A | 220.74084 | -2.8907612 | 1.08304227 |
| 17015 | NR_046343.1 | FAM90A7P | 39.7557176 | 10.2883129 | 3.85475067 |
| 17014 | NM_001307928.1 | SERPINB12 | 39.7557176 | 10.2883129 | 3.85475067 |
| 17016 | NR_117086.1 | ZNF714 | 17.4034548 | -5.3211765 | 1.99387623 |
| 17017 | NM_003862.2 | #N/A | 55.6989541 | -3.0174744 | 1.13084535 |
| 17018 | NR_073199.1 | #N/A | 35.4520178 | 2.87686036 | 1.07833649 |
| 17019 | NM_203341.2 | SELENOF | 23.3467659 | -5.7592689 | 2.15936567 |
| 17020 | NM_138430.4 | #N/A | 10.1093006 | -4.5548189 | 1.70787212 |
| 17021 | NR_045555.1 | #N/A | 10.2490976 | -4.5742559 | 1.71518688 |
| 17022 | NM_001184782.1 | RADX | 14.5518346 | -5.061959 | 1.89818918 |
| 17023 | NM_001098424.1 | DLG1 | 48.1267385 | -6.8004841 | 2.55015305 |
| 17024 | NM_145011.3 | #N/A | 61.6553121 | -7.156386 | 2.6837705 |
| 17025 | NM_014553.2 | #N/A | 29.667291 | 4.17807483 | 1.56688104 |
| 17026 | NR_027133.1 | LINC00924 | 47.8727041 | 2.22948778 | 0.83622371 |
| 17027 | NM_001135669.1 | #N/A | 66.9248778 | -7.2740282 | 2.7285354 |

| | | | | | |
|---|---|---|---|---|---|
| 17029 | NR_046312.1 | KIAA1656 | 38.0515612 | -2.1458677 | 0.80496387 |
| 17028 | NM_138802.2 | #N/A | 15.4355578 | -5.1559405 | 1.9341178 |
| 17030 | NR_126015.1 | LYRM4-AS1 | 15.5312765 | 4.20046709 | 1.57589067 |
| 17031 | NM_001243425.1 | #N/A | 11.1771596 | -4.690252 | 1.75966833 |
| 17032 | NM_001159703.1 | #N/A | 14.1615356 | -5.0443791 | 1.89306261 |
| 17033 | NR_004404.1 | #N/A | 15.8655972 | 5.47685506 | 2.05553049 |
| 17035 | NM_005793.4 | NME6 | 48.1498731 | -6.8012407 | 2.55277148 |
| 17034 | NM_001278284.1 | #N/A | 51.2995107 | -6.8929595 | 2.58720863 |
| 17036 | NR_144347.1 | #N/A | 30.6289928 | 9.9210664 | 3.72375935 |
| 17037 | NM_030824.2 | #N/A | 10.2995532 | -4.5650022 | 1.71365837 |
| 17038 | NR_110090.1 | SIRPG-AS1 | 33.6067179 | 1.81469434 | 0.68129735 |
| 17039 | NM_001130100.1 | #N/A | 58.9022699 | -7.0911358 | 2.66262155 |
| 17040 | NM_001281740.2 | #N/A | 12.4610469 | -4.8533435 | 1.82237327 |
| 17041 | NM_001303052.1 | #N/A | 55.3715494 | -7.0027551 | 2.62960132 |
| 17042 | NR_121676.1 | LINC01179 | 26.4121127 | 2.30551372 | 0.86576506 |
| 17043 | NM_001320598.1 | #N/A | 14.6752436 | -5.0734536 | 1.90530604 |
| 17044 | NM_003311.3 | #N/A | 18.6310915 | -4.1419635 | 1.55553879 |
| 17045 | NM_001177382.1 | #N/A | 59.7522727 | -7.1120781 | 2.67128724 |
| 17046 | NM_001286779.1 | GARNL3 | 49.0512604 | -6.8280436 | 2.56472295 |
| 17047 | NM_033408.3 | TMEM205 | 11.287852 | -4.703735 | 1.76681045 |
| 17048 | NM_030783.2 | #N/A | 55.0694952 | -6.9948749 | 2.62780609 |
| 17049 | NM_004945.3 | DNM2 | 63.7335402 | -7.2038275 | 2.70632576 |
| 17050 | NM_001347969.1 | #N/A | 50.6706189 | -6.8749333 | 2.58353638 |
| 17051 | NM_001252052.1 | PCMT1 | 54.371833 | -6.9764496 | 2.62180637 |
| 17052 | NM_016587.3 | #N/A | 620.618448 | -3.9209349 | 1.47354388 |
| 17053 | NM_001100400.1 | PDS5A | 11.8036984 | -4.7611855 | 1.78947214 |
| 17054 | NM_001324229.1 | #N/A | 11.2695159 | -4.7035779 | 1.76782917 |
| 17055 | NM_020883.1 | #N/A | 175.253824 | -2.9031237 | 1.09112747 |
| 17056 | NM_001320437.1 | #N/A | 46.868733 | -6.7624335 | 2.54177723 |
| 17057 | NM_053285.1 | #N/A | 22.3080764 | -5.6942304 | 2.14034988 |
| 17058 | NM_001130041.1 | #N/A | 68.4353308 | -7.3069119 | 2.74666201 |
| 17059 | NM_015686.2 | FAM155B | 72.7674028 | -3.6382943 | 1.3677396 |
| 17060 | NM_001323516.1 | #N/A | 47.8054302 | -6.7909132 | 2.55295458 |
| 17061 | NM_001293197.1 | #N/A | 13.2863095 | -4.9362224 | 1.85581552 |
| 17062 | NM_001242798.1 | ZNF322 | 62.7195134 | -7.1805376 | 2.69979011 |
| 17063 | NM_001135214.2 | #N/A | 50.862066 | -6.8803813 | 2.58696009 |
| 17064 | NR_117101.1 | CASC11 | 14.3809208 | 8.06635522 | 3.03328571 |
| 17065 | NM_001290043.1 | TAP2 | 47.8473202 | -6.7923241 | 2.55422964 |
| 17066 | NR_036586.1 | LINC02245 | 64.7015034 | 2.53495103 | 0.95330843 |
| 17067 | NM_024775.9 | #N/A | 18.2490578 | -4.1145691 | 1.54736682 |
| 17068 | NM_001329755.1 | #N/A | 10.6716841 | -4.6320827 | 1.7421437 |
| 17069 | NM_001256755.1 | #N/A | 9.00553555 | 8.12538143 | 3.05633601 |
| 17070 | NM_001185092.1 | #N/A | 28.3246131 | -2.2923747 | 0.86234911 |

| | | | | | |
|---|---|---|---|---|---|
| 17071 | NR_103772.1 | LOC100288162 | 34.7121367 | -3.4686066 | 1.30487587 |
| 17072 | NM_001100626.1 | #N/A | 58.5583833 | -7.0825127 | 2.66499843 |
| 17073 | NM_001281535.1 | EPB41L3 | 54.4481857 | -6.9788707 | 2.62645844 |
| 17074 | NM_001114380.1 | #N/A | 17.7854864 | 2.53960579 | 0.95580158 |
| 17075 | NM_012311.3 | #N/A | 253.152896 | 1.03771237 | 0.39060101 |
| 17076 | NR_136606.1 | #N/A | 56.0573743 | -7.0201231 | 2.6426187 |
| 17077 | NM_001173480.1 | #N/A | 60.4385511 | -7.1278346 | 2.68327925 |
| 17078 | NR_026995.1 | RAPGEF4-AS1 | 15.2391819 | 5.1390565 | 1.93469719 |
| 17079 | NM_001294139.1 | PRAMEF1 | 10.1264907 | 5.29061113 | 1.9922358 |
| 17081 | NR_126411.1 | SGMS1-AS1 | 41.6062331 | 3.09581214 | 1.16576634 |
| 17080 | NR_125352.1 | #N/A | 11.4638498 | -4.7356118 | 1.78326385 |
| 17082 | NM_001126337.1 | #N/A | 121.338482 | -2.5920357 | 0.97625096 |
| 17083 | NR_001575.1 | RNF138P1 | 43.996138 | 1.67476225 | 0.63078187 |
| 17084 | NM_001198799.2 | #N/A | 19.1881451 | -3.9174246 | 1.47549469 |
| 17085 | NM_001145009.1 | #N/A | 13.4934562 | -4.9615703 | 1.86881295 |
| 17086 | NM_016120.3 | #N/A | 341.241706 | -1.6952285 | 0.63852862 |
| 17087 | NM_021179.2 | CCDC181 | 19.6472229 | -5.4960188 | 2.07022186 |
| 17089 | NM_020808.4 | SIPA1L2 | 281.506013 | -2.5311033 | 0.95369825 |
| 17088 | NM_001142450.1 | #N/A | 49.0731116 | -6.8288611 | 2.57304304 |
| 17090 | NM_139312.2 | #N/A | 14.4857868 | -5.0713423 | 1.91101312 |
| 17091 | NM_032142.3 | #N/A | 119.258041 | -2.3824831 | 0.89781305 |
| 17092 | NR_102327.1 | HCG18 | 23.6594872 | -3.1631905 | 1.192106 |
| 17093 | NM_001130841.1 | RTCA | 11.9277046 | -4.789736 | 1.80528398 |
| 17094 | NM_001345844.1 | C19orf57 | 11.4961496 | -4.7176281 | 1.77815804 |
| 17095 | NM_024056.3 | TMEM106C | 60.5083957 | -7.1296475 | 2.68818976 |
| 17096 | NM_001287192.1 | #N/A | 155.264518 | -3.15589 | 1.19028556 |
| 17097 | NR_028106.1 | DCAF8 | 11.6844254 | -4.7638156 | 1.79687386 |
| 17098 | NM_207034.2 | #N/A | 13.5579331 | -4.9606164 | 1.8711263 |
| 17099 | NM_001160367.1 | #N/A | 62.338244 | -7.1719603 | 2.70537503 |
| 17100 | NM_206914.1 | #N/A | 11.1666392 | -4.6954047 | 1.77122835 |
| 17101 | NR_003545.1 | #N/A | 14.5787903 | -5.0687398 | 1.91211272 |
| 17102 | NM_182975.3 | ZNF326 | 9.80265852 | -4.5133911 | 1.70274222 |
| 17103 | NM_022160.2 | #N/A | 12.7281443 | -4.8899668 | 1.84494285 |
| 17104 | NM_003749.2 | #N/A | 367.352858 | -3.0231843 | 1.14078853 |
| 17105 | NM_024050.5 | #N/A | 286.244008 | -1.6469635 | 0.62149521 |
| 17106 | NM_080750.4 | DPH3P1 | 71.5535191 | 1.58557594 | 0.5983388 |
| 17107 | NM_001114089.3 | ENTPD6 | 65.8751735 | -7.2515459 | 2.73650382 |
| 17108 | NR_103790.1 | LINC00581 | 22.0471093 | 3.08541086 | 1.16460728 |
| 17109 | NM_017453.3 | STAU1 | 60.6483568 | -7.1338325 | 2.6929017 |
| 17110 | NR_111941.1 | MISP3 | 14.5775257 | -5.0758137 | 1.91626235 |
| 17111 | NR_003136.1 | #N/A | 11.8834115 | -4.7853312 | 1.80679023 |
| 17112 | NM_021012.4 | #N/A | 39.9771723 | -2.8148442 | 1.06283992 |
| 17113 | NM_001287339.1 | CAB39L | 38.5509989 | 10.2438306 | 3.86798212 |

| | | | | | |
|---|---|---|---|---|---|
| 17114 | NR_135124.1 | GPRACR | 70.9287038 | 3.9599842 | 1.49530654 |
| 17115 | NM_001003962.2 | #N/A | 61.3109654 | -7.1488545 | 2.69956373 |
| 17116 | NM_001321785.1 | TASOR2 | 61.2765048 | -7.1485252 | 2.69957051 |
| 17117 | NM_006729.4 | #N/A | 11.4776896 | -4.7375756 | 1.78919773 |
| 17118 | NM_001134285.2 | #N/A | 17.2782826 | -3.2155837 | 1.2146788 |
| 17119 | NM_178523.4 | #N/A | 16.3391283 | -5.2461444 | 1.98184226 |
| 17120 | NM_001172220.1 | #N/A | 59.5777927 | -7.1069904 | 2.68488219 |
| 17122 | NR_077247.1 | RPS18P9 | 9.70475308 | -4.4873053 | 1.69535319 |
| 17121 | NM_173598.4 | #N/A | 46.1103806 | -2.862444 | 1.08145894 |
| 17123 | NM_001287442.1 | JADE1 | 48.98093 | -6.8258611 | 2.57920963 |
| 17124 | NM_019601.3 | #N/A | 9.75503443 | -4.5048256 | 1.70220084 |
| 17125 | NM_001330301.1 | #N/A | 62.0430068 | -7.1653091 | 2.70755219 |
| 17126 | NM_020155.3 | #N/A | 10.3990753 | -4.5999117 | 1.73827715 |
| 17127 | NR_002318.2 | CATSPER2P1 | 9.58845687 | -4.4641182 | 1.68729004 |
| 17128 | NM_001329237.1 | #N/A | 64.9537711 | -7.2330666 | 2.7339656 |
| 17129 | NM_014400.2 | #N/A | 10.5329066 | -4.6006397 | 1.73899367 |
| 17130 | NM_213654.3 | ARMC8 | 15.1457556 | -5.1331454 | 1.94051603 |
| 17131 | NM_153277.2 | #N/A | 19.3688854 | 8.4649924 | 3.20039715 |
| 17133 | NR_110812.1 | TMEM108-AS1 | 28.1135782 | 5.04098738 | 1.90617332 |
| 17132 | NM_001025234.1 | TSPAN4 | 17.1012566 | -5.3144627 | 2.00959212 |
| 17134 | NM_001304794.1 | #N/A | 13.7659377 | -5.0050939 | 1.89277766 |
| 17135 | NM_001321732.1 | #N/A | 14.1242325 | -5.0394071 | 1.9062454 |
| 17136 | NM_032453.1 | #N/A | 198.489914 | 1.28872249 | 0.48753303 |
| 17137 | NM_201517.2 | H2AZ2 | 17.616498 | -5.3555289 | 2.02627724 |
| 17138 | NM_139314.2 | #N/A | 20.9240091 | -5.5995671 | 2.11873237 |
| 17139 | NR_038242.1 | VEZT | 50.60956 | -6.8733553 | 2.60102359 |
| 17140 | NM_001282723.1 | LDAH | 11.3619889 | -4.7096318 | 1.7824478 |
| 17141 | NM_002878.3 | RAD51D | 17.3609329 | -5.3224279 | 2.01449826 |
| 17142 | NM_001282524.1 | NPIPB6 | 10.7037509 | -4.6310253 | 1.75286912 |
| 17143 | NM_000783.3 | #N/A | 14.0347404 | -5.0252958 | 1.90218421 |
| 17144 | NM_080413.2 | VPS16 | 17.3423043 | -5.3209501 | 2.01424814 |
| 17145 | NM_001042493.2 | #N/A | 20.6460156 | -5.5731727 | 2.10980328 |
| 17146 | NM_024503.4 | #N/A | 59.3714259 | -7.1018175 | 2.68865066 |
| 17147 | NM_001319062.1 | GUCA1A | 14.9853387 | -5.1064633 | 1.93419496 |
| 17148 | NM_001256092.1 | #N/A | 47.2726027 | -6.7749312 | 2.56630799 |
| 17149 | NM_001261408.1 | #N/A | 15.965409 | -5.2058218 | 1.97221898 |
| 17150 | NR_033945.1 | LINC01192 | 14.2571398 | 3.98130469 | 1.50843731 |
| 17151 | NM_001145785.1 | #N/A | 14.18371 | -3.730111 | 1.4133479 |
| 17152 | NR_110220.1 | LINC01237 | 12.7582629 | 5.01565988 | 1.90052099 |
| 17153 | NM_022111.3 | #N/A | 58.3250722 | -7.0764073 | 2.68159438 |
| 17154 | NM_001185063.2 | ZNF75D | 12.9180767 | -4.896499 | 1.85555484 |
| 17155 | NM_020728.2 | #N/A | 376.062049 | -1.7316121 | 0.6563915 |
| 17156 | NR_135566.1 | LINC01857 | 23.8338568 | 2.94565107 | 1.11662474 |

| | | | | | |
|---|---|---|---|---|---|
| 17158 | NM_007350.3 | PHLDA1 | 540.894061 | -1.7700413 | 0.67100003 |
| 17157 | NM_001243270.1 | #N/A | 13.0096053 | -4.9044805 | 1.85920369 |
| 17159 | NM_024926.3 | #N/A | 57.3586897 | -7.0525978 | 2.67353891 |
| 17160 | NR_015430.1 | #N/A | 17.4410015 | 1.97623832 | 0.74930921 |
| 17161 | NM_022045.4 | #N/A | 18.3952684 | -3.3258158 | 1.26104823 |
| 17162 | NM_006298.3 | #N/A | 179.504583 | -2.5894266 | 0.98184212 |
| 17163 | NM_145663.2 | #N/A | 10.3148568 | -4.5758887 | 1.7351162 |
| 17164 | NM_024311.3 | #N/A | 18.6148672 | -5.4209718 | 2.05563828 |
| 17165 | NM_198474.3 | #N/A | 19.696485 | -2.8565827 | 1.08349612 |
| 17166 | NM_001204344.1 | TNFSF15 | 26.8611213 | -2.61736 | 0.99278977 |
| 17167 | NM_006291.2 | #N/A | 72.1901734 | -3.312135 | 1.25642525 |
| 17168 | NM_004018.2 | #N/A | 45.3174158 | -6.7139704 | 2.54694681 |
| 17169 | NM_001318136.1 | #N/A | 12.55007 | -4.8664543 | 1.8461719 |
| 17170 | NM_015440.4 | MTHFD1L | 61.6895982 | -7.1577198 | 2.71545728 |
| 17171 | NM_018931.2 | #N/A | 45.325217 | 2.04223927 | 0.77480148 |
| 17172 | NR_037929.1 | SLMO2-ATP5E | 12.6348375 | -4.8778113 | 1.85060249 |
| 17173 | NM_001281447.1 | GTF2IRD2 | 11.1107402 | -4.6883409 | 1.77874648 |
| 17174 | NM_025198.4 | MTERF2 | 19.7949625 | -5.5179138 | 2.09354345 |
| 17175 | NM_001346749.1 | #N/A | 29.4839837 | 9.86623755 | 3.74364937 |
| 17176 | NM_032504.1 | UNC80 | 13.7314291 | -4.9813594 | 1.89019397 |
| 17177 | NR_120551.1 | LOC101928535 | 20.9105435 | 4.04596211 | 1.5353462 |
| 17178 | NM_001206947.1 | #N/A | 72.8268199 | -7.3957324 | 2.80652947 |
| 17179 | NM_001199295.1 | #N/A | 44.7311497 | -6.6952233 | 2.54121451 |
| 17180 | NR_135657.1 | LOC101928000 | 14.1924326 | -5.035616 | 1.91167 |
| 17181 | NM_001144869.2 | #N/A | 11.3331204 | -4.7029682 | 1.78557611 |
| 17182 | NM_031936.4 | #N/A | 10.3695801 | -4.5863246 | 1.74138855 |
| 17183 | NM_001243224.1 | RAD9A | 14.3051859 | -5.0581084 | 1.92080105 |
| 17184 | NR_138037.1 | HCG20 | 30.4943665 | 1.78902681 | 0.67939192 |
| 17185 | NM_019035.4 | #N/A | 50.6260046 | -6.8738842 | 2.61042687 |
| 17186 | NM_053055.4 | #N/A | 201.424589 | 1.60844045 | 0.610912 |
| 17187 | NR_024597.1 | #N/A | 11.23399 | -4.694801 | 1.78329011 |
| 17188 | NM_001317851.1 | HEMK1 | 45.8918729 | -6.7318856 | 2.55717623 |
| 17189 | NM_024429.1 | PRKAG2 | 57.6956987 | -7.0605607 | 2.68203444 |
| 17190 | NR_145416.1 | ZNRD1ASP | 11.5217655 | -4.7273841 | 1.79579617 |
| 17191 | NM_001318485.1 | #N/A | 10.0910674 | -4.5541568 | 1.73007649 |
| 17192 | NM_153027.1 | #N/A | 12.7081715 | -4.872678 | 1.85116975 |
| 17193 | NR_049896.1 | C18orf61 | 38.6347125 | 2.7124152 | 1.03050049 |
| 17194 | NM_001170570.1 | #N/A | 58.5350359 | 6.45007895 | 2.45062151 |
| 17195 | NM_001321123.1 | #N/A | 60.6989307 | -7.1340636 | 2.71049114 |
| 17196 | NM_138463.3 | #N/A | 9.41537102 | -4.4532238 | 1.69214262 |
| 17197 | NM_006434.2 | SORBS1 | 49.0726713 | -6.8283854 | 2.5948105 |
| 17198 | NM_001242861.2 | PRKACB | 61.7523162 | -7.1579611 | 2.72028911 |
| 17199 | NM_001220.4 | #N/A | 16.1962578 | -5.2264973 | 1.98635207 |

| | | | | | |
|---|---|---|---|---|---|
| 17200 | NM_001130919.1 | #N/A | 46.6563812 | -6.7556298 | 2.56755978 |
| 17201 | NM_015297.2 | #N/A | 63.3420306 | -7.1954159 | 2.73489586 |
| 17202 | NM_000834.3 | #N/A | 13.9554326 | -5.0026789 | 1.90163931 |
| 17203 | NM_001204453.1 | PSMC2 | 10.0094655 | -4.5237392 | 1.71974352 |
| 17204 | NR_033992.2 | #N/A | 10.103234 | -4.538526 | 1.72564192 |
| 17205 | NM_005990.3 | #N/A | 111.568532 | -3.0108718 | 1.14502253 |
| 17206 | NM_001135806.1 | #N/A | 60.2970585 | -7.1244506 | 2.70949918 |
| 17207 | NM_001185012.1 | NDUFA2 | 10.2281487 | -4.5568553 | 1.73326839 |
| 17208 | NM_001253908.1 | AKR1C3 | 29.1977315 | 9.85219897 | 3.74883599 |
| 17209 | NM_001300861.1 | #N/A | 16.3294977 | -5.2468435 | 1.99659592 |
| 17210 | NR_037583.1 | #N/A | 11.2062213 | -4.6840499 | 1.78247188 |
| 17211 | NM_001023570.3 | #N/A | 46.1973508 | -6.7419556 | 2.5656515 |
| 17212 | NR_073470.1 | #N/A | 14.5636209 | -5.0698244 | 1.92937704 |
| 17213 | NM_001301701.1 | ERI3 | 13.2821617 | -4.9486594 | 1.88328274 |
| 17214 | NM_001085429.1 | #N/A | 66.0205281 | 2.9071749 | 1.10639042 |
| 17215 | NM_001166247.1 | GRIK2 | 15.8242079 | -5.1889475 | 1.97478871 |
| 17216 | NM_001134439.1 | #N/A | 47.273995 | -6.7751951 | 2.57858395 |
| 17217 | NM_000898.4 | #N/A | 13.7051429 | -4.9951258 | 1.90112685 |
| 17218 | NM_024014.3 | #N/A | 12.9469177 | -4.911568 | 1.86956416 |
| 17219 | NM_001009877.2 | BRD9 | 11.4312119 | -4.7200619 | 1.7967038 |
| 17220 | NM_001077197.1 | PDE11A | 37.3462802 | 10.1979306 | 3.88197124 |
| 17221 | NM_147195.3 | #N/A | 37.3462802 | 10.1979306 | 3.88197124 |
| 17222 | NM_001244767.1 | #N/A | 15.7716012 | -3.8957961 | 1.48313809 |
| 17223 | NM_197977.2 | #N/A | 12.9703777 | -4.9077578 | 1.86846487 |
| 17224 | NM_001297734.1 | #N/A | 47.1375572 | -6.7705094 | 2.57772075 |
| 17225 | NR_144321.1 | TRIM8 | 15.0499896 | -5.1297762 | 1.95330505 |
| 17226 | NM_001160233.1 | ATP1A1 | 18.2349504 | -5.3953191 | 2.05447603 |
| 17227 | NR_135542.1 | #N/A | 72.8191445 | -7.3961841 | 2.81703072 |
| 17228 | NM_001278194.1 | GANAB | 74.0419698 | -7.4217189 | 2.82736321 |
| 17229 | NM_152534.4 | NEK10 | 14.1639181 | -5.0413421 | 1.9206608 |
| 17230 | NM_005429.4 | #N/A | 10.0314351 | -4.5471416 | 1.73259566 |
| 17231 | NM_182920.1 | #N/A | 44.7458609 | -6.6956908 | 2.55135558 |
| 17232 | NR_144634.1 | LOC728715 | 12.4163364 | -4.8281686 | 1.83976558 |
| 17233 | NM_001345871.1 | #N/A | 47.3897916 | -6.7784007 | 2.58343753 |
| 17234 | NM_001288747.1 | DKC1 | 10.3861513 | -4.5986824 | 1.75275402 |
| 17235 | NR_104101.1 | VPS29 | 9.16162966 | -4.406934 | 1.67971217 |
| 17236 | NM_020395.3 | #N/A | 14.0800245 | -5.0249346 | 1.91533401 |
| 17237 | NM_014876.5 | #N/A | 324.978617 | -2.5138054 | 0.95833812 |
| 17238 | NM_017899.3 | #N/A | 14.4539881 | -5.0705858 | 1.93307876 |
| 17239 | NM_001024372.1 | #N/A | 19.3186385 | -5.4867471 | 2.09185656 |
| 17240 | NM_001033506.1 | CSTF3 | 10.8256536 | -4.6494926 | 1.77273215 |
| 17241 | NM_020405.4 | #N/A | 144.857111 | 1.63793044 | 0.62460229 |
| 17242 | NM_001001662.2 | #N/A | 12.2184535 | -4.8325125 | 1.84293096 |

| | | | | | |
|---|---|---|---|---|---|
| 17243 | NM_001320899.1 | #N/A | 13.3426258 | -4.9573111 | 1.89060666 |
| 17244 | NM_001526.4 | HCRTR2 | 13.9961857 | -5.0159029 | 1.9131299 |
| 17245 | NR_046359.1 | #N/A | 14.0623364 | -5.0184001 | 1.91411493 |
| 17246 | NM_001320512.1 | NMNAT3 | 11.7169113 | -4.7461563 | 1.81034406 |
| 17247 | NM_178824.3 | #N/A | 13.5288339 | -4.9768875 | 1.89840739 |
| 17248 | NM_001322029.1 | GPATCH2L | 10.7789553 | -4.6528441 | 1.7748378 |
| 17249 | NM_021013.3 | KRT34 | 21.8766792 | 8.63587667 | 3.2947278 |
| 17250 | NR_126330.1 | #N/A | 11.5327648 | -4.744121 | 1.8099727 |
| 17251 | NM_001031709.2 | #N/A | 9.48718473 | -4.4521242 | 1.69906239 |
| 17252 | NM_001423.2 | #N/A | 153.057475 | -2.5930385 | 0.98978638 |
| 17253 | NM_001319052.1 | GALNT15 | 18.1316677 | -3.8368137 | 1.46458039 |
| 17255 | NR_110839.1 | LINC02087 | 14.6806984 | 4.85534931 | 1.85344244 |
| 17254 | NM_001323534.1 | #N/A | 44.0297569 | -6.6722843 | 2.54704391 |
| 17256 | NM_001278662.1 | #N/A | 21.2687569 | -3.5539838 | 1.35698176 |
| 17257 | NM_001282709.1 | EXOSC2 | 61.8420483 | 4.96624447 | 1.89633446 |
| 17258 | NM_001288616.1 | #N/A | 11.0932477 | -4.6891569 | 1.79055983 |
| 17259 | NM_001330233.1 | MPP3 | 22.712821 | -5.709762 | 2.18054688 |
| 17260 | NM_001556.2 | #N/A | 47.1794665 | -6.7716013 | 2.58634918 |
| 17261 | NM_001718.5 | #N/A | 9.46692282 | -4.4612892 | 1.70394619 |
| 17262 | NR_138051.1 | #N/A | 46.1935978 | -6.7412449 | 2.57494751 |
| 17263 | NR_109827.1 | #N/A | 33.637689 | 4.37325454 | 1.6705927 |
| 17264 | NM_001256547.1 | #N/A | 38.5506028 | 5.26963585 | 2.01338667 |
| 17265 | NM_001009923.1 | #N/A | 9.73356035 | -4.4933758 | 1.71682808 |
| 17266 | NM_004036.4 | #N/A | 187.821001 | -2.356932 | 0.90067636 |
| 17267 | NM_001317721.1 | LRWD1 | 12.3201873 | -4.8192479 | 1.84166381 |
| 17268 | NM_001287062.1 | #N/A | 15.7747458 | -5.181992 | 1.98058551 |
| 17269 | NM_001166263.1 | MTIF3 | 11.2451288 | -4.6918086 | 1.79335658 |
| 17270 | NM_006425.4 | #N/A | 234.273402 | -1.7697149 | 0.67651932 |
| 17271 | NM_001134296.1 | AP3M2 | 14.4758445 | -5.0653021 | 1.93643593 |
| 17272 | NM_018327.2 | #N/A | 16.1331146 | -5.2290359 | 1.99915158 |
| 17273 | NM_145297.3 | ZNF626 | 9.81003814 | -4.5005931 | 1.72075666 |
| 17274 | NR_073516.1 | #N/A | 19.4939586 | -3.1476591 | 1.20355301 |
| 17275 | NM_006906.1 | #N/A | 58.8515145 | -7.0895154 | 2.71108278 |
| 17276 | NM_001318253.1 | #N/A | 19.489478 | -5.4876602 | 2.09856155 |
| 17277 | NM_001320591.1 | #N/A | 56.4344243 | -7.0281157 | 2.68766573 |
| 17278 | NM_002191.3 | #N/A | 9.42577744 | -4.4477158 | 1.70126137 |
| 17279 | NM_004602.3 | STAU1 | 85.9362466 | -5.7244856 | 2.18969369 |
| 17280 | NM_007204.4 | #N/A | 123.131894 | -1.6963203 | 0.64888975 |
| 17281 | NM_201436.2 | H2AZ2 | 10.4409151 | -4.5889338 | 1.75546586 |
| 17282 | NM_001142614.1 | #N/A | 49.1108758 | -6.8293981 | 2.61347136 |
| 17283 | NM_001318492.1 | #N/A | 11.3375535 | -4.6969112 | 1.79743187 |
| 17285 | NM_001330370.1 | LRRTM4 | 28.6252269 | 9.82370371 | 3.75948187 |
| 17284 | NM_001289953.1 | ZNF761 | 28.6252269 | 9.82370371 | 3.75948187 |

| | | | | | |
|---|---|---|---|---|---|
| 17286 | NM_001837.3 | #N/A | 28.6252269 | 9.82370371 | 3.75948187 |
| 17287 | NM_004375.4 | #N/A | 45.4540801 | -6.7185154 | 2.57162839 |
| 17288 | NR_037575.1 | #N/A | 14.2848928 | -5.0523815 | 1.93390016 |
| 17289 | NM_001321006.1 | #N/A | 74.7165105 | 1.8170365 | 0.69563176 |
| 17290 | NM_001042595.2 | TMEM91 | 9.10650196 | -4.4010278 | 1.68492377 |
| 17291 | NM_001007189.1 | #N/A | 9.88430105 | -4.5192789 | 1.73021197 |
| 17292 | NM_001301867.1 | #N/A | 10.0656805 | -4.5533176 | 1.74325761 |
| 17293 | NR_126005.1 | KRT73-AS1 | 25.5235116 | 2.78010867 | 1.06438784 |
| 17294 | NM_000787.3 | #N/A | 10.6260946 | -4.6313082 | 1.77329971 |
| 17295 | NM_001190716.1 | #N/A | 63.6712386 | -7.2035084 | 2.7583443 |
| 17296 | NR_073472.1 | #N/A | 14.5790075 | -5.0633569 | 1.93907918 |
| 17297 | NM_002644.3 | #N/A | 45.3272962 | 2.11594076 | 0.81041512 |
| 17298 | NM_021072.3 | #N/A | 26.0284064 | -2.6519737 | 1.01574982 |
| 17299 | NM_001079804.2 | #N/A | 60.1841542 | -7.1229209 | 2.72843298 |
| 17300 | NM_032898.4 | #N/A | 81.6229538 | -1.8486437 | 0.70812619 |
| 17301 | NM_001077498.2 | #N/A | 55.4777584 | -7.0042431 | 2.68311302 |
| 17302 | NM_001145290.1 | #N/A | 80.8851114 | 1.80024002 | 0.68964188 |
| 17303 | NM_015042.1 | #N/A | 381.694064 | -3.6186767 | 1.38628127 |
| 17304 | NM_001306219.2 | #N/A | 43.3336704 | -6.6495536 | 2.54800937 |
| 17305 | NM_153645.2 | NUP50 | 37.650439 | 3.70186478 | 1.41852326 |
| 17306 | NR_111951.1 | LINC00869 | 12.0777445 | -4.8018197 | 1.84010812 |
| 17307 | NR_036551.1 | RDM1P5 | 10.168243 | -4.5485748 | 1.74324549 |
| 17308 | NM_001243253.1 | #N/A | 18.0166677 | -3.2823915 | 1.25818787 |
| 17309 | NM_198401.3 | #N/A | 20.0506092 | -5.5330138 | 2.12088732 |
| 17310 | NM_001076682.3 | #N/A | 10.9777302 | -4.6655899 | 1.78842884 |
| 17311 | NM_181304.2 | #N/A | 11.460597 | -4.7160407 | 1.80777364 |
| 17312 | NR_126342.1 | LOC171391 | 9.35209146 | -4.4286346 | 1.69766633 |
| 17313 | NM_003608.3 | #N/A | 15.0439189 | 3.74420796 | 1.43554444 |
| 17314 | NM_001318868.1 | DOK1 | 13.3362059 | -4.949711 | 1.89780855 |
| 17316 | NR_026933.2 | C20orf173 | 9.91429358 | 6.50635216 | 2.49503231 |
| 17315 | NM_001322189.1 | #N/A | 13.2691208 | -4.9386695 | 1.89383856 |
| 17317 | NR_045039.1 | #N/A | 9.47341453 | -4.458154 | 1.70958606 |
| 17318 | NR_040034.1 | LOC339298 | 20.0506203 | 5.44052 | 2.08683061 |
| 17319 | NM_001033551.2 | #N/A | 55.6544064 | -7.0098756 | 2.68934696 |
| 17320 | NM_001257379.1 | SEC14L3 | 13.5472427 | 7.97748438 | 3.06106257 |
| 17321 | NM_001248.3 | #N/A | 9.7571484 | -4.4977778 | 1.72588616 |
| 17322 | NM_173084.2 | #N/A | 83.0318107 | -1.5199033 | 0.58329838 |
| 17323 | NM_001034116.1 | EIF2B4 | 44.6564857 | -6.6929465 | 2.56874044 |
| 17324 | NM_001305203.1 | #N/A | 51.2491223 | -6.891209 | 2.64486465 |
| 17325 | NM_025133.4 | FBXO11 | 28.3389747 | 9.80924166 | 3.76494593 |
| 17326 | NM_145284.5 | FAM122B | 42.4477811 | -6.6196765 | 2.54083635 |
| 17327 | NM_001174117.1 | #N/A | 59.9404121 | -7.1164878 | 2.73163906 |
| 17328 | NM_001025161.2 | #N/A | 15.7959325 | -5.1908262 | 1.99258179 |

| | | | | | |
|---|---|---|---|---|---|
| 17329 | NM_001289132.1 | CCDC32 | 33.9870641 | 1.7688991 | 0.67905359 |
| 17330 | NM_001282763.1 | CDV3 | 53.3872395 | -6.949655 | 2.66798049 |
| 17331 | NR_024528.1 | RPL23AP7 | 9.97229448 | -4.5400378 | 1.74323927 |
| 17332 | NR_121582.1 | LOC101928786 | 58.2463663 | 3.04653538 | 1.16984778 |
| 17333 | NM_001005361.2 | DNM2 | 79.8965568 | -7.5293601 | 2.89162193 |
| 17334 | NM_001204140.1 | #N/A | 54.9540363 | -6.9901851 | 2.68518654 |
| 17335 | NM_014900.4 | #N/A | 12.6923471 | -4.8836207 | 1.87604615 |
| 17336 | NM_100486.3 | #N/A | 59.5948862 | -3.0427103 | 1.16911806 |
| 17337 | NR_040599.1 | CARD8-AS1 | 67.885825 | 2.90383614 | 1.11583992 |
| 17338 | NM_001306094.1 | NPIPB15 | 10.6325168 | -4.6241482 | 1.77716229 |
| 17340 | NR_110554.1 | LINC02288 | 40.1765554 | 2.68912952 | 1.03359058 |
| 17339 | NM_001059.2 | #N/A | 13.3369368 | -4.9541658 | 1.90418413 |
| 17341 | NR_045789.1 | NSUN4 | 10.6910054 | -4.6381811 | 1.78282143 |
| 17342 | NR_038457.1 | #N/A | 13.7770285 | -4.9921903 | 1.91897145 |
| 17343 | NM_153442.3 | #N/A | 19.3716283 | -4.1937548 | 1.61220393 |
| 17344 | NM_001220474.2 | #N/A | 57.1722 | -7.0473272 | 2.70928102 |
| 17345 | NM_021147.4 | #N/A | 23.3634517 | -5.7604762 | 2.21471312 |
| 17346 | NM_001206769.1 | DLG2 | 13.7475646 | -4.9912006 | 1.91928774 |
| 17347 | NR_003109.1 | TRNAU1AP | 12.8149444 | -4.8866439 | 1.87917484 |
| 17348 | NM_001130923.1 | #N/A | 45.0725738 | -6.7061137 | 2.57939218 |
| 17349 | NR_117094.1 | PAUPAR | 35.0460055 | 3.40066454 | 1.30803534 |
| 17350 | NM_003506.3 | #N/A | 42.3010219 | -6.6147846 | 2.54436966 |
| 17351 | NR_125440.1 | #N/A | 17.36326 | -5.3255593 | 2.04852946 |
| 17352 | NM_001311312.1 | #N/A | 10.7209325 | -4.6372422 | 1.78385698 |
| 17353 | NM_001345881.1 | #N/A | 252.239377 | 1.69798832 | 0.65321244 |
| 17354 | NR_033344.1 | #N/A | 9.49885598 | -4.4601068 | 1.71603671 |
| 17355 | NM_001294314.1 | TMEM116 | 10.5522687 | -4.6177709 | 1.77687376 |
| 17356 | NR_135086.1 | PPP1R14B-AS1 | 13.093684 | -4.9184421 | 1.89298053 |
| 17357 | NM_001005465.1 | OR10G3 | 28.0527224 | 9.7946329 | 3.77050741 |
| 17358 | NM_001142311.1 | #N/A | 58.8042178 | -7.0886106 | 2.72884594 |
| 17359 | NM_152622.4 | MIER3 | 42.1193054 | -6.6085795 | 2.54428506 |
| 17360 | NM_001198629.1 | #N/A | 54.7915181 | -6.9860476 | 2.68999614 |
| 17363 | NR_120320.1 | LINC01581 | 13.4135992 | 7.11630635 | 2.74047374 |
| 17361 | NM_001330509.1 | SYT17 | 14.1102389 | -5.0186689 | 1.93267372 |
| 17362 | NR_027019.1 | #N/A | 13.4135992 | 7.11630635 | 2.74047374 |
| 17364 | NM_001270891.1 | #N/A | 9.29925823 | -4.4339174 | 1.70757011 |
| 17365 | NM_172000.3 | #N/A | 38.0105901 | 3.92305468 | 1.51083866 |
| 17366 | NM_001330230.1 | #N/A | 13.6868661 | -4.9935141 | 1.92313625 |
| 17367 | NM_000231.2 | SGCG | 12.231062 | -4.8233915 | 1.85767011 |
| 17368 | NM_001142676.1 | #N/A | 16.5654264 | -3.9825526 | 1.53384278 |
| 17369 | NM_001330489.1 | #N/A | 41.8932857 | -6.6007523 | 2.54219808 |
| 17370 | NM_001007471.2 | #N/A | 13.7159297 | -4.9969539 | 1.92459133 |
| 17371 | NR_023351.2 | #N/A | 31.6176422 | 6.08286002 | 2.34313284 |

| | | | | | |
|---|---|---|---|---|---|
| 17372 | NR_049784.1 | #N/A | 11.1764699 | -4.7042374 | 1.81208917 |
| 17373 | NM_001308050.1 | PRUNE2 | 45.0986881 | -6.7072355 | 2.58442188 |
| 17374 | NM_001190983.1 | #N/A | 24.4375417 | -3.2339043 | 1.24618556 |
| 17375 | NR_026598.1 | #N/A | 15.9092602 | -5.2035832 | 2.00549278 |
| 17376 | NR_038958.1 | LOC100507412 | 456.69299 | 2.20931395 | 0.85164027 |
| 17377 | NM_001300885.1 | #N/A | 46.9124485 | -6.7635568 | 2.60723633 |
| 17378 | NM_004586.2 | #N/A | 317.510925 | -1.4004361 | 0.53985336 |
| 17379 | NM_001037164.2 | ARL4A | 43.3486162 | -6.6499642 | 2.56375694 |
| 17380 | NM_005873.2 | #N/A | 15.4652512 | -5.1554507 | 1.98763746 |
| 17381 | NM_001184702.1 | GYG2 | 42.8971721 | -6.6346104 | 2.55801095 |
| 17382 | NR_126525.1 | C5orf64 | 13.794084 | 6.19988603 | 2.39055327 |
| 17383 | NR_132750.1 | #N/A | 30.5024703 | -2.3250883 | 0.89652032 |
| 17384 | NM_001303429.1 | #N/A | 57.501244 | -7.0549999 | 2.7203983 |
| 17385 | NM_001301839.1 | SAP30BP | 22.0008295 | -5.6716868 | 2.18709963 |
| 17386 | NM_001329866.1 | #N/A | 37.3901013 | 6.33940811 | 2.44467284 |
| 17387 | NR_001587.1 | AURKAP1 | 10.9380775 | -4.6584233 | 1.79658116 |
| 17388 | NM_001271893.3 | #N/A | 10.3354329 | -4.5818366 | 1.76707844 |
| 17389 | NM_014311.2 | SMUG1 | 15.066436 | -5.1193072 | 1.97480794 |
| 17390 | NR_104169.2 | NDUFA5 | 9.1854278 | -4.4044112 | 1.6991069 |
| 17391 | NR_015378.3 | ZNF674-AS1 | 38.825277 | -1.8822967 | 0.72621947 |
| 17392 | NM_001257320.2 | APBB1 | 24.6039749 | -5.8256983 | 2.2477917 |
| 17393 | NM_001135861.2 | #N/A | 14.3921133 | -5.0445963 | 1.94662091 |
| 17394 | NM_001145795.1 | SH2B1 | 55.076337 | -6.9934256 | 2.6986726 |
| 17395 | NM_014468.3 | #N/A | 49.5244879 | 1.69870839 | 0.65559467 |
| 17396 | NM_180990.3 | #N/A | 20.7376577 | 4.20124471 | 1.62143648 |
| 17397 | NM_001010861.2 | #N/A | 55.4632582 | -7.0031221 | 2.70292102 |
| 17398 | NM_001289975.1 | BAZ2B | 55.8744427 | -7.0137336 | 2.70711672 |
| 17399 | NM_001099225.1 | CCDC43 | 10.5098639 | -4.6113213 | 1.77992941 |
| 17401 | NM_001134888.2 | RTL1 | 11.6425338 | -4.7632672 | 1.83863117 |
| 17400 | NM_001009936.2 | #N/A | 10.9171951 | -4.6582475 | 1.79810202 |
| 17402 | NM_033430.2 | #N/A | 14.5138888 | -5.0677184 | 1.95613476 |
| 17403 | NM_018260.2 | ZNF701 | 10.7349953 | -4.6362637 | 1.78967025 |
| 17404 | NM_152366.4 | #N/A | 9.9100952 | -4.507408 | 1.7399171 |
| 17405 | NM_198267.1 | #N/A | 10.6156761 | -4.6212082 | 1.78388538 |
| 17406 | NM_001136036.2 | #N/A | 14.7599031 | -5.0896572 | 1.96474638 |
| 17407 | NM_181868.1 | APAF1 | 45.5794096 | -6.7218258 | 2.5950461 |
| 17408 | NM_001305082.1 | #N/A | 9.65560966 | -4.4848799 | 1.73149396 |
| 17409 | NR_138534.1 | #N/A | 12.9013201 | -4.8874675 | 1.88696636 |
| 17410 | NM_001199323.1 | BLOC1S5 | 9.92400199 | 8.26824643 | 3.19249531 |
| 17412 | NR_027279.1 | USP17L6P | 27.7664701 | 9.77987443 | 3.77616893 |
| 17411 | NM_152638.3 | #N/A | 27.7664701 | 9.77987443 | 3.77616893 |
| 17413 | NM_014447.3 | ARFIP1 | 11.9547057 | -4.7834398 | 1.8470181 |
| 17414 | NM_001128635.1 | RIMBP3B | 9.96046467 | -4.5132741 | 1.74283485 |

| | | | | | |
|---|---|---|---|---|---|
| 17415 | NM_001320782.1 | ZNF544 | 13.9072337 | -4.9987378 | 1.93048029 |
| 17416 | NM_144588.6 | #N/A | 57.8957713 | -7.0658918 | 2.72892044 |
| 17417 | NM_001167989.1 | PQBP1 | 19.5753208 | -5.5076745 | 2.12747786 |
| 17418 | NM_001080497.2 | #N/A | 356.29038 | -2.7804548 | 1.07403048 |
| 17419 | NM_001040662.1 | #N/A | 13.2501632 | -4.9285133 | 1.90393423 |
| 17420 | NM_001304952.1 | #N/A | 11.0278252 | -4.6565994 | 1.79897754 |
| 17421 | NM_018279.3 | #N/A | 185.418941 | -2.6486408 | 1.02329669 |
| 17422 | NM_152592.4 | #N/A | 57.2510426 | 1.12080509 | 0.43312897 |
| 17423 | NM_001142602.1 | #N/A | 9.64637347 | -4.4736506 | 1.72886913 |
| 17424 | NM_001080436.1 | #N/A | 107.97265 | -3.3227084 | 1.28414298 |
| 17425 | NM_004525.2 | #N/A | 64.7679333 | -1.9828236 | 0.76633376 |
| 17426 | NM_001346007.1 | C4orf47 | 12.2188559 | -4.8278575 | 1.86595114 |
| 17427 | NM_001271186.1 | #N/A | 42.941673 | -6.6365201 | 2.56512777 |
| 17428 | NM_001130523.2 | #N/A | 65.6500183 | -7.2467719 | 2.80107157 |
| 17429 | NM_019863.2 | F8 | 10.2342182 | -4.5615673 | 1.76321428 |
| 17430 | NM_139285.3 | #N/A | 15.6008198 | -5.1789753 | 2.00208433 |
| 17431 | NM_138434.2 | ZBED6CL | 12.4165236 | -4.8366608 | 1.86991369 |
| 17432 | NM_153050.2 | MTMR3 | 60.7091147 | -7.1353847 | 2.75906317 |
| 17433 | NM_001174127.1 | SLC11A2 | 50.5368393 | -6.8695315 | 2.65632101 |
| 17434 | NM_001040648.1 | CRCP | 58.3983798 | 4.84822972 | 1.87481814 |
| 17435 | NM_001122870.2 | #N/A | 12.5977983 | -4.8582115 | 1.87876509 |
| 17436 | NM_173465.3 | #N/A | 11.3213137 | -4.7188452 | 1.82493425 |
| 17437 | NM_001171806.1 | #N/A | 16.2263282 | -5.2319364 | 2.0236887 |
| 17438 | NM_199361.2 | #N/A | 52.1721947 | -6.9153596 | 2.6752131 |
| 17439 | NM_018689.2 | #N/A | 111.594604 | -6.0711025 | 2.34865484 |
| 17440 | NM_001042574.2 | #N/A | 48.3325572 | -6.8062604 | 2.6331951 |
| 17441 | NM_001346409.1 | #N/A | 59.9752275 | -7.1172095 | 2.75354261 |
| 17442 | NM_001330246.1 | #N/A | 53.2848171 | -6.9456112 | 2.68803928 |
| 17443 | NM_001321007.1 | #N/A | 17.934924 | -5.3659902 | 2.07711669 |
| 17444 | NM_002772.2 | #N/A | 21.405265 | 6.33823689 | 2.45372928 |
| 17445 | NM_001004303.4 | #N/A | 18.4199501 | -5.4173785 | 2.09744586 |
| 17446 | NM_014063.6 | #N/A | 41.7800798 | -6.5968972 | 2.55423717 |
| 17447 | NM_001163629.1 | MROH9 | 33.918896 | 8.32948886 | 3.22513893 |
| 17448 | NM_001206987.2 | METTL23 | 10.024325 | -4.5464346 | 1.76044001 |
| 17449 | NM_001184797.1 | #N/A | 13.5145704 | -4.9632239 | 1.92204644 |
| 17451 | NM_003015.3 | SFRP5 | 11.3655473 | -4.730646 | 1.83201327 |
| 17450 | NM_001320410.1 | #N/A | 24.8324687 | 3.88562569 | 1.5047657 |
| 17452 | NR_027691.1 | #N/A | 11.0424616 | -4.6759521 | 1.81104292 |
| 17453 | NM_001319078.1 | #N/A | 55.6420129 | -7.0086462 | 2.71514942 |
| 17454 | NM_006635.3 | #N/A | 10.7707934 | -4.6327055 | 1.79483376 |
| 17455 | NM_001172674.1 | ZNF347 | 115.633634 | 1.46197468 | 0.5664795 |
| 17456 | NM_018194.5 | #N/A | 14.4973185 | -5.0693681 | 1.96467859 |
| 17457 | NR_103862.1 | LOC100507283 | 9.13904458 | -4.4111627 | 1.70966753 |

| | | | | | |
|---|---|---|---|---|---|
| 17458 | NM_001282234.1 | PSMA6 | 10.2459737 | -4.5811684 | 1.77562643 |
| 17459 | NR_138482.1 | ZNF518A | 20.494553 | 5.22939131 | 2.02701411 |
| 17460 | NM_001288802.1 | #N/A | 56.8077426 | -7.0380489 | 2.72822608 |
| 17461 | NM_001143823.2 | #N/A | 13.1460546 | -4.9271039 | 1.91003143 |
| 17462 | NM_201428.2 | #N/A | 52.4714813 | -6.9236159 | 2.68420356 |
| 17463 | NM_001347821.1 | #N/A | 12.6544393 | -4.8760156 | 1.89044259 |
| 17464 | NM_001727.1 | #N/A | 12.6402017 | 7.96215192 | 3.08704985 |
| 17465 | NM_153635.2 | #N/A | 9.94386388 | -4.5194588 | 1.75230711 |
| 17466 | NM_138964.3 | PROKR1 | 19.0885635 | 3.43956136 | 1.33364493 |
| 17467 | NM_030961.2 | #N/A | 93.2476209 | -2.0088029 | 0.77889533 |
| 17468 | NM_001282865.1 | TMEM108 | 27.25523 | 5.16674061 | 2.00351544 |
| 17469 | NM_001171934.1 | CDH23 | 25.4643889 | -4.2416006 | 1.64521745 |
| 17470 | NM_015296.2 | #N/A | 56.1814887 | -7.0217568 | 2.72362661 |
| 17471 | NR_110514.1 | EP300-AS1 | 158.311911 | 1.4596743 | 0.56629576 |
| 17472 | NM_006414.4 | #N/A | 13.6406618 | -4.9706505 | 1.92852062 |
| 17473 | NM_181829.2 | #N/A | 49.4168798 | -6.8379147 | 2.65320855 |
| 17474 | NM_001316975.1 | #N/A | 12.5605181 | -4.8633123 | 1.88728565 |
| 17475 | NR_131961.1 | LINC02476 | 18.8296475 | 2.17002416 | 0.84215486 |
| 17476 | NM_001316898.1 | #N/A | 52.4302992 | -6.922698 | 2.68672623 |
| 17477 | NM_001323311.1 | PURG | 9.92002607 | -4.5276123 | 1.75739924 |
| 17478 | NM_173619.3 | #N/A | 16.0707573 | -5.2238005 | 2.02773428 |
| 17479 | NM_001042633.2 | #N/A | 41.7963444 | -6.5971649 | 2.56090326 |
| 17480 | NR_104612.1 | TRMT10B | 13.9028967 | -5.0011501 | 1.94142899 |
| 17481 | NM_001308267.1 | #N/A | 52.6070504 | -6.9271529 | 2.6893648 |
| 17482 | NM_178820.4 | #N/A | 133.729355 | 1.26806931 | 0.49231127 |
| 17483 | NM_001201552.1 | #N/A | 52.0797599 | -6.9129661 | 2.68453223 |
| 17484 | NM_001166108.1 | #N/A | 9.75785405 | -4.5049273 | 1.74950297 |
| 17485 | NM_001127372.2 | #N/A | 49.4777738 | -6.840569 | 2.65660259 |
| 17486 | NM_148896.4 | #N/A | 12.7750148 | -4.877891 | 1.89446115 |
| 17487 | NM_001032280.2 | #N/A | 9.44202554 | -4.4562231 | 1.73078488 |
| 17490 | NR_001560.1 | CYCSP52 | 16.9111638 | 8.35230783 | 3.24449143 |
| 17489 | NM_001199824.2 | SBK3 | 16.9111638 | 8.35230783 | 3.24449143 |
| 17488 | NM_001014291.3 | #N/A | 16.9111638 | 8.35230783 | 3.24449143 |
| 17491 | NM_153620.2 | #N/A | 15.1826048 | -5.1421901 | 1.99757567 |
| 17492 | NM_001256356.1 | UBE2L3 | 14.0431609 | -5.0307132 | 1.95436725 |
| 17493 | NM_212540.2 | #N/A | 27.1939656 | 9.74989225 | 3.78780313 |
| 17494 | NM_001164178.1 | ENTPD1 | 51.0506878 | -6.8857919 | 2.67515442 |
| 17495 | NM_173822.4 | #N/A | 59.3228212 | -7.1003286 | 2.75897444 |
| 17496 | NM_017698.2 | TMEM164 | 103.36476 | -3.2629622 | 1.26801717 |
| 17497 | NR_047653.1 | #N/A | 60.5120508 | -7.1287104 | 2.77046083 |
| 17498 | NR_126399.1 | C8orf31 | 23.5274169 | 3.55895057 | 1.38336277 |
| 17499 | NM_001322331.1 | #N/A | 13.4908981 | -4.9549548 | 1.92603998 |
| 17500 | NR_037658.1 | BLOC1S1-RDH5 | 13.0097719 | -4.9014402 | 1.90525606 |

| | | | | | |
|---|---|---|---|---|---|
| 17501 | NR_135140.1 | YAF2 | 14.6232857 | -5.0778099 | 1.97416234 |
| 17502 | NM_001077663.2 | URGCP | 14.6219754 | -5.0728221 | 1.97225404 |
| 17503 | NM_001317101.1 | DNAJB4 | 15.6411419 | -5.1761091 | 2.0124868 |
| 17504 | NM_001033056.3 | GLUL | 288.015502 | 1.48537228 | 0.57753881 |
| 17505 | NM_001199779.1 | #N/A | 40.5615682 | -6.554248 | 2.5484036 |
| 17506 | NM_018378.2 | #N/A | 9.50522281 | -4.4644953 | 1.73600123 |
| 17507 | NM_024824.4 | #N/A | 40.1226785 | -6.5385394 | 2.54260917 |
| 17508 | NM_201626.2 | #N/A | 11.6372877 | -4.7483107 | 1.84648197 |
| 17509 | NM_173078.4 | #N/A | 52.6855502 | -6.9292221 | 2.69484581 |
| 17510 | NM_001165994.1 | #N/A | 15.9225693 | 5.27728703 | 2.05258287 |
| 17511 | NR_130790.1 | MIEF1 | 9.15288518 | -4.3939923 | 1.70929196 |
| 17512 | NR_134285.1 | ZNF517 | 9.02277707 | -4.3807424 | 1.70412852 |
| 17513 | NR_138070.1 | #N/A | 11.610972 | -4.7453478 | 1.84600721 |
| 17514 | NM_001001712.2 | #N/A | 9.70709505 | 5.01985334 | 1.95296091 |
| 17515 | NM_000545.6 | HNF1A | 10.9307117 | -4.6561857 | 1.81187987 |
| 17516 | NR_110399.2 | LOC101928437 | 8.41299288 | 4.75848794 | 1.85191022 |
| 17517 | NM_001766.3 | CD1D | 26.4856156 | 3.76222587 | 1.46432024 |
| 17518 | NM_024917.5 | #N/A | 13.2315742 | -4.9242026 | 1.91692076 |
| 17519 | NR_135074.1 | #N/A | 16.8758738 | -5.2774233 | 2.05464481 |
| 17520 | NR_136155.1 | #N/A | 40.0006823 | -6.5340456 | 2.54401524 |
| 17521 | NM_018073.7 | #N/A | 87.071516 | -3.3866168 | 1.31864323 |
| 17522 | NM_001270497.1 | #N/A | 11.9299105 | -4.7876571 | 1.8642137 |
| 17523 | NM_001271863.1 | KLHDC8A | 14.9851197 | -5.1196107 | 1.99352882 |
| 17524 | NM_172037.4 | #N/A | 139.74228 | -1.8178849 | 0.70786973 |
| 17525 | NM_001303487.1 | #N/A | 48.9110913 | -6.8223144 | 2.65664701 |
| 17526 | NM_001290223.1 | DOCK1 | 42.2912456 | -6.6140693 | 2.57573279 |
| 17527 | NM_018676.3 | #N/A | 10.4452531 | -4.598335 | 1.79122211 |
| 17528 | NM_033142.1 | CGB7 | 22.9095572 | 4.8805763 | 1.90132959 |
| 17529 | NR_135292.1 | LINC00863 | 49.6919244 | -2.3762401 | 0.92580336 |
| 17530 | NM_001330508.1 | #N/A | 85.6586836 | 2.02063101 | 0.78738786 |
| 17531 | NM_198867.1 | ALKBH6 | 12.6590857 | -4.8792875 | 1.90143291 |
| 17532 | NM_001195302.1 | #N/A | 15.5372142 | -5.1731001 | 2.01608473 |
| 17533 | NM_001282802.1 | #N/A | 52.9763823 | -6.9371135 | 2.70379082 |
| 17534 | NM_001243156.1 | #N/A | 59.6128345 | -7.1070609 | 2.77006144 |
| 17535 | NM_001286515.1 | #N/A | 9.67407063 | -4.4774262 | 1.74518893 |
| 17536 | NM_001271522.1 | #N/A | 10.5144614 | -4.6020908 | 1.79396892 |
| 17537 | NM_013239.4 | #N/A | 157.237982 | -2.5206347 | 0.98264925 |
| 17538 | NM_024888.2 | PLPPR3 | 19.8747998 | -5.5263663 | 2.15454136 |
| 17539 | NM_003574.5 | #N/A | 16.4321889 | -5.2458995 | 2.04546466 |
| 17540 | NM_001206671.3 | #N/A | 53.1032428 | -6.940435 | 2.70629891 |
| 17541 | NM_001013627.2 | NHSL2 | 8.93406613 | -4.3701484 | 1.70422807 |
| 17542 | NM_001135254.1 | #N/A | 12.1609393 | -4.8209506 | 1.880172 |
| 17543 | NR_135264.1 | C1orf61 | 19.4283829 | -5.4850304 | 2.13923266 |

| | | | | | |
|---|---|---|---|---|---|
| 17544 | NM_203472.2 | #N/A | 11.7942811 | -4.7752931 | 1.86248265 |
| 17545 | NM_000905.3 | #N/A | 17.7656973 | -5.3572228 | 2.08958753 |
| 17546 | NM_001243646.1 | CD2BP2 | 10.0375046 | -4.5496841 | 1.77478106 |
| 17547 | NR_109763.1 | #N/A | 9.65322998 | -4.4744309 | 1.74541831 |
| 17548 | NM_000273.2 | #N/A | 16.9188236 | -5.2965295 | 2.06614638 |
| 17549 | NM_178457.2 | ZNF831 | 6.40323184 | 6.8227073 | 2.66218801 |
| 17550 | NM_031492.3 | #N/A | 144.220167 | -2.4960519 | 0.97397562 |
| 17551 | NM_020452.3 | #N/A | 287.203851 | -2.1966604 | 0.85725335 |
| 17552 | NM_001012514.2 | #N/A | 47.5201995 | -6.7825078 | 2.64736071 |
| 17553 | NM_013250.2 | #N/A | 10.0871841 | -4.5440922 | 1.77400076 |
| 17554 | NR_110742.1 | ZNF451-AS1 | 7.55517721 | 2.81600041 | 1.09942004 |
| 17555 | NM_001384.4 | #N/A | 42.7429836 | -6.6296409 | 2.58882321 |
| 17556 | NR_027441.1 | LINC02447 | 22.292344 | 2.38120964 | 0.9299457 |
| 17557 | NM_001272020.1 | #N/A | 39.4763212 | -6.5150486 | 2.54466725 |
| 17558 | NM_001271985.1 | PUS3 | 14.8570919 | -5.1007978 | 1.9924163 |
| 17559 | NM_012210.3 | TRIM32 | 51.6388877 | -6.9002126 | 2.69591109 |
| 17560 | NM_001318085.1 | SEC24B | 39.7516248 | -6.5250746 | 2.54953229 |
| 17561 | NM_005996.3 | #N/A | 41.1275077 | -6.5742568 | 2.56887701 |
| 17562 | NM_001347890.1 | TMX2 | 10.8073747 | -4.6454998 | 1.81526272 |
| 17563 | NM_018097.2 | #N/A | 56.1968906 | -7.0221247 | 2.74398506 |
| 17564 | NM_001164712.1 | #N/A | 9.44390072 | -4.4604828 | 1.74303642 |
| 17565 | NM_001320037.1 | LOC730098 | 14.1828881 | -5.0334193 | 1.96710711 |
| 17566 | NM_001256721.1 | CCT4 | 39.8835507 | -6.5299418 | 2.55201232 |
| 17567 | NR_136747.1 | #N/A | 10.5454261 | -4.6063879 | 1.80034181 |
| 17568 | NM_002998.3 | #N/A | 848.258774 | -3.8568178 | 1.5075259 |
| 17569 | NR_024321.1 | LINC00115 | 9.30062397 | -4.4398087 | 1.73544301 |
| 17570 | NR_110010.1 | #N/A | 24.0545678 | -4.1613357 | 1.62674078 |
| 17572 | NR_134329.1 | LINC02624 | 26.6214611 | 9.71927938 | 3.79987145 |
| 17571 | NM_001257138.2 | #N/A | 26.6214611 | 9.71927938 | 3.79987145 |
| 17573 | NM_001347923.1 | SNX19 | 17.1497998 | -5.3035895 | 2.07358258 |
| 17574 | NM_001204055.1 | #N/A | 33.732124 | 10.0507756 | 3.92969588 |
| 17575 | NR_038392.1 | #N/A | 13.7117204 | -4.9946866 | 1.95293456 |
| 17576 | NM_001289922.2 | #N/A | 9.95939538 | 6.22973035 | 2.43629249 |
| 17577 | NM_001331070.1 | #N/A | 39.2232651 | -6.5059386 | 2.54433006 |
| 17578 | NM_015224.3 | TASOR | 11.3144177 | -4.7196757 | 1.84591525 |
| 17579 | NM_001288999.1 | #N/A | 43.5068672 | -6.6553662 | 2.60315947 |
| 17580 | NR_039989.1 | MIR4458HG | 9.47113013 | -4.4615961 | 1.74530622 |
| 17581 | NM_176890.2 | TAS2R50 | 7.25856599 | 5.11037935 | 1.99931449 |
| 17582 | NR_136599.1 | #N/A | 65.7905326 | -7.2496973 | 2.83636912 |
| 17584 | NM_201444.2 | DGKA | 17.9484459 | -5.3815643 | 2.10555428 |
| 17583 | NM_138789.3 | PIH1D2 | 9.93008873 | -4.53006 | 1.77239927 |
| 17585 | NM_001142279.2 | RNASEH2B | 9.67402711 | -4.4726681 | 1.75055256 |
| 17586 | NM_017413.4 | #N/A | 13.7341531 | -3.6991039 | 1.44792043 |

| | | | | | |
|---|---|---|---|---|---|
| 17587 | NR_135006.1 | LOC105372582 | 33.1595519 | 1.74917943 | 0.68468513 |
| 17588 | NM_001317921.1 | #N/A | 39.3625179 | -6.5110573 | 2.54893951 |
| 17589 | NM_173791.4 | #N/A | 368.800102 | -1.2505802 | 0.48961182 |
| 17590 | NM_018218.2 | #N/A | 155.601348 | -2.0013816 | 0.78358646 |
| 17591 | NM_201380.3 | #N/A | 39.1723447 | -6.5041005 | 2.54677971 |
| 17592 | NM_145805.2 | #N/A | 12.9769954 | -4.9117931 | 1.92348718 |
| 17593 | NR_135259.1 | #N/A | 12.1442915 | -4.7997334 | 1.87966643 |
| 17594 | NM_001005376.2 | PLAUR | 49.6142964 | 3.71618105 | 1.45553605 |
| 17595 | NR_145432.1 | #N/A | 10.8969045 | -4.6648308 | 1.82723441 |
| 17596 | NM_000829.3 | #N/A | 45.3833909 | -6.7161554 | 2.6308049 |
| 17597 | NM_001011645.2 | #N/A | 20.2363357 | -3.9567741 | 1.54991327 |
| 17598 | NM_001330311.1 | #N/A | 289.34224 | -3.6151392 | 1.41622265 |
| 17599 | NM_001100621.2 | #N/A | 39.4087476 | -6.5127498 | 2.55144017 |
| 17600 | NR_040772.1 | DCST1-AS1 | 8.72120153 | -4.3294135 | 1.69621655 |
| 17601 | NM_001256404.1 | #N/A | 10.1140023 | -4.5489258 | 1.7823486 |
| 17602 | NM_170771.2 | #N/A | 25.071834 | 4.85803379 | 1.90345131 |
| 17603 | NM_016631.3 | #N/A | 52.1847237 | -6.9152397 | 2.70958334 |
| 17604 | NM_153604.3 | MYOCD | 30.2833159 | 4.40875735 | 1.72768612 |
| 17605 | NM_021923.3 | FGFRL1 | 47.8082341 | -6.7900044 | 2.66098751 |
| 17606 | NM_024841.4 | PRR5L | 12.348834 | -4.8345024 | 1.89485294 |
| 17607 | NM_000531.5 | #N/A | 16.9343693 | 8.27062123 | 3.24174112 |
| 17608 | NM_000481.3 | #N/A | 40.7235965 | -6.5597062 | 2.57129098 |
| 17609 | NM_001033522.1 | #N/A | 16.5546993 | -5.263832 | 2.0633206 |
| 17610 | NM_001257330.1 | #N/A | 12.5918241 | -4.8620331 | 1.90582506 |
| 17611 | NR_033267.1 | LOC100335030 | 58.0202546 | 2.33216119 | 0.91418691 |
| 17612 | NM_001267611.1 | GPR161 | 39.420367 | -6.5131171 | 2.55358056 |
| 17613 | NM_001004065.4 | PALM2AKAP2 | 51.727354 | -6.9038096 | 2.70684025 |
| 17614 | NM_152666.2 | #N/A | 13.8003167 | -5.0070205 | 1.96315211 |
| 17615 | NM_005446.3 | #N/A | 10.8102849 | -4.65699 | 1.82593824 |
| 17616 | NM_001142885.1 | #N/A | 98.8527544 | -5.9377052 | 2.32878232 |
| 17618 | NR_110934.1 | LINC02125 | 22.0934715 | 6.61926626 | 2.5962592 |
| 17617 | NM_001042365.3 | PTPN20 | 26.3352088 | 9.70372457 | 3.8060761 |
| 17619 | NM_001308236.1 | DOK3 | 12.7330243 | -4.8804823 | 1.91451351 |
| 17620 | NM_001243013.1 | #N/A | 13.3497932 | -4.9574181 | 1.94472863 |
| 17621 | NM_001184704.1 | GYG2 | 39.6260363 | -6.5206555 | 2.55800855 |
| 17622 | NM_001320818.1 | COQ10B | 52.0788733 | 6.12203722 | 2.40175707 |
| 17623 | NM_001018037.1 | #N/A | 49.8505934 | -6.8499031 | 2.68739145 |
| 17624 | NM_177405.2 | #N/A | 139.263019 | -2.3942927 | 0.93935586 |
| 17625 | NR_040100.1 | ENTPD3-AS1 | 78.7700105 | 2.52971759 | 0.99250124 |
| 17626 | NM_012418.3 | #N/A | 9.12818815 | -4.4107271 | 1.73056528 |
| 17627 | NM_001317138.1 | #N/A | 13.0208229 | -4.9150396 | 1.92899318 |
| 17628 | NM_001346228.1 | WDR48 | 14.2010008 | -5.0340407 | 1.97574623 |
| 17629 | NR_110833.1 | LINC01448 | 48.2166953 | 6.64195343 | 2.60698938 |

| | | | | | |
|---|---|---|---|---|---|
| 17630 | NM_005298.3 | #N/A | 6.64283282 | 5.17992676 | 2.03327929 |
| 17631 | NM_001207000.1 | HNRNPDL | 14.247163 | -5.0381405 | 1.97766214 |
| 17632 | NM_001123065.1 | C7orf65 | 29.6321502 | 3.65067794 | 1.43305638 |
| 17633 | NM_000403.3 | #N/A | 13.6563482 | -4.98605 | 1.95758119 |
| 17634 | NM_012465.3 | #N/A | 18.753621 | -3.8647552 | 1.51745608 |
| 17635 | NR_027254.1 | LINC00671 | 43.3435283 | 3.3785268 | 1.32660746 |
| 17636 | NM_001300932.1 | #N/A | 12.1699787 | -4.8139334 | 1.8902775 |
| 17637 | NR_134916.1 | LINC01842 | 19.2120618 | 2.88323556 | 1.13216451 |
| 17638 | NM_001171946.1 | #N/A | 8.81538934 | -4.3528541 | 1.70945028 |
| 17639 | NM_001326358.1 | #N/A | 44.6427438 | -6.6917621 | 2.62814378 |
| 17640 | NM_002651.3 | #N/A | 40.7168 | -6.5596272 | 2.57625788 |
| 17641 | NR_133002.1 | MIR9-3HG | 50.9380088 | -6.8805564 | 2.70273664 |
| 17642 | NM_001127890.4 | #N/A | 12.0532298 | -4.7993538 | 1.88552531 |
| 17643 | NM_001281438.1 | ADRM1 | 13.7125101 | -4.9914957 | 1.96108326 |
| 17644 | NR_037577.1 | #N/A | 11.9197716 | -4.7819066 | 1.87878891 |
| 17645 | NM_006181.2 | #N/A | 9.67114115 | -4.4879325 | 1.76349303 |
| 17646 | NM_001145465.1 | NANOGNB | 16.040226 | 3.55233063 | 1.39594769 |
| 17647 | NM_006202.2 | #N/A | 50.7089145 | -6.8737964 | 2.70160984 |
| 17648 | NM_172071.3 | #N/A | 60.9258246 | -7.1382397 | 2.80583448 |
| 17649 | NM_032776.2 | #N/A | 52.1952635 | -6.9150406 | 2.71826192 |
| 17650 | NR_034082.1 | #N/A | 8.82461343 | -4.3437229 | 1.70747932 |
| 17651 | NM_001048171.1 | MUTYH | 11.4949146 | -4.7231196 | 1.85665499 |
| 17652 | NR_110084.1 | LOC101927391 | 12.9626255 | -3.6024556 | 1.41621293 |
| 17653 | NM_001164579.1 | #N/A | 83.6741145 | -3.3312984 | 1.30969288 |
| 17654 | NM_020950.1 | #N/A | 61.4167971 | 2.19508932 | 0.8630028 |
| 17655 | NM_021231.1 | CACTIN | 13.8371742 | -4.9862705 | 1.96056768 |
| 17656 | NR_134658.1 | #N/A | 12.4780285 | -4.8636108 | 1.9124901 |
| 17657 | NM_001282973.1 | SEPTIN2 | 14.7278564 | -5.0848335 | 2.00012625 |
| 17658 | NM_080630.3 | #N/A | 53.006239 | -6.9397456 | 2.72973805 |
| 17659 | NR_103764.1 | ECI2-DT | 8.89987575 | 5.19140641 | 2.04215289 |
| 17660 | NM_000761.4 | #N/A | 113.512407 | 2.43402566 | 0.95749075 |
| 17661 | NM_001199381.1 | #N/A | 11.4517714 | -4.7365229 | 1.86333955 |
| 17662 | NM_001130955.1 | #N/A | 11.8200477 | -4.7823491 | 1.88144242 |
| 17663 | NM_001195243.1 | #N/A | 9.36335824 | -4.4424027 | 1.74773652 |
| 17664 | NM_001166240.1 | CGREF1 | 10.7823685 | -4.6392915 | 1.82538898 |
| 17665 | NM_001247994.1 | #N/A | 43.5990518 | -6.6579222 | 2.61982992 |
| 17666 | NM_001318324.1 | #N/A | 39.4467094 | -6.5138362 | 2.56327873 |
| 17667 | NR_073202.1 | #N/A | 26.0489565 | 9.68799964 | 3.81239872 |
| 17668 | NR_003500.1 | RP9P | 72.7922335 | -2.7235549 | 1.07186511 |
| 17669 | NM_016328.2 | #N/A | 38.4922548 | -6.478889 | 2.54981617 |
| 17672 | NR_110077.1 | CACNA2D1-AS1 | 33.588575 | 3.65897926 | 1.44029454 |
| 17671 | NM_016826.2 | OGG1 | 9.25788976 | -4.4249864 | 1.74184572 |
| 17670 | NM_012459.2 | #N/A | 128.72811 | -2.7980545 | 1.10141451 |

| | | | | | |
|---|---|---|---|---|---|
| 17673 | NM_152796.2 | HIF3A | 52.5671237 | 2.02078311 | 0.79551114 |
| 17674 | NM_001008708.3 | #N/A | 9.7020651 | -4.4810996 | 1.764099 |
| 17676 | NM_024061.3 | ZNF655 | 12.1110831 | -4.8030811 | 1.8911061 |
| 17675 | NM_001005782.1 | #N/A | 16.6562962 | -5.2690636 | 2.0745828 |
| 17677 | NR_002157.1 | OR2A9P | 22.7040713 | 5.10146364 | 2.00868185 |
| 17678 | NM_053016.5 | #N/A | 49.9122012 | -6.8520545 | 2.6980133 |
| 17679 | NM_001304763.1 | #N/A | 128.659374 | -2.30233 | 0.90657345 |
| 17680 | NR_120563.1 | LOC101928847 | 53.2211316 | 2.35967087 | 0.92919538 |
| 17681 | NM_001314033.2 | #N/A | 11.9034779 | -4.7731871 | 1.87961579 |
| 17682 | NM_198965.1 | #N/A | 11.0324337 | -4.6735641 | 1.84072863 |
| 17683 | NM_001324220.1 | #N/A | 49.6133901 | -6.8431848 | 2.69528126 |
| 17684 | NR_046311.1 | #N/A | 9.70959855 | -4.4930829 | 1.7698422 |
| 17685 | NM_001286770.1 | ZNF34 | 8.9337401 | -4.37646 | 1.72397421 |
| 17686 | NR_038942.1 | PARTICL | 17.4749207 | -4.0469007 | 1.59441608 |
| 17687 | NM_000676.2 | ADORA2B | 20.6165681 | -2.6953542 | 1.06200093 |
| 17688 | NM_001320458.1 | #N/A | 42.6545385 | 2.39558001 | 0.9439262 |
| 17689 | NM_207345.3 | #N/A | 12.3771549 | 7.84997672 | 3.09338599 |
| 17690 | NR_048561.1 | PPM1E | 58.4920203 | -7.0795964 | 2.79014227 |
| 17691 | NM_001142327.1 | DMTF1 | 38.9678742 | -6.4963865 | 2.56039744 |
| 17692 | NR_052019.1 | SNAPIN | 13.0321229 | -4.9187633 | 1.93915952 |
| 17693 | NR_119378.1 | LOC102723373 | 34.3762788 | 6.65755729 | 2.62485695 |
| 17694 | NR_027271.1 | CIRBP-AS1 | 8.68719609 | -4.3347819 | 1.70910619 |
| 17695 | NM_001005408.1 | #N/A | 57.7226802 | -7.0621401 | 2.78446422 |
| 17696 | NM_014220.2 | #N/A | 10.5599925 | -4.6203558 | 1.82176481 |
| 17697 | NM_001103148.1 | #N/A | 38.8658697 | -6.4926702 | 2.56007567 |
| 17698 | NR_033915.1 | #N/A | 9.93139796 | -4.5296856 | 1.78606931 |
| 17699 | NM_152504.3 | #N/A | 11.0945746 | -4.6870615 | 1.84816703 |
| 17700 | NM_012176.2 | #N/A | 9.35258499 | -4.439219 | 1.75059395 |
| 17701 | NM_001193282.2 | #N/A | 18.9249957 | -5.4570524 | 2.15216024 |
| 17702 | NR_073133.1 | CFAP221 | 9.77794262 | -4.5065715 | 1.77733945 |
| 17703 | NM_012100.3 | #N/A | 50.8922453 | -6.8786525 | 2.71297055 |
| 17704 | NM_006307.4 | #N/A | 40.4045875 | -6.5486371 | 2.582843 |
| 17705 | NR_104143.1 | LINC00682 | 12.300862 | -4.832862 | 1.90614459 |
| 17706 | NM_001079881.1 | PRKD2 | 14.1967238 | -5.0330658 | 1.9852798 |
| 17707 | NM_181306.2 | #N/A | 10.2388802 | -4.5639141 | 1.80026356 |
| 17709 | NR_120424.1 | LOC101929259 | 81.383736 | 1.49285133 | 0.58892262 |
| 17708 | NM_001172779.1 | #N/A | 10.0993709 | -4.5520862 | 1.79576853 |
| 17710 | NM_021722.5 | #N/A | 43.0843565 | -6.6410448 | 2.61996127 |
| 17711 | NM_182506.3 | MAGEB10 | 63.2463992 | 3.08919106 | 1.21899491 |
| 17712 | NM_001100910.1 | FAM72B | 10.3902523 | -4.5901457 | 1.81184133 |
| 17713 | NM_004941.2 | #N/A | 38.1032619 | -6.4639377 | 2.55178939 |
| 17714 | NM_001145265.1 | #N/A | 17.2272398 | -5.3150092 | 2.09833666 |
| 17720 | NR_125387.1 | LOC100129636 | 15.9464716 | 7.16431464 | 2.82856998 |

| | | | | | |
|---|---|---|---|---|---|
| 17717 | NM_002494.3 | NDUFC1 | 11.8791708 | -4.787173 | 1.89000206 |
| 17719 | NR_002183.1 | ST13P4 | 15.9464716 | 7.16431464 | 2.82856998 |
| 17715 | NM_001164269.1 | #N/A | 32.5274053 | 9.99819121 | 3.94736036 |
| 17716 | NM_001199268.1 | #N/A | 32.5274053 | 9.99819121 | 3.94736036 |
| 17718 | NM_080681.2 | #N/A | 45.0089553 | -6.7026586 | 2.64631144 |
| 17721 | NM_138448.3 | #N/A | 14.1393485 | -5.0316017 | 1.98668067 |
| 17722 | NM_001172648.3 | #N/A | 26.7849747 | -4.6817969 | 1.84867856 |
| 17723 | NM_002240.4 | #N/A | 11.938892 | -4.7841058 | 1.88911671 |
| 17724 | NM_018189.3 | DPPA4 | 48.287982 | 2.45185395 | 0.96818375 |
| 17725 | NM_001330334.1 | EED | 10.4341382 | -4.6032955 | 1.81797218 |
| 17726 | NM_001889.3 | #N/A | 37.7940266 | -6.4523357 | 2.54834949 |
| 17727 | NR_038459.1 | GFOD1 | 33.937046 | 8.14939194 | 3.21882597 |
| 17728 | NM_147190.3 | #N/A | 50.645763 | -6.8728865 | 2.71493793 |
| 17729 | NM_172003.3 | CBWD2 | 49.213877 | -6.8310084 | 2.69845635 |
| 17730 | NM_201612.2 | #N/A | 10.0136213 | -4.5483188 | 1.79691944 |
| 17731 | NR_136256.1 | LOC102724738 | 45.794738 | 3.75524877 | 1.48363784 |
| 17732 | NM_000773.3 | #N/A | 10.2214679 | -4.5716302 | 1.80636486 |
| 17733 | NR_073475.1 | #N/A | 39.4175849 | -6.5129709 | 2.57341792 |
| 17734 | NM_199229.2 | #N/A | 48.3616266 | -6.8060362 | 2.68980041 |
| 17735 | NM_001322242.1 | #N/A | 38.6937162 | -6.4861653 | 2.56359183 |
| 17736 | NM_001100817.1 | #N/A | 8.17357445 | 6.35389534 | 2.51161032 |
| 17737 | NR_026887.1 | #N/A | 11.4381824 | -4.7195939 | 1.86561588 |
| 17738 | NM_001286495.1 | #N/A | 10.9728811 | -4.6753927 | 1.84821937 |
| 17739 | NM_001146019.1 | #N/A | 44.921963 | -6.7001587 | 2.64893277 |
| 17740 | NR_046236.1 | #N/A | 10.1854355 | -4.5542806 | 1.80097738 |
| 17741 | NM_001345970.1 | #N/A | 38.9909991 | -6.4975687 | 2.56975944 |
| 17742 | NM_001320670.1 | #N/A | 49.3153048 | -6.8348005 | 2.70344587 |
| 17743 | NM_001128626.1 | #N/A | 49.1197975 | -6.8289053 | 2.70272103 |
| 17744 | NM_006522.3 | #N/A | 8.45422316 | -4.2874974 | 1.69702974 |
| 17745 | NR_033893.1 | LINC01299 | 39.4363131 | 3.03791233 | 1.20247546 |
| 17746 | NM_001303542.2 | #N/A | 65.9678448 | -7.2546601 | 2.87177497 |
| 17747 | NR_040017.1 | RNF157-AS1 | 8.84035411 | -4.3613259 | 1.72658601 |
| 17748 | NM_001330140.1 | #N/A | 9.22729069 | -4.4058418 | 1.7443433 |
| 17749 | NM_014140.3 | #N/A | 37.5442258 | -6.442887 | 2.55104604 |
| 17750 | NM_019041.6 | #N/A | 48.4567696 | -6.8083895 | 2.69584542 |
| 17751 | NM_021709.2 | #N/A | 10.4425048 | -4.5797255 | 1.81342053 |
| 17752 | NM_001243996.2 | #N/A | 38.1438604 | -6.4656695 | 2.56038369 |
| 17753 | NM_176794.1 | MRPL43 | 9.90752138 | -4.529892 | 1.79385435 |
| 17754 | NM_002502.5 | #N/A | 38.4326268 | -6.476179 | 2.56464152 |
| 17755 | NM_001282681.1 | #N/A | 48.4928462 | -6.809425 | 2.69684245 |
| 17756 | NM_182854.3 | #N/A | 13.4713851 | 4.08925467 | 1.61968513 |
| 17757 | NM_015685.5 | SDCBP2 | 8.9562012 | -4.3609256 | 1.72755591 |
| 17758 | NR_144951.1 | #N/A | 37.6163109 | -6.4454658 | 2.55416505 |

| | | | | | |
|---|---|---|---|---|---|
| 17759 | NM_000678.3 | #N/A | 12.1712039 | -4.8215659 | 1.91073799 |
| 17760 | NM_001291937.1 | NTRK2 | 40.1100436 | -6.5381798 | 2.59134934 |
| 17762 | NM_001320707.1 | PKN2 | 39.6118652 | -6.5199481 | 2.58421469 |
| 17761 | NM_001282758.1 | TRA2A | 10.0545406 | -4.5399235 | 1.79940953 |
| 17763 | NR_135845.1 | FAM219B | 48.7390483 | -6.816368 | 2.70176337 |
| 17764 | NR_028077.1 | #N/A | 13.9306575 | -5.0017366 | 1.98275484 |
| 17765 | NM_001165924.1 | EMX2 | 22.0608641 | 6.0532041 | 2.39983061 |
| 17766 | NM_001256164.1 | TMCO1 | 43.3214239 | 6.05291708 | 2.40001548 |
| 17767 | NM_001129731.2 | #N/A | 16.5797914 | -5.2588689 | 2.08529002 |
| 17768 | NR_003110.2 | CCT6P1 | 33.068355 | -3.358128 | 1.33163725 |
| 17769 | NM_001024647.3 | RAB3IP | 97.7673559 | 3.38412929 | 1.3419919 |
| 17770 | NM_001077239.1 | RNF214 | 39.4058448 | -6.5122232 | 2.58267967 |
| 17771 | NM_133464.3 | #N/A | 10.4627185 | -4.6023946 | 1.82525875 |
| 17772 | NM_001039380.3 | #N/A | 17.9031923 | -3.7951393 | 1.50513679 |
| 17773 | NM_024555.5 | #N/A | 19.9118601 | -5.5225219 | 2.19028384 |
| 17774 | NM_001881.3 | CREM | 11.8252311 | -4.774304 | 1.89357134 |
| 17775 | NM_001199694.2 | SLC4A2 | 13.4404986 | -4.9703912 | 1.97143385 |
| 17778 | NM_003573.2 | LTBP4 | 58.7216803 | -7.0848622 | 2.81032601 |
| 17776 | NM_001144769.2 | #N/A | 48.3905872 | -6.8072906 | 2.700194 |
| 17777 | NM_001300780.1 | #N/A | 37.0155287 | -6.4222676 | 2.5474972 |
| 17779 | NM_017805.2 | #N/A | 9.58740327 | -4.4695681 | 1.77288888 |
| 17780 | NR_036446.1 | ACTG1P17 | 49.504668 | 2.41147601 | 0.95658116 |
| 17782 | NM_177534.2 | SULT1A1 | 10.4794364 | -4.6122269 | 1.82984513 |
| 17781 | NM_032411.2 | #N/A | 11.2533869 | -4.7077186 | 1.86772249 |
| 17783 | NM_003627.5 | #N/A | 8.96283373 | -4.3857032 | 1.74011119 |
| 17784 | NM_001199280.1 | #N/A | 44.1289901 | -6.6742297 | 2.6481704 |
| 17785 | NM_001256706.1 | #N/A | 9.67954839 | -4.4719148 | 1.77447211 |
| 17786 | NM_001244735.1 | TMEM126A | 11.8140089 | -4.7793965 | 1.89686311 |
| 17787 | NR_024007.1 | SMCR5 | 9.88450513 | 4.17912406 | 1.6587246 |
| 17788 | NR_024210.1 | #N/A | 11.9070668 | -4.7733012 | 1.89466526 |
| 17789 | NR_120420.1 | LOC100507006 | 10.9936455 | -4.6733321 | 1.85514719 |
| 17790 | NM_004016.2 | #N/A | 66.9791112 | -7.27612 | 2.88840021 |
| 17791 | NM_006350.3 | #N/A | 15.1815879 | -5.1349991 | 2.03870916 |
| 17792 | NM_004430.2 | #N/A | 8.63147665 | -4.3127329 | 1.71227874 |
| 17794 | NR_027902.1 | LINC00303 | 12.0909026 | 7.81622728 | 3.10332082 |
| 17793 | NM_001048181.2 | OPN1MW2 | 12.0909026 | 7.81622728 | 3.10332082 |
| 17795 | NM_153499.2 | CAMKK2 | 46.398673 | -6.7467108 | 2.67871736 |
| 17796 | NR_033354.1 | #N/A | 10.4134013 | -4.5890941 | 1.8221073 |
| 17798 | NM_178237.2 | EXOC1 | 47.2060227 | -6.7713092 | 2.68935425 |
| 17797 | NM_001317803.1 | #N/A | 14.5832954 | -5.0867292 | 2.02028898 |
| 17799 | NM_182932.2 | #N/A | 47.2644272 | -6.7729783 | 2.69062726 |
| 17800 | NM_001080400.1 | #N/A | 10.3706704 | -4.586162 | 1.82192581 |
| 17801 | NM_201648.2 | #N/A | 14.685316 | 6.75783859 | 2.68481634 |

| | | | | | |
|---|---|---|---|---|---|
| 17802 | NR_046215.1 | LINC00536 | 8.90820838 | 3.78092905 | 1.50237102 |
| 17803 | NM_015554.2 | #N/A | 36.6054897 | -6.4063842 | 2.54573081 |
| 17804 | NR_051960.1 | FALEC | 20.3795082 | 2.11679533 | 0.84118998 |
| 17807 | NR_033874.1 | FLJ20021 | 9.95056951 | -4.5241539 | 1.79849105 |
| 17805 | NM_002195.1 | #N/A | 25.1901997 | 9.63976703 | 3.83210921 |
| 17806 | NM_139240.3 | #N/A | 25.1901997 | 9.63976703 | 3.83210921 |
| 17808 | NR_039994.2 | TMEM161B-AS1 | 9.30056836 | -4.4162111 | 1.7560609 |
| 17809 | NM_005574.3 | #N/A | 9.96821003 | -4.5285093 | 1.80091829 |
| 17810 | NM_001242409.1 | #N/A | 47.0357358 | -6.7659716 | 2.69082285 |
| 17811 | NM_152228.2 | #N/A | 9.34699623 | -4.4378624 | 1.76511989 |
| 17812 | NM_013334.3 | GMPPB | 8.7624996 | -4.3499888 | 1.73020787 |
| 17813 | NM_033429.2 | CALML4 | 41.8386859 | -6.5987107 | 2.62466272 |
| 17815 | NR_027440.1 | LOC100272217 | 9.24608978 | -4.411187 | 1.7550195 |
| 17814 | NM_001302622.1 | #N/A | 49.1025961 | -6.8287442 | 2.71686918 |
| 17816 | NM_001190919.1 | THRA | 46.9963207 | -6.7644791 | 2.69164281 |
| 17817 | NM_004561.3 | #N/A | 14.2896685 | 5.07951998 | 2.02122298 |
| 17818 | NM_024007.4 | #N/A | 36.6485664 | -6.4078357 | 2.54980818 |
| 17819 | NM_183397.2 | #N/A | 291.191793 | 3.79709063 | 1.51096697 |
| 17820 | NM_012318.2 | #N/A | 441.369791 | -1.4568532 | 0.57975397 |
| 17821 | NM_001017420.2 | #N/A | 14.1354832 | -3.7229112 | 1.48174815 |
| 17822 | NM_001204747.1 | #N/A | 36.5093353 | -6.4024244 | 2.54893208 |
| 17823 | NM_001330303.1 | #N/A | 39.207463 | -6.5047577 | 2.5900891 |
| 17824 | NR_105045.1 | LINC01843 | 11.6094943 | 4.78240565 | 1.90432159 |
| 17825 | NM_001257976.1 | #N/A | 50.3348373 | -6.8646263 | 2.733677 |
| 17826 | NM_016014.3 | ABHD17B | 9.49757747 | -4.4592041 | 1.77609972 |
| 17827 | NM_007189.2 | #N/A | 238.29995 | -1.2925307 | 0.51491387 |
| 17828 | NM_001001415.3 | #N/A | 40.3727969 | -6.5474151 | 2.60844973 |
| 17829 | NM_018836.3 | #N/A | 10.7055714 | -4.635234 | 1.84677454 |
| 17830 | NM_002342.2 | #N/A | 8.96015875 | -4.3789146 | 1.74480938 |
| 17831 | NM_001317333.1 | #N/A | 229.865202 | -1.8493155 | 0.73694468 |
| 17832 | NM_001271785.1 | #N/A | 51.4878034 | -6.8958965 | 2.7481803 |
| 17833 | NM_001324506.1 | KMT5A | 19.1125242 | -4.1806463 | 1.66610729 |
| 17834 | NM_001270470.1 | SOCS2 | 13.6847114 | -4.992663 | 1.99020887 |
| 17835 | NM_032997.2 | ZWINT | 11.1606848 | -4.7002173 | 1.87400282 |
| 17836 | NM_001288709.1 | DBNDD1 | 16.0497073 | -5.2204104 | 2.08170166 |
| 17837 | NM_001004746.1 | OR5T2 | 31.3226866 | 9.94362062 | 3.96622086 |
| 17838 | NM_080423.2 | PTPN2 | 31.3226866 | 9.94362062 | 3.96622086 |
| 17839 | NR_037878.1 | LINC02275 | 73.5020989 | 3.73577355 | 1.49012833 |
| 17840 | NM_006340.2 | #N/A | 10.0360942 | -4.5247165 | 1.80595088 |
| 17842 | NM_001281484.1 | ARFGAP1 | 38.0723413 | -6.4631573 | 2.57972891 |
| 17841 | NM_001199671.1 | #N/A | 43.8500479 | -6.6646134 | 2.66014215 |
| 17843 | NR_040059.1 | SRP14-AS1 | 16.3122856 | -5.2351685 | 2.08993062 |
| 17844 | NM_033244.3 | #N/A | 14.2426484 | -5.03185 | 2.00893203 |

| | | | | | |
|---|---|---|---|---|---|
| 17845 | NM_001331071.1 | #N/A | 46.2023774 | -6.7400404 | 2.69135915 |
| 17846 | NR_048575.2 | GLUD1P3 | 8.79938117 | -4.3549363 | 1.73911374 |
| 17847 | NM_175902.4 | #N/A | 37.0850784 | -6.4251683 | 2.56595555 |
| 17848 | NM_001329474.1 | TMEM243 | 9.93233435 | -4.5202042 | 1.80536279 |
| 17849 | NM_014056.3 | HIGD1A | 436.06749 | -1.7693316 | 0.7067183 |
| 17850 | NR_002912.1 | SNORA67 | 9.32912473 | -4.4157688 | 1.76378622 |
| 17851 | NR_136151.1 | SNTG2-AS1 | 10.406225 | -4.5920506 | 1.83417954 |
| 17852 | NM_003348.3 | #N/A | 506.348873 | -3.709746 | 1.48190372 |
| 17853 | NM_014010.4 | #N/A | 259.650342 | -3.0425194 | 1.21555388 |
| 17854 | NR_135592.1 | #N/A | 9.20078887 | -4.4066705 | 1.76069075 |
| 17855 | NM_001144765.1 | #N/A | 152.240171 | 1.54304283 | 0.61655344 |
| 17856 | NM_001004441.2 | #N/A | 21.949031 | 4.74764612 | 1.8970466 |
| 17857 | NR_073585.1 | #N/A | 13.3827995 | -4.9598639 | 1.98185214 |
| 17858 | NM_024119.2 | #N/A | 16.609561 | -5.2695551 | 2.10578398 |
| 17859 | NM_001244755.1 | #N/A | 11.1627146 | 6.70162278 | 2.67822977 |
| 17860 | NR_029388.2 | PARGP1 | 8.95420316 | -4.3694618 | 1.74636485 |
| 17861 | NM_022128.2 | #N/A | 9.5738929 | -4.4729408 | 1.7879732 |
| 17862 | NM_001093772.1 | KIT | 35.8773462 | -6.3772316 | 2.54930489 |
| 17863 | NM_001172517.1 | SATB2 | 48.9898868 | -6.8237502 | 2.72821588 |
| 17864 | NM_001267561.1 | #N/A | 27.5221126 | -5.9957863 | 2.39717043 |
| 17865 | NR_048576.1 | EPN2-AS1 | 12.5906262 | 2.69424233 | 1.07723757 |
| 17866 | NR_135583.1 | #N/A | 19.8256749 | -5.5135563 | 2.20476542 |
| 17867 | NM_000276.3 | #N/A | 52.2360929 | -6.9167083 | 2.76612045 |
| 17868 | NR_104187.1 | APOBEC3B-AS1 | 7.88732218 | 6.30298081 | 2.52102191 |
| 17869 | NM_001101376.2 | #N/A | 14.3537299 | -5.0609919 | 2.02440122 |
| 17870 | NM_000613.2 | #N/A | 8.95155598 | -4.3753976 | 1.75028218 |
| 17871 | NM_000296.3 | #N/A | 564.495528 | -2.7644901 | 1.10590167 |
| 17872 | NM_031934.5 | #N/A | 44.162589 | -6.6747956 | 2.67022987 |
| 17873 | NM_001290327.1 | EFCAB2 | 11.7414639 | -4.7629501 | 1.90545712 |
| 17874 | NM_001199900.1 | PDK2 | 10.1672488 | -4.5487858 | 1.8201764 |
| 17875 | NM_015914.6 | #N/A | 36.0455837 | -6.3838711 | 2.55459515 |
| 17876 | NM_024341.2 | #N/A | 48.3573651 | -6.805982 | 2.72376716 |
| 17877 | NM_004035.6 | #N/A | 250.076032 | -1.9173611 | 0.76738965 |
| 17878 | NM_001243279.2 | #N/A | 46.1104948 | -6.7368532 | 2.69634568 |
| 17879 | NM_001242839.1 | GUK1 | 8.64265822 | -4.3269448 | 1.73220092 |
| 17881 | NM_001329122.1 | FCGR3A | 24.6176952 | 9.60668606 | 3.84590558 |
| 17880 | NM_001270768.1 | STRIP1 | 24.6176952 | 9.60668606 | 3.84590558 |
| 17882 | NM_018712.3 | #N/A | 45.4435016 | -6.7161744 | 2.68880258 |
| 17883 | NR_134495.1 | HINT1 | 52.856918 | 1.42166372 | 0.56918773 |
| 17884 | NM_001134387.1 | ZDHHC4 | 13.9310042 | -5.0077631 | 2.00502511 |
| 17885 | NM_001038705.1 | #N/A | 25.0312462 | 5.71394638 | 2.28792588 |
| 17886 | NM_003684.5 | #N/A | 19.2496116 | -3.8726475 | 1.55063856 |
| 17887 | NR_111930.1 | LOC100132057 | 8.93588278 | -4.3632158 | 1.74720961 |

| | | | | | |
|---|---|---|---|---|---|
| 17888 | NM_001008486.1 | SLC41A3 | 16.1757164 | -5.2327716 | 2.09563871 |
| 17889 | NM_001013660.2 | #N/A | 46.2174649 | 2.65186818 | 1.06218787 |
| 17890 | NM_013374.5 | #N/A | 311.73492 | -3.4412161 | 1.37844488 |
| 17891 | NM_001166284.1 | CCT7 | 10.7152305 | -4.6310803 | 1.8551474 |
| 17892 | NM_001306141.1 | SPDYE5 | 41.714548 | 3.70899026 | 1.48583306 |
| 17893 | NM_001277129.1 | NLRP12 | 69.0366021 | 2.79477438 | 1.1196343 |
| 17894 | NM_001286704.1 | #N/A | 11.3828558 | -4.7321181 | 1.89577813 |
| 17895 | NR_120425.1 | MAP3K4 | 18.60139 | -5.4173588 | 2.17042594 |
| 17896 | NM_001258024.1 | #N/A | 16.2536128 | -3.6722445 | 1.47131148 |
| 17897 | NM_152792.2 | ASPRV1 | 17.5877613 | -3.7665308 | 1.50912105 |
| 17898 | NM_003890.2 | FCGBP | 20.9012241 | -3.9696929 | 1.59070045 |
| 17899 | NM_001322056.1 | PDE1C | 49.4781498 | -6.8385814 | 2.74049271 |
| 17900 | NM_001127493.1 | #N/A | 233.462854 | -3.6450704 | 1.46076537 |
| 17901 | NM_012069.4 | #N/A | 32.9730854 | 2.46052973 | 0.98606595 |
| 17902 | NM_015431.3 | #N/A | 165.039172 | 1.43905131 | 0.57688972 |
| 17903 | NM_002281.3 | #N/A | 9.91606966 | -4.5303312 | 1.81626805 |
| 17904 | NM_000574.4 | #N/A | 12.6628966 | -4.8837568 | 1.95831117 |
| 17905 | NM_001330537.1 | SDCBP | 9.11869835 | -4.3993257 | 1.76424932 |
| 17906 | NM_182985.4 | #N/A | 10.6436697 | -4.633239 | 1.85807028 |
| 17907 | NR_026732.1 | LINC01551 | 25.259467 | -5.8663884 | 2.35282423 |
| 17909 | NM_023001.2 | ARID4A | 52.4884272 | 3.70859896 | 1.48758616 |
| 17908 | NM_002468.4 | #N/A | 45.7392278 | -6.7252204 | 2.69762476 |
| 17910 | NM_001318063.1 | #N/A | 17.0524824 | -3.7289678 | 1.49578108 |
| 17911 | NM_001330499.1 | #N/A | 8.2387966 | -4.2542693 | 1.70659262 |
| 17912 | NM_152611.4 | #N/A | 15.7342877 | 4.46611062 | 1.79162518 |
| 17913 | NM_001290331.1 | #N/A | 10.3186568 | -4.5883248 | 1.84073293 |
| 17915 | NR_125852.1 | CT69 | 18.0345527 | 2.45178731 | 0.98364922 |
| 17914 | NM_004671.4 | #N/A | 36.1662665 | -6.3886738 | 2.56311225 |
| 17916 | NM_005246.3 | #N/A | 47.119845 | -6.769801 | 2.7161883 |
| 17917 | NM_001286609.1 | ARMC2 | 21.980381 | 6.01749511 | 2.41506761 |
| 17918 | NM_198681.3 | PLEKHG5 | 40.0971225 | -6.5374409 | 2.62380376 |
| 17919 | NM_024627.5 | #N/A | 227.017202 | -1.3002841 | 0.52194378 |
| 17920 | NM_058219.2 | #N/A | 154.259391 | -1.362543 | 0.5469326 |
| 17921 | NM_145301.2 | TVP23C | 10.0403915 | -4.5248558 | 1.81636804 |
| 17922 | NR_049718.1 | COX10-AS1 | 67.7238142 | 1.44158287 | 0.5787762 |
| 17923 | NM_001272087.1 | #N/A | 8.62915348 | -4.3076503 | 1.72968173 |
| 17924 | NM_001278212.1 | #N/A | 36.3928644 | -6.3982053 | 2.56957594 |
| 17925 | NM_001297701.1 | #N/A | 13.2845931 | -4.9527872 | 1.98932272 |
| 17926 | NR_136304.1 | #N/A | 29.2223923 | -1.8681076 | 0.75033794 |
| 17927 | NM_001318829.1 | TSC2 | 45.64175 | -6.7229182 | 2.70067953 |
| 17928 | NM_001290018.1 | PRMT7 | 10.0793386 | -4.5272138 | 1.81873183 |
| 17929 | NM_033066.2 | #N/A | 17.6468314 | 5.3510855 | 2.14999774 |
| 17930 | NM_001253793.1 | #N/A | 11.5996708 | 4.04537493 | 1.62540278 |

| | | | | | |
|---|---|---|---|---|---|
| 17931 | NR_125819.1 | LOC101929550 | 41.8945856 | 3.72674575 | 1.49742058 |
| 17932 | NR_033694.1 | #N/A | 30.2335513 | 3.91065754 | 1.57167205 |
| 17933 | NR_038874.1 | LOC286177 | 20.0463603 | 3.48763526 | 1.4016934 |
| 17934 | NM_001704.2 | #N/A | 151.619846 | -3.0209075 | 1.21431034 |
| 17935 | NM_005311.4 | #N/A | 49.5205369 | -6.8398409 | 2.74978397 |
| 17936 | NM_001261403.2 | #N/A | 10.7190514 | -4.6237754 | 1.85896133 |
| 17937 | NM_001134666.2 | #N/A | 46.6737881 | 5.28292317 | 2.12411275 |
| 17938 | NM_004543.4 | #N/A | 42.1400574 | 4.03466067 | 1.62222046 |
| 17939 | NM_001242849.1 | RNF146 | 36.1716656 | -6.3894158 | 2.56912861 |
| 17940 | NM_001278718.1 | #N/A | 13.6169919 | -4.9829591 | 2.00374101 |
| 17941 | NM_017709.3 | #N/A | 15.2874869 | -5.1507842 | 2.07132 |
| 17942 | NR_045418.1 | #N/A | 44.8927577 | -6.698391 | 2.69365764 |
| 17943 | NM_024786.2 | ZDHHC11 | 94.8628419 | -2.2091964 | 0.88840707 |
| 17944 | NM_000502.5 | #N/A | 8.37698909 | -4.2848716 | 1.72329505 |
| 17945 | NM_001008405.2 | #N/A | 10.658972 | -4.6067523 | 1.85280387 |
| 17946 | NM_001319224.1 | EXO1 | 8.59493592 | -4.3139921 | 1.73523971 |
| 17947 | NM_001190882.2 | ORC4 | 54.2843602 | -6.9732516 | 2.80492392 |
| 17948 | NM_152995.5 | NFXL1 | 37.2953968 | -6.433011 | 2.58766186 |
| 17949 | NM_001079684.2 | KIAA1191 | 35.0005192 | -6.3416837 | 2.55109673 |
| 17950 | NM_001242617.1 | #N/A | 11.9446206 | -4.784631 | 1.92497785 |
| 17951 | NR_125718.1 | POT1-AS1 | 23.015933 | 1.6403438 | 0.66001309 |
| 17952 | NM_005640.2 | #N/A | 11.279086 | -4.7135881 | 1.89689453 |
| 17953 | NM_018559.3 | #N/A | 8.41024922 | -4.2804321 | 1.72258375 |
| 17954 | NM_001322436.1 | HNRNPH3 | 11.3840674 | -4.7064278 | 1.89418181 |
| 17955 | NR_135742.1 | #N/A | 9.86482298 | -4.5011081 | 1.81159222 |
| 17956 | NM_001258280.1 | #N/A | 17.1661091 | -5.3068995 | 2.1360466 |
| 17957 | NM_001270552.1 | #N/A | 37.7093539 | -6.4489994 | 2.59577681 |
| 17958 | NM_181846.2 | #N/A | 36.8583387 | -6.4158545 | 2.58262974 |
| 17959 | NM_138639.1 | BCL2L12 | 34.8517921 | -6.3354843 | 2.55040133 |
| 17960 | NM_148179.2 | #N/A | 8.46213712 | -4.3004733 | 1.73118139 |
| 17961 | NR_026968.1 | RPL34-AS1 | 40.624927 | 2.84764332 | 1.14644973 |
| 17962 | NM_004767.3 | #N/A | 22.906227 | 2.8584894 | 1.1509061 |
| 17963 | NR_026835.1 | FLJ37201 | 40.5389464 | 4.5398813 | 1.82825397 |
| 17964 | NM_001330426.1 | #N/A | 99.9352011 | -2.6912702 | 1.08382209 |
| 17965 | NM_178815.3 | #N/A | 123.250838 | -3.1987866 | 1.2882359 |
| 17967 | NM_001346471.1 | AMZ2 | 46.1056678 | -6.7370098 | 2.71361037 |
| 17966 | NM_001242789.1 | BRF1 | 36.1913594 | -6.3900938 | 2.5738854 |
| 17968 | NR_024418.1 | #N/A | 9.53455522 | -4.4625173 | 1.79797563 |
| 17969 | NR_024240.1 | HLA-J | 8.33780208 | -4.2709613 | 1.7208747 |
| 17972 | NM_001256503.1 | CADM2 | 8.71928328 | 8.07975784 | 3.2560631 |
| 17970 | NM_001127615.2 | ENOX1 | 8.71928328 | 8.07975784 | 3.2560631 |
| 17971 | NM_001199881.1 | #N/A | 8.71928328 | 8.07975784 | 3.2560631 |
| 17973 | NM_001286048.1 | #N/A | 8.71928328 | 8.07975784 | 3.2560631 |

| | | | | | |
|---|---|---|---|---|---|
| 17974 | NM_001286767.1 | #N/A | 8.71928328 | 8.07975784 | 3.2560631 |
| 17975 | NR_038923.1 | ZNRD2-AS1 | 8.91103103 | -4.3689747 | 1.76077643 |
| 17976 | NM_018132.3 | #N/A | 18.9621594 | -3.0839245 | 1.24312769 |
| 17977 | NM_183005.4 | #N/A | 10.6561127 | -4.610922 | 1.8586635 |
| 17978 | NM_001039395.3 | #N/A | 12.9782465 | -4.9154778 | 1.98160462 |
| 17979 | NM_024017.4 | #N/A | 258.861119 | -3.6114926 | 1.45604723 |
| 17980 | NM_001172420.1 | #N/A | 32.9843221 | 5.53109235 | 2.23002262 |
| 17981 | NR_103795.1 | NFXL1 | 30.1179679 | 9.88690542 | 3.98639987 |
| 17982 | NM_004334.2 | #N/A | 9.46912201 | -4.4549402 | 1.79632324 |
| 17983 | NM_181774.3 | #N/A | 24.0451906 | 9.57283532 | 3.86026435 |
| 17984 | NM_133264.4 | #N/A | 274.501673 | -1.4491913 | 0.58446587 |
| 17985 | NM_001300969.1 | CCDC181 | 36.212597 | -6.3908876 | 2.57816944 |
| 17986 | NM_001320703.1 | #N/A | 44.6169264 | -6.689342 | 2.69864484 |
| 17987 | NM_001135208.1 | FKBP2 | 12.6608866 | -4.8784553 | 1.96815292 |
| 17988 | NR_110466.1 | HAGLR | 47.355389 | -6.7765291 | 2.73410429 |
| 17989 | NM_001291315.1 | #N/A | 10.2124201 | -4.560696 | 1.84020369 |
| 17990 | NM_001508.2 | #N/A | 13.2371697 | -4.9271318 | 1.98818929 |
| 17991 | NM_152365.2 | #N/A | 11.42367 | 4.30898173 | 1.73875859 |
| 17992 | NM_001258320.1 | #N/A | 11.3231519 | -4.7006271 | 1.89698571 |
| 17993 | NM_001330082.1 | NRXN1 | 42.3987292 | -6.6172618 | 2.67061369 |
| 17994 | NM_001330588.1 | #N/A | 9.34481093 | -4.4284967 | 1.78746337 |
| 17995 | NM_014880.4 | #N/A | 34.3709849 | -6.3154733 | 2.54913329 |
| 17996 | NR_136748.1 | PAK3 | 47.3758285 | -6.77526 | 2.73474069 |
| 17997 | NM_199289.2 | #N/A | 57.8408973 | -2.9579112 | 1.19400186 |
| 17998 | NR_038864.1 | LOC401324 | 41.215773 | 3.59004828 | 1.44934333 |
| 17999 | NR_004401.2 | SEC1P | 18.1254203 | 2.33197326 | 0.94147536 |
| 18001 | NR_110723.1 | ZNF793-AS1 | 16.7970928 | -3.1991316 | 1.29170665 |
| 18000 | NM_020927.2 | #N/A | 754.376991 | -3.682215 | 1.48675808 |
| 18002 | NM_001304532.1 | #N/A | 37.2028945 | -6.4297763 | 2.59632217 |
| 18003 | NM_182523.1 | #N/A | 12.6024781 | -4.8600119 | 1.96264623 |
| 18004 | NR_026693.1 | #N/A | 11.2229286 | -4.7082011 | 1.90162028 |
| 18005 | NM_197956.3 | #N/A | 69.2519433 | -1.6752747 | 0.67667595 |
| 18006 | NM_004230.3 | #N/A | 167.454834 | -0.8250049 | 0.33324684 |
| 18007 | NM_001243099.1 | IL5RA | 32.685008 | 4.72829507 | 1.91040598 |
| 18008 | NR_037158.1 | IL21R-AS1 | 36.9622982 | 1.71274063 | 0.6920474 |
| 18009 | NM_014358.3 | #N/A | 17.04736 | 4.45218734 | 1.79897651 |
| 18010 | NM_001281715.1 | PHB | 13.9975753 | -3.7292721 | 1.50720833 |
| 18011 | NM_144600.3 | #N/A | 59.6194302 | -5.8546635 | 2.36628961 |
| 18012 | NM_144582.2 | #N/A | 253.619339 | -3.4211675 | 1.38282936 |
| 18013 | NM_001193557.1 | #N/A | 14.3674345 | -5.0635079 | 2.04670986 |
| 18014 | NM_001324021.1 | PDXDC1 | 34.9466413 | -6.339694 | 2.56270915 |
| 18015 | NR_073092.1 | #N/A | 70.7632312 | 7.72862811 | 3.12447148 |
| 18016 | NR_036494.1 | LOC643201 | 24.1873166 | 3.26073119 | 1.31844453 |

| | | | | | |
|---|---|---|---|---|---|
| 18017 | NM_017771.4 | #N/A | 34.6934509 | -6.3291314 | 2.55917017 |
| 18018 | NM_173163.2 | #N/A | 11.2551244 | -4.7097742 | 1.90448336 |
| 18019 | NM_001024847.2 | TGFBR2 | 18.6454709 | -5.4325371 | 2.19680282 |
| 18020 | NM_001142274.1 | #N/A | 286.369995 | -2.9701247 | 1.20107944 |
| 18021 | NM_001347729.1 | PITRM1 | 9.05505107 | -4.3863049 | 1.77380786 |
| 18022 | NR_026761.2 | LINC00467 | 8.51588609 | -4.297605 | 1.73796444 |
| 18023 | NM_001329848.1 | MYT1L | 11.1674734 | -4.6833951 | 1.89411583 |
| 18024 | NR_036512.1 | LOC100129917 | 8.82286628 | -4.3599915 | 1.7634093 |
| 18027 | NR_028323.1 | SLC5A6 | 27.5987685 | -4.3089937 | 1.74288414 |
| 18025 | NM_001313972.1 | #N/A | 11.4120664 | -4.7246296 | 1.91100436 |
| 18026 | NM_016390.3 | #N/A | 175.214199 | -2.8531264 | 1.15401178 |
| 18028 | NM_001199636.1 | #N/A | 16.0786528 | -5.208332 | 2.10713712 |
| 18029 | NM_001199052.1 | #N/A | 10.5285827 | -4.608235 | 1.864807 |
| 18030 | NM_001289807.1 | CRYAB | 36.0080161 | -6.3824402 | 2.5834104 |
| 18032 | NR_027141.2 | DENND10P1 | 9.33739252 | -4.431768 | 1.79409567 |
| 18031 | NM_013310.3 | #N/A | 10.3998847 | -4.5911466 | 1.85860734 |
| 18033 | NM_194302.3 | #N/A | 21.7419289 | -5.6565678 | 2.29021074 |
| 18034 | NM_001004346.3 | #N/A | 10.6205733 | -4.6292689 | 1.87434035 |
| 18035 | NM_001320675.1 | #N/A | 42.5938695 | 6.0413007 | 2.44606125 |
| 18036 | NM_002600.3 | #N/A | 12.9902566 | -4.9204331 | 1.99220234 |
| 18037 | NM_153834.3 | #N/A | 12.6067818 | 2.71633192 | 1.10006817 |
| 18038 | NM_001319167.1 | #N/A | 12.9698052 | -4.9045342 | 1.98639867 |
| 18039 | NM_001195203.2 | #N/A | 45.007765 | -6.7042374 | 2.71533249 |
| 18040 | NR_104485.1 | #N/A | 9.22129454 | -4.4220176 | 1.79103696 |
| 18041 | NM_001256310.1 | #N/A | 167.698868 | -3.3820473 | 1.36986033 |
| 18042 | NM_006896.3 | #N/A | 14.936589 | -3.8255697 | 1.54954566 |
| 18043 | NM_022476.3 | #N/A | 34.3717206 | -6.3157949 | 2.55825644 |
| 18044 | NR_031650.1 | MIR1248 | 25.0563249 | 5.67646293 | 2.29945391 |
| 18045 | NM_175634.2 | #N/A | 45.7199901 | -6.7248175 | 2.72421543 |
| 18046 | NM_001127582.1 | #N/A | 34.8441378 | -6.3351923 | 2.56646979 |
| 18047 | NR_137639.1 | #N/A | 14.6259787 | -5.0866136 | 2.06066 |
| 18048 | NM_001318042.1 | #N/A | 48.1923918 | -6.8021807 | 2.75642228 |
| 18049 | NM_020169.3 | #N/A | 10.1682897 | -4.5699366 | 1.85189978 |
| 18050 | NM_001321857.1 | #N/A | 35.2400417 | -6.3518342 | 2.57410825 |
| 18051 | NM_001278255.1 | POLE3 | 103.842853 | -5.2960916 | 2.14631849 |
| 18052 | NM_001202521.1 | DDR1 | 35.9110373 | -6.37826 | 2.58533287 |
| 18053 | NM_012095.5 | #N/A | 42.793394 | -6.6293475 | 2.68710395 |
| 18054 | NM_001282742.1 | OSBP2 | 36.7790726 | -6.4130328 | 2.59972926 |
| 18055 | NM_004135.3 | #N/A | 195.290874 | -3.3903908 | 1.3744166 |
| 18056 | NM_005734.4 | #N/A | 33.7607305 | -6.2896824 | 2.54985676 |
| 18057 | NM_001123395.1 | #N/A | 47.9374899 | -6.7946217 | 2.75460769 |
| 18058 | NM_001284330.1 | #N/A | 8.33881601 | -4.2768226 | 1.7338769 |
| 18059 | NR_126580.1 | AGAP7P | 10.6207447 | -4.6216275 | 1.87370957 |

| | | | | | |
|---|---|---|---|---|---|
| 18060 | NM_014964.4 | EPN2 | 14.4334092 | -5.0703101 | 2.05572978 |
| 18061 | NM_001700.4 | #N/A | 8.34884192 | -4.2754691 | 1.73362284 |
| 18062 | NM_001243089.1 | #N/A | 37.8301036 | -6.4533578 | 2.61710749 |
| 18063 | NM_021629.3 | #N/A | 350.063155 | -1.5568234 | 0.63136773 |
| 18064 | NR_125421.1 | LOC101927657 | 21.3674679 | 5.44313995 | 2.20748396 |
| 18065 | NM_001135248.1 | MFAP2 | 35.3642337 | -6.3561384 | 2.5784864 |
| 18066 | NM_001101357.1 | #N/A | 12.4441371 | -4.855588 | 1.97000818 |
| 18067 | NM_197958.2 | #N/A | 8.96211712 | -4.3798157 | 1.77731244 |
| 18068 | NM_001322963.1 | #N/A | 12.1428038 | -4.8127584 | 1.95340669 |
| 18069 | NM_024421.2 | DSC1 | 8.00374042 | -4.2127838 | 1.71001969 |
| 18070 | NM_032843.4 | #N/A | 36.9628456 | -6.4201076 | 2.60610279 |
| 18071 | NR_138153.1 | XKR6 | 10.3170549 | -4.5599329 | 1.85116955 |
| 18072 | NM_005406.2 | #N/A | 204.396448 | -3.2741168 | 1.32933295 |
| 18073 | NM_002463.1 | #N/A | 9.15993878 | -4.3899837 | 1.78249556 |
| 18074 | NM_024611.5 | #N/A | 35.9071395 | -6.3784587 | 2.59031724 |
| 18075 | NM_001110822.1 | #N/A | 14.5017264 | 8.12850926 | 3.30112443 |
| 18076 | NM_000247.2 | #N/A | 11.025959 | -4.6791557 | 1.90045907 |
| 18077 | NM_001301110.1 | #N/A | 42.169446 | 3.83704126 | 1.55843587 |
| 18078 | NM_148169.2 | FBXO17 | 42.7217676 | 5.00882451 | 2.03444509 |
| 18079 | NM_030643.4 | APOL4 | 18.3740287 | -5.4128258 | 2.19856642 |
| 18080 | NM_002497.3 | #N/A | 17.2892135 | -3.7316195 | 1.51572614 |
| 18081 | NM_144718.3 | #N/A | 12.10698 | -4.8057138 | 1.95202635 |
| 18082 | NM_001321383.1 | ABHD11 | 8.39617966 | -4.2817772 | 1.73926782 |
| 18083 | NM_017868.3 | #N/A | 37.9431105 | -3.4730379 | 1.41081732 |
| 18084 | NM_001287740.1 | #N/A | 43.1973797 | -2.4265756 | 0.98575482 |
| 18085 | NM_001308357.1 | ZNF433 | 12.1430426 | -4.8089308 | 1.95358835 |
| 18086 | NM_001297609.1 | #N/A | 42.6663553 | -6.625248 | 2.69153753 |
| 18087 | NM_173211.1 | TGIF1 | 23.4726861 | 9.53816716 | 3.87522091 |
| 18088 | NM_024586.5 | #N/A | 33.8479468 | -6.2932033 | 2.55714697 |
| 18089 | NM_181830.2 | #N/A | 46.7578607 | -6.7562224 | 2.74574349 |
| 18090 | NR_144356.1 | PPIEL | 11.1223281 | -4.6749322 | 1.89994029 |
| 18091 | NM_001304968.1 | TSGA13 | 20.8853845 | 2.81899219 | 1.14580824 |
| 18092 | NM_175888.3 | #N/A | 10.0956878 | -4.5557783 | 1.85183882 |
| 18093 | NM_176871.3 | #N/A | 9.35432651 | -4.4438195 | 1.80631703 |
| 18094 | NM_001014980.2 | #N/A | 10.7403542 | -4.6387161 | 1.88591856 |
| 18095 | NM_001258326.1 | #N/A | 34.5652419 | -6.3239128 | 2.57149469 |
| 18096 | NR_073595.1 | TYSND1 | 18.0490283 | -3.7852136 | 1.5392174 |
| 18098 | NM_201440.1 | PPHLN1 | 12.4600327 | -4.8385807 | 1.96778432 |
| 18097 | NM_001145714.1 | #N/A | 37.5395766 | -6.4427363 | 2.62019277 |
| 18099 | NM_207393.1 | #N/A | 19.7150391 | 4.21201449 | 1.71298887 |
| 18100 | NR_046504.1 | NLGN4Y-AS1 | 5.8753044 | 3.4593156 | 1.4068865 |
| 18101 | NM_001145030.1 | #N/A | 11.2321458 | 7.70994376 | 3.13562825 |
| 18103 | NR_024603.1 | MPDU1 | 8.85676255 | -4.3576801 | 1.77238631 |

| | | | | | |
|---|---|---|---|---|---|
| 18102 | NM_017839.4 | #N/A | 70.1468086 | -1.5003463 | 0.6102285 |
| 18104 | NM_138363.2 | #N/A | 36.8969677 | -6.4179933 | 2.6104146 |
| 18105 | NM_001308170.1 | AMPD2 | 42.5509064 | -6.6214466 | 2.69323327 |
| 18106 | NR_026742.1 | USP27X-AS1 | 8.08317279 | -4.2318868 | 1.72136524 |
| 18107 | NM_001324244.1 | #N/A | 33.5721031 | -6.2815733 | 2.55527289 |
| 18108 | NM_001346877.1 | #N/A | 11.0450478 | -4.6893439 | 1.90780455 |
| 18109 | NR_024460.1 | #N/A | 33.5167047 | -6.2790755 | 2.55460635 |
| 18110 | NM_001161013.2 | ARMC10 | 31.9786712 | 4.62535333 | 1.88186988 |
| 18111 | NR_130151.1 | #N/A | 9.0759186 | -4.3756298 | 1.78029639 |
| 18112 | NM_001290072.1 | #N/A | 9.45657267 | -4.4363074 | 1.80544194 |
| 18113 | NM_001291594.1 | #N/A | 20.004567 | -4.2586533 | 1.73317315 |
| 18114 | NR_104587.1 | #N/A | 39.507007 | -6.5145034 | 2.65127456 |
| 18115 | NR_027005.2 | KHDC1 | 8.07028849 | -4.2192755 | 1.71762504 |
| 18116 | NR_028045.1 | KLRA1P | 9.80884282 | -4.4990709 | 1.83177978 |
| 18117 | NM_182639.3 | HPS1 | 9.98736114 | -4.5304025 | 1.8446033 |
| 18118 | NR_135133.1 | LOC101928932 | 11.2572245 | 7.70984995 | 3.13932402 |
| 18120 | NM_001300919.1 | HMGA2 | 38.9506018 | 1.61999377 | 0.65967605 |
| 18119 | NM_001128160.1 | UBP1 | 137.507604 | -3.3437748 | 1.36161339 |
| 18121 | NM_001141970.1 | #N/A | 33.9710611 | -6.2988896 | 2.56515763 |
| 18122 | NM_002917.1 | #N/A | 211.235218 | -2.6470631 | 1.07799385 |
| 18123 | NM_001001479.3 | #N/A | 48.0764894 | -1.9131804 | 0.7791576 |
| 18125 | NM_001258299.1 | SHTN1 | 10.7198777 | -4.6271999 | 1.88453474 |
| 18124 | NM_001039375.2 | #N/A | 10.2346404 | -4.5580277 | 1.85635997 |
| 18126 | NM_171846.3 | #N/A | 32.9930378 | -6.2566019 | 2.54851527 |
| 18127 | NM_001257137.2 | #N/A | 34.1287968 | -6.305171 | 2.56897065 |
| 18128 | NM_181578.4 | RFC5 | 10.5466055 | -4.5911628 | 1.87071814 |
| 18129 | NM_001193304.2 | #N/A | 133.907713 | -4.8129852 | 1.96117113 |
| 18130 | NM_005144.4 | HR | 45.5061163 | -6.7173016 | 2.73732426 |
| 18131 | NM_007298.3 | BRCA1 | 55.6546791 | 3.14784729 | 1.28301719 |
| 18132 | NM_001145310.3 | LRTOMT | 9.01108922 | -4.3962274 | 1.79190737 |
| 18133 | NR_028582.1 | #N/A | 14.9795271 | -5.1159461 | 2.08555176 |
| 18134 | NM_001256526.1 | TMEM250 | 10.1094008 | -4.5565561 | 1.85769322 |
| 18135 | NM_001282747.1 | ARMC3 | 12.3591196 | -4.8368116 | 1.97200163 |
| 18136 | NM_015715.4 | #N/A | 9.49684786 | -4.4543523 | 1.81615699 |
| 18137 | NM_004224.3 | GPR50 | 8.89384858 | -4.3766315 | 1.78454939 |
| 18140 | NM_001294337.1 | C5orf38 | 11.3139458 | -4.7190083 | 1.9244899 |
| 18139 | NM_001199112.1 | MDH1 | 28.9132492 | 9.82787563 | 4.0080377 |
| 18138 | NM_001168410.1 | #N/A | 28.9132492 | 9.82787563 | 4.0080377 |
| 18141 | NR_131338.1 | #N/A | 28.9132492 | 9.82787563 | 4.0080377 |
| 18142 | NM_012155.2 | #N/A | 33.9479798 | -6.2978488 | 2.56862173 |
| 18143 | NM_006267.4 | #N/A | 472.530206 | -3.5149104 | 1.43390233 |
| 18144 | NM_198970.1 | #N/A | 35.5235295 | -6.362657 | 2.59574805 |
| 18145 | NM_001039199.2 | #N/A | 54.3879847 | -6.9761886 | 2.84701501 |

| | | | | | |
|---|---|---|---|---|---|
| 18146 | NM_021625.4 | #N/A | 61.8206972 | -7.1612295 | 2.92251498 |
| 18147 | NM_001270400.1 | TUBA1A | 8.26943825 | -4.2502706 | 1.73459144 |
| 18148 | NM_001101653.1 | #N/A | 72.3550949 | 4.84712954 | 1.97827627 |
| 18149 | NM_017925.6 | #N/A | 37.8883312 | -6.455407 | 2.63471144 |
| 18150 | NM_001012756.2 | #N/A | 113.106369 | -2.2159141 | 0.90443043 |
| 18151 | NM_022168.3 | #N/A | 17.3988302 | -5.3360845 | 2.17807829 |
| 18152 | NM_001308402.1 | #N/A | 11.2241166 | -4.6809642 | 1.9107545 |
| 18153 | NM_053284.2 | #N/A | 9.73917692 | -4.5026935 | 1.83807179 |
| 18154 | NM_145286.2 | #N/A | 10.0849098 | -4.5550862 | 1.85970827 |
| 18155 | NM_001135914.1 | #N/A | 8.73521369 | -4.3409512 | 1.77234362 |
| 18156 | NM_001177665.1 | #N/A | 33.8047536 | -6.2912699 | 2.56864116 |
| 18157 | NR_002808.2 | ITPK1-AS1 | 27.065427 | 3.50119403 | 1.42952783 |
| 18158 | NM_002506.2 | #N/A | 8.08683391 | -4.2303647 | 1.72730521 |
| 18159 | NM_001257207.1 | FGF1 | 13.0974723 | -4.9283452 | 2.01237612 |
| 18160 | NR_015436.1 | #N/A | 11.8329457 | -4.7653448 | 1.9458587 |
| 18161 | NM_020826.2 | #N/A | 404.409572 | -2.0292732 | 0.8286927 |
| 18162 | NM_205548.2 | #N/A | 8.67183503 | -4.3325813 | 1.76947992 |
| 18163 | NM_001206833.1 | #N/A | 42.2449664 | -6.6113677 | 2.70023348 |
| 18164 | NM_001316339.1 | #N/A | 17.1643699 | 6.70568289 | 2.73903441 |
| 18165 | NM_178813.5 | #N/A | 8.44699914 | -4.2932296 | 1.75370625 |
| 18166 | NM_001242878.2 | #N/A | 6.35725956 | 7.6696694 | 3.1329531 |
| 18167 | NM_001308056.1 | STARD4 | 48.0465751 | -6.7957781 | 2.77608044 |
| 18168 | NM_001164832.2 | #N/A | 11.2782337 | -4.7131993 | 1.92549687 |
| 18169 | NR_024420.1 | LINC00937 | 8.92650601 | -4.3662002 | 1.78382206 |
| 18170 | NM_171999.3 | SALL3 | 367.892918 | -3.5565328 | 1.45318519 |
| 18171 | NM_147148.2 | #N/A | 10.4610296 | -4.5943279 | 1.8773632 |
| 18172 | NM_001319170.1 | #N/A | 12.4049237 | -4.8408649 | 1.97820756 |
| 18173 | NM_001126051.1 | HDGF | 8.58125093 | -4.3208876 | 1.76598797 |
| 18174 | NR_125966.1 | LINC01732 | 25.0676129 | 4.61698702 | 1.88739655 |
| 18175 | NM_001265610.1 | TRAK1 | 18.599636 | -4.1410122 | 1.69325762 |
| 18176 | NM_001042428.1 | #N/A | 42.4137996 | -6.6172636 | 2.70596542 |
| 18177 | NM_017986.3 | #N/A | 42.9809996 | 2.42503442 | 0.99175569 |
| 18178 | NM_001257396.1 | #N/A | 10.2143773 | -4.5487388 | 1.86045988 |
| 18180 | NR_027468.3 | LINC01138 | 101.385947 | 1.13080309 | 0.46256163 |
| 18179 | NM_001271867.1 | #N/A | 12.0602296 | -4.7930552 | 1.96062956 |
| 18181 | NM_173206.3 | #N/A | 32.5135931 | -6.2353941 | 2.55080744 |
| 18182 | NM_001261411.1 | #N/A | 9.5851487 | -4.4844107 | 1.83453387 |
| 18183 | NM_001270374.1 | TMEM254 | 12.0300117 | -4.8051768 | 1.96598964 |
| 18184 | NR_033201.2 | HOXB-AS3 | 10.8592667 | -4.6604557 | 1.9068814 |
| 18185 | NM_001197221.1 | #N/A | 32.6455367 | -6.2413299 | 2.55409528 |
| 18186 | NM_022153.1 | #N/A | 9.50246646 | -4.4677522 | 1.82851611 |
| 18187 | NM_025204.3 | #N/A | 32.7035652 | -6.2437257 | 2.55551496 |
| 18188 | NM_001317950.1 | #N/A | 34.1587284 | -6.3069652 | 2.58147336 |

| | | | | | |
|---|---|---|---|---|---|
| 18189 | NM_182678.2 | UBE2E3 | 33.9636491 | -6.2981028 | 2.57826892 |
| 18190 | NM_001170414.2 | TJP2 | 16.8255305 | -3.696699 | 1.51342849 |
| 18191 | NM_001040097.1 | #N/A | 10.8910747 | -4.6473235 | 1.90278061 |
| 18192 | NM_001166694.1 | #N/A | 22.9001816 | 9.50264293 | 3.89081366 |
| 18193 | NM_014503.2 | #N/A | 102.852197 | -2.3616292 | 0.9670211 |
| 18194 | NM_001135673.3 | #N/A | 33.8883537 | -6.2948017 | 2.57781585 |
| 18195 | NM_018304.3 | #N/A | 485.181306 | 1.01350828 | 0.41504644 |
| 18196 | NM_001025070.1 | #N/A | 11.5718269 | -4.7520154 | 1.94619512 |
| 18197 | NM_001347829.1 | #N/A | 38.0270992 | -6.4610582 | 2.64624331 |
| 18198 | NM_000534.4 | PMS1 | 33.9949091 | -6.2993471 | 2.58015908 |
| 18199 | NR_027413.2 | LINC00910 | 59.2024942 | 2.62767039 | 1.0766058 |
| 18200 | NM_015017.4 | USP33 | 364.757159 | 0.6641014 | 0.27215096 |
| 18201 | NR_027882.1 | #N/A | 11.7941465 | -3.4658011 | 1.42040967 |
| 18202 | NM_001077242.1 | #N/A | 9.24579448 | -4.4164028 | 1.81016533 |
| 18203 | NM_001197295.1 | ECM2 | 8.69023588 | -4.3430378 | 1.78012469 |
| 18204 | NR_038293.1 | LINC01012 | 7.99184135 | -4.2141661 | 1.72738812 |
| 18205 | NM_001270524.1 | #N/A | 18.1937346 | -5.3997436 | 2.21340412 |
| 18208 | NR_031680.1 | MIR548H4 | 14.5249319 | 8.04606171 | 3.29821875 |
| 18206 | NM_001128225.2 | #N/A | 43.6683978 | -6.65974 | 2.72996087 |
| 18207 | NM_152418.3 | #N/A | 14.5249319 | 8.04606171 | 3.29821875 |
| 18209 | NM_173176.2 | #N/A | 42.8528313 | -6.6329572 | 2.71901918 |
| 18210 | NM_001146289.1 | P3H1 | 45.1145524 | -6.7072045 | 2.74948899 |
| 18211 | NM_000360.3 | #N/A | 14.8720977 | -5.1117367 | 2.09550512 |
| 18212 | NR_040662.1 | HCP5 | 8.66392107 | -4.3217569 | 1.77197601 |
| 18213 | NM_001293172.1 | LOC102724957 | 10.6364358 | 7.71478754 | 3.16328785 |
| 18214 | NR_110104.1 | VPS33B-DT | 73.2996739 | 4.22635578 | 1.73311086 |
| 18215 | NM_001078650.2 | #N/A | 9.84421514 | -4.5172939 | 1.85262332 |
| 18216 | NM_175569.2 | XG | 11.5348641 | 5.18988733 | 2.12862804 |
| 18217 | NM_000135.2 | #N/A | 39.8384186 | -6.5268451 | 2.67703819 |
| 18218 | NM_018250.3 | #N/A | 42.0222502 | -6.6028996 | 2.70854498 |
| 18219 | NM_001130713.2 | LEF1 | 32.991194 | -6.2561838 | 2.56637626 |
| 18220 | NM_001166137.1 | SPATA31C2 | 10.9458936 | 7.67268585 | 3.14760775 |
| 18221 | NM_001324132.1 | #N/A | 34.836217 | -6.3345275 | 2.59864142 |
| 18222 | NR_015421.1 | LOC154761 | 8.82424863 | -4.3425358 | 1.78155866 |
| 18223 | NM_001137610.1 | #N/A | 8.36434471 | -4.2854344 | 1.75818963 |
| 18224 | NM_001306076.1 | CCDC18 | 10.0343065 | -4.5506561 | 1.86706669 |
| 18225 | NM_001136538.1 | #N/A | 17.714611 | -5.3586676 | 2.19909491 |
| 18226 | NM_001348120.1 | TRIM16 | 41.6036502 | -6.5889134 | 2.7042073 |
| 18227 | NM_001282410.1 | PPP1R7 | 40.9563723 | -6.5662609 | 2.69495295 |
| 18228 | NM_001242848.1 | RNF146 | 34.2187304 | -6.3089297 | 2.58967885 |
| 18229 | NM_001319840.1 | #N/A | 43.1089206 | -6.6403899 | 2.72581048 |
| 18230 | NM_017748.4 | #N/A | 168.88173 | -1.1204042 | 0.45993338 |
| 18231 | NM_001323645.1 | #N/A | 32.5744295 | -6.2381976 | 2.56099537 |

| | | | | | |
|---|---|---|---|---|---|
| 18232 | NR_033982.1 | LINC01801 | 8.97418458 | -4.3773217 | 1.79715793 |
| 18233 | NM_001157.2 | #N/A | 32.0775249 | -6.2159691 | 2.55222714 |
| 18234 | NR_034169.1 | FAM133DP | 9.98491195 | -4.5183445 | 1.85535898 |
| 18235 | NM_001261835.1 | #N/A | 40.0866311 | -6.5365516 | 2.68413577 |
| 18236 | NM_173179.3 | #N/A | 31.990143 | -6.2121377 | 2.55122254 |
| 18237 | NM_001303416.1 | #N/A | 12.0807998 | -4.7907158 | 1.96752829 |
| 18238 | NM_003844.3 | #N/A | 28.0731977 | 3.66409195 | 1.50490122 |
| 18240 | NM_001271750.1 | C5AR2 | 8.56364186 | 2.68411811 | 1.10244263 |
| 18239 | NM_001037813.3 | #N/A | 45.2188269 | -1.7266334 | 0.70917848 |
| 18241 | NM_022903.3 | #N/A | 68.2202746 | -2.2454331 | 0.92232393 |
| 18242 | NM_001193427.1 | SUV39H2 | 5.61792203 | 7.45652228 | 3.06288145 |
| 18243 | NR_029378.1 | LOC100289511 | 37.7754065 | 1.68652196 | 0.69278826 |
| 18244 | NM_016131.4 | #N/A | 668.551116 | -3.7043237 | 1.52168405 |
| 18245 | NM_001316978.1 | #N/A | 8.00160066 | -4.2118547 | 1.73033371 |
| 18246 | NM_198715.2 | #N/A | 22.5153349 | -4.0481966 | 1.66316547 |
| 18247 | NM_001286645.1 | ADD1 | 16.7225998 | -5.2727152 | 2.16633199 |
| 18248 | NM_133379.4 | #N/A | 32.8257735 | 5.3738173 | 2.2080158 |
| 18249 | NM_000856.5 | #N/A | 11.9444251 | -4.7773126 | 1.96295456 |
| 18250 | NM_001347680.1 | LOC389831 | 47.1473163 | -6.7703417 | 2.78211181 |
| 18251 | NM_001330.3 | CTF1 | 16.5641469 | -3.6794641 | 1.51251107 |
| 18253 | NM_033661.4 | WDR4 | 9.662344 | -4.4842174 | 1.84336152 |
| 18252 | NM_001199144.1 | #N/A | 22.6139293 | 9.48454662 | 3.89886155 |
| 18254 | NM_001256835.1 | #N/A | 33.2147452 | -6.2662359 | 2.57600251 |
| 18255 | NM_213647.2 | #N/A | 13.9616147 | -5.0203574 | 2.06385902 |
| 18256 | NM_001318532.1 | #N/A | 10.0902497 | -4.5543335 | 1.87232281 |
| 18257 | NM_020899.3 | #N/A | 32.4203687 | -6.2311823 | 2.5617231 |
| 18258 | NM_001002249.2 | ANAPC11 | 12.0505004 | -4.8111944 | 1.97807525 |
| 18259 | NM_001321113.1 | #N/A | 11.4564622 | -4.7342848 | 1.94673735 |
| 18260 | NR_023391.1 | CYP4F30P | 18.8857989 | 3.02756887 | 1.24506209 |
| 18261 | NM_001321759.1 | #N/A | 43.0287934 | -6.637044 | 2.72948394 |
| 18263 | NM_001134373.2 | NT5C2 | 303.824794 | -2.8192857 | 1.15960275 |
| 18262 | NM_001126181.1 | #N/A | 13.0292242 | -4.9204955 | 2.02383945 |
| 18264 | NR_037648.1 | #N/A | 33.0689026 | -6.2594784 | 2.57461811 |
| 18265 | NM_017890.4 | VPS13B | 51.7352735 | -6.9021303 | 2.83910504 |
| 18266 | NM_001276.2 | #N/A | 9.86625111 | -4.5226558 | 1.86059332 |
| 18267 | NM_000106.5 | #N/A | 11.1391697 | -4.6897855 | 1.92943643 |
| 18268 | NR_024349.1 | #N/A | 217.176034 | 1.77408218 | 0.72990273 |
| 18271 | NR_131786.1 | LINC01621 | 12.4290387 | 4.17983494 | 1.71992019 |
| 18269 | NM_001145668.1 | #N/A | 35.6990968 | 4.05578355 | 1.66885472 |
| 18270 | NM_145244.3 | #N/A | 20.0995558 | -3.9114762 | 1.60948349 |
| 18272 | NM_001282714.1 | #N/A | 16.4058344 | -5.2517981 | 2.16105472 |
| 18273 | NM_001173516.1 | MAP7D3 | 5.55821239 | 7.45947738 | 3.06979798 |
| 18274 | NM_001321980.1 | GLRX3 | 8.88034473 | -4.3446238 | 1.78803018 |

| | | | | | |
|---|---|---|---|---|---|
| 18275 | NM_001172440.1 | #N/A | 20.1564131 | 5.44900071 | 2.24267931 |
| 18276 | NM_024029.4 | #N/A | 9.74942171 | -4.5020647 | 1.85323748 |
| 18277 | NR_024066.2 | CLRN1-AS1 | 20.0295354 | 1.87508747 | 0.77191584 |
| 18278 | NM_001136558.1 | GPR107 | 36.5566097 | -6.4037658 | 2.63631562 |
| 18279 | NM_001100592.2 | ATP6V0E2 | 40.3810704 | -6.5475211 | 2.69603628 |
| 18280 | NM_001257335.1 | ATP5F1A | 9.06901882 | -4.4051831 | 1.81411166 |
| 18281 | NM_001288622.1 | #N/A | 32.9444283 | -6.254454 | 2.57605416 |
| 18282 | NR_033827.1 | KDM4A-AS1 | 85.9672912 | 1.6785822 | 0.69143215 |
| 18283 | NM_001256015.1 | PSMC3IP | 7.88762763 | -4.1902046 | 1.72622091 |
| 18284 | NM_001267572.1 | TBC1D2 | 9.23438983 | -4.4105786 | 1.81706167 |
| 18285 | NM_007197.3 | FZD10 | 8.77372084 | -4.3515708 | 1.79293817 |
| 18286 | NM_030877.4 | #N/A | 182.76982 | -3.2930996 | 1.35696777 |
| 18287 | NM_004196.4 | #N/A | 8.70037659 | -4.3207846 | 1.7805324 |
| 18288 | NM_203487.2 | #N/A | 34.3308596 | -6.3133627 | 2.60174226 |
| 18289 | NM_001243144.1 | #N/A | 10.493453 | -4.5945599 | 1.89367852 |
| 18290 | NM_001301824.1 | #N/A | 8.26205863 | -4.265867 | 1.75868074 |
| 18291 | NM_152882.3 | #N/A | 38.4833024 | -6.4775986 | 2.67083624 |
| 18292 | NM_001289010.1 | MAN1C1 | 38.8969584 | 6.19593927 | 2.55485206 |
| 18293 | NM_001177693.1 | ARHGEF28 | 9.8967185 | -4.5121612 | 1.86060108 |
| 18294 | NM_001346031.1 | #N/A | 31.5076618 | -6.190195 | 2.55277153 |
| 18295 | NM_172193.2 | #N/A | 11.9313537 | -3.4784415 | 1.4344971 |
| 18296 | NM_001324212.1 | #N/A | 43.0088068 | -6.6382448 | 2.73785508 |
| 18297 | NR_028594.1 | ZNF833P | 9.20314459 | -4.4182216 | 1.82257132 |
| 18298 | NM_001135147.1 | SLC39A8 | 9.42142955 | -4.4564769 | 1.83860326 |
| 18299 | NM_001291416.1 | KDM6A | 31.5971241 | -6.1944568 | 2.55571668 |
| 18300 | NM_024702.2 | #N/A | 10.2693842 | 3.78131375 | 1.56015644 |
| 18301 | NM_001330203.1 | EIF4E2 | 44.3703041 | -6.6815661 | 2.75685116 |
| 18302 | NM_001289398.1 | #N/A | 14.176241 | -5.0346785 | 2.07786577 |
| 18303 | NR_135258.1 | LOC105370489 | 22.327677 | 9.46622 | 3.90708447 |
| 18304 | NR_024004.1 | #N/A | 63.6554247 | 3.52229532 | 1.45395507 |
| 18305 | NM_153693.4 | #N/A | 13.4392474 | -4.9535094 | 2.04482604 |
| 18306 | NR_126055.1 | THCAT155 | 21.3108788 | -3.9740542 | 1.64059978 |
| 18307 | NM_001040159.1 | #N/A | 39.8946568 | -6.5281449 | 2.69515114 |
| 18308 | NM_000064.3 | #N/A | 41.7116728 | -3.559048 | 1.46957668 |
| 18309 | NM_001321877.1 | #N/A | 34.3768887 | -6.3160723 | 2.60800042 |
| 18310 | NM_001199455.1 | BRD2 | 133.542748 | -2.3775343 | 0.98176732 |
| 18311 | NM_001271734.1 | #N/A | 11.6978544 | -4.7654159 | 1.96811937 |
| 18312 | NM_001324305.1 | RPN2 | 14.7969109 | -5.1032605 | 2.10768816 |
| 18313 | NM_001167972.1 | TRMT2B | 108.348656 | 1.53787205 | 0.63517292 |
| 18314 | NM_022648.5 | #N/A | 31.7695102 | -6.202217 | 2.56188204 |
| 18315 | NM_001282685.1 | KCTD17 | 9.70707715 | -4.4795083 | 1.85039159 |
| 18316 | NM_017583.5 | #N/A | 528.298172 | -3.5089301 | 1.44955878 |
| 18317 | NM_001142477.1 | NREP | 38.7951965 | -6.487212 | 2.6800418 |

| | | | | | |
|---|---|---|---|---|---|
| 18318 | NR_038897.1 | LINC01305 | 7.63689501 | 4.86247965 | 2.00887287 |
| 18319 | NM_014591.4 | #N/A | 8.39927887 | -4.2853434 | 1.77045093 |
| 18321 | NM_181539.4 | KRT26 | 17.4068342 | 3.45285433 | 1.42654356 |
| 18320 | NM_181333.3 | #N/A | 12.5211344 | -4.8659617 | 2.01038024 |
| 18322 | NM_001195308.1 | CGGBP1 | 12.6835883 | -4.8755574 | 2.01445469 |
| 18323 | NM_001256183.1 | #N/A | 496.750943 | -3.5866978 | 1.48199825 |
| 18324 | NR_134296.1 | BRD9 | 9.20674919 | -4.412318 | 1.82341167 |
| 18326 | NR_122070.1 | LINC02158 | 11.208167 | 2.13699737 | 0.88316556 |
| 18325 | NM_001301015.2 | #N/A | 12.4509524 | -4.8444426 | 2.00207892 |
| 18327 | NR_003142.2 | SNHG9 | 10.3350968 | -4.5768377 | 1.8920232 |
| 18328 | NM_001040424.2 | PRDM15 | 31.7452082 | -6.2008626 | 2.56345406 |
| 18329 | NM_005015.3 | #N/A | 288.000955 | -3.3218261 | 1.37328454 |
| 18330 | NR_033980.1 | LINC01185 | 8.03973879 | 4.63842162 | 1.91786014 |
| 18331 | NM_001012267.2 | #N/A | 11.0658922 | -4.6786845 | 1.93452498 |
| 18332 | NM_001322335.1 | #N/A | 70.1535084 | 6.26783689 | 2.59166662 |
| 18333 | NM_000603.4 | #N/A | 8.56049144 | -4.3090958 | 1.78188997 |
| 18334 | NR_108097.1 | LINC01108 | 8.78328425 | 5.67839532 | 2.34835373 |
| 18335 | NM_001321207.1 | #N/A | 44.1017446 | -6.6731972 | 2.7598079 |
| 18336 | NR_036505.1 | LOC100128361 | 8.32114477 | -4.271333 | 1.76649357 |
| 18338 | NR_125389.1 | PKIA-AS1 | 10.3865404 | 4.70691474 | 1.94703314 |
| 18337 | NM_020771.3 | #N/A | 34.4729879 | -6.3201093 | 2.6143274 |
| 18339 | NM_001146267.1 | GPR85 | 220.383068 | 2.13624506 | 0.88367853 |
| 18340 | NM_001258207.1 | #N/A | 23.3245781 | 6.65315043 | 2.75232341 |
| 18341 | NM_004673.3 | #N/A | 7.92954703 | -4.1988593 | 1.73700281 |
| 18342 | NM_001320568.1 | #N/A | 39.689862 | -6.5210972 | 2.69789568 |
| 18344 | NM_001320108.1 | RDH16 | 62.8206624 | 6.36973478 | 2.63572622 |
| 18343 | NM_001286680.1 | #N/A | 10.2714359 | -4.5722225 | 1.89192744 |
| 18345 | NM_001304564.1 | MIEF1 | 29.228062 | 4.71358317 | 1.95059414 |
| 18346 | NM_001033677.1 | CABP1 | 15.2665156 | -5.1428492 | 2.1284001 |
| 18347 | NR_138258.1 | LINC02210 | 9.28717866 | -4.4192602 | 1.82907051 |
| 18348 | NR_027673.1 | ADARB1 | 9.93150382 | -4.5200434 | 1.8708825 |
| 18349 | NM_152888.2 | #N/A | 8.88583934 | -4.3526891 | 1.80180903 |
| 18350 | NM_001098819.3 | PDE4C | 36.6023742 | -6.4059884 | 2.65194427 |
| 18351 | NM_001242857.2 | PRKACB | 44.2558272 | -6.6793557 | 2.76544762 |
| 18352 | NR_130897.1 | CZIB | 8.10236539 | -4.234315 | 1.75317608 |
| 18354 | NR_145451.1 | LINC02012 | 52.3424765 | 2.47943591 | 1.02662111 |
| 18353 | NM_199184.1 | #N/A | 8.81130895 | -4.3483773 | 1.80046924 |
| 18355 | NM_199349.2 | #N/A | 41.9699056 | 2.74517734 | 1.13703209 |
| 18356 | NR_120400.1 | LINC02333 | 16.8602194 | 5.83381382 | 2.41683488 |
| 18357 | NM_022659.3 | #N/A | 11.5903346 | -3.4424915 | 1.42623811 |
| 18358 | NM_001345975.1 | #N/A | 43.0296072 | -6.6361757 | 2.74996584 |
| 18359 | NM_001330317.1 | #N/A | 11.7376181 | -4.7590263 | 1.9722029 |
| 18361 | NM_001256737.1 | GDAP1L1 | 10.2378912 | -4.5509261 | 1.88605705 |

| | | | | | |
|---|---|---|---|---|---|
| 18364 | NR_109986.1 | LOC101928882 | 22.0414247 | 9.44765329 | 3.9154883 |
| 18360 | NM_001171606.1 | PREPL | 31.0183051 | -6.1674815 | 2.55606113 |
| 18362 | NM_001284268.1 | #N/A | 22.0414247 | 9.44765329 | 3.9154883 |
| 18363 | NM_173589.3 | #N/A | 9.43582108 | -4.4459039 | 1.8425283 |
| 18365 | NM_001142483.1 | NREP | 10.3008644 | -4.5716007 | 1.8946842 |
| 18366 | NM_001015072.3 | #N/A | 7.76740371 | -4.1622704 | 1.72510331 |
| 18367 | NM_001282956.1 | #N/A | 9.4045581 | -4.4537146 | 1.84605277 |
| 18368 | NM_003006.4 | SELPLG | 10.5344291 | -4.6131569 | 1.91216868 |
| 18369 | NM_001297421.1 | RAD52 | 22.5939086 | 5.74204265 | 2.38013967 |
| 18370 | NR_126431.1 | #N/A | 13.5297782 | -4.9767633 | 2.06295981 |
| 18371 | NM_198839.2 | ACACA | 177.289552 | -3.2490251 | 1.34687807 |
| 18372 | NR_103490.1 | #N/A | 9.93598246 | -4.5195469 | 1.87384328 |
| 18373 | NM_058192.2 | #N/A | 33.5376981 | -6.2800912 | 2.60395331 |
| 18374 | NM_001058.3 | #N/A | 10.7043604 | -4.635593 | 1.92248848 |
| 18375 | NM_001113491.1 | #N/A | 36.701064 | -6.4096227 | 2.65834106 |
| 18376 | NM_001198910.1 | #N/A | 10.6760298 | -4.6285214 | 1.92015819 |
| 18377 | NM_133180.2 | #N/A | 14.9732594 | -5.1123326 | 2.12110852 |
| 18378 | NM_001286769.1 | ZNF34 | 31.5503934 | -6.1917896 | 2.56940791 |
| 18379 | NM_000095.2 | #N/A | 12.0702291 | -4.8094482 | 1.9959324 |
| 18380 | NM_032153.5 | ZIC4 | 36.3744165 | -6.3971694 | 2.65508778 |
| 18381 | NM_199443.2 | #N/A | 37.5574472 | -6.4413063 | 2.67347702 |
| 18382 | NR_144327.1 | PIGG | 12.6351837 | -4.8811586 | 2.02598532 |
| 18383 | NM_004405.3 | #N/A | 8.14980607 | -4.2398582 | 1.75983671 |
| 18384 | NM_001007027.2 | ALG8 | 15.490586 | -3.8770055 | 1.60926627 |
| 18385 | NM_014289.3 | #N/A | 106.932664 | -2.7479675 | 1.1407041 |
| 18386 | NR_109927.1 | #N/A | 24.9855159 | 4.40559062 | 1.82925607 |
| 18387 | NR_135568.1 | #N/A | 68.8708856 | 6.05602092 | 2.51451795 |
| 18388 | NM_001166161.1 | PPP1R9A | 30.8253497 | -6.1585003 | 2.55739121 |
| 18389 | NM_001308177.1 | #N/A | 42.6064998 | -6.6226608 | 2.75028645 |
| 18390 | NM_032334.2 | #N/A | 101.040475 | -1.8126055 | 0.75295534 |
| 18391 | NM_201382.3 | #N/A | 30.8833351 | -6.1615878 | 2.55975019 |
| 18392 | NM_181646.3 | #N/A | 11.6265374 | 4.03614402 | 1.67681771 |
| 18393 | NM_001318766.1 | #N/A | 12.021113 | -4.7983829 | 1.99391621 |
| 18394 | NM_001171091.1 | #N/A | 30.7548241 | -6.1551891 | 2.55784791 |
| 18395 | NM_001313741.1 | ATP8A2 | 98.731069 | -5.8594499 | 2.43504013 |
| 18396 | NR_048568.1 | #N/A | 14.1905435 | -5.029824 | 2.0902513 |
| 18397 | NM_001242900.1 | #N/A | 13.8075389 | -5.0026639 | 2.07952084 |
| 18398 | NM_001307975.1 | #N/A | 39.372971 | -6.5093417 | 2.70636091 |
| 18399 | NM_001346766.1 | #N/A | 42.2141389 | -6.6094083 | 2.74810052 |
| 18400 | NM_001163436.2 | #N/A | 10.763296 | -4.6353803 | 1.92744893 |
| 18401 | NM_001258384.2 | IDH3B | 9.24510455 | -4.4049735 | 1.83179656 |
| 18402 | NR_144465.1 | LINC01149 | 19.9141802 | 4.24064929 | 1.76358397 |
| 18403 | NM_174855.3 | #N/A | 13.0019205 | -4.9168585 | 2.04482006 |

| | | | | | |
|---|---|---|---|---|---|
| 18404 | NR_131972.1 | GBAT2 | 8.32790307 | -4.268803 | 1.77537541 |
| 18407 | NR_038405.1 | FGF13-AS1 | 22.7847702 | 3.73129308 | 1.55196151 |
| 18405 | NM_001256006.1 | SYT14 | 12.6916033 | -4.8617837 | 2.02216137 |
| 18406 | NM_032780.3 | #N/A | 39.7328961 | -6.5219595 | 2.71269392 |
| 18408 | NR_026890.1 | RPSAP9 | 7.80557678 | -4.1710263 | 1.73494238 |
| 18409 | NM_006279.3 | #N/A | 44.3394846 | -6.6820793 | 2.77951599 |
| 18410 | NM_001330563.1 | #N/A | 9.40504957 | -4.4252861 | 1.84087027 |
| 18411 | NM_001034024.1 | #N/A | 41.0232907 | -6.567468 | 2.7321641 |
| 18412 | NM_001018112.1 | #N/A | 9.36916 | -4.4285907 | 1.84243695 |
| 18413 | NM_007219.4 | #N/A | 39.2224222 | -6.5031644 | 2.70566096 |
| 18414 | NM_178821.2 | #N/A | 16.1117624 | -5.2250911 | 2.17398344 |
| 18415 | NM_001317996.1 | PPIL2 | 126.221947 | -1.5533639 | 0.64635081 |
| 18417 | NR_132317.1 | LOC105373609 | 64.0550013 | 2.41593425 | 1.0054011 |
| 18416 | NM_001146171.1 | TATDN3 | 12.6137337 | -4.8675303 | 2.02565897 |
| 18418 | NM_015230.3 | #N/A | 8.48013439 | -4.2979116 | 1.78866608 |
| 18419 | NM_001012508.3 | SPRN | 107.146226 | -1.4688045 | 0.61128339 |
| 18420 | NR_145417.1 | ZNRD1ASP | 21.7551725 | 9.42885007 | 3.92407906 |
| 18421 | NM_007152.4 | #N/A | 34.3276205 | -6.313781 | 2.62783467 |
| 18422 | NM_018169.3 | #N/A | 122.176419 | -1.6892671 | 0.70307885 |
| 18423 | NM_016496.4 | #N/A | 43.0985474 | -1.9214564 | 0.79985487 |
| 18424 | NM_001142857.1 | EFCAB1 | 14.5319666 | -5.0778554 | 2.11444393 |
| 18425 | NR_026723.1 | SARNP | 9.25022961 | -4.4116025 | 1.8370167 |
| 18426 | NM_001198773.2 | #N/A | 33.5650735 | -6.2807348 | 2.61543247 |
| 18427 | NM_018003.2 | #N/A | 77.0521399 | -3.1940527 | 1.33008036 |
| 18428 | NM_001326421.1 | PISD | 11.6771979 | -4.7598856 | 1.98218779 |
| 18429 | NR_136299.1 | LOC101929583 | 20.2338072 | 2.76152502 | 1.1500244 |
| 18430 | NM_203351.1 | #N/A | 18.5079668 | -4.1385478 | 1.72362571 |
| 18431 | NR_003675.2 | GUSBP5 | 27.1360901 | 3.37118194 | 1.40417646 |
| 18432 | NM_001297436.1 | #N/A | 9.2405277 | -4.4039871 | 1.83479191 |
| 18433 | NM_001198784.1 | #N/A | 8.08376542 | -4.2249234 | 1.76043671 |
| 18435 | NR_033733.1 | CRYBB2P1 | 46.0169535 | -2.270342 | 0.94617693 |
| 18434 | NM_001017981.1 | #N/A | 127.052462 | -3.2277867 | 1.34519591 |
| 18436 | NR_136547.1 | MIR99AHG | 11.2922723 | -4.6985934 | 1.95818357 |
| 18437 | NM_001271944.1 | #N/A | 9.67150955 | -4.4726395 | 1.86407267 |
| 18438 | NM_001321982.1 | #N/A | 14.2746566 | -5.0442794 | 2.10243702 |
| 18439 | NM_005169.3 | #N/A | 12.0591457 | -4.8057399 | 2.00306812 |
| 18440 | NM_001128428.2 | FSTL5 | 40.0613126 | -6.5334557 | 2.72332857 |
| 18441 | NM_001314049.1 | SIRT1 | 15.3812485 | -5.1495662 | 2.14652461 |
| 18442 | NM_001178044.1 | SLC44A4 | 46.3822627 | 3.83972673 | 1.60062254 |
| 18443 | NR_046430.1 | #N/A | 62.2576983 | 3.4908808 | 1.45521233 |
| 18444 | NM_001287511.1 | PPCS | 8.28627808 | -4.2742649 | 1.78198518 |
| 18445 | NM_052817.2 | #N/A | 35.9516394 | -6.3786734 | 2.66002739 |
| 18446 | NM_001318891.1 | ZNF184 | 37.5828595 | -6.4413774 | 2.68629325 |

| | | | | | |
|---|---|---|---|---|---|
| 18447 | NR_046567.1 | SHANK2-AS1 | 63.2029138 | 2.89230667 | 1.20624227 |
| 18448 | NR_120595.1 | LINC01315 | 9.41620446 | -4.4418314 | 1.85252284 |
| 18449 | NM_001330999.1 | #N/A | 31.0285347 | -6.1683876 | 2.57271025 |
| 18450 | NM_021806.3 | #N/A | 39.4065848 | -6.5098018 | 2.71527809 |
| 18451 | NM_001242877.2 | #N/A | 33.1547904 | -6.2636261 | 2.61307932 |
| 18452 | NM_001010892.2 | #N/A | 36.2924468 | -6.3937661 | 2.66747966 |
| 18453 | NR_103549.1 | LUCAT1 | 74.9397983 | 3.67209568 | 1.53206091 |
| 18454 | NM_138576.3 | #N/A | 36.0868662 | -6.3852578 | 2.66423306 |
| 18455 | NR_027347.1 | ZNRF2P2 | 9.24790644 | -4.4080967 | 1.83941367 |
| 18456 | NM_001142286.1 | #N/A | 30.0537333 | -6.1222563 | 2.55476523 |
| 18457 | NM_012486.2 | PSEN2 | 43.5667887 | -6.6539851 | 2.77681327 |
| 18458 | NM_005608.2 | #N/A | 7.57884377 | -4.1285539 | 1.72328765 |
| 18459 | NR_121184.1 | GNA14-AS1 | 15.8530856 | 8.16253001 | 3.4071717 |
| 18460 | NM_024993.5 | #N/A | 38.0871638 | -6.4612212 | 2.69793539 |
| 18461 | NM_032979.4 | #N/A | 39.9960543 | -6.5311981 | 2.7272872 |
| 18462 | NM_178868.4 | #N/A | 8.54594722 | -4.3192118 | 1.80361508 |
| 18463 | NM_181515.1 | #N/A | 13.7405028 | -4.9914211 | 2.08440382 |
| 18464 | NM_001029997.3 | #N/A | 30.7997623 | -6.1572398 | 2.5717885 |
| 18465 | NM_016170.4 | #N/A | 9.01951149 | -4.384424 | 1.83134502 |
| 18466 | NM_130810.3 | #N/A | 12.9892597 | -4.9185453 | 2.05449561 |
| 18467 | NM_002652.2 | #N/A | 10.0639312 | 7.63549327 | 3.18944493 |
| 18468 | NM_001321918.1 | #N/A | 11.4250146 | -4.7107128 | 1.96813905 |
| 18469 | NM_001143992.1 | #N/A | 30.7108679 | -6.1530651 | 2.57092996 |
| 18470 | NM_001253914.1 | TBC1D1 | 21.4689202 | 9.40979713 | 3.93286321 |
| 18471 | NM_001323326.1 | #N/A | 39.6364204 | -6.5180416 | 2.72425568 |
| 18472 | NM_033172.2 | #N/A | 21.4689202 | 9.40979713 | 3.93286321 |
| 18473 | NM_001134403.1 | CYB561D1 | 36.1879703 | -6.3901156 | 2.67083411 |
| 18474 | NR_103871.1 | #N/A | 38.4213136 | -6.4742153 | 2.70596961 |
| 18475 | NM_001256668.1 | FAM193A | 38.0969363 | -6.4620246 | 2.70111951 |
| 18478 | NR_135085.1 | LINC01696 | 26.5038117 | 9.7020354 | 4.05612343 |
| 18477 | NR_121182.1 | LOC101927358 | 26.5038117 | 9.7020354 | 4.05612343 |
| 18476 | NM_001284216.1 | #N/A | 26.5038117 | 9.7020354 | 4.05612343 |
| 18479 | NM_144702.2 | #N/A | 16.8425558 | -5.28868 | 2.21117804 |
| 18480 | NR_110183.1 | LOC101928254 | 49.9344537 | 5.34664078 | 2.2354483 |
| 18482 | NR_040015.1 | LOC100631378 | 14.4162244 | 3.62228039 | 1.51459007 |
| 18481 | NM_001195752.1 | THAP3 | 38.2518386 | -6.4673644 | 2.7042087 |
| 18483 | NM_001256496.1 | MAN2C1 | 30.3239833 | -6.1352584 | 2.56569407 |
| 18484 | NM_003638.2 | #N/A | 29.8660346 | -6.1132645 | 2.55659054 |
| 18485 | NM_001322012.1 | #N/A | 31.0006481 | -6.1667665 | 2.57900108 |
| 18486 | NM_001206850.1 | #N/A | 14.3620843 | -5.053051 | 2.11349369 |
| 18487 | NR_034118.2 | NDUFA6-DT | 86.1683265 | -2.2722894 | 0.95047906 |
| 18488 | NM_178812.3 | #N/A | 583.856227 | -2.2274431 | 0.93174282 |
| 18489 | NM_001300900.1 | #N/A | 10.9106064 | -4.6672611 | 1.95238358 |

| | | | | | |
|---|---|---|---|---|---|
| 18490 | NM_000618.4 | #N/A | 36.4712448 | -6.400534 | 2.67775445 |
| 18491 | NM_001319105.1 | OPCML | 37.989123 | -6.4580931 | 2.70201276 |
| 18492 | NM_024504.3 | #N/A | 17.1088105 | 2.76500819 | 1.15697737 |
| 18493 | NM_001011703.2 | #N/A | 10.0631531 | 4.01021872 | 1.67808595 |
| 18494 | NM_001195430.1 | #N/A | 114.900159 | -3.3706792 | 1.41058793 |
| 18495 | NM_001170407.1 | CDK1 | 8.09370612 | -4.2222222 | 1.7670327 |
| 18496 | NM_001257134.1 | #N/A | 10.2605664 | -4.5709472 | 1.91310545 |
| 18497 | NM_001195101.1 | #N/A | 14.3912741 | -5.0564345 | 2.11639422 |
| 18498 | NM_139281.2 | WDR36 | 146.040636 | -2.7451781 | 1.14928086 |
| 18499 | NM_001178108.1 | ZNF185 | 30.754357 | -6.1554847 | 2.57718824 |
| 18500 | NM_001318763.1 | #N/A | 35.6172663 | -6.3649931 | 2.66521098 |
| 18501 | NM_145720.3 | #N/A | 8.62365798 | -4.3192202 | 1.8087909 |
| 18502 | NM_001750.6 | #N/A | 38.0708449 | -6.4612397 | 2.70600822 |
| 18503 | NM_001042417.1 | RUFY2 | 37.6883746 | -6.4461652 | 2.69978711 |
| 18504 | NM_181552.3 | #N/A | 127.701976 | -4.7048668 | 1.97052382 |
| 18505 | NM_001346946.1 | EXO5 | 8.4904139 | -4.2963494 | 1.79952847 |
| 18506 | NR_033202.2 | HOXB-AS3 | 12.2552696 | -4.8332559 | 2.02458177 |
| 18507 | NM_001009555.3 | SH3D19 | 29.5437718 | -6.0974695 | 2.5545035 |
| 18508 | NM_001286783.1 | #N/A | 12.086574 | -4.7952996 | 2.00907811 |
| 18509 | NM_000440.2 | #N/A | 86.7686267 | 1.9446483 | 0.81480396 |
| 18510 | NM_001329964.1 | TP63 | 13.7863523 | 3.79080029 | 1.5884275 |
| 18511 | NM_206840.2 | NVL | 38.3732548 | -6.472847 | 2.71271523 |
| 18512 | NM_001099337.1 | #N/A | 11.6483148 | -4.7637592 | 1.99654818 |
| 18513 | NM_001166359.1 | SHMT2 | 32.5239307 | -6.2352909 | 2.61395413 |
| 18514 | NM_001142555.2 | #N/A | 38.8159826 | -6.4878122 | 2.71994013 |
| 18515 | NM_001305156.1 | PPP2R3C | 30.4579972 | -6.1415567 | 2.57532733 |
| 18516 | NM_012456.2 | #N/A | 58.6349869 | -3.2985042 | 1.38314971 |
| 18517 | NM_016827.2 | OGG1 | 10.9171833 | -4.6669839 | 1.95715762 |
| 18518 | NR_125367.1 | MYHAS | 9.51516328 | 3.90325035 | 1.6373418 |
| 18519 | NM_001320134.1 | NTRK3 | 37.6376358 | -6.444175 | 2.70323564 |
| 18520 | NM_001329544.1 | #N/A | 16.5734866 | -5.2553623 | 2.20463985 |
| 18521 | NM_000697.2 | #N/A | 8.47588155 | -4.286386 | 1.7983871 |
| 18522 | NM_003790.2 | TNFRSF25 | 30.3176814 | -6.1344302 | 2.57406822 |
| 18523 | NM_001112706.2 | #N/A | 16.2537598 | -5.2393855 | 2.19865924 |
| 18524 | NM_207386.3 | #N/A | 48.5150732 | -6.8108655 | 2.85824243 |
| 18525 | NR_144512.1 | SELENOF | 9.95044079 | -4.5205658 | 1.89737008 |
| 18526 | NR_033842.1 | TMEM72-AS1 | 33.0109882 | 4.35430125 | 1.82768579 |
| 18529 | NM_153444.1 | OR5P2 | 21.1826679 | 9.39048802 | 3.94184742 |
| 18527 | NM_001326341.1 | #N/A | 21.1826679 | 9.39048802 | 3.94184742 |
| 18528 | NM_152631.2 | #N/A | 21.1826679 | 9.39048802 | 3.94184742 |
| 18530 | NM_001346875.1 | #N/A | 9.80336121 | -4.4954713 | 1.88724537 |
| 18531 | NM_001330320.1 | SNX11 | 8.19996342 | -4.2481406 | 1.78349678 |
| 18532 | NM_001191005.2 | #N/A | 113.229856 | -2.126728 | 0.89296339 |

| | | | | | |
|---|---|---|---|---|---|
| 18533 | NM_001282279.1 | #N/A | 41.2174912 | -6.5763132 | 2.76124773 |
| 18534 | NM_001018159.1 | NAE1 | 9.20460854 | -4.4242206 | 1.85780465 |
| 18535 | NM_001320970.1 | #N/A | 49.1727232 | -6.8285356 | 2.86797002 |
| 18536 | NM_001171.5 | ABCC6 | 10.0808682 | -4.5558614 | 1.91433314 |
| 18537 | NM_198189.2 | COPS8 | 11.1194246 | -4.6985657 | 1.97441391 |
| 18538 | NM_000312.3 | PROC | 13.2602875 | 3.74079138 | 1.5720667 |
| 18539 | NM_001007024.1 | #N/A | 36.7185377 | -6.4099581 | 2.69410619 |
| 18540 | NM_001286491.1 | #N/A | 14.9347585 | -5.1078952 | 2.14688496 |
| 18541 | NR_039999.1 | LINC01569 | 41.1240962 | 1.60935861 | 0.67653597 |
| 18542 | NM_001178020.2 | #N/A | 8.05829324 | -4.223306 | 1.77572379 |
| 18543 | NM_005919.3 | #N/A | 7.78072504 | -4.179013 | 1.75745728 |
| 18544 | NM_000722.3 | #N/A | 202.688803 | -1.8997838 | 0.79894933 |
| 18545 | NM_017798.3 | #N/A | 385.457712 | -2.7306941 | 1.14845597 |
| 18546 | NM_001130071.1 | EPN1 | 8.78949788 | -4.3457558 | 1.82825826 |
| 18547 | NM_145047.4 | #N/A | 29.4559719 | -6.0933423 | 2.56348475 |
| 18549 | NR_002332.2 | ST7-OT3 | 17.4313857 | 4.02172263 | 1.69208699 |
| 18548 | NM_004750.4 | #N/A | 9.34360374 | -4.4411047 | 1.86853312 |
| 18550 | NM_194328.2 | RNF38 | 41.2495171 | -6.5777172 | 2.76774265 |
| 18551 | NM_005420.2 | #N/A | 14.5383711 | -5.0766831 | 2.13672711 |
| 18552 | NM_001282908.1 | CCT8 | 10.7999424 | -4.6343661 | 1.95085921 |
| 18553 | NR_073546.1 | #N/A | 39.2777014 | -6.5059131 | 2.73882164 |
| 18554 | NM_014959.3 | #N/A | 31.8034419 | -6.2034365 | 2.61152422 |
| 18555 | NM_001085480.2 | #N/A | 5.49782374 | 4.90386126 | 2.06483922 |
| 18556 | NM_002985.2 | #N/A | 589.482528 | 2.05068032 | 0.86358941 |
| 18557 | NM_001039584.1 | PICK1 | 29.8724035 | -6.1136896 | 2.57524828 |
| 18558 | NR_110906.1 | #N/A | 293.761827 | 3.52067719 | 1.48299338 |
| 18559 | NR_024456.1 | LOC100190986 | 75.8980037 | -1.4029294 | 0.59096464 |
| 18560 | NM_001323615.1 | SMG5 | 29.5901474 | -6.0997768 | 2.5696012 |
| 18561 | NM_001159749.1 | #N/A | 42.4105214 | -1.9461327 | 0.81999234 |
| 18562 | NM_001321922.1 | #N/A | 29.172517 | -6.0790835 | 2.56145268 |
| 18563 | NM_001307931.1 | #N/A | 67.4183671 | 2.1782235 | 0.91781363 |
| 18564 | NM_014568.2 | #N/A | 162.431035 | 1.84419118 | 0.77710165 |
| 18565 | NR_039891.2 | MIR3064 | 7.76472783 | -4.1774711 | 1.76062414 |
| 18566 | NM_020663.4 | #N/A | 119.417958 | 1.40487733 | 0.59212179 |
| 18567 | NR_144328.1 | PIGG | 9.48569119 | -4.4628266 | 1.88145198 |
| 18568 | NM_001005356.2 | #N/A | 20.8964157 | 9.37091611 | 3.95103871 |
| 18569 | NM_001198863.1 | #N/A | 20.8964157 | 9.37091611 | 3.95103871 |
| 18570 | NM_001320901.1 | #N/A | 20.8964157 | 9.37091611 | 3.95103871 |
| 18571 | NM_017506.1 | #N/A | 20.8964157 | 9.37091611 | 3.95103871 |
| 18572 | NM_001242883.1 | NAXD | 37.7806412 | -6.4496734 | 2.71960861 |
| 18573 | NM_014467.2 | #N/A | 9.32877317 | -4.4454455 | 1.87449164 |
| 18574 | NM_001018137.2 | #N/A | 9.91215765 | -4.5246773 | 1.90794265 |
| 18575 | NM_001206789.1 | #N/A | 16.0727966 | -5.2111328 | 2.1975538 |

| | | | | | |
|---|---|---|---|---|---|
| 18576 | NM_014350.3 | #N/A | 11.5445968 | -4.7499461 | 2.00334232 |
| 18577 | NM_005516.5 | #N/A | 281.977975 | -2.3120929 | 0.9752857 |
| 18578 | NM_022487.3 | #N/A | 29.987362 | -6.1184702 | 2.58104492 |
| 18579 | NM_001907.2 | #N/A | 10.6976296 | -4.6240536 | 1.95072289 |
| 18580 | NM_006164.4 | #N/A | 164.231693 | -3.1367734 | 1.32389127 |
| 18581 | NM_001278387.1 | #N/A | 14.6076774 | -5.0794694 | 2.14480877 |
| 18582 | NM_001297767.1 | MRPS18C | 11.0152293 | -4.6784909 | 1.97556386 |
| 18583 | NM_001258031.1 | SERINC4 | 11.0841852 | -4.6874154 | 1.97955839 |
| 18585 | NR_135474.1 | LOC101928725 | 47.0541015 | 3.40738745 | 1.43911169 |
| 18584 | NR_126168.1 | LOC221946 | 54.1477948 | 1.50674518 | 0.63637173 |
| 18586 | NM_201553.1 | FGL1 | 15.5942372 | 7.4914664 | 3.16409368 |
| 18587 | NR_138250.1 | #N/A | 43.5820679 | 2.95895333 | 1.24989177 |
| 18588 | NR_037159.1 | ZNF582-AS1 | 8.84366815 | -4.3545905 | 1.83944845 |
| 18589 | NM_001195798.1 | #N/A | 30.9827109 | -6.1656117 | 2.60447901 |
| 18590 | NM_001306210.1 | #N/A | 40.4309692 | -6.5474103 | 2.76598743 |
| 18591 | NM_198490.2 | RAB43 | 30.5447144 | -6.1449634 | 2.59625618 |
| 18592 | NR_136661.1 | TIAL1 | 90.2491081 | -2.1344288 | 0.90182455 |
| 18593 | NM_001308240.1 | MICOS13 | 9.90035678 | -4.525638 | 1.91252249 |
| 18594 | NM_206827.1 | #N/A | 8.25471062 | -4.2590812 | 1.80002182 |
| 18595 | NR_110172.1 | #N/A | 10.3125236 | -4.5671353 | 1.93031881 |
| 18596 | NM_001683.3 | #N/A | 42.8882848 | -6.6310713 | 2.80268525 |
| 18598 | NM_001164181.1 | ENTPD1 | 378.074541 | 1.50627151 | 0.63672609 |
| 18599 | NR_004405.1 | LAMB2P1 | 8.26721328 | -4.2495918 | 1.7963619 |
| 18597 | NM_001008226.1 | #N/A | 11.210534 | -4.700841 | 1.98709759 |
| 18600 | NM_001145320.1 | ADAMTSL2 | 30.499052 | -2.3742032 | 1.00370014 |
| 18601 | NR_024276.1 | LINC00635 | 16.9562656 | 7.89677114 | 3.33837446 |
| 18602 | NM_001276289.1 | PRKAR1A | 12.5470697 | -4.8607577 | 2.0552198 |
| 18603 | NM_001139517.1 | PRKRA | 8.11605241 | -4.2317229 | 1.78935728 |
| 18604 | NR_045669.1 | PSMC3IP | 8.29754575 | -4.2692059 | 1.80526032 |
| 18605 | NR_047666.1 | EPS15L1 | 30.4520757 | -6.140665 | 2.59667069 |
| 18606 | NM_001329921.1 | CALM3 | 15.8442597 | -5.2026742 | 2.20055959 |
| 18607 | NM_003833.4 | MATN4 | 8.90541244 | -4.3537102 | 1.84157064 |
| 18608 | NM_173855.4 | #N/A | 7.72575091 | -4.1546689 | 1.75737686 |
| 18609 | NM_001088.2 | #N/A | 10.6961304 | 5.00738672 | 2.1182656 |
| 18610 | NM_001278277.1 | E2F6 | 13.1934058 | -4.922645 | 2.08266474 |
| 18611 | NM_020749.4 | MTUS1 | 9.07331607 | -4.4033031 | 1.86322857 |
| 18612 | NR_131223.1 | H19 | 30.8260914 | -6.1590511 | 2.60629853 |
| 18613 | NM_001197114.1 | IQCJ-SCHIP1 | 36.0401924 | -6.3815872 | 2.70052361 |
| 18614 | NR_136893.1 | #N/A | 11.1848861 | -4.6912425 | 1.98547403 |
| 18615 | NM_004249.3 | #N/A | 8.36113361 | -4.2887934 | 1.815241 |
| 18616 | NM_022467.3 | CHST8 | 10.8068435 | -4.6465352 | 1.96668183 |
| 18617 | NM_001347702.1 | SYNE1 | 36.0248879 | -6.3810816 | 2.70086873 |
| 18618 | NM_001081004.2 | #N/A | 12.48896 | -4.858187 | 2.05632461 |

| | | | | | |
|---|---|---|---|---|---|
| 18619 | NM_001242354.1 | ODF2 | 28.4701346 | -6.0442535 | 2.55892186 |
| 18620 | NM_002135.4 | NR4A1 | 9.13542224 | -4.4145343 | 1.86930853 |
| 18621 | NM_001267571.1 | #N/A | 14.0395292 | -5.0146336 | 2.12374133 |
| 18622 | NM_138957.3 | MAPK1 | 10.9227872 | -4.6645248 | 1.97556497 |
| 18623 | NR_110278.1 | LOC101927881 | 13.0655729 | 3.94501052 | 1.67099371 |
| 18624 | NM_181809.3 | #N/A | 93.5301107 | 1.63233339 | 0.69142484 |
| 18625 | NM_001289003.1 | PHLPP2 | 144.176326 | 1.30342995 | 0.55215021 |
| 18626 | NM_001169118.1 | #N/A | 44.6236159 | -6.691144 | 2.83502129 |
| 18627 | NM_006183.4 | #N/A | 13.4902628 | -4.9539808 | 2.09908464 |
| 18628 | NR_136658.1 | DYNC1LI2 | 32.9790446 | -6.2561028 | 2.65091741 |
| 18629 | NR_131761.1 | SNCAIP | 8.14500229 | -4.2281786 | 1.79169947 |
| 18634 | NR_144459.1 | ARSD-AS1 | 25.299093 | 9.63475442 | 4.0835958 |
| 18633 | NR_126025.1 | LOC100128317 | 25.299093 | 9.63475442 | 4.0835958 |
| 18632 | NR_109866.1 | LOC101929413 | 25.299093 | 9.63475442 | 4.0835958 |
| 18630 | NM_001164530.1 | ZNF135 | 25.299093 | 9.63475442 | 4.0835958 |
| 18631 | NM_207365.3 | #N/A | 25.299093 | 9.63475442 | 4.0835958 |
| 18635 | NR_034114.1 | #N/A | 7.30689506 | -4.0824933 | 1.73038954 |
| 18636 | NM_000820.3 | #N/A | 418.122277 | -2.7469371 | 1.16442746 |
| 18637 | NM_000337.5 | SGCD | 29.9815535 | -6.1186821 | 2.59386 |
| 18638 | NM_001031692.2 | #N/A | 29.2235301 | -6.0817043 | 2.57860061 |
| 18639 | NR_047576.2 | #N/A | 8.33769913 | -4.2715976 | 1.81112023 |
| 18640 | NR_046681.1 | PDZRN3-AS1 | 7.92481458 | -4.2068781 | 1.78398463 |
| 18641 | NM_001320709.1 | #N/A | 35.5846101 | -6.3647402 | 2.69923151 |
| 18642 | NM_001330210.1 | OFD1 | 9.31116916 | -4.4173081 | 1.87361493 |
| 18643 | NM_001200001.1 | NOTCH2 | 7.51456456 | 7.86314197 | 3.33569274 |
| 18644 | NM_001159936.1 | EBNA1BP2 | 7.48423376 | -4.1129632 | 1.74481817 |
| 18645 | NM_001284277.1 | PITPNB | 42.671961 | -6.6268005 | 2.81163825 |
| 18646 | NM_001307937.1 | #N/A | 12.6127197 | -4.8663156 | 2.06476898 |
| 18647 | NM_001318053.1 | #N/A | 195.517617 | 1.09979796 | 0.46666012 |
| 18648 | NR_046322.1 | STARD7-AS1 | 30.317991 | -3.1915563 | 1.35427828 |
| 18649 | NM_025003.3 | #N/A | 7.55075312 | -4.128075 | 1.75240052 |
| 18650 | NM_001146028.1 | #N/A | 578.543394 | -3.6401585 | 1.54531451 |
| 18651 | NM_005228.4 | #N/A | 31.9679392 | -6.2111347 | 2.63676147 |
| 18653 | NM_144497.2 | AKAP12 | 58.3231817 | -3.329561 | 1.41351429 |
| 18652 | NM_001099771.2 | POTEF | 11.6664252 | -4.7641189 | 2.02254068 |
| 18654 | NR_136593.1 | LOC100287497 | 8.06656997 | -4.2329476 | 1.79732487 |
| 18655 | NR_002181.1 | PPY2P | 5.78475503 | 7.53319277 | 3.1991615 |
| 18656 | NM_001102592.1 | #N/A | 8.34353073 | -4.2706732 | 1.81367851 |
| 18657 | NM_001320137.1 | #N/A | 11.5098436 | -4.7385748 | 2.01269278 |
| 18658 | NM_014929.3 | FASTKD2 | 616.154188 | 3.95623707 | 1.68042411 |
| 18659 | NM_024783.3 | #N/A | 15.3778432 | -5.1588427 | 2.19129179 |
| 18660 | NM_001330583.1 | LMO7 | 35.5818087 | -6.3630387 | 2.70297809 |
| 18661 | NR_038902.1 | JAKMIP2-AS1 | 8.09983212 | -4.2352334 | 1.79921504 |

| | | | | | | |
|---|---|---|---|---|---|---|
| 18662 | NM_032756.2 | | #N/A | 9.17120645 | -4.3905803 | 1.86532621 |
| 18663 | NR_103525.1 | | #N/A | 10.2932626 | -4.5716066 | 1.94298193 |
| 18664 | NR_037927.1 | SSR4 | | 9.14846779 | -4.4013574 | 1.8707951 |
| 18665 | NM_001282302.1 | | #N/A | 7.84831574 | -4.1852575 | 1.77899288 |
| 18666 | NM_032028.3 | | #N/A | 8.81944186 | 6.12080801 | 2.60174549 |
| 18667 | NM_001267595.1 | | #N/A | 37.3759896 | -6.4337811 | 2.73481342 |
| 18668 | NM_000327.3 | ROM1 | | 9.24367962 | -4.4328367 | 1.88448558 |
| 18669 | NM_001560.2 | | #N/A | 164.362668 | -3.1363593 | 1.33333561 |
| 18670 | NR_134934.1 | EEF1AKMT1 | | 8.08718661 | -4.2120198 | 1.79085178 |
| 18671 | NM_001112812.1 | | #N/A | 29.5369003 | -6.0971174 | 2.59259319 |
| 18672 | NM_001198900.1 | YY1AP1 | | 37.9043095 | -6.4555812 | 2.74535978 |
| 18673 | NM_002017.4 | | #N/A | 17.3476099 | -3.7045438 | 1.57541784 |
| 18674 | NM_001083606.1 | | #N/A | 31.1103783 | -6.1722201 | 2.62501361 |
| 18675 | NM_001877.4 | | #N/A | 10.1014121 | 5.36395554 | 2.28129122 |
| 18676 | NR_073443.1 | HDAC2 | | 40.3945439 | 2.51567812 | 1.06992756 |
| 18677 | NM_016052.3 | | #N/A | 158.094908 | -1.1641294 | 0.49514149 |
| 18678 | NM_139170.2 | | #N/A | 8.84649109 | -4.3658094 | 1.85712601 |
| 18679 | NM_001261443.1 | GNAL | | 34.3118092 | -6.3119963 | 2.68514973 |
| 18680 | NM_153368.2 | | #N/A | 14.519565 | 2.98332926 | 1.26911004 |
| 18681 | NM_014728.3 | FRMPD4 | | 16.9642223 | -3.6732621 | 1.56272669 |
| 18682 | NM_001242851.1 | RNF146 | | 20.3239111 | 9.33095513 | 3.97011282 |
| 18683 | NR_001543.3 | | #N/A | 15.4392364 | 1.77524568 | 0.75535889 |
| 18684 | NM_001243079.1 | | #N/A | 8.29470035 | -4.2601382 | 1.81294807 |
| 18685 | NM_001276418.1 | SEC16A | | 167.558858 | -2.5365653 | 1.07948411 |
| 18686 | NM_015551.1 | | #N/A | 67.0438199 | -3.0071314 | 1.27981596 |
| 18687 | NM_001199850.1 | DAP3 | | 28.4568902 | -6.0432409 | 2.57227586 |
| 18688 | NM_001330316.1 | | #N/A | 185.179144 | 3.41252158 | 1.45258892 |
| 18689 | NM_001282461.1 | | #N/A | 8.83282358 | -4.337269 | 1.84627008 |
| 18690 | NM_020128.2 | | #N/A | 16.3480533 | -3.6249841 | 1.54316786 |
| 18691 | NM_001289119.1 | ZCCHC7 | | 39.3135904 | -6.5065222 | 2.77015823 |
| 18692 | NR_038323.1 | LOC100507156 | | 47.6106737 | 1.74882979 | 0.74460024 |
| 18693 | NM_001278210.1 | NUP153 | | 33.7077347 | -6.2845982 | 2.676316 |
| 18694 | NR_026828.1 | LINC00661 | | 9.80088447 | 7.5132326 | 3.19978619 |
| 18695 | NM_203415.1 | | #N/A | 35.0362028 | -6.3415893 | 2.70084274 |
| 18696 | NM_001370.1 | | #N/A | 28.2391018 | -4.2933047 | 1.82869396 |
| 18697 | NM_003898.3 | | #N/A | 28.7261624 | -6.0572086 | 2.58001556 |
| 18698 | NM_194296.1 | SPATA24 | | 7.24495254 | -4.0682798 | 1.73297141 |
| 18699 | NM_001326411.1 | PISD | | 33.5871799 | -6.2797019 | 2.67528512 |
| 18700 | NM_001166695.2 | | #N/A | 38.6401656 | -6.4815099 | 2.76166757 |
| 18701 | NM_001308189.1 | | #N/A | 34.1944923 | -6.3085276 | 2.68802606 |
| 18702 | NM_001244883.1 | | #N/A | 7.30005155 | -4.0841904 | 1.74034274 |
| 18703 | NM_004322.3 | BAD | | 7.94536284 | -4.2011592 | 1.79027404 |
| 18704 | NM_020698.3 | | #N/A | 15.8619605 | -3.5922978 | 1.5308351 |

| | | | | | |
|---|---|---|---|---|---|
| 18705 | NM_001206480.2 | ELMO1 | 11.2221017 | -4.6808679 | 1.99478752 |
| 18706 | NM_001007259.2 | SETD4 | 10.0516147 | -4.5328916 | 1.9318367 |
| 18707 | NR_027266.1 | LINC00310 | 20.8501708 | 3.56738809 | 1.52050279 |
| 18708 | NM_001034853.1 | #N/A | 14.5149268 | -5.0759881 | 2.16363125 |
| 18709 | NM_001006630.1 | CHRM2 | 31.0975438 | -6.1709911 | 2.63076377 |
| 18711 | NM_139266.2 | STAT1 | 26.5560387 | -4.6752482 | 1.99324842 |
| 18710 | NM_001198850.1 | #N/A | 8.98197876 | -4.374406 | 1.86498793 |
| 18712 | NM_001010844.3 | #N/A | 73.5523591 | 2.10348371 | 0.89682326 |
| 18713 | NR_045492.1 | LSM1 | 10.3167883 | -4.5632282 | 1.94559213 |
| 18714 | NR_040115.1 | ANKRD62P1-PARP4P3 | 17.278343 | 6.25909885 | 2.66909392 |
| 18715 | NM_004688.2 | #N/A | 8.50201091 | -4.2992495 | 1.8335582 |
| 18716 | NM_018349.3 | MCTP2 | 7.26545437 | -4.071409 | 1.73651309 |
| 18717 | NM_001112741.1 | KCNC1 | 14.5498376 | -3.7826568 | 1.61365376 |
| 18718 | NM_001024940.2 | TRIM17 | 10.309838 | -4.5779909 | 1.95299433 |
| 18719 | NM_001308155.1 | #N/A | 45.418792 | -6.714394 | 2.86474737 |
| 18722 | NR_136577.1 | LOC105373156 | 11.9105901 | -4.7781124 | 2.03905286 |
| 18721 | NR_110698.1 | ZPLD2P | 27.9472735 | -2.7489571 | 1.17311336 |
| 18720 | NM_006398.3 | #N/A | 10.285141 | -4.5829848 | 1.95579122 |
| 18723 | NM_001194937.1 | #N/A | 32.1108066 | -6.2180037 | 2.65357015 |
| 18724 | NM_001261444.1 | #N/A | 131.992379 | 6.33911015 | 2.70534825 |
| 18725 | NM_001134396.1 | C7orf50 | 9.16702335 | -4.4157136 | 1.8847417 |
| 18726 | NM_152503.5 | #N/A | 14.4376874 | -2.9728365 | 1.26906909 |
| 18727 | NM_004437.3 | #N/A | 10.9997664 | -4.6685013 | 1.99296356 |
| 18728 | NM_003505.1 | #N/A | 241.4134 | -2.7953946 | 1.19335727 |
| 18729 | NM_001286660.1 | #N/A | 7.547429 | -4.1169136 | 1.75757737 |
| 18730 | NM_017794.4 | FOCAD | 235.920694 | -1.9003145 | 0.81131215 |
| 18731 | NM_001146285.1 | TCF7L2 | 11.6728856 | -4.7435708 | 2.02548905 |
| 18732 | NR_133933.1 | PACRGL | 7.42907735 | -4.1133041 | 1.75642343 |
| 18733 | NM_001199189.1 | CCNH | 8.70240447 | -4.3305468 | 1.84940255 |
| 18734 | NM_001324330.1 | #N/A | 46.3291659 | -6.7453053 | 2.88072779 |
| 18735 | NM_001184776.1 | #N/A | 36.3953679 | -6.3958616 | 2.73159357 |
| 18736 | NR_122113.1 | DUXAP8 | 8.16233743 | -4.2242249 | 1.8042112 |
| 18737 | NM_173588.3 | #N/A | 7.90866469 | -4.1997137 | 1.79424863 |
| 18738 | NR_122112.1 | DUXAP9 | 8.24645967 | -4.257287 | 1.81890972 |
| 18739 | NM_015283.1 | DPY19L1 | 255.90271 | -3.2783174 | 1.40089498 |
| 18740 | NM_199512.2 | CCDC80 | 32.9182299 | -6.2533895 | 2.6724084 |
| 18741 | NR_037161.1 | ZNF582-AS1 | 9.48034414 | -4.4463855 | 1.90019902 |
| 18742 | NM_001488.4 | TADA2A | 8.67708384 | -4.3136901 | 1.84369975 |
| 18743 | NM_198938.2 | PTGES2 | 12.6129449 | -4.8662007 | 2.07988037 |
| 18744 | NR_027054.1 | #N/A | 26.7293207 | 7.63141522 | 3.26183567 |
| 18745 | NM_001345917.1 | #N/A | 41.5724733 | 1.50263831 | 0.642301 |
| 18746 | NM_001003942.1 | BMF | 24.0943743 | 9.56411543 | 4.08837523 |
| 18747 | NM_001204388.1 | DLG1 | 24.0943743 | 9.56411543 | 4.08837523 |

| | | | | | |
|---|---|---|---|---|---|
| 18748 | NM_001282301.2 | NPAP1L | 21.4718558 | 4.08959483 | 1.7485589 |
| 18750 | NR_120686.1 | LINC01503 | 77.1740851 | 2.23113789 | 0.95410377 |
| 18749 | NM_001278293.2 | #N/A | 28.9363704 | -6.0673161 | 2.59456642 |
| 18751 | NM_001330143.1 | #N/A | 25.725327 | 5.87892004 | 2.5140424 |
| 18752 | NM_001271824.1 | SERPINB6 | 12.8188345 | -4.8911837 | 2.09175223 |
| 18753 | NM_001145645.2 | TNFSF13B | 61.0580274 | 3.99343474 | 1.70801176 |
| 18754 | NM_005587.3 | #N/A | 33.1870734 | -6.2642839 | 2.67932074 |
| 18755 | NM_181492.2 | TCF20 | 27.6156041 | -6.0004988 | 2.56663371 |
| 18756 | NM_001199919.1 | #N/A | 27.2753905 | -5.9823841 | 2.55893662 |
| 18757 | NM_001256010.2 | #N/A | 31.9382422 | -6.2088707 | 2.6558325 |
| 18758 | NM_001271329.1 | #N/A | 97.5352089 | 2.90308919 | 1.24177755 |
| 18759 | NM_001289098.1 | PXK | 36.9564996 | -6.4166007 | 2.74488925 |
| 18760 | NR_102401.1 | DNAH17-AS1 | 51.3620097 | 1.52306567 | 0.65165945 |
| 18761 | NM_001197248.2 | BTN3A2 | 101.884164 | 3.53054224 | 1.51087373 |
| 18762 | NM_001202470.2 | RPS10-NUDT3 | 10.6862104 | -4.6323421 | 1.98260355 |
| 18763 | NM_004977.2 | #N/A | 43.5998508 | -1.748657 | 0.74841528 |
| 18764 | NM_001329909.1 | GPI | 53.4741192 | -6.9518101 | 2.97570415 |
| 18765 | NR_038965.1 | LOC100506497 | 9.39423673 | 3.60344408 | 1.54245035 |
| 18766 | NM_021569.3 | #N/A | 35.4482321 | -6.3575956 | 2.72148249 |
| 18767 | NM_001297778.1 | NMNAT1 | 28.1391191 | -6.0268365 | 2.5799896 |
| 18768 | NM_022066.3 | #N/A | 431.273758 | -2.766064 | 1.18422307 |
| 18769 | NM_032144.2 | #N/A | 9.11206582 | -4.3829152 | 1.87643781 |
| 18770 | NR_024111.1 | SBDSP1 | 35.3175837 | -6.3531761 | 2.72023688 |
| 18771 | NM_001134776.1 | #N/A | 15.3847361 | -5.1573897 | 2.20834808 |
| 18772 | NM_001145412.3 | NFE2L2 | 27.6019813 | -5.999648 | 2.56917366 |
| 18773 | NM_001166305.1 | #N/A | 8.91977641 | -4.364066 | 1.86888168 |
| 18774 | NR_132741.1 | #N/A | 35.0920765 | -6.3430801 | 2.71656293 |
| 18775 | NM_000110.3 | #N/A | 32.4658612 | 1.59014634 | 0.68106059 |
| 18776 | NM_001256672.1 | #N/A | 34.405716 | -6.3148653 | 2.70499136 |
| 18777 | NM_001286586.1 | #N/A | 27.7956663 | -6.009832 | 2.57443196 |
| 18778 | NM_001002255.1 | #N/A | 12.2959017 | 6.3537475 | 2.72232032 |
| 18779 | NM_001083961.1 | #N/A | 28.2269053 | -6.0313758 | 2.58470461 |
| 18780 | NM_178120.4 | #N/A | 8.4670678 | -4.3046171 | 1.84481917 |
| 18781 | NM_001330209.1 | OFD1 | 35.1802172 | -6.347457 | 2.72084436 |
| 18782 | NR_109914.1 | SLC35B3 | 8.42447057 | -4.3002258 | 1.84379528 |
| 18783 | NM_015595.3 | #N/A | 32.2726867 | -6.2223285 | 2.6679906 |
| 18784 | NM_001256932.1 | #N/A | 7.68778906 | -4.1663675 | 1.78684505 |
| 18785 | NM_001136052.2 | CPT1C | 27.5325775 | -5.9956861 | 2.57149468 |
| 18787 | NR_135539.1 | NDUFA7 | 8.59800733 | -4.3067165 | 1.84709662 |
| 18786 | NM_001242450.1 | PHF21B | 29.0295122 | -6.0724909 | 2.60441468 |
| 18788 | NR_104598.1 | #N/A | 9.90713079 | -4.5211696 | 1.93930956 |
| 18789 | NM_001287336.1 | KIAA1191 | 28.1277164 | -6.0262303 | 2.58512127 |
| 18790 | NM_001329976.1 | #N/A | 28.5082783 | -6.0465029 | 2.59384709 |

| | | | | | |
|---|---|---|---|---|---|
| 18791 | NM_000741.3 | #N/A | 7.88609531 | -4.1957772 | 1.8000641 |
| 18792 | NM_001329916.1 | PNO1 | 24.9861206 | 4.61500351 | 1.98027848 |
| 18793 | NM_001330503.1 | #N/A | 14.1733405 | -5.0421114 | 2.16360581 |
| 18794 | NR_045787.1 | CERS6-AS1 | 34.4706743 | -6.3177666 | 2.71118259 |
| 18795 | NR_135524.1 | LOC105373878 | 9.02680007 | -4.3928967 | 1.88526332 |
| 18797 | NM_025262.3 | LY6G5C | 9.44163608 | -4.4602823 | 1.9144955 |
| 18796 | NM_001282280.1 | PYCR1 | 37.2073743 | -6.4276456 | 2.75896908 |
| 18798 | NR_003713.1 | LOC728613 | 58.7836273 | -1.321773 | 0.56742194 |
| 18799 | NR_110802.1 | RUNDC3A-AS1 | 203.732239 | 1.51864428 | 0.65195568 |
| 18800 | NR_135752.1 | CKMT1B | 34.2108653 | -6.3087027 | 2.70849723 |
| 18801 | NM_001242875.2 | #N/A | 27.192149 | -5.9777195 | 2.56662719 |
| 18802 | NM_001317744.1 | #N/A | 10.2856464 | -4.5779919 | 1.96570598 |
| 18803 | NM_001301302.1 | OAZ2 | 10.9043236 | -4.6566496 | 1.99956091 |
| 18806 | NR_002798.1 | NAPSB | 10.57013 | 4.34587757 | 1.86643182 |
| 18804 | NM_001290114.1 | #N/A | 35.4609846 | -6.3579634 | 2.73058136 |
| 18805 | NM_031471.5 | #N/A | 7.50559481 | -4.1097429 | 1.76502205 |
| 18807 | NM_001282806.1 | MARCHF7 | 11.7127361 | -4.7696268 | 2.04854675 |
| 18808 | NR_026885.1 | LINC00653 | 7.13154343 | -4.0447326 | 1.73724627 |
| 18809 | NR_131751.1 | PRUNE2 | 19.7514066 | 9.28984998 | 3.99004925 |
| 18810 | NM_003740.3 | #N/A | 8.04843301 | -4.2299233 | 1.8169043 |
| 18811 | NM_001014797.2 | #N/A | 37.588362 | -6.4426104 | 2.76736082 |
| 18812 | NR_108080.1 | CD81-AS1 | 13.6928017 | 3.46935434 | 1.49025848 |
| 18814 | NM_001319103.1 | OPCML | 37.1541939 | -6.4268636 | 2.76076943 |
| 18813 | NM_001194.3 | #N/A | 179.240089 | -2.8432468 | 1.22136187 |
| 18815 | NM_020741.2 | #N/A | 7.46501246 | -4.1171459 | 1.76861705 |
| 18816 | NM_015711.3 | BICRA | 138.747851 | -3.1017332 | 1.33248633 |
| 18817 | NM_001171795.1 | #N/A | 36.8364909 | -6.4139179 | 2.75542026 |
| 18819 | NR_135304.1 | BOLA2 | 8.96721348 | -4.3880556 | 1.88520964 |
| 18818 | NM_001136509.1 | #N/A | 7.60044475 | -4.1401209 | 1.77868603 |
| 18820 | NM_001145343.1 | ZNF566 | 27.9088408 | -6.0158214 | 2.58457678 |
| 18821 | NM_001320666.1 | #N/A | 31.975384 | -6.2098324 | 2.66799069 |
| 18822 | NM_198941.2 | SERINC3 | 29.2634897 | -6.0836858 | 2.61390541 |
| 18823 | NM_001277372.2 | #N/A | 12.501022 | -4.8599337 | 2.08819104 |
| 18824 | NM_001194991.1 | #N/A | 33.4613947 | -6.2738178 | 2.69597254 |
| 18825 | NM_198977.1 | #N/A | 27.8766596 | -6.0136284 | 2.58419861 |
| 18826 | NM_001270424.1 | DHRS12 | 7.51142349 | -4.1258157 | 1.77302528 |
| 18827 | NM_138481.1 | #N/A | 9.05802427 | -4.3792857 | 1.88200973 |
| 18828 | NM_001113203.2 | #N/A | 7.30811908 | -4.0801218 | 1.75374493 |
| 18829 | NM_001317900.1 | #N/A | 29.9986329 | -6.1196886 | 2.63061971 |
| 18830 | NM_001198786.1 | #N/A | 53.8219418 | -5.7078796 | 2.45398102 |
| 18831 | NM_017516.2 | #N/A | 12.3278653 | -3.5272382 | 1.51656901 |
| 18832 | NM_001012751.2 | ABI1 | 39.6806604 | -6.5192248 | 2.80331032 |
| 18833 | NM_001168466.1 | #N/A | 34.5523356 | -6.3200121 | 2.71776109 |

| | | | | | |
|---|---|---|---|---|---|
| 18834 | NM_001288759.2 | #N/A | 9.33151628 | -4.4148297 | 1.89859722 |
| 18835 | NM_000043.5 | #N/A | 34.2331387 | -6.3080158 | 2.71278649 |
| 18836 | NM_001277090.1 | #N/A | 75.5427033 | -1.9957711 | 0.8583682 |
| 18837 | NM_001172557.1 | #N/A | 9.08080669 | -4.4003709 | 1.89289335 |
| 18838 | NM_001242476.1 | ZNF345 | 7.32093389 | -4.081627 | 1.75582883 |
| 18839 | NM_001005389.1 | #N/A | 9.99175659 | -4.5316902 | 1.94954362 |
| 18840 | NR_038945.1 | HTR5A-AS1 | 26.2420085 | 1.75643231 | 0.75562885 |
| 18842 | NR_073064.1 | EXOC2 | 13.3899347 | -4.9536786 | 2.13149156 |
| 18841 | NM_001098670.1 | #N/A | 14.4313479 | -5.0564035 | 2.17567846 |
| 18843 | NM_001256435.2 | #N/A | 8.38522794 | -4.2669762 | 1.83604142 |
| 18844 | NR_026845.1 | URB1-AS1 | 8.52223607 | -4.2893075 | 1.84594862 |
| 18845 | NM_018911.2 | #N/A | 9.64329017 | -4.4678557 | 1.92296732 |
| 18846 | NM_001348122.1 | TRIM16 | 27.8271275 | -6.0113487 | 2.58736448 |
| 18847 | NM_001318747.1 | MSANTD4 | 26.9357134 | -5.9645891 | 2.56728131 |
| 18849 | NM_001256718.1 | SNAP91 | 34.9650941 | -6.3368969 | 2.72757653 |
| 18848 | NM_001242547.1 | #N/A | 33.760715 | -6.2876051 | 2.70636855 |
| 18850 | NM_001084393.1 | #N/A | 7.90397867 | -4.1989744 | 1.80738415 |
| 18851 | NM_004458.2 | #N/A | 103.880141 | -5.8976268 | 2.53858719 |
| 18852 | NM_138959.2 | #N/A | 32.2677691 | -6.222888 | 2.67920799 |
| 18853 | NM_001008388.4 | #N/A | 207.038074 | -2.5121826 | 1.08174528 |
| 18854 | NM_006335.2 | #N/A | 249.582544 | -2.6012459 | 1.1201114 |
| 18855 | NM_001318539.1 | BEND6 | 9.07168373 | -4.3976539 | 1.89436807 |
| 18856 | NM_004923.3 | TESMIN | 7.11702589 | -4.0472294 | 1.74343869 |
| 18857 | NR_024421.1 | ZNF503-AS2 | 7.96532945 | -4.1979371 | 1.80841802 |
| 18858 | NM_001320557.1 | #N/A | 13.4446139 | -4.9525985 | 2.13379427 |
| 18859 | NM_001324100.1 | KLF8 | 22.8896556 | 9.4898288 | 4.08879095 |
| 18860 | NR_030329.1 | MIR599 | 22.8896556 | 9.4898288 | 4.08879095 |
| 18861 | NM_005562.2 | #N/A | 7.57498354 | -4.1342073 | 1.78128989 |
| 18862 | NR_026938.2 | ADCY10P1 | 7.79851019 | -4.1783958 | 1.80041094 |
| 18863 | NM_001130005.1 | #N/A | 28.1585215 | -6.0286967 | 2.59798034 |
| 18864 | NM_020348.2 | #N/A | 34.2612703 | -6.3080652 | 2.71857676 |
| 18865 | NR_040054.1 | IQCH-AS1 | 7.18714985 | -4.0507341 | 1.7459607 |
| 18866 | NM_001199214.1 | #N/A | 8.91104314 | -4.3811881 | 1.88854748 |
| 18867 | NM_001278498.1 | ADORA2A | 19.0975717 | 7.33057815 | 3.15996748 |
| 18868 | NM_001286580.1 | KCTD20 | 36.9179936 | -6.4170605 | 2.7663985 |
| 18869 | NR_024412.1 | #N/A | 8.58487034 | -4.3214469 | 1.86298749 |
| 18870 | NM_172169.2 | #N/A | 37.0116242 | -6.4194375 | 2.7676 |
| 18871 | NR_135175.1 | LOC105371049 | 7.0787676 | -4.0358084 | 1.73997082 |
| 18872 | NM_001321679.1 | #N/A | 30.2946668 | -6.1328211 | 2.64461076 |
| 18873 | NR_136339.1 | LOC105371414 | 7.22906925 | -4.0604385 | 1.75107362 |
| 18874 | NM_001286409.1 | #N/A | 8.55956676 | -4.3136068 | 1.86043098 |
| 18875 | NM_001330145.1 | #N/A | 33.6078837 | -6.2803682 | 2.70867334 |
| 18876 | NM_001172219.1 | #N/A | 27.0714191 | -5.9717812 | 2.57572805 |

| | | | | | |
|---|---|---|---|---|---|
| 18877 | NM_018908.2 | #N/A | 8.74465883 | -4.3274765 | 1.86669503 |
| 18878 | NM_001001502.2 | #N/A | 15.952641 | -5.1985136 | 2.24269532 |
| 18879 | NM_002773.4 | #N/A | 9.13944703 | -4.4009066 | 1.89867823 |
| 18880 | NR_144372.1 | #N/A | 7.87744903 | -4.1859125 | 1.80603462 |
| 18881 | NM_001284243.1 | CCSER2 | 32.0419267 | -6.2136971 | 2.68096611 |
| 18882 | NM_001130859.2 | PPIP5K1 | 35.7368358 | -6.3716192 | 2.74929984 |
| 18883 | NM_001300988.1 | PTBP2 | 36.2430236 | -6.3883495 | 2.75683128 |
| 18884 | NM_145236.2 | #N/A | 13.1746786 | -3.6385225 | 1.57021729 |
| 18885 | NM_198527.3 | #N/A | 7.67559558 | -4.1519301 | 1.79189886 |
| 18886 | NM_001203246.1 | GPBP1 | 30.6950365 | -6.1516981 | 2.65505971 |
| 18889 | NR_120490.1 | LOC102724421 | 19.4651543 | 9.26885278 | 4.00038699 |
| 18887 | NM_002597.4 | #N/A | 19.4651543 | 9.26885278 | 4.00038699 |
| 18888 | NR_073580.1 | #N/A | 19.4651543 | 9.26885278 | 4.00038699 |
| 18890 | NM_001282484.1 | GOLGA8R | 102.786358 | 3.56765294 | 1.53987231 |
| 18891 | NM_004636.3 | SEMA3B | 35.7575819 | -6.3691884 | 2.7494888 |
| 18892 | NM_001282530.1 | UBAP2 | 7.9798174 | -4.2210407 | 1.8224879 |
| 18893 | NR_104487.1 | #N/A | 33.4191463 | -6.2742521 | 2.70915605 |
| 18894 | NM_005435.3 | #N/A | 12.4492601 | -4.8552008 | 2.09674548 |
| 18895 | NR_103738.1 | SOX9-AS1 | 8.0644598 | -4.2078587 | 1.81736065 |
| 18896 | NM_001001560.2 | GGA1 | 33.4226301 | -6.2729596 | 2.70957196 |
| 18897 | NM_033631.3 | #N/A | 26.850729 | -5.9600312 | 2.57442138 |
| 18898 | NR_135064.1 | LOC101928510 | 39.0701469 | 3.68477186 | 1.59165854 |
| 18899 | NM_153456.3 | HS6ST3 | 105.251574 | -1.9492396 | 0.84200089 |
| 18900 | NM_004722.3 | #N/A | 91.5243033 | -2.2291089 | 0.96298656 |
| 18901 | NM_001097599.1 | SLC35G2 | 10.3589864 | -4.5789406 | 1.9782776 |
| 18902 | NM_198437.2 | #N/A | 39.9111647 | 3.33988007 | 1.44322073 |
| 18903 | NM_024790.6 | CSPP1 | 165.07426 | -2.3046667 | 0.99594215 |
| 18904 | NR_136749.1 | SHOC2 | 8.05864504 | -4.2248903 | 1.82605903 |
| 18905 | NM_018355.3 | ZNF415 | 14.6062919 | -5.0712137 | 2.19211899 |
| 18906 | NM_001145.4 | ANG | 7.08879351 | -4.033718 | 1.74369536 |
| 18907 | NM_001306087.1 | #N/A | 10.0882555 | -4.5416675 | 1.96326142 |
| 18908 | NM_002697.3 | #N/A | 85.9896656 | -5.6540831 | 2.44430896 |
| 18909 | NR_136705.1 | KLF8 | 50.5792346 | 2.33817598 | 1.01087082 |
| 18910 | NM_001287436.2 | COBL | 11.1897981 | -4.7009099 | 2.03241855 |
| 18911 | NM_016426.6 | #N/A | 85.6330999 | -1.7035554 | 0.73663987 |
| 18913 | NM_001320445.1 | MORC3 | 39.3633156 | -6.5070404 | 2.81379684 |
| 18912 | NM_001278694.1 | #N/A | 9.30295609 | -4.4332726 | 1.9170447 |
| 18914 | NM_153610.4 | #N/A | 34.8741384 | 2.75592785 | 1.19174635 |
| 18915 | NR_037869.1 | LOC653160 | 7.56364839 | -4.1355681 | 1.78861099 |
| 18916 | NM_178234.2 | TUSC3 | 238.446792 | -2.7965155 | 1.20965611 |
| 18917 | NM_033497.2 | HK1 | 28.1026281 | -6.025359 | 2.60660211 |
| 18918 | NM_001322165.1 | #N/A | 12.4997967 | -4.8473517 | 2.09704524 |
| 18920 | NM_001317188.1 | BRINP3 | 9.63830683 | -4.4678102 | 1.93304242 |

| | | | | | |
|---|---|---|---|---|---|
| 18919 | NM_000530.7 | #N/A | 9.43709396 | -4.4588399 | 1.92915328 |
| 18921 | NM_001135644.1 | DCTN4 | 8.74349516 | -4.3447039 | 1.87986569 |
| 18922 | NM_001242314.1 | #N/A | 37.0149889 | -6.4186103 | 2.77752028 |
| 18923 | NM_001321240.1 | SLC25A39 | 30.4274885 | -6.140247 | 2.65727728 |
| 18924 | NR_027285.1 | FLVCR1-DT | 8.55266674 | -4.3086122 | 1.86492475 |
| 18925 | NM_001282729.1 | SUPT7L | 30.1588568 | -6.1264559 | 2.65178029 |
| 18926 | NM_001320924.2 | NAA38 | 11.6785222 | -4.7610161 | 2.06087294 |
| 18927 | NR_131238.1 | RHOXF1-AS1 | 34.6504488 | 3.38649154 | 1.4658813 |
| 18929 | NR_073431.1 | TMEM9B-AS1 | 7.99883755 | -4.2168027 | 1.82540669 |
| 18928 | NM_001002020.2 | #N/A | 7.89792216 | -4.194885 | 1.81590526 |
| 18930 | NM_001199893.1 | #N/A | 9.55859511 | -4.4747549 | 1.93724001 |
| 18931 | NR_122104.1 | LINC02072 | 7.42649854 | -4.1013855 | 1.77586419 |
| 18932 | NM_001099407.1 | DZANK1 | 116.913185 | 0.98530387 | 0.426668 |
| 18933 | NM_001321726.1 | LOC101928120 | 7.48976803 | -4.1119949 | 1.78067571 |
| 18934 | NM_001318765.1 | #N/A | 29.6737749 | -6.1030161 | 2.64290515 |
| 18935 | NM_001323067.1 | PFKP | 44.6625956 | -6.6925082 | 2.89824597 |
| 18936 | NM_032279.3 | #N/A | 69.8624478 | 1.83728128 | 0.79567416 |
| 18937 | NM_003583.3 | #N/A | 61.2246119 | -3.9650801 | 1.71720007 |
| 18938 | NR_136605.1 | #N/A | 32.9255343 | -6.2509212 | 2.70760388 |
| 18939 | NM_001321975.1 | #N/A | 13.0706326 | 5.64652206 | 2.44593448 |
| 18940 | NR_046477.1 | #N/A | 15.864458 | -3.585638 | 1.55345298 |
| 18941 | NM_025149.5 | ACSF2 | 27.1290179 | -5.9750434 | 2.5888459 |
| 18942 | NR_144524.1 | #N/A | 8.57905377 | -4.3261962 | 1.87468069 |
| 18943 | NM_021173.4 | #N/A | 35.2526616 | -6.3504773 | 2.75250049 |
| 18944 | NM_001166208.1 | #N/A | 25.9116813 | -5.9084967 | 2.56110914 |
| 18945 | NM_001286537.1 | #N/A | 36.6484783 | -2.6572622 | 1.15185736 |
| 18946 | NM_181702.2 | #N/A | 30.441801 | -6.1402682 | 2.66173759 |
| 18947 | NR_028093.1 | LPAL2 | 29.6104145 | 1.97569369 | 0.85650198 |
| 18948 | NM_023932.3 | #N/A | 17.1903183 | -5.3097716 | 2.30200096 |
| 18949 | NM_198700.2 | CELF1 | 15.9167681 | -5.1992245 | 2.25435393 |
| 18950 | NM_001270960.1 | #N/A | 201.443894 | -3.2390329 | 1.40455472 |
| 18951 | NM_001307977.1 | WDR74 | 33.2523681 | -6.2650507 | 2.71695392 |
| 18952 | NM_001128208.1 | #N/A | 7.38263851 | -4.087213 | 1.7725358 |
| 18953 | NR_047558.1 | ZNF32-AS2 | 7.32168123 | -4.0876543 | 1.77291848 |
| 18954 | NM_015531.5 | #N/A | 19.1789021 | 9.24754364 | 4.01098332 |
| 18955 | NM_152599.3 | #N/A | 19.1789021 | 9.24754364 | 4.01098332 |
| 18956 | NM_001321524.1 | ZNF845 | 27.6402192 | -6.0010648 | 2.60363037 |
| 18957 | NM_005402.3 | #N/A | 288.112013 | -2.7174227 | 1.17905985 |
| 18958 | NM_001300950.1 | YEATS4 | 8.00348953 | -4.2048758 | 1.82449483 |
| 18959 | NM_024820.2 | #N/A | 9.26768582 | -4.4258948 | 1.92047313 |
| 18960 | NM_016239.3 | #N/A | 27.8693798 | -3.0802708 | 1.33686632 |
| 18961 | NR_003505.3 | PPP4R1L | 13.7970254 | -3.4108632 | 1.48047864 |
| 18962 | NM_005824.2 | #N/A | 13.4224571 | -4.9424244 | 2.1453815 |

| | | | | | |
|---|---|---|---|---|---|
| 18963 | NM_001161584.1 | TEX10 | 9.02269993 | 5.60880584 | 2.43503959 |
| 18964 | NM_001267619.1 | ARPP21 | 36.7052469 | -6.4063834 | 2.78156696 |
| 18965 | NM_001202559.1 | CHURC1-FNTB | 33.2461789 | -6.2659719 | 2.720795 |
| 18966 | NM_001012752.2 | ABI1 | 29.3251525 | -6.0872912 | 2.64329008 |
| 18967 | NM_001270409.1 | LIN9 | 8.83732618 | -4.3363516 | 1.88303687 |
| 18968 | NM_001195610.1 | DCDC2 | 30.0222309 | -3.6285053 | 1.57571562 |
| 18969 | NM_001135024.1 | PDHX | 35.1149475 | -6.3445027 | 2.75536501 |
| 18970 | NM_024894.3 | #N/A | 35.4393993 | -6.3569637 | 2.76081769 |
| 18971 | NR_027423.1 | #N/A | 7.99118214 | -4.1964162 | 1.82257043 |
| 18978 | NR_132311.1 | ATXN2 | 21.6849369 | 9.41150441 | 4.08925292 |
| 18972 | NM_001025596.2 | CELF1 | 21.6849369 | 9.41150441 | 4.08925292 |
| 18974 | NM_030932.3 | DIAPH3 | 21.6849369 | 9.41150441 | 4.08925292 |
| 18975 | NM_152453.3 | TMCO5A | 21.6849369 | 9.41150441 | 4.08925292 |
| 18973 | NM_001318394.1 | #N/A | 21.6849369 | 9.41150441 | 4.08925292 |
| 18976 | NR_026787.1 | #N/A | 21.6849369 | 9.41150441 | 4.08925292 |
| 18977 | NR_104199.1 | #N/A | 21.6849369 | 9.41150441 | 4.08925292 |
| 18979 | NR_109821.1 | NR2F1-AS1 | 8.8282003 | -4.3432793 | 1.88723913 |
| 18980 | NM_001145157.1 | #N/A | 9.1800601 | -4.3997289 | 1.91186892 |
| 18981 | NM_001308258.1 | #N/A | 10.6859996 | -4.6358248 | 2.01466016 |
| 18982 | NM_001267050.1 | #N/A | 27.5454946 | -5.9965382 | 2.60606578 |
| 18983 | NM_021088.3 | #N/A | 27.4870147 | -5.9939725 | 2.60505332 |
| 18984 | NM_001278221.2 | #N/A | 15.3844307 | -3.5353142 | 1.5369822 |
| 18985 | NM_001282797.1 | ZNF7 | 32.9857725 | -6.2540714 | 2.71933121 |
| 18986 | NM_080631.3 | VPS41 | 4.98570786 | 7.29838859 | 3.1735259 |
| 18987 | NM_033544.2 | RCCD1 | 11.5029265 | -4.7330889 | 2.05813104 |
| 18988 | NM_001734.4 | #N/A | 32.8644708 | -6.2488935 | 2.71738486 |
| 18989 | NM_014659.5 | PPIP5K1 | 31.3089192 | -6.1780464 | 2.6867082 |
| 18990 | NM_178517.4 | PIGW | 9.83819514 | -4.4956111 | 1.95520029 |
| 18991 | NM_001322219.1 | #N/A | 8.09212939 | -4.2393558 | 1.84402562 |
| 18992 | NR_033489.1 | #N/A | 29.7662048 | -6.1078356 | 2.65677744 |
| 18993 | NM_001077445.2 | JADE3 | 32.9803629 | -6.2531496 | 2.72007698 |
| 18994 | NM_001085372.2 | #N/A | 87.4106782 | -1.546406 | 0.67274297 |
| 18995 | NM_198428.2 | #N/A | 36.0412577 | -6.3818953 | 2.77640676 |
| 18996 | NR_073537.1 | CEP76 | 9.55668915 | -4.4473496 | 1.93494161 |
| 18997 | NR_026815.1 | TMEM191A | 9.18248622 | -4.4023821 | 1.91565639 |
| 18998 | NM_001272011.1 | #N/A | 25.9473975 | -5.9104081 | 2.57194546 |
| 18999 | NM_001301063.1 | #N/A | 79.2671605 | -5.5471424 | 2.41392519 |
| 19000 | NR_027354.2 | LSP1P5 | 31.2708567 | 3.67860458 | 1.60083837 |
| 19001 | NM_005155.6 | #N/A | 26.6316244 | -5.9482763 | 2.58859014 |
| 19002 | NM_002076.3 | #N/A | 718.248659 | -2.3412133 | 1.01891357 |
| 19003 | NM_001242834.1 | NDRG4 | 26.3550607 | -5.9328347 | 2.58216487 |
| 19004 | NM_001308257.1 | ABHD3 | 7.14942679 | -4.0485393 | 1.76241182 |
| 19005 | NM_001346304.1 | #N/A | 20.9722819 | -3.1667561 | 1.37861517 |

| | | | | | |
|---|---|---|---|---|---|
| 19006 | NM_013956.4 | #N/A | 27.3039059 | -5.9836485 | 2.60494678 |
| 19007 | NM_020340.4 | #N/A | 290.7549 | -1.2395517 | 0.53965284 |
| 19009 | NM_014885.4 | ANAPC10 | 9.2670692 | -4.4024392 | 1.91698629 |
| 19008 | NM_001301020.1 | OAZ1 | 9.80699248 | -4.4877904 | 1.95415504 |
| 19010 | NM_001291921.1 | #N/A | 9.45286003 | -4.4578477 | 1.94115771 |
| 19011 | NM_001256760.2 | HTR2C | 47.8679661 | -6.7924113 | 2.95807128 |
| 19012 | NR_024607.1 | MIR503HG | 9.41723411 | -4.4415094 | 1.93465922 |
| 19013 | NR_135685.1 | LINC02259 | 6.16980856 | 5.95195913 | 2.59264659 |
| 19014 | NM_017673.6 | #N/A | 26.9876294 | -5.9668496 | 2.59918487 |
| 19015 | NM_031889.2 | #N/A | 18.2073954 | 7.69064522 | 3.35005664 |
| 19016 | NM_001329456.1 | ZNF304 | 12.0825265 | -4.7926417 | 2.08788508 |
| 19017 | NM_001037339.2 | PDE4B | 13.3135295 | -4.9364039 | 2.15054506 |
| 19018 | NM_001144770.1 | DST | 41.9141897 | -6.5976153 | 2.87458269 |
| 19019 | NM_175922.3 | #N/A | 11.65744 | -4.7613771 | 2.07459245 |
| 19020 | NM_001290.4 | #N/A | 28.3070039 | -6.0356958 | 2.62988407 |
| 19021 | NR_103839.1 | FRY-AS1 | 40.125532 | 1.55020559 | 0.67560453 |
| 19022 | NM_001193487.2 | GTF2A1L | 18.8926498 | 9.22591372 | 4.02184799 |
| 19023 | NM_020710.2 | #N/A | 396.325243 | -2.745177 | 1.1967071 |
| 19024 | NM_022573.2 | #N/A | 18.8926498 | 9.22591372 | 4.02184799 |
| 19025 | NM_001320453.1 | C1orf61 | 16.2755043 | -5.2253207 | 2.27790882 |
| 19026 | NM_020354.3 | #N/A | 130.069819 | -2.2196666 | 0.96776561 |
| 19027 | NM_001281458.1 | SLC35C2 | 27.0456006 | -5.9696449 | 2.60306926 |
| 19028 | NM_001009186.1 | #N/A | 26.7777527 | -5.9563099 | 2.59744338 |
| 19030 | NR_002971.1 | SNORA38 | 10.7830763 | 7.63045967 | 3.32764099 |
| 19029 | NM_001316903.1 | #N/A | 10.7830763 | 7.63045967 | 3.32764099 |
| 19031 | NM_145254.2 | #N/A | 25.4000123 | -5.879751 | 2.5642187 |
| 19032 | NM_001321788.1 | #N/A | 32.9782091 | -6.2537789 | 2.72761087 |
| 19034 | NR_027253.1 | BASP1-AS1 | 60.0134484 | 1.363896 | 0.594943 |
| 19033 | NM_001307955.1 | DYNAP | 11.0151199 | 3.01920341 | 1.31701189 |
| 19035 | NM_017785.4 | #N/A | 32.4435634 | -6.229259 | 2.71748202 |
| 19036 | NM_032037.3 | #N/A | 7.74618526 | -4.1548281 | 1.81261821 |
| 19037 | NM_001178110.1 | ZNF185 | 25.3448014 | -5.876778 | 2.56449523 |
| 19038 | NM_003867.3 | #N/A | 13.0416402 | -4.9056385 | 2.14077622 |
| 19039 | NM_018690.3 | #N/A | 14.2335445 | 3.13616405 | 1.36866375 |
| 19040 | NM_001199729.2 | #N/A | 19.9890171 | -5.5261095 | 2.41180854 |
| 19041 | NM_001318918.1 | GLIS2 | 26.8455883 | -5.9588489 | 2.60079012 |
| 19042 | NM_001031726.3 | C19orf12 | 33.519101 | -6.2781393 | 2.74017482 |
| 19043 | NM_001042517.1 | #N/A | 32.2270743 | -6.2205487 | 2.7150673 |
| 19044 | NR_136566.1 | PHTF1 | 28.5381617 | -6.0470244 | 2.63986598 |
| 19045 | NM_003323.2 | #N/A | 11.8846099 | 4.42891036 | 1.93367956 |
| 19046 | NM_007250.5 | KLF8 | 33.049304 | -6.2555362 | 2.7312786 |
| 19047 | NM_153638.3 | PANK2 | 38.6105178 | -6.4804837 | 2.83026795 |
| 19048 | NM_001284368.1 | NDUFB8 | 13.0398807 | 3.38687831 | 1.47921324 |

| | | | | | |
|---|---|---|---|---|---|
| 19049 | NM_001288760.2 | #N/A | 8.70100888 | -4.3128099 | 1.88393644 |
| 19050 | NM_001040161.2 | #N/A | 10.1093044 | -4.537549 | 1.98219 |
| 19051 | NM_197961.3 | #N/A | 32.847621 | -6.2467674 | 2.72900394 |
| 19052 | NM_001042351.2 | #N/A | 30.4391394 | -6.1404265 | 2.68267264 |
| 19053 | NM_001039670.2 | #N/A | 32.0275728 | -6.2116134 | 2.71389613 |
| 19054 | NM_003298.4 | #N/A | 34.0094634 | -6.2992006 | 2.75234667 |
| 19055 | NM_001039569.1 | #N/A | 32.6029193 | -6.2356042 | 2.72470173 |
| 19056 | NR_046099.1 | MIR646HG | 32.7071946 | 2.88024026 | 1.25873998 |
| 19057 | NM_006237.3 | #N/A | 18.6031405 | -5.4320455 | 2.37407935 |
| 19058 | NM_022059.3 | #N/A | 34.2901963 | -1.8148162 | 0.79318597 |
| 19059 | NR_135490.1 | #N/A | 8.52161859 | -4.309836 | 1.88372466 |
| 19060 | NM_001252607.1 | THBS3 | 40.7329245 | -6.5581262 | 2.86651315 |
| 19061 | NM_001271560.1 | LOC389895 | 9.02637071 | -4.3797484 | 1.91444462 |
| 19062 | NM_007139.3 | #N/A | 15.2788009 | -5.1331655 | 2.24383076 |
| 19063 | NM_001282880.1 | ZNF502 | 11.473207 | -3.4399063 | 1.50369771 |
| 19064 | NM_198855.3 | #N/A | 35.4337875 | -6.3572586 | 2.77944418 |
| 19065 | NM_001321391.1 | #N/A | 10.8298508 | -4.6440664 | 2.03050816 |
| 19066 | NM_001321026.1 | #N/A | 10.9181118 | -4.6533549 | 2.03480676 |
| 19067 | NM_016623.4 | #N/A | 28.5156879 | -6.0461047 | 2.64386269 |
| 19068 | NM_001321993.1 | #N/A | 30.4188638 | -6.1369625 | 2.68374455 |
| 19069 | NM_001310319.1 | #N/A | 31.8609916 | -6.2042541 | 2.71327254 |
| 19070 | NM_001305655.1 | #N/A | 26.0427139 | -5.9157135 | 2.58718322 |
| 19071 | NR_027355.2 | LSP1P5 | 9.91560754 | -4.5162738 | 1.97522956 |
| 19072 | NM_181465.2 | MRPL55 | 7.80562232 | -4.1826369 | 1.82943986 |
| 19073 | NM_207371.3 | #N/A | 61.5737466 | -1.9897342 | 0.87033263 |
| 19074 | NM_001304533.1 | #N/A | 8.73223041 | -4.3377437 | 1.8974513 |
| 19075 | NM_177969.3 | PPM1B | 11.5079555 | 4.92149243 | 2.1529414 |
| 19076 | NM_001270629.1 | MED4 | 14.5141215 | -5.074619 | 2.22004584 |
| 19077 | NM_015602.3 | TOR1AIP1 | 174.546373 | -0.9371769 | 0.40999752 |
| 19078 | NM_015912.3 | #N/A | 8.73859521 | -4.3381225 | 1.89784523 |
| 19079 | NM_198585.2 | #N/A | 14.8579519 | 4.6755849 | 2.04567748 |
| 19082 | NR_121608.1 | LOC100507291 | 7.07549999 | -4.03382 | 1.76524053 |
| 19081 | NM_199189.2 | MATR3 | 33.7708199 | -6.2886759 | 2.75200035 |
| 19080 | NM_020367.5 | #N/A | 31.699633 | -6.1960457 | 2.71144023 |
| 19083 | NM_001301224.1 | #N/A | 11.249028 | -4.6965828 | 2.05542661 |
| 19085 | NR_038386.1 | C2orf92 | 7.35500883 | -4.08782 | 1.78908354 |
| 19084 | NM_001042496.1 | #N/A | 34.7598332 | -6.3295991 | 2.77020278 |
| 19086 | NM_016476.11 | ANAPC11 | 11.7999699 | -4.765512 | 2.08574894 |
| 19087 | NM_001171166.1 | #N/A | 30.8742611 | -6.1607937 | 2.69650781 |
| 19088 | NM_001287335.1 | KIAA1191 | 31.7787403 | -6.1995135 | 2.71364528 |
| 19089 | NR_109995.1 | LOC100506476 | 7.41024956 | -4.1127177 | 1.80026463 |
| 19090 | NM_020725.1 | ATXN7L1 | 26.404305 | -5.9352202 | 2.5981093 |
| 19091 | NM_001128620.1 | PAK1 | 10.0319165 | -4.5384584 | 1.98679837 |

| | | | | | |
|---|---|---|---|---|---|
| 19092 | NR_072989.1 | #N/A | 8.22827424 | -4.256932 | 1.86365628 |
| 19093 | NM_001080392.1 | #N/A | 197.611954 | -2.0929522 | 0.91631948 |
| 19094 | NM_133370.3 | #N/A | 86.4328468 | -2.4643832 | 1.07895357 |
| 19095 | NM_000080.3 | #N/A | 6.91528634 | -4.0039803 | 1.75344003 |
| 19096 | NR_136541.1 | MIR99AHG | 7.78823068 | -4.1795929 | 1.83042167 |
| 19097 | NM_000222.2 | KIT | 37.7149018 | -6.4453409 | 2.82287169 |
| 19098 | NM_001330643.1 | CCDC149 | 33.4741539 | -6.2763241 | 2.74915308 |
| 19099 | NR_002605.2 | DLEU1 | 35.7024233 | 2.72376844 | 1.19316056 |
| 19100 | NM_181674.2 | PPP2R2B | 9.82787483 | -4.4892478 | 1.96693156 |
| 19101 | NM_139166.4 | #N/A | 7.98455458 | 4.9983994 | 2.19006117 |
| 19106 | NR_134275.1 | LINC02217 | 18.6063975 | 9.20395356 | 4.03299127 |
| 19103 | NM_001204141.1 | MBD1 | 18.6063975 | 9.20395356 | 4.03299127 |
| 19105 | NR_077228.1 | MTHFD2P1 | 52.7801724 | 1.87660773 | 0.82226766 |
| 19104 | NM_001271805.1 | PFKFB1 | 18.9025983 | 4.00205651 | 1.75357806 |
| 19102 | NM_001190801.1 | PLK4 | 18.6063975 | 9.20395356 | 4.03299127 |
| 19107 | NM_001099791.2 | #N/A | 17.5909943 | -2.7257841 | 1.19446068 |
| 19108 | NM_001330727.1 | #N/A | 10.3439422 | -4.5807715 | 2.00770126 |
| 19109 | NM_001242830.1 | ELOVL5 | 10.9109454 | -4.6642009 | 2.04428866 |
| 19110 | NM_001139.2 | #N/A | 8.07828854 | -4.2153322 | 1.84757911 |
| 19111 | NM_178493.5 | #N/A | 7.80441016 | -4.1616588 | 1.82409471 |
| 19112 | NM_001242764.1 | #N/A | 28.1837122 | -6.0289637 | 2.64258332 |
| 19113 | NR_038862.1 | FLJ42102 | 27.753662 | 1.92569273 | 0.84411293 |
| 19115 | NM_001322329.1 | C2orf76 | 20.4802182 | 9.32868469 | 4.08976925 |
| 19116 | NR_036054.1 | MIR1972-1 | 20.4802182 | 9.32868469 | 4.08976925 |
| 19114 | NM_001005280.1 | OR10A7 | 20.4802182 | 9.32868469 | 4.08976925 |
| 19117 | NR_120615.1 | PLCE1-AS2 | 20.4802182 | 9.32868469 | 4.08976925 |
| 19118 | NR_135779.1 | #N/A | 20.4802182 | 9.32868469 | 4.08976925 |
| 19119 | NM_022783.3 | #N/A | 8.97311616 | -4.3725733 | 1.91741925 |
| 19120 | NM_001304941.1 | #N/A | 35.1653872 | -6.3466353 | 2.78313221 |
| 19121 | NR_038272.1 | TTN-AS1 | 7.62249461 | -4.1290929 | 1.81072589 |
| 19122 | NR_045562.1 | #N/A | 26.3610014 | -5.9328239 | 2.60189682 |
| 19123 | NM_001278206.1 | #N/A | 19.1181588 | 7.15693793 | 3.13880322 |
| 19124 | NM_000307.4 | #N/A | 8.59576936 | -4.304862 | 1.88799645 |
| 19125 | NM_001347953.1 | RPH3A | 31.6789512 | -6.1961428 | 2.71766809 |
| 19127 | NR_034171.1 | EIF3J-DT | 10.1182312 | -4.5352357 | 1.98931325 |
| 19126 | NM_001320849.1 | FAHD2B | 7.40950514 | -4.0909044 | 1.79440358 |
| 19128 | NM_000865.2 | #N/A | 7.58664382 | -4.1463997 | 1.81900702 |
| 19129 | NM_199340.3 | #N/A | 75.6752101 | 2.14672781 | 0.94188271 |
| 19130 | NM_001042702.3 | #N/A | 7.77022041 | -4.1791358 | 1.83394993 |
| 19131 | NM_001317061.1 | #N/A | 16.3572019 | -3.6079526 | 1.58332803 |
| 19132 | NM_001205019.1 | GK | 26.5203346 | -5.9416183 | 2.60753029 |
| 19133 | NM_005681.3 | #N/A | 9.21890005 | -4.4066224 | 1.93421542 |
| 19134 | NM_001195000.1 | MR1 | 25.5972248 | -5.8909839 | 2.58636564 |

| | | | | | |
|---|---|---|---|---|---|
| 19135 | NM_153373.3 | #N/A | 25.2508475 | -5.8711354 | 2.57767511 |
| 19136 | NM_001308069.1 | XXYLT1 | 29.7937741 | -6.108103 | 2.68174099 |
| 19137 | NM_001017388.2 | TLR10 | 31.7964378 | 2.97326358 | 1.30553545 |
| 19138 | NM_004067.3 | CHN2 | 25.744406 | -5.8991166 | 2.59055885 |
| 19139 | NM_175077.2 | #N/A | 29.0887459 | 2.50584893 | 1.10050355 |
| 19140 | NM_032885.5 | #N/A | 25.6427821 | -5.8934622 | 2.58829507 |
| 19141 | NM_001301035.1 | ERF | 16.5363617 | -3.6264732 | 1.59274849 |
| 19142 | NM_001282980.1 | #N/A | 31.7288149 | -6.1996674 | 2.7230415 |
| 19143 | NR_027467.2 | #N/A | 40.4895592 | 1.29722337 | 0.56983102 |
| 19144 | NM_005314.2 | #N/A | 12.4930148 | 3.5138946 | 1.54378525 |
| 19145 | NR_026759.1 | FAM95B1 | 8.25590684 | -4.2467443 | 1.86580886 |
| 19146 | NM_001142546.1 | LUZP1 | 171.527877 | -1.5101182 | 0.66353083 |
| 19147 | NM_001014990.2 | MRPL22 | 12.1309751 | -4.8095787 | 2.11357511 |
| 19148 | NR_103508.1 | PHYKPL | 8.26468002 | -4.2523335 | 1.8687174 |
| 19149 | NM_001204137.1 | #N/A | 26.844157 | -5.9592112 | 2.61893223 |
| 19150 | NM_006974.2 | #N/A | 35.3591616 | -6.3545912 | 2.7927052 |
| 19151 | NM_033124.4 | #N/A | 26.6061018 | -5.946517 | 2.61337872 |
| 19152 | NR_022011.1 | PWARSN | 74.9939595 | 1.39330401 | 0.61234611 |
| 19153 | NM_001172745.1 | #N/A | 7.67031694 | -4.1568642 | 1.82706144 |
| 19154 | NM_152559.2 | #N/A | 7.98767282 | -4.2194008 | 1.85458504 |
| 19155 | NM_001329454.1 | #N/A | 7.24195153 | -4.0671691 | 1.78776059 |
| 19156 | NR_120591.1 | #N/A | 9.48490912 | -4.4597018 | 1.96029022 |
| 19157 | NM_001329556.1 | #N/A | 13.3122403 | -4.9444406 | 2.17344491 |
| 19158 | NR_030684.1 | #N/A | 10.5185866 | -4.6135599 | 2.02830173 |
| 19159 | NM_016179.2 | #N/A | 8.88168765 | -4.3700013 | 1.92146767 |
| 19160 | NM_001252156.1 | GEMIN5 | 39.5368159 | -6.513588 | 2.86402719 |
| 19161 | NR_134475.1 | #N/A | 8.0135394 | -4.2026246 | 1.84791502 |
| 19162 | NM_001330402.1 | #N/A | 31.0393275 | -6.1658433 | 2.71140805 |
| 19163 | NM_001256675.1 | #N/A | 9.16587243 | -4.4051752 | 1.93719802 |
| 19165 | NM_181093.3 | SCYL3 | 8.62153709 | -4.3328041 | 1.90548903 |
| 19164 | NM_001301188.1 | TRIB3 | 31.0015074 | -6.1644792 | 2.71104621 |
| 19166 | NM_001308046.1 | TNK2 | 24.9750393 | -5.8554804 | 2.57546515 |
| 19167 | NM_001135676.1 | #N/A | 8.90781406 | -4.3589101 | 1.91723486 |
| 19168 | NM_001145354.1 | #N/A | 31.9719074 | -6.2072825 | 2.73045992 |
| 19169 | NM_014979.3 | #N/A | 309.811561 | -2.1403329 | 0.94152342 |
| 19170 | NM_001288723.1 | #N/A | 26.4775281 | -5.9396964 | 2.6129158 |
| 19171 | NM_001304563.1 | #N/A | 12.1488535 | -4.8026757 | 2.11291118 |
| 19172 | NM_001318792.1 | TMEM63B | 29.1990949 | -6.0789053 | 2.67494907 |
| 19173 | NM_001288757.1 | BABAM1 | 12.3382462 | -4.8356893 | 2.12792224 |
| 19174 | NR_136695.1 | #N/A | 7.76781292 | -4.1539775 | 1.82810306 |
| 19175 | NM_173653.3 | #N/A | 7.58274481 | -4.1406362 | 1.8223921 |
| 19176 | NM_001257972.1 | #N/A | 7.93093051 | -4.2029405 | 1.84986997 |
| 19177 | NM_001242786.1 | BRF1 | 10.4356465 | -4.5809681 | 2.01637956 |

| | | | | | |
|---|---|---|---|---|---|
| 19178 | NR_003003.2 | SCARNA17 | 7.14811755 | -4.0487616 | 1.78233386 |
| 19179 | NM_001005169.1 | OR52I1 | 8.96720634 | 7.38075651 | 3.2493574 |
| 19180 | NM_001042430.2 | #N/A | 7.92930913 | -4.1926138 | 1.84606601 |
| 19181 | NR_038391.1 | #N/A | 8.67917296 | -4.325189 | 1.90459486 |
| 19182 | NM_053042.2 | #N/A | 155.620863 | -1.8807523 | 0.82828964 |
| 19184 | NM_001256702.1 | USP3 | 18.3201452 | 9.18165313 | 4.04442397 |
| 19183 | NM_001012415.2 | #N/A | 18.3201452 | 9.18165313 | 4.04442397 |
| 19185 | NM_001105198.1 | TMEM177 | 9.76184005 | -4.4877071 | 1.97686215 |
| 19186 | NM_001319847.1 | #N/A | 8.54569519 | -4.2941496 | 1.89162085 |
| 19187 | NM_014405.3 | #N/A | 691.84951 | -3.4872214 | 1.5361483 |
| 19188 | NR_026800.2 | FAM30A | 8.63266988 | 7.4156308 | 3.26702972 |
| 19189 | NR_033879.1 | SPATA3-AS1 | 8.30019187 | 6.26070066 | 2.75820805 |
| 19190 | NM_001032392.2 | PLGLB1 | 31.5580892 | 4.73381125 | 2.08575461 |
| 19191 | NM_001102416.2 | #N/A | 7.93740914 | 2.47617769 | 1.09129665 |
| 19192 | NR_125344.1 | IGFL4 | 42.3836045 | 2.27479701 | 1.00265691 |
| 19193 | NM_001278412.1 | AMN1 | 9.33755732 | -4.4450947 | 1.9593152 |
| 19194 | NM_003167.3 | #N/A | 14.561739 | -3.4609028 | 1.52560856 |
| 19195 | NM_012248.3 | #N/A | 240.143841 | -3.1814273 | 1.40242313 |
| 19196 | NM_001080779.1 | #N/A | 7.54724558 | -4.1136872 | 1.81363628 |
| 19197 | NM_001258461.1 | FASTK | 27.5349736 | -5.9964029 | 2.64384015 |
| 19198 | NR_037852.1 | #N/A | 33.8719943 | -6.2939801 | 2.77530574 |
| 19199 | NM_001321626.1 | #N/A | 36.5920764 | -6.4024412 | 2.82326082 |
| 19200 | NR_126032.1 | #N/A | 32.2018579 | -6.2179037 | 2.7421994 |
| 19201 | NR_125766.1 | BBOX1-AS1 | 8.22851595 | -4.2353663 | 1.86789937 |
| 19202 | NM_001130112.3 | RFC5 | 9.77658068 | -4.4858175 | 1.97841386 |
| 19203 | NM_001197051.1 | #N/A | 32.2152104 | -6.2181375 | 2.74252426 |
| 19204 | NM_001321082.1 | #N/A | 30.7315462 | -6.1504748 | 2.7127444 |
| 19205 | NM_001024935.1 | WASF1 | 24.9764747 | -5.8555628 | 2.58293952 |
| 19206 | NM_001079866.1 | #N/A | 30.6850078 | -6.1497248 | 2.71287861 |
| 19207 | NM_012144.3 | #N/A | 35.295479 | -6.3534243 | 2.80292646 |
| 19209 | NR_134854.1 | SLC39A13 | 7.01995097 | -4.0105931 | 1.76938236 |
| 19208 | NR_104185.1 | #N/A | 41.7850044 | -1.7270049 | 0.76191933 |
| 19210 | NM_001286707.1 | RMDN1 | 59.4279212 | 4.70727403 | 2.0777794 |
| 19211 | NM_203330.2 | CD59 | 24.305023 | -5.816362 | 2.5677091 |
| 19213 | NR_046598.1 | FGF12-AS2 | 8.59472431 | 6.45027045 | 2.84768795 |
| 19212 | NM_001097594.2 | #N/A | 8.59472431 | 6.45027045 | 2.84768795 |
| 19214 | NM_007031.1 | #N/A | 29.8326448 | 3.36249245 | 1.48456522 |
| 19215 | NM_001324214.1 | #N/A | 12.88909 | -4.8855896 | 2.15705567 |
| 19216 | NM_018254.3 | #N/A | 24.295677 | -5.8159044 | 2.56797251 |
| 19217 | NM_001346873.1 | RUBCN | 24.2672482 | -5.813919 | 2.56715166 |
| 19218 | NM_139278.2 | #N/A | 8.62790278 | -4.3206815 | 1.9078095 |
| 19219 | NM_001316893.1 | #N/A | 8.69668767 | -4.3155532 | 1.90565686 |
| 19220 | NM_153221.2 | CILP2 | 37.8054598 | -1.9086605 | 0.84284467 |

| | | | | | |
|---|---|---|---|---|---|
| 19221 | NM_001039350.2 | #N/A | 11.2882475 | -4.7062658 | 2.07840791 |
| 19222 | NM_001278425.1 | #N/A | 10.2431631 | -4.5712009 | 2.01889544 |
| 19223 | NM_015394.4 | #N/A | 90.7987681 | -2.1624499 | 0.95508095 |
| 19224 | NM_001310127.1 | ZNF888 | 13.6615673 | -3.3826335 | 1.49413543 |
| 19225 | NM_001300734.1 | DDB2 | 12.5718136 | -4.8651207 | 2.14905144 |
| 19226 | NM_001323614.1 | SMG5 | 28.0251876 | -6.0210327 | 2.66064082 |
| 19227 | NM_031205.3 | #N/A | 10.6001574 | -4.6031758 | 2.03423384 |
| 19228 | NM_032489.2 | #N/A | 7.05713792 | -4.0314262 | 1.78170727 |
| 19229 | NM_001164749.1 | #N/A | 36.7546161 | -6.4082413 | 2.83289123 |
| 19230 | NM_001006641.3 | SLC25A25 | 35.7418946 | -6.3693464 | 2.81604873 |
| 19231 | NM_001282342.1 | #N/A | 25.1120521 | -5.8631533 | 2.59222997 |
| 19232 | NM_005373.2 | #N/A | 97.9710525 | 1.09660506 | 0.48487481 |
| 19233 | NM_001346354.1 | ATG13 | 33.8233296 | -6.291949 | 2.78210967 |
| 19235 | NM_001282063.1 | DOC2A | 12.5499841 | -4.86577 | 2.15158772 |
| 19236 | NM_018443.3 | ZNF302 | 34.7601275 | -6.3280466 | 2.79820755 |
| 19234 | NM_001258454.1 | #N/A | 27.6169189 | -5.999692 | 2.65303488 |
| 19240 | NR_036476.1 | TMEM198B | 24.9589616 | -5.8544155 | 2.58890998 |
| 19237 | NM_001079675.2 | #N/A | 26.2444869 | -5.9265393 | 2.62083835 |
| 19238 | NM_001167941.1 | #N/A | 64.0416432 | 6.00609435 | 2.6560157 |
| 19239 | NM_001271762.1 | #N/A | 10.4091626 | -4.5939298 | 2.03149589 |
| 19241 | NM_181814.1 | ABHD12B | 9.73973776 | 4.33559232 | 1.91735855 |
| 19242 | NM_022477.3 | #N/A | 31.6743381 | -6.1957619 | 2.74124087 |
| 19243 | NM_001042402.1 | #N/A | 9.85935619 | -4.5068468 | 1.9941242 |
| 19244 | NM_001321065.1 | #N/A | 11.1645161 | -4.6856158 | 2.07333869 |
| 19245 | NR_131903.1 | LINC00566 | 21.6243352 | 5.11469988 | 2.26341577 |
| 19252 | NR_024447.1 | LOC100128288 | 42.5310066 | -1.9252876 | 0.85218572 |
| 19254 | NR_133639.1 | MIR194-2HG | 19.2754994 | 9.24081972 | 4.09035017 |
| 19246 | NM_001009611.4 | PRAMEF4 | 19.2754994 | 9.24081972 | 4.09035017 |
| 19253 | NR_122122.1 | TSL | 19.2754994 | 9.24081972 | 4.09035017 |
| 19250 | NM_001320964.1 | TSPY1 | 19.2754994 | 9.24081972 | 4.09035017 |
| 19247 | NM_001220479.2 | #N/A | 19.2754994 | 9.24081972 | 4.09035017 |
| 19248 | NM_001278607.1 | #N/A | 19.2754994 | 9.24081972 | 4.09035017 |
| 19249 | NM_001301029.1 | #N/A | 33.2443447 | -6.2646207 | 2.77296031 |
| 19251 | NM_021574.2 | #N/A | 19.2754994 | 9.24081972 | 4.09035017 |
| 19255 | NM_018914.2 | #N/A | 30.3924431 | -6.1353521 | 2.71584434 |
| 19256 | NM_001242778.1 | #N/A | 32.4065854 | -6.2275768 | 2.75672038 |
| 19257 | NM_001139443.1 | #N/A | 8.6418851 | -4.3166274 | 1.91084445 |
| 19258 | NM_001330324.1 | #N/A | 27.9188121 | -6.0163116 | 2.66348239 |
| 19259 | NR_003291.1 | ANKRD18DP | 7.44402874 | 4.57524085 | 2.02556776 |
| 19260 | NM_173508.3 | #N/A | 9.51932106 | -4.4597886 | 1.97482248 |
| 19261 | NM_001320664.1 | SHROOM2 | 25.8639159 | -5.9063483 | 2.61549969 |
| 19265 | NR_027921.3 | CCL15-CCL14 | 18.033893 | 9.15900178 | 4.05615751 |
| 19262 | NM_001278392.1 | CEACAM18 | 18.033893 | 9.15900178 | 4.05615751 |

| | | | | | |
|---|---|---|---|---|---|
| 19268 | NR_126377.1 | LINC00437 | 18.033893 | 9.15900178 | 4.05615751 |
| 19266 | NR_031579.1 | MIR2113 | 18.033893 | 9.15900178 | 4.05615751 |
| 19263 | NM_001330062.1 | PIP4K2A | 18.033893 | 9.15900178 | 4.05615751 |
| 19264 | NM_014489.3 | #N/A | 18.033893 | 9.15900178 | 4.05615751 |
| 19267 | NR_033557.1 | #N/A | 18.033893 | 9.15900178 | 4.05615751 |
| 19269 | NR_104269.1 | RALGAPA1P1 | 34.1680261 | 3.47334043 | 1.53825267 |
| 19270 | NM_005391.4 | PDK3 | 25.9422279 | 3.65394099 | 1.61827753 |
| 19271 | NM_000369.2 | #N/A | 7.83041756 | -4.1705276 | 1.84716648 |
| 19272 | NM_153809.2 | TAF1L | 22.8307778 | 3.4232807 | 1.51631586 |
| 19273 | NM_015352.1 | #N/A | 308.082947 | -2.5750539 | 1.14061958 |
| 19274 | NM_001319302.1 | RILPL1 | 8.08762476 | -4.2372321 | 1.87707466 |
| 19275 | NM_001130726.3 | CCDC149 | 32.7067718 | -6.2412723 | 2.76536817 |
| 19276 | NM_001321985.1 | #N/A | 25.7554712 | -5.8990228 | 2.61383373 |
| 19277 | NR_135073.1 | DEPDC4 | 45.9146385 | 2.80505454 | 1.24307515 |
| 19278 | NR_047655.1 | #N/A | 31.7397968 | -6.1995281 | 2.74742067 |
| 19279 | NM_052988.4 | #N/A | 29.3299375 | -6.086782 | 2.69761687 |
| 19280 | NM_001171133.2 | #N/A | 6.86694564 | -3.9832025 | 1.7654888 |
| 19281 | NM_001320365.1 | #N/A | 26.1438074 | -5.9209418 | 2.6243795 |
| 19282 | NM_138815.3 | #N/A | 5.88355629 | 5.8839489 | 2.60793893 |
| 19283 | NM_001282401.1 | #N/A | 8.73368023 | -4.328012 | 1.91884916 |
| 19284 | NM_017929.5 | #N/A | 28.3666178 | -6.0364785 | 2.67668053 |
| 19285 | NM_001037171.1 | #N/A | 9.42441486 | -4.4576761 | 1.97669474 |
| 19286 | NR_134583.1 | SETD6 | 24.4950266 | -5.8270379 | 2.58433197 |
| 19287 | NR_027392.2 | INTS4P2 | 15.8234346 | 2.49849471 | 1.10819245 |
| 19288 | NM_001291501.1 | #N/A | 19.0100045 | 5.30788097 | 2.35459677 |
| 19289 | NM_001048212.3 | CEMP1 | 49.7275188 | -2.1891294 | 0.97126824 |
| 19290 | NM_001347872.1 | #N/A | 8.37467599 | -4.2889115 | 1.90295717 |
| 19291 | NM_001330694.1 | #N/A | 30.1732989 | -6.1248618 | 2.71820065 |
| 19292 | NM_144669.2 | #N/A | 7.83358697 | -4.1694323 | 1.85039793 |
| 19293 | NR_034179.1 | LOC653712 | 6.89838911 | -3.9880838 | 1.76993367 |
| 19294 | NR_110085.1 | LOC101927446 | 11.2724132 | 2.59551365 | 1.15194564 |
| 19295 | NM_001288777.1 | #N/A | 26.6024052 | -5.9469197 | 2.6394457 |
| 19296 | NM_001303024.1 | ZNRD2 | 7.94646199 | -4.2066865 | 1.86713728 |
| 19297 | NR_040001.2 | LINC01116 | 8.61477968 | -4.321449 | 1.91814326 |
| 19298 | NM_001017395.3 | #N/A | 132.631078 | -2.4741837 | 1.09835269 |
| 19299 | NM_025217.3 | #N/A | 8.27166525 | -4.2587187 | 1.8907976 |
| 19300 | NM_001256964.1 | #N/A | 9.20686731 | -4.4076047 | 1.95721786 |
| 19302 | NM_015429.3 | ABI3BP | 7.92688888 | -4.2067318 | 1.86816366 |
| 19301 | NM_003318.4 | #N/A | 29.0914247 | -6.072154 | 2.69658226 |
| 19303 | NM_001323031.2 | SV2B | 33.5991654 | -6.2797159 | 2.78889824 |
| 19304 | NM_001330166.1 | #N/A | 8.52206362 | -4.2854736 | 1.90324473 |
| 19305 | NR_125822.1 | LOC101929709 | 7.18640454 | -4.0521506 | 1.79964219 |
| 19306 | NM_001145115.2 | #N/A | 8.49742107 | -4.2971001 | 1.90864505 |

| | | | | | |
|---|---|---|---|---|---|
| 19307 | NM_001167869.1 | #N/A | 31.6143271 | -6.192958 | 2.75079937 |
| 19308 | NM_020429.2 | #N/A | 32.9620381 | -6.2543677 | 2.77823586 |
| 19309 | NM_144584.2 | HENMT1 | 7.61227432 | -4.1301003 | 1.83505819 |
| 19310 | NM_001015056.1 | RTKN | 23.9350263 | -5.7943638 | 2.5745991 |
| 19311 | NM_001286807.1 | #N/A | 9.30770919 | -4.4208888 | 1.96434614 |
| 19312 | NR_038951.1 | INKA2-AS1 | 8.89247276 | -4.3591573 | 1.93696969 |
| 19313 | NM_015433.2 | #N/A | 23.369814 | -1.9171427 | 0.85192479 |
| 19314 | NM_006458.3 | TRIM3 | 8.83443039 | -4.3657203 | 1.94008653 |
| 19315 | NM_001282208.1 | SIGMAR1 | 13.6527775 | -4.9757079 | 2.21119263 |
| 19316 | NM_001311311.1 | VKORC1 | 7.32894403 | -4.0922572 | 1.8186638 |
| 19317 | NM_172341.3 | #N/A | 144.715316 | -2.2208743 | 0.98722311 |
| 19318 | NM_198504.2 | #N/A | 7.06067415 | -4.0403949 | 1.7963557 |
| 19319 | NM_001037582.2 | #N/A | 646.140645 | -1.3521721 | 0.6012347 |
| 19320 | NM_001199184.1 | #N/A | 25.9298409 | -5.9088328 | 2.62754593 |
| 19321 | NM_001291822.1 | #N/A | 13.304219 | 7.26159991 | 3.22920803 |
| 19322 | NM_031901.5 | MRPS21 | 97.817255 | -2.2419189 | 0.99713755 |
| 19323 | NR_144447.1 | ZNF8-ERVK3-1 | 9.613047 | -4.4849595 | 1.99476514 |
| 19324 | NM_001160223.1 | RNF170 | 9.14863529 | -4.4108036 | 1.96197622 |
| 19325 | NR_134473.1 | #N/A | 8.50213159 | -4.2970618 | 1.91168866 |
| 19326 | NM_000845.2 | #N/A | 28.9564093 | -6.0682042 | 2.69979848 |
| 19327 | NM_001347712.1 | FRA10AC1 | 7.54354007 | -4.1291046 | 1.83738945 |
| 19328 | NR_120459.1 | CNOT2-DT | 6.79355407 | -3.9772713 | 1.76987454 |
| 19329 | NM_030587.2 | B4GALT2 | 29.8973108 | -6.1125666 | 2.72010866 |
| 19330 | NM_001318249.1 | QPRT | 12.1944331 | 4.76838591 | 2.12216038 |
| 19331 | NR_073084.1 | CRIP2 | 8.41969934 | -4.2964504 | 1.91225134 |
| 19332 | NR_110228.1 | LINC01250 | 14.8206214 | 6.18518039 | 2.75311536 |
| 19333 | NM_001007074.1 | RPL32 | 7.15817127 | -4.063333 | 1.80873085 |
| 19334 | NM_001389.3 | #N/A | 28.9910706 | -6.0695999 | 2.70176668 |
| 19335 | NM_182965.2 | #N/A | 7.87618331 | -4.1915301 | 1.86585724 |
| 19336 | NM_001143676.1 | SGK1 | 11.4731593 | -4.7301388 | 2.10569459 |
| 19337 | NM_001282490.1 | GOLGA8H | 8.19724674 | -4.2457263 | 1.89042509 |
| 19338 | NM_001319107.1 | IQCB1 | 41.7447937 | -6.5924932 | 2.93542644 |
| 19339 | NR_028388.2 | FUS | 42.3929755 | 2.24245836 | 0.99850873 |
| 19340 | NM_001177716.1 | KLF11 | 17.7476407 | 9.13598314 | 4.06820397 |
| 19343 | NR_107013.1 | MIR7973-2 | 17.7476407 | 9.13598314 | 4.06820397 |
| 19341 | NM_001318840.1 | #N/A | 17.7476407 | 9.13598314 | 4.06820397 |
| 19342 | NM_020432.4 | #N/A | 17.7476407 | 9.13598314 | 4.06820397 |
| 19344 | NM_001139499.1 | #N/A | 7.21652104 | -4.0600196 | 1.80823108 |
| 19345 | NM_018143.2 | #N/A | 133.703801 | -1.7388369 | 0.77448271 |
| 19346 | NM_138350.3 | THAP3 | 7.8435001 | -4.1943433 | 1.86834141 |
| 19347 | NR_137287.1 | #N/A | 30.9721251 | -6.1654191 | 2.74643435 |
| 19348 | NM_001303268.1 | POLR2B | 30.2233328 | -6.1284891 | 2.73021764 |
| 19349 | NM_003919.2 | #N/A | 26.4768795 | -5.9397702 | 2.64662798 |

| | | | | | |
|---|---|---|---|---|---|
| 19350 | NR_073132.1 | #N/A | 8.69080875 | -4.3338419 | 1.9312173 |
| 19351 | NM_001098632.2 | AKT1S1 | 24.3131171 | -5.8167701 | 2.59228146 |
| 19353 | NM_207171.2 | POGZ | 226.251467 | -2.6863583 | 1.19721837 |
| 19352 | NM_001146683.1 | TMEM253 | 16.4091943 | 3.57818361 | 1.59467328 |
| 19354 | NR_047678.1 | TARS1 | 29.5161758 | -6.09327 | 2.71559973 |
| 19356 | NM_182910.2 | SYNE2 | 24.1340308 | -5.8056277 | 2.58767094 |
| 19355 | NM_032438.3 | #N/A | 24.507703 | -5.8278837 | 2.59758626 |
| 19357 | NR_024496.1 | PTGER4P2-CDK2AP2P2 | 60.5463252 | 3.28374272 | 1.46411098 |
| 19358 | NM_001257135.1 | #N/A | 25.6244821 | -5.8919926 | 2.62709104 |
| 19359 | NM_001270436.1 | #N/A | 28.2387132 | -6.0310001 | 2.68904932 |
| 19360 | NR_138133.1 | #N/A | 10.443945 | -4.6049257 | 2.05371197 |
| 19361 | NM_015172.3 | #N/A | 968.785531 | -2.9576781 | 1.31918641 |
| 19362 | NM_173054.2 | RELN | 35.8116977 | -6.3716031 | 2.84219494 |
| 19363 | NM_000397.3 | #N/A | 8.54475661 | 3.81240869 | 1.70074306 |
| 19364 | NM_145172.4 | #N/A | 29.7016063 | -6.1047803 | 2.72377766 |
| 19365 | NM_003610.3 | #N/A | 183.889213 | -3.1028201 | 1.38445334 |
| 19366 | NM_001195125.1 | #N/A | 8.01524688 | -4.1980236 | 1.87322009 |
| 19367 | NM_001256394.1 | NOL8 | 13.1242415 | -4.9157636 | 2.19388594 |
| 19368 | NM_018309.4 | #N/A | 27.6424874 | -5.9986914 | 2.67733564 |
| 19369 | NM_021958.3 | #N/A | 8.89368493 | -4.371287 | 1.95102211 |
| 19370 | NM_001324062.1 | MIA3 | 24.3744465 | -5.8208635 | 2.59805224 |
| 19371 | NM_001243423.1 | #N/A | 7.47146247 | -4.1176583 | 1.83793007 |
| 19372 | NM_001345998.1 | #N/A | 10.8207682 | -4.6354418 | 2.06944973 |
| 19373 | NM_004626.2 | #N/A | 29.520973 | -3.1107993 | 1.38878577 |
| 19374 | NM_000163.4 | #N/A | 25.3049128 | -5.8739995 | 2.62247523 |
| 19375 | NM_000629.2 | #N/A | 325.300091 | -1.013037 | 0.45232859 |
| 19376 | NM_001258275.2 | #N/A | 8.66849994 | -4.3273181 | 1.93217016 |
| 19377 | NM_177542.3 | SNRPD2 | 8.57903782 | -4.3014322 | 1.92071572 |
| 19378 | NM_001320868.1 | #N/A | 7.20975177 | -4.0671489 | 1.81625119 |
| 19379 | NR_036487.1 | FGF14-AS2 | 6.98501681 | -4.0117384 | 1.79168582 |
| 19380 | NR_073030.1 | TIMM23 | 25.7595904 | -5.8997512 | 2.63553017 |
| 19381 | NM_001244871.1 | DAB2 | 30.6823373 | -6.14956 | 2.74765985 |
| 19382 | NM_024312.4 | #N/A | 405.672114 | -1.783368 | 0.79692253 |
| 19383 | NM_001161577.1 | #N/A | 7.45443562 | -4.1179468 | 1.84034265 |
| 19384 | NM_030567.4 | #N/A | 10.6421501 | -4.6177996 | 2.06394116 |
| 19385 | NM_001320298.1 | CBS | 28.1132541 | -6.0237729 | 2.69250475 |
| 19386 | NM_001242546.1 | ANAPC16 | 378.840822 | 0.90934718 | 0.40666238 |
| 19389 | NM_033397.3 | ITPRIP | 18.0707807 | 9.1472549 | 4.09100858 |
| 19388 | NM_005549.2 | KCNA10 | 18.0707807 | 9.1472549 | 4.09100858 |
| 19391 | NR_031663.1 | MIR1261 | 18.0707807 | 9.1472549 | 4.09100858 |
| 19387 | NM_001146017.1 | TJAP1 | 18.0707807 | 9.1472549 | 4.09100858 |
| 19390 | NM_201569.2 | #N/A | 18.0707807 | 9.1472549 | 4.09100858 |
| 19392 | NR_073124.1 | #N/A | 18.0707807 | 9.1472549 | 4.09100858 |

| | | | | | |
|---|---|---|---|---|---|
| 19393 | NR_111984.1 | #N/A | 18.0707807 | 9.1472549 | 4.09100858 |
| 19394 | NM_001322311.1 | PHACTR1 | 20.5116735 | -3.5403901 | 1.58343321 |
| 19395 | NM_001261415.2 | SOX5 | 34.731102 | -6.3262683 | 2.83006691 |
| 19396 | NR_135282.1 | CARNMT1 | 23.0637062 | -5.7408165 | 2.56850944 |
| 19399 | NM_182944.2 | NIN | 6.89210953 | -3.9906104 | 1.78554306 |
| 19397 | NM_001301135.1 | #N/A | 7.922184 | -4.1876211 | 1.87366619 |
| 19398 | NM_003356.3 | #N/A | 7.92559151 | -4.1874526 | 1.87362016 |
| 19400 | NM_181354.4 | OXR1 | 10.5163636 | -4.6108837 | 2.06313213 |
| 19401 | NM_017541.3 | #N/A | 7.70908264 | -4.1593067 | 1.86155298 |
| 19402 | NM_001207008.1 | AFDN | 196.954732 | -3.0386494 | 1.36002581 |
| 19403 | NM_001276501.1 | #N/A | 7.15822867 | -4.0491263 | 1.81246476 |
| 19404 | NR_036515.1 | #N/A | 117.855176 | 1.5334316 | 0.68639916 |
| 19405 | NM_014258.3 | #N/A | 28.3710768 | 3.29847027 | 1.47664116 |
| 19406 | NM_053044.4 | #N/A | 10.3326492 | -4.5882917 | 2.05418023 |
| 19407 | NM_015687.4 | #N/A | 8.82146892 | -4.3639361 | 1.95380161 |
| 19408 | NM_015354.2 | #N/A | 294.121574 | -2.5606858 | 1.14648261 |
| 19409 | NM_001004459.1 | OR1S2 | 17.4613884 | 9.11259888 | 4.08057596 |
| 19410 | NM_001283057.1 | #N/A | 17.4613884 | 9.11259888 | 4.08057596 |
| 19411 | NM_001308134.1 | FANCM | 32.5279482 | 5.99900178 | 2.68654798 |
| 19412 | NR_015375.2 | VLDLR-AS1 | 6.80422911 | -3.9778414 | 1.78140002 |
| 19413 | NR_037593.1 | CAHM | 7.82615286 | -4.1799225 | 1.87218214 |
| 19414 | NM_001243737.1 | TRAPPC13 | 28.9552275 | -6.0658289 | 2.71694236 |
| 19415 | NM_021979.3 | HSPA2 | 9.57894554 | -4.4782509 | 2.00592017 |
| 19416 | NM_001329641.1 | #N/A | 23.0098203 | -5.7373976 | 2.57009637 |
| 19417 | NM_001177637.2 | DAG1 | 27.393475 | -5.9869102 | 2.68215438 |
| 19419 | NM_052869.1 | TTYH2 | 9.42250691 | -4.4531188 | 1.99519027 |
| 19418 | NM_001207073.1 | #N/A | 7.62024479 | -4.1486099 | 1.85874687 |
| 19420 | NM_182911.3 | #N/A | 10.1556235 | -4.5621744 | 2.04403784 |
| 19421 | NM_198066.3 | #N/A | 111.390517 | -2.5239774 | 1.13086779 |
| 19422 | NM_198489.2 | #N/A | 9.50745069 | -4.456681 | 1.9971144 |
| 19423 | NM_001271162.1 | #N/A | 10.9677898 | 7.29327154 | 3.26870468 |
| 19424 | NM_030984.3 | #N/A | 10.9677898 | 7.29327154 | 3.26870468 |
| 19425 | NM_001316345.1 | #N/A | 9.1812492 | -4.4033258 | 1.97380579 |
| 19426 | NM_020150.4 | #N/A | 353.92686 | -2.6700317 | 1.19685296 |
| 19427 | NM_001143788.1 | PPHLN1 | 29.2316205 | -6.0790586 | 2.72506458 |
| 19428 | NM_001204836.1 | #N/A | 23.2313975 | -5.7509258 | 2.57801752 |
| 19429 | NM_006456.2 | #N/A | 13.8008868 | -5.0020641 | 2.24291479 |
| 19431 | NM_133374.2 | ZNF618 | 59.4136786 | -5.1618838 | 2.31465572 |
| 19430 | NM_002239.3 | #N/A | 29.0963495 | -6.0727227 | 2.72309506 |
| 19432 | NR_144455.1 | BARX1-DT | 10.9107758 | 7.62646837 | 3.42028838 |
| 19433 | NM_002737.2 | #N/A | 238.469725 | -2.5320834 | 1.13564153 |
| 19434 | NM_005293.2 | #N/A | 9.08311708 | -4.3983241 | 1.97287076 |
| 19435 | NM_024076.2 | #N/A | 19.9038984 | -2.4965592 | 1.11986059 |

| | | | | | |
|---|---|---|---|---|---|
| 19436 | NR_040081.1 | LOC100506804 | 18.988999 | 3.97840883 | 1.78464725 |
| 19437 | NM_001145314.1 | CEP44 | 7.37124685 | -4.0814834 | 1.83093446 |
| 19438 | NM_001278451.1 | SP140 | 23.2158941 | 7.04606356 | 3.16116628 |
| 19439 | NM_198427.1 | #N/A | 9.63148817 | -4.4596552 | 2.0010214 |
| 19440 | NM_175571.3 | #N/A | 7.85685034 | 4.48320132 | 2.01172008 |
| 19441 | NM_024744.15 | #N/A | 7.27038505 | -4.0806873 | 1.83125035 |
| 19442 | NM_001303131.1 | GAS2L3 | 10.5483423 | -4.6166931 | 2.07189869 |
| 19444 | NR_024394.1 | LINC00689 | 21.6429259 | 4.00162738 | 1.79599197 |
| 19443 | NM_018910.2 | #N/A | 24.4195838 | -5.8233562 | 2.61362323 |
| 19445 | NM_016319.3 | #N/A | 23.188522 | -5.7488803 | 2.58022214 |
| 19446 | NR_033658.1 | YY1P2 | 10.9358544 | 7.62647528 | 3.4233173 |
| 19447 | NM_001318748.1 | MSANTD4 | 28.7121733 | -6.0538886 | 2.71778871 |
| 19448 | NM_001300888.1 | PBDC1 | 10.7853656 | -4.6358479 | 2.08163119 |
| 19449 | NM_001040452.2 | #N/A | 27.7672927 | -6.0078175 | 2.69771669 |
| 19450 | NM_001281476.2 | FASTKD1 | 29.1658752 | -6.0756626 | 2.72854575 |
| 19451 | NM_001081753.2 | #N/A | 31.7240558 | -6.2000685 | 2.78523985 |
| 19452 | NM_015314.2 | #N/A | 27.6473111 | -5.9989325 | 2.69510754 |
| 19453 | NM_001346092.1 | #N/A | 116.418136 | -1.0709422 | 0.48114865 |
| 19454 | NM_017433.4 | #N/A | 10.5145757 | -4.6110523 | 2.07185225 |
| 19455 | NM_138347.4 | #N/A | 26.8542147 | -5.9578921 | 2.67716453 |
| 19456 | NM_152569.2 | #N/A | 39.0326147 | 1.43424266 | 0.64456988 |
| 19457 | NM_032329.5 | #N/A | 172.476598 | -1.6035643 | 0.7206739 |
| 19458 | NM_001288614.1 | #N/A | 7.98984128 | -4.2145358 | 1.89417061 |
| 19459 | NM_020630.4 | #N/A | 102.447717 | -3.1891466 | 1.43334459 |
| 19460 | NR_046848.1 | FAM155A-IT1 | 13.1059085 | 3.93661253 | 1.76939511 |
| 19461 | NR_026936.1 | LINC01554 | 16.4846602 | 2.21315819 | 0.99476494 |
| 19462 | NM_148894.2 | #N/A | 482.974474 | -2.8752993 | 1.2924647 |
| 19463 | NR_046473.1 | MEG3 | 22.8891391 | 1.94853748 | 0.87589916 |
| 19464 | NR_125888.1 | LOC101928052 | 11.8094311 | 2.43470944 | 1.09449004 |
| 19465 | NM_152386.3 | #N/A | 39.5315127 | -6.5164138 | 2.92961506 |
| 19466 | NR_027163.1 | SNHG29 | 31.0996289 | -6.1677641 | 2.77291465 |
| 19467 | NR_026564.1 | SLC25A51P4 | 7.41307812 | -4.1041909 | 1.84523186 |
| 19468 | NM_001173543.1 | BANP | 15.8264611 | -3.1288936 | 1.40679636 |
| 19469 | NM_001303473.1 | #N/A | 6.96383714 | -4.0139001 | 1.80476521 |
| 19470 | NR_030725.1 | #N/A | 28.6204454 | -6.0486543 | 2.7201675 |
| 19471 | NR_040087.2 | #N/A | 22.8804835 | -5.7295689 | 2.57680213 |
| 19472 | NM_001083608.1 | #N/A | 28.6534002 | -6.050883 | 2.72137402 |
| 19473 | NM_130446.2 | #N/A | 35.0881431 | 1.83045403 | 0.8233553 |
| 19474 | NM_018271.4 | THNSL2 | 11.3272971 | -4.721662 | 2.12394485 |
| 19475 | NM_000831.3 | #N/A | 504.685164 | -3.3384832 | 1.50207341 |
| 19476 | NM_001282915.1 | #N/A | 22.8000221 | -5.7240277 | 2.57536854 |
| 19477 | NM_001318955.1 | #N/A | 7.32000719 | -4.0832882 | 1.8371486 |
| 19478 | NM_005914.3 | #N/A | 7.06699341 | -4.0264956 | 1.81163086 |

| | | | | | |
|---|---|---|---|---|---|
| 19479 | NM_002299.3 | #N/A | 3.10582192 | 4.76657663 | 2.14479124 |
| 19480 | NM_001288799.1 | #N/A | 15.1176818 | -5.1318694 | 2.30930469 |
| 19481 | NM_022096.5 | #N/A | 9.85545807 | -4.4968516 | 2.02380017 |
| 19482 | NM_005545.3 | #N/A | 23.4227934 | -5.763138 | 2.59399139 |
| 19484 | NR_038838.1 | LINC02431 | 17.1751362 | 9.0888342 | 4.09144168 |
| 19483 | NM_001256403.1 | #N/A | 17.1751362 | 9.0888342 | 4.09144168 |
| 19485 | NM_001014449.2 | #N/A | 22.5988308 | -5.7114981 | 2.57149158 |
| 19486 | NM_175737.3 | #N/A | 74.705741 | 1.2113304 | 0.54537372 |
| 19487 | NM_003063.2 | #N/A | 9.43556748 | -4.4470388 | 2.00223936 |
| 19488 | NM_001242817.1 | #N/A | 29.8657754 | -6.1101967 | 2.7512995 |
| 19489 | NM_014708.4 | #N/A | 90.6464018 | -2.1064154 | 0.94858765 |
| 19490 | NM_001330712.1 | ZNF131 | 28.3247441 | -6.0340539 | 2.71769886 |
| 19491 | NM_017436.5 | #N/A | 9.18882948 | -4.4212191 | 1.99134381 |
| 19492 | NM_001242539.2 | NCOA6 | 11.0138826 | -4.676917 | 2.10659042 |
| 19493 | NM_080654.2 | CCDC34 | 9.08865022 | -4.3858281 | 1.97571235 |
| 19494 | NM_004188.6 | #N/A | 24.837484 | 3.12772636 | 1.40904314 |
| 19495 | NR_026879.1 | FAM185A | 9.41147675 | -4.4445755 | 2.00242371 |
| 19496 | NM_001015882.2 | #N/A | 19.9639065 | -3.830291 | 1.7258536 |
| 19497 | NM_000692.4 | #N/A | 151.836514 | -1.9004342 | 0.85632439 |
| 19498 | NM_018223.2 | CHFR | 10.4188071 | -4.6014741 | 2.073452 |
| 19499 | NM_006794.3 | #N/A | 6.6782718 | -3.9520073 | 1.78082328 |
| 19500 | NR_002171.2 | OR7E156P | 24.6463607 | 3.84015187 | 1.73046635 |
| 19501 | NM_001308421.1 | #N/A | 7.93213883 | -4.2069317 | 1.8960497 |
| 19502 | NM_024422.4 | #N/A | 26.6409021 | -5.9462842 | 2.68000927 |
| 19503 | NM_001277403.1 | #N/A | 14.701978 | -3.4533847 | 1.55646628 |
| 19504 | NM_001265579.1 | FAM13A | 29.6716398 | -6.103471 | 2.75156367 |
| 19505 | NM_001330673.1 | PSMA4 | 7.97141376 | -4.2130659 | 1.89950532 |
| 19506 | NM_014574.3 | #N/A | 127.444508 | -2.3193921 | 1.04586285 |
| 19507 | NM_025236.3 | RNF39 | 7.74859276 | -4.1783355 | 1.88416531 |
| 19508 | NM_015328.3 | #N/A | 31.994169 | -6.211567 | 2.80121132 |
| 19509 | NM_001348094.1 | #N/A | 29.2307877 | -6.0781901 | 2.741526 |
| 19510 | NM_000890.3 | #N/A | 51.5343919 | 1.89304322 | 0.85386324 |
| 19512 | NM_001286555.1 | DUSP22 | 29.5458755 | -6.0961959 | 2.74979218 |
| 19511 | NM_001204083.1 | #N/A | 7.22098398 | -4.0666184 | 1.8343206 |
| 19513 | NM_001321058.1 | #N/A | 27.2116361 | -5.9789335 | 2.69689755 |
| 19514 | NM_001142405.1 | #N/A | 32.034528 | -6.2096495 | 2.8010161 |
| 19515 | NM_001282886.1 | #N/A | 24.2238893 | -5.8108982 | 2.62121515 |
| 19516 | NR_120653.1 | LOC101929279 | 29.5809542 | 1.61348474 | 0.72790887 |
| 19517 | NM_001032394.2 | ADGRG6 | 30.6455449 | -6.1477799 | 2.77376309 |
| 19519 | NR_132988.1 | LOC106660606 | 13.6174794 | 2.02082573 | 0.91177918 |
| 19518 | NM_001305108.1 | PTOV1 | 30.6766067 | -6.1489185 | 2.7743233 |
| 19520 | NM_001198902.1 | YY1AP1 | 30.764518 | -6.1539565 | 2.77678164 |
| 19521 | NM_014433.2 | #N/A | 6.45284802 | -3.9026211 | 1.76100749 |

| | | | | | |
|---|---|---|---|---|---|
| 19522 | NM_001162495.2 | #N/A | 28.3411088 | -6.0347846 | 2.72315348 |
| 19523 | NM_030953.3 | #N/A | 7.7404817 | -4.1751795 | 1.88406589 |
| 19524 | NM_001271733.1 | MST1L | 6.72271349 | -3.9647941 | 1.78938228 |
| 19525 | NM_002216.2 | #N/A | 7.20815618 | -4.0632518 | 1.83395141 |
| 19527 | NM_203506.2 | GRB2 | 16.8888839 | 9.06471412 | 4.09162563 |
| 19526 | NM_144499.2 | #N/A | 16.8888839 | 9.06471412 | 4.09162563 |
| 19528 | NM_001199039.1 | #N/A | 9.21763546 | -4.422094 | 1.9962026 |
| 19529 | NM_001201479.1 | #N/A | 7.89578127 | -4.1733209 | 1.88392322 |
| 19530 | NR_110758.1 | LINC01756 | 5.59730403 | 5.81250606 | 2.6245199 |
| 19531 | NM_001025591.3 | SCGB2B2 | 29.0708733 | 3.32402679 | 1.5010743 |
| 19532 | NM_172070.3 | #N/A | 293.676929 | -3.1018624 | 1.40078495 |
| 19533 | NR_046706.1 | MID1IP1-AS1 | 7.60208896 | -4.1252069 | 1.86310286 |
| 19534 | NM_002661.4 | #N/A | 10.1945041 | -3.2553667 | 1.47027721 |
| 19535 | NM_001079842.2 | OCIAD1 | 28.1422395 | -6.024358 | 2.72120382 |
| 19536 | NM_001286792.1 | #N/A | 27.8786275 | -6.0137444 | 2.71642696 |
| 19537 | NM_144781.2 | #N/A | 28.5829099 | -6.0479853 | 2.73195052 |
| 19538 | NM_001293236.1 | #N/A | 23.0976223 | -5.7426785 | 2.59407082 |
| 19539 | NM_002401.3 | #N/A | 22.4764364 | -5.7035547 | 2.57641744 |
| 19540 | NM_000257.3 | #N/A | 16.7534472 | -3.6173903 | 1.63413659 |
| 19541 | NM_005135.2 | SLC12A6 | 68.0603059 | 5.87079602 | 2.65216093 |
| 19542 | NM_001308192.1 | CPEB4 | 27.6462096 | -6.0015179 | 2.71148209 |
| 19543 | NM_001286279.1 | TUBGCP3 | 7.14758144 | -4.0630342 | 1.83610887 |
| 19544 | NM_001254732.2 | SPECC1L | 28.6521639 | -6.0524872 | 2.73643806 |
| 19545 | NM_001039573.2 | #N/A | 22.8827488 | -5.7290043 | 2.59031581 |
| 19546 | NM_001144918.1 | #N/A | 28.7136976 | -6.0545617 | 2.73766472 |
| 19547 | NM_001135652.2 | EIF2AK2 | 16.866062 | 9.04719952 | 4.09176112 |
| 19552 | NR_104294.1 | LOC100422212 | 16.866062 | 9.04719952 | 4.09176112 |
| 19550 | NM_030905.2 | OR2J2 | 16.866062 | 9.04719952 | 4.09176112 |
| 19548 | NM_001173552.1 | TMPRSS4 | 16.866062 | 9.04719952 | 4.09176112 |
| 19549 | NM_001199087.1 | #N/A | 16.866062 | 9.04719952 | 4.09176112 |
| 19551 | NR_073104.1 | #N/A | 16.866062 | 9.04719952 | 4.09176112 |
| 19553 | NM_018656.2 | #N/A | 195.056001 | 1.20507332 | 0.54506726 |
| 19554 | NM_004133.4 | #N/A | 16.6185337 | 5.88588918 | 2.66239739 |
| 19555 | NM_001199514.1 | #N/A | 9.68591027 | -4.4846773 | 2.02863935 |
| 19556 | NM_001303229.1 | #N/A | 29.5422215 | -6.0933477 | 2.75638193 |
| 19557 | NM_003059.2 | #N/A | 7.50000516 | -4.1285778 | 1.86785999 |
| 19558 | NR_028058.1 | PMS2P9 | 33.2003733 | -1.74065 | 0.78751979 |
| 19559 | NR_024422.1 | ZNF503-AS2 | 9.35107549 | -4.4280322 | 2.00350675 |
| 19560 | NM_001130725.1 | PSMB5 | 26.0101173 | -5.9143626 | 2.6763009 |
| 19562 | NR_047470.1 | P4HA2-AS1 | 6.70766072 | -3.9579194 | 1.79102564 |
| 19561 | NM_001024465.2 | #N/A | 22.2200676 | -5.6871621 | 2.57351797 |
| 19563 | NM_001007088.1 | #N/A | 26.2420969 | -5.9239983 | 2.68077374 |
| 19564 | NM_001297608.1 | #N/A | 29.8889402 | -6.1136878 | 2.7666387 |

| | | | | | |
|---|---|---|---|---|---|
| 19565 | NM_007287.2 | #N/A | 27.9920231 | -6.0167337 | 2.72297587 |
| 19566 | NM_052838.4 | #N/A | 6.85980683 | -3.9925916 | 1.80691932 |
| 19570 | NM_002173.3 | IFNA16 | 16.6026316 | 9.04018343 | 4.09181585 |
| 19567 | NM_001256468.1 | KAT6B | 23.248262 | -5.7523079 | 2.60365449 |
| 19568 | NM_001291491.1 | #N/A | 16.6026316 | 9.04018343 | 4.09181585 |
| 19569 | NM_001329057.1 | #N/A | 16.6026316 | 9.04018343 | 4.09181585 |
| 19572 | NR_028415.1 | LINC00942 | 17.5199975 | 2.4792807 | 1.12223251 |
| 19571 | NM_001282656.1 | ZNF143 | 24.6271734 | -5.8350128 | 2.64116566 |
| 19573 | NM_001195071.1 | TMEM184B | 23.255323 | -5.7531948 | 2.60421883 |
| 19574 | NM_080704.3 | #N/A | 29.6349989 | -6.1016056 | 2.76208759 |
| 19575 | NM_001161012.2 | #N/A | 27.6086549 | -5.9989957 | 2.71574738 |
| 19576 | NM_080865.3 | #N/A | 6.96559728 | -3.9950937 | 1.80886642 |
| 19578 | NR_024505.1 | LINC00598 | 12.3705738 | 6.13285091 | 2.77703638 |
| 19577 | NM_020403.4 | #N/A | 34.4752866 | -6.3170459 | 2.86046707 |
| 19579 | NM_001278712.1 | #N/A | 7.38092006 | -4.0988073 | 1.85619148 |
| 19580 | NM_001008401.3 | #N/A | 30.5396345 | -6.1417584 | 2.78145338 |
| 19581 | NM_005556.3 | #N/A | 8.44990398 | -4.2936184 | 1.94456424 |
| 19582 | NM_001291006.1 | #N/A | 72.6453891 | 3.16383657 | 1.43297923 |
| 19583 | NM_001164239.1 | DHX16 | 28.6556763 | -6.0492538 | 2.73996807 |
| 19584 | NM_014791.3 | #N/A | 13.9257753 | -5.0032817 | 2.26653917 |
| 19585 | NM_024622.5 | #N/A | 22.1059112 | -5.6795941 | 2.57289691 |
| 19586 | NR_045104.1 | #N/A | 22.1833695 | -5.6849489 | 2.57536515 |
| 19588 | NM_001111045.1 | CCNA1 | 22.075777 | -5.677664 | 2.57226199 |
| 19587 | NM_001024212.1 | S100A13 | 8.05633576 | -4.2067028 | 1.90582702 |
| 19589 | NM_001256484.1 | #N/A | 26.8246446 | -5.9581275 | 2.70008647 |
| 19590 | NM_012182.2 | #N/A | 7.05023993 | -4.0298126 | 1.82623542 |
| 19591 | NM_001081003.2 | #N/A | 7.4057023 | -4.0841236 | 1.85096521 |
| 19592 | NM_004164.2 | #N/A | 5.8323602 | 4.82876688 | 2.18842796 |
| 19593 | NM_001301089.1 | SNX19 | 22.275659 | -5.6904059 | 2.57900992 |
| 19594 | NM_020653.3 | #N/A | 22.6776516 | -5.7165275 | 2.59089156 |
| 19595 | NM_181808.3 | #N/A | 7.06251951 | -4.0240783 | 1.82393433 |
| 19596 | NM_025170.5 | #N/A | 12.4825966 | 5.69174827 | 2.58025233 |
| 19597 | NM_001199782.1 | POC1B-GALNT4 | 101.020398 | 3.5722244 | 1.61957446 |
| 19598 | NM_001205319.1 | NXN | 64.4546037 | 5.79378563 | 2.62718191 |
| 19600 | NM_001017398.1 | TRIM36 | 8.51238172 | -4.3085587 | 1.95395719 |
| 19599 | NM_001001483.2 | ZDHHC13 | 28.3597963 | -6.0367491 | 2.73768583 |
| 19601 | NR_137280.1 | #N/A | 8.728598 | -4.3346524 | 1.96611849 |
| 19602 | NR_026781.1 | LINC00242 | 7.93964718 | -4.2020864 | 1.90619311 |
| 19603 | NM_001286462.1 | C21orf58 | 6.38772311 | -3.886684 | 1.76330691 |
| 19604 | NM_001321183.1 | FAM156B | 11.0463142 | -4.6664848 | 2.11710172 |
| 19605 | NM_001330059.1 | #N/A | 24.4152771 | -5.8220617 | 2.6414693 |
| 19606 | NM_001303625.1 | RPL10 | 29.9294711 | -6.112604 | 2.77373123 |
| 19607 | NM_175873.5 | #N/A | 7.80815648 | -4.1641777 | 1.88967778 |

| | | | | | |
|---|---|---|---|---|---|
| 19608 | NR_024540.1 | WASH7P | 318.169396 | -3.1132941 | 1.41308402 |
| 19609 | NM_001321201.1 | COA1 | 16.3163794 | 9.01522816 | 4.09201269 |
| 19611 | NR_102749.1 | EGFLAM-AS2 | 16.3163794 | 9.01522816 | 4.09201269 |
| 19610 | NM_005390.4 | #N/A | 16.3163794 | 9.01522816 | 4.09201269 |
| 19612 | NR_045128.1 | TXLNGY | 6.81976059 | -3.9829096 | 1.80788023 |
| 19613 | NM_001277120.1 | #N/A | 28.3539225 | -6.0349986 | 2.73953906 |
| 19614 | NM_032970.3 | #N/A | 189.863803 | -1.9932864 | 0.90484907 |
| 19615 | NM_024949.5 | #N/A | 136.252061 | -2.156596 | 0.97903362 |
| 19616 | NM_001282593.1 | SLAMF7 | 21.938519 | 3.32430462 | 1.50934918 |
| 19617 | NM_001320597.1 | #N/A | 24.9921657 | -5.8560979 | 2.65899211 |
| 19618 | NM_001302652.1 | #N/A | 30.6919382 | -6.1478812 | 2.79179188 |
| 19619 | NR_037897.1 | SAP30L-AS1 | 8.59277754 | -4.3258816 | 1.96449854 |
| 19620 | NM_001199398.1 | #N/A | 30.7646411 | -6.1518588 | 2.79401152 |
| 19621 | NM_002181.3 | #N/A | 11.2870267 | 4.38362787 | 1.99092938 |
| 19622 | NM_080722.4 | ADAMTS14 | 31.8966475 | -6.2042933 | 2.81855244 |
| 19623 | NM_005511.1 | #N/A | 55.9625148 | 1.53654238 | 0.69808894 |
| 19624 | NM_033150.2 | #N/A | 26.2434342 | -5.9253375 | 2.69228728 |
| 19625 | NM_001258325.1 | SFI1 | 31.0920243 | -6.1675965 | 2.80257246 |
| 19626 | NM_053046.3 | #N/A | 10.2552133 | -4.5554921 | 2.07052814 |
| 19627 | NM_001278176.1 | #N/A | 22.3051822 | -5.6929691 | 2.58771605 |
| 19628 | NM_001292033.1 | #N/A | 41.776689 | 1.20332659 | 0.54698835 |
| 19629 | NM_001190918.1 | #N/A | 28.5655214 | -6.0459583 | 2.74851711 |
| 19630 | NM_001290185.1 | #N/A | 9.85204478 | -4.511858 | 2.05112881 |
| 19631 | NM_001324237.1 | #N/A | 8.17495514 | -4.2358922 | 1.925676 |
| 19632 | NM_017580.2 | #N/A | 318.93606 | -3.1166124 | 1.41680244 |
| 19633 | NM_001005353.2 | AK4 | 24.0674903 | -5.8019056 | 2.63767689 |
| 19634 | NM_003979.3 | #N/A | 37.1600307 | 3.78335242 | 1.72007082 |
| 19635 | NM_001145454.2 | #N/A | 99.8595748 | 1.42345907 | 0.64719997 |
| 19636 | NM_001323294.1 | #N/A | 28.9041162 | -6.0645929 | 2.75822415 |
| 19637 | NR_024053.2 | HCG18 | 21.7716682 | -5.6577373 | 2.57343854 |
| 19638 | NR_036433.2 | FAM172BP | 20.9126059 | 1.96487844 | 0.89389121 |
| 19639 | NM_001135256.1 | RBBP4 | 23.1667095 | 5.57056733 | 2.53436323 |
| 19640 | NM_001006937.2 | TCEAL4 | 27.9445851 | -6.0141849 | 2.7362524 |
| 19641 | NM_001286517.1 | #N/A | 22.2123956 | -5.6868322 | 2.58792552 |
| 19642 | NM_001006620.1 | MAPKAP1 | 25.4600183 | -5.8822067 | 2.67689057 |
| 19643 | NR_134582.1 | LOC101927447 | 28.5546233 | 1.36179624 | 0.61976418 |
| 19644 | NM_018917.3 | #N/A | 8.21912821 | -4.2415284 | 1.9304856 |
| 19646 | NM_001286702.1 | ARMC1 | 16.0301271 | 8.9898335 | 4.09221648 |
| 19649 | NR_038372.1 | CSMD2-AS1 | 16.0301271 | 8.9898335 | 4.09221648 |
| 19650 | NR_132986.1 | MIR3150BHG | 16.0301271 | 8.9898335 | 4.09221648 |
| 19645 | NM_001005243.1 | OR9K2 | 16.0301271 | 8.9898335 | 4.09221648 |
| 19647 | NM_001315508.1 | PCM1 | 68.2168458 | -1.40362 | 0.6389086 |
| 19648 | NR_038327.2 | TEKT4P2 | 16.0301271 | 8.9898335 | 4.09221648 |

| | | | | | |
|---|---|---|---|---|---|
| 19651 | NR_135668.1 | LHX1-DT | 6.43517678 | -3.8957195 | 1.7737945 |
| 19652 | NM_001015887.1 | #N/A | 7.73773632 | -4.1773096 | 1.90228536 |
| 19653 | NM_173607.3 | #N/A | 109.276474 | -1.3450662 | 0.6125259 |
| 19654 | NM_003742.2 | #N/A | 4.09801095 | 4.42050072 | 2.01312018 |
| 19655 | NM_001555.4 | #N/A | 25.9591678 | -5.9082477 | 2.69119501 |
| 19656 | NM_001318853.1 | #N/A | 26.0831625 | -5.9177903 | 2.69582456 |
| 19657 | NM_004274.4 | #N/A | 480.484617 | -2.8238619 | 1.28640552 |
| 19658 | NM_018939.3 | PCDHB6 | 8.17480043 | -4.2287057 | 1.92660549 |
| 19659 | NM_001205296.1 | TTF1 | 22.0708851 | 6.97320265 | 3.17710472 |
| 19660 | NR_110585.1 | LINC01819 | 20.4436203 | 1.5577437 | 0.70986526 |
| 19661 | NM_001728.3 | #N/A | 8.24184736 | -4.2584041 | 1.94094124 |
| 19663 | NR_120462.1 | LINC01486 | 10.3321376 | 3.35649662 | 1.52996805 |
| 19662 | NM_001080505.1 | #N/A | 19.3744836 | 3.62236347 | 1.65114407 |
| 19664 | NM_001161403.1 | #N/A | 22.0793105 | -5.678351 | 2.58837037 |
| 19665 | NM_001214907.1 | ZNF48 | 23.1359651 | -5.7455134 | 2.61940168 |
| 19666 | NM_001301860.1 | #N/A | 27.7531368 | -6.0058571 | 2.73837362 |
| 19667 | NM_001278422.1 | DYNC1I1 | 8.40143547 | -4.2945649 | 1.95815155 |
| 19668 | NM_018248.2 | #N/A | 24.1757965 | 1.60526003 | 0.73199252 |
| 19669 | NR_046058.1 | #N/A | 21.7428609 | -5.6557528 | 2.57914906 |
| 19670 | NM_001195304.1 | BBIP1 | 7.01182693 | -4.0065601 | 1.82712091 |
| 19671 | NM_032034.3 | SLC4A11 | 9.15375028 | -4.4074555 | 2.01005213 |
| 19672 | NM_001161563.2 | #N/A | 26.8494324 | -5.9564735 | 2.71686572 |
| 19673 | NR_122079.1 | LOC439933 | 18.8634914 | 3.09262963 | 1.41062971 |
| 19674 | NM_001082970.1 | #N/A | 36.3782585 | -6.3930834 | 2.91610318 |
| 19675 | NR_133664.1 | LINC00449 | 6.89078256 | -3.9942076 | 1.82195856 |
| 19676 | NM_001167931.1 | #N/A | 8.8480609 | -4.3606369 | 1.98915347 |
| 19677 | NM_138455.3 | #N/A | 15.5601193 | -3.5215766 | 1.6064286 |
| 19679 | NM_032328.3 | EFCAB2 | 10.3905071 | -4.5963624 | 2.09697574 |
| 19678 | NM_031496.1 | #N/A | 13.6034737 | -4.9664055 | 2.26581746 |
| 19680 | NM_001201335.1 | #N/A | 7.57828188 | -4.134214 | 1.88623612 |
| 19681 | NM_001199830.1 | HPSE | 54.2910627 | 5.83392435 | 2.66220974 |
| 19682 | NM_015514.1 | #N/A | 6.994564 | -4.012178 | 1.83089878 |
| 19683 | NM_001198827.1 | C8orf58 | 27.7616278 | -6.0043934 | 2.74005636 |
| 19684 | NM_001039651.1 | SAPCD1 | 6.88464469 | -4.0066075 | 1.82853526 |
| 19685 | NM_007351.2 | #N/A | 12.5322377 | -4.8638296 | 2.21976428 |
| 19686 | NR_037573.1 | #N/A | 147.32516 | -2.5227227 | 1.15131028 |
| 19687 | NM_198834.2 | #N/A | 31.6857307 | -6.1979948 | 2.82886689 |
| 19688 | NM_001287260.1 | #N/A | 220.229407 | 4.68383719 | 2.13787806 |
| 19689 | NM_199464.2 | #N/A | 6.42916379 | -3.8991138 | 1.77971746 |
| 19690 | NM_001291328.1 | #N/A | 7.78323252 | -4.1683078 | 1.9028371 |
| 19693 | NM_005959.3 | MTNR1B | 15.7438748 | 8.96397664 | 4.09242767 |
| 19691 | NM_001251825.1 | #N/A | 15.7438748 | 8.96397664 | 4.09242767 |
| 19692 | NM_001401.3 | #N/A | 15.7438748 | 8.96397664 | 4.09242767 |

| | | | | | |
|---|---|---|---|---|---|
| 19694 | NM_173059.2 | #N/A | 8.33714092 | 5.58054461 | 2.54806373 |
| 19695 | NM_004031.2 | #N/A | 6.64026914 | -3.9460291 | 1.80178257 |
| 19696 | NM_001161425.1 | #N/A | 27.2453535 | -5.9771809 | 2.72989173 |
| 19697 | NM_001164392.1 | MTF2 | 22.4858336 | -5.7035886 | 2.60505287 |
| 19698 | NM_001127174.1 | #N/A | 51.4420297 | 3.2742749 | 1.49557496 |
| 19699 | NM_012163.2 | LRRC29 | 7.05476026 | -4.0227569 | 1.83766875 |
| 19700 | NM_138432.3 | SDSL | 31.0393338 | -1.7960942 | 0.82062265 |
| 19701 | NM_001323606.1 | SYNDIG1 | 7.46998484 | -4.1191612 | 1.88207232 |
| 19703 | NM_207168.1 | ENSA | 10.414007 | -4.588606 | 2.09671347 |
| 19702 | NM_001002843.2 | ZNF280D | 28.3015639 | -6.03148 | 2.75599955 |
| 19704 | NM_001145462.1 | #N/A | 7.18888129 | -4.042613 | 1.84724695 |
| 19705 | NM_001002243.2 | #N/A | 22.0961314 | -5.6793651 | 2.59544178 |
| 19706 | NM_033213.4 | #N/A | 23.5583886 | -5.771065 | 2.63736042 |
| 19708 | NM_001289968.1 | PPP3CB | 27.1474713 | -5.9752364 | 2.73173279 |
| 19707 | NM_001288998.1 | #N/A | 28.3081083 | -6.0315795 | 2.75752728 |
| 19709 | NM_017646.5 | #N/A | 22.2138624 | -5.6864384 | 2.59973908 |
| 19710 | NR_138537.1 | #N/A | 27.5531466 | -5.992454 | 2.73986643 |
| 19711 | NR_033234.1 | #N/A | 22.9290316 | -5.7316514 | 2.62096009 |
| 19712 | NM_003417.4 | #N/A | 376.76694 | 0.95935609 | 0.43870158 |
| 19713 | NM_178837.4 | DNAAF3 | 28.0294603 | -6.0212963 | 2.75352581 |
| 19714 | NM_014761.3 | #N/A | 26.9159197 | -5.9588936 | 2.72510148 |
| 19715 | NR_024616.1 | INE1 | 48.8765304 | 1.32625588 | 0.60654279 |
| 19716 | NM_001647.3 | #N/A | 10.1846994 | -4.5680384 | 2.08924469 |
| 19717 | NM_139320.1 | CHRFAM7A | 7.01146123 | -4.0254465 | 1.84118527 |
| 19718 | NM_183356.3 | ASNS | 27.1779749 | -5.9736688 | 2.73251545 |
| 19719 | NM_133174.2 | APBB3 | 21.9048972 | -5.6662525 | 2.59221325 |
| 19720 | NM_001195087.1 | GGACT | 10.4303334 | -3.3014368 | 1.51037296 |
| 19721 | NM_001166133.1 | FRAS1 | 9.90966974 | 5.93652158 | 2.71648043 |
| 19722 | NM_001292023.1 | #N/A | 25.2024567 | 1.74146795 | 0.79694728 |
| 19723 | NR_038978.1 | MORF4L2-AS1 | 7.46563289 | -4.1224066 | 1.88658608 |
| 19724 | NM_005519.1 | HMX2 | 7.88741843 | -4.1804122 | 1.9132068 |
| 19725 | NM_001320541.1 | #N/A | 23.0457929 | -5.7394324 | 2.62678775 |
| 19726 | NM_001204189.1 | #N/A | 15.6613433 | 8.93968642 | 4.0926295 |
| 19727 | NM_001321909.1 | #N/A | 15.6613433 | 8.93968642 | 4.0926295 |
| 19728 | NM_002170.3 | #N/A | 15.6613433 | 8.93968642 | 4.0926295 |
| 19729 | NM_002685.3 | #N/A | 15.6613433 | 8.93968642 | 4.0926295 |
| 19730 | NM_001040708.1 | HEY1 | 26.3325929 | -5.9318565 | 2.71573026 |
| 19731 | NM_001145143.1 | HTR3D | 15.4576225 | 8.93765925 | 4.0926465 |
| 19732 | NM_001316963.1 | #N/A | 7.4349457 | -4.1138588 | 1.88379978 |
| 19733 | NR_037164.1 | #N/A | 9.41113231 | -4.4536294 | 2.03971548 |
| 19734 | NM_001195259.1 | TGFBR3L | 6.79907535 | -3.9738052 | 1.82097006 |
| 19735 | NM_000209.3 | #N/A | 17.3848521 | 6.11502227 | 2.80249067 |
| 19736 | NM_021223.2 | #N/A | 7.55010601 | -4.1306758 | 1.89322611 |

| | | | | | |
|---|---|---|---|---|---|
| 19737 | NR_002788.2 | CYP4Z2P | 12.1592099 | 7.13124617 | 3.26905037 |
| 19738 | NR_038993.1 | ZBTB20-AS1 | 8.63664371 | 6.06733654 | 2.78148495 |
| 19739 | NM_001206648.1 | #N/A | 74.5213499 | 3.63509195 | 1.66677007 |
| 19744 | NR_033934.1 | MRPL45P2 | 6.27300476 | -3.8603193 | 1.77014058 |
| 19740 | NM_001256214.1 | #N/A | 30.7980687 | -6.1566871 | 2.82312246 |
| 19741 | NM_005930.3 | #N/A | 27.5144495 | -5.9916454 | 2.74739567 |
| 19742 | NM_080603.4 | #N/A | 114.153051 | -0.823504 | 0.3775998 |
| 19743 | NM_178130.3 | #N/A | 8.38798048 | -4.2863062 | 1.96546113 |
| 19745 | NR_046478.1 | #N/A | 11.8395679 | -4.7728452 | 2.18853466 |
| 19746 | NM_020185.4 | DUSP22 | 7.3466043 | -4.0996685 | 1.8801677 |
| 19747 | NM_017733.4 | PIGG | 24.4496512 | -5.825413 | 2.67181262 |
| 19748 | NR_102368.1 | SNORA11E | 6.61921638 | -3.9440819 | 1.80903238 |
| 19749 | NM_001139488.1 | #N/A | 21.2946093 | -5.6259936 | 2.58056931 |
| 19750 | NM_198456.2 | #N/A | 7.88012491 | -4.1771434 | 1.91611775 |
| 19751 | NR_026928.1 | N4BP2L2-IT2 | 6.45843881 | -3.912034 | 1.79471491 |
| 19752 | NM_001039592.1 | #N/A | 27.6555516 | -6.0019053 | 2.75383859 |
| 19753 | NM_001330522.1 | FAM124A | 8.39060863 | -4.2785074 | 1.96372991 |
| 19754 | NM_001145160.1 | #N/A | 38.1278578 | 1.44977837 | 0.66542429 |
| 19755 | NM_020834.2 | #N/A | 115.733075 | 0.92885106 | 0.42632883 |
| 19756 | NM_001003699.3 | #N/A | 22.2952743 | -5.6925064 | 2.61292705 |
| 19757 | NM_021038.4 | #N/A | 28.5845297 | -6.0488052 | 2.77651796 |
| 19758 | NR_120565.1 | LOC101928894 | 12.4325163 | 5.83578022 | 2.67878823 |
| 19760 | NR_104331.1 | CTBP1-AS | 32.3389286 | -1.7404152 | 0.79896721 |
| 19759 | NM_001171083.2 | SMPD4 | 28.2381811 | -6.0278554 | 2.76720323 |
| 19761 | NM_001193636.1 | DHRS4L2 | 10.1847792 | -4.5459051 | 2.0871045 |
| 19762 | NM_001281773.2 | ZMYND8 | 27.2240642 | -5.9773017 | 2.74425501 |
| 19763 | NM_002563.4 | #N/A | 13.4408739 | -3.3273259 | 1.52762824 |
| 19764 | NM_001286985.1 | #N/A | 16.5418883 | 6.94908637 | 3.19055608 |
| 19766 | NR_033434.1 | UMPS | 35.2633402 | 6.26007706 | 2.87445601 |
| 19765 | NM_005447.3 | #N/A | 6.25607883 | -3.8522082 | 1.76884211 |
| 19767 | NR_135261.1 | C1orf61 | 13.8298171 | -4.9933372 | 2.29304489 |
| 19768 | NM_001172663.1 | RAB40C | 8.90250049 | -4.3654176 | 2.00476911 |
| 19769 | NM_181864.2 | #N/A | 11.0686522 | -4.6700215 | 2.14479517 |
| 19770 | NM_001348010.1 | #N/A | 16.1523586 | -3.9461678 | 1.81243449 |
| 19772 | NM_020433.4 | JPH2 | 18.1889859 | -3.7006835 | 1.69973044 |
| 19771 | NM_001301024.1 | #N/A | 26.892509 | 5.52101857 | 2.53581202 |
| 19773 | NM_001007534.2 | C3orf56 | 15.1713703 | 8.91085059 | 4.09287351 |
| 19774 | NM_001101362.2 | KBTBD13 | 15.1713703 | 8.91085059 | 4.09287351 |
| 19775 | NM_001256293.1 | #N/A | 7.99565989 | -4.2125987 | 1.93500203 |
| 19776 | NM_013389.2 | #N/A | 7.06413704 | -4.0198437 | 1.84666422 |
| 19777 | NM_152464.2 | #N/A | 138.260299 | -1.6442602 | 0.75539616 |
| 19778 | NM_001012716.2 | #N/A | 6.76853279 | -3.9560541 | 1.81750259 |
| 19779 | NM_001004316.2 | LEKR1 | 7.3868232 | 6.13288879 | 2.81778547 |

| | | | | | |
|---|---|---|---|---|---|
| 19781 | NR_110139.1 | LOC101929718 | 6.81428283 | -3.9927594 | 1.83454415 |
| 19780 | NM_001167929.1 | #N/A | 27.3094274 | -5.982495 | 2.74877456 |
| 19783 | NR_026965.1 | METTL21EP | 19.6103414 | 2.76598468 | 1.27100479 |
| 19782 | NM_001043353.1 | #N/A | 27.6929637 | -6.0000304 | 2.75707155 |
| 19784 | NM_001040197.1 | AGTRAP | 7.37797442 | -4.081325 | 1.8754784 |
| 19785 | NM_001297738.1 | #N/A | 6.82670028 | -3.9786071 | 1.8285445 |
| 19786 | NM_020815.2 | #N/A | 100.939199 | -2.3098129 | 1.06179412 |
| 19787 | NM_015953.4 | NOSIP | 8.29823569 | -4.2773446 | 1.96636102 |
| 19788 | NM_001315531.1 | #N/A | 8.27871998 | -4.2694339 | 1.96294587 |
| 19789 | NM_001318758.1 | #N/A | 27.9331181 | -6.0147019 | 2.76582725 |
| 19790 | NR_047004.1 | LINC00379 | 11.3544282 | 5.45517636 | 2.50874332 |
| 19791 | NM_001256168.1 | ZNF620 | 83.9362677 | 4.03915771 | 1.85759203 |
| 19792 | NM_175709.4 | #N/A | 24.714082 | -5.8374417 | 2.68463068 |
| 19793 | NM_014969.5 | WDR47 | 27.6143455 | -5.9962353 | 2.75822716 |
| 19794 | NM_001042573.2 | ENGASE | 195.596771 | -2.3090501 | 1.06219186 |
| 19795 | NR_002814.2 | LOC374443 | 8.03650702 | -4.203956 | 1.93389051 |
| 19797 | NR_034007.1 | LINC00880 | 9.52700023 | -4.4705389 | 2.0567396 |
| 19796 | NM_021785.4 | #N/A | 23.4082519 | -5.7625031 | 2.65112407 |
| 19798 | NM_001317741.1 | SH3BGR | 6.58764689 | -3.9155981 | 1.80148413 |
| 19799 | NR_134315.1 | LINC02679 | 14.2737049 | 4.20388294 | 1.93439287 |
| 19800 | NR_046455.1 | #N/A | 19.3598958 | 2.41250965 | 1.11014043 |
| 19801 | NM_001291956.1 | #N/A | 45.0080873 | 1.46737329 | 0.6752506 |
| 19802 | NM_000435.2 | #N/A | 705.665437 | -2.8229684 | 1.29909648 |
| 19803 | NM_001012981.4 | #N/A | 169.048749 | -2.9760856 | 1.36959171 |
| 19804 | NM_001168467.1 | #N/A | 25.7197296 | -5.8975317 | 2.71409369 |
| 19805 | NM_145288.2 | #N/A | 10.1682568 | -3.2359922 | 1.48926532 |
| 19806 | NM_198089.2 | #N/A | 21.0109199 | -5.606341 | 2.58029976 |
| 19807 | NM_017490.3 | #N/A | 21.2285598 | -5.6217105 | 2.587432 |
| 19808 | NM_003809.2 | #N/A | 7.2166518 | -4.0726858 | 1.87454035 |
| 19809 | NM_001146191.1 | #N/A | 20.3887242 | -3.8295765 | 1.76278063 |
| 19810 | NM_001286829.1 | #N/A | 37.0078676 | -6.418565 | 2.95452134 |
| 19811 | NR_047527.1 | FGF7P6 | 9.81751484 | -4.4903906 | 2.06719037 |
| 19812 | NM_014046.3 | #N/A | 150.277154 | -2.9896402 | 1.37633203 |
| 19813 | NR_103459.1 | #N/A | 18.8770083 | 6.9283035 | 3.18992506 |
| 19814 | NM_005156.6 | #N/A | 33.2410556 | -6.2660051 | 2.88506802 |
| 19815 | NM_001286798.1 | NTMT1 | 7.9291228 | -4.2009157 | 1.9344759 |
| 19816 | NM_001297655.1 | #N/A | 26.1101792 | -5.9162002 | 2.72505787 |
| 19817 | NM_181305.2 | #N/A | 25.4856022 | -5.8844655 | 2.71086817 |
| 19821 | NM_001160234.1 | ATP1A1 | 14.885118 | 8.88353292 | 4.09310917 |
| 19818 | NM_000498.3 | CYP11B2 | 14.885118 | 8.88353292 | 4.09310917 |
| 19823 | NR_104229.1 | NME7 | 14.885118 | 8.88353292 | 4.09310917 |
| 19819 | NM_001029939.3 | #N/A | 14.885118 | 8.88353292 | 4.09310917 |
| 19820 | NM_001086.2 | #N/A | 14.885118 | 8.88353292 | 4.09310917 |

| | | | | | |
|---|---|---|---|---|---|
| 19822 | NM_001346268.1 | #N/A | 14.885118 | 8.88353292 | 4.09310917 |
| 19824 | NR_022006.1 | KIAA0087 | 45.1082968 | 3.14424238 | 1.44886938 |
| 19825 | NM_001301782.1 | #N/A | 16.0858863 | -3.54214 | 1.63231429 |
| 19826 | NM_001130524.1 | #N/A | 6.67085339 | -3.9575555 | 1.82378779 |
| 19827 | NM_001034833.1 | NHP2 | 26.0969721 | -5.9160676 | 2.72670019 |
| 19828 | NR_104183.1 | VASH1-AS1 | 6.44478048 | -3.9065482 | 1.80084784 |
| 19829 | NR_033984.1 | LINC02139 | 62.9347937 | 1.80360174 | 0.83147187 |
| 19830 | NM_007037.5 | #N/A | 41.4709723 | 1.9675516 | 0.90710886 |
| 19831 | NM_001321115.1 | #N/A | 32.0064915 | 3.6677272 | 1.69128337 |
| 19832 | NM_032184.1 | #N/A | 25.9285095 | -2.9438946 | 1.35752781 |
| 19833 | NR_132738.1 | LNCTAM34A | 6.98083257 | -4.0292138 | 1.85832544 |
| 19834 | NM_133338.2 | #N/A | 10.3542297 | -4.5691725 | 2.10740778 |
| 19835 | NR_028100.1 | DCAF11 | 23.8038026 | -5.7867651 | 2.66941337 |
| 19836 | NR_038352.1 | #N/A | 40.1404015 | -5.2795988 | 2.43550908 |
| 19837 | NM_133342.2 | #N/A | 21.6599842 | -5.6499222 | 2.60647712 |
| 19839 | NM_006883.2 | SHOX | 72.5141398 | 1.96725525 | 0.90760387 |
| 19838 | NM_001301009.1 | VSTM2A | 27.0501766 | -5.9682111 | 2.75348626 |
| 19840 | NM_018842.4 | #N/A | 23.3283528 | -3.2513736 | 1.50010135 |
| 19843 | NR_037615.1 | CNOT2 | 8.57609421 | -4.2951035 | 1.98176073 |
| 19841 | NM_001329135.1 | DYNC1LI1 | 10.3977737 | -4.5880932 | 2.11692406 |
| 19842 | NM_001330600.1 | #N/A | 10.3023602 | -4.5804712 | 2.11344231 |
| 19849 | NR_125804.1 | FOXCUT | 8.474962 | -4.3067237 | 1.98723753 |
| 19848 | NR_026825.2 | RPSAP52 | 16.5681662 | 5.09548088 | 2.35122033 |
| 19847 | NR_015379.3 | UCA1 | 7.45039447 | 2.984228 | 1.37697837 |
| 19844 | NM_001251888.1 | #N/A | 8.15191328 | -4.2331181 | 1.95322933 |
| 19845 | NM_001321032.1 | #N/A | 25.3757752 | -5.8771534 | 2.71185146 |
| 19846 | NM_178505.7 | #N/A | 7.98570551 | -4.2219364 | 1.94813154 |
| 19850 | NR_046762.1 | ALMS1-IT1 | 10.4388566 | -3.2825435 | 1.51472344 |
| 19851 | NM_001142468.2 | #N/A | 8.88937443 | -4.3478581 | 2.00646601 |
| 19852 | NR_033698.1 | FNTA | 22.2663327 | -5.6898733 | 2.62591097 |
| 19853 | NM_001199480.1 | #N/A | 26.7953614 | -5.9527054 | 2.74744325 |
| 19854 | NM_006495.3 | #N/A | 9.68285152 | 7.53915329 | 3.47980891 |
| 19855 | NM_001286678.1 | PRMT2 | 8.2513457 | -4.2426634 | 1.9585556 |
| 19856 | NR_110564.1 | LINC01828 | 5.86013264 | 3.42776099 | 1.58238607 |
| 19858 | NR_109795.1 | LINC01508 | 8.00829149 | -4.2074297 | 1.94301233 |
| 19857 | NR_046282.1 | #N/A | 20.9046386 | -5.5988681 | 2.58557486 |
| 19859 | NM_001170938.1 | #N/A | 20.8660713 | -5.5967831 | 2.58484081 |
| 19860 | NM_001178138.1 | #N/A | 23.9097915 | -5.791825 | 2.67496313 |
| 19861 | NM_001134771.1 | #N/A | 105.980611 | -1.9779811 | 0.91369857 |
| 19862 | NM_152333.3 | #N/A | 25.9108916 | -5.9053842 | 2.72839293 |
| 19863 | NM_021570.3 | #N/A | 6.54920632 | -3.9324281 | 1.81705344 |
| 19864 | NM_001102450.2 | RGS8 | 20.7552807 | -5.5890941 | 2.58265529 |
| 19865 | NM_001243789.1 | #N/A | 12.5082399 | -2.746647 | 1.269224 |

| | | | | | |
|---|---|---|---|---|---|
| 19866 | NM_183241.2 | #N/A | 22.6969628 | -5.7173474 | 2.64212774 |
| 19867 | NM_001166237.1 | GSDMD | 6.68103199 | -3.9625475 | 1.83127313 |
| 19868 | NM_003787.4 | NOL4 | 30.7682572 | -6.1519634 | 2.84315852 |
| 19869 | NM_015576.2 | #N/A | 28.8164369 | -6.0611067 | 2.80131199 |
| 19870 | NR_036549.1 | CCNT2-AS1 | 9.02056983 | -4.3804508 | 2.02458205 |
| 19871 | NM_032364.5 | DNAJC14 | 205.447657 | -2.3393906 | 1.08127578 |
| 19874 | NM_001313913.1 | F9 | 14.5988657 | 8.85568723 | 4.09335397 |
| 19873 | NM_001256305.1 | HPGD | 14.5988657 | 8.85568723 | 4.09335397 |
| 19872 | NM_001201404.2 | #N/A | 14.5988657 | 8.85568723 | 4.09335397 |
| 19875 | NM_001320296.1 | #N/A | 6.73640051 | -3.9615015 | 1.83106409 |
| 19876 | NM_014614.2 | #N/A | 285.230018 | -3.0560539 | 1.41260068 |
| 19877 | NR_110938.1 | LOC101928446 | 28.8608202 | 1.81137584 | 0.83731132 |
| 19878 | NM_001252151.1 | #N/A | 21.0095792 | -5.6067646 | 2.59186639 |
| 19879 | NM_001297761.1 | #N/A | 8.30561418 | -4.2541804 | 1.96677924 |
| 19880 | NM_001288821.1 | OLFM3 | 9.74503697 | -4.4810431 | 2.0717063 |
| 19881 | NM_001303282.1 | ZNF18 | 25.8899252 | -5.9048169 | 2.73002772 |
| 19882 | NR_027291.1 | NAP1L6P | 7.00968425 | -4.0225528 | 1.85992862 |
| 19883 | NM_001301778.1 | #N/A | 21.6088176 | -5.6474702 | 2.61135528 |
| 19884 | NM_001199216.1 | MAG | 16.0962659 | 6.10238341 | 2.82208569 |
| 19885 | NM_005069.4 | #N/A | 6.9728047 | -4.0214408 | 1.85978201 |
| 19886 | NM_001033559.2 | DNAAF4 | 7.9954498 | -4.2171098 | 1.95060638 |
| 19887 | NR_102701.1 | LINC00871 | 5.34324953 | 4.63618158 | 2.14459226 |
| 19890 | NR_125896.1 | LINC02268 | 34.6949393 | 2.41699902 | 1.1184601 |
| 19888 | NM_001324025.1 | #N/A | 25.9857906 | -5.9097682 | 2.73472986 |
| 19889 | NM_016111.3 | #N/A | 237.030731 | -3.0214192 | 1.39815822 |
| 19891 | NR_109921.1 | TMEM9 | 23.8577281 | -5.7901642 | 2.67971712 |
| 19892 | NM_015973.4 | #N/A | 7.42443303 | -4.1056423 | 1.90037248 |
| 19893 | NM_016821.2 | OGG1 | 26.166641 | -5.9187991 | 2.73980904 |
| 19894 | NM_001242757.1 | #N/A | 6.36553153 | -3.8894489 | 1.80046027 |
| 19895 | NR_103861.1 | LINC02408 | 20.2729317 | 2.97205674 | 1.37602032 |
| 19896 | NR_120612.1 | NUTM2B-AS1 | 7.48848749 | -4.107785 | 1.90199269 |
| 19897 | NR_046253.1 | FGD5-AS1 | 23.4335297 | -5.7639081 | 2.66887743 |

| stat | pvalue | padj | Mock1 | Mock 2 | Mock 3 | Mock 4 |
|---|---|---|---|---|---|---|
| 21.7052809 | 1.83E-104 | 5.91E-100 | 301.235252 | 325.815378 | 279.560321 | 309.85472 |
| 17.4451118 | 3.75E-68 | 6.05E-64 | 486.484685 | 557.458205 | 584.43063 | 570.401636 |
| 16.4227669 | 1.31E-60 | 1.41E-56 | 1168.05506 | 938.478182 | 1042.31132 | 1120.6319 |
| 14.7392466 | 3.61E-49 | 2.91E-45 | 127.461447 | 134.493557 | 131.151755 | 129.9933 |
| 14.7001514 | 6.43E-49 | 4.15E-45 | 14.7543797 | 11.9068743 | 12.6549939 | 12.8872668 |
| -13.624258 | 2.87E-42 | 1.55E-38 | 26202.1389 | 62638.8183 | 52625.2171 | 59728.5595 |
| 13.2157381 | 7.12E-40 | 3.28E-36 | 0 | 0 | 0 | 0 |
| -13.129179 | 2.24E-39 | 9.04E-36 | 5176.73805 | 6357.72965 | 5685.54365 | 5982.49338 |
| -12.881834 | 5.70E-38 | 2.04E-34 | 2716.44524 | 4656.39968 | 4070.30624 | 4776.69346 |
| 12.0547234 | 1.83E-33 | 5.91E-30 | 95.0837801 | 91.7370541 | 106.992222 | 92.4521315 |
| -12.012291 | 3.06E-33 | 8.99E-30 | 13727.7208 | 31575.6775 | 16197.2418 | 23004.3316 |
| 11.8579694 | 1.96E-32 | 5.26E-29 | 26.6398522 | 17.5897006 | 17.2568099 | 21.292006 |
| -11.590529 | 4.60E-31 | 1.14E-27 | 2236.10821 | 3604.53558 | 3157.99622 | 3497.49215 |
| -11.466869 | 1.94E-30 | 4.46E-27 | 2060.28518 | 3962.01242 | 2772.59413 | 3346.20685 |
| 11.3557779 | 6.94E-30 | 1.49E-26 | 450.828268 | 325.274157 | 373.897549 | 379.333897 |
| 11.3044351 | 1.25E-29 | 2.52E-26 | 381.974496 | 268.175282 | 266.905327 | 248.780281 |
| -11.263568 | 1.99E-29 | 3.77E-26 | 3961.55094 | 9676.50024 | 7721.84722 | 8568.9118 |
| -11.148374 | 7.29E-29 | 1.31E-25 | 6635.78226 | 12647.8066 | 11581.6204 | 12858.1304 |
| 11.1094572 | 1.13E-28 | 1.92E-25 | 993.051721 | 823.739212 | 882.398214 | 893.703938 |
| -11.059219 | 1.98E-28 | 3.19E-25 | 1172.56334 | 1160.64963 | 947.974092 | 1107.18431 |
| 10.8865109 | 1.34E-27 | 2.06E-24 | 44.263139 | 56.2870421 | 66.7263317 | 78.4442328 |
| -10.804117 | 3.29E-27 | 4.83E-24 | 1498.79907 | 1631.24178 | 1421.96114 | 1492.12137 |
| 10.5665819 | 4.26E-26 | 5.98E-23 | 22.1315695 | 21.6488623 | 21.8586259 | 21.852322 |
| -10.526096 | 6.55E-26 | 8.81E-23 | 3242.68478 | 7283.48912 | 7674.6786 | 7356.38809 |
| 10.485493 | 1.01E-25 | 1.30E-22 | 12.7051603 | 16.2366467 | 14.9559019 | 13.4475828 |
| 10.37552 | 3.20E-25 | 3.98E-22 | 9.01656536 | 15.966036 | 12.6549939 | 19.0507423 |
| 10.3310533 | 5.10E-25 | 6.10E-22 | 213.938505 | 319.320719 | 299.118039 | 364.765683 |
| 10.2745178 | 9.18E-25 | 1.06E-21 | 39.3450125 | 23.272527 | 26.4604419 | 21.292006 |
| 10.2555686 | 1.12E-24 | 1.24E-21 | 550.010487 | 610.227307 | 640.802875 | 697.03304 |
| -10.105227 | 5.24E-24 | 5.64E-21 | 1322.97604 | 1884.26286 | 1732.58372 | 1899.47107 |
| 10.0986525 | 5.60E-24 | 5.83E-21 | 0 | 0 | 0 | 0 |
| 10.0323424 | 1.10E-23 | 1.11E-20 | 0 | 0 | 0 | 0 |
| -10.029311 | 1.13E-23 | 1.11E-20 | 1429.53545 | 2250.39924 | 2230.7303 | 2294.49381 |
| -9.9424532 | 2.72E-23 | 2.58E-20 | 10426.0185 | 31577.5718 | 25957.6935 | 28382.8044 |
| 9.92234258 | 3.33E-23 | 3.07E-20 | 0 | 0 | 0 | 0 |
| -9.9012381 | 4.11E-23 | 3.59E-20 | 655.340364 | 950.385056 | 906.557748 | 876.334144 |
| 9.90298347 | 4.04E-23 | 3.59E-20 | 197.954594 | 248.961917 | 220.887167 | 243.177122 |
| -9.6714375 | 3.99E-22 | 3.39E-19 | 587.716124 | 720.095283 | 613.19198 | 593.934906 |
| 9.63553447 | 5.66E-22 | 4.69E-19 | 0.40984388 | 0.81183234 | 1.150454 | 0.56031595 |
| -9.5991223 | 8.06E-22 | 6.51E-19 | 1912.33154 | 4430.71029 | 4041.54489 | 4267.92658 |
| 9.57513205 | 1.02E-21 | 7.82E-19 | 0 | 0 | 0 | 0 |
| -9.5754142 | 1.01E-21 | 7.82E-19 | 53811.6817 | 13047.7693 | 11605.7799 | 13566.9301 |

| | | | | | | |
|---|---|---|---|---|---|---|
| 9.53362089 | 1.52E-21 | 1.13E-18 | 168.445835 | 113.385916 | 108.142676 | 121.028245 |
| -9.5319039 | 1.54E-21 | 1.13E-18 | 9639.52806 | 28360.5509 | 24271.1279 | 24727.3031 |
| -9.5012754 | 2.07E-21 | 1.49E-18 | 9064.92694 | 20076.8843 | 17069.2859 | 19329.7796 |
| 9.31135203 | 1.26E-20 | 8.86E-18 | 238.938982 | 153.977533 | 193.276271 | 155.207518 |
| -9.235642 | 2.57E-20 | 1.76E-17 | 2044.71112 | 9410.76046 | 7081.04434 | 7884.76603 |
| -9.2249678 | 2.84E-20 | 1.91E-17 | 7213.66213 | 32626.1886 | 29373.3914 | 30911.5103 |
| -9.1639221 | 5.00E-20 | 3.30E-17 | 1687.32725 | 6194.01012 | 5004.47488 | 5405.92827 |
| 9.12101046 | 7.44E-20 | 4.81E-17 | 0 | 0 | 0 | 0 |
| 9.10220478 | 8.85E-20 | 5.60E-17 | 74.5915861 | 64.946587 | 71.3281477 | 61.6347543 |
| 9.07998107 | 1.09E-19 | 6.74E-17 | 56.1486116 | 35.7206228 | 28.7613499 | 33.058641 |
| 9.07173126 | 1.17E-19 | 7.14E-17 | 37.705637 | 27.3316887 | 34.5136199 | 24.0935858 |
| -9.049979 | 1.43E-19 | 8.48E-17 | 477.877964 | 661.643355 | 610.891072 | 652.207764 |
| 9.04892488 | 1.44E-19 | 8.48E-17 | 0 | 0 | 0 | 0 |
| -9.0336319 | 1.66E-19 | 9.57E-17 | 5509.53128 | 9380.72266 | 8008.31026 | 8643.43382 |
| 9.02346779 | 1.82E-19 | 1.03E-16 | 9.42640924 | 11.6362635 | 13.8054479 | 12.3269509 |
| -9.0186339 | 1.90E-19 | 1.06E-16 | 5404.61124 | 11159.7179 | 9540.71498 | 9998.27779 |
| -8.9802748 | 2.70E-19 | 1.48E-16 | 914.361696 | 2150.00264 | 1840.72639 | 1961.10582 |
| -8.966446 | 3.06E-19 | 1.65E-16 | 5747.65057 | 15412.9075 | 10863.7371 | 12834.5971 |
| -8.9596698 | 3.26E-19 | 1.72E-16 | 6001.75378 | 19288.5951 | 10388.5996 | 13449.824 |
| -8.9559955 | 3.37E-19 | 1.75E-16 | 5196.00071 | 14989.6723 | 11788.7021 | 13602.7903 |
| -8.9470974 | 3.65E-19 | 1.87E-16 | 5226.32916 | 11775.6281 | 8698.58266 | 11451.177 |
| -8.9453941 | 3.71E-19 | 1.87E-16 | 3832.85997 | 10636.3567 | 9069.02885 | 9876.68923 |
| 8.90159903 | 5.50E-19 | 2.73E-16 | 72.5423667 | 49.2511618 | 48.3190678 | 35.2999048 |
| -8.8995023 | 5.61E-19 | 2.74E-16 | 1063.13502 | 3666.77606 | 3042.95082 | 3199.40407 |
| -8.8945456 | 5.87E-19 | 2.83E-16 | 3993.10892 | 13158.9904 | 13685.8007 | 13579.257 |
| -8.875743 | 6.95E-19 | 3.30E-16 | 1578.30878 | 1569.0013 | 1953.47088 | 1746.50481 |
| 8.85443223 | 8.41E-19 | 3.94E-16 | 26.2300083 | 26.7904671 | 34.5136199 | 21.292006 |
| -8.8104577 | 1.25E-18 | 5.75E-16 | 880.344654 | 2886.33457 | 2911.79906 | 2296.73507 |
| -8.7939324 | 1.44E-18 | 6.51E-16 | 4597.2188 | 11463.8844 | 17375.3067 | 14013.5019 |
| -8.7932928 | 1.45E-18 | 6.51E-16 | 4251.72041 | 9659.99298 | 8375.30509 | 9011.00109 |
| -8.7617695 | 1.92E-18 | 8.50E-16 | 4086.55333 | 9212.94398 | 8483.44776 | 8867.5602 |
| -8.7548596 | 2.04E-18 | 8.92E-16 | 1679.95006 | 4967.06085 | 4238.27252 | 4665.7509 |
| 8.73327865 | 2.47E-18 | 1.07E-15 | 25.0004767 | 27.0610779 | 26.4604419 | 22.4126379 |
| -8.718164 | 2.83E-18 | 1.20E-15 | 606.978786 | 947.949559 | 893.902754 | 940.210162 |
| 8.70990601 | 3.04E-18 | 1.28E-15 | 34.4268859 | 26.7904671 | 29.9118039 | 28.5761134 |
| -8.6800858 | 3.95E-18 | 1.64E-15 | 1160.26802 | 4114.36629 | 3649.24007 | 3814.63098 |
| -8.6706442 | 4.30E-18 | 1.76E-15 | 3091.04254 | 8275.27763 | 7702.2895 | 8599.72918 |
| -8.6623892 | 4.62E-18 | 1.86E-15 | 2547.9994 | 4178.50104 | 3939.15448 | 4138.4936 |
| -8.6595151 | 4.74E-18 | 1.89E-15 | 3775.48182 | 5732.07752 | 6710.59816 | 7255.53122 |
| -8.6455852 | 5.35E-18 | 2.11E-15 | 3148.01084 | 9439.71581 | 8684.77721 | 8933.67748 |
| -8.6267562 | 6.31E-18 | 2.46E-15 | 657.389583 | 1426.66003 | 1459.92612 | 1549.2736 |
| 8.61919994 | 6.74E-18 | 2.59E-15 | 17.6232868 | 20.0251977 | 16.1063559 | 19.6110582 |
| -8.6132246 | 7.10E-18 | 2.70E-15 | 3913.18937 | 8169.46881 | 5250.67203 | 6071.58362 |

| | | | | | | |
|---|---|---|---|---|---|---|
| -8.5913258 | 8.60E-18 | 3.23E-15 | 3427.52437 | 9530.09981 | 9639.65403 | 8905.66169 |
| -8.5554148 | 1.17E-17 | 4.36E-15 | 2333.24121 | 3846.191 | 3324.81205 | 3510.93973 |
| 8.53890857 | 1.35E-17 | 4.93E-15 | 4.91812656 | 4.32977247 | 5.75226998 | 3.92221164 |
| -8.5384813 | 1.36E-17 | 4.93E-15 | 2013.97283 | 5624.37443 | 5011.3776 | 5429.46154 |
| 8.49788104 | 1.93E-17 | 6.93E-15 | 0 | 0 | 0 | 0 |
| 8.48949999 | 2.08E-17 | 7.33E-15 | 36.4761053 | 24.8961917 | 19.5577179 | 25.2142177 |
| -8.4888035 | 2.09E-17 | 7.33E-15 | 3422.60624 | 3864.32193 | 3816.0559 | 4865.22338 |
| -8.4689694 | 2.48E-17 | 8.60E-15 | 62591.7672 | 14925.2669 | 13700.7566 | 15569.4993 |
| -8.4596911 | 2.68E-17 | 9.21E-15 | 3957.86235 | 3466.79469 | 3314.45796 | 3516.54289 |
| -8.4520479 | 2.86E-17 | 9.73E-15 | 574.60112 | 526.067355 | 650.006507 | 606.261856 |
| -8.4440555 | 3.07E-17 | 1.03E-14 | 2500.86736 | 3293.06257 | 2252.58892 | 2619.47706 |
| -8.4234456 | 3.66E-17 | 1.22E-14 | 2993.90954 | 7445.58498 | 5341.5579 | 6890.76554 |
| -8.3983774 | 4.53E-17 | 1.49E-14 | 2971.77797 | 5480.4095 | 3994.37627 | 4670.23343 |
| -8.3630615 | 6.11E-17 | 1.99E-14 | 597.552377 | 1569.27191 | 1323.02209 | 1399.66924 |
| -8.3540302 | 6.60E-17 | 2.13E-14 | 2946.36765 | 15898.3833 | 9228.94195 | 11085.851 |
| -8.3481449 | 6.93E-17 | 2.22E-14 | 2938.99046 | 15366.3625 | 8889.55802 | 10490.2352 |
| -8.3384021 | 7.53E-17 | 2.38E-14 | 1906.59373 | 2417.90731 | 2175.50851 | 2302.89855 |
| 8.33357616 | 7.84E-17 | 2.46E-14 | 60.6568942 | 44.109557 | 43.7172518 | 47.0665397 |
| -8.3317873 | 7.96E-17 | 2.47E-14 | 2584.47551 | 6329.0449 | 5792.53587 | 6253.6863 |
| 8.31270792 | 9.35E-17 | 2.88E-14 | 49.1812656 | 43.5683354 | 52.9208838 | 37.5411686 |
| 8.30388082 | 1.01E-16 | 3.07E-14 | 13.1150042 | 10.8244312 | 16.1063559 | 11.7666349 |
| -8.271776 | 1.32E-16 | 3.98E-14 | 2664.80491 | 4599.84202 | 3305.25433 | 3757.47875 |
| -8.27061 | 1.33E-16 | 3.98E-14 | 2435.70218 | 8979.40687 | 9392.30642 | 10408.9894 |
| -8.2688272 | 1.35E-16 | 4.01E-14 | 4789.84542 | 3097.14037 | 3275.34252 | 3008.89664 |
| -8.2671521 | 1.37E-16 | 4.03E-14 | 415.991538 | 546.092552 | 484.341132 | 503.724038 |
| -8.2615223 | 1.44E-16 | 4.18E-14 | 2479.96532 | 4235.87053 | 5243.76931 | 5468.68366 |
| -8.2575423 | 1.49E-16 | 4.29E-14 | 2650.87022 | 6113.90933 | 3890.83541 | 4669.67312 |
| -8.2480398 | 1.61E-16 | 4.60E-14 | 2729.56024 | 8645.47317 | 4762.87954 | 6030.12024 |
| -8.2444586 | 1.66E-16 | 4.70E-14 | 405.745441 | 322.297438 | 340.534383 | 383.256109 |
| -8.2338072 | 1.81E-16 | 5.09E-14 | 945.919675 | 621.86357 | 571.775636 | 667.336295 |
| -8.2247368 | 1.96E-16 | 5.44E-14 | 812.31057 | 1336.00542 | 1035.4086 | 1174.42223 |
| -8.2157509 | 2.11E-16 | 5.71E-14 | 2251.27243 | 4501.88092 | 2419.40475 | 3256.55629 |
| 8.21503617 | 2.12E-16 | 5.71E-14 | 22.9512573 | 22.7313054 | 27.6108959 | 22.4126379 |
| -8.2160058 | 2.10E-16 | 5.71E-14 | 1188.54725 | 5342.12739 | 2804.80684 | 3483.48425 |
| -8.2174529 | 2.08E-16 | 5.71E-14 | 2277.50244 | 10261.0195 | 9777.70851 | 10006.1222 |
| 8.20819806 | 2.25E-16 | 5.99E-14 | 0 | 0 | 0 | 0 |
| -8.198215 | 2.44E-16 | 6.46E-14 | 3267.27541 | 3879.47613 | 2509.14016 | 2838.5606 |
| -8.1926757 | 2.55E-16 | 6.71E-14 | 1809.46073 | 2213.32556 | 2850.825 | 3009.45696 |
| -8.1905009 | 2.60E-16 | 6.77E-14 | 2215.20617 | 7558.15906 | 7621.75772 | 8237.20476 |
| -8.1887486 | 2.64E-16 | 6.82E-14 | 2275.86307 | 3387.50573 | 3710.21414 | 3781.57234 |
| -8.1777536 | 2.89E-16 | 7.41E-14 | 2251.27243 | 4613.91378 | 3758.5332 | 4090.30642 |
| -8.1704853 | 3.07E-16 | 7.81E-14 | 2420.12811 | 7909.41185 | 4698.45412 | 5644.62287 |
| -8.1678979 | 3.14E-16 | 7.92E-14 | 1958.6439 | 5630.59848 | 5954.74988 | 6223.42924 |

| | | | | | | |
|---|---|---|---|---|---|---|
| -8.1650943 | 3.21E-16 | 8.04E-14 | 1719.29508 | 3268.16638 | 3458.26471 | 3533.35237 |
| -8.1411901 | 3.91E-16 | 9.72E-14 | 2129.5488 | 3684.36576 | 3781.54228 | 3838.16425 |
| -8.1348388 | 4.12E-16 | 1.02E-13 | 1199.20319 | 4952.98909 | 7035.02618 | 6124.81363 |
| 8.13076832 | 4.27E-16 | 1.04E-13 | 0 | 0 | 0 | 0 |
| -8.1222759 | 4.58E-16 | 1.11E-13 | 2057.41628 | 5644.12902 | 5712.00409 | 6181.40555 |
| -8.1162992 | 4.81E-16 | 1.16E-13 | 15899.0736 | 4829.86119 | 4203.7589 | 5115.68461 |
| -8.108763 | 5.11E-16 | 1.22E-13 | 2241.43618 | 3352.32633 | 2596.57467 | 2965.192 |
| -8.104292 | 5.31E-16 | 1.26E-13 | 586.896436 | 1178.23933 | 1058.41768 | 1155.9318 |
| -8.0999836 | 5.50E-16 | 1.30E-13 | 2009.0547 | 5174.88993 | 5110.31665 | 5547.12789 |
| -8.0934996 | 5.80E-16 | 1.36E-13 | 415.991538 | 558.270037 | 537.262016 | 623.631651 |
| -8.0903658 | 5.95E-16 | 1.38E-13 | 833.212608 | 1276.47105 | 1130.89628 | 1197.9555 |
| -8.0822518 | 6.36E-16 | 1.47E-13 | 1971.34906 | 5889.30239 | 5271.38021 | 5598.67696 |
| -8.0749203 | 6.75E-16 | 1.55E-13 | 1974.62781 | 6520.09611 | 5325.45154 | 6061.49793 |
| -8.06671 | 7.22E-16 | 1.64E-13 | 1932.41389 | 6381.00217 | 6557.58777 | 7116.57286 |
| -8.0589815 | 7.69E-16 | 1.74E-13 | 414.762006 | 610.768529 | 509.65112 | 568.720688 |
| -8.0578231 | 7.77E-16 | 1.74E-13 | 1924.21702 | 7691.02895 | 6566.79141 | 6631.89957 |
| -8.0547172 | 7.97E-16 | 1.77E-13 | 2221.76367 | 2566.20202 | 2198.51759 | 2511.33608 |
| -8.0507181 | 8.23E-16 | 1.82E-13 | 2350.45465 | 2213.86678 | 2478.07791 | 3045.31718 |
| -8.0460401 | 8.55E-16 | 1.87E-13 | 1908.64295 | 5453.34842 | 4655.88732 | 5296.10635 |
| 8.04677261 | 8.50E-16 | 1.87E-13 | 12.2953164 | 6.22404792 | 9.20363196 | 7.28410733 |
| -8.0317118 | 9.61E-16 | 2.08E-13 | 1987.33297 | 3172.64077 | 2618.43329 | 2974.71737 |
| -8.0274221 | 9.95E-16 | 2.14E-13 | 1892.24919 | 5586.75954 | 4403.93789 | 5015.94837 |
| 8.02461882 | 1.02E-15 | 2.18E-13 | 0 | 0 | 0 | 0 |
| -8.0195881 | 1.06E-15 | 2.25E-13 | 1553.71815 | 3316.06449 | 2533.2997 | 2878.90334 |
| -8.0180667 | 1.07E-15 | 2.27E-13 | 1847.16637 | 4247.7774 | 3963.31401 | 4193.96488 |
| -8.0119099 | 1.13E-15 | 2.37E-13 | 1836.10058 | 5480.13889 | 4668.54231 | 5169.47494 |
| -8.009678 | 1.15E-15 | 2.40E-13 | 1825.85449 | 4329.77247 | 4128.97939 | 4193.40456 |
| 8.00863275 | 1.16E-15 | 2.40E-13 | 263.939459 | 257.350851 | 210.533081 | 240.935858 |
| -7.9907704 | 1.34E-15 | 2.76E-13 | 1784.8701 | 10065.6385 | 7782.82128 | 8717.39553 |
| -7.9824016 | 1.44E-15 | 2.93E-13 | 1763.14837 | 5288.81707 | 5432.44377 | 5887.79999 |
| -7.9689667 | 1.60E-15 | 3.25E-13 | 2219.30461 | 4010.45175 | 2267.54482 | 2913.08262 |
| -7.9637787 | 1.67E-15 | 3.37E-13 | 1730.36086 | 4652.88174 | 4438.45151 | 4731.86819 |
| -7.9553601 | 1.79E-15 | 3.58E-13 | 1429.53545 | 2025.25107 | 1681.96374 | 1534.70538 |
| -7.9538377 | 1.81E-15 | 3.60E-13 | 1883.23263 | 3917.36164 | 2672.50463 | 3259.91819 |
| -7.9433935 | 1.97E-15 | 3.90E-13 | 2319.30652 | 2576.21462 | 1907.45272 | 2109.02923 |
| 7.93915835 | 2.04E-15 | 4.01E-13 | 32.7875104 | 17.3190899 | 26.4604419 | 25.2142177 |
| -7.9317173 | 2.16E-15 | 4.23E-13 | 1742.24633 | 2789.45591 | 2603.47739 | 2839.68123 |
| 7.92443038 | 2.29E-15 | 4.46E-13 | 4.0984388 | 2.97671857 | 3.45136199 | 3.92221164 |
| 7.92160187 | 2.34E-15 | 4.53E-13 | 9.42640924 | 4.05916169 | 4.60181598 | 3.92221164 |
| -7.9175091 | 2.42E-15 | 4.66E-13 | 1647.98224 | 4777.3627 | 4336.06111 | 4814.79495 |
| 7.91448258 | 2.48E-15 | 4.72E-13 | 0 | 0 | 0 | 0 |
| -7.9141933 | 2.49E-15 | 4.72E-13 | 1652.08068 | 4883.71273 | 4158.89119 | 4444.98642 |
| -7.9135849 | 2.50E-15 | 4.72E-13 | 1667.65475 | 3895.17156 | 3348.97158 | 3735.62643 |

| | | | | | | |
|---|---|---|---|---|---|---|
| -7.9130199 | 2.51E-15 | 4.72E-13 | 1681.58944 | 3528.22334 | 3016.49038 | 3389.91149 |
| -7.9019673 | 2.75E-15 | 5.12E-13 | 1600.8502 | 5661.44811 | 5851.20902 | 6046.92972 |
| -7.8990172 | 2.81E-15 | 5.22E-13 | 1604.94863 | 4712.1455 | 4882.52676 | 4721.7825 |
| -7.8981141 | 2.83E-15 | 5.22E-13 | 1298.79526 | 2494.49016 | 1853.38139 | 2124.15776 |
| -7.8955926 | 2.89E-15 | 5.30E-13 | 1613.55536 | 4185.80753 | 4820.40224 | 4485.88948 |
| -7.8908496 | 3.00E-15 | 5.47E-13 | 474.599213 | 672.197175 | 704.077845 | 804.613702 |
| -7.8893872 | 3.04E-15 | 5.51E-13 | 1591.42379 | 4441.80533 | 4714.56047 | 4747.55703 |
| 7.88290589 | 3.20E-15 | 5.77E-13 | 6.96734596 | 6.22404792 | 8.05317797 | 10.0856871 |
| -7.8753028 | 3.40E-15 | 6.08E-13 | 1606.99785 | 3767.44327 | 3172.95212 | 3438.09866 |
| -7.8748689 | 3.41E-15 | 6.08E-13 | 1245.10571 | 3716.02722 | 3400.74201 | 4107.1159 |
| -7.8742395 | 3.43E-15 | 6.08E-13 | 1850.85496 | 2538.59972 | 1890.19591 | 2254.71138 |
| -7.8688878 | 3.58E-15 | 6.31E-13 | 1837.73996 | 4671.55388 | 2619.58375 | 3416.80665 |
| -7.8673776 | 3.62E-15 | 6.35E-13 | 1741.0168 | 2469.59397 | 1996.03768 | 2303.45886 |
| -7.8612215 | 3.80E-15 | 6.64E-13 | 1695.52413 | 2930.71474 | 2222.67712 | 2528.70588 |
| -7.8532879 | 4.05E-15 | 7.03E-13 | 3373.83482 | 2986.46056 | 1722.22963 | 2259.75422 |
| 7.84919046 | 4.19E-15 | 7.19E-13 | 74.5915861 | 73.3355211 | 87.4345036 | 114.864769 |
| -7.8493717 | 4.18E-15 | 7.19E-13 | 1727.08211 | 2143.23737 | 1884.44364 | 2139.84661 |
| -7.8483198 | 4.22E-15 | 7.20E-13 | 1563.96425 | 3109.85907 | 2831.26728 | 2941.65873 |
| 7.8470449 | 4.26E-15 | 7.24E-13 | 0 | 0 | 0 | 0 |
| -7.8408575 | 4.47E-15 | 7.56E-13 | 1975.85735 | 5630.32787 | 2559.76014 | 3752.43591 |
| -7.824631 | 5.09E-15 | 8.56E-13 | 1547.16065 | 3424.85002 | 2792.15185 | 2908.03977 |
| -7.8127861 | 5.59E-15 | 9.36E-13 | 1534.04564 | 3339.87824 | 2640.29192 | 3024.02517 |
| -7.8058996 | 5.91E-15 | 9.83E-13 | 1554.94768 | 3970.94257 | 2766.84186 | 3537.8349 |
| -7.8014694 | 6.12E-15 | 1.01E-12 | 1494.29079 | 2575.6734 | 2707.01825 | 2709.68793 |
| -7.7990999 | 6.24E-15 | 1.03E-12 | 1565.60362 | 2966.97658 | 2238.78348 | 2544.39472 |
| -7.7910704 | 6.64E-15 | 1.09E-12 | 1406.17435 | 5642.50536 | 5611.91459 | 5584.66906 |
| -7.7896867 | 6.72E-15 | 1.09E-12 | 688.127874 | 888.685799 | 960.629086 | 1015.2925 |
| -7.7896161 | 6.72E-15 | 1.09E-12 | 1459.45406 | 3405.63666 | 2976.22449 | 3126.56299 |
| -7.7787364 | 7.33E-15 | 1.18E-12 | 1431.17483 | 3242.99958 | 3097.02216 | 3353.49095 |
| -7.775909 | 7.49E-15 | 1.20E-12 | 1416.42045 | 3492.50272 | 3678.00142 | 3463.87319 |
| -7.7747387 | 7.56E-15 | 1.21E-12 | 1439.78155 | 3693.0253 | 4908.9872 | 4391.75641 |
| -7.7735274 | 7.63E-15 | 1.21E-12 | 1404.53498 | 4956.50703 | 4238.27252 | 4483.64822 |
| -7.7720989 | 7.72E-15 | 1.22E-12 | 1406.17435 | 5055.00935 | 4219.86526 | 4484.20854 |
| -7.769992 | 7.85E-15 | 1.23E-12 | 1399.61685 | 4265.63771 | 3871.27769 | 4292.58048 |
| -7.7698988 | 7.85E-15 | 1.23E-12 | 1583.63675 | 6527.67321 | 3575.61102 | 4411.36746 |
| 7.76699959 | 8.04E-15 | 1.25E-12 | 0 | 0 | 0 | 0 |
| 7.76466782 | 8.19E-15 | 1.27E-12 | 0.81968776 | 1.3530539 | 2.30090799 | 1.1206319 |
| -7.7639613 | 8.23E-15 | 1.27E-12 | 883.213561 | 4826.88447 | 3951.80947 | 4233.74731 |
| -7.7600799 | 8.49E-15 | 1.30E-12 | 1562.73471 | 5028.21889 | 2860.02863 | 3927.25448 |
| -7.7528448 | 8.99E-15 | 1.37E-12 | 1707.81945 | 3056.00753 | 1818.86777 | 2338.19845 |
| 7.75227953 | 9.03E-15 | 1.37E-12 | 18.8528185 | 16.7778683 | 20.7081719 | 26.8951655 |
| -7.7504158 | 9.16E-15 | 1.39E-12 | 475.828745 | 713.871235 | 720.184201 | 774.356641 |
| -7.748789 | 9.28E-15 | 1.40E-12 | 2221.76367 | 2762.39483 | 1545.05972 | 1971.19151 |

| | | | | | | |
|---|---|---|---|---|---|---|
| -7.7453041 | 9.54E-15 | 1.43E-12 | 1531.17674 | 2295.59124 | 1806.21277 | 2073.16901 |
| -7.7451222 | 9.55E-15 | 1.43E-12 | 1400.02669 | 2762.66544 | 2572.41513 | 2677.1896 |
| -7.7366175 | 1.02E-14 | 1.52E-12 | 1345.9273 | 6357.72965 | 4944.65127 | 5472.04555 |
| 7.73481977 | 1.04E-14 | 1.53E-12 | 0 | 0 | 0 | 0 |
| 7.72809628 | 1.09E-14 | 1.60E-12 | 151.642236 | 86.5954493 | 92.0363196 | 79.0045488 |
| -7.7281401 | 1.09E-14 | 1.60E-12 | 1327.89417 | 4482.93817 | 4116.3244 | 4285.29638 |
| -7.7261371 | 1.11E-14 | 1.62E-12 | 765.178524 | 1803.89145 | 1234.43714 | 1466.90715 |
| -7.7254039 | 1.11E-14 | 1.62E-12 | 1347.56668 | 3440.81606 | 3141.88986 | 3552.96343 |
| -7.7178939 | 1.18E-14 | 1.71E-12 | 746.325705 | 2020.10947 | 1706.12328 | 1858.00769 |
| -7.7144181 | 1.22E-14 | 1.75E-12 | 1324.61542 | 3299.28662 | 3269.59025 | 3443.70182 |
| -7.7113516 | 1.24E-14 | 1.78E-12 | 360.662614 | 460.579546 | 484.341132 | 568.720688 |
| -7.7108953 | 1.25E-14 | 1.78E-12 | 368.039804 | 421.070372 | 479.739316 | 480.751084 |
| -7.7069259 | 1.29E-14 | 1.83E-12 | 1370.51793 | 2518.57452 | 3279.94434 | 3168.58669 |
| -7.7066808 | 1.29E-14 | 1.83E-12 | 271.726492 | 293.071474 | 271.507143 | 307.05314 |
| -7.7051174 | 1.31E-14 | 1.84E-12 | 1260.67977 | 7009.08979 | 6187.14159 | 6378.63676 |
| -7.6923641 | 1.44E-14 | 2.03E-12 | 1313.13979 | 6652.42478 | 4582.25826 | 5139.21788 |
| -7.6908065 | 1.46E-14 | 2.04E-12 | 1306.58229 | 3634.30276 | 3007.28674 | 3442.02087 |
| -7.6857377 | 1.52E-14 | 2.12E-12 | 1170.92396 | 3669.75278 | 2100.729 | 2562.88515 |
| -7.68341 | 1.55E-14 | 2.15E-12 | 1205.35085 | 3670.294 | 3052.15445 | 3515.98258 |
| -7.6797834 | 1.59E-14 | 2.19E-12 | 1281.17197 | 3134.75527 | 2999.23357 | 2997.13001 |
| -7.6797622 | 1.59E-14 | 2.19E-12 | 1286.49994 | 3645.66842 | 3062.50854 | 3343.40527 |
| -7.6778665 | 1.62E-14 | 2.21E-12 | 341.809796 | 370.195546 | 394.60572 | 426.400437 |
| -7.6766296 | 1.63E-14 | 2.23E-12 | 1269.2865 | 3828.06008 | 3365.07794 | 3531.67142 |
| 7.67490621 | 1.66E-14 | 2.25E-12 | 0.40984388 | 0.27061078 | 1.150454 | 0.56031595 |
| -7.6702195 | 1.72E-14 | 2.32E-12 | 1417.24014 | 2223.87938 | 1821.16867 | 2603.2279 |
| -7.663073 | 1.82E-14 | 2.44E-12 | 503.698128 | 1179.05116 | 716.732839 | 965.984695 |
| -7.6587013 | 1.88E-14 | 2.52E-12 | 1399.20701 | 1920.5247 | 1637.09604 | 1743.70323 |
| -7.657481 | 1.90E-14 | 2.53E-12 | 1378.71481 | 3233.25759 | 2116.83535 | 2573.53115 |
| -7.6555283 | 1.93E-14 | 2.56E-12 | 1194.69491 | 3952.27043 | 3068.26081 | 3428.01297 |
| 7.65082739 | 2.00E-14 | 2.64E-12 | 0 | 0 | 0 | 0 |
| 7.6375137 | 2.21E-14 | 2.92E-12 | 33.6071982 | 16.2366467 | 19.5577179 | 14.0078987 |
| 7.63554983 | 2.25E-14 | 2.95E-12 | 0 | 0 | 0 | 0 |
| -7.6311595 | 2.33E-14 | 3.04E-12 | 1203.30163 | 3856.74482 | 3371.98066 | 3606.75376 |
| -7.622027 | 2.50E-14 | 3.23E-12 | 1227.07258 | 2818.95249 | 3728.6214 | 3397.1956 |
| -7.622127 | 2.50E-14 | 3.23E-12 | 1190.59647 | 5141.6048 | 5111.4671 | 5176.19873 |
| -7.6225681 | 2.49E-14 | 3.23E-12 | 381.154808 | 912.499547 | 867.442313 | 957.01964 |
| -7.6151455 | 2.63E-14 | 3.39E-12 | 1191.41616 | 2898.24144 | 3176.40348 | 3111.43446 |
| -7.6141343 | 2.65E-14 | 3.40E-12 | 1317.23823 | 1686.17576 | 1607.18423 | 1810.38083 |
| 7.61238875 | 2.69E-14 | 3.43E-12 | 43.4434513 | 45.4626109 | 40.2658898 | 41.4633802 |
| -7.6101445 | 2.74E-14 | 3.48E-12 | 1216.00679 | 2937.2094 | 2398.69658 | 2680.5515 |
| -7.6077509 | 2.79E-14 | 3.53E-12 | 2078.72816 | 7304.86737 | 6380.41786 | 7666.80312 |
| -7.6044663 | 2.86E-14 | 3.61E-12 | 1271.33572 | 2799.73912 | 1987.9845 | 2425.04743 |
| -7.6016713 | 2.92E-14 | 3.67E-12 | 1176.66178 | 3572.3329 | 2988.87948 | 3422.97013 |

| | | | | | | |
|---|---|---|---|---|---|---|
| -7.5997248 | 2.97E-14 | 3.71E-12 | 1509.86485 | 6080.89482 | 6692.19089 | 7013.47473 |
| 7.59791395 | 3.01E-14 | 3.75E-12 | 0 | 0 | 0 | 0 |
| -7.5946755 | 3.09E-14 | 3.83E-12 | 1157.80896 | 3408.07215 | 3145.34122 | 3566.97133 |
| -7.5929665 | 3.13E-14 | 3.85E-12 | 1120.92301 | 5414.92169 | 4830.75633 | 4836.64727 |
| -7.5932647 | 3.12E-14 | 3.85E-12 | 1531.17674 | 1593.35627 | 1317.26982 | 1524.6197 |
| -7.5924373 | 3.14E-14 | 3.85E-12 | 1300.43463 | 3492.77333 | 2129.49035 | 2661.50076 |
| -7.5849297 | 3.33E-14 | 4.05E-12 | 1147.15302 | 3166.14612 | 2980.8263 | 3204.44691 |
| -7.5853991 | 3.31E-14 | 4.05E-12 | 1135.67739 | 5275.55714 | 4249.77706 | 4261.76311 |
| -7.5835335 | 3.36E-14 | 4.07E-12 | 514.354069 | 1484.84135 | 1352.9339 | 1524.6197 |
| -7.5838435 | 3.35E-14 | 4.07E-12 | 1237.72852 | 2698.26008 | 2036.30357 | 2110.71018 |
| -7.580795 | 3.43E-14 | 4.14E-12 | 1149.61208 | 5208.98689 | 4378.62791 | 4622.04626 |
| -7.5781257 | 3.51E-14 | 4.21E-12 | 1312.3201 | 3546.89548 | 2098.42809 | 2465.39017 |
| -7.5743674 | 3.61E-14 | 4.32E-12 | 1339.3698 | 2581.89744 | 1650.90148 | 1952.70108 |
| -7.5730747 | 3.64E-14 | 4.34E-12 | 1105.75879 | 4480.23206 | 4062.25306 | 4271.84879 |
| -7.5712278 | 3.70E-14 | 4.39E-12 | 1163.54678 | 3042.20638 | 2434.36065 | 2696.24034 |
| -7.5695557 | 3.75E-14 | 4.41E-12 | 1081.98784 | 3159.11024 | 2420.55521 | 2683.91339 |
| 7.56957676 | 3.74E-14 | 4.41E-12 | 37.705637 | 23.8137486 | 31.0622579 | 32.498325 |
| -7.5643846 | 3.90E-14 | 4.55E-12 | 585.666904 | 1351.15962 | 1285.05711 | 1335.23291 |
| -7.5651251 | 3.87E-14 | 4.55E-12 | 1383.22309 | 2516.13902 | 1577.27243 | 1784.6063 |
| -7.5642205 | 3.90E-14 | 4.55E-12 | 1159.03849 | 3014.33347 | 2391.79386 | 2606.58979 |
| -7.5632956 | 3.93E-14 | 4.56E-12 | 1159.03849 | 3779.89136 | 2721.97415 | 3224.05797 |
| -7.55533 | 4.18E-14 | 4.83E-12 | 1174.20272 | 1938.65562 | 1828.0714 | 1915.15991 |
| -7.5538182 | 4.23E-14 | 4.87E-12 | 1190.59647 | 1699.16508 | 1872.9391 | 1931.40907 |
| -7.5527252 | 4.26E-14 | 4.90E-12 | 1087.31581 | 3867.83987 | 3669.94825 | 4005.1384 |
| -7.5516786 | 4.30E-14 | 4.92E-12 | 597.962221 | 1529.49212 | 926.115466 | 1073.00504 |
| -7.5485777 | 4.40E-14 | 5.02E-12 | 1207.80991 | 1959.22204 | 1604.88332 | 1833.9141 |
| -7.5477739 | 4.43E-14 | 5.03E-12 | 1120.92301 | 2385.70463 | 2455.06883 | 2470.99333 |
| -7.544021 | 4.56E-14 | 5.14E-12 | 1131.9888 | 2585.41538 | 2195.06622 | 2374.61899 |
| -7.5441652 | 4.55E-14 | 5.14E-12 | 1154.94005 | 2485.01878 | 1989.13496 | 2427.28869 |
| 7.54202643 | 4.63E-14 | 5.19E-12 | 11.4756286 | 4.60038325 | 3.45136199 | 2.80157974 |
| -7.5423673 | 4.62E-14 | 5.19E-12 | 1194.69491 | 2083.97361 | 1657.80421 | 1873.13622 |
| -7.5404315 | 4.68E-14 | 5.20E-12 | 1124.20176 | 2516.13902 | 2221.52667 | 2296.17476 |
| -7.5408987 | 4.67E-14 | 5.20E-12 | 1536.50471 | 1981.68274 | 1248.24258 | 1606.98614 |
| -7.5404371 | 4.68E-14 | 5.20E-12 | 1259.45024 | 2154.33241 | 1535.85608 | 1752.10797 |
| -7.5395814 | 4.71E-14 | 5.21E-12 | 1223.38398 | 1659.3853 | 1463.37748 | 1669.18121 |
| -7.5376868 | 4.78E-14 | 5.27E-12 | 964.362649 | 1608.23986 | 1359.83662 | 1548.71328 |
| -7.5352762 | 4.87E-14 | 5.35E-12 | 1246.74508 | 1898.064 | 1455.3243 | 1654.05268 |
| -7.5333844 | 4.94E-14 | 5.41E-12 | 347.54761 | 561.246756 | 594.784716 | 622.511019 |
| -7.5262686 | 5.22E-14 | 5.70E-12 | 1110.26707 | 2397.07028 | 2109.93263 | 2386.94594 |
| -7.5249898 | 5.27E-14 | 5.73E-12 | 1211.49851 | 1554.38832 | 1453.0234 | 1483.71663 |
| 7.5244699 | 5.29E-14 | 5.74E-12 | 0 | 0 | 0 | 0 |
| -7.5224879 | 5.37E-14 | 5.80E-12 | 472.14015 | 874.343427 | 732.839195 | 854.481822 |
| 7.52104625 | 5.43E-14 | 5.85E-12 | 0 | 0 | 0 | 0 |

| | | | | | | |
|---|---|---|---|---|---|---|
| -7.5035073 | 6.21E-14 | 6.66E-12 | 1210.26898 | 1357.38367 | 1686.56556 | 1678.14627 |
| -7.499256 | 6.42E-14 | 6.86E-12 | 1222.56429 | 2367.3031 | 1550.81199 | 1802.53641 |
| -7.4984904 | 6.46E-14 | 6.88E-12 | 1176.25194 | 1449.93255 | 1742.9378 | 1882.10127 |
| -7.4933924 | 6.71E-14 | 7.13E-12 | 961.083898 | 2091.55071 | 2024.79903 | 2311.8636 |
| -7.4921045 | 6.78E-14 | 7.17E-12 | 1122.97223 | 1636.65399 | 1642.84831 | 1573.9275 |
| -7.4906187 | 6.85E-14 | 7.23E-12 | 1046.33143 | 2450.65122 | 2498.78608 | 2642.45001 |
| -7.4871292 | 7.04E-14 | 7.40E-12 | 1160.26802 | 2340.24202 | 1607.18423 | 1923.00434 |
| -7.4830323 | 7.26E-14 | 7.61E-12 | 840.999642 | 1207.19469 | 1166.56035 | 1199.63645 |
| -7.4825008 | 7.29E-14 | 7.62E-12 | 1103.29972 | 2000.35488 | 1648.60058 | 1821.02683 |
| -7.4809865 | 7.38E-14 | 7.68E-12 | 1180.76022 | 2502.33787 | 2413.65248 | 2654.21665 |
| -7.4794548 | 7.46E-14 | 7.75E-12 | 1357.81277 | 1466.43981 | 1205.67579 | 1290.96795 |
| -7.475881 | 7.67E-14 | 7.94E-12 | 1020.9211 | 4018.57007 | 3171.80167 | 3261.03882 |
| -7.4740998 | 7.77E-14 | 8.00E-12 | 1018.87189 | 2915.01931 | 2628.78738 | 2925.40957 |
| -7.4739908 | 7.78E-14 | 8.00E-12 | 1003.70766 | 3115.81251 | 3122.33214 | 3208.36912 |
| -7.4733473 | 7.82E-14 | 8.01E-12 | 1290.18853 | 1426.11881 | 2480.37881 | 2622.27864 |
| -7.47169 | 7.92E-14 | 8.09E-12 | 1013.95376 | 2972.93002 | 2719.67325 | 2763.47826 |
| -7.4680541 | 8.14E-14 | 8.29E-12 | 1184.85866 | 2143.50798 | 1454.17385 | 1657.41458 |
| -7.4667464 | 8.22E-14 | 8.35E-12 | 1000.42891 | 3107.15297 | 2969.32176 | 3357.97348 |
| -7.463285 | 8.44E-14 | 8.54E-12 | 1245.92539 | 4466.97213 | 3967.91583 | 4448.90863 |
| -7.4622458 | 8.51E-14 | 8.58E-12 | 1141.41521 | 2745.34635 | 1714.17645 | 2113.51176 |
| -7.4602965 | 8.63E-14 | 8.68E-12 | 467.222023 | 1063.50036 | 609.740618 | 792.286751 |
| -7.4587451 | 8.74E-14 | 8.76E-12 | 1227.89226 | 1872.35598 | 1282.7562 | 1513.41338 |
| -7.4575643 | 8.81E-14 | 8.81E-12 | 1056.98737 | 1922.68959 | 1690.01692 | 1870.33464 |
| -7.4548284 | 9.00E-14 | 8.97E-12 | 1056.57752 | 2018.4858 | 1726.83145 | 1793.57135 |
| -7.452403 | 9.17E-14 | 9.10E-12 | 992.641877 | 2631.96044 | 2708.16871 | 2816.14796 |
| -7.4520127 | 9.19E-14 | 9.10E-12 | 1045.51174 | 2194.38281 | 1797.00914 | 2014.89615 |
| 7.45039744 | 9.31E-14 | 9.19E-12 | 55.3289238 | 76.8534613 | 70.1776937 | 99.7362389 |
| -7.4462928 | 9.60E-14 | 9.42E-12 | 780.342747 | 893.286182 | 803.016889 | 901.548361 |
| -7.4462927 | 9.60E-14 | 9.42E-12 | 1030.34751 | 2833.29486 | 2142.14534 | 2325.31119 |
| -7.4456092 | 9.65E-14 | 9.44E-12 | 2104.13848 | 1475.91119 | 1040.01041 | 1286.48542 |
| -7.443905 | 9.78E-14 | 9.54E-12 | 957.395304 | 3489.79661 | 3763.13502 | 3945.18459 |
| -7.4391494 | 1.01E-13 | 9.85E-12 | 1155.75974 | 1493.7715 | 1234.43714 | 1376.69629 |
| -7.437911 | 1.02E-13 | 9.92E-12 | 444.68061 | 935.230853 | 893.902754 | 979.432278 |
| -7.4329274 | 1.06E-13 | 1.02E-11 | 979.117029 | 2613.28829 | 2501.08699 | 2394.23005 |
| -7.4332636 | 1.06E-13 | 1.02E-11 | 901.656536 | 2036.0755 | 1992.58632 | 2044.03258 |
| -7.4317046 | 1.07E-13 | 1.03E-11 | 261.890239 | 293.883306 | 357.791193 | 323.862618 |
| -7.4305031 | 1.08E-13 | 1.04E-11 | 979.936717 | 2329.95881 | 2356.12978 | 2493.40597 |
| -7.4235916 | 1.14E-13 | 1.09E-11 | 907.804194 | 1954.89227 | 1221.78214 | 1514.53401 |
| -7.4214729 | 1.16E-13 | 1.10E-11 | 943.050768 | 3129.61366 | 3069.41126 | 3219.01512 |
| 7.42063697 | 1.17E-13 | 1.11E-11 | 0 | 0 | 0 | 0 |
| 7.41815069 | 1.19E-13 | 1.12E-11 | 0 | 0 | 0 | 0 |
| -7.4180145 | 1.19E-13 | 1.12E-11 | 1137.31677 | 1674.81011 | 1252.8444 | 1404.15177 |
| -7.4173991 | 1.19E-13 | 1.12E-11 | 941.001548 | 3022.45179 | 2816.31138 | 3129.36457 |

| | | | | | | |
|---|---|---|---|---|---|---|
| 7.41536981 | 1.21E-13 | 1.14E-11 | 9.83625312 | 13.8011497 | 12.6549939 | 19.0507423 |
| -7.4139527 | 1.23E-13 | 1.15E-11 | 10214.1292 | 17280.3925 | 8780.26489 | 12454.7029 |
| -7.4135284 | 1.23E-13 | 1.15E-11 | 885.262781 | 4886.41884 | 2616.13239 | 3416.24634 |
| -7.4130827 | 1.23E-13 | 1.15E-11 | 731.161482 | 894.368625 | 860.539589 | 940.770478 |
| -7.4113185 | 1.25E-13 | 1.16E-11 | 937.722797 | 4926.46923 | 3589.41647 | 3888.59268 |
| -7.4093311 | 1.27E-13 | 1.17E-11 | 920.509354 | 3556.36686 | 3452.51244 | 3928.93543 |
| -7.4087928 | 1.27E-13 | 1.18E-11 | 996.330472 | 1870.19109 | 1676.21147 | 1850.72358 |
| -7.4036527 | 1.32E-13 | 1.22E-11 | 521.731259 | 2302.0859 | 1906.30227 | 2013.2152 |
| 7.39984256 | 1.36E-13 | 1.25E-11 | 13.524848 | 15.4248144 | 6.90272397 | 12.8872668 |
| -7.397578 | 1.39E-13 | 1.27E-11 | 931.165295 | 2610.58219 | 2631.08829 | 2706.88635 |
| -7.3954559 | 1.41E-13 | 1.29E-11 | 1026.24908 | 1712.69562 | 1421.96114 | 1515.09433 |
| -7.3908912 | 1.46E-13 | 1.33E-11 | 278.283994 | 434.330301 | 372.747095 | 407.349695 |
| -7.3891592 | 1.48E-13 | 1.34E-11 | 1017.64235 | 1440.19057 | 1648.60058 | 1436.08978 |
| -7.380342 | 1.58E-13 | 1.43E-11 | 264.349303 | 313.367282 | 384.251634 | 377.092633 |
| -7.3779486 | 1.61E-13 | 1.44E-11 | 984.445 | 2637.91388 | 1818.86777 | 2157.77672 |
| -7.3778414 | 1.61E-13 | 1.44E-11 | 872.967464 | 3913.8437 | 3996.67718 | 4439.38326 |
| -7.3789154 | 1.60E-13 | 1.44E-11 | 987.723751 | 1912.9476 | 1546.21017 | 1663.01774 |
| -7.3783415 | 1.60E-13 | 1.44E-11 | 1026.65892 | 1320.85121 | 1428.86386 | 1612.02898 |
| -7.3769689 | 1.62E-13 | 1.44E-11 | 962.31343 | 2288.01414 | 1778.60188 | 2046.27384 |
| -7.3759394 | 1.63E-13 | 1.45E-11 | 465.172804 | 1702.1418 | 1347.18163 | 1465.22621 |
| 7.37517127 | 1.64E-13 | 1.45E-11 | 27.45954 | 14.8835929 | 13.8054479 | 14.0078987 |
| -7.3755067 | 1.64E-13 | 1.45E-11 | 834.44214 | 1409.61155 | 1470.28021 | 1573.36718 |
| -7.3737743 | 1.66E-13 | 1.46E-11 | 964.362649 | 2569.71996 | 1870.6382 | 2144.88945 |
| -7.3727172 | 1.67E-13 | 1.47E-11 | 891.820283 | 3242.45836 | 3034.89764 | 2991.52685 |
| -7.3686135 | 1.72E-13 | 1.51E-11 | 967.641401 | 1999.54305 | 1618.68877 | 1716.24775 |
| -7.3656326 | 1.76E-13 | 1.54E-11 | 961.083898 | 1939.46745 | 1655.5033 | 1641.72573 |
| 7.36183946 | 1.81E-13 | 1.58E-11 | 13.524848 | 11.9068743 | 16.1063559 | 15.6888466 |
| -7.3607915 | 1.83E-13 | 1.59E-11 | 874.60684 | 3287.37975 | 3083.21671 | 3254.31503 |
| -7.359689 | 1.84E-13 | 1.60E-11 | 1251.25337 | 1428.82491 | 1205.67579 | 1043.86861 |
| 7.35932824 | 1.85E-13 | 1.60E-11 | 0 | 0 | 0 | 0 |
| -7.3540527 | 1.92E-13 | 1.66E-11 | 977.477654 | 1815.52772 | 1433.46568 | 1556.55771 |
| -7.3542698 | 1.92E-13 | 1.66E-11 | 820.097604 | 1635.57155 | 1738.33599 | 1949.8995 |
| -7.3521797 | 1.95E-13 | 1.67E-11 | 933.624358 | 1701.05936 | 1693.46828 | 1655.73363 |
| -7.3507665 | 1.97E-13 | 1.69E-11 | 885.672625 | 2810.02233 | 2458.52019 | 2736.58309 |
| -7.3480311 | 2.01E-13 | 1.72E-11 | 952.477177 | 1605.53375 | 1446.12067 | 1541.42917 |
| -7.3471416 | 2.02E-13 | 1.73E-11 | 937.722797 | 2316.96949 | 1730.28281 | 2036.74847 |
| -7.3441292 | 2.07E-13 | 1.76E-11 | 1051.6594 | 3221.89194 | 1693.46828 | 2195.8782 |
| -7.3437233 | 2.08E-13 | 1.76E-11 | 2130.36849 | 1522.99747 | 1092.9313 | 1280.88226 |
| -7.3423597 | 2.10E-13 | 1.77E-11 | 876.246215 | 1873.16781 | 1297.71211 | 1491.00074 |
| -7.3405601 | 2.13E-13 | 1.79E-11 | 913.132164 | 1954.89227 | 1734.88463 | 1851.28389 |
| -7.3383805 | 2.16E-13 | 1.82E-11 | 879.93481 | 2250.66985 | 2457.36973 | 2412.72047 |
| -7.3366055 | 2.19E-13 | 1.84E-11 | 920.919198 | 2087.49155 | 1669.30875 | 1952.14076 |
| -7.3351422 | 2.21E-13 | 1.85E-11 | 853.704802 | 3200.24307 | 2876.13499 | 3146.73437 |

| | | | | | | |
|---|---|---|---|---|---|---|
| -7.3335225 | 2.24E-13 | 1.87E-11 | 870.918245 | 4743.80696 | 3107.37624 | 4173.7935 |
| -7.3335648 | 2.24E-13 | 1.87E-11 | 941.411392 | 1854.49567 | 1484.08565 | 1615.9512 |
| -7.3302777 | 2.30E-13 | 1.91E-11 | 895.918722 | 2045.54688 | 1823.46958 | 1914.03928 |
| -7.3263461 | 2.37E-13 | 1.96E-11 | 824.196043 | 4163.61745 | 3941.45539 | 3570.89354 |
| -7.325065 | 2.39E-13 | 1.97E-11 | 886.082468 | 2032.01634 | 1938.51498 | 1878.17906 |
| -7.3226887 | 2.43E-13 | 2.00E-11 | 941.001548 | 2889.85251 | 1837.27503 | 2177.38778 |
| -7.3217195 | 2.45E-13 | 2.01E-11 | 858.622928 | 2515.5978 | 2353.82887 | 2443.53785 |
| -7.3212672 | 2.46E-13 | 2.01E-11 | 1281.99166 | 2588.93332 | 1182.66671 | 1594.65919 |
| -7.3195631 | 2.49E-13 | 2.03E-11 | 453.287331 | 654.878086 | 679.918311 | 691.429881 |
| -7.3181362 | 2.51E-13 | 2.05E-11 | 240.168514 | 234.348935 | 257.701695 | 244.297754 |
| -7.3151903 | 2.57E-13 | 2.09E-11 | 900.836848 | 1550.59976 | 1641.69785 | 1669.18121 |
| -7.312407 | 2.62E-13 | 2.12E-11 | 926.247169 | 2262.57672 | 1591.07788 | 1836.71568 |
| -7.3126643 | 2.62E-13 | 2.12E-11 | 833.212608 | 3270.33127 | 2864.63045 | 2966.87295 |
| 7.3117785 | 2.64E-13 | 2.13E-11 | 172.954117 | 163.990132 | 171.417645 | 185.464579 |
| -7.31103 | 2.65E-13 | 2.13E-11 | 974.198903 | 1470.49897 | 1172.31262 | 1357.64554 |
| -7.3109264 | 2.65E-13 | 2.13E-11 | 393.450125 | 672.738397 | 541.863832 | 576.565111 |
| -7.3084183 | 2.70E-13 | 2.17E-11 | 846.327612 | 2347.54851 | 2562.06105 | 2716.41172 |
| -7.3069811 | 2.73E-13 | 2.18E-11 | 1245.92539 | 1164.43818 | 1027.35542 | 1057.87651 |
| 7.30626039 | 2.75E-13 | 2.19E-11 | 22.5414134 | 19.2133653 | 24.1595339 | 7.84442328 |
| 7.30316328 | 2.81E-13 | 2.24E-11 | 117.625194 | 85.5130062 | 78.2308717 | 72.8410733 |
| -7.2989274 | 2.90E-13 | 2.30E-11 | 921.738886 | 3396.16528 | 2091.52536 | 2365.65394 |
| -7.2955581 | 2.97E-13 | 2.35E-11 | 915.591228 | 1985.7419 | 1450.72249 | 1655.17331 |
| -7.2951658 | 2.98E-13 | 2.35E-11 | 825.01573 | 2668.22228 | 2513.74198 | 2720.89425 |
| -7.293132 | 3.03E-13 | 2.38E-11 | 824.196043 | 1629.88872 | 1165.4099 | 1368.85186 |
| -7.2916586 | 3.06E-13 | 2.41E-11 | 888.951376 | 1575.76657 | 2113.38399 | 1991.36288 |
| -7.2891932 | 3.12E-13 | 2.44E-11 | 813.949946 | 2799.1979 | 2771.44367 | 3161.8629 |
| -7.2879401 | 3.15E-13 | 2.46E-11 | 843.458705 | 2007.66137 | 2327.36843 | 2441.8569 |
| -7.2845595 | 3.23E-13 | 2.52E-11 | 284.021809 | 365.324552 | 341.684837 | 383.816425 |
| -7.2838879 | 3.24E-13 | 2.52E-11 | 1033.21642 | 1029.67401 | 976.735442 | 1053.9543 |
| -7.2811016 | 3.31E-13 | 2.57E-11 | 803.294005 | 3272.49615 | 2874.98453 | 2858.73197 |
| -7.2798233 | 3.34E-13 | 2.59E-11 | 843.048861 | 1808.49184 | 1970.72769 | 2044.03258 |
| -7.2754613 | 3.45E-13 | 2.67E-11 | 307.792754 | 687.351379 | 532.6602 | 614.10628 |
| -7.2745773 | 3.48E-13 | 2.68E-11 | 863.541055 | 1508.38448 | 1531.25427 | 1618.19246 |
| -7.2731887 | 3.51E-13 | 2.70E-11 | 802.474317 | 2665.24556 | 2563.2115 | 2854.24944 |
| -7.2724041 | 3.53E-13 | 2.71E-11 | 534.436419 | 1695.37653 | 1526.65245 | 1616.51151 |
| -7.2708347 | 3.57E-13 | 2.73E-11 | 925.017637 | 1824.18726 | 1272.40212 | 1505.56895 |
| -7.266603 | 3.69E-13 | 2.81E-11 | 913.951852 | 1334.92297 | 1170.01171 | 1224.85066 |
| -7.2661233 | 3.70E-13 | 2.82E-11 | 827.884637 | 1868.83804 | 1952.32043 | 2140.40692 |
| -7.2632762 | 3.78E-13 | 2.87E-11 | 918.050291 | 1207.4653 | 1141.25036 | 1180.5857 |
| -7.2621378 | 3.81E-13 | 2.89E-11 | 819.277916 | 1993.319 | 2149.04806 | 1933.65034 |
| -7.2603493 | 3.86E-13 | 2.92E-11 | 909.853413 | 1198.26453 | 1302.31392 | 1158.17307 |
| -7.2568138 | 3.96E-13 | 2.99E-11 | 808.212131 | 2346.46607 | 2032.85221 | 2264.79706 |
| -7.2545992 | 4.03E-13 | 3.03E-11 | 845.507924 | 2088.03277 | 1585.32561 | 1906.75517 |

| | | | | | | |
|---|---|---|---|---|---|---|
| -7.2530083 | 4.08E-13 | 3.05E-11 | 854.934334 | 1669.12729 | 1404.70433 | 1507.81022 |
| -7.2530019 | 4.08E-13 | 3.05E-11 | 445.910141 | 828.339595 | 577.527906 | 646.604605 |
| -7.2516386 | 4.12E-13 | 3.08E-11 | 830.753545 | 1593.35627 | 1637.09604 | 1697.19701 |
| -7.2498793 | 4.17E-13 | 3.11E-11 | 1038.13455 | 3174.26444 | 1427.71341 | 2014.33584 |
| -7.2479756 | 4.23E-13 | 3.15E-11 | 805.753068 | 2195.46525 | 1961.52406 | 1977.9153 |
| -7.2473214 | 4.25E-13 | 3.16E-11 | 784.441186 | 2596.51043 | 2352.67842 | 2486.12186 |
| -7.2462457 | 4.28E-13 | 3.17E-11 | 809.851507 | 1848.54223 | 2229.57984 | 2153.29419 |
| -7.2441182 | 4.35E-13 | 3.19E-11 | 1182.39959 | 1301.63785 | 1007.7977 | 957.01964 |
| -7.2439582 | 4.36E-13 | 3.19E-11 | 898.377785 | 2307.76872 | 1428.86386 | 1758.83176 |
| -7.2442916 | 4.35E-13 | 3.19E-11 | 1107.39816 | 2106.16369 | 1060.71858 | 1450.09768 |
| -7.2446682 | 4.33E-13 | 3.19E-11 | 870.918245 | 1921.33653 | 1375.94298 | 1596.90045 |
| -7.2441346 | 4.35E-13 | 3.19E-11 | 816.409009 | 1908.34721 | 1680.81329 | 1806.45862 |
| -7.2424391 | 4.41E-13 | 3.22E-11 | 1961.10297 | 1801.45596 | 853.636865 | 1426.56441 |
| -7.2415244 | 4.44E-13 | 3.23E-11 | 801.244785 | 2758.87689 | 2130.6408 | 2268.71928 |
| -7.2409779 | 4.45E-13 | 3.24E-11 | 798.785722 | 2376.23325 | 1964.97542 | 2195.8782 |
| -7.2403392 | 4.48E-13 | 3.25E-11 | 870.098557 | 1791.44336 | 1504.79383 | 1318.98374 |
| 7.23708963 | 4.58E-13 | 3.32E-11 | 0 | 0 | 0 | 0 |
| -7.2352754 | 4.65E-13 | 3.35E-11 | 810.671195 | 1707.28341 | 1652.05194 | 1838.39663 |
| -7.2355041 | 4.64E-13 | 3.35E-11 | 854.114646 | 1763.84106 | 1351.78344 | 1501.64674 |
| -7.2327628 | 4.73E-13 | 3.40E-11 | 883.623405 | 3929.80973 | 1977.63042 | 2803.26069 |
| -7.2307394 | 4.80E-13 | 3.45E-11 | 1020.10142 | 1854.76628 | 1067.62131 | 1322.34564 |
| -7.228276 | 4.89E-13 | 3.49E-11 | 1158.2188 | 1762.75862 | 929.566828 | 1319.54406 |
| -7.2283071 | 4.89E-13 | 3.49E-11 | 766.817899 | 2085.86789 | 1807.36323 | 1911.2377 |
| -7.2268697 | 4.94E-13 | 3.52E-11 | 854.52449 | 1335.73481 | 1275.85348 | 1202.99834 |
| -7.2264317 | 4.96E-13 | 3.53E-11 | 759.850553 | 2469.86458 | 2598.87558 | 2677.1896 |
| -7.2249772 | 5.01E-13 | 3.56E-11 | 775.014777 | 2560.24858 | 2122.58762 | 2426.16806 |
| -7.2220803 | 5.12E-13 | 3.63E-11 | 893.049814 | 1528.68029 | 1187.26852 | 1176.10318 |
| -7.220506 | 5.18E-13 | 3.65E-11 | 947.55905 | 1574.95473 | 1054.96631 | 1227.65224 |
| -7.2206023 | 5.18E-13 | 3.65E-11 | 764.76868 | 2218.19656 | 2292.85481 | 2347.16351 |
| -7.2202471 | 5.19E-13 | 3.65E-11 | 2010.69407 | 3644.58597 | 2810.55911 | 3250.39282 |
| -7.2194739 | 5.22E-13 | 3.66E-11 | 779.52306 | 2123.75339 | 1880.99228 | 2126.39902 |
| 7.21758538 | 5.29E-13 | 3.71E-11 | 0 | 0 | 0 | 0 |
| -7.2146578 | 5.41E-13 | 3.78E-11 | 794.277439 | 1857.47239 | 1631.34377 | 1667.50026 |
| -7.2113577 | 5.54E-13 | 3.86E-11 | 881.98403 | 1541.66961 | 1130.89628 | 1258.46962 |
| -7.2109191 | 5.56E-13 | 3.86E-11 | 820.097604 | 1461.29821 | 2285.95209 | 2072.60869 |
| -7.2105981 | 5.57E-13 | 3.86E-11 | 779.113216 | 2014.96786 | 1757.8937 | 1895.54885 |
| -7.2104323 | 5.58E-13 | 3.86E-11 | 852.885114 | 2570.8024 | 1543.90926 | 1937.57255 |
| 7.20974199 | 5.61E-13 | 3.88E-11 | 0 | 0.81183234 | 0 | 1.68094785 |
| -7.2087426 | 5.65E-13 | 3.90E-11 | 783.621498 | 3136.10832 | 3176.40348 | 3227.98018 |
| -7.2077336 | 5.69E-13 | 3.92E-11 | 759.850553 | 3288.7328 | 2368.78478 | 2771.32268 |
| -7.2066483 | 5.73E-13 | 3.94E-11 | 747.555237 | 2393.55234 | 2394.09476 | 2677.1896 |
| -7.1986214 | 6.08E-13 | 4.17E-11 | 643.864735 | 790.183475 | 689.121943 | 747.461475 |
| -7.1973872 | 6.14E-13 | 4.19E-11 | 732.391013 | 2621.40662 | 2552.85742 | 2512.45671 |

| | | | | | | |
|---|---|---|---|---|---|---|
| -7.1973213 | 6.14E-13 | 4.19E-11 | 736.079608 | 2372.71531 | 2582.76922 | 2598.18505 |
| -7.196942 | 6.16E-13 | 4.19E-11 | 754.932427 | 2049.33543 | 2334.27116 | 2576.89305 |
| -7.1955688 | 6.22E-13 | 4.22E-11 | 775.014777 | 3004.32087 | 2002.94041 | 2581.37558 |
| -7.1955827 | 6.22E-13 | 4.22E-11 | 403.696222 | 931.983523 | 820.273699 | 847.197714 |
| -7.1930747 | 6.33E-13 | 4.29E-11 | 875.836371 | 1197.4527 | 1023.90406 | 1067.40188 |
| -7.1910926 | 6.43E-13 | 4.34E-11 | 1084.03706 | 1468.87531 | 884.699122 | 1136.88106 |
| -7.1870197 | 6.62E-13 | 4.46E-11 | 746.325705 | 2182.74654 | 1953.47088 | 2281.60654 |
| -7.1841087 | 6.76E-13 | 4.55E-11 | 914.361696 | 1290.27219 | 947.974092 | 1119.51127 |
| -7.1825825 | 6.84E-13 | 4.59E-11 | 1515.19282 | 1516.77342 | 878.946852 | 1040.50672 |
| -7.1824981 | 6.84E-13 | 4.59E-11 | 709.8496 | 1327.07526 | 1115.94038 | 1215.88561 |
| -7.1819931 | 6.87E-13 | 4.59E-11 | 428.286855 | 1722.97883 | 1528.95336 | 1753.2286 |
| -7.1813612 | 6.90E-13 | 4.60E-11 | 845.917768 | 1068.91258 | 1057.26722 | 1116.70969 |
| -7.179955 | 6.97E-13 | 4.64E-11 | 1838.96949 | 1545.72877 | 851.335957 | 1175.54286 |
| -7.1767435 | 7.14E-13 | 4.74E-11 | 799.195566 | 1738.67426 | 1309.21665 | 1485.9579 |
| -7.1766283 | 7.15E-13 | 4.74E-11 | 1228.30211 | 1262.6699 | 840.981871 | 1031.54166 |
| -7.1760029 | 7.18E-13 | 4.75E-11 | 775.834465 | 1741.10975 | 1416.20887 | 1611.46867 |
| -7.1756851 | 7.19E-13 | 4.75E-11 | 734.850077 | 2081.2675 | 1968.42679 | 2042.91195 |
| 7.17390983 | 7.29E-13 | 4.80E-11 | 9.42640924 | 7.57710182 | 6.90272397 | 6.72379138 |
| -7.1728923 | 7.34E-13 | 4.83E-11 | 772.965558 | 1760.05251 | 1451.87294 | 1522.93875 |
| -7.171724 | 7.41E-13 | 4.85E-11 | 694.685376 | 2974.28307 | 2919.85224 | 3063.24729 |
| -7.1715064 | 7.42E-13 | 4.85E-11 | 808.212131 | 1644.77232 | 1235.58759 | 1364.36933 |
| -7.1714798 | 7.42E-13 | 4.85E-11 | 770.09665 | 2039.32283 | 1943.1168 | 1893.30759 |
| -7.1705183 | 7.47E-13 | 4.87E-11 | 775.834465 | 1353.32451 | 1364.43844 | 1392.94545 |
| -7.1695463 | 7.52E-13 | 4.90E-11 | 756.981646 | 1752.47541 | 1518.59927 | 1722.41123 |
| -7.1691286 | 7.55E-13 | 4.90E-11 | 734.440233 | 1891.02812 | 1885.5941 | 1966.70898 |
| 7.16842752 | 7.59E-13 | 4.92E-11 | 0 | 0 | 0 | 0 |
| -7.1679678 | 7.61E-13 | 4.93E-11 | 701.652722 | 1349.26534 | 1138.94946 | 1217.56656 |
| -7.1655134 | 7.75E-13 | 5.00E-11 | 1112.31629 | 2589.20393 | 992.841798 | 2010.41362 |
| -7.1648726 | 7.79E-13 | 5.02E-11 | 747.145393 | 1579.55512 | 1719.92872 | 1716.80807 |
| -7.1644297 | 7.81E-13 | 5.02E-11 | 669.6849 | 1032.65073 | 870.893675 | 954.778376 |
| -7.1637247 | 7.85E-13 | 5.04E-11 | 743.046954 | 2025.52168 | 1685.4151 | 1822.14746 |
| -7.1631865 | 7.88E-13 | 5.05E-11 | 695.09522 | 2710.97879 | 2712.77052 | 2921.48736 |
| -7.1625087 | 7.92E-13 | 5.06E-11 | 803.703849 | 1173.36834 | 1110.18811 | 1216.44592 |
| -7.1624157 | 7.93E-13 | 5.06E-11 | 729.112262 | 1909.97088 | 1946.56816 | 1852.40453 |
| -7.1602354 | 8.05E-13 | 5.13E-11 | 741.407579 | 2720.72077 | 2424.00657 | 2525.34398 |
| -7.1546376 | 8.39E-13 | 5.31E-11 | 795.916815 | 1708.63646 | 1220.63169 | 1451.21831 |
| 7.15453896 | 8.40E-13 | 5.31E-11 | 0 | 0 | 0 | 0 |
| -7.1545941 | 8.39E-13 | 5.31E-11 | 288.120248 | 518.760864 | 411.86253 | 485.233612 |
| -7.1535756 | 8.45E-13 | 5.34E-11 | 451.238112 | 887.062134 | 798.415073 | 795.088331 |
| -7.1526395 | 8.51E-13 | 5.37E-11 | 755.752115 | 4543.55498 | 2424.00657 | 3097.42656 |
| -7.1514285 | 8.59E-13 | 5.41E-11 | 737.30914 | 2497.46688 | 1820.01822 | 2021.61994 |
| 7.15090377 | 8.62E-13 | 5.42E-11 | 29.9186032 | 11.0950419 | 13.8054479 | 14.0078987 |
| -7.1504409 | 8.65E-13 | 5.42E-11 | 480.337027 | 958.232769 | 670.714679 | 828.707288 |

| | | | | | | |
|---|---|---|---|---|---|---|
| -7.1492083 | 8.73E-13 | 5.46E-11 | 1013.13407 | 1754.64029 | 929.566828 | 1211.9634 |
| -7.147496 | 8.84E-13 | 5.52E-11 | 858.622928 | 995.847667 | 1111.33856 | 933.48637 |
| -7.1461769 | 8.92E-13 | 5.56E-11 | 774.604933 | 2232.26832 | 3195.9612 | 3138.88994 |
| -7.1419564 | 9.20E-13 | 5.69E-11 | 744.68633 | 1376.86764 | 1421.96114 | 1454.01989 |
| -7.1418362 | 9.21E-13 | 5.69E-11 | 634.438326 | 1176.61567 | 1043.46177 | 1118.39063 |
| -7.1424011 | 9.17E-13 | 5.69E-11 | 746.325705 | 2662.26885 | 1947.71861 | 2306.82076 |
| -7.1424594 | 9.17E-13 | 5.69E-11 | 496.320939 | 1970.31708 | 1753.29189 | 2059.72143 |
| 7.1419635 | 9.20E-13 | 5.69E-11 | 0 | 0 | 0 | 0 |
| 7.13980848 | 9.35E-13 | 5.76E-11 | 0 | 0 | 0 | 0 |
| -7.1364222 | 9.58E-13 | 5.87E-11 | 748.784769 | 1550.87038 | 1296.56165 | 1491.00074 |
| -7.1368166 | 9.55E-13 | 5.87E-11 | 716.407102 | 1694.29409 | 1914.35545 | 2021.61994 |
| -7.1364506 | 9.58E-13 | 5.87E-11 | 714.357883 | 2237.68053 | 2000.6395 | 2049.63574 |
| -7.1362446 | 9.59E-13 | 5.87E-11 | 690.177094 | 2530.21079 | 2205.42031 | 2331.47466 |
| -7.1357427 | 9.63E-13 | 5.88E-11 | 478.697652 | 1522.72685 | 1562.31653 | 1443.9342 |
| -7.12988 | 1.00E-12 | 6.12E-11 | 772.14587 | 1586.32039 | 1167.71081 | 1450.09768 |
| -7.1276887 | 1.02E-12 | 6.21E-11 | 505.747348 | 1659.92652 | 1317.26982 | 1454.5802 |
| -7.1272578 | 1.02E-12 | 6.21E-11 | 779.52306 | 2256.35268 | 1408.15569 | 1630.51941 |
| -7.1263042 | 1.03E-12 | 6.25E-11 | 758.621022 | 1156.59047 | 1378.24389 | 1287.04573 |
| -7.1253968 | 1.04E-12 | 6.27E-11 | 958.214991 | 1492.14784 | 863.990951 | 1164.89686 |
| -7.1240681 | 1.05E-12 | 6.32E-11 | 748.784769 | 1438.5669 | 1290.80938 | 1224.85066 |
| -7.1238108 | 1.05E-12 | 6.32E-11 | 645.094267 | 375.337151 | 390.003904 | 369.24821 |
| -7.1224836 | 1.06E-12 | 6.37E-11 | 768.867119 | 1932.70218 | 1279.30484 | 1567.76402 |
| -7.1219566 | 1.06E-12 | 6.39E-11 | 668.865212 | 2894.7235 | 2427.45793 | 2646.37223 |
| -7.1212574 | 1.07E-12 | 6.41E-11 | 806.572756 | 1660.19713 | 1074.52403 | 1392.38513 |
| -7.1203627 | 1.08E-12 | 6.44E-11 | 663.537242 | 2814.62271 | 2535.60061 | 2599.866 |
| -7.1199506 | 1.08E-12 | 6.44E-11 | 734.440233 | 1525.16235 | 1296.56165 | 1397.98829 |
| -7.1184253 | 1.09E-12 | 6.50E-11 | 15892.926 | 4016.94641 | 1305.76528 | 3089.58214 |
| -7.117256 | 1.10E-12 | 6.55E-11 | 811.490882 | 1806.86817 | 1102.13493 | 1447.85641 |
| -7.1148214 | 1.12E-12 | 6.64E-11 | 675.832558 | 2500.98482 | 2114.53444 | 2310.18266 |
| -7.1150487 | 1.12E-12 | 6.64E-11 | 803.294005 | 1194.74659 | 967.53181 | 1036.02419 |
| -7.113834 | 1.13E-12 | 6.67E-11 | 642.22536 | 3008.92125 | 2995.7822 | 3199.40407 |
| 7.11283225 | 1.14E-12 | 6.71E-11 | 0 | 0 | 0 | 0 |
| -7.1124135 | 1.14E-12 | 6.72E-11 | 672.143963 | 2171.92211 | 2177.80941 | 2226.13526 |
| -7.1119786 | 1.14E-12 | 6.73E-11 | 737.718984 | 2025.52168 | 1416.20887 | 1607.54646 |
| 7.11059509 | 1.16E-12 | 6.78E-11 | 0 | 0 | 0 | 0 |
| -7.109418 | 1.17E-12 | 6.83E-11 | 770.506494 | 1040.22784 | 1037.7095 | 1081.40978 |
| -7.1085499 | 1.17E-12 | 6.86E-11 | 702.47241 | 1672.64523 | 1573.82107 | 1529.66254 |
| -7.1080873 | 1.18E-12 | 6.87E-11 | 706.980693 | 1697.0002 | 1742.9378 | 1464.66589 |
| -7.1071419 | 1.18E-12 | 6.90E-11 | 733.210701 | 1321.12182 | 1194.17125 | 1272.47752 |
| -7.1048934 | 1.20E-12 | 7.01E-11 | 679.521153 | 2814.89332 | 2073.1181 | 2314.10487 |
| -7.1030456 | 1.22E-12 | 7.09E-11 | 689.76725 | 1918.90103 | 1639.39694 | 1872.01558 |
| -7.1015902 | 1.23E-12 | 7.15E-11 | 709.029912 | 2196.27708 | 1571.52016 | 1814.30304 |
| -7.1012819 | 1.24E-12 | 7.15E-11 | 708.210225 | 1480.51157 | 1380.54479 | 1415.9184 |

| | | | | | | |
|---|---|---|---|---|---|---|
| -7.0994005 | 1.25E-12 | 7.24E-11 | 738.128828 | 1515.42036 | 1168.86126 | 1346.99954 |
| -7.0983857 | 1.26E-12 | 7.26E-11 | 675.832558 | 2185.72326 | 1861.43456 | 1945.41697 |
| -7.0985504 | 1.26E-12 | 7.26E-11 | 749.604456 | 1512.71426 | 1125.14401 | 1304.97584 |
| -7.095755 | 1.29E-12 | 7.38E-11 | 705.751161 | 1351.15962 | 1418.50978 | 1399.10892 |
| -7.0957946 | 1.29E-12 | 7.38E-11 | 652.061613 | 1277.01227 | 1239.03895 | 1260.15057 |
| -7.0951312 | 1.29E-12 | 7.39E-11 | 251.644142 | 363.159666 | 425.667978 | 394.462428 |
| -7.0950868 | 1.29E-12 | 7.39E-11 | 698.373971 | 1494.85394 | 1432.31522 | 1602.50361 |
| 7.09255985 | 1.32E-12 | 7.51E-11 | 30.3284471 | 17.5897006 | 25.3099879 | 18.4904263 |
| -7.0918953 | 1.32E-12 | 7.52E-11 | 667.63568 | 1928.1018 | 2444.71474 | 2240.70348 |
| -7.0919129 | 1.32E-12 | 7.52E-11 | 351.236205 | 530.667738 | 552.217918 | 599.538065 |
| -7.0912693 | 1.33E-12 | 7.54E-11 | 670.914431 | 1886.15713 | 1851.08048 | 1801.97609 |
| -7.0899894 | 1.34E-12 | 7.60E-11 | 1008.21594 | 1604.1807 | 737.441011 | 1163.77623 |
| -7.0882379 | 1.36E-12 | 7.68E-11 | 700.013347 | 1450.20317 | 1360.98708 | 1363.80902 |
| -7.0865847 | 1.37E-12 | 7.76E-11 | 556.977833 | 2339.43019 | 1348.33208 | 1646.20826 |
| -7.0849811 | 1.39E-12 | 7.84E-11 | 681.570372 | 2268.53016 | 1672.76011 | 2017.13742 |
| -7.0845568 | 1.39E-12 | 7.85E-11 | 724.60398 | 1528.13907 | 1164.25944 | 1428.80567 |
| -7.083911 | 1.40E-12 | 7.87E-11 | 937.312953 | 1605.53375 | 876.645944 | 1097.65894 |
| -7.0811873 | 1.43E-12 | 8.01E-11 | 669.275056 | 1956.78654 | 1684.26465 | 1819.9062 |
| -7.0804544 | 1.44E-12 | 8.04E-11 | 423.368728 | 1465.89859 | 1172.31262 | 1255.10772 |
| -7.0788924 | 1.45E-12 | 8.12E-11 | 743.866642 | 1034.2744 | 1010.09861 | 1119.51127 |
| -7.0763828 | 1.48E-12 | 8.25E-11 | 626.651292 | 3360.17404 | 2667.90282 | 2772.44331 |
| -7.0749258 | 1.50E-12 | 8.32E-11 | 695.09522 | 1991.69533 | 1405.85478 | 1601.38298 |
| 7.07419061 | 1.50E-12 | 8.34E-11 | 12.7051603 | 4.32977247 | 4.60181598 | 6.72379138 |
| -7.0742174 | 1.50E-12 | 8.34E-11 | 761.489929 | 1246.16264 | 961.77954 | 1066.84157 |
| -7.0732594 | 1.51E-12 | 8.38E-11 | 736.899296 | 1914.30065 | 1212.57851 | 1552.07518 |
| -7.0723404 | 1.52E-12 | 8.41E-11 | 677.471934 | 2141.3431 | 1573.82107 | 1939.2535 |
| -7.0723342 | 1.52E-12 | 8.41E-11 | 854.52449 | 828.339595 | 852.486411 | 892.583306 |
| -7.0699265 | 1.55E-12 | 8.54E-11 | 621.323322 | 2961.29376 | 2520.6447 | 2719.2133 |
| -7.0682253 | 1.57E-12 | 8.63E-11 | 695.914908 | 1480.78218 | 1230.98578 | 1359.88681 |
| -7.0674405 | 1.58E-12 | 8.66E-11 | 748.374925 | 1008.02515 | 939.920914 | 1012.49092 |
| -7.0673393 | 1.58E-12 | 8.66E-11 | 668.865212 | 1491.87723 | 1633.64467 | 1663.01774 |
| -7.0630072 | 1.63E-12 | 8.92E-11 | 686.898343 | 2068.00757 | 1473.73157 | 1633.32099 |
| -7.0616023 | 1.65E-12 | 8.99E-11 | 663.537242 | 2068.00757 | 1614.08696 | 1775.64124 |
| -7.0597614 | 1.67E-12 | 9.10E-11 | 435.2542 | 922.241535 | 528.058384 | 688.628301 |
| -7.0575618 | 1.69E-12 | 9.23E-11 | 702.882254 | 1836.36475 | 1274.70303 | 1470.26905 |
| -7.0563632 | 1.71E-12 | 9.29E-11 | 723.374448 | 1479.42913 | 1057.26722 | 1353.16302 |
| -7.0536271 | 1.74E-12 | 9.46E-11 | 321.317602 | 537.974229 | 633.900151 | 629.795126 |
| -7.0524021 | 1.76E-12 | 9.53E-11 | 963.952806 | 1405.55239 | 765.051907 | 1078.04789 |
| -7.0514577 | 1.77E-12 | 9.57E-11 | 239.75867 | 320.132552 | 381.950726 | 376.532317 |
| -7.0513186 | 1.77E-12 | 9.57E-11 | 712.308663 | 1281.07143 | 1033.10769 | 1158.73338 |
| -7.0495825 | 1.79E-12 | 9.67E-11 | 705.751161 | 1289.46036 | 1052.66541 | 1180.02539 |
| 7.04926951 | 1.80E-12 | 9.68E-11 | 0 | 0 | 0 | 0 |
| -7.0483042 | 1.81E-12 | 9.73E-11 | 660.258491 | 1570.08374 | 1418.50978 | 1471.38968 |

| | | | | | | |
|---|---|---|---|---|---|---|
| 7.04774558 | 1.82E-12 | 9.75E-11 | 47.1320462 | 32.4732935 | 35.6640739 | 34.1792729 |
| -7.0459787 | 1.84E-12 | 9.86E-11 | 670.914431 | 1356.03061 | 1340.2789 | 1590.17666 |
| -7.0437565 | 1.87E-12 | 1.00E-10 | 912.312477 | 1410.69399 | 778.857355 | 1098.21926 |
| -7.0413278 | 1.90E-12 | 1.02E-10 | 754.522583 | 1034.2744 | 853.636865 | 966.545011 |
| -7.0411571 | 1.91E-12 | 1.02E-10 | 658.209271 | 1693.48226 | 1394.35024 | 1531.9038 |
| 7.04022779 | 1.92E-12 | 1.02E-10 | 0 | 0 | 0 | 0 |
| -7.0364522 | 1.97E-12 | 1.05E-10 | 797.146346 | 908.169775 | 1035.4086 | 800.131175 |
| -7.0361086 | 1.98E-12 | 1.05E-10 | 645.913955 | 1770.60633 | 1508.24519 | 1576.72908 |
| -7.0347533 | 2.00E-12 | 1.06E-10 | 707.390537 | 1488.08867 | 1059.56813 | 1238.85856 |
| -7.0316262 | 2.04E-12 | 1.08E-10 | 638.536765 | 1631.783 | 1519.74973 | 1702.23985 |
| -7.0312746 | 2.05E-12 | 1.08E-10 | 296.726969 | 871.366709 | 922.664104 | 998.48302 |
| -7.0271088 | 2.11E-12 | 1.11E-10 | 624.602073 | 1796.85557 | 1694.61874 | 1784.6063 |
| -7.0273604 | 2.10E-12 | 1.11E-10 | 840.999642 | 783.418206 | 853.636865 | 809.656546 |
| -7.0273266 | 2.11E-12 | 1.11E-10 | 745.096174 | 1411.77643 | 933.01819 | 1129.03664 |
| -7.0263944 | 2.12E-12 | 1.11E-10 | 705.341317 | 1721.89639 | 1100.98447 | 1486.51821 |
| 7.02505695 | 2.14E-12 | 1.12E-10 | 0 | 0 | 0 | 0 |
| -7.0243342 | 2.15E-12 | 1.12E-10 | 779.113216 | 2037.42856 | 1033.10769 | 1488.75948 |
| -7.0245041 | 2.15E-12 | 1.12E-10 | 679.930997 | 1143.87176 | 1049.21404 | 1167.69844 |
| -7.0241517 | 2.15E-12 | 1.12E-10 | 670.094744 | 1314.08594 | 1126.29446 | 1214.20466 |
| -7.0239998 | 2.16E-12 | 1.12E-10 | 358.203551 | 472.757031 | 418.765254 | 479.630452 |
| -7.023498 | 2.16E-12 | 1.12E-10 | 654.110832 | 1630.15933 | 1308.06619 | 1418.71998 |
| -7.0226265 | 2.18E-12 | 1.13E-10 | 622.14301 | 1674.81011 | 1761.34507 | 1814.86336 |
| 7.02209018 | 2.19E-12 | 1.13E-10 | 0 | 0 | 0 | 0 |
| -7.0216933 | 2.19E-12 | 1.13E-10 | 585.666904 | 2761.04178 | 2419.40475 | 2701.28319 |
| -7.0211354 | 2.20E-12 | 1.14E-10 | 611.077225 | 1949.20944 | 1869.48774 | 1976.23435 |
| -7.0200441 | 2.22E-12 | 1.14E-10 | 665.176617 | 1284.04815 | 1140.09991 | 1186.74918 |
| -7.0191378 | 2.23E-12 | 1.15E-10 | 570.912525 | 2884.44029 | 2815.16093 | 2973.03642 |
| -7.0178456 | 2.25E-12 | 1.16E-10 | 710.259444 | 1520.56197 | 1012.39952 | 1246.14267 |
| -7.0170395 | 2.27E-12 | 1.16E-10 | 594.273626 | 2604.89936 | 2185.86259 | 2331.47466 |
| -7.0170326 | 2.27E-12 | 1.16E-10 | 714.357883 | 1044.01639 | 934.168644 | 902.108677 |
| -7.0176877 | 2.26E-12 | 1.16E-10 | 616.815039 | 1797.12618 | 1715.32691 | 1767.2365 |
| -7.0170549 | 2.27E-12 | 1.16E-10 | 423.778572 | 794.242637 | 660.360593 | 685.826721 |
| -7.014779 | 2.30E-12 | 1.17E-10 | 987.723751 | 784.500649 | 791.512349 | 844.396135 |
| -7.0130562 | 2.33E-12 | 1.18E-10 | 307.792754 | 868.119379 | 842.132325 | 758.107478 |
| -7.0130257 | 2.33E-12 | 1.18E-10 | 756.161958 | 1554.1177 | 921.51365 | 1206.36024 |
| -7.0128288 | 2.34E-12 | 1.18E-10 | 670.504588 | 1290.00158 | 1066.47085 | 1171.62065 |
| -7.0120648 | 2.35E-12 | 1.19E-10 | 769.686807 | 1564.67153 | 908.858656 | 1172.18096 |
| -7.012252 | 2.35E-12 | 1.19E-10 | 446.729829 | 1738.40365 | 1470.28021 | 1851.28389 |
| 7.01195075 | 2.35E-12 | 1.19E-10 | 0 | 0 | 0 | 0 |
| -7.0110254 | 2.37E-12 | 1.19E-10 | 408.204504 | 854.859451 | 770.804177 | 837.672343 |
| -7.0105609 | 2.37E-12 | 1.19E-10 | 251.644142 | 336.639809 | 339.383929 | 389.9799 |
| -7.0106871 | 2.37E-12 | 1.19E-10 | 640.585984 | 1900.4995 | 1448.42158 | 1515.65464 |
| -7.0090881 | 2.40E-12 | 1.20E-10 | 2724.23227 | 1466.71042 | 827.176423 | 1192.91265 |

| | | | | | | |
|---|---|---|---|---|---|---|
| 7.00916133 | 2.40E-12 | 1.20E-10 | 0 | 0 | 0 | 0 |
| 7.00679069 | 2.44E-12 | 1.22E-10 | 20.0823501 | 12.9893174 | 13.8054479 | 19.0507423 |
| -7.0053769 | 2.46E-12 | 1.23E-10 | 730.341794 | 1113.29275 | 893.902754 | 880.816671 |
| -7.0048401 | 2.47E-12 | 1.23E-10 | 628.700512 | 1433.4253 | 1373.64207 | 1441.69294 |
| -7.0039398 | 2.49E-12 | 1.24E-10 | 638.536765 | 1570.35435 | 1268.95076 | 1503.32769 |
| -7.0033891 | 2.50E-12 | 1.24E-10 | 635.258014 | 1307.8619 | 1320.72119 | 1312.25995 |
| -7.0029794 | 2.51E-12 | 1.24E-10 | 623.782385 | 1789.00786 | 1478.33338 | 1547.03233 |
| -7.0000763 | 2.56E-12 | 1.27E-10 | 554.108926 | 1376.59703 | 1135.49809 | 1219.80782 |
| -6.9991707 | 2.57E-12 | 1.27E-10 | 299.595876 | 464.909319 | 413.012984 | 423.038541 |
| -6.9987565 | 2.58E-12 | 1.27E-10 | 617.634727 | 2445.239 | 1696.91964 | 1957.18361 |
| -6.9983874 | 2.59E-12 | 1.28E-10 | 670.094744 | 1122.49351 | 977.885896 | 1084.21136 |
| -6.9973888 | 2.61E-12 | 1.28E-10 | 570.502681 | 2860.35594 | 2404.44885 | 3198.28343 |
| -6.9954601 | 2.64E-12 | 1.30E-10 | 693.455845 | 906.54611 | 1140.09991 | 989.517965 |
| -6.9953111 | 2.65E-12 | 1.30E-10 | 228.692885 | 409.97533 | 358.941647 | 404.548115 |
| -6.9904051 | 2.74E-12 | 1.34E-10 | 653.700988 | 1205.30041 | 1031.95723 | 1161.53496 |
| 6.98997436 | 2.75E-12 | 1.34E-10 | 0 | 0 | 0 | 0 |
| -6.9885852 | 2.78E-12 | 1.36E-10 | 959.854367 | 3000.26171 | 1444.97022 | 1976.79467 |
| -6.9880292 | 2.79E-12 | 1.36E-10 | 663.537242 | 1191.22865 | 976.735442 | 1108.86526 |
| -6.9861152 | 2.83E-12 | 1.38E-10 | 658.209271 | 1016.41409 | 1007.7977 | 1045.54956 |
| -6.9855405 | 2.84E-12 | 1.38E-10 | 628.700512 | 1309.48556 | 1201.07397 | 1263.51246 |
| 6.98391684 | 2.87E-12 | 1.39E-10 | 19.2626624 | 16.2366467 | 16.1063559 | 17.3697944 |
| 6.98258884 | 2.90E-12 | 1.40E-10 | 0 | 0 | 0 | 0 |
| -6.9824606 | 2.90E-12 | 1.40E-10 | 639.766297 | 1231.82027 | 1103.28538 | 1129.03664 |
| -6.9816207 | 2.92E-12 | 1.41E-10 | 546.321892 | 1617.71124 | 1250.54349 | 1339.15512 |
| -6.9815313 | 2.92E-12 | 1.41E-10 | 670.504588 | 631.876169 | 635.050605 | 792.286751 |
| -6.9802089 | 2.95E-12 | 1.42E-10 | 555.748301 | 2490.97222 | 2461.97155 | 2782.529 |
| -6.9797677 | 2.96E-12 | 1.42E-10 | 602.470503 | 1874.52087 | 1639.39694 | 1520.69748 |
| -6.9780251 | 2.99E-12 | 1.44E-10 | 647.963174 | 1235.60882 | 1127.44492 | 1542.54981 |
| -6.9778311 | 3.00E-12 | 1.44E-10 | 661.488022 | 964.186206 | 967.53181 | 972.148171 |
| -6.9753247 | 3.05E-12 | 1.46E-10 | 606.568942 | 1513.7967 | 1373.64207 | 1417.59935 |
| -6.9749176 | 3.06E-12 | 1.46E-10 | 392.630437 | 611.30975 | 505.049304 | 547.428682 |
| -6.9747423 | 3.06E-12 | 1.46E-10 | 626.241449 | 1288.64853 | 1136.64855 | 1282.00289 |
| -6.9737135 | 3.09E-12 | 1.47E-10 | 243.857109 | 478.981079 | 445.225696 | 449.373391 |
| -6.9736945 | 3.09E-12 | 1.47E-10 | 695.09522 | 924.135811 | 811.070067 | 920.599104 |
| -6.9732309 | 3.10E-12 | 1.47E-10 | 546.321892 | 2981.86018 | 2549.40605 | 2646.37223 |
| -6.9721209 | 3.12E-12 | 1.48E-10 | 594.273626 | 1202.32369 | 1230.98578 | 1358.20586 |
| -6.9723161 | 3.12E-12 | 1.48E-10 | 391.810749 | 709.270852 | 706.378753 | 746.90116 |
| -6.971432 | 3.14E-12 | 1.49E-10 | 625.831605 | 1418.27109 | 1143.55127 | 1325.70753 |
| -6.9704413 | 3.16E-12 | 1.49E-10 | 674.603026 | 1061.33548 | 862.840497 | 1010.24966 |
| -6.96937 | 3.18E-12 | 1.50E-10 | 644.274579 | 1241.56225 | 997.443614 | 1171.06033 |
| -6.9670258 | 3.24E-12 | 1.53E-10 | 712.308663 | 1252.92791 | 829.477331 | 1044.98924 |
| -6.9662892 | 3.25E-12 | 1.53E-10 | 722.144916 | 839.705248 | 760.450091 | 911.634048 |
| -6.9663547 | 3.25E-12 | 1.53E-10 | 534.846263 | 1181.75727 | 1057.26722 | 1095.41768 |

| | | | | | | |
|---|---|---|---|---|---|---|
| -6.9664718 | 3.25E-12 | 1.53E-10 | 264.759146 | 689.786876 | 716.732839 | 658.931556 |
| -6.9655736 | 3.27E-12 | 1.53E-10 | 581.568466 | 1778.99526 | 1630.19331 | 1727.45407 |
| -6.9629764 | 3.33E-12 | 1.56E-10 | 541.813609 | 2766.454 | 2461.97155 | 2550.5582 |
| -6.9629733 | 3.33E-12 | 1.56E-10 | 588.125968 | 1616.89941 | 1563.46698 | 1496.04358 |
| -6.9618091 | 3.36E-12 | 1.57E-10 | 631.159575 | 1351.43023 | 1064.16995 | 1196.27455 |
| -6.9612743 | 3.37E-12 | 1.57E-10 | 678.701465 | 924.135811 | 1141.25036 | 884.178567 |
| -6.9599353 | 3.40E-12 | 1.58E-10 | 648.782862 | 1235.87943 | 967.53181 | 1055.07493 |
| -6.9589942 | 3.43E-12 | 1.59E-10 | 687.308187 | 2224.96183 | 1151.60445 | 1555.99739 |
| -6.9583496 | 3.44E-12 | 1.60E-10 | 636.897389 | 1313.0035 | 1012.39952 | 1167.69844 |
| -6.957526 | 3.46E-12 | 1.60E-10 | 584.437373 | 1635.30094 | 1478.33338 | 1674.78437 |
| -6.9567614 | 3.48E-12 | 1.61E-10 | 607.798474 | 1821.48115 | 1476.03248 | 1500.52611 |
| -6.9561477 | 3.50E-12 | 1.62E-10 | 576.650339 | 1842.04757 | 1687.71601 | 1616.51151 |
| -6.9557597 | 3.51E-12 | 1.62E-10 | 636.077702 | 1437.48446 | 1059.56813 | 1180.02539 |
| -6.9554302 | 3.51E-12 | 1.62E-10 | 647.55333 | 1648.01964 | 1118.24128 | 1202.43803 |
| 6.95456198 | 3.54E-12 | 1.63E-10 | 92.214873 | 94.4431619 | 142.656295 | 172.577312 |
| -6.9522381 | 3.60E-12 | 1.65E-10 | 613.946132 | 1127.90573 | 1119.39174 | 1160.41433 |
| -6.948442 | 3.69E-12 | 1.69E-10 | 339.760576 | 644.865487 | 473.987046 | 557.514369 |
| -6.9477644 | 3.71E-12 | 1.70E-10 | 563.125491 | 775.029271 | 675.316495 | 774.916957 |
| -6.9466302 | 3.74E-12 | 1.71E-10 | 746.735549 | 1230.46721 | 759.299637 | 970.467223 |
| -6.9460862 | 3.76E-12 | 1.72E-10 | 594.273626 | 1342.77069 | 1252.8444 | 1293.76953 |
| -6.9453833 | 3.77E-12 | 1.72E-10 | 594.273626 | 1379.84436 | 1234.43714 | 1332.99164 |
| -6.9455091 | 3.77E-12 | 1.72E-10 | 620.09379 | 1084.87861 | 1018.15179 | 1067.9622 |
| -6.9455289 | 3.77E-12 | 1.72E-10 | 525.419854 | 2636.29021 | 2646.04419 | 2598.74537 |
| -6.9450406 | 3.78E-12 | 1.72E-10 | 654.93052 | 866.766326 | 878.946852 | 898.186466 |
| -6.9448041 | 3.79E-12 | 1.72E-10 | 644.684423 | 1208.27713 | 990.54089 | 948.614901 |
| -6.9406115 | 3.90E-12 | 1.77E-10 | 582.388153 | 1384.44475 | 1365.58889 | 1395.18671 |
| -6.9399219 | 3.92E-12 | 1.77E-10 | 709.439756 | 885.70908 | 724.786017 | 855.602454 |
| -6.9395737 | 3.93E-12 | 1.78E-10 | 632.798951 | 1004.77782 | 914.610926 | 980.55291 |
| -6.9390861 | 3.95E-12 | 1.78E-10 | 559.436896 | 1953.2686 | 2282.50073 | 1764.43492 |
| -6.9368181 | 4.01E-12 | 1.80E-10 | 591.404719 | 1225.32561 | 1297.71211 | 1224.85066 |
| -6.9369138 | 4.01E-12 | 1.80E-10 | 444.270766 | 752.027355 | 696.024667 | 808.535914 |
| -6.9361838 | 4.03E-12 | 1.81E-10 | 634.028482 | 1320.5806 | 968.682264 | 1097.09863 |
| 6.93579165 | 4.04E-12 | 1.81E-10 | 1.22953164 | 0 | 0 | 0 |
| -6.9355073 | 4.05E-12 | 1.81E-10 | 627.880824 | 986.646901 | 1132.04673 | 953.097429 |
| -6.9354905 | 4.05E-12 | 1.81E-10 | 637.717077 | 917.911763 | 903.106386 | 909.953101 |
| -6.9342001 | 4.09E-12 | 1.82E-10 | 564.355023 | 1651.53759 | 2124.88853 | 1757.15081 |
| -6.9316021 | 4.16E-12 | 1.86E-10 | 587.716124 | 940.643068 | 1112.48901 | 1245.02204 |
| -6.9313551 | 4.17E-12 | 1.86E-10 | 542.223453 | 1915.38309 | 2027.09994 | 2126.39902 |
| -6.9302193 | 4.20E-12 | 1.87E-10 | 633.618638 | 1190.14621 | 1042.31132 | 1186.74918 |
| -6.9282344 | 4.26E-12 | 1.89E-10 | 586.076748 | 906.54611 | 1473.73157 | 1223.73003 |
| -6.9250424 | 4.36E-12 | 1.93E-10 | 590.175187 | 1318.14511 | 1122.8431 | 1284.24415 |
| -6.9248512 | 4.36E-12 | 1.93E-10 | 640.585984 | 850.259068 | 983.638166 | 1090.37484 |
| -6.9234726 | 4.41E-12 | 1.95E-10 | 630.339887 | 1011.54309 | 870.893675 | 939.08953 |

| | | | | | | |
|---|---|---|---|---|---|---|
| -6.9231397 | 4.42E-12 | 1.95E-10 | 582.388153 | 1472.66386 | 1217.18033 | 1315.62185 |
| 6.92288483 | 4.43E-12 | 1.95E-10 | 0 | 0 | 0 | 0 |
| -6.9219601 | 4.45E-12 | 1.96E-10 | 614.76582 | 1068.10075 | 934.168644 | 1006.88776 |
| -6.9220387 | 4.45E-12 | 1.96E-10 | 578.289715 | 1530.57457 | 1285.05711 | 1353.16302 |
| -6.9206584 | 4.50E-12 | 1.97E-10 | 592.224407 | 1159.29658 | 1084.87812 | 1127.35569 |
| -6.919596 | 4.53E-12 | 1.99E-10 | 586.486592 | 1843.40063 | 1295.4112 | 1459.62305 |
| -6.9168854 | 4.62E-12 | 2.02E-10 | 550.830175 | 1820.66932 | 1662.40602 | 2176.82746 |
| -6.9159718 | 4.65E-12 | 2.03E-10 | 592.63425 | 1162.2733 | 1040.01041 | 1183.38728 |
| -6.9158553 | 4.65E-12 | 2.03E-10 | 552.059706 | 1602.01581 | 1557.71471 | 1577.84971 |
| -6.9153823 | 4.67E-12 | 2.04E-10 | 621.323322 | 958.232769 | 880.097306 | 922.840367 |
| -6.9134945 | 4.73E-12 | 2.06E-10 | 597.142533 | 1021.55569 | 1023.90406 | 1069.64315 |
| -6.9122407 | 4.77E-12 | 2.08E-10 | 514.354069 | 2328.06453 | 2254.88983 | 2431.2109 |
| -6.9093913 | 4.87E-12 | 2.11E-10 | 534.436419 | 1871.81476 | 1784.35415 | 1839.51726 |
| -6.9095112 | 4.86E-12 | 2.11E-10 | 803.294005 | 826.44532 | 651.156961 | 774.356641 |
| -6.9065519 | 4.97E-12 | 2.15E-10 | 579.519246 | 1207.4653 | 1092.9313 | 1188.43013 |
| -6.9060045 | 4.98E-12 | 2.16E-10 | 1838.96949 | 634.311666 | 1178.06489 | 1132.39853 |
| -6.905157 | 5.01E-12 | 2.17E-10 | 559.027052 | 1562.77725 | 1337.978 | 1406.39303 |
| -6.9045244 | 5.04E-12 | 2.17E-10 | 672.553807 | 1131.42367 | 753.547367 | 1043.3083 |
| -6.9041511 | 5.05E-12 | 2.18E-10 | 594.68347 | 1049.15799 | 965.230902 | 1090.93515 |
| -6.9029006 | 5.10E-12 | 2.19E-10 | 1239.36789 | 6280.87618 | 4974.56308 | 5413.7727 |
| -6.9014235 | 5.15E-12 | 2.21E-10 | 310.251817 | 676.256337 | 490.093402 | 545.187418 |
| -6.9004636 | 5.18E-12 | 2.22E-10 | 527.05923 | 2053.12398 | 1775.15051 | 1965.58835 |
| -6.9004873 | 5.18E-12 | 2.22E-10 | 548.371111 | 1548.16427 | 1417.35932 | 1504.44832 |
| -6.9006909 | 5.18E-12 | 2.22E-10 | 647.55333 | 852.423954 | 759.299637 | 807.415282 |
| -6.8999864 | 5.20E-12 | 2.22E-10 | 588.535812 | 995.847667 | 1170.01171 | 1073.56536 |
| -6.8996471 | 5.21E-12 | 2.23E-10 | 343.039328 | 837.540361 | 704.077845 | 774.356641 |
| -6.8986022 | 5.25E-12 | 2.24E-10 | 553.699082 | 1474.28752 | 1328.77436 | 1395.74703 |
| -6.8975504 | 5.29E-12 | 2.25E-10 | 1318.87761 | 989.623619 | 670.714679 | 812.458125 |
| -6.8961705 | 5.34E-12 | 2.27E-10 | 572.142056 | 1231.54966 | 1193.02079 | 1091.49547 |
| -6.8962204 | 5.34E-12 | 2.27E-10 | 590.585031 | 1044.287 | 953.726362 | 1025.37819 |
| -6.8963699 | 5.33E-12 | 2.27E-10 | 626.241449 | 873.260984 | 803.016889 | 950.856165 |
| -6.8956997 | 5.36E-12 | 2.27E-10 | 204.512096 | 271.152001 | 232.391707 | 252.142177 |
| 6.89377099 | 5.43E-12 | 2.30E-10 | 70.4931473 | 46.8156648 | 70.1776937 | 39.2221164 |
| -6.8935388 | 5.44E-12 | 2.30E-10 | 592.224407 | 1243.99775 | 961.77954 | 1120.6319 |
| -6.8931932 | 5.46E-12 | 2.30E-10 | 607.798474 | 1245.62142 | 899.655024 | 1134.07948 |
| -6.8933509 | 5.45E-12 | 2.30E-10 | 545.09236 | 1545.18755 | 1439.21795 | 1415.35809 |
| -6.8929313 | 5.47E-12 | 2.30E-10 | 561.486116 | 1336.00542 | 1213.72897 | 1211.40308 |
| -6.8918044 | 5.51E-12 | 2.32E-10 | 589.765343 | 965.809871 | 958.328178 | 1027.61945 |
| -6.8910187 | 5.54E-12 | 2.33E-10 | 623.372541 | 907.357942 | 817.972791 | 816.380337 |
| -6.8885934 | 5.63E-12 | 2.36E-10 | 549.600643 | 1492.95967 | 1281.60575 | 1435.52946 |
| -6.8881779 | 5.65E-12 | 2.37E-10 | 543.452985 | 1469.95775 | 1410.4566 | 1383.98039 |
| -6.8851504 | 5.77E-12 | 2.41E-10 | 595.503158 | 876.508314 | 638.501967 | 716.644098 |
| -6.8851384 | 5.77E-12 | 2.41E-10 | 524.600166 | 1868.83804 | 1656.65375 | 1717.9287 |

| | | | | | | |
|---|---|---|---|---|---|---|
| -6.8845919 | 5.80E-12 | 2.41E-10 | 620.09379 | 1090.29083 | 866.291859 | 1302.73458 |
| -6.8847246 | 5.79E-12 | 2.41E-10 | 572.142056 | 1063.77097 | 1040.01041 | 1144.16517 |
| 6.88371294 | 5.83E-12 | 2.43E-10 | 0 | 0 | 0 | 0 |
| -6.8818555 | 5.91E-12 | 2.45E-10 | 566.814086 | 1261.31684 | 1057.26722 | 1177.22381 |
| -6.8819413 | 5.90E-12 | 2.45E-10 | 540.584078 | 1343.58252 | 1436.91704 | 1451.77862 |
| -6.8813828 | 5.93E-12 | 2.46E-10 | 561.076272 | 1340.6058 | 1106.73674 | 1266.31404 |
| -6.879473 | 6.01E-12 | 2.49E-10 | 556.977833 | 1143.87176 | 1191.87034 | 1168.25875 |
| -6.8778234 | 6.08E-12 | 2.51E-10 | 295.497437 | 249.232528 | 318.675757 | 270.632603 |
| -6.8779908 | 6.07E-12 | 2.51E-10 | 545.502204 | 1531.3864 | 2466.57337 | 2437.93469 |
| -6.8761116 | 6.15E-12 | 2.53E-10 | 569.682993 | 1243.45653 | 1019.30224 | 1106.624 |
| -6.8761756 | 6.15E-12 | 2.53E-10 | 554.51877 | 1184.73399 | 1137.799 | 1196.83487 |
| -6.8763845 | 6.14E-12 | 2.53E-10 | 548.780955 | 1317.60388 | 1234.43714 | 1209.72213 |
| -6.8748648 | 6.20E-12 | 2.55E-10 | 611.896913 | 822.927379 | 803.016889 | 817.500969 |
| -6.8742775 | 6.23E-12 | 2.56E-10 | 943.050768 | 1016.14348 | 606.289256 | 802.372438 |
| -6.8729726 | 6.29E-12 | 2.58E-10 | 1239.77774 | 831.316314 | 725.936471 | 734.574209 |
| -6.871536 | 6.35E-12 | 2.60E-10 | 813.540102 | 772.864385 | 640.802875 | 710.480623 |
| -6.8697283 | 6.43E-12 | 2.63E-10 | 639.766297 | 1712.69562 | 949.124546 | 1144.16517 |
| -6.8692365 | 6.45E-12 | 2.63E-10 | 333.203074 | 876.508314 | 737.441011 | 762.590006 |
| -6.8692598 | 6.45E-12 | 2.63E-10 | 590.994875 | 1066.20647 | 851.335957 | 988.397333 |
| -6.8683318 | 6.50E-12 | 2.64E-10 | 626.241449 | 1340.6058 | 839.831417 | 1048.91146 |
| -6.8667011 | 6.57E-12 | 2.67E-10 | 600.421284 | 1031.56829 | 814.521429 | 918.35784 |
| -6.8659069 | 6.61E-12 | 2.68E-10 | 534.026576 | 1319.49816 | 1312.66801 | 1405.2724 |
| -6.8635816 | 6.72E-12 | 2.72E-10 | 551.649862 | 1576.03718 | 1159.65763 | 1324.5869 |
| -6.8629263 | 6.75E-12 | 2.73E-10 | 263.119771 | 676.797559 | 551.067464 | 639.880813 |
| -6.8619982 | 6.79E-12 | 2.75E-10 | 561.486116 | 1195.28781 | 983.638166 | 1172.18096 |
| -6.8613163 | 6.82E-12 | 2.76E-10 | 529.108449 | 1456.1566 | 1310.3671 | 1400.22956 |
| -6.8592203 | 6.92E-12 | 2.79E-10 | 529.518293 | 1500.80738 | 1328.77436 | 1332.99164 |
| -6.8590831 | 6.93E-12 | 2.79E-10 | 561.486116 | 1190.41682 | 984.78862 | 1064.6003 |
| -6.8570057 | 7.03E-12 | 2.83E-10 | 513.124538 | 2001.43732 | 1540.4579 | 1832.79347 |
| -6.8563606 | 7.06E-12 | 2.84E-10 | 644.684423 | 970.410254 | 694.874213 | 856.723085 |
| -6.8556739 | 7.10E-12 | 2.85E-10 | 542.633297 | 1407.98788 | 1144.70173 | 1227.09193 |
| -6.8554232 | 7.11E-12 | 2.85E-10 | 1709.45882 | 5124.28571 | 4934.29719 | 5746.04005 |
| -6.8525823 | 7.25E-12 | 2.91E-10 | 513.534382 | 1544.10511 | 1511.69655 | 1480.35474 |
| -6.8515763 | 7.30E-12 | 2.92E-10 | 565.584554 | 977.716745 | 903.106386 | 966.545011 |
| -6.8488797 | 7.44E-12 | 2.97E-10 | 605.339411 | 759.063236 | 766.202361 | 855.042138 |
| -6.8486832 | 7.45E-12 | 2.97E-10 | 538.944702 | 1670.75095 | 1191.87034 | 1427.12472 |
| -6.8475616 | 7.51E-12 | 2.99E-10 | 510.665474 | 1448.85011 | 1551.96244 | 1564.40213 |
| -6.8474984 | 7.52E-12 | 2.99E-10 | 615.585508 | 732.002158 | 794.963711 | 922.840367 |
| 6.84728664 | 7.53E-12 | 2.99E-10 | 0 | 0 | 0 | 0 |
| -6.8466828 | 7.56E-12 | 3.00E-10 | 558.207364 | 1009.91943 | 947.974092 | 950.295849 |
| -6.8451646 | 7.64E-12 | 3.03E-10 | 367.62996 | 395.632959 | 390.003904 | 449.933707 |
| -6.8440288 | 7.70E-12 | 3.05E-10 | 520.911571 | 1401.76384 | 1270.10121 | 1340.83607 |
| -6.8437673 | 7.71E-12 | 3.05E-10 | 708.210225 | 1013.97859 | 681.068765 | 725.048838 |

| | | | | | | |
|---|---|---|---|---|---|---|
| -6.8430904 | 7.75E-12 | 3.06E-10 | 595.503158 | 819.409439 | 752.396913 | 802.932754 |
| -6.8423833 | 7.79E-12 | 3.06E-10 | 579.109402 | 2193.84159 | 1149.30354 | 1613.14962 |
| -6.842347 | 7.79E-12 | 3.06E-10 | 593.863782 | 1126.82328 | 791.512349 | 1060.11777 |
| -6.842579 | 7.78E-12 | 3.06E-10 | 557.387677 | 1030.75646 | 938.77046 | 937.968898 |
| -6.8429834 | 7.76E-12 | 3.06E-10 | 475.828745 | 1012.89615 | 976.735442 | 1001.84492 |
| -6.8427534 | 7.77E-12 | 3.06E-10 | 522.141103 | 1287.56609 | 1262.04803 | 1302.73458 |
| -6.8397723 | 7.93E-12 | 3.11E-10 | 529.108449 | 1178.78055 | 1148.15309 | 1253.98709 |
| -6.8385722 | 8.00E-12 | 3.13E-10 | 559.436896 | 984.482015 | 1457.62521 | 1391.82482 |
| -6.8382961 | 8.01E-12 | 3.14E-10 | 522.550947 | 1921.60714 | 1320.72119 | 1664.13837 |
| -6.8379841 | 8.03E-12 | 3.14E-10 | 563.125491 | 889.768242 | 976.735442 | 880.256355 |
| -6.8370121 | 8.09E-12 | 3.16E-10 | 577.470027 | 825.092266 | 870.893675 | 1020.89566 |
| -6.8361648 | 8.13E-12 | 3.17E-10 | 554.108926 | 936.313296 | 918.062288 | 989.517965 |
| -6.8356492 | 8.16E-12 | 3.18E-10 | 443.860922 | 2775.65476 | 2888.78998 | 2801.01943 |
| -6.8335723 | 8.28E-12 | 3.22E-10 | 497.960314 | 1674.26889 | 1535.85608 | 1559.9196 |
| -6.8326588 | 8.34E-12 | 3.24E-10 | 529.518293 | 1360.08978 | 1103.28538 | 1219.2475 |
| -6.8319573 | 8.38E-12 | 3.25E-10 | 547.551424 | 1354.67756 | 1023.90406 | 1080.84946 |
| -6.8315698 | 8.40E-12 | 3.26E-10 | 590.585031 | 1091.10266 | 777.706901 | 927.883211 |
| -6.8300656 | 8.49E-12 | 3.29E-10 | 543.862829 | 1202.05308 | 966.381356 | 1070.76378 |
| -6.8277267 | 8.63E-12 | 3.34E-10 | 613.126444 | 849.447236 | 676.466949 | 823.664444 |
| -6.8267361 | 8.69E-12 | 3.35E-10 | 943.460612 | 886.791523 | 576.377452 | 763.150322 |
| -6.8261346 | 8.72E-12 | 3.36E-10 | 513.534382 | 1490.79478 | 1228.68487 | 1347.55986 |
| -6.8258163 | 8.74E-12 | 3.37E-10 | 538.944702 | 1084.06678 | 956.02727 | 1039.38608 |
| -6.8248711 | 8.80E-12 | 3.38E-10 | 535.256107 | 1103.55076 | 1083.72766 | 975.510067 |
| 6.82496912 | 8.79E-12 | 3.38E-10 | 0 | 0 | 0 | 0 |
| -6.8247452 | 8.81E-12 | 3.38E-10 | 508.616255 | 1432.07224 | 1257.44622 | 1484.27695 |
| -6.8237997 | 8.87E-12 | 3.40E-10 | 640.995828 | 754.192241 | 965.230902 | 682.464825 |
| -6.8217551 | 8.99E-12 | 3.44E-10 | 578.699558 | 764.20484 | 776.556447 | 860.084981 |
| -6.8202055 | 9.09E-12 | 3.48E-10 | 581.97831 | 900.322062 | 732.839195 | 842.715187 |
| -6.8196563 | 9.13E-12 | 3.49E-10 | 513.534382 | 2283.14314 | 1444.97022 | 1682.62879 |
| -6.8189563 | 9.17E-12 | 3.50E-10 | 1223.79383 | 831.045703 | 600.536986 | 844.956451 |
| -6.8179824 | 9.23E-12 | 3.52E-10 | 833.622452 | 729.837271 | 617.793796 | 669.017243 |
| -6.8177009 | 9.25E-12 | 3.52E-10 | 562.305803 | 1011.8137 | 799.565527 | 951.416481 |
| -6.8168729 | 9.30E-12 | 3.54E-10 | 3323.83387 | 1501.3486 | 814.521429 | 987.837017 |
| -6.816531 | 9.33E-12 | 3.54E-10 | 548.780955 | 848.906014 | 927.26592 | 944.132373 |
| -6.8161461 | 9.35E-12 | 3.55E-10 | 532.3872 | 1047.80494 | 957.177724 | 1022.57661 |
| -6.8147332 | 9.44E-12 | 3.58E-10 | 590.585031 | 994.494613 | 753.547367 | 1145.84611 |
| -6.8144819 | 9.46E-12 | 3.58E-10 | 1025.01954 | 970.139643 | 577.527906 | 748.582107 |
| -6.8141667 | 9.48E-12 | 3.58E-10 | 476.238588 | 1673.72767 | 2066.21538 | 1891.62664 |
| -6.8138033 | 9.51E-12 | 3.59E-10 | 483.615778 | 1632.05361 | 1630.19331 | 1553.75613 |
| -6.8139006 | 9.50E-12 | 3.59E-10 | 572.961744 | 1008.29576 | 832.928693 | 955.338692 |
| -6.8136161 | 9.52E-12 | 3.59E-10 | 533.616732 | 930.089248 | 1036.55905 | 1019.77503 |
| -6.8118573 | 9.63E-12 | 3.63E-10 | 547.961267 | 1127.90573 | 865.141405 | 1004.08618 |
| 6.81117582 | 9.68E-12 | 3.64E-10 | 347.957454 | 241.114204 | 274.958505 | 265.029444 |

| | | | | | | |
|---|---|---|---|---|---|---|
| -6.808781 | 9.84E-12 | 3.68E-10 | 453.697175 | 2204.66602 | 2081.17128 | 2196.43852 |
| -6.809194 | 9.81E-12 | 3.68E-10 | 530.337981 | 1004.77782 | 944.52273 | 983.914806 |
| -6.8089225 | 9.83E-12 | 3.68E-10 | 638.126921 | 842.681966 | 625.846973 | 762.02969 |
| -6.808914 | 9.83E-12 | 3.68E-10 | 382.794184 | 888.956409 | 625.846973 | 889.22141 |
| -6.807746 | 9.91E-12 | 3.70E-10 | 482.386247 | 1728.12044 | 1481.78475 | 1683.74943 |
| -6.8065636 | 1.00E-11 | 3.73E-10 | 507.386723 | 1434.77835 | 1186.11807 | 1223.73003 |
| -6.8055045 | 1.01E-11 | 3.75E-10 | 562.715647 | 817.244553 | 759.299637 | 802.932754 |
| -6.8044584 | 1.01E-11 | 3.77E-10 | 771.326182 | 1340.33519 | 794.963711 | 898.746782 |
| -6.8043515 | 1.02E-11 | 3.77E-10 | 545.09236 | 904.922445 | 828.326877 | 877.454776 |
| -6.8046791 | 1.01E-11 | 3.77E-10 | 517.63282 | 1095.70304 | 1042.31132 | 1039.9464 |
| -6.8043999 | 1.01E-11 | 3.77E-10 | 577.879871 | 832.398757 | 738.591465 | 728.971049 |
| -6.8039487 | 1.02E-11 | 3.77E-10 | 503.698128 | 1242.10348 | 1155.05581 | 1290.96795 |
| -6.8038149 | 1.02E-11 | 3.77E-10 | 556.977833 | 1058.08815 | 850.185503 | 836.551711 |
| -6.8029901 | 1.02E-11 | 3.79E-10 | 500.419377 | 1190.95804 | 1268.95076 | 1273.03784 |
| -6.8028092 | 1.03E-11 | 3.79E-10 | 566.404242 | 771.511331 | 746.644643 | 834.870763 |
| -6.8017687 | 1.03E-11 | 3.81E-10 | 659.438803 | 654.878086 | 667.263317 | 630.915758 |
| -6.8010263 | 1.04E-11 | 3.83E-10 | 549.600643 | 790.454086 | 893.902754 | 937.408582 |
| -6.8005756 | 1.04E-11 | 3.84E-10 | 609.028006 | 650.818924 | 697.175121 | 751.383687 |
| -6.7999035 | 1.05E-11 | 3.85E-10 | 529.108449 | 946.325895 | 908.858656 | 963.183116 |
| -6.7996244 | 1.05E-11 | 3.85E-10 | 467.222023 | 1954.08044 | 1678.51238 | 1796.93325 |
| -6.7968243 | 1.07E-11 | 3.92E-10 | 463.123584 | 1858.01361 | 1717.62782 | 1853.52516 |
| -6.7964133 | 1.07E-11 | 3.93E-10 | 558.207364 | 1086.23167 | 574.076544 | 786.683592 |
| -6.793836 | 1.09E-11 | 4.00E-10 | 494.271719 | 1394.45734 | 1201.07397 | 1286.48542 |
| -6.792723 | 1.10E-11 | 4.02E-10 | 503.698128 | 1269.16455 | 1074.52403 | 1210.84276 |
| -6.7924612 | 1.10E-11 | 4.03E-10 | 473.369681 | 2028.4984 | 1563.46698 | 1682.06848 |
| -6.7909518 | 1.11E-11 | 4.06E-10 | 493.042188 | 1322.20427 | 1237.8885 | 1199.63645 |
| -6.7900074 | 1.12E-11 | 4.09E-10 | 482.386247 | 1662.90324 | 1363.28798 | 1440.5723 |
| -6.7893093 | 1.13E-11 | 4.10E-10 | 4201.30961 | 1217.4779 | 933.01819 | 1219.80782 |
| -6.7886949 | 1.13E-11 | 4.11E-10 | 1309.4512 | 846.470517 | 635.050605 | 743.539264 |
| -6.7875249 | 1.14E-11 | 4.14E-10 | 498.370158 | 1236.42065 | 1148.15309 | 1112.22716 |
| -6.7872528 | 1.14E-11 | 4.15E-10 | 588.945655 | 1206.65346 | 737.441011 | 949.175217 |
| -6.7870619 | 1.14E-11 | 4.15E-10 | 490.17328 | 1322.20427 | 1183.81716 | 1359.32649 |
| -6.785782 | 1.15E-11 | 4.18E-10 | 471.730306 | 680.315499 | 586.731538 | 655.009344 |
| -6.785573 | 1.16E-11 | 4.18E-10 | 693.455845 | 695.469702 | 582.129722 | 680.783878 |
| -6.7837152 | 1.17E-11 | 4.23E-10 | 458.615302 | 1719.7315 | 1677.36193 | 1862.49021 |
| -6.7827853 | 1.18E-11 | 4.25E-10 | 600.01144 | 754.192241 | 629.298335 | 711.040939 |
| 6.78261145 | 1.18E-11 | 4.25E-10 | 10.6559409 | 9.20076649 | 9.20363196 | 10.646003 |
| 6.78064671 | 1.20E-11 | 4.31E-10 | 134.018949 | 105.808815 | 97.7885896 | 114.304454 |
| -6.7798664 | 1.20E-11 | 4.32E-10 | 620.503634 | 620.239906 | 758.149183 | 889.22141 |
| -6.7799214 | 1.20E-11 | 4.32E-10 | 565.584554 | 858.918613 | 692.573305 | 773.236009 |
| -6.7786255 | 1.21E-11 | 4.35E-10 | 525.419854 | 964.186206 | 855.937773 | 1090.93515 |
| -6.7786918 | 1.21E-11 | 4.35E-10 | 505.747348 | 1027.77974 | 1337.978 | 1125.11442 |
| -6.7778463 | 1.22E-11 | 4.36E-10 | 525.829698 | 853.235787 | 852.486411 | 945.253005 |

| | | | | | | |
|---|---|---|---|---|---|---|
| -6.777959 | 1.22E-11 | 4.36E-10 | 510.255631 | 1336.81725 | 1004.34634 | 1098.77958 |
| -6.7773451 | 1.22E-11 | 4.37E-10 | 493.042188 | 1550.32915 | 1193.02079 | 1235.49667 |
| -6.7773101 | 1.22E-11 | 4.37E-10 | 550.010487 | 809.938062 | 717.883293 | 824.785076 |
| -6.7768624 | 1.23E-11 | 4.38E-10 | 615.995352 | 927.38314 | 620.094704 | 815.259705 |
| -6.776032 | 1.24E-11 | 4.40E-10 | 670.914431 | 604.003259 | 632.749697 | 661.733135 |
| -6.7756991 | 1.24E-11 | 4.40E-10 | 220.086164 | 372.360432 | 316.374849 | 319.380091 |
| -6.775816 | 1.24E-11 | 4.40E-10 | 543.043141 | 1133.58855 | 789.211441 | 1016.41313 |
| -6.7741757 | 1.25E-11 | 4.44E-10 | 524.190322 | 921.159092 | 819.123245 | 954.21806 |
| -6.7728054 | 1.26E-11 | 4.48E-10 | 494.271719 | 1450.20317 | 1120.54219 | 1190.67139 |
| -6.7717807 | 1.27E-11 | 4.50E-10 | 548.371111 | 1034.2744 | 911.159564 | 745.220212 |
| -6.7704879 | 1.28E-11 | 4.54E-10 | 474.189369 | 1518.9383 | 1285.05711 | 1367.73123 |
| -6.7702967 | 1.29E-11 | 4.54E-10 | 600.01144 | 687.351379 | 615.492888 | 740.737684 |
| -6.7700843 | 1.29E-11 | 4.54E-10 | 453.287331 | 1866.67315 | 1576.12197 | 1846.80137 |
| -6.7693849 | 1.29E-11 | 4.56E-10 | 532.3872 | 1009.37821 | 783.459171 | 903.789625 |
| 6.76700433 | 1.31E-11 | 4.63E-10 | 0.40984388 | 1.08244312 | 0 | 2.24126379 |
| -6.7662647 | 1.32E-11 | 4.65E-10 | 525.01001 | 939.019404 | 800.715981 | 851.119926 |
| -6.7655481 | 1.33E-11 | 4.67E-10 | 479.51734 | 1669.93912 | 1228.68487 | 1414.79777 |
| -6.7635737 | 1.35E-11 | 4.72E-10 | 453.287331 | 1634.75972 | 1531.25427 | 1684.87006 |
| 6.76323276 | 1.35E-11 | 4.73E-10 | 30.3284471 | 22.4606947 | 23.0090799 | 21.852322 |
| -6.7629117 | 1.35E-11 | 4.74E-10 | 559.436896 | 891.933128 | 675.316495 | 806.29465 |
| -6.7620666 | 1.36E-11 | 4.76E-10 | 497.960314 | 1246.43325 | 968.682264 | 1133.51916 |
| -6.7618944 | 1.36E-11 | 4.76E-10 | 588.535812 | 772.593774 | 1400.10251 | 953.097429 |
| -6.759981 | 1.38E-11 | 4.82E-10 | 861.491836 | 1008.56637 | 536.111562 | 752.504319 |
| -6.7591655 | 1.39E-11 | 4.84E-10 | 486.074842 | 1070.80685 | 1064.16995 | 1109.98589 |
| -6.7587737 | 1.39E-11 | 4.84E-10 | 468.861399 | 1236.42065 | 1362.13753 | 1413.67714 |
| -6.7589073 | 1.39E-11 | 4.84E-10 | 388.531998 | 391.032576 | 283.011683 | 307.613456 |
| -6.7584954 | 1.39E-11 | 4.85E-10 | 548.371111 | 691.951762 | 733.989649 | 724.488522 |
| -6.7571521 | 1.41E-11 | 4.89E-10 | 500.009534 | 1768.44144 | 1769.39824 | 1739.2207 |
| -6.7555164 | 1.42E-11 | 4.94E-10 | 486.484685 | 1103.82137 | 1010.09861 | 1104.38273 |
| -6.7550892 | 1.43E-11 | 4.95E-10 | 518.452508 | 900.592673 | 778.857355 | 883.057935 |
| -6.7545794 | 1.43E-11 | 4.96E-10 | 506.976879 | 1232.90271 | 899.655024 | 1251.18551 |
| -6.7530763 | 1.45E-11 | 5.00E-10 | 475.009057 | 1294.06075 | 1115.94038 | 1241.09983 |
| -6.7526725 | 1.45E-11 | 5.01E-10 | 440.582171 | 1064.31219 | 922.664104 | 1122.87316 |
| -6.751342 | 1.46E-11 | 5.05E-10 | 506.567036 | 1017.49653 | 840.981871 | 936.848266 |
| -6.7512503 | 1.47E-11 | 5.05E-10 | 538.534858 | 757.980792 | 699.476029 | 740.737684 |
| -6.7493914 | 1.48E-11 | 5.11E-10 | 523.370635 | 1291.08403 | 839.831417 | 1058.99714 |
| -6.749349 | 1.49E-11 | 5.11E-10 | 521.321415 | 934.41902 | 762.750999 | 840.473923 |
| -6.7490784 | 1.49E-11 | 5.11E-10 | 1388.96091 | 4384.16523 | 3581.36329 | 4033.1542 |
| -6.7470496 | 1.51E-11 | 5.18E-10 | 529.108449 | 992.600338 | 775.405993 | 793.967699 |
| -6.7459809 | 1.52E-11 | 5.20E-10 | 1037.31486 | 545.821942 | 712.131023 | 860.084981 |
| -6.7460772 | 1.52E-11 | 5.20E-10 | 517.222976 | 890.850685 | 811.070067 | 770.434429 |
| -6.7455583 | 1.52E-11 | 5.21E-10 | 584.847217 | 774.758661 | 609.740618 | 675.741034 |
| -6.7444281 | 1.54E-11 | 5.25E-10 | 448.369205 | 2683.6471 | 1723.38009 | 1865.85211 |

| | | | | | | |
|---|---|---|---|---|---|---|
| -6.7442096 | 1.54E-11 | 5.25E-10 | 518.862352 | 763.122397 | 785.760079 | 782.201064 |
| -6.7429382 | 1.55E-11 | 5.29E-10 | 457.795614 | 1371.72604 | 1275.85348 | 1332.43133 |
| -6.7424914 | 1.56E-11 | 5.30E-10 | 485.255154 | 1185.54582 | 964.080448 | 1070.76378 |
| -6.7422207 | 1.56E-11 | 5.31E-10 | 355.744488 | 625.922732 | 436.022064 | 511.568461 |
| -6.7418732 | 1.56E-11 | 5.31E-10 | 425.008103 | 1829.32887 | 1914.35545 | 2070.36743 |
| -6.7410484 | 1.57E-11 | 5.33E-10 | 469.271243 | 1431.53102 | 1184.96762 | 1174.42223 |
| -6.7411074 | 1.57E-11 | 5.33E-10 | 470.910618 | 1122.2229 | 1103.28538 | 1179.46507 |
| -6.7389136 | 1.60E-11 | 5.41E-10 | 557.387677 | 922.512146 | 646.555145 | 824.22476 |
| -6.7384633 | 1.60E-11 | 5.41E-10 | 572.142056 | 657.584193 | 737.441011 | 615.226912 |
| -6.7384966 | 1.60E-11 | 5.41E-10 | 551.240019 | 799.925463 | 636.201059 | 760.909058 |
| -6.737922 | 1.61E-11 | 5.43E-10 | 654.520676 | 765.557894 | 562.572004 | 641.001445 |
| -6.7364048 | 1.62E-11 | 5.48E-10 | 472.959837 | 1688.61126 | 1133.19719 | 1581.21161 |
| 6.73584069 | 1.63E-11 | 5.49E-10 | 141.805982 | 110.138587 | 98.9390436 | 94.6933953 |
| -6.7354146 | 1.63E-11 | 5.50E-10 | 494.681563 | 1167.14429 | 893.902754 | 977.75133 |
| -6.7334305 | 1.66E-11 | 5.57E-10 | 535.665951 | 820.762493 | 967.53181 | 695.352092 |
| -6.7330122 | 1.66E-11 | 5.58E-10 | 490.583124 | 1352.51267 | 1089.47993 | 1240.53951 |
| -6.7331202 | 1.66E-11 | 5.58E-10 | 556.977833 | 717.659786 | 624.696519 | 667.896611 |
| -6.7311614 | 1.68E-11 | 5.64E-10 | 414.352163 | 2024.16863 | 1908.60318 | 2175.70683 |
| -6.7301055 | 1.70E-11 | 5.68E-10 | 276.644619 | 299.566133 | 246.197155 | 291.364293 |
| -6.7294864 | 1.70E-11 | 5.70E-10 | 485.255154 | 1026.6973 | 1040.01041 | 879.135723 |
| -6.7290489 | 1.71E-11 | 5.71E-10 | 503.288285 | 970.680865 | 774.255539 | 931.245107 |
| -6.7284282 | 1.72E-11 | 5.73E-10 | 510.665474 | 851.0709 | 769.653723 | 750.823371 |
| -6.7279965 | 1.72E-11 | 5.74E-10 | 1492.24157 | 956.338493 | 540.713378 | 866.808773 |
| -6.7277183 | 1.72E-11 | 5.74E-10 | 468.451555 | 721.718948 | 725.936471 | 695.352092 |
| -6.7268519 | 1.73E-11 | 5.77E-10 | 405.335597 | 977.987356 | 987.089528 | 981.673542 |
| -6.7261518 | 1.74E-11 | 5.79E-10 | 495.911095 | 895.99229 | 1199.92352 | 1151.44927 |
| -6.7256328 | 1.75E-11 | 5.80E-10 | 421.729352 | 1035.35684 | 1013.54997 | 1043.3083 |
| -6.7255621 | 1.75E-11 | 5.80E-10 | 457.38577 | 1228.84355 | 1136.64855 | 1217.00624 |
| -6.7250978 | 1.75E-11 | 5.82E-10 | 461.074365 | 1194.20537 | 1091.78084 | 1153.69054 |
| -6.7228648 | 1.78E-11 | 5.90E-10 | 559.027052 | 642.159379 | 629.298335 | 764.83127 |
| -6.722407 | 1.79E-11 | 5.91E-10 | 419.270289 | 1778.18343 | 1736.03508 | 1918.52181 |
| -6.7190204 | 1.83E-11 | 6.05E-10 | 541.403765 | 896.262901 | 685.670581 | 692.550512 |
| -6.7148169 | 1.88E-11 | 6.21E-10 | 512.714694 | 936.313296 | 725.936471 | 779.9598 |
| -6.7144801 | 1.89E-11 | 6.21E-10 | 501.239065 | 801.819739 | 733.989649 | 849.438978 |
| -6.7145564 | 1.89E-11 | 6.21E-10 | 509.845787 | 1328.42831 | 845.583687 | 968.225959 |
| -6.7147473 | 1.88E-11 | 6.21E-10 | 451.647956 | 1312.73289 | 1120.54219 | 1342.51701 |
| -6.7123491 | 1.92E-11 | 6.30E-10 | 426.647479 | 1562.77725 | 1505.94428 | 1546.47202 |
| -6.7122356 | 1.92E-11 | 6.30E-10 | 458.205458 | 1252.11608 | 1331.07527 | 1044.42893 |
| 6.71007179 | 1.95E-11 | 6.38E-10 | 24.5906328 | 13.2599282 | 21.8586259 | 13.4475828 |
| -6.7100656 | 1.95E-11 | 6.38E-10 | 449.598736 | 873.802206 | 651.156961 | 741.858316 |
| -6.7092503 | 1.96E-11 | 6.41E-10 | 459.025146 | 1245.08019 | 1044.61223 | 1076.36694 |
| -6.7090909 | 1.96E-11 | 6.41E-10 | 534.436419 | 657.313583 | 630.448789 | 709.920307 |
| -6.7082426 | 1.97E-11 | 6.44E-10 | 452.467643 | 1162.00269 | 1074.52403 | 1151.44927 |

| | | | | | | |
|---|---|---|---|---|---|---|
| -6.7080056 | 1.97E-11 | 6.44E-10 | 604.519723 | 752.027355 | 861.690043 | 552.471525 |
| -6.707414 | 1.98E-11 | 6.46E-10 | 640.995828 | 532.832624 | 586.731538 | 489.155823 |
| -6.7064787 | 1.99E-11 | 6.50E-10 | 424.188416 | 2113.47019 | 1600.28151 | 1617.63214 |
| -6.7058178 | 2.00E-11 | 6.52E-10 | 421.729352 | 878.943811 | 857.088227 | 847.75803 |
| -6.7043081 | 2.02E-11 | 6.58E-10 | 547.961267 | 975.281248 | 625.846973 | 845.516766 |
| -6.7008311 | 2.07E-11 | 6.73E-10 | 490.583124 | 1261.85806 | 897.354116 | 1006.88776 |
| -6.6994764 | 2.09E-11 | 6.79E-10 | 491.812656 | 818.326996 | 761.600545 | 743.539264 |
| -6.699265 | 2.09E-11 | 6.79E-10 | 421.319509 | 1663.98568 | 1433.46568 | 1568.88466 |
| -6.6989918 | 2.10E-11 | 6.80E-10 | 454.107019 | 993.41217 | 1134.34764 | 1085.89231 |
| -6.6988383 | 2.10E-11 | 6.80E-10 | 525.419854 | 846.199906 | 637.351513 | 736.255156 |
| -6.6977778 | 2.12E-11 | 6.84E-10 | 472.549994 | 940.913679 | 830.627785 | 913.314996 |
| -6.6972909 | 2.12E-11 | 6.85E-10 | 442.63139 | 1337.35847 | 1140.09991 | 1185.06823 |
| -6.6973733 | 2.12E-11 | 6.85E-10 | 458.205458 | 1211.79507 | 965.230902 | 1260.71088 |
| -6.6965384 | 2.13E-11 | 6.88E-10 | 503.698128 | 713.600625 | 930.717282 | 1032.10198 |
| -6.6962543 | 2.14E-11 | 6.88E-10 | 509.845787 | 671.926565 | 920.363196 | 851.119926 |
| -6.6962351 | 2.14E-11 | 6.88E-10 | 438.942795 | 1146.30726 | 1394.35024 | 1385.10102 |
| -6.695761 | 2.15E-11 | 6.89E-10 | 458.615302 | 966.351092 | 953.726362 | 1014.17187 |
| -6.6946018 | 2.16E-11 | 6.94E-10 | 466.402335 | 873.531595 | 976.735442 | 908.832469 |
| -6.6938974 | 2.17E-11 | 6.97E-10 | 466.812179 | 771.511331 | 637.351513 | 689.188617 |
| -6.6935222 | 2.18E-11 | 6.98E-10 | 487.714217 | 941.18429 | 746.644643 | 834.870763 |
| -6.6923843 | 2.20E-11 | 7.02E-10 | 486.074842 | 937.66635 | 747.795097 | 840.473923 |
| -6.6923454 | 2.20E-11 | 7.02E-10 | 484.025622 | 834.022421 | 740.892373 | 847.197714 |
| 6.69195718 | 2.20E-11 | 7.03E-10 | 0 | 0 | 0 | 0 |
| -6.6872142 | 2.27E-11 | 7.26E-10 | 561.895959 | 953.902996 | 633.900151 | 670.137875 |
| -6.6860618 | 2.29E-11 | 7.30E-10 | 417.630914 | 1860.1785 | 1407.00524 | 1606.98614 |
| -6.6860354 | 2.29E-11 | 7.30E-10 | 447.549517 | 1264.83478 | 1080.2763 | 1025.9385 |
| -6.6857621 | 2.30E-11 | 7.31E-10 | 520.911571 | 1237.23248 | 745.494189 | 866.248457 |
| 6.68551038 | 2.30E-11 | 7.31E-10 | 0 | 0 | 0 | 0 |
| -6.6830791 | 2.34E-11 | 7.43E-10 | 585.257061 | 1006.6721 | 587.881992 | 735.134525 |
| -6.6828336 | 2.34E-11 | 7.43E-10 | 497.55047 | 805.60829 | 693.723759 | 693.671144 |
| -6.6825293 | 2.35E-11 | 7.44E-10 | 453.697175 | 559.08187 | 534.961108 | 614.666596 |
| -6.6820903 | 2.36E-11 | 7.46E-10 | 404.925753 | 1649.91392 | 1589.92742 | 1640.6051 |
| -6.681944 | 2.36E-11 | 7.46E-10 | 494.681563 | 701.42314 | 712.131023 | 835.991395 |
| -6.6813729 | 2.37E-11 | 7.48E-10 | 477.46812 | 846.741128 | 750.096005 | 788.36454 |
| -6.68129 | 2.37E-11 | 7.48E-10 | 529.108449 | 594.261271 | 468.234776 | 573.763531 |
| -6.6809325 | 2.37E-11 | 7.49E-10 | 499.59969 | 677.068169 | 669.564225 | 739.056736 |
| -6.6805419 | 2.38E-11 | 7.50E-10 | 484.84531 | 768.805224 | 706.378753 | 814.699389 |
| -6.680124 | 2.39E-11 | 7.51E-10 | 445.910141 | 1126.01145 | 962.929994 | 1142.48422 |
| -6.6797366 | 2.39E-11 | 7.53E-10 | 454.107019 | 1128.71756 | 900.805478 | 1038.82577 |
| -6.6769352 | 2.44E-11 | 7.66E-10 | 442.221546 | 1098.40915 | 984.78862 | 1075.80662 |
| -6.6761308 | 2.45E-11 | 7.70E-10 | 468.451555 | 797.219355 | 807.618705 | 907.151521 |
| -6.6756189 | 2.46E-11 | 7.72E-10 | 452.0578 | 894.368625 | 939.920914 | 985.595754 |
| -6.6745949 | 2.48E-11 | 7.76E-10 | 601.650816 | 874.884649 | 532.6602 | 715.523466 |

| | | | | | | |
|---|---|---|---|---|---|---|
| -6.6743723 | 2.48E-11 | 7.77E-10 | 456.975926 | 839.975858 | 930.717282 | 962.6228 |
| -6.6738571 | 2.49E-11 | 7.79E-10 | 492.632344 | 903.569392 | 721.334655 | 714.402834 |
| -6.6737074 | 2.49E-11 | 7.79E-10 | 529.518293 | 906.816721 | 609.740618 | 744.09958 |
| -6.6734407 | 2.50E-11 | 7.79E-10 | 483.615778 | 880.296865 | 838.680963 | 694.23146 |
| -6.6734699 | 2.50E-11 | 7.79E-10 | 485.255154 | 1080.81945 | 753.547367 | 860.645297 |
| -6.6727228 | 2.51E-11 | 7.81E-10 | 490.17328 | 665.431906 | 837.530509 | 825.905708 |
| -6.6727319 | 2.51E-11 | 7.81E-10 | 553.289238 | 970.680865 | 666.112863 | 637.079234 |
| -6.6725801 | 2.51E-11 | 7.81E-10 | 463.533428 | 1657.76163 | 1446.12067 | 1510.6118 |
| -6.6719975 | 2.52E-11 | 7.83E-10 | 445.500297 | 1005.86027 | 935.319098 | 981.673542 |
| -6.6718054 | 2.53E-11 | 7.84E-10 | 541.403765 | 585.872337 | 577.527906 | 638.760181 |
| -6.6715006 | 2.53E-11 | 7.85E-10 | 395.499344 | 1814.44527 | 1631.34377 | 2091.65944 |
| -6.6713315 | 2.53E-11 | 7.85E-10 | 482.796091 | 761.228122 | 686.821035 | 756.986847 |
| -6.6701715 | 2.56E-11 | 7.90E-10 | 446.319985 | 1157.4023 | 924.965012 | 1042.74798 |
| -6.6694418 | 2.57E-11 | 7.93E-10 | 471.730306 | 765.016673 | 769.653723 | 754.185267 |
| -6.6691395 | 2.57E-11 | 7.94E-10 | 489.763437 | 741.744146 | 658.059685 | 712.161571 |
| -6.6687812 | 2.58E-11 | 7.95E-10 | 444.270766 | 999.636218 | 935.319098 | 964.303748 |
| -6.6688725 | 2.58E-11 | 7.95E-10 | 443.041234 | 1062.14731 | 943.372276 | 988.957649 |
| -6.668264 | 2.59E-11 | 7.97E-10 | 451.238112 | 1014.79042 | 869.743221 | 926.762579 |
| -6.6675703 | 2.60E-11 | 8.00E-10 | 452.0578 | 937.395739 | 1214.87942 | 954.778376 |
| -6.6669028 | 2.61E-11 | 8.03E-10 | 444.270766 | 1026.15607 | 959.478632 | 1224.29035 |
| -6.6667193 | 2.62E-11 | 8.03E-10 | 619.683946 | 667.867403 | 506.199758 | 628.674494 |
| -6.665845 | 2.63E-11 | 8.06E-10 | 452.877487 | 1120.32863 | 877.796398 | 944.692689 |
| -6.6658615 | 2.63E-11 | 8.06E-10 | 527.469073 | 570.176912 | 765.051907 | 765.391586 |
| -6.6647896 | 2.65E-11 | 8.11E-10 | 461.894053 | 1056.73509 | 819.123245 | 874.653196 |
| -6.6631097 | 2.68E-11 | 8.20E-10 | 1375.02622 | 866.766326 | 575.226998 | 664.534715 |
| -6.6625401 | 2.69E-11 | 8.22E-10 | 537.305327 | 983.399571 | 623.546066 | 713.842519 |
| -6.6624626 | 2.69E-11 | 8.22E-10 | 554.108926 | 1092.1851 | 1011.24906 | 1033.78293 |
| -6.6622929 | 2.70E-11 | 8.22E-10 | 395.0895 | 1583.88489 | 1580.72379 | 1728.01439 |
| -6.6620497 | 2.70E-11 | 8.23E-10 | 504.517816 | 396.986013 | 380.800272 | 414.073486 |
| -6.6611584 | 2.72E-11 | 8.25E-10 | 491.812656 | 776.652936 | 643.103783 | 694.23146 |
| -6.6611203 | 2.72E-11 | 8.25E-10 | 404.925753 | 1064.04158 | 862.840497 | 1014.73218 |
| -6.6611497 | 2.72E-11 | 8.25E-10 | 330.744011 | 427.565031 | 332.481205 | 358.602207 |
| -6.6608876 | 2.72E-11 | 8.26E-10 | 310.661661 | 590.47272 | 517.704298 | 553.592157 |
| -6.6602426 | 2.73E-11 | 8.28E-10 | 284.021809 | 614.286469 | 554.518826 | 685.266405 |
| -6.6603518 | 2.73E-11 | 8.28E-10 | 679.930997 | 643.512433 | 568.324274 | 521.093832 |
| -6.6589113 | 2.76E-11 | 8.35E-10 | 463.943272 | 810.749894 | 737.441011 | 829.82792 |
| -6.6577976 | 2.78E-11 | 8.40E-10 | 426.647479 | 1031.56829 | 1172.31262 | 1186.74918 |
| -6.6575654 | 2.78E-11 | 8.41E-10 | 417.630914 | 1195.28781 | 1233.28668 | 1176.66349 |
| -6.6556724 | 2.82E-11 | 8.51E-10 | 414.762006 | 691.13993 | 656.909231 | 697.593356 |
| -6.6546459 | 2.84E-11 | 8.56E-10 | 442.63139 | 813.456002 | 746.644643 | 732.332945 |
| -6.6537939 | 2.86E-11 | 8.60E-10 | 467.222023 | 746.344529 | 714.431931 | 791.726435 |
| -6.6520154 | 2.89E-11 | 8.69E-10 | 429.92623 | 1335.19358 | 1040.01041 | 1083.09073 |
| 6.65210643 | 2.89E-11 | 8.69E-10 | 25.8201644 | 15.966036 | 16.1063559 | 18.4904263 |

| | | | | | | |
|---|---|---|---|---|---|---|
| -6.6518701 | 2.89E-11 | 8.69E-10 | 418.040758 | 1839.07086 | 1248.24258 | 1396.30734 |
| -6.6478723 | 2.97E-11 | 8.92E-10 | 586.076748 | 660.560912 | 561.42155 | 526.696992 |
| -6.6471395 | 2.99E-11 | 8.96E-10 | 452.877487 | 838.081583 | 759.299637 | 846.637398 |
| -6.6468687 | 2.99E-11 | 8.97E-10 | 474.599213 | 742.555978 | 658.059685 | 712.721887 |
| -6.6459886 | 3.01E-11 | 9.01E-10 | 407.794661 | 1319.76877 | 1274.70303 | 1205.79992 |
| -6.6452497 | 3.03E-11 | 9.05E-10 | 397.13872 | 1344.93557 | 1792.40732 | 1733.61755 |
| -6.6449548 | 3.03E-11 | 9.06E-10 | 461.484209 | 1420.16537 | 852.486411 | 1098.77958 |
| -6.6446318 | 3.04E-11 | 9.07E-10 | 585.257061 | 624.028457 | 600.536986 | 506.525618 |
| -6.6426093 | 3.08E-11 | 9.19E-10 | 432.385293 | 1240.2092 | 939.920914 | 1047.23051 |
| -6.6423676 | 3.09E-11 | 9.19E-10 | 450.418424 | 969.327811 | 771.954631 | 871.2913 |
| -6.6419503 | 3.10E-11 | 9.20E-10 | 479.51734 | 844.305631 | 646.555145 | 736.255156 |
| -6.6418783 | 3.10E-11 | 9.20E-10 | 423.368728 | 1074.5954 | 993.992252 | 1024.81787 |
| -6.6418062 | 3.10E-11 | 9.20E-10 | 466.812179 | 789.642254 | 694.874213 | 906.030889 |
| -6.6419347 | 3.10E-11 | 9.20E-10 | 433.614825 | 1060.25303 | 899.655024 | 944.692689 |
| -6.6404302 | 3.13E-11 | 9.27E-10 | 525.829698 | 582.895618 | 559.120642 | 575.444479 |
| -6.6402446 | 3.13E-11 | 9.28E-10 | 656.569896 | 634.852888 | 486.64204 | 612.425332 |
| -6.6401191 | 3.13E-11 | 9.28E-10 | 413.532475 | 1221.26645 | 1076.82494 | 1186.74918 |
| -6.6393686 | 3.15E-11 | 9.31E-10 | 479.927183 | 859.730445 | 636.201059 | 781.640748 |
| -6.6390551 | 3.16E-11 | 9.33E-10 | 538.125014 | 972.304529 | 590.1829 | 704.317147 |
| -6.6385618 | 3.17E-11 | 9.35E-10 | 516.813133 | 796.407523 | 570.625182 | 657.250608 |
| -6.6378702 | 3.18E-11 | 9.38E-10 | 398.778095 | 1127.90573 | 885.849576 | 922.840367 |
| -6.6366732 | 3.21E-11 | 9.45E-10 | 449.598736 | 764.475451 | 768.503269 | 802.932754 |
| -6.6355701 | 3.23E-11 | 9.51E-10 | 427.467167 | 1223.97255 | 944.52273 | 1037.70514 |
| -6.6354246 | 3.24E-11 | 9.51E-10 | 430.745918 | 936.854517 | 907.708202 | 894.82457 |
| -6.6353645 | 3.24E-11 | 9.51E-10 | 220.086164 | 568.553247 | 544.16474 | 499.24151 |
| -6.6348798 | 3.25E-11 | 9.53E-10 | 428.696698 | 1106.52748 | 949.124546 | 926.202263 |
| -6.6329832 | 3.29E-11 | 9.65E-10 | 550.010487 | 529.585295 | 844.433233 | 701.515568 |
| 6.63197499 | 3.31E-11 | 9.70E-10 | 22.9512573 | 21.6488623 | 17.2568099 | 12.3269509 |
| -6.6316097 | 3.32E-11 | 9.72E-10 | 569.682993 | 552.3166 | 546.465648 | 537.903311 |
| -6.6305666 | 3.34E-11 | 9.78E-10 | 434.024669 | 1680.76355 | 1046.91314 | 1169.37938 |
| -6.6298052 | 3.36E-11 | 9.82E-10 | 301.645096 | 572.071187 | 569.474728 | 571.522268 |
| -6.6296559 | 3.36E-11 | 9.82E-10 | 463.123584 | 832.128146 | 693.723759 | 699.83462 |
| -6.6289751 | 3.38E-11 | 9.86E-10 | 470.500774 | 735.520098 | 648.856053 | 849.438978 |
| -6.6288378 | 3.38E-11 | 9.86E-10 | 477.877964 | 645.136097 | 929.566828 | 776.037589 |
| 6.6286655 | 3.39E-11 | 9.86E-10 | 0 | 0 | 0 | 0 |
| -6.6282059 | 3.40E-11 | 9.88E-10 | 445.090454 | 956.609104 | 803.016889 | 779.9598 |
| 6.62797553 | 3.40E-11 | 9.89E-10 | 4.50828268 | 2.16488623 | 1.150454 | 4.48252759 |
| -6.6270133 | 3.43E-11 | 9.95E-10 | 409.434036 | 1127.36451 | 1061.86904 | 1099.90021 |
| -6.6261707 | 3.45E-11 | 9.99E-10 | 404.925753 | 1113.02213 | 1168.86126 | 1302.73458 |
| -6.6258152 | 3.45E-11 | 1.00E-09 | 459.434989 | 665.702517 | 713.281477 | 752.504319 |
| -6.6254975 | 3.46E-11 | 1.00E-09 | 814.359789 | 762.581176 | 488.942948 | 583.849218 |
| -6.6250959 | 3.47E-11 | 1.00E-09 | 416.401382 | 1167.95612 | 956.02727 | 1101.02084 |
| -6.6231043 | 3.52E-11 | 1.02E-09 | 460.254677 | 728.213607 | 659.210139 | 692.550512 |

| | | | | | | |
|---|---|---|---|---|---|---|
| -6.6223632 | 3.54E-11 | 1.02E-09 | 568.863305 | 829.692649 | 517.704298 | 637.079234 |
| -6.6221547 | 3.54E-11 | 1.02E-09 | 322.956977 | 328.521486 | 411.86253 | 410.151274 |
| -6.6218593 | 3.55E-11 | 1.02E-09 | 416.811226 | 969.0572 | 1347.18163 | 1227.65224 |
| -6.6210064 | 3.57E-11 | 1.03E-09 | 433.614825 | 913.852601 | 813.370975 | 818.061285 |
| -6.6208351 | 3.57E-11 | 1.03E-09 | 473.369681 | 787.206757 | 610.891072 | 714.402834 |
| -6.6192426 | 3.61E-11 | 1.04E-09 | 440.992015 | 844.846852 | 739.741919 | 787.804224 |
| -6.6187845 | 3.62E-11 | 1.04E-09 | 411.893099 | 1110.85725 | 992.841798 | 1007.44808 |
| -6.618713 | 3.62E-11 | 1.04E-09 | 455.336551 | 689.245654 | 692.573305 | 685.826721 |
| -6.6172731 | 3.66E-11 | 1.05E-09 | 407.384817 | 1070.26563 | 1571.52016 | 1524.05938 |
| -6.6166589 | 3.67E-11 | 1.05E-09 | 446.319985 | 1174.72139 | 826.025969 | 849.999294 |
| -6.6152299 | 3.71E-11 | 1.06E-09 | 438.123108 | 812.914781 | 743.193281 | 783.321696 |
| -6.6134789 | 3.75E-11 | 1.07E-09 | 426.647479 | 853.506397 | 816.822337 | 903.229309 |
| -6.612375 | 3.78E-11 | 1.08E-09 | 462.71374 | 622.946014 | 662.661501 | 692.550512 |
| -6.6124087 | 3.78E-11 | 1.08E-09 | 585.666904 | 515.513534 | 532.6602 | 559.195317 |
| -6.6119972 | 3.79E-11 | 1.08E-09 | 402.056846 | 824.551044 | 669.564225 | 700.394936 |
| -6.6105472 | 3.83E-11 | 1.09E-09 | 438.532952 | 942.537344 | 733.989649 | 842.154871 |
| -6.6103694 | 3.83E-11 | 1.09E-09 | 490.583124 | 717.118565 | 557.970188 | 632.03639 |
| -6.6095147 | 3.86E-11 | 1.10E-09 | 444.270766 | 694.65787 | 547.616102 | 669.577559 |
| -6.6090203 | 3.87E-11 | 1.10E-09 | 408.204504 | 991.517895 | 983.638166 | 1002.40523 |
| -6.6062936 | 3.94E-11 | 1.12E-09 | 416.811226 | 1081.36067 | 911.159564 | 900.988045 |
| -6.6055822 | 3.96E-11 | 1.12E-09 | 410.253724 | 939.019404 | 982.487712 | 943.572057 |
| -6.60491 | 3.98E-11 | 1.13E-09 | 458.205458 | 732.543379 | 625.846973 | 666.215663 |
| -6.6042042 | 4.00E-11 | 1.13E-09 | 431.975449 | 755.815906 | 761.600545 | 828.146972 |
| -6.6028595 | 4.03E-11 | 1.14E-09 | 417.22107 | 1058.35876 | 849.035049 | 950.856165 |
| -6.602191 | 4.05E-11 | 1.15E-09 | 386.072935 | 745.262086 | 762.750999 | 795.648647 |
| -6.6020016 | 4.06E-11 | 1.15E-09 | 483.205934 | 837.269751 | 577.527906 | 702.6362 |
| 6.60171586 | 4.06E-11 | 1.15E-09 | 0 | 0 | 0 | 0 |
| -6.6015731 | 4.07E-11 | 1.15E-09 | 397.958407 | 1222.07828 | 1045.76268 | 1085.33199 |
| -6.6004052 | 4.10E-11 | 1.15E-09 | 458.205458 | 842.952577 | 623.546066 | 753.064635 |
| -6.6004313 | 4.10E-11 | 1.15E-09 | 478.287808 | 559.893702 | 739.741919 | 729.531365 |
| -6.5999662 | 4.11E-11 | 1.16E-09 | 503.288285 | 561.787978 | 905.407294 | 716.083782 |
| -6.5973316 | 4.19E-11 | 1.18E-09 | 445.500297 | 671.385343 | 663.811955 | 727.850417 |
| -6.5966164 | 4.21E-11 | 1.18E-09 | 432.385293 | 1226.67866 | 999.744522 | 1134.07948 |
| -6.5966201 | 4.21E-11 | 1.18E-09 | 422.958884 | 859.189224 | 793.813257 | 787.243908 |
| -6.5948955 | 4.26E-11 | 1.19E-09 | 413.122631 | 1216.39545 | 907.708202 | 968.786275 |
| -6.5944125 | 4.27E-11 | 1.20E-09 | 426.237635 | 905.193056 | 743.193281 | 828.707288 |
| -6.5942589 | 4.27E-11 | 1.20E-09 | 423.368728 | 1097.0561 | 805.317797 | 907.151521 |
| -6.5930241 | 4.31E-11 | 1.20E-09 | 418.040758 | 1499.45433 | 934.168644 | 1134.6398 |
| -6.5927138 | 4.32E-11 | 1.21E-09 | 386.072935 | 1313.27411 | 1118.24128 | 1210.28245 |
| -6.5922838 | 4.33E-11 | 1.21E-09 | 400.417471 | 1126.28206 | 981.337258 | 978.871962 |
| -6.5920115 | 4.34E-11 | 1.21E-09 | 471.320462 | 933.065966 | 631.599243 | 693.671144 |
| -6.590795 | 4.37E-11 | 1.22E-09 | 694.275533 | 716.306732 | 452.12842 | 599.538065 |
| -6.5902352 | 4.39E-11 | 1.22E-09 | 407.384817 | 965.809871 | 875.49549 | 907.711837 |

| | | | | | | |
|---|---|---|---|---|---|---|
| -6.5903032 | 4.39E-11 | 1.22E-09 | 418.860445 | 815.350278 | 797.264619 | 800.131175 |
| -6.5898724 | 4.40E-11 | 1.22E-09 | 498.370158 | 694.116649 | 684.520127 | 513.249409 |
| -6.5892917 | 4.42E-11 | 1.23E-09 | 544.272673 | 758.522014 | 677.617403 | 482.432032 |
| -6.5881359 | 4.45E-11 | 1.24E-09 | 427.877011 | 727.942996 | 735.140103 | 748.021791 |
| -6.5879649 | 4.46E-11 | 1.24E-09 | 440.992015 | 832.128146 | 673.015587 | 699.83462 |
| -6.5877257 | 4.47E-11 | 1.24E-09 | 452.877487 | 614.82769 | 736.290557 | 643.242709 |
| -6.5868614 | 4.49E-11 | 1.24E-09 | 436.893576 | 715.765511 | 876.645944 | 700.394936 |
| -6.5865572 | 4.50E-11 | 1.24E-09 | 445.910141 | 694.65787 | 626.997427 | 771.555061 |
| -6.5831707 | 4.61E-11 | 1.27E-09 | 784.031342 | 768.534613 | 441.774334 | 622.511019 |
| -6.5828506 | 4.62E-11 | 1.27E-09 | 403.696222 | 1032.10951 | 874.345036 | 930.124475 |
| -6.5827278 | 4.62E-11 | 1.27E-09 | 423.778572 | 954.98544 | 724.786017 | 888.100779 |
| -6.5820519 | 4.64E-11 | 1.28E-09 | 413.532475 | 877.590757 | 788.060987 | 819.742233 |
| -6.5816331 | 4.65E-11 | 1.28E-09 | 401.647002 | 1037.52173 | 895.053208 | 925.641947 |
| -6.5814713 | 4.66E-11 | 1.28E-09 | 557.387677 | 668.138014 | 464.783414 | 611.865016 |
| -6.5812666 | 4.66E-11 | 1.28E-09 | 331.973543 | 325.815378 | 300.268493 | 337.870517 |
| -6.5806552 | 4.68E-11 | 1.29E-09 | 745.096174 | 446.237175 | 448.677058 | 375.411686 |
| -6.5791073 | 4.73E-11 | 1.30E-09 | 427.057323 | 812.102948 | 686.821035 | 750.263055 |
| -6.5790937 | 4.73E-11 | 1.30E-09 | 383.204028 | 1207.73591 | 1071.07267 | 1129.59695 |
| 6.57849654 | 4.75E-11 | 1.30E-09 | 315.579788 | 156.954252 | 194.426725 | 139.518671 |
| -6.5781436 | 4.76E-11 | 1.30E-09 | 444.270766 | 1215.85423 | 728.237379 | 904.910257 |
| -6.5777393 | 4.78E-11 | 1.30E-09 | 417.630914 | 858.106781 | 738.591465 | 793.967699 |
| -6.577821 | 4.77E-11 | 1.30E-09 | 391.400905 | 998.824386 | 977.885896 | 1049.47177 |
| -6.5773448 | 4.79E-11 | 1.31E-09 | 445.500297 | 721.718948 | 610.891072 | 648.845868 |
| -6.5772766 | 4.79E-11 | 1.31E-09 | 447.959361 | 602.920816 | 635.050605 | 629.795126 |
| -6.5757417 | 4.84E-11 | 1.32E-09 | 423.778572 | 917.641152 | 729.387833 | 760.909058 |
| -6.5758318 | 4.84E-11 | 1.32E-09 | 369.269336 | 1437.21385 | 1242.49032 | 1309.45837 |
| -6.5749023 | 4.87E-11 | 1.32E-09 | 390.991061 | 1175.26261 | 1076.82494 | 950.295849 |
| -6.5733037 | 4.92E-11 | 1.34E-09 | 385.663091 | 1080.27823 | 1054.96631 | 1010.24966 |
| 6.57291659 | 4.93E-11 | 1.34E-09 | 44.6729829 | 15.4248144 | 14.9559019 | 21.292006 |
| -6.5726952 | 4.94E-11 | 1.34E-09 | 359.433083 | 1408.79972 | 1405.85478 | 1458.50241 |
| -6.5718351 | 4.97E-11 | 1.35E-09 | 462.303897 | 708.188409 | 553.368372 | 631.476074 |
| -6.571839 | 4.97E-11 | 1.35E-09 | 555.338457 | 874.614038 | 500.447488 | 623.631651 |
| -6.5713469 | 4.99E-11 | 1.35E-09 | 3424.65546 | 10366.0165 | 9439.47503 | 10247.6184 |
| -6.5707092 | 5.01E-11 | 1.35E-09 | 394.679656 | 942.807955 | 997.443614 | 893.703938 |
| -6.5700603 | 5.03E-11 | 1.36E-09 | 481.976403 | 513.619259 | 699.476029 | 649.9665 |
| -6.5699854 | 5.03E-11 | 1.36E-09 | 354.105112 | 1439.37873 | 1500.19201 | 1647.8892 |
| -6.56996 | 5.03E-11 | 1.36E-09 | 635.258014 | 656.772361 | 444.075242 | 587.211114 |
| -6.569474 | 5.05E-11 | 1.36E-09 | 434.844357 | 707.917798 | 671.865133 | 624.191967 |
| -6.5694244 | 5.05E-11 | 1.36E-09 | 402.46669 | 2449.29816 | 1174.61353 | 1630.51941 |
| -6.5696717 | 5.04E-11 | 1.36E-09 | 389.76153 | 1099.22098 | 931.867736 | 1027.05913 |
| -6.5687092 | 5.08E-11 | 1.36E-09 | 385.663091 | 1846.37735 | 1236.73805 | 1211.9634 |
| -6.5687907 | 5.07E-11 | 1.36E-09 | 336.071982 | 656.772361 | 563.722458 | 616.347543 |
| -6.5681718 | 5.09E-11 | 1.37E-09 | 395.0895 | 1127.0939 | 875.49549 | 1103.2621 |

| | | | | | | |
|---|---|---|---|---|---|---|
| -6.5679904 | 5.10E-11 | 1.37E-09 | 390.581218 | 974.469416 | 943.372276 | 947.494269 |
| -6.5672194 | 5.13E-11 | 1.37E-09 | 412.302943 | 797.489966 | 759.299637 | 746.90116 |
| -6.5670905 | 5.13E-11 | 1.37E-09 | 374.187462 | 1130.61184 | 828.326877 | 976.070382 |
| -6.5661442 | 5.16E-11 | 1.38E-09 | 402.46669 | 836.457918 | 829.477331 | 827.02634 |
| -6.5658508 | 5.17E-11 | 1.38E-09 | 445.910141 | 711.435738 | 609.740618 | 601.779329 |
| -6.5651304 | 5.20E-11 | 1.39E-09 | 380.744964 | 1267.8115 | 1043.46177 | 1079.72883 |
| -6.5639686 | 5.24E-11 | 1.40E-09 | 477.058276 | 661.643355 | 514.252936 | 597.296801 |
| -6.5640187 | 5.24E-11 | 1.40E-09 | 409.024192 | 874.343427 | 763.901453 | 775.477273 |
| -6.5633432 | 5.26E-11 | 1.40E-09 | 426.237635 | 638.641439 | 729.387833 | 730.091681 |
| -6.5631066 | 5.27E-11 | 1.40E-09 | 369.269336 | 1419.08293 | 1179.21535 | 1210.28245 |
| -6.5629346 | 5.28E-11 | 1.40E-09 | 391.810749 | 907.087332 | 912.310018 | 1036.5845 |
| -6.5620733 | 5.31E-11 | 1.41E-09 | 778.293528 | 529.855906 | 514.252936 | 553.031841 |
| -6.5619254 | 5.31E-11 | 1.41E-09 | 338.940889 | 1692.94103 | 1698.0701 | 1706.72238 |
| -6.5612127 | 5.34E-11 | 1.42E-09 | 511.895006 | 771.781942 | 579.828814 | 1163.21591 |
| -6.5600383 | 5.38E-11 | 1.43E-09 | 388.531998 | 988.270565 | 892.7523 | 978.871962 |
| -6.5594959 | 5.40E-11 | 1.43E-09 | 495.501251 | 947.137727 | 545.315194 | 699.83462 |
| -6.5586057 | 5.43E-11 | 1.44E-09 | 393.859969 | 1041.58089 | 836.380055 | 990.638597 |
| -6.5586146 | 5.43E-11 | 1.44E-09 | 416.401382 | 771.781942 | 691.422851 | 711.040939 |
| -6.558522 | 5.43E-11 | 1.44E-09 | 392.220593 | 1034.00379 | 861.690043 | 931.805423 |
| -6.5578111 | 5.46E-11 | 1.44E-09 | 406.155285 | 831.045703 | 797.264619 | 739.056736 |
| -6.5561226 | 5.52E-11 | 1.46E-09 | 473.369681 | 1009.91943 | 586.731538 | 739.617052 |
| -6.5561117 | 5.52E-11 | 1.46E-09 | 621.733166 | 752.839188 | 427.968886 | 671.818822 |
| -6.5554916 | 5.55E-11 | 1.46E-09 | 528.698605 | 875.425871 | 491.243856 | 674.060086 |
| -6.5546071 | 5.58E-11 | 1.47E-09 | 386.072935 | 1620.95857 | 1034.25814 | 1205.79992 |
| -6.5545112 | 5.58E-11 | 1.47E-09 | 482.386247 | 579.648289 | 567.17382 | 494.198667 |
| -6.5544366 | 5.59E-11 | 1.47E-09 | 381.564652 | 1218.56034 | 941.071368 | 1134.07948 |
| -6.553966 | 5.60E-11 | 1.47E-09 | 390.991061 | 1886.42774 | 1033.10769 | 1471.95 |
| -6.5522412 | 5.67E-11 | 1.49E-09 | 472.959837 | 657.313583 | 507.350212 | 588.331746 |
| -6.551997 | 5.68E-11 | 1.49E-09 | 461.484209 | 606.709367 | 516.553844 | 623.071335 |
| -6.5512655 | 5.71E-11 | 1.49E-09 | 372.548087 | 1263.75234 | 1073.37358 | 1067.40188 |
| -6.5511677 | 5.71E-11 | 1.49E-09 | 359.433083 | 1726.76738 | 1247.09213 | 1400.22956 |
| -6.5509777 | 5.72E-11 | 1.49E-09 | 402.056846 | 888.415188 | 750.096005 | 805.174018 |
| -6.5510947 | 5.71E-11 | 1.49E-09 | 411.483255 | 1179.59239 | 774.255539 | 916.116576 |
| -6.550617 | 5.73E-11 | 1.50E-09 | 2772.184 | 841.599523 | 655.758777 | 950.856165 |
| -6.5499832 | 5.75E-11 | 1.50E-09 | 471.320462 | 505.771546 | 568.324274 | 619.709439 |
| -6.5491559 | 5.79E-11 | 1.51E-09 | 392.630437 | 952.549943 | 804.167343 | 901.548361 |
| -6.548962 | 5.79E-11 | 1.51E-09 | 411.483255 | 993.141559 | 762.750999 | 754.185267 |
| -6.5487145 | 5.80E-11 | 1.51E-09 | 435.664044 | 602.650205 | 586.731538 | 625.312599 |
| -6.5485004 | 5.81E-11 | 1.51E-09 | 364.351209 | 1203.40613 | 1127.44492 | 1173.86191 |
| -6.5470878 | 5.87E-11 | 1.52E-09 | 404.925753 | 799.113631 | 704.077845 | 809.656546 |
| -6.5460637 | 5.91E-11 | 1.53E-09 | 398.368251 | 769.075834 | 790.361895 | 892.02299 |
| -6.5458539 | 5.92E-11 | 1.53E-09 | 412.712787 | 809.39684 | 668.413771 | 729.531365 |
| -6.5450035 | 5.95E-11 | 1.54E-09 | 409.024192 | 806.961343 | 678.767857 | 753.064635 |

| | | | | | | |
|---|---|---|---|---|---|---|
| -6.544796 | 5.96E-11 | 1.54E-09 | 362.711834 | 1110.04542 | 1201.07397 | 1214.76498 |
| -6.5443503 | 5.98E-11 | 1.55E-09 | 747.145393 | 551.504768 | 760.450091 | 444.890863 |
| -6.5421589 | 6.06E-11 | 1.57E-09 | 435.664044 | 854.31823 | 597.085624 | 706.558411 |
| -6.5415802 | 6.09E-11 | 1.57E-09 | 346.318079 | 1690.77615 | 1431.16477 | 1352.6027 |
| -6.5415509 | 6.09E-11 | 1.57E-09 | 470.09093 | 813.997224 | 525.757476 | 858.404033 |
| -6.5412281 | 6.10E-11 | 1.57E-09 | 556.158145 | 610.497918 | 436.022064 | 615.226912 |
| -6.5403434 | 6.14E-11 | 1.58E-09 | 392.220593 | 795.054469 | 826.025969 | 801.812122 |
| -6.5395695 | 6.17E-11 | 1.59E-09 | 382.794184 | 930.630469 | 857.088227 | 905.470573 |
| -6.5390975 | 6.19E-11 | 1.59E-09 | 370.908711 | 1359.54855 | 998.594068 | 1173.86191 |
| -6.5389321 | 6.20E-11 | 1.59E-09 | 367.62996 | 1065.66525 | 1031.95723 | 1072.44473 |
| -6.5387146 | 6.20E-11 | 1.59E-09 | 244.266952 | 311.202396 | 256.551241 | 313.776931 |
| -6.5384783 | 6.21E-11 | 1.60E-09 | 345.498391 | 1294.60197 | 1589.92742 | 1720.16996 |
| -6.5383192 | 6.22E-11 | 1.60E-09 | 430.745918 | 1262.39928 | 698.325575 | 963.183116 |
| -6.5380338 | 6.23E-11 | 1.60E-09 | 433.614825 | 579.9189 | 595.93517 | 584.96985 |
| -6.5357796 | 6.33E-11 | 1.62E-09 | 429.92623 | 732.81399 | 624.696519 | 587.77143 |
| -6.5350725 | 6.36E-11 | 1.63E-09 | 363.941365 | 1130.07061 | 1111.33856 | 1035.46387 |
| -6.5349676 | 6.36E-11 | 1.63E-09 | 370.089024 | 1035.62745 | 980.186804 | 999.603652 |
| -6.5344853 | 6.38E-11 | 1.63E-09 | 341.809796 | 1357.65428 | 1432.31522 | 1465.22621 |
| -6.5337964 | 6.41E-11 | 1.64E-09 | 389.351686 | 1962.19876 | 1060.71858 | 1270.23626 |
| -6.5337898 | 6.41E-11 | 1.64E-09 | 455.336551 | 539.056672 | 668.413771 | 543.50647 |
| -6.532532 | 6.47E-11 | 1.65E-09 | 289.759623 | 890.309463 | 794.963711 | 874.09288 |
| -6.5323837 | 6.47E-11 | 1.65E-09 | 398.368251 | 1011.00187 | 1125.14401 | 750.823371 |
| -6.5314307 | 6.51E-11 | 1.65E-09 | 410.663568 | 1240.2092 | 737.441011 | 993.440177 |
| -6.5314324 | 6.51E-11 | 1.65E-09 | 361.072458 | 1036.16867 | 1201.07397 | 1164.33654 |
| -6.5313516 | 6.52E-11 | 1.65E-09 | 662.30771 | 1074.05418 | 418.765254 | 991.759229 |
| -6.5314234 | 6.51E-11 | 1.65E-09 | 376.236682 | 865.954493 | 998.594068 | 1002.96555 |
| -6.5298351 | 6.58E-11 | 1.67E-09 | 404.925753 | 813.185391 | 655.758777 | 763.150322 |
| -6.5298281 | 6.58E-11 | 1.67E-09 | 381.974496 | 925.759475 | 808.769159 | 907.711837 |
| -6.5285719 | 6.64E-11 | 1.68E-09 | 373.367775 | 929.277416 | 931.867736 | 1093.73673 |
| -6.5278072 | 6.67E-11 | 1.69E-09 | 390.581218 | 834.834254 | 737.441011 | 800.691491 |
| -6.5275145 | 6.69E-11 | 1.69E-09 | 388.122154 | 751.215523 | 896.203662 | 848.878662 |
| -6.5272899 | 6.70E-11 | 1.69E-09 | 400.417471 | 1216.39545 | 1239.03895 | 1253.42678 |
| -6.5273485 | 6.69E-11 | 1.69E-09 | 408.614348 | 674.632672 | 652.307415 | 650.526816 |
| -6.5271085 | 6.71E-11 | 1.69E-09 | 399.187939 | 770.428888 | 692.573305 | 709.920307 |
| -6.526734 | 6.72E-11 | 1.69E-09 | 431.155762 | 705.482301 | 553.368372 | 634.277654 |
| -6.5265324 | 6.73E-11 | 1.70E-09 | 437.30342 | 554.752097 | 549.91701 | 572.642899 |
| -6.5259695 | 6.76E-11 | 1.70E-09 | 401.237158 | 1402.03445 | 1453.0234 | 1535.2657 |
| -6.5254251 | 6.78E-11 | 1.71E-09 | 382.794184 | 1141.16566 | 821.424153 | 985.595754 |
| -6.5252703 | 6.79E-11 | 1.71E-09 | 395.909188 | 1090.29083 | 745.494189 | 911.073732 |
| -6.5249647 | 6.80E-11 | 1.71E-09 | 345.908235 | 1243.18592 | 1260.89758 | 1321.22501 |
| -6.5244632 | 6.82E-11 | 1.71E-09 | 386.892623 | 817.785775 | 783.459171 | 767.63285 |
| -6.5233378 | 6.88E-11 | 1.72E-09 | 382.38434 | 857.83617 | 783.459171 | 852.240558 |
| -6.5231112 | 6.89E-11 | 1.72E-09 | 407.794661 | 1452.63866 | 788.060987 | 1090.93515 |

| | | | | | | |
|---|---|---|---|---|---|---|
| -6.5232152 | 6.88E-11 | 1.72E-09 | 400.417471 | 1179.863 | 1040.01041 | 1117.83032 |
| -6.5215744 | 6.96E-11 | 1.74E-09 | 1029.52783 | 739.038038 | 836.380055 | 394.462428 |
| -6.5215319 | 6.96E-11 | 1.74E-09 | 393.040281 | 1220.184 | 840.981871 | 860.084981 |
| -6.5208808 | 6.99E-11 | 1.74E-09 | 389.351686 | 1335.4642 | 819.123245 | 1101.02084 |
| -6.5209962 | 6.98E-11 | 1.74E-09 | 739.768203 | 1436.40202 | 1291.95984 | 1256.22836 |
| -6.5204238 | 7.01E-11 | 1.75E-09 | 410.253724 | 1087.85533 | 715.582385 | 785.56296 |
| -6.5197138 | 7.04E-11 | 1.75E-09 | 376.646526 | 1019.66142 | 816.822337 | 949.735533 |
| -6.5197487 | 7.04E-11 | 1.75E-09 | 647.55333 | 831.586924 | 467.084322 | 514.370041 |
| -6.5188483 | 7.08E-11 | 1.76E-09 | 374.597306 | 1121.95229 | 885.849576 | 909.953101 |
| -6.5172057 | 7.16E-11 | 1.78E-09 | 342.629484 | 1249.68058 | 1263.19849 | 1275.83941 |
| -6.5160136 | 7.22E-11 | 1.79E-09 | 352.875581 | 1100.57404 | 1079.12585 | 1201.31739 |
| -6.5158186 | 7.23E-11 | 1.79E-09 | 386.072935 | 798.301798 | 729.387833 | 778.278853 |
| -6.5157697 | 7.23E-11 | 1.79E-09 | 375.416994 | 924.406422 | 1336.82754 | 989.517965 |
| -6.5145255 | 7.29E-11 | 1.81E-09 | 390.581218 | 869.472433 | 708.679661 | 758.667794 |
| -6.5141333 | 7.31E-11 | 1.81E-09 | 439.762483 | 729.025439 | 523.456568 | 629.23481 |
| -6.5137748 | 7.33E-11 | 1.81E-09 | 329.924323 | 1804.43268 | 1372.49162 | 1569.44497 |
| -6.5129059 | 7.37E-11 | 1.82E-09 | 219.67632 | 653.795642 | 570.625182 | 725.609153 |
| -6.5128433 | 7.37E-11 | 1.82E-09 | 425.417947 | 535.538732 | 620.094704 | 615.226912 |
| -6.513022 | 7.37E-11 | 1.82E-09 | 428.286855 | 694.387259 | 566.023366 | 569.281004 |
| -6.5125029 | 7.39E-11 | 1.82E-09 | 477.46812 | 652.171978 | 423.36707 | 567.600056 |
| -6.5124965 | 7.39E-11 | 1.82E-09 | 402.46669 | 632.417391 | 858.238681 | 806.854966 |
| -6.5125417 | 7.39E-11 | 1.82E-09 | 375.826838 | 853.777008 | 838.680963 | 1035.46387 |
| -6.5108082 | 7.47E-11 | 1.84E-09 | 711.488976 | 473.298253 | 990.54089 | 533.420783 |
| -6.5109262 | 7.47E-11 | 1.84E-09 | 393.859969 | 696.822756 | 678.767857 | 693.671144 |
| -6.5108696 | 7.47E-11 | 1.84E-09 | 401.237158 | 847.823571 | 635.050605 | 763.150322 |
| -6.5104596 | 7.49E-11 | 1.84E-09 | 442.221546 | 499.276888 | 624.696519 | 655.56966 |
| 6.50909077 | 7.56E-11 | 1.85E-09 | 71.722679 | 69.0057487 | 65.5758777 | 65.556966 |
| -6.5091604 | 7.56E-11 | 1.85E-09 | 369.269336 | 963.103763 | 829.477331 | 951.416481 |
| -6.5089607 | 7.57E-11 | 1.85E-09 | 426.647479 | 601.838373 | 556.819734 | 541.265206 |
| -6.5077234 | 7.63E-11 | 1.87E-09 | 402.876534 | 706.294134 | 615.492888 | 646.604605 |
| -6.5074518 | 7.64E-11 | 1.87E-09 | 377.05637 | 884.897248 | 756.998729 | 858.964349 |
| -6.5058634 | 7.72E-11 | 1.89E-09 | 377.466213 | 758.251403 | 803.016889 | 820.302549 |
| -6.5053349 | 7.75E-11 | 1.89E-09 | 1641.0149 | 571.259355 | 666.112863 | 696.472724 |
| -6.5051933 | 7.76E-11 | 1.89E-09 | 457.795614 | 949.843835 | 557.970188 | 651.087132 |
| -6.504888 | 7.78E-11 | 1.90E-09 | 491.402812 | 661.643355 | 448.677058 | 572.642899 |
| -6.5042629 | 7.81E-11 | 1.90E-09 | 377.466213 | 829.422038 | 771.954631 | 766.512218 |
| -6.5024141 | 7.90E-11 | 1.92E-09 | 333.612918 | 1484.57073 | 1228.68487 | 1363.2487 |
| -6.5020406 | 7.92E-11 | 1.93E-09 | 509.845787 | 609.956696 | 441.774334 | 735.134525 |
| -6.5013212 | 7.96E-11 | 1.94E-09 | 462.71374 | 774.758661 | 495.845672 | 602.899961 |
| -6.4979374 | 8.14E-11 | 1.98E-09 | 377.876057 | 754.192241 | 732.839195 | 786.683592 |
| -6.4970203 | 8.19E-11 | 1.99E-09 | 368.449648 | 881.379308 | 788.060987 | 874.653196 |
| -6.4968811 | 8.20E-11 | 1.99E-09 | 366.810273 | 903.840002 | 806.468251 | 982.794174 |
| -6.4967888 | 8.21E-11 | 1.99E-09 | 361.072458 | 972.033919 | 852.486411 | 961.502168 |

| | | | | | | |
|---|---|---|---|---|---|---|
| -6.4961245 | 8.24E-11 | 2.00E-09 | 361.072458 | 954.444218 | 873.194582 | 892.02299 |
| -6.495931 | 8.25E-11 | 2.00E-09 | 399.187939 | 727.942996 | 708.679661 | 591.693642 |
| -6.4948045 | 8.31E-11 | 2.01E-09 | 365.990585 | 1009.1076 | 812.220521 | 902.108677 |
| -6.4944679 | 8.33E-11 | 2.01E-09 | 413.942319 | 616.451355 | 554.518826 | 717.204414 |
| -6.4938388 | 8.37E-11 | 2.02E-09 | 350.416517 | 953.632386 | 1161.95854 | 1079.72883 |
| -6.4935269 | 8.38E-11 | 2.02E-09 | 367.220116 | 963.915595 | 794.963711 | 867.929404 |
| -6.4932022 | 8.40E-11 | 2.03E-09 | 411.073412 | 678.962445 | 545.315194 | 637.079234 |
| -6.4926666 | 8.43E-11 | 2.03E-09 | 392.220593 | 677.609391 | 624.696519 | 658.37124 |
| -6.4918267 | 8.48E-11 | 2.04E-09 | 439.352639 | 787.747978 | 543.014286 | 573.763531 |
| -6.4916861 | 8.49E-11 | 2.04E-09 | 339.350733 | 1078.38395 | 1481.78475 | 1396.86766 |
| -6.4913669 | 8.51E-11 | 2.04E-09 | 446.729829 | 633.770445 | 437.172518 | 512.689093 |
| -6.4908044 | 8.54E-11 | 2.05E-09 | 376.236682 | 749.862469 | 713.281477 | 792.847067 |
| -6.4908513 | 8.54E-11 | 2.05E-09 | 365.170897 | 905.193056 | 786.910533 | 907.711837 |
| -6.4906453 | 8.55E-11 | 2.05E-09 | 350.826361 | 1159.02597 | 965.230902 | 997.922704 |
| -6.489922 | 8.59E-11 | 2.06E-09 | 377.466213 | 839.705248 | 702.927391 | 759.22811 |
| -6.4896233 | 8.61E-11 | 2.06E-09 | 404.515909 | 1261.04623 | 686.821035 | 979.432278 |
| -6.4895067 | 8.61E-11 | 2.06E-09 | 375.00715 | 778.547212 | 716.732839 | 757.547163 |
| -6.4893002 | 8.62E-11 | 2.06E-09 | 428.286855 | 650.277702 | 525.757476 | 535.662047 |
| -6.4889271 | 8.64E-11 | 2.07E-09 | 372.957931 | 860.271667 | 742.042827 | 763.710638 |
| -6.4887386 | 8.66E-11 | 2.07E-09 | 393.040281 | 640.535714 | 625.846973 | 628.674494 |
| -6.4879261 | 8.70E-11 | 2.08E-09 | 297.546657 | 1782.5132 | 1923.55908 | 1840.63789 |
| -6.4868079 | 8.77E-11 | 2.09E-09 | 387.302467 | 740.120481 | 626.997427 | 697.593356 |
| -6.4855275 | 8.84E-11 | 2.11E-09 | 311.481349 | 1496.207 | 1786.65505 | 1540.86886 |
| -6.485463 | 8.85E-11 | 2.11E-09 | 380.335121 | 769.346445 | 678.767857 | 694.23146 |
| -6.4848056 | 8.88E-11 | 2.11E-09 | 381.564652 | 658.396026 | 700.626483 | 707.679043 |
| -6.4841218 | 8.93E-11 | 2.12E-09 | 327.46526 | 1270.51761 | 1252.8444 | 1263.51246 |
| -6.4836816 | 8.95E-11 | 2.13E-09 | 399.597783 | 605.085702 | 607.43971 | 573.763531 |
| -6.4832182 | 8.98E-11 | 2.13E-09 | 429.92623 | 779.088433 | 446.37615 | 606.822172 |
| -6.4828305 | 9.00E-11 | 2.14E-09 | 422.54904 | 517.949031 | 582.129722 | 520.533516 |
| -6.4817017 | 9.07E-11 | 2.15E-09 | 384.843403 | 690.328098 | 632.749697 | 666.215663 |
| -6.4799922 | 9.17E-11 | 2.17E-09 | 353.695268 | 868.38999 | 956.02727 | 892.583306 |
| -6.4796415 | 9.19E-11 | 2.18E-09 | 360.662614 | 895.180457 | 785.760079 | 857.283401 |
| -6.4792619 | 9.22E-11 | 2.18E-09 | 429.92623 | 509.560097 | 570.625182 | 709.359991 |
| -6.4784515 | 9.27E-11 | 2.19E-09 | 395.0895 | 663.53763 | 568.324274 | 651.087132 |
| -6.4784107 | 9.27E-11 | 2.19E-09 | 379.105589 | 813.185391 | 651.156961 | 750.823371 |
| -6.4785891 | 9.26E-11 | 2.19E-09 | 479.927183 | 467.886037 | 829.477331 | 620.830071 |
| -6.4773259 | 9.34E-11 | 2.20E-09 | 331.973543 | 1193.66415 | 1107.8872 | 1172.74128 |
| -6.4768473 | 9.37E-11 | 2.21E-09 | 652.061613 | 577.754013 | 433.721156 | 477.949504 |
| -6.4766717 | 9.38E-11 | 2.21E-09 | 352.465737 | 1079.4664 | 858.238681 | 962.062484 |
| -6.4766065 | 9.38E-11 | 2.21E-09 | 775.834465 | 494.947115 | 482.040224 | 516.611305 |
| -6.4759881 | 9.42E-11 | 2.22E-09 | 402.876534 | 536.079953 | 620.094704 | 638.760181 |
| -6.4753901 | 9.46E-11 | 2.22E-09 | 398.778095 | 663.26702 | 553.368372 | 615.787228 |
| -6.47536 | 9.46E-11 | 2.22E-09 | 389.351686 | 830.504481 | 618.94425 | 689.748933 |

| | | | | | | |
|---|---|---|---|---|---|---|
| -6.4748373 | 9.49E-11 | 2.23E-09 | 368.859492 | 722.26017 | 768.503269 | 727.850417 |
| -6.4746551 | 9.50E-11 | 2.23E-09 | 487.304373 | 559.893702 | 455.579782 | 460.57971 |
| -6.4738163 | 9.56E-11 | 2.24E-09 | 361.482302 | 793.972026 | 793.813257 | 780.520116 |
| -6.4732509 | 9.59E-11 | 2.25E-09 | 421.729352 | 773.676218 | 509.65112 | 701.515568 |
| -6.4730602 | 9.60E-11 | 2.25E-09 | 361.482302 | 900.322062 | 747.795097 | 920.599104 |
| -6.4728721 | 9.62E-11 | 2.25E-09 | 389.351686 | 624.569678 | 593.634262 | 618.588807 |
| -6.4722349 | 9.66E-11 | 2.26E-09 | 376.236682 | 732.002158 | 644.254237 | 721.126626 |
| -6.4720671 | 9.67E-11 | 2.26E-09 | 393.450125 | 663.26702 | 561.42155 | 643.803025 |
| -6.4713709 | 9.71E-11 | 2.26E-09 | 396.728876 | 806.961343 | 594.784716 | 639.880813 |
| -6.4713589 | 9.71E-11 | 2.26E-09 | 384.433559 | 736.873152 | 608.590164 | 664.534715 |
| -6.4702196 | 9.79E-11 | 2.28E-09 | 349.186986 | 687.080768 | 620.094704 | 611.3047 |
| -6.4701664 | 9.79E-11 | 2.28E-09 | 352.055893 | 1008.83698 | 877.796398 | 847.75803 |
| -6.4692086 | 9.85E-11 | 2.29E-09 | 398.778095 | 571.259355 | 566.023366 | 559.195317 |
| -6.4688933 | 9.87E-11 | 2.29E-09 | 342.21964 | 655.689918 | 530.359292 | 605.141225 |
| -6.4674878 | 9.96E-11 | 2.31E-09 | 331.153855 | 1082.98434 | 1091.78084 | 1124.55411 |
| -6.467325 | 9.98E-11 | 2.32E-09 | 409.434036 | 583.978061 | 511.952028 | 546.30805 |
| -6.4669492 | 1.00E-10 | 2.32E-09 | 423.778572 | 537.974229 | 835.229601 | 612.985648 |
| -6.4666904 | 1.00E-10 | 2.32E-09 | 372.957931 | 875.696481 | 674.166041 | 744.09958 |
| -6.4655061 | 1.01E-10 | 2.34E-09 | 359.433083 | 1007.48393 | 761.600545 | 869.050036 |
| -6.4652954 | 1.01E-10 | 2.34E-09 | 311.891193 | 1584.69672 | 1340.2789 | 1376.13597 |
| -6.4643617 | 1.02E-10 | 2.35E-09 | 1326.2548 | 2636.0196 | 2613.83148 | 2702.96414 |
| -6.4641669 | 1.02E-10 | 2.35E-09 | 398.368251 | 750.944912 | 551.067464 | 642.682393 |
| -6.4639692 | 1.02E-10 | 2.36E-09 | 405.335597 | 949.032002 | 631.599243 | 1212.52371 |
| -6.4629227 | 1.03E-10 | 2.37E-09 | 365.580741 | 684.645271 | 750.096005 | 782.201064 |
| 6.46280626 | 1.03E-10 | 2.37E-09 | 0 | 0 | 0 | 0 |
| -6.4612091 | 1.04E-10 | 2.39E-09 | 493.042188 | 464.368097 | 615.492888 | 449.373391 |
| -6.4608687 | 1.04E-10 | 2.40E-09 | 392.220593 | 562.329199 | 577.527906 | 654.449028 |
| -6.4607321 | 1.04E-10 | 2.40E-09 | 1050.02002 | 532.291403 | 461.332052 | 667.336295 |
| -6.4605853 | 1.04E-10 | 2.40E-09 | 364.351209 | 812.914781 | 683.369673 | 775.477273 |
| -6.4599986 | 1.05E-10 | 2.41E-09 | 840.179954 | 608.333032 | 442.924788 | 499.801826 |
| -6.4595766 | 1.05E-10 | 2.41E-09 | 389.76153 | 809.12623 | 582.129722 | 672.939454 |
| -6.459138 | 1.05E-10 | 2.41E-09 | 368.449648 | 1023.17936 | 692.573305 | 904.349941 |
| -6.4592143 | 1.05E-10 | 2.41E-09 | 411.073412 | 540.139115 | 507.350212 | 513.249409 |
| -6.4589518 | 1.05E-10 | 2.41E-09 | 348.367298 | 1376.32642 | 897.354116 | 1065.16062 |
| -6.4591118 | 1.05E-10 | 2.41E-09 | 377.05637 | 1274.84738 | 1191.87034 | 1329.62975 |
| -6.4590427 | 1.05E-10 | 2.41E-09 | 322.547134 | 1227.21988 | 1112.48901 | 1217.56656 |
| -6.458623 | 1.06E-10 | 2.42E-09 | 404.515909 | 605.356313 | 560.271096 | 508.206565 |
| -6.4584814 | 1.06E-10 | 2.42E-09 | 379.515433 | 723.342613 | 597.085624 | 661.172819 |
| 6.45758848 | 1.06E-10 | 2.43E-09 | 0.81968776 | 0.27061078 | 1.150454 | 0.56031595 |
| -6.4573507 | 1.07E-10 | 2.43E-09 | 350.006673 | 854.588841 | 807.618705 | 822.543813 |
| -6.4572306 | 1.07E-10 | 2.43E-09 | 422.958884 | 862.165942 | 549.91701 | 596.176169 |
| -6.457036 | 1.07E-10 | 2.43E-09 | 227.053509 | 772.052553 | 744.343735 | 661.172819 |
| -6.4564328 | 1.07E-10 | 2.44E-09 | 418.040758 | 747.156361 | 808.769159 | 504.284354 |

| | | | | | | |
|---|---|---|---|---|---|---|
| -6.4563709 | 1.07E-10 | 2.44E-09 | 427.057323 | 635.123499 | 468.234776 | 553.592157 |
| -6.45515 | 1.08E-10 | 2.46E-09 | 375.416994 | 720.907116 | 609.740618 | 660.052187 |
| -6.4549723 | 1.08E-10 | 2.46E-09 | 378.695745 | 613.474636 | 623.546066 | 609.063436 |
| -6.4534704 | 1.09E-10 | 2.48E-09 | 376.646526 | 809.938062 | 602.837894 | 773.236009 |
| -6.4533054 | 1.09E-10 | 2.48E-09 | 260.250864 | 890.580074 | 774.255539 | 760.348742 |
| -6.4525776 | 1.10E-10 | 2.49E-09 | 362.30199 | 918.723595 | 689.121943 | 812.458125 |
| -6.4520205 | 1.10E-10 | 2.50E-09 | 425.417947 | 1058.62937 | 559.120642 | 741.858316 |
| -6.4517558 | 1.11E-10 | 2.50E-09 | 460.664521 | 638.641439 | 425.667978 | 577.125427 |
| -6.451316 | 1.11E-10 | 2.51E-09 | 356.974019 | 791.265918 | 722.485109 | 906.591205 |
| -6.4507691 | 1.11E-10 | 2.52E-09 | 393.040281 | 605.356313 | 617.793796 | 521.093832 |
| -6.4506212 | 1.11E-10 | 2.52E-09 | 365.170897 | 644.053654 | 771.954631 | 773.236009 |
| -6.4503437 | 1.12E-10 | 2.52E-09 | 512.30485 | 612.121582 | 403.809352 | 523.895412 |
| -6.4501176 | 1.12E-10 | 2.52E-09 | 343.859015 | 875.425871 | 838.680963 | 848.878662 |
| -6.4494051 | 1.12E-10 | 2.53E-09 | 393.450125 | 519.843307 | 660.360593 | 647.725237 |
| -6.4490043 | 1.13E-10 | 2.54E-09 | 448.779049 | 432.977247 | 635.050605 | 582.168271 |
| -6.4474089 | 1.14E-10 | 2.56E-09 | 384.433559 | 654.066253 | 551.067464 | 602.899961 |
| -6.4472914 | 1.14E-10 | 2.56E-09 | 360.25277 | 730.378493 | 671.865133 | 772.115377 |
| -6.4461501 | 1.15E-10 | 2.58E-09 | 259.021332 | 228.666108 | 193.276271 | 209.558165 |
| -6.4458136 | 1.15E-10 | 2.58E-09 | 373.367775 | 711.435738 | 587.881992 | 718.325046 |
| -6.4459408 | 1.15E-10 | 2.58E-09 | 406.155285 | 531.208959 | 551.067464 | 599.538065 |
| -6.4457522 | 1.15E-10 | 2.58E-09 | 380.744964 | 565.035307 | 689.121943 | 618.588807 |
| -6.4454971 | 1.15E-10 | 2.58E-09 | 475.828745 | 718.471619 | 422.216616 | 597.857117 |
| -6.4442359 | 1.16E-10 | 2.60E-09 | 418.860445 | 509.289486 | 521.15566 | 461.700342 |
| -6.4428257 | 1.17E-10 | 2.63E-09 | 540.993922 | 409.434109 | 546.465648 | 471.786029 |
| -6.4422312 | 1.18E-10 | 2.63E-09 | 354.105112 | 719.824673 | 719.033747 | 785.56296 |
| -6.4422675 | 1.18E-10 | 2.63E-09 | 316.809319 | 1177.4275 | 1128.59537 | 1134.07948 |
| -6.4421824 | 1.18E-10 | 2.63E-09 | 362.711834 | 787.747978 | 661.511047 | 681.344194 |
| -6.4423723 | 1.18E-10 | 2.63E-09 | 372.138243 | 612.121582 | 618.94425 | 615.787228 |
| -6.4409253 | 1.19E-10 | 2.65E-09 | 454.926707 | 985.293847 | 924.965012 | 963.743432 |
| -6.4409024 | 1.19E-10 | 2.65E-09 | 372.138243 | 802.631571 | 595.93517 | 730.091681 |
| -6.4400401 | 1.19E-10 | 2.66E-09 | 365.990585 | 784.77126 | 625.846973 | 691.990197 |
| -6.4395423 | 1.20E-10 | 2.67E-09 | 341.809796 | 828.610206 | 801.866435 | 883.618251 |
| -6.4387672 | 1.20E-10 | 2.68E-09 | 331.973543 | 986.917512 | 885.849576 | 957.579956 |
| -6.4388486 | 1.20E-10 | 2.68E-09 | 352.465737 | 814.809056 | 715.582385 | 731.212313 |
| -6.437817 | 1.21E-10 | 2.69E-09 | 384.023715 | 575.589127 | 533.810654 | 583.849218 |
| -6.437841 | 1.21E-10 | 2.69E-09 | 352.055893 | 800.737295 | 733.989649 | 709.920307 |
| -6.4382481 | 1.21E-10 | 2.69E-09 | 345.908235 | 988.270565 | 765.051907 | 853.921506 |
| -6.4380122 | 1.21E-10 | 2.69E-09 | 381.974496 | 629.711283 | 537.262016 | 666.775979 |
| -6.4377346 | 1.21E-10 | 2.69E-09 | 443.451078 | 571.259355 | 447.526604 | 714.402834 |
| -6.4379089 | 1.21E-10 | 2.69E-09 | 192.21678 | 173.190899 | 172.568099 | 208.997849 |
| 6.43724904 | 1.22E-10 | 2.69E-09 | 117.21535 | 86.8660601 | 94.3372276 | 104.779082 |
| -6.4372306 | 1.22E-10 | 2.69E-09 | 166.806459 | 198.08709 | 216.285351 | 197.231214 |
| -6.4368683 | 1.22E-10 | 2.69E-09 | 411.483255 | 542.304001 | 478.588862 | 486.354243 |

| | | | | | | |
|---|---|---|---|---|---|---|
| -6.4367934 | 1.22E-10 | 2.69E-09 | 331.973543 | 965.53926 | 900.805478 | 906.030889 |
| -6.4360064 | 1.23E-10 | 2.71E-09 | 342.21964 | 786.124313 | 842.132325 | 964.864064 |
| -6.4356204 | 1.23E-10 | 2.71E-09 | 363.531521 | 639.723882 | 644.254237 | 648.285553 |
| 6.43354038 | 1.25E-10 | 2.75E-09 | 7.78703372 | 10.2832096 | 6.90272397 | 6.72379138 |
| -6.4333928 | 1.25E-10 | 2.75E-09 | 529.928137 | 634.852888 | 426.818432 | 451.054339 |
| -6.4334773 | 1.25E-10 | 2.75E-09 | 385.253247 | 839.975858 | 563.722458 | 670.137875 |
| -6.4331552 | 1.25E-10 | 2.75E-09 | 805.343224 | 2649.00892 | 2449.31656 | 2430.09027 |
| -6.4327758 | 1.25E-10 | 2.75E-09 | 390.581218 | 644.324265 | 501.597942 | 613.545964 |
| -6.4323091 | 1.26E-10 | 2.76E-09 | 347.54761 | 735.249487 | 733.989649 | 788.36454 |
| -6.432007 | 1.26E-10 | 2.76E-09 | 440.582171 | 500.629941 | 483.190678 | 428.641701 |
| -6.4317905 | 1.26E-10 | 2.77E-09 | 737.718984 | 415.658157 | 647.705599 | 471.225713 |
| -6.4315203 | 1.26E-10 | 2.77E-09 | 374.597306 | 616.721966 | 568.324274 | 713.842519 |
| -6.4311386 | 1.27E-10 | 2.77E-09 | 314.350256 | 1119.51679 | 1072.22312 | 1177.22381 |
| -6.430451 | 1.27E-10 | 2.78E-09 | 362.30199 | 955.526661 | 656.909231 | 773.796325 |
| -6.4302028 | 1.27E-10 | 2.79E-09 | 366.810273 | 619.428073 | 911.159564 | 778.278853 |
| -6.4301196 | 1.28E-10 | 2.79E-09 | 347.957454 | 826.44532 | 698.325575 | 785.56296 |
| -6.4300657 | 1.28E-10 | 2.79E-09 | 382.38434 | 572.071187 | 551.067464 | 531.739835 |
| -6.4299189 | 1.28E-10 | 2.79E-09 | 332.793231 | 1394.18673 | 923.814558 | 1053.39398 |
| -6.4294112 | 1.28E-10 | 2.79E-09 | 261.070552 | 391.303187 | 440.62388 | 439.287704 |
| -6.4290441 | 1.28E-10 | 2.80E-09 | 477.877964 | 494.135283 | 426.818432 | 440.968652 |
| -6.4284764 | 1.29E-10 | 2.80E-09 | 340.990108 | 1750.31052 | 956.02727 | 1181.14602 |
| -6.428769 | 1.29E-10 | 2.80E-09 | 338.121201 | 807.502565 | 816.822337 | 816.940653 |
| -6.4287238 | 1.29E-10 | 2.80E-09 | 373.777618 | 783.688816 | 569.474728 | 679.10293 |
| -6.4285318 | 1.29E-10 | 2.80E-09 | 344.268859 | 612.121582 | 517.704298 | 602.899961 |
| -6.428549 | 1.29E-10 | 2.80E-09 | 388.941842 | 660.290301 | 508.500666 | 581.607955 |
| -6.4280863 | 1.29E-10 | 2.81E-09 | 542.223453 | 536.079953 | 410.712076 | 449.373391 |
| -6.4277288 | 1.30E-10 | 2.81E-09 | 342.21964 | 791.265918 | 769.653723 | 760.348742 |
| -6.4276154 | 1.30E-10 | 2.81E-09 | 377.05637 | 896.804122 | 586.731538 | 720.005994 |
| -6.4275185 | 1.30E-10 | 2.81E-09 | 284.431653 | 1500.80738 | 1594.52924 | 1725.21281 |
| -6.4268512 | 1.30E-10 | 2.82E-09 | 393.859969 | 734.437655 | 511.952028 | 602.899961 |
| -6.4254808 | 1.31E-10 | 2.84E-09 | 342.629484 | 844.305631 | 730.538287 | 784.442328 |
| -6.4246782 | 1.32E-10 | 2.86E-09 | 368.859492 | 586.413558 | 602.837894 | 580.487323 |
| -6.4246214 | 1.32E-10 | 2.86E-09 | 341.399952 | 832.128146 | 728.237379 | 824.785076 |
| -6.4243975 | 1.32E-10 | 2.86E-09 | 300.825408 | 1359.81917 | 1213.72897 | 1295.45047 |
| -6.4239534 | 1.33E-10 | 2.86E-09 | 360.25277 | 685.457104 | 608.590164 | 644.363341 |
| -6.4239627 | 1.33E-10 | 2.86E-09 | 347.957454 | 750.944912 | 699.476029 | 704.877463 |
| -6.4237391 | 1.33E-10 | 2.87E-09 | 384.023715 | 709.541463 | 520.005206 | 616.907859 |
| -6.4233854 | 1.33E-10 | 2.87E-09 | 346.727922 | 759.875068 | 686.821035 | 810.216862 |
| -6.4234467 | 1.33E-10 | 2.87E-09 | 340.580264 | 865.413272 | 730.538287 | 832.6295 |
| -6.4230586 | 1.34E-10 | 2.87E-09 | 332.383387 | 884.085415 | 837.530509 | 833.189816 |
| -6.4230771 | 1.34E-10 | 2.87E-09 | 330.334167 | 971.222086 | 989.390436 | 818.061285 |
| -6.4216304 | 1.35E-10 | 2.89E-09 | 354.514956 | 1552.76465 | 816.822337 | 1081.9701 |
| -6.4217455 | 1.35E-10 | 2.89E-09 | 345.908235 | 870.554877 | 710.980569 | 744.659896 |

| | | | | | | |
|---|---|---|---|---|---|---|
| -6.4212566 | 1.35E-10 | 2.89E-09 | 292.218686 | 410.245941 | 302.569401 | 248.780281 |
| -6.4215448 | 1.35E-10 | 2.89E-09 | 370.089024 | 611.580361 | 553.368372 | 596.176169 |
| -6.4215972 | 1.35E-10 | 2.89E-09 | 370.089024 | 632.958612 | 557.970188 | 587.211114 |
| -6.4212738 | 1.35E-10 | 2.89E-09 | 370.498867 | 589.931499 | 559.120642 | 581.607955 |
| -6.4217085 | 1.35E-10 | 2.89E-09 | 328.284948 | 832.128146 | 1026.20496 | 1023.13692 |
| -6.421212 | 1.35E-10 | 2.89E-09 | 340.990108 | 983.670182 | 776.556447 | 787.243908 |
| -6.4211424 | 1.35E-10 | 2.89E-09 | 351.646049 | 706.294134 | 646.555145 | 715.523466 |
| -6.4197911 | 1.36E-10 | 2.91E-09 | 353.285425 | 718.201008 | 628.147881 | 709.920307 |
| -6.4189259 | 1.37E-10 | 2.93E-09 | 336.071982 | 964.186206 | 771.954631 | 839.353291 |
| -6.4167096 | 1.39E-10 | 2.97E-09 | 336.891669 | 846.470517 | 745.494189 | 796.769279 |
| -6.4167503 | 1.39E-10 | 2.97E-09 | 344.268859 | 1041.8515 | 712.131023 | 934.607002 |
| -6.4163596 | 1.40E-10 | 2.97E-09 | 348.777142 | 813.185391 | 651.156961 | 752.504319 |
| -6.4164499 | 1.39E-10 | 2.97E-09 | 384.023715 | 505.500935 | 530.359292 | 525.57636 |
| -6.4163794 | 1.40E-10 | 2.97E-09 | 351.236205 | 771.781942 | 653.457869 | 674.620402 |
| -6.4161026 | 1.40E-10 | 2.97E-09 | 351.646049 | 665.161295 | 660.360593 | 782.76138 |
| -6.4158454 | 1.40E-10 | 2.97E-09 | 374.597306 | 530.397127 | 557.970188 | 611.865016 |
| -6.4155785 | 1.40E-10 | 2.98E-09 | 350.826361 | 766.369727 | 630.448789 | 743.539264 |
| -6.4154578 | 1.40E-10 | 2.98E-09 | 362.30199 | 596.967379 | 587.881992 | 663.414083 |
| -6.4147581 | 1.41E-10 | 2.99E-09 | 342.21964 | 686.268936 | 778.857355 | 785.002644 |
| -6.4145255 | 1.41E-10 | 2.99E-09 | 372.548087 | 541.492169 | 548.766556 | 594.495221 |
| -6.4133117 | 1.42E-10 | 3.01E-09 | 327.875104 | 941.996122 | 844.433233 | 833.189816 |
| -6.4129127 | 1.43E-10 | 3.02E-09 | 362.711834 | 598.861654 | 608.590164 | 573.203215 |
| -6.4128402 | 1.43E-10 | 3.02E-09 | 390.991061 | 1257.52829 | 601.68744 | 1115.02874 |
| -6.4129491 | 1.43E-10 | 3.02E-09 | 303.694315 | 1298.66113 | 1122.8431 | 1146.40643 |
| -6.4126374 | 1.43E-10 | 3.02E-09 | 367.220116 | 549.881103 | 582.129722 | 592.253958 |
| -6.4124874 | 1.43E-10 | 3.02E-09 | 457.795614 | 523.631858 | 395.756174 | 540.70489 |
| -6.4120054 | 1.44E-10 | 3.03E-09 | 347.957454 | 736.602541 | 637.351513 | 750.823371 |
| -6.4113421 | 1.44E-10 | 3.04E-09 | 326.235728 | 984.752625 | 823.725061 | 892.02299 |
| -6.4112112 | 1.44E-10 | 3.04E-09 | 310.251817 | 1064.31219 | 1110.18811 | 1010.24966 |
| -6.4104874 | 1.45E-10 | 3.05E-09 | 346.318079 | 840.787691 | 654.608323 | 876.334144 |
| -6.4094398 | 1.46E-10 | 3.07E-09 | 390.581218 | 585.872337 | 1160.80808 | 1143.60485 |
| -6.4092019 | 1.46E-10 | 3.07E-09 | 360.662614 | 484.663905 | 490.093402 | 488.035191 |
| -6.4087579 | 1.47E-10 | 3.08E-09 | 331.153855 | 795.32508 | 791.512349 | 801.251807 |
| -6.4080745 | 1.47E-10 | 3.09E-09 | 355.334644 | 834.022421 | 744.343735 | 604.020593 |
| -6.4072201 | 1.48E-10 | 3.11E-09 | 358.203551 | 661.372744 | 580.979268 | 607.382488 |
| -6.4069067 | 1.49E-10 | 3.11E-09 | 330.334167 | 761.228122 | 887.00003 | 971.027539 |
| -6.4046253 | 1.51E-10 | 3.16E-09 | 316.399475 | 1456.42721 | 944.52273 | 1174.98254 |
| -6.4042788 | 1.51E-10 | 3.16E-09 | 341.809796 | 517.949031 | 473.987046 | 536.222363 |
| -6.4038737 | 1.51E-10 | 3.17E-09 | 273.775712 | 890.038853 | 1365.58889 | 1140.24296 |
| -6.4031554 | 1.52E-10 | 3.18E-09 | 326.235728 | 602.108984 | 719.033747 | 656.690292 |
| -6.4030213 | 1.52E-10 | 3.18E-09 | 328.284948 | 976.904913 | 768.503269 | 876.334144 |
| -6.4023107 | 1.53E-10 | 3.20E-09 | 329.924323 | 744.179643 | 852.486411 | 819.181917 |
| -6.4014774 | 1.54E-10 | 3.21E-09 | 456.156238 | 719.283451 | 829.477331 | 717.76473 |

| | | | | | | |
|---|---|---|---|---|---|---|
| -6.4010262 | 1.54E-10 | 3.22E-09 | 338.531045 | 812.373559 | 681.068765 | 731.212313 |
| -6.4008659 | 1.54E-10 | 3.22E-09 | 344.268859 | 684.37466 | 654.608323 | 655.56966 |
| -6.3991704 | 1.56E-10 | 3.25E-09 | 344.268859 | 681.397942 | 679.918311 | 632.596706 |
| -6.3984404 | 1.57E-10 | 3.27E-09 | 438.942795 | 944.70223 | 895.053208 | 897.62615 |
| -6.3982035 | 1.57E-10 | 3.27E-09 | 311.481349 | 971.492697 | 1090.63039 | 932.926054 |
| -6.3956751 | 1.60E-10 | 3.32E-09 | 716.816946 | 440.283738 | 483.190678 | 455.536866 |
| -6.3949582 | 1.61E-10 | 3.34E-09 | 313.120724 | 991.247284 | 906.557748 | 917.797524 |
| -6.3921859 | 1.64E-10 | 3.40E-09 | 386.072935 | 752.839188 | 507.350212 | 561.996896 |
| -6.391425 | 1.64E-10 | 3.41E-09 | 911.082945 | 407.539833 | 507.350212 | 613.545964 |
| -6.3902422 | 1.66E-10 | 3.43E-09 | 381.564652 | 486.016959 | 479.739316 | 552.471525 |
| -6.3901375 | 1.66E-10 | 3.43E-09 | 438.532952 | 624.840289 | 501.597942 | 410.151274 |
| -6.3899645 | 1.66E-10 | 3.44E-09 | 366.810273 | 566.11775 | 517.704298 | 522.214464 |
| -6.3891087 | 1.67E-10 | 3.45E-09 | 338.121201 | 662.725798 | 673.015587 | 667.336295 |
| -6.3891108 | 1.67E-10 | 3.45E-09 | 308.612442 | 1040.22784 | 939.920914 | 932.926054 |
| -6.3887704 | 1.67E-10 | 3.46E-09 | 367.62996 | 553.669654 | 496.996126 | 579.927007 |
| -6.3884606 | 1.68E-10 | 3.46E-09 | 193.856155 | 191.05121 | 297.967585 | 252.142177 |
| -6.3879785 | 1.68E-10 | 3.47E-09 | 328.284948 | 745.262086 | 743.193281 | 732.893261 |
| -6.3873298 | 1.69E-10 | 3.48E-09 | 347.957454 | 675.985726 | 580.979268 | 621.390387 |
| -6.387218 | 1.69E-10 | 3.48E-09 | 422.139196 | 570.176912 | 408.411168 | 499.801826 |
| -6.3868999 | 1.69E-10 | 3.49E-09 | 310.661661 | 1256.17524 | 910.00911 | 1015.85281 |
| -6.3866184 | 1.70E-10 | 3.49E-09 | 346.727922 | 609.415475 | 626.997427 | 593.37459 |
| 6.38649231 | 1.70E-10 | 3.49E-09 | 0 | 0 | 0 | 0 |
| -6.3864087 | 1.70E-10 | 3.49E-09 | 358.613395 | 539.056672 | 579.828814 | 546.868366 |
| -6.3859975 | 1.70E-10 | 3.50E-09 | 315.579788 | 948.490781 | 842.132325 | 846.637398 |
| -6.3858969 | 1.70E-10 | 3.50E-09 | 321.727446 | 2045.27627 | 1006.64725 | 1310.579 |
| -6.3848984 | 1.72E-10 | 3.52E-09 | 317.219163 | 1018.57897 | 819.123245 | 864.007193 |
| -6.3836596 | 1.73E-10 | 3.55E-09 | 193.446311 | 190.780599 | 199.028541 | 186.024895 |
| -6.3833785 | 1.73E-10 | 3.55E-09 | 350.826361 | 1289.46036 | 1150.454 | 1191.23171 |
| -6.3832445 | 1.73E-10 | 3.55E-09 | 353.695268 | 632.14678 | 538.41247 | 599.538065 |
| -6.3826473 | 1.74E-10 | 3.56E-09 | 320.08807 | 757.16896 | 842.132325 | 834.310447 |
| -6.3824379 | 1.74E-10 | 3.57E-09 | 290.989155 | 1250.2218 | 1191.87034 | 1093.17642 |
| -6.3822876 | 1.74E-10 | 3.57E-09 | 352.875581 | 607.79181 | 613.19198 | 537.903311 |
| -6.3809684 | 1.76E-10 | 3.60E-09 | 297.136813 | 1233.71454 | 1057.26722 | 1034.34324 |
| 6.38061102 | 1.76E-10 | 3.60E-09 | 17.6232868 | 11.6362635 | 6.90272397 | 10.0856871 |
| -6.3797757 | 1.77E-10 | 3.62E-09 | 561.076272 | 606.709367 | 383.10118 | 447.132127 |
| -6.3796148 | 1.78E-10 | 3.62E-09 | 346.318079 | 602.379594 | 785.760079 | 631.476074 |
| -6.378591 | 1.79E-10 | 3.64E-09 | 286.071028 | 1249.40997 | 1295.4112 | 1138.56201 |
| -6.3783678 | 1.79E-10 | 3.65E-09 | 368.859492 | 610.768529 | 478.588862 | 564.23816 |
| -6.3781082 | 1.79E-10 | 3.65E-09 | 441.401859 | 570.447522 | 401.508444 | 458.898762 |
| 6.37779857 | 1.80E-10 | 3.65E-09 | 34.017042 | 20.2958084 | 25.3099879 | 27.4554815 |
| -6.3774193 | 1.80E-10 | 3.66E-09 | 2661.52616 | 7449.37353 | 6091.65391 | 6986.57956 |
| -6.3773952 | 1.80E-10 | 3.66E-09 | 236.479919 | 645.94793 | 579.828814 | 592.814274 |
| -6.3771948 | 1.80E-10 | 3.66E-09 | 368.449648 | 492.511618 | 496.996126 | 550.230262 |

| | | | | | | |
|---|---|---|---|---|---|---|
| -6.3768037 | 1.81E-10 | 3.67E-09 | 450.418424 | 460.038325 | 434.87161 | 397.824324 |
| -6.3767331 | 1.81E-10 | 3.67E-09 | 450.828268 | 525.255522 | 363.543463 | 439.84802 |
| -6.3767492 | 1.81E-10 | 3.67E-09 | 314.350256 | 830.23387 | 840.981871 | 813.578757 |
| -6.3761408 | 1.82E-10 | 3.68E-09 | 375.416994 | 561.787978 | 459.031144 | 519.412884 |
| -6.3759453 | 1.82E-10 | 3.68E-09 | 356.974019 | 846.470517 | 548.766556 | 833.750132 |
| -6.3755666 | 1.82E-10 | 3.69E-09 | 312.71088 | 1062.41792 | 801.866435 | 959.82122 |
| -6.3739853 | 1.84E-10 | 3.72E-09 | 345.498391 | 574.777295 | 618.94425 | 727.850417 |
| -6.3736614 | 1.85E-10 | 3.73E-09 | 527.05923 | 424.047091 | 417.6148 | 429.762333 |
| -6.3734424 | 1.85E-10 | 3.73E-09 | 426.237635 | 549.610492 | 400.35799 | 466.182869 |
| -6.3733187 | 1.85E-10 | 3.73E-09 | 368.859492 | 518.490253 | 473.987046 | 511.008145 |
| -6.3730885 | 1.85E-10 | 3.74E-09 | 295.497437 | 1453.17988 | 1038.85996 | 1123.99379 |
| -6.3727858 | 1.86E-10 | 3.74E-09 | 365.990585 | 540.950948 | 492.39431 | 544.066786 |
| -6.3726653 | 1.86E-10 | 3.74E-09 | 316.399475 | 809.39684 | 770.804177 | 796.769279 |
| -6.3724396 | 1.86E-10 | 3.74E-09 | 292.218686 | 1037.52173 | 1687.71601 | 1524.05938 |
| -6.3723608 | 1.86E-10 | 3.74E-09 | 298.366345 | 1237.50309 | 945.673184 | 1087.01294 |
| -6.3723922 | 1.86E-10 | 3.74E-09 | 360.662614 | 598.861654 | 529.208838 | 504.284354 |
| -6.3722064 | 1.86E-10 | 3.74E-09 | 346.318079 | 589.660888 | 553.368372 | 602.339645 |
| -6.3712562 | 1.87E-10 | 3.76E-09 | 314.7601 | 1110.31603 | 853.636865 | 829.82792 |
| -6.3711641 | 1.88E-10 | 3.76E-09 | 327.875104 | 810.479284 | 663.811955 | 718.885362 |
| -6.370797 | 1.88E-10 | 3.77E-09 | 384.843403 | 1047.53433 | 523.456568 | 815.259705 |
| -6.3706255 | 1.88E-10 | 3.77E-09 | 418.860445 | 481.687187 | 393.455266 | 500.922458 |
| -6.3690032 | 1.90E-10 | 3.81E-09 | 310.661661 | 932.524745 | 804.167343 | 846.077082 |
| -6.3688855 | 1.90E-10 | 3.81E-09 | 372.138243 | 756.898349 | 484.341132 | 648.285553 |
| 6.36820602 | 1.91E-10 | 3.83E-09 | 0 | 0 | 0 | 0 |
| -6.3673831 | 1.92E-10 | 3.84E-09 | 333.203074 | 704.399858 | 608.590164 | 663.414083 |
| -6.3666632 | 1.93E-10 | 3.86E-09 | 408.614348 | 418.093654 | 473.987046 | 427.521069 |
| -6.366349 | 1.94E-10 | 3.87E-09 | 343.039328 | 623.487235 | 556.819734 | 587.77143 |
| -6.3654189 | 1.95E-10 | 3.89E-09 | 311.071505 | 638.370828 | 626.997427 | 641.001445 |
| -6.3651532 | 1.95E-10 | 3.89E-09 | 577.470027 | 467.886037 | 402.658898 | 422.478225 |
| -6.3643394 | 1.96E-10 | 3.91E-09 | 466.812179 | 427.29442 | 399.207536 | 428.081385 |
| -6.3631753 | 1.98E-10 | 3.94E-09 | 368.449648 | 693.575427 | 473.987046 | 603.460277 |
| -6.3630738 | 1.98E-10 | 3.94E-09 | 315.989631 | 1003.69538 | 747.795097 | 834.870763 |
| -6.3626269 | 1.98E-10 | 3.95E-09 | 356.564176 | 516.866588 | 531.509746 | 646.604605 |
| -6.3621981 | 1.99E-10 | 3.95E-09 | 309.432129 | 898.969008 | 792.662803 | 819.181917 |
| -6.3619997 | 1.99E-10 | 3.96E-09 | 304.104159 | 925.218254 | 827.176423 | 930.684791 |
| -6.3611512 | 2.00E-10 | 3.98E-09 | 320.08807 | 736.873152 | 702.927391 | 706.558411 |
| -6.3599011 | 2.02E-10 | 4.01E-09 | 338.531045 | 653.525032 | 572.92609 | 586.650798 |
| -6.359307 | 2.03E-10 | 4.02E-09 | 381.154808 | 465.991762 | 432.570702 | 505.404986 |
| -6.3588682 | 2.03E-10 | 4.02E-09 | 333.203074 | 739.038038 | 678.767857 | 586.090482 |
| -6.3588425 | 2.03E-10 | 4.02E-09 | 336.891669 | 654.066253 | 572.92609 | 600.098381 |
| -6.3590255 | 2.03E-10 | 4.02E-09 | 340.17042 | 578.565846 | 559.120642 | 595.615853 |
| -6.3578105 | 2.05E-10 | 4.05E-09 | 1101.66035 | 733.355211 | 362.393009 | 593.934906 |
| -6.3575215 | 2.05E-10 | 4.05E-09 | 362.711834 | 508.207043 | 468.234776 | 503.163722 |

| | | | | | | |
|---|---|---|---|---|---|---|
| -6.356419 | 2.07E-10 | 4.07E-09 | 270.906805 | 1338.98214 | 1259.74712 | 1332.43133 |
| -6.356369 | 2.07E-10 | 4.07E-09 | 302.05494 | 1131.96489 | 807.618705 | 875.773828 |
| -6.3565219 | 2.06E-10 | 4.07E-09 | 335.252294 | 680.85672 | 576.377452 | 765.391586 |
| -6.3563941 | 2.07E-10 | 4.07E-09 | 411.893099 | 764.20484 | 417.6148 | 742.418632 |
| -6.3552914 | 2.08E-10 | 4.10E-09 | 543.452985 | 470.050923 | 346.286653 | 410.151274 |
| -6.3553784 | 2.08E-10 | 4.10E-09 | 425.417947 | 472.48642 | 419.915708 | 397.824324 |
| -6.3545318 | 2.09E-10 | 4.12E-09 | 306.153378 | 961.480098 | 774.255539 | 891.462674 |
| -6.3542591 | 2.09E-10 | 4.12E-09 | 307.792754 | 892.203739 | 782.308717 | 797.329595 |
| 6.35391361 | 2.10E-10 | 4.12E-09 | 15.1642236 | 13.530539 | 14.9559019 | 14.0078987 |
| -6.3539079 | 2.10E-10 | 4.12E-09 | 279.513526 | 380.478755 | 358.941647 | 403.427483 |
| -6.3540117 | 2.10E-10 | 4.12E-09 | 295.907281 | 468.697869 | 306.020763 | 327.78483 |
| -6.3527896 | 2.11E-10 | 4.15E-09 | 310.251817 | 750.13308 | 767.352815 | 835.991395 |
| -6.3527681 | 2.11E-10 | 4.15E-09 | 277.054463 | 1211.25385 | 1145.85218 | 1207.48087 |
| -6.3512811 | 2.14E-10 | 4.19E-09 | 404.106066 | 512.536816 | 421.066162 | 418.556014 |
| -6.3509745 | 2.14E-10 | 4.19E-09 | 311.891193 | 1161.73207 | 756.998729 | 959.82122 |
| -6.3503458 | 2.15E-10 | 4.21E-09 | 320.497914 | 724.425056 | 669.564225 | 658.931556 |
| -6.3501078 | 2.15E-10 | 4.21E-09 | 320.08807 | 677.609391 | 658.059685 | 706.558411 |
| -6.3500534 | 2.15E-10 | 4.21E-09 | 340.580264 | 662.184577 | 529.208838 | 605.141225 |
| -6.3497967 | 2.16E-10 | 4.21E-09 | 254.103206 | 1425.30697 | 1695.76919 | 1738.66039 |
| -6.349817 | 2.16E-10 | 4.21E-09 | 301.235252 | 800.737295 | 926.115466 | 985.595754 |
| -6.3496729 | 2.16E-10 | 4.21E-09 | 414.762006 | 410.787163 | 413.012984 | 429.762333 |
| -6.3480633 | 2.18E-10 | 4.25E-09 | 358.613395 | 614.557079 | 499.297034 | 490.276455 |
| -6.3476635 | 2.19E-10 | 4.26E-09 | 450.828268 | 428.647474 | 381.950726 | 439.287704 |
| -6.3472008 | 2.19E-10 | 4.27E-09 | 450.828268 | 413.493271 | 984.78862 | 725.048838 |
| -6.3464376 | 2.20E-10 | 4.29E-09 | 525.829698 | 562.59981 | 347.437107 | 491.957403 |
| -6.3464312 | 2.20E-10 | 4.29E-09 | 452.0578 | 530.938349 | 379.649818 | 430.322649 |
| -6.3462858 | 2.21E-10 | 4.29E-09 | 320.907758 | 721.448337 | 625.846973 | 693.110828 |
| -6.3461156 | 2.21E-10 | 4.29E-09 | 251.234298 | 356.935618 | 327.879389 | 308.734088 |
| -6.3459975 | 2.21E-10 | 4.29E-09 | 286.480872 | 1365.7726 | 1022.7536 | 1049.47177 |
| -6.3456805 | 2.21E-10 | 4.30E-09 | 296.726969 | 888.685799 | 875.49549 | 879.135723 |
| -6.3444829 | 2.23E-10 | 4.33E-09 | 299.186032 | 813.726613 | 887.00003 | 872.411932 |
| -6.3443065 | 2.23E-10 | 4.33E-09 | 459.844833 | 601.567762 | 360.092101 | 517.731937 |
| -6.3440725 | 2.24E-10 | 4.33E-09 | 309.432129 | 786.665535 | 712.131023 | 768.193166 |
| -6.3441214 | 2.24E-10 | 4.33E-09 | 441.401859 | 817.515164 | 406.11026 | 570.961952 |
| -6.3435904 | 2.24E-10 | 4.35E-09 | 344.268859 | 710.082684 | 529.208838 | 574.884163 |
| -6.3421744 | 2.27E-10 | 4.38E-09 | 384.433559 | 537.433007 | 446.37615 | 717.76473 |
| -6.3419773 | 2.27E-10 | 4.39E-09 | 310.251817 | 905.734278 | 730.538287 | 745.780528 |
| -6.3411175 | 2.28E-10 | 4.41E-09 | 326.645572 | 581.001343 | 625.846973 | 627.553862 |
| -6.3405872 | 2.29E-10 | 4.42E-09 | 343.039328 | 556.375762 | 502.748396 | 536.222363 |
| -6.3404538 | 2.29E-10 | 4.42E-09 | 315.579788 | 725.236888 | 674.166041 | 665.095031 |
| -6.3403032 | 2.29E-10 | 4.42E-09 | 467.222023 | 795.595691 | 907.708202 | 885.299199 |
| -6.3398652 | 2.30E-10 | 4.43E-09 | 650.012394 | 516.325367 | 379.649818 | 433.684544 |
| -6.3395124 | 2.30E-10 | 4.44E-09 | 318.448695 | 685.186493 | 643.103783 | 648.845868 |

| | | | | | | |
|---|---|---|---|---|---|---|
| -6.3394205 | 2.31E-10 | 4.44E-09 | 286.480872 | 922.782757 | 1026.20496 | 1113.90811 |
| -6.3391681 | 2.31E-10 | 4.44E-09 | 321.727446 | 616.992576 | 628.147881 | 659.491872 |
| -6.3386909 | 2.32E-10 | 4.46E-09 | 302.05494 | 861.624721 | 755.848275 | 824.785076 |
| -6.3384512 | 2.32E-10 | 4.46E-09 | 296.317125 | 879.755643 | 814.521429 | 918.35784 |
| -6.3380561 | 2.33E-10 | 4.47E-09 | 334.022762 | 394.821127 | 386.552542 | 402.306851 |
| -6.3378124 | 2.33E-10 | 4.47E-09 | 313.120724 | 768.805224 | 651.156961 | 761.469374 |
| -6.3374596 | 2.34E-10 | 4.48E-09 | 356.974019 | 530.397127 | 475.1375 | 463.38129 |
| -6.3373077 | 2.34E-10 | 4.48E-09 | 289.349779 | 919.264817 | 952.575908 | 921.15942 |
| -6.3361754 | 2.36E-10 | 4.51E-09 | 368.859492 | 454.89672 | 455.579782 | 435.925808 |
| -6.3360659 | 2.36E-10 | 4.51E-09 | 350.826361 | 532.020792 | 465.933868 | 511.008145 |
| -6.3353307 | 2.37E-10 | 4.53E-09 | 304.923847 | 713.871235 | 952.575908 | 933.48637 |
| -6.3346821 | 2.38E-10 | 4.55E-09 | 304.923847 | 772.593774 | 758.149183 | 728.971049 |
| -6.3345189 | 2.38E-10 | 4.55E-09 | 398.368251 | 750.944912 | 413.012984 | 629.23481 |
| -6.3345902 | 2.38E-10 | 4.55E-09 | 283.202121 | 977.175523 | 1121.69265 | 978.871962 |
| -6.3334916 | 2.40E-10 | 4.58E-09 | 315.169944 | 633.499834 | 693.723759 | 678.542614 |
| -6.3315848 | 2.43E-10 | 4.63E-09 | 344.268859 | 493.864672 | 496.996126 | 541.825522 |
| -6.3297545 | 2.46E-10 | 4.68E-09 | 377.05637 | 497.653223 | 576.377452 | 411.832222 |
| -6.3296939 | 2.46E-10 | 4.68E-09 | 445.500297 | 521.737582 | 368.145279 | 433.124228 |
| -6.3294164 | 2.46E-10 | 4.69E-09 | 365.580741 | 482.499019 | 358.941647 | 418.556014 |
| -6.3292328 | 2.46E-10 | 4.69E-09 | 275.005243 | 1178.50994 | 1059.56813 | 1055.63525 |
| -6.3289326 | 2.47E-10 | 4.70E-09 | 381.974496 | 503.065438 | 501.597942 | 394.462428 |
| -6.327639 | 2.49E-10 | 4.73E-09 | 320.497914 | 650.548313 | 575.226998 | 669.017243 |
| -6.3275162 | 2.49E-10 | 4.73E-09 | 327.055416 | 675.444505 | 546.465648 | 609.623752 |
| -6.3271087 | 2.50E-10 | 4.74E-09 | 351.646049 | 551.234157 | 446.37615 | 544.066786 |
| -6.3260053 | 2.52E-10 | 4.78E-09 | 392.220593 | 791.265918 | 637.351513 | 422.478225 |
| -6.3256154 | 2.52E-10 | 4.78E-09 | 347.957454 | 1185.54582 | 607.43971 | 774.916957 |
| -6.3251504 | 2.53E-10 | 4.80E-09 | 314.7601 | 625.381511 | 639.652421 | 723.36789 |
| -6.3247074 | 2.54E-10 | 4.81E-09 | 308.202598 | 820.221272 | 654.608323 | 748.021791 |
| -6.3243136 | 2.54E-10 | 4.82E-09 | 299.595876 | 806.961343 | 724.786017 | 795.648647 |
| -6.3235385 | 2.56E-10 | 4.84E-09 | 306.563222 | 867.036936 | 666.112863 | 780.520116 |
| -6.3235836 | 2.56E-10 | 4.84E-09 | 275.415087 | 764.475451 | 883.548668 | 848.878662 |
| -6.3234089 | 2.56E-10 | 4.84E-09 | 317.219163 | 744.179643 | 607.43971 | 635.398286 |
| -6.3228241 | 2.57E-10 | 4.85E-09 | 356.154332 | 537.974229 | 407.260714 | 501.482774 |
| -6.322614 | 2.57E-10 | 4.85E-09 | 298.776188 | 776.111715 | 776.556447 | 741.298 |
| -6.3226545 | 2.57E-10 | 4.85E-09 | 322.956977 | 600.214708 | 562.572004 | 586.090482 |
| -6.3226065 | 2.57E-10 | 4.85E-09 | 273.775712 | 1000.98927 | 1310.3671 | 1514.53401 |
| -6.322048 | 2.58E-10 | 4.86E-09 | 308.202598 | 653.525032 | 691.422851 | 702.6362 |
| -6.3219829 | 2.58E-10 | 4.86E-09 | 359.023239 | 439.742516 | 579.828814 | 652.207764 |
| -6.3211431 | 2.60E-10 | 4.89E-09 | 350.826361 | 488.181846 | 514.252936 | 444.890863 |
| -6.3183083 | 2.64E-10 | 4.98E-09 | 351.646049 | 470.050923 | 444.075242 | 490.836771 |
| -6.3177756 | 2.65E-10 | 4.99E-09 | 370.908711 | 612.933415 | 410.712076 | 545.747734 |
| -6.317203 | 2.66E-10 | 5.01E-09 | 307.38291 | 787.206757 | 641.953329 | 706.558411 |
| -6.3169884 | 2.67E-10 | 5.01E-09 | 299.595876 | 949.302613 | 720.184201 | 794.528015 |

| | | | | | | |
|---|---|---|---|---|---|---|
| -6.3170003 | 2.67E-10 | 5.01E-09 | 286.890716 | 969.598422 | 813.370975 | 926.762579 |
| -6.3164304 | 2.68E-10 | 5.02E-09 | 346.318079 | 569.63569 | 447.526604 | 542.946154 |
| -6.3163645 | 2.68E-10 | 5.02E-09 | 419.270289 | 367.218827 | 471.686138 | 429.202017 |
| -6.3161108 | 2.68E-10 | 5.03E-09 | 510.665474 | 865.142661 | 347.437107 | 643.242709 |
| -6.3145927 | 2.71E-10 | 5.07E-09 | 317.219163 | 681.127331 | 597.085624 | 582.168271 |
| -6.3134202 | 2.73E-10 | 5.11E-09 | 352.875581 | 826.174709 | 484.341132 | 636.518918 |
| -6.3134652 | 2.73E-10 | 5.11E-09 | 308.612442 | 621.86357 | 660.360593 | 695.352092 |
| -6.3131441 | 2.73E-10 | 5.11E-09 | 333.203074 | 1048.07555 | 849.035049 | 938.529214 |
| -6.3129424 | 2.74E-10 | 5.11E-09 | 331.563699 | 693.034205 | 585.581084 | 647.725237 |
| -6.3129793 | 2.74E-10 | 5.11E-09 | 276.644619 | 921.429703 | 1007.7977 | 1034.90356 |
| -6.3125218 | 2.75E-10 | 5.12E-09 | 311.481349 | 693.575427 | 592.483808 | 663.414083 |
| -6.3124174 | 2.75E-10 | 5.12E-09 | 309.841973 | 705.752912 | 600.536986 | 689.188617 |
| -6.3106532 | 2.78E-10 | 5.18E-09 | 324.596353 | 680.85672 | 528.058384 | 593.37459 |
| -6.3106557 | 2.78E-10 | 5.18E-09 | 466.812179 | 603.191427 | 335.932567 | 587.77143 |
| -6.3101595 | 2.79E-10 | 5.19E-09 | 287.710404 | 834.563643 | 849.035049 | 802.372438 |
| -6.3097243 | 2.80E-10 | 5.20E-09 | 157.789894 | 185.097773 | 181.771731 | 178.180472 |
| -6.3095525 | 2.80E-10 | 5.20E-09 | 318.858539 | 638.100217 | 563.722458 | 565.358792 |
| -6.3094992 | 2.80E-10 | 5.20E-09 | 519.272196 | 389.408911 | 414.163438 | 399.505271 |
| -6.3084736 | 2.82E-10 | 5.24E-09 | 325.416041 | 545.28072 | 556.819734 | 516.050989 |
| -6.3082206 | 2.82E-10 | 5.24E-09 | 431.155762 | 419.446708 | 378.499364 | 394.462428 |
| -6.3078996 | 2.83E-10 | 5.25E-09 | 468.041711 | 1098.67976 | 1193.02079 | 1187.3095 |
| -6.3074411 | 2.84E-10 | 5.26E-09 | 320.08807 | 578.024624 | 540.713378 | 569.84132 |
| -6.3070852 | 2.84E-10 | 5.26E-09 | 332.793231 | 555.83454 | 498.14658 | 493.078035 |
| -6.307337 | 2.84E-10 | 5.26E-09 | 358.613395 | 795.595691 | 469.38523 | 582.728587 |
| -6.3071239 | 2.84E-10 | 5.26E-09 | 164.347396 | 273.587498 | 263.453965 | 278.477026 |
| -6.3071556 | 2.84E-10 | 5.26E-09 | 345.908235 | 517.40781 | 453.278874 | 458.898762 |
| -6.3064104 | 2.86E-10 | 5.28E-09 | 308.202598 | 688.975044 | 597.085624 | 667.896611 |
| -6.3063091 | 2.86E-10 | 5.28E-09 | 317.629007 | 650.277702 | 540.713378 | 615.787228 |
| -6.3059258 | 2.86E-10 | 5.29E-09 | 404.925753 | 419.717318 | 371.596641 | 423.598857 |
| -6.3055928 | 2.87E-10 | 5.30E-09 | 281.972589 | 880.296865 | 845.583687 | 895.384886 |
| -6.3047941 | 2.89E-10 | 5.32E-09 | 331.973543 | 495.758947 | 505.049304 | 498.681194 |
| -6.3042595 | 2.90E-10 | 5.34E-09 | 279.513526 | 989.082398 | 944.52273 | 841.594555 |
| -6.3033482 | 2.91E-10 | 5.37E-09 | 278.693838 | 977.987356 | 1029.65633 | 841.034239 |
| -6.3030709 | 2.92E-10 | 5.37E-09 | 416.811226 | 434.05969 | 372.747095 | 397.264008 |
| -6.3023673 | 2.93E-10 | 5.40E-09 | 326.645572 | 647.030373 | 493.544764 | 607.382488 |
| -6.3021925 | 2.93E-10 | 5.40E-09 | 407.794661 | 448.13145 | 366.994825 | 544.627102 |
| -6.3008095 | 2.96E-10 | 5.44E-09 | 331.153855 | 497.923834 | 483.190678 | 518.852568 |
| -6.3004485 | 2.97E-10 | 5.45E-09 | 312.301037 | 663.808241 | 891.601846 | 623.631651 |
| -6.3001866 | 2.97E-10 | 5.46E-09 | 350.006673 | 484.934516 | 511.952028 | 417.995698 |
| -6.2992635 | 2.99E-10 | 5.49E-09 | 316.399475 | 529.314684 | 616.643342 | 604.580909 |
| -6.2981328 | 3.01E-10 | 5.53E-09 | 298.366345 | 814.538445 | 663.811955 | 700.394936 |
| -6.2970907 | 3.03E-10 | 5.56E-09 | 510.665474 | 517.40781 | 335.932567 | 444.330547 |
| -6.2971428 | 3.03E-10 | 5.56E-09 | 345.498391 | 453.273055 | 499.297034 | 434.805176 |

| | | | | | | |
|---|---|---|---|---|---|---|
| -6.2968354 | 3.04E-10 | 5.56E-09 | 373.367775 | 386.161582 | 517.704298 | 452.174971 |
| -6.2963036 | 3.05E-10 | 5.58E-09 | 352.875581 | 625.652121 | 441.774334 | 484.673296 |
| -6.2962137 | 3.05E-10 | 5.58E-09 | 339.350733 | 516.866588 | 445.225696 | 476.268556 |
| -6.2957001 | 3.06E-10 | 5.59E-09 | 309.432129 | 781.794541 | 647.705599 | 587.77143 |
| -6.2957271 | 3.06E-10 | 5.59E-09 | 265.16899 | 996.93011 | 1194.17125 | 1044.98924 |
| -6.2953837 | 3.07E-10 | 5.60E-09 | 490.992968 | 395.632959 | 363.543463 | 439.84802 |
| -6.2951298 | 3.07E-10 | 5.61E-09 | 325.825885 | 566.11775 | 486.64204 | 523.895412 |
| -6.2945381 | 3.08E-10 | 5.62E-09 | 274.5954 | 889.22702 | 1202.22443 | 971.027539 |
| -6.2944627 | 3.08E-10 | 5.62E-09 | 283.202121 | 862.436553 | 812.220521 | 782.76138 |
| -6.2942168 | 3.09E-10 | 5.63E-09 | 313.940412 | 582.354397 | 537.262016 | 592.253958 |
| -6.2937103 | 3.10E-10 | 5.64E-09 | 332.793231 | 1143.60115 | 850.185503 | 941.89111 |
| -6.2926887 | 3.12E-10 | 5.68E-09 | 305.333691 | 668.408624 | 610.891072 | 820.862865 |
| -6.2923026 | 3.13E-10 | 5.69E-09 | 292.218686 | 671.385343 | 765.051907 | 786.123276 |
| -6.2915772 | 3.14E-10 | 5.71E-09 | 334.84245 | 1042.66333 | 559.120642 | 716.083782 |
| -6.2909263 | 3.16E-10 | 5.73E-09 | 350.006673 | 506.042157 | 413.012984 | 458.338446 |
| -6.2909884 | 3.15E-10 | 5.73E-09 | 311.071505 | 548.79866 | 599.386532 | 669.577559 |
| -6.2908008 | 3.16E-10 | 5.73E-09 | 173.363961 | 259.515737 | 260.002603 | 263.348496 |
| -6.2905812 | 3.16E-10 | 5.74E-09 | 259.021332 | 490.346732 | 327.879389 | 393.902112 |
| -6.2899291 | 3.18E-10 | 5.76E-09 | 260.660708 | 995.577056 | 1217.18033 | 1144.72548 |
| -6.2897493 | 3.18E-10 | 5.76E-09 | 1045.10189 | 651.089535 | 631.599243 | 670.69819 |
| -6.289716 | 3.18E-10 | 5.76E-09 | 311.071505 | 796.136912 | 521.15566 | 808.535914 |
| -6.2889508 | 3.20E-10 | 5.78E-09 | 306.973066 | 699.258253 | 595.93517 | 584.409534 |
| -6.2885001 | 3.21E-10 | 5.80E-09 | 687.718031 | 1067.0183 | 970.983172 | 1076.36694 |
| -6.2884071 | 3.21E-10 | 5.80E-09 | 252.873674 | 1156.86108 | 1253.99485 | 1152.56991 |
| 6.2879896 | 3.22E-10 | 5.81E-09 | 24.5906328 | 25.4374132 | 21.8586259 | 25.7745336 |
| -6.2879326 | 3.22E-10 | 5.81E-09 | 321.317602 | 567.470804 | 1038.85996 | 978.871962 |
| -6.2876774 | 3.22E-10 | 5.81E-09 | 284.021809 | 835.646086 | 746.644643 | 778.839169 |
| -6.2874789 | 3.23E-10 | 5.81E-09 | 314.350256 | 967.704146 | 574.076544 | 921.15942 |
| -6.2874783 | 3.23E-10 | 5.81E-09 | 290.989155 | 690.869319 | 702.927391 | 735.134525 |
| -6.2876085 | 3.22E-10 | 5.81E-09 | 324.186509 | 620.510517 | 829.477331 | 534.541415 |
| -6.2877978 | 3.22E-10 | 5.81E-09 | 280.333214 | 791.265918 | 832.928693 | 819.742233 |
| -6.2872926 | 3.23E-10 | 5.81E-09 | 462.71374 | 380.478755 | 418.765254 | 373.170422 |
| -6.2871975 | 3.23E-10 | 5.81E-09 | 313.940412 | 579.9189 | 520.005206 | 572.642899 |
| -6.2869178 | 3.24E-10 | 5.82E-09 | 1187.31772 | 482.76963 | 533.810654 | 462.260658 |
| -6.2858683 | 3.26E-10 | 5.86E-09 | 350.006673 | 652.98381 | 421.066162 | 570.401636 |
| -6.2857625 | 3.26E-10 | 5.86E-09 | 289.759623 | 798.031188 | 678.767857 | 738.49642 |
| -6.2854952 | 3.27E-10 | 5.87E-09 | 279.92337 | 782.606373 | 849.035049 | 810.216862 |
| -6.2853918 | 3.27E-10 | 5.87E-09 | 293.038374 | 1569.0013 | 813.370975 | 1046.67019 |
| -6.2843455 | 3.29E-10 | 5.90E-09 | 491.812656 | 385.62036 | 372.747095 | 414.073486 |
| -6.2841146 | 3.30E-10 | 5.91E-09 | 352.465737 | 513.88987 | 742.042827 | 462.820974 |
| -6.2837032 | 3.31E-10 | 5.92E-09 | 347.54761 | 451.37878 | 411.86253 | 441.528968 |
| -6.2830504 | 3.32E-10 | 5.94E-09 | 314.7601 | 571.529966 | 549.91701 | 509.327197 |
| -6.282939 | 3.32E-10 | 5.94E-09 | 298.776188 | 686.268936 | 624.696519 | 615.226912 |

| | | | | | | |
|---|---|---|---|---|---|---|
| -6.28223 | 3.34E-10 | 5.97E-09 | 344.678703 | 495.488337 | 411.86253 | 459.459078 |
| -6.2814552 | 3.35E-10 | 5.99E-09 | 453.697175 | 346.652408 | 456.730236 | 617.468175 |
| -6.2810653 | 3.36E-10 | 6.00E-09 | 301.645096 | 762.310565 | 593.634262 | 642.682393 |
| -6.2804079 | 3.38E-10 | 6.02E-09 | 1039.36408 | 2848.99028 | 2187.01305 | 2716.41172 |
| -6.2802944 | 3.38E-10 | 6.02E-09 | 286.890716 | 755.545295 | 685.670581 | 746.340844 |
| -6.280565 | 3.37E-10 | 6.02E-09 | 258.611488 | 1159.56719 | 1034.25814 | 1023.13692 |
| -6.280489 | 3.38E-10 | 6.02E-09 | 311.481349 | 565.305918 | 593.634262 | 518.292252 |
| -6.2802919 | 3.38E-10 | 6.02E-09 | 331.563699 | 580.460121 | 441.774334 | 532.300151 |
| -6.2800794 | 3.38E-10 | 6.02E-09 | 247.13586 | 1252.92791 | 1253.99485 | 1172.18096 |
| -6.279451 | 3.40E-10 | 6.04E-09 | 423.778572 | 611.580361 | 339.383929 | 577.125427 |
| -6.27948 | 3.40E-10 | 6.04E-09 | 392.630437 | 441.907402 | 543.014286 | 364.205367 |
| -6.2794375 | 3.40E-10 | 6.04E-09 | 236.889763 | 773.676218 | 868.592767 | 958.700588 |
| -6.27931 | 3.40E-10 | 6.04E-09 | 412.712787 | 430.271139 | 352.038923 | 409.030642 |
| -6.2786587 | 3.42E-10 | 6.06E-09 | 375.416994 | 610.497918 | 554.518826 | 520.533516 |
| -6.2777302 | 3.44E-10 | 6.09E-09 | 309.841973 | 559.08187 | 534.961108 | 546.868366 |
| -6.2769199 | 3.45E-10 | 6.12E-09 | 317.219163 | 737.414373 | 508.500666 | 616.347543 |
| -6.2767314 | 3.46E-10 | 6.13E-09 | 292.62853 | 629.711283 | 675.316495 | 690.869565 |
| -6.2764877 | 3.46E-10 | 6.13E-09 | 332.793231 | 604.544481 | 469.38523 | 471.225713 |
| -6.2760478 | 3.47E-10 | 6.15E-09 | 247.545703 | 1103.00954 | 1571.52016 | 1334.67259 |
| -6.2758818 | 3.48E-10 | 6.15E-09 | 320.08807 | 467.344816 | 543.014286 | 544.627102 |
| -6.2756326 | 3.48E-10 | 6.15E-09 | 261.070552 | 1125.19962 | 975.584988 | 971.027539 |
| -6.2756878 | 3.48E-10 | 6.15E-09 | 350.416517 | 418.634875 | 404.959806 | 434.24486 |
| -6.2753754 | 3.49E-10 | 6.16E-09 | 273.775712 | 959.315212 | 820.273699 | 825.345392 |
| -6.2754029 | 3.49E-10 | 6.16E-09 | 312.71088 | 627.817008 | 510.801574 | 551.350893 |
| -6.2743632 | 3.51E-10 | 6.19E-09 | 300.415564 | 594.531882 | 586.731538 | 597.857117 |
| -6.2741698 | 3.52E-10 | 6.20E-09 | 284.021809 | 733.896433 | 698.325575 | 753.064635 |
| -6.2741118 | 3.52E-10 | 6.20E-09 | 314.350256 | 1187.4401 | 626.997427 | 803.49307 |
| -6.2738469 | 3.52E-10 | 6.20E-09 | 316.809319 | 579.9189 | 479.739316 | 543.50647 |
| -6.2734973 | 3.53E-10 | 6.21E-09 | 289.759623 | 755.545295 | 676.466949 | 650.526816 |
| -6.273547 | 3.53E-10 | 6.21E-09 | 258.201644 | 980.152242 | 1037.7095 | 1064.6003 |
| -6.2728241 | 3.55E-10 | 6.24E-09 | 290.169467 | 720.636505 | 658.059685 | 651.087132 |
| -6.2725218 | 3.55E-10 | 6.24E-09 | 302.874627 | 705.752912 | 579.828814 | 578.806375 |
| -6.2725257 | 3.55E-10 | 6.24E-09 | 292.218686 | 611.850972 | 698.325575 | 692.550512 |
| -6.2723312 | 3.56E-10 | 6.24E-09 | 313.530568 | 618.886852 | 508.500666 | 530.058887 |
| -6.2722223 | 3.56E-10 | 6.25E-09 | 347.957454 | 492.241007 | 563.722458 | 400.625903 |
| -6.2714755 | 3.58E-10 | 6.27E-09 | 260.660708 | 903.298781 | 1133.19719 | 1082.53041 |
| -6.271351 | 3.58E-10 | 6.27E-09 | 299.186032 | 589.931499 | 626.997427 | 579.927007 |
| -6.270201 | 3.61E-10 | 6.32E-09 | 280.743058 | 736.061319 | 742.042827 | 739.056736 |
| -6.2698273 | 3.61E-10 | 6.33E-09 | 322.547134 | 618.616241 | 482.040224 | 497.560562 |
| -6.2697194 | 3.62E-10 | 6.33E-09 | 300.00572 | 664.890684 | 555.66928 | 620.830071 |
| -6.26957 | 3.62E-10 | 6.33E-09 | 315.989631 | 637.558996 | 586.731538 | 637.63955 |
| -6.2693058 | 3.63E-10 | 6.34E-09 | 288.120248 | 677.609391 | 650.006507 | 694.23146 |
| -6.2688842 | 3.64E-10 | 6.35E-09 | 240.988201 | 1273.49433 | 1233.28668 | 1253.42678 |

| | | | | | | |
|---|---|---|---|---|---|---|
| -6.2685268 | 3.64E-10 | 6.36E-09 | 288.120248 | 661.372744 | 664.962409 | 675.741034 |
| 6.26818837 | 3.65E-10 | 6.37E-09 | 0 | 0 | 0 | 0 |
| -6.2677168 | 3.66E-10 | 6.39E-09 | 335.662138 | 487.640624 | 454.429328 | 415.754434 |
| -6.2671373 | 3.68E-10 | 6.41E-09 | 301.235252 | 612.392193 | 546.465648 | 584.96985 |
| -6.2665944 | 3.69E-10 | 6.42E-09 | 422.54904 | 421.882205 | 337.083021 | 469.544765 |
| -6.266516 | 3.69E-10 | 6.42E-09 | 312.71088 | 576.40096 | 536.111562 | 485.233612 |
| -6.2667481 | 3.69E-10 | 6.42E-09 | 284.841497 | 811.561727 | 691.422851 | 681.904509 |
| -6.2665225 | 3.69E-10 | 6.42E-09 | 302.464783 | 602.379594 | 792.662803 | 587.211114 |
| -6.2657397 | 3.71E-10 | 6.44E-09 | 306.153378 | 519.843307 | 546.465648 | 585.530166 |
| -6.2659475 | 3.71E-10 | 6.44E-09 | 301.235252 | 702.776193 | 538.41247 | 689.748933 |
| -6.2658659 | 3.71E-10 | 6.44E-09 | 367.220116 | 456.249774 | 363.543463 | 432.563912 |
| -6.2657734 | 3.71E-10 | 6.44E-09 | 253.283518 | 1981.41212 | 1110.18811 | 1312.82027 |
| -6.2650511 | 3.73E-10 | 6.47E-09 | 295.087594 | 758.792625 | 579.828814 | 677.982298 |
| -6.263299 | 3.77E-10 | 6.54E-09 | 287.30056 | 733.625822 | 654.608323 | 644.363341 |
| -6.2630872 | 3.77E-10 | 6.54E-09 | 258.201644 | 992.329727 | 923.814558 | 1027.05913 |
| -6.2622711 | 3.79E-10 | 6.57E-09 | 273.365868 | 833.210589 | 804.167343 | 750.263055 |
| -6.2616624 | 3.81E-10 | 6.59E-09 | 310.661661 | 499.006277 | 517.704298 | 509.327197 |
| -6.2613219 | 3.82E-10 | 6.60E-09 | 272.136336 | 747.156361 | 1043.46177 | 970.467223 |
| -6.2614002 | 3.82E-10 | 6.60E-09 | 313.940412 | 708.188409 | 494.695218 | 582.728587 |
| -6.2613476 | 3.82E-10 | 6.60E-09 | 381.974496 | 383.996696 | 364.693917 | 430.322649 |
| -6.2605687 | 3.84E-10 | 6.63E-09 | 179.921463 | 496.57078 | 453.278874 | 471.225713 |
| -6.2604945 | 3.84E-10 | 6.63E-09 | 290.579311 | 641.618157 | 597.085624 | 679.663246 |
| -6.2604031 | 3.84E-10 | 6.63E-09 | 268.857585 | 1007.48393 | 797.264619 | 860.084981 |
| -6.2593919 | 3.86E-10 | 6.67E-09 | 257.7918 | 910.875883 | 1007.7977 | 985.035438 |
| -6.2587451 | 3.88E-10 | 6.69E-09 | 320.497914 | 456.520384 | 543.014286 | 637.079234 |
| -6.2572113 | 3.92E-10 | 6.75E-09 | 336.071982 | 415.928768 | 518.854752 | 613.545964 |
| -6.256474 | 3.94E-10 | 6.78E-09 | 349.186986 | 473.568864 | 383.10118 | 422.478225 |
| -6.2562021 | 3.94E-10 | 6.79E-09 | 283.202121 | 719.283451 | 631.599243 | 732.332945 |
| -6.2557097 | 3.96E-10 | 6.81E-09 | 279.103682 | 745.262086 | 682.219219 | 706.558411 |
| -6.2548251 | 3.98E-10 | 6.84E-09 | 262.709927 | 1682.65782 | 935.319098 | 1162.09528 |
| -6.2547671 | 3.98E-10 | 6.84E-09 | 279.92337 | 798.301798 | 686.821035 | 686.387037 |
| -6.2543684 | 3.99E-10 | 6.85E-09 | 327.055416 | 1072.97174 | 537.262016 | 723.928206 |
| -6.2534138 | 4.02E-10 | 6.89E-09 | 313.940412 | 474.921917 | 586.731538 | 688.628301 |
| -6.2534468 | 4.01E-10 | 6.89E-09 | 237.70945 | 264.11612 | 272.657597 | 279.597658 |
| -6.2532621 | 4.02E-10 | 6.89E-09 | 304.514003 | 1317.33327 | 1370.19071 | 1306.09648 |
| -6.2531579 | 4.02E-10 | 6.89E-09 | 272.956024 | 785.583092 | 988.239982 | 754.185267 |
| -6.2529839 | 4.03E-10 | 6.90E-09 | 282.792277 | 743.909032 | 676.466949 | 637.63955 |
| -6.2528691 | 4.03E-10 | 6.90E-09 | 289.759623 | 678.691834 | 583.280176 | 626.433231 |
| -6.2520011 | 4.05E-10 | 6.93E-09 | 315.579788 | 489.264289 | 462.482506 | 468.984449 |
| -6.2517613 | 4.06E-10 | 6.94E-09 | 474.599213 | 479.522301 | 335.932567 | 389.419584 |
| -6.2515399 | 4.06E-10 | 6.95E-09 | 303.284471 | 651.360145 | 505.049304 | 573.763531 |
| -6.2492288 | 4.12E-10 | 7.05E-09 | 265.988678 | 857.024338 | 863.990951 | 768.193166 |
| 6.24860504 | 4.14E-10 | 7.07E-09 | 0 | 0 | 0 | 0 |

| | | | | | | |
|---|---|---|---|---|---|---|
| -6.248587 | 4.14E-10 | 7.07E-09 | 302.05494 | 505.500935 | 408.411168 | 479.630452 |
| -6.2482478 | 4.15E-10 | 7.08E-09 | 271.316649 | 889.22702 | 719.033747 | 788.36454 |
| -6.2480132 | 4.16E-10 | 7.09E-09 | 565.584554 | 523.902468 | 326.728935 | 410.71159 |
| -6.2472866 | 4.18E-10 | 7.12E-09 | 293.448218 | 590.202109 | 562.572004 | 704.317147 |
| -6.2465683 | 4.20E-10 | 7.14E-09 | 289.349779 | 720.636505 | 593.634262 | 600.098381 |
| -6.2466459 | 4.19E-10 | 7.14E-09 | 306.153378 | 507.1246 | 485.491586 | 565.919108 |
| -6.2464399 | 4.20E-10 | 7.14E-09 | 301.645096 | 569.906301 | 494.695218 | 550.790577 |
| -6.2457017 | 4.22E-10 | 7.17E-09 | 293.448218 | 593.99066 | 626.997427 | 834.310447 |
| -6.245326 | 4.23E-10 | 7.19E-09 | 297.136813 | 534.185678 | 781.158263 | 677.421982 |
| -6.2451019 | 4.24E-10 | 7.19E-09 | 606.568942 | 566.11775 | 327.879389 | 411.271906 |
| -6.2439887 | 4.27E-10 | 7.24E-09 | 312.301037 | 633.770445 | 464.783414 | 528.938255 |
| -6.2436279 | 4.28E-10 | 7.25E-09 | 275.824931 | 1476.99363 | 792.662803 | 1061.23841 |
| -6.2435355 | 4.28E-10 | 7.25E-09 | 281.152902 | 769.887667 | 666.112863 | 629.23481 |
| -6.24299 | 4.29E-10 | 7.27E-09 | 296.317125 | 582.895618 | 517.704298 | 560.315949 |
| -6.2429473 | 4.29E-10 | 7.27E-09 | 313.120724 | 530.667738 | 452.12842 | 474.587608 |
| -6.2430748 | 4.29E-10 | 7.27E-09 | 300.415564 | 577.212792 | 532.6602 | 500.362142 |
| -6.2428327 | 4.30E-10 | 7.27E-09 | 290.989155 | 645.136097 | 540.713378 | 629.23481 |
| -6.2424408 | 4.31E-10 | 7.29E-09 | 297.546657 | 586.95478 | 503.89885 | 582.728587 |
| -6.2414046 | 4.34E-10 | 7.33E-09 | 299.186032 | 629.170062 | 511.952028 | 541.265206 |
| -6.2412655 | 4.34E-10 | 7.33E-09 | 317.219163 | 550.692936 | 433.721156 | 485.233612 |
| -6.2411301 | 4.34E-10 | 7.34E-09 | 320.497914 | 424.858923 | 450.977966 | 500.362142 |
| -6.2409773 | 4.35E-10 | 7.34E-09 | 333.203074 | 556.105151 | 399.207536 | 473.466977 |
| -6.2403954 | 4.36E-10 | 7.36E-09 | 354.514956 | 487.370013 | 462.482506 | 414.633802 |
| -6.2399004 | 4.38E-10 | 7.38E-09 | 326.235728 | 975.010637 | 494.695218 | 697.03304 |
| -6.2373196 | 4.45E-10 | 7.50E-09 | 275.415087 | 929.006805 | 662.661501 | 769.313797 |
| -6.2373988 | 4.45E-10 | 7.50E-09 | 281.152902 | 733.084601 | 590.1829 | 718.325046 |
| -6.2371235 | 4.46E-10 | 7.50E-09 | 261.890239 | 762.851786 | 960.629086 | 866.808773 |
| -6.2358901 | 4.49E-10 | 7.56E-09 | 331.973543 | 522.008193 | 387.702996 | 475.147924 |
| -6.2352473 | 4.51E-10 | 7.58E-09 | 398.778095 | 411.869606 | 331.330751 | 350.197468 |
| -6.2352668 | 4.51E-10 | 7.58E-09 | 243.037421 | 1018.84958 | 1128.59537 | 1015.85281 |
| -6.2350574 | 4.52E-10 | 7.59E-09 | 263.939459 | 821.033104 | 740.892373 | 811.89781 |
| -6.2338469 | 4.55E-10 | 7.64E-09 | 638.536765 | 2122.12973 | 1946.56816 | 2009.85331 |
| -6.2330347 | 4.57E-10 | 7.68E-09 | 302.464783 | 478.710468 | 493.544764 | 551.350893 |
| -6.2323597 | 4.59E-10 | 7.71E-09 | 331.563699 | 415.928768 | 394.60572 | 480.190768 |
| -6.2318009 | 4.61E-10 | 7.72E-09 | 359.433083 | 536.621175 | 350.888469 | 543.50647 |
| -6.2318096 | 4.61E-10 | 7.72E-09 | 283.202121 | 630.523115 | 568.324274 | 630.355442 |
| -6.2319556 | 4.61E-10 | 7.72E-09 | 309.022285 | 569.906301 | 446.37615 | 502.603406 |
| -6.2311047 | 4.63E-10 | 7.75E-09 | 300.415564 | 469.239091 | 602.837894 | 590.57301 |
| -6.2307267 | 4.64E-10 | 7.75E-09 | 251.234298 | 911.687715 | 895.053208 | 920.038788 |
| -6.2307806 | 4.64E-10 | 7.75E-09 | 284.021809 | 800.737295 | 607.43971 | 619.709439 |
| -6.2308815 | 4.64E-10 | 7.75E-09 | 307.792754 | 714.141846 | 607.43971 | 659.491872 |
| -6.2309693 | 4.64E-10 | 7.75E-09 | 265.16899 | 779.629655 | 755.848275 | 728.410733 |
| -6.2304635 | 4.65E-10 | 7.76E-09 | 267.628054 | 684.915882 | 798.415073 | 758.667794 |

| | | | | | | |
|---|---|---|---|---|---|---|
| -6.2303121 | 4.66E-10 | 7.76E-09 | 331.973543 | 579.377678 | 388.85345 | 522.77478 |
| -6.230411 | 4.65E-10 | 7.76E-09 | 250.414611 | 1014.2492 | 1048.06359 | 848.878662 |
| -6.2303712 | 4.65E-10 | 7.76E-09 | 296.726969 | 522.008193 | 495.845672 | 529.498571 |
| -6.2299504 | 4.67E-10 | 7.77E-09 | 313.940412 | 510.371929 | 440.62388 | 440.968652 |
| -6.2299281 | 4.67E-10 | 7.77E-09 | 281.152902 | 631.876169 | 586.731538 | 616.347543 |
| -6.2297545 | 4.67E-10 | 7.77E-09 | 295.087594 | 541.76278 | 495.845672 | 551.911209 |
| -6.2292668 | 4.69E-10 | 7.79E-09 | 311.481349 | 651.630756 | 584.43063 | 441.528968 |
| -6.229277 | 4.69E-10 | 7.79E-09 | 690.996782 | 395.091738 | 354.339831 | 488.035191 |
| -6.2293999 | 4.68E-10 | 7.79E-09 | 340.990108 | 382.373031 | 511.952028 | 607.382488 |
| -6.2290831 | 4.69E-10 | 7.79E-09 | 364.761053 | 396.715402 | 358.941647 | 371.489474 |
| -6.228698 | 4.70E-10 | 7.81E-09 | 324.596353 | 432.436025 | 395.756174 | 442.649599 |
| -6.2283243 | 4.71E-10 | 7.82E-09 | 286.480872 | 1190.95804 | 646.555145 | 847.197714 |
| -6.2277691 | 4.73E-10 | 7.84E-09 | 293.038374 | 575.859738 | 541.863832 | 503.724038 |
| 6.22771037 | 4.73E-10 | 7.84E-09 | 85.247527 | 63.0523115 | 72.4786017 | 63.8760181 |
| -6.2274833 | 4.74E-10 | 7.85E-09 | 286.071028 | 660.01969 | 533.810654 | 671.818822 |
| -6.2274466 | 4.74E-10 | 7.85E-09 | 290.169467 | 604.815091 | 520.005206 | 556.954053 |
| -6.2269785 | 4.76E-10 | 7.87E-09 | 370.498867 | 347.19363 | 393.455266 | 386.057689 |
| -6.2260991 | 4.78E-10 | 7.91E-09 | 325.825885 | 396.986013 | 486.64204 | 561.996896 |
| -6.2252381 | 4.81E-10 | 7.95E-09 | 324.596353 | 477.357414 | 416.464346 | 402.306851 |
| -6.225154 | 4.81E-10 | 7.95E-09 | 391.400905 | 356.394396 | 393.455266 | 557.514369 |
| -6.2245566 | 4.83E-10 | 7.98E-09 | 268.447741 | 727.131164 | 674.166041 | 702.075884 |
| -6.2240425 | 4.85E-10 | 8.00E-09 | 259.431176 | 908.710996 | 730.538287 | 878.575407 |
| -6.2233704 | 4.87E-10 | 8.03E-09 | 263.119771 | 854.859451 | 715.582385 | 758.667794 |
| -6.2230433 | 4.88E-10 | 8.04E-09 | 304.923847 | 870.554877 | 625.846973 | 693.110828 |
| -6.222401 | 4.90E-10 | 8.07E-09 | 267.628054 | 703.588026 | 683.369673 | 703.756831 |
| -6.2222681 | 4.90E-10 | 8.07E-09 | 291.808843 | 663.808241 | 532.6602 | 530.619203 |
| -6.2221634 | 4.90E-10 | 8.07E-09 | 337.711357 | 546.904385 | 892.7523 | 471.225713 |
| -6.2210223 | 4.94E-10 | 8.13E-09 | 289.759623 | 650.818924 | 501.597942 | 619.709439 |
| -6.2206959 | 4.95E-10 | 8.14E-09 | 308.202598 | 451.37878 | 465.933868 | 435.925808 |
| -6.2205718 | 4.95E-10 | 8.14E-09 | 478.697652 | 482.76963 | 297.967585 | 480.190768 |
| -6.2198091 | 4.98E-10 | 8.18E-09 | 288.530091 | 691.410541 | 516.553844 | 602.899961 |
| -6.2196451 | 4.98E-10 | 8.18E-09 | 324.186509 | 517.678421 | 391.154358 | 449.933707 |
| -6.2190737 | 5.00E-10 | 8.21E-09 | 603.700035 | 527.420409 | 324.428027 | 395.022744 |
| -6.217544 | 5.05E-10 | 8.28E-09 | 283.611965 | 661.643355 | 526.90793 | 627.553862 |
| -6.2172732 | 5.06E-10 | 8.30E-09 | 438.123108 | 401.045175 | 325.578481 | 387.17832 |
| -6.2167516 | 5.08E-10 | 8.32E-09 | 238.529138 | 986.646901 | 993.992252 | 1149.76833 |
| -6.216604 | 5.08E-10 | 8.32E-09 | 260.660708 | 783.147595 | 728.237379 | 737.375788 |
| -6.2163721 | 5.09E-10 | 8.33E-09 | 267.21821 | 642.159379 | 731.688741 | 722.807574 |
| -6.2161791 | 5.09E-10 | 8.34E-09 | 289.759623 | 585.331115 | 499.297034 | 533.981099 |
| -6.215673 | 5.11E-10 | 8.35E-09 | 324.596353 | 503.877271 | 384.251634 | 444.330547 |
| -6.2157442 | 5.11E-10 | 8.35E-09 | 303.694315 | 528.232241 | 430.269794 | 530.619203 |
| -6.215287 | 5.12E-10 | 8.37E-09 | 298.776188 | 484.393295 | 480.88977 | 597.857117 |
| -6.2151793 | 5.13E-10 | 8.37E-09 | 270.087117 | 650.277702 | 632.749697 | 702.6362 |

| | | | | | | |
|---|---|---|---|---|---|---|
| -6.2145297 | 5.15E-10 | 8.40E-09 | 335.662138 | 474.380696 | 370.446187 | 406.229063 |
| -6.2143344 | 5.15E-10 | 8.41E-09 | 288.939935 | 664.890684 | 495.845672 | 615.787228 |
| -6.2142853 | 5.16E-10 | 8.41E-09 | 366.810273 | 507.1246 | 540.713378 | 333.948305 |
| -6.2135464 | 5.18E-10 | 8.44E-09 | 279.103682 | 657.042972 | 693.723759 | 544.627102 |
| -6.2133353 | 5.19E-10 | 8.45E-09 | 293.858062 | 523.631858 | 482.040224 | 490.276455 |
| -6.213321 | 5.19E-10 | 8.45E-09 | 320.08807 | 674.903283 | 1030.80678 | 847.197714 |
| -6.2127522 | 5.21E-10 | 8.47E-09 | 303.694315 | 479.25169 | 456.730236 | 437.04644 |
| -6.2126545 | 5.21E-10 | 8.47E-09 | 295.497437 | 709.270852 | 511.952028 | 530.058887 |
| -6.2104239 | 5.28E-10 | 8.59E-09 | 322.547134 | 420.799762 | 402.658898 | 388.298952 |
| -6.209645 | 5.31E-10 | 8.63E-09 | 286.480872 | 600.485319 | 496.996126 | 558.635001 |
| -6.2095257 | 5.31E-10 | 8.63E-09 | 278.283994 | 885.167859 | 582.129722 | 705.998095 |
| -6.2078751 | 5.37E-10 | 8.72E-09 | 288.939935 | 597.23799 | 484.341132 | 540.144574 |
| -6.2072104 | 5.39E-10 | 8.74E-09 | 1147.97271 | 3546.35426 | 3325.9625 | 3702.00747 |
| -6.2071865 | 5.39E-10 | 8.74E-09 | 229.102729 | 647.571594 | 487.792494 | 531.179519 |
| -6.2072457 | 5.39E-10 | 8.74E-09 | 281.152902 | 588.578445 | 521.15566 | 578.806375 |
| 6.20736789 | 5.39E-10 | 8.74E-09 | 34.8367298 | 24.0843593 | 28.7613499 | 29.1364293 |
| -6.2069223 | 5.40E-10 | 8.75E-09 | 260.250864 | 709.812074 | 696.024667 | 755.305899 |
| -6.2066074 | 5.41E-10 | 8.76E-09 | 278.283994 | 662.455187 | 544.16474 | 590.57301 |
| -6.2065371 | 5.42E-10 | 8.76E-09 | 263.119771 | 676.797559 | 933.01819 | 750.263055 |
| -6.2066765 | 5.41E-10 | 8.76E-09 | 295.907281 | 448.943283 | 587.881992 | 540.70489 |
| -6.2063312 | 5.42E-10 | 8.76E-09 | 297.546657 | 465.721151 | 453.278874 | 518.292252 |
| -6.2062425 | 5.43E-10 | 8.76E-09 | 264.759146 | 754.192241 | 640.802875 | 688.067985 |
| 6.20606469 | 5.43E-10 | 8.77E-09 | 58.6076748 | 39.7797845 | 41.4163438 | 42.5840121 |
| -6.2055913 | 5.45E-10 | 8.79E-09 | 291.808843 | 528.502852 | 820.273699 | 593.37459 |
| -6.2056432 | 5.45E-10 | 8.79E-09 | 302.05494 | 447.049007 | 483.190678 | 434.24486 |
| -6.2052498 | 5.46E-10 | 8.80E-09 | 360.662614 | 538.24484 | 334.782113 | 471.225713 |
| -6.2052064 | 5.46E-10 | 8.80E-09 | 294.267906 | 573.694852 | 452.12842 | 532.300151 |
| -6.2045352 | 5.49E-10 | 8.83E-09 | 314.350256 | 471.133366 | 387.702996 | 500.362142 |
| -6.2046037 | 5.48E-10 | 8.83E-09 | 283.202121 | 560.976145 | 506.199758 | 553.031841 |
| -6.2023335 | 5.56E-10 | 8.95E-09 | 293.448218 | 501.171163 | 465.933868 | 470.105081 |
| -6.2021427 | 5.57E-10 | 8.96E-09 | 318.038851 | 376.419594 | 487.792494 | 484.11298 |
| -6.2020036 | 5.57E-10 | 8.96E-09 | 279.103682 | 544.198277 | 538.41247 | 577.685743 |
| -6.2015542 | 5.59E-10 | 8.98E-09 | 174.593493 | 289.282923 | 274.958505 | 276.235763 |
| -6.2005395 | 5.63E-10 | 9.03E-09 | 276.234775 | 606.438756 | 586.731538 | 537.342995 |
| -6.2003274 | 5.63E-10 | 9.04E-09 | 279.92337 | 843.493799 | 571.775636 | 624.752283 |
| -6.1998882 | 5.65E-10 | 9.06E-09 | 351.646049 | 336.639809 | 482.040224 | 521.093832 |
| -6.1988972 | 5.69E-10 | 9.11E-09 | 240.578358 | 843.493799 | 1066.47085 | 964.303748 |
| -6.1989253 | 5.69E-10 | 9.11E-09 | 276.644619 | 557.999427 | 586.731538 | 541.825522 |
| -6.198283 | 5.71E-10 | 9.14E-09 | 240.988201 | 907.899164 | 857.088227 | 959.260904 |
| -6.1982224 | 5.71E-10 | 9.14E-09 | 262.709927 | 721.177726 | 628.147881 | 691.990197 |
| -6.1976999 | 5.73E-10 | 9.17E-09 | 349.186986 | 335.286755 | 357.791193 | 330.026094 |
| -6.1975008 | 5.74E-10 | 9.17E-09 | 261.890239 | 641.888768 | 692.573305 | 683.585457 |
| -6.1975298 | 5.74E-10 | 9.17E-09 | 275.415087 | 856.212505 | 566.023366 | 694.23146 |

| | | | | | | |
|---|---|---|---|---|---|---|
| -6.1972107 | 5.75E-10 | 9.18E-09 | 268.037897 | 665.431906 | 587.881992 | 631.476074 |
| -6.1965729 | 5.77E-10 | 9.21E-09 | 201.233345 | 801.278517 | 793.813257 | 769.313797 |
| -6.1963206 | 5.78E-10 | 9.22E-09 | 295.497437 | 681.668553 | 633.900151 | 665.655347 |
| -6.1961221 | 5.79E-10 | 9.23E-09 | 420.499821 | 620.510517 | 302.569401 | 539.023943 |
| -6.1959271 | 5.79E-10 | 9.24E-09 | 619.683946 | 537.974229 | 315.224395 | 393.341796 |
| -6.1957891 | 5.80E-10 | 9.24E-09 | 268.857585 | 699.258253 | 585.581084 | 616.907859 |
| -6.1954787 | 5.81E-10 | 9.26E-09 | 308.202598 | 518.760864 | 410.712076 | 432.003596 |
| -6.1951036 | 5.82E-10 | 9.27E-09 | 255.742581 | 706.564744 | 709.830115 | 742.418632 |
| -6.194842 | 5.83E-10 | 9.27E-09 | 327.055416 | 446.778396 | 358.941647 | 407.349695 |
| -6.1949562 | 5.83E-10 | 9.27E-09 | 306.153378 | 476.004361 | 666.112863 | 453.855918 |
| -6.1948818 | 5.83E-10 | 9.27E-09 | 299.595876 | 499.818109 | 422.216616 | 454.97655 |
| -6.194455 | 5.85E-10 | 9.29E-09 | 311.071505 | 390.761965 | 423.36707 | 470.105081 |
| -6.1940447 | 5.86E-10 | 9.31E-09 | 305.743534 | 500.359331 | 419.915708 | 428.641701 |
| -6.1939225 | 5.87E-10 | 9.31E-09 | 461.484209 | 777.735379 | 701.776937 | 725.048838 |
| -6.1939948 | 5.87E-10 | 9.31E-09 | 264.349303 | 646.218541 | 610.891072 | 673.49977 |
| -6.1937899 | 5.87E-10 | 9.31E-09 | 281.152902 | 969.598422 | 559.120642 | 741.298 |
| -6.1933234 | 5.89E-10 | 9.34E-09 | 464.76296 | 466.532983 | 288.763953 | 504.284354 |
| -6.1922175 | 5.93E-10 | 9.40E-09 | 288.939935 | 522.008193 | 464.783414 | 612.985648 |
| -6.1917634 | 5.95E-10 | 9.42E-09 | 361.072458 | 647.300984 | 603.988348 | 636.518918 |
| -6.1911785 | 5.97E-10 | 9.45E-09 | 345.498391 | 335.557366 | 409.561622 | 453.855918 |
| -6.1911256 | 5.97E-10 | 9.45E-09 | 281.972589 | 538.786061 | 510.801574 | 493.078035 |
| -6.190326 | 6.00E-10 | 9.49E-09 | 256.972113 | 930.089248 | 739.741919 | 684.706089 |
| -6.1903759 | 6.00E-10 | 9.49E-09 | 250.824455 | 910.605272 | 702.927391 | 833.189816 |
| -6.1901627 | 6.01E-10 | 9.49E-09 | 218.446788 | 1242.6447 | 1089.47993 | 1294.32984 |
| -6.1897577 | 6.03E-10 | 9.51E-09 | 229.102729 | 1003.69538 | 1056.11677 | 996.241757 |
| -6.1893056 | 6.04E-10 | 9.54E-09 | 274.185556 | 616.451355 | 551.067464 | 536.782679 |
| -6.1888121 | 6.06E-10 | 9.56E-09 | 303.284471 | 416.7406 | 429.11934 | 415.194118 |
| -6.1884496 | 6.08E-10 | 9.58E-09 | 252.053986 | 843.493799 | 696.024667 | 707.118727 |
| -6.188425 | 6.08E-10 | 9.58E-09 | 295.497437 | 561.787978 | 453.278874 | 450.494023 |
| -6.1882316 | 6.08E-10 | 9.58E-09 | 236.479919 | 939.560625 | 956.02727 | 865.127825 |
| -6.1874166 | 6.12E-10 | 9.63E-09 | 346.727922 | 512.807426 | 400.35799 | 348.51652 |
| -6.1866826 | 6.14E-10 | 9.67E-09 | 223.774758 | 1583.34367 | 1616.38786 | 1366.6106 |
| -6.1864128 | 6.15E-10 | 9.68E-09 | 268.447741 | 610.768529 | 556.819734 | 604.020593 |
| -6.1857647 | 6.18E-10 | 9.71E-09 | 274.5954 | 652.442589 | 509.65112 | 612.985648 |
| -6.1855045 | 6.19E-10 | 9.73E-09 | 299.595876 | 715.224289 | 448.677058 | 543.50647 |
| -6.1850885 | 6.21E-10 | 9.75E-09 | 276.644619 | 768.805224 | 525.757476 | 626.433231 |
| -6.1847745 | 6.22E-10 | 9.76E-09 | 238.938982 | 926.030086 | 860.539589 | 837.112027 |
| -6.1847061 | 6.22E-10 | 9.76E-09 | 272.54618 | 557.458205 | 579.828814 | 530.058887 |
| -6.1846255 | 6.23E-10 | 9.76E-09 | 285.25134 | 476.816193 | 470.535684 | 497.560562 |
| -6.1845391 | 6.23E-10 | 9.76E-09 | 283.611965 | 581.001343 | 464.783414 | 527.257308 |
| -6.1829522 | 6.29E-10 | 9.86E-09 | 289.349779 | 537.703618 | 528.058384 | 432.003596 |
| -6.1824245 | 6.31E-10 | 9.88E-09 | 244.676796 | 799.384242 | 767.352815 | 802.372438 |
| -6.1823007 | 6.32E-10 | 9.89E-09 | 266.398522 | 591.013942 | 571.775636 | 601.779329 |

| | | | | | | |
|---|---|---|---|---|---|---|
| -6.1819203 | 6.33E-10 | 9.91E-09 | 282.382433 | 528.232241 | 556.819734 | 461.140026 |
| -6.1811528 | 6.36E-10 | 9.94E-09 | 274.5954 | 684.645271 | 507.350212 | 609.623752 |
| -6.1811223 | 6.36E-10 | 9.94E-09 | 297.956501 | 488.181846 | 400.35799 | 496.43993 |
| -6.1811536 | 6.36E-10 | 9.94E-09 | 271.726492 | 624.028457 | 516.553844 | 590.57301 |
| -6.1809072 | 6.37E-10 | 9.95E-09 | 258.611488 | 672.467786 | 612.041526 | 694.791776 |
| -6.1807872 | 6.38E-10 | 9.95E-09 | 309.841973 | 375.337151 | 484.341132 | 437.04644 |
| -6.1804839 | 6.39E-10 | 9.97E-09 | 374.597306 | 682.209774 | 342.835291 | 470.665397 |
| -6.1801363 | 6.40E-10 | 9.97E-09 | 392.220593 | 641.888768 | 675.316495 | 670.69819 |
| -6.1803062 | 6.40E-10 | 9.97E-09 | 411.483255 | 752.027355 | 314.073941 | 578.806375 |
| -6.1801293 | 6.40E-10 | 9.97E-09 | 413.942319 | 750.13308 | 549.91701 | 607.382488 |
| -6.1801109 | 6.41E-10 | 9.97E-09 | 433.614825 | 477.628025 | 304.870309 | 383.256109 |
| -6.1789349 | 6.45E-10 | 1.00E-08 | 288.120248 | 453.543666 | 549.91701 | 659.491872 |
| -6.1783018 | 6.48E-10 | 1.01E-08 | 493.042188 | 1107.33931 | 307.171217 | 1157.61275 |
| -6.1783127 | 6.48E-10 | 1.01E-08 | 285.661184 | 500.629941 | 442.924788 | 494.198667 |
| -6.1777571 | 6.50E-10 | 1.01E-08 | 295.907281 | 471.403977 | 407.260714 | 446.011495 |
| -6.1776053 | 6.51E-10 | 1.01E-08 | 257.381957 | 797.489966 | 618.94425 | 691.990197 |
| -6.1772584 | 6.52E-10 | 1.01E-08 | 382.38434 | 310.931785 | 480.88977 | 403.427483 |
| -6.1773291 | 6.52E-10 | 1.01E-08 | 269.267429 | 639.994493 | 545.315194 | 553.592157 |
| -6.1771046 | 6.53E-10 | 1.01E-08 | 300.825408 | 980.964074 | 483.190678 | 707.118727 |
| -6.1762465 | 6.56E-10 | 1.02E-08 | 322.13729 | 394.821127 | 349.738015 | 413.51317 |
| -6.1754327 | 6.60E-10 | 1.02E-08 | 248.775235 | 808.314397 | 702.927391 | 712.161571 |
| -6.1742678 | 6.65E-10 | 1.03E-08 | 279.513526 | 601.838373 | 528.058384 | 832.6295 |
| -6.1735798 | 6.68E-10 | 1.03E-08 | 268.447741 | 643.241822 | 546.465648 | 547.428682 |
| -6.1733261 | 6.69E-10 | 1.03E-08 | 283.202121 | 454.084887 | 471.686138 | 512.128777 |
| -6.1727169 | 6.71E-10 | 1.04E-08 | 966.002025 | 504.418492 | 356.640739 | 380.454529 |
| -6.1723985 | 6.73E-10 | 1.04E-08 | 247.545703 | 841.328912 | 671.865133 | 800.131175 |
| -6.1717503 | 6.75E-10 | 1.04E-08 | 1218.05601 | 355.041342 | 433.721156 | 411.271906 |
| -6.1718055 | 6.75E-10 | 1.04E-08 | 424.188416 | 544.198277 | 387.702996 | 310.975351 |
| -6.1717848 | 6.75E-10 | 1.04E-08 | 313.940412 | 394.279905 | 375.048002 | 373.730738 |
| -6.1718295 | 6.75E-10 | 1.04E-08 | 271.726492 | 653.795642 | 505.049304 | 574.323847 |
| -6.1712264 | 6.78E-10 | 1.04E-08 | 290.989155 | 512.266205 | 452.12842 | 420.236961 |
| -6.1708133 | 6.79E-10 | 1.05E-08 | 347.54761 | 398.339067 | 330.180297 | 359.162523 |
| -6.17079 | 6.79E-10 | 1.05E-08 | 271.726492 | 587.225391 | 661.511047 | 500.362142 |
| -6.1704114 | 6.81E-10 | 1.05E-08 | 251.644142 | 788.2892 | 661.511047 | 674.620402 |
| -6.1697637 | 6.84E-10 | 1.05E-08 | 284.021809 | 565.305918 | 442.924788 | 494.758983 |
| -6.1697416 | 6.84E-10 | 1.05E-08 | 952.067333 | 290.635977 | 637.351513 | 513.809725 |
| 6.16952319 | 6.85E-10 | 1.05E-08 | 2.86890716 | 2.43549701 | 1.150454 | 2.80157974 |
| -6.1692296 | 6.86E-10 | 1.05E-08 | 322.13729 | 400.503953 | 343.985745 | 393.341796 |
| -6.1692651 | 6.86E-10 | 1.05E-08 | 304.923847 | 459.767714 | 391.154358 | 391.660848 |
| -6.1693053 | 6.86E-10 | 1.05E-08 | 281.562746 | 480.334133 | 449.827512 | 504.84467 |
| -6.1691511 | 6.87E-10 | 1.05E-08 | 227.053509 | 1151.17825 | 962.929994 | 1065.16062 |
| -6.1690641 | 6.87E-10 | 1.05E-08 | 339.350733 | 444.072289 | 471.686138 | 324.422934 |
| -6.1689529 | 6.87E-10 | 1.05E-08 | 297.546657 | 546.904385 | 398.057082 | 473.466977 |

| | | | | | | |
|---|---|---|---|---|---|---|
| -6.1688569 | 6.88E-10 | 1.05E-08 | 381.154808 | 481.957798 | 311.773033 | 386.057689 |
| -6.1681471 | 6.91E-10 | 1.06E-08 | 243.447265 | 914.123212 | 706.378753 | 808.535914 |
| -6.1679217 | 6.92E-10 | 1.06E-08 | 266.398522 | 520.655139 | 606.289256 | 579.927007 |
| -6.1677787 | 6.93E-10 | 1.06E-08 | 260.660708 | 572.071187 | 599.386532 | 609.623752 |
| -6.1675768 | 6.93E-10 | 1.06E-08 | 283.611965 | 455.979163 | 445.225696 | 486.914559 |
| -6.1671347 | 6.95E-10 | 1.06E-08 | 335.252294 | 437.848241 | 333.631659 | 380.454529 |
| -6.1671417 | 6.95E-10 | 1.06E-08 | 275.824931 | 538.51545 | 465.933868 | 526.696992 |
| -6.1668431 | 6.97E-10 | 1.06E-08 | 300.00572 | 1265.376 | 838.680963 | 1030.98135 |
| -6.1668727 | 6.97E-10 | 1.06E-08 | 403.286378 | 601.02654 | 241.595339 | 513.809725 |
| -6.1666997 | 6.97E-10 | 1.06E-08 | 361.482302 | 348.817294 | 342.835291 | 341.792729 |
| -6.165745 | 7.02E-10 | 1.07E-08 | 314.350256 | 450.025726 | 348.587561 | 442.649599 |
| -6.165574 | 7.02E-10 | 1.07E-08 | 289.349779 | 497.923834 | 416.464346 | 446.571811 |
| -6.1654971 | 7.03E-10 | 1.07E-08 | 293.858062 | 458.144049 | 400.35799 | 428.081385 |
| -6.16389 | 7.10E-10 | 1.08E-08 | 215.987725 | 1143.87176 | 1045.76268 | 1079.16852 |
| -6.1635859 | 7.11E-10 | 1.08E-08 | 213.938505 | 407.539833 | 254.250333 | 314.337247 |
| -6.1632973 | 7.12E-10 | 1.08E-08 | 302.05494 | 473.298253 | 415.313892 | 379.333897 |
| -6.1631229 | 7.13E-10 | 1.08E-08 | 345.908235 | 782.606373 | 368.145279 | 505.965302 |
| -6.1618567 | 7.19E-10 | 1.09E-08 | 229.922417 | 992.329727 | 908.858656 | 830.948552 |
| -6.1608839 | 7.23E-10 | 1.10E-08 | 302.464783 | 418.093654 | 384.251634 | 384.376741 |
| -6.160899 | 7.23E-10 | 1.10E-08 | 347.957454 | 336.91042 | 361.242555 | 471.225713 |
| -6.1608547 | 7.24E-10 | 1.10E-08 | 295.087594 | 415.928768 | 423.36707 | 393.902112 |
| -6.1605374 | 7.25E-10 | 1.10E-08 | 292.218686 | 396.174181 | 432.570702 | 455.536866 |
| -6.1603442 | 7.26E-10 | 1.10E-08 | 269.677273 | 547.716217 | 486.64204 | 540.70489 |
| -6.1601256 | 7.27E-10 | 1.10E-08 | 381.564652 | 351.794013 | 368.145279 | 601.779329 |
| -6.1588763 | 7.33E-10 | 1.11E-08 | 261.890239 | 585.872337 | 546.465648 | 555.833421 |
| -6.1588161 | 7.33E-10 | 1.11E-08 | 385.253247 | 394.279905 | 323.277573 | 378.773581 |
| -6.1583428 | 7.35E-10 | 1.11E-08 | 261.070552 | 715.4949 | 532.6602 | 665.655347 |
| -6.158306 | 7.35E-10 | 1.11E-08 | 249.185079 | 761.769343 | 632.749697 | 676.30135 |
| -6.1579011 | 7.37E-10 | 1.11E-08 | 309.432129 | 469.780313 | 355.490285 | 419.11633 |
| -6.157551 | 7.39E-10 | 1.12E-08 | 191.806936 | 891.121296 | 551.067464 | 616.347543 |
| -6.157195 | 7.40E-10 | 1.12E-08 | 297.136813 | 1673.18645 | 1071.07267 | 1135.76043 |
| -6.1571906 | 7.40E-10 | 1.12E-08 | 487.304373 | 376.419594 | 468.234776 | 296.407137 |
| -6.1573094 | 7.40E-10 | 1.12E-08 | 239.348826 | 832.128146 | 751.246459 | 729.531365 |
| -6.155899 | 7.47E-10 | 1.13E-08 | 284.431653 | 524.714301 | 461.332052 | 420.236961 |
| -6.1558885 | 7.47E-10 | 1.13E-08 | 347.957454 | 417.552432 | 319.826211 | 511.568461 |
| -6.1547771 | 7.52E-10 | 1.13E-08 | 261.890239 | 582.625007 | 531.509746 | 551.911209 |
| -6.1545625 | 7.53E-10 | 1.13E-08 | 311.891193 | 674.091451 | 430.269794 | 426.400437 |
| -6.1542942 | 7.54E-10 | 1.14E-08 | 251.234298 | 854.047619 | 633.900151 | 666.215663 |
| -6.1542242 | 7.54E-10 | 1.14E-08 | 305.743534 | 439.471905 | 366.994825 | 383.816425 |
| -6.1540755 | 7.55E-10 | 1.14E-08 | 250.004767 | 731.460936 | 739.741919 | 595.055537 |
| -6.1532752 | 7.59E-10 | 1.14E-08 | 276.644619 | 521.466971 | 437.172518 | 502.04309 |
| -6.1528857 | 7.61E-10 | 1.14E-08 | 218.036944 | 420.799762 | 356.640739 | 377.092633 |
| -6.1526058 | 7.62E-10 | 1.15E-08 | 279.513526 | 430.271139 | 510.801574 | 466.182869 |

| | | | | | | |
|---|---|---|---|---|---|---|
| -6.1524809 | 7.63E-10 | 1.15E-08 | 331.563699 | 494.135283 | 609.740618 | 341.232413 |
| -6.1518878 | 7.66E-10 | 1.15E-08 | 259.84102 | 591.555163 | 525.757476 | 629.795126 |
| 6.15046527 | 7.73E-10 | 1.16E-08 | 0 | 0 | 0 | 0 |
| -6.1498771 | 7.75E-10 | 1.16E-08 | 909.44357 | 2065.57208 | 1837.27503 | 1941.49476 |
| -6.1491351 | 7.79E-10 | 1.17E-08 | 279.92337 | 433.789079 | 441.774334 | 453.855918 |
| -6.1490675 | 7.79E-10 | 1.17E-08 | 273.365868 | 477.357414 | 456.730236 | 533.981099 |
| -6.1485963 | 7.82E-10 | 1.17E-08 | 246.726016 | 743.909032 | 743.193281 | 606.261856 |
| -6.1484752 | 7.82E-10 | 1.17E-08 | 291.398999 | 401.045175 | 403.809352 | 403.427483 |
| 6.14806949 | 7.84E-10 | 1.17E-08 | 0 | 0 | 0 | 0 |
| -6.1474618 | 7.87E-10 | 1.18E-08 | 284.021809 | 416.469989 | 424.517524 | 434.805176 |
| -6.147255 | 7.88E-10 | 1.18E-08 | 277.464307 | 492.782229 | 423.36707 | 495.879615 |
| -6.1473181 | 7.88E-10 | 1.18E-08 | 208.200691 | 1246.97447 | 1142.40082 | 1049.47177 |
| -6.1472851 | 7.88E-10 | 1.18E-08 | 276.234775 | 766.640337 | 777.706901 | 467.303501 |
| -6.1471366 | 7.89E-10 | 1.18E-08 | 252.873674 | 717.930397 | 572.92609 | 624.752283 |
| -6.1470378 | 7.89E-10 | 1.18E-08 | 254.103206 | 592.366996 | 586.731538 | 577.685743 |
| -6.146801 | 7.91E-10 | 1.18E-08 | 397.13872 | 365.053941 | 308.321671 | 354.679995 |
| -6.1455887 | 7.97E-10 | 1.19E-08 | 317.219163 | 552.857822 | 342.835291 | 553.031841 |
| -6.1453165 | 7.98E-10 | 1.19E-08 | 264.349303 | 510.371929 | 506.199758 | 518.292252 |
| -6.145217 | 7.99E-10 | 1.19E-08 | 247.13586 | 721.177726 | 600.536986 | 679.663246 |
| -6.1451231 | 7.99E-10 | 1.19E-08 | 312.301037 | 645.406708 | 700.626483 | 372.04979 |
| -6.1448798 | 8.00E-10 | 1.19E-08 | 253.283518 | 603.732648 | 591.333354 | 762.590006 |
| -6.1446604 | 8.01E-10 | 1.19E-08 | 306.563222 | 382.10242 | 349.738015 | 395.58306 |
| -6.1444269 | 8.03E-10 | 1.19E-08 | 284.021809 | 494.405893 | 406.11026 | 437.04644 |
| -6.1442508 | 8.03E-10 | 1.19E-08 | 234.840543 | 708.45902 | 996.29316 | 870.170668 |
| -6.1442212 | 8.04E-10 | 1.19E-08 | 295.497437 | 542.033391 | 391.154358 | 427.521069 |
| -6.1438604 | 8.05E-10 | 1.20E-08 | 911.902633 | 713.330014 | 513.102482 | 269.511971 |
| -6.1438637 | 8.05E-10 | 1.20E-08 | 328.284948 | 428.376863 | 320.976665 | 385.497373 |
| -6.1429199 | 8.10E-10 | 1.20E-08 | 273.365868 | 874.884649 | 513.102482 | 615.226912 |
| -6.1422237 | 8.14E-10 | 1.21E-08 | 251.644142 | 687.62199 | 560.271096 | 630.915758 |
| -6.1418508 | 8.16E-10 | 1.21E-08 | 270.906805 | 396.715402 | 501.597942 | 485.793927 |
| -6.1418781 | 8.16E-10 | 1.21E-08 | 289.349779 | 387.514636 | 439.473426 | 410.71159 |
| -6.141778 | 8.16E-10 | 1.21E-08 | 248.775235 | 773.676218 | 586.731538 | 671.818822 |
| -6.1412328 | 8.19E-10 | 1.21E-08 | 231.151948 | 936.583907 | 744.343735 | 820.302549 |
| -6.1409797 | 8.20E-10 | 1.21E-08 | 260.250864 | 548.257439 | 514.252936 | 528.37794 |
| -6.1407526 | 8.21E-10 | 1.21E-08 | 234.020855 | 746.61514 | 819.123245 | 729.531365 |
| -6.1407541 | 8.21E-10 | 1.21E-08 | 279.92337 | 463.826875 | 406.11026 | 463.38129 |
| -6.1400567 | 8.25E-10 | 1.22E-08 | 257.7918 | 545.28072 | 529.208838 | 553.592157 |
| -6.1397919 | 8.26E-10 | 1.22E-08 | 246.726016 | 632.958612 | 606.289256 | 647.725237 |
| -6.1398415 | 8.26E-10 | 1.22E-08 | 276.234775 | 469.509702 | 432.570702 | 439.84802 |
| -6.1393947 | 8.28E-10 | 1.22E-08 | 278.283994 | 752.027355 | 462.482506 | 559.755633 |
| -6.1376414 | 8.38E-10 | 1.23E-08 | 253.283518 | 815.620888 | 576.377452 | 628.674494 |
| -6.1375141 | 8.38E-10 | 1.24E-08 | 270.087117 | 458.144049 | 463.63296 | 477.949504 |
| -6.1371285 | 8.40E-10 | 1.24E-08 | 303.694315 | 394.550516 | 343.985745 | 406.229063 |

| | | | | | | |
|---|---|---|---|---|---|---|
| -6.1368382 | 8.42E-10 | 1.24E-08 | 234.430699 | 887.873966 | 684.520127 | 810.777178 |
| -6.1367112 | 8.42E-10 | 1.24E-08 | 293.448218 | 562.59981 | 425.667978 | 397.824324 |
| 6.13632648 | 8.45E-10 | 1.24E-08 | 58.197831 | 34.367569 | 35.6640739 | 31.3776931 |
| -6.1348262 | 8.53E-10 | 1.25E-08 | 256.562269 | 686.810157 | 528.058384 | 569.84132 |
| -6.1345515 | 8.54E-10 | 1.26E-08 | 272.136336 | 534.7269 | 431.420248 | 481.871716 |
| -6.1337282 | 8.58E-10 | 1.26E-08 | 372.548087 | 974.469416 | 984.78862 | 940.210162 |
| -6.1335804 | 8.59E-10 | 1.26E-08 | 295.907281 | 453.543666 | 377.34891 | 375.972002 |
| -6.1335757 | 8.59E-10 | 1.26E-08 | 230.332261 | 757.710182 | 766.202361 | 946.373637 |
| -6.1333296 | 8.61E-10 | 1.26E-08 | 242.627577 | 813.997224 | 614.342434 | 699.274304 |
| -6.1332265 | 8.61E-10 | 1.26E-08 | 289.759623 | 471.945199 | 370.446187 | 428.641701 |
| -6.1329673 | 8.63E-10 | 1.26E-08 | 244.676796 | 636.476553 | 748.945551 | 606.261856 |
| -6.1322062 | 8.67E-10 | 1.27E-08 | 269.267429 | 574.236073 | 436.022064 | 536.222363 |
| -6.1319236 | 8.68E-10 | 1.27E-08 | 383.204028 | 494.947115 | 281.861229 | 479.630452 |
| -6.1316662 | 8.70E-10 | 1.27E-08 | 267.628054 | 657.854804 | 455.579782 | 568.720688 |
| -6.1312375 | 8.72E-10 | 1.28E-08 | 274.5954 | 1103.55076 | 973.28408 | 1022.01629 |
| -6.1307634 | 8.75E-10 | 1.28E-08 | 234.020855 | 733.355211 | 684.520127 | 773.236009 |
| -6.1306414 | 8.75E-10 | 1.28E-08 | 319.268382 | 616.992576 | 340.534383 | 606.822172 |
| -6.1297044 | 8.80E-10 | 1.29E-08 | 195.495531 | 1285.94242 | 1158.50717 | 1303.2949 |
| -6.129292 | 8.83E-10 | 1.29E-08 | 249.185079 | 581.542564 | 1012.39952 | 937.408582 |
| -6.128547 | 8.87E-10 | 1.29E-08 | 271.726492 | 425.670756 | 460.181598 | 456.657498 |
| -6.1284639 | 8.87E-10 | 1.29E-08 | 320.08807 | 319.861941 | 369.295733 | 394.462428 |
| -6.1276302 | 8.92E-10 | 1.30E-08 | 303.694315 | 348.817294 | 399.207536 | 483.552664 |
| -6.1273187 | 8.94E-10 | 1.30E-08 | 277.874151 | 506.583379 | 393.455266 | 501.482774 |
| -6.1271946 | 8.94E-10 | 1.30E-08 | 318.448695 | 726.319331 | 578.67836 | 357.481575 |
| -6.1269266 | 8.96E-10 | 1.30E-08 | 313.940412 | 389.950133 | 326.728935 | 363.084735 |
| -6.1268246 | 8.97E-10 | 1.31E-08 | 249.594923 | 702.234972 | 553.368372 | 591.693642 |
| -6.1265241 | 8.98E-10 | 1.31E-08 | 273.365868 | 414.034492 | 447.526604 | 479.070136 |
| -6.1262299 | 9.00E-10 | 1.31E-08 | 289.759623 | 386.432193 | 376.198456 | 384.937057 |
| -6.1262225 | 9.00E-10 | 1.31E-08 | 278.283994 | 456.249774 | 401.508444 | 418.556014 |
| -6.1261318 | 9.00E-10 | 1.31E-08 | 240.168514 | 409.70472 | 239.294431 | 332.267358 |
| -6.1255284 | 9.04E-10 | 1.31E-08 | 293.038374 | 432.706636 | 360.092101 | 387.17832 |
| -6.1253908 | 9.05E-10 | 1.31E-08 | 240.578358 | 718.201008 | 597.085624 | 688.628301 |
| -6.1244642 | 9.10E-10 | 1.32E-08 | 265.578834 | 523.361247 | 436.022064 | 521.654148 |
| -6.1237217 | 9.14E-10 | 1.33E-08 | 409.434036 | 430.271139 | 281.861229 | 378.213265 |
| -6.1236271 | 9.15E-10 | 1.33E-08 | 233.201168 | 1277.8241 | 738.591465 | 890.342042 |
| 6.12350617 | 9.15E-10 | 1.33E-08 | 20.9020379 | 24.0843593 | 14.9559019 | 17.3697944 |
| -6.123369 | 9.16E-10 | 1.33E-08 | 644.274579 | 453.002444 | 280.710775 | 437.04644 |
| -6.1230468 | 9.18E-10 | 1.33E-08 | 247.545703 | 596.967379 | 545.315194 | 610.184068 |
| -6.1221365 | 9.23E-10 | 1.34E-08 | 264.349303 | 500.08872 | 445.225696 | 481.871716 |
| -6.1218724 | 9.25E-10 | 1.34E-08 | 265.988678 | 607.79181 | 452.12842 | 500.362142 |
| -6.120392 | 9.33E-10 | 1.35E-08 | 488.533905 | 629.440672 | 260.002603 | 781.640748 |
| -6.120253 | 9.34E-10 | 1.35E-08 | 263.529615 | 473.839474 | 485.491586 | 451.054339 |
| -6.12013 | 9.35E-10 | 1.35E-08 | 254.103206 | 533.103235 | 522.306114 | 508.766881 |

| | | | | | | |
|---|---|---|---|---|---|---|
| -6.1198333 | 9.37E-10 | 1.35E-08 | 293.858062 | 447.319618 | 350.888469 | 396.703692 |
| -6.1197372 | 9.37E-10 | 1.35E-08 | 382.38434 | 357.206228 | 302.569401 | 338.991149 |
| -6.11968 | 9.38E-10 | 1.35E-08 | 222.545227 | 432.706636 | 377.34891 | 392.221164 |
| -6.1185722 | 9.44E-10 | 1.36E-08 | 258.201644 | 567.200193 | 463.63296 | 600.658697 |
| -6.117734 | 9.49E-10 | 1.37E-08 | 341.809796 | 350.440959 | 304.870309 | 354.679995 |
| -6.1175725 | 9.50E-10 | 1.37E-08 | 281.972589 | 483.040241 | 378.499364 | 420.797277 |
| -6.1172379 | 9.52E-10 | 1.37E-08 | 220.905851 | 790.454086 | 816.822337 | 853.36119 |
| -6.1171261 | 9.53E-10 | 1.37E-08 | 435.664044 | 290.365366 | 350.888469 | 408.470327 |
| -6.1167385 | 9.55E-10 | 1.37E-08 | 242.627577 | 598.049822 | 575.226998 | 647.725237 |
| -6.1164192 | 9.57E-10 | 1.38E-08 | 236.889763 | 604.544481 | 686.821035 | 727.290101 |
| -6.1162908 | 9.58E-10 | 1.38E-08 | 266.398522 | 634.852888 | 524.607022 | 441.528968 |
| -6.1161093 | 9.59E-10 | 1.38E-08 | 258.201644 | 547.174995 | 476.287954 | 486.354243 |
| -6.1158631 | 9.60E-10 | 1.38E-08 | 218.856632 | 844.846852 | 929.566828 | 794.528015 |
| -6.1156521 | 9.62E-10 | 1.38E-08 | 262.300083 | 501.712385 | 454.429328 | 461.140026 |
| -6.1154473 | 9.63E-10 | 1.38E-08 | 259.021332 | 471.133366 | 488.942948 | 485.233612 |
| -6.1149641 | 9.66E-10 | 1.39E-08 | 271.726492 | 514.16048 | 400.35799 | 474.027293 |
| -6.1139296 | 9.72E-10 | 1.39E-08 | 202.053033 | 1171.47406 | 993.992252 | 1066.28125 |
| -6.1139931 | 9.72E-10 | 1.39E-08 | 252.873674 | 649.46587 | 493.544764 | 567.03974 |
| -6.1137217 | 9.73E-10 | 1.39E-08 | 233.611012 | 663.26702 | 652.307415 | 694.23146 |
| -6.1134669 | 9.75E-10 | 1.40E-08 | 235.250387 | 927.38314 | 621.245158 | 795.088331 |
| -6.1133113 | 9.76E-10 | 1.40E-08 | 234.430699 | 745.262086 | 614.342434 | 713.282203 |
| -6.1125316 | 9.81E-10 | 1.40E-08 | 308.612442 | 342.052025 | 331.330751 | 371.489474 |
| -6.1122823 | 9.82E-10 | 1.40E-08 | 284.021809 | 412.140217 | 487.792494 | 370.368842 |
| -6.1119186 | 9.84E-10 | 1.41E-08 | 557.387677 | 672.197175 | 240.444885 | 626.993547 |
| -6.1115181 | 9.87E-10 | 1.41E-08 | 272.956024 | 486.016959 | 433.721156 | 396.703692 |
| -6.111268 | 9.88E-10 | 1.41E-08 | 529.108449 | 525.796744 | 281.861229 | 367.567262 |
| -6.1111342 | 9.89E-10 | 1.41E-08 | 238.529138 | 1062.14731 | 662.661501 | 730.091681 |
| -6.1109749 | 9.90E-10 | 1.41E-08 | 238.938982 | 636.747163 | 579.828814 | 648.845868 |
| -6.1106167 | 9.92E-10 | 1.42E-08 | 256.562269 | 470.862756 | 562.572004 | 493.638351 |
| -6.1104864 | 9.93E-10 | 1.42E-08 | 279.103682 | 419.717318 | 370.446187 | 436.486124 |
| -6.1095366 | 9.99E-10 | 1.42E-08 | 322.956977 | 451.37878 | 441.774334 | 303.130928 |
| -6.1094016 | 1.00E-09 | 1.42E-08 | 235.250387 | 635.935331 | 670.714679 | 621.950703 |
| -6.1089116 | 1.00E-09 | 1.43E-08 | 255.332737 | 515.784145 | 506.199758 | 427.521069 |
| -6.1087536 | 1.00E-09 | 1.43E-08 | 250.824455 | 491.158564 | 663.811955 | 591.133326 |
| -6.1083394 | 1.01E-09 | 1.43E-08 | 292.62853 | 479.25169 | 362.393009 | 434.805176 |
| -6.1080632 | 1.01E-09 | 1.43E-08 | 270.087117 | 401.857007 | 465.933868 | 427.521069 |
| -6.1078193 | 1.01E-09 | 1.44E-08 | 236.070075 | 589.390277 | 820.273699 | 733.453577 |
| -6.1073826 | 1.01E-09 | 1.44E-08 | 236.479919 | 662.725798 | 621.245158 | 602.899961 |
| -6.1073075 | 1.01E-09 | 1.44E-08 | 293.448218 | 476.274971 | 352.038923 | 387.17832 |
| -6.106748 | 1.02E-09 | 1.44E-08 | 421.319509 | 387.785247 | 325.578481 | 356.360943 |
| -6.1067107 | 1.02E-09 | 1.44E-08 | 217.217256 | 900.051451 | 808.769159 | 796.769279 |
| -6.1065439 | 1.02E-09 | 1.44E-08 | 201.233345 | 1088.39655 | 1000.89498 | 1025.9385 |
| -6.106315 | 1.02E-09 | 1.44E-08 | 243.857109 | 598.861654 | 560.271096 | 541.265206 |

| | | | | | | |
|---|---|---|---|---|---|---|
| -6.1063842 | 1.02E-09 | 1.44E-08 | 223.364915 | 729.025439 | 738.591465 | 807.975598 |
| -6.1059406 | 1.02E-09 | 1.45E-08 | 236.070075 | 614.286469 | 641.953329 | 612.425332 |
| -6.1059821 | 1.02E-09 | 1.45E-08 | 283.202121 | 441.366181 | 355.490285 | 411.832222 |
| -6.105509 | 1.02E-09 | 1.45E-08 | 246.316172 | 560.164313 | 557.970188 | 516.611305 |
| -6.1056159 | 1.02E-09 | 1.45E-08 | 243.037421 | 578.024624 | 585.581084 | 541.265206 |
| -6.1055227 | 1.02E-09 | 1.45E-08 | 242.627577 | 664.078852 | 528.058384 | 674.620402 |
| -6.105395 | 1.03E-09 | 1.45E-08 | 249.185079 | 547.445606 | 510.801574 | 512.689093 |
| -6.1053216 | 1.03E-09 | 1.45E-08 | 242.627577 | 623.216624 | 537.262016 | 587.77143 |
| -6.1050564 | 1.03E-09 | 1.45E-08 | 238.119294 | 635.66472 | 570.625182 | 630.355442 |
| -6.1049112 | 1.03E-09 | 1.45E-08 | 182.79037 | 222.983282 | 181.771731 | 243.737438 |
| -6.1034051 | 1.04E-09 | 1.46E-08 | 249.594923 | 565.847139 | 484.341132 | 542.946154 |
| -6.1034409 | 1.04E-09 | 1.46E-08 | 279.513526 | 397.797845 | 373.897549 | 383.256109 |
| -6.1033235 | 1.04E-09 | 1.46E-08 | 570.502681 | 417.552432 | 302.569401 | 351.3181 |
| -6.1029915 | 1.04E-09 | 1.47E-08 | 240.988201 | 599.944097 | 545.315194 | 670.137875 |
| -6.1029911 | 1.04E-09 | 1.47E-08 | 266.398522 | 423.505869 | 425.667978 | 425.279805 |
| -6.1027029 | 1.04E-09 | 1.47E-08 | 381.154808 | 430.271139 | 285.312591 | 351.878416 |
| -6.1017575 | 1.05E-09 | 1.48E-08 | 295.497437 | 367.489438 | 345.136199 | 452.735286 |
| -6.1015664 | 1.05E-09 | 1.48E-08 | 272.956024 | 783.147595 | 662.661501 | 734.574209 |
| -6.1015508 | 1.05E-09 | 1.48E-08 | 248.365391 | 541.492169 | 498.14658 | 523.895412 |
| -6.1011895 | 1.05E-09 | 1.48E-08 | 238.529138 | 559.623091 | 608.590164 | 619.709439 |
| -6.1012184 | 1.05E-09 | 1.48E-08 | 255.742581 | 525.526133 | 442.924788 | 535.662047 |
| -6.1011401 | 1.05E-09 | 1.48E-08 | 327.46526 | 315.532168 | 334.782113 | 330.58641 |
| -6.100825 | 1.06E-09 | 1.48E-08 | 410.663568 | 334.745534 | 288.763953 | 409.030642 |
| -6.1002964 | 1.06E-09 | 1.48E-08 | 221.725539 | 763.93423 | 766.202361 | 717.76473 |
| -6.1000115 | 1.06E-09 | 1.49E-08 | 245.906328 | 646.218541 | 509.65112 | 559.755633 |
| -6.0992974 | 1.07E-09 | 1.49E-08 | 206.971159 | 927.924362 | 916.911834 | 915.55626 |
| -6.0993435 | 1.07E-09 | 1.49E-08 | 257.381957 | 500.08872 | 403.809352 | 454.97655 |
| -6.0991046 | 1.07E-09 | 1.49E-08 | 317.219163 | 422.423426 | 304.870309 | 373.170422 |
| -6.0990773 | 1.07E-09 | 1.49E-08 | 247.13586 | 508.748265 | 526.90793 | 522.214464 |
| -6.099147 | 1.07E-09 | 1.49E-08 | 270.087117 | 421.611594 | 494.695218 | 396.143376 |
| -6.0988888 | 1.07E-09 | 1.49E-08 | 364.761053 | 401.586396 | 402.658898 | 277.356395 |
| -6.0986776 | 1.07E-09 | 1.49E-08 | 307.38291 | 449.484504 | 334.782113 | 352.999048 |
| -6.0984144 | 1.07E-09 | 1.50E-08 | 270.087117 | 937.936961 | 602.837894 | 490.276455 |
| -6.0970633 | 1.08E-09 | 1.51E-08 | 260.250864 | 468.427259 | 419.915708 | 469.544765 |
| -6.0970881 | 1.08E-09 | 1.51E-08 | 240.168514 | 642.159379 | 540.713378 | 577.685743 |
| -6.0961238 | 1.09E-09 | 1.52E-08 | 242.627577 | 584.519283 | 511.952028 | 584.409534 |
| -6.095933 | 1.09E-09 | 1.52E-08 | 263.119771 | 668.679235 | 432.570702 | 537.903311 |
| -6.0953836 | 1.09E-09 | 1.52E-08 | 279.103682 | 355.311953 | 409.561622 | 396.143376 |
| -6.0947105 | 1.10E-09 | 1.53E-08 | 251.234298 | 574.236073 | 459.031144 | 525.57636 |
| -6.0945402 | 1.10E-09 | 1.53E-08 | 342.21964 | 634.582277 | 656.909231 | 670.137875 |
| -6.0944175 | 1.10E-09 | 1.53E-08 | 256.562269 | 694.928481 | 460.181598 | 551.350893 |
| -6.094332 | 1.10E-09 | 1.53E-08 | 218.036944 | 833.4812 | 708.679661 | 860.084981 |
| -6.0924095 | 1.11E-09 | 1.55E-08 | 246.726016 | 544.198277 | 482.040224 | 605.141225 |

| | | | | | | |
|---|---|---|---|---|---|---|
| -6.092427 | 1.11E-09 | 1.55E-08 | 238.529138 | 526.879187 | 632.749697 | 629.23481 |
| -6.0923476 | 1.11E-09 | 1.55E-08 | 281.972589 | 395.091738 | 376.198456 | 349.637152 |
| -6.09187 | 1.12E-09 | 1.55E-08 | 202.462877 | 896.262901 | 1017.00133 | 980.55291 |
| -6.0918579 | 1.12E-09 | 1.55E-08 | 277.054463 | 594.531882 | 403.809352 | 425.840121 |
| -6.0914952 | 1.12E-09 | 1.55E-08 | 249.594923 | 579.107067 | 456.730236 | 571.522268 |
| -6.0914164 | 1.12E-09 | 1.55E-08 | 288.939935 | 481.145965 | 334.782113 | 430.882964 |
| -6.0912866 | 1.12E-09 | 1.55E-08 | 336.071982 | 380.478755 | 381.950726 | 286.32145 |
| -6.0909261 | 1.12E-09 | 1.56E-08 | 208.610535 | 918.723595 | 873.194582 | 830.388236 |
| -6.0909892 | 1.12E-09 | 1.56E-08 | 258.611488 | 488.452456 | 413.012984 | 499.24151 |
| -6.090746 | 1.12E-09 | 1.56E-08 | 176.232868 | 1241.02103 | 1522.05064 | 1433.2882 |
| -6.0904287 | 1.13E-09 | 1.56E-08 | 294.267906 | 556.916983 | 384.251634 | 370.929158 |
| -6.0900128 | 1.13E-09 | 1.56E-08 | 259.84102 | 445.425342 | 529.208838 | 424.719489 |
| -6.0889244 | 1.14E-09 | 1.57E-08 | 232.38148 | 681.127331 | 820.273699 | 583.288903 |
| -6.0886872 | 1.14E-09 | 1.57E-08 | 279.513526 | 381.290588 | 494.695218 | 376.532317 |
| -6.088179 | 1.14E-09 | 1.58E-08 | 239.348826 | 614.557079 | 528.058384 | 555.273105 |
| -6.087912 | 1.14E-09 | 1.58E-08 | 263.119771 | 416.199378 | 410.712076 | 420.797277 |
| 6.08772995 | 1.15E-09 | 1.58E-08 | 0 | 0 | 0 | 0 |
| -6.0873744 | 1.15E-09 | 1.58E-08 | 281.562746 | 667.596792 | 492.39431 | 532.300151 |
| -6.0871398 | 1.15E-09 | 1.59E-08 | 268.447741 | 383.996696 | 401.508444 | 451.054339 |
| -6.0868518 | 1.15E-09 | 1.59E-08 | 276.644619 | 392.115019 | 353.189377 | 416.875066 |
| -6.0865204 | 1.15E-09 | 1.59E-08 | 286.890716 | 391.573797 | 329.029843 | 386.618005 |
| -6.0863061 | 1.16E-09 | 1.59E-08 | 418.860445 | 534.7269 | 268.055781 | 398.384639 |
| -6.0862605 | 1.16E-09 | 1.59E-08 | 459.025146 | 450.296336 | 286.463045 | 329.465778 |
| -6.0858922 | 1.16E-09 | 1.59E-08 | 394.269812 | 419.446708 | 314.073941 | 296.407137 |
| -6.0857687 | 1.16E-09 | 1.60E-08 | 305.333691 | 636.747163 | 332.481205 | 514.930357 |
| -6.0857101 | 1.16E-09 | 1.60E-08 | 270.906805 | 466.262372 | 371.596641 | 410.71159 |
| -6.0849517 | 1.17E-09 | 1.60E-08 | 282.792277 | 363.159666 | 362.393009 | 348.51652 |
| -6.0845038 | 1.17E-09 | 1.61E-08 | 268.447741 | 418.905486 | 371.596641 | 422.478225 |
| -6.0843375 | 1.17E-09 | 1.61E-08 | 270.087117 | 378.58448 | 386.552542 | 402.867167 |
| -6.0840643 | 1.17E-09 | 1.61E-08 | 448.779049 | 354.770731 | 607.43971 | 291.924609 |
| -6.0839688 | 1.17E-09 | 1.61E-08 | 332.793231 | 394.550516 | 285.312591 | 364.765683 |
| -6.0840125 | 1.17E-09 | 1.61E-08 | 331.153855 | 351.252791 | 294.516223 | 335.068937 |
| -6.0840909 | 1.17E-09 | 1.61E-08 | 205.741628 | 396.444791 | 355.490285 | 389.9799 |
| -6.0820201 | 1.19E-09 | 1.63E-08 | 218.856632 | 732.543379 | 692.573305 | 727.850417 |
| -6.081904 | 1.19E-09 | 1.63E-08 | 262.300083 | 450.025726 | 408.411168 | 401.186219 |
| -6.0817785 | 1.19E-09 | 1.63E-08 | 243.447265 | 572.883019 | 483.190678 | 521.654148 |
| -6.0815821 | 1.19E-09 | 1.63E-08 | 300.00572 | 458.41466 | 309.472125 | 454.97655 |
| -6.0814157 | 1.19E-09 | 1.63E-08 | 273.775712 | 400.774564 | 470.535684 | 423.038541 |
| -6.0813418 | 1.19E-09 | 1.63E-08 | 702.882254 | 688.433822 | 468.234776 | 546.30805 |
| -6.0802657 | 1.20E-09 | 1.64E-08 | 229.922417 | 648.112816 | 585.581084 | 601.219013 |
| -6.0798611 | 1.20E-09 | 1.64E-08 | 238.938982 | 612.121582 | 508.500666 | 549.669946 |
| -6.0792605 | 1.21E-09 | 1.65E-08 | 200.003813 | 869.472433 | 682.219219 | 748.582107 |
| -6.0793202 | 1.21E-09 | 1.65E-08 | 414.762006 | 330.145151 | 284.162137 | 374.291054 |

| | | | | | | |
|---|---|---|---|---|---|---|
| 6.07834052 | 1.21E-09 | 1.66E-08 | 17.213443 | 10.8244312 | 21.8586259 | 12.3269509 |
| -6.0781858 | 1.22E-09 | 1.66E-08 | 255.332737 | 413.493271 | 578.67836 | 574.323847 |
| -6.0783244 | 1.21E-09 | 1.66E-08 | 255.332737 | 531.750181 | 413.012984 | 479.070136 |
| -6.0781243 | 1.22E-09 | 1.66E-08 | 211.479442 | 733.896433 | 930.717282 | 822.543813 |
| -6.0782061 | 1.22E-09 | 1.66E-08 | 252.46383 | 459.226492 | 453.278874 | 434.805176 |
| -6.0782791 | 1.21E-09 | 1.66E-08 | 240.988201 | 605.356313 | 549.91701 | 486.914559 |
| -6.0779213 | 1.22E-09 | 1.66E-08 | 211.069598 | 948.490781 | 867.442313 | 725.609153 |
| -6.0779366 | 1.22E-09 | 1.66E-08 | 241.807889 | 567.470804 | 486.64204 | 519.412884 |
| -6.0775357 | 1.22E-09 | 1.66E-08 | 241.398045 | 525.526133 | 488.942948 | 529.498571 |
| -6.0775707 | 1.22E-09 | 1.66E-08 | 741.817423 | 385.079139 | 327.879389 | 389.419584 |
| -6.0773985 | 1.22E-09 | 1.66E-08 | 284.431653 | 605.356313 | 403.809352 | 388.298952 |
| -6.0767732 | 1.23E-09 | 1.67E-08 | 218.856632 | 865.954493 | 673.015587 | 720.005994 |
| -6.0766137 | 1.23E-09 | 1.67E-08 | 599.191752 | 361.26539 | 368.145279 | 310.975351 |
| -6.0762915 | 1.23E-09 | 1.67E-08 | 275.005243 | 397.256624 | 343.985745 | 415.754434 |
| -6.07593 | 1.23E-09 | 1.67E-08 | 342.629484 | 378.313869 | 283.011683 | 340.672097 |
| -6.0757269 | 1.23E-09 | 1.67E-08 | 280.743058 | 419.176097 | 346.286653 | 373.170422 |
| -6.0750561 | 1.24E-09 | 1.68E-08 | 256.562269 | 707.376577 | 445.225696 | 527.817624 |
| -6.0747636 | 1.24E-09 | 1.68E-08 | 248.775235 | 500.629941 | 434.87161 | 489.716139 |
| -6.0743851 | 1.24E-09 | 1.69E-08 | 245.906328 | 586.95478 | 478.588862 | 477.389188 |
| -6.0740619 | 1.25E-09 | 1.69E-08 | 292.62853 | 320.673773 | 341.684837 | 382.135477 |
| -6.0734654 | 1.25E-09 | 1.69E-08 | 255.742581 | 466.803594 | 399.207536 | 481.3114 |
| -6.0729032 | 1.26E-09 | 1.70E-08 | 372.138243 | 434.05969 | 263.453965 | 403.427483 |
| -6.0730291 | 1.26E-09 | 1.70E-08 | 247.13586 | 441.636792 | 514.252936 | 486.914559 |
| 6.07296439 | 1.26E-09 | 1.70E-08 | 0 | 0 | 0 | 0 |
| -6.0723739 | 1.26E-09 | 1.70E-08 | 218.036944 | 1045.64005 | 671.865133 | 820.862865 |
| -6.0723339 | 1.26E-09 | 1.70E-08 | 203.692408 | 638.91205 | 603.988348 | 590.012694 |
| -6.0719164 | 1.26E-09 | 1.71E-08 | 245.906328 | 515.513534 | 449.827512 | 483.552664 |
| -6.0714873 | 1.27E-09 | 1.71E-08 | 330.334167 | 386.702803 | 287.613499 | 336.189569 |
| -6.0714161 | 1.27E-09 | 1.71E-08 | 257.381957 | 398.609678 | 545.315194 | 474.587608 |
| -6.0710864 | 1.27E-09 | 1.71E-08 | 239.348826 | 573.424241 | 602.837894 | 476.268556 |
| -6.0709155 | 1.27E-09 | 1.71E-08 | 261.480395 | 770.97011 | 804.167343 | 783.882012 |
| -6.0706297 | 1.27E-09 | 1.72E-08 | 238.529138 | 639.18266 | 483.190678 | 590.012694 |
| -6.0705901 | 1.27E-09 | 1.72E-08 | 257.7918 | 477.357414 | 399.207536 | 421.357593 |
| -6.0705024 | 1.28E-09 | 1.72E-08 | 296.726969 | 491.699786 | 314.073941 | 413.51317 |
| -6.0701288 | 1.28E-09 | 1.72E-08 | 307.792754 | 307.955067 | 334.782113 | 324.98325 |
| -6.0697452 | 1.28E-09 | 1.72E-08 | 165.576927 | 375.607761 | 263.453965 | 297.527769 |
| -6.0697213 | 1.28E-09 | 1.72E-08 | 268.447741 | 442.719235 | 402.658898 | 419.11633 |
| -6.0694939 | 1.28E-09 | 1.72E-08 | 317.629007 | 313.908504 | 304.870309 | 339.551465 |
| -6.0693701 | 1.28E-09 | 1.73E-08 | 234.840543 | 649.46587 | 571.775636 | 511.568461 |
| -6.069203 | 1.29E-09 | 1.73E-08 | 269.677273 | 445.695953 | 379.649818 | 364.205367 |
| -6.0690648 | 1.29E-09 | 1.73E-08 | 258.201644 | 448.672672 | 394.60572 | 411.832222 |
| -6.0689143 | 1.29E-09 | 1.73E-08 | 232.38148 | 550.963546 | 592.483808 | 538.463627 |
| -6.0686481 | 1.29E-09 | 1.73E-08 | 243.857109 | 868.119379 | 511.952028 | 625.312599 |

| | | | | | | |
|---|---|---|---|---|---|---|
| -6.0685864 | 1.29E-09 | 1.73E-08 | 252.053986 | 430.54175 | 423.36707 | 458.338446 |
| -6.0684396 | 1.29E-09 | 1.73E-08 | 213.938505 | 1049.15799 | 730.538287 | 765.951902 |
| -6.0680839 | 1.29E-09 | 1.73E-08 | 187.708497 | 1090.56144 | 1029.65633 | 1068.52251 |
| -6.0680045 | 1.30E-09 | 1.73E-08 | 404.515909 | 256.809629 | 614.342434 | 531.179519 |
| -6.0677571 | 1.30E-09 | 1.74E-08 | 243.857109 | 483.852073 | 471.686138 | 468.424133 |
| -6.0675621 | 1.30E-09 | 1.74E-08 | 252.873674 | 404.563115 | 463.63296 | 453.855918 |
| -6.0673502 | 1.30E-09 | 1.74E-08 | 220.905851 | 762.310565 | 693.723759 | 607.942804 |
| -6.0673301 | 1.30E-09 | 1.74E-08 | 246.316172 | 875.15526 | 511.952028 | 600.658697 |
| -6.0673178 | 1.30E-09 | 1.74E-08 | 268.037897 | 401.045175 | 384.251634 | 357.481575 |
| -6.0672012 | 1.30E-09 | 1.74E-08 | 259.431176 | 439.471905 | 384.251634 | 410.71159 |
| -6.0661272 | 1.31E-09 | 1.75E-08 | 296.317125 | 483.581462 | 310.622579 | 409.590958 |
| -6.0652537 | 1.32E-09 | 1.76E-08 | 229.102729 | 798.84302 | 544.16474 | 692.550512 |
| -6.0653778 | 1.32E-09 | 1.76E-08 | 243.857109 | 442.448624 | 502.748396 | 522.214464 |
| -6.0654538 | 1.32E-09 | 1.76E-08 | 256.972113 | 404.021893 | 404.959806 | 414.073486 |
| -6.0653853 | 1.32E-09 | 1.76E-08 | 242.627577 | 573.424241 | 457.88069 | 503.163722 |
| -6.065305 | 1.32E-09 | 1.76E-08 | 248.775235 | 459.767714 | 475.1375 | 422.478225 |
| -6.0650842 | 1.32E-09 | 1.76E-08 | 261.480395 | 437.307019 | 372.747095 | 406.789379 |
| -6.0646751 | 1.32E-09 | 1.76E-08 | 453.697175 | 833.751811 | 250.798971 | 682.464825 |
| -6.0643389 | 1.32E-09 | 1.76E-08 | 303.694315 | 547.445606 | 459.031144 | 308.734088 |
| -6.0639665 | 1.33E-09 | 1.77E-08 | 217.217256 | 775.841104 | 654.608323 | 672.379138 |
| -6.0635815 | 1.33E-09 | 1.77E-08 | 274.5954 | 402.127618 | 353.189377 | 347.395888 |
| -6.0635716 | 1.33E-09 | 1.77E-08 | 331.153855 | 424.047091 | 281.861229 | 354.679995 |
| -6.0631837 | 1.33E-09 | 1.77E-08 | 209.020379 | 812.64417 | 753.547367 | 737.936104 |
| -6.0629613 | 1.34E-09 | 1.77E-08 | 195.905375 | 916.82932 | 899.655024 | 959.260904 |
| -6.0611398 | 1.35E-09 | 1.79E-08 | 217.217256 | 772.323164 | 629.298335 | 693.671144 |
| -6.060816 | 1.35E-09 | 1.80E-08 | 193.856155 | 503.60666 | 461.332052 | 472.906661 |
| -6.0608202 | 1.35E-09 | 1.80E-08 | 522.141103 | 504.147882 | 510.801574 | 235.332698 |
| -6.0608237 | 1.35E-09 | 1.80E-08 | 175.413181 | 1207.4653 | 1604.88332 | 1169.37938 |
| -6.0601794 | 1.36E-09 | 1.80E-08 | 245.496484 | 473.298253 | 461.332052 | 433.684544 |
| -6.0599213 | 1.36E-09 | 1.80E-08 | 281.152902 | 474.380696 | 323.277573 | 485.793927 |
| -6.0597796 | 1.36E-09 | 1.80E-08 | 259.021332 | 746.344529 | 781.158263 | 773.796325 |
| -6.0597346 | 1.36E-09 | 1.80E-08 | 338.531045 | 274.39933 | 477.438408 | 408.470327 |
| -6.0594453 | 1.37E-09 | 1.81E-08 | 219.67632 | 586.95478 | 755.848275 | 745.780528 |
| -6.0592339 | 1.37E-09 | 1.81E-08 | 232.38148 | 704.129247 | 499.297034 | 671.258506 |
| -6.0592585 | 1.37E-09 | 1.81E-08 | 223.774758 | 674.362062 | 571.775636 | 614.10628 |
| -6.059041 | 1.37E-09 | 1.81E-08 | 229.512573 | 595.073103 | 537.262016 | 548.549314 |
| -6.0589217 | 1.37E-09 | 1.81E-08 | 334.022762 | 389.950133 | 442.924788 | 273.994499 |
| -6.0585689 | 1.37E-09 | 1.81E-08 | 209.430223 | 761.228122 | 737.441011 | 725.609153 |
| -6.0579861 | 1.38E-09 | 1.82E-08 | 306.153378 | 719.554062 | 593.634262 | 637.63955 |
| -6.0577874 | 1.38E-09 | 1.82E-08 | 188.938029 | 1074.86601 | 977.885896 | 972.148171 |
| -6.0575829 | 1.38E-09 | 1.82E-08 | 232.791324 | 535.538732 | 580.979268 | 500.922458 |
| -6.057532 | 1.38E-09 | 1.82E-08 | 259.431176 | 970.410254 | 806.468251 | 862.886561 |
| -6.0574249 | 1.38E-09 | 1.82E-08 | 210.659754 | 723.342613 | 696.024667 | 777.158221 |

| | | | | | | |
|---|---|---|---|---|---|---|
| -6.0573515 | 1.38E-09 | 1.82E-08 | 243.037421 | 471.133366 | 439.473426 | 492.517719 |
| -6.0570575 | 1.39E-09 | 1.83E-08 | 276.234775 | 346.923019 | 393.455266 | 335.629253 |
| -6.0569688 | 1.39E-09 | 1.83E-08 | 202.872721 | 844.846852 | 770.804177 | 839.913607 |
| -6.056801 | 1.39E-09 | 1.83E-08 | 198.774282 | 1000.71866 | 876.645944 | 811.89781 |
| -6.0562488 | 1.39E-09 | 1.83E-08 | 415.17185 | 353.688288 | 271.507143 | 345.71494 |
| -6.0559951 | 1.40E-09 | 1.83E-08 | 268.857585 | 355.853175 | 369.295733 | 355.800627 |
| -6.056039 | 1.40E-09 | 1.83E-08 | 197.54475 | 945.784673 | 827.176423 | 889.781726 |
| 6.05590321 | 1.40E-09 | 1.83E-08 | 259.021332 | 209.182132 | 210.533081 | 150.164674 |
| -6.0552252 | 1.40E-09 | 1.84E-08 | 317.629007 | 327.439043 | 384.251634 | 289.123029 |
| -6.0544669 | 1.41E-09 | 1.85E-08 | 226.233822 | 880.838086 | 651.156961 | 638.760181 |
| -6.0538744 | 1.41E-09 | 1.86E-08 | 249.185079 | 400.774564 | 506.199758 | 460.019394 |
| -6.0533378 | 1.42E-09 | 1.86E-08 | 252.873674 | 547.986828 | 460.181598 | 396.703692 |
| -6.0529787 | 1.42E-09 | 1.86E-08 | 265.578834 | 455.708552 | 483.190678 | 345.71494 |
| -6.0525732 | 1.43E-09 | 1.87E-08 | 218.446788 | 622.404792 | 625.846973 | 640.441129 |
| -6.0522541 | 1.43E-09 | 1.87E-08 | 289.349779 | 350.440959 | 304.870309 | 344.033992 |
| -6.051938 | 1.43E-09 | 1.87E-08 | 254.922893 | 504.689103 | 392.304812 | 416.31475 |
| -6.0515073 | 1.43E-09 | 1.88E-08 | 244.676796 | 536.621175 | 462.482506 | 426.400437 |
| -6.0508447 | 1.44E-09 | 1.88E-08 | 245.08664 | 610.768529 | 426.818432 | 513.249409 |
| -6.0507474 | 1.44E-09 | 1.89E-08 | 200.823501 | 921.700314 | 753.547367 | 860.645297 |
| -6.0506335 | 1.44E-09 | 1.89E-08 | 269.677273 | 341.781414 | 356.640739 | 366.44663 |
| -6.0502323 | 1.45E-09 | 1.89E-08 | 270.087117 | 333.39248 | 369.295733 | 394.462428 |
| -6.0501145 | 1.45E-09 | 1.89E-08 | 223.364915 | 593.449439 | 553.368372 | 614.666596 |
| -6.049892 | 1.45E-09 | 1.89E-08 | 245.906328 | 416.199378 | 600.536986 | 525.016044 |
| -6.0495876 | 1.45E-09 | 1.90E-08 | 234.020855 | 508.748265 | 492.39431 | 495.319299 |
| -6.0492974 | 1.45E-09 | 1.90E-08 | 395.909188 | 240.843593 | 272.657597 | 255.504073 |
| -6.0489912 | 1.46E-09 | 1.90E-08 | 243.447265 | 409.434109 | 492.39431 | 429.762333 |
| -6.0484353 | 1.46E-09 | 1.91E-08 | 172.13443 | 1154.96681 | 1256.29576 | 1280.32194 |
| -6.048358 | 1.46E-09 | 1.91E-08 | 248.365391 | 587.225391 | 401.508444 | 548.549314 |
| -6.0476266 | 1.47E-09 | 1.91E-08 | 241.807889 | 490.617343 | 432.570702 | 453.295602 |
| -6.0476191 | 1.47E-09 | 1.91E-08 | 294.67775 | 402.93945 | 310.622579 | 324.98325 |
| -6.0474246 | 1.47E-09 | 1.92E-08 | 211.889286 | 709.812074 | 655.758777 | 682.464825 |
| -6.0471609 | 1.47E-09 | 1.92E-08 | 264.349303 | 372.360432 | 585.581084 | 444.890863 |
| -6.0468731 | 1.48E-09 | 1.92E-08 | 347.137766 | 313.637893 | 280.710775 | 392.221164 |
| -6.0467904 | 1.48E-09 | 1.92E-08 | 211.479442 | 697.363978 | 691.422851 | 659.491872 |
| -6.0466966 | 1.48E-09 | 1.92E-08 | 254.922893 | 428.106253 | 371.596641 | 479.630452 |
| -6.0462477 | 1.48E-09 | 1.93E-08 | 240.168514 | 458.685271 | 456.730236 | 448.813075 |
| -6.0461095 | 1.48E-09 | 1.93E-08 | 284.431653 | 586.95478 | 567.17382 | 325.543566 |
| -6.0458697 | 1.49E-09 | 1.93E-08 | 303.284471 | 452.190612 | 314.073941 | 350.197468 |
| -6.0449482 | 1.49E-09 | 1.94E-08 | 232.791324 | 500.629941 | 554.518826 | 479.070136 |
| -6.0447476 | 1.50E-09 | 1.94E-08 | 233.611012 | 569.63569 | 491.243856 | 481.3114 |
| -6.0443854 | 1.50E-09 | 1.94E-08 | 225.00429 | 741.473535 | 553.368372 | 577.125427 |
| -6.0443186 | 1.50E-09 | 1.94E-08 | 266.808366 | 440.013127 | 340.534383 | 502.603406 |
| -6.0440141 | 1.50E-09 | 1.95E-08 | 261.890239 | 407.269223 | 352.038923 | 370.929158 |

| | | | | | | |
|---|---|---|---|---|---|---|
| -6.0432446 | 1.51E-09 | 1.96E-08 | 181.150995 | 343.67569 | 381.950726 | 387.738636 |
| -6.0426853 | 1.52E-09 | 1.96E-08 | 215.168037 | 744.991475 | 643.103783 | 607.382488 |
| -6.0426074 | 1.52E-09 | 1.96E-08 | 208.610535 | 759.875068 | 687.971489 | 678.542614 |
| -6.0425915 | 1.52E-09 | 1.96E-08 | 289.349779 | 361.536001 | 294.516223 | 381.014845 |
| -6.0421319 | 1.52E-09 | 1.97E-08 | 247.545703 | 490.887953 | 392.304812 | 443.770231 |
| -6.0417796 | 1.52E-09 | 1.97E-08 | 280.743058 | 329.062707 | 322.127119 | 394.462428 |
| -6.0418024 | 1.52E-09 | 1.97E-08 | 352.875581 | 1046.18127 | 1034.25814 | 1112.22716 |
| -6.0418895 | 1.52E-09 | 1.97E-08 | 232.791324 | 497.112001 | 473.987046 | 510.447829 |
| -6.0412372 | 1.53E-09 | 1.97E-08 | 288.120248 | 312.014228 | 315.224395 | 338.430833 |
| -6.0410439 | 1.53E-09 | 1.98E-08 | 245.08664 | 436.765798 | 403.809352 | 439.287704 |
| -6.0404935 | 1.54E-09 | 1.98E-08 | 220.086164 | 663.26702 | 548.766556 | 611.3047 |
| -6.0403604 | 1.54E-09 | 1.98E-08 | 204.512096 | 671.655954 | 924.965012 | 847.75803 |
| -6.039134 | 1.55E-09 | 2.00E-08 | 232.38148 | 553.128433 | 477.438408 | 484.673296 |
| -6.0380139 | 1.56E-09 | 2.01E-08 | 243.037421 | 417.552432 | 430.269794 | 432.003596 |
| -6.0378397 | 1.56E-09 | 2.01E-08 | 239.75867 | 679.233056 | 449.827512 | 524.455728 |
| -6.0371681 | 1.57E-09 | 2.02E-08 | 227.053509 | 484.663905 | 597.085624 | 572.082584 |
| -6.0370559 | 1.57E-09 | 2.02E-08 | 195.495531 | 787.206757 | 872.044129 | 853.36119 |
| -6.0364398 | 1.58E-09 | 2.03E-08 | 1024.6097 | 1601.7452 | 1138.94946 | 1284.80447 |
| -6.0349497 | 1.59E-09 | 2.04E-08 | 245.08664 | 445.966564 | 402.658898 | 409.590958 |
| -6.0348538 | 1.59E-09 | 2.04E-08 | 232.791324 | 540.680337 | 445.225696 | 531.739835 |
| -6.0344696 | 1.59E-09 | 2.05E-08 | 205.331784 | 760.145679 | 681.068765 | 706.558411 |
| -6.03372 | 1.60E-09 | 2.06E-08 | 276.644619 | 662.725798 | 348.587561 | 488.595507 |
| -6.0337181 | 1.60E-09 | 2.06E-08 | 263.119771 | 387.244025 | 331.330751 | 377.092633 |
| -6.0334409 | 1.61E-09 | 2.06E-08 | 219.266476 | 669.220457 | 557.970188 | 570.401636 |
| -6.0330229 | 1.61E-09 | 2.06E-08 | 225.414134 | 528.502852 | 503.89885 | 599.538065 |
| -6.0330326 | 1.61E-09 | 2.06E-08 | 180.741151 | 1064.58281 | 938.77046 | 1102.70179 |
| -6.0327107 | 1.61E-09 | 2.07E-08 | 238.529138 | 472.48642 | 414.163438 | 489.716139 |
| -6.0323955 | 1.62E-09 | 2.07E-08 | 252.46383 | 826.44532 | 415.313892 | 671.818822 |
| -6.0318172 | 1.62E-09 | 2.08E-08 | 345.088547 | 866.766326 | 842.132325 | 893.143622 |
| -6.0311776 | 1.63E-09 | 2.08E-08 | 223.774758 | 519.572696 | 530.359292 | 541.825522 |
| -6.0308777 | 1.63E-09 | 2.09E-08 | 219.266476 | 538.786061 | 638.501967 | 576.004795 |
| -6.030414 | 1.64E-09 | 2.09E-08 | 200.413657 | 800.466685 | 692.573305 | 878.015091 |
| -6.0302098 | 1.64E-09 | 2.09E-08 | 237.299606 | 459.226492 | 482.040224 | 419.676646 |
| -6.0301936 | 1.64E-09 | 2.09E-08 | 268.447741 | 371.277989 | 322.127119 | 346.835572 |
| -6.0299531 | 1.64E-09 | 2.09E-08 | 223.774758 | 572.071187 | 506.199758 | 533.420783 |
| -6.0296788 | 1.64E-09 | 2.10E-08 | 218.856632 | 553.669654 | 576.377452 | 566.479424 |
| -6.0295238 | 1.64E-09 | 2.10E-08 | 366.810273 | 600.485319 | 255.400787 | 498.120878 |
| -6.0283472 | 1.66E-09 | 2.11E-08 | 224.594446 | 525.526133 | 525.757476 | 504.284354 |
| -6.0279457 | 1.66E-09 | 2.12E-08 | 252.46383 | 430.271139 | 376.198456 | 367.006946 |
| -6.0269511 | 1.67E-09 | 2.13E-08 | 215.987725 | 545.010109 | 626.997427 | 686.947353 |
| -6.0268715 | 1.67E-09 | 2.13E-08 | 190.577404 | 364.78333 | 218.586259 | 258.865968 |
| -6.0262095 | 1.68E-09 | 2.14E-08 | 290.579311 | 352.064624 | 431.420248 | 294.165873 |
| -6.0259805 | 1.68E-09 | 2.14E-08 | 212.708974 | 578.024624 | 927.26592 | 795.088331 |

| | | | | | | |
|---|---|---|---|---|---|---|
| -6.0260401 | 1.68E-09 | 2.14E-08 | 298.366345 | 277.91727 | 376.198456 | 358.602207 |
| -6.0260684 | 1.68E-09 | 2.14E-08 | 286.480872 | 786.394924 | 468.234776 | 567.03974 |
| -6.0256426 | 1.68E-09 | 2.14E-08 | 269.267429 | 2123.48278 | 2036.30357 | 1899.47107 |
| -6.0252213 | 1.69E-09 | 2.15E-08 | 216.397569 | 586.142948 | 556.819734 | 586.090482 |
| -6.0249811 | 1.69E-09 | 2.15E-08 | 235.660231 | 598.049822 | 433.721156 | 489.716139 |
| -6.0245541 | 1.70E-09 | 2.15E-08 | 207.790847 | 679.774277 | 612.041526 | 737.375788 |
| -6.0244347 | 1.70E-09 | 2.15E-08 | 234.840543 | 491.429175 | 417.6148 | 488.035191 |
| -6.0239454 | 1.70E-09 | 2.16E-08 | 234.020855 | 597.5086 | 424.517524 | 583.849218 |
| -6.0238158 | 1.70E-09 | 2.16E-08 | 346.318079 | 484.393295 | 272.657597 | 342.913361 |
| -6.0235525 | 1.71E-09 | 2.16E-08 | 266.808366 | 315.261558 | 358.941647 | 373.170422 |
| -6.0233981 | 1.71E-09 | 2.17E-08 | 258.201644 | 545.821942 | 355.490285 | 431.44328 |
| -6.0232472 | 1.71E-09 | 2.17E-08 | 215.168037 | 625.922732 | 551.067464 | 596.176169 |
| -6.0232838 | 1.71E-09 | 2.17E-08 | 217.6271 | 645.677319 | 532.6602 | 577.125427 |
| -6.0228646 | 1.71E-09 | 2.17E-08 | 223.774758 | 627.275786 | 492.39431 | 536.222363 |
| -6.0225953 | 1.72E-09 | 2.17E-08 | 236.070075 | 610.768529 | 456.730236 | 454.416234 |
| -6.0223484 | 1.72E-09 | 2.17E-08 | 204.102252 | 788.55981 | 632.749697 | 734.574209 |
| -6.0217084 | 1.73E-09 | 2.18E-08 | 318.038851 | 540.139115 | 478.588862 | 528.938255 |
| -6.0215627 | 1.73E-09 | 2.18E-08 | 180.331307 | 899.239619 | 1037.7095 | 1043.86861 |
| -6.02162 | 1.73E-09 | 2.18E-08 | 265.16899 | 505.230325 | 784.609625 | 395.58306 |
| -6.0212756 | 1.73E-09 | 2.19E-08 | 223.364915 | 584.248672 | 486.64204 | 528.37794 |
| -6.0209362 | 1.73E-09 | 2.19E-08 | 313.940412 | 506.583379 | 273.808051 | 419.11633 |
| -6.0208927 | 1.73E-09 | 2.19E-08 | 242.217733 | 423.235259 | 385.402088 | 467.863817 |
| -6.0199442 | 1.74E-09 | 2.20E-08 | 371.318555 | 362.347833 | 280.710775 | 289.123029 |
| -6.0197994 | 1.75E-09 | 2.20E-08 | 199.59397 | 697.634589 | 746.644643 | 733.453577 |
| -6.0192945 | 1.75E-09 | 2.21E-08 | 209.430223 | 639.994493 | 651.156961 | 591.693642 |
| 6.01902191 | 1.75E-09 | 2.21E-08 | 6.1476582 | 2.43549701 | 4.60181598 | 1.1206319 |
| -6.0188189 | 1.76E-09 | 2.21E-08 | 307.38291 | 386.432193 | 266.905327 | 372.04979 |
| -6.0180729 | 1.77E-09 | 2.22E-08 | 220.496007 | 639.18266 | 514.252936 | 532.300151 |
| -6.0178016 | 1.77E-09 | 2.22E-08 | 231.971636 | 471.674588 | 425.667978 | 482.432032 |
| -6.0177998 | 1.77E-09 | 2.22E-08 | 253.693362 | 395.091738 | 338.233475 | 392.221164 |
| -6.0175544 | 1.77E-09 | 2.23E-08 | 746.735549 | 530.397127 | 300.268493 | 318.259459 |
| -6.0174107 | 1.77E-09 | 2.23E-08 | 632.389107 | 543.927666 | 296.817131 | 363.084735 |
| -6.0172602 | 1.77E-09 | 2.23E-08 | 227.053509 | 615.639523 | 491.243856 | 477.949504 |
| -6.016982 | 1.78E-09 | 2.23E-08 | 229.512573 | 551.775379 | 471.686138 | 449.933707 |
| -6.0165132 | 1.78E-09 | 2.24E-08 | 334.432606 | 278.729103 | 296.817131 | 379.894213 |
| -6.0159731 | 1.79E-09 | 2.24E-08 | 291.808843 | 375.337151 | 291.064861 | 475.70824 |
| -6.015434 | 1.79E-09 | 2.25E-08 | 251.644142 | 379.125702 | 368.145279 | 350.757784 |
| -6.0153796 | 1.79E-09 | 2.25E-08 | 205.331784 | 667.596792 | 677.617403 | 620.269755 |
| -6.0151605 | 1.80E-09 | 2.25E-08 | 266.808366 | 537.703618 | 323.277573 | 463.38129 |
| -6.0147694 | 1.80E-09 | 2.26E-08 | 235.660231 | 404.292504 | 480.88977 | 440.408336 |
| -6.0147056 | 1.80E-09 | 2.26E-08 | 412.712787 | 352.335234 | 265.754873 | 315.457879 |
| -6.0131542 | 1.82E-09 | 2.28E-08 | 294.67775 | 375.607761 | 276.108959 | 339.551465 |
| -6.012249 | 1.83E-09 | 2.29E-08 | 238.529138 | 406.186779 | 414.163438 | 401.746535 |

| | | | | | | |
|---|---|---|---|---|---|---|
| 6.01214143 | 1.83E-09 | 2.29E-08 | 26.2300083 | 10.8244312 | 18.4072639 | 7.84442328 |
| -6.0117295 | 1.84E-09 | 2.30E-08 | 260.250864 | 469.509702 | 325.578481 | 402.306851 |
| -6.0116349 | 1.84E-09 | 2.30E-08 | 307.792754 | 419.717318 | 264.604419 | 388.298952 |
| -6.0115492 | 1.84E-09 | 2.30E-08 | 247.13586 | 466.262372 | 365.844371 | 389.9799 |
| -6.0112104 | 1.84E-09 | 2.30E-08 | 220.086164 | 576.67157 | 491.243856 | 521.093832 |
| -6.0110841 | 1.84E-09 | 2.30E-08 | 204.92194 | 697.093367 | 609.740618 | 653.888712 |
| -6.0109229 | 1.84E-09 | 2.30E-08 | 232.791324 | 511.995594 | 408.411168 | 471.786029 |
| -6.010261 | 1.85E-09 | 2.31E-08 | 223.774758 | 480.604744 | 514.252936 | 472.906661 |
| 6.00976038 | 1.86E-09 | 2.32E-08 | 0 | 0 | 0 | 0 |
| -6.0097292 | 1.86E-09 | 2.32E-08 | 234.840543 | 414.575714 | 413.012984 | 451.614655 |
| -6.0093234 | 1.86E-09 | 2.32E-08 | 320.497914 | 340.42836 | 260.002603 | 351.3181 |
| -6.0090874 | 1.87E-09 | 2.32E-08 | 247.545703 | 365.865773 | 375.048002 | 364.765683 |
| -6.0089285 | 1.87E-09 | 2.33E-08 | 270.496961 | 298.7543 | 348.587561 | 343.473677 |
| -6.0086974 | 1.87E-09 | 2.33E-08 | 243.447265 | 362.618444 | 400.35799 | 424.159173 |
| -6.0083231 | 1.87E-09 | 2.33E-08 | 464.353116 | 363.430276 | 257.701695 | 339.551465 |
| -6.0080811 | 1.88E-09 | 2.34E-08 | 216.397569 | 566.658972 | 521.15566 | 526.696992 |
| -6.0077276 | 1.88E-09 | 2.34E-08 | 206.151472 | 654.066253 | 629.298335 | 594.495221 |
| -6.0074072 | 1.89E-09 | 2.34E-08 | 256.562269 | 455.437941 | 326.728935 | 400.065587 |
| -6.007452 | 1.88E-09 | 2.34E-08 | 351.236205 | 447.860839 | 270.356689 | 313.216615 |
| -6.006788 | 1.89E-09 | 2.35E-08 | 220.086164 | 517.949031 | 525.757476 | 480.751084 |
| -6.006486 | 1.90E-09 | 2.35E-08 | 234.020855 | 423.77648 | 442.924788 | 404.548115 |
| -6.006561 | 1.89E-09 | 2.35E-08 | 318.038851 | 263.84551 | 366.994825 | 326.664198 |
| -6.0064718 | 1.90E-09 | 2.35E-08 | 301.235252 | 296.860025 | 286.463045 | 309.294404 |
| -6.0063894 | 1.90E-09 | 2.35E-08 | 211.889286 | 664.349463 | 536.111562 | 589.452378 |
| -6.0060939 | 1.90E-09 | 2.36E-08 | 263.529615 | 305.248959 | 407.260714 | 396.703692 |
| -6.005682 | 1.91E-09 | 2.36E-08 | 406.155285 | 760.145679 | 417.6148 | 528.37794 |
| -6.005111 | 1.91E-09 | 2.37E-08 | 272.136336 | 433.247857 | 307.171217 | 319.940407 |
| -6.0044251 | 1.92E-09 | 2.38E-08 | 599.191752 | 1286.21303 | 1259.74712 | 1257.34899 |
| -6.0042539 | 1.92E-09 | 2.38E-08 | 277.054463 | 294.695138 | 315.224395 | 328.345146 |
| -6.0040381 | 1.92E-09 | 2.38E-08 | 215.987725 | 545.010109 | 506.199758 | 541.265206 |
| -6.0038524 | 1.93E-09 | 2.38E-08 | 295.907281 | 375.337151 | 365.844371 | 348.51652 |
| -6.0038386 | 1.93E-09 | 2.38E-08 | 214.758193 | 499.276888 | 601.68744 | 582.168271 |
| -6.0021265 | 1.95E-09 | 2.41E-08 | 201.643189 | 638.370828 | 644.254237 | 663.414083 |
| -6.0020179 | 1.95E-09 | 2.41E-08 | 222.545227 | 473.298253 | 477.438408 | 476.828872 |
| -6.0019091 | 1.95E-09 | 2.41E-08 | 250.414611 | 337.722252 | 491.243856 | 435.365492 |
| -6.0013747 | 1.96E-09 | 2.42E-08 | 267.628054 | 543.115834 | 384.251634 | 442.089283 |
| -6.0012044 | 1.96E-09 | 2.42E-08 | 318.038851 | 493.05284 | 258.852149 | 445.451179 |
| -6.0010579 | 1.96E-09 | 2.42E-08 | 389.351686 | 307.413845 | 265.754873 | 331.707042 |
| -6.0009802 | 1.96E-09 | 2.42E-08 | 216.397569 | 539.056672 | 490.093402 | 550.230262 |
| -6.0007896 | 1.96E-09 | 2.42E-08 | 343.449171 | 334.745534 | 306.020763 | 263.908812 |
| 6.00075132 | 1.96E-09 | 2.42E-08 | 19.6725062 | 7.30649104 | 12.6549939 | 11.7666349 |
| -5.9997202 | 1.98E-09 | 2.43E-08 | 222.955071 | 507.1246 | 442.924788 | 496.43993 |
| -5.9994527 | 1.98E-09 | 2.44E-08 | 238.529138 | 583.707451 | 379.649818 | 585.530166 |

| | | | | | | |
|---|---|---|---|---|---|---|
| -5.9993998 | 1.98E-09 | 2.44E-08 | 215.168037 | 528.502852 | 536.111562 | 507.646249 |
| -5.9990019 | 1.99E-09 | 2.44E-08 | 240.578358 | 450.837558 | 388.85345 | 370.929158 |
| -5.9978847 | 2.00E-09 | 2.46E-08 | 275.005243 | 345.840576 | 288.763953 | 321.621355 |
| -5.9972418 | 2.01E-09 | 2.47E-08 | 233.611012 | 465.179929 | 434.87161 | 384.376741 |
| -5.9970902 | 2.01E-09 | 2.47E-08 | 219.266476 | 258.703905 | 251.949425 | 257.18502 |
| -5.9968209 | 2.01E-09 | 2.47E-08 | 205.331784 | 569.63569 | 628.147881 | 617.468175 |
| -5.9968233 | 2.01E-09 | 2.47E-08 | 223.364915 | 491.158564 | 461.332052 | 447.132127 |
| -5.9965236 | 2.02E-09 | 2.47E-08 | 226.233822 | 502.253606 | 454.429328 | 425.279805 |
| -5.9964738 | 2.02E-09 | 2.47E-08 | 276.234775 | 301.189797 | 299.118039 | 352.438732 |
| -5.9961963 | 2.02E-09 | 2.48E-08 | 223.774758 | 483.581462 | 463.63296 | 628.114178 |
| -5.9960988 | 2.02E-09 | 2.48E-08 | 214.758193 | 541.221558 | 505.049304 | 517.171621 |
| -5.9960934 | 2.02E-09 | 2.48E-08 | 211.479442 | 553.399043 | 529.208838 | 552.471525 |
| -5.9952736 | 2.03E-09 | 2.49E-08 | 237.299606 | 432.977247 | 369.295733 | 405.108431 |
| -5.9948858 | 2.04E-09 | 2.49E-08 | 259.431176 | 332.039426 | 314.073941 | 336.749885 |
| -5.9948066 | 2.04E-09 | 2.49E-08 | 233.201168 | 576.130349 | 400.35799 | 466.743185 |
| -5.9943015 | 2.04E-09 | 2.50E-08 | 224.184602 | 420.25854 | 532.6602 | 528.37794 |
| -5.9931028 | 2.06E-09 | 2.52E-08 | 253.283518 | 535.268121 | 345.136199 | 403.987799 |
| -5.9931541 | 2.06E-09 | 2.52E-08 | 271.726492 | 388.326468 | 299.118039 | 360.283155 |
| -5.9927557 | 2.06E-09 | 2.52E-08 | 285.661184 | 319.050109 | 274.958505 | 350.197468 |
| -5.9922194 | 2.07E-09 | 2.53E-08 | 230.742104 | 541.221558 | 418.765254 | 425.279805 |
| -5.9920997 | 2.07E-09 | 2.53E-08 | 207.381003 | 698.446421 | 537.262016 | 606.822172 |
| -5.991848 | 2.07E-09 | 2.53E-08 | 218.446788 | 464.638708 | 494.695218 | 499.801826 |
| -5.9917051 | 2.08E-09 | 2.53E-08 | 219.266476 | 613.474636 | 459.031144 | 518.852568 |
| -5.9912867 | 2.08E-09 | 2.54E-08 | 172.954117 | 889.497631 | 1017.00133 | 1018.65439 |
| -5.9910453 | 2.08E-09 | 2.54E-08 | 227.873197 | 472.21581 | 402.658898 | 447.132127 |
| -5.9900173 | 2.10E-09 | 2.56E-08 | 213.118818 | 512.536816 | 561.42155 | 502.603406 |
| -5.9897358 | 2.10E-09 | 2.56E-08 | 268.037897 | 373.984097 | 287.613499 | 350.197468 |
| -5.9891622 | 2.11E-09 | 2.57E-08 | 205.331784 | 587.225391 | 691.422851 | 557.514369 |
| -5.9887725 | 2.11E-09 | 2.58E-08 | 293.038374 | 350.170348 | 561.42155 | 312.656299 |
| -5.9886568 | 2.12E-09 | 2.58E-08 | 225.00429 | 415.387546 | 463.63296 | 453.855918 |
| -5.9885168 | 2.12E-09 | 2.58E-08 | 171.314742 | 866.225104 | 1107.8872 | 1293.76953 |
| -5.9881168 | 2.12E-09 | 2.58E-08 | 272.136336 | 352.335234 | 280.710775 | 346.275256 |
| -5.9880469 | 2.12E-09 | 2.58E-08 | 218.856632 | 510.913151 | 442.924788 | 533.981099 |
| 5.98789187 | 2.13E-09 | 2.58E-08 | 0 | 0 | 0 | 0 |
| -5.987341 | 2.13E-09 | 2.59E-08 | 240.168514 | 360.453558 | 364.693917 | 382.135477 |
| -5.9872972 | 2.13E-09 | 2.59E-08 | 221.315695 | 431.082971 | 554.518826 | 613.545964 |
| -5.9872634 | 2.13E-09 | 2.59E-08 | 560.256584 | 305.51957 | 280.710775 | 392.78148 |
| -5.9871258 | 2.14E-09 | 2.59E-08 | 193.036467 | 507.1246 | 427.968886 | 485.233612 |
| -5.9862575 | 2.15E-09 | 2.61E-08 | 257.381957 | 335.557366 | 310.622579 | 328.905462 |
| -5.985116 | 2.16E-09 | 2.62E-08 | 192.626624 | 699.799475 | 678.767857 | 697.593356 |
| -5.9846597 | 2.17E-09 | 2.63E-08 | 219.266476 | 551.775379 | 450.977966 | 471.786029 |
| -5.9845161 | 2.17E-09 | 2.63E-08 | 265.988678 | 914.123212 | 408.411168 | 499.801826 |
| -5.9843067 | 2.17E-09 | 2.63E-08 | 820.507448 | 2918.26664 | 2788.70048 | 3086.78056 |

| | | | | | | |
|---|---|---|---|---|---|---|
| -5.9841607 | 2.18E-09 | 2.63E-08 | 634.028482 | 473.027642 | 254.250333 | 344.594308 |
| 5.98343847 | 2.18E-09 | 2.64E-08 | 11.4756286 | 12.4480958 | 9.20363196 | 11.206319 |
| -5.9830636 | 2.19E-09 | 2.65E-08 | 311.481349 | 402.93945 | 257.701695 | 331.707042 |
| -5.9828504 | 2.19E-09 | 2.65E-08 | 372.548087 | 390.491354 | 233.542161 | 458.898762 |
| -5.9826331 | 2.20E-09 | 2.66E-08 | 222.135383 | 512.266205 | 430.269794 | 446.571811 |
| -5.982142 | 2.20E-09 | 2.66E-08 | 194.675843 | 651.089535 | 733.989649 | 642.122077 |
| -5.9817585 | 2.21E-09 | 2.67E-08 | 302.464783 | 383.726085 | 284.162137 | 284.640502 |
| -5.9816195 | 2.21E-09 | 2.67E-08 | 0 | 615.910133 | 442.924788 | 0 |
| -5.9815865 | 2.21E-09 | 2.67E-08 | 246.726016 | 358.018061 | 337.083021 | 336.189569 |
| -5.9812549 | 2.21E-09 | 2.67E-08 | 409.024192 | 386.702803 | 228.940345 | 472.906661 |
| -5.9812267 | 2.21E-09 | 2.67E-08 | 221.315695 | 809.12623 | 499.297034 | 531.739835 |
| -5.9810038 | 2.22E-09 | 2.67E-08 | 289.349779 | 375.337151 | 271.507143 | 361.964103 |
| -5.9806958 | 2.22E-09 | 2.68E-08 | 292.62853 | 365.053941 | 270.356689 | 305.932508 |
| -5.9800497 | 2.23E-09 | 2.69E-08 | 205.741628 | 523.902468 | 606.289256 | 567.600056 |
| -5.9798485 | 2.23E-09 | 2.69E-08 | 210.659754 | 479.522301 | 574.076544 | 550.790577 |
| -5.9793366 | 2.24E-09 | 2.70E-08 | 217.6271 | 455.167331 | 480.88977 | 460.019394 |
| -5.9793331 | 2.24E-09 | 2.70E-08 | 342.629484 | 932.254134 | 276.108959 | 832.6295 |
| -5.9791739 | 2.24E-09 | 2.70E-08 | 214.348349 | 519.302085 | 461.332052 | 502.04309 |
| -5.9789885 | 2.25E-09 | 2.70E-08 | 277.054463 | 472.48642 | 279.560321 | 408.470327 |
| -5.9789218 | 2.25E-09 | 2.70E-08 | 250.004767 | 428.647474 | 314.073941 | 382.695793 |
| -5.9782022 | 2.26E-09 | 2.71E-08 | 208.200691 | 748.238804 | 508.500666 | 612.985648 |
| -5.9780394 | 2.26E-09 | 2.71E-08 | 280.333214 | 496.300169 | 277.259413 | 460.019394 |
| -5.9777671 | 2.26E-09 | 2.72E-08 | 220.496007 | 430.81236 | 444.075242 | 464.501921 |
| -5.977304 | 2.27E-09 | 2.72E-08 | 215.168037 | 467.074205 | 468.234776 | 526.136676 |
| -5.9763374 | 2.28E-09 | 2.74E-08 | 187.298653 | 860.542278 | 685.670581 | 736.815472 |
| -5.9760799 | 2.29E-09 | 2.74E-08 | 204.92194 | 587.766612 | 564.872912 | 523.335096 |
| -5.9760979 | 2.29E-09 | 2.74E-08 | 208.610535 | 532.562013 | 524.607022 | 507.646249 |
| -5.9760382 | 2.29E-09 | 2.74E-08 | 229.102729 | 396.174181 | 400.35799 | 392.221164 |
| -5.9758771 | 2.29E-09 | 2.74E-08 | 265.16899 | 358.288672 | 284.162137 | 397.264008 |
| -5.9752078 | 2.30E-09 | 2.75E-08 | 223.774758 | 444.072289 | 403.809352 | 509.887513 |
| -5.9749397 | 2.30E-09 | 2.76E-08 | 256.972113 | 316.885222 | 303.719855 | 335.629253 |
| -5.9745114 | 2.31E-09 | 2.76E-08 | 232.38148 | 393.468073 | 392.304812 | 365.325998 |
| -5.9743809 | 2.31E-09 | 2.76E-08 | 193.036467 | 604.544481 | 739.741919 | 714.402834 |
| -5.9737409 | 2.32E-09 | 2.77E-08 | 189.347873 | 645.677319 | 746.644643 | 737.375788 |
| -5.9733994 | 2.32E-09 | 2.78E-08 | 204.102252 | 835.916697 | 549.91701 | 625.872915 |
| -5.9733647 | 2.32E-09 | 2.78E-08 | 243.857109 | 458.41466 | 502.748396 | 485.233612 |
| -5.9727835 | 2.33E-09 | 2.79E-08 | 185.249434 | 965.809871 | 701.776937 | 766.512218 |
| -5.9722873 | 2.34E-09 | 2.79E-08 | 243.447265 | 395.632959 | 337.083021 | 338.991149 |
| -5.9720711 | 2.34E-09 | 2.80E-08 | 177.052556 | 944.431619 | 786.910533 | 838.232659 |
| -5.9721866 | 2.34E-09 | 2.80E-08 | 306.153378 | 274.128719 | 307.171217 | 280.157974 |
| -5.9721302 | 2.34E-09 | 2.80E-08 | 223.774758 | 438.930684 | 434.87161 | 395.58306 |
| -5.9716646 | 2.35E-09 | 2.80E-08 | 245.08664 | 338.534085 | 329.029843 | 342.353045 |
| -5.9703142 | 2.37E-09 | 2.82E-08 | 175.823024 | 826.986541 | 1079.12585 | 824.785076 |

| | | | | | | |
|---|---|---|---|---|---|---|
| -5.9701343 | 2.37E-09 | 2.83E-08 | 0 | 316.344001 | 0 | 428.081385 |
| -5.9700332 | 2.37E-09 | 2.83E-08 | 228.692885 | 453.814277 | 377.34891 | 397.824324 |
| 5.96964927 | 2.38E-09 | 2.83E-08 | 31.5579788 | 15.6954252 | 21.8586259 | 14.0078987 |
| -5.9689421 | 2.39E-09 | 2.84E-08 | 251.234298 | 326.62721 | 311.773033 | 324.98325 |
| -5.9685249 | 2.39E-09 | 2.85E-08 | 221.315695 | 405.104336 | 453.278874 | 430.882964 |
| -5.9674336 | 2.41E-09 | 2.87E-08 | 219.67632 | 616.180744 | 473.987046 | 434.24486 |
| -5.9671702 | 2.41E-09 | 2.87E-08 | 227.463353 | 427.565031 | 439.473426 | 366.44663 |
| -5.9668653 | 2.42E-09 | 2.88E-08 | 243.857109 | 364.242109 | 327.879389 | 331.146726 |
| -5.9667305 | 2.42E-09 | 2.88E-08 | 330.744011 | 248.150084 | 505.049304 | 382.135477 |
| -5.966609 | 2.42E-09 | 2.88E-08 | 248.775235 | 310.119953 | 329.029843 | 355.800627 |
| -5.9663718 | 2.43E-09 | 2.88E-08 | 244.676796 | 484.122684 | 329.029843 | 388.298952 |
| -5.9662983 | 2.43E-09 | 2.88E-08 | 238.938982 | 770.428888 | 377.34891 | 635.398286 |
| -5.966273 | 2.43E-09 | 2.88E-08 | 236.070075 | 371.007378 | 341.684837 | 369.808526 |
| -5.9651482 | 2.44E-09 | 2.90E-08 | 254.922893 | 296.318803 | 419.915708 | 361.403787 |
| -5.9645732 | 2.45E-09 | 2.91E-08 | 202.462877 | 600.75593 | 505.049304 | 595.055537 |
| -5.964423 | 2.46E-09 | 2.91E-08 | 218.036944 | 425.941366 | 429.11934 | 458.338446 |
| -5.9642438 | 2.46E-09 | 2.91E-08 | 334.022762 | 252.209246 | 314.073941 | 307.05314 |
| -5.9641459 | 2.46E-09 | 2.91E-08 | 210.24991 | 484.663905 | 471.686138 | 497.000246 |
| -5.9638206 | 2.46E-09 | 2.92E-08 | 210.24991 | 565.305918 | 463.63296 | 494.198667 |
| -5.9637238 | 2.47E-09 | 2.92E-08 | 495.501251 | 309.308121 | 316.374849 | 280.71829 |
| -5.9637047 | 2.47E-09 | 2.92E-08 | 214.758193 | 538.51545 | 432.570702 | 473.466977 |
| 5.96258826 | 2.48E-09 | 2.94E-08 | 113.116911 | 70.0881918 | 75.9299637 | 56.5919108 |
| -5.9622338 | 2.49E-09 | 2.94E-08 | 294.67775 | 363.430276 | 270.356689 | 286.32145 |
| -5.9620493 | 2.49E-09 | 2.94E-08 | 220.086164 | 523.631858 | 411.86253 | 485.233612 |
| -5.9618892 | 2.49E-09 | 2.95E-08 | 197.54475 | 572.341798 | 564.872912 | 618.028491 |
| -5.9617683 | 2.50E-09 | 2.95E-08 | 335.662138 | 255.997797 | 304.870309 | 299.769033 |
| -5.9616346 | 2.50E-09 | 2.95E-08 | 250.414611 | 687.351379 | 362.393009 | 446.571811 |
| -5.9616586 | 2.50E-09 | 2.95E-08 | 217.6271 | 459.767714 | 410.712076 | 449.373391 |
| -5.9615627 | 2.50E-09 | 2.95E-08 | 261.890239 | 376.690205 | 279.560321 | 410.151274 |
| -5.9611867 | 2.50E-09 | 2.95E-08 | 241.807889 | 329.062707 | 301.418947 | 316.018195 |
| -5.9612314 | 2.50E-09 | 2.95E-08 | 216.807412 | 432.436025 | 572.92609 | 456.657498 |
| -5.9608437 | 2.51E-09 | 2.96E-08 | 229.512573 | 562.329199 | 415.313892 | 389.419584 |
| -5.9605846 | 2.51E-09 | 2.96E-08 | 228.692885 | 377.231426 | 369.295733 | 386.057689 |
| -5.9604174 | 2.52E-09 | 2.96E-08 | 196.725062 | 615.639523 | 607.43971 | 551.911209 |
| -5.9598975 | 2.52E-09 | 2.97E-08 | 214.348349 | 424.317702 | 521.15566 | 471.225713 |
| -5.959695 | 2.53E-09 | 2.97E-08 | 259.84102 | 285.223761 | 337.083021 | 390.540216 |
| -5.9596077 | 2.53E-09 | 2.97E-08 | 202.872721 | 568.282636 | 653.457869 | 498.681194 |
| -5.959666 | 2.53E-09 | 2.97E-08 | 249.594923 | 392.926851 | 295.666677 | 368.687894 |
| -5.959675 | 2.53E-09 | 2.97E-08 | 243.037421 | 386.432193 | 411.86253 | 392.221164 |
| -5.9595272 | 2.53E-09 | 2.97E-08 | 312.71088 | 589.390277 | 348.587561 | 287.442082 |
| -5.959386 | 2.53E-09 | 2.97E-08 | 231.971636 | 402.93945 | 346.286653 | 374.291054 |
| -5.9593049 | 2.53E-09 | 2.97E-08 | 247.955547 | 398.880288 | 297.967585 | 400.065587 |
| -5.9590507 | 2.54E-09 | 2.98E-08 | 222.955071 | 416.7406 | 379.649818 | 435.925808 |

| | | | | | | |
|---|---|---|---|---|---|---|
| -5.9590494 | 2.54E-09 | 2.98E-08 | 308.612442 | 402.668839 | 241.595339 | 398.384639 |
| -5.9581103 | 2.55E-09 | 2.99E-08 | 235.660231 | 358.288672 | 371.596641 | 333.387989 |
| -5.9571005 | 2.57E-09 | 3.01E-08 | 213.938505 | 425.941366 | 621.245158 | 532.860467 |
| -5.9567895 | 2.57E-09 | 3.01E-08 | 266.398522 | 278.187881 | 310.622579 | 362.524419 |
| -5.9566375 | 2.57E-09 | 3.02E-08 | 245.08664 | 394.550516 | 690.272397 | 410.151274 |
| -5.9561724 | 2.58E-09 | 3.02E-08 | 256.972113 | 361.806612 | 279.560321 | 361.403787 |
| -5.9560694 | 2.58E-09 | 3.02E-08 | 227.463353 | 425.400145 | 357.791193 | 391.660848 |
| -5.955685 | 2.59E-09 | 3.03E-08 | 14342.8964 | 16258.025 | 8490.35049 | 11171.5794 |
| -5.9556656 | 2.59E-09 | 3.03E-08 | 247.955547 | 627.275786 | 480.88977 | 493.078035 |
| -5.9555486 | 2.59E-09 | 3.03E-08 | 217.217256 | 460.038325 | 415.313892 | 463.941605 |
| -5.9552184 | 2.60E-09 | 3.04E-08 | 214.348349 | 485.746349 | 480.88977 | 411.271906 |
| -5.9546734 | 2.61E-09 | 3.05E-08 | 166.806459 | 240.302372 | 220.887167 | 230.850171 |
| -5.954194 | 2.61E-09 | 3.05E-08 | 297.136813 | 426.482588 | 253.099879 | 345.71494 |
| -5.9540302 | 2.62E-09 | 3.05E-08 | 224.184602 | 472.21581 | 364.693917 | 433.124228 |
| -5.953877 | 2.62E-09 | 3.06E-08 | 171.724586 | 1122.2229 | 771.954631 | 908.272153 |
| -5.95374 | 2.62E-09 | 3.06E-08 | 228.692885 | 490.346732 | 358.941647 | 406.789379 |
| -5.9534631 | 2.63E-09 | 3.06E-08 | 215.168037 | 452.190612 | 424.517524 | 426.960753 |
| -5.953495 | 2.62E-09 | 3.06E-08 | 198.774282 | 545.28072 | 539.562924 | 561.996896 |
| -5.9531125 | 2.63E-09 | 3.07E-08 | 222.135383 | 390.220744 | 392.304812 | 403.987799 |
| -5.9527209 | 2.64E-09 | 3.07E-08 | 195.495531 | 687.351379 | 528.058384 | 655.56966 |
| -5.9521566 | 2.65E-09 | 3.08E-08 | 311.071505 | 270.88139 | 289.914407 | 275.115131 |
| -5.9522365 | 2.65E-09 | 3.08E-08 | 220.905851 | 403.751282 | 386.552542 | 418.556014 |
| -5.9521595 | 2.65E-09 | 3.08E-08 | 195.085687 | 613.204026 | 561.42155 | 572.642899 |
| -5.9520165 | 2.65E-09 | 3.08E-08 | 216.397569 | 480.334133 | 404.959806 | 433.124228 |
| -5.9518641 | 2.65E-09 | 3.08E-08 | 273.775712 | 313.096671 | 295.666677 | 268.951655 |
| -5.9505902 | 2.67E-09 | 3.11E-08 | 236.479919 | 664.620074 | 598.236078 | 670.69819 |
| -5.9504633 | 2.67E-09 | 3.11E-08 | 176.642712 | 733.355211 | 785.760079 | 791.166119 |
| -5.9499029 | 2.68E-09 | 3.12E-08 | 204.92194 | 572.341798 | 498.14658 | 474.587608 |
| -5.9499216 | 2.68E-09 | 3.12E-08 | 178.691932 | 763.122397 | 717.883293 | 746.340844 |
| -5.9494128 | 2.69E-09 | 3.12E-08 | 166.806459 | 987.999955 | 829.477331 | 908.272153 |
| -5.9488824 | 2.70E-09 | 3.13E-08 | 207.381003 | 459.767714 | 471.686138 | 495.319299 |
| -5.9487897 | 2.70E-09 | 3.13E-08 | 201.643189 | 489.80551 | 529.208838 | 544.066786 |
| -5.9487763 | 2.70E-09 | 3.13E-08 | 231.971636 | 417.281821 | 330.180297 | 386.618005 |
| -5.9483782 | 2.71E-09 | 3.14E-08 | 337.711357 | 1725.68494 | 1374.79252 | 1417.03903 |
| -5.9479069 | 2.72E-09 | 3.15E-08 | 211.889286 | 420.529151 | 491.243856 | 569.281004 |
| -5.9471454 | 2.73E-09 | 3.16E-08 | 248.775235 | 336.098588 | 292.215315 | 321.061039 |
| -5.946819 | 2.73E-09 | 3.17E-08 | 245.08664 | 440.824959 | 408.411168 | 298.648401 |
| -5.946842 | 2.73E-09 | 3.17E-08 | 205.741628 | 453.273055 | 509.65112 | 505.404986 |
| -5.9463908 | 2.74E-09 | 3.17E-08 | 343.039328 | 314.179115 | 246.197155 | 298.088085 |
| -5.9461632 | 2.75E-09 | 3.18E-08 | 157.38005 | 1010.73126 | 982.487712 | 1051.15272 |
| -5.9460268 | 2.75E-09 | 3.18E-08 | 175.413181 | 807.231954 | 893.902754 | 706.558411 |
| -5.9457226 | 2.75E-09 | 3.18E-08 | 245.906328 | 337.992863 | 295.666677 | 378.213265 |
| -5.9455616 | 2.76E-09 | 3.18E-08 | 198.774282 | 656.50175 | 666.112863 | 486.914559 |

| | | | | | | |
|---|---|---|---|---|---|---|
| -5.9454433 | 2.76E-09 | 3.18E-08 | 275.824931 | 457.602828 | 314.073941 | 288.562714 |
| -5.9452918 | 2.76E-09 | 3.19E-08 | 228.283041 | 425.400145 | 510.801574 | 349.637152 |
| -5.9450677 | 2.76E-09 | 3.19E-08 | 251.644142 | 423.77648 | 444.075242 | 337.870517 |
| -5.9445528 | 2.77E-09 | 3.20E-08 | 262.300083 | 459.226492 | 314.073941 | 308.173772 |
| -5.9443529 | 2.78E-09 | 3.20E-08 | 189.347873 | 675.444505 | 574.076544 | 633.717338 |
| -5.9440383 | 2.78E-09 | 3.20E-08 | 215.987725 | 862.977775 | 436.022064 | 685.266405 |
| -5.9440899 | 2.78E-09 | 3.20E-08 | 217.217256 | 414.034492 | 438.322972 | 387.738636 |
| -5.9440959 | 2.78E-09 | 3.20E-08 | 240.168514 | 332.580648 | 314.073941 | 326.103882 |
| -5.9438972 | 2.78E-09 | 3.21E-08 | 184.429746 | 779.359044 | 849.035049 | 801.812122 |
| -5.9436852 | 2.79E-09 | 3.21E-08 | 217.217256 | 449.213893 | 386.552542 | 416.31475 |
| -5.9433938 | 2.79E-09 | 3.21E-08 | 213.528661 | 454.89672 | 410.712076 | 425.279805 |
| -5.9433305 | 2.79E-09 | 3.21E-08 | 289.349779 | 325.274157 | 247.347609 | 357.481575 |
| -5.9431051 | 2.80E-09 | 3.22E-08 | 235.250387 | 374.795929 | 329.029843 | 462.260658 |
| -5.9430015 | 2.80E-09 | 3.22E-08 | 0 | 395.90357 | 536.111562 | 0 |
| -5.9424623 | 2.81E-09 | 3.23E-08 | 204.102252 | 571.259355 | 463.63296 | 495.319299 |
| -5.9422956 | 2.81E-09 | 3.23E-08 | 245.906328 | 288.741701 | 365.844371 | 379.894213 |
| -5.9421501 | 2.81E-09 | 3.23E-08 | 213.528661 | 434.05969 | 409.561622 | 428.641701 |
| -5.9420501 | 2.81E-09 | 3.23E-08 | 201.643189 | 645.136097 | 471.686138 | 558.074685 |
| -5.942012 | 2.82E-09 | 3.23E-08 | 295.497437 | 348.546684 | 269.206235 | 269.511971 |
| -5.9417837 | 2.82E-09 | 3.23E-08 | 226.233822 | 357.74745 | 361.242555 | 369.24821 |
| -5.9417451 | 2.82E-09 | 3.23E-08 | 200.823501 | 585.331115 | 482.040224 | 518.852568 |
| -5.9410098 | 2.83E-09 | 3.25E-08 | 293.858062 | 469.780313 | 247.347609 | 417.995698 |
| -5.9401006 | 2.85E-09 | 3.26E-08 | 221.725539 | 734.708265 | 417.6148 | 502.603406 |
| -5.939195 | 2.86E-09 | 3.28E-08 | 254.103206 | 665.161295 | 318.675757 | 488.595507 |
| -5.9390857 | 2.87E-09 | 3.28E-08 | 250.414611 | 374.795929 | 294.516223 | 308.734088 |
| -5.9380867 | 2.88E-09 | 3.30E-08 | 209.840067 | 964.998038 | 476.287954 | 687.507669 |
| -5.9376411 | 2.89E-09 | 3.31E-08 | 224.594446 | 389.679522 | 370.446187 | 349.076836 |
| 5.9374342 | 2.90E-09 | 3.31E-08 | 0 | 0 | 0 | 0 |
| -5.9372423 | 2.90E-09 | 3.32E-08 | 392.630437 | 475.192528 | 220.887167 | 365.886314 |
| -5.9366965 | 2.91E-09 | 3.32E-08 | 225.00429 | 345.840576 | 371.596641 | 377.652949 |
| -5.9361782 | 2.92E-09 | 3.33E-08 | 238.529138 | 333.121869 | 317.525303 | 313.776931 |
| -5.9358137 | 2.92E-09 | 3.34E-08 | 209.840067 | 532.562013 | 423.36707 | 443.770231 |
| -5.9356178 | 2.93E-09 | 3.34E-08 | 233.611012 | 314.720336 | 448.677058 | 407.349695 |
| -5.9354134 | 2.93E-09 | 3.35E-08 | 251.644142 | 283.600097 | 361.242555 | 310.415036 |
| -5.9353817 | 2.93E-09 | 3.35E-08 | 234.840543 | 317.155833 | 334.782113 | 338.430833 |
| -5.9349474 | 2.94E-09 | 3.35E-08 | 241.807889 | 418.093654 | 294.516223 | 379.894213 |
| -5.9347125 | 2.94E-09 | 3.36E-08 | 165.167084 | 775.029271 | 1463.37748 | 1130.71758 |
| -5.9342199 | 2.95E-09 | 3.37E-08 | 209.430223 | 503.60666 | 884.699122 | 545.747734 |
| -5.9337521 | 2.96E-09 | 3.37E-08 | 259.84102 | 342.863857 | 266.905327 | 312.656299 |
| -5.9336118 | 2.96E-09 | 3.38E-08 | 230.332261 | 569.365079 | 365.844371 | 397.824324 |
| -5.9331049 | 2.97E-09 | 3.38E-08 | 191.397092 | 543.927666 | 609.740618 | 561.436581 |
| -5.9330131 | 2.97E-09 | 3.38E-08 | 211.069598 | 566.658972 | 525.757476 | 396.143376 |
| -5.933061 | 2.97E-09 | 3.38E-08 | 279.92337 | 577.754013 | 330.180297 | 440.968652 |

| | | | | | | |
|---|---|---|---|---|---|---|
| -5.9331061 | 2.97E-09 | 3.38E-08 | 220.496007 | 398.880288 | 407.260714 | 352.999048 |
| -5.932392 | 2.99E-09 | 3.40E-08 | 227.463353 | 491.158564 | 821.424153 | 426.960753 |
| -5.932306 | 2.99E-09 | 3.40E-08 | 217.217256 | 366.136384 | 432.570702 | 478.50982 |
| -5.9319474 | 2.99E-09 | 3.40E-08 | 213.118818 | 429.729917 | 387.702996 | 428.641701 |
| -5.9318511 | 3.00E-09 | 3.40E-08 | 164.75724 | 778.276601 | 1969.57724 | 1498.84516 |
| -5.9308819 | 3.01E-09 | 3.42E-08 | 210.24991 | 460.308935 | 423.36707 | 407.910011 |
| -5.9308343 | 3.01E-09 | 3.42E-08 | 202.053033 | 458.955881 | 521.15566 | 471.786029 |
| -5.9305493 | 3.02E-09 | 3.43E-08 | 205.741628 | 481.957798 | 427.968886 | 460.57971 |
| -5.9305696 | 3.02E-09 | 3.43E-08 | 236.889763 | 340.157749 | 300.268493 | 351.878416 |
| 5.93041652 | 3.02E-09 | 3.43E-08 | 13.1150042 | 16.2366467 | 13.8054479 | 16.8094785 |
| -5.9303964 | 3.02E-09 | 3.43E-08 | 195.085687 | 538.51545 | 502.748396 | 553.592157 |
| -5.9292302 | 3.04E-09 | 3.45E-08 | 236.889763 | 413.22266 | 301.418947 | 366.44663 |
| -5.9290975 | 3.05E-09 | 3.45E-08 | 282.792277 | 268.445893 | 499.297034 | 347.956204 |
| -5.9288774 | 3.05E-09 | 3.45E-08 | 229.512573 | 416.7406 | 315.224395 | 413.51317 |
| -5.9283607 | 3.06E-09 | 3.46E-08 | 303.284471 | 274.39933 | 256.551241 | 290.803977 |
| -5.9278458 | 3.07E-09 | 3.47E-08 | 165.986771 | 766.910948 | 846.734141 | 954.21806 |
| -5.9277237 | 3.07E-09 | 3.47E-08 | 191.806936 | 507.665822 | 690.272397 | 613.545964 |
| -5.927494 | 3.08E-09 | 3.48E-08 | 221.315695 | 379.666923 | 353.189377 | 366.44663 |
| -5.926506 | 3.09E-09 | 3.50E-08 | 287.30056 | 261.410013 | 273.808051 | 286.881766 |
| -5.9262137 | 3.10E-09 | 3.50E-08 | 359.433083 | 370.736767 | 236.993523 | 291.364293 |
| -5.9257062 | 3.11E-09 | 3.51E-08 | 243.037421 | 865.142661 | 574.076544 | 680.783878 |
| -5.9256479 | 3.11E-09 | 3.51E-08 | 212.708974 | 420.529151 | 378.499364 | 442.649599 |
| -5.9251016 | 3.12E-09 | 3.52E-08 | 234.430699 | 299.566133 | 353.189377 | 352.999048 |
| -5.924577 | 3.13E-09 | 3.53E-08 | 245.906328 | 413.493271 | 302.569401 | 313.216615 |
| -5.9241769 | 3.14E-09 | 3.54E-08 | 204.102252 | 457.332217 | 447.526604 | 439.84802 |
| -5.9240639 | 3.14E-09 | 3.54E-08 | 211.069598 | 428.647474 | 414.163438 | 390.540216 |
| -5.9236987 | 3.15E-09 | 3.55E-08 | 252.053986 | 448.13145 | 317.525303 | 301.44998 |
| -5.9234576 | 3.15E-09 | 3.55E-08 | 219.67632 | 366.406995 | 354.339831 | 405.108431 |
| -5.9233217 | 3.16E-09 | 3.55E-08 | 225.823978 | 517.137199 | 353.189377 | 386.057689 |
| -5.9231166 | 3.16E-09 | 3.56E-08 | 216.807412 | 454.89672 | 364.693917 | 391.660848 |
| -5.9230592 | 3.16E-09 | 3.56E-08 | 340.990108 | 448.672672 | 222.037621 | 519.9732 |
| -5.9229347 | 3.16E-09 | 3.56E-08 | 234.020855 | 330.415761 | 303.719855 | 341.792729 |
| -5.9224991 | 3.17E-09 | 3.57E-08 | 281.562746 | 268.445893 | 274.958505 | 272.313551 |
| -5.9218423 | 3.18E-09 | 3.58E-08 | 161.478489 | 211.88824 | 181.771731 | 222.445432 |
| -5.9216619 | 3.19E-09 | 3.58E-08 | 219.266476 | 422.964648 | 362.393009 | 360.283155 |
| 5.92129533 | 3.19E-09 | 3.59E-08 | 81.5589321 | 72.253078 | 86.2840497 | 82.9267604 |
| -5.9210701 | 3.20E-09 | 3.59E-08 | 219.266476 | 344.758133 | 393.455266 | 435.925808 |
| -5.9209321 | 3.20E-09 | 3.60E-08 | 189.757716 | 541.76278 | 570.625182 | 540.70489 |
| -5.9208396 | 3.20E-09 | 3.60E-08 | 223.364915 | 332.580648 | 388.85345 | 357.481575 |
| -5.9203454 | 3.21E-09 | 3.61E-08 | 760.260397 | 712.788792 | 444.075242 | 201.153426 |
| -5.9200816 | 3.22E-09 | 3.61E-08 | 619.683946 | 401.857007 | 263.453965 | 300.889664 |
| -5.9198467 | 3.22E-09 | 3.61E-08 | 194.265999 | 609.144864 | 494.695218 | 513.249409 |
| -5.919593 | 3.23E-09 | 3.62E-08 | 236.479919 | 340.698971 | 306.020763 | 305.372192 |

| | | | | | | |
|---|---|---|---|---|---|---|
| -5.9194405 | 3.23E-09 | 3.62E-08 | 161.478489 | 275.752384 | 266.905327 | 271.753235 |
| -5.919441 | 3.23E-09 | 3.62E-08 | 211.479442 | 420.25854 | 401.508444 | 383.256109 |
| -5.9194011 | 3.23E-09 | 3.62E-08 | 186.888809 | 670.844121 | 531.509746 | 609.623752 |
| -5.9192422 | 3.23E-09 | 3.62E-08 | 265.988678 | 361.26539 | 253.099879 | 322.18167 |
| -5.9190374 | 3.24E-09 | 3.62E-08 | 186.478965 | 980.964074 | 562.572004 | 731.212313 |
| -5.9190963 | 3.24E-09 | 3.62E-08 | 279.103682 | 317.697055 | 552.217918 | 302.010296 |
| -5.9192992 | 3.23E-09 | 3.62E-08 | 264.349303 | 353.147067 | 369.295733 | 709.359991 |
| -5.9190301 | 3.24E-09 | 3.62E-08 | 195.905375 | 577.212792 | 459.031144 | 547.428682 |
| -5.9190714 | 3.24E-09 | 3.62E-08 | 214.348349 | 463.556265 | 369.295733 | 399.505271 |
| -5.9188755 | 3.24E-09 | 3.62E-08 | 286.890716 | 286.035594 | 347.437107 | 249.340597 |
| -5.91866 | 3.25E-09 | 3.62E-08 | 494.681563 | 943.078565 | 760.450091 | 875.213512 |
| -5.9183574 | 3.25E-09 | 3.63E-08 | 161.478489 | 839.705248 | 843.282779 | 877.454776 |
| -5.9183666 | 3.25E-09 | 3.63E-08 | 230.332261 | 423.505869 | 308.321671 | 374.291054 |
| -5.9179926 | 3.26E-09 | 3.63E-08 | 225.414134 | 627.005175 | 354.339831 | 473.466977 |
| -5.9180202 | 3.26E-09 | 3.63E-08 | 236.479919 | 403.210061 | 289.914407 | 397.264008 |
| -5.9175033 | 3.27E-09 | 3.64E-08 | 185.249434 | 616.451355 | 568.324274 | 573.763531 |
| -5.9173445 | 3.27E-09 | 3.65E-08 | 193.036467 | 861.895332 | 522.306114 | 618.028491 |
| -5.917359 | 3.27E-09 | 3.65E-08 | 359.842927 | 281.43521 | 253.099879 | 286.881766 |
| -5.9169419 | 3.28E-09 | 3.65E-08 | 202.462877 | 660.01969 | 422.216616 | 621.390387 |
| 5.91637194 | 3.29E-09 | 3.66E-08 | 4.0984388 | 6.76526948 | 2.30090799 | 3.36189569 |
| -5.9158147 | 3.30E-09 | 3.68E-08 | 245.496484 | 279.540935 | 384.251634 | 438.727388 |
| -5.915428 | 3.31E-09 | 3.68E-08 | 240.988201 | 336.369198 | 278.409867 | 336.189569 |
| -5.9153789 | 3.31E-09 | 3.68E-08 | 197.134906 | 531.750181 | 470.535684 | 468.984449 |
| -5.9150418 | 3.32E-09 | 3.69E-08 | 234.020855 | 346.381797 | 292.215315 | 344.594308 |
| -5.9150454 | 3.32E-09 | 3.69E-08 | 594.68347 | 273.858108 | 399.207536 | 280.157974 |
| -5.9146037 | 3.33E-09 | 3.70E-08 | 222.545227 | 325.544767 | 409.561622 | 451.614655 |
| -5.913914 | 3.34E-09 | 3.71E-08 | 209.020379 | 380.749366 | 417.6148 | 446.011495 |
| -5.9133234 | 3.35E-09 | 3.72E-08 | 188.528185 | 547.445606 | 667.263317 | 526.696992 |
| -5.9132352 | 3.35E-09 | 3.72E-08 | 246.316172 | 350.71157 | 270.356689 | 383.816425 |
| -5.9129143 | 3.36E-09 | 3.73E-08 | 281.562746 | 286.035594 | 280.710775 | 252.142177 |
| -5.9119781 | 3.38E-09 | 3.75E-08 | 192.626624 | 824.821655 | 505.049304 | 611.865016 |
| -5.910686 | 3.41E-09 | 3.78E-08 | 183.200214 | 721.989559 | 540.713378 | 639.320497 |
| -5.9106052 | 3.41E-09 | 3.78E-08 | 223.364915 | 397.256624 | 326.728935 | 345.154624 |
| -5.9103395 | 3.41E-09 | 3.78E-08 | 219.266476 | 385.62036 | 329.029843 | 401.746535 |
| -5.9101516 | 3.42E-09 | 3.79E-08 | 197.134906 | 455.437941 | 482.040224 | 471.786029 |
| -5.909974 | 3.42E-09 | 3.79E-08 | 256.152425 | 649.46587 | 399.207536 | 487.474875 |
| -5.9098812 | 3.42E-09 | 3.79E-08 | 205.331784 | 456.249774 | 392.304812 | 437.04644 |
| -5.9087542 | 3.45E-09 | 3.81E-08 | 228.283041 | 355.041342 | 306.020763 | 330.58641 |
| -5.9085836 | 3.45E-09 | 3.82E-08 | 183.610058 | 543.115834 | 616.643342 | 593.934906 |
| -5.9083265 | 3.46E-09 | 3.82E-08 | 191.806936 | 508.477654 | 483.190678 | 555.833421 |
| -5.9082567 | 3.46E-09 | 3.82E-08 | 213.938505 | 452.461223 | 350.888469 | 403.427483 |
| -5.9078784 | 3.47E-09 | 3.83E-08 | 268.447741 | 330.415761 | 247.347609 | 299.769033 |
| -5.9076315 | 3.47E-09 | 3.83E-08 | 184.83959 | 600.485319 | 735.140103 | 533.981099 |

| | | | | | | |
|---|---|---|---|---|---|---|
| -5.9072214 | 3.48E-09 | 3.84E-08 | 229.512573 | 364.242109 | 295.666677 | 351.878416 |
| -5.9071567 | 3.48E-09 | 3.84E-08 | 179.921463 | 645.94793 | 577.527906 | 618.588807 |
| -5.9067736 | 3.49E-09 | 3.85E-08 | 293.858062 | 362.077222 | 228.940345 | 349.076836 |
| -5.9065356 | 3.49E-09 | 3.85E-08 | 188.528185 | 506.583379 | 538.41247 | 545.747734 |
| -5.9066228 | 3.49E-09 | 3.85E-08 | 200.003813 | 472.21581 | 422.216616 | 453.855918 |
| 5.9065568 | 3.49E-09 | 3.85E-08 | 54.9190799 | 23.5431378 | 20.7081719 | 30.2570612 |
| -5.9064412 | 3.50E-09 | 3.85E-08 | 189.347873 | 486.558181 | 547.616102 | 584.96985 |
| -5.906314 | 3.50E-09 | 3.85E-08 | 269.677273 | 250.856192 | 312.923487 | 382.135477 |
| -5.9062267 | 3.50E-09 | 3.85E-08 | 233.201168 | 396.715402 | 337.083021 | 295.286505 |
| -5.9061319 | 3.50E-09 | 3.86E-08 | 176.642712 | 657.042972 | 597.085624 | 713.282203 |
| -5.906041 | 3.50E-09 | 3.86E-08 | 201.643189 | 565.576528 | 419.915708 | 461.140026 |
| -5.9057213 | 3.51E-09 | 3.86E-08 | 183.200214 | 698.987643 | 520.005206 | 735.134525 |
| -5.9055606 | 3.51E-09 | 3.86E-08 | 223.774758 | 316.07339 | 346.286653 | 341.232413 |
| -5.9053084 | 3.52E-09 | 3.87E-08 | 204.92194 | 381.561199 | 492.39431 | 471.225713 |
| -5.9046831 | 3.53E-09 | 3.88E-08 | 372.957931 | 297.401246 | 534.961108 | 247.659649 |
| -5.9044334 | 3.54E-09 | 3.89E-08 | 265.578834 | 389.679522 | 247.347609 | 342.353045 |
| -5.9037696 | 3.55E-09 | 3.90E-08 | 235.250387 | 296.860025 | 295.666677 | 317.138827 |
| -5.9037702 | 3.55E-09 | 3.90E-08 | 205.331784 | 402.398229 | 414.163438 | 401.746535 |
| -5.903924 | 3.55E-09 | 3.90E-08 | 222.545227 | 471.674588 | 326.728935 | 383.256109 |
| -5.9037793 | 3.55E-09 | 3.90E-08 | 256.562269 | 516.866588 | 268.055781 | 489.155823 |
| -5.9034306 | 3.56E-09 | 3.90E-08 | 247.545703 | 384.537917 | 410.712076 | 265.029444 |
| -5.9034105 | 3.56E-09 | 3.90E-08 | 223.364915 | 416.7406 | 523.456568 | 328.905462 |
| -5.9033245 | 3.56E-09 | 3.90E-08 | 208.610535 | 372.089821 | 413.012984 | 394.462428 |
| -5.9028327 | 3.57E-09 | 3.91E-08 | 208.610535 | 507.1246 | 379.649818 | 413.51317 |
| -5.9024659 | 3.58E-09 | 3.92E-08 | 169.675366 | 252.479857 | 173.718553 | 191.628054 |
| -5.901945 | 3.59E-09 | 3.93E-08 | 178.282088 | 599.673487 | 1251.69395 | 908.832469 |
| -5.9012928 | 3.61E-09 | 3.95E-08 | 255.332737 | 389.408911 | 269.206235 | 299.208717 |
| -5.9006658 | 3.62E-09 | 3.96E-08 | 214.758193 | 432.436025 | 345.136199 | 511.008145 |
| -5.9004536 | 3.63E-09 | 3.96E-08 | 203.282564 | 399.962732 | 438.322972 | 410.71159 |
| -5.9002578 | 3.63E-09 | 3.97E-08 | 210.659754 | 383.455474 | 365.844371 | 375.411686 |
| -5.8999149 | 3.64E-09 | 3.97E-08 | 199.184126 | 511.724983 | 438.322972 | 424.159173 |
| -5.8998678 | 3.64E-09 | 3.97E-08 | 288.530091 | 248.150084 | 317.525303 | 266.150076 |
| -5.8993634 | 3.65E-09 | 3.98E-08 | 310.661661 | 388.326468 | 662.661501 | 261.107232 |
| -5.8993776 | 3.65E-09 | 3.98E-08 | 177.052556 | 672.197175 | 608.590164 | 594.495221 |
| -5.8990329 | 3.66E-09 | 3.99E-08 | 314.7601 | 426.211977 | 224.338529 | 344.594308 |
| -5.8984818 | 3.67E-09 | 4.00E-08 | 234.020855 | 775.570493 | 372.747095 | 453.855918 |
| -5.8983951 | 3.67E-09 | 4.00E-08 | 239.75867 | 562.058588 | 300.268493 | 424.719489 |
| -5.8982559 | 3.67E-09 | 4.01E-08 | 227.873197 | 765.287283 | 600.536986 | 598.417433 |
| -5.8980397 | 3.68E-09 | 4.01E-08 | 206.561315 | 507.665822 | 386.552542 | 408.470327 |
| -5.8979149 | 3.68E-09 | 4.01E-08 | 282.382433 | 271.693222 | 249.648517 | 349.076836 |
| -5.8974744 | 3.69E-09 | 4.02E-08 | 0 | 284.141318 | 551.067464 | 0 |
| -5.8969779 | 3.70E-09 | 4.03E-08 | 180.741151 | 555.293319 | 598.236078 | 575.444479 |
| -5.8969666 | 3.70E-09 | 4.03E-08 | 303.694315 | 240.031761 | 284.162137 | 379.333897 |

| | | | | | | |
|---|---|---|---|---|---|---|
| -5.896311 | 3.72E-09 | 4.05E-08 | 225.414134 | 512.807426 | 320.976665 | 395.58306 |
| -5.8959303 | 3.73E-09 | 4.05E-08 | 237.70945 | 294.153917 | 293.365769 | 286.32145 |
| -5.8958145 | 3.73E-09 | 4.05E-08 | 174.593493 | 623.487235 | 641.953329 | 626.993547 |
| -5.8954038 | 3.74E-09 | 4.06E-08 | 190.987248 | 462.203211 | 513.102482 | 494.198667 |
| -5.8948032 | 3.75E-09 | 4.08E-08 | 210.24991 | 384.267306 | 350.888469 | 388.859268 |
| -5.894713 | 3.75E-09 | 4.08E-08 | 223.774758 | 394.279905 | 337.083021 | 309.294404 |
| -5.8944732 | 3.76E-09 | 4.08E-08 | 186.069121 | 621.322349 | 476.287954 | 639.320497 |
| -5.8944917 | 3.76E-09 | 4.08E-08 | 211.479442 | 374.254708 | 346.286653 | 411.271906 |
| -5.8938251 | 3.77E-09 | 4.10E-08 | 170.904898 | 702.234972 | 625.846973 | 691.429881 |
| -5.893755 | 3.78E-09 | 4.10E-08 | 186.069121 | 532.562013 | 630.448789 | 497.000246 |
| -5.893687 | 3.78E-09 | 4.10E-08 | 229.102729 | 350.71157 | 291.064861 | 318.819775 |
| -5.893277 | 3.79E-09 | 4.10E-08 | 195.495531 | 490.076121 | 498.14658 | 416.875066 |
| -5.8929866 | 3.79E-09 | 4.11E-08 | 209.430223 | 365.324552 | 369.295733 | 374.85137 |
| -5.8929142 | 3.79E-09 | 4.11E-08 | 159.429269 | 756.627738 | 823.725061 | 822.543813 |
| 5.89276266 | 3.80E-09 | 4.11E-08 | 9.01656536 | 8.11832337 | 8.05317797 | 11.206319 |
| -5.8927161 | 3.80E-09 | 4.11E-08 | 215.168037 | 434.05969 | 332.481205 | 372.610106 |
| -5.8922705 | 3.81E-09 | 4.12E-08 | 193.856155 | 581.813175 | 456.730236 | 460.57971 |
| -5.8913357 | 3.83E-09 | 4.14E-08 | 235.660231 | 278.999713 | 308.321671 | 298.088085 |
| -5.8913545 | 3.83E-09 | 4.14E-08 | 275.824931 | 297.401246 | 235.843069 | 310.975351 |
| -5.8914219 | 3.83E-09 | 4.14E-08 | 174.593493 | 782.876984 | 555.66928 | 781.640748 |
| -5.8907821 | 3.84E-09 | 4.15E-08 | 176.642712 | 632.417391 | 579.828814 | 593.37459 |
| -5.8905215 | 3.85E-09 | 4.16E-08 | 197.134906 | 423.77648 | 427.968886 | 496.43993 |
| -5.890061 | 3.86E-09 | 4.17E-08 | 208.200691 | 914.123212 | 463.63296 | 709.920307 |
| -5.8900805 | 3.86E-09 | 4.17E-08 | 188.528185 | 550.422325 | 473.987046 | 492.517719 |
| -5.8897009 | 3.87E-09 | 4.18E-08 | 210.659754 | 452.461223 | 354.339831 | 372.610106 |
| -5.8895133 | 3.87E-09 | 4.18E-08 | 188.938029 | 535.809343 | 453.278874 | 533.981099 |
| -5.8892582 | 3.88E-09 | 4.18E-08 | 177.052556 | 623.216624 | 562.572004 | 601.219013 |
| -5.8891249 | 3.88E-09 | 4.19E-08 | 178.691932 | 633.770445 | 557.970188 | 564.23816 |
| -5.8889823 | 3.89E-09 | 4.19E-08 | 183.200214 | 548.257439 | 521.15566 | 533.420783 |
| -5.8890366 | 3.88E-09 | 4.19E-08 | 196.315218 | 573.15363 | 407.260714 | 515.490673 |
| -5.8889083 | 3.89E-09 | 4.19E-08 | 178.282088 | 612.121582 | 557.970188 | 576.004795 |
| -5.8882598 | 3.90E-09 | 4.20E-08 | 190.577404 | 472.48642 | 462.482506 | 488.035191 |
| -5.8874811 | 3.92E-09 | 4.22E-08 | 240.578358 | 331.498204 | 375.048002 | 264.469128 |
| -5.8869731 | 3.93E-09 | 4.23E-08 | 308.612442 | 274.940552 | 292.215315 | 236.45333 |
| -5.887009 | 3.93E-09 | 4.23E-08 | 206.561315 | 364.242109 | 375.048002 | 395.58306 |
| -5.8867683 | 3.94E-09 | 4.24E-08 | 187.298653 | 469.509702 | 520.005206 | 514.370041 |
| -5.8866409 | 3.94E-09 | 4.24E-08 | 229.512573 | 442.178013 | 380.800272 | 440.408336 |
| -5.8864464 | 3.95E-09 | 4.24E-08 | 217.6271 | 332.580648 | 429.11934 | 543.50647 |
| -5.8857337 | 3.96E-09 | 4.26E-08 | 152.052079 | 212.429462 | 203.630357 | 195.550266 |
| -5.8857113 | 3.96E-09 | 4.26E-08 | 189.757716 | 506.853989 | 493.544764 | 448.252759 |
| -5.8855289 | 3.97E-09 | 4.26E-08 | 169.675366 | 729.566661 | 595.93517 | 739.617052 |
| -5.8853237 | 3.97E-09 | 4.26E-08 | 206.151472 | 375.878372 | 362.393009 | 401.746535 |
| -5.8853087 | 3.97E-09 | 4.26E-08 | 372.548087 | 271.422611 | 274.958505 | 254.383441 |

| | | | | | | |
|---|---|---|---|---|---|---|
| -5.88491 | 3.98E-09 | 4.27E-08 | 225.00429 | 474.380696 | 448.677058 | 481.3114 |
| -5.8846115 | 3.99E-09 | 4.28E-08 | 204.512096 | 611.850972 | 388.85345 | 458.898762 |
| -5.8845938 | 3.99E-09 | 4.28E-08 | 193.856155 | 448.402061 | 553.368372 | 436.486124 |
| -5.8842231 | 4.00E-09 | 4.29E-08 | 188.118341 | 559.08187 | 457.88069 | 499.801826 |
| -5.8840809 | 4.00E-09 | 4.29E-08 | 209.020379 | 378.313869 | 365.844371 | 346.835572 |
| -5.8840256 | 4.00E-09 | 4.29E-08 | 270.087117 | 297.671857 | 372.747095 | 242.616806 |
| -5.8839336 | 4.01E-09 | 4.29E-08 | 226.233822 | 397.256624 | 396.906628 | 409.030642 |
| -5.8832982 | 4.02E-09 | 4.30E-08 | 180.741151 | 528.773462 | 553.368372 | 545.187418 |
| -5.8831903 | 4.02E-09 | 4.31E-08 | 235.250387 | 320.132552 | 269.206235 | 305.932508 |
| -5.8830409 | 4.03E-09 | 4.31E-08 | 194.675843 | 447.860839 | 442.924788 | 420.797277 |
| -5.8828563 | 4.03E-09 | 4.31E-08 | 208.610535 | 354.22951 | 355.490285 | 405.108431 |
| -5.8827039 | 4.04E-09 | 4.31E-08 | 236.479919 | 342.863857 | 277.259413 | 289.683345 |
| -5.8823838 | 4.04E-09 | 4.32E-08 | 171.314742 | 647.571594 | 637.351513 | 609.063436 |
| -5.8822967 | 4.05E-09 | 4.32E-08 | 170.904898 | 711.706349 | 610.891072 | 638.199865 |
| -5.8820838 | 4.05E-09 | 4.33E-08 | 233.611012 | 393.468073 | 270.356689 | 380.454529 |
| 5.88200972 | 4.05E-09 | 4.33E-08 | 35.2465737 | 20.83703 | 12.6549939 | 21.852322 |
| -5.8817596 | 4.06E-09 | 4.33E-08 | 838.950422 | 3528.49395 | 3169.50076 | 3372.54169 |
| -5.8816718 | 4.06E-09 | 4.33E-08 | 230.332261 | 285.494372 | 293.365769 | 318.259459 |
| -5.8815228 | 4.07E-09 | 4.33E-08 | 195.085687 | 592.637606 | 406.11026 | 516.050989 |
| -5.8813331 | 4.07E-09 | 4.34E-08 | 214.758193 | 363.971498 | 319.826211 | 342.913361 |
| -5.8809371 | 4.08E-09 | 4.35E-08 | 202.053033 | 634.582277 | 404.959806 | 456.657498 |
| -5.8803064 | 4.10E-09 | 4.36E-08 | 215.168037 | 310.119953 | 375.048002 | 389.9799 |
| -5.8801612 | 4.10E-09 | 4.36E-08 | 220.905851 | 343.9463 | 296.817131 | 332.827673 |
| -5.8801891 | 4.10E-09 | 4.36E-08 | 218.446788 | 452.461223 | 354.339831 | 321.621355 |
| -5.8800737 | 4.10E-09 | 4.36E-08 | 205.741628 | 368.03066 | 372.747095 | 367.006946 |
| -5.8797332 | 4.11E-09 | 4.37E-08 | 238.938982 | 386.702803 | 424.517524 | 265.029444 |
| -5.8797724 | 4.11E-09 | 4.37E-08 | 164.75724 | 726.589942 | 696.024667 | 667.896611 |
| -5.8797307 | 4.11E-09 | 4.37E-08 | 145.084733 | 952.549943 | 949.124546 | 941.89111 |
| -5.8793424 | 4.12E-09 | 4.38E-08 | 210.24991 | 399.962732 | 326.728935 | 384.376741 |
| -5.878204 | 4.15E-09 | 4.41E-08 | 192.21678 | 551.234157 | 485.491586 | 517.731937 |
| -5.8781409 | 4.15E-09 | 4.41E-08 | 284.021809 | 425.400145 | 242.745793 | 300.329348 |
| -5.8780301 | 4.15E-09 | 4.41E-08 | 181.560839 | 486.28757 | 560.271096 | 573.763531 |
| -5.8779702 | 4.15E-09 | 4.41E-08 | 168.855679 | 644.324265 | 616.643342 | 692.550512 |
| -5.8779271 | 4.15E-09 | 4.41E-08 | 228.692885 | 279.811546 | 330.180297 | 304.811876 |
| -5.8778579 | 4.16E-09 | 4.41E-08 | 502.468597 | 211.617629 | 444.075242 | 329.465778 |
| -5.8775574 | 4.16E-09 | 4.41E-08 | 213.118818 | 415.116935 | 327.879389 | 349.076836 |
| -5.8774451 | 4.17E-09 | 4.42E-08 | 236.070075 | 277.91727 | 279.560321 | 290.803977 |
| -5.8769082 | 4.18E-09 | 4.43E-08 | 381.564652 | 433.247857 | 196.727633 | 430.322649 |
| -5.8765567 | 4.19E-09 | 4.44E-08 | 195.495531 | 424.317702 | 425.667978 | 409.030642 |
| -5.8765127 | 4.19E-09 | 4.44E-08 | 150.00286 | 882.19114 | 858.238681 | 859.524665 |
| -5.8763929 | 4.19E-09 | 4.44E-08 | 349.59683 | 565.576528 | 284.162137 | 322.18167 |
| -5.8762219 | 4.20E-09 | 4.44E-08 | 186.888809 | 583.978061 | 525.757476 | 443.770231 |
| -5.8761005 | 4.20E-09 | 4.44E-08 | 0 | 325.544767 | 430.269794 | 0 |

| | | | | | | |
|---|---|---|---|---|---|---|
| -5.8759195 | 4.21E-09 | 4.45E-08 | 178.691932 | 553.399043 | 546.465648 | 531.179519 |
| -5.8756877 | 4.21E-09 | 4.45E-08 | 232.38148 | 316.885222 | 266.905327 | 331.707042 |
| -5.8754591 | 4.22E-09 | 4.46E-08 | 167.626147 | 727.942996 | 753.547367 | 588.331746 |
| -5.8752976 | 4.22E-09 | 4.46E-08 | 248.365391 | 433.518468 | 271.507143 | 309.85472 |
| -5.8752853 | 4.22E-09 | 4.46E-08 | 172.544273 | 587.225391 | 629.298335 | 589.452378 |
| -5.874772 | 4.23E-09 | 4.47E-08 | 182.380527 | 552.3166 | 496.996126 | 500.922458 |
| -5.8740983 | 4.25E-09 | 4.49E-08 | 171.314742 | 655.960529 | 563.722458 | 656.690292 |
| -5.8739756 | 4.25E-09 | 4.49E-08 | 204.92194 | 361.536001 | 356.640739 | 392.78148 |
| -5.8734138 | 4.27E-09 | 4.50E-08 | 246.726016 | 353.417678 | 301.418947 | 252.702493 |
| -5.8733412 | 4.27E-09 | 4.50E-08 | 216.807412 | 333.663091 | 561.42155 | 410.71159 |
| -5.8730685 | 4.28E-09 | 4.51E-08 | 179.101776 | 618.886852 | 537.262016 | 510.447829 |
| -5.872794 | 4.29E-09 | 4.52E-08 | 212.708974 | 324.191713 | 340.534383 | 335.068937 |
| -5.8727048 | 4.29E-09 | 4.52E-08 | 267.21821 | 270.610779 | 250.798971 | 259.426284 |
| -5.8725253 | 4.29E-09 | 4.52E-08 | 234.840543 | 337.181031 | 318.675757 | 261.107232 |
| -5.8724347 | 4.29E-09 | 4.52E-08 | 234.840543 | 409.70472 | 264.604419 | 389.419584 |
| 5.871995 | 4.31E-09 | 4.53E-08 | 24.5906328 | 35.4500121 | 47.1686138 | 37.5411686 |
| -5.8714544 | 4.32E-09 | 4.54E-08 | 190.16756 | 553.399043 | 493.544764 | 415.754434 |
| 5.87122774 | 4.33E-09 | 4.55E-08 | 0 | 0 | 0 | 0 |
| -5.8711541 | 4.33E-09 | 4.55E-08 | 220.905851 | 423.235259 | 304.870309 | 513.809725 |
| -5.869985 | 4.36E-09 | 4.58E-08 | 236.479919 | 286.306204 | 303.719855 | 262.78818 |
| -5.869787 | 4.36E-09 | 4.58E-08 | 192.626624 | 424.317702 | 411.86253 | 470.665397 |
| -5.8695041 | 4.37E-09 | 4.59E-08 | 218.446788 | 359.371115 | 291.064861 | 379.894213 |
| -5.8695173 | 4.37E-09 | 4.59E-08 | 218.856632 | 318.779498 | 296.817131 | 350.757784 |
| -5.8690874 | 4.38E-09 | 4.60E-08 | 186.069121 | 443.801678 | 488.942948 | 497.000246 |
| -5.8690623 | 4.38E-09 | 4.60E-08 | 192.626624 | 467.074205 | 403.809352 | 434.805176 |
| -5.8685681 | 4.40E-09 | 4.61E-08 | 204.92194 | 382.373031 | 364.693917 | 343.473677 |
| -5.8676178 | 4.42E-09 | 4.64E-08 | 246.316172 | 382.373031 | 249.648517 | 323.862618 |
| -5.8674136 | 4.43E-09 | 4.64E-08 | 222.135383 | 325.544767 | 285.312591 | 312.095983 |
| -5.8674401 | 4.43E-09 | 4.64E-08 | 206.971159 | 339.616528 | 473.987046 | 385.497373 |
| -5.8672944 | 4.43E-09 | 4.64E-08 | 280.333214 | 353.417678 | 226.639437 | 298.648401 |
| -5.866559 | 4.45E-09 | 4.66E-08 | 218.446788 | 471.674588 | 314.073941 | 356.360943 |
| -5.8661333 | 4.46E-09 | 4.67E-08 | 277.874151 | 388.055857 | 240.444885 | 281.838922 |
| -5.8656455 | 4.47E-09 | 4.68E-08 | 223.364915 | 199.981366 | 148.408565 | 183.223315 |
| -5.8650664 | 4.49E-09 | 4.70E-08 | 284.841497 | 335.557366 | 257.701695 | 240.375542 |
| -5.8639002 | 4.52E-09 | 4.73E-08 | 213.118818 | 360.724169 | 311.773033 | 323.862618 |
| -5.8637489 | 4.53E-09 | 4.73E-08 | 201.233345 | 451.108169 | 350.888469 | 395.58306 |
| -5.863464 | 4.53E-09 | 4.74E-08 | 216.397569 | 405.374947 | 292.215315 | 387.17832 |
| -5.8632107 | 4.54E-09 | 4.74E-08 | 184.019902 | 546.904385 | 479.739316 | 452.735286 |
| -5.8632129 | 4.54E-09 | 4.74E-08 | 181.560839 | 478.710468 | 530.359292 | 485.233612 |
| -5.863105 | 4.54E-09 | 4.74E-08 | 188.528185 | 542.845223 | 456.730236 | 424.159173 |
| -5.8630282 | 4.55E-09 | 4.74E-08 | 161.068645 | 329.87454 | 294.516223 | 271.753235 |
| -5.8629207 | 4.55E-09 | 4.75E-08 | 194.675843 | 446.237175 | 378.499364 | 423.598857 |
| -5.8624958 | 4.56E-09 | 4.76E-08 | 226.233822 | 341.510803 | 277.259413 | 297.527769 |

| | | | | | | |
|---|---|---|---|---|---|---|
| -5.8613829 | 4.59E-09 | 4.79E-08 | 204.512096 | 359.641725 | 356.640739 | 340.672097 |
| -5.8613113 | 4.59E-09 | 4.79E-08 | 211.069598 | 341.510803 | 315.224395 | 329.465778 |
| -5.8612049 | 4.60E-09 | 4.79E-08 | 232.38148 | 351.794013 | 260.002603 | 314.897563 |
| -5.8611748 | 4.60E-09 | 4.79E-08 | 181.970683 | 490.346732 | 467.084322 | 505.965302 |
| -5.8605495 | 4.61E-09 | 4.80E-08 | 208.610535 | 339.887139 | 325.578481 | 338.430833 |
| -5.8600269 | 4.63E-09 | 4.82E-08 | 189.757716 | 453.814277 | 409.561622 | 434.24486 |
| -5.8594765 | 4.64E-09 | 4.83E-08 | 212.29913 | 316.344001 | 331.330751 | 315.457879 |
| -5.8595231 | 4.64E-09 | 4.83E-08 | 194.265999 | 386.973414 | 422.216616 | 398.384639 |
| -5.8589893 | 4.66E-09 | 4.84E-08 | 162.298176 | 703.588026 | 670.714679 | 633.717338 |
| -5.8589187 | 4.66E-09 | 4.84E-08 | 210.659754 | 572.883019 | 325.578481 | 487.474875 |
| 5.85880463 | 4.66E-09 | 4.85E-08 | 114.756286 | 50.8748265 | 37.9649818 | 33.6189569 |
| -5.8583731 | 4.67E-09 | 4.86E-08 | 188.938029 | 565.305918 | 398.057082 | 568.160372 |
| -5.8581049 | 4.68E-09 | 4.86E-08 | 154.101299 | 739.84987 | 766.202361 | 762.590006 |
| -5.857878 | 4.69E-09 | 4.87E-08 | 179.511619 | 478.439858 | 499.297034 | 531.179519 |
| -5.8578796 | 4.69E-09 | 4.87E-08 | 503.288285 | 872.719763 | 887.00003 | 821.983497 |
| -5.8576153 | 4.70E-09 | 4.87E-08 | 195.085687 | 407.269223 | 384.251634 | 388.298952 |
| -5.8573154 | 4.70E-09 | 4.88E-08 | 318.038851 | 300.107354 | 217.435805 | 291.364293 |
| -5.8571102 | 4.71E-09 | 4.88E-08 | 181.150995 | 475.73375 | 524.607022 | 470.105081 |
| -5.8571633 | 4.71E-09 | 4.88E-08 | 193.036467 | 420.529151 | 386.552542 | 411.271906 |
| -5.8568204 | 4.72E-09 | 4.89E-08 | 162.298176 | 665.973127 | 641.953329 | 667.336295 |
| -5.8566965 | 4.72E-09 | 4.89E-08 | 201.643189 | 413.22266 | 332.481205 | 412.392538 |
| -5.8562263 | 4.74E-09 | 4.90E-08 | 147.953641 | 338.804695 | 248.498063 | 274.554815 |
| -5.8562092 | 4.74E-09 | 4.90E-08 | 146.724109 | 837.540361 | 812.220521 | 906.030889 |
| -5.8559039 | 4.74E-09 | 4.91E-08 | 305.333691 | 242.196647 | 249.648517 | 274.554815 |
| -5.8557852 | 4.75E-09 | 4.91E-08 | 175.003337 | 582.895618 | 525.757476 | 513.249409 |
| -5.8555286 | 4.75E-09 | 4.92E-08 | 190.577404 | 594.802493 | 400.35799 | 483.552664 |
| -5.8549687 | 4.77E-09 | 4.93E-08 | 197.54475 | 347.734851 | 432.570702 | 427.521069 |
| -5.8548489 | 4.77E-09 | 4.94E-08 | 314.350256 | 486.558181 | 208.232173 | 421.917909 |
| -5.8547746 | 4.78E-09 | 4.94E-08 | 218.036944 | 297.130636 | 293.365769 | 316.578511 |
| -5.8543836 | 4.79E-09 | 4.95E-08 | 170.904898 | 623.487235 | 556.819734 | 546.868366 |
| -5.8539321 | 4.80E-09 | 4.96E-08 | 186.888809 | 534.99751 | 516.553844 | 486.914559 |
| -5.8538006 | 4.80E-09 | 4.96E-08 | 214.348349 | 295.23636 | 320.976665 | 319.940407 |
| -5.8537393 | 4.81E-09 | 4.96E-08 | 218.446788 | 379.125702 | 286.463045 | 322.741986 |
| -5.8530748 | 4.83E-09 | 4.98E-08 | 205.331784 | 483.310852 | 427.968886 | 322.741986 |
| -5.8530955 | 4.83E-09 | 4.98E-08 | 190.16756 | 529.855906 | 392.304812 | 453.295602 |
| -5.8525091 | 4.84E-09 | 4.99E-08 | 299.186032 | 238.949318 | 253.099879 | 364.765683 |
| -5.8522624 | 4.85E-09 | 5.00E-08 | 162.70802 | 647.571594 | 838.680963 | 619.149123 |
| -5.8521475 | 4.85E-09 | 5.00E-08 | 201.643189 | 548.528049 | 343.985745 | 456.657498 |
| -5.8516909 | 4.87E-09 | 5.01E-08 | 195.495531 | 383.184863 | 371.596641 | 389.9799 |
| -5.8502392 | 4.91E-09 | 5.05E-08 | 195.905375 | 717.389176 | 385.402088 | 560.315949 |
| -5.8502811 | 4.91E-09 | 5.05E-08 | 186.478965 | 454.084887 | 408.411168 | 457.217814 |
| -5.8502374 | 4.91E-09 | 5.05E-08 | 284.431653 | 1136.83588 | 1041.16087 | 1013.61155 |
| -5.8497279 | 4.92E-09 | 5.07E-08 | 186.069121 | 548.79866 | 457.88069 | 416.31475 |

| | | | | | | |
|---|---|---|---|---|---|---|
| -5.849631 | 4.93E-09 | 5.07E-08 | 222.955071 | 373.713486 | 337.083021 | 353.559364 |
| -5.8489776 | 4.95E-09 | 5.08E-08 | 420.089977 | 282.517653 | 227.789891 | 306.492824 |
| -5.8489805 | 4.95E-09 | 5.08E-08 | 248.365391 | 700.881918 | 278.409867 | 500.362142 |
| -5.8478229 | 4.98E-09 | 5.12E-08 | 169.675366 | 567.200193 | 538.41247 | 590.57301 |
| -5.8477235 | 4.98E-09 | 5.12E-08 | 232.791324 | 321.214995 | 366.994825 | 252.702493 |
| -5.8475912 | 4.99E-09 | 5.12E-08 | 2627.91896 | 5108.8609 | 3429.50336 | 4170.43161 |
| -5.8468704 | 5.01E-09 | 5.14E-08 | 243.447265 | 264.657342 | 264.604419 | 252.142177 |
| -5.8464924 | 5.02E-09 | 5.15E-08 | 211.889286 | 371.819211 | 292.215315 | 335.068937 |
| -5.8462882 | 5.03E-09 | 5.16E-08 | 180.331307 | 458.41466 | 461.332052 | 495.319299 |
| -5.8461588 | 5.03E-09 | 5.16E-08 | 281.562746 | 529.855906 | 224.338529 | 449.933707 |
| -5.8458434 | 5.04E-09 | 5.17E-08 | 1063.54487 | 294.965749 | 291.064861 | 529.498571 |
| -5.845508 | 5.05E-09 | 5.17E-08 | 217.217256 | 634.311666 | 318.675757 | 459.459078 |
| -5.8454614 | 5.05E-09 | 5.17E-08 | 202.872721 | 369.113103 | 332.481205 | 333.948305 |
| -5.8455082 | 5.05E-09 | 5.17E-08 | 172.954117 | 548.79866 | 492.39431 | 591.133326 |
| -5.8455311 | 5.05E-09 | 5.17E-08 | 200.823501 | 332.039426 | 358.941647 | 363.084735 |
| -5.8452807 | 5.06E-09 | 5.18E-08 | 304.514003 | 624.299067 | 251.949425 | 335.629253 |
| -5.84524 | 5.06E-09 | 5.18E-08 | 168.035991 | 534.99751 | 699.476029 | 594.495221 |
| -5.8450815 | 5.06E-09 | 5.18E-08 | 462.71374 | 284.411929 | 255.400787 | 272.313551 |
| -5.8444225 | 5.08E-09 | 5.20E-08 | 209.430223 | 348.546684 | 295.666677 | 336.749885 |
| -5.8441502 | 5.09E-09 | 5.21E-08 | 259.431176 | 271.422611 | 239.294431 | 255.504073 |
| -5.8439059 | 5.10E-09 | 5.21E-08 | 277.054463 | 262.221845 | 238.143977 | 253.262809 |
| -5.8437257 | 5.10E-09 | 5.22E-08 | 205.331784 | 326.62721 | 318.675757 | 346.275256 |
| -5.843463 | 5.11E-09 | 5.22E-08 | 182.380527 | 481.145965 | 434.87161 | 444.890863 |
| -5.8427374 | 5.13E-09 | 5.24E-08 | 223.774758 | 328.792097 | 332.481205 | 258.865968 |
| -5.8426001 | 5.14E-09 | 5.25E-08 | 205.741628 | 496.841391 | 448.677058 | 523.895412 |
| -5.8425143 | 5.14E-09 | 5.25E-08 | 188.118341 | 400.503953 | 413.012984 | 412.952854 |
| -5.8417981 | 5.16E-09 | 5.27E-08 | 147.953641 | 284.68254 | 271.507143 | 295.846821 |
| -5.8417994 | 5.16E-09 | 5.27E-08 | 192.21678 | 428.106253 | 396.906628 | 364.205367 |
| -5.8415593 | 5.17E-09 | 5.27E-08 | 332.793231 | 308.225677 | 342.835291 | 205.635953 |
| -5.8414785 | 5.17E-09 | 5.27E-08 | 245.906328 | 274.128719 | 236.993523 | 291.924609 |
| -5.8413704 | 5.18E-09 | 5.27E-08 | 250.824455 | 261.410013 | 240.444885 | 268.951655 |
| -5.8408728 | 5.19E-09 | 5.29E-08 | 375.826838 | 256.809629 | 245.046701 | 272.313551 |
| -5.8397573 | 5.23E-09 | 5.32E-08 | 223.774758 | 383.455474 | 261.153057 | 396.703692 |
| -5.8397042 | 5.23E-09 | 5.32E-08 | 243.447265 | 299.295522 | 236.993523 | 279.037342 |
| -5.8395953 | 5.23E-09 | 5.32E-08 | 195.905375 | 362.618444 | 379.649818 | 351.3181 |
| -5.8394753 | 5.24E-09 | 5.33E-08 | 175.823024 | 573.694852 | 459.031144 | 514.370041 |
| -5.8393605 | 5.24E-09 | 5.33E-08 | 199.59397 | 338.804695 | 403.809352 | 343.473677 |
| -5.8393417 | 5.24E-09 | 5.33E-08 | 142.215826 | 219.194731 | 228.940345 | 208.997849 |
| -5.8381689 | 5.28E-09 | 5.36E-08 | 238.938982 | 336.098588 | 238.143977 | 307.05314 |
| -5.8381865 | 5.28E-09 | 5.36E-08 | 204.512096 | 419.176097 | 310.622579 | 359.722839 |
| -5.8380977 | 5.28E-09 | 5.36E-08 | 226.233822 | 340.698971 | 262.303511 | 287.442082 |
| -5.8373399 | 5.30E-09 | 5.39E-08 | 234.840543 | 385.079139 | 274.958505 | 271.753235 |
| -5.8371825 | 5.31E-09 | 5.39E-08 | 172.13443 | 429.188696 | 410.712076 | 365.325998 |

| | | | | | | |
|---|---|---|---|---|---|---|
| -5.8371646 | 5.31E-09 | 5.39E-08 | 203.692408 | 316.344001 | 326.728935 | 347.395888 |
| -5.8372381 | 5.31E-09 | 5.39E-08 | 256.972113 | 281.705821 | 288.763953 | 226.927959 |
| -5.8370544 | 5.31E-09 | 5.39E-08 | 157.38005 | 672.738397 | 1067.62131 | 691.429881 |
| -5.8360977 | 5.34E-09 | 5.41E-08 | 145.494577 | 771.511331 | 926.115466 | 772.115377 |
| -5.8361052 | 5.34E-09 | 5.41E-08 | 197.54475 | 402.127618 | 335.932567 | 358.041891 |
| -5.8361899 | 5.34E-09 | 5.41E-08 | 309.022285 | 241.114204 | 283.011683 | 237.573962 |
| -5.8357504 | 5.35E-09 | 5.42E-08 | 192.626624 | 371.5486 | 513.102482 | 397.824324 |
| -5.8351524 | 5.37E-09 | 5.44E-08 | 239.348826 | 259.515737 | 248.498063 | 270.632603 |
| -5.8350369 | 5.38E-09 | 5.44E-08 | 283.202121 | 411.057774 | 211.683535 | 437.04644 |
| -5.8346413 | 5.39E-09 | 5.45E-08 | 190.16756 | 400.233342 | 480.88977 | 372.04979 |
| -5.8345578 | 5.39E-09 | 5.46E-08 | 200.823501 | 348.817294 | 331.330751 | 328.345146 |
| -5.8343852 | 5.40E-09 | 5.46E-08 | 156.150518 | 701.964361 | 790.361895 | 704.317147 |
| -5.8340674 | 5.41E-09 | 5.47E-08 | 236.889763 | 339.616528 | 258.852149 | 262.227864 |
| -5.8339695 | 5.41E-09 | 5.47E-08 | 163.937552 | 682.750996 | 543.014286 | 609.063436 |
| -5.8338628 | 5.42E-09 | 5.47E-08 | 217.6271 | 424.317702 | 273.808051 | 363.645051 |
| -5.8339069 | 5.41E-09 | 5.47E-08 | 204.92194 | 376.419594 | 304.870309 | 332.827673 |
| -5.8330054 | 5.44E-09 | 5.50E-08 | 186.888809 | 493.594061 | 384.251634 | 420.236961 |
| -5.8329288 | 5.45E-09 | 5.50E-08 | 210.659754 | 319.050109 | 283.011683 | 327.78483 |
| -5.8327509 | 5.45E-09 | 5.50E-08 | 193.446311 | 416.199378 | 342.835291 | 389.419584 |
| -5.832717 | 5.45E-09 | 5.50E-08 | 337.301513 | 417.552432 | 347.437107 | 410.151274 |
| -5.8319915 | 5.48E-09 | 5.52E-08 | 222.955071 | 286.576815 | 271.507143 | 268.391339 |
| -5.8314984 | 5.49E-09 | 5.54E-08 | 178.282088 | 468.427259 | 448.677058 | 597.857117 |
| -5.8313955 | 5.50E-09 | 5.54E-08 | 163.937552 | 704.670469 | 689.121943 | 518.292252 |
| -5.8309454 | 5.51E-09 | 5.55E-08 | 172.954117 | 580.730732 | 486.64204 | 486.354243 |
| -5.8308491 | 5.51E-09 | 5.55E-08 | 221.315695 | 306.060791 | 266.905327 | 374.291054 |
| 5.83083817 | 5.51E-09 | 5.55E-08 | 4.50828268 | 9.20076649 | 2.30090799 | 6.72379138 |
| -5.8306554 | 5.52E-09 | 5.56E-08 | 201.233345 | 399.42151 | 339.383929 | 318.819775 |
| -5.8306309 | 5.52E-09 | 5.56E-08 | 186.888809 | 668.408624 | 407.260714 | 482.992348 |
| -5.8303389 | 5.53E-09 | 5.56E-08 | 245.906328 | 273.858108 | 241.595339 | 252.702493 |
| -5.830037 | 5.54E-09 | 5.57E-08 | 211.889286 | 290.635977 | 339.383929 | 290.803977 |
| -5.8299106 | 5.55E-09 | 5.57E-08 | 214.758193 | 376.960815 | 277.259413 | 316.578511 |
| 5.82962976 | 5.56E-09 | 5.58E-08 | 0 | 0 | 0 | 0 |
| -5.8295698 | 5.56E-09 | 5.58E-08 | 227.053509 | 260.056959 | 264.604419 | 300.889664 |
| -5.8288391 | 5.58E-09 | 5.60E-08 | 166.396615 | 593.178828 | 552.217918 | 523.895412 |
| -5.8287806 | 5.58E-09 | 5.60E-08 | 278.693838 | 453.543666 | 391.154358 | 451.054339 |
| -5.8287479 | 5.58E-09 | 5.60E-08 | 201.643189 | 337.451642 | 308.321671 | 355.240311 |
| -5.8282638 | 5.60E-09 | 5.62E-08 | 234.430699 | 250.04436 | 266.905327 | 262.227864 |
| -5.8279171 | 5.61E-09 | 5.63E-08 | 222.545227 | 396.986013 | 264.604419 | 319.380091 |
| -5.8278436 | 5.61E-09 | 5.63E-08 | 240.988201 | 230.830995 | 306.020763 | 309.85472 |
| -5.8276791 | 5.62E-09 | 5.63E-08 | 184.019902 | 433.789079 | 401.508444 | 405.108431 |
| -5.8266989 | 5.65E-09 | 5.67E-08 | 181.970683 | 453.002444 | 436.022064 | 399.505271 |
| -5.8262382 | 5.67E-09 | 5.68E-08 | 178.282088 | 470.862756 | 431.420248 | 447.132127 |
| -5.8260708 | 5.67E-09 | 5.68E-08 | 211.889286 | 318.779498 | 272.657597 | 333.948305 |

| | | | | | | |
|---|---|---|---|---|---|---|
| -5.8260114 | 5.68E-09 | 5.68E-08 | 179.921463 | 408.351666 | 465.933868 | 456.657498 |
| -5.8251483 | 5.71E-09 | 5.71E-08 | 181.970683 | 502.524217 | 410.712076 | 418.556014 |
| -5.8242106 | 5.74E-09 | 5.74E-08 | 147.543797 | 844.03502 | 715.582385 | 714.96315 |
| -5.8238673 | 5.75E-09 | 5.75E-08 | 235.250387 | 487.099402 | 261.153057 | 346.275256 |
| -5.8237525 | 5.75E-09 | 5.75E-08 | 169.675366 | 481.957798 | 528.058384 | 533.420783 |
| -5.8237561 | 5.75E-09 | 5.75E-08 | 223.774758 | 1167.4149 | 648.856053 | 824.785076 |
| -5.8224869 | 5.80E-09 | 5.79E-08 | 204.92194 | 477.086804 | 327.879389 | 336.189569 |
| -5.8224804 | 5.80E-09 | 5.79E-08 | 158.609582 | 573.694852 | 618.94425 | 713.282203 |
| -5.8223943 | 5.80E-09 | 5.79E-08 | 224.594446 | 489.80551 | 269.206235 | 407.910011 |
| -5.8224134 | 5.80E-09 | 5.79E-08 | 159.839113 | 622.675403 | 580.979268 | 593.934906 |
| -5.8220965 | 5.81E-09 | 5.80E-08 | 189.347873 | 434.05969 | 346.286653 | 407.910011 |
| -5.8204905 | 5.87E-09 | 5.85E-08 | 181.150995 | 422.694037 | 431.420248 | 402.306851 |
| -5.8199879 | 5.89E-09 | 5.86E-08 | 319.268382 | 230.019162 | 280.710775 | 243.177122 |
| -5.820006 | 5.88E-09 | 5.86E-08 | 190.987248 | 534.456289 | 390.003904 | 372.04979 |
| -5.8200692 | 5.88E-09 | 5.86E-08 | 205.741628 | 320.132552 | 287.613499 | 317.699143 |
| -5.820224 | 5.88E-09 | 5.86E-08 | 192.626624 | 360.182947 | 384.251634 | 517.731937 |
| -5.8200919 | 5.88E-09 | 5.86E-08 | 202.053033 | 377.772648 | 381.950726 | 292.484925 |
| -5.8199886 | 5.89E-09 | 5.86E-08 | 296.317125 | 933.607188 | 815.671883 | 940.210162 |
| -5.8197704 | 5.89E-09 | 5.87E-08 | 170.495054 | 595.884936 | 459.031144 | 529.498571 |
| -5.8196529 | 5.90E-09 | 5.87E-08 | 180.331307 | 458.41466 | 391.154358 | 465.622553 |
| -5.819208 | 5.91E-09 | 5.88E-08 | 211.069598 | 309.849342 | 334.782113 | 270.632603 |
| -5.8191504 | 5.91E-09 | 5.88E-08 | 201.233345 | 311.743618 | 310.622579 | 337.310201 |
| 5.81893177 | 5.92E-09 | 5.89E-08 | 42.2139196 | 31.3908504 | 28.7613499 | 29.1364293 |
| -5.8187712 | 5.93E-09 | 5.89E-08 | 199.59397 | 843.223188 | 379.649818 | 537.903311 |
| -5.8186455 | 5.93E-09 | 5.89E-08 | 196.315218 | 380.208145 | 311.773033 | 402.867167 |
| -5.8186413 | 5.93E-09 | 5.89E-08 | 189.757716 | 394.279905 | 340.534383 | 411.271906 |
| -5.818406 | 5.94E-09 | 5.90E-08 | 189.757716 | 529.314684 | 343.985745 | 494.758983 |
| -5.81823 | 5.95E-09 | 5.90E-08 | 237.299606 | 257.350851 | 239.294431 | 289.683345 |
| -5.8179665 | 5.96E-09 | 5.91E-08 | 210.24991 | 536.891786 | 496.996126 | 523.335096 |
| 5.81781795 | 5.96E-09 | 5.92E-08 | 22.5414134 | 13.8011497 | 10.354086 | 14.5682147 |
| -5.8174444 | 5.98E-09 | 5.93E-08 | 188.528185 | 383.184863 | 362.393009 | 365.886314 |
| -5.8163112 | 6.02E-09 | 5.97E-08 | 251.644142 | 309.578731 | 222.037621 | 275.675447 |
| -5.816135 | 6.02E-09 | 5.97E-08 | 173.363961 | 562.058588 | 773.105085 | 460.57971 |
| -5.8158157 | 6.03E-09 | 5.98E-08 | 515.173757 | 1092.45572 | 918.062288 | 1009.68934 |
| -5.8156466 | 6.04E-09 | 5.98E-08 | 241.807889 | 328.250875 | 227.789891 | 373.730738 |
| -5.8156739 | 6.04E-09 | 5.98E-08 | 197.134906 | 306.602013 | 399.207536 | 368.127578 |
| -5.8153636 | 6.05E-09 | 5.99E-08 | 170.08521 | 756.086517 | 570.625182 | 473.466977 |
| -5.8153158 | 6.05E-09 | 5.99E-08 | 0 | 0 | 799.565527 | 193.869318 |
| -5.8153094 | 6.05E-09 | 5.99E-08 | 146.314265 | 798.031188 | 866.291859 | 652.76808 |
| -5.8153132 | 6.05E-09 | 5.99E-08 | 259.021332 | 215.135569 | 308.321671 | 301.44998 |
| -5.8152257 | 6.06E-09 | 5.99E-08 | 193.036467 | 326.897821 | 409.561622 | 362.524419 |
| -5.8149649 | 6.06E-09 | 6.00E-08 | 199.184126 | 301.731019 | 334.782113 | 349.637152 |
| -5.8144593 | 6.08E-09 | 6.01E-08 | 181.970683 | 436.765798 | 373.897549 | 454.416234 |

| | | | | | | |
|---|---|---|---|---|---|---|
| -5.8142653 | 6.09E-09 | 6.02E-08 | 228.692885 | 343.67569 | 251.949425 | 271.192919 |
| -5.8140744 | 6.10E-09 | 6.02E-08 | 234.020855 | 399.692121 | 238.143977 | 336.749885 |
| -5.813644 | 6.11E-09 | 6.03E-08 | 192.626624 | 465.721151 | 546.465648 | 333.948305 |
| -5.8136113 | 6.11E-09 | 6.03E-08 | 192.21678 | 366.136384 | 330.180297 | 357.481575 |
| -5.8132033 | 6.13E-09 | 6.05E-08 | 145.494577 | 885.70908 | 781.158263 | 663.414083 |
| -5.8132027 | 6.13E-09 | 6.05E-08 | 179.511619 | 401.045175 | 438.322972 | 416.875066 |
| -5.8130602 | 6.13E-09 | 6.05E-08 | 189.347873 | 340.698971 | 434.87161 | 380.454529 |
| -5.8127765 | 6.14E-09 | 6.06E-08 | 180.741151 | 423.77648 | 399.207536 | 402.867167 |
| -5.8128048 | 6.14E-09 | 6.06E-08 | 177.872244 | 596.155546 | 398.057082 | 518.852568 |
| -5.8125778 | 6.15E-09 | 6.06E-08 | 164.347396 | 494.135283 | 650.006507 | 802.372438 |
| -5.8125403 | 6.15E-09 | 6.06E-08 | 238.938982 | 400.233342 | 235.843069 | 318.819775 |
| -5.8120296 | 6.17E-09 | 6.08E-08 | 229.512573 | 250.856192 | 356.640739 | 274.554815 |
| -5.8118321 | 6.18E-09 | 6.08E-08 | 350.416517 | 195.110372 | 346.286653 | 316.578511 |
| -5.8118533 | 6.18E-09 | 6.08E-08 | 201.233345 | 433.518468 | 318.675757 | 324.422934 |
| -5.8114039 | 6.20E-09 | 6.09E-08 | 606.978786 | 2626.27761 | 2663.301 | 2894.59219 |
| -5.8112687 | 6.20E-09 | 6.10E-08 | 181.970683 | 388.055857 | 395.756174 | 468.984449 |
| -5.8111189 | 6.21E-09 | 6.10E-08 | 163.527708 | 553.128433 | 506.199758 | 571.522268 |
| -5.8097488 | 6.26E-09 | 6.15E-08 | 218.036944 | 454.084887 | 261.153057 | 444.890863 |
| -5.8096891 | 6.26E-09 | 6.15E-08 | 175.003337 | 592.637606 | 518.854752 | 531.739835 |
| -5.8092881 | 6.27E-09 | 6.16E-08 | 188.528185 | 405.374947 | 333.631659 | 393.902112 |
| -5.8091274 | 6.28E-09 | 6.16E-08 | 194.265999 | 383.726085 | 315.224395 | 347.395888 |
| -5.8091581 | 6.28E-09 | 6.16E-08 | 209.430223 | 266.010396 | 315.224395 | 308.173772 |
| -5.8090761 | 6.28E-09 | 6.16E-08 | 207.790847 | 337.992863 | 278.409867 | 294.165873 |
| -5.8085718 | 6.30E-09 | 6.18E-08 | 193.856155 | 368.571881 | 320.976665 | 442.649599 |
| -5.8081976 | 6.31E-09 | 6.19E-08 | 278.283994 | 327.439043 | 208.232173 | 281.838922 |
| -5.8080503 | 6.32E-09 | 6.20E-08 | 190.577404 | 422.964648 | 331.330751 | 366.44663 |
| -5.8078754 | 6.33E-09 | 6.20E-08 | 223.364915 | 368.03066 | 245.046701 | 399.505271 |
| -5.8076692 | 6.33E-09 | 6.21E-08 | 192.21678 | 340.42836 | 337.083021 | 418.556014 |
| -5.8071897 | 6.35E-09 | 6.22E-08 | 196.725062 | 464.368097 | 323.277573 | 353.559364 |
| -5.8072375 | 6.35E-09 | 6.22E-08 | 247.13586 | 447.319618 | 243.896247 | 302.010296 |
| -5.8069185 | 6.36E-09 | 6.23E-08 | 222.545227 | 263.304288 | 347.437107 | 264.469128 |
| -5.8062731 | 6.39E-09 | 6.25E-08 | 191.397092 | 458.685271 | 330.180297 | 377.092633 |
| -5.8063488 | 6.38E-09 | 6.25E-08 | 200.003813 | 305.248959 | 306.020763 | 365.886314 |
| -5.8062991 | 6.39E-09 | 6.25E-08 | 181.150995 | 473.298253 | 442.924788 | 365.325998 |
| -5.8058847 | 6.40E-09 | 6.26E-08 | 193.446311 | 355.311953 | 312.923487 | 352.438732 |
| -5.8051662 | 6.43E-09 | 6.29E-08 | 172.954117 | 612.933415 | 446.37615 | 462.260658 |
| -5.8046695 | 6.45E-09 | 6.30E-08 | 179.921463 | 503.065438 | 396.906628 | 401.186219 |
| -5.8046064 | 6.45E-09 | 6.30E-08 | 181.560839 | 384.537917 | 379.649818 | 434.805176 |
| -5.8040686 | 6.47E-09 | 6.32E-08 | 180.741151 | 372.089821 | 441.774334 | 405.668747 |
| -5.8034287 | 6.50E-09 | 6.34E-08 | 177.052556 | 432.977247 | 495.845672 | 393.341796 |
| -5.8027878 | 6.52E-09 | 6.36E-08 | 410.253724 | 249.232528 | 278.409867 | 240.375542 |
| -5.8028028 | 6.52E-09 | 6.36E-08 | 234.430699 | 300.648576 | 247.347609 | 242.05649 |
| -5.802425 | 6.54E-09 | 6.37E-08 | 195.905375 | 372.089821 | 295.666677 | 376.532317 |

| | | | | | | |
|---|---|---|---|---|---|---|
| -5.8024017 | 6.54E-09 | 6.37E-08 | 184.429746 | 382.643642 | 368.145279 | 362.524419 |
| -5.8019462 | 6.55E-09 | 6.39E-08 | 194.675843 | 397.256624 | 306.020763 | 349.637152 |
| -5.8013709 | 6.58E-09 | 6.41E-08 | 257.381957 | 428.106253 | 256.551241 | 254.943757 |
| -5.8010183 | 6.59E-09 | 6.42E-08 | 140.576451 | 802.902182 | 797.264619 | 707.118727 |
| -5.8009112 | 6.60E-09 | 6.42E-08 | 291.808843 | 312.826061 | 278.409867 | 204.515321 |
| -5.8005904 | 6.61E-09 | 6.43E-08 | 319.678226 | 284.141318 | 208.232173 | 266.710392 |
| -5.7998857 | 6.64E-09 | 6.46E-08 | 246.316172 | 389.408911 | 218.586259 | 389.9799 |
| -5.7997433 | 6.64E-09 | 6.46E-08 | 201.233345 | 296.318803 | 301.418947 | 370.929158 |
| -5.799716 | 6.64E-09 | 6.46E-08 | 202.872721 | 442.448624 | 316.374849 | 347.395888 |
| -5.7995385 | 6.65E-09 | 6.47E-08 | 177.4624 | 415.387546 | 453.278874 | 386.057689 |
| -5.7993708 | 6.66E-09 | 6.47E-08 | 173.363961 | 462.744432 | 406.11026 | 466.743185 |
| -5.7991483 | 6.67E-09 | 6.48E-08 | 204.92194 | 348.817294 | 266.905327 | 345.154624 |
| -5.7990174 | 6.67E-09 | 6.48E-08 | 168.855679 | 448.402061 | 468.234776 | 499.24151 |
| 5.79873307 | 6.68E-09 | 6.49E-08 | 56.9682993 | 27.6022995 | 33.3631659 | 25.7745336 |
| -5.7981084 | 6.71E-09 | 6.51E-08 | 197.954594 | 303.354683 | 303.719855 | 314.337247 |
| -5.7979744 | 6.71E-09 | 6.52E-08 | 190.16756 | 324.462324 | 363.543463 | 337.310201 |
| -5.7976786 | 6.72E-09 | 6.53E-08 | 146.314265 | 648.654038 | 689.121943 | 717.76473 |
| -5.7974871 | 6.73E-09 | 6.53E-08 | 214.758193 | 336.369198 | 246.197155 | 332.827673 |
| -5.7972564 | 6.74E-09 | 6.54E-08 | 321.727446 | 193.757318 | 345.136199 | 311.535667 |
| -5.797197 | 6.74E-09 | 6.54E-08 | 177.4624 | 411.328384 | 399.207536 | 398.944955 |
| -5.7970054 | 6.75E-09 | 6.54E-08 | 358.203551 | 294.424528 | 219.736713 | 246.539017 |
| -5.7969584 | 6.75E-09 | 6.54E-08 | 222.955071 | 345.299354 | 283.011683 | 244.297754 |
| -5.7967083 | 6.76E-09 | 6.55E-08 | 202.462877 | 338.263474 | 271.507143 | 331.707042 |
| -5.7967087 | 6.76E-09 | 6.55E-08 | 181.560839 | 392.656241 | 358.941647 | 398.384639 |
| -5.7966282 | 6.77E-09 | 6.55E-08 | 202.462877 | 295.777582 | 292.215315 | 287.442082 |
| -5.7963719 | 6.78E-09 | 6.56E-08 | 214.348349 | 268.987114 | 256.551241 | 305.932508 |
| -5.7960833 | 6.79E-09 | 6.56E-08 | 283.611965 | 372.360432 | 207.081719 | 284.080186 |
| -5.7961286 | 6.79E-09 | 6.56E-08 | 167.626147 | 453.273055 | 594.784716 | 482.432032 |
| -5.7961575 | 6.79E-09 | 6.56E-08 | 185.249434 | 357.476839 | 345.136199 | 393.341796 |
| 5.79522186 | 6.82E-09 | 6.60E-08 | 0 | 3.51794013 | 2.30090799 | 2.80157974 |
| -5.7949068 | 6.84E-09 | 6.61E-08 | 160.248957 | 487.911235 | 595.93517 | 573.203215 |
| -5.7944304 | 6.86E-09 | 6.62E-08 | 202.872721 | 326.897821 | 272.657597 | 304.811876 |
| -5.7940736 | 6.87E-09 | 6.64E-08 | 267.628054 | 589.660888 | 272.657597 | 291.364293 |
| -5.7929589 | 6.92E-09 | 6.68E-08 | 206.561315 | 382.10242 | 958.328178 | 500.362142 |
| -5.7924846 | 6.94E-09 | 6.69E-08 | 162.298176 | 503.065438 | 553.368372 | 489.716139 |
| -5.7916858 | 6.97E-09 | 6.72E-08 | 189.347873 | 398.339067 | 332.481205 | 328.345146 |
| -5.7914671 | 6.98E-09 | 6.73E-08 | 183.200214 | 389.1383 | 343.985745 | 372.04979 |
| -5.7912306 | 6.99E-09 | 6.74E-08 | 186.069121 | 367.760049 | 352.038923 | 333.387989 |
| -5.7902301 | 7.03E-09 | 6.77E-08 | 220.496007 | 316.07339 | 249.648517 | 257.745336 |
| -5.7902739 | 7.03E-09 | 6.77E-08 | 196.725062 | 348.276073 | 285.312591 | 324.98325 |
| -5.7897931 | 7.05E-09 | 6.79E-08 | 165.167084 | 517.40781 | 469.38523 | 467.303501 |
| -5.7893649 | 7.07E-09 | 6.80E-08 | 172.954117 | 635.935331 | 392.304812 | 645.483973 |
| -5.7893538 | 7.07E-09 | 6.80E-08 | 170.904898 | 450.296336 | 425.667978 | 430.322649 |

| | | | | | | |
|---|---|---|---|---|---|---|
| -5.7888409 | 7.09E-09 | 6.82E-08 | 192.626624 | 318.508887 | 307.171217 | 365.886314 |
| -5.7886943 | 7.09E-09 | 6.83E-08 | 225.414134 | 538.786061 | 430.269794 | 468.984449 |
| -5.7885114 | 7.10E-09 | 6.83E-08 | 183.200214 | 855.941894 | 377.34891 | 709.920307 |
| -5.7883861 | 7.11E-09 | 6.83E-08 | 208.200691 | 448.13145 | 410.712076 | 263.908812 |
| -5.7881857 | 7.12E-09 | 6.84E-08 | 186.478965 | 326.3566 | 347.437107 | 382.135477 |
| -5.788054 | 7.12E-09 | 6.84E-08 | 155.740674 | 561.246756 | 536.111562 | 589.452378 |
| -5.7878395 | 7.13E-09 | 6.85E-08 | 194.265999 | 380.749366 | 310.622579 | 308.173772 |
| -5.7878053 | 7.13E-09 | 6.85E-08 | 502.878441 | 332.580648 | 216.285351 | 278.477026 |
| -5.7876786 | 7.14E-09 | 6.85E-08 | 263.119771 | 212.429462 | 325.578481 | 423.598857 |
| -5.787628 | 7.14E-09 | 6.85E-08 | 220.496007 | 339.616528 | 707.529207 | 355.240311 |
| -5.7875045 | 7.14E-09 | 6.86E-08 | 140.986295 | 898.157176 | 689.121943 | 699.83462 |
| -5.7873561 | 7.15E-09 | 6.86E-08 | 143.855202 | 737.143762 | 651.156961 | 680.223562 |
| -5.7872753 | 7.15E-09 | 6.86E-08 | 158.199738 | 514.701702 | 576.377452 | 528.37794 |
| -5.7870666 | 7.16E-09 | 6.87E-08 | 211.479442 | 350.440959 | 293.365769 | 254.383441 |
| -5.7868774 | 7.17E-09 | 6.87E-08 | 193.036467 | 392.38563 | 295.666677 | 347.395888 |
| -5.7866502 | 7.18E-09 | 6.88E-08 | 337.711357 | 1051.8641 | 1043.46177 | 1047.79082 |
| -5.7856122 | 7.22E-09 | 6.92E-08 | 174.183649 | 387.785247 | 700.626483 | 635.398286 |
| -5.7854356 | 7.23E-09 | 6.93E-08 | 192.626624 | 390.761965 | 309.472125 | 318.819775 |
| -5.7853843 | 7.23E-09 | 6.93E-08 | 149.183172 | 554.481486 | 1035.4086 | 839.353291 |
| -5.7848707 | 7.26E-09 | 6.95E-08 | 213.118818 | 330.956983 | 277.259413 | 251.581861 |
| -5.7846849 | 7.26E-09 | 6.95E-08 | 261.890239 | 352.876456 | 223.188075 | 249.900913 |
| -5.7847049 | 7.26E-09 | 6.95E-08 | 269.677273 | 275.211162 | 212.833989 | 247.099333 |
| -5.7844824 | 7.27E-09 | 6.96E-08 | 145.494577 | 595.884936 | 865.141405 | 714.402834 |
| -5.7844319 | 7.28E-09 | 6.96E-08 | 204.512096 | 280.893989 | 265.754873 | 303.691244 |
| -5.7841761 | 7.29E-09 | 6.96E-08 | 163.527708 | 509.560097 | 507.350212 | 451.614655 |
| -5.7841567 | 7.29E-09 | 6.96E-08 | 182.380527 | 438.930684 | 356.640739 | 353.559364 |
| -5.784005 | 7.29E-09 | 6.97E-08 | 189.347873 | 376.419594 | 316.374849 | 324.422934 |
| -5.7839755 | 7.30E-09 | 6.97E-08 | 341.809796 | 320.673773 | 186.373547 | 334.508621 |
| -5.7838547 | 7.30E-09 | 6.97E-08 | 192.21678 | 350.982181 | 303.719855 | 312.656299 |
| -5.7838086 | 7.30E-09 | 6.97E-08 | 183.200214 | 403.210061 | 347.437107 | 345.154624 |
| -5.7838979 | 7.30E-09 | 6.97E-08 | 194.265999 | 303.084073 | 303.719855 | 309.294404 |
| -5.7836096 | 7.31E-09 | 6.97E-08 | 238.938982 | 243.27909 | 246.197155 | 231.410487 |
| -5.7836017 | 7.31E-09 | 6.97E-08 | 218.036944 | 378.855091 | 239.294431 | 331.707042 |
| -5.7833667 | 7.32E-09 | 6.98E-08 | 169.265522 | 457.061606 | 431.420248 | 425.279805 |
| -5.7831145 | 7.33E-09 | 6.99E-08 | 336.481825 | 225.418779 | 225.488983 | 325.543566 |
| -5.7830598 | 7.34E-09 | 6.99E-08 | 192.21678 | 342.863857 | 294.516223 | 327.78483 |
| -5.7829588 | 7.34E-09 | 6.99E-08 | 186.888809 | 343.67569 | 335.932567 | 323.862618 |
| -5.7828686 | 7.34E-09 | 6.99E-08 | 231.151948 | 228.666108 | 319.826211 | 390.540216 |
| -5.7824489 | 7.36E-09 | 7.01E-08 | 167.626147 | 649.46587 | 455.579782 | 463.941605 |
| -5.7821684 | 7.37E-09 | 7.02E-08 | 190.577404 | 310.661174 | 421.066162 | 338.430833 |
| -5.7816429 | 7.40E-09 | 7.04E-08 | 188.938029 | 403.751282 | 306.020763 | 354.11968 |
| -5.7810379 | 7.42E-09 | 7.06E-08 | 170.08521 | 449.484504 | 408.411168 | 431.44328 |
| -5.7809731 | 7.43E-09 | 7.06E-08 | 192.21678 | 296.589414 | 326.728935 | 316.578511 |

| | | | | | | |
|---|---|---|---|---|---|---|
| -5.7809104 | 7.43E-09 | 7.06E-08 | 327.875104 | 295.777582 | 379.649818 | 266.710392 |
| -5.7806585 | 7.44E-09 | 7.07E-08 | 370.908711 | 458.955881 | 543.014286 | 514.930357 |
| -5.7804468 | 7.45E-09 | 7.07E-08 | 172.13443 | 390.491354 | 434.87161 | 422.478225 |
| -5.78047 | 7.45E-09 | 7.07E-08 | 223.774758 | 228.666108 | 284.162137 | 268.951655 |
| -5.7793653 | 7.50E-09 | 7.12E-08 | 163.937552 | 527.96163 | 449.827512 | 467.863817 |
| -5.7793739 | 7.50E-09 | 7.12E-08 | 203.282564 | 633.229223 | 523.456568 | 568.160372 |
| -5.7788168 | 7.52E-09 | 7.14E-08 | 176.642712 | 358.018061 | 432.570702 | 396.143376 |
| -5.7783133 | 7.55E-09 | 7.16E-08 | 181.560839 | 391.573797 | 327.879389 | 393.341796 |
| -5.7779613 | 7.56E-09 | 7.17E-08 | 185.249434 | 382.914252 | 320.976665 | 341.232413 |
| -5.7777034 | 7.57E-09 | 7.18E-08 | 177.872244 | 434.600911 | 432.570702 | 447.692443 |
| -5.777006 | 7.60E-09 | 7.20E-08 | 150.412704 | 634.852888 | 635.050605 | 538.463627 |
| -5.776988 | 7.60E-09 | 7.20E-08 | 205.741628 | 260.056959 | 270.356689 | 287.442082 |
| -5.7769084 | 7.61E-09 | 7.21E-08 | 170.904898 | 452.190612 | 402.658898 | 407.349695 |
| -5.7767667 | 7.61E-09 | 7.21E-08 | 185.249434 | 418.093654 | 324.428027 | 346.275256 |
| -5.7756831 | 7.66E-09 | 7.25E-08 | 207.381003 | 252.209246 | 306.020763 | 274.554815 |
| -5.7753863 | 7.68E-09 | 7.27E-08 | 338.940889 | 546.904385 | 508.500666 | 531.179519 |
| -5.7751036 | 7.69E-09 | 7.28E-08 | 186.069121 | 329.062707 | 430.269794 | 335.068937 |
| -5.7748528 | 7.70E-09 | 7.28E-08 | 192.21678 | 283.058875 | 360.092101 | 340.111781 |
| -5.7742994 | 7.73E-09 | 7.31E-08 | 215.987725 | 316.885222 | 234.692615 | 282.959554 |
| -5.7742005 | 7.73E-09 | 7.31E-08 | 189.757716 | 296.589414 | 337.083021 | 321.061039 |
| -5.7738861 | 7.75E-09 | 7.32E-08 | 193.036467 | 378.855091 | 283.011683 | 337.310201 |
| -5.773814 | 7.75E-09 | 7.32E-08 | 206.561315 | 290.094755 | 276.108959 | 251.581861 |
| -5.7738512 | 7.75E-09 | 7.32E-08 | 202.462877 | 342.863857 | 277.259413 | 272.873867 |
| -5.7737257 | 7.75E-09 | 7.32E-08 | 173.363961 | 499.818109 | 391.154358 | 393.902112 |
| -5.7736259 | 7.76E-09 | 7.32E-08 | 305.743534 | 513.619259 | 190.975363 | 396.703692 |
| -5.7734529 | 7.77E-09 | 7.33E-08 | 179.101776 | 404.563115 | 337.083021 | 379.894213 |
| -5.7732787 | 7.77E-09 | 7.33E-08 | 186.478965 | 307.413845 | 476.287954 | 473.466977 |
| -5.7726425 | 7.80E-09 | 7.36E-08 | 156.560362 | 600.485319 | 468.234776 | 595.615853 |
| -5.7725072 | 7.81E-09 | 7.36E-08 | 302.05494 | 504.689103 | 320.976665 | 379.333897 |
| -5.7722596 | 7.82E-09 | 7.37E-08 | 234.020855 | 230.830995 | 241.595339 | 308.734088 |
| -5.7719107 | 7.84E-09 | 7.38E-08 | 189.757716 | 376.960815 | 296.817131 | 325.543566 |
| -5.7719565 | 7.84E-09 | 7.38E-08 | 229.102729 | 315.532168 | 225.488983 | 260.546916 |
| -5.7716141 | 7.85E-09 | 7.39E-08 | 177.052556 | 365.324552 | 358.941647 | 426.960753 |
| -5.7714404 | 7.86E-09 | 7.40E-08 | 171.724586 | 400.774564 | 453.278874 | 383.256109 |
| -5.7713992 | 7.86E-09 | 7.40E-08 | 169.265522 | 491.970396 | 388.85345 | 440.968652 |
| -5.7708005 | 7.89E-09 | 7.42E-08 | 552.46955 | 1530.57457 | 1212.57851 | 1507.81022 |
| -5.7702519 | 7.92E-09 | 7.44E-08 | 164.347396 | 485.746349 | 436.022064 | 443.209915 |
| 5.77028967 | 7.91E-09 | 7.44E-08 | 67.2143963 | 54.1221558 | 54.0713378 | 36.9808526 |
| -5.7702382 | 7.92E-09 | 7.44E-08 | 146.314265 | 607.521199 | 602.837894 | 628.114178 |
| -5.7703102 | 7.91E-09 | 7.44E-08 | 211.479442 | 307.684456 | 289.914407 | 326.103882 |
| -5.769939 | 7.93E-09 | 7.45E-08 | 167.626147 | 419.987929 | 417.6148 | 438.167072 |
| -5.769952 | 7.93E-09 | 7.45E-08 | 199.184126 | 333.121869 | 273.808051 | 280.71829 |
| -5.7694024 | 7.96E-09 | 7.47E-08 | 209.430223 | 409.434109 | 276.108959 | 280.71829 |

| | | | | | | |
|---|---|---|---|---|---|---|
| -5.7690809 | 7.97E-09 | 7.47E-08 | 250.824455 | 411.057774 | 205.931265 | 373.730738 |
| -5.7690896 | 7.97E-09 | 7.47E-08 | 165.576927 | 458.144049 | 411.86253 | 467.863817 |
| -5.7690748 | 7.97E-09 | 7.47E-08 | 168.445835 | 417.552432 | 401.508444 | 451.614655 |
| -5.769152 | 7.97E-09 | 7.47E-08 | 182.79037 | 327.168432 | 341.684837 | 338.991149 |
| -5.7688414 | 7.98E-09 | 7.48E-08 | 291.398999 | 450.566947 | 196.727633 | 326.664198 |
| -5.7684019 | 8.00E-09 | 7.50E-08 | 278.283994 | 289.282923 | 365.844371 | 373.170422 |
| -5.7681876 | 8.01E-09 | 7.51E-08 | 165.986771 | 227.042444 | 215.134897 | 252.142177 |
| -5.7677604 | 8.03E-09 | 7.52E-08 | 187.708497 | 316.614612 | 301.418947 | 352.999048 |
| -5.7676678 | 8.04E-09 | 7.53E-08 | 194.675843 | 335.557366 | 278.409867 | 295.846821 |
| -5.7675982 | 8.04E-09 | 7.53E-08 | 298.776188 | 255.727186 | 234.692615 | 217.962904 |
| -5.7675321 | 8.04E-09 | 7.53E-08 | 170.495054 | 444.342899 | 380.800272 | 410.151274 |
| -5.7674434 | 8.05E-09 | 7.53E-08 | 162.70802 | 543.386445 | 424.517524 | 485.793927 |
| -5.7673645 | 8.05E-09 | 7.53E-08 | 204.92194 | 298.213079 | 257.701695 | 263.348496 |
| -5.767207 | 8.06E-09 | 7.54E-08 | 152.052079 | 624.840289 | 499.297034 | 599.538065 |
| -5.7670542 | 8.07E-09 | 7.54E-08 | 190.577404 | 322.568049 | 289.914407 | 306.492824 |
| -5.766794 | 8.08E-09 | 7.55E-08 | 183.200214 | 397.256624 | 308.321671 | 369.808526 |
| -5.7666168 | 8.09E-09 | 7.56E-08 | 209.020379 | 275.211162 | 240.444885 | 288.002398 |
| -5.7663736 | 8.10E-09 | 7.56E-08 | 261.070552 | 272.505055 | 200.178995 | 296.967453 |
| -5.7661407 | 8.11E-09 | 7.57E-08 | 166.396615 | 468.427259 | 400.35799 | 446.011495 |
| -5.7660506 | 8.12E-09 | 7.57E-08 | 245.08664 | 701.152529 | 242.745793 | 484.673296 |
| -5.7659966 | 8.12E-09 | 7.58E-08 | 179.101776 | 387.785247 | 325.578481 | 381.014845 |
| -5.7658074 | 8.13E-09 | 7.58E-08 | 170.904898 | 386.702803 | 388.85345 | 436.486124 |
| -5.7656974 | 8.13E-09 | 7.58E-08 | 167.216303 | 446.778396 | 413.012984 | 411.832222 |
| -5.7655806 | 8.14E-09 | 7.59E-08 | 166.396615 | 670.844121 | 576.377452 | 585.530166 |
| -5.7653818 | 8.15E-09 | 7.59E-08 | 200.413657 | 326.897821 | 291.064861 | 258.305652 |
| 5.76500831 | 8.17E-09 | 7.60E-08 | 0.40984388 | 0 | 0 | 0 |
| -5.7650009 | 8.17E-09 | 7.60E-08 | 159.019425 | 522.008193 | 473.987046 | 467.863817 |
| -5.7650382 | 8.16E-09 | 7.60E-08 | 191.397092 | 439.201295 | 288.763953 | 360.283155 |
| -5.7648725 | 8.17E-09 | 7.61E-08 | 186.478965 | 315.532168 | 304.870309 | 353.559364 |
| -5.7636643 | 8.23E-09 | 7.66E-08 | 243.857109 | 339.345917 | 629.298335 | 407.349695 |
| -5.7635739 | 8.24E-09 | 7.66E-08 | 154.511143 | 485.205127 | 569.474728 | 532.860467 |
| -5.7632138 | 8.25E-09 | 7.67E-08 | 246.726016 | 323.650492 | 215.134897 | 250.461229 |
| -5.7633043 | 8.25E-09 | 7.67E-08 | 230.332261 | 230.019162 | 460.181598 | 348.51652 |
| -5.7632089 | 8.25E-09 | 7.67E-08 | 200.003813 | 315.802779 | 263.453965 | 275.115131 |
| -5.7631992 | 8.25E-09 | 7.67E-08 | 177.4624 | 355.853175 | 400.35799 | 340.672097 |
| -5.763263 | 8.25E-09 | 7.67E-08 | 194.675843 | 271.963833 | 293.365769 | 313.776931 |
| -5.7630078 | 8.26E-09 | 7.67E-08 | 534.436419 | 1350.34779 | 1157.35672 | 1280.32194 |
| -5.7630443 | 8.26E-09 | 7.67E-08 | 168.445835 | 663.53763 | 396.906628 | 509.887513 |
| -5.7629078 | 8.27E-09 | 7.68E-08 | 178.691932 | 339.345917 | 373.897549 | 342.353045 |
| -5.762859 | 8.27E-09 | 7.68E-08 | 169.265522 | 383.996696 | 429.11934 | 407.349695 |
| -5.7626687 | 8.28E-09 | 7.68E-08 | 180.331307 | 338.263474 | 329.029843 | 383.256109 |
| -5.7626016 | 8.28E-09 | 7.68E-08 | 197.54475 | 312.014228 | 265.754873 | 282.959554 |
| -5.7620016 | 8.31E-09 | 7.71E-08 | 229.102729 | 414.034492 | 220.887167 | 357.481575 |

| | | | | | | |
|---|---|---|---|---|---|---|
| -5.7617104 | 8.33E-09 | 7.72E-08 | 156.970206 | 680.85672 | 486.64204 | 504.84467 |
| -5.7616257 | 8.33E-09 | 7.72E-08 | 154.511143 | 541.76278 | 539.562924 | 482.432032 |
| -5.761514 | 8.34E-09 | 7.72E-08 | 201.643189 | 275.211162 | 257.701695 | 273.434183 |
| -5.7613621 | 8.34E-09 | 7.73E-08 | 166.806459 | 430.54175 | 399.207536 | 426.400437 |
| -5.761023 | 8.36E-09 | 7.74E-08 | 176.232868 | 371.5486 | 340.534383 | 375.411686 |
| -5.7606578 | 8.38E-09 | 7.76E-08 | 173.363961 | 442.719235 | 346.286653 | 412.952854 |
| -5.7605859 | 8.38E-09 | 7.76E-08 | 160.658801 | 444.072289 | 456.730236 | 497.560562 |
| -5.7605059 | 8.39E-09 | 7.76E-08 | 143.035514 | 968.245368 | 589.032446 | 693.110828 |
| 5.76037739 | 8.39E-09 | 7.76E-08 | 85.6573709 | 69.5469702 | 77.0804177 | 81.2458125 |
| -5.7603662 | 8.39E-09 | 7.76E-08 | 156.150518 | 497.923834 | 478.588862 | 524.455728 |
| -5.7600395 | 8.41E-09 | 7.77E-08 | 143.035514 | 678.962445 | 635.050605 | 585.530166 |
| -5.7600757 | 8.41E-09 | 7.77E-08 | 191.397092 | 313.908504 | 274.958505 | 318.819775 |
| -5.7597736 | 8.42E-09 | 7.78E-08 | 169.675366 | 408.892887 | 395.756174 | 383.256109 |
| -5.7597143 | 8.43E-09 | 7.78E-08 | 164.347396 | 465.991762 | 403.809352 | 452.174971 |
| 5.75960254 | 8.43E-09 | 7.78E-08 | 0 | 0 | 0 | 0 |
| -5.7589946 | 8.46E-09 | 7.81E-08 | 231.151948 | 332.580648 | 218.586259 | 267.831023 |
| -5.758931 | 8.46E-09 | 7.81E-08 | 197.134906 | 262.492456 | 324.428027 | 286.881766 |
| -5.7587508 | 8.47E-09 | 7.82E-08 | 174.183649 | 367.760049 | 398.057082 | 351.3181 |
| -5.7586632 | 8.48E-09 | 7.82E-08 | 175.823024 | 419.987929 | 365.844371 | 341.792729 |
| -5.758635 | 8.48E-09 | 7.82E-08 | 193.856155 | 275.752384 | 296.817131 | 284.640502 |
| -5.7581781 | 8.50E-09 | 7.83E-08 | 211.479442 | 265.739785 | 234.692615 | 263.348496 |
| -5.7581864 | 8.50E-09 | 7.83E-08 | 176.642712 | 326.897821 | 385.402088 | 395.58306 |
| -5.7582324 | 8.50E-09 | 7.83E-08 | 208.200691 | 294.965749 | 236.993523 | 273.994499 |
| -5.7582624 | 8.50E-09 | 7.83E-08 | 234.430699 | 208.099689 | 310.622579 | 309.85472 |
| -5.7577109 | 8.53E-09 | 7.85E-08 | 140.576451 | 819.950661 | 606.289256 | 652.76808 |
| -5.7577356 | 8.52E-09 | 7.85E-08 | 162.298176 | 461.661989 | 460.181598 | 419.676646 |
| -5.7573078 | 8.55E-09 | 7.86E-08 | 138.937075 | 755.274685 | 612.041526 | 681.344194 |
| -5.757259 | 8.55E-09 | 7.86E-08 | 239.348826 | 333.933701 | 207.081719 | 288.002398 |
| -5.7571042 | 8.56E-09 | 7.87E-08 | 169.675366 | 439.201295 | 390.003904 | 376.532317 |
| -5.7570872 | 8.56E-09 | 7.87E-08 | 282.382433 | 342.863857 | 195.577179 | 270.072287 |
| -5.7568759 | 8.57E-09 | 7.88E-08 | 177.872244 | 395.362348 | 365.844371 | 322.741986 |
| -5.7556682 | 8.63E-09 | 7.93E-08 | 157.38005 | 464.638708 | 469.38523 | 514.370041 |
| -5.755408 | 8.64E-09 | 7.94E-08 | 135.24848 | 602.108984 | 943.372276 | 870.730984 |
| -5.7549039 | 8.67E-09 | 7.96E-08 | 198.774282 | 292.800863 | 256.551241 | 275.115131 |
| -5.7548842 | 8.67E-09 | 7.96E-08 | 246.316172 | 299.024911 | 222.037621 | 227.488275 |
| -5.754687 | 8.68E-09 | 7.97E-08 | 180.741151 | 333.933701 | 342.835291 | 314.897563 |
| -5.7544521 | 8.69E-09 | 7.98E-08 | 190.577404 | 280.623378 | 293.365769 | 333.948305 |
| -5.7539585 | 8.72E-09 | 8.00E-08 | 190.577404 | 324.191713 | 270.356689 | 320.500723 |
| -5.7537277 | 8.73E-09 | 8.00E-08 | 184.83959 | 345.299354 | 438.322972 | 304.25156 |
| -5.7537174 | 8.73E-09 | 8.00E-08 | 230.742104 | 498.465055 | 230.090799 | 367.567262 |
| -5.752943 | 8.77E-09 | 8.04E-08 | 158.199738 | 490.076121 | 440.62388 | 483.552664 |
| -5.7526055 | 8.79E-09 | 8.05E-08 | 156.150518 | 439.742516 | 577.527906 | 531.739835 |
| -5.7525485 | 8.79E-09 | 8.05E-08 | 171.724586 | 372.089821 | 407.260714 | 356.921259 |

| | | | | | | |
|---|---|---|---|---|---|---|
| -5.7523855 | 8.80E-09 | 8.06E-08 | 367.62996 | 202.146252 | 284.162137 | 259.426284 |
| -5.7524104 | 8.80E-09 | 8.06E-08 | 176.232868 | 359.371115 | 340.534383 | 343.473677 |
| -5.7520784 | 8.82E-09 | 8.07E-08 | 180.331307 | 335.827977 | 327.879389 | 318.819775 |
| -5.7520204 | 8.82E-09 | 8.07E-08 | 170.08521 | 409.70472 | 387.702996 | 362.524419 |
| -5.75192 | 8.82E-09 | 8.07E-08 | 188.528185 | 381.831809 | 277.259413 | 341.232413 |
| -5.7517719 | 8.83E-09 | 8.07E-08 | 203.692408 | 343.9463 | 398.057082 | 374.291054 |
| -5.7518107 | 8.83E-09 | 8.07E-08 | 187.708497 | 294.153917 | 352.038923 | 294.165873 |
| -5.7515767 | 8.84E-09 | 8.08E-08 | 163.937552 | 398.068456 | 476.287954 | 427.521069 |
| -5.7514281 | 8.85E-09 | 8.08E-08 | 549.600643 | 247.879474 | 217.435805 | 412.952854 |
| -5.7513187 | 8.86E-09 | 8.09E-08 | 677.881777 | 307.955067 | 238.143977 | 303.130928 |
| -5.7506447 | 8.89E-09 | 8.12E-08 | 307.792754 | 312.826061 | 188.674455 | 271.753235 |
| -5.7506588 | 8.89E-09 | 8.12E-08 | 162.298176 | 437.307019 | 414.163438 | 446.011495 |
| -5.7504713 | 8.90E-09 | 8.12E-08 | 184.019902 | 324.462324 | 292.215315 | 340.111781 |
| -5.7502628 | 8.91E-09 | 8.13E-08 | 106.969253 | 912.228936 | 1403.55387 | 1311.69964 |
| -5.7501779 | 8.91E-09 | 8.13E-08 | 159.019425 | 467.074205 | 441.774334 | 456.657498 |
| -5.7499562 | 8.93E-09 | 8.14E-08 | 205.331784 | 240.031761 | 265.754873 | 267.831023 |
| -5.7498439 | 8.93E-09 | 8.14E-08 | 177.872244 | 392.115019 | 347.437107 | 319.380091 |
| -5.7497369 | 8.94E-09 | 8.15E-08 | 268.037897 | 552.3166 | 568.324274 | 552.471525 |
| -5.7492205 | 8.97E-09 | 8.17E-08 | 175.413181 | 343.9463 | 356.640739 | 341.232413 |
| -5.7481709 | 9.02E-09 | 8.22E-08 | 326.235728 | 818.868218 | 685.670581 | 756.426531 |
| -5.7479716 | 9.03E-09 | 8.22E-08 | 190.577404 | 417.281821 | 285.312591 | 514.370041 |
| -5.7478446 | 9.04E-09 | 8.23E-08 | 150.00286 | 616.180744 | 480.88977 | 569.84132 |
| -5.7474645 | 9.06E-09 | 8.24E-08 | 170.904898 | 476.274971 | 347.437107 | 409.590958 |
| -5.7472005 | 9.07E-09 | 8.25E-08 | 173.363961 | 362.347833 | 472.836592 | 356.360943 |
| -5.7459781 | 9.14E-09 | 8.31E-08 | 230.742104 | 384.267306 | 220.887167 | 282.399238 |
| -5.745865 | 9.15E-09 | 8.32E-08 | 287.30056 | 274.669941 | 196.727633 | 250.461229 |
| -5.7449805 | 9.19E-09 | 8.36E-08 | 173.773805 | 326.897821 | 386.552542 | 431.44328 |
| -5.7442614 | 9.23E-09 | 8.39E-08 | 189.347873 | 266.281007 | 366.994825 | 378.773581 |
| -5.7442353 | 9.23E-09 | 8.39E-08 | 167.216303 | 383.184863 | 391.154358 | 384.937057 |
| -5.744035 | 9.24E-09 | 8.40E-08 | 134.018949 | 660.560912 | 714.431931 | 688.628301 |
| -5.7438857 | 9.25E-09 | 8.40E-08 | 178.282088 | 352.064624 | 306.020763 | 352.999048 |
| -5.7430821 | 9.30E-09 | 8.44E-08 | 172.954117 | 368.30127 | 341.684837 | 351.878416 |
| -5.7428527 | 9.31E-09 | 8.45E-08 | 609.028006 | 375.06654 | 251.949425 | 313.776931 |
| -5.7421766 | 9.35E-09 | 8.48E-08 | 172.13443 | 481.687187 | 684.520127 | 368.127578 |
| -5.7414607 | 9.39E-09 | 8.51E-08 | 158.199738 | 497.923834 | 450.977966 | 422.478225 |
| -5.7412142 | 9.40E-09 | 8.52E-08 | 163.937552 | 460.038325 | 399.207536 | 391.660848 |
| -5.7409018 | 9.42E-09 | 8.54E-08 | 196.725062 | 316.07339 | 245.046701 | 335.629253 |
| -5.7408424 | 9.42E-09 | 8.54E-08 | 452.467643 | 342.863857 | 200.178995 | 257.745336 |
| 5.7402038 | 9.46E-09 | 8.57E-08 | 0 | 0 | 0 | 0 |
| -5.7399627 | 9.47E-09 | 8.57E-08 | 194.265999 | 343.9463 | 250.798971 | 316.578511 |
| -5.7399524 | 9.47E-09 | 8.57E-08 | 147.543797 | 497.653223 | 548.766556 | 597.857117 |
| -5.7401293 | 9.46E-09 | 8.57E-08 | 246.316172 | 206.205414 | 240.444885 | 259.426284 |
| -5.7400159 | 9.47E-09 | 8.57E-08 | 207.381003 | 497.923834 | 358.941647 | 444.330547 |

| | | | | | | |
|---|---|---|---|---|---|---|
| -5.7399771 | 9.47E-09 | 8.57E-08 | 190.987248 | 300.377965 | 260.002603 | 298.088085 |
| -5.7399018 | 9.47E-09 | 8.57E-08 | 268.857585 | 273.587498 | 258.852149 | 194.98995 |
| -5.7397786 | 9.48E-09 | 8.57E-08 | 192.21678 | 268.445893 | 325.578481 | 273.434183 |
| -5.7395854 | 9.49E-09 | 8.58E-08 | 185.249434 | 279.270324 | 307.171217 | 311.535667 |
| -5.7394903 | 9.50E-09 | 8.58E-08 | 248.365391 | 217.841677 | 345.136199 | 237.013646 |
| -5.7393044 | 9.51E-09 | 8.59E-08 | 207.790847 | 305.248959 | 225.488983 | 285.200818 |
| -5.7383534 | 9.56E-09 | 8.63E-08 | 180.331307 | 323.921103 | 308.321671 | 306.492824 |
| -5.7380383 | 9.58E-09 | 8.65E-08 | 148.363485 | 482.499019 | 552.217918 | 572.082584 |
| -5.7379193 | 9.58E-09 | 8.65E-08 | 390.171374 | 183.744719 | 366.994825 | 278.477026 |
| -5.7379465 | 9.58E-09 | 8.65E-08 | 191.806936 | 327.980264 | 265.754873 | 278.477026 |
| -5.7378104 | 9.59E-09 | 8.65E-08 | 159.019425 | 486.558181 | 399.207536 | 469.544765 |
| -5.7376805 | 9.60E-09 | 8.66E-08 | 170.904898 | 432.977247 | 408.411168 | 328.345146 |
| -5.7372387 | 9.62E-09 | 8.68E-08 | 151.642236 | 562.058588 | 471.686138 | 486.354243 |
| -5.7370998 | 9.63E-09 | 8.68E-08 | 185.249434 | 338.804695 | 296.817131 | 281.838922 |
| -5.7367487 | 9.65E-09 | 8.69E-08 | 161.478489 | 396.715402 | 437.172518 | 416.31475 |
| -5.7367576 | 9.65E-09 | 8.69E-08 | 146.724109 | 543.657055 | 503.89885 | 583.849218 |
| -5.7364995 | 9.67E-09 | 8.70E-08 | 158.609582 | 438.389462 | 432.570702 | 583.849218 |
| -5.7361669 | 9.68E-09 | 8.72E-08 | 220.086164 | 312.284839 | 213.984443 | 269.511971 |
| -5.7360378 | 9.69E-09 | 8.72E-08 | 150.412704 | 507.1246 | 483.190678 | 511.008145 |
| -5.7356293 | 9.72E-09 | 8.74E-08 | 163.937552 | 381.019977 | 406.11026 | 401.746535 |
| -5.7354829 | 9.72E-09 | 8.75E-08 | 200.823501 | 258.703905 | 240.444885 | 284.640502 |
| -5.7351726 | 9.74E-09 | 8.76E-08 | 187.708497 | 295.23636 | 270.356689 | 340.111781 |
| 5.73427566 | 9.79E-09 | 8.80E-08 | 0 | 0 | 0 | 0 |
| -5.7341901 | 9.80E-09 | 8.81E-08 | 167.626147 | 348.546684 | 436.022064 | 386.057689 |
| -5.7341514 | 9.80E-09 | 8.81E-08 | 150.412704 | 571.259355 | 459.031144 | 520.533516 |
| -5.7340821 | 9.80E-09 | 8.81E-08 | 176.642712 | 330.686372 | 323.277573 | 316.018195 |
| -5.7340269 | 9.81E-09 | 8.81E-08 | 164.75724 | 454.626109 | 366.994825 | 399.505271 |
| -5.7336179 | 9.83E-09 | 8.83E-08 | 178.282088 | 552.3166 | 376.198456 | 336.189569 |
| -5.7332925 | 9.85E-09 | 8.84E-08 | 194.675843 | 288.741701 | 249.648517 | 271.753235 |
| -5.7330875 | 9.86E-09 | 8.85E-08 | 193.446311 | 503.065438 | 289.914407 | 333.948305 |
| -5.7327987 | 9.88E-09 | 8.86E-08 | 0 | 92.2782757 | 570.625182 | 0 |
| -5.7327496 | 9.88E-09 | 8.86E-08 | 152.871767 | 575.318516 | 433.721156 | 513.249409 |
| -5.7325968 | 9.89E-09 | 8.87E-08 | 229.922417 | 527.691019 | 224.338529 | 564.798476 |
| -5.7321116 | 9.92E-09 | 8.89E-08 | 217.217256 | 272.775665 | 247.347609 | 216.842272 |
| -5.7320061 | 9.92E-09 | 8.89E-08 | 102.46097 | 1042.66333 | 1298.86256 | 1092.6161 |
| -5.7306013 | 1.00E-08 | 8.97E-08 | 173.363961 | 457.873438 | 320.976665 | 375.411686 |
| -5.7302231 | 1.00E-08 | 8.98E-08 | 166.806459 | 396.444791 | 373.897549 | 352.999048 |
| -5.7301384 | 1.00E-08 | 8.98E-08 | 155.740674 | 486.016959 | 425.667978 | 444.890863 |
| -5.729344 | 1.01E-08 | 9.02E-08 | 179.101776 | 285.494372 | 384.251634 | 356.360943 |
| -5.7290699 | 1.01E-08 | 9.04E-08 | 193.446311 | 286.306204 | 246.197155 | 284.080186 |
| -5.7287249 | 1.01E-08 | 9.05E-08 | 156.560362 | 465.179929 | 417.6148 | 438.727388 |
| -5.7281459 | 1.02E-08 | 9.08E-08 | 165.986771 | 355.041342 | 385.402088 | 381.014845 |
| -5.7277687 | 1.02E-08 | 9.10E-08 | 151.232392 | 467.074205 | 462.482506 | 549.669946 |

| | | | | | | |
|---|---|---|---|---|---|---|
| -5.7277183 | 1.02E-08 | 9.10E-08 | 111.067691 | 940.643068 | 1188.41898 | 1141.36359 |
| -5.7276334 | 1.02E-08 | 9.10E-08 | 178.282088 | 315.802779 | 270.356689 | 281.278606 |
| -5.7273529 | 1.02E-08 | 9.11E-08 | 164.347396 | 384.808528 | 373.897549 | 381.014845 |
| -5.7270819 | 1.02E-08 | 9.13E-08 | 144.67489 | 586.684169 | 502.748396 | 542.946154 |
| -5.7265534 | 1.02E-08 | 9.15E-08 | 184.429746 | 319.59133 | 277.259413 | 284.640502 |
| -5.7261205 | 1.03E-08 | 9.17E-08 | 238.938982 | 221.900839 | 215.134897 | 289.683345 |
| 5.72591848 | 1.03E-08 | 9.18E-08 | 23.770945 | 13.530539 | 23.0090799 | 12.3269509 |
| -5.7255926 | 1.03E-08 | 9.20E-08 | 193.446311 | 263.574899 | 250.798971 | 297.527769 |
| -5.7254306 | 1.03E-08 | 9.20E-08 | 232.38148 | 309.849342 | 204.780811 | 253.262809 |
| -5.7251736 | 1.03E-08 | 9.21E-08 | 0 | 456.249774 | 120.79767 | 0 |
| -5.7248401 | 1.04E-08 | 9.23E-08 | 199.59397 | 271.152001 | 595.93517 | 415.754434 |
| -5.7247876 | 1.04E-08 | 9.23E-08 | 153.281611 | 438.930684 | 448.677058 | 499.24151 |
| -5.7245607 | 1.04E-08 | 9.24E-08 | 164.347396 | 372.631043 | 364.693917 | 405.108431 |
| -5.7240493 | 1.04E-08 | 9.26E-08 | 195.085687 | 412.681438 | 246.197155 | 365.325998 |
| -5.723925 | 1.04E-08 | 9.27E-08 | 220.496007 | 255.997797 | 213.984443 | 234.772382 |
| -5.7239492 | 1.04E-08 | 9.27E-08 | 225.00429 | 226.230611 | 216.285351 | 245.978701 |
| -5.7238249 | 1.04E-08 | 9.27E-08 | 149.593016 | 480.063522 | 465.933868 | 517.171621 |
| -5.7236001 | 1.04E-08 | 9.28E-08 | 148.363485 | 478.169247 | 492.39431 | 514.930357 |
| -5.7232298 | 1.05E-08 | 9.30E-08 | 163.117864 | 367.760049 | 402.658898 | 383.256109 |
| -5.7231286 | 1.05E-08 | 9.30E-08 | 187.298653 | 307.684456 | 261.153057 | 281.278606 |
| -5.7231302 | 1.05E-08 | 9.30E-08 | 171.724586 | 358.559282 | 311.773033 | 361.964103 |
| -5.722709 | 1.05E-08 | 9.32E-08 | 186.888809 | 329.87454 | 536.111562 | 316.578511 |
| -5.7226413 | 1.05E-08 | 9.32E-08 | 225.414134 | 278.999713 | 203.630357 | 255.504073 |
| -5.7224424 | 1.05E-08 | 9.33E-08 | 166.396615 | 375.607761 | 349.738015 | 362.524419 |
| -5.7222508 | 1.05E-08 | 9.33E-08 | 163.937552 | 658.396026 | 398.057082 | 434.805176 |
| -5.7218096 | 1.05E-08 | 9.36E-08 | 165.167084 | 470.321534 | 341.684837 | 415.194118 |
| -5.7209665 | 1.06E-08 | 9.40E-08 | 159.019425 | 600.485319 | 422.216616 | 416.875066 |
| -5.7203737 | 1.06E-08 | 9.43E-08 | 172.544273 | 358.018061 | 303.719855 | 387.17832 |
| -5.7203263 | 1.06E-08 | 9.43E-08 | 138.527231 | 622.134181 | 551.067464 | 585.530166 |
| -5.7198944 | 1.07E-08 | 9.45E-08 | 177.872244 | 385.079139 | 322.127119 | 291.364293 |
| -5.7196193 | 1.07E-08 | 9.46E-08 | 167.626147 | 342.322636 | 342.835291 | 234.772382 |
| -5.7196488 | 1.07E-08 | 9.46E-08 | 208.610535 | 241.384815 | 235.843069 | 232.531119 |
| -5.7193337 | 1.07E-08 | 9.48E-08 | 137.707544 | 541.76278 | 607.43971 | 661.733135 |
| -5.7192624 | 1.07E-08 | 9.48E-08 | 170.904898 | 380.208145 | 308.321671 | 369.24821 |
| -5.7190828 | 1.07E-08 | 9.48E-08 | 122.133476 | 847.823571 | 888.150484 | 680.223562 |
| -5.7189499 | 1.07E-08 | 9.49E-08 | 143.445358 | 525.255522 | 614.342434 | 503.724038 |
| -5.7186501 | 1.07E-08 | 9.50E-08 | 211.479442 | 259.786348 | 212.833989 | 261.107232 |
| -5.7186531 | 1.07E-08 | 9.50E-08 | 200.823501 | 279.270324 | 251.949425 | 230.850171 |
| -5.7186206 | 1.07E-08 | 9.50E-08 | 232.791324 | 216.218013 | 444.075242 | 303.130928 |
| -5.718263 | 1.08E-08 | 9.51E-08 | 202.462877 | 291.71842 | 230.090799 | 250.461229 |
| -5.7182894 | 1.08E-08 | 9.51E-08 | 132.789417 | 703.858637 | 615.492888 | 620.830071 |
| -5.7181213 | 1.08E-08 | 9.52E-08 | 169.265522 | 357.206228 | 332.481205 | 335.629253 |
| -5.7180317 | 1.08E-08 | 9.52E-08 | 197.54475 | 366.948217 | 331.330751 | 347.956204 |

| | | | | | | |
|---|---|---|---|---|---|---|
| -5.717962 | 1.08E-08 | 9.52E-08 | 167.626147 | 429.729917 | 320.976665 | 403.987799 |
| -5.717679 | 1.08E-08 | 9.54E-08 | 172.13443 | 313.637893 | 325.578481 | 368.127578 |
| -5.7167977 | 1.09E-08 | 9.58E-08 | 186.478965 | 273.316887 | 293.365769 | 260.546916 |
| -5.7166291 | 1.09E-08 | 9.59E-08 | 145.494577 | 517.949031 | 538.41247 | 477.389188 |
| -5.7163956 | 1.09E-08 | 9.60E-08 | 210.24991 | 230.289773 | 271.507143 | 230.850171 |
| -5.7164227 | 1.09E-08 | 9.60E-08 | 146.724109 | 454.355498 | 559.120642 | 559.195317 |
| -5.7161433 | 1.09E-08 | 9.61E-08 | 233.611012 | 391.844408 | 355.490285 | 310.415036 |
| -5.7161171 | 1.09E-08 | 9.61E-08 | 260.250864 | 268.175282 | 188.674455 | 257.745336 |
| -5.7155487 | 1.09E-08 | 9.64E-08 | 176.642712 | 395.90357 | 291.064861 | 330.026094 |
| -5.7154705 | 1.09E-08 | 9.64E-08 | 152.052079 | 491.699786 | 842.132325 | 500.922458 |
| -5.7152637 | 1.10E-08 | 9.65E-08 | 165.167084 | 379.666923 | 334.782113 | 402.867167 |
| -5.7152341 | 1.10E-08 | 9.65E-08 | 215.577881 | 248.691306 | 222.037621 | 223.566063 |
| -5.7151388 | 1.10E-08 | 9.65E-08 | 174.183649 | 351.523402 | 291.064861 | 381.014845 |
| -5.7148173 | 1.10E-08 | 9.67E-08 | 163.117864 | 348.005462 | 404.959806 | 477.389188 |
| -5.7141059 | 1.10E-08 | 9.70E-08 | 160.248957 | 458.685271 | 363.543463 | 405.108431 |
| -5.7137924 | 1.10E-08 | 9.72E-08 | 183.610058 | 295.777582 | 262.303511 | 290.243661 |
| -5.7131826 | 1.11E-08 | 9.75E-08 | 167.626147 | 336.098588 | 341.684837 | 348.51652 |
| -5.7132108 | 1.11E-08 | 9.75E-08 | 165.986771 | 368.571881 | 338.233475 | 354.11968 |
| -5.7129386 | 1.11E-08 | 9.76E-08 | 169.265522 | 354.500121 | 311.773033 | 373.730738 |
| -5.7124116 | 1.11E-08 | 9.79E-08 | 196.725062 | 272.775665 | 240.444885 | 243.177122 |
| -5.7112709 | 1.12E-08 | 9.85E-08 | 176.642712 | 281.705821 | 315.224395 | 319.380091 |
| -5.7103559 | 1.13E-08 | 9.90E-08 | 183.610058 | 447.860839 | 333.631659 | 323.302302 |
| 5.71025534 | 1.13E-08 | 9.91E-08 | 97.1329995 | 97.9611021 | 108.142676 | 84.0473923 |
| -5.7101243 | 1.13E-08 | 9.91E-08 | 140.576451 | 721.718948 | 507.350212 | 560.876265 |
| -5.7094132 | 1.13E-08 | 9.95E-08 | 196.315218 | 226.230611 | 289.914407 | 290.803977 |
| -5.7080578 | 1.14E-08 | 1.00E-07 | 254.922893 | 316.344001 | 283.011683 | 338.430833 |
| -5.7074642 | 1.15E-08 | 1.01E-07 | 179.511619 | 328.250875 | 264.604419 | 318.819775 |
| -5.7072085 | 1.15E-08 | 1.01E-07 | 188.118341 | 254.103522 | 256.551241 | 282.399238 |
| -5.7070017 | 1.15E-08 | 1.01E-07 | 206.561315 | 256.539019 | 213.984443 | 253.262809 |
| -5.7067662 | 1.15E-08 | 1.01E-07 | 294.267906 | 201.063809 | 260.002603 | 222.445432 |
| -5.7063911 | 1.15E-08 | 1.01E-07 | 178.282088 | 303.895905 | 279.560321 | 288.002398 |
| -5.7059954 | 1.16E-08 | 1.01E-07 | 168.035991 | 307.143234 | 369.295733 | 363.084735 |
| -5.7058518 | 1.16E-08 | 1.01E-07 | 247.13586 | 193.486707 | 453.278874 | 363.645051 |
| -5.7053266 | 1.16E-08 | 1.02E-07 | 141.805982 | 559.893702 | 493.544764 | 517.171621 |
| -5.7042427 | 1.17E-08 | 1.02E-07 | 188.118341 | 671.114732 | 291.064861 | 454.416234 |
| -5.7039229 | 1.17E-08 | 1.02E-07 | 179.101776 | 271.152001 | 300.268493 | 287.442082 |
| -5.7037057 | 1.17E-08 | 1.03E-07 | 274.5954 | 207.829078 | 251.949425 | 211.799429 |
| -5.7034413 | 1.17E-08 | 1.03E-07 | 152.461923 | 408.892887 | 421.066162 | 463.941605 |
| -5.7034072 | 1.17E-08 | 1.03E-07 | 158.609582 | 357.476839 | 398.057082 | 428.081385 |
| -5.7024166 | 1.18E-08 | 1.03E-07 | 207.381003 | 210.264575 | 265.754873 | 257.18502 |
| -5.7023645 | 1.18E-08 | 1.03E-07 | 211.889286 | 382.914252 | 224.338529 | 272.313551 |
| -5.7021432 | 1.18E-08 | 1.03E-07 | 159.019425 | 412.140217 | 375.048002 | 364.205367 |
| -5.7020635 | 1.18E-08 | 1.03E-07 | 244.676796 | 326.62721 | 187.524001 | 275.675447 |

| | | | | | | |
|---|---|---|---|---|---|---|
| -5.7020172 | 1.18E-08 | 1.03E-07 | 174.183649 | 300.107354 | 299.118039 | 297.527769 |
| -5.7018971 | 1.18E-08 | 1.03E-07 | 312.71088 | 711.165128 | 599.386532 | 661.172819 |
| -5.7019064 | 1.18E-08 | 1.03E-07 | 174.593493 | 346.111187 | 284.162137 | 414.073486 |
| -5.701839 | 1.19E-08 | 1.03E-07 | 125.412227 | 725.236888 | 617.793796 | 776.597905 |
| -5.7017559 | 1.19E-08 | 1.03E-07 | 198.774282 | 352.605845 | 294.516223 | 337.870517 |
| -5.7017215 | 1.19E-08 | 1.03E-07 | 177.052556 | 351.523402 | 274.958505 | 304.25156 |
| -5.7016483 | 1.19E-08 | 1.03E-07 | 170.495054 | 325.815378 | 302.569401 | 321.621355 |
| -5.7012182 | 1.19E-08 | 1.04E-07 | 255.332737 | 347.46424 | 185.223093 | 277.916711 |
| -5.7011786 | 1.19E-08 | 1.04E-07 | 140.166607 | 854.588841 | 514.252936 | 577.685743 |
| -5.7008451 | 1.19E-08 | 1.04E-07 | 163.117864 | 356.665007 | 346.286653 | 349.637152 |
| -5.7004069 | 1.20E-08 | 1.04E-07 | 277.054463 | 356.394396 | 297.967585 | 177.05984 |
| -5.6998628 | 1.20E-08 | 1.04E-07 | 153.691455 | 386.973414 | 472.836592 | 417.995698 |
| -5.6989942 | 1.21E-08 | 1.05E-07 | 213.938505 | 279.270324 | 213.984443 | 228.048591 |
| -5.6988324 | 1.21E-08 | 1.05E-07 | 211.889286 | 209.723354 | 329.029843 | 268.391339 |
| -5.698677 | 1.21E-08 | 1.05E-07 | 183.200214 | 281.43521 | 258.852149 | 265.58976 |
| -5.6986909 | 1.21E-08 | 1.05E-07 | 135.24848 | 587.766612 | 605.138802 | 530.619203 |
| -5.6986588 | 1.21E-08 | 1.05E-07 | 168.035991 | 307.955067 | 343.985745 | 328.345146 |
| -5.6984178 | 1.21E-08 | 1.05E-07 | 189.347873 | 253.291689 | 279.560321 | 241.496174 |
| -5.6979449 | 1.21E-08 | 1.05E-07 | 123.772852 | 696.281535 | 719.033747 | 669.577559 |
| -5.6978021 | 1.21E-08 | 1.05E-07 | 159.429269 | 446.778396 | 340.534383 | 408.470327 |
| -5.6975627 | 1.22E-08 | 1.06E-07 | 210.659754 | 360.453558 | 308.321671 | 322.18167 |
| -5.6974131 | 1.22E-08 | 1.06E-07 | 136.887856 | 666.514349 | 783.459171 | 767.072534 |
| -5.6972938 | 1.22E-08 | 1.06E-07 | 399.597783 | 249.773749 | 227.789891 | 223.005748 |
| -5.6969028 | 1.22E-08 | 1.06E-07 | 325.825885 | 162.637078 | 395.756174 | 372.04979 |
| -5.6967252 | 1.22E-08 | 1.06E-07 | 205.331784 | 307.684456 | 219.736713 | 245.418385 |
| -5.6964689 | 1.22E-08 | 1.06E-07 | 156.150518 | 377.502037 | 410.712076 | 382.695793 |
| -5.6963473 | 1.22E-08 | 1.06E-07 | 157.38005 | 366.677606 | 403.809352 | 377.652949 |
| -5.6960754 | 1.23E-08 | 1.06E-07 | 176.642712 | 362.889055 | 279.560321 | 294.165873 |
| -5.6959966 | 1.23E-08 | 1.06E-07 | 145.084733 | 484.122684 | 459.031144 | 469.544765 |
| -5.695594 | 1.23E-08 | 1.07E-07 | 137.2977 | 527.149798 | 551.067464 | 542.946154 |
| -5.6955333 | 1.23E-08 | 1.07E-07 | 149.593016 | 410.245941 | 483.190678 | 437.606756 |
| -5.694799 | 1.24E-08 | 1.07E-07 | 168.445835 | 318.508887 | 340.534383 | 303.691244 |
| -5.6945407 | 1.24E-08 | 1.07E-07 | 233.201168 | 262.763067 | 188.674455 | 283.51987 |
| -5.6944227 | 1.24E-08 | 1.07E-07 | 158.609582 | 404.021893 | 342.835291 | 398.384639 |
| -5.6943566 | 1.24E-08 | 1.07E-07 | 177.872244 | 296.860025 | 262.303511 | 299.769033 |
| 5.69373446 | 1.24E-08 | 1.07E-07 | 150.412704 | 70.6294134 | 71.3281477 | 69.4791776 |
| -5.6936226 | 1.24E-08 | 1.08E-07 | 161.068645 | 416.7406 | 340.534383 | 530.619203 |
| -5.6936193 | 1.24E-08 | 1.08E-07 | 149.593016 | 422.423426 | 518.854752 | 419.676646 |
| -5.6933996 | 1.25E-08 | 1.08E-07 | 148.363485 | 445.154732 | 429.11934 | 443.209915 |
| -5.6931238 | 1.25E-08 | 1.08E-07 | 177.4624 | 385.62036 | 303.719855 | 275.115131 |
| -5.692887 | 1.25E-08 | 1.08E-07 | 179.101776 | 336.91042 | 257.701695 | 379.894213 |
| -5.6927435 | 1.25E-08 | 1.08E-07 | 414.762006 | 969.327811 | 1105.58629 | 1045.54956 |
| -5.6925789 | 1.25E-08 | 1.08E-07 | 165.167084 | 563.141031 | 332.481205 | 411.271906 |

| | | | | | | |
|---|---|---|---|---|---|---|
| -5.692424 | 1.25E-08 | 1.08E-07 | 146.724109 | 501.712385 | 479.739316 | 415.754434 |
| -5.6921884 | 1.25E-08 | 1.08E-07 | 156.150518 | 377.231426 | 388.85345 | 377.652949 |
| -5.6919836 | 1.26E-08 | 1.08E-07 | 198.364438 | 353.147067 | 220.887167 | 298.648401 |
| -5.6918032 | 1.26E-08 | 1.08E-07 | 186.069121 | 245.443977 | 258.852149 | 262.227864 |
| -5.6916779 | 1.26E-08 | 1.08E-07 | 181.560839 | 343.405079 | 250.798971 | 299.769033 |
| -5.6915513 | 1.26E-08 | 1.09E-07 | 162.70802 | 376.960815 | 318.675757 | 364.205367 |
| -5.6906281 | 1.27E-08 | 1.09E-07 | 153.691455 | 435.412744 | 383.10118 | 390.540216 |
| -5.6898185 | 1.27E-08 | 1.10E-07 | 145.494577 | 513.078037 | 427.968886 | 466.743185 |
| -5.689782 | 1.27E-08 | 1.10E-07 | 168.035991 | 328.792097 | 294.516223 | 337.310201 |
| -5.6894313 | 1.27E-08 | 1.10E-07 | 152.871767 | 408.081055 | 377.34891 | 441.528968 |
| -5.6892672 | 1.28E-08 | 1.10E-07 | 134.018949 | 533.644456 | 563.722458 | 594.495221 |
| -5.6885572 | 1.28E-08 | 1.10E-07 | 225.823978 | 197.545869 | 254.250333 | 231.410487 |
| -5.6881518 | 1.28E-08 | 1.10E-07 | 154.101299 | 374.254708 | 407.260714 | 392.221164 |
| -5.6880283 | 1.29E-08 | 1.11E-07 | 178.282088 | 539.597894 | 279.560321 | 414.073486 |
| -5.6872883 | 1.29E-08 | 1.11E-07 | 182.79037 | 349.629127 | 241.595339 | 327.78483 |
| -5.6872031 | 1.29E-08 | 1.11E-07 | 141.805982 | 249.232528 | 185.223093 | 213.480376 |
| -5.6869773 | 1.29E-08 | 1.11E-07 | 172.954117 | 602.108984 | 309.472125 | 415.194118 |
| -5.6862731 | 1.30E-08 | 1.12E-07 | 186.888809 | 262.492456 | 319.826211 | 268.951655 |
| -5.6860701 | 1.30E-08 | 1.12E-07 | 222.955071 | 309.03751 | 227.789891 | 248.219965 |
| -5.6855816 | 1.30E-08 | 1.12E-07 | 192.626624 | 355.311953 | 318.675757 | 331.707042 |
| -5.6854177 | 1.30E-08 | 1.12E-07 | 168.445835 | 316.885222 | 309.472125 | 296.407137 |
| -5.6852461 | 1.31E-08 | 1.12E-07 | 185.659278 | 280.623378 | 235.843069 | 318.259459 |
| -5.6849045 | 1.31E-08 | 1.12E-07 | 177.052556 | 382.10242 | 266.905327 | 305.372192 |
| -5.6847934 | 1.31E-08 | 1.12E-07 | 146.314265 | 448.943283 | 453.278874 | 424.159173 |
| -5.6848281 | 1.31E-08 | 1.12E-07 | 183.200214 | 266.551617 | 271.507143 | 243.177122 |
| -5.6844988 | 1.31E-08 | 1.12E-07 | 225.823978 | 260.32757 | 195.577179 | 230.289855 |
| -5.6838612 | 1.32E-08 | 1.13E-07 | 139.346919 | 462.203211 | 564.872912 | 514.930357 |
| -5.6828722 | 1.32E-08 | 1.13E-07 | 208.200691 | 296.048192 | 297.967585 | 200.032794 |
| -5.6827967 | 1.33E-08 | 1.14E-07 | 186.888809 | 522.820025 | 453.278874 | 467.303501 |
| -5.6826845 | 1.33E-08 | 1.14E-07 | 159.429269 | 398.068456 | 326.728935 | 365.325998 |
| -5.6823389 | 1.33E-08 | 1.14E-07 | 189.757716 | 278.187881 | 294.516223 | 223.005748 |
| -5.6821389 | 1.33E-08 | 1.14E-07 | 143.445358 | 466.803594 | 438.322972 | 483.552664 |
| -5.6813716 | 1.34E-08 | 1.14E-07 | 180.331307 | 805.337679 | 659.210139 | 638.199865 |
| -5.6813939 | 1.34E-08 | 1.14E-07 | 200.823501 | 267.634061 | 216.285351 | 230.850171 |
| -5.6811678 | 1.34E-08 | 1.14E-07 | 159.429269 | 386.973414 | 327.879389 | 355.800627 |
| -5.6808997 | 1.34E-08 | 1.15E-07 | 183.200214 | 235.160767 | 312.923487 | 325.543566 |
| -5.6808397 | 1.34E-08 | 1.15E-07 | 171.724586 | 341.510803 | 272.657597 | 304.811876 |
| -5.6807412 | 1.34E-08 | 1.15E-07 | 165.167084 | 317.426444 | 333.631659 | 304.811876 |
| -5.6798534 | 1.35E-08 | 1.15E-07 | 155.740674 | 350.982181 | 673.015587 | 699.274304 |
| -5.6792402 | 1.35E-08 | 1.16E-07 | 146.724109 | 397.527235 | 457.88069 | 460.57971 |
| -5.6782784 | 1.36E-08 | 1.16E-07 | 146.314265 | 713.600625 | 410.712076 | 530.058887 |
| -5.6779772 | 1.36E-08 | 1.16E-07 | 286.071028 | 220.277174 | 207.081719 | 217.962904 |
| -5.677582 | 1.37E-08 | 1.17E-07 | 162.298176 | 322.83866 | 343.985745 | 318.819775 |

| | | | | | | |
|---|---|---|---|---|---|---|
| -5.6773905 | 1.37E-08 | 1.17E-07 | 158.199738 | 342.052025 | 437.172518 | 350.197468 |
| -5.6767295 | 1.37E-08 | 1.17E-07 | 164.75724 | 396.986013 | 302.569401 | 328.345146 |
| -5.6764824 | 1.37E-08 | 1.17E-07 | 149.593016 | 792.889583 | 410.712076 | 526.696992 |
| -5.6765171 | 1.37E-08 | 1.17E-07 | 195.905375 | 266.822228 | 287.613499 | 270.632603 |
| -5.6765033 | 1.37E-08 | 1.17E-07 | 151.642236 | 378.043258 | 423.36707 | 381.014845 |
| -5.6765483 | 1.37E-08 | 1.17E-07 | 181.970683 | 280.893989 | 240.444885 | 262.227864 |
| -5.6765072 | 1.37E-08 | 1.17E-07 | 150.822548 | 410.516552 | 292.215315 | 309.294404 |
| -5.6762945 | 1.38E-08 | 1.17E-07 | 195.495531 | 258.162683 | 279.560321 | 212.359745 |
| -5.6756754 | 1.38E-08 | 1.18E-07 | 182.79037 | 329.333318 | 236.993523 | 368.127578 |
| -5.6754287 | 1.38E-08 | 1.18E-07 | 167.216303 | 421.882205 | 280.710775 | 393.341796 |
| -5.6747277 | 1.39E-08 | 1.18E-07 | 200.413657 | 400.503953 | 289.914407 | 340.111781 |
| -5.6745113 | 1.39E-08 | 1.18E-07 | 167.626147 | 339.616528 | 277.259413 | 328.345146 |
| -5.6743037 | 1.39E-08 | 1.18E-07 | 165.167084 | 496.57078 | 310.622579 | 366.44663 |
| -5.6738213 | 1.40E-08 | 1.19E-07 | 173.773805 | 284.95315 | 341.684837 | 265.58976 |
| -5.6733229 | 1.40E-08 | 1.19E-07 | 152.871767 | 451.108169 | 347.437107 | 408.470327 |
| -5.6727749 | 1.41E-08 | 1.19E-07 | 122.953164 | 785.04187 | 670.714679 | 583.288903 |
| -5.6726368 | 1.41E-08 | 1.19E-07 | 154.101299 | 419.446708 | 376.198456 | 346.835572 |
| 5.6724678 | 1.41E-08 | 1.20E-07 | 1.22953164 | 5.1416048 | 3.45136199 | 3.92221164 |
| -5.6721965 | 1.41E-08 | 1.20E-07 | 156.970206 | 371.5486 | 339.383929 | 345.71494 |
| -5.6722064 | 1.41E-08 | 1.20E-07 | 422.54904 | 320.132552 | 179.470823 | 254.943757 |
| -5.6720634 | 1.41E-08 | 1.20E-07 | 179.511619 | 314.179115 | 263.453965 | 249.340597 |
| -5.6721049 | 1.41E-08 | 1.20E-07 | 181.560839 | 317.426444 | 296.817131 | 233.651751 |
| -5.6719674 | 1.41E-08 | 1.20E-07 | 148.773328 | 415.116935 | 390.003904 | 402.867167 |
| -5.6717242 | 1.41E-08 | 1.20E-07 | 175.823024 | 249.503138 | 293.365769 | 285.200818 |
| -5.6715609 | 1.42E-08 | 1.20E-07 | 140.166607 | 508.207043 | 437.172518 | 487.474875 |
| -5.6703664 | 1.42E-08 | 1.21E-07 | 242.217733 | 210.535186 | 418.765254 | 242.616806 |
| -5.6703771 | 1.42E-08 | 1.21E-07 | 153.691455 | 409.70472 | 352.038923 | 365.886314 |
| -5.6698935 | 1.43E-08 | 1.21E-07 | 159.019425 | 372.089821 | 315.224395 | 347.395888 |
| -5.6698463 | 1.43E-08 | 1.21E-07 | 179.511619 | 326.3566 | 266.905327 | 248.780281 |
| -5.6696122 | 1.43E-08 | 1.21E-07 | 231.151948 | 218.382899 | 195.577179 | 230.289855 |
| -5.6692998 | 1.43E-08 | 1.21E-07 | 147.543797 | 440.283738 | 386.552542 | 411.271906 |
| -5.6690913 | 1.44E-08 | 1.21E-07 | 218.036944 | 258.162683 | 189.824909 | 245.978701 |
| -5.6690165 | 1.44E-08 | 1.22E-07 | 187.298653 | 220.277174 | 281.861229 | 299.208717 |
| -5.6688796 | 1.44E-08 | 1.22E-07 | 186.888809 | 477.628025 | 402.658898 | 421.917909 |
| -5.6685143 | 1.44E-08 | 1.22E-07 | 169.265522 | 320.403163 | 272.657597 | 293.045241 |
| 5.66843464 | 1.44E-08 | 1.22E-07 | 20.492194 | 16.5072575 | 23.0090799 | 17.3697944 |
| -5.6680829 | 1.44E-08 | 1.22E-07 | 223.364915 | 292.259641 | 204.780811 | 214.601008 |
| -5.6678696 | 1.45E-08 | 1.22E-07 | 148.363485 | 398.339067 | 386.552542 | 414.073486 |
| -5.6678157 | 1.45E-08 | 1.22E-07 | 197.54475 | 366.948217 | 913.460472 | 358.041891 |
| -5.6674919 | 1.45E-08 | 1.22E-07 | 168.445835 | 319.050109 | 301.418947 | 271.753235 |
| -5.6670153 | 1.45E-08 | 1.23E-07 | 226.643666 | 210.535186 | 199.028541 | 239.815226 |
| -5.6667456 | 1.46E-08 | 1.23E-07 | 166.396615 | 244.902755 | 209.382627 | 237.013646 |
| -5.6666708 | 1.46E-08 | 1.23E-07 | 310.661661 | 261.680623 | 178.320369 | 244.297754 |

| | | | | | | |
|---|---|---|---|---|---|---|
| -5.6662435 | 1.46E-08 | 1.23E-07 | 157.789894 | 419.717318 | 340.534383 | 329.465778 |
| -5.6655821 | 1.47E-08 | 1.24E-07 | 150.822548 | 631.064337 | 464.783414 | 526.696992 |
| -5.6654166 | 1.47E-08 | 1.24E-07 | 190.987248 | 511.995594 | 355.490285 | 245.418385 |
| -5.6653668 | 1.47E-08 | 1.24E-07 | 141.396139 | 425.670756 | 532.6602 | 448.252759 |
| 5.66514018 | 1.47E-08 | 1.24E-07 | 29.9186032 | 22.7313054 | 31.0622579 | 28.5761134 |
| -5.6647333 | 1.47E-08 | 1.24E-07 | 186.069121 | 832.128146 | 274.958505 | 694.791776 |
| 5.66415582 | 1.48E-08 | 1.24E-07 | 123.772852 | 58.9931499 | 65.5758777 | 63.3157022 |
| -5.6640356 | 1.48E-08 | 1.25E-07 | 173.363961 | 851.0709 | 318.675757 | 538.463627 |
| -5.663717 | 1.48E-08 | 1.25E-07 | 235.660231 | 187.262659 | 371.596641 | 275.675447 |
| -5.6634371 | 1.48E-08 | 1.25E-07 | 149.183172 | 451.920001 | 385.402088 | 371.489474 |
| -5.6632575 | 1.49E-08 | 1.25E-07 | 268.037897 | 317.697055 | 193.276271 | 228.048591 |
| -5.6631593 | 1.49E-08 | 1.25E-07 | 135.658324 | 470.592145 | 495.845672 | 629.795126 |
| -5.6627243 | 1.49E-08 | 1.25E-07 | 186.478965 | 284.141318 | 217.435805 | 276.796079 |
| -5.6623565 | 1.49E-08 | 1.26E-07 | 159.429269 | 327.709654 | 315.224395 | 328.905462 |
| -5.6617927 | 1.50E-08 | 1.26E-07 | 165.167084 | 290.635977 | 291.064861 | 314.337247 |
| -5.6597611 | 1.52E-08 | 1.27E-07 | 813.949946 | 390.761965 | 234.692615 | 245.418385 |
| -5.659656 | 1.52E-08 | 1.27E-07 | 234.020855 | 292.259641 | 190.975363 | 221.885116 |
| -5.6595336 | 1.52E-08 | 1.27E-07 | 488.533905 | 1233.44393 | 1494.43974 | 1459.62305 |
| -5.659532 | 1.52E-08 | 1.27E-07 | 224.184602 | 290.906588 | 200.178995 | 214.601008 |
| -5.6592114 | 1.52E-08 | 1.28E-07 | 254.103206 | 302.813462 | 177.169915 | 244.297754 |
| -5.6592762 | 1.52E-08 | 1.28E-07 | 561.895959 | 390.491354 | 165.665375 | 302.570612 |
| -5.6593158 | 1.52E-08 | 1.28E-07 | 180.741151 | 332.039426 | 248.498063 | 252.702493 |
| -5.659214 | 1.52E-08 | 1.28E-07 | 205.331784 | 222.17145 | 209.382627 | 230.289855 |
| -5.6589696 | 1.52E-08 | 1.28E-07 | 161.478489 | 496.841391 | 294.516223 | 430.322649 |
| -5.6588945 | 1.52E-08 | 1.28E-07 | 162.70802 | 349.087905 | 291.064861 | 308.734088 |
| -5.6586968 | 1.53E-08 | 1.28E-07 | 129.92051 | 492.782229 | 629.298335 | 563.117528 |
| -5.6585318 | 1.53E-08 | 1.28E-07 | 171.314742 | 293.612695 | 356.640739 | 333.387989 |
| -5.6583549 | 1.53E-08 | 1.28E-07 | 156.560362 | 323.10927 | 357.791193 | 331.707042 |
| -5.6578279 | 1.53E-08 | 1.28E-07 | 162.70802 | 298.483689 | 362.393009 | 299.208717 |
| -5.6571539 | 1.54E-08 | 1.29E-07 | 137.707544 | 523.902468 | 425.667978 | 495.319299 |
| -5.6566892 | 1.54E-08 | 1.29E-07 | 172.544273 | 290.906588 | 439.473426 | 288.562714 |
| -5.6564687 | 1.55E-08 | 1.29E-07 | 149.183172 | 348.817294 | 402.658898 | 410.71159 |
| -5.6563008 | 1.55E-08 | 1.29E-07 | 125.822071 | 717.930397 | 503.89885 | 667.336295 |
| -5.656266 | 1.55E-08 | 1.29E-07 | 140.576451 | 436.765798 | 436.022064 | 441.528968 |
| -5.6558526 | 1.55E-08 | 1.30E-07 | 139.756763 | 426.753199 | 624.696519 | 468.984449 |
| -5.655584 | 1.55E-08 | 1.30E-07 | 134.428793 | 453.814277 | 538.41247 | 511.568461 |
| -5.6552033 | 1.56E-08 | 1.30E-07 | 180.741151 | 359.371115 | 236.993523 | 283.51987 |
| -5.6548868 | 1.56E-08 | 1.30E-07 | 366.400429 | 674.632672 | 702.927391 | 690.869565 |
| -5.6548678 | 1.56E-08 | 1.30E-07 | 168.855679 | 400.774564 | 261.153057 | 345.154624 |
| -5.6546841 | 1.56E-08 | 1.30E-07 | 389.76153 | 494.947115 | 141.505841 | 444.330547 |
| -5.6546238 | 1.56E-08 | 1.30E-07 | 145.494577 | 457.873438 | 369.295733 | 422.478225 |
| -5.6546127 | 1.56E-08 | 1.30E-07 | 159.429269 | 343.67569 | 296.817131 | 327.224514 |
| -5.6545559 | 1.56E-08 | 1.30E-07 | 311.071505 | 545.821942 | 582.129722 | 629.795126 |

| | | | | | | |
|---|---|---|---|---|---|---|
| -5.6545095 | 1.56E-08 | 1.30E-07 | 563.125491 | 239.49054 | 240.444885 | 259.426284 |
| -5.6544255 | 1.56E-08 | 1.30E-07 | 160.658801 | 337.722252 | 289.914407 | 324.422934 |
| -5.6536121 | 1.57E-08 | 1.31E-07 | 154.511143 | 478.169247 | 337.083021 | 359.722839 |
| -5.6534627 | 1.57E-08 | 1.31E-07 | 257.381957 | 598.320433 | 477.438408 | 572.082584 |
| -5.6534849 | 1.57E-08 | 1.31E-07 | 143.855202 | 439.471905 | 376.198456 | 467.303501 |
| -5.6533632 | 1.57E-08 | 1.31E-07 | 151.232392 | 398.609678 | 334.782113 | 377.652949 |
| -5.6530705 | 1.58E-08 | 1.31E-07 | 186.888809 | 256.539019 | 222.037621 | 230.850171 |
| -5.6528601 | 1.58E-08 | 1.31E-07 | 127.871291 | 699.528864 | 569.474728 | 504.284354 |
| -5.6528152 | 1.58E-08 | 1.31E-07 | 167.216303 | 334.204312 | 257.701695 | 332.827673 |
| -5.6525251 | 1.58E-08 | 1.32E-07 | 210.24991 | 275.752384 | 218.586259 | 241.496174 |
| -5.6519882 | 1.59E-08 | 1.32E-07 | 172.544273 | 297.671857 | 243.896247 | 281.838922 |
| -5.6520183 | 1.59E-08 | 1.32E-07 | 0 | 253.832911 | 407.260714 | 0 |
| -5.6519377 | 1.59E-08 | 1.32E-07 | 205.741628 | 262.763067 | 225.488983 | 198.351846 |
| -5.6515973 | 1.59E-08 | 1.32E-07 | 171.314742 | 269.528336 | 253.099879 | 302.010296 |
| -5.6512623 | 1.59E-08 | 1.32E-07 | 168.445835 | 333.663091 | 254.250333 | 310.415036 |
| -5.6511686 | 1.59E-08 | 1.32E-07 | 157.789894 | 406.998612 | 317.525303 | 320.500723 |
| -5.6511454 | 1.59E-08 | 1.32E-07 | 165.167084 | 388.597079 | 266.905327 | 348.51652 |
| -5.6502997 | 1.60E-08 | 1.33E-07 | 165.167084 | 303.084073 | 266.905327 | 322.741986 |
| -5.6502401 | 1.60E-08 | 1.33E-07 | 146.314265 | 444.61351 | 408.411168 | 358.602207 |
| -5.649998 | 1.60E-08 | 1.33E-07 | 213.938505 | 189.427545 | 241.595339 | 283.51987 |
| -5.6500413 | 1.60E-08 | 1.33E-07 | 189.757716 | 246.255809 | 222.037621 | 261.667548 |
| -5.6487891 | 1.62E-08 | 1.34E-07 | 192.21678 | 248.961917 | 212.833989 | 222.445432 |
| -5.648653 | 1.62E-08 | 1.34E-07 | 138.527231 | 447.319618 | 531.509746 | 418.556014 |
| -5.6482296 | 1.62E-08 | 1.34E-07 | 177.4624 | 575.318516 | 514.252936 | 603.460277 |
| -5.6479288 | 1.62E-08 | 1.35E-07 | 142.215826 | 411.057774 | 408.411168 | 416.875066 |
| 5.64763595 | 1.63E-08 | 1.35E-07 | 10.246097 | 5.68282636 | 6.90272397 | 6.16347543 |
| -5.6470656 | 1.63E-08 | 1.35E-07 | 146.724109 | 425.941366 | 377.34891 | 364.205367 |
| -5.6466116 | 1.64E-08 | 1.36E-07 | 238.119294 | 242.467258 | 186.373547 | 213.480376 |
| -5.6466493 | 1.64E-08 | 1.36E-07 | 193.856155 | 328.521486 | 204.780811 | 356.921259 |
| -5.6460207 | 1.64E-08 | 1.36E-07 | 178.282088 | 282.788264 | 226.639437 | 261.107232 |
| -5.6456532 | 1.65E-08 | 1.36E-07 | 167.626147 | 283.329486 | 258.852149 | 324.422934 |
| -5.6455439 | 1.65E-08 | 1.36E-07 | 163.527708 | 312.014228 | 296.817131 | 271.753235 |
| -5.6451774 | 1.65E-08 | 1.37E-07 | 147.543797 | 376.148983 | 352.038923 | 421.357593 |
| -5.6450111 | 1.65E-08 | 1.37E-07 | 165.576927 | 273.316887 | 300.268493 | 272.873867 |
| -5.6444929 | 1.66E-08 | 1.37E-07 | 116.805506 | 595.343714 | 756.998729 | 664.534715 |
| -5.64429 | 1.66E-08 | 1.37E-07 | 152.871767 | 437.57763 | 314.073941 | 379.333897 |
| -5.6437688 | 1.66E-08 | 1.38E-07 | 173.773805 | 313.637893 | 232.391707 | 312.656299 |
| -5.6436811 | 1.66E-08 | 1.38E-07 | 190.987248 | 293.071474 | 215.134897 | 231.970803 |
| -5.6436198 | 1.67E-08 | 1.38E-07 | 180.741151 | 353.417678 | 228.940345 | 281.278606 |
| -5.6434491 | 1.67E-08 | 1.38E-07 | 244.266952 | 231.913438 | 401.508444 | 200.59311 |
| -5.6434823 | 1.67E-08 | 1.38E-07 | 211.069598 | 307.955067 | 218.586259 | 205.075637 |
| -5.6434853 | 1.67E-08 | 1.38E-07 | 162.70802 | 364.51272 | 331.330751 | 365.325998 |
| -5.6429514 | 1.67E-08 | 1.38E-07 | 168.855679 | 348.276073 | 266.905327 | 270.632603 |

| | | | | | | |
|---|---|---|---|---|---|---|
| -5.6427706 | 1.67E-08 | 1.38E-07 | 181.560839 | 322.568049 | 375.048002 | 339.551465 |
| -5.6425166 | 1.68E-08 | 1.38E-07 | 130.330354 | 609.144864 | 461.332052 | 529.498571 |
| -5.6424704 | 1.68E-08 | 1.38E-07 | 143.855202 | 434.600911 | 600.536986 | 383.816425 |
| -5.6423056 | 1.68E-08 | 1.38E-07 | 157.789894 | 334.474923 | 293.365769 | 317.699143 |
| -5.6419885 | 1.68E-08 | 1.39E-07 | 213.938505 | 255.456576 | 151.859927 | 207.877217 |
| 5.6420244 | 1.68E-08 | 1.39E-07 | 43.8532952 | 41.1328384 | 39.1154358 | 40.3427483 |
| -5.641907 | 1.68E-08 | 1.39E-07 | 1111.90645 | 5266.89759 | 4684.64867 | 4667.43185 |
| -5.6413879 | 1.69E-08 | 1.39E-07 | 126.641759 | 515.784145 | 556.819734 | 542.385838 |
| -5.6412192 | 1.69E-08 | 1.39E-07 | 145.904421 | 401.586396 | 370.446187 | 367.006946 |
| 5.64102513 | 1.69E-08 | 1.39E-07 | 0 | 0 | 0 | 0 |
| -5.6403339 | 1.70E-08 | 1.40E-07 | 157.789894 | 418.364265 | 293.365769 | 342.353045 |
| -5.6399929 | 1.70E-08 | 1.40E-07 | 238.529138 | 294.424528 | 180.621277 | 227.488275 |
| -5.6398583 | 1.70E-08 | 1.40E-07 | 163.117864 | 329.333318 | 423.36707 | 276.796079 |
| -5.6393356 | 1.71E-08 | 1.40E-07 | 187.298653 | 211.88824 | 238.143977 | 230.289855 |
| -5.6394345 | 1.71E-08 | 1.40E-07 | 201.643189 | 203.228695 | 212.833989 | 260.546916 |
| -5.6393095 | 1.71E-08 | 1.40E-07 | 145.084733 | 412.952049 | 349.738015 | 417.435382 |
| -5.6393257 | 1.71E-08 | 1.40E-07 | 154.920987 | 327.709654 | 333.631659 | 306.492824 |
| -5.6387128 | 1.71E-08 | 1.41E-07 | 151.232392 | 388.86769 | 325.578481 | 341.792729 |
| -5.638273 | 1.72E-08 | 1.41E-07 | 171.314742 | 235.972599 | 304.870309 | 285.761134 |
| -5.6378697 | 1.72E-08 | 1.41E-07 | 157.789894 | 324.191713 | 464.783414 | 317.138827 |
| -5.6376059 | 1.72E-08 | 1.41E-07 | 133.199261 | 494.405893 | 445.225696 | 477.949504 |
| -5.6376792 | 1.72E-08 | 1.41E-07 | 168.035991 | 311.473007 | 262.303511 | 259.426284 |
| -5.6376302 | 1.72E-08 | 1.41E-07 | 188.938029 | 418.634875 | 219.736713 | 317.138827 |
| -5.6372628 | 1.73E-08 | 1.42E-07 | 162.70802 | 284.68254 | 271.507143 | 304.25156 |
| -5.6371874 | 1.73E-08 | 1.42E-07 | 264.759146 | 219.194731 | 186.373547 | 219.643852 |
| 5.6369422 | 1.73E-08 | 1.42E-07 | 38.5253247 | 20.2958084 | 18.4072639 | 22.9729539 |
| -5.6364175 | 1.74E-08 | 1.42E-07 | 149.183172 | 365.324552 | 330.180297 | 365.325998 |
| -5.6358229 | 1.74E-08 | 1.43E-07 | 195.905375 | 516.595977 | 227.789891 | 339.551465 |
| -5.6358305 | 1.74E-08 | 1.43E-07 | 161.068645 | 366.136384 | 266.905327 | 335.068937 |
| -5.6355647 | 1.74E-08 | 1.43E-07 | 155.740674 | 425.670756 | 296.817131 | 349.637152 |
| -5.6355204 | 1.75E-08 | 1.43E-07 | 179.511619 | 437.307019 | 409.561622 | 461.140026 |
| -5.6351348 | 1.75E-08 | 1.43E-07 | 154.511143 | 485.205127 | 370.446187 | 310.415036 |
| -5.6350273 | 1.75E-08 | 1.43E-07 | 168.855679 | 248.150084 | 288.763953 | 261.667548 |
| -5.634796 | 1.75E-08 | 1.43E-07 | 177.872244 | 270.340168 | 218.586259 | 266.710392 |
| -5.6347664 | 1.75E-08 | 1.43E-07 | 141.805982 | 361.806612 | 467.084322 | 430.882964 |
| -5.6345767 | 1.75E-08 | 1.44E-07 | 154.920987 | 317.155833 | 301.418947 | 346.275256 |
| -5.6340969 | 1.76E-08 | 1.44E-07 | 135.24848 | 501.982995 | 445.225696 | 423.038541 |
| -5.6341033 | 1.76E-08 | 1.44E-07 | 129.510666 | 531.750181 | 506.199758 | 466.743185 |
| -5.6337381 | 1.76E-08 | 1.44E-07 | 235.250387 | 452.190612 | 232.391707 | 411.832222 |
| -5.6327816 | 1.77E-08 | 1.45E-07 | 169.675366 | 261.951234 | 358.941647 | 263.908812 |
| -5.6326701 | 1.77E-08 | 1.45E-07 | 190.16756 | 205.122971 | 224.338529 | 239.25491 |
| -5.6326472 | 1.77E-08 | 1.45E-07 | 144.67489 | 417.281821 | 356.640739 | 372.610106 |
| -5.6325107 | 1.78E-08 | 1.45E-07 | 319.268382 | 158.036695 | 429.11934 | 299.208717 |

| | | | | | | |
|---|---|---|---|---|---|---|
| -5.6322841 | 1.78E-08 | 1.45E-07 | 164.75724 | 268.716504 | 262.303511 | 299.208717 |
| -5.6322947 | 1.78E-08 | 1.45E-07 | 151.232392 | 454.355498 | 316.374849 | 367.006946 |
| -5.6313251 | 1.79E-08 | 1.46E-07 | 162.298176 | 270.340168 | 278.409867 | 293.045241 |
| -5.630533 | 1.80E-08 | 1.47E-07 | 181.150995 | 250.585581 | 253.099879 | 232.531119 |
| -5.6303207 | 1.80E-08 | 1.47E-07 | 170.904898 | 384.267306 | 238.143977 | 398.944955 |
| -5.6298782 | 1.80E-08 | 1.47E-07 | 140.166607 | 500.900552 | 390.003904 | 397.824324 |
| -5.6296332 | 1.81E-08 | 1.47E-07 | 168.035991 | 329.062707 | 272.657597 | 248.780281 |
| -5.6294425 | 1.81E-08 | 1.47E-07 | 159.839113 | 352.876456 | 270.356689 | 308.734088 |
| -5.629132 | 1.81E-08 | 1.48E-07 | 162.70802 | 257.08024 | 302.569401 | 335.629253 |
| -5.6286913 | 1.82E-08 | 1.48E-07 | 163.527708 | 363.700887 | 266.905327 | 288.002398 |
| -5.6281153 | 1.82E-08 | 1.48E-07 | 304.923847 | 307.955067 | 158.762651 | 266.710392 |
| -5.6276568 | 1.83E-08 | 1.49E-07 | 138.117388 | 353.417678 | 281.861229 | 327.78483 |
| -5.6274121 | 1.83E-08 | 1.49E-07 | 159.839113 | 387.785247 | 268.055781 | 333.387989 |
| -5.6270871 | 1.83E-08 | 1.49E-07 | 144.265046 | 394.009294 | 356.640739 | 363.645051 |
| -5.6268156 | 1.84E-08 | 1.49E-07 | 184.019902 | 308.225677 | 209.382627 | 258.865968 |
| -5.6261868 | 1.84E-08 | 1.50E-07 | 178.691932 | 287.929869 | 217.435805 | 248.780281 |
| -5.6258655 | 1.85E-08 | 1.50E-07 | 151.232392 | 352.335234 | 317.525303 | 317.699143 |
| -5.625753 | 1.85E-08 | 1.50E-07 | 234.430699 | 330.956983 | 178.320369 | 241.496174 |
| -5.625634 | 1.85E-08 | 1.50E-07 | 125.822071 | 493.32345 | 516.553844 | 541.825522 |
| -5.6254304 | 1.85E-08 | 1.50E-07 | 140.576451 | 394.279905 | 370.446187 | 438.727388 |
| -5.6252995 | 1.85E-08 | 1.50E-07 | 144.265046 | 367.218827 | 468.234776 | 356.921259 |
| -5.6251476 | 1.85E-08 | 1.50E-07 | 154.511143 | 323.379881 | 291.064861 | 317.699143 |
| -5.6251536 | 1.85E-08 | 1.50E-07 | 160.658801 | 292.259641 | 280.710775 | 270.632603 |
| -5.6250235 | 1.85E-08 | 1.51E-07 | 396.319032 | 219.465342 | 208.232173 | 233.091435 |
| -5.6247028 | 1.86E-08 | 1.51E-07 | 165.576927 | 252.479857 | 273.808051 | 262.78818 |
| -5.6247262 | 1.86E-08 | 1.51E-07 | 137.2977 | 388.326468 | 453.278874 | 424.719489 |
| -5.6243343 | 1.86E-08 | 1.51E-07 | 150.412704 | 403.480672 | 311.773033 | 339.551465 |
| -5.624224 | 1.86E-08 | 1.51E-07 | 175.823024 | 235.972599 | 261.153057 | 351.878416 |
| -5.6241558 | 1.86E-08 | 1.51E-07 | 346.318079 | 164.531354 | 260.002603 | 277.356395 |
| -5.6237996 | 1.87E-08 | 1.51E-07 | 173.363961 | 273.316887 | 243.896247 | 229.169223 |
| -5.6235292 | 1.87E-08 | 1.52E-07 | 150.412704 | 329.603929 | 312.923487 | 344.033992 |
| -5.6232944 | 1.87E-08 | 1.52E-07 | 112.707067 | 760.6869 | 579.828814 | 697.593356 |
| -5.623266 | 1.87E-08 | 1.52E-07 | 193.036467 | 295.777582 | 285.312591 | 192.18837 |
| -5.622969 | 1.88E-08 | 1.52E-07 | 164.347396 | 319.050109 | 288.763953 | 247.099333 |
| -5.6227311 | 1.88E-08 | 1.52E-07 | 150.00286 | 327.439043 | 320.976665 | 333.948305 |
| -5.622118 | 1.89E-08 | 1.53E-07 | 222.135383 | 305.79018 | 483.190678 | 196.110582 |
| -5.6221805 | 1.89E-08 | 1.53E-07 | 134.018949 | 461.9326 | 448.677058 | 411.271906 |
| -5.6221166 | 1.89E-08 | 1.53E-07 | 130.330354 | 431.082971 | 536.111562 | 484.673296 |
| -5.6217363 | 1.89E-08 | 1.53E-07 | 147.953641 | 336.369198 | 325.578481 | 386.618005 |
| -5.6210712 | 1.90E-08 | 1.53E-07 | 231.971636 | 239.49054 | 180.621277 | 210.118481 |
| -5.6205675 | 1.90E-08 | 1.54E-07 | 174.183649 | 379.937534 | 250.798971 | 259.426284 |
| -5.6202098 | 1.91E-08 | 1.54E-07 | 202.053033 | 257.892073 | 189.824909 | 219.083536 |
| 5.62016132 | 1.91E-08 | 1.54E-07 | 0 | 0 | 0 | 0 |

| | | | | | | |
|---|---|---|---|---|---|---|
| -5.6197651 | 1.91E-08 | 1.54E-07 | 152.052079 | 359.371115 | 294.516223 | 326.103882 |
| -5.6197942 | 1.91E-08 | 1.54E-07 | 140.576451 | 382.643642 | 375.048002 | 389.9799 |
| -5.6197947 | 1.91E-08 | 1.54E-07 | 168.035991 | 276.022995 | 232.391707 | 284.640502 |
| -5.6197198 | 1.91E-08 | 1.54E-07 | 175.823024 | 230.289773 | 280.710775 | 228.608907 |
| -5.6195588 | 1.91E-08 | 1.54E-07 | 113.526755 | 640.535714 | 587.881992 | 869.610352 |
| -5.6194911 | 1.92E-08 | 1.54E-07 | 150.00286 | 375.607761 | 303.719855 | 338.430833 |
| -5.6192657 | 1.92E-08 | 1.55E-07 | 175.413181 | 983.128961 | 631.599243 | 655.009344 |
| -5.619271 | 1.92E-08 | 1.55E-07 | 177.052556 | 212.429462 | 314.073941 | 303.130928 |
| -5.6188696 | 1.92E-08 | 1.55E-07 | 145.084733 | 391.032576 | 348.587561 | 338.991149 |
| -5.6183594 | 1.93E-08 | 1.55E-07 | 214.758193 | 289.282923 | 181.771731 | 230.289855 |
| -5.6183791 | 1.93E-08 | 1.55E-07 | 147.543797 | 330.956983 | 362.393009 | 326.664198 |
| -5.6184182 | 1.93E-08 | 1.55E-07 | 157.38005 | 297.942468 | 288.763953 | 277.356395 |
| -5.617706 | 1.94E-08 | 1.56E-07 | 150.00286 | 304.707737 | 327.879389 | 358.602207 |
| -5.6175424 | 1.94E-08 | 1.56E-07 | 164.75724 | 258.162683 | 262.303511 | 254.383441 |
| -5.617441 | 1.94E-08 | 1.56E-07 | 181.560839 | 207.558468 | 323.277573 | 277.916711 |
| -5.6172299 | 1.94E-08 | 1.56E-07 | 137.707544 | 467.615426 | 415.313892 | 439.287704 |
| -5.6167643 | 1.95E-08 | 1.56E-07 | 251.234298 | 614.82769 | 491.243856 | 501.482774 |
| -5.6167505 | 1.95E-08 | 1.56E-07 | 205.331784 | 232.454659 | 203.630357 | 192.748686 |
| -5.6166721 | 1.95E-08 | 1.56E-07 | 171.314742 | 246.52642 | 238.143977 | 235.332698 |
| -5.6162666 | 1.95E-08 | 1.57E-07 | 203.692408 | 192.133653 | 349.738015 | 259.9866 |
| -5.6162206 | 1.95E-08 | 1.57E-07 | 1359.04231 | 393.738684 | 215.134897 | 374.85137 |
| -5.6161493 | 1.95E-08 | 1.57E-07 | 135.658324 | 406.186779 | 403.809352 | 436.486124 |
| -5.6161068 | 1.95E-08 | 1.57E-07 | 185.249434 | 329.87454 | 210.533081 | 251.581861 |
| -5.6160872 | 1.95E-08 | 1.57E-07 | 136.068168 | 418.364265 | 421.066162 | 394.462428 |
| -5.6158449 | 1.96E-08 | 1.57E-07 | 147.133953 | 361.536001 | 320.976665 | 337.870517 |
| -5.6157406 | 1.96E-08 | 1.57E-07 | 181.150995 | 325.544767 | 300.268493 | 316.018195 |
| -5.6156124 | 1.96E-08 | 1.57E-07 | 2105.77786 | 325.544767 | 357.791193 | 381.014845 |
| -5.614812 | 1.97E-08 | 1.58E-07 | 140.986295 | 443.260456 | 343.985745 | 400.065587 |
| -5.6147459 | 1.97E-08 | 1.58E-07 | 166.806459 | 289.553534 | 236.993523 | 260.546916 |
| -5.6146773 | 1.97E-08 | 1.58E-07 | 202.053033 | 245.714587 | 538.41247 | 272.313551 |
| -5.614642 | 1.97E-08 | 1.58E-07 | 199.184126 | 320.132552 | 189.824909 | 256.064389 |
| -5.6143814 | 1.97E-08 | 1.58E-07 | 155.33083 | 293.342085 | 288.763953 | 291.924609 |
| -5.6141637 | 1.98E-08 | 1.58E-07 | 528.698605 | 1276.47105 | 1077.97539 | 1126.23506 |
| -5.6137301 | 1.98E-08 | 1.59E-07 | 149.183172 | 302.542851 | 339.383929 | 337.870517 |
| -5.6136724 | 1.98E-08 | 1.59E-07 | 150.822548 | 378.58448 | 307.171217 | 310.415036 |
| -5.6136438 | 1.98E-08 | 1.59E-07 | 145.904421 | 676.256337 | 350.888469 | 449.373391 |
| -5.6133879 | 1.98E-08 | 1.59E-07 | 165.167084 | 341.510803 | 272.657597 | 248.219965 |
| -5.6134158 | 1.98E-08 | 1.59E-07 | 189.347873 | 305.79018 | 204.780811 | 233.091435 |
| -5.613268 | 1.99E-08 | 1.59E-07 | 141.805982 | 419.987929 | 339.383929 | 384.376741 |
| -5.6131448 | 1.99E-08 | 1.59E-07 | 155.740674 | 553.399043 | 295.666677 | 394.462428 |
| -5.6128376 | 1.99E-08 | 1.59E-07 | 190.16756 | 527.96163 | 228.940345 | 335.068937 |
| -5.6127092 | 1.99E-08 | 1.59E-07 | 140.576451 | 358.288672 | 395.756174 | 372.610106 |
| -5.6124991 | 1.99E-08 | 1.59E-07 | 161.068645 | 281.1646 | 250.798971 | 291.364293 |

| | | | | | | |
|---|---|---|---|---|---|---|
| -5.6123679 | 2.00E-08 | 1.59E-07 | 136.068168 | 394.279905 | 415.313892 | 404.548115 |
| -5.6124071 | 2.00E-08 | 1.59E-07 | 156.150518 | 300.648576 | 270.356689 | 302.570612 |
| -5.6122304 | 2.00E-08 | 1.59E-07 | 186.069121 | 283.870707 | 288.763953 | 194.98995 |
| -5.6116828 | 2.00E-08 | 1.60E-07 | 222.135383 | 189.156935 | 391.154358 | 252.702493 |
| -5.6113309 | 2.01E-08 | 1.60E-07 | 134.428793 | 456.249774 | 378.499364 | 467.863817 |
| -5.6113387 | 2.01E-08 | 1.60E-07 | 148.363485 | 334.204312 | 318.675757 | 318.259459 |
| -5.6111675 | 2.01E-08 | 1.60E-07 | 141.396139 | 354.22951 | 358.941647 | 421.357593 |
| -5.6110763 | 2.01E-08 | 1.60E-07 | 166.396615 | 612.933415 | 624.696519 | 647.164921 |
| 5.61058244 | 2.02E-08 | 1.61E-07 | 61.0667381 | 38.9679522 | 44.8677058 | 36.9808526 |
| -5.6105178 | 2.02E-08 | 1.61E-07 | 126.641759 | 435.142133 | 733.989649 | 566.479424 |
| -5.6104453 | 2.02E-08 | 1.61E-07 | 150.412704 | 551.234157 | 307.171217 | 411.832222 |
| -5.6101454 | 2.02E-08 | 1.61E-07 | 147.543797 | 303.354683 | 515.40339 | 452.735286 |
| -5.6097827 | 2.03E-08 | 1.61E-07 | 144.265046 | 471.133366 | 312.923487 | 447.692443 |
| -5.609097 | 2.03E-08 | 1.62E-07 | 122.54332 | 544.198277 | 488.942948 | 524.455728 |
| -5.6090892 | 2.03E-08 | 1.62E-07 | 263.939459 | 322.297438 | 161.063559 | 254.943757 |
| -5.6089091 | 2.04E-08 | 1.62E-07 | 117.21535 | 621.86357 | 544.16474 | 561.436581 |
| -5.6087853 | 2.04E-08 | 1.62E-07 | 123.772852 | 686.539547 | 477.438408 | 513.249409 |
| -5.6081879 | 2.04E-08 | 1.63E-07 | 124.182696 | 464.097486 | 560.271096 | 507.085933 |
| -5.6080243 | 2.05E-08 | 1.63E-07 | 219.67632 | 278.729103 | 169.116737 | 263.348496 |
| -5.6078754 | 2.05E-08 | 1.63E-07 | 159.019425 | 256.809629 | 277.259413 | 303.130928 |
| -5.6077462 | 2.05E-08 | 1.63E-07 | 136.478012 | 386.973414 | 381.950726 | 453.295602 |
| -5.6071841 | 2.06E-08 | 1.63E-07 | 175.823024 | 349.358516 | 265.754873 | 222.445432 |
| -5.6070901 | 2.06E-08 | 1.63E-07 | 158.609582 | 368.571881 | 262.303511 | 296.967453 |
| -5.607011 | 2.06E-08 | 1.64E-07 | 187.298653 | 249.503138 | 190.975363 | 251.021545 |
| -5.6068592 | 2.06E-08 | 1.64E-07 | 159.839113 | 259.515737 | 380.800272 | 298.648401 |
| -5.6067059 | 2.06E-08 | 1.64E-07 | 138.527231 | 459.767714 | 409.561622 | 347.956204 |
| -5.6064487 | 2.07E-08 | 1.64E-07 | 131.559885 | 474.651307 | 429.11934 | 410.71159 |
| -5.6056847 | 2.07E-08 | 1.65E-07 | 132.379573 | 440.554348 | 418.765254 | 412.392538 |
| -5.605352 | 2.08E-08 | 1.65E-07 | 175.823024 | 414.034492 | 373.897549 | 392.78148 |
| -5.6052723 | 2.08E-08 | 1.65E-07 | 375.00715 | 292.530252 | 173.718553 | 221.885116 |
| -5.6047843 | 2.09E-08 | 1.65E-07 | 109.428316 | 721.989559 | 724.786017 | 574.323847 |
| -5.6043135 | 2.09E-08 | 1.66E-07 | 148.773328 | 297.130636 | 318.675757 | 340.111781 |
| -5.6042115 | 2.09E-08 | 1.66E-07 | 204.92194 | 233.537102 | 180.621277 | 219.083536 |
| -5.6041583 | 2.09E-08 | 1.66E-07 | 139.346919 | 415.658157 | 353.189377 | 368.687894 |
| -5.603311 | 2.10E-08 | 1.67E-07 | 170.904898 | 246.52642 | 218.586259 | 273.994499 |
| -5.6032811 | 2.10E-08 | 1.67E-07 | 238.119294 | 167.237462 | 238.143977 | 251.021545 |
| -5.6032209 | 2.10E-08 | 1.67E-07 | 151.642236 | 309.849342 | 287.613499 | 299.208717 |
| -5.6028455 | 2.11E-08 | 1.67E-07 | 146.724109 | 335.016145 | 339.383929 | 304.811876 |
| 5.60247794 | 2.11E-08 | 1.67E-07 | 42.6237635 | 27.6022995 | 32.2127119 | 31.3776931 |
| -5.6024393 | 2.11E-08 | 1.67E-07 | 136.887856 | 365.865773 | 401.508444 | 392.221164 |
| -5.6022976 | 2.12E-08 | 1.67E-07 | 149.183172 | 324.462324 | 292.215315 | 321.621355 |
| -5.6018563 | 2.12E-08 | 1.68E-07 | 140.576451 | 365.324552 | 345.136199 | 367.006946 |
| -5.601711 | 2.12E-08 | 1.68E-07 | 177.872244 | 291.177198 | 201.329449 | 277.916711 |

| | | | | | | |
|---|---|---|---|---|---|---|
| -5.6014838 | 2.13E-08 | 1.68E-07 | 169.675366 | 227.583665 | 236.993523 | 282.399238 |
| -5.6015101 | 2.12E-08 | 1.68E-07 | 151.232392 | 300.107354 | 346.286653 | 285.200818 |
| -5.6014782 | 2.13E-08 | 1.68E-07 | 263.939459 | 191.863042 | 223.188075 | 191.067738 |
| -5.6014556 | 2.13E-08 | 1.68E-07 | 140.576451 | 414.305103 | 324.428027 | 414.073486 |
| -5.6010513 | 2.13E-08 | 1.68E-07 | 186.478965 | 206.746635 | 265.754873 | 207.877217 |
| -5.6003152 | 2.14E-08 | 1.69E-07 | 157.789894 | 347.734851 | 247.347609 | 345.71494 |
| -5.5996286 | 2.15E-08 | 1.70E-07 | 148.363485 | 443.801678 | 288.763953 | 380.454529 |
| -5.5989945 | 2.16E-08 | 1.70E-07 | 147.543797 | 334.745534 | 332.481205 | 293.605557 |
| -5.5987763 | 2.16E-08 | 1.70E-07 | 240.988201 | 366.406995 | 295.666677 | 319.380091 |
| -5.5982878 | 2.16E-08 | 1.71E-07 | 149.183172 | 291.447809 | 366.994825 | 463.38129 |
| -5.5982323 | 2.17E-08 | 1.71E-07 | 150.00286 | 326.3566 | 308.321671 | 284.080186 |
| -5.598113 | 2.17E-08 | 1.71E-07 | 144.67489 | 343.405079 | 322.127119 | 323.302302 |
| -5.5980739 | 2.17E-08 | 1.71E-07 | 141.396139 | 425.941366 | 322.127119 | 375.972002 |
| -5.5975795 | 2.17E-08 | 1.71E-07 | 159.839113 | 249.503138 | 262.303511 | 303.691244 |
| -5.5974603 | 2.18E-08 | 1.71E-07 | 203.692408 | 178.332503 | 226.639437 | 237.573962 |
| -5.5973633 | 2.18E-08 | 1.71E-07 | 155.33083 | 348.546684 | 254.250333 | 324.422934 |
| -5.5969025 | 2.18E-08 | 1.72E-07 | 139.346919 | 327.439043 | 408.411168 | 403.987799 |
| -5.596561 | 2.19E-08 | 1.72E-07 | 193.856155 | 311.473007 | 335.932567 | 184.904263 |
| -5.5961684 | 2.19E-08 | 1.72E-07 | 134.838636 | 571.259355 | 696.024667 | 358.041891 |
| -5.5959854 | 2.19E-08 | 1.73E-07 | 139.756763 | 388.86769 | 334.782113 | 365.886314 |
| -5.5956478 | 2.20E-08 | 1.73E-07 | 143.855202 | 320.673773 | 368.145279 | 323.862618 |
| -5.5954976 | 2.20E-08 | 1.73E-07 | 156.150518 | 280.623378 | 265.754873 | 267.831023 |
| -5.5953741 | 2.20E-08 | 1.73E-07 | 126.641759 | 413.22266 | 566.023366 | 741.858316 |
| -5.5951069 | 2.20E-08 | 1.73E-07 | 181.970683 | 195.380983 | 293.365769 | 257.18502 |
| -5.5948548 | 2.21E-08 | 1.74E-07 | 130.330354 | 393.738684 | 524.607022 | 437.04644 |
| -5.5946553 | 2.21E-08 | 1.74E-07 | 152.461923 | 701.152529 | 364.693917 | 360.843471 |
| -5.5945901 | 2.21E-08 | 1.74E-07 | 324.186509 | 1138.45955 | 908.858656 | 1011.37029 |
| -5.5944153 | 2.21E-08 | 1.74E-07 | 86.8869025 | 830.775092 | 1112.48901 | 997.922704 |
| -5.5944008 | 2.21E-08 | 1.74E-07 | 141.805982 | 471.403977 | 332.481205 | 359.162523 |
| -5.594002 | 2.22E-08 | 1.74E-07 | 181.560839 | 225.68939 | 196.727633 | 228.048591 |
| -5.5931019 | 2.23E-08 | 1.75E-07 | 137.2977 | 387.785247 | 341.684837 | 409.590958 |
| -5.59294 | 2.23E-08 | 1.75E-07 | 251.644142 | 223.795114 | 173.718553 | 209.558165 |
| -5.5928218 | 2.23E-08 | 1.75E-07 | 221.315695 | 166.155018 | 338.233475 | 319.940407 |
| -5.5927647 | 2.23E-08 | 1.75E-07 | 170.495054 | 221.900839 | 225.488983 | 237.013646 |
| -5.5927356 | 2.24E-08 | 1.75E-07 | 188.528185 | 265.469174 | 193.276271 | 217.402588 |
| -5.5926778 | 2.24E-08 | 1.75E-07 | 139.756763 | 351.523402 | 335.932567 | 401.746535 |
| 5.59256907 | 2.24E-08 | 1.75E-07 | 0 | 0 | 0 | 0 |
| -5.5923822 | 2.24E-08 | 1.75E-07 | 140.986295 | 419.176097 | 322.127119 | 360.283155 |
| -5.5921451 | 2.24E-08 | 1.76E-07 | 131.150042 | 384.267306 | 448.677058 | 425.840121 |
| -5.5920792 | 2.24E-08 | 1.76E-07 | 175.413181 | 366.136384 | 432.570702 | 373.730738 |
| -5.5918935 | 2.25E-08 | 1.76E-07 | 155.33083 | 296.048192 | 255.400787 | 345.71494 |
| -5.5915551 | 2.25E-08 | 1.76E-07 | 127.871291 | 728.484217 | 395.756174 | 573.763531 |
| -5.5914803 | 2.25E-08 | 1.76E-07 | 154.511143 | 278.187881 | 271.507143 | 270.632603 |

| | | | | | | |
|---|---|---|---|---|---|---|
| -5.5912682 | 2.25E-08 | 1.76E-07 | 191.806936 | 190.509989 | 236.993523 | 206.196269 |
| -5.5912237 | 2.25E-08 | 1.76E-07 | 144.67489 | 320.673773 | 364.693917 | 306.492824 |
| -5.5904379 | 2.26E-08 | 1.77E-07 | 197.54475 | 293.342085 | 192.125817 | 214.601008 |
| -5.5899071 | 2.27E-08 | 1.78E-07 | 141.396139 | 350.170348 | 316.374849 | 377.652949 |
| -5.5894577 | 2.28E-08 | 1.78E-07 | 184.83959 | 193.216096 | 260.002603 | 313.776931 |
| -5.5894722 | 2.28E-08 | 1.78E-07 | 139.756763 | 366.406995 | 324.428027 | 373.730738 |
| -5.5885961 | 2.29E-08 | 1.79E-07 | 781.162435 | 836.728529 | 606.289256 | 253.262809 |
| -5.5884278 | 2.29E-08 | 1.79E-07 | 141.396139 | 317.967665 | 363.543463 | 342.353045 |
| -5.5873825 | 2.31E-08 | 1.80E-07 | 263.939459 | 266.281007 | 178.320369 | 192.748686 |
| -5.5871387 | 2.31E-08 | 1.80E-07 | 184.83959 | 220.006563 | 189.824909 | 230.850171 |
| -5.587055 | 2.31E-08 | 1.80E-07 | 135.658324 | 387.244025 | 377.34891 | 355.240311 |
| -5.5870031 | 2.31E-08 | 1.80E-07 | 165.986771 | 260.868791 | 265.754873 | 216.281956 |
| -5.5864471 | 2.32E-08 | 1.81E-07 | 176.642712 | 472.757031 | 337.083021 | 223.005748 |
| -5.5864802 | 2.32E-08 | 1.81E-07 | 222.135383 | 297.942468 | 179.470823 | 239.25491 |
| -5.5862437 | 2.32E-08 | 1.81E-07 | 183.610058 | 187.803881 | 258.852149 | 244.297754 |
| -5.586196 | 2.32E-08 | 1.81E-07 | 186.069121 | 298.7543 | 192.125817 | 239.25491 |
| -5.5856282 | 2.33E-08 | 1.81E-07 | 193.446311 | 222.17145 | 207.081719 | 346.275256 |
| -5.5854838 | 2.33E-08 | 1.82E-07 | 98.3625312 | 739.84987 | 725.936471 | 776.597905 |
| -5.5854302 | 2.33E-08 | 1.82E-07 | 175.003337 | 351.252791 | 203.630357 | 318.819775 |
| -5.5852954 | 2.33E-08 | 1.82E-07 | 148.363485 | 308.496288 | 289.914407 | 289.683345 |
| 5.58451066 | 2.34E-08 | 1.82E-07 | 39.7548564 | 19.2133653 | 21.8586259 | 16.2491625 |
| -5.5842642 | 2.35E-08 | 1.83E-07 | 142.62567 | 299.024911 | 452.12842 | 386.618005 |
| -5.5835665 | 2.36E-08 | 1.83E-07 | 254.103206 | 185.638994 | 187.524001 | 224.686695 |
| -5.5835853 | 2.36E-08 | 1.83E-07 | 155.33083 | 275.481773 | 194.426725 | 253.262809 |
| -5.5835809 | 2.36E-08 | 1.83E-07 | 154.101299 | 281.1646 | 251.949425 | 304.25156 |
| -5.5827038 | 2.37E-08 | 1.84E-07 | 168.855679 | 284.141318 | 226.639437 | 223.566063 |
| -5.5826171 | 2.37E-08 | 1.84E-07 | 144.265046 | 347.46424 | 288.763953 | 342.913361 |
| -5.5822225 | 2.37E-08 | 1.85E-07 | 227.463353 | 428.647474 | 176.019461 | 277.916711 |
| -5.5816128 | 2.38E-08 | 1.85E-07 | 200.003813 | 229.477941 | 173.718553 | 235.332698 |
| -5.5815774 | 2.38E-08 | 1.85E-07 | 170.08521 | 392.926851 | 215.134897 | 333.387989 |
| -5.5812873 | 2.39E-08 | 1.85E-07 | 146.314265 | 356.123785 | 276.108959 | 338.430833 |
| -5.5808384 | 2.39E-08 | 1.86E-07 | 116.395662 | 613.474636 | 505.049304 | 505.404986 |
| -5.5808687 | 2.39E-08 | 1.86E-07 | 153.691455 | 258.433294 | 303.719855 | 266.150076 |
| -5.5804317 | 2.40E-08 | 1.86E-07 | 143.035514 | 353.958899 | 299.118039 | 325.543566 |
| -5.5800525 | 2.40E-08 | 1.87E-07 | 227.463353 | 662.725798 | 431.420248 | 442.649599 |
| -5.5798704 | 2.41E-08 | 1.87E-07 | 208.610535 | 176.97945 | 207.081719 | 252.142177 |
| -5.5795191 | 2.41E-08 | 1.87E-07 | 161.478489 | 256.809629 | 235.843069 | 236.45333 |
| -5.5781777 | 2.43E-08 | 1.88E-07 | 165.986771 | 226.230611 | 222.037621 | 254.943757 |
| -5.5779865 | 2.43E-08 | 1.89E-07 | 146.314265 | 314.449725 | 280.710775 | 310.415036 |
| -5.5778809 | 2.43E-08 | 1.89E-07 | 146.724109 | 291.71842 | 300.268493 | 292.484925 |
| -5.5776243 | 2.44E-08 | 1.89E-07 | 143.855202 | 373.172264 | 291.064861 | 324.98325 |
| 5.57765113 | 2.44E-08 | 1.89E-07 | 19.2626624 | 18.6721438 | 11.50454 | 19.0507423 |
| -5.5775233 | 2.44E-08 | 1.89E-07 | 128.281134 | 418.364265 | 704.077845 | 449.933707 |

| | | | | | | |
|---|---|---|---|---|---|---|
| -5.5767883 | 2.45E-08 | 1.90E-07 | 177.4624 | 292.259641 | 287.613499 | 265.58976 |
| -5.5767065 | 2.45E-08 | 1.90E-07 | 220.496007 | 358.829893 | 352.038923 | 327.224514 |
| -5.5761604 | 2.46E-08 | 1.90E-07 | 142.62567 | 405.104336 | 285.312591 | 398.944955 |
| -5.5757903 | 2.46E-08 | 1.91E-07 | 145.084733 | 300.377965 | 311.773033 | 293.045241 |
| 5.57573717 | 2.46E-08 | 1.91E-07 | 25.0004767 | 33.0145151 | 27.6108959 | 17.3697944 |
| -5.5756296 | 2.47E-08 | 1.91E-07 | 126.641759 | 483.852073 | 394.60572 | 430.882964 |
| -5.5755461 | 2.47E-08 | 1.91E-07 | 160.658801 | 361.26539 | 233.542161 | 285.200818 |
| -5.5749445 | 2.48E-08 | 1.91E-07 | 148.363485 | 304.437127 | 266.905327 | 301.44998 |
| -5.5744344 | 2.48E-08 | 1.92E-07 | 119.264569 | 853.777008 | 440.62388 | 611.865016 |
| -5.5743409 | 2.48E-08 | 1.92E-07 | 161.888333 | 629.981894 | 574.076544 | 572.642899 |
| -5.5743769 | 2.48E-08 | 1.92E-07 | 171.314742 | 209.182132 | 215.134897 | 232.531119 |
| -5.5740536 | 2.49E-08 | 1.92E-07 | 181.150995 | 312.55545 | 188.674455 | 327.224514 |
| -5.5738435 | 2.49E-08 | 1.92E-07 | 165.986771 | 421.611594 | 424.517524 | 379.894213 |
| -5.5736418 | 2.49E-08 | 1.93E-07 | 229.922417 | 290.365366 | 250.798971 | 159.690045 |
| -5.5734422 | 2.50E-08 | 1.93E-07 | 153.281611 | 263.033677 | 272.657597 | 254.943757 |
| -5.5734156 | 2.50E-08 | 1.93E-07 | 151.232392 | 257.350851 | 283.011683 | 280.157974 |
| -5.5731762 | 2.50E-08 | 1.93E-07 | 177.872244 | 526.608576 | 230.090799 | 327.78483 |
| -5.573168 | 2.50E-08 | 1.93E-07 | 209.020379 | 320.403163 | 180.621277 | 453.855918 |
| -5.5730876 | 2.50E-08 | 1.93E-07 | 148.363485 | 316.07339 | 270.356689 | 285.200818 |
| -5.5730519 | 2.50E-08 | 1.93E-07 | 244.676796 | 471.945199 | 510.801574 | 429.762333 |
| -5.5727517 | 2.51E-08 | 1.93E-07 | 313.120724 | 273.587498 | 147.258111 | 278.477026 |
| -5.572737 | 2.51E-08 | 1.93E-07 | 136.887856 | 368.30127 | 326.728935 | 351.3181 |
| -5.572667 | 2.51E-08 | 1.93E-07 | 139.346919 | 358.829893 | 307.171217 | 346.835572 |
| -5.5725621 | 2.51E-08 | 1.93E-07 | 219.266476 | 208.099689 | 171.417645 | 214.040692 |
| -5.5724898 | 2.51E-08 | 1.93E-07 | 154.920987 | 291.177198 | 239.294431 | 272.873867 |
| -5.5724357 | 2.51E-08 | 1.93E-07 | 127.051603 | 441.09557 | 393.455266 | 419.676646 |
| -5.5721864 | 2.52E-08 | 1.94E-07 | 148.363485 | 269.257725 | 296.817131 | 357.481575 |
| -5.5720464 | 2.52E-08 | 1.94E-07 | 119.264569 | 463.826875 | 516.553844 | 472.346345 |
| -5.5716981 | 2.52E-08 | 1.94E-07 | 268.037897 | 156.14242 | 235.843069 | 275.675447 |
| -5.5715638 | 2.52E-08 | 1.94E-07 | 127.051603 | 468.427259 | 370.446187 | 481.871716 |
| -5.5715663 | 2.52E-08 | 1.94E-07 | 147.543797 | 260.056959 | 357.791193 | 359.722839 |
| -5.5715114 | 2.53E-08 | 1.94E-07 | 118.854725 | 514.431091 | 476.287954 | 467.863817 |
| -5.5710718 | 2.53E-08 | 1.94E-07 | 151.232392 | 280.082156 | 254.250333 | 284.080186 |
| -5.5708084 | 2.54E-08 | 1.95E-07 | 152.461923 | 270.88139 | 257.701695 | 263.348496 |
| -5.5706235 | 2.54E-08 | 1.95E-07 | 179.511619 | 247.608863 | 217.435805 | 210.678797 |
| -5.570383 | 2.54E-08 | 1.95E-07 | 140.576451 | 363.159666 | 303.719855 | 323.302302 |
| -5.5698823 | 2.55E-08 | 1.96E-07 | 133.609105 | 366.948217 | 346.286653 | 381.014845 |
| -5.5698282 | 2.55E-08 | 1.96E-07 | 137.707544 | 356.123785 | 331.330751 | 326.664198 |
| -5.569161 | 2.56E-08 | 1.96E-07 | 136.887856 | 444.342899 | 375.048002 | 315.457879 |
| -5.5691241 | 2.56E-08 | 1.96E-07 | 151.232392 | 266.281007 | 257.701695 | 300.889664 |
| -5.5688962 | 2.56E-08 | 1.97E-07 | 135.24848 | 382.643642 | 329.029843 | 360.283155 |
| -5.5688353 | 2.56E-08 | 1.97E-07 | 122.54332 | 409.163498 | 479.739316 | 481.3114 |
| -5.5686581 | 2.57E-08 | 1.97E-07 | 127.871291 | 442.448624 | 364.693917 | 458.898762 |

| | | | | | | |
|---|---|---|---|---|---|---|
| -5.5685788 | 2.57E-08 | 1.97E-07 | 195.495531 | 372.089821 | 177.169915 | 340.672097 |
| -5.5682884 | 2.57E-08 | 1.97E-07 | 120.494101 | 441.907402 | 521.15566 | 455.536866 |
| -5.5680244 | 2.58E-08 | 1.97E-07 | 135.24848 | 572.341798 | 347.437107 | 396.703692 |
| -5.5679227 | 2.58E-08 | 1.97E-07 | 147.543797 | 304.437127 | 424.517524 | 285.200818 |
| -5.5677593 | 2.58E-08 | 1.98E-07 | 211.069598 | 304.707737 | 219.736713 | 175.378892 |
| -5.5676679 | 2.58E-08 | 1.98E-07 | 160.658801 | 232.184049 | 233.542161 | 238.134278 |
| -5.5675317 | 2.58E-08 | 1.98E-07 | 297.136813 | 227.313054 | 157.612197 | 256.624704 |
| -5.5672539 | 2.59E-08 | 1.98E-07 | 211.069598 | 379.666923 | 192.125817 | 224.126379 |
| -5.5665728 | 2.60E-08 | 1.99E-07 | 236.889763 | 222.983282 | 179.470823 | 201.713742 |
| -5.5661483 | 2.60E-08 | 1.99E-07 | 122.133476 | 439.201295 | 462.482506 | 435.365492 |
| -5.5660878 | 2.61E-08 | 1.99E-07 | 131.150042 | 396.986013 | 345.136199 | 419.11633 |
| -5.5659877 | 2.61E-08 | 1.99E-07 | 144.67489 | 304.978348 | 274.958505 | 328.345146 |
| -5.5658958 | 2.61E-08 | 1.99E-07 | 132.379573 | 410.245941 | 334.782113 | 396.703692 |
| -5.5657339 | 2.61E-08 | 1.99E-07 | 184.83959 | 187.803881 | 274.958505 | 217.402588 |
| -5.56562 | 2.61E-08 | 1.99E-07 | 174.183649 | 314.449725 | 193.276271 | 316.018195 |
| -5.5654685 | 2.61E-08 | 2.00E-07 | 145.494577 | 273.858108 | 432.570702 | 348.51652 |
| -5.5645473 | 2.63E-08 | 2.01E-07 | 638.536765 | 891.933128 | 656.909231 | 789.485172 |
| -5.5641573 | 2.63E-08 | 2.01E-07 | 149.183172 | 298.213079 | 281.861229 | 254.943757 |
| -5.5640435 | 2.64E-08 | 2.01E-07 | 149.593016 | 264.927953 | 262.303511 | 292.484925 |
| -5.5638478 | 2.64E-08 | 2.01E-07 | 151.642236 | 169.943569 | 220.887167 | 187.705843 |
| -5.5636114 | 2.64E-08 | 2.02E-07 | 161.478489 | 214.053126 | 287.613499 | 314.897563 |
| -5.5631363 | 2.65E-08 | 2.02E-07 | 126.231915 | 751.756744 | 411.86253 | 470.665397 |
| -5.5630082 | 2.65E-08 | 2.02E-07 | 163.117864 | 218.382899 | 245.046701 | 224.686695 |
| -5.5628149 | 2.65E-08 | 2.02E-07 | 151.232392 | 245.714587 | 292.215315 | 340.111781 |
| -5.5627592 | 2.66E-08 | 2.02E-07 | 149.593016 | 274.39933 | 292.215315 | 253.823125 |
| -5.5626984 | 2.66E-08 | 2.02E-07 | 135.24848 | 336.369198 | 353.189377 | 337.870517 |
| -5.5619363 | 2.67E-08 | 2.03E-07 | 134.428793 | 638.100217 | 342.835291 | 439.84802 |
| -5.5618558 | 2.67E-08 | 2.03E-07 | 180.741151 | 261.951234 | 180.621277 | 253.823125 |
| -5.5611582 | 2.68E-08 | 2.04E-07 | 138.117388 | 401.315785 | 292.215315 | 387.738636 |
| -5.5610822 | 2.68E-08 | 2.04E-07 | 117.21535 | 461.9326 | 483.190678 | 501.482774 |
| -5.5610196 | 2.68E-08 | 2.04E-07 | 137.707544 | 371.819211 | 312.923487 | 323.302302 |
| -5.5609238 | 2.68E-08 | 2.04E-07 | 126.641759 | 473.568864 | 370.446187 | 416.875066 |
| -5.5606382 | 2.69E-08 | 2.04E-07 | 138.527231 | 345.569965 | 297.967585 | 341.232413 |
| -5.560642 | 2.69E-08 | 2.04E-07 | 318.448695 | 221.900839 | 192.125817 | 188.266159 |
| -5.5602792 | 2.69E-08 | 2.05E-07 | 136.068168 | 486.558181 | 302.569401 | 449.933707 |
| -5.5602535 | 2.69E-08 | 2.05E-07 | 566.814086 | 236.513821 | 220.887167 | 231.970803 |
| -5.5602241 | 2.69E-08 | 2.05E-07 | 163.117864 | 247.067641 | 217.435805 | 223.005748 |
| -5.5597869 | 2.70E-08 | 2.05E-07 | 139.346919 | 362.077222 | 301.418947 | 314.897563 |
| -5.5597776 | 2.70E-08 | 2.05E-07 | 152.871767 | 340.698971 | 498.14658 | 264.469128 |
| -5.5593243 | 2.71E-08 | 2.06E-07 | 369.67918 | 218.112288 | 224.338529 | 242.05649 |
| -5.5591348 | 2.71E-08 | 2.06E-07 | 150.412704 | 358.288672 | 308.321671 | 350.757784 |
| -5.5585129 | 2.72E-08 | 2.06E-07 | 168.445835 | 300.107354 | 291.064861 | 197.231214 |
| -5.55819 | 2.73E-08 | 2.07E-07 | 247.545703 | 864.060218 | 661.511047 | 633.717338 |

| | | | | | | |
|---|---|---|---|---|---|---|
| -5.5580445 | 2.73E-08 | 2.07E-07 | 139.756763 | 330.686372 | 288.763953 | 336.749885 |
| -5.5573864 | 2.74E-08 | 2.08E-07 | 165.986771 | 243.549701 | 200.178995 | 250.461229 |
| -5.5572635 | 2.74E-08 | 2.08E-07 | 152.871767 | 286.847426 | 233.542161 | 321.061039 |
| -5.5573029 | 2.74E-08 | 2.08E-07 | 134.838636 | 347.46424 | 323.277573 | 349.637152 |
| -5.5569527 | 2.75E-08 | 2.08E-07 | 172.13443 | 321.485606 | 193.276271 | 288.562714 |
| -5.5566205 | 2.75E-08 | 2.08E-07 | 186.478965 | 214.053126 | 177.169915 | 234.212067 |
| -5.5557052 | 2.76E-08 | 2.09E-07 | 165.986771 | 247.879474 | 200.178995 | 237.573962 |
| -5.5556739 | 2.77E-08 | 2.09E-07 | 123.772852 | 505.230325 | 366.994825 | 499.24151 |
| -5.5556243 | 2.77E-08 | 2.09E-07 | 149.183172 | 271.152001 | 250.798971 | 277.916711 |
| -5.5553101 | 2.77E-08 | 2.10E-07 | 135.24848 | 354.22951 | 325.578481 | 327.224514 |
| -5.5551864 | 2.77E-08 | 2.10E-07 | 127.051603 | 452.731834 | 421.066162 | 354.11968 |
| -5.5551679 | 2.77E-08 | 2.10E-07 | 170.495054 | 232.995881 | 193.276271 | 259.426284 |
| 5.55430248 | 2.79E-08 | 2.11E-07 | 0 | 0 | 0 | 0 |
| 5.55432951 | 2.79E-08 | 2.11E-07 | 7.78703372 | 3.24732935 | 4.60181598 | 1.68094785 |
| -5.5541358 | 2.79E-08 | 2.11E-07 | 191.397092 | 189.427545 | 188.674455 | 198.351846 |
| -5.553825 | 2.79E-08 | 2.11E-07 | 238.529138 | 315.261558 | 310.622579 | 348.51652 |
| -5.5534251 | 2.80E-08 | 2.12E-07 | 149.183172 | 393.197462 | 260.002603 | 293.605557 |
| -5.5531121 | 2.81E-08 | 2.12E-07 | 148.363485 | 250.585581 | 288.763953 | 270.632603 |
| 5.55218935 | 2.82E-08 | 2.13E-07 | 0 | 0 | 0 | 0 |
| -5.5519902 | 2.82E-08 | 2.13E-07 | 150.822548 | 303.895905 | 251.949425 | 250.461229 |
| -5.5514383 | 2.83E-08 | 2.14E-07 | 154.920987 | 236.513821 | 271.507143 | 234.212067 |
| -5.5506653 | 2.85E-08 | 2.15E-07 | 180.741151 | 186.992048 | 218.586259 | 197.79153 |
| -5.5502706 | 2.85E-08 | 2.15E-07 | 144.265046 | 258.974516 | 337.083021 | 384.376741 |
| -5.5501709 | 2.85E-08 | 2.15E-07 | 121.313788 | 464.909319 | 404.959806 | 428.641701 |
| -5.5496113 | 2.86E-08 | 2.16E-07 | 142.62567 | 285.223761 | 277.259413 | 304.811876 |
| -5.54957 | 2.86E-08 | 2.16E-07 | 147.543797 | 259.515737 | 263.453965 | 272.313551 |
| -5.5494279 | 2.87E-08 | 2.16E-07 | 128.690978 | 418.905486 | 347.437107 | 374.85137 |
| -5.5494057 | 2.87E-08 | 2.16E-07 | 229.102729 | 370.736767 | 356.640739 | 405.668747 |
| -5.5494763 | 2.87E-08 | 2.16E-07 | 164.75724 | 219.194731 | 211.683535 | 214.601008 |
| -5.549303 | 2.87E-08 | 2.16E-07 | 167.216303 | 241.114204 | 220.887167 | 198.351846 |
| 5.54886076 | 2.88E-08 | 2.17E-07 | 6.96734596 | 8.38893415 | 10.354086 | 9.52537113 |
| -5.5488321 | 2.88E-08 | 2.17E-07 | 124.592539 | 416.7406 | 383.10118 | 401.186219 |
| -5.5485353 | 2.88E-08 | 2.17E-07 | 200.823501 | 199.710755 | 176.019461 | 195.550266 |
| -5.5479876 | 2.89E-08 | 2.17E-07 | 135.24848 | 454.084887 | 320.976665 | 336.749885 |
| -5.5476566 | 2.90E-08 | 2.18E-07 | 131.150042 | 369.924935 | 332.481205 | 357.481575 |
| -5.547368 | 2.90E-08 | 2.18E-07 | 146.314265 | 281.43521 | 256.551241 | 272.873867 |
| -5.5470766 | 2.90E-08 | 2.18E-07 | 213.118818 | 159.93097 | 235.843069 | 233.091435 |
| -5.5470653 | 2.91E-08 | 2.18E-07 | 170.08521 | 213.782516 | 193.276271 | 229.729539 |
| -5.5459903 | 2.92E-08 | 2.20E-07 | 145.084733 | 276.564216 | 258.852149 | 295.286505 |
| -5.5459901 | 2.92E-08 | 2.20E-07 | 1006.98641 | 283.058875 | 270.356689 | 253.262809 |
| -5.5458889 | 2.92E-08 | 2.20E-07 | 167.216303 | 250.585581 | 266.905327 | 192.18837 |
| -5.5455514 | 2.93E-08 | 2.20E-07 | 131.969729 | 329.87454 | 342.835291 | 357.481575 |
| -5.5455709 | 2.93E-08 | 2.20E-07 | 156.150518 | 230.560384 | 253.099879 | 225.247011 |

| | | | | | | |
|---|---|---|---|---|---|---|
| -5.5455104 | 2.93E-08 | 2.20E-07 | 173.363961 | 248.150084 | 213.984443 | 214.040692 |
| -5.545409 | 2.93E-08 | 2.20E-07 | 99.1822189 | 897.345344 | 625.846973 | 611.3047 |
| -5.5449898 | 2.94E-08 | 2.21E-07 | 154.511143 | 227.313054 | 248.498063 | 296.967453 |
| -5.5441483 | 2.95E-08 | 2.22E-07 | 309.022285 | 1121.68168 | 967.53181 | 1018.65439 |
| -5.5440597 | 2.96E-08 | 2.22E-07 | 181.970683 | 324.191713 | 177.169915 | 310.415036 |
| -5.5438437 | 2.96E-08 | 2.22E-07 | 146.724109 | 288.20048 | 245.046701 | 284.080186 |
| -5.5438687 | 2.96E-08 | 2.22E-07 | 131.150042 | 337.451642 | 362.393009 | 339.551465 |
| -5.5435713 | 2.96E-08 | 2.22E-07 | 195.495531 | 253.832911 | 163.364467 | 256.624704 |
| -5.5435322 | 2.96E-08 | 2.22E-07 | 156.150518 | 290.906588 | 213.984443 | 268.951655 |
| -5.5434549 | 2.97E-08 | 2.22E-07 | 136.478012 | 318.508887 | 312.923487 | 305.372192 |
| -5.5433692 | 2.97E-08 | 2.22E-07 | 143.445358 | 353.688288 | 264.604419 | 291.924609 |
| -5.5432549 | 2.97E-08 | 2.22E-07 | 157.789894 | 217.841677 | 268.055781 | 230.289855 |
| -5.5431996 | 2.97E-08 | 2.22E-07 | 148.363485 | 619.428073 | 358.941647 | 405.668747 |
| -5.5424634 | 2.98E-08 | 2.23E-07 | 143.035514 | 337.181031 | 255.400787 | 314.337247 |
| -5.5424672 | 2.98E-08 | 2.23E-07 | 129.510666 | 446.507786 | 346.286653 | 350.757784 |
| -5.5424224 | 2.98E-08 | 2.23E-07 | 129.100822 | 363.700887 | 364.693917 | 344.594308 |
| -5.5419113 | 2.99E-08 | 2.24E-07 | 196.315218 | 227.854276 | 164.514921 | 225.247011 |
| -5.5415641 | 3.00E-08 | 2.24E-07 | 200.003813 | 237.325653 | 200.178995 | 169.775732 |
| -5.5413863 | 3.00E-08 | 2.24E-07 | 137.2977 | 305.248959 | 293.365769 | 330.026094 |
| -5.5413617 | 3.00E-08 | 2.24E-07 | 160.248957 | 268.716504 | 266.905327 | 239.815226 |
| -5.5414367 | 3.00E-08 | 2.24E-07 | 198.364438 | 273.858108 | 196.727633 | 282.959554 |
| -5.5412699 | 3.00E-08 | 2.24E-07 | 155.740674 | 294.424528 | 215.134897 | 326.664198 |
| -5.5411151 | 3.01E-08 | 2.25E-07 | 159.019425 | 399.962732 | 222.037621 | 302.570612 |
| -5.5402756 | 3.02E-08 | 2.25E-07 | 141.396139 | 409.434109 | 283.011683 | 302.010296 |
| -5.5402964 | 3.02E-08 | 2.25E-07 | 154.920987 | 246.52642 | 220.887167 | 240.375542 |
| -5.540168 | 3.02E-08 | 2.26E-07 | 125.002383 | 425.400145 | 371.596641 | 374.291054 |
| -5.5400651 | 3.02E-08 | 2.26E-07 | 130.740198 | 355.311953 | 345.136199 | 331.707042 |
| -5.5398471 | 3.03E-08 | 2.26E-07 | 152.052079 | 238.137486 | 234.692615 | 256.064389 |
| -5.5397579 | 3.03E-08 | 2.26E-07 | 175.823024 | 186.450827 | 243.896247 | 204.515321 |
| -5.5396967 | 3.03E-08 | 2.26E-07 | 372.138243 | 220.006563 | 179.470823 | 215.161324 |
| -5.5391327 | 3.04E-08 | 2.27E-07 | 122.953164 | 414.846324 | 394.60572 | 386.057689 |
| -5.5390576 | 3.04E-08 | 2.27E-07 | 127.871291 | 343.9463 | 400.35799 | 360.283155 |
| -5.538815 | 3.05E-08 | 2.27E-07 | 135.658324 | 337.451642 | 288.763953 | 338.991149 |
| -5.538519 | 3.05E-08 | 2.27E-07 | 167.626147 | 285.494372 | 245.046701 | 192.18837 |
| 5.53838134 | 3.05E-08 | 2.27E-07 | 53.6895483 | 27.0610779 | 39.1154358 | 17.9301104 |
| -5.5382868 | 3.05E-08 | 2.27E-07 | 115.985818 | 399.692121 | 750.096005 | 595.055537 |
| -5.538105 | 3.06E-08 | 2.28E-07 | 304.104159 | 370.736767 | 167.966283 | 197.231214 |
| -5.5375062 | 3.07E-08 | 2.28E-07 | 150.412704 | 237.055043 | 260.002603 | 242.616806 |
| -5.5375239 | 3.07E-08 | 2.28E-07 | 150.412704 | 250.856192 | 242.745793 | 240.375542 |
| -5.5373376 | 3.07E-08 | 2.29E-07 | 180.331307 | 247.338252 | 220.887167 | 221.885116 |
| -5.537136 | 3.07E-08 | 2.29E-07 | 166.806459 | 214.594348 | 212.833989 | 301.44998 |
| -5.537048 | 3.08E-08 | 2.29E-07 | 179.101776 | 365.324552 | 260.002603 | 352.438732 |
| -5.5366193 | 3.08E-08 | 2.29E-07 | 231.971636 | 583.43684 | 528.058384 | 541.265206 |

| | | | | | | |
|---|---|---|---|---|---|---|
| -5.536546 | 3.08E-08 | 2.29E-07 | 184.429746 | 287.659258 | 170.267191 | 292.484925 |
| -5.5358082 | 3.10E-08 | 2.30E-07 | 174.183649 | 222.17145 | 195.577179 | 189.947107 |
| -5.5356195 | 3.10E-08 | 2.30E-07 | 156.970206 | 441.907402 | 222.037621 | 446.011495 |
| -5.5353647 | 3.11E-08 | 2.31E-07 | 121.313788 | 531.208959 | 365.844371 | 451.054339 |
| -5.5352154 | 3.11E-08 | 2.31E-07 | 225.414134 | 310.931785 | 173.718553 | 194.429634 |
| -5.5351361 | 3.11E-08 | 2.31E-07 | 98.7723751 | 530.938349 | 943.372276 | 745.780528 |
| -5.5348503 | 3.11E-08 | 2.31E-07 | 123.363008 | 411.328384 | 372.747095 | 388.298952 |
| -5.5347093 | 3.12E-08 | 2.31E-07 | 160.658801 | 199.169533 | 240.444885 | 244.85807 |
| -5.5345221 | 3.12E-08 | 2.32E-07 | 208.610535 | 350.440959 | 342.835291 | 384.937057 |
| -5.5343848 | 3.12E-08 | 2.32E-07 | 122.54332 | 394.550516 | 380.800272 | 409.030642 |
| -5.5341902 | 3.13E-08 | 2.32E-07 | 164.75724 | 198.357701 | 210.533081 | 229.729539 |
| -5.5340876 | 3.13E-08 | 2.32E-07 | 184.429746 | 237.866875 | 179.470823 | 194.429634 |
| -5.5339843 | 3.13E-08 | 2.32E-07 | 121.723632 | 455.708552 | 400.35799 | 377.652949 |
| -5.5340125 | 3.13E-08 | 2.32E-07 | 188.118341 | 273.046276 | 165.665375 | 271.753235 |
| -5.5332566 | 3.14E-08 | 2.33E-07 | 142.215826 | 577.483403 | 295.666677 | 361.964103 |
| -5.5330295 | 3.15E-08 | 2.33E-07 | 111.477535 | 509.289486 | 464.783414 | 493.638351 |
| -5.532905 | 3.15E-08 | 2.33E-07 | 142.215826 | 400.503953 | 274.958505 | 291.364293 |
| -5.532166 | 3.16E-08 | 2.34E-07 | 151.642236 | 350.440959 | 251.949425 | 519.9732 |
| -5.5319531 | 3.17E-08 | 2.34E-07 | 172.544273 | 279.811546 | 182.922185 | 237.013646 |
| -5.5317124 | 3.17E-08 | 2.35E-07 | 145.904421 | 279.811546 | 249.648517 | 252.702493 |
| -5.5316612 | 3.17E-08 | 2.35E-07 | 160.248957 | 202.687474 | 264.604419 | 228.608907 |
| -5.5317106 | 3.17E-08 | 2.35E-07 | 126.641759 | 344.758133 | 381.950726 | 361.403787 |
| -5.5317838 | 3.17E-08 | 2.35E-07 | 136.478012 | 311.743618 | 286.463045 | 307.05314 |
| -5.5316641 | 3.17E-08 | 2.35E-07 | 153.691455 | 299.836743 | 212.833989 | 277.916711 |
| -5.5313785 | 3.18E-08 | 2.35E-07 | 252.46383 | 632.688002 | 544.16474 | 573.763531 |
| -5.530906 | 3.19E-08 | 2.35E-07 | 228.692885 | 279.270324 | 147.258111 | 266.710392 |
| -5.5306933 | 3.19E-08 | 2.36E-07 | 149.593016 | 232.72527 | 243.896247 | 277.356395 |
| -5.5303275 | 3.20E-08 | 2.36E-07 | 140.576451 | 445.425342 | 262.303511 | 368.127578 |
| -5.5296922 | 3.21E-08 | 2.37E-07 | 139.756763 | 286.306204 | 271.507143 | 287.442082 |
| -5.5293292 | 3.21E-08 | 2.37E-07 | 118.854725 | 565.847139 | 436.022064 | 389.419584 |
| -5.528627 | 3.23E-08 | 2.38E-07 | 274.5954 | 185.097773 | 202.479903 | 180.982051 |
| -5.5283827 | 3.23E-08 | 2.38E-07 | 241.807889 | 455.167331 | 146.107657 | 382.695793 |
| -5.5283838 | 3.23E-08 | 2.38E-07 | 142.62567 | 339.075306 | 248.498063 | 296.967453 |
| -5.5284024 | 3.23E-08 | 2.38E-07 | 126.231915 | 386.161582 | 386.552542 | 331.146726 |
| -5.5279092 | 3.24E-08 | 2.39E-07 | 124.182696 | 350.440959 | 433.721156 | 376.532317 |
| -5.5277786 | 3.24E-08 | 2.39E-07 | 178.691932 | 180.768 | 250.798971 | 201.713742 |
| -5.5276538 | 3.25E-08 | 2.39E-07 | 136.068168 | 287.388647 | 332.481205 | 293.605557 |
| -5.5275302 | 3.25E-08 | 2.39E-07 | 158.609582 | 317.426444 | 209.382627 | 249.900913 |
| -5.5274551 | 3.25E-08 | 2.39E-07 | 660.668334 | 1342.22946 | 1133.19719 | 1187.3095 |
| -5.5272935 | 3.25E-08 | 2.40E-07 | 88.5262781 | 665.161295 | 837.530509 | 782.76138 |
| -5.5266255 | 3.26E-08 | 2.40E-07 | 147.133953 | 251.668025 | 235.843069 | 271.192919 |
| -5.5264813 | 3.27E-08 | 2.41E-07 | 127.051603 | 346.381797 | 375.048002 | 338.991149 |
| -5.5261017 | 3.27E-08 | 2.41E-07 | 128.690978 | 354.770731 | 315.224395 | 400.065587 |

| | | | | | | |
|---|---|---|---|---|---|---|
| -5.5256641 | 3.28E-08 | 2.42E-07 | 131.150042 | 333.663091 | 317.525303 | 319.940407 |
| -5.5255777 | 3.28E-08 | 2.42E-07 | 132.379573 | 307.955067 | 318.675757 | 321.061039 |
| -5.5245735 | 3.30E-08 | 2.43E-07 | 122.133476 | 151.812647 | 162.214013 | 138.958355 |
| -5.5241683 | 3.31E-08 | 2.44E-07 | 154.920987 | 250.04436 | 213.984443 | 223.566063 |
| -5.5239193 | 3.32E-08 | 2.44E-07 | 152.871767 | 303.354683 | 220.887167 | 243.737438 |
| -5.5238531 | 3.32E-08 | 2.44E-07 | 148.363485 | 300.377965 | 231.241253 | 252.142177 |
| -5.5237241 | 3.32E-08 | 2.44E-07 | 195.495531 | 233.807713 | 188.674455 | 171.45668 |
| -5.52314 | 3.33E-08 | 2.45E-07 | 352.055893 | 201.875641 | 194.426725 | 196.110582 |
| -5.5230377 | 3.33E-08 | 2.45E-07 | 146.314265 | 266.551617 | 243.896247 | 242.05649 |
| -5.5228812 | 3.33E-08 | 2.45E-07 | 202.462877 | 973.116362 | 713.281477 | 845.516766 |
| -5.5222099 | 3.35E-08 | 2.46E-07 | 110.248004 | 549.610492 | 464.783414 | 463.38129 |
| -5.5220318 | 3.35E-08 | 2.46E-07 | 161.888333 | 404.021893 | 518.854752 | 475.147924 |
| -5.521903 | 3.35E-08 | 2.46E-07 | 143.035514 | 244.090923 | 269.206235 | 292.484925 |
| -5.5216163 | 3.36E-08 | 2.47E-07 | 145.494577 | 539.056672 | 270.356689 | 344.594308 |
| -5.5215489 | 3.36E-08 | 2.47E-07 | 162.298176 | 310.661174 | 208.232173 | 225.807327 |
| -5.5211536 | 3.37E-08 | 2.47E-07 | 145.904421 | 297.671857 | 287.613499 | 227.488275 |
| -5.520801 | 3.37E-08 | 2.48E-07 | 187.708497 | 183.744719 | 177.169915 | 226.927959 |
| -5.5207308 | 3.38E-08 | 2.48E-07 | 203.692408 | 172.920288 | 261.153057 | 186.024895 |
| -5.5203949 | 3.38E-08 | 2.48E-07 | 136.478012 | 359.371115 | 261.153057 | 354.679995 |
| -5.5204568 | 3.38E-08 | 2.48E-07 | 179.921463 | 230.830995 | 190.975363 | 179.301104 |
| 5.52038039 | 3.38E-08 | 2.48E-07 | 47.951734 | 30.0377965 | 35.6640739 | 27.4554815 |
| -5.5197985 | 3.39E-08 | 2.49E-07 | 129.92051 | 419.176097 | 335.932567 | 309.294404 |
| -5.519528 | 3.40E-08 | 2.49E-07 | 140.166607 | 282.247043 | 257.701695 | 341.792729 |
| -5.5183183 | 3.42E-08 | 2.51E-07 | 131.150042 | 448.402061 | 297.967585 | 352.438732 |
| -5.5182947 | 3.42E-08 | 2.51E-07 | 134.838636 | 291.71842 | 310.622579 | 285.761134 |
| -5.5179707 | 3.43E-08 | 2.51E-07 | 166.806459 | 231.642827 | 184.072639 | 212.359745 |
| -5.5177209 | 3.43E-08 | 2.51E-07 | 146.724109 | 276.834827 | 225.488983 | 261.107232 |
| -5.5170819 | 3.45E-08 | 2.52E-07 | 145.084733 | 256.268408 | 245.046701 | 243.177122 |
| -5.5168515 | 3.45E-08 | 2.52E-07 | 161.888333 | 257.08024 | 190.975363 | 225.807327 |
| -5.5165787 | 3.46E-08 | 2.53E-07 | 150.822548 | 212.970683 | 248.498063 | 271.753235 |
| -5.5155967 | 3.48E-08 | 2.54E-07 | 136.887856 | 295.777582 | 425.667978 | 289.123029 |
| -5.515224 | 3.48E-08 | 2.55E-07 | 168.035991 | 305.51957 | 186.373547 | 242.05649 |
| -5.5152591 | 3.48E-08 | 2.55E-07 | 138.937075 | 321.756216 | 253.099879 | 289.123029 |
| -5.5151254 | 3.49E-08 | 2.55E-07 | 129.100822 | 351.252791 | 317.525303 | 314.897563 |
| -5.5146316 | 3.50E-08 | 2.55E-07 | 165.576927 | 245.985198 | 223.188075 | 183.223315 |
| -5.5142393 | 3.50E-08 | 2.56E-07 | 141.805982 | 279.811546 | 250.798971 | 254.943757 |
| -5.5142469 | 3.50E-08 | 2.56E-07 | 217.6271 | 163.990132 | 186.373547 | 242.05649 |
| -5.5140554 | 3.51E-08 | 2.56E-07 | 136.068168 | 328.521486 | 445.225696 | 277.356395 |
| -5.5141692 | 3.50E-08 | 2.56E-07 | 302.05494 | 181.850444 | 192.125817 | 196.110582 |
| -5.5141116 | 3.51E-08 | 2.56E-07 | 153.691455 | 246.797031 | 203.630357 | 240.375542 |
| -5.5140521 | 3.51E-08 | 2.56E-07 | 120.903945 | 358.559282 | 377.34891 | 430.882964 |
| -5.513862 | 3.51E-08 | 2.56E-07 | 124.592539 | 340.157749 | 352.038923 | 364.765683 |
| -5.5135037 | 3.52E-08 | 2.56E-07 | 109.428316 | 427.565031 | 534.961108 | 508.766881 |

| | | | | | | |
|---|---|---|---|---|---|---|
| -5.5133517 | 3.52E-08 | 2.57E-07 | 147.133953 | 221.900839 | 291.064861 | 332.827673 |
| -5.5133853 | 3.52E-08 | 2.57E-07 | 134.018949 | 290.365366 | 299.118039 | 290.803977 |
| -5.5131897 | 3.52E-08 | 2.57E-07 | 154.511143 | 226.501222 | 205.931265 | 227.488275 |
| -5.5130088 | 3.53E-08 | 2.57E-07 | 160.248957 | 191.592432 | 271.507143 | 235.893014 |
| -5.5129486 | 3.53E-08 | 2.57E-07 | 228.283041 | 663.53763 | 670.714679 | 673.49977 |
| -5.5127711 | 3.53E-08 | 2.57E-07 | 124.182696 | 437.036408 | 352.038923 | 339.551465 |
| -5.5126543 | 3.53E-08 | 2.57E-07 | 147.133953 | 266.822228 | 225.488983 | 242.616806 |
| -5.5125089 | 3.54E-08 | 2.57E-07 | 236.889763 | 583.707451 | 429.11934 | 464.501921 |
| -5.5119328 | 3.55E-08 | 2.58E-07 | 168.445835 | 232.72527 | 176.019461 | 234.772382 |
| -5.5118366 | 3.55E-08 | 2.58E-07 | 122.953164 | 459.226492 | 327.879389 | 397.824324 |
| -5.5116783 | 3.55E-08 | 2.59E-07 | 140.576451 | 275.481773 | 250.798971 | 263.348496 |
| -5.5108732 | 3.57E-08 | 2.60E-07 | 241.807889 | 269.528336 | 348.587561 | 143.440883 |
| -5.510635 | 3.58E-08 | 2.60E-07 | 197.54475 | 170.484791 | 180.621277 | 208.437533 |
| -5.5102229 | 3.58E-08 | 2.60E-07 | 129.100822 | 355.041342 | 296.817131 | 331.146726 |
| -5.5099501 | 3.59E-08 | 2.61E-07 | 136.887856 | 334.474923 | 250.798971 | 341.232413 |
| -5.5098875 | 3.59E-08 | 2.61E-07 | 127.871291 | 352.064624 | 299.118039 | 360.843471 |
| -5.5098474 | 3.59E-08 | 2.61E-07 | 172.544273 | 205.122971 | 181.771731 | 189.947107 |
| -5.5093467 | 3.60E-08 | 2.62E-07 | 131.559885 | 314.449725 | 302.569401 | 294.165873 |
| -5.509272 | 3.60E-08 | 2.62E-07 | 173.363961 | 192.404264 | 226.639437 | 179.301104 |
| -5.5088964 | 3.61E-08 | 2.62E-07 | 163.527708 | 219.465342 | 225.488983 | 183.223315 |
| -5.5089207 | 3.61E-08 | 2.62E-07 | 111.067691 | 758.522014 | 436.022064 | 491.397087 |
| -5.5087574 | 3.61E-08 | 2.62E-07 | 163.937552 | 227.313054 | 181.771731 | 223.566063 |
| -5.5085197 | 3.62E-08 | 2.62E-07 | 138.117388 | 259.245126 | 272.657597 | 277.916711 |
| -5.5079639 | 3.63E-08 | 2.63E-07 | 139.346919 | 258.433294 | 257.701695 | 306.492824 |
| -5.5079811 | 3.63E-08 | 2.63E-07 | 175.413181 | 337.181031 | 182.922185 | 240.375542 |
| -5.5077723 | 3.63E-08 | 2.63E-07 | 149.183172 | 269.257725 | 209.382627 | 262.227864 |
| -5.5076782 | 3.64E-08 | 2.64E-07 | 107.379097 | 417.552432 | 735.140103 | 597.296801 |
| -5.5075424 | 3.64E-08 | 2.64E-07 | 133.199261 | 332.310037 | 286.463045 | 284.640502 |
| -5.5073464 | 3.64E-08 | 2.64E-07 | 134.018949 | 300.377965 | 331.330751 | 268.391339 |
| -5.5073183 | 3.64E-08 | 2.64E-07 | 202.872721 | 222.44206 | 166.815829 | 338.430833 |
| -5.5070803 | 3.65E-08 | 2.64E-07 | 115.16613 | 385.890971 | 427.968886 | 443.770231 |
| -5.5070336 | 3.65E-08 | 2.64E-07 | 173.363961 | 174.543953 | 200.178995 | 212.92006 |
| -5.5070482 | 3.65E-08 | 2.64E-07 | 134.018949 | 308.225677 | 272.657597 | 293.605557 |
| -5.5068091 | 3.65E-08 | 2.64E-07 | 165.986771 | 179.685557 | 218.586259 | 215.161324 |
| 5.50660151 | 3.66E-08 | 2.65E-07 | 147.543797 | 68.1939163 | 80.5317797 | 85.7283401 |
| -5.5065932 | 3.66E-08 | 2.65E-07 | 136.068168 | 272.234444 | 223.188075 | 219.643852 |
| -5.5063681 | 3.66E-08 | 2.65E-07 | 153.691455 | 277.376049 | 203.630357 | 235.332698 |
| -5.5063588 | 3.66E-08 | 2.65E-07 | 144.67489 | 283.600097 | 223.188075 | 265.029444 |
| -5.5062149 | 3.67E-08 | 2.65E-07 | 167.216303 | 212.700072 | 194.426725 | 184.904263 |
| -5.5058697 | 3.67E-08 | 2.65E-07 | 150.822548 | 421.882205 | 307.171217 | 225.247011 |
| 5.50590591 | 3.67E-08 | 2.65E-07 | 143.445358 | 99.5847667 | 103.54086 | 57.1522268 |
| -5.5056368 | 3.68E-08 | 2.66E-07 | 132.379573 | 271.963833 | 364.693917 | 320.500723 |
| -5.5055312 | 3.68E-08 | 2.66E-07 | 917.230603 | 1980.8709 | 1782.05324 | 1984.63909 |

| | | | | | | |
|---|---|---|---|---|---|---|
| -5.5054824 | 3.68E-08 | 2.66E-07 | 143.035514 | 246.797031 | 213.984443 | 197.79153 |
| -5.5051808 | 3.69E-08 | 2.66E-07 | 136.478012 | 256.268408 | 285.312591 | 298.088085 |
| -5.5047826 | 3.70E-08 | 2.67E-07 | 142.62567 | 266.010396 | 335.932567 | 238.694594 |
| -5.5047809 | 3.70E-08 | 2.67E-07 | 187.708497 | 382.373031 | 343.985745 | 361.964103 |
| -5.5047692 | 3.70E-08 | 2.67E-07 | 143.035514 | 245.714587 | 250.798971 | 240.375542 |
| -5.5046857 | 3.70E-08 | 2.67E-07 | 190.577404 | 211.88824 | 165.665375 | 188.266159 |
| -5.5046412 | 3.70E-08 | 2.67E-07 | 236.479919 | 187.262659 | 174.869007 | 180.982051 |
| -5.5045574 | 3.70E-08 | 2.67E-07 | 157.789894 | 252.750468 | 272.657597 | 189.386791 |
| -5.5043152 | 3.71E-08 | 2.67E-07 | 168.855679 | 317.967665 | 186.373547 | 234.772382 |
| -5.5037677 | 3.72E-08 | 2.68E-07 | 150.00286 | 221.359617 | 242.745793 | 215.161324 |
| -5.5035088 | 3.72E-08 | 2.68E-07 | 170.495054 | 171.837845 | 230.090799 | 222.445432 |
| -5.5032676 | 3.73E-08 | 2.69E-07 | 134.018949 | 343.9463 | 261.153057 | 312.656299 |
| -5.5027517 | 3.74E-08 | 2.69E-07 | 220.496007 | 509.289486 | 434.87161 | 466.182869 |
| -5.5023831 | 3.75E-08 | 2.70E-07 | 131.559885 | 320.132552 | 279.560321 | 302.570612 |
| -5.502197 | 3.75E-08 | 2.70E-07 | 137.707544 | 253.291689 | 299.118039 | 266.150076 |
| -5.5020229 | 3.75E-08 | 2.70E-07 | 157.789894 | 227.042444 | 240.444885 | 188.826475 |
| -5.5018048 | 3.76E-08 | 2.70E-07 | 142.62567 | 254.644743 | 235.843069 | 248.219965 |
| -5.5017866 | 3.76E-08 | 2.70E-07 | 124.592539 | 309.578731 | 363.543463 | 368.127578 |
| -5.5011779 | 3.77E-08 | 2.71E-07 | 192.21678 | 270.88139 | 346.286653 | 161.370993 |
| -5.500946 | 3.78E-08 | 2.72E-07 | 164.347396 | 190.239378 | 194.426725 | 238.134278 |
| -5.5006368 | 3.78E-08 | 2.72E-07 | 141.396139 | 244.902755 | 294.516223 | 394.462428 |
| -5.5003617 | 3.79E-08 | 2.72E-07 | 154.101299 | 217.029845 | 202.479903 | 256.624704 |
| -5.5003697 | 3.79E-08 | 2.72E-07 | 138.937075 | 245.714587 | 288.763953 | 261.107232 |
| -5.4999482 | 3.80E-08 | 2.73E-07 | 140.986295 | 262.492456 | 240.444885 | 251.581861 |
| -5.4998621 | 3.80E-08 | 2.73E-07 | 297.136813 | 160.742803 | 476.287954 | 219.083536 |
| -5.4996899 | 3.80E-08 | 2.73E-07 | 135.658324 | 275.752384 | 260.002603 | 290.243661 |
| -5.4996174 | 3.81E-08 | 2.73E-07 | 128.690978 | 538.51545 | 347.437107 | 321.061039 |
| -5.4995641 | 3.81E-08 | 2.73E-07 | 108.608628 | 495.758947 | 454.429328 | 439.84802 |
| -5.4994635 | 3.81E-08 | 2.73E-07 | 136.068168 | 306.872624 | 253.099879 | 281.838922 |
| -5.4993484 | 3.81E-08 | 2.73E-07 | 780.342747 | 323.650492 | 177.169915 | 259.9866 |
| -5.4991408 | 3.82E-08 | 2.74E-07 | 141.805982 | 256.268408 | 232.391707 | 292.484925 |
| -5.4990334 | 3.82E-08 | 2.74E-07 | 118.854725 | 357.74745 | 373.897549 | 395.58306 |
| -5.4989687 | 3.82E-08 | 2.74E-07 | 203.282564 | 179.956168 | 165.665375 | 236.45333 |
| 5.49892851 | 3.82E-08 | 2.74E-07 | 19.6725062 | 10.8244312 | 6.90272397 | 10.646003 |
| -5.498827 | 3.82E-08 | 2.74E-07 | 146.724109 | 435.142133 | 231.241253 | 318.259459 |
| -5.4987836 | 3.82E-08 | 2.74E-07 | 177.052556 | 273.587498 | 174.869007 | 205.635953 |
| -5.4985436 | 3.83E-08 | 2.74E-07 | 128.690978 | 329.333318 | 378.499364 | 284.640502 |
| -5.4984698 | 3.83E-08 | 2.74E-07 | 117.21535 | 408.622277 | 388.85345 | 366.44663 |
| -5.498335 | 3.83E-08 | 2.74E-07 | 136.478012 | 251.668025 | 276.108959 | 290.243661 |
| -5.4983148 | 3.83E-08 | 2.74E-07 | 150.822548 | 248.420695 | 204.780811 | 226.927959 |
| -5.4983544 | 3.83E-08 | 2.74E-07 | 101.641282 | 610.768529 | 496.996126 | 513.809725 |
| -5.4981864 | 3.84E-08 | 2.75E-07 | 155.740674 | 235.972599 | 189.824909 | 245.418385 |
| -5.4979119 | 3.84E-08 | 2.75E-07 | 142.62567 | 620.781127 | 249.648517 | 490.836771 |

| | | | | | | |
|---|---|---|---|---|---|---|
| -5.497868 | 3.84E-08 | 2.75E-07 | 197.954594 | 230.019162 | 185.223093 | 163.612257 |
| -5.4977545 | 3.85E-08 | 2.75E-07 | 165.576927 | 243.820312 | 246.197155 | 174.25826 |
| -5.4977397 | 3.85E-08 | 2.75E-07 | 130.330354 | 297.401246 | 287.613499 | 312.095983 |
| -5.4976318 | 3.85E-08 | 2.75E-07 | 147.953641 | 276.293606 | 205.931265 | 272.873867 |
| -5.4975095 | 3.85E-08 | 2.75E-07 | 129.92051 | 303.354683 | 319.826211 | 285.200818 |
| -5.4969102 | 3.87E-08 | 2.76E-07 | 154.511143 | 255.727186 | 195.577179 | 224.686695 |
| -5.4952903 | 3.90E-08 | 2.79E-07 | 140.166607 | 255.185965 | 235.843069 | 278.477026 |
| -5.4952783 | 3.90E-08 | 2.79E-07 | 156.970206 | 232.995881 | 186.373547 | 232.531119 |
| -5.4952356 | 3.90E-08 | 2.79E-07 | 131.150042 | 268.445893 | 338.233475 | 309.85472 |
| -5.4950564 | 3.91E-08 | 2.79E-07 | 127.051603 | 303.625294 | 353.189377 | 307.05314 |
| -5.4946443 | 3.91E-08 | 2.79E-07 | 259.021332 | 168.861126 | 156.461743 | 148.483726 |
| -5.4940656 | 3.93E-08 | 2.80E-07 | 149.183172 | 257.621462 | 203.630357 | 238.694594 |
| -5.4939733 | 3.93E-08 | 2.80E-07 | 187.708497 | 294.153917 | 167.966283 | 208.997849 |
| -5.4939032 | 3.93E-08 | 2.80E-07 | 140.576451 | 365.053941 | 324.428027 | 229.169223 |
| -5.4936877 | 3.94E-08 | 2.81E-07 | 96.3133118 | 519.031474 | 697.175121 | 588.892062 |
| -5.4926428 | 3.96E-08 | 2.82E-07 | 120.084257 | 314.179115 | 502.748396 | 463.38129 |
| -5.492632 | 3.96E-08 | 2.82E-07 | 134.838636 | 311.202396 | 246.197155 | 312.095983 |
| -5.4921881 | 3.97E-08 | 2.83E-07 | 299.595876 | 232.995881 | 166.815829 | 179.301104 |
| -5.4922245 | 3.97E-08 | 2.83E-07 | 139.346919 | 300.107354 | 253.099879 | 240.935858 |
| -5.4920874 | 3.97E-08 | 2.83E-07 | 117.21535 | 525.796744 | 332.481205 | 465.062237 |
| -5.4918522 | 3.98E-08 | 2.83E-07 | 121.313788 | 316.344001 | 372.747095 | 401.186219 |
| -5.4918535 | 3.98E-08 | 2.83E-07 | 129.92051 | 329.603929 | 284.162137 | 289.683345 |
| -5.4919168 | 3.98E-08 | 2.83E-07 | 136.887856 | 239.219929 | 293.365769 | 323.862618 |
| -5.4917774 | 3.98E-08 | 2.83E-07 | 140.576451 | 243.00848 | 236.993523 | 263.908812 |
| -5.4908951 | 4.00E-08 | 2.84E-07 | 109.018472 | 544.739498 | 483.190678 | 393.902112 |
| -5.4904253 | 4.01E-08 | 2.85E-07 | 151.232392 | 235.701989 | 204.780811 | 296.967453 |
| -5.4902381 | 4.01E-08 | 2.85E-07 | 125.412227 | 310.661174 | 309.472125 | 351.3181 |
| 5.49007429 | 4.02E-08 | 2.86E-07 | 6.55750208 | 5.1416048 | 2.30090799 | 3.92221164 |
| -5.489755 | 4.02E-08 | 2.86E-07 | 127.871291 | 327.439043 | 278.409867 | 347.395888 |
| -5.4897732 | 4.02E-08 | 2.86E-07 | 130.740198 | 286.306204 | 316.374849 | 275.115131 |
| -5.4895692 | 4.03E-08 | 2.86E-07 | 183.200214 | 161.284024 | 240.444885 | 205.075637 |
| -5.4892035 | 4.04E-08 | 2.87E-07 | 138.527231 | 266.010396 | 258.852149 | 237.573962 |
| -5.4891793 | 4.04E-08 | 2.87E-07 | 112.297223 | 412.681438 | 467.084322 | 386.057689 |
| -5.48907 | 4.04E-08 | 2.87E-07 | 177.872244 | 164.260743 | 203.630357 | 204.515321 |
| -5.4890616 | 4.04E-08 | 2.87E-07 | 132.789417 | 313.637893 | 255.400787 | 296.967453 |
| -5.4887721 | 4.05E-08 | 2.87E-07 | 169.265522 | 188.345102 | 195.577179 | 177.620156 |
| -5.4886355 | 4.05E-08 | 2.87E-07 | 150.822548 | 232.184049 | 196.727633 | 233.651751 |
| -5.4886235 | 4.05E-08 | 2.87E-07 | 197.134906 | 188.886324 | 172.568099 | 172.577312 |
| -5.4882525 | 4.06E-08 | 2.88E-07 | 19396.6813 | 4125.19072 | 3618.17782 | 4173.7935 |
| -5.4877218 | 4.07E-08 | 2.89E-07 | 156.560362 | 514.16048 | 436.022064 | 408.470327 |
| -5.487391 | 4.08E-08 | 2.89E-07 | 103.280658 | 587.496002 | 419.915708 | 617.468175 |
| -5.4871831 | 4.08E-08 | 2.89E-07 | 181.560839 | 225.418779 | 189.824909 | 165.293205 |
| -5.4872072 | 4.08E-08 | 2.89E-07 | 136.887856 | 282.247043 | 241.595339 | 263.348496 |

| | | | | | | |
|---|---|---|---|---|---|---|
| -5.4870892 | 4.09E-08 | 2.89E-07 | 141.396139 | 230.560384 | 272.657597 | 236.45333 |
| -5.4869381 | 4.09E-08 | 2.90E-07 | 187.708497 | 173.73212 | 171.417645 | 196.670898 |
| -5.4866056 | 4.10E-08 | 2.90E-07 | 184.83959 | 203.499306 | 171.417645 | 174.25826 |
| -5.4862261 | 4.11E-08 | 2.91E-07 | 132.789417 | 264.927953 | 272.657597 | 286.881766 |
| -5.4853238 | 4.13E-08 | 2.92E-07 | 120.903945 | 337.722252 | 372.747095 | 327.78483 |
| -5.4852323 | 4.13E-08 | 2.92E-07 | 156.970206 | 312.014228 | 195.577179 | 233.091435 |
| -5.4852401 | 4.13E-08 | 2.92E-07 | 146.724109 | 254.374132 | 207.081719 | 231.410487 |
| -5.4851367 | 4.13E-08 | 2.92E-07 | 155.740674 | 216.759234 | 222.037621 | 186.024895 |
| -5.4851704 | 4.13E-08 | 2.92E-07 | 115.985818 | 383.184863 | 352.038923 | 423.598857 |
| -5.4846945 | 4.14E-08 | 2.93E-07 | 153.281611 | 211.347019 | 194.426725 | 215.72164 |
| 5.48380422 | 4.16E-08 | 2.94E-07 | 7.37718984 | 6.22404792 | 4.60181598 | 5.04284354 |
| -5.4838294 | 4.16E-08 | 2.94E-07 | 149.593016 | 225.960001 | 297.967585 | 210.118481 |
| -5.4836524 | 4.17E-08 | 2.94E-07 | 129.510666 | 297.130636 | 281.861229 | 285.200818 |
| -5.4836184 | 4.17E-08 | 2.94E-07 | 182.79037 | 158.307306 | 249.648517 | 216.281956 |
| -5.48352 | 4.17E-08 | 2.94E-07 | 119.674413 | 355.311953 | 334.782113 | 355.240311 |
| -5.4827296 | 4.19E-08 | 2.96E-07 | 173.773805 | 254.374132 | 578.67836 | 244.297754 |
| -5.4826515 | 4.19E-08 | 2.96E-07 | 759.44071 | 341.240193 | 166.815829 | 253.823125 |
| -5.4822822 | 4.20E-08 | 2.96E-07 | 132.379573 | 281.705821 | 258.852149 | 281.838922 |
| -5.4820694 | 4.20E-08 | 2.97E-07 | 131.969729 | 294.424528 | 265.754873 | 270.072287 |
| -5.4818785 | 4.21E-08 | 2.97E-07 | 123.363008 | 442.989845 | 331.330751 | 313.216615 |
| -5.4817916 | 4.21E-08 | 2.97E-07 | 121.723632 | 347.734851 | 342.835291 | 313.776931 |
| 5.48147594 | 4.22E-08 | 2.97E-07 | 0.81968776 | 2.16488623 | 1.150454 | 2.24126379 |
| 5.48147594 | 4.22E-08 | 2.97E-07 | 0.81968776 | 2.16488623 | 1.150454 | 2.24126379 |
| -5.4812456 | 4.22E-08 | 2.98E-07 | 144.67489 | 268.175282 | 210.533081 | 237.013646 |
| -5.4809429 | 4.23E-08 | 2.98E-07 | 122.133476 | 447.860839 | 294.516223 | 398.384639 |
| -5.4807946 | 4.23E-08 | 2.98E-07 | 131.150042 | 297.130636 | 256.551241 | 301.44998 |
| -5.4807434 | 4.24E-08 | 2.98E-07 | 225.414134 | 187.803881 | 164.514921 | 179.86142 |
| -5.4799966 | 4.25E-08 | 3.00E-07 | 143.445358 | 304.978348 | 271.507143 | 293.045241 |
| -5.479832 | 4.26E-08 | 3.00E-07 | 126.641759 | 301.460408 | 324.428027 | 285.200818 |
| -5.4796034 | 4.26E-08 | 3.00E-07 | 129.92051 | 301.731019 | 280.710775 | 271.192919 |
| -5.4794394 | 4.27E-08 | 3.00E-07 | 151.642236 | 216.759234 | 149.559019 | 198.912162 |
| -5.4793168 | 4.27E-08 | 3.00E-07 | 129.92051 | 321.485606 | 262.303511 | 294.726189 |
| -5.4791482 | 4.27E-08 | 3.01E-07 | 106.969253 | 138.82333 | 138.054479 | 144.001199 |
| -5.4789265 | 4.28E-08 | 3.01E-07 | 133.609105 | 271.693222 | 249.648517 | 276.796079 |
| -5.4788079 | 4.28E-08 | 3.01E-07 | 119.264569 | 449.755115 | 553.368372 | 311.535667 |
| -5.4786906 | 4.28E-08 | 3.01E-07 | 130.330354 | 422.694037 | 308.321671 | 274.554815 |
| -5.4783615 | 4.29E-08 | 3.02E-07 | 120.494101 | 333.121869 | 339.383929 | 333.387989 |
| -5.4781623 | 4.30E-08 | 3.02E-07 | 134.838636 | 265.469174 | 242.745793 | 302.570612 |
| -5.4780382 | 4.30E-08 | 3.02E-07 | 135.658324 | 231.913438 | 297.967585 | 286.32145 |
| -5.4780749 | 4.30E-08 | 3.02E-07 | 220.905851 | 537.433007 | 517.704298 | 530.619203 |
| -5.4776522 | 4.31E-08 | 3.03E-07 | 119.674413 | 446.778396 | 310.622579 | 378.213265 |
| -5.477366 | 4.32E-08 | 3.03E-07 | 172.954117 | 233.807713 | 166.815829 | 194.98995 |
| -5.4762157 | 4.35E-08 | 3.05E-07 | 142.215826 | 361.26539 | 226.639437 | 266.710392 |

| | | | | | | |
|---|---|---|---|---|---|---|
| -5.4762129 | 4.35E-08 | 3.05E-07 | 140.166607 | 239.76115 | 225.488983 | 242.616806 |
| -5.4758741 | 4.35E-08 | 3.05E-07 | 137.2977 | 250.04436 | 232.391707 | 276.235763 |
| -5.4758929 | 4.35E-08 | 3.05E-07 | 173.363961 | 563.952864 | 327.879389 | 384.937057 |
| -5.475593 | 4.36E-08 | 3.06E-07 | 145.494577 | 551.504768 | 287.613499 | 417.435382 |
| -5.4754716 | 4.36E-08 | 3.06E-07 | 156.560362 | 199.981366 | 199.028541 | 187.705843 |
| -5.4755382 | 4.36E-08 | 3.06E-07 | 152.871767 | 195.651593 | 199.028541 | 212.92006 |
| -5.4754938 | 4.36E-08 | 3.06E-07 | 116.805506 | 335.827977 | 362.393009 | 421.357593 |
| -5.4752834 | 4.37E-08 | 3.06E-07 | 145.904421 | 206.205414 | 223.188075 | 226.367643 |
| -5.475042 | 4.37E-08 | 3.06E-07 | 156.970206 | 393.468073 | 339.383929 | 364.205367 |
| -5.4748 | 4.38E-08 | 3.07E-07 | 202.053033 | 165.884408 | 170.267191 | 198.351846 |
| -5.4745757 | 4.39E-08 | 3.07E-07 | 448.369205 | 372.360432 | 115.0454 | 338.991149 |
| -5.4745679 | 4.39E-08 | 3.07E-07 | 117.625194 | 571.529966 | 335.932567 | 402.306851 |
| -5.4744113 | 4.39E-08 | 3.07E-07 | 138.117388 | 312.55545 | 223.188075 | 276.235763 |
| -5.4741592 | 4.40E-08 | 3.08E-07 | 105.329877 | 486.558181 | 402.658898 | 519.9732 |
| -5.474045 | 4.40E-08 | 3.08E-07 | 137.2977 | 245.714587 | 249.648517 | 236.45333 |
| -5.4739295 | 4.40E-08 | 3.08E-07 | 130.740198 | 277.376049 | 261.153057 | 281.838922 |
| -5.4737435 | 4.41E-08 | 3.08E-07 | 134.428793 | 259.515737 | 276.108959 | 242.05649 |
| -5.4736743 | 4.41E-08 | 3.08E-07 | 135.658324 | 326.897821 | 239.294431 | 265.58976 |
| -5.4735407 | 4.41E-08 | 3.08E-07 | 137.707544 | 239.219929 | 278.409867 | 234.212067 |
| -5.4733884 | 4.42E-08 | 3.09E-07 | 162.70802 | 316.344001 | 173.718553 | 314.897563 |
| -5.4729177 | 4.43E-08 | 3.09E-07 | 216.397569 | 347.734851 | 144.957203 | 448.813075 |
| -5.4729128 | 4.43E-08 | 3.09E-07 | 121.313788 | 340.969582 | 366.994825 | 299.208717 |
| -5.4724307 | 4.44E-08 | 3.10E-07 | 125.822071 | 305.79018 | 307.171217 | 282.399238 |
| -5.4722027 | 4.44E-08 | 3.10E-07 | 149.593016 | 275.481773 | 211.683535 | 205.635953 |
| -5.4720832 | 4.45E-08 | 3.10E-07 | 151.232392 | 206.476024 | 194.426725 | 210.118481 |
| -5.4717284 | 4.46E-08 | 3.11E-07 | 257.7918 | 129.351952 | 300.268493 | 344.033992 |
| -5.4713905 | 4.47E-08 | 3.12E-07 | 107.379097 | 473.298253 | 414.163438 | 414.073486 |
| -5.4713137 | 4.47E-08 | 3.12E-07 | 166.396615 | 262.763067 | 326.728935 | 254.943757 |
| -5.4710718 | 4.47E-08 | 3.12E-07 | 192.21678 | 149.106539 | 251.949425 | 226.927959 |
| 5.47048179 | 4.49E-08 | 3.13E-07 | 28.2792277 | 22.7313054 | 10.354086 | 20.7316901 |
| -5.4702686 | 4.49E-08 | 3.13E-07 | 141.805982 | 258.433294 | 212.833989 | 233.091435 |
| -5.4701531 | 4.50E-08 | 3.13E-07 | 142.215826 | 281.976432 | 216.285351 | 231.970803 |
| -5.4692215 | 4.52E-08 | 3.15E-07 | 123.772852 | 302.542851 | 308.321671 | 302.570612 |
| -5.4690173 | 4.53E-08 | 3.15E-07 | 151.642236 | 236.784432 | 188.674455 | 210.118481 |
| -5.4689216 | 4.53E-08 | 3.15E-07 | 156.150518 | 177.791282 | 234.692615 | 261.107232 |
| -5.4689375 | 4.53E-08 | 3.15E-07 | 135.24848 | 314.449725 | 256.551241 | 238.694594 |
| -5.4688016 | 4.53E-08 | 3.15E-07 | 138.117388 | 281.43521 | 218.586259 | 261.667548 |
| -5.4683335 | 4.54E-08 | 3.16E-07 | 451.238112 | 243.27909 | 199.028541 | 271.192919 |
| -5.4676137 | 4.56E-08 | 3.17E-07 | 144.67489 | 198.628312 | 251.949425 | 274.554815 |
| -5.4674953 | 4.56E-08 | 3.18E-07 | 106.559409 | 386.973414 | 454.429328 | 512.128777 |
| -5.4674723 | 4.56E-08 | 3.18E-07 | 140.986295 | 277.376049 | 211.683535 | 244.297754 |
| -5.4673144 | 4.57E-08 | 3.18E-07 | 185.249434 | 155.059976 | 258.852149 | 209.558165 |
| -5.4670305 | 4.58E-08 | 3.18E-07 | 151.642236 | 244.902755 | 193.276271 | 202.834373 |

| | | | | | | |
|---|---|---|---|---|---|---|
| -5.4668736 | 4.58E-08 | 3.18E-07 | 129.92051 | 333.663091 | 265.754873 | 266.710392 |
| -5.4668567 | 4.58E-08 | 3.18E-07 | 165.576927 | 194.027929 | 170.267191 | 231.970803 |
| -5.4667727 | 4.58E-08 | 3.18E-07 | 176.642712 | 170.21418 | 178.320369 | 181.542367 |
| -5.4662464 | 4.60E-08 | 3.19E-07 | 188.528185 | 204.311138 | 154.160835 | 190.507423 |
| -5.4661879 | 4.60E-08 | 3.19E-07 | 116.395662 | 309.03751 | 483.190678 | 431.44328 |
| -5.4660611 | 4.60E-08 | 3.20E-07 | 98.3625312 | 446.237175 | 617.793796 | 540.144574 |
| -5.4656929 | 4.61E-08 | 3.20E-07 | 141.396139 | 222.17145 | 223.188075 | 223.566063 |
| -5.4652272 | 4.62E-08 | 3.21E-07 | 129.92051 | 299.566133 | 253.099879 | 274.554815 |
| -5.4650031 | 4.63E-08 | 3.21E-07 | 168.445835 | 295.777582 | 227.789891 | 302.010296 |
| -5.4645781 | 4.64E-08 | 3.22E-07 | 131.969729 | 274.39933 | 242.745793 | 270.632603 |
| -5.4642675 | 4.65E-08 | 3.22E-07 | 172.13443 | 288.20048 | 159.913105 | 304.811876 |
| 5.46417572 | 4.65E-08 | 3.22E-07 | 71.3128351 | 36.803066 | 34.5136199 | 39.2221164 |
| -5.4641765 | 4.65E-08 | 3.22E-07 | 137.707544 | 221.359617 | 365.844371 | 304.25156 |
| -5.4635167 | 4.67E-08 | 3.24E-07 | 124.592539 | 567.200193 | 301.418947 | 365.325998 |
| -5.4630184 | 4.68E-08 | 3.24E-07 | 138.117388 | 366.948217 | 257.701695 | 238.134278 |
| -5.4628579 | 4.69E-08 | 3.25E-07 | 165.576927 | 270.610779 | 164.514921 | 237.573962 |
| -5.4627818 | 4.69E-08 | 3.25E-07 | 138.527231 | 260.056959 | 218.586259 | 237.013646 |
| -5.4627019 | 4.69E-08 | 3.25E-07 | 124.182696 | 329.062707 | 278.409867 | 302.570612 |
| -5.4626292 | 4.69E-08 | 3.25E-07 | 107.78894 | 411.869606 | 459.031144 | 386.618005 |
| -5.4624915 | 4.69E-08 | 3.25E-07 | 170.08521 | 160.201581 | 217.435805 | 206.756585 |
| -5.462387 | 4.70E-08 | 3.25E-07 | 143.035514 | 225.960001 | 204.780811 | 229.169223 |
| -5.4622282 | 4.70E-08 | 3.25E-07 | 109.83816 | 376.419594 | 408.411168 | 400.065587 |
| -5.4621196 | 4.70E-08 | 3.26E-07 | 126.231915 | 279.270324 | 278.409867 | 346.275256 |
| -5.4620629 | 4.71E-08 | 3.26E-07 | 145.494577 | 270.069558 | 193.276271 | 250.461229 |
| -5.461668 | 4.72E-08 | 3.26E-07 | 147.543797 | 226.501222 | 219.736713 | 344.033992 |
| -5.4616101 | 4.72E-08 | 3.26E-07 | 151.642236 | 206.476024 | 185.223093 | 210.118481 |
| 5.46113057 | 4.73E-08 | 3.27E-07 | 46.7222023 | 47.8981079 | 40.2658898 | 53.7903311 |
| -5.4608653 | 4.74E-08 | 3.28E-07 | 152.052079 | 455.437941 | 387.702996 | 423.598857 |
| -5.4605428 | 4.75E-08 | 3.28E-07 | 165.576927 | 236.784432 | 453.278874 | 214.601008 |
| -5.4605312 | 4.75E-08 | 3.28E-07 | 147.133953 | 296.318803 | 216.285351 | 208.997849 |
| -5.4605244 | 4.75E-08 | 3.28E-07 | 250.414611 | 306.872624 | 127.700393 | 364.765683 |
| -5.4604735 | 4.75E-08 | 3.28E-07 | 129.92051 | 252.750468 | 264.604419 | 284.640502 |
| -5.4601338 | 4.76E-08 | 3.29E-07 | 133.199261 | 209.723354 | 195.577179 | 250.461229 |
| -5.4599284 | 4.76E-08 | 3.29E-07 | 142.62567 | 234.890156 | 208.232173 | 216.281956 |
| -5.4595145 | 4.77E-08 | 3.29E-07 | 188.938029 | 175.897006 | 446.37615 | 332.827673 |
| -5.4595255 | 4.77E-08 | 3.29E-07 | 134.838636 | 285.764983 | 224.338529 | 262.227864 |
| -5.4586946 | 4.80E-08 | 3.31E-07 | 124.592539 | 307.413845 | 268.055781 | 346.275256 |
| -5.4584204 | 4.80E-08 | 3.31E-07 | 151.232392 | 179.414947 | 253.099879 | 254.943757 |
| -5.458109 | 4.81E-08 | 3.32E-07 | 124.592539 | 440.824959 | 283.011683 | 318.819775 |
| -5.4578373 | 4.82E-08 | 3.32E-07 | 150.00286 | 223.524504 | 197.878087 | 191.067738 |
| -5.4577135 | 4.82E-08 | 3.32E-07 | 123.363008 | 268.445893 | 334.782113 | 331.707042 |
| -5.457582 | 4.83E-08 | 3.33E-07 | 117.21535 | 393.197462 | 296.817131 | 417.435382 |
| -5.4575349 | 4.83E-08 | 3.33E-07 | 127.871291 | 286.576815 | 264.604419 | 267.270707 |

| | | | | | | |
|---|---|---|---|---|---|---|
| -5.4574968 | 4.83E-08 | 3.33E-07 | 131.559885 | 235.701989 | 301.418947 | 354.679995 |
| -5.4573015 | 4.83E-08 | 3.33E-07 | 143.855202 | 253.291689 | 195.577179 | 234.772382 |
| -5.457224 | 4.84E-08 | 3.33E-07 | 132.379573 | 292.259641 | 243.896247 | 247.099333 |
| -5.4570603 | 4.84E-08 | 3.33E-07 | 166.396615 | 393.738684 | 470.535684 | 447.692443 |
| -5.4570007 | 4.84E-08 | 3.33E-07 | 152.461923 | 409.434109 | 368.145279 | 382.135477 |
| -5.4569757 | 4.84E-08 | 3.33E-07 | 169.675366 | 326.085989 | 167.966283 | 241.496174 |
| -5.4568062 | 4.85E-08 | 3.34E-07 | 102.051126 | 681.668553 | 400.35799 | 522.77478 |
| -5.4559675 | 4.87E-08 | 3.35E-07 | 192.626624 | 183.744719 | 157.612197 | 180.421735 |
| -5.4556329 | 4.88E-08 | 3.35E-07 | 216.807412 | 218.92412 | 154.160835 | 173.137628 |
| -5.455636 | 4.88E-08 | 3.35E-07 | 229.922417 | 263.033677 | 134.603117 | 230.850171 |
| -5.4556854 | 4.88E-08 | 3.35E-07 | 106.149565 | 461.120768 | 411.86253 | 393.902112 |
| -5.455694 | 4.88E-08 | 3.35E-07 | 134.428793 | 223.253893 | 261.153057 | 261.667548 |
| -5.4553689 | 4.89E-08 | 3.36E-07 | 120.494101 | 341.240193 | 293.365769 | 313.216615 |
| -5.4553232 | 4.89E-08 | 3.36E-07 | 158.609582 | 189.968767 | 184.072639 | 180.421735 |
| 5.454757 | 4.90E-08 | 3.37E-07 | 19.6725062 | 7.30649104 | 17.2568099 | 7.28410733 |
| 5.4543762 | 4.91E-08 | 3.37E-07 | 0 | 0 | 0 | 0 |
| -5.4542679 | 4.92E-08 | 3.38E-07 | 107.379097 | 417.552432 | 381.950726 | 415.754434 |
| -5.4542168 | 4.92E-08 | 3.38E-07 | 154.920987 | 185.097773 | 184.072639 | 207.877217 |
| -5.4540378 | 4.92E-08 | 3.38E-07 | 117.21535 | 357.74745 | 346.286653 | 305.372192 |
| -5.4539263 | 4.93E-08 | 3.38E-07 | 148.363485 | 225.148168 | 187.524001 | 267.270707 |
| -5.4534124 | 4.94E-08 | 3.39E-07 | 131.559885 | 429.459307 | 311.773033 | 397.824324 |
| -5.4531208 | 4.95E-08 | 3.39E-07 | 139.346919 | 206.476024 | 239.294431 | 239.25491 |
| -5.4513652 | 5.00E-08 | 3.43E-07 | 151.642236 | 188.615713 | 288.763953 | 216.842272 |
| -5.4512854 | 5.00E-08 | 3.43E-07 | 147.953641 | 202.416863 | 202.479903 | 192.748686 |
| -5.4511669 | 5.00E-08 | 3.43E-07 | 140.576451 | 291.177198 | 212.833989 | 227.488275 |
| -5.451074 | 5.01E-08 | 3.43E-07 | 129.510666 | 277.376049 | 241.595339 | 270.072287 |
| -5.4507686 | 5.02E-08 | 3.44E-07 | 128.281134 | 243.00848 | 286.463045 | 321.621355 |
| -5.4496399 | 5.05E-08 | 3.46E-07 | 114.756286 | 345.840576 | 352.038923 | 327.224514 |
| 5.44954965 | 5.05E-08 | 3.46E-07 | 0 | 0 | 0 | 0 |
| -5.4490869 | 5.06E-08 | 3.47E-07 | 147.133953 | 404.833726 | 216.285351 | 256.624704 |
| -5.4489146 | 5.07E-08 | 3.47E-07 | 118.035037 | 496.300169 | 400.35799 | 290.243661 |
| -5.4488826 | 5.07E-08 | 3.47E-07 | 215.577881 | 200.793198 | 195.577179 | 145.682147 |
| -5.4483236 | 5.08E-08 | 3.48E-07 | 134.428793 | 229.20733 | 253.099879 | 230.850171 |
| -5.4480132 | 5.09E-08 | 3.48E-07 | 124.182696 | 310.931785 | 276.108959 | 272.873867 |
| -5.4478845 | 5.10E-08 | 3.49E-07 | 162.298176 | 207.829078 | 162.214013 | 229.729539 |
| -5.4477206 | 5.10E-08 | 3.49E-07 | 482.796091 | 942.807955 | 846.734141 | 923.400683 |
| -5.4476604 | 5.10E-08 | 3.49E-07 | 141.805982 | 238.678707 | 270.356689 | 196.110582 |
| -5.4471016 | 5.12E-08 | 3.50E-07 | 148.773328 | 190.780599 | 192.125817 | 220.764484 |
| -5.4468855 | 5.13E-08 | 3.50E-07 | 109.018472 | 379.666923 | 364.693917 | 400.065587 |
| -5.4455953 | 5.16E-08 | 3.53E-07 | 196.725062 | 259.786348 | 181.771731 | 155.207518 |
| -5.445448 | 5.17E-08 | 3.53E-07 | 367.62996 | 469.509702 | 463.63296 | 519.9732 |
| -5.4448898 | 5.18E-08 | 3.54E-07 | 123.772852 | 284.141318 | 270.356689 | 289.123029 |
| -5.4448173 | 5.19E-08 | 3.54E-07 | 357.793707 | 588.307834 | 537.262016 | 495.879615 |

| | | | | | | |
|---|---|---|---|---|---|---|
| -5.4447623 | 5.19E-08 | 3.54E-07 | 113.116911 | 378.043258 | 381.950726 | 315.457879 |
| -5.4445754 | 5.19E-08 | 3.54E-07 | 129.100822 | 295.23636 | 235.843069 | 270.632603 |
| -5.4438739 | 5.21E-08 | 3.56E-07 | 136.068168 | 302.542851 | 222.037621 | 235.893014 |
| -5.4438225 | 5.21E-08 | 3.56E-07 | 119.264569 | 303.354683 | 295.666677 | 319.380091 |
| -5.4432259 | 5.23E-08 | 3.57E-07 | 203.692408 | 406.45739 | 213.984443 | 474.027293 |
| -5.4432116 | 5.23E-08 | 3.57E-07 | 162.298176 | 223.795114 | 163.364467 | 194.98995 |
| -5.4424347 | 5.26E-08 | 3.58E-07 | 127.461447 | 397.527235 | 318.675757 | 244.297754 |
| -5.4422957 | 5.26E-08 | 3.59E-07 | 211.889286 | 212.158851 | 169.116737 | 154.086886 |
| -5.4422537 | 5.26E-08 | 3.59E-07 | 118.444881 | 320.673773 | 293.365769 | 317.138827 |
| -5.4421619 | 5.26E-08 | 3.59E-07 | 142.215826 | 232.454659 | 210.533081 | 197.231214 |
| -5.4419015 | 5.27E-08 | 3.59E-07 | 133.199261 | 308.225677 | 311.773033 | 314.897563 |
| -5.4416946 | 5.28E-08 | 3.59E-07 | 165.167084 | 204.85236 | 231.241253 | 243.177122 |
| -5.4417717 | 5.28E-08 | 3.59E-07 | 130.740198 | 568.553247 | 266.905327 | 345.154624 |
| -5.4416895 | 5.28E-08 | 3.59E-07 | 143.445358 | 221.359617 | 194.426725 | 206.756585 |
| -5.4418221 | 5.27E-08 | 3.59E-07 | 112.707067 | 356.665007 | 341.684837 | 339.551465 |
| -5.4417451 | 5.28E-08 | 3.59E-07 | 134.838636 | 256.268408 | 213.984443 | 241.496174 |
| -5.4411877 | 5.29E-08 | 3.60E-07 | 144.67489 | 221.900839 | 186.373547 | 247.099333 |
| -5.4409866 | 5.30E-08 | 3.60E-07 | 120.494101 | 323.921103 | 274.958505 | 307.05314 |
| -5.4408329 | 5.30E-08 | 3.61E-07 | 119.674413 | 355.041342 | 287.613499 | 297.527769 |
| -5.4407205 | 5.31E-08 | 3.61E-07 | 125.412227 | 289.824144 | 254.250333 | 272.873867 |
| -5.4407237 | 5.31E-08 | 3.61E-07 | 114.346443 | 380.749366 | 319.826211 | 327.78483 |
| -5.440602 | 5.31E-08 | 3.61E-07 | 119.674413 | 358.018061 | 276.108959 | 317.699143 |
| -5.4401398 | 5.32E-08 | 3.62E-07 | 446.319985 | 252.750468 | 143.806749 | 235.332698 |
| -5.4399117 | 5.33E-08 | 3.62E-07 | 123.772852 | 277.646659 | 264.604419 | 290.243661 |
| -5.4398349 | 5.33E-08 | 3.62E-07 | 125.412227 | 277.376049 | 256.551241 | 271.753235 |
| 5.4391352 | 5.35E-08 | 3.63E-07 | 60.6568942 | 45.1920001 | 58.6731538 | 49.8681194 |
| -5.4391539 | 5.35E-08 | 3.63E-07 | 278.283994 | 132.599282 | 263.453965 | 216.842272 |
| -5.4389245 | 5.36E-08 | 3.64E-07 | 133.609105 | 265.469174 | 283.011683 | 211.799429 |
| -5.43895 | 5.36E-08 | 3.64E-07 | 118.444881 | 294.424528 | 303.719855 | 320.500723 |
| -5.4385857 | 5.37E-08 | 3.64E-07 | 105.329877 | 345.299354 | 515.40339 | 477.949504 |
| -5.4385245 | 5.37E-08 | 3.64E-07 | 530.337981 | 345.569965 | 163.364467 | 187.145527 |
| -5.4383681 | 5.38E-08 | 3.65E-07 | 122.54332 | 315.532168 | 300.268493 | 258.865968 |
| -5.4382624 | 5.38E-08 | 3.65E-07 | 132.379573 | 223.795114 | 239.294431 | 268.951655 |
| -5.4380447 | 5.39E-08 | 3.65E-07 | 122.133476 | 342.593246 | 258.852149 | 309.85472 |
| -5.4370439 | 5.42E-08 | 3.67E-07 | 134.428793 | 216.488623 | 231.241253 | 240.935858 |
| -5.43695 | 5.42E-08 | 3.67E-07 | 90.5754975 | 561.246756 | 612.041526 | 483.552664 |
| -5.4367789 | 5.43E-08 | 3.67E-07 | 118.854725 | 384.267306 | 266.905327 | 383.816425 |
| -5.4367709 | 5.43E-08 | 3.67E-07 | 131.969729 | 388.326468 | 219.736713 | 377.092633 |
| -5.4368158 | 5.42E-08 | 3.67E-07 | 122.953164 | 364.51272 | 250.798971 | 331.146726 |
| -5.4365692 | 5.43E-08 | 3.68E-07 | 171.724586 | 361.536001 | 163.364467 | 256.624704 |
| -5.4365432 | 5.43E-08 | 3.68E-07 | 149.593016 | 340.969582 | 218.586259 | 297.527769 |
| -5.4365679 | 5.43E-08 | 3.68E-07 | 121.313788 | 339.616528 | 260.002603 | 393.341796 |
| -5.4364119 | 5.44E-08 | 3.68E-07 | 377.05637 | 243.27909 | 338.233475 | 167.534469 |

| | | | | | | |
|---|---|---|---|---|---|---|
| -5.4361718 | 5.44E-08 | 3.68E-07 | 111.067691 | 374.254708 | 324.428027 | 374.291054 |
| -5.4359214 | 5.45E-08 | 3.69E-07 | 292.62853 | 245.173366 | 134.603117 | 209.558165 |
| -5.4357125 | 5.46E-08 | 3.69E-07 | 120.494101 | 317.697055 | 268.055781 | 307.613456 |
| -5.4353474 | 5.47E-08 | 3.70E-07 | 136.068168 | 208.911521 | 248.498063 | 224.686695 |
| -5.4353215 | 5.47E-08 | 3.70E-07 | 123.363008 | 281.1646 | 268.055781 | 271.753235 |
| -5.4353014 | 5.47E-08 | 3.70E-07 | 188.528185 | 304.437127 | 150.709473 | 214.040692 |
| -5.435326 | 5.47E-08 | 3.70E-07 | 153.281611 | 304.437127 | 294.516223 | 322.741986 |
| -5.4351534 | 5.47E-08 | 3.70E-07 | 126.231915 | 246.797031 | 261.153057 | 283.51987 |
| -5.4351717 | 5.47E-08 | 3.70E-07 | 131.559885 | 246.52642 | 232.391707 | 232.531119 |
| -5.4349058 | 5.48E-08 | 3.70E-07 | 143.855202 | 210.535186 | 195.577179 | 271.192919 |
| -5.4348817 | 5.48E-08 | 3.70E-07 | 136.478012 | 253.021079 | 225.488983 | 207.316901 |
| -5.4347871 | 5.49E-08 | 3.70E-07 | 186.478965 | 199.169533 | 192.125817 | 147.363094 |
| -5.4346836 | 5.49E-08 | 3.70E-07 | 106.559409 | 389.679522 | 371.596641 | 388.298952 |
| -5.4341643 | 5.51E-08 | 3.71E-07 | 123.772852 | 758.251403 | 269.206235 | 569.84132 |
| -5.4341267 | 5.51E-08 | 3.71E-07 | 100.821594 | 526.879187 | 411.86253 | 417.995698 |
| -5.4338364 | 5.52E-08 | 3.72E-07 | 129.100822 | 263.033677 | 230.090799 | 309.294404 |
| -5.4337854 | 5.52E-08 | 3.72E-07 | 162.298176 | 231.101605 | 208.232173 | 228.048591 |
| -5.433648 | 5.52E-08 | 3.72E-07 | 125.822071 | 284.141318 | 245.046701 | 266.710392 |
| -5.4335814 | 5.52E-08 | 3.72E-07 | 139.346919 | 214.323737 | 201.329449 | 216.842272 |
| -5.4331206 | 5.54E-08 | 3.73E-07 | 152.052079 | 174.002731 | 190.975363 | 214.040692 |
| -5.4330178 | 5.54E-08 | 3.73E-07 | 91.3951852 | 487.640624 | 580.979268 | 520.533516 |
| -5.4329632 | 5.54E-08 | 3.73E-07 | 125.412227 | 309.578731 | 417.6148 | 252.142177 |
| -5.4323769 | 5.56E-08 | 3.74E-07 | 118.854725 | 262.221845 | 256.551241 | 253.262809 |
| -5.4322317 | 5.57E-08 | 3.75E-07 | 120.494101 | 273.046276 | 311.773033 | 285.761134 |
| -5.4322212 | 5.57E-08 | 3.75E-07 | 138.527231 | 222.44206 | 199.028541 | 223.005748 |
| -5.4319939 | 5.57E-08 | 3.75E-07 | 138.937075 | 232.454659 | 217.435805 | 197.79153 |
| -5.4317158 | 5.58E-08 | 3.76E-07 | 132.379573 | 299.295522 | 210.533081 | 284.080186 |
| -5.4316507 | 5.58E-08 | 3.76E-07 | 180.741151 | 152.353869 | 209.382627 | 178.740788 |
| -5.4316067 | 5.58E-08 | 3.76E-07 | 155.740674 | 206.476024 | 163.364467 | 219.083536 |
| -5.4315084 | 5.59E-08 | 3.76E-07 | 102.46097 | 444.884121 | 423.36707 | 385.497373 |
| -5.4313972 | 5.59E-08 | 3.76E-07 | 134.428793 | 242.467258 | 207.081719 | 265.58976 |
| -5.4313504 | 5.59E-08 | 3.76E-07 | 131.150042 | 234.619546 | 280.710775 | 227.488275 |
| -5.4309347 | 5.61E-08 | 3.77E-07 | 172.544273 | 289.282923 | 174.869007 | 180.982051 |
| -5.4308743 | 5.61E-08 | 3.77E-07 | 133.609105 | 221.089007 | 236.993523 | 221.885116 |
| -5.4303723 | 5.62E-08 | 3.78E-07 | 125.002383 | 300.107354 | 247.347609 | 265.029444 |
| -5.4301298 | 5.63E-08 | 3.78E-07 | 89.3459658 | 537.433007 | 567.17382 | 516.611305 |
| -5.4300765 | 5.63E-08 | 3.78E-07 | 597.142533 | 1828.24642 | 1750.99098 | 1830.5522 |
| -5.4299096 | 5.64E-08 | 3.78E-07 | 152.461923 | 183.203497 | 228.940345 | 178.740788 |
| -5.4299324 | 5.64E-08 | 3.78E-07 | 170.08521 | 213.782516 | 157.612197 | 178.180472 |
| -5.4291529 | 5.66E-08 | 3.80E-07 | 159.429269 | 193.216096 | 277.259413 | 175.378892 |
| -5.4285341 | 5.68E-08 | 3.81E-07 | 118.035037 | 322.026827 | 294.516223 | 284.640502 |
| -5.4278547 | 5.70E-08 | 3.82E-07 | 131.150042 | 274.940552 | 232.391707 | 225.247011 |
| -5.4278901 | 5.70E-08 | 3.82E-07 | 122.133476 | 123.127905 | 163.364467 | 123.269509 |

| | | | | | | |
|---|---|---|---|---|---|---|
| -5.4276524 | 5.71E-08 | 3.83E-07 | 163.117864 | 257.08024 | 163.364467 | 196.110582 |
| -5.4275857 | 5.71E-08 | 3.83E-07 | 111.067691 | 330.415761 | 394.60572 | 329.465778 |
| -5.4272884 | 5.72E-08 | 3.83E-07 | 145.494577 | 193.757318 | 187.524001 | 235.893014 |
| -5.4272813 | 5.72E-08 | 3.83E-07 | 148.363485 | 178.061893 | 196.727633 | 231.970803 |
| -5.4272636 | 5.72E-08 | 3.83E-07 | 2010.28423 | 2638.18449 | 2001.78995 | 2596.50411 |
| -5.4270823 | 5.73E-08 | 3.84E-07 | 115.985818 | 321.214995 | 301.418947 | 302.010296 |
| -5.4266311 | 5.74E-08 | 3.85E-07 | 133.199261 | 309.578731 | 211.683535 | 255.504073 |
| -5.4265239 | 5.75E-08 | 3.85E-07 | 104.920033 | 671.926565 | 354.339831 | 466.743185 |
| -5.4262783 | 5.75E-08 | 3.85E-07 | 211.889286 | 332.310037 | 131.151755 | 286.32145 |
| -5.4261381 | 5.76E-08 | 3.85E-07 | 155.740674 | 172.649677 | 178.320369 | 184.343947 |
| -5.4257411 | 5.77E-08 | 3.86E-07 | 145.494577 | 243.27909 | 190.975363 | 191.628054 |
| -5.4254383 | 5.78E-08 | 3.87E-07 | 156.150518 | 169.402348 | 205.931265 | 178.180472 |
| -5.4254643 | 5.78E-08 | 3.87E-07 | 125.822071 | 369.113103 | 365.844371 | 229.729539 |
| -5.4254542 | 5.78E-08 | 3.87E-07 | 122.953164 | 321.756216 | 241.595339 | 354.679995 |
| -5.4245323 | 5.81E-08 | 3.88E-07 | 100.821594 | 404.292504 | 401.508444 | 471.786029 |
| -5.4242682 | 5.82E-08 | 3.89E-07 | 131.969729 | 226.501222 | 232.391707 | 221.885116 |
| -5.4237994 | 5.83E-08 | 3.90E-07 | 118.854725 | 304.437127 | 269.206235 | 300.329348 |
| -5.4238214 | 5.83E-08 | 3.90E-07 | 115.16613 | 348.005462 | 288.763953 | 317.138827 |
| -5.4238108 | 5.83E-08 | 3.90E-07 | 144.67489 | 179.414947 | 205.931265 | 209.558165 |
| -5.4235753 | 5.84E-08 | 3.90E-07 | 146.314265 | 174.814563 | 241.595339 | 216.281956 |
| -5.423033 | 5.86E-08 | 3.91E-07 | 55.3289238 | 867.848769 | 1320.72119 | 1195.71423 |
| -5.4230578 | 5.86E-08 | 3.91E-07 | 105.739721 | 439.471905 | 347.437107 | 381.575161 |
| -5.4226605 | 5.87E-08 | 3.92E-07 | 118.854725 | 272.505055 | 301.418947 | 289.683345 |
| -5.4226612 | 5.87E-08 | 3.92E-07 | 102.051126 | 433.789079 | 385.402088 | 402.867167 |
| -5.4223797 | 5.88E-08 | 3.92E-07 | 135.658324 | 208.640911 | 209.382627 | 233.651751 |
| -5.4222816 | 5.88E-08 | 3.92E-07 | 217.6271 | 195.922204 | 167.966283 | 151.285306 |
| -5.4217804 | 5.90E-08 | 3.93E-07 | 138.117388 | 227.854276 | 201.329449 | 296.407137 |
| -5.4217169 | 5.90E-08 | 3.94E-07 | 117.625194 | 297.942468 | 287.613499 | 289.683345 |
| -5.42144 | 5.91E-08 | 3.94E-07 | 131.969729 | 243.549701 | 208.232173 | 256.624704 |
| -5.4211984 | 5.92E-08 | 3.94E-07 | 165.576927 | 231.642827 | 150.709473 | 253.823125 |
| -5.4211915 | 5.92E-08 | 3.94E-07 | 121.723632 | 322.297438 | 247.347609 | 291.924609 |
| -5.4210519 | 5.92E-08 | 3.95E-07 | 113.526755 | 120.692407 | 139.204933 | 128.312352 |
| -5.4209418 | 5.93E-08 | 3.95E-07 | 163.937552 | 214.864959 | 172.568099 | 163.612257 |
| -5.4207807 | 5.93E-08 | 3.95E-07 | 222.545227 | 159.93097 | 212.833989 | 215.72164 |
| -5.420784 | 5.93E-08 | 3.95E-07 | 152.871767 | 286.035594 | 170.267191 | 224.686695 |
| -5.4208012 | 5.93E-08 | 3.95E-07 | 127.871291 | 267.634061 | 350.888469 | 231.970803 |
| -5.4205745 | 5.94E-08 | 3.95E-07 | 114.756286 | 327.439043 | 292.215315 | 313.776931 |
| -5.4205837 | 5.94E-08 | 3.95E-07 | 268.857585 | 424.047091 | 421.066162 | 402.306851 |
| -5.4200614 | 5.96E-08 | 3.96E-07 | 125.822071 | 235.972599 | 251.949425 | 288.002398 |
| -5.4200111 | 5.96E-08 | 3.96E-07 | 130.740198 | 242.467258 | 271.507143 | 212.92006 |
| -5.4197891 | 5.97E-08 | 3.97E-07 | 100.411751 | 416.469989 | 421.066162 | 403.987799 |
| -5.4197434 | 5.97E-08 | 3.97E-07 | 255.332737 | 261.680623 | 125.399485 | 230.289855 |
| -5.4196754 | 5.97E-08 | 3.97E-07 | 131.150042 | 266.010396 | 225.488983 | 220.204168 |

| | | | | | | |
|---|---|---|---|---|---|---|
| -5.4195491 | 5.97E-08 | 3.97E-07 | 121.723632 | 249.773749 | 444.075242 | 352.438732 |
| -5.4182935 | 6.02E-08 | 4.00E-07 | 206.971159 | 303.084073 | 141.505841 | 204.515321 |
| -5.4181514 | 6.02E-08 | 4.00E-07 | 101.231438 | 392.656241 | 392.304812 | 444.890863 |
| -5.4179982 | 6.03E-08 | 4.00E-07 | 175.003337 | 150.188982 | 323.277573 | 364.205367 |
| -5.4179399 | 6.03E-08 | 4.00E-07 | 221.315695 | 209.993965 | 144.957203 | 172.577312 |
| 5.41773584 | 6.04E-08 | 4.01E-07 | 295.497437 | 134.764168 | 131.151755 | 103.65845 |
| -5.4167392 | 6.07E-08 | 4.03E-07 | 152.052079 | 239.76115 | 390.003904 | 192.748686 |
| -5.4163281 | 6.08E-08 | 4.04E-07 | 122.54332 | 306.872624 | 235.843069 | 307.05314 |
| -5.4162665 | 6.09E-08 | 4.04E-07 | 126.231915 | 356.123785 | 234.692615 | 273.434183 |
| -5.4159507 | 6.10E-08 | 4.04E-07 | 127.871291 | 737.955595 | 701.776937 | 702.6362 |
| -5.4158554 | 6.10E-08 | 4.04E-07 | 178.282088 | 270.340168 | 189.824909 | 154.086886 |
| -5.4158714 | 6.10E-08 | 4.04E-07 | 120.084257 | 316.344001 | 274.958505 | 321.621355 |
| -5.4156386 | 6.11E-08 | 4.05E-07 | 106.969253 | 509.289486 | 309.472125 | 456.097182 |
| -5.4154933 | 6.11E-08 | 4.05E-07 | 292.218686 | 153.165701 | 163.364467 | 278.477026 |
| -5.4155114 | 6.11E-08 | 4.05E-07 | 129.100822 | 253.291689 | 216.285351 | 243.737438 |
| 5.41534402 | 6.12E-08 | 4.05E-07 | 23.770945 | 6.4946587 | 9.20363196 | 11.206319 |
| -5.4149151 | 6.13E-08 | 4.06E-07 | 200.823501 | 302.00163 | 176.019461 | 225.247011 |
| -5.4148383 | 6.13E-08 | 4.06E-07 | 150.00286 | 170.484791 | 185.223093 | 203.394689 |
| -5.4143734 | 6.15E-08 | 4.07E-07 | 210.659754 | 245.985198 | 146.107657 | 172.016996 |
| -5.4142706 | 6.15E-08 | 4.07E-07 | 136.068168 | 306.060791 | 216.285351 | 215.161324 |
| -5.4143062 | 6.15E-08 | 4.07E-07 | 103.280658 | 507.395211 | 358.941647 | 389.419584 |
| -5.4140749 | 6.16E-08 | 4.08E-07 | 143.035514 | 403.480672 | 396.906628 | 417.995698 |
| 5.413895 | 6.17E-08 | 4.08E-07 | 40.1647002 | 27.0610779 | 23.0090799 | 16.2491625 |
| -5.4136788 | 6.17E-08 | 4.08E-07 | 136.887856 | 270.88139 | 187.524001 | 261.667548 |
| -5.4135494 | 6.18E-08 | 4.09E-07 | 133.609105 | 244.361534 | 196.727633 | 246.539017 |
| -5.4131678 | 6.19E-08 | 4.09E-07 | 295.497437 | 234.078324 | 156.461743 | 162.491625 |
| -5.4131898 | 6.19E-08 | 4.09E-07 | 122.54332 | 267.904671 | 263.453965 | 246.539017 |
| -5.4126921 | 6.21E-08 | 4.10E-07 | 77.4604933 | 641.347547 | 663.811955 | 651.087132 |
| -5.4123699 | 6.22E-08 | 4.11E-07 | 126.641759 | 244.902755 | 240.444885 | 231.970803 |
| -5.4120106 | 6.23E-08 | 4.12E-07 | 110.657848 | 313.637893 | 390.003904 | 318.259459 |
| -5.4117836 | 6.24E-08 | 4.12E-07 | 111.477535 | 304.707737 | 347.437107 | 318.819775 |
| -5.4117191 | 6.24E-08 | 4.12E-07 | 98.3625312 | 449.484504 | 437.172518 | 391.100532 |
| -5.4108708 | 6.27E-08 | 4.14E-07 | 187.298653 | 253.291689 | 136.904025 | 221.885116 |
| -5.4107673 | 6.28E-08 | 4.14E-07 | 121.723632 | 247.067641 | 262.303511 | 281.278606 |
| -5.4104954 | 6.29E-08 | 4.15E-07 | 131.559885 | 241.114204 | 210.533081 | 219.083536 |
| -5.4104429 | 6.29E-08 | 4.15E-07 | 142.62567 | 188.345102 | 197.878087 | 258.305652 |
| -5.4099968 | 6.30E-08 | 4.16E-07 | 174.183649 | 162.366467 | 193.276271 | 157.448782 |
| -5.4098536 | 6.31E-08 | 4.16E-07 | 134.428793 | 205.122971 | 201.329449 | 171.45668 |
| -5.4097546 | 6.31E-08 | 4.16E-07 | 141.805982 | 191.863042 | 185.223093 | 225.247011 |
| -5.4096534 | 6.31E-08 | 4.16E-07 | 119.264569 | 327.439043 | 247.347609 | 300.889664 |
| -5.4095353 | 6.32E-08 | 4.16E-07 | 211.069598 | 220.818396 | 140.355387 | 177.620156 |
| -5.4091537 | 6.33E-08 | 4.17E-07 | 121.723632 | 263.033677 | 249.648517 | 263.908812 |
| -5.4089778 | 6.34E-08 | 4.18E-07 | 119.264569 | 273.858108 | 256.551241 | 316.578511 |

| | | | | | | |
|---|---|---|---|---|---|---|
| -5.408871 | 6.34E-08 | 4.18E-07 | 196.315218 | 191.05121 | 154.160835 | 159.129729 |
| -5.4088729 | 6.34E-08 | 4.18E-07 | 117.625194 | 360.994779 | 281.861229 | 271.192919 |
| -5.4085891 | 6.35E-08 | 4.18E-07 | 112.297223 | 367.489438 | 288.763953 | 321.621355 |
| -5.4082703 | 6.36E-08 | 4.19E-07 | 158.609582 | 185.909605 | 156.461743 | 214.040692 |
| -5.4081426 | 6.37E-08 | 4.19E-07 | 134.428793 | 217.300456 | 234.692615 | 195.550266 |
| -5.4079262 | 6.38E-08 | 4.20E-07 | 177.052556 | 164.260743 | 155.311289 | 179.86142 |
| -5.4076508 | 6.39E-08 | 4.20E-07 | 138.937075 | 202.687474 | 187.524001 | 205.075637 |
| -5.4072887 | 6.40E-08 | 4.21E-07 | 117.625194 | 304.437127 | 254.250333 | 311.535667 |
| -5.4069542 | 6.41E-08 | 4.22E-07 | 134.428793 | 234.348935 | 193.276271 | 223.005748 |
| -5.4066474 | 6.42E-08 | 4.22E-07 | 150.822548 | 166.69624 | 185.223093 | 216.842272 |
| -5.40659 | 6.42E-08 | 4.22E-07 | 124.592539 | 293.342085 | 278.409867 | 221.3248 |
| -5.4063346 | 6.43E-08 | 4.23E-07 | 98.3625312 | 529.314684 | 369.295733 | 443.770231 |
| -5.4061921 | 6.44E-08 | 4.23E-07 | 110.248004 | 322.026827 | 308.321671 | 341.232413 |
| -5.4060625 | 6.44E-08 | 4.23E-07 | 122.54332 | 271.963833 | 256.551241 | 238.694594 |
| -5.4059911 | 6.45E-08 | 4.23E-07 | 127.871291 | 209.723354 | 216.285351 | 192.18837 |
| -5.4058834 | 6.45E-08 | 4.23E-07 | 122.953164 | 245.173366 | 247.347609 | 257.18502 |
| -5.4054114 | 6.47E-08 | 4.24E-07 | 106.149565 | 348.276073 | 342.835291 | 356.921259 |
| -5.4053807 | 6.47E-08 | 4.24E-07 | 117.21535 | 367.218827 | 264.604419 | 291.364293 |
| -5.4053411 | 6.47E-08 | 4.24E-07 | 143.855202 | 185.638994 | 283.011683 | 205.075637 |
| -5.4052668 | 6.47E-08 | 4.25E-07 | 129.92051 | 279.540935 | 215.134897 | 225.807327 |
| -5.4051006 | 6.48E-08 | 4.25E-07 | 144.67489 | 251.938635 | 170.267191 | 221.885116 |
| -5.4026657 | 6.57E-08 | 4.31E-07 | 120.903945 | 254.374132 | 265.754873 | 251.021545 |
| -5.4025656 | 6.57E-08 | 4.31E-07 | 225.414134 | 122.857294 | 307.171217 | 287.442082 |
| -5.4025269 | 6.57E-08 | 4.31E-07 | 110.657848 | 348.546684 | 314.073941 | 299.769033 |
| -5.4024375 | 6.57E-08 | 4.31E-07 | 110.657848 | 310.119953 | 329.029843 | 307.613456 |
| -5.4018141 | 6.60E-08 | 4.32E-07 | 188.528185 | 727.401774 | 744.343735 | 154.086886 |
| -5.4016722 | 6.60E-08 | 4.33E-07 | 134.018949 | 212.970683 | 195.577179 | 216.842272 |
| -5.4016168 | 6.60E-08 | 4.33E-07 | 113.526755 | 287.388647 | 288.763953 | 327.78483 |
| -5.4015637 | 6.61E-08 | 4.33E-07 | 112.297223 | 327.709654 | 307.171217 | 287.442082 |
| -5.4012175 | 6.62E-08 | 4.33E-07 | 174.183649 | 272.234444 | 140.355387 | 276.796079 |
| -5.4012037 | 6.62E-08 | 4.33E-07 | 123.772852 | 500.08872 | 326.728935 | 389.419584 |
| -5.4012536 | 6.62E-08 | 4.33E-07 | 118.854725 | 284.95315 | 273.808051 | 250.461229 |
| -5.4010379 | 6.63E-08 | 4.34E-07 | 170.495054 | 196.734036 | 187.524001 | 145.682147 |
| -5.4007909 | 6.63E-08 | 4.34E-07 | 120.084257 | 240.572983 | 331.330751 | 280.157974 |
| -5.4007033 | 6.64E-08 | 4.34E-07 | 124.182696 | 471.674588 | 249.648517 | 305.372192 |
| -5.400646 | 6.64E-08 | 4.34E-07 | 128.690978 | 248.961917 | 392.304812 | 239.25491 |
| -5.4005368 | 6.64E-08 | 4.34E-07 | 123.772852 | 385.62036 | 554.518826 | 243.737438 |
| -5.4004111 | 6.65E-08 | 4.35E-07 | 196.315218 | 678.691834 | 410.712076 | 498.120878 |
| -5.4002449 | 6.65E-08 | 4.35E-07 | 154.920987 | 220.006563 | 162.214013 | 178.740788 |
| -5.3999188 | 6.67E-08 | 4.36E-07 | 165.576927 | 201.33442 | 144.957203 | 147.363094 |
| -5.3999469 | 6.67E-08 | 4.36E-07 | 89.3459658 | 578.836457 | 427.968886 | 539.023943 |
| -5.3995929 | 6.68E-08 | 4.36E-07 | 114.756286 | 277.376049 | 283.011683 | 307.05314 |
| -5.3995045 | 6.68E-08 | 4.36E-07 | 123.363008 | 323.10927 | 227.789891 | 267.270707 |

| | | | | | | |
|---|---|---|---|---|---|---|
| -5.3992864 | 6.69E-08 | 4.37E-07 | 115.16613 | 292.800863 | 266.905327 | 349.637152 |
| -5.399148 | 6.70E-08 | 4.37E-07 | 130.330354 | 412.681438 | 218.586259 | 288.562714 |
| -5.3988528 | 6.71E-08 | 4.38E-07 | 113.936599 | 298.483689 | 294.516223 | 280.71829 |
| -5.3988131 | 6.71E-08 | 4.38E-07 | 123.772852 | 306.872624 | 223.188075 | 265.029444 |
| -5.3982965 | 6.73E-08 | 4.39E-07 | 123.363008 | 245.714587 | 270.356689 | 228.608907 |
| -5.3980426 | 6.74E-08 | 4.39E-07 | 116.395662 | 270.069558 | 372.747095 | 279.597658 |
| -5.3978318 | 6.75E-08 | 4.40E-07 | 119.674413 | 301.460408 | 238.143977 | 278.477026 |
| -5.3975066 | 6.76E-08 | 4.41E-07 | 131.969729 | 491.970396 | 224.338529 | 316.018195 |
| -5.3972729 | 6.77E-08 | 4.41E-07 | 136.478012 | 187.262659 | 207.081719 | 203.394689 |
| -5.3967775 | 6.78E-08 | 4.42E-07 | 307.38291 | 126.645845 | 213.984443 | 255.504073 |
| -5.3965228 | 6.79E-08 | 4.43E-07 | 90.5754975 | 390.491354 | 636.201059 | 602.899961 |
| -5.3963706 | 6.80E-08 | 4.43E-07 | 147.133953 | 180.49739 | 178.320369 | 173.697944 |
| -5.3960628 | 6.81E-08 | 4.44E-07 | 111.477535 | 302.542851 | 299.118039 | 415.194118 |
| -5.3957914 | 6.82E-08 | 4.44E-07 | 115.575974 | 277.105438 | 323.277573 | 265.58976 |
| 5.39553909 | 6.83E-08 | 4.45E-07 | 37.2957931 | 24.8961917 | 29.9118039 | 17.9301104 |
| -5.3955351 | 6.83E-08 | 4.45E-07 | 195.495531 | 183.744719 | 143.806749 | 171.45668 |
| -5.3955706 | 6.83E-08 | 4.45E-07 | 129.100822 | 232.184049 | 204.780811 | 228.048591 |
| -5.3956261 | 6.83E-08 | 4.45E-07 | 93.4444046 | 405.916169 | 470.535684 | 472.346345 |
| -5.3947618 | 6.86E-08 | 4.46E-07 | 398.368251 | 603.732648 | 590.1829 | 590.012694 |
| -5.3946615 | 6.87E-08 | 4.47E-07 | 104.510189 | 363.430276 | 323.277573 | 379.894213 |
| -5.3945383 | 6.87E-08 | 4.47E-07 | 150.00286 | 296.318803 | 178.320369 | 198.912162 |
| -5.3937013 | 6.90E-08 | 4.49E-07 | 108.608628 | 431.894804 | 280.710775 | 494.758983 |
| -5.3933864 | 6.91E-08 | 4.49E-07 | 140.166607 | 212.700072 | 172.568099 | 207.877217 |
| -5.3932325 | 6.92E-08 | 4.50E-07 | 97.1329995 | 533.915067 | 399.207536 | 386.618005 |
| -5.3932208 | 6.92E-08 | 4.50E-07 | 113.936599 | 266.551617 | 245.046701 | 255.504073 |
| 5.39311254 | 6.92E-08 | 4.50E-07 | 31.5579788 | 18.1309222 | 17.2568099 | 22.4126379 |
| -5.3931376 | 6.92E-08 | 4.50E-07 | 122.54332 | 295.23636 | 218.586259 | 288.002398 |
| -5.3929788 | 6.93E-08 | 4.50E-07 | 140.166607 | 183.203497 | 207.081719 | 182.662999 |
| -5.3928548 | 6.93E-08 | 4.50E-07 | 117.625194 | 253.291689 | 295.666677 | 262.78818 |
| -5.3928762 | 6.93E-08 | 4.50E-07 | 138.117388 | 243.27909 | 179.470823 | 208.997849 |
| -5.3927004 | 6.94E-08 | 4.50E-07 | 134.428793 | 225.418779 | 211.683535 | 186.585211 |
| -5.3922773 | 6.96E-08 | 4.51E-07 | 213.118818 | 380.478755 | 291.064861 | 332.267358 |
| -5.3914986 | 6.99E-08 | 4.53E-07 | 272.136336 | 661.913966 | 889.300938 | 704.877463 |
| -5.3913551 | 6.99E-08 | 4.54E-07 | 109.428316 | 347.734851 | 292.215315 | 315.457879 |
| -5.3911324 | 7.00E-08 | 4.54E-07 | 109.018472 | 329.603929 | 292.215315 | 333.387989 |
| -5.3911588 | 7.00E-08 | 4.54E-07 | 98.3625312 | 406.45739 | 390.003904 | 383.256109 |
| -5.3907716 | 7.02E-08 | 4.55E-07 | 157.789894 | 175.897006 | 322.127119 | 195.550266 |
| -5.3907916 | 7.01E-08 | 4.55E-07 | 107.78894 | 330.145151 | 358.941647 | 298.648401 |
| -5.3906051 | 7.02E-08 | 4.55E-07 | 105.739721 | 391.573797 | 322.127119 | 327.78483 |
| -5.3905573 | 7.02E-08 | 4.55E-07 | 151.232392 | 346.652408 | 655.758777 | 207.316901 |
| -5.3904789 | 7.03E-08 | 4.55E-07 | 170.495054 | 319.59133 | 202.479903 | 250.461229 |
| -5.3902484 | 7.04E-08 | 4.56E-07 | 102.46097 | 398.609678 | 324.428027 | 392.78148 |
| -5.3901833 | 7.04E-08 | 4.56E-07 | 181.560839 | 218.382899 | 241.595339 | 135.036144 |

| | | | | | | |
|---|---|---|---|---|---|---|
| -5.3898223 | 7.05E-08 | 4.57E-07 | 115.985818 | 282.517653 | 257.701695 | 277.356395 |
| -5.3896422 | 7.06E-08 | 4.57E-07 | 114.346443 | 289.012312 | 266.905327 | 286.881766 |
| -5.3895661 | 7.06E-08 | 4.57E-07 | 107.78894 | 401.586396 | 326.728935 | 300.329348 |
| -5.3894225 | 7.07E-08 | 4.57E-07 | 127.871291 | 213.511905 | 212.833989 | 235.332698 |
| -5.3891455 | 7.08E-08 | 4.58E-07 | 218.856632 | 247.608863 | 156.461743 | 150.164674 |
| 5.38902497 | 7.08E-08 | 4.58E-07 | 88.9361219 | 63.8641439 | 67.8767857 | 64.4363341 |
| -5.3881812 | 7.12E-08 | 4.60E-07 | 166.806459 | 170.484791 | 148.408565 | 192.18837 |
| -5.3882028 | 7.12E-08 | 4.60E-07 | 113.936599 | 284.95315 | 297.967585 | 266.150076 |
| -5.3880764 | 7.12E-08 | 4.60E-07 | 111.067691 | 310.390564 | 304.870309 | 282.399238 |
| -5.3880648 | 7.12E-08 | 4.60E-07 | 136.887856 | 203.769917 | 243.896247 | 181.542367 |
| -5.3880434 | 7.12E-08 | 4.60E-07 | 146.314265 | 333.663091 | 346.286653 | 421.917909 |
| -5.3876904 | 7.14E-08 | 4.61E-07 | 156.970206 | 220.006563 | 154.160835 | 280.71829 |
| -5.3876323 | 7.14E-08 | 4.61E-07 | 125.822071 | 324.191713 | 219.736713 | 243.737438 |
| -5.3874876 | 7.14E-08 | 4.61E-07 | 106.149565 | 336.369198 | 319.826211 | 330.58641 |
| -5.3870235 | 7.16E-08 | 4.62E-07 | 119.674413 | 304.978348 | 242.745793 | 248.219965 |
| -5.3867944 | 7.17E-08 | 4.63E-07 | 433.204981 | 205.664192 | 155.311289 | 212.92006 |
| -5.3867431 | 7.17E-08 | 4.63E-07 | 88.9361219 | 459.767714 | 546.465648 | 448.252759 |
| -5.386512 | 7.18E-08 | 4.63E-07 | 126.231915 | 261.410013 | 203.630357 | 247.099333 |
| -5.3863831 | 7.19E-08 | 4.64E-07 | 99.1822189 | 645.136097 | 329.029843 | 507.646249 |
| -5.3863963 | 7.19E-08 | 4.64E-07 | 136.478012 | 214.594348 | 177.169915 | 210.678797 |
| -5.386298 | 7.19E-08 | 4.64E-07 | 103.690502 | 441.366181 | 314.073941 | 368.127578 |
| -5.3862543 | 7.19E-08 | 4.64E-07 | 338.940889 | 536.079953 | 500.447488 | 570.401636 |
| -5.3860062 | 7.20E-08 | 4.64E-07 | 190.577404 | 288.471091 | 150.709473 | 175.378892 |
| -5.3860521 | 7.20E-08 | 4.64E-07 | 194.265999 | 176.438228 | 142.656295 | 173.137628 |
| -5.386015 | 7.20E-08 | 4.64E-07 | 154.920987 | 623.216624 | 583.280176 | 600.098381 |
| -5.3856664 | 7.22E-08 | 4.65E-07 | 146.314265 | 504.418492 | 373.897549 | 370.929158 |
| -5.3855904 | 7.22E-08 | 4.65E-07 | 119.674413 | 675.173894 | 291.064861 | 349.076836 |
| -5.3854965 | 7.22E-08 | 4.65E-07 | 164.75724 | 162.637078 | 155.311289 | 173.697944 |
| -5.3852522 | 7.23E-08 | 4.66E-07 | 147.953641 | 162.366467 | 180.621277 | 186.024895 |
| -5.3852803 | 7.23E-08 | 4.66E-07 | 106.149565 | 312.014228 | 338.233475 | 332.267358 |
| -5.3851676 | 7.24E-08 | 4.66E-07 | 124.592539 | 243.820312 | 210.533081 | 250.461229 |
| -5.3850186 | 7.24E-08 | 4.66E-07 | 119.264569 | 431.082971 | 253.099879 | 289.683345 |
| -5.3850528 | 7.24E-08 | 4.66E-07 | 113.116911 | 283.058875 | 269.206235 | 303.691244 |
| -5.3846194 | 7.26E-08 | 4.67E-07 | 116.395662 | 309.849342 | 261.153057 | 254.383441 |
| -5.3844821 | 7.27E-08 | 4.67E-07 | 97.5428434 | 422.152815 | 395.756174 | 368.127578 |
| -5.3844752 | 7.27E-08 | 4.67E-07 | 129.510666 | 300.919186 | 258.852149 | 316.018195 |
| -5.3843737 | 7.27E-08 | 4.67E-07 | 110.248004 | 302.542851 | 280.710775 | 335.629253 |
| -5.3841313 | 7.28E-08 | 4.68E-07 | 105.739721 | 341.240193 | 309.472125 | 341.232413 |
| -5.383773 | 7.29E-08 | 4.68E-07 | 189.757716 | 188.886324 | 155.311289 | 173.697944 |
| -5.3838043 | 7.29E-08 | 4.68E-07 | 120.494101 | 237.866875 | 239.294431 | 255.504073 |
| -5.3838033 | 7.29E-08 | 4.68E-07 | 122.953164 | 309.308121 | 224.338529 | 244.297754 |
| -5.3837278 | 7.30E-08 | 4.68E-07 | 144.67489 | 214.864959 | 173.718553 | 174.25826 |
| -5.3834863 | 7.31E-08 | 4.69E-07 | 130.740198 | 201.063809 | 200.178995 | 230.850171 |

| | | | | | | |
|---|---|---|---|---|---|---|
| -5.3829749 | 7.33E-08 | 4.70E-07 | 122.133476 | 245.985198 | 219.736713 | 269.511971 |
| -5.3828165 | 7.33E-08 | 4.71E-07 | 102.870814 | 383.996696 | 341.684837 | 332.267358 |
| -5.3827534 | 7.34E-08 | 4.71E-07 | 122.54332 | 222.712671 | 239.294431 | 243.177122 |
| -5.3825318 | 7.34E-08 | 4.71E-07 | 152.871767 | 307.684456 | 171.417645 | 201.153426 |
| -5.3823891 | 7.35E-08 | 4.71E-07 | 106.149565 | 304.166516 | 324.428027 | 370.368842 |
| -5.3822896 | 7.35E-08 | 4.72E-07 | 104.100346 | 548.79866 | 291.064861 | 493.638351 |
| -5.3822037 | 7.36E-08 | 4.72E-07 | 110.657848 | 295.23636 | 280.710775 | 313.216615 |
| -5.3821757 | 7.36E-08 | 4.72E-07 | 170.495054 | 217.571066 | 194.426725 | 140.078987 |
| -5.3819985 | 7.37E-08 | 4.72E-07 | 123.363008 | 231.101605 | 230.090799 | 226.367643 |
| -5.3818712 | 7.37E-08 | 4.72E-07 | 122.953164 | 290.906588 | 213.984443 | 254.943757 |
| -5.3814311 | 7.39E-08 | 4.73E-07 | 121.313788 | 235.972599 | 274.958505 | 228.608907 |
| -5.3812585 | 7.40E-08 | 4.73E-07 | 130.330354 | 304.166516 | 203.630357 | 224.686695 |
| -5.3812552 | 7.40E-08 | 4.73E-07 | 121.723632 | 267.092839 | 216.285351 | 261.107232 |
| -5.3812647 | 7.40E-08 | 4.73E-07 | 127.051603 | 229.748551 | 203.630357 | 274.554815 |
| -5.3809711 | 7.41E-08 | 4.74E-07 | 184.019902 | 197.004647 | 166.815829 | 177.05984 |
| -5.3807656 | 7.42E-08 | 4.74E-07 | 390.991061 | 1236.42065 | 896.203662 | 948.054585 |
| -5.3807899 | 7.42E-08 | 4.74E-07 | 160.658801 | 216.488623 | 147.258111 | 183.783631 |
| -5.3806644 | 7.42E-08 | 4.75E-07 | 153.281611 | 165.072575 | 162.214013 | 175.939208 |
| -5.3802951 | 7.44E-08 | 4.76E-07 | 119.674413 | 176.97945 | 133.452663 | 155.767834 |
| -5.3800886 | 7.44E-08 | 4.76E-07 | 98.7723751 | 428.106253 | 361.242555 | 364.205367 |
| -5.38007 | 7.45E-08 | 4.76E-07 | 102.051126 | 430.81236 | 334.782113 | 343.473677 |
| -5.3799699 | 7.45E-08 | 4.76E-07 | 133.609105 | 245.714587 | 186.373547 | 205.075637 |
| -5.3799672 | 7.45E-08 | 4.76E-07 | 120.084257 | 240.843593 | 232.391707 | 276.796079 |
| -5.3799177 | 7.45E-08 | 4.76E-07 | 104.920033 | 344.758133 | 390.003904 | 300.889664 |
| -5.3796288 | 7.46E-08 | 4.77E-07 | 123.363008 | 229.20733 | 217.435805 | 255.504073 |
| -5.3795612 | 7.47E-08 | 4.77E-07 | 117.625194 | 310.119953 | 236.993523 | 265.58976 |
| -5.3790117 | 7.49E-08 | 4.78E-07 | 127.871291 | 192.404264 | 281.861229 | 268.391339 |
| -5.3788602 | 7.50E-08 | 4.78E-07 | 154.101299 | 254.644743 | 155.311289 | 197.231214 |
| -5.3788655 | 7.50E-08 | 4.78E-07 | 95.493624 | 368.571881 | 436.022064 | 420.236961 |
| -5.3787538 | 7.50E-08 | 4.78E-07 | 115.575974 | 340.157749 | 251.949425 | 270.072287 |
| -5.3787618 | 7.50E-08 | 4.78E-07 | 110.248004 | 293.342085 | 288.763953 | 295.286505 |
| -5.3780953 | 7.53E-08 | 4.80E-07 | 134.018949 | 223.524504 | 181.771731 | 201.713742 |
| -5.3774073 | 7.56E-08 | 4.82E-07 | 182.380527 | 162.637078 | 240.444885 | 149.604358 |
| -5.3771893 | 7.57E-08 | 4.82E-07 | 120.903945 | 522.278804 | 242.745793 | 342.353045 |
| -5.377004 | 7.57E-08 | 4.83E-07 | 134.428793 | 178.061893 | 256.551241 | 288.002398 |
| -5.3762264 | 7.61E-08 | 4.85E-07 | 98.7723751 | 482.499019 | 324.428027 | 425.840121 |
| -5.3757026 | 7.63E-08 | 4.86E-07 | 119.264569 | 364.242109 | 302.569401 | 224.126379 |
| -5.3745897 | 7.68E-08 | 4.89E-07 | 151.642236 | 184.827162 | 166.815829 | 269.511971 |
| -5.3745022 | 7.68E-08 | 4.89E-07 | 111.887379 | 304.437127 | 261.153057 | 288.562714 |
| -5.3741555 | 7.69E-08 | 4.90E-07 | 88.9361219 | 508.477654 | 398.057082 | 523.335096 |
| -5.37409 | 7.70E-08 | 4.90E-07 | 234.840543 | 183.744719 | 146.107657 | 163.612257 |
| -5.3737924 | 7.71E-08 | 4.91E-07 | 108.198784 | 333.39248 | 285.312591 | 300.889664 |
| -5.3738061 | 7.71E-08 | 4.91E-07 | 1002.47813 | 1828.24642 | 1046.91314 | 1354.28365 |

| | | | | | | |
|---|---|---|---|---|---|---|
| -5.3736186 | 7.72E-08 | 4.91E-07 | 140.986295 | 355.853175 | 358.941647 | 359.162523 |
| -5.3733851 | 7.73E-08 | 4.92E-07 | 140.986295 | 163.448911 | 209.382627 | 196.670898 |
| -5.3733933 | 7.73E-08 | 4.92E-07 | 134.428793 | 328.792097 | 233.542161 | 273.434183 |
| -5.3731124 | 7.74E-08 | 4.92E-07 | 113.526755 | 317.155833 | 264.604419 | 259.9866 |
| -5.3729722 | 7.74E-08 | 4.92E-07 | 125.822071 | 228.395498 | 199.028541 | 242.05649 |
| -5.3728927 | 7.75E-08 | 4.93E-07 | 102.051126 | 303.895905 | 431.420248 | 379.894213 |
| -5.3727876 | 7.75E-08 | 4.93E-07 | 204.102252 | 461.391378 | 134.603117 | 290.803977 |
| -5.372337 | 7.77E-08 | 4.94E-07 | 111.067691 | 319.320719 | 263.453965 | 289.123029 |
| -5.3717054 | 7.80E-08 | 4.95E-07 | 113.116911 | 292.800863 | 251.949425 | 280.71829 |
| -5.3717291 | 7.80E-08 | 4.95E-07 | 161.478489 | 284.141318 | 197.878087 | 265.58976 |
| -5.3717162 | 7.80E-08 | 4.95E-07 | 111.887379 | 296.048192 | 280.710775 | 262.227864 |
| -5.3716337 | 7.80E-08 | 4.95E-07 | 179.511619 | 159.119138 | 158.762651 | 155.767834 |
| -5.3713758 | 7.81E-08 | 4.96E-07 | 121.723632 | 239.219929 | 316.374849 | 223.566063 |
| -5.3713073 | 7.82E-08 | 4.96E-07 | 127.871291 | 188.886324 | 268.055781 | 246.539017 |
| -5.3712508 | 7.82E-08 | 4.96E-07 | 105.739721 | 408.351666 | 288.763953 | 334.508621 |
| -5.3710859 | 7.83E-08 | 4.97E-07 | 106.149565 | 299.836743 | 309.472125 | 328.345146 |
| -5.3704793 | 7.85E-08 | 4.98E-07 | 114.756286 | 362.618444 | 262.303511 | 260.546916 |
| -5.3704249 | 7.86E-08 | 4.98E-07 | 132.789417 | 297.401246 | 180.621277 | 250.461229 |
| -5.3703453 | 7.86E-08 | 4.98E-07 | 161.478489 | 202.958084 | 140.355387 | 220.764484 |
| -5.3698269 | 7.88E-08 | 5.00E-07 | 145.904421 | 263.304288 | 296.817131 | 288.002398 |
| -5.3698576 | 7.88E-08 | 5.00E-07 | 120.903945 | 220.006563 | 234.692615 | 243.737438 |
| -5.3697873 | 7.88E-08 | 5.00E-07 | 340.580264 | 197.004647 | 134.603117 | 221.885116 |
| -5.3695283 | 7.89E-08 | 5.00E-07 | 109.428316 | 289.824144 | 299.118039 | 279.037342 |
| -5.3694502 | 7.90E-08 | 5.00E-07 | 123.363008 | 217.571066 | 226.639437 | 217.962904 |
| -5.3689162 | 7.92E-08 | 5.02E-07 | 158.199738 | 183.744719 | 158.762651 | 155.207518 |
| -5.3686105 | 7.93E-08 | 5.02E-07 | 129.92051 | 182.662276 | 235.843069 | 222.445432 |
| -5.3686028 | 7.93E-08 | 5.02E-07 | 88.9361219 | 462.473822 | 441.774334 | 435.925808 |
| -5.368322 | 7.95E-08 | 5.03E-07 | 104.920033 | 343.67569 | 309.472125 | 306.492824 |
| -5.3683201 | 7.95E-08 | 5.03E-07 | 113.116911 | 279.540935 | 256.551241 | 267.831023 |
| -5.368204 | 7.95E-08 | 5.03E-07 | 292.62853 | 292.800863 | 109.29313 | 267.270707 |
| -5.3681918 | 7.95E-08 | 5.03E-07 | 126.641759 | 423.77648 | 200.178995 | 383.816425 |
| -5.3682007 | 7.95E-08 | 5.03E-07 | 152.871767 | 170.484791 | 125.399485 | 136.717091 |
| -5.3674243 | 7.99E-08 | 5.05E-07 | 97.9526873 | 369.113103 | 373.897549 | 355.800627 |
| -5.3671865 | 8.00E-08 | 5.06E-07 | 122.953164 | 273.316887 | 219.736713 | 216.842272 |
| -5.3667762 | 8.02E-08 | 5.07E-07 | 117.21535 | 225.148168 | 285.312591 | 262.227864 |
| -5.3666974 | 8.02E-08 | 5.07E-07 | 152.461923 | 181.579833 | 245.046701 | 156.888466 |
| -5.3667207 | 8.02E-08 | 5.07E-07 | 186.069121 | 805.60829 | 153.010381 | 508.206565 |
| -5.3666761 | 8.02E-08 | 5.07E-07 | 111.887379 | 259.515737 | 274.958505 | 340.111781 |
| -5.3662328 | 8.04E-08 | 5.08E-07 | 143.445358 | 199.981366 | 165.665375 | 171.45668 |
| 5.36597363 | 8.05E-08 | 5.08E-07 | 24.5906328 | 15.6954252 | 6.90272397 | 6.72379138 |
| -5.3659445 | 8.05E-08 | 5.08E-07 | 98.7723751 | 351.523402 | 358.941647 | 362.524419 |
| -5.3656913 | 8.06E-08 | 5.09E-07 | 98.3625312 | 391.303187 | 361.242555 | 342.353045 |
| -5.3656404 | 8.07E-08 | 5.09E-07 | 137.2977 | 342.863857 | 172.568099 | 335.629253 |

| | | | | | | |
|---|---|---|---|---|---|---|
| -5.3653093 | 8.08E-08 | 5.10E-07 | 116.805506 | 304.166516 | 228.940345 | 260.546916 |
| -5.3651966 | 8.09E-08 | 5.10E-07 | 170.904898 | 259.786348 | 133.452663 | 261.667548 |
| -5.3647294 | 8.11E-08 | 5.11E-07 | 82.7884637 | 469.239091 | 522.306114 | 528.37794 |
| -5.3647537 | 8.11E-08 | 5.11E-07 | 112.707067 | 325.003546 | 248.498063 | 276.235763 |
| -5.3643802 | 8.12E-08 | 5.12E-07 | 133.199261 | 212.970683 | 173.718553 | 234.772382 |
| -5.3642794 | 8.13E-08 | 5.12E-07 | 122.953164 | 289.012312 | 209.382627 | 232.531119 |
| -5.3642218 | 8.13E-08 | 5.12E-07 | 147.543797 | 414.305103 | 291.064861 | 347.395888 |
| -5.3640519 | 8.14E-08 | 5.13E-07 | 133.199261 | 186.450827 | 274.958505 | 203.955005 |
| -5.3638893 | 8.14E-08 | 5.13E-07 | 124.592539 | 219.735953 | 200.178995 | 233.651751 |
| -5.3639072 | 8.14E-08 | 5.13E-07 | 112.297223 | 280.352767 | 251.949425 | 277.916711 |
| -5.3638848 | 8.15E-08 | 5.13E-07 | 125.002383 | 232.995881 | 194.426725 | 242.05649 |
| -5.3636047 | 8.16E-08 | 5.14E-07 | 127.051603 | 207.558468 | 196.727633 | 211.799429 |
| -5.363558 | 8.16E-08 | 5.14E-07 | 123.363008 | 239.49054 | 216.285351 | 208.437533 |
| -5.3631009 | 8.18E-08 | 5.15E-07 | 102.46097 | 332.851258 | 310.622579 | 346.835572 |
| -5.3629644 | 8.19E-08 | 5.15E-07 | 131.150042 | 257.892073 | 199.028541 | 190.507423 |
| -5.3628366 | 8.19E-08 | 5.15E-07 | 136.068168 | 173.190899 | 189.824909 | 187.145527 |
| -5.3624926 | 8.21E-08 | 5.16E-07 | 103.690502 | 342.593246 | 300.268493 | 323.862618 |
| -5.3624168 | 8.21E-08 | 5.16E-07 | 124.592539 | 398.609678 | 247.347609 | 313.216615 |
| -5.3623038 | 8.22E-08 | 5.17E-07 | 145.904421 | 259.245126 | 202.479903 | 159.129729 |
| -5.3619889 | 8.23E-08 | 5.17E-07 | 1347.56668 | 3198.07819 | 3064.80944 | 3383.1877 |
| -5.3615657 | 8.25E-08 | 5.19E-07 | 171.314742 | 172.379066 | 172.568099 | 186.024895 |
| -5.3611871 | 8.27E-08 | 5.20E-07 | 136.068168 | 194.56915 | 167.966283 | 203.955005 |
| -5.3611271 | 8.27E-08 | 5.20E-07 | 418.450601 | 1095.97366 | 1571.52016 | 1471.95 |
| -5.360672 | 8.29E-08 | 5.21E-07 | 120.084257 | 408.622277 | 219.736713 | 305.372192 |
| -5.3605251 | 8.30E-08 | 5.21E-07 | 120.494101 | 248.691306 | 262.303511 | 206.756585 |
| -5.3602659 | 8.31E-08 | 5.22E-07 | 127.461447 | 197.545869 | 197.878087 | 210.678797 |
| -5.360012 | 8.32E-08 | 5.22E-07 | 617.634727 | 241.384815 | 155.311289 | 229.729539 |
| -5.3598859 | 8.33E-08 | 5.23E-07 | 116.395662 | 268.987114 | 222.037621 | 293.045241 |
| -5.3595597 | 8.34E-08 | 5.23E-07 | 102.051126 | 582.895618 | 294.516223 | 430.882964 |
| -5.3595485 | 8.34E-08 | 5.23E-07 | 148.363485 | 201.063809 | 148.408565 | 193.309002 |
| -5.3594686 | 8.35E-08 | 5.24E-07 | 274.5954 | 163.719521 | 153.010381 | 180.421735 |
| -5.3593646 | 8.35E-08 | 5.24E-07 | 104.920033 | 689.245654 | 284.162137 | 484.673296 |
| 5.35928896 | 8.36E-08 | 5.24E-07 | 4.91812656 | 6.22404792 | 1.150454 | 1.68094785 |
| -5.3592451 | 8.36E-08 | 5.24E-07 | 92.6247169 | 503.60666 | 390.003904 | 379.894213 |
| -5.358855 | 8.38E-08 | 5.25E-07 | 116.805506 | 277.91727 | 224.338529 | 250.461229 |
| -5.3585559 | 8.39E-08 | 5.26E-07 | 108.198784 | 324.732935 | 256.551241 | 331.146726 |
| -5.3585883 | 8.39E-08 | 5.26E-07 | 322.547134 | 186.450827 | 354.339831 | 128.312352 |
| -5.3581004 | 8.41E-08 | 5.27E-07 | 161.888333 | 254.103522 | 139.204933 | 288.562714 |
| -5.3579459 | 8.42E-08 | 5.27E-07 | 122.54332 | 204.85236 | 245.046701 | 219.083536 |
| -5.3576583 | 8.43E-08 | 5.28E-07 | 125.002383 | 207.287857 | 199.028541 | 228.608907 |
| -5.357653 | 8.43E-08 | 5.28E-07 | 116.805506 | 273.587498 | 218.586259 | 266.710392 |
| -5.3571527 | 8.45E-08 | 5.29E-07 | 126.231915 | 211.88824 | 190.975363 | 219.643852 |
| -5.3571119 | 8.46E-08 | 5.29E-07 | 158.199738 | 155.601198 | 164.514921 | 157.448782 |

| | | | | | | |
|---|---|---|---|---|---|---|
| -5.3570592 | 8.46E-08 | 5.29E-07 | 129.510666 | 486.28757 | 281.861229 | 219.643852 |
| -5.3564075 | 8.49E-08 | 5.31E-07 | 106.559409 | 432.706636 | 281.861229 | 312.656299 |
| -5.3561201 | 8.50E-08 | 5.32E-07 | 150.412704 | 303.354683 | 188.674455 | 170.896364 |
| -5.3560541 | 8.51E-08 | 5.32E-07 | 125.002383 | 513.619259 | 297.967585 | 339.551465 |
| -5.3558864 | 8.51E-08 | 5.32E-07 | 140.576451 | 166.425629 | 177.169915 | 221.3248 |
| -5.355571 | 8.53E-08 | 5.33E-07 | 141.396139 | 195.380983 | 199.028541 | 215.161324 |
| -5.3555841 | 8.53E-08 | 5.33E-07 | 143.445358 | 224.065725 | 188.674455 | 156.888466 |
| 5.35539764 | 8.54E-08 | 5.33E-07 | 0 | 0 | 0 | 0 |
| -5.3553881 | 8.54E-08 | 5.33E-07 | 111.477535 | 388.597079 | 327.879389 | 327.224514 |
| -5.3550593 | 8.55E-08 | 5.34E-07 | 107.78894 | 298.213079 | 274.958505 | 284.080186 |
| -5.3545997 | 8.57E-08 | 5.35E-07 | 129.510666 | 286.576815 | 177.169915 | 281.838922 |
| -5.3544999 | 8.58E-08 | 5.36E-07 | 124.592539 | 225.68939 | 192.125817 | 223.005748 |
| -5.3542446 | 8.59E-08 | 5.36E-07 | 121.723632 | 216.218013 | 208.232173 | 235.332698 |
| -5.353964 | 8.60E-08 | 5.37E-07 | 116.805506 | 311.202396 | 249.648517 | 225.247011 |
| -5.353633 | 8.62E-08 | 5.38E-07 | 154.920987 | 243.00848 | 195.577179 | 210.678797 |
| -5.3536185 | 8.62E-08 | 5.38E-07 | 177.052556 | 179.685557 | 135.753571 | 173.137628 |
| -5.3534017 | 8.63E-08 | 5.38E-07 | 111.887379 | 247.608863 | 260.002603 | 300.329348 |
| -5.3530444 | 8.65E-08 | 5.39E-07 | 136.478012 | 206.205414 | 223.188075 | 161.931309 |
| -5.3529048 | 8.66E-08 | 5.40E-07 | 146.724109 | 271.963833 | 156.461743 | 205.635953 |
| -5.3526313 | 8.67E-08 | 5.40E-07 | 123.363008 | 207.017246 | 209.382627 | 211.799429 |
| -5.3521772 | 8.69E-08 | 5.42E-07 | 136.068168 | 181.579833 | 170.267191 | 217.962904 |
| -5.3517214 | 8.71E-08 | 5.43E-07 | 130.740198 | 228.395498 | 171.417645 | 225.247011 |
| -5.3515299 | 8.72E-08 | 5.43E-07 | 169.265522 | 291.989031 | 138.054479 | 215.72164 |
| -5.3514268 | 8.73E-08 | 5.44E-07 | 120.494101 | 453.814277 | 209.382627 | 385.497373 |
| -5.351393 | 8.73E-08 | 5.44E-07 | 111.067691 | 281.43521 | 242.745793 | 285.200818 |
| -5.3513018 | 8.73E-08 | 5.44E-07 | 161.888333 | 268.716504 | 185.223093 | 247.099333 |
| -5.3513312 | 8.73E-08 | 5.44E-07 | 175.003337 | 199.440144 | 136.904025 | 166.974153 |
| -5.3509839 | 8.75E-08 | 5.45E-07 | 100.001907 | 321.214995 | 352.038923 | 326.664198 |
| -5.3508421 | 8.75E-08 | 5.45E-07 | 122.133476 | 255.456576 | 201.329449 | 220.764484 |
| -5.3506022 | 8.77E-08 | 5.45E-07 | 111.477535 | 333.39248 | 236.993523 | 290.243661 |
| -5.3501588 | 8.79E-08 | 5.47E-07 | 107.78894 | 445.695953 | 272.657597 | 309.294404 |
| -5.3495567 | 8.82E-08 | 5.48E-07 | 125.412227 | 421.882205 | 208.232173 | 281.838922 |
| -5.3493539 | 8.83E-08 | 5.49E-07 | 97.1329995 | 382.914252 | 348.587561 | 334.508621 |
| -5.3490185 | 8.84E-08 | 5.50E-07 | 116.395662 | 214.053126 | 277.259413 | 259.426284 |
| -5.348868 | 8.85E-08 | 5.50E-07 | 102.870814 | 322.026827 | 316.374849 | 294.726189 |
| -5.3487932 | 8.85E-08 | 5.50E-07 | 138.527231 | 208.3703 | 156.461743 | 203.955005 |
| -5.348773 | 8.86E-08 | 5.50E-07 | 197.954594 | 132.599282 | 488.942948 | 311.535667 |
| -5.3485148 | 8.87E-08 | 5.51E-07 | 128.281134 | 234.890156 | 199.028541 | 183.783631 |
| -5.3484426 | 8.87E-08 | 5.51E-07 | 102.870814 | 384.808528 | 288.763953 | 316.018195 |
| -5.348397 | 8.87E-08 | 5.51E-07 | 127.461447 | 224.336336 | 184.072639 | 200.59311 |
| -5.3477906 | 8.90E-08 | 5.53E-07 | 136.068168 | 217.571066 | 161.063559 | 200.59311 |
| -5.3477007 | 8.91E-08 | 5.53E-07 | 133.199261 | 197.545869 | 213.984443 | 166.974153 |
| -5.3473249 | 8.93E-08 | 5.54E-07 | 100.001907 | 375.06654 | 304.870309 | 333.387989 |

| | | | | | | |
|---|---|---|---|---|---|---|
| -5.3471694 | 8.93E-08 | 5.54E-07 | 186.478965 | 141.529437 | 231.241253 | 395.58306 |
| -5.3469508 | 8.94E-08 | 5.55E-07 | 113.116911 | 274.39933 | 246.197155 | 238.694594 |
| -5.3467882 | 8.95E-08 | 5.55E-07 | 130.740198 | 197.545869 | 211.683535 | 172.577312 |
| -5.3464759 | 8.97E-08 | 5.56E-07 | 105.329877 | 318.779498 | 269.206235 | 307.05314 |
| -5.346357 | 8.97E-08 | 5.56E-07 | 117.625194 | 250.314971 | 216.285351 | 229.729539 |
| -5.3463589 | 8.97E-08 | 5.56E-07 | 115.985818 | 219.194731 | 242.745793 | 268.951655 |
| -5.3462334 | 8.98E-08 | 5.57E-07 | 105.739721 | 260.59818 | 390.003904 | 326.664198 |
| -5.3459178 | 9.00E-08 | 5.57E-07 | 117.625194 | 268.445893 | 209.382627 | 245.418385 |
| -5.3459304 | 9.00E-08 | 5.57E-07 | 129.100822 | 207.287857 | 176.019461 | 202.834373 |
| -5.3457 | 9.01E-08 | 5.58E-07 | 478.697652 | 770.428888 | 105.841768 | 280.157974 |
| -5.3456145 | 9.01E-08 | 5.58E-07 | 144.67489 | 170.21418 | 157.612197 | 215.161324 |
| -5.345466 | 9.02E-08 | 5.58E-07 | 100.821594 | 409.97533 | 294.516223 | 335.068937 |
| -5.3451921 | 9.03E-08 | 5.59E-07 | 196.315218 | 202.958084 | 127.700393 | 178.740788 |
| -5.3448979 | 9.05E-08 | 5.60E-07 | 137.707544 | 186.721438 | 165.665375 | 239.25491 |
| -5.3448548 | 9.05E-08 | 5.60E-07 | 140.576451 | 193.216096 | 157.612197 | 174.818576 |
| -5.3447771 | 9.05E-08 | 5.60E-07 | 133.199261 | 176.97945 | 178.320369 | 183.223315 |
| -5.3446965 | 9.06E-08 | 5.60E-07 | 85.6573709 | 506.853989 | 552.217918 | 380.454529 |
| -5.3443692 | 9.07E-08 | 5.61E-07 | 115.985818 | 230.289773 | 228.940345 | 239.815226 |
| -5.344201 | 9.08E-08 | 5.62E-07 | 81.5589321 | 427.565031 | 514.252936 | 578.246059 |
| -5.3440946 | 9.09E-08 | 5.62E-07 | 99.1822189 | 338.804695 | 323.277573 | 327.78483 |
| -5.3436452 | 9.11E-08 | 5.63E-07 | 379.925277 | 242.737869 | 149.559019 | 163.612257 |
| -5.3436316 | 9.11E-08 | 5.63E-07 | 150.822548 | 148.024096 | 166.815829 | 191.628054 |
| -5.3432285 | 9.13E-08 | 5.64E-07 | 197.134906 | 141.529437 | 157.612197 | 251.021545 |
| -5.3430331 | 9.14E-08 | 5.65E-07 | 122.953164 | 235.972599 | 199.028541 | 203.394689 |
| -5.342605 | 9.16E-08 | 5.66E-07 | 127.871291 | 264.11612 | 181.771731 | 212.359745 |
| -5.3425733 | 9.16E-08 | 5.66E-07 | 119.674413 | 208.640911 | 292.215315 | 386.057689 |
| -5.3420616 | 9.19E-08 | 5.67E-07 | 103.690502 | 324.191713 | 300.268493 | 280.71829 |
| -5.3419294 | 9.20E-08 | 5.68E-07 | 108.198784 | 302.27224 | 272.657597 | 254.943757 |
| -5.3418016 | 9.20E-08 | 5.68E-07 | 143.035514 | 237.866875 | 149.559019 | 211.799429 |
| -5.341613 | 9.21E-08 | 5.68E-07 | 102.46097 | 310.661174 | 300.268493 | 302.570612 |
| -5.3415631 | 9.21E-08 | 5.69E-07 | 95.493624 | 321.485606 | 375.048002 | 405.668747 |
| -5.3411567 | 9.24E-08 | 5.70E-07 | 108.608628 | 280.352767 | 260.002603 | 263.908812 |
| -5.3408871 | 9.25E-08 | 5.70E-07 | 101.231438 | 396.174181 | 276.108959 | 360.283155 |
| -5.340785 | 9.25E-08 | 5.71E-07 | 117.625194 | 202.146252 | 320.976665 | 303.130928 |
| -5.3406578 | 9.26E-08 | 5.71E-07 | 145.904421 | 268.175282 | 263.453965 | 225.807327 |
| -5.3402255 | 9.28E-08 | 5.72E-07 | 107.379097 | 341.510803 | 246.197155 | 307.613456 |
| -5.3399873 | 9.30E-08 | 5.73E-07 | 119.674413 | 214.323737 | 232.391707 | 206.196269 |
| -5.3399464 | 9.30E-08 | 5.73E-07 | 123.363008 | 183.744719 | 261.153057 | 247.659649 |
| -5.33954 | 9.32E-08 | 5.74E-07 | 161.068645 | 153.165701 | 155.311289 | 155.207518 |
| -5.3395005 | 9.32E-08 | 5.74E-07 | 106.969253 | 281.705821 | 262.303511 | 284.640502 |
| -5.3392072 | 9.34E-08 | 5.75E-07 | 149.183172 | 189.968767 | 157.612197 | 153.52657 |
| -5.3390605 | 9.34E-08 | 5.75E-07 | 121.723632 | 199.981366 | 204.780811 | 230.850171 |
| 5.33890957 | 9.35E-08 | 5.76E-07 | 14.7543797 | 18.1309222 | 11.50454 | 22.4126379 |

| | | | | | | |
|---|---|---|---|---|---|---|
| -5.3386237 | 9.37E-08 | 5.76E-07 | 128.690978 | 237.055043 | 176.019461 | 200.59311 |
| -5.3385233 | 9.37E-08 | 5.77E-07 | 109.428316 | 312.284839 | 243.896247 | 267.831023 |
| -5.3385524 | 9.37E-08 | 5.77E-07 | 101.231438 | 348.005462 | 295.666677 | 305.932508 |
| -5.3384449 | 9.37E-08 | 5.77E-07 | 301.235252 | 246.797031 | 188.674455 | 123.269509 |
| -5.3382404 | 9.39E-08 | 5.77E-07 | 93.4444046 | 416.199378 | 350.888469 | 358.041891 |
| -5.3380638 | 9.39E-08 | 5.78E-07 | 206.971159 | 214.594348 | 128.850847 | 159.690045 |
| -5.3375885 | 9.42E-08 | 5.79E-07 | 94.2640924 | 429.459307 | 496.996126 | 310.415036 |
| 5.33749123 | 9.42E-08 | 5.79E-07 | 398.368251 | 355.853175 | 346.286653 | 345.154624 |
| -5.3374006 | 9.43E-08 | 5.79E-07 | 119.264569 | 206.476024 | 215.134897 | 227.488275 |
| -5.3374218 | 9.43E-08 | 5.79E-07 | 342.629484 | 191.05121 | 159.913105 | 163.051941 |
| -5.3370172 | 9.45E-08 | 5.80E-07 | 108.198784 | 276.293606 | 263.453965 | 258.865968 |
| -5.3370339 | 9.45E-08 | 5.80E-07 | 114.346443 | 220.006563 | 287.613499 | 244.297754 |
| -5.337055 | 9.45E-08 | 5.80E-07 | 335.662138 | 445.425342 | 388.85345 | 420.797277 |
| -5.3369216 | 9.45E-08 | 5.80E-07 | 131.969729 | 316.614612 | 172.568099 | 245.978701 |
| -5.336875 | 9.46E-08 | 5.80E-07 | 147.133953 | 200.251977 | 154.160835 | 160.810677 |
| -5.3368437 | 9.46E-08 | 5.80E-07 | 152.461923 | 147.212264 | 188.674455 | 160.250361 |
| -5.3366041 | 9.47E-08 | 5.81E-07 | 105.739721 | 353.688288 | 251.949425 | 312.656299 |
| -5.3364249 | 9.48E-08 | 5.82E-07 | 195.085687 | 253.832911 | 143.806749 | 187.705843 |
| -5.3362746 | 9.49E-08 | 5.82E-07 | 120.494101 | 225.418779 | 208.232173 | 202.274057 |
| -5.3360794 | 9.50E-08 | 5.82E-07 | 103.280658 | 279.811546 | 384.251634 | 296.967453 |
| -5.335988 | 9.50E-08 | 5.83E-07 | 101.231438 | 303.625294 | 302.569401 | 314.897563 |
| -5.3358636 | 9.51E-08 | 5.83E-07 | 132.379573 | 225.960001 | 169.116737 | 187.145527 |
| -5.3356901 | 9.52E-08 | 5.83E-07 | 102.051126 | 293.612695 | 331.330751 | 291.924609 |
| -5.3355986 | 9.52E-08 | 5.84E-07 | 118.444881 | 232.72527 | 200.178995 | 245.418385 |
| -5.335509 | 9.53E-08 | 5.84E-07 | 128.281134 | 190.780599 | 189.824909 | 179.301104 |
| 5.33501319 | 9.55E-08 | 5.85E-07 | 6.1476582 | 4.60038325 | 3.45136199 | 3.36189569 |
| -5.3347165 | 9.57E-08 | 5.86E-07 | 155.33083 | 227.042444 | 139.204933 | 276.796079 |
| -5.3347461 | 9.57E-08 | 5.86E-07 | 89.3459658 | 441.366181 | 360.092101 | 420.236961 |
| -5.3346824 | 9.57E-08 | 5.86E-07 | 132.789417 | 125.563402 | 111.594038 | 122.148877 |
| -5.3346567 | 9.57E-08 | 5.86E-07 | 249.185079 | 144.235545 | 159.913105 | 177.05984 |
| -5.3345692 | 9.58E-08 | 5.86E-07 | 128.281134 | 192.674875 | 178.320369 | 188.826475 |
| -5.3342971 | 9.59E-08 | 5.87E-07 | 118.035037 | 202.687474 | 234.692615 | 229.169223 |
| -5.3338984 | 9.61E-08 | 5.88E-07 | 103.690502 | 455.167331 | 281.861229 | 312.095983 |
| -5.3338434 | 9.62E-08 | 5.88E-07 | 114.346443 | 309.308121 | 212.833989 | 267.831023 |
| -5.3336932 | 9.62E-08 | 5.88E-07 | 138.937075 | 350.440959 | 299.118039 | 320.500723 |
| -5.333684 | 9.62E-08 | 5.88E-07 | 211.069598 | 214.053126 | 147.258111 | 141.199619 |
| -5.3323279 | 9.70E-08 | 5.93E-07 | 125.412227 | 275.211162 | 176.019461 | 251.581861 |
| -5.3319841 | 9.71E-08 | 5.94E-07 | 107.379097 | 306.602013 | 247.347609 | 274.554815 |
| -5.3319339 | 9.72E-08 | 5.94E-07 | 97.5428434 | 544.468888 | 320.976665 | 351.3181 |
| -5.3318016 | 9.72E-08 | 5.94E-07 | 101.641282 | 259.515737 | 534.961108 | 535.101731 |
| -5.3317347 | 9.73E-08 | 5.94E-07 | 120.903945 | 257.350851 | 204.780811 | 202.274057 |
| -5.3307655 | 9.78E-08 | 5.97E-07 | 256.152425 | 138.011497 | 201.329449 | 158.569413 |
| -5.3304725 | 9.80E-08 | 5.98E-07 | 143.855202 | 152.353869 | 322.127119 | 259.426284 |

| | | | | | | |
|---|---|---|---|---|---|---|
| -5.3302807 | 9.81E-08 | 5.99E-07 | 100.411751 | 367.760049 | 285.312591 | 313.216615 |
| -5.3300558 | 9.82E-08 | 5.99E-07 | 145.494577 | 150.459593 | 162.214013 | 184.343947 |
| -5.3295843 | 9.84E-08 | 6.01E-07 | 109.018472 | 321.756216 | 322.127119 | 226.367643 |
| -5.3295809 | 9.84E-08 | 6.01E-07 | 125.822071 | 222.983282 | 178.320369 | 197.231214 |
| -5.3294957 | 9.85E-08 | 6.01E-07 | 123.363008 | 264.657342 | 227.789891 | 182.102683 |
| -5.3292582 | 9.86E-08 | 6.01E-07 | 110.657848 | 296.048192 | 225.488983 | 267.270707 |
| -5.3288348 | 9.88E-08 | 6.03E-07 | 92.6247169 | 314.179115 | 494.695218 | 421.357593 |
| -5.3288444 | 9.88E-08 | 6.03E-07 | 112.707067 | 255.727186 | 368.145279 | 235.332698 |
| -5.3283953 | 9.91E-08 | 6.04E-07 | 102.870814 | 287.929869 | 283.011683 | 295.846821 |
| -5.328174 | 9.92E-08 | 6.05E-07 | 139.346919 | 170.21418 | 154.160835 | 192.18837 |
| -5.3281011 | 9.92E-08 | 6.05E-07 | 109.428316 | 258.162683 | 249.648517 | 246.539017 |
| -5.3273965 | 9.96E-08 | 6.07E-07 | 125.412227 | 211.88824 | 173.718553 | 224.126379 |
| -5.3273239 | 9.97E-08 | 6.07E-07 | 175.003337 | 125.02218 | 216.285351 | 231.970803 |
| -5.3273294 | 9.97E-08 | 6.07E-07 | 114.756286 | 216.218013 | 260.002603 | 224.686695 |
| -5.3271245 | 9.98E-08 | 6.08E-07 | 111.887379 | 239.76115 | 228.940345 | 251.581861 |
| -5.3270695 | 9.98E-08 | 6.08E-07 | 130.740198 | 235.431378 | 164.514921 | 202.274057 |
| -5.3260972 | 1.00E-07 | 6.11E-07 | 128.281134 | 233.266492 | 181.771731 | 180.982051 |
| 5.32606078 | 1.00E-07 | 6.11E-07 | 96.7231557 | 48.4393295 | 33.3631659 | 46.5062237 |
| -5.3257904 | 1.01E-07 | 6.12E-07 | 110.248004 | 296.048192 | 270.356689 | 223.566063 |
| -5.3256434 | 1.01E-07 | 6.12E-07 | 149.183172 | 240.843593 | 144.957203 | 184.904263 |
| -5.3256302 | 1.01E-07 | 6.12E-07 | 132.379573 | 222.983282 | 162.214013 | 189.947107 |
| -5.3255335 | 1.01E-07 | 6.12E-07 | 96.3133118 | 377.772648 | 656.909231 | 331.707042 |
| -5.3253134 | 1.01E-07 | 6.13E-07 | 116.395662 | 196.463426 | 272.657597 | 263.348496 |
| -5.3249078 | 1.01E-07 | 6.14E-07 | 130.330354 | 265.469174 | 169.116737 | 208.437533 |
| -5.3248091 | 1.01E-07 | 6.14E-07 | 107.379097 | 340.969582 | 239.294431 | 282.959554 |
| -5.3238824 | 1.02E-07 | 6.17E-07 | 115.575974 | 208.911521 | 234.692615 | 223.566063 |
| -5.323045 | 1.02E-07 | 6.20E-07 | 104.510189 | 271.693222 | 276.108959 | 272.873867 |
| -5.3219735 | 1.03E-07 | 6.23E-07 | 150.00286 | 152.624479 | 150.709473 | 195.550266 |
| -5.3214811 | 1.03E-07 | 6.25E-07 | 130.330354 | 189.698156 | 181.771731 | 165.293205 |
| -5.3214115 | 1.03E-07 | 6.25E-07 | 150.822548 | 141.258827 | 170.267191 | 168.655101 |
| -5.3211753 | 1.03E-07 | 6.26E-07 | 114.756286 | 302.542851 | 416.464346 | 277.916711 |
| -5.3206359 | 1.03E-07 | 6.28E-07 | 257.7918 | 566.658972 | 326.728935 | 414.073486 |
| -5.3205102 | 1.03E-07 | 6.28E-07 | 111.887379 | 343.405079 | 217.435805 | 274.554815 |
| -5.3203565 | 1.04E-07 | 6.28E-07 | 118.444881 | 228.124887 | 189.824909 | 237.013646 |
| -5.3202259 | 1.04E-07 | 6.29E-07 | 103.280658 | 262.492456 | 299.118039 | 284.640502 |
| -5.3201645 | 1.04E-07 | 6.29E-07 | 116.805506 | 213.511905 | 203.630357 | 229.169223 |
| -5.3195347 | 1.04E-07 | 6.31E-07 | 191.397092 | 255.185965 | 126.549939 | 178.740788 |
| -5.319392 | 1.04E-07 | 6.31E-07 | 119.674413 | 265.469174 | 192.125817 | 216.281956 |
| -5.3189376 | 1.04E-07 | 6.33E-07 | 108.608628 | 244.632144 | 269.206235 | 239.25491 |
| -5.3189017 | 1.04E-07 | 6.33E-07 | 222.545227 | 421.882205 | 108.142676 | 373.170422 |
| -5.3180971 | 1.05E-07 | 6.35E-07 | 119.264569 | 185.909605 | 236.993523 | 223.005748 |
| -5.318033 | 1.05E-07 | 6.35E-07 | 175.003337 | 196.734036 | 126.549939 | 175.939208 |
| -5.3177772 | 1.05E-07 | 6.36E-07 | 109.018472 | 355.311953 | 236.993523 | 262.78818 |

| | | | | | | |
|---|---|---|---|---|---|---|
| -5.3177295 | 1.05E-07 | 6.36E-07 | 116.805506 | 202.146252 | 211.683535 | 245.418385 |
| -5.3175092 | 1.05E-07 | 6.37E-07 | 104.100346 | 293.342085 | 254.250333 | 280.157974 |
| -5.3171276 | 1.05E-07 | 6.38E-07 | 165.167084 | 236.784432 | 131.151755 | 184.343947 |
| -5.3168875 | 1.06E-07 | 6.39E-07 | 170.904898 | 129.081342 | 176.019461 | 208.997849 |
| -5.3168424 | 1.06E-07 | 6.39E-07 | 97.9526873 | 367.760049 | 347.437107 | 767.63285 |
| -5.3167508 | 1.06E-07 | 6.39E-07 | 133.199261 | 164.801964 | 165.665375 | 195.550266 |
| -5.31619 | 1.06E-07 | 6.41E-07 | 141.396139 | 143.694324 | 208.232173 | 189.947107 |
| -5.3161343 | 1.06E-07 | 6.41E-07 | 104.510189 | 311.743618 | 254.250333 | 267.831023 |
| -5.3158265 | 1.06E-07 | 6.42E-07 | 102.46097 | 284.141318 | 271.507143 | 283.51987 |
| -5.3157646 | 1.06E-07 | 6.42E-07 | 114.756286 | 248.691306 | 202.479903 | 229.169223 |
| -5.3155256 | 1.06E-07 | 6.43E-07 | 159.429269 | 162.095857 | 158.762651 | 139.518671 |
| -5.3155379 | 1.06E-07 | 6.43E-07 | 108.608628 | 299.836743 | 223.188075 | 271.753235 |
| -5.3154846 | 1.06E-07 | 6.43E-07 | 110.248004 | 271.152001 | 225.488983 | 238.694594 |
| -5.3153057 | 1.06E-07 | 6.43E-07 | 146.314265 | 166.69624 | 142.656295 | 170.336048 |
| -5.3151943 | 1.07E-07 | 6.43E-07 | 164.347396 | 211.347019 | 138.054479 | 319.380091 |
| -5.3149537 | 1.07E-07 | 6.44E-07 | 110.657848 | 226.230611 | 242.745793 | 237.573962 |
| -5.3145904 | 1.07E-07 | 6.45E-07 | 126.231915 | 228.936719 | 201.329449 | 168.094785 |
| -5.3141111 | 1.07E-07 | 6.47E-07 | 105.739721 | 282.247043 | 241.595339 | 270.072287 |
| -5.3139905 | 1.07E-07 | 6.47E-07 | 130.740198 | 273.587498 | 158.762651 | 245.978701 |
| -5.3139815 | 1.07E-07 | 6.47E-07 | 108.198784 | 237.055043 | 306.020763 | 245.418385 |
| -5.3138467 | 1.07E-07 | 6.48E-07 | 109.018472 | 310.661174 | 230.090799 | 249.900913 |
| -5.3135874 | 1.07E-07 | 6.48E-07 | 138.117388 | 146.671042 | 188.674455 | 185.464579 |
| -5.3119696 | 1.08E-07 | 6.54E-07 | 165.576927 | 282.517653 | 126.549939 | 242.616806 |
| -5.3119299 | 1.08E-07 | 6.54E-07 | 110.248004 | 297.942468 | 210.533081 | 324.98325 |
| -5.3118902 | 1.08E-07 | 6.54E-07 | 109.428316 | 252.750468 | 257.701695 | 221.3248 |
| -5.3110046 | 1.09E-07 | 6.57E-07 | 101.231438 | 321.756216 | 269.206235 | 280.157974 |
| -5.3107652 | 1.09E-07 | 6.58E-07 | 180.741151 | 130.975617 | 213.984443 | 160.250361 |
| -5.3103437 | 1.09E-07 | 6.59E-07 | 113.936599 | 243.00848 | 199.028541 | 242.616806 |
| -5.3103008 | 1.09E-07 | 6.59E-07 | 205.331784 | 202.958084 | 125.399485 | 163.612257 |
| -5.3101508 | 1.10E-07 | 6.60E-07 | 101.641282 | 317.155833 | 287.613499 | 259.426284 |
| -5.3093971 | 1.10E-07 | 6.62E-07 | 115.16613 | 230.560384 | 228.940345 | 196.110582 |
| -5.3093303 | 1.10E-07 | 6.62E-07 | 94.6739363 | 284.141318 | 430.269794 | 376.532317 |
| -5.3092268 | 1.10E-07 | 6.63E-07 | 93.0345607 | 364.78333 | 325.578481 | 329.465778 |
| -5.3091319 | 1.10E-07 | 6.63E-07 | 109.018472 | 258.162683 | 218.586259 | 267.270707 |
| 5.30872476 | 1.10E-07 | 6.64E-07 | 0 | 0 | 0 | 0 |
| -5.3086772 | 1.10E-07 | 6.64E-07 | 175.003337 | 121.774851 | 222.037621 | 207.877217 |
| -5.3085895 | 1.10E-07 | 6.64E-07 | 102.46097 | 298.213079 | 295.666677 | 249.900913 |
| -5.3081777 | 1.11E-07 | 6.66E-07 | 109.83816 | 289.824144 | 224.338529 | 235.893014 |
| -5.3080044 | 1.11E-07 | 6.66E-07 | 112.297223 | 259.786348 | 318.675757 | 206.196269 |
| -5.3080352 | 1.11E-07 | 6.66E-07 | 105.329877 | 238.137486 | 265.754873 | 286.32145 |
| -5.3078955 | 1.11E-07 | 6.67E-07 | 117.21535 | 199.981366 | 205.931265 | 203.394689 |
| -5.3076355 | 1.11E-07 | 6.67E-07 | 107.78894 | 243.27909 | 245.046701 | 239.815226 |
| -5.3074898 | 1.11E-07 | 6.68E-07 | 86.4770587 | 391.573797 | 393.455266 | 377.652949 |

| | | | | | | |
|---|---|---|---|---|---|---|
| -5.3074738 | 1.11E-07 | 6.68E-07 | 100.001907 | 359.912336 | 261.153057 | 308.173772 |
| -5.3071461 | 1.11E-07 | 6.69E-07 | 327.46526 | 139.364551 | 317.525303 | 149.604358 |
| -5.3069021 | 1.12E-07 | 6.70E-07 | 181.970683 | 180.226779 | 209.382627 | 122.709193 |
| -5.3067474 | 1.12E-07 | 6.70E-07 | 197.54475 | 117.715689 | 245.046701 | 198.912162 |
| -5.3065494 | 1.12E-07 | 6.71E-07 | 159.839113 | 217.841677 | 286.463045 | 132.234564 |
| -5.3055883 | 1.12E-07 | 6.74E-07 | 109.428316 | 284.141318 | 223.188075 | 235.893014 |
| -5.3053102 | 1.12E-07 | 6.75E-07 | 171.724586 | 207.287857 | 139.204933 | 146.802779 |
| -5.3051493 | 1.13E-07 | 6.75E-07 | 321.727446 | 235.972599 | 127.700393 | 169.775732 |
| -5.305086 | 1.13E-07 | 6.75E-07 | 118.444881 | 272.234444 | 181.771731 | 238.134278 |
| -5.3049826 | 1.13E-07 | 6.76E-07 | 100.411751 | 294.965749 | 266.905327 | 290.803977 |
| -5.3047227 | 1.13E-07 | 6.77E-07 | 91.8050291 | 328.250875 | 371.596641 | 333.948305 |
| -5.3045868 | 1.13E-07 | 6.77E-07 | 745.915861 | 826.44532 | 776.556447 | 902.108677 |
| -5.3045236 | 1.13E-07 | 6.77E-07 | 99.5920628 | 279.270324 | 283.011683 | 300.329348 |
| -5.3037581 | 1.13E-07 | 6.80E-07 | 138.937075 | 170.484791 | 743.193281 | 591.133326 |
| -5.3033721 | 1.14E-07 | 6.81E-07 | 136.887856 | 218.65351 | 326.728935 | 161.370993 |
| -5.3029284 | 1.14E-07 | 6.83E-07 | 126.641759 | 314.720336 | 261.153057 | 218.52322 |
| -5.3020521 | 1.15E-07 | 6.86E-07 | 151.642236 | 148.294707 | 154.160835 | 148.483726 |
| -5.3018951 | 1.15E-07 | 6.86E-07 | 130.330354 | 165.613797 | 171.417645 | 223.005748 |
| -5.3016457 | 1.15E-07 | 6.87E-07 | 109.428316 | 312.284839 | 240.444885 | 222.445432 |
| -5.3016424 | 1.15E-07 | 6.87E-07 | 2625.45989 | 4455.87709 | 4314.20248 | 4273.52974 |
| -5.301318 | 1.15E-07 | 6.88E-07 | 115.985818 | 245.714587 | 195.577179 | 206.196269 |
| -5.3012517 | 1.15E-07 | 6.88E-07 | 319.268382 | 243.549701 | 227.789891 | 108.140978 |
| -5.30099 | 1.15E-07 | 6.89E-07 | 97.9526873 | 286.306204 | 283.011683 | 335.629253 |
| -5.3008086 | 1.15E-07 | 6.90E-07 | 116.395662 | 192.945486 | 270.356689 | 214.601008 |
| -5.3004077 | 1.16E-07 | 6.91E-07 | 111.067691 | 239.76115 | 245.046701 | 205.075637 |
| -5.3002763 | 1.16E-07 | 6.91E-07 | 121.313788 | 169.402348 | 226.639437 | 225.247011 |
| -5.3002306 | 1.16E-07 | 6.91E-07 | 94.6739363 | 540.139115 | 288.763953 | 378.213265 |
| -5.2996775 | 1.16E-07 | 6.93E-07 | 109.018472 | 247.608863 | 216.285351 | 242.05649 |
| -5.2996469 | 1.16E-07 | 6.93E-07 | 240.168514 | 180.768 | 217.435805 | 117.666349 |
| -5.2992482 | 1.16E-07 | 6.94E-07 | 122.54332 | 184.285941 | 373.897549 | 260.546916 |
| -5.2992576 | 1.16E-07 | 6.94E-07 | 162.298176 | 507.395211 | 144.957203 | 310.975351 |
| -5.2992745 | 1.16E-07 | 6.94E-07 | 154.511143 | 176.97945 | 130.001301 | 199.472478 |
| -5.2983796 | 1.17E-07 | 6.98E-07 | 112.297223 | 264.927953 | 194.426725 | 265.029444 |
| -5.2972094 | 1.18E-07 | 7.02E-07 | 141.396139 | 150.459593 | 181.771731 | 150.164674 |
| -5.2962963 | 1.18E-07 | 7.05E-07 | 114.346443 | 224.336336 | 193.276271 | 268.391339 |
| -5.2957586 | 1.19E-07 | 7.07E-07 | 154.101299 | 170.755402 | 136.904025 | 152.405938 |
| -5.2953257 | 1.19E-07 | 7.09E-07 | 163.527708 | 129.081342 | 164.514921 | 183.783631 |
| -5.2951515 | 1.19E-07 | 7.09E-07 | 92.214873 | 349.358516 | 295.666677 | 379.333897 |
| 5.29502033 | 1.19E-07 | 7.10E-07 | 0 | 0 | 0 | 0 |
| -5.2948152 | 1.19E-07 | 7.10E-07 | 107.379097 | 256.268408 | 216.285351 | 257.18502 |
| -5.2944718 | 1.19E-07 | 7.12E-07 | 84.8376831 | 467.344816 | 341.684837 | 414.073486 |
| -5.2942476 | 1.20E-07 | 7.12E-07 | 153.281611 | 162.637078 | 133.452663 | 165.853521 |
| -5.2939474 | 1.20E-07 | 7.13E-07 | 101.231438 | 261.951234 | 260.002603 | 336.749885 |

| | | | | | | |
|---|---|---|---|---|---|---|
| -5.2939442 | 1.20E-07 | 7.13E-07 | 100.001907 | 272.775665 | 277.259413 | 271.753235 |
| -5.2938053 | 1.20E-07 | 7.14E-07 | 109.428316 | 231.101605 | 211.683535 | 244.85807 |
| -5.2936944 | 1.20E-07 | 7.14E-07 | 132.379573 | 241.655426 | 163.364467 | 172.016996 |
| -5.2935779 | 1.20E-07 | 7.14E-07 | 81.1490882 | 435.683354 | 433.721156 | 392.221164 |
| -5.2934501 | 1.20E-07 | 7.15E-07 | 99.5920628 | 292.530252 | 262.303511 | 281.278606 |
| -5.2934598 | 1.20E-07 | 7.15E-07 | 110.248004 | 282.247043 | 203.630357 | 244.297754 |
| -5.2932572 | 1.20E-07 | 7.15E-07 | 118.035037 | 204.85236 | 199.028541 | 181.542367 |
| -5.2929696 | 1.20E-07 | 7.16E-07 | 107.379097 | 281.43521 | 209.382627 | 284.080186 |
| -5.2929106 | 1.20E-07 | 7.16E-07 | 102.051126 | 259.786348 | 253.099879 | 275.115131 |
| 5.29271152 | 1.21E-07 | 7.17E-07 | 10.6559409 | 12.7187066 | 12.6549939 | 10.0856871 |
| -5.2923046 | 1.21E-07 | 7.18E-07 | 349.59683 | 211.076408 | 146.107657 | 195.550266 |
| -5.2922874 | 1.21E-07 | 7.18E-07 | 111.477535 | 398.339067 | 200.178995 | 332.267358 |
| -5.2914834 | 1.21E-07 | 7.21E-07 | 111.887379 | 199.981366 | 222.037621 | 226.927959 |
| -5.2914758 | 1.21E-07 | 7.21E-07 | 113.526755 | 216.218013 | 196.727633 | 212.359745 |
| -5.2914318 | 1.21E-07 | 7.21E-07 | 114.756286 | 185.368384 | 222.037621 | 231.970803 |
| -5.2913877 | 1.21E-07 | 7.21E-07 | 131.969729 | 389.408911 | 316.374849 | 388.298952 |
| -5.2910343 | 1.22E-07 | 7.23E-07 | 136.478012 | 161.284024 | 249.648517 | 166.413837 |
| 5.29093132 | 1.22E-07 | 7.23E-07 | 20.492194 | 12.7187066 | 12.6549939 | 14.0078987 |
| -5.2909231 | 1.22E-07 | 7.23E-07 | 110.657848 | 227.854276 | 205.931265 | 226.367643 |
| -5.2906564 | 1.22E-07 | 7.24E-07 | 108.608628 | 217.841677 | 245.046701 | 224.126379 |
| -5.2906323 | 1.22E-07 | 7.24E-07 | 92.6247169 | 311.743618 | 315.224395 | 335.068937 |
| -5.29022 | 1.22E-07 | 7.25E-07 | 98.7723751 | 311.473007 | 286.463045 | 258.865968 |
| -5.2898719 | 1.22E-07 | 7.26E-07 | 117.625194 | 234.890156 | 184.072639 | 195.550266 |
| -5.2895534 | 1.23E-07 | 7.28E-07 | 106.969253 | 246.797031 | 220.887167 | 239.25491 |
| -5.2895054 | 1.23E-07 | 7.28E-07 | 111.067691 | 216.759234 | 205.931265 | 227.488275 |
| -5.2894789 | 1.23E-07 | 7.28E-07 | 114.756286 | 215.135569 | 187.524001 | 187.145527 |
| -5.2893164 | 1.23E-07 | 7.28E-07 | 123.363008 | 169.131737 | 176.019461 | 198.351846 |
| -5.2886156 | 1.23E-07 | 7.31E-07 | 93.4444046 | 374.254708 | 291.064861 | 314.337247 |
| -5.2884624 | 1.23E-07 | 7.31E-07 | 110.657848 | 256.268408 | 202.479903 | 224.126379 |
| -5.2882653 | 1.23E-07 | 7.32E-07 | 124.592539 | 326.085989 | 163.364467 | 304.811876 |
| -5.2881725 | 1.24E-07 | 7.32E-07 | 266.808366 | 159.93097 | 320.976665 | 127.19172 |
| -5.287305 | 1.24E-07 | 7.36E-07 | 112.707067 | 327.439043 | 188.674455 | 300.889664 |
| -5.2872365 | 1.24E-07 | 7.36E-07 | 99.5920628 | 323.921103 | 241.595339 | 308.734088 |
| -5.2870758 | 1.24E-07 | 7.36E-07 | 136.887856 | 167.237462 | 212.833989 | 146.802779 |
| -5.2868243 | 1.24E-07 | 7.37E-07 | 110.248004 | 237.596264 | 224.338529 | 202.834373 |
| -5.2864984 | 1.25E-07 | 7.38E-07 | 75.8211178 | 477.628025 | 480.88977 | 432.563912 |
| -5.2864806 | 1.25E-07 | 7.38E-07 | 92.214873 | 321.756216 | 303.719855 | 447.692443 |
| -5.2865173 | 1.25E-07 | 7.38E-07 | 99.1822189 | 340.157749 | 240.444885 | 325.543566 |
| -5.2865291 | 1.25E-07 | 7.38E-07 | 119.674413 | 205.934803 | 171.417645 | 242.05649 |
| -5.2860579 | 1.25E-07 | 7.39E-07 | 187.708497 | 835.104864 | 730.538287 | 739.617052 |
| -5.2859542 | 1.25E-07 | 7.40E-07 | 92.6247169 | 290.635977 | 419.915708 | 326.664198 |
| -5.2856498 | 1.25E-07 | 7.41E-07 | 154.920987 | 203.228695 | 131.151755 | 163.051941 |
| -5.2856548 | 1.25E-07 | 7.41E-07 | 114.756286 | 214.323737 | 276.108959 | 190.507423 |

| | | | | | | |
|---|---|---|---|---|---|---|
| -5.2855087 | 1.25E-07 | 7.41E-07 | 106.149565 | 345.028743 | 217.435805 | 272.313551 |
| -5.2855376 | 1.25E-07 | 7.41E-07 | 145.494577 | 258.703905 | 132.302209 | 243.177122 |
| -5.2852401 | 1.26E-07 | 7.42E-07 | 127.871291 | 152.353869 | 251.949425 | 259.426284 |
| -5.2850052 | 1.26E-07 | 7.43E-07 | 101.641282 | 226.501222 | 329.029843 | 319.380091 |
| -5.2849456 | 1.26E-07 | 7.43E-07 | 125.412227 | 170.755402 | 164.514921 | 205.635953 |
| -5.2848027 | 1.26E-07 | 7.43E-07 | 125.002383 | 218.382899 | 176.019461 | 165.293205 |
| -5.2847343 | 1.26E-07 | 7.43E-07 | 86.8869025 | 434.05969 | 437.172518 | 305.372192 |
| -5.2846866 | 1.26E-07 | 7.44E-07 | 110.657848 | 235.160767 | 195.577179 | 239.815226 |
| -5.2838799 | 1.26E-07 | 7.47E-07 | 89.3459658 | 350.71157 | 398.057082 | 305.932508 |
| -5.2836901 | 1.27E-07 | 7.47E-07 | 98.7723751 | 289.553534 | 276.108959 | 258.865968 |
| -5.282807 | 1.27E-07 | 7.51E-07 | 104.100346 | 333.933701 | 215.134897 | 367.006946 |
| -5.2824594 | 1.27E-07 | 7.52E-07 | 107.379097 | 238.137486 | 213.984443 | 233.091435 |
| -5.282449 | 1.27E-07 | 7.52E-07 | 116.805506 | 259.245126 | 200.178995 | 185.464579 |
| -5.2820684 | 1.28E-07 | 7.53E-07 | 135.24848 | 178.061893 | 190.975363 | 140.639303 |
| -5.2820895 | 1.28E-07 | 7.53E-07 | 131.969729 | 175.085174 | 162.214013 | 150.164674 |
| -5.2819757 | 1.28E-07 | 7.54E-07 | 96.7231557 | 289.012312 | 274.958505 | 287.442082 |
| -5.281847 | 1.28E-07 | 7.54E-07 | 118.854725 | 186.180216 | 247.347609 | 183.223315 |
| -5.2816355 | 1.28E-07 | 7.55E-07 | 204.512096 | 222.712671 | 117.346308 | 170.336048 |
| -5.2815123 | 1.28E-07 | 7.55E-07 | 143.035514 | 234.348935 | 134.603117 | 269.511971 |
| -5.2815178 | 1.28E-07 | 7.55E-07 | 111.887379 | 455.437941 | 199.028541 | 351.878416 |
| -5.2815005 | 1.28E-07 | 7.55E-07 | 125.002383 | 172.108456 | 169.116737 | 228.048591 |
| -5.2813856 | 1.28E-07 | 7.55E-07 | 109.018472 | 219.735953 | 209.382627 | 229.169223 |
| -5.281242 | 1.28E-07 | 7.55E-07 | 98.3625312 | 255.997797 | 284.162137 | 286.881766 |
| -5.2812589 | 1.28E-07 | 7.55E-07 | 115.16613 | 232.72527 | 177.169915 | 225.247011 |
| -5.2810731 | 1.28E-07 | 7.56E-07 | 231.151948 | 239.76115 | 507.350212 | 120.467929 |
| -5.2807035 | 1.29E-07 | 7.57E-07 | 114.346443 | 180.49739 | 224.338529 | 219.083536 |
| -5.2793789 | 1.30E-07 | 7.63E-07 | 120.903945 | 245.173366 | 171.417645 | 191.628054 |
| -5.2781107 | 1.31E-07 | 7.68E-07 | 134.428793 | 184.285941 | 173.718553 | 141.199619 |
| -5.2779901 | 1.31E-07 | 7.68E-07 | 140.166607 | 238.678707 | 213.984443 | 205.075637 |
| -5.2774157 | 1.31E-07 | 7.70E-07 | 106.969253 | 239.219929 | 212.833989 | 228.608907 |
| -5.2774229 | 1.31E-07 | 7.70E-07 | 73.7718984 | 522.549415 | 616.643342 | 397.824324 |
| -5.2766648 | 1.32E-07 | 7.73E-07 | 161.478489 | 204.581749 | 119.647216 | 219.083536 |
| -5.2765964 | 1.32E-07 | 7.73E-07 | 107.379097 | 233.537102 | 205.931265 | 245.978701 |
| -5.2766218 | 1.32E-07 | 7.73E-07 | 77.8703372 | 436.495187 | 424.517524 | 419.676646 |
| 5.27640511 | 1.32E-07 | 7.74E-07 | 17.6232868 | 13.8011497 | 20.7081719 | 4.48252759 |
| -5.2764118 | 1.32E-07 | 7.74E-07 | 97.1329995 | 305.79018 | 265.754873 | 270.632603 |
| -5.2763188 | 1.32E-07 | 7.74E-07 | 197.54475 | 144.506156 | 131.151755 | 175.378892 |
| -5.2762824 | 1.32E-07 | 7.74E-07 | 113.936599 | 204.581749 | 185.223093 | 205.635953 |
| -5.2761263 | 1.32E-07 | 7.75E-07 | 116.395662 | 189.427545 | 180.621277 | 224.126379 |
| -5.2759128 | 1.32E-07 | 7.75E-07 | 97.5428434 | 278.187881 | 262.303511 | 280.71829 |
| -5.2758316 | 1.32E-07 | 7.76E-07 | 95.493624 | 499.006277 | 304.870309 | 290.243661 |
| -5.2748374 | 1.33E-07 | 7.80E-07 | 149.593016 | 127.728288 | 225.488983 | 194.429634 |
| -5.2747262 | 1.33E-07 | 7.80E-07 | 119.674413 | 179.414947 | 170.267191 | 195.550266 |

| | | | | | | |
|---|---|---|---|---|---|---|
| -5.2743768 | 1.33E-07 | 7.81E-07 | 132.379573 | 488.181846 | 331.330751 | 373.170422 |
| -5.2741863 | 1.33E-07 | 7.82E-07 | 140.576451 | 228.124887 | 231.241253 | 132.79488 |
| -5.2735675 | 1.34E-07 | 7.85E-07 | 109.018472 | 335.557366 | 196.727633 | 273.434183 |
| -5.2735116 | 1.34E-07 | 7.85E-07 | 95.0837801 | 311.473007 | 279.560321 | 277.356395 |
| -5.2731507 | 1.34E-07 | 7.86E-07 | 161.888333 | 200.251977 | 119.647216 | 229.169223 |
| -5.2730069 | 1.34E-07 | 7.87E-07 | 253.283518 | 205.664192 | 121.948124 | 160.250361 |
| -5.2727647 | 1.34E-07 | 7.87E-07 | 94.6739363 | 303.895905 | 274.958505 | 290.803977 |
| -5.2727246 | 1.34E-07 | 7.87E-07 | 102.870814 | 267.904671 | 218.586259 | 269.511971 |
| -5.2726738 | 1.34E-07 | 7.88E-07 | 85.247527 | 369.654324 | 430.269794 | 322.741986 |
| -5.2720863 | 1.35E-07 | 7.90E-07 | 130.330354 | 241.926037 | 143.806749 | 218.52322 |
| -5.2720079 | 1.35E-07 | 7.90E-07 | 150.00286 | 253.021079 | 144.957203 | 159.129729 |
| -5.2715732 | 1.35E-07 | 7.92E-07 | 102.46097 | 248.961917 | 265.754873 | 225.807327 |
| -5.2712976 | 1.35E-07 | 7.93E-07 | 97.9526873 | 255.997797 | 289.914407 | 262.78818 |
| -5.2713176 | 1.35E-07 | 7.93E-07 | 152.052079 | 188.615713 | 124.249032 | 212.92006 |
| -5.2711961 | 1.36E-07 | 7.93E-07 | 96.3133118 | 286.306204 | 293.365769 | 491.397087 |
| -5.2709412 | 1.36E-07 | 7.94E-07 | 659.438803 | 634.311666 | 671.865133 | 677.982298 |
| -5.2705382 | 1.36E-07 | 7.96E-07 | 208.610535 | 188.074492 | 123.098578 | 153.52657 |
| -5.270034 | 1.36E-07 | 7.98E-07 | 104.100346 | 312.284839 | 271.507143 | 210.118481 |
| -5.2697328 | 1.37E-07 | 7.99E-07 | 106.559409 | 243.549701 | 337.083021 | 214.040692 |
| -5.269746 | 1.37E-07 | 7.99E-07 | 151.232392 | 188.345102 | 163.364467 | 128.312352 |
| -5.2696894 | 1.37E-07 | 7.99E-07 | 76.2309617 | 446.778396 | 407.260714 | 456.097182 |
| -5.2694991 | 1.37E-07 | 7.99E-07 | 98.7723751 | 261.680623 | 264.604419 | 253.823125 |
| -5.2693227 | 1.37E-07 | 8.00E-07 | 107.379097 | 240.031761 | 222.037621 | 203.955005 |
| -5.2690256 | 1.37E-07 | 8.01E-07 | 89.3459658 | 287.118037 | 370.446187 | 349.076836 |
| -5.268902 | 1.37E-07 | 8.02E-07 | 116.805506 | 181.850444 | 177.169915 | 194.98995 |
| -5.2685283 | 1.38E-07 | 8.03E-07 | 123.772852 | 175.897006 | 158.762651 | 170.336048 |
| -5.2683531 | 1.38E-07 | 8.04E-07 | 94.2640924 | 282.788264 | 279.560321 | 299.769033 |
| -5.2682813 | 1.38E-07 | 8.04E-07 | 129.100822 | 207.017246 | 182.922185 | 145.121831 |
| -5.2681865 | 1.38E-07 | 8.04E-07 | 141.805982 | 235.431378 | 194.426725 | 202.274057 |
| -5.2681256 | 1.38E-07 | 8.04E-07 | 118.035037 | 199.440144 | 164.514921 | 210.118481 |
| -5.2680647 | 1.38E-07 | 8.04E-07 | 111.887379 | 181.579833 | 226.639437 | 255.504073 |
| -5.2680316 | 1.38E-07 | 8.04E-07 | 881.164342 | 225.960001 | 166.815829 | 238.694594 |
| -5.2678535 | 1.38E-07 | 8.05E-07 | 230.332261 | 458.685271 | 411.86253 | 435.925808 |
| -5.2677175 | 1.38E-07 | 8.05E-07 | 102.46097 | 224.606947 | 254.250333 | 245.978701 |
| -5.2677039 | 1.38E-07 | 8.05E-07 | 109.83816 | 207.287857 | 195.577179 | 229.729539 |
| -5.267484 | 1.38E-07 | 8.06E-07 | 211.479442 | 254.644743 | 113.894946 | 175.939208 |
| -5.2675028 | 1.38E-07 | 8.06E-07 | 132.379573 | 411.057774 | 292.215315 | 387.17832 |
| -5.267389 | 1.38E-07 | 8.06E-07 | 85.247527 | 363.430276 | 331.330751 | 364.205367 |
| -5.2669926 | 1.39E-07 | 8.08E-07 | 107.379097 | 198.357701 | 294.516223 | 244.297754 |
| -5.2669593 | 1.39E-07 | 8.08E-07 | 115.575974 | 221.089007 | 181.771731 | 183.223315 |
| -5.2669768 | 1.39E-07 | 8.08E-07 | 112.707067 | 196.192815 | 194.426725 | 192.748686 |
| -5.2667393 | 1.39E-07 | 8.09E-07 | 335.662138 | 485.475738 | 387.702996 | 432.003596 |
| -5.2665584 | 1.39E-07 | 8.09E-07 | 117.625194 | 227.042444 | 171.417645 | 187.705843 |

| | | | | | | |
|---|---|---|---|---|---|---|
| -5.2664306 | 1.39E-07 | 8.10E-07 | 138.937075 | 217.571066 | 134.603117 | 180.982051 |
| -5.2664572 | 1.39E-07 | 8.10E-07 | 84.0179954 | 383.996696 | 361.242555 | 341.792729 |
| -5.2663077 | 1.39E-07 | 8.10E-07 | 97.9526873 | 311.202396 | 240.444885 | 276.235763 |
| -5.2662153 | 1.39E-07 | 8.10E-07 | 128.690978 | 205.664192 | 154.160835 | 161.370993 |
| -5.2659327 | 1.39E-07 | 8.11E-07 | 103.280658 | 266.822228 | 210.533081 | 267.831023 |
| -5.2658249 | 1.40E-07 | 8.12E-07 | 108.198784 | 216.218013 | 225.488983 | 199.472478 |
| -5.2652683 | 1.40E-07 | 8.14E-07 | 87.2967464 | 320.403163 | 362.393009 | 324.98325 |
| -5.26512 | 1.40E-07 | 8.14E-07 | 98.3625312 | 263.304288 | 273.808051 | 245.418385 |
| -5.264885 | 1.40E-07 | 8.15E-07 | 214.758193 | 193.216096 | 148.408565 | 166.974153 |
| -5.2646329 | 1.40E-07 | 8.16E-07 | 111.887379 | 198.08709 | 213.984443 | 186.585211 |
| -5.264477 | 1.41E-07 | 8.17E-07 | 117.21535 | 266.551617 | 164.514921 | 276.235763 |
| -5.2639871 | 1.41E-07 | 8.19E-07 | 111.067691 | 191.321821 | 200.178995 | 207.877217 |
| -5.2638392 | 1.41E-07 | 8.19E-07 | 93.8542485 | 271.693222 | 295.666677 | 291.364293 |
| -5.2634094 | 1.41E-07 | 8.21E-07 | 102.46097 | 277.105438 | 238.143977 | 222.445432 |
| -5.2633545 | 1.41E-07 | 8.21E-07 | 79.0998688 | 360.453558 | 425.667978 | 435.365492 |
| -5.263228 | 1.42E-07 | 8.22E-07 | 117.625194 | 173.190899 | 231.241253 | 182.102683 |
| -5.2631533 | 1.42E-07 | 8.22E-07 | 75.4112739 | 391.844408 | 609.740618 | 438.167072 |
| -5.2630536 | 1.42E-07 | 8.22E-07 | 76.2309617 | 415.928768 | 442.924788 | 418.556014 |
| -5.2630744 | 1.42E-07 | 8.22E-07 | 117.21535 | 167.237462 | 245.046701 | 203.394689 |
| -5.2630422 | 1.42E-07 | 8.22E-07 | 124.592539 | 258.433294 | 165.665375 | 179.86142 |
| -5.2623593 | 1.42E-07 | 8.25E-07 | 346.318079 | 173.461509 | 155.311289 | 155.207518 |
| -5.2623898 | 1.42E-07 | 8.25E-07 | 108.198784 | 328.792097 | 190.975363 | 368.687894 |
| -5.2622988 | 1.42E-07 | 8.25E-07 | 247.545703 | 121.774851 | 156.461743 | 208.437533 |
| -5.2618815 | 1.43E-07 | 8.27E-07 | 90.5754975 | 327.709654 | 289.914407 | 306.492824 |
| -5.2617457 | 1.43E-07 | 8.27E-07 | 127.871291 | 168.590515 | 151.859927 | 157.448782 |
| -5.2604848 | 1.44E-07 | 8.33E-07 | 96.3133118 | 245.443977 | 333.631659 | 279.037342 |
| -5.2603443 | 1.44E-07 | 8.33E-07 | 167.626147 | 335.557366 | 124.249032 | 468.984449 |
| -5.2599224 | 1.44E-07 | 8.35E-07 | 120.084257 | 206.746635 | 156.461743 | 192.18837 |
| -5.2598585 | 1.44E-07 | 8.35E-07 | 138.117388 | 255.185965 | 138.054479 | 191.628054 |
| -5.2597552 | 1.44E-07 | 8.35E-07 | 105.739721 | 217.841677 | 207.081719 | 250.461229 |
| -5.2597343 | 1.44E-07 | 8.35E-07 | 278.283994 | 176.438228 | 131.151755 | 159.690045 |
| -5.2597041 | 1.44E-07 | 8.35E-07 | 134.018949 | 197.545869 | 134.603117 | 185.464579 |
| -5.259598 | 1.44E-07 | 8.36E-07 | 124.592539 | 162.907689 | 167.966283 | 156.32815 |
| -5.2593008 | 1.45E-07 | 8.37E-07 | 111.477535 | 215.947402 | 199.028541 | 183.783631 |
| -5.2591884 | 1.45E-07 | 8.37E-07 | 117.21535 | 246.255809 | 235.843069 | 242.616806 |
| -5.2589673 | 1.45E-07 | 8.38E-07 | 107.379097 | 331.498204 | 301.418947 | 296.967453 |
| -5.2588771 | 1.45E-07 | 8.38E-07 | 208.610535 | 150.459593 | 136.904025 | 146.802779 |
| -5.2587724 | 1.45E-07 | 8.39E-07 | 76.6408055 | 413.493271 | 414.163438 | 418.556014 |
| -5.2586954 | 1.45E-07 | 8.39E-07 | 109.018472 | 195.922204 | 233.542161 | 200.59311 |
| -5.2583937 | 1.45E-07 | 8.40E-07 | 115.16613 | 211.617629 | 170.267191 | 192.18837 |
| -5.2581411 | 1.46E-07 | 8.41E-07 | 155.740674 | 188.615713 | 197.878087 | 119.907613 |
| -5.257905 | 1.46E-07 | 8.42E-07 | 105.329877 | 294.695138 | 209.382627 | 413.51317 |
| -5.2573468 | 1.46E-07 | 8.44E-07 | 165.986771 | 154.248144 | 128.850847 | 147.92341 |

| | | | | | | |
|---|---|---|---|---|---|---|
| -5.2572311 | 1.46E-07 | 8.45E-07 | 105.329877 | 302.542851 | 343.985745 | 196.110582 |
| -5.2571614 | 1.46E-07 | 8.45E-07 | 111.067691 | 202.146252 | 185.223093 | 244.85807 |
| -5.2566488 | 1.47E-07 | 8.47E-07 | 109.83816 | 195.651593 | 201.329449 | 198.912162 |
| -5.2563282 | 1.47E-07 | 8.48E-07 | 93.0345607 | 266.551617 | 303.719855 | 289.123029 |
| -5.2563042 | 1.47E-07 | 8.48E-07 | 215.168037 | 189.156935 | 157.612197 | 119.347297 |
| -5.2562607 | 1.47E-07 | 8.48E-07 | 114.756286 | 211.617629 | 534.961108 | 285.200818 |
| -5.2560046 | 1.47E-07 | 8.49E-07 | 92.6247169 | 258.974516 | 335.932567 | 303.130928 |
| -5.2557517 | 1.47E-07 | 8.50E-07 | 92.214873 | 287.659258 | 286.463045 | 289.123029 |
| -5.2557164 | 1.47E-07 | 8.50E-07 | 765.998212 | 1468.6047 | 1163.10899 | 1363.2487 |
| -5.2556273 | 1.48E-07 | 8.51E-07 | 634.028482 | 2286.93169 | 1885.5941 | 2035.06753 |
| -5.2554475 | 1.48E-07 | 8.51E-07 | 101.641282 | 265.198564 | 216.285351 | 244.297754 |
| 5.2553794 | 1.48E-07 | 8.52E-07 | 58.197831 | 18.401533 | 25.3099879 | 22.9729539 |
| -5.2550731 | 1.48E-07 | 8.53E-07 | 172.13443 | 290.094755 | 166.815829 | 198.351846 |
| -5.2546574 | 1.48E-07 | 8.54E-07 | 90.5754975 | 350.71157 | 310.622579 | 272.313551 |
| -5.2546575 | 1.48E-07 | 8.54E-07 | 100.821594 | 227.042444 | 247.347609 | 239.815226 |
| -5.2540508 | 1.49E-07 | 8.57E-07 | 112.297223 | 232.995881 | 309.472125 | 180.982051 |
| -5.2538021 | 1.49E-07 | 8.58E-07 | 70.9029912 | 463.015043 | 424.517524 | 585.530166 |
| -5.2536487 | 1.49E-07 | 8.59E-07 | 86.8869025 | 343.67569 | 319.826211 | 312.095983 |
| -5.2534711 | 1.49E-07 | 8.59E-07 | 198.364438 | 321.214995 | 103.54086 | 368.687894 |
| -5.2532578 | 1.49E-07 | 8.60E-07 | 138.117388 | 202.687474 | 157.612197 | 180.421735 |
| -5.2530487 | 1.50E-07 | 8.61E-07 | 85.6573709 | 322.568049 | 409.561622 | 316.018195 |
| -5.2528367 | 1.50E-07 | 8.62E-07 | 101.231438 | 451.920001 | 226.639437 | 304.25156 |
| -5.2519294 | 1.51E-07 | 8.66E-07 | 639.356453 | 2911.23076 | 2575.8665 | 2853.12881 |
| -5.2518783 | 1.51E-07 | 8.66E-07 | 184.83959 | 155.059976 | 324.428027 | 256.624704 |
| -5.2512492 | 1.51E-07 | 8.69E-07 | 172.544273 | 117.445078 | 301.418947 | 211.799429 |
| -5.2511291 | 1.51E-07 | 8.69E-07 | 102.870814 | 223.253893 | 217.435805 | 235.332698 |
| -5.2509977 | 1.51E-07 | 8.70E-07 | 72.1325229 | 445.154732 | 448.677058 | 457.217814 |
| -5.250984 | 1.51E-07 | 8.70E-07 | 124.182696 | 378.043258 | 376.198456 | 344.033992 |
| -5.2508879 | 1.51E-07 | 8.70E-07 | 109.83816 | 204.311138 | 193.276271 | 191.067738 |
| 5.25076948 | 1.51E-07 | 8.70E-07 | 42.6237635 | 13.2599282 | 17.2568099 | 5.04284354 |
| -5.2505524 | 1.52E-07 | 8.71E-07 | 376.236682 | 173.73212 | 205.931265 | 129.432984 |
| -5.2503747 | 1.52E-07 | 8.72E-07 | 136.068168 | 184.285941 | 130.001301 | 194.429634 |
| -5.2502294 | 1.52E-07 | 8.72E-07 | 101.641282 | 249.503138 | 234.692615 | 216.281956 |
| -5.2495298 | 1.52E-07 | 8.76E-07 | 118.854725 | 199.440144 | 166.815829 | 165.293205 |
| -5.2494357 | 1.53E-07 | 8.76E-07 | 172.13443 | 205.934803 | 116.195854 | 167.534469 |
| -5.2492003 | 1.53E-07 | 8.77E-07 | 123.363008 | 163.448911 | 181.771731 | 151.285306 |
| -5.2490002 | 1.53E-07 | 8.78E-07 | 90.5754975 | 281.976432 | 304.870309 | 293.605557 |
| -5.24893 | 1.53E-07 | 8.78E-07 | 230.742104 | 177.520671 | 139.204933 | 504.284354 |
| -5.2487635 | 1.53E-07 | 8.78E-07 | 102.870814 | 220.547785 | 232.391707 | 216.842272 |
| -5.2486402 | 1.53E-07 | 8.79E-07 | 96.7231557 | 267.904671 | 268.055781 | 237.573962 |
| -5.2484087 | 1.53E-07 | 8.80E-07 | 115.16613 | 185.368384 | 271.507143 | 183.783631 |
| -5.2482765 | 1.54E-07 | 8.80E-07 | 209.840067 | 191.05121 | 117.346308 | 155.767834 |
| -5.2482746 | 1.54E-07 | 8.80E-07 | 111.477535 | 182.121054 | 194.426725 | 189.947107 |

| | | | | | | |
|---|---|---|---|---|---|---|
| -5.2481281 | 1.54E-07 | 8.81E-07 | 127.051603 | 210.805797 | 149.559019 | 163.051941 |
| -5.2480593 | 1.54E-07 | 8.81E-07 | 109.018472 | 212.970683 | 184.072639 | 206.756585 |
| -5.2477934 | 1.54E-07 | 8.82E-07 | 96.7231557 | 308.496288 | 226.639437 | 287.442082 |
| -5.2476306 | 1.54E-07 | 8.82E-07 | 134.018949 | 136.658443 | 156.461743 | 164.172573 |
| -5.2476253 | 1.54E-07 | 8.82E-07 | 73.3620545 | 324.462324 | 836.380055 | 758.107478 |
| -5.2474449 | 1.54E-07 | 8.83E-07 | 108.608628 | 205.664192 | 218.586259 | 184.904263 |
| -5.2473918 | 1.54E-07 | 8.83E-07 | 163.937552 | 194.839761 | 113.894946 | 194.429634 |
| -5.2470772 | 1.55E-07 | 8.85E-07 | 93.4444046 | 285.494372 | 271.507143 | 263.348496 |
| 5.24693622 | 1.55E-07 | 8.85E-07 | 36.8859492 | 56.0164313 | 47.1686138 | 50.9887513 |
| -5.2468496 | 1.55E-07 | 8.85E-07 | 89.3459658 | 328.792097 | 294.516223 | 286.32145 |
| -5.2467226 | 1.55E-07 | 8.86E-07 | 101.231438 | 266.010396 | 210.533081 | 243.737438 |
| -5.2463869 | 1.55E-07 | 8.87E-07 | 473.369681 | 569.094469 | 577.527906 | 545.747734 |
| -5.2459848 | 1.55E-07 | 8.89E-07 | 249.185079 | 98.2317128 | 276.108959 | 229.169223 |
| -5.245702 | 1.56E-07 | 8.90E-07 | 103.690502 | 276.293606 | 204.780811 | 225.807327 |
| -5.2454043 | 1.56E-07 | 8.91E-07 | 126.231915 | 155.871809 | 220.887167 | 158.569413 |
| -5.2454274 | 1.56E-07 | 8.91E-07 | 106.559409 | 222.983282 | 188.674455 | 223.566063 |
| -5.2452006 | 1.56E-07 | 8.92E-07 | 105.329877 | 209.993965 | 212.833989 | 205.075637 |
| -5.24499 | 1.56E-07 | 8.93E-07 | 133.199261 | 219.735953 | 228.940345 | 132.234564 |
| -5.2449229 | 1.56E-07 | 8.93E-07 | 111.477535 | 175.085174 | 241.595339 | 200.59311 |
| -5.2447439 | 1.56E-07 | 8.94E-07 | 109.018472 | 198.898923 | 192.125817 | 193.309002 |
| -5.2443283 | 1.57E-07 | 8.96E-07 | 97.5428434 | 260.59818 | 236.993523 | 245.978701 |
| -5.2442568 | 1.57E-07 | 8.96E-07 | 115.16613 | 159.389749 | 246.197155 | 235.332698 |
| -5.2440657 | 1.57E-07 | 8.97E-07 | 92.6247169 | 256.539019 | 287.613499 | 290.803977 |
| -5.2440561 | 1.57E-07 | 8.97E-07 | 100.411751 | 219.465342 | 231.241253 | 261.107232 |
| -5.2435314 | 1.58E-07 | 8.99E-07 | 100.821594 | 217.300456 | 250.798971 | 229.169223 |
| -5.2431848 | 1.58E-07 | 9.01E-07 | 115.575974 | 185.638994 | 164.514921 | 179.86142 |
| -5.2431223 | 1.58E-07 | 9.01E-07 | 93.0345607 | 329.062707 | 247.347609 | 287.442082 |
| -5.2431453 | 1.58E-07 | 9.01E-07 | 106.969253 | 277.105438 | 186.373547 | 230.850171 |
| -5.2429177 | 1.58E-07 | 9.01E-07 | 132.379573 | 245.443977 | 258.852149 | 307.613456 |
| -5.242712 | 1.58E-07 | 9.02E-07 | 94.2640924 | 265.739785 | 250.798971 | 281.838922 |
| -5.2422037 | 1.59E-07 | 9.05E-07 | 112.707067 | 171.567234 | 225.488983 | 188.826475 |
| -5.2418565 | 1.59E-07 | 9.06E-07 | 99.5920628 | 289.553534 | 223.188075 | 234.772382 |
| -5.2415224 | 1.59E-07 | 9.08E-07 | 170.495054 | 266.281007 | 151.859927 | 215.72164 |
| -5.2409619 | 1.60E-07 | 9.10E-07 | 79.9195566 | 340.698971 | 376.198456 | 388.859268 |
| -5.2409629 | 1.60E-07 | 9.10E-07 | 118.444881 | 177.520671 | 161.063559 | 165.293205 |
| -5.2409146 | 1.60E-07 | 9.10E-07 | 293.858062 | 187.262659 | 149.559019 | 132.79488 |
| -5.2403993 | 1.60E-07 | 9.13E-07 | 1125.02145 | 492.241007 | 396.906628 | 75.6426531 |
| -5.2403285 | 1.60E-07 | 9.13E-07 | 104.920033 | 223.795114 | 211.683535 | 197.231214 |
| -5.2398957 | 1.61E-07 | 9.15E-07 | 104.510189 | 190.780599 | 232.391707 | 237.013646 |
| -5.2398328 | 1.61E-07 | 9.15E-07 | 108.608628 | 209.182132 | 201.329449 | 181.542367 |
| -5.2397718 | 1.61E-07 | 9.15E-07 | 108.608628 | 236.24321 | 338.233475 | 190.507423 |
| -5.2395365 | 1.61E-07 | 9.16E-07 | 91.3951852 | 251.397414 | 467.084322 | 342.353045 |
| -5.2394053 | 1.61E-07 | 9.17E-07 | 103.690502 | 543.115834 | 234.692615 | 294.165873 |

| | | | | | | |
|---|---|---|---|---|---|---|
| -5.2392993 | 1.61E-07 | 9.17E-07 | 165.167084 | 195.380983 | 205.931265 | 112.623506 |
| -5.2390425 | 1.61E-07 | 9.18E-07 | 100.001907 | 243.549701 | 261.153057 | 210.678797 |
| -5.2389388 | 1.62E-07 | 9.18E-07 | 119.674413 | 244.090923 | 199.028541 | 245.418385 |
| -5.2386858 | 1.62E-07 | 9.19E-07 | 149.183172 | 235.160767 | 120.79767 | 192.748686 |
| -5.2385617 | 1.62E-07 | 9.20E-07 | 117.21535 | 152.624479 | 231.241253 | 219.083536 |
| -5.2382046 | 1.62E-07 | 9.22E-07 | 164.347396 | 163.719521 | 120.79767 | 151.845622 |
| -5.2380844 | 1.62E-07 | 9.22E-07 | 123.772852 | 253.5623 | 161.063559 | 170.336048 |
| -5.2378797 | 1.62E-07 | 9.23E-07 | 99.1822189 | 262.492456 | 212.833989 | 320.500723 |
| -5.237885 | 1.62E-07 | 9.23E-07 | 113.936599 | 175.626396 | 146.107657 | 181.542367 |
| -5.2365417 | 1.64E-07 | 9.29E-07 | 98.3625312 | 220.006563 | 245.046701 | 255.504073 |
| -5.2360988 | 1.64E-07 | 9.31E-07 | 102.46097 | 244.090923 | 227.789891 | 199.472478 |
| -5.2359099 | 1.64E-07 | 9.32E-07 | 95.9034679 | 250.314971 | 235.843069 | 269.511971 |
| -5.2359419 | 1.64E-07 | 9.32E-07 | 103.690502 | 246.52642 | 203.630357 | 207.316901 |
| -5.2357247 | 1.64E-07 | 9.33E-07 | 197.954594 | 253.5623 | 104.691314 | 200.032794 |
| -5.235235 | 1.65E-07 | 9.35E-07 | 110.248004 | 172.920288 | 308.321671 | 251.021545 |
| -5.2347448 | 1.65E-07 | 9.37E-07 | 102.051126 | 232.995881 | 220.887167 | 205.075637 |
| -5.2346792 | 1.65E-07 | 9.37E-07 | 116.805506 | 157.495473 | 192.125817 | 172.016996 |
| -5.2344517 | 1.65E-07 | 9.38E-07 | 114.756286 | 215.947402 | 163.364467 | 180.421735 |
| -5.2341454 | 1.66E-07 | 9.40E-07 | 106.559409 | 245.985198 | 182.922185 | 214.040692 |
| -5.23399 | 1.66E-07 | 9.40E-07 | 119.264569 | 211.88824 | 166.815829 | 322.741986 |
| -5.2339383 | 1.66E-07 | 9.40E-07 | 89.3459658 | 284.95315 | 286.463045 | 288.002398 |
| 5.23393333 | 1.66E-07 | 9.40E-07 | 318.858539 | 321.756216 | 325.578481 | 309.294404 |
| -5.23372 | 1.66E-07 | 9.41E-07 | 93.4444046 | 396.715402 | 224.338529 | 391.660848 |
| -5.2337185 | 1.66E-07 | 9.41E-07 | 115.16613 | 168.861126 | 188.674455 | 164.732889 |
| -5.2335877 | 1.66E-07 | 9.42E-07 | 108.608628 | 209.993965 | 185.223093 | 186.024895 |
| -5.2333646 | 1.66E-07 | 9.43E-07 | 111.887379 | 194.56915 | 167.966283 | 188.266159 |
| -5.2328294 | 1.67E-07 | 9.45E-07 | 82.7884637 | 431.894804 | 303.719855 | 342.353045 |
| 5.23212465 | 1.68E-07 | 9.49E-07 | 65.9848647 | 47.8981079 | 43.7172518 | 48.7474875 |
| -5.2321534 | 1.68E-07 | 9.49E-07 | 118.035037 | 244.361534 | 153.010381 | 200.59311 |
| -5.2319139 | 1.68E-07 | 9.49E-07 | 130.330354 | 133.411114 | 249.648517 | 249.340597 |
| -5.2318296 | 1.68E-07 | 9.50E-07 | 106.969253 | 198.898923 | 222.037621 | 183.223315 |
| -5.2316714 | 1.68E-07 | 9.50E-07 | 120.903945 | 171.296623 | 253.099879 | 162.491625 |
| -5.2314219 | 1.68E-07 | 9.51E-07 | 109.83816 | 233.537102 | 166.815829 | 246.539017 |
| -5.2311828 | 1.68E-07 | 9.53E-07 | 116.805506 | 201.33442 | 150.709473 | 215.72164 |
| -5.2309515 | 1.69E-07 | 9.54E-07 | 124.592539 | 158.307306 | 142.656295 | 168.655101 |
| -5.2305964 | 1.69E-07 | 9.55E-07 | 105.739721 | 237.325653 | 193.276271 | 196.110582 |
| -5.2302376 | 1.69E-07 | 9.57E-07 | 100.411751 | 291.989031 | 196.727633 | 274.554815 |
| -5.2302498 | 1.69E-07 | 9.57E-07 | 78.2801811 | 361.806612 | 342.835291 | 498.681194 |
| -5.230051 | 1.69E-07 | 9.58E-07 | 134.428793 | 636.476553 | 366.994825 | 354.11968 |
| -5.2295315 | 1.70E-07 | 9.60E-07 | 77.4604933 | 378.855091 | 387.702996 | 356.360943 |
| -5.2292267 | 1.70E-07 | 9.61E-07 | 96.3133118 | 254.374132 | 227.789891 | 248.219965 |
| -5.2286738 | 1.71E-07 | 9.64E-07 | 103.280658 | 220.547785 | 196.727633 | 211.799429 |
| -5.2286324 | 1.71E-07 | 9.64E-07 | 96.7231557 | 242.737869 | 269.206235 | 223.005748 |

| | | | | | | |
|---|---|---|---|---|---|---|
| -5.2284541 | 1.71E-07 | 9.65E-07 | 109.018472 | 188.345102 | 215.134897 | 175.939208 |
| -5.2284462 | 1.71E-07 | 9.65E-07 | 86.8869025 | 331.498204 | 280.710775 | 503.724038 |
| -5.2280582 | 1.71E-07 | 9.67E-07 | 134.018949 | 202.416863 | 128.850847 | 239.815226 |
| -5.2279383 | 1.71E-07 | 9.67E-07 | 100.001907 | 202.146252 | 404.959806 | 481.3114 |
| -5.2277442 | 1.72E-07 | 9.68E-07 | 102.46097 | 220.277174 | 215.134897 | 200.032794 |
| -5.2276817 | 1.72E-07 | 9.68E-07 | 113.116911 | 218.382899 | 158.762651 | 197.79153 |
| -5.2275603 | 1.72E-07 | 9.69E-07 | 182.79037 | 409.163498 | 440.62388 | 425.840121 |
| -5.2275481 | 1.72E-07 | 9.69E-07 | 102.870814 | 209.723354 | 247.347609 | 197.79153 |
| -5.2274522 | 1.72E-07 | 9.69E-07 | 118.444881 | 175.085174 | 155.311289 | 160.250361 |
| -5.2272842 | 1.72E-07 | 9.70E-07 | 84.0179954 | 377.502037 | 293.365769 | 322.18167 |
| -5.2271784 | 1.72E-07 | 9.70E-07 | 134.018949 | 141.529437 | 151.859927 | 218.52322 |
| -5.2270812 | 1.72E-07 | 9.70E-07 | 112.297223 | 350.170348 | 171.417645 | 256.624704 |
| -5.2269473 | 1.72E-07 | 9.71E-07 | 114.346443 | 173.190899 | 164.514921 | 172.577312 |
| -5.2261219 | 1.73E-07 | 9.75E-07 | 136.887856 | 225.418779 | 239.294431 | 124.950457 |
| -5.2257995 | 1.73E-07 | 9.77E-07 | 86.8869025 | 456.790995 | 317.525303 | 276.796079 |
| -5.2257604 | 1.73E-07 | 9.77E-07 | 163.527708 | 208.640911 | 120.79767 | 149.604358 |
| -5.2256363 | 1.74E-07 | 9.77E-07 | 126.641759 | 175.355785 | 135.753571 | 163.051941 |
| -5.225516 | 1.74E-07 | 9.78E-07 | 120.084257 | 193.486707 | 146.107657 | 170.336048 |
| -5.2254735 | 1.74E-07 | 9.78E-07 | 145.494577 | 126.375234 | 171.417645 | 145.121831 |
| -5.2253639 | 1.74E-07 | 9.78E-07 | 80.7392443 | 448.943283 | 312.923487 | 346.835572 |
| -5.2250212 | 1.74E-07 | 9.80E-07 | 106.559409 | 176.438228 | 228.940345 | 268.951655 |
| -5.2247131 | 1.74E-07 | 9.81E-07 | 115.985818 | 309.308121 | 189.824909 | 225.247011 |
| -5.2246747 | 1.74E-07 | 9.81E-07 | 131.150042 | 185.638994 | 187.524001 | 128.312352 |
| -5.2243231 | 1.75E-07 | 9.83E-07 | 98.3625312 | 281.1646 | 215.134897 | 230.289855 |
| -5.224025 | 1.75E-07 | 9.84E-07 | 118.854725 | 197.81648 | 147.258111 | 174.25826 |
| -5.2239112 | 1.75E-07 | 9.84E-07 | 131.559885 | 164.801964 | 148.408565 | 134.475828 |
| -5.2238886 | 1.75E-07 | 9.84E-07 | 288.530091 | 238.408096 | 157.612197 | 114.304454 |
| -5.2238936 | 1.75E-07 | 9.84E-07 | 105.739721 | 192.674875 | 187.524001 | 216.842272 |
| -5.2238407 | 1.75E-07 | 9.85E-07 | 108.198784 | 209.993965 | 195.577179 | 173.137628 |
| -5.2238017 | 1.75E-07 | 9.85E-07 | 101.641282 | 232.995881 | 200.178995 | 213.480376 |
| -5.2235847 | 1.75E-07 | 9.86E-07 | 111.887379 | 220.818396 | 159.913105 | 200.032794 |
| -5.2235601 | 1.76E-07 | 9.86E-07 | 106.969253 | 210.535186 | 180.621277 | 192.18837 |
| -5.223463 | 1.76E-07 | 9.86E-07 | 118.035037 | 189.156935 | 150.709473 | 166.974153 |
| -5.2234602 | 1.76E-07 | 9.86E-07 | 85.6573709 | 279.270324 | 349.738015 | 307.05314 |
| -5.2233512 | 1.76E-07 | 9.86E-07 | 100.001907 | 276.293606 | 200.178995 | 238.134278 |
| -5.2232181 | 1.76E-07 | 9.87E-07 | 159.019425 | 145.047378 | 120.79767 | 166.974153 |
| -5.2232376 | 1.76E-07 | 9.87E-07 | 110.657848 | 237.325653 | 169.116737 | 191.628054 |
| -5.223098 | 1.76E-07 | 9.87E-07 | 167.216303 | 303.354683 | 119.647216 | 187.705843 |
| -5.2228287 | 1.76E-07 | 9.88E-07 | 295.087594 | 215.676791 | 155.311289 | 172.577312 |
| -5.222215 | 1.77E-07 | 9.91E-07 | 111.477535 | 257.621462 | 161.063559 | 222.445432 |
| -5.2222122 | 1.77E-07 | 9.91E-07 | 90.5754975 | 278.999713 | 269.206235 | 259.426284 |
| -5.2222273 | 1.77E-07 | 9.91E-07 | 91.3951852 | 318.508887 | 249.648517 | 262.227864 |
| -5.2221403 | 1.77E-07 | 9.91E-07 | 141.805982 | 145.047378 | 161.063559 | 127.19172 |

| | | | | | | |
|---|---|---|---|---|---|---|
| -5.2220474 | 1.77E-07 | 9.91E-07 | 81.5589321 | 327.439043 | 311.773033 | 379.333897 |
| -5.222036 | 1.77E-07 | 9.91E-07 | 109.018472 | 175.626396 | 218.586259 | 182.102683 |
| -5.2219836 | 1.77E-07 | 9.92E-07 | 119.264569 | 150.730204 | 159.913105 | 165.853521 |
| -5.2217511 | 1.77E-07 | 9.93E-07 | 124.182696 | 197.004647 | 138.054479 | 233.651751 |
| -5.2213474 | 1.78E-07 | 9.94E-07 | 102.46097 | 217.841677 | 201.329449 | 202.834373 |
| -5.2213661 | 1.78E-07 | 9.94E-07 | 129.100822 | 202.146252 | 155.311289 | 138.958355 |
| -5.2212875 | 1.78E-07 | 9.95E-07 | 102.46097 | 279.270324 | 199.028541 | 214.040692 |
| -5.2211439 | 1.78E-07 | 9.95E-07 | 106.559409 | 183.203497 | 189.824909 | 201.713742 |
| -5.2211744 | 1.78E-07 | 9.95E-07 | 106.149565 | 186.450827 | 199.028541 | 190.507423 |
| -5.2210983 | 1.78E-07 | 9.95E-07 | 79.5097127 | 377.502037 | 320.976665 | 356.921259 |
| -5.2210498 | 1.78E-07 | 9.95E-07 | 95.493624 | 262.221845 | 216.285351 | 294.726189 |
| -5.2204074 | 1.79E-07 | 9.98E-07 | 84.8376831 | 285.764983 | 324.428027 | 314.337247 |
| -5.2202906 | 1.79E-07 | 9.99E-07 | 138.527231 | 222.17145 | 130.001301 | 161.931309 |
| -5.2191592 | 1.80E-07 | 1.00E-06 | 127.051603 | 145.85921 | 159.913105 | 228.608907 |
| -5.2189154 | 1.80E-07 | 1.01E-06 | 82.3786199 | 341.510803 | 295.666677 | 345.154624 |
| -5.2189392 | 1.80E-07 | 1.01E-06 | 106.149565 | 191.05121 | 203.630357 | 182.102683 |
| 5.2188734 | 1.80E-07 | 1.01E-06 | 35.2465737 | 18.401533 | 17.2568099 | 17.3697944 |
| -5.218409 | 1.80E-07 | 1.01E-06 | 110.248004 | 174.273342 | 171.417645 | 200.59311 |
| -5.2183743 | 1.81E-07 | 1.01E-06 | 168.855679 | 171.026012 | 120.79767 | 140.078987 |
| -5.2183124 | 1.81E-07 | 1.01E-06 | 104.100346 | 196.734036 | 190.975363 | 208.997849 |
| -5.2183073 | 1.81E-07 | 1.01E-06 | 117.625194 | 172.649677 | 148.408565 | 169.215416 |
| -5.2178996 | 1.81E-07 | 1.01E-06 | 94.6739363 | 268.175282 | 251.949425 | 223.005748 |
| -5.2176147 | 1.81E-07 | 1.01E-06 | 106.969253 | 305.51957 | 190.975363 | 205.635953 |
| -5.2168807 | 1.82E-07 | 1.02E-06 | 106.559409 | 320.944384 | 279.560321 | 172.577312 |
| -5.2168848 | 1.82E-07 | 1.02E-06 | 82.3786199 | 333.663091 | 289.914407 | 424.719489 |
| -5.2166719 | 1.82E-07 | 1.02E-06 | 89.7558097 | 270.340168 | 281.861229 | 257.745336 |
| -5.2165983 | 1.82E-07 | 1.02E-06 | 102.051126 | 199.440144 | 207.081719 | 207.877217 |
| -5.21646 | 1.82E-07 | 1.02E-06 | 110.248004 | 171.296623 | 201.329449 | 172.016996 |
| -5.2162821 | 1.83E-07 | 1.02E-06 | 81.968776 | 294.153917 | 331.330751 | 377.092633 |
| -5.215562 | 1.83E-07 | 1.02E-06 | 173.773805 | 223.253893 | 117.346308 | 150.72499 |
| -5.2154011 | 1.83E-07 | 1.02E-06 | 99.5920628 | 245.173366 | 213.984443 | 205.075637 |
| -5.2149731 | 1.84E-07 | 1.02E-06 | 135.658324 | 142.882491 | 140.355387 | 136.156776 |
| -5.2148454 | 1.84E-07 | 1.03E-06 | 159.019425 | 312.284839 | 169.116737 | 231.970803 |
| -5.2146591 | 1.84E-07 | 1.03E-06 | 175.003337 | 147.753485 | 120.79767 | 148.483726 |
| -5.2144992 | 1.84E-07 | 1.03E-06 | 104.100346 | 365.324552 | 185.223093 | 275.115131 |
| -5.2144936 | 1.84E-07 | 1.03E-06 | 102.46097 | 194.56915 | 201.329449 | 213.480376 |
| -5.2144464 | 1.84E-07 | 1.03E-06 | 110.248004 | 165.884408 | 230.090799 | 186.024895 |
| -5.2143747 | 1.84E-07 | 1.03E-06 | 99.5920628 | 286.576815 | 203.630357 | 224.126379 |
| -5.2143334 | 1.84E-07 | 1.03E-06 | 106.559409 | 209.452743 | 178.320369 | 186.585211 |
| -5.2140492 | 1.85E-07 | 1.03E-06 | 119.264569 | 449.484504 | 224.338529 | 188.826475 |
| -5.2140493 | 1.85E-07 | 1.03E-06 | 308.202598 | 409.163498 | 464.783414 | 456.097182 |
| -5.2138637 | 1.85E-07 | 1.03E-06 | 219.266476 | 294.965749 | 261.153057 | 231.410487 |
| -5.2137546 | 1.85E-07 | 1.03E-06 | 95.9034679 | 514.16048 | 317.525303 | 389.9799 |

| | | | | | | |
|---|---|---|---|---|---|---|
| -5.2133721 | 1.85E-07 | 1.03E-06 | 107.379097 | 211.076408 | 185.223093 | 174.25826 |
| -5.2133401 | 1.85E-07 | 1.03E-06 | 92.6247169 | 261.139402 | 256.551241 | 235.332698 |
| -5.213279 | 1.86E-07 | 1.03E-06 | 119.264569 | 168.319905 | 169.116737 | 142.880567 |
| -5.2131723 | 1.86E-07 | 1.03E-06 | 133.199261 | 285.223761 | 172.568099 | 197.79153 |
| -5.2126425 | 1.86E-07 | 1.03E-06 | 101.641282 | 223.524504 | 202.479903 | 196.110582 |
| -5.2125591 | 1.86E-07 | 1.04E-06 | 87.2967464 | 323.379881 | 251.949425 | 307.05314 |
| -5.2123232 | 1.86E-07 | 1.04E-06 | 99.5920628 | 194.839761 | 398.057082 | 317.699143 |
| -5.2122906 | 1.87E-07 | 1.04E-06 | 153.281611 | 158.036695 | 150.709473 | 184.904263 |
| -5.2121869 | 1.87E-07 | 1.04E-06 | 117.21535 | 169.672959 | 178.320369 | 145.682147 |
| -5.2118766 | 1.87E-07 | 1.04E-06 | 84.0179954 | 288.471091 | 295.666677 | 350.757784 |
| -5.2118104 | 1.87E-07 | 1.04E-06 | 365.990585 | 185.638994 | 190.975363 | 118.786981 |
| -5.2118218 | 1.87E-07 | 1.04E-06 | 124.182696 | 212.970683 | 170.267191 | 201.153426 |
| -5.211837 | 1.87E-07 | 1.04E-06 | 92.214873 | 286.035594 | 234.692615 | 252.702493 |
| -5.2117703 | 1.87E-07 | 1.04E-06 | 70.9029912 | 318.779498 | 799.565527 | 615.226912 |
| -5.2115504 | 1.87E-07 | 1.04E-06 | 100.411751 | 221.089007 | 213.984443 | 336.749885 |
| -5.2115803 | 1.87E-07 | 1.04E-06 | 106.149565 | 221.900839 | 173.718553 | 196.110582 |
| -5.2114853 | 1.87E-07 | 1.04E-06 | 124.182696 | 169.672959 | 139.204933 | 150.164674 |
| -5.2111102 | 1.88E-07 | 1.04E-06 | 134.018949 | 183.474108 | 196.727633 | 186.024895 |
| -5.211079 | 1.88E-07 | 1.04E-06 | 87.7065903 | 288.471091 | 262.303511 | 285.761134 |
| -5.2105154 | 1.88E-07 | 1.04E-06 | 100.821594 | 271.693222 | 188.674455 | 231.970803 |
| -5.2103631 | 1.88E-07 | 1.04E-06 | 102.46097 | 288.471091 | 193.276271 | 214.040692 |
| -5.2101721 | 1.89E-07 | 1.05E-06 | 124.182696 | 197.275258 | 135.753571 | 165.853521 |
| -5.2101526 | 1.89E-07 | 1.05E-06 | 412.302943 | 151.542036 | 156.461743 | 178.740788 |
| -5.2099529 | 1.89E-07 | 1.05E-06 | 97.1329995 | 244.090923 | 208.232173 | 229.729539 |
| -5.2095848 | 1.89E-07 | 1.05E-06 | 77.8703372 | 328.521486 | 353.189377 | 368.127578 |
| -5.2092371 | 1.90E-07 | 1.05E-06 | 122.133476 | 177.791282 | 141.505841 | 152.405938 |
| -5.2091145 | 1.90E-07 | 1.05E-06 | 104.510189 | 493.594061 | 352.038923 | 416.875066 |
| -5.2086573 | 1.90E-07 | 1.05E-06 | 508.616255 | 1424.22453 | 739.741919 | 1065.72093 |
| 5.20861473 | 1.90E-07 | 1.05E-06 | 22.5414134 | 24.6255809 | 23.0090799 | 30.8173772 |
| -5.2084068 | 1.90E-07 | 1.05E-06 | 128.281134 | 198.898923 | 126.549939 | 195.550266 |
| -5.2084203 | 1.90E-07 | 1.05E-06 | 84.4278393 | 309.578731 | 292.215315 | 290.243661 |
| -5.208321 | 1.91E-07 | 1.05E-06 | 142.62567 | 232.184049 | 165.665375 | 202.274057 |
| -5.2081032 | 1.91E-07 | 1.06E-06 | 103.280658 | 209.452743 | 189.824909 | 191.067738 |
| 5.20784121 | 1.91E-07 | 1.06E-06 | 0 | 0 | 0 | 0.56031595 |
| -5.2078473 | 1.91E-07 | 1.06E-06 | 155.33083 | 550.692936 | 473.987046 | 522.214464 |
| -5.2076393 | 1.91E-07 | 1.06E-06 | 83.6081515 | 294.695138 | 306.020763 | 304.25156 |
| -5.2073789 | 1.92E-07 | 1.06E-06 | 87.7065903 | 416.7406 | 295.666677 | 256.064389 |
| -5.2072703 | 1.92E-07 | 1.06E-06 | 100.411751 | 274.39933 | 204.780811 | 204.515321 |
| -5.2069316 | 1.92E-07 | 1.06E-06 | 122.133476 | 213.241294 | 141.505841 | 165.293205 |
| -5.2065897 | 1.92E-07 | 1.06E-06 | 152.461923 | 148.565318 | 217.435805 | 196.670898 |
| -5.2059876 | 1.93E-07 | 1.07E-06 | 75.4112739 | 358.829893 | 369.295733 | 364.765683 |
| -5.2057142 | 1.93E-07 | 1.07E-06 | 163.117864 | 218.112288 | 106.992222 | 255.504073 |
| -5.2056172 | 1.93E-07 | 1.07E-06 | 99.1822189 | 234.890156 | 193.276271 | 225.247011 |

| | | | | | | |
|---|---|---|---|---|---|---|
| -5.2055725 | 1.93E-07 | 1.07E-06 | 99.5920628 | 288.20048 | 215.134897 | 202.834373 |
| -5.2054146 | 1.94E-07 | 1.07E-06 | 130.330354 | 153.165701 | 165.665375 | 290.803977 |
| -5.2053324 | 1.94E-07 | 1.07E-06 | 70.0833035 | 386.161582 | 429.11934 | 430.322649 |
| -5.2052033 | 1.94E-07 | 1.07E-06 | 106.969253 | 196.463426 | 169.116737 | 187.145527 |
| -5.2048812 | 1.94E-07 | 1.07E-06 | 195.085687 | 188.345102 | 292.215315 | 105.899714 |
| -5.2047096 | 1.94E-07 | 1.07E-06 | 140.986295 | 191.05121 | 120.79767 | 156.32815 |
| -5.2044823 | 1.95E-07 | 1.07E-06 | 83.1983076 | 309.578731 | 289.914407 | 307.613456 |
| -5.2043974 | 1.95E-07 | 1.07E-06 | 109.428316 | 255.997797 | 156.461743 | 256.064389 |
| -5.2041313 | 1.95E-07 | 1.07E-06 | 93.8542485 | 219.735953 | 264.604419 | 239.25491 |
| -5.2035662 | 1.96E-07 | 1.08E-06 | 105.329877 | 302.27224 | 176.019461 | 228.048591 |
| 5.20349609 | 1.96E-07 | 1.08E-06 | 7.78703372 | 4.32977247 | 2.30090799 | 2.24126379 |
| -5.2026882 | 1.96E-07 | 1.08E-06 | 114.346443 | 181.038611 | 148.408565 | 169.775732 |
| -5.2022009 | 1.97E-07 | 1.09E-06 | 91.8050291 | 280.082156 | 218.586259 | 282.399238 |
| -5.2021488 | 1.97E-07 | 1.09E-06 | 90.5754975 | 247.879474 | 256.551241 | 250.461229 |
| -5.2017478 | 1.97E-07 | 1.09E-06 | 216.807412 | 106.891258 | 166.815829 | 267.270707 |
| -5.2009197 | 1.98E-07 | 1.09E-06 | 125.412227 | 141.800048 | 138.054479 | 165.293205 |
| -5.2007993 | 1.98E-07 | 1.09E-06 | 111.067691 | 255.185965 | 151.859927 | 235.893014 |
| -5.200797 | 1.98E-07 | 1.09E-06 | 102.051126 | 267.634061 | 184.072639 | 212.359745 |
| -5.2008422 | 1.98E-07 | 1.09E-06 | 111.067691 | 178.332503 | 170.267191 | 158.009098 |
| -5.2003926 | 1.99E-07 | 1.09E-06 | 97.5428434 | 209.723354 | 215.134897 | 216.842272 |
| -5.2001716 | 1.99E-07 | 1.10E-06 | 79.9195566 | 392.38563 | 304.870309 | 313.216615 |
| -5.1995815 | 2.00E-07 | 1.10E-06 | 214.758193 | 145.047378 | 112.744492 | 206.196269 |
| -5.1994387 | 2.00E-07 | 1.10E-06 | 1094.693 | 1447.76767 | 1654.35285 | 1739.2207 |
| -5.1992098 | 2.00E-07 | 1.10E-06 | 94.2640924 | 232.184049 | 216.285351 | 252.142177 |
| -5.1991538 | 2.00E-07 | 1.10E-06 | 91.8050291 | 222.17145 | 269.206235 | 257.18502 |
| -5.1990686 | 2.00E-07 | 1.10E-06 | 117.21535 | 197.81648 | 139.204933 | 181.542367 |
| -5.198107 | 2.01E-07 | 1.11E-06 | 87.2967464 | 307.684456 | 242.745793 | 288.002398 |
| -5.198056 | 2.01E-07 | 1.11E-06 | 237.299606 | 183.203497 | 144.957203 | 165.853521 |
| -5.1980204 | 2.01E-07 | 1.11E-06 | 154.101299 | 217.841677 | 108.142676 | 227.488275 |
| -5.1978635 | 2.02E-07 | 1.11E-06 | 104.920033 | 208.3703 | 170.267191 | 192.748686 |
| -5.1977524 | 2.02E-07 | 1.11E-06 | 107.78894 | 185.909605 | 161.063559 | 203.955005 |
| -5.1976232 | 2.02E-07 | 1.11E-06 | 103.280658 | 230.289773 | 172.568099 | 208.997849 |
| -5.1974791 | 2.02E-07 | 1.11E-06 | 146.724109 | 240.302372 | 158.762651 | 206.196269 |
| -5.1973857 | 2.02E-07 | 1.11E-06 | 83.1983076 | 339.616528 | 287.613499 | 281.838922 |
| -5.1971222 | 2.02E-07 | 1.11E-06 | 71.722679 | 453.814277 | 325.578481 | 483.552664 |
| -5.1968379 | 2.03E-07 | 1.11E-06 | 104.920033 | 251.668025 | 167.966283 | 213.480376 |
| -5.1964156 | 2.03E-07 | 1.11E-06 | 115.985818 | 244.902755 | 156.461743 | 169.775732 |
| -5.1963089 | 2.03E-07 | 1.12E-06 | 241.398045 | 149.918372 | 125.399485 | 151.845622 |
| -5.1958346 | 2.04E-07 | 1.12E-06 | 97.1329995 | 211.617629 | 203.630357 | 249.900913 |
| -5.1958274 | 2.04E-07 | 1.12E-06 | 93.4444046 | 279.811546 | 213.984443 | 240.375542 |
| -5.1952818 | 2.04E-07 | 1.12E-06 | 158.609582 | 134.764168 | 194.426725 | 122.709193 |
| -5.1948024 | 2.05E-07 | 1.12E-06 | 95.9034679 | 410.516552 | 211.683535 | 273.994499 |
| -5.1948305 | 2.05E-07 | 1.12E-06 | 104.510189 | 181.309222 | 178.320369 | 224.126379 |

| | | | | | | |
|---|---|---|---|---|---|---|
| -5.1947358 | 2.05E-07 | 1.12E-06 | 103.690502 | 186.450827 | 212.833989 | 176.499524 |
| 5.19432973 | 2.05E-07 | 1.13E-06 | 37.705637 | 21.6488623 | 25.3099879 | 16.8094785 |
| -5.1942706 | 2.06E-07 | 1.13E-06 | 176.232868 | 288.20048 | 120.79767 | 155.207518 |
| -5.1940301 | 2.06E-07 | 1.13E-06 | 63.1159575 | 432.706636 | 587.881992 | 457.217814 |
| -5.1938199 | 2.06E-07 | 1.13E-06 | 3496.78798 | 468.427259 | 142.656295 | 286.32145 |
| -5.1936678 | 2.06E-07 | 1.13E-06 | 81.5589321 | 271.152001 | 346.286653 | 327.78483 |
| -5.1934735 | 2.06E-07 | 1.13E-06 | 106.149565 | 282.247043 | 241.595339 | 259.9866 |
| -5.1933964 | 2.06E-07 | 1.13E-06 | 100.001907 | 213.782516 | 185.223093 | 210.678797 |
| -5.1933514 | 2.07E-07 | 1.13E-06 | 256.152425 | 377.772648 | 364.693917 | 405.108431 |
| -5.1931703 | 2.07E-07 | 1.13E-06 | 97.9526873 | 264.657342 | 207.081719 | 201.713742 |
| -5.1930963 | 2.07E-07 | 1.13E-06 | 90.9853413 | 238.408096 | 235.843069 | 256.624704 |
| -5.192953 | 2.07E-07 | 1.13E-06 | 78.6900249 | 389.679522 | 331.330751 | 295.286505 |
| -5.192645 | 2.07E-07 | 1.13E-06 | 98.7723751 | 195.922204 | 251.949425 | 202.834373 |
| -5.1925116 | 2.07E-07 | 1.13E-06 | 95.493624 | 245.714587 | 253.099879 | 196.670898 |
| -5.1919634 | 2.08E-07 | 1.14E-06 | 122.54332 | 189.968767 | 131.151755 | 165.853521 |
| -5.1919897 | 2.08E-07 | 1.14E-06 | 96.7231557 | 216.759234 | 228.940345 | 200.032794 |
| -5.191812 | 2.08E-07 | 1.14E-06 | 200.003813 | 147.212264 | 195.577179 | 110.942558 |
| -5.1917985 | 2.08E-07 | 1.14E-06 | 98.3625312 | 325.544767 | 203.630357 | 223.005748 |
| -5.1917035 | 2.08E-07 | 1.14E-06 | 105.739721 | 171.567234 | 177.169915 | 184.904263 |
| -5.1902698 | 2.10E-07 | 1.15E-06 | 108.608628 | 169.402348 | 167.966283 | 166.413837 |
| -5.1901165 | 2.10E-07 | 1.15E-06 | 97.9526873 | 341.510803 | 184.072639 | 302.010296 |
| -5.1897616 | 2.11E-07 | 1.15E-06 | 100.001907 | 206.205414 | 217.435805 | 182.662999 |
| -5.1896395 | 2.11E-07 | 1.15E-06 | 109.018472 | 189.698156 | 153.010381 | 213.480376 |
| -5.189063 | 2.11E-07 | 1.15E-06 | 187.298653 | 335.286755 | 98.9390436 | 246.539017 |
| -5.188959 | 2.11E-07 | 1.15E-06 | 79.9195566 | 310.390564 | 297.967585 | 328.345146 |
| -5.1889372 | 2.11E-07 | 1.15E-06 | 95.0837801 | 248.691306 | 195.577179 | 257.18502 |
| -5.1887578 | 2.12E-07 | 1.16E-06 | 115.16613 | 373.442875 | 149.559019 | 402.306851 |
| -5.1887132 | 2.12E-07 | 1.16E-06 | 158.609582 | 151.542036 | 116.195854 | 143.440883 |
| -5.1884355 | 2.12E-07 | 1.16E-06 | 108.608628 | 201.33442 | 164.514921 | 164.732889 |
| -5.1882151 | 2.12E-07 | 1.16E-06 | 87.2967464 | 296.589414 | 243.896247 | 261.667548 |
| -5.1879636 | 2.13E-07 | 1.16E-06 | 117.625194 | 135.576 | 173.718553 | 169.775732 |
| -5.1876486 | 2.13E-07 | 1.16E-06 | 113.116911 | 171.837845 | 143.806749 | 180.982051 |
| -5.1873015 | 2.13E-07 | 1.16E-06 | 103.280658 | 202.416863 | 169.116737 | 237.573962 |
| -5.1867919 | 2.14E-07 | 1.17E-06 | 102.46097 | 231.642827 | 193.276271 | 175.378892 |
| -5.1866836 | 2.14E-07 | 1.17E-06 | 104.920033 | 252.209246 | 162.214013 | 217.402588 |
| -5.186345 | 2.14E-07 | 1.17E-06 | 279.513526 | 124.210348 | 148.408565 | 170.336048 |
| -5.1860666 | 2.15E-07 | 1.17E-06 | 356.974019 | 185.638994 | 97.7885896 | 276.796079 |
| -5.1858652 | 2.15E-07 | 1.17E-06 | 85.6573709 | 285.764983 | 266.905327 | 257.745336 |
| -5.1856488 | 2.15E-07 | 1.17E-06 | 95.0837801 | 258.703905 | 210.533081 | 210.118481 |
| -5.1855076 | 2.15E-07 | 1.17E-06 | 89.7558097 | 367.760049 | 243.896247 | 245.418385 |
| -5.1854257 | 2.16E-07 | 1.17E-06 | 239.348826 | 213.241294 | 100.089498 | 173.137628 |
| -5.1851463 | 2.16E-07 | 1.17E-06 | 104.920033 | 162.637078 | 196.727633 | 190.507423 |
| -5.1849311 | 2.16E-07 | 1.18E-06 | 92.214873 | 323.10927 | 208.232173 | 262.227864 |

| | | | | | | |
|---|---|---|---|---|---|---|
| -5.1848831 | 2.16E-07 | 1.18E-06 | 108.608628 | 152.083258 | 189.824909 | 182.102683 |
| -5.1848214 | 2.16E-07 | 1.18E-06 | 101.231438 | 205.393581 | 174.869007 | 240.935858 |
| -5.1847215 | 2.16E-07 | 1.18E-06 | 127.871291 | 269.257725 | 130.001301 | 196.670898 |
| -5.1841917 | 2.17E-07 | 1.18E-06 | 115.575974 | 155.601198 | 143.806749 | 156.888466 |
| -5.1840693 | 2.17E-07 | 1.18E-06 | 169.265522 | 231.372216 | 103.54086 | 185.464579 |
| -5.18346 | 2.18E-07 | 1.18E-06 | 121.723632 | 223.253893 | 147.258111 | 149.604358 |
| -5.1830074 | 2.18E-07 | 1.19E-06 | 107.78894 | 189.156935 | 153.010381 | 210.118481 |
| -5.1822075 | 2.19E-07 | 1.19E-06 | 117.625194 | 169.402348 | 147.258111 | 141.199619 |
| -5.1814802 | 2.20E-07 | 1.20E-06 | 88.9361219 | 249.773749 | 283.011683 | 227.488275 |
| -5.1810902 | 2.21E-07 | 1.20E-06 | 82.3786199 | 284.68254 | 273.808051 | 304.811876 |
| -5.1801896 | 2.22E-07 | 1.20E-06 | 134.428793 | 143.694324 | 120.79767 | 160.810677 |
| -5.1801095 | 2.22E-07 | 1.20E-06 | 115.985818 | 160.742803 | 257.701695 | 160.250361 |
| -5.179899 | 2.22E-07 | 1.21E-06 | 94.6739363 | 220.547785 | 204.780811 | 289.683345 |
| -5.1793943 | 2.23E-07 | 1.21E-06 | 96.3133118 | 213.782516 | 212.833989 | 195.550266 |
| -5.1791718 | 2.23E-07 | 1.21E-06 | 103.690502 | 179.685557 | 176.019461 | 175.939208 |
| -5.1789461 | 2.23E-07 | 1.21E-06 | 88.1164342 | 316.885222 | 215.134897 | 309.294404 |
| -5.1787958 | 2.23E-07 | 1.21E-06 | 91.3951852 | 226.771833 | 225.488983 | 236.45333 |
| 5.17855239 | 2.24E-07 | 1.21E-06 | 190.16756 | 243.820312 | 222.037621 | 210.678797 |
| -5.1784127 | 2.24E-07 | 1.21E-06 | 67.6242402 | 390.761965 | 407.260714 | 415.754434 |
| -5.1780365 | 2.24E-07 | 1.22E-06 | 95.0837801 | 304.166516 | 240.444885 | 194.429634 |
| -5.1777912 | 2.25E-07 | 1.22E-06 | 193.036467 | 218.65351 | 96.6381356 | 302.570612 |
| -5.1776924 | 2.25E-07 | 1.22E-06 | 119.264569 | 323.10927 | 136.904025 | 268.391339 |
| -5.1774588 | 2.25E-07 | 1.22E-06 | 81.5589321 | 316.07339 | 265.754873 | 299.769033 |
| -5.1772865 | 2.25E-07 | 1.22E-06 | 100.411751 | 194.027929 | 376.198456 | 233.091435 |
| -5.1772623 | 2.25E-07 | 1.22E-06 | 114.346443 | 170.755402 | 293.365769 | 168.094785 |
| -5.1771039 | 2.25E-07 | 1.22E-06 | 60.6568942 | 375.337151 | 617.793796 | 609.063436 |
| -5.1768925 | 2.26E-07 | 1.22E-06 | 112.707067 | 147.482875 | 171.417645 | 152.405938 |
| -5.1769315 | 2.26E-07 | 1.22E-06 | 88.1164342 | 278.729103 | 222.037621 | 270.072287 |
| -5.1768925 | 2.26E-07 | 1.22E-06 | 316.809319 | 151.812647 | 124.249032 | 177.05984 |
| -5.177015 | 2.25E-07 | 1.22E-06 | 91.3951852 | 242.737869 | 223.188075 | 223.005748 |
| -5.1769166 | 2.26E-07 | 1.22E-06 | 100.001907 | 193.216096 | 247.347609 | 182.102683 |
| -5.1767737 | 2.26E-07 | 1.22E-06 | 153.691455 | 150.188982 | 144.957203 | 114.864769 |
| -5.176516 | 2.26E-07 | 1.22E-06 | 89.3459658 | 248.961917 | 240.444885 | 229.729539 |
| -5.1760174 | 2.27E-07 | 1.23E-06 | 102.051126 | 219.465342 | 219.736713 | 163.051941 |
| -5.1757119 | 2.27E-07 | 1.23E-06 | 99.1822189 | 235.160767 | 171.417645 | 240.935858 |
| -5.1755547 | 2.27E-07 | 1.23E-06 | 102.46097 | 166.425629 | 209.382627 | 254.943757 |
| -5.1752115 | 2.28E-07 | 1.23E-06 | 93.4444046 | 193.486707 | 401.508444 | 331.707042 |
| -5.173617 | 2.30E-07 | 1.24E-06 | 147.953641 | 188.345102 | 347.437107 | 127.752036 |
| -5.1735447 | 2.30E-07 | 1.24E-06 | 99.1822189 | 570.447522 | 589.032446 | 584.409534 |
| -5.1735517 | 2.30E-07 | 1.24E-06 | 100.411751 | 177.520671 | 224.338529 | 186.585211 |
| -5.1735074 | 2.30E-07 | 1.24E-06 | 111.067691 | 146.941653 | 161.063559 | 187.705843 |
| -5.1730625 | 2.30E-07 | 1.24E-06 | 91.8050291 | 313.367282 | 194.426725 | 308.173772 |
| -5.173078 | 2.30E-07 | 1.24E-06 | 122.54332 | 169.943569 | 124.249032 | 169.215416 |

| | | | | | | |
|---|---|---|---|---|---|---|
| -5.1727513 | 2.31E-07 | 1.25E-06 | 93.0345607 | 208.099689 | 220.887167 | 223.566063 |
| -5.172666 | 2.31E-07 | 1.25E-06 | 112.297223 | 151.542036 | 147.258111 | 169.215416 |
| -5.172661 | 2.31E-07 | 1.25E-06 | 91.3951852 | 209.452743 | 307.171217 | 239.25491 |
| -5.1724707 | 2.31E-07 | 1.25E-06 | 112.707067 | 199.169533 | 171.417645 | 140.639303 |
| -5.1723616 | 2.31E-07 | 1.25E-06 | 103.690502 | 175.355785 | 187.524001 | 165.853521 |
| -5.1723356 | 2.31E-07 | 1.25E-06 | 91.3951852 | 232.72527 | 213.984443 | 308.734088 |
| -5.1722439 | 2.31E-07 | 1.25E-06 | 88.9361219 | 256.268408 | 353.189377 | 223.566063 |
| -5.1719838 | 2.32E-07 | 1.25E-06 | 73.3620545 | 386.973414 | 355.490285 | 316.018195 |
| -5.1717446 | 2.32E-07 | 1.25E-06 | 224.594446 | 152.89509 | 104.691314 | 248.780281 |
| -5.1714284 | 2.32E-07 | 1.25E-06 | 70.0833035 | 345.028743 | 407.260714 | 381.575161 |
| -5.1714077 | 2.32E-07 | 1.25E-06 | 174.593493 | 174.543953 | 165.665375 | 104.218766 |
| -5.1713408 | 2.32E-07 | 1.25E-06 | 90.5754975 | 221.089007 | 239.294431 | 226.927959 |
| -5.1710273 | 2.33E-07 | 1.26E-06 | 100.411751 | 188.345102 | 173.718553 | 206.196269 |
| -5.170861 | 2.33E-07 | 1.26E-06 | 97.1329995 | 300.648576 | 179.470823 | 247.659649 |
| 5.17062139 | 2.33E-07 | 1.26E-06 | 0 | 0 | 0 | 0 |
| -5.1705418 | 2.33E-07 | 1.26E-06 | 100.821594 | 200.793198 | 172.568099 | 186.585211 |
| -5.1701436 | 2.34E-07 | 1.26E-06 | 90.5754975 | 238.137486 | 210.533081 | 250.461229 |
| -5.1693543 | 2.35E-07 | 1.27E-06 | 100.001907 | 181.309222 | 180.621277 | 220.204168 |
| -5.1693598 | 2.35E-07 | 1.27E-06 | 80.7392443 | 294.424528 | 280.710775 | 285.200818 |
| 5.16906895 | 2.35E-07 | 1.27E-06 | 103.690502 | 52.4984912 | 58.6731538 | 77.3236009 |
| -5.168636 | 2.36E-07 | 1.27E-06 | 285.661184 | 683.833439 | 639.652421 | 670.69819 |
| -5.1685232 | 2.36E-07 | 1.27E-06 | 106.149565 | 172.920288 | 164.514921 | 160.250361 |
| -5.1683907 | 2.36E-07 | 1.27E-06 | 102.870814 | 216.759234 | 258.852149 | 160.250361 |
| -5.1682964 | 2.36E-07 | 1.27E-06 | 119.674413 | 138.011497 | 139.204933 | 146.242463 |
| -5.1681554 | 2.36E-07 | 1.27E-06 | 126.641759 | 138.552719 | 348.587561 | 212.359745 |
| -5.1678927 | 2.37E-07 | 1.27E-06 | 86.0672148 | 270.069558 | 236.993523 | 254.943757 |
| -5.1679071 | 2.37E-07 | 1.27E-06 | 122.953164 | 138.282108 | 136.904025 | 137.277407 |
| -5.1678531 | 2.37E-07 | 1.27E-06 | 109.83816 | 214.594348 | 210.533081 | 219.083536 |
| -5.1677464 | 2.37E-07 | 1.27E-06 | 91.3951852 | 194.027929 | 281.861229 | 296.967453 |
| -5.1677218 | 2.37E-07 | 1.27E-06 | 102.051126 | 243.27909 | 158.762651 | 248.780281 |
| -5.1674108 | 2.37E-07 | 1.28E-06 | 87.7065903 | 484.663905 | 238.143977 | 291.364293 |
| -5.1673831 | 2.37E-07 | 1.28E-06 | 81.1490882 | 272.234444 | 287.613499 | 287.442082 |
| -5.1672314 | 2.38E-07 | 1.28E-06 | 71.722679 | 311.743618 | 415.313892 | 381.575161 |
| -5.1671714 | 2.38E-07 | 1.28E-06 | 99.1822189 | 178.603114 | 205.931265 | 186.024895 |
| -5.1670536 | 2.38E-07 | 1.28E-06 | 134.018949 | 147.482875 | 119.647216 | 141.199619 |
| -5.1668398 | 2.38E-07 | 1.28E-06 | 97.9526873 | 229.20733 | 179.470823 | 197.79153 |
| -5.166842 | 2.38E-07 | 1.28E-06 | 105.739721 | 153.977533 | 180.621277 | 174.818576 |
| -5.1667481 | 2.38E-07 | 1.28E-06 | 126.231915 | 240.302372 | 126.549939 | 309.85472 |
| -5.1666584 | 2.38E-07 | 1.28E-06 | 101.231438 | 173.461509 | 189.824909 | 177.620156 |
| -5.1660579 | 2.39E-07 | 1.28E-06 | 125.002383 | 242.196647 | 127.700393 | 177.620156 |
| -5.1658268 | 2.39E-07 | 1.28E-06 | 97.1329995 | 270.88139 | 189.824909 | 200.59311 |
| -5.165315 | 2.40E-07 | 1.29E-06 | 101.641282 | 173.461509 | 209.382627 | 172.577312 |
| -5.1645161 | 2.41E-07 | 1.29E-06 | 142.62567 | 110.138587 | 274.958505 | 257.18502 |

| | | | | | | |
|---|---|---|---|---|---|---|
| -5.1645164 | 2.41E-07 | 1.29E-06 | 130.740198 | 177.791282 | 148.408565 | 151.845622 |
| -5.1642102 | 2.41E-07 | 1.29E-06 | 94.2640924 | 280.893989 | 189.824909 | 226.927959 |
| -5.1636202 | 2.42E-07 | 1.30E-06 | 76.6408055 | 289.282923 | 332.481205 | 321.621355 |
| -5.16278 | 2.43E-07 | 1.30E-06 | 106.149565 | 226.230611 | 155.311289 | 177.620156 |
| -5.1625037 | 2.44E-07 | 1.31E-06 | 101.641282 | 185.638994 | 167.966283 | 178.180472 |
| -5.1624647 | 2.44E-07 | 1.31E-06 | 174.183649 | 144.776767 | 119.647216 | 130.553616 |
| -5.1624174 | 2.44E-07 | 1.31E-06 | 136.478012 | 313.367282 | 115.0454 | 275.675447 |
| -5.1621829 | 2.44E-07 | 1.31E-06 | 654.520676 | 266.551617 | 103.54086 | 226.367643 |
| -5.1619376 | 2.44E-07 | 1.31E-06 | 115.575974 | 238.137486 | 131.151755 | 237.573962 |
| -5.1619358 | 2.44E-07 | 1.31E-06 | 100.411751 | 209.993965 | 167.966283 | 186.585211 |
| -5.1611895 | 2.45E-07 | 1.31E-06 | 111.067691 | 366.136384 | 154.160835 | 244.297754 |
| -5.1610077 | 2.46E-07 | 1.31E-06 | 93.4444046 | 200.251977 | 219.736713 | 205.075637 |
| -5.1607398 | 2.46E-07 | 1.32E-06 | 98.7723751 | 248.691306 | 228.940345 | 281.838922 |
| -5.1605897 | 2.46E-07 | 1.32E-06 | 109.428316 | 182.391665 | 150.709473 | 151.845622 |
| -5.1605617 | 2.46E-07 | 1.32E-06 | 96.3133118 | 222.983282 | 241.595339 | 174.818576 |
| -5.1602949 | 2.47E-07 | 1.32E-06 | 82.3786199 | 250.314971 | 323.277573 | 267.270707 |
| -5.1598487 | 2.47E-07 | 1.32E-06 | 88.1164342 | 283.058875 | 208.232173 | 261.107232 |
| -5.1596307 | 2.47E-07 | 1.32E-06 | 133.199261 | 130.705006 | 130.001301 | 131.674248 |
| -5.1595525 | 2.48E-07 | 1.32E-06 | 89.3459658 | 235.431378 | 219.736713 | 225.807327 |
| -5.1593389 | 2.48E-07 | 1.32E-06 | 82.7884637 | 244.902755 | 327.879389 | 267.831023 |
| -5.1592932 | 2.48E-07 | 1.33E-06 | 95.0837801 | 327.709654 | 188.674455 | 237.573962 |
| -5.1592157 | 2.48E-07 | 1.33E-06 | 79.0998688 | 313.637893 | 264.604419 | 306.492824 |
| -5.1592177 | 2.48E-07 | 1.33E-06 | 97.9526873 | 244.902755 | 218.586259 | 169.215416 |
| -5.1591677 | 2.48E-07 | 1.33E-06 | 88.5262781 | 247.067641 | 212.833989 | 246.539017 |
| -5.1590912 | 2.48E-07 | 1.33E-06 | 147.953641 | 215.135569 | 118.496762 | 361.403787 |
| -5.1590424 | 2.48E-07 | 1.33E-06 | 91.3951852 | 238.408096 | 194.426725 | 256.624704 |
| -5.1586311 | 2.49E-07 | 1.33E-06 | 73.3620545 | 430.271139 | 302.569401 | 335.068937 |
| -5.1584151 | 2.49E-07 | 1.33E-06 | 102.46097 | 177.791282 | 159.913105 | 194.429634 |
| -5.1582423 | 2.49E-07 | 1.33E-06 | 104.100346 | 176.97945 | 225.488983 | 158.569413 |
| -5.1580777 | 2.49E-07 | 1.33E-06 | 98.3625312 | 194.298539 | 194.426725 | 173.697944 |
| -5.1577555 | 2.50E-07 | 1.33E-06 | 100.411751 | 160.472192 | 204.780811 | 207.877217 |
| -5.1574432 | 2.50E-07 | 1.34E-06 | 167.626147 | 156.14242 | 104.691314 | 161.370993 |
| -5.1573602 | 2.50E-07 | 1.34E-06 | 103.690502 | 188.886324 | 164.514921 | 161.931309 |
| -5.1572273 | 2.51E-07 | 1.34E-06 | 118.854725 | 190.780599 | 210.533081 | 124.950457 |
| -5.1572332 | 2.51E-07 | 1.34E-06 | 88.5262781 | 246.797031 | 213.984443 | 237.573962 |
| -5.1567379 | 2.51E-07 | 1.34E-06 | 95.493624 | 240.843593 | 185.223093 | 199.472478 |
| -5.1565508 | 2.52E-07 | 1.34E-06 | 83.6081515 | 271.693222 | 238.143977 | 273.434183 |
| -5.1561182 | 2.52E-07 | 1.34E-06 | 105.739721 | 171.837845 | 154.160835 | 161.931309 |
| -5.1560051 | 2.52E-07 | 1.34E-06 | 73.3620545 | 464.638708 | 283.011683 | 370.929158 |
| -5.1559169 | 2.52E-07 | 1.34E-06 | 152.461923 | 135.576 | 140.355387 | 116.545717 |
| -5.1559363 | 2.52E-07 | 1.34E-06 | 100.001907 | 203.769917 | 161.063559 | 214.040692 |
| -5.1557847 | 2.53E-07 | 1.35E-06 | 195.905375 | 133.411114 | 124.249032 | 137.277407 |
| 5.15567892 | 2.53E-07 | 1.35E-06 | 105.739721 | 62.7817008 | 57.5226998 | 58.8331746 |

| | | | | | | |
|---|---|---|---|---|---|---|
| -5.1554596 | 2.53E-07 | 1.35E-06 | 100.821594 | 167.237462 | 181.771731 | 193.309002 |
| -5.1553244 | 2.53E-07 | 1.35E-06 | 75.00143 | 319.59133 | 316.374849 | 310.415036 |
| -5.155153 | 2.53E-07 | 1.35E-06 | 136.478012 | 201.33442 | 131.151755 | 128.312352 |
| -5.1549365 | 2.54E-07 | 1.35E-06 | 68.8537718 | 360.994779 | 357.791193 | 381.575161 |
| -5.1547509 | 2.54E-07 | 1.35E-06 | 252.46383 | 162.637078 | 118.496762 | 141.199619 |
| -5.154521 | 2.54E-07 | 1.35E-06 | 86.0672148 | 266.281007 | 216.285351 | 273.994499 |
| -5.1537463 | 2.55E-07 | 1.36E-06 | 121.723632 | 139.09394 | 194.426725 | 133.915512 |
| -5.1527378 | 2.57E-07 | 1.37E-06 | 157.38005 | 146.941653 | 118.496762 | 126.631404 |
| -5.1523786 | 2.57E-07 | 1.37E-06 | 102.46097 | 156.954252 | 184.072639 | 175.939208 |
| -5.1522664 | 2.57E-07 | 1.37E-06 | 90.1656536 | 197.275258 | 235.843069 | 273.994499 |
| -5.152235 | 2.57E-07 | 1.37E-06 | 81.5589321 | 236.24321 | 319.826211 | 292.484925 |
| -5.1520697 | 2.58E-07 | 1.37E-06 | 72.5423667 | 326.085989 | 327.879389 | 342.353045 |
| -5.1515836 | 2.58E-07 | 1.37E-06 | 929.52592 | 1913.48882 | 1629.04286 | 1856.32674 |
| -5.1511002 | 2.59E-07 | 1.38E-06 | 102.46097 | 214.864959 | 150.709473 | 223.566063 |
| -5.1508883 | 2.59E-07 | 1.38E-06 | 88.1164342 | 251.397414 | 302.569401 | 204.515321 |
| -5.1509059 | 2.59E-07 | 1.38E-06 | 131.969729 | 247.067641 | 234.692615 | 113.183822 |
| -5.1502498 | 2.60E-07 | 1.38E-06 | 99.5920628 | 182.662276 | 174.869007 | 172.577312 |
| -5.1499433 | 2.61E-07 | 1.38E-06 | 96.3133118 | 189.427545 | 179.470823 | 226.927959 |
| -5.1499439 | 2.61E-07 | 1.38E-06 | 80.3294005 | 238.408096 | 353.189377 | 307.05314 |
| -5.1496782 | 2.61E-07 | 1.39E-06 | 125.822071 | 227.042444 | 233.542161 | 117.106033 |
| -5.1494426 | 2.61E-07 | 1.39E-06 | 95.0837801 | 298.483689 | 210.533081 | 188.826475 |
| -5.1488671 | 2.62E-07 | 1.39E-06 | 365.170897 | 197.81648 | 97.7885896 | 207.877217 |
| -5.1484803 | 2.63E-07 | 1.39E-06 | 91.3951852 | 225.68939 | 194.426725 | 217.962904 |
| -5.1484935 | 2.63E-07 | 1.39E-06 | 87.7065903 | 241.655426 | 253.099879 | 207.877217 |
| -5.1481686 | 2.63E-07 | 1.40E-06 | 75.8211178 | 345.299354 | 362.393009 | 262.227864 |
| -5.1476626 | 2.64E-07 | 1.40E-06 | 71.3128351 | 379.125702 | 312.923487 | 340.111781 |
| -5.1474335 | 2.64E-07 | 1.40E-06 | 84.4278393 | 211.88824 | 315.224395 | 300.329348 |
| -5.1474465 | 2.64E-07 | 1.40E-06 | 76.6408055 | 329.603929 | 307.171217 | 271.192919 |
| -5.1473443 | 2.64E-07 | 1.40E-06 | 102.46097 | 319.59133 | 165.665375 | 216.281956 |
| -5.1470714 | 2.65E-07 | 1.40E-06 | 100.411751 | 173.73212 | 219.736713 | 166.413837 |
| -5.1468877 | 2.65E-07 | 1.40E-06 | 112.297223 | 140.446994 | 165.665375 | 141.199619 |
| -5.1466845 | 2.65E-07 | 1.40E-06 | 568.043618 | 166.425629 | 143.806749 | 197.231214 |
| -5.1466022 | 2.65E-07 | 1.41E-06 | 105.329877 | 149.106539 | 163.364467 | 168.094785 |
| 5.14632686 | 2.66E-07 | 1.41E-06 | 6.1476582 | 7.8477126 | 9.20363196 | 10.646003 |
| -5.1463709 | 2.66E-07 | 1.41E-06 | 95.493624 | 185.097773 | 201.329449 | 185.464579 |
| -5.1463138 | 2.66E-07 | 1.41E-06 | 161.478489 | 166.425629 | 112.744492 | 129.432984 |
| -5.1460393 | 2.66E-07 | 1.41E-06 | 81.5589321 | 232.454659 | 317.525303 | 284.640502 |
| -5.145868 | 2.66E-07 | 1.41E-06 | 109.428316 | 235.160767 | 148.408565 | 338.430833 |
| -5.1455448 | 2.67E-07 | 1.41E-06 | 97.9526873 | 203.499306 | 173.718553 | 177.05984 |
| -5.1454283 | 2.67E-07 | 1.41E-06 | 114.346443 | 153.977533 | 154.160835 | 129.9933 |
| -5.1454426 | 2.67E-07 | 1.41E-06 | 105.739721 | 176.97945 | 215.134897 | 145.682147 |
| -5.1453863 | 2.67E-07 | 1.41E-06 | 95.0837801 | 175.355785 | 202.479903 | 216.281956 |
| -5.144776 | 2.68E-07 | 1.42E-06 | 92.214873 | 226.230611 | 190.975363 | 205.075637 |

| | | | | | | |
|---|---|---|---|---|---|---|
| -5.1446177 | 2.68E-07 | 1.42E-06 | 96.7231557 | 205.934803 | 166.815829 | 211.239113 |
| -5.1444632 | 2.68E-07 | 1.42E-06 | 191.806936 | 148.294707 | 124.249032 | 122.148877 |
| -5.1444592 | 2.68E-07 | 1.42E-06 | 102.051126 | 178.332503 | 157.612197 | 168.655101 |
| -5.1441883 | 2.69E-07 | 1.42E-06 | 111.887379 | 180.226779 | 132.302209 | 160.250361 |
| -5.1442104 | 2.69E-07 | 1.42E-06 | 91.8050291 | 215.676791 | 195.577179 | 205.075637 |
| -5.1436479 | 2.69E-07 | 1.42E-06 | 108.198784 | 188.615713 | 144.957203 | 153.52657 |
| -5.1436585 | 2.69E-07 | 1.42E-06 | 108.198784 | 184.827162 | 141.505841 | 158.009098 |
| -5.1432288 | 2.70E-07 | 1.43E-06 | 101.641282 | 170.755402 | 158.762651 | 173.697944 |
| -5.1427186 | 2.71E-07 | 1.43E-06 | 120.494101 | 206.746635 | 161.063559 | 168.094785 |
| -5.1424973 | 2.71E-07 | 1.43E-06 | 89.3459658 | 208.640911 | 209.382627 | 234.212067 |
| -5.1424845 | 2.71E-07 | 1.43E-06 | 35.6564176 | 930.089248 | 670.714679 | 963.183116 |
| -5.1424853 | 2.71E-07 | 1.43E-06 | 135.658324 | 191.05121 | 180.621277 | 108.140978 |
| -5.1419098 | 2.72E-07 | 1.44E-06 | 771.326182 | 230.289773 | 131.151755 | 200.59311 |
| -5.1413088 | 2.73E-07 | 1.44E-06 | 116.805506 | 124.751569 | 156.461743 | 175.378892 |
| -5.1412401 | 2.73E-07 | 1.44E-06 | 118.444881 | 127.728288 | 138.054479 | 151.285306 |
| -5.1411825 | 2.73E-07 | 1.44E-06 | 70.4931473 | 332.580648 | 394.60572 | 314.897563 |
| -5.1410664 | 2.73E-07 | 1.44E-06 | 81.968776 | 305.51957 | 248.498063 | 243.737438 |
| -5.1408339 | 2.74E-07 | 1.44E-06 | 100.821594 | 153.977533 | 186.373547 | 184.343947 |
| -5.1408078 | 2.74E-07 | 1.44E-06 | 122.54332 | 198.357701 | 148.408565 | 124.950457 |
| -5.1407805 | 2.74E-07 | 1.44E-06 | 86.0672148 | 222.44206 | 240.444885 | 230.850171 |
| -5.1404081 | 2.74E-07 | 1.44E-06 | 113.116911 | 135.846611 | 141.505841 | 169.215416 |
| -5.1399558 | 2.75E-07 | 1.45E-06 | 97.1329995 | 202.687474 | 166.815829 | 188.826475 |
| -5.1394612 | 2.76E-07 | 1.45E-06 | 108.198784 | 162.095857 | 140.355387 | 199.472478 |
| -5.1392195 | 2.76E-07 | 1.45E-06 | 104.100346 | 162.907689 | 148.408565 | 176.499524 |
| -5.1383548 | 2.77E-07 | 1.46E-06 | 95.0837801 | 318.779498 | 281.861229 | 268.951655 |
| -5.1380627 | 2.78E-07 | 1.46E-06 | 125.412227 | 235.160767 | 185.223093 | 209.558165 |
| -5.1377222 | 2.78E-07 | 1.46E-06 | 112.707067 | 137.199665 | 218.586259 | 158.569413 |
| -5.1374759 | 2.78E-07 | 1.47E-06 | 77.0506494 | 274.669941 | 285.312591 | 294.165873 |
| -5.137352 | 2.79E-07 | 1.47E-06 | 98.7723751 | 158.036695 | 188.674455 | 201.713742 |
| -5.1370875 | 2.79E-07 | 1.47E-06 | 398.368251 | 198.357701 | 135.753571 | 135.036144 |
| -5.1370924 | 2.79E-07 | 1.47E-06 | 90.1656536 | 227.583665 | 216.285351 | 192.18837 |
| -5.1367722 | 2.79E-07 | 1.47E-06 | 105.739721 | 221.359617 | 182.922185 | 142.320251 |
| -5.1365571 | 2.80E-07 | 1.47E-06 | 132.789417 | 195.922204 | 158.762651 | 219.083536 |
| -5.1364427 | 2.80E-07 | 1.47E-06 | 163.527708 | 135.034779 | 133.452663 | 115.425085 |
| -5.136209 | 2.80E-07 | 1.47E-06 | 97.1329995 | 194.56915 | 165.665375 | 186.585211 |
| -5.1358448 | 2.81E-07 | 1.48E-06 | 80.3294005 | 256.539019 | 253.099879 | 277.356395 |
| -5.1353948 | 2.82E-07 | 1.48E-06 | 107.78894 | 162.907689 | 148.408565 | 142.320251 |
| -5.1353061 | 2.82E-07 | 1.48E-06 | 271.316649 | 624.840289 | 585.581084 | 647.164921 |
| -5.1351855 | 2.82E-07 | 1.48E-06 | 117.21535 | 174.002731 | 197.878087 | 121.588561 |
| -5.1348054 | 2.82E-07 | 1.48E-06 | 136.887856 | 216.218013 | 234.692615 | 106.46003 |
| -5.1343099 | 2.83E-07 | 1.49E-06 | 211.889286 | 134.493557 | 125.399485 | 132.79488 |
| -5.1342205 | 2.83E-07 | 1.49E-06 | 128.281134 | 178.061893 | 132.302209 | 122.148877 |
| -5.133762 | 2.84E-07 | 1.49E-06 | 79.0998688 | 267.634061 | 253.099879 | 289.683345 |

| | | | | | | |
|---|---|---|---|---|---|---|
| -5.1336712 | 2.84E-07 | 1.49E-06 | 110.657848 | 139.09394 | 162.214013 | 139.518671 |
| -5.1335222 | 2.84E-07 | 1.49E-06 | 202.462877 | 136.117222 | 123.098578 | 130.553616 |
| -5.1330419 | 2.85E-07 | 1.50E-06 | 96.3133118 | 181.850444 | 174.869007 | 180.421735 |
| -5.1327957 | 2.85E-07 | 1.50E-06 | 119.264569 | 148.565318 | 121.948124 | 142.320251 |
| -5.1326429 | 2.86E-07 | 1.50E-06 | 80.7392443 | 273.587498 | 248.498063 | 250.461229 |
| -5.1326177 | 2.86E-07 | 1.50E-06 | 102.46097 | 278.999713 | 154.160835 | 203.394689 |
| -5.1324306 | 2.86E-07 | 1.50E-06 | 181.150995 | 117.174467 | 135.753571 | 132.79488 |
| -5.1323902 | 2.86E-07 | 1.50E-06 | 152.871767 | 139.09394 | 125.399485 | 117.106033 |
| -5.1323567 | 2.86E-07 | 1.50E-06 | 111.067691 | 140.176384 | 139.204933 | 146.802779 |
| -5.1321583 | 2.86E-07 | 1.50E-06 | 106.149565 | 207.017246 | 142.656295 | 165.293205 |
| -5.1320384 | 2.87E-07 | 1.50E-06 | 114.346443 | 359.371115 | 302.569401 | 289.123029 |
| -5.1319231 | 2.87E-07 | 1.50E-06 | 97.5428434 | 211.617629 | 156.461743 | 228.608907 |
| -5.1318354 | 2.87E-07 | 1.50E-06 | 85.247527 | 323.650492 | 203.630357 | 266.710392 |
| -5.1317119 | 2.87E-07 | 1.50E-06 | 100.821594 | 169.672959 | 155.311289 | 205.635953 |
| -5.1310897 | 2.88E-07 | 1.51E-06 | 88.9361219 | 204.581749 | 240.444885 | 201.153426 |
| -5.1309584 | 2.88E-07 | 1.51E-06 | 147.133953 | 148.565318 | 222.037621 | 110.382242 |
| -5.1307325 | 2.89E-07 | 1.51E-06 | 99.5920628 | 173.190899 | 159.913105 | 170.336048 |
| -5.1306376 | 2.89E-07 | 1.51E-06 | 122.133476 | 125.834012 | 133.452663 | 133.355196 |
| -5.1301052 | 2.90E-07 | 1.52E-06 | 104.100346 | 147.753485 | 178.320369 | 154.086886 |
| -5.1301078 | 2.90E-07 | 1.52E-06 | 104.510189 | 233.537102 | 149.559019 | 172.016996 |
| -5.1301339 | 2.90E-07 | 1.52E-06 | 262.709927 | 122.586683 | 144.957203 | 144.561515 |
| -5.1298754 | 2.90E-07 | 1.52E-06 | 86.8869025 | 234.348935 | 199.028541 | 249.340597 |
| -5.129663 | 2.90E-07 | 1.52E-06 | 99.1822189 | 208.640911 | 161.063559 | 170.336048 |
| -5.1295805 | 2.90E-07 | 1.52E-06 | 102.051126 | 237.055043 | 217.435805 | 255.504073 |
| -5.1294006 | 2.91E-07 | 1.52E-06 | 102.46097 | 179.414947 | 153.010381 | 156.32815 |
| -5.1290704 | 2.91E-07 | 1.52E-06 | 95.9034679 | 241.926037 | 162.214013 | 208.997849 |
| -5.128664 | 2.92E-07 | 1.53E-06 | 122.953164 | 201.875641 | 260.002603 | 118.786981 |
| -5.1286244 | 2.92E-07 | 1.53E-06 | 83.1983076 | 361.26539 | 234.692615 | 240.935858 |
| -5.128224 | 2.92E-07 | 1.53E-06 | 107.78894 | 142.34127 | 144.957203 | 151.845622 |
| -5.128157 | 2.93E-07 | 1.53E-06 | 109.018472 | 277.376049 | 133.452663 | 248.780281 |
| -5.1280434 | 2.93E-07 | 1.53E-06 | 100.001907 | 243.00848 | 151.859927 | 200.032794 |
| 5.1280231 | 2.93E-07 | 1.53E-06 | 0 | 0 | 0 | 0 |
| -5.1279074 | 2.93E-07 | 1.53E-06 | 104.100346 | 385.62036 | 484.341132 | 513.809725 |
| -5.1276889 | 2.93E-07 | 1.53E-06 | 138.117388 | 171.296623 | 118.496762 | 124.390141 |
| -5.1275229 | 2.94E-07 | 1.53E-06 | 128.281134 | 199.710755 | 113.894946 | 155.207518 |
| -5.1274897 | 2.94E-07 | 1.53E-06 | 121.313788 | 146.129821 | 120.79767 | 135.59646 |
| -5.1272181 | 2.94E-07 | 1.54E-06 | 103.690502 | 271.152001 | 156.461743 | 181.542367 |
| -5.1270786 | 2.94E-07 | 1.54E-06 | 102.051126 | 163.719521 | 158.762651 | 153.52657 |
| -5.1268895 | 2.95E-07 | 1.54E-06 | 107.78894 | 144.776767 | 161.063559 | 139.518671 |
| -5.1264081 | 2.95E-07 | 1.54E-06 | 95.0837801 | 231.101605 | 163.364467 | 275.675447 |
| -5.1262051 | 2.96E-07 | 1.54E-06 | 117.625194 | 229.20733 | 236.993523 | 238.134278 |
| -5.1258671 | 2.96E-07 | 1.54E-06 | 75.4112739 | 339.345917 | 245.046701 | 347.395888 |
| -5.1257406 | 2.96E-07 | 1.55E-06 | 99.1822189 | 178.873725 | 153.010381 | 183.783631 |

| | | | | | | |
|---|---|---|---|---|---|---|
| -5.1253802 | 2.97E-07 | 1.55E-06 | 93.0345607 | 180.768 | 201.329449 | 185.464579 |
| -5.1252775 | 2.97E-07 | 1.55E-06 | 99.1822189 | 169.672959 | 158.762651 | 170.896364 |
| -5.1251796 | 2.97E-07 | 1.55E-06 | 97.5428434 | 168.590515 | 171.417645 | 171.45668 |
| -5.1249721 | 2.98E-07 | 1.55E-06 | 289.759623 | 701.152529 | 62.1245158 | 615.226912 |
| -5.1247823 | 2.98E-07 | 1.55E-06 | 137.2977 | 104.455761 | 188.674455 | 201.713742 |
| -5.1246953 | 2.98E-07 | 1.55E-06 | 161.068645 | 133.681725 | 274.958505 | 122.709193 |
| -5.1244716 | 2.98E-07 | 1.55E-06 | 102.870814 | 161.825246 | 149.559019 | 156.32815 |
| -5.1243927 | 2.98E-07 | 1.55E-06 | 120.903945 | 229.20733 | 132.302209 | 145.682147 |
| -5.1235937 | 3.00E-07 | 1.56E-06 | 128.690978 | 198.628312 | 165.665375 | 166.974153 |
| -5.1235406 | 3.00E-07 | 1.56E-06 | 84.8376831 | 289.282923 | 195.577179 | 281.838922 |
| -5.1234816 | 3.00E-07 | 1.56E-06 | 83.6081515 | 225.68939 | 223.188075 | 265.029444 |
| -5.1233874 | 3.00E-07 | 1.56E-06 | 71.722679 | 330.415761 | 324.428027 | 288.002398 |
| -5.1228728 | 3.01E-07 | 1.57E-06 | 100.411751 | 155.059976 | 159.913105 | 173.697944 |
| -5.1228 | 3.01E-07 | 1.57E-06 | 124.592539 | 172.649677 | 181.771731 | 173.697944 |
| -5.1227845 | 3.01E-07 | 1.57E-06 | 97.5428434 | 176.97945 | 159.913105 | 178.740788 |
| -5.1227503 | 3.01E-07 | 1.57E-06 | 99.1822189 | 164.260743 | 276.108959 | 187.705843 |
| -5.1222503 | 3.02E-07 | 1.57E-06 | 81.1490882 | 267.634061 | 231.241253 | 247.099333 |
| -5.1221813 | 3.02E-07 | 1.57E-06 | 128.281134 | 224.877557 | 219.736713 | 109.26161 |
| -5.1219925 | 3.02E-07 | 1.57E-06 | 75.4112739 | 334.204312 | 243.896247 | 332.827673 |
| -5.1217935 | 3.03E-07 | 1.57E-06 | 108.198784 | 162.095857 | 134.603117 | 147.363094 |
| -5.1216758 | 3.03E-07 | 1.57E-06 | 93.0345607 | 239.49054 | 177.169915 | 189.947107 |
| -5.1215309 | 3.03E-07 | 1.57E-06 | 115.575974 | 214.864959 | 216.285351 | 119.907613 |
| -5.1215284 | 3.03E-07 | 1.57E-06 | 168.445835 | 527.149798 | 494.695218 | 451.054339 |
| -5.1214743 | 3.03E-07 | 1.57E-06 | 95.0837801 | 225.960001 | 233.542161 | 158.009098 |
| -5.1212778 | 3.03E-07 | 1.58E-06 | 64.3454891 | 379.666923 | 368.145279 | 365.325998 |
| -5.1212731 | 3.03E-07 | 1.58E-06 | 150.00286 | 113.115306 | 128.850847 | 138.398039 |
| -5.1210531 | 3.04E-07 | 1.58E-06 | 88.1164342 | 264.11612 | 223.188075 | 188.826475 |
| -5.1206473 | 3.04E-07 | 1.58E-06 | 143.445358 | 172.379066 | 135.753571 | 132.79488 |
| -5.1204509 | 3.05E-07 | 1.58E-06 | 93.4444046 | 206.205414 | 170.267191 | 191.067738 |
| -5.1198832 | 3.06E-07 | 1.59E-06 | 103.690502 | 137.470276 | 177.169915 | 174.25826 |
| -5.1198309 | 3.06E-07 | 1.59E-06 | 88.9361219 | 194.839761 | 205.931265 | 206.196269 |
| -5.119798 | 3.06E-07 | 1.59E-06 | 80.7392443 | 254.103522 | 234.692615 | 251.581861 |
| -5.1197365 | 3.06E-07 | 1.59E-06 | 93.0345607 | 190.509989 | 180.621277 | 182.662999 |
| -5.1194069 | 3.06E-07 | 1.59E-06 | 143.855202 | 165.613797 | 112.744492 | 125.510772 |
| -5.1191758 | 3.07E-07 | 1.59E-06 | 83.1983076 | 223.795114 | 230.090799 | 240.935858 |
| -5.119111 | 3.07E-07 | 1.59E-06 | 85.247527 | 395.632959 | 204.780811 | 272.313551 |
| -5.1188074 | 3.07E-07 | 1.59E-06 | 195.085687 | 208.640911 | 102.390406 | 137.277407 |
| -5.118496 | 3.08E-07 | 1.60E-06 | 125.002383 | 172.920288 | 113.894946 | 143.440883 |
| -5.1182011 | 3.08E-07 | 1.60E-06 | 88.1164342 | 248.961917 | 185.223093 | 314.337247 |
| -5.1180823 | 3.09E-07 | 1.60E-06 | 92.214873 | 213.511905 | 170.267191 | 256.064389 |
| -5.1179041 | 3.09E-07 | 1.60E-06 | 106.969253 | 131.516839 | 173.718553 | 160.810677 |
| -5.117318 | 3.10E-07 | 1.61E-06 | 208.610535 | 281.705821 | 87.4345036 | 496.43993 |
| -5.1172009 | 3.10E-07 | 1.61E-06 | 81.1490882 | 376.960815 | 218.586259 | 277.356395 |

| | | | | | | |
|---|---|---|---|---|---|---|
| -5.1169579 | 3.11E-07 | 1.61E-06 | 129.100822 | 119.068743 | 134.603117 | 193.309002 |
| -5.1165891 | 3.11E-07 | 1.61E-06 | 91.3951852 | 228.936719 | 169.116737 | 222.445432 |
| -5.1164191 | 3.11E-07 | 1.61E-06 | 108.198784 | 222.17145 | 162.214013 | 422.478225 |
| -5.1161393 | 3.12E-07 | 1.61E-06 | 84.8376831 | 251.938635 | 233.542161 | 200.59311 |
| -5.1159145 | 3.12E-07 | 1.62E-06 | 140.986295 | 137.199665 | 106.992222 | 149.604358 |
| -5.1156468 | 3.13E-07 | 1.62E-06 | 88.9361219 | 232.454659 | 179.470823 | 224.126379 |
| -5.1152996 | 3.13E-07 | 1.62E-06 | 95.0837801 | 267.904671 | 266.905327 | 255.504073 |
| -5.1152136 | 3.13E-07 | 1.62E-06 | 103.690502 | 146.129821 | 178.320369 | 146.802779 |
| -5.1151425 | 3.14E-07 | 1.62E-06 | 103.690502 | 165.343186 | 140.355387 | 155.207518 |
| -5.1149592 | 3.14E-07 | 1.62E-06 | 99.5920628 | 185.368384 | 223.188075 | 147.363094 |
| -5.1147084 | 3.14E-07 | 1.62E-06 | 109.428316 | 207.558468 | 132.302209 | 159.129729 |
| -5.1143002 | 3.15E-07 | 1.63E-06 | 111.887379 | 151.542036 | 153.010381 | 123.829825 |
| -5.1141238 | 3.15E-07 | 1.63E-06 | 235.250387 | 334.745534 | 309.472125 | 325.543566 |
| -5.114066 | 3.15E-07 | 1.63E-06 | 77.0506494 | 288.20048 | 423.36707 | 243.737438 |
| -5.1140432 | 3.15E-07 | 1.63E-06 | 102.051126 | 197.275258 | 146.107657 | 159.690045 |
| -5.1139028 | 3.16E-07 | 1.63E-06 | 88.5262781 | 264.11612 | 225.488983 | 180.982051 |
| -5.113808 | 3.16E-07 | 1.63E-06 | 88.5262781 | 225.68939 | 196.727633 | 192.748686 |
| -5.1134885 | 3.16E-07 | 1.63E-06 | 76.2309617 | 309.03751 | 253.099879 | 267.831023 |
| -5.1132369 | 3.17E-07 | 1.64E-06 | 104.100346 | 192.945486 | 146.107657 | 149.044042 |
| -5.1130629 | 3.17E-07 | 1.64E-06 | 104.920033 | 349.629127 | 370.446187 | 140.078987 |
| -5.1130045 | 3.17E-07 | 1.64E-06 | 151.232392 | 163.719521 | 111.594038 | 122.709193 |
| -5.1124042 | 3.18E-07 | 1.64E-06 | 119.264569 | 203.499306 | 196.727633 | 210.678797 |
| -5.1120506 | 3.19E-07 | 1.64E-06 | 308.202598 | 166.69624 | 105.841768 | 166.413837 |
| -5.1105588 | 3.21E-07 | 1.66E-06 | 106.969253 | 228.395498 | 212.833989 | 128.872668 |
| -5.1105259 | 3.21E-07 | 1.66E-06 | 104.510189 | 166.966851 | 133.452663 | 186.024895 |
| -5.110368 | 3.22E-07 | 1.66E-06 | 69.6734596 | 295.23636 | 309.472125 | 361.964103 |
| -5.1100842 | 3.22E-07 | 1.66E-06 | 145.904421 | 135.034779 | 187.524001 | 107.580662 |
| -5.1095436 | 3.23E-07 | 1.67E-06 | 357.793707 | 744.450253 | 716.732839 | 744.659896 |
| -5.1094176 | 3.23E-07 | 1.67E-06 | 89.3459658 | 211.617629 | 190.975363 | 187.145527 |
| -5.1090353 | 3.24E-07 | 1.67E-06 | 98.7723751 | 195.651593 | 147.258111 | 238.694594 |
| -5.1090556 | 3.24E-07 | 1.67E-06 | 136.068168 | 135.30539 | 108.142676 | 144.001199 |
| -5.1089481 | 3.24E-07 | 1.67E-06 | 111.477535 | 168.861126 | 127.700393 | 139.518671 |
| -5.1088569 | 3.24E-07 | 1.67E-06 | 175.413181 | 119.068743 | 118.496762 | 140.078987 |
| -5.108265 | 3.25E-07 | 1.67E-06 | 91.3951852 | 172.379066 | 223.188075 | 189.947107 |
| -5.1080365 | 3.26E-07 | 1.68E-06 | 131.150042 | 135.576 | 111.594038 | 133.915512 |
| -5.107695 | 3.26E-07 | 1.68E-06 | 70.4931473 | 301.731019 | 377.34891 | 283.51987 |
| -5.1076689 | 3.26E-07 | 1.68E-06 | 89.3459658 | 230.019162 | 182.922185 | 193.869318 |
| -5.1075977 | 3.26E-07 | 1.68E-06 | 341.809796 | 454.626109 | 403.809352 | 430.882964 |
| -5.1072457 | 3.27E-07 | 1.68E-06 | 72.1325229 | 341.240193 | 271.507143 | 293.605557 |
| -5.1071244 | 3.27E-07 | 1.68E-06 | 93.0345607 | 193.216096 | 169.116737 | 178.740788 |
| -5.1070814 | 3.27E-07 | 1.68E-06 | 114.756286 | 227.313054 | 127.700393 | 157.448782 |
| -5.1069297 | 3.27E-07 | 1.68E-06 | 183.200214 | 108.514922 | 135.753571 | 140.639303 |
| -5.1068118 | 3.28E-07 | 1.69E-06 | 121.723632 | 187.803881 | 110.443584 | 168.094785 |

| | | | | | | |
|---|---|---|---|---|---|---|
| -5.1063066 | 3.29E-07 | 1.69E-06 | 86.8869025 | 230.560384 | 200.178995 | 194.98995 |
| -5.1058747 | 3.29E-07 | 1.69E-06 | 100.411751 | 139.09394 | 205.931265 | 187.705843 |
| -5.1058638 | 3.29E-07 | 1.69E-06 | 97.1329995 | 169.672959 | 161.063559 | 159.690045 |
| -5.1055086 | 3.30E-07 | 1.70E-06 | 192.626624 | 182.121054 | 115.0454 | 115.425085 |
| -5.1053805 | 3.30E-07 | 1.70E-06 | 115.575974 | 126.375234 | 133.452663 | 130.553616 |
| -5.1047472 | 3.31E-07 | 1.70E-06 | 93.8542485 | 169.943569 | 167.966283 | 196.110582 |
| -5.1047068 | 3.31E-07 | 1.70E-06 | 112.707067 | 207.558468 | 120.79767 | 172.016996 |
| -5.1045851 | 3.32E-07 | 1.70E-06 | 93.4444046 | 169.672959 | 186.373547 | 172.016996 |
| -5.1045336 | 3.32E-07 | 1.70E-06 | 79.0998688 | 266.822228 | 247.347609 | 229.169223 |
| -5.1043975 | 3.32E-07 | 1.70E-06 | 102.051126 | 147.212264 | 149.559019 | 150.72499 |
| -5.1036457 | 3.33E-07 | 1.71E-06 | 489.763437 | 176.708839 | 121.948124 | 179.301104 |
| -5.1033128 | 3.34E-07 | 1.71E-06 | 93.0345607 | 180.768 | 165.665375 | 217.962904 |
| -5.1029336 | 3.34E-07 | 1.72E-06 | 327.46526 | 180.49739 | 207.081719 | 92.4521315 |
| -5.1028868 | 3.35E-07 | 1.72E-06 | 97.5428434 | 206.746635 | 200.178995 | 144.001199 |
| -5.102713 | 3.35E-07 | 1.72E-06 | 111.477535 | 122.857294 | 142.656295 | 153.52657 |
| -5.1023414 | 3.35E-07 | 1.72E-06 | 116.395662 | 156.954252 | 113.894946 | 164.732889 |
| -5.1023357 | 3.35E-07 | 1.72E-06 | 79.9195566 | 242.467258 | 223.188075 | 256.064389 |
| -5.1021497 | 3.36E-07 | 1.72E-06 | 98.7723751 | 181.579833 | 142.656295 | 176.499524 |
| -5.1020169 | 3.36E-07 | 1.72E-06 | 946.329519 | 1556.28259 | 1531.25427 | 1657.41458 |
| -5.1020246 | 3.36E-07 | 1.72E-06 | 706.161005 | 1049.96982 | 1066.47085 | 1081.40978 |
| -5.1017828 | 3.36E-07 | 1.72E-06 | 50.8206411 | 370.195546 | 761.600545 | 629.23481 |
| -5.1017308 | 3.37E-07 | 1.73E-06 | 93.0345607 | 163.448911 | 182.922185 | 187.145527 |
| -5.1016548 | 3.37E-07 | 1.73E-06 | 140.986295 | 225.960001 | 97.7885896 | 217.962904 |
| -5.1016473 | 3.37E-07 | 1.73E-06 | 97.9526873 | 167.778683 | 148.408565 | 169.775732 |
| -5.1015762 | 3.37E-07 | 1.73E-06 | 88.1164342 | 244.090923 | 184.072639 | 198.351846 |
| -5.1015416 | 3.37E-07 | 1.73E-06 | 84.4278393 | 207.829078 | 238.143977 | 203.955005 |
| -5.101404 | 3.37E-07 | 1.73E-06 | 200.413657 | 495.488337 | 272.657597 | 330.026094 |
| -5.1012437 | 3.37E-07 | 1.73E-06 | 131.150042 | 140.988216 | 127.700393 | 112.06319 |
| -5.1009697 | 3.38E-07 | 1.73E-06 | 131.969729 | 104.996982 | 199.028541 | 240.375542 |
| -5.1008667 | 3.38E-07 | 1.73E-06 | 113.526755 | 138.282108 | 124.249032 | 133.355196 |
| -5.1007891 | 3.38E-07 | 1.73E-06 | 110.657848 | 195.651593 | 260.002603 | 126.631404 |
| -5.1007347 | 3.38E-07 | 1.73E-06 | 124.592539 | 151.542036 | 146.107657 | 109.26161 |
| -5.1006759 | 3.38E-07 | 1.73E-06 | 92.214873 | 167.237462 | 182.922185 | 187.705843 |
| -5.1002739 | 3.39E-07 | 1.74E-06 | 188.938029 | 154.518755 | 105.841768 | 129.9933 |
| -5.0994696 | 3.41E-07 | 1.74E-06 | 100.821594 | 155.601198 | 148.408565 | 147.92341 |
| -5.0989484 | 3.42E-07 | 1.75E-06 | 96.7231557 | 186.180216 | 146.107657 | 186.024895 |
| -5.0988594 | 3.42E-07 | 1.75E-06 | 86.0672148 | 207.287857 | 201.329449 | 198.912162 |
| -5.0985051 | 3.42E-07 | 1.75E-06 | 93.4444046 | 207.287857 | 177.169915 | 161.370993 |
| -5.0980935 | 3.43E-07 | 1.75E-06 | 76.2309617 | 249.773749 | 314.073941 | 244.85807 |
| -5.0979939 | 3.43E-07 | 1.75E-06 | 89.3459658 | 174.273342 | 211.683535 | 192.748686 |
| -5.0979314 | 3.43E-07 | 1.75E-06 | 519.68204 | 1384.71536 | 972.133626 | 1152.56991 |
| -5.0978332 | 3.44E-07 | 1.76E-06 | 107.78894 | 145.317988 | 133.452663 | 134.475828 |
| -5.097831 | 3.44E-07 | 1.76E-06 | 101.641282 | 221.359617 | 138.054479 | 177.620156 |

| | | | | | | |
|---|---|---|---|---|---|---|
| -5.0977271 | 3.44E-07 | 1.76E-06 | 306.153378 | 149.918372 | 144.957203 | 121.588561 |
| -5.0972281 | 3.45E-07 | 1.76E-06 | 84.8376831 | 326.085989 | 197.878087 | 228.048591 |
| -5.0970993 | 3.45E-07 | 1.76E-06 | 107.379097 | 375.337151 | 329.029843 | 316.578511 |
| -5.0969062 | 3.45E-07 | 1.76E-06 | 84.8376831 | 228.395498 | 188.674455 | 226.927959 |
| -5.0968747 | 3.45E-07 | 1.76E-06 | 88.1164342 | 203.499306 | 200.178995 | 179.86142 |
| -5.0963575 | 3.46E-07 | 1.77E-06 | 75.00143 | 434.05969 | 243.896247 | 288.562714 |
| -5.0962801 | 3.46E-07 | 1.77E-06 | 116.805506 | 119.339354 | 189.824909 | 256.624704 |
| -5.0956694 | 3.48E-07 | 1.77E-06 | 83.6081515 | 391.573797 | 189.824909 | 295.846821 |
| -5.0956213 | 3.48E-07 | 1.77E-06 | 201.233345 | 158.577917 | 100.089498 | 141.759935 |
| -5.0954996 | 3.48E-07 | 1.77E-06 | 119.674413 | 151.812647 | 123.098578 | 121.028245 |
| -5.095399 | 3.48E-07 | 1.77E-06 | 93.4444046 | 206.476024 | 154.160835 | 192.18837 |
| -5.0953105 | 3.48E-07 | 1.78E-06 | 101.231438 | 228.124887 | 144.957203 | 165.293205 |
| -5.0953291 | 3.48E-07 | 1.78E-06 | 52.8698605 | 367.760049 | 624.696519 | 533.420783 |
| -5.0952416 | 3.48E-07 | 1.78E-06 | 151.642236 | 221.630228 | 416.464346 | 108.701294 |
| -5.0952155 | 3.48E-07 | 1.78E-06 | 81.5589321 | 215.40618 | 288.763953 | 218.52322 |
| -5.0944155 | 3.50E-07 | 1.78E-06 | 137.2977 | 188.615713 | 100.089498 | 230.850171 |
| -5.0943922 | 3.50E-07 | 1.78E-06 | 102.870814 | 355.582564 | 411.86253 | 410.71159 |
| -5.0939446 | 3.51E-07 | 1.79E-06 | 39.3450125 | 456.249774 | 1127.44492 | 842.715187 |
| -5.0928454 | 3.53E-07 | 1.80E-06 | 106.149565 | 149.37715 | 140.355387 | 130.553616 |
| -5.0927222 | 3.53E-07 | 1.80E-06 | 94.6739363 | 215.135569 | 158.762651 | 169.215416 |
| -5.0923016 | 3.54E-07 | 1.80E-06 | 117.21535 | 178.061893 | 128.850847 | 296.967453 |
| -5.0921201 | 3.54E-07 | 1.80E-06 | 89.7558097 | 237.596264 | 161.063559 | 251.021545 |
| -5.0917306 | 3.55E-07 | 1.81E-06 | 85.247527 | 186.721438 | 230.090799 | 208.997849 |
| -5.0917754 | 3.55E-07 | 1.81E-06 | 107.78894 | 161.825246 | 146.107657 | 124.390141 |
| -5.0917313 | 3.55E-07 | 1.81E-06 | 92.6247169 | 179.144336 | 166.815829 | 169.775732 |
| -5.0915806 | 3.55E-07 | 1.81E-06 | 82.3786199 | 412.952049 | 232.391707 | 228.048591 |
| -5.091597 | 3.55E-07 | 1.81E-06 | 81.5589321 | 205.934803 | 254.250333 | 223.005748 |
| -5.0915097 | 3.55E-07 | 1.81E-06 | 83.6081515 | 209.993965 | 199.028541 | 232.531119 |
| -5.0912991 | 3.56E-07 | 1.81E-06 | 96.3133118 | 155.059976 | 186.373547 | 155.207518 |
| -5.0906873 | 3.57E-07 | 1.81E-06 | 100.821594 | 213.241294 | 208.232173 | 210.118481 |
| -5.090604 | 3.57E-07 | 1.81E-06 | 97.5428434 | 169.672959 | 148.408565 | 155.767834 |
| -5.0903232 | 3.57E-07 | 1.82E-06 | 106.149565 | 189.427545 | 125.399485 | 161.931309 |
| -5.090108 | 3.58E-07 | 1.82E-06 | 127.871291 | 133.681725 | 155.311289 | 108.140978 |
| -5.0897638 | 3.59E-07 | 1.82E-06 | 178.282088 | 116.092024 | 169.116737 | 111.502874 |
| -5.0897333 | 3.59E-07 | 1.82E-06 | 356.564176 | 687.080768 | 615.492888 | 669.577559 |
| -5.089607 | 3.59E-07 | 1.82E-06 | 92.214873 | 188.074492 | 157.612197 | 188.266159 |
| -5.089312 | 3.59E-07 | 1.83E-06 | 110.657848 | 173.73212 | 131.151755 | 129.432984 |
| -5.0888961 | 3.60E-07 | 1.83E-06 | 144.67489 | 97.4198805 | 169.116737 | 201.713742 |
| -5.0885615 | 3.61E-07 | 1.83E-06 | 77.4604933 | 233.807713 | 265.754873 | 235.893014 |
| -5.0882669 | 3.61E-07 | 1.83E-06 | 101.641282 | 151.000815 | 163.364467 | 134.475828 |
| -5.0881796 | 3.62E-07 | 1.83E-06 | 168.855679 | 151.812647 | 121.948124 | 107.580662 |
| 5.08798881 | 3.62E-07 | 1.84E-06 | 18.4429746 | 18.1309222 | 9.20363196 | 19.0507423 |
| -5.0878714 | 3.62E-07 | 1.84E-06 | 88.9361219 | 286.847426 | 186.373547 | 190.507423 |

| | | | | | | |
|---|---|---|---|---|---|---|
| -5.0875263 | 3.63E-07 | 1.84E-06 | 90.9853413 | 183.474108 | 162.214013 | 201.153426 |
| -5.0872746 | 3.63E-07 | 1.84E-06 | 88.1164342 | 170.21418 | 201.329449 | 230.289855 |
| -5.0870374 | 3.64E-07 | 1.84E-06 | 82.7884637 | 201.875641 | 224.338529 | 211.799429 |
| -5.0869464 | 3.64E-07 | 1.85E-06 | 79.9195566 | 253.5623 | 228.940345 | 212.359745 |
| -5.0864949 | 3.65E-07 | 1.85E-06 | 103.690502 | 162.095857 | 134.603117 | 139.518671 |
| -5.0864653 | 3.65E-07 | 1.85E-06 | 95.9034679 | 183.744719 | 186.373547 | 142.880567 |
| -5.0862984 | 3.65E-07 | 1.85E-06 | 104.510189 | 139.905773 | 149.559019 | 131.674248 |
| -5.0862286 | 3.65E-07 | 1.85E-06 | 91.8050291 | 195.380983 | 239.294431 | 157.448782 |
| -5.0862187 | 3.65E-07 | 1.85E-06 | 83.6081515 | 216.218013 | 192.125817 | 222.445432 |
| -5.0861845 | 3.65E-07 | 1.85E-06 | 76.6408055 | 243.00848 | 276.108959 | 231.970803 |
| -5.0861285 | 3.65E-07 | 1.85E-06 | 109.018472 | 118.25691 | 176.019461 | 186.024895 |
| -5.0861161 | 3.65E-07 | 1.85E-06 | 171.314742 | 207.829078 | 319.826211 | 142.320251 |
| -5.0857023 | 3.66E-07 | 1.85E-06 | 107.78894 | 164.531354 | 140.355387 | 124.950457 |
| -5.085713 | 3.66E-07 | 1.85E-06 | 241.398045 | 132.05806 | 130.001301 | 125.510772 |
| -5.0855607 | 3.67E-07 | 1.86E-06 | 61.0667381 | 320.944384 | 401.508444 | 400.065587 |
| -5.0851664 | 3.67E-07 | 1.86E-06 | 92.214873 | 202.687474 | 180.621277 | 156.32815 |
| -5.0849151 | 3.68E-07 | 1.86E-06 | 328.284948 | 247.879474 | 123.098578 | 112.06319 |
| -5.0847859 | 3.68E-07 | 1.86E-06 | 212.708974 | 171.296623 | 100.089498 | 135.036144 |
| -5.0847435 | 3.68E-07 | 1.86E-06 | 111.067691 | 144.776767 | 164.514921 | 117.666349 |
| -5.0846822 | 3.68E-07 | 1.86E-06 | 572.961744 | 825.633487 | 944.52273 | 890.902358 |
| -5.0843167 | 3.69E-07 | 1.87E-06 | 98.3625312 | 150.188982 | 147.258111 | 154.647202 |
| -5.0842837 | 3.69E-07 | 1.87E-06 | 111.067691 | 125.563402 | 127.700393 | 156.888466 |
| -5.0840023 | 3.70E-07 | 1.87E-06 | 146.314265 | 197.004647 | 102.390406 | 137.837723 |
| -5.0837909 | 3.70E-07 | 1.87E-06 | 109.428316 | 186.180216 | 119.647216 | 155.767834 |
| -5.0834786 | 3.71E-07 | 1.87E-06 | 326.645572 | 505.500935 | 392.304812 | 423.598857 |
| -5.0830126 | 3.71E-07 | 1.88E-06 | 47.951734 | 457.873438 | 652.307415 | 484.673296 |
| -5.0827823 | 3.72E-07 | 1.88E-06 | 107.78894 | 155.871809 | 120.79767 | 146.802779 |
| -5.0826048 | 3.72E-07 | 1.88E-06 | 84.4278393 | 180.226779 | 247.347609 | 314.897563 |
| -5.081982 | 3.74E-07 | 1.89E-06 | 77.4604933 | 356.123785 | 253.099879 | 220.764484 |
| -5.0818054 | 3.74E-07 | 1.89E-06 | 95.9034679 | 239.49054 | 144.957203 | 296.967453 |
| 5.08174635 | 3.74E-07 | 1.89E-06 | 0 | 3.78855091 | 0 | 7.28410733 |
| -5.0815924 | 3.74E-07 | 1.89E-06 | 87.2967464 | 138.011497 | 144.957203 | 138.958355 |
| -5.081514 | 3.74E-07 | 1.89E-06 | 74.1817423 | 303.895905 | 300.268493 | 223.566063 |
| -5.0814467 | 3.75E-07 | 1.89E-06 | 94.6739363 | 164.801964 | 164.514921 | 152.405938 |
| -5.0814418 | 3.75E-07 | 1.89E-06 | 106.969253 | 136.658443 | 125.399485 | 155.207518 |
| -5.0812135 | 3.75E-07 | 1.89E-06 | 270.906805 | 218.65351 | 219.736713 | 79.0045488 |
| -5.0806963 | 3.76E-07 | 1.90E-06 | 163.527708 | 222.44206 | 98.9390436 | 141.759935 |
| -5.0804649 | 3.77E-07 | 1.90E-06 | 99.1822189 | 146.129821 | 142.656295 | 177.05984 |
| -5.0804135 | 3.77E-07 | 1.90E-06 | 86.8869025 | 194.027929 | 176.019461 | 202.834373 |
| -5.0804024 | 3.77E-07 | 1.90E-06 | 161.478489 | 156.41303 | 153.010381 | 95.2537113 |
| -5.0802519 | 3.77E-07 | 1.90E-06 | 156.560362 | 175.085174 | 126.549939 | 146.802779 |
| -5.0801749 | 3.77E-07 | 1.90E-06 | 94.6739363 | 156.954252 | 157.612197 | 162.491625 |
| -5.0799128 | 3.78E-07 | 1.90E-06 | 90.5754975 | 163.1783 | 174.869007 | 201.153426 |

| | | | | | | |
|---|---|---|---|---|---|---|
| -5.0797498 | 3.78E-07 | 1.90E-06 | 122.953164 | 148.565318 | 110.443584 | 127.752036 |
| -5.0790657 | 3.79E-07 | 1.91E-06 | 80.7392443 | 269.528336 | 202.479903 | 220.204168 |
| -5.0788768 | 3.80E-07 | 1.91E-06 | 214.348349 | 180.49739 | 106.992222 | 120.467929 |
| -5.0784256 | 3.81E-07 | 1.92E-06 | 143.855202 | 222.44206 | 132.302209 | 141.199619 |
| -5.0784177 | 3.81E-07 | 1.92E-06 | 74.5915861 | 400.503953 | 236.993523 | 267.270707 |
| -5.0782606 | 3.81E-07 | 1.92E-06 | 234.020855 | 192.674875 | 112.744492 | 114.864769 |
| -5.0782714 | 3.81E-07 | 1.92E-06 | 106.149565 | 165.072575 | 124.249032 | 142.320251 |
| -5.0777723 | 3.82E-07 | 1.92E-06 | 73.3620545 | 270.88139 | 262.303511 | 249.340597 |
| -5.0776762 | 3.82E-07 | 1.92E-06 | 122.54332 | 123.669126 | 199.028541 | 123.829825 |
| -5.0775997 | 3.82E-07 | 1.92E-06 | 95.0837801 | 166.966851 | 154.160835 | 152.966254 |
| -5.0775152 | 3.82E-07 | 1.92E-06 | 107.78894 | 259.786348 | 133.452663 | 166.974153 |
| -5.0774735 | 3.82E-07 | 1.92E-06 | 112.297223 | 130.975617 | 118.496762 | 155.207518 |
| -5.0772049 | 3.83E-07 | 1.93E-06 | 200.413657 | 419.446708 | 409.561622 | 421.917909 |
| -5.0771365 | 3.83E-07 | 1.93E-06 | 104.510189 | 164.801964 | 133.452663 | 133.355196 |
| -5.0769822 | 3.83E-07 | 1.93E-06 | 94.2640924 | 158.036695 | 159.913105 | 158.009098 |
| -5.0768304 | 3.84E-07 | 1.93E-06 | 88.5262781 | 236.24321 | 161.063559 | 285.761134 |
| -5.0766916 | 3.84E-07 | 1.93E-06 | 76.6408055 | 235.431378 | 257.701695 | 228.608907 |
| -5.0764005 | 3.85E-07 | 1.93E-06 | 89.3459658 | 161.013414 | 187.524001 | 205.075637 |
| -5.0763109 | 3.85E-07 | 1.93E-06 | 106.149565 | 139.635162 | 126.549939 | 136.717091 |
| -5.0761443 | 3.85E-07 | 1.93E-06 | 81.5589321 | 232.184049 | 197.878087 | 213.480376 |
| -5.0761126 | 3.85E-07 | 1.93E-06 | 134.838636 | 173.461509 | 186.373547 | 97.4949751 |
| -5.0761138 | 3.85E-07 | 1.93E-06 | 108.608628 | 132.869893 | 128.850847 | 128.312352 |
| -5.0756899 | 3.86E-07 | 1.94E-06 | 95.493624 | 320.132552 | 150.709473 | 223.566063 |
| -5.0757004 | 3.86E-07 | 1.94E-06 | 104.100346 | 136.387833 | 130.001301 | 159.129729 |
| -5.0751952 | 3.87E-07 | 1.94E-06 | 70.4931473 | 348.005462 | 261.153057 | 271.192919 |
| -5.0751038 | 3.87E-07 | 1.94E-06 | 82.3786199 | 284.141318 | 185.223093 | 232.531119 |
| -5.0748424 | 3.88E-07 | 1.95E-06 | 93.8542485 | 192.674875 | 148.408565 | 246.539017 |
| -5.0748039 | 3.88E-07 | 1.95E-06 | 84.4278393 | 207.829078 | 203.630357 | 183.223315 |
| -5.0746993 | 3.88E-07 | 1.95E-06 | 113.936599 | 209.452743 | 209.382627 | 211.799429 |
| -5.0746749 | 3.88E-07 | 1.95E-06 | 122.133476 | 110.679809 | 151.859927 | 127.19172 |
| 5.07445391 | 3.89E-07 | 1.95E-06 | 107.78894 | 96.6080482 | 105.841768 | 70.5998095 |
| -5.0742039 | 3.89E-07 | 1.95E-06 | 92.214873 | 179.414947 | 157.612197 | 163.612257 |
| -5.0741505 | 3.89E-07 | 1.95E-06 | 200.003813 | 116.903857 | 112.744492 | 151.285306 |
| -5.0738357 | 3.90E-07 | 1.95E-06 | 97.1329995 | 222.983282 | 211.683535 | 227.488275 |
| -5.0737085 | 3.90E-07 | 1.96E-06 | 658.619115 | 1211.79507 | 1035.4086 | 1160.41433 |
| -5.073339 | 3.91E-07 | 1.96E-06 | 197.134906 | 563.952864 | 452.12842 | 505.404986 |
| -5.0731633 | 3.91E-07 | 1.96E-06 | 70.0833035 | 278.729103 | 274.958505 | 282.399238 |
| -5.0730013 | 3.92E-07 | 1.96E-06 | 534.436419 | 408.622277 | 48.3190678 | 555.833421 |
| -5.0727254 | 3.92E-07 | 1.96E-06 | 80.3294005 | 217.300456 | 210.533081 | 216.281956 |
| -5.0726075 | 3.92E-07 | 1.97E-06 | 80.3294005 | 352.605845 | 193.276271 | 262.227864 |
| -5.0712374 | 3.95E-07 | 1.98E-06 | 159.839113 | 128.54012 | 104.691314 | 131.674248 |
| -5.0710067 | 3.96E-07 | 1.98E-06 | 111.477535 | 234.348935 | 116.195854 | 277.916711 |
| -5.0709319 | 3.96E-07 | 1.98E-06 | 91.3951852 | 155.330587 | 189.824909 | 168.655101 |

| | | | | | | |
|---|---|---|---|---|---|---|
| -5.0704499 | 3.97E-07 | 1.99E-06 | 92.214873 | 172.920288 | 150.709473 | 177.05984 |
| 5.06987726 | 3.98E-07 | 1.99E-06 | 152.871767 | 99.0435452 | 131.151755 | 129.9933 |
| -5.0696889 | 3.98E-07 | 1.99E-06 | 77.0506494 | 218.92412 | 234.692615 | 244.297754 |
| -5.0695246 | 3.99E-07 | 2.00E-06 | 150.00286 | 312.826061 | 90.8858656 | 260.546916 |
| -5.0683105 | 4.01E-07 | 2.01E-06 | 102.870814 | 161.013414 | 130.001301 | 136.717091 |
| -5.0681068 | 4.02E-07 | 2.01E-06 | 91.3951852 | 191.863042 | 166.815829 | 155.207518 |
| -5.0678542 | 4.02E-07 | 2.01E-06 | 115.985818 | 250.585581 | 111.594038 | 193.869318 |
| -5.0677187 | 4.03E-07 | 2.01E-06 | 87.2967464 | 182.121054 | 213.984443 | 168.094785 |
| -5.0674427 | 4.03E-07 | 2.02E-06 | 112.707067 | 155.330587 | 113.894946 | 133.915512 |
| -5.0674101 | 4.03E-07 | 2.02E-06 | 88.9361219 | 286.576815 | 158.762651 | 222.445432 |
| -5.0672834 | 4.04E-07 | 2.02E-06 | 86.0672148 | 182.662276 | 176.019461 | 203.394689 |
| -5.0670168 | 4.04E-07 | 2.02E-06 | 100.001907 | 192.404264 | 205.931265 | 126.631404 |
| -5.0660404 | 4.06E-07 | 2.03E-06 | 87.7065903 | 174.002731 | 172.568099 | 223.566063 |
| -5.0659587 | 4.06E-07 | 2.03E-06 | 83.1983076 | 266.551617 | 172.568099 | 239.25491 |
| -5.0658954 | 4.06E-07 | 2.03E-06 | 101.231438 | 133.140503 | 136.904025 | 161.370993 |
| -5.0656113 | 4.07E-07 | 2.03E-06 | 93.4444046 | 187.262659 | 277.259413 | 154.647202 |
| -5.0654518 | 4.07E-07 | 2.03E-06 | 142.215826 | 267.36345 | 96.6381356 | 188.826475 |
| -5.0649889 | 4.08E-07 | 2.04E-06 | 100.821594 | 189.968767 | 143.806749 | 136.156776 |
| -5.064636 | 4.09E-07 | 2.04E-06 | 80.7392443 | 188.074492 | 234.692615 | 223.005748 |
| -5.0644468 | 4.10E-07 | 2.04E-06 | 81.968776 | 225.148168 | 182.922185 | 217.962904 |
| -5.0644597 | 4.10E-07 | 2.04E-06 | 74.5915861 | 251.938635 | 227.789891 | 252.142177 |
| -5.0643947 | 4.10E-07 | 2.04E-06 | 65.5750208 | 346.923019 | 303.719855 | 284.080186 |
| -5.0640434 | 4.10E-07 | 2.05E-06 | 81.5589321 | 202.146252 | 192.125817 | 231.970803 |
| -5.0640174 | 4.11E-07 | 2.05E-06 | 99.1822189 | 180.49739 | 151.859927 | 132.234564 |
| -5.0632724 | 4.12E-07 | 2.06E-06 | 84.8376831 | 239.49054 | 178.320369 | 189.386791 |
| -5.0632682 | 4.12E-07 | 2.06E-06 | 82.7884637 | 207.558468 | 212.833989 | 181.542367 |
| -5.0631644 | 4.12E-07 | 2.06E-06 | 90.9853413 | 160.742803 | 162.214013 | 166.413837 |
| -5.0631053 | 4.12E-07 | 2.06E-06 | 96.3133118 | 196.463426 | 132.302209 | 206.756585 |
| -5.0621582 | 4.15E-07 | 2.07E-06 | 91.3951852 | 177.791282 | 147.258111 | 184.904263 |
| -5.0620212 | 4.15E-07 | 2.07E-06 | 91.8050291 | 156.41303 | 185.223093 | 155.207518 |
| -5.0619402 | 4.15E-07 | 2.07E-06 | 92.6247169 | 203.769917 | 140.355387 | 208.997849 |
| -5.0618781 | 4.15E-07 | 2.07E-06 | 148.773328 | 805.8789 | 659.210139 | 711.040939 |
| -5.0617904 | 4.15E-07 | 2.07E-06 | 85.6573709 | 174.543953 | 181.771731 | 201.153426 |
| -5.061467 | 4.16E-07 | 2.07E-06 | 111.067691 | 271.422611 | 227.789891 | 237.013646 |
| -5.0611409 | 4.17E-07 | 2.08E-06 | 74.5915861 | 256.268408 | 242.745793 | 226.367643 |
| -5.0609176 | 4.17E-07 | 2.08E-06 | 75.00143 | 245.985198 | 232.391707 | 233.651751 |
| -5.0607401 | 4.18E-07 | 2.08E-06 | 79.0998688 | 235.972599 | 190.975363 | 279.597658 |
| -5.0603156 | 4.19E-07 | 2.08E-06 | 73.3620545 | 221.089007 | 261.153057 | 303.130928 |
| -5.059908 | 4.19E-07 | 2.09E-06 | 113.116911 | 113.927138 | 180.621277 | 138.398039 |
| -5.0597268 | 4.20E-07 | 2.09E-06 | 80.7392443 | 235.431378 | 186.373547 | 216.842272 |
| -5.0595448 | 4.20E-07 | 2.09E-06 | 105.329877 | 139.635162 | 124.249032 | 183.223315 |
| -5.0591511 | 4.21E-07 | 2.09E-06 | 86.8869025 | 199.981366 | 162.214013 | 187.145527 |
| -5.0590254 | 4.21E-07 | 2.10E-06 | 88.1164342 | 224.606947 | 157.612197 | 188.266159 |

| | | | | | | |
|---|---|---|---|---|---|---|
| -5.0586992 | 4.22E-07 | 2.10E-06 | 80.3294005 | 190.780599 | 249.648517 | 210.118481 |
| -5.0586255 | 4.22E-07 | 2.10E-06 | 89.3459658 | 189.968767 | 161.063559 | 164.172573 |
| -5.0580839 | 4.23E-07 | 2.11E-06 | 1308.22166 | 5005.21697 | 4226.76798 | 4364.30092 |
| 5.05799639 | 4.24E-07 | 2.11E-06 | 50.8206411 | 44.109557 | 23.0090799 | 43.704644 |
| -5.0577833 | 4.24E-07 | 2.11E-06 | 137.707544 | 232.184049 | 138.054479 | 193.869318 |
| -5.0573783 | 4.25E-07 | 2.11E-06 | 434.024669 | 210.535186 | 121.948124 | 128.872668 |
| 5.05730289 | 4.25E-07 | 2.11E-06 | 0 | 0 | 0 | 0 |
| -5.0570201 | 4.26E-07 | 2.12E-06 | 88.1164342 | 186.721438 | 193.276271 | 155.207518 |
| -5.056167 | 4.28E-07 | 2.12E-06 | 74.1817423 | 236.513821 | 236.993523 | 241.496174 |
| -5.0560226 | 4.28E-07 | 2.13E-06 | 95.0837801 | 174.273342 | 133.452663 | 182.102683 |
| -5.0559384 | 4.28E-07 | 2.13E-06 | 72.9522106 | 240.572983 | 322.127119 | 233.651751 |
| -5.0557798 | 4.29E-07 | 2.13E-06 | 87.7065903 | 181.038611 | 192.125817 | 158.009098 |
| -5.055655 | 4.29E-07 | 2.13E-06 | 88.1164342 | 184.556551 | 165.665375 | 165.853521 |
| -5.0549892 | 4.30E-07 | 2.14E-06 | 113.936599 | 125.563402 | 112.744492 | 133.355196 |
| -5.0548489 | 4.31E-07 | 2.14E-06 | 86.0672148 | 263.84551 | 166.815829 | 197.79153 |
| -5.0547559 | 4.31E-07 | 2.14E-06 | 91.3951852 | 161.825246 | 151.859927 | 205.075637 |
| -5.0545563 | 4.31E-07 | 2.14E-06 | 86.0672148 | 168.049294 | 181.771731 | 186.024895 |
| -5.0543683 | 4.32E-07 | 2.14E-06 | 77.8703372 | 220.277174 | 215.134897 | 212.359745 |
| -5.0542611 | 4.32E-07 | 2.14E-06 | 143.445358 | 240.302372 | 220.887167 | 271.753235 |
| -5.0541535 | 4.32E-07 | 2.14E-06 | 155.740674 | 567.741415 | 560.271096 | 537.342995 |
| -5.0541687 | 4.32E-07 | 2.14E-06 | 87.2967464 | 172.649677 | 163.364467 | 197.79153 |
| -5.0540969 | 4.32E-07 | 2.14E-06 | 114.346443 | 156.954252 | 106.992222 | 141.759935 |
| -5.0540536 | 4.33E-07 | 2.14E-06 | 101.231438 | 146.671042 | 123.098578 | 153.52657 |
| -5.0537992 | 4.33E-07 | 2.15E-06 | 49.1812656 | 447.319618 | 490.093402 | 432.563912 |
| -5.0537564 | 4.33E-07 | 2.15E-06 | 72.1325229 | 264.657342 | 232.391707 | 260.546916 |
| -5.0535691 | 4.34E-07 | 2.15E-06 | 90.1656536 | 174.273342 | 149.559019 | 174.25826 |
| -5.053238 | 4.34E-07 | 2.15E-06 | 35.2465737 | 539.327283 | 1004.34634 | 668.456927 |
| -5.0532019 | 4.34E-07 | 2.15E-06 | 86.4770587 | 229.20733 | 192.125817 | 160.250361 |
| -5.0531637 | 4.35E-07 | 2.15E-06 | 172.544273 | 266.010396 | 97.7885896 | 141.759935 |
| -5.0528878 | 4.35E-07 | 2.15E-06 | 106.559409 | 171.296623 | 112.744492 | 162.491625 |
| -5.0527892 | 4.35E-07 | 2.16E-06 | 86.4770587 | 195.110372 | 217.435805 | 157.448782 |
| -5.0525614 | 4.36E-07 | 2.16E-06 | 113.116911 | 107.973701 | 210.533081 | 182.102683 |
| -5.0520863 | 4.37E-07 | 2.16E-06 | 70.4931473 | 394.009294 | 223.188075 | 304.811876 |
| -5.0520154 | 4.37E-07 | 2.16E-06 | 166.806459 | 307.684456 | 284.162137 | 276.235763 |
| -5.0519281 | 4.37E-07 | 2.16E-06 | 91.8050291 | 139.364551 | 192.125817 | 187.705843 |
| -5.0516807 | 4.38E-07 | 2.17E-06 | 94.6739363 | 219.194731 | 140.355387 | 166.974153 |
| -5.050571 | 4.40E-07 | 2.18E-06 | 164.347396 | 140.717605 | 113.894946 | 107.580662 |
| -5.0505891 | 4.40E-07 | 2.18E-06 | 159.019425 | 133.140503 | 102.390406 | 123.829825 |
| -5.0501966 | 4.41E-07 | 2.18E-06 | 125.412227 | 158.307306 | 97.7885896 | 155.767834 |
| -5.0499993 | 4.42E-07 | 2.18E-06 | 87.2967464 | 173.461509 | 159.913105 | 188.826475 |
| -5.049915 | 4.42E-07 | 2.18E-06 | 62.2962697 | 446.237175 | 318.675757 | 289.123029 |
| -5.0497869 | 4.42E-07 | 2.19E-06 | 85.6573709 | 241.926037 | 161.063559 | 198.912162 |
| -5.0495034 | 4.43E-07 | 2.19E-06 | 228.283041 | 181.309222 | 123.098578 | 101.417187 |

| | | | | | | |
|---|---|---|---|---|---|---|
| -5.0493642 | 4.43E-07 | 2.19E-06 | 231.151948 | 154.789366 | 104.691314 | 126.631404 |
| -5.0486045 | 4.45E-07 | 2.20E-06 | 178.282088 | 230.019162 | 116.195854 | 188.826475 |
| -5.0483645 | 4.46E-07 | 2.20E-06 | 127.051603 | 97.9611021 | 226.639437 | 192.18837 |
| -5.0482471 | 4.46E-07 | 2.20E-06 | 126.231915 | 279.540935 | 102.390406 | 199.472478 |
| -5.048129 | 4.46E-07 | 2.20E-06 | 83.1983076 | 237.866875 | 186.373547 | 178.180472 |
| -5.0479258 | 4.47E-07 | 2.20E-06 | 63.5258014 | 282.788264 | 343.985745 | 310.415036 |
| -5.0475142 | 4.48E-07 | 2.21E-06 | 77.0506494 | 247.338252 | 197.878087 | 225.247011 |
| -5.0470685 | 4.49E-07 | 2.21E-06 | 77.4604933 | 186.450827 | 381.950726 | 290.803977 |
| -5.0467377 | 4.49E-07 | 2.22E-06 | 80.3294005 | 209.723354 | 349.738015 | 203.394689 |
| -5.0462205 | 4.51E-07 | 2.22E-06 | 68.8537718 | 279.540935 | 236.993523 | 332.267358 |
| -5.0457857 | 4.52E-07 | 2.23E-06 | 118.035037 | 116.633246 | 190.975363 | 123.829825 |
| -5.0457376 | 4.52E-07 | 2.23E-06 | 111.067691 | 200.522587 | 159.913105 | 110.382242 |
| -5.0451038 | 4.53E-07 | 2.24E-06 | 79.5097127 | 213.241294 | 190.975363 | 205.635953 |
| -5.0450233 | 4.53E-07 | 2.24E-06 | 81.1490882 | 328.792097 | 190.975363 | 211.799429 |
| -5.0450132 | 4.53E-07 | 2.24E-06 | 103.280658 | 120.151186 | 162.214013 | 206.756585 |
| -5.0433138 | 4.58E-07 | 2.26E-06 | 78.2801811 | 293.342085 | 204.780811 | 203.394689 |
| -5.0427553 | 4.59E-07 | 2.26E-06 | 84.8376831 | 191.592432 | 173.718553 | 169.775732 |
| -5.0425779 | 4.59E-07 | 2.26E-06 | 126.231915 | 110.950419 | 149.559019 | 112.06319 |
| -5.0422848 | 4.60E-07 | 2.27E-06 | 84.8376831 | 338.804695 | 156.461743 | 284.640502 |
| -5.0422025 | 4.60E-07 | 2.27E-06 | 113.526755 | 137.470276 | 106.992222 | 132.79488 |
| 5.04206872 | 4.61E-07 | 2.27E-06 | 0 | 0 | 0 | 0 |
| -5.0418477 | 4.61E-07 | 2.27E-06 | 78.2801811 | 214.323737 | 190.975363 | 223.566063 |
| -5.0418003 | 4.61E-07 | 2.27E-06 | 79.9195566 | 358.018061 | 219.736713 | 197.231214 |
| -5.0416504 | 4.62E-07 | 2.27E-06 | 97.9526873 | 139.09394 | 165.665375 | 129.432984 |
| -5.0412339 | 4.63E-07 | 2.28E-06 | 160.248957 | 132.599282 | 96.6381356 | 135.59646 |
| -5.0410514 | 4.63E-07 | 2.28E-06 | 89.7558097 | 154.518755 | 153.010381 | 169.215416 |
| 5.04093331 | 4.63E-07 | 2.28E-06 | 0.40984388 | 0.27061078 | 0 | 0 |
| -5.0406596 | 4.64E-07 | 2.28E-06 | 71.722679 | 230.019162 | 247.347609 | 254.383441 |
| -5.040329 | 4.65E-07 | 2.29E-06 | 198.774282 | 488.181846 | 418.765254 | 443.770231 |
| -5.0402576 | 4.65E-07 | 2.29E-06 | 88.1164342 | 211.617629 | 159.913105 | 161.370993 |
| -5.0399587 | 4.66E-07 | 2.29E-06 | 142.215826 | 103.914539 | 117.346308 | 173.697944 |
| -5.0391303 | 4.68E-07 | 2.30E-06 | 90.5754975 | 222.44206 | 136.904025 | 222.445432 |
| -5.0381007 | 4.70E-07 | 2.31E-06 | 65.1651769 | 421.340983 | 248.498063 | 313.216615 |
| -5.0380018 | 4.70E-07 | 2.31E-06 | 86.0672148 | 171.567234 | 176.019461 | 161.370993 |
| -5.0376417 | 4.71E-07 | 2.32E-06 | 81.5589321 | 198.08709 | 174.869007 | 197.79153 |
| -5.0375691 | 4.71E-07 | 2.32E-06 | 78.6900249 | 218.112288 | 187.524001 | 207.877217 |
| -5.0373384 | 4.72E-07 | 2.32E-06 | 96.7231557 | 163.990132 | 138.054479 | 132.234564 |
| -5.0370637 | 4.73E-07 | 2.32E-06 | 130.740198 | 132.869893 | 100.089498 | 126.631404 |
| -5.0367217 | 4.74E-07 | 2.33E-06 | 86.0672148 | 150.459593 | 195.577179 | 199.472478 |
| -5.0364343 | 4.74E-07 | 2.33E-06 | 95.0837801 | 144.776767 | 132.302209 | 161.370993 |
| -5.0362941 | 4.75E-07 | 2.33E-06 | 129.510666 | 100.937821 | 162.214013 | 241.496174 |
| -5.0356982 | 4.76E-07 | 2.34E-06 | 88.9361219 | 182.121054 | 155.311289 | 152.405938 |
| -5.035695 | 4.76E-07 | 2.34E-06 | 85.247527 | 181.038611 | 157.612197 | 212.359745 |

| | | | | | | |
|---|---|---|---|---|---|---|
| 5.03551256 | 4.77E-07 | 2.34E-06 | 68.4439279 | 36.5324552 | 34.5136199 | 29.1364293 |
| -5.035465 | 4.77E-07 | 2.34E-06 | 65.5750208 | 335.016145 | 256.551241 | 282.959554 |
| -5.0353493 | 4.77E-07 | 2.34E-06 | 95.9034679 | 125.834012 | 158.762651 | 160.250361 |
| -5.0350984 | 4.78E-07 | 2.34E-06 | 94.6739363 | 130.434396 | 154.160835 | 177.05984 |
| -5.0350617 | 4.78E-07 | 2.34E-06 | 79.9195566 | 169.402348 | 233.542161 | 253.262809 |
| -5.0349968 | 4.78E-07 | 2.34E-06 | 798.785722 | 1475.64058 | 985.939074 | 1217.56656 |
| -5.034963 | 4.78E-07 | 2.34E-06 | 88.5262781 | 145.047378 | 179.470823 | 212.359745 |
| -5.0344747 | 4.79E-07 | 2.35E-06 | 235.250387 | 239.76115 | 166.815829 | 200.032794 |
| -5.0345303 | 4.79E-07 | 2.35E-06 | 135.24848 | 109.867976 | 140.355387 | 108.140978 |
| -5.0345006 | 4.79E-07 | 2.35E-06 | 72.1325229 | 234.348935 | 230.090799 | 243.177122 |
| -5.0338097 | 4.81E-07 | 2.36E-06 | 66.8045524 | 254.644743 | 277.259413 | 284.080186 |
| -5.0337626 | 4.81E-07 | 2.36E-06 | 82.7884637 | 291.177198 | 158.762651 | 247.099333 |
| -5.0337289 | 4.81E-07 | 2.36E-06 | 234.840543 | 142.611881 | 106.992222 | 126.631404 |
| -5.033088 | 4.83E-07 | 2.36E-06 | 581.97831 | 980.693464 | 775.405993 | 841.594555 |
| -5.0325357 | 4.84E-07 | 2.37E-06 | 93.0345607 | 203.499306 | 202.479903 | 168.655101 |
| -5.0323362 | 4.85E-07 | 2.37E-06 | 66.3947085 | 291.71842 | 273.808051 | 257.18502 |
| -5.032235 | 4.85E-07 | 2.37E-06 | 82.7884637 | 175.897006 | 172.568099 | 215.72164 |
| -5.0321077 | 4.85E-07 | 2.37E-06 | 83.1983076 | 201.875641 | 208.232173 | 158.569413 |
| -5.0320155 | 4.85E-07 | 2.38E-06 | 93.0345607 | 180.49739 | 146.107657 | 138.958355 |
| -5.0317488 | 4.86E-07 | 2.38E-06 | 85.247527 | 149.647761 | 207.081719 | 207.316901 |
| -5.0317689 | 4.86E-07 | 2.38E-06 | 96.7231557 | 176.438228 | 127.700393 | 145.121831 |
| -5.0314715 | 4.87E-07 | 2.38E-06 | 77.4604933 | 199.710755 | 200.178995 | 212.359745 |
| -5.0313146 | 4.87E-07 | 2.38E-06 | 87.7065903 | 178.873725 | 150.709473 | 162.491625 |
| 5.03101052 | 4.88E-07 | 2.39E-06 | 18.8528185 | 11.3656527 | 10.354086 | 12.8872668 |
| -5.0309457 | 4.88E-07 | 2.39E-06 | 87.7065903 | 184.556551 | 146.107657 | 172.577312 |
| -5.030259 | 4.90E-07 | 2.39E-06 | 317.629007 | 639.994493 | 54.0713378 | 393.902112 |
| -5.0299847 | 4.91E-07 | 2.40E-06 | 81.1490882 | 176.438228 | 194.426725 | 187.145527 |
| -5.0295139 | 4.92E-07 | 2.40E-06 | 81.1490882 | 194.027929 | 177.169915 | 186.024895 |
| -5.0294937 | 4.92E-07 | 2.40E-06 | 78.2801811 | 196.463426 | 202.479903 | 195.550266 |
| -5.0294527 | 4.92E-07 | 2.40E-06 | 84.8376831 | 204.581749 | 158.762651 | 173.697944 |
| -5.0292354 | 4.92E-07 | 2.41E-06 | 116.395662 | 281.1646 | 105.841768 | 203.394689 |
| -5.0286938 | 4.94E-07 | 2.41E-06 | 95.0837801 | 166.155018 | 124.249032 | 171.45668 |
| -5.0286579 | 4.94E-07 | 2.41E-06 | 104.920033 | 162.637078 | 135.753571 | 129.432984 |
| -5.0285307 | 4.94E-07 | 2.41E-06 | 95.493624 | 132.869893 | 159.913105 | 135.036144 |
| -5.0284706 | 4.94E-07 | 2.41E-06 | 79.5097127 | 211.88824 | 192.125817 | 181.542367 |
| -5.0283914 | 4.95E-07 | 2.41E-06 | 73.3620545 | 239.76115 | 209.382627 | 231.410487 |
| -5.028145 | 4.95E-07 | 2.42E-06 | 84.8376831 | 164.801964 | 184.072639 | 163.612257 |
| -5.0279668 | 4.96E-07 | 2.42E-06 | 344.268859 | 304.978348 | 309.472125 | 365.325998 |
| -5.0277359 | 4.96E-07 | 2.42E-06 | 104.920033 | 117.445078 | 132.302209 | 123.269509 |
| -5.0277065 | 4.96E-07 | 2.42E-06 | 87.2967464 | 149.106539 | 166.815829 | 170.896364 |
| -5.027663 | 4.96E-07 | 2.42E-06 | 97.1329995 | 196.463426 | 127.700393 | 149.604358 |
| -5.0271383 | 4.98E-07 | 2.43E-06 | 83.1983076 | 153.436312 | 239.294431 | 267.270707 |
| -5.0269767 | 4.98E-07 | 2.43E-06 | 78.2801811 | 192.674875 | 188.674455 | 225.807327 |

| | | | | | | |
|---|---|---|---|---|---|---|
| -5.0267442 | 4.99E-07 | 2.43E-06 | 87.7065903 | 160.472192 | 173.718553 | 148.483726 |
| -5.0266938 | 4.99E-07 | 2.43E-06 | 263.529615 | 213.782516 | 115.0454 | 180.421735 |
| -5.0265532 | 4.99E-07 | 2.43E-06 | 75.4112739 | 215.676791 | 195.577179 | 245.978701 |
| -5.0262757 | 5.00E-07 | 2.44E-06 | 85.247527 | 153.977533 | 181.771731 | 226.367643 |
| -5.026301 | 5.00E-07 | 2.44E-06 | 70.9029912 | 266.010396 | 222.037621 | 238.694594 |
| -5.0261966 | 5.00E-07 | 2.44E-06 | 95.0837801 | 204.85236 | 123.098578 | 205.075637 |
| -5.0261375 | 5.00E-07 | 2.44E-06 | 79.5097127 | 214.864959 | 173.718553 | 203.394689 |
| -5.0260387 | 5.01E-07 | 2.44E-06 | 54.0993922 | 329.333318 | 469.38523 | 368.687894 |
| -5.0255829 | 5.02E-07 | 2.44E-06 | 96.3133118 | 174.273342 | 123.098578 | 152.966254 |
| -5.0255607 | 5.02E-07 | 2.44E-06 | 63.1159575 | 277.646659 | 284.162137 | 321.061039 |
| -5.0254395 | 5.02E-07 | 2.45E-06 | 116.805506 | 210.264575 | 98.9390436 | 221.885116 |
| -5.0251316 | 5.03E-07 | 2.45E-06 | 801.244785 | 273.046276 | 235.843069 | 306.492824 |
| -5.024883 | 5.04E-07 | 2.45E-06 | 79.9195566 | 242.737869 | 163.364467 | 268.951655 |
| -5.0245791 | 5.05E-07 | 2.46E-06 | 131.559885 | 102.290875 | 184.072639 | 123.269509 |
| -5.024549 | 5.05E-07 | 2.46E-06 | 76.2309617 | 269.528336 | 232.391707 | 182.102683 |
| -5.0240217 | 5.06E-07 | 2.46E-06 | 109.83816 | 155.059976 | 116.195854 | 116.545717 |
| -5.0236983 | 5.07E-07 | 2.47E-06 | 85.247527 | 181.038611 | 157.612197 | 164.732889 |
| -5.0235642 | 5.07E-07 | 2.47E-06 | 76.2309617 | 230.019162 | 187.524001 | 216.842272 |
| -5.0229706 | 5.09E-07 | 2.47E-06 | 94.6739363 | 133.140503 | 138.054479 | 141.199619 |
| -5.0229117 | 5.09E-07 | 2.47E-06 | 93.0345607 | 180.49739 | 130.001301 | 155.207518 |
| -5.0222869 | 5.11E-07 | 2.48E-06 | 160.658801 | 273.316887 | 111.594038 | 406.229063 |
| -5.0219942 | 5.11E-07 | 2.49E-06 | 92.6247169 | 170.755402 | 172.568099 | 127.19172 |
| -5.0219763 | 5.11E-07 | 2.49E-06 | 92.6247169 | 146.941653 | 132.302209 | 158.569413 |
| -5.0217454 | 5.12E-07 | 2.49E-06 | 95.0837801 | 200.793198 | 133.452663 | 146.242463 |
| -5.0215732 | 5.12E-07 | 2.49E-06 | 81.5589321 | 209.182132 | 192.125817 | 162.491625 |
| -5.021587 | 5.12E-07 | 2.49E-06 | 78.6900249 | 230.019162 | 190.975363 | 181.542367 |
| -5.0215948 | 5.12E-07 | 2.49E-06 | 85.247527 | 169.131737 | 154.160835 | 205.635953 |
| -5.0214614 | 5.13E-07 | 2.49E-06 | 102.46097 | 154.248144 | 110.443584 | 157.448782 |
| -5.0213902 | 5.13E-07 | 2.49E-06 | 84.0179954 | 175.897006 | 163.364467 | 168.094785 |
| -5.0212022 | 5.13E-07 | 2.49E-06 | 98.7723751 | 123.669126 | 128.850847 | 141.759935 |
| -5.0211747 | 5.14E-07 | 2.49E-06 | 243.037421 | 223.253893 | 182.922185 | 229.169223 |
| -5.0208092 | 5.15E-07 | 2.50E-06 | 89.7558097 | 163.448911 | 142.656295 | 148.483726 |
| -5.0207687 | 5.15E-07 | 2.50E-06 | 102.870814 | 169.402348 | 111.594038 | 146.802779 |
| -5.0203865 | 5.16E-07 | 2.50E-06 | 91.8050291 | 149.106539 | 227.789891 | 148.483726 |
| -5.019354 | 5.18E-07 | 2.51E-06 | 1092.23394 | 1526.24479 | 970.983172 | 1224.85066 |
| -5.0192968 | 5.19E-07 | 2.52E-06 | 564.764867 | 139.09394 | 274.958505 | 104.779082 |
| -5.0192498 | 5.19E-07 | 2.52E-06 | 143.855202 | 159.93097 | 90.8858656 | 135.036144 |
| -5.0191374 | 5.19E-07 | 2.52E-06 | 119.674413 | 145.85921 | 97.7885896 | 135.036144 |
| -5.0189619 | 5.20E-07 | 2.52E-06 | 82.7884637 | 169.672959 | 202.479903 | 164.732889 |
| -5.018976 | 5.19E-07 | 2.52E-06 | 72.5423667 | 242.737869 | 203.630357 | 236.45333 |
| -5.0189416 | 5.20E-07 | 2.52E-06 | 206.151472 | 105.538204 | 158.762651 | 111.502874 |
| -5.0188463 | 5.20E-07 | 2.52E-06 | 85.247527 | 184.285941 | 162.214013 | 155.207518 |
| -5.018541 | 5.21E-07 | 2.52E-06 | 78.2801811 | 220.006563 | 172.568099 | 211.799429 |

| | | | | | | |
|---|---|---|---|---|---|---|
| -5.0184626 | 5.21E-07 | 2.52E-06 | 98.3625312 | 139.635162 | 97.7885896 | 132.79488 |
| -5.0181305 | 5.22E-07 | 2.53E-06 | 88.1164342 | 175.626396 | 164.514921 | 140.078987 |
| -5.0180549 | 5.22E-07 | 2.53E-06 | 81.1490882 | 167.508072 | 181.771731 | 196.670898 |
| -5.0179315 | 5.22E-07 | 2.53E-06 | 85.6573709 | 171.026012 | 155.311289 | 234.212067 |
| -5.0174136 | 5.24E-07 | 2.54E-06 | 97.1329995 | 136.387833 | 128.850847 | 128.312352 |
| -5.0173447 | 5.24E-07 | 2.54E-06 | 99.5920628 | 118.798132 | 132.302209 | 154.086886 |
| -5.0169827 | 5.25E-07 | 2.54E-06 | 81.1490882 | 165.884408 | 188.674455 | 187.145527 |
| -5.0169873 | 5.25E-07 | 2.54E-06 | 307.38291 | 136.658443 | 118.496762 | 133.915512 |
| -5.0169327 | 5.25E-07 | 2.54E-06 | 56.9682993 | 397.256624 | 528.058384 | 270.072287 |
| -5.015921 | 5.28E-07 | 2.55E-06 | 74.5915861 | 254.374132 | 235.843069 | 184.343947 |
| -5.0159192 | 5.28E-07 | 2.55E-06 | 83.6081515 | 165.884408 | 171.417645 | 164.732889 |
| -5.0159454 | 5.28E-07 | 2.55E-06 | 71.722679 | 240.302372 | 211.683535 | 230.850171 |
| -5.0156326 | 5.29E-07 | 2.56E-06 | 87.2967464 | 234.890156 | 149.559019 | 169.215416 |
| -5.015527 | 5.29E-07 | 2.56E-06 | 75.8211178 | 199.169533 | 227.789891 | 192.748686 |
| -5.0155532 | 5.29E-07 | 2.56E-06 | 115.985818 | 294.965749 | 307.171217 | 292.484925 |
| -5.015257 | 5.30E-07 | 2.56E-06 | 81.968776 | 212.429462 | 153.010381 | 222.445432 |
| -5.0152343 | 5.30E-07 | 2.56E-06 | 85.247527 | 274.128719 | 158.762651 | 183.783631 |
| -5.01489 | 5.31E-07 | 2.56E-06 | 84.4278393 | 186.450827 | 153.010381 | 168.655101 |
| -5.0147638 | 5.31E-07 | 2.57E-06 | 118.444881 | 139.09394 | 108.142676 | 110.942558 |
| -5.0145997 | 5.31E-07 | 2.57E-06 | 152.052079 | 272.775665 | 139.204933 | 168.094785 |
| -5.0145488 | 5.32E-07 | 2.57E-06 | 88.5262781 | 158.577917 | 169.116737 | 138.398039 |
| -5.0145269 | 5.32E-07 | 2.57E-06 | 231.151948 | 575.859738 | 521.15566 | 591.693642 |
| -5.0143856 | 5.32E-07 | 2.57E-06 | 241.807889 | 156.14242 | 102.390406 | 122.148877 |
| -5.0141528 | 5.33E-07 | 2.57E-06 | 84.4278393 | 340.42836 | 165.665375 | 206.756585 |
| -5.0138531 | 5.34E-07 | 2.57E-06 | 65.5750208 | 276.022995 | 253.099879 | 263.908812 |
| -5.0138601 | 5.33E-07 | 2.57E-06 | 82.7884637 | 177.791282 | 156.461743 | 202.274057 |
| -5.0136139 | 5.34E-07 | 2.58E-06 | 68.4439279 | 233.266492 | 257.701695 | 244.297754 |
| -5.0132552 | 5.35E-07 | 2.58E-06 | 158.609582 | 335.286755 | 364.693917 | 361.403787 |
| -5.013202 | 5.35E-07 | 2.58E-06 | 53.2797044 | 342.593246 | 423.36707 | 351.3181 |
| -5.0125457 | 5.37E-07 | 2.59E-06 | 133.609105 | 140.176384 | 128.850847 | 94.6933953 |
| 5.01250667 | 5.37E-07 | 2.59E-06 | 0 | 0 | 0 | 0 |
| -5.0122569 | 5.38E-07 | 2.59E-06 | 121.313788 | 182.932887 | 104.691314 | 123.269509 |
| -5.0119722 | 5.39E-07 | 2.60E-06 | 68.8537718 | 247.338252 | 224.338529 | 256.624704 |
| -5.011916 | 5.39E-07 | 2.60E-06 | 78.6900249 | 523.631858 | 653.457869 | 612.985648 |
| -5.0117304 | 5.39E-07 | 2.60E-06 | 92.214873 | 137.470276 | 134.603117 | 165.293205 |
| -5.0116892 | 5.40E-07 | 2.60E-06 | 83.6081515 | 181.579833 | 162.214013 | 160.250361 |
| -5.0116597 | 5.40E-07 | 2.60E-06 | 112.297223 | 228.395498 | 157.612197 | 198.351846 |
| -5.0116188 | 5.40E-07 | 2.60E-06 | 35.2465737 | 429.729917 | 865.141405 | 754.745583 |
| -5.0113517 | 5.40E-07 | 2.60E-06 | 64.3454891 | 233.807713 | 306.020763 | 315.457879 |
| -5.0112532 | 5.41E-07 | 2.60E-06 | 142.215826 | 183.474108 | 192.125817 | 85.1680242 |
| 5.01122947 | 5.41E-07 | 2.60E-06 | 127.461447 | 71.9824673 | 90.8858656 | 79.0045488 |
| -5.0112452 | 5.41E-07 | 2.60E-06 | 103.690502 | 271.963833 | 163.364467 | 191.067738 |
| -5.0111511 | 5.41E-07 | 2.60E-06 | 89.7558097 | 199.710755 | 227.789891 | 130.553616 |

| | | | | | | |
|---|---|---|---|---|---|---|
| -5.0108405 | 5.42E-07 | 2.61E-06 | 89.3459658 | 233.266492 | 200.178995 | 132.234564 |
| -5.0107887 | 5.42E-07 | 2.61E-06 | 74.5915861 | 201.063809 | 202.479903 | 216.281956 |
| -5.0095861 | 5.45E-07 | 2.62E-06 | 75.00143 | 191.592432 | 223.188075 | 203.394689 |
| -5.0094873 | 5.46E-07 | 2.63E-06 | 73.3620545 | 297.130636 | 186.373547 | 243.177122 |
| -5.0094087 | 5.46E-07 | 2.63E-06 | 94.6739363 | 150.730204 | 196.727633 | 125.510772 |
| -5.0092485 | 5.46E-07 | 2.63E-06 | 92.214873 | 139.635162 | 134.603117 | 179.86142 |
| -5.009165 | 5.47E-07 | 2.63E-06 | 116.805506 | 159.66036 | 130.001301 | 100.296555 |
| -5.0090399 | 5.47E-07 | 2.63E-06 | 79.5097127 | 195.380983 | 181.771731 | 172.016996 |
| -5.0090517 | 5.47E-07 | 2.63E-06 | 79.5097127 | 172.649677 | 205.931265 | 178.180472 |
| -5.0088578 | 5.48E-07 | 2.63E-06 | 111.067691 | 189.427545 | 269.206235 | 107.020346 |
| -5.0088043 | 5.48E-07 | 2.63E-06 | 78.2801811 | 216.218013 | 201.329449 | 168.655101 |
| -5.0085492 | 5.48E-07 | 2.64E-06 | 91.8050291 | 169.131737 | 139.204933 | 133.915512 |
| -5.0083679 | 5.49E-07 | 2.64E-06 | 223.774758 | 542.033391 | 822.574607 | 593.934906 |
| -5.0080426 | 5.50E-07 | 2.64E-06 | 90.5754975 | 293.883306 | 130.001301 | 254.383441 |
| -5.007822 | 5.50E-07 | 2.64E-06 | 94.6739363 | 152.624479 | 120.79767 | 146.802779 |
| -5.0074156 | 5.52E-07 | 2.65E-06 | 103.690502 | 135.846611 | 108.142676 | 135.59646 |
| -5.0071609 | 5.52E-07 | 2.65E-06 | 118.854725 | 109.326755 | 132.302209 | 107.580662 |
| -5.0069729 | 5.53E-07 | 2.65E-06 | 94.6739363 | 119.068743 | 157.612197 | 156.888466 |
| -5.0069204 | 5.53E-07 | 2.65E-06 | 75.00143 | 526.608576 | 164.514921 | 553.592157 |
| -5.0066634 | 5.54E-07 | 2.66E-06 | 80.7392443 | 167.237462 | 243.896247 | 179.301104 |
| -5.0065085 | 5.54E-07 | 2.66E-06 | 120.903945 | 258.162683 | 103.54086 | 160.250361 |
| -5.0065151 | 5.54E-07 | 2.66E-06 | 102.870814 | 111.491641 | 104.691314 | 130.553616 |
| -5.0062065 | 5.55E-07 | 2.66E-06 | 172.13443 | 131.246228 | 90.8858656 | 231.410487 |
| -5.006049 | 5.56E-07 | 2.66E-06 | 86.0672148 | 186.721438 | 142.656295 | 163.051941 |
| -5.0060728 | 5.56E-07 | 2.66E-06 | 62.2962697 | 276.022995 | 274.958505 | 294.165873 |
| -5.0060844 | 5.55E-07 | 2.66E-06 | 168.855679 | 138.282108 | 86.2840497 | 202.834373 |
| -5.0058713 | 5.56E-07 | 2.67E-06 | 96.3133118 | 218.382899 | 116.195854 | 218.52322 |
| -5.0056593 | 5.57E-07 | 2.67E-06 | 170.495054 | 143.423713 | 117.346308 | 96.9346591 |
| -5.0055657 | 5.57E-07 | 2.67E-06 | 92.6247169 | 190.780599 | 209.382627 | 216.842272 |
| -5.0052986 | 5.58E-07 | 2.67E-06 | 95.493624 | 152.083258 | 117.346308 | 154.086886 |
| -5.0051247 | 5.58E-07 | 2.68E-06 | 89.3459658 | 212.700072 | 220.887167 | 223.005748 |
| -5.004965 | 5.59E-07 | 2.68E-06 | 84.8376831 | 171.837845 | 146.107657 | 168.094785 |
| -5.0045433 | 5.60E-07 | 2.68E-06 | 86.0672148 | 160.472192 | 154.160835 | 147.92341 |
| -5.0041418 | 5.61E-07 | 2.69E-06 | 70.4931473 | 217.029845 | 232.391707 | 225.807327 |
| -5.0039074 | 5.62E-07 | 2.69E-06 | 116.805506 | 250.585581 | 182.922185 | 224.126379 |
| -5.0037019 | 5.62E-07 | 2.69E-06 | 86.0672148 | 178.061893 | 138.054479 | 175.939208 |
| -5.003506 | 5.63E-07 | 2.70E-06 | 55.3289238 | 347.46424 | 303.719855 | 378.213265 |
| -5.0034013 | 5.63E-07 | 2.70E-06 | 61.8864259 | 333.121869 | 278.409867 | 261.667548 |
| -5.0033034 | 5.64E-07 | 2.70E-06 | 131.559885 | 146.129821 | 93.1867736 | 126.631404 |
| -5.0033068 | 5.64E-07 | 2.70E-06 | 81.1490882 | 283.058875 | 156.461743 | 213.480376 |
| -5.003331 | 5.63E-07 | 2.70E-06 | 62.2962697 | 284.68254 | 257.701695 | 307.05314 |
| -5.0029844 | 5.64E-07 | 2.70E-06 | 622.14301 | 189.698156 | 110.443584 | 173.697944 |
| -5.0027935 | 5.65E-07 | 2.70E-06 | 69.2636157 | 247.879474 | 307.171217 | 205.635953 |

| | | | | | | |
|---|---|---|---|---|---|---|
| -5.0025777 | 5.66E-07 | 2.71E-06 | 95.493624 | 120.692407 | 158.762651 | 135.59646 |
| -5.002185 | 5.67E-07 | 2.71E-06 | 135.658324 | 225.418779 | 85.1335957 | 210.118481 |
| -5.0020037 | 5.67E-07 | 2.71E-06 | 112.297223 | 107.432479 | 123.098578 | 114.864769 |
| -5.0014737 | 5.69E-07 | 2.72E-06 | 93.4444046 | 371.277989 | 285.312591 | 127.752036 |
| -5.0013214 | 5.69E-07 | 2.72E-06 | 95.493624 | 126.645845 | 128.850847 | 112.06319 |
| -5.001255 | 5.70E-07 | 2.72E-06 | 108.608628 | 113.656527 | 110.443584 | 141.199619 |
| -5.0011887 | 5.70E-07 | 2.72E-06 | 89.7558097 | 144.776767 | 138.054479 | 191.628054 |
| -5.0011587 | 5.70E-07 | 2.72E-06 | 80.3294005 | 167.778683 | 192.125817 | 168.655101 |
| -5.0008614 | 5.71E-07 | 2.73E-06 | 80.3294005 | 155.059976 | 200.178995 | 195.550266 |
| -5.0008072 | 5.71E-07 | 2.73E-06 | 73.3620545 | 199.440144 | 199.028541 | 250.461229 |
| -5.0006749 | 5.71E-07 | 2.73E-06 | 67.6242402 | 226.501222 | 240.444885 | 261.107232 |
| -5.0003247 | 5.72E-07 | 2.73E-06 | 97.9526873 | 130.163785 | 116.195854 | 139.518671 |
| -5.0000273 | 5.73E-07 | 2.74E-06 | 100.001907 | 116.362635 | 127.700393 | 124.950457 |
| -4.999968 | 5.73E-07 | 2.74E-06 | 108.608628 | 120.151186 | 106.992222 | 151.845622 |
| 4.99976788 | 5.74E-07 | 2.74E-06 | 27.45954 | 21.9194731 | 23.0090799 | 22.9729539 |
| -4.9996798 | 5.74E-07 | 2.74E-06 | 77.0506494 | 175.355785 | 190.975363 | 203.394689 |
| -4.9994545 | 5.75E-07 | 2.74E-06 | 166.806459 | 130.705006 | 87.4345036 | 186.024895 |
| -4.9992881 | 5.75E-07 | 2.75E-06 | 113.116911 | 139.905773 | 97.7885896 | 136.156776 |
| -4.9991068 | 5.76E-07 | 2.75E-06 | 151.232392 | 447.860839 | 459.031144 | 461.700342 |
| -4.9990117 | 5.76E-07 | 2.75E-06 | 71.722679 | 190.239378 | 245.046701 | 238.134278 |
| -4.9983749 | 5.78E-07 | 2.76E-06 | 689.76725 | 1154.69619 | 1076.82494 | 1162.09528 |
| -4.9981995 | 5.79E-07 | 2.76E-06 | 78.2801811 | 189.968767 | 186.373547 | 170.336048 |
| -4.9977402 | 5.80E-07 | 2.77E-06 | 313.940412 | 119.068743 | 123.098578 | 144.561515 |
| -4.9970165 | 5.82E-07 | 2.78E-06 | 182.79037 | 253.291689 | 85.1335957 | 143.440883 |
| -4.9968799 | 5.83E-07 | 2.78E-06 | 118.444881 | 169.943569 | 118.496762 | 148.483726 |
| -4.9968807 | 5.83E-07 | 2.78E-06 | 74.1817423 | 185.909605 | 223.188075 | 202.834373 |
| -4.9967958 | 5.83E-07 | 2.78E-06 | 82.7884637 | 200.251977 | 146.107657 | 179.301104 |
| -4.9967671 | 5.83E-07 | 2.78E-06 | 68.0340841 | 288.471091 | 308.321671 | 197.79153 |
| -4.9963528 | 5.84E-07 | 2.78E-06 | 80.7392443 | 207.287857 | 192.125817 | 151.285306 |
| -4.9960841 | 5.85E-07 | 2.79E-06 | 88.9361219 | 158.036695 | 131.151755 | 150.164674 |
| -4.9958587 | 5.86E-07 | 2.79E-06 | 71.3128351 | 205.122971 | 256.551241 | 207.877217 |
| -4.9957503 | 5.86E-07 | 2.79E-06 | 111.887379 | 98.5023236 | 180.621277 | 151.845622 |
| -4.9956508 | 5.86E-07 | 2.79E-06 | 88.5262781 | 222.44206 | 205.931265 | 231.410487 |
| -4.9952285 | 5.88E-07 | 2.80E-06 | 107.78894 | 119.068743 | 109.29313 | 117.106033 |
| -4.9949152 | 5.89E-07 | 2.80E-06 | 111.477535 | 227.583665 | 149.559019 | 107.020346 |
| -4.9945969 | 5.90E-07 | 2.81E-06 | 279.92337 | 323.650492 | 363.543463 | 373.730738 |
| 4.99400171 | 5.91E-07 | 2.81E-06 | 16.3937552 | 6.76526948 | 3.45136199 | 11.7666349 |
| -4.9940828 | 5.91E-07 | 2.81E-06 | 100.821594 | 139.09394 | 151.859927 | 107.580662 |
| -4.9940475 | 5.91E-07 | 2.81E-06 | 81.1490882 | 167.508072 | 159.913105 | 174.25826 |
| -4.9940272 | 5.91E-07 | 2.81E-06 | 73.3620545 | 186.180216 | 247.347609 | 206.756585 |
| -4.993937 | 5.92E-07 | 2.81E-06 | 77.4604933 | 175.085174 | 195.577179 | 179.86142 |
| -4.9938052 | 5.92E-07 | 2.81E-06 | 73.3620545 | 249.773749 | 181.771731 | 216.842272 |
| -4.99374 | 5.92E-07 | 2.81E-06 | 51.6403289 | 393.197462 | 366.994825 | 335.068937 |

| | | | | | | |
|---|---|---|---|---|---|---|
| -4.9934528 | 5.93E-07 | 2.82E-06 | 54.0993922 | 311.202396 | 346.286653 | 368.687894 |
| -4.9934106 | 5.93E-07 | 2.82E-06 | 82.7884637 | 166.966851 | 234.692615 | 155.767834 |
| -4.9932186 | 5.94E-07 | 2.82E-06 | 87.2967464 | 151.812647 | 136.904025 | 151.845622 |
| 4.99304495 | 5.94E-07 | 2.82E-06 | 35.2465737 | 27.3316887 | 39.1154358 | 47.6268556 |
| -4.9928478 | 5.95E-07 | 2.83E-06 | 88.1164342 | 161.013414 | 130.001301 | 159.129729 |
| -4.9923269 | 5.97E-07 | 2.83E-06 | 79.9195566 | 220.818396 | 158.762651 | 178.740788 |
| -4.9921092 | 5.97E-07 | 2.84E-06 | 337.711357 | 105.267593 | 136.904025 | 158.009098 |
| -4.9915846 | 5.99E-07 | 2.84E-06 | 117.625194 | 121.50424 | 284.162137 | 135.036144 |
| -4.9915767 | 5.99E-07 | 2.84E-06 | 83.6081515 | 181.579833 | 140.355387 | 206.196269 |
| -4.9914316 | 5.99E-07 | 2.84E-06 | 103.690502 | 123.669126 | 106.992222 | 132.79488 |
| -4.991413 | 5.99E-07 | 2.84E-06 | 71.722679 | 215.135569 | 196.727633 | 223.005748 |
| -4.9913101 | 6.00E-07 | 2.85E-06 | 82.7884637 | 165.343186 | 151.859927 | 162.491625 |
| -4.9911498 | 6.00E-07 | 2.85E-06 | 101.641282 | 105.808815 | 149.559019 | 140.639303 |
| -4.99052 | 6.02E-07 | 2.86E-06 | 250.004767 | 412.952049 | 341.684837 | 522.214464 |
| -4.9902184 | 6.03E-07 | 2.86E-06 | 117.625194 | 156.14242 | 238.143977 | 120.467929 |
| -4.9902168 | 6.03E-07 | 2.86E-06 | 82.7884637 | 158.307306 | 182.922185 | 150.164674 |
| -4.9900335 | 6.04E-07 | 2.86E-06 | 72.9522106 | 197.545869 | 203.630357 | 204.515321 |
| -4.9899003 | 6.04E-07 | 2.86E-06 | 51.6403289 | 277.105438 | 690.272397 | 453.295602 |
| -4.9898742 | 6.04E-07 | 2.86E-06 | 72.9522106 | 250.04436 | 272.657597 | 173.137628 |
| -4.9898149 | 6.04E-07 | 2.86E-06 | 112.297223 | 99.5847667 | 220.887167 | 166.974153 |
| 4.9894676 | 6.05E-07 | 2.87E-06 | 45.4926707 | 57.0988744 | 32.2127119 | 49.8681194 |
| -4.9891289 | 6.07E-07 | 2.87E-06 | 76.2309617 | 199.169533 | 169.116737 | 203.394689 |
| -4.9887015 | 6.08E-07 | 2.88E-06 | 126.641759 | 135.846611 | 109.29313 | 206.196269 |
| -4.9886716 | 6.08E-07 | 2.88E-06 | 107.379097 | 202.958084 | 106.992222 | 141.199619 |
| -4.9881473 | 6.10E-07 | 2.89E-06 | 79.0998688 | 209.182132 | 196.727633 | 152.966254 |
| -4.9878419 | 6.11E-07 | 2.89E-06 | 74.5915861 | 235.972599 | 170.267191 | 304.811876 |
| -4.9878138 | 6.11E-07 | 2.89E-06 | 77.4604933 | 173.461509 | 178.320369 | 185.464579 |
| -4.9876222 | 6.11E-07 | 2.89E-06 | 77.0506494 | 216.218013 | 159.913105 | 208.437533 |
| -4.9876273 | 6.11E-07 | 2.89E-06 | 68.8537718 | 226.230611 | 224.338529 | 219.643852 |
| -4.9875272 | 6.12E-07 | 2.89E-06 | 85.247527 | 145.317988 | 148.408565 | 191.067738 |
| -4.9868761 | 6.14E-07 | 2.90E-06 | 65.5750208 | 444.072289 | 333.631659 | 201.153426 |
| -4.9865492 | 6.15E-07 | 2.91E-06 | 257.7918 | 127.457677 | 105.841768 | 134.475828 |
| -4.9863497 | 6.15E-07 | 2.91E-06 | 88.9361219 | 165.343186 | 128.850847 | 144.561515 |
| -4.986009 | 6.16E-07 | 2.92E-06 | 84.4278393 | 137.199665 | 177.169915 | 165.293205 |
| -4.9859061 | 6.17E-07 | 2.92E-06 | 1088.95519 | 202.146252 | 153.010381 | 159.690045 |
| -4.985453 | 6.18E-07 | 2.92E-06 | 81.5589321 | 236.513821 | 150.709473 | 178.740788 |
| -4.9848467 | 6.20E-07 | 2.93E-06 | 121.723632 | 193.486707 | 88.5849576 | 193.869318 |
| -4.9843597 | 6.22E-07 | 2.94E-06 | 134.018949 | 176.97945 | 83.9831417 | 169.215416 |
| -4.9843813 | 6.22E-07 | 2.94E-06 | 47.5418901 | 397.527235 | 469.38523 | 352.438732 |
| -4.9841988 | 6.22E-07 | 2.94E-06 | 93.8542485 | 186.992048 | 118.496762 | 147.92341 |
| -4.9841158 | 6.22E-07 | 2.94E-06 | 79.9195566 | 238.949318 | 186.373547 | 153.52657 |
| -4.9840946 | 6.23E-07 | 2.94E-06 | 93.4444046 | 147.482875 | 117.346308 | 137.837723 |
| -4.9839755 | 6.23E-07 | 2.94E-06 | 69.2636157 | 203.228695 | 226.639437 | 234.772382 |

| | | | | | | |
|---|---|---|---|---|---|---|
| -4.9838236 | 6.23E-07 | 2.94E-06 | 118.035037 | 188.074492 | 94.3372276 | 144.001199 |
| -4.9837019 | 6.24E-07 | 2.95E-06 | 93.4444046 | 145.047378 | 116.195854 | 165.293205 |
| -4.9836017 | 6.24E-07 | 2.95E-06 | 86.0672148 | 154.518755 | 141.505841 | 142.320251 |
| -4.9832256 | 6.25E-07 | 2.95E-06 | 90.9853413 | 136.387833 | 125.399485 | 139.518671 |
| -4.9831289 | 6.26E-07 | 2.95E-06 | 88.1164342 | 182.121054 | 352.038923 | 156.888466 |
| -4.9829486 | 6.26E-07 | 2.96E-06 | 105.329877 | 139.635162 | 102.390406 | 126.071088 |
| -4.982604 | 6.27E-07 | 2.96E-06 | 58.197831 | 269.528336 | 331.330751 | 296.967453 |
| -4.9822066 | 6.29E-07 | 2.97E-06 | 112.707067 | 113.927138 | 106.992222 | 111.502874 |
| -4.9819621 | 6.29E-07 | 2.97E-06 | 97.5428434 | 140.176384 | 115.0454 | 121.588561 |
| -4.9819136 | 6.30E-07 | 2.97E-06 | 262.709927 | 465.991762 | 74.7795097 | 180.982051 |
| -4.9817904 | 6.30E-07 | 2.97E-06 | 72.5423667 | 192.133653 | 215.134897 | 195.550266 |
| -4.9817064 | 6.30E-07 | 2.97E-06 | 127.871291 | 107.432479 | 109.29313 | 109.26161 |
| 4.98155303 | 6.31E-07 | 2.97E-06 | 81.1490882 | 28.9553534 | 21.8586259 | 29.6967453 |
| -4.9813036 | 6.32E-07 | 2.98E-06 | 97.9526873 | 166.425629 | 119.647216 | 121.588561 |
| -4.9807932 | 6.33E-07 | 2.98E-06 | 77.4604933 | 160.201581 | 203.630357 | 183.783631 |
| 4.9807583 | 6.33E-07 | 2.98E-06 | 3.68859492 | 10.0125988 | 6.90272397 | 8.40473923 |
| -4.980653 | 6.34E-07 | 2.99E-06 | 52.8698605 | 339.887139 | 356.640739 | 320.500723 |
| -4.9805668 | 6.34E-07 | 2.99E-06 | 70.0833035 | 217.841677 | 201.329449 | 219.643852 |
| -4.9802088 | 6.35E-07 | 2.99E-06 | 61.0667381 | 311.473007 | 250.798971 | 267.831023 |
| -4.9800552 | 6.36E-07 | 2.99E-06 | 70.9029912 | 210.805797 | 204.780811 | 205.075637 |
| -4.9800104 | 6.36E-07 | 2.99E-06 | 234.430699 | 174.002731 | 117.346308 | 94.6933953 |
| -4.9800386 | 6.36E-07 | 2.99E-06 | 81.968776 | 180.768 | 149.559019 | 156.32815 |
| -4.9797198 | 6.37E-07 | 3.00E-06 | 95.0837801 | 139.635162 | 119.647216 | 122.148877 |
| -4.9796647 | 6.37E-07 | 3.00E-06 | 475.009057 | 925.759475 | 1022.7536 | 1103.82242 |
| -4.9791959 | 6.38E-07 | 3.00E-06 | 109.018472 | 112.844695 | 142.656295 | 104.779082 |
| -4.9788217 | 6.40E-07 | 3.01E-06 | 92.214873 | 244.902755 | 253.099879 | 117.106033 |
| -4.9785504 | 6.41E-07 | 3.01E-06 | 75.8211178 | 186.450827 | 230.090799 | 168.094785 |
| -4.9783855 | 6.41E-07 | 3.02E-06 | 73.3620545 | 203.228695 | 186.373547 | 192.18837 |
| -4.9783047 | 6.41E-07 | 3.02E-06 | 88.1164342 | 232.995881 | 123.098578 | 219.643852 |
| -4.9779174 | 6.43E-07 | 3.02E-06 | 86.0672148 | 172.108456 | 128.850847 | 163.051941 |
| -4.9777197 | 6.43E-07 | 3.03E-06 | 87.7065903 | 192.945486 | 205.931265 | 202.274057 |
| -4.9774402 | 6.44E-07 | 3.03E-06 | 63.9356453 | 247.338252 | 239.294431 | 258.305652 |
| -4.9772562 | 6.45E-07 | 3.03E-06 | 75.8211178 | 209.723354 | 166.815829 | 186.585211 |
| -4.9771755 | 6.45E-07 | 3.03E-06 | 105.329877 | 174.273342 | 116.195854 | 113.744138 |
| -4.9771839 | 6.45E-07 | 3.03E-06 | 100.411751 | 113.927138 | 133.452663 | 193.309002 |
| -4.9766862 | 6.47E-07 | 3.04E-06 | 87.2967464 | 149.918372 | 164.514921 | 127.19172 |
| 4.97655303 | 6.47E-07 | 3.04E-06 | 12.7051603 | 19.2133653 | 39.1154358 | 22.4126379 |
| -4.9762073 | 6.48E-07 | 3.05E-06 | 70.9029912 | 223.795114 | 184.072639 | 228.608907 |
| -4.9760648 | 6.49E-07 | 3.05E-06 | 92.214873 | 125.02218 | 126.549939 | 132.234564 |
| -4.9758752 | 6.50E-07 | 3.05E-06 | 64.755333 | 231.101605 | 234.692615 | 268.391339 |
| -4.9757526 | 6.50E-07 | 3.05E-06 | 147.543797 | 201.33442 | 93.1867736 | 119.347297 |
| -4.9756214 | 6.50E-07 | 3.05E-06 | 74.1817423 | 231.101605 | 163.364467 | 273.434183 |
| -4.9755606 | 6.51E-07 | 3.05E-06 | 83.1983076 | 152.353869 | 339.383929 | 203.955005 |

| | | | | | | |
|---|---|---|---|---|---|---|
| -4.9753485 | 6.51E-07 | 3.06E-06 | 101.641282 | 103.373318 | 135.753571 | 133.915512 |
| 4.97519392 | 6.52E-07 | 3.06E-06 | 0 | 0 | 0 | 0 |
| -4.9751086 | 6.52E-07 | 3.06E-06 | 82.7884637 | 200.251977 | 136.904025 | 175.378892 |
| -4.9749278 | 6.53E-07 | 3.06E-06 | 82.3786199 | 188.345102 | 135.753571 | 198.912162 |
| -4.9747452 | 6.53E-07 | 3.06E-06 | 94.6739363 | 109.867976 | 179.470823 | 177.620156 |
| -4.9744417 | 6.54E-07 | 3.07E-06 | 85.6573709 | 164.531354 | 130.001301 | 156.888466 |
| -4.9744505 | 6.54E-07 | 3.07E-06 | 93.0345607 | 121.233629 | 141.505841 | 123.829825 |
| -4.9743814 | 6.55E-07 | 3.07E-06 | 134.838636 | 149.106539 | 83.9831417 | 167.534469 |
| -4.9732503 | 6.58E-07 | 3.09E-06 | 55.3289238 | 372.631043 | 260.002603 | 355.800627 |
| -4.9731355 | 6.59E-07 | 3.09E-06 | 86.4770587 | 132.599282 | 151.859927 | 141.199619 |
| -4.972494 | 6.61E-07 | 3.10E-06 | 107.78894 | 197.275258 | 165.665375 | 194.429634 |
| -4.9723961 | 6.61E-07 | 3.10E-06 | 75.4112739 | 171.567234 | 179.470823 | 194.429634 |
| -4.9721838 | 6.62E-07 | 3.10E-06 | 118.854725 | 107.161869 | 105.841768 | 112.06319 |
| -4.9718177 | 6.63E-07 | 3.11E-06 | 91.8050291 | 130.434396 | 119.647216 | 149.044042 |
| -4.9716071 | 6.64E-07 | 3.11E-06 | 104.510189 | 172.649677 | 97.7885896 | 180.421735 |
| -4.9715426 | 6.64E-07 | 3.11E-06 | 184.019902 | 93.090108 | 115.0454 | 150.164674 |
| -4.9715265 | 6.64E-07 | 3.11E-06 | 84.0179954 | 145.047378 | 151.859927 | 142.880567 |
| -4.9714288 | 6.65E-07 | 3.11E-06 | 90.5754975 | 184.827162 | 125.399485 | 138.398039 |
| -4.9713536 | 6.65E-07 | 3.11E-06 | 112.707067 | 92.8194972 | 167.966283 | 184.904263 |
| 4.97086224 | 6.67E-07 | 3.12E-06 | 11.0657848 | 6.4946587 | 6.90272397 | 8.96505518 |
| -4.9703042 | 6.68E-07 | 3.13E-06 | 86.4770587 | 134.764168 | 162.214013 | 135.59646 |
| -4.9703052 | 6.68E-07 | 3.13E-06 | 93.4444046 | 127.998899 | 117.346308 | 130.553616 |
| -4.9700716 | 6.69E-07 | 3.13E-06 | 87.2967464 | 159.389749 | 141.505841 | 127.752036 |
| -4.969509 | 6.71E-07 | 3.14E-06 | 106.559409 | 111.491641 | 121.948124 | 105.339398 |
| -4.9692545 | 6.72E-07 | 3.14E-06 | 81.968776 | 167.778683 | 138.054479 | 175.378892 |
| -4.9687419 | 6.74E-07 | 3.15E-06 | 82.7884637 | 193.216096 | 138.054479 | 160.810677 |
| -4.9682583 | 6.76E-07 | 3.16E-06 | 77.4604933 | 169.402348 | 165.665375 | 170.896364 |
| -4.9682079 | 6.76E-07 | 3.16E-06 | 90.9853413 | 143.694324 | 116.195854 | 145.121831 |
| -4.9680377 | 6.76E-07 | 3.16E-06 | 61.476582 | 243.820312 | 333.631659 | 247.099333 |
| -4.9678312 | 6.77E-07 | 3.16E-06 | 78.2801811 | 190.239378 | 149.559019 | 180.982051 |
| -4.9674179 | 6.79E-07 | 3.17E-06 | 84.8376831 | 444.342899 | 429.11934 | 401.746535 |
| -4.9673218 | 6.79E-07 | 3.17E-06 | 84.4278393 | 139.905773 | 141.505841 | 177.620156 |
| -4.9671887 | 6.79E-07 | 3.17E-06 | 77.8703372 | 234.078324 | 148.408565 | 298.648401 |
| -4.9670981 | 6.80E-07 | 3.17E-06 | 108.198784 | 130.434396 | 149.559019 | 131.674248 |
| -4.9668306 | 6.81E-07 | 3.18E-06 | 79.0998688 | 204.581749 | 148.408565 | 172.577312 |
| -4.9665811 | 6.81E-07 | 3.18E-06 | 95.0837801 | 254.103522 | 111.594038 | 205.075637 |
| -4.9665108 | 6.82E-07 | 3.18E-06 | 56.9682993 | 318.508887 | 261.153057 | 309.294404 |
| -4.9663149 | 6.82E-07 | 3.19E-06 | 77.8703372 | 192.133653 | 150.709473 | 180.421735 |
| -4.966276 | 6.83E-07 | 3.19E-06 | 58.197831 | 277.91727 | 306.020763 | 265.58976 |
| -4.9662388 | 6.83E-07 | 3.19E-06 | 74.1817423 | 174.814563 | 202.479903 | 178.740788 |
| -4.9659643 | 6.84E-07 | 3.19E-06 | 646.323799 | 173.190899 | 123.098578 | 155.207518 |
| -4.9658692 | 6.84E-07 | 3.19E-06 | 84.4278393 | 134.493557 | 148.408565 | 179.86142 |
| -4.9654733 | 6.85E-07 | 3.20E-06 | 95.493624 | 155.330587 | 181.771731 | 178.740788 |

| | | | | | | |
|---|---|---|---|---|---|---|
| -4.9654269 | 6.86E-07 | 3.20E-06 | 85.6573709 | 131.787449 | 147.258111 | 142.880567 |
| -4.9653415 | 6.86E-07 | 3.20E-06 | 86.8869025 | 185.097773 | 120.79767 | 189.947107 |
| -4.9651675 | 6.86E-07 | 3.20E-06 | 79.0998688 | 204.040527 | 146.107657 | 175.939208 |
| -4.9651816 | 6.86E-07 | 3.20E-06 | 72.1325229 | 200.793198 | 178.320369 | 204.515321 |
| -4.9650304 | 6.87E-07 | 3.20E-06 | 188.528185 | 124.751569 | 207.081719 | 85.1680242 |
| -4.9649647 | 6.87E-07 | 3.20E-06 | 80.7392443 | 148.835929 | 156.461743 | 159.690045 |
| -4.9647143 | 6.88E-07 | 3.21E-06 | 81.1490882 | 154.789366 | 149.559019 | 154.647202 |
| -4.9647206 | 6.88E-07 | 3.21E-06 | 92.214873 | 140.446994 | 127.700393 | 117.106033 |
| -4.9646708 | 6.88E-07 | 3.21E-06 | 82.7884637 | 143.423713 | 143.806749 | 168.655101 |
| -4.9645512 | 6.89E-07 | 3.21E-06 | 61.476582 | 203.769917 | 402.658898 | 348.51652 |
| -4.9642523 | 6.90E-07 | 3.21E-06 | 77.4604933 | 154.789366 | 188.674455 | 171.45668 |
| -4.9641419 | 6.90E-07 | 3.21E-06 | 78.2801811 | 193.216096 | 159.913105 | 158.009098 |
| -4.9641087 | 6.90E-07 | 3.21E-06 | 102.870814 | 103.643928 | 188.674455 | 140.078987 |
| -4.963389 | 6.93E-07 | 3.22E-06 | 95.9034679 | 151.271426 | 240.444885 | 119.907613 |
| -4.9632542 | 6.93E-07 | 3.23E-06 | 64.3454891 | 329.333318 | 212.833989 | 244.297754 |
| -4.9628659 | 6.95E-07 | 3.23E-06 | 108.608628 | 140.176384 | 96.6381356 | 122.709193 |
| -4.9626745 | 6.95E-07 | 3.24E-06 | 104.510189 | 115.280192 | 117.346308 | 192.18837 |
| -4.9622036 | 6.97E-07 | 3.24E-06 | 84.0179954 | 154.248144 | 133.452663 | 149.044042 |
| -4.9621486 | 6.97E-07 | 3.24E-06 | 79.5097127 | 176.438228 | 163.364467 | 146.802779 |
| -4.9621203 | 6.97E-07 | 3.24E-06 | 73.3620545 | 187.803881 | 177.169915 | 188.266159 |
| -4.9619116 | 6.98E-07 | 3.25E-06 | 64.3454891 | 221.900839 | 232.391707 | 256.624704 |
| -4.9617545 | 6.99E-07 | 3.25E-06 | 74.1817423 | 215.676791 | 164.514921 | 188.826475 |
| -4.9616787 | 6.99E-07 | 3.25E-06 | 92.6247169 | 245.443977 | 115.0454 | 188.266159 |
| -4.9614619 | 7.00E-07 | 3.25E-06 | 237.70945 | 260.59818 | 69.0272397 | 174.25826 |
| -4.9611667 | 7.01E-07 | 3.26E-06 | 94.2640924 | 136.929054 | 148.408565 | 108.701294 |
| -4.9610094 | 7.01E-07 | 3.26E-06 | 70.9029912 | 194.027929 | 186.373547 | 214.601008 |
| -4.9600516 | 7.05E-07 | 3.27E-06 | 95.9034679 | 378.58448 | 278.409867 | 296.407137 |
| -4.9600298 | 7.05E-07 | 3.27E-06 | 93.4444046 | 113.385916 | 138.054479 | 124.950457 |
| -4.9599702 | 7.05E-07 | 3.27E-06 | 81.5589321 | 138.011497 | 185.223093 | 158.009098 |
| -4.9599631 | 7.05E-07 | 3.27E-06 | 105.329877 | 164.260743 | 102.390406 | 122.148877 |
| -4.9594419 | 7.07E-07 | 3.28E-06 | 95.9034679 | 222.17145 | 110.443584 | 165.293205 |
| -4.9592591 | 7.08E-07 | 3.29E-06 | 81.968776 | 159.389749 | 138.054479 | 155.207518 |
| -4.9590237 | 7.08E-07 | 3.29E-06 | 80.3294005 | 177.791282 | 153.010381 | 146.242463 |
| -4.9589203 | 7.09E-07 | 3.29E-06 | 68.8537718 | 218.112288 | 222.037621 | 188.266159 |
| -4.9588509 | 7.09E-07 | 3.29E-06 | 69.6734596 | 224.606947 | 177.169915 | 269.511971 |
| -4.9588626 | 7.09E-07 | 3.29E-06 | 101.231438 | 103.643928 | 102.390406 | 97.4949751 |
| 4.95873619 | 7.10E-07 | 3.29E-06 | 13.524848 | 4.32977247 | 5.75226998 | 1.1206319 |
| -4.9581953 | 7.12E-07 | 3.30E-06 | 64.755333 | 242.467258 | 308.321671 | 203.955005 |
| -4.9577193 | 7.13E-07 | 3.31E-06 | 135.24848 | 80.912623 | 291.064861 | 240.935858 |
| -4.9568459 | 7.16E-07 | 3.32E-06 | 84.8376831 | 130.434396 | 174.869007 | 142.880567 |
| -4.9567234 | 7.17E-07 | 3.32E-06 | 71.722679 | 192.133653 | 202.479903 | 178.180472 |
| -4.9565004 | 7.18E-07 | 3.33E-06 | 78.6900249 | 189.968767 | 142.656295 | 175.939208 |
| -4.9562007 | 7.19E-07 | 3.33E-06 | 79.9195566 | 154.518755 | 147.258111 | 195.550266 |

| | | | | | | |
|---|---|---|---|---|---|---|
| -4.956123 | 7.19E-07 | 3.33E-06 | 120.084257 | 137.740887 | 94.3372276 | 111.502874 |
| -4.9553782 | 7.22E-07 | 3.35E-06 | 91.8050291 | 179.956168 | 109.29313 | 170.336048 |
| -4.9548056 | 7.24E-07 | 3.35E-06 | 104.510189 | 154.518755 | 279.560321 | 113.744138 |
| -4.9546436 | 7.25E-07 | 3.36E-06 | 72.1325229 | 171.296623 | 216.285351 | 190.507423 |
| -4.9543548 | 7.26E-07 | 3.36E-06 | 72.9522106 | 221.630228 | 173.718553 | 178.740788 |
| -4.9543221 | 7.26E-07 | 3.36E-06 | 68.0340841 | 234.348935 | 204.780811 | 196.110582 |
| -4.9542697 | 7.26E-07 | 3.36E-06 | 77.0506494 | 215.135569 | 156.461743 | 165.853521 |
| 4.95378402 | 7.28E-07 | 3.37E-06 | 12.7051603 | 12.9893174 | 14.9559019 | 15.6888466 |
| -4.9534849 | 7.29E-07 | 3.37E-06 | 91.3951852 | 124.480958 | 116.195854 | 139.518671 |
| -4.9534601 | 7.29E-07 | 3.37E-06 | 111.477535 | 124.210348 | 249.648517 | 237.013646 |
| -4.9533159 | 7.30E-07 | 3.38E-06 | 125.002383 | 146.941653 | 116.195854 | 117.106033 |
| -4.9527063 | 7.32E-07 | 3.39E-06 | 94.6739363 | 104.996982 | 148.408565 | 149.604358 |
| -4.9524886 | 7.33E-07 | 3.39E-06 | 103.280658 | 143.964935 | 149.559019 | 131.113932 |
| -4.9524412 | 7.33E-07 | 3.39E-06 | 270.087117 | 112.844695 | 119.647216 | 126.071088 |
| -4.9524253 | 7.33E-07 | 3.39E-06 | 75.8211178 | 213.241294 | 146.107657 | 225.807327 |
| -4.9523599 | 7.33E-07 | 3.39E-06 | 83.1983076 | 136.658443 | 158.762651 | 138.398039 |
| -4.9522091 | 7.34E-07 | 3.39E-06 | 87.2967464 | 158.577917 | 556.819734 | 236.45333 |
| -4.9520357 | 7.34E-07 | 3.40E-06 | 71.722679 | 182.662276 | 195.577179 | 182.662999 |
| -4.9518794 | 7.35E-07 | 3.40E-06 | 70.0833035 | 215.135569 | 225.488983 | 173.137628 |
| -4.9518954 | 7.35E-07 | 3.40E-06 | 71.3128351 | 202.687474 | 193.276271 | 176.499524 |
| -4.9517047 | 7.36E-07 | 3.40E-06 | 76.2309617 | 148.565318 | 192.125817 | 185.464579 |
| 4.95166681 | 7.36E-07 | 3.40E-06 | 92.6247169 | 74.688575 | 73.6290557 | 66.6775979 |
| -4.9515962 | 7.36E-07 | 3.40E-06 | 84.0179954 | 147.753485 | 142.656295 | 131.113932 |
| -4.9515676 | 7.36E-07 | 3.40E-06 | 78.6900249 | 175.897006 | 215.134897 | 141.759935 |
| -4.9512381 | 7.37E-07 | 3.41E-06 | 68.8537718 | 315.261558 | 187.524001 | 215.161324 |
| -4.9501118 | 7.42E-07 | 3.43E-06 | 61.476582 | 305.79018 | 215.134897 | 262.227864 |
| -4.9500867 | 7.42E-07 | 3.43E-06 | 70.0833035 | 165.343186 | 242.745793 | 256.064389 |
| -4.9500166 | 7.42E-07 | 3.43E-06 | 84.4278393 | 192.945486 | 127.700393 | 153.52657 |
| -4.9498569 | 7.43E-07 | 3.43E-06 | 69.6734596 | 239.49054 | 186.373547 | 191.628054 |
| -4.949569 | 7.44E-07 | 3.43E-06 | 572.142056 | 1121.41107 | 1217.18033 | 1187.86981 |
| -4.9495453 | 7.44E-07 | 3.43E-06 | 73.7718984 | 186.450827 | 169.116737 | 174.25826 |
| -4.9495368 | 7.44E-07 | 3.43E-06 | 113.936599 | 104.18515 | 112.744492 | 103.65845 |
| -4.949162 | 7.45E-07 | 3.44E-06 | 109.018472 | 124.210348 | 94.3372276 | 122.709193 |
| -4.9489867 | 7.46E-07 | 3.44E-06 | 96.7231557 | 169.943569 | 187.524001 | 101.977503 |
| -4.9489042 | 7.46E-07 | 3.44E-06 | 84.0179954 | 160.201581 | 125.399485 | 189.947107 |
| -4.9484628 | 7.48E-07 | 3.45E-06 | 77.8703372 | 152.083258 | 182.922185 | 154.086886 |
| -4.9483939 | 7.48E-07 | 3.45E-06 | 130.740198 | 173.190899 | 118.496762 | 140.639303 |
| -4.9479424 | 7.50E-07 | 3.46E-06 | 66.8045524 | 209.993965 | 195.577179 | 256.064389 |
| -4.9478587 | 7.50E-07 | 3.46E-06 | 98.3625312 | 162.637078 | 178.320369 | 99.7362389 |
| -4.9476994 | 7.51E-07 | 3.46E-06 | 109.018472 | 250.856192 | 93.1867736 | 204.515321 |
| -4.9477179 | 7.51E-07 | 3.46E-06 | 81.968776 | 132.869893 | 151.859927 | 179.301104 |
| -4.9475639 | 7.51E-07 | 3.46E-06 | 81.1490882 | 131.516839 | 174.869007 | 163.051941 |
| -4.9474467 | 7.52E-07 | 3.46E-06 | 116.395662 | 162.907689 | 103.54086 | 104.218766 |

| | | | | | | |
|---|---|---|---|---|---|---|
| -4.9467787 | 7.55E-07 | 3.48E-06 | 86.0672148 | 164.531354 | 135.753571 | 124.950457 |
| -4.9467379 | 7.55E-07 | 3.48E-06 | 63.5258014 | 379.666923 | 195.577179 | 278.477026 |
| -4.9459362 | 7.58E-07 | 3.49E-06 | 31.5579788 | 528.502852 | 577.527906 | 611.3047 |
| 4.94575493 | 7.58E-07 | 3.49E-06 | 65.5750208 | 35.7206228 | 34.5136199 | 48.1871716 |
| -4.945157 | 7.61E-07 | 3.50E-06 | 86.8869025 | 141.800048 | 119.647216 | 138.398039 |
| -4.9450091 | 7.61E-07 | 3.50E-06 | 484.84531 | 197.275258 | 200.178995 | 76.763285 |
| -4.9449231 | 7.62E-07 | 3.51E-06 | 71.3128351 | 177.25006 | 226.639437 | 179.301104 |
| -4.9448734 | 7.62E-07 | 3.51E-06 | 88.9361219 | 116.633246 | 130.001301 | 138.958355 |
| -4.9447356 | 7.62E-07 | 3.51E-06 | 102.46097 | 142.34127 | 103.54086 | 111.502874 |
| -4.9446854 | 7.63E-07 | 3.51E-06 | 69.2636157 | 179.414947 | 216.285351 | 201.713742 |
| -4.9444127 | 7.64E-07 | 3.51E-06 | 103.690502 | 292.530252 | 196.727633 | 238.694594 |
| -4.9443173 | 7.64E-07 | 3.51E-06 | 87.7065903 | 158.307306 | 117.346308 | 137.837723 |
| -4.9441304 | 7.65E-07 | 3.52E-06 | 74.1817423 | 208.3703 | 190.975363 | 152.405938 |
| -4.9437447 | 7.66E-07 | 3.52E-06 | 305.333691 | 142.34127 | 164.514921 | 168.094785 |
| -4.9437327 | 7.66E-07 | 3.52E-06 | 80.7392443 | 152.624479 | 135.753571 | 183.223315 |
| -4.9435719 | 7.67E-07 | 3.53E-06 | 81.5589321 | 156.954252 | 130.001301 | 169.775732 |
| -4.9433108 | 7.68E-07 | 3.53E-06 | 57.3781432 | 257.621462 | 273.808051 | 277.916711 |
| -4.9432283 | 7.68E-07 | 3.53E-06 | 119.674413 | 106.350036 | 103.54086 | 104.779082 |
| -4.9428013 | 7.70E-07 | 3.54E-06 | 123.363008 | 107.973701 | 95.4876816 | 115.985401 |
| 4.94269001 | 7.71E-07 | 3.54E-06 | 2.86890716 | 0.81183234 | 1.150454 | 0.56031595 |
| 4.94255591 | 7.71E-07 | 3.54E-06 | 16.3937552 | 11.9068743 | 9.20363196 | 7.28410733 |
| -4.942449 | 7.71E-07 | 3.54E-06 | 64.3454891 | 203.769917 | 257.701695 | 223.566063 |
| -4.9424239 | 7.72E-07 | 3.54E-06 | 82.7884637 | 162.907689 | 151.859927 | 126.631404 |
| -4.9421696 | 7.73E-07 | 3.55E-06 | 84.8376831 | 135.846611 | 127.700393 | 138.398039 |
| -4.9420928 | 7.73E-07 | 3.55E-06 | 63.9356453 | 249.773749 | 210.533081 | 222.445432 |
| -4.9419771 | 7.73E-07 | 3.55E-06 | 64.3454891 | 254.915354 | 193.276271 | 251.581861 |
| -4.9408976 | 7.78E-07 | 3.57E-06 | 72.9522106 | 177.25006 | 251.949425 | 168.655101 |
| -4.9400739 | 7.81E-07 | 3.58E-06 | 138.117388 | 92.0076649 | 123.098578 | 110.942558 |
| -4.9400431 | 7.81E-07 | 3.58E-06 | 96.3133118 | 151.542036 | 124.249032 | 104.779082 |
| -4.939822 | 7.82E-07 | 3.59E-06 | 3175.06054 | 16147.3452 | 28648.6054 | 14021.3463 |
| -4.9390786 | 7.85E-07 | 3.60E-06 | 68.4439279 | 186.450827 | 195.577179 | 270.632603 |
| -4.9388471 | 7.86E-07 | 3.60E-06 | 58.197831 | 257.892073 | 272.657597 | 251.021545 |
| -4.9387765 | 7.86E-07 | 3.60E-06 | 79.0998688 | 159.66036 | 144.957203 | 146.802779 |
| -4.9385298 | 7.87E-07 | 3.61E-06 | 76.2309617 | 415.658157 | 139.204933 | 344.594308 |
| -4.9382541 | 7.88E-07 | 3.61E-06 | 86.8869025 | 124.480958 | 173.718553 | 129.432984 |
| -4.9381951 | 7.88E-07 | 3.61E-06 | 84.0179954 | 165.884408 | 150.709473 | 121.588561 |
| -4.9380469 | 7.89E-07 | 3.62E-06 | 82.3786199 | 126.375234 | 178.320369 | 152.966254 |
| -4.9379787 | 7.89E-07 | 3.62E-06 | 66.3947085 | 195.922204 | 231.241253 | 203.394689 |
| -4.9377793 | 7.90E-07 | 3.62E-06 | 92.6247169 | 178.332503 | 104.691314 | 158.569413 |
| -4.9377448 | 7.90E-07 | 3.62E-06 | 88.1164342 | 171.026012 | 124.249032 | 125.510772 |
| -4.9376536 | 7.91E-07 | 3.62E-06 | 91.3951852 | 264.657342 | 130.001301 | 147.363094 |
| -4.937603 | 7.91E-07 | 3.62E-06 | 70.4931473 | 195.380983 | 184.072639 | 177.620156 |
| -4.9375177 | 7.91E-07 | 3.62E-06 | 98.3625312 | 153.165701 | 247.347609 | 110.382242 |

| | | | | | | |
|---|---|---|---|---|---|---|
| -4.9374955 | 7.91E-07 | 3.62E-06 | 80.7392443 | 159.93097 | 130.001301 | 161.931309 |
| -4.9372024 | 7.93E-07 | 3.63E-06 | 102.051126 | 236.24321 | 184.072639 | 227.488275 |
| -4.9369184 | 7.94E-07 | 3.63E-06 | 75.4112739 | 181.850444 | 151.859927 | 165.293205 |
| -4.9368191 | 7.94E-07 | 3.63E-06 | 79.9195566 | 193.757318 | 140.355387 | 148.483726 |
| -4.9366486 | 7.95E-07 | 3.64E-06 | 63.5258014 | 246.255809 | 216.285351 | 214.601008 |
| -4.936524 | 7.95E-07 | 3.64E-06 | 83.1983076 | 132.869893 | 169.116737 | 252.702493 |
| -4.9364802 | 7.95E-07 | 3.64E-06 | 33.1973543 | 378.58448 | 876.645944 | 629.23481 |
| -4.936366 | 7.96E-07 | 3.64E-06 | 69.2636157 | 182.932887 | 187.524001 | 205.075637 |
| -4.9361754 | 7.97E-07 | 3.64E-06 | 63.9356453 | 305.248959 | 197.878087 | 231.410487 |
| -4.9361025 | 7.97E-07 | 3.64E-06 | 84.4278393 | 135.576 | 139.204933 | 125.510772 |
| -4.9359126 | 7.98E-07 | 3.65E-06 | 66.8045524 | 441.366181 | 186.373547 | 262.227864 |
| -4.9353938 | 8.00E-07 | 3.66E-06 | 82.7884637 | 142.34127 | 127.700393 | 146.242463 |
| -4.9352574 | 8.00E-07 | 3.66E-06 | 77.4604933 | 174.002731 | 138.054479 | 175.939208 |
| -4.9352243 | 8.01E-07 | 3.66E-06 | 73.3620545 | 174.002731 | 161.063559 | 179.301104 |
| -4.9351431 | 8.01E-07 | 3.66E-06 | 89.3459658 | 153.436312 | 184.072639 | 110.382242 |
| -4.9350859 | 8.01E-07 | 3.66E-06 | 64.3454891 | 369.924935 | 204.780811 | 229.729539 |
| -4.9341641 | 8.05E-07 | 3.68E-06 | 195.085687 | 303.895905 | 284.162137 | 397.264008 |
| -4.9340774 | 8.05E-07 | 3.68E-06 | 78.2801811 | 310.661174 | 133.452663 | 256.064389 |
| -4.9338125 | 8.06E-07 | 3.68E-06 | 95.9034679 | 413.763881 | 104.691314 | 398.944955 |
| -4.9334144 | 8.08E-07 | 3.69E-06 | 82.3786199 | 129.622563 | 153.010381 | 137.277407 |
| -4.9329424 | 8.10E-07 | 3.70E-06 | 56.5584554 | 313.367282 | 289.914407 | 233.651751 |
| -4.9317783 | 8.15E-07 | 3.72E-06 | 73.3620545 | 157.224863 | 171.417645 | 188.266159 |
| -4.9309798 | 8.18E-07 | 3.73E-06 | 84.0179954 | 151.542036 | 121.948124 | 138.398039 |
| -4.930716 | 8.19E-07 | 3.74E-06 | 157.38005 | 330.686372 | 232.391707 | 71.7204414 |
| -4.9303378 | 8.21E-07 | 3.75E-06 | 79.9195566 | 159.93097 | 132.302209 | 150.72499 |
| -4.9300674 | 8.22E-07 | 3.75E-06 | 93.8542485 | 137.470276 | 104.691314 | 121.588561 |
| -4.9300756 | 8.22E-07 | 3.75E-06 | 67.2143963 | 197.81648 | 200.178995 | 193.869318 |
| -4.9299115 | 8.23E-07 | 3.75E-06 | 93.4444046 | 250.585581 | 170.267191 | 110.942558 |
| -4.9293808 | 8.25E-07 | 3.76E-06 | 109.428316 | 201.33442 | 96.6381356 | 131.113932 |
| -4.9293361 | 8.25E-07 | 3.76E-06 | 78.2801811 | 175.626396 | 245.046701 | 138.958355 |
| -4.9292732 | 8.25E-07 | 3.76E-06 | 94.6739363 | 113.115306 | 109.29313 | 117.106033 |
| -4.9289484 | 8.27E-07 | 3.77E-06 | 92.214873 | 208.099689 | 150.709473 | 109.821926 |
| -4.9285125 | 8.29E-07 | 3.78E-06 | 79.9195566 | 127.457677 | 165.665375 | 166.974153 |
| -4.9281893 | 8.30E-07 | 3.78E-06 | 271.316649 | 147.753485 | 89.7354116 | 130.553616 |
| -4.9275623 | 8.33E-07 | 3.79E-06 | 80.3294005 | 153.706923 | 130.001301 | 150.164674 |
| -4.9275479 | 8.33E-07 | 3.79E-06 | 97.1329995 | 106.620647 | 111.594038 | 113.183822 |
| -4.9274623 | 8.33E-07 | 3.79E-06 | 96.7231557 | 102.020264 | 117.346308 | 122.709193 |
| -4.9270203 | 8.35E-07 | 3.80E-06 | 198.774282 | 65.4878086 | 294.516223 | 201.153426 |
| -4.926477 | 8.37E-07 | 3.81E-06 | 77.8703372 | 143.964935 | 169.116737 | 144.001199 |
| -4.9264156 | 8.38E-07 | 3.81E-06 | 66.3947085 | 182.121054 | 350.888469 | 230.289855 |
| -4.9257723 | 8.40E-07 | 3.83E-06 | 99.5920628 | 114.46836 | 101.239952 | 149.604358 |
| -4.9256928 | 8.41E-07 | 3.83E-06 | 75.4112739 | 177.25006 | 147.258111 | 159.690045 |
| -4.9256954 | 8.41E-07 | 3.83E-06 | 74.1817423 | 159.119138 | 162.214013 | 229.169223 |

| | | | | | | |
|---|---|---|---|---|---|---|
| -4.9252645 | 8.42E-07 | 3.83E-06 | 81.5589321 | 155.059976 | 228.940345 | 133.355196 |
| -4.9251465 | 8.43E-07 | 3.84E-06 | 135.658324 | 107.161869 | 95.4876816 | 109.26161 |
| -4.9250048 | 8.44E-07 | 3.84E-06 | 47.5418901 | 274.128719 | 501.597942 | 375.411686 |
| -4.9249768 | 8.44E-07 | 3.84E-06 | 77.8703372 | 146.671042 | 142.656295 | 156.888466 |
| -4.9248606 | 8.44E-07 | 3.84E-06 | 98.3625312 | 100.66721 | 215.134897 | 156.32815 |
| -4.9246046 | 8.45E-07 | 3.84E-06 | 63.9356453 | 197.81648 | 245.046701 | 212.359745 |
| -4.9244837 | 8.46E-07 | 3.85E-06 | 73.3620545 | 170.484791 | 157.612197 | 168.655101 |
| -4.9241295 | 8.47E-07 | 3.85E-06 | 66.8045524 | 236.513821 | 167.966283 | 271.192919 |
| -4.9238645 | 8.49E-07 | 3.86E-06 | 305.333691 | 235.972599 | 83.9831417 | 119.347297 |
| -4.9235574 | 8.50E-07 | 3.86E-06 | 99.5920628 | 155.601198 | 105.841768 | 109.821926 |
| 4.92336102 | 8.51E-07 | 3.87E-06 | 10.6559409 | 9.20076649 | 5.75226998 | 7.28410733 |
| -4.9231828 | 8.51E-07 | 3.87E-06 | 184.429746 | 171.567234 | 172.568099 | 231.410487 |
| -4.9230813 | 8.52E-07 | 3.87E-06 | 69.6734596 | 199.710755 | 166.815829 | 188.826475 |
| -4.9229108 | 8.53E-07 | 3.87E-06 | 81.968776 | 157.224863 | 121.948124 | 148.483726 |
| -4.9225654 | 8.54E-07 | 3.88E-06 | 130.330354 | 98.7729344 | 121.948124 | 96.9346591 |
| -4.9220618 | 8.56E-07 | 3.89E-06 | 85.6573709 | 152.624479 | 111.594038 | 158.569413 |
| -4.9219985 | 8.57E-07 | 3.89E-06 | 83.1983076 | 170.21418 | 189.824909 | 116.545717 |
| -4.9218465 | 8.57E-07 | 3.89E-06 | 81.5589321 | 165.072575 | 133.452663 | 131.113932 |
| -4.9216837 | 8.58E-07 | 3.89E-06 | 214.758193 | 371.5486 | 212.833989 | 257.18502 |
| -4.9216298 | 8.58E-07 | 3.90E-06 | 117.21535 | 217.300456 | 83.9831417 | 170.336048 |
| -4.9212942 | 8.60E-07 | 3.90E-06 | 79.5097127 | 201.063809 | 130.001301 | 159.690045 |
| -4.9212805 | 8.60E-07 | 3.90E-06 | 88.9361219 | 140.446994 | 128.850847 | 108.701294 |
| -4.920902 | 8.61E-07 | 3.91E-06 | 66.8045524 | 192.404264 | 215.134897 | 184.343947 |
| -4.9205693 | 8.63E-07 | 3.91E-06 | 97.9526873 | 152.89509 | 96.6381356 | 129.432984 |
| -4.9204905 | 8.63E-07 | 3.92E-06 | 58.197831 | 240.843593 | 257.701695 | 243.177122 |
| -4.9202185 | 8.64E-07 | 3.92E-06 | 675.01287 | 264.927953 | 314.073941 | 314.337247 |
| -4.9200575 | 8.65E-07 | 3.92E-06 | 78.6900249 | 220.547785 | 132.302209 | 168.655101 |
| 4.92006453 | 8.65E-07 | 3.92E-06 | 54.509236 | 53.0397127 | 63.2749697 | 61.0744384 |
| -4.9198868 | 8.66E-07 | 3.93E-06 | 153.281611 | 199.169533 | 241.595339 | 215.161324 |
| -4.9196393 | 8.67E-07 | 3.93E-06 | 81.5589321 | 154.248144 | 120.79767 | 158.009098 |
| 4.9195222 | 8.68E-07 | 3.93E-06 | 64.3454891 | 134.222946 | 136.904025 | 200.032794 |
| -4.9193189 | 8.68E-07 | 3.93E-06 | 86.0672148 | 167.508072 | 112.744492 | 141.759935 |
| 4.91890395 | 8.70E-07 | 3.94E-06 | 42.6237635 | 38.6973414 | 37.9649818 | 39.2221164 |
| -4.9187822 | 8.71E-07 | 3.94E-06 | 161.888333 | 93.3607188 | 108.142676 | 119.347297 |
| -4.9185081 | 8.72E-07 | 3.95E-06 | 93.4444046 | 116.633246 | 123.098578 | 104.218766 |
| -4.9182249 | 8.73E-07 | 3.95E-06 | 65.9848647 | 182.391665 | 205.931265 | 212.359745 |
| -4.9180445 | 8.74E-07 | 3.96E-06 | 93.4444046 | 117.174467 | 103.54086 | 135.036144 |
| -4.9178394 | 8.75E-07 | 3.96E-06 | 56.5584554 | 235.972599 | 265.754873 | 277.916711 |
| -4.917548 | 8.76E-07 | 3.97E-06 | 93.0345607 | 197.004647 | 101.239952 | 163.051941 |
| -4.9172606 | 8.78E-07 | 3.97E-06 | 72.9522106 | 144.235545 | 192.125817 | 190.507423 |
| -4.9168337 | 8.80E-07 | 3.98E-06 | 90.1656536 | 165.613797 | 102.390406 | 171.45668 |
| -4.9167476 | 8.80E-07 | 3.98E-06 | 147.133953 | 139.09394 | 93.1867736 | 98.615607 |
| -4.9165539 | 8.81E-07 | 3.98E-06 | 75.00143 | 161.013414 | 169.116737 | 143.440883 |

| | | | | | | |
|---|---|---|---|---|---|---|
| -4.9164033 | 8.81E-07 | 3.99E-06 | 65.5750208 | 203.499306 | 190.975363 | 202.274057 |
| -4.9161781 | 8.83E-07 | 3.99E-06 | 101.231438 | 175.626396 | 178.320369 | 204.515321 |
| -4.9157878 | 8.84E-07 | 4.00E-06 | 100.821594 | 149.37715 | 92.0363196 | 174.818576 |
| -4.915428 | 8.86E-07 | 4.01E-06 | 105.739721 | 145.85921 | 95.4876816 | 221.885116 |
| -4.9152576 | 8.87E-07 | 4.01E-06 | 78.6900249 | 152.083258 | 180.621277 | 129.432984 |
| -4.9150923 | 8.87E-07 | 4.01E-06 | 77.8703372 | 140.717605 | 157.612197 | 138.958355 |
| -4.9147166 | 8.89E-07 | 4.02E-06 | 60.6568942 | 180.49739 | 423.36707 | 335.629253 |
| -4.9145011 | 8.90E-07 | 4.02E-06 | 69.2636157 | 206.476024 | 173.718553 | 171.45668 |
| -4.9142756 | 8.91E-07 | 4.03E-06 | 139.346919 | 137.470276 | 78.2308717 | 138.958355 |
| -4.9142363 | 8.91E-07 | 4.03E-06 | 108.608628 | 108.244312 | 109.29313 | 95.8140272 |
| -4.9141832 | 8.92E-07 | 4.03E-06 | 77.0506494 | 196.734036 | 134.603117 | 159.690045 |
| -4.9141157 | 8.92E-07 | 4.03E-06 | 72.9522106 | 302.542851 | 158.762651 | 187.705843 |
| -4.9140494 | 8.92E-07 | 4.03E-06 | 89.3459658 | 156.14242 | 103.54086 | 145.121831 |
| -4.9138522 | 8.93E-07 | 4.03E-06 | 95.0837801 | 115.821413 | 115.0454 | 102.537819 |
| -4.9137445 | 8.94E-07 | 4.03E-06 | 96.7231557 | 110.409198 | 241.595339 | 141.759935 |
| -4.9136006 | 8.94E-07 | 4.04E-06 | 77.4604933 | 147.753485 | 135.753571 | 158.569413 |
| -4.9134631 | 8.95E-07 | 4.04E-06 | 690.177094 | 1412.85888 | 733.989649 | 1043.3083 |
| -4.9133953 | 8.95E-07 | 4.04E-06 | 517.63282 | 936.042685 | 893.902754 | 952.537113 |
| -4.9128284 | 8.98E-07 | 4.05E-06 | 59.0175187 | 358.829893 | 211.683535 | 248.219965 |
| -4.9127841 | 8.98E-07 | 4.05E-06 | 65.1651769 | 232.184049 | 182.922185 | 202.274057 |
| -4.9123158 | 9.00E-07 | 4.06E-06 | 84.4278393 | 138.552719 | 121.948124 | 121.588561 |
| -4.9122794 | 9.00E-07 | 4.06E-06 | 70.9029912 | 186.992048 | 155.311289 | 180.982051 |
| -4.9113134 | 9.05E-07 | 4.08E-06 | 54.9190799 | 318.508887 | 454.429328 | 216.281956 |
| -4.9111726 | 9.05E-07 | 4.08E-06 | 68.0340841 | 172.920288 | 192.125817 | 186.585211 |
| -4.9110788 | 9.06E-07 | 4.08E-06 | 47.5418901 | 307.684456 | 362.393009 | 316.578511 |
| 4.91098175 | 9.06E-07 | 4.08E-06 | 84.4278393 | 25.4374132 | 37.9649818 | 16.8094785 |
| -4.9109727 | 9.06E-07 | 4.08E-06 | 84.4278393 | 212.700072 | 115.0454 | 164.732889 |
| -4.9108945 | 9.07E-07 | 4.09E-06 | 148.773328 | 143.694324 | 80.5317797 | 119.907613 |
| -4.9107585 | 9.07E-07 | 4.09E-06 | 86.4770587 | 137.470276 | 115.0454 | 119.907613 |
| -4.9106456 | 9.08E-07 | 4.09E-06 | 111.477535 | 231.372216 | 83.9831417 | 212.359745 |
| -4.9102892 | 9.09E-07 | 4.10E-06 | 90.5754975 | 103.643928 | 126.549939 | 126.631404 |
| -4.9097664 | 9.12E-07 | 4.11E-06 | 124.592539 | 108.244312 | 285.312591 | 118.226665 |
| -4.9097745 | 9.12E-07 | 4.11E-06 | 1909.05279 | 4338.43201 | 3776.94047 | 4063.97158 |
| -4.9096018 | 9.13E-07 | 4.11E-06 | 80.7392443 | 139.635162 | 148.408565 | 126.631404 |
| -4.9095315 | 9.13E-07 | 4.11E-06 | 55.3289238 | 263.574899 | 240.444885 | 285.200818 |
| -4.9093056 | 9.14E-07 | 4.11E-06 | 59.0175187 | 351.523402 | 185.223093 | 333.387989 |
| -4.9090998 | 9.15E-07 | 4.12E-06 | 55.7387677 | 263.033677 | 228.940345 | 313.216615 |
| -4.909091 | 9.15E-07 | 4.12E-06 | 77.4604933 | 184.285941 | 142.656295 | 138.398039 |
| -4.908787 | 9.16E-07 | 4.12E-06 | 575.830651 | 1135.75344 | 901.955932 | 1076.92725 |
| -4.9085077 | 9.18E-07 | 4.13E-06 | 65.5750208 | 205.664192 | 182.922185 | 198.351846 |
| -4.9082079 | 9.19E-07 | 4.13E-06 | 64.3454891 | 221.359617 | 189.824909 | 199.472478 |
| -4.907802 | 9.21E-07 | 4.14E-06 | 128.690978 | 247.608863 | 82.8326877 | 152.966254 |
| -4.9075121 | 9.22E-07 | 4.15E-06 | 117.625194 | 125.563402 | 100.089498 | 93.5727634 |

| | | | | | | |
|---|---|---|---|---|---|---|
| -4.9071943 | 9.24E-07 | 4.15E-06 | 154.511143 | 190.239378 | 69.0272397 | 189.947107 |
| -4.9064449 | 9.27E-07 | 4.17E-06 | 93.4444046 | 112.032863 | 119.647216 | 103.098135 |
| -4.9061075 | 9.29E-07 | 4.18E-06 | 102.051126 | 185.097773 | 88.5849576 | 186.585211 |
| -4.9058357 | 9.30E-07 | 4.18E-06 | 90.5754975 | 135.846611 | 101.239952 | 128.872668 |
| 4.90576151 | 9.31E-07 | 4.18E-06 | 6.96734596 | 4.60038325 | 2.30090799 | 2.80157974 |
| -4.90568 | 9.31E-07 | 4.18E-06 | 70.9029912 | 165.613797 | 172.568099 | 161.931309 |
| -4.9056072 | 9.31E-07 | 4.18E-06 | 81.968776 | 130.163785 | 121.948124 | 138.398039 |
| -4.9056196 | 9.31E-07 | 4.18E-06 | 153.691455 | 159.119138 | 77.0804177 | 126.631404 |
| -4.904964 | 9.34E-07 | 4.20E-06 | 75.8211178 | 223.524504 | 136.904025 | 166.413837 |
| -4.904916 | 9.35E-07 | 4.20E-06 | 74.1817423 | 195.110372 | 138.054479 | 168.094785 |
| -4.9048295 | 9.35E-07 | 4.20E-06 | 107.78894 | 140.446994 | 157.612197 | 85.7283401 |
| -4.9046751 | 9.36E-07 | 4.20E-06 | 69.2636157 | 273.316887 | 158.762651 | 196.670898 |
| -4.9044744 | 9.37E-07 | 4.20E-06 | 69.2636157 | 175.085174 | 185.223093 | 164.172573 |
| 4.90414185 | 9.38E-07 | 4.21E-06 | 41.3942319 | 21.6488623 | 21.8586259 | 17.9301104 |
| -4.9036949 | 9.41E-07 | 4.22E-06 | 112.297223 | 159.66036 | 83.9831417 | 133.355196 |
| -4.9036599 | 9.41E-07 | 4.22E-06 | 100.001907 | 113.385916 | 157.612197 | 98.055291 |
| -4.9034605 | 9.42E-07 | 4.22E-06 | 107.78894 | 95.7962158 | 102.390406 | 113.744138 |
| -4.9033963 | 9.42E-07 | 4.23E-06 | 72.9522106 | 157.224863 | 185.223093 | 147.363094 |
| -4.903048 | 9.44E-07 | 4.23E-06 | 181.970683 | 323.650492 | 62.1245158 | 233.651751 |
| -4.9019841 | 9.49E-07 | 4.25E-06 | 79.5097127 | 126.375234 | 154.160835 | 134.475828 |
| -4.9017056 | 9.50E-07 | 4.26E-06 | 73.7718984 | 188.074492 | 140.355387 | 161.370993 |
| -4.9016642 | 9.50E-07 | 4.26E-06 | 77.4604933 | 130.705006 | 149.559019 | 145.682147 |
| -4.9011297 | 9.53E-07 | 4.27E-06 | 78.2801811 | 130.705006 | 139.204933 | 174.25826 |
| -4.9010427 | 9.53E-07 | 4.27E-06 | 63.1159575 | 175.626396 | 432.570702 | 279.037342 |
| -4.9006075 | 9.55E-07 | 4.28E-06 | 66.8045524 | 184.827162 | 182.922185 | 181.542367 |
| -4.9002162 | 9.57E-07 | 4.29E-06 | 69.6734596 | 241.114204 | 162.214013 | 172.577312 |
| -4.9000796 | 9.58E-07 | 4.29E-06 | 223.364915 | 728.213607 | 571.775636 | 662.293451 |
| -4.8999543 | 9.59E-07 | 4.29E-06 | 97.5428434 | 215.947402 | 230.090799 | 93.0124475 |
| -4.8996633 | 9.60E-07 | 4.30E-06 | 67.6242402 | 188.074492 | 173.718553 | 176.499524 |
| -4.8994481 | 9.61E-07 | 4.30E-06 | 61.476582 | 379.396312 | 181.771731 | 263.908812 |
| -4.8993539 | 9.62E-07 | 4.31E-06 | 79.0998688 | 142.882491 | 126.549939 | 137.837723 |
| -4.8987467 | 9.64E-07 | 4.32E-06 | 65.5750208 | 221.630228 | 163.364467 | 229.169223 |
| -4.8976398 | 9.70E-07 | 4.34E-06 | 65.9848647 | 186.992048 | 184.072639 | 186.024895 |
| 4.897597 | 9.70E-07 | 4.34E-06 | 0 | 0 | 0 | 0 |
| -4.897427 | 9.71E-07 | 4.35E-06 | 39.7548564 | 372.089821 | 440.62388 | 379.894213 |
| -4.8972529 | 9.72E-07 | 4.35E-06 | 64.3454891 | 260.056959 | 173.718553 | 208.437533 |
| -4.8966599 | 9.75E-07 | 4.36E-06 | 168.855679 | 83.6187308 | 133.452663 | 114.864769 |
| -4.8965136 | 9.76E-07 | 4.36E-06 | 88.5262781 | 102.832096 | 227.789891 | 196.110582 |
| -4.8962142 | 9.77E-07 | 4.37E-06 | 76.2309617 | 378.855091 | 171.417645 | 169.775732 |
| -4.8958915 | 9.79E-07 | 4.38E-06 | 59.0175187 | 237.055043 | 210.533081 | 233.091435 |
| -4.8958361 | 9.79E-07 | 4.38E-06 | 59.8372065 | 214.323737 | 231.241253 | 215.72164 |
| -4.8955537 | 9.80E-07 | 4.38E-06 | 65.9848647 | 198.628312 | 177.169915 | 183.223315 |
| -4.8953243 | 9.81E-07 | 4.39E-06 | 83.6081515 | 135.576 | 119.647216 | 117.666349 |

| | | | | | | |
|---|---|---|---|---|---|---|
| -4.8949485 | 9.83E-07 | 4.39E-06 | 146.724109 | 97.4198805 | 92.0363196 | 127.19172 |
| -4.8944032 | 9.86E-07 | 4.41E-06 | 39.3450125 | 335.286755 | 537.262016 | 398.384639 |
| -4.8941418 | 9.87E-07 | 4.41E-06 | 62.7061136 | 210.535186 | 223.188075 | 184.343947 |
| -4.8940488 | 9.88E-07 | 4.41E-06 | 45.9025146 | 254.644743 | 491.243856 | 388.859268 |
| -4.8937292 | 9.89E-07 | 4.42E-06 | 89.7558097 | 114.46836 | 132.302209 | 103.65845 |
| -4.893672 | 9.90E-07 | 4.42E-06 | 69.6734596 | 163.1783 | 169.116737 | 165.853521 |
| -4.8934853 | 9.91E-07 | 4.42E-06 | 52.0501728 | 346.923019 | 352.038923 | 218.52322 |
| -4.8934206 | 9.91E-07 | 4.42E-06 | 105.329877 | 294.424528 | 222.037621 | 89.0902358 |
| -4.8934009 | 9.91E-07 | 4.42E-06 | 74.1817423 | 217.841677 | 128.850847 | 198.912162 |
| -4.8931851 | 9.92E-07 | 4.43E-06 | 74.1817423 | 156.41303 | 141.505841 | 148.483726 |
| -4.8931464 | 9.92E-07 | 4.43E-06 | 91.3951852 | 249.773749 | 335.932567 | 265.029444 |
| -4.8930752 | 9.93E-07 | 4.43E-06 | 84.8376831 | 120.692407 | 125.399485 | 188.266159 |
| -4.8929278 | 9.93E-07 | 4.43E-06 | 95.0837801 | 137.199665 | 139.204933 | 166.974153 |
| -4.892936 | 9.93E-07 | 4.43E-06 | 66.3947085 | 233.266492 | 243.896247 | 153.52657 |
| -4.8928334 | 9.94E-07 | 4.43E-06 | 97.5428434 | 222.983282 | 184.072639 | 197.79153 |
| -4.8928561 | 9.94E-07 | 4.43E-06 | 76.6408055 | 131.787449 | 144.957203 | 186.585211 |
| -4.8926156 | 9.95E-07 | 4.44E-06 | 82.3786199 | 114.73897 | 130.001301 | 134.475828 |
| -4.8925998 | 9.95E-07 | 4.44E-06 | 62.7061136 | 223.795114 | 195.577179 | 192.748686 |
| -4.8925024 | 9.96E-07 | 4.44E-06 | 71.722679 | 232.995881 | 135.753571 | 208.997849 |
| -4.8925213 | 9.96E-07 | 4.44E-06 | 84.8376831 | 128.810731 | 108.142676 | 136.156776 |
| -4.8924105 | 9.96E-07 | 4.44E-06 | 95.9034679 | 109.056144 | 102.390406 | 104.779082 |
| -4.8922447 | 9.97E-07 | 4.44E-06 | 54.9190799 | 254.644743 | 261.153057 | 238.134278 |
| -4.8921647 | 9.97E-07 | 4.44E-06 | 91.8050291 | 191.321821 | 104.691314 | 131.113932 |
| -4.8921083 | 9.98E-07 | 4.44E-06 | 65.9848647 | 175.897006 | 180.621277 | 196.670898 |
| -4.8920851 | 9.98E-07 | 4.44E-06 | 65.9848647 | 169.943569 | 184.072639 | 224.686695 |
| -4.8919009 | 9.99E-07 | 4.45E-06 | 89.3459658 | 125.292791 | 105.841768 | 111.502874 |
| -4.8916509 | 1.00E-06 | 4.45E-06 | 61.0667381 | 217.029845 | 199.028541 | 216.281956 |
| -4.8915383 | 1.00E-06 | 4.46E-06 | 104.100346 | 101.208431 | 208.232173 | 118.786981 |
| -4.8915064 | 1.00E-06 | 4.46E-06 | 74.1817423 | 142.34127 | 297.967585 | 187.705843 |
| -4.891248 | 1.00E-06 | 4.46E-06 | 52.8698605 | 268.987114 | 284.162137 | 247.659649 |
| -4.8910823 | 1.00E-06 | 4.46E-06 | 67.2143963 | 218.112288 | 158.762651 | 188.266159 |
| -4.8906787 | 1.00E-06 | 4.47E-06 | 90.1656536 | 159.66036 | 151.859927 | 152.966254 |
| 4.89010693 | 1.01E-06 | 4.48E-06 | 17.213443 | 13.8011497 | 14.9559019 | 8.40473923 |
| -4.8899861 | 1.01E-06 | 4.49E-06 | 93.8542485 | 118.798132 | 101.239952 | 105.899714 |
| -4.8899253 | 1.01E-06 | 4.49E-06 | 65.1651769 | 203.499306 | 185.223093 | 176.499524 |
| -4.8894712 | 1.01E-06 | 4.50E-06 | 69.6734596 | 168.590515 | 173.718553 | 154.647202 |
| -4.8894331 | 1.01E-06 | 4.50E-06 | 85.247527 | 123.939737 | 125.399485 | 108.701294 |
| -4.8893333 | 1.01E-06 | 4.50E-06 | 80.7392443 | 131.787449 | 117.346308 | 141.759935 |
| -4.8891046 | 1.01E-06 | 4.50E-06 | 77.4604933 | 143.423713 | 125.399485 | 146.242463 |
| -4.8890989 | 1.01E-06 | 4.50E-06 | 85.6573709 | 118.527521 | 110.443584 | 122.148877 |
| -4.8889899 | 1.01E-06 | 4.51E-06 | 66.3947085 | 269.528336 | 154.160835 | 223.005748 |
| -4.8885584 | 1.02E-06 | 4.51E-06 | 109.428316 | 207.558468 | 463.63296 | 103.65845 |
| -4.8883608 | 1.02E-06 | 4.52E-06 | 68.0340841 | 189.968767 | 162.214013 | 170.336048 |

| | | | | | | |
|---|---|---|---|---|---|---|
| -4.8881225 | 1.02E-06 | 4.52E-06 | 90.9853413 | 198.08709 | 228.940345 | 215.161324 |
| -4.8879479 | 1.02E-06 | 4.53E-06 | 81.5589321 | 120.151186 | 312.923487 | 206.756585 |
| -4.8879371 | 1.02E-06 | 4.53E-06 | 115.16613 | 191.321821 | 86.2840497 | 125.510772 |
| -4.8877945 | 1.02E-06 | 4.53E-06 | 69.2636157 | 170.21418 | 155.311289 | 175.378892 |
| -4.887536 | 1.02E-06 | 4.53E-06 | 68.8537718 | 187.53327 | 158.762651 | 165.293205 |
| -4.8873067 | 1.02E-06 | 4.54E-06 | 115.16613 | 184.827162 | 119.647216 | 87.9696039 |
| -4.8867915 | 1.02E-06 | 4.55E-06 | 74.1817423 | 139.905773 | 151.859927 | 145.121831 |
| -4.8867566 | 1.03E-06 | 4.55E-06 | 72.5423667 | 143.964935 | 169.116737 | 149.604358 |
| -4.8866723 | 1.03E-06 | 4.55E-06 | 61.0667381 | 208.640911 | 211.683535 | 200.59311 |
| -4.8865385 | 1.03E-06 | 4.55E-06 | 177.052556 | 80.3714014 | 189.824909 | 112.06319 |
| -4.8860838 | 1.03E-06 | 4.56E-06 | 107.78894 | 234.078324 | 179.470823 | 149.044042 |
| -4.8855945 | 1.03E-06 | 4.57E-06 | 90.9853413 | 107.161869 | 108.142676 | 108.701294 |
| -4.8852592 | 1.03E-06 | 4.58E-06 | 89.7558097 | 122.045461 | 188.674455 | 109.821926 |
| -4.8850584 | 1.03E-06 | 4.59E-06 | 111.477535 | 167.778683 | 170.267191 | 80.1251807 |
| -4.8848342 | 1.04E-06 | 4.59E-06 | 297.136813 | 118.798132 | 105.841768 | 123.269509 |
| -4.8845789 | 1.04E-06 | 4.60E-06 | 532.3872 | 376.419594 | 66.7263317 | 296.967453 |
| -4.8845724 | 1.04E-06 | 4.60E-06 | 65.5750208 | 175.626396 | 177.169915 | 194.429634 |
| 4.88450384 | 1.04E-06 | 4.60E-06 | 22.1315695 | 16.7778683 | 17.2568099 | 12.8872668 |
| -4.8845012 | 1.04E-06 | 4.60E-06 | 61.476582 | 228.395498 | 187.524001 | 207.316901 |
| -4.8839674 | 1.04E-06 | 4.61E-06 | 122.54332 | 102.561485 | 87.4345036 | 149.044042 |
| -4.8837715 | 1.04E-06 | 4.61E-06 | 130.740198 | 170.484791 | 73.6290557 | 222.445432 |
| -4.8838456 | 1.04E-06 | 4.61E-06 | 131.150042 | 80.1007906 | 121.948124 | 128.312352 |
| -4.883818 | 1.04E-06 | 4.61E-06 | 74.5915861 | 125.563402 | 222.037621 | 189.947107 |
| -4.8837917 | 1.04E-06 | 4.61E-06 | 72.1325229 | 155.330587 | 146.107657 | 154.086886 |
| -4.8836659 | 1.04E-06 | 4.61E-06 | 70.9029912 | 183.744719 | 146.107657 | 158.569413 |
| -4.8834753 | 1.04E-06 | 4.62E-06 | 82.3786199 | 128.54012 | 119.647216 | 117.106033 |
| -4.8833129 | 1.04E-06 | 4.62E-06 | 63.9356453 | 193.486707 | 204.780811 | 175.939208 |
| -4.8832352 | 1.04E-06 | 4.62E-06 | 72.1325229 | 159.389749 | 139.204933 | 168.094785 |
| -4.8831662 | 1.04E-06 | 4.62E-06 | 65.5750208 | 158.577917 | 255.400787 | 204.515321 |
| -4.8830937 | 1.04E-06 | 4.62E-06 | 138.937075 | 109.867976 | 85.1335957 | 113.744138 |
| -4.8830879 | 1.04E-06 | 4.62E-06 | 79.9195566 | 188.074492 | 111.594038 | 178.740788 |
| -4.8826495 | 1.05E-06 | 4.63E-06 | 70.0833035 | 187.803881 | 153.010381 | 155.767834 |
| -4.8820706 | 1.05E-06 | 4.64E-06 | 69.6734596 | 220.547785 | 151.859927 | 165.293205 |
| -4.8820814 | 1.05E-06 | 4.64E-06 | 139.756763 | 112.574084 | 92.0363196 | 98.615607 |
| -4.8818714 | 1.05E-06 | 4.65E-06 | 140.576451 | 450.566947 | 366.994825 | 397.264008 |
| -4.8818955 | 1.05E-06 | 4.65E-06 | 71.722679 | 143.694324 | 156.461743 | 203.394689 |
| -4.8817254 | 1.05E-06 | 4.65E-06 | 181.150995 | 102.561485 | 102.390406 | 106.46003 |
| -4.8815852 | 1.05E-06 | 4.65E-06 | 69.6734596 | 166.69624 | 155.311289 | 160.810677 |
| -4.8812028 | 1.05E-06 | 4.66E-06 | 159.839113 | 86.3248385 | 103.54086 | 131.674248 |
| -4.8809504 | 1.06E-06 | 4.67E-06 | 69.6734596 | 184.556551 | 159.913105 | 151.285306 |
| -4.8808646 | 1.06E-06 | 4.67E-06 | 76.2309617 | 142.611881 | 146.107657 | 126.631404 |
| -4.880685 | 1.06E-06 | 4.67E-06 | 73.3620545 | 150.459593 | 140.355387 | 147.363094 |
| -4.8803858 | 1.06E-06 | 4.68E-06 | 90.9853413 | 133.681725 | 134.603117 | 94.6933953 |

| | | | | | | |
|---|---|---|---|---|---|---|
| -4.8802984 | 1.06E-06 | 4.68E-06 | 57.3781432 | 192.945486 | 268.055781 | 260.546916 |
| -4.8801322 | 1.06E-06 | 4.68E-06 | 60.2470503 | 203.769917 | 240.444885 | 195.550266 |
| -4.8794897 | 1.06E-06 | 4.70E-06 | 68.4439279 | 273.587498 | 138.054479 | 296.967453 |
| -4.8792831 | 1.06E-06 | 4.70E-06 | 206.971159 | 501.441774 | 348.587561 | 418.556014 |
| -4.87919 | 1.07E-06 | 4.70E-06 | 82.7884637 | 109.597366 | 142.656295 | 125.510772 |
| -4.8787617 | 1.07E-06 | 4.71E-06 | 76.2309617 | 192.945486 | 126.549939 | 148.483726 |
| -4.8785924 | 1.07E-06 | 4.72E-06 | 62.2962697 | 185.909605 | 195.577179 | 207.877217 |
| -4.8784867 | 1.07E-06 | 4.72E-06 | 88.9361219 | 188.886324 | 138.054479 | 102.537819 |
| 4.8784302 | 1.07E-06 | 4.72E-06 | 69.2636157 | 38.9679522 | 35.6640739 | 39.7824324 |
| -4.8780137 | 1.07E-06 | 4.73E-06 | 80.7392443 | 145.317988 | 142.656295 | 110.382242 |
| -4.8780057 | 1.07E-06 | 4.73E-06 | 68.8537718 | 168.049294 | 157.612197 | 162.491625 |
| -4.8776947 | 1.07E-06 | 4.73E-06 | 88.5262781 | 135.30539 | 97.7885896 | 126.631404 |
| -4.8763316 | 1.08E-06 | 4.77E-06 | 70.4931473 | 159.93097 | 151.859927 | 152.966254 |
| -4.87627 | 1.08E-06 | 4.77E-06 | 64.3454891 | 188.886324 | 172.568099 | 191.628054 |
| -4.8762769 | 1.08E-06 | 4.77E-06 | 1888.5606 | 8286.91389 | 7026.973 | 7579.39384 |
| -4.8761298 | 1.08E-06 | 4.77E-06 | 77.0506494 | 119.880575 | 146.107657 | 146.802779 |
| -4.8760685 | 1.08E-06 | 4.77E-06 | 77.4604933 | 130.163785 | 125.399485 | 140.639303 |
| -4.8760022 | 1.08E-06 | 4.77E-06 | 69.2636157 | 238.408096 | 143.806749 | 184.343947 |
| -4.8759068 | 1.08E-06 | 4.77E-06 | 60.2470503 | 209.182132 | 188.674455 | 234.772382 |
| -4.8756605 | 1.08E-06 | 4.78E-06 | 49.1812656 | 293.612695 | 273.808051 | 281.278606 |
| -4.8756659 | 1.08E-06 | 4.78E-06 | 74.1817423 | 145.047378 | 132.302209 | 150.72499 |
| -4.8756678 | 1.08E-06 | 4.78E-06 | 54.0993922 | 258.162683 | 246.197155 | 233.651751 |
| -4.8756891 | 1.08E-06 | 4.78E-06 | 64.3454891 | 190.509989 | 197.878087 | 168.655101 |
| -4.8750767 | 1.09E-06 | 4.79E-06 | 65.5750208 | 188.074492 | 158.762651 | 205.075637 |
| -4.8746689 | 1.09E-06 | 4.80E-06 | 65.1651769 | 180.226779 | 166.815829 | 195.550266 |
| -4.8745516 | 1.09E-06 | 4.80E-06 | 94.6739363 | 134.764168 | 92.0363196 | 117.106033 |
| -4.8743192 | 1.09E-06 | 4.81E-06 | 69.6734596 | 148.835929 | 164.514921 | 161.370993 |
| -4.8742474 | 1.09E-06 | 4.81E-06 | 130.330354 | 196.463426 | 234.692615 | 188.826475 |
| 4.87402732 | 1.09E-06 | 4.81E-06 | 60.2470503 | 40.5916169 | 33.3631659 | 36.9808526 |
| -4.8739996 | 1.09E-06 | 4.81E-06 | 65.5750208 | 185.097773 | 173.718553 | 170.336048 |
| -4.873674 | 1.10E-06 | 4.82E-06 | 61.476582 | 233.266492 | 185.223093 | 193.869318 |
| -4.8735479 | 1.10E-06 | 4.82E-06 | 70.0833035 | 228.395498 | 132.302209 | 241.496174 |
| -4.8732765 | 1.10E-06 | 4.83E-06 | 81.968776 | 171.837845 | 115.0454 | 126.631404 |
| -4.8732297 | 1.10E-06 | 4.83E-06 | 58.197831 | 197.004647 | 322.127119 | 221.885116 |
| -4.8730162 | 1.10E-06 | 4.83E-06 | 88.1164342 | 278.458492 | 250.798971 | 270.632603 |
| -4.8727855 | 1.10E-06 | 4.84E-06 | 65.9848647 | 190.509989 | 162.214013 | 175.378892 |
| -4.8726456 | 1.10E-06 | 4.84E-06 | 216.807412 | 113.656527 | 269.206235 | 79.0045488 |
| -4.8725335 | 1.10E-06 | 4.84E-06 | 127.051603 | 73.6061319 | 162.214013 | 165.853521 |
| -4.8722441 | 1.10E-06 | 4.85E-06 | 93.8542485 | 300.377965 | 217.435805 | 226.927959 |
| -4.8722581 | 1.10E-06 | 4.85E-06 | 77.8703372 | 143.964935 | 130.001301 | 121.028245 |
| -4.872094 | 1.10E-06 | 4.85E-06 | 84.8376831 | 107.973701 | 117.346308 | 116.545717 |
| -4.8720777 | 1.10E-06 | 4.85E-06 | 84.4278393 | 105.538204 | 119.647216 | 132.234564 |
| -4.8717282 | 1.11E-06 | 4.86E-06 | 61.0667381 | 206.476024 | 196.727633 | 192.748686 |

| | | | | | | |
|---|---|---|---|---|---|---|
| -4.8714667 | 1.11E-06 | 4.86E-06 | 80.3294005 | 131.787449 | 113.894946 | 127.19172 |
| -4.8713631 | 1.11E-06 | 4.87E-06 | 122.953164 | 117.174467 | 205.931265 | 158.009098 |
| -4.8709879 | 1.11E-06 | 4.87E-06 | 75.4112739 | 192.945486 | 123.098578 | 155.767834 |
| -4.8703053 | 1.11E-06 | 4.89E-06 | 83.6081515 | 126.104623 | 108.142676 | 117.666349 |
| -4.8702562 | 1.11E-06 | 4.89E-06 | 104.510189 | 94.9843835 | 95.4876816 | 114.304454 |
| -4.8702166 | 1.11E-06 | 4.89E-06 | 66.8045524 | 159.93097 | 196.727633 | 164.172573 |
| -4.869697 | 1.12E-06 | 4.90E-06 | 68.0340841 | 179.144336 | 150.709473 | 166.974153 |
| -4.8695013 | 1.12E-06 | 4.91E-06 | 110.657848 | 113.385916 | 82.8326877 | 122.148877 |
| -4.8695078 | 1.12E-06 | 4.91E-06 | 384.023715 | 65.2171978 | 184.072639 | 194.429634 |
| -4.8694578 | 1.12E-06 | 4.91E-06 | 56.1486116 | 237.325653 | 224.338529 | 220.764484 |
| -4.8691623 | 1.12E-06 | 4.91E-06 | 75.4112739 | 132.328671 | 130.001301 | 144.561515 |
| -4.8686004 | 1.12E-06 | 4.93E-06 | 73.7718984 | 215.947402 | 128.850847 | 162.491625 |
| -4.8684453 | 1.12E-06 | 4.93E-06 | 93.0345607 | 185.097773 | 94.3372276 | 137.837723 |
| -4.8683706 | 1.13E-06 | 4.93E-06 | 1146.74318 | 6651.61295 | 3796.49818 | 4707.7746 |
| -4.867978 | 1.13E-06 | 4.94E-06 | 51.6403289 | 290.094755 | 236.993523 | 259.9866 |
| -4.8679801 | 1.13E-06 | 4.94E-06 | 101.231438 | 129.081342 | 250.798971 | 103.098135 |
| -4.8678277 | 1.13E-06 | 4.94E-06 | 139.346919 | 436.495187 | 391.154358 | 533.981099 |
| -4.8669962 | 1.13E-06 | 4.96E-06 | 65.9848647 | 168.049294 | 169.116737 | 173.137628 |
| -4.8668556 | 1.13E-06 | 4.97E-06 | 52.0501728 | 225.148168 | 268.055781 | 291.364293 |
| -4.8668441 | 1.13E-06 | 4.97E-06 | 96.7231557 | 149.918372 | 94.3372276 | 109.26161 |
| -4.8666284 | 1.14E-06 | 4.97E-06 | 162.70802 | 106.620647 | 87.4345036 | 113.183822 |
| -4.8663132 | 1.14E-06 | 4.98E-06 | 61.476582 | 293.612695 | 169.116737 | 219.643852 |
| -4.866102 | 1.14E-06 | 4.98E-06 | 67.2143963 | 172.108456 | 155.311289 | 168.655101 |
| -4.8658104 | 1.14E-06 | 4.99E-06 | 83.6081515 | 188.074492 | 101.239952 | 170.336048 |
| -4.8657514 | 1.14E-06 | 4.99E-06 | 53.2797044 | 205.122971 | 330.180297 | 273.434183 |
| -4.8652914 | 1.14E-06 | 5.00E-06 | 122.133476 | 193.486707 | 83.9831417 | 115.985401 |
| -4.8649927 | 1.14E-06 | 5.01E-06 | 85.6573709 | 114.73897 | 138.054479 | 103.098135 |
| -4.8648035 | 1.15E-06 | 5.01E-06 | 50.4107972 | 273.316887 | 260.002603 | 261.667548 |
| -4.8644942 | 1.15E-06 | 5.02E-06 | 69.2636157 | 155.871809 | 272.657597 | 164.732889 |
| 4.8645048 | 1.15E-06 | 5.02E-06 | 15.9839113 | 35.1794013 | 54.0713378 | 51.5490673 |
| -4.8644785 | 1.15E-06 | 5.02E-06 | 920.919198 | 1322.47488 | 989.390436 | 1211.40308 |
| -4.8643602 | 1.15E-06 | 5.02E-06 | 95.9034679 | 213.782516 | 89.7354116 | 165.293205 |
| -4.8642769 | 1.15E-06 | 5.02E-06 | 78.6900249 | 114.73897 | 131.151755 | 132.79488 |
| -4.8642013 | 1.15E-06 | 5.02E-06 | 18.4429746 | 775.029271 | 890.451392 | 631.476074 |
| -4.8641424 | 1.15E-06 | 5.02E-06 | 151.232392 | 114.197749 | 88.5849576 | 100.296555 |
| -4.8640651 | 1.15E-06 | 5.03E-06 | 82.7884637 | 179.414947 | 166.815829 | 101.977503 |
| -4.8638864 | 1.15E-06 | 5.03E-06 | 86.0672148 | 232.184049 | 104.691314 | 160.250361 |
| -4.8638705 | 1.15E-06 | 5.03E-06 | 100.821594 | 115.821413 | 93.1867736 | 98.055291 |
| -4.8632029 | 1.16E-06 | 5.05E-06 | 45.4926707 | 277.91727 | 369.295733 | 296.967453 |
| -4.8630884 | 1.16E-06 | 5.05E-06 | 4063.19223 | 11660.0773 | 10466.8305 | 11676.9844 |
| -4.8625698 | 1.16E-06 | 5.06E-06 | 76.6408055 | 135.30539 | 134.603117 | 120.467929 |
| -4.8618706 | 1.16E-06 | 5.08E-06 | 67.2143963 | 171.837845 | 155.311289 | 162.491625 |
| -4.8616686 | 1.16E-06 | 5.08E-06 | 87.7065903 | 108.244312 | 212.833989 | 131.113932 |

| | | | | | | |
|---|---|---|---|---|---|---|
| -4.861091 | 1.17E-06 | 5.10E-06 | 72.9522106 | 132.599282 | 162.214013 | 137.837723 |
| -4.8610531 | 1.17E-06 | 5.10E-06 | 106.559409 | 185.909605 | 79.3813257 | 164.172573 |
| -4.8609657 | 1.17E-06 | 5.10E-06 | 86.4770587 | 102.020264 | 120.79767 | 126.631404 |
| -4.8609341 | 1.17E-06 | 5.10E-06 | 90.9853413 | 230.830995 | 209.382627 | 235.893014 |
| -4.8608144 | 1.17E-06 | 5.10E-06 | 173.773805 | 205.122971 | 98.9390436 | 85.1680242 |
| -4.8605766 | 1.17E-06 | 5.11E-06 | 89.3459658 | 121.233629 | 94.3372276 | 141.199619 |
| -4.8604445 | 1.17E-06 | 5.11E-06 | 55.7387677 | 347.46424 | 268.055781 | 186.024895 |
| -4.8600897 | 1.17E-06 | 5.12E-06 | 141.805982 | 103.102707 | 158.762651 | 81.8061285 |
| -4.8600731 | 1.17E-06 | 5.12E-06 | 145.494577 | 193.216096 | 200.178995 | 212.92006 |
| -4.8599091 | 1.17E-06 | 5.12E-06 | 84.8376831 | 139.635162 | 98.9390436 | 128.872668 |
| -4.8596314 | 1.18E-06 | 5.13E-06 | 36.0662614 | 492.782229 | 425.667978 | 334.508621 |
| -4.8593791 | 1.18E-06 | 5.13E-06 | 93.8542485 | 139.905773 | 128.850847 | 88.5299199 |
| -4.8593998 | 1.18E-06 | 5.13E-06 | 67.2143963 | 144.776767 | 208.232173 | 173.697944 |
| -4.8593191 | 1.18E-06 | 5.13E-06 | 70.9029912 | 150.188982 | 138.054479 | 153.52657 |
| -4.8592843 | 1.18E-06 | 5.13E-06 | 51.6403289 | 232.995881 | 257.701695 | 272.313551 |
| -4.8589581 | 1.18E-06 | 5.14E-06 | 76.6408055 | 112.844695 | 169.116737 | 190.507423 |
| -4.8588197 | 1.18E-06 | 5.14E-06 | 63.5258014 | 166.966851 | 209.382627 | 176.499524 |
| -4.8586254 | 1.18E-06 | 5.15E-06 | 68.4439279 | 145.588599 | 158.762651 | 168.655101 |
| -4.8580581 | 1.19E-06 | 5.16E-06 | 78.6900249 | 169.131737 | 396.906628 | 149.044042 |
| -4.8578851 | 1.19E-06 | 5.17E-06 | 127.871291 | 147.212264 | 93.1867736 | 89.0902358 |
| -4.8576543 | 1.19E-06 | 5.17E-06 | 1197.97366 | 1846.91857 | 1470.28021 | 1670.86216 |
| -4.8573431 | 1.19E-06 | 5.18E-06 | 62.2962697 | 183.474108 | 181.771731 | 185.464579 |
| -4.8565867 | 1.19E-06 | 5.20E-06 | 51.230485 | 292.530252 | 273.808051 | 220.764484 |
| -4.8565343 | 1.19E-06 | 5.20E-06 | 62.7061136 | 176.167617 | 187.524001 | 180.421735 |
| -4.8563656 | 1.20E-06 | 5.20E-06 | 74.5915861 | 131.246228 | 130.001301 | 135.59646 |
| -4.856229 | 1.20E-06 | 5.21E-06 | 97.9526873 | 112.844695 | 88.5849576 | 146.242463 |
| -4.8558512 | 1.20E-06 | 5.21E-06 | 133.609105 | 130.434396 | 85.1335957 | 96.3743432 |
| -4.855545 | 1.20E-06 | 5.22E-06 | 70.0833035 | 153.165701 | 178.320369 | 135.59646 |
| -4.8553968 | 1.20E-06 | 5.23E-06 | 76.6408055 | 155.330587 | 124.249032 | 122.709193 |
| -4.8552121 | 1.20E-06 | 5.23E-06 | 90.5754975 | 169.402348 | 89.7354116 | 172.577312 |
| -4.8551098 | 1.20E-06 | 5.23E-06 | 79.5097127 | 191.05121 | 115.0454 | 135.036144 |
| -4.8547865 | 1.21E-06 | 5.24E-06 | 49.5911095 | 226.230611 | 301.418947 | 298.648401 |
| -4.8545253 | 1.21E-06 | 5.25E-06 | 96.3133118 | 103.914539 | 128.850847 | 91.3314996 |
| -4.8542819 | 1.21E-06 | 5.25E-06 | 72.5423667 | 186.450827 | 127.700393 | 149.044042 |
| -4.8541557 | 1.21E-06 | 5.25E-06 | 100.821594 | 140.446994 | 81.6822337 | 154.647202 |
| -4.8540586 | 1.21E-06 | 5.26E-06 | 2059.4655 | 5267.98004 | 4952.70445 | 5155.46704 |
| -4.8537634 | 1.21E-06 | 5.26E-06 | 74.1817423 | 129.893174 | 139.204933 | 129.9933 |
| -4.8536731 | 1.21E-06 | 5.26E-06 | 74.1817423 | 133.681725 | 125.399485 | 146.802779 |
| -4.8535622 | 1.21E-06 | 5.27E-06 | 79.0998688 | 150.459593 | 111.594038 | 125.510772 |
| -4.8534192 | 1.21E-06 | 5.27E-06 | 103.690502 | 160.742803 | 187.524001 | 185.464579 |
| -4.8533718 | 1.21E-06 | 5.27E-06 | 58.6076748 | 227.583665 | 179.470823 | 221.3248 |
| -4.8532547 | 1.21E-06 | 5.27E-06 | 59.8372065 | 222.712671 | 196.727633 | 180.421735 |
| -4.8531282 | 1.22E-06 | 5.28E-06 | 55.7387677 | 214.053126 | 205.931265 | 259.426284 |

| | | | | | | |
|---|---|---|---|---|---|---|
| 4.85291708 | 1.22E-06 | 5.28E-06 | 0 | 0 | 0 | 0 |
| -4.8521653 | 1.22E-06 | 5.30E-06 | 69.6734596 | 156.683641 | 135.753571 | 162.491625 |
| -4.8520719 | 1.22E-06 | 5.30E-06 | 173.363961 | 92.5488865 | 95.4876816 | 119.907613 |
| -4.8520127 | 1.22E-06 | 5.30E-06 | 75.4112739 | 123.398515 | 138.054479 | 127.19172 |
| -4.8518869 | 1.22E-06 | 5.30E-06 | 157.789894 | 117.174467 | 90.8858656 | 94.6933953 |
| -4.8518751 | 1.22E-06 | 5.30E-06 | 75.4112739 | 192.945486 | 121.948124 | 141.759935 |
| -4.8512591 | 1.23E-06 | 5.32E-06 | 85.6573709 | 258.433294 | 309.472125 | 225.807327 |
| -4.8510502 | 1.23E-06 | 5.33E-06 | 78.2801811 | 120.963018 | 126.549939 | 115.985401 |
| -4.850902 | 1.23E-06 | 5.33E-06 | 140.166607 | 199.440144 | 66.7263317 | 168.655101 |
| -4.850801 | 1.23E-06 | 5.33E-06 | 96.3133118 | 92.8194972 | 103.54086 | 104.218766 |
| -4.8505014 | 1.23E-06 | 5.34E-06 | 77.0506494 | 226.771833 | 136.904025 | 129.432984 |
| -4.8503746 | 1.23E-06 | 5.34E-06 | 67.2143963 | 197.545869 | 146.107657 | 160.250361 |
| -4.8503125 | 1.23E-06 | 5.34E-06 | 65.5750208 | 169.672959 | 159.913105 | 163.612257 |
| -4.8502416 | 1.23E-06 | 5.34E-06 | 74.5915861 | 175.897006 | 127.700393 | 130.553616 |
| -4.8500137 | 1.23E-06 | 5.35E-06 | 70.0833035 | 143.153102 | 141.505841 | 152.966254 |
| -4.849792 | 1.24E-06 | 5.35E-06 | 55.7387677 | 294.424528 | 189.824909 | 233.091435 |
| -4.8496748 | 1.24E-06 | 5.36E-06 | 47.1320462 | 266.281007 | 263.453965 | 488.595507 |
| -4.8494462 | 1.24E-06 | 5.36E-06 | 79.9195566 | 112.574084 | 116.195854 | 122.709193 |
| -4.848866 | 1.24E-06 | 5.38E-06 | 59.4273626 | 233.537102 | 184.072639 | 189.947107 |
| -4.8486965 | 1.24E-06 | 5.38E-06 | 63.5258014 | 165.613797 | 186.373547 | 171.45668 |
| -4.8483516 | 1.24E-06 | 5.39E-06 | 113.936599 | 141.800048 | 92.0363196 | 91.3314996 |
| -4.8483204 | 1.25E-06 | 5.39E-06 | 50.8206411 | 342.052025 | 231.241253 | 241.496174 |
| -4.8482535 | 1.25E-06 | 5.39E-06 | 71.722679 | 158.036695 | 133.452663 | 136.156776 |
| -4.8480898 | 1.25E-06 | 5.39E-06 | 69.2636157 | 158.036695 | 148.408565 | 140.639303 |
| -4.8480204 | 1.25E-06 | 5.39E-06 | 72.9522106 | 151.542036 | 132.302209 | 129.432984 |
| -4.8475916 | 1.25E-06 | 5.40E-06 | 72.5423667 | 140.988216 | 151.859927 | 125.510772 |
| -4.8476447 | 1.25E-06 | 5.40E-06 | 77.0506494 | 135.576 | 112.744492 | 131.674248 |
| -4.8476085 | 1.25E-06 | 5.40E-06 | 69.6734596 | 207.829078 | 178.320369 | 128.312352 |
| 4.84744789 | 1.25E-06 | 5.41E-06 | 293.858062 | 104.726372 | 110.443584 | 90.2108677 |
| -4.8472396 | 1.25E-06 | 5.41E-06 | 92.214873 | 231.913438 | 192.125817 | 202.834373 |
| -4.8471955 | 1.25E-06 | 5.41E-06 | 91.3951852 | 149.918372 | 87.4345036 | 142.880567 |
| -4.8469868 | 1.25E-06 | 5.42E-06 | 89.3459658 | 150.459593 | 92.0363196 | 188.826475 |
| -4.8468041 | 1.25E-06 | 5.42E-06 | 61.0667381 | 214.053126 | 158.762651 | 249.340597 |
| -4.8463254 | 1.26E-06 | 5.43E-06 | 75.4112739 | 117.174467 | 141.505841 | 131.674248 |
| -4.8459778 | 1.26E-06 | 5.44E-06 | 46.3123584 | 267.36345 | 309.472125 | 282.399238 |
| -4.8457744 | 1.26E-06 | 5.45E-06 | 72.5423667 | 151.000815 | 130.001301 | 132.79488 |
| -4.8457279 | 1.26E-06 | 5.45E-06 | 71.3128351 | 161.284024 | 177.169915 | 123.269509 |
| -4.845634 | 1.26E-06 | 5.45E-06 | 65.5750208 | 203.769917 | 207.081719 | 138.958355 |
| -4.845597 | 1.26E-06 | 5.45E-06 | 69.2636157 | 145.047378 | 150.709473 | 144.001199 |
| -4.8454703 | 1.26E-06 | 5.45E-06 | 41.8040758 | 329.062707 | 294.516223 | 365.886314 |
| -4.8451418 | 1.27E-06 | 5.46E-06 | 62.2962697 | 200.251977 | 156.461743 | 204.515321 |
| -4.8450872 | 1.27E-06 | 5.46E-06 | 129.510666 | 149.647761 | 70.1776937 | 191.067738 |
| -4.8449735 | 1.27E-06 | 5.47E-06 | 70.9029912 | 162.907689 | 140.355387 | 132.79488 |

| | | | | | | |
|---|---|---|---|---|---|---|
| -4.8449024 | 1.27E-06 | 5.47E-06 | 139.346919 | 107.432479 | 86.2840497 | 100.296555 |
| -4.8445594 | 1.27E-06 | 5.48E-06 | 79.5097127 | 119.880575 | 111.594038 | 156.888466 |
| -4.8445047 | 1.27E-06 | 5.48E-06 | 134.428793 | 114.197749 | 74.7795097 | 146.802779 |
| -4.8444903 | 1.27E-06 | 5.48E-06 | 69.6734596 | 152.353869 | 139.204933 | 144.561515 |
| -4.8444064 | 1.27E-06 | 5.48E-06 | 69.6734596 | 133.681725 | 153.010381 | 155.207518 |
| -4.8443793 | 1.27E-06 | 5.48E-06 | 65.9848647 | 157.224863 | 158.762651 | 162.491625 |
| -4.8443401 | 1.27E-06 | 5.48E-06 | 74.1817423 | 119.339354 | 144.957203 | 135.59646 |
| -4.8442492 | 1.27E-06 | 5.48E-06 | 82.7884637 | 115.821413 | 159.913105 | 285.761134 |
| -4.8439311 | 1.27E-06 | 5.49E-06 | 65.9848647 | 156.954252 | 171.417645 | 153.52657 |
| -4.8437924 | 1.27E-06 | 5.49E-06 | 133.609105 | 544.468888 | 416.464346 | 471.225713 |
| -4.8435652 | 1.28E-06 | 5.50E-06 | 109.83816 | 90.3840002 | 93.1867736 | 108.701294 |
| -4.8435055 | 1.28E-06 | 5.50E-06 | 88.9361219 | 110.138587 | 93.1867736 | 118.786981 |
| -4.8434058 | 1.28E-06 | 5.50E-06 | 134.838636 | 197.545869 | 174.869007 | 185.464579 |
| -4.8425568 | 1.28E-06 | 5.52E-06 | 74.5915861 | 127.187066 | 132.302209 | 123.829825 |
| -4.8424745 | 1.28E-06 | 5.52E-06 | 77.8703372 | 143.153102 | 111.594038 | 122.709193 |
| -4.8422833 | 1.28E-06 | 5.53E-06 | 56.9682993 | 233.807713 | 194.426725 | 201.153426 |
| -4.8422242 | 1.28E-06 | 5.53E-06 | 73.3620545 | 145.047378 | 118.496762 | 163.612257 |
| -4.8418389 | 1.29E-06 | 5.54E-06 | 113.936599 | 94.7137727 | 106.992222 | 219.643852 |
| -4.841626 | 1.29E-06 | 5.54E-06 | 98.3625312 | 82.5362876 | 156.461743 | 137.277407 |
| -4.8410482 | 1.29E-06 | 5.56E-06 | 108.608628 | 196.192815 | 236.993523 | 76.763285 |
| -4.8409523 | 1.29E-06 | 5.56E-06 | 80.7392443 | 108.785533 | 111.594038 | 121.588561 |
| -4.840863 | 1.29E-06 | 5.56E-06 | 69.2636157 | 139.905773 | 146.107657 | 148.483726 |
| -4.840724 | 1.29E-06 | 5.57E-06 | 88.1164342 | 128.269509 | 208.232173 | 102.537819 |
| -4.8405132 | 1.30E-06 | 5.57E-06 | 62.7061136 | 214.594348 | 150.709473 | 198.351846 |
| -4.8404795 | 1.30E-06 | 5.57E-06 | 77.4604933 | 192.945486 | 105.841768 | 172.577312 |
| -4.8402866 | 1.30E-06 | 5.58E-06 | 76.6408055 | 125.02218 | 192.125817 | 124.390141 |
| -4.8400365 | 1.30E-06 | 5.58E-06 | 37.705637 | 306.602013 | 485.491586 | 368.687894 |
| -4.8400221 | 1.30E-06 | 5.58E-06 | 79.9195566 | 118.527521 | 110.443584 | 114.304454 |
| 4.83948165 | 1.30E-06 | 5.60E-06 | 43.8532952 | 15.6954252 | 19.5577179 | 15.6888466 |
| -4.8394932 | 1.30E-06 | 5.60E-06 | 86.0672148 | 125.292791 | 95.4876816 | 112.623506 |
| -4.8393668 | 1.30E-06 | 5.60E-06 | 90.5754975 | 96.3374374 | 103.54086 | 103.098135 |
| -4.8393473 | 1.30E-06 | 5.60E-06 | 69.6734596 | 140.988216 | 154.160835 | 136.717091 |
| -4.8391145 | 1.30E-06 | 5.60E-06 | 119.674413 | 121.233629 | 79.3813257 | 105.339398 |
| -4.8390725 | 1.30E-06 | 5.60E-06 | 84.0179954 | 111.22103 | 100.089498 | 115.985401 |
| -4.8390407 | 1.30E-06 | 5.60E-06 | 74.1817423 | 133.411114 | 118.496762 | 158.569413 |
| -4.8389883 | 1.31E-06 | 5.61E-06 | 62.7061136 | 170.484791 | 166.815829 | 182.102683 |
| -4.8389232 | 1.31E-06 | 5.61E-06 | 77.4604933 | 108.785533 | 131.151755 | 130.553616 |
| -4.8378828 | 1.31E-06 | 5.63E-06 | 63.1159575 | 218.65351 | 146.107657 | 202.834373 |
| -4.837869 | 1.31E-06 | 5.63E-06 | 62.2962697 | 179.414947 | 165.665375 | 178.740788 |
| -4.8378085 | 1.31E-06 | 5.64E-06 | 75.8211178 | 116.633246 | 131.151755 | 123.829825 |
| -4.8375515 | 1.31E-06 | 5.64E-06 | 209.020379 | 93.090108 | 111.594038 | 107.020346 |
| -4.8373402 | 1.32E-06 | 5.65E-06 | 163.937552 | 176.438228 | 109.29313 | 75.0823371 |
| -4.8370676 | 1.32E-06 | 5.65E-06 | 84.4278393 | 146.129821 | 94.3372276 | 131.113932 |

| | | | | | | |
|---|---|---|---|---|---|---|
| -4.8368981 | 1.32E-06 | 5.66E-06 | 70.0833035 | 172.649677 | 194.426725 | 122.709193 |
| -4.8366215 | 1.32E-06 | 5.67E-06 | 89.7558097 | 103.914539 | 93.1867736 | 116.545717 |
| -4.8361683 | 1.32E-06 | 5.68E-06 | 88.5262781 | 209.182132 | 89.7354116 | 187.145527 |
| -4.835775 | 1.33E-06 | 5.69E-06 | 67.2143963 | 158.848527 | 140.355387 | 159.129729 |
| -4.8356524 | 1.33E-06 | 5.69E-06 | 98.3625312 | 120.421797 | 106.992222 | 85.1680242 |
| -4.8350759 | 1.33E-06 | 5.71E-06 | 77.0506494 | 148.294707 | 116.195854 | 117.106033 |
| -4.8347119 | 1.33E-06 | 5.72E-06 | 205.741628 | 277.646659 | 96.6381356 | 85.7283401 |
| -4.8345172 | 1.33E-06 | 5.72E-06 | 57.7879871 | 218.65351 | 182.922185 | 197.79153 |
| -4.8344466 | 1.34E-06 | 5.72E-06 | 100.411751 | 79.8301798 | 173.718553 | 155.767834 |
| -4.8344709 | 1.34E-06 | 5.72E-06 | 86.0672148 | 111.762252 | 95.4876816 | 112.06319 |
| -4.8343764 | 1.34E-06 | 5.72E-06 | 86.0672148 | 115.280192 | 95.4876816 | 109.821926 |
| -4.8340055 | 1.34E-06 | 5.73E-06 | 42.6237635 | 318.779498 | 307.171217 | 298.088085 |
| -4.8339289 | 1.34E-06 | 5.73E-06 | 104.510189 | 113.115306 | 108.142676 | 82.3664444 |
| -4.833792 | 1.34E-06 | 5.74E-06 | 66.3947085 | 144.506156 | 209.382627 | 155.767834 |
| -4.833798 | 1.34E-06 | 5.74E-06 | 286.480872 | 518.760864 | 388.85345 | 470.665397 |
| -4.8337591 | 1.34E-06 | 5.74E-06 | 134.018949 | 123.398515 | 294.516223 | 177.620156 |
| -4.8335312 | 1.34E-06 | 5.74E-06 | 62.2962697 | 175.085174 | 174.869007 | 165.853521 |
| -4.8334343 | 1.34E-06 | 5.74E-06 | 79.9195566 | 100.66721 | 146.107657 | 132.234564 |
| -4.8333539 | 1.34E-06 | 5.75E-06 | 76.2309617 | 119.339354 | 116.195854 | 148.483726 |
| -4.8333205 | 1.34E-06 | 5.75E-06 | 52.0501728 | 212.700072 | 248.498063 | 242.616806 |
| -4.8328042 | 1.35E-06 | 5.76E-06 | 81.968776 | 109.867976 | 128.850847 | 102.537819 |
| -4.8324315 | 1.35E-06 | 5.77E-06 | 66.8045524 | 166.69624 | 134.603117 | 186.024895 |
| -4.8322951 | 1.35E-06 | 5.77E-06 | 112.297223 | 190.780599 | 72.4786017 | 210.118481 |
| -4.832037 | 1.35E-06 | 5.78E-06 | 94.2640924 | 136.929054 | 89.7354116 | 104.218766 |
| -4.8316917 | 1.35E-06 | 5.79E-06 | 71.722679 | 136.658443 | 127.700393 | 134.475828 |
| -4.8316167 | 1.35E-06 | 5.79E-06 | 52.4600166 | 215.947402 | 241.595339 | 229.729539 |
| -4.8315296 | 1.35E-06 | 5.79E-06 | 144.265046 | 97.9611021 | 86.2840497 | 107.580662 |
| -4.8313829 | 1.36E-06 | 5.80E-06 | 88.9361219 | 93.3607188 | 110.443584 | 104.779082 |
| 4.8312779 | 1.36E-06 | 5.80E-06 | 109.018472 | 114.197749 | 138.054479 | 104.779082 |
| -4.8310559 | 1.36E-06 | 5.80E-06 | 51.6403289 | 273.316887 | 224.338529 | 220.764484 |
| 4.83090081 | 1.36E-06 | 5.81E-06 | 15.5740674 | 8.38893415 | 12.6549939 | 11.7666349 |
| -4.8305087 | 1.36E-06 | 5.82E-06 | 89.7558097 | 104.455761 | 136.904025 | 93.5727634 |
| 4.83039973 | 1.36E-06 | 5.82E-06 | 13.9346919 | 8.38893415 | 5.75226998 | 9.52537113 |
| -4.829821 | 1.37E-06 | 5.84E-06 | 70.0833035 | 145.588599 | 171.417645 | 125.510772 |
| -4.8296627 | 1.37E-06 | 5.84E-06 | 68.8537718 | 139.09394 | 171.417645 | 135.59646 |
| -4.8283732 | 1.38E-06 | 5.88E-06 | 67.6242402 | 148.835929 | 135.753571 | 177.620156 |
| -4.8283754 | 1.38E-06 | 5.88E-06 | 71.722679 | 129.351952 | 134.603117 | 129.9933 |
| -4.8281308 | 1.38E-06 | 5.88E-06 | 107.379097 | 77.9359044 | 126.549939 | 119.907613 |
| -4.8276462 | 1.38E-06 | 5.90E-06 | 73.7718984 | 143.423713 | 128.850847 | 116.545717 |
| -4.8275539 | 1.38E-06 | 5.90E-06 | 85.247527 | 102.290875 | 98.9390436 | 115.425085 |
| -4.8273864 | 1.38E-06 | 5.90E-06 | 67.6242402 | 155.059976 | 146.107657 | 138.958355 |
| -4.8271031 | 1.39E-06 | 5.91E-06 | 79.9195566 | 100.66721 | 161.063559 | 132.234564 |
| -4.8268952 | 1.39E-06 | 5.92E-06 | 118.854725 | 96.6080482 | 87.4345036 | 99.1759229 |

| | | | | | | |
|---|---|---|---|---|---|---|
| -4.8266302 | 1.39E-06 | 5.92E-06 | 84.8376831 | 156.683641 | 92.0363196 | 135.036144 |
| -4.826426 | 1.39E-06 | 5.93E-06 | 60.6568942 | 309.308121 | 158.762651 | 206.196269 |
| -4.8259789 | 1.39E-06 | 5.94E-06 | 49.5911095 | 290.635977 | 219.736713 | 248.780281 |
| -4.8253255 | 1.40E-06 | 5.96E-06 | 52.4600166 | 207.558468 | 222.037621 | 259.426284 |
| -4.8253148 | 1.40E-06 | 5.96E-06 | 74.5915861 | 126.375234 | 117.346308 | 123.829825 |
| -4.8251916 | 1.40E-06 | 5.96E-06 | 142.215826 | 148.835929 | 65.5758777 | 194.98995 |
| -4.8248956 | 1.40E-06 | 5.97E-06 | 88.9361219 | 98.7729344 | 103.54086 | 96.9346591 |
| -4.8246134 | 1.40E-06 | 5.98E-06 | 67.6242402 | 159.119138 | 134.603117 | 148.483726 |
| -4.8245386 | 1.40E-06 | 5.98E-06 | 89.3459658 | 154.789366 | 92.0363196 | 113.744138 |
| -4.8245393 | 1.40E-06 | 5.98E-06 | 61.8864259 | 308.766899 | 187.524001 | 161.931309 |
| -4.8243922 | 1.40E-06 | 5.98E-06 | 45.9025146 | 234.890156 | 292.215315 | 320.500723 |
| -4.8244042 | 1.40E-06 | 5.98E-06 | 72.1325229 | 120.421797 | 136.904025 | 174.818576 |
| -4.8241994 | 1.41E-06 | 5.99E-06 | 78.6900249 | 146.400432 | 116.195854 | 107.020346 |
| -4.823956 | 1.41E-06 | 5.99E-06 | 46.3123584 | 264.657342 | 407.260714 | 239.25491 |
| -4.8237422 | 1.41E-06 | 6.00E-06 | 53.2797044 | 207.287857 | 234.692615 | 215.72164 |
| -4.8231207 | 1.41E-06 | 6.02E-06 | 57.7879871 | 268.445893 | 179.470823 | 187.705843 |
| -4.8231202 | 1.41E-06 | 6.02E-06 | 157.789894 | 142.882491 | 67.8767857 | 129.432984 |
| -4.8230817 | 1.41E-06 | 6.02E-06 | 52.8698605 | 233.807713 | 217.435805 | 211.799429 |
| -4.8222393 | 1.42E-06 | 6.04E-06 | 71.722679 | 178.332503 | 115.0454 | 221.3248 |
| -4.8219659 | 1.42E-06 | 6.05E-06 | 64.3454891 | 192.945486 | 178.320369 | 137.837723 |
| -4.8219478 | 1.42E-06 | 6.05E-06 | 76.2309617 | 137.470276 | 121.948124 | 108.701294 |
| -4.8219205 | 1.42E-06 | 6.05E-06 | 75.00143 | 147.753485 | 109.29313 | 131.113932 |
| -4.8218748 | 1.42E-06 | 6.05E-06 | 67.2143963 | 155.330587 | 131.151755 | 171.45668 |
| -4.8217621 | 1.42E-06 | 6.05E-06 | 111.067691 | 210.264575 | 271.507143 | 245.978701 |
| -4.8217344 | 1.42E-06 | 6.05E-06 | 82.7884637 | 112.032863 | 104.691314 | 101.417187 |
| -4.8211949 | 1.43E-06 | 6.07E-06 | 85.247527 | 126.645845 | 94.3372276 | 170.336048 |
| -4.8208125 | 1.43E-06 | 6.08E-06 | 128.690978 | 130.434396 | 77.0804177 | 101.977503 |
| -4.820743 | 1.43E-06 | 6.08E-06 | 63.1159575 | 151.000815 | 184.072639 | 162.491625 |
| -4.8204822 | 1.43E-06 | 6.09E-06 | 68.4439279 | 147.753485 | 157.612197 | 127.752036 |
| -4.8204686 | 1.43E-06 | 6.09E-06 | 73.3620545 | 132.328671 | 116.195854 | 170.336048 |
| -4.820493 | 1.43E-06 | 6.09E-06 | 84.8376831 | 143.153102 | 92.0363196 | 121.028245 |
| -4.820198 | 1.43E-06 | 6.09E-06 | 68.4439279 | 187.53327 | 144.957203 | 131.113932 |
| -4.820146 | 1.43E-06 | 6.09E-06 | 82.7884637 | 210.535186 | 233.542161 | 210.118481 |
| -4.8198556 | 1.44E-06 | 6.10E-06 | 111.887379 | 178.061893 | 72.4786017 | 147.363094 |
| -4.8198472 | 1.44E-06 | 6.10E-06 | 75.00143 | 132.869893 | 115.0454 | 117.666349 |
| -4.819323 | 1.44E-06 | 6.12E-06 | 204.92194 | 113.115306 | 88.5849576 | 105.339398 |
| -4.8189339 | 1.44E-06 | 6.13E-06 | 49.1812656 | 258.433294 | 235.843069 | 237.573962 |
| -4.8188569 | 1.44E-06 | 6.13E-06 | 89.7558097 | 133.681725 | 94.3372276 | 99.1759229 |
| -4.8187541 | 1.44E-06 | 6.13E-06 | 69.6734596 | 118.25691 | 196.727633 | 169.775732 |
| -4.8184113 | 1.45E-06 | 6.14E-06 | 69.2636157 | 125.563402 | 148.408565 | 142.880567 |
| -4.8183526 | 1.45E-06 | 6.14E-06 | 92.214873 | 85.5130062 | 139.204933 | 178.740788 |
| -4.8179483 | 1.45E-06 | 6.15E-06 | 200.413657 | 58.9931499 | 399.207536 | 182.102683 |
| -4.8177408 | 1.45E-06 | 6.16E-06 | 59.8372065 | 159.389749 | 189.824909 | 193.869318 |

| | | | | | | |
|---|---|---|---|---|---|---|
| -4.8176817 | 1.45E-06 | 6.16E-06 | 46.7222023 | 214.323737 | 330.180297 | 300.329348 |
| -4.8176378 | 1.45E-06 | 6.16E-06 | 76.2309617 | 112.032863 | 125.399485 | 113.744138 |
| -4.8175915 | 1.45E-06 | 6.16E-06 | 538.534858 | 1564.40091 | 1510.5461 | 1598.5814 |
| -4.8173226 | 1.45E-06 | 6.17E-06 | 68.8537718 | 218.112288 | 127.700393 | 156.888466 |
| -4.8173262 | 1.45E-06 | 6.17E-06 | 66.3947085 | 172.920288 | 128.850847 | 172.016996 |
| -4.8172667 | 1.46E-06 | 6.17E-06 | 56.9682993 | 199.710755 | 207.081719 | 176.499524 |
| -4.8169529 | 1.46E-06 | 6.18E-06 | 70.9029912 | 136.117222 | 130.001301 | 125.510772 |
| -4.8167308 | 1.46E-06 | 6.18E-06 | 50.0009534 | 265.198564 | 272.657597 | 202.274057 |
| -4.8166627 | 1.46E-06 | 6.19E-06 | 117.625194 | 86.8660601 | 231.241253 | 112.06319 |
| -4.8159211 | 1.47E-06 | 6.21E-06 | 573.781432 | 1479.69974 | 934.168644 | 1178.34444 |
| -4.8159464 | 1.47E-06 | 6.21E-06 | 307.38291 | 935.230853 | 1045.76268 | 969.906907 |
| -4.8157747 | 1.47E-06 | 6.21E-06 | 57.7879871 | 172.379066 | 187.524001 | 257.745336 |
| -4.8152664 | 1.47E-06 | 6.23E-06 | 73.3620545 | 119.609964 | 121.948124 | 162.491625 |
| -4.8149652 | 1.47E-06 | 6.23E-06 | 104.920033 | 87.4072817 | 103.54086 | 93.0124475 |
| -4.8147931 | 1.47E-06 | 6.24E-06 | 86.0672148 | 230.289773 | 112.744492 | 118.226665 |
| -4.8142621 | 1.48E-06 | 6.26E-06 | 206.971159 | 145.047378 | 85.1335957 | 94.1330794 |
| -4.8141873 | 1.48E-06 | 6.26E-06 | 97.1329995 | 106.891258 | 82.8326877 | 135.59646 |
| -4.8141193 | 1.48E-06 | 6.26E-06 | 64.755333 | 181.850444 | 146.107657 | 147.363094 |
| -4.8137501 | 1.48E-06 | 6.27E-06 | 87.7065903 | 134.764168 | 92.0363196 | 104.779082 |
| -4.8135971 | 1.48E-06 | 6.27E-06 | 97.9526873 | 89.8427787 | 158.762651 | 100.856871 |
| -4.813449 | 1.48E-06 | 6.28E-06 | 39.3450125 | 194.027929 | 676.466949 | 688.067985 |
| -4.8133281 | 1.48E-06 | 6.28E-06 | 82.7884637 | 166.69624 | 103.54086 | 278.477026 |
| -4.8133317 | 1.48E-06 | 6.28E-06 | 86.0672148 | 104.996982 | 115.0454 | 91.8918156 |
| -4.8131334 | 1.49E-06 | 6.28E-06 | 61.0667381 | 180.226779 | 162.214013 | 165.293205 |
| -4.8129537 | 1.49E-06 | 6.29E-06 | 63.5258014 | 195.380983 | 134.603117 | 194.98995 |
| -4.8125385 | 1.49E-06 | 6.30E-06 | 56.5584554 | 222.712671 | 181.771731 | 186.024895 |
| -4.8124574 | 1.49E-06 | 6.30E-06 | 78.6900249 | 123.669126 | 106.992222 | 106.46003 |
| -4.8122632 | 1.49E-06 | 6.31E-06 | 75.4112739 | 158.307306 | 118.496762 | 113.183822 |
| -4.8121208 | 1.49E-06 | 6.31E-06 | 89.7558097 | 324.462324 | 105.841768 | 155.767834 |
| -4.8121359 | 1.49E-06 | 6.31E-06 | 88.1164342 | 185.909605 | 108.142676 | 103.098135 |
| -4.8121044 | 1.49E-06 | 6.31E-06 | 78.6900249 | 116.633246 | 182.922185 | 112.06319 |
| -4.8118047 | 1.50E-06 | 6.32E-06 | 94.6739363 | 84.9717847 | 105.841768 | 133.355196 |
| -4.8117017 | 1.50E-06 | 6.32E-06 | 70.9029912 | 116.633246 | 158.762651 | 137.837723 |
| -4.8116041 | 1.50E-06 | 6.32E-06 | 75.8211178 | 316.07339 | 334.782113 | 351.878416 |
| -4.8113313 | 1.50E-06 | 6.33E-06 | 81.968776 | 119.880575 | 93.1867736 | 131.674248 |
| -4.8105104 | 1.51E-06 | 6.36E-06 | 68.4439279 | 131.787449 | 169.116737 | 132.234564 |
| -4.8103761 | 1.51E-06 | 6.36E-06 | 70.0833035 | 125.834012 | 128.850847 | 139.518671 |
| -4.8102156 | 1.51E-06 | 6.36E-06 | 82.7884637 | 207.558468 | 93.1867736 | 164.172573 |
| -4.810237 | 1.51E-06 | 6.36E-06 | 125.002383 | 87.9485032 | 108.142676 | 88.5299199 |
| -4.8099508 | 1.51E-06 | 6.37E-06 | 35.2465737 | 444.884121 | 307.171217 | 382.695793 |
| -4.8090539 | 1.52E-06 | 6.40E-06 | 41.8040758 | 364.78333 | 258.852149 | 302.010296 |
| -4.8090433 | 1.52E-06 | 6.40E-06 | 55.3289238 | 190.509989 | 185.223093 | 254.943757 |
| -4.8090618 | 1.52E-06 | 6.40E-06 | 62.2962697 | 172.379066 | 147.258111 | 170.896364 |

| | | | | | | |
|---|---|---|---|---|---|---|
| -4.8089988 | 1.52E-06 | 6.40E-06 | 77.4604933 | 124.480958 | 100.089498 | 135.59646 |
| -4.8089068 | 1.52E-06 | 6.40E-06 | 568.453461 | 1030.21524 | 758.149183 | 918.918156 |
| -4.8088772 | 1.52E-06 | 6.40E-06 | 58.6076748 | 154.518755 | 251.949425 | 201.713742 |
| -4.8086451 | 1.52E-06 | 6.41E-06 | 57.3781432 | 192.674875 | 263.453965 | 166.974153 |
| -4.8086564 | 1.52E-06 | 6.41E-06 | 68.0340841 | 125.292791 | 211.683535 | 156.888466 |
| -4.8085051 | 1.52E-06 | 6.41E-06 | 139.346919 | 66.0290301 | 185.223093 | 141.759935 |
| -4.8083724 | 1.52E-06 | 6.41E-06 | 81.968776 | 167.508072 | 95.4876816 | 127.752036 |
| -4.8079277 | 1.53E-06 | 6.43E-06 | 81.968776 | 204.85236 | 97.7885896 | 142.880567 |
| -4.8078092 | 1.53E-06 | 6.43E-06 | 105.329877 | 99.8553775 | 80.5317797 | 109.821926 |
| -4.8074506 | 1.53E-06 | 6.44E-06 | 98.7723751 | 89.0309463 | 102.390406 | 91.8918156 |
| -4.8072097 | 1.53E-06 | 6.45E-06 | 91.8050291 | 131.787449 | 81.6822337 | 121.588561 |
| -4.807006 | 1.53E-06 | 6.45E-06 | 199.184126 | 211.347019 | 51.7704298 | 238.134278 |
| -4.8068506 | 1.53E-06 | 6.46E-06 | 142.62567 | 126.375234 | 131.151755 | 142.320251 |
| -4.806635 | 1.53E-06 | 6.46E-06 | 94.2640924 | 116.092024 | 81.6822337 | 109.821926 |
| -4.8061019 | 1.54E-06 | 6.48E-06 | 57.7879871 | 340.698971 | 340.534383 | 140.639303 |
| -4.8060301 | 1.54E-06 | 6.48E-06 | 72.1325229 | 159.93097 | 121.948124 | 119.907613 |
| 4.80595162 | 1.54E-06 | 6.48E-06 | 25.0004767 | 12.1774851 | 21.8586259 | 11.206319 |
| -4.8058652 | 1.54E-06 | 6.49E-06 | 97.5428434 | 89.8427787 | 93.1867736 | 98.055291 |
| -4.8056649 | 1.54E-06 | 6.49E-06 | 65.1651769 | 188.074492 | 132.302209 | 158.009098 |
| -4.8056017 | 1.54E-06 | 6.49E-06 | 64.3454891 | 203.228695 | 130.001301 | 180.421735 |
| -4.8053333 | 1.54E-06 | 6.50E-06 | 99.1822189 | 104.996982 | 101.239952 | 82.9267604 |
| -4.8051413 | 1.55E-06 | 6.51E-06 | 48.7714217 | 222.44206 | 240.444885 | 253.262809 |
| -4.8049378 | 1.55E-06 | 6.51E-06 | 87.2967464 | 107.161869 | 88.5849576 | 107.580662 |
| -4.8046805 | 1.55E-06 | 6.52E-06 | 115.985818 | 136.117222 | 108.142676 | 75.0823371 |
| -4.8044785 | 1.55E-06 | 6.52E-06 | 29.5087594 | 398.609678 | 486.64204 | 426.400437 |
| -4.8042573 | 1.55E-06 | 6.53E-06 | 69.2636157 | 118.25691 | 179.470823 | 146.802779 |
| -4.8038298 | 1.56E-06 | 6.54E-06 | 75.8211178 | 309.849342 | 163.364467 | 121.028245 |
| -4.8035247 | 1.56E-06 | 6.55E-06 | 73.3620545 | 186.450827 | 109.29313 | 142.880567 |
| -4.8032722 | 1.56E-06 | 6.56E-06 | 60.6568942 | 152.89509 | 173.718553 | 176.499524 |
| -4.8031462 | 1.56E-06 | 6.56E-06 | 68.4439279 | 114.73897 | 274.958505 | 237.573962 |
| -4.8031153 | 1.56E-06 | 6.56E-06 | 146.724109 | 111.762252 | 95.4876816 | 82.3664444 |
| -4.8030252 | 1.56E-06 | 6.57E-06 | 66.3947085 | 138.552719 | 134.603117 | 156.32815 |
| -4.8028007 | 1.56E-06 | 6.57E-06 | 100.411751 | 96.0668266 | 82.8326877 | 120.467929 |
| -4.8027683 | 1.56E-06 | 6.57E-06 | 69.6734596 | 119.880575 | 133.452663 | 158.569413 |
| -4.802336 | 1.57E-06 | 6.59E-06 | 77.0506494 | 119.880575 | 125.399485 | 100.296555 |
| -4.8021106 | 1.57E-06 | 6.59E-06 | 69.2636157 | 151.812647 | 119.647216 | 136.717091 |
| -4.8019263 | 1.57E-06 | 6.60E-06 | 279.92337 | 363.430276 | 329.029843 | 365.325998 |
| -4.8018037 | 1.57E-06 | 6.60E-06 | 50.8206411 | 208.3703 | 212.833989 | 260.546916 |
| -4.8015098 | 1.57E-06 | 6.61E-06 | 70.4931473 | 160.201581 | 163.364467 | 110.382242 |
| -4.801171 | 1.58E-06 | 6.62E-06 | 56.1486116 | 235.972599 | 166.815829 | 198.351846 |
| -4.8011564 | 1.58E-06 | 6.62E-06 | 128.690978 | 70.9000241 | 119.647216 | 129.432984 |
| -4.8008651 | 1.58E-06 | 6.63E-06 | 77.8703372 | 103.643928 | 231.241253 | 155.207518 |
| -4.8007237 | 1.58E-06 | 6.63E-06 | 73.7718984 | 128.810731 | 123.098578 | 108.140978 |

| | | | | | | |
|---|---|---|---|---|---|---|
| -4.8003005 | 1.58E-06 | 6.65E-06 | 86.8869025 | 106.079425 | 87.4345036 | 116.545717 |
| -4.799814 | 1.59E-06 | 6.66E-06 | 120.084257 | 205.934803 | 106.992222 | 178.180472 |
| -4.7996998 | 1.59E-06 | 6.66E-06 | 77.0506494 | 107.70309 | 120.79767 | 105.339398 |
| -4.7993708 | 1.59E-06 | 6.67E-06 | 46.3123584 | 282.788264 | 614.342434 | 222.445432 |
| -4.7991935 | 1.59E-06 | 6.68E-06 | 64.755333 | 148.835929 | 162.214013 | 135.036144 |
| -4.7992035 | 1.59E-06 | 6.68E-06 | 61.476582 | 165.613797 | 165.665375 | 150.72499 |
| -4.7991416 | 1.59E-06 | 6.68E-06 | 56.9682993 | 197.004647 | 173.718553 | 180.982051 |
| -4.7989448 | 1.60E-06 | 6.68E-06 | 98.3625312 | 171.026012 | 147.258111 | 129.432984 |
| -4.7988165 | 1.60E-06 | 6.69E-06 | 65.5750208 | 142.070659 | 151.859927 | 135.59646 |
| -4.7986184 | 1.60E-06 | 6.69E-06 | 72.9522106 | 116.362635 | 113.894946 | 138.398039 |
| -4.7985673 | 1.60E-06 | 6.69E-06 | 82.3786199 | 135.846611 | 88.5849576 | 145.121831 |
| -4.7984385 | 1.60E-06 | 6.70E-06 | 58.6076748 | 192.945486 | 157.612197 | 182.662999 |
| -4.7982758 | 1.60E-06 | 6.70E-06 | 220.496007 | 126.104623 | 112.744492 | 108.701294 |
| -4.7980171 | 1.60E-06 | 6.71E-06 | 62.2962697 | 177.791282 | 144.957203 | 156.888466 |
| -4.7978612 | 1.60E-06 | 6.71E-06 | 66.8045524 | 115.280192 | 297.967585 | 291.924609 |
| -4.7978802 | 1.60E-06 | 6.71E-06 | 163.527708 | 77.9359044 | 97.7885896 | 208.997849 |
| -4.7977554 | 1.60E-06 | 6.72E-06 | 115.985818 | 67.382084 | 205.931265 | 196.110582 |
| -4.7977322 | 1.60E-06 | 6.72E-06 | 99.1822189 | 107.432479 | 103.54086 | 80.6854966 |
| -4.79721 | 1.61E-06 | 6.73E-06 | 69.6734596 | 139.364551 | 131.151755 | 233.651751 |
| 4.79700734 | 1.61E-06 | 6.74E-06 | 47.951734 | 103.914539 | 78.2308717 | 85.1680242 |
| -4.7968594 | 1.61E-06 | 6.74E-06 | 54.509236 | 216.759234 | 189.824909 | 186.585211 |
| 4.79672344 | 1.61E-06 | 6.75E-06 | 61.8864259 | 30.3084073 | 24.1595339 | 35.2999048 |
| -4.7967306 | 1.61E-06 | 6.75E-06 | 47.1320462 | 274.940552 | 222.037621 | 248.219965 |
| -4.7964424 | 1.62E-06 | 6.75E-06 | 94.2640924 | 147.753485 | 146.107657 | 139.518671 |
| -4.7950928 | 1.63E-06 | 6.80E-06 | 65.5750208 | 162.637078 | 128.850847 | 147.363094 |
| 4.79493976 | 1.63E-06 | 6.80E-06 | 0 | 0 | 0 | 0 |
| -4.7946781 | 1.63E-06 | 6.81E-06 | 58.197831 | 159.119138 | 178.320369 | 196.670898 |
| -4.7944158 | 1.63E-06 | 6.82E-06 | 68.8537718 | 132.599282 | 464.783414 | 224.686695 |
| -4.7943337 | 1.63E-06 | 6.82E-06 | 84.8376831 | 121.233629 | 136.904025 | 86.2886561 |
| -4.7942697 | 1.63E-06 | 6.82E-06 | 60.2470503 | 171.837845 | 210.533081 | 146.802779 |
| -4.7941269 | 1.63E-06 | 6.83E-06 | 100.411751 | 74.1473535 | 201.329449 | 199.472478 |
| -4.7940756 | 1.63E-06 | 6.83E-06 | 61.476582 | 244.361534 | 158.762651 | 154.086886 |
| -4.793995 | 1.63E-06 | 6.83E-06 | 81.1490882 | 111.762252 | 97.7885896 | 100.856871 |
| -4.7937872 | 1.64E-06 | 6.84E-06 | 63.9356453 | 160.742803 | 135.753571 | 152.405938 |
| -4.7936266 | 1.64E-06 | 6.84E-06 | 118.444881 | 89.0309463 | 93.1867736 | 91.3314996 |
| -4.7930763 | 1.64E-06 | 6.86E-06 | 90.1656536 | 83.34812 | 195.577179 | 149.044042 |
| -4.7930721 | 1.64E-06 | 6.86E-06 | 86.0672148 | 137.199665 | 101.239952 | 256.624704 |
| -4.7929536 | 1.64E-06 | 6.86E-06 | 61.8864259 | 157.766084 | 149.559019 | 158.569413 |
| -4.7928246 | 1.64E-06 | 6.86E-06 | 68.0340841 | 147.212264 | 118.496762 | 144.561515 |
| -4.792766 | 1.64E-06 | 6.87E-06 | 80.3294005 | 94.1725511 | 215.134897 | 162.491625 |
| -4.7921215 | 1.65E-06 | 6.89E-06 | 100.411751 | 89.3015571 | 88.5849576 | 137.277407 |
| -4.7920858 | 1.65E-06 | 6.89E-06 | 72.9522106 | 119.339354 | 199.028541 | 124.390141 |
| -4.7920424 | 1.65E-06 | 6.89E-06 | 199.59397 | 536.079953 | 392.304812 | 366.44663 |

| | | | | | | |
|---|---|---|---|---|---|---|
| -4.7919431 | 1.65E-06 | 6.89E-06 | 70.0833035 | 197.275258 | 148.408565 | 114.304454 |
| -4.7919102 | 1.65E-06 | 6.89E-06 | 62.2962697 | 160.472192 | 153.010381 | 146.802779 |
| -4.7917525 | 1.65E-06 | 6.89E-06 | 99.1822189 | 95.7962158 | 88.5849576 | 90.7711837 |
| -4.7916152 | 1.65E-06 | 6.90E-06 | 220.086164 | 201.60503 | 62.1245158 | 127.19172 |
| -4.7916408 | 1.65E-06 | 6.90E-06 | 58.197831 | 171.837845 | 177.169915 | 167.534469 |
| -4.7914 | 1.66E-06 | 6.90E-06 | 98.7723751 | 88.7603356 | 105.841768 | 87.409288 |
| -4.791105 | 1.66E-06 | 6.91E-06 | 109.018472 | 77.9359044 | 162.214013 | 107.020346 |
| -4.7907784 | 1.66E-06 | 6.92E-06 | 70.9029912 | 112.303473 | 132.302209 | 127.752036 |
| -4.7907095 | 1.66E-06 | 6.92E-06 | 54.9190799 | 169.402348 | 217.435805 | 200.032794 |
| -4.7907223 | 1.66E-06 | 6.92E-06 | 89.3459658 | 94.9843835 | 90.8858656 | 98.055291 |
| -4.7903834 | 1.66E-06 | 6.93E-06 | 177.052556 | 88.7603356 | 118.496762 | 89.0902358 |
| -4.7902595 | 1.67E-06 | 6.94E-06 | 63.1159575 | 164.260743 | 151.859927 | 138.398039 |
| -4.7897795 | 1.67E-06 | 6.95E-06 | 62.2962697 | 160.742803 | 139.204933 | 166.413837 |
| -4.7898044 | 1.67E-06 | 6.95E-06 | 63.1159575 | 160.742803 | 150.709473 | 139.518671 |
| -4.7896964 | 1.67E-06 | 6.96E-06 | 77.8703372 | 123.398515 | 234.692615 | 117.666349 |
| -4.7896554 | 1.67E-06 | 6.96E-06 | 86.4770587 | 131.516839 | 88.5849576 | 103.098135 |
| -4.7881551 | 1.68E-06 | 7.01E-06 | 56.1486116 | 189.156935 | 166.815829 | 196.110582 |
| -4.7879771 | 1.68E-06 | 7.01E-06 | 74.1817423 | 118.798132 | 103.54086 | 126.631404 |
| -4.7878417 | 1.69E-06 | 7.02E-06 | 189.757716 | 153.165701 | 131.151755 | 137.837723 |
| -4.7874414 | 1.69E-06 | 7.03E-06 | 51.230485 | 250.314971 | 172.568099 | 322.741986 |
| -4.7873366 | 1.69E-06 | 7.03E-06 | 198.774282 | 103.373318 | 83.9831417 | 111.502874 |
| -4.7872229 | 1.69E-06 | 7.04E-06 | 78.6900249 | 110.950419 | 95.4876816 | 114.864769 |
| -4.7871609 | 1.69E-06 | 7.04E-06 | 63.1159575 | 156.954252 | 142.656295 | 145.121831 |
| -4.7871131 | 1.69E-06 | 7.04E-06 | 70.0833035 | 149.647761 | 108.142676 | 164.172573 |
| -4.7865676 | 1.70E-06 | 7.05E-06 | 67.6242402 | 147.753485 | 142.656295 | 117.666349 |
| -4.7865866 | 1.70E-06 | 7.05E-06 | 78.2801811 | 131.516839 | 102.390406 | 102.537819 |
| -4.7864053 | 1.70E-06 | 7.06E-06 | 72.9522106 | 172.649677 | 119.647216 | 322.741986 |
| -4.785958 | 1.70E-06 | 7.07E-06 | 68.0340841 | 467.615426 | 121.948124 | 260.546916 |
| -4.7859346 | 1.70E-06 | 7.07E-06 | 66.3947085 | 121.774851 | 166.815829 | 142.880567 |
| -4.7859067 | 1.70E-06 | 7.07E-06 | 61.476582 | 155.871809 | 148.408565 | 158.009098 |
| -4.7856168 | 1.70E-06 | 7.08E-06 | 143.445358 | 209.182132 | 178.320369 | 59.3934906 |
| -4.7855716 | 1.71E-06 | 7.08E-06 | 139.756763 | 134.222946 | 67.8767857 | 115.425085 |
| -4.7852301 | 1.71E-06 | 7.10E-06 | 85.6573709 | 230.560384 | 194.426725 | 210.118481 |
| -4.785012 | 1.71E-06 | 7.10E-06 | 45.0828268 | 193.757318 | 360.092101 | 326.103882 |
| -4.7847045 | 1.71E-06 | 7.11E-06 | 59.0175187 | 186.721438 | 194.426725 | 144.001199 |
| -4.7845823 | 1.71E-06 | 7.12E-06 | 65.5750208 | 142.611881 | 125.399485 | 158.009098 |
| -4.784465 | 1.71E-06 | 7.12E-06 | 66.8045524 | 142.34127 | 121.948124 | 141.199619 |
| -4.784459 | 1.71E-06 | 7.12E-06 | 209.840067 | 723.072002 | 851.335957 | 944.692689 |
| -4.7844663 | 1.71E-06 | 7.12E-06 | 48.7714217 | 225.960001 | 231.241253 | 216.842272 |
| -4.7840333 | 1.72E-06 | 7.13E-06 | 88.5262781 | 88.7603356 | 127.700393 | 95.8140272 |
| -4.78348 | 1.72E-06 | 7.15E-06 | 63.1159575 | 154.789366 | 135.753571 | 152.966254 |
| -4.783446 | 1.72E-06 | 7.15E-06 | 67.2143963 | 220.006563 | 128.850847 | 141.759935 |
| -4.7832743 | 1.72E-06 | 7.15E-06 | 75.8211178 | 313.637893 | 426.818432 | 290.243661 |

| | | | | | | |
|---|---|---|---|---|---|---|
| -4.7832515 | 1.72E-06 | 7.15E-06 | 181.560839 | 44.109557 | 801.866435 | 427.521069 |
| -4.7832502 | 1.72E-06 | 7.15E-06 | 54.9190799 | 174.814563 | 182.922185 | 208.997849 |
| -4.7831982 | 1.73E-06 | 7.16E-06 | 59.0175187 | 214.594348 | 149.559019 | 169.775732 |
| -4.7830769 | 1.73E-06 | 7.16E-06 | 65.5750208 | 146.129821 | 128.850847 | 137.837723 |
| -4.7827311 | 1.73E-06 | 7.17E-06 | 69.2636157 | 200.251977 | 140.355387 | 117.666349 |
| -4.7825379 | 1.73E-06 | 7.18E-06 | 58.6076748 | 170.484791 | 163.364467 | 163.612257 |
| -4.7825037 | 1.73E-06 | 7.18E-06 | 89.7558097 | 100.66721 | 87.4345036 | 94.6933953 |
| -4.7822365 | 1.73E-06 | 7.19E-06 | 95.9034679 | 152.624479 | 126.549939 | 142.320251 |
| -4.782143 | 1.73E-06 | 7.19E-06 | 61.8864259 | 144.235545 | 182.922185 | 145.682147 |
| -4.7821123 | 1.73E-06 | 7.19E-06 | 181.150995 | 218.112288 | 55.2217918 | 163.612257 |
| -4.7816701 | 1.74E-06 | 7.20E-06 | 68.0340841 | 171.837845 | 112.744492 | 150.164674 |
| -4.7815269 | 1.74E-06 | 7.21E-06 | 66.3947085 | 131.787449 | 135.753571 | 129.9933 |
| -4.7813959 | 1.74E-06 | 7.21E-06 | 78.6900249 | 103.914539 | 176.019461 | 112.06319 |
| -4.7810076 | 1.74E-06 | 7.22E-06 | 66.8045524 | 126.104623 | 134.603117 | 132.79488 |
| -4.7807024 | 1.75E-06 | 7.23E-06 | 74.5915861 | 110.950419 | 135.753571 | 103.65845 |
| -4.7806548 | 1.75E-06 | 7.23E-06 | 64.3454891 | 144.506156 | 149.559019 | 130.553616 |
| -4.7804295 | 1.75E-06 | 7.24E-06 | 65.1651769 | 135.034779 | 130.001301 | 155.767834 |
| -4.7798996 | 1.75E-06 | 7.26E-06 | 117.625194 | 97.6904913 | 210.533081 | 128.312352 |
| -4.7795381 | 1.76E-06 | 7.27E-06 | 85.6573709 | 118.527521 | 88.5849576 | 97.4949751 |
| -4.7795278 | 1.76E-06 | 7.27E-06 | 60.6568942 | 171.567234 | 162.214013 | 141.759935 |
| -4.7795938 | 1.76E-06 | 7.27E-06 | 73.7718984 | 124.480958 | 105.841768 | 110.942558 |
| -4.779556 | 1.76E-06 | 7.27E-06 | 131.969729 | 91.1958326 | 81.6822337 | 104.779082 |
| -4.7795321 | 1.76E-06 | 7.27E-06 | 65.9848647 | 147.212264 | 121.948124 | 142.880567 |
| -4.7793504 | 1.76E-06 | 7.27E-06 | 90.1656536 | 165.613797 | 153.010381 | 166.974153 |
| -4.7789911 | 1.76E-06 | 7.29E-06 | 91.3951852 | 205.122971 | 162.214013 | 160.810677 |
| -4.7789541 | 1.76E-06 | 7.29E-06 | 61.0667381 | 211.88824 | 154.160835 | 144.561515 |
| -4.7789217 | 1.76E-06 | 7.29E-06 | 138.937075 | 76.5828505 | 103.54086 | 105.339398 |
| 4.77871573 | 1.76E-06 | 7.29E-06 | 20.0823501 | 17.5897006 | 13.8054479 | 16.8094785 |
| -4.7787137 | 1.76E-06 | 7.29E-06 | 68.8537718 | 143.964935 | 110.443584 | 142.320251 |
| -4.7782572 | 1.77E-06 | 7.31E-06 | 129.100822 | 150.188982 | 71.3281477 | 309.294404 |
| -4.7782139 | 1.77E-06 | 7.31E-06 | 110.657848 | 186.180216 | 83.9831417 | 98.055291 |
| -4.7777252 | 1.77E-06 | 7.33E-06 | 63.1159575 | 162.907689 | 148.408565 | 132.234564 |
| -4.7776487 | 1.77E-06 | 7.33E-06 | 93.0345607 | 148.565318 | 155.311289 | 135.036144 |
| -4.7775628 | 1.77E-06 | 7.33E-06 | 68.8537718 | 145.588599 | 117.346308 | 122.709193 |
| -4.7774843 | 1.78E-06 | 7.33E-06 | 64.3454891 | 176.708839 | 123.098578 | 155.767834 |
| -4.7774583 | 1.78E-06 | 7.33E-06 | 55.3289238 | 171.837845 | 201.329449 | 175.939208 |
| -4.7774807 | 1.78E-06 | 7.33E-06 | 62.2962697 | 148.835929 | 140.355387 | 152.966254 |
| -4.7770603 | 1.78E-06 | 7.34E-06 | 72.5423667 | 112.303473 | 108.142676 | 131.674248 |
| -4.7770094 | 1.78E-06 | 7.35E-06 | 61.476582 | 153.977533 | 147.258111 | 149.044042 |
| -4.7768874 | 1.78E-06 | 7.35E-06 | 58.6076748 | 163.719521 | 153.010381 | 181.542367 |
| -4.7762842 | 1.79E-06 | 7.37E-06 | 73.3620545 | 156.954252 | 207.081719 | 212.359745 |
| -4.7762587 | 1.79E-06 | 7.37E-06 | 125.822071 | 168.319905 | 62.1245158 | 169.775732 |
| -4.7761551 | 1.79E-06 | 7.37E-06 | 202.053033 | 551.775379 | 633.900151 | 769.313797 |

| | | | | | | |
|---|---|---|---|---|---|---|
| -4.7757984 | 1.79E-06 | 7.38E-06 | 74.1817423 | 151.542036 | 96.6381356 | 138.398039 |
| -4.7754993 | 1.79E-06 | 7.39E-06 | 198.774282 | 114.73897 | 65.5758777 | 253.823125 |
| -4.7752956 | 1.79E-06 | 7.40E-06 | 99.5920628 | 113.115306 | 77.0804177 | 100.296555 |
| -4.7749045 | 1.80E-06 | 7.41E-06 | 65.5750208 | 173.73212 | 133.452663 | 127.19172 |
| -4.7748768 | 1.80E-06 | 7.41E-06 | 75.4112739 | 101.208431 | 111.594038 | 109.26161 |
| -4.7747624 | 1.80E-06 | 7.42E-06 | 87.2967464 | 81.1832337 | 235.843069 | 234.212067 |
| -4.7747408 | 1.80E-06 | 7.42E-06 | 62.2962697 | 124.480958 | 188.674455 | 217.402588 |
| -4.7746688 | 1.80E-06 | 7.42E-06 | 61.0667381 | 127.187066 | 234.692615 | 202.834373 |
| -4.774508 | 1.80E-06 | 7.42E-06 | 176.232868 | 109.867976 | 82.8326877 | 96.3743432 |
| -4.7742261 | 1.80E-06 | 7.43E-06 | 358.613395 | 581.001343 | 562.572004 | 623.071335 |
| -4.7732512 | 1.81E-06 | 7.47E-06 | 93.8542485 | 89.8427787 | 97.7885896 | 86.848972 |
| -4.7732436 | 1.81E-06 | 7.47E-06 | 46.7222023 | 224.877557 | 256.551241 | 221.885116 |
| -4.7732176 | 1.81E-06 | 7.47E-06 | 74.5915861 | 141.800048 | 127.700393 | 96.3743432 |
| -4.7731022 | 1.81E-06 | 7.47E-06 | 64.3454891 | 126.645845 | 140.355387 | 159.690045 |
| -4.7731125 | 1.81E-06 | 7.47E-06 | 75.8211178 | 135.30539 | 102.390406 | 105.339398 |
| -4.7726708 | 1.82E-06 | 7.49E-06 | 58.197831 | 177.791282 | 324.428027 | 160.810677 |
| -4.7724306 | 1.82E-06 | 7.49E-06 | 41.8040758 | 300.377965 | 280.710775 | 242.05649 |
| -4.7724449 | 1.82E-06 | 7.49E-06 | 59.0175187 | 165.072575 | 155.311289 | 158.009098 |
| -4.771824 | 1.83E-06 | 7.51E-06 | 74.5915861 | 126.104623 | 104.691314 | 104.779082 |
| -4.7717925 | 1.83E-06 | 7.51E-06 | 63.9356453 | 154.518755 | 125.399485 | 148.483726 |
| -4.7717273 | 1.83E-06 | 7.52E-06 | 59.8372065 | 163.448911 | 166.815829 | 142.880567 |
| -4.771106 | 1.83E-06 | 7.54E-06 | 77.4604933 | 177.25006 | 90.8858656 | 151.285306 |
| -4.7709906 | 1.83E-06 | 7.54E-06 | 73.3620545 | 127.998899 | 105.841768 | 187.145527 |
| -4.7708517 | 1.83E-06 | 7.55E-06 | 1133.62817 | 2225.23244 | 1937.36453 | 2169.54335 |
| -4.7707789 | 1.84E-06 | 7.55E-06 | 67.2143963 | 149.918372 | 117.346308 | 129.432984 |
| -4.7707046 | 1.84E-06 | 7.55E-06 | 38.5253247 | 301.731019 | 414.163438 | 253.262809 |
| -4.7705761 | 1.84E-06 | 7.55E-06 | 100.001907 | 135.846611 | 71.3281477 | 132.79488 |
| -4.7704968 | 1.84E-06 | 7.56E-06 | 43.4434513 | 273.858108 | 255.400787 | 240.375542 |
| -4.7704706 | 1.84E-06 | 7.56E-06 | 135.658324 | 192.404264 | 200.178995 | 201.153426 |
| -4.7701299 | 1.84E-06 | 7.57E-06 | 61.8864259 | 141.258827 | 140.355387 | 164.172573 |
| -4.7700946 | 1.84E-06 | 7.57E-06 | 373.777618 | 153.977533 | 81.6822337 | 115.985401 |
| -4.7699637 | 1.84E-06 | 7.57E-06 | 145.494577 | 84.4305631 | 85.1335957 | 113.744138 |
| -4.7699 | 1.84E-06 | 7.57E-06 | 53.2797044 | 364.242109 | 181.771731 | 196.110582 |
| -4.7699138 | 1.84E-06 | 7.57E-06 | 87.7065903 | 235.431378 | 105.841768 | 110.382242 |
| -4.7698677 | 1.84E-06 | 7.57E-06 | 65.9848647 | 145.047378 | 116.195854 | 166.974153 |
| -4.7686005 | 1.86E-06 | 7.62E-06 | 113.526755 | 85.5130062 | 85.1335957 | 99.1759229 |
| -4.7681599 | 1.86E-06 | 7.64E-06 | 62.2962697 | 175.085174 | 148.408565 | 131.113932 |
| -4.7680232 | 1.86E-06 | 7.64E-06 | 39.3450125 | 256.268408 | 349.738015 | 287.442082 |
| -4.7677147 | 1.86E-06 | 7.65E-06 | 67.2143963 | 154.248144 | 128.850847 | 116.545717 |
| -4.7671782 | 1.87E-06 | 7.67E-06 | 55.7387677 | 250.314971 | 219.736713 | 143.440883 |
| -4.766863 | 1.87E-06 | 7.68E-06 | 66.8045524 | 167.237462 | 110.443584 | 154.647202 |
| -4.7667966 | 1.87E-06 | 7.68E-06 | 101.641282 | 128.810731 | 108.142676 | 73.9617052 |
| -4.7667042 | 1.87E-06 | 7.68E-06 | 78.6900249 | 119.609964 | 93.1867736 | 102.537819 |

| | | | | | | |
|---|---|---|---|---|---|---|
| -4.7667229 | 1.87E-06 | 7.68E-06 | 70.4931473 | 104.18515 | 170.267191 | 137.277407 |
| -4.7667446 | 1.87E-06 | 7.68E-06 | 222.955071 | 396.715402 | 333.631659 | 363.084735 |
| -4.7656563 | 1.88E-06 | 7.72E-06 | 83.6081515 | 150.730204 | 92.0363196 | 249.900913 |
| 4.76529558 | 1.89E-06 | 7.73E-06 | 27.8693838 | 11.0950419 | 9.20363196 | 13.4475828 |
| -4.7652692 | 1.89E-06 | 7.73E-06 | 49.5911095 | 214.864959 | 222.037621 | 192.748686 |
| 4.76522358 | 1.89E-06 | 7.74E-06 | 34.4268859 | 22.4606947 | 36.8145279 | 13.4475828 |
| -4.7649064 | 1.89E-06 | 7.75E-06 | 98.3625312 | 107.161869 | 80.5317797 | 90.7711837 |
| -4.7648508 | 1.89E-06 | 7.75E-06 | 93.8542485 | 76.5828505 | 117.346308 | 109.821926 |
| -4.7648167 | 1.89E-06 | 7.75E-06 | 155.740674 | 356.123785 | 311.773033 | 373.170422 |
| -4.7645519 | 1.89E-06 | 7.76E-06 | 57.7879871 | 227.313054 | 144.957203 | 173.697944 |
| -4.7645128 | 1.89E-06 | 7.76E-06 | 881.574186 | 163.448911 | 201.329449 | 74.5220212 |
| -4.7644312 | 1.89E-06 | 7.76E-06 | 135.658324 | 217.571066 | 247.347609 | 228.048591 |
| -4.7639729 | 1.90E-06 | 7.78E-06 | 79.9195566 | 101.479042 | 104.691314 | 108.140978 |
| -4.7638271 | 1.90E-06 | 7.78E-06 | 31.1481349 | 382.914252 | 361.242555 | 369.808526 |
| -4.7637873 | 1.90E-06 | 7.78E-06 | 181.560839 | 155.601198 | 56.3722458 | 161.931309 |
| -4.7632308 | 1.91E-06 | 7.80E-06 | 56.5584554 | 180.226779 | 151.859927 | 180.421735 |
| -4.763206 | 1.91E-06 | 7.80E-06 | 61.0667381 | 163.1783 | 142.656295 | 140.639303 |
| -4.7631824 | 1.91E-06 | 7.80E-06 | 72.5423667 | 115.821413 | 108.142676 | 106.46003 |
| -4.762875 | 1.91E-06 | 7.81E-06 | 70.4931473 | 124.751569 | 103.54086 | 135.59646 |
| -4.7620299 | 1.92E-06 | 7.84E-06 | 61.8864259 | 178.603114 | 123.098578 | 173.137628 |
| -4.7619402 | 1.92E-06 | 7.85E-06 | 40.5745441 | 226.771833 | 315.224395 | 318.819775 |
| -4.7618797 | 1.92E-06 | 7.85E-06 | 58.197831 | 199.440144 | 338.233475 | 145.121831 |
| -4.7611022 | 1.93E-06 | 7.88E-06 | 101.641282 | 116.092024 | 73.6290557 | 103.098135 |
| -4.7606758 | 1.93E-06 | 7.89E-06 | 79.0998688 | 152.89509 | 88.5849576 | 122.148877 |
| -4.760561 | 1.93E-06 | 7.90E-06 | 64.3454891 | 116.092024 | 155.311289 | 168.094785 |
| -4.7602005 | 1.93E-06 | 7.91E-06 | 118.854725 | 139.635162 | 69.0272397 | 107.020346 |
| -4.7597952 | 1.94E-06 | 7.93E-06 | 51.6403289 | 253.5623 | 179.470823 | 187.145527 |
| -4.7597056 | 1.94E-06 | 7.93E-06 | 76.6408055 | 121.50424 | 93.1867736 | 107.020346 |
| -4.7596928 | 1.94E-06 | 7.93E-06 | 65.5750208 | 128.269509 | 121.948124 | 166.974153 |
| -4.7596422 | 1.94E-06 | 7.93E-06 | 60.2470503 | 136.929054 | 161.063559 | 154.086886 |
| -4.7594811 | 1.94E-06 | 7.93E-06 | 42.2139196 | 295.23636 | 243.896247 | 247.099333 |
| -4.7593697 | 1.94E-06 | 7.94E-06 | 103.280658 | 91.7370541 | 102.390406 | 79.0045488 |
| -4.7593079 | 1.94E-06 | 7.94E-06 | 79.0998688 | 110.409198 | 92.0363196 | 99.1759229 |
| -4.7592988 | 1.94E-06 | 7.94E-06 | 70.0833035 | 242.737869 | 200.178995 | 168.655101 |
| -4.7591967 | 1.94E-06 | 7.94E-06 | 84.4278393 | 203.499306 | 186.373547 | 192.18837 |
| -4.757851 | 1.96E-06 | 7.99E-06 | 59.0175187 | 155.330587 | 143.806749 | 164.732889 |
| -4.7573957 | 1.96E-06 | 8.01E-06 | 161.068645 | 87.9485032 | 81.6822337 | 115.425085 |
| -4.7572262 | 1.96E-06 | 8.02E-06 | 62.2962697 | 158.307306 | 155.311289 | 123.829825 |
| -4.7569997 | 1.96E-06 | 8.02E-06 | 61.8864259 | 216.218013 | 117.346308 | 238.694594 |
| -4.7569992 | 1.96E-06 | 8.02E-06 | 68.0340841 | 136.658443 | 234.692615 | 122.148877 |
| -4.7565907 | 1.97E-06 | 8.04E-06 | 322.956977 | 75.2297966 | 125.399485 | 138.398039 |
| -4.7564549 | 1.97E-06 | 8.04E-06 | 99.1822189 | 132.328671 | 140.355387 | 70.5998095 |
| -4.7564588 | 1.97E-06 | 8.04E-06 | 112.707067 | 83.6187308 | 89.7354116 | 91.3314996 |

| | | | | | | |
|---|---|---|---|---|---|---|
| -4.7564487 | 1.97E-06 | 8.04E-06 | 59.0175187 | 145.588599 | 162.214013 | 151.845622 |
| -4.7562507 | 1.97E-06 | 8.05E-06 | 66.8045524 | 175.897006 | 116.195854 | 129.9933 |
| -4.7557397 | 1.98E-06 | 8.07E-06 | 61.8864259 | 157.766084 | 125.399485 | 155.207518 |
| -4.7555226 | 1.98E-06 | 8.07E-06 | 58.6076748 | 154.248144 | 169.116737 | 144.561515 |
| -4.7551756 | 1.98E-06 | 8.09E-06 | 73.7718984 | 143.423713 | 155.311289 | 149.604358 |
| -4.7551143 | 1.98E-06 | 8.09E-06 | 51.230485 | 193.486707 | 179.470823 | 211.799429 |
| -4.7548601 | 1.99E-06 | 8.10E-06 | 73.3620545 | 114.46836 | 159.913105 | 101.417187 |
| -4.7548453 | 1.99E-06 | 8.10E-06 | 78.6900249 | 91.7370541 | 134.603117 | 103.098135 |
| -4.7546862 | 1.99E-06 | 8.10E-06 | 69.2636157 | 105.808815 | 133.452663 | 120.467929 |
| -4.7545881 | 1.99E-06 | 8.11E-06 | 78.2801811 | 113.927138 | 111.594038 | 87.9696039 |
| -4.7542059 | 1.99E-06 | 8.12E-06 | 128.690978 | 287.118037 | 306.020763 | 325.543566 |
| 4.75393561 | 1.99E-06 | 8.13E-06 | 2.86890716 | 2.70610779 | 3.45136199 | 2.24126379 |
| -4.7537756 | 2.00E-06 | 8.13E-06 | 46.7222023 | 242.737869 | 243.896247 | 193.869318 |
| -4.7537331 | 2.00E-06 | 8.13E-06 | 96.3133118 | 85.2423954 | 101.239952 | 179.86142 |
| -4.7537303 | 2.00E-06 | 8.13E-06 | 63.1159575 | 151.000815 | 124.249032 | 138.958355 |
| -4.7534445 | 2.00E-06 | 8.15E-06 | 69.6734596 | 110.950419 | 123.098578 | 110.382242 |
| -4.7532577 | 2.00E-06 | 8.15E-06 | 67.6242402 | 120.692407 | 136.904025 | 111.502874 |
| -4.7532273 | 2.00E-06 | 8.15E-06 | 104.920033 | 134.222946 | 131.151755 | 102.537819 |
| -4.7530351 | 2.00E-06 | 8.16E-06 | 104.100346 | 92.8194972 | 81.6822337 | 161.931309 |
| -4.7528647 | 2.01E-06 | 8.16E-06 | 55.7387677 | 164.801964 | 159.913105 | 177.620156 |
| -4.7522155 | 2.01E-06 | 8.19E-06 | 46.3123584 | 257.621462 | 202.479903 | 224.126379 |
| -4.75206 | 2.01E-06 | 8.20E-06 | 66.8045524 | 141.258827 | 108.142676 | 140.078987 |
| -4.751932 | 2.01E-06 | 8.20E-06 | 62.2962697 | 183.744719 | 118.496762 | 232.531119 |
| -4.7515232 | 2.02E-06 | 8.21E-06 | 75.8211178 | 111.491641 | 98.9390436 | 97.4949751 |
| -4.7513074 | 2.02E-06 | 8.22E-06 | 56.5584554 | 190.239378 | 173.718553 | 144.001199 |
| -4.7510533 | 2.02E-06 | 8.23E-06 | 59.0175187 | 150.188982 | 153.010381 | 146.802779 |
| -4.7503693 | 2.03E-06 | 8.26E-06 | 93.0345607 | 104.996982 | 87.4345036 | 82.9267604 |
| -4.7501677 | 2.03E-06 | 8.27E-06 | 54.9190799 | 179.414947 | 165.665375 | 165.293205 |
| -4.74971 | 2.04E-06 | 8.28E-06 | 77.4604933 | 87.4072817 | 116.195854 | 117.106033 |
| -4.7496118 | 2.04E-06 | 8.29E-06 | 59.0175187 | 139.364551 | 185.223093 | 149.044042 |
| -4.7494463 | 2.04E-06 | 8.29E-06 | 66.3947085 | 122.316072 | 119.647216 | 122.709193 |
| -4.7493889 | 2.04E-06 | 8.29E-06 | 76.6408055 | 98.7729344 | 94.3372276 | 113.183822 |
| -4.7493207 | 2.04E-06 | 8.29E-06 | 90.5754975 | 105.538204 | 93.1867736 | 80.6854966 |
| -4.7493233 | 2.04E-06 | 8.29E-06 | 56.1486116 | 163.448911 | 159.913105 | 164.732889 |
| -4.7491179 | 2.04E-06 | 8.30E-06 | 63.1159575 | 161.554635 | 164.514921 | 115.425085 |
| -4.7486519 | 2.05E-06 | 8.32E-06 | 51.230485 | 166.69624 | 201.329449 | 224.126379 |
| -4.7480163 | 2.05E-06 | 8.34E-06 | 60.2470503 | 156.683641 | 131.151755 | 153.52657 |
| -4.7479756 | 2.05E-06 | 8.35E-06 | 88.1164342 | 96.0668266 | 87.4345036 | 87.409288 |
| -4.747795 | 2.06E-06 | 8.35E-06 | 66.8045524 | 239.219929 | 218.586259 | 208.997849 |
| 4.747755 | 2.06E-06 | 8.35E-06 | 12.7051603 | 9.20076649 | 9.20363196 | 14.0078987 |
| -4.7475305 | 2.06E-06 | 8.36E-06 | 38.5253247 | 337.722252 | 620.094704 | 474.027293 |
| -4.747552 | 2.06E-06 | 8.36E-06 | 36.8859492 | 271.152001 | 322.127119 | 303.130928 |
| -4.7474394 | 2.06E-06 | 8.36E-06 | 85.247527 | 217.841677 | 200.178995 | 78.4442328 |

| | | | | | | |
|---|---|---|---|---|---|---|
| -4.7471175 | 2.06E-06 | 8.37E-06 | 72.9522106 | 102.561485 | 101.239952 | 121.028245 |
| -4.7470113 | 2.06E-06 | 8.38E-06 | 61.0667381 | 144.506156 | 287.613499 | 153.52657 |
| -4.746955 | 2.07E-06 | 8.38E-06 | 70.4931473 | 119.068743 | 100.089498 | 135.59646 |
| -4.7468457 | 2.07E-06 | 8.38E-06 | 70.9029912 | 123.669126 | 98.9390436 | 149.604358 |
| -4.7467823 | 2.07E-06 | 8.38E-06 | 47.5418901 | 206.746635 | 210.533081 | 215.72164 |
| -4.7467797 | 2.07E-06 | 8.38E-06 | 101.231438 | 106.620647 | 81.6822337 | 84.0473923 |
| -4.746715 | 2.07E-06 | 8.38E-06 | 100.001907 | 82.2656769 | 98.9390436 | 85.7283401 |
| -4.7465014 | 2.07E-06 | 8.39E-06 | 110.248004 | 95.7962158 | 80.5317797 | 87.409288 |
| -4.7464234 | 2.07E-06 | 8.39E-06 | 79.5097127 | 97.4198805 | 97.7885896 | 91.3314996 |
| -4.7459398 | 2.08E-06 | 8.41E-06 | 64.3454891 | 136.929054 | 140.355387 | 115.985401 |
| 4.74559797 | 2.08E-06 | 8.43E-06 | 14.3445358 | 16.5072575 | 17.2568099 | 13.4475828 |
| -4.7452368 | 2.08E-06 | 8.44E-06 | 109.83816 | 161.554635 | 64.4254237 | 202.274057 |
| -4.7448221 | 2.09E-06 | 8.46E-06 | 275.005243 | 667.596792 | 594.784716 | 606.822172 |
| -4.7442253 | 2.09E-06 | 8.48E-06 | 85.247527 | 97.1492697 | 81.6822337 | 117.106033 |
| -4.7441045 | 2.09E-06 | 8.49E-06 | 49.5911095 | 168.861126 | 225.488983 | 225.247011 |
| -4.7437652 | 2.10E-06 | 8.50E-06 | 65.1651769 | 151.000815 | 127.700393 | 114.864769 |
| -4.7437505 | 2.10E-06 | 8.50E-06 | 96.7231557 | 118.527521 | 89.7354116 | 79.0045488 |
| 4.74366097 | 2.10E-06 | 8.50E-06 | 6.1476582 | 5.68282636 | 2.30090799 | 3.92221164 |
| -4.7434476 | 2.10E-06 | 8.51E-06 | 59.8372065 | 164.801964 | 150.709473 | 130.553616 |
| -4.7434293 | 2.10E-06 | 8.51E-06 | 65.5750208 | 151.812647 | 116.195854 | 122.709193 |
| 4.74332624 | 2.10E-06 | 8.51E-06 | 11.8854725 | 5.1416048 | 0 | 5.04284354 |
| -4.7431766 | 2.10E-06 | 8.52E-06 | 109.018472 | 104.996982 | 73.6290557 | 94.6933953 |
| -4.7431046 | 2.10E-06 | 8.52E-06 | 82.3786199 | 124.210348 | 93.1867736 | 88.5299199 |
| -4.7429332 | 2.11E-06 | 8.52E-06 | 66.8045524 | 119.068743 | 113.894946 | 123.829825 |
| -4.7428967 | 2.11E-06 | 8.53E-06 | 55.7387677 | 195.922204 | 140.355387 | 184.904263 |
| -4.7426951 | 2.11E-06 | 8.53E-06 | 102.870814 | 105.808815 | 112.744492 | 119.907613 |
| -4.7423731 | 2.11E-06 | 8.55E-06 | 57.3781432 | 158.307306 | 144.957203 | 161.931309 |
| -4.7417647 | 2.12E-06 | 8.57E-06 | 60.6568942 | 129.081342 | 155.311289 | 142.880567 |
| -4.7414533 | 2.12E-06 | 8.58E-06 | 69.2636157 | 121.233629 | 101.239952 | 127.19172 |
| -4.7413621 | 2.12E-06 | 8.58E-06 | 63.9356453 | 124.210348 | 135.753571 | 207.316901 |
| -4.7412071 | 2.12E-06 | 8.59E-06 | 102.46097 | 127.457677 | 74.7795097 | 93.5727634 |
| -4.7411602 | 2.12E-06 | 8.59E-06 | 312.71088 | 438.660073 | 342.835291 | 400.065587 |
| -4.7409696 | 2.13E-06 | 8.60E-06 | 60.6568942 | 217.571066 | 123.098578 | 166.413837 |
| -4.7409731 | 2.13E-06 | 8.60E-06 | 64.755333 | 163.448911 | 108.142676 | 154.647202 |
| -4.7408936 | 2.13E-06 | 8.60E-06 | 55.7387677 | 182.391665 | 163.364467 | 149.044042 |
| -4.7408369 | 2.13E-06 | 8.60E-06 | 58.6076748 | 159.66036 | 134.603117 | 219.643852 |
| -4.7408424 | 2.13E-06 | 8.60E-06 | 63.5258014 | 173.461509 | 117.346308 | 138.398039 |
| -4.7405984 | 2.13E-06 | 8.61E-06 | 75.00143 | 114.46836 | 90.8858656 | 108.140978 |
| -4.7403853 | 2.13E-06 | 8.62E-06 | 57.3781432 | 155.059976 | 144.957203 | 162.491625 |
| -4.7402452 | 2.13E-06 | 8.62E-06 | 58.6076748 | 220.547785 | 155.311289 | 137.277407 |
| -4.7401793 | 2.14E-06 | 8.62E-06 | 67.6242402 | 109.867976 | 117.346308 | 120.467929 |
| -4.7400028 | 2.14E-06 | 8.63E-06 | 88.9361219 | 88.7603356 | 132.302209 | 85.7283401 |
| -4.7399585 | 2.14E-06 | 8.63E-06 | 93.8542485 | 162.095857 | 71.3281477 | 135.59646 |

| | | | | | | |
|---|---|---|---|---|---|---|
| -4.739648 | 2.14E-06 | 8.64E-06 | 69.6734596 | 120.692407 | 100.089498 | 122.148877 |
| -4.739234 | 2.15E-06 | 8.66E-06 | 1047.15111 | 147.212264 | 98.9390436 | 160.810677 |
| -4.7391563 | 2.15E-06 | 8.66E-06 | 66.8045524 | 242.467258 | 104.691314 | 182.662999 |
| -4.7384703 | 2.15E-06 | 8.69E-06 | 99.5920628 | 113.927138 | 74.7795097 | 93.0124475 |
| -4.7382404 | 2.16E-06 | 8.70E-06 | 38.9351686 | 282.517653 | 248.498063 | 289.123029 |
| -4.738274 | 2.16E-06 | 8.70E-06 | 56.1486116 | 199.440144 | 134.603117 | 189.386791 |
| 4.73824619 | 2.16E-06 | 8.70E-06 | 69.6734596 | 35.9912336 | 50.6199758 | 38.1014845 |
| -4.7381184 | 2.16E-06 | 8.70E-06 | 70.0833035 | 173.73212 | 185.223093 | 169.775732 |
| -4.7380941 | 2.16E-06 | 8.70E-06 | 67.6242402 | 128.810731 | 104.691314 | 123.829825 |
| -4.7380439 | 2.16E-06 | 8.70E-06 | 54.0993922 | 192.674875 | 149.559019 | 181.542367 |
| -4.7380192 | 2.16E-06 | 8.70E-06 | 82.3786199 | 102.290875 | 89.7354116 | 87.9696039 |
| -4.7375245 | 2.16E-06 | 8.72E-06 | 120.903945 | 178.603114 | 157.612197 | 60.5141225 |
| -4.7375457 | 2.16E-06 | 8.72E-06 | 55.7387677 | 158.036695 | 159.913105 | 160.250361 |
| -4.737235 | 2.17E-06 | 8.73E-06 | 88.9361219 | 166.425629 | 75.9299637 | 126.071088 |
| 4.73710368 | 2.17E-06 | 8.74E-06 | 9.83625312 | 11.6362635 | 8.05317797 | 9.52537113 |
| -4.737121 | 2.17E-06 | 8.74E-06 | 72.1325229 | 122.045461 | 101.239952 | 101.417187 |
| -4.7368993 | 2.17E-06 | 8.74E-06 | 59.0175187 | 149.918372 | 155.311289 | 133.915512 |
| -4.7368046 | 2.17E-06 | 8.75E-06 | 87.2967464 | 109.326755 | 75.9299637 | 110.382242 |
| -4.7366312 | 2.17E-06 | 8.75E-06 | 84.8376831 | 116.633246 | 90.8858656 | 84.6077082 |
| -4.7364286 | 2.18E-06 | 8.76E-06 | 52.0501728 | 185.368384 | 176.019461 | 174.25826 |
| -4.7363341 | 2.18E-06 | 8.76E-06 | 55.7387677 | 145.047378 | 164.514921 | 181.542367 |
| -4.7358769 | 2.18E-06 | 8.78E-06 | 63.1159575 | 150.730204 | 131.151755 | 118.786981 |
| -4.7358934 | 2.18E-06 | 8.78E-06 | 65.5750208 | 122.857294 | 128.850847 | 112.623506 |
| -4.7354402 | 2.19E-06 | 8.80E-06 | 150.00286 | 175.355785 | 93.1867736 | 124.390141 |
| -4.7353635 | 2.19E-06 | 8.80E-06 | 76.6408055 | 142.070659 | 249.648517 | 256.624704 |
| -4.7352255 | 2.19E-06 | 8.81E-06 | 109.83816 | 134.222946 | 75.9299637 | 87.9696039 |
| -4.7351971 | 2.19E-06 | 8.81E-06 | 82.3786199 | 99.0435452 | 81.6822337 | 111.502874 |
| -4.7351698 | 2.19E-06 | 8.81E-06 | 134.428793 | 88.219114 | 192.125817 | 77.8839169 |
| -4.7351083 | 2.19E-06 | 8.81E-06 | 75.4112739 | 133.411114 | 89.7354116 | 109.26161 |
| -4.7348126 | 2.19E-06 | 8.82E-06 | 50.8206411 | 199.440144 | 174.869007 | 183.783631 |
| -4.7339152 | 2.20E-06 | 8.86E-06 | 97.1329995 | 101.749653 | 77.0804177 | 89.6505518 |
| -4.7339304 | 2.20E-06 | 8.86E-06 | 63.1159575 | 127.457677 | 125.399485 | 128.872668 |
| -4.7337567 | 2.20E-06 | 8.86E-06 | 63.1159575 | 109.597366 | 203.630357 | 174.25826 |
| -4.7336725 | 2.20E-06 | 8.86E-06 | 59.4273626 | 164.801964 | 133.452663 | 140.078987 |
| -4.7333161 | 2.21E-06 | 8.88E-06 | 58.197831 | 169.672959 | 178.320369 | 127.19172 |
| -4.7330403 | 2.21E-06 | 8.89E-06 | 54.509236 | 168.590515 | 148.408565 | 218.52322 |
| -4.7329521 | 2.21E-06 | 8.89E-06 | 75.00143 | 172.920288 | 197.878087 | 86.848972 |
| -4.732883 | 2.21E-06 | 8.89E-06 | 68.4439279 | 124.751569 | 102.390406 | 117.106033 |
| -4.7327601 | 2.21E-06 | 8.90E-06 | 68.4439279 | 129.351952 | 116.195854 | 102.537819 |
| -4.7326525 | 2.22E-06 | 8.90E-06 | 64.3454891 | 125.292791 | 123.098578 | 119.907613 |
| -4.7321261 | 2.22E-06 | 8.92E-06 | 315.579788 | 640.806325 | 838.680963 | 819.181917 |
| -4.7320613 | 2.22E-06 | 8.93E-06 | 81.968776 | 102.561485 | 143.806749 | 84.6077082 |
| -4.7317384 | 2.23E-06 | 8.94E-06 | 80.3294005 | 95.7962158 | 103.54086 | 107.580662 |

| | | | | | | |
|---|---|---|---|---|---|---|
| -4.7315371 | 2.23E-06 | 8.95E-06 | 67.2143963 | 125.02218 | 110.443584 | 111.502874 |
| -4.7310564 | 2.23E-06 | 8.97E-06 | 62.2962697 | 127.457677 | 126.549939 | 134.475828 |
| -4.73086 | 2.24E-06 | 8.97E-06 | 81.1490882 | 80.1007906 | 111.594038 | 117.106033 |
| -4.7308286 | 2.24E-06 | 8.98E-06 | 57.3781432 | 204.581749 | 144.957203 | 144.561515 |
| -4.7307262 | 2.24E-06 | 8.98E-06 | 122.54332 | 117.9863 | 72.4786017 | 89.6505518 |
| 4.73022754 | 2.24E-06 | 9.00E-06 | 554.928613 | 416.469989 | 425.667978 | 401.746535 |
| -4.729788 | 2.25E-06 | 9.02E-06 | 77.0506494 | 129.622563 | 101.239952 | 89.0902358 |
| -4.7297247 | 2.25E-06 | 9.02E-06 | 73.7718984 | 117.174467 | 88.5849576 | 118.226665 |
| -4.7293134 | 2.25E-06 | 9.04E-06 | 60.2470503 | 194.56915 | 115.0454 | 188.266159 |
| -4.7292459 | 2.25E-06 | 9.04E-06 | 60.6568942 | 130.434396 | 134.603117 | 140.078987 |
| -4.7289401 | 2.26E-06 | 9.05E-06 | 63.5258014 | 134.764168 | 113.894946 | 172.016996 |
| -4.7288393 | 2.26E-06 | 9.05E-06 | 130.740198 | 90.9252218 | 149.559019 | 71.7204414 |
| -4.7288228 | 2.26E-06 | 9.05E-06 | 58.197831 | 162.095857 | 154.160835 | 131.113932 |
| -4.7287201 | 2.26E-06 | 9.06E-06 | 63.1159575 | 116.903857 | 136.904025 | 129.432984 |
| -4.7287043 | 2.26E-06 | 9.06E-06 | 63.9356453 | 105.538204 | 171.417645 | 185.464579 |
| -4.7286156 | 2.26E-06 | 9.06E-06 | 72.1325229 | 98.5023236 | 101.239952 | 108.701294 |
| -4.7284072 | 2.26E-06 | 9.07E-06 | 68.4439279 | 102.290875 | 132.302209 | 112.623506 |
| -4.7272935 | 2.28E-06 | 9.12E-06 | 236.070075 | 76.3122397 | 110.443584 | 114.304454 |
| -4.7268927 | 2.28E-06 | 9.13E-06 | 51.230485 | 178.603114 | 173.718553 | 182.662999 |
| -4.7267553 | 2.28E-06 | 9.14E-06 | 68.0340841 | 114.197749 | 119.647216 | 103.098135 |
| -4.7265707 | 2.28E-06 | 9.15E-06 | 87.2967464 | 89.0309463 | 80.5317797 | 106.46003 |
| -4.726413 | 2.29E-06 | 9.15E-06 | 330.744011 | 58.7225391 | 358.941647 | 111.502874 |
| -4.7263129 | 2.29E-06 | 9.15E-06 | 59.0175187 | 119.068743 | 179.470823 | 187.145527 |
| -4.7263137 | 2.29E-06 | 9.15E-06 | 162.70802 | 68.7351379 | 121.948124 | 101.977503 |
| 4.72626978 | 2.29E-06 | 9.15E-06 | 4.50828268 | 3.24732935 | 2.30090799 | 4.48252759 |
| -4.7261208 | 2.29E-06 | 9.16E-06 | 93.0345607 | 95.7962158 | 74.7795097 | 101.417187 |
| -4.7259765 | 2.29E-06 | 9.17E-06 | 59.4273626 | 148.294707 | 127.700393 | 146.802779 |
| -4.7258415 | 2.29E-06 | 9.17E-06 | 97.5428434 | 151.812647 | 83.9831417 | 86.2886561 |
| -4.725594 | 2.29E-06 | 9.18E-06 | 394.679656 | 436.765798 | 584.43063 | 510.447829 |
| -4.7255458 | 2.29E-06 | 9.18E-06 | 63.9356453 | 146.129821 | 110.443584 | 131.113932 |
| -4.72524 | 2.30E-06 | 9.19E-06 | 49.5911095 | 237.866875 | 218.586259 | 156.32815 |
| 4.72504049 | 2.30E-06 | 9.20E-06 | 0 | 0 | 0 | 0 |
| -4.7246023 | 2.31E-06 | 9.22E-06 | 57.3781432 | 156.41303 | 132.302209 | 166.413837 |
| -4.7245025 | 2.31E-06 | 9.22E-06 | 98.3625312 | 87.6778924 | 75.9299637 | 107.580662 |
| -4.7244635 | 2.31E-06 | 9.23E-06 | 96.7231557 | 137.470276 | 69.0272397 | 115.425085 |
| -4.7241598 | 2.31E-06 | 9.24E-06 | 38.5253247 | 258.162683 | 250.798971 | 294.165873 |
| -4.7240848 | 2.31E-06 | 9.24E-06 | 96.3133118 | 71.1706349 | 109.29313 | 108.701294 |
| -4.723772 | 2.32E-06 | 9.25E-06 | 45.0828268 | 180.49739 | 234.692615 | 289.123029 |
| -4.7236957 | 2.32E-06 | 9.26E-06 | 83.1983076 | 101.479042 | 151.859927 | 84.0473923 |
| -4.7236599 | 2.32E-06 | 9.26E-06 | 63.5258014 | 113.656527 | 127.700393 | 133.355196 |
| -4.7234845 | 2.32E-06 | 9.26E-06 | 57.3781432 | 178.603114 | 190.975363 | 124.390141 |
| -4.7229258 | 2.32E-06 | 9.29E-06 | 15.5740674 | 422.694037 | 754.697821 | 849.999294 |
| -4.7228357 | 2.33E-06 | 9.29E-06 | 71.3128351 | 90.9252218 | 124.249032 | 138.398039 |

| | | | | | | |
|---|---|---|---|---|---|---|
| -4.7222589 | 2.33E-06 | 9.32E-06 | 56.5584554 | 137.199665 | 158.762651 | 159.129729 |
| -4.7221347 | 2.33E-06 | 9.32E-06 | 144.265046 | 97.4198805 | 165.665375 | 67.2379138 |
| -4.7221493 | 2.33E-06 | 9.32E-06 | 56.1486116 | 286.035594 | 149.559019 | 160.810677 |
| -4.7219996 | 2.34E-06 | 9.32E-06 | 61.0667381 | 136.929054 | 134.603117 | 123.269509 |
| -4.7220084 | 2.34E-06 | 9.32E-06 | 86.4770587 | 112.844695 | 77.0804177 | 96.3743432 |
| -4.7217404 | 2.34E-06 | 9.33E-06 | 62.2962697 | 128.269509 | 124.249032 | 125.510772 |
| -4.7212464 | 2.34E-06 | 9.36E-06 | 68.4439279 | 274.39933 | 142.656295 | 115.425085 |
| -4.7211278 | 2.35E-06 | 9.36E-06 | 58.197831 | 154.248144 | 217.435805 | 129.432984 |
| -4.7208766 | 2.35E-06 | 9.37E-06 | 59.8372065 | 143.423713 | 121.948124 | 158.569413 |
| -4.7208033 | 2.35E-06 | 9.37E-06 | 76.6408055 | 95.2549943 | 89.7354116 | 98.615607 |
| -4.7206034 | 2.35E-06 | 9.38E-06 | 63.9356453 | 202.687474 | 117.346308 | 132.234564 |
| -4.7204636 | 2.35E-06 | 9.39E-06 | 56.1486116 | 125.02218 | 224.338529 | 198.351846 |
| -4.7199151 | 2.36E-06 | 9.41E-06 | 64.755333 | 111.22103 | 120.79767 | 157.448782 |
| -4.7196465 | 2.36E-06 | 9.42E-06 | 57.7879871 | 181.309222 | 131.151755 | 146.802779 |
| -4.7193534 | 2.37E-06 | 9.43E-06 | 59.8372065 | 156.683641 | 155.311289 | 118.226665 |
| -4.7193045 | 2.37E-06 | 9.43E-06 | 50.8206411 | 156.14242 | 256.551241 | 187.145527 |
| -4.7192379 | 2.37E-06 | 9.44E-06 | 63.1159575 | 123.939737 | 117.346308 | 127.752036 |
| -4.7191991 | 2.37E-06 | 9.44E-06 | 200.823501 | 516.595977 | 459.031144 | 500.362142 |
| -4.7184997 | 2.38E-06 | 9.47E-06 | 75.8211178 | 172.108456 | 154.160835 | 82.9267604 |
| -4.7184014 | 2.38E-06 | 9.47E-06 | 63.9356453 | 134.764168 | 119.647216 | 113.183822 |
| -4.718321 | 2.38E-06 | 9.47E-06 | 157.38005 | 101.749653 | 73.6290557 | 98.055291 |
| -4.7181476 | 2.38E-06 | 9.48E-06 | 45.4926707 | 249.503138 | 196.727633 | 200.032794 |
| -4.7178388 | 2.38E-06 | 9.49E-06 | 94.6739363 | 136.929054 | 85.1335957 | 130.553616 |
| -4.7178068 | 2.38E-06 | 9.49E-06 | 30.738291 | 326.3566 | 390.003904 | 311.535667 |
| -4.7175896 | 2.39E-06 | 9.50E-06 | 52.0501728 | 179.144336 | 161.063559 | 170.896364 |
| -4.7175196 | 2.39E-06 | 9.51E-06 | 45.0828268 | 229.748551 | 311.773033 | 177.620156 |
| -4.7171848 | 2.39E-06 | 9.52E-06 | 123.363008 | 222.17145 | 208.232173 | 247.099333 |
| -4.7171087 | 2.39E-06 | 9.52E-06 | 48.7714217 | 185.368384 | 174.869007 | 215.161324 |
| -4.7165345 | 2.40E-06 | 9.55E-06 | 77.8703372 | 133.952336 | 153.010381 | 127.19172 |
| -4.7159716 | 2.41E-06 | 9.57E-06 | 62.2962697 | 142.882491 | 110.443584 | 142.880567 |
| -4.7158558 | 2.41E-06 | 9.58E-06 | 61.0667381 | 179.144336 | 113.894946 | 147.92341 |
| -4.7156772 | 2.41E-06 | 9.58E-06 | 95.9034679 | 89.3015571 | 74.7795097 | 101.977503 |
| -4.7156357 | 2.41E-06 | 9.59E-06 | 57.3781432 | 161.554635 | 132.302209 | 146.242463 |
| -4.7153927 | 2.41E-06 | 9.60E-06 | 75.4112739 | 236.513821 | 239.294431 | 245.978701 |
| -4.7148549 | 2.42E-06 | 9.62E-06 | 72.5423667 | 105.538204 | 95.4876816 | 96.9346591 |
| -4.7147327 | 2.42E-06 | 9.62E-06 | 169.675366 | 372.089821 | 371.596641 | 328.905462 |
| -4.7146306 | 2.42E-06 | 9.63E-06 | 53.2797044 | 271.693222 | 232.391707 | 129.9933 |
| -4.7146133 | 2.42E-06 | 9.63E-06 | 47.5418901 | 186.992048 | 200.178995 | 194.98995 |
| -4.7143629 | 2.42E-06 | 9.64E-06 | 155.740674 | 92.8194972 | 79.3813257 | 94.6933953 |
| -4.7139236 | 2.43E-06 | 9.66E-06 | 75.8211178 | 164.801964 | 149.559019 | 151.285306 |
| -4.7136197 | 2.43E-06 | 9.67E-06 | 86.4770587 | 274.669941 | 276.108959 | 321.621355 |
| -4.7127549 | 2.44E-06 | 9.71E-06 | 74.1817423 | 101.479042 | 88.5849576 | 102.537819 |
| -4.7126814 | 2.44E-06 | 9.71E-06 | 56.9682993 | 130.975617 | 158.762651 | 151.285306 |

| | | | | | | |
|---|---|---|---|---|---|---|
| -4.7122822 | 2.45E-06 | 9.73E-06 | 80.3294005 | 94.1725511 | 97.7885896 | 82.3664444 |
| -4.7122624 | 2.45E-06 | 9.73E-06 | 385.663091 | 1109.23358 | 945.673184 | 1070.76378 |
| -4.7120563 | 2.45E-06 | 9.74E-06 | 218.856632 | 644.324265 | 626.997427 | 632.03639 |
| -4.7120431 | 2.45E-06 | 9.74E-06 | 58.197831 | 132.05806 | 134.603117 | 161.931309 |
| -4.7119498 | 2.45E-06 | 9.74E-06 | 72.1325229 | 147.212264 | 102.390406 | 97.4949751 |
| -4.7116802 | 2.46E-06 | 9.75E-06 | 51.230485 | 225.68939 | 142.656295 | 206.196269 |
| -4.7116904 | 2.46E-06 | 9.75E-06 | 52.4600166 | 240.843593 | 147.258111 | 174.818576 |
| -4.7116471 | 2.46E-06 | 9.75E-06 | 81.1490882 | 83.0775092 | 89.7354116 | 101.977503 |
| -4.7115825 | 2.46E-06 | 9.76E-06 | 70.0833035 | 122.316072 | 90.8858656 | 140.078987 |
| -4.7111193 | 2.46E-06 | 9.78E-06 | 65.1651769 | 103.643928 | 146.107657 | 122.148877 |
| -4.710801 | 2.47E-06 | 9.79E-06 | 40.984388 | 266.281007 | 220.887167 | 233.091435 |
| -4.7101895 | 2.47E-06 | 9.82E-06 | 87.7065903 | 125.563402 | 83.9831417 | 238.694594 |
| -4.7102117 | 2.47E-06 | 9.82E-06 | 62.2962697 | 160.742803 | 189.824909 | 105.899714 |
| -4.70994 | 2.48E-06 | 9.83E-06 | 92.214873 | 109.597366 | 71.3281477 | 99.7362389 |
| 4.70922819 | 2.49E-06 | 9.86E-06 | 9.01656536 | 6.76526948 | 6.90272397 | 5.04284354 |
| -4.7088723 | 2.49E-06 | 9.88E-06 | 71.3128351 | 178.873725 | 151.859927 | 152.966254 |
| -4.7086124 | 2.49E-06 | 9.89E-06 | 28.2792277 | 252.209246 | 507.350212 | 479.070136 |
| -4.7081798 | 2.50E-06 | 9.91E-06 | 60.2470503 | 157.224863 | 111.594038 | 161.931309 |
| -4.7080687 | 2.50E-06 | 9.91E-06 | 55.3289238 | 169.402348 | 139.204933 | 149.604358 |
| -4.7080088 | 2.50E-06 | 9.91E-06 | 65.9848647 | 124.480958 | 100.089498 | 122.709193 |
| -4.7078455 | 2.50E-06 | 9.92E-06 | 52.4600166 | 179.144336 | 161.063559 | 154.647202 |
| -4.7077897 | 2.50E-06 | 9.92E-06 | 65.5750208 | 318.508887 | 150.709473 | 122.709193 |
| -4.707748 | 2.50E-06 | 9.92E-06 | 69.6734596 | 118.527521 | 98.9390436 | 99.7362389 |
| -4.7074147 | 2.51E-06 | 9.94E-06 | 77.8703372 | 209.452743 | 128.850847 | 154.086886 |
| -4.7071959 | 2.51E-06 | 9.95E-06 | 107.78894 | 66.2996409 | 121.948124 | 105.899714 |
| -4.7068655 | 2.52E-06 | 9.96E-06 | 74.1817423 | 83.6187308 | 199.028541 | 227.488275 |
| -4.7064555 | 2.52E-06 | 9.98E-06 | 66.3947085 | 117.445078 | 109.29313 | 104.218766 |
| -4.7062018 | 2.52E-06 | 9.99E-06 | 63.1159575 | 103.914539 | 141.505841 | 145.121831 |
| -4.7061811 | 2.52E-06 | 9.99E-06 | 228.692885 | 1187.4401 | 1840.72639 | 1547.59265 |
| -4.7060754 | 2.53E-06 | 1.00E-05 | 238.119294 | 607.79181 | 761.600545 | 705.437779 |
| -4.7059416 | 2.53E-06 | 1.00E-05 | 61.0667381 | 150.188982 | 118.496762 | 124.390141 |
| -4.705964 | 2.53E-06 | 1.00E-05 | 161.888333 | 101.208431 | 211.683535 | 64.4363341 |
| -4.7058859 | 2.53E-06 | 1.00E-05 | 61.0667381 | 112.844695 | 139.204933 | 137.277407 |
| 4.70561128 | 2.53E-06 | 1.00E-05 | 0 | 0 | 0 | 0 |
| 4.70551271 | 2.53E-06 | 1.00E-05 | 42.2139196 | 27.8729103 | 17.2568099 | 17.9301104 |
| -4.7053466 | 2.53E-06 | 1.00E-05 | 49.1812656 | 165.884408 | 189.824909 | 188.266159 |
| -4.705306 | 2.53E-06 | 1.00E-05 | 59.4273626 | 154.789366 | 142.656295 | 117.106033 |
| -4.7050444 | 2.54E-06 | 1.00E-05 | 79.0998688 | 105.267593 | 78.2308717 | 117.106033 |
| 4.70484319 | 2.54E-06 | 1.00E-05 | 37.705637 | 23.8137486 | 32.2127119 | 24.0935858 |
| -4.7047974 | 2.54E-06 | 1.00E-05 | 74.5915861 | 102.020264 | 89.7354116 | 93.0124475 |
| -4.7045253 | 2.54E-06 | 1.01E-05 | 55.7387677 | 153.706923 | 131.151755 | 187.705843 |
| -4.7044437 | 2.55E-06 | 1.01E-05 | 82.7884637 | 107.973701 | 74.7795097 | 124.950457 |
| -4.7038379 | 2.55E-06 | 1.01E-05 | 73.3620545 | 91.7370541 | 101.239952 | 94.1330794 |

| | | | | | | |
|---|---|---|---|---|---|---|
| -4.7038566 | 2.55E-06 | 1.01E-05 | 55.3289238 | 124.751569 | 239.294431 | 187.145527 |
| -4.7036988 | 2.55E-06 | 1.01E-05 | 76.6408055 | 87.9485032 | 93.1867736 | 93.0124475 |
| -4.7036369 | 2.56E-06 | 1.01E-05 | 50.0009534 | 133.681725 | 272.657597 | 307.05314 |
| -4.7033446 | 2.56E-06 | 1.01E-05 | 99.5920628 | 140.176384 | 66.7263317 | 109.821926 |
| -4.7030328 | 2.56E-06 | 1.01E-05 | 74.1817423 | 94.4431619 | 101.239952 | 89.0902358 |
| -4.702778 | 2.57E-06 | 1.01E-05 | 54.509236 | 122.586683 | 217.435805 | 248.780281 |
| -4.7027365 | 2.57E-06 | 1.01E-05 | 112.707067 | 77.9359044 | 78.2308717 | 123.829825 |
| -4.7027079 | 2.57E-06 | 1.01E-05 | 57.7879871 | 202.687474 | 123.098578 | 151.285306 |
| -4.7026855 | 2.57E-06 | 1.01E-05 | 53.2797044 | 266.822228 | 157.612197 | 153.52657 |
| -4.7021428 | 2.57E-06 | 1.02E-05 | 93.0345607 | 80.6420122 | 89.7354116 | 154.647202 |
| -4.7019442 | 2.58E-06 | 1.02E-05 | 68.0340841 | 111.491641 | 95.4876816 | 115.425085 |
| -4.7018255 | 2.58E-06 | 1.02E-05 | 63.9356453 | 137.740887 | 109.29313 | 113.744138 |
| -4.7015309 | 2.58E-06 | 1.02E-05 | 72.5423667 | 85.2423954 | 112.744492 | 115.985401 |
| -4.7014994 | 2.58E-06 | 1.02E-05 | 258.611488 | 57.0988744 | 184.072639 | 121.028245 |
| -4.7010262 | 2.59E-06 | 1.02E-05 | 99.1822189 | 73.8767427 | 133.452663 | 89.0902358 |
| -4.7004687 | 2.60E-06 | 1.02E-05 | 40.984388 | 220.818396 | 245.046701 | 228.608907 |
| -4.7001981 | 2.60E-06 | 1.03E-05 | 65.9848647 | 202.958084 | 121.948124 | 110.942558 |
| 4.70019615 | 2.60E-06 | 1.03E-05 | 0 | 0 | 0 | 0 |
| -4.7001099 | 2.60E-06 | 1.03E-05 | 57.7879871 | 155.059976 | 123.098578 | 142.880567 |
| -4.6995462 | 2.61E-06 | 1.03E-05 | 47.5418901 | 319.050109 | 172.568099 | 188.826475 |
| -4.6994873 | 2.61E-06 | 1.03E-05 | 92.6247169 | 358.559282 | 222.037621 | 265.58976 |
| -4.6992703 | 2.61E-06 | 1.03E-05 | 97.9526873 | 136.117222 | 103.54086 | 149.604358 |
| -4.6990931 | 2.61E-06 | 1.03E-05 | 94.6739363 | 86.0542278 | 131.151755 | 76.763285 |
| -4.6987946 | 2.62E-06 | 1.03E-05 | 43.4434513 | 230.830995 | 220.887167 | 193.309002 |
| -4.6985139 | 2.62E-06 | 1.03E-05 | 195.495531 | 101.749653 | 73.6290557 | 105.339398 |
| -4.698407 | 2.62E-06 | 1.03E-05 | 82.3786199 | 126.104623 | 75.9299637 | 101.977503 |
| -4.6983943 | 2.62E-06 | 1.03E-05 | 46.7222023 | 196.463426 | 187.524001 | 188.266159 |
| -4.6982362 | 2.62E-06 | 1.03E-05 | 77.0506494 | 102.561485 | 95.4876816 | 82.3664444 |
| -4.6982216 | 2.62E-06 | 1.03E-05 | 57.7879871 | 192.674875 | 134.603117 | 129.9933 |
| -4.6982693 | 2.62E-06 | 1.03E-05 | 120.084257 | 281.976432 | 55.2217918 | 208.437533 |
| 4.69825276 | 2.62E-06 | 1.03E-05 | 0.81968776 | 0.54122156 | 0 | 1.68094785 |
| -4.6980606 | 2.63E-06 | 1.03E-05 | 82.3786199 | 191.592432 | 104.691314 | 88.5299199 |
| -4.6976263 | 2.63E-06 | 1.04E-05 | 122.133476 | 147.482875 | 56.3722458 | 165.853521 |
| -4.6974064 | 2.63E-06 | 1.04E-05 | 132.789417 | 89.3015571 | 98.9390436 | 73.4013893 |
| -4.6972115 | 2.64E-06 | 1.04E-05 | 53.2797044 | 199.981366 | 226.639437 | 127.19172 |
| -4.697175 | 2.64E-06 | 1.04E-05 | 80.3294005 | 132.599282 | 142.656295 | 74.5220212 |
| -4.6970178 | 2.64E-06 | 1.04E-05 | 59.4273626 | 171.026012 | 139.204933 | 115.425085 |
| -4.6968633 | 2.64E-06 | 1.04E-05 | 46.3123584 | 175.085174 | 250.798971 | 189.386791 |
| -4.6968184 | 2.64E-06 | 1.04E-05 | 213.118818 | 1160.37902 | 834.079147 | 939.08953 |
| -4.6967144 | 2.64E-06 | 1.04E-05 | 54.9190799 | 158.577917 | 203.630357 | 129.9933 |
| -4.6965836 | 2.65E-06 | 1.04E-05 | 102.870814 | 73.8767427 | 92.0363196 | 156.888466 |
| -4.6964066 | 2.65E-06 | 1.04E-05 | 48.3615778 | 225.960001 | 156.461743 | 193.309002 |
| -4.6963091 | 2.65E-06 | 1.04E-05 | 49.5911095 | 162.637078 | 194.426725 | 170.336048 |

| | | | | | | |
|---|---|---|---|---|---|---|
| -4.696143 | 2.65E-06 | 1.04E-05 | 60.6568942 | 199.169533 | 208.232173 | 215.72164 |
| -4.6959914 | 2.65E-06 | 1.04E-05 | 48.7714217 | 238.408096 | 151.859927 | 195.550266 |
| -4.6958183 | 2.66E-06 | 1.04E-05 | 91.8050291 | 126.104623 | 112.744492 | 118.786981 |
| -4.6953418 | 2.66E-06 | 1.05E-05 | 79.0998688 | 132.599282 | 87.4345036 | 89.0902358 |
| -4.6952373 | 2.66E-06 | 1.05E-05 | 163.937552 | 113.385916 | 72.4786017 | 89.0902358 |
| -4.6949413 | 2.67E-06 | 1.05E-05 | 61.8864259 | 123.398515 | 113.894946 | 120.467929 |
| -4.6948537 | 2.67E-06 | 1.05E-05 | 133.609105 | 101.749653 | 82.8326877 | 76.202969 |
| -4.6947072 | 2.67E-06 | 1.05E-05 | 48.7714217 | 207.017246 | 161.063559 | 179.301104 |
| -4.6944531 | 2.67E-06 | 1.05E-05 | 74.1817423 | 91.1958326 | 96.6381356 | 141.759935 |
| -4.6943645 | 2.67E-06 | 1.05E-05 | 538.125014 | 80.1007906 | 150.709473 | 122.709193 |
| -4.6943746 | 2.67E-06 | 1.05E-05 | 58.6076748 | 139.09394 | 133.452663 | 122.148877 |
| -4.6943471 | 2.67E-06 | 1.05E-05 | 47.1320462 | 216.218013 | 156.461743 | 228.048591 |
| -4.6941863 | 2.68E-06 | 1.05E-05 | 47.1320462 | 211.347019 | 178.320369 | 178.740788 |
| -4.6937855 | 2.68E-06 | 1.05E-05 | 63.9356453 | 97.9611021 | 156.461743 | 141.199619 |
| -4.6934679 | 2.69E-06 | 1.05E-05 | 56.5584554 | 134.222946 | 132.302209 | 157.448782 |
| -4.6932076 | 2.69E-06 | 1.06E-05 | 60.6568942 | 124.480958 | 126.549939 | 117.106033 |
| -4.6930821 | 2.69E-06 | 1.06E-05 | 57.3781432 | 134.222946 | 149.559019 | 126.631404 |
| -4.6930249 | 2.69E-06 | 1.06E-05 | 45.4926707 | 205.393581 | 217.435805 | 175.939208 |
| -4.6929313 | 2.69E-06 | 1.06E-05 | 61.0667381 | 134.764168 | 109.29313 | 132.79488 |
| 4.69284422 | 2.69E-06 | 1.06E-05 | 6.96734596 | 4.32977247 | 0 | 1.1206319 |
| -4.6928436 | 2.69E-06 | 1.06E-05 | 54.9190799 | 156.41303 | 132.302209 | 155.207518 |
| -4.6928642 | 2.69E-06 | 1.06E-05 | 50.0009534 | 214.323737 | 201.329449 | 141.759935 |
| -4.6927676 | 2.70E-06 | 1.06E-05 | 99.5920628 | 70.9000241 | 101.239952 | 91.8918156 |
| -4.6927322 | 2.70E-06 | 1.06E-05 | 64.3454891 | 130.705006 | 117.346308 | 101.417187 |
| 4.69270164 | 2.70E-06 | 1.06E-05 | 22.9512573 | 17.8603114 | 17.2568099 | 14.0078987 |
| -4.6924355 | 2.70E-06 | 1.06E-05 | 68.4439279 | 110.138587 | 98.9390436 | 98.055291 |
| -4.6924084 | 2.70E-06 | 1.06E-05 | 80.7392443 | 74.4179643 | 117.346308 | 113.183822 |
| -4.6923773 | 2.70E-06 | 1.06E-05 | 85.247527 | 102.020264 | 72.4786017 | 101.977503 |
| -4.69224 | 2.70E-06 | 1.06E-05 | 45.9025146 | 162.907689 | 223.188075 | 226.367643 |
| -4.6921888 | 2.70E-06 | 1.06E-05 | 59.8372065 | 126.104623 | 119.647216 | 174.818576 |
| -4.6915734 | 2.71E-06 | 1.06E-05 | 25.0004767 | 495.758947 | 300.268493 | 448.252759 |
| -4.6915141 | 2.71E-06 | 1.06E-05 | 69.2636157 | 96.6080482 | 97.7885896 | 126.071088 |
| -4.6912201 | 2.72E-06 | 1.06E-05 | 64.3454891 | 95.2549943 | 169.116737 | 185.464579 |
| -4.6911457 | 2.72E-06 | 1.06E-05 | 68.4439279 | 102.561485 | 135.753571 | 96.3743432 |
| -4.6910675 | 2.72E-06 | 1.06E-05 | 71.722679 | 129.081342 | 82.8326877 | 127.19172 |
| -4.6908736 | 2.72E-06 | 1.06E-05 | 63.1159575 | 129.351952 | 101.239952 | 137.837723 |
| -4.6900412 | 2.73E-06 | 1.07E-05 | 53.6895483 | 162.366467 | 136.904025 | 157.448782 |
| -4.6900024 | 2.73E-06 | 1.07E-05 | 56.1486116 | 122.586683 | 151.859927 | 183.223315 |
| 4.6898545 | 2.73E-06 | 1.07E-05 | 31.1481349 | 12.7187066 | 23.0090799 | 12.8872668 |
| -4.6891048 | 2.74E-06 | 1.07E-05 | 61.8864259 | 111.491641 | 116.195854 | 145.682147 |
| -4.6884989 | 2.75E-06 | 1.08E-05 | 83.6081515 | 180.226779 | 87.4345036 | 96.9346591 |
| -4.68757 | 2.76E-06 | 1.08E-05 | 125.822071 | 257.892073 | 208.232173 | 223.005748 |
| -4.6874846 | 2.77E-06 | 1.08E-05 | 193.036467 | 270.340168 | 316.374849 | 293.045241 |

| | | | | | | |
|---|---|---|---|---|---|---|
| -4.6874141 | 2.77E-06 | 1.08E-05 | 61.0667381 | 154.789366 | 118.496762 | 114.304454 |
| -4.6867391 | 2.78E-06 | 1.09E-05 | 72.5423667 | 168.590515 | 190.975363 | 161.931309 |
| -4.6866097 | 2.78E-06 | 1.09E-05 | 59.4273626 | 143.694324 | 150.709473 | 109.26161 |
| -4.6859984 | 2.79E-06 | 1.09E-05 | 61.476582 | 129.081342 | 109.29313 | 121.028245 |
| -4.685631 | 2.79E-06 | 1.09E-05 | 52.0501728 | 173.190899 | 149.559019 | 150.72499 |
| -4.6855448 | 2.79E-06 | 1.09E-05 | 75.4112739 | 97.1492697 | 83.9831417 | 91.8918156 |
| -4.6853762 | 2.79E-06 | 1.09E-05 | 70.9029912 | 125.834012 | 95.4876816 | 92.4521315 |
| -4.6852431 | 2.80E-06 | 1.09E-05 | 1302.89369 | 158.307306 | 78.2308717 | 175.939208 |
| 4.68525812 | 2.80E-06 | 1.09E-05 | 31.5579788 | 20.83703 | 21.8586259 | 19.0507423 |
| -4.684977 | 2.80E-06 | 1.09E-05 | 51.230485 | 199.710755 | 132.302209 | 206.196269 |
| -4.6848333 | 2.80E-06 | 1.09E-05 | 65.9848647 | 146.400432 | 93.1867736 | 124.950457 |
| -4.6847618 | 2.80E-06 | 1.09E-05 | 61.476582 | 115.280192 | 163.364467 | 114.864769 |
| -4.684623 | 2.80E-06 | 1.10E-05 | 77.0506494 | 81.9950661 | 101.239952 | 89.6505518 |
| -4.6843046 | 2.81E-06 | 1.10E-05 | 95.9034679 | 122.586683 | 65.5758777 | 107.020346 |
| -4.6840551 | 2.81E-06 | 1.10E-05 | 43.0336074 | 216.488623 | 200.178995 | 205.635953 |
| -4.6839369 | 2.81E-06 | 1.10E-05 | 3846.79466 | 3645.93903 | 3078.61489 | 3534.473 |
| -4.6835855 | 2.82E-06 | 1.10E-05 | 169.265522 | 84.9717847 | 87.4345036 | 90.2108677 |
| -4.6828248 | 2.83E-06 | 1.10E-05 | 52.0501728 | 187.262659 | 142.656295 | 154.086886 |
| -4.6828011 | 2.83E-06 | 1.10E-05 | 100.821594 | 84.7011739 | 71.3281477 | 105.899714 |
| -4.6827288 | 2.83E-06 | 1.10E-05 | 45.9025146 | 195.380983 | 196.727633 | 175.378892 |
| -4.6827167 | 2.83E-06 | 1.10E-05 | 40.984388 | 281.43521 | 200.178995 | 214.601008 |
| -4.6826099 | 2.83E-06 | 1.10E-05 | 201.233345 | 115.009581 | 74.7795097 | 90.7711837 |
| -4.6825307 | 2.83E-06 | 1.11E-05 | 4.91812656 | 677.880002 | 1724.53054 | 1422.64219 |
| -4.6824081 | 2.84E-06 | 1.11E-05 | 56.9682993 | 209.182132 | 110.443584 | 187.145527 |
| -4.6822497 | 2.84E-06 | 1.11E-05 | 56.9682993 | 125.834012 | 138.054479 | 134.475828 |
| -4.6816877 | 2.85E-06 | 1.11E-05 | 173.773805 | 662.184577 | 721.334655 | 697.593356 |
| -4.6814343 | 2.85E-06 | 1.11E-05 | 47.5418901 | 156.683641 | 192.125817 | 201.713742 |
| -4.6813302 | 2.85E-06 | 1.11E-05 | 67.2143963 | 136.658443 | 90.8858656 | 115.985401 |
| -4.6812758 | 2.85E-06 | 1.11E-05 | 61.476582 | 116.362635 | 120.79767 | 111.502874 |
| -4.6812135 | 2.85E-06 | 1.11E-05 | 112.297223 | 140.988216 | 78.2308717 | 76.202969 |
| -4.6808971 | 2.86E-06 | 1.11E-05 | 75.4112739 | 90.654611 | 96.6381356 | 83.4870763 |
| -4.6807938 | 2.86E-06 | 1.11E-05 | 37.2957931 | 255.727186 | 246.197155 | 242.05649 |
| -4.6807938 | 2.86E-06 | 1.11E-05 | 68.4439279 | 98.7729344 | 93.1867736 | 117.106033 |
| -4.680661 | 2.86E-06 | 1.11E-05 | 93.4444046 | 330.145151 | 164.514921 | 238.694594 |
| -4.6804487 | 2.86E-06 | 1.11E-05 | 42.6237635 | 195.110372 | 272.657597 | 194.98995 |
| -4.6802936 | 2.86E-06 | 1.12E-05 | 85.247527 | 115.009581 | 70.1776937 | 103.65845 |
| -4.6801172 | 2.87E-06 | 1.12E-05 | 153.281611 | 93.9019404 | 65.5758777 | 119.347297 |
| -4.6799017 | 2.87E-06 | 1.12E-05 | 70.9029912 | 95.2549943 | 93.1867736 | 93.5727634 |
| -4.6795693 | 2.87E-06 | 1.12E-05 | 81.1490882 | 180.49739 | 153.010381 | 180.982051 |
| -4.6795306 | 2.88E-06 | 1.12E-05 | 111.067691 | 311.743618 | 54.0713378 | 271.192919 |
| -4.6794652 | 2.88E-06 | 1.12E-05 | 51.6403289 | 138.282108 | 162.214013 | 191.628054 |
| -4.6793711 | 2.88E-06 | 1.12E-05 | 76.2309617 | 104.18515 | 124.249032 | 77.8839169 |
| -4.679308 | 2.88E-06 | 1.12E-05 | 131.150042 | 234.348935 | 49.4695218 | 217.402588 |

| | | | | | | |
|---|---|---|---|---|---|---|
| -4.6792964 | 2.88E-06 | 1.12E-05 | 68.4439279 | 107.973701 | 94.3372276 | 96.9346591 |
| -4.6791423 | 2.88E-06 | 1.12E-05 | 55.3289238 | 159.389749 | 146.107657 | 124.390141 |
| -4.6788289 | 2.89E-06 | 1.12E-05 | 64.755333 | 121.774851 | 103.54086 | 102.537819 |
| -4.6786588 | 2.89E-06 | 1.12E-05 | 256.152425 | 79.0183475 | 105.841768 | 104.779082 |
| -4.6783846 | 2.89E-06 | 1.12E-05 | 54.9190799 | 142.882491 | 135.753571 | 139.518671 |
| 4.67811339 | 2.90E-06 | 1.13E-05 | 63.9356453 | 34.9087905 | 46.0181598 | 22.4126379 |
| -4.6776842 | 2.90E-06 | 1.13E-05 | 83.1983076 | 115.009581 | 72.4786017 | 97.4949751 |
| 4.67733303 | 2.91E-06 | 1.13E-05 | 0 | 0 | 0 | 0 |
| -4.6769029 | 2.91E-06 | 1.13E-05 | 411.893099 | 333.663091 | 287.613499 | 194.429634 |
| -4.6764398 | 2.92E-06 | 1.13E-05 | 56.5584554 | 131.787449 | 126.549939 | 138.958355 |
| -4.6762347 | 2.92E-06 | 1.14E-05 | 71.722679 | 179.144336 | 98.9390436 | 101.417187 |
| -4.6761292 | 2.92E-06 | 1.14E-05 | 75.00143 | 94.4431619 | 90.8858656 | 83.4870763 |
| 4.67596823 | 2.93E-06 | 1.14E-05 | 47.951734 | 55.7458205 | 58.6731538 | 50.4284354 |
| -4.6758937 | 2.93E-06 | 1.14E-05 | 38.9351686 | 247.879474 | 227.789891 | 223.005748 |
| -4.6751125 | 2.94E-06 | 1.14E-05 | 63.9356453 | 178.061893 | 163.364467 | 92.4521315 |
| -4.6750552 | 2.94E-06 | 1.14E-05 | 89.3459658 | 123.127905 | 67.8767857 | 103.098135 |
| -4.6750245 | 2.94E-06 | 1.14E-05 | 135.658324 | 206.746635 | 341.684837 | 279.597658 |
| -4.6745516 | 2.95E-06 | 1.14E-05 | 118.035037 | 98.7729344 | 117.346308 | 63.3157022 |
| -4.6745535 | 2.95E-06 | 1.14E-05 | 85.247527 | 163.1783 | 133.452663 | 186.585211 |
| -4.6741924 | 2.95E-06 | 1.15E-05 | 37.2957931 | 221.089007 | 251.949425 | 267.831023 |
| -4.6738404 | 2.96E-06 | 1.15E-05 | 63.5258014 | 182.662276 | 96.6381356 | 140.639303 |
| -4.673451 | 2.96E-06 | 1.15E-05 | 52.8698605 | 168.319905 | 136.904025 | 146.242463 |
| -4.6732478 | 2.96E-06 | 1.15E-05 | 240.168514 | 568.012025 | 465.933868 | 488.035191 |
| -4.672842 | 2.97E-06 | 1.15E-05 | 480.337027 | 756.086517 | 608.590164 | 743.539264 |
| -4.6727191 | 2.97E-06 | 1.15E-05 | 45.9025146 | 198.898923 | 187.524001 | 169.215416 |
| -4.6726967 | 2.97E-06 | 1.15E-05 | 58.197831 | 125.292791 | 116.195854 | 155.207518 |
| -4.6724778 | 2.98E-06 | 1.15E-05 | 82.7884637 | 96.6080482 | 74.7795097 | 135.036144 |
| 4.6723789 | 2.98E-06 | 1.15E-05 | 0 | 2.16488623 | 0 | 0 |
| -4.6723706 | 2.98E-06 | 1.15E-05 | 67.2143963 | 101.208431 | 126.549939 | 93.0124475 |
| -4.6719795 | 2.98E-06 | 1.16E-05 | 47.951734 | 189.427545 | 165.665375 | 164.172573 |
| -4.6714708 | 2.99E-06 | 1.16E-05 | 56.1486116 | 156.14242 | 120.79767 | 135.59646 |
| -4.6713401 | 2.99E-06 | 1.16E-05 | 247.955547 | 101.208431 | 71.3281477 | 121.028245 |
| -4.6712737 | 2.99E-06 | 1.16E-05 | 508.616255 | 137.740887 | 90.8858656 | 104.779082 |
| -4.6711857 | 2.99E-06 | 1.16E-05 | 104.510189 | 90.654611 | 208.232173 | 79.0045488 |
| -4.6708629 | 3.00E-06 | 1.16E-05 | 53.6895483 | 169.943569 | 143.806749 | 130.553616 |
| -4.6708346 | 3.00E-06 | 1.16E-05 | 101.641282 | 98.5023236 | 72.4786017 | 81.2458125 |
| 4.67042751 | 3.01E-06 | 1.16E-05 | 0 | 0 | 0 | 0 |
| 4.67031804 | 3.01E-06 | 1.16E-05 | 0 | 0 | 0 | 0 |
| -4.6701557 | 3.01E-06 | 1.17E-05 | 74.1817423 | 103.373318 | 104.691314 | 78.4442328 |
| -4.6700701 | 3.01E-06 | 1.17E-05 | 63.1159575 | 99.3141559 | 112.744492 | 126.071088 |
| -4.6697882 | 3.02E-06 | 1.17E-05 | 72.9522106 | 94.4431619 | 97.7885896 | 82.9267604 |
| -4.6697302 | 3.02E-06 | 1.17E-05 | 134.018949 | 168.319905 | 157.612197 | 158.569413 |
| -4.6695943 | 3.02E-06 | 1.17E-05 | 97.9526873 | 121.50424 | 65.5758777 | 96.9346591 |

| | | | | | | |
|---|---|---|---|---|---|---|
| -4.6696178 | 3.02E-06 | 1.17E-05 | 55.7387677 | 178.603114 | 115.0454 | 154.647202 |
| -4.6687954 | 3.03E-06 | 1.17E-05 | 53.6895483 | 146.129821 | 135.753571 | 142.320251 |
| -4.6686648 | 3.03E-06 | 1.17E-05 | 95.493624 | 80.3714014 | 80.5317797 | 81.8061285 |
| -4.6685977 | 3.03E-06 | 1.17E-05 | 59.4273626 | 133.140503 | 126.549939 | 108.140978 |
| -4.6680412 | 3.04E-06 | 1.18E-05 | 54.509236 | 122.857294 | 167.966283 | 145.121831 |
| -4.6678278 | 3.04E-06 | 1.18E-05 | 54.9190799 | 201.875641 | 228.940345 | 110.942558 |
| -4.6676996 | 3.05E-06 | 1.18E-05 | 76.2309617 | 84.4305631 | 94.3372276 | 83.4870763 |
| -4.6677164 | 3.05E-06 | 1.18E-05 | 119.674413 | 158.848527 | 89.7354116 | 122.148877 |
| -4.6676117 | 3.05E-06 | 1.18E-05 | 58.197831 | 128.810731 | 111.594038 | 150.72499 |
| -4.6674953 | 3.05E-06 | 1.18E-05 | 52.4600166 | 291.177198 | 119.647216 | 215.161324 |
| -4.667486 | 3.05E-06 | 1.18E-05 | 55.7387677 | 138.282108 | 155.311289 | 119.347297 |
| -4.6672495 | 3.05E-06 | 1.18E-05 | 61.0667381 | 103.643928 | 234.692615 | 155.767834 |
| -4.667088 | 3.05E-06 | 1.18E-05 | 62.7061136 | 259.786348 | 166.815829 | 190.507423 |
| -4.6667686 | 3.06E-06 | 1.18E-05 | 49.1812656 | 172.920288 | 157.612197 | 158.009098 |
| -4.6663842 | 3.07E-06 | 1.18E-05 | 37.705637 | 232.72527 | 272.657597 | 216.281956 |
| -4.6660593 | 3.07E-06 | 1.19E-05 | 52.4600166 | 211.076408 | 142.656295 | 140.078987 |
| -4.6658275 | 3.07E-06 | 1.19E-05 | 59.0175187 | 146.400432 | 103.54086 | 170.336048 |
| -4.6658351 | 3.07E-06 | 1.19E-05 | 62.7061136 | 109.056144 | 101.239952 | 125.510772 |
| -4.6658028 | 3.07E-06 | 1.19E-05 | 62.2962697 | 96.8786589 | 142.656295 | 181.542367 |
| -4.6656917 | 3.08E-06 | 1.19E-05 | 62.7061136 | 120.151186 | 98.9390436 | 114.864769 |
| -4.6653434 | 3.08E-06 | 1.19E-05 | 335.252294 | 454.355498 | 437.172518 | 489.155823 |
| -4.6653502 | 3.08E-06 | 1.19E-05 | 66.8045524 | 155.871809 | 86.2840497 | 134.475828 |
| -4.6653012 | 3.08E-06 | 1.19E-05 | 51.230485 | 152.083258 | 148.408565 | 152.405938 |
| -4.6652592 | 3.08E-06 | 1.19E-05 | 55.3289238 | 131.787449 | 147.258111 | 125.510772 |
| -4.6649187 | 3.09E-06 | 1.19E-05 | 63.9356453 | 112.303473 | 95.4876816 | 116.545717 |
| -4.664878 | 3.09E-06 | 1.19E-05 | 52.4600166 | 136.117222 | 143.806749 | 165.853521 |
| -4.6647638 | 3.09E-06 | 1.19E-05 | 81.1490882 | 235.701989 | 153.010381 | 186.585211 |
| -4.6647819 | 3.09E-06 | 1.19E-05 | 68.4439279 | 127.187066 | 82.8326877 | 129.432984 |
| -4.664689 | 3.09E-06 | 1.19E-05 | 39.7548564 | 233.266492 | 273.808051 | 187.145527 |
| 4.66451566 | 3.09E-06 | 1.19E-05 | 21.7217256 | 10.2832096 | 8.05317797 | 7.84442328 |
| 4.66427397 | 3.10E-06 | 1.19E-05 | 0 | 0 | 0 | 0 |
| -4.6641522 | 3.10E-06 | 1.19E-05 | 41.3942319 | 199.169533 | 226.639437 | 201.713742 |
| -4.6639143 | 3.10E-06 | 1.20E-05 | 90.9853413 | 120.421797 | 67.8767857 | 94.6933953 |
| -4.6634077 | 3.11E-06 | 1.20E-05 | 53.2797044 | 133.952336 | 136.904025 | 179.301104 |
| -4.6632018 | 3.11E-06 | 1.20E-05 | 70.9029912 | 156.954252 | 171.417645 | 164.732889 |
| 4.66300711 | 3.12E-06 | 1.20E-05 | 21.3118818 | 15.6954252 | 12.6549939 | 7.84442328 |
| -4.6629728 | 3.12E-06 | 1.20E-05 | 50.4107972 | 245.173366 | 133.452663 | 176.499524 |
| -4.6629023 | 3.12E-06 | 1.20E-05 | 43.0336074 | 173.190899 | 215.134897 | 214.601008 |
| -4.6624769 | 3.12E-06 | 1.20E-05 | 41.8040758 | 301.189797 | 169.116737 | 224.686695 |
| -4.6622835 | 3.13E-06 | 1.20E-05 | 45.0828268 | 238.949318 | 143.806749 | 271.192919 |
| -4.6617885 | 3.13E-06 | 1.21E-05 | 43.4434513 | 249.773749 | 173.718553 | 191.628054 |
| -4.6606759 | 3.15E-06 | 1.21E-05 | 237.299606 | 403.210061 | 269.206235 | 310.975351 |
| -4.6604955 | 3.15E-06 | 1.21E-05 | 69.6734596 | 270.88139 | 79.3813257 | 290.803977 |

| | | | | | | |
|---|---|---|---|---|---|---|
| -4.6602562 | 3.16E-06 | 1.22E-05 | 70.4931473 | 149.106539 | 96.6381356 | 91.8918156 |
| -4.6602023 | 3.16E-06 | 1.22E-05 | 77.0506494 | 210.535186 | 205.931265 | 188.266159 |
| -4.6600792 | 3.16E-06 | 1.22E-05 | 67.6242402 | 118.798132 | 90.8858656 | 96.3743432 |
| -4.6598274 | 3.16E-06 | 1.22E-05 | 63.1159575 | 114.73897 | 196.727633 | 108.701294 |
| -4.6597954 | 3.17E-06 | 1.22E-05 | 67.6242402 | 133.952336 | 178.320369 | 86.2886561 |
| -4.659291 | 3.17E-06 | 1.22E-05 | 105.329877 | 88.219114 | 80.5317797 | 74.5220212 |
| -4.6592289 | 3.17E-06 | 1.22E-05 | 773.375401 | 1700.51814 | 1601.43196 | 1572.24655 |
| -4.6592407 | 3.17E-06 | 1.22E-05 | 57.3781432 | 115.009581 | 142.656295 | 122.709193 |
| -4.6590841 | 3.18E-06 | 1.22E-05 | 47.951734 | 151.542036 | 167.966283 | 186.585211 |
| -4.6586343 | 3.18E-06 | 1.22E-05 | 67.6242402 | 91.1958326 | 155.311289 | 107.020346 |
| -4.6583595 | 3.19E-06 | 1.23E-05 | 37.2957931 | 240.031761 | 253.099879 | 216.281956 |
| -4.6581705 | 3.19E-06 | 1.23E-05 | 68.0340841 | 199.440144 | 94.3372276 | 119.907613 |
| -4.6579282 | 3.19E-06 | 1.23E-05 | 87.7065903 | 130.434396 | 112.744492 | 121.588561 |
| -4.6579251 | 3.19E-06 | 1.23E-05 | 48.7714217 | 160.472192 | 158.762651 | 161.370993 |
| -4.657702 | 3.20E-06 | 1.23E-05 | 44.6729829 | 186.721438 | 173.718553 | 191.628054 |
| -4.6576628 | 3.20E-06 | 1.23E-05 | 59.4273626 | 132.869893 | 102.390406 | 133.355196 |
| -4.6575435 | 3.20E-06 | 1.23E-05 | 71.3128351 | 91.1958326 | 92.0363196 | 86.2886561 |
| -4.6574368 | 3.20E-06 | 1.23E-05 | 106.149565 | 89.3015571 | 98.9390436 | 80.1251807 |
| -4.6573984 | 3.20E-06 | 1.23E-05 | 59.0175187 | 117.715689 | 110.443584 | 125.510772 |
| -4.6573854 | 3.20E-06 | 1.23E-05 | 56.1486116 | 126.104623 | 120.79767 | 137.837723 |
| -4.6573085 | 3.20E-06 | 1.23E-05 | 53.6895483 | 158.577917 | 139.204933 | 126.071088 |
| -4.6572779 | 3.20E-06 | 1.23E-05 | 80.3294005 | 85.2423954 | 195.577179 | 101.417187 |
| -4.6572455 | 3.20E-06 | 1.23E-05 | 69.2636157 | 98.5023236 | 158.762651 | 94.1330794 |
| -4.657025 | 3.21E-06 | 1.23E-05 | 79.0998688 | 97.6904913 | 72.4786017 | 116.545717 |
| -4.65676 | 3.21E-06 | 1.23E-05 | 65.1651769 | 100.125988 | 96.6381356 | 133.915512 |
| -4.6566549 | 3.21E-06 | 1.23E-05 | 46.7222023 | 149.647761 | 220.887167 | 179.301104 |
| -4.6563895 | 3.22E-06 | 1.24E-05 | 84.4278393 | 112.032863 | 92.0363196 | 93.5727634 |
| 4.65617151 | 3.22E-06 | 1.24E-05 | 8.60672148 | 7.03588026 | 13.8054479 | 12.8872668 |
| -4.6559125 | 3.23E-06 | 1.24E-05 | 72.1325229 | 103.643928 | 82.8326877 | 89.6505518 |
| -4.6554778 | 3.23E-06 | 1.24E-05 | 61.8864259 | 121.233629 | 101.239952 | 108.140978 |
| -4.6552088 | 3.24E-06 | 1.24E-05 | 67.2143963 | 113.385916 | 93.1867736 | 92.4521315 |
| -4.6546879 | 3.24E-06 | 1.24E-05 | 65.1651769 | 86.5954493 | 130.001301 | 133.355196 |
| -4.6542777 | 3.25E-06 | 1.25E-05 | 68.0340841 | 83.34812 | 241.595339 | 187.705843 |
| -4.6539962 | 3.26E-06 | 1.25E-05 | 140.576451 | 162.637078 | 52.9208838 | 122.709193 |
| -4.6538834 | 3.26E-06 | 1.25E-05 | 56.1486116 | 125.02218 | 127.700393 | 123.829825 |
| -4.6533127 | 3.27E-06 | 1.25E-05 | 65.1651769 | 214.594348 | 89.7354116 | 154.086886 |
| -4.653245 | 3.27E-06 | 1.25E-05 | 58.197831 | 129.081342 | 197.878087 | 112.623506 |
| -4.6532056 | 3.27E-06 | 1.25E-05 | 50.4107972 | 140.988216 | 159.913105 | 152.966254 |
| -4.6529833 | 3.27E-06 | 1.25E-05 | 69.2636157 | 194.56915 | 80.5317797 | 160.810677 |
| -4.6526497 | 3.28E-06 | 1.26E-05 | 54.9190799 | 126.916455 | 242.745793 | 141.199619 |
| 4.65238372 | 3.28E-06 | 1.26E-05 | 0.40984388 | 0 | 0 | 0 |
| -4.652303 | 3.28E-06 | 1.26E-05 | 58.6076748 | 110.409198 | 119.647216 | 121.028245 |
| -4.6520645 | 3.29E-06 | 1.26E-05 | 297.546657 | 104.996982 | 90.8858656 | 92.4521315 |

| | | | | | | |
|---|---|---|---|---|---|---|
| 4.65138664 | 3.30E-06 | 1.26E-05 | 34.017042 | 20.5664192 | 20.7081719 | 20.7316901 |
| -4.6513037 | 3.30E-06 | 1.26E-05 | 56.1486116 | 278.458492 | 340.534383 | 367.567262 |
| -4.6507712 | 3.31E-06 | 1.27E-05 | 193.856155 | 111.762252 | 71.3281477 | 89.0902358 |
| -4.6506372 | 3.31E-06 | 1.27E-05 | 50.4107972 | 166.155018 | 128.850847 | 199.472478 |
| 4.65035457 | 3.31E-06 | 1.27E-05 | 0 | 0 | 0 | 0 |
| 4.65033852 | 3.31E-06 | 1.27E-05 | 0 | 0 | 0 | 0 |
| -4.6496719 | 3.32E-06 | 1.27E-05 | 56.9682993 | 167.508072 | 104.691314 | 147.92341 |
| -4.6495292 | 3.33E-06 | 1.27E-05 | 59.4273626 | 256.809629 | 132.302209 | 119.347297 |
| -4.6491925 | 3.33E-06 | 1.28E-05 | 67.6242402 | 111.491641 | 180.621277 | 92.4521315 |
| -4.6490436 | 3.33E-06 | 1.28E-05 | 75.8211178 | 73.0649104 | 105.841768 | 109.26161 |
| -4.6489945 | 3.34E-06 | 1.28E-05 | 61.8864259 | 119.880575 | 94.3372276 | 123.829825 |
| -4.6488314 | 3.34E-06 | 1.28E-05 | 60.6568942 | 138.552719 | 100.089498 | 115.425085 |
| -4.6486553 | 3.34E-06 | 1.28E-05 | 75.8211178 | 122.586683 | 73.6290557 | 102.537819 |
| -4.6485267 | 3.34E-06 | 1.28E-05 | 56.9682993 | 102.290875 | 178.320369 | 162.491625 |
| -4.6473049 | 3.36E-06 | 1.29E-05 | 82.7884637 | 122.857294 | 119.647216 | 66.6775979 |
| -4.6472505 | 3.36E-06 | 1.29E-05 | 89.7558097 | 136.929054 | 62.1245158 | 129.432984 |
| -4.6468408 | 3.37E-06 | 1.29E-05 | 65.1651769 | 116.362635 | 97.7885896 | 92.4521315 |
| -4.646604 | 3.37E-06 | 1.29E-05 | 26.6398522 | 286.306204 | 456.730236 | 313.216615 |
| -4.6465496 | 3.38E-06 | 1.29E-05 | 62.7061136 | 93.3607188 | 124.249032 | 110.382242 |
| -4.6461919 | 3.38E-06 | 1.29E-05 | 69.2636157 | 141.529437 | 92.0363196 | 92.4521315 |
| -4.6458999 | 3.39E-06 | 1.29E-05 | 135.658324 | 87.9485032 | 70.1776937 | 91.3314996 |
| -4.645916 | 3.39E-06 | 1.29E-05 | 84.8376831 | 112.032863 | 111.594038 | 109.26161 |
| -4.6458204 | 3.39E-06 | 1.29E-05 | 67.2143963 | 98.2317128 | 86.2840497 | 103.098135 |
| -4.6452175 | 3.40E-06 | 1.30E-05 | 50.0009534 | 144.776767 | 195.577179 | 140.078987 |
| -4.6447001 | 3.41E-06 | 1.30E-05 | 152.871767 | 274.128719 | 277.259413 | 271.192919 |
| -4.6442469 | 3.41E-06 | 1.30E-05 | 54.9190799 | 126.645845 | 120.79767 | 138.398039 |
| -4.6442427 | 3.41E-06 | 1.30E-05 | 220.086164 | 95.525605 | 83.9831417 | 89.0902358 |
| -4.6439271 | 3.42E-06 | 1.31E-05 | 56.9682993 | 225.960001 | 97.7885896 | 223.566063 |
| -4.6436894 | 3.42E-06 | 1.31E-05 | 56.1486116 | 162.366467 | 108.142676 | 137.277407 |
| -4.643528 | 3.43E-06 | 1.31E-05 | 55.7387677 | 133.952336 | 111.594038 | 139.518671 |
| -4.6434915 | 3.43E-06 | 1.31E-05 | 94.6739363 | 67.6526948 | 94.3372276 | 89.0902358 |
| -4.6434505 | 3.43E-06 | 1.31E-05 | 181.560839 | 67.382084 | 101.239952 | 100.296555 |
| -4.6433646 | 3.43E-06 | 1.31E-05 | 61.8864259 | 92.8194972 | 159.913105 | 127.752036 |
| -4.6433444 | 3.43E-06 | 1.31E-05 | 90.9853413 | 110.138587 | 67.8767857 | 86.2886561 |
| -4.6431507 | 3.43E-06 | 1.31E-05 | 35.2465737 | 309.03751 | 230.090799 | 221.3248 |
| -4.6428557 | 3.44E-06 | 1.31E-05 | 44.263139 | 209.452743 | 157.612197 | 189.386791 |
| -4.6424487 | 3.44E-06 | 1.31E-05 | 66.3947085 | 103.373318 | 89.7354116 | 94.1330794 |
| -4.6424286 | 3.44E-06 | 1.31E-05 | 68.4439279 | 94.1725511 | 85.1335957 | 96.9346591 |
| -4.6423609 | 3.44E-06 | 1.31E-05 | 52.4600166 | 120.963018 | 149.559019 | 158.569413 |
| -4.6422204 | 3.45E-06 | 1.31E-05 | 31.9678226 | 309.578731 | 264.604419 | 249.340597 |
| -4.6421821 | 3.45E-06 | 1.31E-05 | 76.2309617 | 100.66721 | 199.028541 | 88.5299199 |
| -4.6421803 | 3.45E-06 | 1.31E-05 | 88.5262781 | 81.9950661 | 77.0804177 | 79.0045488 |
| -4.6421361 | 3.45E-06 | 1.31E-05 | 140.166607 | 257.621462 | 301.418947 | 304.25156 |

| | | | | | | |
|---|---|---|---|---|---|---|
| -4.6420705 | 3.45E-06 | 1.31E-05 | 91.3951852 | 161.284024 | 144.957203 | 61.0744384 |
| -4.6419899 | 3.45E-06 | 1.32E-05 | 87.2967464 | 86.0542278 | 70.1776937 | 89.6505518 |
| -4.6416469 | 3.46E-06 | 1.32E-05 | 300.825408 | 719.824673 | 674.166041 | 673.49977 |
| -4.6413615 | 3.46E-06 | 1.32E-05 | 62.2962697 | 110.409198 | 94.3372276 | 111.502874 |
| -4.6404435 | 3.48E-06 | 1.32E-05 | 70.4931473 | 279.270324 | 234.692615 | 212.92006 |
| -4.6403911 | 3.48E-06 | 1.32E-05 | 64.755333 | 99.0435452 | 117.346308 | 90.2108677 |
| -4.6396667 | 3.49E-06 | 1.33E-05 | 76.2309617 | 129.351952 | 71.3281477 | 108.701294 |
| -4.6394655 | 3.49E-06 | 1.33E-05 | 40.984388 | 245.173366 | 165.665375 | 221.3248 |
| -4.6393805 | 3.49E-06 | 1.33E-05 | 43.4434513 | 264.386731 | 189.824909 | 158.569413 |
| -4.639337 | 3.50E-06 | 1.33E-05 | 62.7061136 | 120.421797 | 92.0363196 | 108.701294 |
| -4.6392969 | 3.50E-06 | 1.33E-05 | 72.1325229 | 115.009581 | 74.7795097 | 135.036144 |
| -4.6390687 | 3.50E-06 | 1.33E-05 | 60.2470503 | 151.271426 | 102.390406 | 111.502874 |
| -4.6385539 | 3.51E-06 | 1.34E-05 | 2.45906328 | 899.239619 | 2088.074 | 1736.41912 |
| -4.6385111 | 3.51E-06 | 1.34E-05 | 63.1159575 | 171.296623 | 93.1867736 | 121.028245 |
| -4.6384615 | 3.51E-06 | 1.34E-05 | 81.968776 | 71.7118565 | 90.8858656 | 87.9696039 |
| -4.6380825 | 3.52E-06 | 1.34E-05 | 84.8376831 | 118.527521 | 190.975363 | 70.0394936 |
| -4.6379626 | 3.52E-06 | 1.34E-05 | 241.807889 | 434.330301 | 281.861229 | 352.999048 |
| -4.637814 | 3.52E-06 | 1.34E-05 | 51.6403289 | 126.375234 | 144.957203 | 153.52657 |
| 4.6377433 | 3.52E-06 | 1.34E-05 | 40.5745441 | 20.83703 | 25.3099879 | 21.292006 |
| -4.6376304 | 3.52E-06 | 1.34E-05 | 208.200691 | 283.329486 | 337.083021 | 349.637152 |
| -4.637575 | 3.53E-06 | 1.34E-05 | 76.6408055 | 83.8893415 | 81.6822337 | 81.2458125 |
| -4.6374268 | 3.53E-06 | 1.34E-05 | 65.1651769 | 102.832096 | 104.691314 | 87.409288 |
| -4.637179 | 3.53E-06 | 1.34E-05 | 69.6734596 | 123.669126 | 138.054479 | 76.763285 |
| -4.6371911 | 3.53E-06 | 1.34E-05 | 61.0667381 | 93.090108 | 119.647216 | 130.553616 |
| -4.6370617 | 3.53E-06 | 1.34E-05 | 93.0345607 | 74.688575 | 81.6822337 | 147.363094 |
| -4.6362899 | 3.55E-06 | 1.35E-05 | 86.8869025 | 121.50424 | 64.4254237 | 162.491625 |
| -4.6357393 | 3.56E-06 | 1.35E-05 | 41.3942319 | 201.063809 | 179.470823 | 203.394689 |
| -4.6356812 | 3.56E-06 | 1.35E-05 | 99.1822189 | 102.290875 | 116.195854 | 61.0744384 |
| -4.6351604 | 3.57E-06 | 1.35E-05 | 33.1973543 | 279.540935 | 277.259413 | 223.005748 |
| -4.6351009 | 3.57E-06 | 1.35E-05 | 53.2797044 | 177.520671 | 123.098578 | 130.553616 |
| -4.6351111 | 3.57E-06 | 1.35E-05 | 95.493624 | 60.8874253 | 112.744492 | 123.269509 |
| -4.6346702 | 3.58E-06 | 1.36E-05 | 48.7714217 | 123.669126 | 196.727633 | 196.670898 |
| -4.6340719 | 3.59E-06 | 1.36E-05 | 73.7718984 | 73.8767427 | 98.9390436 | 112.623506 |
| -4.6339217 | 3.59E-06 | 1.36E-05 | 66.3947085 | 81.1832337 | 116.195854 | 123.269509 |
| -4.6336088 | 3.59E-06 | 1.36E-05 | 60.6568942 | 97.6904913 | 116.195854 | 107.580662 |
| -4.6335005 | 3.60E-06 | 1.36E-05 | 41.8040758 | 173.461509 | 212.833989 | 191.628054 |
| -4.6333843 | 3.60E-06 | 1.37E-05 | 58.197831 | 131.246228 | 98.9390436 | 131.674248 |
| -4.6332643 | 3.60E-06 | 1.37E-05 | 58.6076748 | 114.197749 | 108.142676 | 110.382242 |
| -4.6332288 | 3.60E-06 | 1.37E-05 | 71.3128351 | 93.6313296 | 88.5849576 | 80.1251807 |
| -4.6331761 | 3.60E-06 | 1.37E-05 | 53.2797044 | 131.787449 | 121.948124 | 140.639303 |
| -4.6329204 | 3.61E-06 | 1.37E-05 | 62.2962697 | 115.821413 | 93.1867736 | 104.218766 |
| -4.6329368 | 3.61E-06 | 1.37E-05 | 54.9190799 | 116.092024 | 126.549939 | 130.553616 |
| -4.6328604 | 3.61E-06 | 1.37E-05 | 65.9848647 | 93.090108 | 88.5849576 | 115.425085 |

| | | | | | | |
|---|---|---|---|---|---|---|
| -4.6327233 | 3.61E-06 | 1.37E-05 | 47.951734 | 289.282923 | 132.302209 | 186.024895 |
| -4.6326497 | 3.61E-06 | 1.37E-05 | 52.0501728 | 140.176384 | 126.549939 | 141.199619 |
| -4.6325929 | 3.61E-06 | 1.37E-05 | 60.6568942 | 98.5023236 | 116.195854 | 105.339398 |
| -4.6324353 | 3.61E-06 | 1.37E-05 | 74.1817423 | 72.253078 | 118.496762 | 105.339398 |
| -4.6321182 | 3.62E-06 | 1.37E-05 | 126.641759 | 94.7137727 | 118.496762 | 59.9538065 |
| -4.6319705 | 3.62E-06 | 1.37E-05 | 68.4439279 | 159.119138 | 77.0804177 | 178.180472 |
| -4.6315043 | 3.63E-06 | 1.38E-05 | 63.1159575 | 110.679809 | 197.878087 | 103.098135 |
| -4.6313407 | 3.63E-06 | 1.38E-05 | 95.9034679 | 100.396599 | 111.594038 | 145.121831 |
| -4.6310289 | 3.64E-06 | 1.38E-05 | 52.0501728 | 211.347019 | 130.001301 | 135.036144 |
| -4.6307003 | 3.64E-06 | 1.38E-05 | 60.6568942 | 115.280192 | 113.894946 | 94.6933953 |
| -4.6305829 | 3.65E-06 | 1.38E-05 | 61.0667381 | 216.759234 | 154.160835 | 187.705843 |
| -4.6303291 | 3.65E-06 | 1.38E-05 | 58.197831 | 135.30539 | 96.6381356 | 145.682147 |
| -4.6300934 | 3.66E-06 | 1.38E-05 | 48.7714217 | 139.635162 | 153.010381 | 157.448782 |
| -4.6298184 | 3.66E-06 | 1.39E-05 | 59.8372065 | 150.730204 | 93.1867736 | 129.9933 |
| -4.6296868 | 3.66E-06 | 1.39E-05 | 138.937075 | 79.2889583 | 66.7263317 | 121.028245 |
| -4.6297102 | 3.66E-06 | 1.39E-05 | 70.0833035 | 109.056144 | 83.9831417 | 85.1680242 |
| -4.6294637 | 3.67E-06 | 1.39E-05 | 53.2797044 | 126.375234 | 158.762651 | 121.028245 |
| -4.6293433 | 3.67E-06 | 1.39E-05 | 71.3128351 | 84.7011739 | 291.064861 | 155.767834 |
| -4.6290536 | 3.67E-06 | 1.39E-05 | 30.3284471 | 248.420695 | 260.002603 | 382.135477 |
| -4.6287035 | 3.68E-06 | 1.39E-05 | 77.8703372 | 88.4897248 | 72.4786017 | 115.425085 |
| -4.6286398 | 3.68E-06 | 1.39E-05 | 163.937552 | 126.375234 | 54.0713378 | 112.623506 |
| -4.6283984 | 3.69E-06 | 1.39E-05 | 43.8532952 | 175.355785 | 192.125817 | 162.491625 |
| -4.6283388 | 3.69E-06 | 1.39E-05 | 66.3947085 | 151.812647 | 79.3813257 | 136.156776 |
| -4.6282678 | 3.69E-06 | 1.39E-05 | 59.0175187 | 103.914539 | 119.647216 | 105.899714 |
| -4.6276912 | 3.70E-06 | 1.40E-05 | 44.263139 | 176.97945 | 164.514921 | 175.378892 |
| -4.627714 | 3.70E-06 | 1.40E-05 | 190.577404 | 544.198277 | 393.455266 | 453.855918 |
| -4.6273697 | 3.70E-06 | 1.40E-05 | 259.431176 | 310.931785 | 277.259413 | 276.235763 |
| -4.6268967 | 3.71E-06 | 1.40E-05 | 84.4278393 | 117.715689 | 120.79767 | 115.425085 |
| -4.6268551 | 3.71E-06 | 1.40E-05 | 66.3947085 | 80.3714014 | 111.594038 | 122.148877 |
| -4.6267172 | 3.72E-06 | 1.40E-05 | 47.951734 | 152.624479 | 164.514921 | 142.320251 |
| -4.6267101 | 3.72E-06 | 1.40E-05 | 44.6729829 | 136.387833 | 211.683535 | 228.608907 |
| -4.6264224 | 3.72E-06 | 1.41E-05 | 36.8859492 | 310.390564 | 355.490285 | 157.448782 |
| -4.625988 | 3.73E-06 | 1.41E-05 | 66.3947085 | 245.443977 | 147.258111 | 154.647202 |
| -4.6257977 | 3.73E-06 | 1.41E-05 | 120.494101 | 184.827162 | 48.3190678 | 222.445432 |
| -4.6257996 | 3.73E-06 | 1.41E-05 | 68.4439279 | 175.626396 | 97.7885896 | 94.1330794 |
| -4.625735 | 3.73E-06 | 1.41E-05 | 51.6403289 | 201.063809 | 117.346308 | 152.966254 |
| -4.6255978 | 3.74E-06 | 1.41E-05 | 108.198784 | 198.357701 | 58.6731538 | 117.106033 |
| -4.6252562 | 3.74E-06 | 1.41E-05 | 45.9025146 | 159.93097 | 151.859927 | 180.982051 |
| -4.6250914 | 3.74E-06 | 1.41E-05 | 64.755333 | 100.125988 | 96.6381356 | 87.9696039 |
| -4.6246107 | 3.75E-06 | 1.42E-05 | 126.641759 | 107.973701 | 60.9740618 | 90.7711837 |
| -4.6245082 | 3.75E-06 | 1.42E-05 | 119.264569 | 80.1007906 | 73.6290557 | 83.4870763 |
| -4.6244481 | 3.76E-06 | 1.42E-05 | 65.5750208 | 91.1958326 | 89.7354116 | 121.028245 |
| -4.6241475 | 3.76E-06 | 1.42E-05 | 68.8537718 | 85.2423954 | 83.9831417 | 100.296555 |

| | | | | | | |
|---|---|---|---|---|---|---|
| 4.62380493 | 3.77E-06 | 1.42E-05 | 144.67489 | 107.432479 | 119.647216 | 89.0902358 |
| -4.6237947 | 3.77E-06 | 1.42E-05 | 55.7387677 | 120.963018 | 171.417645 | 110.942558 |
| -4.6236645 | 3.77E-06 | 1.42E-05 | 107.379097 | 162.907689 | 158.762651 | 140.078987 |
| -4.6235726 | 3.77E-06 | 1.42E-05 | 59.0175187 | 104.996982 | 102.390406 | 128.312352 |
| -4.623466 | 3.77E-06 | 1.42E-05 | 374.597306 | 93.6313296 | 70.1776937 | 158.569413 |
| -4.6233274 | 3.78E-06 | 1.42E-05 | 49.1812656 | 147.212264 | 185.223093 | 130.553616 |
| -4.6228236 | 3.79E-06 | 1.43E-05 | 75.4112739 | 295.506971 | 66.7263317 | 302.570612 |
| -4.6227023 | 3.79E-06 | 1.43E-05 | 41.3942319 | 206.476024 | 163.364467 | 208.437533 |
| -4.6221029 | 3.80E-06 | 1.43E-05 | 60.2470503 | 147.753485 | 88.5849576 | 142.880567 |
| -4.6214875 | 3.81E-06 | 1.44E-05 | 50.8206411 | 120.421797 | 161.063559 | 147.363094 |
| -4.6214088 | 3.81E-06 | 1.44E-05 | 200.003813 | 344.758133 | 308.321671 | 356.360943 |
| -4.6207785 | 3.82E-06 | 1.44E-05 | 56.5584554 | 146.941653 | 111.594038 | 109.821926 |
| -4.6206273 | 3.83E-06 | 1.44E-05 | 45.0828268 | 191.592432 | 157.612197 | 156.32815 |
| -4.6202926 | 3.83E-06 | 1.44E-05 | 43.0336074 | 189.968767 | 165.665375 | 175.939208 |
| -4.6200312 | 3.84E-06 | 1.45E-05 | 29.0989155 | 283.058875 | 287.613499 | 281.838922 |
| -4.6197115 | 3.84E-06 | 1.45E-05 | 99.1822189 | 92.8194972 | 72.4786017 | 73.4013893 |
| -4.6193566 | 3.85E-06 | 1.45E-05 | 47.1320462 | 193.757318 | 127.700393 | 183.783631 |
| -4.6193754 | 3.85E-06 | 1.45E-05 | 82.3786199 | 243.549701 | 101.239952 | 86.848972 |
| -4.6191263 | 3.85E-06 | 1.45E-05 | 99.5920628 | 74.9591858 | 133.452663 | 72.2807574 |
| -4.6190548 | 3.85E-06 | 1.45E-05 | 50.0009534 | 164.260743 | 128.850847 | 138.958355 |
| -4.6182224 | 3.87E-06 | 1.46E-05 | 54.9190799 | 149.647761 | 124.249032 | 108.140978 |
| -4.6180494 | 3.87E-06 | 1.46E-05 | 71.722679 | 89.8427787 | 102.390406 | 75.0823371 |
| 4.61779546 | 3.88E-06 | 1.46E-05 | 25.4103206 | 57.0988744 | 47.1686138 | 35.8602207 |
| -4.6175885 | 3.88E-06 | 1.46E-05 | 100.001907 | 116.362635 | 58.6731538 | 101.417187 |
| -4.6169094 | 3.89E-06 | 1.47E-05 | 46.3123584 | 173.461509 | 146.107657 | 155.767834 |
| -4.6169311 | 3.89E-06 | 1.47E-05 | 62.7061136 | 86.8660601 | 111.594038 | 104.218766 |
| -4.6165214 | 3.90E-06 | 1.47E-05 | 48.7714217 | 118.798132 | 180.621277 | 221.885116 |
| -4.6160682 | 3.91E-06 | 1.47E-05 | 42.2139196 | 166.155018 | 185.223093 | 188.826475 |
| -4.6159584 | 3.91E-06 | 1.47E-05 | 52.4600166 | 135.034779 | 131.151755 | 119.347297 |
| 4.61574552 | 3.92E-06 | 1.47E-05 | 0.81968776 | 2.70610779 | 1.150454 | 0 |
| -4.6155612 | 3.92E-06 | 1.47E-05 | 59.0175187 | 97.1492697 | 117.346308 | 107.580662 |
| -4.6154304 | 3.92E-06 | 1.47E-05 | 56.1486116 | 163.719521 | 103.54086 | 124.390141 |
| -4.6153318 | 3.92E-06 | 1.48E-05 | 77.4604933 | 93.3607188 | 170.267191 | 79.5648647 |
| 4.61517344 | 3.93E-06 | 1.48E-05 | 53.2797044 | 33.2851258 | 31.0622579 | 35.2999048 |
| -4.6148376 | 3.93E-06 | 1.48E-05 | 68.8537718 | 126.645845 | 81.6822337 | 91.3314996 |
| -4.6143875 | 3.94E-06 | 1.48E-05 | 58.197831 | 286.035594 | 89.7354116 | 221.885116 |
| -4.6141094 | 3.95E-06 | 1.48E-05 | 506.157192 | 2147.02592 | 1133.19719 | 1480.35474 |
| -4.6137583 | 3.95E-06 | 1.49E-05 | 70.9029912 | 80.3714014 | 95.4876816 | 81.2458125 |
| -4.6137591 | 3.95E-06 | 1.49E-05 | 54.0993922 | 103.373318 | 144.957203 | 154.647202 |
| -4.6136112 | 3.96E-06 | 1.49E-05 | 275.005243 | 150.188982 | 283.011683 | 38.6618005 |
| -4.6133855 | 3.96E-06 | 1.49E-05 | 77.0506494 | 83.0775092 | 238.143977 | 112.623506 |
| -4.6127795 | 3.97E-06 | 1.49E-05 | 61.8864259 | 181.038611 | 178.320369 | 177.05984 |
| -4.612727 | 3.97E-06 | 1.49E-05 | 142.215826 | 56.2870421 | 134.603117 | 98.055291 |

| | | | | | | |
|---|---|---|---|---|---|---|
| -4.6122693 | 3.98E-06 | 1.50E-05 | 63.9356453 | 90.9252218 | 97.7885896 | 90.2108677 |
| -4.6117141 | 3.99E-06 | 1.50E-05 | 85.247527 | 143.153102 | 269.206235 | 68.9188617 |
| -4.6113178 | 4.00E-06 | 1.50E-05 | 61.0667381 | 112.303473 | 124.249032 | 87.409288 |
| -4.6110345 | 4.01E-06 | 1.50E-05 | 50.0009534 | 121.774851 | 144.957203 | 156.32815 |
| -4.6110032 | 4.01E-06 | 1.50E-05 | 65.1651769 | 90.3840002 | 133.452663 | 90.2108677 |
| -4.6108816 | 4.01E-06 | 1.50E-05 | 224.184602 | 118.527521 | 96.6381356 | 100.296555 |
| -4.6107543 | 4.01E-06 | 1.51E-05 | 151.642236 | 96.8786589 | 64.4254237 | 90.7711837 |
| -4.6104903 | 4.02E-06 | 1.51E-05 | 53.6895483 | 139.09394 | 117.346308 | 115.985401 |
| -4.6099701 | 4.03E-06 | 1.51E-05 | 45.9025146 | 177.25006 | 132.302209 | 189.947107 |
| -4.609882 | 4.03E-06 | 1.51E-05 | 333.203074 | 486.558181 | 461.332052 | 580.487323 |
| -4.6096671 | 4.03E-06 | 1.51E-05 | 47.951734 | 192.133653 | 123.098578 | 166.974153 |
| -4.6096825 | 4.03E-06 | 1.51E-05 | 186.478965 | 116.362635 | 69.0272397 | 80.6854966 |
| -4.6093105 | 4.04E-06 | 1.51E-05 | 144.265046 | 128.810731 | 297.967585 | 52.1093832 |
| -4.6091739 | 4.04E-06 | 1.52E-05 | 202.462877 | 115.550803 | 66.7263317 | 87.409288 |
| -4.6091443 | 4.04E-06 | 1.52E-05 | 60.6568942 | 152.353869 | 87.4345036 | 124.950457 |
| -4.6086789 | 4.05E-06 | 1.52E-05 | 240.578358 | 299.024911 | 247.347609 | 243.737438 |
| -4.6080209 | 4.07E-06 | 1.52E-05 | 304.923847 | 518.760864 | 491.243856 | 495.319299 |
| -4.6078816 | 4.07E-06 | 1.52E-05 | 198.364438 | 58.4519283 | 127.700393 | 99.1759229 |
| -4.6075935 | 4.07E-06 | 1.53E-05 | 55.3289238 | 113.927138 | 109.29313 | 121.028245 |
| -4.6069612 | 4.09E-06 | 1.53E-05 | 53.2797044 | 128.810731 | 113.894946 | 124.390141 |
| -4.6068285 | 4.09E-06 | 1.53E-05 | 68.0340841 | 97.4198805 | 78.2308717 | 90.2108677 |
| -4.6066247 | 4.09E-06 | 1.53E-05 | 41.8040758 | 212.700072 | 257.701695 | 141.759935 |
| -4.6060344 | 4.10E-06 | 1.54E-05 | 59.8372065 | 116.903857 | 88.5849576 | 121.028245 |
| -4.6059586 | 4.11E-06 | 1.54E-05 | 52.0501728 | 123.398515 | 123.098578 | 131.113932 |
| -4.605916 | 4.11E-06 | 1.54E-05 | 137.707544 | 122.316072 | 71.3281477 | 70.0394936 |
| -4.6056196 | 4.11E-06 | 1.54E-05 | 59.0175187 | 92.0076649 | 132.302209 | 179.86142 |
| -4.6048752 | 4.13E-06 | 1.54E-05 | 108.608628 | 117.9863 | 57.5226998 | 95.2537113 |
| -4.6048753 | 4.13E-06 | 1.54E-05 | 103.690502 | 143.694324 | 75.9299637 | 110.942558 |
| -4.6047001 | 4.13E-06 | 1.55E-05 | 104.100346 | 144.235545 | 58.6731538 | 96.9346591 |
| -4.6046461 | 4.13E-06 | 1.55E-05 | 83.1983076 | 80.3714014 | 77.0804177 | 73.9617052 |
| -4.6043512 | 4.14E-06 | 1.55E-05 | 60.6568942 | 98.2317128 | 106.992222 | 92.4521315 |
| -4.60425 | 4.14E-06 | 1.55E-05 | 49.5911095 | 110.138587 | 188.674455 | 192.748686 |
| -4.6039918 | 4.14E-06 | 1.55E-05 | 41.3942319 | 188.074492 | 157.612197 | 205.635953 |
| -4.6039856 | 4.14E-06 | 1.55E-05 | 59.0175187 | 112.844695 | 95.4876816 | 103.65845 |
| -4.6038989 | 4.15E-06 | 1.55E-05 | 65.1651769 | 83.8893415 | 98.9390436 | 137.277407 |
| -4.603659 | 4.15E-06 | 1.55E-05 | 43.4434513 | 164.531354 | 162.214013 | 172.016996 |
| -4.6034788 | 4.15E-06 | 1.55E-05 | 59.4273626 | 102.832096 | 104.691314 | 95.8140272 |
| -4.6032344 | 4.16E-06 | 1.56E-05 | 50.8206411 | 153.436312 | 115.0454 | 141.199619 |
| -4.6030103 | 4.16E-06 | 1.56E-05 | 52.8698605 | 143.153102 | 113.894946 | 121.028245 |
| -4.6029288 | 4.17E-06 | 1.56E-05 | 84.4278393 | 81.1832337 | 136.904025 | 72.2807574 |
| -4.6018465 | 4.19E-06 | 1.56E-05 | 65.9848647 | 76.8534613 | 112.744492 | 130.553616 |
| -4.6016969 | 4.19E-06 | 1.57E-05 | 56.1486116 | 98.5023236 | 133.452663 | 183.783631 |
| -4.6016904 | 4.19E-06 | 1.57E-05 | 83.6081515 | 124.751569 | 82.8326877 | 111.502874 |

| | | | | | | |
|---|---|---|---|---|---|---|
| -4.6016104 | 4.19E-06 | 1.57E-05 | 41.8040758 | 175.355785 | 185.223093 | 165.853521 |
| 4.60150853 | 4.19E-06 | 1.57E-05 | 54.9190799 | 24.8961917 | 23.0090799 | 21.292006 |
| -4.6012523 | 4.20E-06 | 1.57E-05 | 47.1320462 | 158.577917 | 138.054479 | 145.121831 |
| -4.6005969 | 4.21E-06 | 1.57E-05 | 117.21535 | 230.560384 | 325.578481 | 45.9459078 |
| -4.5999489 | 4.23E-06 | 1.58E-05 | 40.984388 | 250.856192 | 170.267191 | 166.413837 |
| -4.5999309 | 4.23E-06 | 1.58E-05 | 63.9356453 | 129.351952 | 86.2840497 | 94.6933953 |
| -4.5996391 | 4.23E-06 | 1.58E-05 | 57.7879871 | 110.409198 | 106.992222 | 98.055291 |
| -4.5994901 | 4.24E-06 | 1.58E-05 | 53.6895483 | 108.244312 | 123.098578 | 126.631404 |
| -4.5993789 | 4.24E-06 | 1.58E-05 | 56.1486116 | 116.903857 | 104.691314 | 108.140978 |
| -4.5993886 | 4.24E-06 | 1.58E-05 | 63.1159575 | 93.3607188 | 87.4345036 | 95.2537113 |
| 4.5994212 | 4.24E-06 | 1.58E-05 | 33.6071982 | 22.7313054 | 27.6108959 | 28.0157974 |
| -4.5992066 | 4.24E-06 | 1.58E-05 | 68.0340841 | 150.730204 | 117.346308 | 75.0823371 |
| -4.5992083 | 4.24E-06 | 1.58E-05 | 114.346443 | 70.0881918 | 144.957203 | 72.2807574 |
| -4.599121 | 4.24E-06 | 1.58E-05 | 53.6895483 | 133.681725 | 103.54086 | 135.036144 |
| -4.598442 | 4.26E-06 | 1.59E-05 | 79.9195566 | 123.398515 | 85.1335957 | 70.0394936 |
| -4.5983149 | 4.26E-06 | 1.59E-05 | 77.8703372 | 167.778683 | 81.6822337 | 85.7283401 |
| -4.5982956 | 4.26E-06 | 1.59E-05 | 38.5253247 | 166.425629 | 337.083021 | 201.713742 |
| -4.5981691 | 4.26E-06 | 1.59E-05 | 143.035514 | 87.6778924 | 105.841768 | 112.623506 |
| -4.5976577 | 4.27E-06 | 1.59E-05 | 64.3454891 | 80.3714014 | 102.390406 | 98.055291 |
| -4.5974797 | 4.28E-06 | 1.59E-05 | 49.5911095 | 138.82333 | 132.302209 | 128.872668 |
| -4.5967156 | 4.29E-06 | 1.60E-05 | 63.5258014 | 95.7962158 | 90.8858656 | 86.2886561 |
| -4.5965126 | 4.30E-06 | 1.60E-05 | 52.8698605 | 206.476024 | 111.594038 | 133.355196 |
| -4.5961086 | 4.30E-06 | 1.60E-05 | 61.476582 | 88.219114 | 95.4876816 | 107.020346 |
| -4.5959886 | 4.31E-06 | 1.60E-05 | 57.3781432 | 114.46836 | 97.7885896 | 106.46003 |
| -4.5959233 | 4.31E-06 | 1.60E-05 | 55.3289238 | 124.210348 | 190.975363 | 104.218766 |
| -4.5954912 | 4.32E-06 | 1.61E-05 | 159.019425 | 64.946587 | 510.801574 | 109.821926 |
| -4.5954547 | 4.32E-06 | 1.61E-05 | 57.3781432 | 152.624479 | 94.3372276 | 118.226665 |
| -4.5953352 | 4.32E-06 | 1.61E-05 | 42.6237635 | 163.719521 | 154.160835 | 203.394689 |
| -4.5953343 | 4.32E-06 | 1.61E-05 | 52.4600166 | 150.730204 | 120.79767 | 112.623506 |
| -4.5949467 | 4.33E-06 | 1.61E-05 | 51.230485 | 152.353869 | 154.160835 | 107.020346 |
| -4.5945145 | 4.34E-06 | 1.61E-05 | 677.471934 | 135.576 | 70.1776937 | 131.674248 |
| -4.5945317 | 4.34E-06 | 1.61E-05 | 132.789417 | 93.3607188 | 67.8767857 | 78.4442328 |
| -4.5942676 | 4.34E-06 | 1.62E-05 | 63.9356453 | 124.480958 | 82.8326877 | 96.3743432 |
| -4.594199 | 4.34E-06 | 1.62E-05 | 57.7879871 | 132.869893 | 110.443584 | 93.5727634 |
| -4.5941762 | 4.34E-06 | 1.62E-05 | 93.4444046 | 122.045461 | 66.7263317 | 77.8839169 |
| -4.5942235 | 4.34E-06 | 1.62E-05 | 60.2470503 | 147.482875 | 92.0363196 | 104.218766 |
| -4.5937422 | 4.35E-06 | 1.62E-05 | 41.3942319 | 197.004647 | 176.019461 | 158.569413 |
| -4.5936818 | 4.35E-06 | 1.62E-05 | 61.476582 | 89.8427787 | 94.3372276 | 123.829825 |
| -4.5936155 | 4.36E-06 | 1.62E-05 | 62.2962697 | 171.026012 | 83.9831417 | 122.709193 |
| -4.5931912 | 4.37E-06 | 1.62E-05 | 52.8698605 | 124.480958 | 124.249032 | 110.382242 |
| -4.5931254 | 4.37E-06 | 1.62E-05 | 181.150995 | 96.3374374 | 96.6381356 | 64.4363341 |
| -4.5922616 | 4.38E-06 | 1.63E-05 | 105.329877 | 174.543953 | 50.6199758 | 151.285306 |
| -4.5922083 | 4.39E-06 | 1.63E-05 | 63.1159575 | 86.3248385 | 124.249032 | 91.8918156 |

| | | | | | | |
|---|---|---|---|---|---|---|
| -4.5921059 | 4.39E-06 | 1.63E-05 | 78.6900249 | 151.542036 | 65.5758777 | 228.048591 |
| 4.59194586 | 4.39E-06 | 1.63E-05 | 0.81968776 | 0.27061078 | 2.30090799 | 0.56031595 |
| -4.5919408 | 4.39E-06 | 1.63E-05 | 56.1486116 | 116.903857 | 97.7885896 | 114.864769 |
| -4.5919082 | 4.39E-06 | 1.63E-05 | 112.297223 | 67.382084 | 75.9299637 | 93.0124475 |
| -4.5915548 | 4.40E-06 | 1.63E-05 | 68.0340841 | 127.187066 | 77.0804177 | 93.5727634 |
| -4.5914952 | 4.40E-06 | 1.63E-05 | 64.755333 | 83.8893415 | 96.6381356 | 85.7283401 |
| -4.5914308 | 4.40E-06 | 1.64E-05 | 50.8206411 | 174.002731 | 312.923487 | 257.18502 |
| -4.5909954 | 4.41E-06 | 1.64E-05 | 77.4604933 | 110.679809 | 65.5758777 | 149.604358 |
| -4.5908108 | 4.42E-06 | 1.64E-05 | 58.197831 | 97.1492697 | 98.9390436 | 109.821926 |
| -4.5905049 | 4.42E-06 | 1.64E-05 | 77.4604933 | 71.7118565 | 79.3813257 | 83.4870763 |
| 4.59048494 | 4.42E-06 | 1.64E-05 | 61.8864259 | 71.9824673 | 62.1245158 | 53.7903311 |
| -4.590451 | 4.42E-06 | 1.64E-05 | 1409.86295 | 4436.93433 | 4306.1493 | 4825.44095 |
| -4.5899271 | 4.43E-06 | 1.65E-05 | 60.2470503 | 84.7011739 | 153.010381 | 121.588561 |
| -4.5897355 | 4.44E-06 | 1.65E-05 | 66.3947085 | 78.7477367 | 103.54086 | 86.848972 |
| -4.5895656 | 4.44E-06 | 1.65E-05 | 45.0828268 | 161.013414 | 164.514921 | 137.277407 |
| -4.5892365 | 4.45E-06 | 1.65E-05 | 52.8698605 | 110.409198 | 125.399485 | 118.786981 |
| -4.5891115 | 4.45E-06 | 1.65E-05 | 66.3947085 | 79.0183475 | 104.691314 | 86.2886561 |
| 4.58888644 | 4.46E-06 | 1.65E-05 | 0 | 0 | 0 | 0 |
| 4.58888644 | 4.46E-06 | 1.65E-05 | 0 | 0 | 0 | 0 |
| -4.5885672 | 4.46E-06 | 1.66E-05 | 64.755333 | 226.771833 | 192.125817 | 188.826475 |
| -4.5883273 | 4.47E-06 | 1.66E-05 | 47.5418901 | 142.070659 | 130.001301 | 149.044042 |
| -4.5882458 | 4.47E-06 | 1.66E-05 | 45.4926707 | 150.459593 | 143.806749 | 152.966254 |
| -4.588224 | 4.47E-06 | 1.66E-05 | 132.379573 | 58.4519283 | 87.4345036 | 119.907613 |
| -4.5879778 | 4.48E-06 | 1.66E-05 | 38.9351686 | 311.202396 | 314.073941 | 284.640502 |
| 4.5876435 | 4.48E-06 | 1.66E-05 | 20.9020379 | 24.8961917 | 24.1595339 | 25.7745336 |
| -4.5875733 | 4.48E-06 | 1.66E-05 | 56.1486116 | 117.174467 | 110.443584 | 97.4949751 |
| 4.58746468 | 4.49E-06 | 1.66E-05 | 0 | 0 | 0 | 0 |
| -4.5874286 | 4.49E-06 | 1.66E-05 | 60.6568942 | 176.97945 | 118.496762 | 88.5299199 |
| 4.58692616 | 4.50E-06 | 1.67E-05 | 4.50828268 | 14.3423713 | 3.45136199 | 5.04284354 |
| -4.5869028 | 4.50E-06 | 1.67E-05 | 63.5258014 | 91.7370541 | 81.6822337 | 103.098135 |
| -4.586698 | 4.50E-06 | 1.67E-05 | 64.755333 | 107.161869 | 77.0804177 | 98.055291 |
| -4.5866484 | 4.50E-06 | 1.67E-05 | 63.1159575 | 100.396599 | 82.8326877 | 91.8918156 |
| -4.5862017 | 4.51E-06 | 1.67E-05 | 86.0672148 | 152.353869 | 66.7263317 | 90.2108677 |
| -4.5861349 | 4.52E-06 | 1.67E-05 | 100.821594 | 92.2782757 | 62.1245158 | 82.9267604 |
| -4.5857535 | 4.52E-06 | 1.67E-05 | 120.903945 | 103.643928 | 71.3281477 | 68.3585457 |
| -4.5854539 | 4.53E-06 | 1.68E-05 | 46.3123584 | 156.683641 | 132.302209 | 149.044042 |
| -4.5852791 | 4.53E-06 | 1.68E-05 | 99.5920628 | 229.20733 | 113.894946 | 155.207518 |
| -4.5852213 | 4.54E-06 | 1.68E-05 | 59.8372065 | 119.339354 | 92.0363196 | 94.1330794 |
| -4.5850816 | 4.54E-06 | 1.68E-05 | 84.4278393 | 361.806612 | 211.683535 | 278.477026 |
| -4.5848231 | 4.54E-06 | 1.68E-05 | 53.6895483 | 124.210348 | 112.744492 | 107.020346 |
| -4.5847694 | 4.54E-06 | 1.68E-05 | 47.951734 | 119.609964 | 165.665375 | 145.682147 |
| -4.5847881 | 4.54E-06 | 1.68E-05 | 59.8372065 | 98.2317128 | 118.496762 | 219.083536 |
| -4.5845677 | 4.55E-06 | 1.68E-05 | 57.7879871 | 112.844695 | 121.948124 | 89.0902358 |

| | | | | | | |
|---|---|---|---|---|---|---|
| -4.5845181 | 4.55E-06 | 1.68E-05 | 107.78894 | 107.432479 | 59.8236078 | 83.4870763 |
| -4.58448 | 4.55E-06 | 1.68E-05 | 52.4600166 | 132.05806 | 116.195854 | 110.942558 |
| -4.5844267 | 4.55E-06 | 1.68E-05 | 66.8045524 | 150.459593 | 71.3281477 | 161.931309 |
| -4.5837425 | 4.57E-06 | 1.69E-05 | 48.7714217 | 145.047378 | 130.001301 | 125.510772 |
| -4.5837527 | 4.57E-06 | 1.69E-05 | 64.755333 | 106.079425 | 95.4876816 | 77.3236009 |
| -4.583648 | 4.57E-06 | 1.69E-05 | 52.0501728 | 148.565318 | 120.79767 | 109.26161 |
| -4.58291 | 4.59E-06 | 1.69E-05 | 61.0667381 | 82.5362876 | 105.841768 | 104.779082 |
| -4.5828008 | 4.59E-06 | 1.70E-05 | 45.9025146 | 166.155018 | 151.859927 | 129.9933 |
| -4.5827539 | 4.59E-06 | 1.70E-05 | 67.2143963 | 70.3588026 | 224.338529 | 221.885116 |
| -4.5826199 | 4.59E-06 | 1.70E-05 | 59.0175187 | 135.30539 | 86.2840497 | 112.623506 |
| 4.58212067 | 4.60E-06 | 1.70E-05 | 6.55750208 | 1.62366467 | 0 | 3.36189569 |
| -4.5820625 | 4.60E-06 | 1.70E-05 | 50.4107972 | 141.529437 | 121.948124 | 118.226665 |
| 4.58177086 | 4.61E-06 | 1.70E-05 | 14.7543797 | 42.4858923 | 31.0622579 | 16.8094785 |
| -4.5803209 | 4.64E-06 | 1.71E-05 | 48.3615778 | 122.857294 | 144.957203 | 135.036144 |
| -4.5802384 | 4.64E-06 | 1.72E-05 | 46.3123584 | 187.53327 | 125.399485 | 149.604358 |
| -4.5799158 | 4.65E-06 | 1.72E-05 | 61.8864259 | 142.070659 | 105.841768 | 82.9267604 |
| -4.579847 | 4.65E-06 | 1.72E-05 | 106.149565 | 108.785533 | 85.1335957 | 59.3934906 |
| -4.5797992 | 4.65E-06 | 1.72E-05 | 71.722679 | 107.161869 | 75.9299637 | 77.8839169 |
| -4.5797984 | 4.65E-06 | 1.72E-05 | 100.001907 | 95.525605 | 60.9740618 | 82.9267604 |
| -4.5795479 | 4.66E-06 | 1.72E-05 | 63.1159575 | 108.244312 | 89.7354116 | 82.3664444 |
| -4.5795179 | 4.66E-06 | 1.72E-05 | 150.00286 | 180.226779 | 182.922185 | 197.231214 |
| -4.579391 | 4.66E-06 | 1.72E-05 | 101.641282 | 150.188982 | 50.6199758 | 152.405938 |
| -4.5793982 | 4.66E-06 | 1.72E-05 | 83.6081515 | 66.5702517 | 97.7885896 | 77.8839169 |
| -4.5789437 | 4.67E-06 | 1.72E-05 | 47.1320462 | 117.174467 | 158.762651 | 170.896364 |
| -4.5789594 | 4.67E-06 | 1.72E-05 | 62.2962697 | 144.506156 | 85.1335957 | 99.7362389 |
| -4.578906 | 4.67E-06 | 1.72E-05 | 838.950422 | 1439.37873 | 1180.3658 | 1366.05028 |
| -4.5787364 | 4.68E-06 | 1.73E-05 | 57.3781432 | 96.8786589 | 98.9390436 | 105.339398 |
| -4.5784586 | 4.68E-06 | 1.73E-05 | 53.6895483 | 136.117222 | 105.841768 | 109.26161 |
| -4.5783087 | 4.69E-06 | 1.73E-05 | 64.3454891 | 92.8194972 | 81.6822337 | 85.7283401 |
| -4.5782828 | 4.69E-06 | 1.73E-05 | 44.6729829 | 142.611881 | 141.505841 | 183.223315 |
| 4.57798328 | 4.69E-06 | 1.73E-05 | 0 | 0 | 0 | 0 |
| -4.5778323 | 4.70E-06 | 1.73E-05 | 41.8040758 | 184.827162 | 155.311289 | 160.810677 |
| -4.5777988 | 4.70E-06 | 1.73E-05 | 45.0828268 | 146.671042 | 135.753571 | 170.336048 |
| -4.5776342 | 4.70E-06 | 1.73E-05 | 118.444881 | 107.432479 | 83.9831417 | 82.9267604 |
| -4.5774121 | 4.71E-06 | 1.73E-05 | 213.938505 | 215.676791 | 446.37615 | 31.3776931 |
| -4.5770497 | 4.72E-06 | 1.74E-05 | 117.21535 | 192.945486 | 199.028541 | 194.98995 |
| -4.5766776 | 4.72E-06 | 1.74E-05 | 52.8698605 | 124.210348 | 105.841768 | 115.985401 |
| -4.5764413 | 4.73E-06 | 1.74E-05 | 88.1164342 | 79.5595691 | 65.5758777 | 82.9267604 |
| -4.5760607 | 4.74E-06 | 1.74E-05 | 46.7222023 | 149.37715 | 128.850847 | 141.759935 |
| -4.5759164 | 4.74E-06 | 1.75E-05 | 48.7714217 | 178.873725 | 165.665375 | 107.020346 |
| -4.5758298 | 4.74E-06 | 1.75E-05 | 66.8045524 | 118.25691 | 79.3813257 | 84.6077082 |
| -4.5754936 | 4.75E-06 | 1.75E-05 | 53.2797044 | 187.262659 | 96.6381356 | 144.001199 |
| -4.5752781 | 4.76E-06 | 1.75E-05 | 57.7879871 | 90.1133895 | 102.390406 | 119.347297 |

| | | | | | | |
|---|---|---|---|---|---|---|
| -4.5752336 | 4.76E-06 | 1.75E-05 | 54.0993922 | 157.495473 | 94.3372276 | 132.79488 |
| -4.5751689 | 4.76E-06 | 1.75E-05 | 56.5584554 | 143.423713 | 86.2840497 | 158.569413 |
| -4.5749703 | 4.76E-06 | 1.75E-05 | 370.089024 | 76.5828505 | 220.887167 | 64.4363341 |
| -4.5749028 | 4.76E-06 | 1.75E-05 | 44.6729829 | 129.893174 | 185.223093 | 155.767834 |
| -4.574705 | 4.77E-06 | 1.75E-05 | 43.8532952 | 197.275258 | 127.700393 | 174.25826 |
| -4.5745683 | 4.77E-06 | 1.76E-05 | 103.690502 | 558.540648 | 388.85345 | 435.925808 |
| -4.5744351 | 4.78E-06 | 1.76E-05 | 53.6895483 | 144.235545 | 115.0454 | 100.856871 |
| -4.5744088 | 4.78E-06 | 1.76E-05 | 75.8211178 | 106.350036 | 78.2308717 | 70.5998095 |
| -4.5740386 | 4.78E-06 | 1.76E-05 | 65.1651769 | 88.219114 | 83.9831417 | 80.6854966 |
| -4.5734419 | 4.80E-06 | 1.76E-05 | 581.158622 | 70.0881918 | 127.700393 | 122.148877 |
| -4.5734134 | 4.80E-06 | 1.76E-05 | 985.264687 | 2701.2368 | 2326.21798 | 2609.39137 |
| -4.5733847 | 4.80E-06 | 1.76E-05 | 77.4604933 | 135.846611 | 131.151755 | 123.269509 |
| -4.5727389 | 4.81E-06 | 1.77E-05 | 86.4770587 | 82.8068984 | 62.1245158 | 95.8140272 |
| -4.5722091 | 4.83E-06 | 1.77E-05 | 63.9356453 | 131.516839 | 73.6290557 | 119.907613 |
| -4.5721445 | 4.83E-06 | 1.77E-05 | 98.7723751 | 90.1133895 | 113.894946 | 57.7125427 |
| -4.5720788 | 4.83E-06 | 1.77E-05 | 48.7714217 | 138.011497 | 124.249032 | 124.390141 |
| -4.5717936 | 4.84E-06 | 1.78E-05 | 65.1651769 | 113.927138 | 89.7354116 | 77.3236009 |
| -4.5717648 | 4.84E-06 | 1.78E-05 | 40.984388 | 167.778683 | 177.169915 | 157.448782 |
| -4.5711664 | 4.85E-06 | 1.78E-05 | 78.2801811 | 100.937821 | 92.0363196 | 104.779082 |
| -4.5700506 | 4.88E-06 | 1.79E-05 | 67.6242402 | 107.432479 | 71.3281477 | 92.4521315 |
| -4.5694447 | 4.89E-06 | 1.80E-05 | 179.511619 | 553.399043 | 409.561622 | 438.167072 |
| -4.56933 | 4.89E-06 | 1.80E-05 | 45.4926707 | 143.423713 | 130.001301 | 163.612257 |
| -4.5692674 | 4.89E-06 | 1.80E-05 | 938.542485 | 2481.50084 | 2505.6888 | 2489.48376 |
| -4.5689201 | 4.90E-06 | 1.80E-05 | 45.9025146 | 128.269509 | 154.160835 | 143.440883 |
| -4.5674561 | 4.94E-06 | 1.81E-05 | 41.3942319 | 158.577917 | 169.116737 | 160.250361 |
| -4.5674776 | 4.94E-06 | 1.81E-05 | 58.197831 | 81.1832337 | 147.258111 | 149.604358 |
| -4.5671317 | 4.94E-06 | 1.81E-05 | 43.0336074 | 132.328671 | 164.514921 | 202.274057 |
| -4.5665642 | 4.96E-06 | 1.82E-05 | 43.0336074 | 135.034779 | 258.852149 | 163.612257 |
| -4.5660308 | 4.97E-06 | 1.82E-05 | 70.4931473 | 95.7962158 | 66.7263317 | 101.417187 |
| -4.5660303 | 4.97E-06 | 1.82E-05 | 57.7879871 | 106.620647 | 88.5849576 | 100.296555 |
| -4.5660423 | 4.97E-06 | 1.82E-05 | 50.4107972 | 133.952336 | 120.79767 | 110.942558 |
| -4.5657093 | 4.98E-06 | 1.83E-05 | 54.509236 | 95.525605 | 260.002603 | 156.888466 |
| -4.5654377 | 4.98E-06 | 1.83E-05 | 46.7222023 | 112.574084 | 195.577179 | 160.810677 |
| -4.5654006 | 4.99E-06 | 1.83E-05 | 80.7392443 | 128.269509 | 58.6731538 | 114.304454 |
| -4.5653892 | 4.99E-06 | 1.83E-05 | 60.6568942 | 119.609964 | 79.3813257 | 104.779082 |
| -4.5652985 | 4.99E-06 | 1.83E-05 | 34.8367298 | 194.839761 | 204.780811 | 211.799429 |
| -4.5646885 | 5.00E-06 | 1.83E-05 | 151.642236 | 205.934803 | 36.8145279 | 276.796079 |
| -4.5646164 | 5.00E-06 | 1.83E-05 | 1277.89322 | 5922.85812 | 5695.89773 | 6374.71455 |
| -4.5645085 | 5.01E-06 | 1.83E-05 | 49.5911095 | 106.891258 | 134.603117 | 154.647202 |
| -4.5645295 | 5.01E-06 | 1.83E-05 | 35.2465737 | 230.560384 | 199.028541 | 182.662999 |
| -4.5632011 | 5.04E-06 | 1.85E-05 | 218.856632 | 75.2297966 | 96.6381356 | 81.8061285 |
| -4.5631117 | 5.04E-06 | 1.85E-05 | 106.559409 | 66.2996409 | 81.6822337 | 77.3236009 |
| 4.56280768 | 5.05E-06 | 1.85E-05 | 63.9356453 | 47.8981079 | 37.9649818 | 45.3855918 |

| | | | | | | |
|---|---|---|---|---|---|---|
| -4.5626383 | 5.05E-06 | 1.85E-05 | 56.5584554 | 117.715689 | 90.8858656 | 101.417187 |
| -4.5623314 | 5.06E-06 | 1.85E-05 | 65.5750208 | 138.552719 | 72.4786017 | 107.580662 |
| -4.5620243 | 5.07E-06 | 1.86E-05 | 52.4600166 | 117.715689 | 113.894946 | 103.65845 |
| -4.5620302 | 5.07E-06 | 1.86E-05 | 59.0175187 | 138.552719 | 86.2840497 | 101.417187 |
| 4.56156152 | 5.08E-06 | 1.86E-05 | 120.494101 | 58.9931499 | 88.5849576 | 83.4870763 |
| -4.5613346 | 5.08E-06 | 1.86E-05 | 65.1651769 | 103.914539 | 73.6290557 | 90.2108677 |
| 4.56127909 | 5.08E-06 | 1.86E-05 | 12.7051603 | 13.8011497 | 11.50454 | 20.7316901 |
| -4.5609556 | 5.09E-06 | 1.86E-05 | 47.1320462 | 238.678707 | 104.691314 | 192.18837 |
| -4.5609263 | 5.09E-06 | 1.86E-05 | 90.1656536 | 91.4664434 | 57.5226998 | 115.425085 |
| -4.5605555 | 5.10E-06 | 1.87E-05 | 39.7548564 | 199.981366 | 140.355387 | 202.834373 |
| -4.5605231 | 5.10E-06 | 1.87E-05 | 50.8206411 | 106.891258 | 157.612197 | 118.786981 |
| 4.55953284 | 5.13E-06 | 1.88E-05 | 75.00143 | 58.7225391 | 72.4786017 | 41.4633802 |
| -4.5592926 | 5.13E-06 | 1.88E-05 | 42.2139196 | 164.531354 | 163.364467 | 141.759935 |
| -4.5591285 | 5.14E-06 | 1.88E-05 | 53.6895483 | 156.683641 | 89.7354116 | 141.199619 |
| -4.5588861 | 5.14E-06 | 1.88E-05 | 48.3615778 | 119.609964 | 139.204933 | 122.148877 |
| -4.5588685 | 5.14E-06 | 1.88E-05 | 89.7558097 | 136.658443 | 96.6381356 | 58.2728587 |
| -4.5586272 | 5.15E-06 | 1.88E-05 | 63.5258014 | 75.2297966 | 170.267191 | 123.269509 |
| -4.5585847 | 5.15E-06 | 1.88E-05 | 76.2309617 | 80.3714014 | 72.4786017 | 72.2807574 |
| -4.5581758 | 5.16E-06 | 1.89E-05 | 72.5423667 | 168.861126 | 62.1245158 | 175.939208 |
| -4.5581358 | 5.16E-06 | 1.89E-05 | 64.755333 | 254.103522 | 224.338529 | 438.727388 |
| -4.5579258 | 5.17E-06 | 1.89E-05 | 59.0175187 | 145.047378 | 162.214013 | 156.32815 |
| -4.5578535 | 5.17E-06 | 1.89E-05 | 48.7714217 | 139.09394 | 130.001301 | 111.502874 |
| -4.5576167 | 5.17E-06 | 1.89E-05 | 92.214873 | 101.208431 | 56.3722458 | 94.1330794 |
| -4.5574009 | 5.18E-06 | 1.89E-05 | 47.5418901 | 158.577917 | 119.647216 | 126.631404 |
| -4.5573162 | 5.18E-06 | 1.89E-05 | 52.0501728 | 121.50424 | 105.841768 | 109.26161 |
| -4.5566787 | 5.20E-06 | 1.90E-05 | 42.6237635 | 160.201581 | 148.408565 | 147.363094 |
| -4.5565795 | 5.20E-06 | 1.90E-05 | 54.509236 | 100.66721 | 101.239952 | 105.899714 |
| -4.5564247 | 5.20E-06 | 1.90E-05 | 65.9848647 | 102.290875 | 155.311289 | 75.0823371 |
| -4.5562722 | 5.21E-06 | 1.90E-05 | 77.4604933 | 127.998899 | 209.382627 | 170.896364 |
| -4.5550619 | 5.24E-06 | 1.91E-05 | 47.951734 | 135.034779 | 123.098578 | 121.588561 |
| 4.55498401 | 5.24E-06 | 1.91E-05 | 59.0175187 | 24.6255809 | 29.9118039 | 25.2142177 |
| -4.5548758 | 5.24E-06 | 1.91E-05 | 45.0828268 | 120.963018 | 148.408565 | 178.180472 |
| -4.5545462 | 5.25E-06 | 1.92E-05 | 44.263139 | 127.457677 | 161.063559 | 150.164674 |
| 4.554337 | 5.26E-06 | 1.92E-05 | 4.50828268 | 2.16488623 | 3.45136199 | 3.36189569 |
| -4.5541834 | 5.26E-06 | 1.92E-05 | 110.248004 | 81.4538445 | 174.869007 | 149.044042 |
| 4.5541153 | 5.26E-06 | 1.92E-05 | 0.40984388 | 3.24732935 | 0 | 0.56031595 |
| -4.5539816 | 5.26E-06 | 1.92E-05 | 58.6076748 | 99.8553775 | 90.8858656 | 87.409288 |
| -4.5538061 | 5.27E-06 | 1.92E-05 | 55.7387677 | 85.5130062 | 120.79767 | 123.269509 |
| -4.5537648 | 5.27E-06 | 1.92E-05 | 566.814086 | 77.1240721 | 115.0454 | 113.183822 |
| -4.5536677 | 5.27E-06 | 1.92E-05 | 54.9190799 | 105.267593 | 97.7885896 | 99.7362389 |
| -4.5536291 | 5.27E-06 | 1.92E-05 | 75.4112739 | 89.3015571 | 93.1867736 | 63.3157022 |
| -4.5534387 | 5.28E-06 | 1.92E-05 | 44.6729829 | 204.85236 | 119.647216 | 156.888466 |
| -4.5532711 | 5.28E-06 | 1.93E-05 | 36.4761053 | 246.52642 | 202.479903 | 155.207518 |

| | | | | | | |
|---|---|---|---|---|---|---|
| -4.5529815 | 5.29E-06 | 1.93E-05 | 45.4926707 | 130.163785 | 148.408565 | 132.79488 |
| -4.5525703 | 5.30E-06 | 1.93E-05 | 63.5258014 | 87.6778924 | 83.9831417 | 78.4442328 |
| -4.5518456 | 5.32E-06 | 1.94E-05 | 55.3289238 | 102.832096 | 94.3372276 | 101.417187 |
| -4.5512071 | 5.33E-06 | 1.94E-05 | 124.592539 | 184.285941 | 90.8858656 | 132.79488 |
| -4.5512249 | 5.33E-06 | 1.94E-05 | 55.7387677 | 113.927138 | 92.0363196 | 98.055291 |
| -4.5510598 | 5.34E-06 | 1.94E-05 | 95.0837801 | 84.4305631 | 59.8236078 | 84.6077082 |
| -4.5510421 | 5.34E-06 | 1.94E-05 | 40.1647002 | 147.212264 | 177.169915 | 171.45668 |
| -4.5508278 | 5.34E-06 | 1.95E-05 | 70.9029912 | 71.7118565 | 108.142676 | 77.3236009 |
| -4.5507826 | 5.34E-06 | 1.95E-05 | 104.920033 | 413.22266 | 334.782113 | 389.419584 |
| -4.5506405 | 5.35E-06 | 1.95E-05 | 295.907281 | 349.899737 | 307.171217 | 316.578511 |
| -4.5501154 | 5.36E-06 | 1.95E-05 | 27.45954 | 306.060791 | 343.985745 | 194.429634 |
| -4.5495971 | 5.37E-06 | 1.96E-05 | 47.1320462 | 113.385916 | 170.267191 | 133.915512 |
| -4.5495001 | 5.38E-06 | 1.96E-05 | 40.5745441 | 188.345102 | 151.859927 | 152.405938 |
| -4.5483984 | 5.41E-06 | 1.97E-05 | 49.5911095 | 181.309222 | 101.239952 | 139.518671 |
| 4.54833713 | 5.41E-06 | 1.97E-05 | 4.0984388 | 7.57710182 | 9.20363196 | 6.72379138 |
| -4.5481526 | 5.41E-06 | 1.97E-05 | 53.2797044 | 119.339354 | 92.0363196 | 146.802779 |
| -4.5480074 | 5.42E-06 | 1.97E-05 | 381.564652 | 824.280433 | 776.556447 | 786.123276 |
| -4.5479279 | 5.42E-06 | 1.97E-05 | 56.5584554 | 176.167617 | 83.9831417 | 131.113932 |
| -4.5477949 | 5.42E-06 | 1.97E-05 | 75.4112739 | 78.7477367 | 77.0804177 | 68.3585457 |
| -4.5473625 | 5.43E-06 | 1.98E-05 | 54.509236 | 116.903857 | 90.8858656 | 107.020346 |
| -4.5472609 | 5.43E-06 | 1.98E-05 | 59.0175187 | 103.643928 | 81.6822337 | 94.1330794 |
| -4.5468459 | 5.45E-06 | 1.98E-05 | 41.3942319 | 156.41303 | 149.559019 | 156.888466 |
| -4.5468039 | 5.45E-06 | 1.98E-05 | 54.9190799 | 139.635162 | 85.1335957 | 174.25826 |
| -4.5466927 | 5.45E-06 | 1.98E-05 | 56.5584554 | 102.561485 | 101.239952 | 86.848972 |
| -4.5462607 | 5.46E-06 | 1.98E-05 | 56.9682993 | 329.062707 | 209.382627 | 253.262809 |
| -4.5462516 | 5.46E-06 | 1.98E-05 | 60.6568942 | 79.0183475 | 93.1867736 | 91.8918156 |
| -4.5461838 | 5.46E-06 | 1.99E-05 | 51.6403289 | 115.280192 | 109.29313 | 104.218766 |
| -4.5460776 | 5.47E-06 | 1.99E-05 | 71.3128351 | 175.897006 | 510.801574 | 80.1251807 |
| -4.5459835 | 5.47E-06 | 1.99E-05 | 79.9195566 | 154.518755 | 78.2308717 | 73.4013893 |
| -4.5458987 | 5.47E-06 | 1.99E-05 | 179.921463 | 233.266492 | 32.2127119 | 231.970803 |
| 4.54586669 | 5.47E-06 | 1.99E-05 | 0 | 0 | 0 | 0 |
| -4.5457105 | 5.48E-06 | 1.99E-05 | 143.855202 | 69.2763595 | 74.7795097 | 86.848972 |
| -4.5454616 | 5.48E-06 | 1.99E-05 | 53.2797044 | 116.362635 | 92.0363196 | 144.001199 |
| -4.5452526 | 5.49E-06 | 1.99E-05 | 57.7879871 | 97.4198805 | 124.249032 | 84.0473923 |
| -4.5451233 | 5.49E-06 | 1.99E-05 | 97.9526873 | 101.749653 | 60.9740618 | 74.5220212 |
| -4.5449616 | 5.49E-06 | 1.99E-05 | 229.102729 | 342.052025 | 318.675757 | 322.741986 |
| -4.5449247 | 5.50E-06 | 1.99E-05 | 50.8206411 | 118.798132 | 104.691314 | 114.304454 |
| -4.5447646 | 5.50E-06 | 2.00E-05 | 71.3128351 | 125.02218 | 128.850847 | 62.1950703 |
| -4.544698 | 5.50E-06 | 2.00E-05 | 53.2797044 | 192.945486 | 151.859927 | 87.9696039 |
| -4.5444972 | 5.51E-06 | 2.00E-05 | 82.7884637 | 99.5847667 | 108.142676 | 100.856871 |
| -4.5443026 | 5.51E-06 | 2.00E-05 | 49.5911095 | 143.694324 | 108.142676 | 115.425085 |
| -4.5440226 | 5.52E-06 | 2.00E-05 | 172.544273 | 69.2763595 | 90.8858656 | 79.0045488 |
| -4.5439619 | 5.52E-06 | 2.00E-05 | 42.2139196 | 154.789366 | 143.806749 | 148.483726 |

| | | | | | | |
|---|---|---|---|---|---|---|
| -4.5438418 | 5.52E-06 | 2.00E-05 | 45.4926707 | 126.104623 | 133.452663 | 140.639303 |
| -4.5435203 | 5.53E-06 | 2.01E-05 | 38.1154808 | 149.918372 | 187.524001 | 192.748686 |
| -4.5434827 | 5.53E-06 | 2.01E-05 | 85.6573709 | 78.477126 | 64.4254237 | 76.763285 |
| -4.5433376 | 5.54E-06 | 2.01E-05 | 66.8045524 | 71.1706349 | 81.6822337 | 93.5727634 |
| -4.5432634 | 5.54E-06 | 2.01E-05 | 47.951734 | 119.880575 | 127.700393 | 118.786981 |
| -4.5432592 | 5.54E-06 | 2.01E-05 | 62.7061136 | 70.6294134 | 147.258111 | 134.475828 |
| -4.5432329 | 5.54E-06 | 2.01E-05 | 34.017042 | 155.601198 | 299.118039 | 228.048591 |
| -4.5430203 | 5.55E-06 | 2.01E-05 | 95.9034679 | 72.253078 | 69.0272397 | 75.0823371 |
| -4.5429496 | 5.55E-06 | 2.01E-05 | 46.7222023 | 137.199665 | 113.894946 | 141.199619 |
| -4.542686 | 5.55E-06 | 2.01E-05 | 82.3786199 | 70.6294134 | 66.7263317 | 87.409288 |
| 4.54260761 | 5.56E-06 | 2.01E-05 | 13.524848 | 25.4374132 | 14.9559019 | 27.4554815 |
| -4.5425385 | 5.56E-06 | 2.01E-05 | 50.8206411 | 115.550803 | 113.894946 | 104.779082 |
| -4.5424798 | 5.56E-06 | 2.01E-05 | 50.4107972 | 128.810731 | 132.302209 | 98.055291 |
| -4.5424253 | 5.56E-06 | 2.01E-05 | 40.5745441 | 197.81648 | 235.843069 | 123.829825 |
| -4.5422675 | 5.57E-06 | 2.02E-05 | 56.1486116 | 115.821413 | 83.9831417 | 147.92341 |
| -4.5422293 | 5.57E-06 | 2.02E-05 | 106.969253 | 74.9591858 | 66.7263317 | 77.3236009 |
| -4.5421424 | 5.57E-06 | 2.02E-05 | 64.3454891 | 155.601198 | 67.8767857 | 149.044042 |
| -4.5418083 | 5.58E-06 | 2.02E-05 | 59.8372065 | 83.0775092 | 98.9390436 | 84.6077082 |
| -4.5418164 | 5.58E-06 | 2.02E-05 | 68.8537718 | 164.260743 | 176.019461 | 174.818576 |
| -4.541676 | 5.58E-06 | 2.02E-05 | 39.3450125 | 162.366467 | 189.824909 | 150.164674 |
| -4.5416539 | 5.58E-06 | 2.02E-05 | 62.7061136 | 171.296623 | 71.3281477 | 137.277407 |
| -4.5413722 | 5.59E-06 | 2.02E-05 | 509.435943 | 496.57078 | 391.154358 | 443.209915 |
| -4.541259 | 5.59E-06 | 2.02E-05 | 47.951734 | 135.034779 | 200.178995 | 108.701294 |
| -4.5404919 | 5.61E-06 | 2.03E-05 | 44.6729829 | 132.869893 | 156.461743 | 126.631404 |
| -4.5403672 | 5.62E-06 | 2.03E-05 | 180.331307 | 73.8767427 | 62.1245158 | 128.312352 |
| 4.53995431 | 5.63E-06 | 2.04E-05 | 30.3284471 | 29.2259641 | 26.4604419 | 26.8951655 |
| -4.5393744 | 5.64E-06 | 2.04E-05 | 45.0828268 | 232.454659 | 132.302209 | 129.9933 |
| -4.5390533 | 5.65E-06 | 2.04E-05 | 48.7714217 | 114.73897 | 126.549939 | 113.183822 |
| -4.5388726 | 5.66E-06 | 2.05E-05 | 50.8206411 | 119.068743 | 102.390406 | 112.623506 |
| -4.538779 | 5.66E-06 | 2.05E-05 | 52.0501728 | 117.174467 | 109.29313 | 97.4949751 |
| -4.5385871 | 5.66E-06 | 2.05E-05 | 46.7222023 | 121.233629 | 139.204933 | 121.028245 |
| -4.5385267 | 5.66E-06 | 2.05E-05 | 67.2143963 | 92.8194972 | 69.0272397 | 82.9267604 |
| -4.5380984 | 5.68E-06 | 2.05E-05 | 55.3289238 | 83.0775092 | 130.001301 | 117.666349 |
| -4.53783 | 5.68E-06 | 2.05E-05 | 51.6403289 | 150.730204 | 90.8858656 | 134.475828 |
| -4.5378522 | 5.68E-06 | 2.05E-05 | 53.2797044 | 114.197749 | 96.6381356 | 100.296555 |
| -4.5374834 | 5.69E-06 | 2.06E-05 | 88.5262781 | 117.445078 | 98.9390436 | 90.7711837 |
| -4.5373941 | 5.70E-06 | 2.06E-05 | 54.0993922 | 133.681725 | 93.1867736 | 101.417187 |
| -4.5369465 | 5.71E-06 | 2.06E-05 | 72.9522106 | 179.144336 | 141.505841 | 142.880567 |
| -4.536759 | 5.71E-06 | 2.06E-05 | 54.0993922 | 94.9843835 | 97.7885896 | 105.899714 |
| -4.5364591 | 5.72E-06 | 2.07E-05 | 57.3781432 | 136.929054 | 79.3813257 | 114.304454 |
| -4.5360722 | 5.73E-06 | 2.07E-05 | 54.0993922 | 151.542036 | 112.744492 | 89.6505518 |
| -4.5359518 | 5.73E-06 | 2.07E-05 | 102.051126 | 235.701989 | 49.4695218 | 141.759935 |
| -4.5356984 | 5.74E-06 | 2.07E-05 | 56.9682993 | 119.609964 | 88.5849576 | 90.2108677 |

| | | | | | | |
|---|---|---|---|---|---|---|
| -4.5350317 | 5.76E-06 | 2.08E-05 | 179.101776 | 243.820312 | 205.931265 | 238.134278 |
| -4.5347829 | 5.77E-06 | 2.08E-05 | 54.0993922 | 144.506156 | 188.674455 | 84.6077082 |
| -4.534572 | 5.77E-06 | 2.08E-05 | 56.1486116 | 110.950419 | 92.0363196 | 88.5299199 |
| -4.5345323 | 5.77E-06 | 2.08E-05 | 49.1812656 | 104.726372 | 118.496762 | 128.872668 |
| -4.5344955 | 5.77E-06 | 2.08E-05 | 39.7548564 | 161.013414 | 174.869007 | 145.682147 |
| -4.5344604 | 5.78E-06 | 2.08E-05 | 43.8532952 | 152.353869 | 132.302209 | 132.234564 |
| -4.5344293 | 5.78E-06 | 2.08E-05 | 59.0175187 | 89.5721679 | 89.7354116 | 82.3664444 |
| -4.5343261 | 5.78E-06 | 2.08E-05 | 40.984388 | 192.674875 | 120.79767 | 209.558165 |
| -4.5340361 | 5.79E-06 | 2.09E-05 | 69.6734596 | 71.4412457 | 77.0804177 | 76.763285 |
| -4.5338357 | 5.79E-06 | 2.09E-05 | 53.2797044 | 168.319905 | 110.443584 | 95.8140272 |
| -4.533239 | 5.81E-06 | 2.09E-05 | 195.085687 | 77.9359044 | 73.6290557 | 89.0902358 |
| 4.53276514 | 5.82E-06 | 2.10E-05 | 0 | 0 | 0 | 0 |
| -4.5324873 | 5.83E-06 | 2.10E-05 | 45.4926707 | 121.50424 | 147.258111 | 126.071088 |
| -4.5322628 | 5.84E-06 | 2.10E-05 | 38.5253247 | 171.567234 | 153.010381 | 172.577312 |
| -4.5322413 | 5.84E-06 | 2.10E-05 | 77.0506494 | 122.586683 | 219.736713 | 63.8760181 |
| -4.5320721 | 5.84E-06 | 2.10E-05 | 63.5258014 | 86.5954493 | 92.0363196 | 71.7204414 |
| -4.5320442 | 5.84E-06 | 2.10E-05 | 95.9034679 | 190.509989 | 69.0272397 | 214.040692 |
| -4.531993 | 5.84E-06 | 2.11E-05 | 76.6408055 | 69.817581 | 80.5317797 | 69.4791776 |
| -4.5319317 | 5.84E-06 | 2.11E-05 | 57.7879871 | 80.1007906 | 96.6381356 | 111.502874 |
| -4.5313929 | 5.86E-06 | 2.11E-05 | 41.3942319 | 205.934803 | 133.452663 | 149.044042 |
| -4.5307764 | 5.88E-06 | 2.12E-05 | 179.921463 | 140.446994 | 39.1154358 | 161.931309 |
| -4.5307652 | 5.88E-06 | 2.12E-05 | 78.6900249 | 96.3374374 | 60.9740618 | 80.1251807 |
| -4.5305695 | 5.88E-06 | 2.12E-05 | 46.3123584 | 199.169533 | 143.806749 | 108.140978 |
| -4.5302009 | 5.89E-06 | 2.12E-05 | 38.9351686 | 261.951234 | 146.107657 | 160.250361 |
| -4.5301576 | 5.89E-06 | 2.12E-05 | 31.1481349 | 264.657342 | 173.718553 | 241.496174 |
| -4.5299159 | 5.90E-06 | 2.12E-05 | 52.0501728 | 93.9019404 | 207.081719 | 132.234564 |
| -4.5292562 | 5.92E-06 | 2.13E-05 | 44.6729829 | 139.635162 | 189.824909 | 115.985401 |
| -4.5292699 | 5.92E-06 | 2.13E-05 | 39.3450125 | 211.617629 | 140.355387 | 160.810677 |
| -4.5289671 | 5.93E-06 | 2.13E-05 | 56.9682993 | 86.5954493 | 88.5849576 | 114.304454 |
| -4.5284928 | 5.94E-06 | 2.14E-05 | 49.5911095 | 319.59133 | 349.738015 | 360.283155 |
| -4.5282895 | 5.95E-06 | 2.14E-05 | 67.2143963 | 152.083258 | 63.2749697 | 140.078987 |
| -4.5282456 | 5.95E-06 | 2.14E-05 | 43.0336074 | 139.905773 | 144.957203 | 132.79488 |
| -4.5281255 | 5.95E-06 | 2.14E-05 | 55.7387677 | 115.821413 | 80.5317797 | 135.036144 |
| 4.52755869 | 5.97E-06 | 2.15E-05 | 33.6071982 | 18.401533 | 31.0622579 | 21.292006 |
| -4.5274346 | 5.97E-06 | 2.15E-05 | 54.9190799 | 97.9611021 | 93.1867736 | 92.4521315 |
| -4.5267527 | 5.99E-06 | 2.15E-05 | 95.493624 | 65.4878086 | 80.5317797 | 70.5998095 |
| -4.5267434 | 5.99E-06 | 2.15E-05 | 1283.22119 | 99.8553775 | 51.7704298 | 249.900913 |
| -4.526599 | 5.99E-06 | 2.15E-05 | 55.3289238 | 97.9611021 | 85.1335957 | 114.304454 |
| -4.5265021 | 6.00E-06 | 2.16E-05 | 60.6568942 | 104.455761 | 109.29313 | 71.7204414 |
| -4.5258626 | 6.01E-06 | 2.16E-05 | 58.197831 | 87.1366709 | 81.6822337 | 97.4949751 |
| -4.5254759 | 6.03E-06 | 2.17E-05 | 39.3450125 | 231.913438 | 134.603117 | 166.974153 |
| -4.5253358 | 6.03E-06 | 2.17E-05 | 58.6076748 | 94.7137727 | 78.2308717 | 93.0124475 |
| -4.5251043 | 6.04E-06 | 2.17E-05 | 314.7601 | 526.067355 | 415.313892 | 489.155823 |

| | | | | | | |
|---|---|---|---|---|---|---|
| -4.5250391 | 6.04E-06 | 2.17E-05 | 45.4926707 | 356.123785 | 418.765254 | 390.540216 |
| -4.5249563 | 6.04E-06 | 2.17E-05 | 142.215826 | 73.8767427 | 83.9831417 | 68.3585457 |
| -4.5247299 | 6.05E-06 | 2.17E-05 | 54.9190799 | 165.884408 | 78.2308717 | 176.499524 |
| -4.5247463 | 6.05E-06 | 2.17E-05 | 50.8206411 | 110.679809 | 117.346308 | 96.9346591 |
| -4.5246703 | 6.05E-06 | 2.17E-05 | 54.509236 | 114.73897 | 123.098578 | 82.3664444 |
| -4.5246638 | 6.05E-06 | 2.17E-05 | 138.937075 | 215.676791 | 98.9390436 | 49.3078035 |
| -4.5245586 | 6.05E-06 | 2.17E-05 | 67.6242402 | 70.6294134 | 74.7795097 | 86.848972 |
| -4.5244223 | 6.06E-06 | 2.17E-05 | 60.6568942 | 113.656527 | 103.54086 | 71.7204414 |
| -4.5243475 | 6.06E-06 | 2.17E-05 | 30.738291 | 222.44206 | 190.975363 | 238.694594 |
| -4.5241614 | 6.06E-06 | 2.18E-05 | 51.230485 | 99.8553775 | 108.142676 | 161.370993 |
| -4.523341 | 6.09E-06 | 2.18E-05 | 48.7714217 | 150.459593 | 95.4876816 | 138.398039 |
| -4.5228107 | 6.10E-06 | 2.19E-05 | 63.9356453 | 208.3703 | 153.010381 | 216.281956 |
| -4.5227284 | 6.10E-06 | 2.19E-05 | 50.8206411 | 158.036695 | 98.9390436 | 110.382242 |
| -4.5227356 | 6.10E-06 | 2.19E-05 | 47.951734 | 145.047378 | 121.948124 | 103.65845 |
| 4.52204161 | 6.12E-06 | 2.20E-05 | 0 | 0 | 0 | 0 |
| -4.5213427 | 6.14E-06 | 2.20E-05 | 65.1651769 | 70.6294134 | 90.8858656 | 78.4442328 |
| -4.5212152 | 6.15E-06 | 2.21E-05 | 173.773805 | 182.391665 | 202.479903 | 228.608907 |
| -4.521147 | 6.15E-06 | 2.21E-05 | 43.4434513 | 200.522587 | 116.195854 | 147.92341 |
| -4.5210415 | 6.15E-06 | 2.21E-05 | 121.313788 | 110.950419 | 46.0181598 | 131.674248 |
| -4.5209144 | 6.16E-06 | 2.21E-05 | 41.8040758 | 198.898923 | 116.195854 | 172.016996 |
| -4.5207462 | 6.16E-06 | 2.21E-05 | 64.3454891 | 88.4897248 | 67.8767857 | 98.615607 |
| -4.5207618 | 6.16E-06 | 2.21E-05 | 55.3289238 | 97.6904913 | 92.0363196 | 87.9696039 |
| -4.5202159 | 6.18E-06 | 2.21E-05 | 358.203551 | 118.25691 | 64.4254237 | 96.3743432 |
| -4.5201076 | 6.18E-06 | 2.21E-05 | 53.2797044 | 127.457677 | 82.8326877 | 135.59646 |
| -4.5200342 | 6.18E-06 | 2.22E-05 | 52.4600166 | 90.1133895 | 124.249032 | 104.779082 |
| -4.5197826 | 6.19E-06 | 2.22E-05 | 43.4434513 | 124.480958 | 131.151755 | 165.293205 |
| -4.5193479 | 6.20E-06 | 2.22E-05 | 162.298176 | 89.3015571 | 63.2749697 | 80.6854966 |
| -4.5193707 | 6.20E-06 | 2.22E-05 | 45.9025146 | 197.81648 | 211.683535 | 225.247011 |
| -4.5188561 | 6.22E-06 | 2.23E-05 | 56.9682993 | 107.432479 | 81.6822337 | 90.2108677 |
| -4.518459 | 6.23E-06 | 2.23E-05 | 66.3947085 | 67.6526948 | 108.142676 | 84.0473923 |
| -4.5183652 | 6.23E-06 | 2.23E-05 | 52.0501728 | 92.5488865 | 103.54086 | 112.06319 |
| -4.5182447 | 6.24E-06 | 2.23E-05 | 358.203551 | 433.247857 | 361.242555 | 405.668747 |
| 4.51807744 | 6.24E-06 | 2.23E-05 | 24.5906328 | 17.3190899 | 19.5577179 | 8.40473923 |
| -4.5178188 | 6.25E-06 | 2.24E-05 | 49.5911095 | 88.219114 | 167.966283 | 173.137628 |
| -4.5177524 | 6.25E-06 | 2.24E-05 | 45.9025146 | 106.079425 | 141.505841 | 140.639303 |
| -4.5177264 | 6.25E-06 | 2.24E-05 | 1370.10809 | 4671.28327 | 4307.29976 | 4768.28872 |
| -4.5175542 | 6.26E-06 | 2.24E-05 | 117.625194 | 96.0668266 | 62.1245158 | 69.4791776 |
| -4.5175364 | 6.26E-06 | 2.24E-05 | 54.9190799 | 100.396599 | 83.9831417 | 100.856871 |
| -4.517135 | 6.27E-06 | 2.24E-05 | 326.235728 | 112.844695 | 58.6731538 | 106.46003 |
| -4.5171318 | 6.27E-06 | 2.24E-05 | 65.9848647 | 206.476024 | 74.7795097 | 581.607955 |
| -4.5169975 | 6.27E-06 | 2.24E-05 | 59.4273626 | 134.222946 | 72.4786017 | 175.378892 |
| -4.5169661 | 6.27E-06 | 2.24E-05 | 111.887379 | 166.425629 | 48.3190678 | 104.779082 |
| -4.5167178 | 6.28E-06 | 2.25E-05 | 125.412227 | 74.688575 | 101.239952 | 58.8331746 |

| | | | | | | |
|---|---|---|---|---|---|---|
| -4.5164006 | 6.29E-06 | 2.25E-05 | 68.4439279 | 103.373318 | 65.5758777 | 81.8061285 |
| -4.516237 | 6.29E-06 | 2.25E-05 | 63.5258014 | 113.656527 | 69.0272397 | 90.7711837 |
| -4.5161997 | 6.30E-06 | 2.25E-05 | 119.674413 | 108.785533 | 81.6822337 | 93.0124475 |
| -4.5161717 | 6.30E-06 | 2.25E-05 | 47.951734 | 116.362635 | 115.0454 | 109.821926 |
| -4.5161401 | 6.30E-06 | 2.25E-05 | 52.0501728 | 108.514922 | 94.3372276 | 100.296555 |
| -4.5154815 | 6.32E-06 | 2.26E-05 | 43.0336074 | 174.273342 | 153.010381 | 113.744138 |
| -4.5151757 | 6.33E-06 | 2.26E-05 | 43.4434513 | 225.960001 | 118.496762 | 143.440883 |
| -4.514909 | 6.33E-06 | 2.26E-05 | 118.854725 | 66.2996409 | 77.0804177 | 73.9617052 |
| -4.5147282 | 6.34E-06 | 2.26E-05 | 43.8532952 | 168.861126 | 131.151755 | 118.786981 |
| -4.5147241 | 6.34E-06 | 2.26E-05 | 101.231438 | 69.2763595 | 72.4786017 | 70.0394936 |
| -4.5144275 | 6.35E-06 | 2.27E-05 | 45.4926707 | 128.269509 | 115.0454 | 206.756585 |
| -4.5143999 | 6.35E-06 | 2.27E-05 | 51.230485 | 107.161869 | 96.6381356 | 103.65845 |
| -4.5141581 | 6.36E-06 | 2.27E-05 | 57.3781432 | 99.3141559 | 80.5317797 | 87.409288 |
| -4.5139656 | 6.36E-06 | 2.27E-05 | 58.6076748 | 141.800048 | 77.0804177 | 101.417187 |
| -4.5138522 | 6.37E-06 | 2.27E-05 | 45.4926707 | 135.846611 | 112.744492 | 127.19172 |
| -4.5137945 | 6.37E-06 | 2.27E-05 | 34.8367298 | 207.017246 | 195.577179 | 154.647202 |
| -4.5137249 | 6.37E-06 | 2.27E-05 | 22.1315695 | 425.670756 | 295.666677 | 231.410487 |
| -4.5136149 | 6.37E-06 | 2.27E-05 | 59.8372065 | 79.2889583 | 79.3813257 | 99.1759229 |
| -4.5134027 | 6.38E-06 | 2.28E-05 | 117.625194 | 153.977533 | 182.922185 | 189.386791 |
| -4.5133178 | 6.38E-06 | 2.28E-05 | 48.3615778 | 116.092024 | 110.443584 | 108.140978 |
| -4.5126471 | 6.40E-06 | 2.28E-05 | 51.6403289 | 145.85921 | 94.3372276 | 104.218766 |
| -4.5124756 | 6.41E-06 | 2.29E-05 | 90.5754975 | 152.353869 | 105.841768 | 52.6696992 |
| -4.5124276 | 6.41E-06 | 2.29E-05 | 38.9351686 | 201.60503 | 136.904025 | 156.32815 |
| -4.5124365 | 6.41E-06 | 2.29E-05 | 61.0667381 | 106.350036 | 70.1776937 | 94.1330794 |
| -4.5123793 | 6.41E-06 | 2.29E-05 | 38.5253247 | 155.059976 | 159.913105 | 154.647202 |
| -4.512365 | 6.41E-06 | 2.29E-05 | 54.0993922 | 84.7011739 | 100.089498 | 122.709193 |
| -4.512252 | 6.41E-06 | 2.29E-05 | 0 | 0 | 79.3813257 | 0 |
| -4.511353 | 6.44E-06 | 2.30E-05 | 34.8367298 | 155.601198 | 178.320369 | 251.581861 |
| -4.510662 | 6.46E-06 | 2.30E-05 | 52.0501728 | 96.6080482 | 180.621277 | 107.580662 |
| -4.5098258 | 6.49E-06 | 2.31E-05 | 54.509236 | 106.891258 | 80.5317797 | 122.709193 |
| -4.5095744 | 6.50E-06 | 2.31E-05 | 38.5253247 | 186.180216 | 148.408565 | 146.802779 |
| -4.509519 | 6.50E-06 | 2.32E-05 | 44.6729829 | 123.127905 | 130.001301 | 122.148877 |
| -4.5094264 | 6.50E-06 | 2.32E-05 | 184.429746 | 124.480958 | 92.0363196 | 94.6933953 |
| -4.509432 | 6.50E-06 | 2.32E-05 | 72.5423667 | 70.3588026 | 403.809352 | 158.569413 |
| 4.50929097 | 6.50E-06 | 2.32E-05 | 255.742581 | 115.009581 | 136.904025 | 126.631404 |
| -4.5092142 | 6.51E-06 | 2.32E-05 | 44.263139 | 142.611881 | 121.948124 | 122.148877 |
| -4.5091557 | 6.51E-06 | 2.32E-05 | 36.0662614 | 216.488623 | 226.639437 | 132.234564 |
| -4.5080844 | 6.54E-06 | 2.33E-05 | 197.54475 | 97.9611021 | 69.0272397 | 73.4013893 |
| 4.50790909 | 6.55E-06 | 2.33E-05 | 20.492194 | 12.7187066 | 17.2568099 | 14.0078987 |
| -4.507867 | 6.55E-06 | 2.33E-05 | 0 | 0 | 78.2308717 | 0 |
| -4.507867 | 6.55E-06 | 2.33E-05 | 0 | 0 | 78.2308717 | 0 |
| -4.5078915 | 6.55E-06 | 2.33E-05 | 63.9356453 | 64.6759762 | 278.409867 | 198.351846 |
| -4.5072899 | 6.57E-06 | 2.34E-05 | 54.509236 | 160.201581 | 79.3813257 | 124.950457 |

| | | | | | | |
|---|---|---|---|---|---|---|
| -4.5070957 | 6.57E-06 | 2.34E-05 | 58.6076748 | 104.726372 | 74.7795097 | 89.6505518 |
| -4.5069625 | 6.58E-06 | 2.34E-05 | 52.4600166 | 112.032863 | 88.5849576 | 99.7362389 |
| -4.5068852 | 6.58E-06 | 2.34E-05 | 141.396139 | 150.730204 | 37.9649818 | 185.464579 |
| -4.5064226 | 6.59E-06 | 2.35E-05 | 331.153855 | 97.9611021 | 51.7704298 | 147.363094 |
| -4.5060549 | 6.60E-06 | 2.35E-05 | 212.29913 | 179.685557 | 31.0622579 | 209.558165 |
| -4.5060165 | 6.61E-06 | 2.35E-05 | 180.741151 | 437.307019 | 347.437107 | 413.51317 |
| -4.5059879 | 6.61E-06 | 2.35E-05 | 50.4107972 | 107.432479 | 97.7885896 | 103.098135 |
| -4.505773 | 6.61E-06 | 2.35E-05 | 66.8045524 | 169.672959 | 158.762651 | 172.016996 |
| -4.5057483 | 6.61E-06 | 2.35E-05 | 48.3615778 | 104.18515 | 106.992222 | 122.148877 |
| 4.50548309 | 6.62E-06 | 2.35E-05 | 134.428793 | 85.2423954 | 80.5317797 | 81.2458125 |
| -4.5044892 | 6.65E-06 | 2.37E-05 | 63.5258014 | 77.9359044 | 87.4345036 | 70.0394936 |
| 4.50432484 | 6.66E-06 | 2.37E-05 | 26.2300083 | 21.9194731 | 10.354086 | 19.6110582 |
| -4.5043448 | 6.66E-06 | 2.37E-05 | 93.8542485 | 139.09394 | 64.4254237 | 68.9188617 |
| -4.5037373 | 6.68E-06 | 2.37E-05 | 45.4926707 | 125.292791 | 106.992222 | 149.604358 |
| -4.5036927 | 6.68E-06 | 2.37E-05 | 88.1164342 | 77.6652936 | 67.8767857 | 64.99665 |
| -4.5036518 | 6.68E-06 | 2.37E-05 | 77.4604933 | 68.7351379 | 71.3281477 | 69.4791776 |
| -4.502743 | 6.71E-06 | 2.38E-05 | 45.0828268 | 125.02218 | 132.302209 | 111.502874 |
| -4.5024137 | 6.72E-06 | 2.39E-05 | 42.6237635 | 130.705006 | 132.302209 | 132.234564 |
| -4.5023323 | 6.72E-06 | 2.39E-05 | 881.98403 | 3930.89218 | 3215.51892 | 3438.65898 |
| -4.5022617 | 6.72E-06 | 2.39E-05 | 41.3942319 | 135.576 | 128.850847 | 157.448782 |
| -4.501984 | 6.73E-06 | 2.39E-05 | 54.0993922 | 90.3840002 | 87.4345036 | 96.3743432 |
| -4.5017888 | 6.74E-06 | 2.39E-05 | 61.476582 | 155.871809 | 70.1776937 | 109.26161 |
| -4.5017355 | 6.74E-06 | 2.39E-05 | 65.1651769 | 94.9843835 | 65.5758777 | 128.312352 |
| -4.5015294 | 6.75E-06 | 2.39E-05 | 115.575974 | 119.609964 | 110.443584 | 74.5220212 |
| -4.5013611 | 6.75E-06 | 2.40E-05 | 127.461447 | 143.423713 | 40.2658898 | 154.086886 |
| -4.5013083 | 6.75E-06 | 2.40E-05 | 42.6237635 | 151.542036 | 120.79767 | 132.79488 |
| -4.5012449 | 6.76E-06 | 2.40E-05 | 39.7548564 | 172.108456 | 128.850847 | 152.966254 |
| 4.50120979 | 6.76E-06 | 2.40E-05 | 0 | 0 | 0 | 0 |
| -4.50102 | 6.76E-06 | 2.40E-05 | 50.4107972 | 97.4198805 | 97.7885896 | 126.631404 |
| -4.5009453 | 6.77E-06 | 2.40E-05 | 50.8206411 | 93.3607188 | 112.744492 | 99.7362389 |
| -4.5000247 | 6.79E-06 | 2.41E-05 | 49.5911095 | 125.02218 | 95.4876816 | 202.834373 |
| -4.500034 | 6.79E-06 | 2.41E-05 | 68.8537718 | 66.0290301 | 73.6290557 | 84.0473923 |
| -4.4992448 | 6.82E-06 | 2.42E-05 | 634.438326 | 88.7603356 | 65.5758777 | 160.250361 |
| -4.4989188 | 6.83E-06 | 2.42E-05 | 56.1486116 | 79.8301798 | 104.691314 | 86.848972 |
| -4.4989217 | 6.83E-06 | 2.42E-05 | 55.3289238 | 177.25006 | 174.869007 | 168.655101 |
| -4.4986576 | 6.84E-06 | 2.42E-05 | 55.3289238 | 117.174467 | 79.3813257 | 95.2537113 |
| -4.4983151 | 6.85E-06 | 2.43E-05 | 58.197831 | 76.0416289 | 86.2840497 | 86.848972 |
| -4.4981822 | 6.85E-06 | 2.43E-05 | 54.9190799 | 266.281007 | 396.906628 | 70.0394936 |
| -4.4976603 | 6.87E-06 | 2.43E-05 | 134.838636 | 84.9717847 | 49.4695218 | 141.759935 |
| -4.4974237 | 6.88E-06 | 2.44E-05 | 55.3289238 | 79.5595691 | 96.6381356 | 124.950457 |
| -4.4971399 | 6.89E-06 | 2.44E-05 | 57.7879871 | 131.246228 | 130.001301 | 70.0394936 |
| -4.4968351 | 6.90E-06 | 2.44E-05 | 136.478012 | 102.561485 | 58.6731538 | 71.7204414 |
| -4.4965479 | 6.91E-06 | 2.45E-05 | 32.3776665 | 165.884408 | 234.692615 | 188.826475 |

| | | | | | | |
|---|---|---|---|---|---|---|
| -4.4964383 | 6.91E-06 | 2.45E-05 | 52.8698605 | 84.1599523 | 101.239952 | 127.752036 |
| -4.4959867 | 6.92E-06 | 2.45E-05 | 66.3947085 | 68.4645271 | 75.9299637 | 75.6426531 |
| -4.4958505 | 6.93E-06 | 2.45E-05 | 445.090454 | 696.010924 | 492.39431 | 568.720688 |
| -4.4956101 | 6.94E-06 | 2.46E-05 | 83.1983076 | 57.640096 | 79.3813257 | 112.623506 |
| -4.4952343 | 6.95E-06 | 2.46E-05 | 30.3284471 | 213.241294 | 173.718553 | 244.297754 |
| -4.4951667 | 6.95E-06 | 2.46E-05 | 45.9025146 | 111.491641 | 110.443584 | 133.915512 |
| -4.4950012 | 6.96E-06 | 2.46E-05 | 47.951734 | 102.290875 | 108.142676 | 114.864769 |
| -4.4950167 | 6.96E-06 | 2.46E-05 | 48.3615778 | 92.0076649 | 132.302209 | 179.86142 |
| -4.4947265 | 6.97E-06 | 2.47E-05 | 112.297223 | 181.038611 | 172.568099 | 174.25826 |
| -4.4945701 | 6.97E-06 | 2.47E-05 | 40.984388 | 139.364551 | 131.151755 | 141.759935 |
| -4.4942773 | 6.98E-06 | 2.47E-05 | 56.5584554 | 132.869893 | 79.3813257 | 93.0124475 |
| -4.4937582 | 7.00E-06 | 2.47E-05 | 773.375401 | 1282.69509 | 1157.35672 | 1261.2712 |
| -4.4937444 | 7.00E-06 | 2.47E-05 | 44.6729829 | 133.952336 | 136.904025 | 105.339398 |
| -4.4937084 | 7.00E-06 | 2.47E-05 | 106.969253 | 102.290875 | 55.2217918 | 74.5220212 |
| -4.4937306 | 7.00E-06 | 2.47E-05 | 47.1320462 | 124.210348 | 124.249032 | 97.4949751 |
| -4.4937087 | 7.00E-06 | 2.47E-05 | 63.9356453 | 67.1114732 | 80.5317797 | 84.0473923 |
| -4.4935188 | 7.01E-06 | 2.48E-05 | 49.1812656 | 94.7137727 | 108.142676 | 112.06319 |
| -4.4932063 | 7.02E-06 | 2.48E-05 | 41.8040758 | 140.717605 | 130.001301 | 128.872668 |
| -4.4929182 | 7.03E-06 | 2.48E-05 | 62.2962697 | 100.66721 | 136.904025 | 67.2379138 |
| -4.4927703 | 7.03E-06 | 2.48E-05 | 45.4926707 | 126.375234 | 103.54086 | 132.79488 |
| -4.4926887 | 7.03E-06 | 2.49E-05 | 35.6564176 | 177.791282 | 190.975363 | 144.001199 |
| -4.4925444 | 7.04E-06 | 2.49E-05 | 182.380527 | 144.776767 | 142.656295 | 138.398039 |
| -4.4925037 | 7.04E-06 | 2.49E-05 | 49.1812656 | 93.6313296 | 106.992222 | 119.347297 |
| -4.4920079 | 7.06E-06 | 2.49E-05 | 50.8206411 | 87.1366709 | 124.249032 | 105.339398 |
| -4.4915862 | 7.07E-06 | 2.50E-05 | 56.5584554 | 83.0775092 | 92.0363196 | 149.044042 |
| -4.4916175 | 7.07E-06 | 2.50E-05 | 47.1320462 | 97.9611021 | 203.630357 | 131.113932 |
| -4.4916419 | 7.07E-06 | 2.50E-05 | 94.2640924 | 102.832096 | 50.6199758 | 167.534469 |
| -4.4916071 | 7.07E-06 | 2.50E-05 | 46.3123584 | 109.056144 | 141.505841 | 107.020346 |
| -4.4913744 | 7.08E-06 | 2.50E-05 | 69.6734596 | 79.0183475 | 63.2749697 | 76.202969 |
| -4.4909043 | 7.09E-06 | 2.50E-05 | 54.509236 | 86.8660601 | 83.9831417 | 94.6933953 |
| -4.4908691 | 7.09E-06 | 2.50E-05 | 61.0667381 | 124.480958 | 116.195854 | 127.752036 |
| -4.4907882 | 7.10E-06 | 2.50E-05 | 59.4273626 | 108.514922 | 123.098578 | 67.7982298 |
| -4.4904623 | 7.11E-06 | 2.51E-05 | 55.7387677 | 87.9485032 | 87.4345036 | 80.6854966 |
| -4.4899199 | 7.12E-06 | 2.51E-05 | 118.444881 | 70.0881918 | 78.2308717 | 64.99665 |
| -4.4898784 | 7.13E-06 | 2.51E-05 | 50.8206411 | 106.350036 | 94.3372276 | 94.1330794 |
| 4.48980394 | 7.13E-06 | 2.52E-05 | 65.9848647 | 49.7923834 | 60.9740618 | 53.2300151 |
| -4.4894577 | 7.14E-06 | 2.52E-05 | 77.0506494 | 104.726372 | 94.3372276 | 80.1251807 |
| -4.4893752 | 7.14E-06 | 2.52E-05 | 43.0336074 | 140.176384 | 117.346308 | 126.071088 |
| 4.48933989 | 7.14E-06 | 2.52E-05 | 0.81968776 | 0.27061078 | 0 | 0.56031595 |
| -4.4891713 | 7.15E-06 | 2.52E-05 | 63.5258014 | 140.446994 | 65.5758777 | 101.977503 |
| -4.4891209 | 7.15E-06 | 2.52E-05 | 88.1164342 | 52.2278804 | 96.6381356 | 96.9346591 |
| -4.4882095 | 7.18E-06 | 2.53E-05 | 47.5418901 | 191.863042 | 88.5849576 | 152.966254 |
| -4.4881816 | 7.18E-06 | 2.53E-05 | 29.5087594 | 242.196647 | 197.878087 | 190.507423 |

| | | | | | | |
|---|---|---|---|---|---|---|
| -4.4878685 | 7.19E-06 | 2.54E-05 | 46.3123584 | 138.552719 | 104.691314 | 110.382242 |
| -4.4877828 | 7.20E-06 | 2.54E-05 | 152.052079 | 173.73212 | 235.843069 | 223.566063 |
| -4.4872819 | 7.21E-06 | 2.54E-05 | 279.92337 | 506.312768 | 553.368372 | 578.246059 |
| -4.4872143 | 7.22E-06 | 2.54E-05 | 34.4268859 | 186.992048 | 161.063559 | 169.215416 |
| -4.4869505 | 7.22E-06 | 2.55E-05 | 42.2139196 | 129.081342 | 119.647216 | 144.001199 |
| -4.486937 | 7.23E-06 | 2.55E-05 | 49.5911095 | 101.208431 | 101.239952 | 97.4949751 |
| -4.4865384 | 7.24E-06 | 2.55E-05 | 256.972113 | 670.573511 | 571.775636 | 612.985648 |
| -4.4863052 | 7.25E-06 | 2.55E-05 | 76.2309617 | 99.8553775 | 113.894946 | 106.46003 |
| -4.4862406 | 7.25E-06 | 2.55E-05 | 40.5745441 | 191.592432 | 113.894946 | 155.207518 |
| -4.4862121 | 7.25E-06 | 2.55E-05 | 79.0998688 | 62.7817008 | 90.8858656 | 67.2379138 |
| -4.4861445 | 7.25E-06 | 2.55E-05 | 35.6564176 | 198.628312 | 171.417645 | 141.199619 |
| -4.4856912 | 7.27E-06 | 2.56E-05 | 55.7387677 | 289.824144 | 66.7263317 | 253.823125 |
| -4.4852143 | 7.28E-06 | 2.56E-05 | 35.2465737 | 162.366467 | 161.063559 | 172.577312 |
| -4.4852566 | 7.28E-06 | 2.56E-05 | 52.0501728 | 92.5488865 | 87.4345036 | 101.977503 |
| -4.4852351 | 7.28E-06 | 2.56E-05 | 83.1983076 | 81.9950661 | 105.841768 | 54.910963 |
| -4.4852495 | 7.28E-06 | 2.56E-05 | 51.6403289 | 97.6904913 | 88.5849576 | 96.3743432 |
| -4.4851546 | 7.29E-06 | 2.56E-05 | 89.3459658 | 162.366467 | 49.4695218 | 112.623506 |
| -4.4847117 | 7.30E-06 | 2.57E-05 | 39.7548564 | 119.068743 | 232.391707 | 154.647202 |
| -4.4837457 | 7.33E-06 | 2.58E-05 | 52.4600166 | 96.0668266 | 82.8326877 | 107.580662 |
| -4.4837091 | 7.34E-06 | 2.58E-05 | 47.951734 | 90.9252218 | 188.674455 | 132.234564 |
| -4.4834029 | 7.35E-06 | 2.58E-05 | 43.0336074 | 157.766084 | 116.195854 | 120.467929 |
| -4.4830842 | 7.36E-06 | 2.59E-05 | 27.8693838 | 161.825246 | 341.684837 | 251.021545 |
| -4.4828965 | 7.36E-06 | 2.59E-05 | 45.0828268 | 113.115306 | 113.894946 | 118.226665 |
| -4.4826993 | 7.37E-06 | 2.59E-05 | 59.4273626 | 84.9717847 | 109.29313 | 70.5998095 |
| -4.4821478 | 7.39E-06 | 2.60E-05 | 40.984388 | 153.165701 | 149.559019 | 114.864769 |
| -4.4820986 | 7.39E-06 | 2.60E-05 | 46.7222023 | 110.409198 | 103.54086 | 113.183822 |
| -4.4816511 | 7.41E-06 | 2.60E-05 | 58.6076748 | 94.9843835 | 74.7795097 | 77.8839169 |
| -4.4815627 | 7.41E-06 | 2.60E-05 | 43.0336074 | 154.518755 | 123.098578 | 113.183822 |
| -4.4815063 | 7.41E-06 | 2.60E-05 | 68.4439279 | 104.455761 | 57.5226998 | 104.779082 |
| -4.481345 | 7.42E-06 | 2.61E-05 | 53.2797044 | 115.009581 | 75.9299637 | 120.467929 |
| -4.4811955 | 7.42E-06 | 2.61E-05 | 122.953164 | 162.095857 | 131.151755 | 156.32815 |
| -4.4811223 | 7.43E-06 | 2.61E-05 | 24.1807889 | 268.175282 | 258.852149 | 226.367643 |
| -4.4809872 | 7.43E-06 | 2.61E-05 | 33.6071982 | 168.590515 | 165.665375 | 191.067738 |
| -4.4809428 | 7.43E-06 | 2.61E-05 | 45.4926707 | 108.785533 | 121.948124 | 110.382242 |
| -4.480948 | 7.43E-06 | 2.61E-05 | 52.8698605 | 143.964935 | 96.6381356 | 85.7283401 |
| -4.4805977 | 7.44E-06 | 2.61E-05 | 50.0009534 | 101.479042 | 100.089498 | 91.8918156 |
| -4.4805538 | 7.44E-06 | 2.61E-05 | 44.6729829 | 138.011497 | 115.0454 | 108.140978 |
| -4.4797565 | 7.47E-06 | 2.62E-05 | 65.5750208 | 145.85921 | 62.1245158 | 105.339398 |
| -4.4797484 | 7.47E-06 | 2.62E-05 | 100.411751 | 73.0649104 | 59.8236078 | 75.6426531 |
| -4.4796641 | 7.48E-06 | 2.62E-05 | 40.1647002 | 167.237462 | 112.744492 | 164.732889 |
| -4.4796215 | 7.48E-06 | 2.62E-05 | 35.2465737 | 173.461509 | 165.665375 | 152.405938 |
| -4.4795919 | 7.48E-06 | 2.62E-05 | 54.509236 | 103.102707 | 178.320369 | 84.0473923 |
| -4.479438 | 7.48E-06 | 2.63E-05 | 54.9190799 | 99.8553775 | 79.3813257 | 85.1680242 |

| | | | | | | |
|---|---|---|---|---|---|---|
| -4.4791475 | 7.49E-06 | 2.63E-05 | 31.1481349 | 195.380983 | 232.391707 | 163.612257 |
| -4.4787178 | 7.51E-06 | 2.63E-05 | 54.509236 | 84.9717847 | 80.5317797 | 101.977503 |
| -4.4786572 | 7.51E-06 | 2.63E-05 | 42.2139196 | 158.848527 | 110.443584 | 134.475828 |
| -4.4783398 | 7.52E-06 | 2.64E-05 | 100.411751 | 60.075593 | 109.29313 | 65.556966 |
| -4.4782386 | 7.53E-06 | 2.64E-05 | 39.7548564 | 164.260743 | 150.709473 | 118.786981 |
| -4.4782445 | 7.53E-06 | 2.64E-05 | 25.0004767 | 171.296623 | 376.198456 | 286.881766 |
| -4.4780626 | 7.53E-06 | 2.64E-05 | 44.263139 | 107.973701 | 139.204933 | 116.545717 |
| 4.4779595 | 7.54E-06 | 2.64E-05 | 0 | 0 | 0 | 0 |
| -4.4774043 | 7.56E-06 | 2.65E-05 | 60.6568942 | 70.6294134 | 139.204933 | 91.3314996 |
| -4.4773666 | 7.56E-06 | 2.65E-05 | 49.5911095 | 86.8660601 | 360.092101 | 180.421735 |
| 4.47733954 | 7.56E-06 | 2.65E-05 | 5.32797044 | 6.76526948 | 0 | 5.60315949 |
| -4.4770421 | 7.57E-06 | 2.65E-05 | 51.230485 | 102.832096 | 82.8326877 | 108.140978 |
| -4.4768194 | 7.58E-06 | 2.65E-05 | 51.230485 | 97.1492697 | 89.7354116 | 92.4521315 |
| -4.4767408 | 7.58E-06 | 2.66E-05 | 240.578358 | 333.663091 | 207.081719 | 245.978701 |
| -4.4767029 | 7.58E-06 | 2.66E-05 | 45.4926707 | 113.385916 | 110.443584 | 110.942558 |
| -4.4758816 | 7.61E-06 | 2.67E-05 | 395.909188 | 1802.26779 | 3101.62397 | 2437.37438 |
| -4.4758389 | 7.61E-06 | 2.67E-05 | 132.379573 | 51.9572696 | 93.1867736 | 87.409288 |
| -4.4755691 | 7.62E-06 | 2.67E-05 | 38.9351686 | 173.190899 | 119.647216 | 156.888466 |
| -4.4755409 | 7.62E-06 | 2.67E-05 | 40.984388 | 118.25691 | 135.753571 | 146.802779 |
| -4.475469 | 7.62E-06 | 2.67E-05 | 176.642712 | 120.421797 | 43.7172518 | 105.899714 |
| -4.4753579 | 7.63E-06 | 2.67E-05 | 51.6403289 | 113.385916 | 80.5317797 | 104.218766 |
| -4.4751064 | 7.64E-06 | 2.67E-05 | 47.951734 | 186.721438 | 108.142676 | 101.417187 |
| -4.4747567 | 7.65E-06 | 2.68E-05 | 53.2797044 | 79.0183475 | 95.4876816 | 95.8140272 |
| -4.474733 | 7.65E-06 | 2.68E-05 | 59.8372065 | 78.2065152 | 73.6290557 | 76.202969 |
| -4.4747085 | 7.65E-06 | 2.68E-05 | 49.1812656 | 95.7962158 | 103.54086 | 96.3743432 |
| -4.4743424 | 7.66E-06 | 2.68E-05 | 35.2465737 | 135.30539 | 205.931265 | 173.697944 |
| -4.4741107 | 7.67E-06 | 2.68E-05 | 45.4926707 | 107.432479 | 124.249032 | 106.46003 |
| -4.4740879 | 7.67E-06 | 2.68E-05 | 96.7231557 | 147.212264 | 111.594038 | 117.666349 |
| -4.4740348 | 7.68E-06 | 2.69E-05 | 49.1812656 | 90.3840002 | 117.346308 | 99.7362389 |
| -4.4737262 | 7.69E-06 | 2.69E-05 | 75.00143 | 61.9698684 | 70.1776937 | 76.763285 |
| -4.4737081 | 7.69E-06 | 2.69E-05 | 43.8532952 | 118.527521 | 111.594038 | 123.829825 |
| -4.4729555 | 7.71E-06 | 2.70E-05 | 247.545703 | 503.60666 | 442.924788 | 487.474875 |
| -4.472917 | 7.72E-06 | 2.70E-05 | 34.4268859 | 177.791282 | 149.559019 | 174.818576 |
| -4.4728352 | 7.72E-06 | 2.70E-05 | 43.8532952 | 108.244312 | 133.452663 | 117.666349 |
| -4.472707 | 7.72E-06 | 2.70E-05 | 71.722679 | 103.914539 | 62.1245158 | 72.2807574 |
| -4.4727031 | 7.72E-06 | 2.70E-05 | 47.5418901 | 126.916455 | 88.5849576 | 124.390141 |
| -4.4724271 | 7.73E-06 | 2.70E-05 | 67.2143963 | 94.7137727 | 58.6731538 | 89.0902358 |
| -4.4723072 | 7.74E-06 | 2.70E-05 | 48.7714217 | 115.821413 | 138.054479 | 85.7283401 |
| -4.4715834 | 7.76E-06 | 2.71E-05 | 109.83816 | 78.477126 | 63.2749697 | 66.6775979 |
| -4.4713764 | 7.77E-06 | 2.72E-05 | 40.1647002 | 200.251977 | 120.79767 | 134.475828 |
| -4.4713863 | 7.77E-06 | 2.72E-05 | 95.0837801 | 69.0057487 | 323.277573 | 90.2108677 |
| -4.4708295 | 7.79E-06 | 2.72E-05 | 48.3615778 | 109.597366 | 96.6381356 | 97.4949751 |
| -4.4706845 | 7.80E-06 | 2.72E-05 | 57.7879871 | 66.5702517 | 125.399485 | 117.106033 |

| | | | | | | |
|---|---|---|---|---|---|---|
| -4.4706446 | 7.80E-06 | 2.72E-05 | 44.263139 | 133.952336 | 98.9390436 | 135.036144 |
| 4.47043976 | 7.81E-06 | 2.73E-05 | 2.86890716 | 4.87099402 | 1.150454 | 2.80157974 |
| -4.4700473 | 7.82E-06 | 2.73E-05 | 97.5428434 | 60.3462037 | 74.7795097 | 178.180472 |
| -4.4699672 | 7.82E-06 | 2.73E-05 | 35.6564176 | 212.429462 | 130.001301 | 172.577312 |
| -4.4699816 | 7.82E-06 | 2.73E-05 | 47.1320462 | 101.749653 | 101.239952 | 109.821926 |
| -4.4699335 | 7.82E-06 | 2.73E-05 | 97.1329995 | 122.857294 | 44.8677058 | 173.137628 |
| -4.4697609 | 7.83E-06 | 2.73E-05 | 47.5418901 | 117.174467 | 95.4876816 | 103.098135 |
| -4.4693903 | 7.84E-06 | 2.74E-05 | 44.6729829 | 99.8553775 | 139.204933 | 119.907613 |
| -4.4690351 | 7.86E-06 | 2.74E-05 | 48.3615778 | 102.290875 | 132.302209 | 91.3314996 |
| -4.4688424 | 7.86E-06 | 2.74E-05 | 45.0828268 | 102.832096 | 254.250333 | 135.59646 |
| -4.4687407 | 7.87E-06 | 2.74E-05 | 45.9025146 | 95.525605 | 125.399485 | 117.666349 |
| -4.4680895 | 7.89E-06 | 2.75E-05 | 61.476582 | 94.9843835 | 64.4254237 | 84.0473923 |
| -4.4680385 | 7.89E-06 | 2.75E-05 | 186.069121 | 290.365366 | 411.86253 | 300.329348 |
| -4.467902 | 7.90E-06 | 2.75E-05 | 71.722679 | 74.9591858 | 111.594038 | 61.0744384 |
| -4.4676042 | 7.91E-06 | 2.76E-05 | 95.493624 | 80.6420122 | 52.9208838 | 142.320251 |
| -4.4668326 | 7.94E-06 | 2.77E-05 | 215.168037 | 78.477126 | 222.037621 | 107.020346 |
| -4.46678 | 7.94E-06 | 2.77E-05 | 54.9190799 | 197.004647 | 69.0272397 | 162.491625 |
| -4.4662793 | 7.96E-06 | 2.77E-05 | 88.1164342 | 92.2782757 | 56.3722458 | 70.0394936 |
| -4.4657974 | 7.98E-06 | 2.78E-05 | 80.3294005 | 95.2549943 | 57.5226998 | 70.5998095 |
| -4.4657969 | 7.98E-06 | 2.78E-05 | 50.8206411 | 80.6420122 | 103.54086 | 115.425085 |
| -4.4650249 | 8.01E-06 | 2.79E-05 | 42.6237635 | 118.25691 | 116.195854 | 126.631404 |
| -4.4647742 | 8.02E-06 | 2.79E-05 | 54.509236 | 92.5488865 | 85.1335957 | 76.763285 |
| -4.4644113 | 8.03E-06 | 2.80E-05 | 101.231438 | 297.942468 | 261.153057 | 305.372192 |
| -4.4643378 | 8.03E-06 | 2.80E-05 | 40.5745441 | 165.613797 | 103.54086 | 224.126379 |
| -4.464067 | 8.04E-06 | 2.80E-05 | 42.2139196 | 131.787449 | 118.496762 | 117.106033 |
| 4.46303868 | 8.08E-06 | 2.81E-05 | 0 | 0 | 0 | 0 |
| -4.4629738 | 8.08E-06 | 2.81E-05 | 44.6729829 | 108.244312 | 115.0454 | 109.821926 |
| -4.462918 | 8.09E-06 | 2.82E-05 | 63.5258014 | 139.635162 | 102.390406 | 120.467929 |
| -4.4624402 | 8.10E-06 | 2.82E-05 | 85.6573709 | 89.0309463 | 86.2840497 | 52.1093832 |
| -4.4624534 | 8.10E-06 | 2.82E-05 | 63.1159575 | 78.477126 | 65.5758777 | 74.5220212 |
| -4.4621405 | 8.11E-06 | 2.82E-05 | 175.413181 | 148.294707 | 226.639437 | 32.498325 |
| -4.461477 | 8.14E-06 | 2.83E-05 | 75.00143 | 99.0435452 | 79.3813257 | 56.0315949 |
| -4.4613989 | 8.14E-06 | 2.83E-05 | 120.494101 | 81.4538445 | 60.9740618 | 68.3585457 |
| -4.461233 | 8.15E-06 | 2.84E-05 | 102.46097 | 243.27909 | 266.905327 | 266.710392 |
| -4.4611283 | 8.15E-06 | 2.84E-05 | 53.6895483 | 71.4412457 | 161.063559 | 131.674248 |
| -4.4610627 | 8.16E-06 | 2.84E-05 | 43.4434513 | 111.762252 | 109.29313 | 136.717091 |
| -4.4609291 | 8.16E-06 | 2.84E-05 | 102.870814 | 193.486707 | 40.2658898 | 156.32815 |
| -4.4607334 | 8.17E-06 | 2.84E-05 | 44.6729829 | 101.749653 | 112.744492 | 123.269509 |
| -4.4601313 | 8.19E-06 | 2.85E-05 | 116.395662 | 198.628312 | 242.745793 | 160.810677 |
| -4.4601043 | 8.19E-06 | 2.85E-05 | 47.1320462 | 86.5954493 | 121.948124 | 129.432984 |
| -4.4600655 | 8.19E-06 | 2.85E-05 | 38.1154808 | 166.425629 | 131.151755 | 134.475828 |
| 4.45992905 | 8.20E-06 | 2.85E-05 | 6.55750208 | 1.89427545 | 3.45136199 | 4.48252759 |
| -4.4599261 | 8.20E-06 | 2.85E-05 | 82.3786199 | 70.6294134 | 153.010381 | 64.4363341 |

| | | | | | | |
|---|---|---|---|---|---|---|
| -4.4594943 | 8.22E-06 | 2.86E-05 | 49.1812656 | 107.161869 | 87.4345036 | 97.4949751 |
| -4.4594445 | 8.22E-06 | 2.86E-05 | 129.100822 | 100.937821 | 50.6199758 | 79.5648647 |
| -4.4594373 | 8.22E-06 | 2.86E-05 | 40.1647002 | 145.047378 | 199.028541 | 109.26161 |
| -4.4592739 | 8.22E-06 | 2.86E-05 | 49.1812656 | 115.280192 | 199.028541 | 89.6505518 |
| -4.4589938 | 8.23E-06 | 2.86E-05 | 33.6071982 | 141.258827 | 202.479903 | 173.697944 |
| -4.4590088 | 8.23E-06 | 2.86E-05 | 72.1325229 | 265.469174 | 48.3190678 | 275.675447 |
| -4.4584754 | 8.25E-06 | 2.87E-05 | 35.6564176 | 267.36345 | 134.603117 | 155.207518 |
| -4.4584197 | 8.26E-06 | 2.87E-05 | 53.2797044 | 85.5130062 | 83.9831417 | 82.3664444 |
| -4.4583681 | 8.26E-06 | 2.87E-05 | 43.0336074 | 126.104623 | 123.098578 | 105.899714 |
| 4.45809848 | 8.27E-06 | 2.87E-05 | 36.8859492 | 41.67406 | 31.0622579 | 33.058641 |
| -4.4578751 | 8.28E-06 | 2.87E-05 | 69.2636157 | 174.002731 | 100.089498 | 61.6347543 |
| -4.4577645 | 8.28E-06 | 2.88E-05 | 38.1154808 | 146.941653 | 167.966283 | 121.028245 |
| -4.4571678 | 8.30E-06 | 2.88E-05 | 37.705637 | 228.936719 | 118.496762 | 155.767834 |
| -4.4567737 | 8.32E-06 | 2.89E-05 | 47.951734 | 93.6313296 | 104.691314 | 96.9346591 |
| -4.4566845 | 8.32E-06 | 2.89E-05 | 52.8698605 | 144.776767 | 79.3813257 | 97.4949751 |
| -4.4566062 | 8.33E-06 | 2.89E-05 | 65.5750208 | 137.740887 | 56.3722458 | 126.631404 |
| -4.456501 | 8.33E-06 | 2.89E-05 | 50.4107972 | 84.4305631 | 95.4876816 | 94.6933953 |
| -4.4560811 | 8.35E-06 | 2.90E-05 | 245.906328 | 528.502852 | 320.976665 | 423.038541 |
| -4.4554564 | 8.37E-06 | 2.90E-05 | 25.4103206 | 275.481773 | 240.444885 | 184.904263 |
| -4.4553903 | 8.37E-06 | 2.91E-05 | 46.3123584 | 88.7603356 | 131.151755 | 118.226665 |
| 4.455225 | 8.38E-06 | 2.91E-05 | 67.2143963 | 55.2045989 | 48.3190678 | 23.5332698 |
| -4.4551901 | 8.38E-06 | 2.91E-05 | 113.936599 | 113.385916 | 123.098578 | 42.5840121 |
| -4.455119 | 8.38E-06 | 2.91E-05 | 243.037421 | 96.3374374 | 77.0804177 | 112.06319 |
| -4.4546779 | 8.40E-06 | 2.91E-05 | 51.6403289 | 116.903857 | 75.9299637 | 103.65845 |
| -4.4545859 | 8.41E-06 | 2.91E-05 | 32.3776665 | 193.486707 | 150.709473 | 179.86142 |
| -4.4542791 | 8.42E-06 | 2.92E-05 | 45.4926707 | 109.326755 | 105.841768 | 101.977503 |
| -4.454197 | 8.42E-06 | 2.92E-05 | 52.8698605 | 73.6061319 | 100.089498 | 103.098135 |
| -4.4540208 | 8.43E-06 | 2.92E-05 | 42.6237635 | 144.235545 | 116.195854 | 107.580662 |
| -4.4539368 | 8.43E-06 | 2.92E-05 | 43.4434513 | 107.432479 | 111.594038 | 125.510772 |
| -4.4538922 | 8.43E-06 | 2.92E-05 | 47.951734 | 119.609964 | 157.612197 | 84.0473923 |
| -4.4535497 | 8.45E-06 | 2.93E-05 | 45.4926707 | 120.963018 | 100.089498 | 103.098135 |
| -4.4530592 | 8.47E-06 | 2.93E-05 | 68.0340841 | 137.199665 | 126.549939 | 128.872668 |
| -4.4528816 | 8.47E-06 | 2.94E-05 | 37.705637 | 123.127905 | 353.189377 | 161.931309 |
| -4.452718 | 8.48E-06 | 2.94E-05 | 54.509236 | 148.294707 | 72.4786017 | 105.899714 |
| -4.4527355 | 8.48E-06 | 2.94E-05 | 50.0009534 | 112.032863 | 82.8326877 | 95.2537113 |
| -4.4526604 | 8.48E-06 | 2.94E-05 | 52.0501728 | 105.538204 | 83.9831417 | 82.3664444 |
| -4.4518391 | 8.51E-06 | 2.95E-05 | 63.9356453 | 70.6294134 | 81.6822337 | 64.4363341 |
| -4.4511537 | 8.54E-06 | 2.96E-05 | 40.1647002 | 123.939737 | 138.054479 | 121.028245 |
| 4.45094614 | 8.55E-06 | 2.96E-05 | 58.6076748 | 135.846611 | 81.6822337 | 95.8140272 |
| -4.4506485 | 8.56E-06 | 2.96E-05 | 45.4926707 | 127.998899 | 96.6381356 | 105.339398 |
| -4.4498462 | 8.59E-06 | 2.97E-05 | 43.4434513 | 214.864959 | 92.0363196 | 153.52657 |
| -4.4494769 | 8.61E-06 | 2.98E-05 | 42.2139196 | 135.034779 | 178.320369 | 99.1759229 |
| -4.4494474 | 8.61E-06 | 2.98E-05 | 55.3289238 | 79.0183475 | 93.1867736 | 73.4013893 |

| | | | | | | |
|---|---|---|---|---|---|---|
| -4.4492182 | 8.62E-06 | 2.98E-05 | 42.6237635 | 111.22103 | 130.001301 | 110.382242 |
| -4.4491392 | 8.62E-06 | 2.98E-05 | 63.1159575 | 109.867976 | 60.9740618 | 85.7283401 |
| -4.4489504 | 8.63E-06 | 2.99E-05 | 38.5253247 | 132.869893 | 127.700393 | 142.880567 |
| -4.4485079 | 8.65E-06 | 2.99E-05 | 30.3284471 | 180.768 | 155.311289 | 273.994499 |
| -4.4477537 | 8.68E-06 | 3.00E-05 | 133.199261 | 431.894804 | 365.844371 | 362.524419 |
| -4.4472094 | 8.70E-06 | 3.01E-05 | 57.7879871 | 86.8660601 | 92.0363196 | 65.556966 |
| -4.4466112 | 8.72E-06 | 3.02E-05 | 39.7548564 | 124.480958 | 120.79767 | 140.639303 |
| -4.4461493 | 8.74E-06 | 3.02E-05 | 45.9025146 | 152.083258 | 83.9831417 | 139.518671 |
| -4.4459388 | 8.75E-06 | 3.03E-05 | 45.0828268 | 125.834012 | 101.239952 | 100.296555 |
| -4.4454605 | 8.77E-06 | 3.03E-05 | 111.067691 | 59.5343714 | 60.9740618 | 117.106033 |
| -4.4451457 | 8.78E-06 | 3.04E-05 | 117.21535 | 120.692407 | 55.2217918 | 65.556966 |
| -4.4447803 | 8.80E-06 | 3.04E-05 | 45.4926707 | 112.032863 | 102.390406 | 98.055291 |
| -4.4447451 | 8.80E-06 | 3.04E-05 | 309.432129 | 207.558468 | 35.6640739 | 242.616806 |
| -4.4446406 | 8.80E-06 | 3.04E-05 | 955.755928 | 1303.53212 | 701.776937 | 900.427729 |
| -4.4445691 | 8.81E-06 | 3.04E-05 | 54.509236 | 71.4412457 | 93.1867736 | 85.1680242 |
| -4.4444942 | 8.81E-06 | 3.04E-05 | 46.7222023 | 102.020264 | 101.239952 | 94.1330794 |
| -4.4443085 | 8.82E-06 | 3.05E-05 | 61.8864259 | 59.2637606 | 134.603117 | 113.744138 |
| -4.4440509 | 8.83E-06 | 3.05E-05 | 32.7875104 | 167.508072 | 158.762651 | 165.853521 |
| -4.4439804 | 8.83E-06 | 3.05E-05 | 40.1647002 | 127.457677 | 127.700393 | 118.786981 |
| -4.4436022 | 8.85E-06 | 3.05E-05 | 52.0501728 | 77.1240721 | 95.4876816 | 86.848972 |
| -4.4433605 | 8.86E-06 | 3.06E-05 | 98.7723751 | 77.1240721 | 60.9740618 | 63.8760181 |
| -4.4431671 | 8.86E-06 | 3.06E-05 | 46.3123584 | 135.034779 | 82.8326877 | 159.129729 |
| -4.4429526 | 8.87E-06 | 3.06E-05 | 50.0009534 | 120.963018 | 75.9299637 | 112.623506 |
| -4.4428071 | 8.88E-06 | 3.06E-05 | 88.9361219 | 98.5023236 | 66.7263317 | 82.3664444 |
| -4.4426757 | 8.88E-06 | 3.07E-05 | 75.4112739 | 103.102707 | 55.2217918 | 75.0823371 |
| 4.44267222 | 8.88E-06 | 3.07E-05 | 81.5589321 | 46.0038325 | 49.4695218 | 50.9887513 |
| -4.442559 | 8.89E-06 | 3.07E-05 | 53.6895483 | 106.891258 | 118.496762 | 68.9188617 |
| -4.4424927 | 8.89E-06 | 3.07E-05 | 771.326182 | 1304.61457 | 1000.89498 | 963.743432 |
| -4.4420995 | 8.91E-06 | 3.07E-05 | 48.3615778 | 94.7137727 | 88.5849576 | 97.4949751 |
| -4.4419933 | 8.91E-06 | 3.07E-05 | 53.2797044 | 96.8786589 | 78.2308717 | 79.0045488 |
| -4.441443 | 8.94E-06 | 3.08E-05 | 68.0340841 | 93.090108 | 77.0804177 | 57.7125427 |
| -4.4414497 | 8.94E-06 | 3.08E-05 | 47.1320462 | 125.563402 | 83.9831417 | 111.502874 |
| -4.4413898 | 8.94E-06 | 3.08E-05 | 35.2465737 | 142.611881 | 153.010381 | 150.164674 |
| -4.4408562 | 8.96E-06 | 3.09E-05 | 38.1154808 | 129.081342 | 143.806749 | 128.872668 |
| -4.4404712 | 8.98E-06 | 3.09E-05 | 122.953164 | 67.1114732 | 65.5758777 | 71.7204414 |
| -4.4400058 | 9.00E-06 | 3.10E-05 | 48.7714217 | 81.9950661 | 104.691314 | 99.1759229 |
| -4.4399766 | 9.00E-06 | 3.10E-05 | 45.4926707 | 104.18515 | 103.54086 | 98.055291 |
| -4.4399503 | 9.00E-06 | 3.10E-05 | 49.1812656 | 102.290875 | 80.5317797 | 101.417187 |
| -4.4398903 | 9.00E-06 | 3.10E-05 | 20.9020379 | 345.028743 | 209.382627 | 270.632603 |
| -4.4398818 | 9.00E-06 | 3.10E-05 | 61.476582 | 87.9485032 | 64.4254237 | 72.2807574 |
| -4.439422 | 9.02E-06 | 3.11E-05 | 40.1647002 | 126.916455 | 109.29313 | 158.569413 |
| -4.4388446 | 9.04E-06 | 3.12E-05 | 56.9682993 | 97.1492697 | 188.674455 | 76.202969 |
| -4.4387939 | 9.05E-06 | 3.12E-05 | 47.1320462 | 113.115306 | 82.8326877 | 129.9933 |

| | | | | | | |
|---|---|---|---|---|---|---|
| -4.4386915 | 9.05E-06 | 3.12E-05 | 244.676796 | 205.934803 | 159.913105 | 196.110582 |
| -4.4379268 | 9.08E-06 | 3.13E-05 | 63.9356453 | 131.246228 | 78.2308717 | 66.1172819 |
| -4.4379007 | 9.08E-06 | 3.13E-05 | 54.509236 | 91.1958326 | 77.0804177 | 74.5220212 |
| -4.4374488 | 9.10E-06 | 3.13E-05 | 48.3615778 | 86.0542278 | 93.1867736 | 101.977503 |
| -4.4373373 | 9.11E-06 | 3.14E-05 | 52.0501728 | 82.2656769 | 80.5317797 | 85.7283401 |
| -4.4372787 | 9.11E-06 | 3.14E-05 | 44.263139 | 145.047378 | 88.5849576 | 129.432984 |
| -4.437188 | 9.11E-06 | 3.14E-05 | 73.7718984 | 99.3141559 | 108.142676 | 96.3743432 |
| -4.437067 | 9.12E-06 | 3.14E-05 | 42.6237635 | 225.418779 | 112.744492 | 114.304454 |
| -4.4366838 | 9.14E-06 | 3.14E-05 | 59.8372065 | 92.8194972 | 90.8858656 | 61.0744384 |
| -4.4366196 | 9.14E-06 | 3.14E-05 | 641.405672 | 103.914539 | 57.5226998 | 126.631404 |
| -4.4364832 | 9.14E-06 | 3.15E-05 | 51.6403289 | 149.37715 | 71.3281477 | 121.588561 |
| -4.4362984 | 9.15E-06 | 3.15E-05 | 67.6242402 | 90.3840002 | 86.2840497 | 55.4712789 |
| -4.4362194 | 9.16E-06 | 3.15E-05 | 28.2792277 | 158.036695 | 235.843069 | 208.997849 |
| -4.4360827 | 9.16E-06 | 3.15E-05 | 58.197831 | 75.2297966 | 66.7263317 | 88.5299199 |
| -4.4353309 | 9.19E-06 | 3.16E-05 | 200.823501 | 69.5469702 | 67.8767857 | 87.409288 |
| -4.4353039 | 9.19E-06 | 3.16E-05 | 80.3294005 | 85.5130062 | 60.9740618 | 61.0744384 |
| -4.43509 | 9.20E-06 | 3.16E-05 | 33.1973543 | 157.495473 | 176.019461 | 146.242463 |
| -4.434959 | 9.21E-06 | 3.17E-05 | 71.3128351 | 65.4878086 | 65.5758777 | 118.786981 |
| -4.4348785 | 9.21E-06 | 3.17E-05 | 267.628054 | 143.423713 | 26.4604419 | 291.364293 |
| -4.4347655 | 9.22E-06 | 3.17E-05 | 46.3123584 | 105.538204 | 87.4345036 | 134.475828 |
| -4.4345879 | 9.22E-06 | 3.17E-05 | 37.705637 | 152.353869 | 115.0454 | 149.044042 |
| -4.4345565 | 9.23E-06 | 3.17E-05 | 49.1812656 | 87.1366709 | 89.7354116 | 90.2108677 |
| -4.4344574 | 9.23E-06 | 3.17E-05 | 43.8532952 | 107.161869 | 101.239952 | 112.06319 |
| -4.4343432 | 9.24E-06 | 3.17E-05 | 53.6895483 | 95.7962158 | 87.4345036 | 70.5998095 |
| -4.4342307 | 9.24E-06 | 3.17E-05 | 34.8367298 | 205.122971 | 142.656295 | 135.036144 |
| -4.4340819 | 9.25E-06 | 3.18E-05 | 59.0175187 | 70.0881918 | 81.6822337 | 68.3585457 |
| -4.4340984 | 9.25E-06 | 3.18E-05 | 36.0662614 | 114.46836 | 180.621277 | 170.336048 |
| -4.4340492 | 9.25E-06 | 3.18E-05 | 48.7714217 | 157.224863 | 154.160835 | 161.370993 |
| 4.43389741 | 9.25E-06 | 3.18E-05 | 0.40984388 | 0.27061078 | 2.30090799 | 0.56031595 |
| -4.4338094 | 9.26E-06 | 3.18E-05 | 37.705637 | 181.038611 | 143.806749 | 113.183822 |
| -4.4337799 | 9.26E-06 | 3.18E-05 | 57.7879871 | 77.6652936 | 70.1776937 | 72.2807574 |
| -4.4337568 | 9.26E-06 | 3.18E-05 | 58.6076748 | 130.705006 | 103.54086 | 163.051941 |
| -4.433742 | 9.26E-06 | 3.18E-05 | 45.9025146 | 86.0542278 | 111.594038 | 128.312352 |
| -4.4335047 | 9.27E-06 | 3.18E-05 | 50.8206411 | 83.34812 | 95.4876816 | 80.6854966 |
| -4.4334254 | 9.27E-06 | 3.18E-05 | 61.8864259 | 98.5023236 | 165.665375 | 118.786981 |
| -4.4332858 | 9.28E-06 | 3.18E-05 | 45.0828268 | 178.603114 | 85.1335957 | 135.036144 |
| -4.4332935 | 9.28E-06 | 3.18E-05 | 52.0501728 | 96.0668266 | 82.8326877 | 76.763285 |
| -4.4329253 | 9.30E-06 | 3.19E-05 | 43.8532952 | 100.125988 | 108.142676 | 110.942558 |
| -4.4327109 | 9.31E-06 | 3.19E-05 | 45.9025146 | 107.70309 | 98.9390436 | 93.0124475 |
| -4.4321894 | 9.33E-06 | 3.20E-05 | 72.9522106 | 95.525605 | 55.2217918 | 73.4013893 |
| -4.4319409 | 9.34E-06 | 3.20E-05 | 49.1812656 | 75.5004074 | 242.745793 | 157.448782 |
| 4.43174206 | 9.35E-06 | 3.21E-05 | 72.5423667 | 29.7671857 | 43.7172518 | 20.7316901 |
| -4.4317159 | 9.35E-06 | 3.21E-05 | 52.4600166 | 81.1832337 | 77.0804177 | 86.2886561 |

| | | | | | | |
|---|---|---|---|---|---|---|
| -4.4310831 | 9.38E-06 | 3.21E-05 | 77.4604933 | 73.6061319 | 120.79767 | 56.0315949 |
| -4.4306413 | 9.40E-06 | 3.22E-05 | 45.0828268 | 101.208431 | 100.089498 | 100.856871 |
| -4.4306057 | 9.40E-06 | 3.22E-05 | 96.7231557 | 96.3374374 | 44.8677058 | 103.65845 |
| -4.4304376 | 9.40E-06 | 3.22E-05 | 53.2797044 | 131.246228 | 71.3281477 | 96.3743432 |
| -4.4289005 | 9.47E-06 | 3.25E-05 | 143.855202 | 79.2889583 | 72.4786017 | 59.3934906 |
| -4.4287552 | 9.48E-06 | 3.25E-05 | 88.9361219 | 120.963018 | 43.7172518 | 162.491625 |
| 4.42871626 | 9.48E-06 | 3.25E-05 | 8.1968776 | 1.89427545 | 0 | 1.1206319 |
| -4.4286155 | 9.48E-06 | 3.25E-05 | 53.2797044 | 68.1939163 | 143.806749 | 113.183822 |
| -4.4285777 | 9.49E-06 | 3.25E-05 | 58.6076748 | 95.2549943 | 88.5849576 | 61.6347543 |
| -4.4283335 | 9.50E-06 | 3.25E-05 | 55.7387677 | 66.0290301 | 95.4876816 | 85.1680242 |
| -4.4282489 | 9.50E-06 | 3.25E-05 | 43.8532952 | 87.4072817 | 142.656295 | 129.432984 |
| -4.4281709 | 9.50E-06 | 3.25E-05 | 28.6890716 | 163.990132 | 209.382627 | 191.628054 |
| -4.4274178 | 9.54E-06 | 3.27E-05 | 22.1315695 | 202.146252 | 538.41247 | 215.161324 |
| -4.4266718 | 9.57E-06 | 3.28E-05 | 31.1481349 | 143.153102 | 218.586259 | 172.016996 |
| -4.4261389 | 9.59E-06 | 3.28E-05 | 26.6398522 | 217.841677 | 202.479903 | 183.223315 |
| -4.4261398 | 9.59E-06 | 3.28E-05 | 102.46097 | 102.290875 | 75.9299637 | 49.3078035 |
| -4.4261822 | 9.59E-06 | 3.28E-05 | 46.7222023 | 112.844695 | 90.8858656 | 91.3314996 |
| -4.4260499 | 9.60E-06 | 3.28E-05 | 53.6895483 | 71.9824673 | 79.3813257 | 91.3314996 |
| -4.4259481 | 9.60E-06 | 3.29E-05 | 59.4273626 | 67.9233056 | 87.4345036 | 67.7982298 |
| -4.4255991 | 9.62E-06 | 3.29E-05 | 46.7222023 | 98.2317128 | 87.4345036 | 101.417187 |
| -4.4255522 | 9.62E-06 | 3.29E-05 | 114.346443 | 64.1347547 | 150.709473 | 56.0315949 |
| -4.4254574 | 9.62E-06 | 3.29E-05 | 50.8206411 | 213.782516 | 326.728935 | 273.994499 |
| -4.4253659 | 9.63E-06 | 3.29E-05 | 66.3947085 | 121.50424 | 173.718553 | 178.180472 |
| -4.4252838 | 9.63E-06 | 3.29E-05 | 60.6568942 | 86.5954493 | 63.2749697 | 71.7204414 |
| -4.4249053 | 9.65E-06 | 3.30E-05 | 93.0345607 | 67.382084 | 109.29313 | 53.7903311 |
| -4.4247833 | 9.65E-06 | 3.30E-05 | 39.3450125 | 127.187066 | 156.461743 | 107.580662 |
| -4.4242252 | 9.68E-06 | 3.31E-05 | 47.1320462 | 80.3714014 | 108.142676 | 124.950457 |
| -4.4241743 | 9.68E-06 | 3.31E-05 | 63.9356453 | 125.834012 | 54.0713378 | 127.19172 |
| -4.4236854 | 9.70E-06 | 3.32E-05 | 365.170897 | 105.538204 | 44.8677058 | 130.553616 |
| -4.4231215 | 9.73E-06 | 3.33E-05 | 32.7875104 | 131.516839 | 165.665375 | 203.955005 |
| -4.4230658 | 9.73E-06 | 3.33E-05 | 35.2465737 | 143.694324 | 131.151755 | 156.32815 |
| -4.4230405 | 9.73E-06 | 3.33E-05 | 28.2792277 | 149.37715 | 234.692615 | 209.558165 |
| -4.4223928 | 9.76E-06 | 3.34E-05 | 206.561315 | 77.1240721 | 63.2749697 | 82.9267604 |
| -4.4222601 | 9.77E-06 | 3.34E-05 | 40.5745441 | 133.952336 | 134.603117 | 99.7362389 |
| -4.4215135 | 9.80E-06 | 3.35E-05 | 59.8372065 | 185.097773 | 64.4254237 | 108.140978 |
| -4.4213349 | 9.81E-06 | 3.35E-05 | 59.8372065 | 73.8767427 | 63.2749697 | 76.202969 |
| -4.4210462 | 9.82E-06 | 3.35E-05 | 32.3776665 | 220.006563 | 142.656295 | 150.72499 |
| -4.4205815 | 9.84E-06 | 3.36E-05 | 37.705637 | 130.975617 | 139.204933 | 119.347297 |
| -4.420458 | 9.85E-06 | 3.36E-05 | 78.6900249 | 59.8049822 | 65.5758777 | 68.9188617 |
| -4.4204209 | 9.85E-06 | 3.36E-05 | 131.559885 | 139.635162 | 46.0181598 | 77.3236009 |
| -4.4193364 | 9.90E-06 | 3.38E-05 | 47.5418901 | 111.491641 | 78.2308717 | 108.140978 |
| -4.4191701 | 9.91E-06 | 3.38E-05 | 56.9682993 | 83.34812 | 67.8767857 | 71.7204414 |
| -4.4190568 | 9.91E-06 | 3.38E-05 | 85.247527 | 139.09394 | 58.6731538 | 66.6775979 |

| | | | | | | |
|---|---|---|---|---|---|---|
| -4.418897 | 9.92E-06 | 3.39E-05 | 72.9522106 | 187.53327 | 56.3722458 | 97.4949751 |
| -4.4187605 | 9.93E-06 | 3.39E-05 | 73.7718984 | 117.715689 | 93.1867736 | 134.475828 |
| -4.4184575 | 9.94E-06 | 3.39E-05 | 47.5418901 | 126.645845 | 78.2308717 | 107.020346 |
| -4.418341 | 9.95E-06 | 3.39E-05 | 93.4444046 | 129.081342 | 67.8767857 | 96.3743432 |
| -4.4176706 | 9.98E-06 | 3.40E-05 | 52.0501728 | 77.3946828 | 83.9831417 | 77.3236009 |
| -4.4175612 | 9.98E-06 | 3.41E-05 | 59.0175187 | 72.253078 | 65.5758777 | 72.8410733 |
| -4.4173117 | 9.99E-06 | 3.41E-05 | 134.428793 | 334.745534 | 306.020763 | 309.294404 |
| -4.4170725 | 1.00E-05 | 3.41E-05 | 36.0662614 | 111.491641 | 155.311289 | 182.102683 |
| -4.4170351 | 1.00E-05 | 3.41E-05 | 54.0993922 | 72.7942996 | 155.311289 | 93.0124475 |
| -4.4165592 | 1.00E-05 | 3.42E-05 | 40.984388 | 214.594348 | 97.7885896 | 133.915512 |
| -4.4164478 | 1.00E-05 | 3.42E-05 | 52.4600166 | 76.0416289 | 75.9299637 | 86.2886561 |
| -4.4163116 | 1.00E-05 | 3.42E-05 | 68.4439279 | 117.715689 | 95.4876816 | 52.6696992 |
| -4.4162609 | 1.00E-05 | 3.42E-05 | 110.657848 | 83.0775092 | 44.8677058 | 136.717091 |
| -4.4161262 | 1.00E-05 | 3.42E-05 | 55.3289238 | 68.4645271 | 77.0804177 | 78.4442328 |
| -4.416036 | 1.01E-05 | 3.43E-05 | 41.3942319 | 82.8068984 | 301.418947 | 205.075637 |
| -4.4156955 | 1.01E-05 | 3.43E-05 | 32.7875104 | 119.068743 | 202.479903 | 193.869318 |
| -4.4154467 | 1.01E-05 | 3.43E-05 | 71.3128351 | 61.1580361 | 104.691314 | 63.8760181 |
| -4.4153786 | 1.01E-05 | 3.44E-05 | 39.7548564 | 123.669126 | 104.691314 | 137.837723 |
| -4.4152062 | 1.01E-05 | 3.44E-05 | 34.017042 | 153.706923 | 146.107657 | 139.518671 |
| -4.4149837 | 1.01E-05 | 3.44E-05 | 54.509236 | 71.4412457 | 105.841768 | 75.6426531 |
| -4.4149535 | 1.01E-05 | 3.44E-05 | 45.0828268 | 97.9611021 | 134.603117 | 90.2108677 |
| -4.41482 | 1.01E-05 | 3.44E-05 | 48.7714217 | 136.658443 | 72.4786017 | 120.467929 |
| -4.4147674 | 1.01E-05 | 3.44E-05 | 110.248004 | 52.7691019 | 97.7885896 | 67.2379138 |
| -4.4142508 | 1.01E-05 | 3.45E-05 | 42.6237635 | 97.1492697 | 106.992222 | 121.028245 |
| -4.4142583 | 1.01E-05 | 3.45E-05 | 54.509236 | 83.0775092 | 73.6290557 | 71.1601255 |
| -4.4141718 | 1.01E-05 | 3.45E-05 | 152.461923 | 461.120768 | 430.269794 | 405.668747 |
| -4.4141241 | 1.01E-05 | 3.45E-05 | 43.0336074 | 106.620647 | 101.239952 | 104.218766 |
| -4.4140409 | 1.01E-05 | 3.45E-05 | 95.0837801 | 390.491354 | 31.0622579 | 336.189569 |
| -4.41384 | 1.02E-05 | 3.46E-05 | 32.3776665 | 209.993965 | 141.505841 | 147.363094 |
| -4.413787 | 1.02E-05 | 3.46E-05 | 37.2957931 | 127.998899 | 138.054479 | 121.588561 |
| 4.41367336 | 1.02E-05 | 3.46E-05 | 55.3289238 | 44.3801678 | 49.4695218 | 40.3427483 |
| -4.413513 | 1.02E-05 | 3.46E-05 | 50.0009534 | 79.0183475 | 90.8858656 | 81.8061285 |
| -4.4132798 | 1.02E-05 | 3.46E-05 | 50.0009534 | 81.7244553 | 90.8858656 | 79.0045488 |
| -4.4132484 | 1.02E-05 | 3.46E-05 | 77.0506494 | 93.3607188 | 163.364467 | 52.1093832 |
| -4.4131542 | 1.02E-05 | 3.46E-05 | 86.4770587 | 70.3588026 | 123.098578 | 53.2300151 |
| -4.4131521 | 1.02E-05 | 3.46E-05 | 59.4273626 | 70.6294134 | 65.5758777 | 71.1601255 |
| -4.4124912 | 1.02E-05 | 3.47E-05 | 37.2957931 | 176.97945 | 127.700393 | 115.425085 |
| -4.4122843 | 1.02E-05 | 3.48E-05 | 43.0336074 | 115.821413 | 89.7354116 | 129.9933 |
| -4.4122711 | 1.02E-05 | 3.48E-05 | 50.4107972 | 116.633246 | 71.3281477 | 98.615607 |
| -4.4121229 | 1.02E-05 | 3.48E-05 | 37.2957931 | 115.009581 | 158.762651 | 128.312352 |
| -4.4120758 | 1.02E-05 | 3.48E-05 | 38.1154808 | 127.728288 | 194.426725 | 110.382242 |
| -4.4120569 | 1.02E-05 | 3.48E-05 | 238.119294 | 306.060791 | 320.976665 | 330.026094 |
| -4.4118876 | 1.02E-05 | 3.48E-05 | 47.951734 | 99.0435452 | 81.6822337 | 89.0902358 |

| | | | | | | |
|---|---|---|---|---|---|---|
| -4.411553 | 1.03E-05 | 3.49E-05 | 44.263139 | 151.000815 | 96.6381356 | 97.4949751 |
| -4.4111679 | 1.03E-05 | 3.49E-05 | 63.1159575 | 68.7351379 | 60.9740618 | 93.5727634 |
| 4.41112478 | 1.03E-05 | 3.49E-05 | 2.0492194 | 3.24732935 | 0 | 1.68094785 |
| -4.410393 | 1.03E-05 | 3.50E-05 | 76.6408055 | 61.1580361 | 77.0804177 | 176.499524 |
| -4.4102554 | 1.03E-05 | 3.51E-05 | 639.766297 | 91.1958326 | 65.5758777 | 114.304454 |
| -4.4100861 | 1.03E-05 | 3.51E-05 | 36.8859492 | 139.905773 | 136.904025 | 117.106033 |
| -4.4099141 | 1.03E-05 | 3.51E-05 | 40.5745441 | 120.151186 | 106.992222 | 112.623506 |
| -4.4095517 | 1.04E-05 | 3.52E-05 | 34.8367298 | 103.102707 | 246.197155 | 194.98995 |
| -4.4095031 | 1.04E-05 | 3.52E-05 | 81.968776 | 56.2870421 | 63.2749697 | 98.055291 |
| -4.4094228 | 1.04E-05 | 3.52E-05 | 108.198784 | 196.463426 | 37.9649818 | 126.631404 |
| -4.409249 | 1.04E-05 | 3.52E-05 | 49.1812656 | 108.514922 | 75.9299637 | 90.2108677 |
| -4.4091041 | 1.04E-05 | 3.52E-05 | 54.509236 | 67.1114732 | 80.5317797 | 101.417187 |
| -4.4086406 | 1.04E-05 | 3.53E-05 | 46.7222023 | 85.2423954 | 89.7354116 | 113.744138 |
| -4.4085795 | 1.04E-05 | 3.53E-05 | 159.839113 | 347.734851 | 329.029843 | 342.353045 |
| -4.4085605 | 1.04E-05 | 3.53E-05 | 56.9682993 | 81.1832337 | 75.9299637 | 63.8760181 |
| -4.4085114 | 1.04E-05 | 3.53E-05 | 52.4600166 | 129.622563 | 131.151755 | 64.4363341 |
| -4.4077389 | 1.04E-05 | 3.54E-05 | 53.2797044 | 240.843593 | 64.4254237 | 159.690045 |
| -4.407448 | 1.05E-05 | 3.55E-05 | 43.4434513 | 134.493557 | 104.691314 | 91.3314996 |
| -4.4065881 | 1.05E-05 | 3.56E-05 | 59.4273626 | 178.061893 | 207.081719 | 154.647202 |
| -4.406324 | 1.05E-05 | 3.57E-05 | 53.2797044 | 82.2656769 | 90.8858656 | 67.2379138 |
| -4.4061329 | 1.05E-05 | 3.57E-05 | 49.5911095 | 71.9824673 | 123.098578 | 99.7362389 |
| -4.4056004 | 1.05E-05 | 3.58E-05 | 40.1647002 | 122.586683 | 195.577179 | 100.856871 |
| -4.4056151 | 1.05E-05 | 3.58E-05 | 38.1154808 | 164.801964 | 184.072639 | 95.2537113 |
| -4.4054297 | 1.06E-05 | 3.58E-05 | 40.984388 | 114.197749 | 150.709473 | 97.4949751 |
| -4.4048136 | 1.06E-05 | 3.59E-05 | 29.0989155 | 215.40618 | 402.658898 | 305.372192 |
| -4.4048331 | 1.06E-05 | 3.59E-05 | 34.4268859 | 118.25691 | 164.514921 | 159.129729 |
| -4.4048082 | 1.06E-05 | 3.59E-05 | 59.4273626 | 100.937821 | 64.4254237 | 70.5998095 |
| -4.4045195 | 1.06E-05 | 3.59E-05 | 32.3776665 | 206.476024 | 159.913105 | 127.752036 |
| -4.4045366 | 1.06E-05 | 3.59E-05 | 200.003813 | 539.597894 | 514.252936 | 522.214464 |
| -4.4044246 | 1.06E-05 | 3.59E-05 | 63.9356453 | 72.5236888 | 62.1245158 | 64.4363341 |
| -4.404319 | 1.06E-05 | 3.59E-05 | 43.4434513 | 117.445078 | 121.948124 | 85.7283401 |
| -4.403289 | 1.07E-05 | 3.61E-05 | 52.0501728 | 100.396599 | 67.8767857 | 87.9696039 |
| -4.4028707 | 1.07E-05 | 3.62E-05 | 41.8040758 | 109.597366 | 112.744492 | 98.055291 |
| -4.4025907 | 1.07E-05 | 3.62E-05 | 43.0336074 | 96.6080482 | 104.691314 | 102.537819 |
| -4.4016481 | 1.07E-05 | 3.64E-05 | 28.2792277 | 120.151186 | 240.444885 | 449.933707 |
| -4.4014866 | 1.08E-05 | 3.64E-05 | 51.230485 | 83.0775092 | 71.3281477 | 86.848972 |
| -4.4012659 | 1.08E-05 | 3.64E-05 | 63.1159575 | 196.463426 | 128.850847 | 196.670898 |
| -4.4006732 | 1.08E-05 | 3.65E-05 | 64.3454891 | 100.396599 | 52.9208838 | 90.7711837 |
| -4.4006016 | 1.08E-05 | 3.65E-05 | 75.00143 | 78.7477367 | 55.2217918 | 64.99665 |
| -4.400605 | 1.08E-05 | 3.65E-05 | 50.0009534 | 85.5130062 | 75.9299637 | 82.3664444 |
| -4.4005855 | 1.08E-05 | 3.65E-05 | 115.985818 | 252.479857 | 226.639437 | 220.764484 |
| -4.4004132 | 1.08E-05 | 3.66E-05 | 50.4107972 | 119.339354 | 66.7263317 | 124.390141 |
| -4.4002941 | 1.08E-05 | 3.66E-05 | 42.6237635 | 227.313054 | 128.850847 | 91.3314996 |

| | | | | | | |
|---|---|---|---|---|---|---|
| -4.4003087 | 1.08E-05 | 3.66E-05 | 732.391013 | 1104.6332 | 957.177724 | 1021.45597 |
| -4.3997674 | 1.08E-05 | 3.66E-05 | 153.281611 | 526.067355 | 416.464346 | 501.482774 |
| -4.3997064 | 1.08E-05 | 3.67E-05 | 77.4604933 | 43.0271139 | 182.922185 | 190.507423 |
| -4.3996235 | 1.08E-05 | 3.67E-05 | 41.8040758 | 126.375234 | 104.691314 | 96.9346591 |
| -4.3996227 | 1.08E-05 | 3.67E-05 | 54.0993922 | 76.0416289 | 67.8767857 | 78.4442328 |
| -4.3995456 | 1.08E-05 | 3.67E-05 | 27.8693838 | 121.233629 | 280.710775 | 266.710392 |
| -4.3994361 | 1.09E-05 | 3.67E-05 | 29.9186032 | 130.705006 | 200.178995 | 203.394689 |
| -4.3991861 | 1.09E-05 | 3.67E-05 | 56.1486116 | 180.768 | 194.426725 | 191.067738 |
| -4.3990143 | 1.09E-05 | 3.67E-05 | 56.5584554 | 115.821413 | 110.443584 | 111.502874 |
| -4.3989818 | 1.09E-05 | 3.68E-05 | 54.0993922 | 70.3588026 | 82.8326877 | 70.5998095 |
| 4.398763 | 1.09E-05 | 3.68E-05 | 22.1315695 | 14.3423713 | 31.0622579 | 30.8173772 |
| -4.3986884 | 1.09E-05 | 3.68E-05 | 56.9682993 | 94.9843835 | 59.8236078 | 85.1680242 |
| -4.398499 | 1.09E-05 | 3.68E-05 | 148.773328 | 334.474923 | 386.552542 | 390.540216 |
| -4.398379 | 1.09E-05 | 3.68E-05 | 64.3454891 | 175.355785 | 50.6199758 | 152.966254 |
| -4.3982752 | 1.09E-05 | 3.69E-05 | 68.8537718 | 66.8408624 | 59.8236078 | 64.99665 |
| -4.397958 | 1.09E-05 | 3.69E-05 | 101.641282 | 59.5343714 | 60.9740618 | 74.5220212 |
| -4.3976789 | 1.09E-05 | 3.69E-05 | 65.5750208 | 94.4431619 | 81.6822337 | 103.098135 |
| -4.3976058 | 1.09E-05 | 3.69E-05 | 47.1320462 | 95.7962158 | 77.0804177 | 107.020346 |
| -4.397606 | 1.09E-05 | 3.69E-05 | 51.230485 | 70.0881918 | 88.5849576 | 82.9267604 |
| -4.3974772 | 1.10E-05 | 3.70E-05 | 95.9034679 | 415.658157 | 276.108959 | 312.656299 |
| -4.3973124 | 1.10E-05 | 3.70E-05 | 36.4761053 | 134.493557 | 116.195854 | 136.156776 |
| -4.39729 | 1.10E-05 | 3.70E-05 | 51.230485 | 79.2889583 | 83.9831417 | 72.2807574 |
| -4.3972635 | 1.10E-05 | 3.70E-05 | 41.3942319 | 139.09394 | 94.3372276 | 110.942558 |
| -4.3971594 | 1.10E-05 | 3.70E-05 | 84.4278393 | 66.2996409 | 60.9740618 | 61.6347543 |
| -4.3968229 | 1.10E-05 | 3.71E-05 | 52.8698605 | 109.056144 | 135.753571 | 63.8760181 |
| -4.3964235 | 1.10E-05 | 3.71E-05 | 415.991538 | 105.808815 | 67.8767857 | 76.763285 |
| -4.3960521 | 1.10E-05 | 3.72E-05 | 39.7548564 | 156.14242 | 151.859927 | 89.6505518 |
| -4.3959793 | 1.10E-05 | 3.72E-05 | 41.8040758 | 84.4305631 | 172.568099 | 138.958355 |
| -4.3949257 | 1.11E-05 | 3.74E-05 | 36.0662614 | 181.850444 | 110.443584 | 136.156776 |
| -4.3949328 | 1.11E-05 | 3.74E-05 | 35.2465737 | 205.934803 | 97.7885896 | 201.153426 |
| -4.3944597 | 1.11E-05 | 3.74E-05 | 63.5258014 | 80.3714014 | 62.1245158 | 62.1950703 |
| -4.3943202 | 1.11E-05 | 3.75E-05 | 38.9351686 | 101.208431 | 121.948124 | 139.518671 |
| -4.394206 | 1.11E-05 | 3.75E-05 | 30.738291 | 209.182132 | 134.603117 | 162.491625 |
| -4.3940073 | 1.11E-05 | 3.75E-05 | 329.104636 | 252.479857 | 31.0622579 | 100.856871 |
| -4.3938949 | 1.11E-05 | 3.75E-05 | 46.7222023 | 75.2297966 | 108.142676 | 105.339398 |
| -4.3937233 | 1.11E-05 | 3.75E-05 | 71.722679 | 109.326755 | 47.1686138 | 122.148877 |
| -4.3936817 | 1.11E-05 | 3.76E-05 | 47.1320462 | 168.049294 | 157.612197 | 69.4791776 |
| -4.3928548 | 1.12E-05 | 3.77E-05 | 88.5262781 | 146.400432 | 169.116737 | 178.180472 |
| -4.3924751 | 1.12E-05 | 3.78E-05 | 22.9512573 | 214.864959 | 205.931265 | 225.247011 |
| -4.3922178 | 1.12E-05 | 3.78E-05 | 330.744011 | 644.865487 | 610.891072 | 625.312599 |
| -4.3916636 | 1.12E-05 | 3.79E-05 | 114.756286 | 195.922204 | 185.223093 | 207.316901 |
| -4.391282 | 1.13E-05 | 3.79E-05 | 64.3454891 | 55.4752097 | 73.6290557 | 80.1251807 |
| -4.3911739 | 1.13E-05 | 3.80E-05 | 311.071505 | 508.748265 | 838.680963 | 570.401636 |

| | | | | | | |
|---|---|---|---|---|---|---|
| -4.3908255 | 1.13E-05 | 3.80E-05 | 79.0998688 | 51.416048 | 72.4786017 | 110.942558 |
| -4.3905139 | 1.13E-05 | 3.81E-05 | 1818.88714 | 4445.05266 | 3574.46056 | 3992.25113 |
| -4.3904782 | 1.13E-05 | 3.81E-05 | 40.984388 | 145.85921 | 155.311289 | 84.6077082 |
| -4.3904658 | 1.13E-05 | 3.81E-05 | 61.476582 | 63.5935331 | 75.9299637 | 62.1950703 |
| -4.3903386 | 1.13E-05 | 3.81E-05 | 57.7879871 | 73.0649104 | 64.4254237 | 67.7982298 |
| -4.3903303 | 1.13E-05 | 3.81E-05 | 54.0993922 | 69.2763595 | 70.1776937 | 78.4442328 |
| -4.3901059 | 1.13E-05 | 3.81E-05 | 36.8859492 | 125.292791 | 111.594038 | 142.320251 |
| -4.3900629 | 1.13E-05 | 3.81E-05 | 43.4434513 | 95.2549943 | 92.0363196 | 105.899714 |
| -4.3898094 | 1.13E-05 | 3.82E-05 | 53.6895483 | 80.1007906 | 64.4254237 | 95.2537113 |
| -4.3894456 | 1.14E-05 | 3.82E-05 | 36.0662614 | 118.798132 | 170.267191 | 119.907613 |
| -4.3891523 | 1.14E-05 | 3.83E-05 | 62.7061136 | 59.5343714 | 70.1776937 | 67.7982298 |
| -4.3887104 | 1.14E-05 | 3.83E-05 | 45.0828268 | 119.339354 | 77.0804177 | 113.744138 |
| -4.388545 | 1.14E-05 | 3.84E-05 | 56.9682993 | 83.8893415 | 87.4345036 | 58.8331746 |
| -4.3881837 | 1.14E-05 | 3.84E-05 | 45.4926707 | 81.7244553 | 94.3372276 | 123.269509 |
| -4.3879598 | 1.14E-05 | 3.85E-05 | 40.984388 | 125.563402 | 90.8858656 | 118.226665 |
| -4.3875637 | 1.15E-05 | 3.85E-05 | 46.3123584 | 91.1958326 | 101.239952 | 79.0045488 |
| -4.3874626 | 1.15E-05 | 3.85E-05 | 47.5418901 | 85.783617 | 81.6822337 | 84.0473923 |
| -4.3874527 | 1.15E-05 | 3.85E-05 | 41.8040758 | 116.362635 | 97.7885896 | 98.055291 |
| -4.3861201 | 1.15E-05 | 3.88E-05 | 120.903945 | 103.102707 | 58.6731538 | 55.4712789 |
| -4.3859943 | 1.15E-05 | 3.88E-05 | 466.402335 | 845.658685 | 814.521429 | 745.780528 |
| -4.3858486 | 1.16E-05 | 3.88E-05 | 52.8698605 | 90.3840002 | 64.4254237 | 82.3664444 |
| 4.38517833 | 1.16E-05 | 3.89E-05 | 126.231915 | 19.7545869 | 28.7613499 | 27.4554815 |
| -4.3851505 | 1.16E-05 | 3.89E-05 | 41.3942319 | 126.916455 | 123.098578 | 86.848972 |
| -4.3850007 | 1.16E-05 | 3.90E-05 | 45.9025146 | 84.1599523 | 85.1335957 | 101.977503 |
| -4.3845382 | 1.16E-05 | 3.90E-05 | 59.8372065 | 75.5004074 | 65.5758777 | 61.6347543 |
| -4.3845138 | 1.16E-05 | 3.90E-05 | 66.3947085 | 227.313054 | 124.249032 | 155.767834 |
| 4.38432927 | 1.16E-05 | 3.91E-05 | 29.0989155 | 7.57710182 | 4.60181598 | 7.84442328 |
| -4.3843029 | 1.16E-05 | 3.91E-05 | 59.0175187 | 75.7710182 | 60.9740618 | 67.7982298 |
| -4.3841727 | 1.16E-05 | 3.91E-05 | 29.9186032 | 120.421797 | 208.232173 | 210.678797 |
| -4.383881 | 1.17E-05 | 3.91E-05 | 43.4434513 | 162.366467 | 162.214013 | 75.0823371 |
| 4.38380889 | 1.17E-05 | 3.91E-05 | 156.970206 | 86.8660601 | 78.2308717 | 79.0045488 |
| -4.3833489 | 1.17E-05 | 3.92E-05 | 97.9526873 | 79.8301798 | 49.4695218 | 71.1601255 |
| -4.3829343 | 1.17E-05 | 3.93E-05 | 36.4761053 | 119.339354 | 124.249032 | 124.950457 |
| -4.3827144 | 1.17E-05 | 3.93E-05 | 31.9678226 | 142.882491 | 181.771731 | 132.234564 |
| -4.3824222 | 1.17E-05 | 3.94E-05 | 43.4434513 | 92.5488865 | 90.8858656 | 104.779082 |
| -4.3820048 | 1.18E-05 | 3.95E-05 | 43.0336074 | 107.70309 | 109.29313 | 84.6077082 |
| -4.3817784 | 1.18E-05 | 3.95E-05 | 83.1983076 | 213.782516 | 51.7704298 | 90.7711837 |
| 4.38073015 | 1.18E-05 | 3.97E-05 | 11.8854725 | 8.11832337 | 17.2568099 | 9.52537113 |
| -4.3806769 | 1.18E-05 | 3.97E-05 | 38.5253247 | 145.588599 | 94.3372276 | 132.79488 |
| -4.3806953 | 1.18E-05 | 3.97E-05 | 35.6564176 | 151.000815 | 105.841768 | 146.242463 |
| -4.3805156 | 1.18E-05 | 3.97E-05 | 41.3942319 | 113.115306 | 92.0363196 | 107.020346 |
| -4.3803946 | 1.18E-05 | 3.97E-05 | 45.0828268 | 107.70309 | 86.2840497 | 85.7283401 |
| -4.3798207 | 1.19E-05 | 3.98E-05 | 51.6403289 | 127.728288 | 97.7885896 | 64.99665 |

| | | | | | | |
|---|---|---|---|---|---|---|
| 4.37980157 | 1.19E-05 | 3.98E-05 | 518.042664 | 871.63732 | 806.468251 | 773.236009 |
| -4.3797676 | 1.19E-05 | 3.98E-05 | 78.6900249 | 75.5004074 | 50.6199758 | 70.5998095 |
| -4.378858 | 1.19E-05 | 4.00E-05 | 33.1973543 | 128.54012 | 132.302209 | 168.094785 |
| -4.3785269 | 1.19E-05 | 4.00E-05 | 54.9190799 | 99.3141559 | 140.355387 | 129.9933 |
| -4.3783731 | 1.20E-05 | 4.01E-05 | 52.4600166 | 104.726372 | 171.417645 | 66.6775979 |
| -4.3776766 | 1.20E-05 | 4.02E-05 | 72.9522106 | 64.4053654 | 55.2217918 | 69.4791776 |
| -4.3776201 | 1.20E-05 | 4.02E-05 | 54.509236 | 126.645845 | 112.744492 | 109.26161 |
| -4.3772039 | 1.20E-05 | 4.03E-05 | 63.5258014 | 70.3588026 | 67.8767857 | 57.1522268 |
| -4.3770919 | 1.20E-05 | 4.03E-05 | 32.3776665 | 175.626396 | 134.603117 | 132.79488 |
| 4.37700541 | 1.20E-05 | 4.03E-05 | 0 | 0 | 0 | 0 |
| -4.3768629 | 1.20E-05 | 4.03E-05 | 41.8040758 | 96.8786589 | 95.4876816 | 107.580662 |
| -4.3765574 | 1.21E-05 | 4.04E-05 | 35.2465737 | 105.808815 | 142.656295 | 163.612257 |
| -4.3762776 | 1.21E-05 | 4.04E-05 | 81.1490882 | 83.0775092 | 50.6199758 | 66.6775979 |
| -4.3758631 | 1.21E-05 | 4.05E-05 | 53.2797044 | 58.4519283 | 212.833989 | 163.612257 |
| -4.3753805 | 1.21E-05 | 4.06E-05 | 59.4273626 | 74.1473535 | 56.3722458 | 83.4870763 |
| -4.3752458 | 1.21E-05 | 4.06E-05 | 170.904898 | 103.373318 | 41.4163438 | 92.4521315 |
| -4.3752199 | 1.21E-05 | 4.06E-05 | 44.263139 | 84.9717847 | 95.4876816 | 93.0124475 |
| -4.3750985 | 1.21E-05 | 4.06E-05 | 42.2139196 | 100.396599 | 98.9390436 | 93.0124475 |
| -4.3750516 | 1.21E-05 | 4.06E-05 | 194.675843 | 580.18951 | 517.704298 | 414.073486 |
| -4.3748236 | 1.22E-05 | 4.07E-05 | 54.9190799 | 63.8641439 | 82.8326877 | 124.390141 |
| -4.3746287 | 1.22E-05 | 4.07E-05 | 84.8376831 | 81.9950661 | 48.3190678 | 71.7204414 |
| -4.3746023 | 1.22E-05 | 4.07E-05 | 57.7879871 | 190.780599 | 154.160835 | 164.732889 |
| -4.3744195 | 1.22E-05 | 4.07E-05 | 38.5253247 | 106.350036 | 118.496762 | 110.382242 |
| -4.3735009 | 1.22E-05 | 4.09E-05 | 45.9025146 | 97.1492697 | 74.7795097 | 104.218766 |
| -4.3726608 | 1.23E-05 | 4.10E-05 | 40.984388 | 103.102707 | 100.089498 | 99.7362389 |
| -4.3723494 | 1.23E-05 | 4.11E-05 | 37.2957931 | 121.774851 | 119.647216 | 108.701294 |
| -4.3721153 | 1.23E-05 | 4.11E-05 | 115.575974 | 130.434396 | 43.7172518 | 75.0823371 |
| -4.3717145 | 1.23E-05 | 4.12E-05 | 52.4600166 | 78.2065152 | 72.4786017 | 67.2379138 |
| -4.3715876 | 1.23E-05 | 4.12E-05 | 50.8206411 | 166.155018 | 103.54086 | 68.3585457 |
| -4.3707856 | 1.24E-05 | 4.14E-05 | 36.4761053 | 153.165701 | 112.744492 | 113.183822 |
| -4.3704858 | 1.24E-05 | 4.14E-05 | 112.707067 | 100.937821 | 37.9649818 | 118.226665 |
| -4.3701345 | 1.24E-05 | 4.15E-05 | 40.1647002 | 113.115306 | 90.8858656 | 121.588561 |
| -4.3699819 | 1.24E-05 | 4.15E-05 | 55.3289238 | 98.7729344 | 59.8236078 | 144.001199 |
| -4.3699271 | 1.24E-05 | 4.15E-05 | 38.5253247 | 122.857294 | 96.6381356 | 123.269509 |
| -4.3698635 | 1.24E-05 | 4.15E-05 | 34.8367298 | 170.755402 | 132.302209 | 109.821926 |
| -4.3691656 | 1.25E-05 | 4.17E-05 | 427.057323 | 117.715689 | 77.0804177 | 58.2728587 |
| -4.3690415 | 1.25E-05 | 4.17E-05 | 33.6071982 | 145.588599 | 148.408565 | 115.985401 |
| -4.3689789 | 1.25E-05 | 4.17E-05 | 65.5750208 | 52.4984912 | 89.7354116 | 73.4013893 |
| -4.3682436 | 1.25E-05 | 4.18E-05 | 53.6895483 | 66.0290301 | 70.1776937 | 73.4013893 |
| -4.367939 | 1.25E-05 | 4.19E-05 | 54.509236 | 113.385916 | 57.5226998 | 100.856871 |
| -4.3679194 | 1.25E-05 | 4.19E-05 | 59.4273626 | 151.542036 | 103.54086 | 110.382242 |
| -4.3678791 | 1.25E-05 | 4.19E-05 | 62.2962697 | 120.151186 | 110.443584 | 111.502874 |
| -4.3675908 | 1.26E-05 | 4.19E-05 | 29.9186032 | 145.317988 | 199.028541 | 139.518671 |

| | | | | | | |
|---|---|---|---|---|---|---|
| -4.3675313 | 1.26E-05 | 4.19E-05 | 56.9682993 | 160.201581 | 197.878087 | 294.726189 |
| -4.3675073 | 1.26E-05 | 4.19E-05 | 56.5584554 | 80.6420122 | 124.249032 | 61.0744384 |
| -4.3674416 | 1.26E-05 | 4.19E-05 | 98.7723751 | 64.946587 | 103.54086 | 49.3078035 |
| -4.3674396 | 1.26E-05 | 4.19E-05 | 76.2309617 | 85.783617 | 52.9208838 | 62.1950703 |
| -4.3673599 | 1.26E-05 | 4.19E-05 | 35.2465737 | 138.552719 | 116.195854 | 122.709193 |
| -4.3673803 | 1.26E-05 | 4.19E-05 | 54.509236 | 105.267593 | 172.568099 | 62.1950703 |
| -4.3670871 | 1.26E-05 | 4.20E-05 | 64.3454891 | 202.958084 | 148.408565 | 48.7474875 |
| -4.3669313 | 1.26E-05 | 4.20E-05 | 40.1647002 | 87.9485032 | 123.098578 | 118.226665 |
| -4.3668902 | 1.26E-05 | 4.20E-05 | 45.4926707 | 87.6778924 | 78.2308717 | 98.055291 |
| -4.3660656 | 1.27E-05 | 4.22E-05 | 72.9522106 | 63.5935331 | 57.5226998 | 62.7553862 |
| -4.3658089 | 1.27E-05 | 4.22E-05 | 49.5911095 | 136.117222 | 141.505841 | 62.1950703 |
| -4.3653369 | 1.27E-05 | 4.23E-05 | 60.2470503 | 57.640096 | 71.3281477 | 68.3585457 |
| -4.3650696 | 1.27E-05 | 4.24E-05 | 47.5418901 | 86.5954493 | 77.0804177 | 77.8839169 |
| -4.3650415 | 1.27E-05 | 4.24E-05 | 74.5915861 | 99.8553775 | 95.4876816 | 45.3855918 |
| -4.3648324 | 1.27E-05 | 4.24E-05 | 35.6564176 | 125.292791 | 118.496762 | 120.467929 |
| -4.3644896 | 1.27E-05 | 4.25E-05 | 40.5745441 | 127.187066 | 89.7354116 | 105.339398 |
| -4.3643307 | 1.28E-05 | 4.25E-05 | 75.00143 | 89.0309463 | 80.5317797 | 93.5727634 |
| -4.3641074 | 1.28E-05 | 4.25E-05 | 59.8372065 | 79.0183475 | 64.4254237 | 158.569413 |
| -4.3640604 | 1.28E-05 | 4.25E-05 | 46.7222023 | 97.9611021 | 70.1776937 | 106.46003 |
| -4.3635366 | 1.28E-05 | 4.26E-05 | 38.1154808 | 133.411114 | 101.239952 | 109.26161 |
| -4.3630183 | 1.28E-05 | 4.27E-05 | 39.7548564 | 117.174467 | 121.948124 | 89.0902358 |
| -4.3629054 | 1.28E-05 | 4.27E-05 | 43.8532952 | 71.1706349 | 202.479903 | 154.647202 |
| -4.3628825 | 1.28E-05 | 4.27E-05 | 31.9678226 | 192.133653 | 133.452663 | 127.19172 |
| 4.36281918 | 1.28E-05 | 4.27E-05 | 45.9025146 | 43.5683354 | 54.0713378 | 37.5411686 |
| -4.362826 | 1.28E-05 | 4.27E-05 | 43.8532952 | 82.5362876 | 127.700393 | 91.3314996 |
| -4.3627265 | 1.28E-05 | 4.28E-05 | 364.351209 | 92.8194972 | 50.6199758 | 103.65845 |
| -4.3623969 | 1.29E-05 | 4.28E-05 | 38.5253247 | 119.609964 | 98.9390436 | 111.502874 |
| -4.3623076 | 1.29E-05 | 4.28E-05 | 56.1486116 | 62.2404792 | 86.2840497 | 66.1172819 |
| -4.3622531 | 1.29E-05 | 4.28E-05 | 44.263139 | 112.844695 | 73.6290557 | 128.872668 |
| -4.3622678 | 1.29E-05 | 4.28E-05 | 63.9356453 | 111.22103 | 143.806749 | 160.250361 |
| -4.3619951 | 1.29E-05 | 4.29E-05 | 83.6081515 | 49.2511618 | 69.0272397 | 81.2458125 |
| -4.3619487 | 1.29E-05 | 4.29E-05 | 20.0823501 | 174.002731 | 312.923487 | 249.900913 |
| -4.3617219 | 1.29E-05 | 4.29E-05 | 44.6729829 | 111.491641 | 94.3372276 | 76.763285 |
| -4.3616162 | 1.29E-05 | 4.29E-05 | 42.2139196 | 98.2317128 | 92.0363196 | 92.4521315 |
| -4.3609298 | 1.30E-05 | 4.31E-05 | 34.8367298 | 131.516839 | 133.452663 | 112.623506 |
| -4.3607656 | 1.30E-05 | 4.31E-05 | 56.1486116 | 143.964935 | 54.0713378 | 126.631404 |
| -4.3607367 | 1.30E-05 | 4.31E-05 | 57.3781432 | 82.2656769 | 57.5226998 | 117.106033 |
| -4.3603005 | 1.30E-05 | 4.32E-05 | 61.476582 | 64.6759762 | 59.8236078 | 64.4363341 |
| -4.3598019 | 1.30E-05 | 4.33E-05 | 50.0009534 | 80.1007906 | 128.850847 | 71.7204414 |
| -4.3597391 | 1.30E-05 | 4.33E-05 | 47.5418901 | 119.609964 | 69.0272397 | 94.6933953 |
| -4.3595473 | 1.30E-05 | 4.33E-05 | 45.4926707 | 80.912623 | 80.5317797 | 99.1759229 |
| -4.3587951 | 1.31E-05 | 4.35E-05 | 71.722679 | 84.4305631 | 69.0272397 | 50.4284354 |
| -4.3587104 | 1.31E-05 | 4.35E-05 | 190.16756 | 125.292791 | 46.0181598 | 70.0394936 |

| | | | | | | |
|---|---|---|---|---|---|---|
| -4.358214 | 1.31E-05 | 4.36E-05 | 94.2640924 | 192.133653 | 36.8145279 | 129.432984 |
| -4.3579403 | 1.31E-05 | 4.36E-05 | 43.0336074 | 86.3248385 | 92.0363196 | 94.1330794 |
| -4.3576025 | 1.31E-05 | 4.37E-05 | 74.1817423 | 65.4878086 | 50.6199758 | 89.6505518 |
| -4.3573926 | 1.32E-05 | 4.37E-05 | 38.1154808 | 128.269509 | 104.691314 | 102.537819 |
| -4.3571745 | 1.32E-05 | 4.38E-05 | 37.705637 | 104.996982 | 117.346308 | 109.26161 |
| -4.3567239 | 1.32E-05 | 4.38E-05 | 270.087117 | 100.66721 | 57.5226998 | 70.5998095 |
| -4.356727 | 1.32E-05 | 4.38E-05 | 47.1320462 | 77.9359044 | 80.5317797 | 78.4442328 |
| -4.3565171 | 1.32E-05 | 4.39E-05 | 44.6729829 | 85.2423954 | 85.1335957 | 85.1680242 |
| -4.3562723 | 1.32E-05 | 4.39E-05 | 44.6729829 | 75.5004074 | 111.594038 | 93.0124475 |
| -4.3562353 | 1.32E-05 | 4.39E-05 | 49.5911095 | 91.1958326 | 73.6290557 | 70.0394936 |
| -4.3561738 | 1.32E-05 | 4.39E-05 | 34.8367298 | 154.518755 | 111.594038 | 121.028245 |
| -4.3560243 | 1.32E-05 | 4.39E-05 | 229.922417 | 181.309222 | 29.9118039 | 110.382242 |
| -4.3560411 | 1.32E-05 | 4.39E-05 | 45.9025146 | 83.8893415 | 100.089498 | 75.0823371 |
| -4.3558217 | 1.33E-05 | 4.40E-05 | 33.6071982 | 163.1783 | 121.948124 | 120.467929 |
| -4.3555131 | 1.33E-05 | 4.40E-05 | 50.4107972 | 211.88824 | 225.488983 | 197.79153 |
| -4.3554514 | 1.33E-05 | 4.41E-05 | 85.247527 | 148.294707 | 58.6731538 | 59.3934906 |
| -4.3553563 | 1.33E-05 | 4.41E-05 | 41.8040758 | 91.1958326 | 96.6381356 | 94.1330794 |
| -4.3551946 | 1.33E-05 | 4.41E-05 | 44.263139 | 111.762252 | 75.9299637 | 95.8140272 |
| -4.3549343 | 1.33E-05 | 4.41E-05 | 56.5584554 | 94.7137727 | 55.2217918 | 82.9267604 |
| -4.3545046 | 1.33E-05 | 4.42E-05 | 65.9848647 | 80.912623 | 87.4345036 | 49.3078035 |
| -4.3544895 | 1.33E-05 | 4.42E-05 | 41.8040758 | 103.102707 | 83.9831417 | 103.65845 |
| -4.3544093 | 1.33E-05 | 4.42E-05 | 54.0993922 | 69.2763595 | 74.7795097 | 62.1950703 |
| -4.3544254 | 1.33E-05 | 4.42E-05 | 14.3445358 | 192.404264 | 383.10118 | 418.556014 |
| -4.3541045 | 1.34E-05 | 4.43E-05 | 54.0993922 | 120.421797 | 59.8236078 | 86.2886561 |
| -4.3540802 | 1.34E-05 | 4.43E-05 | 52.4600166 | 106.350036 | 79.3813257 | 62.7553862 |
| -4.3539311 | 1.34E-05 | 4.43E-05 | 47.951734 | 114.46836 | 165.665375 | 66.6775979 |
| -4.3536056 | 1.34E-05 | 4.44E-05 | 54.9190799 | 82.5362876 | 59.8236078 | 70.5998095 |
| -4.3536156 | 1.34E-05 | 4.44E-05 | 327.875104 | 92.0076649 | 62.1245158 | 76.202969 |
| -4.3535035 | 1.34E-05 | 4.44E-05 | 58.197831 | 57.0988744 | 69.0272397 | 84.0473923 |
| 4.35349977 | 1.34E-05 | 4.44E-05 | 30.738291 | 23.5431378 | 26.4604419 | 22.9729539 |
| -4.3531311 | 1.34E-05 | 4.45E-05 | 64.3454891 | 49.5217726 | 87.4345036 | 92.4521315 |
| -4.3529989 | 1.34E-05 | 4.45E-05 | 37.705637 | 144.506156 | 97.7885896 | 110.942558 |
| -4.3528203 | 1.34E-05 | 4.45E-05 | 42.2139196 | 89.8427787 | 108.142676 | 86.848972 |
| -4.3527913 | 1.34E-05 | 4.45E-05 | 42.6237635 | 125.834012 | 78.2308717 | 104.779082 |
| -4.3527576 | 1.34E-05 | 4.45E-05 | 40.5745441 | 106.891258 | 85.1335957 | 125.510772 |
| -4.3521159 | 1.35E-05 | 4.46E-05 | 48.7714217 | 71.1706349 | 73.6290557 | 87.9696039 |
| -4.3520714 | 1.35E-05 | 4.46E-05 | 109.83816 | 139.905773 | 202.479903 | 67.2379138 |
| -4.3516464 | 1.35E-05 | 4.47E-05 | 76.6408055 | 78.477126 | 126.549939 | 128.312352 |
| -4.3514821 | 1.35E-05 | 4.48E-05 | 36.0662614 | 103.373318 | 120.79767 | 132.234564 |
| -4.3509015 | 1.36E-05 | 4.49E-05 | 150.822548 | 151.000815 | 57.5226998 | 50.9887513 |
| -4.3507488 | 1.36E-05 | 4.49E-05 | 45.0828268 | 94.7137727 | 74.7795097 | 89.6505518 |
| -4.3505895 | 1.36E-05 | 4.49E-05 | 34.4268859 | 167.778683 | 103.54086 | 134.475828 |
| -4.3503531 | 1.36E-05 | 4.50E-05 | 39.7548564 | 90.3840002 | 124.249032 | 101.977503 |

| | | | | | | |
|---|---|---|---|---|---|---|
| -4.3503282 | 1.36E-05 | 4.50E-05 | 42.6237635 | 145.047378 | 74.7795097 | 119.907613 |
| -4.3501663 | 1.36E-05 | 4.50E-05 | 45.4926707 | 101.208431 | 128.850847 | 70.5998095 |
| -4.3500992 | 1.36E-05 | 4.50E-05 | 37.2957931 | 120.421797 | 102.390406 | 110.942558 |
| -4.3499298 | 1.36E-05 | 4.50E-05 | 28.6890716 | 113.115306 | 204.780811 | 224.126379 |
| -4.3498127 | 1.36E-05 | 4.51E-05 | 63.1159575 | 85.783617 | 55.2217918 | 63.8760181 |
| 4.34971867 | 1.36E-05 | 4.51E-05 | 0 | 0.81183234 | 0 | 0.56031595 |
| -4.3494768 | 1.36E-05 | 4.51E-05 | 41.8040758 | 124.210348 | 78.2308717 | 112.06319 |
| -4.349467 | 1.36E-05 | 4.51E-05 | 62.7061136 | 319.050109 | 247.347609 | 256.624704 |
| -4.3492208 | 1.37E-05 | 4.52E-05 | 45.4926707 | 126.916455 | 72.4786017 | 95.2537113 |
| -4.3489737 | 1.37E-05 | 4.52E-05 | 47.5418901 | 73.8767427 | 77.0804177 | 80.1251807 |
| -4.3487601 | 1.37E-05 | 4.52E-05 | 62.2962697 | 79.5595691 | 119.647216 | 53.2300151 |
| -4.3486531 | 1.37E-05 | 4.53E-05 | 51.230485 | 65.2171978 | 73.6290557 | 74.5220212 |
| -4.3482324 | 1.37E-05 | 4.53E-05 | 56.9682993 | 76.5828505 | 55.2217918 | 93.0124475 |
| -4.3479027 | 1.37E-05 | 4.54E-05 | 115.985818 | 224.606947 | 204.780811 | 28.0157974 |
| -4.3479032 | 1.37E-05 | 4.54E-05 | 61.0667381 | 79.2889583 | 63.2749697 | 56.5919108 |
| -4.347791 | 1.38E-05 | 4.54E-05 | 92.214873 | 48.4393295 | 66.7263317 | 82.9267604 |
| -4.3477358 | 1.38E-05 | 4.54E-05 | 318.448695 | 539.327283 | 434.87161 | 476.268556 |
| -4.3476876 | 1.38E-05 | 4.54E-05 | 64.755333 | 97.4198805 | 153.010381 | 50.4284354 |
| -4.3473924 | 1.38E-05 | 4.55E-05 | 31.9678226 | 165.343186 | 163.364467 | 108.701294 |
| -4.3470668 | 1.38E-05 | 4.55E-05 | 54.0993922 | 102.832096 | 55.2217918 | 103.65845 |
| -4.3468971 | 1.38E-05 | 4.56E-05 | 49.5911095 | 75.5004074 | 78.2308717 | 67.2379138 |
| -4.346488 | 1.38E-05 | 4.57E-05 | 37.2957931 | 121.774851 | 108.142676 | 101.977503 |
| -4.346413 | 1.38E-05 | 4.57E-05 | 106.969253 | 71.4412457 | 60.9740618 | 55.4712789 |
| -4.3461251 | 1.39E-05 | 4.57E-05 | 46.3123584 | 78.7477367 | 75.9299637 | 84.0473923 |
| -4.3455839 | 1.39E-05 | 4.58E-05 | 288.530091 | 125.563402 | 56.3722458 | 61.6347543 |
| -4.345495 | 1.39E-05 | 4.58E-05 | 162.298176 | 76.0416289 | 50.6199758 | 79.0045488 |
| -4.3450893 | 1.39E-05 | 4.59E-05 | 40.1647002 | 90.654611 | 103.54086 | 100.296555 |
| -4.3449925 | 1.39E-05 | 4.59E-05 | 51.6403289 | 81.7244553 | 62.1245158 | 74.5220212 |
| -4.3448533 | 1.39E-05 | 4.60E-05 | 51.6403289 | 100.937821 | 100.089498 | 58.2728587 |
| -4.3447613 | 1.39E-05 | 4.60E-05 | 1502.89751 | 4095.42353 | 3304.10387 | 3757.47875 |
| 4.34473034 | 1.39E-05 | 4.60E-05 | 0 | 0 | 0 | 0 |
| -4.3446913 | 1.39E-05 | 4.60E-05 | 60.2470503 | 54.6633774 | 70.1776937 | 71.1601255 |
| 4.34421029 | 1.40E-05 | 4.61E-05 | 6.1476582 | 7.8477126 | 6.90272397 | 5.04284354 |
| -4.3436819 | 1.40E-05 | 4.62E-05 | 53.6895483 | 271.152001 | 155.311289 | 225.247011 |
| -4.3436177 | 1.40E-05 | 4.62E-05 | 42.2139196 | 77.9359044 | 104.691314 | 135.59646 |
| 4.34345431 | 1.40E-05 | 4.62E-05 | 171.724586 | 148.565318 | 147.258111 | 169.775732 |
| -4.3431262 | 1.40E-05 | 4.63E-05 | 126.231915 | 314.179115 | 212.833989 | 385.497373 |
| -4.3427587 | 1.41E-05 | 4.64E-05 | 40.1647002 | 141.529437 | 92.0363196 | 95.2537113 |
| -4.342585 | 1.41E-05 | 4.64E-05 | 41.8040758 | 77.1240721 | 121.948124 | 108.701294 |
| -4.3419319 | 1.41E-05 | 4.65E-05 | 46.7222023 | 114.197749 | 73.6290557 | 80.1251807 |
| -4.3418655 | 1.41E-05 | 4.65E-05 | 40.984388 | 102.290875 | 83.9831417 | 148.483726 |
| -4.3415453 | 1.41E-05 | 4.66E-05 | 45.0828268 | 81.7244553 | 80.5317797 | 81.2458125 |
| -4.3414787 | 1.42E-05 | 4.66E-05 | 75.8211178 | 135.034779 | 40.2658898 | 136.156776 |

| | | | | | | |
|---|---|---|---|---|---|---|
| -4.3413458 | 1.42E-05 | 4.66E-05 | 50.0009534 | 79.5595691 | 87.4345036 | 62.7553862 |
| -4.3409797 | 1.42E-05 | 4.67E-05 | 47.5418901 | 67.9233056 | 81.6822337 | 86.848972 |
| -4.3409911 | 1.42E-05 | 4.67E-05 | 40.984388 | 84.4305631 | 109.29313 | 96.9346591 |
| -4.3407377 | 1.42E-05 | 4.68E-05 | 36.4761053 | 126.916455 | 117.346308 | 98.615607 |
| -4.3403771 | 1.42E-05 | 4.68E-05 | 47.5418901 | 75.5004074 | 80.5317797 | 71.7204414 |
| -4.3402949 | 1.42E-05 | 4.68E-05 | 35.6564176 | 104.455761 | 133.452663 | 114.304454 |
| -4.3394712 | 1.43E-05 | 4.70E-05 | 31.9678226 | 128.269509 | 280.710775 | 123.829825 |
| -4.3390973 | 1.43E-05 | 4.71E-05 | 74.1817423 | 89.3015571 | 51.7704298 | 60.5141225 |
| -4.3384003 | 1.44E-05 | 4.72E-05 | 57.7879871 | 65.4878086 | 58.6731538 | 65.556966 |
| -4.3382663 | 1.44E-05 | 4.73E-05 | 42.2139196 | 99.3141559 | 101.239952 | 78.4442328 |
| -4.3379745 | 1.44E-05 | 4.73E-05 | 69.2636157 | 88.7603356 | 47.1686138 | 75.0823371 |
| -4.3379913 | 1.44E-05 | 4.73E-05 | 40.5745441 | 192.133653 | 83.9831417 | 114.304454 |
| 4.33776829 | 1.44E-05 | 4.73E-05 | 33.1973543 | 26.5198564 | 11.50454 | 17.9301104 |
| 4.33725194 | 1.44E-05 | 4.75E-05 | 34.017042 | 30.579018 | 26.4604419 | 26.3348496 |
| -4.3366044 | 1.45E-05 | 4.76E-05 | 184.83959 | 71.7118565 | 73.6290557 | 58.2728587 |
| -4.336455 | 1.45E-05 | 4.76E-05 | 121.723632 | 88.7603356 | 64.4254237 | 48.7474875 |
| -4.3354564 | 1.45E-05 | 4.78E-05 | 65.1651769 | 103.373318 | 173.718553 | 164.172573 |
| -4.3354442 | 1.45E-05 | 4.78E-05 | 44.6729829 | 171.296623 | 205.931265 | 63.3157022 |
| -4.3352314 | 1.46E-05 | 4.79E-05 | 68.8537718 | 79.8301798 | 56.3722458 | 56.0315949 |
| -4.3351163 | 1.46E-05 | 4.79E-05 | 44.6729829 | 73.6061319 | 230.090799 | 122.709193 |
| -4.3350966 | 1.46E-05 | 4.79E-05 | 34.4268859 | 112.574084 | 115.0454 | 132.79488 |
| -4.3349941 | 1.46E-05 | 4.79E-05 | 65.1651769 | 68.7351379 | 81.6822337 | 50.4284354 |
| -4.3349618 | 1.46E-05 | 4.79E-05 | 84.4278393 | 92.0076649 | 47.1686138 | 64.99665 |
| -4.3348421 | 1.46E-05 | 4.79E-05 | 47.5418901 | 90.1133895 | 64.4254237 | 93.5727634 |
| 4.33480125 | 1.46E-05 | 4.79E-05 | 123.772852 | 61.6992576 | 59.8236078 | 48.1871716 |
| 4.33360549 | 1.47E-05 | 4.82E-05 | 0.40984388 | 1.89427545 | 1.150454 | 1.1206319 |
| -4.3335973 | 1.47E-05 | 4.82E-05 | 38.9351686 | 110.409198 | 112.744492 | 86.848972 |
| -4.3336342 | 1.47E-05 | 4.82E-05 | 106.559409 | 80.3714014 | 48.3190678 | 64.99665 |
| -4.3334652 | 1.47E-05 | 4.82E-05 | 82.7884637 | 36.2618444 | 167.966283 | 184.904263 |
| -4.3329005 | 1.47E-05 | 4.83E-05 | 7.78703372 | 296.860025 | 496.996126 | 531.179519 |
| -4.3325314 | 1.47E-05 | 4.84E-05 | 46.7222023 | 74.9591858 | 73.6290557 | 79.5648647 |
| -4.3321012 | 1.48E-05 | 4.85E-05 | 42.6237635 | 103.373318 | 79.3813257 | 87.9696039 |
| -4.3317442 | 1.48E-05 | 4.86E-05 | 72.1325229 | 67.382084 | 49.4695218 | 107.580662 |
| 4.33127942 | 1.48E-05 | 4.87E-05 | 14.3445358 | 14.6129821 | 23.0090799 | 17.9301104 |
| -4.3311313 | 1.48E-05 | 4.87E-05 | 52.0501728 | 68.1939163 | 82.8326877 | 61.6347543 |
| -4.3309052 | 1.48E-05 | 4.87E-05 | 64.3454891 | 85.5130062 | 47.1686138 | 101.977503 |
| -4.3308453 | 1.49E-05 | 4.87E-05 | 44.6729829 | 80.912623 | 75.9299637 | 86.848972 |
| -4.3305968 | 1.49E-05 | 4.88E-05 | 33.1973543 | 211.617629 | 93.1867736 | 163.612257 |
| -4.3300869 | 1.49E-05 | 4.89E-05 | 59.4273626 | 101.479042 | 59.8236078 | 62.1950703 |
| -4.3300471 | 1.49E-05 | 4.89E-05 | 46.3123584 | 76.5828505 | 80.5317797 | 72.8410733 |
| -4.3299841 | 1.49E-05 | 4.89E-05 | 35.2465737 | 148.565318 | 100.089498 | 116.545717 |
| -4.3296777 | 1.49E-05 | 4.90E-05 | 62.7061136 | 152.353869 | 50.6199758 | 94.1330794 |
| -4.3296 | 1.49E-05 | 4.90E-05 | 34.8367298 | 120.151186 | 108.142676 | 118.786981 |

| | | | | | | |
|---|---|---|---|---|---|---|
| -4.3289896 | 1.50E-05 | 4.91E-05 | 34.4268859 | 124.210348 | 133.452663 | 103.098135 |
| -4.3289124 | 1.50E-05 | 4.91E-05 | 30.738291 | 112.303473 | 202.479903 | 144.001199 |
| -4.3288627 | 1.50E-05 | 4.91E-05 | 93.8542485 | 287.118037 | 216.285351 | 250.461229 |
| -4.328823 | 1.50E-05 | 4.91E-05 | 148.363485 | 34.0969582 | 97.7885896 | 140.639303 |
| -4.3282803 | 1.50E-05 | 4.93E-05 | 162.298176 | 72.7942996 | 55.2217918 | 70.0394936 |
| -4.3281299 | 1.50E-05 | 4.93E-05 | 36.4761053 | 176.97945 | 90.8858656 | 122.709193 |
| -4.3279227 | 1.51E-05 | 4.93E-05 | 51.6403289 | 292.530252 | 55.2217918 | 169.775732 |
| -4.3277762 | 1.51E-05 | 4.94E-05 | 77.0506494 | 84.7011739 | 80.5317797 | 44.2649599 |
| -4.3276699 | 1.51E-05 | 4.94E-05 | 69.2636157 | 68.7351379 | 51.7704298 | 62.1950703 |
| -4.3274903 | 1.51E-05 | 4.94E-05 | 54.9190799 | 81.9950661 | 257.701695 | 80.6854966 |
| -4.3272558 | 1.51E-05 | 4.95E-05 | 43.4434513 | 81.9950661 | 266.905327 | 114.864769 |
| -4.3272193 | 1.51E-05 | 4.95E-05 | 41.8040758 | 92.8194972 | 83.9831417 | 87.9696039 |
| -4.3269106 | 1.51E-05 | 4.95E-05 | 1395.92826 | 601.567762 | 511.952028 | 577.685743 |
| -4.3267525 | 1.51E-05 | 4.96E-05 | 72.9522106 | 85.5130062 | 46.0181598 | 71.7204414 |
| -4.3266907 | 1.51E-05 | 4.96E-05 | 32.3776665 | 159.389749 | 118.496762 | 118.786981 |
| -4.32654 | 1.51E-05 | 4.96E-05 | 38.1154808 | 118.527521 | 87.4345036 | 111.502874 |
| -4.32566 | 1.52E-05 | 4.98E-05 | 33.6071982 | 136.387833 | 109.29313 | 121.028245 |
| -4.3255392 | 1.52E-05 | 4.98E-05 | 36.4761053 | 113.927138 | 106.992222 | 101.417187 |
| -4.3252386 | 1.52E-05 | 4.99E-05 | 313.940412 | 119.339354 | 242.745793 | 165.293205 |
| -4.3250253 | 1.53E-05 | 4.99E-05 | 23.3611012 | 157.766084 | 202.479903 | 196.670898 |
| -4.3250342 | 1.53E-05 | 4.99E-05 | 43.0336074 | 112.032863 | 120.79767 | 69.4791776 |
| -4.324852 | 1.53E-05 | 4.99E-05 | 82.3786199 | 74.688575 | 87.4345036 | 74.5220212 |
| -4.3246703 | 1.53E-05 | 5.00E-05 | 91.3951852 | 93.3607188 | 46.0181598 | 64.4363341 |
| -4.3244756 | 1.53E-05 | 5.00E-05 | 174.183649 | 483.310852 | 518.854752 | 542.946154 |
| -4.3242273 | 1.53E-05 | 5.01E-05 | 38.5253247 | 94.7137727 | 111.594038 | 94.1330794 |
| -4.3241325 | 1.53E-05 | 5.01E-05 | 42.2139196 | 94.4431619 | 78.2308717 | 91.3314996 |
| -4.3240295 | 1.53E-05 | 5.01E-05 | 44.263139 | 78.7477367 | 131.151755 | 81.2458125 |
| -4.3240311 | 1.53E-05 | 5.01E-05 | 30.3284471 | 140.988216 | 140.355387 | 128.312352 |
| -4.3235786 | 1.54E-05 | 5.02E-05 | 29.0989155 | 125.834012 | 177.169915 | 143.440883 |
| -4.3235265 | 1.54E-05 | 5.02E-05 | 56.9682993 | 110.950419 | 50.6199758 | 144.001199 |
| -4.3233064 | 1.54E-05 | 5.02E-05 | 38.5253247 | 98.7729344 | 94.3372276 | 101.977503 |
| -4.3233272 | 1.54E-05 | 5.02E-05 | 42.2139196 | 106.620647 | 95.4876816 | 75.0823371 |
| -4.3232579 | 1.54E-05 | 5.02E-05 | 58.197831 | 77.6652936 | 60.9740618 | 57.1522268 |
| -4.3232245 | 1.54E-05 | 5.02E-05 | 36.0662614 | 115.550803 | 95.4876816 | 123.829825 |
| -4.3231646 | 1.54E-05 | 5.03E-05 | 202.462877 | 396.715402 | 215.134897 | 252.142177 |
| -4.3230419 | 1.54E-05 | 5.03E-05 | 44.263139 | 75.5004074 | 80.5317797 | 86.2886561 |
| -4.3224548 | 1.54E-05 | 5.04E-05 | 82.3786199 | 41.4034492 | 88.5849576 | 101.977503 |
| -4.321773 | 1.55E-05 | 5.06E-05 | 53.2797044 | 69.2763595 | 59.8236078 | 67.2379138 |
| -4.3216604 | 1.55E-05 | 5.06E-05 | 49.1812656 | 136.117222 | 57.5226998 | 180.982051 |
| -4.3211557 | 1.55E-05 | 5.07E-05 | 55.3289238 | 64.4053654 | 57.5226998 | 72.2807574 |
| -4.3210526 | 1.55E-05 | 5.07E-05 | 57.7879871 | 50.8748265 | 88.5849576 | 105.899714 |
| -4.3208639 | 1.55E-05 | 5.07E-05 | 34.8367298 | 125.834012 | 126.549939 | 98.055291 |
| -4.3202036 | 1.56E-05 | 5.09E-05 | 81.5589321 | 40.5916169 | 202.479903 | 112.623506 |

| | | | | | | |
|---|---|---|---|---|---|---|
| -4.3198499 | 1.56E-05 | 5.10E-05 | 69.6734596 | 117.715689 | 89.7354116 | 154.086886 |
| -4.3197148 | 1.56E-05 | 5.10E-05 | 55.7387677 | 103.102707 | 52.9208838 | 81.8061285 |
| -4.3197349 | 1.56E-05 | 5.10E-05 | 34.4268859 | 116.362635 | 130.001301 | 103.65845 |
| -4.319552 | 1.56E-05 | 5.10E-05 | 36.8859492 | 96.0668266 | 212.833989 | 111.502874 |
| -4.3192426 | 1.57E-05 | 5.11E-05 | 40.984388 | 121.774851 | 72.4786017 | 149.604358 |
| -4.3190985 | 1.57E-05 | 5.11E-05 | 33.6071982 | 157.495473 | 125.399485 | 100.856871 |
| -4.3190736 | 1.57E-05 | 5.11E-05 | 35.6564176 | 111.491641 | 98.9390436 | 122.148877 |
| -4.3190506 | 1.57E-05 | 5.11E-05 | 115.575974 | 191.863042 | 263.453965 | 26.3348496 |
| -4.3187962 | 1.57E-05 | 5.12E-05 | 33.1973543 | 93.6313296 | 158.762651 | 158.009098 |
| -4.3183023 | 1.57E-05 | 5.13E-05 | 52.8698605 | 100.66721 | 111.594038 | 104.218766 |
| -4.3183145 | 1.57E-05 | 5.13E-05 | 51.230485 | 71.9824673 | 109.29313 | 62.7553862 |
| -4.3181262 | 1.57E-05 | 5.13E-05 | 41.3942319 | 86.5954493 | 83.9831417 | 91.8918156 |
| -4.3179408 | 1.57E-05 | 5.14E-05 | 37.705637 | 102.290875 | 115.0454 | 90.7711837 |
| -4.3174408 | 1.58E-05 | 5.15E-05 | 31.1481349 | 255.185965 | 98.9390436 | 162.491625 |
| -4.3173909 | 1.58E-05 | 5.15E-05 | 249.185079 | 465.45054 | 422.216616 | 453.855918 |
| -4.3171774 | 1.58E-05 | 5.15E-05 | 55.3289238 | 76.5828505 | 60.9740618 | 59.3934906 |
| -4.3167953 | 1.58E-05 | 5.16E-05 | 50.4107972 | 79.8301798 | 62.1245158 | 121.028245 |
| -4.3165859 | 1.58E-05 | 5.16E-05 | 37.705637 | 108.514922 | 86.2840497 | 128.312352 |
| -4.316468 | 1.59E-05 | 5.17E-05 | 133.199261 | 110.138587 | 55.2217918 | 85.1680242 |
| -4.3163302 | 1.59E-05 | 5.17E-05 | 298.366345 | 584.248672 | 576.377452 | 533.420783 |
| -4.3159206 | 1.59E-05 | 5.18E-05 | 47.1320462 | 233.807713 | 192.125817 | 235.893014 |
| -4.3159063 | 1.59E-05 | 5.18E-05 | 1266.41759 | 3440.54545 | 1886.74455 | 2550.5582 |
| -4.3153109 | 1.59E-05 | 5.19E-05 | 33.1973543 | 147.212264 | 100.089498 | 132.234564 |
| -4.3152247 | 1.59E-05 | 5.19E-05 | 42.2139196 | 104.18515 | 128.850847 | 71.1601255 |
| -4.3148586 | 1.60E-05 | 5.20E-05 | 34.017042 | 114.46836 | 109.29313 | 122.148877 |
| -4.3146488 | 1.60E-05 | 5.21E-05 | 65.5750208 | 83.0775092 | 147.258111 | 49.8681194 |
| -4.3143298 | 1.60E-05 | 5.21E-05 | 51.230485 | 73.6061319 | 98.9390436 | 59.3934906 |
| -4.3143353 | 1.60E-05 | 5.21E-05 | 184.019902 | 154.789366 | 27.6108959 | 122.148877 |
| -4.3141843 | 1.60E-05 | 5.21E-05 | 30.738291 | 111.762252 | 134.603117 | 212.92006 |
| -4.3140797 | 1.60E-05 | 5.22E-05 | 31.9678226 | 106.350036 | 272.657597 | 136.717091 |
| -4.3133944 | 1.61E-05 | 5.23E-05 | 52.4600166 | 73.3355211 | 56.3722458 | 86.2886561 |
| -4.3130738 | 1.61E-05 | 5.24E-05 | 46.7222023 | 66.2996409 | 82.8326877 | 115.985401 |
| -4.3130544 | 1.61E-05 | 5.24E-05 | 44.6729829 | 91.1958326 | 66.7263317 | 97.4949751 |
| -4.3129933 | 1.61E-05 | 5.24E-05 | 111.067691 | 74.688575 | 112.744492 | 40.3427483 |
| -4.312783 | 1.61E-05 | 5.24E-05 | 35.2465737 | 94.7137727 | 123.098578 | 120.467929 |
| -4.3124631 | 1.61E-05 | 5.25E-05 | 113.526755 | 53.5809343 | 52.9208838 | 94.1330794 |
| -4.3122577 | 1.62E-05 | 5.26E-05 | 35.6564176 | 90.9252218 | 124.249032 | 123.829825 |
| -4.3122422 | 1.62E-05 | 5.26E-05 | 54.509236 | 126.104623 | 72.4786017 | 61.0744384 |
| -4.3121135 | 1.62E-05 | 5.26E-05 | 43.4434513 | 76.0416289 | 95.4876816 | 77.8839169 |
| -4.3118844 | 1.62E-05 | 5.26E-05 | 49.5911095 | 139.09394 | 55.2217918 | 128.872668 |
| -4.311687 | 1.62E-05 | 5.27E-05 | 72.9522106 | 84.4305631 | 44.8677058 | 71.7204414 |
| -4.3113234 | 1.62E-05 | 5.28E-05 | 88.5262781 | 36.5324552 | 132.302209 | 113.744138 |
| -4.3112462 | 1.62E-05 | 5.28E-05 | 166.806459 | 81.4538445 | 47.1686138 | 75.0823371 |

| | | | | | | |
|---|---|---|---|---|---|---|
| -4.311194 | 1.62E-05 | 5.28E-05 | 32.7875104 | 123.939737 | 119.647216 | 115.425085 |
| 4.31103398 | 1.62E-05 | 5.28E-05 | 0 | 0 | 0 | 0 |
| -4.3110386 | 1.62E-05 | 5.28E-05 | 37.705637 | 125.292791 | 147.258111 | 79.5648647 |
| -4.3109928 | 1.63E-05 | 5.28E-05 | 32.7875104 | 143.423713 | 121.948124 | 106.46003 |
| -4.310967 | 1.63E-05 | 5.28E-05 | 106.559409 | 81.9950661 | 46.0181598 | 64.99665 |
| -4.3107369 | 1.63E-05 | 5.29E-05 | 61.476582 | 75.7710182 | 55.2217918 | 57.7125427 |
| -4.3106325 | 1.63E-05 | 5.29E-05 | 75.8211178 | 71.4412457 | 47.1686138 | 132.79488 |
| -4.3104811 | 1.63E-05 | 5.29E-05 | 47.951734 | 61.6992576 | 80.5317797 | 82.9267604 |
| -4.3104163 | 1.63E-05 | 5.29E-05 | 495.091407 | 953.632386 | 936.469552 | 983.35449 |
| -4.3099384 | 1.63E-05 | 5.30E-05 | 35.2465737 | 120.692407 | 108.142676 | 100.296555 |
| -4.3095871 | 1.64E-05 | 5.31E-05 | 33.6071982 | 113.656527 | 108.142676 | 128.312352 |
| -4.3095175 | 1.64E-05 | 5.31E-05 | 46.7222023 | 74.1473535 | 69.0272397 | 73.9617052 |
| -4.3087167 | 1.64E-05 | 5.33E-05 | 61.0667381 | 240.572983 | 125.399485 | 152.966254 |
| -4.3084022 | 1.64E-05 | 5.34E-05 | 124.592539 | 397.527235 | 376.198456 | 427.521069 |
| -4.3080135 | 1.65E-05 | 5.35E-05 | 87.2967464 | 143.964935 | 200.178995 | 34.1792729 |
| -4.3079032 | 1.65E-05 | 5.35E-05 | 38.1154808 | 106.350036 | 92.0363196 | 93.5727634 |
| -4.3078396 | 1.65E-05 | 5.35E-05 | 42.2139196 | 80.3714014 | 82.8326877 | 84.6077082 |
| -4.307547 | 1.65E-05 | 5.36E-05 | 29.5087594 | 156.41303 | 140.355387 | 119.907613 |
| -4.3075095 | 1.65E-05 | 5.36E-05 | 251.234298 | 627.817008 | 595.93517 | 638.199865 |
| -4.3074597 | 1.65E-05 | 5.36E-05 | 42.6237635 | 68.1939163 | 106.992222 | 126.071088 |
| -4.3073517 | 1.65E-05 | 5.36E-05 | 59.8372065 | 87.1366709 | 54.0713378 | 61.6347543 |
| -4.3071971 | 1.65E-05 | 5.36E-05 | 704.931473 | 1301.36724 | 1392.04933 | 1407.51366 |
| -4.3070328 | 1.65E-05 | 5.37E-05 | 64.3454891 | 61.6992576 | 50.6199758 | 73.9617052 |
| -4.3068722 | 1.66E-05 | 5.37E-05 | 165.576927 | 475.463139 | 59.8236078 | 186.585211 |
| -4.3065741 | 1.66E-05 | 5.38E-05 | 27.8693838 | 104.996982 | 219.736713 | 191.067738 |
| -4.30628 | 1.66E-05 | 5.38E-05 | 29.5087594 | 153.977533 | 117.346308 | 145.682147 |
| -4.3049106 | 1.67E-05 | 5.42E-05 | 41.3942319 | 106.350036 | 121.948124 | 70.5998095 |
| -4.3047152 | 1.67E-05 | 5.42E-05 | 95.9034679 | 97.1492697 | 36.8145279 | 178.180472 |
| -4.3046165 | 1.67E-05 | 5.42E-05 | 50.8206411 | 83.34812 | 63.2749697 | 63.3157022 |
| -4.3045941 | 1.67E-05 | 5.42E-05 | 50.0009534 | 57.9107067 | 77.0804177 | 82.9267604 |
| -4.3043749 | 1.67E-05 | 5.43E-05 | 349.59683 | 85.783617 | 46.0181598 | 103.65845 |
| -4.3038707 | 1.68E-05 | 5.44E-05 | 47.1320462 | 138.82333 | 162.214013 | 57.1522268 |
| -4.3034311 | 1.68E-05 | 5.45E-05 | 560.666428 | 1216.39545 | 632.749697 | 856.162769 |
| -4.303007 | 1.68E-05 | 5.46E-05 | 121.723632 | 211.347019 | 193.276271 | 201.713742 |
| -4.3024964 | 1.69E-05 | 5.47E-05 | 25.4103206 | 160.201581 | 161.063559 | 155.207518 |
| -4.3012464 | 1.70E-05 | 5.50E-05 | 52.0501728 | 194.56915 | 56.3722458 | 113.183822 |
| -4.3006103 | 1.70E-05 | 5.52E-05 | 42.6237635 | 97.1492697 | 136.904025 | 70.0394936 |
| -4.3003862 | 1.71E-05 | 5.52E-05 | 61.476582 | 114.197749 | 48.3190678 | 79.5648647 |
| -4.2999932 | 1.71E-05 | 5.53E-05 | 60.6568942 | 172.920288 | 42.5667978 | 187.705843 |
| -4.2998128 | 1.71E-05 | 5.53E-05 | 36.4761053 | 93.090108 | 174.869007 | 102.537819 |
| -4.2995304 | 1.71E-05 | 5.54E-05 | 63.1159575 | 63.5935331 | 127.700393 | 56.5919108 |
| -4.29938 | 1.71E-05 | 5.54E-05 | 67.6242402 | 97.4198805 | 74.7795097 | 173.697944 |
| -4.2980975 | 1.72E-05 | 5.58E-05 | 34.017042 | 170.21418 | 86.2840497 | 141.759935 |

| | | | | | | |
|---|---|---|---|---|---|---|
| -4.2979909 | 1.72E-05 | 5.58E-05 | 47.951734 | 102.832096 | 149.559019 | 59.3934906 |
| -4.2978385 | 1.72E-05 | 5.58E-05 | 34.017042 | 145.588599 | 97.7885896 | 112.06319 |
| -4.2973832 | 1.73E-05 | 5.59E-05 | 33.6071982 | 135.034779 | 110.443584 | 102.537819 |
| -4.2970254 | 1.73E-05 | 5.60E-05 | 49.1812656 | 62.51109 | 67.8767857 | 70.5998095 |
| -4.2966137 | 1.73E-05 | 5.61E-05 | 38.1154808 | 91.7370541 | 93.1867736 | 95.2537113 |
| -4.2963538 | 1.74E-05 | 5.62E-05 | 43.8532952 | 129.081342 | 115.0454 | 62.7553862 |
| -4.296341 | 1.74E-05 | 5.62E-05 | 24.5906328 | 174.543953 | 125.399485 | 435.925808 |
| -4.2963152 | 1.74E-05 | 5.62E-05 | 40.5745441 | 92.2782757 | 79.3813257 | 87.409288 |
| -4.2962133 | 1.74E-05 | 5.62E-05 | 42.6237635 | 114.197749 | 80.5317797 | 75.0823371 |
| -4.2954868 | 1.74E-05 | 5.64E-05 | 141.805982 | 138.282108 | 37.9649818 | 72.2807574 |
| -4.2951766 | 1.75E-05 | 5.64E-05 | 64.3454891 | 90.9252218 | 95.4876816 | 101.417187 |
| -4.2950025 | 1.75E-05 | 5.65E-05 | 113.116911 | 136.929054 | 172.568099 | 164.732889 |
| -4.2943407 | 1.75E-05 | 5.66E-05 | 31.9678226 | 144.776767 | 103.54086 | 124.390141 |
| -4.2943529 | 1.75E-05 | 5.66E-05 | 41.3942319 | 141.800048 | 69.0272397 | 109.26161 |
| -4.2941195 | 1.75E-05 | 5.67E-05 | 42.2139196 | 99.5847667 | 78.2308717 | 76.202969 |
| -4.2940944 | 1.75E-05 | 5.67E-05 | 47.1320462 | 72.7942996 | 64.4254237 | 72.8410733 |
| -4.2940277 | 1.75E-05 | 5.67E-05 | 47.1320462 | 62.51109 | 74.7795097 | 77.3236009 |
| -4.2938361 | 1.76E-05 | 5.67E-05 | 63.5258014 | 94.1725511 | 43.7172518 | 104.779082 |
| -4.2937256 | 1.76E-05 | 5.68E-05 | 35.2465737 | 116.633246 | 135.753571 | 87.409288 |
| 4.29343119 | 1.76E-05 | 5.68E-05 | 160.248957 | 171.837845 | 195.577179 | 178.740788 |
| -4.2932411 | 1.76E-05 | 5.69E-05 | 40.1647002 | 96.0668266 | 121.948124 | 74.5220212 |
| -4.2931366 | 1.76E-05 | 5.69E-05 | 40.5745441 | 92.2782757 | 78.2308717 | 87.409288 |
| -4.2930849 | 1.76E-05 | 5.69E-05 | 71.3128351 | 56.8282636 | 78.2308717 | 50.4284354 |
| -4.2929216 | 1.76E-05 | 5.69E-05 | 29.5087594 | 120.963018 | 132.302209 | 143.440883 |
| -4.2928964 | 1.76E-05 | 5.69E-05 | 654.520676 | 145.588599 | 23.0090799 | 160.250361 |
| -4.2924818 | 1.77E-05 | 5.70E-05 | 38.1154808 | 114.46836 | 111.594038 | 77.8839169 |
| -4.2923879 | 1.77E-05 | 5.71E-05 | 79.9195566 | 62.51109 | 52.9208838 | 57.1522268 |
| -4.2924076 | 1.77E-05 | 5.71E-05 | 38.1154808 | 131.787449 | 81.6822337 | 99.1759229 |
| -4.2922773 | 1.77E-05 | 5.71E-05 | 58.6076748 | 72.5236888 | 56.3722458 | 56.0315949 |
| -4.292268 | 1.77E-05 | 5.71E-05 | 116.395662 | 101.208431 | 32.2127119 | 187.145527 |
| -4.2920148 | 1.77E-05 | 5.71E-05 | 51.230485 | 151.812647 | 51.7704298 | 243.177122 |
| -4.2919832 | 1.77E-05 | 5.71E-05 | 75.4112739 | 38.9679522 | 108.142676 | 132.234564 |
| 4.29185826 | 1.77E-05 | 5.72E-05 | 15.9839113 | 9.20076649 | 5.75226998 | 7.28410733 |
| -4.2916549 | 1.77E-05 | 5.72E-05 | 72.9522106 | 92.0076649 | 72.4786017 | 95.8140272 |
| -4.2915856 | 1.77E-05 | 5.72E-05 | 58.197831 | 70.3588026 | 89.7354116 | 49.8681194 |
| -4.291408 | 1.78E-05 | 5.73E-05 | 775.834465 | 1615.54635 | 1465.67839 | 1627.15751 |
| -4.2913209 | 1.78E-05 | 5.73E-05 | 47.1320462 | 75.5004074 | 62.1245158 | 75.6426531 |
| -4.291274 | 1.78E-05 | 5.73E-05 | 70.0833035 | 151.542036 | 39.1154358 | 124.950457 |
| -4.2911463 | 1.78E-05 | 5.73E-05 | 40.984388 | 122.316072 | 77.0804177 | 85.7283401 |
| -4.290984 | 1.78E-05 | 5.73E-05 | 233.611012 | 133.952336 | 88.5849576 | 78.4442328 |
| -4.2908771 | 1.78E-05 | 5.74E-05 | 36.0662614 | 113.385916 | 89.7354116 | 104.779082 |
| -4.2908506 | 1.78E-05 | 5.74E-05 | 39.7548564 | 90.9252218 | 85.1335957 | 84.6077082 |
| -4.2906543 | 1.78E-05 | 5.74E-05 | 47.5418901 | 139.364551 | 127.700393 | 55.4712789 |

| | | | | | | |
|---|---|---|---|---|---|---|
| 4.2903663 | 1.78E-05 | 5.75E-05 | 15.1642236 | 46.8156648 | 32.2127119 | 33.6189569 |
| -4.290126 | 1.79E-05 | 5.75E-05 | 39.3450125 | 75.5004074 | 104.691314 | 101.977503 |
| -4.2897606 | 1.79E-05 | 5.76E-05 | 43.0336074 | 121.50424 | 66.7263317 | 94.1330794 |
| -4.2888913 | 1.80E-05 | 5.78E-05 | 94.2640924 | 115.280192 | 144.957203 | 121.028245 |
| -4.2879004 | 1.80E-05 | 5.81E-05 | 233.611012 | 102.832096 | 82.8326877 | 99.7362389 |
| -4.2878798 | 1.80E-05 | 5.81E-05 | 41.8040758 | 94.4431619 | 79.3813257 | 75.0823371 |
| -4.2878273 | 1.80E-05 | 5.81E-05 | 86.0672148 | 89.8427787 | 41.4163438 | 70.5998095 |
| -4.2876879 | 1.81E-05 | 5.81E-05 | 49.1812656 | 102.832096 | 54.0713378 | 116.545717 |
| -4.2875614 | 1.81E-05 | 5.82E-05 | 31.5579788 | 102.561485 | 141.505841 | 127.19172 |
| -4.2875548 | 1.81E-05 | 5.82E-05 | 62.2962697 | 62.2404792 | 50.6199758 | 64.4363341 |
| -4.2874572 | 1.81E-05 | 5.82E-05 | 33.6071982 | 89.0309463 | 231.241253 | 134.475828 |
| -4.2874725 | 1.81E-05 | 5.82E-05 | 42.2139196 | 135.576 | 95.4876816 | 69.4791776 |
| 4.28720682 | 1.81E-05 | 5.82E-05 | 475.418901 | 213.241294 | 295.666677 | 245.418385 |
| -4.2871695 | 1.81E-05 | 5.82E-05 | 47.1320462 | 72.253078 | 88.5849576 | 61.6347543 |
| -4.2871258 | 1.81E-05 | 5.82E-05 | 50.0009534 | 58.1813175 | 78.2308717 | 67.2379138 |
| -4.2867023 | 1.81E-05 | 5.84E-05 | 68.4439279 | 156.41303 | 79.3813257 | 48.1871716 |
| -4.2864105 | 1.82E-05 | 5.84E-05 | 209.020379 | 179.685557 | 94.3372276 | 29.1364293 |
| -4.2863487 | 1.82E-05 | 5.84E-05 | 96.7231557 | 131.246228 | 133.452663 | 138.398039 |
| -4.2862139 | 1.82E-05 | 5.85E-05 | 45.9025146 | 124.210348 | 95.4876816 | 60.5141225 |
| -4.2861512 | 1.82E-05 | 5.85E-05 | 41.8040758 | 77.9359044 | 77.0804177 | 112.623506 |
| 4.28605695 | 1.82E-05 | 5.85E-05 | 31.5579788 | 30.0377965 | 34.5136199 | 29.6967453 |
| -4.2853479 | 1.82E-05 | 5.87E-05 | 52.4600166 | 79.5595691 | 59.8236078 | 59.3934906 |
| -4.2844308 | 1.83E-05 | 5.89E-05 | 54.9190799 | 74.9591858 | 50.6199758 | 94.6933953 |
| -4.2841945 | 1.83E-05 | 5.90E-05 | 46.7222023 | 78.7477367 | 71.3281477 | 62.7553862 |
| -4.2840512 | 1.84E-05 | 5.90E-05 | 110.248004 | 64.4053654 | 44.8677058 | 79.5648647 |
| -4.283904 | 1.84E-05 | 5.90E-05 | 68.8537718 | 63.5935331 | 222.037621 | 66.6775979 |
| -4.2838004 | 1.84E-05 | 5.91E-05 | 31.9678226 | 101.479042 | 124.249032 | 132.234564 |
| -4.2828744 | 1.84E-05 | 5.93E-05 | 35.6564176 | 94.7137727 | 106.992222 | 98.615607 |
| -4.2826375 | 1.85E-05 | 5.94E-05 | 95.493624 | 162.907689 | 182.922185 | 172.016996 |
| -4.2822576 | 1.85E-05 | 5.94E-05 | 40.1647002 | 89.0309463 | 116.195854 | 74.5220212 |
| -4.2820002 | 1.85E-05 | 5.95E-05 | 116.805506 | 79.0183475 | 36.8145279 | 104.218766 |
| -4.2819763 | 1.85E-05 | 5.95E-05 | 29.9186032 | 304.978348 | 356.640739 | 394.462428 |
| -4.2818698 | 1.85E-05 | 5.95E-05 | 81.968776 | 97.6904913 | 36.8145279 | 132.79488 |
| -4.2817378 | 1.85E-05 | 5.96E-05 | 79.5097127 | 81.4538445 | 44.8677058 | 62.1950703 |
| -4.280834 | 1.86E-05 | 5.98E-05 | 26.2300083 | 142.070659 | 157.612197 | 140.639303 |
| -4.2805812 | 1.86E-05 | 5.99E-05 | 36.0662614 | 93.090108 | 93.1867736 | 113.744138 |
| 4.28025386 | 1.87E-05 | 5.99E-05 | 77.0506494 | 83.34812 | 121.948124 | 86.2886561 |
| -4.2802093 | 1.87E-05 | 5.99E-05 | 14.7543797 | 276.564216 | 261.153057 | 212.359745 |
| -4.2797086 | 1.87E-05 | 6.01E-05 | 47.1320462 | 77.9359044 | 59.8236078 | 74.5220212 |
| -4.2796341 | 1.87E-05 | 6.01E-05 | 49.1812656 | 77.1240721 | 96.6381356 | 56.0315949 |
| -4.2791424 | 1.88E-05 | 6.02E-05 | 38.5253247 | 76.0416289 | 117.346308 | 98.055291 |
| -4.278815 | 1.88E-05 | 6.03E-05 | 39.3450125 | 107.973701 | 73.6290557 | 97.4949751 |
| -4.2788291 | 1.88E-05 | 6.03E-05 | 57.7879871 | 49.5217726 | 72.4786017 | 68.3585457 |

| | | | | | | |
|---|---|---|---|---|---|---|
| -4.278612 | 1.88E-05 | 6.03E-05 | 43.4434513 | 105.267593 | 64.4254237 | 87.409288 |
| -4.2784346 | 1.88E-05 | 6.04E-05 | 37.2957931 | 80.912623 | 100.089498 | 136.717091 |
| -4.2778054 | 1.89E-05 | 6.06E-05 | 37.2957931 | 105.538204 | 78.2308717 | 120.467929 |
| -4.277686 | 1.89E-05 | 6.06E-05 | 39.7548564 | 142.070659 | 65.5758777 | 186.024895 |
| -4.2772354 | 1.89E-05 | 6.07E-05 | 59.8372065 | 46.2744432 | 178.320369 | 110.942558 |
| -4.2772121 | 1.89E-05 | 6.07E-05 | 273.365868 | 530.938349 | 333.631659 | 416.31475 |
| -4.2763874 | 1.90E-05 | 6.09E-05 | 39.7548564 | 81.1832337 | 80.5317797 | 116.545717 |
| -4.2760545 | 1.90E-05 | 6.10E-05 | 36.0662614 | 98.7729344 | 90.8858656 | 100.856871 |
| -4.2759299 | 1.90E-05 | 6.10E-05 | 24.1807889 | 131.516839 | 185.223093 | 174.25826 |
| -4.2753753 | 1.91E-05 | 6.12E-05 | 34.4268859 | 113.656527 | 90.8858656 | 112.623506 |
| -4.2752173 | 1.91E-05 | 6.12E-05 | 42.2139196 | 84.9717847 | 78.2308717 | 71.1601255 |
| -4.2747831 | 1.91E-05 | 6.13E-05 | 38.5253247 | 95.2549943 | 78.2308717 | 96.3743432 |
| -4.2742708 | 1.92E-05 | 6.15E-05 | 50.8206411 | 78.477126 | 56.3722458 | 122.709193 |
| -4.274175 | 1.92E-05 | 6.15E-05 | 54.0993922 | 64.1347547 | 55.2217918 | 61.0744384 |
| -4.2739465 | 1.92E-05 | 6.15E-05 | 40.5745441 | 101.479042 | 88.5849576 | 71.1601255 |
| -4.2738031 | 1.92E-05 | 6.16E-05 | 44.263139 | 114.197749 | 60.9740618 | 93.5727634 |
| -4.2735141 | 1.92E-05 | 6.16E-05 | 47.951734 | 62.2404792 | 67.8767857 | 66.1172819 |
| 4.27331927 | 1.93E-05 | 6.17E-05 | 45.4926707 | 16.5072575 | 19.5577179 | 13.4475828 |
| -4.2732475 | 1.93E-05 | 6.17E-05 | 45.9025146 | 98.7729344 | 58.6731538 | 136.156776 |
| -4.273054 | 1.93E-05 | 6.18E-05 | 76.6408055 | 152.89509 | 149.559019 | 134.475828 |
| -4.2730328 | 1.93E-05 | 6.18E-05 | 45.4926707 | 166.155018 | 111.594038 | 59.3934906 |
| -4.2725948 | 1.93E-05 | 6.19E-05 | 28.2792277 | 108.244312 | 158.762651 | 150.164674 |
| -4.2712775 | 1.94E-05 | 6.22E-05 | 32.7875104 | 103.102707 | 120.79767 | 105.899714 |
| -4.2712096 | 1.94E-05 | 6.22E-05 | 56.5584554 | 120.151186 | 73.6290557 | 52.1093832 |
| -4.2710523 | 1.95E-05 | 6.23E-05 | 34.8367298 | 91.1958326 | 108.142676 | 104.779082 |
| -4.2710255 | 1.95E-05 | 6.23E-05 | 286.071028 | 51.9572696 | 86.2840497 | 73.9617052 |
| -4.2706232 | 1.95E-05 | 6.24E-05 | 35.2465737 | 117.715689 | 132.302209 | 80.6854966 |
| -4.2702865 | 1.95E-05 | 6.25E-05 | 35.6564176 | 99.3141559 | 92.0363196 | 99.1759229 |
| -4.2702167 | 1.95E-05 | 6.25E-05 | 73.7718984 | 57.0988744 | 134.603117 | 53.2300151 |
| -4.2701105 | 1.95E-05 | 6.25E-05 | 26.2300083 | 136.929054 | 132.302209 | 168.094785 |
| -4.2700829 | 1.95E-05 | 6.25E-05 | 59.8372065 | 65.7584193 | 48.3190678 | 66.6775979 |
| -4.2699783 | 1.95E-05 | 6.25E-05 | 55.3289238 | 60.3462037 | 52.9208838 | 67.7982298 |
| -4.269825 | 1.96E-05 | 6.26E-05 | 37.2957931 | 84.4305631 | 106.992222 | 88.5299199 |
| -4.2692732 | 1.96E-05 | 6.27E-05 | 45.0828268 | 130.705006 | 121.948124 | 115.985401 |
| -4.2689652 | 1.96E-05 | 6.28E-05 | 42.6237635 | 75.5004074 | 69.0272397 | 86.2886561 |
| -4.2683847 | 1.97E-05 | 6.30E-05 | 78.2801811 | 69.817581 | 75.9299637 | 42.5840121 |
| -4.2679471 | 1.97E-05 | 6.31E-05 | 38.1154808 | 97.4198805 | 80.5317797 | 89.0902358 |
| -4.267848 | 1.97E-05 | 6.31E-05 | 43.0336074 | 106.079425 | 69.0272397 | 75.6426531 |
| -4.2673955 | 1.98E-05 | 6.32E-05 | 64.3454891 | 71.4412457 | 120.79767 | 47.0665397 |
| -4.2672633 | 1.98E-05 | 6.33E-05 | 37.705637 | 94.1725511 | 87.4345036 | 85.1680242 |
| 4.26705303 | 1.98E-05 | 6.33E-05 | 30.738291 | 10.5538204 | 12.6549939 | 24.0935858 |
| -4.266871 | 1.98E-05 | 6.34E-05 | 161.478489 | 47.6274971 | 113.894946 | 54.910963 |
| 4.26650039 | 1.99E-05 | 6.35E-05 | 3.27875104 | 1.89427545 | 0 | 2.80157974 |

| | | | | | | |
|---|---|---|---|---|---|---|
| -4.2663765 | 1.99E-05 | 6.35E-05 | 30.3284471 | 186.721438 | 98.9390436 | 128.312352 |
| -4.266334 | 1.99E-05 | 6.35E-05 | 48.7714217 | 101.479042 | 95.4876816 | 52.6696992 |
| -4.2662913 | 1.99E-05 | 6.35E-05 | 484.84531 | 545.821942 | 915.76138 | 700.955252 |
| -4.2660981 | 1.99E-05 | 6.35E-05 | 33.1973543 | 95.525605 | 124.249032 | 106.46003 |
| -4.2657807 | 1.99E-05 | 6.36E-05 | 128.281134 | 122.586683 | 121.948124 | 29.1364293 |
| 4.26572569 | 1.99E-05 | 6.36E-05 | 0 | 0 | 0 | 0 |
| 4.26572569 | 1.99E-05 | 6.36E-05 | 0 | 0 | 0 | 0 |
| -4.2657158 | 1.99E-05 | 6.36E-05 | 45.4926707 | 66.5702517 | 65.5758777 | 74.5220212 |
| -4.2656289 | 1.99E-05 | 6.36E-05 | 47.1320462 | 61.1580361 | 66.7263317 | 71.1601255 |
| -4.2656223 | 1.99E-05 | 6.36E-05 | 18.8528185 | 179.414947 | 162.214013 | 336.749885 |
| -4.2655331 | 1.99E-05 | 6.37E-05 | 68.4439279 | 93.3607188 | 81.6822337 | 41.4633802 |
| 4.26539758 | 2.00E-05 | 6.37E-05 | 5.73781432 | 4.87099402 | 4.60181598 | 2.80157974 |
| -4.2653718 | 2.00E-05 | 6.37E-05 | 41.8040758 | 74.1473535 | 72.4786017 | 98.055291 |
| -4.2652739 | 2.00E-05 | 6.37E-05 | 65.1651769 | 100.125988 | 42.5667978 | 163.612257 |
| -4.2652355 | 2.00E-05 | 6.37E-05 | 30.738291 | 125.563402 | 209.382627 | 98.055291 |
| -4.2647074 | 2.00E-05 | 6.39E-05 | 169.265522 | 87.4072817 | 36.8145279 | 90.7711837 |
| -4.2646506 | 2.00E-05 | 6.39E-05 | 32.3776665 | 153.977533 | 85.1335957 | 140.078987 |
| -4.2640873 | 2.01E-05 | 6.40E-05 | 58.6076748 | 55.2045989 | 65.5758777 | 53.7903311 |
| -4.2637559 | 2.01E-05 | 6.41E-05 | 33.1973543 | 108.244312 | 139.204933 | 91.8918156 |
| -4.2632056 | 2.02E-05 | 6.43E-05 | 107.379097 | 139.364551 | 130.001301 | 155.767834 |
| -4.2631005 | 2.02E-05 | 6.43E-05 | 46.7222023 | 54.9339882 | 110.443584 | 96.9346591 |
| -4.2630972 | 2.02E-05 | 6.43E-05 | 150.822548 | 71.1706349 | 48.3190678 | 68.3585457 |
| -4.2629553 | 2.02E-05 | 6.43E-05 | 56.1486116 | 84.9717847 | 55.2217918 | 56.5919108 |
| -4.2629459 | 2.02E-05 | 6.43E-05 | 52.8698605 | 89.5721679 | 202.479903 | 58.8331746 |
| -4.2624781 | 2.02E-05 | 6.44E-05 | 36.8859492 | 127.728288 | 207.081719 | 73.4013893 |
| -4.2624704 | 2.02E-05 | 6.44E-05 | 500.829221 | 993.41217 | 773.105085 | 1058.43683 |
| -4.2624888 | 2.02E-05 | 6.44E-05 | 37.2957931 | 76.0416289 | 100.089498 | 117.106033 |
| -4.2620207 | 2.03E-05 | 6.46E-05 | 35.2465737 | 103.914539 | 90.8858656 | 96.3743432 |
| -4.2618521 | 2.03E-05 | 6.46E-05 | 20.0823501 | 202.416863 | 157.612197 | 201.153426 |
| -4.2617434 | 2.03E-05 | 6.46E-05 | 179.921463 | 74.1473535 | 42.5667978 | 86.2886561 |
| 4.26141326 | 2.03E-05 | 6.47E-05 | 0 | 0 | 0 | 0 |
| -4.2613885 | 2.03E-05 | 6.47E-05 | 29.0989155 | 119.880575 | 133.452663 | 119.907613 |
| -4.26126 | 2.03E-05 | 6.47E-05 | 176.232868 | 428.647474 | 437.172518 | 381.575161 |
| -4.2609665 | 2.04E-05 | 6.48E-05 | 41.3942319 | 89.3015571 | 65.5758777 | 99.1759229 |
| -4.2605826 | 2.04E-05 | 6.49E-05 | 35.6564176 | 96.0668266 | 93.1867736 | 93.5727634 |
| -4.2605279 | 2.04E-05 | 6.49E-05 | 32.7875104 | 115.009581 | 96.6381356 | 110.942558 |
| -4.2604464 | 2.04E-05 | 6.50E-05 | 38.9351686 | 82.5362876 | 79.3813257 | 89.0902358 |
| 4.2602858 | 2.04E-05 | 6.50E-05 | 22.9512573 | 9.20076649 | 0 | 12.8872668 |
| -4.2601279 | 2.04E-05 | 6.50E-05 | 35.6564176 | 90.9252218 | 96.6381356 | 95.2537113 |
| -4.2600826 | 2.04E-05 | 6.50E-05 | 48.3615778 | 60.075593 | 62.1245158 | 80.1251807 |
| -4.259611 | 2.05E-05 | 6.52E-05 | 45.4926707 | 96.8786589 | 162.214013 | 151.285306 |
| -4.2593316 | 2.05E-05 | 6.52E-05 | 36.8859492 | 106.079425 | 110.443584 | 75.6426531 |
| -4.2588471 | 2.05E-05 | 6.54E-05 | 45.4926707 | 113.385916 | 56.3722458 | 95.2537113 |

| | | | | | | |
|---|---|---|---|---|---|---|
| -4.2587625 | 2.06E-05 | 6.54E-05 | 60.6568942 | 102.020264 | 94.3372276 | 105.899714 |
| -4.2584849 | 2.06E-05 | 6.55E-05 | 32.3776665 | 99.3141559 | 140.355387 | 102.537819 |
| -4.2583715 | 2.06E-05 | 6.55E-05 | 36.8859492 | 77.6652936 | 108.142676 | 98.055291 |
| 4.25818176 | 2.06E-05 | 6.56E-05 | 2.86890716 | 1.89427545 | 1.150454 | 3.36189569 |
| -4.2581177 | 2.06E-05 | 6.56E-05 | 47.1320462 | 73.3355211 | 65.5758777 | 59.9538065 |
| -4.2580684 | 2.06E-05 | 6.56E-05 | 48.7714217 | 56.8282636 | 83.9831417 | 65.556966 |
| 4.25809037 | 2.06E-05 | 6.56E-05 | 33.6071982 | 23.272527 | 18.4072639 | 21.852322 |
| -4.2576465 | 2.07E-05 | 6.57E-05 | 87.2967464 | 71.4412457 | 44.8677058 | 60.5141225 |
| -4.2575092 | 2.07E-05 | 6.57E-05 | 114.756286 | 78.477126 | 64.4254237 | 69.4791776 |
| -4.2574975 | 2.07E-05 | 6.57E-05 | 64.3454891 | 125.292791 | 83.9831417 | 97.4949751 |
| -4.2574654 | 2.07E-05 | 6.57E-05 | 54.9190799 | 142.611881 | 124.249032 | 129.432984 |
| -4.2572917 | 2.07E-05 | 6.58E-05 | 72.5423667 | 95.525605 | 40.2658898 | 75.0823371 |
| -4.2572996 | 2.07E-05 | 6.58E-05 | 141.805982 | 54.6633774 | 70.1776937 | 57.7125427 |
| -4.2569802 | 2.07E-05 | 6.58E-05 | 27.8693838 | 140.176384 | 117.346308 | 134.475828 |
| -4.2565701 | 2.08E-05 | 6.60E-05 | 51.6403289 | 67.1114732 | 56.3722458 | 58.8331746 |
| -4.2563476 | 2.08E-05 | 6.60E-05 | 39.7548564 | 85.783617 | 119.647216 | 71.7204414 |
| -4.2560695 | 2.08E-05 | 6.61E-05 | 39.7548564 | 71.1706349 | 88.5849576 | 87.9696039 |
| -4.255903 | 2.08E-05 | 6.61E-05 | 31.5579788 | 154.518755 | 95.4876816 | 114.304454 |
| -4.2559181 | 2.08E-05 | 6.61E-05 | 49.1812656 | 109.326755 | 63.2749697 | 63.3157022 |
| -4.2557478 | 2.08E-05 | 6.62E-05 | 49.5911095 | 135.846611 | 54.0713378 | 88.5299199 |
| -4.2551032 | 2.09E-05 | 6.63E-05 | 21.3118818 | 165.343186 | 182.922185 | 166.413837 |
| -4.25512 | 2.09E-05 | 6.63E-05 | 43.4434513 | 89.5721679 | 63.2749697 | 75.6426531 |
| -4.2550265 | 2.09E-05 | 6.64E-05 | 40.984388 | 103.914539 | 75.9299637 | 72.2807574 |
| -4.2549166 | 2.09E-05 | 6.64E-05 | 45.0828268 | 142.070659 | 56.3722458 | 107.020346 |
| -4.2548079 | 2.09E-05 | 6.64E-05 | 31.1481349 | 134.493557 | 109.29313 | 103.098135 |
| -4.2547789 | 2.09E-05 | 6.64E-05 | 36.0662614 | 77.3946828 | 108.142676 | 110.382242 |
| -4.25474 | 2.09E-05 | 6.64E-05 | 53.2797044 | 52.7691019 | 123.098578 | 75.0823371 |
| -4.2547493 | 2.09E-05 | 6.64E-05 | 66.3947085 | 147.212264 | 82.8326877 | 44.2649599 |
| -4.2546378 | 2.09E-05 | 6.64E-05 | 14.3445358 | 153.977533 | 318.675757 | 345.154624 |
| -4.2542076 | 2.10E-05 | 6.66E-05 | 41.8040758 | 61.9698684 | 108.142676 | 96.9346591 |
| -4.2541664 | 2.10E-05 | 6.66E-05 | 94.6739363 | 61.9698684 | 60.9740618 | 48.7474875 |
| -4.2539228 | 2.10E-05 | 6.66E-05 | 40.5745441 | 79.8301798 | 138.054479 | 75.6426531 |
| -4.2538874 | 2.10E-05 | 6.66E-05 | 14.7543797 | 260.868791 | 222.037621 | 220.764484 |
| -4.2538564 | 2.10E-05 | 6.66E-05 | 38.1154808 | 97.9611021 | 124.249032 | 71.7204414 |
| -4.2536544 | 2.10E-05 | 6.67E-05 | 44.6729829 | 117.445078 | 55.2217918 | 115.425085 |
| -4.2534868 | 2.10E-05 | 6.67E-05 | 61.476582 | 76.8534613 | 81.6822337 | 43.704644 |
| -4.2533854 | 2.11E-05 | 6.68E-05 | 34.8367298 | 106.079425 | 94.3372276 | 90.7711837 |
| -4.2533635 | 2.11E-05 | 6.68E-05 | 86.4770587 | 57.640096 | 47.1686138 | 67.2379138 |
| -4.2529786 | 2.11E-05 | 6.69E-05 | 42.6237635 | 148.024096 | 67.8767857 | 86.2886561 |
| -4.2529702 | 2.11E-05 | 6.69E-05 | 146.314265 | 66.8408624 | 59.8236078 | 54.910963 |
| -4.2527158 | 2.11E-05 | 6.69E-05 | 59.8372065 | 96.0668266 | 52.9208838 | 56.5919108 |
| -4.2526685 | 2.11E-05 | 6.69E-05 | 36.8859492 | 133.411114 | 74.7795097 | 102.537819 |
| -4.2522027 | 2.12E-05 | 6.71E-05 | 32.3776665 | 104.996982 | 106.992222 | 104.779082 |

| | | | | | | |
|---|---|---|---|---|---|---|
| -4.2520175 | 2.12E-05 | 6.71E-05 | 103.280658 | 74.4179643 | 47.1686138 | 56.5919108 |
| -4.25168 | 2.12E-05 | 6.72E-05 | 57.7879871 | 58.7225391 | 52.9208838 | 57.7125427 |
| -4.2514652 | 2.12E-05 | 6.73E-05 | 31.1481349 | 109.056144 | 101.239952 | 136.156776 |
| -4.2514005 | 2.12E-05 | 6.73E-05 | 77.8703372 | 120.421797 | 43.7172518 | 63.8760181 |
| -4.2513056 | 2.13E-05 | 6.73E-05 | 56.5584554 | 53.0397127 | 56.3722458 | 63.3157022 |
| -4.2512782 | 2.13E-05 | 6.73E-05 | 102.870814 | 97.6904913 | 57.5226998 | 43.704644 |
| -4.2511778 | 2.13E-05 | 6.73E-05 | 57.7879871 | 104.455761 | 70.1776937 | 122.148877 |
| -4.2505042 | 2.13E-05 | 6.75E-05 | 41.8040758 | 92.2782757 | 62.1245158 | 97.4949751 |
| -4.2505167 | 2.13E-05 | 6.75E-05 | 33.6071982 | 103.643928 | 106.992222 | 92.4521315 |
| -4.2503888 | 2.13E-05 | 6.75E-05 | 59.8372065 | 165.343186 | 106.992222 | 115.985401 |
| -4.2498021 | 2.14E-05 | 6.77E-05 | 22.5414134 | 142.882491 | 172.568099 | 170.336048 |
| -4.249489 | 2.14E-05 | 6.78E-05 | 35.2465737 | 102.832096 | 85.1335957 | 97.4949751 |
| 4.24922121 | 2.15E-05 | 6.79E-05 | 4.50828268 | 2.43549701 | 2.30090799 | 3.36189569 |
| -4.2489728 | 2.15E-05 | 6.79E-05 | 52.0501728 | 74.688575 | 56.3722458 | 56.5919108 |
| -4.2489655 | 2.15E-05 | 6.79E-05 | 36.4761053 | 86.8660601 | 130.001301 | 83.4870763 |
| -4.2489048 | 2.15E-05 | 6.80E-05 | 45.9025146 | 74.1473535 | 151.859927 | 68.3585457 |
| -4.2486499 | 2.15E-05 | 6.80E-05 | 24.5906328 | 216.759234 | 105.841768 | 185.464579 |
| -4.2483399 | 2.15E-05 | 6.81E-05 | 36.4761053 | 90.3840002 | 80.5317797 | 105.899714 |
| -4.2482739 | 2.15E-05 | 6.81E-05 | 36.4761053 | 86.0542278 | 89.7354116 | 90.2108677 |
| -4.2479651 | 2.16E-05 | 6.82E-05 | 59.0175187 | 93.3607188 | 52.9208838 | 56.0315949 |
| -4.2472604 | 2.16E-05 | 6.84E-05 | 34.8367298 | 101.479042 | 113.894946 | 82.3664444 |
| -4.2469946 | 2.17E-05 | 6.85E-05 | 36.8859492 | 122.045461 | 73.6290557 | 100.856871 |
| -4.2468792 | 2.17E-05 | 6.85E-05 | 20.492194 | 176.167617 | 158.762651 | 193.309002 |
| -4.2468002 | 2.17E-05 | 6.85E-05 | 26.6398522 | 138.552719 | 140.355387 | 122.709193 |
| -4.2466761 | 2.17E-05 | 6.86E-05 | 38.5253247 | 97.1492697 | 69.0272397 | 114.864769 |
| -4.2463697 | 2.17E-05 | 6.87E-05 | 228.692885 | 515.242923 | 388.85345 | 425.279805 |
| -4.2458837 | 2.18E-05 | 6.88E-05 | 29.5087594 | 188.886324 | 102.390406 | 118.226665 |
| -4.2453673 | 2.18E-05 | 6.90E-05 | 41.3942319 | 133.681725 | 72.4786017 | 77.8839169 |
| -4.2452762 | 2.18E-05 | 6.90E-05 | 37.705637 | 88.219114 | 78.2308717 | 87.9696039 |
| -4.2449822 | 2.19E-05 | 6.91E-05 | 30.3284471 | 112.574084 | 123.098578 | 106.46003 |
| -4.2444364 | 2.19E-05 | 6.92E-05 | 22.5414134 | 131.787449 | 196.727633 | 166.974153 |
| -4.244186 | 2.19E-05 | 6.93E-05 | 33.6071982 | 94.1725511 | 190.975363 | 98.055291 |
| -4.2441843 | 2.19E-05 | 6.93E-05 | 48.7714217 | 70.6294134 | 54.0713378 | 67.2379138 |
| -4.2440619 | 2.20E-05 | 6.93E-05 | 74.5915861 | 70.9000241 | 87.4345036 | 40.3427483 |
| 4.24387247 | 2.20E-05 | 6.94E-05 | 0 | 0 | 0 | 0 |
| -4.2437289 | 2.20E-05 | 6.94E-05 | 52.4600166 | 73.6061319 | 106.992222 | 50.4284354 |
| 4.24349192 | 2.20E-05 | 6.95E-05 | 72.1325229 | 37.6148983 | 48.3190678 | 45.3855918 |
| -4.2434055 | 2.20E-05 | 6.95E-05 | 427.877011 | 306.331402 | 309.472125 | 321.061039 |
| -4.2433417 | 2.20E-05 | 6.95E-05 | 24.5906328 | 160.201581 | 117.346308 | 180.421735 |
| -4.243357 | 2.20E-05 | 6.95E-05 | 32.3776665 | 102.290875 | 109.29313 | 99.7362389 |
| -4.2431382 | 2.20E-05 | 6.96E-05 | 38.5253247 | 139.09394 | 71.3281477 | 94.1330794 |
| 4.24300464 | 2.21E-05 | 6.96E-05 | 84.4278393 | 28.4141318 | 46.0181598 | 17.9301104 |
| -4.2421073 | 2.21E-05 | 6.99E-05 | 30.3284471 | 105.538204 | 108.142676 | 131.674248 |

| | | | | | | |
|---|---|---|---|---|---|---|
| -4.2420167 | 2.22E-05 | 6.99E-05 | 29.5087594 | 135.30539 | 112.744492 | 108.701294 |
| -4.2419506 | 2.22E-05 | 6.99E-05 | 40.5745441 | 80.3714014 | 92.0363196 | 66.6775979 |
| -4.241736 | 2.22E-05 | 7.00E-05 | 35.2465737 | 108.244312 | 81.6822337 | 96.3743432 |
| -4.2417021 | 2.22E-05 | 7.00E-05 | 60.2470503 | 54.6633774 | 50.6199758 | 82.3664444 |
| -4.2416695 | 2.22E-05 | 7.00E-05 | 44.6729829 | 81.7244553 | 80.5317797 | 57.1522268 |
| -4.241203 | 2.22E-05 | 7.01E-05 | 89.3459658 | 75.2297966 | 43.7172518 | 58.8331746 |
| -4.2412224 | 2.22E-05 | 7.01E-05 | 290.579311 | 438.930684 | 393.455266 | 393.902112 |
| -4.2411153 | 2.22E-05 | 7.01E-05 | 43.4434513 | 60.075593 | 79.3813257 | 79.0045488 |
| -4.2410132 | 2.23E-05 | 7.01E-05 | 72.1325229 | 223.795114 | 94.3372276 | 117.106033 |
| -4.2410111 | 2.23E-05 | 7.01E-05 | 36.4761053 | 80.912623 | 94.3372276 | 89.0902358 |
| -4.2409152 | 2.23E-05 | 7.02E-05 | 62.2962697 | 262.221845 | 41.4163438 | 129.9933 |
| -4.2408398 | 2.23E-05 | 7.02E-05 | 29.9186032 | 177.25006 | 92.0363196 | 129.432984 |
| -4.2407958 | 2.23E-05 | 7.02E-05 | 32.7875104 | 110.138587 | 102.390406 | 94.1330794 |
| -4.2407325 | 2.23E-05 | 7.02E-05 | 13.524848 | 267.36345 | 213.984443 | 252.702493 |
| -4.2403973 | 2.23E-05 | 7.03E-05 | 53.6895483 | 49.5217726 | 71.3281477 | 63.8760181 |
| -4.2403926 | 2.23E-05 | 7.03E-05 | 71.722679 | 70.0881918 | 222.037621 | 52.6696992 |
| -4.2403782 | 2.23E-05 | 7.03E-05 | 716.407102 | 2554.02453 | 2518.3438 | 2732.66088 |
| -4.2401975 | 2.23E-05 | 7.03E-05 | 40.5745441 | 91.1958326 | 65.5758777 | 83.4870763 |
| -4.240215 | 2.23E-05 | 7.03E-05 | 81.968776 | 31.3908504 | 142.656295 | 165.293205 |
| -4.2400557 | 2.23E-05 | 7.04E-05 | 35.6564176 | 80.912623 | 110.443584 | 91.3314996 |
| -4.239974 | 2.24E-05 | 7.04E-05 | 31.9678226 | 136.117222 | 95.4876816 | 101.977503 |
| -4.2399311 | 2.24E-05 | 7.04E-05 | 222.955071 | 213.241294 | 126.549939 | 234.772382 |
| -4.2399012 | 2.24E-05 | 7.04E-05 | 64.755333 | 84.4305631 | 89.7354116 | 103.098135 |
| -4.2396758 | 2.24E-05 | 7.05E-05 | 97.9526873 | 87.1366709 | 77.0804177 | 36.9808526 |
| -4.2394722 | 2.24E-05 | 7.05E-05 | 36.0662614 | 77.9359044 | 93.1867736 | 110.382242 |
| -4.2394244 | 2.24E-05 | 7.05E-05 | 45.0828268 | 72.7942996 | 57.5226998 | 85.1680242 |
| -4.2392446 | 2.24E-05 | 7.06E-05 | 40.5745441 | 103.643928 | 63.2749697 | 91.3314996 |
| -4.2391301 | 2.24E-05 | 7.06E-05 | 124.182696 | 120.692407 | 27.6108959 | 121.028245 |
| -4.2390206 | 2.24E-05 | 7.06E-05 | 33.6071982 | 132.05806 | 77.0804177 | 130.553616 |
| -4.2388962 | 2.25E-05 | 7.07E-05 | 68.4439279 | 99.8553775 | 140.355387 | 80.1251807 |
| -4.2383728 | 2.25E-05 | 7.08E-05 | 37.2957931 | 84.1599523 | 77.0804177 | 104.218766 |
| -4.2383443 | 2.25E-05 | 7.08E-05 | 334.84245 | 88.4897248 | 33.3631659 | 122.148877 |
| 4.23828459 | 2.25E-05 | 7.08E-05 | 76.2309617 | 30.579018 | 37.9649818 | 33.058641 |
| -4.2382997 | 2.25E-05 | 7.08E-05 | 49.5911095 | 56.0164313 | 104.691314 | 66.1172819 |
| -4.237994 | 2.26E-05 | 7.09E-05 | 231.561792 | 109.867976 | 223.188075 | 21.852322 |
| -4.2379698 | 2.26E-05 | 7.09E-05 | 40.5745441 | 75.5004074 | 71.3281477 | 76.763285 |
| -4.2375724 | 2.26E-05 | 7.10E-05 | 61.476582 | 71.4412457 | 44.8677058 | 63.3157022 |
| -4.2374373 | 2.26E-05 | 7.11E-05 | 187.298653 | 163.719521 | 104.691314 | 122.148877 |
| -4.2371036 | 2.26E-05 | 7.12E-05 | 29.5087594 | 185.638994 | 82.8326877 | 168.094785 |
| -4.2370758 | 2.26E-05 | 7.12E-05 | 37.2957931 | 82.8068984 | 110.443584 | 77.3236009 |
| -4.2368656 | 2.27E-05 | 7.12E-05 | 64.3454891 | 101.749653 | 47.1686138 | 59.9538065 |
| -4.2368316 | 2.27E-05 | 7.12E-05 | 56.9682993 | 113.115306 | 105.841768 | 132.234564 |
| 4.23659692 | 2.27E-05 | 7.13E-05 | 21.7217256 | 16.2366467 | 17.2568099 | 12.8872668 |

| | | | | | | |
|---|---|---|---|---|---|---|
| -4.2365416 | 2.27E-05 | 7.13E-05 | 91.8050291 | 32.2026827 | 115.0454 | 109.821926 |
| -4.2364677 | 2.27E-05 | 7.13E-05 | 26.6398522 | 104.455761 | 190.975363 | 141.199619 |
| -4.236438 | 2.27E-05 | 7.13E-05 | 36.8859492 | 100.937821 | 71.3281477 | 110.382242 |
| -4.2364101 | 2.27E-05 | 7.13E-05 | 63.1159575 | 102.020264 | 41.4163438 | 78.4442328 |
| -4.236279 | 2.27E-05 | 7.14E-05 | 45.4926707 | 102.561485 | 144.957203 | 54.350647 |
| -4.2360496 | 2.27E-05 | 7.14E-05 | 40.5745441 | 72.253078 | 73.6290557 | 76.202969 |
| -4.2360473 | 2.27E-05 | 7.14E-05 | 53.6895483 | 93.3607188 | 44.8677058 | 98.615607 |
| -4.2359583 | 2.28E-05 | 7.14E-05 | 50.8206411 | 64.6759762 | 56.3722458 | 56.5919108 |
| -4.2357655 | 2.28E-05 | 7.15E-05 | 40.5745441 | 96.0668266 | 65.5758777 | 80.6854966 |
| -4.2357718 | 2.28E-05 | 7.15E-05 | 52.0501728 | 57.0988744 | 54.0713378 | 64.4363341 |
| -4.2351305 | 2.28E-05 | 7.17E-05 | 37.2957931 | 74.688575 | 88.5849576 | 94.1330794 |
| -4.2350342 | 2.29E-05 | 7.17E-05 | 24.5906328 | 182.932887 | 124.249032 | 138.958355 |
| 4.23487372 | 2.29E-05 | 7.18E-05 | 193.446311 | 147.753485 | 128.850847 | 129.432984 |
| -4.2348347 | 2.29E-05 | 7.18E-05 | 28.6890716 | 132.869893 | 100.089498 | 138.958355 |
| -4.2344848 | 2.29E-05 | 7.19E-05 | 43.8532952 | 57.3694852 | 86.2840497 | 109.26161 |
| -4.234212 | 2.29E-05 | 7.20E-05 | 55.7387677 | 47.8981079 | 101.239952 | 68.9188617 |
| -4.233937 | 2.30E-05 | 7.20E-05 | 52.0501728 | 76.8534613 | 52.9208838 | 58.2728587 |
| -4.2338256 | 2.30E-05 | 7.21E-05 | 14.3445358 | 169.943569 | 378.499364 | 228.608907 |
| -4.2337251 | 2.30E-05 | 7.21E-05 | 56.5584554 | 104.996982 | 59.8236078 | 52.6696992 |
| -4.2336187 | 2.30E-05 | 7.21E-05 | 292.218686 | 55.4752097 | 69.0272397 | 77.3236009 |
| -4.2334088 | 2.30E-05 | 7.22E-05 | 47.1320462 | 55.4752097 | 72.4786017 | 65.556966 |
| -4.2330872 | 2.31E-05 | 7.23E-05 | 38.9351686 | 97.6904913 | 127.700393 | 65.556966 |
| -4.2328474 | 2.31E-05 | 7.23E-05 | 90.5754975 | 148.835929 | 164.514921 | 28.5761134 |
| -4.2327287 | 2.31E-05 | 7.24E-05 | 29.0989155 | 157.766084 | 97.7885896 | 122.709193 |
| -4.2327062 | 2.31E-05 | 7.24E-05 | 50.0009534 | 109.326755 | 138.054479 | 105.899714 |
| -4.232602 | 2.31E-05 | 7.24E-05 | 50.8206411 | 61.6992576 | 60.9740618 | 53.7903311 |
| -4.2317263 | 2.32E-05 | 7.27E-05 | 54.9190799 | 45.1920001 | 86.2840497 | 79.5648647 |
| -4.2314411 | 2.32E-05 | 7.28E-05 | 25.8201644 | 171.026012 | 157.612197 | 104.779082 |
| -4.2314127 | 2.32E-05 | 7.28E-05 | 90.9853413 | 177.520671 | 92.0363196 | 34.1792729 |
| -4.2313741 | 2.32E-05 | 7.28E-05 | 36.0662614 | 89.3015571 | 81.6822337 | 90.2108677 |
| -4.2313502 | 2.32E-05 | 7.28E-05 | 283.202121 | 912.228936 | 873.194582 | 862.886561 |
| -4.2307652 | 2.33E-05 | 7.29E-05 | 53.6895483 | 85.5130062 | 49.4695218 | 62.7553862 |
| -4.2304497 | 2.33E-05 | 7.30E-05 | 54.9190799 | 103.643928 | 81.6822337 | 45.3855918 |
| -4.2303584 | 2.33E-05 | 7.31E-05 | 28.6890716 | 122.316072 | 124.249032 | 108.140978 |
| -4.2300412 | 2.34E-05 | 7.32E-05 | 41.3942319 | 76.8534613 | 102.390406 | 63.8760181 |
| -4.230011 | 2.34E-05 | 7.32E-05 | 50.4107972 | 89.3015571 | 49.4695218 | 72.2807574 |
| -4.2294891 | 2.34E-05 | 7.33E-05 | 31.9678226 | 108.785533 | 87.4345036 | 132.79488 |
| -4.229349 | 2.34E-05 | 7.34E-05 | 39.7548564 | 90.1133895 | 64.4254237 | 90.7711837 |
| -4.2286413 | 2.35E-05 | 7.36E-05 | 89.7558097 | 37.8855091 | 200.178995 | 81.2458125 |
| -4.2284086 | 2.35E-05 | 7.37E-05 | 47.951734 | 51.416048 | 92.0363196 | 76.202969 |
| -4.2283796 | 2.35E-05 | 7.37E-05 | 40.984388 | 63.8641439 | 146.107657 | 91.8918156 |
| -4.2283556 | 2.35E-05 | 7.37E-05 | 68.8537718 | 80.912623 | 41.4163438 | 64.99665 |
| -4.2282613 | 2.36E-05 | 7.37E-05 | 160.658801 | 60.6168145 | 112.744492 | 40.3427483 |

| | | | | | | |
|---|---|---|---|---|---|---|
| 4.22818423 | 2.36E-05 | 7.37E-05 | 0.40984388 | 1.08244312 | 0 | 0.56031595 |
| -4.2281938 | 2.36E-05 | 7.37E-05 | 49.1812656 | 49.7923834 | 77.0804177 | 85.7283401 |
| -4.2281861 | 2.36E-05 | 7.37E-05 | 48.7714217 | 73.3355211 | 56.3722458 | 128.872668 |
| -4.2280709 | 2.36E-05 | 7.37E-05 | 84.8376831 | 62.51109 | 40.2658898 | 83.4870763 |
| -4.2279557 | 2.36E-05 | 7.38E-05 | 73.3620545 | 64.4053654 | 51.7704298 | 49.3078035 |
| -4.2269991 | 2.37E-05 | 7.41E-05 | 36.0662614 | 112.032863 | 70.1776937 | 111.502874 |
| -4.2267565 | 2.37E-05 | 7.41E-05 | 38.1154808 | 90.9252218 | 67.8767857 | 101.977503 |
| -4.2262953 | 2.38E-05 | 7.43E-05 | 38.9351686 | 69.2763595 | 81.6822337 | 111.502874 |
| -4.2255823 | 2.38E-05 | 7.45E-05 | 50.8206411 | 133.952336 | 150.709473 | 149.604358 |
| 4.2253414 | 2.39E-05 | 7.46E-05 | 46.7222023 | 31.3908504 | 26.4604419 | 24.6539017 |
| -4.2253457 | 2.39E-05 | 7.46E-05 | 29.9186032 | 175.085174 | 192.125817 | 205.075637 |
| -4.2252233 | 2.39E-05 | 7.46E-05 | 38.5253247 | 131.787449 | 66.7263317 | 94.6933953 |
| -4.2251008 | 2.39E-05 | 7.46E-05 | 37.705637 | 74.1473535 | 88.5849576 | 81.2458125 |
| -4.2249172 | 2.39E-05 | 7.47E-05 | 85.6573709 | 90.654611 | 65.5758777 | 39.2221164 |
| -4.2247561 | 2.39E-05 | 7.47E-05 | 36.8859492 | 86.0542278 | 78.2308717 | 84.6077082 |
| -4.2242706 | 2.40E-05 | 7.49E-05 | 51.230485 | 61.6992576 | 50.6199758 | 79.0045488 |
| -4.2238526 | 2.40E-05 | 7.50E-05 | 78.6900249 | 73.6061319 | 156.461743 | 40.3427483 |
| -4.2237794 | 2.40E-05 | 7.50E-05 | 21.7217256 | 138.282108 | 212.833989 | 147.92341 |
| -4.2237303 | 2.40E-05 | 7.50E-05 | 60.2470503 | 89.0309463 | 46.0181598 | 60.5141225 |
| -4.2235654 | 2.40E-05 | 7.51E-05 | 40.5745441 | 100.937821 | 65.5758777 | 76.202969 |
| -4.2232923 | 2.41E-05 | 7.52E-05 | 28.2792277 | 126.645845 | 108.142676 | 121.028245 |
| -4.2230605 | 2.41E-05 | 7.52E-05 | 87.2967464 | 92.2782757 | 59.8236078 | 99.1759229 |
| -4.2229107 | 2.41E-05 | 7.53E-05 | 73.7718984 | 52.2278804 | 140.355387 | 53.2300151 |
| -4.2225982 | 2.42E-05 | 7.54E-05 | 441.401859 | 643.783044 | 571.775636 | 630.915758 |
| -4.2225643 | 2.42E-05 | 7.54E-05 | 27.0496961 | 124.480958 | 142.656295 | 111.502874 |
| -4.2221346 | 2.42E-05 | 7.55E-05 | 36.0662614 | 115.280192 | 81.6822337 | 80.6854966 |
| -4.2219119 | 2.42E-05 | 7.56E-05 | 40.5745441 | 80.912623 | 74.7795097 | 65.556966 |
| -4.2219329 | 2.42E-05 | 7.56E-05 | 172.13443 | 62.7817008 | 49.4695218 | 72.2807574 |
| -4.2215272 | 2.43E-05 | 7.57E-05 | 33.1973543 | 171.026012 | 87.4345036 | 97.4949751 |
| 4.22145409 | 2.43E-05 | 7.57E-05 | 27.0496961 | 27.3316887 | 33.3631659 | 50.9887513 |
| 4.22142416 | 2.43E-05 | 7.57E-05 | 15.9839113 | 10.8244312 | 8.05317797 | 16.2491625 |
| -4.2214231 | 2.43E-05 | 7.57E-05 | 48.7714217 | 173.190899 | 95.4876816 | 103.098135 |
| -4.2204944 | 2.44E-05 | 7.60E-05 | 36.4761053 | 81.1832337 | 116.195854 | 77.8839169 |
| -4.2203147 | 2.44E-05 | 7.61E-05 | 32.7875104 | 125.563402 | 86.2840497 | 97.4949751 |
| -4.2201169 | 2.44E-05 | 7.61E-05 | 42.2139196 | 70.3588026 | 63.2749697 | 72.2807574 |
| -4.2199083 | 2.44E-05 | 7.62E-05 | 85.247527 | 78.477126 | 43.7172518 | 54.910963 |
| -4.2198602 | 2.44E-05 | 7.62E-05 | 45.0828268 | 92.2782757 | 52.9208838 | 82.9267604 |
| -4.2198184 | 2.44E-05 | 7.62E-05 | 45.9025146 | 86.8660601 | 51.7704298 | 106.46003 |
| -4.2196643 | 2.45E-05 | 7.63E-05 | 36.8859492 | 112.032863 | 65.5758777 | 122.709193 |
| -4.2193361 | 2.45E-05 | 7.64E-05 | 262.709927 | 365.865773 | 332.481205 | 382.135477 |
| 4.21911465 | 2.45E-05 | 7.64E-05 | 9.01656536 | 14.8835929 | 0 | 14.5682147 |
| -4.2190285 | 2.45E-05 | 7.65E-05 | 242.217733 | 78.2065152 | 60.9740618 | 105.899714 |
| 4.21886965 | 2.46E-05 | 7.65E-05 | 9.83625312 | 8.93015571 | 8.05317797 | 9.52537113 |

| | | | | | | |
|---|---|---|---|---|---|---|
| -4.2185339 | 2.46E-05 | 7.66E-05 | 38.9351686 | 79.5595691 | 69.0272397 | 81.8061285 |
| -4.2183288 | 2.46E-05 | 7.67E-05 | 33.1973543 | 138.011497 | 108.142676 | 78.4442328 |
| -4.2182242 | 2.46E-05 | 7.67E-05 | 36.4761053 | 77.9359044 | 89.7354116 | 82.3664444 |
| -4.2181725 | 2.46E-05 | 7.67E-05 | 19.2626624 | 192.133653 | 192.125817 | 147.92341 |
| -4.2181406 | 2.46E-05 | 7.67E-05 | 34.8367298 | 88.7603356 | 82.8326877 | 95.2537113 |
| -4.217986 | 2.46E-05 | 7.68E-05 | 35.2465737 | 73.3355211 | 112.744492 | 98.615607 |
| -4.217982 | 2.46E-05 | 7.68E-05 | 66.3947085 | 68.7351379 | 71.3281477 | 41.4633802 |
| -4.217202 | 2.47E-05 | 7.70E-05 | 48.3615778 | 52.7691019 | 64.4254237 | 64.99665 |
| -4.2171003 | 2.47E-05 | 7.70E-05 | 41.8040758 | 70.9000241 | 62.1245158 | 80.6854966 |
| -4.216823 | 2.48E-05 | 7.71E-05 | 84.0179954 | 88.7603356 | 33.3631659 | 124.950457 |
| -4.2165955 | 2.48E-05 | 7.72E-05 | 64.755333 | 65.2171978 | 56.3722458 | 45.9459078 |
| -4.2159907 | 2.49E-05 | 7.74E-05 | 35.2465737 | 87.6778924 | 81.6822337 | 90.7711837 |
| -4.2156131 | 2.49E-05 | 7.75E-05 | 53.6895483 | 99.0435452 | 72.4786017 | 99.1759229 |
| -4.215536 | 2.49E-05 | 7.75E-05 | 86.0672148 | 95.7962158 | 32.2127119 | 110.942558 |
| -4.2153445 | 2.49E-05 | 7.76E-05 | 58.6076748 | 57.640096 | 69.0272397 | 46.5062237 |
| -4.2153166 | 2.49E-05 | 7.76E-05 | 29.5087594 | 104.455761 | 165.665375 | 99.7362389 |
| -4.2151341 | 2.50E-05 | 7.77E-05 | 45.0828268 | 164.531354 | 49.4695218 | 129.9933 |
| -4.2148898 | 2.50E-05 | 7.77E-05 | 65.1651769 | 63.0523115 | 42.5667978 | 67.2379138 |
| -4.2147702 | 2.50E-05 | 7.78E-05 | 38.9351686 | 93.090108 | 63.2749697 | 93.0124475 |
| -4.2145104 | 2.50E-05 | 7.78E-05 | 33.1973543 | 111.22103 | 89.7354116 | 88.5299199 |
| -4.2143617 | 2.50E-05 | 7.79E-05 | 38.9351686 | 98.2317128 | 69.0272397 | 75.6426531 |
| -4.2142377 | 2.51E-05 | 7.79E-05 | 27.45954 | 105.538204 | 124.249032 | 136.156776 |
| -4.2142063 | 2.51E-05 | 7.79E-05 | 54.509236 | 52.4984912 | 54.0713378 | 84.6077082 |
| -4.2141311 | 2.51E-05 | 7.79E-05 | 183.610058 | 89.0309463 | 248.498063 | 102.537819 |
| -4.2139698 | 2.51E-05 | 7.80E-05 | 34.4268859 | 155.059976 | 64.4254237 | 191.067738 |
| -4.213639 | 2.51E-05 | 7.81E-05 | 38.1154808 | 80.1007906 | 103.54086 | 68.9188617 |
| -4.2134261 | 2.52E-05 | 7.82E-05 | 306.563222 | 438.930684 | 403.809352 | 436.486124 |
| -4.2129042 | 2.52E-05 | 7.83E-05 | 50.0009534 | 110.679809 | 64.4254237 | 54.910963 |
| -4.2127336 | 2.52E-05 | 7.84E-05 | 36.8859492 | 90.9252218 | 73.6290557 | 82.3664444 |
| -4.2124368 | 2.53E-05 | 7.85E-05 | 27.8693838 | 124.751569 | 106.992222 | 121.588561 |
| -4.2121408 | 2.53E-05 | 7.86E-05 | 40.984388 | 147.482875 | 73.6290557 | 72.8410733 |
| -4.212154 | 2.53E-05 | 7.86E-05 | 29.0989155 | 126.375234 | 108.142676 | 103.098135 |
| -4.2112843 | 2.54E-05 | 7.89E-05 | 53.6895483 | 227.042444 | 212.833989 | 36.9808526 |
| -4.2107428 | 2.55E-05 | 7.91E-05 | 59.8372065 | 105.808815 | 87.4345036 | 40.3427483 |
| -4.2104329 | 2.55E-05 | 7.92E-05 | 32.3776665 | 74.9591858 | 124.249032 | 160.250361 |
| -4.2101125 | 2.55E-05 | 7.93E-05 | 31.9678226 | 104.726372 | 123.098578 | 83.4870763 |
| -4.2100323 | 2.55E-05 | 7.93E-05 | 94.6739363 | 73.0649104 | 44.8677058 | 53.7903311 |
| -4.209933 | 2.55E-05 | 7.93E-05 | 572.5519 | 94.4431619 | 43.7172518 | 84.0473923 |
| -4.2093784 | 2.56E-05 | 7.95E-05 | 423.368728 | 589.119666 | 439.473426 | 478.50982 |
| -4.2092406 | 2.56E-05 | 7.95E-05 | 38.9351686 | 131.516839 | 131.151755 | 57.7125427 |
| -4.2084357 | 2.57E-05 | 7.98E-05 | 27.0496961 | 133.140503 | 115.0454 | 114.864769 |
| -4.2082504 | 2.57E-05 | 7.99E-05 | 29.5087594 | 119.339354 | 94.3372276 | 118.226665 |
| -4.2081804 | 2.57E-05 | 7.99E-05 | 37.2957931 | 84.1599523 | 73.6290557 | 79.0045488 |

| | | | | | | |
|---|---|---|---|---|---|---|
| -4.208153 | 2.57E-05 | 7.99E-05 | 29.9186032 | 90.3840002 | 118.496762 | 121.588561 |
| -4.2077656 | 2.58E-05 | 8.00E-05 | 52.0501728 | 49.7923834 | 64.4254237 | 58.2728587 |
| -4.2075797 | 2.58E-05 | 8.01E-05 | 30.738291 | 90.3840002 | 150.709473 | 101.977503 |
| -4.2072845 | 2.58E-05 | 8.02E-05 | 68.0340841 | 54.3927666 | 47.1686138 | 58.8331746 |
| -4.2073005 | 2.58E-05 | 8.02E-05 | 220.905851 | 66.2996409 | 63.2749697 | 57.1522268 |
| -4.2070656 | 2.59E-05 | 8.02E-05 | 144.265046 | 61.1580361 | 48.3190678 | 66.6775979 |
| -4.2070091 | 2.59E-05 | 8.02E-05 | 60.2470503 | 53.851545 | 72.4786017 | 47.0665397 |
| -4.2066351 | 2.59E-05 | 8.04E-05 | 68.8537718 | 77.3946828 | 37.9649818 | 77.3236009 |
| -4.2066347 | 2.59E-05 | 8.04E-05 | 29.0989155 | 214.864959 | 79.3813257 | 150.72499 |
| -4.2066347 | 2.59E-05 | 8.04E-05 | 51.6403289 | 218.92412 | 140.355387 | 183.783631 |
| -4.2061091 | 2.60E-05 | 8.05E-05 | 32.3776665 | 90.9252218 | 106.992222 | 90.7711837 |
| -4.2060337 | 2.60E-05 | 8.06E-05 | 65.5750208 | 122.586683 | 130.001301 | 69.4791776 |
| -4.2056777 | 2.60E-05 | 8.07E-05 | 42.2139196 | 99.5847667 | 133.452663 | 54.910963 |
| -4.2050779 | 2.61E-05 | 8.09E-05 | 27.45954 | 126.916455 | 95.4876816 | 166.974153 |
| -4.2048031 | 2.61E-05 | 8.10E-05 | 21.7217256 | 148.565318 | 171.417645 | 137.837723 |
| -4.2045062 | 2.62E-05 | 8.11E-05 | 23.770945 | 305.51957 | 103.54086 | 153.52657 |
| -4.2044948 | 2.62E-05 | 8.11E-05 | 33.6071982 | 88.7603356 | 93.1867736 | 87.409288 |
| -4.2037576 | 2.63E-05 | 8.13E-05 | 71.3128351 | 84.1599523 | 35.6640739 | 99.7362389 |
| -4.2037221 | 2.63E-05 | 8.13E-05 | 67.6242402 | 138.011497 | 93.1867736 | 117.666349 |
| -4.2036762 | 2.63E-05 | 8.13E-05 | 20.9020379 | 179.956168 | 111.594038 | 275.675447 |
| -4.2033513 | 2.63E-05 | 8.14E-05 | 93.8542485 | 145.317988 | 200.178995 | 212.359745 |
| -4.20332 | 2.63E-05 | 8.14E-05 | 33.1973543 | 105.808815 | 83.9831417 | 90.7711837 |
| -4.2031688 | 2.63E-05 | 8.15E-05 | 269.267429 | 72.7942996 | 57.5226998 | 62.1950703 |
| 4.20294047 | 2.63E-05 | 8.16E-05 | 39.3450125 | 26.2492456 | 34.5136199 | 31.3776931 |
| -4.2028463 | 2.64E-05 | 8.16E-05 | 55.7387677 | 66.0290301 | 65.5758777 | 44.8252759 |
| -4.202755 | 2.64E-05 | 8.16E-05 | 57.3781432 | 54.3927666 | 48.3190678 | 60.5141225 |
| -4.2026054 | 2.64E-05 | 8.17E-05 | 31.5579788 | 95.2549943 | 142.656295 | 88.5299199 |
| -4.2025025 | 2.64E-05 | 8.17E-05 | 53.6895483 | 115.009581 | 41.4163438 | 106.46003 |
| -4.2022227 | 2.64E-05 | 8.18E-05 | 40.984388 | 63.0523115 | 69.0272397 | 91.3314996 |
| -4.2021206 | 2.64E-05 | 8.18E-05 | 101.231438 | 153.436312 | 25.3099879 | 142.320251 |
| -4.2019458 | 2.65E-05 | 8.19E-05 | 40.1647002 | 95.525605 | 72.4786017 | 63.8760181 |
| -4.2017518 | 2.65E-05 | 8.19E-05 | 47.5418901 | 66.8408624 | 51.7704298 | 62.7553862 |
| -4.2015569 | 2.65E-05 | 8.20E-05 | 57.3781432 | 64.946587 | 93.1867736 | 44.2649599 |
| -4.2011606 | 2.66E-05 | 8.21E-05 | 35.2465737 | 90.3840002 | 72.4786017 | 105.899714 |
| -4.2005535 | 2.66E-05 | 8.23E-05 | 34.4268859 | 76.8534613 | 96.6381356 | 91.3314996 |
| -4.2004142 | 2.66E-05 | 8.24E-05 | 39.3450125 | 121.50424 | 66.7263317 | 77.3236009 |
| -4.1999387 | 2.67E-05 | 8.26E-05 | 43.4434513 | 76.8534613 | 58.6731538 | 63.3157022 |
| -4.1996 | 2.67E-05 | 8.27E-05 | 36.0662614 | 71.1706349 | 87.4345036 | 91.3314996 |
| -4.1992981 | 2.68E-05 | 8.28E-05 | 30.738291 | 145.047378 | 104.691314 | 86.2886561 |
| -4.1993108 | 2.68E-05 | 8.28E-05 | 47.1320462 | 79.5595691 | 78.2308717 | 48.7474875 |
| -4.199041 | 2.68E-05 | 8.28E-05 | 37.2957931 | 74.4179643 | 75.9299637 | 123.269509 |
| -4.1990502 | 2.68E-05 | 8.28E-05 | 40.1647002 | 70.6294134 | 66.7263317 | 110.382242 |
| -4.1989184 | 2.68E-05 | 8.29E-05 | 36.4761053 | 115.821413 | 64.4254237 | 103.098135 |

| | | | | | | |
|---|---|---|---|---|---|---|
| -4.1982278 | 2.69E-05 | 8.31E-05 | 20.0823501 | 197.81648 | 128.850847 | 177.620156 |
| -4.1977119 | 2.70E-05 | 8.33E-05 | 48.3615778 | 72.5236888 | 96.6381356 | 48.7474875 |
| -4.1975836 | 2.70E-05 | 8.33E-05 | 60.2470503 | 119.068743 | 88.5849576 | 90.7711837 |
| -4.1974845 | 2.70E-05 | 8.34E-05 | 56.5584554 | 142.882491 | 39.1154358 | 113.744138 |
| -4.1972736 | 2.70E-05 | 8.34E-05 | 37.2957931 | 82.8068984 | 101.239952 | 66.6775979 |
| -4.1971982 | 2.70E-05 | 8.35E-05 | 29.5087594 | 116.903857 | 96.6381356 | 105.899714 |
| -4.197055 | 2.70E-05 | 8.35E-05 | 43.8532952 | 53.851545 | 77.0804177 | 71.7204414 |
| -4.19675 | 2.71E-05 | 8.36E-05 | 179.101776 | 75.7710182 | 44.8677058 | 64.4363341 |
| -4.1966676 | 2.71E-05 | 8.36E-05 | 134.018949 | 290.906588 | 289.914407 | 313.776931 |
| -4.1962705 | 2.71E-05 | 8.38E-05 | 20.492194 | 132.328671 | 189.824909 | 163.051941 |
| -4.196195 | 2.71E-05 | 8.38E-05 | 39.3450125 | 132.328671 | 138.054479 | 54.910963 |
| -4.1962108 | 2.71E-05 | 8.38E-05 | 288.530091 | 117.174467 | 69.0272397 | 109.26161 |
| -4.1960652 | 2.72E-05 | 8.38E-05 | 49.5911095 | 118.527521 | 43.7172518 | 111.502874 |
| -4.1955386 | 2.72E-05 | 8.40E-05 | 103.690502 | 115.280192 | 31.0622579 | 81.8061285 |
| -4.1952078 | 2.73E-05 | 8.41E-05 | 39.3450125 | 73.0649104 | 69.0272397 | 69.4791776 |
| -4.1950112 | 2.73E-05 | 8.42E-05 | 42.6237635 | 99.3141559 | 57.5226998 | 71.7204414 |
| -4.194736 | 2.73E-05 | 8.43E-05 | 35.2465737 | 110.950419 | 80.5317797 | 76.763285 |
| -4.1944628 | 2.74E-05 | 8.44E-05 | 40.984388 | 70.3588026 | 64.4254237 | 66.1172819 |
| -4.1942984 | 2.74E-05 | 8.44E-05 | 38.1154808 | 68.7351379 | 172.568099 | 89.6505518 |
| -4.1941895 | 2.74E-05 | 8.45E-05 | 32.3776665 | 102.290875 | 119.647216 | 77.8839169 |
| -4.1938812 | 2.74E-05 | 8.46E-05 | 38.9351686 | 120.151186 | 118.496762 | 56.5919108 |
| -4.1938523 | 2.74E-05 | 8.46E-05 | 39.3450125 | 63.5935331 | 82.8326877 | 72.8410733 |
| -4.1932298 | 2.75E-05 | 8.48E-05 | 31.9678226 | 83.8893415 | 117.346308 | 94.1330794 |
| -4.1928481 | 2.75E-05 | 8.49E-05 | 44.6729829 | 56.0164313 | 75.9299637 | 60.5141225 |
| -4.1925611 | 2.76E-05 | 8.50E-05 | 31.9678226 | 76.3122397 | 126.549939 | 110.382242 |
| -4.1925808 | 2.76E-05 | 8.50E-05 | 166.806459 | 70.0881918 | 43.7172518 | 68.3585457 |
| -4.1922261 | 2.76E-05 | 8.51E-05 | 67.2143963 | 48.980551 | 55.2217918 | 52.6696992 |
| -4.1917787 | 2.77E-05 | 8.53E-05 | 92.214873 | 44.9213893 | 54.0713378 | 67.2379138 |
| -4.191535 | 2.77E-05 | 8.54E-05 | 34.8367298 | 168.319905 | 142.656295 | 140.078987 |
| -4.191446 | 2.77E-05 | 8.54E-05 | 17.213443 | 118.798132 | 268.055781 | 244.85807 |
| -4.1907615 | 2.78E-05 | 8.57E-05 | 29.0989155 | 260.59818 | 147.258111 | 191.067738 |
| -4.190694 | 2.78E-05 | 8.57E-05 | 281.152902 | 326.3566 | 303.719855 | 373.170422 |
| -4.190683 | 2.78E-05 | 8.57E-05 | 58.6076748 | 47.0862756 | 55.2217918 | 58.8331746 |
| -4.1899594 | 2.79E-05 | 8.59E-05 | 56.5584554 | 42.2152815 | 87.4345036 | 70.5998095 |
| -4.1897266 | 2.79E-05 | 8.60E-05 | 35.2465737 | 75.7710182 | 79.3813257 | 99.1759229 |
| -4.1894927 | 2.80E-05 | 8.61E-05 | 28.2792277 | 150.730204 | 90.8858656 | 116.545717 |
| 4.18921622 | 2.80E-05 | 8.62E-05 | 0 | 0 | 0 | 0 |
| -4.1891367 | 2.80E-05 | 8.62E-05 | 61.0667381 | 57.9107067 | 49.4695218 | 50.4284354 |
| -4.1887627 | 2.80E-05 | 8.64E-05 | 37.705637 | 101.749653 | 60.9740618 | 95.2537113 |
| -4.1887211 | 2.81E-05 | 8.64E-05 | 38.9351686 | 78.477126 | 62.1245158 | 81.8061285 |
| -4.1883116 | 2.81E-05 | 8.65E-05 | 44.6729829 | 57.9107067 | 70.1776937 | 57.1522268 |
| -4.1877671 | 2.82E-05 | 8.67E-05 | 22.1315695 | 167.778683 | 153.010381 | 120.467929 |
| -4.1861688 | 2.84E-05 | 8.73E-05 | 49.1812656 | 50.0629941 | 60.9740618 | 86.2886561 |

| | | | | | | |
|---|---|---|---|---|---|---|
| -4.1860011 | 2.84E-05 | 8.74E-05 | 47.1320462 | 90.3840002 | 50.6199758 | 65.556966 |
| -4.1859954 | 2.84E-05 | 8.74E-05 | 35.2465737 | 85.783617 | 94.3372276 | 71.1601255 |
| -4.1859767 | 2.84E-05 | 8.74E-05 | 26.6398522 | 98.7729344 | 136.904025 | 125.510772 |
| -4.1854362 | 2.85E-05 | 8.76E-05 | 47.1320462 | 97.9611021 | 77.0804177 | 48.1871716 |
| -4.1854498 | 2.85E-05 | 8.76E-05 | 42.6237635 | 66.2996409 | 60.9740618 | 61.0744384 |
| -4.1853974 | 2.85E-05 | 8.76E-05 | 97.1329995 | 163.990132 | 31.0622579 | 83.4870763 |
| -4.185317 | 2.85E-05 | 8.76E-05 | 47.5418901 | 74.4179643 | 47.1686138 | 74.5220212 |
| 4.18525638 | 2.85E-05 | 8.76E-05 | 0 | 0.27061078 | 0 | 0 |
| -4.1846919 | 2.86E-05 | 8.78E-05 | 24.5906328 | 126.916455 | 105.841768 | 214.601008 |
| -4.1846783 | 2.86E-05 | 8.78E-05 | 37.2957931 | 83.0775092 | 72.4786017 | 70.5998095 |
| 4.1843475 | 2.86E-05 | 8.79E-05 | 21.7217256 | 14.6129821 | 21.8586259 | 12.3269509 |
| -4.1840208 | 2.86E-05 | 8.80E-05 | 39.7548564 | 59.5343714 | 77.0804177 | 78.4442328 |
| -4.1840297 | 2.86E-05 | 8.80E-05 | 37.2957931 | 100.396599 | 63.2749697 | 85.7283401 |
| -4.1836003 | 2.87E-05 | 8.82E-05 | 75.00143 | 97.1492697 | 32.2127119 | 124.390141 |
| -4.1831547 | 2.87E-05 | 8.84E-05 | 100.411751 | 104.18515 | 32.2127119 | 75.0823371 |
| -4.1830067 | 2.88E-05 | 8.84E-05 | 388.122154 | 763.93423 | 799.565527 | 772.115377 |
| -4.1828865 | 2.88E-05 | 8.84E-05 | 41.8040758 | 64.946587 | 60.9740618 | 97.4949751 |
| -4.1825036 | 2.88E-05 | 8.86E-05 | 40.1647002 | 129.351952 | 58.6731538 | 85.7283401 |
| -4.1824582 | 2.88E-05 | 8.86E-05 | 22.9512573 | 106.891258 | 211.683535 | 144.561515 |
| -4.1821592 | 2.89E-05 | 8.87E-05 | 54.9190799 | 149.918372 | 39.1154358 | 379.894213 |
| -4.1821191 | 2.89E-05 | 8.87E-05 | 45.9025146 | 60.3462037 | 59.8236078 | 54.350647 |
| -4.182102 | 2.89E-05 | 8.87E-05 | 49.1812656 | 66.2996409 | 66.7263317 | 47.0665397 |
| -4.1820281 | 2.89E-05 | 8.87E-05 | 40.984388 | 66.5702517 | 59.8236078 | 85.1680242 |
| 4.1812305 | 2.90E-05 | 8.90E-05 | 69.6734596 | 58.7225391 | 43.7172518 | 49.8681194 |
| -4.1811958 | 2.90E-05 | 8.90E-05 | 26.6398522 | 115.009581 | 150.709473 | 99.7362389 |
| 4.18100496 | 2.90E-05 | 8.91E-05 | 5.32797044 | 4.60038325 | 0 | 6.72379138 |
| -4.180489 | 2.91E-05 | 8.93E-05 | 59.4273626 | 62.51109 | 43.7172518 | 57.1522268 |
| -4.1804814 | 2.91E-05 | 8.93E-05 | 202.462877 | 459.767714 | 418.765254 | 461.140026 |
| -4.180361 | 2.91E-05 | 8.93E-05 | 44.6729829 | 64.1347547 | 55.2217918 | 59.9538065 |
| -4.1801288 | 2.91E-05 | 8.94E-05 | 209.840067 | 271.152001 | 194.426725 | 283.51987 |
| -4.179813 | 2.92E-05 | 8.95E-05 | 30.3284471 | 72.7942996 | 174.869007 | 133.915512 |
| -4.1787427 | 2.93E-05 | 8.99E-05 | 27.8693838 | 108.244312 | 101.239952 | 117.106033 |
| -4.1785375 | 2.93E-05 | 9.00E-05 | 90.1656536 | 68.1939163 | 41.4163438 | 56.5919108 |
| -4.1785089 | 2.93E-05 | 9.00E-05 | 20.9020379 | 85.2423954 | 260.002603 | 273.994499 |
| -4.1782803 | 2.94E-05 | 9.01E-05 | 46.7222023 | 77.3946828 | 50.6199758 | 61.0744384 |
| -4.1777478 | 2.94E-05 | 9.03E-05 | 109.428316 | 112.574084 | 57.5226998 | 37.5411686 |
| -4.1776794 | 2.94E-05 | 9.03E-05 | 31.1481349 | 82.5362876 | 109.29313 | 98.055291 |
| -4.1774002 | 2.95E-05 | 9.04E-05 | 39.7548564 | 110.138587 | 54.0713378 | 102.537819 |
| -4.1770368 | 2.95E-05 | 9.06E-05 | 29.9186032 | 138.82333 | 94.3372276 | 90.7711837 |
| -4.1768853 | 2.96E-05 | 9.06E-05 | 29.5087594 | 115.009581 | 81.6822337 | 135.036144 |
| -4.1767177 | 2.96E-05 | 9.07E-05 | 88.5262781 | 114.197749 | 100.089498 | 108.701294 |
| -4.1766198 | 2.96E-05 | 9.07E-05 | 64.3454891 | 67.6526948 | 63.2749697 | 66.6775979 |
| -4.1764454 | 2.96E-05 | 9.08E-05 | 115.985818 | 46.0038325 | 50.6199758 | 75.6426531 |

| | | | | | | |
|---|---|---|---|---|---|---|
| -4.1759829 | 2.97E-05 | 9.09E-05 | 25.8201644 | 147.212264 | 94.3372276 | 137.837723 |
| -4.175829 | 2.97E-05 | 9.10E-05 | 38.1154808 | 69.5469702 | 71.3281477 | 70.0394936 |
| -4.1756898 | 2.97E-05 | 9.10E-05 | 36.8859492 | 62.2404792 | 95.4876816 | 87.9696039 |
| -4.1751379 | 2.98E-05 | 9.12E-05 | 52.4600166 | 138.552719 | 85.1335957 | 44.2649599 |
| -4.1750928 | 2.98E-05 | 9.13E-05 | 43.4434513 | 54.6633774 | 66.7263317 | 63.3157022 |
| -4.1750199 | 2.98E-05 | 9.13E-05 | 36.8859492 | 111.22103 | 60.9740618 | 93.5727634 |
| -4.1749645 | 2.98E-05 | 9.13E-05 | 40.984388 | 56.2870421 | 73.6290557 | 73.4013893 |
| -4.1749025 | 2.98E-05 | 9.13E-05 | 45.0828268 | 51.9572696 | 64.4254237 | 85.1680242 |
| -4.1747508 | 2.98E-05 | 9.14E-05 | 39.3450125 | 93.3607188 | 64.4254237 | 66.6775979 |
| -4.1746896 | 2.98E-05 | 9.14E-05 | 69.6734596 | 329.333318 | 314.073941 | 316.578511 |
| -4.1743665 | 2.99E-05 | 9.15E-05 | 36.4761053 | 110.679809 | 88.5849576 | 63.3157022 |
| -4.1742843 | 2.99E-05 | 9.15E-05 | 37.2957931 | 79.8301798 | 66.7263317 | 74.5220212 |
| -4.1740875 | 2.99E-05 | 9.16E-05 | 34.8367298 | 68.1939163 | 134.603117 | 91.8918156 |
| -4.1741085 | 2.99E-05 | 9.16E-05 | 40.984388 | 70.0881918 | 65.5758777 | 59.3934906 |
| -4.1740797 | 2.99E-05 | 9.16E-05 | 82.3786199 | 155.059976 | 106.992222 | 29.6967453 |
| -4.1740531 | 2.99E-05 | 9.16E-05 | 85.6573709 | 26.2492456 | 155.311289 | 159.129729 |
| -4.1739356 | 2.99E-05 | 9.16E-05 | 39.7548564 | 62.51109 | 92.0363196 | 66.1172819 |
| -4.1738764 | 2.99E-05 | 9.16E-05 | 35.2465737 | 79.5595691 | 75.9299637 | 79.0045488 |
| -4.1737188 | 3.00E-05 | 9.17E-05 | 61.0667381 | 48.1687187 | 48.3190678 | 77.8839169 |
| -4.1736434 | 3.00E-05 | 9.17E-05 | 39.7548564 | 113.927138 | 90.8858656 | 55.4712789 |
| -4.1735998 | 3.00E-05 | 9.17E-05 | 125.822071 | 208.099689 | 319.826211 | 333.948305 |
| -4.173548 | 3.00E-05 | 9.17E-05 | 70.9029912 | 69.817581 | 57.5226998 | 73.9617052 |
| -4.1735173 | 3.00E-05 | 9.17E-05 | 29.9186032 | 89.5721679 | 103.54086 | 102.537819 |
| -4.1734512 | 3.00E-05 | 9.17E-05 | 98.3625312 | 34.0969582 | 118.496762 | 67.7982298 |
| -4.1725921 | 3.01E-05 | 9.21E-05 | 38.1154808 | 77.3946828 | 64.4254237 | 72.2807574 |
| -4.1724189 | 3.01E-05 | 9.21E-05 | 393.040281 | 947.137727 | 712.131023 | 742.978948 |
| -4.1724198 | 3.01E-05 | 9.21E-05 | 51.230485 | 103.643928 | 89.7354116 | 85.1680242 |
| -4.1719764 | 3.02E-05 | 9.23E-05 | 42.6237635 | 96.3374374 | 55.2217918 | 68.3585457 |
| -4.1718905 | 3.02E-05 | 9.23E-05 | 52.0501728 | 48.980551 | 57.5226998 | 54.350647 |
| -4.171848 | 3.02E-05 | 9.23E-05 | 48.7714217 | 77.1240721 | 50.6199758 | 55.4712789 |
| 4.17106635 | 3.03E-05 | 9.27E-05 | 79.5097127 | 44.109557 | 43.7172518 | 45.3855918 |
| -4.1709601 | 3.03E-05 | 9.27E-05 | 33.1973543 | 115.550803 | 128.850847 | 66.1172819 |
| -4.1709392 | 3.03E-05 | 9.27E-05 | 36.0662614 | 75.7710182 | 75.9299637 | 145.121831 |
| -4.1709079 | 3.03E-05 | 9.27E-05 | 50.4107972 | 56.8282636 | 47.1686138 | 69.4791776 |
| -4.170852 | 3.03E-05 | 9.27E-05 | 101.641282 | 135.30539 | 24.1595339 | 146.802779 |
| -4.1708532 | 3.03E-05 | 9.27E-05 | 47.951734 | 82.2656769 | 52.9208838 | 54.910963 |
| -4.1702057 | 3.04E-05 | 9.29E-05 | 25.8201644 | 115.821413 | 115.0454 | 117.106033 |
| -4.1701526 | 3.04E-05 | 9.30E-05 | 36.4761053 | 86.0542278 | 69.0272397 | 73.4013893 |
| -4.1700409 | 3.05E-05 | 9.30E-05 | 37.705637 | 76.5828505 | 64.4254237 | 75.0823371 |
| -4.1695361 | 3.05E-05 | 9.32E-05 | 49.1812656 | 113.927138 | 115.0454 | 112.623506 |
| -4.1691635 | 3.06E-05 | 9.33E-05 | 263.939459 | 525.255522 | 437.172518 | 505.965302 |
| -4.1689416 | 3.06E-05 | 9.34E-05 | 50.8206411 | 174.273342 | 121.948124 | 124.390141 |
| -4.1689613 | 3.06E-05 | 9.34E-05 | 34.4268859 | 83.6187308 | 80.5317797 | 76.202969 |

| | | | | | | |
|---|---|---|---|---|---|---|
| 4.16873772 | 3.06E-05 | 9.35E-05 | 104.100346 | 100.396599 | 95.4876816 | 98.055291 |
| -4.1686274 | 3.06E-05 | 9.35E-05 | 36.4761053 | 90.3840002 | 71.3281477 | 70.0394936 |
| -4.1682731 | 3.07E-05 | 9.37E-05 | 247.955547 | 465.991762 | 460.181598 | 458.898762 |
| -4.1681868 | 3.07E-05 | 9.37E-05 | 47.1320462 | 142.882491 | 46.0181598 | 95.2537113 |
| 4.16785408 | 3.07E-05 | 9.38E-05 | 80.7392443 | 46.2744432 | 60.9740618 | 36.9808526 |
| -4.1677286 | 3.08E-05 | 9.39E-05 | 30.3284471 | 92.5488865 | 86.2840497 | 143.440883 |
| -4.167626 | 3.08E-05 | 9.39E-05 | 16.8035991 | 157.766084 | 301.418947 | 146.802779 |
| -4.1675591 | 3.08E-05 | 9.39E-05 | 42.6237635 | 119.339354 | 49.4695218 | 198.912162 |
| -4.167482 | 3.08E-05 | 9.39E-05 | 44.263139 | 77.9359044 | 56.3722458 | 56.5919108 |
| -4.1672383 | 3.08E-05 | 9.40E-05 | 27.0496961 | 107.432479 | 127.700393 | 99.7362389 |
| -4.1671699 | 3.08E-05 | 9.40E-05 | 11.0657848 | 304.978348 | 159.913105 | 327.78483 |
| -4.166996 | 3.09E-05 | 9.41E-05 | 35.6564176 | 73.8767427 | 117.346308 | 72.8410733 |
| -4.1664748 | 3.09E-05 | 9.43E-05 | 52.8698605 | 83.8893415 | 47.1686138 | 55.4712789 |
| -4.1662233 | 3.10E-05 | 9.44E-05 | 40.1647002 | 102.832096 | 57.5226998 | 73.9617052 |
| -4.1661159 | 3.10E-05 | 9.44E-05 | 34.8367298 | 67.6526948 | 127.700393 | 87.9696039 |
| -4.166046 | 3.10E-05 | 9.45E-05 | 51.230485 | 67.6526948 | 64.4254237 | 44.2649599 |
| -4.16578 | 3.10E-05 | 9.46E-05 | 59.8372065 | 74.688575 | 40.2658898 | 61.6347543 |
| -4.1651646 | 3.11E-05 | 9.48E-05 | 31.1481349 | 88.4897248 | 85.1335957 | 104.779082 |
| -4.164707 | 3.12E-05 | 9.50E-05 | 27.0496961 | 113.385916 | 90.8858656 | 170.896364 |
| -4.1645977 | 3.12E-05 | 9.50E-05 | 36.0662614 | 87.1366709 | 70.1776937 | 72.2807574 |
| -4.164527 | 3.12E-05 | 9.51E-05 | 469.681086 | 667.326181 | 793.813257 | 700.955252 |
| 4.16409315 | 3.13E-05 | 9.52E-05 | 12.7051603 | 10.8244312 | 9.20363196 | 15.6888466 |
| -4.1639722 | 3.13E-05 | 9.53E-05 | 35.6564176 | 103.643928 | 66.7263317 | 80.1251807 |
| -4.1638982 | 3.13E-05 | 9.53E-05 | 32.7875104 | 83.8893415 | 97.7885896 | 77.8839169 |
| -4.1631646 | 3.14E-05 | 9.56E-05 | 59.8372065 | 171.296623 | 96.6381356 | 101.977503 |
| -4.1628741 | 3.14E-05 | 9.57E-05 | 37.2957931 | 77.6652936 | 66.7263317 | 70.0394936 |
| -4.1625956 | 3.15E-05 | 9.58E-05 | 38.1154808 | 104.726372 | 66.7263317 | 67.7982298 |
| -4.1622497 | 3.15E-05 | 9.59E-05 | 127.461447 | 155.330587 | 79.3813257 | 28.0157974 |
| -4.1621919 | 3.15E-05 | 9.60E-05 | 38.1154808 | 74.4179643 | 67.8767857 | 64.99665 |
| -4.1618096 | 3.16E-05 | 9.61E-05 | 35.2465737 | 90.1133895 | 72.4786017 | 73.4013893 |
| -4.1616459 | 3.16E-05 | 9.62E-05 | 423.368728 | 1541.94022 | 1268.95076 | 1179.46507 |
| -4.1609478 | 3.17E-05 | 9.65E-05 | 31.1481349 | 139.364551 | 89.7354116 | 81.8061285 |
| -4.1603962 | 3.18E-05 | 9.67E-05 | 44.6729829 | 70.3588026 | 51.7704298 | 112.06319 |
| -4.160207 | 3.18E-05 | 9.67E-05 | 30.738291 | 78.7477367 | 97.7885896 | 114.304454 |
| -4.1600389 | 3.18E-05 | 9.68E-05 | 32.7875104 | 130.975617 | 64.4254237 | 121.588561 |
| -4.1600422 | 3.18E-05 | 9.68E-05 | 231.561792 | 78.477126 | 42.5667978 | 67.2379138 |
| -4.1598748 | 3.18E-05 | 9.69E-05 | 36.0662614 | 67.382084 | 75.9299637 | 80.1251807 |
| -4.1598945 | 3.18E-05 | 9.69E-05 | 120.084257 | 41.4034492 | 72.4786017 | 57.7125427 |
| -4.1594055 | 3.19E-05 | 9.70E-05 | 69.6734596 | 48.7099402 | 58.6731538 | 47.0665397 |
| -4.1593165 | 3.19E-05 | 9.71E-05 | 43.8532952 | 52.7691019 | 59.8236078 | 68.3585457 |
| -4.1591994 | 3.19E-05 | 9.71E-05 | 59.0175187 | 65.4878086 | 40.2658898 | 62.7553862 |
| -4.1586721 | 3.20E-05 | 9.73E-05 | 38.1154808 | 62.2404792 | 90.8858656 | 68.9188617 |
| -4.1582465 | 3.21E-05 | 9.75E-05 | 26.6398522 | 93.6313296 | 126.549939 | 117.666349 |

| | | | | | | |
|---|---|---|---|---|---|---|
| -4.1580374 | 3.21E-05 | 9.76E-05 | 34.4268859 | 97.9611021 | 98.9390436 | 64.4363341 |
| -4.1576503 | 3.22E-05 | 9.77E-05 | 36.8859492 | 83.34812 | 96.6381356 | 59.9538065 |
| -4.1573149 | 3.22E-05 | 9.79E-05 | 109.018472 | 44.6507786 | 44.8677058 | 123.269509 |
| -4.1572217 | 3.22E-05 | 9.79E-05 | 47.5418901 | 45.1920001 | 72.4786017 | 85.7283401 |
| -4.1571757 | 3.22E-05 | 9.79E-05 | 61.0667381 | 59.5343714 | 42.5667978 | 108.701294 |
| -4.156957 | 3.23E-05 | 9.80E-05 | 56.1486116 | 87.9485032 | 66.7263317 | 82.3664444 |
| -4.1566116 | 3.23E-05 | 9.81E-05 | 327.46526 | 584.519283 | 485.491586 | 485.233612 |
| 4.15631688 | 3.23E-05 | 9.83E-05 | 4.91812656 | 5.95343714 | 0 | 7.28410733 |
| -4.1563 | 3.23E-05 | 9.83E-05 | 38.5253247 | 78.2065152 | 63.2749697 | 64.99665 |
| -4.1561014 | 3.24E-05 | 9.83E-05 | 35.6564176 | 107.70309 | 60.9740618 | 93.0124475 |
| 4.15553147 | 3.25E-05 | 9.86E-05 | 0 | 4.60038325 | 0 | 3.36189569 |
| -4.1554599 | 3.25E-05 | 9.86E-05 | 37.705637 | 191.321821 | 48.3190678 | 173.137628 |
| -4.1552058 | 3.25E-05 | 9.87E-05 | 35.6564176 | 69.5469702 | 82.8326877 | 72.2807574 |
| -4.1551974 | 3.25E-05 | 9.87E-05 | 159.019425 | 82.5362876 | 34.5136199 | 72.2807574 |
| -4.1549803 | 3.25E-05 | 9.88E-05 | 35.2465737 | 80.3714014 | 94.3372276 | 65.556966 |
| -4.1549284 | 3.25E-05 | 9.88E-05 | 22.5414134 | 138.82333 | 125.399485 | 127.752036 |
| -4.1545886 | 3.26E-05 | 9.89E-05 | 30.738291 | 94.1725511 | 110.443584 | 79.0045488 |
| 4.1544312 | 3.26E-05 | 9.90E-05 | 8.60672148 | 4.05916169 | 8.05317797 | 5.04284354 |
| -4.1544344 | 3.26E-05 | 9.90E-05 | 36.4761053 | 69.817581 | 67.8767857 | 79.5648647 |
| -4.1543102 | 3.26E-05 | 9.90E-05 | 32.3776665 | 79.0183475 | 80.5317797 | 108.140978 |
| -4.1541207 | 3.27E-05 | 9.91E-05 | 39.7548564 | 63.8641439 | 75.9299637 | 58.8331746 |
| -4.1540712 | 3.27E-05 | 9.91E-05 | 35.2465737 | 84.7011739 | 70.1776937 | 73.9617052 |
| -4.1540082 | 3.27E-05 | 9.91E-05 | 29.9186032 | 86.3248385 | 138.054479 | 89.0902358 |
| -4.1538867 | 3.27E-05 | 9.92E-05 | 28.6890716 | 171.296623 | 72.4786017 | 136.156776 |
| -4.1533983 | 3.28E-05 | 9.94E-05 | 65.5750208 | 380.478755 | 63.2749697 | 61.6347543 |
| -4.1533068 | 3.28E-05 | 9.94E-05 | 50.8206411 | 44.9213893 | 58.6731538 | 67.7982298 |
| -4.153232 | 3.28E-05 | 9.94E-05 | 164.347396 | 76.0416289 | 35.6640739 | 74.5220212 |
| -4.1532108 | 3.28E-05 | 9.94E-05 | 36.8859492 | 78.2065152 | 70.1776937 | 65.556966 |
| -4.152709 | 3.29E-05 | 9.96E-05 | 29.0989155 | 129.622563 | 78.2308717 | 112.623506 |
| -4.1523685 | 3.29E-05 | 9.98E-05 | 70.4931473 | 71.9824673 | 44.8677058 | 47.0665397 |
| -4.1515493 | 3.30E-05 | 0.00010011 | 33.6071982 | 119.880575 | 70.1776937 | 84.6077082 |
| -4.1514216 | 3.30E-05 | 0.00010015 | 200.823501 | 45.7332217 | 74.7795097 | 59.9538065 |
| -4.1512932 | 3.31E-05 | 0.0001002 | 32.7875104 | 84.7011739 | 101.239952 | 72.8410733 |
| 4.1512385 | 3.31E-05 | 0.00010021 | 0 | 0 | 0 | 0 |
| -4.1510461 | 3.31E-05 | 0.00010029 | 67.6242402 | 65.2171978 | 55.2217918 | 40.9030642 |
| -4.1507839 | 3.31E-05 | 0.00010037 | 41.8040758 | 50.3336049 | 96.6381356 | 79.0045488 |
| -4.1507846 | 3.31E-05 | 0.00010037 | 31.5579788 | 85.783617 | 113.894946 | 77.8839169 |
| -4.1507903 | 3.31E-05 | 0.00010037 | 21.3118818 | 112.303473 | 186.373547 | 140.639303 |
| -4.150544 | 3.32E-05 | 0.00010047 | 56.9682993 | 68.4645271 | 39.1154358 | 69.4791776 |
| -4.1504313 | 3.32E-05 | 0.00010051 | 27.0496961 | 104.18515 | 97.7885896 | 117.106033 |
| 4.14987142 | 3.33E-05 | 0.00010074 | 0.40984388 | 1.3530539 | 1.150454 | 1.1206319 |
| -4.1498843 | 3.33E-05 | 0.00010074 | 51.6403289 | 68.4645271 | 52.9208838 | 46.5062237 |
| -4.1496686 | 3.33E-05 | 0.00010082 | 51.6403289 | 104.455761 | 101.239952 | 39.2221164 |

| | | | | | | |
|---|---|---|---|---|---|---|
| -4.1495558 | 3.33E-05 | 0.00010086 | 15.1642236 | 169.131737 | 177.169915 | 214.040692 |
| -4.1488843 | 3.34E-05 | 0.00010114 | 19.6725062 | 138.282108 | 195.577179 | 129.9933 |
| -4.1488708 | 3.34E-05 | 0.00010114 | 41.3942319 | 89.5721679 | 49.4695218 | 86.848972 |
| -4.1487416 | 3.34E-05 | 0.00010119 | 30.738291 | 117.174467 | 78.2308717 | 91.8918156 |
| 4.14799359 | 3.35E-05 | 0.00010151 | 108.608628 | 75.2297966 | 70.1776937 | 75.0823371 |
| -4.1476431 | 3.36E-05 | 0.00010166 | 52.4600166 | 86.8660601 | 108.142676 | 66.6775979 |
| -4.1470781 | 3.37E-05 | 0.0001019 | 52.0501728 | 40.8622277 | 83.9831417 | 73.4013893 |
| -4.1467504 | 3.37E-05 | 0.00010203 | 21.7217256 | 126.375234 | 163.364467 | 121.028245 |
| -4.1463707 | 3.38E-05 | 0.00010219 | 44.263139 | 63.0523115 | 49.4695218 | 63.8760181 |
| -4.1462375 | 3.38E-05 | 0.00010224 | 28.6890716 | 110.950419 | 88.5849576 | 96.9346591 |
| -4.1460935 | 3.38E-05 | 0.0001023 | 111.887379 | 138.011497 | 25.3099879 | 95.2537113 |
| -4.1458536 | 3.39E-05 | 0.0001024 | 41.8040758 | 47.8981079 | 104.691314 | 97.4949751 |
| -4.1457619 | 3.39E-05 | 0.00010243 | 50.4107972 | 60.3462037 | 47.1686138 | 52.6696992 |
| -4.1455996 | 3.39E-05 | 0.00010249 | 51.6403289 | 93.090108 | 39.1154358 | 114.304454 |
| -4.1455149 | 3.39E-05 | 0.00010252 | 61.0667381 | 58.9931499 | 40.2658898 | 61.0744384 |
| -4.145084 | 3.40E-05 | 0.0001027 | 52.4600166 | 63.8641439 | 47.1686138 | 126.631404 |
| -4.1448055 | 3.40E-05 | 0.00010282 | 56.9682993 | 42.4858923 | 71.3281477 | 54.910963 |
| -4.1447876 | 3.40E-05 | 0.00010282 | 31.1481349 | 87.6778924 | 79.3813257 | 98.055291 |
| -4.1446444 | 3.40E-05 | 0.00010287 | 511.485162 | 857.294948 | 714.431931 | 815.820021 |
| -4.1442629 | 3.41E-05 | 0.00010303 | 33.1973543 | 82.8068984 | 70.1776937 | 105.899714 |
| -4.1438437 | 3.42E-05 | 0.00010321 | 64.755333 | 143.423713 | 197.878087 | 207.316901 |
| -4.1430637 | 3.43E-05 | 0.00010355 | 48.3615778 | 71.4412457 | 43.7172518 | 64.99665 |
| -4.1430293 | 3.43E-05 | 0.00010356 | 41.3942319 | 56.5576528 | 58.6731538 | 65.556966 |
| -4.1427697 | 3.43E-05 | 0.00010367 | 36.4761053 | 72.7942996 | 65.5758777 | 71.1601255 |
| -4.1423789 | 3.44E-05 | 0.00010383 | 81.1490882 | 86.8660601 | 44.8677058 | 44.8252759 |
| -4.1422923 | 3.44E-05 | 0.00010386 | 34.017042 | 74.4179643 | 78.2308717 | 76.202969 |
| -4.1422541 | 3.44E-05 | 0.00010387 | 50.8206411 | 137.470276 | 174.869007 | 36.9808526 |
| -4.1421373 | 3.44E-05 | 0.00010391 | 34.8367298 | 106.079425 | 71.3281477 | 71.1601255 |
| -4.1420641 | 3.44E-05 | 0.00010394 | 92.214873 | 120.151186 | 25.3099879 | 126.071088 |
| -4.1418859 | 3.44E-05 | 0.00010401 | 67.2143963 | 124.751569 | 29.9118039 | 140.639303 |
| -4.1415299 | 3.45E-05 | 0.00010416 | 27.0496961 | 91.7370541 | 132.302209 | 101.417187 |
| -4.1410948 | 3.46E-05 | 0.00010435 | 23.3611012 | 122.857294 | 112.744492 | 132.234564 |
| -4.1407824 | 3.46E-05 | 0.00010448 | 33.6071982 | 88.219114 | 74.7795097 | 73.9617052 |
| -4.1403132 | 3.47E-05 | 0.00010469 | 41.3942319 | 59.2637606 | 57.5226998 | 60.5141225 |
| -4.1400581 | 3.47E-05 | 0.00010479 | 32.3776665 | 78.7477367 | 86.2840497 | 79.5648647 |
| -4.1399373 | 3.47E-05 | 0.00010484 | 90.9853413 | 77.1240721 | 63.2749697 | 35.2999048 |
| -4.1395295 | 3.48E-05 | 0.00010501 | 28.2792277 | 102.020264 | 109.29313 | 86.2886561 |
| -4.1395224 | 3.48E-05 | 0.00010501 | 46.3123584 | 84.7011739 | 48.3190678 | 59.3934906 |
| -4.1393207 | 3.48E-05 | 0.00010509 | 40.5745441 | 94.4431619 | 51.7704298 | 74.5220212 |
| 4.1392195 | 3.48E-05 | 0.00010512 | 4.0984388 | 2.70610779 | 1.150454 | 5.60315949 |
| -4.1392396 | 3.48E-05 | 0.00010512 | 34.4268859 | 111.491641 | 187.524001 | 146.242463 |
| -4.1391772 | 3.49E-05 | 0.00010513 | 52.0501728 | 78.477126 | 40.2658898 | 68.9188617 |
| -4.1391544 | 3.49E-05 | 0.00010513 | 162.298176 | 424.858923 | 452.12842 | 414.073486 |

| | | | | | | |
|---|---|---|---|---|---|---|
| -4.1388315 | 3.49E-05 | 0.00010526 | 38.1154808 | 114.197749 | 58.6731538 | 75.0823371 |
| -4.1387674 | 3.49E-05 | 0.00010528 | 31.5579788 | 80.912623 | 86.2840497 | 84.6077082 |
| -4.1385853 | 3.49E-05 | 0.00010536 | 35.2465737 | 74.1473535 | 66.7263317 | 78.4442328 |
| -4.1382874 | 3.50E-05 | 0.00010549 | 26.2300083 | 108.514922 | 164.514921 | 89.6505518 |
| -4.1379578 | 3.50E-05 | 0.00010561 | 30.3284471 | 105.538204 | 103.54086 | 73.9617052 |
| -4.1379525 | 3.50E-05 | 0.00010561 | 30.738291 | 68.7351379 | 119.647216 | 111.502874 |
| -4.1379413 | 3.50E-05 | 0.00010561 | 28.6890716 | 120.692407 | 86.2840497 | 93.0124475 |
| -4.1377678 | 3.51E-05 | 0.00010568 | 593.044094 | 421.340983 | 440.62388 | 508.766881 |
| -4.1373844 | 3.51E-05 | 0.00010584 | 430.336074 | 1524.62113 | 2166.30487 | 1954.94234 |
| -4.1374037 | 3.51E-05 | 0.00010584 | 45.4926707 | 100.396599 | 42.5667978 | 109.26161 |
| -4.1372562 | 3.51E-05 | 0.00010589 | 350.826361 | 625.652121 | 500.447488 | 473.466977 |
| -4.1371122 | 3.52E-05 | 0.00010595 | 44.263139 | 58.7225391 | 49.4695218 | 64.4363341 |
| -4.1370381 | 3.52E-05 | 0.00010597 | 313.530568 | 60.075593 | 52.9208838 | 72.2807574 |
| -4.1368211 | 3.52E-05 | 0.00010605 | 177.052556 | 52.4984912 | 57.5226998 | 59.3934906 |
| -4.1368322 | 3.52E-05 | 0.00010605 | 46.7222023 | 65.4878086 | 47.1686138 | 56.5919108 |
| -4.136621 | 3.52E-05 | 0.00010614 | 101.231438 | 51.416048 | 44.8677058 | 61.0744384 |
| -4.1364134 | 3.53E-05 | 0.00010622 | 34.017042 | 89.5721679 | 77.0804177 | 68.3585457 |
| -4.1361487 | 3.53E-05 | 0.00010633 | 38.5253247 | 123.127905 | 52.9208838 | 91.3314996 |
| -4.1360124 | 3.53E-05 | 0.00010639 | 163.937552 | 59.8049822 | 44.8677058 | 65.556966 |
| -4.135686 | 3.54E-05 | 0.00010653 | 28.6890716 | 169.131737 | 178.320369 | 63.8760181 |
| 4.135062 | 3.55E-05 | 0.00010678 | 33.1973543 | 18.6721438 | 19.5577179 | 22.9729539 |
| -4.1350556 | 3.55E-05 | 0.00010678 | 36.0662614 | 80.1007906 | 64.4254237 | 69.4791776 |
| -4.1350725 | 3.55E-05 | 0.00010678 | 26.6398522 | 173.461509 | 239.294431 | 66.6775979 |
| -4.1350654 | 3.55E-05 | 0.00010678 | 38.5253247 | 58.7225391 | 74.7795097 | 63.8760181 |
| -4.1350242 | 3.55E-05 | 0.00010679 | 32.7875104 | 99.3141559 | 69.0272397 | 84.6077082 |
| -4.1349168 | 3.55E-05 | 0.00010683 | 51.6403289 | 71.7118565 | 49.4695218 | 47.6268556 |
| -4.1344438 | 3.56E-05 | 0.00010704 | 64.755333 | 120.421797 | 146.107657 | 30.8173772 |
| -4.1340754 | 3.56E-05 | 0.0001072 | 41.8040758 | 61.6992576 | 51.7704298 | 75.0823371 |
| -4.133792 | 3.57E-05 | 0.00010732 | 54.0993922 | 53.3103235 | 48.3190678 | 107.020346 |
| -4.1337113 | 3.57E-05 | 0.00010734 | 38.1154808 | 62.2404792 | 66.7263317 | 63.8760181 |
| -4.1337252 | 3.57E-05 | 0.00010734 | 32.3776665 | 87.9485032 | 85.1335957 | 72.2807574 |
| -4.1332577 | 3.58E-05 | 0.00010754 | 63.5258014 | 76.8534613 | 77.0804177 | 35.2999048 |
| -4.1329046 | 3.58E-05 | 0.0001077 | 26.6398522 | 125.292791 | 130.001301 | 81.8061285 |
| -4.1328803 | 3.58E-05 | 0.0001077 | 27.0496961 | 94.7137727 | 105.841768 | 103.65845 |
| -4.132774 | 3.58E-05 | 0.00010774 | 28.2792277 | 99.0435452 | 87.4345036 | 103.65845 |
| -4.1324832 | 3.59E-05 | 0.00010786 | 97.9526873 | 61.1580361 | 47.1686138 | 47.6268556 |
| -4.1322727 | 3.59E-05 | 0.00010795 | 251.234298 | 757.980792 | 739.741919 | 736.815472 |
| -4.1321264 | 3.59E-05 | 0.00010801 | 38.9351686 | 59.8049822 | 67.8767857 | 61.0744384 |
| -4.1320711 | 3.60E-05 | 0.00010803 | 64.755333 | 46.545054 | 46.0181598 | 58.8331746 |
| -4.1319163 | 3.60E-05 | 0.00010809 | 29.5087594 | 80.1007906 | 105.841768 | 93.5727634 |
| -4.1318305 | 3.60E-05 | 0.00010812 | 46.3123584 | 73.0649104 | 43.7172518 | 70.0394936 |
| -4.1317732 | 3.60E-05 | 0.00010814 | 148.773328 | 354.500121 | 317.525303 | 341.792729 |
| -4.1315664 | 3.60E-05 | 0.00010822 | 45.4926707 | 122.316072 | 47.1686138 | 75.0823371 |

| | | | | | | |
|---|---|---|---|---|---|---|
| -4.1311816 | 3.61E-05 | 0.00010839 | 40.984388 | 77.6652936 | 55.2217918 | 58.8331746 |
| -4.1311769 | 3.61E-05 | 0.00010839 | 19.6725062 | 119.068743 | 156.461743 | 161.370993 |
| -4.1308885 | 3.61E-05 | 0.00010851 | 43.8532952 | 72.253078 | 80.5317797 | 45.9459078 |
| -4.130776 | 3.62E-05 | 0.00010856 | 39.3450125 | 74.9591858 | 86.2840497 | 52.1093832 |
| -4.1307384 | 3.62E-05 | 0.00010856 | 65.9848647 | 56.5576528 | 46.0181598 | 47.0665397 |
| -4.1302802 | 3.62E-05 | 0.00010877 | 34.8367298 | 127.998899 | 54.0713378 | 137.837723 |
| -4.1301794 | 3.62E-05 | 0.00010881 | 13.1150042 | 160.201581 | 346.286653 | 174.25826 |
| -4.1294614 | 3.64E-05 | 0.00010914 | 236.479919 | 428.918085 | 404.959806 | 415.194118 |
| -4.1291233 | 3.64E-05 | 0.00010929 | 52.4600166 | 53.3103235 | 58.6731538 | 44.2649599 |
| -4.1288679 | 3.65E-05 | 0.0001094 | 22.1315695 | 69.817581 | 372.747095 | 211.239113 |
| -4.1284393 | 3.65E-05 | 0.00010959 | 63.9356453 | 151.000815 | 82.8326877 | 258.305652 |
| -4.1281812 | 3.66E-05 | 0.00010971 | 153.691455 | 56.5576528 | 37.9649818 | 86.2886561 |
| -4.128142 | 3.66E-05 | 0.00010972 | 52.4600166 | 41.1328384 | 66.7263317 | 62.1950703 |
| -4.1281091 | 3.66E-05 | 0.00010972 | 36.8859492 | 73.6061319 | 58.6731538 | 75.0823371 |
| -4.1277356 | 3.66E-05 | 0.00010989 | 324.186509 | 1481.59402 | 1747.53962 | 1702.23985 |
| -4.1275678 | 3.67E-05 | 0.00010996 | 27.0496961 | 162.637078 | 112.744492 | 80.1251807 |
| -4.1274331 | 3.67E-05 | 0.00011001 | 36.4761053 | 63.8641439 | 97.7885896 | 66.1172819 |
| -4.1273002 | 3.67E-05 | 0.00011007 | 34.017042 | 113.385916 | 71.3281477 | 71.7204414 |
| -4.1270362 | 3.67E-05 | 0.00011018 | 31.1481349 | 94.7137727 | 73.6290557 | 89.0902358 |
| -4.1269139 | 3.68E-05 | 0.00011023 | 41.8040758 | 98.7729344 | 48.3190678 | 76.202969 |
| -4.1267396 | 3.68E-05 | 0.0001103 | 61.0667381 | 64.6759762 | 39.1154358 | 56.5919108 |
| -4.1265439 | 3.68E-05 | 0.00011039 | 47.951734 | 132.599282 | 119.647216 | 122.148877 |
| -4.1263944 | 3.68E-05 | 0.00011045 | 38.9351686 | 75.5004074 | 57.5226998 | 61.6347543 |
| -4.1263644 | 3.69E-05 | 0.00011045 | 33.6071982 | 70.9000241 | 80.5317797 | 74.5220212 |
| -4.1260101 | 3.69E-05 | 0.00011061 | 27.0496961 | 96.0668266 | 95.4876816 | 109.26161 |
| -4.1259314 | 3.69E-05 | 0.00011064 | 31.1481349 | 70.9000241 | 166.815829 | 95.8140272 |
| -4.1255959 | 3.70E-05 | 0.00011079 | 28.2792277 | 94.7137727 | 128.850847 | 82.3664444 |
| 4.12544392 | 3.70E-05 | 0.00011086 | 0 | 0 | 0 | 0 |
| 4.12466809 | 3.71E-05 | 0.00011122 | 34.017042 | 15.1542036 | 12.6549939 | 15.1285306 |
| -4.1241035 | 3.72E-05 | 0.00011148 | 172.13443 | 47.8981079 | 81.6822337 | 47.0665397 |
| -4.1240346 | 3.72E-05 | 0.0001115 | 54.9190799 | 61.1580361 | 39.1154358 | 72.8410733 |
| -4.1235217 | 3.73E-05 | 0.00011174 | 37.705637 | 70.3588026 | 58.6731538 | 66.1172819 |
| -4.1234909 | 3.73E-05 | 0.00011175 | 68.8537718 | 86.8660601 | 32.2127119 | 80.1251807 |
| 4.1232109 | 3.74E-05 | 0.00011187 | 0 | 0 | 0 | 0 |
| -4.1227421 | 3.74E-05 | 0.00011209 | 45.4926707 | 59.2637606 | 58.6731538 | 47.0665397 |
| -4.1225621 | 3.75E-05 | 0.00011217 | 30.3284471 | 80.6420122 | 93.1867736 | 84.0473923 |
| -4.1222716 | 3.75E-05 | 0.0001123 | 22.1315695 | 150.188982 | 140.355387 | 100.296555 |
| -4.1218489 | 3.76E-05 | 0.0001125 | 29.5087594 | 83.0775092 | 95.4876816 | 87.9696039 |
| -4.1217922 | 3.76E-05 | 0.00011251 | 36.4761053 | 74.9591858 | 58.6731538 | 73.9617052 |
| -4.1217198 | 3.76E-05 | 0.00011254 | 26.6398522 | 83.34812 | 148.408565 | 106.46003 |
| -4.1214504 | 3.76E-05 | 0.00011266 | 47.951734 | 68.1939163 | 47.1686138 | 51.5490673 |
| -4.1213201 | 3.77E-05 | 0.00011271 | 19.6725062 | 125.02218 | 134.603117 | 166.413837 |
| -4.1202609 | 3.78E-05 | 0.00011321 | 15.9839113 | 113.385916 | 226.639437 | 221.885116 |

| | | | | | | |
|---|---|---|---|---|---|---|
| -4.1202709 | 3.78E-05 | 0.00011321 | 30.3284471 | 51.1454373 | 211.683535 | 242.05649 |
| -4.1198965 | 3.79E-05 | 0.00011338 | 33.1973543 | 115.550803 | 63.2749697 | 87.409288 |
| -4.1184864 | 3.81E-05 | 0.00011406 | 31.5579788 | 133.681725 | 60.9740618 | 117.106033 |
| -4.1183298 | 3.82E-05 | 0.00011413 | 50.4107972 | 61.6992576 | 147.258111 | 50.4284354 |
| -4.1182154 | 3.82E-05 | 0.00011418 | 31.9678226 | 93.3607188 | 69.0272397 | 84.6077082 |
| -4.1180925 | 3.82E-05 | 0.00011423 | 50.8206411 | 82.8068984 | 69.0272397 | 74.5220212 |
| -4.1177462 | 3.83E-05 | 0.00011439 | 36.4761053 | 77.3946828 | 82.8326877 | 56.0315949 |
| -4.1175779 | 3.83E-05 | 0.00011446 | 54.9190799 | 102.832096 | 59.8236078 | 42.0236961 |
| -4.1171312 | 3.84E-05 | 0.00011466 | 102.051126 | 82.5362876 | 59.8236078 | 86.2886561 |
| -4.1171451 | 3.84E-05 | 0.00011466 | 161.888333 | 271.963833 | 262.303511 | 281.838922 |
| -4.1168351 | 3.84E-05 | 0.0001148 | 38.9351686 | 70.6294134 | 64.4254237 | 54.350647 |
| -4.1166982 | 3.84E-05 | 0.00011486 | 46.3123584 | 82.2656769 | 41.4163438 | 79.5648647 |
| -4.1162453 | 3.85E-05 | 0.00011507 | 31.9678226 | 102.561485 | 65.5758777 | 90.7711837 |
| -4.116159 | 3.85E-05 | 0.0001151 | 37.705637 | 60.6168145 | 66.7263317 | 61.6347543 |
| -4.1160253 | 3.85E-05 | 0.00011516 | 31.9678226 | 97.9611021 | 66.7263317 | 87.9696039 |
| -4.1159469 | 3.86E-05 | 0.00011519 | 151.642236 | 77.6652936 | 41.4163438 | 52.6696992 |
| -4.1157394 | 3.86E-05 | 0.00011528 | 166.396615 | 258.162683 | 279.560321 | 296.407137 |
| -4.1157185 | 3.86E-05 | 0.00011528 | 160.248957 | 66.2996409 | 36.8145279 | 71.7204414 |
| -4.1155054 | 3.86E-05 | 0.00011538 | 44.6729829 | 65.4878086 | 44.8677058 | 64.99665 |
| -4.1154021 | 3.87E-05 | 0.00011542 | 19.2626624 | 106.891258 | 146.107657 | 237.013646 |
| -4.1143959 | 3.88E-05 | 0.00011591 | 32.3776665 | 86.8660601 | 81.6822337 | 68.3585457 |
| 4.11388707 | 3.89E-05 | 0.00011616 | 47.1320462 | 21.3782516 | 23.0090799 | 16.2491625 |
| -4.1138483 | 3.89E-05 | 0.00011617 | 36.4761053 | 125.563402 | 49.4695218 | 130.553616 |
| -4.1138184 | 3.89E-05 | 0.00011617 | 36.4761053 | 110.409198 | 54.0713378 | 86.2886561 |
| -4.1134283 | 3.90E-05 | 0.00011636 | 32.3776665 | 91.4664434 | 65.5758777 | 86.848972 |
| -4.113388 | 3.90E-05 | 0.00011637 | 25.4103206 | 95.7962158 | 120.79767 | 101.977503 |
| -4.1131238 | 3.90E-05 | 0.00011649 | 274.185556 | 546.092552 | 483.190678 | 518.292252 |
| 4.11267605 | 3.91E-05 | 0.0001167 | 4.50828268 | 5.95343714 | 4.60181598 | 5.60315949 |
| -4.1126142 | 3.91E-05 | 0.00011671 | 34.4268859 | 69.5469702 | 66.7263317 | 76.202969 |
| -4.1126196 | 3.91E-05 | 0.00011671 | 50.8206411 | 99.3141559 | 86.2840497 | 102.537819 |
| -4.1122252 | 3.92E-05 | 0.0001169 | 40.984388 | 50.3336049 | 66.7263317 | 61.6347543 |
| -4.1115155 | 3.93E-05 | 0.00011725 | 29.9186032 | 108.785533 | 83.9831417 | 76.763285 |
| -4.111456 | 3.93E-05 | 0.00011727 | 61.476582 | 57.0988744 | 37.9649818 | 63.3157022 |
| -4.1113974 | 3.93E-05 | 0.00011728 | 34.8367298 | 86.0542278 | 63.2749697 | 69.4791776 |
| -4.1114162 | 3.93E-05 | 0.00011728 | 23.770945 | 106.079425 | 98.9390436 | 171.45668 |
| -4.1110558 | 3.94E-05 | 0.00011744 | 46.3123584 | 60.8874253 | 65.5758777 | 43.704644 |
| -4.1101421 | 3.95E-05 | 0.00011789 | 175.823024 | 485.746349 | 437.172518 | 474.027293 |
| -4.1091164 | 3.97E-05 | 0.00011841 | 34.8367298 | 65.7584193 | 66.7263317 | 75.6426531 |
| -4.1088147 | 3.98E-05 | 0.00011855 | 22.1315695 | 117.174467 | 150.709473 | 106.46003 |
| -4.1086691 | 3.98E-05 | 0.00011862 | 49.1812656 | 122.316072 | 158.762651 | 104.779082 |
| -4.1086126 | 3.98E-05 | 0.00011862 | 22.5414134 | 129.351952 | 144.957203 | 96.9346591 |
| 4.10861181 | 3.98E-05 | 0.00011862 | 0 | 1.89427545 | 0 | 0 |
| -4.1084203 | 3.98E-05 | 0.00011871 | 47.5418901 | 48.7099402 | 48.3190678 | 59.3934906 |

| | | | | | | |
|---|---|---|---|---|---|---|
| -4.1080242 | 3.99E-05 | 0.0001189 | 29.0989155 | 86.5954493 | 97.7885896 | 80.6854966 |
| -4.1077535 | 4.00E-05 | 0.00011901 | 40.5745441 | 74.4179643 | 48.3190678 | 68.9188617 |
| -4.107745 | 4.00E-05 | 0.00011901 | 70.4931473 | 64.946587 | 52.9208838 | 38.6618005 |
| -4.1077711 | 3.99E-05 | 0.00011901 | 31.5579788 | 106.079425 | 93.1867736 | 64.4363341 |
| -4.1077166 | 4.00E-05 | 0.00011902 | 25.8201644 | 103.102707 | 93.1867736 | 109.26161 |
| -4.1075454 | 4.00E-05 | 0.00011909 | 30.738291 | 106.350036 | 67.8767857 | 91.8918156 |
| -4.1075284 | 4.00E-05 | 0.00011909 | 57.7879871 | 49.7923834 | 46.0181598 | 107.580662 |
| -4.1070891 | 4.01E-05 | 0.00011931 | 43.0336074 | 79.5595691 | 71.3281477 | 44.8252759 |
| -4.1070483 | 4.01E-05 | 0.00011932 | 27.45954 | 96.0668266 | 93.1867736 | 91.3314996 |
| -4.1069554 | 4.01E-05 | 0.00011936 | 31.1481349 | 98.2317128 | 70.1776937 | 82.9267604 |
| -4.1065137 | 4.02E-05 | 0.00011957 | 94.2640924 | 114.197749 | 36.8145279 | 50.4284354 |
| -4.1058595 | 4.03E-05 | 0.0001199 | 85.6573709 | 40.3210061 | 93.1867736 | 46.5062237 |
| -4.1056644 | 4.03E-05 | 0.00011999 | 29.0989155 | 77.3946828 | 90.8858656 | 97.4949751 |
| -4.1054772 | 4.03E-05 | 0.00012008 | 43.8532952 | 70.3588026 | 138.054479 | 49.8681194 |
| -4.1050973 | 4.04E-05 | 0.00012026 | 34.8367298 | 92.8194972 | 62.1245158 | 69.4791776 |
| -4.1049844 | 4.04E-05 | 0.0001203 | 27.45954 | 87.9485032 | 86.2840497 | 126.071088 |
| -4.1049931 | 4.04E-05 | 0.0001203 | 26.6398522 | 182.662276 | 78.2308717 | 108.701294 |
| -4.104478 | 4.05E-05 | 0.00012055 | 36.8859492 | 62.51109 | 60.9740618 | 96.3743432 |
| -4.1043208 | 4.06E-05 | 0.00012062 | 64.3454891 | 64.4053654 | 55.2217918 | 38.1014845 |
| -4.104253 | 4.06E-05 | 0.00012065 | 117.21535 | 212.700072 | 154.160835 | 183.223315 |
| -4.1041962 | 4.06E-05 | 0.00012067 | 34.4268859 | 87.1366709 | 70.1776937 | 62.7553862 |
| -4.1039654 | 4.06E-05 | 0.00012078 | 30.3284471 | 103.102707 | 209.382627 | 68.3585457 |
| -4.103589 | 4.07E-05 | 0.00012095 | 33.6071982 | 81.1832337 | 89.7354116 | 60.5141225 |
| -4.1035956 | 4.07E-05 | 0.00012095 | 105.739721 | 253.5623 | 181.771731 | 225.247011 |
| -4.1035652 | 4.07E-05 | 0.00012095 | 29.0989155 | 80.3714014 | 94.3372276 | 86.2886561 |
| -4.1032454 | 4.07E-05 | 0.00012111 | 22.5414134 | 180.49739 | 89.7354116 | 130.553616 |
| -4.1030948 | 4.08E-05 | 0.00012118 | 40.5745441 | 90.654611 | 50.6199758 | 63.8760181 |
| 4.10281578 | 4.08E-05 | 0.00012131 | 15.5740674 | 9.47137727 | 5.75226998 | 6.16347543 |
| -4.1026127 | 4.09E-05 | 0.00012141 | 51.230485 | 171.837845 | 88.5849576 | 136.717091 |
| -4.1025848 | 4.09E-05 | 0.00012141 | 51.230485 | 99.0435452 | 94.3372276 | 103.65845 |
| -4.1025636 | 4.09E-05 | 0.00012141 | 35.2465737 | 60.075593 | 96.6381356 | 69.4791776 |
| -4.1022733 | 4.09E-05 | 0.00012155 | 45.9025146 | 46.8156648 | 89.7354116 | 56.5919108 |
| -4.1020661 | 4.09E-05 | 0.00012164 | 25.0004767 | 104.726372 | 116.195854 | 94.1330794 |
| -4.1020759 | 4.09E-05 | 0.00012164 | 31.5579788 | 68.1939163 | 146.107657 | 85.1680242 |
| -4.1018857 | 4.10E-05 | 0.00012172 | 25.8201644 | 86.0542278 | 208.232173 | 101.977503 |
| -4.1016115 | 4.10E-05 | 0.00012185 | 27.8693838 | 87.1366709 | 93.1867736 | 91.8918156 |
| -4.1011493 | 4.11E-05 | 0.00012209 | 44.263139 | 79.2889583 | 44.8677058 | 61.6347543 |
| -4.1001588 | 4.13E-05 | 0.0001226 | 31.5579788 | 171.837845 | 115.0454 | 58.8331746 |
| -4.0999723 | 4.13E-05 | 0.00012269 | 29.9186032 | 90.9252218 | 77.0804177 | 82.3664444 |
| -4.0997554 | 4.14E-05 | 0.00012279 | 32.3776665 | 146.400432 | 52.9208838 | 147.363094 |
| -4.0990488 | 4.15E-05 | 0.00012315 | 29.5087594 | 103.914539 | 72.4786017 | 89.6505518 |
| -4.0986927 | 4.15E-05 | 0.00012333 | 35.6564176 | 51.9572696 | 112.744492 | 95.8140272 |
| -4.0986655 | 4.16E-05 | 0.00012334 | 32.3776665 | 79.8301798 | 64.4254237 | 108.140978 |

| | | | | | | |
|---|---|---|---|---|---|---|
| -4.0982621 | 4.16E-05 | 0.00012354 | 29.5087594 | 96.8786589 | 70.1776937 | 100.856871 |
| -4.0980476 | 4.17E-05 | 0.00012364 | 50.4107972 | 58.9931499 | 51.7704298 | 43.144328 |
| -4.0978549 | 4.17E-05 | 0.00012373 | 89.7558097 | 67.1114732 | 29.9118039 | 119.347297 |
| 4.0973475 | 4.18E-05 | 0.00012399 | 329.104636 | 176.708839 | 165.665375 | 146.242463 |
| -4.0971836 | 4.18E-05 | 0.00012407 | 38.9351686 | 52.7691019 | 102.390406 | 66.6775979 |
| 4.09701491 | 4.19E-05 | 0.00012415 | 24.1807889 | 8.11832337 | 9.20363196 | 11.206319 |
| -4.0967452 | 4.19E-05 | 0.00012427 | 31.1481349 | 172.108456 | 65.5758777 | 95.8140272 |
| -4.0967592 | 4.19E-05 | 0.00012427 | 26.6398522 | 100.125988 | 85.1335957 | 104.218766 |
| -4.0967126 | 4.19E-05 | 0.00012428 | 38.9351686 | 140.176384 | 42.5667978 | 186.585211 |
| -4.0964617 | 4.20E-05 | 0.0001244 | 22.5414134 | 149.918372 | 83.9831417 | 159.129729 |
| -4.0961336 | 4.20E-05 | 0.00012457 | 1714.37695 | 1670.75095 | 949.124546 | 1279.20131 |
| -4.0959994 | 4.20E-05 | 0.00012463 | 47.951734 | 95.525605 | 63.2749697 | 42.5840121 |
| -4.0958611 | 4.21E-05 | 0.00012469 | 31.5579788 | 79.8301798 | 71.3281477 | 78.4442328 |
| -4.0957316 | 4.21E-05 | 0.00012475 | 60.2470503 | 34.6381797 | 169.116737 | 86.848972 |
| -4.0957108 | 4.21E-05 | 0.00012475 | 23.770945 | 102.561485 | 113.894946 | 108.701294 |
| -4.0956028 | 4.21E-05 | 0.00012479 | 30.738291 | 95.2549943 | 81.6822337 | 70.0394936 |
| -4.0953587 | 4.22E-05 | 0.00012491 | 26.2300083 | 94.9843835 | 100.089498 | 94.1330794 |
| -4.0953178 | 4.22E-05 | 0.00012493 | 44.263139 | 43.0271139 | 67.8767857 | 73.9617052 |
| -4.0951686 | 4.22E-05 | 0.00012499 | 25.0004767 | 105.538204 | 109.29313 | 93.5727634 |
| -4.0950721 | 4.22E-05 | 0.00012503 | 90.5754975 | 31.6614612 | 65.5758777 | 84.6077082 |
| -4.0945942 | 4.23E-05 | 0.00012528 | 43.8532952 | 83.34812 | 70.1776937 | 43.144328 |
| 4.09392215 | 4.24E-05 | 0.00012563 | 28.2792277 | 18.6721438 | 12.6549939 | 12.3269509 |
| -4.0937864 | 4.24E-05 | 0.0001257 | 132.379573 | 33.5557366 | 126.549939 | 52.1093832 |
| -4.0933078 | 4.25E-05 | 0.00012594 | 61.8864259 | 91.7370541 | 117.346308 | 31.3776931 |
| -4.0927246 | 4.26E-05 | 0.00012625 | 39.7548564 | 93.3607188 | 108.142676 | 101.417187 |
| -4.0924922 | 4.27E-05 | 0.00012637 | 105.329877 | 46.2744432 | 43.7172518 | 65.556966 |
| -4.0922937 | 4.27E-05 | 0.00012645 | 56.1486116 | 53.0397127 | 41.4163438 | 52.1093832 |
| -4.0922937 | 4.27E-05 | 0.00012645 | 36.0662614 | 71.9824673 | 57.5226998 | 65.556966 |
| -4.0916271 | 4.28E-05 | 0.0001268 | 47.1320462 | 54.6633774 | 81.6822337 | 44.2649599 |
| -4.0913279 | 4.29E-05 | 0.00012695 | 35.2465737 | 69.2763595 | 62.1245158 | 64.4363341 |
| -4.0909932 | 4.30E-05 | 0.00012713 | 38.5253247 | 48.980551 | 75.9299637 | 73.9617052 |
| -4.0906017 | 4.30E-05 | 0.00012732 | 75.8211178 | 58.1813175 | 180.621277 | 38.6618005 |
| -4.0905892 | 4.30E-05 | 0.00012732 | 27.8693838 | 127.998899 | 172.568099 | 63.8760181 |
| -4.0905735 | 4.30E-05 | 0.00012732 | 38.5253247 | 50.8748265 | 65.5758777 | 68.9188617 |
| -4.0905421 | 4.30E-05 | 0.00012733 | 35.6564176 | 67.382084 | 58.6731538 | 68.3585457 |
| 4.09051802 | 4.30E-05 | 0.00012733 | 21.7217256 | 39.5091738 | 35.6640739 | 34.1792729 |
| -4.090064 | 4.31E-05 | 0.00012757 | 34.017042 | 69.817581 | 63.2749697 | 72.2807574 |
| -4.0897625 | 4.32E-05 | 0.00012772 | 55.7387677 | 108.514922 | 93.1867736 | 75.0823371 |
| -4.0892681 | 4.33E-05 | 0.00012798 | 34.8367298 | 93.9019404 | 55.2217918 | 79.0045488 |
| -4.0890858 | 4.33E-05 | 0.00012807 | 752.06352 | 61.4286469 | 33.3631659 | 109.821926 |
| -4.0887297 | 4.34E-05 | 0.00012826 | 16.8035991 | 175.085174 | 204.780811 | 109.821926 |
| -4.0886314 | 4.34E-05 | 0.0001283 | 26.2300083 | 140.176384 | 80.5317797 | 98.615607 |
| 4.08850721 | 4.34E-05 | 0.00012836 | 39.3450125 | 43.2977247 | 43.7172518 | 38.6618005 |

| | | | | | | |
|---|---|---|---|---|---|---|
| -4.0884552 | 4.34E-05 | 0.00012837 | 32.7875104 | 102.832096 | 251.949425 | 61.6347543 |
| -4.0883412 | 4.34E-05 | 0.00012841 | 43.8532952 | 84.7011739 | 41.4163438 | 76.763285 |
| -4.0883454 | 4.34E-05 | 0.00012841 | 28.6890716 | 71.1706349 | 133.452663 | 92.4521315 |
| -4.0883193 | 4.35E-05 | 0.00012841 | 39.7548564 | 213.782516 | 44.8677058 | 115.425085 |
| -4.0883033 | 4.35E-05 | 0.00012841 | 29.9186032 | 97.1492697 | 75.9299637 | 76.763285 |
| -4.0882025 | 4.35E-05 | 0.00012845 | 46.3123584 | 87.6778924 | 64.4254237 | 42.0236961 |
| -4.0881725 | 4.35E-05 | 0.00012846 | 45.9025146 | 74.688575 | 42.5667978 | 58.8331746 |
| -4.0877361 | 4.36E-05 | 0.00012868 | 41.8040758 | 50.6042157 | 78.2308717 | 167.534469 |
| -4.0877281 | 4.36E-05 | 0.00012868 | 41.3942319 | 79.2889583 | 157.612197 | 49.3078035 |
| -4.0876798 | 4.36E-05 | 0.00012868 | 33.6071982 | 80.6420122 | 89.7354116 | 57.7125427 |
| -4.087681 | 4.36E-05 | 0.00012868 | 59.4273626 | 29.2259641 | 182.922185 | 129.9933 |
| -4.0876138 | 4.36E-05 | 0.00012871 | 49.1812656 | 86.3248385 | 102.390406 | 36.9808526 |
| -4.0874378 | 4.36E-05 | 0.00012879 | 37.705637 | 66.8408624 | 51.7704298 | 85.7283401 |
| -4.0870738 | 4.37E-05 | 0.00012899 | 46.7222023 | 67.1114732 | 46.0181598 | 49.3078035 |
| -4.0868959 | 4.37E-05 | 0.00012907 | 45.9025146 | 43.0271139 | 66.7263317 | 56.5919108 |
| -4.0868466 | 4.37E-05 | 0.00012909 | 21.3118818 | 150.188982 | 123.098578 | 100.856871 |
| -4.0867232 | 4.38E-05 | 0.00012914 | 42.2139196 | 53.3103235 | 55.2217918 | 50.4284354 |
| -4.0866801 | 4.38E-05 | 0.00012916 | 40.1647002 | 53.3103235 | 113.894946 | 62.7553862 |
| -4.0864946 | 4.38E-05 | 0.00012925 | 27.0496961 | 79.0183475 | 97.7885896 | 105.339398 |
| -4.086046 | 4.39E-05 | 0.00012949 | 39.7548564 | 77.3946828 | 63.2749697 | 48.7474875 |
| -4.0860244 | 4.39E-05 | 0.00012949 | 82.3786199 | 58.7225391 | 39.1154358 | 51.5490673 |
| -4.0859237 | 4.39E-05 | 0.00012953 | 47.951734 | 40.8622277 | 70.1776937 | 107.020346 |
| 4.08581562 | 4.39E-05 | 0.00012958 | 7.37718984 | 5.41221558 | 6.90272397 | 4.48252759 |
| -4.0855882 | 4.40E-05 | 0.00012969 | 60.2470503 | 79.2889583 | 33.3631659 | 70.0394936 |
| -4.0854846 | 4.40E-05 | 0.00012974 | 52.4600166 | 106.620647 | 36.8145279 | 72.8410733 |
| -4.0850012 | 4.41E-05 | 0.00013 | 27.8693838 | 91.4664434 | 131.151755 | 74.5220212 |
| -4.0847896 | 4.41E-05 | 0.00013011 | 22.5414134 | 169.943569 | 111.594038 | 94.1330794 |
| -4.0840539 | 4.43E-05 | 0.00013051 | 23.770945 | 110.679809 | 117.346308 | 93.5727634 |
| -4.0831919 | 4.44E-05 | 0.00013098 | 43.4434513 | 57.640096 | 46.0181598 | 55.4712789 |
| -4.0831343 | 4.44E-05 | 0.000131 | 104.100346 | 54.3927666 | 48.3190678 | 46.5062237 |
| -4.0831053 | 4.44E-05 | 0.000131 | 52.0501728 | 75.7710182 | 50.6199758 | 42.0236961 |
| -4.0829203 | 4.45E-05 | 0.0001311 | 29.0989155 | 84.9717847 | 87.4345036 | 76.202969 |
| -4.0827047 | 4.45E-05 | 0.00013121 | 27.45954 | 108.514922 | 73.6290557 | 101.417187 |
| -4.0818325 | 4.47E-05 | 0.00013169 | 92.6247169 | 115.280192 | 21.8586259 | 161.931309 |
| -4.0816606 | 4.47E-05 | 0.00013176 | 38.1154808 | 70.0881918 | 67.8767857 | 49.8681194 |
| -4.0816741 | 4.47E-05 | 0.00013176 | 49.1812656 | 77.1240721 | 47.1686138 | 46.5062237 |
| -4.0815741 | 4.47E-05 | 0.0001318 | 241.398045 | 394.550516 | 299.118039 | 397.824324 |
| -4.0808564 | 4.49E-05 | 0.00013219 | 46.7222023 | 75.5004074 | 43.7172518 | 53.2300151 |
| -4.0806247 | 4.49E-05 | 0.00013231 | 18.0331307 | 102.561485 | 195.577179 | 158.569413 |
| -4.0800306 | 4.50E-05 | 0.00013264 | 31.5579788 | 82.5362876 | 67.8767857 | 73.9617052 |
| -4.0795723 | 4.51E-05 | 0.00013289 | 25.4103206 | 112.574084 | 86.2840497 | 98.615607 |
| -4.0795473 | 4.51E-05 | 0.00013289 | 53.6895483 | 84.9717847 | 42.5667978 | 49.3078035 |
| -4.0793421 | 4.52E-05 | 0.000133 | 359.433083 | 308.766899 | 304.870309 | 298.648401 |

| | | | | | | |
|---|---|---|---|---|---|---|
| 4.07926206 | 4.52E-05 | 0.00013303 | 9.42640924 | 4.60038325 | 3.45136199 | 3.92221164 |
| -4.0792031 | 4.52E-05 | 0.00013305 | 27.0496961 | 154.518755 | 59.8236078 | 177.05984 |
| 4.07907116 | 4.52E-05 | 0.00013312 | 0 | 0 | 0 | 0 |
| -4.0789523 | 4.52E-05 | 0.00013317 | 39.7548564 | 79.8301798 | 44.8677058 | 81.8061285 |
| -4.0786394 | 4.53E-05 | 0.00013334 | 49.1812656 | 59.2637606 | 39.1154358 | 64.4363341 |
| -4.0784913 | 4.53E-05 | 0.00013341 | 38.9351686 | 90.654611 | 46.0181598 | 132.234564 |
| -4.0783081 | 4.54E-05 | 0.00013351 | 47.5418901 | 75.7710182 | 158.762651 | 43.144328 |
| -4.0781031 | 4.54E-05 | 0.00013361 | 29.0989155 | 129.081342 | 74.7795097 | 80.1251807 |
| -4.0780773 | 4.54E-05 | 0.00013361 | 40.984388 | 61.1580361 | 46.0181598 | 72.2807574 |
| -4.0777858 | 4.55E-05 | 0.00013377 | 41.8040758 | 46.0038325 | 75.9299637 | 58.8331746 |
| -4.0774972 | 4.55E-05 | 0.00013391 | 37.2957931 | 97.4198805 | 109.29313 | 46.5062237 |
| -4.0774954 | 4.55E-05 | 0.00013391 | 35.2465737 | 74.688575 | 63.2749697 | 58.2728587 |
| -4.0774632 | 4.55E-05 | 0.00013392 | 36.8859492 | 54.9339882 | 62.1245158 | 63.8760181 |
| -4.0774031 | 4.55E-05 | 0.00013394 | 59.0175187 | 86.0542278 | 135.753571 | 32.498325 |
| -4.0773229 | 4.56E-05 | 0.00013397 | 22.1315695 | 132.599282 | 155.311289 | 85.7283401 |
| -4.0767992 | 4.57E-05 | 0.00013426 | 52.8698605 | 71.4412457 | 35.6640739 | 68.9188617 |
| -4.0767004 | 4.57E-05 | 0.00013431 | 28.2792277 | 101.479042 | 70.1776937 | 98.055291 |
| -4.076017 | 4.58E-05 | 0.00013469 | 39.7548564 | 71.4412457 | 46.0181598 | 91.8918156 |
| -4.0755203 | 4.59E-05 | 0.00013497 | 19.2626624 | 99.5847667 | 133.452663 | 215.161324 |
| -4.0754447 | 4.59E-05 | 0.000135 | 113.526755 | 77.3946828 | 24.1595339 | 168.094785 |
| -4.0754137 | 4.59E-05 | 0.000135 | 47.951734 | 133.952336 | 87.4345036 | 38.1014845 |
| -4.0750763 | 4.60E-05 | 0.00013519 | 26.2300083 | 130.434396 | 81.6822337 | 90.7711837 |
| -4.07496 | 4.60E-05 | 0.00013524 | 42.6237635 | 59.5343714 | 54.0713378 | 46.5062237 |
| -4.0748527 | 4.60E-05 | 0.00013529 | 173.773805 | 222.983282 | 222.037621 | 225.807327 |
| 4.07478357 | 4.61E-05 | 0.00013532 | 25.8201644 | 17.8603114 | 16.1063559 | 12.8872668 |
| -4.0745922 | 4.61E-05 | 0.00013542 | 36.4761053 | 50.8748265 | 73.6290557 | 75.0823371 |
| -4.0741178 | 4.62E-05 | 0.00013568 | 30.3284471 | 72.253078 | 90.8858656 | 72.8410733 |
| -4.0740478 | 4.62E-05 | 0.00013571 | 32.7875104 | 92.8194972 | 62.1245158 | 70.5998095 |
| -4.0738878 | 4.62E-05 | 0.00013579 | 34.8367298 | 52.2278804 | 100.089498 | 82.3664444 |
| -4.0736205 | 4.63E-05 | 0.00013594 | 42.6237635 | 47.8981079 | 94.3372276 | 57.1522268 |
| -4.0732684 | 4.64E-05 | 0.00013613 | 11.4756286 | 186.450827 | 205.931265 | 190.507423 |
| -4.0730146 | 4.64E-05 | 0.00013627 | 48.3615778 | 106.079425 | 124.249032 | 98.615607 |
| -4.0725412 | 4.65E-05 | 0.00013653 | 34.017042 | 70.3588026 | 75.9299637 | 57.7125427 |
| -4.0722679 | 4.66E-05 | 0.00013668 | 27.0496961 | 102.561485 | 81.6822337 | 86.2886561 |
| -4.0720686 | 4.66E-05 | 0.00013678 | 300.00572 | 597.779211 | 531.509746 | 589.452378 |
| -4.072021 | 4.66E-05 | 0.0001368 | 89.3459658 | 55.4752097 | 43.7172518 | 46.5062237 |
| -4.0719245 | 4.66E-05 | 0.00013684 | 38.5253247 | 78.477126 | 64.4254237 | 48.7474875 |
| -4.0717838 | 4.67E-05 | 0.00013691 | 27.0496961 | 134.764168 | 69.0272397 | 106.46003 |
| -4.0713802 | 4.67E-05 | 0.00013714 | 67.2143963 | 26.5198564 | 234.692615 | 121.028245 |
| -4.0710374 | 4.68E-05 | 0.00013732 | 65.5750208 | 80.1007906 | 36.8145279 | 50.4284354 |
| -4.0710326 | 4.68E-05 | 0.00013732 | 29.9186032 | 103.914539 | 74.7795097 | 71.7204414 |
| -4.0707922 | 4.69E-05 | 0.00013745 | 30.3284471 | 120.421797 | 58.6731538 | 104.218766 |
| -4.0707222 | 4.69E-05 | 0.00013748 | 6.96734596 | 266.822228 | 277.259413 | 247.099333 |

| | | | | | | |
|---|---|---|---|---|---|---|
| -4.0705537 | 4.69E-05 | 0.00013756 | 38.1154808 | 89.5721679 | 64.4254237 | 50.4284354 |
| -4.0701561 | 4.70E-05 | 0.00013779 | 24.1807889 | 114.197749 | 101.239952 | 90.7711837 |
| -4.0699048 | 4.70E-05 | 0.00013792 | 23.3611012 | 169.672959 | 80.5317797 | 118.226665 |
| -4.0696608 | 4.71E-05 | 0.00013806 | 25.0004767 | 100.937821 | 87.4345036 | 104.218766 |
| -4.0696042 | 4.71E-05 | 0.00013808 | 94.6739363 | 94.9843835 | 51.7704298 | 79.5648647 |
| -4.0687533 | 4.73E-05 | 0.00013857 | 78.2801811 | 47.0862756 | 56.3722458 | 41.4633802 |
| -4.0687334 | 4.73E-05 | 0.00013857 | 38.9351686 | 98.2317128 | 48.3190678 | 67.7982298 |
| -4.0683933 | 4.73E-05 | 0.00013875 | 6.96734596 | 153.706923 | 327.879389 | 419.676646 |
| -4.0683982 | 4.73E-05 | 0.00013875 | 42.2139196 | 51.6866588 | 101.239952 | 53.7903311 |
| -4.068141 | 4.74E-05 | 0.00013888 | 38.1154808 | 79.0183475 | 47.1686138 | 71.1601255 |
| -4.0679128 | 4.74E-05 | 0.00013901 | 29.0989155 | 125.563402 | 63.2749697 | 98.055291 |
| -4.0678767 | 4.74E-05 | 0.00013902 | 24.5906328 | 73.6061319 | 119.647216 | 162.491625 |
| -4.0678141 | 4.75E-05 | 0.00013904 | 18.4429746 | 99.0435452 | 156.461743 | 173.697944 |
| -4.0675164 | 4.75E-05 | 0.0001392 | 32.7875104 | 72.5236888 | 70.1776937 | 62.7553862 |
| -4.0671874 | 4.76E-05 | 0.00013939 | 216.807412 | 70.0881918 | 52.9208838 | 44.8252759 |
| 4.06690084 | 4.76E-05 | 0.00013955 | 127.461447 | 137.740887 | 132.302209 | 113.744138 |
| -4.066854 | 4.77E-05 | 0.00013956 | 77.0506494 | 197.004647 | 179.470823 | 187.705843 |
| -4.0667868 | 4.77E-05 | 0.00013959 | 34.4268859 | 94.1725511 | 100.089498 | 50.4284354 |
| -4.0667386 | 4.77E-05 | 0.00013961 | 29.0989155 | 79.5595691 | 94.3372276 | 71.7204414 |
| -4.0664643 | 4.77E-05 | 0.00013976 | 20.9020379 | 102.832096 | 119.647216 | 133.355196 |
| -4.065928 | 4.78E-05 | 0.00014007 | 38.5253247 | 49.7923834 | 60.9740618 | 60.5141225 |
| -4.0658627 | 4.79E-05 | 0.00014009 | 36.0662614 | 100.125988 | 49.4695218 | 79.5648647 |
| -4.0658024 | 4.79E-05 | 0.00014012 | 90.5754975 | 32.4732935 | 77.0804177 | 58.8331746 |
| -4.0657346 | 4.79E-05 | 0.00014015 | 22.9512573 | 95.7962158 | 115.0454 | 108.140978 |
| -4.0652468 | 4.80E-05 | 0.00014043 | 61.8864259 | 40.5916169 | 64.4254237 | 45.3855918 |
| -4.0651553 | 4.80E-05 | 0.00014047 | 41.3942319 | 86.5954493 | 90.8858656 | 88.5299199 |
| -4.0649454 | 4.80E-05 | 0.00014057 | 28.2792277 | 94.4431619 | 80.5317797 | 76.202969 |
| 4.06495634 | 4.80E-05 | 0.00014057 | 5.73781432 | 4.60038325 | 6.90272397 | 5.04284354 |
| -4.064748 | 4.81E-05 | 0.00014068 | 18.0331307 | 99.3141559 | 182.922185 | 156.888466 |
| -4.0642158 | 4.82E-05 | 0.00014096 | 28.2792277 | 114.73897 | 103.54086 | 64.4363341 |
| -4.0642421 | 4.82E-05 | 0.00014096 | 36.8859492 | 66.8408624 | 50.6199758 | 66.6775979 |
| -4.0642152 | 4.82E-05 | 0.00014096 | 112.297223 | 144.235545 | 155.311289 | 17.9301104 |
| -4.0641625 | 4.82E-05 | 0.00014098 | 33.1973543 | 56.2870421 | 79.3813257 | 78.4442328 |
| -4.0640387 | 4.82E-05 | 0.00014104 | 33.1973543 | 64.4053654 | 69.0272397 | 64.99665 |
| -4.0639687 | 4.82E-05 | 0.00014107 | 40.1647002 | 38.9679522 | 142.656295 | 139.518671 |
| -4.0637544 | 4.83E-05 | 0.00014119 | 57.7879871 | 81.4538445 | 358.941647 | 41.4633802 |
| -4.0636794 | 4.83E-05 | 0.00014122 | 38.9351686 | 51.6866588 | 96.6381356 | 58.8331746 |
| -4.063558 | 4.83E-05 | 0.00014128 | 189.347873 | 329.87454 | 387.702996 | 382.695793 |
| -4.0631994 | 4.84E-05 | 0.00014148 | 45.9025146 | 108.785533 | 35.6640739 | 114.864769 |
| -4.0631314 | 4.84E-05 | 0.00014151 | 29.0989155 | 79.5595691 | 77.0804177 | 79.0045488 |
| -4.0627591 | 4.85E-05 | 0.00014173 | 35.6564176 | 53.851545 | 65.5758777 | 66.6775979 |
| -4.0626987 | 4.85E-05 | 0.00014175 | 66.8045524 | 60.8874253 | 33.3631659 | 63.8760181 |
| -4.0618714 | 4.87E-05 | 0.00014224 | 39.3450125 | 44.3801678 | 75.9299637 | 77.8839169 |

| | | | | | | |
|---|---|---|---|---|---|---|
| -4.0616896 | 4.87E-05 | 0.00014234 | 38.1154808 | 45.7332217 | 88.5849576 | 77.3236009 |
| -4.0614414 | 4.88E-05 | 0.00014248 | 25.4103206 | 88.219114 | 96.6381356 | 94.1330794 |
| -4.0612827 | 4.88E-05 | 0.00014256 | 1904.54451 | 1917.54798 | 1397.8016 | 1927.48686 |
| -4.0611301 | 4.88E-05 | 0.00014264 | 18.0331307 | 110.950419 | 148.408565 | 149.604358 |
| -4.0608797 | 4.89E-05 | 0.00014278 | 392.630437 | 43.8389462 | 88.5849576 | 50.9887513 |
| -4.0607804 | 4.89E-05 | 0.00014283 | 170.495054 | 182.662276 | 174.869007 | 261.667548 |
| -4.0606862 | 4.89E-05 | 0.00014286 | 70.0833035 | 84.4305631 | 50.6199758 | 60.5141225 |
| -4.0606915 | 4.89E-05 | 0.00014286 | 87.7065903 | 42.4858923 | 40.2658898 | 78.4442328 |
| -4.0604378 | 4.90E-05 | 0.00014299 | 39.3450125 | 46.545054 | 67.8767857 | 61.0744384 |
| -4.0604375 | 4.90E-05 | 0.00014299 | 40.984388 | 45.7332217 | 77.0804177 | 57.7125427 |
| -4.0600481 | 4.91E-05 | 0.00014321 | 36.0662614 | 71.1706349 | 57.5226998 | 56.0315949 |
| -4.0585014 | 4.94E-05 | 0.00014415 | 30.3284471 | 61.6992576 | 88.5849576 | 87.9696039 |
| -4.0584431 | 4.94E-05 | 0.00014418 | 40.984388 | 55.4752097 | 72.4786017 | 45.9459078 |
| -4.0583636 | 4.94E-05 | 0.00014421 | 52.8698605 | 58.9931499 | 77.0804177 | 36.4205367 |
| -4.0567943 | 4.98E-05 | 0.00014517 | 21.3118818 | 152.353869 | 95.4876816 | 110.942558 |
| -4.0567534 | 4.98E-05 | 0.00014518 | 75.8211178 | 71.9824673 | 41.4163438 | 41.4633802 |
| -4.0565452 | 4.98E-05 | 0.0001453 | 34.4268859 | 83.0775092 | 51.7704298 | 77.3236009 |
| -4.0565064 | 4.98E-05 | 0.00014531 | 33.6071982 | 75.5004074 | 55.2217918 | 73.9617052 |
| -4.0564552 | 4.98E-05 | 0.00014533 | 61.0667381 | 277.646659 | 194.426725 | 21.852322 |
| -4.0563745 | 4.98E-05 | 0.00014537 | 26.2300083 | 97.9611021 | 96.6381356 | 77.3236009 |
| -4.0555903 | 5.00E-05 | 0.00014584 | 34.017042 | 74.9591858 | 54.0713378 | 73.4013893 |
| -4.0553818 | 5.01E-05 | 0.00014596 | 93.4444046 | 58.4519283 | 35.6640739 | 56.5919108 |
| -4.0553577 | 5.01E-05 | 0.00014596 | 23.770945 | 123.127905 | 89.7354116 | 95.8140272 |
| -4.0551452 | 5.01E-05 | 0.00014608 | 78.2801811 | 109.867976 | 25.3099879 | 91.8918156 |
| -4.0543102 | 5.03E-05 | 0.00014659 | 33.6071982 | 68.7351379 | 59.8236078 | 64.4363341 |
| -4.054115 | 5.03E-05 | 0.0001467 | 29.0989155 | 71.9824673 | 90.8858656 | 74.5220212 |
| -4.0539539 | 5.04E-05 | 0.00014679 | 31.1481349 | 81.4538445 | 87.4345036 | 59.3934906 |
| -4.0538644 | 5.04E-05 | 0.00014683 | 30.738291 | 116.633246 | 64.4254237 | 75.0823371 |
| -4.0536718 | 5.04E-05 | 0.00014694 | 35.2465737 | 60.6168145 | 63.2749697 | 56.5919108 |
| -4.0534583 | 5.05E-05 | 0.00014706 | 65.5750208 | 169.943569 | 146.107657 | 155.767834 |
| -4.053327 | 5.05E-05 | 0.00014713 | 54.0993922 | 60.075593 | 43.7172518 | 42.5840121 |
| -4.0532779 | 5.05E-05 | 0.00014714 | 39.7548564 | 67.382084 | 43.7172518 | 76.763285 |
| -4.0531937 | 5.05E-05 | 0.00014718 | 26.2300083 | 104.996982 | 86.2840497 | 80.1251807 |
| -4.0529798 | 5.06E-05 | 0.00014731 | 40.1647002 | 46.0038325 | 57.5226998 | 60.5141225 |
| -4.0527698 | 5.06E-05 | 0.00014742 | 33.6071982 | 73.8767427 | 92.0363196 | 53.7903311 |
| -4.0525425 | 5.07E-05 | 0.00014755 | 27.8693838 | 112.032863 | 67.8767857 | 89.6505518 |
| -4.051982 | 5.08E-05 | 0.00014788 | 38.5253247 | 46.0038325 | 74.7795097 | 64.4363341 |
| -4.0519951 | 5.08E-05 | 0.00014788 | 31.9678226 | 68.1939163 | 66.7263317 | 67.7982298 |
| 4.05162767 | 5.09E-05 | 0.00014809 | 5.32797044 | 1.62366467 | 4.60181598 | 3.36189569 |
| -4.0512909 | 5.09E-05 | 0.00014829 | 124.592539 | 26.7904671 | 63.2749697 | 124.390141 |
| -4.0511484 | 5.10E-05 | 0.00014837 | 979.117029 | 3357.19733 | 1754.44234 | 3235.8246 |
| -4.0511171 | 5.10E-05 | 0.00014838 | 1097.97175 | 4213.13922 | 2747.28414 | 3227.98018 |
| -4.0510889 | 5.10E-05 | 0.00014838 | 34.017042 | 69.817581 | 77.0804177 | 53.7903311 |

| | | | | | | |
|---|---|---|---|---|---|---|
| -4.0508159 | 5.10E-05 | 0.00014854 | 29.9186032 | 79.8301798 | 146.107657 | 66.1172819 |
| -4.0502815 | 5.12E-05 | 0.00014887 | 21.3118818 | 105.538204 | 119.647216 | 107.020346 |
| -4.0500896 | 5.12E-05 | 0.00014898 | 29.0989155 | 67.9233056 | 89.7354116 | 79.0045488 |
| -4.0499429 | 5.12E-05 | 0.00014906 | 25.4103206 | 90.654611 | 87.4345036 | 93.5727634 |
| -4.0498349 | 5.13E-05 | 0.00014911 | 13.524848 | 169.943569 | 208.232173 | 134.475828 |
| -4.0497874 | 5.13E-05 | 0.00014913 | 48.3615778 | 49.2511618 | 42.5667978 | 50.4284354 |
| -4.0493362 | 5.14E-05 | 0.0001494 | 251.234298 | 240.302372 | 402.658898 | 372.610106 |
| -4.0493104 | 5.14E-05 | 0.0001494 | 56.9682993 | 90.654611 | 146.107657 | 79.0045488 |
| -4.0492508 | 5.14E-05 | 0.00014943 | 36.4761053 | 69.817581 | 51.7704298 | 58.8331746 |
| -4.0486657 | 5.15E-05 | 0.00014979 | 357.383863 | 675.444505 | 645.404691 | 704.877463 |
| -4.0483608 | 5.16E-05 | 0.00014997 | 41.8040758 | 47.3568864 | 49.4695218 | 56.0315949 |
| -4.0482391 | 5.16E-05 | 0.00015004 | 25.4103206 | 94.1725511 | 117.346308 | 76.763285 |
| -4.0479756 | 5.17E-05 | 0.00015019 | 16.8035991 | 113.115306 | 151.859927 | 160.810677 |
| -4.0479347 | 5.17E-05 | 0.0001502 | 4147.62006 | 1578.20206 | 1137.799 | 1469.70873 |
| -4.0476754 | 5.17E-05 | 0.00015036 | 15.1642236 | 85.2423954 | 253.099879 | 237.013646 |
| -4.0473367 | 5.18E-05 | 0.00015056 | 30.738291 | 64.6759762 | 70.1776937 | 98.055291 |
| -4.0471602 | 5.18E-05 | 0.00015066 | 44.263139 | 59.5343714 | 40.2658898 | 59.9538065 |
| -4.046488 | 5.20E-05 | 0.00015108 | 35.2465737 | 59.2637606 | 79.3813257 | 54.350647 |
| -4.0457594 | 5.22E-05 | 0.00015154 | 27.0496961 | 111.762252 | 87.4345036 | 71.1601255 |
| -4.0455398 | 5.22E-05 | 0.00015167 | 39.7548564 | 70.3588026 | 43.7172518 | 63.3157022 |
| -4.0454507 | 5.22E-05 | 0.00015171 | 53.2797044 | 82.2656769 | 32.2127119 | 84.0473923 |
| -4.0453915 | 5.22E-05 | 0.00015174 | 41.3942319 | 73.3355211 | 44.8677058 | 54.350647 |
| -4.0453643 | 5.22E-05 | 0.00015174 | 57.3781432 | 70.9000241 | 33.3631659 | 61.0744384 |
| -4.0452261 | 5.23E-05 | 0.00015181 | 34.017042 | 79.8301798 | 62.1245158 | 56.5919108 |
| -4.0450485 | 5.23E-05 | 0.00015192 | 32.3776665 | 69.817581 | 60.9740618 | 67.7982298 |
| -4.0449691 | 5.23E-05 | 0.00015195 | 139.346919 | 57.0988744 | 39.1154358 | 58.2728587 |
| -4.0448547 | 5.24E-05 | 0.00015201 | 25.0004767 | 96.8786589 | 77.0804177 | 117.666349 |
| -4.0446736 | 5.24E-05 | 0.00015212 | 27.45954 | 89.3015571 | 75.9299637 | 81.2458125 |
| -4.0442397 | 5.25E-05 | 0.00015239 | 19.6725062 | 103.643928 | 146.107657 | 115.985401 |
| -4.043867 | 5.26E-05 | 0.00015262 | 27.8693838 | 83.8893415 | 81.6822337 | 74.5220212 |
| -4.0436192 | 5.26E-05 | 0.00015276 | 413.942319 | 888.144577 | 615.492888 | 763.150322 |
| -4.0435322 | 5.27E-05 | 0.00015281 | 29.0989155 | 76.8534613 | 73.6290557 | 75.0823371 |
| -4.0431849 | 5.27E-05 | 0.00015302 | 50.4107972 | 52.4984912 | 56.3722458 | 38.6618005 |
| -4.0431079 | 5.27E-05 | 0.00015306 | 27.0496961 | 117.715689 | 121.948124 | 61.6347543 |
| -4.0429809 | 5.28E-05 | 0.00015313 | 19.2626624 | 106.079425 | 158.762651 | 113.744138 |
| -4.0428575 | 5.28E-05 | 0.00015318 | 34.017042 | 74.4179643 | 59.8236078 | 57.7125427 |
| -4.0428712 | 5.28E-05 | 0.00015318 | 38.5253247 | 53.3103235 | 72.4786017 | 49.3078035 |
| -4.0424353 | 5.29E-05 | 0.00015344 | 22.9512573 | 135.034779 | 85.1335957 | 101.977503 |
| -4.042412 | 5.29E-05 | 0.00015344 | 61.476582 | 64.6759762 | 77.0804177 | 31.9380091 |
| -4.0422048 | 5.30E-05 | 0.00015356 | 26.2300083 | 87.4072817 | 85.1335957 | 84.0473923 |
| -4.0416073 | 5.31E-05 | 0.00015394 | 49.5911095 | 70.0881918 | 70.1776937 | 35.2999048 |
| -4.0412547 | 5.32E-05 | 0.00015416 | 38.5253247 | 46.0038325 | 101.239952 | 67.7982298 |
| -4.0406127 | 5.33E-05 | 0.00015457 | 20.492194 | 108.785533 | 184.072639 | 94.1330794 |

| | | | | | | |
|---|---|---|---|---|---|---|
| -4.0405644 | 5.33E-05 | 0.00015459 | 23.770945 | 224.877557 | 120.79767 | 70.0394936 |
| -4.0402817 | 5.34E-05 | 0.00015476 | 43.8532952 | 132.328671 | 40.2658898 | 76.763285 |
| -4.0402397 | 5.34E-05 | 0.00015477 | 31.5579788 | 73.3355211 | 66.7263317 | 63.3157022 |
| -4.0399723 | 5.35E-05 | 0.00015494 | 32.7875104 | 75.7710182 | 57.5226998 | 66.1172819 |
| -4.0395501 | 5.36E-05 | 0.0001552 | 29.5087594 | 63.5935331 | 82.8326877 | 81.8061285 |
| -4.0390216 | 5.37E-05 | 0.00015554 | 463.123584 | 405.374947 | 486.64204 | 458.338446 |
| -4.0382634 | 5.38E-05 | 0.00015603 | 35.2465737 | 49.7923834 | 131.151755 | 79.5648647 |
| -4.0381823 | 5.39E-05 | 0.00015607 | 76.6408055 | 37.6148983 | 65.5758777 | 45.9459078 |
| -4.037717 | 5.40E-05 | 0.00015636 | 45.9025146 | 63.5935331 | 71.3281477 | 37.5411686 |
| -4.0373236 | 5.41E-05 | 0.00015659 | 30.3284471 | 88.219114 | 65.5758777 | 68.3585457 |
| -4.0373103 | 5.41E-05 | 0.00015659 | 26.2300083 | 73.8767427 | 102.390406 | 87.9696039 |
| -4.0373479 | 5.41E-05 | 0.00015659 | 54.509236 | 36.2618444 | 87.4345036 | 55.4712789 |
| -4.0371702 | 5.41E-05 | 0.00015667 | 33.1973543 | 72.7942996 | 72.4786017 | 53.7903311 |
| 4.03687566 | 5.42E-05 | 0.00015685 | 242.217733 | 227.042444 | 169.116737 | 207.316901 |
| -4.0367934 | 5.42E-05 | 0.00015689 | 32.3776665 | 63.3229223 | 69.0272397 | 61.6347543 |
| -4.0364471 | 5.43E-05 | 0.00015711 | 45.0828268 | 229.20733 | 26.4604419 | 230.289855 |
| -4.0358041 | 5.44E-05 | 0.00015753 | 32.3776665 | 68.7351379 | 67.8767857 | 58.8331746 |
| -4.035732 | 5.44E-05 | 0.00015756 | 130.740198 | 65.4878086 | 35.6640739 | 54.910963 |
| -4.0356276 | 5.45E-05 | 0.00015762 | 40.5745441 | 53.5809343 | 46.0181598 | 53.2300151 |
| -4.0355067 | 5.45E-05 | 0.00015769 | 26.2300083 | 70.9000241 | 92.0363196 | 107.020346 |
| -4.0350872 | 5.46E-05 | 0.00015795 | 32.7875104 | 50.6042157 | 98.9390436 | 86.2886561 |
| -4.0348997 | 5.46E-05 | 0.00015807 | 39.3450125 | 90.1133895 | 85.1335957 | 40.9030642 |
| -4.0348298 | 5.46E-05 | 0.0001581 | 29.0989155 | 71.1706349 | 72.4786017 | 78.4442328 |
| 4.03454078 | 5.47E-05 | 0.00015828 | 6.55750208 | 3.51794013 | 3.45136199 | 1.1206319 |
| -4.034485 | 5.47E-05 | 0.0001583 | 21.7217256 | 104.996982 | 120.79767 | 93.5727634 |
| -4.0340817 | 5.48E-05 | 0.00015856 | 1052.47908 | 420.25854 | 340.534383 | 485.233612 |
| -4.0338049 | 5.49E-05 | 0.00015873 | 29.5087594 | 99.8553775 | 67.8767857 | 69.4791776 |
| -4.0335939 | 5.49E-05 | 0.00015886 | 26.6398522 | 93.9019404 | 83.9831417 | 74.5220212 |
| -4.0334648 | 5.50E-05 | 0.00015893 | 29.5087594 | 107.70309 | 71.3281477 | 66.6775979 |
| -4.0333333 | 5.50E-05 | 0.00015901 | 38.5253247 | 49.5217726 | 55.2217918 | 52.6696992 |
| -4.0332057 | 5.50E-05 | 0.00015908 | 29.9186032 | 60.3462037 | 80.5317797 | 85.1680242 |
| -4.0330517 | 5.51E-05 | 0.00015917 | 65.1651769 | 62.2404792 | 63.2749697 | 50.9887513 |
| -4.0329862 | 5.51E-05 | 0.0001592 | 40.1647002 | 42.7565031 | 119.647216 | 75.0823371 |
| -4.0329475 | 5.51E-05 | 0.00015921 | 85.6573709 | 69.5469702 | 28.7613499 | 68.3585457 |
| -4.0326504 | 5.52E-05 | 0.0001594 | 45.9025146 | 200.522587 | 98.9390436 | 141.199619 |
| -4.0325249 | 5.52E-05 | 0.00015947 | 84.8376831 | 44.6507786 | 44.8677058 | 49.3078035 |
| -4.0320048 | 5.53E-05 | 0.00015981 | 50.8206411 | 66.5702517 | 51.7704298 | 37.5411686 |
| -4.0317651 | 5.54E-05 | 0.00015996 | 27.0496961 | 60.8874253 | 120.79767 | 109.26161 |
| -4.0315879 | 5.54E-05 | 0.00016006 | 29.0989155 | 135.034779 | 69.0272397 | 73.4013893 |
| -4.031569 | 5.54E-05 | 0.00016006 | 55.3289238 | 73.8767427 | 105.841768 | 31.9380091 |
| -4.030877 | 5.56E-05 | 0.00016052 | 34.8367298 | 54.9339882 | 56.3722458 | 68.3585457 |
| -4.0307072 | 5.56E-05 | 0.00016062 | 35.6564176 | 54.6633774 | 67.8767857 | 53.2300151 |
| -4.0297366 | 5.58E-05 | 0.00016127 | 66.8045524 | 359.912336 | 242.745793 | 256.064389 |

| | | | | | | |
|---|---|---|---|---|---|---|
| -4.0293327 | 5.59E-05 | 0.00016152 | 26.2300083 | 90.9252218 | 86.2840497 | 75.6426531 |
| -4.0293415 | 5.59E-05 | 0.00016152 | 73.3620545 | 72.253078 | 66.7263317 | 29.6967453 |
| -4.0290239 | 5.60E-05 | 0.00016172 | 29.0989155 | 86.3248385 | 67.8767857 | 71.1601255 |
| 4.02899166 | 5.60E-05 | 0.00016173 | 2.86890716 | 11.9068743 | 1.150454 | 6.16347543 |
| -4.0285766 | 5.61E-05 | 0.000162 | 31.5579788 | 72.7942996 | 56.3722458 | 98.615607 |
| -4.0285398 | 5.61E-05 | 0.00016201 | 28.6890716 | 65.4878086 | 75.9299637 | 98.615607 |
| -4.028053 | 5.62E-05 | 0.00016233 | 42.2139196 | 116.092024 | 37.9649818 | 84.6077082 |
| -4.0280217 | 5.62E-05 | 0.00016234 | 28.2792277 | 83.6187308 | 63.2749697 | 103.098135 |
| -4.0274726 | 5.64E-05 | 0.0001627 | 22.1315695 | 107.432479 | 118.496762 | 86.2886561 |
| -4.027369 | 5.64E-05 | 0.00016276 | 27.8693838 | 84.1599523 | 66.7263317 | 87.9696039 |
| -4.026802 | 5.65E-05 | 0.00016314 | 27.8693838 | 107.432479 | 65.5758777 | 81.8061285 |
| -4.0266115 | 5.66E-05 | 0.00016326 | 34.4268859 | 90.1133895 | 62.1245158 | 53.2300151 |
| -4.0264909 | 5.66E-05 | 0.00016333 | 32.3776665 | 66.8408624 | 58.6731538 | 64.99665 |
| -4.0264111 | 5.66E-05 | 0.00016337 | 20.492194 | 96.0668266 | 126.549939 | 111.502874 |
| -4.0263277 | 5.67E-05 | 0.00016341 | 33.6071982 | 64.1347547 | 70.1776937 | 53.2300151 |
| -4.0262776 | 5.67E-05 | 0.00016343 | 39.3450125 | 46.2744432 | 51.7704298 | 63.8760181 |
| -4.0257121 | 5.68E-05 | 0.00016381 | 50.4107972 | 42.4858923 | 42.5667978 | 59.3934906 |
| -4.0256609 | 5.68E-05 | 0.00016383 | 17.213443 | 135.034779 | 111.594038 | 146.242463 |
| -4.0256319 | 5.68E-05 | 0.00016384 | 487.304373 | 850.529679 | 773.105085 | 869.610352 |
| -4.0255314 | 5.68E-05 | 0.00016389 | 69.2636157 | 47.8981079 | 40.2658898 | 47.6268556 |
| -4.0254011 | 5.69E-05 | 0.00016397 | 32.7875104 | 106.620647 | 52.9208838 | 73.9617052 |
| -4.0253183 | 5.69E-05 | 0.00016401 | 26.2300083 | 93.9019404 | 81.6822337 | 76.202969 |
| -4.0251407 | 5.69E-05 | 0.00016412 | 53.6895483 | 55.2045989 | 48.3190678 | 38.1014845 |
| -4.0250798 | 5.70E-05 | 0.00016415 | 46.3123584 | 83.8893415 | 89.7354116 | 73.4013893 |
| -4.0250514 | 5.70E-05 | 0.00016415 | 32.7875104 | 54.3927666 | 66.7263317 | 73.4013893 |
| -4.0246298 | 5.71E-05 | 0.00016443 | 37.2957931 | 51.416048 | 50.6199758 | 67.7982298 |
| 4.02457452 | 5.71E-05 | 0.00016446 | 0 | 0 | 0 | 0 |
| -4.0243546 | 5.71E-05 | 0.00016459 | 26.2300083 | 64.4053654 | 113.894946 | 101.977503 |
| -4.0241973 | 5.72E-05 | 0.00016469 | 31.5579788 | 86.8660601 | 59.8236078 | 64.4363341 |
| -4.0237764 | 5.73E-05 | 0.00016496 | 54.0993922 | 36.5324552 | 54.0713378 | 54.350647 |
| -4.0237712 | 5.73E-05 | 0.00016496 | 87.7065903 | 44.9213893 | 69.0272397 | 36.9808526 |
| -4.02349 | 5.73E-05 | 0.00016514 | 40.5745441 | 145.85921 | 130.001301 | 128.872668 |
| -4.0234541 | 5.74E-05 | 0.00016515 | 51.230485 | 82.5362876 | 36.8145279 | 53.2300151 |
| -4.0228599 | 5.75E-05 | 0.00016555 | 37.2957931 | 44.3801678 | 66.7263317 | 75.0823371 |
| -4.0224025 | 5.76E-05 | 0.00016586 | 13.9346919 | 160.742803 | 123.098578 | 191.067738 |
| -4.0221791 | 5.77E-05 | 0.000166 | 61.8864259 | 49.2511618 | 35.6640739 | 57.1522268 |
| -4.0219975 | 5.77E-05 | 0.00016612 | 35.6564176 | 58.7225391 | 74.7795097 | 49.3078035 |
| -4.0218111 | 5.78E-05 | 0.00016623 | 51.6403289 | 80.912623 | 35.6640739 | 197.79153 |
| -4.0209885 | 5.80E-05 | 0.0001668 | 29.9186032 | 81.4538445 | 59.8236078 | 76.202969 |
| -4.0208568 | 5.80E-05 | 0.00016688 | 29.5087594 | 77.9359044 | 60.9740618 | 81.2458125 |
| 4.02078827 | 5.80E-05 | 0.00016691 | 76.2309617 | 39.238563 | 36.8145279 | 38.1014845 |
| -4.0206502 | 5.80E-05 | 0.000167 | 31.5579788 | 99.5847667 | 57.5226998 | 68.3585457 |
| -4.0203911 | 5.81E-05 | 0.00016717 | 40.5745441 | 56.8282636 | 47.1686138 | 47.0665397 |

| | | | | | | |
|---|---|---|---|---|---|---|
| -4.0197427 | 5.83E-05 | 0.00016761 | 27.8693838 | 70.9000241 | 78.2308717 | 77.8839169 |
| -4.0196774 | 5.83E-05 | 0.00016764 | 49.1812656 | 88.219114 | 39.1154358 | 51.5490673 |
| -4.0196044 | 5.83E-05 | 0.00016768 | 50.4107972 | 45.1920001 | 64.4254237 | 39.7824324 |
| -4.0195311 | 5.83E-05 | 0.00016772 | 22.9512573 | 97.4198805 | 120.79767 | 81.8061285 |
| -4.0188865 | 5.85E-05 | 0.00016816 | 810.261351 | 605.356313 | 549.91701 | 564.23816 |
| -4.0181917 | 5.86E-05 | 0.00016864 | 46.3123584 | 85.2423954 | 85.1335957 | 34.1792729 |
| -4.0180823 | 5.87E-05 | 0.00016871 | 52.4600166 | 35.9912336 | 70.1776937 | 52.1093832 |
| -4.0179618 | 5.87E-05 | 0.00016878 | 26.6398522 | 96.8786589 | 65.5758777 | 94.1330794 |
| -4.017928 | 5.87E-05 | 0.00016879 | 31.5579788 | 122.586683 | 142.656295 | 45.9459078 |
| -4.0175671 | 5.88E-05 | 0.00016903 | 62.2962697 | 64.6759762 | 57.5226998 | 32.498325 |
| -4.0175313 | 5.88E-05 | 0.00016904 | 31.1481349 | 66.2996409 | 58.6731538 | 78.4442328 |
| -4.0173649 | 5.89E-05 | 0.00016914 | 25.8201644 | 96.0668266 | 87.4345036 | 72.2807574 |
| -4.0172381 | 5.89E-05 | 0.00016922 | 35.6564176 | 64.6759762 | 66.7263317 | 47.0665397 |
| -4.0171149 | 5.89E-05 | 0.00016929 | 70.4931473 | 26.7904671 | 80.5317797 | 96.9346591 |
| -4.0169698 | 5.90E-05 | 0.00016938 | 67.6242402 | 93.090108 | 63.2749697 | 29.6967453 |
| -4.016831 | 5.90E-05 | 0.00016947 | 62.2962697 | 73.0649104 | 28.7613499 | 110.382242 |
| -4.0167958 | 5.90E-05 | 0.00016948 | 1055.75783 | 1658.30285 | 1369.04025 | 1526.30064 |
| -4.0166951 | 5.90E-05 | 0.00016954 | 37.2957931 | 100.66721 | 57.5226998 | 50.4284354 |
| -4.0164056 | 5.91E-05 | 0.00016973 | 31.9678226 | 60.8874253 | 63.2749697 | 63.3157022 |
| -4.0157361 | 5.93E-05 | 0.0001702 | 836.491359 | 1501.3486 | 1382.8457 | 1562.16086 |
| -4.0156823 | 5.93E-05 | 0.00017022 | 28.6890716 | 93.3607188 | 77.0804177 | 62.1950703 |
| -4.0151074 | 5.94E-05 | 0.00017062 | 122.54332 | 43.8389462 | 69.0272397 | 39.7824324 |
| -4.0150385 | 5.94E-05 | 0.00017065 | 47.5418901 | 37.3442875 | 57.5226998 | 54.910963 |
| -4.0146973 | 5.95E-05 | 0.00017089 | 25.8201644 | 92.2782757 | 101.239952 | 68.3585457 |
| -4.0144623 | 5.96E-05 | 0.00017104 | 28.6890716 | 77.9359044 | 65.5758777 | 75.0823371 |
| -4.0144219 | 5.96E-05 | 0.00017106 | 40.5745441 | 150.459593 | 32.2127119 | 198.351846 |
| -4.0143467 | 5.96E-05 | 0.0001711 | 32.7875104 | 71.7118565 | 52.9208838 | 114.864769 |
| -4.0139548 | 5.97E-05 | 0.00017136 | 197.134906 | 285.494372 | 317.525303 | 266.150076 |
| -4.0138153 | 5.97E-05 | 0.00017145 | 34.017042 | 80.912623 | 49.4695218 | 66.1172819 |
| -4.013722 | 5.98E-05 | 0.0001715 | 152.871767 | 469.780313 | 638.501967 | 635.958602 |
| -4.013192 | 5.99E-05 | 0.00017187 | 29.5087594 | 200.793198 | 57.5226998 | 93.5727634 |
| -4.0130542 | 5.99E-05 | 0.00017196 | 17.213443 | 145.588599 | 98.9390436 | 150.72499 |
| -4.0130066 | 6.00E-05 | 0.00017198 | 59.0175187 | 62.51109 | 32.2127119 | 58.2728587 |
| -4.0126073 | 6.01E-05 | 0.00017225 | 11.8854725 | 132.328671 | 185.223093 | 206.756585 |
| -4.0125298 | 6.01E-05 | 0.00017229 | 19.6725062 | 153.706923 | 88.5849576 | 117.106033 |
| -4.0125174 | 6.01E-05 | 0.00017229 | 86.4770587 | 34.0969582 | 86.2840497 | 45.9459078 |
| -4.0123682 | 6.01E-05 | 0.00017238 | 29.0989155 | 76.3122397 | 72.4786017 | 64.4363341 |
| -4.0122363 | 6.01E-05 | 0.00017246 | 92.6247169 | 191.05121 | 143.806749 | 150.72499 |
| -4.0113759 | 6.04E-05 | 0.00017308 | 27.8693838 | 64.6759762 | 126.549939 | 77.8839169 |
| -4.011052 | 6.04E-05 | 0.0001733 | 94.2640924 | 94.1725511 | 28.7613499 | 54.910963 |
| -4.0109065 | 6.05E-05 | 0.00017339 | 28.6890716 | 70.0881918 | 67.8767857 | 77.3236009 |
| -4.0106595 | 6.05E-05 | 0.00017355 | 60.2470503 | 60.6168145 | 42.5667978 | 38.6618005 |
| -4.0106476 | 6.06E-05 | 0.00017355 | 102.051126 | 307.413845 | 297.967585 | 299.208717 |

| | | | | | | |
|---|---|---|---|---|---|---|
| -4.0106263 | 6.06E-05 | 0.00017355 | 32.7875104 | 54.1221558 | 72.4786017 | 61.0744384 |
| -4.010437 | 6.06E-05 | 0.00017368 | 31.5579788 | 56.2870421 | 169.116737 | 80.1251807 |
| -4.0102738 | 6.06E-05 | 0.00017378 | 90.9853413 | 38.9679522 | 46.0181598 | 56.0315949 |
| -4.0100803 | 6.07E-05 | 0.00017391 | 14.7543797 | 208.3703 | 87.4345036 | 217.402588 |
| -4.0098918 | 6.07E-05 | 0.00017403 | 14.3445358 | 115.821413 | 136.904025 | 238.134278 |
| -4.0097509 | 6.08E-05 | 0.00017412 | 41.3942319 | 57.3694852 | 40.2658898 | 57.1522268 |
| -4.009626 | 6.08E-05 | 0.0001742 | 203.282564 | 197.004647 | 216.285351 | 254.943757 |
| -4.0095369 | 6.08E-05 | 0.00017425 | 72.9522106 | 48.1687187 | 40.2658898 | 45.9459078 |
| -4.0081324 | 6.12E-05 | 0.00017527 | 32.3776665 | 61.6992576 | 63.2749697 | 57.1522268 |
| -4.008055 | 6.12E-05 | 0.00017531 | 17.213443 | 150.188982 | 102.390406 | 133.355196 |
| -4.0079968 | 6.12E-05 | 0.00017534 | 18.8528185 | 97.1492697 | 178.320369 | 107.020346 |
| -4.0077223 | 6.13E-05 | 0.00017553 | 37.2957931 | 60.8874253 | 43.7172518 | 67.2379138 |
| -4.0075904 | 6.13E-05 | 0.00017561 | 29.9186032 | 64.6759762 | 119.647216 | 65.556966 |
| -4.0073886 | 6.14E-05 | 0.00017574 | 30.738291 | 63.3229223 | 63.2749697 | 67.2379138 |
| 4.00693306 | 6.15E-05 | 0.00017607 | 6.1476582 | 5.1416048 | 8.05317797 | 5.60315949 |
| -4.00688 | 6.15E-05 | 0.00017609 | 19.6725062 | 126.645845 | 101.239952 | 104.779082 |
| -4.0066684 | 6.16E-05 | 0.00017623 | 38.1154808 | 59.8049822 | 60.9740618 | 43.144328 |
| -4.0066593 | 6.16E-05 | 0.00017623 | 47.5418901 | 63.5935331 | 36.8145279 | 51.5490673 |
| -4.0062165 | 6.17E-05 | 0.00017654 | 29.0989155 | 83.8893415 | 56.3722458 | 95.2537113 |
| -4.006038 | 6.17E-05 | 0.00017666 | 876.246215 | 2988.89606 | 3241.97936 | 3674.55199 |
| -4.0059583 | 6.18E-05 | 0.0001767 | 42.6237635 | 59.5343714 | 37.9649818 | 72.2807574 |
| -4.0054385 | 6.19E-05 | 0.00017708 | 31.5579788 | 96.0668266 | 50.6199758 | 79.5648647 |
| -4.0053335 | 6.19E-05 | 0.00017714 | 33.1973543 | 120.421797 | 51.7704298 | 71.1601255 |
| -4.0047549 | 6.21E-05 | 0.00017756 | 127.051603 | 319.320719 | 223.188075 | 266.150076 |
| -4.0043326 | 6.22E-05 | 0.00017784 | 53.6895483 | 38.6973414 | 46.0181598 | 50.9887513 |
| -4.0043368 | 6.22E-05 | 0.00017784 | 48.3615778 | 138.552719 | 28.7613499 | 127.752036 |
| -4.0042303 | 6.22E-05 | 0.0001779 | 33.6071982 | 75.5004074 | 50.6199758 | 61.6347543 |
| -4.0029346 | 6.26E-05 | 0.00017887 | 86.8869025 | 17.5897006 | 131.151755 | 238.134278 |
| -4.0027498 | 6.26E-05 | 0.00017899 | 33.6071982 | 74.1473535 | 94.3372276 | 47.0665397 |
| -4.0026905 | 6.26E-05 | 0.00017902 | 31.5579788 | 69.2763595 | 58.6731538 | 60.5141225 |
| -4.0021835 | 6.28E-05 | 0.00017939 | 19.6725062 | 77.6652936 | 204.780811 | 124.950457 |
| -4.0020047 | 6.28E-05 | 0.00017951 | 27.0496961 | 96.8786589 | 58.6731538 | 108.701294 |
| 4.00184854 | 6.28E-05 | 0.00017961 | 0 | 1.89427545 | 0 | 2.80157974 |
| -4.001661 | 6.29E-05 | 0.00017974 | 27.45954 | 67.382084 | 88.5849576 | 72.8410733 |
| -4.0014165 | 6.30E-05 | 0.00017991 | 35.2465737 | 70.3588026 | 51.7704298 | 52.1093832 |
| -4.0012548 | 6.30E-05 | 0.00018001 | 44.6729829 | 38.4267306 | 72.4786017 | 52.6696992 |
| -4.0011687 | 6.30E-05 | 0.00018006 | 47.1320462 | 63.5935331 | 35.6640739 | 54.910963 |
| -4.000477 | 6.32E-05 | 0.00018057 | 50.8206411 | 51.416048 | 56.3722458 | 35.8602207 |
| -3.9997552 | 6.34E-05 | 0.00018111 | 25.0004767 | 85.783617 | 79.3813257 | 82.9267604 |
| -3.9996742 | 6.34E-05 | 0.00018115 | 26.2300083 | 107.432479 | 66.7263317 | 81.8061285 |
| -3.9993945 | 6.35E-05 | 0.00018135 | 21.7217256 | 93.9019404 | 92.0363196 | 107.020346 |
| -3.9991119 | 6.36E-05 | 0.00018155 | 44.6729829 | 78.477126 | 35.6640739 | 62.7553862 |
| -3.9989778 | 6.36E-05 | 0.00018164 | 23.3611012 | 109.867976 | 95.4876816 | 75.6426531 |

| | | | | | | |
|---|---|---|---|---|---|---|
| -3.9987348 | 6.37E-05 | 0.00018181 | 39.3450125 | 50.6042157 | 43.7172518 | 55.4712789 |
| -3.998681 | 6.37E-05 | 0.00018184 | 266.398522 | 57.3694852 | 36.8145279 | 70.0394936 |
| -3.9980683 | 6.39E-05 | 0.00018229 | 26.6398522 | 80.912623 | 79.3813257 | 70.0394936 |
| -3.9980088 | 6.39E-05 | 0.00018232 | 29.9186032 | 74.688575 | 66.7263317 | 59.9538065 |
| -3.9979781 | 6.39E-05 | 0.00018233 | 43.4434513 | 62.51109 | 37.9649818 | 53.7903311 |
| -3.9977837 | 6.39E-05 | 0.00018246 | 51.230485 | 28.4141318 | 154.160835 | 109.821926 |
| -3.9976416 | 6.40E-05 | 0.00018256 | 22.9512573 | 80.1007906 | 110.443584 | 91.3314996 |
| -3.9970931 | 6.41E-05 | 0.00018296 | 52.4600166 | 44.9213893 | 41.4163438 | 44.8252759 |
| -3.9970373 | 6.41E-05 | 0.00018299 | 52.4600166 | 89.5721679 | 58.6731538 | 33.6189569 |
| -3.9969324 | 6.42E-05 | 0.00018305 | 36.0662614 | 47.6274971 | 54.0713378 | 72.2807574 |
| -3.9966346 | 6.42E-05 | 0.00018327 | 35.2465737 | 44.3801678 | 69.0272397 | 73.9617052 |
| -3.9963657 | 6.43E-05 | 0.00018346 | 22.9512573 | 87.1366709 | 139.204933 | 79.5648647 |
| -3.9960841 | 6.44E-05 | 0.00018366 | 26.2300083 | 45.7332217 | 158.762651 | 291.924609 |
| -3.9959979 | 6.44E-05 | 0.00018371 | 27.45954 | 71.4412457 | 66.7263317 | 110.942558 |
| -3.9956688 | 6.45E-05 | 0.00018395 | 48.7714217 | 40.5916169 | 42.5667978 | 56.0315949 |
| -3.9953446 | 6.46E-05 | 0.00018419 | 55.7387677 | 53.5809343 | 37.9649818 | 43.704644 |
| -3.9951617 | 6.47E-05 | 0.00018431 | 388.941842 | 854.859451 | 715.582385 | 767.072534 |
| -3.9948314 | 6.47E-05 | 0.00018454 | 94.2640924 | 51.6866588 | 35.6640739 | 53.2300151 |
| -3.9948249 | 6.47E-05 | 0.00018454 | 106.149565 | 76.8534613 | 148.408565 | 21.292006 |
| -3.994779 | 6.48E-05 | 0.00018456 | 43.4434513 | 45.7332217 | 41.4163438 | 54.350647 |
| -3.9946353 | 6.48E-05 | 0.00018465 | 27.8693838 | 67.9233056 | 70.1776937 | 77.3236009 |
| -3.9946268 | 6.48E-05 | 0.00018465 | 36.0662614 | 103.643928 | 41.4163438 | 81.8061285 |
| -3.9940622 | 6.50E-05 | 0.00018507 | 37.705637 | 54.6633774 | 47.1686138 | 49.3078035 |
| -3.9940186 | 6.50E-05 | 0.00018509 | 32.7875104 | 96.3374374 | 46.0181598 | 82.9267604 |
| -3.9936961 | 6.51E-05 | 0.00018533 | 31.9678226 | 59.8049822 | 73.6290557 | 54.350647 |
| -3.993613 | 6.51E-05 | 0.00018538 | 28.6890716 | 70.6294134 | 72.4786017 | 63.8760181 |
| -3.9935869 | 6.51E-05 | 0.00018538 | 47.5418901 | 52.2278804 | 96.6381356 | 38.6618005 |
| -3.993336 | 6.52E-05 | 0.00018556 | 51.6403289 | 73.3355211 | 71.3281477 | 81.8061285 |
| -3.9932686 | 6.52E-05 | 0.0001856 | 31.5579788 | 59.2637606 | 67.8767857 | 57.1522268 |
| -3.992691 | 6.53E-05 | 0.00018603 | 56.5584554 | 100.125988 | 29.9118039 | 67.2379138 |
| -3.9926553 | 6.53E-05 | 0.00018604 | 33.1973543 | 50.6042157 | 59.8236078 | 71.1601255 |
| -3.9926343 | 6.53E-05 | 0.00018604 | 37.705637 | 57.3694852 | 46.0181598 | 49.8681194 |
| -3.9925713 | 6.54E-05 | 0.00018608 | 73.7718984 | 66.8408624 | 26.4604419 | 85.7283401 |
| -3.992538 | 6.54E-05 | 0.00018609 | 58.197831 | 27.8729103 | 110.443584 | 82.9267604 |
| -3.9920716 | 6.55E-05 | 0.00018644 | 36.4761053 | 75.7710182 | 41.4163438 | 72.8410733 |
| -3.991952 | 6.55E-05 | 0.00018651 | 21.3118818 | 86.0542278 | 103.54086 | 107.020346 |
| -3.9916844 | 6.56E-05 | 0.00018671 | 25.4103206 | 100.66721 | 71.3281477 | 79.0045488 |
| -3.990926 | 6.58E-05 | 0.00018729 | 102.051126 | 391.303187 | 352.038923 | 364.205367 |
| -3.990871 | 6.58E-05 | 0.00018732 | 45.4926707 | 48.980551 | 37.9649818 | 54.350647 |
| -3.9905932 | 6.59E-05 | 0.00018752 | 27.8693838 | 56.0164313 | 88.5849576 | 110.382242 |
| -3.990474 | 6.59E-05 | 0.0001876 | 85.6573709 | 55.2045989 | 28.7613499 | 76.763285 |
| -3.9895929 | 6.62E-05 | 0.00018828 | 179.921463 | 621.322349 | 521.15566 | 537.342995 |
| -3.9894811 | 6.62E-05 | 0.00018835 | 26.6398522 | 69.817581 | 80.5317797 | 75.0823371 |

| | | | | | | |
|---|---|---|---|---|---|---|
| -3.9894518 | 6.62E-05 | 0.00018836 | 32.3776665 | 126.375234 | 55.2217918 | 65.556966 |
| 3.98927286 | 6.63E-05 | 0.00018848 | 1.63937552 | 0.27061078 | 0 | 0.56031595 |
| -3.9892001 | 6.63E-05 | 0.00018853 | 34.4268859 | 61.4286469 | 54.0713378 | 50.4284354 |
| -3.9888201 | 6.64E-05 | 0.00018879 | 31.9678226 | 73.6061319 | 64.4254237 | 51.5490673 |
| -3.9888337 | 6.64E-05 | 0.00018879 | 11.8854725 | 140.988216 | 171.417645 | 173.697944 |
| -3.9885562 | 6.65E-05 | 0.00018899 | 25.0004767 | 103.643928 | 102.390406 | 63.3157022 |
| -3.9882481 | 6.66E-05 | 0.00018922 | 56.9682993 | 122.045461 | 64.4254237 | 156.888466 |
| -3.988203 | 6.66E-05 | 0.00018924 | 33.6071982 | 159.119138 | 41.4163438 | 107.580662 |
| -3.9875175 | 6.68E-05 | 0.00018976 | 29.9186032 | 84.4305631 | 58.6731538 | 63.8760181 |
| -3.9875097 | 6.68E-05 | 0.00018976 | 35.6564176 | 46.545054 | 57.5226998 | 56.0315949 |
| -3.9872787 | 6.68E-05 | 0.00018992 | 160.658801 | 210.264575 | 167.966283 | 184.343947 |
| -3.9872426 | 6.68E-05 | 0.00018994 | 26.6398522 | 69.817581 | 78.2308717 | 75.6426531 |
| -3.9867094 | 6.70E-05 | 0.00019035 | 27.0496961 | 79.5595691 | 69.0272397 | 71.7204414 |
| 3.98665667 | 6.70E-05 | 0.00019037 | 2.86890716 | 2.70610779 | 1.150454 | 6.72379138 |
| -3.9864963 | 6.71E-05 | 0.00019048 | 129.100822 | 40.8622277 | 43.7172518 | 61.0744384 |
| -3.9863611 | 6.71E-05 | 0.00019058 | 26.6398522 | 77.1240721 | 67.8767857 | 80.1251807 |
| -3.9862257 | 6.71E-05 | 0.00019067 | 24.1807889 | 90.1133895 | 90.8858656 | 73.9617052 |
| -3.9858194 | 6.72E-05 | 0.00019098 | 89.7558097 | 54.9339882 | 39.1154358 | 43.144328 |
| 3.98577915 | 6.73E-05 | 0.00019099 | 65.5750208 | 57.0988744 | 63.2749697 | 49.3078035 |
| -3.9856322 | 6.73E-05 | 0.00019109 | 29.0989155 | 114.197749 | 50.6199758 | 94.6933953 |
| -3.9849757 | 6.75E-05 | 0.00019161 | 36.4761053 | 52.2278804 | 63.2749697 | 45.3855918 |
| -3.9847802 | 6.75E-05 | 0.00019175 | 26.6398522 | 70.6294134 | 70.1776937 | 85.7283401 |
| -3.9844493 | 6.76E-05 | 0.000192 | 140.986295 | 117.445078 | 176.019461 | 14.0078987 |
| -3.9842282 | 6.77E-05 | 0.00019216 | 36.8859492 | 64.1347547 | 41.4163438 | 66.1172819 |
| -3.9841641 | 6.77E-05 | 0.0001922 | 45.4926707 | 43.2977247 | 52.9208838 | 40.9030642 |
| -3.9833525 | 6.79E-05 | 0.00019284 | 52.4600166 | 47.8981079 | 92.0363196 | 36.9808526 |
| -3.9831241 | 6.80E-05 | 0.000193 | 37.2957931 | 47.0862756 | 48.3190678 | 54.350647 |
| -3.9829288 | 6.81E-05 | 0.00019313 | 66.8045524 | 43.0271139 | 47.1686138 | 40.3427483 |
| -3.9829414 | 6.81E-05 | 0.00019313 | 74.1817423 | 155.330587 | 140.355387 | 132.234564 |
| -3.9828458 | 6.81E-05 | 0.00019318 | 45.9025146 | 42.4858923 | 44.8677058 | 44.8252759 |
| -3.9826727 | 6.81E-05 | 0.0001933 | 43.0336074 | 53.0397127 | 44.8677058 | 41.4633802 |
| -3.9826311 | 6.82E-05 | 0.00019332 | 44.6729829 | 166.966851 | 26.4604419 | 183.223315 |
| -3.9821907 | 6.83E-05 | 0.00019366 | 42.6237635 | 52.2278804 | 37.9649818 | 56.0315949 |
| -3.9818685 | 6.84E-05 | 0.00019391 | 26.2300083 | 54.6633774 | 376.198456 | 113.183822 |
| -3.981257 | 6.86E-05 | 0.00019438 | 11.4756286 | 138.282108 | 262.303511 | 137.277407 |
| -3.9812518 | 6.86E-05 | 0.00019438 | 778.703372 | 1007.21332 | 796.114165 | 894.264254 |
| -3.9806087 | 6.87E-05 | 0.00019489 | 29.9186032 | 65.7584193 | 60.9740618 | 62.7553862 |
| -3.9800342 | 6.89E-05 | 0.00019534 | 27.45954 | 71.9824673 | 62.1245158 | 101.417187 |
| 3.97999474 | 6.89E-05 | 0.00019536 | 0.81968776 | 0.54122156 | 0 | 0.56031595 |
| -3.9799191 | 6.89E-05 | 0.0001954 | 25.0004767 | 139.09394 | 81.6822337 | 68.3585457 |
| 3.97956644 | 6.90E-05 | 0.00019567 | 25.4103206 | 18.1309222 | 12.6549939 | 19.6110582 |
| -3.9794268 | 6.91E-05 | 0.00019577 | 44.263139 | 37.0736767 | 95.4876816 | 58.8331746 |
| -3.9794152 | 6.91E-05 | 0.00019577 | 43.8532952 | 105.538204 | 100.089498 | 99.1759229 |

| | | | | | | |
|---|---|---|---|---|---|---|
| -3.9792334 | 6.91E-05 | 0.0001959 | 31.5579788 | 102.020264 | 74.7795097 | 48.7474875 |
| -3.9790485 | 6.92E-05 | 0.00019603 | 74.1817423 | 69.817581 | 25.3099879 | 86.2886561 |
| -3.9790251 | 6.92E-05 | 0.00019603 | 36.0662614 | 76.0416289 | 40.2658898 | 85.7283401 |
| -3.9788581 | 6.92E-05 | 0.00019616 | 86.8869025 | 48.7099402 | 31.0622579 | 71.7204414 |
| -3.9787013 | 6.93E-05 | 0.00019627 | 33.1973543 | 99.8553775 | 103.54086 | 42.0236961 |
| -3.9781399 | 6.95E-05 | 0.00019671 | 35.6564176 | 54.6633774 | 46.0181598 | 77.3236009 |
| -3.9779263 | 6.95E-05 | 0.00019687 | 31.1481349 | 66.8408624 | 65.5758777 | 52.6696992 |
| -3.9777443 | 6.96E-05 | 0.00019701 | 23.770945 | 69.0057487 | 105.841768 | 92.4521315 |
| -3.9776286 | 6.96E-05 | 0.00019709 | 65.1651769 | 150.459593 | 67.8767857 | 28.5761134 |
| -3.977327 | 6.97E-05 | 0.00019732 | 23.770945 | 87.4072817 | 86.2840497 | 77.8839169 |
| -3.9772468 | 6.97E-05 | 0.00019737 | 25.8201644 | 114.197749 | 142.656295 | 165.293205 |
| -3.9771903 | 6.97E-05 | 0.0001974 | 39.3450125 | 63.3229223 | 109.29313 | 40.9030642 |
| -3.9766409 | 6.99E-05 | 0.00019784 | 21.3118818 | 95.525605 | 79.3813257 | 131.113932 |
| -3.9765951 | 6.99E-05 | 0.00019786 | 215.168037 | 58.4519283 | 51.7704298 | 42.0236961 |
| -3.9760386 | 7.01E-05 | 0.0001983 | 34.8367298 | 105.808815 | 44.8677058 | 67.7982298 |
| -3.9758661 | 7.01E-05 | 0.00019843 | 30.738291 | 70.0881918 | 51.7704298 | 70.5998095 |
| -3.9755857 | 7.02E-05 | 0.00019865 | 315.579788 | 901.133895 | 677.617403 | 700.955252 |
| -3.9755143 | 7.02E-05 | 0.00019869 | 21.7217256 | 89.5721679 | 96.6381356 | 90.2108677 |
| -3.9753585 | 7.03E-05 | 0.0001988 | 31.5579788 | 56.5576528 | 67.8767857 | 54.910963 |
| -3.9748254 | 7.04E-05 | 0.00019923 | 45.9025146 | 30.3084073 | 126.549939 | 100.856871 |
| -3.9747605 | 7.05E-05 | 0.00019927 | 27.45954 | 67.382084 | 120.79767 | 65.556966 |
| -3.974583 | 7.05E-05 | 0.0001994 | 37.705637 | 43.2977247 | 60.9740618 | 49.8681194 |
| -3.9744899 | 7.05E-05 | 0.00019943 | 31.5579788 | 60.8874253 | 51.7704298 | 75.0823371 |
| -3.9744788 | 7.05E-05 | 0.00019943 | 23.770945 | 76.8534613 | 81.6822337 | 94.1330794 |
| -3.9744976 | 7.05E-05 | 0.00019943 | 24.5906328 | 74.1473535 | 81.6822337 | 84.6077082 |
| -3.974274 | 7.06E-05 | 0.00019959 | 27.0496961 | 72.7942996 | 71.3281477 | 68.3585457 |
| -3.9741146 | 7.06E-05 | 0.0001997 | 54.509236 | 27.8729103 | 143.806749 | 87.9696039 |
| -3.9738888 | 7.07E-05 | 0.00019987 | 25.4103206 | 96.0668266 | 67.8767857 | 77.3236009 |
| -3.9737183 | 7.08E-05 | 0.0002 | 31.5579788 | 48.980551 | 75.9299637 | 67.7982298 |
| -3.9733856 | 7.09E-05 | 0.00020026 | 26.2300083 | 79.2889583 | 62.1245158 | 99.1759229 |
| -3.972917 | 7.10E-05 | 0.00020064 | 60.6568942 | 50.6042157 | 37.9649818 | 42.0236961 |
| -3.9728794 | 7.10E-05 | 0.00020065 | 75.8211178 | 136.117222 | 81.6822337 | 104.779082 |
| -3.9728092 | 7.10E-05 | 0.0002007 | 41.3942319 | 54.9339882 | 40.2658898 | 47.0665397 |
| -3.9726099 | 7.11E-05 | 0.00020085 | 28.2792277 | 61.4286469 | 67.8767857 | 73.4013893 |
| -3.9724262 | 7.11E-05 | 0.00020098 | 28.2792277 | 69.5469702 | 65.5758777 | 64.99665 |
| -3.9722099 | 7.12E-05 | 0.00020115 | 13.1150042 | 162.366467 | 119.647216 | 161.370993 |
| -3.9715615 | 7.14E-05 | 0.00020168 | 34.017042 | 45.1920001 | 69.0272397 | 103.65845 |
| -3.9712496 | 7.15E-05 | 0.00020193 | 42.2139196 | 99.0435452 | 120.79767 | 112.623506 |
| -3.9711763 | 7.15E-05 | 0.00020197 | 70.4931473 | 106.079425 | 21.8586259 | 107.580662 |
| -3.9711423 | 7.15E-05 | 0.00020198 | 31.1481349 | 51.6866588 | 66.7263317 | 64.4363341 |
| -3.9711126 | 7.15E-05 | 0.00020199 | 27.0496961 | 69.5469702 | 75.9299637 | 66.6775979 |
| -3.970893 | 7.16E-05 | 0.00020216 | 31.1481349 | 92.2782757 | 47.1686138 | 78.4442328 |
| -3.9708007 | 7.16E-05 | 0.00020222 | 447.959361 | 427.023809 | 355.490285 | 363.645051 |

| | | | | | | |
|---|---|---|---|---|---|---|
| -3.9707095 | 7.17E-05 | 0.00020228 | 48.3615778 | 55.4752097 | 37.9649818 | 43.144328 |
| -3.9704687 | 7.17E-05 | 0.00020246 | 39.7548564 | 63.8641439 | 37.9649818 | 57.7125427 |
| -3.970093 | 7.18E-05 | 0.00020277 | 25.0004767 | 82.5362876 | 75.9299637 | 74.5220212 |
| 3.96995237 | 7.19E-05 | 0.00020285 | 28.6890716 | 18.1309222 | 18.4072639 | 25.2142177 |
| -3.9699528 | 7.19E-05 | 0.00020285 | 27.0496961 | 59.5343714 | 75.9299637 | 101.977503 |
| -3.9694094 | 7.21E-05 | 0.0002033 | 27.45954 | 61.6992576 | 72.4786017 | 76.202969 |
| -3.969301 | 7.21E-05 | 0.00020337 | 190.16756 | 150.188982 | 11.50454 | 135.036144 |
| -3.9688929 | 7.22E-05 | 0.0002037 | 337.711357 | 58.9931499 | 26.4604419 | 95.8140272 |
| -3.9688224 | 7.22E-05 | 0.00020374 | 42.6237635 | 57.0988744 | 35.6640739 | 61.0744384 |
| -3.968788 | 7.22E-05 | 0.00020376 | 71.3128351 | 136.929054 | 72.4786017 | 25.2142177 |
| 3.96848592 | 7.23E-05 | 0.000204 | 145.084733 | 206.205414 | 173.718553 | 177.05984 |
| 3.96764319 | 7.26E-05 | 0.0002047 | 13.1150042 | 10.0125988 | 6.90272397 | 10.646003 |
| -3.9675735 | 7.26E-05 | 0.00020474 | 22.5414134 | 192.133653 | 56.3722458 | 133.355196 |
| -3.9673762 | 7.27E-05 | 0.00020489 | 38.1154808 | 66.5702517 | 42.5667978 | 132.79488 |
| -3.9672617 | 7.27E-05 | 0.00020497 | 29.5087594 | 63.8641439 | 58.6731538 | 65.556966 |
| -3.9670624 | 7.28E-05 | 0.00020513 | 27.0496961 | 79.5595691 | 63.2749697 | 71.1601255 |
| -3.9669833 | 7.28E-05 | 0.00020516 | 42.2139196 | 79.5595691 | 33.3631659 | 81.8061285 |
| 3.9669992 | 7.28E-05 | 0.00020516 | 36.8859492 | 25.4374132 | 24.1595339 | 29.6967453 |
| -3.9658562 | 7.31E-05 | 0.00020611 | 36.8859492 | 33.2851258 | 171.417645 | 131.113932 |
| -3.9655986 | 7.32E-05 | 0.00020632 | 128.281134 | 51.6866588 | 63.2749697 | 33.058641 |
| 3.96554121 | 7.32E-05 | 0.00020633 | 0 | 0 | 0 | 0 |
| -3.965545 | 7.32E-05 | 0.00020633 | 26.2300083 | 65.4878086 | 73.6290557 | 81.8061285 |
| 3.96526038 | 7.33E-05 | 0.00020656 | 2.0492194 | 1.89427545 | 3.45136199 | 2.24126379 |
| -3.9650835 | 7.34E-05 | 0.00020669 | 27.45954 | 90.654611 | 52.9208838 | 100.856871 |
| -3.9649488 | 7.34E-05 | 0.00020679 | 330.334167 | 92.2782757 | 23.0090799 | 70.5998095 |
| -3.9645699 | 7.35E-05 | 0.0002071 | 43.0336074 | 47.6274971 | 39.1154358 | 48.1871716 |
| -3.9644155 | 7.36E-05 | 0.00020722 | 366.810273 | 805.067068 | 591.333354 | 649.9665 |
| -3.963598 | 7.38E-05 | 0.00020791 | 28.6890716 | 89.0309463 | 58.6731538 | 63.8760181 |
| -3.9631602 | 7.40E-05 | 0.00020827 | 38.5253247 | 41.67406 | 67.8767857 | 49.3078035 |
| -3.9630514 | 7.40E-05 | 0.00020835 | 82.7884637 | 43.8389462 | 32.2127119 | 76.763285 |
| -3.9629343 | 7.40E-05 | 0.00020844 | 97.5428434 | 91.1958326 | 100.089498 | 89.6505518 |
| -3.9625942 | 7.41E-05 | 0.00020871 | 28.6890716 | 63.0523115 | 60.9740618 | 69.4791776 |
| -3.962468 | 7.42E-05 | 0.00020881 | 55.3289238 | 142.882491 | 21.8586259 | 179.301104 |
| -3.9616463 | 7.44E-05 | 0.00020951 | 42.6237635 | 38.1561199 | 48.3190678 | 53.7903311 |
| -3.9612018 | 7.46E-05 | 0.00020988 | 30.738291 | 51.9572696 | 63.2749697 | 65.556966 |
| -3.9611321 | 7.46E-05 | 0.00020992 | 39.3450125 | 45.1920001 | 44.8677058 | 75.6426531 |
| -3.9611138 | 7.46E-05 | 0.00020992 | 91.8050291 | 66.2996409 | 58.6731538 | 28.0157974 |
| -3.9610211 | 7.46E-05 | 0.00020995 | 31.5579788 | 111.762252 | 41.4163438 | 112.06319 |
| -3.961009 | 7.46E-05 | 0.00020995 | 27.45954 | 72.253078 | 59.8236078 | 76.202969 |
| -3.9610147 | 7.46E-05 | 0.00020995 | 26.2300083 | 112.574084 | 51.7704298 | 114.864769 |
| -3.9610001 | 7.46E-05 | 0.00020995 | 25.4103206 | 86.0542278 | 62.1245158 | 88.5299199 |
| -3.9606539 | 7.47E-05 | 0.00021023 | 28.2792277 | 65.2171978 | 63.2749697 | 66.1172819 |
| -3.9598561 | 7.50E-05 | 0.00021092 | 68.4439279 | 49.7923834 | 32.2127119 | 53.7903311 |

| | | | | | | |
|---|---|---|---|---|---|---|
| -3.9597752 | 7.50E-05 | 0.00021097 | 19.2626624 | 179.956168 | 77.0804177 | 108.701294 |
| -3.9596458 | 7.51E-05 | 0.00021107 | 18.8528185 | 71.7118565 | 173.718553 | 128.312352 |
| -3.9595609 | 7.51E-05 | 0.00021113 | 27.45954 | 69.5469702 | 59.8236078 | 80.6854966 |
| -3.9595227 | 7.51E-05 | 0.00021114 | 38.1154808 | 42.2152815 | 48.3190678 | 54.910963 |
| -3.9593933 | 7.51E-05 | 0.00021124 | 39.3450125 | 71.1706349 | 35.6640739 | 80.1251807 |
| -3.9586029 | 7.54E-05 | 0.00021192 | 46.3123584 | 30.8496288 | 112.744492 | 81.2458125 |
| -3.9585333 | 7.54E-05 | 0.00021196 | 27.8693838 | 69.2763595 | 86.2840497 | 57.1522268 |
| -3.958371 | 7.55E-05 | 0.00021209 | 29.9186032 | 57.0988744 | 57.5226998 | 73.9617052 |
| -3.9581007 | 7.55E-05 | 0.00021231 | 34.4268859 | 48.1687187 | 52.9208838 | 52.1093832 |
| -3.9580447 | 7.56E-05 | 0.00021234 | 31.9678226 | 86.8660601 | 50.6199758 | 58.2728587 |
| -3.9569047 | 7.59E-05 | 0.00021333 | 149.593016 | 97.6904913 | 96.6381356 | 105.899714 |
| -3.9568876 | 7.59E-05 | 0.00021333 | 21.3118818 | 105.267593 | 85.1335957 | 85.7283401 |
| -3.9568506 | 7.59E-05 | 0.00021335 | 27.8693838 | 94.1725511 | 60.9740618 | 63.8760181 |
| -3.9565041 | 7.61E-05 | 0.00021364 | 292.218686 | 61.6992576 | 44.8677058 | 47.0665397 |
| -3.9563239 | 7.61E-05 | 0.00021378 | 27.8693838 | 59.8049822 | 73.6290557 | 66.6775979 |
| 3.95622848 | 7.61E-05 | 0.00021385 | 4.0984388 | 1.62366467 | 0 | 15.1285306 |
| -3.9556774 | 7.63E-05 | 0.00021432 | 33.1973543 | 66.8408624 | 43.7172518 | 84.0473923 |
| -3.9556544 | 7.63E-05 | 0.00021433 | 25.8201644 | 92.8194972 | 71.3281477 | 64.99665 |
| -3.9554021 | 7.64E-05 | 0.00021453 | 21.7217256 | 96.0668266 | 72.4786017 | 113.744138 |
| -3.9548559 | 7.66E-05 | 0.00021501 | 30.738291 | 99.3141559 | 49.4695218 | 67.7982298 |
| -3.9547712 | 7.66E-05 | 0.00021506 | 27.45954 | 95.7962158 | 127.700393 | 49.8681194 |
| -3.9546094 | 7.67E-05 | 0.00021519 | 2921.77702 | 675.985726 | 773.105085 | 900.988045 |
| -3.9543741 | 7.67E-05 | 0.00021538 | 43.0336074 | 109.326755 | 55.2217918 | 39.2221164 |
| -3.9540372 | 7.68E-05 | 0.00021567 | 25.8201644 | 108.244312 | 71.3281477 | 64.4363341 |
| -3.9539894 | 7.69E-05 | 0.00021569 | 34.017042 | 59.2637606 | 64.4254237 | 43.704644 |
| -3.9539661 | 7.69E-05 | 0.00021569 | 59.8372065 | 42.2152815 | 44.8677058 | 39.2221164 |
| -3.9538888 | 7.69E-05 | 0.00021575 | 68.8537718 | 57.3694852 | 28.7613499 | 59.3934906 |
| -3.953255 | 7.71E-05 | 0.0002163 | 38.1154808 | 153.706923 | 134.603117 | 109.26161 |
| -3.9530476 | 7.72E-05 | 0.00021647 | 73.7718984 | 112.032863 | 105.841768 | 98.615607 |
| -3.9525742 | 7.73E-05 | 0.00021688 | 35.2465737 | 58.4519283 | 46.0181598 | 48.7474875 |
| -3.9523208 | 7.74E-05 | 0.00021709 | 42.2139196 | 33.0145151 | 77.0804177 | 89.0902358 |
| -3.9521013 | 7.75E-05 | 0.00021727 | 48.3615778 | 78.7477367 | 39.1154358 | 42.0236961 |
| -3.9519292 | 7.75E-05 | 0.00021741 | 73.7718984 | 56.0164313 | 29.9118039 | 54.350647 |
| 3.95149674 | 7.77E-05 | 0.00021778 | 0 | 0 | 0 | 0 |
| -3.9513153 | 7.77E-05 | 0.00021793 | 600.831128 | 942.537344 | 869.743221 | 930.684791 |
| -3.9507343 | 7.79E-05 | 0.00021844 | 28.2792277 | 73.8767427 | 52.9208838 | 82.9267604 |
| -3.9501675 | 7.81E-05 | 0.00021894 | 39.7548564 | 56.5576528 | 40.2658898 | 45.9459078 |
| -3.9498959 | 7.82E-05 | 0.00021915 | 31.9678226 | 48.4393295 | 100.089498 | 62.1950703 |
| -3.9498984 | 7.82E-05 | 0.00021915 | 20.492194 | 90.654611 | 106.992222 | 85.1680242 |
| -3.949601 | 7.83E-05 | 0.0002194 | 21.3118818 | 125.292791 | 74.7795097 | 90.7711837 |
| -3.9494781 | 7.83E-05 | 0.00021949 | 31.1481349 | 50.8748265 | 92.0363196 | 59.3934906 |
| -3.9491026 | 7.84E-05 | 0.00021982 | 44.263139 | 97.9611021 | 85.1335957 | 114.304454 |
| -3.9488144 | 7.85E-05 | 0.00022006 | 48.7714217 | 31.9320719 | 58.6731538 | 61.6347543 |

| | | | | | | |
|---|---|---|---|---|---|---|
| -3.948458 | 7.87E-05 | 0.00022037 | 19.6725062 | 90.9252218 | 111.594038 | 91.3314996 |
| -3.9484309 | 7.87E-05 | 0.00022038 | 34.4268859 | 76.0416289 | 43.7172518 | 56.5919108 |
| -3.948377 | 7.87E-05 | 0.00022041 | 11.4756286 | 113.115306 | 241.595339 | 151.285306 |
| -3.9483015 | 7.87E-05 | 0.00022046 | 291.808843 | 45.7332217 | 39.1154358 | 72.2807574 |
| -3.9481865 | 7.87E-05 | 0.00022055 | 55.3289238 | 38.9679522 | 64.4254237 | 37.5411686 |
| -3.9479957 | 7.88E-05 | 0.0002207 | 55.7387677 | 43.8389462 | 41.4163438 | 39.7824324 |
| -3.9479408 | 7.88E-05 | 0.00022073 | 40.5745441 | 100.396599 | 54.0713378 | 40.3427483 |
| -3.9477306 | 7.89E-05 | 0.00022091 | 23.3611012 | 83.8893415 | 78.2308717 | 77.8839169 |
| -3.9476743 | 7.89E-05 | 0.00022094 | 90.1656536 | 76.8534613 | 253.099879 | 20.7316901 |
| -3.9474522 | 7.90E-05 | 0.00022113 | 109.018472 | 310.931785 | 277.259413 | 284.640502 |
| -3.9472657 | 7.90E-05 | 0.00022128 | 32.7875104 | 65.7584193 | 55.2217918 | 47.0665397 |
| -3.9470696 | 7.91E-05 | 0.00022144 | 16.8035991 | 192.945486 | 79.3813257 | 127.752036 |
| -3.9467916 | 7.92E-05 | 0.00022168 | 34.017042 | 102.832096 | 40.2658898 | 77.3236009 |
| -3.9467095 | 7.92E-05 | 0.00022174 | 8.60672148 | 365.595163 | 193.276271 | 112.623506 |
| 3.94666086 | 7.92E-05 | 0.00022176 | 0.40984388 | 0 | 1.150454 | 0 |
| -3.9464639 | 7.93E-05 | 0.00022193 | 51.6403289 | 36.5324552 | 85.1335957 | 43.704644 |
| -3.9461707 | 7.94E-05 | 0.00022216 | 38.5253247 | 70.9000241 | 82.8326877 | 80.1251807 |
| -3.9461846 | 7.94E-05 | 0.00022216 | 25.0004767 | 86.5954493 | 77.0804177 | 64.4363341 |
| -3.9460439 | 7.95E-05 | 0.00022226 | 59.8372065 | 94.9843835 | 118.496762 | 101.417187 |
| -3.9459307 | 7.95E-05 | 0.00022234 | 28.6890716 | 71.9824673 | 62.1245158 | 56.5919108 |
| -3.9456374 | 7.96E-05 | 0.0002226 | 51.230485 | 62.7817008 | 309.472125 | 296.407137 |
| -3.9452258 | 7.97E-05 | 0.00022296 | 36.8859492 | 52.2278804 | 41.4163438 | 51.5490673 |
| -3.9446568 | 7.99E-05 | 0.00022347 | 37.705637 | 57.0988744 | 37.9649818 | 66.6775979 |
| -3.9445125 | 8.00E-05 | 0.00022359 | 38.1154808 | 42.2152815 | 81.6822337 | 49.3078035 |
| -3.9444397 | 8.00E-05 | 0.00022363 | 332.793231 | 760.957511 | 743.193281 | 670.137875 |
| -3.9441803 | 8.01E-05 | 0.00022386 | 241.398045 | 538.24484 | 526.90793 | 461.140026 |
| -3.943822 | 8.02E-05 | 0.00022417 | 33.1973543 | 38.1561199 | 110.443584 | 106.46003 |
| -3.9437849 | 8.02E-05 | 0.00022419 | 25.4103206 | 70.6294134 | 69.0272397 | 75.6426531 |
| -3.9437382 | 8.02E-05 | 0.00022421 | 65.5750208 | 42.2152815 | 35.6640739 | 50.9887513 |
| -3.9433158 | 8.04E-05 | 0.00022457 | 295.087594 | 805.8789 | 763.901453 | 804.053386 |
| -3.9433303 | 8.04E-05 | 0.00022457 | 57.7879871 | 50.6042157 | 95.4876816 | 31.3776931 |
| -3.9432369 | 8.04E-05 | 0.00022462 | 77.0506494 | 39.238563 | 50.6199758 | 39.7824324 |
| -3.9432132 | 8.04E-05 | 0.00022463 | 52.8698605 | 81.1832337 | 27.6108959 | 71.1601255 |
| -3.9431262 | 8.04E-05 | 0.00022469 | 37.2957931 | 40.8622277 | 49.4695218 | 59.3934906 |
| -3.9423532 | 8.07E-05 | 0.00022539 | 31.5579788 | 53.3103235 | 55.2217918 | 54.350647 |
| -3.9422589 | 8.07E-05 | 0.00022546 | 240.578358 | 680.315499 | 511.952028 | 587.77143 |
| -3.9421228 | 8.08E-05 | 0.00022557 | 25.0004767 | 63.0523115 | 80.5317797 | 81.8061285 |
| -3.9417607 | 8.09E-05 | 0.00022589 | 54.0993922 | 32.2026827 | 49.4695218 | 58.8331746 |
| -3.9416216 | 8.09E-05 | 0.00022601 | 25.8201644 | 74.1473535 | 308.321671 | 70.0394936 |
| -3.9413564 | 8.10E-05 | 0.00022624 | 9.01656536 | 346.923019 | 113.894946 | 178.180472 |
| -3.9411345 | 8.11E-05 | 0.00022643 | 36.0662614 | 50.8748265 | 74.7795097 | 42.5840121 |
| -3.9406628 | 8.13E-05 | 0.00022685 | 43.4434513 | 34.0969582 | 63.2749697 | 54.910963 |
| -3.9406346 | 8.13E-05 | 0.00022686 | 34.8367298 | 45.4626109 | 50.6199758 | 76.763285 |

| | | | | | | |
|---|---|---|---|---|---|---|
| -3.9405207 | 8.13E-05 | 0.00022693 | 291.808843 | 275.481773 | 182.922185 | 235.893014 |
| -3.9404928 | 8.13E-05 | 0.00022693 | 33.1973543 | 105.267593 | 37.9649818 | 103.098135 |
| -3.9404955 | 8.13E-05 | 0.00022693 | 41.3942319 | 105.808815 | 29.9118039 | 131.113932 |
| -3.9402558 | 8.14E-05 | 0.00022714 | 31.9678226 | 80.1007906 | 101.239952 | 42.0236961 |
| 3.94006508 | 8.15E-05 | 0.0002273 | 0 | 0 | 0 | 0 |
| -3.9399754 | 8.15E-05 | 0.00022736 | 25.4103206 | 69.817581 | 73.6290557 | 69.4791776 |
| -3.9399099 | 8.15E-05 | 0.00022741 | 42.6237635 | 46.545054 | 117.346308 | 44.8252759 |
| 3.93958196 | 8.16E-05 | 0.0002277 | 20.9020379 | 11.6362635 | 14.9559019 | 6.72379138 |
| -3.9395323 | 8.16E-05 | 0.00022773 | 21.7217256 | 91.1958326 | 89.7354116 | 77.3236009 |
| -3.9392783 | 8.17E-05 | 0.00022795 | 33.1973543 | 120.692407 | 94.3372276 | 38.6618005 |
| -3.9390746 | 8.18E-05 | 0.00022812 | 54.0993922 | 140.176384 | 60.9740618 | 124.390141 |
| -3.9389936 | 8.18E-05 | 0.00022818 | 84.0179954 | 66.5702517 | 27.6108959 | 53.7903311 |
| -3.9389152 | 8.19E-05 | 0.00022821 | 60.6568942 | 42.2152815 | 43.7172518 | 38.6618005 |
| -3.9389219 | 8.18E-05 | 0.00022821 | 22.1315695 | 90.654611 | 75.9299637 | 86.2886561 |
| -3.938821 | 8.19E-05 | 0.00022828 | 32.3776665 | 49.7923834 | 120.79767 | 60.5141225 |
| -3.9386431 | 8.19E-05 | 0.00022843 | 31.1481349 | 67.1114732 | 46.0181598 | 68.3585457 |
| -3.9381323 | 8.21E-05 | 0.0002289 | 33.1973543 | 73.0649104 | 57.5226998 | 44.2649599 |
| -3.9381088 | 8.21E-05 | 0.0002289 | 78.6900249 | 51.1454373 | 60.9740618 | 29.6967453 |
| -3.938042 | 8.21E-05 | 0.00022895 | 107.78894 | 323.379881 | 406.11026 | 330.58641 |
| -3.9369203 | 8.25E-05 | 0.00023 | 38.5253247 | 52.2278804 | 39.1154358 | 79.5648647 |
| -3.936868 | 8.26E-05 | 0.00023003 | 8.60672148 | 80.3714014 | 406.11026 | 265.58976 |
| -3.9368429 | 8.26E-05 | 0.00023003 | 22.9512573 | 67.6526948 | 103.54086 | 82.9267604 |
| -3.9367836 | 8.26E-05 | 0.00023007 | 38.1154808 | 61.9698684 | 44.8677058 | 42.0236961 |
| -3.9367576 | 8.26E-05 | 0.00023008 | 49.5911095 | 48.1687187 | 36.8145279 | 42.0236961 |
| -3.9364875 | 8.27E-05 | 0.00023032 | 30.738291 | 64.1347547 | 65.5758777 | 47.6268556 |
| -3.9363073 | 8.27E-05 | 0.00023047 | 33.1973543 | 128.810731 | 67.8767857 | 45.3855918 |
| -3.9361679 | 8.28E-05 | 0.00023058 | 30.738291 | 57.640096 | 59.8236078 | 50.9887513 |
| -3.9356951 | 8.30E-05 | 0.00023102 | 16.8035991 | 182.662276 | 79.3813257 | 122.148877 |
| -3.9355504 | 8.30E-05 | 0.00023114 | 47.5418901 | 55.7458205 | 39.1154358 | 38.6618005 |
| -3.935467 | 8.30E-05 | 0.0002312 | 37.2957931 | 44.3801678 | 63.2749697 | 43.144328 |
| -3.9350175 | 8.32E-05 | 0.00023161 | 163.527708 | 107.161869 | 18.4072639 | 74.5220212 |
| 3.93489694 | 8.32E-05 | 0.00023171 | 46.3123584 | 37.0736767 | 60.9740618 | 36.4205367 |
| -3.9346196 | 8.33E-05 | 0.00023195 | 29.5087594 | 52.2278804 | 64.4254237 | 61.6347543 |
| -3.9343927 | 8.34E-05 | 0.00023215 | 38.9351686 | 59.8049822 | 36.8145279 | 90.2108677 |
| -3.9342009 | 8.35E-05 | 0.00023232 | 20.0823501 | 60.6168145 | 164.514921 | 130.553616 |
| -3.9341489 | 8.35E-05 | 0.00023233 | 24.5906328 | 107.161869 | 56.3722458 | 91.8918156 |
| -3.9341657 | 8.35E-05 | 0.00023233 | 26.6398522 | 130.975617 | 44.8677058 | 126.071088 |
| -3.9339983 | 8.35E-05 | 0.00023245 | 31.5579788 | 50.0629941 | 52.9208838 | 73.9617052 |
| -3.933753 | 8.36E-05 | 0.00023267 | 24.1807889 | 63.3229223 | 88.5849576 | 81.2458125 |
| -3.9333229 | 8.38E-05 | 0.00023307 | 97.9526873 | 47.0862756 | 33.3631659 | 54.910963 |
| -3.9328863 | 8.39E-05 | 0.00023347 | 33.1973543 | 41.4034492 | 90.8858656 | 68.3585457 |
| 3.93281313 | 8.40E-05 | 0.00023352 | 0 | 1.62366467 | 5.75226998 | 0 |
| -3.932553 | 8.40E-05 | 0.00023376 | 31.9678226 | 64.1347547 | 54.0713378 | 47.6268556 |

| | | | | | | |
|---|---|---|---|---|---|---|
| -3.932432 | 8.41E-05 | 0.00023385 | 28.6890716 | 86.8660601 | 46.0181598 | 115.425085 |
| -3.9324055 | 8.41E-05 | 0.00023386 | 21.3118818 | 94.1725511 | 117.346308 | 68.9188617 |
| -3.9322782 | 8.41E-05 | 0.00023396 | 36.4761053 | 41.9446708 | 133.452663 | 64.4363341 |
| -3.9321521 | 8.42E-05 | 0.00023407 | 25.8201644 | 71.4412457 | 70.1776937 | 64.4363341 |
| -3.9319252 | 8.43E-05 | 0.00023427 | 43.8532952 | 31.6614612 | 89.7354116 | 68.3585457 |
| -3.9318177 | 8.43E-05 | 0.00023435 | 30.738291 | 63.0523115 | 49.4695218 | 57.7125427 |
| -3.9317621 | 8.43E-05 | 0.00023439 | 105.739721 | 50.3336049 | 34.5136199 | 49.8681194 |
| -3.9314897 | 8.44E-05 | 0.00023462 | 34.8367298 | 43.2977247 | 80.5317797 | 52.6696992 |
| -3.9314789 | 8.44E-05 | 0.00023462 | 33.1973543 | 69.5469702 | 41.4163438 | 64.99665 |
| 3.93112281 | 8.46E-05 | 0.00023495 | 78.6900249 | 63.8641439 | 58.6731538 | 63.8760181 |
| -3.9310336 | 8.46E-05 | 0.00023502 | 21.7217256 | 74.9591858 | 89.7354116 | 89.6505518 |
| -3.9309436 | 8.46E-05 | 0.00023508 | 33.1973543 | 42.7565031 | 59.8236078 | 62.1950703 |
| -3.9308213 | 8.47E-05 | 0.00023518 | 40.1647002 | 90.654611 | 80.5317797 | 92.4521315 |
| -3.9307574 | 8.47E-05 | 0.0002352 | 23.770945 | 68.7351379 | 78.2308717 | 81.8061285 |
| -3.9307778 | 8.47E-05 | 0.0002352 | 15.9839113 | 87.1366709 | 150.709473 | 126.071088 |
| -3.9306464 | 8.47E-05 | 0.00023529 | 28.6890716 | 58.9931499 | 55.2217918 | 71.1601255 |
| -3.9301076 | 8.49E-05 | 0.0002358 | 36.8859492 | 52.2278804 | 40.2658898 | 49.8681194 |
| -3.9293108 | 8.52E-05 | 0.00023656 | 49.1812656 | 81.7244553 | 55.2217918 | 73.4013893 |
| -3.9291074 | 8.53E-05 | 0.00023674 | 38.9351686 | 91.1958326 | 37.9649818 | 56.0315949 |
| -3.9289578 | 8.53E-05 | 0.00023687 | 27.0496961 | 64.4053654 | 78.2308717 | 57.7125427 |
| -3.9288432 | 8.54E-05 | 0.00023696 | 61.476582 | 260.59818 | 236.993523 | 216.281956 |
| -3.9286748 | 8.54E-05 | 0.00023711 | 23.3611012 | 57.3694852 | 104.691314 | 106.46003 |
| -3.9285717 | 8.55E-05 | 0.00023719 | 22.1315695 | 111.491641 | 57.5226998 | 140.639303 |
| -3.9284523 | 8.55E-05 | 0.00023729 | 75.4112739 | 177.791282 | 128.850847 | 151.845622 |
| -3.9282315 | 8.56E-05 | 0.00023748 | 38.5253247 | 88.219114 | 37.9649818 | 56.0315949 |
| -3.9278186 | 8.57E-05 | 0.00023787 | 28.2792277 | 64.946587 | 113.894946 | 54.350647 |
| -3.9276257 | 8.58E-05 | 0.00023804 | 16.3937552 | 115.550803 | 102.390406 | 113.183822 |
| -3.9272659 | 8.59E-05 | 0.00023838 | 28.6890716 | 54.1221558 | 63.2749697 | 62.7553862 |
| -3.9266892 | 8.61E-05 | 0.00023893 | 30.3284471 | 70.0881918 | 73.6290557 | 45.3855918 |
| -3.9266136 | 8.62E-05 | 0.00023895 | 44.263139 | 90.1133895 | 32.2127119 | 60.5141225 |
| -3.9266215 | 8.61E-05 | 0.00023895 | 18.0331307 | 87.4072817 | 116.195854 | 104.218766 |
| -3.9266027 | 8.62E-05 | 0.00023895 | 28.6890716 | 87.4072817 | 54.0713378 | 59.9538065 |
| -3.9261611 | 8.63E-05 | 0.00023937 | 34.4268859 | 67.9233056 | 65.5758777 | 39.2221164 |
| -3.9259999 | 8.64E-05 | 0.00023951 | 22.1315695 | 90.3840002 | 119.647216 | 64.4363341 |
| -3.9259222 | 8.64E-05 | 0.00023957 | 20.0823501 | 129.893174 | 75.9299637 | 90.2108677 |
| 3.92539682 | 8.66E-05 | 0.00024007 | 34.4268859 | 23.8137486 | 23.0090799 | 26.3348496 |
| -3.9251923 | 8.67E-05 | 0.00024025 | 9.42640924 | 94.9843835 | 276.108959 | 221.885116 |
| -3.925118 | 8.67E-05 | 0.00024031 | 31.1481349 | 60.3462037 | 47.1686138 | 58.8331746 |
| -3.9249554 | 8.67E-05 | 0.00024045 | 45.4926707 | 51.9572696 | 44.8677058 | 35.2999048 |
| -3.9248804 | 8.68E-05 | 0.00024048 | 16.8035991 | 116.362635 | 92.0363196 | 118.786981 |
| -3.9248963 | 8.68E-05 | 0.00024048 | 29.5087594 | 47.0862756 | 90.8858656 | 68.9188617 |
| -3.9248282 | 8.68E-05 | 0.00024052 | 18.0331307 | 86.0542278 | 120.79767 | 102.537819 |
| -3.9246539 | 8.69E-05 | 0.00024067 | 19.6725062 | 106.350036 | 79.3813257 | 96.9346591 |

| | | | | | | |
|---|---|---|---|---|---|---|
| -3.9242797 | 8.70E-05 | 0.00024102 | 28.6890716 | 66.2996409 | 50.6199758 | 70.5998095 |
| -3.9239839 | 8.71E-05 | 0.0002413 | 53.2797044 | 45.1920001 | 33.3631659 | 49.3078035 |
| -3.9235728 | 8.72E-05 | 0.00024169 | 31.1481349 | 55.7458205 | 50.6199758 | 53.7903311 |
| -3.922991 | 8.75E-05 | 0.00024225 | 14.3445358 | 93.090108 | 125.399485 | 188.826475 |
| -3.9229378 | 8.75E-05 | 0.00024229 | 18.4429746 | 152.89509 | 80.5317797 | 96.3743432 |
| -3.922909 | 8.75E-05 | 0.00024229 | 52.4600166 | 44.9213893 | 42.5667978 | 36.4205367 |
| -3.9222299 | 8.77E-05 | 0.00024296 | 77.0506494 | 112.303473 | 17.2568099 | 151.285306 |
| -3.9221899 | 8.77E-05 | 0.00024298 | 32.3776665 | 58.1813175 | 43.7172518 | 61.0744384 |
| -3.9220678 | 8.78E-05 | 0.00024308 | 20.9020379 | 83.34812 | 89.7354116 | 84.0473923 |
| -3.9218104 | 8.79E-05 | 0.00024332 | 42.6237635 | 37.0736767 | 49.4695218 | 44.2649599 |
| -3.9213943 | 8.80E-05 | 0.00024371 | 71.722679 | 60.6168145 | 28.7613499 | 49.8681194 |
| -3.9213833 | 8.80E-05 | 0.00024371 | 26.6398522 | 55.7458205 | 70.1776937 | 75.0823371 |
| -3.9210963 | 8.81E-05 | 0.00024398 | 35.6564176 | 55.2045989 | 41.4163438 | 92.4521315 |
| -3.9210673 | 8.82E-05 | 0.00024399 | 48.7714217 | 54.6633774 | 33.3631659 | 44.8252759 |
| -3.9206251 | 8.83E-05 | 0.00024441 | 22.5414134 | 63.0523115 | 88.5849576 | 108.140978 |
| 3.9205089 | 8.84E-05 | 0.00024451 | 0 | 0 | 0 | 0 |
| -3.9202866 | 8.84E-05 | 0.00024472 | 27.8693838 | 178.332503 | 41.4163438 | 114.304454 |
| -3.9201091 | 8.85E-05 | 0.00024487 | 34.4268859 | 42.7565031 | 49.4695218 | 58.2728587 |
| -3.9191912 | 8.88E-05 | 0.00024579 | 35.6564176 | 43.0271139 | 46.0181598 | 51.5490673 |
| -3.9190697 | 8.89E-05 | 0.00024589 | 29.5087594 | 79.5595691 | 164.514921 | 47.0665397 |
| -3.918888 | 8.90E-05 | 0.00024606 | 16.8035991 | 115.009581 | 150.709473 | 81.2458125 |
| -3.918829 | 8.90E-05 | 0.00024609 | 52.8698605 | 61.1580361 | 46.0181598 | 31.9380091 |
| -3.9184982 | 8.91E-05 | 0.00024641 | 29.9186032 | 55.2045989 | 50.6199758 | 70.5998095 |
| -3.918282 | 8.92E-05 | 0.00024661 | 24.1807889 | 69.5469702 | 87.4345036 | 66.1172819 |
| -3.9180901 | 8.93E-05 | 0.00024679 | 26.6398522 | 112.574084 | 64.4254237 | 58.8331746 |
| -3.9177651 | 8.94E-05 | 0.0002471 | 102.870814 | 63.3229223 | 29.9118039 | 47.6268556 |
| -3.9174932 | 8.95E-05 | 0.00024736 | 64.755333 | 39.7797845 | 36.8145279 | 47.6268556 |
| -3.9174575 | 8.95E-05 | 0.00024737 | 184.429746 | 393.468073 | 338.233475 | 368.127578 |
| 3.91702134 | 8.96E-05 | 0.0002478 | 34.4268859 | 21.9194731 | 8.05317797 | 11.206319 |
| -3.9169838 | 8.97E-05 | 0.00024781 | 20.492194 | 63.0523115 | 117.346308 | 118.226665 |
| -3.9168963 | 8.97E-05 | 0.00024788 | 29.0989155 | 52.7691019 | 86.2840497 | 56.5919108 |
| -3.9166934 | 8.98E-05 | 0.00024807 | 36.4761053 | 68.7351379 | 90.8858656 | 35.8602207 |
| -3.9161151 | 9.00E-05 | 0.00024865 | 79.5097127 | 29.7671857 | 65.5758777 | 46.5062237 |
| -3.9160529 | 9.00E-05 | 0.00024869 | 35.6564176 | 40.5916169 | 51.7704298 | 75.0823371 |
| -3.9155438 | 9.02E-05 | 0.00024919 | 20.492194 | 87.1366709 | 77.0804177 | 100.296555 |
| -3.915163 | 9.03E-05 | 0.00024956 | 1004.93719 | 2026.0629 | 1210.2776 | 1520.13717 |
| -3.9150116 | 9.04E-05 | 0.0002497 | 48.3615778 | 124.210348 | 89.7354116 | 89.6505518 |
| -3.9148126 | 9.05E-05 | 0.00024988 | 21.3118818 | 82.5362876 | 123.098578 | 70.0394936 |
| -3.9141674 | 9.07E-05 | 0.00025053 | 43.4434513 | 61.1580361 | 35.6640739 | 43.704644 |
| -3.9141145 | 9.07E-05 | 0.00025057 | 52.4600166 | 68.7351379 | 26.4604419 | 86.848972 |
| -3.91407 | 9.08E-05 | 0.00025059 | 37.2957931 | 41.9446708 | 42.5667978 | 57.1522268 |
| -3.9138739 | 9.08E-05 | 0.00025077 | 43.8532952 | 41.4034492 | 37.9649818 | 45.9459078 |
| -3.9134904 | 9.10E-05 | 0.00025115 | 25.4103206 | 58.1813175 | 77.0804177 | 72.8410733 |

| | | | | | | |
|---|---|---|---|---|---|---|
| -3.9123988 | 9.14E-05 | 0.00025227 | 76.2309617 | 229.477941 | 136.904025 | 157.448782 |
| -3.9123492 | 9.14E-05 | 0.0002523 | 27.8693838 | 59.5343714 | 54.0713378 | 80.1251807 |
| -3.9119841 | 9.15E-05 | 0.00025266 | 23.770945 | 63.5935331 | 73.6290557 | 91.3314996 |
| -3.9118333 | 9.16E-05 | 0.00025279 | 11.4756286 | 550.692936 | 744.343735 | 695.912408 |
| -3.9115497 | 9.17E-05 | 0.00025307 | 35.2465737 | 44.3801678 | 43.7172518 | 53.2300151 |
| -3.9115146 | 9.17E-05 | 0.00025308 | 30.3284471 | 50.6042157 | 51.7704298 | 68.3585457 |
| -3.9112624 | 9.18E-05 | 0.00025333 | 10.6559409 | 153.977533 | 222.037621 | 109.821926 |
| -3.9101737 | 9.22E-05 | 0.00025445 | 29.0989155 | 133.681725 | 74.7795097 | 47.0665397 |
| -3.9101058 | 9.23E-05 | 0.0002545 | 11.8854725 | 127.998899 | 186.373547 | 117.666349 |
| -3.9097526 | 9.24E-05 | 0.00025485 | 30.738291 | 71.4412457 | 51.7704298 | 49.3078035 |
| -3.9094659 | 9.25E-05 | 0.00025513 | 29.5087594 | 77.1240721 | 74.7795097 | 44.2649599 |
| -3.9093134 | 9.26E-05 | 0.00025527 | 37.2957931 | 82.8068984 | 77.0804177 | 33.6189569 |
| -3.908901 | 9.27E-05 | 0.00025566 | 24.5906328 | 126.645845 | 48.3190678 | 108.701294 |
| -3.9089062 | 9.27E-05 | 0.00025566 | 32.7875104 | 49.2511618 | 44.8677058 | 57.7125427 |
| -3.9087314 | 9.28E-05 | 0.00025582 | 29.0989155 | 200.793198 | 102.390406 | 40.9030642 |
| -3.9083679 | 9.29E-05 | 0.00025618 | 33.1973543 | 56.0164313 | 63.2749697 | 40.9030642 |
| -3.9081635 | 9.30E-05 | 0.00025638 | 19.6725062 | 59.5343714 | 115.0454 | 217.402588 |
| -3.9080875 | 9.30E-05 | 0.00025642 | 32.3776665 | 54.9339882 | 43.7172518 | 79.5648647 |
| -3.9080912 | 9.30E-05 | 0.00025642 | 90.1656536 | 22.4606947 | 89.7354116 | 63.8760181 |
| 3.90754489 | 9.32E-05 | 0.00025697 | 0 | 0 | 0 | 0 |
| -3.9070249 | 9.34E-05 | 0.0002575 | 28.2792277 | 55.7458205 | 151.859927 | 63.8760181 |
| -3.906951 | 9.35E-05 | 0.00025756 | 21.7217256 | 72.5236888 | 77.0804177 | 96.3743432 |
| -3.9063434 | 9.37E-05 | 0.00025818 | 44.263139 | 49.7923834 | 32.2127119 | 57.7125427 |
| -3.9059395 | 9.39E-05 | 0.00025859 | 31.1481349 | 41.1328384 | 72.4786017 | 89.6505518 |
| 3.90586218 | 9.39E-05 | 0.00025865 | 38.9351686 | 17.3190899 | 20.7081719 | 17.3697944 |
| -3.9056433 | 9.40E-05 | 0.00025887 | 30.3284471 | 39.7797845 | 104.691314 | 90.7711837 |
| -3.905607 | 9.40E-05 | 0.00025888 | 38.1154808 | 56.5576528 | 35.6640739 | 52.1093832 |
| -3.9050009 | 9.42E-05 | 0.00025951 | 90.9853413 | 353.147067 | 253.099879 | 273.994499 |
| -3.9049336 | 9.43E-05 | 0.00025956 | 22.9512573 | 113.656527 | 60.9740618 | 80.6854966 |
| -3.9045383 | 9.44E-05 | 0.00025996 | 77.0506494 | 44.6507786 | 29.9118039 | 61.0744384 |
| -3.9043849 | 9.45E-05 | 0.00026011 | 48.3615778 | 60.6168145 | 162.214013 | 33.058641 |
| -3.904335 | 9.45E-05 | 0.00026012 | 25.4103206 | 66.2996409 | 64.4254237 | 66.1172819 |
| -3.9043268 | 9.45E-05 | 0.00026012 | 33.6071982 | 114.46836 | 105.841768 | 121.028245 |
| -3.9039133 | 9.46E-05 | 0.00026055 | 33.6071982 | 142.882491 | 31.0622579 | 138.398039 |
| -3.9032734 | 9.49E-05 | 0.00026121 | 31.1481349 | 50.0629941 | 48.3190678 | 57.7125427 |
| -3.9028117 | 9.51E-05 | 0.00026169 | 44.6729829 | 62.2404792 | 41.4163438 | 35.8602207 |
| -3.9023191 | 9.53E-05 | 0.0002622 | 23.770945 | 96.0668266 | 72.4786017 | 61.6347543 |
| -3.9022798 | 9.53E-05 | 0.00026222 | 18.0331307 | 84.1599523 | 104.691314 | 101.977503 |
| -3.9020387 | 9.54E-05 | 0.00026246 | 41.3942319 | 44.3801678 | 36.8145279 | 45.3855918 |
| -3.9019603 | 9.54E-05 | 0.00026252 | 29.5087594 | 51.416048 | 58.6731538 | 53.7903311 |
| -3.9017886 | 9.55E-05 | 0.00026269 | 29.9186032 | 55.7458205 | 50.6199758 | 54.350647 |
| -3.9015578 | 9.56E-05 | 0.00026292 | 57.7879871 | 81.9950661 | 80.5317797 | 79.5648647 |
| -3.9009236 | 9.58E-05 | 0.00026358 | 22.5414134 | 99.0435452 | 67.8767857 | 72.8410733 |

| | | | | | | |
|---|---|---|---|---|---|---|
| -3.9006648 | 9.59E-05 | 0.00026384 | 26.2300083 | 79.2889583 | 55.2217918 | 64.99665 |
| -3.9006173 | 9.59E-05 | 0.00026387 | 16.3937552 | 131.787449 | 109.29313 | 85.7283401 |
| -3.9005506 | 9.60E-05 | 0.00026392 | 43.0336074 | 85.2423954 | 86.2840497 | 84.6077082 |
| -3.9003353 | 9.61E-05 | 0.00026414 | 21.7217256 | 76.5828505 | 71.3281477 | 94.1330794 |
| -3.9002241 | 9.61E-05 | 0.00026423 | 25.4103206 | 71.4412457 | 65.5758777 | 60.5141225 |
| -3.9000909 | 9.62E-05 | 0.00026433 | 19.6725062 | 72.5236888 | 95.4876816 | 104.779082 |
| -3.9000775 | 9.62E-05 | 0.00026433 | 25.0004767 | 51.416048 | 97.7885896 | 85.1680242 |
| -3.9001029 | 9.62E-05 | 0.00026433 | 49.5911095 | 30.8496288 | 74.7795097 | 49.3078035 |
| -3.9000167 | 9.62E-05 | 0.00026437 | 38.1154808 | 36.5324552 | 77.0804177 | 51.5490673 |
| 3.89985937 | 9.62E-05 | 0.00026452 | 18.0331307 | 13.530539 | 14.9559019 | 11.7666349 |
| -3.8995288 | 9.64E-05 | 0.00026484 | 57.7879871 | 185.368384 | 23.0090799 | 85.1680242 |
| -3.8995298 | 9.64E-05 | 0.00026484 | 22.1315695 | 74.9591858 | 69.0272397 | 95.2537113 |
| -3.899487 | 9.64E-05 | 0.00026486 | 17.213443 | 95.525605 | 94.3372276 | 110.382242 |
| -3.8991516 | 9.65E-05 | 0.0002652 | 24.5906328 | 73.6061319 | 63.2749697 | 67.7982298 |
| -3.8989487 | 9.66E-05 | 0.0002654 | 29.0989155 | 48.4393295 | 58.6731538 | 81.8061285 |
| -3.8985858 | 9.68E-05 | 0.00026578 | 20.492194 | 79.0183475 | 79.3813257 | 94.6933953 |
| -3.8981076 | 9.69E-05 | 0.00026628 | 25.4103206 | 92.5488865 | 70.1776937 | 54.910963 |
| -3.8979104 | 9.70E-05 | 0.00026648 | 29.9186032 | 74.1473535 | 59.8236078 | 45.3855918 |
| -3.8978859 | 9.70E-05 | 0.00026648 | 24.5906328 | 87.9485032 | 54.0713378 | 82.3664444 |
| -3.8975359 | 9.72E-05 | 0.00026684 | 0 | 2403.29433 | 1325.323 | 1322.90595 |
| -3.897467 | 9.72E-05 | 0.0002669 | 28.2792277 | 55.7458205 | 69.0272397 | 52.1093832 |
| 3.89739679 | 9.72E-05 | 0.00026695 | 54.509236 | 46.2744432 | 37.9649818 | 46.5062237 |
| -3.8973028 | 9.73E-05 | 0.00026703 | 40.1647002 | 38.6973414 | 48.3190678 | 40.9030642 |
| -3.897046 | 9.74E-05 | 0.00026724 | 17.6232868 | 98.2317128 | 87.4345036 | 107.580662 |
| -3.8970692 | 9.74E-05 | 0.00026724 | 109.428316 | 75.7710182 | 44.8677058 | 160.250361 |
| -3.8970279 | 9.74E-05 | 0.00026724 | 979.526873 | 3457.0527 | 4995.27125 | 4381.67072 |
| -3.8970581 | 9.74E-05 | 0.00026724 | 24.1807889 | 159.119138 | 40.2658898 | 165.853521 |
| -3.8967938 | 9.75E-05 | 0.00026748 | 173.363961 | 261.680623 | 246.197155 | 238.134278 |
| -3.8966761 | 9.75E-05 | 0.00026759 | 33.1973543 | 138.552719 | 143.806749 | 132.79488 |
| -3.896139 | 9.77E-05 | 0.00026816 | 34.4268859 | 43.2977247 | 46.0181598 | 48.1871716 |
| -3.8959625 | 9.78E-05 | 0.00026833 | 28.6890716 | 46.545054 | 64.4254237 | 79.5648647 |
| -3.8959131 | 9.78E-05 | 0.00026836 | 205.741628 | 58.1813175 | 83.9831417 | 74.5220212 |
| -3.8958201 | 9.79E-05 | 0.00026844 | 118.035037 | 230.019162 | 261.153057 | 201.713742 |
| -3.8957194 | 9.79E-05 | 0.00026853 | 33.1973543 | 40.0503953 | 79.3813257 | 57.1522268 |
| -3.8956462 | 9.79E-05 | 0.00026859 | 36.4761053 | 39.5091738 | 55.2217918 | 44.2649599 |
| -3.8954249 | 9.80E-05 | 0.00026877 | 20.9020379 | 115.821413 | 87.4345036 | 65.556966 |
| -3.8954436 | 9.80E-05 | 0.00026877 | 80.3294005 | 132.05806 | 92.0363196 | 93.5727634 |
| -3.8954557 | 9.80E-05 | 0.00026877 | 767.637587 | 1455.88599 | 1317.26982 | 1366.6106 |
| -3.8954016 | 9.80E-05 | 0.00026877 | 29.5087594 | 67.6526948 | 46.0181598 | 59.9538065 |
| -3.8950425 | 9.82E-05 | 0.00026914 | 20.492194 | 95.525605 | 79.3813257 | 77.3236009 |
| -3.8948171 | 9.83E-05 | 0.00026937 | 24.5906328 | 69.0057487 | 58.6731538 | 84.6077082 |
| -3.8947198 | 9.83E-05 | 0.00026946 | 62.2962697 | 62.51109 | 24.1595339 | 94.1330794 |
| -3.8946505 | 9.83E-05 | 0.00026951 | 46.3123584 | 98.7729344 | 28.7613499 | 64.4363341 |

| | | | | | | |
|---|---|---|---|---|---|---|
| -3.8946241 | 9.84E-05 | 0.00026952 | 38.5253247 | 52.4984912 | 41.4163438 | 39.2221164 |
| -3.8940522 | 9.86E-05 | 0.00027012 | 36.8859492 | 115.550803 | 29.9118039 | 110.382242 |
| 3.8940433 | 9.86E-05 | 0.00027012 | 0 | 0 | 0 | 0 |
| -3.8938644 | 9.87E-05 | 0.0002703 | 38.1154808 | 78.477126 | 65.5758777 | 33.058641 |
| -3.8931728 | 9.89E-05 | 0.00027104 | 27.45954 | 63.0523115 | 59.8236078 | 53.7903311 |
| -3.8931101 | 9.90E-05 | 0.00027109 | 14.3445358 | 78.7477367 | 142.656295 | 170.336048 |
| -3.8928654 | 9.91E-05 | 0.00027134 | 18.4429746 | 105.538204 | 81.6822337 | 93.0124475 |
| -3.892702 | 9.91E-05 | 0.0002715 | 26.6398522 | 51.416048 | 81.6822337 | 65.556966 |
| -3.8924539 | 9.92E-05 | 0.00027176 | 26.2300083 | 68.7351379 | 65.5758777 | 54.910963 |
| -3.8921743 | 9.93E-05 | 0.00027205 | 43.4434513 | 128.269509 | 59.8236078 | 33.6189569 |
| -3.8919259 | 9.95E-05 | 0.0002723 | 240.988201 | 56.5576528 | 35.6640739 | 50.9887513 |
| -3.891848 | 9.95E-05 | 0.00027237 | 31.1481349 | 81.1832337 | 125.399485 | 92.4521315 |
| -3.8912307 | 9.97E-05 | 0.00027304 | 45.4926707 | 66.5702517 | 28.7613499 | 105.899714 |
| -3.8910213 | 9.98E-05 | 0.00027325 | 40.5745441 | 45.4626109 | 35.6640739 | 46.5062237 |
| 3.89096254 | 9.98E-05 | 0.00027329 | 0 | 0 | 0 | 0 |
| -3.8907815 | 9.99E-05 | 0.00027347 | 35.2465737 | 43.5683354 | 41.4163438 | 53.7903311 |
| -3.8902828 | 0.00010013 | 0.00027401 | 18.4429746 | 80.1007906 | 104.691314 | 94.1330794 |
| -3.8898863 | 0.00010029 | 0.00027444 | 26.6398522 | 119.609964 | 232.391707 | 38.6618005 |
| -3.8897278 | 0.00010036 | 0.00027459 | 21.7217256 | 82.2656769 | 81.6822337 | 68.9188617 |
| -3.8893711 | 0.0001005 | 0.00027498 | 48.3615778 | 43.0271139 | 32.2127119 | 54.910963 |
| -3.8893406 | 0.00010052 | 0.00027499 | 36.0662614 | 36.5324552 | 52.9208838 | 56.5919108 |
| -3.888913 | 0.00010069 | 0.00027545 | 44.263139 | 46.545054 | 73.6290557 | 33.058641 |
| -3.8888282 | 0.00010073 | 0.00027552 | 45.9025146 | 88.7603356 | 26.4604419 | 81.2458125 |
| 3.88875561 | 0.00010076 | 0.00027558 | 17.213443 | 8.65954493 | 6.90272397 | 10.0856871 |
| -3.8887292 | 0.00010077 | 0.00027559 | 52.0501728 | 85.5130062 | 67.8767857 | 43.704644 |
| -3.8883198 | 0.00010094 | 0.00027603 | 39.7548564 | 103.914539 | 44.8677058 | 42.0236961 |
| -3.8882573 | 0.00010097 | 0.00027605 | 74.1817423 | 106.891258 | 16.1063559 | 206.756585 |
| -3.888269 | 0.00010096 | 0.00027605 | 41.3942319 | 52.7691019 | 36.8145279 | 40.3427483 |
| -3.888225 | 0.00010098 | 0.00027607 | 25.8201644 | 62.2404792 | 88.5849576 | 55.4712789 |
| -3.8877854 | 0.00010116 | 0.00027654 | 17.6232868 | 109.056144 | 101.239952 | 81.2458125 |
| -3.8874684 | 0.0001013 | 0.00027688 | 20.0823501 | 94.1725511 | 78.2308717 | 80.1251807 |
| -3.8871376 | 0.00010143 | 0.00027723 | 27.0496961 | 51.6866588 | 69.0272397 | 61.6347543 |
| -3.8870872 | 0.00010145 | 0.00027727 | 36.0662614 | 56.2870421 | 35.6640739 | 55.4712789 |
| -3.8865854 | 0.00010166 | 0.00027782 | 64.755333 | 70.0881918 | 24.1595339 | 61.6347543 |
| -3.8865157 | 0.00010169 | 0.00027788 | 36.0662614 | 50.8748265 | 46.0181598 | 38.6618005 |
| -3.8864535 | 0.00010172 | 0.00027792 | 121.313788 | 34.6381797 | 56.3722458 | 42.5840121 |
| -3.8860035 | 0.00010191 | 0.00027841 | 25.4103206 | 58.1813175 | 67.8767857 | 64.99665 |
| -3.8859295 | 0.00010194 | 0.00027848 | 8506.30973 | 49488.2168 | 89123.3701 | 46244.5562 |
| -3.8854604 | 0.00010214 | 0.00027899 | 43.0336074 | 47.0862756 | 71.3281477 | 33.058641 |
| -3.8852758 | 0.00010221 | 0.00027918 | 45.9025146 | 243.820312 | 523.456568 | 541.825522 |
| -3.8849107 | 0.00010237 | 0.00027958 | 29.5087594 | 60.075593 | 44.8677058 | 63.3157022 |
| -3.8845702 | 0.00010251 | 0.00027994 | 48.7714217 | 62.7817008 | 52.9208838 | 29.1364293 |
| -3.8845023 | 0.00010254 | 0.00028 | 32.7875104 | 53.5809343 | 55.2217918 | 40.3427483 |

| | | | | | | |
|---|---|---|---|---|---|---|
| -3.8844529 | 0.00010256 | 0.00028003 | 50.4107972 | 326.085989 | 284.162137 | 299.769033 |
| -3.8842005 | 0.00010267 | 0.0002803 | 65.5750208 | 65.4878086 | 63.2749697 | 24.6539017 |
| -3.8837556 | 0.00010286 | 0.00028079 | 684.029436 | 1207.4653 | 1074.52403 | 1079.16852 |
| -3.883413 | 0.000103 | 0.00028116 | 22.9512573 | 55.4752097 | 96.6381356 | 86.848972 |
| -3.8833004 | 0.00010305 | 0.00028127 | 34.4268859 | 61.9698684 | 56.3722458 | 36.9808526 |
| -3.8825803 | 0.00010335 | 0.00028208 | 44.263139 | 29.4965749 | 60.9740618 | 77.3236009 |
| -3.882342 | 0.00010346 | 0.00028233 | 68.8537718 | 141.800048 | 16.1063559 | 133.355196 |
| -3.8821471 | 0.00010354 | 0.00028253 | 26.6398522 | 61.4286469 | 58.6731538 | 57.1522268 |
| 3.88196842 | 0.00010361 | 0.00028272 | 21.3118818 | 11.6362635 | 12.6549939 | 17.9301104 |
| 3.88159512 | 0.00010377 | 0.00028313 | 6.96734596 | 5.68282636 | 3.45136199 | 3.92221164 |
| -3.8815456 | 0.00010379 | 0.00028316 | 70.0833035 | 27.8729103 | 46.0181598 | 71.1601255 |
| -3.8812893 | 0.0001039 | 0.00028344 | 27.8693838 | 72.5236888 | 62.1245158 | 47.6268556 |
| 3.88125778 | 0.00010392 | 0.00028344 | 24.1807889 | 9.74198805 | 18.4072639 | 14.5682147 |
| 3.88124291 | 0.00010392 | 0.00028344 | 0.81968776 | 1.08244312 | 3.45136199 | 0 |
| -3.8812131 | 0.00010394 | 0.00028345 | 42.2139196 | 90.1133895 | 83.9831417 | 83.4870763 |
| -3.8807859 | 0.00010412 | 0.00028393 | 16.8035991 | 123.398515 | 88.5849576 | 91.8918156 |
| -3.8805454 | 0.00010422 | 0.00028418 | 27.8693838 | 51.416048 | 56.3722458 | 61.6347543 |
| -3.880521 | 0.00010423 | 0.00028419 | 37.2957931 | 42.7565031 | 41.4163438 | 42.0236961 |
| -3.880339 | 0.00010431 | 0.00028438 | 34.017042 | 59.8049822 | 42.5667978 | 43.704644 |
| 3.87999545 | 0.00010446 | 0.00028476 | 227.463353 | 248.420695 | 199.028541 | 233.091435 |
| 3.87995341 | 0.00010448 | 0.00028478 | 2.86890716 | 0.81183234 | 1.150454 | 1.68094785 |
| -3.879936 | 0.00010448 | 0.00028478 | 40.5745441 | 47.3568864 | 48.3190678 | 34.1792729 |
| -3.8795143 | 0.00010467 | 0.00028525 | 21.3118818 | 74.688575 | 77.0804177 | 77.8839169 |
| -3.8792496 | 0.00010478 | 0.00028553 | 93.4444046 | 237.325653 | 136.904025 | 151.845622 |
| -3.8787293 | 0.000105 | 0.00028612 | 57.7879871 | 64.1347547 | 34.5136199 | 35.8602207 |
| -3.8783867 | 0.00010515 | 0.0002865 | 25.8201644 | 55.7458205 | 60.9740618 | 70.5998095 |
| -3.8778255 | 0.00010539 | 0.00028714 | 34.4268859 | 33.0145151 | 87.4345036 | 113.183822 |
| -3.8775201 | 0.00010553 | 0.00028747 | 42.6237635 | 53.5809343 | 59.8236078 | 30.8173772 |
| -3.8774178 | 0.00010557 | 0.00028756 | 18.0331307 | 97.1492697 | 74.7795097 | 114.304454 |
| -3.8774023 | 0.00010558 | 0.00028756 | 86.8869025 | 47.8981079 | 49.4695218 | 30.8173772 |
| -3.876763 | 0.00010586 | 0.0002883 | 4.0984388 | 196.734036 | 254.250333 | 274.554815 |
| -3.8763993 | 0.00010601 | 0.0002887 | 27.0496961 | 67.9233056 | 58.6731538 | 51.5490673 |
| -3.8762435 | 0.00010608 | 0.00028886 | 27.8693838 | 79.8301798 | 44.8677058 | 68.3585457 |
| 3.87621204 | 0.0001061 | 0.00028888 | 1.63937552 | 1.89427545 | 0 | 2.24126379 |
| 3.87613221 | 0.00010613 | 0.00028895 | 0.40984388 | 0.27061078 | 0 | 0 |
| -3.8760865 | 0.00010615 | 0.00028898 | 28.2792277 | 77.9359044 | 48.3190678 | 56.5919108 |
| -3.8757589 | 0.00010629 | 0.00028934 | 86.4770587 | 49.7923834 | 31.0622579 | 45.9459078 |
| -3.8754896 | 0.00010641 | 0.00028964 | 29.5087594 | 49.5217726 | 50.6199758 | 82.9267604 |
| -3.8754509 | 0.00010643 | 0.00028966 | 89.7558097 | 148.294707 | 46.0181598 | 25.2142177 |
| -3.8753874 | 0.00010646 | 0.00028971 | 45.4926707 | 41.9446708 | 42.5667978 | 35.2999048 |
| -3.8753636 | 0.00010647 | 0.00028971 | 93.0345607 | 40.8622277 | 33.3631659 | 53.2300151 |
| -3.8752732 | 0.00010651 | 0.0002898 | 26.6398522 | 48.4393295 | 73.6290557 | 67.7982298 |
| 3.87469938 | 0.00010676 | 0.00029046 | 38.1154808 | 30.579018 | 46.0181598 | 42.5840121 |

| | | | | | | |
|---|---|---|---|---|---|---|
| -3.8743985 | 0.00010689 | 0.00029079 | 292.218686 | 44.3801678 | 48.3190678 | 46.5062237 |
| -3.8741759 | 0.00010699 | 0.00029103 | 149.593016 | 280.893989 | 212.833989 | 245.418385 |
| -3.8738844 | 0.00010711 | 0.00029136 | 25.4103206 | 61.9698684 | 57.5226998 | 66.6775979 |
| -3.8735525 | 0.00010726 | 0.00029173 | 19.6725062 | 81.9950661 | 74.7795097 | 95.2537113 |
| -3.8733662 | 0.00010734 | 0.00029193 | 67.2143963 | 23.272527 | 67.8767857 | 71.1601255 |
| -3.8730193 | 0.0001075 | 0.00029232 | 12.7051603 | 155.871809 | 93.1867736 | 136.717091 |
| -3.8728981 | 0.00010755 | 0.00029244 | 36.8859492 | 56.2870421 | 37.9649818 | 42.5840121 |
| -3.872132 | 0.00010789 | 0.00029334 | 35.2465737 | 45.1920001 | 39.1154358 | 47.6268556 |
| -3.872049 | 0.00010792 | 0.00029341 | 26.2300083 | 92.8194972 | 121.948124 | 119.907613 |
| -3.871254 | 0.00010828 | 0.00029435 | 137.2977 | 116.903857 | 13.8054479 | 91.3314996 |
| -3.871075 | 0.00010836 | 0.00029454 | 29.5087594 | 55.4752097 | 108.142676 | 48.1871716 |
| -3.8710371 | 0.00010837 | 0.00029456 | 42.6237635 | 38.9679522 | 36.8145279 | 44.8252759 |
| -3.8709704 | 0.0001084 | 0.00029461 | 81.968776 | 27.6022995 | 46.0181598 | 64.99665 |
| -3.8708793 | 0.00010844 | 0.0002947 | 34.8367298 | 38.1561199 | 64.4254237 | 47.0665397 |
| -3.8705657 | 0.00010858 | 0.00029505 | 267.21821 | 450.025726 | 478.588862 | 417.995698 |
| -3.870207 | 0.00010874 | 0.00029546 | 37.705637 | 35.4500121 | 46.0181598 | 53.2300151 |
| -3.8696354 | 0.000109 | 0.00029613 | 26.6398522 | 96.3374374 | 46.0181598 | 72.2807574 |
| -3.8695501 | 0.00010904 | 0.00029621 | 20.492194 | 151.812647 | 58.6731538 | 91.8918156 |
| -3.8693948 | 0.00010911 | 0.00029637 | 39.3450125 | 102.290875 | 95.4876816 | 92.4521315 |
| -3.869269 | 0.00010916 | 0.00029648 | 24.5906328 | 53.3103235 | 81.6822337 | 69.4791776 |
| 3.8692677 | 0.00010916 | 0.00029648 | 19.6725062 | 12.7187066 | 12.6549939 | 8.40473923 |
| -3.8689284 | 0.00010931 | 0.00029687 | 506.157192 | 1800.1029 | 1817.71731 | 1824.38873 |
| -3.8688407 | 0.00010935 | 0.00029693 | 73.3620545 | 22.7313054 | 97.7885896 | 58.8331746 |
| -3.8688379 | 0.00010936 | 0.00029693 | 35.2465737 | 36.5324552 | 63.2749697 | 49.3078035 |
| -3.8687637 | 0.00010939 | 0.00029699 | 33.1973543 | 45.4626109 | 44.8677058 | 84.0473923 |
| -3.8687329 | 0.0001094 | 0.000297 | 180.741151 | 41.9446708 | 44.8677058 | 46.5062237 |
| -3.868154 | 0.00010966 | 0.00029769 | 38.5253247 | 88.7603356 | 55.2217918 | 84.0473923 |
| -3.8680466 | 0.00010971 | 0.00029779 | 27.8693838 | 82.8068984 | 46.0181598 | 61.6347543 |
| -3.8678883 | 0.00010978 | 0.00029796 | 24.5906328 | 47.3568864 | 123.098578 | 85.7283401 |
| -3.8677405 | 0.00010985 | 0.00029812 | 29.9186032 | 33.0145151 | 125.399485 | 139.518671 |
| -3.8676879 | 0.00010987 | 0.00029815 | 20.9020379 | 95.525605 | 59.8236078 | 94.6933953 |
| -3.8676301 | 0.0001099 | 0.0002982 | 50.0009534 | 58.4519283 | 37.9649818 | 33.6189569 |
| -3.8674019 | 0.00011 | 0.00029845 | 33.6071982 | 240.843593 | 34.5136199 | 89.0902358 |
| -3.8672123 | 0.00011009 | 0.00029866 | 45.0828268 | 108.514922 | 82.8326877 | 25.7745336 |
| 3.86707645 | 0.00011015 | 0.0002988 | 0 | 0 | 0 | 0 |
| -3.8669242 | 0.00011022 | 0.00029896 | 18.8528185 | 98.2317128 | 83.9831417 | 77.3236009 |
| -3.8662004 | 0.00011054 | 0.00029983 | 19.6725062 | 66.5702517 | 93.1867736 | 95.8140272 |
| -3.8659962 | 0.00011064 | 0.00030005 | 27.0496961 | 62.51109 | 51.7704298 | 56.5919108 |
| -3.8657356 | 0.00011075 | 0.00030035 | 21.7217256 | 88.219114 | 85.1335957 | 59.3934906 |
| -3.8653866 | 0.00011091 | 0.00030075 | 77.0506494 | 50.6042157 | 31.0622579 | 42.5840121 |
| -3.8647515 | 0.0001112 | 0.0003015 | 26.6398522 | 55.2045989 | 55.2217918 | 61.0744384 |
| -3.8647434 | 0.00011121 | 0.0003015 | 35.6564176 | 51.1454373 | 37.9649818 | 43.704644 |
| -3.8641747 | 0.00011147 | 0.00030217 | 71.722679 | 89.0309463 | 81.6822337 | 77.3236009 |

| | | | | | | |
|---|---|---|---|---|---|---|
| -3.8640966 | 0.0001115 | 0.00030224 | 25.8201644 | 63.8641439 | 60.9740618 | 54.350647 |
| -3.8640422 | 0.00011153 | 0.00030229 | 47.1320462 | 32.7439043 | 60.9740618 | 40.3427483 |
| -3.8638067 | 0.00011163 | 0.00030255 | 23.770945 | 69.817581 | 71.3281477 | 58.2728587 |
| -3.863689 | 0.00011169 | 0.00030267 | 45.0828268 | 71.7118565 | 27.6108959 | 59.9538065 |
| -3.8634831 | 0.00011178 | 0.0003029 | 36.8859492 | 74.9591858 | 74.7795097 | 62.7553862 |
| -3.8631945 | 0.00011191 | 0.00030324 | 356.974019 | 52.2278804 | 34.5136199 | 54.350647 |
| -3.8630486 | 0.00011198 | 0.00030339 | 29.9186032 | 87.4072817 | 43.7172518 | 56.0315949 |
| -3.8630257 | 0.00011199 | 0.00030339 | 47.951734 | 42.4858923 | 31.0622579 | 53.2300151 |
| -3.862916 | 0.00011204 | 0.00030349 | 40.984388 | 32.4732935 | 78.2308717 | 49.8681194 |
| -3.8628905 | 0.00011205 | 0.00030349 | 21.3118818 | 68.1939163 | 83.9831417 | 72.8410733 |
| -3.8628871 | 0.00011205 | 0.00030349 | 396.319032 | 615.368912 | 433.721156 | 519.9732 |
| -3.8620511 | 0.00011244 | 0.00030451 | 70.0833035 | 69.2763595 | 63.2749697 | 22.9729539 |
| 3.86161385 | 0.00011264 | 0.00030503 | 46.3123584 | 13.8011497 | 13.8054479 | 11.7666349 |
| -3.8614616 | 0.00011271 | 0.00030519 | 24.1807889 | 94.9843835 | 108.142676 | 45.9459078 |
| -3.8611133 | 0.00011287 | 0.0003056 | 23.770945 | 63.3229223 | 62.1245158 | 70.5998095 |
| -3.8609108 | 0.00011297 | 0.00030583 | 305.333691 | 171.026012 | 132.302209 | 158.569413 |
| -3.8606886 | 0.00011307 | 0.00030608 | 29.9186032 | 38.4267306 | 97.7885896 | 73.9617052 |
| -3.8604164 | 0.00011319 | 0.0003064 | 19.2626624 | 84.7011739 | 86.2840497 | 76.202969 |
| -3.8602589 | 0.00011327 | 0.00030657 | 11.8854725 | 113.927138 | 136.904025 | 126.071088 |
| -3.8601514 | 0.00011332 | 0.00030668 | 39.7548564 | 61.4286469 | 57.5226998 | 30.8173772 |
| -3.8598112 | 0.00011347 | 0.00030705 | 35.6564176 | 46.0038325 | 36.8145279 | 48.1871716 |
| -3.8598195 | 0.00011347 | 0.00030705 | 27.45954 | 69.0057487 | 49.4695218 | 54.350647 |
| -3.8597476 | 0.0001135 | 0.00030708 | 28.6890716 | 73.6061319 | 57.5226998 | 44.2649599 |
| -3.8597479 | 0.0001135 | 0.00030708 | 102.46097 | 45.7332217 | 57.5226998 | 29.1364293 |
| -3.8593885 | 0.00011367 | 0.00030746 | 14.7543797 | 95.7962158 | 139.204933 | 94.6933953 |
| -3.8594054 | 0.00011366 | 0.00030746 | 40.1647002 | 57.640096 | 49.4695218 | 31.9380091 |
| -3.8594232 | 0.00011365 | 0.00030746 | 19.6725062 | 80.912623 | 92.0363196 | 71.7204414 |
| -3.8592093 | 0.00011375 | 0.00030766 | 25.8201644 | 86.8660601 | 50.6199758 | 62.1950703 |
| -3.8588016 | 0.00011394 | 0.00030814 | 23.3611012 | 107.973701 | 57.5226998 | 67.2379138 |
| -3.8584741 | 0.0001141 | 0.0003085 | 33.1973543 | 44.3801678 | 44.8677058 | 43.144328 |
| -3.8584812 | 0.00011409 | 0.0003085 | 30.3284471 | 45.1920001 | 56.3722458 | 47.0665397 |
| -3.8583661 | 0.00011415 | 0.00030862 | 23.770945 | 76.0416289 | 90.8858656 | 51.5490673 |
| -3.8581236 | 0.00011426 | 0.0003089 | 27.8693838 | 49.5217726 | 51.7704298 | 65.556966 |
| -3.8576494 | 0.00011448 | 0.00030947 | 109.428316 | 53.5809343 | 33.3631659 | 40.9030642 |
| -3.8574602 | 0.00011457 | 0.00030968 | 26.2300083 | 56.5576528 | 69.0272397 | 52.1093832 |
| -3.857283 | 0.00011465 | 0.00030988 | 30.3284471 | 102.020264 | 41.4163438 | 61.0744384 |
| 3.8571136 | 0.00011473 | 0.00031007 | 0.40984388 | 0.54122156 | 0 | 0.56031595 |
| 3.85667672 | 0.00011494 | 0.0003106 | 141.396139 | 87.9485032 | 106.992222 | 89.6505518 |
| -3.8561334 | 0.00011519 | 0.00031126 | 38.1154808 | 57.0988744 | 40.2658898 | 36.9808526 |
| -3.8560526 | 0.00011523 | 0.00031134 | 28.2792277 | 308.496288 | 427.968886 | 403.427483 |
| -3.8559553 | 0.00011528 | 0.00031144 | 47.5418901 | 39.5091738 | 41.4163438 | 35.2999048 |
| -3.8558429 | 0.00011533 | 0.00031156 | 34.017042 | 54.1221558 | 35.6640739 | 55.4712789 |
| -3.8556439 | 0.00011543 | 0.00031178 | 31.5579788 | 60.075593 | 65.5758777 | 37.5411686 |

| | | | | | | |
|---|---|---|---|---|---|---|
| -3.8555439 | 0.00011547 | 0.00031188 | 31.5579788 | 97.6904913 | 39.1154358 | 60.5141225 |
| -3.8551687 | 0.00011565 | 0.00031234 | 25.8201644 | 53.0397127 | 57.5226998 | 69.4791776 |
| 3.85469034 | 0.00011588 | 0.00031292 | 58.197831 | 21.1076408 | 37.9649818 | 26.8951655 |
| -3.8546671 | 0.00011589 | 0.00031293 | 43.0336074 | 40.0503953 | 33.3631659 | 53.2300151 |
| -3.8542876 | 0.00011607 | 0.00031339 | 29.9186032 | 54.6633774 | 46.0181598 | 47.6268556 |
| 3.85385039 | 0.00011627 | 0.00031392 | 0.40984388 | 0 | 0 | 0 |
| -3.8537705 | 0.00011631 | 0.00031397 | 49.1812656 | 46.2744432 | 33.3631659 | 38.6618005 |
| -3.8537708 | 0.00011631 | 0.00031397 | 18.8528185 | 66.8408624 | 93.1867736 | 102.537819 |
| -3.8536535 | 0.00011637 | 0.00031409 | 32.7875104 | 63.5935331 | 67.8767857 | 35.2999048 |
| -3.8529032 | 0.00011673 | 0.00031503 | 28.6890716 | 58.4519283 | 95.4876816 | 44.2649599 |
| -3.852862 | 0.00011675 | 0.00031506 | 71.722679 | 135.034779 | 150.709473 | 119.347297 |
| -3.8527552 | 0.0001168 | 0.00031517 | 28.2792277 | 57.3694852 | 44.8677058 | 60.5141225 |
| -3.8523357 | 0.000117 | 0.00031568 | 31.9678226 | 39.7797845 | 51.7704298 | 49.8681194 |
| -3.8522988 | 0.00011701 | 0.0003157 | 163.117864 | 50.3336049 | 34.5136199 | 47.0665397 |
| -3.8522571 | 0.00011703 | 0.00031573 | 54.509236 | 38.4267306 | 130.001301 | 36.4205367 |
| -3.8521887 | 0.00011707 | 0.00031579 | 42.6237635 | 31.6614612 | 49.4695218 | 46.5062237 |
| -3.8518068 | 0.00011725 | 0.00031626 | 35.2465737 | 81.9950661 | 46.0181598 | 39.2221164 |
| -3.8517127 | 0.00011729 | 0.00031636 | 28.6890716 | 53.0397127 | 44.8677058 | 64.99665 |
| -3.8514433 | 0.00011742 | 0.00031668 | 29.0989155 | 98.2317128 | 39.1154358 | 72.2807574 |
| -3.8506193 | 0.00011782 | 0.00031772 | 47.1320462 | 42.7565031 | 31.0622579 | 74.5220212 |
| -3.8505845 | 0.00011784 | 0.00031774 | 16.3937552 | 75.2297966 | 119.647216 | 103.098135 |
| -3.8504718 | 0.00011789 | 0.00031786 | 25.8201644 | 58.7225391 | 66.7263317 | 51.5490673 |
| -3.8499152 | 0.00011816 | 0.00031855 | 75.00143 | 39.5091738 | 35.6640739 | 42.5840121 |
| 3.84881295 | 0.00011869 | 0.00031996 | 0 | 0 | 0 | 0 |
| -3.8487766 | 0.00011871 | 0.00031998 | 22.1315695 | 61.4286469 | 75.9299637 | 72.2807574 |
| 3.84859159 | 0.0001188 | 0.0003202 | 320.907758 | 371.5486 | 500.447488 | 383.256109 |
| -3.8482651 | 0.00011896 | 0.0003206 | 101.641282 | 51.6866588 | 28.7613499 | 48.1871716 |
| -3.8481088 | 0.00011903 | 0.00032078 | 35.2465737 | 63.5935331 | 46.0181598 | 36.4205367 |
| -3.8479773 | 0.0001191 | 0.00032092 | 25.0004767 | 60.075593 | 56.3722458 | 62.1950703 |
| -3.8478691 | 0.00011915 | 0.00032104 | 23.770945 | 66.8408624 | 57.5226998 | 67.2379138 |
| -3.8475467 | 0.00011931 | 0.00032143 | 63.5258014 | 121.774851 | 148.408565 | 16.2491625 |
| -3.8469561 | 0.00011959 | 0.00032218 | 38.1154808 | 54.6633774 | 31.0622579 | 59.3934906 |
| -3.8469071 | 0.00011962 | 0.00032222 | 20.0823501 | 138.282108 | 80.5317797 | 61.6347543 |
| -3.8468658 | 0.00011964 | 0.00032225 | 29.9186032 | 52.2278804 | 44.8677058 | 48.1871716 |
| -3.8467579 | 0.00011969 | 0.00032236 | 27.8693838 | 44.9213893 | 74.7795097 | 56.0315949 |
| -3.846601 | 0.00011977 | 0.00032254 | 645.094267 | 1426.11881 | 1213.72897 | 1274.15847 |
| -3.8465265 | 0.0001198 | 0.00032261 | 31.5579788 | 52.7691019 | 47.1686138 | 40.9030642 |
| -3.8464837 | 0.00011983 | 0.00032264 | 29.5087594 | 54.6633774 | 71.3281477 | 41.4633802 |
| -3.846247 | 0.00011994 | 0.00032293 | 36.8859492 | 36.2618444 | 49.4695218 | 41.4633802 |
| -3.846168 | 0.00011998 | 0.000323 | 24.1807889 | 52.4984912 | 95.4876816 | 64.99665 |
| -3.8460274 | 0.00012005 | 0.00032316 | 61.8864259 | 89.0309463 | 28.7613499 | 40.9030642 |
| -3.8455656 | 0.00012027 | 0.00032374 | 23.770945 | 69.817581 | 64.4254237 | 57.1522268 |
| -3.8452758 | 0.00012042 | 0.0003241 | 53.6895483 | 40.8622277 | 36.8145279 | 36.4205367 |

| | | | | | | |
|---|---|---|---|---|---|---|
| -3.8448164 | 0.00012064 | 0.00032468 | 10.246097 | 124.210348 | 144.957203 | 136.156776 |
| -3.8443699 | 0.00012086 | 0.00032519 | 14.3445358 | 104.996982 | 95.4876816 | 115.425085 |
| -3.8443904 | 0.00012085 | 0.00032519 | 27.8693838 | 54.3927666 | 49.4695218 | 51.5490673 |
| -3.8443811 | 0.00012086 | 0.00032519 | 28.6890716 | 43.8389462 | 55.2217918 | 56.5919108 |
| -3.8439772 | 0.00012106 | 0.00032568 | 31.5579788 | 216.488623 | 111.594038 | 29.6967453 |
| -3.8437939 | 0.00012115 | 0.0003259 | 31.5579788 | 47.3568864 | 42.5667978 | 80.6854966 |
| -3.8431448 | 0.00012147 | 0.00032674 | 25.0004767 | 69.817581 | 49.4695218 | 68.3585457 |
| -3.8430833 | 0.0001215 | 0.00032679 | 33.1973543 | 88.7603356 | 51.7704298 | 38.6618005 |
| -3.8428452 | 0.00012162 | 0.00032708 | 19.6725062 | 109.597366 | 74.7795097 | 67.7982298 |
| -3.8425412 | 0.00012177 | 0.00032746 | 36.8859492 | 94.9843835 | 28.7613499 | 81.2458125 |
| -3.842374 | 0.00012185 | 0.00032766 | 229.512573 | 765.828505 | 620.094704 | 616.347543 |
| -3.8422024 | 0.00012194 | 0.00032786 | 143.855202 | 46.2744432 | 36.8145279 | 44.2649599 |
| -3.8421799 | 0.00012195 | 0.00032786 | 38.1154808 | 56.0164313 | 36.8145279 | 38.6618005 |
| -3.8420025 | 0.00012203 | 0.00032807 | 33.1973543 | 193.216096 | 103.54086 | 137.837723 |
| -3.8419495 | 0.00012206 | 0.00032811 | 63.9356453 | 32.4732935 | 70.1776937 | 34.7395888 |
| -3.8417537 | 0.00012216 | 0.00032835 | 13.9346919 | 110.409198 | 124.249032 | 91.3314996 |
| -3.8417068 | 0.00012218 | 0.00032838 | 66.8045524 | 46.545054 | 32.2127119 | 39.2221164 |
| -3.8413122 | 0.00012238 | 0.00032889 | 327.055416 | 555.022708 | 294.516223 | 564.798476 |
| -3.8412169 | 0.00012243 | 0.00032899 | 28.6890716 | 42.7565031 | 60.9740618 | 54.910963 |
| -3.8411904 | 0.00012244 | 0.00032899 | 22.1315695 | 65.4878086 | 62.1245158 | 82.3664444 |
| -3.8406518 | 0.00012271 | 0.00032969 | 38.5253247 | 98.5023236 | 85.1335957 | 110.382242 |
| -3.840466 | 0.0001228 | 0.00032991 | 40.984388 | 62.2404792 | 31.0622579 | 45.9459078 |
| -3.8403555 | 0.00012286 | 0.00033001 | 72.5423667 | 48.980551 | 103.54086 | 75.6426531 |
| -3.8403541 | 0.00012286 | 0.00033001 | 34.4268859 | 57.9107067 | 34.5136199 | 92.4521315 |
| -3.8403262 | 0.00012287 | 0.00033002 | 25.8201644 | 59.5343714 | 51.7704298 | 58.8331746 |
| -3.8398966 | 0.00012309 | 0.00033057 | 85.247527 | 64.1347547 | 20.7081719 | 68.9188617 |
| -3.8398635 | 0.0001231 | 0.00033058 | 32.3776665 | 60.8874253 | 40.2658898 | 44.2649599 |
| -3.839786 | 0.00012314 | 0.00033065 | 23.3611012 | 54.1221558 | 73.6290557 | 70.0394936 |
| -3.839772 | 0.00012315 | 0.00033065 | 25.8201644 | 60.075593 | 54.0713378 | 54.910963 |
| 3.83967393 | 0.0001232 | 0.00033076 | 8.1968776 | 8.11832337 | 8.05317797 | 5.60315949 |
| -3.8396512 | 0.00012321 | 0.00033076 | 43.0336074 | 50.8748265 | 31.0622579 | 43.144328 |
| -3.8394851 | 0.00012329 | 0.00033096 | 15.9839113 | 76.8534613 | 119.647216 | 99.7362389 |
| -3.8394575 | 0.00012331 | 0.00033097 | 176.642712 | 93.9019404 | 51.7704298 | 67.2379138 |
| -3.8389049 | 0.00012358 | 0.00033168 | 194.675843 | 386.702803 | 265.754873 | 468.424133 |
| -3.8387678 | 0.00012365 | 0.00033184 | 23.770945 | 105.808815 | 56.3722458 | 61.0744384 |
| -3.8386522 | 0.00012371 | 0.00033197 | 18.0331307 | 94.9843835 | 86.2840497 | 74.5220212 |
| -3.8385148 | 0.00012378 | 0.00033213 | 65.1651769 | 45.1920001 | 55.2217918 | 27.4554815 |
| -3.8381781 | 0.00012395 | 0.00033256 | 22.5414134 | 123.669126 | 139.204933 | 41.4633802 |
| -3.8380219 | 0.00012403 | 0.00033274 | 23.3611012 | 59.2637606 | 63.2749697 | 67.7982298 |
| -3.8371938 | 0.00012445 | 0.00033384 | 25.8201644 | 54.1221558 | 56.3722458 | 56.5919108 |
| -3.8369574 | 0.00012457 | 0.00033413 | 70.9029912 | 38.1561199 | 43.7172518 | 34.7395888 |
| -3.8369174 | 0.00012459 | 0.00033416 | 83.6081515 | 51.9572696 | 51.7704298 | 26.3348496 |
| -3.8368811 | 0.00012461 | 0.00033418 | 31.9678226 | 87.6778924 | 33.3631659 | 75.0823371 |

| | | | | | | |
|---|---|---|---|---|---|---|
| 3.83567671 | 0.00012522 | 0.00033579 | 106.969253 | 59.2637606 | 58.6731538 | 62.7553862 |
| -3.8355859 | 0.00012527 | 0.00033589 | 24.5906328 | 49.7923834 | 64.4254237 | 80.1251807 |
| -3.8352757 | 0.00012542 | 0.00033629 | 65.9848647 | 61.4286469 | 43.7172518 | 47.0665397 |
| -3.8351504 | 0.00012549 | 0.00033643 | 14.7543797 | 97.4198805 | 93.1867736 | 112.623506 |
| -3.8350756 | 0.00012553 | 0.0003365 | 19.2626624 | 97.1492697 | 66.7263317 | 80.1251807 |
| -3.8347318 | 0.0001257 | 0.00033695 | 49.1812656 | 28.143521 | 67.8767857 | 46.5062237 |
| -3.8347009 | 0.00012572 | 0.00033696 | 25.4103206 | 67.1114732 | 58.6731538 | 50.4284354 |
| -3.8346246 | 0.00012576 | 0.00033704 | 188.118341 | 278.187881 | 255.400787 | 272.873867 |
| -3.8345203 | 0.00012581 | 0.00033712 | 22.9512573 | 88.4897248 | 59.8236078 | 59.9538065 |
| -3.8345009 | 0.00012582 | 0.00033712 | 32.3776665 | 56.8282636 | 36.8145279 | 49.8681194 |
| -3.834503 | 0.00012582 | 0.00033712 | 86.4770587 | 41.1328384 | 32.2127119 | 47.0665397 |
| -3.8343752 | 0.00012588 | 0.00033727 | 34.4268859 | 100.396599 | 125.399485 | 123.829825 |
| -3.8340651 | 0.00012604 | 0.00033767 | 28.2792277 | 83.0775092 | 77.0804177 | 38.1014845 |
| -3.8337977 | 0.00012618 | 0.000338 | 23.770945 | 105.267593 | 60.9740618 | 55.4712789 |
| -3.8337245 | 0.00012622 | 0.00033808 | 20.9020379 | 89.0309463 | 75.9299637 | 59.9538065 |
| -3.8332989 | 0.00012644 | 0.00033863 | 92.214873 | 39.5091738 | 36.8145279 | 42.5840121 |
| -3.8331974 | 0.00012649 | 0.00033875 | 56.5584554 | 77.3946828 | 23.0090799 | 59.9538065 |
| -3.8318928 | 0.00012716 | 0.00034052 | 120.494101 | 43.2977247 | 40.2658898 | 39.2221164 |
| -3.8316585 | 0.00012728 | 0.00034082 | 49.1812656 | 72.5236888 | 26.4604419 | 50.4284354 |
| -3.831567 | 0.00012733 | 0.00034091 | 32.3776665 | 40.3210061 | 42.5667978 | 58.8331746 |
| -3.8315266 | 0.00012735 | 0.00034094 | 41.8040758 | 83.34812 | 59.8236078 | 72.8410733 |
| -3.8311639 | 0.00012754 | 0.00034142 | 47.951734 | 27.0610779 | 54.0713378 | 57.1522268 |
| -3.8308522 | 0.0001277 | 0.00034182 | 28.6890716 | 58.1813175 | 46.0181598 | 47.0665397 |
| -3.8306379 | 0.00012781 | 0.00034209 | 27.45954 | 64.4053654 | 43.7172518 | 57.1522268 |
| 3.83028017 | 0.000128 | 0.00034256 | 12.7051603 | 12.7187066 | 10.354086 | 15.1285306 |
| -3.8301337 | 0.00012807 | 0.00034274 | 23.770945 | 42.2152815 | 111.594038 | 100.296555 |
| -3.8300104 | 0.00012814 | 0.00034288 | 30.738291 | 37.3442875 | 77.0804177 | 57.1522268 |
| -3.82963 | 0.00012834 | 0.00034338 | 94.2640924 | 36.2618444 | 109.29313 | 27.4554815 |
| -3.8291753 | 0.00012857 | 0.00034399 | 36.8859492 | 46.2744432 | 33.3631659 | 68.9188617 |
| 3.8289922 | 0.00012867 | 0.00034422 | 24.1807889 | 17.0484791 | 20.7081719 | 16.8094785 |
| -3.8287308 | 0.00012881 | 0.00034455 | 25.0004767 | 81.9950661 | 47.1686138 | 64.99665 |
| -3.8286402 | 0.00012885 | 0.00034465 | 46.7222023 | 71.1706349 | 24.1595339 | 75.0823371 |
| -3.8278058 | 0.00012929 | 0.00034579 | 26.2300083 | 37.6148983 | 153.010381 | 94.6933953 |
| -3.8269895 | 0.00012972 | 0.00034691 | 30.738291 | 42.2152815 | 43.7172518 | 53.7903311 |
| -3.8262211 | 0.00013013 | 0.00034797 | 25.0004767 | 79.8301798 | 43.7172518 | 79.5648647 |
| -3.8257953 | 0.00013035 | 0.00034854 | 28.6890716 | 51.416048 | 42.5667978 | 56.5919108 |
| 3.82572971 | 0.00013039 | 0.0003486 | 5.32797044 | 5.41221558 | 5.75226998 | 5.60315949 |
| -3.8256056 | 0.00013045 | 0.00034872 | 27.0496961 | 83.0775092 | 97.7885896 | 38.1014845 |
| -3.8256197 | 0.00013044 | 0.00034872 | 9.42640924 | 193.486707 | 123.098578 | 117.666349 |
| -3.8253213 | 0.0001306 | 0.0003491 | 49.1812656 | 43.2977247 | 28.7613499 | 49.8681194 |
| -3.8253001 | 0.00013061 | 0.0003491 | 32.7875104 | 77.6652936 | 77.0804177 | 31.9380091 |
| -3.8250445 | 0.00013075 | 0.00034943 | 27.8693838 | 36.5324552 | 116.195854 | 84.6077082 |
| -3.8246129 | 0.00013098 | 0.00035001 | 165.576927 | 63.8641439 | 128.850847 | 14.5682147 |

| | | | | | | |
|---|---|---|---|---|---|---|
| -3.8243176 | 0.00013113 | 0.00035041 | 56.5584554 | 109.597366 | 20.7081719 | 72.8410733 |
| -3.8240771 | 0.00013126 | 0.00035072 | 13.9346919 | 105.538204 | 96.6381356 | 107.020346 |
| -3.8238128 | 0.0001314 | 0.00035107 | 114.346443 | 28.6847426 | 50.6199758 | 48.7474875 |
| -3.8236952 | 0.00013147 | 0.0003512 | 41.3942319 | 71.9824673 | 75.9299637 | 26.3348496 |
| -3.8233546 | 0.00013165 | 0.00035166 | 29.5087594 | 65.7584193 | 67.8767857 | 36.9808526 |
| -3.8224922 | 0.00013211 | 0.00035286 | 36.4761053 | 104.455761 | 93.1867736 | 102.537819 |
| -3.8221373 | 0.0001323 | 0.00035334 | 36.4761053 | 102.832096 | 26.4604419 | 99.1759229 |
| -3.8217874 | 0.00013249 | 0.00035382 | 31.1481349 | 115.550803 | 143.806749 | 28.5761134 |
| 3.82168144 | 0.00013254 | 0.00035394 | 33.6071982 | 18.6721438 | 17.2568099 | 11.7666349 |
| -3.8210051 | 0.00013291 | 0.00035488 | 20.492194 | 61.1580361 | 80.5317797 | 76.202969 |
| -3.820666 | 0.00013309 | 0.00035534 | 55.3289238 | 34.367569 | 35.6640739 | 43.144328 |
| -3.8203708 | 0.00013325 | 0.00035574 | 22.1315695 | 82.8068984 | 48.3190678 | 114.304454 |
| -3.8202142 | 0.00013334 | 0.00035593 | 15.5740674 | 101.749653 | 103.54086 | 79.0045488 |
| 3.82012518 | 0.00013338 | 0.00035603 | 153.691455 | 183.474108 | 112.744492 | 100.296555 |
| -3.8197133 | 0.00013361 | 0.0003566 | 21.3118818 | 66.2996409 | 75.9299637 | 62.7553862 |
| -3.8196891 | 0.00013362 | 0.0003566 | 964.772493 | 3701.14363 | 3102.77443 | 3307.54504 |
| -3.8193806 | 0.00013379 | 0.00035702 | 46.7222023 | 45.1920001 | 31.0622579 | 39.7824324 |
| -3.8193218 | 0.00013382 | 0.00035708 | 26.2300083 | 64.1347547 | 50.6199758 | 49.8681194 |
| -3.8189681 | 0.00013401 | 0.00035756 | 18.4429746 | 64.946587 | 82.8326877 | 104.218766 |
| -3.8184802 | 0.00013428 | 0.00035821 | 42.2139196 | 165.613797 | 190.975363 | 175.939208 |
| -3.8184772 | 0.00013428 | 0.00035821 | 16.3937552 | 100.125988 | 83.9831417 | 83.4870763 |
| -3.8180747 | 0.0001345 | 0.00035871 | 27.8693838 | 71.7118565 | 37.9649818 | 76.763285 |
| -3.8181095 | 0.00013448 | 0.00035871 | 63.1159575 | 28.6847426 | 41.4163438 | 51.5490673 |
| -3.8180802 | 0.00013449 | 0.00035871 | 52.4600166 | 85.2423954 | 131.151755 | 54.910963 |
| -3.8175031 | 0.00013481 | 0.00035951 | 45.9025146 | 80.3714014 | 29.9118039 | 43.144328 |
| -3.8170543 | 0.00013505 | 0.00036013 | 55.3289238 | 50.0629941 | 94.3372276 | 80.6854966 |
| -3.8163339 | 0.00013545 | 0.00036116 | 113.936599 | 113.656527 | 11.50454 | 116.545717 |
| -3.8159168 | 0.00013568 | 0.00036174 | 374.597306 | 882.461751 | 762.750999 | 809.656546 |
| -3.815653 | 0.00013582 | 0.00036209 | 56.5584554 | 52.7691019 | 25.3099879 | 50.4284354 |
| -3.8155427 | 0.00013588 | 0.00036223 | 33.6071982 | 46.2744432 | 88.5849576 | 38.1014845 |
| -3.8151013 | 0.00013613 | 0.00036284 | 111.067691 | 53.5809343 | 28.7613499 | 43.704644 |
| 3.81494004 | 0.00013622 | 0.00036305 | 0 | 0 | 0 | 0 |
| -3.8147844 | 0.0001363 | 0.00036325 | 26.6398522 | 56.2870421 | 58.6731538 | 44.8252759 |
| -3.8145905 | 0.00013641 | 0.00036351 | 32.3776665 | 36.5324552 | 77.0804177 | 49.3078035 |
| -3.8144177 | 0.0001365 | 0.00036373 | 28.6890716 | 55.7458205 | 46.0181598 | 44.2649599 |
| 3.81426387 | 0.00013659 | 0.00036393 | 0 | 0 | 0 | 0 |
| -3.8140234 | 0.00013672 | 0.00036425 | 445.090454 | 1031.29768 | 956.02727 | 987.276701 |
| -3.8138087 | 0.00013684 | 0.00036451 | 24.5906328 | 46.0038325 | 86.2840497 | 64.99665 |
| -3.8138168 | 0.00013684 | 0.00036451 | 61.0667381 | 65.2171978 | 44.8677058 | 75.6426531 |
| -3.8135439 | 0.00013699 | 0.00036487 | 18.4429746 | 74.4179643 | 70.1776937 | 98.615607 |
| -3.8132995 | 0.00013712 | 0.0003652 | 19.2626624 | 95.525605 | 59.8236078 | 84.0473923 |
| -3.8130208 | 0.00013728 | 0.00036558 | 28.2792277 | 64.946587 | 40.2658898 | 54.910963 |
| -3.8128828 | 0.00013736 | 0.00036576 | 56.5584554 | 46.545054 | 35.6640739 | 32.498325 |

| | | | | | | |
|---|---|---|---|---|---|---|
| -3.8124579 | 0.00013759 | 0.00036632 | 86.8869025 | 47.3568864 | 87.4345036 | 48.1871716 |
| -3.8124684 | 0.00013759 | 0.00036632 | 80.3294005 | 39.5091738 | 29.9118039 | 50.4284354 |
| -3.8122781 | 0.00013769 | 0.00036656 | 25.8201644 | 42.7565031 | 71.3281477 | 66.1172819 |
| -3.8121776 | 0.00013775 | 0.00036668 | 120.903945 | 186.180216 | 147.258111 | 177.620156 |
| -3.8121036 | 0.00013779 | 0.00036676 | 54.509236 | 207.287857 | 225.488983 | 248.219965 |
| -3.8118346 | 0.00013794 | 0.00036713 | 132.379573 | 40.3210061 | 89.7354116 | 77.8839169 |
| -3.8111161 | 0.00013834 | 0.00036817 | 18.4429746 | 122.045461 | 104.691314 | 53.7903311 |
| -3.8108488 | 0.00013849 | 0.00036854 | 40.5745441 | 58.1813175 | 29.9118039 | 44.2649599 |
| 3.81073561 | 0.00013855 | 0.00036867 | 2.45906328 | 2.97671857 | 0 | 0 |
| -3.8106646 | 0.00013859 | 0.00036875 | 34.8367298 | 95.7962158 | 121.948124 | 106.46003 |
| -3.8102539 | 0.00013882 | 0.00036933 | 31.9678226 | 69.817581 | 32.2127119 | 70.5998095 |
| -3.8100785 | 0.00013892 | 0.00036956 | 23.3611012 | 65.4878086 | 52.9208838 | 63.8760181 |
| -3.809864 | 0.00013904 | 0.00036985 | 64.755333 | 37.3442875 | 29.9118039 | 51.5490673 |
| -3.8096851 | 0.00013914 | 0.00037009 | 18.0331307 | 60.3462037 | 164.514921 | 84.6077082 |
| -3.8094748 | 0.00013926 | 0.00037038 | 29.9186032 | 73.6061319 | 35.6640739 | 61.0744384 |
| -3.8093272 | 0.00013935 | 0.00037057 | 121.313788 | 220.547785 | 169.116737 | 186.024895 |
| -3.809272 | 0.00013938 | 0.00037059 | 27.8693838 | 61.9698684 | 41.4163438 | 53.2300151 |
| -3.8092773 | 0.00013937 | 0.00037059 | 17.213443 | 74.1473535 | 86.2840497 | 90.2108677 |
| -3.8092196 | 0.00013941 | 0.00037064 | 27.8693838 | 35.9912336 | 93.1867736 | 79.0045488 |
| -3.8091088 | 0.00013947 | 0.00037077 | 102.46097 | 216.759234 | 232.391707 | 234.212067 |
| -3.8090083 | 0.00013953 | 0.00037089 | 24.5906328 | 76.0416289 | 48.3190678 | 58.8331746 |
| -3.8088906 | 0.00013959 | 0.00037104 | 20.492194 | 135.30539 | 48.3190678 | 93.0124475 |
| -3.8086503 | 0.00013973 | 0.00037137 | 19.2626624 | 126.375234 | 50.6199758 | 101.417187 |
| -3.8082323 | 0.00013996 | 0.00037197 | 36.8859492 | 28.6847426 | 65.5758777 | 76.763285 |
| -3.8079948 | 0.0001401 | 0.00037229 | 66.8045524 | 68.7351379 | 66.7263317 | 52.6696992 |
| -3.8076005 | 0.00014032 | 0.00037286 | 26.6398522 | 58.9931499 | 43.7172518 | 56.5919108 |
| -3.807541 | 0.00014036 | 0.00037291 | 31.5579788 | 40.8622277 | 40.2658898 | 49.8681194 |
| -3.8074071 | 0.00014043 | 0.00037309 | 15.9839113 | 96.8786589 | 108.142676 | 71.7204414 |
| -3.8072768 | 0.00014051 | 0.00037325 | 19.2626624 | 100.125988 | 58.6731538 | 81.8061285 |
| -3.8064306 | 0.00014099 | 0.0003745 | 197.954594 | 383.184863 | 437.172518 | 438.167072 |
| -3.8059657 | 0.00014125 | 0.00037517 | 60.6568942 | 53.5809343 | 27.6108959 | 40.3427483 |
| 3.80581679 | 0.00014134 | 0.00037537 | 408.614348 | 350.170348 | 403.809352 | 374.291054 |
| -3.8045187 | 0.00014208 | 0.00037731 | 41.8040758 | 80.3714014 | 98.9390436 | 24.0935858 |
| -3.804471 | 0.00014211 | 0.00037735 | 278.693838 | 621.59296 | 510.801574 | 572.642899 |
| -3.8042683 | 0.00014222 | 0.00037763 | 68.8537718 | 138.011497 | 135.753571 | 116.545717 |
| -3.8039219 | 0.00014242 | 0.00037813 | 25.0004767 | 54.6633774 | 49.4695218 | 61.6347543 |
| -3.803659 | 0.00014257 | 0.0003785 | 22.9512573 | 59.2637606 | 54.0713378 | 104.218766 |
| -3.8031504 | 0.00014287 | 0.00037925 | 9.83625312 | 242.196647 | 141.505841 | 80.1251807 |
| 3.80264582 | 0.00014316 | 0.00037999 | 5.73781432 | 10.2832096 | 12.6549939 | 15.6888466 |
| 3.80255748 | 0.00014321 | 0.00038009 | 15.1642236 | 20.83703 | 25.3099879 | 30.8173772 |
| 3.80228368 | 0.00014337 | 0.00038048 | 0 | 0.54122156 | 0 | 0 |
| -3.8021456 | 0.00014345 | 0.00038066 | 36.8859492 | 65.2171978 | 29.9118039 | 50.9887513 |
| -3.8018926 | 0.00014359 | 0.00038102 | 18.0331307 | 89.0309463 | 74.7795097 | 72.8410733 |

| | | | | | | |
|---|---|---|---|---|---|---|
| -3.8018644 | 0.00014361 | 0.00038103 | 28.6890716 | 54.1221558 | 41.4163438 | 47.6268556 |
| -3.8018385 | 0.00014363 | 0.00038104 | 32.7875104 | 33.5557366 | 51.7704298 | 65.556966 |
| -3.8017783 | 0.00014366 | 0.0003811 | 27.8693838 | 60.6168145 | 54.0713378 | 40.3427483 |
| -3.8016208 | 0.00014375 | 0.00038131 | 88.1164342 | 60.075593 | 36.8145279 | 29.1364293 |
| 3.80113851 | 0.00014403 | 0.00038199 | 180.331307 | 88.4897248 | 100.089498 | 90.2108677 |
| -3.8011494 | 0.00014403 | 0.00038199 | 30.3284471 | 59.8049822 | 97.7885896 | 35.8602207 |
| -3.800517 | 0.00014439 | 0.00038292 | 45.4926707 | 58.7225391 | 29.9118039 | 38.6618005 |
| -3.8002982 | 0.00014452 | 0.00038323 | 56.9682993 | 34.9087905 | 37.9649818 | 36.9808526 |
| -3.8000486 | 0.00014467 | 0.00038358 | 27.0496961 | 48.4393295 | 44.8677058 | 62.1950703 |
| -3.7986502 | 0.00014549 | 0.00038572 | 45.9025146 | 44.9213893 | 63.2749697 | 52.6696992 |
| -3.7984337 | 0.00014561 | 0.000386 | 26.6398522 | 56.2870421 | 54.0713378 | 43.704644 |
| -3.7984293 | 0.00014562 | 0.000386 | 17.6232868 | 75.5004074 | 82.8326877 | 79.0045488 |
| -3.7983648 | 0.00014565 | 0.00038607 | 65.5750208 | 119.880575 | 14.9559019 | 111.502874 |
| -3.798339 | 0.00014567 | 0.00038608 | 15.5740674 | 106.079425 | 89.7354116 | 77.3236009 |
| -3.7980686 | 0.00014583 | 0.00038647 | 30.738291 | 46.2744432 | 42.5667978 | 40.3427483 |
| 3.79786131 | 0.00014595 | 0.0003867 | 130.740198 | 32.2026827 | 35.6640739 | 28.5761134 |
| -3.7978577 | 0.00014595 | 0.0003867 | 59.8372065 | 76.3122397 | 103.54086 | 59.3934906 |
| -3.7978661 | 0.00014595 | 0.0003867 | 21.3118818 | 48.7099402 | 98.9390436 | 80.6854966 |
| -3.7974432 | 0.0001462 | 0.00038732 | 36.8859492 | 76.8534613 | 31.0622579 | 48.7474875 |
| -3.796963 | 0.00014648 | 0.00038804 | 26.2300083 | 49.5217726 | 52.9208838 | 48.7474875 |
| -3.7965894 | 0.0001467 | 0.00038859 | 34.017042 | 25.4374132 | 167.966283 | 103.65845 |
| -3.7965097 | 0.00014675 | 0.00038868 | 14.7543797 | 87.1366709 | 97.7885896 | 94.1330794 |
| -3.7963859 | 0.00014682 | 0.00038884 | 142.215826 | 143.423713 | 87.4345036 | 128.312352 |
| -3.79628 | 0.00014688 | 0.00038891 | 18.0331307 | 141.529437 | 55.2217918 | 91.8918156 |
| -3.7963037 | 0.00014687 | 0.00038891 | 70.9029912 | 19.4839761 | 119.647216 | 59.9538065 |
| 3.79629693 | 0.00014687 | 0.00038891 | 0 | 0 | 0 | 0 |
| -3.7962626 | 0.00014689 | 0.00038891 | 42.2139196 | 36.2618444 | 37.9649818 | 35.8602207 |
| -3.7957608 | 0.00014719 | 0.00038967 | 20.492194 | 86.3248385 | 74.7795097 | 54.910963 |
| -3.7950693 | 0.0001476 | 0.00039072 | 49.5911095 | 133.140503 | 127.700393 | 17.3697944 |
| -3.7945772 | 0.0001479 | 0.00039147 | 21.3118818 | 96.3374374 | 49.4695218 | 77.3236009 |
| -3.7944496 | 0.00014797 | 0.00039163 | 36.0662614 | 26.2492456 | 96.6381356 | 88.5299199 |
| 3.79430359 | 0.00014806 | 0.00039183 | 220.086164 | 122.316072 | 117.346308 | 124.390141 |
| -3.7942233 | 0.00014811 | 0.00039193 | 18.8528185 | 72.253078 | 79.3813257 | 68.9188617 |
| -3.7941202 | 0.00014817 | 0.00039203 | 181.560839 | 80.1007906 | 54.0713378 | 67.7982298 |
| -3.7941188 | 0.00014817 | 0.00039203 | 124.182696 | 58.1813175 | 143.806749 | 15.1285306 |
| -3.7939192 | 0.00014829 | 0.00039228 | 89.7558097 | 44.6507786 | 26.4604419 | 51.5490673 |
| -3.7939382 | 0.00014828 | 0.00039228 | 48.3615778 | 138.282108 | 23.0090799 | 66.6775979 |
| -3.7938596 | 0.00014832 | 0.00039234 | 29.0989155 | 47.8981079 | 42.5667978 | 44.2649599 |
| -3.7936073 | 0.00014847 | 0.00039271 | 28.6890716 | 150.730204 | 40.2658898 | 62.1950703 |
| -3.7932335 | 0.0001487 | 0.00039327 | 12.7051603 | 86.5954493 | 217.435805 | 87.409288 |
| -3.7930916 | 0.00014878 | 0.00039346 | 28.6890716 | 58.4519283 | 41.4163438 | 45.3855918 |
| -3.7929108 | 0.00014889 | 0.00039372 | 32.7875104 | 44.3801678 | 34.5136199 | 57.1522268 |
| -3.7926162 | 0.00014907 | 0.00039415 | 16.8035991 | 137.740887 | 67.8767857 | 79.0045488 |

| | | | | | | |
|---|---|---|---|---|---|---|
| -3.7923899 | 0.0001492 | 0.00039448 | 74.1817423 | 92.2782757 | 18.4072639 | 62.1950703 |
| -3.7920497 | 0.00014941 | 0.00039499 | 17.213443 | 79.2889583 | 72.4786017 | 91.3314996 |
| -3.7915566 | 0.00014971 | 0.00039574 | 30.738291 | 72.253078 | 35.6640739 | 50.9887513 |
| -3.7907387 | 0.0001502 | 0.00039701 | 296.317125 | 38.1561199 | 32.2127119 | 62.7553862 |
| -3.7903544 | 0.00015043 | 0.00039759 | 35.6564176 | 35.7206228 | 36.8145279 | 48.7474875 |
| -3.7891113 | 0.00015119 | 0.00039956 | 29.9186032 | 41.67406 | 41.4163438 | 47.6268556 |
| -3.7890889 | 0.0001512 | 0.00039956 | 23.770945 | 56.8282636 | 56.3722458 | 53.7903311 |
| -3.7887242 | 0.00015142 | 0.00040011 | 28.6890716 | 174.002731 | 132.302209 | 119.907613 |
| -3.7886465 | 0.00015147 | 0.00040021 | 42.6237635 | 29.7671857 | 40.2658898 | 54.350647 |
| -3.7883914 | 0.00015163 | 0.00040058 | 26.2300083 | 58.4519283 | 44.8677058 | 50.4284354 |
| -3.7883384 | 0.00015166 | 0.00040064 | 29.9186032 | 37.3442875 | 63.2749697 | 47.0665397 |
| -3.7877974 | 0.00015199 | 0.00040148 | 47.5418901 | 35.1794013 | 49.4695218 | 31.3776931 |
| -3.787606 | 0.00015211 | 0.00040175 | 31.5579788 | 69.5469702 | 31.0622579 | 70.5998095 |
| -3.7872227 | 0.00015234 | 0.00040234 | 38.1154808 | 76.0416289 | 26.4604419 | 63.8760181 |
| -3.7870642 | 0.00015244 | 0.00040256 | 23.770945 | 86.3248385 | 69.0272397 | 43.704644 |
| -3.7868372 | 0.00015258 | 0.0004029 | 225.00429 | 237.596264 | 167.966283 | 214.040692 |
| -3.7866456 | 0.00015269 | 0.00040318 | 25.0004767 | 63.0523115 | 79.3813257 | 42.0236961 |
| -3.7861445 | 0.000153 | 0.00040396 | 18.0331307 | 73.3355211 | 86.2840497 | 69.4791776 |
| -3.7858645 | 0.00015318 | 0.00040438 | 16.8035991 | 76.5828505 | 82.8326877 | 82.9267604 |
| -3.7847036 | 0.00015389 | 0.00040624 | 11.0657848 | 149.918372 | 79.3813257 | 142.320251 |
| -3.7846183 | 0.00015394 | 0.00040634 | 57.3781432 | 35.7206228 | 34.5136199 | 38.1014845 |
| -3.7844752 | 0.00015403 | 0.00040655 | 94.2640924 | 204.581749 | 138.054479 | 177.05984 |
| -3.7843175 | 0.00015413 | 0.00040677 | 65.9848647 | 60.075593 | 26.4604419 | 38.6618005 |
| -3.7842672 | 0.00015416 | 0.00040682 | 60.6568942 | 68.4645271 | 98.9390436 | 85.7283401 |
| -3.7839832 | 0.00015434 | 0.00040725 | 21.7217256 | 72.5236888 | 49.4695218 | 77.8839169 |
| -3.7837812 | 0.00015446 | 0.00040755 | 120.084257 | 33.8263474 | 35.6640739 | 51.5490673 |
| -3.7836577 | 0.00015454 | 0.00040772 | 21.7217256 | 51.1454373 | 82.8326877 | 65.556966 |
| -3.7834098 | 0.00015469 | 0.00040809 | 39.3450125 | 76.8534613 | 27.6108959 | 52.1093832 |
| -3.7833505 | 0.00015473 | 0.00040815 | 29.9186032 | 46.8156648 | 36.8145279 | 59.3934906 |
| -3.7832317 | 0.00015481 | 0.00040832 | 21.3118818 | 114.197749 | 73.6290557 | 48.7474875 |
| -3.7826182 | 0.00015519 | 0.00040929 | 24.1807889 | 54.3927666 | 73.6290557 | 47.6268556 |
| -3.7825393 | 0.00015524 | 0.00040939 | 17.213443 | 102.290875 | 73.6290557 | 70.5998095 |
| -3.7822288 | 0.00015543 | 0.00040986 | 39.7548564 | 55.2045989 | 26.4604419 | 73.4013893 |
| 3.78217412 | 0.00015546 | 0.00040992 | 12.2953164 | 5.41221558 | 17.2568099 | 7.84442328 |
| -3.7818983 | 0.00015564 | 0.00041034 | 63.1159575 | 52.7691019 | 47.1686138 | 24.6539017 |
| -3.7818022 | 0.0001557 | 0.00041047 | 27.45954 | 52.2278804 | 64.4254237 | 39.2221164 |
| -3.7812507 | 0.00015604 | 0.00041134 | 177.4624 | 45.7332217 | 41.4163438 | 35.8602207 |
| -3.7811808 | 0.00015609 | 0.00041143 | 43.8532952 | 34.0969582 | 35.6640739 | 38.6618005 |
| -3.780957 | 0.00015623 | 0.00041176 | 205.741628 | 154.518755 | 59.8236078 | 280.71829 |
| -3.7809197 | 0.00015625 | 0.00041179 | 28.2792277 | 62.7817008 | 35.6640739 | 63.3157022 |
| -3.7806155 | 0.00015644 | 0.00041226 | 13.524848 | 179.144336 | 50.6199758 | 171.45668 |
| -3.7804919 | 0.00015652 | 0.00041243 | 40.5745441 | 84.4305631 | 65.5758777 | 89.0902358 |
| -3.7798295 | 0.00015694 | 0.0004135 | 28.6890716 | 37.0736767 | 79.3813257 | 53.7903311 |

| | | | | | | |
|---|---|---|---|---|---|---|
| 3.77962559 | 0.00015706 | 0.0004138 | 48.7714217 | 31.3908504 | 46.0181598 | 30.8173772 |
| -3.7795761 | 0.0001571 | 0.00041385 | 88.9361219 | 75.7710182 | 41.4163438 | 23.5332698 |
| -3.7795419 | 0.00015712 | 0.00041387 | 21.3118818 | 91.7370541 | 54.0713378 | 62.1950703 |
| -3.7781574 | 0.00015799 | 0.00041615 | 43.4434513 | 24.8961917 | 58.6731538 | 67.2379138 |
| -3.7781361 | 0.00015801 | 0.00041615 | 22.9512573 | 62.7817008 | 58.6731538 | 51.5490673 |
| -3.7776432 | 0.00015832 | 0.00041694 | 310.251817 | 509.018876 | 534.961108 | 535.101731 |
| -3.7775759 | 0.00015836 | 0.00041698 | 29.0989155 | 58.9931499 | 36.8145279 | 112.623506 |
| -3.777593 | 0.00015835 | 0.00041698 | 73.7718984 | 129.893174 | 26.4604419 | 89.6505518 |
| -3.7770277 | 0.00015871 | 0.00041787 | 11.4756286 | 96.0668266 | 104.691314 | 135.036144 |
| -3.7769178 | 0.00015878 | 0.00041802 | 73.3620545 | 28.4141318 | 56.3722458 | 36.4205367 |
| -3.7766818 | 0.00015893 | 0.00041838 | 29.0989155 | 72.5236888 | 36.8145279 | 50.9887513 |
| -3.7765361 | 0.00015902 | 0.00041859 | 22.5414134 | 120.151186 | 59.8236078 | 51.5490673 |
| -3.7757584 | 0.00015952 | 0.00041986 | 29.0989155 | 41.1328384 | 41.4163438 | 51.5490673 |
| -3.7756741 | 0.00015958 | 0.00041997 | 19.6725062 | 60.3462037 | 81.6822337 | 66.1172819 |
| -3.775496 | 0.00015969 | 0.00042024 | 72.5423667 | 47.6274971 | 33.3631659 | 32.498325 |
| 3.77539112 | 0.00015976 | 0.00042038 | 4.0984388 | 1.89427545 | 3.45136199 | 1.68094785 |
| -3.7747203 | 0.00016019 | 0.00042148 | 24.1807889 | 43.2977247 | 64.4254237 | 66.6775979 |
| -3.7746719 | 0.00016022 | 0.00042153 | 173.773805 | 45.7332217 | 26.4604419 | 57.1522268 |
| -3.7745121 | 0.00016032 | 0.00042176 | 72.5423667 | 45.7332217 | 27.6108959 | 41.4633802 |
| -3.7739179 | 0.0001607 | 0.00042273 | 9.83625312 | 122.586683 | 116.195854 | 124.390141 |
| -3.7732266 | 0.00016115 | 0.00042387 | 43.0336074 | 44.3801678 | 28.7613499 | 91.8918156 |
| -3.7730304 | 0.00016128 | 0.00042417 | 243.447265 | 431.624193 | 361.242555 | 396.703692 |
| -3.7728646 | 0.00016138 | 0.00042442 | 52.4600166 | 33.0145151 | 32.2127119 | 87.409288 |
| -3.7722507 | 0.00016178 | 0.00042543 | 30.738291 | 46.2744432 | 37.9649818 | 40.9030642 |
| -3.7721548 | 0.00016184 | 0.00042556 | 175.823024 | 348.817294 | 319.826211 | 317.138827 |
| -3.7721296 | 0.00016186 | 0.00042557 | 15.9839113 | 74.1473535 | 74.7795097 | 115.425085 |
| -3.771402 | 0.00016233 | 0.00042678 | 27.45954 | 40.0503953 | 52.9208838 | 48.7474875 |
| -3.7713484 | 0.00016237 | 0.00042683 | 13.524848 | 61.6992576 | 176.019461 | 114.304454 |
| -3.7712704 | 0.00016242 | 0.00042693 | 13.1150042 | 67.1114732 | 124.249032 | 135.59646 |
| -3.7708972 | 0.00016266 | 0.00042754 | 24.5906328 | 50.0629941 | 57.5226998 | 48.7474875 |
| -3.7705311 | 0.0001629 | 0.00042813 | 36.4761053 | 71.7118565 | 36.8145279 | 35.8602207 |
| -3.7704527 | 0.00016295 | 0.00042823 | 62.2962697 | 58.1813175 | 27.6108959 | 35.8602207 |
| -3.7703203 | 0.00016304 | 0.00042842 | 54.509236 | 33.5557366 | 29.9118039 | 56.5919108 |
| -3.7702868 | 0.00016306 | 0.00042844 | 24.1807889 | 46.545054 | 58.6731538 | 56.0315949 |
| -3.7699077 | 0.00016331 | 0.00042906 | 25.0004767 | 40.3210061 | 71.3281477 | 63.3157022 |
| -3.769508 | 0.00016357 | 0.00042971 | 31.9678226 | 70.0881918 | 41.4163438 | 36.9808526 |
| -3.7694555 | 0.0001636 | 0.00042977 | 34.017042 | 42.2152815 | 65.5758777 | 33.058641 |
| -3.7689135 | 0.00016396 | 0.00043067 | 21.3118818 | 65.4878086 | 52.9208838 | 68.9188617 |
| -3.7683701 | 0.00016432 | 0.00043157 | 23.3611012 | 86.8660601 | 50.6199758 | 52.6696992 |
| -3.7681588 | 0.00016446 | 0.0004319 | 28.6890716 | 35.9912336 | 54.0713378 | 53.2300151 |
| -3.7678635 | 0.00016465 | 0.00043235 | 22.9512573 | 105.538204 | 123.098578 | 35.8602207 |
| -3.7678603 | 0.00016465 | 0.00043235 | 10.246097 | 92.0076649 | 111.594038 | 172.577312 |
| 3.76745655 | 0.00016492 | 0.00043301 | 10.6559409 | 9.74198805 | 6.90272397 | 6.16347543 |

| | | | | | | |
|---|---|---|---|---|---|---|
| -3.7672165 | 0.00016508 | 0.00043339 | 25.0004767 | 44.109557 | 67.8767857 | 51.5490673 |
| -3.7670031 | 0.00016522 | 0.00043373 | 88.5262781 | 248.691306 | 248.498063 | 299.769033 |
| -3.7660922 | 0.00016582 | 0.00043528 | 26.6398522 | 42.4858923 | 51.7704298 | 47.6268556 |
| 3.76586054 | 0.00016598 | 0.00043565 | 65.9848647 | 46.0038325 | 37.9649818 | 45.3855918 |
| -3.7657448 | 0.00016605 | 0.00043581 | 592.224407 | 1278.63593 | 1623.29059 | 1503.32769 |
| -3.7656439 | 0.00016612 | 0.00043595 | 31.5579788 | 129.081342 | 25.3099879 | 105.339398 |
| -3.7651291 | 0.00016646 | 0.00043682 | 25.0004767 | 57.640096 | 52.9208838 | 44.2649599 |
| -3.7648922 | 0.00016662 | 0.0004372 | 79.5097127 | 38.1561199 | 32.2127119 | 40.3427483 |
| -3.7644585 | 0.00016691 | 0.00043792 | 40.984388 | 40.8622277 | 29.9118039 | 40.3427483 |
| -3.7640971 | 0.00016715 | 0.00043852 | 25.0004767 | 54.3927666 | 44.8677058 | 53.2300151 |
| -3.7639383 | 0.00016726 | 0.00043876 | 259.431176 | 459.226492 | 368.145279 | 373.170422 |
| -3.7638825 | 0.0001673 | 0.00043882 | 24.5906328 | 42.2152815 | 81.6822337 | 57.7125427 |
| -3.7638487 | 0.00016732 | 0.00043885 | 474.189369 | 938.748793 | 972.133626 | 953.657745 |
| -3.7634384 | 0.00016759 | 0.00043953 | 134.838636 | 31.1202396 | 31.0622579 | 68.9188617 |
| -3.7629405 | 0.00016793 | 0.00044037 | 47.5418901 | 41.1328384 | 34.5136199 | 31.9380091 |
| -3.762492 | 0.00016823 | 0.00044113 | 36.4761053 | 59.8049822 | 51.7704298 | 28.0157974 |
| -3.7624265 | 0.00016827 | 0.00044121 | 27.8693838 | 48.7099402 | 41.4163438 | 43.704644 |
| -3.7621937 | 0.00016843 | 0.00044158 | 16.3937552 | 77.3946828 | 86.2840497 | 73.9617052 |
| -3.7616396 | 0.0001688 | 0.00044253 | 17.6232868 | 68.4645271 | 87.4345036 | 68.9188617 |
| -3.761526 | 0.00016888 | 0.00044269 | 28.6890716 | 67.1114732 | 37.9649818 | 45.9459078 |
| -3.7610305 | 0.00016921 | 0.00044353 | 13.524848 | 94.9843835 | 90.8858656 | 93.5727634 |
| -3.7610058 | 0.00016923 | 0.00044354 | 48.3615778 | 44.9213893 | 29.9118039 | 35.2999048 |
| -3.7607222 | 0.00016942 | 0.00044401 | 41.3942319 | 31.9320719 | 34.5136199 | 45.9459078 |
| -3.7606361 | 0.00016948 | 0.00044412 | 35.2465737 | 49.7923834 | 32.2127119 | 38.6618005 |
| -3.760609 | 0.0001695 | 0.00044414 | 35.6564176 | 30.3084073 | 231.241253 | 64.4363341 |
| -3.7605058 | 0.00016957 | 0.00044428 | 28.6890716 | 47.3568864 | 39.1154358 | 43.704644 |
| -3.7602858 | 0.00016972 | 0.00044464 | 20.492194 | 67.1114732 | 58.6731538 | 62.1950703 |
| -3.7599619 | 0.00016994 | 0.00044518 | 43.4434513 | 33.2851258 | 42.5667978 | 33.058641 |
| -3.759551 | 0.00017022 | 0.00044587 | 31.1481349 | 46.545054 | 58.6731538 | 33.058641 |
| -3.7594246 | 0.0001703 | 0.00044606 | 120.903945 | 46.2744432 | 28.7613499 | 43.704644 |
| -3.7590236 | 0.00017058 | 0.00044674 | 117.625194 | 191.863042 | 203.630357 | 160.810677 |
| -3.7588646 | 0.00017069 | 0.00044699 | 26.6398522 | 46.0038325 | 44.8677058 | 46.5062237 |
| -3.7585717 | 0.00017089 | 0.00044744 | 32.3776665 | 53.5809343 | 37.9649818 | 35.8602207 |
| -3.7585768 | 0.00017088 | 0.00044744 | 251.234298 | 1379.57375 | 1224.08305 | 1278.08068 |
| -3.7580873 | 0.00017122 | 0.00044827 | 25.4103206 | 61.9698684 | 47.1686138 | 44.8252759 |
| -3.7574969 | 0.00017162 | 0.00044926 | 29.5087594 | 39.5091738 | 40.2658898 | 45.3855918 |
| -3.7575066 | 0.00017161 | 0.00044926 | 107.78894 | 100.937821 | 12.6549939 | 81.8061285 |
| -3.7573054 | 0.00017175 | 0.00044956 | 29.5087594 | 49.7923834 | 36.8145279 | 43.144328 |
| -3.7572357 | 0.0001718 | 0.00044965 | 21.3118818 | 71.4412457 | 51.7704298 | 61.6347543 |
| -3.7572073 | 0.00017182 | 0.00044967 | 56.5584554 | 43.2977247 | 25.3099879 | 47.0665397 |
| -3.7569618 | 0.00017199 | 0.00045007 | 50.8206411 | 20.5664192 | 132.302209 | 66.6775979 |
| -3.7561003 | 0.00017258 | 0.00045159 | 84.4278393 | 216.488623 | 196.727633 | 238.694594 |
| -3.7558672 | 0.00017274 | 0.00045193 | 30.738291 | 40.8622277 | 35.6640739 | 49.3078035 |

| | | | | | | |
|---|---|---|---|---|---|---|
| -3.7558858 | 0.00017273 | 0.00045193 | 62.7061136 | 48.980551 | 24.1595339 | 45.3855918 |
| -3.7555178 | 0.00017298 | 0.00045249 | 22.5414134 | 69.2763595 | 52.9208838 | 51.5490673 |
| -3.7555349 | 0.00017297 | 0.00045249 | 32.3776665 | 37.0736767 | 37.9649818 | 40.3427483 |
| -3.7553456 | 0.0001731 | 0.00045277 | 120.084257 | 70.0881918 | 24.1595339 | 38.6618005 |
| 3.75515411 | 0.00017323 | 0.00045308 | 15.1642236 | 20.5664192 | 25.3099879 | 19.6110582 |
| -3.7548678 | 0.00017343 | 0.00045356 | 62.7061136 | 25.4374132 | 195.577179 | 42.5840121 |
| -3.7545339 | 0.00017366 | 0.00045413 | 26.6398522 | 41.9446708 | 44.8677058 | 60.5141225 |
| 3.75444725 | 0.00017372 | 0.00045425 | 8.60672148 | 1.89427545 | 2.30090799 | 8.96505518 |
| -3.7543047 | 0.00017382 | 0.00045447 | 64.755333 | 45.4626109 | 26.4604419 | 40.9030642 |
| -3.7539182 | 0.00017409 | 0.00045511 | 27.8693838 | 63.8641439 | 41.4163438 | 42.0236961 |
| -3.7539114 | 0.0001741 | 0.00045511 | 50.4107972 | 31.6614612 | 42.5667978 | 33.6189569 |
| -3.7538825 | 0.00017412 | 0.00045512 | 25.0004767 | 57.0988744 | 43.7172518 | 128.872668 |
| -3.7528974 | 0.0001748 | 0.00045688 | 22.9512573 | 54.9339882 | 48.3190678 | 63.3157022 |
| 3.75282323 | 0.00017485 | 0.00045698 | 8.1968776 | 4.32977247 | 2.30090799 | 5.60315949 |
| -3.7527975 | 0.00017487 | 0.00045699 | 21.3118818 | 94.1725511 | 58.6731538 | 51.5490673 |
| -3.7526096 | 0.000175 | 0.00045729 | 104.100346 | 224.606947 | 174.869007 | 206.196269 |
| -3.7524967 | 0.00017508 | 0.00045746 | 12.2953164 | 209.993965 | 72.4786017 | 94.1330794 |
| -3.7522346 | 0.00017527 | 0.00045791 | 19.2626624 | 43.5683354 | 127.700393 | 95.8140272 |
| -3.7520821 | 0.00017537 | 0.00045815 | 25.4103206 | 42.4858923 | 55.2217918 | 49.8681194 |
| -3.7517605 | 0.0001756 | 0.0004587 | 27.8693838 | 42.4858923 | 73.6290557 | 41.4633802 |
| -3.751355 | 0.00017588 | 0.0004594 | 30.738291 | 62.2404792 | 81.6822337 | 30.2570612 |
| -3.7510506 | 0.0001761 | 0.00045992 | 44.6729829 | 44.9213893 | 28.7613499 | 128.312352 |
| -3.7509752 | 0.00017615 | 0.00046003 | 55.3289238 | 67.9233056 | 23.0090799 | 44.8252759 |
| -3.7509303 | 0.00017618 | 0.00046007 | 37.705637 | 53.851545 | 41.4163438 | 29.1364293 |
| -3.7506973 | 0.00017634 | 0.00046046 | 23.3611012 | 46.545054 | 65.5758777 | 53.2300151 |
| -3.7504634 | 0.00017651 | 0.00046085 | 28.6890716 | 73.0649104 | 34.5136199 | 52.6696992 |
| -3.7502058 | 0.00017669 | 0.00046129 | 27.45954 | 110.138587 | 71.3281477 | 33.058641 |
| 3.75013586 | 0.00017674 | 0.00046138 | 72.9522106 | 32.4732935 | 37.9649818 | 35.8602207 |
| -3.7499092 | 0.0001769 | 0.00046176 | 21.7217256 | 83.0775092 | 69.0272397 | 44.8252759 |
| -3.7498748 | 0.00017692 | 0.00046179 | 19.6725062 | 67.382084 | 62.1245158 | 62.1950703 |
| -3.7496952 | 0.00017705 | 0.00046208 | 18.8528185 | 60.6168145 | 70.1776937 | 71.1601255 |
| -3.7494648 | 0.00017721 | 0.00046247 | 40.984388 | 34.6381797 | 32.2127119 | 69.4791776 |
| 3.74929728 | 0.00017733 | 0.00046274 | 0 | 0 | 0 | 0 |
| -3.7486204 | 0.00017781 | 0.00046395 | 30.738291 | 38.9679522 | 39.1154358 | 39.7824324 |
| -3.7482323 | 0.00017809 | 0.00046463 | 95.0837801 | 34.6381797 | 33.3631659 | 44.2649599 |
| -3.7480867 | 0.00017819 | 0.00046485 | 28.6890716 | 47.3568864 | 43.7172518 | 37.5411686 |
| -3.7480755 | 0.0001782 | 0.00046485 | 13.524848 | 66.5702517 | 123.098578 | 107.020346 |
| -3.747883 | 0.00017833 | 0.00046517 | 30.738291 | 69.5469702 | 35.6640739 | 42.5840121 |
| -3.7476328 | 0.00017851 | 0.00046559 | 48.3615778 | 191.592432 | 82.8326877 | 124.950457 |
| 3.74753383 | 0.00017858 | 0.00046574 | 52.8698605 | 52.2278804 | 63.2749697 | 66.1172819 |
| -3.7473854 | 0.00017869 | 0.00046598 | 6.96734596 | 105.808815 | 187.524001 | 154.086886 |
| -3.7472329 | 0.0001788 | 0.00046622 | 18.4429746 | 111.22103 | 50.6199758 | 80.6854966 |
| -3.7466786 | 0.00017919 | 0.0004672 | 20.492194 | 60.075593 | 69.0272397 | 55.4712789 |

| | | | | | | |
|---|---|---|---|---|---|---|
| -3.7466672 | 0.0001792 | 0.0004672 | 30.3284471 | 35.7206228 | 46.0181598 | 40.9030642 |
| -3.7464924 | 0.00017932 | 0.00046749 | 33.1973543 | 91.7370541 | 31.0622579 | 50.9887513 |
| -3.7462925 | 0.00017947 | 0.00046782 | 101.231438 | 171.026012 | 186.373547 | 156.32815 |
| -3.7460884 | 0.00017961 | 0.00046817 | 19.2626624 | 96.8786589 | 48.3190678 | 80.6854966 |
| -3.7460392 | 0.00017965 | 0.00046822 | 19.6725062 | 72.253078 | 54.0713378 | 70.0394936 |
| -3.7459862 | 0.00017969 | 0.00046827 | 61.476582 | 33.0145151 | 29.9118039 | 48.1871716 |
| -3.7459743 | 0.0001797 | 0.00046827 | 20.0823501 | 59.5343714 | 81.6822337 | 55.4712789 |
| -3.7456786 | 0.00017991 | 0.00046878 | 103.690502 | 49.7923834 | 18.4072639 | 79.0045488 |
| -3.7453916 | 0.00018011 | 0.00046928 | 61.0667381 | 37.8855091 | 34.5136199 | 33.058641 |
| -3.7450285 | 0.00018037 | 0.00046992 | 23.3611012 | 56.2870421 | 49.4695218 | 50.9887513 |
| 3.74474622 | 0.00018058 | 0.00047041 | 6.55750208 | 5.1416048 | 0 | 6.16347543 |
| -3.7443269 | 0.00018088 | 0.00047116 | 49.1812656 | 41.1328384 | 32.2127119 | 32.498325 |
| -3.743835 | 0.00018123 | 0.00047204 | 36.0662614 | 73.0649104 | 39.1154358 | 32.498325 |
| 3.7437659 | 0.00018128 | 0.00047214 | 195.495531 | 172.649677 | 157.612197 | 152.405938 |
| -3.7437287 | 0.00018131 | 0.00047217 | 49.5911095 | 41.67406 | 27.6108959 | 39.2221164 |
| -3.743644 | 0.00018137 | 0.00047229 | 25.4103206 | 47.0862756 | 42.5667978 | 66.6775979 |
| -3.7430219 | 0.00018182 | 0.00047342 | 23.770945 | 76.3122397 | 37.9649818 | 75.6426531 |
| -3.7429797 | 0.00018185 | 0.00047346 | 19.2626624 | 85.783617 | 51.7704298 | 72.2807574 |
| -3.7428431 | 0.00018195 | 0.00047368 | 21.7217256 | 40.3210061 | 93.1867736 | 82.9267604 |
| 3.74276259 | 0.00018201 | 0.00047376 | 100.821594 | 35.7206228 | 66.7263317 | 33.6189569 |
| -3.742771 | 0.000182 | 0.00047376 | 14.7543797 | 78.2065152 | 70.1776937 | 130.553616 |
| -3.7425302 | 0.00018218 | 0.00047416 | 25.0004767 | 41.1328384 | 83.9831417 | 52.6696992 |
| -3.7421268 | 0.00018247 | 0.00047488 | 59.0175187 | 96.6080482 | 14.9559019 | 111.502874 |
| -3.7419899 | 0.00018257 | 0.0004751 | 27.45954 | 57.3694852 | 37.9649818 | 45.3855918 |
| -3.7416166 | 0.00018284 | 0.00047577 | 28.2792277 | 39.7797845 | 47.1686138 | 99.7362389 |
| -3.7414688 | 0.00018295 | 0.00047601 | 29.0989155 | 35.7206228 | 48.3190678 | 43.704644 |
| -3.7409819 | 0.0001833 | 0.0004769 | 30.738291 | 46.545054 | 36.8145279 | 37.5411686 |
| -3.7408638 | 0.00018339 | 0.00047708 | 208.610535 | 36.2618444 | 27.6108959 | 64.99665 |
| -3.740539 | 0.00018363 | 0.00047766 | 25.8201644 | 42.2152815 | 46.0181598 | 52.1093832 |
| -3.7404488 | 0.00018369 | 0.00047779 | 49.1812656 | 45.7332217 | 24.1595339 | 49.3078035 |
| -3.7403578 | 0.00018376 | 0.00047793 | 106.969253 | 230.830995 | 201.329449 | 221.3248 |
| -3.7401834 | 0.00018389 | 0.00047822 | 22.5414134 | 51.1454373 | 65.5758777 | 49.8681194 |
| -3.739543 | 0.00018436 | 0.0004794 | 27.0496961 | 35.7206228 | 63.2749697 | 52.6696992 |
| -3.7395115 | 0.00018438 | 0.00047942 | 30.738291 | 45.1920001 | 36.8145279 | 37.5411686 |
| 3.73935142 | 0.0001845 | 0.00047969 | 0 | 0 | 0 | 0 |
| -3.7389222 | 0.00018481 | 0.00048047 | 27.0496961 | 58.7225391 | 37.9649818 | 46.5062237 |
| -3.7384334 | 0.00018517 | 0.00048137 | 52.8698605 | 42.4858923 | 24.1595339 | 52.1093832 |
| -3.7383343 | 0.00018524 | 0.00048152 | 325.416041 | 734.167044 | 669.564225 | 757.547163 |
| -3.7379174 | 0.00018555 | 0.00048228 | 40.1647002 | 40.8622277 | 32.2127119 | 33.6189569 |
| -3.7377402 | 0.00018568 | 0.00048258 | 36.0662614 | 29.7671857 | 40.2658898 | 48.7474875 |
| -3.7373571 | 0.00018596 | 0.00048328 | 33.1973543 | 53.5809343 | 37.9649818 | 33.058641 |
| 3.73719664 | 0.00018608 | 0.00048355 | 36.4761053 | 24.0843593 | 17.2568099 | 35.2999048 |
| -3.7371356 | 0.00018613 | 0.00048363 | 36.8859492 | 42.7565031 | 33.3631659 | 33.6189569 |

| | | | | | | |
|---|---|---|---|---|---|---|
| -3.7370411 | 0.0001862 | 0.00048374 | 17.6232868 | 59.2637606 | 86.2840497 | 72.2807574 |
| -3.7370347 | 0.0001862 | 0.00048374 | 27.45954 | 47.0862756 | 37.9649818 | 46.5062237 |
| -3.7366943 | 0.00018646 | 0.00048436 | 43.0336074 | 46.2744432 | 35.6640739 | 29.1364293 |
| 3.73618449 | 0.00018683 | 0.00048522 | 7.37718984 | 3.51794013 | 4.60181598 | 4.48252759 |
| -3.7361949 | 0.00018683 | 0.00048522 | 90.5754975 | 55.2045989 | 36.8145279 | 52.6696992 |
| -3.7361909 | 0.00018683 | 0.00048522 | 35.6564176 | 33.2851258 | 35.6640739 | 39.7824324 |
| 3.73598307 | 0.00018698 | 0.00048557 | 5.32797044 | 5.68282636 | 4.60181598 | 4.48252759 |
| -3.7357426 | 0.00018716 | 0.000486 | 169.675366 | 199.981366 | 167.966283 | 191.628054 |
| -3.7352597 | 0.00018752 | 0.00048689 | 41.8040758 | 40.8622277 | 49.4695218 | 26.8951655 |
| -3.734953 | 0.00018775 | 0.00048745 | 22.9512573 | 81.1832337 | 39.1154358 | 71.7204414 |
| -3.7347787 | 0.00018788 | 0.00048775 | 334.84245 | 810.749894 | 694.874213 | 730.651997 |
| -3.7346988 | 0.00018794 | 0.00048786 | 23.3611012 | 48.1687187 | 59.8236078 | 48.1871716 |
| -3.7346095 | 0.00018801 | 0.000488 | 38.9351686 | 88.4897248 | 35.6640739 | 34.1792729 |
| -3.7345718 | 0.00018803 | 0.00048803 | 26.6398522 | 67.6526948 | 37.9649818 | 48.1871716 |
| -3.7341462 | 0.00018835 | 0.00048882 | 18.0331307 | 62.2404792 | 67.8767857 | 75.0823371 |
| -3.7339881 | 0.00018847 | 0.00048908 | 49.5911095 | 29.4965749 | 33.3631659 | 47.0665397 |
| -3.7338353 | 0.00018859 | 0.00048934 | 76.6408055 | 41.67406 | 24.1595339 | 50.9887513 |
| -3.7334026 | 0.00018891 | 0.00049014 | 62.7061136 | 41.9446708 | 56.3722458 | 42.0236961 |
| -3.7332614 | 0.00018902 | 0.00049038 | 28.6890716 | 47.8981079 | 34.5136199 | 71.1601255 |
| -3.7331405 | 0.00018911 | 0.00049058 | 70.4931473 | 16.7778683 | 67.8767857 | 84.0473923 |
| -3.7327142 | 0.00018943 | 0.00049137 | 31.5579788 | 69.2763595 | 81.6822337 | 27.4554815 |
| -3.7322918 | 0.00018975 | 0.00049215 | 24.1807889 | 49.7923834 | 49.4695218 | 142.880567 |
| -3.7322446 | 0.00018978 | 0.00049221 | 181.970683 | 43.2977247 | 25.3099879 | 56.0315949 |
| -3.7321664 | 0.00018984 | 0.00049232 | 25.8201644 | 48.980551 | 44.8677058 | 42.5840121 |
| -3.731961 | 0.00018999 | 0.0004926 | 29.9186032 | 120.421797 | 135.753571 | 152.966254 |
| -3.7319733 | 0.00018999 | 0.0004926 | 27.45954 | 36.2618444 | 54.0713378 | 46.5062237 |
| -3.7319814 | 0.00018998 | 0.0004926 | 530.747825 | 854.588841 | 767.352815 | 750.823371 |
| -3.7315237 | 0.00019033 | 0.00049342 | 27.8693838 | 33.0145151 | 79.3813257 | 57.7125427 |
| -3.7313486 | 0.00019046 | 0.00049372 | 20.0823501 | 73.3355211 | 47.1686138 | 77.8839169 |
| -3.7307859 | 0.00019088 | 0.00049479 | 19.2626624 | 60.3462037 | 56.3722458 | 82.3664444 |
| -3.7306514 | 0.00019099 | 0.00049501 | 26.2300083 | 38.6973414 | 49.4695218 | 49.3078035 |
| -3.7306166 | 0.00019101 | 0.00049504 | 238.119294 | 538.786061 | 339.383929 | 421.917909 |
| -3.7305973 | 0.00019103 | 0.00049504 | 15.9839113 | 70.3588026 | 73.6290557 | 85.7283401 |
| -3.7305558 | 0.00019106 | 0.00049508 | 32.7875104 | 44.6507786 | 33.3631659 | 36.9808526 |
| -3.7302451 | 0.00019129 | 0.00049565 | 22.5414134 | 41.67406 | 79.3813257 | 62.7553862 |
| -3.7299631 | 0.00019151 | 0.00049617 | 22.1315695 | 55.4752097 | 49.4695218 | 57.1522268 |
| -3.7298385 | 0.0001916 | 0.00049637 | 21.7217256 | 94.1725511 | 48.3190678 | 55.4712789 |
| -3.729375 | 0.00019196 | 0.00049724 | 22.5414134 | 64.6759762 | 48.3190678 | 50.9887513 |
| -3.7291722 | 0.00019211 | 0.00049761 | 342.21964 | 434.05969 | 454.429328 | 477.389188 |
| -3.7291227 | 0.00019215 | 0.00049766 | 26.6398522 | 132.05806 | 172.568099 | 146.802779 |
| -3.7290966 | 0.00019217 | 0.00049767 | 103.280658 | 57.640096 | 24.1595339 | 39.7824324 |
| -3.7290084 | 0.00019223 | 0.00049781 | 53.6895483 | 134.222946 | 70.1776937 | 17.9301104 |
| -3.7286748 | 0.00019249 | 0.00049843 | 9.01656536 | 71.9824673 | 220.887167 | 132.79488 |

| | | | | | | |
|---|---|---|---|---|---|---|
| -3.7279282 | 0.00019306 | 0.00049987 | 180.331307 | 7.03588026 | 286.463045 | 66.6775979 |
| -3.7277816 | 0.00019317 | 0.00050012 | 23.770945 | 66.0290301 | 39.1154358 | 62.7553862 |
| -3.7276575 | 0.00019327 | 0.00050032 | 81.5589321 | 45.4626109 | 32.2127119 | 31.9380091 |
| -3.7272725 | 0.00019356 | 0.00050105 | 18.4429746 | 149.918372 | 108.142676 | 104.779082 |
| -3.7271852 | 0.00019363 | 0.00050118 | 4.91812656 | 120.963018 | 211.683535 | 183.783631 |
| -3.7266682 | 0.00019403 | 0.00050217 | 26.2300083 | 32.4732935 | 73.6290557 | 79.5648647 |
| -3.726577 | 0.0001941 | 0.00050231 | 20.9020379 | 86.3248385 | 394.60572 | 33.6189569 |
| -3.7261529 | 0.00019442 | 0.00050312 | 21.3118818 | 50.6042157 | 58.6731538 | 58.8331746 |
| -3.7260076 | 0.00019454 | 0.00050337 | 25.4103206 | 49.7923834 | 41.4163438 | 47.0665397 |
| -3.7257067 | 0.00019477 | 0.00050393 | 31.9678226 | 66.8408624 | 57.5226998 | 73.9617052 |
| -3.7255547 | 0.00019489 | 0.00050419 | 20.9020379 | 62.7817008 | 75.9299637 | 46.5062237 |
| -3.7246091 | 0.00019562 | 0.00050604 | 32.7875104 | 66.8408624 | 29.9118039 | 45.3855918 |
| -3.7243538 | 0.00019582 | 0.00050651 | 15.9839113 | 60.8874253 | 94.3372276 | 80.6854966 |
| -3.7242972 | 0.00019586 | 0.00050659 | 48.7714217 | 30.579018 | 52.9208838 | 30.8173772 |
| -3.7242635 | 0.00019589 | 0.0005066 | 7.37718984 | 102.290875 | 209.382627 | 119.907613 |
| -3.7242519 | 0.0001959 | 0.0005066 | 40.984388 | 89.5721679 | 155.311289 | 19.0507423 |
| -3.7242181 | 0.00019592 | 0.00050662 | 40.1647002 | 51.1454373 | 26.4604419 | 40.9030642 |
| -3.7241689 | 0.00019596 | 0.00050668 | 18.0331307 | 68.4645271 | 65.5758777 | 64.99665 |
| 3.72400104 | 0.00019609 | 0.00050698 | 186.069121 | 97.1492697 | 105.841768 | 61.6347543 |
| -3.7235049 | 0.00019648 | 0.00050794 | 29.0989155 | 37.3442875 | 39.1154358 | 43.144328 |
| -3.7230083 | 0.00019686 | 0.0005089 | 30.3284471 | 59.8049822 | 32.2127119 | 44.2649599 |
| -3.7225142 | 0.00019725 | 0.00050985 | 50.8206411 | 41.67406 | 26.4604419 | 39.2221164 |
| -3.7222763 | 0.00019743 | 0.00051025 | 33.6071982 | 50.3336049 | 39.1154358 | 30.8173772 |
| -3.7222782 | 0.00019743 | 0.00051025 | 30.3284471 | 38.4267306 | 34.5136199 | 49.3078035 |
| -3.7215031 | 0.00019804 | 0.00051178 | 29.0989155 | 58.1813175 | 86.2840497 | 30.8173772 |
| -3.7208425 | 0.00019856 | 0.00051308 | 31.9678226 | 43.8389462 | 50.6199758 | 30.2570612 |
| -3.720318 | 0.00019897 | 0.0005141 | 18.4429746 | 60.3462037 | 59.8236078 | 79.5648647 |
| -3.7202638 | 0.00019901 | 0.00051417 | 25.0004767 | 38.4267306 | 58.6731538 | 51.5490673 |
| -3.719422 | 0.00019968 | 0.00051585 | 25.8201644 | 69.2763595 | 44.8677058 | 40.3427483 |
| -3.7192348 | 0.00019983 | 0.00051619 | 23.3611012 | 44.9213893 | 48.3190678 | 63.3157022 |
| -3.7187908 | 0.00020018 | 0.00051701 | 26.6398522 | 44.6507786 | 40.2658898 | 43.144328 |
| -3.7187938 | 0.00020018 | 0.00051701 | 97.1329995 | 131.246228 | 90.8858656 | 94.6933953 |
| -3.7187325 | 0.00020022 | 0.00051709 | 18.8528185 | 68.1939163 | 62.1245158 | 58.8331746 |
| -3.7186455 | 0.00020029 | 0.00051723 | 30.3284471 | 58.9931499 | 48.3190678 | 30.8173772 |
| -3.7184765 | 0.00020043 | 0.00051753 | 43.0336074 | 85.5130062 | 18.4072639 | 128.872668 |
| -3.7184114 | 0.00020048 | 0.00051762 | 29.0989155 | 68.1939163 | 34.5136199 | 43.704644 |
| -3.7183174 | 0.00020055 | 0.00051777 | 26.6398522 | 45.7332217 | 36.8145279 | 56.5919108 |
| -3.7182607 | 0.0002006 | 0.00051785 | 23.3611012 | 43.2977247 | 88.5849576 | 51.5490673 |
| -3.7179845 | 0.00020082 | 0.00051837 | 29.9186032 | 46.545054 | 57.5226998 | 31.3776931 |
| -3.7175298 | 0.00020118 | 0.00051927 | 29.5087594 | 51.9572696 | 32.2127119 | 45.3855918 |
| -3.7174305 | 0.00020126 | 0.00051943 | 205.741628 | 167.237462 | 178.320369 | 133.915512 |
| -3.7172319 | 0.00020142 | 0.0005198 | 28.2792277 | 58.4519283 | 31.0622579 | 79.0045488 |
| -3.7166096 | 0.00020191 | 0.00052104 | 74.5915861 | 79.8301798 | 39.1154358 | 21.852322 |

| | | | | | | |
|---|---|---|---|---|---|---|
| -3.7163785 | 0.0002021 | 0.00052147 | 28.2792277 | 41.9446708 | 40.2658898 | 38.1014845 |
| -3.7162102 | 0.00020223 | 0.00052178 | 24.1807889 | 71.9824673 | 37.9649818 | 57.1522268 |
| -3.7158686 | 0.00020251 | 0.00052244 | 45.9025146 | 223.795114 | 238.143977 | 270.072287 |
| -3.7156948 | 0.00020265 | 0.00052276 | 23.770945 | 65.4878086 | 46.0181598 | 44.8252759 |
| -3.7154336 | 0.00020286 | 0.00052326 | 59.8372065 | 368.842492 | 506.199758 | 553.031841 |
| -3.7152248 | 0.00020302 | 0.00052365 | 22.9512573 | 53.851545 | 48.3190678 | 48.1871716 |
| -3.715053 | 0.00020316 | 0.00052396 | 83.1983076 | 45.7332217 | 48.3190678 | 76.202969 |
| -3.7147806 | 0.00020338 | 0.00052444 | 37.705637 | 54.3927666 | 24.1595339 | 62.7553862 |
| -3.7147861 | 0.00020338 | 0.00052444 | 299.186032 | 783.688816 | 646.555145 | 974.949751 |
| -3.7146342 | 0.0002035 | 0.0005247 | 195.495531 | 443.531067 | 432.570702 | 413.51317 |
| 3.71449304 | 0.00020361 | 0.00052495 | 0 | 0 | 0 | 0 |
| 3.71440903 | 0.00020368 | 0.00052509 | 313.530568 | 239.76115 | 224.338529 | 220.204168 |
| -3.7142673 | 0.00020379 | 0.00052534 | 22.9512573 | 45.7332217 | 78.2308717 | 48.1871716 |
| -3.7139525 | 0.00020405 | 0.00052595 | 26.6398522 | 42.7565031 | 41.4163438 | 42.0236961 |
| -3.7138716 | 0.00020411 | 0.00052608 | 22.9512573 | 46.2744432 | 67.8767857 | 47.0665397 |
| -3.7136867 | 0.00020426 | 0.00052642 | 34.4268859 | 27.6022995 | 60.9740618 | 46.5062237 |
| 3.7135394 | 0.00020438 | 0.00052669 | 27.8693838 | 24.0843593 | 28.7613499 | 16.2491625 |
| -3.7128439 | 0.00020494 | 0.00052809 | 23.770945 | 56.2870421 | 46.0181598 | 44.8252759 |
| -3.7127715 | 0.000205 | 0.00052816 | 22.1315695 | 53.851545 | 47.1686138 | 56.0315949 |
| -3.7127775 | 0.000205 | 0.00052816 | 47.5418901 | 50.3336049 | 54.0713378 | 53.7903311 |
| -3.7127086 | 0.00020505 | 0.00052825 | 28.2792277 | 38.4267306 | 222.037621 | 53.7903311 |
| -3.7124498 | 0.00020526 | 0.00052871 | 40.5745441 | 37.3442875 | 27.6108959 | 43.144328 |
| -3.71246 | 0.00020525 | 0.00052871 | 70.4931473 | 13.8011497 | 98.9390436 | 88.5299199 |
| -3.7122911 | 0.00020539 | 0.00052899 | 54.509236 | 149.106539 | 111.594038 | 95.2537113 |
| -3.7118327 | 0.00020576 | 0.00052991 | 45.4926707 | 42.7565031 | 26.4604419 | 38.1014845 |
| -3.7117694 | 0.00020582 | 0.00053 | 19.2626624 | 120.421797 | 41.4163438 | 84.0473923 |
| -3.7116371 | 0.00020592 | 0.00053024 | 57.3781432 | 45.1920001 | 42.5667978 | 44.8252759 |
| -3.7112283 | 0.00020626 | 0.00053105 | 61.8864259 | 24.6255809 | 41.4163438 | 45.9459078 |
| -3.7110629 | 0.00020639 | 0.00053136 | 48.7714217 | 32.4732935 | 71.3281477 | 28.5761134 |
| -3.7105298 | 0.00020683 | 0.00053243 | 9.01656536 | 134.764168 | 140.355387 | 84.0473923 |
| -3.7103461 | 0.00020698 | 0.00053278 | 55.7387677 | 70.9000241 | 49.4695218 | 42.5840121 |
| -3.710095 | 0.00020718 | 0.00053326 | 47.1320462 | 21.1076408 | 60.9740618 | 61.0744384 |
| -3.7100421 | 0.00020722 | 0.00053333 | 26.6398522 | 41.67406 | 39.1154358 | 46.5062237 |
| -3.7087887 | 0.00020825 | 0.00053594 | 15.1642236 | 84.7011739 | 70.1776937 | 75.6426531 |
| 3.70873279 | 0.0002083 | 0.00053601 | 0 | 0 | 0 | 0 |
| -3.7085707 | 0.00020843 | 0.00053631 | 238.119294 | 1597.95665 | 798.415073 | 1063.47967 |
| -3.7081154 | 0.00020881 | 0.00053724 | 18.4429746 | 48.7099402 | 77.0804177 | 80.1251807 |
| -3.7080908 | 0.00020883 | 0.00053724 | 20.9020379 | 46.0038325 | 64.4254237 | 61.6347543 |
| 3.70803473 | 0.00020887 | 0.00053728 | 0.40984388 | 0.54122156 | 0 | 0 |
| -3.7080517 | 0.00020886 | 0.00053728 | 29.9186032 | 57.0988744 | 28.7613499 | 73.4013893 |
| -3.707875 | 0.00020901 | 0.00053757 | 29.5087594 | 67.9233056 | 48.3190678 | 31.3776931 |
| -3.7077869 | 0.00020908 | 0.00053772 | 133.609105 | 52.7691019 | 28.7613499 | 34.7395888 |
| -3.7076053 | 0.00020923 | 0.00053806 | 53.2797044 | 29.4965749 | 32.2127119 | 43.144328 |

| | | | | | | |
|---|---|---|---|---|---|---|
| -3.7072129 | 0.00020955 | 0.00053877 | 19.2626624 | 84.1599523 | 42.5667978 | 91.3314996 |
| -3.7072249 | 0.00020954 | 0.00053877 | 13.524848 | 85.2423954 | 83.9831417 | 82.3664444 |
| -3.7072424 | 0.00020953 | 0.00053877 | 26.2300083 | 36.5324552 | 46.0181598 | 54.910963 |
| -3.7069402 | 0.00020978 | 0.0005393 | 40.5745441 | 32.2026827 | 35.6640739 | 33.6189569 |
| -3.7062744 | 0.00021033 | 0.00054068 | 44.263139 | 35.7206228 | 34.5136199 | 30.8173772 |
| 3.7060393 | 0.00021053 | 0.00054114 | 0 | 0 | 0 | 0 |
| -3.7058778 | 0.00021066 | 0.00054144 | 144.67489 | 129.893174 | 256.551241 | 23.5332698 |
| -3.705531 | 0.00021095 | 0.00054214 | 43.4434513 | 36.803066 | 29.9118039 | 34.7395888 |
| -3.7054856 | 0.00021099 | 0.00054219 | 42.2139196 | 83.6187308 | 73.6290557 | 20.1713742 |
| -3.7054606 | 0.00021101 | 0.0005422 | 17.213443 | 103.643928 | 44.8677058 | 99.7362389 |
| -3.7054024 | 0.00021106 | 0.00054228 | 22.5414134 | 42.2152815 | 63.2749697 | 54.350647 |
| -3.705333 | 0.00021111 | 0.00054239 | 29.5087594 | 42.2152815 | 32.2127119 | 50.9887513 |
| -3.7049192 | 0.00021146 | 0.00054323 | 21.7217256 | 55.2045989 | 63.2749697 | 44.8252759 |
| -3.7048929 | 0.00021148 | 0.00054325 | 109.018472 | 49.5217726 | 35.6640739 | 28.0157974 |
| -3.7047145 | 0.00021163 | 0.00054359 | 18.8528185 | 99.8553775 | 52.9208838 | 59.3934906 |
| -3.7043002 | 0.00021198 | 0.00054443 | 25.0004767 | 45.1920001 | 40.2658898 | 50.4284354 |
| -3.7041703 | 0.00021208 | 0.00054467 | 16.3937552 | 39.5091738 | 143.806749 | 123.829825 |
| -3.7038884 | 0.00021232 | 0.00054523 | 29.0989155 | 40.3210061 | 34.5136199 | 63.3157022 |
| -3.7033269 | 0.00021279 | 0.00054635 | 169.675366 | 180.226779 | 304.870309 | 115.425085 |
| -3.7033423 | 0.00021278 | 0.00054635 | 676.242402 | 401.586396 | 403.809352 | 403.987799 |
| -3.7030179 | 0.00021305 | 0.00054697 | 25.4103206 | 85.2423954 | 213.984443 | 28.5761134 |
| -3.7028988 | 0.00021315 | 0.00054719 | 417.630914 | 71.9824673 | 11.50454 | 65.556966 |
| 3.70278484 | 0.00021325 | 0.00054735 | 116.805506 | 65.7584193 | 94.3372276 | 67.7982298 |
| -3.7028047 | 0.00021323 | 0.00054735 | 102.46097 | 41.67406 | 27.6108959 | 40.9030642 |
| -3.7027077 | 0.00021331 | 0.00054747 | 19.6725062 | 87.6778924 | 42.5667978 | 76.763285 |
| -3.7025718 | 0.00021342 | 0.00054772 | 80.3294005 | 31.1202396 | 51.7704298 | 29.1364293 |
| -3.7016565 | 0.0002142 | 0.00054966 | 30.3284471 | 33.8263474 | 36.8145279 | 48.1871716 |
| -3.7014234 | 0.00021439 | 0.00055012 | 78.2801811 | 30.3084073 | 35.6640739 | 39.2221164 |
| -3.7011639 | 0.00021461 | 0.00055064 | 50.4107972 | 37.3442875 | 27.6108959 | 37.5411686 |
| -3.7010253 | 0.00021473 | 0.00055089 | 45.4926707 | 28.143521 | 71.3281477 | 33.6189569 |
| 3.70097436 | 0.00021477 | 0.00055096 | 98.7723751 | 189.156935 | 161.063559 | 178.740788 |
| 3.7009393 | 0.0002148 | 0.00055099 | 1.22953164 | 2.97671857 | 1.150454 | 3.36189569 |
| -3.7008891 | 0.00021485 | 0.00055106 | 29.9186032 | 65.2171978 | 29.9118039 | 48.7474875 |
| -3.7008025 | 0.00021492 | 0.0005512 | 16.8035991 | 45.1920001 | 116.195854 | 94.6933953 |
| -3.7005719 | 0.00021511 | 0.00055166 | 20.9020379 | 51.6866588 | 51.7704298 | 134.475828 |
| -3.7004091 | 0.00021525 | 0.00055193 | 22.5414134 | 57.640096 | 46.0181598 | 48.1871716 |
| -3.7004054 | 0.00021526 | 0.00055193 | 38.5253247 | 87.9485032 | 21.8586259 | 64.4363341 |
| -3.6998213 | 0.00021575 | 0.00055316 | 9.83625312 | 88.4897248 | 133.452663 | 103.65845 |
| 3.69965761 | 0.00021589 | 0.00055347 | 29.9186032 | 14.6129821 | 16.1063559 | 22.9729539 |
| -3.6988986 | 0.00021654 | 0.00055506 | 13.1150042 | 81.1832337 | 85.1335957 | 87.9696039 |
| -3.6988913 | 0.00021654 | 0.00055506 | 20.492194 | 58.1813175 | 56.3722458 | 50.9887513 |
| -3.6988709 | 0.00021656 | 0.00055506 | 20.0823501 | 46.0038325 | 73.6290557 | 62.1950703 |
| -3.6985835 | 0.00021681 | 0.00055565 | 21.7217256 | 58.1813175 | 69.0272397 | 42.0236961 |

| | | | | | | |
|---|---|---|---|---|---|---|
| 3.69854032 | 0.00021684 | 0.0005557 | 40.1647002 | 20.0251977 | 11.50454 | 14.0078987 |
| -3.6984824 | 0.00021689 | 0.00055578 | 65.1651769 | 139.364551 | 106.992222 | 118.786981 |
| -3.698415 | 0.00021695 | 0.00055588 | 45.4926707 | 36.803066 | 27.6108959 | 38.1014845 |
| -3.6983238 | 0.00021703 | 0.00055604 | 25.4103206 | 49.7923834 | 47.1686138 | 37.5411686 |
| -3.6979609 | 0.00021734 | 0.00055679 | 13.524848 | 67.382084 | 95.4876816 | 92.4521315 |
| -3.6979318 | 0.00021736 | 0.00055681 | 112.297223 | 164.531354 | 88.5849576 | 77.3236009 |
| -3.6978221 | 0.00021746 | 0.000557 | 29.5087594 | 94.4431619 | 41.4163438 | 36.9808526 |
| -3.6976688 | 0.00021759 | 0.0005573 | 24.5906328 | 62.7817008 | 40.2658898 | 44.8252759 |
| -3.6975547 | 0.00021769 | 0.0005575 | 22.5414134 | 50.6042157 | 48.3190678 | 48.1871716 |
| -3.6972702 | 0.00021793 | 0.00055808 | 19.2626624 | 67.1114732 | 58.6731538 | 53.2300151 |
| -3.6972113 | 0.00021798 | 0.00055817 | 39.7548564 | 37.6148983 | 28.7613499 | 36.4205367 |
| -3.6970683 | 0.0002181 | 0.00055844 | 49.1812656 | 71.7118565 | 25.3099879 | 35.8602207 |
| -3.6969215 | 0.00021823 | 0.00055872 | 18.8528185 | 50.8748265 | 74.7795097 | 62.7553862 |
| -3.6967502 | 0.00021838 | 0.00055905 | 65.5750208 | 142.34127 | 125.399485 | 103.098135 |
| -3.6961922 | 0.00021886 | 0.00056024 | 54.509236 | 79.8301798 | 18.4072639 | 56.0315949 |
| 3.69610577 | 0.00021893 | 0.00056038 | 42.2139196 | 23.272527 | 32.2127119 | 18.4904263 |
| -3.6960685 | 0.00021896 | 0.00056042 | 20.0823501 | 120.692407 | 40.2658898 | 73.9617052 |
| -3.69576 | 0.00021923 | 0.00056106 | 94.6739363 | 33.5557366 | 39.1154358 | 33.6189569 |
| -3.6956401 | 0.00021933 | 0.00056128 | 14.7543797 | 69.5469702 | 82.8326877 | 77.3236009 |
| -3.6955013 | 0.00021945 | 0.00056154 | 45.0828268 | 83.34812 | 17.2568099 | 105.899714 |
| 3.69530205 | 0.00021963 | 0.00056193 | 115.985818 | 128.54012 | 142.656295 | 129.9933 |
| -3.6949 | 0.00021997 | 0.00056278 | 18.0331307 | 66.0290301 | 80.5317797 | 52.1093832 |
| -3.6938264 | 0.0002209 | 0.00056512 | 23.770945 | 37.6148983 | 66.7263317 | 53.2300151 |
| -3.6931652 | 0.00022148 | 0.00056654 | 21.3118818 | 97.6904913 | 69.0272397 | 38.6618005 |
| -3.6930225 | 0.0002216 | 0.00056682 | 71.3128351 | 49.7923834 | 72.4786017 | 45.9459078 |
| -3.6928144 | 0.00022179 | 0.00056719 | 14.7543797 | 148.294707 | 85.1335957 | 53.2300151 |
| -3.6928231 | 0.00022178 | 0.00056719 | 14.3445358 | 54.3927666 | 113.894946 | 96.3743432 |
| -3.6925701 | 0.000222 | 0.00056769 | 58.197831 | 46.8156648 | 21.8586259 | 45.9459078 |
| -3.6924627 | 0.00022209 | 0.00056789 | 25.8201644 | 44.6507786 | 36.8145279 | 66.1172819 |
| -3.6922951 | 0.00022224 | 0.00056819 | 38.5253247 | 63.8641439 | 27.6108959 | 36.9808526 |
| -3.6922712 | 0.00022226 | 0.00056819 | 14.7543797 | 110.679809 | 69.0272397 | 66.6775979 |
| -3.6922683 | 0.00022226 | 0.00056819 | 21.3118818 | 69.817581 | 98.9390436 | 38.1014845 |
| -3.691933 | 0.00022256 | 0.00056889 | 0 | 1195.55842 | 791.512349 | 1020.89566 |
| -3.6917411 | 0.00022272 | 0.00056923 | 43.8532952 | 58.1813175 | 25.3099879 | 36.9808526 |
| 3.69175493 | 0.00022271 | 0.00056923 | 0.40984388 | 0.54122156 | 0 | 0 |
| 3.69157819 | 0.00022287 | 0.00056955 | 6.96734596 | 11.9068743 | 3.45136199 | 0 |
| -3.6914338 | 0.00022299 | 0.00056983 | 11.8854725 | 68.1939163 | 111.594038 | 107.020346 |
| -3.6912242 | 0.00022318 | 0.00057025 | 33.6071982 | 90.9252218 | 75.9299637 | 96.9346591 |
| -3.6906925 | 0.00022364 | 0.0005714 | 40.1647002 | 37.6148983 | 27.6108959 | 38.1014845 |
| -3.6906416 | 0.00022369 | 0.00057147 | 19.2626624 | 62.51109 | 64.4254237 | 50.4284354 |
| -3.6899664 | 0.00022428 | 0.00057294 | 15.1642236 | 64.6759762 | 96.6381356 | 69.4791776 |
| -3.6898834 | 0.00022436 | 0.00057309 | 25.4103206 | 99.8553775 | 27.6108959 | 96.9346591 |
| -3.6895003 | 0.00022469 | 0.00057386 | 48.7714217 | 30.3084073 | 80.5317797 | 29.6967453 |

| | | | | | | |
|---|---|---|---|---|---|---|
| -3.6895041 | 0.00022469 | 0.00057386 | 39.3450125 | 48.980551 | 27.6108959 | 34.7395888 |
| -3.6894445 | 0.00022474 | 0.00057394 | 113.116911 | 83.6187308 | 93.1867736 | 80.6854966 |
| -3.6885401 | 0.00022554 | 0.00057594 | 25.0004767 | 75.2297966 | 44.8677058 | 39.2221164 |
| -3.6881583 | 0.00022588 | 0.00057676 | 15.5740674 | 89.0309463 | 57.5226998 | 79.0045488 |
| -3.6880856 | 0.00022595 | 0.00057688 | 264.759146 | 679.503666 | 438.322972 | 502.603406 |
| 3.687657 | 0.00022633 | 0.0005778 | 0.81968776 | 0 | 0 | 0.56031595 |
| -3.6872488 | 0.00022669 | 0.00057868 | 23.770945 | 44.6507786 | 43.7172518 | 47.6268556 |
| -3.6868617 | 0.00022704 | 0.00057952 | 84.4278393 | 50.8748265 | 71.3281477 | 59.9538065 |
| -3.6868015 | 0.00022709 | 0.00057961 | 56.5584554 | 29.2259641 | 31.0622579 | 43.144328 |
| -3.6862998 | 0.00022754 | 0.00058071 | 57.7879871 | 165.072575 | 245.046701 | 174.25826 |
| -3.6860815 | 0.00022773 | 0.00058116 | 27.45954 | 47.8981079 | 33.3631659 | 43.704644 |
| -3.6859887 | 0.00022782 | 0.00058133 | 445.910141 | 659.207858 | 533.810654 | 574.323847 |
| -3.6854587 | 0.00022829 | 0.00058249 | 49.5911095 | 81.4538445 | 63.2749697 | 48.1871716 |
| -3.6854075 | 0.00022834 | 0.00058256 | 23.770945 | 45.7332217 | 41.4163438 | 50.4284354 |
| -3.6852218 | 0.0002285 | 0.00058294 | 18.4429746 | 67.382084 | 54.0713378 | 61.0744384 |
| 3.68510885 | 0.0002286 | 0.00058315 | 2.0492194 | 1.89427545 | 1.150454 | 0.56031595 |
| -3.684881 | 0.00022881 | 0.00058363 | 1025.01954 | 297.942468 | 691.422851 | 774.916957 |
| 3.68448498 | 0.00022917 | 0.00058448 | 0.81968776 | 1.3530539 | 0 | 1.1206319 |
| -3.6844704 | 0.00022918 | 0.00058448 | 455.336551 | 57.640096 | 23.0090799 | 35.2999048 |
| -3.6841651 | 0.00022945 | 0.00058513 | 23.3611012 | 63.0523115 | 40.2658898 | 47.6268556 |
| -3.6840601 | 0.00022955 | 0.00058533 | 230.742104 | 263.304288 | 234.692615 | 262.227864 |
| -3.6839389 | 0.00022966 | 0.00058556 | 46.7222023 | 46.545054 | 25.3099879 | 35.8602207 |
| -3.6836413 | 0.00022993 | 0.0005862 | 66.8045524 | 28.4141318 | 39.1154358 | 35.2999048 |
| -3.683584 | 0.00022998 | 0.00058628 | 43.0336074 | 75.2297966 | 57.5226998 | 37.5411686 |
| -3.6833799 | 0.00023016 | 0.00058671 | 29.0989155 | 35.1794013 | 71.3281477 | 38.1014845 |
| -3.6833258 | 0.00023021 | 0.00058679 | 17.6232868 | 60.8874253 | 58.6731538 | 70.0394936 |
| -3.6831812 | 0.00023034 | 0.00058707 | 25.8201644 | 45.7332217 | 44.8677058 | 36.4205367 |
| 3.68314739 | 0.00023037 | 0.0005871 | 68.0340841 | 44.109557 | 47.1686138 | 37.5411686 |
| -3.6830776 | 0.00023043 | 0.00058722 | 13.9346919 | 102.561485 | 69.0272397 | 73.4013893 |
| -3.6824204 | 0.00023103 | 0.00058869 | 13.9346919 | 76.8534613 | 111.594038 | 63.8760181 |
| -3.6822895 | 0.00023115 | 0.00058894 | 36.4761053 | 30.8496288 | 35.6640739 | 34.7395888 |
| -3.681968 | 0.00023144 | 0.00058964 | 254.922893 | 382.914252 | 410.712076 | 411.832222 |
| -3.6814097 | 0.00023195 | 0.00059089 | 31.5579788 | 106.350036 | 86.2840497 | 90.7711837 |
| -3.6805699 | 0.00023271 | 0.00059279 | 29.9186032 | 52.2278804 | 28.7613499 | 47.0665397 |
| -3.6803552 | 0.00023291 | 0.00059324 | 27.8693838 | 38.1561199 | 75.9299637 | 37.5411686 |
| -3.6802775 | 0.00023298 | 0.00059338 | 25.8201644 | 33.2851258 | 50.6199758 | 60.5141225 |
| -3.6795032 | 0.00023369 | 0.00059513 | 217.217256 | 553.669654 | 582.129722 | 582.728587 |
| -3.6791483 | 0.00023401 | 0.00059587 | 25.8201644 | 51.416048 | 36.8145279 | 41.4633802 |
| -3.6791467 | 0.00023402 | 0.00059587 | 18.4429746 | 55.7458205 | 58.6731538 | 63.8760181 |
| -3.6786742 | 0.00023445 | 0.00059693 | 65.9848647 | 32.2026827 | 39.1154358 | 30.2570612 |
| 3.67860382 | 0.00023451 | 0.00059705 | 0 | 3.78855091 | 0 | 0 |
| -3.6785845 | 0.00023453 | 0.00059705 | 47.1320462 | 39.5091738 | 27.6108959 | 33.6189569 |
| -3.6782443 | 0.00023485 | 0.0005978 | 27.0496961 | 44.109557 | 35.6640739 | 39.7824324 |

| | | | | | | |
|---|---|---|---|---|---|---|
| -3.6782167 | 0.00023487 | 0.00059781 | 23.3611012 | 57.640096 | 36.8145279 | 56.5919108 |
| -3.6781753 | 0.00023491 | 0.00059786 | 24.1807889 | 38.4267306 | 59.8236078 | 44.8252759 |
| 3.6777301 | 0.00023532 | 0.00059886 | 6.96734596 | 3.78855091 | 3.45136199 | 2.24126379 |
| -3.6775378 | 0.0002355 | 0.00059926 | 24.1807889 | 52.4984912 | 36.8145279 | 50.4284354 |
| -3.6773394 | 0.00023568 | 0.00059968 | 102.051126 | 31.6614612 | 37.9649818 | 35.8602207 |
| -3.6772388 | 0.00023577 | 0.00059987 | 22.1315695 | 41.67406 | 82.8326877 | 50.4284354 |
| -3.6770379 | 0.00023596 | 0.0006003 | 30.738291 | 26.2492456 | 70.1776937 | 58.8331746 |
| -3.6769818 | 0.00023601 | 0.00060038 | 275.824931 | 441.09557 | 394.60572 | 425.840121 |
| -3.6769231 | 0.00023606 | 0.00060047 | 29.5087594 | 61.9698684 | 27.6108959 | 54.350647 |
| -3.6767182 | 0.00023625 | 0.00060091 | 19.6725062 | 69.0057487 | 57.5226998 | 47.0665397 |
| -3.6766961 | 0.00023627 | 0.00060091 | 29.9186032 | 44.3801678 | 56.3722458 | 29.1364293 |
| -3.6761972 | 0.00023674 | 0.00060203 | 23.3611012 | 34.9087905 | 60.9740618 | 70.0394936 |
| -3.6761824 | 0.00023675 | 0.00060203 | 20.9020379 | 48.7099402 | 48.3190678 | 99.1759229 |
| -3.6758115 | 0.00023709 | 0.00060286 | 20.9020379 | 50.3336049 | 77.0804177 | 45.3855918 |
| -3.675737 | 0.00023716 | 0.00060298 | 109.018472 | 595.614325 | 599.386532 | 587.77143 |
| -3.6756582 | 0.00023724 | 0.00060312 | 34.4268859 | 71.4412457 | 21.8586259 | 91.8918156 |
| -3.6754859 | 0.0002374 | 0.00060348 | 31.5579788 | 35.1794013 | 88.5849576 | 36.4205367 |
| -3.6754189 | 0.00023746 | 0.00060359 | 25.8201644 | 47.3568864 | 40.2658898 | 37.5411686 |
| -3.6753365 | 0.00023754 | 0.00060374 | 234.840543 | 417.823043 | 357.791193 | 381.575161 |
| -3.6750457 | 0.00023781 | 0.00060438 | 30.3284471 | 43.8389462 | 29.9118039 | 40.9030642 |
| -3.6749552 | 0.00023789 | 0.00060455 | 93.4444046 | 40.0503953 | 27.6108959 | 38.1014845 |
| -3.6744871 | 0.00023833 | 0.00060561 | 34.4268859 | 36.2618444 | 28.7613499 | 40.3427483 |
| -3.674328 | 0.00023848 | 0.00060594 | 45.0828268 | 17.0484791 | 82.8326877 | 100.856871 |
| 3.67397128 | 0.00023881 | 0.00060674 | 0.40984388 | 0 | 0 | 0.56031595 |
| -3.673552 | 0.0002392 | 0.00060769 | 44.6729829 | 101.208431 | 43.7172518 | 23.5332698 |
| 3.67329125 | 0.00023945 | 0.00060821 | 23.770945 | 14.8835929 | 16.1063559 | 10.646003 |
| -3.6733064 | 0.00023943 | 0.00060821 | 19.6725062 | 66.5702517 | 52.9208838 | 49.8681194 |
| -3.6732517 | 0.00023948 | 0.00060826 | 10.246097 | 112.574084 | 103.54086 | 81.8061285 |
| -3.6731965 | 0.00023954 | 0.00060834 | 21.7217256 | 65.4878086 | 37.9649818 | 64.4363341 |
| -3.6730278 | 0.00023969 | 0.0006087 | 20.0823501 | 43.2977247 | 58.6731538 | 87.9696039 |
| 3.67262198 | 0.00024007 | 0.00060961 | 34.8367298 | 16.5072575 | 16.1063559 | 16.2491625 |
| -3.6725869 | 0.00024011 | 0.00060961 | 46.7222023 | 33.8263474 | 25.3099879 | 56.5919108 |
| -3.6726067 | 0.00024009 | 0.00060961 | 93.0345607 | 145.588599 | 109.29313 | 109.821926 |
| 3.67164945 | 0.00024099 | 0.0006118 | 116.805506 | 66.2996409 | 69.0272397 | 47.0665397 |
| -3.6714691 | 0.00024116 | 0.00061218 | 62.2962697 | 38.4267306 | 28.7613499 | 33.058641 |
| -3.6711597 | 0.00024145 | 0.00061288 | 15.5740674 | 101.749653 | 52.9208838 | 76.202969 |
| -3.6709814 | 0.00024162 | 0.00061325 | 76.6408055 | 24.3549701 | 287.613499 | 28.0157974 |
| -3.6709581 | 0.00024164 | 0.00061326 | 8.1968776 | 81.4538445 | 144.957203 | 129.9933 |
| -3.670645 | 0.00024194 | 0.00061387 | 13.1150042 | 81.7244553 | 69.0272397 | 95.8140272 |
| -3.6706564 | 0.00024193 | 0.00061387 | 20.0823501 | 50.8748265 | 48.3190678 | 68.9188617 |
| -3.6706818 | 0.0002419 | 0.00061387 | 30.738291 | 24.6255809 | 74.7795097 | 72.8410733 |
| -3.6706224 | 0.00024196 | 0.00061388 | 22.1315695 | 80.3714014 | 41.4163438 | 49.8681194 |
| -3.6705936 | 0.00024199 | 0.0006139 | 18.4429746 | 76.3122397 | 51.7704298 | 56.5919108 |

| | | | | | | |
|---|---|---|---|---|---|---|
| -3.6705598 | 0.00024202 | 0.00061393 | 19.6725062 | 53.851545 | 55.2217918 | 53.2300151 |
| -3.6703418 | 0.00024223 | 0.00061436 | 13.1150042 | 86.3248385 | 98.9390436 | 64.99665 |
| -3.6703503 | 0.00024222 | 0.00061436 | 36.4761053 | 31.6614612 | 31.0622579 | 38.1014845 |
| -3.6688819 | 0.00024361 | 0.00061783 | 10.6559409 | 153.706923 | 94.3372276 | 69.4791776 |
| -3.6688019 | 0.00024369 | 0.00061797 | 33.1973543 | 35.4500121 | 34.5136199 | 31.9380091 |
| -3.6682283 | 0.00024424 | 0.00061931 | 57.7879871 | 20.5664192 | 46.0181598 | 54.350647 |
| -3.6668741 | 0.00024553 | 0.00062255 | 44.263139 | 59.8049822 | 52.9208838 | 49.3078035 |
| -3.6665164 | 0.00024588 | 0.00062337 | 43.4434513 | 103.914539 | 17.2568099 | 75.6426531 |
| -3.6664853 | 0.00024591 | 0.0006234 | 165.576927 | 42.2152815 | 26.4604419 | 43.144328 |
| -3.6664341 | 0.00024596 | 0.00062347 | 13.9346919 | 65.4878086 | 86.2840497 | 79.0045488 |
| -3.6661495 | 0.00024623 | 0.00062412 | 6.96734596 | 96.6080482 | 117.346308 | 174.818576 |
| -3.6656919 | 0.00024667 | 0.00062519 | 24.5906328 | 43.8389462 | 36.8145279 | 52.1093832 |
| 3.66562044 | 0.00024674 | 0.00062531 | 0 | 0.27061078 | 0 | 0 |
| -3.6650558 | 0.00024728 | 0.00062664 | 16.3937552 | 68.4645271 | 51.7704298 | 96.3743432 |
| -3.6649873 | 0.00024735 | 0.00062676 | 22.5414134 | 42.7565031 | 44.8677058 | 53.7903311 |
| -3.6648288 | 0.0002475 | 0.0006271 | 29.0989155 | 32.4732935 | 36.8145279 | 43.144328 |
| -3.664767 | 0.00024756 | 0.0006272 | 28.2792277 | 43.0271139 | 34.5136199 | 35.8602207 |
| -3.6644493 | 0.00024787 | 0.00062793 | 27.0496961 | 58.9931499 | 28.7613499 | 84.6077082 |
| -3.6643538 | 0.00024796 | 0.00062812 | 14.7543797 | 90.1133895 | 62.1245158 | 70.5998095 |
| -3.664011 | 0.0002483 | 0.00062891 | 95.493624 | 61.6992576 | 66.7263317 | 74.5220212 |
| -3.6639691 | 0.00024834 | 0.00062897 | 44.6729829 | 76.8534613 | 23.0090799 | 40.3427483 |
| -3.6637287 | 0.00024857 | 0.00062951 | 24.5906328 | 53.0397127 | 47.1686138 | 35.2999048 |
| -3.6636134 | 0.00024868 | 0.00062974 | 51.6403289 | 33.0145151 | 41.4163438 | 26.8951655 |
| -3.6635343 | 0.00024876 | 0.00062989 | 12.7051603 | 130.705006 | 47.1686138 | 130.553616 |
| -3.6634464 | 0.00024884 | 0.00063005 | 20.0823501 | 55.4752097 | 43.7172518 | 94.6933953 |
| -3.6633144 | 0.00024897 | 0.00063033 | 24.1807889 | 35.7206228 | 67.8767857 | 47.6268556 |
| -3.6630838 | 0.0002492 | 0.00063085 | 23.3611012 | 62.51109 | 85.1335957 | 33.058641 |
| -3.6628403 | 0.00024943 | 0.0006314 | 21.7217256 | 43.8389462 | 46.0181598 | 65.556966 |
| -3.6622902 | 0.00024997 | 0.00063271 | 33.1973543 | 35.9912336 | 43.7172518 | 28.5761134 |
| -3.662127 | 0.00025013 | 0.00063304 | 18.4429746 | 58.9931499 | 51.7704298 | 62.7553862 |
| -3.6621166 | 0.00025014 | 0.00063304 | 15.1642236 | 78.7477367 | 62.1245158 | 70.5998095 |
| -3.6619857 | 0.00025027 | 0.00063331 | 20.9020379 | 53.851545 | 42.5667978 | 60.5141225 |
| -3.6616953 | 0.00025055 | 0.00063398 | 33.1973543 | 30.3084073 | 72.4786017 | 36.4205367 |
| -3.6616566 | 0.00025059 | 0.00063402 | 50.4107972 | 27.0610779 | 297.967585 | 35.2999048 |
| 3.6603501 | 0.00025187 | 0.00063722 | 48.7714217 | 23.5431378 | 36.8145279 | 21.852322 |
| -3.6602069 | 0.00025201 | 0.00063752 | 110.657848 | 281.976432 | 238.143977 | 385.497373 |
| -3.6599587 | 0.00025226 | 0.00063809 | 21.3118818 | 101.479042 | 33.3631659 | 76.763285 |
| -3.6598665 | 0.00025235 | 0.00063827 | 15.1642236 | 76.5828505 | 70.1776937 | 62.7553862 |
| -3.659597 | 0.00025261 | 0.00063889 | 48.3615778 | 56.0164313 | 81.6822337 | 18.4904263 |
| -3.6591166 | 0.00025309 | 0.00064004 | 13.524848 | 71.9824673 | 89.7354116 | 71.1601255 |
| -3.6590365 | 0.00025317 | 0.00064019 | 20.492194 | 70.3588026 | 66.7263317 | 38.6618005 |
| -3.6584214 | 0.00025377 | 0.00064165 | 19.6725062 | 51.9572696 | 50.6199758 | 56.5919108 |
| -3.6584126 | 0.00025378 | 0.00064165 | 87.7065903 | 50.0629941 | 133.452663 | 14.0078987 |

| | | | | | | |
|---|---|---|---|---|---|---|
| -3.6582767 | 0.00025392 | 0.0006419 | 39.3450125 | 32.7439043 | 40.2658898 | 28.0157974 |
| 3.65827327 | 0.00025392 | 0.0006419 | 9.01656536 | 1.08244312 | 4.60181598 | 1.1206319 |
| 3.65822436 | 0.00025397 | 0.00064197 | 4.50828268 | 2.43549701 | 3.45136199 | 5.60315949 |
| -3.658033 | 0.00025416 | 0.0006424 | 132.379573 | 50.8748265 | 27.6108959 | 32.498325 |
| -3.6577621 | 0.00025443 | 0.00064303 | 60.6568942 | 40.0503953 | 29.9118039 | 29.6967453 |
| -3.6572628 | 0.00025492 | 0.00064423 | 22.9512573 | 40.0503953 | 149.559019 | 52.1093832 |
| -3.6572049 | 0.00025498 | 0.00064433 | 36.4761053 | 25.9786348 | 40.2658898 | 41.4633802 |
| 3.6571748 | 0.00025501 | 0.00064435 | 18.0331307 | 7.57710182 | 9.20363196 | 13.4475828 |
| -3.6567681 | 0.00025542 | 0.00064532 | 12.7051603 | 94.1725511 | 134.603117 | 54.350647 |
| -3.6565611 | 0.00025562 | 0.00064579 | 166.396615 | 42.4858923 | 29.9118039 | 36.4205367 |
| -3.656528 | 0.00025565 | 0.00064583 | 33.6071982 | 35.4500121 | 31.0622579 | 33.6189569 |
| -3.6561935 | 0.00025599 | 0.00064662 | 12.7051603 | 76.0416289 | 77.0804177 | 88.5299199 |
| -3.6556079 | 0.00025657 | 0.00064801 | 19.2626624 | 45.4626109 | 86.2840497 | 54.910963 |
| -3.6556017 | 0.00025658 | 0.00064801 | 93.4444046 | 163.1783 | 151.859927 | 99.1759229 |
| 3.65533765 | 0.00025684 | 0.00064863 | 20.0823501 | 8.11832337 | 17.2568099 | 7.84442328 |
| -3.6544917 | 0.00025769 | 0.00065072 | 43.0336074 | 57.640096 | 20.7081719 | 48.7474875 |
| -3.6543336 | 0.00025785 | 0.00065107 | 27.45954 | 31.1202396 | 42.5667978 | 49.3078035 |
| 3.65427229 | 0.00025791 | 0.00065117 | 39.7548564 | 24.3549701 | 31.0622579 | 25.2142177 |
| -3.6534592 | 0.00025873 | 0.00065319 | 18.0331307 | 62.2404792 | 67.8767857 | 48.7474875 |
| -3.6533831 | 0.00025881 | 0.00065333 | 25.8201644 | 50.3336049 | 46.0181598 | 32.498325 |
| -3.6531612 | 0.00025903 | 0.00065385 | 48.7714217 | 216.488623 | 171.417645 | 198.912162 |
| -3.6531094 | 0.00025908 | 0.00065393 | 57.3781432 | 84.9717847 | 14.9559019 | 67.7982298 |
| -3.6530338 | 0.00025916 | 0.00065407 | 35.6564176 | 63.8641439 | 75.9299637 | 73.4013893 |
| 3.6529718 | 0.00025922 | 0.00065418 | 0.81968776 | 3.24732935 | 2.30090799 | 3.36189569 |
| 3.65278558 | 0.00025941 | 0.00065458 | 24.5906328 | 12.1774851 | 8.05317797 | 20.1713742 |
| -3.6527719 | 0.00025942 | 0.00065458 | 26.2300083 | 66.8408624 | 27.6108959 | 79.0045488 |
| -3.6526767 | 0.00025952 | 0.00065477 | 82.7884637 | 148.024096 | 157.612197 | 163.051941 |
| -3.6525889 | 0.00025961 | 0.00065495 | 245.906328 | 390.491354 | 349.738015 | 346.275256 |
| -3.6522537 | 0.00025995 | 0.00065575 | 46.3123584 | 34.9087905 | 25.3099879 | 40.3427483 |
| -3.6520658 | 0.00026014 | 0.00065618 | 30.738291 | 64.1347547 | 31.0622579 | 36.4205367 |
| -3.6517843 | 0.00026042 | 0.00065685 | 39.7548564 | 56.0164313 | 98.9390436 | 20.7316901 |
| -3.6515463 | 0.00026067 | 0.00065741 | 15.5740674 | 110.138587 | 50.6199758 | 71.7204414 |
| -3.6515239 | 0.00026069 | 0.00065741 | 20.9020379 | 39.5091738 | 59.8236078 | 59.3934906 |
| -3.6514015 | 0.00026081 | 0.00065768 | 233.201168 | 40.0503953 | 27.6108959 | 42.5840121 |
| -3.6510714 | 0.00026115 | 0.00065847 | 15.5740674 | 101.749653 | 55.2217918 | 64.99665 |
| -3.6506373 | 0.00026159 | 0.00065953 | 25.0004767 | 69.817581 | 46.0181598 | 34.1792729 |
| -3.6505841 | 0.00026164 | 0.00065962 | 40.5745441 | 34.0969582 | 31.0622579 | 30.8173772 |
| -3.650473 | 0.00026176 | 0.00065985 | 19.6725062 | 92.0076649 | 116.195854 | 32.498325 |
| -3.6503379 | 0.0002619 | 0.00066015 | 61.0667381 | 174.543953 | 184.072639 | 164.732889 |
| -3.6501909 | 0.00026205 | 0.00066047 | 20.0823501 | 53.3103235 | 50.6199758 | 48.7474875 |
| -3.6501252 | 0.00026211 | 0.00066059 | 20.0823501 | 53.0397127 | 44.8677058 | 58.8331746 |
| -3.6499999 | 0.00026224 | 0.00066081 | 295.087594 | 277.646659 | 174.869007 | 253.262809 |
| -3.6500019 | 0.00026224 | 0.00066081 | 24.1807889 | 56.5576528 | 32.2127119 | 60.5141225 |

| | | | | | | |
|---|---|---|---|---|---|---|
| -3.6499356 | 0.00026231 | 0.00066087 | 22.1315695 | 61.1580361 | 104.691314 | 34.7395888 |
| -3.6499426 | 0.0002623 | 0.00066087 | 32.7875104 | 49.7923834 | 26.4604419 | 39.7824324 |
| -3.64951 | 0.00026274 | 0.00066192 | 28.2792277 | 41.9446708 | 34.5136199 | 34.1792729 |
| -3.6493814 | 0.00026287 | 0.0006622 | 13.9346919 | 122.857294 | 43.7172518 | 114.304454 |
| -3.6482372 | 0.00026405 | 0.0006651 | 22.5414134 | 48.980551 | 39.1154358 | 49.3078035 |
| -3.6481532 | 0.00026413 | 0.00066527 | 390.171374 | 970.680865 | 785.760079 | 932.926054 |
| -3.6480447 | 0.00026424 | 0.00066549 | 27.45954 | 53.0397127 | 33.3631659 | 36.9808526 |
| -3.6477651 | 0.00026453 | 0.00066617 | 33.6071982 | 37.8855091 | 26.4604419 | 52.1093832 |
| -3.6468315 | 0.00026549 | 0.00066854 | 43.0336074 | 44.109557 | 29.9118039 | 28.0157974 |
| -3.646571 | 0.00026576 | 0.00066916 | 18.0331307 | 46.0038325 | 62.1245158 | 76.763285 |
| -3.6464508 | 0.00026589 | 0.00066943 | 159.019425 | 334.474923 | 265.754873 | 329.465778 |
| -3.6463745 | 0.00026597 | 0.00066957 | 20.492194 | 89.0309463 | 41.4163438 | 53.2300151 |
| -3.6462368 | 0.00026611 | 0.00066988 | 27.0496961 | 39.7797845 | 33.3631659 | 39.7824324 |
| -3.6461921 | 0.00026616 | 0.00066994 | 16.3937552 | 47.8981079 | 83.9831417 | 73.4013893 |
| -3.646093 | 0.00026626 | 0.00067015 | 16.3937552 | 81.7244553 | 75.9299637 | 47.6268556 |
| -3.6455442 | 0.00026683 | 0.00067153 | 22.5414134 | 54.3927666 | 41.4163438 | 42.5840121 |
| -3.6454621 | 0.00026691 | 0.00067169 | 98.3625312 | 64.4053654 | 43.7172518 | 51.5490673 |
| -3.6451117 | 0.00026728 | 0.0006725 | 20.9020379 | 40.8622277 | 63.2749697 | 50.9887513 |
| -3.6451149 | 0.00026727 | 0.0006725 | 15.1642236 | 53.5809343 | 96.6381356 | 70.0394936 |
| -3.6450876 | 0.0002673 | 0.00067251 | 42.2139196 | 32.2026827 | 34.5136199 | 29.1364293 |
| -3.6449112 | 0.00026748 | 0.00067292 | 53.2797044 | 68.4645271 | 19.5577179 | 42.5840121 |
| -3.6446173 | 0.00026779 | 0.00067364 | 27.0496961 | 35.9912336 | 66.7263317 | 35.8602207 |
| -3.6444103 | 0.00026801 | 0.00067413 | 14.7543797 | 65.2171978 | 66.7263317 | 75.6426531 |
| -3.643895 | 0.00026854 | 0.00067543 | 20.9020379 | 43.5683354 | 56.3722458 | 47.6268556 |
| -3.6436849 | 0.00026876 | 0.00067593 | 21.3118818 | 55.4752097 | 40.2658898 | 51.5490673 |
| -3.6436188 | 0.00026883 | 0.00067605 | 11.8854725 | 61.1580361 | 106.992222 | 96.9346591 |
| -3.64348 | 0.00026898 | 0.00067636 | 38.9351686 | 30.8496288 | 28.7613499 | 38.6618005 |
| -3.643447 | 0.00026901 | 0.00067639 | 19.6725062 | 46.8156648 | 58.6731538 | 50.9887513 |
| -3.6432277 | 0.00026924 | 0.00067692 | 21.3118818 | 54.6633774 | 39.1154358 | 56.0315949 |
| -3.6429336 | 0.00026955 | 0.00067764 | 113.936599 | 26.7904671 | 34.5136199 | 45.3855918 |
| -3.6429035 | 0.00026958 | 0.00067766 | 51.6403289 | 44.9213893 | 20.7081719 | 45.3855918 |
| -3.6427433 | 0.00026975 | 0.00067798 | 229.922417 | 27.8729103 | 32.2127119 | 51.5490673 |
| -3.6427491 | 0.00026974 | 0.00067798 | 20.9020379 | 54.1221558 | 41.4163438 | 53.2300151 |
| -3.6424246 | 0.00027008 | 0.00067877 | 10.246097 | 80.3714014 | 95.4876816 | 103.098135 |
| -3.6423661 | 0.00027014 | 0.00067887 | 48.7714217 | 25.4374132 | 39.1154358 | 35.2999048 |
| -3.6418243 | 0.00027071 | 0.00068025 | 43.0336074 | 35.4500121 | 62.1245158 | 24.0935858 |
| -3.6414406 | 0.00027112 | 0.00068121 | 9.01656536 | 78.7477367 | 156.461743 | 93.0124475 |
| -3.6413648 | 0.0002712 | 0.00068136 | 69.6734596 | 20.5664192 | 34.5136199 | 168.094785 |
| -3.6401032 | 0.00027253 | 0.00068465 | 22.5414134 | 43.0271139 | 44.8677058 | 43.144328 |
| -3.6398958 | 0.00027275 | 0.00068515 | 27.0496961 | 33.8263474 | 43.7172518 | 35.8602207 |
| -3.6395638 | 0.0002731 | 0.00068592 | 65.1651769 | 158.848527 | 71.3281477 | 71.7204414 |
| -3.6395474 | 0.00027312 | 0.00068592 | 15.9839113 | 63.5935331 | 54.0713378 | 81.2458125 |
| -3.6395642 | 0.0002731 | 0.00068592 | 19.6725062 | 44.3801678 | 51.7704298 | 65.556966 |

| | | | | | | |
|---|---|---|---|---|---|---|
| -3.639437 | 0.00027323 | 0.0006861 | 42.6237635 | 152.353869 | 44.8677058 | 228.608907 |
| -3.6394421 | 0.00027323 | 0.0006861 | 61.0667381 | 116.903857 | 118.496762 | 109.26161 |
| -3.6390318 | 0.00027366 | 0.00068713 | 15.1642236 | 68.1939163 | 60.9740618 | 71.1601255 |
| -3.6388574 | 0.00027385 | 0.00068754 | 17.213443 | 79.8301798 | 62.1245158 | 48.1871716 |
| -3.6384907 | 0.00027424 | 0.00068847 | 35.6564176 | 34.367569 | 80.5317797 | 29.1364293 |
| -3.6384448 | 0.00027429 | 0.00068854 | 72.5423667 | 56.5576528 | 25.3099879 | 29.1364293 |
| -3.6380419 | 0.00027472 | 0.00068956 | 9.01656536 | 63.0523115 | 158.762651 | 119.907613 |
| -3.6380011 | 0.00027476 | 0.00068962 | 17.213443 | 57.3694852 | 57.5226998 | 60.5141225 |
| -3.6373528 | 0.00027545 | 0.0006913 | 26.6398522 | 74.1473535 | 39.1154358 | 34.7395888 |
| -3.6372828 | 0.00027553 | 0.00069144 | 21.3118818 | 50.8748265 | 54.0713378 | 39.7824324 |
| -3.6372235 | 0.00027559 | 0.00069154 | 15.9839113 | 88.7603356 | 50.6199758 | 66.6775979 |
| -3.6369904 | 0.00027584 | 0.00069211 | 21.3118818 | 48.4393295 | 41.4163438 | 52.1093832 |
| -3.6366771 | 0.00027618 | 0.0006929 | 44.6729829 | 36.5324552 | 46.0181598 | 23.5332698 |
| -3.6364259 | 0.00027645 | 0.00069352 | 22.9512573 | 40.8622277 | 65.5758777 | 39.7824324 |
| -3.6353752 | 0.00027758 | 0.0006963 | 23.3611012 | 45.4626109 | 37.9649818 | 43.704644 |
| -3.6350289 | 0.00027795 | 0.00069718 | 20.492194 | 88.7603356 | 28.7613499 | 163.612257 |
| -3.6349115 | 0.00027808 | 0.00069745 | 45.0828268 | 18.1309222 | 73.6290557 | 60.5141225 |
| -3.6348296 | 0.00027816 | 0.00069761 | 56.1486116 | 25.9786348 | 35.6640739 | 36.4205367 |
| 3.63396507 | 0.0002791 | 0.0006999 | 85.247527 | 76.3122397 | 79.3813257 | 67.2379138 |
| -3.6337292 | 0.00027935 | 0.00070049 | 273.775712 | 1653.16125 | 885.849576 | 1165.45717 |
| -3.6335456 | 0.00027955 | 0.00070093 | 38.5253247 | 38.9679522 | 24.1595339 | 40.3427483 |
| -3.6333267 | 0.00027979 | 0.00070147 | 18.0331307 | 66.5702517 | 41.4163438 | 88.5299199 |
| -3.633304 | 0.00027982 | 0.00070148 | 13.9346919 | 58.7225391 | 108.142676 | 67.7982298 |
| 3.63324588 | 0.00027988 | 0.00070159 | 35.2465737 | 12.1774851 | 13.8054479 | 11.7666349 |
| -3.6330083 | 0.00028014 | 0.00070213 | 26.6398522 | 44.109557 | 31.0622579 | 73.9617052 |
| -3.6330076 | 0.00028014 | 0.00070213 | 155.740674 | 305.51957 | 238.143977 | 273.994499 |
| -3.6328298 | 0.00028033 | 0.00070255 | 21.7217256 | 41.1328384 | 43.7172518 | 58.8331746 |
| -3.6327228 | 0.00028045 | 0.00070279 | 34.4268859 | 55.2045989 | 27.6108959 | 33.6189569 |
| -3.6325054 | 0.00028068 | 0.00070333 | 27.45954 | 35.1794013 | 33.3631659 | 40.9030642 |
| -3.6324217 | 0.00028077 | 0.00070345 | 17.213443 | 43.0271139 | 92.0363196 | 68.9188617 |
| -3.6324313 | 0.00028076 | 0.00070345 | 18.8528185 | 56.2870421 | 44.8677058 | 63.3157022 |
| -3.6323572 | 0.00028084 | 0.00070357 | 13.9346919 | 50.6042157 | 78.2308717 | 128.872668 |
| -3.6323349 | 0.00028087 | 0.00070358 | 45.0828268 | 40.8622277 | 48.3190678 | 21.852322 |
| -3.6319897 | 0.00028124 | 0.00070446 | 41.3942319 | 43.8389462 | 24.1595339 | 35.2999048 |
| -3.6316422 | 0.00028162 | 0.00070536 | 32.3776665 | 27.6022995 | 37.9649818 | 64.99665 |
| -3.6315574 | 0.00028172 | 0.00070554 | 11.8854725 | 83.0775092 | 82.8326877 | 76.202969 |
| -3.6313326 | 0.00028196 | 0.0007061 | 22.9512573 | 82.8068984 | 71.3281477 | 86.848972 |
| -3.6312178 | 0.00028209 | 0.00070636 | 259.84102 | 873.531595 | 0 | 1009.68934 |
| -3.6310808 | 0.00028224 | 0.00070668 | 39.7548564 | 61.6992576 | 50.6199758 | 69.4791776 |
| -3.6307754 | 0.00028257 | 0.00070746 | 19.2626624 | 38.6973414 | 85.1335957 | 62.7553862 |
| -3.63074 | 0.00028261 | 0.0007075 | 50.8206411 | 43.5683354 | 66.7263317 | 56.0315949 |
| 3.62854518 | 0.00028502 | 0.00071349 | 29.9186032 | 21.3782516 | 13.8054479 | 17.3697944 |
| 3.62817426 | 0.00028543 | 0.00071446 | 45.0828268 | 26.7904671 | 29.9118039 | 20.7316901 |

| | | | | | | |
|---|---|---|---|---|---|---|
| -3.6280854 | 0.00028553 | 0.00071465 | 50.0009534 | 51.9572696 | 51.7704298 | 33.058641 |
| 3.6280493 | 0.00028557 | 0.00071469 | 19.2626624 | 13.530539 | 14.9559019 | 13.4475828 |
| -3.6277398 | 0.00028591 | 0.00071549 | 140.576451 | 34.0969582 | 33.3631659 | 35.8602207 |
| -3.6277095 | 0.00028595 | 0.00071552 | 12.2953164 | 61.6992576 | 155.311289 | 70.5998095 |
| -3.6276815 | 0.00028598 | 0.00071554 | 54.0993922 | 153.436312 | 139.204933 | 149.044042 |
| -3.6274144 | 0.00028627 | 0.00071621 | 20.0823501 | 65.2171978 | 39.1154358 | 56.5919108 |
| -3.6273997 | 0.00028629 | 0.00071621 | 17.213443 | 54.3927666 | 58.6731538 | 58.8331746 |
| -3.6273783 | 0.00028631 | 0.00071622 | 100.411751 | 47.3568864 | 51.7704298 | 53.2300151 |
| 3.6269609 | 0.00028678 | 0.00071732 | 22.1315695 | 19.2133653 | 25.3099879 | 16.8094785 |
| -3.6269342 | 0.00028681 | 0.00071734 | 13.524848 | 89.0309463 | 82.8326877 | 56.0315949 |
| -3.6268104 | 0.00028694 | 0.00071763 | 19.6725062 | 57.9107067 | 40.2658898 | 62.7553862 |
| -3.6261041 | 0.00028773 | 0.00071954 | 29.0989155 | 32.2026827 | 35.6640739 | 34.7395888 |
| -3.6259916 | 0.00028785 | 0.00071979 | 24.1807889 | 44.3801678 | 33.3631659 | 51.5490673 |
| -3.6258072 | 0.00028806 | 0.00072025 | 280.743058 | 811.291116 | 0 | 967.105327 |
| -3.6256023 | 0.00028829 | 0.00072077 | 21.3118818 | 36.5324552 | 87.4345036 | 53.2300151 |
| -3.6254362 | 0.00028847 | 0.00072117 | 31.1481349 | 51.1454373 | 24.1595339 | 52.6696992 |
| -3.6252124 | 0.00028872 | 0.00072174 | 14.7543797 | 88.7603356 | 57.5226998 | 63.8760181 |
| -3.6250959 | 0.00028885 | 0.00072201 | 45.0828268 | 15.1542036 | 73.6290557 | 160.250361 |
| -3.6248545 | 0.00028912 | 0.00072263 | 9.42640924 | 114.46836 | 88.5849576 | 85.1680242 |
| 3.62443975 | 0.00028959 | 0.00072374 | 44.6729829 | 14.8835929 | 34.5136199 | 20.1713742 |
| -3.6243274 | 0.00028971 | 0.00072399 | 22.5414134 | 43.2977247 | 49.4695218 | 38.1014845 |
| -3.6242633 | 0.00028979 | 0.00072412 | 17.6232868 | 74.1473535 | 44.8677058 | 61.6347543 |
| -3.6234579 | 0.00029069 | 0.00072632 | 61.0667381 | 124.480958 | 109.29313 | 107.020346 |
| 3.62342955 | 0.00029072 | 0.00072634 | 0.40984388 | 0.54122156 | 0 | 0.56031595 |
| -3.6227185 | 0.00029152 | 0.00072829 | 29.5087594 | 34.0969582 | 31.0622579 | 36.9808526 |
| -3.6224308 | 0.00029185 | 0.00072904 | 18.0331307 | 53.5809343 | 56.3722458 | 52.1093832 |
| -3.6222541 | 0.00029205 | 0.00072948 | 49.5911095 | 100.937821 | 50.6199758 | 87.9696039 |
| -3.6221786 | 0.00029213 | 0.00072964 | 31.5579788 | 43.8389462 | 25.3099879 | 70.0394936 |
| -3.6220909 | 0.00029223 | 0.00072983 | 26.2300083 | 33.2851258 | 48.3190678 | 35.8602207 |
| -3.621427 | 0.00029298 | 0.00073165 | 20.0823501 | 34.6381797 | 70.1776937 | 76.202969 |
| -3.6212733 | 0.00029316 | 0.00073197 | 20.492194 | 43.8389462 | 63.2749697 | 43.704644 |
| -3.6212864 | 0.00029314 | 0.00073197 | 46.7222023 | 61.9698684 | 17.2568099 | 60.5141225 |
| 3.6209229 | 0.00029355 | 0.00073291 | 0 | 0 | 0 | 0 |
| -3.6207125 | 0.00029379 | 0.00073339 | 16.3937552 | 73.0649104 | 52.9208838 | 58.2728587 |
| -3.6207145 | 0.00029379 | 0.00073339 | 175.003337 | 62.2404792 | 49.4695218 | 58.2728587 |
| -3.6206176 | 0.0002939 | 0.0007336 | 39.3450125 | 27.0610779 | 32.2127119 | 73.9617052 |
| -3.6203224 | 0.00029424 | 0.00073436 | 125.002383 | 29.4965749 | 32.2127119 | 41.4633802 |
| -3.6203104 | 0.00029425 | 0.00073436 | 27.45954 | 28.9553534 | 43.7172518 | 65.556966 |
| -3.6200962 | 0.00029449 | 0.00073491 | 23.770945 | 45.7332217 | 40.2658898 | 36.4205367 |
| 3.62007529 | 0.00029452 | 0.00073492 | 56.9682993 | 35.1794013 | 40.2658898 | 35.2999048 |
| -3.6198797 | 0.00029474 | 0.00073541 | 68.8537718 | 35.1794013 | 19.5577179 | 73.4013893 |
| -3.6198368 | 0.00029479 | 0.00073548 | 29.9186032 | 39.7797845 | 36.8145279 | 28.5761134 |
| -3.6197485 | 0.00029489 | 0.00073567 | 22.9512573 | 48.7099402 | 44.8677058 | 35.8602207 |

| | | | | | | |
|---|---|---|---|---|---|---|
| -3.6194152 | 0.00029527 | 0.00073657 | 21.7217256 | 49.2511618 | 42.5667978 | 41.4633802 |
| -3.6193794 | 0.00029531 | 0.00073661 | 6.1476582 | 110.950419 | 136.904025 | 117.666349 |
| -3.6192257 | 0.00029549 | 0.00073699 | 39.7548564 | 27.6022995 | 49.4695218 | 29.1364293 |
| -3.6191857 | 0.00029553 | 0.00073705 | 16.8035991 | 115.009581 | 34.5136199 | 98.615607 |
| 3.61902882 | 0.00029571 | 0.00073744 | 0 | 0.81183234 | 2.30090799 | 0.56031595 |
| -3.6185451 | 0.00029626 | 0.00073876 | 20.0823501 | 57.9107067 | 102.390406 | 36.9808526 |
| -3.6179716 | 0.00029692 | 0.00074034 | 20.9020379 | 43.8389462 | 67.8767857 | 41.4633802 |
| -3.617735 | 0.00029719 | 0.00074096 | 37.705637 | 19.4839761 | 67.8767857 | 64.4363341 |
| -3.6171397 | 0.00029788 | 0.00074261 | 23.3611012 | 46.2744432 | 33.3631659 | 60.5141225 |
| -3.6163434 | 0.00029879 | 0.00074484 | 34.8367298 | 86.5954493 | 36.8145279 | 27.4554815 |
| 3.61630763 | 0.00029884 | 0.00074488 | 3.68859492 | 3.51794013 | 1.150454 | 2.24126379 |
| -3.6159642 | 0.00029923 | 0.00074576 | 34.017042 | 76.0416289 | 18.4072639 | 120.467929 |
| -3.615971 | 0.00029922 | 0.00074576 | 29.9186032 | 35.1794013 | 31.0622579 | 33.6189569 |
| -3.6159268 | 0.00029927 | 0.00074581 | 39.7548564 | 66.2996409 | 167.966283 | 17.3697944 |
| -3.6154864 | 0.00029978 | 0.00074702 | 24.5906328 | 48.1687187 | 47.1686138 | 31.3776931 |
| -3.615212 | 0.0003001 | 0.0007477 | 16.3937552 | 49.5217726 | 94.3372276 | 57.1522268 |
| -3.6152251 | 0.00030009 | 0.0007477 | 60.2470503 | 41.9446708 | 77.0804177 | 17.9301104 |
| -3.6151549 | 0.00030017 | 0.0007478 | 22.9512573 | 84.9717847 | 34.5136199 | 48.1871716 |
| -3.6150699 | 0.00030027 | 0.00074799 | 14.3445358 | 76.0416289 | 56.3722458 | 72.8410733 |
| -3.6149098 | 0.00030045 | 0.0007484 | 25.8201644 | 62.7817008 | 102.390406 | 26.3348496 |
| -3.6148879 | 0.00030048 | 0.0007484 | 776.654152 | 783.688816 | 905.407294 | 907.711837 |
| -3.6147556 | 0.00030063 | 0.00074873 | 22.9512573 | 49.7923834 | 39.1154358 | 38.6618005 |
| -3.6147218 | 0.00030067 | 0.00074877 | 27.8693838 | 48.1687187 | 31.0622579 | 34.7395888 |
| -3.614262 | 0.0003012 | 0.00075004 | 19.6725062 | 67.6526948 | 35.6640739 | 71.1601255 |
| -3.6140108 | 0.0003015 | 0.00075071 | 16.3937552 | 45.1920001 | 93.1867736 | 64.4363341 |
| -3.6138682 | 0.00030166 | 0.00075106 | 57.7879871 | 53.0397127 | 24.1595339 | 29.6967453 |
| 3.61369381 | 0.00030187 | 0.00075151 | 8.1968776 | 1.3530539 | 1.150454 | 1.1206319 |
| -3.6131638 | 0.00030248 | 0.00075299 | 31.5579788 | 39.7797845 | 25.3099879 | 45.3855918 |
| -3.6130902 | 0.00030257 | 0.00075315 | 21.3118818 | 41.1328384 | 42.5667978 | 53.2300151 |
| -3.6128766 | 0.00030282 | 0.00075371 | 38.9351686 | 24.6255809 | 44.8677058 | 34.1792729 |
| -3.6127152 | 0.00030301 | 0.00075412 | 332.383387 | 658.666636 | 661.511047 | 652.207764 |
| 3.6124071 | 0.00030337 | 0.00075496 | 4.91812656 | 0 | 0 | 0 |
| 3.61231927 | 0.00030347 | 0.00075516 | 10.246097 | 1.89427545 | 1.150454 | 1.68094785 |
| -3.6121726 | 0.00030364 | 0.00075553 | 251.234298 | 43.0271139 | 17.2568099 | 57.1522268 |
| -3.6121105 | 0.00030372 | 0.00075556 | 31.9678226 | 84.4305631 | 24.1595339 | 47.6268556 |
| -3.6121008 | 0.00030373 | 0.00075556 | 16.8035991 | 81.9950661 | 35.6640739 | 142.320251 |
| -3.6121094 | 0.00030372 | 0.00075556 | 33.6071982 | 67.1114732 | 29.9118039 | 31.3776931 |
| -3.6119625 | 0.00030389 | 0.00075591 | 93.8542485 | 24.8961917 | 24.1595339 | 85.1680242 |
| -3.6118094 | 0.00030407 | 0.00075629 | 27.0496961 | 61.1580361 | 28.7613499 | 42.5840121 |
| -3.6116413 | 0.00030427 | 0.00075673 | 20.9020379 | 89.8427787 | 88.5849576 | 29.1364293 |
| -3.6111872 | 0.0003048 | 0.00075794 | 36.8859492 | 69.2763595 | 59.8236078 | 57.1522268 |
| -3.6112049 | 0.00030478 | 0.00075794 | 41.3942319 | 49.2511618 | 32.2127119 | 24.6539017 |
| -3.6109605 | 0.00030507 | 0.00075854 | 25.8201644 | 35.7206228 | 33.3631659 | 50.9887513 |

| | | | | | | |
|---|---|---|---|---|---|---|
| -3.6108267 | 0.00030522 | 0.00075887 | 25.8201644 | 44.9213893 | 77.0804177 | 30.2570612 |
| -3.6106693 | 0.00030541 | 0.00075927 | 22.5414134 | 50.8748265 | 34.5136199 | 48.1871716 |
| -3.6105909 | 0.0003055 | 0.00075945 | 89.3459658 | 28.6847426 | 59.8236078 | 24.0935858 |
| -3.6105048 | 0.0003056 | 0.00075964 | 26.2300083 | 46.0038325 | 50.6199758 | 28.5761134 |
| -3.6101285 | 0.00030605 | 0.00076068 | 21.3118818 | 63.8641439 | 41.4163438 | 41.4633802 |
| -3.6099713 | 0.00030623 | 0.00076109 | 88.1164342 | 244.361534 | 211.683535 | 230.850171 |
| 3.60950288 | 0.00030678 | 0.0007624 | 163.117864 | 75.7710182 | 98.9390436 | 97.4949751 |
| -3.6091561 | 0.00030719 | 0.00076336 | 68.4439279 | 37.6148983 | 95.4876816 | 17.9301104 |
| -3.6087111 | 0.00030772 | 0.00076462 | 15.5740674 | 47.3568864 | 79.3813257 | 70.5998095 |
| -3.6086357 | 0.00030781 | 0.00076478 | 20.0823501 | 42.7565031 | 48.3190678 | 50.4284354 |
| 3.60767535 | 0.00030895 | 0.00076756 | 63.1159575 | 17.3190899 | 24.1595339 | 19.0507423 |
| -3.6074042 | 0.00030928 | 0.0007683 | 218.036944 | 45.7332217 | 26.4604419 | 33.6189569 |
| 3.60719805 | 0.00030952 | 0.00076885 | 400.827315 | 115.821413 | 185.223093 | 160.250361 |
| -3.6071594 | 0.00030957 | 0.00076891 | 103.690502 | 26.5198564 | 33.3631659 | 41.4633802 |
| -3.6069263 | 0.00030985 | 0.00076948 | 57.3781432 | 34.9087905 | 25.3099879 | 34.7395888 |
| -3.6069318 | 0.00030984 | 0.00076948 | 22.5414134 | 51.416048 | 75.9299637 | 32.498325 |
| -3.6067335 | 0.00031008 | 0.00076999 | 14.7543797 | 61.6992576 | 66.7263317 | 63.3157022 |
| -3.606137 | 0.00031079 | 0.0007717 | 24.5906328 | 65.2171978 | 26.4604419 | 95.8140272 |
| -3.6059535 | 0.00031101 | 0.00077211 | 40.1647002 | 76.8534613 | 17.2568099 | 66.1172819 |
| -3.6059552 | 0.00031101 | 0.00077211 | 23.3611012 | 31.1202396 | 71.3281477 | 50.9887513 |
| 3.60594093 | 0.00031102 | 0.00077211 | 7.78703372 | 72.253078 | 73.6290557 | 34.7395888 |
| 3.60571284 | 0.0003113 | 0.00077267 | 0 | 0 | 0 | 0 |
| -3.6057246 | 0.00031128 | 0.00077267 | 22.9512573 | 38.4267306 | 40.2658898 | 42.0236961 |
| -3.6055141 | 0.00031154 | 0.0007732 | 71.722679 | 39.7797845 | 29.9118039 | 27.4554815 |
| -3.6051131 | 0.00031202 | 0.00077433 | 59.4273626 | 29.2259641 | 31.0622579 | 33.058641 |
| -3.6043584 | 0.00031292 | 0.00077651 | 35.6564176 | 53.5809343 | 39.1154358 | 59.9538065 |
| -3.6043433 | 0.00031294 | 0.00077651 | 166.806459 | 246.255809 | 215.134897 | 243.177122 |
| -3.6042985 | 0.000313 | 0.00077659 | 26.2300083 | 37.6148983 | 33.3631659 | 35.8602207 |
| -3.6041697 | 0.00031315 | 0.00077691 | 27.8693838 | 28.143521 | 52.9208838 | 39.2221164 |
| -3.6034697 | 0.000314 | 0.00077895 | 250.004767 | 354.770731 | 317.525303 | 336.749885 |
| -3.6033171 | 0.00031418 | 0.00077935 | 215.987725 | 331.227594 | 339.383929 | 180.421735 |
| -3.6032951 | 0.00031421 | 0.00077935 | 22.1315695 | 55.7458205 | 33.3631659 | 51.5490673 |
| -3.6032725 | 0.00031424 | 0.00077936 | 129.510666 | 66.8408624 | 17.2568099 | 38.1014845 |
| -3.6032134 | 0.00031431 | 0.00077948 | 19.6725062 | 44.3801678 | 88.5849576 | 44.2649599 |
| -3.6025027 | 0.00031517 | 0.00078155 | 21.3118818 | 40.3210061 | 41.4163438 | 53.7903311 |
| -3.602173 | 0.00031557 | 0.00078249 | 14.3445358 | 64.1347547 | 74.7795097 | 58.8331746 |
| -3.6016324 | 0.00031623 | 0.00078406 | 16.8035991 | 49.7923834 | 75.9299637 | 52.6696992 |
| -3.6014844 | 0.00031641 | 0.00078442 | 22.5414134 | 40.3210061 | 46.0181598 | 37.5411686 |
| -3.6014702 | 0.00031642 | 0.00078442 | 42.6237635 | 39.7797845 | 23.0090799 | 35.8602207 |
| -3.60095 | 0.00031706 | 0.00078594 | 8.60672148 | 103.102707 | 101.239952 | 85.1680242 |
| -3.600909 | 0.00031711 | 0.000786 | 35.2465737 | 70.3588026 | 71.3281477 | 57.1522268 |
| -3.6008596 | 0.00031717 | 0.00078609 | 26.2300083 | 40.5916169 | 29.9118039 | 41.4633802 |
| -3.6004438 | 0.00031767 | 0.00078729 | 33.1973543 | 33.8263474 | 36.8145279 | 26.8951655 |

| | | | | | | |
|---|---|---|---|---|---|---|
| -3.6003805 | 0.00031775 | 0.00078742 | 18.0331307 | 105.267593 | 223.188075 | 193.309002 |
| -3.6000661 | 0.00031814 | 0.00078831 | 343.859015 | 51.6866588 | 18.4072639 | 38.6618005 |
| -3.5999367 | 0.00031829 | 0.00078864 | 16.3937552 | 38.9679522 | 112.744492 | 73.4013893 |
| -3.5997911 | 0.00031847 | 0.00078902 | 23.3611012 | 51.6866588 | 34.5136199 | 40.3427483 |
| 3.59951658 | 0.00031881 | 0.00078979 | 1.22953164 | 1.62366467 | 3.45136199 | 1.1206319 |
| -3.5991709 | 0.00031923 | 0.00079078 | 42.2139196 | 43.0271139 | 24.1595339 | 31.9380091 |
| -3.5984351 | 0.00032014 | 0.00079296 | 11.8854725 | 114.197749 | 60.9740618 | 73.4013893 |
| -3.5983969 | 0.00032018 | 0.00079302 | 69.2636157 | 28.4141318 | 35.6640739 | 30.8173772 |
| -3.5982478 | 0.00032037 | 0.00079341 | 18.0331307 | 54.9339882 | 50.6199758 | 49.3078035 |
| -3.5982163 | 0.00032041 | 0.00079345 | 20.0823501 | 43.2977247 | 43.7172518 | 53.2300151 |
| -3.5981448 | 0.0003205 | 0.00079361 | 21.3118818 | 42.4858923 | 41.4163438 | 44.8252759 |
| -3.5975352 | 0.00032125 | 0.00079541 | 32.7875104 | 30.8496288 | 32.2127119 | 30.8173772 |
| -3.5972735 | 0.00032157 | 0.00079615 | 19.2626624 | 81.9950661 | 40.2658898 | 50.4284354 |
| -3.5972404 | 0.00032161 | 0.00079619 | 49.1812656 | 44.109557 | 79.3813257 | 49.3078035 |
| 3.59704973 | 0.00032185 | 0.00079671 | 1.22953164 | 0.27061078 | 0 | 0.56031595 |
| -3.5964349 | 0.00032261 | 0.00079853 | 54.509236 | 47.6274971 | 18.4072639 | 45.3855918 |
| -3.5960969 | 0.00032303 | 0.00079951 | 35.6564176 | 67.1114732 | 21.8586259 | 43.144328 |
| -3.5960439 | 0.00032309 | 0.00079961 | 4.0984388 | 112.844695 | 153.010381 | 170.336048 |
| -3.5954348 | 0.00032385 | 0.00080136 | 16.8035991 | 49.2511618 | 55.2217918 | 64.4363341 |
| -3.5954384 | 0.00032385 | 0.00080136 | 132.789417 | 119.339354 | 155.311289 | 122.709193 |
| -3.5953987 | 0.0003239 | 0.00080141 | 22.9512573 | 39.7797845 | 35.6640739 | 47.0665397 |
| -3.5951049 | 0.00032426 | 0.00080225 | 45.0828268 | 59.8049822 | 67.8767857 | 16.8094785 |
| -3.595064 | 0.00032431 | 0.00080232 | 29.0989155 | 34.367569 | 28.7613499 | 38.1014845 |
| -3.5948809 | 0.00032454 | 0.00080282 | 17.213443 | 53.3103235 | 46.0181598 | 76.763285 |
| -3.5940999 | 0.00032551 | 0.00080517 | 16.3937552 | 47.6274971 | 60.9740618 | 66.1172819 |
| -3.5938707 | 0.0003258 | 0.00080576 | 22.5414134 | 54.1221558 | 31.0622579 | 57.7125427 |
| -3.5938903 | 0.00032578 | 0.00080576 | 137.2977 | 27.0610779 | 33.3631659 | 41.4633802 |
| -3.5936574 | 0.00032607 | 0.00080636 | 11.0657848 | 75.2297966 | 80.5317797 | 82.3664444 |
| 3.59363335 | 0.0003261 | 0.00080637 | 2.0492194 | 3.24732935 | 2.30090799 | 1.1206319 |
| -3.593241 | 0.00032659 | 0.00080752 | 22.9512573 | 42.4858923 | 49.4695218 | 33.6189569 |
| -3.5929879 | 0.00032691 | 0.00080825 | 38.1154808 | 88.4897248 | 49.4695218 | 73.9617052 |
| -3.5925936 | 0.0003274 | 0.00080941 | 27.8693838 | 114.73897 | 23.0090799 | 63.3157022 |
| -3.5924917 | 0.00032753 | 0.00080966 | 43.8532952 | 36.5324552 | 31.0622579 | 25.7745336 |
| -3.5921241 | 0.00032799 | 0.00081074 | 70.4931473 | 47.0862756 | 20.7081719 | 35.8602207 |
| -3.5921004 | 0.00032802 | 0.00081075 | 31.9678226 | 36.803066 | 90.8858656 | 28.0157974 |
| -3.5920799 | 0.00032805 | 0.00081076 | 26.6398522 | 27.8729103 | 43.7172518 | 44.8252759 |
| -3.592044 | 0.00032809 | 0.00081081 | 64.3454891 | 347.46424 | 294.516223 | 314.337247 |
| -3.5918551 | 0.00032833 | 0.00081133 | 27.45954 | 27.0610779 | 42.5667978 | 47.0665397 |
| -3.5916233 | 0.00032862 | 0.00081199 | 14.3445358 | 71.7118565 | 90.8858656 | 48.1871716 |
| -3.5913558 | 0.00032896 | 0.00081276 | 22.1315695 | 53.0397127 | 47.1686138 | 33.6189569 |
| -3.5912478 | 0.0003291 | 0.00081304 | 70.4931473 | 45.1920001 | 21.8586259 | 34.1792729 |
| -3.5894442 | 0.00033138 | 0.00081862 | 14.3445358 | 44.109557 | 86.2840497 | 87.9696039 |
| -3.5893983 | 0.00033144 | 0.0008187 | 19.6725062 | 44.9213893 | 43.7172518 | 50.4284354 |

| | | | | | | |
|---|---|---|---|---|---|---|
| -3.5891837 | 0.00033171 | 0.00081931 | 23.770945 | 38.1561199 | 56.3722458 | 33.6189569 |
| -3.5890651 | 0.00033187 | 0.00081963 | 13.524848 | 87.4072817 | 47.1686138 | 90.2108677 |
| -3.5888191 | 0.00033218 | 0.00082034 | 122.133476 | 15.1542036 | 60.9740618 | 47.6268556 |
| -3.5886367 | 0.00033241 | 0.00082078 | 20.9020379 | 74.688575 | 156.461743 | 27.4554815 |
| 3.58865461 | 0.00033239 | 0.00082078 | 31.9678226 | 13.8011497 | 14.9559019 | 15.1285306 |
| -3.5878554 | 0.00033341 | 0.00082318 | 291.398999 | 52.7691019 | 70.1776937 | 57.1522268 |
| -3.5877233 | 0.00033358 | 0.00082354 | 15.5740674 | 49.2511618 | 57.5226998 | 86.848972 |
| -3.5876007 | 0.00033373 | 0.00082386 | 68.4439279 | 35.7206228 | 19.5577179 | 53.2300151 |
| -3.587572 | 0.00033377 | 0.00082389 | 26.2300083 | 51.416048 | 27.6108959 | 43.144328 |
| -3.5873972 | 0.000334 | 0.00082438 | 55.3289238 | 39.5091738 | 31.0622579 | 24.6539017 |
| -3.5872734 | 0.00033415 | 0.00082471 | 27.8693838 | 67.6526948 | 28.7613499 | 38.1014845 |
| -3.5872254 | 0.00033422 | 0.0008248 | 15.1642236 | 78.2065152 | 52.9208838 | 57.7125427 |
| -3.5869641 | 0.00033455 | 0.0008255 | 25.8201644 | 32.4732935 | 40.2658898 | 34.1792729 |
| -3.5869767 | 0.00033453 | 0.0008255 | 70.4931473 | 38.9679522 | 20.7081719 | 41.4633802 |
| -3.5866774 | 0.00033492 | 0.0008263 | 31.5579788 | 22.7313054 | 62.1245158 | 47.6268556 |
| -3.5866707 | 0.00033493 | 0.0008263 | 31.1481349 | 48.980551 | 44.8677058 | 23.5332698 |
| 3.58660408 | 0.00033501 | 0.00082645 | 15.9839113 | 28.4141318 | 27.6108959 | 33.058641 |
| -3.5865391 | 0.0003351 | 0.00082659 | 217.217256 | 224.065725 | 180.621277 | 220.204168 |
| -3.5859277 | 0.00033588 | 0.00082847 | 17.6232868 | 36.803066 | 81.6822337 | 68.9188617 |
| -3.5858947 | 0.00033592 | 0.00082851 | 32.7875104 | 33.0145151 | 35.6640739 | 26.8951655 |
| -3.5857251 | 0.00033614 | 0.00082898 | 163.527708 | 40.0503953 | 23.0090799 | 40.9030642 |
| -3.5849378 | 0.00033716 | 0.00083142 | 20.0823501 | 62.51109 | 34.5136199 | 57.1522268 |
| 3.5842807 | 0.00033801 | 0.00083346 | 3.68859492 | 3.51794013 | 6.90272397 | 6.72379138 |
| -3.5841702 | 0.00033815 | 0.00083375 | 14.3445358 | 52.7691019 | 69.0272397 | 71.7204414 |
| -3.5840972 | 0.00033825 | 0.00083392 | 43.4434513 | 82.5362876 | 43.7172518 | 60.5141225 |
| -3.5839511 | 0.00033844 | 0.00083432 | 33.6071982 | 33.2851258 | 72.4786017 | 26.8951655 |
| -3.5837642 | 0.00033868 | 0.00083485 | 24.5906328 | 34.6381797 | 36.8145279 | 36.9808526 |
| -3.5831006 | 0.00033954 | 0.00083691 | 12.7051603 | 68.4645271 | 60.9740618 | 85.1680242 |
| 3.58299369 | 0.00033968 | 0.00083719 | 112.297223 | 153.436312 | 140.355387 | 103.65845 |
| -3.5826734 | 0.0003401 | 0.00083816 | 35.6564176 | 66.0290301 | 54.0713378 | 70.5998095 |
| -3.582588 | 0.00034021 | 0.00083837 | 18.8528185 | 62.2404792 | 34.5136199 | 96.3743432 |
| -3.5825299 | 0.00034028 | 0.00083849 | 47.1320462 | 39.7797845 | 21.8586259 | 35.8602207 |
| -3.5821335 | 0.0003408 | 0.0008397 | 526.239542 | 22.4606947 | 14.9559019 | 79.0045488 |
| -3.5820251 | 0.00034094 | 0.00083998 | 25.4103206 | 76.5828505 | 31.0622579 | 40.3427483 |
| -3.5817475 | 0.0003413 | 0.00084081 | 14.7543797 | 94.1725511 | 52.9208838 | 56.5919108 |
| -3.5816754 | 0.0003414 | 0.00084098 | 524.190322 | 1423.95392 | 1157.35672 | 1327.38848 |
| -3.5815183 | 0.0003416 | 0.00084142 | 112.707067 | 265.739785 | 93.1867736 | 191.628054 |
| 3.58149938 | 0.00034163 | 0.00084142 | 0.40984388 | 0.27061078 | 0 | 0.56031595 |
| -3.5814389 | 0.00034171 | 0.00084155 | 34.4268859 | 143.153102 | 18.4072639 | 63.3157022 |
| -3.5812236 | 0.00034199 | 0.00084218 | 18.8528185 | 33.8263474 | 65.5758777 | 86.848972 |
| -3.5808886 | 0.00034243 | 0.00084319 | 51.6403289 | 10.0125988 | 135.753571 | 107.580662 |
| -3.5807424 | 0.00034262 | 0.0008436 | 26.6398522 | 60.6168145 | 26.4604419 | 44.8252759 |
| -3.579497 | 0.00034426 | 0.00084757 | 40.5745441 | 31.1202396 | 24.1595339 | 40.9030642 |

| | | | | | | |
|---|---|---|---|---|---|---|
| -3.5791199 | 0.00034475 | 0.00084873 | 17.213443 | 65.7584193 | 44.8677058 | 53.2300151 |
| -3.5789508 | 0.00034498 | 0.00084921 | 17.213443 | 50.3336049 | 59.8236078 | 48.1871716 |
| -3.5786078 | 0.00034543 | 0.00085026 | 22.5414134 | 39.7797845 | 35.6640739 | 43.144328 |
| -3.5785793 | 0.00034547 | 0.00085029 | 25.0004767 | 42.7565031 | 36.8145279 | 31.3776931 |
| 3.57848599 | 0.00034559 | 0.00085053 | 4.0984388 | 2.70610779 | 4.60181598 | 4.48252759 |
| -3.5782157 | 0.00034595 | 0.00085134 | 19.2626624 | 43.5683354 | 44.8677058 | 50.9887513 |
| -3.5780918 | 0.00034611 | 0.00085168 | 58.6076748 | 147.482875 | 151.859927 | 173.137628 |
| -3.5778248 | 0.00034647 | 0.00085249 | 49.5911095 | 78.7477367 | 12.6549939 | 94.1330794 |
| -3.577272 | 0.0003472 | 0.00085416 | 6.55750208 | 71.7118565 | 151.859927 | 128.872668 |
| -3.5772772 | 0.00034719 | 0.00085416 | 12.7051603 | 59.8049822 | 92.0363196 | 64.99665 |
| -3.5771569 | 0.00034735 | 0.00085447 | 19.6725062 | 54.1221558 | 49.4695218 | 38.1014845 |
| -3.5767807 | 0.00034785 | 0.00085564 | 27.0496961 | 28.4141318 | 44.8677058 | 35.8602207 |
| -3.5760971 | 0.00034876 | 0.00085781 | 378.285901 | 555.56393 | 0 | 745.780528 |
| -3.5755931 | 0.00034943 | 0.00085937 | 67.2143963 | 141.529437 | 78.2308717 | 115.425085 |
| -3.5755815 | 0.00034945 | 0.00085937 | 542.633297 | 639.453271 | 598.236078 | 604.020593 |
| -3.5755269 | 0.00034952 | 0.00085949 | 13.9346919 | 59.5343714 | 81.6822337 | 56.5919108 |
| -3.5753046 | 0.00034982 | 0.00086015 | 15.5740674 | 79.8301798 | 54.0713378 | 50.4284354 |
| -3.5750186 | 0.0003502 | 0.00086103 | 573.781432 | 406.45739 | 0 | 665.095031 |
| -3.5749165 | 0.00035034 | 0.0008613 | 21.3118818 | 50.8748265 | 32.2127119 | 73.9617052 |
| -3.5748364 | 0.00035045 | 0.0008615 | 32.3776665 | 43.0271139 | 58.6731538 | 22.4126379 |
| -3.5745593 | 0.00035082 | 0.00086234 | 40.1647002 | 54.1221558 | 33.3631659 | 80.6854966 |
| -3.5737113 | 0.00035196 | 0.00086508 | 20.492194 | 73.8767427 | 42.5667978 | 38.6618005 |
| 3.57342105 | 0.00035235 | 0.00086597 | 2.86890716 | 1.08244312 | 1.150454 | 1.68094785 |
| -3.5731429 | 0.00035272 | 0.00086682 | 42.2139196 | 42.2152815 | 21.8586259 | 34.7395888 |
| -3.5726413 | 0.0003534 | 0.00086842 | 13.524848 | 77.1240721 | 55.2217918 | 67.2379138 |
| -3.5724873 | 0.00035361 | 0.00086887 | 27.45954 | 51.416048 | 24.1595339 | 53.2300151 |
| -3.5722527 | 0.00035392 | 0.00086958 | 18.0331307 | 55.4752097 | 41.4163438 | 55.4712789 |
| -3.5712129 | 0.00035533 | 0.00087291 | 29.9186032 | 29.2259641 | 32.2127119 | 33.058641 |
| -3.5712197 | 0.00035532 | 0.00087291 | 95.9034679 | 34.0969582 | 23.0090799 | 42.0236961 |
| -3.5709757 | 0.00035565 | 0.00087363 | 27.45954 | 27.6022995 | 41.4163438 | 36.4205367 |
| -3.570822 | 0.00035586 | 0.00087408 | 29.0989155 | 30.0377965 | 31.0622579 | 34.7395888 |
| -3.5704448 | 0.00035638 | 0.00087527 | 12.2953164 | 75.2297966 | 59.8236078 | 78.4442328 |
| -3.5703835 | 0.00035646 | 0.00087541 | 22.9512573 | 63.5935331 | 62.1245158 | 27.4554815 |
| -3.5700961 | 0.00035685 | 0.0008763 | 20.492194 | 38.6973414 | 67.8767857 | 40.9030642 |
| -3.5700383 | 0.00035693 | 0.00087643 | 16.3937552 | 64.946587 | 40.2658898 | 77.8839169 |
| 3.56997292 | 0.00035702 | 0.00087658 | 0 | 0 | 0 | 0 |
| -3.5697774 | 0.00035728 | 0.00087717 | 31.5579788 | 62.2404792 | 19.5577179 | 77.3236009 |
| 3.56973001 | 0.00035735 | 0.00087726 | 84.0179954 | 52.2278804 | 42.5667978 | 24.0935858 |
| -3.5694846 | 0.00035768 | 0.00087802 | 24.1807889 | 31.1202396 | 42.5667978 | 38.6618005 |
| -3.5693518 | 0.00035787 | 0.00087839 | 38.9351686 | 43.2977247 | 19.5577179 | 75.6426531 |
| -3.568906 | 0.00035847 | 0.00087982 | 22.9512573 | 98.5023236 | 131.151755 | 116.545717 |
| -3.5688427 | 0.00035856 | 0.00087997 | 31.1481349 | 29.4965749 | 28.7613499 | 36.4205367 |
| -3.5684831 | 0.00035905 | 0.00088104 | 37.705637 | 33.8263474 | 24.1595339 | 34.7395888 |

| | | | | | | |
|---|---|---|---|---|---|---|
| -3.5684962 | 0.00035904 | 0.00088104 | 36.0662614 | 52.2278804 | 26.4604419 | 28.5761134 |
| -3.5684589 | 0.00035909 | 0.00088106 | 13.524848 | 61.9698684 | 260.002603 | 43.704644 |
| -3.5683693 | 0.00035921 | 0.00088129 | 39.3450125 | 28.4141318 | 26.4604419 | 36.9808526 |
| 3.5680941 | 0.00035959 | 0.00088215 | 0 | 0.27061078 | 0 | 0.56031595 |
| -3.5678751 | 0.00035989 | 0.00088282 | 18.8528185 | 52.4984912 | 40.2658898 | 49.8681194 |
| -3.5674358 | 0.00036049 | 0.00088424 | 78.6900249 | 55.2045989 | 94.3372276 | 11.7666349 |
| -3.5672466 | 0.00036075 | 0.00088481 | 17.6232868 | 38.9679522 | 104.691314 | 54.910963 |
| -3.5672121 | 0.0003608 | 0.00088486 | 20.0823501 | 38.4267306 | 43.7172518 | 51.5490673 |
| -3.566963 | 0.00036114 | 0.00088563 | 17.213443 | 75.2297966 | 54.0713378 | 41.4633802 |
| -3.5667378 | 0.00036145 | 0.00088632 | 20.9020379 | 35.9912336 | 42.5667978 | 53.7903311 |
| -3.5665473 | 0.00036172 | 0.0008869 | 14.7543797 | 54.9339882 | 56.3722458 | 68.9188617 |
| 3.56652539 | 0.00036175 | 0.00088691 | 34.017042 | 20.0251977 | 21.8586259 | 15.1285306 |
| 3.56642706 | 0.00036188 | 0.00088717 | 48.7714217 | 28.9553534 | 32.2127119 | 33.6189569 |
| -3.5663781 | 0.00036195 | 0.00088727 | 19.2626624 | 47.6274971 | 39.1154358 | 53.2300151 |
| -3.5654552 | 0.00036323 | 0.00089033 | 226.643666 | 493.05284 | 482.040224 | 495.879615 |
| -3.5653932 | 0.00036331 | 0.00089047 | 22.1315695 | 165.072575 | 26.4604419 | 64.99665 |
| -3.5652994 | 0.00036344 | 0.00089073 | 17.6232868 | 54.3927666 | 55.2217918 | 42.5840121 |
| -3.5652058 | 0.00036357 | 0.00089098 | 22.5414134 | 36.2618444 | 35.6640739 | 48.1871716 |
| -3.5642653 | 0.00036488 | 0.00089411 | 31.9678226 | 41.1328384 | 25.3099879 | 32.498325 |
| -3.5634552 | 0.00036601 | 0.00089681 | 134.428793 | 227.313054 | 139.204933 | 188.826475 |
| -3.5633323 | 0.00036618 | 0.00089716 | 20.492194 | 39.5091738 | 47.1686138 | 40.3427483 |
| -3.5632253 | 0.00036633 | 0.00089746 | 21.3118818 | 39.238563 | 42.5667978 | 38.6618005 |
| -3.5624946 | 0.00036735 | 0.00089989 | 13.9346919 | 91.4664434 | 58.6731538 | 52.6696992 |
| -3.5617625 | 0.00036837 | 0.00090233 | 18.4429746 | 38.9679522 | 49.4695218 | 67.2379138 |
| -3.561221 | 0.00036913 | 0.00090413 | 38.9351686 | 48.7099402 | 34.5136199 | 50.4284354 |
| -3.5611372 | 0.00036925 | 0.00090435 | 157.38005 | 221.359617 | 213.984443 | 212.359745 |
| -3.5610903 | 0.00036932 | 0.00090444 | 27.0496961 | 83.8893415 | 104.691314 | 86.848972 |
| -3.5608474 | 0.00036966 | 0.00090521 | 28.2792277 | 69.0057487 | 24.1595339 | 44.8252759 |
| -3.5608247 | 0.00036969 | 0.00090522 | 43.0336074 | 30.579018 | 25.3099879 | 33.6189569 |
| -3.5603797 | 0.00037032 | 0.00090669 | 25.4103206 | 27.6022995 | 44.8677058 | 41.4633802 |
| -3.5600504 | 0.00037078 | 0.00090776 | 22.9512573 | 40.5916169 | 33.3631659 | 82.3664444 |
| -3.5597591 | 0.0003712 | 0.00090869 | 57.7879871 | 113.927138 | 101.239952 | 127.19172 |
| -3.5597196 | 0.00037125 | 0.00090876 | 26.6398522 | 49.7923834 | 24.1595339 | 63.3157022 |
| -3.5593012 | 0.00037184 | 0.00091014 | 18.4429746 | 56.0164313 | 42.5667978 | 45.3855918 |
| -3.5592426 | 0.00037193 | 0.00091028 | 26.2300083 | 37.0736767 | 29.9118039 | 34.1792729 |
| -3.5592213 | 0.00037196 | 0.00091028 | 40.984388 | 24.8961917 | 28.7613499 | 66.6775979 |
| -3.5588154 | 0.00037253 | 0.00091162 | 11.4756286 | 47.0862756 | 96.6381356 | 106.46003 |
| -3.558667 | 0.00037274 | 0.00091205 | 119.264569 | 33.0145151 | 31.0622579 | 31.3776931 |
| -3.558652 | 0.00037276 | 0.00091205 | 23.3611012 | 36.2618444 | 40.2658898 | 33.6189569 |
| -3.5584866 | 0.000373 | 0.00091255 | 15.9839113 | 55.2045989 | 54.0713378 | 52.1093832 |
| -3.5583042 | 0.00037326 | 0.00091312 | 25.8201644 | 125.292791 | 123.098578 | 74.5220212 |
| -3.5576466 | 0.00037419 | 0.00091534 | 48.7714217 | 107.70309 | 18.4072639 | 38.6618005 |
| -3.5575334 | 0.00037435 | 0.00091566 | 9.83625312 | 70.9000241 | 102.390406 | 76.202969 |

| | | | | | | |
|---|---|---|---|---|---|---|
| -3.5571027 | 0.00037497 | 0.0009171 | 11.0657848 | 46.545054 | 116.195854 | 99.1759229 |
| -3.5570657 | 0.00037502 | 0.00091716 | 559.027052 | 811.561727 | 746.644643 | 867.929404 |
| -3.5565359 | 0.00037578 | 0.00091894 | 20.9020379 | 43.0271139 | 37.9649818 | 40.9030642 |
| 3.55585552 | 0.00037675 | 0.00092125 | 2.45906328 | 2.43549701 | 1.150454 | 1.1206319 |
| -3.5556322 | 0.00037707 | 0.00092196 | 156.970206 | 261.680623 | 227.789891 | 225.807327 |
| -3.5550278 | 0.00037794 | 0.00092401 | 39.3450125 | 122.045461 | 19.5577179 | 44.2649599 |
| -3.5548631 | 0.00037818 | 0.00092452 | 13.1150042 | 134.493557 | 28.7613499 | 151.285306 |
| -3.554606 | 0.00037855 | 0.00092536 | 23.770945 | 43.2977247 | 28.7613499 | 47.0665397 |
| -3.5544501 | 0.00037877 | 0.00092584 | 14.7543797 | 74.9591858 | 48.3190678 | 59.3934906 |
| -3.5543338 | 0.00037894 | 0.00092617 | 20.9020379 | 48.980551 | 40.2658898 | 36.4205367 |
| -3.5543093 | 0.00037897 | 0.00092619 | 22.1315695 | 99.3141559 | 41.4163438 | 35.2999048 |
| -3.5538982 | 0.00037957 | 0.00092751 | 12.2953164 | 61.6992576 | 75.9299637 | 67.2379138 |
| -3.5538963 | 0.00037957 | 0.00092751 | 26.2300083 | 39.7797845 | 41.4163438 | 26.8951655 |
| -3.5538377 | 0.00037965 | 0.00092764 | 23.770945 | 50.8748265 | 43.7172518 | 28.5761134 |
| -3.5532539 | 0.0003805 | 0.00092963 | 14.7543797 | 54.6633774 | 78.2308717 | 50.4284354 |
| -3.5531307 | 0.00038068 | 0.00093 | 33.1973543 | 49.5217726 | 21.8586259 | 38.6618005 |
| -3.5526983 | 0.0003813 | 0.00093146 | 14.3445358 | 71.7118565 | 56.3722458 | 53.7903311 |
| -3.552594 | 0.00038145 | 0.00093176 | 29.0989155 | 45.1920001 | 40.2658898 | 24.0935858 |
| -3.5523046 | 0.00038187 | 0.00093271 | 20.0823501 | 61.6992576 | 35.6640739 | 44.2649599 |
| -3.5517358 | 0.0003827 | 0.00093466 | 0 | 826.986541 | 471.686138 | 941.330794 |
| -3.5515529 | 0.00038297 | 0.00093524 | 52.8698605 | 30.8496288 | 27.6108959 | 29.6967453 |
| -3.5513654 | 0.00038324 | 0.00093583 | 29.9186032 | 55.7458205 | 20.7081719 | 60.5141225 |
| -3.5510074 | 0.00038376 | 0.00093697 | 31.9678226 | 62.51109 | 26.4604419 | 31.3776931 |
| 3.55101493 | 0.00038375 | 0.00093697 | 76.2309617 | 76.8534613 | 56.3722458 | 80.6854966 |
| -3.5506842 | 0.00038423 | 0.00093805 | 229.512573 | 54.3927666 | 17.2568099 | 36.4205367 |
| -3.550571 | 0.0003844 | 0.00093838 | 27.0496961 | 23.5431378 | 55.2217918 | 144.561515 |
| -3.5505407 | 0.00038444 | 0.00093842 | 17.213443 | 94.4431619 | 50.6199758 | 40.9030642 |
| -3.5494137 | 0.00038609 | 0.00094237 | 25.0004767 | 55.7458205 | 24.1595339 | 68.9188617 |
| -3.5493718 | 0.00038615 | 0.00094245 | 41.8040758 | 24.6255809 | 31.0622579 | 34.1792729 |
| -3.5492653 | 0.00038631 | 0.00094276 | 11.4756286 | 71.1706349 | 77.0804177 | 65.556966 |
| -3.5492191 | 0.00038638 | 0.00094285 | 31.1481349 | 31.6614612 | 100.089498 | 30.2570612 |
| -3.5489764 | 0.00038673 | 0.00094365 | 25.4103206 | 30.579018 | 75.9299637 | 36.4205367 |
| -3.5485302 | 0.00038739 | 0.00094518 | 36.8859492 | 32.2026827 | 24.1595339 | 34.1792729 |
| -3.5485053 | 0.00038742 | 0.0009452 | 62.7061136 | 29.7671857 | 27.6108959 | 31.9380091 |
| -3.5481666 | 0.00038792 | 0.00094634 | 9.42640924 | 59.5343714 | 94.3372276 | 108.701294 |
| -3.5480092 | 0.00038815 | 0.00094684 | 28.2792277 | 25.1668025 | 47.1686138 | 36.9808526 |
| -3.5476782 | 0.00038864 | 0.00094796 | 22.5414134 | 48.980551 | 40.2658898 | 31.3776931 |
| -3.5475768 | 0.00038879 | 0.00094821 | 33.1973543 | 75.5004074 | 210.533081 | 14.5682147 |
| -3.5475691 | 0.0003888 | 0.00094821 | 90.9853413 | 37.0736767 | 21.8586259 | 37.5411686 |
| -3.5474795 | 0.00038894 | 0.00094846 | 62.7061136 | 34.0969582 | 21.8586259 | 37.5411686 |
| -3.5473054 | 0.00038919 | 0.00094901 | 22.9512573 | 43.2977247 | 46.0181598 | 29.6967453 |
| -3.5471128 | 0.00038948 | 0.00094964 | 29.0989155 | 34.6381797 | 26.4604419 | 34.1792729 |
| -3.5466886 | 0.00039011 | 0.00095109 | 12.7051603 | 56.0164313 | 79.3813257 | 64.99665 |

| | | | | | | |
|---|---|---|---|---|---|---|
| 3.54652948 | 0.00039034 | 0.0009516 | 7.78703372 | 10.0125988 | 0 | 5.04284354 |
| -3.5457046 | 0.00039156 | 0.00095451 | 15.9839113 | 61.9698684 | 40.2658898 | 72.2807574 |
| -3.5452559 | 0.00039223 | 0.00095606 | 51.230485 | 28.9553534 | 29.9118039 | 28.5761134 |
| 3.5450295 | 0.00039257 | 0.00095681 | 90.9853413 | 42.7565031 | 32.2127119 | 38.6618005 |
| -3.5449823 | 0.00039264 | 0.00095691 | 19.6725062 | 66.8408624 | 41.4163438 | 38.1014845 |
| -3.5443841 | 0.00039353 | 0.00095894 | 12.7051603 | 63.3229223 | 56.3722458 | 80.6854966 |
| 3.544393 | 0.00039352 | 0.00095894 | 34.4268859 | 13.2599282 | 28.7613499 | 15.1285306 |
| -3.5441505 | 0.00039388 | 0.00095972 | 14.7543797 | 57.0988744 | 57.5226998 | 54.910963 |
| -3.5437468 | 0.00039448 | 0.00096112 | 11.0657848 | 60.6168145 | 70.1776937 | 96.3743432 |
| -3.5434937 | 0.00039486 | 0.00096197 | 13.9346919 | 73.3355211 | 49.4695218 | 63.3157022 |
| -3.5433571 | 0.00039507 | 0.00096239 | 52.0501728 | 34.6381797 | 48.3190678 | 19.6110582 |
| -3.5414409 | 0.00039795 | 0.00096934 | 13.1150042 | 146.129821 | 41.4163438 | 71.7204414 |
| -3.5409582 | 0.00039868 | 0.00097104 | 36.0662614 | 72.253078 | 14.9559019 | 105.339398 |
| -3.540875 | 0.0003988 | 0.00097127 | 39.7548564 | 40.5916169 | 36.8145279 | 66.6775979 |
| -3.5406673 | 0.00039912 | 0.00097182 | 195.495531 | 172.108456 | 165.665375 | 183.223315 |
| -3.5406799 | 0.0003991 | 0.00097182 | 20.0823501 | 41.1328384 | 39.1154358 | 42.5840121 |
| -3.5406752 | 0.0003991 | 0.00097182 | 153.281611 | 419.987929 | 476.287954 | 498.681194 |
| -3.5404192 | 0.00039949 | 0.00097266 | 19.6725062 | 36.5324552 | 44.8677058 | 47.6268556 |
| -3.5401237 | 0.00039994 | 0.00097367 | 115.575974 | 24.6255809 | 85.1335957 | 19.6110582 |
| -3.5399527 | 0.0004002 | 0.00097423 | 22.9512573 | 32.7439043 | 40.2658898 | 35.8602207 |
| -3.5399205 | 0.00040025 | 0.00097428 | 36.4761053 | 40.3210061 | 40.2658898 | 64.4363341 |
| -3.5392208 | 0.00040131 | 0.00097679 | 32.7875104 | 27.0610779 | 28.7613499 | 34.1792729 |
| -3.5391064 | 0.00040148 | 0.00097714 | 15.9839113 | 74.4179643 | 37.9649818 | 66.1172819 |
| -3.5388482 | 0.00040188 | 0.00097802 | 19.2626624 | 41.9446708 | 37.9649818 | 61.6347543 |
| -3.5387317 | 0.00040205 | 0.00097838 | 29.5087594 | 80.1007906 | 19.5577179 | 59.9538065 |
| -3.538573 | 0.0004023 | 0.00097889 | 68.0340841 | 13.8011497 | 57.5226998 | 56.0315949 |
| -3.5384184 | 0.00040253 | 0.00097939 | 23.3611012 | 37.6148983 | 31.0622579 | 64.99665 |
| 3.53811106 | 0.000403 | 0.00098046 | 0.40984388 | 0.54122156 | 0 | 0 |
| -3.5374125 | 0.00040407 | 0.00098294 | 24.1807889 | 28.4141318 | 43.7172518 | 38.6618005 |
| -3.5374051 | 0.00040408 | 0.00098294 | 18.0331307 | 57.640096 | 34.5136199 | 66.1172819 |
| -3.5367126 | 0.00040514 | 0.00098544 | 137.707544 | 415.928768 | 444.075242 | 435.925808 |
| -3.5355239 | 0.00040697 | 0.00098981 | 22.1315695 | 51.1454373 | 55.2217918 | 28.0157974 |
| -3.5349447 | 0.00040786 | 0.00099186 | 31.9678226 | 64.6759762 | 35.6640739 | 23.5332698 |
| -3.5349384 | 0.00040787 | 0.00099186 | 47.5418901 | 19.7545869 | 55.2217918 | 33.6189569 |
| -3.5349001 | 0.00040793 | 0.00099193 | 39.7548564 | 20.83703 | 34.5136199 | 50.9887513 |
| -3.5338864 | 0.0004095 | 0.00099566 | 17.6232868 | 52.2278804 | 36.8145279 | 69.4791776 |
| -3.5338308 | 0.00040958 | 0.0009958 | 44.263139 | 92.0076649 | 14.9559019 | 52.6696992 |
| -3.5330848 | 0.00041074 | 0.00099854 | 23.770945 | 48.980551 | 31.0622579 | 34.1792729 |
| -3.5325717 | 0.00041154 | 0.0010004 | 10.6559409 | 51.6866588 | 74.7795097 | 132.79488 |
| -3.5325199 | 0.00041162 | 0.00100052 | 34.8367298 | 26.2492456 | 27.6108959 | 36.4205367 |
| -3.5322462 | 0.00041205 | 0.00100148 | 25.8201644 | 30.8496288 | 41.4163438 | 29.6967453 |
| -3.5322251 | 0.00041208 | 0.00100149 | 28.2792277 | 62.2404792 | 62.1245158 | 43.704644 |
| -3.5319776 | 0.00041246 | 0.00100235 | 20.492194 | 40.3210061 | 35.6640739 | 79.0045488 |

| | | | | | | |
|---|---|---|---|---|---|---|
| -3.5314521 | 0.00041328 | 0.00100427 | 20.9020379 | 33.0145151 | 41.4163438 | 63.3157022 |
| 3.53112194 | 0.0004138 | 0.00100545 | 14.3445358 | 36.5324552 | 55.2217918 | 21.852322 |
| -3.5308316 | 0.00041426 | 0.00100648 | 17.213443 | 57.9107067 | 42.5667978 | 46.5062237 |
| -3.5307901 | 0.00041432 | 0.00100653 | 349.59683 | 641.618157 | 605.138802 | 626.993547 |
| 3.53077702 | 0.00041434 | 0.00100653 | 0 | 0 | 0 | 0 |
| -3.5306256 | 0.00041458 | 0.00100703 | 133.609105 | 36.5324552 | 20.7081719 | 41.4633802 |
| -3.5304502 | 0.00041485 | 0.00100763 | 14.3445358 | 63.0523115 | 51.7704298 | 56.5919108 |
| -3.5302508 | 0.00041517 | 0.00100831 | 13.9346919 | 50.6042157 | 67.8767857 | 59.3934906 |
| -3.5300722 | 0.00041545 | 0.00100891 | 15.5740674 | 53.851545 | 58.6731538 | 46.5062237 |
| -3.5299799 | 0.00041559 | 0.00100919 | 30.738291 | 28.4141318 | 28.7613499 | 31.9380091 |
| -3.5298706 | 0.00041576 | 0.00100953 | 30.738291 | 39.238563 | 21.8586259 | 47.0665397 |
| -3.5296571 | 0.0004161 | 0.00101021 | 29.9186032 | 37.8855091 | 31.0622579 | 25.7745336 |
| -3.5296539 | 0.0004161 | 0.00101021 | 35.6564176 | 30.0377965 | 54.0713378 | 23.5332698 |
| -3.5289035 | 0.00041729 | 0.001013 | 66.3947085 | 128.810731 | 128.850847 | 102.537819 |
| -3.528799 | 0.00041745 | 0.00101326 | 23.3611012 | 62.7817008 | 42.5667978 | 28.0157974 |
| -3.5287949 | 0.00041746 | 0.00101326 | 18.0331307 | 49.7923834 | 37.9649818 | 52.6696992 |
| -3.5287564 | 0.00041752 | 0.00101333 | 22.5414134 | 38.1561199 | 31.0622579 | 42.0236961 |
| -3.5283974 | 0.00041808 | 0.00101463 | 26.6398522 | 23.272527 | 62.1245158 | 47.0665397 |
| -3.5280181 | 0.00041868 | 0.00101601 | 8.60672148 | 86.5954493 | 154.160835 | 56.0315949 |
| -3.5274041 | 0.00041966 | 0.00101829 | 32.7875104 | 27.8729103 | 26.4604419 | 35.8602207 |
| -3.5270948 | 0.00042015 | 0.00101941 | 38.5253247 | 26.2492456 | 26.4604419 | 71.7204414 |
| -3.5270709 | 0.00042018 | 0.00101942 | 24.1807889 | 146.129821 | 81.6822337 | 101.977503 |
| -3.5270346 | 0.00042024 | 0.00101949 | 32.7875104 | 45.1920001 | 19.5577179 | 57.1522268 |
| -3.5268961 | 0.00042046 | 0.00101994 | 16.3937552 | 47.8981079 | 49.4695218 | 50.4284354 |
| -3.5265697 | 0.00042098 | 0.00102112 | 109.018472 | 41.1328384 | 41.4163438 | 18.4904263 |
| -3.5263916 | 0.00042126 | 0.00102173 | 17.6232868 | 35.4500121 | 54.0713378 | 67.7982298 |
| -3.5261149 | 0.0004217 | 0.00102273 | 27.8693838 | 26.2492456 | 43.7172518 | 31.9380091 |
| 3.52567251 | 0.00042241 | 0.00102436 | 48.7714217 | 33.8263474 | 25.3099879 | 23.5332698 |
| -3.5255468 | 0.00042261 | 0.00102477 | 26.6398522 | 26.5198564 | 34.5136199 | 39.2221164 |
| -3.5248054 | 0.00042379 | 0.00102756 | 31.1481349 | 47.0862756 | 36.8145279 | 22.4126379 |
| -3.5245036 | 0.00042428 | 0.00102866 | 34.4268859 | 41.1328384 | 19.5577179 | 57.7125427 |
| -3.5239675 | 0.00042514 | 0.00103066 | 287.710404 | 476.004361 | 556.819734 | 494.198667 |
| -3.523613 | 0.00042571 | 0.00103197 | 25.4103206 | 45.4626109 | 24.1595339 | 52.1093832 |
| 3.52316678 | 0.00042642 | 0.00103363 | 18.0331307 | 6.22404792 | 6.90272397 | 5.60315949 |
| -3.5231321 | 0.00042648 | 0.00103368 | 27.45954 | 34.9087905 | 25.3099879 | 53.7903311 |
| -3.5224431 | 0.00042759 | 0.0010363 | 25.0004767 | 46.2744432 | 35.6640739 | 27.4554815 |
| -3.5222417 | 0.00042791 | 0.00103701 | 42.6237635 | 104.455761 | 110.443584 | 10.646003 |
| -3.5221776 | 0.00042802 | 0.00103718 | 22.5414134 | 66.0290301 | 24.1595339 | 67.2379138 |
| 3.52203872 | 0.00042824 | 0.00103765 | 14.3445358 | 8.11832337 | 12.6549939 | 9.52537113 |
| -3.5218598 | 0.00042853 | 0.00103827 | 12.2953164 | 68.1939163 | 106.992222 | 46.5062237 |
| -3.5211604 | 0.00042966 | 0.00104093 | 16.3937552 | 62.7817008 | 58.6731538 | 38.1014845 |
| 3.5207834 | 0.00043027 | 0.00104234 | 0 | 0 | 0 | 0 |
| -3.5199713 | 0.00043159 | 0.00104545 | 19.2626624 | 70.9000241 | 29.9118039 | 57.1522268 |

| | | | | | | |
|---|---|---|---|---|---|---|
| 3.51974878 | 0.00043196 | 0.00104619 | 0.81968776 | 0.54122156 | 0 | 0.56031595 |
| -3.5197454 | 0.00043196 | 0.00104619 | 21.3118818 | 41.67406 | 35.6640739 | 35.2999048 |
| -3.5196587 | 0.0004321 | 0.00104645 | 21.7217256 | 34.367569 | 34.5136199 | 43.704644 |
| -3.5193393 | 0.00043262 | 0.00104763 | 18.0331307 | 42.7565031 | 39.1154358 | 65.556966 |
| -3.5192211 | 0.00043282 | 0.00104802 | 12.7051603 | 46.545054 | 72.4786017 | 80.1251807 |
| -3.518437 | 0.0004341 | 0.00105105 | 20.0823501 | 43.5683354 | 36.8145279 | 38.6618005 |
| -3.5175113 | 0.00043561 | 0.00105464 | 20.9020379 | 35.1794013 | 35.6640739 | 48.7474875 |
| -3.5174609 | 0.0004357 | 0.00105476 | 25.4103206 | 39.238563 | 35.6640739 | 26.8951655 |
| -3.5173282 | 0.00043591 | 0.00105516 | 11.0657848 | 106.079425 | 62.1245158 | 58.2728587 |
| -3.51732 | 0.00043593 | 0.00105516 | 30.3284471 | 75.5004074 | 40.2658898 | 22.4126379 |
| -3.5170042 | 0.00043645 | 0.00105634 | 117.21535 | 55.4752097 | 39.1154358 | 54.350647 |
| -3.5169585 | 0.00043652 | 0.00105644 | 13.9346919 | 41.4034492 | 90.8858656 | 64.4363341 |
| -3.5168559 | 0.00043669 | 0.00105677 | 35.2465737 | 21.1076408 | 51.7704298 | 34.7395888 |
| -3.5167967 | 0.00043679 | 0.00105693 | 20.0823501 | 38.4267306 | 37.9649818 | 40.9030642 |
| -3.5167302 | 0.0004369 | 0.00105711 | 18.0331307 | 57.640096 | 34.5136199 | 53.2300151 |
| -3.5165385 | 0.00043721 | 0.0010578 | 22.1315695 | 34.0969582 | 56.3722458 | 33.058641 |
| -3.5163295 | 0.00043756 | 0.00105855 | 21.7217256 | 40.5916169 | 34.5136199 | 34.7395888 |
| -3.5158165 | 0.0004384 | 0.00106052 | 26.6398522 | 44.3801678 | 27.6108959 | 30.8173772 |
| -3.5157126 | 0.00043858 | 0.00106085 | 108.198784 | 35.4500121 | 19.5577179 | 42.5840121 |
| 3.5155197 | 0.00043889 | 0.00106155 | 22.9512573 | 23.0019162 | 18.4072639 | 14.0078987 |
| -3.5153009 | 0.00043926 | 0.00106226 | 44.6729829 | 22.7313054 | 28.7613499 | 39.7824324 |
| -3.5153021 | 0.00043925 | 0.00106226 | 53.6895483 | 29.4965749 | 23.0090799 | 35.8602207 |
| -3.5152543 | 0.00043933 | 0.00106237 | 235.660231 | 31.3908504 | 25.3099879 | 36.4205367 |
| -3.5149443 | 0.00043985 | 0.00106353 | 21.3118818 | 37.0736767 | 36.8145279 | 35.8602207 |
| -3.514852 | 0.00044 | 0.00106382 | 31.9678226 | 60.6168145 | 17.2568099 | 75.6426531 |
| -3.5147852 | 0.00044011 | 0.00106401 | 26.6398522 | 25.1668025 | 39.1154358 | 36.9808526 |
| -3.514346 | 0.00044084 | 0.00106569 | 16.3937552 | 41.4034492 | 52.9208838 | 53.7903311 |
| -3.5142003 | 0.00044108 | 0.00106619 | 1.63937552 | 132.05806 | 234.692615 | 187.145527 |
| -3.5138762 | 0.00044162 | 0.00106742 | 88.1164342 | 31.3908504 | 24.1595339 | 35.2999048 |
| -3.5136861 | 0.00044193 | 0.0010681 | 0 | 633.499834 | 586.731538 | 771.555061 |
| -3.5136537 | 0.00044199 | 0.00106815 | 21.3118818 | 42.4858923 | 52.9208838 | 29.6967453 |
| -3.513551 | 0.00044216 | 0.00106848 | 209.430223 | 33.2851258 | 33.3631659 | 26.3348496 |
| -3.5134971 | 0.00044225 | 0.00106862 | 30.738291 | 24.8961917 | 34.5136199 | 31.3776931 |
| -3.5130317 | 0.00044302 | 0.00107041 | 8.1968776 | 53.3103235 | 146.107657 | 95.2537113 |
| -3.5126817 | 0.00044361 | 0.00107174 | 56.5584554 | 97.1492697 | 105.841768 | 110.382242 |
| -3.5123728 | 0.00044412 | 0.00107291 | 769.276963 | 1461.29821 | 1166.56035 | 1251.18551 |
| -3.51231 | 0.00044423 | 0.00107308 | 70.4931473 | 23.0019162 | 33.3631659 | 33.6189569 |
| -3.5122715 | 0.00044429 | 0.00107316 | 24.1807889 | 33.0145151 | 32.2127119 | 75.6426531 |
| -3.5121773 | 0.00044445 | 0.00107328 | 42.2139196 | 28.143521 | 24.1595339 | 34.1792729 |
| -3.5121616 | 0.00044448 | 0.00107328 | 23.3611012 | 72.5236888 | 28.7613499 | 39.2221164 |
| -3.5122118 | 0.00044439 | 0.00107328 | 27.45954 | 50.0629941 | 23.0090799 | 39.7824324 |
| -3.5121703 | 0.00044446 | 0.00107328 | 18.0331307 | 35.7206228 | 48.3190678 | 56.0315949 |
| -3.511559 | 0.00044549 | 0.00107564 | 50.8206411 | 98.5023236 | 131.151755 | 100.296555 |

| | | | | | | |
|---|---|---|---|---|---|---|
| -3.5113106 | 0.0004459 | 0.00107656 | 25.0004767 | 31.9320719 | 28.7613499 | 38.1014845 |
| -3.5112674 | 0.00044598 | 0.00107666 | 22.1315695 | 38.4267306 | 35.6640739 | 32.498325 |
| -3.5112181 | 0.00044606 | 0.00107678 | 80.7392443 | 34.367569 | 13.8054479 | 101.977503 |
| -3.510799 | 0.00044676 | 0.00107839 | 15.5740674 | 49.2511618 | 49.4695218 | 52.1093832 |
| -3.5096187 | 0.00044875 | 0.00108311 | 12.7051603 | 117.174467 | 35.6640739 | 87.409288 |
| -3.5095573 | 0.00044885 | 0.00108328 | 21.3118818 | 44.9213893 | 59.8236078 | 28.5761134 |
| -3.509457 | 0.00044902 | 0.00108361 | 49.1812656 | 24.3549701 | 24.1595339 | 49.8681194 |
| -3.5093918 | 0.00044913 | 0.00108379 | 38.5253247 | 54.1221558 | 16.1063559 | 56.5919108 |
| -3.5089152 | 0.00044994 | 0.00108566 | 22.9512573 | 45.7332217 | 127.700393 | 28.0157974 |
| -3.5088081 | 0.00045012 | 0.00108601 | 49.5911095 | 15.966036 | 67.8767857 | 41.4633802 |
| -3.5087775 | 0.00045017 | 0.00108606 | 22.5414134 | 30.8496288 | 47.1686138 | 34.1792729 |
| -3.5087567 | 0.00045021 | 0.00108606 | 11.4756286 | 59.5343714 | 63.2749697 | 77.8839169 |
| -3.5083392 | 0.00045091 | 0.00108768 | 24.1807889 | 35.9912336 | 27.6108959 | 56.0315949 |
| -3.5082955 | 0.00045099 | 0.00108778 | 229.512573 | 511.183762 | 296.817131 | 333.948305 |
| -3.5081712 | 0.0004512 | 0.00108821 | 17.213443 | 47.0862756 | 37.9649818 | 62.7553862 |
| -3.5078904 | 0.00045168 | 0.00108928 | 158.199738 | 105.538204 | 249.648517 | 132.234564 |
| -3.5078213 | 0.00045179 | 0.00108948 | 25.0004767 | 38.9679522 | 31.0622579 | 29.1364293 |
| -3.5076444 | 0.00045209 | 0.00109012 | 22.5414134 | 36.5324552 | 31.0622579 | 36.9808526 |
| -3.5075316 | 0.00045228 | 0.0010905 | 88.1164342 | 16.2366467 | 25.3099879 | 144.561515 |
| -3.507498 | 0.00045234 | 0.00109056 | 38.1154808 | 101.479042 | 88.5849576 | 12.3269509 |
| -3.5071729 | 0.00045289 | 0.00109181 | 156.560362 | 30.8496288 | 25.3099879 | 36.9808526 |
| -3.5070451 | 0.00045311 | 0.00109225 | 17.6232868 | 38.9679522 | 57.5226998 | 43.144328 |
| 3.50669096 | 0.00045372 | 0.00109363 | 1.63937552 | 0.81183234 | 0 | 0.56031595 |
| -3.5066142 | 0.00045385 | 0.00109386 | 15.5740674 | 142.882491 | 134.603117 | 162.491625 |
| -3.5064531 | 0.00045412 | 0.00109444 | 13.524848 | 56.8282636 | 48.3190678 | 74.5220212 |
| -3.5061814 | 0.00045459 | 0.00109548 | 131.969729 | 267.904671 | 269.206235 | 262.78818 |
| -3.5058691 | 0.00045512 | 0.00109668 | 22.1315695 | 26.7904671 | 65.5758777 | 46.5062237 |
| -3.5046772 | 0.00045716 | 0.00110152 | 25.0004767 | 37.8855091 | 25.3099879 | 51.5490673 |
| 3.50464974 | 0.00045721 | 0.00110155 | 145.904421 | 62.7817008 | 56.3722458 | 58.8331746 |
| -3.504139 | 0.00045809 | 0.00110358 | 34.4268859 | 29.2259641 | 34.5136199 | 24.0935858 |
| -3.5038933 | 0.00045851 | 0.00110452 | 42.2139196 | 62.51109 | 25.3099879 | 24.0935858 |
| -3.5035588 | 0.00045909 | 0.00110583 | 16.8035991 | 54.1221558 | 40.2658898 | 47.6268556 |
| -3.5034745 | 0.00045923 | 0.00110601 | 32.3776665 | 35.4500121 | 43.7172518 | 21.292006 |
| -3.5034876 | 0.00045921 | 0.00110601 | 99.5920628 | 33.5557366 | 27.6108959 | 28.5761134 |
| 3.50343333 | 0.0004593 | 0.0011061 | 4.50828268 | 0 | 2.30090799 | 5.04284354 |
| -3.503217 | 0.00045967 | 0.00110692 | 29.0989155 | 28.4141318 | 29.9118039 | 29.1364293 |
| -3.5031307 | 0.00045982 | 0.00110719 | 47.951734 | 35.7206228 | 52.9208838 | 58.2728587 |
| -3.5028164 | 0.00046037 | 0.00110842 | 286.890716 | 435.412744 | 502.748396 | 460.019394 |
| -3.5024874 | 0.00046094 | 0.00110962 | 25.4103206 | 69.817581 | 21.8586259 | 52.1093832 |
| -3.5025021 | 0.00046091 | 0.00110962 | 25.0004767 | 31.9320719 | 29.9118039 | 32.498325 |
| -3.501692 | 0.00046231 | 0.00111285 | 229.512573 | 588.849055 | 547.616102 | 579.927007 |
| -3.5013361 | 0.00046293 | 0.00111426 | 30.3284471 | 23.8137486 | 64.4254237 | 33.058641 |
| -3.5012856 | 0.00046302 | 0.00111439 | 21.3118818 | 52.7691019 | 26.4604419 | 120.467929 |

| | | | | | | |
|---|---|---|---|---|---|---|
| -3.5009513 | 0.0004636 | 0.0011157 | 6.96734596 | 62.2404792 | 123.098578 | 108.701294 |
| -3.5008651 | 0.00046375 | 0.00111598 | 134.018949 | 46.0038325 | 10.354086 | 74.5220212 |
| -3.5004797 | 0.00046442 | 0.00111751 | 25.0004767 | 26.5198564 | 39.1154358 | 34.7395888 |
| -3.500321 | 0.0004647 | 0.00111804 | 17.213443 | 61.4286469 | 35.6640739 | 50.4284354 |
| -3.5003152 | 0.00046471 | 0.00111804 | 20.492194 | 54.1221558 | 28.7613499 | 48.1871716 |
| -3.5001534 | 0.00046499 | 0.00111863 | 16.8035991 | 119.880575 | 35.6640739 | 50.9887513 |
| -3.5000733 | 0.00046513 | 0.00111886 | 8.1968776 | 69.2763595 | 135.753571 | 68.9188617 |
| -3.5000585 | 0.00046516 | 0.00111886 | 21.3118818 | 56.0164313 | 27.6108959 | 46.5062237 |
| -3.5000068 | 0.00046525 | 0.001119 | 16.3937552 | 69.2763595 | 41.4163438 | 44.8252759 |
| -3.4998644 | 0.00046549 | 0.00111951 | 186.069121 | 72.253078 | 11.50454 | 38.1014845 |
| -3.4997931 | 0.00046562 | 0.00111973 | 18.8528185 | 41.67406 | 43.7172518 | 36.9808526 |
| -3.4997357 | 0.00046572 | 0.00111988 | 13.9346919 | 58.9931499 | 47.1686138 | 61.6347543 |
| -3.4996356 | 0.00046589 | 0.00112022 | 54.0993922 | 67.382084 | 43.7172518 | 14.5682147 |
| -3.4995237 | 0.00046609 | 0.00112061 | 23.3611012 | 34.9087905 | 169.116737 | 34.7395888 |
| -3.499289 | 0.0004665 | 0.00112151 | 16.8035991 | 90.654611 | 36.8145279 | 48.7474875 |
| 3.49917736 | 0.0004667 | 0.0011219 | 20.492194 | 6.4946587 | 5.75226998 | 6.72379138 |
| -3.4990499 | 0.00046692 | 0.00112227 | 12.2953164 | 55.2045989 | 75.9299637 | 57.1522268 |
| -3.4990497 | 0.00046692 | 0.00112227 | 48.3615778 | 78.477126 | 29.9118039 | 19.6110582 |
| -3.4989504 | 0.00046709 | 0.0011226 | 12.2953164 | 48.1687187 | 79.3813257 | 66.1172819 |
| -3.4988164 | 0.00046733 | 0.00112308 | 9.01656536 | 85.2423954 | 65.5758777 | 84.0473923 |
| -3.4982204 | 0.00046837 | 0.00112551 | 22.1315695 | 41.67406 | 28.7613499 | 39.7824324 |
| -3.4981004 | 0.00046858 | 0.00112594 | 17.213443 | 62.51109 | 36.8145279 | 47.0665397 |
| 3.49759186 | 0.00046948 | 0.001128 | 10.6559409 | 6.76526948 | 5.75226998 | 4.48252759 |
| 3.49739471 | 0.00046983 | 0.00112875 | 5.32797044 | 4.05916169 | 0 | 2.80157974 |
| -3.4970909 | 0.00047036 | 0.00112995 | 52.8698605 | 92.8194972 | 71.3281477 | 10.646003 |
| -3.4968431 | 0.0004708 | 0.00113092 | 9.42640924 | 61.1580361 | 140.355387 | 62.1950703 |
| -3.4967167 | 0.00047102 | 0.00113137 | 136.478012 | 38.4267306 | 20.7081719 | 35.2999048 |
| -3.4966792 | 0.00047109 | 0.00113145 | 13.9346919 | 52.7691019 | 121.948124 | 43.144328 |
| -3.4963425 | 0.00047168 | 0.00113279 | 43.8532952 | 43.5683354 | 113.894946 | 15.6888466 |
| -3.4952541 | 0.00047361 | 0.00113733 | 19.2626624 | 34.0969582 | 46.0181598 | 41.4633802 |
| -3.4949762 | 0.0004741 | 0.00113843 | 85.6573709 | 245.985198 | 216.285351 | 99.7362389 |
| -3.494631 | 0.00047472 | 0.00113982 | 41.8040758 | 51.9572696 | 17.2568099 | 38.6618005 |
| -3.4941605 | 0.00047555 | 0.00114175 | 19.6725062 | 38.4267306 | 37.9649818 | 38.1014845 |
| -3.4941337 | 0.0004756 | 0.00114178 | 56.1486116 | 22.7313054 | 33.3631659 | 30.8173772 |
| -3.4941001 | 0.00047566 | 0.00114184 | 23.3611012 | 43.0271139 | 32.2127119 | 29.6967453 |
| 3.49344807 | 0.00047683 | 0.00114454 | 11.8854725 | 7.8477126 | 9.20363196 | 5.04284354 |
| -3.4933301 | 0.00047704 | 0.00114497 | 15.9839113 | 46.8156648 | 46.0181598 | 49.3078035 |
| 3.49328012 | 0.00047713 | 0.00114509 | 21.3118818 | 8.65954493 | 19.5577179 | 11.7666349 |
| -3.4931766 | 0.00047731 | 0.00114545 | 55.3289238 | 71.1706349 | 9.20363196 | 123.269509 |
| -3.4928431 | 0.00047791 | 0.0011468 | 16.8035991 | 46.545054 | 58.6731538 | 38.1014845 |
| -3.492286 | 0.00047891 | 0.00114911 | 18.8528185 | 36.5324552 | 44.8677058 | 39.7824324 |
| -3.492234 | 0.000479 | 0.00114925 | 17.6232868 | 29.7671857 | 72.4786017 | 62.7553862 |
| -3.4921969 | 0.00047906 | 0.00114932 | 43.0336074 | 79.8301798 | 104.691314 | 89.0902358 |

| | | | | | | |
|---|---|---|---|---|---|---|
| -3.4920374 | 0.00047935 | 0.00114992 | 17.6232868 | 46.545054 | 43.7172518 | 39.2221164 |
| -3.4906881 | 0.00048178 | 0.00115566 | 15.1642236 | 75.5004074 | 50.6199758 | 40.9030642 |
| -3.490524 | 0.00048207 | 0.00115628 | 25.4103206 | 115.009581 | 17.2568099 | 76.202969 |
| -3.4903637 | 0.00048236 | 0.00115689 | 22.1315695 | 56.2870421 | 47.1686138 | 25.7745336 |
| -3.4903041 | 0.00048247 | 0.00115706 | 18.8528185 | 43.5683354 | 41.4163438 | 35.8602207 |
| -3.4900704 | 0.00048289 | 0.00115799 | 22.1315695 | 75.7710182 | 132.302209 | 131.113932 |
| -3.4899341 | 0.00048314 | 0.00115849 | 22.1315695 | 41.67406 | 36.8145279 | 29.1364293 |
| -3.4898836 | 0.00048323 | 0.00115863 | 12.7051603 | 61.4286469 | 50.6199758 | 66.1172819 |
| -3.48938 | 0.00048414 | 0.00116073 | 38.1154808 | 30.3084073 | 21.8586259 | 84.0473923 |
| -3.4890733 | 0.0004847 | 0.00116197 | 16.3937552 | 62.51109 | 36.8145279 | 50.9887513 |
| 3.48893405 | 0.00048495 | 0.00116249 | 121.313788 | 51.9572696 | 41.4163438 | 58.8331746 |
| -3.4888798 | 0.00048505 | 0.00116264 | 32.7875104 | 23.272527 | 32.2127119 | 79.0045488 |
| -3.4888192 | 0.00048516 | 0.00116282 | 18.8528185 | 41.1328384 | 36.8145279 | 41.4633802 |
| -3.4886951 | 0.00048538 | 0.00116327 | 81.968776 | 54.1221558 | 14.9559019 | 37.5411686 |
| -3.4885647 | 0.00048562 | 0.00116375 | 18.0331307 | 62.51109 | 29.9118039 | 59.9538065 |
| -3.4883285 | 0.00048605 | 0.0011647 | 5.32797044 | 116.092024 | 85.1335957 | 113.744138 |
| -3.4881896 | 0.0004863 | 0.00116521 | 13.524848 | 78.477126 | 40.2658898 | 66.1172819 |
| -3.4878248 | 0.00048697 | 0.00116672 | 29.5087594 | 40.8622277 | 21.8586259 | 35.2999048 |
| -3.4870705 | 0.00048834 | 0.00116993 | 26.6398522 | 22.1900839 | 42.5667978 | 51.5490673 |
| -3.4865312 | 0.00048933 | 0.0011722 | 51.6403289 | 77.6652936 | 10.354086 | 79.5648647 |
| -3.4863795 | 0.00048961 | 0.00117278 | 49.5911095 | 30.3084073 | 24.1595339 | 29.1364293 |
| -3.4860791 | 0.00049016 | 0.00117401 | 23.770945 | 28.6847426 | 40.2658898 | 31.3776931 |
| -3.4859803 | 0.00049034 | 0.00117436 | 13.9346919 | 92.2782757 | 41.4163438 | 55.4712789 |
| -3.4851978 | 0.00049177 | 0.00117771 | 761.899773 | 840.787691 | 647.705599 | 843.835819 |
| -3.4848755 | 0.00049237 | 0.00117904 | 13.9346919 | 80.3714014 | 49.4695218 | 46.5062237 |
| -3.4847656 | 0.00049257 | 0.00117942 | 40.1647002 | 166.425629 | 189.824909 | 192.18837 |
| -3.4847493 | 0.0004926 | 0.00117942 | 522.141103 | 1485.65318 | 1382.8457 | 1459.62305 |
| -3.4846329 | 0.00049281 | 0.00117985 | 52.8698605 | 47.6274971 | 44.8677058 | 42.0236961 |
| -3.4842796 | 0.00049346 | 0.00118132 | 16.8035991 | 34.0969582 | 52.9208838 | 63.3157022 |
| -3.4840674 | 0.00049386 | 0.00118217 | 29.0989155 | 41.1328384 | 34.5136199 | 22.4126379 |
| -3.4839395 | 0.00049409 | 0.00118265 | 278.693838 | 600.75593 | 461.332052 | 529.498571 |
| -3.4835422 | 0.00049483 | 0.00118432 | 17.6232868 | 46.2744432 | 42.5667978 | 38.6618005 |
| -3.483211 | 0.00049544 | 0.00118569 | 27.0496961 | 41.4034492 | 23.0090799 | 35.8602207 |
| -3.4831695 | 0.00049551 | 0.00118579 | 30.3284471 | 41.67406 | 20.7081719 | 100.856871 |
| -3.4829997 | 0.00049583 | 0.00118645 | 15.9839113 | 60.075593 | 37.9649818 | 50.9887513 |
| -3.4818925 | 0.00049788 | 0.00119128 | 18.8528185 | 58.1813175 | 29.9118039 | 49.8681194 |
| -3.4814494 | 0.00049871 | 0.00119317 | 21.3118818 | 30.579018 | 49.4695218 | 34.7395888 |
| 3.48118913 | 0.00049919 | 0.00119424 | 42.6237635 | 28.4141318 | 26.4604419 | 17.9301104 |
| -3.480868 | 0.00049979 | 0.00119558 | 12.7051603 | 55.7458205 | 66.7263317 | 52.1093832 |
| -3.4806072 | 0.00050028 | 0.00119666 | 159.429269 | 95.525605 | 67.8767857 | 79.5648647 |
| -3.4802692 | 0.00050091 | 0.001198 | 27.8693838 | 37.8855091 | 23.0090799 | 34.1792729 |
| 3.48026741 | 0.00050091 | 0.001198 | 8.60672148 | 8.38893415 | 11.50454 | 4.48252759 |
| -3.4801623 | 0.00050111 | 0.00119838 | 62.7061136 | 38.9679522 | 32.2127119 | 19.6110582 |

| | | | | | | |
|---|---|---|---|---|---|---|
| -3.4799205 | 0.00050156 | 0.00119937 | 27.8693838 | 18.6721438 | 77.0804177 | 56.5919108 |
| -3.4797444 | 0.00050189 | 0.00120006 | 3.68859492 | 72.7942996 | 200.178995 | 133.915512 |
| -3.4797277 | 0.00050192 | 0.00120006 | 23.770945 | 42.2152815 | 55.2217918 | 24.0935858 |
| -3.479702 | 0.00050197 | 0.00120009 | 121.723632 | 516.595977 | 248.498063 | 345.71494 |
| -3.479463 | 0.00050242 | 0.00120107 | 26.6398522 | 20.0251977 | 55.2217918 | 60.5141225 |
| 3.47943457 | 0.00050247 | 0.00120111 | 0 | 0 | 0 | 0 |
| -3.4793825 | 0.00050257 | 0.00120125 | 13.524848 | 58.4519283 | 52.9208838 | 51.5490673 |
| -3.4792933 | 0.00050274 | 0.00120156 | 18.0331307 | 47.3568864 | 55.2217918 | 32.498325 |
| 3.47926331 | 0.00050279 | 0.0012016 | 0.40984388 | 0.27061078 | 0 | 0 |
| -3.4792254 | 0.00050287 | 0.0012016 | 17.213443 | 27.8729103 | 66.7263317 | 89.6505518 |
| -3.4792348 | 0.00050285 | 0.0012016 | 15.9839113 | 71.7118565 | 36.8145279 | 49.8681194 |
| -3.4789618 | 0.00050336 | 0.00120269 | 15.9839113 | 52.7691019 | 123.098578 | 34.1792729 |
| -3.4786352 | 0.00050397 | 0.00120407 | 14.7543797 | 79.5595691 | 32.2127119 | 77.3236009 |
| -3.4785678 | 0.0005041 | 0.00120428 | 16.8035991 | 76.8534613 | 34.5136199 | 48.7474875 |
| -3.4784284 | 0.00050436 | 0.00120482 | 24.5906328 | 31.3908504 | 35.6640739 | 27.4554815 |
| -3.4780143 | 0.00050514 | 0.0012066 | 10.6559409 | 48.4393295 | 141.505841 | 61.6347543 |
| -3.4778749 | 0.00050541 | 0.00120713 | 27.0496961 | 40.5916169 | 21.8586259 | 41.4633802 |
| 3.4775467 | 0.00050602 | 0.00120852 | 0 | 0 | 0 | 0 |
| -3.4774196 | 0.00050626 | 0.00120894 | 315.169944 | 495.758947 | 0 | 530.619203 |
| -3.4774137 | 0.00050628 | 0.00120894 | 14.7543797 | 50.8748265 | 147.258111 | 38.1014845 |
| -3.4771583 | 0.00050676 | 0.00121001 | 6.1476582 | 120.963018 | 75.9299637 | 93.5727634 |
| -3.4763518 | 0.00050829 | 0.00121356 | 16.8035991 | 68.1939163 | 47.1686138 | 35.2999048 |
| -3.4760528 | 0.00050885 | 0.00121483 | 11.8854725 | 73.6061319 | 73.6290557 | 45.3855918 |
| -3.475987 | 0.00050898 | 0.00121503 | 24.5906328 | 54.9339882 | 20.7081719 | 71.1601255 |
| -3.4758697 | 0.0005092 | 0.00121548 | 36.8859492 | 35.9912336 | 18.4072639 | 45.9459078 |
| -3.4758115 | 0.00050931 | 0.00121565 | 50.8206411 | 64.1347547 | 59.8236078 | 57.1522268 |
| -3.4756905 | 0.00050954 | 0.00121611 | 831.983076 | 732.272768 | 543.014286 | 666.775979 |
| -3.4748324 | 0.00051117 | 0.00121991 | 24.1807889 | 26.7904671 | 35.6640739 | 33.6189569 |
| -3.4746357 | 0.00051155 | 0.00122072 | 115.985818 | 18.401533 | 98.9390436 | 20.1713742 |
| -3.4743366 | 0.00051212 | 0.00122199 | 70.4931473 | 51.6866588 | 14.9559019 | 37.5411686 |
| -3.4742064 | 0.00051237 | 0.00122249 | 34.017042 | 42.2152815 | 20.7081719 | 31.3776931 |
| -3.4736018 | 0.00051352 | 0.00122516 | 157.38005 | 272.775665 | 249.648517 | 260.546916 |
| -3.4734541 | 0.00051381 | 0.00122574 | 18.4429746 | 34.0969582 | 66.7263317 | 39.7824324 |
| -3.4733269 | 0.00051405 | 0.00122624 | 115.575974 | 318.779498 | 500.447488 | 358.041891 |
| 3.47329336 | 0.00051411 | 0.0012263 | 58.6076748 | 89.8427787 | 69.0272397 | 68.3585457 |
| -3.4732523 | 0.00051419 | 0.00122639 | 14.7543797 | 76.3122397 | 42.5667978 | 47.0665397 |
| -3.4731987 | 0.00051429 | 0.00122655 | 17.6232868 | 39.238563 | 40.2658898 | 44.8252759 |
| -3.4724796 | 0.00051567 | 0.00122967 | 36.8859492 | 50.6042157 | 28.7613499 | 21.292006 |
| -3.4724773 | 0.00051568 | 0.00122967 | 20.9020379 | 38.6973414 | 28.7613499 | 47.6268556 |
| -3.4720901 | 0.00051642 | 0.00123135 | 24.5906328 | 47.8981079 | 71.3281477 | 21.852322 |
| -3.4719184 | 0.00051675 | 0.00123205 | 245.496484 | 315.261558 | 170.267191 | 151.285306 |
| -3.4717095 | 0.00051716 | 0.00123292 | 20.0823501 | 46.0038325 | 27.6108959 | 68.9188617 |
| -3.4714653 | 0.00051763 | 0.00123395 | 19.6725062 | 33.5557366 | 35.6640739 | 46.5062237 |

| | | | | | | |
|---|---|---|---|---|---|---|
| -3.4713124 | 0.00051792 | 0.00123456 | 307.792754 | 401.857007 | 408.411168 | 429.762333 |
| -3.4710663 | 0.0005184 | 0.0012356 | 20.0823501 | 52.2278804 | 39.1154358 | 30.2570612 |
| -3.471038 | 0.00051845 | 0.00123564 | 25.0004767 | 26.7904671 | 32.2127119 | 33.058641 |
| -3.4710137 | 0.0005185 | 0.00123566 | 18.4429746 | 63.0523115 | 34.5136199 | 39.2221164 |
| -3.4705707 | 0.00051935 | 0.00123761 | 35.6564176 | 67.382084 | 17.2568099 | 40.9030642 |
| -3.4702078 | 0.00052006 | 0.00123919 | 9.83625312 | 40.5916169 | 97.7885896 | 110.942558 |
| -3.4700464 | 0.00052037 | 0.00123984 | 18.8528185 | 32.4732935 | 41.4163438 | 47.0665397 |
| -3.4698845 | 0.00052068 | 0.0012405 | 20.492194 | 50.3336049 | 63.2749697 | 25.7745336 |
| -3.4695349 | 0.00052136 | 0.00124202 | 9.42640924 | 76.5828505 | 75.9299637 | 62.1950703 |
| -3.4683632 | 0.00052364 | 0.00124736 | 227.053509 | 452.190612 | 473.987046 | 491.957403 |
| -3.4683352 | 0.00052369 | 0.0012474 | 33.6071982 | 84.7011739 | 13.8054479 | 70.5998095 |
| -3.4682671 | 0.00052383 | 0.00124762 | 0 | 582.625007 | 675.316495 | 539.584259 |
| -3.4680652 | 0.00052422 | 0.00124847 | 21.7217256 | 117.9863 | 81.6822337 | 109.821926 |
| -3.4679485 | 0.00052445 | 0.00124884 | 57.7879871 | 73.3355211 | 9.20363196 | 84.6077082 |
| -3.4679448 | 0.00052446 | 0.00124884 | 19.6725062 | 41.67406 | 39.1154358 | 31.9380091 |
| -3.4679102 | 0.00052452 | 0.00124891 | 25.4103206 | 101.749653 | 70.1776937 | 85.1680242 |
| 3.46747472 | 0.00052537 | 0.00125077 | 41.8040758 | 73.6061319 | 95.4876816 | 53.7903311 |
| -3.4674705 | 0.00052538 | 0.00125077 | 14.7543797 | 61.9698684 | 48.3190678 | 42.5840121 |
| -3.4671042 | 0.0005261 | 0.00125229 | 20.492194 | 27.6022995 | 44.8677058 | 47.0665397 |
| -3.4671065 | 0.00052609 | 0.00125229 | 243.857109 | 795.595691 | 704.077845 | 737.936104 |
| -3.4667481 | 0.0005268 | 0.00125386 | 11.8854725 | 60.3462037 | 80.5317797 | 47.6268556 |
| -3.4664396 | 0.0005274 | 0.00125521 | 18.0331307 | 37.0736767 | 37.9649818 | 47.6268556 |
| -3.4663914 | 0.00052749 | 0.00125534 | 15.1642236 | 62.7817008 | 47.1686138 | 40.9030642 |
| -3.4659587 | 0.00052834 | 0.00125727 | 22.1315695 | 45.7332217 | 39.1154358 | 26.3348496 |
| -3.4654737 | 0.0005293 | 0.00125945 | 23.770945 | 30.0377965 | 28.7613499 | 35.2999048 |
| -3.4654281 | 0.00052939 | 0.00125957 | 36.0662614 | 33.5557366 | 28.7613499 | 22.4126379 |
| -3.4651655 | 0.00052991 | 0.00126071 | 90.5754975 | 181.309222 | 215.134897 | 136.156776 |
| 3.4650065 | 0.00053022 | 0.00126127 | 47.951734 | 35.7206228 | 26.4604419 | 21.292006 |
| -3.4650117 | 0.00053021 | 0.00126127 | 46.3123584 | 41.9446708 | 59.8236078 | 44.8252759 |
| 3.46484271 | 0.00053054 | 0.00126194 | 13.524848 | 11.3656527 | 13.8054479 | 9.52537113 |
| -3.4648019 | 0.00053062 | 0.00126204 | 17.6232868 | 35.9912336 | 40.2658898 | 52.1093832 |
| 3.46467782 | 0.00053087 | 0.00126253 | 145.494577 | 143.694324 | 169.116737 | 177.05984 |
| -3.4646069 | 0.00053101 | 0.00126277 | 22.9512573 | 85.2423954 | 85.1335957 | 176.499524 |
| -3.4645486 | 0.00053112 | 0.00126295 | 34.4268859 | 43.5683354 | 19.5577179 | 33.058641 |
| -3.4644857 | 0.00053125 | 0.00126315 | 9.01656536 | 34.9087905 | 131.151755 | 125.510772 |
| -3.4639241 | 0.00053236 | 0.0012657 | 2.0492194 | 132.869893 | 155.311289 | 166.413837 |
| -3.46351 | 0.00053318 | 0.00126756 | 45.9025146 | 76.0416289 | 9.20363196 | 130.553616 |
| -3.4633461 | 0.0005335 | 0.00126824 | 27.45954 | 78.7477367 | 23.0090799 | 36.4205367 |
| -3.463254 | 0.00053368 | 0.00126858 | 119.674413 | 33.2851258 | 19.5577179 | 38.6618005 |
| -3.4629622 | 0.00053426 | 0.00126986 | 40.5745441 | 32.7439043 | 18.4072639 | 43.144328 |
| -3.4628543 | 0.00053448 | 0.00127027 | 19.2626624 | 37.0736767 | 39.1154358 | 34.7395888 |
| -3.4628275 | 0.00053453 | 0.00127031 | 95.493624 | 42.7565031 | 19.5577179 | 29.6967453 |
| -3.4626488 | 0.00053489 | 0.00127106 | 69.2636157 | 26.2492456 | 37.9649818 | 22.9729539 |

| | | | | | | |
|---|---|---|---|---|---|---|
| -3.4625554 | 0.00053507 | 0.00127141 | 20.0823501 | 46.0038325 | 62.1245158 | 26.8951655 |
| -3.4621462 | 0.00053589 | 0.00127325 | 18.0331307 | 32.7439043 | 50.6199758 | 43.144328 |
| -3.4619406 | 0.0005363 | 0.00127413 | 21.3118818 | 39.7797845 | 35.6640739 | 29.1364293 |
| -3.46162 | 0.00053693 | 0.00127555 | 17.6232868 | 71.9824673 | 32.2127119 | 45.3855918 |
| -3.4613245 | 0.00053752 | 0.00127686 | 13.524848 | 69.5469702 | 50.6199758 | 45.3855918 |
| -3.4612852 | 0.0005376 | 0.00127695 | 41.3942319 | 24.8961917 | 31.0622579 | 25.7745336 |
| -3.4612111 | 0.00053775 | 0.00127721 | 26.2300083 | 64.4053654 | 19.5577179 | 50.4284354 |
| -3.4610044 | 0.00053816 | 0.0012781 | 40.5745441 | 51.6866588 | 37.9649818 | 64.4363341 |
| -3.4606189 | 0.00053894 | 0.00127983 | 67.6242402 | 53.851545 | 16.1063559 | 31.9380091 |
| -3.4598895 | 0.0005404 | 0.00128321 | 35.2465737 | 55.7458205 | 19.5577179 | 31.9380091 |
| -3.4597603 | 0.00054066 | 0.00128373 | 13.9346919 | 75.5004074 | 35.6640739 | 65.556966 |
| -3.4592626 | 0.00054166 | 0.00128595 | 16.8035991 | 59.2637606 | 37.9649818 | 40.3427483 |
| -3.4592557 | 0.00054167 | 0.00128595 | 29.5087594 | 25.1668025 | 31.0622579 | 28.0157974 |
| -3.4591183 | 0.00054195 | 0.00128651 | 25.8201644 | 47.0862756 | 20.7081719 | 47.6268556 |
| 3.45861588 | 0.00054296 | 0.00128882 | 0 | 0 | 0 | 0 |
| -3.4582851 | 0.00054363 | 0.00129028 | 14.7543797 | 48.7099402 | 48.3190678 | 46.5062237 |
| -3.4582704 | 0.00054366 | 0.00129028 | 13.9346919 | 39.7797845 | 88.5849576 | 51.5490673 |
| -3.4579742 | 0.00054425 | 0.00129151 | 40.5745441 | 53.851545 | 31.0622579 | 38.6618005 |
| -3.4579837 | 0.00054423 | 0.00129151 | 13.9346919 | 70.0881918 | 32.2127119 | 95.2537113 |
| 3.45789125 | 0.00054442 | 0.00129174 | 343.859015 | 658.937247 | 337.083021 | 537.342995 |
| -3.4578875 | 0.00054443 | 0.00129174 | 18.4429746 | 64.4053654 | 25.3099879 | 72.8410733 |
| -3.4574458 | 0.00054532 | 0.00129376 | 36.4761053 | 69.2763595 | 40.2658898 | 77.8839169 |
| -3.4570189 | 0.00054619 | 0.00129572 | 18.0331307 | 34.0969582 | 48.3190678 | 39.7824324 |
| 3.45691695 | 0.00054639 | 0.00129611 | 59.4273626 | 43.0271139 | 50.6199758 | 32.498325 |
| 3.45682643 | 0.00054658 | 0.00129645 | 9.83625312 | 6.76526948 | 6.90272397 | 6.72379138 |
| 3.45651454 | 0.00054721 | 0.00129783 | 7.78703372 | 10.5538204 | 9.20363196 | 12.8872668 |
| -3.4565011 | 0.00054724 | 0.00129783 | 22.1315695 | 37.0736767 | 33.3631659 | 28.5761134 |
| -3.4560972 | 0.00054806 | 0.00129968 | 33.1973543 | 126.645845 | 31.0622579 | 23.5332698 |
| 3.45554989 | 0.00054917 | 0.00130223 | 239.75867 | 213.782516 | 187.524001 | 187.145527 |
| -3.4555256 | 0.00054922 | 0.00130225 | 16.3937552 | 73.8767427 | 80.5317797 | 26.8951655 |
| -3.4554979 | 0.00054928 | 0.00130229 | 87.7065903 | 17.0484791 | 40.2658898 | 36.9808526 |
| -3.4551772 | 0.00054993 | 0.00130374 | 36.8859492 | 39.5091738 | 46.0181598 | 17.3697944 |
| -3.4550063 | 0.00055028 | 0.00130447 | 14.3445358 | 38.9679522 | 64.4254237 | 53.7903311 |
| -3.4547081 | 0.00055089 | 0.00130582 | 13.9346919 | 50.8748265 | 40.2658898 | 82.3664444 |
| -3.4545284 | 0.00055126 | 0.00130659 | 11.4756286 | 47.6274971 | 80.5317797 | 61.0744384 |
| -3.4542622 | 0.0005518 | 0.00130779 | 34.017042 | 35.1794013 | 70.1776937 | 19.0507423 |
| -3.454127 | 0.00055208 | 0.00130835 | 430.336074 | 900.322062 | 751.246459 | 788.924856 |
| -3.4538686 | 0.00055261 | 0.00130951 | 79.9195566 | 220.547785 | 346.286653 | 401.186219 |
| -3.4536423 | 0.00055307 | 0.00131051 | 15.5740674 | 47.8981079 | 40.2658898 | 48.7474875 |
| -3.4535965 | 0.00055316 | 0.00131063 | 18.8528185 | 95.2549943 | 134.603117 | 114.304454 |
| -3.4532016 | 0.00055397 | 0.00131246 | 15.1642236 | 51.1454373 | 44.8677058 | 43.704644 |
| -3.4528538 | 0.00055469 | 0.00131406 | 24.1807889 | 32.2026827 | 25.3099879 | 46.5062237 |
| -3.452461 | 0.0005555 | 0.00131578 | 25.8201644 | 27.8729103 | 33.3631659 | 26.8951655 |

| | | | | | | |
|---|---|---|---|---|---|---|
| -3.4524606 | 0.0005555 | 0.00131578 | 28.6890716 | 36.2618444 | 41.4163438 | 20.7316901 |
| -3.4524269 | 0.00055557 | 0.00131585 | 27.45954 | 57.9107067 | 54.0713378 | 18.4904263 |
| -3.4523819 | 0.00055566 | 0.00131597 | 29.5087594 | 65.7584193 | 69.0272397 | 68.9188617 |
| -3.4522427 | 0.00055595 | 0.00131655 | 32.3776665 | 51.6866588 | 16.1063559 | 59.3934906 |
| -3.4519223 | 0.00055661 | 0.00131802 | 11.0657848 | 61.4286469 | 96.6381356 | 46.5062237 |
| -3.4510956 | 0.00055832 | 0.00132197 | 22.1315695 | 29.4965749 | 32.2127119 | 61.6347543 |
| -3.4507716 | 0.00055899 | 0.00132346 | 60.2470503 | 50.6042157 | 144.957203 | 9.52537113 |
| -3.4507036 | 0.00055913 | 0.00132369 | 14.7543797 | 45.1920001 | 47.1686138 | 49.3078035 |
| -3.4505083 | 0.00055953 | 0.00132456 | 16.8035991 | 47.8981079 | 34.5136199 | 48.1871716 |
| -3.4504498 | 0.00055965 | 0.00132474 | 20.0823501 | 34.0969582 | 37.9649818 | 32.498325 |
| -3.4498712 | 0.00056085 | 0.00132749 | 14.7543797 | 63.8641439 | 40.2658898 | 47.0665397 |
| 3.44975669 | 0.00056109 | 0.00132796 | 18.8528185 | 11.3656527 | 8.05317797 | 12.3269509 |
| -3.4494642 | 0.0005617 | 0.0013293 | 21.3118818 | 30.0377965 | 33.3631659 | 35.8602207 |
| -3.4490881 | 0.00056248 | 0.00133105 | 23.3611012 | 53.0397127 | 58.6731538 | 55.4712789 |
| -3.4489399 | 0.00056279 | 0.00133169 | 24.5906328 | 46.545054 | 21.8586259 | 42.0236961 |
| -3.4488482 | 0.00056298 | 0.00133204 | 14.3445358 | 33.8263474 | 102.390406 | 57.1522268 |
| -3.4485375 | 0.00056363 | 0.00133348 | 20.9020379 | 34.9087905 | 34.5136199 | 30.8173772 |
| 3.4483134 | 0.0005641 | 0.00133449 | 13.524848 | 8.93015571 | 10.354086 | 5.60315949 |
| -3.4482802 | 0.00056417 | 0.00133455 | 8.1968776 | 75.2297966 | 63.2749697 | 91.8918156 |
| -3.4481565 | 0.00056443 | 0.00133507 | 9.83625312 | 52.7691019 | 69.0272397 | 85.7283401 |
| -3.4471586 | 0.00056652 | 0.00133991 | 31.9678226 | 41.4034492 | 66.7263317 | 36.9808526 |
| -3.4470588 | 0.00056673 | 0.00134031 | 177.872244 | 203.228695 | 187.524001 | 203.955005 |
| -3.4470075 | 0.00056683 | 0.00134046 | 18.0331307 | 62.2404792 | 70.1776937 | 25.7745336 |
| -3.4468909 | 0.00056708 | 0.00134094 | 18.8528185 | 33.0145151 | 55.2217918 | 35.2999048 |
| -3.4468158 | 0.00056724 | 0.00134122 | 17.213443 | 27.8729103 | 67.8767857 | 59.3934906 |
| -3.446531 | 0.00056783 | 0.00134253 | 29.9186032 | 36.5324552 | 20.7081719 | 33.6189569 |
| -3.4464507 | 0.000568 | 0.00134283 | 4.91812656 | 138.82333 | 59.8236078 | 132.79488 |
| -3.4463624 | 0.00056819 | 0.00134317 | 17.213443 | 39.238563 | 46.0181598 | 36.9808526 |
| -3.446315 | 0.00056829 | 0.00134331 | 65.1651769 | 35.7206228 | 18.4072639 | 33.6189569 |
| -3.4460416 | 0.00056886 | 0.00134457 | 509.026099 | 556.105151 | 0 | 240.935858 |
| 3.44570891 | 0.00056956 | 0.00134613 | 46.3123584 | 36.803066 | 31.0622579 | 31.3776931 |
| 3.44521646 | 0.0005706 | 0.00134849 | 28.2792277 | 21.6488623 | 37.9649818 | 21.852322 |
| -3.4448533 | 0.00057137 | 0.0013502 | 18.4429746 | 47.6274971 | 63.2749697 | 28.0157974 |
| -3.4447897 | 0.0005715 | 0.00135042 | 40.984388 | 65.2171978 | 13.8054479 | 48.1871716 |
| -3.4447692 | 0.00057155 | 0.00135042 | 50.4107972 | 97.6904913 | 144.957203 | 147.363094 |
| -3.4447406 | 0.00057161 | 0.00135047 | 14.3445358 | 55.2045989 | 37.9649818 | 60.5141225 |
| -3.4444573 | 0.00057221 | 0.00135179 | 24.1807889 | 23.0019162 | 42.5667978 | 38.6618005 |
| -3.444429 | 0.00057227 | 0.00135183 | 35.2465737 | 34.367569 | 21.8586259 | 27.4554815 |
| -3.444068 | 0.00057303 | 0.00135353 | 11.8854725 | 49.2511618 | 70.1776937 | 55.4712789 |
| -3.4439112 | 0.00057336 | 0.00135422 | 30.3284471 | 27.3316887 | 23.0090799 | 48.1871716 |
| -3.4437047 | 0.0005738 | 0.00135516 | 26.6398522 | 40.8622277 | 20.7081719 | 40.3427483 |
| -3.4432483 | 0.00057477 | 0.00135735 | 21.7217256 | 54.9339882 | 47.1686138 | 23.5332698 |
| -3.4432052 | 0.00057486 | 0.00135736 | 15.1642236 | 61.9698684 | 36.8145279 | 48.7474875 |

| | | | | | | |
|---|---|---|---|---|---|---|
| -3.4432185 | 0.00057483 | 0.00135736 | 16.3937552 | 48.980551 | 35.6640739 | 45.9459078 |
| -3.4429165 | 0.00057548 | 0.00135871 | 24.5906328 | 37.0736767 | 23.0090799 | 62.1950703 |
| -3.4425013 | 0.00057636 | 0.0013607 | 18.4429746 | 42.4858923 | 36.8145279 | 34.1792729 |
| -3.4419788 | 0.00057748 | 0.00136323 | 31.5579788 | 26.5198564 | 25.3099879 | 79.5648647 |
| -3.4416127 | 0.00057826 | 0.00136498 | 19.2626624 | 55.2045989 | 69.0272397 | 24.6539017 |
| -3.4415092 | 0.00057848 | 0.0013654 | 32.3776665 | 63.3229223 | 14.9559019 | 59.3934906 |
| -3.4414443 | 0.00057862 | 0.00136563 | 28.2792277 | 21.1076408 | 112.744492 | 39.2221164 |
| -3.4409536 | 0.00057967 | 0.00136791 | 17.6232868 | 37.0736767 | 49.4695218 | 35.2999048 |
| -3.4409652 | 0.00057964 | 0.00136791 | 20.9020379 | 20.0251977 | 67.8767857 | 97.4949751 |
| -3.4403898 | 0.00058088 | 0.00137066 | 768.047431 | 1668.04484 | 1302.31392 | 1400.78987 |
| -3.4398521 | 0.00058203 | 0.00137329 | 173.773805 | 494.135283 | 342.835291 | 383.256109 |
| -3.4397015 | 0.00058236 | 0.00137395 | 25.0004767 | 37.6148983 | 24.1595339 | 31.3776931 |
| -3.4396149 | 0.00058254 | 0.00137429 | 15.9839113 | 30.3084073 | 56.3722458 | 72.8410733 |
| -3.4395867 | 0.0005826 | 0.00137433 | 29.0989155 | 14.6129821 | 94.3372276 | 69.4791776 |
| -3.4391789 | 0.00058348 | 0.00137631 | 15.5740674 | 41.1328384 | 69.0272397 | 38.6618005 |
| -3.4381528 | 0.0005857 | 0.00138143 | 313.940412 | 1409.61155 | 699.476029 | 921.719735 |
| -3.4374745 | 0.00058717 | 0.00138479 | 16.8035991 | 43.2977247 | 35.6640739 | 44.8252759 |
| -3.4374148 | 0.0005873 | 0.001385 | 7.37718984 | 48.7099402 | 100.089498 | 117.666349 |
| -3.4372654 | 0.00058762 | 0.00138566 | 26.2300083 | 26.2492456 | 26.4604419 | 35.2999048 |
| -3.4372418 | 0.00058767 | 0.00138568 | 21.7217256 | 29.7671857 | 29.9118039 | 48.1871716 |
| -3.4368334 | 0.00058856 | 0.00138767 | 76.2309617 | 32.2026827 | 35.6640739 | 43.704644 |
| -3.4364799 | 0.00058933 | 0.00138938 | 66.8045524 | 36.803066 | 42.5667978 | 15.6888466 |
| -3.4360348 | 0.0005903 | 0.00139156 | 11.4756286 | 68.1939163 | 48.3190678 | 60.5141225 |
| 3.43580335 | 0.0005908 | 0.00139265 | 0 | 0 | 0 | 0 |
| -3.4356542 | 0.00059112 | 0.00139332 | 18.0331307 | 28.6847426 | 50.6199758 | 49.8681194 |
| -3.4353787 | 0.00059173 | 0.00139463 | 15.9839113 | 38.4267306 | 44.8677058 | 44.8252759 |
| -3.4345188 | 0.00059361 | 0.00139896 | 27.0496961 | 24.0843593 | 51.7704298 | 28.5761134 |
| -3.4342421 | 0.00059421 | 0.00140029 | 34.017042 | 18.9427545 | 36.8145279 | 36.4205367 |
| -3.4341709 | 0.00059437 | 0.00140055 | 13.524848 | 38.9679522 | 56.3722458 | 63.8760181 |
| -3.4333543 | 0.00059616 | 0.00140468 | 29.0989155 | 35.9912336 | 47.1686138 | 19.6110582 |
| -3.433101 | 0.00059672 | 0.00140589 | 6.55750208 | 65.2171978 | 106.992222 | 87.9696039 |
| -3.4326166 | 0.00059779 | 0.0014083 | 158.609582 | 458.41466 | 387.702996 | 470.665397 |
| -3.4324485 | 0.00059816 | 0.00140907 | 11.4756286 | 56.8282636 | 58.6731538 | 55.4712789 |
| -3.4323205 | 0.00059844 | 0.00140963 | 16.8035991 | 46.545054 | 66.7263317 | 30.8173772 |
| -3.4320778 | 0.00059898 | 0.00141079 | 34.4268859 | 40.8622277 | 72.4786017 | 52.1093832 |
| -3.4319481 | 0.00059926 | 0.00141136 | 292.218686 | 459.226492 | 543.014286 | 600.098381 |
| -3.4305932 | 0.00060226 | 0.00141833 | 16.8035991 | 64.4053654 | 71.3281477 | 26.3348496 |
| -3.430387 | 0.00060272 | 0.0014193 | 562.305803 | 387.514636 | 427.968886 | 442.089283 |
| -3.4300607 | 0.00060345 | 0.00142091 | 32.3776665 | 114.46836 | 48.3190678 | 69.4791776 |
| 3.4297488 | 0.00060414 | 0.00142244 | 0.81968776 | 0.27061078 | 0 | 0.56031595 |
| -3.4295521 | 0.00060458 | 0.00142336 | 9.83625312 | 87.9485032 | 51.7704298 | 63.3157022 |
| -3.4291786 | 0.00060541 | 0.00142522 | 52.8698605 | 36.803066 | 34.5136199 | 34.7395888 |
| -3.4291229 | 0.00060554 | 0.00142527 | 10.246097 | 63.5935331 | 51.7704298 | 76.202969 |

| | | | | | | |
|---|---|---|---|---|---|---|
| -3.4291106 | 0.00060556 | 0.00142527 | 15.1642236 | 34.9087905 | 51.7704298 | 55.4712789 |
| -3.4291499 | 0.00060547 | 0.00142527 | 8.60672148 | 60.8874253 | 60.9740618 | 103.65845 |
| -3.4289094 | 0.00060601 | 0.00142622 | 15.9839113 | 35.7206228 | 62.1245158 | 41.4633802 |
| -3.4288143 | 0.00060622 | 0.00142661 | 19.2626624 | 74.9591858 | 24.1595339 | 52.6696992 |
| -3.4287955 | 0.00060627 | 0.00142661 | 19.2626624 | 94.7137727 | 28.7613499 | 40.9030642 |
| 3.4279198 | 0.00060823 | 0.00143112 | 43.8532952 | 18.401533 | 27.6108959 | 21.852322 |
| -3.4273568 | 0.00060949 | 0.00143398 | 15.5740674 | 35.9912336 | 69.0272397 | 42.5840121 |
| -3.426569 | 0.00061126 | 0.00143804 | 18.4429746 | 42.4858923 | 46.0181598 | 29.1364293 |
| -3.4261469 | 0.00061221 | 0.00144017 | 29.9186032 | 35.9912336 | 44.8677058 | 19.0507423 |
| -3.4255946 | 0.00061346 | 0.001443 | 13.524848 | 56.2870421 | 48.3190678 | 44.2649599 |
| -3.4254432 | 0.0006138 | 0.0014437 | 40.1647002 | 51.9572696 | 21.8586259 | 23.5332698 |
| 3.42521229 | 0.00061432 | 0.00144482 | 13.524848 | 6.22404792 | 9.20363196 | 7.84442328 |
| -3.425038 | 0.00061471 | 0.00144564 | 22.9512573 | 48.1687187 | 36.8145279 | 23.5332698 |
| -3.4245975 | 0.00061571 | 0.00144788 | 138.937075 | 35.4500121 | 17.2568099 | 36.9808526 |
| -3.424323 | 0.00061633 | 0.00144924 | 8.1968776 | 82.8068984 | 94.3372276 | 52.1093832 |
| -3.4241215 | 0.00061679 | 0.00145021 | 25.8201644 | 37.8855091 | 21.8586259 | 33.6189569 |
| -3.423737 | 0.00061766 | 0.00145216 | 36.8859492 | 55.7458205 | 26.4604419 | 61.0744384 |
| -3.4233768 | 0.00061848 | 0.00145398 | 24.1807889 | 26.7904671 | 29.9118039 | 29.6967453 |
| -3.4233457 | 0.00061855 | 0.00145404 | 79.5097127 | 24.3549701 | 25.3099879 | 33.058641 |
| -3.4232459 | 0.00061878 | 0.00145447 | 349.59683 | 698.987643 | 553.368372 | 608.50312 |
| -3.4231964 | 0.00061889 | 0.00145463 | 396.728876 | 385.62036 | 0 | 384.376741 |
| -3.422895 | 0.00061958 | 0.00145613 | 17.213443 | 40.3210061 | 32.2127119 | 54.350647 |
| -3.4226613 | 0.00062011 | 0.00145728 | 24.1807889 | 25.1668025 | 50.6199758 | 29.6967453 |
| 3.42217519 | 0.00062122 | 0.00145978 | 16.3937552 | 9.47137727 | 2.30090799 | 7.28410733 |
| -3.4210243 | 0.00062386 | 0.00146587 | 31.9678226 | 21.3782516 | 71.3281477 | 29.1364293 |
| -3.420737 | 0.00062452 | 0.00146731 | 15.9839113 | 41.67406 | 54.0713378 | 35.2999048 |
| -3.4204588 | 0.00062516 | 0.00146871 | 14.3445358 | 47.6274971 | 44.8677058 | 44.8252759 |
| -3.4195669 | 0.00062721 | 0.00147342 | 38.5253247 | 69.5469702 | 72.4786017 | 68.9188617 |
| -3.419523 | 0.00062731 | 0.00147355 | 23.3611012 | 68.4645271 | 20.7081719 | 45.3855918 |
| -3.4192551 | 0.00062793 | 0.0014749 | 65.1651769 | 31.1202396 | 16.1063559 | 45.3855918 |
| 3.419041 | 0.00062842 | 0.00147584 | 38.1154808 | 19.4839761 | 26.4604419 | 22.9729539 |
| -3.4190602 | 0.00062838 | 0.00147584 | 18.0331307 | 48.4393295 | 27.6108959 | 50.4284354 |
| -3.4183574 | 0.00063 | 0.00147945 | 29.0989155 | 43.0271139 | 130.001301 | 17.9301104 |
| -3.4182687 | 0.00063021 | 0.00147982 | 21.7217256 | 39.7797845 | 29.9118039 | 28.0157974 |
| -3.4179688 | 0.0006309 | 0.00148134 | 332.793231 | 340.969582 | 0 | 502.603406 |
| 3.41720317 | 0.00063268 | 0.00148541 | 0.40984388 | 1.89427545 | 3.45136199 | 2.24126379 |
| -3.4168784 | 0.00063344 | 0.00148707 | 20.492194 | 31.3908504 | 29.9118039 | 36.4205367 |
| -3.4167293 | 0.00063378 | 0.00148778 | 24.5906328 | 70.6294134 | 36.8145279 | 22.4126379 |
| 3.41667541 | 0.00063391 | 0.00148797 | 25.0004767 | 5.41221558 | 0 | 8.40473923 |
| -3.4162419 | 0.00063492 | 0.00149023 | 137.2977 | 33.5557366 | 18.4072639 | 35.2999048 |
| -3.416041 | 0.00063539 | 0.00149122 | 25.4103206 | 58.9931499 | 18.4072639 | 49.3078035 |
| -3.4157201 | 0.00063614 | 0.00149287 | 17.6232868 | 38.1561199 | 39.1154358 | 34.1792729 |
| -3.4156079 | 0.0006364 | 0.00149338 | 45.4926707 | 35.4500121 | 20.7081719 | 25.7745336 |

| | | | | | | |
|---|---|---|---|---|---|---|
| -3.414954 | 0.00063793 | 0.00149686 | 11.0657848 | 58.1813175 | 41.4163438 | 105.339398 |
| -3.4146361 | 0.00063867 | 0.0014985 | 0 | 606.979978 | 571.775636 | 429.762333 |
| -3.4145498 | 0.00063888 | 0.00149886 | 31.9678226 | 53.0397127 | 26.4604419 | 21.852322 |
| -3.4144707 | 0.00063906 | 0.00149919 | 22.1315695 | 51.416048 | 28.7613499 | 28.5761134 |
| -3.4143364 | 0.00063938 | 0.00149982 | 28.2792277 | 94.1725511 | 14.9559019 | 58.2728587 |
| -3.4143103 | 0.00063944 | 0.00149985 | 25.0004767 | 191.321821 | 62.1245158 | 89.6505518 |
| -3.4136175 | 0.00064107 | 0.00150356 | 17.6232868 | 37.6148983 | 55.2217918 | 31.3776931 |
| -3.4135095 | 0.00064132 | 0.00150394 | 13.9346919 | 67.382084 | 50.6199758 | 36.4205367 |
| -3.41352 | 0.00064129 | 0.00150394 | 41.3942319 | 31.3908504 | 17.2568099 | 40.9030642 |
| -3.4132399 | 0.00064195 | 0.00150532 | 276.644619 | 669.220457 | 514.252936 | 530.619203 |
| -3.4130674 | 0.00064236 | 0.00150617 | 17.6232868 | 28.6847426 | 48.3190678 | 46.5062237 |
| -3.4130411 | 0.00064242 | 0.0015062 | 200.003813 | 343.67569 | 324.428027 | 319.380091 |
| -3.412641 | 0.00064337 | 0.0015083 | 9.01656536 | 66.2996409 | 63.2749697 | 68.3585457 |
| -3.4123934 | 0.00064395 | 0.00150957 | 35.2465737 | 44.109557 | 58.6731538 | 15.1285306 |
| -3.4122019 | 0.0006444 | 0.00151052 | 13.9346919 | 61.6992576 | 31.0622579 | 80.1251807 |
| -3.411493 | 0.00064608 | 0.00151434 | 23.770945 | 22.4606947 | 41.4163438 | 35.2999048 |
| 3.41134308 | 0.00064644 | 0.00151506 | 0.40984388 | 0.81183234 | 1.150454 | 1.1206319 |
| -3.4110499 | 0.00064713 | 0.00151658 | 13.9346919 | 54.9339882 | 44.8677058 | 42.0236961 |
| -3.4110195 | 0.0006472 | 0.00151664 | 18.4429746 | 28.143521 | 44.8677058 | 42.0236961 |
| -3.4109654 | 0.00064733 | 0.00151681 | 18.8528185 | 58.4519283 | 26.4604419 | 42.5840121 |
| -3.4109493 | 0.00064737 | 0.00151681 | 16.3937552 | 43.5683354 | 40.2658898 | 35.8602207 |
| -3.4105198 | 0.00064839 | 0.00151909 | 13.1150042 | 52.7691019 | 48.3190678 | 45.9459078 |
| -3.4101755 | 0.00064921 | 0.0015209 | 36.4761053 | 58.1813175 | 44.8677058 | 68.3585457 |
| -3.4101115 | 0.00064936 | 0.00152115 | 47.5418901 | 19.4839761 | 24.1595339 | 58.2728587 |
| -3.4095775 | 0.00065064 | 0.00152402 | 48.7714217 | 98.7729344 | 49.4695218 | 142.320251 |
| -3.4094399 | 0.00065096 | 0.00152457 | 18.4429746 | 43.5683354 | 27.6108959 | 45.3855918 |
| -3.4094509 | 0.00065094 | 0.00152457 | 16.3937552 | 47.6274971 | 29.9118039 | 58.8331746 |
| -3.4094175 | 0.00065102 | 0.00152458 | 19.2626624 | 56.8282636 | 27.6108959 | 37.5411686 |
| -3.4092715 | 0.00065137 | 0.00152529 | 4.50828268 | 115.009581 | 55.2217918 | 166.974153 |
| -3.409241 | 0.00065144 | 0.00152535 | 98.3625312 | 23.5431378 | 19.5577179 | 48.1871716 |
| -3.4091419 | 0.00065168 | 0.00152579 | 24.5906328 | 44.3801678 | 19.5577179 | 82.9267604 |
| -3.4088552 | 0.00065236 | 0.00152728 | 21.3118818 | 47.0862756 | 21.8586259 | 97.4949751 |
| -3.4085949 | 0.00065298 | 0.00152863 | 12.7051603 | 71.4412457 | 40.2658898 | 51.5490673 |
| -3.4080399 | 0.00065431 | 0.00153163 | 15.1642236 | 36.5324552 | 54.0713378 | 42.5840121 |
| -3.4078305 | 0.00065482 | 0.0015327 | 40.1647002 | 32.7439043 | 16.1063559 | 73.4013893 |
| -3.4077005 | 0.00065513 | 0.00153332 | 12.2953164 | 53.0397127 | 50.6199758 | 50.4284354 |
| 3.40649802 | 0.00065802 | 0.00153998 | 382.38434 | 539.597894 | 472.836592 | 439.84802 |
| -3.4061875 | 0.00065877 | 0.00154162 | 20.9020379 | 25.9786348 | 35.6640739 | 36.9808526 |
| -3.4057651 | 0.00065979 | 0.00154389 | 20.492194 | 38.4267306 | 41.4163438 | 25.2142177 |
| 3.40559816 | 0.00066019 | 0.00154472 | 159.839113 | 112.303473 | 126.549939 | 128.872668 |
| -3.405546 | 0.00066032 | 0.00154491 | 12.2953164 | 40.8622277 | 79.3813257 | 50.9887513 |
| 3.40522468 | 0.0006611 | 0.00154661 | 53.2797044 | 58.7225391 | 69.0272397 | 66.1172819 |
| -3.4040716 | 0.00066389 | 0.00155305 | 17.213443 | 34.0969582 | 37.9649818 | 39.7824324 |

| | | | | | | |
|---|---|---|---|---|---|---|
| -3.4038078 | 0.00066453 | 0.00155443 | 25.8201644 | 34.0969582 | 20.7081719 | 38.6618005 |
| -3.4035812 | 0.00066509 | 0.00155561 | 26.6398522 | 31.3908504 | 23.0090799 | 28.5761134 |
| -3.4035469 | 0.00066517 | 0.00155569 | 13.524848 | 57.0988744 | 35.6640739 | 59.9538065 |
| -3.4034644 | 0.00066537 | 0.00155605 | 27.45954 | 30.579018 | 26.4604419 | 24.0935858 |
| -3.4034422 | 0.00066542 | 0.00155606 | 255.742581 | 354.22951 | 284.162137 | 333.387989 |
| -3.4033985 | 0.00066553 | 0.0015562 | 23.770945 | 26.7904671 | 32.2127119 | 26.8951655 |
| -3.4033451 | 0.00066566 | 0.00155639 | 6.1476582 | 92.8194972 | 74.7795097 | 81.2458125 |
| -3.4030335 | 0.00066642 | 0.00155805 | 39.3450125 | 58.1813175 | 17.2568099 | 29.6967453 |
| 3.40300604 | 0.00066649 | 0.0015581 | 0.81968776 | 0 | 1.150454 | 0 |
| -3.4029178 | 0.0006667 | 0.00155849 | 19.2626624 | 33.5557366 | 29.9118039 | 38.1014845 |
| -3.4026369 | 0.00066739 | 0.00155998 | 19.6725062 | 37.0736767 | 26.4604419 | 57.1522268 |
| -3.4022032 | 0.00066845 | 0.00156234 | 15.9839113 | 40.5916169 | 65.5758777 | 33.058641 |
| -3.4017315 | 0.0006696 | 0.00156493 | 15.9839113 | 30.0377965 | 49.4695218 | 56.5919108 |
| -3.401601 | 0.00066992 | 0.00156556 | 43.0336074 | 66.0290301 | 17.2568099 | 28.0157974 |
| 3.40144751 | 0.0006703 | 0.00156633 | 1.63937552 | 3.51794013 | 0 | 2.24126379 |
| 3.40064611 | 0.00067227 | 0.00157074 | 42.6237635 | 24.6255809 | 21.8586259 | 17.9301104 |
| -3.40064 | 0.00067228 | 0.00157074 | 19.2626624 | 38.4267306 | 35.6640739 | 28.5761134 |
| -3.3997656 | 0.00067444 | 0.00157565 | 28.6890716 | 29.4965749 | 81.6822337 | 22.4126379 |
| -3.3997373 | 0.00067451 | 0.0015757 | 10.246097 | 96.0668266 | 36.8145279 | 76.763285 |
| -3.399675 | 0.00067466 | 0.00157595 | 12.2953164 | 48.1687187 | 50.6199758 | 53.2300151 |
| -3.3996438 | 0.00067474 | 0.00157601 | 8.1968776 | 56.2870421 | 86.2840497 | 70.0394936 |
| -3.3992816 | 0.00067563 | 0.00157799 | 20.492194 | 44.3801678 | 44.8677058 | 23.5332698 |
| -3.3989396 | 0.00067648 | 0.00157985 | 15.1642236 | 65.4878086 | 34.5136199 | 42.5840121 |
| 3.39879321 | 0.00067684 | 0.00158058 | 0 | 3.51794013 | 0 | 0 |
| -3.3987447 | 0.00067696 | 0.00158075 | 21.3118818 | 21.3782516 | 48.3190678 | 58.2728587 |
| -3.3983826 | 0.00067786 | 0.00158272 | 172.544273 | 391.032576 | 394.60572 | 379.894213 |
| -3.3982971 | 0.00067807 | 0.00158311 | 16.8035991 | 36.5324552 | 36.8145279 | 39.2221164 |
| -3.3981798 | 0.00067836 | 0.00158367 | 12.2953164 | 54.3927666 | 43.7172518 | 56.0315949 |
| -3.3979748 | 0.00067887 | 0.00158474 | 36.8859492 | 24.8961917 | 20.7081719 | 37.5411686 |
| 3.39770109 | 0.00067955 | 0.00158621 | 9.42640924 | 6.22404792 | 5.75226998 | 1.68094785 |
| -3.3972658 | 0.00068063 | 0.00158862 | 258.611488 | 33.8263474 | 16.1063559 | 32.498325 |
| -3.3962693 | 0.00068311 | 0.0015943 | 20.492194 | 26.5198564 | 33.3631659 | 38.6618005 |
| -3.3961131 | 0.0006835 | 0.0015951 | 15.5740674 | 40.5916169 | 35.6640739 | 48.7474875 |
| 3.39608558 | 0.00068357 | 0.00159514 | 6.55750208 | 3.24732935 | 2.30090799 | 1.1206319 |
| 3.3958033 | 0.00068428 | 0.00159667 | 0 | 3.51794013 | 8.05317797 | 6.16347543 |
| -3.3954015 | 0.00068528 | 0.0015989 | 16.8035991 | 43.5683354 | 36.8145279 | 34.1792729 |
| -3.3952868 | 0.00068557 | 0.00159946 | 19.6725062 | 27.0610779 | 34.5136199 | 46.5062237 |
| -3.394951 | 0.00068641 | 0.00160131 | 160.658801 | 615.368912 | 0 | 498.120878 |
| -3.3947209 | 0.00068699 | 0.00160254 | 14.7543797 | 47.6274971 | 37.9649818 | 43.704644 |
| -3.3939136 | 0.00068901 | 0.00160715 | 71.3128351 | 27.3316887 | 29.9118039 | 22.4126379 |
| -3.3938265 | 0.00068923 | 0.00160755 | 70.9029912 | 238.137486 | 218.586259 | 179.86142 |
| 3.39344062 | 0.00069021 | 0.0016097 | 23.3611012 | 14.0717605 | 12.6549939 | 14.0078987 |
| -3.3933724 | 0.00069038 | 0.00160998 | 13.524848 | 63.3229223 | 43.7172518 | 40.3427483 |

| | | | | | | |
|---|---|---|---|---|---|---|
| -3.3931097 | 0.00069104 | 0.00161141 | 74.5915861 | 186.992048 | 178.320369 | 182.662999 |
| 3.39222264 | 0.00069328 | 0.00161652 | 0 | 0 | 0 | 0 |
| 3.39218115 | 0.00069339 | 0.00161665 | 0.40984388 | 0.54122156 | 0 | 0 |
| -3.3918222 | 0.00069429 | 0.00161865 | 9.42640924 | 57.640096 | 86.2840497 | 51.5490673 |
| -3.3917861 | 0.00069439 | 0.00161869 | 10.6559409 | 38.4267306 | 67.8767857 | 85.1680242 |
| -3.3917317 | 0.00069452 | 0.00161869 | 20.0823501 | 27.3316887 | 32.2127119 | 40.9030642 |
| -3.3917715 | 0.00069442 | 0.00161869 | 24.5906328 | 38.1561199 | 25.3099879 | 25.7745336 |
| -3.3917384 | 0.00069451 | 0.00161869 | 17.6232868 | 60.3462037 | 25.3099879 | 49.8681194 |
| -3.3917174 | 0.00069456 | 0.00161869 | 66.3947085 | 138.82333 | 111.594038 | 108.140978 |
| -3.3916587 | 0.00069471 | 0.00161892 | 29.0989155 | 31.1202396 | 56.3722458 | 19.6110582 |
| -3.3914974 | 0.00069512 | 0.00161975 | 71.3128351 | 123.669126 | 125.399485 | 126.631404 |
| 3.3912788 | 0.00069567 | 0.00162081 | 19.2626624 | 14.3423713 | 17.2568099 | 10.646003 |
| -3.3912884 | 0.00069565 | 0.00162081 | 20.492194 | 41.67406 | 23.0090799 | 50.9887513 |
| -3.3911809 | 0.00069592 | 0.00162127 | 24.5906328 | 55.4752097 | 17.2568099 | 59.3934906 |
| -3.3911255 | 0.00069606 | 0.00162148 | 20.9020379 | 29.2259641 | 28.7613499 | 33.6189569 |
| -3.3910827 | 0.00069617 | 0.00162162 | 25.0004767 | 23.8137486 | 41.4163438 | 26.8951655 |
| -3.3909774 | 0.00069644 | 0.00162213 | 20.492194 | 30.579018 | 27.6108959 | 38.1014845 |
| -3.3904732 | 0.00069772 | 0.001625 | 8.1968776 | 59.8049822 | 70.1776937 | 74.5220212 |
| -3.3902424 | 0.00069831 | 0.00162625 | 16.3937552 | 62.2404792 | 25.3099879 | 62.1950703 |
| 3.39019623 | 0.00069843 | 0.0016264 | 0 | 0.54122156 | 0 | 0.56031595 |
| 3.39015124 | 0.00069854 | 0.00162655 | 0 | 0 | 0 | 1.1206319 |
| -3.3899453 | 0.00069907 | 0.00162766 | 20.9020379 | 44.6507786 | 23.0090799 | 40.3427483 |
| -3.3899164 | 0.00069914 | 0.00162771 | 27.8693838 | 70.6294134 | 17.2568099 | 41.4633802 |
| 3.38938825 | 0.00070049 | 0.00163073 | 0 | 0 | 0 | 0 |
| -3.3893477 | 0.00070059 | 0.00163086 | 16.8035991 | 27.3316887 | 55.2217918 | 48.7474875 |
| -3.3889767 | 0.00070154 | 0.00163295 | 157.789894 | 144.776767 | 90.8858656 | 87.9696039 |
| 3.38889955 | 0.00070174 | 0.00163329 | 27.45954 | 18.6721438 | 14.9559019 | 19.0507423 |
| -3.3888566 | 0.00070185 | 0.00163343 | 18.8528185 | 72.5236888 | 19.5577179 | 75.0823371 |
| -3.3887985 | 0.000702 | 0.00163366 | 40.984388 | 57.9107067 | 24.1595339 | 19.6110582 |
| -3.3886139 | 0.00070247 | 0.00163464 | 12.7051603 | 31.1202396 | 97.7885896 | 61.0744384 |
| -3.3883036 | 0.00070326 | 0.00163637 | 15.5740674 | 68.4645271 | 31.0622579 | 44.2649599 |
| 3.38822082 | 0.00070348 | 0.00163675 | 20.0823501 | 12.1774851 | 13.8054479 | 11.206319 |
| -3.3877563 | 0.00070467 | 0.00163933 | 13.9346919 | 40.3210061 | 43.7172518 | 50.4284354 |
| -3.3877487 | 0.00070469 | 0.00163933 | 564.764867 | 1290.27219 | 974.434534 | 1079.72883 |
| 3.38752003 | 0.00070528 | 0.00164058 | 0 | 0 | 0 | 0 |
| -3.386615 | 0.00070761 | 0.00164588 | 23.3611012 | 20.83703 | 48.3190678 | 38.1014845 |
| -3.3862172 | 0.00070863 | 0.00164815 | 18.0331307 | 36.2618444 | 28.7613499 | 51.5490673 |
| -3.3860096 | 0.00070917 | 0.00164928 | 18.0331307 | 42.4858923 | 27.6108959 | 41.4633802 |
| -3.3858897 | 0.00070948 | 0.00164988 | 19.6725062 | 33.5557366 | 27.6108959 | 36.4205367 |
| -3.3858506 | 0.00070958 | 0.00165 | 16.3937552 | 41.9446708 | 44.8677058 | 31.3776931 |
| -3.3853107 | 0.00071098 | 0.00165313 | 25.4103206 | 18.9427545 | 41.4163438 | 47.0665397 |
| -3.3849217 | 0.00071199 | 0.00165526 | 19.6725062 | 88.7603356 | 143.806749 | 81.2458125 |
| -3.384917 | 0.000712 | 0.00165526 | 28.2792277 | 17.0484791 | 51.7704298 | 43.144328 |

| | | | | | | |
|---|---|---|---|---|---|---|
| -3.3847915 | 0.00071232 | 0.00165586 | 47.951734 | 71.9824673 | 54.0713378 | 68.3585457 |
| 3.38477842 | 0.00071236 | 0.00165586 | 25.4103206 | 12.9893174 | 13.8054479 | 12.8872668 |
| -3.3846691 | 0.00071264 | 0.0016564 | 20.492194 | 34.6381797 | 26.4604419 | 33.058641 |
| -3.3840178 | 0.00071433 | 0.00166021 | 12.7051603 | 72.5236888 | 35.6640739 | 53.7903311 |
| -3.3837903 | 0.00071493 | 0.00166147 | 132.789417 | 196.734036 | 220.887167 | 189.947107 |
| -3.3835357 | 0.00071559 | 0.00166289 | 25.0004767 | 509.018876 | 1098.68357 | 1071.88441 |
| -3.3833505 | 0.00071607 | 0.00166389 | 17.6232868 | 27.6022995 | 42.5667978 | 45.3855918 |
| -3.3833183 | 0.00071616 | 0.00166397 | 12.2953164 | 63.0523115 | 37.9649818 | 56.5919108 |
| -3.3829704 | 0.00071706 | 0.00166596 | 17.213443 | 41.9446708 | 27.6108959 | 64.4363341 |
| -3.3829023 | 0.00071724 | 0.00166625 | 15.1642236 | 68.1939163 | 27.6108959 | 56.5919108 |
| -3.3825197 | 0.00071824 | 0.00166845 | 8.1968776 | 69.5469702 | 78.2308717 | 56.0315949 |
| -3.3823818 | 0.0007186 | 0.00166917 | 189.347873 | 360.453558 | 292.215315 | 304.25156 |
| -3.3820645 | 0.00071943 | 0.00167098 | 2.86890716 | 79.0183475 | 162.214013 | 120.467929 |
| -3.3817599 | 0.00072023 | 0.00167272 | 49.1812656 | 60.075593 | 14.9559019 | 29.6967453 |
| -3.3805457 | 0.00072342 | 0.00167996 | 17.6232868 | 20.5664192 | 101.239952 | 63.3157022 |
| -3.3805329 | 0.00072345 | 0.00167996 | 16.3937552 | 55.7458205 | 28.7613499 | 44.2649599 |
| -3.379602 | 0.00072591 | 0.00168554 | 21.7217256 | 34.367569 | 23.0090799 | 40.3427483 |
| -3.3789008 | 0.00072776 | 0.00168972 | 39.7548564 | 44.9213893 | 17.2568099 | 28.0157974 |
| -3.3788521 | 0.00072789 | 0.0016899 | 340.17042 | 385.890971 | 0 | 340.672097 |
| -3.3787709 | 0.00072811 | 0.00169028 | 44.263139 | 42.2152815 | 12.6549939 | 53.7903311 |
| -3.3785568 | 0.00072867 | 0.00169147 | 152.052079 | 351.523402 | 355.490285 | 363.084735 |
| -3.3781847 | 0.00072966 | 0.00169364 | 14.3445358 | 45.1920001 | 34.5136199 | 56.0315949 |
| -3.3780136 | 0.00073011 | 0.00169457 | 19.2626624 | 23.8137486 | 42.5667978 | 55.4712789 |
| -3.3777185 | 0.0007309 | 0.00169624 | 706.161005 | 228.936719 | 0 | 273.994499 |
| -3.3776841 | 0.00073099 | 0.00169624 | 328.694792 | 373.172264 | 0 | 361.964103 |
| -3.3777006 | 0.00073095 | 0.00169624 | 21.3118818 | 33.5557366 | 27.6108959 | 28.0157974 |
| -3.3776528 | 0.00073107 | 0.00169631 | 11.4756286 | 70.3588026 | 41.4163438 | 53.2300151 |
| -3.3775728 | 0.00073129 | 0.00169668 | 26.2300083 | 49.5217726 | 17.2568099 | 44.8252759 |
| -3.3771642 | 0.00073237 | 0.00169908 | 18.8528185 | 63.8641439 | 24.1595339 | 42.5840121 |
| -3.3770068 | 0.00073279 | 0.00169993 | 12.7051603 | 40.0503953 | 55.2217918 | 49.8681194 |
| -3.376954 | 0.00073293 | 0.00170014 | 41.3942319 | 66.0290301 | 34.5136199 | 14.5682147 |
| -3.3764998 | 0.00073414 | 0.0017027 | 18.4429746 | 47.0862756 | 29.9118039 | 31.9380091 |
| -3.3765094 | 0.00073412 | 0.0017027 | 263.529615 | 461.391378 | 0 | 363.084735 |
| -3.3759252 | 0.00073568 | 0.00170614 | 23.3611012 | 126.375234 | 64.4254237 | 79.0045488 |
| -3.3755003 | 0.00073682 | 0.00170865 | 25.4103206 | 45.7332217 | 17.2568099 | 56.5919108 |
| -3.3753099 | 0.00073733 | 0.00170971 | 15.1642236 | 47.8981079 | 36.8145279 | 38.1014845 |
| -3.3750962 | 0.0007379 | 0.00171092 | 22.9512573 | 36.5324552 | 21.8586259 | 34.1792729 |
| -3.375055 | 0.00073801 | 0.00171105 | 17.6232868 | 29.7671857 | 42.5667978 | 35.2999048 |
| -3.3745507 | 0.00073936 | 0.00171407 | 11.4756286 | 74.4179643 | 36.8145279 | 60.5141225 |
| -3.374253 | 0.00074016 | 0.0017158 | 19.2626624 | 36.803066 | 26.4604419 | 36.4205367 |
| -3.3739385 | 0.00074101 | 0.00171764 | 300.00572 | 471.674588 | 0 | 307.05314 |
| -3.3738245 | 0.00074132 | 0.00171822 | 63.1159575 | 25.708024 | 27.6108959 | 23.5332698 |
| -3.3734381 | 0.00074236 | 0.00172051 | 11.0657848 | 62.51109 | 43.7172518 | 56.5919108 |

| | | | | | | |
|---|---|---|---|---|---|---|
| -3.373142 | 0.00074316 | 0.00172224 | 26.6398522 | 29.7671857 | 21.8586259 | 28.5761134 |
| -3.372958 | 0.00074365 | 0.00172327 | 12.7051603 | 66.2996409 | 35.6640739 | 52.6696992 |
| -3.3726401 | 0.00074451 | 0.00172514 | 23.3611012 | 42.4858923 | 44.8677058 | 19.6110582 |
| -3.3717252 | 0.00074699 | 0.00173075 | 5.73781432 | 112.303473 | 67.8767857 | 73.4013893 |
| -3.3711987 | 0.00074842 | 0.00173394 | 36.8859492 | 52.7691019 | 54.0713378 | 12.8872668 |
| -3.3709288 | 0.00074915 | 0.00173552 | 24.1807889 | 59.2637606 | 20.7081719 | 34.1792729 |
| -3.3708742 | 0.0007493 | 0.00173573 | 14.7543797 | 41.4034492 | 40.2658898 | 39.7824324 |
| -3.370373 | 0.00075066 | 0.00173877 | 15.5740674 | 50.8748265 | 111.594038 | 25.7745336 |
| -3.3701633 | 0.00075124 | 0.00173997 | 14.7543797 | 45.4626109 | 33.3631659 | 48.7474875 |
| -3.3695544 | 0.0007529 | 0.00174369 | 18.0331307 | 37.0736767 | 57.5226998 | 26.8951655 |
| -3.369353 | 0.00075345 | 0.00174484 | 20.0823501 | 29.4965749 | 27.6108959 | 36.4205367 |
| -3.3688991 | 0.00075469 | 0.00174759 | 55.7387677 | 38.1561199 | 14.9559019 | 33.6189569 |
| -3.3687488 | 0.0007551 | 0.00174842 | 18.4429746 | 36.2618444 | 26.4604419 | 50.9887513 |
| -3.3683581 | 0.00075617 | 0.00175077 | 22.1315695 | 34.9087905 | 23.0090799 | 32.498325 |
| -3.3680564 | 0.000757 | 0.00175257 | 13.524848 | 69.5469702 | 36.8145279 | 42.5840121 |
| -3.367816 | 0.00075766 | 0.00175397 | 16.3937552 | 32.7439043 | 128.850847 | 35.2999048 |
| -3.3672218 | 0.0007593 | 0.00175763 | 0 | 750.944912 | 301.418947 | 474.027293 |
| -3.3671769 | 0.00075942 | 0.00175779 | 25.0004767 | 14.6129821 | 94.3372276 | 61.0744384 |
| -3.3667138 | 0.0007607 | 0.00176062 | 8.60672148 | 36.2618444 | 138.054479 | 72.2807574 |
| -3.3666408 | 0.0007609 | 0.00176096 | 6.96734596 | 60.3462037 | 136.904025 | 54.910963 |
| -3.3664603 | 0.0007614 | 0.00176198 | 15.5740674 | 37.6148983 | 41.4163438 | 35.8602207 |
| 3.3663491 | 0.0007617 | 0.00176257 | 10.246097 | 18.401533 | 11.50454 | 12.8872668 |
| -3.366178 | 0.00076218 | 0.00176353 | 31.9678226 | 25.9786348 | 42.5667978 | 19.6110582 |
| -3.3661191 | 0.00076234 | 0.00176378 | 17.6232868 | 28.6847426 | 73.6290557 | 35.8602207 |
| -3.3656629 | 0.0007636 | 0.00176658 | 17.213443 | 51.1454373 | 34.5136199 | 30.2570612 |
| -3.3648675 | 0.0007658 | 0.00177155 | 31.9678226 | 31.9320719 | 18.4072639 | 30.8173772 |
| -3.364759 | 0.00076611 | 0.00177209 | 15.5740674 | 30.579018 | 44.8677058 | 46.5062237 |
| -3.364744 | 0.00076615 | 0.00177209 | 44.6729829 | 42.7565031 | 31.0622579 | 15.6888466 |
| -3.3646614 | 0.00076638 | 0.00177249 | 13.524848 | 31.3908504 | 67.8767857 | 52.6696992 |
| -3.3643597 | 0.00076721 | 0.00177409 | 2.45906328 | 124.751569 | 108.142676 | 122.148877 |
| -3.3643531 | 0.00076723 | 0.00177409 | 9.01656536 | 47.3568864 | 57.5226998 | 95.2537113 |
| -3.3643889 | 0.00076713 | 0.00177409 | 13.9346919 | 55.4752097 | 41.4163438 | 36.9808526 |
| -3.3641857 | 0.0007677 | 0.00177504 | 18.8528185 | 33.8263474 | 29.9118039 | 31.3776931 |
| -3.3637649 | 0.00076887 | 0.0017775 | 32.7875104 | 41.67406 | 18.4072639 | 27.4554815 |
| -3.3637844 | 0.00076882 | 0.0017775 | 50.8206411 | 90.3840002 | 104.691314 | 93.0124475 |
| -3.3628186 | 0.00077151 | 0.00178347 | 38.9351686 | 33.0145151 | 18.4072639 | 27.4554815 |
| -3.3625889 | 0.00077215 | 0.00178483 | 22.9512573 | 35.7206228 | 25.3099879 | 25.7745336 |
| -3.3621831 | 0.00077329 | 0.00178733 | 22.9512573 | 33.5557366 | 29.9118039 | 22.9729539 |
| -3.361988 | 0.00077383 | 0.00178846 | 27.8693838 | 19.2133653 | 41.4163438 | 30.2570612 |
| -3.3617101 | 0.00077461 | 0.00179013 | 58.6076748 | 29.4965749 | 20.7081719 | 26.3348496 |
| -3.3614416 | 0.00077537 | 0.00179175 | 15.5740674 | 41.1328384 | 33.3631659 | 40.9030642 |
| -3.3613966 | 0.00077549 | 0.00179191 | 13.524848 | 60.8874253 | 36.8145279 | 42.5840121 |
| -3.3607922 | 0.00077719 | 0.00179559 | 18.8528185 | 37.0736767 | 29.9118039 | 29.6967453 |

| | | | | | | |
|---|---|---|---|---|---|---|
| -3.3607831 | 0.00077722 | 0.00179559 | 8.1968776 | 51.9572696 | 78.2308717 | 67.7982298 |
| -3.3607708 | 0.00077725 | 0.00179559 | 17.6232868 | 41.67406 | 26.4604419 | 42.5840121 |
| -3.3605471 | 0.00077788 | 0.00179675 | 18.0331307 | 26.5198564 | 40.2658898 | 39.2221164 |
| -3.3605335 | 0.00077792 | 0.00179675 | 61.8864259 | 25.9786348 | 20.7081719 | 30.2570612 |
| -3.3605548 | 0.00077786 | 0.00179675 | 27.0496961 | 57.0988744 | 63.2749697 | 57.7125427 |
| -3.3602546 | 0.00077871 | 0.00179843 | 352.465737 | 307.955067 | 393.455266 | 350.757784 |
| 3.35973086 | 0.00078018 | 0.00180172 | 35.6564176 | 31.3908504 | 80.5317797 | 7.84442328 |
| -3.3593547 | 0.00078125 | 0.00180404 | 13.9346919 | 48.1687187 | 56.3722458 | 32.498325 |
| 3.35914577 | 0.00078184 | 0.00180528 | 4.50828268 | 3.78855091 | 1.150454 | 1.1206319 |
| 3.35877451 | 0.00078289 | 0.00180748 | 45.0828268 | 18.1309222 | 19.5577179 | 18.4904263 |
| -3.3587693 | 0.0007829 | 0.00180748 | 38.1154808 | 51.1454373 | 39.1154358 | 14.0078987 |
| 3.35815752 | 0.00078464 | 0.00181136 | 9.83625312 | 13.2599282 | 13.8054479 | 10.0856871 |
| -3.3577622 | 0.00078576 | 0.00181382 | 18.0331307 | 31.1202396 | 36.8145279 | 30.8173772 |
| -3.3575483 | 0.00078637 | 0.00181509 | 6.96734596 | 31.1202396 | 162.214013 | 101.417187 |
| -3.3572639 | 0.00078718 | 0.00181683 | 24.1807889 | 53.3103235 | 37.9649818 | 54.910963 |
| -3.356961 | 0.00078804 | 0.00181869 | 39.7548564 | 199.440144 | 118.496762 | 163.051941 |
| -3.3560545 | 0.00079063 | 0.00182453 | 11.4756286 | 55.7458205 | 36.8145279 | 71.1601255 |
| -3.3558964 | 0.00079108 | 0.00182545 | 10.6559409 | 45.7332217 | 65.5758777 | 50.9887513 |
| -3.3558171 | 0.00079131 | 0.00182584 | 21.7217256 | 33.5557366 | 23.0090799 | 32.498325 |
| -3.3556594 | 0.00079176 | 0.00182675 | 24.5906328 | 33.0145151 | 19.5577179 | 44.8252759 |
| -3.3555487 | 0.00079208 | 0.00182735 | 12.7051603 | 69.817581 | 37.9649818 | 43.704644 |
| 3.35538573 | 0.00079254 | 0.0018283 | 27.45954 | 9.74198805 | 11.50454 | 11.7666349 |
| -3.3552067 | 0.00079306 | 0.00182935 | 70.0833035 | 20.0251977 | 27.6108959 | 30.2570612 |
| -3.3549928 | 0.00079367 | 0.00183064 | 15.5740674 | 44.6507786 | 85.1335957 | 27.4554815 |
| -3.3547921 | 0.00079425 | 0.00183183 | 10.6559409 | 51.416048 | 56.3722458 | 49.8681194 |
| -3.3541804 | 0.000796 | 0.00183576 | 19.2626624 | 38.9679522 | 25.3099879 | 34.1792729 |
| -3.3539763 | 0.00079659 | 0.00183698 | 29.5087594 | 24.0843593 | 23.0090799 | 28.0157974 |
| -3.3534548 | 0.00079809 | 0.00184031 | 40.1647002 | 32.4732935 | 16.1063559 | 34.1792729 |
| -3.3532715 | 0.00079862 | 0.0018414 | 17.213443 | 31.9320719 | 31.0622579 | 57.7125427 |
| -3.352649 | 0.00080042 | 0.00184542 | 56.1486116 | 16.2366467 | 26.4604419 | 45.9459078 |
| -3.3523288 | 0.00080135 | 0.00184742 | 2.45906328 | 82.5362876 | 150.709473 | 126.631404 |
| -3.352213 | 0.00080168 | 0.00184806 | 31.5579788 | 14.8835929 | 41.4163438 | 58.8331746 |
| -3.3521682 | 0.00080181 | 0.00184823 | 28.6890716 | 36.5324552 | 18.4072639 | 30.2570612 |
| -3.352085 | 0.00080205 | 0.00184865 | 23.3611012 | 49.2511618 | 57.5226998 | 17.3697944 |
| -3.3520198 | 0.00080224 | 0.00184896 | 6.96734596 | 69.817581 | 57.5226998 | 89.0902358 |
| -3.351854 | 0.00080272 | 0.00184993 | 21.3118818 | 46.0038325 | 20.7081719 | 39.2221164 |
| -3.3514342 | 0.00080394 | 0.00185261 | 2.86890716 | 74.4179643 | 179.470823 | 100.296555 |
| -3.3508075 | 0.00080576 | 0.00185667 | 261.480395 | 319.320719 | 0 | 451.614655 |
| 3.35059212 | 0.00080639 | 0.00185798 | 37.705637 | 54.6633774 | 55.2217918 | 46.5062237 |
| -3.3492809 | 0.00081022 | 0.00186667 | 35.2465737 | 51.9572696 | 12.6549939 | 49.8681194 |
| -3.3491725 | 0.00081053 | 0.00186726 | 29.5087594 | 22.4606947 | 24.1595339 | 68.9188617 |
| 3.3488785 | 0.00081139 | 0.00186911 | 73.7718984 | 47.8981079 | 47.1686138 | 48.7474875 |
| -3.3487245 | 0.00081184 | 0.00187002 | 40.984388 | 25.1668025 | 26.4604419 | 21.852322 |

| | | | | | | |
|---|---|---|---|---|---|---|
| 3.3483781 | 0.00081286 | 0.00187222 | 0.40984388 | 0 | 2.30090799 | 0 |
| -3.3482593 | 0.00081321 | 0.00187289 | 10.6559409 | 37.6148983 | 82.8326877 | 56.0315949 |
| -3.3482111 | 0.00081335 | 0.00187308 | 115.985818 | 304.166516 | 286.463045 | 300.329348 |
| 3.34818495 | 0.00081343 | 0.00187313 | 0 | 0.27061078 | 0 | 0 |
| -3.3480957 | 0.00081369 | 0.0018736 | 20.0823501 | 35.9912336 | 23.0090799 | 39.2221164 |
| -3.3479928 | 0.00081399 | 0.00187416 | 13.524848 | 48.980551 | 35.6640739 | 45.9459078 |
| -3.3477075 | 0.00081483 | 0.00187596 | 18.0331307 | 53.0397127 | 31.0622579 | 29.1364293 |
| -3.3475527 | 0.00081528 | 0.00187687 | 68.4439279 | 26.7904671 | 16.1063559 | 41.4633802 |
| -3.347076 | 0.00081669 | 0.00187997 | 13.9346919 | 57.640096 | 27.6108959 | 65.556966 |
| -3.3468373 | 0.00081739 | 0.00188145 | 36.0662614 | 27.8729103 | 19.5577179 | 27.4554815 |
| -3.3467675 | 0.0008176 | 0.00188179 | 7.37718984 | 34.6381797 | 101.239952 | 113.744138 |
| -3.3463171 | 0.00081893 | 0.00188472 | 25.0004767 | 51.9572696 | 14.9559019 | 84.6077082 |
| -3.3462432 | 0.00081915 | 0.00188508 | 77.0506494 | 228.124887 | 196.727633 | 206.756585 |
| -3.3456838 | 0.0008208 | 0.00188876 | 40.984388 | 21.9194731 | 27.6108959 | 24.0935858 |
| -3.345498 | 0.00082135 | 0.00188989 | 15.9839113 | 39.238563 | 29.9118039 | 43.704644 |
| -3.3454703 | 0.00082143 | 0.00188994 | 20.492194 | 38.6973414 | 21.8586259 | 39.2221164 |
| -3.345347 | 0.0008218 | 0.00189065 | 33.6071982 | 56.2870421 | 80.5317797 | 72.8410733 |
| -3.3452355 | 0.00082213 | 0.00189127 | 448.779049 | 662.996409 | 495.845672 | 584.96985 |
| -3.3447224 | 0.00082365 | 0.00189464 | 195.905375 | 178.873725 | 181.771731 | 160.810677 |
| -3.3445642 | 0.00082412 | 0.00189559 | 15.1642236 | 34.0969582 | 43.7172518 | 36.9808526 |
| -3.3439969 | 0.00082581 | 0.00189925 | 19.6725062 | 29.2259641 | 26.4604419 | 36.4205367 |
| -3.3439886 | 0.00082583 | 0.00189925 | 28.6890716 | 19.4839761 | 35.6640739 | 27.4554815 |
| -3.343952 | 0.00082594 | 0.00189937 | 25.4103206 | 24.3549701 | 27.6108959 | 24.6539017 |
| -3.3437203 | 0.00082663 | 0.00190082 | 230.742104 | 264.657342 | 207.081719 | 326.664198 |
| -3.3433185 | 0.00082783 | 0.00190344 | 71.3128351 | 26.5198564 | 18.4072639 | 33.6189569 |
| -3.3430173 | 0.00082873 | 0.00190537 | 22.5414134 | 33.5557366 | 69.0272397 | 21.292006 |
| -3.3428178 | 0.00082932 | 0.00190661 | 24.5906328 | 17.3190899 | 43.7172518 | 51.5490673 |
| -3.3427228 | 0.00082961 | 0.00190712 | 13.1150042 | 115.550803 | 28.7613499 | 50.4284354 |
| -3.3425621 | 0.00083009 | 0.00190809 | 21.3118818 | 33.2851258 | 32.2127119 | 22.9729539 |
| -3.3424612 | 0.00083039 | 0.00190865 | 31.5579788 | 36.2618444 | 24.1595339 | 20.1713742 |
| -3.342397 | 0.00083058 | 0.00190895 | 31.5579788 | 61.1580361 | 12.6549939 | 53.2300151 |
| -3.3421363 | 0.00083136 | 0.00191048 | 23.3611012 | 61.4286469 | 82.8326877 | 73.9617052 |
| -3.3421461 | 0.00083133 | 0.00191048 | 18.4429746 | 26.5198564 | 36.8145279 | 33.6189569 |
| -3.342099 | 0.00083147 | 0.0019106 | 180.331307 | 519.302085 | 488.942948 | 430.882964 |
| -3.3419988 | 0.00083177 | 0.00191115 | 47.5418901 | 111.491641 | 93.1867736 | 110.942558 |
| -3.3417926 | 0.00083239 | 0.00191243 | 18.0331307 | 68.7351379 | 16.1063559 | 122.709193 |
| -3.3416805 | 0.00083273 | 0.00191307 | 16.8035991 | 41.4034492 | 36.8145279 | 28.5761134 |
| -3.3416161 | 0.00083292 | 0.00191338 | 17.6232868 | 61.6992576 | 21.8586259 | 50.9887513 |
| -3.3415449 | 0.00083314 | 0.00191373 | 10.6559409 | 44.6507786 | 64.4254237 | 49.3078035 |
| -3.3413923 | 0.00083359 | 0.00191465 | 72.5423667 | 13.2599282 | 71.3281477 | 25.7745336 |
| -3.3396615 | 0.00083881 | 0.00192648 | 95.493624 | 29.7671857 | 25.3099879 | 21.852322 |
| -3.3393976 | 0.0008396 | 0.00192818 | 11.8854725 | 42.7565031 | 55.2217918 | 44.2649599 |
| -3.3389857 | 0.00084085 | 0.0019309 | 15.9839113 | 56.2870421 | 25.3099879 | 48.1871716 |

| | | | | | | |
|---|---|---|---|---|---|---|
| 3.33856197 | 0.00084213 | 0.00193362 | 20.0823501 | 13.8011497 | 16.1063559 | 15.6888466 |
| -3.3385549 | 0.00084215 | 0.00193362 | 0 | 2263.92978 | 2628.78738 | 3340.60369 |
| -3.3384624 | 0.00084243 | 0.00193413 | 8.1968776 | 74.688575 | 72.4786017 | 46.5062237 |
| -3.3383982 | 0.00084263 | 0.00193444 | 17.213443 | 53.851545 | 27.6108959 | 34.7395888 |
| -3.3379711 | 0.00084393 | 0.00193728 | 36.8859492 | 92.0076649 | 48.3190678 | 11.206319 |
| -3.3379407 | 0.00084402 | 0.00193735 | 38.5253247 | 14.8835929 | 111.594038 | 30.8173772 |
| -3.3378137 | 0.0008444 | 0.0019381 | 9.83625312 | 57.0988744 | 50.6199758 | 54.910963 |
| -3.3374837 | 0.00084541 | 0.00194026 | 92.214873 | 27.8729103 | 23.0090799 | 25.2142177 |
| -3.3373459 | 0.00084583 | 0.00194109 | 10.6559409 | 51.9572696 | 57.5226998 | 44.8252759 |
| -3.3372097 | 0.00084624 | 0.0019419 | 13.524848 | 38.1561199 | 43.7172518 | 42.5840121 |
| -3.3369783 | 0.00084695 | 0.00194338 | 16.3937552 | 32.7439043 | 32.2127119 | 40.3427483 |
| -3.3368798 | 0.00084725 | 0.00194393 | 15.1642236 | 69.0057487 | 35.6640739 | 32.498325 |
| -3.3366606 | 0.00084791 | 0.00194533 | 65.9848647 | 135.846611 | 144.957203 | 128.312352 |
| -3.336178 | 0.00084939 | 0.00194857 | 19.2626624 | 42.4858923 | 44.8677058 | 21.852322 |
| -3.3360462 | 0.00084979 | 0.00194936 | 55.3289238 | 18.1309222 | 42.5667978 | 22.9729539 |
| -3.3359935 | 0.00084995 | 0.00194959 | 15.9839113 | 34.0969582 | 37.9649818 | 33.6189569 |
| -3.3355466 | 0.00085132 | 0.00195258 | 17.213443 | 28.143521 | 34.5136199 | 39.7824324 |
| -3.3353233 | 0.000852 | 0.00195401 | 13.9346919 | 58.4519283 | 37.9649818 | 35.2999048 |
| -3.3351854 | 0.00085243 | 0.00195484 | 15.1642236 | 38.1561199 | 31.0622579 | 50.4284354 |
| -3.3346882 | 0.00085395 | 0.0019582 | 6.55750208 | 56.0164313 | 117.346308 | 60.5141225 |
| -3.3345819 | 0.00085428 | 0.00195881 | 24.1807889 | 115.821413 | 10.354086 | 87.9696039 |
| -3.3345234 | 0.00085446 | 0.00195909 | 15.1642236 | 40.5916169 | 36.8145279 | 34.1792729 |
| -3.334482 | 0.00085458 | 0.0019592 | 23.3611012 | 33.2851258 | 25.3099879 | 23.5332698 |
| -3.3344678 | 0.00085463 | 0.0019592 | 18.0331307 | 31.3908504 | 27.6108959 | 39.7824324 |
| -3.3342677 | 0.00085524 | 0.00196047 | 20.0823501 | 68.4645271 | 23.0090799 | 34.7395888 |
| -3.3341272 | 0.00085568 | 0.00196132 | 133.199261 | 33.8263474 | 20.7081719 | 23.5332698 |
| -3.334096 | 0.00085577 | 0.0019614 | 119.674413 | 184.556551 | 260.002603 | 285.761134 |
| -3.3339715 | 0.00085615 | 0.00196214 | 14.3445358 | 70.6294134 | 26.4604419 | 52.1093832 |
| 3.3332818 | 0.00085828 | 0.00196687 | 84.0179954 | 61.9698684 | 56.3722458 | 57.7125427 |
| -3.3332372 | 0.00085842 | 0.00196705 | 15.1642236 | 121.233629 | 89.7354116 | 126.071088 |
| -3.3332014 | 0.00085853 | 0.00196716 | 18.0331307 | 55.4752097 | 35.6640739 | 25.2142177 |
| -3.3331433 | 0.00085871 | 0.00196743 | 69.2636157 | 51.9572696 | 14.9559019 | 24.6539017 |
| -3.3329517 | 0.0008593 | 0.00196865 | 656.97974 | 793.701415 | 760.450091 | 833.750132 |
| -3.332828 | 0.00085968 | 0.00196938 | 11.8854725 | 46.8156648 | 42.5667978 | 49.3078035 |
| -3.3326759 | 0.00086015 | 0.00197032 | 40.984388 | 17.0484791 | 78.2308717 | 24.6539017 |
| -3.3323462 | 0.00086117 | 0.00197252 | 16.3937552 | 72.7942996 | 79.3813257 | 19.0507423 |
| -3.3322621 | 0.00086143 | 0.00197297 | 11.8854725 | 41.67406 | 48.3190678 | 48.7474875 |
| -3.332062 | 0.00086205 | 0.00197425 | 9.83625312 | 56.2870421 | 62.1245158 | 45.3855918 |
| 3.33174393 | 0.00086304 | 0.00197637 | 73.7718984 | 30.3084073 | 27.6108959 | 20.7316901 |
| -3.3314539 | 0.00086394 | 0.00197829 | 18.0331307 | 35.9912336 | 47.1686138 | 24.6539017 |
| -3.3313746 | 0.00086418 | 0.00197871 | 12.2953164 | 31.1202396 | 52.9208838 | 77.3236009 |
| 3.33124793 | 0.00086458 | 0.0019794 | 12.2953164 | 4.87099402 | 0 | 7.84442328 |
| -3.3312383 | 0.00086461 | 0.0019794 | 28.2792277 | 23.272527 | 79.3813257 | 23.5332698 |

| | | | | | | |
|---|---|---|---|---|---|---|
| -3.331072 | 0.00086512 | 0.00198045 | 23.770945 | 25.708024 | 23.0090799 | 30.2570612 |
| -3.3309193 | 0.0008656 | 0.00198139 | 242.217733 | 800.466685 | 0 | 166.974153 |
| -3.3301352 | 0.00086804 | 0.00198684 | 10.6559409 | 47.8981079 | 54.0713378 | 48.7474875 |
| -3.3299413 | 0.00086864 | 0.00198808 | 10.246097 | 47.8981079 | 49.4695218 | 59.3934906 |
| -3.3298854 | 0.00086882 | 0.00198834 | 15.1642236 | 34.9087905 | 33.3631659 | 45.9459078 |
| -3.3295557 | 0.00086985 | 0.00199055 | 24.5906328 | 24.3549701 | 25.3099879 | 72.8410733 |
| -3.3293543 | 0.00087048 | 0.00199185 | 49.5911095 | 46.545054 | 12.6549939 | 34.7395888 |
| -3.3292887 | 0.00087068 | 0.00199218 | 19.6725062 | 21.6488623 | 42.5667978 | 39.7824324 |
| -3.3286669 | 0.00087263 | 0.00199649 | 44.6729829 | 52.2278804 | 31.0622579 | 43.144328 |
| -3.3284917 | 0.00087318 | 0.00199761 | 15.5740674 | 28.9553534 | 42.5667978 | 41.4633802 |
| -3.3283295 | 0.00087368 | 0.00199863 | 17.213443 | 68.7351379 | 25.3099879 | 38.1014845 |
| -3.3279238 | 0.00087496 | 0.0020014 | 15.1642236 | 36.803066 | 34.5136199 | 38.1014845 |
| -3.3275413 | 0.00087616 | 0.00200401 | 16.8035991 | 53.5809343 | 28.7613499 | 33.058641 |
| -3.3271537 | 0.00087738 | 0.00200666 | 27.8693838 | 85.2423954 | 44.8677058 | 67.2379138 |
| -3.3270168 | 0.00087781 | 0.0020075 | 29.0989155 | 15.966036 | 36.8145279 | 46.5062237 |
| -3.3269298 | 0.00087808 | 0.00200798 | 20.9020379 | 31.9320719 | 24.1595339 | 28.5761134 |
| -3.3267197 | 0.00087875 | 0.00200936 | 15.5740674 | 24.3549701 | 57.5226998 | 52.6696992 |
| -3.326639 | 0.000879 | 0.0020098 | 12.2953164 | 76.3122397 | 37.9649818 | 40.3427483 |
| -3.3265379 | 0.00087932 | 0.00201038 | 13.9346919 | 75.2297966 | 66.7263317 | 23.5332698 |
| -3.326106 | 0.00088068 | 0.00201336 | 121.723632 | 31.9320719 | 23.0090799 | 21.852322 |
| -3.3260631 | 0.00088082 | 0.00201353 | 22.1315695 | 32.4732935 | 27.6108959 | 22.9729539 |
| -3.3257302 | 0.00088187 | 0.00201579 | 16.8035991 | 33.0145151 | 33.3631659 | 31.9380091 |
| -3.3255202 | 0.00088254 | 0.00201716 | 218.856632 | 531.47957 | 467.084322 | 492.517719 |
| -3.3253751 | 0.000883 | 0.00201807 | 35.2465737 | 23.8137486 | 43.7172518 | 18.4904263 |
| -3.3248414 | 0.00088469 | 0.0020218 | 21.3118818 | 23.272527 | 34.5136199 | 28.0157974 |
| -3.3245561 | 0.00088559 | 0.00202372 | 28.2792277 | 27.3316887 | 18.4072639 | 44.2649599 |
| -3.324432 | 0.00088599 | 0.00202448 | 28.6890716 | 29.4965749 | 35.6640739 | 17.9301104 |
| -3.3242462 | 0.00088658 | 0.00202554 | 210.659754 | 415.387546 | 539.562924 | 495.319299 |
| -3.3242579 | 0.00088654 | 0.00202554 | 15.1642236 | 56.8282636 | 28.7613499 | 39.7824324 |
| 3.32374624 | 0.00088817 | 0.00202903 | 0.81968776 | 1.3530539 | 2.30090799 | 2.24126379 |
| 3.32366485 | 0.00088843 | 0.00202948 | 0.81968776 | 0.54122156 | 0 | 0 |
| -3.323641 | 0.00088851 | 0.00202951 | 36.0662614 | 61.4286469 | 33.3631659 | 53.7903311 |
| -3.3234342 | 0.00088916 | 0.00203087 | 36.0662614 | 36.2618444 | 18.4072639 | 24.0935858 |
| -3.322924 | 0.00089079 | 0.00203445 | 12.2953164 | 55.4752097 | 39.1154358 | 42.0236961 |
| -3.3228277 | 0.0008911 | 0.00203501 | 17.6232868 | 31.6614612 | 46.0181598 | 26.8951655 |
| -3.3227392 | 0.00089138 | 0.00203551 | 13.1150042 | 40.0503953 | 41.4163438 | 42.5840121 |
| -3.3223777 | 0.00089254 | 0.002038 | 507.386723 | 469.239091 | 491.243856 | 522.214464 |
| -3.3222633 | 0.0008929 | 0.0020387 | 212.708974 | 263.574899 | 236.993523 | 266.150076 |
| -3.3221448 | 0.00089328 | 0.00203927 | 89.3459658 | 182.932887 | 158.762651 | 148.483726 |
| -3.3221448 | 0.00089328 | 0.00203927 | 25.8201644 | 51.416048 | 70.1776937 | 63.3157022 |
| -3.3216732 | 0.00089479 | 0.00204258 | 20.0823501 | 88.4897248 | 55.2217918 | 16.8094785 |
| -3.3215199 | 0.00089529 | 0.00204356 | 11.8854725 | 59.8049822 | 33.3631659 | 54.910963 |
| -3.3212186 | 0.00089625 | 0.00204562 | 29.5087594 | 37.8855091 | 14.9559019 | 46.5062237 |

| | | | | | | |
|---|---|---|---|---|---|---|
| 3.32070007 | 0.00089792 | 0.00204928 | 0 | 0 | 0 | 0 |
| -3.3203327 | 0.0008991 | 0.00205183 | 61.476582 | 35.9912336 | 21.8586259 | 19.6110582 |
| -3.3200264 | 0.00090009 | 0.00205394 | 29.0989155 | 56.8282636 | 14.9559019 | 37.5411686 |
| -3.3196457 | 0.00090132 | 0.00205659 | 6.96734596 | 106.079425 | 55.2217918 | 56.0315949 |
| -3.3196275 | 0.00090138 | 0.00205659 | 47.5418901 | 24.8961917 | 17.2568099 | 35.8602207 |
| -3.3195768 | 0.00090154 | 0.00205681 | 14.3445358 | 43.8389462 | 31.0622579 | 44.2649599 |
| -3.3193622 | 0.00090223 | 0.00205822 | 112.707067 | 598.320433 | 740.892373 | 661.172819 |
| -3.3193458 | 0.00090229 | 0.00205822 | 21.3118818 | 27.6022995 | 27.6108959 | 25.2142177 |
| 3.31921972 | 0.00090269 | 0.00205886 | 2.45906328 | 1.3530539 | 2.30090799 | 3.36189569 |
| -3.3192312 | 0.00090266 | 0.00205886 | 14.7543797 | 45.4626109 | 29.9118039 | 42.0236961 |
| -3.3190996 | 0.00090308 | 0.0020596 | 15.9839113 | 33.8263474 | 37.9649818 | 31.3776931 |
| -3.3187261 | 0.00090429 | 0.00206221 | 183.200214 | 414.575714 | 0 | 407.910011 |
| 3.31855179 | 0.00090486 | 0.00206336 | 9.83625312 | 14.6129821 | 12.6549939 | 17.3697944 |
| -3.3185192 | 0.00090496 | 0.00206345 | 20.0823501 | 58.9931499 | 42.5667978 | 19.6110582 |
| -3.3182504 | 0.00090583 | 0.00206529 | 292.218686 | 656.772361 | 648.856053 | 660.612503 |
| -3.3182025 | 0.00090599 | 0.0020655 | 154.920987 | 243.27909 | 296.817131 | 262.227864 |
| -3.3180815 | 0.00090638 | 0.00206625 | 186.069121 | 346.381797 | 310.622579 | 360.283155 |
| -3.317755 | 0.00090744 | 0.00206852 | 46.3123584 | 22.7313054 | 23.0090799 | 25.7745336 |
| -3.3176174 | 0.00090789 | 0.00206939 | 13.524848 | 41.1328384 | 39.1154358 | 39.2221164 |
| -3.3172296 | 0.00090915 | 0.00207212 | 25.8201644 | 37.3442875 | 42.5667978 | 16.8094785 |
| -3.3169157 | 0.00091017 | 0.00207431 | 15.1642236 | 19.2133653 | 103.54086 | 66.1172819 |
| -3.3168807 | 0.00091028 | 0.00207442 | 41.8040758 | 20.83703 | 24.1595339 | 26.8951655 |
| -3.3165117 | 0.00091149 | 0.00207701 | 17.213443 | 30.0377965 | 36.8145279 | 29.6967453 |
| -3.3162363 | 0.00091239 | 0.00207877 | 25.4103206 | 52.2278804 | 65.5758777 | 66.6775979 |
| -3.3162449 | 0.00091236 | 0.00207877 | 36.8859492 | 11.9068743 | 47.1686138 | 56.5919108 |
| -3.3158715 | 0.00091358 | 0.00208134 | 311.071505 | 522.278804 | 408.411168 | 453.855918 |
| -3.3153689 | 0.00091522 | 0.00208493 | 29.0989155 | 56.5576528 | 12.6549939 | 56.0315949 |
| -3.3153501 | 0.00091528 | 0.00208493 | 9.83625312 | 60.6168145 | 24.1595339 | 214.040692 |
| -3.3152874 | 0.00091549 | 0.00208525 | 35.6564176 | 44.9213893 | 33.3631659 | 14.5682147 |
| 3.31496055 | 0.00091656 | 0.00208754 | 18.4429746 | 8.11832337 | 4.60181598 | 8.40473923 |
| -3.3147132 | 0.00091737 | 0.00208924 | 16.8035991 | 33.0145151 | 35.6640739 | 29.1364293 |
| -3.314564 | 0.00091786 | 0.00209021 | 31.5579788 | 18.1309222 | 57.5226998 | 25.7745336 |
| -3.3140537 | 0.00091954 | 0.00209374 | 22.5414134 | 57.9107067 | 17.2568099 | 42.0236961 |
| -3.314053 | 0.00091954 | 0.00209374 | 313.940412 | 360.182947 | 0 | 266.710392 |
| -3.3138007 | 0.00092037 | 0.00209548 | 15.1642236 | 33.8263474 | 32.2127119 | 45.9459078 |
| -3.313593 | 0.00092105 | 0.00209689 | 32.3776665 | 82.5362876 | 102.390406 | 76.763285 |
| -3.3134033 | 0.00092168 | 0.00209816 | 11.0657848 | 56.0164313 | 60.9740618 | 35.2999048 |
| -3.3130475 | 0.00092285 | 0.00210069 | 173.363961 | 184.556551 | 182.922185 | 177.05984 |
| 3.31229503 | 0.00092534 | 0.0021062 | 87.7065903 | 38.9679522 | 16.1063559 | 47.6268556 |
| -3.3115172 | 0.00092792 | 0.00211191 | 27.45954 | 51.416048 | 44.8677058 | 55.4712789 |
| -3.3110106 | 0.0009296 | 0.00211559 | 0 | 302.813462 | 1050.3645 | 212.92006 |
| 3.31090036 | 0.00092996 | 0.00211628 | 24.1807889 | 14.6129821 | 13.8054479 | 11.206319 |
| -3.3108426 | 0.00093015 | 0.00211656 | 14.3445358 | 44.6507786 | 28.7613499 | 49.3078035 |

| | | | | | | |
|---|---|---|---|---|---|---|
| -3.3106759 | 0.00093071 | 0.00211768 | 22.1315695 | 65.7584193 | 13.8054479 | 77.8839169 |
| -3.310405 | 0.00093161 | 0.00211958 | 38.9351686 | 8.65954493 | 78.2308717 | 69.4791776 |
| -3.3097721 | 0.00093372 | 0.00212423 | 148.363485 | 135.034779 | 192.125817 | 156.32815 |
| -3.3096686 | 0.00093406 | 0.00212473 | 28.6890716 | 50.8748265 | 18.4072639 | 26.3348496 |
| 3.309647 | 0.00093414 | 0.00212473 | 3.27875104 | 3.24732935 | 0 | 0 |
| -3.3096486 | 0.00093413 | 0.00212473 | 16.8035991 | 102.561485 | 77.0804177 | 122.148877 |
| -3.3095109 | 0.00093459 | 0.00212561 | 9.83625312 | 136.929054 | 14.9559019 | 149.604358 |
| -3.3094185 | 0.0009349 | 0.00212616 | 0 | 469.509702 | 300.268493 | 534.541415 |
| -3.3091274 | 0.00093587 | 0.00212822 | 19.2626624 | 37.6148983 | 21.8586259 | 37.5411686 |
| -3.308369 | 0.00093841 | 0.00213385 | 22.1315695 | 76.5828505 | 17.2568099 | 42.0236961 |
| -3.3071342 | 0.00094256 | 0.00214312 | 26.6398522 | 31.6614612 | 17.2568099 | 35.8602207 |
| 3.30692711 | 0.00094325 | 0.00214456 | 146.314265 | 70.3588026 | 62.1245158 | 61.0744384 |
| -3.3068355 | 0.00094356 | 0.00214511 | 11.4756286 | 39.7797845 | 63.2749697 | 41.4633802 |
| -3.3063945 | 0.00094505 | 0.00214834 | 18.8528185 | 40.8622277 | 29.9118039 | 24.6539017 |
| 3.30589535 | 0.00094673 | 0.00215201 | 39.3450125 | 21.1076408 | 16.1063559 | 8.96505518 |
| 3.30537244 | 0.0009485 | 0.00215575 | 2.45906328 | 3.51794013 | 1.150454 | 2.24126379 |
| -3.30537 | 0.00094851 | 0.00215575 | 36.4761053 | 39.5091738 | 93.1867736 | 71.7204414 |
| -3.3052664 | 0.00094886 | 0.00215639 | 15.1642236 | 42.2152815 | 29.9118039 | 36.9808526 |
| -3.3052424 | 0.00094894 | 0.00215643 | 10.6559409 | 40.5916169 | 47.1686138 | 62.1950703 |
| -3.3051513 | 0.00094925 | 0.00215698 | 32.3776665 | 46.2744432 | 44.8677058 | 38.6618005 |
| -3.3051134 | 0.00094938 | 0.00215712 | 16.8035991 | 29.2259641 | 31.0622579 | 36.4205367 |
| -3.3044126 | 0.00095176 | 0.00216236 | 17.213443 | 23.8137486 | 41.4163438 | 40.3427483 |
| -3.3042142 | 0.00095243 | 0.00216371 | 20.0823501 | 34.0969582 | 29.9118039 | 22.9729539 |
| -3.3041991 | 0.00095248 | 0.00216371 | 19.2626624 | 31.9320719 | 33.3631659 | 23.5332698 |
| -3.3040985 | 0.00095282 | 0.00216433 | 17.6232868 | 29.2259641 | 27.6108959 | 53.2300151 |
| -3.3038869 | 0.00095354 | 0.00216581 | 20.0823501 | 66.5702517 | 17.2568099 | 49.8681194 |
| -3.3036399 | 0.00095438 | 0.00216757 | 13.524848 | 67.6526948 | 54.0713378 | 25.7745336 |
| 3.30343268 | 0.00095509 | 0.00216902 | 0 | 0 | 0 | 0 |
| -3.3034057 | 0.00095518 | 0.00216908 | 13.524848 | 42.4858923 | 74.7795097 | 29.1364293 |
| -3.3033393 | 0.00095541 | 0.00216944 | 150.412704 | 411.598995 | 346.286653 | 383.256109 |
| -3.3030964 | 0.00095624 | 0.00217116 | 15.9839113 | 47.3568864 | 41.4163438 | 25.2142177 |
| -3.3027362 | 0.00095746 | 0.0021738 | 25.0004767 | 45.7332217 | 29.9118039 | 18.4904263 |
| -3.3026743 | 0.00095768 | 0.00217413 | 45.0828268 | 30.579018 | 16.1063559 | 29.1364293 |
| -3.3025097 | 0.00095824 | 0.00217513 | 12.2953164 | 49.2511618 | 34.5136199 | 48.1871716 |
| -3.302506 | 0.00095825 | 0.00217513 | 37.2957931 | 23.272527 | 17.2568099 | 58.8331746 |
| -3.3023998 | 0.00095861 | 0.0021758 | 95.0837801 | 24.6255809 | 23.0090799 | 25.7745336 |
| -3.3019879 | 0.00096002 | 0.00217884 | 1000.01907 | 282.247043 | 172.568099 | 0 |
| -3.3007143 | 0.00096439 | 0.0021886 | 8.60672148 | 44.6507786 | 69.0272397 | 59.3934906 |
| 3.3006702 | 0.00096454 | 0.00218879 | 2.0492194 | 1.62366467 | 0 | 1.1206319 |
| -3.3002932 | 0.00096584 | 0.00219158 | 69.2636157 | 20.0251977 | 16.1063559 | 57.1522268 |
| -3.3002573 | 0.00096596 | 0.00219171 | 15.5740674 | 42.4858923 | 40.2658898 | 26.8951655 |
| -3.2999192 | 0.00096713 | 0.0021942 | 22.9512573 | 20.0251977 | 39.1154358 | 28.5761134 |
| -3.2996622 | 0.00096801 | 0.00219605 | 10.6559409 | 67.9233056 | 35.6640739 | 52.1093832 |

| | | | | | | |
|---|---|---|---|---|---|---|
| -3.2995093 | 0.00096854 | 0.0021971 | 7.37718984 | 44.109557 | 77.0804177 | 75.0823371 |
| -3.2985296 | 0.00097193 | 0.00220462 | 19.2626624 | 84.9717847 | 28.7613499 | 26.3348496 |
| -3.2982841 | 0.00097278 | 0.0022064 | 35.2465737 | 89.5721679 | 8.05317797 | 72.8410733 |
| -3.2981968 | 0.00097308 | 0.00220662 | 6.96734596 | 58.7225391 | 65.5758777 | 67.2379138 |
| -3.2982291 | 0.00097297 | 0.00220662 | 27.45954 | 24.3549701 | 25.3099879 | 21.852322 |
| -3.2982099 | 0.00097303 | 0.00220662 | 13.9346919 | 71.4412457 | 23.0090799 | 59.3934906 |
| -3.297993 | 0.00097379 | 0.00220806 | 13.524848 | 29.4965749 | 49.4695218 | 46.5062237 |
| -3.2973292 | 0.00097609 | 0.00221313 | 60.2470503 | 18.401533 | 34.5136199 | 22.9729539 |
| -3.2966765 | 0.00097836 | 0.00221813 | 20.0823501 | 21.1076408 | 33.3631659 | 53.7903311 |
| -3.2965827 | 0.00097869 | 0.00221871 | 29.5087594 | 25.1668025 | 18.4072639 | 32.498325 |
| -3.2965522 | 0.00097879 | 0.0022188 | 21.3118818 | 47.0862756 | 21.8586259 | 28.0157974 |
| 3.29644942 | 0.00097915 | 0.00221945 | 0.40984388 | 1.08244312 | 1.150454 | 0 |
| -3.2962915 | 0.0009797 | 0.00222055 | 9.83625312 | 54.6633774 | 47.1686138 | 49.3078035 |
| -3.2958056 | 0.0009814 | 0.00222423 | 88.9361219 | 66.2996409 | 16.1063559 | 17.3697944 |
| -3.2956344 | 0.000982 | 0.00222543 | 21.3118818 | 32.2026827 | 29.9118039 | 21.292006 |
| 3.29551554 | 0.00098241 | 0.00222622 | 2.45906328 | 1.08244312 | 0 | 1.1206319 |
| -3.2953186 | 0.0009831 | 0.00222762 | 13.9346919 | 31.1202396 | 40.2658898 | 44.8252759 |
| 3.29515032 | 0.00098369 | 0.0022288 | 0.40984388 | 1.3530539 | 0 | 1.68094785 |
| -3.2950919 | 0.0009839 | 0.00222911 | 47.951734 | 54.6633774 | 13.8054479 | 26.3348496 |
| -3.2948747 | 0.00098466 | 0.00223067 | 266.398522 | 287.388647 | 314.073941 | 317.699143 |
| -3.2946736 | 0.00098536 | 0.00223212 | 8.60672148 | 43.0271139 | 94.3372276 | 50.4284354 |
| -3.2946167 | 0.00098556 | 0.00223241 | 5.73781432 | 37.6148983 | 133.452663 | 87.409288 |
| -3.2945234 | 0.00098589 | 0.002233 | 172.13443 | 265.198564 | 235.843069 | 269.511971 |
| -3.2941079 | 0.00098735 | 0.00223614 | 19.6725062 | 31.9320719 | 24.1595339 | 28.0157974 |
| -3.2940072 | 0.0009877 | 0.00223679 | 22.5414134 | 77.1240721 | 20.7081719 | 30.2570612 |
| -3.2936012 | 0.00098913 | 0.00223986 | 13.1150042 | 38.4267306 | 36.8145279 | 43.144328 |
| 3.29342041 | 0.00098976 | 0.00224115 | 353.695268 | 117.174467 | 81.6822337 | 179.86142 |
| -3.293188 | 0.00099058 | 0.00224284 | 1934.87296 | 2701.77802 | 2647.19464 | 2845.28439 |
| -3.2929688 | 0.00099135 | 0.0022444 | 31.1481349 | 38.1561199 | 14.9559019 | 33.058641 |
| -3.2929532 | 0.00099141 | 0.0022444 | 59.0175187 | 12.9893174 | 37.9649818 | 34.1792729 |
| -3.2926441 | 0.0009925 | 0.00224671 | 18.8528185 | 24.3549701 | 28.7613499 | 38.1014845 |
| 3.29193884 | 0.00099499 | 0.0022522 | 186.478965 | 241.926037 | 187.524001 | 214.040692 |
| -3.291902 | 0.00099512 | 0.00225233 | 31.9678226 | 23.5431378 | 20.7081719 | 25.7745336 |
| -3.2918782 | 0.00099521 | 0.00225237 | 26.2300083 | 28.9553534 | 24.1595339 | 20.7316901 |
| -3.2917991 | 0.00099549 | 0.00225284 | 60.6568942 | 199.169533 | 315.224395 | 155.207518 |
| -3.2912985 | 0.00099726 | 0.0022567 | 33.1973543 | 39.5091738 | 13.8054479 | 36.4205367 |
| -3.2910187 | 0.00099825 | 0.00225874 | 16.3937552 | 40.3210061 | 37.9649818 | 25.2142177 |
| -3.2910048 | 0.0009983 | 0.00225874 | 16.8035991 | 32.7439043 | 37.9649818 | 26.3348496 |
| 3.29097527 | 0.00099841 | 0.00225882 | 44.263139 | 13.2599282 | 37.9649818 | 19.6110582 |
| -3.2908709 | 0.00099878 | 0.0022595 | 19.2626624 | 29.2259641 | 25.3099879 | 28.5761134 |
| -3.2903512 | 0.00100062 | 0.00226352 | 21.3118818 | 27.6022995 | 39.1154358 | 21.292006 |
| -3.289504 | 0.00100364 | 0.00227018 | 11.8854725 | 50.6042157 | 35.6640739 | 45.3855918 |
| -3.2893164 | 0.00100431 | 0.00227154 | 20.492194 | 31.9320719 | 32.2127119 | 21.292006 |

| | | | | | | |
|---|---|---|---|---|---|---|
| -3.2889732 | 0.00100554 | 0.00227415 | 20.9020379 | 15.4248144 | 72.4786017 | 54.350647 |
| -3.2887523 | 0.00100633 | 0.00227577 | 17.213443 | 43.2977247 | 25.3099879 | 31.9380091 |
| -3.2885158 | 0.00100717 | 0.00227737 | 246.726016 | 197.004647 | 124.249032 | 187.145527 |
| -3.2885167 | 0.00100717 | 0.00227737 | 19.6725062 | 46.2744432 | 26.4604419 | 24.6539017 |
| -3.2882408 | 0.00100816 | 0.00227927 | 24.5906328 | 19.7545869 | 32.2127119 | 26.3348496 |
| -3.2882468 | 0.00100813 | 0.00227927 | 13.524848 | 52.7691019 | 31.0622579 | 39.2221164 |
| -3.2875573 | 0.00101061 | 0.00228466 | 173.363961 | 262.221845 | 248.498063 | 285.200818 |
| 3.28741769 | 0.00101111 | 0.00228563 | 0.40984388 | 0.27061078 | 1.150454 | 1.1206319 |
| 3.28687022 | 0.00101308 | 0.00228992 | 2.86890716 | 9.47137727 | 8.05317797 | 8.40473923 |
| -3.2863028 | 0.00101512 | 0.00229437 | 27.0496961 | 58.9931499 | 11.50454 | 72.2807574 |
| -3.2848727 | 0.00102028 | 0.00230589 | 27.0496961 | 25.1668025 | 18.4072639 | 38.6618005 |
| -3.2848468 | 0.00102038 | 0.00230594 | 309.022285 | 306.060791 | 0 | 266.150076 |
| -3.2846951 | 0.00102093 | 0.00230702 | 12.2953164 | 29.4965749 | 62.1245158 | 48.7474875 |
| -3.284393 | 0.00102202 | 0.00230933 | 5.73781432 | 47.8981079 | 60.9740618 | 140.639303 |
| 3.28392306 | 0.00102373 | 0.00231303 | 1.63937552 | 0.81183234 | 2.30090799 | 1.68094785 |
| -3.2838599 | 0.00102396 | 0.00231338 | 14.3445358 | 40.3210061 | 29.9118039 | 39.7824324 |
| -3.2837166 | 0.00102448 | 0.0023144 | 11.0657848 | 41.67406 | 46.0181598 | 46.5062237 |
| -3.2835468 | 0.0010251 | 0.00231563 | 82.3786199 | 50.6042157 | 8.05317797 | 45.3855918 |
| -3.2835045 | 0.00102525 | 0.00231581 | 97.1329995 | 114.46836 | 124.249032 | 111.502874 |
| -3.2834789 | 0.00102534 | 0.00231586 | 11.4756286 | 39.238563 | 51.7704298 | 41.4633802 |
| -3.2833514 | 0.00102581 | 0.00231675 | 11.8854725 | 58.7225391 | 26.4604419 | 73.4013893 |
| -3.2832012 | 0.00102635 | 0.00231782 | 289.349779 | 418.093654 | 406.11026 | 0 |
| -3.2830483 | 0.00102691 | 0.00231892 | 5.73781432 | 64.946587 | 92.0363196 | 58.2728587 |
| -3.282936 | 0.00102732 | 0.00231968 | 14.3445358 | 34.6381797 | 51.7704298 | 30.2570612 |
| -3.2829123 | 0.00102741 | 0.00231971 | 9.01656536 | 47.8981079 | 52.9208838 | 56.5919108 |
| -3.2824977 | 0.00102892 | 0.00232296 | 42.6237635 | 36.803066 | 34.5136199 | 39.2221164 |
| -3.2823201 | 0.00102957 | 0.00232426 | 414.352163 | 626.734564 | 509.65112 | 565.358792 |
| 3.2822035 | 0.00102999 | 0.00232506 | 1.63937552 | 0.54122156 | 3.45136199 | 0.56031595 |
| -3.2821801 | 0.00103008 | 0.00232509 | 152.871767 | 30.579018 | 21.8586259 | 20.1713742 |
| -3.2820706 | 0.00103048 | 0.00232583 | 13.9346919 | 66.2996409 | 33.3631659 | 32.498325 |
| 3.28201603 | 0.00103068 | 0.00232612 | 11.0657848 | 5.95343714 | 5.75226998 | 9.52537113 |
| -3.2814978 | 0.00103257 | 0.00233024 | 25.0004767 | 30.8496288 | 23.0090799 | 21.292006 |
| -3.2813798 | 0.00103301 | 0.00233089 | 28.2792277 | 16.7778683 | 28.7613499 | 40.3427483 |
| -3.2813844 | 0.00103299 | 0.00233089 | 17.213443 | 41.67406 | 23.0090799 | 36.9808526 |
| -3.2810768 | 0.00103412 | 0.00233323 | 20.9020379 | 43.2977247 | 17.2568099 | 47.6268556 |
| -3.2809856 | 0.00103445 | 0.00233382 | 477.877964 | 1697.81203 | 1356.38526 | 1380.05818 |
| 3.28078481 | 0.00103519 | 0.00233532 | 4.0984388 | 2.43549701 | 2.30090799 | 5.60315949 |
| -3.2804711 | 0.00103634 | 0.00233775 | 11.8854725 | 79.2889583 | 24.1595339 | 65.556966 |
| -3.2804134 | 0.00103655 | 0.00233807 | 16.8035991 | 31.1202396 | 26.4604419 | 37.5411686 |
| -3.2802681 | 0.00103708 | 0.00233911 | 16.3937552 | 28.6847426 | 32.2127119 | 32.498325 |
| -3.2801958 | 0.00103735 | 0.00233955 | 32.7875104 | 34.6381797 | 21.8586259 | 19.0507423 |
| -3.2801415 | 0.00103755 | 0.00233972 | 34.4268859 | 21.6488623 | 19.5577179 | 30.8173772 |
| -3.2801349 | 0.00103757 | 0.00233972 | 11.4756286 | 51.1454373 | 33.3631659 | 52.6696992 |

| | | | | | | |
|---|---|---|---|---|---|---|
| -3.2799049 | 0.00103842 | 0.00234147 | 14.7543797 | 41.67406 | 26.4604419 | 45.3855918 |
| -3.2798743 | 0.00103853 | 0.00234156 | 15.1642236 | 35.4500121 | 56.3722458 | 26.8951655 |
| -3.2797648 | 0.00103894 | 0.0023423 | 38.5253247 | 34.367569 | 13.8054479 | 33.6189569 |
| -3.278766 | 0.00104262 | 0.00235045 | 295.497437 | 333.933701 | 0 | 245.978701 |
| -3.2786013 | 0.00104323 | 0.00235165 | 175.413181 | 17.0484791 | 32.2127119 | 23.5332698 |
| -3.278539 | 0.00104346 | 0.00235201 | 191.806936 | 444.342899 | 425.667978 | 414.073486 |
| -3.2782043 | 0.0010447 | 0.00235464 | 16.3937552 | 50.8748265 | 59.8236078 | 20.1713742 |
| -3.2781825 | 0.00104478 | 0.00235465 | 23.3611012 | 39.238563 | 20.7081719 | 24.6539017 |
| -3.2780418 | 0.0010453 | 0.00235566 | 58.6076748 | 87.1366709 | 79.3813257 | 5.04284354 |
| -3.2775828 | 0.001047 | 0.00235933 | 34.8367298 | 62.2404792 | 8.05317797 | 99.1759229 |
| -3.2775218 | 0.00104723 | 0.00235968 | 22.9512573 | 29.2259641 | 23.0090799 | 22.9729539 |
| -3.2774757 | 0.0010474 | 0.0023599 | 32.7875104 | 34.6381797 | 31.0622579 | 32.498325 |
| -3.2772501 | 0.00104823 | 0.00236162 | 220.086164 | 438.930684 | 404.959806 | 433.684544 |
| -3.2771036 | 0.00104878 | 0.00236268 | 23.770945 | 24.6255809 | 21.8586259 | 26.3348496 |
| -3.2770657 | 0.00104892 | 0.00236283 | 12.7051603 | 49.2511618 | 42.5667978 | 31.9380091 |
| -3.2770053 | 0.00104914 | 0.00236317 | 27.0496961 | 27.8729103 | 23.0090799 | 20.7316901 |
| -3.2769488 | 0.00104935 | 0.00236348 | 172.544273 | 601.567762 | 644.254237 | 708.239359 |
| -3.2768224 | 0.00104982 | 0.00236437 | 14.7543797 | 27.8729103 | 54.0713378 | 35.2999048 |
| -3.2763448 | 0.0010516 | 0.00236821 | 138.937075 | 23.0019162 | 25.3099879 | 22.9729539 |
| -3.276291 | 0.0010518 | 0.0023685 | 231.561792 | 158.307306 | 174.869007 | 169.215416 |
| -3.2760616 | 0.00105266 | 0.00237026 | 16.8035991 | 46.0038325 | 28.7613499 | 27.4554815 |
| -3.2760214 | 0.00105281 | 0.00237043 | 5.73781432 | 69.0057487 | 82.8326877 | 57.7125427 |
| -3.2758306 | 0.00105352 | 0.0023717 | 18.8528185 | 38.4267306 | 23.0090799 | 29.1364293 |
| -3.2758466 | 0.00105346 | 0.0023717 | 380.335121 | 380.749366 | 419.915708 | 361.403787 |
| -3.2755714 | 0.00105448 | 0.00237371 | 11.8854725 | 41.67406 | 41.4163438 | 40.9030642 |
| -3.2753013 | 0.00105549 | 0.00237582 | 12.7051603 | 31.9320719 | 42.5667978 | 48.1871716 |
| -3.2750493 | 0.00105644 | 0.00237777 | 18.8528185 | 31.1202396 | 24.1595339 | 28.5761134 |
| 3.27502763 | 0.00105652 | 0.00237779 | 50.0009534 | 32.4732935 | 25.3099879 | 19.0507423 |
| -3.2749927 | 0.00105665 | 0.00237792 | 18.4429746 | 33.2851258 | 24.1595339 | 29.1364293 |
| -3.2747082 | 0.00105771 | 0.00238015 | 11.8854725 | 47.3568864 | 63.2749697 | 30.2570612 |
| -3.2740468 | 0.00106019 | 0.00238556 | 20.9020379 | 20.5664192 | 42.5667978 | 28.5761134 |
| -3.2739024 | 0.00106073 | 0.00238661 | 326.645572 | 770.428888 | 717.883293 | 764.83127 |
| -3.2737083 | 0.00106146 | 0.00238808 | 12.7051603 | 30.3084073 | 41.4163438 | 61.6347543 |
| -3.2732585 | 0.00106315 | 0.00239172 | 19.6725062 | 23.5431378 | 35.6640739 | 25.7745336 |
| -3.2730999 | 0.00106375 | 0.0023929 | 13.9346919 | 43.5683354 | 32.2127119 | 34.7395888 |
| -3.2725797 | 0.00106571 | 0.00239714 | 227.053509 | 338.534085 | 0 | 304.25156 |
| -3.2722883 | 0.00106681 | 0.00239944 | 22.1315695 | 43.8389462 | 39.1154358 | 49.8681194 |
| -3.2716971 | 0.00106904 | 0.0024043 | 112.297223 | 341.240193 | 544.16474 | 457.77813 |
| -3.2715992 | 0.00106941 | 0.00240496 | 85.6573709 | 23.0019162 | 18.4072639 | 31.9380091 |
| -3.2714975 | 0.0010698 | 0.00240566 | 11.8854725 | 36.5324552 | 44.8677058 | 43.144328 |
| -3.2710968 | 0.00107131 | 0.00240876 | 15.1642236 | 46.8156648 | 33.3631659 | 28.0157974 |
| -3.2710935 | 0.00107132 | 0.00240876 | 14.7543797 | 35.1794013 | 35.6640739 | 30.8173772 |
| -3.2708888 | 0.0010721 | 0.00241034 | 6.1476582 | 47.0862756 | 92.0363196 | 70.0394936 |

| | | | | | | |
|---|---|---|---|---|---|---|
| -3.2705083 | 0.00107354 | 0.00241342 | 14.3445358 | 38.1561199 | 34.5136199 | 31.9380091 |
| -3.2704764 | 0.00107367 | 0.00241352 | 11.4756286 | 54.1221558 | 40.2658898 | 37.5411686 |
| -3.2703056 | 0.00107431 | 0.00241481 | 31.5579788 | 24.3549701 | 17.2568099 | 33.6189569 |
| -3.2702062 | 0.00107469 | 0.00241549 | 49.1812656 | 95.2549943 | 82.8326877 | 81.2458125 |
| -3.2699307 | 0.00107574 | 0.00241768 | 21.7217256 | 43.2977247 | 23.0090799 | 23.5332698 |
| 3.26950114 | 0.00107737 | 0.00242118 | 5.73781432 | 0 | 0 | 0 |
| -3.2692997 | 0.00107814 | 0.00242274 | 36.8859492 | 27.8729103 | 63.2749697 | 14.0078987 |
| -3.268877 | 0.00107975 | 0.00242619 | 18.0331307 | 28.9553534 | 24.1595339 | 35.8602207 |
| -3.2688483 | 0.00107986 | 0.00242627 | 31.5579788 | 29.4965749 | 17.2568099 | 25.7745336 |
| -3.2686761 | 0.00108052 | 0.00242758 | 25.8201644 | 26.5198564 | 35.6640739 | 17.9301104 |
| -3.2683818 | 0.00108164 | 0.00242993 | 16.8035991 | 36.2618444 | 32.2127119 | 25.2142177 |
| -3.268308 | 0.00108193 | 0.0024304 | 41.8040758 | 20.0251977 | 21.8586259 | 27.4554815 |
| -3.2680544 | 0.0010829 | 0.00243241 | 27.8693838 | 50.0629941 | 50.6199758 | 43.704644 |
| -3.2676444 | 0.00108447 | 0.00243577 | 56.1486116 | 26.2492456 | 14.9559019 | 34.7395888 |
| -3.2676104 | 0.0010846 | 0.00243589 | 21.7217256 | 30.8496288 | 19.5577179 | 30.2570612 |
| -3.2674929 | 0.00108505 | 0.00243662 | 9.83625312 | 65.2171978 | 50.6199758 | 36.9808526 |
| -3.2674861 | 0.00108507 | 0.00243662 | 22.1315695 | 62.51109 | 59.8236078 | 87.9696039 |
| -3.2674309 | 0.00108528 | 0.00243693 | 9.42640924 | 69.0057487 | 37.9649818 | 51.5490673 |
| -3.2673072 | 0.00108576 | 0.00243782 | 11.0657848 | 54.1221558 | 39.1154358 | 40.9030642 |
| -3.2672427 | 0.00108601 | 0.00243821 | 168.855679 | 327.439043 | 279.560321 | 249.340597 |
| -3.2671884 | 0.00108621 | 0.00243851 | 17.213443 | 30.579018 | 26.4604419 | 30.8173772 |
| -3.2670165 | 0.00108687 | 0.00243982 | 11.8854725 | 50.8748265 | 32.2127119 | 46.5062237 |
| -3.2666095 | 0.00108844 | 0.00244316 | 20.0823501 | 19.2133653 | 34.5136199 | 49.3078035 |
| -3.2663945 | 0.00108926 | 0.00244484 | 10.6559409 | 66.5702517 | 34.5136199 | 46.5062237 |
| -3.2662044 | 0.00109 | 0.00244632 | 18.8528185 | 70.9000241 | 14.9559019 | 62.1950703 |
| -3.2661267 | 0.00109029 | 0.00244682 | 25.4103206 | 25.4374132 | 23.0090799 | 21.852322 |
| 3.26592234 | 0.00109108 | 0.00244833 | 2.0492194 | 0.27061078 | 0 | 1.1206319 |
| -3.2659124 | 0.00109112 | 0.00244833 | 12.2953164 | 30.3084073 | 66.7263317 | 41.4633802 |
| -3.2649043 | 0.00109501 | 0.00245689 | 7.78703372 | 48.980551 | 51.7704298 | 72.8410733 |
| -3.2647816 | 0.00109549 | 0.00245778 | 13.1150042 | 35.4500121 | 33.3631659 | 47.6268556 |
| 3.26349944 | 0.00110045 | 0.00246876 | 24.5906328 | 13.8011497 | 0 | 0 |
| 3.26279076 | 0.00110321 | 0.00247477 | 0.40984388 | 0 | 1.150454 | 0 |
| -3.2626234 | 0.00110386 | 0.00247606 | 13.9346919 | 27.3316887 | 46.0181598 | 40.9030642 |
| -3.2624041 | 0.00110472 | 0.0024778 | 13.1150042 | 30.3084073 | 48.3190678 | 39.2221164 |
| -3.2622882 | 0.00110517 | 0.00247864 | 18.8528185 | 34.0969582 | 23.0090799 | 28.0157974 |
| -3.2620197 | 0.00110621 | 0.00248082 | 17.6232868 | 33.0145151 | 25.3099879 | 28.5761134 |
| -3.2613035 | 0.00110901 | 0.00248692 | 31.5579788 | 28.4141318 | 17.2568099 | 25.7745336 |
| -3.2608981 | 0.0011106 | 0.00249031 | 336.481825 | 661.102133 | 607.43971 | 729.531365 |
| -3.2602961 | 0.00111296 | 0.00249543 | 19.6725062 | 92.2782757 | 11.50454 | 79.5648647 |
| -3.2601659 | 0.00111347 | 0.0024964 | 18.4429746 | 45.4626109 | 29.9118039 | 22.4126379 |
| -3.2599775 | 0.00111421 | 0.00249788 | 21.3118818 | 74.4179643 | 14.9559019 | 45.9459078 |
| -3.2594048 | 0.00111646 | 0.00250276 | 10.6559409 | 96.8786589 | 17.2568099 | 113.183822 |
| 3.2592306 | 0.00111715 | 0.00250412 | 0.40984388 | 0 | 0 | 0 |

| | | | | | | |
|---|---|---|---|---|---|---|
| -3.2592073 | 0.00111724 | 0.00250415 | 29.5087594 | 30.3084073 | 27.6108959 | 16.8094785 |
| -3.25903 | 0.00111794 | 0.00250546 | 2.45906328 | 63.0523115 | 126.549939 | 123.269509 |
| -3.2589843 | 0.00111812 | 0.00250546 | 13.1150042 | 34.367569 | 35.6640739 | 41.4633802 |
| -3.2589828 | 0.00111812 | 0.00250546 | 31.9678226 | 22.1900839 | 18.4072639 | 31.3776931 |
| -3.2589805 | 0.00111813 | 0.00250546 | 18.8528185 | 31.1202396 | 23.0090799 | 28.5761134 |
| -3.2586301 | 0.00111952 | 0.00250838 | 22.5414134 | 20.83703 | 29.9118039 | 24.6539017 |
| -3.2585744 | 0.00111974 | 0.0025087 | 21.3118818 | 34.367569 | 19.5577179 | 28.5761134 |
| -3.2584424 | 0.00112026 | 0.00250969 | 20.0823501 | 24.3549701 | 31.0622579 | 23.5332698 |
| -3.2583877 | 0.00112047 | 0.00251 | 31.1481349 | 29.4965749 | 24.1595339 | 17.9301104 |
| -3.2580632 | 0.00112175 | 0.0025127 | 15.9839113 | 32.7439043 | 41.4163438 | 25.2142177 |
| -3.2574686 | 0.00112411 | 0.0025178 | 420.089977 | 384.808528 | 396.906628 | 470.665397 |
| -3.2572858 | 0.00112483 | 0.00251924 | 11.8854725 | 47.3568864 | 31.0622579 | 49.8681194 |
| 3.25612327 | 0.00112945 | 0.00252941 | 0 | 0.27061078 | 0 | 0.56031595 |
| 3.25604826 | 0.00112975 | 0.0025299 | 4.91812656 | 6.76526948 | 11.50454 | 7.28410733 |
| -3.2556473 | 0.00113134 | 0.0025333 | 12.7051603 | 42.7565031 | 42.5667978 | 31.3776931 |
| 3.25558138 | 0.0011316 | 0.00253371 | 0 | 0.54122156 | 2.30090799 | 1.68094785 |
| -3.255258 | 0.00113289 | 0.00253642 | 38.5253247 | 16.5072575 | 26.4604419 | 29.1364293 |
| -3.255011 | 0.00113388 | 0.00253845 | 26.2300083 | 29.4965749 | 59.8236078 | 16.2491625 |
| -3.2545357 | 0.00113578 | 0.00254253 | 22.9512573 | 25.9786348 | 39.1154358 | 19.0507423 |
| -3.2545045 | 0.0011359 | 0.00254263 | 17.213443 | 42.4858923 | 19.5577179 | 53.2300151 |
| -3.2543915 | 0.00113636 | 0.00254347 | 13.9346919 | 40.0503953 | 33.3631659 | 31.9380091 |
| -3.2535976 | 0.00113954 | 0.00255041 | 5.73781432 | 91.4664434 | 50.6199758 | 66.1172819 |
| -3.2527372 | 0.00114299 | 0.00255797 | 60.2470503 | 60.8874253 | 8.05317797 | 40.9030642 |
| -3.2526948 | 0.00114316 | 0.00255799 | 15.5740674 | 29.4965749 | 35.6640739 | 29.1364293 |
| -3.2527142 | 0.00114308 | 0.00255799 | 18.0331307 | 32.2026827 | 24.1595339 | 28.0157974 |
| -3.2520871 | 0.00114561 | 0.00256326 | 33.1973543 | 43.2977247 | 13.8054479 | 29.6967453 |
| 3.25207058 | 0.00114568 | 0.00256326 | 30.3284471 | 18.6721438 | 5.75226998 | 15.1285306 |
| -3.2519749 | 0.00114606 | 0.00256395 | 66.8045524 | 33.8263474 | 16.1063559 | 23.5332698 |
| -3.2518023 | 0.00114676 | 0.00256533 | 13.524848 | 28.6847426 | 44.8677058 | 38.6618005 |
| -3.2516309 | 0.00114745 | 0.0025667 | 0.40984388 | 96.0668266 | 239.294431 | 221.3248 |
| 3.25129156 | 0.00114882 | 0.00256958 | 0.40984388 | 0.54122156 | 0 | 0 |
| -3.2510015 | 0.00114999 | 0.00257203 | 22.1315695 | 15.4248144 | 48.3190678 | 44.8252759 |
| -3.2509709 | 0.00115012 | 0.00257213 | 122.133476 | 13.2599282 | 26.4604419 | 37.5411686 |
| -3.2508456 | 0.00115062 | 0.00257308 | 66.3947085 | 12.9893174 | 33.3631659 | 32.498325 |
| -3.2508035 | 0.00115079 | 0.00257329 | 8.1968776 | 141.529437 | 24.1595339 | 72.2807574 |
| -3.2504305 | 0.0011523 | 0.00257631 | 9.01656536 | 38.9679522 | 55.2217918 | 59.3934906 |
| -3.2504477 | 0.00115223 | 0.00257631 | 30.3284471 | 27.6022995 | 14.9559019 | 48.1871716 |
| -3.2502833 | 0.0011529 | 0.00257746 | 11.4756286 | 34.9087905 | 46.0181598 | 43.704644 |
| -3.2501226 | 0.00115355 | 0.00257874 | 30.3284471 | 34.9087905 | 58.6731538 | 57.1522268 |
| -3.2498839 | 0.00115452 | 0.00258073 | 18.8528185 | 28.9553534 | 21.8586259 | 33.058641 |
| -3.249728 | 0.00115515 | 0.00258196 | 8.60672148 | 53.5809343 | 48.3190678 | 50.9887513 |
| -3.2494318 | 0.00115636 | 0.00258448 | 108.608628 | 206.746635 | 246.197155 | 286.881766 |
| -3.2492574 | 0.00115707 | 0.00258588 | 0 | 450.837558 | 339.383929 | 341.232413 |

| | | | | | | |
|---|---|---|---|---|---|---|
| -3.2489907 | 0.00115815 | 0.00258813 | 29.0989155 | 24.6255809 | 23.0090799 | 20.1713742 |
| -3.2485149 | 0.00116009 | 0.00259228 | 5.32797044 | 106.891258 | 31.0622579 | 119.347297 |
| -3.2481488 | 0.00116159 | 0.00259544 | 26.6398522 | 16.7778683 | 32.2127119 | 28.5761134 |
| -3.2479965 | 0.00116221 | 0.00259665 | 8.60672148 | 47.6274971 | 47.1686138 | 59.3934906 |
| -3.2477082 | 0.00116339 | 0.0025991 | 37.705637 | 21.9194731 | 20.7081719 | 24.0935858 |
| -3.2475326 | 0.0011641 | 0.00260035 | 31.1481349 | 27.8729103 | 16.1063559 | 28.5761134 |
| -3.2475329 | 0.0011641 | 0.00260035 | 112.707067 | 63.8641439 | 70.1776937 | 62.1950703 |
| 3.24749846 | 0.00116424 | 0.00260048 | 23.3611012 | 15.4248144 | 21.8586259 | 15.1285306 |
| -3.2472664 | 0.00116519 | 0.00260242 | 20.9020379 | 35.4500121 | 19.5577179 | 28.0157974 |
| -3.246504 | 0.00116832 | 0.00260922 | 17.213443 | 33.2851258 | 25.3099879 | 28.0157974 |
| -3.2463375 | 0.001169 | 0.00261057 | 131.969729 | 25.1668025 | 16.1063559 | 30.2570612 |
| -3.245147 | 0.0011739 | 0.0026213 | 20.9020379 | 21.3782516 | 44.8677058 | 24.6539017 |
| -3.2451298 | 0.00117397 | 0.0026213 | 41.3942319 | 15.6954252 | 27.6108959 | 29.1364293 |
| -3.2448296 | 0.00117521 | 0.00262388 | 14.3445358 | 51.416048 | 25.3099879 | 38.1014845 |
| -3.244776 | 0.00117543 | 0.0026242 | 10.6559409 | 48.980551 | 40.2658898 | 40.9030642 |
| -3.2443323 | 0.00117726 | 0.00262811 | 259.431176 | 159.93097 | 0 | 474.587608 |
| -3.2442645 | 0.00117754 | 0.00262855 | 106.969253 | 26.2492456 | 13.8054479 | 35.2999048 |
| -3.2435735 | 0.0011804 | 0.00263475 | 9.83625312 | 43.2977247 | 41.4163438 | 54.910963 |
| 3.24331114 | 0.00118149 | 0.002637 | 2.45906328 | 3.24732935 | 4.60181598 | 2.80157974 |
| -3.2431992 | 0.00118196 | 0.00263785 | 11.0657848 | 33.0145151 | 60.9740618 | 42.0236961 |
| -3.2426923 | 0.00118406 | 0.00264237 | 69.6734596 | 24.3549701 | 24.1595339 | 20.1713742 |
| 3.24196367 | 0.00118709 | 0.0026489 | 0 | 0 | 0 | 0 |
| -3.2419496 | 0.00118715 | 0.0026489 | 12.7051603 | 45.4626109 | 36.8145279 | 31.9380091 |
| 3.24122958 | 0.00119015 | 0.00265541 | 1.22953164 | 1.3530539 | 0 | 1.1206319 |
| -3.2412001 | 0.00119028 | 0.0026555 | 9.83625312 | 37.8855091 | 56.3722458 | 46.5062237 |
| -3.2402278 | 0.00119434 | 0.00266439 | 25.4103206 | 33.2851258 | 14.9559019 | 40.3427483 |
| 3.24011042 | 0.00119483 | 0.0026653 | 15.1642236 | 12.9893174 | 10.354086 | 9.52537113 |
| 3.24004232 | 0.00119512 | 0.00266576 | 1.63937552 | 0.54122156 | 1.150454 | 2.24126379 |
| -3.2398437 | 0.00119595 | 0.00266738 | 6.1476582 | 67.382084 | 138.054479 | 127.19172 |
| -3.2398293 | 0.00119601 | 0.00266738 | 522.141103 | 1099.76221 | 1035.4086 | 1046.10988 |
| -3.2397685 | 0.00119627 | 0.00266758 | 14.3445358 | 27.8729103 | 54.0713378 | 31.9380091 |
| -3.2397869 | 0.00119619 | 0.00266758 | 74.1817423 | 51.6866588 | 42.5667978 | 7.84442328 |
| -3.2396485 | 0.00119677 | 0.00266815 | 31.5579788 | 40.8622277 | 36.8145279 | 42.5840121 |
| -3.2396747 | 0.00119666 | 0.00266815 | 6.1476582 | 91.7370541 | 24.1595339 | 154.647202 |
| -3.2396555 | 0.00119674 | 0.00266815 | 9.42640924 | 75.2297966 | 40.2658898 | 40.9030642 |
| -3.2395067 | 0.00119737 | 0.00266911 | 16.8035991 | 50.8748265 | 42.5667978 | 19.6110582 |
| -3.2395176 | 0.00119732 | 0.00266911 | 10.246097 | 43.8389462 | 62.1245158 | 35.2999048 |
| 3.23907722 | 0.00119917 | 0.00267288 | 18.0331307 | 14.0717605 | 18.4072639 | 10.0856871 |
| -3.2390646 | 0.00119922 | 0.00267288 | 9.42640924 | 68.4645271 | 35.6640739 | 48.7474875 |
| -3.2385863 | 0.00120124 | 0.00267718 | 23.770945 | 44.6507786 | 136.904025 | 11.7666349 |
| -3.2385216 | 0.00120151 | 0.0026776 | 34.4268859 | 28.6847426 | 28.7613499 | 43.704644 |
| -3.2383471 | 0.00120224 | 0.00267906 | 159.019425 | 461.120768 | 431.420248 | 451.614655 |
| -3.2382518 | 0.00120265 | 0.00267942 | 62.2962697 | 14.6129821 | 56.3722458 | 19.6110582 |

| | | | | | | |
|---|---|---|---|---|---|---|
| -3.2382848 | 0.00120251 | 0.00267942 | 95.9034679 | 21.3782516 | 21.8586259 | 25.7745336 |
| 3.23824963 | 0.00120266 | 0.00267942 | 0.40984388 | 0 | 0 | 0 |
| 3.23820392 | 0.00120285 | 0.00267966 | 0.40984388 | 1.89427545 | 0 | 1.1206319 |
| -3.2381469 | 0.00120309 | 0.00268001 | 308.612442 | 524.173079 | 496.996126 | 502.04309 |
| -3.2377714 | 0.00120467 | 0.00268336 | 32.3776665 | 21.1076408 | 19.5577179 | 26.8951655 |
| 3.2364603 | 0.00121022 | 0.00269553 | 0.81968776 | 0.54122156 | 0 | 0.56031595 |
| -3.2360454 | 0.00121198 | 0.00269926 | 14.7543797 | 30.3084073 | 31.0622579 | 33.6189569 |
| -3.2359099 | 0.00121256 | 0.00270036 | 24.1807889 | 51.6866588 | 16.1063559 | 30.2570612 |
| -3.2357101 | 0.00121341 | 0.00270206 | 26.6398522 | 22.1900839 | 27.6108959 | 19.6110582 |
| -3.2354088 | 0.00121469 | 0.00270473 | 4.0984388 | 74.688575 | 78.2308717 | 75.0823371 |
| -3.2353522 | 0.00121493 | 0.00270508 | 17.213443 | 25.1668025 | 31.0622579 | 27.4554815 |
| -3.2351684 | 0.00121571 | 0.00270664 | 31.9678226 | 21.1076408 | 23.0090799 | 21.852322 |
| -3.2347697 | 0.00121741 | 0.00271023 | 11.8854725 | 48.7099402 | 35.6640739 | 35.2999048 |
| -3.234519 | 0.00121848 | 0.00271242 | 9.42640924 | 38.9679522 | 63.2749697 | 44.2649599 |
| -3.2339545 | 0.00122089 | 0.0027176 | 18.0331307 | 44.6507786 | 33.3631659 | 20.1713742 |
| -3.2335548 | 0.0012226 | 0.00272122 | 14.3445358 | 35.7206228 | 25.3099879 | 52.6696992 |
| -3.2334638 | 0.00122299 | 0.0027219 | 5.32797044 | 63.8641439 | 64.4254237 | 71.1601255 |
| -3.2333529 | 0.00122346 | 0.00272277 | 4.91812656 | 45.7332217 | 94.3372276 | 82.9267604 |
| -3.2332993 | 0.00122369 | 0.00272309 | 11.8854725 | 36.2618444 | 47.1686138 | 34.7395888 |
| -3.233047 | 0.00122477 | 0.00272531 | 40.984388 | 39.5091738 | 27.6108959 | 13.4475828 |
| -3.2327948 | 0.00122586 | 0.00272753 | 10.6559409 | 42.2152815 | 37.9649818 | 47.0665397 |
| 3.23271537 | 0.0012262 | 0.0027281 | 40.1647002 | 53.0397127 | 28.7613499 | 35.2999048 |
| -3.2326325 | 0.00122655 | 0.00272871 | 11.8854725 | 40.5916169 | 29.9118039 | 57.1522268 |
| -3.2324376 | 0.00122739 | 0.00273038 | 3.27875104 | 53.851545 | 86.2840497 | 133.355196 |
| -3.2323749 | 0.00122766 | 0.00273079 | 59.4273626 | 30.579018 | 46.0181598 | 52.1093832 |
| -3.2320565 | 0.00122903 | 0.00273365 | 11.8854725 | 36.803066 | 39.1154358 | 38.6618005 |
| -3.2318259 | 0.00123002 | 0.00273567 | 22.5414134 | 20.5664192 | 25.3099879 | 25.2142177 |
| -3.2314816 | 0.0012315 | 0.00273877 | 138.117388 | 503.877271 | 0 | 259.9866 |
| 3.23143318 | 0.00123171 | 0.00273905 | 0 | 4.87099402 | 8.05317797 | 5.60315949 |
| -3.2312798 | 0.00123237 | 0.00274033 | 9.01656536 | 47.0862756 | 63.2749697 | 39.2221164 |
| -3.2309399 | 0.00123384 | 0.0027434 | 21.3118818 | 42.2152815 | 14.9559019 | 52.6696992 |
| -3.230692 | 0.00123491 | 0.00274559 | 22.5414134 | 31.9320719 | 18.4072639 | 26.3348496 |
| -3.2305963 | 0.00123532 | 0.00274632 | 11.8854725 | 63.8641439 | 27.6108959 | 43.704644 |
| -3.2305083 | 0.0012357 | 0.00274698 | 13.524848 | 36.5324552 | 42.5667978 | 28.0157974 |
| -3.2303626 | 0.00123633 | 0.00274819 | 7.78703372 | 71.7118565 | 60.9740618 | 37.5411686 |
| -3.2300454 | 0.00123771 | 0.00275105 | 96.7231557 | 138.82333 | 153.010381 | 138.958355 |
| -3.2297958 | 0.00123879 | 0.00275327 | 89.3459658 | 73.8767427 | 79.3813257 | 49.3078035 |
| -3.2289869 | 0.0012423 | 0.00276088 | 12.7051603 | 35.1794013 | 35.6640739 | 36.4205367 |
| -3.2285462 | 0.00124421 | 0.00276494 | 128.690978 | 27.6022995 | 21.8586259 | 19.0507423 |
| -3.2285184 | 0.00124433 | 0.00276502 | 43.4434513 | 24.3549701 | 23.0090799 | 19.0507423 |
| -3.228404 | 0.00124483 | 0.00276594 | 14.3445358 | 39.5091738 | 33.3631659 | 27.4554815 |
| -3.2282453 | 0.00124552 | 0.00276728 | 131.559885 | 21.6488623 | 34.5136199 | 15.6888466 |
| -3.227748 | 0.00124769 | 0.00277191 | 11.8854725 | 30.0377965 | 43.7172518 | 46.5062237 |

| | | | | | | |
|---|---|---|---|---|---|---|
| -3.2273884 | 0.00124926 | 0.0027752 | 54.509236 | 16.2366467 | 20.7081719 | 37.5411686 |
| 3.22699567 | 0.00125097 | 0.00277882 | 15.9839113 | 10.0125988 | 0 | 0 |
| -3.2265365 | 0.00125298 | 0.00278309 | 17.6232868 | 80.912623 | 69.0272397 | 89.0902358 |
| -3.2262704 | 0.00125415 | 0.00278549 | 23.770945 | 31.9320719 | 16.1063559 | 32.498325 |
| -3.2261088 | 0.00125486 | 0.00278687 | 10.6559409 | 35.4500121 | 49.4695218 | 42.5840121 |
| -3.2258045 | 0.00125619 | 0.00278964 | 15.9839113 | 34.367569 | 41.4163438 | 22.4126379 |
| -3.2257311 | 0.00125651 | 0.00279017 | 179.101776 | 276.293606 | 0 | 357.481575 |
| -3.2240059 | 0.00126411 | 0.00280684 | 9.01656536 | 85.783617 | 39.1154358 | 40.3427483 |
| -3.2229037 | 0.00126898 | 0.00281747 | 23.3611012 | 32.2026827 | 21.8586259 | 20.1713742 |
| -3.2228691 | 0.00126914 | 0.00281761 | 33.1973543 | 11.6362635 | 41.4163438 | 43.144328 |
| -3.2227791 | 0.00126953 | 0.00281831 | 315.169944 | 730.378493 | 676.466949 | 735.134525 |
| -3.2226573 | 0.00127007 | 0.00281931 | 333.203074 | 197.275258 | 0 | 265.029444 |
| -3.2223931 | 0.00127125 | 0.00282172 | 8.1968776 | 50.8748265 | 41.4163438 | 63.3157022 |
| -3.2223509 | 0.00127143 | 0.00282194 | 277.464307 | 545.821942 | 509.65112 | 516.050989 |
| -3.22228 | 0.00127175 | 0.00282244 | 13.9346919 | 70.6294134 | 16.1063559 | 81.2458125 |
| -3.2217623 | 0.00127405 | 0.00282735 | 16.3937552 | 26.2492456 | 31.0622579 | 27.4554815 |
| -3.2216265 | 0.00127465 | 0.0028285 | 233.611012 | 483.310852 | 517.704298 | 469.544765 |
| -3.221366 | 0.00127581 | 0.00283088 | 150.00286 | 73.3355211 | 71.3281477 | 108.701294 |
| -3.2212631 | 0.00127627 | 0.0028317 | 12.2953164 | 35.1794013 | 33.3631659 | 42.5840121 |
| -3.2211712 | 0.00127668 | 0.00283241 | 8.60672148 | 55.2045989 | 65.5758777 | 35.2999048 |
| -3.2211478 | 0.00127678 | 0.00283245 | 12.2953164 | 87.6778924 | 16.1063559 | 84.6077082 |
| -3.2206661 | 0.00127893 | 0.00283702 | 18.0331307 | 24.8961917 | 26.4604419 | 26.3348496 |
| -3.2205283 | 0.00127955 | 0.00283819 | 10.246097 | 36.5324552 | 59.8236078 | 39.2221164 |
| -3.2202833 | 0.00128064 | 0.00284042 | 12.7051603 | 31.9320719 | 33.3631659 | 44.8252759 |
| -3.220075 | 0.00128157 | 0.0028421 | 21.3118818 | 34.9087905 | 23.0090799 | 20.7316901 |
| -3.2200906 | 0.0012815 | 0.0028421 | 11.4756286 | 25.1668025 | 109.29313 | 43.144328 |
| -3.2198943 | 0.00128238 | 0.00284369 | 23.770945 | 33.2851258 | 43.7172518 | 15.1285306 |
| -3.2198482 | 0.00128258 | 0.00284395 | 47.1320462 | 19.2133653 | 20.7081719 | 26.8951655 |
| -3.2197568 | 0.00128299 | 0.00284447 | 12.7051603 | 50.8748265 | 27.6108959 | 38.6618005 |
| -3.2197617 | 0.00128297 | 0.00284447 | 71.722679 | 21.1076408 | 21.8586259 | 24.0935858 |
| -3.2194794 | 0.00128424 | 0.00284703 | 45.0828268 | 25.9786348 | 14.9559019 | 29.1364293 |
| -3.2192467 | 0.00128528 | 0.00284914 | 12.7051603 | 37.8855091 | 41.4163438 | 29.6967453 |
| -3.2190487 | 0.00128617 | 0.00285092 | 12.2953164 | 33.0145151 | 32.2127119 | 56.0315949 |
| -3.2183341 | 0.00128938 | 0.00285783 | 9.42640924 | 54.1221558 | 34.5136199 | 53.7903311 |
| -3.2182491 | 0.00128976 | 0.00285832 | 7.37718984 | 57.3694852 | 36.8145279 | 85.1680242 |
| -3.2182459 | 0.00128977 | 0.00285832 | 146.724109 | 25.708024 | 20.7081719 | 20.1713742 |
| -3.217692 | 0.00129227 | 0.00286365 | 20.9020379 | 34.9087905 | 48.3190678 | 16.2491625 |
| -3.2175134 | 0.00129307 | 0.00286524 | 27.45954 | 14.8835929 | 28.7613499 | 63.8760181 |
| -3.2174561 | 0.00129333 | 0.00286561 | 33.1973543 | 17.3190899 | 43.7172518 | 20.1713742 |
| -3.2173936 | 0.00129361 | 0.00286604 | 7.37718984 | 39.238563 | 64.4254237 | 64.99665 |
| -3.2169556 | 0.00129559 | 0.00287022 | 13.9346919 | 35.7206228 | 29.9118039 | 31.9380091 |
| -3.2166942 | 0.00129677 | 0.00287264 | 20.0823501 | 36.2618444 | 32.2127119 | 17.9301104 |
| -3.2165552 | 0.0012974 | 0.00287383 | 13.9346919 | 29.2259641 | 33.3631659 | 34.7395888 |

| | | | | | | |
|---|---|---|---|---|---|---|
| -3.2163389 | 0.00129837 | 0.0028758 | 19.2626624 | 20.2958084 | 28.7613499 | 29.6967453 |
| -3.216244 | 0.0012988 | 0.00287656 | 6.55750208 | 54.6633774 | 98.9390436 | 40.9030642 |
| -3.2160994 | 0.00129946 | 0.00287781 | 15.1642236 | 97.4198805 | 18.4072639 | 44.2649599 |
| -3.2159664 | 0.00130006 | 0.00287895 | 18.4429746 | 83.6187308 | 98.9390436 | 10.0856871 |
| -3.2156823 | 0.00130135 | 0.0028816 | 17.6232868 | 26.7904671 | 25.3099879 | 26.3348496 |
| -3.215187 | 0.0013036 | 0.00288638 | 134.018949 | 316.614612 | 330.180297 | 294.726189 |
| -3.2148294 | 0.00130522 | 0.00288978 | 13.1150042 | 40.8622277 | 26.4604419 | 42.5840121 |
| -3.2148006 | 0.00130535 | 0.00288987 | 139.756763 | 167.508072 | 164.514921 | 183.783631 |
| -3.2143758 | 0.00130728 | 0.00289395 | 17.213443 | 29.2259641 | 24.1595339 | 27.4554815 |
| -3.214293 | 0.00130766 | 0.00289459 | 237.70945 | 534.185678 | 448.677058 | 532.300151 |
| -3.2140987 | 0.00130855 | 0.00289635 | 25.8201644 | 24.0843593 | 19.5577179 | 22.9729539 |
| -3.2140004 | 0.00130899 | 0.00289714 | 15.9839113 | 32.2026827 | 25.3099879 | 29.1364293 |
| -3.213801 | 0.0013099 | 0.00289896 | 273.775712 | 331.498204 | 0 | 389.9799 |
| -3.2134476 | 0.00131152 | 0.00290233 | 111.067691 | 799.925463 | 200.178995 | 763.710638 |
| -3.2131089 | 0.00131306 | 0.00290555 | 136.068168 | 24.8961917 | 16.1063559 | 26.8951655 |
| -3.2130739 | 0.00131323 | 0.00290571 | 11.4756286 | 45.1920001 | 50.6199758 | 28.5761134 |
| -3.2123497 | 0.00131654 | 0.00291284 | 16.3937552 | 29.4965749 | 23.0090799 | 37.5411686 |
| -3.2119648 | 0.00131831 | 0.00291655 | 12.2953164 | 36.2618444 | 33.3631659 | 38.1014845 |
| 3.2118865 | 0.00131866 | 0.00291714 | 0 | 0 | 0 | 0 |
| -3.2116129 | 0.00131992 | 0.00291972 | 19.2626624 | 22.7313054 | 24.1595339 | 27.4554815 |
| 3.21085524 | 0.00132341 | 0.00292723 | 11.4756286 | 6.76526948 | 5.75226998 | 6.72379138 |
| -3.210696 | 0.00132414 | 0.00292865 | 17.6232868 | 15.4248144 | 69.0272397 | 49.8681194 |
| -3.2102236 | 0.00132632 | 0.00293327 | 211.069598 | 296.318803 | 0 | 261.107232 |
| -3.209757 | 0.00132847 | 0.00293783 | 147.543797 | 200.522587 | 135.753571 | 192.18837 |
| -3.2095659 | 0.00132936 | 0.00293959 | 29.5087594 | 22.7313054 | 21.8586259 | 20.1713742 |
| -3.2094114 | 0.00133007 | 0.00294097 | 43.8532952 | 20.0251977 | 23.0090799 | 21.852322 |
| -3.2092125 | 0.00133099 | 0.0029428 | 36.8859492 | 110.950419 | 28.7613499 | 59.3934906 |
| -3.2087068 | 0.00133333 | 0.00294778 | 90.1656536 | 465.991762 | 0 | 367.006946 |
| -3.2085066 | 0.00133426 | 0.00294963 | 0 | 133.411114 | 801.866435 | 326.664198 |
| -3.2083161 | 0.00133515 | 0.00295138 | 0 | 226.771833 | 366.994825 | 477.949504 |
| -3.2080386 | 0.00133644 | 0.00295403 | 4.0984388 | 88.219114 | 90.8858656 | 49.8681194 |
| -3.2079775 | 0.00133672 | 0.00295445 | 65.5750208 | 25.708024 | 21.8586259 | 19.0507423 |
| -3.2076503 | 0.00133824 | 0.00295761 | 20.9020379 | 38.1561199 | 16.1063559 | 35.2999048 |
| -3.2073731 | 0.00133953 | 0.00296026 | 49.1812656 | 63.3229223 | 52.9208838 | 63.3157022 |
| -3.2067805 | 0.00134229 | 0.00296616 | 21.3118818 | 78.7477367 | 13.8054479 | 40.9030642 |
| 3.20593056 | 0.00134626 | 0.00297474 | 4.0984388 | 4.05916169 | 10.354086 | 7.28410733 |
| -3.205779 | 0.00134697 | 0.0029761 | 32.3776665 | 21.1076408 | 26.4604419 | 18.4904263 |
| -3.2049426 | 0.00135089 | 0.00298456 | 12.2953164 | 71.1706349 | 17.2568099 | 81.8061285 |
| -3.2040036 | 0.00135531 | 0.0029941 | 12.7051603 | 25.708024 | 40.2658898 | 49.8681194 |
| -3.2039583 | 0.00135552 | 0.00299437 | 149.183172 | 287.388647 | 258.852149 | 304.811876 |
| -3.2037962 | 0.00135628 | 0.00299585 | 24.1807889 | 50.3336049 | 51.7704298 | 50.9887513 |
| -3.2037343 | 0.00135658 | 0.00299629 | 25.8201644 | 19.7545869 | 43.7172518 | 19.6110582 |
| -3.2036801 | 0.00135683 | 0.00299665 | 5.73781432 | 50.3336049 | 83.9831417 | 55.4712789 |

| | | | | | | |
|---|---|---|---|---|---|---|
| -3.20366 | 0.00135693 | 0.00299665 | 20.0823501 | 27.3316887 | 40.2658898 | 18.4904263 |
| -3.2018195 | 0.00136563 | 0.00301566 | 151.642236 | 282.517653 | 0 | 357.481575 |
| -3.2017639 | 0.00136589 | 0.00301603 | 20.0823501 | 32.4732935 | 55.2217918 | 16.8094785 |
| -3.2015009 | 0.00136714 | 0.00301858 | 21.7217256 | 28.4141318 | 20.7081719 | 21.852322 |
| -3.2013974 | 0.00136763 | 0.00301946 | 15.5740674 | 31.1202396 | 39.1154358 | 22.9729539 |
| -3.201343 | 0.00136789 | 0.00301962 | 16.3937552 | 25.1668025 | 25.3099879 | 36.9808526 |
| -3.2013432 | 0.00136789 | 0.00301962 | 17.6232868 | 25.708024 | 29.9118039 | 22.9729539 |
| -3.2011705 | 0.00136871 | 0.00302122 | 50.8206411 | 348.276073 | 204.780811 | 355.800627 |
| -3.2008503 | 0.00137023 | 0.00302437 | 154.101299 | 313.367282 | 0 | 316.018195 |
| -3.2008261 | 0.00137034 | 0.00302442 | 67.6242402 | 79.8301798 | 70.1776937 | 76.763285 |
| -3.2006928 | 0.00137098 | 0.00302561 | 20.9020379 | 47.6274971 | 16.1063559 | 32.498325 |
| -3.1999969 | 0.00137429 | 0.00303272 | 33.6071982 | 25.4374132 | 16.1063559 | 25.2142177 |
| -3.1999668 | 0.00137443 | 0.00303283 | 61.476582 | 51.6866588 | 88.5849576 | 64.4363341 |
| -3.1998709 | 0.00137489 | 0.00303363 | 15.1642236 | 28.143521 | 29.9118039 | 28.5761134 |
| -3.1996129 | 0.00137612 | 0.00303614 | 15.5740674 | 26.2492456 | 26.4604419 | 36.4205367 |
| -3.1988988 | 0.00137954 | 0.00304346 | 24.5906328 | 44.109557 | 16.1063559 | 25.7745336 |
| -3.1987362 | 0.00138031 | 0.00304497 | 737.718984 | 1481.59402 | 1319.57073 | 1475.87221 |
| -3.1986345 | 0.0013808 | 0.00304584 | 56.9682993 | 22.4606947 | 18.4072639 | 24.6539017 |
| -3.1979219 | 0.00138422 | 0.00305317 | 10.246097 | 35.9912336 | 58.6731538 | 36.4205367 |
| -3.1977327 | 0.00138513 | 0.00305496 | 32.3776665 | 15.6954252 | 26.4604419 | 26.8951655 |
| -3.1976485 | 0.00138553 | 0.00305564 | 9.83625312 | 49.7923834 | 42.5667978 | 36.4205367 |
| -3.1974895 | 0.00138629 | 0.00305712 | 11.4756286 | 41.9446708 | 73.6290557 | 25.7745336 |
| -3.1968676 | 0.00138929 | 0.00306351 | 8.60672148 | 41.4034492 | 40.2658898 | 66.6775979 |
| -3.1963739 | 0.00139167 | 0.00306855 | 18.4429746 | 24.3549701 | 24.1595339 | 25.2142177 |
| -3.1963438 | 0.00139181 | 0.00306866 | 109.018472 | 15.966036 | 17.2568099 | 40.3427483 |
| -3.1962151 | 0.00139243 | 0.00306982 | 21.3118818 | 20.83703 | 31.0622579 | 21.292006 |
| -3.1961597 | 0.0013927 | 0.0030702 | 26.6398522 | 15.6954252 | 25.3099879 | 39.7824324 |
| -3.1960025 | 0.00139346 | 0.00307166 | 169.265522 | 622.675403 | 561.42155 | 564.23816 |
| -3.1958314 | 0.00139429 | 0.00307327 | 19.2626624 | 43.5683354 | 14.9559019 | 51.5490673 |
| -3.1956568 | 0.00139513 | 0.00307492 | 17.6232868 | 30.0377965 | 25.3099879 | 22.9729539 |
| -3.1956109 | 0.00139535 | 0.00307517 | 50.8206411 | 97.4198805 | 18.4072639 | 91.3314996 |
| -3.1955944 | 0.00139543 | 0.00307517 | 12.7051603 | 30.3084073 | 34.5136199 | 37.5411686 |
| 3.19537671 | 0.00139648 | 0.00307728 | 13.1150042 | 6.76526948 | 4.60181598 | 5.04284354 |
| -3.1952293 | 0.0013972 | 0.00307864 | 13.9346919 | 35.4500121 | 23.0090799 | 74.5220212 |
| -3.1950699 | 0.00139797 | 0.00308013 | 13.1150042 | 35.7206228 | 35.6640739 | 28.5761134 |
| 3.19504424 | 0.00139809 | 0.0030802 | 0.40984388 | 1.62366467 | 0 | 1.68094785 |
| -3.1941048 | 0.00140265 | 0.00309003 | 149.183172 | 306.872624 | 301.418947 | 338.991149 |
| -3.1935707 | 0.00140525 | 0.00309554 | 177.052556 | 241.655426 | 0 | 342.913361 |
| -3.193548 | 0.00140536 | 0.00309557 | 0 | 514.431091 | 189.824909 | 364.765683 |
| -3.1934505 | 0.00140583 | 0.0030964 | 24.5906328 | 43.0271139 | 52.9208838 | 60.5141225 |
| -3.1933606 | 0.00140627 | 0.00309716 | 20.9020379 | 37.6148983 | 25.3099879 | 18.4904263 |
| -3.1932682 | 0.00140672 | 0.00309794 | 18.8528185 | 40.0503953 | 24.1595339 | 21.292006 |
| -3.1924532 | 0.0014107 | 0.00310648 | 1.63937552 | 61.1580361 | 148.408565 | 119.347297 |

| | | | | | | |
|---|---|---|---|---|---|---|
| -3.1915133 | 0.0014153 | 0.00311639 | 150.00286 | 31.3908504 | 14.9559019 | 20.7316901 |
| -3.1903184 | 0.00142116 | 0.00312909 | 13.524848 | 33.2851258 | 27.6108959 | 35.8602207 |
| -3.1898899 | 0.00142327 | 0.00313352 | 19.2626624 | 23.5431378 | 23.0090799 | 24.6539017 |
| -3.1895203 | 0.00142509 | 0.00313732 | 18.4429746 | 36.2618444 | 17.2568099 | 38.1014845 |
| -3.1894741 | 0.00142532 | 0.00313761 | 128.690978 | 56.0164313 | 27.6108959 | 39.7824324 |
| -3.1888914 | 0.0014282 | 0.00314372 | 0 | 322.568049 | 294.516223 | 379.894213 |
| -3.1887012 | 0.00142913 | 0.00314558 | 30.738291 | 23.8137486 | 54.0713378 | 14.5682147 |
| -3.1885677 | 0.0014298 | 0.00314682 | 168.035991 | 483.310852 | 315.224395 | 381.575161 |
| 3.18793923 | 0.00143291 | 0.00315345 | 22.9512573 | 14.6129821 | 14.9559019 | 20.7316901 |
| -3.1877097 | 0.00143404 | 0.00315574 | 22.1315695 | 36.803066 | 28.7613499 | 16.2491625 |
| -3.18742 | 0.00143548 | 0.00315869 | 55.3289238 | 43.2977247 | 6.90272397 | 57.7125427 |
| -3.187201 | 0.00143657 | 0.00316086 | 14.3445358 | 67.9233056 | 40.2658898 | 20.1713742 |
| -3.1869916 | 0.00143761 | 0.00316294 | 7.78703372 | 70.0881918 | 49.4695218 | 37.5411686 |
| -3.1868418 | 0.00143835 | 0.00316436 | 9.42640924 | 69.5469702 | 26.4604419 | 57.1522268 |
| -3.1866204 | 0.00143946 | 0.00316657 | 14.3445358 | 25.4374132 | 47.1686138 | 29.1364293 |
| -3.186233 | 0.00144138 | 0.00317059 | 11.4756286 | 61.4286469 | 36.8145279 | 28.5761134 |
| -3.1855333 | 0.00144487 | 0.00317806 | 40.1647002 | 58.7225391 | 9.20363196 | 35.2999048 |
| 3.18463259 | 0.00144938 | 0.00318775 | 31.5579788 | 25.4374132 | 27.6108959 | 17.3697944 |
| -3.1844242 | 0.00145042 | 0.00318982 | 25.0004767 | 15.6954252 | 32.2127119 | 26.8951655 |
| -3.1843794 | 0.00145065 | 0.0031901 | 4.91812656 | 45.4626109 | 180.621277 | 40.9030642 |
| 3.183783 | 0.00145364 | 0.00319646 | 12.7051603 | 10.5538204 | 14.9559019 | 14.5682147 |
| -3.1836387 | 0.00145436 | 0.00319784 | 15.9839113 | 31.3908504 | 21.8586259 | 33.058641 |
| -3.1833739 | 0.0014557 | 0.00320055 | 23.770945 | 22.1900839 | 25.3099879 | 19.0507423 |
| -3.1832163 | 0.00145649 | 0.00320207 | 22.5414134 | 21.6488623 | 29.9118039 | 19.0507423 |
| -3.1830629 | 0.00145726 | 0.00320333 | 13.524848 | 28.9553534 | 28.7613499 | 40.3427483 |
| -3.1830771 | 0.00145719 | 0.00320333 | 73.7718984 | 349.629127 | 360.092101 | 348.51652 |
| -3.1827191 | 0.00145899 | 0.00320692 | 24.1807889 | 20.2958084 | 69.0272397 | 20.1713742 |
| -3.1826631 | 0.00145927 | 0.00320732 | 9.01656536 | 28.9553534 | 58.6731538 | 60.5141225 |
| -3.1824603 | 0.0014603 | 0.00320913 | 8.1968776 | 162.637078 | 24.1595339 | 48.1871716 |
| -3.1824672 | 0.00146026 | 0.00320913 | 284.431653 | 171.567234 | 0 | 284.080186 |
| -3.1824279 | 0.00146046 | 0.00320927 | 31.5579788 | 29.7671857 | 16.1063559 | 21.852322 |
| 3.18208025 | 0.00146221 | 0.00321291 | 0 | 0 | 0 | 0 |
| -3.1818606 | 0.00146332 | 0.00321513 | 20.9020379 | 25.708024 | 19.5577179 | 24.0935858 |
| -3.1817468 | 0.0014639 | 0.00321617 | 120.903945 | 25.708024 | 14.9559019 | 25.7745336 |
| -3.1816092 | 0.00146459 | 0.00321748 | 9.83625312 | 37.3442875 | 40.2658898 | 45.9459078 |
| -3.1815474 | 0.00146491 | 0.00321773 | 16.8035991 | 28.6847426 | 23.0090799 | 26.8951655 |
| -3.1815619 | 0.00146483 | 0.00321773 | 16.3937552 | 30.0377965 | 20.7081719 | 40.3427483 |
| -3.1811287 | 0.00146702 | 0.00322216 | 31.9678226 | 23.272527 | 16.1063559 | 26.3348496 |
| -3.1808114 | 0.00146863 | 0.00322548 | 54.9190799 | 30.0377965 | 35.6640739 | 39.2221164 |
| -3.180748 | 0.00146895 | 0.00322596 | 797.146346 | 1376.05581 | 1405.85478 | 1440.5723 |
| -3.1806397 | 0.0014695 | 0.00322695 | 27.0496961 | 47.0862756 | 14.9559019 | 24.6539017 |
| -3.1803311 | 0.00147107 | 0.00323017 | 14.7543797 | 33.0145151 | 29.9118039 | 25.2142177 |
| -3.180299 | 0.00147123 | 0.00323031 | 15.1642236 | 34.0969582 | 27.6108959 | 25.2142177 |

| | | | | | | |
|---|---|---|---|---|---|---|
| -3.1802405 | 0.00147153 | 0.00323074 | 15.1642236 | 30.579018 | 26.4604419 | 27.4554815 |
| -3.179664 | 0.00147446 | 0.00323695 | 12.7051603 | 27.3316887 | 37.9649818 | 36.4205367 |
| -3.1794827 | 0.00147538 | 0.00323876 | 20.492194 | 25.1668025 | 20.7081719 | 22.9729539 |
| -3.1791816 | 0.00147692 | 0.0032419 | 297.546657 | 374.254708 | 366.994825 | 276.235763 |
| -3.1779879 | 0.00148301 | 0.00325506 | 23.770945 | 75.5004074 | 12.6549939 | 35.8602207 |
| -3.1779105 | 0.00148341 | 0.00325571 | 15.1642236 | 28.143521 | 25.3099879 | 31.3776931 |
| -3.1778057 | 0.00148394 | 0.00325666 | 58.6076748 | 28.6847426 | 12.6549939 | 29.1364293 |
| -3.1777224 | 0.00148437 | 0.00325738 | 13.1150042 | 43.2977247 | 75.9299637 | 20.1713742 |
| -3.176943 | 0.00148836 | 0.00326592 | 15.1642236 | 25.4374132 | 26.4604419 | 35.8602207 |
| 3.17661225 | 0.00149006 | 0.00326943 | 0.81968776 | 0.54122156 | 1.150454 | 0.56031595 |
| -3.1759621 | 0.0014934 | 0.00327654 | 254.513049 | 458.41466 | 517.704298 | 470.665397 |
| 3.17523199 | 0.00149717 | 0.00328457 | 27.0496961 | 10.5538204 | 18.4072639 | 6.72379138 |
| -3.1750751 | 0.00149798 | 0.00328612 | 68.0340841 | 67.382084 | 56.3722458 | 75.0823371 |
| -3.1748851 | 0.00149896 | 0.00328805 | 59.0175187 | 29.7671857 | 11.50454 | 31.9380091 |
| -3.1747853 | 0.00149947 | 0.00328896 | 87.7065903 | 65.7584193 | 18.4072639 | 10.646003 |
| -3.1743761 | 0.00150159 | 0.00329338 | 6.1476582 | 66.0290301 | 57.5226998 | 45.9459078 |
| -3.1740248 | 0.00150341 | 0.00329714 | 9.01656536 | 29.2259641 | 83.9831417 | 45.3855918 |
| 3.17371673 | 0.001505 | 0.00330042 | 30.738291 | 11.6362635 | 4.60181598 | 14.5682147 |
| -3.1734145 | 0.00150657 | 0.00330363 | 2.86890716 | 112.032863 | 57.5226998 | 82.3664444 |
| -3.1733436 | 0.00150694 | 0.00330422 | 11.4756286 | 53.5809343 | 25.3099879 | 43.144328 |
| -3.1724466 | 0.0015116 | 0.00331422 | 38.5253247 | 19.2133653 | 25.3099879 | 19.0507423 |
| -3.1720081 | 0.00151389 | 0.003319 | 218.856632 | 354.22951 | 413.012984 | 358.602207 |
| -3.1713549 | 0.0015173 | 0.00332625 | 53.6895483 | 60.075593 | 8.05317797 | 32.498325 |
| 3.17129718 | 0.0015176 | 0.00332668 | 0 | 0 | 0 | 0 |
| 3.17126777 | 0.00151775 | 0.00332679 | 0.40984388 | 0.27061078 | 1.150454 | 0 |
| -3.1710562 | 0.00151886 | 0.00332899 | 11.4756286 | 33.5557366 | 44.8677058 | 31.3776931 |
| -3.1709438 | 0.00151945 | 0.00333005 | 24.1807889 | 44.9213893 | 47.1686138 | 61.0744384 |
| -3.1706548 | 0.00152096 | 0.00333314 | 274.185556 | 295.777582 | 307.171217 | 0 |
| -3.1705298 | 0.00152161 | 0.00333412 | 32.7875104 | 20.0251977 | 34.5136199 | 16.2491625 |
| -3.1705356 | 0.00152158 | 0.00333412 | 27.8693838 | 29.7671857 | 13.8054479 | 29.6967453 |
| -3.1704591 | 0.00152198 | 0.00333471 | 78.6900249 | 36.2618444 | 9.20363196 | 33.6189569 |
| -3.1703717 | 0.00152244 | 0.00333548 | 13.9346919 | 27.8729103 | 34.5136199 | 28.0157974 |
| -3.1701661 | 0.00152352 | 0.00333762 | 11.8854725 | 40.8622277 | 33.3631659 | 30.8173772 |
| -3.1701104 | 0.00152381 | 0.00333803 | 228.283041 | 406.186779 | 312.923487 | 349.637152 |
| -3.1697591 | 0.00152565 | 0.00334184 | 21.7217256 | 23.0019162 | 54.0713378 | 18.4904263 |
| -3.1693292 | 0.00152791 | 0.00334636 | 36.4761053 | 38.9679522 | 24.1595339 | 29.1364293 |
| -3.1693276 | 0.00152792 | 0.00334636 | 32.7875104 | 45.1920001 | 8.05317797 | 77.8839169 |
| -3.1691219 | 0.001529 | 0.0033485 | 7.78703372 | 66.5702517 | 35.6640739 | 49.8681194 |
| -3.1690844 | 0.0015292 | 0.0033487 | 36.0662614 | 18.1309222 | 17.2568099 | 34.1792729 |
| -3.1688609 | 0.00153038 | 0.00335105 | 92.214873 | 23.0019162 | 16.1063559 | 26.3348496 |
| -3.1685522 | 0.001532 | 0.00335439 | 27.0496961 | 47.3568864 | 9.20363196 | 75.6426531 |
| -3.1675024 | 0.00153754 | 0.00336629 | 4.91812656 | 122.045461 | 96.6381356 | 91.8918156 |
| -3.1671553 | 0.00153938 | 0.00337008 | 16.8035991 | 25.708024 | 26.4604419 | 23.5332698 |

| | | | | | | |
|---|---|---|---|---|---|---|
| -3.1668966 | 0.00154075 | 0.00337285 | 61.8864259 | 31.9320719 | 44.8677058 | 30.8173772 |
| -3.166679 | 0.0015419 | 0.00337515 | 8.60672148 | 56.0164313 | 47.1686138 | 34.7395888 |
| -3.1666481 | 0.00154207 | 0.00337528 | 13.1150042 | 42.7565031 | 23.0090799 | 39.7824324 |
| 3.16540137 | 0.00154869 | 0.00338955 | 2.0492194 | 0.54122156 | 3.45136199 | 1.68094785 |
| -3.1653714 | 0.00154885 | 0.00338967 | 6.55750208 | 30.3084073 | 88.5849576 | 66.6775979 |
| -3.165337 | 0.00154903 | 0.00338984 | 27.45954 | 25.9786348 | 13.8054479 | 69.4791776 |
| -3.1645716 | 0.00155311 | 0.00339854 | 10.6559409 | 44.109557 | 39.1154358 | 31.3776931 |
| -3.1645089 | 0.00155345 | 0.00339904 | 9.42640924 | 75.5004074 | 24.1595339 | 55.4712789 |
| 3.16385852 | 0.00155692 | 0.00340631 | 88.9361219 | 130.705006 | 121.948124 | 129.9933 |
| -3.1638474 | 0.00155698 | 0.00340631 | 11.8854725 | 51.9572696 | 24.1595339 | 41.4633802 |
| -3.1636352 | 0.00155812 | 0.00340857 | 9.83625312 | 39.7797845 | 44.8677058 | 35.2999048 |
| -3.1635663 | 0.00155849 | 0.00340914 | 8.1968776 | 44.3801678 | 56.3722458 | 38.6618005 |
| -3.163478 | 0.00155896 | 0.00340994 | 11.4756286 | 38.9679522 | 42.5667978 | 28.0157974 |
| -3.1633838 | 0.00155947 | 0.00341082 | 20.9020379 | 23.0019162 | 26.4604419 | 19.0507423 |
| 3.16323411 | 0.00156027 | 0.00341234 | 5.73781432 | 2.16488623 | 0 | 0 |
| -3.1628566 | 0.00156229 | 0.00341654 | 31.1481349 | 21.6488623 | 23.0090799 | 17.9301104 |
| 3.16246597 | 0.00156439 | 0.00342089 | 8.1968776 | 6.4946587 | 2.30090799 | 8.40473923 |
| -3.1624437 | 0.00156451 | 0.00342092 | 15.9839113 | 17.0484791 | 54.0713378 | 40.9030642 |
| -3.1623384 | 0.00156508 | 0.00342193 | 397.13872 | 175.626396 | 0 | 173.697944 |
| -3.1623033 | 0.00156526 | 0.00342211 | 20.9020379 | 25.708024 | 17.2568099 | 29.1364293 |
| -3.1620689 | 0.00156652 | 0.00342463 | 13.524848 | 28.143521 | 26.4604419 | 56.0315949 |
| 3.16204662 | 0.00156664 | 0.00342466 | 4.91812656 | 11.0950419 | 1.150454 | 4.48252759 |
| -3.1619556 | 0.00156713 | 0.0034255 | 15.1642236 | 129.351952 | 10.354086 | 64.99665 |
| -3.1618429 | 0.00156774 | 0.0034266 | 4.50828268 | 83.6187308 | 44.8677058 | 73.9617052 |
| -3.1617134 | 0.00156844 | 0.00342789 | 13.524848 | 44.3801678 | 25.3099879 | 30.2570612 |
| -3.1612776 | 0.00157079 | 0.00343256 | 6.55750208 | 73.0649104 | 52.9208838 | 39.7824324 |
| -3.1612839 | 0.00157075 | 0.00343256 | 144.67489 | 283.600097 | 0 | 295.846821 |
| -3.1611721 | 0.00157136 | 0.00343357 | 18.8528185 | 17.8603114 | 28.7613499 | 32.498325 |
| -3.1605977 | 0.00157446 | 0.00344011 | 148.363485 | 269.798947 | 203.630357 | 186.024895 |
| -3.1598 | 0.00157877 | 0.00344931 | 13.9346919 | 23.5431378 | 32.2127119 | 38.6618005 |
| -3.1591276 | 0.00158242 | 0.00345704 | 7.78703372 | 81.9950661 | 28.7613499 | 55.4712789 |
| -3.1590868 | 0.00158264 | 0.00345729 | 19.2626624 | 100.66721 | 26.4604419 | 18.4904263 |
| -3.1590123 | 0.00158305 | 0.00345794 | 13.1150042 | 33.5557366 | 29.9118039 | 29.1364293 |
| -3.1587706 | 0.00158436 | 0.00346058 | 15.1642236 | 73.8767427 | 14.9559019 | 50.4284354 |
| -3.1586261 | 0.00158515 | 0.00346206 | 24.1807889 | 16.5072575 | 23.0090799 | 32.498325 |
| -3.1583632 | 0.00158658 | 0.00346495 | 565.584554 | 82.8068984 | 0 | 236.45333 |
| -3.1583207 | 0.00158681 | 0.00346522 | 75.00143 | 25.9786348 | 13.8054479 | 26.8951655 |
| -3.1582004 | 0.00158746 | 0.00346642 | 14.3445358 | 37.3442875 | 21.8586259 | 34.7395888 |
| -3.1581535 | 0.00158772 | 0.00346674 | 22.9512573 | 27.3316887 | 16.1063559 | 26.3348496 |
| -3.1579351 | 0.00158891 | 0.00346864 | 15.5740674 | 28.6847426 | 33.3631659 | 21.852322 |
| 3.15797022 | 0.00158872 | 0.00346864 | 1.22953164 | 0.27061078 | 0 | 0.56031595 |
| -3.1579402 | 0.00158888 | 0.00346864 | 25.0004767 | 33.5557366 | 12.6549939 | 40.3427483 |
| -3.1573469 | 0.00159212 | 0.0034754 | 0 | 270.610779 | 391.154358 | 280.157974 |

| | | | | | | |
|---|---|---|---|---|---|---|
| -3.1571294 | 0.00159331 | 0.00347776 | 17.213443 | 22.4606947 | 26.4604419 | 24.6539017 |
| -3.1570137 | 0.00159394 | 0.00347891 | 9.01656536 | 39.238563 | 47.1686138 | 39.7824324 |
| -3.1568983 | 0.00159457 | 0.00348005 | 9.42640924 | 66.0290301 | 33.3631659 | 36.9808526 |
| -3.1566109 | 0.00159614 | 0.00348325 | 11.0657848 | 32.4732935 | 40.2658898 | 34.7395888 |
| -3.1557222 | 0.00160101 | 0.00349364 | 7.37718984 | 126.645845 | 20.7081719 | 67.7982298 |
| -3.1555357 | 0.00160204 | 0.00349564 | 8.60672148 | 23.5431378 | 81.6822337 | 61.6347543 |
| -3.155389 | 0.00160284 | 0.00349716 | 18.4429746 | 20.0251977 | 28.7613499 | 24.0935858 |
| -3.1550074 | 0.00160494 | 0.0035015 | 23.770945 | 29.2259641 | 14.9559019 | 28.0157974 |
| -3.1546058 | 0.00160715 | 0.00350609 | 25.8201644 | 16.2366467 | 33.3631659 | 21.852322 |
| -3.1544423 | 0.00160805 | 0.00350781 | 142.62567 | 288.471091 | 250.798971 | 273.434183 |
| -3.1542963 | 0.00160886 | 0.00350933 | 215.577881 | 217.029845 | 0 | 251.021545 |
| -3.1541598 | 0.00160961 | 0.00351074 | 34.017042 | 112.574084 | 32.2127119 | 8.96505518 |
| 3.15397029 | 0.00161065 | 0.00351278 | 132.379573 | 84.9717847 | 88.5849576 | 83.4870763 |
| -3.1539203 | 0.00161093 | 0.00351315 | 0 | 692.492984 | 202.479903 | 184.343947 |
| -3.153668 | 0.00161232 | 0.00351595 | 47.1320462 | 48.1687187 | 8.05317797 | 37.5411686 |
| -3.1535129 | 0.00161318 | 0.00351758 | 13.524848 | 53.5809343 | 16.1063559 | 72.2807574 |
| -3.1533747 | 0.00161395 | 0.00351901 | 22.5414134 | 63.3229223 | 47.1686138 | 50.4284354 |
| -3.1532725 | 0.00161451 | 0.00352 | 10.246097 | 27.0610779 | 42.5667978 | 58.8331746 |
| -3.1531687 | 0.00161508 | 0.00352101 | 7.37718984 | 44.3801678 | 79.3813257 | 36.4205367 |
| -3.1529861 | 0.00161609 | 0.00352298 | 14.3445358 | 25.4374132 | 46.0181598 | 25.7745336 |
| -3.1528984 | 0.00161658 | 0.0035238 | 165.576927 | 385.890971 | 340.534383 | 0 |
| -3.1527361 | 0.00161748 | 0.00352552 | 26.6398522 | 30.579018 | 12.6549939 | 37.5411686 |
| -3.1525122 | 0.00161872 | 0.00352799 | 12.7051603 | 41.4034492 | 23.0090799 | 40.9030642 |
| -3.1522937 | 0.00161993 | 0.00353039 | 11.0657848 | 54.1221558 | 28.7613499 | 34.7395888 |
| 3.15217898 | 0.00162057 | 0.00353154 | 139.346919 | 172.379066 | 196.727633 | 172.016996 |
| -3.1510987 | 0.00162658 | 0.00354439 | 16.8035991 | 67.9233056 | 16.1063559 | 36.9808526 |
| -3.1509557 | 0.00162737 | 0.00354589 | 22.5414134 | 41.9446708 | 13.8054479 | 31.3776931 |
| -3.1505162 | 0.00162982 | 0.00355099 | 39.7548564 | 105.808815 | 92.0363196 | 93.0124475 |
| -3.1502789 | 0.00163115 | 0.00355363 | 41.3942319 | 192.674875 | 73.6290557 | 133.915512 |
| -3.1502323 | 0.00163141 | 0.00355396 | 148.773328 | 262.221845 | 209.382627 | 227.488275 |
| -3.1501667 | 0.00163177 | 0.0035541 | 26.6398522 | 39.238563 | 40.2658898 | 45.3855918 |
| -3.1501619 | 0.0016318 | 0.0035541 | 58.6076748 | 26.2492456 | 40.2658898 | 10.646003 |
| -3.1501872 | 0.00163166 | 0.0035541 | 7.37718984 | 67.9233056 | 47.1686138 | 36.9808526 |
| 3.14995569 | 0.00163295 | 0.00355637 | 2.45906328 | 3.24732935 | 0 | 1.1206319 |
| -3.1495191 | 0.00163539 | 0.00356144 | 18.8528185 | 37.6148983 | 16.1063559 | 31.9380091 |
| -3.1491437 | 0.0016375 | 0.00356578 | 20.0823501 | 29.2259641 | 32.2127119 | 16.2491625 |
| -3.1487078 | 0.00163994 | 0.00357086 | 32.3776665 | 22.1900839 | 14.9559019 | 28.5761134 |
| -3.1484399 | 0.00164144 | 0.0035739 | 232.791324 | 31.9320719 | 17.2568099 | 11.206319 |
| -3.1481056 | 0.00164332 | 0.00357775 | 17.213443 | 31.6614612 | 20.7081719 | 24.6539017 |
| -3.1475877 | 0.00164624 | 0.00358385 | 93.8542485 | 434.871522 | 375.048002 | 352.999048 |
| -3.14747 | 0.0016469 | 0.00358505 | 286.890716 | 263.033677 | 0 | 149.604358 |
| -3.1469883 | 0.00164962 | 0.00359072 | 234.020855 | 176.167617 | 80.5317797 | 113.744138 |
| 3.14695614 | 0.0016498 | 0.00359087 | 0.40984388 | 1.3530539 | 2.30090799 | 0.56031595 |

| | | | | | | |
|---|---|---|---|---|---|---|
| -3.1464988 | 0.00165238 | 0.00359625 | 16.3937552 | 31.6614612 | 23.0090799 | 23.5332698 |
| 3.14633989 | 0.00165328 | 0.00359796 | 2.0492194 | 1.62366467 | 1.150454 | 1.1206319 |
| -3.1462542 | 0.00165376 | 0.00359877 | 16.8035991 | 17.3190899 | 52.9208838 | 33.058641 |
| 3.1461234 | 0.0016545 | 0.00360014 | 2.0492194 | 0.54122156 | 0 | 0.56031595 |
| -3.145991 | 0.00165525 | 0.00360153 | 10.6559409 | 56.5576528 | 37.9649818 | 27.4554815 |
| -3.1457334 | 0.00165671 | 0.00360446 | 34.4268859 | 78.477126 | 52.9208838 | 64.4363341 |
| -3.1456449 | 0.00165721 | 0.00360531 | 62.2962697 | 5.95343714 | 242.745793 | 15.6888466 |
| 3.14553417 | 0.00165784 | 0.00360643 | 0.40984388 | 0.81183234 | 1.150454 | 0 |
| 3.14533292 | 0.00165898 | 0.00360867 | 0 | 0 | 1.150454 | 0 |
| -3.1451862 | 0.00165981 | 0.00361024 | 6.96734596 | 41.4034492 | 63.2749697 | 46.5062237 |
| -3.1450831 | 0.0016604 | 0.00361126 | 34.8367298 | 48.1687187 | 27.6108959 | 77.8839169 |
| -3.1444636 | 0.00166392 | 0.00361868 | 19.6725062 | 29.4965749 | 17.2568099 | 26.3348496 |
| 3.14427437 | 0.00166499 | 0.00362077 | 6.96734596 | 8.65954493 | 8.05317797 | 10.0856871 |
| -3.1441919 | 0.00166546 | 0.00362155 | 13.524848 | 34.367569 | 25.3099879 | 30.2570612 |
| -3.1438211 | 0.00166757 | 0.0036259 | 18.4429746 | 25.4374132 | 31.0622579 | 18.4904263 |
| -3.1432644 | 0.00167075 | 0.00363256 | 148.363485 | 234.078324 | 124.249032 | 169.775732 |
| -3.1432195 | 0.001671 | 0.00363284 | 5.73781432 | 78.7477367 | 58.6731538 | 38.6618005 |
| 3.14320234 | 0.0016711 | 0.00363284 | 28.2792277 | 10.2832096 | 11.50454 | 15.6888466 |
| -3.1430197 | 0.00167215 | 0.00363486 | 872.55762 | 326.897821 | 179.470823 | 275.115131 |
| -3.142996 | 0.00167228 | 0.00363491 | 9.01656536 | 34.0969582 | 49.4695218 | 42.0236961 |
| -3.1427116 | 0.00167391 | 0.0036382 | 11.0657848 | 13.2599282 | 109.29313 | 73.9617052 |
| -3.1426206 | 0.00167443 | 0.00363908 | 79.0998688 | 46.545054 | 43.7172518 | 24.0935858 |
| -3.1423061 | 0.00167623 | 0.00364275 | 261.890239 | 315.802779 | 278.409867 | 333.387989 |
| -3.1422543 | 0.00167652 | 0.00364315 | 15.1642236 | 27.8729103 | 23.0090799 | 29.6967453 |
| -3.1417319 | 0.00167952 | 0.00364916 | 25.0004767 | 46.0038325 | 39.1154358 | 54.350647 |
| -3.1417438 | 0.00167945 | 0.00364916 | 6.1476582 | 40.3210061 | 77.0804177 | 50.4284354 |
| -3.1415593 | 0.00168051 | 0.00365107 | 43.0336074 | 45.7332217 | 78.2308717 | 81.8061285 |
| -3.1413698 | 0.0016816 | 0.00365319 | 6.55750208 | 58.1813175 | 56.3722458 | 40.3427483 |
| -3.1405099 | 0.00168654 | 0.00366368 | 662.30771 | 1666.69179 | 1251.69395 | 1495.48327 |
| -3.1402291 | 0.00168816 | 0.00366695 | 17.6232868 | 23.0019162 | 20.7081719 | 30.8173772 |
| -3.1399507 | 0.00168976 | 0.00367019 | 11.4756286 | 27.3316887 | 42.5667978 | 35.2999048 |
| -3.1394669 | 0.00169256 | 0.00367601 | 13.524848 | 36.2618444 | 24.1595339 | 30.8173772 |
| -3.139131 | 0.0016945 | 0.00367998 | 6.1476582 | 80.912623 | 36.8145279 | 53.7903311 |
| -3.1387792 | 0.00169653 | 0.00368415 | 12.2953164 | 47.0862756 | 19.5577179 | 57.1522268 |
| -3.1387419 | 0.00169675 | 0.00368437 | 16.8035991 | 71.4412457 | 13.8054479 | 44.2649599 |
| -3.1385838 | 0.00169766 | 0.00368611 | 8.1968776 | 44.6507786 | 26.4604419 | 109.26161 |
| -3.1381829 | 0.00169999 | 0.00369091 | 71.722679 | 18.1309222 | 14.9559019 | 34.7395888 |
| -3.1380191 | 0.00170094 | 0.00369272 | 21.3118818 | 34.0969582 | 14.9559019 | 28.5761134 |
| -3.1377801 | 0.00170233 | 0.00369549 | 5.32797044 | 36.2618444 | 92.0363196 | 61.0744384 |
| -3.1372626 | 0.00170533 | 0.00370177 | 15.9839113 | 20.0251977 | 39.1154358 | 27.4554815 |
| -3.1371081 | 0.00170623 | 0.00370347 | 13.1150042 | 63.3229223 | 21.8586259 | 33.6189569 |
| -3.1369895 | 0.00170692 | 0.00370472 | 322.547134 | 414.846324 | 308.321671 | 389.9799 |
| -3.1367487 | 0.00170833 | 0.00370751 | 53.2797044 | 30.8496288 | 10.354086 | 31.9380091 |

| | | | | | | |
|---|---|---|---|---|---|---|
| -3.1364222 | 0.00171023 | 0.0037114 | 133.609105 | 282.247043 | 0 | 280.157974 |
| -3.1359929 | 0.00171273 | 0.00371658 | 123.772852 | 215.40618 | 253.099879 | 276.796079 |
| -3.135866 | 0.00171347 | 0.00371794 | 8.60672148 | 67.9233056 | 95.4876816 | 19.0507423 |
| -3.1355288 | 0.00171545 | 0.00372197 | 6.96734596 | 60.075593 | 70.1776937 | 30.8173772 |
| -3.1352943 | 0.00171682 | 0.00372469 | 5.73781432 | 72.7942996 | 72.4786017 | 33.6189569 |
| -3.1351401 | 0.00171772 | 0.0037264 | 19.6725062 | 29.2259641 | 16.1063559 | 30.2570612 |
| -3.1349541 | 0.00171881 | 0.00372851 | 29.9186032 | 14.0717605 | 29.9118039 | 24.6539017 |
| -3.1348212 | 0.00171959 | 0.00372995 | 0 | 224.877557 | 447.526604 | 252.142177 |
| -3.1346058 | 0.00172085 | 0.00373244 | 165.986771 | 172.108456 | 0 | 362.524419 |
| -3.1336664 | 0.00172637 | 0.00374416 | 19.2626624 | 36.2618444 | 14.9559019 | 34.1792729 |
| -3.1333521 | 0.00172822 | 0.00374792 | 28.6890716 | 25.1668025 | 12.6549939 | 50.9887513 |
| -3.1332952 | 0.00172856 | 0.0037484 | 7.37718984 | 60.6168145 | 69.0272397 | 28.5761134 |
| -3.1332259 | 0.00172896 | 0.00374903 | 45.4926707 | 14.3423713 | 14.9559019 | 88.5299199 |
| -3.1330553 | 0.00172997 | 0.00375096 | 13.524848 | 33.5557366 | 40.2658898 | 21.852322 |
| -3.1330262 | 0.00173014 | 0.00375108 | 42.2139196 | 62.2404792 | 13.8054479 | 17.3697944 |
| -3.1328509 | 0.00173117 | 0.00375307 | 659.028959 | 139.905773 | 94.3372276 | 117.666349 |
| -3.1327603 | 0.00173171 | 0.00375397 | 29.0989155 | 142.070659 | 131.151755 | 150.164674 |
| -3.1323966 | 0.00173386 | 0.00375837 | 306.153378 | 185.097773 | 0 | 182.662999 |
| 3.13216819 | 0.0017352 | 0.00376105 | 8.60672148 | 4.87099402 | 3.45136199 | 4.48252759 |
| -3.1319985 | 0.00173621 | 0.00376297 | 20.0823501 | 24.0843593 | 24.1595339 | 18.4904263 |
| -3.1317217 | 0.00173785 | 0.00376626 | 6.55750208 | 31.9320719 | 69.0272397 | 63.3157022 |
| -3.1309914 | 0.00174217 | 0.00377539 | 11.4756286 | 41.67406 | 29.9118039 | 30.8173772 |
| -3.1307653 | 0.00174351 | 0.00377804 | 17.6232868 | 20.2958084 | 35.6640739 | 22.4126379 |
| 3.13069622 | 0.00174392 | 0.00377868 | 0 | 2.16488623 | 0 | 0 |
| -3.1306132 | 0.00174442 | 0.00377949 | 0 | 342.863857 | 356.640739 | 204.515321 |
| -3.1304215 | 0.00174556 | 0.00378171 | 13.524848 | 22.1900839 | 34.5136199 | 35.8602207 |
| -3.1303374 | 0.00174606 | 0.00378254 | 140.986295 | 24.8961917 | 10.354086 | 30.8173772 |
| -3.1301153 | 0.00174738 | 0.00378514 | 13.1150042 | 26.7904671 | 32.2127119 | 29.1364293 |
| -3.1298001 | 0.00174925 | 0.00378895 | 18.0331307 | 52.7691019 | 10.354086 | 95.2537113 |
| -3.1296742 | 0.00175 | 0.00379032 | 20.492194 | 35.1794013 | 13.8054479 | 37.5411686 |
| -3.1295225 | 0.00175091 | 0.00379203 | 27.0496961 | 20.2958084 | 35.6640739 | 15.6888466 |
| -3.1293866 | 0.00175172 | 0.00379353 | 13.9346919 | 25.1668025 | 42.5667978 | 25.2142177 |
| -3.1293662 | 0.00175184 | 0.00379354 | 48.7714217 | 22.4606947 | 23.0090799 | 16.2491625 |
| -3.1290649 | 0.00175364 | 0.00379717 | 25.4103206 | 27.3316887 | 14.9559019 | 24.0935858 |
| -3.1285202 | 0.00175689 | 0.00380396 | 43.0336074 | 100.396599 | 139.204933 | 117.666349 |
| -3.1283451 | 0.00175794 | 0.00380597 | 47.1320462 | 49.5217726 | 40.2658898 | 50.9887513 |
| -3.1281454 | 0.00175913 | 0.00380831 | 13.9346919 | 32.2026827 | 46.0181598 | 20.7316901 |
| -3.1280513 | 0.00175969 | 0.00380927 | 159.839113 | 235.701989 | 0 | 267.270707 |
| -3.1278999 | 0.0017606 | 0.00381098 | 17.213443 | 26.7904671 | 20.7081719 | 24.0935858 |
| -3.127846 | 0.00176092 | 0.00381142 | 10.246097 | 34.6381797 | 43.7172518 | 31.9380091 |
| -3.1277174 | 0.00176169 | 0.00381283 | 16.8035991 | 35.4500121 | 18.4072639 | 28.0157974 |
| -3.127587 | 0.00176248 | 0.00381427 | 353.695268 | 263.304288 | 204.780811 | 0 |
| -3.1273585 | 0.00176385 | 0.00381672 | 13.1150042 | 48.4393295 | 18.4072639 | 47.6268556 |

| | | | | | | |
|---|---|---|---|---|---|---|
| -3.1273621 | 0.00176383 | 0.00381672 | 36.0662614 | 23.5431378 | 11.50454 | 52.1093832 |
| -3.1271965 | 0.00176482 | 0.00381857 | 19.6725062 | 19.2133653 | 48.3190678 | 21.292006 |
| -3.1271434 | 0.00176514 | 0.00381901 | 20.492194 | 33.5557366 | 19.5577179 | 19.6110582 |
| -3.1271163 | 0.0017653 | 0.0038191 | 65.1651769 | 112.844695 | 46.0181598 | 87.9696039 |
| -3.1269294 | 0.00176642 | 0.00382127 | 21.3118818 | 28.143521 | 70.1776937 | 14.5682147 |
| -3.1268258 | 0.00176705 | 0.00382236 | 10.246097 | 33.8263474 | 33.3631659 | 42.0236961 |
| -3.1266629 | 0.00176803 | 0.00382397 | 18.4429746 | 52.4984912 | 12.6549939 | 47.0665397 |
| -3.1266808 | 0.00176792 | 0.00382397 | 19.6725062 | 107.161869 | 55.2217918 | 70.5998095 |
| -3.1260453 | 0.00177174 | 0.00383175 | 19.2626624 | 23.8137486 | 18.4072639 | 25.7745336 |
| -3.1259075 | 0.00177257 | 0.00383329 | 27.8693838 | 47.6274971 | 35.6640739 | 10.646003 |
| -3.1256562 | 0.00177409 | 0.00383606 | 58.197831 | 21.6488623 | 18.4072639 | 20.7316901 |
| -3.1256586 | 0.00177407 | 0.00383606 | 42.2139196 | 38.6973414 | 16.1063559 | 16.2491625 |
| -3.1254746 | 0.00177518 | 0.00383817 | 25.4103206 | 36.5324552 | 46.0181598 | 57.7125427 |
| -3.1252504 | 0.00177654 | 0.00384058 | 9.83625312 | 37.0736767 | 39.1154358 | 34.7395888 |
| -3.1252594 | 0.00177648 | 0.00384058 | 32.3776665 | 56.8282636 | 13.8054479 | 20.1713742 |
| -3.124138 | 0.00178327 | 0.00385487 | 113.936599 | 157.495473 | 97.7885896 | 114.864769 |
| -3.1239199 | 0.00178459 | 0.00385747 | 48.7714217 | 63.0523115 | 88.5849576 | 3.92221164 |
| -3.1235288 | 0.00178696 | 0.00386234 | 17.213443 | 58.9931499 | 80.5317797 | 10.646003 |
| -3.1234507 | 0.00178744 | 0.00386285 | 0 | 456.790995 | 201.329449 | 255.504073 |
| -3.1234592 | 0.00178739 | 0.00386285 | 12.2953164 | 34.6381797 | 42.5667978 | 23.5332698 |
| -3.1232262 | 0.0017888 | 0.00386554 | 18.8528185 | 23.5431378 | 18.4072639 | 27.4554815 |
| 3.12311728 | 0.00178946 | 0.00386671 | 6.1476582 | 9.47137727 | 6.90272397 | 4.48252759 |
| 3.12248095 | 0.00179334 | 0.00387482 | 13.9346919 | 13.530539 | 18.4072639 | 8.96505518 |
| -3.1223555 | 0.0017941 | 0.00387622 | 25.4103206 | 18.6721438 | 17.2568099 | 31.9380091 |
| -3.1219179 | 0.00179677 | 0.00388172 | 19.6725062 | 22.4606947 | 23.0090799 | 19.6110582 |
| -3.1218672 | 0.00179708 | 0.00388213 | 11.4756286 | 32.2026827 | 35.6640739 | 29.6967453 |
| -3.1217734 | 0.00179765 | 0.00388311 | 7.78703372 | 34.367569 | 44.8677058 | 58.2728587 |
| 3.12167468 | 0.00179826 | 0.00388415 | 0 | 0 | 11.50454 | 10.646003 |
| -3.1216291 | 0.00179853 | 0.00388449 | 22.5414134 | 26.2492456 | 16.1063559 | 23.5332698 |
| -3.1215311 | 0.00179913 | 0.00388552 | 7.37718984 | 32.2026827 | 59.8236078 | 53.2300151 |
| -3.121157 | 0.00180142 | 0.00388994 | 8.60672148 | 74.4179643 | 17.2568099 | 91.8918156 |
| -3.1211684 | 0.00180135 | 0.00388994 | 14.7543797 | 21.9194731 | 26.4604419 | 39.7824324 |
| -3.1208431 | 0.00180334 | 0.00389383 | 20.492194 | 34.0969582 | 13.8054479 | 35.8602207 |
| -3.1203156 | 0.00180657 | 0.00390055 | 73.3620545 | 170.755402 | 156.461743 | 165.853521 |
| -3.1200396 | 0.00180827 | 0.00390395 | 110.248004 | 158.848527 | 105.841768 | 141.759935 |
| -3.1198967 | 0.00180914 | 0.00390558 | 119.264569 | 281.1646 | 0 | 283.51987 |
| -3.1196642 | 0.00181057 | 0.0039084 | 13.9346919 | 43.8389462 | 33.3631659 | 20.1713742 |
| -3.1194972 | 0.0018116 | 0.00391036 | 8.1968776 | 35.9912336 | 47.1686138 | 44.2649599 |
| -3.1193717 | 0.00181237 | 0.00391176 | 11.8854725 | 44.109557 | 26.4604419 | 30.8173772 |
| -3.1191629 | 0.00181366 | 0.00391427 | 14.7543797 | 23.0019162 | 27.6108959 | 28.0157974 |
| -3.1190686 | 0.00181424 | 0.00391526 | 70.0833035 | 31.9320719 | 6.90272397 | 49.8681194 |
| -3.1190053 | 0.00181463 | 0.00391584 | 11.0657848 | 53.0397127 | 54.0713378 | 20.1713742 |
| 3.11894876 | 0.00181498 | 0.00391633 | 46.3123584 | 22.4606947 | 28.7613499 | 31.9380091 |

| | | | | | | |
|---|---|---|---|---|---|---|
| -3.1188693 | 0.00181546 | 0.00391713 | 22.9512573 | 23.0019162 | 17.2568099 | 22.4126379 |
| -3.1185418 | 0.00181748 | 0.00392112 | 8.60672148 | 40.5916169 | 47.1686138 | 35.2999048 |
| -3.1185296 | 0.00181756 | 0.00392112 | 6.55750208 | 39.7797845 | 50.6199758 | 58.2728587 |
| -3.1183799 | 0.00181848 | 0.00392285 | 0 | 249.503138 | 326.728935 | 289.123029 |
| -3.1182752 | 0.00181913 | 0.00392399 | 18.0331307 | 27.6022995 | 20.7081719 | 21.292006 |
| -3.1182398 | 0.00181935 | 0.00392419 | 9.42640924 | 33.5557366 | 39.1154358 | 41.4633802 |
| -3.1182037 | 0.00181957 | 0.00392441 | 22.9512573 | 20.83703 | 19.5577179 | 20.7316901 |
| -3.1178145 | 0.00182197 | 0.00392934 | 20.492194 | 20.83703 | 67.8767857 | 19.0507423 |
| -3.1174742 | 0.00182408 | 0.00393361 | 13.524848 | 24.6255809 | 28.7613499 | 31.3776931 |
| -3.1166381 | 0.00182926 | 0.00394452 | 11.8854725 | 35.1794013 | 27.6108959 | 31.9380091 |
| 3.11628165 | 0.00183147 | 0.00394903 | 3.68859492 | 0 | 0 | 0 |
| -3.1160641 | 0.00183282 | 0.00395168 | 4.0984388 | 48.4393295 | 74.7795097 | 67.7982298 |
| -3.1155205 | 0.00183621 | 0.00395871 | 26.2300083 | 41.4034492 | 42.5667978 | 34.7395888 |
| 3.11521325 | 0.00183812 | 0.00396257 | 0 | 0 | 0 | 0 |
| -3.1151852 | 0.00183829 | 0.00396268 | 15.1642236 | 65.2171978 | 57.5226998 | 12.8872668 |
| 3.11495181 | 0.00183975 | 0.00396555 | 0 | 0 | 0 | 0 |
| -3.1148137 | 0.00184061 | 0.00396714 | 17.213443 | 28.143521 | 20.7081719 | 22.4126379 |
| -3.114263 | 0.00184405 | 0.00397429 | 0 | 253.5623 | 354.339831 | 254.943757 |
| -3.1142201 | 0.00184432 | 0.00397452 | 51.230485 | 73.0649104 | 57.5226998 | 73.4013893 |
| -3.1142068 | 0.0018444 | 0.00397452 | 8.60672148 | 31.3908504 | 57.5226998 | 40.3427483 |
| -3.1140228 | 0.00184555 | 0.00397667 | 8.60672148 | 69.2763595 | 21.8586259 | 62.1950703 |
| 3.114008 | 0.00184564 | 0.00397667 | 4.50828268 | 7.8477126 | 14.9559019 | 2.24126379 |
| -3.1135274 | 0.00184865 | 0.00398288 | 18.0331307 | 51.6866588 | 12.6549939 | 45.9459078 |
| 3.11338155 | 0.00184957 | 0.00398459 | 0 | 0 | 0 | 0 |
| -3.1132193 | 0.00185058 | 0.00398651 | 22.5414134 | 39.5091738 | 42.5667978 | 11.7666349 |
| -3.1131144 | 0.00185124 | 0.0039874 | 145.904421 | 221.359617 | 508.500666 | 0 |
| -3.1131189 | 0.00185121 | 0.0039874 | 32.3776665 | 9.47137727 | 35.6640739 | 49.3078035 |
| -3.1130896 | 0.0018514 | 0.00398747 | 18.0331307 | 12.4480958 | 42.5667978 | 72.8410733 |
| -3.1129027 | 0.00185257 | 0.00398973 | 36.4761053 | 21.3782516 | 105.841768 | 30.2570612 |
| -3.1119488 | 0.00185857 | 0.00400237 | 10.246097 | 46.545054 | 54.0713378 | 22.9729539 |
| -3.1114909 | 0.00186145 | 0.00400832 | 6.55750208 | 31.6614612 | 103.54086 | 44.2649599 |
| -3.1114445 | 0.00186174 | 0.00400868 | 4.91812656 | 50.3336049 | 56.3722458 | 63.3157022 |
| -3.111369 | 0.00186222 | 0.00400944 | 14.3445358 | 35.1794013 | 28.7613499 | 21.292006 |
| -3.1111449 | 0.00186363 | 0.00401221 | 30.738291 | 35.7206228 | 21.8586259 | 34.1792729 |
| -3.1108015 | 0.0018658 | 0.00401661 | 174.183649 | 325.544767 | 325.578481 | 412.392538 |
| 3.11078117 | 0.00186593 | 0.00401662 | 0 | 1.08244312 | 0 | 0 |
| 3.11055923 | 0.00186733 | 0.00401938 | 26.6398522 | 40.8622277 | 40.2658898 | 47.0665397 |
| -3.1104567 | 0.00186798 | 0.00402049 | 166.396615 | 111.22103 | 86.2840497 | 146.242463 |
| -3.1104383 | 0.0018681 | 0.00402049 | 0 | 234.348935 | 426.818432 | 219.083536 |
| -3.1100129 | 0.00187079 | 0.00402601 | 387.302467 | 617.263187 | 564.872912 | 624.191967 |
| -3.1099543 | 0.00187116 | 0.00402654 | 30.738291 | 86.8660601 | 26.4604419 | 10.646003 |
| 3.10982833 | 0.00187196 | 0.00402772 | 10.246097 | 6.76526948 | 6.90272397 | 11.206319 |
| -3.1098477 | 0.00187184 | 0.00402772 | 182.380527 | 384.808528 | 354.339831 | 332.827673 |

| | | | | | | |
|---|---|---|---|---|---|---|
| -3.1098075 | 0.00187209 | 0.00402774 | 7.78703372 | 42.7565031 | 111.594038 | 25.7745336 |
| -3.10894 | 0.0018776 | 0.00403931 | 93.0345607 | 19.7545869 | 18.4072639 | 21.852322 |
| -3.1084134 | 0.00188095 | 0.00404598 | 30.3284471 | 41.67406 | 10.354086 | 31.3776931 |
| -3.1084231 | 0.00188089 | 0.00404598 | 161.888333 | 243.27909 | 253.099879 | 254.383441 |
| -3.1081591 | 0.00188257 | 0.00404919 | 25.8201644 | 22.4606947 | 18.4072639 | 19.0507423 |
| -3.1076752 | 0.00188565 | 0.00405556 | 51.6403289 | 29.4965749 | 11.50454 | 25.7745336 |
| -3.1074367 | 0.00188717 | 0.00405856 | 32.3776665 | 37.6148983 | 31.0622579 | 10.646003 |
| -3.1072495 | 0.00188837 | 0.00406086 | 11.0657848 | 84.1599523 | 73.6290557 | 13.4475828 |
| -3.1070575 | 0.0018896 | 0.00406323 | 23.770945 | 42.4858923 | 46.0181598 | 43.144328 |
| -3.1068928 | 0.00189065 | 0.00406522 | 16.3937552 | 24.8961917 | 23.0090799 | 22.4126379 |
| -3.1068475 | 0.00189094 | 0.00406558 | 14.3445358 | 27.3316887 | 25.3099879 | 25.2142177 |
| -3.1068056 | 0.00189121 | 0.00406588 | 35.2465737 | 66.0290301 | 10.354086 | 24.6539017 |
| -3.1061494 | 0.00189541 | 0.00407437 | 283.202121 | 388.055857 | 357.791193 | 385.497373 |
| -3.1061574 | 0.00189536 | 0.00407437 | 690.177094 | 1990.8835 | 1519.74973 | 1705.60175 |
| -3.1059459 | 0.00189671 | 0.00407691 | 9.01656536 | 32.4732935 | 48.3190678 | 37.5411686 |
| -3.1047537 | 0.00190438 | 0.0040931 | 19.6725062 | 70.6294134 | 72.4786017 | 8.40473923 |
| -3.1042606 | 0.00190755 | 0.00409966 | 8.1968776 | 32.7439043 | 54.0713378 | 41.4633802 |
| -3.1040753 | 0.00190875 | 0.00410195 | 11.0657848 | 23.5431378 | 39.1154358 | 45.9459078 |
| -3.1039352 | 0.00190965 | 0.00410362 | 29.9186032 | 4.05916169 | 94.3372276 | 93.5727634 |
| -3.1036908 | 0.00191123 | 0.00410674 | 14.3445358 | 66.5702517 | 13.8054479 | 50.4284354 |
| -3.1030693 | 0.00191525 | 0.0041151 | 7.78703372 | 52.4984912 | 39.1154358 | 38.1014845 |
| -3.1028717 | 0.00191653 | 0.00411757 | 11.4756286 | 37.3442875 | 24.1595339 | 39.2221164 |
| -3.1026791 | 0.00191777 | 0.00411998 | 9.42640924 | 66.8408624 | 32.2127119 | 30.8173772 |
| -3.102654 | 0.00191794 | 0.00412006 | 11.4756286 | 46.545054 | 25.3099879 | 31.9380091 |
| -3.1026304 | 0.00191809 | 0.00412011 | 23.770945 | 25.708024 | 40.2658898 | 13.4475828 |
| -3.1024049 | 0.00191955 | 0.00412298 | 156.970206 | 189.427545 | 0 | 291.924609 |
| 3.10228361 | 0.00192034 | 0.00412438 | 44.263139 | 31.1202396 | 29.9118039 | 27.4554815 |
| -3.1022644 | 0.00192046 | 0.00412438 | 18.4429746 | 24.0843593 | 17.2568099 | 30.2570612 |
| -3.1020823 | 0.00192165 | 0.00412665 | 151.232392 | 161.825246 | 166.815829 | 163.612257 |
| -3.1019329 | 0.00192262 | 0.00412846 | 400.007627 | 600.214708 | 632.749697 | 655.009344 |
| -3.1019082 | 0.00192278 | 0.00412853 | 79.0998688 | 20.2958084 | 17.2568099 | 22.4126379 |
| -3.1015498 | 0.0019251 | 0.00413325 | 5.32797044 | 60.3462037 | 51.7704298 | 48.1871716 |
| -3.101524 | 0.00192527 | 0.00413334 | 7.78703372 | 16.7778683 | 87.4345036 | 85.7283401 |
| -3.1014852 | 0.00192552 | 0.00413361 | 792.22822 | 868.931212 | 759.299637 | 853.921506 |
| -3.1014187 | 0.00192596 | 0.00413426 | 41.8040758 | 90.1133895 | 109.29313 | 93.0124475 |
| -3.1012936 | 0.00192677 | 0.00413573 | 14.7543797 | 33.8263474 | 50.6199758 | 17.3697944 |
| -3.1011143 | 0.00192794 | 0.00413796 | 13.524848 | 31.3908504 | 20.7081719 | 39.7824324 |
| 3.10101839 | 0.00192856 | 0.00413903 | 0.40984388 | 0.27061078 | 0 | 0 |
| -3.1007835 | 0.00193009 | 0.00414204 | 39.3450125 | 88.4897248 | 18.4072639 | 11.7666349 |
| -3.1005856 | 0.00193138 | 0.00414453 | 15.1642236 | 33.0145151 | 18.4072639 | 32.498325 |
| -3.100328 | 0.00193306 | 0.00414771 | 14.7543797 | 29.2259641 | 20.7081719 | 29.1364293 |
| -3.1003189 | 0.00193312 | 0.00414771 | 22.1315695 | 38.9679522 | 118.496762 | 59.9538065 |
| -3.1002769 | 0.0019334 | 0.00414802 | 0 | 295.777582 | 199.028541 | 354.679995 |

| | | | | | | |
|---|---|---|---|---|---|---|
| -3.1002393 | 0.00193364 | 0.00414827 | 8.60672148 | 39.238563 | 43.7172518 | 35.2999048 |
| -3.0999742 | 0.00193537 | 0.00415171 | 23.770945 | 30.8496288 | 17.2568099 | 18.4904263 |
| -3.0983827 | 0.0019458 | 0.0041738 | 10.6559409 | 50.8748265 | 17.2568099 | 81.8061285 |
| -3.0983006 | 0.00194634 | 0.00417468 | 12.2953164 | 40.3210061 | 23.0090799 | 32.498325 |
| -3.0978806 | 0.0019491 | 0.00418032 | 16.8035991 | 56.8282636 | 13.8054479 | 38.6618005 |
| -3.0976854 | 0.00195038 | 0.0041828 | 107.78894 | 273.858108 | 0 | 282.399238 |
| -3.0976569 | 0.00195057 | 0.00418292 | 14.3445358 | 32.7439043 | 26.4604419 | 21.852322 |
| -3.0975119 | 0.00195153 | 0.00418469 | 11.0657848 | 33.5557366 | 29.9118039 | 32.498325 |
| -3.0973256 | 0.00195275 | 0.00418704 | 7.37718984 | 62.7817008 | 50.6199758 | 29.6967453 |
| -3.0972232 | 0.00195343 | 0.00418794 | 543.862829 | 1085.96106 | 745.494189 | 753.624951 |
| -3.0972349 | 0.00195335 | 0.00418794 | 12.2953164 | 30.8496288 | 27.6108959 | 29.1364293 |
| 3.09705124 | 0.00195456 | 0.00419009 | 0.40984388 | 0.54122156 | 3.45136199 | 1.68094785 |
| -3.096824 | 0.00195606 | 0.00419302 | 15.9839113 | 25.708024 | 20.7081719 | 25.2142177 |
| -3.0964675 | 0.00195841 | 0.00419779 | 11.4756286 | 23.0019162 | 43.7172518 | 36.4205367 |
| -3.0961389 | 0.00196058 | 0.00420216 | 19.2626624 | 20.0251977 | 21.8586259 | 21.292006 |
| 3.09580021 | 0.00196283 | 0.00420648 | 10.246097 | 14.3423713 | 14.9559019 | 16.2491625 |
| -3.0957955 | 0.00196286 | 0.00420648 | 57.7879871 | 24.8961917 | 17.2568099 | 17.9301104 |
| -3.0955463 | 0.00196451 | 0.00420973 | 23.3611012 | 15.6954252 | 20.7081719 | 39.2221164 |
| -3.0954648 | 0.00196505 | 0.00421061 | 11.0657848 | 47.6274971 | 47.1686138 | 20.7316901 |
| -3.0951696 | 0.001967 | 0.00421452 | 6.55750208 | 99.0435452 | 18.4072639 | 82.3664444 |
| -3.0944099 | 0.00197205 | 0.00422505 | 43.0336074 | 33.8263474 | 9.20363196 | 32.498325 |
| -3.0941838 | 0.00197355 | 0.00422799 | 11.0657848 | 28.9553534 | 50.6199758 | 27.4554815 |
| -3.0940367 | 0.00197453 | 0.00422981 | 126.231915 | 344.216911 | 358.941647 | 335.629253 |
| -3.0938981 | 0.00197545 | 0.0042315 | 25.0004767 | 29.2259641 | 24.1595339 | 14.0078987 |
| 3.09344507 | 0.00197847 | 0.00423769 | 0 | 0.54122156 | 0 | 1.1206319 |
| -3.0932685 | 0.00197965 | 0.00423993 | 194.675843 | 97.4198805 | 0 | 417.435382 |
| -3.0929509 | 0.00198177 | 0.00424419 | 12.2953164 | 33.5557366 | 23.0090799 | 35.8602207 |
| -3.0928443 | 0.00198248 | 0.00424543 | 27.0496961 | 71.4412457 | 29.9118039 | 54.910963 |
| 3.09263433 | 0.00198388 | 0.00424815 | 0.40984388 | 0.81183234 | 0 | 0 |
| -3.0922506 | 0.00198645 | 0.00425337 | 11.8854725 | 32.4732935 | 29.9118039 | 27.4554815 |
| -3.0920052 | 0.00198809 | 0.0042566 | 84.8376831 | 323.10927 | 343.985745 | 359.162523 |
| -3.0919502 | 0.00198846 | 0.00425711 | 11.0657848 | 28.9553534 | 31.0622579 | 35.8602207 |
| -3.0918315 | 0.00198926 | 0.00425853 | 30.738291 | 18.6721438 | 19.5577179 | 19.6110582 |
| -3.0917797 | 0.0019896 | 0.00425899 | 161.068645 | 47.8981079 | 23.0090799 | 45.3855918 |
| -3.0916621 | 0.00199039 | 0.0042604 | 208.200691 | 306.060791 | 274.958505 | 336.189569 |
| -3.0911793 | 0.00199363 | 0.00426705 | 138.937075 | 168.861126 | 120.79767 | 98.615607 |
| -3.0911464 | 0.00199385 | 0.00426724 | 7.37718984 | 35.9912336 | 42.5667978 | 53.7903311 |
| -3.0901371 | 0.00200064 | 0.00428148 | 32.3776665 | 23.5431378 | 14.9559019 | 21.292006 |
| -3.0897401 | 0.00200332 | 0.00428692 | 334.84245 | 113.927138 | 0 | 208.437533 |
| -3.0896733 | 0.00200377 | 0.0042876 | 6.96734596 | 43.8389462 | 44.8677058 | 44.2649599 |
| 3.08948803 | 0.00200502 | 0.00428971 | 1.22953164 | 1.62366467 | 2.30090799 | 2.24126379 |
| -3.0894974 | 0.00200495 | 0.00428971 | 20.492194 | 18.9427545 | 20.7081719 | 21.852322 |
| -3.0893537 | 0.00200592 | 0.00429137 | 15.5740674 | 47.3568864 | 86.2840497 | 11.7666349 |

| | | | | | | |
|---|---|---|---|---|---|---|
| -3.089154 | 0.00200727 | 0.00429397 | 18.4429746 | 24.0843593 | 21.8586259 | 19.0507423 |
| -3.0889073 | 0.00200894 | 0.00429725 | 24.1807889 | 16.7778683 | 19.5577179 | 25.2142177 |
| -3.0887415 | 0.00201006 | 0.00429936 | 6.55750208 | 31.9320719 | 48.3190678 | 73.4013893 |
| -3.0885493 | 0.00201136 | 0.00430137 | 17.213443 | 33.0145151 | 24.1595339 | 17.9301104 |
| -3.0885435 | 0.0020114 | 0.00430137 | 29.5087594 | 18.1309222 | 25.3099879 | 16.8094785 |
| -3.0885597 | 0.00201129 | 0.00430137 | 15.9839113 | 19.7545869 | 24.1595339 | 29.6967453 |
| -3.0884366 | 0.00201213 | 0.00430235 | 17.213443 | 18.9427545 | 24.1595339 | 25.2142177 |
| -3.0884378 | 0.00201212 | 0.00430235 | 17.6232868 | 13.2599282 | 54.0713378 | 38.1014845 |
| -3.0878919 | 0.00201582 | 0.00430996 | 9.42640924 | 41.9446708 | 31.0622579 | 36.4205367 |
| -3.0876288 | 0.0020176 | 0.00431349 | 11.0657848 | 27.8729103 | 29.9118039 | 40.3427483 |
| -3.0867718 | 0.00202343 | 0.00432566 | 282.792277 | 417.281821 | 381.950726 | 447.132127 |
| -3.0867358 | 0.00202367 | 0.0043259 | 39.7548564 | 20.83703 | 32.2127119 | 12.8872668 |
| -3.0865923 | 0.00202465 | 0.0043277 | 102.870814 | 261.410013 | 0 | 290.243661 |
| -3.0860822 | 0.00202813 | 0.00433484 | 25.8201644 | 35.1794013 | 11.50454 | 29.6967453 |
| -3.0859217 | 0.00202922 | 0.0043369 | 122.133476 | 265.739785 | 0 | 243.177122 |
| -3.0857705 | 0.00203025 | 0.00433882 | 25.0004767 | 36.803066 | 12.6549939 | 25.2142177 |
| -3.0855556 | 0.00203172 | 0.00434138 | 76.2309617 | 37.3442875 | 8.05317797 | 28.5761134 |
| -3.0855561 | 0.00203172 | 0.00434138 | 8.60672148 | 44.9213893 | 51.7704298 | 27.4554815 |
| -3.0852624 | 0.00203373 | 0.00434537 | 19.6725062 | 43.0271139 | 12.6549939 | 34.7395888 |
| -3.0852242 | 0.00203399 | 0.00434564 | 8.60672148 | 36.803066 | 35.6640739 | 42.5840121 |
| -3.0848917 | 0.00203626 | 0.00435022 | 14.3445358 | 29.2259641 | 19.5577179 | 34.1792729 |
| -3.0848576 | 0.0020365 | 0.00435043 | 21.3118818 | 33.2851258 | 47.1686138 | 29.1364293 |
| -3.0848301 | 0.00203668 | 0.00435054 | 4.0984388 | 44.3801678 | 83.9831417 | 57.7125427 |
| -3.0847367 | 0.00203732 | 0.00435162 | 226.643666 | 166.155018 | 173.718553 | 153.52657 |
| -3.084619 | 0.00203813 | 0.00435305 | 26.6398522 | 11.9068743 | 32.2127119 | 30.2570612 |
| -3.0845367 | 0.00203869 | 0.00435397 | 6.55750208 | 76.5828505 | 92.0363196 | 19.6110582 |
| 3.08422183 | 0.00204085 | 0.00435829 | 158.199738 | 27.3316887 | 39.1154358 | 39.2221164 |
| 3.08381436 | 0.00204365 | 0.00436398 | 0 | 0 | 0 | 0 |
| -3.0837661 | 0.00204398 | 0.0043644 | 13.524848 | 27.8729103 | 21.8586259 | 33.6189569 |
| -3.083624 | 0.00204496 | 0.00436619 | 14.3445358 | 28.6847426 | 25.3099879 | 22.4126379 |
| -3.0836001 | 0.00204512 | 0.00436626 | 147.543797 | 21.9194731 | 9.20363196 | 31.9380091 |
| -3.0833958 | 0.00204653 | 0.00436897 | 9.01656536 | 75.5004074 | 14.9559019 | 83.4870763 |
| -3.0829811 | 0.00204938 | 0.00437477 | 16.8035991 | 24.0843593 | 44.8677058 | 17.9301104 |
| -3.0829393 | 0.00204967 | 0.00437509 | 11.4756286 | 48.1687187 | 92.0363196 | 15.1285306 |
| -3.0827104 | 0.00205125 | 0.00437817 | 58.197831 | 21.9194731 | 12.6549939 | 28.0157974 |
| -3.0816401 | 0.00205864 | 0.00439365 | 13.9346919 | 21.6488623 | 39.1154358 | 25.2142177 |
| -3.0813014 | 0.00206098 | 0.00439836 | 8.1968776 | 49.2511618 | 18.4072639 | 122.148877 |
| -3.0812229 | 0.00206152 | 0.00439923 | 12.7051603 | 35.1794013 | 23.0090799 | 29.1364293 |
| -3.0809702 | 0.00206327 | 0.00440268 | 4.0984388 | 41.67406 | 81.6822337 | 62.1950703 |
| -3.0806032 | 0.00206582 | 0.00440781 | 19.6725062 | 25.4374132 | 31.0622579 | 15.1285306 |
| -3.0804058 | 0.00206719 | 0.00441045 | 0 | 233.537102 | 243.896247 | 328.905462 |
| 3.08034173 | 0.00206763 | 0.0044111 | 18.8528185 | 39.5091738 | 29.9118039 | 28.0157974 |
| -3.0803023 | 0.00206791 | 0.0044114 | 132.789417 | 479.25169 | 347.437107 | 581.047639 |

| | | | | | | |
|---|---|---|---|---|---|---|
| -3.0801879 | 0.0020687 | 0.0044128 | 70.0833035 | 337.181031 | 0 | 305.932508 |
| -3.0793225 | 0.00207472 | 0.00442535 | 34.017042 | 40.3210061 | 74.7795097 | 6.72379138 |
| -3.0790458 | 0.00207665 | 0.00442917 | 18.0331307 | 18.1309222 | 24.1595339 | 23.5332698 |
| 3.07897361 | 0.00207715 | 0.00442995 | 7.78703372 | 9.74198805 | 2.30090799 | 6.16347543 |
| -3.0787513 | 0.0020787 | 0.00443296 | 221.725539 | 679.503666 | 482.040224 | 543.50647 |
| -3.0785918 | 0.00207981 | 0.00443504 | 150.412704 | 18.6721438 | 18.4072639 | 17.9301104 |
| -3.0781813 | 0.00208268 | 0.00444086 | 8.1968776 | 59.8049822 | 27.6108959 | 43.144328 |
| -3.0768922 | 0.00209171 | 0.00445982 | 10.246097 | 28.9553534 | 37.9649818 | 33.058641 |
| -3.0757579 | 0.00209968 | 0.00447652 | 8.60672148 | 43.0271139 | 33.3631659 | 37.5411686 |
| -3.0755024 | 0.00210148 | 0.00448006 | 10.246097 | 26.5198564 | 39.1154358 | 36.4205367 |
| -3.0752065 | 0.00210357 | 0.00448421 | 11.8854725 | 33.2851258 | 37.9649818 | 22.4126379 |
| -3.0751502 | 0.00210397 | 0.00448454 | 27.0496961 | 20.83703 | 16.1063559 | 21.292006 |
| -3.0751457 | 0.002104 | 0.00448454 | 41.3942319 | 14.6129821 | 25.3099879 | 20.1713742 |
| -3.0748643 | 0.00210598 | 0.00448847 | 20.9020379 | 25.9786348 | 13.8054479 | 33.6189569 |
| -3.0746168 | 0.00210773 | 0.0044919 | 5.73781432 | 37.6148983 | 72.4786017 | 45.9459078 |
| -3.0744271 | 0.00210907 | 0.00449446 | 16.8035991 | 23.8137486 | 55.2217918 | 17.9301104 |
| 3.07409535 | 0.00211142 | 0.00449917 | 6.96734596 | 6.4946587 | 4.60181598 | 6.72379138 |
| -3.0738317 | 0.00211329 | 0.00450285 | 19.6725062 | 196.192815 | 232.391707 | 143.440883 |
| -3.0735398 | 0.00211535 | 0.00450696 | 32.3776665 | 18.9427545 | 16.1063559 | 22.9729539 |
| -3.0735142 | 0.00211554 | 0.00450705 | 3.68859492 | 63.8641439 | 39.1154358 | 96.9346591 |
| -3.0734868 | 0.00211573 | 0.00450716 | 19.2626624 | 7.03588026 | 106.992222 | 59.9538065 |
| -3.0733292 | 0.00211685 | 0.00450925 | 13.524848 | 58.9931499 | 12.6549939 | 62.1950703 |
| -3.0732747 | 0.00211723 | 0.00450977 | 259.84102 | 52.2278804 | 0 | 475.147924 |
| -3.0732028 | 0.00211775 | 0.00451056 | 13.524848 | 24.8961917 | 24.1595339 | 29.1364293 |
| -3.0727701 | 0.00212082 | 0.00451681 | 12.7051603 | 27.3316887 | 27.6108959 | 26.3348496 |
| -3.0726445 | 0.00212171 | 0.00451841 | 162.70802 | 299.295522 | 274.958505 | 284.640502 |
| -3.0726234 | 0.00212186 | 0.00451844 | 184.019902 | 396.174181 | 375.048002 | 377.652949 |
| -3.0724343 | 0.00212321 | 0.004521 | 235.250387 | 1005.58966 | 733.989649 | 897.065834 |
| -3.0723746 | 0.00212363 | 0.00452161 | 0 | 264.657342 | 201.329449 | 333.387989 |
| -3.0723225 | 0.002124 | 0.0045221 | 20.492194 | 35.9912336 | 26.4604419 | 14.0078987 |
| -3.0721087 | 0.00212552 | 0.00452504 | 13.9346919 | 20.5664192 | 48.3190678 | 25.7745336 |
| -3.0719049 | 0.00212697 | 0.00452783 | 6.1476582 | 46.0038325 | 55.2217918 | 40.3427483 |
| -3.0717685 | 0.00212795 | 0.0045296 | 101.231438 | 71.9824673 | 75.9299637 | 72.2807574 |
| -3.0716544 | 0.00212876 | 0.00453104 | 54.0993922 | 16.2366467 | 14.9559019 | 31.9380091 |
| -3.0711665 | 0.00213224 | 0.00453815 | 12.2953164 | 31.1202396 | 23.0090799 | 33.058641 |
| -3.0710047 | 0.0021334 | 0.00454031 | 7.78703372 | 27.8729103 | 52.9208838 | 49.3078035 |
| -3.0709266 | 0.00213396 | 0.0045412 | 83.1983076 | 24.6255809 | 17.2568099 | 16.8094785 |
| -3.0709063 | 0.0021341 | 0.00454121 | 120.903945 | 162.095857 | 148.408565 | 169.215416 |
| -3.0704611 | 0.00213729 | 0.00454768 | 31.5579788 | 22.7313054 | 14.9559019 | 20.7316901 |
| -3.0703681 | 0.00213795 | 0.0045488 | 9.42640924 | 25.9786348 | 50.6199758 | 37.5411686 |
| -3.069812 | 0.00214194 | 0.00455698 | 286.071028 | 220.547785 | 0 | 113.744138 |
| -3.0692438 | 0.00214601 | 0.00456535 | 12.7051603 | 33.0145151 | 21.8586259 | 30.8173772 |
| -3.0689047 | 0.00214845 | 0.00457024 | 36.0662614 | 122.586683 | 127.700393 | 81.2458125 |

| | | | | | | |
|---|---|---|---|---|---|---|
| -3.0686649 | 0.00215018 | 0.0045736 | 282.382433 | 517.949031 | 557.970188 | 558.074685 |
| -3.0684392 | 0.0021518 | 0.00457676 | 15.9839113 | 22.4606947 | 23.0090799 | 21.852322 |
| 3.06807915 | 0.0021544 | 0.00458146 | 4.50828268 | 1.08244312 | 2.30090799 | 3.36189569 |
| -3.0680734 | 0.00215444 | 0.00458146 | 18.8528185 | 39.5091738 | 13.8054479 | 29.1364293 |
| -3.0680833 | 0.00215437 | 0.00458146 | 108.608628 | 17.5897006 | 14.9559019 | 25.7745336 |
| -3.0679316 | 0.00215546 | 0.00458333 | 124.182696 | 261.139402 | 0 | 219.643852 |
| -3.0667289 | 0.00216415 | 0.00460151 | 27.8693838 | 19.2133653 | 26.4604419 | 15.1285306 |
| -3.0665237 | 0.00216564 | 0.00460437 | 11.0657848 | 32.2026827 | 25.3099879 | 36.4205367 |
| -3.0664879 | 0.0021659 | 0.00460461 | 15.9839113 | 19.4839761 | 25.3099879 | 24.0935858 |
| 3.06607528 | 0.00216889 | 0.00461067 | 49.5911095 | 27.8729103 | 33.3631659 | 24.6539017 |
| -3.0658785 | 0.00217031 | 0.0046134 | 12.7051603 | 29.2259641 | 24.1595339 | 28.0157974 |
| -3.0656158 | 0.00217222 | 0.00461715 | 16.8035991 | 40.3210061 | 19.5577179 | 20.7316901 |
| -3.0655086 | 0.002173 | 0.0046185 | 26.6398522 | 58.4519283 | 40.2658898 | 32.498325 |
| -3.0650877 | 0.00217606 | 0.0046247 | 9.01656536 | 32.7439043 | 35.6640739 | 38.6618005 |
| -3.0648982 | 0.00217744 | 0.00462733 | 41.8040758 | 41.9446708 | 12.6549939 | 17.3697944 |
| -3.0647441 | 0.00217856 | 0.00462941 | 6.1476582 | 40.3210061 | 56.3722458 | 43.704644 |
| -3.0645428 | 0.00218003 | 0.00463222 | 674.603026 | 974.740026 | 1097.53311 | 1132.95885 |
| -3.0645128 | 0.00218025 | 0.00463238 | 2.45906328 | 63.5935331 | 66.7263317 | 85.1680242 |
| -3.0643178 | 0.00218167 | 0.0046351 | 105.329877 | 170.484791 | 197.878087 | 179.301104 |
| -3.0637273 | 0.00218598 | 0.00464395 | 11.8854725 | 42.7565031 | 24.1595339 | 27.4554815 |
| -3.0633814 | 0.00218851 | 0.00464902 | 21.3118818 | 48.4393295 | 11.50454 | 31.9380091 |
| -3.0632497 | 0.00218947 | 0.00465076 | 6.96734596 | 47.0862756 | 37.9649818 | 43.144328 |
| -3.0631782 | 0.00219 | 0.00465156 | 9.83625312 | 37.8855091 | 36.8145279 | 27.4554815 |
| -3.0620569 | 0.00219822 | 0.00466872 | 9.01656536 | 30.0377965 | 49.4695218 | 33.058641 |
| -3.0619743 | 0.00219882 | 0.0046697 | 62.7061136 | 21.9194731 | 19.5577179 | 16.2491625 |
| -3.0618151 | 0.00219999 | 0.00467187 | 35.6564176 | 23.0019162 | 35.6640739 | 11.206319 |
| -3.0615704 | 0.00220179 | 0.00467539 | 20.0823501 | 57.9107067 | 52.9208838 | 68.9188617 |
| -3.0609501 | 0.00220636 | 0.00468478 | 376.646526 | 38.6973414 | 20.7081719 | 2.24126379 |
| -3.060892 | 0.00220679 | 0.00468538 | 25.0004767 | 63.8641439 | 6.90272397 | 56.0315949 |
| 3.06074303 | 0.00220788 | 0.0046874 | 3.27875104 | 1.08244312 | 1.150454 | 1.68094785 |
| -3.0607063 | 0.00220816 | 0.00468767 | 13.1150042 | 31.9320719 | 19.5577179 | 35.2999048 |
| 3.06049064 | 0.00220975 | 0.00469074 | 0 | 0 | 0 | 0 |
| -3.0603329 | 0.00221091 | 0.0046929 | 434.024669 | 2282.33131 | 1252.8444 | 1585.69413 |
| -3.0602285 | 0.00221168 | 0.00469423 | 81.5589321 | 427.835642 | 368.145279 | 232.531119 |
| -3.0599692 | 0.0022136 | 0.00469798 | 122.54332 | 193.757318 | 0 | 285.761134 |
| 3.05972998 | 0.00221537 | 0.00470143 | 0.40984388 | 0.27061078 | 0 | 0 |
| -3.059305 | 0.00221851 | 0.00470748 | 9.42640924 | 54.6633774 | 29.9118039 | 29.6967453 |
| -3.0593192 | 0.00221841 | 0.00470748 | 777.063996 | 1933.51402 | 1493.28929 | 1669.18121 |
| -3.0589123 | 0.00222142 | 0.00471335 | 32.3776665 | 18.9427545 | 33.3631659 | 100.296555 |
| -3.0587129 | 0.0022229 | 0.00471587 | 21.3118818 | 21.3782516 | 16.1063559 | 22.9729539 |
| 3.05873085 | 0.00222277 | 0.00471587 | 154.920987 | 67.382084 | 57.5226998 | 80.6854966 |
| -3.0586887 | 0.00222308 | 0.00471594 | 8.60672148 | 54.1221558 | 16.1063559 | 104.218766 |
| -3.0585384 | 0.0022242 | 0.004718 | 760.670241 | 1202.32369 | 1203.37488 | 1177.22381 |

| | | | | | | |
|---|---|---|---|---|---|---|
| -3.0584277 | 0.00222502 | 0.00471943 | 11.0657848 | 27.6022995 | 27.6108959 | 36.9808526 |
| -3.0582925 | 0.00222602 | 0.00472125 | 14.7543797 | 34.0969582 | 44.8677058 | 15.6888466 |
| -3.0577255 | 0.00223024 | 0.00472988 | 15.1642236 | 30.8496288 | 82.8326877 | 15.1285306 |
| -3.0571415 | 0.00223459 | 0.00473879 | 523.780479 | 1011.54309 | 970.983172 | 1026.49882 |
| -3.056573 | 0.00223883 | 0.00474748 | 19.6725062 | 10.8244312 | 36.8145279 | 69.4791776 |
| -3.0565352 | 0.00223911 | 0.00474776 | 13.524848 | 27.3316887 | 24.1595339 | 24.0935858 |
| -3.0563366 | 0.0022406 | 0.0047506 | 0 | 205.122971 | 320.976665 | 245.978701 |
| -3.0561992 | 0.00224162 | 0.00475246 | 17.213443 | 28.143521 | 16.1063559 | 26.3348496 |
| 3.05588866 | 0.00224395 | 0.00475708 | 4.50828268 | 9.74198805 | 5.75226998 | 11.7666349 |
| -3.0558646 | 0.00224413 | 0.00475715 | 15.9839113 | 47.6274971 | 31.0622579 | 15.1285306 |
| -3.0551362 | 0.00224958 | 0.0047684 | 23.3611012 | 41.4034492 | 37.9649818 | 45.9459078 |
| -3.0547732 | 0.00225231 | 0.00477386 | 303.284471 | 870.825487 | 783.459171 | 860.645297 |
| -3.0546792 | 0.00225301 | 0.00477505 | 11.4756286 | 38.1561199 | 24.1595339 | 29.1364293 |
| -3.0544479 | 0.00225475 | 0.00477842 | 14.7543797 | 29.7671857 | 20.7081719 | 22.9729539 |
| -3.0543354 | 0.0022556 | 0.00477957 | 772.14587 | 45.1920001 | 0 | 131.674248 |
| -3.0543477 | 0.00225551 | 0.00477957 | 31.1481349 | 22.1900839 | 16.1063559 | 18.4904263 |
| -3.0543162 | 0.00225574 | 0.00477957 | 13.1150042 | 24.6255809 | 25.3099879 | 26.8951655 |
| -3.0538893 | 0.00225895 | 0.00478607 | 11.4756286 | 23.8137486 | 35.6640739 | 30.2570612 |
| -3.0538355 | 0.00225936 | 0.00478661 | 64.3454891 | 16.5072575 | 17.2568099 | 24.0935858 |
| 3.0536402 | 0.00226083 | 0.00478941 | 7.37718984 | 8.11832337 | 5.75226998 | 11.7666349 |
| -3.0535762 | 0.00226131 | 0.00479012 | 12.2953164 | 22.7313054 | 37.9649818 | 26.8951655 |
| -3.0530298 | 0.00226543 | 0.00479854 | 103.280658 | 5.1416048 | 24.1595339 | 63.3157022 |
| -3.0529669 | 0.00226591 | 0.00479923 | 93.4444046 | 18.9427545 | 11.50454 | 31.3776931 |
| -3.0528038 | 0.00226714 | 0.00480152 | 13.524848 | 35.1794013 | 29.9118039 | 19.0507423 |
| -3.0526389 | 0.00226839 | 0.00480369 | 67.6242402 | 17.5897006 | 19.5577179 | 19.6110582 |
| -3.0526287 | 0.00226846 | 0.00480369 | 15.9839113 | 14.3423713 | 39.1154358 | 36.9808526 |
| -3.0523616 | 0.00227048 | 0.00480765 | 218.036944 | 470.862756 | 400.35799 | 416.875066 |
| -3.0521612 | 0.002272 | 0.00481055 | 11.4756286 | 24.0843593 | 33.3631659 | 30.8173772 |
| -3.0519882 | 0.00227331 | 0.00481301 | 12.7051603 | 33.0145151 | 24.1595339 | 24.6539017 |
| -3.0510895 | 0.00228013 | 0.00482712 | 10.6559409 | 36.5324552 | 41.4163438 | 22.4126379 |
| -3.0506307 | 0.00228361 | 0.00483419 | 47.1320462 | 26.2492456 | 13.8054479 | 19.0507423 |
| -3.0504462 | 0.00228502 | 0.00483684 | 10.6559409 | 39.238563 | 21.8586259 | 40.3427483 |
| 3.05034677 | 0.00228577 | 0.00483812 | 63.5258014 | 113.656527 | 144.957203 | 131.674248 |
| -3.0501812 | 0.00228703 | 0.00484048 | 17.6232868 | 37.0736767 | 12.6549939 | 40.3427483 |
| -3.049756 | 0.00229027 | 0.00484702 | 18.0331307 | 23.5431378 | 24.1595339 | 16.8094785 |
| -3.0496666 | 0.00229096 | 0.00484814 | 72.1325229 | 16.7778683 | 13.8054479 | 30.2570612 |
| -3.0493043 | 0.00229372 | 0.00485335 | 20.0823501 | 20.2958084 | 26.4604419 | 16.2491625 |
| -3.0493213 | 0.00229359 | 0.00485335 | 3.68859492 | 61.6992576 | 46.0181598 | 71.7204414 |
| -3.0485931 | 0.00229916 | 0.00486454 | 738.128828 | 184.827162 | 111.594038 | 144.001199 |
| -3.048482 | 0.00230001 | 0.00486583 | 0 | 195.110372 | 318.675757 | 248.219965 |
| -3.0484543 | 0.00230022 | 0.00486583 | 32.7875104 | 35.4500121 | 63.2749697 | 7.28410733 |
| 3.04847283 | 0.00230008 | 0.00486583 | 27.45954 | 11.3656527 | 0 | 0 |
| 3.0483813 | 0.00230078 | 0.00486669 | 53.6895483 | 42.4858923 | 27.6108959 | 33.058641 |

| | | | | | | |
|---|---|---|---|---|---|---|
| -3.047755 | 0.00230558 | 0.00487653 | 45.9025146 | 21.9194731 | 10.354086 | 35.2999048 |
| -3.0476828 | 0.00230613 | 0.00487738 | 63.1159575 | 15.6954252 | 21.8586259 | 19.6110582 |
| -3.0471879 | 0.00230993 | 0.0048851 | 6.96734596 | 187.53327 | 56.3722458 | 11.7666349 |
| -3.0460087 | 0.00231901 | 0.00490398 | 10.6559409 | 35.7206228 | 26.4604419 | 30.2570612 |
| -3.0458626 | 0.00232014 | 0.00490604 | 275.005243 | 546.633774 | 395.756174 | 526.136676 |
| -3.0456448 | 0.00232182 | 0.00490927 | 35.2465737 | 31.6614612 | 13.8054479 | 17.3697944 |
| -3.0451802 | 0.00232541 | 0.00491654 | 15.5740674 | 34.9087905 | 16.1063559 | 28.5761134 |
| -3.0451212 | 0.00232587 | 0.00491718 | 11.0657848 | 37.8855091 | 24.1595339 | 30.2570612 |
| -3.0443054 | 0.00233218 | 0.00493022 | 14.7543797 | 51.9572696 | 33.3631659 | 15.1285306 |
| -3.0442594 | 0.00233254 | 0.00493065 | 1.63937552 | 48.980551 | 109.29313 | 95.8140272 |
| -3.044021 | 0.00233439 | 0.00493391 | 8.1968776 | 35.4500121 | 34.5136199 | 40.9030642 |
| -3.044025 | 0.00233436 | 0.00493391 | 152.461923 | 204.040527 | 0 | 201.713742 |
| -3.0438762 | 0.00233551 | 0.00493596 | 40.5745441 | 28.6847426 | 20.7081719 | 12.3269509 |
| -3.043852 | 0.0023357 | 0.00493599 | 42.2139196 | 15.966036 | 19.5577179 | 20.7316901 |
| -3.0438352 | 0.00233583 | 0.00493599 | 326.235728 | 180.226779 | 0 | 104.218766 |
| -3.043754 | 0.00233646 | 0.004937 | 11.8854725 | 80.1007906 | 39.1154358 | 15.6888466 |
| -3.0437341 | 0.00233662 | 0.004937 | 16.8035991 | 20.0251977 | 21.8586259 | 21.292006 |
| -3.0434667 | 0.00233869 | 0.00494107 | 13.1150042 | 21.6488623 | 26.4604419 | 29.6967453 |
| -3.0432485 | 0.00234039 | 0.00494433 | 27.8693838 | 59.2637606 | 50.6199758 | 6.72379138 |
| -3.0430306 | 0.00234209 | 0.00494759 | 0 | 235.160767 | 220.887167 | 290.243661 |
| -3.0430052 | 0.00234228 | 0.00494768 | 13.1150042 | 29.2259641 | 23.0090799 | 24.6539017 |
| 3.0429644 | 0.0023426 | 0.00494803 | 19.2626624 | 10.5538204 | 21.8586259 | 18.4904263 |
| -3.0428347 | 0.00234361 | 0.00494984 | 43.4434513 | 17.3190899 | 27.6108959 | 14.5682147 |
| -3.0421271 | 0.00234913 | 0.00496116 | 31.9678226 | 18.1309222 | 21.8586259 | 16.2491625 |
| -3.0420314 | 0.00234987 | 0.00496242 | 6.55750208 | 38.1561199 | 41.4163438 | 49.3078035 |
| -3.0418347 | 0.00235141 | 0.00496533 | 12.7051603 | 46.2744432 | 16.1063559 | 40.9030642 |
| -3.0418102 | 0.0023516 | 0.00496541 | 12.7051603 | 23.0019162 | 35.6640739 | 24.0935858 |
| -3.0415521 | 0.00235362 | 0.00496935 | 15.1642236 | 54.1221558 | 112.744492 | 61.6347543 |
| -3.0414637 | 0.00235431 | 0.00497048 | 11.4756286 | 38.1561199 | 18.4072639 | 48.7474875 |
| -3.041407 | 0.00235475 | 0.00497109 | 16.8035991 | 31.1202396 | 14.9559019 | 29.1364293 |
| -3.0409783 | 0.00235811 | 0.00497753 | 18.0331307 | 21.9194731 | 16.1063559 | 42.5840121 |
| -3.0409807 | 0.00235809 | 0.00497753 | 16.3937552 | 11.9068743 | 29.9118039 | 129.432984 |
| -3.0403065 | 0.00236337 | 0.00498832 | 243.037421 | 249.503138 | 167.966283 | 225.807327 |
| -3.0401122 | 0.0023649 | 0.00499121 | 27.8693838 | 21.6488623 | 14.9559019 | 20.1713742 |
| -3.0396611 | 0.00236844 | 0.00499836 | 66.8045524 | 287.659258 | 0 | 299.208717 |
| 3.0394167 | 0.00237037 | 0.0050021 | 52.4600166 | 34.6381797 | 50.6199758 | 33.6189569 |
| -3.0393539 | 0.00237086 | 0.00500281 | 14.3445358 | 47.0862756 | 14.9559019 | 35.2999048 |
| -3.0391947 | 0.00237212 | 0.00500513 | 10.246097 | 31.9320719 | 31.0622579 | 29.1364293 |
| -3.0388056 | 0.00237518 | 0.00501109 | 13.1150042 | 26.5198564 | 132.302209 | 16.2491625 |
| 3.03879685 | 0.00237525 | 0.00501109 | 159.019425 | 84.9717847 | 78.2308717 | 80.6854966 |
| -3.0386843 | 0.00237614 | 0.00501263 | 13.1150042 | 21.3782516 | 25.3099879 | 32.498325 |
| -3.0384647 | 0.00237787 | 0.00501596 | 18.8528185 | 33.0145151 | 13.8054479 | 26.3348496 |
| -3.0382247 | 0.00237976 | 0.00501963 | 25.0004767 | 17.0484791 | 20.7081719 | 18.4904263 |

| | | | | | | |
|---|---|---|---|---|---|---|
| -3.0378882 | 0.00238242 | 0.00502466 | 43.0336074 | 8.93015571 | 32.2127119 | 30.8173772 |
| -3.0378834 | 0.00238246 | 0.00502466 | 6.1476582 | 68.4645271 | 28.7613499 | 49.8681194 |
| -3.0378599 | 0.00238265 | 0.00502472 | 18.0331307 | 27.0610779 | 16.1063559 | 22.4126379 |
| -3.037596 | 0.00238473 | 0.0050288 | 9.42640924 | 36.2618444 | 35.6640739 | 28.0157974 |
| -3.0375428 | 0.00238516 | 0.00502936 | 10.6559409 | 38.6973414 | 24.1595339 | 31.3776931 |
| -3.0370682 | 0.00238891 | 0.00503695 | 4.50828268 | 44.109557 | 55.2217918 | 58.2728587 |
| -3.0369769 | 0.00238964 | 0.00503815 | 23.3611012 | 107.161869 | 36.8145279 | 64.4363341 |
| -3.0366967 | 0.00239186 | 0.00504251 | 60.6568942 | 30.3084073 | 35.6640739 | 29.1364293 |
| -3.0365389 | 0.00239311 | 0.00504482 | 0 | 145.047378 | 360.092101 | 266.710392 |
| -3.0362715 | 0.00239524 | 0.00504884 | 17.213443 | 35.9912336 | 14.9559019 | 25.7745336 |
| -3.0362592 | 0.00239533 | 0.00504884 | 2.86890716 | 53.0397127 | 82.8326877 | 59.9538065 |
| -3.0359052 | 0.00239815 | 0.00505444 | 26.6398522 | 17.0484791 | 63.2749697 | 14.5682147 |
| -3.0357004 | 0.00239978 | 0.00505755 | 10.6559409 | 30.0377965 | 24.1595339 | 41.4633802 |
| -3.0355782 | 0.00240075 | 0.00505927 | 39.7548564 | 19.7545869 | 14.9559019 | 21.292006 |
| -3.0354492 | 0.00240178 | 0.0050611 | 22.1315695 | 57.0988744 | 17.2568099 | 16.8094785 |
| -3.0353065 | 0.00240291 | 0.00506316 | 9.42640924 | 50.6042157 | 24.1595339 | 35.2999048 |
| -3.0349058 | 0.00240611 | 0.00506957 | 9.42640924 | 29.2259641 | 37.9649818 | 31.9380091 |
| -3.0348456 | 0.00240659 | 0.00507025 | 37.705637 | 18.6721438 | 17.2568099 | 19.0507423 |
| -3.0347452 | 0.00240739 | 0.0050716 | 59.4273626 | 23.272527 | 13.8054479 | 20.1713742 |
| -3.0345912 | 0.00240862 | 0.00507386 | 21.3118818 | 31.1202396 | 44.8677058 | 26.8951655 |
| -3.0345617 | 0.00240886 | 0.00507403 | 13.1150042 | 37.6148983 | 17.2568099 | 34.7395888 |
| -3.034467 | 0.00240961 | 0.00507529 | 72.1325229 | 26.7904671 | 16.1063559 | 15.1285306 |
| -3.0341815 | 0.00241189 | 0.00507976 | 8.60672148 | 32.7439043 | 37.9649818 | 34.1792729 |
| -3.0341599 | 0.00241207 | 0.0050798 | 110.248004 | 203.228695 | 0 | 262.227864 |
| -3.0336808 | 0.0024159 | 0.00508754 | 8.60672148 | 29.4965749 | 35.6640739 | 42.0236961 |
| -3.0332167 | 0.00241962 | 0.00509504 | 131.559885 | 204.311138 | 0 | 219.083536 |
| -3.0328976 | 0.00242218 | 0.0051001 | 22.9512573 | 11.0950419 | 42.5667978 | 29.1364293 |
| -3.0325641 | 0.00242486 | 0.0051054 | 8.60672148 | 19.4839761 | 85.1335957 | 42.5840121 |
| -3.0325104 | 0.00242529 | 0.00510598 | 17.213443 | 20.5664192 | 28.7613499 | 17.3697944 |
| -3.0324583 | 0.00242571 | 0.00510653 | 10.6559409 | 31.9320719 | 29.9118039 | 26.8951655 |
| -3.0318042 | 0.00243097 | 0.00511727 | 7.37718984 | 33.5557366 | 40.2658898 | 43.704644 |
| -3.0315664 | 0.00243288 | 0.00512097 | 9.83625312 | 35.7206228 | 24.1595339 | 40.3427483 |
| -3.0307659 | 0.00243934 | 0.00513423 | 13.1150042 | 34.367569 | 23.0090799 | 22.4126379 |
| -3.0306884 | 0.00243997 | 0.00513521 | 9.83625312 | 37.0736767 | 21.8586259 | 49.3078035 |
| -3.0306671 | 0.00244014 | 0.00513524 | 120.494101 | 15.6954252 | 8.05317797 | 47.6268556 |
| 3.03064525 | 0.00244032 | 0.00513528 | 3.68859492 | 7.30649104 | 4.60181598 | 3.36189569 |
| -3.0304388 | 0.00244199 | 0.00513845 | 12.7051603 | 25.9786348 | 26.4604419 | 23.5332698 |
| -3.0299446 | 0.00244599 | 0.00514653 | 343.039328 | 379.666923 | 383.10118 | 427.521069 |
| -3.0293078 | 0.00245115 | 0.00515706 | 13.524848 | 20.2958084 | 27.6108959 | 26.3348496 |
| -3.0292536 | 0.00245159 | 0.00515765 | 71.722679 | 33.2851258 | 60.9740618 | 4.48252759 |
| -3.0291451 | 0.00245247 | 0.00515916 | 8.60672148 | 31.1202396 | 35.6640739 | 37.5411686 |
| -3.0290571 | 0.00245318 | 0.00516033 | 6.96734596 | 18.1309222 | 173.718553 | 35.8602207 |
| -3.0280953 | 0.002461 | 0.00517645 | 27.45954 | 54.3927666 | 5.75226998 | 65.556966 |

| | | | | | | |
|---|---|---|---|---|---|---|
| -3.0280378 | 0.00246147 | 0.00517709 | 1.22953164 | 55.2045989 | 112.744492 | 100.296555 |
| -3.0280155 | 0.00246165 | 0.00517714 | 16.8035991 | 38.4267306 | 27.6108959 | 14.5682147 |
| -3.0275323 | 0.00246559 | 0.00518509 | 27.0496961 | 21.1076408 | 12.6549939 | 28.0157974 |
| -3.0273184 | 0.00246734 | 0.00518842 | 5.73781432 | 33.2851258 | 40.2658898 | 79.0045488 |
| -3.0270991 | 0.00246913 | 0.00519163 | 20.0823501 | 45.1920001 | 46.0181598 | 49.8681194 |
| -3.0270922 | 0.00246919 | 0.00519163 | 14.3445358 | 65.4878086 | 74.7795097 | 8.96505518 |
| -3.0265587 | 0.00247355 | 0.00520046 | 29.5087594 | 36.5324552 | 9.20363196 | 30.2570612 |
| -3.0262351 | 0.0024762 | 0.00520569 | 57.7879871 | 75.7710182 | 5.75226998 | 24.0935858 |
| -3.0262044 | 0.00247645 | 0.00520588 | 13.524848 | 54.6633774 | 29.9118039 | 16.8094785 |
| -3.02583 | 0.00247952 | 0.00521199 | 18.4429746 | 16.2366467 | 41.4163438 | 20.1713742 |
| 3.02541376 | 0.00248293 | 0.00521883 | 6.96734596 | 3.24732935 | 3.45136199 | 2.24126379 |
| -3.0250359 | 0.00248604 | 0.00522501 | 341.809796 | 834.022421 | 600.536986 | 728.410733 |
| -3.0249171 | 0.00248701 | 0.00522673 | 52.4600166 | 166.425629 | 0 | 532.860467 |
| -3.0244848 | 0.00249057 | 0.00523386 | 17.6232868 | 52.7691019 | 11.50454 | 35.2999048 |
| -3.0244257 | 0.00249106 | 0.00523454 | 12.7051603 | 22.4606947 | 23.0090799 | 40.9030642 |
| -3.02374 | 0.00249671 | 0.00524574 | 22.5414134 | 17.8603114 | 24.1595339 | 16.2491625 |
| -3.0237451 | 0.00249667 | 0.00524574 | 13.1150042 | 26.2492456 | 23.0090799 | 24.0935858 |
| -3.0234732 | 0.00249891 | 0.00525002 | 79.5097127 | 47.3568864 | 179.470823 | 100.856871 |
| -3.023362 | 0.00249983 | 0.00525161 | 68.8537718 | 75.7710182 | 82.8326877 | 66.6775979 |
| -3.0232675 | 0.00250061 | 0.00525291 | 13.524848 | 25.708024 | 27.6108959 | 20.1713742 |
| -3.0232322 | 0.0025009 | 0.00525318 | 17.213443 | 15.1542036 | 35.6640739 | 24.0935858 |
| -3.0230911 | 0.00250207 | 0.00525529 | 8.60672148 | 45.1920001 | 31.0622579 | 30.8173772 |
| -3.0229489 | 0.00250324 | 0.00525742 | 65.1651769 | 53.851545 | 36.8145279 | 117.666349 |
| -3.0228584 | 0.00250399 | 0.00525865 | 6.55750208 | 49.7923834 | 112.744492 | 19.0507423 |
| -3.0227859 | 0.00250459 | 0.00525956 | 9.01656536 | 31.9320719 | 33.3631659 | 33.6189569 |
| 3.02248225 | 0.00250711 | 0.0052645 | 1.63937552 | 2.43549701 | 1.150454 | 2.24126379 |
| -3.0224426 | 0.00250744 | 0.00526485 | 13.524848 | 28.143521 | 19.5577179 | 26.8951655 |
| -3.0218834 | 0.00251207 | 0.00527424 | 140.166607 | 829.422038 | 633.900151 | 936.28795 |
| -3.0218169 | 0.00251263 | 0.00527506 | 26.2300083 | 25.1668025 | 13.8054479 | 19.6110582 |
| -3.0213266 | 0.0025167 | 0.00528326 | 29.5087594 | 23.8137486 | 20.7081719 | 13.4475828 |
| -3.0209451 | 0.00251987 | 0.00528958 | 33.6071982 | 17.8603114 | 14.9559019 | 22.9729539 |
| -3.020712 | 0.00252181 | 0.00529331 | 12.2953164 | 30.8496288 | 19.5577179 | 33.6189569 |
| -3.0204765 | 0.00252377 | 0.00529708 | 146.724109 | 0 | 347.437107 | 336.749885 |
| -3.0202665 | 0.00252552 | 0.00530042 | 28.6890716 | 35.1794013 | 29.9118039 | 33.6189569 |
| -3.0201067 | 0.00252686 | 0.00530287 | 9.01656536 | 26.5198564 | 42.5667978 | 34.1792729 |
| -3.0197111 | 0.00253016 | 0.00530945 | 12.2953164 | 27.8729103 | 19.5577179 | 44.2649599 |
| -3.0196446 | 0.00253071 | 0.00531027 | 14.7543797 | 32.7439043 | 14.9559019 | 36.4205367 |
| -3.0196019 | 0.00253107 | 0.00531068 | 20.9020379 | 14.8835929 | 19.5577179 | 37.5411686 |
| 3.01944333 | 0.0025324 | 0.00531311 | 0.81968776 | 1.62366467 | 1.150454 | 2.24126379 |
| -3.0191534 | 0.00253482 | 0.00531785 | 11.8854725 | 38.4267306 | 16.1063559 | 53.2300151 |
| -3.0187313 | 0.00253836 | 0.00532492 | 9.01656536 | 24.0843593 | 50.6199758 | 35.2999048 |
| -3.0186721 | 0.00253885 | 0.00532562 | 18.4429746 | 14.3423713 | 90.8858656 | 22.4126379 |
| -3.0182827 | 0.00254212 | 0.00533212 | 22.5414134 | 27.3316887 | 31.0622579 | 11.7666349 |

| | | | | | | |
|---|---|---|---|---|---|---|
| -3.0182446 | 0.00254244 | 0.00533244 | 49.5911095 | 21.3782516 | 13.8054479 | 20.7316901 |
| 3.01738072 | 0.00254969 | 0.00534732 | 1.63937552 | 2.16488623 | 2.30090799 | 2.80157974 |
| -3.0171223 | 0.00255187 | 0.00535153 | 137.707544 | 14.0717605 | 17.2568099 | 20.7316901 |
| -3.0164226 | 0.00255776 | 0.00536355 | 39.7548564 | 38.6973414 | 31.0622579 | 44.8252759 |
| -3.0162392 | 0.00255931 | 0.00536645 | 20.492194 | 24.3549701 | 13.8054479 | 24.0935858 |
| -3.0160032 | 0.00256131 | 0.00537028 | 22.9512573 | 24.8961917 | 11.50454 | 36.4205367 |
| -3.015875 | 0.00256239 | 0.0053722 | 11.8854725 | 29.2259641 | 51.7704298 | 19.0507423 |
| -3.0155719 | 0.00256495 | 0.00537722 | 34.4268859 | 24.0843593 | 19.5577179 | 13.4475828 |
| -3.0153627 | 0.00256672 | 0.00538058 | 11.0657848 | 33.8263474 | 39.1154358 | 20.1713742 |
| -3.0151838 | 0.00256824 | 0.00538341 | 9.42640924 | 34.367569 | 28.7613499 | 31.9380091 |
| -3.015089 | 0.00256904 | 0.00538474 | 4.0984388 | 52.2278804 | 66.7263317 | 42.5840121 |
| 3.01458134 | 0.00257334 | 0.00539341 | 0.40984388 | 0.54122156 | 0 | 0 |
| -3.0143692 | 0.00257514 | 0.00539683 | 99.1822189 | 161.554635 | 0 | 322.18167 |
| -3.0140217 | 0.00257809 | 0.00540267 | 8.1968776 | 37.3442875 | 44.8677058 | 27.4554815 |
| -3.0139273 | 0.0025789 | 0.005404 | 24.1807889 | 52.2278804 | 49.4695218 | 47.6268556 |
| -3.0138624 | 0.00257945 | 0.0054048 | 53.6895483 | 46.545054 | 19.5577179 | 42.0236961 |
| 3.01381911 | 0.00257982 | 0.00540522 | 0 | 0 | 0 | 0 |
| -3.0137316 | 0.00258056 | 0.00540643 | 74.1817423 | 93.090108 | 90.8858656 | 103.098135 |
| -3.0133848 | 0.00258351 | 0.00541226 | 11.8854725 | 44.109557 | 17.2568099 | 35.8602207 |
| -3.012938 | 0.00258732 | 0.00541954 | 15.1642236 | 41.1328384 | 13.8054479 | 33.6189569 |
| -3.0129465 | 0.00258725 | 0.00541954 | 224.184602 | 294.695138 | 311.773033 | 289.123029 |
| -3.0124878 | 0.00259116 | 0.00542723 | 6.96734596 | 31.9320719 | 48.3190678 | 39.7824324 |
| -3.0124252 | 0.00259169 | 0.005428 | 11.4756286 | 18.9427545 | 57.5226998 | 30.2570612 |
| -3.0123768 | 0.00259211 | 0.00542816 | 10.246097 | 32.2026827 | 28.7613499 | 27.4554815 |
| -3.0123828 | 0.00259205 | 0.00542816 | 21.7217256 | 20.83703 | 13.8054479 | 26.8951655 |
| 3.01227629 | 0.00259296 | 0.0054296 | 16.3937552 | 13.8011497 | 14.9559019 | 15.1285306 |
| -3.0117324 | 0.00259762 | 0.00543899 | 33.6071982 | 21.6488623 | 16.1063559 | 16.8094785 |
| -3.01166 | 0.00259823 | 0.00543993 | 72.1325229 | 36.5324552 | 24.1595339 | 34.7395888 |
| -3.0114675 | 0.00259988 | 0.00544303 | 127.461447 | 0 | 123.098578 | 765.391586 |
| -3.0114389 | 0.00260013 | 0.00544319 | 28.2792277 | 10.8244312 | 29.9118039 | 26.3348496 |
| 3.01124732 | 0.00260177 | 0.00544627 | 114.346443 | 135.034779 | 112.744492 | 97.4949751 |
| -3.0107898 | 0.00260569 | 0.00545413 | 11.4756286 | 33.8263474 | 19.5577179 | 35.8602207 |
| -3.0105314 | 0.00260791 | 0.00545842 | 276.234775 | 126.916455 | 0 | 146.802779 |
| -3.0105084 | 0.00260811 | 0.00545848 | 17.213443 | 18.1309222 | 23.0090799 | 19.6110582 |
| -3.0100392 | 0.00261214 | 0.00546618 | 20.492194 | 23.272527 | 20.7081719 | 15.1285306 |
| -3.0100489 | 0.00261206 | 0.00546618 | 0 | 197.545869 | 317.525303 | 195.550266 |
| -3.0100209 | 0.0026123 | 0.00546618 | 0 | 214.323737 | 248.498063 | 234.212067 |
| -3.0090708 | 0.00262048 | 0.00548295 | 88.5262781 | 238.137486 | 0 | 238.694594 |
| -3.009014 | 0.00262097 | 0.00548362 | 62.2962697 | 56.8282636 | 11.50454 | 12.3269509 |
| -3.0086787 | 0.00262386 | 0.00548932 | 9.01656536 | 26.2492456 | 50.6199758 | 30.2570612 |
| -3.008571 | 0.00262479 | 0.00549091 | 16.3937552 | 25.708024 | 18.4072639 | 19.6110582 |
| 3.00819485 | 0.00262805 | 0.00549736 | 1.22953164 | 5.1416048 | 1.150454 | 1.68094785 |
| -3.0079057 | 0.00263055 | 0.00550223 | 16.8035991 | 20.2958084 | 21.8586259 | 18.4904263 |

| | | | | | | |
|---|---|---|---|---|---|---|
| -3.0078101 | 0.00263138 | 0.00550361 | 13.9346919 | 21.3782516 | 23.0090799 | 23.5332698 |
| -3.0076999 | 0.00263233 | 0.00550513 | 16.8035991 | 22.4606947 | 21.8586259 | 17.3697944 |
| 3.00768669 | 0.00263244 | 0.00550513 | 377.466213 | 332.310037 | 311.773033 | 333.387989 |
| -3.007522 | 0.00263387 | 0.00550776 | 46.7222023 | 13.8011497 | 41.4163438 | 13.4475828 |
| 3.00750148 | 0.00263405 | 0.00550777 | 0 | 0 | 0 | 0 |
| -3.0070883 | 0.00263763 | 0.00551491 | 9.42640924 | 32.4732935 | 26.4604419 | 36.4205367 |
| -3.0066499 | 0.00264144 | 0.00552237 | 12.7051603 | 25.708024 | 34.5136199 | 19.6110582 |
| -3.0066378 | 0.00264154 | 0.00552237 | 122.54332 | 21.1076408 | 10.354086 | 23.5332698 |
| -3.0065705 | 0.00264213 | 0.00552324 | 13.1150042 | 31.3908504 | 18.4072639 | 28.0157974 |
| -3.0063359 | 0.00264417 | 0.00552714 | 15.9839113 | 33.2851258 | 12.6549939 | 45.9459078 |
| -3.0061946 | 0.0026454 | 0.00552935 | 438.532952 | 1549.24671 | 970.983172 | 994.000493 |
| 3.00615402 | 0.00264575 | 0.00552973 | 2.0492194 | 1.89427545 | 2.30090799 | 2.24126379 |
| -3.0058109 | 0.00264874 | 0.00553562 | 11.4756286 | 24.3549701 | 66.7263317 | 21.852322 |
| -3.0057636 | 0.00264915 | 0.00553612 | 16.3937552 | 21.1076408 | 19.5577179 | 20.1713742 |
| 3.00562562 | 0.00265035 | 0.00553792 | 10.6559409 | 4.87099402 | 8.05317797 | 11.206319 |
| -3.0056323 | 0.00265029 | 0.00553792 | 24.5906328 | 24.3549701 | 11.50454 | 28.5761134 |
| -3.0055524 | 0.00265099 | 0.00553853 | 20.9020379 | 21.3782516 | 16.1063559 | 19.0507423 |
| -3.0055571 | 0.00265095 | 0.00553853 | 309.841973 | 425.941366 | 381.950726 | 449.933707 |
| -3.0052814 | 0.00265335 | 0.00554311 | 15.1642236 | 15.1542036 | 29.9118039 | 31.9380091 |
| -3.0051975 | 0.00265408 | 0.00554428 | 22.1315695 | 20.5664192 | 13.8054479 | 25.2142177 |
| -3.0049183 | 0.00265652 | 0.00554902 | 23.3611012 | 27.8729103 | 10.354086 | 38.1014845 |
| -3.004726 | 0.0026582 | 0.00555217 | 9.83625312 | 25.1668025 | 28.7613499 | 42.5840121 |
| -3.0046174 | 0.00265915 | 0.00555379 | 16.3937552 | 98.2317128 | 47.1686138 | 56.0315949 |
| -3.0045469 | 0.00265977 | 0.00555472 | 33.1973543 | 33.5557366 | 9.20363196 | 26.3348496 |
| -3.0043109 | 0.00266183 | 0.00555831 | 38.5253247 | 18.6721438 | 11.50454 | 31.9380091 |
| -3.0043272 | 0.00266169 | 0.00555831 | 11.0657848 | 27.3316887 | 24.1595339 | 31.3776931 |
| -3.0038394 | 0.00266596 | 0.00556657 | 19.2626624 | 15.966036 | 21.8586259 | 20.7316901 |
| -3.0037434 | 0.0026668 | 0.00556796 | 0 | 224.877557 | 379.649818 | 132.234564 |
| -3.0036347 | 0.00266775 | 0.00556959 | 13.1150042 | 30.0377965 | 33.3631659 | 17.3697944 |
| -3.0033492 | 0.00267026 | 0.00557374 | 9.01656536 | 35.1794013 | 35.6640739 | 26.8951655 |
| -3.0033709 | 0.00267007 | 0.00557374 | 9.42640924 | 28.9553534 | 54.0713378 | 24.6539017 |
| 3.00336812 | 0.00267009 | 0.00557374 | 8.1968776 | 5.68282636 | 4.60181598 | 3.92221164 |
| -3.0032711 | 0.00267094 | 0.00557481 | 20.492194 | 34.9087905 | 21.8586259 | 13.4475828 |
| -3.0027548 | 0.00267548 | 0.00558392 | 9.01656536 | 29.4965749 | 31.0622579 | 35.8602207 |
| -3.0023945 | 0.00267865 | 0.00559017 | 15.5740674 | 23.272527 | 17.2568099 | 24.6539017 |
| 3.00221247 | 0.00268025 | 0.00559315 | 0 | 0.54122156 | 1.150454 | 0 |
| -3.0021085 | 0.00268117 | 0.0055947 | 12.7051603 | 50.6042157 | 21.8586259 | 21.852322 |
| -3.0018281 | 0.00268364 | 0.00559949 | 13.524848 | 44.9213893 | 20.7081719 | 21.292006 |
| -3.0011939 | 0.00268923 | 0.00561081 | 89.3459658 | 109.056144 | 144.957203 | 130.553616 |
| -3.0010302 | 0.00269068 | 0.00561346 | 70.0833035 | 94.1725511 | 81.6822337 | 80.1251807 |
| -3.0001056 | 0.00269886 | 0.00563017 | 15.5740674 | 26.7904671 | 19.5577179 | 19.0507423 |
| -2.9999927 | 0.00269986 | 0.00563153 | 14.7543797 | 29.7671857 | 19.5577179 | 20.1713742 |
| -2.9999943 | 0.00269985 | 0.00563153 | 6.1476582 | 26.7904671 | 78.2308717 | 41.4633802 |

| | | | | | | |
|---|---|---|---|---|---|---|
| -2.9998592 | 0.00270104 | 0.00563327 | 7.78703372 | 41.1328384 | 35.6640739 | 30.8173772 |
| -2.9998628 | 0.00270101 | 0.00563327 | 11.8854725 | 24.8961917 | 23.0090799 | 29.6967453 |
| 2.99977904 | 0.00270176 | 0.00563439 | 11.0657848 | 5.95343714 | 0 | 2.24126379 |
| 2.99963269 | 0.00270305 | 0.00563673 | 0 | 24.3549701 | 28.7613499 | 31.3776931 |
| -2.9994742 | 0.00270446 | 0.0056393 | 11.8854725 | 24.3549701 | 42.5667978 | 21.292006 |
| -2.9991067 | 0.00270772 | 0.00564574 | 18.0331307 | 18.1309222 | 18.4072639 | 21.852322 |
| -2.9990807 | 0.00270796 | 0.00564586 | 7.37718984 | 47.3568864 | 42.5667978 | 26.8951655 |
| -2.998756 | 0.00271084 | 0.00565151 | 10.246097 | 53.851545 | 19.5577179 | 33.6189569 |
| 2.99835711 | 0.00271439 | 0.00565855 | 0 | 0 | 0 | 0 |
| -2.9976751 | 0.00272047 | 0.00567086 | 0 | 212.429462 | 363.543463 | 142.880567 |
| -2.9973092 | 0.00272374 | 0.00567731 | 12.2953164 | 24.6255809 | 23.0090799 | 26.3348496 |
| -2.997222 | 0.00272452 | 0.0056782 | 21.3118818 | 23.8137486 | 17.2568099 | 16.2491625 |
| 2.99723221 | 0.00272443 | 0.0056782 | 0.81968776 | 1.08244312 | 1.150454 | 0 |
| -2.9967792 | 0.00272848 | 0.00568608 | 50.8206411 | 14.6129821 | 19.5577179 | 20.1713742 |
| -2.9965932 | 0.00273015 | 0.00568919 | 9.83625312 | 25.1668025 | 31.0622579 | 34.1792729 |
| -2.9963856 | 0.00273201 | 0.0056927 | 0 | 288.471091 | 392.304812 | 89.6505518 |
| 2.99636352 | 0.00273221 | 0.00569274 | 5.32797044 | 4.05916169 | 1.150454 | 5.04284354 |
| 2.99627754 | 0.00273298 | 0.00569373 | 4.50828268 | 2.97671857 | 3.45136199 | 2.80157974 |
| 2.99627105 | 0.00273303 | 0.00569373 | 22.9512573 | 15.4248144 | 10.354086 | 8.96505518 |
| 2.99596873 | 0.00273574 | 0.00569901 | 17.6232868 | 5.95343714 | 12.6549939 | 9.52537113 |
| -2.9955449 | 0.00273955 | 0.00570657 | 9.01656536 | 24.8961917 | 35.6640739 | 38.1014845 |
| -2.9952048 | 0.00274261 | 0.00571257 | 5.73781432 | 44.3801678 | 31.0622579 | 61.0744384 |
| -2.9951157 | 0.00274341 | 0.00571387 | 23.770945 | 25.1668025 | 42.5667978 | 10.646003 |
| -2.9945381 | 0.00274861 | 0.00572433 | 90.1656536 | 310.931785 | 0 | 164.172573 |
| -2.9942769 | 0.00275096 | 0.00572886 | 109.83816 | 242.737869 | 0 | 176.499524 |
| -2.9935836 | 0.00275722 | 0.00574153 | 8.1968776 | 31.6614612 | 39.1154358 | 31.3776931 |
| -2.9935266 | 0.00275773 | 0.00574223 | 10.6559409 | 66.2996409 | 20.7081719 | 27.4554815 |
| -2.9933795 | 0.00275906 | 0.00574463 | 709.8496 | 2659.83335 | 1870.6382 | 2075.97059 |
| -2.9932384 | 0.00276034 | 0.00574691 | 17.213443 | 31.9320719 | 12.6549939 | 31.3776931 |
| -2.9929217 | 0.00276321 | 0.00575251 | 165.167084 | 216.759234 | 182.922185 | 216.842272 |
| -2.9921174 | 0.0027705 | 0.00576694 | 29.5087594 | 22.7313054 | 32.2127119 | 10.646003 |
| -2.9921348 | 0.00277034 | 0.00576694 | 159.429269 | 305.79018 | 304.870309 | 316.018195 |
| -2.9920721 | 0.00277091 | 0.00576743 | 9.83625312 | 28.143521 | 28.7613499 | 30.8173772 |
| -2.9917664 | 0.00277368 | 0.00577283 | 24.5906328 | 14.0717605 | 20.7081719 | 21.292006 |
| 2.99115164 | 0.00277927 | 0.00578409 | 3.27875104 | 4.60038325 | 3.45136199 | 7.28410733 |
| -2.9909844 | 0.0027808 | 0.00578689 | 11.8854725 | 24.8961917 | 20.7081719 | 39.2221164 |
| -2.9907528 | 0.00278291 | 0.00579091 | 8.1968776 | 51.416048 | 23.0090799 | 39.2221164 |
| -2.9906273 | 0.00278405 | 0.00579291 | 45.9025146 | 38.9679522 | 6.90272397 | 29.1364293 |
| -2.9902204 | 0.00278776 | 0.00580026 | 5.73781432 | 17.3190899 | 95.4876816 | 63.8760181 |
| -2.9899718 | 0.00279003 | 0.00580461 | 19.2626624 | 33.5557366 | 11.50454 | 29.6967453 |
| -2.9899477 | 0.00279025 | 0.0058047 | 20.9020379 | 21.6488623 | 12.6549939 | 35.2999048 |
| 2.98990709 | 0.00279062 | 0.00580509 | 54.509236 | 23.5431378 | 34.5136199 | 24.6539017 |
| -2.9895768 | 0.00279364 | 0.00581091 | 94.6739363 | 15.966036 | 18.4072639 | 17.9301104 |

| | | | | | | |
|---|---|---|---|---|---|---|
| -2.9895616 | 0.00279378 | 0.00581091 | 6.96734596 | 46.545054 | 39.1154358 | 30.2570612 |
| -2.9894358 | 0.00279493 | 0.00581293 | 6.96734596 | 57.3694852 | 27.6108959 | 38.1014845 |
| -2.9893401 | 0.00279581 | 0.005814 | 18.0331307 | 28.143521 | 29.9118039 | 12.8872668 |
| -2.9893494 | 0.00279572 | 0.005814 | 472.14015 | 1015.33164 | 1221.78214 | 1113.34779 |
| -2.9890351 | 0.0027986 | 0.00581944 | 11.0657848 | 27.0610779 | 54.0713378 | 20.1713742 |
| -2.9888753 | 0.00280006 | 0.00582211 | 14.3445358 | 17.5897006 | 33.3631659 | 22.4126379 |
| -2.9888461 | 0.00280033 | 0.00582229 | 29.9186032 | 10.2832096 | 21.8586259 | 42.0236961 |
| -2.9886021 | 0.00280257 | 0.00582656 | 14.7543797 | 29.2259641 | 14.9559019 | 31.3776931 |
| -2.9882237 | 0.00280604 | 0.00583341 | 13.9346919 | 55.2045989 | 11.50454 | 43.704644 |
| -2.9881823 | 0.00280642 | 0.00583382 | 10.6559409 | 48.980551 | 20.7081719 | 28.5761134 |
| -2.988127 | 0.00280693 | 0.0058345 | 5.32797044 | 34.9087905 | 46.0181598 | 54.350647 |
| -2.9873633 | 0.00281395 | 0.00584872 | 771.736026 | 151.542036 | 134.603117 | 227.488275 |
| -2.9872986 | 0.00281455 | 0.00584923 | 9.83625312 | 24.3549701 | 32.2127119 | 32.498325 |
| 2.98729722 | 0.00281456 | 0.00584923 | 0 | 0.27061078 | 0 | 0 |
| 2.9871652 | 0.00281578 | 0.00585138 | 4.0984388 | 5.68282636 | 8.05317797 | 7.28410733 |
| -2.9870558 | 0.00281678 | 0.0058531 | 0 | 243.549701 | 174.869007 | 254.383441 |
| -2.9863225 | 0.00282355 | 0.00586678 | 268.857585 | 123.669126 | 247.347609 | 0 |
| -2.9859201 | 0.00282726 | 0.00587412 | 52.0501728 | 60.075593 | 132.302209 | 114.864769 |
| -2.9857901 | 0.00282847 | 0.00587625 | 98.3625312 | 194.56915 | 399.207536 | 0 |
| 2.98553104 | 0.00283086 | 0.00588085 | 0 | 0 | 0 | 0 |
| -2.9852328 | 0.00283363 | 0.00588582 | 18.8528185 | 21.1076408 | 13.8054479 | 41.4633802 |
| -2.985236 | 0.0028336 | 0.00588582 | 10.246097 | 27.6022995 | 31.0622579 | 25.7745336 |
| -2.9851262 | 0.00283461 | 0.0058875 | 12.7051603 | 18.1309222 | 46.0181598 | 25.2142177 |
| -2.9840893 | 0.00284424 | 0.00590711 | 31.1481349 | 28.9553534 | 8.05317797 | 39.7824324 |
| -2.9836189 | 0.00284861 | 0.00591581 | 114.346443 | 221.359617 | 0 | 175.939208 |
| 2.98339657 | 0.00285068 | 0.00591973 | 31.5579788 | 13.8011497 | 11.50454 | 14.5682147 |
| -2.9829079 | 0.00285524 | 0.00592881 | 104.920033 | 139.905773 | 0 | 295.846821 |
| -2.9827282 | 0.00285692 | 0.00593191 | 20.0823501 | 36.2618444 | 54.0713378 | 44.8252759 |
| -2.9825038 | 0.00285901 | 0.00593588 | 33.6071982 | 15.1542036 | 24.1595339 | 15.6888466 |
| 2.98244942 | 0.00285952 | 0.00593655 | 0 | 0.27061078 | 1.150454 | 0 |
| -2.9823557 | 0.00286039 | 0.00593799 | 2.0492194 | 79.5595691 | 59.8236078 | 62.1950703 |
| 2.98220851 | 0.00286177 | 0.00594046 | 52.8698605 | 28.9553534 | 19.5577179 | 19.6110582 |
| 2.98192106 | 0.00286446 | 0.00594566 | 23.770945 | 11.3656527 | 16.1063559 | 11.7666349 |
| -2.9810697 | 0.00287243 | 0.00596183 | 61.8864259 | 14.0717605 | 16.1063559 | 24.6539017 |
| -2.9797515 | 0.00288482 | 0.00598716 | 28.6890716 | 18.9427545 | 17.2568099 | 16.2491625 |
| 2.97945611 | 0.00288761 | 0.00599255 | 5.32797044 | 4.60038325 | 2.30090799 | 1.68094785 |
| -2.978831 | 0.0028935 | 0.00600441 | 0 | 282.247043 | 467.084322 | 62.1950703 |
| 2.97880718 | 0.00289373 | 0.00600449 | 50.0009534 | 13.2599282 | 17.2568099 | 15.6888466 |
| -2.9785506 | 0.00289615 | 0.00600913 | 141.396139 | 13.2599282 | 13.8054479 | 22.9729539 |
| -2.9782611 | 0.00289889 | 0.00601442 | 94.2640924 | 229.477941 | 0 | 198.351846 |
| -2.9777517 | 0.00290371 | 0.00602404 | 31.5579788 | 24.3549701 | 35.6640739 | 40.9030642 |
| -2.9775103 | 0.002906 | 0.0060284 | 54.509236 | 16.7778683 | 17.2568099 | 18.4904263 |
| -2.9774327 | 0.00290673 | 0.00602954 | 8.1968776 | 27.3316887 | 48.3190678 | 30.2570612 |

| | | | | | | |
|---|---|---|---|---|---|---|
| 2.97733814 | 0.00290763 | 0.00603101 | 1.63937552 | 1.08244312 | 1.150454 | 2.80157974 |
| -2.9770061 | 0.00291078 | 0.00603716 | 13.524848 | 24.6255809 | 31.0622579 | 17.3697944 |
| -2.976512 | 0.00291548 | 0.0060465 | 10.246097 | 56.5576528 | 26.4604419 | 21.852322 |
| -2.9763098 | 0.0029174 | 0.00605011 | 140.986295 | 146.671042 | 0 | 207.877217 |
| -2.9758583 | 0.0029217 | 0.00605863 | 25.0004767 | 16.7778683 | 16.1063559 | 20.1713742 |
| 2.97550746 | 0.00292504 | 0.00606479 | 1.63937552 | 1.62366467 | 1.150454 | 2.80157974 |
| -2.9755082 | 0.00292504 | 0.00606479 | 16.8035991 | 43.8389462 | 11.50454 | 31.9380091 |
| -2.9754713 | 0.00292539 | 0.00606511 | 121.723632 | 290.635977 | 0 | 118.226665 |
| -2.9751548 | 0.00292841 | 0.00607098 | 271.316649 | 84.9717847 | 0 | 179.86142 |
| -2.9750561 | 0.00292935 | 0.00607251 | 14.3445358 | 25.4374132 | 20.7081719 | 19.0507423 |
| -2.9750385 | 0.00292952 | 0.00607251 | 12.7051603 | 31.1202396 | 17.2568099 | 29.1364293 |
| 2.974458 | 0.00293507 | 0.00608362 | 0.81968776 | 0.54122156 | 2.30090799 | 2.24126379 |
| -2.9743516 | 0.00293608 | 0.00608534 | 8.1968776 | 41.67406 | 24.1595339 | 38.6618005 |
| -2.9741364 | 0.00293814 | 0.00608921 | 45.4926707 | 25.708024 | 13.8054479 | 15.6888466 |
| -2.9739537 | 0.00293989 | 0.00609245 | 15.5740674 | 27.3316887 | 13.8054479 | 33.6189569 |
| -2.9732703 | 0.00294645 | 0.00610564 | 43.0336074 | 18.6721438 | 16.1063559 | 17.3697944 |
| -2.9732488 | 0.00294666 | 0.00610568 | 5.73781432 | 47.8981079 | 70.1776937 | 24.6539017 |
| -2.9731105 | 0.00294798 | 0.00610804 | 12.7051603 | 48.7099402 | 95.4876816 | 8.96505518 |
| -2.9730105 | 0.00294894 | 0.00610925 | 23.770945 | 24.3549701 | 73.6290557 | 9.52537113 |
| -2.9730102 | 0.00294895 | 0.00610925 | 13.524848 | 24.0843593 | 27.6108959 | 17.9301104 |
| -2.9728658 | 0.00295033 | 0.00611173 | 14.3445358 | 30.3084073 | 13.8054479 | 40.9030642 |
| -2.9727832 | 0.00295113 | 0.00611298 | 18.0331307 | 17.8603114 | 54.0713378 | 16.2491625 |
| -2.9726225 | 0.00295267 | 0.00611579 | 17.213443 | 16.2366467 | 28.7613499 | 18.4904263 |
| -2.9723353 | 0.00295544 | 0.00612113 | 0 | 384.808528 | 113.894946 | 209.558165 |
| -2.9721842 | 0.00295689 | 0.00612374 | 258.611488 | 138.282108 | 0 | 115.985401 |
| -2.9721069 | 0.00295764 | 0.00612489 | 0 | 302.27224 | 196.727633 | 164.732889 |
| -2.9720186 | 0.00295849 | 0.00612626 | 10.6559409 | 27.0610779 | 28.7613499 | 24.0935858 |
| -2.971973 | 0.00295893 | 0.00612678 | 0 | 264.927953 | 112.744492 | 314.337247 |
| -2.9717555 | 0.00296102 | 0.00613073 | 243.037421 | 108.785533 | 0 | 156.888466 |
| -2.9717048 | 0.00296151 | 0.00613135 | 13.9346919 | 20.0251977 | 19.5577179 | 28.5761134 |
| -2.971552 | 0.00296299 | 0.00613361 | 182.79037 | 264.927953 | 0 | 84.0473923 |
| -2.9715562 | 0.00296295 | 0.00613361 | 68.4439279 | 16.2366467 | 20.7081719 | 15.6888466 |
| -2.9702793 | 0.00297529 | 0.00615869 | 177.4624 | 123.669126 | 334.782113 | 0 |
| -2.970166 | 0.00297639 | 0.00616057 | 6.55750208 | 75.7710182 | 21.8586259 | 43.144328 |
| -2.9700029 | 0.00297797 | 0.00616344 | 31.9678226 | 34.6381797 | 18.4072639 | 11.206319 |
| 2.96988265 | 0.00297914 | 0.00616546 | 78.2801811 | 188.345102 | 227.789891 | 162.491625 |
| -2.9698157 | 0.00297978 | 0.00616641 | 24.5906328 | 35.1794013 | 13.8054479 | 16.2491625 |
| -2.9694846 | 0.002983 | 0.00617238 | 14.3445358 | 36.2618444 | 13.8054479 | 31.3776931 |
| -2.9694791 | 0.00298305 | 0.00617238 | 0 | 209.182132 | 205.931265 | 229.169223 |
| -2.9692933 | 0.00298486 | 0.00617571 | 23.3611012 | 25.4374132 | 16.1063559 | 15.1285306 |
| 2.96926129 | 0.00298517 | 0.00617596 | 18.0331307 | 6.22404792 | 12.6549939 | 12.8872668 |
| -2.9688103 | 0.00298955 | 0.00618464 | 7.37718984 | 39.238563 | 32.2127119 | 33.6189569 |
| -2.9687481 | 0.00299016 | 0.00618549 | 37.705637 | 37.3442875 | 10.354086 | 17.9301104 |

| | | | | | | |
|---|---|---|---|---|---|---|
| 2.96871027 | 0.00299052 | 0.00618586 | 2.45906328 | 2.16488623 | 0 | 3.36189569 |
| -2.9683981 | 0.00299356 | 0.00619175 | 7.78703372 | 70.9000241 | 109.29313 | 9.52537113 |
| -2.9681465 | 0.00299601 | 0.00619642 | 29.0989155 | 20.83703 | 18.4072639 | 14.0078987 |
| -2.9677732 | 0.00299966 | 0.00620355 | 13.9346919 | 20.83703 | 24.1595339 | 19.6110582 |
| -2.9669184 | 0.00300801 | 0.00622043 | 7.78703372 | 51.9572696 | 28.7613499 | 29.1364293 |
| -2.9666571 | 0.00301056 | 0.00622532 | 11.0657848 | 33.2851258 | 25.3099879 | 21.852322 |
| -2.9658049 | 0.00301892 | 0.00624207 | 0 | 233.807713 | 208.232173 | 198.351846 |
| -2.9657915 | 0.00301905 | 0.00624207 | 22.5414134 | 95.7962158 | 80.5317797 | 68.9188617 |
| -2.9655618 | 0.00302131 | 0.00624633 | 13.524848 | 26.5198564 | 17.2568099 | 25.7745336 |
| -2.9652034 | 0.00302483 | 0.00625321 | 97.5428434 | 23.0019162 | 9.20363196 | 22.9729539 |
| -2.9642821 | 0.0030339 | 0.00627156 | 25.4103206 | 21.3782516 | 14.9559019 | 16.8094785 |
| -2.9642312 | 0.0030344 | 0.0062722 | 9.83625312 | 12.1774851 | 62.1245158 | 61.0744384 |
| -2.9639885 | 0.0030368 | 0.00627594 | 15.5740674 | 17.3190899 | 19.5577179 | 25.2142177 |
| -2.9639925 | 0.00303676 | 0.00627594 | 137.2977 | 151.542036 | 0 | 193.309002 |
| -2.9640092 | 0.00303659 | 0.00627594 | 727.472887 | 651.630756 | 340.534383 | 482.992348 |
| -2.9639496 | 0.00303718 | 0.00627633 | 13.524848 | 23.8137486 | 28.7613499 | 17.3697944 |
| -2.96385 | 0.00303816 | 0.00627756 | 22.1315695 | 24.3549701 | 10.354086 | 44.8252759 |
| -2.9638661 | 0.003038 | 0.00627756 | 22.9512573 | 74.688575 | 52.9208838 | 28.0157974 |
| -2.9632454 | 0.00304414 | 0.0062895 | 14.7543797 | 30.8496288 | 16.1063559 | 22.4126379 |
| 2.96300719 | 0.00304649 | 0.00629397 | 0 | 0.27061078 | 0 | 0 |
| -2.9628741 | 0.00304781 | 0.00629629 | 9.42640924 | 42.7565031 | 18.4072639 | 41.4633802 |
| 2.96278059 | 0.00304874 | 0.0062978 | 25.8201644 | 18.401533 | 23.0090799 | 17.9301104 |
| 2.9625551 | 0.00305097 | 0.00630201 | 2.86890716 | 0.81183234 | 2.30090799 | 3.36189569 |
| 2.96201274 | 0.00305635 | 0.00631272 | 3.68859492 | 13.8011497 | 5.75226998 | 10.0856871 |
| 2.96145338 | 0.00306191 | 0.00632339 | 0 | 0 | 0 | 0 |
| -2.9614593 | 0.00306185 | 0.00632339 | 13.9346919 | 21.9194731 | 24.1595339 | 18.4904263 |
| -2.9611901 | 0.00306453 | 0.00632839 | 30.738291 | 13.530539 | 26.4604419 | 16.2491625 |
| -2.9610597 | 0.00306583 | 0.00633067 | 4.91812656 | 40.3210061 | 37.9649818 | 56.5919108 |
| -2.9609791 | 0.00306663 | 0.00633192 | 122.54332 | 103.914539 | 0 | 299.769033 |
| -2.9607312 | 0.0030691 | 0.00633661 | 4.0984388 | 72.7942996 | 100.089498 | 18.4904263 |
| -2.9600434 | 0.00307596 | 0.00635037 | 9.01656536 | 50.8748265 | 24.1595339 | 27.4554815 |
| 2.95947795 | 0.00308161 | 0.00636163 | 0 | 3.24732935 | 0 | 0 |
| -2.9589412 | 0.00308698 | 0.00637231 | 2.45906328 | 67.6526948 | 43.7172518 | 73.4013893 |
| -2.9585702 | 0.0030907 | 0.00637958 | 94.6739363 | 467.074205 | 271.507143 | 333.948305 |
| -2.95842 | 0.00309221 | 0.00638228 | 0 | 132.05806 | 239.294431 | 282.959554 |
| -2.9581483 | 0.00309493 | 0.0063875 | 13.9346919 | 22.4606947 | 23.0090799 | 18.4904263 |
| -2.9576191 | 0.00310025 | 0.00639766 | 54.0993922 | 122.586683 | 133.452663 | 150.72499 |
| -2.9576385 | 0.00310005 | 0.00639766 | 10.6559409 | 41.1328384 | 16.1063559 | 40.3427483 |
| -2.9573494 | 0.00310296 | 0.00640285 | 53.6895483 | 7.30649104 | 26.4604419 | 33.6189569 |
| -2.9569501 | 0.00310698 | 0.00641074 | 11.4756286 | 25.4374132 | 21.8586259 | 26.3348496 |
| -2.9568887 | 0.0031076 | 0.0064116 | 8.60672148 | 29.2259641 | 29.9118039 | 32.498325 |
| -2.9567332 | 0.00310917 | 0.00641443 | 5.73781432 | 33.5557366 | 31.0622579 | 72.2807574 |
| 2.95626512 | 0.00311389 | 0.00642376 | 0 | 0 | 0 | 0.56031595 |

| | | | | | | |
|---|---|---|---|---|---|---|
| -2.9561914 | 0.00311464 | 0.00642489 | 9.42640924 | 40.8622277 | 23.0090799 | 28.5761134 |
| -2.955999 | 0.00311658 | 0.00642848 | 20.0823501 | 12.1774851 | 21.8586259 | 42.0236961 |
| -2.9554296 | 0.00312234 | 0.00643995 | 8.1968776 | 24.0843593 | 43.7172518 | 33.6189569 |
| -2.9553635 | 0.00312301 | 0.00644092 | 20.0823501 | 24.6255809 | 11.50454 | 29.1364293 |
| -2.9550892 | 0.00312579 | 0.00644624 | 140.986295 | 340.969582 | 281.861229 | 371.489474 |
| -2.954985 | 0.00312684 | 0.006448 | 13.1150042 | 27.0610779 | 14.9559019 | 55.4712789 |
| -2.9545674 | 0.00313108 | 0.00645632 | 0 | 192.133653 | 225.488983 | 208.997849 |
| 2.95437357 | 0.00313305 | 0.00645997 | 13.524848 | 7.8477126 | 4.60181598 | 8.40473923 |
| -2.9532089 | 0.00314489 | 0.00648398 | 6.55750208 | 32.7439043 | 44.8677058 | 34.1792729 |
| -2.9528961 | 0.00314808 | 0.00649014 | 64.755333 | 15.1542036 | 32.2127119 | 11.206319 |
| -2.9526367 | 0.00315073 | 0.00649476 | 181.150995 | 411.869606 | 196.727633 | 333.948305 |
| -2.9526542 | 0.00315055 | 0.00649476 | 13.524848 | 23.272527 | 20.7081719 | 20.1713742 |
| -2.9523599 | 0.00315355 | 0.00650017 | 13.9346919 | 30.8496288 | 14.9559019 | 27.4554815 |
| 2.9519715 | 0.00315752 | 0.00650794 | 11.0657848 | 3.51794013 | 0 | 4.48252759 |
| -2.9519252 | 0.00315799 | 0.00650808 | 11.8854725 | 18.9427545 | 25.3099879 | 31.3776931 |
| -2.9519378 | 0.00315787 | 0.00650808 | 22.9512573 | 54.6633774 | 70.1776937 | 60.5141225 |
| -2.9518709 | 0.00315855 | 0.00650881 | 61.8864259 | 23.5431378 | 31.0622579 | 7.84442328 |
| -2.9516167 | 0.00316115 | 0.00651376 | 24.1807889 | 25.708024 | 11.50454 | 21.292006 |
| -2.9514819 | 0.00316253 | 0.00651619 | 68.0340841 | 18.6721438 | 9.20363196 | 30.8173772 |
| 2.95130736 | 0.00316432 | 0.00651945 | 12.7051603 | 12.9893174 | 14.9559019 | 7.28410733 |
| -2.95127 | 0.0031647 | 0.00651983 | 19.2626624 | 21.3782516 | 52.9208838 | 12.3269509 |
| 2.95115514 | 0.00316588 | 0.00652184 | 56.9682993 | 50.0629941 | 54.0713378 | 35.2999048 |
| -2.9510453 | 0.003167 | 0.00652374 | 278.283994 | 125.834012 | 0 | 104.218766 |
| 2.95095938 | 0.00316789 | 0.00652472 | 214.758193 | 247.608863 | 226.639437 | 234.772382 |
| -2.9509704 | 0.00316777 | 0.00652472 | 76.6408055 | 199.710755 | 0 | 237.013646 |
| -2.9506505 | 0.00317106 | 0.00653083 | 3.27875104 | 41.4034492 | 124.249032 | 31.3776931 |
| 2.9502636 | 0.00317503 | 0.0065386 | 0 | 0.27061078 | 1.150454 | 0 |
| -2.9502381 | 0.00317529 | 0.00653872 | 59.4273626 | 28.9553534 | 9.20363196 | 20.1713742 |
| -2.950174 | 0.00317595 | 0.00653966 | 6.96734596 | 38.9679522 | 31.0622579 | 35.8602207 |
| -2.9500417 | 0.00317731 | 0.00654205 | 165.167084 | 168.049294 | 148.408565 | 149.044042 |
| -2.9499295 | 0.00317846 | 0.00654401 | 6.1476582 | 39.238563 | 41.4163438 | 33.6189569 |
| -2.9496783 | 0.00318105 | 0.0065484 | 68.0340841 | 187.262659 | 0 | 277.356395 |
| -2.9496736 | 0.0031811 | 0.0065484 | 0 | 219.194731 | 394.60572 | 89.0902358 |
| -2.9496631 | 0.00318121 | 0.0065484 | 10.6559409 | 32.7439043 | 19.5577179 | 30.2570612 |
| 2.94949594 | 0.00318293 | 0.00655152 | 0.40984388 | 0 | 0 | 0 |
| -2.9491364 | 0.00318663 | 0.00655873 | 98.3625312 | 191.321821 | 0 | 194.98995 |
| -2.9486708 | 0.00319144 | 0.0065682 | 27.45954 | 38.6973414 | 44.8677058 | 6.72379138 |
| -2.9483134 | 0.00319513 | 0.00657538 | 63.5258014 | 45.1920001 | 55.2217918 | 60.5141225 |
| -2.9481808 | 0.0031965 | 0.00657778 | 14.3445358 | 26.7904671 | 17.2568099 | 20.7316901 |
| -2.9479764 | 0.00319861 | 0.00658171 | 266.808366 | 302.542851 | 75.9299637 | 0 |
| -2.9474557 | 0.00320401 | 0.00659239 | 8.1968776 | 24.0843593 | 34.5136199 | 39.7824324 |
| -2.9472942 | 0.00320568 | 0.00659541 | 16.8035991 | 28.143521 | 16.1063559 | 17.9301104 |
| 2.94724113 | 0.00320623 | 0.00659612 | 0.81968776 | 0.54122156 | 0 | 0.56031595 |

| | | | | | | |
|---|---|---|---|---|---|---|
| 2.94708616 | 0.00320784 | 0.00659901 | 184.429746 | 145.85921 | 97.7885896 | 123.829825 |
| -2.9469434 | 0.00320932 | 0.00660164 | 13.9346919 | 12.1774851 | 44.8677058 | 35.8602207 |
| -2.9468214 | 0.00321059 | 0.00660382 | 40.1647002 | 29.7671857 | 37.9649818 | 6.72379138 |
| 2.94652567 | 0.00321366 | 0.00660972 | 0 | 0 | 1.150454 | 0 |
| 2.94622542 | 0.00321678 | 0.00661572 | 0 | 2.43549701 | 0 | 0 |
| -2.9461428 | 0.00321764 | 0.00661706 | 25.4103206 | 6.4946587 | 55.2217918 | 39.7824324 |
| -2.94592 | 0.00321996 | 0.00662141 | 13.9346919 | 17.5897006 | 20.7081719 | 28.5761134 |
| -2.9457171 | 0.00322207 | 0.00662533 | 11.4756286 | 23.272527 | 28.7613499 | 21.292006 |
| -2.9455545 | 0.00322376 | 0.00662839 | 14.3445358 | 26.2492456 | 13.8054479 | 44.8252759 |
| 2.94544667 | 0.00322489 | 0.00663028 | 1.22953164 | 0.27061078 | 0 | 0.56031595 |
| -2.9452415 | 0.00322703 | 0.00663426 | 12.2953164 | 23.8137486 | 29.9118039 | 18.4904263 |
| -2.9451129 | 0.00322837 | 0.00663659 | 199.59397 | 294.695138 | 246.197155 | 275.115131 |
| -2.944935 | 0.00323023 | 0.00663999 | 9.83625312 | 22.7313054 | 27.6108959 | 33.6189569 |
| -2.944831 | 0.00323131 | 0.0066418 | 56.9682993 | 20.0251977 | 14.9559019 | 16.2491625 |
| -2.9448028 | 0.00323161 | 0.00664198 | 12.2953164 | 11.6362635 | 48.3190678 | 48.7474875 |
| -2.9447309 | 0.00323236 | 0.0066431 | 265.578834 | 902.216338 | 851.335957 | 911.634048 |
| -2.9446779 | 0.00323291 | 0.00664352 | 796.326659 | 512.266205 | 463.63296 | 446.571811 |
| -2.944672 | 0.00323297 | 0.00664352 | 15.9839113 | 18.6721438 | 16.1063559 | 29.1364293 |
| -2.9445147 | 0.00323462 | 0.00664647 | 13.524848 | 14.3423713 | 24.1595339 | 86.848972 |
| -2.9439201 | 0.00324084 | 0.00665883 | 36.0662614 | 34.6381797 | 9.20363196 | 20.1713742 |
| -2.9438158 | 0.00324193 | 0.00666065 | 15.9839113 | 40.8622277 | 10.354086 | 38.6618005 |
| -2.9436922 | 0.00324322 | 0.00666289 | 22.9512573 | 26.5198564 | 10.354086 | 26.3348496 |
| -2.9434186 | 0.00324609 | 0.00666836 | 11.8854725 | 36.803066 | 43.7172518 | 14.0078987 |
| 2.94272564 | 0.00325337 | 0.00668287 | 8.60672148 | 30.8496288 | 41.4163438 | 30.8173772 |
| -2.9425933 | 0.00325476 | 0.00668531 | 14.3445358 | 33.0145151 | 13.8054479 | 28.0157974 |
| -2.9420687 | 0.00326028 | 0.00669622 | 6.55750208 | 22.1900839 | 54.0713378 | 46.5062237 |
| -2.9418927 | 0.00326213 | 0.00669875 | 402.46669 | 536.079953 | 536.111562 | 571.522268 |
| 2.94190283 | 0.00326202 | 0.00669875 | 3.27875104 | 7.8477126 | 1.150454 | 6.16347543 |
| -2.9419033 | 0.00326202 | 0.00669875 | 450.00858 | 783.147595 | 715.582385 | 728.410733 |
| -2.94175 | 0.00326363 | 0.00670141 | 8.60672148 | 29.7671857 | 54.0713378 | 21.852322 |
| -2.9416521 | 0.00326466 | 0.0067031 | 13.9346919 | 17.8603114 | 26.4604419 | 20.1713742 |
| -2.9412792 | 0.0032686 | 0.00671075 | 36.0662614 | 16.2366467 | 86.2840497 | 8.40473923 |
| -2.9407587 | 0.0032741 | 0.00672161 | 343.449171 | 500.08872 | 626.997427 | 638.199865 |
| -2.9406316 | 0.00327544 | 0.00672394 | 210.24991 | 154.518755 | 0 | 108.140978 |
| -2.9405951 | 0.00327582 | 0.0067243 | 31.1481349 | 101.208431 | 50.6199758 | 20.7316901 |
| -2.9399021 | 0.00328316 | 0.00673893 | 84.0179954 | 100.125988 | 566.023366 | 0 |
| -2.9396784 | 0.00328553 | 0.00674337 | 45.9025146 | 18.401533 | 89.7354116 | 35.8602207 |
| -2.9394684 | 0.00328776 | 0.00674751 | 28.6890716 | 24.6255809 | 20.7081719 | 11.206319 |
| -2.9390024 | 0.00329271 | 0.00675681 | 9.42640924 | 40.5916169 | 25.3099879 | 24.0935858 |
| -2.9390145 | 0.00329258 | 0.00675681 | 0 | 254.915354 | 92.0363196 | 320.500723 |
| -2.938434 | 0.00329875 | 0.00676878 | 114.346443 | 8.93015571 | 42.5667978 | 11.206319 |
| -2.9380326 | 0.00330302 | 0.00677712 | 90.9853413 | 14.3423713 | 18.4072639 | 16.8094785 |
| -2.9380052 | 0.00330331 | 0.00677728 | 16.3937552 | 13.2599282 | 32.2127119 | 23.5332698 |

| | | | | | | |
|---|---|---|---|---|---|---|
| -2.9378275 | 0.00330521 | 0.00678074 | 18.0331307 | 24.0843593 | 17.2568099 | 15.6888466 |
| -2.9377723 | 0.0033058 | 0.00678152 | 28.2792277 | 46.545054 | 14.9559019 | 38.1014845 |
| -2.9375789 | 0.00330786 | 0.00678532 | 16.8035991 | 11.0950419 | 37.9649818 | 31.3776931 |
| -2.9374946 | 0.00330876 | 0.00678673 | 184.019902 | 122.586683 | 0 | 154.086886 |
| -2.9372455 | 0.00331142 | 0.00679175 | 21.7217256 | 16.7778683 | 19.5577179 | 15.6888466 |
| 2.93713633 | 0.00331258 | 0.00679371 | 0 | 15.6954252 | 0 | 5.60315949 |
| 2.93699322 | 0.00331411 | 0.00679642 | 135.24848 | 181.038611 | 179.470823 | 138.958355 |
| -2.93694 | 0.00331468 | 0.00679715 | 115.985818 | 169.131737 | 0 | 176.499524 |
| -2.936894 | 0.00331517 | 0.00679773 | 6.1476582 | 30.3084073 | 31.0622579 | 63.3157022 |
| -2.9367617 | 0.00331659 | 0.0068002 | 15.1642236 | 16.2366467 | 35.6640739 | 19.0507423 |
| -2.9361943 | 0.00332266 | 0.00681221 | 14.3445358 | 14.8835929 | 41.4163438 | 22.9729539 |
| -2.9359015 | 0.0033258 | 0.00681822 | 41.3942319 | 20.83703 | 6.90272397 | 56.0315949 |
| -2.9358113 | 0.00332677 | 0.00681977 | 4.91812656 | 36.2618444 | 47.1686138 | 43.704644 |
| -2.9356666 | 0.00332832 | 0.00682251 | 13.9346919 | 19.2133653 | 18.4072639 | 26.3348496 |
| -2.9355747 | 0.0033293 | 0.0068241 | 274.5954 | 87.4072817 | 0 | 138.398039 |
| -2.9351069 | 0.00333433 | 0.00683396 | 264.759146 | 84.9717847 | 0 | 147.92341 |
| -2.9350863 | 0.00333455 | 0.00683398 | 31.5579788 | 7.03588026 | 34.5136199 | 36.4205367 |
| -2.9345018 | 0.00334084 | 0.00684556 | 11.0657848 | 24.8961917 | 89.7354116 | 16.2491625 |
| -2.9345138 | 0.00334071 | 0.00684556 | 6.96734596 | 36.2618444 | 31.0622579 | 35.2999048 |
| -2.9345127 | 0.00334072 | 0.00684556 | 11.4756286 | 16.7778683 | 32.2127119 | 31.3776931 |
| -2.9342319 | 0.00334374 | 0.00685109 | 18.8528185 | 18.401533 | 16.1063559 | 18.4904263 |
| 2.93392536 | 0.00334705 | 0.00685742 | 78.2801811 | 41.1328384 | 44.8677058 | 48.1871716 |
| -2.933795 | 0.00334845 | 0.00685987 | 9.42640924 | 34.367569 | 23.0090799 | 28.0157974 |
| -2.9336346 | 0.00335018 | 0.00686272 | 27.8693838 | 24.3549701 | 20.7081719 | 11.206319 |
| -2.9336266 | 0.00335027 | 0.00686272 | 8.1968776 | 35.9912336 | 19.5577179 | 54.350647 |
| -2.9327156 | 0.00336012 | 0.00688245 | 5.73781432 | 53.0397127 | 29.9118039 | 38.1014845 |
| -2.9325397 | 0.00336202 | 0.00688591 | 5.73781432 | 38.6973414 | 43.7172518 | 33.6189569 |
| -2.9324458 | 0.00336304 | 0.00688756 | 25.4103206 | 30.8496288 | 31.0622579 | 33.058641 |
| -2.9322413 | 0.00336525 | 0.00689166 | 12.2953164 | 32.4732935 | 13.8054479 | 44.2649599 |
| -2.9319937 | 0.00336794 | 0.00689628 | 103.280658 | 169.943569 | 0 | 191.067738 |
| -2.9320063 | 0.0033678 | 0.00689628 | 12.2953164 | 20.0251977 | 25.3099879 | 21.852322 |
| -2.9314705 | 0.00337361 | 0.00690747 | 6.1476582 | 27.3316887 | 65.5758777 | 33.058641 |
| -2.931322 | 0.00337523 | 0.00691033 | 26.6398522 | 36.803066 | 13.8054479 | 14.0078987 |
| -2.931183 | 0.00337674 | 0.00691299 | 14.7543797 | 23.0019162 | 56.3722458 | 14.0078987 |
| -2.9311547 | 0.00337705 | 0.00691318 | 128.281134 | 656.50175 | 330.180297 | 435.925808 |
| -2.930593 | 0.00338316 | 0.00692525 | 35.2465737 | 23.5431378 | 17.2568099 | 12.3269509 |
| -2.9302437 | 0.00338696 | 0.0069326 | 13.524848 | 13.530539 | 28.7613499 | 44.2649599 |
| -2.9299701 | 0.00338995 | 0.00693827 | 0 | 196.192815 | 201.329449 | 199.472478 |
| -2.9297829 | 0.00339199 | 0.00694201 | 103.280658 | 169.943569 | 0 | 188.826475 |
| -2.9295431 | 0.00339461 | 0.00694693 | 50.4107972 | 24.6255809 | 26.4604419 | 38.1014845 |
| -2.9293219 | 0.00339702 | 0.00695143 | 0 | 43.2977247 | 543.014286 | 224.126379 |
| -2.9291094 | 0.00339935 | 0.00695575 | 17.6232868 | 12.9893174 | 43.7172518 | 20.7316901 |
| -2.9287232 | 0.00340357 | 0.00696395 | 20.0823501 | 16.7778683 | 14.9559019 | 21.852322 |

| | | | | | | |
|---|---|---|---|---|---|---|
| -2.9283128 | 0.00340807 | 0.00697271 | 31.1481349 | 20.83703 | 11.50454 | 20.7316901 |
| -2.9279972 | 0.00341153 | 0.00697935 | 16.8035991 | 18.9427545 | 16.1063559 | 20.1713742 |
| -2.9276375 | 0.00341548 | 0.00698654 | 154.101299 | 220.818396 | 177.169915 | 0 |
| -2.9276537 | 0.0034153 | 0.00698654 | 299.595876 | 103.914539 | 0 | 100.856871 |
| -2.9275506 | 0.00341643 | 0.00698805 | 24.5906328 | 18.1309222 | 17.2568099 | 15.1285306 |
| -2.9271861 | 0.00342044 | 0.00699581 | 5.73781432 | 30.8496288 | 46.0181598 | 39.7824324 |
| -2.92705 | 0.00342194 | 0.00699843 | 75.4112739 | 211.88824 | 0 | 200.032794 |
| 2.9269969 | 0.00342252 | 0.00699873 | 3.27875104 | 1.08244312 | 4.60181598 | 0 |
| -2.9269976 | 0.00342252 | 0.00699873 | 18.8528185 | 20.2958084 | 20.7081719 | 14.0078987 |
| 2.92687565 | 0.00342386 | 0.00700102 | 173.363961 | 99.3141559 | 108.142676 | 105.899714 |
| -2.9268087 | 0.00342459 | 0.00700208 | 13.9346919 | 16.7778683 | 20.7081719 | 27.4554815 |
| -2.9264887 | 0.00342812 | 0.00700885 | 8.60672148 | 34.6381797 | 20.7081719 | 40.3427483 |
| -2.9264668 | 0.00342836 | 0.0070089 | 22.1315695 | 4.32977247 | 83.9831417 | 55.4712789 |
| -2.9263675 | 0.00342946 | 0.00701069 | 36.4761053 | 20.0251977 | 12.6549939 | 18.4904263 |
| -2.9263425 | 0.00342973 | 0.00701081 | 13.524848 | 13.530539 | 34.5136199 | 30.8173772 |
| -2.9262746 | 0.00343048 | 0.0070119 | 11.0657848 | 20.0251977 | 27.6108959 | 25.7745336 |
| -2.9260024 | 0.00343348 | 0.00701759 | 16.3937552 | 16.7778683 | 17.2568099 | 22.9729539 |
| -2.9258481 | 0.00343519 | 0.00702063 | 18.0331307 | 29.7671857 | 11.50454 | 25.2142177 |
| -2.9255295 | 0.00343871 | 0.00702738 | 17.6232868 | 27.8729103 | 41.4163438 | 10.646003 |
| -2.9252651 | 0.00344163 | 0.0070329 | 8.1968776 | 46.545054 | 18.4072639 | 42.0236961 |
| -2.9247236 | 0.00344762 | 0.00704471 | 61.476582 | 14.0717605 | 13.8054479 | 24.0935858 |
| -2.9245374 | 0.00344969 | 0.00704848 | 262.300083 | 590.743331 | 322.127119 | 453.855918 |
| -2.9243299 | 0.00345199 | 0.00705273 | 8.60672148 | 96.3374374 | 9.20363196 | 58.8331746 |
| -2.9242983 | 0.00345234 | 0.007053 | 61.8864259 | 115.821413 | 83.9831417 | 101.977503 |
| -2.9241997 | 0.00345343 | 0.00705479 | 13.1150042 | 24.6255809 | 16.1063559 | 27.4554815 |
| -2.9241098 | 0.00345443 | 0.00705638 | 62.7061136 | 40.8622277 | 5.75226998 | 23.5332698 |
| -2.9238873 | 0.0034569 | 0.00706098 | 21.3118818 | 17.8603114 | 14.9559019 | 18.4904263 |
| -2.9233626 | 0.00346273 | 0.00707244 | 10.6559409 | 28.4141318 | 23.0090799 | 22.9729539 |
| -2.9233388 | 0.00346299 | 0.00707254 | 175.823024 | 114.197749 | 119.647216 | 129.432984 |
| 2.92316628 | 0.00346491 | 0.00707601 | 0 | 1.3530539 | 0 | 1.68094785 |
| -2.9227509 | 0.00346954 | 0.00708501 | 34.8367298 | 33.5557366 | 28.7613499 | 24.6539017 |
| -2.9227059 | 0.00347004 | 0.00708558 | 659.028959 | 1099.76221 | 960.629086 | 1185.62855 |
| 2.92222248 | 0.00347543 | 0.00709569 | 295.907281 | 180.226779 | 233.542161 | 162.491625 |
| -2.9222287 | 0.00347536 | 0.00709569 | 48.7714217 | 27.3316887 | 18.4072639 | 10.0856871 |
| 2.92217406 | 0.00347597 | 0.0070959 | 0.81968776 | 1.08244312 | 1.150454 | 0 |
| -2.9221758 | 0.00347595 | 0.0070959 | 44.6729829 | 23.272527 | 10.354086 | 20.1713742 |
| -2.9218884 | 0.00347916 | 0.00710151 | 16.8035991 | 20.5664192 | 31.0622579 | 13.4475828 |
| 2.92189686 | 0.00347907 | 0.00710151 | 136.478012 | 106.620647 | 106.992222 | 83.4870763 |
| -2.9218586 | 0.00347949 | 0.00710174 | 16.3937552 | 24.0843593 | 24.1595339 | 13.4475828 |
| -2.9214539 | 0.00348402 | 0.00711052 | 15.1642236 | 5.1416048 | 65.5758777 | 87.9696039 |
| -2.9209374 | 0.0034898 | 0.00712187 | 182.79037 | 121.774851 | 106.992222 | 114.304454 |
| -2.9208474 | 0.00349081 | 0.00712305 | 10.246097 | 54.1221558 | 20.7081719 | 22.9729539 |
| -2.9208466 | 0.00349082 | 0.00712305 | 0 | 189.968767 | 189.824909 | 207.316901 |

| | | | | | | |
|---|---|---|---|---|---|---|
| 2.92060869 | 0.00349348 | 0.00712804 | 0.40984388 | 0.27061078 | 1.150454 | 0 |
| -2.9204996 | 0.00349471 | 0.00712963 | 5.73781432 | 33.8263474 | 29.9118039 | 58.2728587 |
| -2.9205151 | 0.00349453 | 0.00712963 | 4.50828268 | 43.2977247 | 50.6199758 | 36.4205367 |
| 2.92030985 | 0.00349684 | 0.00713353 | 6.96734596 | 14.6129821 | 18.4072639 | 21.852322 |
| -2.9200352 | 0.00349992 | 0.00713937 | 49.1812656 | 21.1076408 | 13.8054479 | 15.6888466 |
| -2.9199848 | 0.00350049 | 0.00713962 | 15.1642236 | 36.5324552 | 13.8054479 | 22.4126379 |
| -2.91999 | 0.00350043 | 0.00713962 | 8.60672148 | 22.7313054 | 46.0181598 | 27.4554815 |
| -2.9190381 | 0.00351113 | 0.00716088 | 212.708974 | 264.11612 | 242.745793 | 245.418385 |
| -2.918988 | 0.0035117 | 0.00716158 | 103.280658 | 107.161869 | 104.691314 | 98.055291 |
| -2.918954 | 0.00351208 | 0.00716191 | 8.60672148 | 53.5809343 | 21.8586259 | 27.4554815 |
| -2.918443 | 0.00351784 | 0.00717321 | 0 | 125.834012 | 293.365769 | 191.067738 |
| -2.9183932 | 0.0035184 | 0.0071739 | 15.9839113 | 27.3316887 | 35.6640739 | 11.7666349 |
| 2.91735084 | 0.00353018 | 0.00719747 | 8.1968776 | 7.57710182 | 4.60181598 | 6.72379138 |
| -2.9171316 | 0.00353267 | 0.00720207 | 44.6729829 | 89.0309463 | 90.8858656 | 103.098135 |
| -2.916974 | 0.00353445 | 0.00720526 | 115.985818 | 14.6129821 | 11.50454 | 21.852322 |
| -2.9168897 | 0.00353541 | 0.00720675 | 4.0984388 | 64.946587 | 26.4604419 | 57.1522268 |
| -2.9167678 | 0.00353679 | 0.00720911 | 16.8035991 | 32.2026827 | 9.20363196 | 57.1522268 |
| -2.916369 | 0.00354131 | 0.00721788 | 41.3942319 | 23.0019162 | 43.7172518 | 34.7395888 |
| -2.9160161 | 0.00354532 | 0.00722559 | 19.6725062 | 8.65954493 | 43.7172518 | 33.6189569 |
| -2.9158395 | 0.00354733 | 0.00722923 | 2.86890716 | 82.8068984 | 63.2749697 | 29.1364293 |
| 2.91562849 | 0.00354973 | 0.00723366 | 0 | 8.65954493 | 0 | 0 |
| -2.9155617 | 0.00355049 | 0.00723475 | 39.7548564 | 56.5576528 | 8.05317797 | 16.8094785 |
| -2.9151516 | 0.00355516 | 0.00724381 | 6.1476582 | 54.1221558 | 26.4604419 | 35.8602207 |
| -2.9146809 | 0.00356052 | 0.00725428 | 143.035514 | 138.011497 | 0 | 156.32815 |
| -2.9143373 | 0.00356445 | 0.00726182 | 0 | 221.900839 | 151.859927 | 211.799429 |
| -2.9143168 | 0.00356468 | 0.00726183 | 14.3445358 | 27.3316887 | 12.6549939 | 37.5411686 |
| 2.91412866 | 0.00356683 | 0.00726575 | 0.40984388 | 0.54122156 | 0 | 0 |
| -2.9138891 | 0.00356957 | 0.00727087 | 21.7217256 | 27.8729103 | 26.4604419 | 10.0856871 |
| -2.9138143 | 0.00357042 | 0.00727216 | 11.8854725 | 22.1900839 | 24.1595339 | 20.1713742 |
| -2.9137353 | 0.00357133 | 0.00727354 | 5.32797044 | 25.708024 | 60.9740618 | 42.0236961 |
| -2.9136629 | 0.00357215 | 0.00727477 | 16.3937552 | 15.6954252 | 17.2568099 | 26.8951655 |
| -2.9136113 | 0.00357274 | 0.00727551 | 127.871291 | 254.103522 | 169.116737 | 0 |
| -2.9135163 | 0.00357383 | 0.00727727 | 41.8040758 | 33.2851258 | 23.0090799 | 30.2570612 |
| -2.9133915 | 0.00357526 | 0.00727971 | 13.524848 | 25.9786348 | 16.1063559 | 22.4126379 |
| -2.9132613 | 0.00357675 | 0.00728183 | 10.6559409 | 76.0416289 | 70.1776937 | 64.99665 |
| -2.9132574 | 0.0035768 | 0.00728183 | 279.92337 | 454.355498 | 682.219219 | 0 |
| 2.91324154 | 0.00357698 | 0.00728183 | 0 | 0 | 0 | 0 |
| -2.9132196 | 0.00357723 | 0.00728189 | 4.0984388 | 43.0271139 | 60.9740618 | 34.7395888 |
| -2.9130512 | 0.00357916 | 0.00728536 | 12.2953164 | 20.0251977 | 25.3099879 | 20.1713742 |
| -2.9129143 | 0.00358073 | 0.00728809 | 14.3445358 | 20.2958084 | 16.1063559 | 24.6539017 |
| -2.9127579 | 0.00358252 | 0.00729128 | 29.9186032 | 29.4965749 | 12.6549939 | 14.5682147 |
| 2.91271492 | 0.00358302 | 0.00729183 | 5.32797044 | 10.8244312 | 4.60181598 | 5.60315949 |
| -2.9122445 | 0.00358842 | 0.00730236 | 330.334167 | 285.494372 | 0 | 24.0935858 |

| | | | | | | |
|---|---|---|---|---|---|---|
| -2.911843 | 0.00359303 | 0.00731129 | 31.1481349 | 40.3210061 | 6.90272397 | 27.4554815 |
| -2.9116655 | 0.00359507 | 0.00731498 | 5.32797044 | 65.4878086 | 23.0090799 | 45.3855918 |
| -2.9114379 | 0.00359769 | 0.00731985 | 9.83625312 | 24.6255809 | 24.1595339 | 28.0157974 |
| 2.91134391 | 0.00359878 | 0.00732159 | 31.1481349 | 20.2958084 | 0 | 30.2570612 |
| -2.9111814 | 0.00360065 | 0.00732494 | 13.9346919 | 23.8137486 | 25.3099879 | 15.1285306 |
| -2.9108123 | 0.00360491 | 0.00733314 | 20.9020379 | 26.5198564 | 9.20363196 | 38.1014845 |
| -2.9106584 | 0.00360668 | 0.00733629 | 100.001907 | 174.814563 | 0 | 170.896364 |
| -2.9103438 | 0.00361031 | 0.00734322 | 32.7875104 | 15.6954252 | 50.6199758 | 28.0157974 |
| -2.9101529 | 0.00361252 | 0.00734724 | 15.5740674 | 50.6042157 | 12.6549939 | 22.9729539 |
| -2.9099591 | 0.00361476 | 0.00735133 | 7.78703372 | 44.3801678 | 26.4604419 | 26.8951655 |
| -2.9098764 | 0.00361572 | 0.00735282 | 0 | 232.454659 | 128.850847 | 228.608907 |
| -2.9097895 | 0.00361672 | 0.0073544 | 54.0993922 | 114.73897 | 56.3722458 | 79.5648647 |
| -2.9097633 | 0.00361703 | 0.00735455 | 11.4756286 | 21.9194731 | 39.1154358 | 18.4904263 |
| 2.90933294 | 0.00362201 | 0.00736422 | 0.81968776 | 1.62366467 | 0 | 1.1206319 |
| -2.9091633 | 0.00362398 | 0.00736775 | 0 | 476.004361 | 23.0090799 | 360.843471 |
| -2.9090948 | 0.00362477 | 0.00736859 | 21.7217256 | 15.6954252 | 16.1063559 | 18.4904263 |
| -2.9090885 | 0.00362484 | 0.00736859 | 11.0657848 | 40.5916169 | 18.4072639 | 23.5332698 |
| -2.9089592 | 0.00362634 | 0.00737117 | 7.37718984 | 40.8622277 | 28.7613499 | 28.0157974 |
| -2.9089252 | 0.00362674 | 0.00737151 | 13.9346919 | 14.3423713 | 26.4604419 | 26.3348496 |
| -2.9086612 | 0.0036298 | 0.00737727 | 20.9020379 | 30.8496288 | 8.05317797 | 45.9459078 |
| -2.908563 | 0.00363094 | 0.00737912 | 0 | 152.89509 | 88.5849576 | 429.202017 |
| -2.9085175 | 0.00363147 | 0.00737973 | 15.5740674 | 14.3423713 | 49.4695218 | 19.6110582 |
| -2.9074854 | 0.00364347 | 0.00740367 | 230.742104 | 1199.61758 | 1125.14401 | 982.794174 |
| -2.9069451 | 0.00364977 | 0.007416 | 17.213443 | 38.4267306 | 10.354086 | 28.0157974 |
| -2.9068099 | 0.00365135 | 0.00741874 | 17.213443 | 13.530539 | 20.7081719 | 23.5332698 |
| 2.90677927 | 0.00365171 | 0.007419 | 27.0496961 | 31.6614612 | 47.1686138 | 76.763285 |
| -2.9066892 | 0.00365276 | 0.00742066 | 8.60672148 | 29.2259641 | 36.8145279 | 22.4126379 |
| 2.90654396 | 0.00365446 | 0.00742333 | 2.86890716 | 2.97671857 | 0 | 4.48252759 |
| 2.90653753 | 0.00365453 | 0.00742333 | 35.6564176 | 7.30649104 | 19.5577179 | 12.8872668 |
| -2.9065027 | 0.00365494 | 0.00742369 | 22.9512573 | 69.0057487 | 31.0622579 | 50.4284354 |
| -2.9060045 | 0.00366076 | 0.00743505 | 83.1983076 | 375.607761 | 312.923487 | 329.465778 |
| -2.9057553 | 0.00366368 | 0.00744051 | 36.4761053 | 413.493271 | 316.374849 | 394.462428 |
| -2.9057092 | 0.00366422 | 0.00744114 | 9.83625312 | 23.5431378 | 28.7613499 | 24.0935858 |
| 2.90511367 | 0.0036712 | 0.00745484 | 423.368728 | 293.883306 | 104.691314 | 237.013646 |
| -2.904712 | 0.00367591 | 0.00746394 | 118.444881 | 84.7011739 | 109.29313 | 28.0157974 |
| -2.9044972 | 0.00367843 | 0.0074686 | 38.9351686 | 10.2832096 | 14.9559019 | 38.6618005 |
| -2.9040214 | 0.00368403 | 0.00747949 | 96.7231557 | 143.694324 | 0 | 206.196269 |
| 2.9038752 | 0.00368575 | 0.00748251 | 0.40984388 | 0.27061078 | 1.150454 | 0 |
| 2.9037263 | 0.0036875 | 0.0074856 | 89.3459658 | 120.692407 | 106.992222 | 127.19172 |
| -2.9036477 | 0.00368843 | 0.00748701 | 473.779525 | 750.13308 | 698.325575 | 592.814274 |
| -2.9033684 | 0.00369172 | 0.00749322 | 7.78703372 | 25.708024 | 35.6640739 | 30.2570612 |
| -2.9031039 | 0.00369484 | 0.00749908 | 13.1150042 | 25.9786348 | 24.1595339 | 15.6888466 |
| -2.9025486 | 0.0037014 | 0.00751191 | 0 | 205.934803 | 89.7354116 | 328.345146 |

| | | | | | | |
|---|---|---|---|---|---|---|
| -2.9020353 | 0.00370747 | 0.00752376 | 27.0496961 | 11.6362635 | 35.6640739 | 15.6888466 |
| -2.9019765 | 0.00370816 | 0.0075247 | 9.01656536 | 47.8981079 | 14.9559019 | 42.5840121 |
| -2.90173 | 0.00371108 | 0.00753015 | 102.051126 | 138.282108 | 0 | 201.153426 |
| 2.90161689 | 0.00371242 | 0.00753239 | 1.22953164 | 5.1416048 | 3.45136199 | 2.24126379 |
| -2.9015549 | 0.00371316 | 0.00753341 | 9.83625312 | 24.0843593 | 34.5136199 | 21.292006 |
| -2.9012959 | 0.00371623 | 0.00753917 | 5.73781432 | 47.3568864 | 21.8586259 | 54.910963 |
| -2.9008862 | 0.00372109 | 0.00754856 | 0 | 187.53327 | 187.524001 | 189.947107 |
| -2.9006684 | 0.00372368 | 0.00755333 | 238.529138 | 485.475738 | 439.473426 | 469.544765 |
| -2.9004676 | 0.00372606 | 0.0075577 | 11.8854725 | 75.2297966 | 6.90272397 | 61.6347543 |
| -2.9002674 | 0.00372844 | 0.00756205 | 13.524848 | 33.8263474 | 12.6549939 | 30.2570612 |
| -2.9002144 | 0.00372907 | 0.00756285 | 10.246097 | 18.9427545 | 25.3099879 | 37.5411686 |
| -2.9000617 | 0.00373089 | 0.00756559 | 120.084257 | 212.429462 | 200.178995 | 217.402588 |
| -2.9000717 | 0.00373077 | 0.00756559 | 52.8698605 | 44.9213893 | 4.60181598 | 28.5761134 |
| -2.8998069 | 0.00373393 | 0.00757127 | 8.60672148 | 23.5431378 | 27.6108959 | 34.7395888 |
| -2.8997422 | 0.0037347 | 0.00757235 | 106.149565 | 14.8835929 | 12.6549939 | 19.0507423 |
| -2.899655 | 0.00373574 | 0.00757398 | 19.2626624 | 54.9339882 | 6.90272397 | 39.2221164 |
| -2.8996309 | 0.00373602 | 0.00757409 | 25.0004767 | 41.67406 | 25.3099879 | 36.9808526 |
| -2.8995144 | 0.00373741 | 0.00757643 | 22.9512573 | 50.6042157 | 8.05317797 | 25.7745336 |
| -2.8992133 | 0.003741 | 0.00758324 | 3.27875104 | 87.9485032 | 40.2658898 | 34.1792729 |
| -2.8983216 | 0.00375166 | 0.00760435 | 11.8854725 | 23.272527 | 24.1595339 | 18.4904263 |
| 2.89823188 | 0.00375273 | 0.00760605 | 27.8693838 | 13.2599282 | 10.354086 | 14.5682147 |
| -2.8979443 | 0.00375617 | 0.00761255 | 9.83625312 | 13.8011497 | 65.5758777 | 34.1792729 |
| -2.8976987 | 0.00375912 | 0.00761804 | 0 | 137.740887 | 204.780811 | 228.608907 |
| -2.8966892 | 0.00377123 | 0.00764211 | 11.0657848 | 32.7439043 | 17.2568099 | 25.7745336 |
| -2.8961525 | 0.00377769 | 0.00765471 | 9.42640924 | 23.8137486 | 23.0090799 | 33.6189569 |
| -2.8960191 | 0.00377929 | 0.00765749 | 7.37718984 | 37.3442875 | 27.6108959 | 29.1364293 |
| -2.8954652 | 0.00378597 | 0.00767053 | 19.6725062 | 42.4858923 | 89.7354116 | 5.04284354 |
| -2.8953151 | 0.00378778 | 0.00767372 | 71.3128351 | 163.990132 | 0 | 228.048591 |
| -2.8949705 | 0.00379194 | 0.00768167 | 113.936599 | 165.343186 | 0 | 146.802779 |
| -2.8949354 | 0.00379237 | 0.00768205 | 11.0657848 | 17.8603114 | 24.1595339 | 33.058641 |
| -2.8948871 | 0.00379295 | 0.00768275 | 9.42640924 | 31.6614612 | 20.7081719 | 28.0157974 |
| -2.8948061 | 0.00379393 | 0.00768424 | 15.5740674 | 25.9786348 | 12.6549939 | 24.6539017 |
| -2.8946439 | 0.00379589 | 0.00768725 | 17.213443 | 37.6148983 | 12.6549939 | 19.6110582 |
| -2.8946635 | 0.00379565 | 0.00768725 | 22.5414134 | 18.401533 | 12.6549939 | 20.1713742 |
| -2.8945345 | 0.00379721 | 0.00768945 | 0 | 65.4878086 | 224.338529 | 363.645051 |
| -2.8943493 | 0.00379945 | 0.0076935 | 22.9512573 | 18.9427545 | 16.1063559 | 14.5682147 |
| -2.8939607 | 0.00380416 | 0.00770255 | 22.5414134 | 14.3423713 | 19.5577179 | 16.2491625 |
| 2.89391647 | 0.00380469 | 0.00770315 | 1.22953164 | 2.16488623 | 2.30090799 | 1.68094785 |
| -2.8937011 | 0.0038073 | 0.0077076 | 11.8854725 | 29.4965749 | 19.5577179 | 19.6110582 |
| -2.8936958 | 0.00380737 | 0.0077076 | 102.46097 | 17.5897006 | 10.354086 | 19.6110582 |
| -2.8929681 | 0.0038162 | 0.00772499 | 159.839113 | 222.44206 | 162.214013 | 186.585211 |
| 2.89251732 | 0.00382168 | 0.0077356 | 7.37718984 | 8.65954493 | 3.45136199 | 19.0507423 |
| -2.892278 | 0.00382459 | 0.00774053 | 19.6725062 | 30.3084073 | 25.3099879 | 26.3348496 |

| | | | | | | |
|---|---|---|---|---|---|---|
| -2.8922967 | 0.00382437 | 0.00774053 | 12.7051603 | 30.8496288 | 20.7081719 | 16.8094785 |
| -2.8921711 | 0.00382589 | 0.00774268 | 11.8854725 | 35.1794013 | 14.9559019 | 26.8951655 |
| -2.8918893 | 0.00382933 | 0.00774914 | 153.691455 | 82.8068984 | 0 | 208.997849 |
| -2.8918427 | 0.0038299 | 0.0077498 | 10.246097 | 45.1920001 | 14.9559019 | 32.498325 |
| -2.8917768 | 0.0038307 | 0.00775094 | 9.83625312 | 31.3908504 | 23.0090799 | 22.4126379 |
| -2.8915938 | 0.00383293 | 0.00775497 | 29.0989155 | 15.6954252 | 13.8054479 | 19.6110582 |
| -2.8908523 | 0.00384199 | 0.00777281 | 3.68859492 | 13.8011497 | 85.1335957 | 92.4521315 |
| -2.8906188 | 0.00384484 | 0.00777809 | 12.2953164 | 20.0251977 | 20.7081719 | 21.292006 |
| -2.890377 | 0.0038478 | 0.00778359 | 17.213443 | 21.6488623 | 13.8054479 | 19.0507423 |
| -2.88982 | 0.00385462 | 0.00779691 | 11.4756286 | 34.367569 | 17.2568099 | 22.9729539 |
| -2.8897055 | 0.00385603 | 0.00779926 | 62.2962697 | 175.355785 | 0 | 233.091435 |
| -2.8896686 | 0.00385648 | 0.00779969 | 0 | 204.581749 | 235.843069 | 126.631404 |
| 2.88924498 | 0.00386168 | 0.00780971 | 9.01656536 | 4.60038325 | 3.45136199 | 6.16347543 |
| -2.8892151 | 0.00386205 | 0.00780997 | 0 | 95.525605 | 370.446187 | 154.647202 |
| -2.8891939 | 0.00386231 | 0.00781 | 5.73781432 | 19.2133653 | 48.3190678 | 61.6347543 |
| -2.8891515 | 0.00386283 | 0.00781057 | 15.1642236 | 43.0271139 | 64.4254237 | 49.8681194 |
| 2.8888639 | 0.00386636 | 0.00781723 | 1.63937552 | 0 | 0 | 0 |
| -2.8888395 | 0.00386666 | 0.00781734 | 7.37718984 | 27.3316887 | 37.9649818 | 28.0157974 |
| 2.88881975 | 0.00386691 | 0.00781734 | 0 | 0 | 0 | 0 |
| 2.88854524 | 0.00387028 | 0.00782368 | 18.8528185 | 21.6488623 | 17.2568099 | 14.5682147 |
| 2.88830835 | 0.0038732 | 0.00782909 | 0 | 0 | 0 | 0 |
| -2.8882403 | 0.00387404 | 0.00783029 | 18.4429746 | 46.545054 | 6.90272397 | 44.8252759 |
| -2.8880179 | 0.00387678 | 0.00783534 | 15.1642236 | 25.9786348 | 12.6549939 | 25.2142177 |
| -2.8874684 | 0.00388356 | 0.00784855 | 5.73781432 | 32.4732935 | 29.9118039 | 49.3078035 |
| -2.8872724 | 0.00388598 | 0.00785295 | 9.01656536 | 27.0610779 | 20.7081719 | 36.9808526 |
| -2.8869394 | 0.00389009 | 0.00786077 | 46.7222023 | 74.1473535 | 28.7613499 | 51.5490673 |
| -2.8866246 | 0.00389398 | 0.00786814 | 6.55750208 | 33.2851258 | 33.3631659 | 31.3776931 |
| 2.88652205 | 0.00389525 | 0.00787022 | 0 | 4.87099402 | 0 | 0 |
| -2.88621 | 0.00389912 | 0.00787753 | 3.27875104 | 32.7439043 | 55.2217918 | 59.9538065 |
| -2.886064 | 0.00390093 | 0.0078807 | 10.6559409 | 25.708024 | 19.5577179 | 25.2142177 |
| -2.8860091 | 0.00390161 | 0.00788158 | 11.4756286 | 88.4897248 | 10.354086 | 32.498325 |
| -2.8858871 | 0.00390312 | 0.00788414 | 10.6559409 | 28.6847426 | 20.7081719 | 21.852322 |
| -2.8852938 | 0.00391049 | 0.00789852 | 251.644142 | 407.269223 | 441.774334 | 458.898762 |
| -2.8847694 | 0.003917 | 0.00791119 | 15.5740674 | 15.4248144 | 19.5577179 | 19.6110582 |
| 2.88469755 | 0.0039179 | 0.0079125 | 8.1968776 | 6.76526948 | 5.75226998 | 7.84442328 |
| -2.8845398 | 0.00391986 | 0.00791597 | 39.7548564 | 11.0950419 | 14.9559019 | 28.5761134 |
| -2.8844827 | 0.00392057 | 0.00791691 | 9.83625312 | 21.1076408 | 24.1595339 | 30.2570612 |
| -2.8840977 | 0.00392537 | 0.0079261 | 15.1642236 | 19.4839761 | 16.1063559 | 19.0507423 |
| 2.88395287 | 0.00392718 | 0.00792925 | 4.0984388 | 5.95343714 | 2.30090799 | 6.16347543 |
| -2.883755 | 0.00392964 | 0.00793374 | 9.42640924 | 20.2958084 | 27.6108959 | 30.8173772 |
| -2.8834414 | 0.00393356 | 0.00794115 | 10.6559409 | 18.1309222 | 25.3099879 | 29.6967453 |
| -2.88327 | 0.0039357 | 0.00794497 | 18.4429746 | 21.9194731 | 80.5317797 | 9.52537113 |
| -2.8831884 | 0.00393672 | 0.00794654 | 98.7723751 | 14.8835929 | 12.6549939 | 18.4904263 |

| | | | | | | |
|---|---|---|---|---|---|---|
| -2.8831651 | 0.00393701 | 0.00794663 | 15.5740674 | 31.3908504 | 17.2568099 | 15.1285306 |
| -2.8827012 | 0.00394281 | 0.00795784 | 181.970683 | 364.51272 | 314.073941 | 363.645051 |
| -2.8821653 | 0.00394952 | 0.00797089 | 22.1315695 | 20.0251977 | 12.6549939 | 17.9301104 |
| -2.882028 | 0.00395125 | 0.00797387 | 13.1150042 | 28.9553534 | 14.9559019 | 22.4126379 |
| -2.8815847 | 0.00395681 | 0.00798459 | 13.1150042 | 25.708024 | 16.1063559 | 20.7316901 |
| -2.8809617 | 0.00396464 | 0.00799989 | 125.412227 | 136.929054 | 0 | 149.604358 |
| 2.8808878 | 0.00396557 | 0.00800127 | 0.40984388 | 0 | 0 | 0 |
| -2.8807968 | 0.00396671 | 0.00800308 | 18.4429746 | 18.1309222 | 17.2568099 | 15.1285306 |
| -2.8806632 | 0.00396839 | 0.00800497 | 2.86890716 | 41.4034492 | 59.8236078 | 49.3078035 |
| 2.88066934 | 0.00396832 | 0.00800497 | 157.38005 | 201.063809 | 210.533081 | 136.717091 |
| 2.88068034 | 0.00396818 | 0.00800497 | 38.1154808 | 30.0377965 | 26.4604419 | 19.6110582 |
| -2.8801753 | 0.00397454 | 0.00801687 | 284.021809 | 858.106781 | 881.24776 | 831.508868 |
| -2.8800538 | 0.00397607 | 0.00801946 | 29.5087594 | 36.5324552 | 9.20363196 | 17.9301104 |
| -2.8797227 | 0.00398025 | 0.00802738 | 21.3118818 | 19.2133653 | 26.4604419 | 11.206319 |
| 2.87963479 | 0.00398136 | 0.00802912 | 13.9346919 | 6.76526948 | 5.75226998 | 6.16347543 |
| -2.8793024 | 0.00398556 | 0.00803708 | 96.7231557 | 167.508072 | 147.258111 | 173.137628 |
| -2.8792311 | 0.00398646 | 0.0080384 | 0 | 286.847426 | 58.6731538 | 290.243661 |
| -2.8791248 | 0.00398781 | 0.00804061 | 9.42640924 | 25.9786348 | 24.1595339 | 24.6539017 |
| 2.87906216 | 0.0039886 | 0.0080417 | 38.5253247 | 21.1076408 | 12.6549939 | 25.7745336 |
| -2.8783209 | 0.00399798 | 0.00806012 | 15.1642236 | 12.9893174 | 21.8586259 | 30.8173772 |
| -2.8782809 | 0.00399849 | 0.00806064 | 7.78703372 | 21.6488623 | 41.4163438 | 30.2570612 |
| -2.8782371 | 0.00399905 | 0.00806126 | 14.3445358 | 35.9912336 | 16.1063559 | 17.3697944 |
| -2.8778721 | 0.00400367 | 0.00807009 | 10.246097 | 23.0019162 | 27.6108959 | 20.7316901 |
| 2.87780704 | 0.0040045 | 0.00807125 | 44.263139 | 25.708024 | 29.9118039 | 20.1713742 |
| -2.8776631 | 0.00400633 | 0.00807443 | 12.2953164 | 27.8729103 | 14.9559019 | 25.7745336 |
| 2.87759416 | 0.0040072 | 0.00807569 | 0 | 0 | 0 | 0 |
| -2.8775065 | 0.00400832 | 0.00807692 | 204.102252 | 554.752097 | 557.970188 | 564.23816 |
| -2.8775167 | 0.00400819 | 0.00807692 | 30.3284471 | 20.0251977 | 12.6549939 | 16.2491625 |
| -2.8774579 | 0.00400893 | 0.00807766 | 12.2953164 | 27.8729103 | 16.1063559 | 22.4126379 |
| -2.877096 | 0.00401353 | 0.00808595 | 9.01656536 | 35.9912336 | 13.8054479 | 61.6347543 |
| -2.877095 | 0.00401355 | 0.00808595 | 11.4756286 | 17.0484791 | 24.1595339 | 26.8951655 |
| 2.87702466 | 0.00401444 | 0.00808725 | 1.63937552 | 2.97671857 | 2.30090799 | 0.56031595 |
| 2.87699033 | 0.00401488 | 0.00808762 | 185.249434 | 54.3927666 | 71.3281477 | 40.9030642 |
| -2.8767913 | 0.00401741 | 0.00809222 | 157.789894 | 243.820312 | 115.0454 | 0 |
| -2.876455 | 0.0040217 | 0.00810034 | 111.477535 | 121.233629 | 0 | 184.343947 |
| -2.8762884 | 0.00402382 | 0.00810411 | 216.807412 | 156.41303 | 0 | 71.1601255 |
| -2.8761733 | 0.00402529 | 0.00810656 | 54.509236 | 23.5431378 | 10.354086 | 16.8094785 |
| -2.8758571 | 0.00402932 | 0.00811418 | 20.492194 | 47.3568864 | 28.7613499 | 41.4633802 |
| -2.8758189 | 0.00402981 | 0.00811466 | 54.509236 | 16.7778683 | 18.4072639 | 12.8872668 |
| -2.875756 | 0.00403061 | 0.00811577 | 69.2636157 | 225.148168 | 0 | 153.52657 |
| -2.8755886 | 0.00403275 | 0.00811957 | 78.2801811 | 126.375234 | 239.294431 | 188.266159 |
| -2.8754027 | 0.00403512 | 0.00812384 | 18.4429746 | 24.8961917 | 44.8677058 | 9.52537113 |
| 2.875293 | 0.00403653 | 0.00812616 | 25.8201644 | 16.5072575 | 14.9559019 | 14.0078987 |

| | | | | | | |
|---|---|---|---|---|---|---|
| -2.8752728 | 0.00403679 | 0.00812617 | 5.32797044 | 35.4500121 | 33.3631659 | 40.9030642 |
| -2.8750031 | 0.00404023 | 0.00813261 | 10.246097 | 16.7778683 | 36.8145279 | 26.8951655 |
| -2.8746241 | 0.00404509 | 0.00814187 | 6.96734596 | 28.6847426 | 34.5136199 | 29.6967453 |
| -2.8739719 | 0.00405345 | 0.0081582 | 1477.48719 | 3995.29754 | 3580.21283 | 3744.03117 |
| -2.8738768 | 0.00405467 | 0.00816014 | 72.1325229 | 22.1900839 | 4.60181598 | 42.5840121 |
| -2.8735541 | 0.00405882 | 0.00816797 | 18.8528185 | 19.7545869 | 25.3099879 | 11.7666349 |
| -2.8731621 | 0.00406386 | 0.00817761 | 6.96734596 | 25.708024 | 50.6199758 | 26.3348496 |
| 2.87309804 | 0.00406468 | 0.00817876 | 2.0492194 | 1.3530539 | 0 | 1.1206319 |
| -2.8728096 | 0.00406839 | 0.00818572 | 32.3776665 | 15.6954252 | 17.2568099 | 14.5682147 |
| -2.872544 | 0.00407181 | 0.00819209 | 12.2953164 | 15.6954252 | 28.7613499 | 22.4126379 |
| -2.8723584 | 0.00407421 | 0.00819639 | 6.55750208 | 42.7565031 | 34.5136199 | 24.0935858 |
| 2.87212029 | 0.00407728 | 0.0082017 | 0 | 0 | 0 | 0 |
| -2.8721143 | 0.00407735 | 0.0082017 | 15.5740674 | 25.9786348 | 12.6549939 | 21.852322 |
| -2.8716003 | 0.00408399 | 0.00821454 | 406.155285 | 829.151427 | 796.114165 | 867.369088 |
| -2.8710552 | 0.00409104 | 0.00822821 | 83.6081515 | 15.6954252 | 12.6549939 | 17.9301104 |
| -2.8703429 | 0.00410027 | 0.00824626 | 9.83625312 | 25.708024 | 24.1595339 | 21.852322 |
| 2.87029458 | 0.0041009 | 0.008247 | 100.821594 | 103.643928 | 105.841768 | 95.8140272 |
| -2.8698476 | 0.0041067 | 0.00825815 | 82.3786199 | 142.070659 | 119.647216 | 127.752036 |
| -2.8696911 | 0.00410873 | 0.00826173 | 20.9020379 | 28.6847426 | 28.7613499 | 8.96505518 |
| -2.8695884 | 0.00411006 | 0.0082639 | 2.86890716 | 76.3122397 | 72.4786017 | 22.4126379 |
| -2.869488 | 0.00411137 | 0.00826601 | 11.0657848 | 13.530539 | 33.3631659 | 37.5411686 |
| -2.8694354 | 0.00411205 | 0.00826687 | 21.7217256 | 32.2026827 | 32.2127119 | 35.2999048 |
| -2.8688721 | 0.00411938 | 0.00828109 | 12.2953164 | 28.143521 | 28.7613499 | 14.0078987 |
| 2.86879856 | 0.00412034 | 0.0082825 | 0.40984388 | 0 | 0 | 0 |
| -2.8687644 | 0.00412079 | 0.00828288 | 90.5754975 | 247.879474 | 281.861229 | 255.504073 |
| -2.8684242 | 0.00412522 | 0.00829127 | 6.55750208 | 37.8855091 | 27.6108959 | 31.3776931 |
| -2.8681721 | 0.00412851 | 0.00829737 | 11.8854725 | 37.6148983 | 13.8054479 | 26.8951655 |
| -2.8680969 | 0.00412949 | 0.00829882 | 22.1315695 | 17.3190899 | 19.5577179 | 12.8872668 |
| -2.8680512 | 0.00413009 | 0.0082995 | 15.5740674 | 24.8961917 | 34.5136199 | 11.206319 |
| -2.8677757 | 0.00413368 | 0.00830622 | 5.73781432 | 20.5664192 | 77.0804177 | 34.1792729 |
| -2.8677268 | 0.00413432 | 0.00830698 | 15.1642236 | 23.0019162 | 14.9559019 | 17.9301104 |
| -2.8675831 | 0.0041362 | 0.00831024 | 20.0823501 | 67.6526948 | 31.0622579 | 39.2221164 |
| -2.8675451 | 0.0041367 | 0.00831072 | 12.7051603 | 21.1076408 | 26.4604419 | 15.6888466 |
| -2.867319 | 0.00413965 | 0.00831614 | 0 | 106.891258 | 247.347609 | 199.472478 |
| -2.8671968 | 0.00414125 | 0.00831884 | 15.9839113 | 13.530539 | 24.1595339 | 19.0507423 |
| 2.86706446 | 0.00414299 | 0.0083218 | 38.9351686 | 21.9194731 | 18.4072639 | 17.9301104 |
| -2.8669806 | 0.00414408 | 0.00832349 | 20.9020379 | 11.6362635 | 35.6640739 | 16.8094785 |
| -2.8669312 | 0.00414473 | 0.00832427 | 23.3611012 | 14.8835929 | 17.2568099 | 15.6888466 |
| -2.8662894 | 0.00415314 | 0.00834064 | 12.2953164 | 22.4606947 | 14.9559019 | 34.1792729 |
| -2.8660245 | 0.00415662 | 0.00834711 | 48.3615778 | 19.2133653 | 13.8054479 | 14.5682147 |
| -2.8658293 | 0.00415918 | 0.00835174 | 0 | 211.347019 | 171.417645 | 150.164674 |
| -2.865808 | 0.00415946 | 0.00835178 | 7.78703372 | 27.0610779 | 27.6108959 | 29.6967453 |
| -2.8657302 | 0.00416049 | 0.00835331 | 23.3611012 | 17.3190899 | 13.8054479 | 16.8094785 |

| | | | | | | |
|---|---|---|---|---|---|---|
| -2.8650961 | 0.00416883 | 0.00836909 | 8.60672148 | 33.0145151 | 19.5577179 | 31.3776931 |
| -2.8650737 | 0.00416912 | 0.00836909 | 5.73781432 | 61.1580361 | 16.1063559 | 54.350647 |
| -2.8650818 | 0.00416902 | 0.00836909 | 63.1159575 | 224.606947 | 0 | 158.009098 |
| -2.8642561 | 0.0041799 | 0.0083902 | 12.7051603 | 33.5557366 | 13.8054479 | 24.0935858 |
| -2.8640459 | 0.00418267 | 0.00839525 | 98.7723751 | 201.875641 | 205.931265 | 0 |
| -2.8639979 | 0.00418331 | 0.008396 | 5.73781432 | 17.8603114 | 55.2217918 | 51.5490673 |
| 2.86341435 | 0.00419102 | 0.00841096 | 15.5740674 | 18.9427545 | 23.0090799 | 16.8094785 |
| 2.86337155 | 0.00419159 | 0.00841157 | 21.7217256 | 10.0125988 | 16.1063559 | 11.7666349 |
| -2.8630005 | 0.0041965 | 0.0084209 | 9.42640924 | 47.6274971 | 14.9559019 | 31.9380091 |
| 2.86290944 | 0.00419771 | 0.0084228 | 179.511619 | 171.567234 | 196.727633 | 198.351846 |
| -2.8622974 | 0.00420582 | 0.00843856 | 9.01656536 | 26.7904671 | 28.7613499 | 21.292006 |
| -2.8618736 | 0.00421145 | 0.00844933 | 12.2953164 | 22.1900839 | 14.9559019 | 33.058641 |
| -2.8616533 | 0.00421438 | 0.00845468 | 12.2953164 | 47.8981079 | 13.8054479 | 23.5332698 |
| -2.8615681 | 0.00421551 | 0.00845643 | 15.1642236 | 15.966036 | 17.2568099 | 20.1713742 |
| -2.8615155 | 0.00421621 | 0.0084573 | 0 | 207.558468 | 161.063559 | 159.129729 |
| -2.8614904 | 0.00421654 | 0.00845745 | 21.3118818 | 18.1309222 | 14.9559019 | 15.1285306 |
| -2.8614376 | 0.00421725 | 0.00845833 | 0 | 295.777582 | 230.090799 | 67.7982298 |
| -2.8609944 | 0.00422314 | 0.00846964 | 17.6232868 | 22.7313054 | 14.9559019 | 15.1285306 |
| -2.8607487 | 0.00422642 | 0.00847568 | 118.444881 | 98.7729344 | 0 | 194.429634 |
| -2.8603616 | 0.00423158 | 0.0084855 | 11.8854725 | 20.0251977 | 21.8586259 | 19.0507423 |
| -2.8603154 | 0.0042322 | 0.00848621 | 444.68061 | 1135.75344 | 729.387833 | 936.848266 |
| -2.8602205 | 0.00423347 | 0.00848823 | 25.8201644 | 23.0019162 | 51.7704298 | 7.28410733 |
| -2.8601192 | 0.00423482 | 0.00849041 | 93.0345607 | 135.846611 | 0 | 179.86142 |
| -2.859815 | 0.00423888 | 0.00849803 | 32.3776665 | 14.3423713 | 13.8054479 | 19.6110582 |
| -2.8596729 | 0.00424078 | 0.00850131 | 9.83625312 | 19.7545869 | 31.0622579 | 22.9729539 |
| -2.8594001 | 0.00424443 | 0.0085081 | 32.7875104 | 25.1668025 | 28.7613499 | 27.4554815 |
| -2.8593464 | 0.00424515 | 0.00850901 | 15.5740674 | 17.8603114 | 14.9559019 | 20.1713742 |
| 2.85926297 | 0.00424627 | 0.00851019 | 50.0009534 | 25.4374132 | 23.0090799 | 35.8602207 |
| -2.8592791 | 0.00424605 | 0.00851019 | 6.1476582 | 45.7332217 | 31.0622579 | 26.3348496 |
| -2.8591238 | 0.00424813 | 0.0085134 | 79.5097127 | 15.6954252 | 8.05317797 | 28.5761134 |
| -2.8591014 | 0.00424843 | 0.00851347 | 33.1973543 | 40.0503953 | 47.1686138 | 40.3427483 |
| -2.8586896 | 0.00425395 | 0.008524 | 400.417471 | 719.283451 | 767.352815 | 760.909058 |
| -2.85822 | 0.00426025 | 0.0085361 | 20.0823501 | 19.4839761 | 12.6549939 | 17.9301104 |
| -2.8579273 | 0.00426418 | 0.00854344 | 5.73781432 | 38.1561199 | 23.0090799 | 49.8681194 |
| -2.8578289 | 0.0042655 | 0.00854556 | 10.246097 | 18.9427545 | 28.7613499 | 22.9729539 |
| -2.8577704 | 0.00426629 | 0.00854661 | 40.5745441 | 76.0416289 | 109.29313 | 49.3078035 |
| -2.8576518 | 0.00426788 | 0.00854927 | 2.86890716 | 49.5217726 | 52.9208838 | 41.4633802 |
| -2.8569806 | 0.00427692 | 0.00856684 | 37.2957931 | 10.0125988 | 23.0090799 | 18.4904263 |
| 2.85684374 | 0.00427876 | 0.00857 | 6.96734596 | 7.30649104 | 9.20363196 | 14.0078987 |
| -2.8566821 | 0.00428094 | 0.00857384 | 0 | 158.307306 | 208.232173 | 156.888466 |
| -2.8565077 | 0.0042833 | 0.00857802 | 12.7051603 | 19.2133653 | 21.8586259 | 17.3697944 |
| -2.8564266 | 0.00428439 | 0.00857968 | 18.8528185 | 16.7778683 | 16.1063559 | 15.6888466 |
| -2.8555359 | 0.00429643 | 0.00860324 | 24.5906328 | 16.2366467 | 14.9559019 | 15.6888466 |

| | | | | | | |
|---|---|---|---|---|---|---|
| 2.85489083 | 0.00430516 | 0.0086202 | 151.642236 | 135.30539 | 130.001301 | 131.674248 |
| -2.8548462 | 0.00430577 | 0.00862088 | 21.3118818 | 18.6721438 | 20.7081719 | 11.7666349 |
| 2.85466234 | 0.00430826 | 0.00862534 | 0 | 0.27061078 | 0 | 0 |
| -2.8541885 | 0.00431469 | 0.00863768 | 8.60672148 | 34.6381797 | 20.7081719 | 26.3348496 |
| 2.85410147 | 0.00431587 | 0.00863951 | 14.3445358 | 13.530539 | 13.8054479 | 10.646003 |
| 2.85399663 | 0.0043173 | 0.00864182 | 8.1968776 | 4.05916169 | 4.60181598 | 2.80157974 |
| 2.85374041 | 0.00432078 | 0.00864826 | 21.7217256 | 8.11832337 | 4.60181598 | 14.0078987 |
| 2.85364768 | 0.00432204 | 0.00865025 | 60.6568942 | 44.3801678 | 55.2217918 | 40.9030642 |
| -2.8534139 | 0.00432523 | 0.00865608 | 5.73781432 | 63.0523115 | 14.9559019 | 54.910963 |
| 2.85298372 | 0.00433108 | 0.00866727 | 2.86890716 | 2.70610779 | 2.30090799 | 2.24126379 |
| -2.8529431 | 0.00433164 | 0.0086673 | 0 | 129.081342 | 227.789891 | 169.775732 |
| -2.8529598 | 0.00433141 | 0.0086673 | 388.531998 | 428.376863 | 447.526604 | 472.346345 |
| -2.8528195 | 0.00433332 | 0.00867014 | 18.0331307 | 25.1668025 | 11.50454 | 19.6110582 |
| -2.8527838 | 0.00433381 | 0.00867057 | 32.7875104 | 25.4374132 | 6.90272397 | 29.6967453 |
| -2.8525315 | 0.00433725 | 0.00867692 | 25.4103206 | 18.6721438 | 40.2658898 | 8.96505518 |
| -2.8524939 | 0.00433776 | 0.00867741 | 29.9186032 | 15.1542036 | 12.6549939 | 20.1713742 |
| 2.85184991 | 0.00434656 | 0.00869447 | 0 | 0 | 0 | 2.24126379 |
| -2.8513884 | 0.00435288 | 0.00870656 | 4.0984388 | 28.6847426 | 49.4695218 | 47.6268556 |
| -2.8510593 | 0.00435738 | 0.00871504 | 11.4756286 | 46.2744432 | 16.1063559 | 20.7316901 |
| -2.8509235 | 0.00435924 | 0.00871822 | 151.232392 | 200.251977 | 202.479903 | 197.231214 |
| -2.8508903 | 0.0043597 | 0.00871859 | 15.5740674 | 10.0125988 | 35.6640739 | 29.1364293 |
| -2.8507219 | 0.00436201 | 0.00872267 | 19.2626624 | 27.3316887 | 9.20363196 | 26.8951655 |
| -2.8506192 | 0.00436342 | 0.00872495 | 42.2139196 | 28.9553534 | 32.2127119 | 21.852322 |
| -2.8505675 | 0.00436413 | 0.00872582 | 22.1315695 | 38.9679522 | 19.5577179 | 25.2142177 |
| -2.8505166 | 0.00436483 | 0.00872668 | 6.96734596 | 31.6614612 | 27.6108959 | 29.6967453 |
| -2.8504213 | 0.00436614 | 0.00872823 | 33.1973543 | 18.1309222 | 12.6549939 | 16.2491625 |
| 2.85042071 | 0.00436614 | 0.00872823 | 0.40984388 | 0 | 1.150454 | 0.56031595 |
| -2.8501562 | 0.00436978 | 0.00873495 | 36.8859492 | 22.1900839 | 54.0713378 | 26.3348496 |
| 2.84925199 | 0.00438222 | 0.00875928 | 9.01656536 | 2.70610779 | 1.150454 | 3.36189569 |
| -2.8490404 | 0.00438513 | 0.00876456 | 12.7051603 | 32.7439043 | 51.7704298 | 10.0856871 |
| -2.8487536 | 0.00438909 | 0.00877192 | 4.0984388 | 39.238563 | 44.8677058 | 36.4205367 |
| -2.8486828 | 0.00439006 | 0.00877333 | 55.3289238 | 130.705006 | 0 | 271.753235 |
| -2.8483486 | 0.00439468 | 0.00878201 | 262.709927 | 195.110372 | 212.833989 | 238.694594 |
| -2.8477676 | 0.00440271 | 0.00879751 | 0 | 309.578731 | 62.1245158 | 216.842272 |
| -2.8473388 | 0.00440864 | 0.00880882 | 50.0009534 | 30.579018 | 4.60181598 | 33.6189569 |
| -2.8463007 | 0.00442304 | 0.00883705 | 55.7387677 | 179.414947 | 0 | 201.153426 |
| -2.8456344 | 0.00443231 | 0.00885501 | 12.2953164 | 18.1309222 | 18.4072639 | 22.4126379 |
| -2.8454413 | 0.00443499 | 0.00885983 | 28.6890716 | 23.5431378 | 9.20363196 | 20.1713742 |
| 2.84539815 | 0.00443559 | 0.00886049 | 0 | 0 | 0 | 0 |
| -2.8452138 | 0.00443816 | 0.00886507 | 105.739721 | 115.009581 | 0 | 173.137628 |
| -2.8445236 | 0.00444779 | 0.00888375 | 45.4926707 | 16.5072575 | 78.2308717 | 5.04284354 |
| -2.8444112 | 0.00444936 | 0.00888633 | 0 | 270.069558 | 90.8858656 | 183.223315 |
| -2.8443651 | 0.00445 | 0.00888707 | 10.246097 | 17.3190899 | 29.9118039 | 24.0935858 |

| | | | | | | |
|---|---|---|---|---|---|---|
| -2.8436789 | 0.0044596 | 0.00890568 | 34.017042 | 15.6954252 | 36.8145279 | 8.96505518 |
| -2.8436221 | 0.00446039 | 0.00890671 | 11.4756286 | 40.5916169 | 11.50454 | 34.1792729 |
| 2.84351984 | 0.00446182 | 0.00890902 | 0 | 0.27061078 | 0 | 0.56031595 |
| -2.8433187 | 0.00446464 | 0.0089141 | 39.7548564 | 19.4839761 | 9.20363196 | 21.852322 |
| -2.8432328 | 0.00446584 | 0.00891595 | 227.463353 | 258.433294 | 250.798971 | 256.624704 |
| -2.8429859 | 0.0044693 | 0.00892231 | 0 | 141.529437 | 196.727633 | 172.016996 |
| -2.842814 | 0.00447172 | 0.00892657 | 12.7051603 | 15.966036 | 35.6640739 | 17.9301104 |
| -2.8427704 | 0.00447233 | 0.00892724 | 9.01656536 | 27.0610779 | 27.6108959 | 20.1713742 |
| -2.8423078 | 0.00447882 | 0.00893914 | 11.4756286 | 51.1454373 | 11.50454 | 30.2570612 |
| -2.8423064 | 0.00447884 | 0.00893914 | 7.37718984 | 56.8282636 | 56.3722458 | 42.0236961 |
| -2.8421647 | 0.00448083 | 0.00894256 | 6.1476582 | 50.6042157 | 16.1063559 | 51.5490673 |
| -2.8418563 | 0.00448517 | 0.00895066 | 19.6725062 | 77.3946828 | 10.354086 | 16.2491625 |
| -2.8418178 | 0.00448571 | 0.00895119 | 10.6559409 | 19.2133653 | 25.3099879 | 20.7316901 |
| -2.841762 | 0.0044865 | 0.0089522 | 11.0657848 | 26.5198564 | 20.7081719 | 17.9301104 |
| 2.84171434 | 0.00448717 | 0.00895298 | 0 | 0.27061078 | 0 | 0 |
| -2.8415637 | 0.00448929 | 0.00895666 | 13.1150042 | 29.2259641 | 14.9559019 | 19.0507423 |
| -2.841454 | 0.00449083 | 0.00895919 | 9.42640924 | 31.6614612 | 37.9649818 | 15.1285306 |
| -2.8412606 | 0.00449356 | 0.00896407 | 0 | 25.708024 | 332.481205 | 344.033992 |
| -2.8411539 | 0.00449506 | 0.00896652 | 345.498391 | 341.510803 | 257.701695 | 327.224514 |
| -2.8411301 | 0.0044954 | 0.00896663 | 13.524848 | 11.3656527 | 32.2127119 | 29.1364293 |
| -2.8405582 | 0.00450346 | 0.00898217 | 23.3611012 | 20.83703 | 14.9559019 | 12.8872668 |
| 2.84043788 | 0.00450516 | 0.00898501 | 23.3611012 | 11.3656527 | 13.8054479 | 14.5682147 |
| -2.8402369 | 0.004508 | 0.00899011 | 36.8859492 | 14.3423713 | 17.2568099 | 14.5682147 |
| -2.8401593 | 0.0045091 | 0.00899174 | 10.6559409 | 19.7545869 | 23.0090799 | 21.292006 |
| -2.8400224 | 0.00451104 | 0.00899505 | 9.01656536 | 15.6954252 | 35.6640739 | 34.1792729 |
| -2.8399558 | 0.00451198 | 0.00899637 | 5.73781432 | 51.6866588 | 23.0090799 | 34.1792729 |
| -2.8397996 | 0.00451419 | 0.00900022 | 586.486592 | 33.2851258 | 0 | 56.5919108 |
| 2.83934201 | 0.00452067 | 0.00901258 | 1.22953164 | 6.22404792 | 3.45136199 | 2.24126379 |
| 2.83923291 | 0.00452221 | 0.00901511 | 0.40984388 | 0.54122156 | 0 | 0 |
| -2.8390497 | 0.00452481 | 0.00901973 | 13.1150042 | 19.4839761 | 18.4072639 | 17.3697944 |
| -2.8384698 | 0.00453304 | 0.00903558 | 73.3620545 | 226.771833 | 0 | 117.666349 |
| 2.83815941 | 0.00453745 | 0.00904381 | 9.01656536 | 8.11832337 | 10.354086 | 12.3269509 |
| -2.8377199 | 0.0045437 | 0.00905484 | 13.9346919 | 11.9068743 | 32.2127119 | 24.0935858 |
| -2.8377313 | 0.00454354 | 0.00905484 | 5.73781432 | 25.708024 | 47.1686138 | 31.3776931 |
| -2.8377113 | 0.00454383 | 0.00905484 | 33.1973543 | 21.1076408 | 17.2568099 | 10.646003 |
| -2.8375758 | 0.00454576 | 0.00905813 | 296.317125 | 475.463139 | 507.350212 | 481.3114 |
| -2.8374801 | 0.00454712 | 0.00906028 | 0 | 120.692407 | 185.223093 | 205.075637 |
| -2.8372651 | 0.00455018 | 0.00906583 | 26.6398522 | 14.3423713 | 19.5577179 | 13.4475828 |
| -2.8369689 | 0.00455441 | 0.00907368 | 17.213443 | 32.4732935 | 35.6640739 | 8.40473923 |
| -2.8368985 | 0.00455541 | 0.00907512 | 5.32797044 | 55.4752097 | 49.4695218 | 17.9301104 |
| -2.8368256 | 0.00455645 | 0.00907663 | 221.315695 | 476.816193 | 522.306114 | 498.681194 |
| -2.8364472 | 0.00456185 | 0.00908684 | 0 | 79.8301798 | 247.347609 | 214.601008 |
| -2.8361927 | 0.00456549 | 0.00909336 | 17.213443 | 18.401533 | 18.4072639 | 13.4475828 |

| | | | | | | |
|---|---|---|---|---|---|---|
| -2.8361787 | 0.00456569 | 0.00909336 | 218.856632 | 576.40096 | 429.11934 | 563.677844 |
| -2.8360634 | 0.00456734 | 0.00909608 | 7.78703372 | 19.7545869 | 31.0622579 | 35.2999048 |
| -2.8356957 | 0.0045726 | 0.009106 | 28.6890716 | 24.3549701 | 20.7081719 | 8.96505518 |
| -2.8352836 | 0.0045785 | 0.00911719 | 9.01656536 | 15.6954252 | 34.5136199 | 34.1792729 |
| -2.8351537 | 0.00458036 | 0.00912033 | 15.9839113 | 12.1774851 | 28.7613499 | 19.0507423 |
| -2.8350401 | 0.00458199 | 0.00912271 | 90.1656536 | 135.034779 | 276.108959 | 0 |
| -2.8350313 | 0.00458212 | 0.00912271 | 61.8864259 | 48.7099402 | 65.5758777 | 63.8760181 |
| 2.83417497 | 0.00459442 | 0.00914662 | 22.5414134 | 13.8011497 | 21.8586259 | 9.52537113 |
| -2.8340361 | 0.00459641 | 0.00915004 | 0 | 128.269509 | 142.656295 | 242.616806 |
| -2.8338703 | 0.0045988 | 0.00915422 | 105.329877 | 194.298539 | 172.568099 | 0 |
| -2.8334667 | 0.00460461 | 0.00916522 | 28.2792277 | 13.530539 | 19.5577179 | 14.0078987 |
| -2.8332816 | 0.00460728 | 0.00916997 | 10.6559409 | 18.401533 | 34.5136199 | 19.0507423 |
| -2.8330318 | 0.00461088 | 0.00917657 | 161.068645 | 0 | 387.702996 | 80.1251807 |
| 2.83285057 | 0.00461349 | 0.00918121 | 0.81968776 | 2.43549701 | 0 | 2.24126379 |
| -2.8319671 | 0.00462626 | 0.00920604 | 146.314265 | 156.954252 | 156.461743 | 0 |
| -2.8318337 | 0.00462819 | 0.00920932 | 0 | 226.771833 | 170.267191 | 113.744138 |
| -2.8307244 | 0.00464427 | 0.00924075 | 13.9346919 | 16.5072575 | 16.1063559 | 21.852322 |
| -2.8305164 | 0.00464729 | 0.00924619 | 160.248957 | 116.903857 | 0 | 104.218766 |
| -2.830362 | 0.00464954 | 0.00925008 | 17.6232868 | 18.401533 | 11.50454 | 25.2142177 |
| -2.8301884 | 0.00465206 | 0.00925453 | 106.559409 | 13.530539 | 12.6549939 | 15.6888466 |
| -2.8296508 | 0.00465988 | 0.00926953 | 28.2792277 | 30.3084073 | 21.8586259 | 7.84442328 |
| -2.8294081 | 0.00466342 | 0.00927599 | 0 | 125.563402 | 180.621277 | 194.429634 |
| 2.82934324 | 0.00466436 | 0.0092773 | 52.0501728 | 41.4034492 | 59.8236078 | 58.8331746 |
| -2.8292076 | 0.00466634 | 0.00928066 | 14.3445358 | 26.2492456 | 10.354086 | 37.5411686 |
| -2.8291139 | 0.00466771 | 0.0092828 | 5.32797044 | 46.8156648 | 26.4604419 | 33.058641 |
| -2.8290574 | 0.00466853 | 0.00928387 | 6.96734596 | 40.5916169 | 14.9559019 | 54.910963 |
| -2.8288815 | 0.0046711 | 0.0092884 | 0 | 183.744719 | 148.408565 | 163.051941 |
| -2.8288159 | 0.00467206 | 0.00928973 | 37.705637 | 15.6954252 | 9.20363196 | 28.0157974 |
| -2.8287239 | 0.0046734 | 0.00929183 | 278.693838 | 787.477367 | 702.927391 | 773.796325 |
| -2.8286149 | 0.00467499 | 0.00929442 | 18.4429746 | 30.8496288 | 26.4604419 | 8.96505518 |
| 2.82853675 | 0.00467613 | 0.00929555 | 0 | 0.27061078 | 0 | 0 |
| 2.82853675 | 0.00467613 | 0.00929555 | 0 | 0.27061078 | 0 | 0 |
| -2.8281785 | 0.00468137 | 0.00930539 | 14.3445358 | 19.4839761 | 13.8054479 | 21.292006 |
| -2.8280999 | 0.00468252 | 0.0093071 | 8.60672148 | 33.8263474 | 23.0090799 | 21.292006 |
| -2.8278696 | 0.00468589 | 0.00931322 | 36.0662614 | 59.2637606 | 69.0272397 | 65.556966 |
| -2.8276292 | 0.00468941 | 0.00931907 | 282.792277 | 498.194444 | 485.491586 | 545.187418 |
| -2.8276442 | 0.00468919 | 0.00931907 | 8.1968776 | 36.2618444 | 21.8586259 | 23.5332698 |
| 2.82732764 | 0.00469383 | 0.00932728 | 0 | 0 | 0 | 0 |
| 2.82720903 | 0.00469557 | 0.00933016 | 0 | 3.51794013 | 4.60181598 | 2.24126379 |
| -2.8270851 | 0.00469738 | 0.0093332 | 0 | 159.119138 | 130.001301 | 210.118481 |
| -2.8268571 | 0.00470073 | 0.00933927 | 80.7392443 | 107.161869 | 97.7885896 | 101.417187 |
| -2.8264743 | 0.00470635 | 0.00934986 | 0 | 81.1832337 | 279.560321 | 174.818576 |
| -2.8260981 | 0.00471188 | 0.00936028 | 0 | 199.710755 | 169.116737 | 128.312352 |

| | | | | | | |
|---|---|---|---|---|---|---|
| -2.8260536 | 0.00471254 | 0.009361 | 209.430223 | 294.153917 | 309.472125 | 314.337247 |
| -2.8260239 | 0.00471297 | 0.00936129 | 11.0657848 | 23.0019162 | 23.0090799 | 16.8094785 |
| -2.825935 | 0.00471428 | 0.00936332 | 65.5750208 | 237.866875 | 0 | 115.425085 |
| 2.8255262 | 0.0047203 | 0.00937469 | 7.37718984 | 6.76526948 | 6.90272397 | 3.92221164 |
| -2.8254894 | 0.00472084 | 0.00937519 | 58.197831 | 435.953965 | 498.14658 | 490.276455 |
| 2.82527671 | 0.00472398 | 0.00938084 | 6.1476582 | 0 | 0 | 0 |
| -2.8252555 | 0.00472429 | 0.00938089 | 12.2953164 | 19.2133653 | 17.2568099 | 20.1713742 |
| -2.825115 | 0.00472637 | 0.00938443 | 15.9839113 | 16.2366467 | 16.1063559 | 16.8094785 |
| -2.8242975 | 0.00473844 | 0.00940782 | 14.3445358 | 32.7439043 | 11.50454 | 22.4126379 |
| -2.8239298 | 0.00474388 | 0.00941804 | 22.5414134 | 22.7313054 | 25.3099879 | 8.96505518 |
| -2.8238208 | 0.00474549 | 0.00942008 | 11.8854725 | 16.2366467 | 23.0090799 | 20.1713742 |
| -2.8238317 | 0.00474533 | 0.00942008 | 109.018472 | 40.0503953 | 16.1063559 | 36.4205367 |
| 2.82367786 | 0.00474761 | 0.00942371 | 3.68859492 | 0 | 0 | 0.56031595 |
| -2.8230507 | 0.00475691 | 0.00944158 | 50.4107972 | 50.8748265 | 24.1595339 | 39.7824324 |
| -2.8230027 | 0.00475762 | 0.00944241 | 27.45954 | 14.3423713 | 18.4072639 | 13.4475828 |
| -2.8228326 | 0.00476014 | 0.00944684 | 26.6398522 | 30.3084073 | 43.7172518 | 5.60315949 |
| -2.8225907 | 0.00476373 | 0.00945281 | 31.1481349 | 36.2618444 | 6.90272397 | 20.7316901 |
| -2.8225982 | 0.00476362 | 0.00945281 | 18.4429746 | 25.708024 | 10.354086 | 20.1713742 |
| -2.8218768 | 0.00477435 | 0.0094733 | 24.1807889 | 37.0736767 | 8.05317797 | 20.1713742 |
| -2.8208616 | 0.00478949 | 0.00950274 | 0 | 423.235259 | 224.338529 | 27.4554815 |
| -2.8204905 | 0.00479503 | 0.00951316 | 11.4756286 | 18.1309222 | 18.4072639 | 23.5332698 |
| 2.82040917 | 0.00479625 | 0.00951498 | 0 | 20.2958084 | 9.20363196 | 8.96505518 |
| -2.8203819 | 0.00479665 | 0.00951521 | 20.9020379 | 9.74198805 | 40.2658898 | 17.9301104 |
| -2.8202662 | 0.00479838 | 0.00951805 | 9.01656536 | 25.1668025 | 17.2568099 | 38.1014845 |
| -2.8202028 | 0.00479933 | 0.00951935 | 11.8854725 | 13.2599282 | 24.1595339 | 33.6189569 |
| -2.8200016 | 0.00480234 | 0.00952474 | 115.575974 | 0 | 224.338529 | 211.239113 |
| -2.8195067 | 0.00480975 | 0.00953885 | 14.7543797 | 17.5897006 | 16.1063559 | 16.8094785 |
| 2.81936508 | 0.00481188 | 0.00954247 | 6.55750208 | 1.89427545 | 3.45136199 | 2.24126379 |
| -2.8182561 | 0.00482853 | 0.00957491 | 0 | 144.235545 | 166.815829 | 174.25826 |
| -2.8176568 | 0.00483755 | 0.00959221 | 27.8693838 | 18.1309222 | 27.6108959 | 8.96505518 |
| -2.8166391 | 0.0048529 | 0.00962206 | 157.789894 | 308.225677 | 291.064861 | 307.05314 |
| -2.8165815 | 0.00485377 | 0.0096232 | 80.3294005 | 104.455761 | 0 | 209.558165 |
| -2.8165373 | 0.00485444 | 0.00962393 | 428.286855 | 936.583907 | 915.76138 | 899.307098 |
| 2.81633931 | 0.00485743 | 0.00962927 | 136.068168 | 63.8641439 | 132.302209 | 61.6347543 |
| -2.8162573 | 0.00485867 | 0.00963114 | 118.035037 | 90.654611 | 0 | 168.094785 |
| -2.8162189 | 0.00485925 | 0.0096317 | 10.246097 | 14.8835929 | 34.5136199 | 25.2142177 |
| -2.8161543 | 0.00486023 | 0.00963305 | 19.6725062 | 31.3908504 | 36.8145279 | 33.6189569 |
| -2.8159394 | 0.00486348 | 0.00963835 | 32.3776665 | 20.5664192 | 9.20363196 | 19.6110582 |
| -2.8159184 | 0.0048638 | 0.00963835 | 4.91812656 | 45.7332217 | 23.0090799 | 42.5840121 |
| -2.8159358 | 0.00486354 | 0.00963835 | 7.37718984 | 26.5198564 | 27.6108959 | 26.3348496 |
| -2.8156516 | 0.00486784 | 0.00964535 | 19.2626624 | 18.9427545 | 17.2568099 | 12.3269509 |
| -2.8156455 | 0.00486793 | 0.00964535 | 32.3776665 | 18.401533 | 12.6549939 | 14.5682147 |
| -2.8155405 | 0.00486953 | 0.00964791 | 18.8528185 | 17.5897006 | 13.8054479 | 15.6888466 |

| | | | | | | |
|---|---|---|---|---|---|---|
| -2.8153999 | 0.00487166 | 0.00965154 | 7.37718984 | 17.8603114 | 39.1154358 | 33.6189569 |
| 2.81512251 | 0.00487586 | 0.00965929 | 0 | 0.27061078 | 0 | 0 |
| -2.8147509 | 0.00488151 | 0.00966987 | 29.9186032 | 19.4839761 | 37.9649818 | 36.9808526 |
| 2.81456551 | 0.00488432 | 0.00967485 | 0 | 0 | 0 | 0.56031595 |
| -2.8143919 | 0.00488696 | 0.00967949 | 84.4278393 | 14.0717605 | 9.20363196 | 22.4126379 |
| -2.814332 | 0.00488787 | 0.0096807 | 29.5087594 | 31.6614612 | 28.7613499 | 39.7824324 |
| -2.8142278 | 0.00488946 | 0.00968265 | 8.60672148 | 27.3316887 | 20.7081719 | 25.2142177 |
| -2.8142371 | 0.00488932 | 0.00968265 | 12.2953164 | 55.7458205 | 31.0622579 | 9.52537113 |
| -2.8141388 | 0.00489081 | 0.00968474 | 68.0340841 | 180.226779 | 240.444885 | 0 |
| 2.81377385 | 0.00489637 | 0.00969514 | 247.955547 | 254.644743 | 233.542161 | 253.262809 |
| -2.8131839 | 0.00490536 | 0.00971235 | 251.644142 | 633.229223 | 590.1829 | 628.114178 |
| -2.8128375 | 0.00491065 | 0.00972222 | 87.2967464 | 92.0076649 | 0 | 214.040692 |
| 2.81251622 | 0.00491555 | 0.00973134 | 1.63937552 | 1.08244312 | 2.30090799 | 0.56031595 |
| -2.8123408 | 0.00491824 | 0.00973606 | 22.1315695 | 12.7187066 | 40.2658898 | 12.3269509 |
| -2.8123093 | 0.00491872 | 0.00973641 | 161.888333 | 82.8068984 | 0 | 131.113932 |
| -2.8122757 | 0.00491923 | 0.00973683 | 35.2465737 | 17.0484791 | 10.354086 | 19.6110582 |
| -2.8122438 | 0.00491972 | 0.0097372 | 142.62567 | 467.344816 | 384.251634 | 416.31475 |
| 2.81215497 | 0.00492108 | 0.0097393 | 3.68859492 | 4.05916169 | 4.60181598 | 3.36189569 |
| -2.812085 | 0.00492215 | 0.00974082 | 90.1656536 | 86.0542278 | 0 | 219.643852 |
| -2.811821 | 0.00492619 | 0.00974822 | 15.1642236 | 30.0377965 | 9.20363196 | 31.9380091 |
| 2.81154933 | 0.00493035 | 0.00975586 | 0 | 0.54122156 | 0 | 0 |
| -2.8112881 | 0.00493436 | 0.0097628 | 81.968776 | 138.011497 | 0 | 154.647202 |
| -2.811281 | 0.00493447 | 0.0097628 | 120.903945 | 0 | 236.993523 | 182.662999 |
| -2.8112348 | 0.00493518 | 0.0097636 | 8.60672148 | 49.2511618 | 44.8677058 | 11.206319 |
| -2.8108211 | 0.00494153 | 0.00977557 | 100.411751 | 0 | 380.800272 | 116.545717 |
| -2.8106801 | 0.00494369 | 0.00977925 | 15.9839113 | 43.0271139 | 28.7613499 | 8.40473923 |
| -2.8101961 | 0.00495113 | 0.00979337 | 346.727922 | 757.16896 | 331.330751 | 548.549314 |
| 2.81003815 | 0.00495356 | 0.00979758 | 1.63937552 | 1.08244312 | 1.150454 | 1.1206319 |
| 2.80971209 | 0.00495858 | 0.00980691 | 16.3937552 | 16.5072575 | 33.3631659 | 13.4475828 |
| 2.80959597 | 0.00496037 | 0.00980985 | 6.1476582 | 0.54122156 | 1.150454 | 1.1206319 |
| -2.8094969 | 0.0049619 | 0.00981226 | 15.9839113 | 17.0484791 | 13.8054479 | 18.4904263 |
| -2.8089013 | 0.00497109 | 0.00982984 | 73.3620545 | 206.746635 | 0 | 110.942558 |
| -2.8086383 | 0.00497515 | 0.00983726 | 4.50828268 | 24.6255809 | 78.2308717 | 28.5761134 |
| -2.8083149 | 0.00498015 | 0.00984655 | 15.5740674 | 28.6847426 | 8.05317797 | 48.7474875 |
| -2.8081404 | 0.00498285 | 0.00985128 | 11.0657848 | 26.2492456 | 16.1063559 | 20.7316901 |
| -2.8072927 | 0.00499598 | 0.00987664 | 9.83625312 | 22.7313054 | 18.4072639 | 24.6539017 |
| -2.8072333 | 0.0049969 | 0.00987786 | 9.42640924 | 46.8156648 | 27.6108959 | 14.0078987 |
| -2.8071577 | 0.00499808 | 0.00987957 | 307.792754 | 515.784145 | 521.15566 | 507.646249 |
| -2.8064004 | 0.00500984 | 0.00990222 | 12.7051603 | 23.8137486 | 11.50454 | 39.7824324 |
| -2.8059796 | 0.00501639 | 0.00991455 | 6.96734596 | 28.9553534 | 31.0622579 | 23.5332698 |
| -2.8059106 | 0.00501746 | 0.00991607 | 12.2953164 | 20.0251977 | 14.9559019 | 22.4126379 |
| -2.8054437 | 0.00502474 | 0.00992983 | 8.1968776 | 29.2259641 | 17.2568099 | 35.2999048 |
| -2.8053881 | 0.0050256 | 0.00993094 | 6.1476582 | 19.7545869 | 43.7172518 | 35.2999048 |

| | | | | | | |
|---|---|---|---|---|---|---|
| -2.8048723 | 0.00503365 | 0.00994624 | 0 | 165.343186 | 150.709473 | 155.767834 |
| -2.8047175 | 0.00503607 | 0.00995041 | 11.0657848 | 24.6255809 | 18.4072639 | 17.9301104 |
| 2.80455622 | 0.00503859 | 0.00995478 | 0.40984388 | 0.54122156 | 0 | 0 |
| -2.8045112 | 0.00503929 | 0.00995556 | 4.91812656 | 37.6148983 | 29.9118039 | 34.7395888 |
| -2.8043213 | 0.00504226 | 0.00996081 | 8.1968776 | 29.4965749 | 18.4072639 | 30.2570612 |
| 2.80427073 | 0.00504305 | 0.00996177 | 13.524848 | 6.4946587 | 8.05317797 | 7.28410733 |
| -2.8036654 | 0.00505253 | 0.00997988 | 30.738291 | 21.6488623 | 10.354086 | 15.6888466 |
| 2.80352495 | 0.00505473 | 0.00998361 | 0 | 0 | 0 | 0 |
| 2.80336664 | 0.00505721 | 0.00998791 | 0.81968776 | 0.54122156 | 4.60181598 | 0.56031595 |
| -2.8032503 | 0.00505904 | 0.0099909 | 9.42640924 | 34.6381797 | 37.9649818 | 12.8872668 |
| -2.8031774 | 0.00506018 | 0.00999255 | 21.3118818 | 20.0251977 | 17.2568099 | 11.206319 |
| -2.8028918 | 0.00506467 | 0.00999956 | 15.1642236 | 59.2637606 | 8.05317797 | 26.8951655 |
| -2.8029032 | 0.00506449 | 0.00999956 | 12.7051603 | 18.9427545 | 13.8054479 | 30.8173772 |
| -2.8029302 | 0.00506406 | 0.00999956 | 18.0331307 | 22.7313054 | 12.6549939 | 15.1285306 |
| -2.8026076 | 0.00506913 | 0.01000777 | 15.1642236 | 21.3782516 | 12.6549939 | 18.4904263 |
| -2.8025819 | 0.00506953 | 0.01000795 | 9.01656536 | 20.0251977 | 21.8586259 | 28.5761134 |
| -2.8025164 | 0.00507056 | 0.01000937 | 60.6568942 | 42.4858923 | 6.90272397 | 12.3269509 |
| -2.8023454 | 0.00507325 | 0.01001407 | 43.4434513 | 56.8282636 | 19.5577179 | 40.9030642 |
| -2.8021409 | 0.00507647 | 0.01001919 | 5.32797044 | 31.6614612 | 24.1595339 | 48.1871716 |
| -2.8021571 | 0.00507622 | 0.01001919 | 11.8854725 | 22.7313054 | 20.7081719 | 15.1285306 |
| -2.8020886 | 0.00507729 | 0.0100202 | 11.4756286 | 32.7439043 | 6.90272397 | 96.9346591 |
| -2.80197 | 0.00507916 | 0.01002328 | 11.8854725 | 17.8603114 | 42.5667978 | 15.1285306 |
| -2.80188 | 0.00508058 | 0.01002546 | 75.4112739 | 153.706923 | 0 | 142.880567 |
| 2.80158317 | 0.00508525 | 0.01003407 | 3.27875104 | 3.51794013 | 2.30090799 | 1.68094785 |
| 2.80142889 | 0.00508769 | 0.01003826 | 0 | 0 | 0 | 0.56031595 |
| 2.80088811 | 0.00509622 | 0.01005448 | 3.68859492 | 5.68282636 | 2.30090799 | 3.36189569 |
| -2.8008251 | 0.00509721 | 0.01005583 | 65.1651769 | 172.920288 | 0 | 144.001199 |
| -2.8004734 | 0.00510277 | 0.01006557 | 104.920033 | 0 | 318.675757 | 135.59646 |
| -2.8004871 | 0.00510256 | 0.01006557 | 102.46097 | 125.02218 | 0 | 130.553616 |
| -2.8001389 | 0.00510806 | 0.01007539 | 0 | 75.5004074 | 250.798971 | 181.542367 |
| -2.7997921 | 0.00511355 | 0.0100856 | 110.248004 | 119.339354 | 219.736713 | 0 |
| -2.7991783 | 0.00512328 | 0.01010418 | 3.68859492 | 59.2637606 | 52.9208838 | 20.7316901 |
| 2.79905392 | 0.00512526 | 0.01010745 | 63.5258014 | 28.6847426 | 13.8054479 | 31.9380091 |
| -2.7986735 | 0.0051313 | 0.01011875 | 13.1150042 | 19.7545869 | 13.8054479 | 21.852322 |
| 2.79834064 | 0.00513659 | 0.01012856 | 49.1812656 | 19.4839761 | 6.90272397 | 21.292006 |
| 2.7982696 | 0.00513772 | 0.01013017 | 13.524848 | 47.0862756 | 49.4695218 | 0 |
| -2.7981789 | 0.00513916 | 0.0101324 | 54.9190799 | 83.34812 | 128.850847 | 135.036144 |
| -2.7977874 | 0.0051454 | 0.01014407 | 139.756763 | 103.102707 | 0 | 114.304454 |
| -2.7975629 | 0.00514897 | 0.0101505 | 25.0004767 | 205.664192 | 0 | 250.461229 |
| -2.7975424 | 0.0051493 | 0.01015053 | 5.32797044 | 26.5198564 | 35.6640739 | 35.2999048 |
| 2.79743424 | 0.00515103 | 0.01015281 | 0.81968776 | 0.27061078 | 0 | 1.68094785 |
| -2.7974304 | 0.00515109 | 0.01015281 | 31.5579788 | 25.4374132 | 5.75226998 | 33.058641 |
| -2.7971441 | 0.00515565 | 0.01016119 | 22.5414134 | 21.6488623 | 8.05317797 | 28.5761134 |

| | | | | | | |
|---|---|---|---|---|---|---|
| -2.7970726 | 0.00515679 | 0.01016282 | 7.78703372 | 28.4141318 | 19.5577179 | 30.8173772 |
| 2.79692987 | 0.00515907 | 0.01016669 | 0.40984388 | 4.32977247 | 1.150454 | 6.16347543 |
| -2.7966481 | 0.00516357 | 0.01017494 | 149.593016 | 90.3840002 | 0 | 120.467929 |
| -2.7964643 | 0.00516651 | 0.01017959 | 5.32797044 | 51.6866588 | 13.8054479 | 61.0744384 |
| 2.79646077 | 0.00516657 | 0.01017959 | 6.1476582 | 5.1416048 | 4.60181598 | 8.96505518 |
| 2.79628431 | 0.00516939 | 0.01018453 | 37.2957931 | 23.0019162 | 26.4604419 | 14.5682147 |
| -2.7959092 | 0.00517539 | 0.01019574 | 18.0331307 | 11.0950419 | 25.3099879 | 17.3697944 |
| -2.7957245 | 0.00517835 | 0.01020094 | 29.0989155 | 16.7778683 | 11.50454 | 16.8094785 |
| -2.7955352 | 0.00518139 | 0.0102063 | 9.42640924 | 25.9786348 | 16.1063559 | 29.1364293 |
| -2.7947467 | 0.00519404 | 0.0102306 | 55.3289238 | 124.751569 | 111.594038 | 99.1759229 |
| -2.7944995 | 0.00519801 | 0.0102378 | 3.68859492 | 56.5576528 | 62.1245158 | 18.4904263 |
| -2.7944496 | 0.00519882 | 0.01023875 | 16.3937552 | 22.7313054 | 9.20363196 | 35.8602207 |
| -2.7943239 | 0.00520084 | 0.01024211 | 6.55750208 | 24.6255809 | 31.0622579 | 29.1364293 |
| -2.7940532 | 0.00520519 | 0.01025006 | 61.0667381 | 4.05916169 | 18.4072639 | 51.5490673 |
| -2.793771 | 0.00520973 | 0.01025838 | 62.2962697 | 155.601198 | 195.577179 | 156.32815 |
| -2.7937264 | 0.00521045 | 0.01025917 | 9.01656536 | 44.9213893 | 18.4072639 | 20.1713742 |
| -2.7934659 | 0.00521465 | 0.01026643 | 5.73781432 | 41.1328384 | 20.7081719 | 36.9808526 |
| -2.7934582 | 0.00521478 | 0.01026643 | 205.331784 | 86.0542278 | 0 | 87.9696039 |
| -2.7932046 | 0.00521887 | 0.01027386 | 163.117864 | 63.8641439 | 248.498063 | 0 |
| -2.7930105 | 0.005222 | 0.0102794 | 77.8703372 | 15.4248144 | 12.6549939 | 14.0078987 |
| 2.79279757 | 0.00522544 | 0.01028491 | 50.8206411 | 23.272527 | 18.4072639 | 33.6189569 |
| -2.7927977 | 0.00522544 | 0.01028491 | 15.9839113 | 8.11832337 | 32.2127119 | 35.2999048 |
| -2.7927502 | 0.0052262 | 0.01028579 | 13.9346919 | 12.7187066 | 23.0090799 | 19.6110582 |
| -2.7926739 | 0.00522744 | 0.01028759 | 5.32797044 | 47.0862756 | 110.443584 | 8.96505518 |
| -2.7918709 | 0.00524043 | 0.01031253 | 0 | 160.472192 | 147.258111 | 152.966254 |
| -2.7915421 | 0.00524575 | 0.01032238 | 175.003337 | 82.2656769 | 189.824909 | 0 |
| -2.7914177 | 0.00524777 | 0.01032572 | 132.379573 | 123.398515 | 0 | 97.4949751 |
| -2.7911021 | 0.00525289 | 0.01033516 | 11.0657848 | 24.3549701 | 20.7081719 | 15.6888466 |
| -2.7907842 | 0.00525805 | 0.01034469 | 175.823024 | 206.746635 | 243.896247 | 227.488275 |
| -2.7907187 | 0.00525912 | 0.01034615 | 295.907281 | 563.682253 | 591.333354 | 602.899961 |
| 2.79060299 | 0.005261 | 0.01034922 | 59.8372065 | 259.245126 | 195.577179 | 146.242463 |
| 2.78992559 | 0.00527202 | 0.01037027 | 0.81968776 | 0.81183234 | 1.150454 | 0.56031595 |
| -2.7894184 | 0.00528028 | 0.01038589 | 13.9346919 | 23.5431378 | 12.6549939 | 19.0507423 |
| -2.7892494 | 0.00528304 | 0.01039068 | 4.50828268 | 43.2977247 | 26.4604419 | 37.5411686 |
| -2.7890991 | 0.00528549 | 0.01039487 | 22.9512573 | 14.3423713 | 19.5577179 | 12.3269509 |
| -2.7889438 | 0.00528802 | 0.01039922 | 4.91812656 | 12.7187066 | 44.8677058 | 85.7283401 |
| -2.7886827 | 0.00529229 | 0.01040698 | 10.246097 | 32.7439043 | 11.50454 | 38.1014845 |
| -2.787894 | 0.00530519 | 0.01043171 | 77.8703372 | 64.946587 | 56.3722458 | 70.0394936 |
| 2.78755319 | 0.00531077 | 0.01044205 | 0 | 0.27061078 | 0 | 0 |
| -2.787486 | 0.00531187 | 0.01044358 | 24.5906328 | 10.0125988 | 44.8677058 | 13.4475828 |
| -2.7873522 | 0.00531407 | 0.01044726 | 15.1642236 | 17.8603114 | 11.50454 | 34.7395888 |
| -2.787306 | 0.00531483 | 0.01044811 | 75.00143 | 81.1832337 | 33.3631659 | 71.7204414 |
| -2.7859472 | 0.00533716 | 0.01049137 | 49.5911095 | 60.8874253 | 12.6549939 | 64.99665 |

| | | | | | | |
|---|---|---|---|---|---|---|
| -2.7858869 | 0.00533815 | 0.01049268 | 9.01656536 | 29.7671857 | 14.9559019 | 31.3776931 |
| -2.7858038 | 0.00533952 | 0.01049474 | 11.0657848 | 24.3549701 | 17.2568099 | 17.9301104 |
| -2.7856349 | 0.0053423 | 0.01049893 | 126.641759 | 115.280192 | 0 | 105.899714 |
| -2.785649 | 0.00534207 | 0.01049893 | 68.4439279 | 135.576 | 0 | 162.491625 |
| -2.7855341 | 0.00534396 | 0.01050155 | 17.6232868 | 22.7313054 | 9.20363196 | 25.7745336 |
| -2.7853221 | 0.00534746 | 0.01050778 | 9.83625312 | 30.8496288 | 43.7172518 | 11.7666349 |
| -2.785194 | 0.00534957 | 0.01051129 | 28.6890716 | 43.0271139 | 56.3722458 | 24.6539017 |
| -2.7848434 | 0.00535536 | 0.01052202 | 2.0492194 | 43.8389462 | 50.6199758 | 50.9887513 |
| 2.78469152 | 0.00535787 | 0.01052631 | 8.60672148 | 5.41221558 | 4.60181598 | 8.40473923 |
| -2.7846256 | 0.00535896 | 0.01052781 | 0 | 192.674875 | 139.204933 | 127.752036 |
| -2.7845378 | 0.00536041 | 0.01053002 | 337.301513 | 839.705248 | 755.848275 | 824.785076 |
| 2.78440932 | 0.00536253 | 0.01053356 | 8.60672148 | 8.11832337 | 4.60181598 | 10.0856871 |
| 2.78405827 | 0.00536834 | 0.01054432 | 22.9512573 | 15.4248144 | 13.8054479 | 10.646003 |
| -2.7840041 | 0.00536924 | 0.01054544 | 10.6559409 | 31.9320719 | 16.1063559 | 19.0507423 |
| -2.7839138 | 0.00537073 | 0.01054766 | 184.019902 | 353.958899 | 345.136199 | 377.652949 |
| -2.7838964 | 0.00537102 | 0.01054766 | 20.9020379 | 20.2958084 | 21.8586259 | 9.52537113 |
| -2.7837543 | 0.00537337 | 0.01055164 | 0 | 149.647761 | 162.214013 | 142.320251 |
| -2.7835822 | 0.00537622 | 0.0105566 | 3.68859492 | 43.5683354 | 23.0090799 | 58.8331746 |
| -2.7834918 | 0.00537772 | 0.0105589 | 2.0492194 | 46.2744432 | 63.2749697 | 38.6618005 |
| 2.78333276 | 0.00538036 | 0.01056343 | 24.1807889 | 9.47137727 | 2.30090799 | 11.7666349 |
| -2.7829608 | 0.00538653 | 0.01057491 | 15.9839113 | 18.6721438 | 16.1063559 | 13.4475828 |
| -2.7826899 | 0.00539103 | 0.0105831 | 20.492194 | 12.1774851 | 14.9559019 | 19.6110582 |
| -2.7823095 | 0.00539735 | 0.01059487 | 9.01656536 | 24.6255809 | 18.4072639 | 24.6539017 |
| -2.7821527 | 0.00539996 | 0.01059867 | 47.5418901 | 15.1542036 | 9.20363196 | 22.4126379 |
| -2.7821338 | 0.00540028 | 0.01059867 | 11.4756286 | 31.9320719 | 34.5136199 | 10.646003 |
| -2.7821389 | 0.00540019 | 0.01059867 | 103.690502 | 15.966036 | 10.354086 | 13.4475828 |
| -2.7818361 | 0.00540523 | 0.01060775 | 194.265999 | 101.208431 | 0 | 74.5220212 |
| -2.7818028 | 0.00540579 | 0.0106082 | 17.6232868 | 33.8263474 | 33.3631659 | 44.2649599 |
| -2.781626 | 0.00540873 | 0.0106114 | 7.37718984 | 26.7904671 | 24.1595339 | 25.2142177 |
| -2.7816285 | 0.00540869 | 0.0106114 | 46.7222023 | 14.8835929 | 9.20363196 | 22.9729539 |
| -2.7816259 | 0.00540874 | 0.0106114 | 100.411751 | 100.66721 | 0 | 149.044042 |
| -2.7816829 | 0.00540778 | 0.0106114 | 15.1642236 | 12.7187066 | 14.9559019 | 57.7125427 |
| -2.7814455 | 0.00541174 | 0.01061665 | 57.7879871 | 59.8049822 | 57.5226998 | 57.1522268 |
| -2.7803734 | 0.00542964 | 0.01065113 | 0 | 123.127905 | 161.063559 | 169.775732 |
| -2.7801467 | 0.00543344 | 0.01065792 | 234.430699 | 258.974516 | 0 | 394.462428 |
| -2.7799477 | 0.00543676 | 0.0106638 | 113.936599 | 121.50424 | 0 | 252.142177 |
| -2.7798966 | 0.00543762 | 0.01066419 | 9.42640924 | 19.7545869 | 36.8145279 | 17.3697944 |
| -2.7799077 | 0.00543743 | 0.01066419 | 6.1476582 | 45.1920001 | 57.5226998 | 12.8872668 |
| 2.77971269 | 0.0054407 | 0.01066893 | 16.8035991 | 8.93015571 | 6.90272397 | 9.52537113 |
| -2.7797127 | 0.0054407 | 0.01066893 | 8.60672148 | 50.3336049 | 40.2658898 | 10.646003 |
| -2.7792712 | 0.0054481 | 0.01068279 | 2.0492194 | 37.3442875 | 41.4163438 | 72.2807574 |
| -2.7784852 | 0.0054613 | 0.01070802 | 113.116911 | 181.309222 | 128.850847 | 0 |
| 2.77840469 | 0.00546265 | 0.01071003 | 28.6890716 | 49.5217726 | 0 | 10.0856871 |

| | | | | | | |
|---|---|---|---|---|---|---|
| -2.7782203 | 0.00546576 | 0.01071546 | 72.9522106 | 124.751569 | 0 | 160.250361 |
| -2.7773679 | 0.00548011 | 0.01074295 | 8.1968776 | 27.6022995 | 21.8586259 | 21.852322 |
| -2.7773327 | 0.0054807 | 0.01074346 | 7.78703372 | 36.5324552 | 18.4072639 | 25.2142177 |
| -2.7771414 | 0.00548393 | 0.01074913 | 27.0496961 | 66.2996409 | 8.05317797 | 50.9887513 |
| -2.7769143 | 0.00548776 | 0.01075599 | 15.9839113 | 41.1328384 | 35.6640739 | 42.0236961 |
| 2.77678537 | 0.00548994 | 0.01075961 | 41.3942319 | 51.1454373 | 63.2749697 | 52.6696992 |
| -2.7765605 | 0.00549374 | 0.0107664 | 20.0823501 | 28.9553534 | 19.5577179 | 8.96505518 |
| 2.77605152 | 0.00550235 | 0.01078196 | 14.3445358 | 11.6362635 | 14.9559019 | 8.40473923 |
| -2.7760688 | 0.00550206 | 0.01078196 | 11.0657848 | 40.0503953 | 14.9559019 | 18.4904263 |
| 2.77592092 | 0.00550456 | 0.01078564 | 0.81968776 | 3.51794013 | 0 | 4.48252759 |
| -2.7757908 | 0.00550676 | 0.0107893 | 61.0667381 | 257.621462 | 0 | 84.0473923 |
| -2.7754947 | 0.00551178 | 0.01079847 | 4.91812656 | 30.8496288 | 26.4604419 | 43.144328 |
| -2.7753269 | 0.00551463 | 0.01080339 | 22.9512573 | 32.7439043 | 5.75226998 | 32.498325 |
| -2.774962 | 0.00552082 | 0.01081486 | 31.1481349 | 20.2958084 | 20.7081719 | 8.40473923 |
| -2.7743956 | 0.00553044 | 0.01083306 | 0 | 89.0309463 | 185.223093 | 190.507423 |
| -2.7740746 | 0.0055359 | 0.01084309 | 13.1150042 | 21.3782516 | 14.9559019 | 16.2491625 |
| -2.7737977 | 0.00554061 | 0.01085107 | 5.73781432 | 27.8729103 | 28.7613499 | 31.9380091 |
| -2.7737957 | 0.00554065 | 0.01085107 | 76.6408055 | 155.059976 | 0 | 120.467929 |
| -2.7737028 | 0.00554223 | 0.01085352 | 13.1150042 | 18.1309222 | 12.6549939 | 33.058641 |
| -2.7735252 | 0.00554526 | 0.01085879 | 6.1476582 | 20.0251977 | 35.6640739 | 34.7395888 |
| 2.77343866 | 0.00554673 | 0.01086101 | 265.578834 | 398.068456 | 391.154358 | 361.964103 |
| -2.7731311 | 0.00555198 | 0.01087063 | 27.0496961 | 230.289773 | 0 | 187.145527 |
| -2.7730654 | 0.0055531 | 0.01087216 | 310.661661 | 697.093367 | 663.811955 | 702.6362 |
| -2.7729334 | 0.00555535 | 0.01087591 | 13.1150042 | 29.2259641 | 10.354086 | 26.3348496 |
| -2.7728125 | 0.00555741 | 0.01087929 | 18.4429746 | 12.4480958 | 20.7081719 | 14.5682147 |
| 2.77262857 | 0.00556055 | 0.01088478 | 236.070075 | 236.513821 | 216.285351 | 220.204168 |
| -2.7725061 | 0.00556265 | 0.01088769 | 132.789417 | 149.918372 | 0 | 71.1601255 |
| -2.7725021 | 0.00556272 | 0.01088769 | 44.263139 | 12.9893174 | 13.8054479 | 16.2491625 |
| -2.7722067 | 0.00556777 | 0.01089692 | 26.2300083 | 9.74198805 | 72.4786017 | 10.646003 |
| 2.77212159 | 0.00556922 | 0.01089881 | 39.7548564 | 19.7545869 | 32.2127119 | 19.0507423 |
| -2.7721109 | 0.00556941 | 0.01089881 | 90.1656536 | 68.7351379 | 0 | 218.52322 |
| -2.7719215 | 0.00557265 | 0.01090449 | 6.55750208 | 26.2492456 | 29.9118039 | 25.2142177 |
| -2.7715093 | 0.00557971 | 0.01091764 | 87.7065903 | 94.9843835 | 73.6290557 | 112.623506 |
| -2.7713285 | 0.00558281 | 0.01092305 | 107.379097 | 103.102707 | 0 | 129.432984 |
| -2.7712089 | 0.00558486 | 0.0109264 | 25.4103206 | 11.3656527 | 16.1063559 | 17.3697944 |
| -2.7711774 | 0.0055854 | 0.01092679 | 29.9186032 | 24.8961917 | 6.90272397 | 22.4126379 |
| -2.7711228 | 0.00558634 | 0.01092796 | 41.3942319 | 23.272527 | 10.354086 | 12.8872668 |
| -2.7710296 | 0.00558793 | 0.01093043 | 13.524848 | 26.7904671 | 11.50454 | 20.7316901 |
| -2.7708129 | 0.00559166 | 0.01093638 | 30.3284471 | 9.74198805 | 17.2568099 | 19.6110582 |
| -2.7708156 | 0.00559161 | 0.01093638 | 17.213443 | 57.3694852 | 39.1154358 | 29.6967453 |
| -2.7707319 | 0.00559305 | 0.01093844 | 8.60672148 | 22.7313054 | 18.4072639 | 29.1364293 |
| -2.7705733 | 0.00559577 | 0.0109429 | 8.60672148 | 32.2026827 | 14.9559019 | 29.6967453 |
| -2.7705598 | 0.005596 | 0.0109429 | 28.2792277 | 24.8961917 | 4.60181598 | 54.350647 |

| | | | | | | |
|---|---|---|---|---|---|---|
| -2.7702969 | 0.00560052 | 0.01095107 | 5.32797044 | 32.2026827 | 29.9118039 | 29.6967453 |
| -2.7698216 | 0.0056087 | 0.0109664 | 6.55750208 | 55.7458205 | 24.1595339 | 19.6110582 |
| -2.7692727 | 0.00561816 | 0.01098423 | 12.7051603 | 18.9427545 | 13.8054479 | 20.7316901 |
| -2.7691928 | 0.00561954 | 0.01098625 | 18.0331307 | 32.2026827 | 9.20363196 | 19.0507423 |
| 2.76860603 | 0.00562967 | 0.01100539 | 0.40984388 | 0.54122156 | 1.150454 | 0 |
| -2.7683895 | 0.00563341 | 0.01101204 | 29.9186032 | 18.1309222 | 6.90272397 | 34.1792729 |
| -2.7683384 | 0.00563429 | 0.0110131 | 10.6559409 | 18.6721438 | 23.0090799 | 16.8094785 |
| -2.7680411 | 0.00563943 | 0.01102205 | 148.773328 | 87.4072817 | 0 | 107.580662 |
| -2.7680343 | 0.00563955 | 0.01102205 | 11.8854725 | 25.708024 | 14.9559019 | 17.3697944 |
| -2.7679864 | 0.00564038 | 0.011023 | 157.789894 | 43.2977247 | 0 | 188.266159 |
| -2.7677068 | 0.00564522 | 0.01103133 | 65.5750208 | 126.104623 | 0 | 165.293205 |
| -2.7677008 | 0.00564532 | 0.01103133 | 267.21821 | 333.121869 | 210.533081 | 265.029444 |
| -2.7676738 | 0.00564579 | 0.01103157 | 14.7543797 | 13.8011497 | 18.4072639 | 16.2491625 |
| -2.7676304 | 0.00564655 | 0.01103238 | 4.0984388 | 57.640096 | 14.9559019 | 59.9538065 |
| -2.767543 | 0.00564806 | 0.01103467 | 8.1968776 | 21.3782516 | 26.4604419 | 21.852322 |
| -2.7674165 | 0.00565025 | 0.01103828 | 137.707544 | 280.082156 | 258.852149 | 267.270707 |
| -2.7671972 | 0.00565405 | 0.01104501 | 40.984388 | 10.5538204 | 37.9649818 | 9.52537113 |
| -2.7671784 | 0.00565438 | 0.01104501 | 14.7543797 | 35.7206228 | 48.3190678 | 44.8252759 |
| -2.7671563 | 0.00565476 | 0.01104509 | 19.2626624 | 14.3423713 | 13.8054479 | 16.2491625 |
| -2.7669769 | 0.00565788 | 0.0110505 | 103.280658 | 211.88824 | 138.054479 | 187.145527 |
| -2.766937 | 0.00565857 | 0.01105119 | 0 | 100.937821 | 193.276271 | 157.448782 |
| -2.7669096 | 0.00565904 | 0.01105145 | 55.3289238 | 10.8244312 | 19.5577179 | 13.4475828 |
| -2.7664934 | 0.00566627 | 0.0110649 | 13.524848 | 23.5431378 | 10.354086 | 29.6967453 |
| -2.7664168 | 0.0056676 | 0.01106682 | 8.60672148 | 18.6721438 | 27.6108959 | 22.4126379 |
| -2.7659033 | 0.00567654 | 0.0110836 | 0 | 224.606947 | 147.258111 | 89.6505518 |
| 2.76574715 | 0.00567926 | 0.01108756 | 33.6071982 | 104.455761 | 47.1686138 | 115.985401 |
| -2.7657659 | 0.00567893 | 0.01108756 | 9.01656536 | 24.0843593 | 21.8586259 | 19.0507423 |
| -2.7657071 | 0.00567995 | 0.01108825 | 18.0331307 | 18.1309222 | 20.7081719 | 10.646003 |
| -2.7651392 | 0.00568985 | 0.01110691 | 6.1476582 | 24.8961917 | 29.9118039 | 29.1364293 |
| -2.7651156 | 0.00569026 | 0.01110704 | 22.5414134 | 16.7778683 | 14.9559019 | 12.3269509 |
| 2.76476035 | 0.00569646 | 0.01111847 | 85.247527 | 228.124887 | 225.488983 | 223.005748 |
| 2.76439784 | 0.0057028 | 0.01113016 | 1.22953164 | 0.27061078 | 0 | 0 |
| -2.7641767 | 0.00570666 | 0.01113703 | 43.4434513 | 256.809629 | 0 | 107.580662 |
| -2.7640678 | 0.00570857 | 0.01114007 | 62.2962697 | 184.285941 | 0 | 115.425085 |
| -2.764013 | 0.00570953 | 0.01114127 | 88.1164342 | 11.9068743 | 13.8054479 | 14.0078987 |
| -2.7639825 | 0.00571006 | 0.01114164 | 9.83625312 | 20.5664192 | 54.0713378 | 14.0078987 |
| 2.76370847 | 0.00571486 | 0.01115033 | 92.214873 | 63.5935331 | 56.3722458 | 60.5141225 |
| -2.7635829 | 0.00571706 | 0.01115394 | 34.8367298 | 49.7923834 | 3.45136199 | 29.6967453 |
| -2.7631036 | 0.00572546 | 0.01116966 | 9.42640924 | 18.1309222 | 26.4604419 | 19.6110582 |
| -2.7628089 | 0.00573063 | 0.01117907 | 81.1490882 | 113.115306 | 0 | 147.92341 |
| -2.7627465 | 0.00573173 | 0.01118054 | 0 | 97.9611021 | 185.223093 | 165.293205 |
| -2.7626955 | 0.00573262 | 0.01118161 | 4.50828268 | 112.032863 | 19.5577179 | 21.852322 |
| -2.7621163 | 0.0057428 | 0.01120079 | 43.8532952 | 19.2133653 | 40.2658898 | 5.04284354 |

| | | | | | | |
|---|---|---|---|---|---|---|
| -2.7616542 | 0.00575093 | 0.01121597 | 23.770945 | 21.1076408 | 11.50454 | 13.4475828 |
| 2.761151 | 0.0057598 | 0.01123259 | 0 | 2.43549701 | 17.2568099 | 0 |
| -2.7608611 | 0.00576492 | 0.01124189 | 4.91812656 | 34.6381797 | 26.4604419 | 34.7395888 |
| -2.7607856 | 0.00576625 | 0.01124381 | 0 | 117.445078 | 227.789891 | 108.701294 |
| -2.7604352 | 0.00577244 | 0.0112552 | 6.96734596 | 14.6129821 | 32.2127119 | 49.8681194 |
| 2.76032443 | 0.0057744 | 0.01125789 | 0.40984388 | 0.54122156 | 0 | 2.80157974 |
| -2.7603177 | 0.00577452 | 0.01125789 | 0 | 174.273342 | 101.239952 | 168.094785 |
| -2.7600122 | 0.00577992 | 0.01126774 | 27.0496961 | 11.3656527 | 37.9649818 | 10.646003 |
| -2.7599424 | 0.00578116 | 0.01126947 | 47.951734 | 23.8137486 | 21.8586259 | 6.16347543 |
| -2.7598336 | 0.00578308 | 0.01127254 | 4.91812656 | 31.1202396 | 41.4163438 | 25.2142177 |
| -2.7596399 | 0.00578651 | 0.01127854 | 17.6232868 | 27.6022995 | 56.3722458 | 6.16347543 |
| -2.7593566 | 0.00579153 | 0.01128764 | 5.73781432 | 17.3190899 | 50.6199758 | 33.6189569 |
| -2.7583676 | 0.00580908 | 0.01132117 | 21.3118818 | 8.65954493 | 23.0090799 | 21.292006 |
| -2.7583295 | 0.00580976 | 0.01132181 | 0 | 193.216096 | 302.569401 | 38.1014845 |
| -2.7582972 | 0.00581033 | 0.01132224 | 12.2953164 | 17.3190899 | 14.9559019 | 20.1713742 |
| -2.758096 | 0.00581391 | 0.01132853 | 21.7217256 | 52.2278804 | 8.05317797 | 16.2491625 |
| 2.75783784 | 0.00581851 | 0.0113368 | 1.63937552 | 1.62366467 | 1.150454 | 0.56031595 |
| -2.757301 | 0.00582807 | 0.01135475 | 20.9020379 | 22.1900839 | 5.75226998 | 66.1172819 |
| -2.7571449 | 0.00583085 | 0.01135948 | 15.1642236 | 10.2832096 | 24.1595339 | 21.852322 |
| -2.7567543 | 0.00583782 | 0.01137237 | 11.4756286 | 18.1309222 | 18.4072639 | 16.8094785 |
| -2.7567152 | 0.00583852 | 0.01137305 | 71.722679 | 131.787449 | 0 | 138.398039 |
| -2.7564424 | 0.00584339 | 0.01138185 | 4.50828268 | 33.8263474 | 74.7795097 | 17.3697944 |
| -2.7562725 | 0.00584643 | 0.01138708 | 18.8528185 | 43.0271139 | 4.60181598 | 44.8252759 |
| 2.75606263 | 0.00585018 | 0.0113937 | 23.3611012 | 29.4965749 | 25.3099879 | 19.6110582 |
| -2.7559497 | 0.0058522 | 0.01139695 | 71.3128351 | 15.4248144 | 8.05317797 | 20.1713742 |
| -2.7558114 | 0.00585467 | 0.01140108 | 11.0657848 | 43.2977247 | 18.4072639 | 14.0078987 |
| -2.7557882 | 0.00585509 | 0.0114012 | 200.823501 | 323.921103 | 234.692615 | 326.103882 |
| -2.7553622 | 0.00586272 | 0.01141537 | 9.01656536 | 37.8855091 | 24.1595339 | 14.5682147 |
| -2.7550824 | 0.00586774 | 0.01142445 | 22.9512573 | 23.0019162 | 9.20363196 | 16.8094785 |
| -2.7548809 | 0.00587135 | 0.01143079 | 13.9346919 | 21.1076408 | 11.50454 | 20.7316901 |
| -2.7548411 | 0.00587206 | 0.01143149 | 10.6559409 | 24.0843593 | 23.0090799 | 14.0078987 |
| -2.7546983 | 0.00587463 | 0.0114358 | 14.3445358 | 27.8729103 | 29.9118039 | 8.96505518 |
| -2.75446 | 0.00587891 | 0.01144344 | 93.8542485 | 18.9427545 | 6.90272397 | 16.2491625 |
| -2.7542665 | 0.00588239 | 0.01144952 | 9.01656536 | 26.7904671 | 13.8054479 | 34.7395888 |
| -2.7541516 | 0.00588445 | 0.01145285 | 11.0657848 | 17.5897006 | 19.5577179 | 17.3697944 |
| -2.7540159 | 0.00588689 | 0.0114569 | 8.1968776 | 15.6954252 | 50.6199758 | 22.4126379 |
| -2.7537521 | 0.00589164 | 0.01146545 | 29.0989155 | 155.871809 | 0 | 236.45333 |
| -2.7537044 | 0.0058925 | 0.01146643 | 34.8367298 | 29.2259641 | 6.90272397 | 17.3697944 |
| -2.7530755 | 0.00590383 | 0.01148779 | 36.4761053 | 165.343186 | 0 | 191.067738 |
| -2.7529207 | 0.00590662 | 0.01149253 | 8.1968776 | 22.4606947 | 19.5577179 | 26.8951655 |
| 2.75265771 | 0.00591137 | 0.01150107 | 0.40984388 | 3.78855091 | 3.45136199 | 5.04284354 |
| -2.7526151 | 0.00591214 | 0.01150188 | 9.01656536 | 14.6129821 | 27.6108959 | 28.5761134 |
| 2.75248881 | 0.00591442 | 0.01150562 | 4.91812656 | 3.78855091 | 5.75226998 | 2.80157974 |

| | | | | | | |
|---|---|---|---|---|---|---|
| -2.7523382 | 0.00591714 | 0.01151022 | 16.8035991 | 28.9553534 | 54.0713378 | 6.16347543 |
| -2.75222 | 0.00591927 | 0.01151368 | 11.4756286 | 15.966036 | 19.5577179 | 17.9301104 |
| -2.7514968 | 0.00593236 | 0.01153844 | 30.738291 | 142.34127 | 0 | 244.85807 |
| -2.7512482 | 0.00593686 | 0.01154633 | 9.83625312 | 20.5664192 | 16.1063559 | 25.2142177 |
| -2.7512335 | 0.00593713 | 0.01154633 | 83.1983076 | 136.117222 | 0 | 113.183822 |
| -2.7511388 | 0.00593885 | 0.01154897 | 7.78703372 | 21.9194731 | 26.4604419 | 21.852322 |
| -2.7509641 | 0.00594202 | 0.01155444 | 8.1968776 | 24.3549701 | 17.2568099 | 30.2570612 |
| -2.7507388 | 0.0059461 | 0.01156169 | 0 | 173.461509 | 87.4345036 | 182.662999 |
| -2.7506348 | 0.00594799 | 0.01156466 | 168.445835 | 222.44206 | 203.630357 | 219.083536 |
| -2.7503779 | 0.00595266 | 0.01157304 | 37.705637 | 18.9427545 | 12.6549939 | 11.7666349 |
| 2.75001131 | 0.00595932 | 0.0115853 | 0 | 0 | 0 | 0 |
| -2.7499903 | 0.0059597 | 0.01158534 | 21.3118818 | 16.7778683 | 13.8054479 | 12.8872668 |
| -2.749628 | 0.0059663 | 0.01159746 | 7.78703372 | 28.4141318 | 17.2568099 | 28.5761134 |
| -2.7491708 | 0.00597462 | 0.01161295 | 5.73781432 | 15.966036 | 57.5226998 | 32.498325 |
| 2.74871816 | 0.00598288 | 0.0116283 | 0 | 26.2492456 | 23.0090799 | 22.4126379 |
| 2.74865205 | 0.00598409 | 0.01162994 | 54.509236 | 20.5664192 | 42.5667978 | 36.4205367 |
| -2.7484211 | 0.00598831 | 0.01163744 | 23.770945 | 28.9553534 | 8.05317797 | 17.3697944 |
| -2.7480546 | 0.005995 | 0.01164975 | 5.32797044 | 36.5324552 | 28.7613499 | 25.2142177 |
| -2.7479762 | 0.00599644 | 0.01165184 | 10.6559409 | 26.7904671 | 42.5667978 | 10.646003 |
| -2.747749 | 0.00600059 | 0.01165921 | 6.55750208 | 24.0843593 | 27.6108959 | 26.3348496 |
| -2.7477054 | 0.00600139 | 0.01166006 | 24.1807889 | 8.38893415 | 19.5577179 | 22.9729539 |
| 2.74752994 | 0.0060046 | 0.0116656 | 106.149565 | 24.8961917 | 18.4072639 | 24.6539017 |
| -2.7473701 | 0.00600753 | 0.01167058 | 43.8532952 | 221.630228 | 120.79767 | 157.448782 |
| -2.7458672 | 0.00603512 | 0.01172347 | 5.32797044 | 28.143521 | 28.7613499 | 31.3776931 |
| -2.7457964 | 0.00603642 | 0.0117246 | 8.1968776 | 40.0503953 | 12.6549939 | 32.498325 |
| -2.745803 | 0.0060363 | 0.0117246 | 18.0331307 | 38.4267306 | 35.6640739 | 28.5761134 |
| -2.7455534 | 0.00604089 | 0.01173257 | 10.6559409 | 20.2958084 | 14.9559019 | 22.4126379 |
| -2.7451671 | 0.00604801 | 0.01174569 | 155.740674 | 97.4198805 | 0 | 81.2458125 |
| 2.74497453 | 0.00605156 | 0.01175188 | 0 | 0 | 0 | 0 |
| 2.74487166 | 0.00605346 | 0.01175485 | 16.3937552 | 10.5538204 | 11.50454 | 8.40473923 |
| -2.743996 | 0.00606963 | 0.01178555 | 232.791324 | 336.369198 | 357.791193 | 344.033992 |
| 2.74378533 | 0.00607352 | 0.01179142 | 0 | 0 | 0 | 0 |
| -2.7437733 | 0.00607375 | 0.01179142 | 130.740198 | 199.981366 | 0 | 43.144328 |
| -2.743778 | 0.00607366 | 0.01179142 | 25.0004767 | 84.7011739 | 88.5849576 | 84.0473923 |
| -2.742991 | 0.00608824 | 0.01181884 | 9.83625312 | 29.2259641 | 14.9559019 | 20.7316901 |
| 2.74261952 | 0.00609513 | 0.0118315 | 63.5258014 | 17.0484791 | 18.4072639 | 20.7316901 |
| 2.74253385 | 0.00609672 | 0.01183388 | 1.63937552 | 0.81183234 | 0 | 0.56031595 |
| -2.7422252 | 0.00610245 | 0.01184429 | 381.154808 | 536.079953 | 498.14658 | 527.257308 |
| -2.742153 | 0.00610379 | 0.01184547 | 35.6564176 | 152.89509 | 184.072639 | 154.647202 |
| -2.7421635 | 0.0061036 | 0.01184547 | 12.7051603 | 14.6129821 | 20.7081719 | 15.6888466 |
| -2.7418503 | 0.00610942 | 0.01185568 | 34.4268859 | 31.3908504 | 21.8586259 | 5.60315949 |
| -2.7415074 | 0.0061158 | 0.01186735 | 52.4600166 | 18.6721438 | 10.354086 | 13.4475828 |
| -2.741382 | 0.00611813 | 0.01187116 | 13.9346919 | 16.5072575 | 16.1063559 | 14.5682147 |

| | | | | | | |
|---|---|---|---|---|---|---|
| -2.7412376 | 0.00612082 | 0.01187567 | 226.233822 | 139.09394 | 0 | 276.235763 |
| -2.741051 | 0.0061243 | 0.0118817 | 8.60672148 | 20.0251977 | 26.4604419 | 19.0507423 |
| 2.74095177 | 0.00612615 | 0.01188458 | 0 | 1.3530539 | 0 | 0 |
| -2.7405664 | 0.00613334 | 0.01189781 | 50.8206411 | 18.9427545 | 13.8054479 | 10.0856871 |
| -2.7405403 | 0.00613383 | 0.01189804 | 37.705637 | 18.9427545 | 10.354086 | 14.0078987 |
| -2.7404631 | 0.00613527 | 0.01190012 | 15.5740674 | 21.1076408 | 9.20363196 | 28.0157974 |
| -2.7403412 | 0.00613754 | 0.01190382 | 16.3937552 | 19.2133653 | 16.1063559 | 11.7666349 |
| -2.7402757 | 0.00613877 | 0.01190548 | 54.0993922 | 20.0251977 | 8.05317797 | 16.2491625 |
| -2.7399404 | 0.00614503 | 0.01191627 | 4.91812656 | 30.3084073 | 33.3631659 | 28.0157974 |
| -2.7399382 | 0.00614507 | 0.01191627 | 57.7879871 | 20.2958084 | 8.05317797 | 15.6888466 |
| -2.7399071 | 0.00614566 | 0.01191669 | 81.968776 | 134.222946 | 0 | 109.821926 |
| -2.7398397 | 0.00614692 | 0.01191842 | 248.775235 | 171.026012 | 209.382627 | 182.662999 |
| -2.7395281 | 0.00615275 | 0.011929 | 5.32797044 | 37.6148983 | 21.8586259 | 31.9380091 |
| -2.7394877 | 0.0061535 | 0.01192975 | 5.73781432 | 34.367569 | 24.1595339 | 26.8951655 |
| -2.7392369 | 0.0061582 | 0.01193814 | 7.37718984 | 25.9786348 | 18.4072639 | 30.2570612 |
| -2.7391592 | 0.00615965 | 0.01194025 | 19.2626624 | 11.0950419 | 24.1595339 | 14.0078987 |
| -2.738543 | 0.00617121 | 0.01196193 | 60.2470503 | 215.135569 | 0 | 86.2886561 |
| 2.73842033 | 0.00617351 | 0.01196567 | 9.42640924 | 26.5198564 | 18.4072639 | 7.28410733 |
| -2.7381431 | 0.00617872 | 0.01197505 | 9.01656536 | 49.2511618 | 20.7081719 | 14.5682147 |
| -2.7380499 | 0.00618047 | 0.01197772 | 398.368251 | 555.56393 | 776.556447 | 725.048838 |
| -2.7379446 | 0.00618245 | 0.01198084 | 53.2797044 | 189.427545 | 63.2749697 | 191.067738 |
| -2.7378102 | 0.00618498 | 0.01198358 | 30.3284471 | 53.3103235 | 50.6199758 | 40.9030642 |
| -2.7378294 | 0.00618461 | 0.01198358 | 7.37718984 | 21.1076408 | 20.7081719 | 33.6189569 |
| -2.7378125 | 0.00618493 | 0.01198358 | 131.559885 | 101.749653 | 123.098578 | 113.744138 |
| -2.7370504 | 0.00619928 | 0.01201057 | 24.1807889 | 30.8496288 | 44.8677058 | 33.058641 |
| -2.7367503 | 0.00620494 | 0.01202081 | 19.2626624 | 8.38893415 | 37.9649818 | 17.9301104 |
| 2.73658183 | 0.00620812 | 0.01202624 | 20.492194 | 11.6362635 | 8.05317797 | 6.16347543 |
| -2.736516 | 0.00620936 | 0.01202793 | 36.8859492 | 26.2492456 | 33.3631659 | 4.48252759 |
| -2.7364483 | 0.00621064 | 0.01202968 | 4.50828268 | 34.9087905 | 32.2127119 | 28.5761134 |
| -2.7364007 | 0.00621153 | 0.0120307 | 18.0331307 | 10.5538204 | 16.1063559 | 22.4126379 |
| 2.7360835 | 0.00621752 | 0.01204158 | 0.81968776 | 0.27061078 | 0 | 0 |
| -2.73534 | 0.00623159 | 0.0120681 | 9.83625312 | 36.2618444 | 11.50454 | 27.4554815 |
| -2.7350413 | 0.00623725 | 0.01207833 | 2.86890716 | 54.9339882 | 79.3813257 | 64.4363341 |
| -2.7348212 | 0.00624142 | 0.01208563 | 9.01656536 | 24.6255809 | 32.2127119 | 14.0078987 |
| -2.7348027 | 0.00624177 | 0.01208563 | 19.6725062 | 11.9068743 | 16.1063559 | 15.6888466 |
| -2.7347233 | 0.00624327 | 0.01208592 | 31.1481349 | 19.7545869 | 9.20363196 | 15.6888466 |
| -2.7347092 | 0.00624354 | 0.01208592 | 40.984388 | 15.1542036 | 10.354086 | 16.8094785 |
| 2.73469623 | 0.00624379 | 0.01208592 | 0 | 1.62366467 | 1.150454 | 0.56031595 |
| -2.7347585 | 0.00624261 | 0.01208592 | 6.1476582 | 22.4606947 | 33.3631659 | 25.7745336 |
| -2.7347093 | 0.00624354 | 0.01208592 | 57.7879871 | 140.176384 | 0 | 141.199619 |
| -2.7344773 | 0.00624794 | 0.01209323 | 13.9346919 | 12.9893174 | 17.2568099 | 17.9301104 |
| -2.7336499 | 0.00626366 | 0.01212294 | 5.32797044 | 49.7923834 | 31.0622579 | 18.4904263 |
| 2.7331322 | 0.00627351 | 0.01214128 | 8.1968776 | 17.0484791 | 0 | 20.7316901 |

| | | | | | | |
|---|---|---|---|---|---|---|
| -2.7329027 | 0.00627789 | 0.01214901 | 104.920033 | 201.60503 | 179.470823 | 205.635953 |
| -2.7325877 | 0.00628389 | 0.01215991 | 0 | 83.6187308 | 172.568099 | 172.577312 |
| -2.732403 | 0.00628742 | 0.012166 | 13.9346919 | 12.4480958 | 24.1595339 | 15.6888466 |
| -2.7322575 | 0.0062902 | 0.01217064 | 64.3454891 | 6.4946587 | 16.1063559 | 24.0935858 |
| -2.7319791 | 0.00629551 | 0.0121802 | 8.1968776 | 26.5198564 | 28.7613499 | 15.6888466 |
| -2.7319232 | 0.00629658 | 0.01218154 | 4.0984388 | 49.2511618 | 44.8677058 | 18.4904263 |
| -2.7317935 | 0.00629906 | 0.01218561 | 0 | 123.398515 | 187.524001 | 109.821926 |
| -2.7316819 | 0.0063012 | 0.01218901 | 15.5740674 | 22.1900839 | 13.8054479 | 12.8872668 |
| -2.7311618 | 0.00631115 | 0.01220342 | 110.248004 | 192.674875 | 0 | 50.9887513 |
| -2.7311604 | 0.00631118 | 0.01220342 | 11.4756286 | 15.966036 | 42.5667978 | 14.0078987 |
| -2.7311991 | 0.00631043 | 0.01220342 | 5.73781432 | 21.1076408 | 36.8145279 | 28.5761134 |
| 2.73115436 | 0.00631129 | 0.01220342 | 0 | 0 | 0 | 0 |
| 2.73115436 | 0.00631129 | 0.01220342 | 0 | 0 | 0 | 0 |
| -2.731231 | 0.00630982 | 0.01220342 | 27.8693838 | 14.3423713 | 12.6549939 | 14.5682147 |
| -2.731215 | 0.00631013 | 0.01220342 | 61.0667381 | 18.1309222 | 20.7081719 | 6.72379138 |
| -2.731032 | 0.00631363 | 0.01220722 | 0 | 121.233629 | 173.718553 | 121.588561 |
| -2.7308569 | 0.00631699 | 0.01221297 | 11.4756286 | 11.9068743 | 34.5136199 | 20.1713742 |
| -2.7308214 | 0.00631767 | 0.01221356 | 9.83625312 | 19.2133653 | 16.1063559 | 24.0935858 |
| -2.7302288 | 0.00632904 | 0.01223427 | 42.6237635 | 24.8961917 | 8.05317797 | 14.0078987 |
| -2.7302233 | 0.00632914 | 0.01223427 | 0 | 120.692407 | 231.241253 | 85.7283401 |
| -2.7300991 | 0.00633153 | 0.01223815 | 8.1968776 | 37.8855091 | 19.5577179 | 17.9301104 |
| -2.7298838 | 0.00633567 | 0.01224541 | 14.3445358 | 11.3656527 | 21.8586259 | 17.9301104 |
| -2.7296768 | 0.00633964 | 0.01225237 | 30.738291 | 175.626396 | 0 | 182.102683 |
| -2.7293146 | 0.00634661 | 0.0122651 | 0 | 232.995881 | 94.3372276 | 108.701294 |
| -2.7289897 | 0.00635287 | 0.01227646 | 87.7065903 | 28.6847426 | 25.3099879 | 22.9729539 |
| -2.7288865 | 0.00635486 | 0.01227957 | 15.9839113 | 12.9893174 | 16.1063559 | 15.6888466 |
| -2.7287287 | 0.0063579 | 0.01228471 | 21.7217256 | 24.0843593 | 5.75226998 | 40.3427483 |
| -2.7281366 | 0.00636932 | 0.01230531 | 6.1476582 | 16.5072575 | 36.8145279 | 35.2999048 |
| -2.7281448 | 0.00636916 | 0.01230531 | 4.0984388 | 24.6255809 | 31.0622579 | 49.8681194 |
| -2.7280425 | 0.00637114 | 0.01230669 | 7.37718984 | 30.0377965 | 14.9559019 | 35.2999048 |
| -2.7280404 | 0.00637118 | 0.01230669 | 308.612442 | 610.227307 | 449.827512 | 498.120878 |
| -2.7280426 | 0.00637114 | 0.01230669 | 6.96734596 | 23.5431378 | 25.3099879 | 23.5332698 |
| -2.7279903 | 0.00637215 | 0.01230782 | 247.955547 | 486.28757 | 524.607022 | 416.31475 |
| -2.7277424 | 0.00637694 | 0.01231634 | 12.7051603 | 22.1900839 | 13.8054479 | 15.6888466 |
| -2.7276813 | 0.00637812 | 0.01231788 | 15.5740674 | 14.8835929 | 14.9559019 | 14.5682147 |
| -2.727454 | 0.00638251 | 0.01232563 | 11.8854725 | 14.0717605 | 28.7613499 | 15.6888466 |
| -2.7273431 | 0.00638466 | 0.01232904 | 13.9346919 | 25.4374132 | 19.5577179 | 10.646003 |
| -2.7271706 | 0.006388 | 0.01233475 | 6.96734596 | 53.3103235 | 11.50454 | 35.8602207 |
| -2.7270836 | 0.00638968 | 0.01233726 | 11.0657848 | 7.57710182 | 49.4695218 | 35.8602207 |
| 2.72688754 | 0.00639348 | 0.01234386 | 0 | 0 | 0 | 0 |
| -2.7268335 | 0.00639453 | 0.01234514 | 97.5428434 | 115.009581 | 0 | 101.417187 |
| -2.7262808 | 0.00640525 | 0.0123651 | 54.509236 | 9.47137727 | 8.05317797 | 36.9808526 |
| 2.72621631 | 0.0064065 | 0.01236677 | 68.0340841 | 46.0038325 | 65.5758777 | 75.6426531 |

| | | | | | | |
|---|---|---|---|---|---|---|
| -2.7259999 | 0.0064107 | 0.01237414 | 18.0331307 | 12.9893174 | 11.50454 | 28.0157974 |
| -2.7253699 | 0.00642295 | 0.01239704 | 9.01656536 | 27.0610779 | 21.8586259 | 15.6888466 |
| -2.725323 | 0.00642386 | 0.01239806 | 100.411751 | 99.8553775 | 0 | 112.623506 |
| -2.7249052 | 0.00643199 | 0.01241228 | 121.313788 | 110.679809 | 148.408565 | 0 |
| -2.7249072 | 0.00643196 | 0.01241228 | 93.8542485 | 12.1774851 | 18.4072639 | 8.40473923 |
| 2.72474741 | 0.00643507 | 0.01241701 | 0 | 0 | 0 | 0 |
| -2.72474 | 0.00643521 | 0.01241701 | 116.805506 | 139.364551 | 0 | 67.7982298 |
| -2.7244968 | 0.00643996 | 0.01242541 | 107.78894 | 79.8301798 | 47.1686138 | 61.6347543 |
| -2.724421 | 0.00644143 | 0.01242752 | 6.55750208 | 41.67406 | 16.1063559 | 29.1364293 |
| -2.7242776 | 0.00644423 | 0.01243218 | 27.45954 | 11.0950419 | 10.354086 | 30.8173772 |
| -2.7240085 | 0.00644948 | 0.01244157 | 0 | 132.599282 | 171.417645 | 108.140978 |
| -2.7239648 | 0.00645034 | 0.01244247 | 49.5911095 | 131.787449 | 0 | 161.931309 |
| -2.7237009 | 0.00645549 | 0.01245167 | 9.42640924 | 31.3908504 | 13.8054479 | 22.4126379 |
| 2.72327216 | 0.00646388 | 0.0124671 | 19.2626624 | 25.9786348 | 11.50454 | 21.292006 |
| -2.7232458 | 0.0064644 | 0.01246735 | 14.7543797 | 12.9893174 | 19.5577179 | 14.5682147 |
| -2.7231317 | 0.00646663 | 0.01247091 | 14.7543797 | 43.8389462 | 19.5577179 | 29.6967453 |
| 2.72280448 | 0.00647304 | 0.01248178 | 12.7051603 | 7.30649104 | 12.6549939 | 5.60315949 |
| -2.722818 | 0.00647277 | 0.01248178 | 299.595876 | 55.7458205 | 0 | 52.6696992 |
| -2.7221539 | 0.00648579 | 0.01250488 | 14.3445358 | 18.1309222 | 14.9559019 | 13.4475828 |
| -2.7221649 | 0.00648558 | 0.01250488 | 7.78703372 | 20.83703 | 20.7081719 | 25.7745336 |
| -2.7219414 | 0.00648997 | 0.01251218 | 34.8367298 | 202.146252 | 32.2127119 | 145.121831 |
| -2.7215419 | 0.00649781 | 0.01252657 | 9.42640924 | 31.3908504 | 20.7081719 | 14.5682147 |
| -2.7213545 | 0.0065015 | 0.01253292 | 5.73781432 | 40.5916169 | 24.1595339 | 22.4126379 |
| -2.7211853 | 0.00650483 | 0.01253859 | 104.920033 | 94.4431619 | 0 | 111.502874 |
| 2.72069238 | 0.00651454 | 0.01255655 | 0 | 0 | 0 | 0 |
| -2.7206222 | 0.00651592 | 0.01255847 | 77.4604933 | 64.4053654 | 57.5226998 | 72.8410733 |
| -2.7201175 | 0.00652587 | 0.0125769 | 7.37718984 | 24.8961917 | 17.2568099 | 31.9380091 |
| -2.7197395 | 0.00653334 | 0.01259054 | 11.8854725 | 18.401533 | 18.4072639 | 14.0078987 |
| -2.7197085 | 0.00653395 | 0.01259097 | 56.1486116 | 179.414947 | 0 | 103.098135 |
| -2.7188979 | 0.00654998 | 0.01262111 | 13.9346919 | 37.0736767 | 51.7704298 | 38.1014845 |
| -2.7188484 | 0.00655096 | 0.01262224 | 18.4429746 | 9.47137727 | 23.0090799 | 17.3697944 |
| 2.71827712 | 0.00656229 | 0.0126433 | 0 | 0 | 0 | 0 |
| -2.7181546 | 0.00656472 | 0.01264723 | 133.609105 | 107.161869 | 134.603117 | 0 |
| -2.7177494 | 0.00657276 | 0.01266198 | 0 | 64.1347547 | 170.267191 | 200.032794 |
| -2.7176788 | 0.00657416 | 0.01266392 | 15.1642236 | 19.4839761 | 14.9559019 | 12.3269509 |
| -2.7173815 | 0.00658007 | 0.01267455 | 8.1968776 | 25.4374132 | 24.1595339 | 16.8094785 |
| -2.7169316 | 0.00658902 | 0.01269103 | 4.0984388 | 22.4606947 | 31.0622579 | 53.2300151 |
| -2.7165662 | 0.0065963 | 0.01270429 | 86.4770587 | 97.1492697 | 0 | 127.752036 |
| -2.7165347 | 0.00659693 | 0.01270474 | 16.3937552 | 11.6362635 | 20.7081719 | 14.5682147 |
| -2.7159448 | 0.00660869 | 0.01272664 | 19.6725062 | 17.3190899 | 12.6549939 | 12.8872668 |
| -2.715746 | 0.00661266 | 0.01273353 | 54.9190799 | 161.825246 | 120.79767 | 142.320251 |
| -2.7155669 | 0.00661624 | 0.01273966 | 18.4429746 | 7.30649104 | 57.5226998 | 17.9301104 |
| -2.7154498 | 0.00661858 | 0.0127434 | 5.73781432 | 35.4500121 | 21.8586259 | 26.3348496 |

| | | | | | | |
|---|---|---|---|---|---|---|
| -2.7151668 | 0.00662424 | 0.01275354 | 15.5740674 | 14.0717605 | 13.8054479 | 16.2491625 |
| -2.7149566 | 0.00662845 | 0.01276087 | 20.9020379 | 26.7904671 | 25.3099879 | 6.72379138 |
| -2.7146678 | 0.00663423 | 0.01277124 | 9.42640924 | 18.9427545 | 17.2568099 | 21.852322 |
| -2.7145859 | 0.00663587 | 0.01277329 | 17.213443 | 10.5538204 | 16.1063559 | 20.7316901 |
| -2.7145754 | 0.00663608 | 0.01277329 | 7.37718984 | 15.6954252 | 34.5136199 | 26.3348496 |
| 2.71448931 | 0.0066378 | 0.01277584 | 6.1476582 | 2.16488623 | 3.45136199 | 4.48252759 |
| -2.7144404 | 0.00663878 | 0.01277697 | 180.741151 | 46.0038325 | 0 | 118.786981 |
| -2.7142893 | 0.00664181 | 0.01278204 | 0 | 136.658443 | 233.542161 | 66.1172819 |
| 2.71402968 | 0.00664702 | 0.0127913 | 6.55750208 | 8.93015571 | 0 | 3.92221164 |
| -2.7138633 | 0.00665036 | 0.01279696 | 4.91812656 | 53.5809343 | 17.2568099 | 32.498325 |
| -2.7135504 | 0.00665664 | 0.01280829 | 23.770945 | 56.5576528 | 36.8145279 | 42.0236961 |
| -2.7135238 | 0.00665718 | 0.01280855 | 6.96734596 | 38.6973414 | 32.2127119 | 14.0078987 |
| -2.7133978 | 0.00665971 | 0.01281266 | 26.2300083 | 16.5072575 | 12.6549939 | 12.3269509 |
| -2.713358 | 0.00666051 | 0.01281344 | 83.1983076 | 116.903857 | 0 | 108.701294 |
| -2.7133159 | 0.00666136 | 0.0128143 | 0 | 322.568049 | 381.950726 | 299.208717 |
| -2.7131346 | 0.006665 | 0.01282055 | 56.5584554 | 8.93015571 | 9.20363196 | 30.8173772 |
| -2.712818 | 0.00667137 | 0.01283204 | 100.411751 | 99.5847667 | 0 | 105.899714 |
| -2.7127378 | 0.00667299 | 0.01283438 | 13.1150042 | 25.708024 | 11.50454 | 17.3697944 |
| -2.7126355 | 0.00667505 | 0.01283758 | 45.4926707 | 110.679809 | 0 | 192.18837 |
| -2.712503 | 0.00667772 | 0.01284195 | 30.738291 | 17.5897006 | 17.2568099 | 8.96505518 |
| -2.7123916 | 0.00667996 | 0.0128455 | 97.1329995 | 98.7729344 | 187.524001 | 0 |
| 2.71233655 | 0.00668107 | 0.01284687 | 6.96734596 | 1.89427545 | 3.45136199 | 5.04284354 |
| -2.7122338 | 0.00668314 | 0.01284984 | 10.246097 | 17.5897006 | 17.2568099 | 19.0507423 |
| -2.7122203 | 0.00668342 | 0.01284984 | 54.509236 | 37.6148983 | 467.084322 | 0 |
| -2.7118055 | 0.00669178 | 0.01286517 | 4.91812656 | 40.3210061 | 27.6108959 | 23.5332698 |
| -2.7116396 | 0.00669513 | 0.01287084 | 11.0657848 | 14.6129821 | 17.2568099 | 21.292006 |
| -2.7114573 | 0.00669882 | 0.01287639 | 13.1150042 | 10.8244312 | 26.4604419 | 19.0507423 |
| 2.71146429 | 0.00669868 | 0.01287639 | 136.068168 | 142.611881 | 128.850847 | 113.744138 |
| -2.7113736 | 0.00670051 | 0.01287887 | 8.60672148 | 30.0377965 | 19.5577179 | 16.8094785 |
| -2.7113282 | 0.00670143 | 0.01287987 | 6.96734596 | 43.2977247 | 14.9559019 | 26.8951655 |
| -2.7110815 | 0.00670641 | 0.01288869 | 0 | 322.568049 | 100.089498 | 54.910963 |
| -2.7109903 | 0.00670826 | 0.01289147 | 14.7543797 | 9.47137727 | 20.7081719 | 26.3348496 |
| -2.7109424 | 0.00670923 | 0.01289256 | 78.6900249 | 135.30539 | 167.966283 | 0 |
| -2.7109101 | 0.00670988 | 0.01289305 | 6.96734596 | 23.8137486 | 24.1595339 | 22.4126379 |
| -2.7101949 | 0.00672437 | 0.0129195 | 14.3445358 | 23.8137486 | 13.8054479 | 12.8872668 |
| 2.71019107 | 0.00672445 | 0.0129195 | 0 | 1.08244312 | 6.90272397 | 2.80157974 |
| -2.7081823 | 0.00676529 | 0.01299706 | 25.4103206 | 38.6973414 | 4.60181598 | 26.3348496 |
| -2.7081658 | 0.00676562 | 0.01299706 | 57.3781432 | 8.38893415 | 21.8586259 | 12.8872668 |
| 2.70811287 | 0.0067667 | 0.01299836 | 5.32797044 | 2.16488623 | 1.150454 | 5.60315949 |
| -2.7080613 | 0.00676775 | 0.01299961 | 20.0823501 | 14.8835929 | 20.7081719 | 10.0856871 |
| -2.70797 | 0.00676961 | 0.01300241 | 49.5911095 | 14.8835929 | 10.354086 | 15.1285306 |
| -2.7077978 | 0.00677313 | 0.01300761 | 0 | 108.785533 | 83.9831417 | 229.729539 |
| -2.7078135 | 0.00677281 | 0.01300761 | 8.1968776 | 12.4480958 | 21.8586259 | 64.99665 |

| | | | | | | |
|---|---|---|---|---|---|---|
| -2.7076366 | 0.00677642 | 0.01301316 | 65.9848647 | 102.561485 | 94.3372276 | 106.46003 |
| -2.7071578 | 0.0067862 | 0.01303117 | 8.60672148 | 22.4606947 | 17.2568099 | 21.852322 |
| -2.706876 | 0.00679196 | 0.01304146 | 33.1973543 | 3.78855091 | 32.2127119 | 34.1792729 |
| 2.70672047 | 0.00679514 | 0.01304679 | 9.83625312 | 8.93015571 | 11.50454 | 12.3269509 |
| -2.7066947 | 0.00679567 | 0.01304703 | 67.2143963 | 0 | 247.347609 | 158.009098 |
| -2.7065481 | 0.00679867 | 0.01305201 | 175.003337 | 404.021893 | 264.604419 | 303.130928 |
| 2.70651366 | 0.00679938 | 0.01305259 | 0 | 1.08244312 | 0 | 0.56031595 |
| -2.7061135 | 0.00680758 | 0.01306755 | 32.7875104 | 21.6488623 | 10.354086 | 11.7666349 |
| -2.7060332 | 0.00680922 | 0.01306994 | 84.8376831 | 82.5362876 | 16.1063559 | 85.7283401 |
| -2.7059885 | 0.00681014 | 0.01307073 | 84.0179954 | 86.0542278 | 0 | 139.518671 |
| -2.7059735 | 0.00681045 | 0.01307073 | 19.6725062 | 21.6488623 | 6.90272397 | 29.6967453 |
| -2.7058125 | 0.00681375 | 0.01307629 | 21.3118818 | 8.38893415 | 14.9559019 | 36.9808526 |
| -2.705523 | 0.00681969 | 0.01308692 | 0 | 161.284024 | 121.948124 | 113.744138 |
| -2.7052683 | 0.00682493 | 0.01309618 | 73.7718984 | 108.785533 | 0 | 125.510772 |
| 2.70514807 | 0.00682739 | 0.01309974 | 0 | 0 | 0 | 0 |
| -2.7051386 | 0.00682759 | 0.01309974 | 0 | 172.649677 | 112.744492 | 113.744138 |
| -2.7050991 | 0.0068284 | 0.01310052 | 5.73781432 | 25.708024 | 21.8586259 | 34.1792729 |
| -2.7050079 | 0.00683028 | 0.01310333 | 54.509236 | 88.7603356 | 0 | 194.98995 |
| 2.70497667 | 0.00683092 | 0.01310379 | 1.22953164 | 0.27061078 | 0 | 0 |
| -2.7047442 | 0.0068357 | 0.01311215 | 8.1968776 | 23.0019162 | 20.7081719 | 19.0507423 |
| -2.7047252 | 0.00683609 | 0.01311215 | 18.0331307 | 25.4374132 | 10.354086 | 14.0078987 |
| -2.7044562 | 0.00684163 | 0.013122 | 48.7714217 | 51.416048 | 59.8236078 | 67.7982298 |
| -2.7043875 | 0.00684304 | 0.01312328 | 75.4112739 | 146.129821 | 156.461743 | 0 |
| -2.7043843 | 0.00684311 | 0.01312328 | 13.1150042 | 42.4858923 | 6.90272397 | 30.8173772 |
| -2.7036824 | 0.00685758 | 0.01314869 | 77.4604933 | 141.258827 | 157.612197 | 0 |
| -2.7036938 | 0.00685734 | 0.01314869 | 52.4600166 | 10.0125988 | 10.354086 | 22.9729539 |
| -2.7037198 | 0.00685681 | 0.01314869 | 11.0657848 | 23.272527 | 13.8054479 | 17.3697944 |
| -2.7034826 | 0.0068617 | 0.01315581 | 9.01656536 | 23.272527 | 16.1063559 | 20.7316901 |
| -2.7030116 | 0.00687144 | 0.01317369 | 0 | 132.05806 | 172.568099 | 95.2537113 |
| -2.7028472 | 0.00687483 | 0.01317942 | 3.68859492 | 39.238563 | 41.4163438 | 23.5332698 |
| 2.7027201 | 0.00687746 | 0.01318356 | 0.81968776 | 0 | 0 | 1.1206319 |
| -2.7027032 | 0.00687781 | 0.01318356 | 9.83625312 | 17.5897006 | 18.4072639 | 18.4904263 |
| -2.7022619 | 0.00688695 | 0.01320029 | 12.7051603 | 21.3782516 | 46.0181598 | 8.96505518 |
| -2.7021125 | 0.00689004 | 0.01320544 | 26.6398522 | 10.2832096 | 16.1063559 | 15.6888466 |
| -2.7020161 | 0.00689204 | 0.01320849 | 0 | 139.635162 | 124.249032 | 128.312352 |
| -2.7013587 | 0.00690568 | 0.01323384 | 12.2953164 | 15.4248144 | 13.8054479 | 20.1713742 |
| -2.701114 | 0.00691076 | 0.01324279 | 4.0984388 | 26.2492456 | 32.2127119 | 37.5411686 |
| -2.7010787 | 0.0069115 | 0.01324341 | 22.5414134 | 10.2832096 | 36.8145279 | 11.206319 |
| -2.7009986 | 0.00691316 | 0.01324582 | 66.3947085 | 141.529437 | 0 | 103.65845 |
| -2.7009632 | 0.0069139 | 0.01324644 | 19.2626624 | 38.6973414 | 26.4604419 | 29.1364293 |
| -2.7007974 | 0.00691735 | 0.01325226 | 95.0837801 | 131.516839 | 134.603117 | 132.79488 |
| -2.7001931 | 0.00692992 | 0.01327557 | 161.888333 | 276.022995 | 255.400787 | 249.340597 |
| -2.6999154 | 0.00693571 | 0.01328587 | 229.102729 | 380.208145 | 303.719855 | 317.138827 |

| | | | | | | |
|---|---|---|---|---|---|---|
| -2.6996318 | 0.00694162 | 0.01329641 | 0 | 197.545869 | 86.2840497 | 121.588561 |
| -2.6995956 | 0.00694238 | 0.01329707 | 6.55750208 | 32.7439043 | 19.5577179 | 23.5332698 |
| -2.6993663 | 0.00694717 | 0.01330544 | 10.246097 | 14.0717605 | 18.4072639 | 25.2142177 |
| -2.6989185 | 0.00695652 | 0.01332257 | 284.431653 | 396.715402 | 770.804177 | 503.163722 |
| -2.6988779 | 0.00695737 | 0.0133234 | 15.9839113 | 20.0251977 | 29.9118039 | 8.40473923 |
| 2.69864239 | 0.00696229 | 0.01333204 | 0 | 0 | 0 | 0 |
| -2.6982378 | 0.00697076 | 0.01334747 | 10.246097 | 48.7099402 | 51.7704298 | 52.1093832 |
| -2.6979187 | 0.00697745 | 0.01335948 | 147.133953 | 102.290875 | 0 | 63.3157022 |
| -2.6977387 | 0.00698122 | 0.01336591 | 0 | 125.02218 | 166.815829 | 101.977503 |
| 2.69757474 | 0.00698466 | 0.0133717 | 26.2300083 | 18.6721438 | 12.6549939 | 7.84442328 |
| -2.6973675 | 0.00698901 | 0.01337923 | 184.83959 | 556.105151 | 288.763953 | 417.995698 |
| -2.6972109 | 0.0069923 | 0.01338473 | 18.0331307 | 14.0717605 | 37.9649818 | 9.52537113 |
| -2.6970327 | 0.00699604 | 0.01339111 | 170.08521 | 420.529151 | 427.968886 | 416.31475 |
| -2.6962089 | 0.00701337 | 0.01342347 | 102.870814 | 87.6778924 | 0 | 107.580662 |
| -2.695988 | 0.00701802 | 0.01343158 | 11.8854725 | 19.2133653 | 16.1063559 | 14.0078987 |
| -2.6959333 | 0.00701917 | 0.01343299 | 17.213443 | 90.9252218 | 135.753571 | 115.985401 |
| -2.6956852 | 0.0070244 | 0.01344141 | 155.740674 | 566.11775 | 706.378753 | 612.425332 |
| -2.6956977 | 0.00702414 | 0.01344141 | 10.246097 | 28.6847426 | 11.50454 | 23.5332698 |
| -2.6951925 | 0.0070348 | 0.0134605 | 0 | 75.5004074 | 199.028541 | 132.234564 |
| -2.6949105 | 0.00704075 | 0.0134711 | 312.71088 | 28.9553534 | 0 | 75.6426531 |
| -2.6947522 | 0.0070441 | 0.0134767 | 0 | 140.446994 | 141.505841 | 107.020346 |
| -2.6947155 | 0.00704488 | 0.01347739 | 0.81968776 | 27.0610779 | 92.0363196 | 66.6775979 |
| -2.694486 | 0.00704973 | 0.01348587 | 14.7543797 | 12.4480958 | 19.5577179 | 14.0078987 |
| -2.694359 | 0.00705241 | 0.01349021 | 170.495054 | 223.524504 | 120.79767 | 130.553616 |
| -2.6943275 | 0.00705308 | 0.01349069 | 309.022285 | 162.366467 | 289.914407 | 107.020346 |
| -2.6941775 | 0.00705626 | 0.01349597 | 1.63937552 | 35.7206228 | 46.0181598 | 56.5919108 |
| -2.6939137 | 0.00706185 | 0.01350585 | 69.2636157 | 188.615713 | 0 | 68.9188617 |
| -2.6938528 | 0.00706314 | 0.01350752 | 12.2953164 | 29.4965749 | 19.5577179 | 10.646003 |
| -2.6936076 | 0.00706833 | 0.01351666 | 28.6890716 | 13.2599282 | 11.50454 | 15.6888466 |
| -2.6935835 | 0.00706884 | 0.01351684 | 243.037421 | 458.955881 | 373.897549 | 497.560562 |
| 2.69308787 | 0.00707936 | 0.01353614 | 3.68859492 | 3.24732935 | 4.60181598 | 1.68094785 |
| -2.6927847 | 0.0070858 | 0.01354766 | 177.872244 | 238.949318 | 299.118039 | 271.753235 |
| -2.6920147 | 0.00710218 | 0.01357817 | 6.96734596 | 21.3782516 | 29.9118039 | 19.6110582 |
| -2.6919935 | 0.00710263 | 0.01357823 | 9.01656536 | 28.143521 | 17.2568099 | 16.8094785 |
| -2.6917049 | 0.00710878 | 0.01358896 | 18.4429746 | 20.83703 | 24.1595339 | 7.84442328 |
| -2.6916732 | 0.00710946 | 0.01358896 | 0 | 161.284024 | 100.089498 | 127.752036 |
| -2.6916707 | 0.00710951 | 0.01358896 | 115.985818 | 250.856192 | 194.426725 | 237.013646 |
| -2.6915596 | 0.00711188 | 0.01359269 | 4.50828268 | 35.1794013 | 27.6108959 | 26.8951655 |
| -2.6908783 | 0.00712642 | 0.01361886 | 9.83625312 | 26.5198564 | 14.9559019 | 17.3697944 |
| -2.6908803 | 0.00712638 | 0.01361886 | 288.120248 | 399.150899 | 356.640739 | 363.645051 |
| -2.6902928 | 0.00713894 | 0.01364197 | 0 | 112.303473 | 143.806749 | 128.872668 |
| 2.69022816 | 0.00714032 | 0.01364381 | 1.22953164 | 0 | 2.30090799 | 0.56031595 |
| -2.6896755 | 0.00715215 | 0.01366561 | 9.01656536 | 24.0843593 | 16.1063559 | 19.0507423 |

| | | | | | | |
|---|---|---|---|---|---|---|
| -2.6895703 | 0.00715441 | 0.01366912 | 192.626624 | 315.532168 | 248.498063 | 295.846821 |
| -2.6892445 | 0.00716139 | 0.01368152 | 3.27875104 | 39.238563 | 24.1595339 | 44.2649599 |
| -2.6891884 | 0.0071626 | 0.01368152 | 10.6559409 | 22.1900839 | 21.8586259 | 12.3269509 |
| -2.6891893 | 0.00716258 | 0.01368152 | 11.4756286 | 14.3423713 | 14.9559019 | 22.9729539 |
| -2.689224 | 0.00716184 | 0.01368152 | 422.139196 | 555.293319 | 511.952028 | 522.214464 |
| -2.6891258 | 0.00716394 | 0.01368328 | 11.8854725 | 11.9068743 | 29.9118039 | 16.8094785 |
| -2.6890088 | 0.00716645 | 0.01368727 | 6.96734596 | 26.2492456 | 20.7081719 | 21.852322 |
| -2.688988 | 0.0071669 | 0.01368731 | 72.1325229 | 103.643928 | 0 | 123.829825 |
| 2.68893357 | 0.00716807 | 0.01368792 | 0 | 0 | 0 | 0 |
| 2.68893357 | 0.00716807 | 0.01368792 | 0 | 0 | 0 | 0 |
| -2.6886649 | 0.00717384 | 0.01369813 | 34.8367298 | 18.401533 | 11.50454 | 11.206319 |
| -2.6886272 | 0.00717465 | 0.01369887 | 193.446311 | 214.323737 | 201.329449 | 202.274057 |
| -2.6884213 | 0.00717908 | 0.01370651 | 5.32797044 | 34.367569 | 27.6108959 | 21.292006 |
| -2.6882327 | 0.00718313 | 0.01371345 | 15.1642236 | 16.5072575 | 14.9559019 | 12.3269509 |
| -2.6882052 | 0.00718372 | 0.01371376 | 13.9346919 | 18.9427545 | 10.354086 | 20.1713742 |
| -2.6876293 | 0.00719612 | 0.01373662 | 6.55750208 | 35.7206228 | 33.3631659 | 14.0078987 |
| 2.68751168 | 0.00719866 | 0.01374065 | 1.63937552 | 1.3530539 | 0 | 2.80157974 |
| -2.6874031 | 0.007201 | 0.0137443 | 72.5423667 | 122.857294 | 0 | 103.098135 |
| -2.6872407 | 0.0072045 | 0.01375018 | 74.5915861 | 83.8893415 | 0 | 145.121831 |
| -2.6866773 | 0.00721666 | 0.01377257 | 120.903945 | 214.594348 | 255.400787 | 272.313551 |
| -2.6865498 | 0.00721942 | 0.01377702 | 39.3450125 | 54.6633774 | 59.8236078 | 42.5840121 |
| -2.6863107 | 0.00722459 | 0.01378607 | 12.2953164 | 28.4141318 | 26.4604419 | 8.96505518 |
| -2.6861794 | 0.00722743 | 0.01379067 | 71.722679 | 94.1725511 | 0 | 134.475828 |
| -2.6861497 | 0.00722807 | 0.01379108 | 45.9025146 | 39.7797845 | 10.354086 | 6.72379138 |
| -2.6857501 | 0.00723672 | 0.01380677 | 0 | 125.02218 | 127.700393 | 127.752036 |
| 2.68562285 | 0.00723948 | 0.01381121 | 0 | 0 | 0 | 0 |
| -2.6855678 | 0.00724067 | 0.01381267 | 48.7714217 | 26.2492456 | 35.6640739 | 20.7316901 |
| -2.6854917 | 0.00724232 | 0.013815 | 6.1476582 | 25.4374132 | 26.4604419 | 21.852322 |
| -2.6854245 | 0.00724377 | 0.01381696 | 18.8528185 | 29.2259641 | 8.05317797 | 16.8094785 |
| -2.6847884 | 0.00725757 | 0.01384247 | 167.216303 | 328.792097 | 375.048002 | 349.637152 |
| 2.68403825 | 0.00727388 | 0.01387275 | 1.22953164 | 2.70610779 | 2.30090799 | 1.68094785 |
| 2.68389968 | 0.00727689 | 0.01387768 | 12.2953164 | 15.1542036 | 11.50454 | 11.206319 |
| -2.6838167 | 0.0072787 | 0.0138803 | 62.7061136 | 14.6129821 | 6.90272397 | 20.1713742 |
| -2.6835618 | 0.00728425 | 0.01389007 | 60.6568942 | 175.897006 | 0 | 80.6854966 |
| -2.6833691 | 0.00728845 | 0.01389725 | 7.37718984 | 19.2133653 | 18.4072639 | 33.6189569 |
| -2.6833048 | 0.00728985 | 0.0138991 | 0 | 199.710755 | 173.718553 | 49.3078035 |
| -2.6830834 | 0.00729468 | 0.01390749 | 71.722679 | 14.3423713 | 3.45136199 | 39.7824324 |
| -2.6828786 | 0.00729915 | 0.01391519 | 0 | 113.927138 | 174.869007 | 97.4949751 |
| -2.682629 | 0.0073046 | 0.01392475 | 0 | 73.8767427 | 247.347609 | 95.2537113 |
| 2.68218711 | 0.00731425 | 0.01394234 | 168.035991 | 145.85921 | 173.718553 | 107.020346 |
| -2.6820968 | 0.00731623 | 0.01394528 | 588.125968 | 536.891786 | 556.819734 | 602.899961 |
| -2.6820471 | 0.00731732 | 0.01394653 | 197.134906 | 334.204312 | 299.118039 | 307.05314 |
| -2.6817993 | 0.00732274 | 0.01395604 | 129.510666 | 232.184049 | 222.037621 | 201.153426 |

| | | | | | | |
|---|---|---|---|---|---|---|
| -2.6815694 | 0.00732777 | 0.01396481 | 5.32797044 | 21.1076408 | 25.3099879 | 37.5411686 |
| -2.6813771 | 0.00733198 | 0.01397201 | 7.37718984 | 26.7904671 | 16.1063559 | 25.7745336 |
| -2.6811871 | 0.00733615 | 0.01397912 | 271.316649 | 53.851545 | 87.4345036 | 0 |
| -2.6809954 | 0.00734035 | 0.01398631 | 27.45954 | 18.401533 | 5.75226998 | 31.9380091 |
| -2.6806042 | 0.00734894 | 0.01400101 | 288.120248 | 255.185965 | 240.444885 | 369.24821 |
| -2.6806184 | 0.00734863 | 0.01400101 | 113.526755 | 0 | 113.894946 | 195.550266 |
| -2.6804978 | 0.00735127 | 0.01400464 | 18.0331307 | 24.8961917 | 27.6108959 | 17.3697944 |
| -2.6802986 | 0.00735565 | 0.01401215 | 60.2470503 | 127.457677 | 0 | 113.744138 |
| -2.6802566 | 0.00735657 | 0.01401308 | 40.1647002 | 11.9068743 | 13.8054479 | 13.4475828 |
| -2.680162 | 0.00735865 | 0.01401622 | 9.42640924 | 29.2259641 | 9.20363196 | 42.0236961 |
| -2.6795991 | 0.00737104 | 0.01403898 | 75.4112739 | 93.3607188 | 205.931265 | 0 |
| -2.6794732 | 0.00737381 | 0.01404343 | 24.1807889 | 23.0019162 | 5.75226998 | 26.3348496 |
| -2.67918 | 0.00738027 | 0.0140549 | 11.4756286 | 9.20076649 | 24.1595339 | 35.8602207 |
| -2.6790706 | 0.00738268 | 0.01405867 | 22.1315695 | 31.6614612 | 3.45136199 | 51.5490673 |
| 2.6789809 | 0.00738466 | 0.0140616 | 0 | 21.1076408 | 0 | 36.4205367 |
| 2.67892416 | 0.00738591 | 0.01406316 | 143.035514 | 111.22103 | 135.753571 | 128.312352 |
| -2.6787761 | 0.00738918 | 0.01406855 | 23.3611012 | 22.7313054 | 8.05317797 | 15.6888466 |
| -2.6786597 | 0.00739175 | 0.01407261 | 9.42640924 | 23.8137486 | 14.9559019 | 18.4904263 |
| -2.6784924 | 0.00739544 | 0.01407881 | 2.0492194 | 45.4626109 | 55.2217918 | 27.4554815 |
| -2.6784188 | 0.00739707 | 0.01408107 | 723.784292 | 1212.6069 | 919.212742 | 749.142423 |
| -2.6782002 | 0.0074019 | 0.01408944 | 40.5745441 | 121.774851 | 0 | 164.172573 |
| -2.6778066 | 0.0074106 | 0.01410517 | 0 | 74.9591858 | 136.904025 | 178.740788 |
| -2.6776819 | 0.00741336 | 0.01410959 | 9.42640924 | 15.1542036 | 17.2568099 | 26.8951655 |
| 2.67739044 | 0.00741981 | 0.01412104 | 46.7222023 | 40.0503953 | 26.4604419 | 30.2570612 |
| -2.6771645 | 0.00742482 | 0.01412974 | 9.83625312 | 11.0950419 | 37.9649818 | 21.852322 |
| -2.6770544 | 0.00742726 | 0.01413354 | 0 | 150.459593 | 186.373547 | 62.7553862 |
| -2.6768775 | 0.00743118 | 0.01414018 | 9.01656536 | 27.3316887 | 21.8586259 | 13.4475828 |
| -2.6767221 | 0.00743463 | 0.0141459 | 89.7558097 | 126.916455 | 134.603117 | 0 |
| -2.6766764 | 0.00743564 | 0.014147 | 86.0672148 | 112.844695 | 0 | 90.2108677 |
| -2.6765541 | 0.00743836 | 0.01415133 | 75.8211178 | 192.133653 | 148.408565 | 171.45668 |
| -2.6764933 | 0.00743971 | 0.01415306 | 72.9522106 | 113.385916 | 177.169915 | 0 |
| 2.67640373 | 0.00744169 | 0.01415535 | 777.883684 | 278.187881 | 136.904025 | 286.881766 |
| 2.67639954 | 0.00744179 | 0.01415535 | 0 | 8.11832337 | 0 | 1.68094785 |
| -2.6752234 | 0.00746795 | 0.01420428 | 15.9839113 | 31.1202396 | 6.90272397 | 24.0935858 |
| 2.67449331 | 0.00748423 | 0.0142344 | 40.984388 | 40.8622277 | 43.7172518 | 32.498325 |
| -2.6743593 | 0.00748722 | 0.01423926 | 65.5750208 | 186.180216 | 128.850847 | 151.285306 |
| -2.6740261 | 0.00749466 | 0.01425257 | 0 | 89.8427787 | 159.913105 | 129.9933 |
| -2.6738183 | 0.00749931 | 0.01426056 | 14.7543797 | 23.0019162 | 6.90272397 | 41.4633802 |
| -2.6737605 | 0.0075006 | 0.01426218 | 0 | 110.950419 | 253.099879 | 56.5919108 |
| -2.6735009 | 0.00750641 | 0.01427238 | 0 | 123.127905 | 133.452663 | 115.985401 |
| 2.67343975 | 0.00750778 | 0.01427415 | 16.8035991 | 11.9068743 | 14.9559019 | 5.60315949 |
| 2.67341399 | 0.00750835 | 0.0142744 | 77.0506494 | 85.5130062 | 78.2308717 | 80.1251807 |
| -2.6730128 | 0.00751734 | 0.01429064 | 9.01656536 | 15.966036 | 17.2568099 | 26.3348496 |

| | | | | | | |
|---|---|---|---|---|---|---|
| -2.6729159 | 0.00751951 | 0.01429393 | 0 | 149.918372 | 196.727633 | 57.1522268 |
| -2.672716 | 0.00752399 | 0.01430161 | 12.7051603 | 11.0950419 | 24.1595339 | 16.8094785 |
| -2.6726313 | 0.00752589 | 0.01430438 | 13.524848 | 21.1076408 | 16.1063559 | 11.206319 |
| -2.6725028 | 0.00752877 | 0.01430842 | 20.9020379 | 19.2133653 | 8.05317797 | 18.4904263 |
| -2.6724971 | 0.0075289 | 0.01430842 | 12.2953164 | 8.65954493 | 43.7172518 | 21.292006 |
| -2.6720547 | 0.00753884 | 0.01432645 | 86.0672148 | 15.4248144 | 11.50454 | 8.96505518 |
| -2.6717653 | 0.00754534 | 0.01433797 | 100.411751 | 267.092839 | 286.463045 | 259.426284 |
| -2.671566 | 0.00754982 | 0.01434564 | 18.8528185 | 11.0950419 | 12.6549939 | 19.6110582 |
| 2.67130552 | 0.00755568 | 0.0143551 | 2.45906328 | 3.51794013 | 3.45136199 | 4.48252759 |
| -2.6713052 | 0.00755569 | 0.0143551 | 11.0657848 | 26.7904671 | 17.2568099 | 12.3269509 |
| 2.67127688 | 0.00755633 | 0.01435547 | 0 | 0 | 0 | 0 |
| -2.6710349 | 0.00756178 | 0.01436498 | 9.83625312 | 30.0377965 | 65.5758777 | 6.72379138 |
| 2.67095902 | 0.00756349 | 0.01436653 | 0 | 0.27061078 | 0 | 0 |
| 2.67095902 | 0.00756349 | 0.01436653 | 0 | 0.27061078 | 0 | 0 |
| -2.6708227 | 0.00756656 | 0.01437153 | 47.5418901 | 90.3840002 | 104.691314 | 108.701294 |
| -2.6707584 | 0.00756801 | 0.01437343 | 5.32797044 | 23.272527 | 32.2127119 | 24.0935858 |
| -2.6701204 | 0.0075824 | 0.01439992 | 9.01656536 | 24.6255809 | 17.2568099 | 16.2491625 |
| -2.6700514 | 0.00758396 | 0.01440204 | 0 | 158.036695 | 120.79767 | 96.3743432 |
| 2.66985093 | 0.00758849 | 0.01440979 | 2.0492194 | 2.16488623 | 6.90272397 | 2.24126379 |
| -2.6697471 | 0.00759084 | 0.0144134 | 17.6232868 | 15.1542036 | 11.50454 | 14.5682147 |
| -2.6697001 | 0.0075919 | 0.01441411 | 79.5097127 | 103.102707 | 0 | 103.098135 |
| -2.6696911 | 0.00759211 | 0.01441411 | 77.0506494 | 17.5897006 | 6.90272397 | 14.0078987 |
| -2.669643 | 0.00759319 | 0.01441532 | 300.825408 | 155.871809 | 0 | 275.675447 |
| -2.6696137 | 0.00759386 | 0.01441574 | 13.1150042 | 22.7313054 | 10.354086 | 17.9301104 |
| -2.6695473 | 0.00759536 | 0.01441774 | 9.01656536 | 23.8137486 | 23.0090799 | 13.4475828 |
| -2.6693736 | 0.00759929 | 0.01442435 | 10.6559409 | 20.5664192 | 11.50454 | 24.6539017 |
| -2.6693444 | 0.00759995 | 0.01442475 | 7.78703372 | 23.272527 | 25.3099879 | 15.6888466 |
| -2.6691512 | 0.00760432 | 0.01443221 | 125.412227 | 94.9843835 | 0 | 70.0394936 |
| -2.6691121 | 0.00760521 | 0.01443304 | 195.085687 | 318.779498 | 361.242555 | 359.162523 |
| 2.66899569 | 0.00760784 | 0.01443635 | 0 | 0 | 0 | 0 |
| 2.66899569 | 0.00760784 | 0.01443635 | 0 | 0 | 0 | 0 |
| -2.6687597 | 0.00761319 | 0.01444564 | 6.96734596 | 23.0019162 | 62.1245158 | 12.3269509 |
| -2.6683352 | 0.00762282 | 0.01446306 | 28.2792277 | 132.869893 | 70.1776937 | 87.409288 |
| 2.66786888 | 0.0076334 | 0.0144823 | 27.0496961 | 5.95343714 | 5.75226998 | 7.28410733 |
| -2.6671115 | 0.00765063 | 0.01451412 | 2.86890716 | 74.688575 | 12.6549939 | 49.8681194 |
| -2.6669555 | 0.00765418 | 0.01452001 | 20.492194 | 13.530539 | 11.50454 | 15.1285306 |
| -2.666914 | 0.00765513 | 0.01452095 | 21.3118818 | 12.9893174 | 11.50454 | 15.6888466 |
| -2.6667305 | 0.00765931 | 0.01452802 | 8.1968776 | 21.9194731 | 44.8677058 | 12.3269509 |
| -2.6666965 | 0.00766008 | 0.01452864 | 90.1656536 | 72.5236888 | 0 | 126.071088 |
| -2.6665417 | 0.00766361 | 0.01453448 | 0 | 102.832096 | 123.098578 | 144.001199 |
| 2.66649141 | 0.00766476 | 0.0145358 | 13.9346919 | 0.54122156 | 2.30090799 | 0.56031595 |
| 2.66613796 | 0.00767282 | 0.01455024 | 38.9351686 | 15.1542036 | 10.354086 | 22.4126379 |
| -2.6659094 | 0.00767804 | 0.01455928 | 0 | 84.4305631 | 236.993523 | 80.1251807 |

| | | | | | | |
|---|---|---|---|---|---|---|
| -2.6657938 | 0.00768068 | 0.01456299 | 68.8537718 | 49.2511618 | 32.2127119 | 53.2300151 |
| -2.6657841 | 0.0076809 | 0.01456299 | 4.50828268 | 23.5431378 | 26.4604419 | 38.1014845 |
| 2.6654559 | 0.0076884 | 0.01457636 | 4.0984388 | 1.62366467 | 0 | 2.80157974 |
| -2.6654182 | 0.00768926 | 0.01457714 | 8.1968776 | 21.3782516 | 19.5577179 | 17.9301104 |
| -2.6646657 | 0.00770649 | 0.01460894 | 15.9839113 | 46.2744432 | 11.50454 | 11.206319 |
| 2.66444847 | 0.00771147 | 0.01461752 | 0 | 1.89427545 | 1.150454 | 1.1206319 |
| -2.6642575 | 0.00771585 | 0.01462376 | 121.723632 | 65.7584193 | 0 | 101.417187 |
| -2.6642457 | 0.00771612 | 0.01462376 | 65.1651769 | 174.273342 | 0 | 68.3585457 |
| 2.66426089 | 0.00771577 | 0.01462376 | 0 | 0 | 0 | 0 |
| -2.6638928 | 0.00772422 | 0.01463825 | 10.246097 | 15.1542036 | 20.7081719 | 15.6888466 |
| 2.66358638 | 0.00773126 | 0.01465073 | 27.8693838 | 18.1309222 | 11.50454 | 16.2491625 |
| -2.6632158 | 0.00773978 | 0.01466586 | 81.1490882 | 175.626396 | 96.6381356 | 0 |
| -2.6631995 | 0.00774015 | 0.01466586 | 8.1968776 | 32.4732935 | 18.4072639 | 15.6888466 |
| -2.6630482 | 0.00774363 | 0.01467159 | 31.9678226 | 160.742803 | 0 | 139.518671 |
| 2.66297847 | 0.00774524 | 0.01467377 | 18.0331307 | 10.2832096 | 13.8054479 | 3.92221164 |
| -2.6628025 | 0.00774929 | 0.01468059 | 8.60672148 | 10.0125988 | 41.4163438 | 28.0157974 |
| -2.6627195 | 0.0077512 | 0.01468335 | 9.42640924 | 35.9912336 | 29.9118039 | 34.7395888 |
| -2.6624161 | 0.00775819 | 0.01469573 | 98.7723751 | 187.262659 | 72.4786017 | 0 |
| -2.6622929 | 0.00776103 | 0.01470017 | 55.7387677 | 114.73897 | 0 | 123.829825 |
| -2.6622749 | 0.00776145 | 0.01470017 | 8.1968776 | 17.5897006 | 18.4072639 | 23.5332698 |
| -2.6618688 | 0.00777082 | 0.01471706 | 32.3776665 | 160.201581 | 0 | 137.837723 |
| -2.6618479 | 0.0077713 | 0.0147171 | 0 | 116.903857 | 190.975363 | 74.5220212 |
| -2.6610553 | 0.00778962 | 0.01475093 | 46.7222023 | 139.635162 | 0 | 117.666349 |
| -2.6609324 | 0.00779246 | 0.01475545 | 33.6071982 | 146.941653 | 0 | 145.682147 |
| -2.6608878 | 0.00779349 | 0.01475654 | 372.548087 | 1086.77289 | 1058.41768 | 1083.09073 |
| -2.6606648 | 0.00779866 | 0.01476428 | 8.1968776 | 14.3423713 | 25.3099879 | 22.9729539 |
| -2.6606518 | 0.00779896 | 0.01476428 | 10.246097 | 24.8961917 | 20.7081719 | 11.7666349 |
| -2.6606641 | 0.00779867 | 0.01476428 | 95.9034679 | 340.157749 | 262.303511 | 293.045241 |
| -2.6605139 | 0.00780215 | 0.01476947 | 82.3786199 | 89.5721679 | 0 | 109.26161 |
| -2.6604203 | 0.00780432 | 0.01477271 | 97.5428434 | 15.4248144 | 5.75226998 | 15.1285306 |
| -2.6602879 | 0.00780739 | 0.01477765 | 186.888809 | 0 | 135.753571 | 87.9696039 |
| -2.6600782 | 0.00781225 | 0.01478599 | 36.8859492 | 142.34127 | 121.948124 | 113.744138 |
| -2.6600211 | 0.00781357 | 0.01478762 | 66.3947085 | 96.6080482 | 0 | 123.829825 |
| -2.659867 | 0.00781715 | 0.01479353 | 9.83625312 | 28.143521 | 32.2127119 | 9.52537113 |
| -2.6596651 | 0.00782184 | 0.01480153 | 52.8698605 | 107.161869 | 216.285351 | 0 |
| -2.6596395 | 0.00782243 | 0.01480179 | 45.4926707 | 143.694324 | 0 | 115.985401 |
| 2.65927973 | 0.00783079 | 0.01481673 | 0.40984388 | 0 | 0 | 0 |
| -2.6592457 | 0.00783158 | 0.01481737 | 63.5258014 | 119.339354 | 0 | 104.218766 |
| 2.65910901 | 0.00783476 | 0.01482251 | 43.8532952 | 17.0484791 | 24.1595339 | 16.2491625 |
| -2.659078 | 0.00783548 | 0.014823 | 10.6559409 | 35.4500121 | 23.0090799 | 38.6618005 |
| -2.6588408 | 0.007841 | 0.01483258 | 23.3611012 | 16.7778683 | 13.8054479 | 10.0856871 |
| 2.65853669 | 0.00784808 | 0.0148451 | 0 | 0 | 0 | 0 |
| -2.6582908 | 0.00785381 | 0.01485507 | 38.9351686 | 34.0969582 | 42.5667978 | 34.7395888 |

| | | | | | | |
|---|---|---|---|---|---|---|
| -2.658189 | 0.00785618 | 0.01485868 | 37.705637 | 43.0271139 | 60.9740618 | 52.1093832 |
| -2.6576048 | 0.00786981 | 0.01488359 | 70.0833035 | 169.672959 | 111.594038 | 0 |
| -2.6571411 | 0.00788064 | 0.0149032 | 65.1651769 | 210.535186 | 0 | 50.9887513 |
| 2.65704288 | 0.00788294 | 0.01490667 | 9.42640924 | 7.30649104 | 8.05317797 | 3.92221164 |
| 2.65670681 | 0.0078908 | 0.01492067 | 201.233345 | 188.615713 | 194.426725 | 158.009098 |
| -2.6565025 | 0.00789559 | 0.01492884 | 141.805982 | 28.6847426 | 0 | 165.853521 |
| -2.6563894 | 0.00789823 | 0.01493297 | 0 | 120.692407 | 103.54086 | 138.398039 |
| 2.65625883 | 0.00790129 | 0.01493788 | 0.81968776 | 2.16488623 | 1.150454 | 0.56031595 |
| 2.65561493 | 0.0079164 | 0.01496505 | 0.40984388 | 0.81183234 | 1.150454 | 0.56031595 |
| 2.65560261 | 0.00791668 | 0.01496505 | 16.8035991 | 12.1774851 | 11.50454 | 7.84442328 |
| -2.6555867 | 0.00791706 | 0.01496505 | 18.0331307 | 13.530539 | 9.20363196 | 28.0157974 |
| -2.6550916 | 0.00792869 | 0.01498616 | 102.46097 | 217.571066 | 146.107657 | 198.912162 |
| 2.65505768 | 0.00792948 | 0.01498679 | 30.738291 | 20.0251977 | 26.4604419 | 21.292006 |
| -2.6549906 | 0.00793106 | 0.01498889 | 21.3118818 | 27.3316887 | 37.9649818 | 21.292006 |
| -2.6549314 | 0.00793245 | 0.01499064 | 22.9512573 | 5.95343714 | 19.5577179 | 32.498325 |
| -2.6548982 | 0.00793323 | 0.01499124 | 367.62996 | 513.348648 | 398.057082 | 504.84467 |
| -2.654797 | 0.00793561 | 0.01499486 | 6.96734596 | 9.74198805 | 77.0804177 | 24.0935858 |
| -2.6539876 | 0.00795468 | 0.01502912 | 192.626624 | 466.803594 | 455.579782 | 446.571811 |
| -2.6540019 | 0.00795434 | 0.01502912 | 54.0993922 | 143.964935 | 0 | 96.3743432 |
| -2.6537454 | 0.00796039 | 0.01503903 | 13.9346919 | 17.8603114 | 8.05317797 | 47.0665397 |
| -2.6536517 | 0.0079626 | 0.01504232 | 132.789417 | 177.520671 | 176.019461 | 174.25826 |
| -2.6534473 | 0.00796742 | 0.01505056 | 19.2626624 | 37.3442875 | 46.0181598 | 34.1792729 |
| -2.653176 | 0.00797383 | 0.01506178 | 28.6890716 | 8.93015571 | 12.6549939 | 21.292006 |
| -2.6530983 | 0.00797566 | 0.01506436 | 13.9346919 | 9.47137727 | 28.7613499 | 16.8094785 |
| -2.6522114 | 0.00799665 | 0.01510311 | 0 | 130.434396 | 89.7354116 | 142.880567 |
| -2.6513721 | 0.00801655 | 0.01513981 | 51.230485 | 251.668025 | 331.330751 | 252.702493 |
| -2.6511686 | 0.00802138 | 0.01514805 | 22.9512573 | 24.3549701 | 14.9559019 | 7.84442328 |
| -2.6511393 | 0.00802208 | 0.01514848 | 10.246097 | 22.4606947 | 39.1154358 | 9.52537113 |
| -2.6510041 | 0.00802529 | 0.01515366 | 0 | 123.398515 | 179.470823 | 71.1601255 |
| -2.6509313 | 0.00802702 | 0.01515604 | 11.4756286 | 27.0610779 | 17.2568099 | 11.206319 |
| -2.6508583 | 0.00802875 | 0.01515843 | 4.91812656 | 20.0251977 | 34.5136199 | 28.0157974 |
| -2.6506602 | 0.00803346 | 0.01516643 | 15.1642236 | 17.0484791 | 10.354086 | 16.2491625 |
| -2.6504706 | 0.00803797 | 0.01517406 | 32.7875104 | 16.5072575 | 6.90272397 | 20.1713742 |
| -2.650083 | 0.0080472 | 0.01519059 | 279.103682 | 670.573511 | 544.16474 | 588.331746 |
| -2.650002 | 0.00804913 | 0.01519335 | 226.643666 | 467.615426 | 377.34891 | 417.995698 |
| 2.64996344 | 0.00805005 | 0.0151942 | 63.9356453 | 33.8263474 | 31.0622579 | 39.7824324 |
| -2.6499309 | 0.00805082 | 0.01519477 | 0 | 137.740887 | 207.081719 | 50.4284354 |
| 2.64931442 | 0.00806553 | 0.01522163 | 5.73781432 | 5.41221558 | 6.90272397 | 6.16347543 |
| -2.6491247 | 0.00807005 | 0.01522929 | 19.6725062 | 161.554635 | 0 | 182.662999 |
| -2.6488094 | 0.00807759 | 0.01524261 | 4.91812656 | 25.9786348 | 20.7081719 | 35.8602207 |
| -2.6485262 | 0.00808436 | 0.0152545 | 8.60672148 | 30.3084073 | 17.2568099 | 15.1285306 |
| -2.6484178 | 0.00808695 | 0.0152585 | 26.2300083 | 81.1832337 | 63.2749697 | 57.1522268 |
| 2.64836555 | 0.0080882 | 0.01525996 | 0 | 0 | 0 | 0 |

| | | | | | | |
|---|---|---|---|---|---|---|
| 2.64827586 | 0.00809035 | 0.01526312 | 18.0331307 | 8.38893415 | 10.354086 | 11.206319 |
| -2.6481518 | 0.00809332 | 0.01526783 | 0 | 148.024096 | 72.4786017 | 147.363094 |
| -2.6480231 | 0.0080964 | 0.01527275 | 79.0998688 | 222.983282 | 65.5758777 | 0 |
| -2.647877 | 0.0080999 | 0.01527846 | 22.5414134 | 11.9068743 | 9.20363196 | 25.2142177 |
| -2.6472708 | 0.00811443 | 0.01530499 | 23.770945 | 20.83703 | 27.6108959 | 28.0157974 |
| -2.6471049 | 0.00811842 | 0.0153116 | 5.32797044 | 51.416048 | 14.9559019 | 26.3348496 |
| -2.6470399 | 0.00811998 | 0.01531365 | 0 | 133.952336 | 127.700393 | 95.8140272 |
| -2.6468262 | 0.00812511 | 0.01532154 | 11.0657848 | 17.0484791 | 16.1063559 | 14.0078987 |
| -2.6468356 | 0.00812489 | 0.01532154 | 25.4103206 | 72.253078 | 85.1335957 | 80.1251807 |
| -2.6464933 | 0.00813311 | 0.01533573 | 48.7714217 | 103.914539 | 0 | 141.199619 |
| -2.6464713 | 0.00813364 | 0.01533584 | 18.4429746 | 15.6954252 | 11.50454 | 12.8872668 |
| -2.6464159 | 0.00813497 | 0.01533745 | 0 | 145.047378 | 162.214013 | 64.99665 |
| -2.6462476 | 0.00813902 | 0.01534419 | 21.3118818 | 18.401533 | 10.354086 | 12.3269509 |
| -2.6457325 | 0.00815142 | 0.01536667 | 11.8854725 | 13.2599282 | 17.2568099 | 15.1285306 |
| -2.6456319 | 0.00815385 | 0.01537035 | 148.363485 | 226.230611 | 0 | 15.1285306 |
| -2.6455759 | 0.0081552 | 0.015372 | 9.42640924 | 14.0717605 | 18.4072639 | 21.292006 |
| -2.6452476 | 0.00816312 | 0.01538602 | 4.91812656 | 30.0377965 | 17.2568099 | 38.6618005 |
| 2.64498186 | 0.00816953 | 0.01539721 | 0 | 0 | 0 | 0.56031595 |
| 2.64455878 | 0.00817975 | 0.01541517 | 0.40984388 | 2.43549701 | 3.45136199 | 3.36189569 |
| -2.644548 | 0.00818001 | 0.01541517 | 31.9678226 | 54.1221558 | 10.354086 | 6.16347543 |
| -2.6443116 | 0.00818573 | 0.01542503 | 27.8693838 | 33.2851258 | 5.75226998 | 15.6888466 |
| -2.6436298 | 0.00820223 | 0.01545523 | 29.9186032 | 21.6488623 | 4.60181598 | 28.5761134 |
| 2.64335424 | 0.00820891 | 0.01546692 | 141.396139 | 130.434396 | 139.204933 | 131.674248 |
| -2.6430386 | 0.00821657 | 0.01548044 | 9.42640924 | 46.8156648 | 5.75226998 | 43.704644 |
| -2.6428855 | 0.00822028 | 0.01548654 | 66.8045524 | 30.0377965 | 26.4604419 | 2.24126379 |
| -2.6425578 | 0.00822824 | 0.01550063 | 45.4926707 | 170.755402 | 0 | 87.409288 |
| -2.6422271 | 0.00823628 | 0.01551487 | 8.1968776 | 14.6129821 | 20.7081719 | 24.6539017 |
| -2.6420613 | 0.00824031 | 0.01552156 | 4.0984388 | 34.9087905 | 49.4695218 | 15.6888466 |
| -2.6419687 | 0.00824257 | 0.0155249 | 11.0657848 | 23.5431378 | 18.4072639 | 11.206319 |
| -2.6418555 | 0.00824532 | 0.01552918 | 5.73781432 | 34.9087905 | 16.1063559 | 27.4554815 |
| -2.6416557 | 0.00825019 | 0.01553744 | 3.68859492 | 33.5557366 | 48.3190678 | 18.4904263 |
| -2.6415604 | 0.00825251 | 0.0155409 | 2.0492194 | 28.4141318 | 58.6731538 | 34.7395888 |
| -2.6414058 | 0.00825628 | 0.01554709 | 0 | 87.9485032 | 125.399485 | 142.880567 |
| -2.6400975 | 0.00828822 | 0.01560633 | 4.91812656 | 19.4839761 | 41.4163438 | 24.0935858 |
| -2.6399525 | 0.00829176 | 0.0156121 | 54.509236 | 128.269509 | 0 | 100.856871 |
| -2.639576 | 0.00830098 | 0.01562854 | 9.83625312 | 43.5683354 | 35.6640739 | 6.72379138 |
| 2.63935708 | 0.00830634 | 0.01563772 | 4.91812656 | 1.89427545 | 0 | 2.80157974 |
| -2.6392023 | 0.00831014 | 0.01564395 | 15.9839113 | 21.9194731 | 25.3099879 | 20.1713742 |
| 2.63909734 | 0.00831271 | 0.01564788 | 0.81968776 | 1.3530539 | 0 | 2.24126379 |
| -2.6388806 | 0.00831803 | 0.01565698 | 0 | 119.068743 | 109.29313 | 121.588561 |
| -2.6388328 | 0.0083192 | 0.01565827 | 6.55750208 | 27.0610779 | 28.7613499 | 15.1285306 |
| -2.6380782 | 0.00833774 | 0.01569224 | 600.831128 | 464.097486 | 421.066162 | 464.501921 |
| 2.63799552 | 0.00833977 | 0.01569516 | 8.1968776 | 5.1416048 | 11.50454 | 6.16347543 |

| | | | | | | |
|---|---|---|---|---|---|---|
| -2.6379155 | 0.00834174 | 0.01569612 | 262.709927 | 681.939163 | 1025.05451 | 863.446877 |
| -2.6379468 | 0.00834097 | 0.01569612 | 5.73781432 | 20.0251977 | 28.7613499 | 23.5332698 |
| -2.637926 | 0.00834148 | 0.01569612 | 180.741151 | 67.9233056 | 95.4876816 | 0 |
| 2.63741364 | 0.00835409 | 0.01571844 | 8.1968776 | 9.20076649 | 5.75226998 | 10.646003 |
| -2.6373423 | 0.00835585 | 0.01572083 | 18.0331307 | 34.367569 | 28.7613499 | 25.7745336 |
| -2.6373147 | 0.00835653 | 0.0157212 | 201.233345 | 301.460408 | 284.162137 | 214.601008 |
| -2.6372232 | 0.00835878 | 0.01572452 | 15.5740674 | 18.9427545 | 18.4072639 | 8.96505518 |
| -2.6371234 | 0.00836124 | 0.01572823 | 4.91812656 | 30.579018 | 64.4254237 | 11.7666349 |
| -2.6364494 | 0.00837787 | 0.0157586 | 25.4103206 | 30.3084073 | 27.6108959 | 29.6967453 |
| -2.6363688 | 0.00837986 | 0.01576142 | 47.5418901 | 32.7439043 | 43.7172518 | 28.5761134 |
| -2.6361576 | 0.00838508 | 0.01577032 | 247.13586 | 60.6168145 | 46.0181598 | 62.1950703 |
| -2.6360858 | 0.00838685 | 0.01577273 | 114.346443 | 76.3122397 | 0 | 81.2458125 |
| -2.6359703 | 0.00838971 | 0.01577718 | 8.1968776 | 28.143521 | 11.50454 | 27.4554815 |
| -2.6359169 | 0.00839103 | 0.01577875 | 0 | 123.669126 | 62.1245158 | 184.343947 |
| 2.63582261 | 0.00839336 | 0.01578221 | 18.8528185 | 23.272527 | 20.7081719 | 22.9729539 |
| -2.6357963 | 0.00839401 | 0.01578252 | 8.60672148 | 33.2851258 | 11.50454 | 22.4126379 |
| -2.6357555 | 0.00839502 | 0.0157835 | 10.6559409 | 17.0484791 | 11.50454 | 27.4554815 |
| -2.6356815 | 0.00839685 | 0.01578602 | 3.68859492 | 23.272527 | 18.4072639 | 73.4013893 |
| 2.63545983 | 0.00840234 | 0.01579542 | 0 | 0 | 0 | 0 |
| -2.6353694 | 0.00840457 | 0.0157987 | 14.3445358 | 6.22404792 | 28.7613499 | 33.058641 |
| 2.63521159 | 0.00840848 | 0.01580513 | 6.55750208 | 2.16488623 | 2.30090799 | 2.80157974 |
| -2.6351879 | 0.00840907 | 0.01580532 | 0 | 64.4053654 | 322.127119 | 50.4284354 |
| -2.6346549 | 0.00842229 | 0.01582923 | 78.6900249 | 94.4431619 | 0 | 95.2537113 |
| -2.634145 | 0.00843495 | 0.0158521 | 4.91812656 | 33.0145151 | 27.6108959 | 19.6110582 |
| -2.633866 | 0.00844188 | 0.01586421 | 8.60672148 | 19.4839761 | 26.4604419 | 13.4475828 |
| -2.633717 | 0.00844558 | 0.01587025 | 9.01656536 | 19.7545869 | 14.9559019 | 18.4904263 |
| -2.6333328 | 0.00845515 | 0.01588729 | 36.0662614 | 22.1900839 | 4.60181598 | 22.9729539 |
| 2.63327657 | 0.00845655 | 0.01588899 | 22.1315695 | 12.7187066 | 19.5577179 | 15.1285306 |
| -2.6332414 | 0.00845742 | 0.01588971 | 47.1320462 | 182.662276 | 0 | 73.9617052 |
| 2.63285128 | 0.00846714 | 0.01590705 | 84.4278393 | 161.554635 | 109.29313 | 118.786981 |
| -2.6326625 | 0.00847185 | 0.01591497 | 8.1968776 | 22.1900839 | 23.0090799 | 14.0078987 |
| -2.6325466 | 0.00847474 | 0.0159189 | 80.7392443 | 66.2996409 | 0 | 128.312352 |
| -2.6325392 | 0.00847492 | 0.0159189 | 0 | 112.574084 | 127.700393 | 105.899714 |
| -2.6324726 | 0.00847659 | 0.01592109 | 7.78703372 | 19.7545869 | 26.4604419 | 15.1285306 |
| -2.6323442 | 0.00847979 | 0.01592618 | 13.9346919 | 22.1900839 | 12.6549939 | 11.7666349 |
| -2.6322156 | 0.008483 | 0.01593128 | 5.73781432 | 21.1076408 | 25.3099879 | 24.0935858 |
| 2.63213382 | 0.00848504 | 0.01593419 | 16.3937552 | 13.2599282 | 18.4072639 | 7.84442328 |
| 2.63201761 | 0.00848795 | 0.01593779 | 0 | 0 | 0 | 7.84442328 |
| -2.6320188 | 0.00848792 | 0.01593779 | 39.7548564 | 182.932887 | 141.505841 | 0 |
| -2.6317071 | 0.00849571 | 0.01595143 | 15.5740674 | 15.966036 | 11.50454 | 13.4475828 |
| -2.6315546 | 0.00849952 | 0.01595767 | 48.7714217 | 80.3714014 | 0 | 165.293205 |
| -2.6313237 | 0.0085053 | 0.01596759 | 0 | 64.4053654 | 201.329449 | 104.779082 |
| -2.6312039 | 0.0085083 | 0.01597229 | 4.50828268 | 43.0271139 | 34.5136199 | 15.1285306 |

| | | | | | | |
|---|---|---|---|---|---|---|
| -2.631148 | 0.0085097 | 0.01597399 | 77.8703372 | 61.4286469 | 0 | 140.639303 |
| -2.6309652 | 0.00851427 | 0.01598165 | 0 | 179.414947 | 153.010381 | 47.6268556 |
| -2.6307191 | 0.00852044 | 0.0159923 | 5.73781432 | 15.966036 | 36.8145279 | 25.2142177 |
| -2.6304732 | 0.00852661 | 0.01600294 | 15.9839113 | 12.7187066 | 20.7081719 | 10.646003 |
| -2.6300508 | 0.00853721 | 0.01602191 | 10.246097 | 16.7778683 | 20.7081719 | 12.8872668 |
| -2.6295306 | 0.00855028 | 0.01604551 | 71.3128351 | 195.651593 | 189.824909 | 167.534469 |
| -2.6294345 | 0.0085527 | 0.01604911 | 0 | 84.4305631 | 86.2840497 | 191.067738 |
| -2.6290535 | 0.00856229 | 0.01606617 | 9.01656536 | 15.4248144 | 19.5577179 | 17.3697944 |
| 2.62806882 | 0.00858711 | 0.01611182 | 0 | 0 | 0 | 0 |
| -2.6278945 | 0.00859152 | 0.01611914 | 10.246097 | 42.7565031 | 5.75226998 | 39.2221164 |
| -2.6278394 | 0.00859291 | 0.01612081 | 12.2953164 | 8.93015571 | 24.1595339 | 21.852322 |
| -2.6277753 | 0.00859453 | 0.01612291 | 89.3459658 | 131.246228 | 0 | 56.5919108 |
| -2.6277002 | 0.00859642 | 0.01612554 | 8.60672148 | 33.5557366 | 36.8145279 | 8.40473923 |
| -2.6276774 | 0.008597 | 0.01612568 | 9.01656536 | 28.4141318 | 9.20363196 | 33.058641 |
| 2.62762119 | 0.00859842 | 0.01612741 | 32.3776665 | 19.2133653 | 17.2568099 | 15.6888466 |
| -2.6275963 | 0.00859905 | 0.01612765 | 13.1150042 | 5.41221558 | 29.9118039 | 46.5062237 |
| -2.6274867 | 0.00860182 | 0.01613191 | 81.5589321 | 148.294707 | 0 | 53.7903311 |
| -2.6274553 | 0.00860261 | 0.01613246 | 41.8040758 | 21.1076408 | 10.354086 | 8.96505518 |
| -2.6271193 | 0.00861111 | 0.01614746 | 7.37718984 | 33.5557366 | 12.6549939 | 24.0935858 |
| -2.6270674 | 0.00861243 | 0.01614899 | 9.42640924 | 12.1774851 | 18.4072639 | 28.5761134 |
| 2.62699798 | 0.00861418 | 0.01615041 | 0 | 0 | 0 | 0 |
| 2.62699798 | 0.00861418 | 0.01615041 | 0 | 0 | 0 | 0 |
| -2.6267252 | 0.00862109 | 0.01616242 | 33.6071982 | 21.1076408 | 25.3099879 | 12.8872668 |
| -2.6266257 | 0.00862361 | 0.01616621 | 6.1476582 | 21.9194731 | 17.2568099 | 32.498325 |
| -2.6265488 | 0.00862556 | 0.01616892 | 50.4107972 | 90.654611 | 0 | 141.759935 |
| -2.6262033 | 0.00863432 | 0.01618441 | 34.4268859 | 37.6148983 | 12.6549939 | 5.60315949 |
| -2.626129 | 0.00863621 | 0.016187 | 2.45906328 | 34.0969582 | 43.7172518 | 29.1364293 |
| -2.6255248 | 0.00865155 | 0.01621481 | 0 | 22.7313054 | 177.169915 | 237.013646 |
| -2.6249613 | 0.00866588 | 0.01624073 | 67.2143963 | 348.276073 | 28.7613499 | 0 |
| -2.6247956 | 0.0086701 | 0.01624769 | 47.951734 | 13.2599282 | 9.20363196 | 14.5682147 |
| -2.6244678 | 0.00867845 | 0.0162624 | 14.3445358 | 17.5897006 | 9.20363196 | 19.0507423 |
| -2.624366 | 0.00868104 | 0.01626632 | 119.264569 | 74.688575 | 0 | 74.5220212 |
| -2.624339 | 0.00868173 | 0.01626666 | 9.83625312 | 12.7187066 | 36.8145279 | 15.1285306 |
| -2.6237912 | 0.00869571 | 0.0162919 | 43.8532952 | 123.939737 | 0 | 116.545717 |
| -2.6236895 | 0.0086983 | 0.01629582 | 17.6232868 | 17.5897006 | 8.05317797 | 19.0507423 |
| -2.6236245 | 0.00869996 | 0.01629798 | 13.9346919 | 14.3423713 | 13.8054479 | 12.8872668 |
| -2.6235292 | 0.0087024 | 0.0163016 | 4.50828268 | 30.0377965 | 24.1595339 | 25.7745336 |
| -2.6230882 | 0.00871367 | 0.01632177 | 257.7918 | 502.253606 | 544.16474 | 495.879615 |
| -2.6230622 | 0.00871434 | 0.01632207 | 50.0009534 | 14.0717605 | 9.20363196 | 13.4475828 |
| -2.6229079 | 0.00871828 | 0.01632851 | 4.0984388 | 54.3927666 | 10.354086 | 47.0665397 |
| -2.6227835 | 0.00872147 | 0.01633353 | 8.1968776 | 22.1900839 | 14.9559019 | 19.6110582 |
| 2.62235739 | 0.00873238 | 0.01635302 | 84.0179954 | 95.7962158 | 93.1867736 | 71.7204414 |
| -2.6221886 | 0.00873671 | 0.01636017 | 25.4103206 | 20.2958084 | 5.75226998 | 21.852322 |

| | | | | | | |
|---|---|---|---|---|---|---|
| -2.6220743 | 0.00873964 | 0.01636471 | 40.984388 | 18.1309222 | 8.05317797 | 12.8872668 |
| -2.6218308 | 0.00874588 | 0.01637545 | 4.50828268 | 28.4141318 | 24.1595339 | 26.8951655 |
| -2.6217862 | 0.00874703 | 0.01637665 | 4.91812656 | 27.6022995 | 33.3631659 | 18.4904263 |
| -2.6216875 | 0.00874956 | 0.01638044 | 15.5740674 | 12.9893174 | 32.2127119 | 9.52537113 |
| -2.621612 | 0.0087515 | 0.01638312 | 42.6237635 | 17.0484791 | 8.05317797 | 13.4475828 |
| -2.6215602 | 0.00875283 | 0.01638466 | 23.3611012 | 18.1309222 | 8.05317797 | 15.1285306 |
| 2.62111992 | 0.00876414 | 0.01640485 | 0 | 0 | 1.150454 | 0 |
| -2.6211008 | 0.00876463 | 0.01640485 | 8.1968776 | 28.143521 | 13.8054479 | 19.0507423 |
| -2.6203418 | 0.00878417 | 0.01644046 | 15.1642236 | 10.5538204 | 14.9559019 | 16.2491625 |
| -2.6197961 | 0.00879824 | 0.01646584 | 166.396615 | 306.602013 | 159.913105 | 221.885116 |
| -2.6197358 | 0.00879979 | 0.01646779 | 23.770945 | 19.2133653 | 33.3631659 | 25.2142177 |
| 2.61963858 | 0.0088023 | 0.01647153 | 2.0492194 | 1.3530539 | 0 | 2.24126379 |
| -2.6196189 | 0.00880281 | 0.01647153 | 79.9195566 | 73.8767427 | 0 | 110.382242 |
| -2.6190358 | 0.00881787 | 0.01649876 | 23.3611012 | 15.6954252 | 41.4163438 | 43.704644 |
| 2.61886527 | 0.00882228 | 0.01650605 | 0 | 12.9893174 | 0 | 8.96505518 |
| -2.6188217 | 0.00882341 | 0.0165072 | 13.9346919 | 28.143521 | 14.9559019 | 9.52537113 |
| -2.6185 | 0.00883173 | 0.01652182 | 2.45906328 | 33.5557366 | 85.1335957 | 15.1285306 |
| -2.6182085 | 0.00883928 | 0.01653402 | 81.1490882 | 49.5217726 | 0 | 152.405938 |
| -2.6182101 | 0.00883924 | 0.01653402 | 14.7543797 | 14.8835929 | 10.354086 | 16.8094785 |
| -2.6180126 | 0.00884435 | 0.01654256 | 62.7061136 | 67.6526948 | 0 | 146.802779 |
| 2.61778622 | 0.00885022 | 0.01655258 | 0.81968776 | 8.11832337 | 5.75226998 | 3.36189569 |
| 2.61729946 | 0.00886285 | 0.01657524 | 2.86890716 | 5.1416048 | 3.45136199 | 0 |
| -2.6172544 | 0.00886403 | 0.01657647 | 16.8035991 | 15.966036 | 16.1063559 | 9.52537113 |
| -2.6168467 | 0.00887462 | 0.01659532 | 126.641759 | 317.967665 | 293.365769 | 295.846821 |
| -2.6167902 | 0.00887609 | 0.01659711 | 19.2626624 | 7.8477126 | 33.3631659 | 13.4475828 |
| -2.616394 | 0.0088864 | 0.01661542 | 18.0331307 | 23.8137486 | 48.3190678 | 4.48252759 |
| -2.6162162 | 0.00889102 | 0.01662311 | 10.6559409 | 9.74198805 | 21.8586259 | 25.2142177 |
| -2.6159119 | 0.00889895 | 0.01663697 | 218.446788 | 312.014228 | 337.083021 | 348.51652 |
| -2.6157861 | 0.00890223 | 0.01664214 | 41.3942319 | 5.1416048 | 20.7081719 | 19.6110582 |
| -2.6156275 | 0.00890637 | 0.0166489 | 59.4273626 | 18.6721438 | 8.05317797 | 10.646003 |
| -2.6154733 | 0.00891039 | 0.01665546 | 9.42640924 | 17.0484791 | 17.2568099 | 15.1285306 |
| -2.6153057 | 0.00891476 | 0.01666267 | 20.0823501 | 30.0377965 | 28.7613499 | 29.1364293 |
| -2.6150125 | 0.00892242 | 0.01667601 | 0 | 171.296623 | 118.496762 | 63.3157022 |
| -2.6149627 | 0.00892372 | 0.01667706 | 4.50828268 | 8.93015571 | 58.6731538 | 44.8252759 |
| -2.6149516 | 0.00892401 | 0.01667706 | 100.411751 | 54.1221558 | 184.072639 | 0 |
| -2.6143636 | 0.00893938 | 0.01670482 | 15.1642236 | 11.3656527 | 12.6549939 | 17.3697944 |
| -2.614286 | 0.00894142 | 0.01670765 | 216.397569 | 128.810731 | 0 | 166.974153 |
| -2.6141888 | 0.00894396 | 0.01671144 | 103.280658 | 167.237462 | 189.824909 | 171.45668 |
| -2.6140832 | 0.00894672 | 0.01671564 | 9.42640924 | 12.9893174 | 17.2568099 | 22.9729539 |
| -2.6131521 | 0.00897114 | 0.01676028 | 47.1320462 | 67.1114732 | 0 | 180.421735 |
| -2.6131234 | 0.00897189 | 0.01676071 | 14.3445358 | 9.20076649 | 29.9118039 | 14.5682147 |
| 2.61304724 | 0.00897389 | 0.01676154 | 0 | 0 | 0 | 0 |
| 2.61304724 | 0.00897389 | 0.01676154 | 0 | 0 | 0 | 0 |

| | | | | | | |
|---|---|---|---|---|---|---|
| 2.61304724 | 0.00897389 | 0.01676154 | 0 | 0 | 0 | 0 |
| -2.612553 | 0.00898688 | 0.01678475 | 56.5584554 | 138.282108 | 0 | 77.8839169 |
| -2.6125348 | 0.00898735 | 0.01678475 | 7.78703372 | 29.4965749 | 9.20363196 | 39.2221164 |
| 2.61206663 | 0.00899967 | 0.01680678 | 45.4926707 | 28.143521 | 67.8767857 | 23.5332698 |
| -2.6120041 | 0.00900132 | 0.01680888 | 14.7543797 | 14.3423713 | 12.6549939 | 12.8872668 |
| -2.6119799 | 0.00900195 | 0.0168091 | 11.4756286 | 21.1076408 | 14.9559019 | 11.7666349 |
| -2.6119591 | 0.0090025 | 0.01680915 | 13.524848 | 20.2958084 | 9.20363196 | 17.3697944 |
| 2.61193201 | 0.00900322 | 0.01680951 | 10.246097 | 7.57710182 | 9.20363196 | 8.96505518 |
| -2.6116895 | 0.0090096 | 0.01682046 | 12.2953164 | 25.9786348 | 10.354086 | 15.1285306 |
| -2.6115334 | 0.00901372 | 0.01682717 | 0 | 135.034779 | 39.1154358 | 207.877217 |
| -2.6112172 | 0.00902206 | 0.01684177 | 14.7543797 | 15.966036 | 49.4695218 | 7.28410733 |
| 2.61093444 | 0.00902952 | 0.01685472 | 33.6071982 | 18.6721438 | 20.7081719 | 15.1285306 |
| -2.6108532 | 0.00903166 | 0.01685775 | 22.1315695 | 38.4267306 | 33.3631659 | 49.8681194 |
| -2.610627 | 0.00903764 | 0.01686763 | 15.1642236 | 215.947402 | 0 | 129.9933 |
| -2.6106134 | 0.009038 | 0.01686763 | 81.968776 | 114.73897 | 116.195854 | 124.950457 |
| -2.6104912 | 0.00904123 | 0.01687267 | 0 | 121.774851 | 102.390406 | 108.701294 |
| 2.61039834 | 0.00904368 | 0.01687628 | 69.6734596 | 37.0736767 | 29.9118039 | 36.9808526 |
| -2.6103481 | 0.00904501 | 0.01687779 | 381.974496 | 633.770445 | 556.819734 | 623.631651 |
| -2.6097053 | 0.00906202 | 0.01690855 | 106.149565 | 86.0542278 | 0 | 67.7982298 |
| 2.60966098 | 0.0090632 | 0.01690977 | 4.0984388 | 11.3656527 | 0 | 14.5682147 |
| -2.6095313 | 0.00906663 | 0.0169152 | 6.96734596 | 27.3316887 | 24.1595339 | 14.0078987 |
| -2.6092566 | 0.00907392 | 0.01692781 | 19.6725062 | 10.0125988 | 19.5577179 | 11.7666349 |
| -2.6088246 | 0.00908538 | 0.01694745 | 19.6725062 | 24.0843593 | 36.8145279 | 24.0935858 |
| -2.6088202 | 0.0090855 | 0.01694745 | 2.0492194 | 45.1920001 | 51.7704298 | 21.292006 |
| -2.6087647 | 0.00908697 | 0.01694866 | 7.78703372 | 17.3190899 | 16.1063559 | 24.6539017 |
| -2.6087562 | 0.0090872 | 0.01694866 | 7.78703372 | 15.966036 | 28.7613499 | 16.2491625 |
| -2.6086602 | 0.00908975 | 0.01695244 | 11.0657848 | 14.3423713 | 17.2568099 | 13.4475828 |
| 2.60821459 | 0.00910159 | 0.01697354 | 0.81968776 | 2.97671857 | 1.150454 | 6.16347543 |
| -2.6081192 | 0.00910412 | 0.01697729 | 7.78703372 | 18.6721438 | 12.6549939 | 40.9030642 |
| 2.60772261 | 0.00911468 | 0.01699403 | 0.81968776 | 0.27061078 | 0 | 0 |
| -2.6077563 | 0.00911378 | 0.01699403 | 4.91812656 | 28.6847426 | 24.1595339 | 21.852322 |
| -2.6077389 | 0.00911424 | 0.01699403 | 11.4756286 | 18.6721438 | 13.8054479 | 12.8872668 |
| 2.60707313 | 0.00913199 | 0.01702532 | 3.27875104 | 0.27061078 | 0 | 1.68094785 |
| -2.6065345 | 0.00914636 | 0.01705113 | 19.2626624 | 133.681725 | 0 | 180.982051 |
| 2.60611608 | 0.00915754 | 0.01707099 | 0 | 0 | 0 | 0.56031595 |
| -2.6060686 | 0.00915881 | 0.01707237 | 11.4756286 | 19.7545869 | 16.1063559 | 11.206319 |
| -2.6057046 | 0.00916855 | 0.01708954 | 77.8703372 | 110.138587 | 132.302209 | 109.821926 |
| -2.6055363 | 0.00917306 | 0.01709695 | 53.6895483 | 129.081342 | 0 | 85.1680242 |
| -2.6055054 | 0.00917388 | 0.0170975 | 26.2300083 | 151.271426 | 0 | 129.9933 |
| 2.60541369 | 0.00917634 | 0.01710109 | 0 | 0 | 0 | 0 |
| -2.605314 | 0.00917901 | 0.01710508 | 86.8869025 | 79.8301798 | 0 | 87.9696039 |
| -2.6052079 | 0.00918185 | 0.01710939 | 0 | 163.990132 | 47.1686138 | 148.483726 |
| -2.6050756 | 0.0091854 | 0.01711501 | 4.0984388 | 42.2152815 | 32.2127119 | 16.2491625 |

| | | | | | | |
|---|---|---|---|---|---|---|
| 2.60494773 | 0.00918883 | 0.01712042 | 20.0823501 | 21.9194731 | 10.354086 | 20.1713742 |
| -2.6048373 | 0.00919179 | 0.01712495 | 116.805506 | 143.694324 | 59.8236078 | 0 |
| -2.6043687 | 0.00920437 | 0.01714739 | 13.524848 | 20.2958084 | 9.20363196 | 16.8094785 |
| 2.60421521 | 0.00920849 | 0.01715408 | 16.8035991 | 20.5664192 | 16.1063559 | 15.6888466 |
| -2.6038536 | 0.00921821 | 0.0171712 | 0 | 27.0610779 | 394.60572 | 57.7125427 |
| -2.60324 | 0.00923473 | 0.01720097 | 0 | 94.4431619 | 127.700393 | 107.580662 |
| -2.6031453 | 0.00923728 | 0.01720473 | 34.8367298 | 21.3782516 | 12.6549939 | 7.28410733 |
| -2.6025689 | 0.00925282 | 0.01723269 | 89.7558097 | 59.8049822 | 170.267191 | 18.4904263 |
| 2.602377 | 0.009258 | 0.01724134 | 31.5579788 | 20.83703 | 21.8586259 | 12.8872668 |
| -2.6019842 | 0.00926861 | 0.0172601 | 8.1968776 | 23.272527 | 14.9559019 | 17.3697944 |
| 2.6017357 | 0.00927533 | 0.0172711 | 21.3118818 | 14.8835929 | 9.20363196 | 12.8872668 |
| -2.6017262 | 0.00927559 | 0.0172711 | 42.6237635 | 10.8244312 | 9.20363196 | 17.3697944 |
| -2.601596 | 0.00927911 | 0.01727666 | 22.9512573 | 13.530539 | 8.05317797 | 19.6110582 |
| -2.6014927 | 0.0092819 | 0.01728087 | 4.91812656 | 20.0251977 | 21.8586259 | 35.8602207 |
| -2.6012558 | 0.00928832 | 0.01729181 | 8.1968776 | 31.6614612 | 48.3190678 | 26.3348496 |
| -2.6011799 | 0.00929037 | 0.01729464 | 0 | 64.4053654 | 113.894946 | 164.732889 |
| -2.6010033 | 0.00929515 | 0.01730255 | 107.78894 | 14.3423713 | 4.60181598 | 13.4475828 |
| -2.6005483 | 0.00930749 | 0.01732451 | 10.246097 | 37.3442875 | 27.6108959 | 7.28410733 |
| -2.60042 | 0.00931097 | 0.01732999 | 10.246097 | 30.0377965 | 28.7613499 | 7.84442328 |
| -2.5998814 | 0.0093256 | 0.01735622 | 145.904421 | 59.5343714 | 0 | 64.99665 |
| 2.59982619 | 0.0093271 | 0.01735801 | 20.492194 | 3.78855091 | 8.05317797 | 1.68094785 |
| -2.5997734 | 0.00932853 | 0.01735968 | 97.1329995 | 85.5130062 | 0 | 71.1601255 |
| -2.5996986 | 0.00933057 | 0.01736246 | 2.45906328 | 35.1794013 | 37.9649818 | 28.5761134 |
| -2.5995594 | 0.00933435 | 0.0173685 | 8.1968776 | 23.272527 | 13.8054479 | 19.0507423 |
| 2.59944269 | 0.00933753 | 0.01737341 | 137.2977 | 148.835929 | 219.736713 | 75.0823371 |
| -2.5990743 | 0.00934755 | 0.01739106 | 19.2626624 | 12.7187066 | 13.8054479 | 11.206319 |
| -2.5988177 | 0.00935454 | 0.01740306 | 11.8854725 | 18.1309222 | 9.20363196 | 24.0935858 |
| -2.5982529 | 0.00936995 | 0.01743071 | 4.91812656 | 27.6022995 | 25.3099879 | 20.7316901 |
| 2.59769624 | 0.00938515 | 0.01745799 | 0 | 0 | 0 | 0 |
| -2.5976588 | 0.00938617 | 0.01745888 | 0 | 196.734036 | 98.9390436 | 57.1522268 |
| -2.597421 | 0.00939267 | 0.01746997 | 96.3133118 | 85.2423954 | 0 | 71.1601255 |
| -2.5970475 | 0.00940289 | 0.01748797 | 0 | 83.8893415 | 108.142676 | 136.717091 |
| 2.59674313 | 0.00941123 | 0.01750045 | 0.40984388 | 0 | 0 | 0.56031595 |
| -2.5967492 | 0.00941106 | 0.01750045 | 5.32797044 | 21.1076408 | 41.4163438 | 16.8094785 |
| 2.59674313 | 0.00941123 | 0.01750045 | 0.40984388 | 0 | 0 | 0.56031595 |
| -2.5966239 | 0.00941449 | 0.01750535 | 14.3445358 | 17.5897006 | 12.6549939 | 11.206319 |
| 2.59660729 | 0.00941495 | 0.01750535 | 9.42640924 | 5.68282636 | 9.20363196 | 2.80157974 |
| -2.5965472 | 0.0094166 | 0.01750741 | 43.8532952 | 16.2366467 | 6.90272397 | 15.1285306 |
| -2.5964736 | 0.00941861 | 0.01750912 | 5.73781432 | 25.708024 | 18.4072639 | 23.5332698 |
| -2.5964543 | 0.00941914 | 0.01750912 | 31.9678226 | 36.803066 | 12.6549939 | 16.2491625 |
| -2.5964744 | 0.00941859 | 0.01750912 | 74.5915861 | 87.6778924 | 0 | 89.0902358 |
| -2.5963714 | 0.00942142 | 0.01751233 | 10.246097 | 35.4500121 | 6.90272397 | 29.6967453 |
| 2.5960372 | 0.00943059 | 0.01752774 | 0 | 5.1416048 | 0 | 0 |

| | | | | | | |
|---|---|---|---|---|---|---|
| -2.5960297 | 0.00943079 | 0.01752774 | 12.2953164 | 25.9786348 | 8.05317797 | 20.7316901 |
| -2.5952557 | 0.00945206 | 0.01756626 | 51.6403289 | 146.671042 | 0 | 72.2807574 |
| -2.5950423 | 0.00945793 | 0.01757615 | 23.770945 | 24.3549701 | 34.5136199 | 58.8331746 |
| -2.5946656 | 0.0094683 | 0.01759441 | 3.27875104 | 40.5916169 | 23.0090799 | 28.5761134 |
| 2.5941868 | 0.0094815 | 0.01761792 | 479.107496 | 113.656527 | 120.79767 | 112.06319 |
| -2.594148 | 0.00948257 | 0.0176189 | 36.8859492 | 94.9843835 | 0 | 149.604358 |
| -2.5941046 | 0.00948376 | 0.01762011 | 390.581218 | 466.803594 | 355.490285 | 400.065587 |
| -2.5938357 | 0.00949118 | 0.01763288 | 127.461447 | 68.1939163 | 0 | 64.4363341 |
| -2.5937581 | 0.00949333 | 0.01763585 | 3.68859492 | 29.2259641 | 40.2658898 | 19.6110582 |
| -2.5936599 | 0.00949604 | 0.01763987 | 79.0998688 | 60.6168145 | 0 | 117.666349 |
| 2.5934942 | 0.00950061 | 0.01764736 | 1.22953164 | 0.81183234 | 0 | 0 |
| -2.5934586 | 0.0095016 | 0.01764817 | 35.6564176 | 39.5091738 | 43.7172518 | 52.1093832 |
| -2.5933702 | 0.00950404 | 0.01765169 | 0 | 61.6992576 | 186.373547 | 96.9346591 |
| -2.5932457 | 0.00950748 | 0.01765706 | 2.86890716 | 82.2656769 | 35.6640739 | 11.206319 |
| 2.59315194 | 0.00951007 | 0.01766086 | 0 | 5.1416048 | 0 | 0 |
| -2.5929379 | 0.009516 | 0.01767084 | 6.96734596 | 18.401533 | 18.4072639 | 21.852322 |
| -2.5928881 | 0.00951737 | 0.01767239 | 8.60672148 | 22.7313054 | 16.1063559 | 14.5682147 |
| -2.5923064 | 0.00953348 | 0.01770128 | 4.50828268 | 15.6954252 | 27.6108959 | 42.5840121 |
| -2.5921919 | 0.00953666 | 0.01770615 | 10.6559409 | 14.3423713 | 16.1063559 | 14.0078987 |
| -2.5919117 | 0.00954443 | 0.01771956 | 52.0501728 | 55.4752097 | 37.9649818 | 55.4712789 |
| -2.591743 | 0.00954911 | 0.01772723 | 2.0492194 | 10.0125988 | 85.1335957 | 50.4284354 |
| -2.5914631 | 0.00955688 | 0.01774064 | 10.6559409 | 6.76526948 | 43.7172518 | 25.2142177 |
| -2.5914317 | 0.00955775 | 0.01774124 | 0 | 71.4412457 | 113.894946 | 145.121831 |
| 2.59109547 | 0.00956709 | 0.01775756 | 40.984388 | 20.5664192 | 35.6640739 | 17.3697944 |
| 2.5910634 | 0.00956799 | 0.0177582 | 1.22953164 | 1.3530539 | 4.60181598 | 5.60315949 |
| -2.5909459 | 0.00957125 | 0.01776324 | 0 | 82.5362876 | 164.514921 | 85.7283401 |
| -2.5908501 | 0.00957392 | 0.01776716 | 0 | 68.1939163 | 164.514921 | 102.537819 |
| -2.5907327 | 0.00957719 | 0.01777221 | 26.2300083 | 11.9068743 | 10.354086 | 14.5682147 |
| -2.5906595 | 0.00957922 | 0.01777357 | 26.2300083 | 24.8961917 | 9.20363196 | 9.52537113 |
| -2.5906469 | 0.00957957 | 0.01777357 | 6.96734596 | 21.1076408 | 17.2568099 | 20.1713742 |
| -2.5906796 | 0.00957866 | 0.01777357 | 42.2139196 | 11.0950419 | 27.6108959 | 6.16347543 |
| -2.5905687 | 0.00958175 | 0.01777557 | 14.3445358 | 23.8137486 | 18.4072639 | 7.84442328 |
| -2.5905878 | 0.00958122 | 0.01777557 | 16.3937552 | 9.47137727 | 20.7081719 | 12.8872668 |
| -2.5905298 | 0.00958283 | 0.01777655 | 7.78703372 | 22.4606947 | 14.9559019 | 18.4904263 |
| -2.5904907 | 0.00958392 | 0.01777755 | 3.68859492 | 22.4606947 | 29.9118039 | 32.498325 |
| -2.5902529 | 0.00959054 | 0.01778881 | 61.8864259 | 53.5809343 | 0 | 158.009098 |
| -2.5901793 | 0.00959259 | 0.0177916 | 11.4756286 | 20.2958084 | 14.9559019 | 11.206319 |
| -2.5901191 | 0.00959427 | 0.01779369 | 6.1476582 | 15.4248144 | 35.6640739 | 20.1713742 |
| 2.58990088 | 0.00960036 | 0.0178023 | 0 | 0 | 0 | 0 |
| 2.58989325 | 0.00960057 | 0.0178023 | 0 | 0 | 0 | 0 |
| 2.58989325 | 0.00960057 | 0.0178023 | 0 | 0 | 0 | 0 |
| -2.5898175 | 0.00960268 | 0.01780519 | 6.1476582 | 16.2366467 | 23.0090799 | 26.3348496 |
| -2.5896166 | 0.00960829 | 0.01781456 | 11.4756286 | 14.0717605 | 23.0090799 | 11.206319 |

| | | | | | | |
|---|---|---|---|---|---|---|
| -2.5893752 | 0.00961503 | 0.01782603 | 4.91812656 | 15.966036 | 35.6640739 | 26.8951655 |
| -2.5892627 | 0.00961817 | 0.01783084 | 0 | 176.708839 | 120.79767 | 49.8681194 |
| -2.5888281 | 0.00963032 | 0.01785233 | 6.96734596 | 72.7942996 | 5.75226998 | 31.9380091 |
| -2.5888044 | 0.00963098 | 0.01785253 | 257.7918 | 572.883019 | 593.634262 | 561.996896 |
| -2.5885943 | 0.00963686 | 0.0178624 | 8.1968776 | 23.8137486 | 37.9649818 | 9.52537113 |
| -2.58847 | 0.00964033 | 0.01786782 | 13.9346919 | 10.5538204 | 29.9118039 | 11.7666349 |
| -2.588341 | 0.00964394 | 0.01787349 | 153.281611 | 286.306204 | 304.870309 | 292.484925 |
| 2.58769365 | 0.00966209 | 0.01790608 | 41.8040758 | 46.545054 | 37.9649818 | 39.7824324 |
| -2.5876167 | 0.00966425 | 0.01790906 | 11.0657848 | 11.0950419 | 16.1063559 | 19.6110582 |
| -2.587491 | 0.00966777 | 0.01791457 | 126.641759 | 205.393581 | 110.443584 | 169.215416 |
| -2.5874152 | 0.0096699 | 0.01791748 | 108.608628 | 111.491641 | 57.5226998 | 72.8410733 |
| -2.587344 | 0.0096719 | 0.01792015 | 37.2957931 | 11.0950419 | 10.354086 | 14.5682147 |
| -2.5872084 | 0.00967571 | 0.01792618 | 54.9190799 | 122.045461 | 0 | 80.6854966 |
| -2.5871427 | 0.00967755 | 0.01792857 | 17.213443 | 170.755402 | 205.931265 | 0 |
| -2.5870748 | 0.00967946 | 0.01793107 | 7.78703372 | 18.401533 | 18.4072639 | 16.8094785 |
| -2.5867918 | 0.00968741 | 0.01794477 | 59.8372065 | 12.1774851 | 11.50454 | 10.0856871 |
| -2.586569 | 0.00969368 | 0.01795535 | 5.73781432 | 15.966036 | 26.4604419 | 26.3348496 |
| -2.5861621 | 0.00970513 | 0.01797553 | 11.4756286 | 212.700072 | 0 | 140.078987 |
| -2.5861074 | 0.00970667 | 0.01797735 | 66.8045524 | 114.46836 | 121.948124 | 0 |
| 2.58597334 | 0.00971044 | 0.01798331 | 7.37718984 | 14.8835929 | 0 | 0 |
| -2.5858536 | 0.00971382 | 0.01798853 | 12.2953164 | 24.3549701 | 32.2127119 | 6.72379138 |
| -2.5857617 | 0.00971641 | 0.01799229 | 31.9678226 | 12.7187066 | 10.354086 | 12.8872668 |
| -2.5853465 | 0.00972812 | 0.01801295 | 4.50828268 | 23.5431378 | 14.9559019 | 54.350647 |
| -2.5849752 | 0.0097386 | 0.01803132 | 52.4600166 | 123.669126 | 136.904025 | 0 |
| -2.5849275 | 0.00973995 | 0.01803278 | 102.051126 | 302.542851 | 0 | 257.745336 |
| -2.5847915 | 0.00974379 | 0.01803886 | 48.3615778 | 51.6866588 | 0 | 189.947107 |
| -2.5847464 | 0.00974506 | 0.01804018 | 0 | 121.233629 | 40.2658898 | 198.351846 |
| -2.583895 | 0.00976915 | 0.01808374 | 0 | 94.9843835 | 131.151755 | 93.5727634 |
| -2.5833841 | 0.00978363 | 0.01810951 | 29.0989155 | 4.05916169 | 62.1245158 | 12.3269509 |
| 2.58310358 | 0.00979159 | 0.0181232 | 0 | 0.81183234 | 0 | 2.24126379 |
| -2.5828454 | 0.00979892 | 0.01813573 | 78.2801811 | 7.8477126 | 11.50454 | 12.8872668 |
| -2.582727 | 0.00980229 | 0.01814092 | 58.197831 | 102.832096 | 0 | 89.6505518 |
| 2.58267599 | 0.00980373 | 0.01814256 | 0 | 0 | 0 | 1.1206319 |
| -2.5825558 | 0.00980715 | 0.01814784 | 11.8854725 | 20.5664192 | 9.20363196 | 18.4904263 |
| -2.5822601 | 0.00981556 | 0.01816235 | 4.91812656 | 17.8603114 | 23.0090799 | 35.2999048 |
| -2.5822117 | 0.00981694 | 0.01816282 | 20.492194 | 26.7904671 | 6.90272397 | 14.0078987 |
| 2.5822131 | 0.00981689 | 0.01816282 | 6.55750208 | 2.70610779 | 2.30090799 | 5.60315949 |
| -2.5819113 | 0.00982548 | 0.0181776 | 8.1968776 | 16.2366467 | 13.8054479 | 28.0157974 |
| -2.5813114 | 0.00984257 | 0.01820817 | 0 | 86.0542278 | 70.1776937 | 177.620156 |
| -2.5811335 | 0.00984765 | 0.01821652 | 6.96734596 | 16.7778683 | 20.7081719 | 20.1713742 |
| 2.58080775 | 0.00985695 | 0.01823267 | 64.755333 | 76.8534613 | 79.3813257 | 76.202969 |
| -2.5802531 | 0.00987279 | 0.01826093 | 43.0336074 | 4.87099402 | 16.1063559 | 22.9729539 |
| -2.5801289 | 0.00987634 | 0.01826645 | 13.1150042 | 16.2366467 | 10.354086 | 15.1285306 |

| | | | | | | |
|---|---|---|---|---|---|---|
| -2.5800294 | 0.00987919 | 0.01827067 | 16.3937552 | 22.4606947 | 8.05317797 | 14.5682147 |
| 2.57984949 | 0.00988434 | 0.01827915 | 2.0492194 | 2.97671857 | 1.150454 | 0 |
| -2.5797162 | 0.00988816 | 0.01828516 | 31.1481349 | 142.882491 | 166.815829 | 0 |
| -2.5795931 | 0.00989168 | 0.01829063 | 36.8859492 | 5.68282636 | 17.2568099 | 19.0507423 |
| -2.579393 | 0.00989741 | 0.01830018 | 0 | 100.66721 | 112.744492 | 101.417187 |
| -2.5792984 | 0.00990012 | 0.01830414 | 5.73781432 | 30.579018 | 14.9559019 | 24.6539017 |
| 2.57921067 | 0.00990264 | 0.01830774 | 0 | 0.27061078 | 0 | 0 |
| -2.5791477 | 0.00990444 | 0.01831003 | 10.246097 | 19.2133653 | 19.5577179 | 10.646003 |
| 2.57906831 | 0.00990672 | 0.01831319 | 2.86890716 | 8.93015571 | 1.150454 | 5.04284354 |
| -2.5790409 | 0.00990751 | 0.0183136 | 216.397569 | 103.643928 | 66.7263317 | 107.020346 |
| 2.57883743 | 0.00991334 | 0.01832334 | 2.45906328 | 1.89427545 | 4.60181598 | 0 |
| -2.5781398 | 0.00993338 | 0.01835932 | 62.7061136 | 13.8011497 | 48.3190678 | 20.7316901 |
| -2.5780908 | 0.00993479 | 0.01836088 | 0 | 49.2511618 | 144.957203 | 142.880567 |
| 2.57758297 | 0.0099494 | 0.01838683 | 153.691455 | 82.8068984 | 82.8326877 | 76.763285 |
| -2.5774422 | 0.00995345 | 0.01839326 | 5.73781432 | 23.5431378 | 39.1154358 | 13.4475828 |
| -2.5772247 | 0.00995972 | 0.01840379 | 94.2640924 | 127.457677 | 74.7795097 | 0 |
| -2.5768819 | 0.0099696 | 0.018421 | 5.32797044 | 28.6847426 | 17.2568099 | 24.0935858 |
| 2.57675192 | 0.00997335 | 0.01842687 | 11.0657848 | 5.1416048 | 13.8054479 | 5.60315949 |
| -2.5766295 | 0.00997688 | 0.01843234 | 0 | 100.396599 | 92.0363196 | 122.148877 |
| -2.576314 | 0.00998599 | 0.01844811 | 9.83625312 | 22.4606947 | 12.6549939 | 14.5682147 |
| -2.5761761 | 0.00998998 | 0.01845442 | 10.6559409 | 41.9446708 | 4.60181598 | 39.2221164 |
| -2.5761086 | 0.00999193 | 0.01845697 | 97.1329995 | 52.2278804 | 0 | 101.417187 |
| -2.576015 | 0.00999463 | 0.01846091 | 4.0984388 | 23.5431378 | 33.3631659 | 22.4126379 |
| -2.575758 | 0.01000206 | 0.0184731 | 0 | 98.2317128 | 130.001301 | 87.409288 |
| 2.57574709 | 0.01000238 | 0.0184731 | 114.756286 | 105.808815 | 59.8236078 | 85.7283401 |
| -2.5751101 | 0.01002082 | 0.0185061 | 0 | 112.032863 | 102.390406 | 98.055291 |
| -2.5749755 | 0.01002472 | 0.01851224 | 10.246097 | 21.1076408 | 11.50454 | 15.6888466 |
| -2.5749312 | 0.010026 | 0.01851355 | 23.770945 | 160.472192 | 0 | 112.623506 |
| -2.5748171 | 0.01002931 | 0.0185186 | 5.32797044 | 15.6954252 | 27.6108959 | 28.0157974 |
| -2.5746834 | 0.01003319 | 0.01852471 | 19.2626624 | 11.9068743 | 10.354086 | 15.1285306 |
| 2.57430417 | 0.01004419 | 0.01854184 | 0 | 0.27061078 | 0 | 0 |
| 2.57430417 | 0.01004419 | 0.01854184 | 0 | 0.27061078 | 0 | 0 |
| 2.57430417 | 0.01004419 | 0.01854184 | 0 | 0.27061078 | 0 | 0 |
| -2.5742154 | 0.01004677 | 0.01854554 | 6.55750208 | 48.4393295 | 9.20363196 | 26.8951655 |
| -2.5740879 | 0.01005047 | 0.01855131 | 8.1968776 | 41.67406 | 8.05317797 | 26.3348496 |
| 2.57402297 | 0.01005236 | 0.01855374 | 0 | 0 | 0 | 0 |
| -2.5739792 | 0.01005363 | 0.01855502 | 23.3611012 | 191.05121 | 0 | 91.8918156 |
| -2.5735391 | 0.01006643 | 0.01857758 | 63.1159575 | 0 | 142.656295 | 150.164674 |
| -2.5732792 | 0.01007399 | 0.01859047 | 126.231915 | 181.309222 | 118.496762 | 158.009098 |
| -2.5731136 | 0.01007881 | 0.01859831 | 105.329877 | 0 | 194.426725 | 63.3157022 |
| 2.5726806 | 0.01009143 | 0.01862053 | 6.96734596 | 5.41221558 | 8.05317797 | 0 |
| -2.5726127 | 0.01009341 | 0.01862312 | 5.73781432 | 12.1774851 | 32.2127119 | 30.8173772 |
| -2.5725887 | 0.01009411 | 0.01862334 | 6.1476582 | 21.6488623 | 36.8145279 | 13.4475828 |

| | | | | | | |
|---|---|---|---|---|---|---|
| -2.5721339 | 0.01010738 | 0.01864676 | 4.0984388 | 35.1794013 | 32.2127119 | 16.2491625 |
| -2.5720936 | 0.01010855 | 0.01864786 | 4.0984388 | 25.4374132 | 40.2658898 | 17.9301104 |
| -2.5719965 | 0.01011139 | 0.01865203 | 2.86890716 | 30.3084073 | 27.6108959 | 33.058641 |
| 2.57190036 | 0.0101142 | 0.01865508 | 244.676796 | 147.212264 | 199.028541 | 133.915512 |
| -2.5719035 | 0.01011411 | 0.01865508 | 63.5258014 | 95.7962158 | 0 | 84.0473923 |
| -2.571712 | 0.0101197 | 0.01866416 | 13.1150042 | 20.0251977 | 13.8054479 | 10.0856871 |
| -2.5715865 | 0.01012337 | 0.01866987 | 86.8869025 | 78.2065152 | 0 | 75.6426531 |
| -2.5715446 | 0.0101246 | 0.01867106 | 6.55750208 | 26.2492456 | 23.0090799 | 14.0078987 |
| -2.571287 | 0.01013213 | 0.01868388 | 0 | 99.5847667 | 139.204933 | 77.3236009 |
| 2.571047 | 0.01013916 | 0.01869577 | 0.40984388 | 2.97671857 | 0 | 1.1206319 |
| -2.5706505 | 0.01015077 | 0.01871505 | 12.2953164 | 12.4480958 | 18.4072639 | 11.7666349 |
| -2.5706643 | 0.01015037 | 0.01871505 | 11.8854725 | 11.6362635 | 13.8054479 | 16.8094785 |
| -2.5706009 | 0.01015223 | 0.01871666 | 5.73781432 | 20.83703 | 19.5577179 | 23.5332698 |
| 2.57038085 | 0.01015868 | 0.01872749 | 1.63937552 | 1.08244312 | 0 | 0.56031595 |
| -2.5698093 | 0.01017545 | 0.01875734 | 6.96734596 | 22.1900839 | 21.8586259 | 14.5682147 |
| 2.5695025 | 0.01018447 | 0.01877288 | 0 | 0.81183234 | 2.30090799 | 0.56031595 |
| 2.5692644 | 0.01019147 | 0.01878472 | 2.86890716 | 6.22404792 | 4.60181598 | 6.16347543 |
| -2.5688086 | 0.01020488 | 0.01880837 | 11.4756286 | 19.2133653 | 41.4163438 | 7.28410733 |
| -2.5685331 | 0.01021299 | 0.01882225 | 15.1642236 | 19.4839761 | 62.1245158 | 4.48252759 |
| -2.5683988 | 0.01021696 | 0.01882847 | 83.6081515 | 69.5469702 | 0 | 86.848972 |
| -2.5682586 | 0.01022109 | 0.01883501 | 57.3781432 | 140.717605 | 156.461743 | 138.958355 |
| -2.5681911 | 0.01022308 | 0.0188376 | 7.37718984 | 16.7778683 | 13.8054479 | 33.6189569 |
| -2.5681147 | 0.01022533 | 0.01884005 | 5.73781432 | 27.8729103 | 10.354086 | 45.9459078 |
| -2.5681066 | 0.01022557 | 0.01884005 | 288.120248 | 174.002731 | 131.151755 | 147.92341 |
| -2.5680169 | 0.01022822 | 0.01884385 | 65.5750208 | 108.244312 | 119.647216 | 0 |
| -2.5678398 | 0.01023344 | 0.0188524 | 51.230485 | 72.5236888 | 0 | 129.9933 |
| -2.5671495 | 0.01025384 | 0.0188889 | 9.42640924 | 15.966036 | 11.50454 | 25.7745336 |
| 2.56692807 | 0.01026039 | 0.01889989 | 4.50828268 | 1.89427545 | 0 | 2.24126379 |
| -2.5666791 | 0.01026776 | 0.01891238 | 40.5745441 | 74.688575 | 78.2308717 | 84.0473923 |
| 2.56624609 | 0.01028059 | 0.01893493 | 57.7879871 | 35.9912336 | 35.6640739 | 36.9808526 |
| -2.5661108 | 0.0102846 | 0.01894123 | 6.55750208 | 35.4500121 | 12.6549939 | 21.292006 |
| -2.565914 | 0.01029043 | 0.0189509 | 20.492194 | 49.2511618 | 19.5577179 | 3.92221164 |
| -2.565699 | 0.01029681 | 0.01896157 | 39.7548564 | 114.73897 | 163.364467 | 0 |
| -2.5656691 | 0.0102977 | 0.01896213 | 97.1329995 | 0 | 194.426725 | 66.1172819 |
| -2.5655825 | 0.01030027 | 0.01896578 | 12.2953164 | 9.47137727 | 16.1063559 | 20.1713742 |
| -2.5653125 | 0.01030829 | 0.01897946 | 6.96734596 | 18.1309222 | 17.2568099 | 20.7316901 |
| -2.5651419 | 0.01031336 | 0.01898771 | 144.265046 | 236.784432 | 239.294431 | 254.383441 |
| -2.564985 | 0.01031803 | 0.01899522 | 2.0492194 | 56.8282636 | 16.1063559 | 44.2649599 |
| -2.5646493 | 0.01032801 | 0.01901252 | 2.45906328 | 33.0145151 | 35.6640739 | 27.4554815 |
| -2.5645486 | 0.01033101 | 0.01901695 | 0 | 61.1580361 | 140.355387 | 117.106033 |
| -2.5642978 | 0.01033848 | 0.01902961 | 10.6559409 | 15.6954252 | 13.8054479 | 13.4475828 |
| -2.5641009 | 0.01034435 | 0.01903933 | 38.1154808 | 13.2599282 | 11.50454 | 10.0856871 |
| -2.5640177 | 0.01034683 | 0.01904281 | 3.27875104 | 9.47137727 | 49.4695218 | 54.350647 |

| | | | | | | |
|---|---|---|---|---|---|---|
| -2.5639396 | 0.01034915 | 0.01904601 | 8.1968776 | 20.83703 | 10.354086 | 31.3776931 |
| -2.5637705 | 0.0103542 | 0.01905421 | 2.0492194 | 31.6614612 | 44.8677058 | 28.0157974 |
| -2.5635185 | 0.01036172 | 0.01906587 | 13.1150042 | 20.5664192 | 8.05317797 | 18.4904263 |
| -2.5635292 | 0.0103614 | 0.01906587 | 9.83625312 | 17.3190899 | 19.5577179 | 11.206319 |
| -2.5634822 | 0.0103628 | 0.01906677 | 68.4439279 | 12.7187066 | 6.90272397 | 13.4475828 |
| 2.56281948 | 0.0103826 | 0.01910212 | 0 | 0 | 0 | 0.56031595 |
| -2.5627458 | 0.01038481 | 0.01910509 | 136.887856 | 242.737869 | 181.771731 | 240.375542 |
| -2.5624401 | 0.01039395 | 0.01912082 | 281.152902 | 463.826875 | 375.048002 | 425.279805 |
| -2.5619885 | 0.01040748 | 0.01914461 | 27.45954 | 169.131737 | 0 | 88.5299199 |
| -2.5614939 | 0.01042231 | 0.0191708 | 7.78703372 | 17.0484791 | 14.9559019 | 20.7316901 |
| 2.56135081 | 0.0104266 | 0.0191776 | 4.0984388 | 1.62366467 | 2.30090799 | 2.24126379 |
| -2.5608705 | 0.01044103 | 0.01920305 | 38.5253247 | 112.032863 | 0 | 105.899714 |
| 2.56058998 | 0.01044946 | 0.01921746 | 13.524848 | 9.74198805 | 6.90272397 | 8.40473923 |
| -2.5602752 | 0.01045893 | 0.01923378 | 70.0833035 | 77.1240721 | 0 | 89.6505518 |
| -2.5601064 | 0.01046401 | 0.01924203 | 3.27875104 | 23.5431378 | 26.4604419 | 35.8602207 |
| -2.5595105 | 0.01048197 | 0.01927395 | 42.6237635 | 109.597366 | 157.612197 | 0 |
| -2.5593222 | 0.01048765 | 0.0192833 | 61.0667381 | 84.4305631 | 0 | 93.0124475 |
| -2.5591949 | 0.01049149 | 0.01928927 | 48.7714217 | 113.385916 | 0 | 84.6077082 |
| -2.5591336 | 0.01049334 | 0.01929157 | 16.3937552 | 8.93015571 | 11.50454 | 28.0157974 |
| -2.5590973 | 0.01049444 | 0.01929248 | 0 | 113.385916 | 181.771731 | 42.0236961 |
| -2.5590302 | 0.01049646 | 0.01929511 | 12.2953164 | 20.5664192 | 10.354086 | 13.4475828 |
| -2.5587927 | 0.01050364 | 0.0193072 | 3.68859492 | 21.9194731 | 25.3099879 | 34.1792729 |
| -2.5587423 | 0.01050516 | 0.0193089 | 58.6076748 | 95.525605 | 0 | 85.1680242 |
| -2.5586185 | 0.0105089 | 0.01931467 | 7.37718984 | 15.6954252 | 16.1063559 | 24.0935858 |
| -2.5583758 | 0.01051624 | 0.01932706 | 667.225837 | 1855.57811 | 1065.3204 | 1327.9488 |
| -2.5583143 | 0.0105181 | 0.01932938 | 16.3937552 | 17.3190899 | 9.20363196 | 12.8872668 |
| -2.5580816 | 0.01052514 | 0.01934122 | 35.6564176 | 15.6954252 | 65.5758777 | 20.1713742 |
| 2.55779163 | 0.01053392 | 0.01935515 | 0 | 0 | 0 | 0 |
| 2.55779163 | 0.01053392 | 0.01935515 | 0 | 0 | 0 | 0 |
| -2.5576939 | 0.01053688 | 0.01935949 | 3.68859492 | 31.6614612 | 55.2217918 | 12.3269509 |
| 2.55764719 | 0.01053829 | 0.01936098 | 0 | 0 | 0 | 0 |
| -2.5575289 | 0.01054188 | 0.01936647 | 11.0657848 | 26.7904671 | 5.75226998 | 38.6618005 |
| 2.55705354 | 0.0105563 | 0.01939183 | 0.81968776 | 0.54122156 | 0 | 0 |
| -2.5570341 | 0.01055689 | 0.01939183 | 76.2309617 | 87.9485032 | 0 | 71.1601255 |
| -2.5568214 | 0.01056334 | 0.01940258 | 19.2626624 | 28.6847426 | 12.6549939 | 7.28410733 |
| -2.5566494 | 0.01056857 | 0.01941108 | 37.705637 | 141.529437 | 0 | 81.8061285 |
| -2.55634 | 0.01057797 | 0.01942724 | 20.0823501 | 15.1542036 | 11.50454 | 10.0856871 |
| 2.55606578 | 0.01058631 | 0.01944145 | 1.22953164 | 0.54122156 | 0 | 0.56031595 |
| -2.5559781 | 0.01058898 | 0.01944525 | 12.7051603 | 154.248144 | 227.789891 | 0 |
| -2.5558896 | 0.01059167 | 0.01944798 | 3.68859492 | 62.2404792 | 11.50454 | 30.2570612 |
| -2.5558913 | 0.01059162 | 0.01944798 | 24.1807889 | 12.7187066 | 10.354086 | 12.3269509 |
| -2.5550036 | 0.01061867 | 0.01949645 | 10.6559409 | 14.8835929 | 21.8586259 | 10.646003 |
| -2.5547701 | 0.01062579 | 0.01950842 | 28.2792277 | 18.401533 | 14.9559019 | 19.0507423 |

| | | | | | | |
|---|---|---|---|---|---|---|
| 2.55472091 | 0.0106273 | 0.01951007 | 18.4429746 | 20.83703 | 18.4072639 | 19.6110582 |
| -2.5544181 | 0.01063654 | 0.01952593 | 64.755333 | 96.3374374 | 0 | 75.0823371 |
| -2.554228 | 0.01064235 | 0.01953549 | 553.289238 | 399.692121 | 429.11934 | 457.77813 |
| -2.5541299 | 0.01064535 | 0.01953988 | 207.381003 | 220.547785 | 200.178995 | 210.678797 |
| -2.5538528 | 0.01065383 | 0.01955433 | 82.3786199 | 88.219114 | 0 | 64.4363341 |
| -2.5535876 | 0.01066194 | 0.01956811 | 4.91812656 | 32.7439043 | 16.1063559 | 24.0935858 |
| -2.5535028 | 0.01066454 | 0.01957177 | 6.1476582 | 15.4248144 | 33.3631659 | 17.9301104 |
| 2.55313569 | 0.01067579 | 0.01959129 | 7.78703372 | 9.47137727 | 14.9559019 | 7.84442328 |
| -2.552946 | 0.01068161 | 0.01960085 | 8.60672148 | 26.2492456 | 10.354086 | 20.1713742 |
| -2.5528899 | 0.01068332 | 0.01960178 | 29.0989155 | 150.188982 | 0 | 93.0124475 |
| -2.5529003 | 0.01068301 | 0.01960178 | 14.3445358 | 31.3908504 | 25.3099879 | 43.144328 |
| -2.552663 | 0.01069029 | 0.01961344 | 167.626147 | 422.152815 | 556.819734 | 517.171621 |
| -2.5525779 | 0.0106929 | 0.01961712 | 61.476582 | 98.7729344 | 0 | 76.202969 |
| -2.5523943 | 0.01069854 | 0.01962634 | 11.4756286 | 12.4480958 | 14.9559019 | 13.4475828 |
| -2.5522088 | 0.01070423 | 0.01963457 | 23.3611012 | 8.38893415 | 13.8054479 | 15.1285306 |
| 2.55222383 | 0.01070377 | 0.01963457 | 4.91812656 | 2.43549701 | 0 | 2.24126379 |
| -2.5521413 | 0.01070631 | 0.01963725 | 0 | 93.3607188 | 155.311289 | 64.4363341 |
| 2.55182773 | 0.01071595 | 0.01965382 | 4.50828268 | 2.97671857 | 4.60181598 | 3.36189569 |
| -2.5516859 | 0.01072031 | 0.0196607 | 68.0340841 | 133.681725 | 85.1335957 | 0 |
| -2.5513866 | 0.01072952 | 0.01967648 | 6.1476582 | 30.3084073 | 25.3099879 | 12.3269509 |
| 2.55128985 | 0.0107325 | 0.01968083 | 0.40984388 | 0 | 0 | 0 |
| -2.5511333 | 0.01073732 | 0.01968631 | 110.657848 | 46.2744432 | 0 | 87.409288 |
| -2.5511459 | 0.01073694 | 0.01968631 | 4.0984388 | 40.0503953 | 10.354086 | 44.8252759 |
| -2.5511434 | 0.01073701 | 0.01968631 | 23.770945 | 20.83703 | 26.4604419 | 4.48252759 |
| 2.551077 | 0.01073906 | 0.01968838 | 27.8693838 | 23.272527 | 23.0090799 | 22.4126379 |
| -2.5505822 | 0.01075432 | 0.01971523 | 56.9682993 | 96.0668266 | 0 | 83.4870763 |
| -2.550505 | 0.0107567 | 0.01971761 | 43.8532952 | 180.226779 | 86.2840497 | 0 |
| -2.5505005 | 0.01075684 | 0.01971761 | 10.246097 | 44.9213893 | 6.90272397 | 20.7316901 |
| -2.5504641 | 0.01075796 | 0.01971855 | 25.8201644 | 18.1309222 | 6.90272397 | 14.0078987 |
| -2.5497038 | 0.01078145 | 0.01976048 | 0 | 184.827162 | 141.505841 | 263.348496 |
| 2.54953983 | 0.01078652 | 0.0197678 | 1.63937552 | 0.81183234 | 0 | 0 |
| 2.54953509 | 0.01078667 | 0.0197678 | 0 | 0 | 0 | 0 |
| -2.5492024 | 0.01079696 | 0.01978555 | 4.91812656 | 31.6614612 | 23.0090799 | 16.8094785 |
| -2.5491567 | 0.01079838 | 0.01978701 | 7.78703372 | 36.803066 | 8.05317797 | 27.4554815 |
| -2.549114 | 0.0107997 | 0.01978831 | 54.509236 | 102.561485 | 0 | 80.6854966 |
| 2.5489827 | 0.01080377 | 0.01979464 | 0 | 8.38893415 | 0 | 0 |
| -2.5489041 | 0.0108062 | 0.01979798 | 0 | 112.844695 | 95.4876816 | 90.7711837 |
| -2.5488665 | 0.01080737 | 0.01979899 | 211.889286 | 215.40618 | 155.311289 | 187.705843 |
| 2.54883067 | 0.01080848 | 0.0197999 | 31.5579788 | 26.2492456 | 37.9649818 | 24.0935858 |
| -2.5487204 | 0.01081189 | 0.01980504 | 12.2953164 | 18.6721438 | 10.354086 | 13.4475828 |
| -2.5479818 | 0.01083481 | 0.01984589 | 15.1642236 | 34.9087905 | 23.0090799 | 5.04284354 |
| -2.5479187 | 0.01083677 | 0.01984836 | 16.3937552 | 29.2259641 | 35.6640739 | 3.92221164 |
| 2.54774856 | 0.01084206 | 0.01985691 | 2.45906328 | 0 | 3.45136199 | 0 |

| | | | | | | |
|---|---|---|---|---|---|---|
| 2.54757268 | 0.01084753 | 0.0198658 | 0.40984388 | 0.54122156 | 0 | 1.1206319 |
| -2.5475234 | 0.01084906 | 0.01986748 | 6.1476582 | 33.5557366 | 36.8145279 | 8.96505518 |
| 2.54747684 | 0.01085051 | 0.019869 | 9.01656536 | 5.68282636 | 5.75226998 | 4.48252759 |
| -2.5470463 | 0.0108639 | 0.0198924 | 9.83625312 | 44.109557 | 20.7081719 | 7.28410733 |
| -2.5468646 | 0.01086956 | 0.01990164 | 16.3937552 | 37.3442875 | 21.8586259 | 29.6967453 |
| 2.54674191 | 0.01087338 | 0.01990751 | 23.770945 | 9.74198805 | 0 | 7.84442328 |
| -2.5466808 | 0.01087529 | 0.01990986 | 4.91812656 | 21.6488623 | 19.5577179 | 26.8951655 |
| 2.54665778 | 0.010876 | 0.01991005 | 10.246097 | 5.68282636 | 5.75226998 | 3.92221164 |
| -2.5463473 | 0.01088568 | 0.01992664 | 15.1642236 | 11.0950419 | 11.50454 | 15.1285306 |
| -2.5461933 | 0.01089049 | 0.01993369 | 74.5915861 | 33.0145151 | 0 | 160.250361 |
| -2.5461842 | 0.01089077 | 0.01993369 | 127.461447 | 55.2045989 | 0 | 61.6347543 |
| -2.5457739 | 0.01090358 | 0.019956 | 0 | 62.2404792 | 116.195854 | 127.19172 |
| -2.5453669 | 0.0109163 | 0.01997815 | 5.73781432 | 28.4141318 | 24.1595339 | 14.0078987 |
| -2.5452747 | 0.01091918 | 0.01998229 | 9.83625312 | 17.3190899 | 8.05317797 | 47.0665397 |
| -2.5452069 | 0.0109213 | 0.01998504 | 5.73781432 | 15.966036 | 19.5577179 | 30.2570612 |
| -2.5449108 | 0.01093056 | 0.02000086 | 11.0657848 | 21.9194731 | 14.9559019 | 10.0856871 |
| 2.54474479 | 0.01093576 | 0.02000924 | 5.73781432 | 1.89427545 | 2.30090799 | 3.36189569 |
| -2.5443335 | 0.01094865 | 0.02003168 | 0 | 74.688575 | 144.957203 | 84.6077082 |
| -2.5440701 | 0.01095691 | 0.02004565 | 22.5414134 | 79.5595691 | 263.453965 | 0 |
| -2.5439199 | 0.01096162 | 0.020052 | 57.7879871 | 55.2045989 | 200.178995 | 0 |
| -2.5439388 | 0.01096103 | 0.020052 | 10.6559409 | 12.1774851 | 16.1063559 | 14.0078987 |
| -2.5438865 | 0.01096267 | 0.02005278 | 6.55750208 | 12.4480958 | 25.3099879 | 24.6539017 |
| -2.5437246 | 0.01096775 | 0.02006095 | 15.1642236 | 19.4839761 | 19.5577179 | 21.852322 |
| -2.5435722 | 0.01097254 | 0.02006856 | 60.6568942 | 154.518755 | 116.195854 | 141.759935 |
| 2.54354833 | 0.01097329 | 0.0200688 | 30.738291 | 24.8961917 | 26.4604419 | 32.498325 |
| -2.5432789 | 0.01098175 | 0.02008314 | 10.246097 | 14.3423713 | 49.4695218 | 8.96505518 |
| -2.5430776 | 0.01098808 | 0.02009358 | 14.3445358 | 35.7206228 | 5.75226998 | 19.0507423 |
| -2.5422563 | 0.01101394 | 0.02013858 | 23.3611012 | 21.3782516 | 40.2658898 | 3.36189569 |
| -2.5422753 | 0.01101334 | 0.02013858 | 13.1150042 | 171.567234 | 0 | 133.355196 |
| 2.54212426 | 0.0110181 | 0.02014505 | 0.81968776 | 0.27061078 | 1.150454 | 0.56031595 |
| 2.54208791 | 0.01101925 | 0.020146 | 70.9029912 | 34.367569 | 44.8677058 | 35.8602207 |
| -2.5419537 | 0.01102348 | 0.0201526 | 9.42640924 | 24.8961917 | 8.05317797 | 26.3348496 |
| -2.5418525 | 0.01102667 | 0.02015729 | 14.3445358 | 20.5664192 | 5.75226998 | 30.2570612 |
| -2.5418034 | 0.01102822 | 0.02015898 | 9.42640924 | 17.3190899 | 11.50454 | 17.9301104 |
| -2.5415358 | 0.01103666 | 0.02017328 | 6.1476582 | 21.6488623 | 18.4072639 | 18.4904263 |
| -2.5413566 | 0.01104232 | 0.02018248 | 31.5579788 | 86.5954493 | 0 | 143.440883 |
| -2.5412126 | 0.01104687 | 0.02018965 | 103.690502 | 50.0629941 | 0 | 82.9267604 |
| 2.54118216 | 0.01104783 | 0.02019027 | 0 | 0 | 0 | 0 |
| -2.5409493 | 0.0110552 | 0.02020258 | 36.0662614 | 137.199665 | 113.894946 | 123.829825 |
| -2.540924 | 0.011056 | 0.02020289 | 81.968776 | 89.0309463 | 0 | 59.9538065 |
| 2.54043819 | 0.01107137 | 0.0202297 | 4.91812656 | 7.30649104 | 5.75226998 | 10.0856871 |
| -2.5404009 | 0.01107255 | 0.0202297 | 13.1150042 | 11.9068743 | 10.354086 | 20.1713742 |
| -2.5404191 | 0.01107197 | 0.0202297 | 61.476582 | 246.52642 | 194.426725 | 223.566063 |

| | | | | | | |
|---|---|---|---|---|---|---|
| 2.54023233 | 0.01107789 | 0.02023831 | 37.705637 | 24.8961917 | 13.8054479 | 24.6539017 |
| -2.5401633 | 0.01108007 | 0.02024116 | 8.60672148 | 12.1774851 | 17.2568099 | 20.1713742 |
| -2.5398264 | 0.01109075 | 0.02025884 | 17.213443 | 21.6488623 | 28.7613499 | 5.04284354 |
| -2.5398184 | 0.011091 | 0.02025884 | 4.91812656 | 16.7778683 | 13.8054479 | 64.4363341 |
| 2.53970715 | 0.01109453 | 0.02026414 | 4.91812656 | 0.81183234 | 0 | 1.68094785 |
| -2.5396667 | 0.01109582 | 0.02026534 | 0 | 133.952336 | 73.6290557 | 91.8918156 |
| -2.5395956 | 0.01109807 | 0.02026831 | 185.659278 | 151.000815 | 164.514921 | 191.628054 |
| 2.53947763 | 0.01110181 | 0.020274 | 29.0989155 | 23.272527 | 19.5577179 | 19.6110582 |
| -2.5394483 | 0.01110275 | 0.02027455 | 6.1476582 | 12.4480958 | 27.6108959 | 25.2142177 |
| -2.5389751 | 0.01111777 | 0.02030085 | 15.1642236 | 8.11832337 | 12.6549939 | 30.2570612 |
| -2.5389502 | 0.01111857 | 0.02030114 | 0 | 97.4198805 | 73.6290557 | 126.631404 |
| -2.5386912 | 0.0111268 | 0.02031502 | 8.1968776 | 20.5664192 | 13.8054479 | 15.6888466 |
| -2.5385878 | 0.01113009 | 0.02031988 | 17.6232868 | 12.1774851 | 10.354086 | 13.4475828 |
| -2.538171 | 0.01114335 | 0.02034295 | 7.78703372 | 27.3316887 | 29.9118039 | 38.1014845 |
| -2.5379961 | 0.01114892 | 0.02035197 | 32.3776665 | 33.5557366 | 23.0090799 | 24.0935858 |
| 2.53788911 | 0.01115233 | 0.02035704 | 22.1315695 | 11.9068743 | 25.3099879 | 13.4475828 |
| 2.53766479 | 0.01115948 | 0.02036894 | 0.40984388 | 0 | 0 | 0 |
| -2.5373604 | 0.01116919 | 0.02038551 | 0 | 95.7962158 | 224.338529 | 30.8173772 |
| -2.5372571 | 0.01117249 | 0.02039038 | 50.4107972 | 90.3840002 | 0 | 93.0124475 |
| -2.5365439 | 0.01119527 | 0.0204308 | 14.3445358 | 13.8011497 | 6.90272397 | 43.144328 |
| 2.53635052 | 0.01120146 | 0.02044093 | 0 | 1.62366467 | 0 | 2.24126379 |
| -2.5362859 | 0.01120352 | 0.02044355 | 11.4756286 | 15.6954252 | 11.50454 | 13.4475828 |
| -2.5362654 | 0.01120418 | 0.02044359 | 12.7051603 | 72.5236888 | 0 | 261.107232 |
| -2.5361977 | 0.01120634 | 0.02044639 | 27.45954 | 12.7187066 | 8.05317797 | 15.1285306 |
| -2.5361243 | 0.01120869 | 0.02044863 | 50.8206411 | 95.525605 | 0 | 86.848972 |
| -2.5361197 | 0.01120884 | 0.02044863 | 25.0004767 | 11.9068743 | 10.354086 | 12.3269509 |
| -2.5360595 | 0.01121077 | 0.02045099 | 6.96734596 | 28.9553534 | 14.9559019 | 15.6888466 |
| -2.5358359 | 0.01121793 | 0.0204629 | 13.9346919 | 11.0950419 | 10.354086 | 20.7316901 |
| -2.5356162 | 0.01122497 | 0.02047458 | 84.0179954 | 9.74198805 | 6.90272397 | 12.8872668 |
| -2.5355717 | 0.0112264 | 0.02047603 | 23.770945 | 11.0950419 | 10.354086 | 13.4475828 |
| -2.5354689 | 0.01122969 | 0.02048088 | 47.1320462 | 69.5469702 | 188.674455 | 0 |
| -2.5354376 | 0.01123069 | 0.02048155 | 40.1647002 | 111.491641 | 0 | 90.7711837 |
| -2.5354121 | 0.01123151 | 0.02048189 | 4.91812656 | 30.3084073 | 18.4072639 | 20.1713742 |
| -2.5351922 | 0.01123857 | 0.02049359 | 20.492194 | 3.51794013 | 25.3099879 | 35.8602207 |
| -2.5351366 | 0.01124035 | 0.02049569 | 6.96734596 | 16.5072575 | 16.1063559 | 21.852322 |
| 2.53488535 | 0.01124841 | 0.02050808 | 81.1490882 | 42.4858923 | 43.7172518 | 36.9808526 |
| -2.5348958 | 0.01124808 | 0.02050808 | 26.2300083 | 11.0950419 | 11.50454 | 11.7666349 |
| -2.5347874 | 0.01125156 | 0.02051265 | 28.6890716 | 116.633246 | 0 | 113.183822 |
| 2.53421162 | 0.01127006 | 0.02054523 | 24.1807889 | 10.5538204 | 21.8586259 | 16.8094785 |
| -2.5334148 | 0.01129572 | 0.02059084 | 6.96734596 | 20.83703 | 13.8054479 | 20.7316901 |
| -2.5331 | 0.01130587 | 0.02060818 | 61.8864259 | 70.3588026 | 0 | 96.3743432 |
| -2.532963 | 0.01131029 | 0.02061507 | 2.0492194 | 24.8961917 | 29.9118039 | 46.5062237 |
| 2.53283981 | 0.01131426 | 0.02061587 | 0 | 0 | 1.150454 | 0.56031595 |

| | | | | | | |
|---|---|---|---|---|---|---|
| -2.532893 | 0.01131255 | 0.02061587 | 12.7051603 | 35.1794013 | 16.1063559 | 7.28410733 |
| 2.53283981 | 0.01131426 | 0.02061587 | 0 | 0 | 1.150454 | 0.56031595 |
| 2.53288028 | 0.01131296 | 0.02061587 | 0 | 0 | 0 | 0 |
| 2.53288028 | 0.01131296 | 0.02061587 | 0 | 0 | 0 | 0 |
| -2.5328306 | 0.01131456 | 0.02061587 | 94.2640924 | 86.0542278 | 89.7354116 | 0 |
| -2.5326675 | 0.01131983 | 0.0206243 | 3.27875104 | 32.7439043 | 24.1595339 | 24.6539017 |
| -2.53251 | 0.01132491 | 0.02063241 | 2.45906328 | 48.4393295 | 41.4163438 | 66.6775979 |
| -2.5324564 | 0.01132664 | 0.02063439 | 4.91812656 | 24.6255809 | 24.1595339 | 17.9301104 |
| 2.53242627 | 0.01132762 | 0.02063501 | 27.45954 | 23.8137486 | 14.9559019 | 12.8872668 |
| -2.5321045 | 0.01133802 | 0.02065279 | 22.5414134 | 12.9893174 | 6.90272397 | 20.1713742 |
| -2.5319666 | 0.01134248 | 0.02065975 | 90.9853413 | 66.2996409 | 0 | 69.4791776 |
| 2.53179016 | 0.01134819 | 0.02066898 | 1.22953164 | 0 | 0 | 0 |
| -2.5315078 | 0.01135733 | 0.02068446 | 0 | 147.753485 | 100.089498 | 56.0315949 |
| -2.5314504 | 0.01135919 | 0.02068668 | 0 | 67.382084 | 102.390406 | 125.510772 |
| -2.5311757 | 0.01136809 | 0.02070172 | 17.213443 | 20.83703 | 6.90272397 | 15.1285306 |
| 2.53110878 | 0.01137026 | 0.0207045 | 13.524848 | 4.87099402 | 10.354086 | 7.28410733 |
| -2.5308454 | 0.0113788 | 0.02071772 | 27.0496961 | 12.7187066 | 10.354086 | 11.206319 |
| -2.530864 | 0.01137819 | 0.02071772 | 43.4434513 | 102.290875 | 0 | 90.7711837 |
| -2.5303127 | 0.01139609 | 0.02074803 | 0 | 108.244312 | 101.239952 | 80.6854966 |
| -2.5301084 | 0.01140273 | 0.02075894 | 48.3615778 | 86.8660601 | 0 | 96.9346591 |
| 2.52980938 | 0.01141245 | 0.02077547 | 0.40984388 | 0.54122156 | 0 | 0 |
| -2.5297779 | 0.01141347 | 0.02077616 | 5.32797044 | 17.3190899 | 23.0090799 | 22.9729539 |
| -2.5296741 | 0.01141685 | 0.02078114 | 17.213443 | 28.6847426 | 12.6549939 | 7.28410733 |
| -2.5293804 | 0.01142641 | 0.02079737 | 96.7231557 | 96.8786589 | 75.9299637 | 0 |
| -2.5287828 | 0.01144588 | 0.02083164 | 22.5414134 | 7.30649104 | 17.2568099 | 14.0078987 |
| -2.5284735 | 0.01145597 | 0.02084882 | 65.1651769 | 115.550803 | 0 | 53.2300151 |
| -2.528181 | 0.01146552 | 0.02086503 | 0 | 133.411114 | 64.4254237 | 98.055291 |
| -2.5266778 | 0.01151471 | 0.02095335 | 0 | 139.635162 | 81.6822337 | 73.4013893 |
| -2.5264716 | 0.01152147 | 0.02096448 | 12.7051603 | 12.4480958 | 13.8054479 | 11.7666349 |
| 2.52638197 | 0.01152441 | 0.02096865 | 25.8201644 | 8.11832337 | 5.75226998 | 7.28410733 |
| -2.5261938 | 0.01153059 | 0.0209787 | 286.890716 | 0 | 29.9118039 | 79.0045488 |
| -2.5259824 | 0.01153753 | 0.02099014 | 17.6232868 | 12.1774851 | 10.354086 | 12.8872668 |
| -2.5257882 | 0.0115439 | 0.02100056 | 9.42640924 | 14.6129821 | 19.5577179 | 11.7666349 |
| -2.5255863 | 0.01155054 | 0.02101145 | 59.8372065 | 91.4664434 | 0 | 73.9617052 |
| -2.5255118 | 0.01155299 | 0.02101473 | 41.3942319 | 100.937821 | 148.408565 | 0 |
| -2.5254625 | 0.01155461 | 0.02101649 | 7.78703372 | 12.7187066 | 26.4604419 | 15.6888466 |
| -2.5252736 | 0.01156082 | 0.0210266 | 50.4107972 | 95.525605 | 0 | 82.9267604 |
| -2.5252284 | 0.01156231 | 0.02102813 | 15.5740674 | 16.2366467 | 6.90272397 | 20.7316901 |
| -2.5251791 | 0.01156394 | 0.0210299 | 80.7392443 | 48.4393295 | 0 | 101.417187 |
| -2.5249621 | 0.01157108 | 0.0210417 | 0 | 73.6061319 | 127.700393 | 89.6505518 |
| 2.52472199 | 0.01157899 | 0.02105489 | 4.50828268 | 1.3530539 | 1.150454 | 1.68094785 |
| -2.5243325 | 0.01159182 | 0.02107705 | 10.246097 | 11.6362635 | 18.4072639 | 13.4475828 |
| -2.5235119 | 0.01161891 | 0.02112512 | 55.3289238 | 78.477126 | 0 | 91.8918156 |

| | | | | | | |
|---|---|---|---|---|---|---|
| -2.523405 | 0.01162244 | 0.02113035 | 5.73781432 | 31.6614612 | 12.6549939 | 22.9729539 |
| -2.5230793 | 0.01163322 | 0.02114874 | 123.772852 | 77.6652936 | 0 | 39.2221164 |
| -2.5229901 | 0.01163617 | 0.02115172 | 38.5253247 | 96.6080482 | 0 | 102.537819 |
| -2.5230074 | 0.0116356 | 0.02115172 | 11.8854725 | 22.1900839 | 18.4072639 | 7.84442328 |
| -2.522933 | 0.01163806 | 0.02115397 | 47.951734 | 73.0649104 | 171.417645 | 0 |
| -2.5226198 | 0.01164843 | 0.02117163 | 14.3445358 | 22.1900839 | 42.5667978 | 4.48252759 |
| 2.5223464 | 0.01165748 | 0.0211869 | 3.68859492 | 0 | 0 | 0 |
| 2.52203252 | 0.01166789 | 0.02120462 | 0 | 7.30649104 | 0 | 0 |
| -2.5218885 | 0.01167267 | 0.02121211 | 3.68859492 | 12.1774851 | 26.4604419 | 57.1522268 |
| -2.5218039 | 0.01167548 | 0.02121602 | 30.738291 | 56.8282636 | 43.7172518 | 52.6696992 |
| 2.5217211 | 0.01167823 | 0.02121981 | 25.0004767 | 30.8496288 | 0 | 38.1014845 |
| -2.5214986 | 0.01168561 | 0.02123085 | 43.4434513 | 72.5236888 | 0 | 120.467929 |
| -2.5215026 | 0.01168548 | 0.02123085 | 28.6890716 | 11.9068743 | 12.6549939 | 9.52537113 |
| -2.5214581 | 0.01168696 | 0.0212321 | 17.6232868 | 20.83703 | 25.3099879 | 36.4205367 |
| -2.5213727 | 0.0116898 | 0.02123606 | 6.55750208 | 3.78855091 | 37.9649818 | 71.1601255 |
| -2.5213225 | 0.01169146 | 0.02123789 | 4.91812656 | 23.0019162 | 17.2568099 | 25.7745336 |
| -2.5212062 | 0.01169533 | 0.02124372 | 17.6232868 | 44.3801678 | 5.75226998 | 12.8872668 |
| -2.5210108 | 0.01170183 | 0.02125074 | 0 | 50.3336049 | 258.852149 | 43.144328 |
| -2.5210376 | 0.01170093 | 0.02125074 | 0 | 100.937821 | 65.5758777 | 123.829825 |
| -2.5210107 | 0.01170183 | 0.02125074 | 81.968776 | 62.2404792 | 0 | 77.8839169 |
| -2.521065 | 0.01170002 | 0.02125074 | 8.60672148 | 13.2599282 | 13.8054479 | 21.852322 |
| 2.52093196 | 0.01170445 | 0.0212543 | 31.5579788 | 16.2366467 | 17.2568099 | 17.9301104 |
| -2.5205559 | 0.01171696 | 0.02127464 | 20.0823501 | 24.6255809 | 9.20363196 | 8.96505518 |
| -2.5205664 | 0.01171661 | 0.02127464 | 31.9678226 | 8.38893415 | 10.354086 | 16.8094785 |
| -2.520358 | 0.01172355 | 0.0212854 | 12.7051603 | 17.8603114 | 9.20363196 | 14.0078987 |
| -2.5203173 | 0.01172491 | 0.02128667 | 79.9195566 | 97.4198805 | 87.4345036 | 0 |
| -2.5201381 | 0.01173088 | 0.02129631 | 9.42640924 | 10.0125988 | 20.7081719 | 17.9301104 |
| -2.5196318 | 0.01174777 | 0.02132577 | 7.37718984 | 25.1668025 | 9.20363196 | 29.1364293 |
| 2.51948037 | 0.01175282 | 0.02133374 | 2.45906328 | 1.89427545 | 1.150454 | 0.56031595 |
| -2.5193375 | 0.01175759 | 0.0213412 | 5.73781432 | 19.4839761 | 32.2127119 | 14.0078987 |
| -2.5191166 | 0.01176497 | 0.02135339 | 32.3776665 | 10.2832096 | 12.6549939 | 10.646003 |
| -2.5190831 | 0.01176609 | 0.02135423 | 267.21821 | 0 | 115.0454 | 19.6110582 |
| -2.5187502 | 0.01177722 | 0.02137322 | 3.68859492 | 45.1920001 | 28.7613499 | 13.4475828 |
| -2.5187096 | 0.01177858 | 0.02137449 | 10.6559409 | 13.2599282 | 16.1063559 | 11.7666349 |
| 2.51866556 | 0.01178005 | 0.02137476 | 0.40984388 | 0 | 0 | 0 |
| 2.51866556 | 0.01178005 | 0.02137476 | 0.40984388 | 0 | 0 | 0 |
| -2.5186348 | 0.01178108 | 0.02137542 | 145.904421 | 56.5576528 | 75.9299637 | 0 |
| -2.5185642 | 0.01178344 | 0.02137851 | 6.1476582 | 20.0251977 | 17.2568099 | 19.0507423 |
| -2.5178197 | 0.01180837 | 0.02142134 | 0 | 100.66721 | 82.8326877 | 99.7362389 |
| -2.5178226 | 0.01180828 | 0.02142134 | 14.3445358 | 50.0629941 | 4.60181598 | 18.4904263 |
| -2.5172488 | 0.01182753 | 0.02145488 | 0 | 107.973701 | 95.4876816 | 80.1251807 |
| -2.5172057 | 0.01182897 | 0.0214563 | 27.45954 | 10.2832096 | 14.9559019 | 9.52537113 |
| 2.51705805 | 0.01183393 | 0.02146409 | 0 | 0.81183234 | 0 | 0.56031595 |

| | | | | | | |
|---|---|---|---|---|---|---|
| 2.51664137 | 0.01184794 | 0.02148828 | 1.22953164 | 3.24732935 | 0 | 2.24126379 |
| -2.5165207 | 0.01185199 | 0.02149443 | 70.9029912 | 80.3714014 | 0 | 68.3585457 |
| 2.51642957 | 0.01185506 | 0.02149878 | 10.246097 | 10.8244312 | 13.8054479 | 5.60315949 |
| -2.5155276 | 0.01188544 | 0.02155039 | 6.96734596 | 17.0484791 | 14.9559019 | 20.7316901 |
| 2.51552514 | 0.01188552 | 0.02155039 | 0 | 0 | 0 | 0 |
| 2.51552514 | 0.01188552 | 0.02155039 | 0 | 0 | 0 | 0 |
| -2.5148394 | 0.01190866 | 0.02159114 | 9.83625312 | 14.6129821 | 20.7081719 | 10.646003 |
| -2.5145556 | 0.01191825 | 0.02160731 | 17.213443 | 8.11832337 | 11.50454 | 22.9729539 |
| -2.5144619 | 0.01192142 | 0.02161184 | 0 | 107.161869 | 95.4876816 | 79.5648647 |
| -2.5141988 | 0.01193031 | 0.02162675 | 8.1968776 | 17.8603114 | 12.6549939 | 17.3697944 |
| -2.5141423 | 0.01193223 | 0.02162901 | 17.213443 | 13.8011497 | 10.354086 | 11.206319 |
| -2.5141176 | 0.01193306 | 0.02162931 | 157.789894 | 41.1328384 | 0 | 52.1093832 |
| -2.5134689 | 0.01195503 | 0.02166719 | 8.1968776 | 15.4248144 | 18.4072639 | 13.4475828 |
| -2.5134608 | 0.01195531 | 0.02166719 | 0 | 146.941653 | 55.2217918 | 92.4521315 |
| -2.5131414 | 0.01196614 | 0.0216856 | 0 | 93.090108 | 110.443584 | 78.4442328 |
| 2.51309233 | 0.0119678 | 0.0216874 | 0.81968776 | 1.89427545 | 2.30090799 | 0 |
| -2.5130658 | 0.0119687 | 0.02168781 | 70.4931473 | 61.4286469 | 0 | 87.9696039 |
| 2.51302026 | 0.01197025 | 0.02168939 | 60.2470503 | 41.1328384 | 0 | 118.226665 |
| -2.5128817 | 0.01197495 | 0.0216967 | 352.875581 | 665.431906 | 635.050605 | 596.176169 |
| -2.5125128 | 0.01198747 | 0.02171817 | 11.8854725 | 20.5664192 | 29.9118039 | 20.7316901 |
| -2.5118066 | 0.01201149 | 0.02176046 | 61.8864259 | 71.4412457 | 0 | 85.7283401 |
| -2.5114031 | 0.01202523 | 0.02178413 | 64.755333 | 43.2977247 | 0 | 127.19172 |
| 2.5113435 | 0.01202726 | 0.02178659 | 1.22953164 | 0.54122156 | 1.150454 | 1.68094785 |
| -2.5111329 | 0.01203444 | 0.02179837 | 0 | 151.271426 | 43.7172518 | 107.020346 |
| -2.5106721 | 0.01205016 | 0.02182561 | 10.246097 | 21.1076408 | 16.1063559 | 9.52537113 |
| -2.5101881 | 0.01206669 | 0.02185433 | 172.954117 | 359.371115 | 322.127119 | 336.749885 |
| -2.5100791 | 0.01207041 | 0.02185985 | 30.738291 | 111.762252 | 0 | 99.7362389 |
| -2.5099079 | 0.01207627 | 0.02186922 | 6.96734596 | 21.1076408 | 11.50454 | 24.6539017 |
| -2.5096808 | 0.01208403 | 0.02188206 | 9.83625312 | 18.9427545 | 12.6549939 | 12.3269509 |
| -2.509436 | 0.01209241 | 0.02189601 | 263.119771 | 323.10927 | 243.896247 | 389.9799 |
| -2.5092591 | 0.01209847 | 0.02190575 | 0 | 112.032863 | 159.913105 | 36.9808526 |
| -2.5092299 | 0.01209947 | 0.02190633 | 5.73781432 | 38.9679522 | 39.1154358 | 29.1364293 |
| -2.5086125 | 0.01212063 | 0.02194342 | 6.1476582 | 47.0862756 | 11.50454 | 17.3697944 |
| -2.5081164 | 0.01213767 | 0.02197302 | 9.01656536 | 33.5557366 | 11.50454 | 12.8872668 |
| -2.5077611 | 0.01214988 | 0.0219939 | 11.0657848 | 32.4732935 | 3.45136199 | 49.3078035 |
| 2.50707688 | 0.01217342 | 0.02203405 | 0 | 0 | 0 | 0 |
| 2.50707688 | 0.01217342 | 0.02203405 | 0 | 0 | 0 | 0 |
| 2.50701465 | 0.01217557 | 0.02203669 | 20.0823501 | 15.1542036 | 13.8054479 | 8.96505518 |
| -2.5054483 | 0.01222963 | 0.0221333 | 10.246097 | 15.6954252 | 25.3099879 | 8.96505518 |
| -2.505363 | 0.01223258 | 0.02213635 | 47.1320462 | 117.9863 | 0 | 63.3157022 |
| -2.5053599 | 0.01223269 | 0.02213635 | 59.8372065 | 92.8194972 | 110.443584 | 0 |
| -2.5049485 | 0.01224692 | 0.02216087 | 38.1154808 | 18.401533 | 39.1154358 | 2.24126379 |
| -2.5047388 | 0.01225419 | 0.02217277 | 26.2300083 | 5.95343714 | 44.8677058 | 8.40473923 |

| | | | | | | |
|---|---|---|---|---|---|---|
| -2.5043259 | 0.0122685 | 0.02219742 | 0 | 78.7477367 | 90.8858656 | 107.580662 |
| -2.5041124 | 0.0122759 | 0.02220957 | 10.246097 | 16.5072575 | 10.354086 | 15.6888466 |
| -2.5040061 | 0.01227959 | 0.02221501 | 102.051126 | 71.7118565 | 0 | 48.7474875 |
| -2.503765 | 0.01228796 | 0.0222289 | 6.55750208 | 17.3190899 | 16.1063559 | 19.6110582 |
| -2.5035882 | 0.0122941 | 0.02223719 | 290.169467 | 662.184577 | 662.661501 | 658.37124 |
| -2.5035737 | 0.01229461 | 0.02223719 | 10.6559409 | 10.0125988 | 21.8586259 | 13.4475828 |
| -2.5035993 | 0.01229372 | 0.02223719 | 27.45954 | 12.1774851 | 14.9559019 | 7.84442328 |
| -2.503365 | 0.01230186 | 0.02224905 | 271.316649 | 934.689631 | 850.185503 | 866.248457 |
| -2.5029902 | 0.0123149 | 0.02227139 | 109.018472 | 588.849055 | 361.242555 | 414.633802 |
| -2.5028078 | 0.01232124 | 0.02228162 | 11.8854725 | 16.5072575 | 18.4072639 | 8.40473923 |
| 2.50269115 | 0.0123253 | 0.02228771 | 129.92051 | 87.4072817 | 87.4345036 | 71.1601255 |
| 2.50265129 | 0.01232669 | 0.02228839 | 4.91812656 | 1.62366467 | 1.150454 | 1.1206319 |
| -2.5026408 | 0.01232706 | 0.02228839 | 11.8854725 | 25.708024 | 4.60181598 | 38.1014845 |
| -2.5024196 | 0.01233476 | 0.02230107 | 67.6242402 | 9.47137727 | 5.75226998 | 16.8094785 |
| 2.50225834 | 0.01234038 | 0.02230998 | 0 | 0 | 1.150454 | 0.56031595 |
| -2.5020326 | 0.01234826 | 0.02232296 | 8.1968776 | 14.8835929 | 14.9559019 | 15.6888466 |
| -2.5016823 | 0.01236048 | 0.02234381 | 21.7217256 | 14.0717605 | 13.8054479 | 7.84442328 |
| -2.5015571 | 0.01236485 | 0.02235046 | 63.5258014 | 66.5702517 | 0 | 85.1680242 |
| -2.5011768 | 0.01237814 | 0.02237197 | 0 | 71.7118565 | 164.514921 | 57.7125427 |
| -2.5011932 | 0.01237756 | 0.02237197 | 139.756763 | 12.9893174 | 2.30090799 | 10.0856871 |
| 2.50106606 | 0.01238201 | 0.02237772 | 7.78703372 | 2.70610779 | 2.30090799 | 3.92221164 |
| -2.5007451 | 0.01239324 | 0.02239675 | 1.63937552 | 29.7671857 | 70.1776937 | 17.3697944 |
| -2.5005087 | 0.01240151 | 0.02241045 | 0 | 30.0377965 | 124.249032 | 159.129729 |
| 2.50016899 | 0.01241341 | 0.0224307 | 0.40984388 | 0.54122156 | 0 | 0 |
| -2.4999945 | 0.01241952 | 0.02244049 | 52.8698605 | 14.6129821 | 5.75226998 | 12.8872668 |
| -2.4998241 | 0.0124255 | 0.02245003 | 8.60672148 | 17.3190899 | 12.6549939 | 15.1285306 |
| -2.4997612 | 0.0124277 | 0.02245276 | 127.461447 | 725.507499 | 1619.83923 | 677.421982 |
| -2.4997082 | 0.01242956 | 0.02245486 | 107.379097 | 50.6042157 | 106.992222 | 0 |
| -2.4996365 | 0.01243208 | 0.02245815 | 31.1481349 | 12.9893174 | 20.7081719 | 5.60315949 |
| -2.4990906 | 0.01245125 | 0.02249152 | 6.1476582 | 16.7778683 | 20.7081719 | 17.3697944 |
| -2.4989757 | 0.01245529 | 0.02249756 | 74.1817423 | 54.1221558 | 0 | 87.9696039 |
| -2.4987385 | 0.01246362 | 0.02251136 | 29.5087594 | 140.988216 | 119.647216 | 0 |
| -2.4985496 | 0.01247027 | 0.0225221 | 278.693838 | 314.449725 | 356.640739 | 365.886314 |
| -2.4985124 | 0.01247158 | 0.0225232 | 0 | 68.1939163 | 109.29313 | 99.1759229 |
| -2.4979463 | 0.01249151 | 0.02255794 | 9.83625312 | 16.5072575 | 11.50454 | 14.0078987 |
| 2.49789961 | 0.01249316 | 0.02255839 | 0 | 0 | 0 | 0 |
| 2.49789961 | 0.01249316 | 0.02255839 | 0 | 0 | 0 | 0 |
| -2.497831 | 0.01249558 | 0.0225615 | 0 | 90.3840002 | 93.1867736 | 89.0902358 |
| 2.4977062 | 0.01249997 | 0.02256817 | 48.3615778 | 30.3084073 | 26.4604419 | 28.0157974 |
| -2.4976062 | 0.0125035 | 0.02257328 | 2.86890716 | 27.3316887 | 27.6108959 | 25.7745336 |
| 2.49743509 | 0.01250954 | 0.02258165 | 1.63937552 | 0 | 2.30090799 | 1.68094785 |
| -2.4974534 | 0.01250889 | 0.02258165 | 21.3118818 | 14.3423713 | 34.5136199 | 38.1014845 |
| -2.4972481 | 0.01251614 | 0.0225923 | 8.60672148 | 15.4248144 | 17.2568099 | 12.3269509 |

| | | | | | | |
|---|---|---|---|---|---|---|
| -2.4969818 | 0.01252554 | 0.02260801 | 15.9839113 | 38.4267306 | 2.30090799 | 40.3427483 |
| 2.49660937 | 0.0125387 | 0.0226305 | 18.4429746 | 22.1900839 | 9.20363196 | 15.1285306 |
| -2.4964481 | 0.0125444 | 0.02263952 | 325.416041 | 462.744432 | 425.667978 | 569.84132 |
| -2.4963409 | 0.01254819 | 0.0226451 | 10.246097 | 10.0125988 | 12.6549939 | 31.3776931 |
| 2.49623619 | 0.0125519 | 0.02265052 | 4.50828268 | 10.5538204 | 12.6549939 | 6.16347543 |
| 2.4961493 | 0.01255497 | 0.02265446 | 36.0662614 | 18.1309222 | 24.1595339 | 15.6888466 |
| -2.496135 | 0.01255548 | 0.02265446 | 14.3445358 | 31.3908504 | 5.75226998 | 16.8094785 |
| -2.4959888 | 0.01256066 | 0.02266253 | 4.0984388 | 13.530539 | 80.5317797 | 13.4475828 |
| -2.4958988 | 0.01256384 | 0.02266701 | 29.0989155 | 19.7545869 | 20.7081719 | 20.7316901 |
| -2.4958441 | 0.01256578 | 0.02266924 | 14.3445358 | 30.0377965 | 26.4604419 | 27.4554815 |
| -2.4955628 | 0.01257575 | 0.02268595 | 22.9512573 | 13.2599282 | 49.4695218 | 32.498325 |
| -2.4953839 | 0.01258209 | 0.02269613 | 0 | 115.550803 | 142.656295 | 38.6618005 |
| -2.4953155 | 0.01258452 | 0.022699 | 85.6573709 | 471.133366 | 433.721156 | 352.438732 |
| 2.49529936 | 0.01258509 | 0.022699 | 19.2626624 | 10.8244312 | 14.9559019 | 9.52537113 |
| 2.49449985 | 0.01261348 | 0.02274893 | 87.2967464 | 141.800048 | 83.9831417 | 101.977503 |
| -2.4943076 | 0.01262031 | 0.02275999 | 9.42640924 | 25.708024 | 10.354086 | 14.0078987 |
| -2.4938615 | 0.01263618 | 0.02278734 | 23.3611012 | 27.8729103 | 3.45136199 | 21.292006 |
| -2.4935964 | 0.01264562 | 0.02280309 | 8.1968776 | 18.6721438 | 14.9559019 | 12.8872668 |
| -2.4935757 | 0.01264636 | 0.02280314 | 9.42640924 | 27.3316887 | 8.05317797 | 19.0507423 |
| -2.493339 | 0.0126548 | 0.02281708 | 0.40984388 | 31.6614612 | 70.1776937 | 49.3078035 |
| 2.49303138 | 0.01266577 | 0.02283533 | 8.60672148 | 3.24732935 | 0 | 29.6967453 |
| -2.4930155 | 0.01266633 | 0.02283533 | 0 | 88.7603356 | 116.195854 | 69.4791776 |
| -2.4929904 | 0.01266723 | 0.02283567 | 16.3937552 | 24.8961917 | 21.8586259 | 31.9380091 |
| -2.4928441 | 0.01267245 | 0.0228438 | 13.1150042 | 11.3656527 | 11.50454 | 13.4475828 |
| 2.49277062 | 0.01267507 | 0.02284725 | 1.22953164 | 1.89427545 | 2.30090799 | 2.24126379 |
| -2.4926619 | 0.01267895 | 0.02285298 | 26.6398522 | 13.8011497 | 6.90272397 | 14.5682147 |
| 2.49254232 | 0.01268322 | 0.02285812 | 12.2953164 | 4.87099402 | 5.75226998 | 7.28410733 |
| -2.4925455 | 0.01268311 | 0.02285812 | 88.9361219 | 46.8156648 | 0 | 81.2458125 |
| -2.4923902 | 0.01268866 | 0.02286664 | 13.9346919 | 107.973701 | 0 | 160.810677 |
| 2.49164665 | 0.01271525 | 0.02291328 | 0 | 0 | 0 | 3.92221164 |
| -2.4915891 | 0.01271731 | 0.02291571 | 150.412704 | 40.8622277 | 0 | 49.3078035 |
| -2.4912341 | 0.01273002 | 0.02293606 | 189.347873 | 315.532168 | 307.171217 | 328.345146 |
| -2.4912448 | 0.01272964 | 0.02293606 | 156.970206 | 199.710755 | 211.683535 | 216.281956 |
| -2.4911558 | 0.01273283 | 0.02293984 | 7.37718984 | 11.3656527 | 23.0090799 | 18.4904263 |
| 2.49074319 | 0.01274762 | 0.02296521 | 30.738291 | 50.6042157 | 43.7172518 | 50.9887513 |
| -2.4904295 | 0.01275888 | 0.02298421 | 10.246097 | 10.8244312 | 17.2568099 | 13.4475828 |
| -2.4899849 | 0.01277485 | 0.0230117 | 65.9848647 | 104.18515 | 0 | 48.1871716 |
| -2.4896851 | 0.01278563 | 0.02302854 | 19.2626624 | 36.2618444 | 3.45136199 | 20.7316901 |
| -2.4896884 | 0.01278551 | 0.02302854 | 36.4761053 | 45.1920001 | 39.1154358 | 35.2999048 |
| -2.4893432 | 0.01279794 | 0.02304942 | 0 | 123.398515 | 75.9299637 | 74.5220212 |
| -2.4892146 | 0.01280257 | 0.02305648 | 9.42640924 | 8.93015571 | 25.3099879 | 16.8094785 |
| 2.48887959 | 0.01281464 | 0.02307692 | 0.81968776 | 1.62366467 | 0 | 2.24126379 |
| 2.4888446 | 0.0128159 | 0.02307791 | 0.81968776 | 2.43549701 | 1.150454 | 3.36189569 |

| | | | | | | |
|---|---|---|---|---|---|---|
| 2.4887769 | 0.01281834 | 0.02308101 | 11.0657848 | 9.20076649 | 3.45136199 | 9.52537113 |
| 2.48821472 | 0.01283862 | 0.02311624 | 7.78703372 | 5.68282636 | 0 | 7.84442328 |
| 2.48815845 | 0.01284065 | 0.02311861 | 3.27875104 | 4.87099402 | 5.75226998 | 5.04284354 |
| -2.4877557 | 0.0128552 | 0.02314352 | 46.7222023 | 242.467258 | 316.374849 | 256.064389 |
| -2.4874103 | 0.01286769 | 0.02316471 | 70.9029912 | 0 | 102.390406 | 123.829825 |
| -2.4872897 | 0.01287205 | 0.02317128 | 23.3611012 | 8.38893415 | 24.1595339 | 8.40473923 |
| 2.48711994 | 0.0128782 | 0.02317975 | 7.78703372 | 5.68282636 | 0 | 0 |
| 2.4871223 | 0.01287811 | 0.02317975 | 10.6559409 | 7.30649104 | 0 | 9.52537113 |
| -2.4869973 | 0.01288264 | 0.02318645 | 62.7061136 | 104.455761 | 0 | 49.8681194 |
| -2.4868279 | 0.01288877 | 0.0231962 | 51.230485 | 9.47137727 | 10.354086 | 10.646003 |
| -2.4867158 | 0.01289283 | 0.02320092 | 61.476582 | 13.2599282 | 6.90272397 | 10.0856871 |
| -2.4867269 | 0.01289243 | 0.02320092 | 0 | 90.1133895 | 105.841768 | 73.4013893 |
| -2.4866939 | 0.01289363 | 0.02320106 | 161.478489 | 121.233629 | 117.346308 | 121.028245 |
| -2.4864411 | 0.01290279 | 0.02321625 | 11.8854725 | 15.6954252 | 10.354086 | 12.3269509 |
| -2.486368 | 0.01290544 | 0.02321973 | 11.0657848 | 7.8477126 | 29.9118039 | 15.1285306 |
| -2.4861073 | 0.0129149 | 0.02323545 | 14.3445358 | 9.47137727 | 11.50454 | 16.2491625 |
| -2.4860751 | 0.01291607 | 0.02323626 | 139.346919 | 0 | 34.5136199 | 151.845622 |
| -2.4860323 | 0.01291762 | 0.02323776 | 36.4761053 | 83.0775092 | 0 | 104.218766 |
| -2.4858656 | 0.01292368 | 0.02324735 | 84.4278393 | 61.6992576 | 0 | 63.8760181 |
| -2.4855512 | 0.0129351 | 0.0232666 | 6.96734596 | 28.6847426 | 27.6108959 | 8.40473923 |
| 2.48532009 | 0.0129435 | 0.02328042 | 15.1642236 | 14.6129821 | 14.9559019 | 11.206319 |
| -2.4848973 | 0.01295888 | 0.02330593 | 31.1481349 | 10.0125988 | 6.90272397 | 19.6110582 |
| -2.4848906 | 0.01295913 | 0.02330593 | 9.42640924 | 14.3423713 | 13.8054479 | 12.8872668 |
| -2.4846759 | 0.01296694 | 0.02331869 | 6.55750208 | 10.0125988 | 28.7613499 | 22.9729539 |
| -2.4846144 | 0.01296918 | 0.02332142 | 18.0331307 | 9.74198805 | 23.0090799 | 8.40473923 |
| -2.4844493 | 0.0129752 | 0.02333093 | 1.63937552 | 28.4141318 | 47.1686138 | 25.7745336 |
| -2.4844198 | 0.01297627 | 0.02333156 | 33.1973543 | 94.4431619 | 0 | 98.615607 |
| -2.4842332 | 0.01298308 | 0.0233425 | 93.0345607 | 37.3442875 | 0 | 90.7711837 |
| -2.4841127 | 0.01298747 | 0.0233478 | 58.6076748 | 69.817581 | 0 | 80.6854966 |
| -2.4841263 | 0.01298697 | 0.0233478 | 12.7051603 | 16.5072575 | 10.354086 | 11.206319 |
| 2.4838798 | 0.01299597 | 0.02336177 | 9.42640924 | 15.4248144 | 18.4072639 | 11.206319 |
| 2.48368603 | 0.01300304 | 0.02337318 | 10.6559409 | 11.9068743 | 1.150454 | 6.16347543 |
| 2.48317868 | 0.01302158 | 0.0234052 | 1.22953164 | 4.32977247 | 5.75226998 | 7.84442328 |
| -2.4831291 | 0.01302339 | 0.02340715 | 65.1651769 | 228.124887 | 92.0363196 | 176.499524 |
| -2.4830752 | 0.01302536 | 0.0234094 | 128.690978 | 176.167617 | 201.329449 | 189.947107 |
| -2.482674 | 0.01304004 | 0.02343379 | 31.5579788 | 123.127905 | 121.948124 | 0 |
| -2.4826645 | 0.01304039 | 0.02343379 | 45.4926707 | 105.538204 | 0 | 66.1172819 |
| -2.4819676 | 0.01306591 | 0.02347836 | 8.1968776 | 21.1076408 | 17.2568099 | 10.646003 |
| -2.4818549 | 0.01307005 | 0.02348448 | 10.6559409 | 12.1774851 | 11.50454 | 15.6888466 |
| 2.48145002 | 0.01308491 | 0.02350464 | 0 | 0 | 0 | 0 |
| 2.48145002 | 0.01308491 | 0.02350464 | 0 | 0 | 0 | 0 |
| 2.48145002 | 0.01308491 | 0.02350464 | 0 | 0 | 0 | 0 |
| 2.48145002 | 0.01308491 | 0.02350464 | 0 | 0 | 0 | 0 |

| | | | | | | |
|---|---|---|---|---|---|---|
| 2.48145002 | 0.01308491 | 0.02350464 | 0 | 0 | 0 | 0 |
| -2.4812774 | 0.01309125 | 0.02351472 | 8.60672148 | 15.4248144 | 11.50454 | 17.9301104 |
| -2.4807785 | 0.01310958 | 0.0235454 | 17.213443 | 28.6847426 | 25.3099879 | 33.6189569 |
| -2.4807729 | 0.01310979 | 0.0235454 | 15.5740674 | 5.68282636 | 25.3099879 | 17.3697944 |
| -2.4805543 | 0.01311783 | 0.02355853 | 46.7222023 | 12.4480958 | 8.05317797 | 10.646003 |
| -2.4803403 | 0.01312571 | 0.02357137 | 109.018472 | 631.334948 | 314.073941 | 433.684544 |
| 2.48028531 | 0.01312773 | 0.02357369 | 2.86890716 | 0 | 5.75226998 | 0 |
| 2.48015898 | 0.01313238 | 0.02358074 | 0 | 0 | 0 | 0 |
| -2.4800326 | 0.01313704 | 0.02358778 | 6.96734596 | 18.6721438 | 13.8054479 | 17.3697944 |
| 2.47983932 | 0.01314416 | 0.02359926 | 0 | 0 | 0 | 0 |
| -2.4795139 | 0.01315616 | 0.02361949 | 280.743058 | 369.924935 | 357.791193 | 372.610106 |
| -2.478847 | 0.01318078 | 0.02366238 | 104.100346 | 71.7118565 | 0 | 41.4633802 |
| -2.4787782 | 0.01318333 | 0.02366563 | 0 | 63.8641439 | 103.54086 | 100.296555 |
| -2.4786973 | 0.01318632 | 0.02366968 | 6.1476582 | 41.9446708 | 16.1063559 | 11.7666349 |
| -2.4785189 | 0.01319291 | 0.02368021 | 0 | 123.398515 | 40.2658898 | 120.467929 |
| -2.4783648 | 0.01319861 | 0.02368912 | 6.96734596 | 23.5431378 | 19.5577179 | 11.206319 |
| -2.4782006 | 0.01320469 | 0.02369781 | 3.68859492 | 16.5072575 | 36.8145279 | 22.4126379 |
| 2.47819436 | 0.01320492 | 0.02369781 | 2.45906328 | 0.27061078 | 2.30090799 | 1.68094785 |
| -2.4779455 | 0.01321413 | 0.02371303 | 5.32797044 | 18.1309222 | 29.9118039 | 14.5682147 |
| -2.4778057 | 0.01321931 | 0.023721 | 57.7879871 | 25.708024 | 0 | 170.896364 |
| -2.4775315 | 0.01322947 | 0.02373792 | 10.246097 | 17.8603114 | 19.5577179 | 8.40473923 |
| -2.4774982 | 0.01323071 | 0.02373881 | 75.8211178 | 59.8049822 | 0 | 70.5998095 |
| -2.477478 | 0.01323146 | 0.02373883 | 0 | 48.1687187 | 158.762651 | 77.3236009 |
| -2.4773087 | 0.01323774 | 0.02374878 | 180.741151 | 43.5683354 | 51.7704298 | 53.7903311 |
| 2.47701715 | 0.01324855 | 0.02376687 | 9.01656536 | 6.4946587 | 13.8054479 | 6.72379138 |
| 2.476935 | 0.0132516 | 0.02377102 | 12.2953164 | 5.95343714 | 8.05317797 | 6.16347543 |
| -2.4766704 | 0.01326143 | 0.023786 | 15.1642236 | 30.0377965 | 23.0090799 | 29.1364293 |
| -2.476674 | 0.0132613 | 0.023786 | 1470.11 | 1204.75919 | 725.936471 | 951.976797 |
| -2.476494 | 0.01326799 | 0.02379644 | 35.6564176 | 115.280192 | 0 | 72.2807574 |
| -2.4762547 | 0.01327688 | 0.02381107 | 3.68859492 | 24.0843593 | 29.9118039 | 17.9301104 |
| -2.4758892 | 0.01329048 | 0.02383414 | 34.017042 | 13.530539 | 6.90272397 | 12.8872668 |
| -2.4757414 | 0.01329598 | 0.02384268 | 80.7392443 | 104.726372 | 88.5849576 | 93.0124475 |
| -2.4756571 | 0.01329912 | 0.02384699 | 201.643189 | 212.700072 | 186.373547 | 186.024895 |
| 2.47502108 | 0.01332283 | 0.02388817 | 0 | 4.60038325 | 4.60181598 | 4.48252759 |
| 2.47488918 | 0.01332775 | 0.02389567 | 27.0496961 | 16.5072575 | 20.7081719 | 16.8094785 |
| 2.47484461 | 0.01332942 | 0.02389732 | 2.45906328 | 2.43549701 | 2.30090799 | 1.1206319 |
| -2.4742911 | 0.01335009 | 0.02393305 | 13.1150042 | 27.3316887 | 13.8054479 | 28.0157974 |
| -2.4741957 | 0.01335365 | 0.02393812 | 0 | 106.620647 | 131.151755 | 118.226665 |
| -2.4740345 | 0.01335968 | 0.02394759 | 201.233345 | 491.970396 | 338.233475 | 417.995698 |
| -2.4739744 | 0.01336193 | 0.02395029 | 11.8854725 | 40.0503953 | 3.45136199 | 30.8173772 |
| -2.473825 | 0.01336752 | 0.02395897 | 54.509236 | 96.3374374 | 0 | 58.8331746 |
| 2.47357934 | 0.01337671 | 0.02397412 | 0 | 3.78855091 | 0 | 0 |
| 2.47316524 | 0.01339222 | 0.02400059 | 10.246097 | 3.24732935 | 6.90272397 | 3.92221164 |

| | | | | | | |
|---|---|---|---|---|---|---|
| -2.4731186 | 0.01339397 | 0.02400239 | 51.6403289 | 89.8427787 | 0 | 66.6775979 |
| -2.4729931 | 0.01339867 | 0.02400949 | 12.2953164 | 33.5557366 | 14.9559019 | 6.72379138 |
| -2.4729288 | 0.01340108 | 0.02401247 | 78.6900249 | 4.60038325 | 21.8586259 | 6.72379138 |
| -2.4728795 | 0.01340293 | 0.02401445 | 201.233345 | 563.952864 | 630.448789 | 224.686695 |
| -2.4728185 | 0.01340522 | 0.02401722 | 13.1150042 | 8.38893415 | 12.6549939 | 20.1713742 |
| -2.4727807 | 0.01340664 | 0.02401843 | 10.246097 | 10.8244312 | 12.6549939 | 17.3697944 |
| -2.4726023 | 0.01341333 | 0.02402909 | 6.55750208 | 10.0125988 | 21.8586259 | 28.5761134 |
| -2.4724784 | 0.01341798 | 0.02403608 | 13.524848 | 18.9427545 | 11.50454 | 8.96505518 |
| -2.4723351 | 0.01342336 | 0.0240422 | 95.493624 | 16.5072575 | 16.1063559 | 30.2570612 |
| -2.472328 | 0.01342363 | 0.0240422 | 26.6398522 | 18.9427545 | 18.4072639 | 4.48252759 |
| -2.4723546 | 0.01342263 | 0.0240422 | 160.248957 | 289.553534 | 268.055781 | 259.426284 |
| -2.4717575 | 0.01344507 | 0.02407926 | 4.91812656 | 6.22404792 | 49.4695218 | 35.8602207 |
| -2.4711592 | 0.01346758 | 0.02411825 | 8.1968776 | 27.6022995 | 18.4072639 | 8.96505518 |
| -2.4705483 | 0.01349061 | 0.02415814 | 36.8859492 | 83.34812 | 0 | 95.8140272 |
| -2.470196 | 0.0135039 | 0.02417927 | 6.96734596 | 18.9427545 | 16.1063559 | 14.0078987 |
| -2.470208 | 0.01350345 | 0.02417927 | 6.1476582 | 18.9427545 | 12.6549939 | 24.6539017 |
| -2.4698897 | 0.01351547 | 0.02419864 | 104.100346 | 10.0125988 | 3.45136199 | 12.8872668 |
| -2.4698123 | 0.01351839 | 0.02420017 | 13.524848 | 30.8496288 | 11.50454 | 7.84442328 |
| 2.46980761 | 0.01351857 | 0.02420017 | 7.37718984 | 0 | 0 | 0 |
| -2.4698461 | 0.01351712 | 0.02420017 | 11.8854725 | 47.3568864 | 8.05317797 | 10.646003 |
| 2.46923964 | 0.01354005 | 0.02423727 | 9.01656536 | 2.97671857 | 3.45136199 | 2.80157974 |
| -2.4690583 | 0.01354691 | 0.02424821 | 17.213443 | 27.3316887 | 29.9118039 | 3.36189569 |
| -2.46903 | 0.01354799 | 0.02424878 | 16.8035991 | 183.203497 | 0 | 70.0394936 |
| -2.4689706 | 0.01355023 | 0.02425146 | 9.42640924 | 15.966036 | 9.20363196 | 20.7316901 |
| -2.4688994 | 0.01355293 | 0.02425495 | 102.051126 | 278.458492 | 288.763953 | 286.32145 |
| -2.4688332 | 0.01355544 | 0.02425809 | 11.0657848 | 32.7439043 | 14.9559019 | 29.1364293 |
| -2.4687888 | 0.01355712 | 0.02425975 | 61.476582 | 89.8427787 | 0 | 54.910963 |
| 2.46861349 | 0.01356376 | 0.0242703 | 1.22953164 | 0 | 2.30090799 | 2.24126379 |
| -2.4685337 | 0.01356679 | 0.02427437 | 0 | 115.009581 | 115.0454 | 44.2649599 |
| -2.4684461 | 0.01357011 | 0.02427809 | 43.8532952 | 81.7244553 | 0 | 83.4870763 |
| -2.4684391 | 0.01357038 | 0.02427809 | 3.68859492 | 48.4393295 | 12.6549939 | 22.9729539 |
| -2.4677571 | 0.01359625 | 0.02432304 | 10.246097 | 106.891258 | 0 | 172.016996 |
| -2.4677019 | 0.01359835 | 0.02432544 | 14.3445358 | 23.5431378 | 5.75226998 | 17.3697944 |
| -2.4675863 | 0.01360274 | 0.02433196 | 70.0833035 | 98.7729344 | 0 | 42.5840121 |
| -2.4675236 | 0.01360512 | 0.02433487 | 98.7723751 | 207.017246 | 308.321671 | 0 |
| -2.4670943 | 0.01362145 | 0.02436136 | 81.1490882 | 36.2618444 | 0 | 98.055291 |
| -2.4670975 | 0.01362133 | 0.02436136 | 40.5745441 | 102.290875 | 113.894946 | 0 |
| -2.466808 | 0.01363234 | 0.0243795 | 126.231915 | 46.8156648 | 0 | 47.6268556 |
| -2.4667854 | 0.0136332 | 0.02437968 | 126.641759 | 328.250875 | 327.879389 | 331.146726 |
| -2.4666807 | 0.01363719 | 0.02438547 | 59.0175187 | 68.4645271 | 0 | 75.0823371 |
| -2.4666386 | 0.01363879 | 0.02438662 | 214.758193 | 15.1542036 | 0 | 57.7125427 |
| -2.4666241 | 0.01363934 | 0.02438662 | 10.246097 | 15.4248144 | 10.354086 | 14.0078987 |
| -2.4665656 | 0.01364158 | 0.02438925 | 6.55750208 | 28.143521 | 17.2568099 | 11.7666349 |

| | | | | | | |
|---|---|---|---|---|---|---|
| -2.4664283 | 0.0136468 | 0.02439725 | 26.6398522 | 46.2744432 | 9.20363196 | 4.48252759 |
| -2.4662049 | 0.01365532 | 0.02441112 | 9.42640924 | 15.1542036 | 11.50454 | 14.0078987 |
| -2.465836 | 0.01366939 | 0.02443493 | 29.0989155 | 76.8534613 | 0 | 121.028245 |
| -2.4657949 | 0.01367096 | 0.02443637 | 252.873674 | 512.807426 | 499.297034 | 521.654148 |
| 2.46576648 | 0.01367204 | 0.02443696 | 2.86890716 | 1.89427545 | 0 | 0.56031595 |
| -2.4650657 | 0.01369881 | 0.02448345 | 61.8864259 | 43.8389462 | 0 | 106.46003 |
| -2.4647553 | 0.01371069 | 0.02450332 | 43.0336074 | 14.6129821 | 8.05317797 | 8.96505518 |
| -2.4642913 | 0.01372845 | 0.0245337 | 8.1968776 | 18.9427545 | 11.50454 | 15.1285306 |
| -2.463777 | 0.01374816 | 0.02456757 | 4.50828268 | 19.2133653 | 14.9559019 | 34.1792729 |
| -2.4635879 | 0.01375542 | 0.02457918 | 13.1150042 | 11.6362635 | 11.50454 | 11.7666349 |
| -2.4634898 | 0.01375918 | 0.02458454 | 29.9186032 | 88.7603356 | 0 | 103.098135 |
| -2.4632713 | 0.01376757 | 0.02459817 | 8.60672148 | 12.1774851 | 29.9118039 | 11.206319 |
| -2.4629773 | 0.01377887 | 0.02461699 | 233.611012 | 297.942468 | 323.277573 | 306.492824 |
| -2.4628301 | 0.01378452 | 0.02462573 | 13.524848 | 8.93015571 | 21.8586259 | 10.646003 |
| -2.4624238 | 0.01380015 | 0.02465229 | 34.017042 | 90.654611 | 0 | 90.7711837 |
| 2.46234562 | 0.01380316 | 0.0246563 | 0 | 0.27061078 | 0 | 0 |
| -2.4621187 | 0.0138119 | 0.02466974 | 5.73781432 | 27.0610779 | 11.50454 | 21.852322 |
| 2.46211046 | 0.01381221 | 0.02466974 | 0 | 14.6129821 | 3.45136199 | 12.8872668 |
| 2.46201018 | 0.01381607 | 0.02467527 | 0 | 9.20076649 | 0 | 5.60315949 |
| -2.4619797 | 0.01381725 | 0.02467601 | 82.3786199 | 7.30649104 | 14.9559019 | 5.60315949 |
| -2.4619352 | 0.01381896 | 0.0246777 | 13.9346919 | 24.0843593 | 29.9118039 | 28.5761134 |
| -2.4619103 | 0.01381992 | 0.02467805 | 34.017042 | 10.0125988 | 23.0090799 | 5.60315949 |
| -2.4618274 | 0.01382312 | 0.02468239 | 16.8035991 | 9.74198805 | 11.50454 | 12.3269509 |
| -2.4617205 | 0.01382724 | 0.02468838 | 95.9034679 | 35.4500121 | 44.8677058 | 36.9808526 |
| -2.4616421 | 0.01383026 | 0.02469241 | 29.5087594 | 83.8893415 | 49.4695218 | 69.4791776 |
| -2.4615886 | 0.01383232 | 0.02469473 | 4.50828268 | 14.6129821 | 19.5577179 | 34.1792729 |
| -2.4615105 | 0.01383533 | 0.02469874 | 0 | 87.4072817 | 83.9831417 | 84.6077082 |
| 2.46132218 | 0.0138426 | 0.02471035 | 0 | 0 | 0 | 0 |
| -2.4610253 | 0.01385406 | 0.02472944 | 67.6242402 | 51.416048 | 0 | 84.0473923 |
| -2.4606167 | 0.01386984 | 0.02475624 | 0 | 57.640096 | 169.116737 | 53.7903311 |
| -2.460568 | 0.01387173 | 0.02475824 | 4.91812656 | 15.1542036 | 27.6108959 | 19.0507423 |
| 2.46026525 | 0.01388344 | 0.02477776 | 13.1150042 | 7.30649104 | 5.75226998 | 2.24126379 |
| -2.4601376 | 0.01388838 | 0.02478384 | 15.5740674 | 15.1542036 | 5.75226998 | 24.0935858 |
| -2.4601548 | 0.01388771 | 0.02478384 | 23.770945 | 10.8244312 | 9.20363196 | 12.3269509 |
| -2.4596588 | 0.01390692 | 0.02481555 | 31.9678226 | 10.8244312 | 13.8054479 | 7.84442328 |
| -2.4592362 | 0.0139233 | 0.02484341 | 49.5911095 | 101.208431 | 0 | 56.5919108 |
| -2.4591806 | 0.01392545 | 0.02484589 | 15.1642236 | 34.6381797 | 32.2127119 | 19.0507423 |
| -2.4588979 | 0.01393643 | 0.02486337 | 4.0984388 | 16.5072575 | 35.6640739 | 18.4904263 |
| -2.4588787 | 0.01393717 | 0.02486337 | 28.6890716 | 122.586683 | 0 | 73.9617052 |
| 2.45886857 | 0.01393756 | 0.02486337 | 0.40984388 | 5.1416048 | 3.45136199 | 2.80157974 |
| 2.45884484 | 0.01393848 | 0.02486364 | 0.40984388 | 1.89427545 | 1.150454 | 2.24126379 |
| 2.45881946 | 0.01393947 | 0.02486402 | 0.40984388 | 0 | 0 | 0 |
| -2.4586514 | 0.01394599 | 0.02487291 | 9.01656536 | 12.4480958 | 11.50454 | 20.1713742 |

| | | | | | | |
|---|---|---|---|---|---|---|
| -2.4586632 | 0.01394554 | 0.02487291 | 114.756286 | 104.726372 | 71.3281477 | 70.5998095 |
| -2.4586107 | 0.01394758 | 0.02487436 | 98.3625312 | 94.4431619 | 0 | 28.5761134 |
| -2.4585492 | 0.01394997 | 0.02487725 | 0 | 85.2423954 | 75.9299637 | 94.1330794 |
| -2.4584479 | 0.0139539 | 0.02488289 | 11.4756286 | 14.3423713 | 10.354086 | 12.3269509 |
| -2.4582789 | 0.01396047 | 0.02489323 | 52.0501728 | 69.817581 | 0 | 79.5648647 |
| -2.4579792 | 0.01397213 | 0.02491264 | 18.4429746 | 30.3084073 | 5.75226998 | 11.7666349 |
| -2.4579425 | 0.01397356 | 0.02491381 | 61.8864259 | 56.2870421 | 0 | 82.9267604 |
| 2.45784971 | 0.01397717 | 0.02491887 | 0 | 7.30649104 | 0 | 6.72379138 |
| -2.4578097 | 0.01397872 | 0.02492027 | 9.42640924 | 10.2832096 | 21.8586259 | 12.8872668 |
| -2.4571864 | 0.014003 | 0.02496217 | 11.8854725 | 7.30649104 | 21.8586259 | 15.6888466 |
| -2.4571424 | 0.01400472 | 0.02496385 | 6.96734596 | 54.3927666 | 13.8054479 | 43.144328 |
| -2.4571214 | 0.01400554 | 0.02496394 | 80.7392443 | 75.7710182 | 80.5317797 | 0 |
| -2.456459 | 0.01403138 | 0.02500862 | 11.4756286 | 10.2832096 | 12.6549939 | 14.0078987 |
| -2.45612 | 0.01404463 | 0.02503085 | 5.73781432 | 16.7778683 | 18.4072639 | 17.9301104 |
| -2.4560308 | 0.01404811 | 0.02503568 | 8.1968776 | 24.3549701 | 18.4072639 | 8.96505518 |
| 2.45589493 | 0.01405342 | 0.02504376 | 0 | 0 | 0 | 0.56031595 |
| 2.45574137 | 0.01405943 | 0.0250517 | 35.2465737 | 18.6721438 | 19.5577179 | 16.8094785 |
| -2.4557446 | 0.0140593 | 0.0250517 | 91.8050291 | 295.506971 | 169.116737 | 225.247011 |
| -2.4555566 | 0.01406666 | 0.02506258 | 50.8206411 | 94.1725511 | 0 | 58.8331746 |
| -2.4555457 | 0.01406709 | 0.02506258 | 163.117864 | 394.279905 | 299.118039 | 323.302302 |
| -2.4554473 | 0.01407094 | 0.02506806 | 38.5253247 | 80.1007906 | 72.4786017 | 67.7982298 |
| -2.455354 | 0.01407459 | 0.0250718 | 5.73781432 | 12.7187066 | 18.4072639 | 27.4554815 |
| -2.4553577 | 0.01407445 | 0.0250718 | 5.73781432 | 19.2133653 | 23.0090799 | 13.4475828 |
| -2.4549988 | 0.01408851 | 0.0250952 | 68.0340841 | 65.4878086 | 0 | 64.4363341 |
| -2.454357 | 0.01411368 | 0.02513865 | 93.4444046 | 44.6507786 | 0 | 66.6775979 |
| -2.4542248 | 0.01411887 | 0.02514651 | 8.1968776 | 10.0125988 | 31.0622579 | 14.0078987 |
| -2.4541383 | 0.01412227 | 0.02515117 | 73.7718984 | 355.853175 | 0 | 359.722839 |
| -2.4539663 | 0.01412902 | 0.02516181 | 0 | 38.6973414 | 134.603117 | 99.7362389 |
| 2.45347243 | 0.01414844 | 0.02519501 | 22.9512573 | 20.0251977 | 0 | 17.9301104 |
| -2.4533787 | 0.01415212 | 0.02520018 | 14.7543797 | 18.401533 | 6.90272397 | 14.0078987 |
| -2.453042 | 0.01416538 | 0.02522239 | 2.0492194 | 36.803066 | 23.0090799 | 28.0157974 |
| -2.4528033 | 0.01417478 | 0.02523774 | 13.1150042 | 27.6022995 | 12.6549939 | 7.28410733 |
| -2.4527422 | 0.01417719 | 0.02524064 | 32.3776665 | 3.78855091 | 17.2568099 | 20.7316901 |
| -2.4526252 | 0.0141818 | 0.02524746 | 6.1476582 | 16.2366467 | 16.1063559 | 18.4904263 |
| -2.4525135 | 0.0141862 | 0.0252539 | 16.8035991 | 17.8603114 | 8.05317797 | 10.646003 |
| -2.4520827 | 0.0142032 | 0.02528146 | 6.55750208 | 31.3908504 | 9.20363196 | 20.7316901 |
| 2.45204171 | 0.01420482 | 0.02528146 | 0 | 0 | 0 | 0 |
| 2.45204171 | 0.01420482 | 0.02528146 | 0 | 0 | 0 | 0 |
| 2.45204171 | 0.01420482 | 0.02528146 | 0 | 0 | 0 | 0 |
| -2.4518397 | 0.01421279 | 0.02529426 | 90.1656536 | 69.817581 | 0 | 43.704644 |
| -2.45129 | 0.01423452 | 0.02533153 | 435.664044 | 811.291116 | 717.883293 | 747.461475 |
| -2.4511843 | 0.0142387 | 0.02533757 | 36.8859492 | 62.51109 | 0 | 113.744138 |
| -2.4503519 | 0.01427167 | 0.02539344 | 222.955071 | 0 | 32.2127119 | 71.1601255 |

| | | | | | | |
|---|---|---|---|---|---|---|
| -2.4503654 | 0.01427113 | 0.02539344 | 322.547134 | 31.1202396 | 17.2568099 | 0 |
| -2.4503007 | 0.0142737 | 0.02539565 | 9.01656536 | 11.6362635 | 14.9559019 | 14.0078987 |
| 2.45017828 | 0.01427855 | 0.02540288 | 18.0331307 | 9.74198805 | 0 | 0 |
| -2.4501382 | 0.01428014 | 0.02540431 | 27.8693838 | 63.8641439 | 0 | 135.59646 |
| -2.4500659 | 0.01428301 | 0.02540802 | 164.75724 | 161.013414 | 141.505841 | 152.966254 |
| -2.4499048 | 0.0142894 | 0.02541798 | 78.2801811 | 11.3656527 | 6.90272397 | 7.84442328 |
| -2.4497989 | 0.0142936 | 0.02542406 | 10.6559409 | 7.57710182 | 35.6640739 | 13.4475828 |
| -2.4496832 | 0.0142982 | 0.02543083 | 25.8201644 | 10.5538204 | 8.05317797 | 14.0078987 |
| -2.4493552 | 0.01431122 | 0.0254526 | 27.45954 | 13.2599282 | 6.90272397 | 12.8872668 |
| -2.4492718 | 0.01431454 | 0.02545653 | 8.1968776 | 18.6721438 | 12.6549939 | 12.8872668 |
| -2.4492599 | 0.01431501 | 0.02545653 | 51.230485 | 54.6633774 | 0 | 96.9346591 |
| 2.44919613 | 0.01431755 | 0.02545964 | 9.01656536 | 4.05916169 | 5.75226998 | 6.16347543 |
| -2.4491125 | 0.01432087 | 0.02546415 | 10.6559409 | 15.1542036 | 11.50454 | 11.206319 |
| -2.4490179 | 0.01432463 | 0.02546943 | 5.32797044 | 46.0038325 | 13.8054479 | 13.4475828 |
| -2.4489675 | 0.01432664 | 0.02547159 | 4.0984388 | 15.6954252 | 29.9118039 | 21.292006 |
| -2.4487644 | 0.01433472 | 0.02548456 | 120.494101 | 549.069271 | 811.070067 | 682.464825 |
| -2.4485055 | 0.01434503 | 0.02550148 | 10.246097 | 13.530539 | 9.20363196 | 19.0507423 |
| -2.448443 | 0.01434751 | 0.0255045 | 0 | 104.996982 | 64.4254237 | 84.0473923 |
| 2.44819228 | 0.0143575 | 0.02552085 | 1.22953164 | 0 | 0 | 0.56031595 |
| -2.4480894 | 0.0143616 | 0.02552673 | 13.1150042 | 16.5072575 | 12.6549939 | 8.40473923 |
| 2.44806391 | 0.01436262 | 0.02552713 | 0 | 0 | 0 | 0 |
| -2.4479759 | 0.01436613 | 0.02553196 | 0 | 91.7370541 | 167.966283 | 28.5761134 |
| -2.4477834 | 0.0143738 | 0.0255442 | 7.78703372 | 35.9912336 | 13.8054479 | 10.0856871 |
| -2.4476658 | 0.0143785 | 0.02555113 | 6.96734596 | 16.5072575 | 14.9559019 | 15.1285306 |
| -2.4474051 | 0.0143889 | 0.02556822 | 225.414134 | 823.468601 | 493.544764 | 570.961952 |
| -2.4472238 | 0.01439614 | 0.02557967 | 22.1315695 | 14.8835929 | 20.7081719 | 5.04284354 |
| -2.4470966 | 0.01440122 | 0.0255873 | 6.55750208 | 9.74198805 | 16.1063559 | 42.0236961 |
| -2.4467254 | 0.01441606 | 0.02561225 | 9.83625312 | 17.3190899 | 9.20363196 | 15.1285306 |
| 2.44621991 | 0.01443629 | 0.02564678 | 4.0984388 | 2.70610779 | 0 | 4.48252759 |
| -2.445589 | 0.01446158 | 0.02569028 | 4.91812656 | 28.9553534 | 35.6640739 | 40.3427483 |
| -2.4454354 | 0.01446774 | 0.02569981 | 0 | 111.762252 | 58.6731538 | 84.0473923 |
| 2.44519335 | 0.01447745 | 0.02571565 | 22.9512573 | 18.6721438 | 13.8054479 | 16.2491625 |
| -2.444954 | 0.01448706 | 0.02573131 | 6.1476582 | 12.9893174 | 26.4604419 | 15.6888466 |
| 2.44465387 | 0.01449912 | 0.02575009 | 96.7231557 | 77.6652936 | 55.2217918 | 59.9538065 |
| -2.4446511 | 0.01449923 | 0.02575009 | 4.50828268 | 11.6362635 | 28.7613499 | 27.4554815 |
| -2.4444786 | 0.01450617 | 0.025761 | 62.7061136 | 59.5343714 | 0 | 72.8410733 |
| -2.4444415 | 0.01450766 | 0.02576223 | 14.3445358 | 19.4839761 | 5.75226998 | 17.9301104 |
| -2.4441517 | 0.01451932 | 0.02578152 | 41.8040758 | 12.1774851 | 10.354086 | 7.84442328 |
| -2.4440197 | 0.01452463 | 0.02578953 | 7.37718984 | 27.8729103 | 8.05317797 | 21.852322 |
| -2.4436559 | 0.01453929 | 0.02581413 | 52.4600166 | 72.253078 | 0 | 71.1601255 |
| -2.4433759 | 0.01455057 | 0.02583275 | 9.01656536 | 19.2133653 | 8.05317797 | 20.7316901 |
| -2.4432358 | 0.01455622 | 0.02584136 | 54.9190799 | 60.6168145 | 0 | 80.6854966 |
| -2.4431649 | 0.01455908 | 0.02584501 | 60.6568942 | 102.832096 | 0 | 41.4633802 |

| | | | | | | |
|---|---|---|---|---|---|---|
| -2.4427641 | 0.01457526 | 0.02587231 | 39.7548564 | 67.6526948 | 0 | 96.3743432 |
| -2.442599 | 0.01458193 | 0.02588272 | 32.3776665 | 15.4248144 | 20.7081719 | 25.2142177 |
| -2.4423853 | 0.01459056 | 0.02589663 | 4.50828268 | 16.5072575 | 24.1595339 | 20.1713742 |
| 2.44232795 | 0.01459288 | 0.02589932 | 0 | 0 | 0 | 0 |
| -2.4421692 | 0.0145993 | 0.02590929 | 81.1490882 | 151.812647 | 167.966283 | 168.094785 |
| -2.4419129 | 0.01460967 | 0.02592484 | 82.7884637 | 37.6148983 | 0 | 82.9267604 |
| 2.44191538 | 0.01460957 | 0.02592484 | 315.579788 | 402.127618 | 385.402088 | 337.870517 |
| -2.4416953 | 0.01461848 | 0.02593905 | 6.55750208 | 35.1794013 | 9.20363196 | 18.4904263 |
| -2.4415964 | 0.01462248 | 0.02594472 | 150.822548 | 39.238563 | 0 | 38.1014845 |
| -2.4414568 | 0.01462814 | 0.02595333 | 41.3942319 | 61.1580361 | 0 | 101.417187 |
| 2.44069872 | 0.01465888 | 0.02600644 | 31.9678226 | 22.1900839 | 19.5577179 | 14.5682147 |
| 2.44019495 | 0.01467934 | 0.02604131 | 281.562746 | 326.3566 | 314.073941 | 305.932508 |
| -2.440001 | 0.01468722 | 0.02605386 | 16.8035991 | 15.966036 | 16.1063559 | 20.1713742 |
| -2.4397787 | 0.01469626 | 0.02606847 | 6.96734596 | 19.4839761 | 17.2568099 | 11.7666349 |
| -2.439738 | 0.01469792 | 0.02606997 | 16.3937552 | 17.0484791 | 8.05317797 | 10.646003 |
| -2.4396174 | 0.01470282 | 0.02607724 | 14.7543797 | 11.6362635 | 10.354086 | 11.206319 |
| -2.4395652 | 0.01470495 | 0.02607958 | 84.8376831 | 7.30649104 | 8.05317797 | 8.96505518 |
| 2.4395173 | 0.0147069 | 0.02607997 | 0.40984388 | 0 | 0 | 0 |
| -2.4395001 | 0.0147076 | 0.02607997 | 20.9020379 | 42.7565031 | 0 | 198.351846 |
| 2.4395173 | 0.0147069 | 0.02607997 | 0.40984388 | 0 | 0 | 0 |
| -2.4394669 | 0.01470895 | 0.02608095 | 174.183649 | 15.6954252 | 0 | 67.2379138 |
| -2.4394367 | 0.01471018 | 0.02608169 | 9.83625312 | 126.375234 | 0 | 134.475828 |
| -2.4393816 | 0.01471242 | 0.02608423 | 54.9190799 | 14.6129821 | 3.45136199 | 16.2491625 |
| -2.4389478 | 0.0147301 | 0.02611414 | 10.246097 | 9.47137727 | 12.6549939 | 19.6110582 |
| 2.43885094 | 0.01473404 | 0.0261197 | 0 | 0.27061078 | 0 | 0 |
| 2.43859517 | 0.01474448 | 0.02613676 | 6.96734596 | 9.20076649 | 11.50454 | 14.5682147 |
| -2.4383229 | 0.01475559 | 0.02615501 | 6.96734596 | 23.8137486 | 10.354086 | 17.9301104 |
| 2.43813726 | 0.01476317 | 0.02616702 | 0 | 1.08244312 | 2.30090799 | 0.56031595 |
| -2.4380844 | 0.01476532 | 0.0261694 | 48.3615778 | 114.73897 | 75.9299637 | 0 |
| -2.4378032 | 0.01477682 | 0.02618834 | 0 | 88.4897248 | 109.29313 | 54.350647 |
| -2.43775 | 0.01477899 | 0.02619075 | 71.722679 | 43.2977247 | 0 | 82.9267604 |
| 2.43762453 | 0.01478412 | 0.02619697 | 0.40984388 | 0 | 0 | 0 |
| -2.4376305 | 0.01478388 | 0.02619697 | 99.1822189 | 31.3908504 | 0 | 78.4442328 |
| -2.4374924 | 0.01478952 | 0.0262051 | 8.60672148 | 9.74198805 | 16.1063559 | 18.4904263 |
| -2.4374131 | 0.01479277 | 0.02620942 | 11.0657848 | 17.5897006 | 9.20363196 | 12.3269509 |
| -2.4373292 | 0.0147962 | 0.02621406 | 14.7543797 | 27.8729103 | 8.05317797 | 9.52537113 |
| -2.4367605 | 0.01481949 | 0.02625388 | 27.45954 | 2.97671857 | 8.05317797 | 67.7982298 |
| -2.4365415 | 0.01482847 | 0.02626834 | 0 | 103.914539 | 87.4345036 | 58.2728587 |
| -2.436503 | 0.01483005 | 0.0262697 | 0 | 90.654611 | 81.6822337 | 73.4013893 |
| -2.4361823 | 0.0148432 | 0.02629156 | 34.017042 | 76.0416289 | 0 | 95.2537113 |
| -2.4361158 | 0.01484593 | 0.02629495 | 0 | 63.3229223 | 56.3722458 | 138.958355 |
| -2.436014 | 0.01485011 | 0.0263009 | 198.774282 | 200.793198 | 211.683535 | 201.153426 |
| -2.4358489 | 0.01485689 | 0.02631147 | 46.3123584 | 76.8534613 | 0 | 72.2807574 |

| | | | | | | |
|---|---|---|---|---|---|---|
| -2.4356912 | 0.01486337 | 0.0263215 | 11.0657848 | 10.0125988 | 10.354086 | 22.4126379 |
| -2.4355078 | 0.0148709 | 0.0263334 | 71.722679 | 55.7458205 | 0 | 64.99665 |
| -2.435294 | 0.0148797 | 0.02634752 | 5.73781432 | 14.8835929 | 24.1595339 | 15.1285306 |
| -2.4352537 | 0.01488135 | 0.02634901 | 84.8376831 | 113.115306 | 42.5667978 | 0 |
| -2.434965 | 0.01489323 | 0.02636859 | 57.3781432 | 69.0057487 | 0 | 65.556966 |
| -2.4348904 | 0.0148963 | 0.02637258 | 5.73781432 | 9.20076649 | 36.8145279 | 20.7316901 |
| 2.4347724 | 0.01490116 | 0.02637974 | 8.60672148 | 6.76526948 | 2.30090799 | 4.48252759 |
| 2.43470095 | 0.0149041 | 0.02638249 | 2.86890716 | 2.43549701 | 3.45136199 | 4.48252759 |
| -2.4346951 | 0.01490434 | 0.02638249 | 47.5418901 | 71.1706349 | 49.4695218 | 62.1950703 |
| -2.4345385 | 0.01491079 | 0.02639246 | 58.6076748 | 122.857294 | 95.4876816 | 99.7362389 |
| 2.43447956 | 0.01491322 | 0.02639531 | 0 | 0 | 0 | 0 |
| 2.43439743 | 0.01491661 | 0.02639985 | 33.6071982 | 20.0251977 | 17.2568099 | 17.9301104 |
| -2.434358 | 0.01491823 | 0.02640128 | 347.137766 | 1247.7863 | 1115.94038 | 1148.64769 |
| -2.4341286 | 0.01492769 | 0.02641657 | 10.6559409 | 14.6129821 | 12.6549939 | 10.0856871 |
| -2.4340312 | 0.01493171 | 0.02642223 | 9.42640924 | 56.5576528 | 31.0622579 | 30.8173772 |
| -2.4339368 | 0.0149356 | 0.02642768 | 1.22953164 | 32.7439043 | 31.0622579 | 35.2999048 |
| 2.43377665 | 0.01494221 | 0.02643792 | 2.0492194 | 3.51794013 | 3.45136199 | 0 |
| -2.4337357 | 0.0149439 | 0.02643946 | 4.0984388 | 15.1542036 | 33.3631659 | 19.0507423 |
| -2.4335261 | 0.01495256 | 0.02645333 | 0 | 123.127905 | 24.1595339 | 135.59646 |
| -2.4326858 | 0.0149873 | 0.02651334 | 20.9020379 | 33.8263474 | 17.2568099 | 20.1713742 |
| -2.4326305 | 0.01498959 | 0.02651448 | 6.1476582 | 17.3190899 | 12.6549939 | 21.852322 |
| 2.43264515 | 0.01498898 | 0.02651448 | 0 | 0 | 0 | 0 |
| -2.4325426 | 0.01499323 | 0.02651947 | 99.1822189 | 53.0397127 | 0 | 47.0665397 |
| -2.4325098 | 0.01499458 | 0.02652041 | 6.55750208 | 53.851545 | 14.9559019 | 8.40473923 |
| -2.432451 | 0.01499702 | 0.02652327 | 22.1315695 | 11.3656527 | 5.75226998 | 21.292006 |
| -2.4324183 | 0.01499837 | 0.02652421 | 80.7392443 | 46.545054 | 0 | 67.2379138 |
| -2.4322605 | 0.01500491 | 0.02653432 | 10.246097 | 31.1202396 | 4.60181598 | 26.3348496 |
| -2.4319073 | 0.01501955 | 0.02655875 | 6.55750208 | 22.1900839 | 8.05317797 | 31.9380091 |
| 2.43166096 | 0.01502977 | 0.02657536 | 6.55750208 | 4.60038325 | 6.90272397 | 3.36189569 |
| -2.4316113 | 0.01503183 | 0.02657755 | 0 | 98.7729344 | 118.496762 | 40.9030642 |
| -2.4312513 | 0.01504677 | 0.02660106 | 200.823501 | 517.40781 | 437.172518 | 552.471525 |
| -2.4312677 | 0.01504609 | 0.02660106 | 4.91812656 | 33.8263474 | 8.05317797 | 31.3776931 |
| -2.431226 | 0.01504783 | 0.02660146 | 52.4600166 | 47.8981079 | 0 | 98.055291 |
| -2.4310937 | 0.01505332 | 0.02660973 | 72.5423667 | 0 | 205.931265 | 31.9380091 |
| -2.43076 | 0.01506719 | 0.02663278 | 13.1150042 | 16.2366467 | 5.75226998 | 24.0935858 |
| -2.4306505 | 0.01507175 | 0.02663938 | 14.7543797 | 7.57710182 | 9.20363196 | 35.2999048 |
| 2.43057344 | 0.01507495 | 0.02664358 | 176.232868 | 84.7011739 | 130.001301 | 93.5727634 |
| 2.43024936 | 0.01508844 | 0.02666304 | 4.50828268 | 0.81183234 | 1.150454 | 1.1206319 |
| 2.43027958 | 0.01508718 | 0.02666304 | 0.81968776 | 5.68282636 | 9.20363196 | 7.84442328 |
| -2.430268 | 0.01508766 | 0.02666304 | 43.8532952 | 17.0484791 | 13.8054479 | 38.6618005 |
| -2.4302014 | 0.01509044 | 0.02666511 | 70.0833035 | 9.74198805 | 4.60181598 | 14.0078987 |
| 2.4299571 | 0.01510061 | 0.02668163 | 0 | 0 | 0 | 0 |
| -2.4298381 | 0.01510557 | 0.02668893 | 11.8854725 | 10.0125988 | 21.8586259 | 9.52537113 |

| | | | | | | |
|---|---|---|---|---|---|---|
| 2.42968341 | 0.01511202 | 0.02669886 | 0.81968776 | 2.16488623 | 2.30090799 | 0 |
| -2.4292972 | 0.01512812 | 0.02672586 | 7.37718984 | 18.9427545 | 9.20363196 | 22.9729539 |
| 2.42913459 | 0.01513491 | 0.02673639 | 15.5740674 | 8.93015571 | 6.90272397 | 11.7666349 |
| -2.4290589 | 0.01513807 | 0.02674051 | 29.0989155 | 55.2045989 | 0 | 135.036144 |
| -2.4285731 | 0.01515837 | 0.02677489 | 14.3445358 | 114.197749 | 0 | 113.744138 |
| -2.4282866 | 0.01517035 | 0.02679459 | 13.524848 | 22.1900839 | 8.05317797 | 10.646003 |
| -2.4279202 | 0.01518568 | 0.0268202 | 38.1154808 | 83.34812 | 0 | 76.202969 |
| 2.42768896 | 0.01519537 | 0.02683584 | 62.7061136 | 46.8156648 | 42.5667978 | 42.0236961 |
| -2.4273861 | 0.01520806 | 0.02685679 | 11.8854725 | 13.2599282 | 12.6549939 | 9.52537113 |
| -2.4273137 | 0.0152111 | 0.02686068 | 6.1476582 | 14.0717605 | 17.2568099 | 17.9301104 |
| -2.4270613 | 0.01522169 | 0.02687791 | 20.0823501 | 13.8011497 | 10.354086 | 8.40473923 |
| -2.4268075 | 0.01523234 | 0.02689524 | 136.887856 | 299.566133 | 302.569401 | 299.769033 |
| -2.4266813 | 0.01523764 | 0.02690313 | 7.37718984 | 13.530539 | 18.4072639 | 12.8872668 |
| -2.4265904 | 0.01524145 | 0.0269084 | 36.4761053 | 55.2045989 | 0 | 114.304454 |
| -2.4262618 | 0.01525526 | 0.0269313 | 4.91812656 | 13.8011497 | 20.7081719 | 23.5332698 |
| -2.4256062 | 0.01528284 | 0.02697852 | 14.7543797 | 11.6362635 | 8.05317797 | 15.1285306 |
| -2.4253073 | 0.01529543 | 0.02699927 | 30.3284471 | 40.3210061 | 0 | 160.250361 |
| 2.42516558 | 0.01530141 | 0.02700834 | 0 | 5.95343714 | 0 | 0 |
| -2.4251094 | 0.01530377 | 0.02701104 | 12.7051603 | 19.2133653 | 21.8586259 | 5.60315949 |
| -2.4248919 | 0.01531295 | 0.02702575 | 71.722679 | 57.3694852 | 0 | 59.9538065 |
| -2.4248508 | 0.01531468 | 0.02702733 | 13.9346919 | 16.2366467 | 18.4072639 | 21.292006 |
| -2.4246151 | 0.01532462 | 0.0270434 | 10.6559409 | 110.679809 | 0 | 136.717091 |
| -2.4241694 | 0.01534345 | 0.02707514 | 8.1968776 | 15.966036 | 9.20363196 | 21.852322 |
| -2.4238383 | 0.01535744 | 0.02709836 | 8.60672148 | 21.3782516 | 8.05317797 | 18.4904263 |
| -2.4237651 | 0.01536054 | 0.02710234 | 56.9682993 | 76.5828505 | 0 | 56.0315949 |
| 2.42367602 | 0.01536431 | 0.0271075 | 2.45906328 | 0.81183234 | 2.30090799 | 2.80157974 |
| -2.4236223 | 0.01536658 | 0.02711003 | 69.2636157 | 0 | 75.9299637 | 121.028245 |
| -2.4230047 | 0.01539273 | 0.02715468 | 2.0492194 | 33.0145151 | 26.4604419 | 23.5332698 |
| 2.42283474 | 0.01539993 | 0.0271659 | 0 | 0 | 0 | 0 |
| 2.42256132 | 0.01541152 | 0.02718487 | 13.1150042 | 15.1542036 | 18.4072639 | 10.646003 |
| -2.42246 | 0.01541582 | 0.02719096 | 2.86890716 | 28.6847426 | 34.5136199 | 14.5682147 |
| -2.4223179 | 0.01542185 | 0.02720012 | 8.60672148 | 37.6148983 | 39.1154358 | 35.2999048 |
| -2.4221814 | 0.01542765 | 0.02720885 | 0 | 75.2297966 | 85.1335957 | 79.0045488 |
| -2.4218185 | 0.01544306 | 0.02723396 | 32.7875104 | 57.0988744 | 31.0622579 | 114.864769 |
| -2.4218065 | 0.01544357 | 0.02723396 | 89.7558097 | 88.4897248 | 0 | 28.0157974 |
| -2.4216882 | 0.0154486 | 0.02724134 | 225.823978 | 183.744719 | 208.232173 | 119.907613 |
| -2.4213043 | 0.01546492 | 0.02726864 | 8.60672148 | 21.6488623 | 5.75226998 | 34.1792729 |
| -2.4212598 | 0.01546682 | 0.02727049 | 3.27875104 | 44.9213893 | 9.20363196 | 31.3776931 |
| 2.42118643 | 0.01546994 | 0.0272745 | 73.3620545 | 39.5091738 | 78.2308717 | 70.5998095 |
| -2.4209612 | 0.01547953 | 0.02728992 | 46.3123584 | 78.7477367 | 0 | 65.556966 |
| -2.4208434 | 0.01548455 | 0.02729727 | 6.1476582 | 17.3190899 | 23.0090799 | 11.7666349 |
| -2.4206884 | 0.01549115 | 0.02730742 | 503.698128 | 959.856434 | 756.998729 | 811.89781 |
| -2.4205637 | 0.01549646 | 0.0273153 | 57.3781432 | 55.7458205 | 119.647216 | 0 |

| | | | | | | |
|---|---|---|---|---|---|---|
| 2.42050143 | 0.01549912 | 0.02731849 | 0.40984388 | 1.3530539 | 0 | 1.1206319 |
| -2.4204813 | 0.01549998 | 0.02731851 | 11.4756286 | 17.8603114 | 12.6549939 | 8.40473923 |
| 2.42043387 | 0.015502 | 0.02732024 | 5.32797044 | 2.43549701 | 4.60181598 | 4.48252759 |
| -2.4204186 | 0.01550265 | 0.02732024 | 37.705637 | 24.3549701 | 6.90272397 | 6.16347543 |
| -2.4202865 | 0.01550828 | 0.02732867 | 2.86890716 | 26.2492456 | 18.4072639 | 28.5761134 |
| -2.4201768 | 0.01551296 | 0.02733543 | 624.602073 | 548.257439 | 1271.25166 | 413.51317 |
| -2.4198145 | 0.01552843 | 0.02736118 | 6.1476582 | 16.2366467 | 13.8054479 | 19.0507423 |
| 2.41970187 | 0.01553324 | 0.02736667 | 5.73781432 | 6.4946587 | 4.60181598 | 5.04284354 |
| -2.4197061 | 0.01553305 | 0.02736667 | 2.86890716 | 24.0843593 | 26.4604419 | 21.292006 |
| -2.4190177 | 0.01556248 | 0.0274167 | 4.50828268 | 19.4839761 | 18.4072639 | 19.6110582 |
| -2.4189482 | 0.01556546 | 0.02742044 | 45.9025146 | 63.3229223 | 0 | 81.2458125 |
| -2.4188914 | 0.01556789 | 0.02742323 | 320.08807 | 485.746349 | 410.712076 | 424.719489 |
| 2.41854008 | 0.01558293 | 0.02744823 | 0.81968776 | 1.08244312 | 0 | 1.68094785 |
| -2.4185185 | 0.01558385 | 0.02744836 | 4.0984388 | 26.5198564 | 18.4072639 | 17.3697944 |
| 2.41845801 | 0.01558644 | 0.02745143 | 0 | 0 | 0 | 10.646003 |
| -2.4182727 | 0.01559439 | 0.02746392 | 18.8528185 | 8.11832337 | 11.50454 | 12.8872668 |
| 2.41803236 | 0.01560469 | 0.02748056 | 0.40984388 | 0.54122156 | 1.150454 | 0 |
| -2.4179934 | 0.01560636 | 0.027482 | 115.16613 | 0 | 58.6731538 | 90.7711837 |
| -2.4179726 | 0.01560725 | 0.02748208 | 8.1968776 | 16.7778683 | 11.50454 | 13.4475828 |
| 2.41748054 | 0.01562837 | 0.02751626 | 0.81968776 | 0.81183234 | 2.30090799 | 0.56031595 |
| -2.4174896 | 0.01562798 | 0.02751626 | 34.017042 | 120.151186 | 0 | 52.6696992 |
| 2.41744592 | 0.01562986 | 0.02751738 | 75.00143 | 92.8194972 | 131.151755 | 112.06319 |
| 2.41728512 | 0.01563676 | 0.02752653 | 2.0492194 | 0 | 0 | 0.56031595 |
| -2.4173014 | 0.01563607 | 0.02752653 | 11.8854725 | 14.0717605 | 12.6549939 | 8.96505518 |
| -2.417105 | 0.0156445 | 0.02753866 | 0 | 92.8194972 | 72.4786017 | 72.8410733 |
| 2.4166906 | 0.01566232 | 0.02756702 | 0 | 8.65954493 | 0 | 0 |
| -2.4167008 | 0.01566189 | 0.02756702 | 4.91812656 | 24.3549701 | 16.1063559 | 16.2491625 |
| 2.41648587 | 0.01567113 | 0.02758103 | 0 | 9.74198805 | 0 | 2.24126379 |
| -2.4162982 | 0.01567922 | 0.02759375 | 1.63937552 | 31.1202396 | 21.8586259 | 36.9808526 |
| -2.4161235 | 0.01568674 | 0.02760548 | 6.55750208 | 18.401533 | 19.5577179 | 11.206319 |
| -2.4159953 | 0.01569226 | 0.02761369 | 4.91812656 | 19.4839761 | 17.2568099 | 17.9301104 |
| -2.4157327 | 0.01570358 | 0.02763211 | 6.55750208 | 14.3423713 | 18.4072639 | 14.0078987 |
| -2.4155818 | 0.01571009 | 0.02764206 | 146.314265 | 34.6381797 | 0 | 38.6618005 |
| -2.4152892 | 0.01572272 | 0.02766277 | 33.6071982 | 185.909605 | 46.0181598 | 0 |
| -2.4152252 | 0.01572548 | 0.02766612 | 10.246097 | 16.7778683 | 11.50454 | 10.0856871 |
| 2.41514216 | 0.01572907 | 0.02766992 | 24.5906328 | 17.0484791 | 26.4604419 | 16.8094785 |
| -2.4151356 | 0.01572935 | 0.02766992 | 6.55750208 | 15.6954252 | 13.8054479 | 16.8094785 |
| 2.41433586 | 0.01576392 | 0.02772922 | 15.1642236 | 19.4839761 | 13.8054479 | 10.646003 |
| 2.41382391 | 0.01578609 | 0.0277667 | 1.63937552 | 1.62366467 | 0 | 0 |
| -2.4136864 | 0.01579205 | 0.02777566 | 20.0823501 | 16.5072575 | 16.1063559 | 15.1285306 |
| -2.4131848 | 0.0158138 | 0.02781241 | 0 | 46.2744432 | 155.311289 | 56.5919108 |
| -2.413051 | 0.0158196 | 0.0278211 | 3.27875104 | 19.4839761 | 23.0090799 | 24.6539017 |
| -2.4129313 | 0.0158248 | 0.0278262 | 7.37718984 | 14.6129821 | 29.9118039 | 9.52537113 |

| | | | | | | |
|---|---|---|---|---|---|---|
| 2.41289274 | 0.01582648 | 0.0278262 | 0 | 0 | 0 | 0 |
| -2.412885 | 0.01582681 | 0.0278262 | 64.3454891 | 50.6042157 | 0 | 71.1601255 |
| 2.41289274 | 0.01582648 | 0.0278262 | 0 | 0 | 0 | 0 |
| -2.4129366 | 0.01582457 | 0.0278262 | 6.55750208 | 12.7187066 | 13.8054479 | 23.5332698 |
| -2.4128563 | 0.01582806 | 0.02782687 | 4.91812656 | 22.7313054 | 20.7081719 | 13.4475828 |
| -2.4127659 | 0.01583199 | 0.02783226 | 10.246097 | 11.9068743 | 13.8054479 | 10.646003 |
| -2.4125608 | 0.01584089 | 0.02784641 | 16.3937552 | 12.9893174 | 5.75226998 | 21.292006 |
| -2.4125261 | 0.0158424 | 0.02784754 | 5.32797044 | 28.4141318 | 12.6549939 | 16.8094785 |
| 2.41248138 | 0.01584435 | 0.02784944 | 0 | 4.60038325 | 0 | 0 |
| -2.4124383 | 0.01584622 | 0.02785122 | 9.83625312 | 37.0736767 | 3.45136199 | 30.8173772 |
| -2.4122637 | 0.01585381 | 0.02786304 | 372.138243 | 177.791282 | 211.683535 | 244.297754 |
| -2.411913 | 0.01586907 | 0.02788834 | 6.1476582 | 12.1774851 | 14.9559019 | 26.3348496 |
| -2.4117526 | 0.01587605 | 0.02789909 | 28.6890716 | 88.4897248 | 0 | 84.0473923 |
| -2.4112462 | 0.01589811 | 0.02793635 | 4.91812656 | 27.6022995 | 12.6549939 | 19.0507423 |
| -2.4111363 | 0.0159029 | 0.02794324 | 130.740198 | 21.1076408 | 0 | 68.3585457 |
| -2.4104896 | 0.01593112 | 0.0279913 | 4.50828268 | 21.6488623 | 13.8054479 | 24.0935858 |
| -2.4102174 | 0.01594302 | 0.02801068 | 1.63937552 | 33.0145151 | 32.2127119 | 22.9729539 |
| -2.4098118 | 0.01596075 | 0.0280403 | 55.3289238 | 46.0038325 | 0 | 87.9696039 |
| -2.4096248 | 0.01596893 | 0.02805315 | 4.0984388 | 33.2851258 | 11.50454 | 23.5332698 |
| -2.4094003 | 0.01597876 | 0.02806889 | 138.117388 | 55.4752097 | 0 | 24.6539017 |
| -2.4093367 | 0.01598155 | 0.02807226 | 45.9025146 | 92.0076649 | 87.4345036 | 0 |
| -2.4092764 | 0.01598419 | 0.02807537 | 23.770945 | 36.2618444 | 3.45136199 | 12.3269509 |
| -2.4092339 | 0.01598605 | 0.02807711 | 9.01656536 | 12.7187066 | 10.354086 | 16.8094785 |
| -2.4091759 | 0.01598859 | 0.02808005 | 27.8693838 | 32.4732935 | 24.1595339 | 6.72379138 |
| -2.4090099 | 0.01599586 | 0.02809129 | 52.8698605 | 268.175282 | 120.79767 | 160.250361 |
| 2.40840563 | 0.01602237 | 0.02813478 | 1.22953164 | 5.68282636 | 2.30090799 | 3.36189569 |
| 2.40842223 | 0.01602164 | 0.02813478 | 11.8854725 | 0 | 0 | 0 |
| -2.4081182 | 0.01603499 | 0.02815541 | 68.4439279 | 48.7099402 | 0 | 67.7982298 |
| -2.4079895 | 0.01604064 | 0.02816381 | 21.7217256 | 99.3141559 | 134.603117 | 0 |
| -2.4073214 | 0.01607002 | 0.02821385 | 100.821594 | 135.846611 | 159.913105 | 142.320251 |
| -2.4071051 | 0.01607954 | 0.02822903 | 63.5258014 | 74.1473535 | 0 | 47.6268556 |
| 2.40702612 | 0.01608302 | 0.0282336 | 0.40984388 | 2.97671857 | 2.30090799 | 2.24126379 |
| -2.4065118 | 0.01610568 | 0.02827185 | 13.524848 | 6.22404792 | 10.354086 | 42.0236961 |
| -2.4063937 | 0.01611089 | 0.02827946 | 52.8698605 | 56.0164313 | 0 | 75.6426531 |
| -2.4063053 | 0.01611479 | 0.02828323 | 0 | 227.583665 | 219.736713 | 137.837723 |
| -2.406325 | 0.01611392 | 0.02828323 | 2.86890716 | 26.7904671 | 43.7172518 | 11.7666349 |
| -2.4056811 | 0.01614235 | 0.02833006 | 5.73781432 | 51.416048 | 18.4072639 | 7.28410733 |
| -2.4052009 | 0.01616358 | 0.02836577 | 31.5579788 | 106.891258 | 97.7885896 | 0 |
| -2.4050824 | 0.01616882 | 0.02837342 | 76.6408055 | 0 | 92.0363196 | 84.6077082 |
| -2.4049303 | 0.01617555 | 0.0283837 | 17.6232868 | 20.0251977 | 23.0090799 | 3.92221164 |
| -2.4047286 | 0.01618448 | 0.02839782 | 6.55750208 | 11.3656527 | 21.8586259 | 15.6888466 |
| 2.40456329 | 0.0161918 | 0.02840912 | 2.45906328 | 4.05916169 | 2.30090799 | 2.24126379 |
| -2.4045434 | 0.01619269 | 0.02840913 | 3.68859492 | 37.0736767 | 10.354086 | 26.8951655 |

| | | | | | | |
|---|---|---|---|---|---|---|
| -2.4044509 | 0.01619678 | 0.02841478 | 8.1968776 | 17.3190899 | 13.8054479 | 10.646003 |
| 2.40424331 | 0.01620598 | 0.02842676 | 8.1968776 | 4.87099402 | 3.45136199 | 1.68094785 |
| -2.4042511 | 0.01620564 | 0.02842676 | 17.213443 | 6.22404792 | 44.8677058 | 7.84442328 |
| -2.4042372 | 0.01620626 | 0.02842676 | 30.3284471 | 89.5721679 | 118.496762 | 0 |
| -2.4041296 | 0.01621103 | 0.02843358 | 11.8854725 | 12.1774851 | 13.8054479 | 8.96505518 |
| -2.4040442 | 0.01621481 | 0.02843868 | 29.5087594 | 190.509989 | 46.0181598 | 0 |
| -2.4039098 | 0.01622077 | 0.02844759 | 10.6559409 | 7.57710182 | 25.3099879 | 12.8872668 |
| -2.4037605 | 0.0162274 | 0.02845767 | 0 | 55.4752097 | 135.753571 | 54.910963 |
| -2.4036593 | 0.01623189 | 0.028464 | 5.73781432 | 12.9893174 | 19.5577179 | 17.9301104 |
| -2.40354 | 0.01623719 | 0.02847174 | 0 | 58.4519283 | 101.239952 | 75.6426531 |
| -2.4034641 | 0.01624056 | 0.02847611 | 72.1325229 | 9.20076649 | 8.05317797 | 7.28410733 |
| -2.4032829 | 0.01624861 | 0.02848867 | 75.4112739 | 182.121054 | 204.780811 | 186.024895 |
| 2.40295566 | 0.01626316 | 0.02851256 | 34.8367298 | 25.708024 | 31.0622579 | 15.1285306 |
| -2.4029367 | 0.016264 | 0.02851256 | 6.55750208 | 10.5538204 | 11.50454 | 47.0665397 |
| -2.4028586 | 0.01626748 | 0.02851711 | 7.78703372 | 18.6721438 | 12.6549939 | 11.7666349 |
| -2.4028208 | 0.01626916 | 0.02851713 | 73.7718984 | 172.649677 | 154.160835 | 145.682147 |
| 2.40281858 | 0.01626926 | 0.02851713 | 0 | 0 | 0 | 0 |
| -2.4026553 | 0.01627652 | 0.02852676 | 24.5906328 | 107.973701 | 0 | 73.4013893 |
| -2.4026708 | 0.01627583 | 0.02852676 | 111.887379 | 167.508072 | 172.568099 | 191.067738 |
| -2.4022563 | 0.01629429 | 0.02855635 | 34.4268859 | 69.817581 | 44.8677058 | 79.5648647 |
| -2.4015086 | 0.01632762 | 0.02861205 | 58.6076748 | 11.0950419 | 4.60181598 | 12.8872668 |
| -2.4015037 | 0.01632784 | 0.02861205 | 6.55750208 | 17.5897006 | 20.7081719 | 10.646003 |
| -2.4014135 | 0.01633187 | 0.02861719 | 35.2465737 | 47.3568864 | 0 | 118.786981 |
| -2.4013983 | 0.01633255 | 0.02861719 | 179.101776 | 72.253078 | 105.841768 | 76.202969 |
| -2.4013293 | 0.01633563 | 0.02862104 | 9.01656536 | 13.8011497 | 6.90272397 | 40.3427483 |
| 2.40127515 | 0.01633805 | 0.02862372 | 6.1476582 | 10.2832096 | 4.60181598 | 5.04284354 |
| -2.4010711 | 0.01634716 | 0.02863813 | 4.50828268 | 46.2744432 | 32.2127119 | 26.3348496 |
| 2.400825 | 0.01635816 | 0.02865584 | 4.50828268 | 7.8477126 | 8.05317797 | 6.72379138 |
| -2.4002651 | 0.0163832 | 0.02869815 | 6.55750208 | 12.9893174 | 23.0090799 | 12.8872668 |
| -2.3999292 | 0.01639824 | 0.02872295 | 7.78703372 | 14.6129821 | 12.6549939 | 13.4475828 |
| -2.3994899 | 0.01641793 | 0.02875431 | 48.3615778 | 77.1240721 | 43.7172518 | 86.2886561 |
| -2.3994919 | 0.01641784 | 0.02875431 | 91.3951852 | 250.585581 | 167.966283 | 208.997849 |
| -2.3994652 | 0.01641904 | 0.02875469 | 4.0984388 | 20.2958084 | 29.9118039 | 13.4475828 |
| -2.3993912 | 0.01642236 | 0.02875895 | 6.55750208 | 9.47137727 | 20.7081719 | 21.292006 |
| -2.3992535 | 0.01642854 | 0.02876821 | 2.0492194 | 20.83703 | 21.8586259 | 40.9030642 |
| -2.3991894 | 0.01643141 | 0.02877168 | 3.68859492 | 35.7206228 | 17.2568099 | 15.6888466 |
| -2.3990699 | 0.01643677 | 0.02877951 | 0 | 66.8408624 | 123.098578 | 50.4284354 |
| -2.399025 | 0.01643879 | 0.02878148 | 2.0492194 | 13.2599282 | 29.9118039 | 47.0665397 |
| 2.39889583 | 0.01644459 | 0.0287897 | 19.2626624 | 3.78855091 | 5.75226998 | 5.60315949 |
| 2.39888071 | 0.01644527 | 0.0287897 | 22.1315695 | 12.4480958 | 21.8586259 | 0 |
| -2.3985973 | 0.016458 | 0.02881043 | 14.7543797 | 14.6129821 | 6.90272397 | 13.4475828 |
| -2.3979728 | 0.01648609 | 0.02885803 | 86.0672148 | 65.2171978 | 64.4254237 | 0 |
| -2.3978683 | 0.01649079 | 0.0288647 | 46.7222023 | 61.4286469 | 117.346308 | 0 |

| | | | | | | |
|---|---|---|---|---|---|---|
| 2.39778256 | 0.01649465 | 0.02886989 | 28.6890716 | 14.3423713 | 23.0090799 | 15.6888466 |
| -2.3977202 | 0.01649746 | 0.02887324 | 5.73781432 | 20.0251977 | 18.4072639 | 12.3269509 |
| -2.3976224 | 0.01650187 | 0.02887939 | 73.3620545 | 74.1473535 | 0 | 38.6618005 |
| -2.3974715 | 0.01650867 | 0.02888972 | 0 | 68.1939163 | 113.894946 | 54.350647 |
| -2.3970287 | 0.01652863 | 0.02892309 | 26.6398522 | 95.525605 | 0 | 76.763285 |
| -2.3969315 | 0.01653301 | 0.02892919 | 149.593016 | 37.3442875 | 0 | 30.8173772 |
| 2.396834 | 0.01653741 | 0.02893532 | 26.6398522 | 8.11832337 | 14.9559019 | 11.7666349 |
| -2.3966589 | 0.01654532 | 0.02894758 | 18.8528185 | 74.9591858 | 0 | 122.709193 |
| -2.3964684 | 0.01655392 | 0.02896106 | 6.1476582 | 15.1542036 | 21.8586259 | 12.3269509 |
| -2.3964066 | 0.01655671 | 0.02896438 | 62.7061136 | 64.946587 | 0 | 52.6696992 |
| -2.3962667 | 0.01656304 | 0.02897387 | 0 | 29.7671857 | 159.913105 | 71.7204414 |
| -2.3957428 | 0.01658673 | 0.02901374 | 10.246097 | 11.3656527 | 12.6549939 | 11.206319 |
| 2.39569083 | 0.01658908 | 0.02901628 | 0 | 0 | 1.150454 | 0 |
| -2.3948762 | 0.01662598 | 0.02907925 | 0 | 69.5469702 | 80.5317797 | 78.4442328 |
| -2.3947599 | 0.01663125 | 0.02908591 | 0 | 40.5916169 | 104.691314 | 94.6933953 |
| -2.3947525 | 0.01663159 | 0.02908591 | 14.7543797 | 18.9427545 | 8.05317797 | 9.52537113 |
| -2.3946517 | 0.01663616 | 0.02909233 | 2.0492194 | 30.579018 | 19.5577179 | 30.2570612 |
| -2.394147 | 0.01665907 | 0.02913081 | 40.984388 | 60.8874253 | 0 | 82.9267604 |
| -2.3941005 | 0.01666118 | 0.02913293 | 6.55750208 | 20.83703 | 14.9559019 | 11.7666349 |
| -2.3940403 | 0.01666392 | 0.02913614 | 41.3942319 | 24.6255809 | 5.75226998 | 6.16347543 |
| 2.39398812 | 0.01666629 | 0.02913871 | 0 | 0.27061078 | 0 | 0 |
| -2.3934858 | 0.01668913 | 0.02917706 | 4.50828268 | 12.7187066 | 34.5136199 | 16.8094785 |
| -2.3933227 | 0.01669655 | 0.02918845 | 41.8040758 | 59.5343714 | 0 | 82.9267604 |
| 2.39260728 | 0.01672914 | 0.02924162 | 0 | 0 | 0 | 0 |
| -2.3925954 | 0.01672968 | 0.02924162 | 0 | 43.2977247 | 111.594038 | 82.9267604 |
| 2.39260728 | 0.01672914 | 0.02924162 | 0 | 0 | 0 | 0 |
| -2.3925543 | 0.01673155 | 0.02924172 | 26.2300083 | 147.753485 | 0 | 43.144328 |
| -2.3925676 | 0.01673094 | 0.02924172 | 0 | 69.817581 | 60.9740618 | 99.7362389 |
| -2.3923505 | 0.01674085 | 0.02925639 | 0 | 91.1958326 | 69.0272397 | 68.3585457 |
| 2.39194777 | 0.01675923 | 0.02928375 | 0 | 0 | 0 | 0 |
| 2.39194777 | 0.01675923 | 0.02928375 | 0 | 0 | 0 | 0 |
| 2.39194777 | 0.01675923 | 0.02928375 | 0 | 0 | 0 | 0 |
| -2.3917929 | 0.0167663 | 0.02929452 | 77.8703372 | 7.8477126 | 8.05317797 | 7.28410733 |
| 2.39175327 | 0.01676811 | 0.0292961 | 0 | 8.38893415 | 5.75226998 | 0 |
| 2.39159127 | 0.01677551 | 0.02930586 | 5.32797044 | 1.62366467 | 3.45136199 | 2.24126379 |
| -2.3915922 | 0.01677547 | 0.02930586 | 0 | 80.912623 | 90.8858656 | 57.7125427 |
| -2.3912665 | 0.01679036 | 0.02933021 | 48.3615778 | 80.912623 | 0 | 52.6696992 |
| -2.3911786 | 0.01679438 | 0.02933565 | 56.1486116 | 69.817581 | 0 | 53.2300151 |
| -2.3911454 | 0.0167959 | 0.02933672 | 92.6247169 | 45.1920001 | 0 | 48.1871716 |
| -2.3908522 | 0.01680932 | 0.02935857 | 1.63937552 | 29.4965749 | 26.4604419 | 28.5761134 |
| -2.390678 | 0.01681729 | 0.02937091 | 72.9522106 | 130.434396 | 134.603117 | 123.269509 |
| -2.3906201 | 0.01681995 | 0.02937395 | 505.747348 | 899.51023 | 852.486411 | 899.867413 |
| -2.3905452 | 0.01682338 | 0.02937836 | 10.6559409 | 22.1900839 | 4.60181598 | 28.0157974 |

| | | | | | | |
|---|---|---|---|---|---|---|
| -2.3902617 | 0.01683637 | 0.02939945 | 134.018949 | 18.1309222 | 0 | 66.6775979 |
| -2.3901046 | 0.01684358 | 0.02941044 | 34.017042 | 113.385916 | 80.5317797 | 0 |
| 2.38985503 | 0.01685502 | 0.02942884 | 4.91812656 | 8.93015571 | 3.45136199 | 4.48252759 |
| 2.38975764 | 0.01685949 | 0.02943506 | 0.40984388 | 1.62366467 | 2.30090799 | 2.80157974 |
| -2.3895563 | 0.01686874 | 0.0294496 | 51.230485 | 251.668025 | 157.612197 | 192.748686 |
| -2.3894421 | 0.01687399 | 0.02945717 | 7.78703372 | 14.6129821 | 14.9559019 | 11.206319 |
| -2.3892814 | 0.01688137 | 0.02946847 | 4.50828268 | 20.83703 | 13.8054479 | 22.4126379 |
| -2.3891742 | 0.01688629 | 0.02947547 | 1.63937552 | 32.4732935 | 26.4604419 | 25.7745336 |
| -2.3886051 | 0.01691247 | 0.02951956 | 125.822071 | 208.640911 | 232.391707 | 242.616806 |
| -2.3884498 | 0.01691962 | 0.02953045 | 85.247527 | 60.6168145 | 0 | 38.6618005 |
| -2.3881761 | 0.01693223 | 0.02955086 | 52.8698605 | 86.0542278 | 74.7795097 | 0 |
| -2.3879046 | 0.01694474 | 0.02957109 | 20.9020379 | 8.65954493 | 12.6549939 | 9.52537113 |
| -2.3877384 | 0.0169524 | 0.02958287 | 0 | 98.5023236 | 62.1245158 | 67.7982298 |
| -2.3876569 | 0.01695617 | 0.02958784 | 0 | 67.9233056 | 72.4786017 | 85.7283401 |
| -2.3876224 | 0.01695776 | 0.02958901 | 218.446788 | 290.094755 | 0 | 241.496174 |
| -2.3874862 | 0.01696404 | 0.02959837 | 10.6559409 | 9.20076649 | 10.354086 | 20.7316901 |
| -2.3872861 | 0.01697328 | 0.02961289 | 5.32797044 | 33.2851258 | 6.90272397 | 28.0157974 |
| -2.386949 | 0.01698885 | 0.02963846 | 64.3454891 | 55.2045989 | 0 | 57.7125427 |
| -2.3868159 | 0.016995 | 0.02964759 | 3.27875104 | 16.2366467 | 34.5136199 | 18.4904263 |
| 2.38664563 | 0.01700287 | 0.02965972 | 65.5750208 | 35.9912336 | 36.8145279 | 36.4205367 |
| 2.38651137 | 0.01700908 | 0.02966895 | 2.0492194 | 5.1416048 | 1.150454 | 2.24126379 |
| -2.3861137 | 0.01702748 | 0.02969944 | 0 | 93.3607188 | 49.4695218 | 87.409288 |
| -2.3859976 | 0.01703286 | 0.02970722 | 10.246097 | 35.9912336 | 4.60181598 | 19.0507423 |
| -2.3853865 | 0.01706119 | 0.02975501 | 66.3947085 | 27.3316887 | 0 | 101.417187 |
| -2.3852776 | 0.01706624 | 0.02976222 | 0 | 49.5217726 | 113.894946 | 69.4791776 |
| -2.3847674 | 0.01708992 | 0.0298003 | 39.7548564 | 84.1599523 | 0 | 58.8331746 |
| -2.3847774 | 0.01708946 | 0.0298003 | 29.9186032 | 107.432479 | 87.4345036 | 105.339398 |
| -2.3845723 | 0.01709899 | 0.02981449 | 6.55750208 | 25.708024 | 6.90272397 | 26.3348496 |
| 2.38389464 | 0.01713051 | 0.02986784 | 1.63937552 | 0.54122156 | 2.30090799 | 2.80157974 |
| -2.3838747 | 0.01713144 | 0.02986785 | 0 | 84.1599523 | 82.8326877 | 58.8331746 |
| -2.3837736 | 0.01713614 | 0.02987444 | 4.50828268 | 34.6381797 | 56.3722458 | 3.92221164 |
| -2.3834613 | 0.01715069 | 0.02989819 | 6.96734596 | 15.4248144 | 17.2568099 | 11.206319 |
| -2.3831653 | 0.01716448 | 0.02992062 | 41.8040758 | 54.9339882 | 0 | 85.1680242 |
| -2.3829911 | 0.01717261 | 0.02993316 | 64.3454891 | 23.0019162 | 3.45136199 | 6.72379138 |
| -2.3828859 | 0.01717752 | 0.02994011 | 0 | 168.319905 | 110.443584 | 12.3269509 |
| -2.382543 | 0.01719352 | 0.02996639 | 6.1476582 | 22.7313054 | 21.8586259 | 8.96505518 |
| 2.38241238 | 0.01719962 | 0.0299754 | 1.22953164 | 5.95343714 | 3.45136199 | 6.72379138 |
| 2.38225558 | 0.01720695 | 0.02998331 | 0 | 0 | 0 | 0 |
| 2.38225558 | 0.01720695 | 0.02998331 | 0 | 0 | 0 | 0 |
| 2.38225558 | 0.01720695 | 0.02998331 | 0 | 0 | 0 | 0 |
| -2.382028 | 0.01721759 | 0.03000023 | 10.6559409 | 18.401533 | 24.1595339 | 5.60315949 |
| -2.3819166 | 0.0172228 | 0.03000769 | 8.1968776 | 12.7187066 | 10.354086 | 17.9301104 |
| -2.3816519 | 0.01723518 | 0.03002655 | 41.8040758 | 144.776767 | 159.913105 | 264.469128 |

| | | | | | | |
|---|---|---|---|---|---|---|
| -2.3816455 | 0.01723548 | 0.03002655 | 0 | 94.7137727 | 28.7613499 | 123.829825 |
| -2.381424 | 0.01724585 | 0.03004299 | 9.01656536 | 21.3782516 | 6.90272397 | 17.9301104 |
| -2.3809648 | 0.01726736 | 0.03007884 | 13.9346919 | 48.7099402 | 232.391707 | 0 |
| -2.3798686 | 0.01731881 | 0.03016684 | 27.8693838 | 16.7778683 | 5.75226998 | 10.0856871 |
| -2.3797268 | 0.01732548 | 0.03017682 | 17.6232868 | 33.2851258 | 4.60181598 | 11.206319 |
| 2.37953732 | 0.01733439 | 0.03019071 | 0.81968776 | 3.78855091 | 3.45136199 | 2.80157974 |
| -2.3792522 | 0.0173478 | 0.03021244 | 58.6076748 | 117.9863 | 43.7172518 | 0 |
| -2.3792123 | 0.01734968 | 0.03021408 | 2.86890716 | 39.7797845 | 39.1154358 | 7.84442328 |
| 2.37882194 | 0.01736806 | 0.03024447 | 34.4268859 | 21.6488623 | 28.7613499 | 15.1285306 |
| -2.3783575 | 0.01738996 | 0.03028096 | 8.1968776 | 12.9893174 | 10.354086 | 16.8094785 |
| -2.3778746 | 0.01741275 | 0.03031901 | 11.8854725 | 12.1774851 | 8.05317797 | 14.5682147 |
| -2.3778526 | 0.01741378 | 0.03031918 | 108.198784 | 250.585581 | 376.198456 | 331.146726 |
| -2.377709 | 0.01742057 | 0.03032935 | 407.384817 | 577.754013 | 571.775636 | 599.538065 |
| -2.3769923 | 0.01745445 | 0.03038639 | 6.1476582 | 14.8835929 | 12.6549939 | 19.0507423 |
| -2.3769762 | 0.01745521 | 0.03038639 | 71.3128351 | 61.1580361 | 0 | 44.2649599 |
| 2.37678243 | 0.01746439 | 0.03039908 | 0.40984388 | 4.05916169 | 3.45136199 | 3.92221164 |
| -2.3767867 | 0.01746418 | 0.03039908 | 6.1476582 | 22.1900839 | 10.354086 | 17.3697944 |
| -2.3765639 | 0.01747473 | 0.03041546 | 9.83625312 | 69.0057487 | 0 | 168.655101 |
| -2.3759155 | 0.01750547 | 0.03046731 | 62.2962697 | 6.76526948 | 9.20363196 | 8.96505518 |
| -2.3755513 | 0.01752276 | 0.03049576 | 4.0984388 | 12.4480958 | 24.1595339 | 24.0935858 |
| -2.3754424 | 0.01752793 | 0.03050311 | 0 | 111.22103 | 88.5849576 | 35.8602207 |
| -2.3754084 | 0.01752954 | 0.03050428 | 26.2300083 | 75.5004074 | 0 | 89.0902358 |
| 2.37493613 | 0.01755199 | 0.03054169 | 0.40984388 | 0.54122156 | 0 | 1.1206319 |
| 2.37460104 | 0.01756793 | 0.03056778 | 652.471457 | 165.343186 | 142.656295 | 192.18837 |
| -2.3740195 | 0.01759562 | 0.03061266 | 43.8532952 | 85.5130062 | 0 | 49.8681194 |
| 2.37403433 | 0.01759491 | 0.03061266 | 100.001907 | 66.0290301 | 0 | 95.2537113 |
| -2.3739651 | 0.01759822 | 0.03061553 | 96.7231557 | 90.3840002 | 110.443584 | 93.5727634 |
| -2.3738224 | 0.01760502 | 0.03062571 | 80.7392443 | 51.416048 | 0 | 45.3855918 |
| -2.3733548 | 0.01762732 | 0.03066286 | 50.0009534 | 61.6992576 | 63.2749697 | 43.704644 |
| -2.3732953 | 0.01763016 | 0.03066615 | 48.3615778 | 54.3927666 | 0 | 72.2807574 |
| 2.3732743 | 0.01763116 | 0.03066624 | 49.5911095 | 17.5897006 | 28.7613499 | 30.2570612 |
| 2.37316594 | 0.01763634 | 0.03067359 | 177.052556 | 60.075593 | 35.6640739 | 74.5220212 |
| -2.3727217 | 0.01765756 | 0.03070885 | 12.7051603 | 15.1542036 | 9.20363196 | 9.52537113 |
| 2.37261547 | 0.01766264 | 0.03071602 | 61.476582 | 103.102707 | 71.3281477 | 76.763285 |
| -2.3720119 | 0.01769152 | 0.03076459 | 21.7217256 | 8.11832337 | 6.90272397 | 20.1713742 |
| 2.37176014 | 0.01770358 | 0.03077893 | 0 | 0 | 0 | 0 |
| 2.37176014 | 0.01770358 | 0.03077893 | 0 | 0 | 0 | 0 |
| 2.37176014 | 0.01770358 | 0.03077893 | 0 | 0 | 0 | 0 |
| 2.37176014 | 0.01770358 | 0.03077893 | 0 | 0 | 0 | 0 |
| -2.3715447 | 0.0177139 | 0.03079356 | 0 | 94.9843835 | 87.4345036 | 44.2649599 |
| -2.3715473 | 0.01771378 | 0.03079356 | 12.7051603 | 26.2492456 | 8.05317797 | 8.96505518 |
| -2.3714955 | 0.01771626 | 0.03079601 | 5.32797044 | 21.3782516 | 10.354086 | 22.4126379 |
| -2.3713335 | 0.01772403 | 0.03080785 | 1.22953164 | 18.401533 | 44.8677058 | 31.9380091 |

| | | | | | | |
|---|---|---|---|---|---|---|
| -2.3710107 | 0.01773952 | 0.03083311 | 35.6564176 | 19.4839761 | 4.60181598 | 9.52537113 |
| -2.3706827 | 0.01775527 | 0.03085882 | 397.958407 | 479.25169 | 294.516223 | 358.041891 |
| -2.37054 | 0.01776212 | 0.03086907 | 54.9190799 | 38.1561199 | 0 | 86.848972 |
| -2.3704308 | 0.01776737 | 0.03087653 | 3.68859492 | 15.6954252 | 20.7081719 | 24.0935858 |
| -2.3693588 | 0.01781896 | 0.03096451 | 182.380527 | 275.211162 | 254.250333 | 215.72164 |
| -2.3682622 | 0.01787187 | 0.03105479 | 6.1476582 | 8.38893415 | 10.354086 | 62.7553862 |
| -2.3681801 | 0.01787583 | 0.03106001 | 11.8854725 | 15.4248144 | 4.60181598 | 34.1792729 |
| -2.3679096 | 0.0178889 | 0.03108105 | 36.4761053 | 14.0717605 | 10.354086 | 5.60315949 |
| 2.36770187 | 0.01789895 | 0.03109516 | 9.42640924 | 13.2599282 | 8.05317797 | 14.5682147 |
| 2.36771233 | 0.01789845 | 0.03109516 | 52.0501728 | 32.2026827 | 24.1595339 | 23.5332698 |
| 2.36765 | 0.01790146 | 0.03109784 | 0.81968776 | 0 | 0 | 0 |
| 2.36736764 | 0.01791513 | 0.03111991 | 20.492194 | 12.4480958 | 9.20363196 | 10.0856871 |
| -2.3673349 | 0.01791671 | 0.03112098 | 6.96734596 | 12.1774851 | 11.50454 | 22.4126379 |
| -2.367311 | 0.01791787 | 0.03112132 | 92.214873 | 28.6847426 | 0 | 64.99665 |
| -2.367115 | 0.01792736 | 0.03113613 | 0 | 110.409198 | 105.841768 | 26.3348496 |
| -2.3668556 | 0.01793993 | 0.03115629 | 37.2957931 | 47.3568864 | 0 | 98.615607 |
| -2.3667894 | 0.01794314 | 0.03116019 | 94.2640924 | 127.998899 | 127.700393 | 143.440883 |
| -2.3663188 | 0.01796596 | 0.03119814 | 5.73781432 | 27.0610779 | 10.354086 | 16.2491625 |
| -2.3661275 | 0.01797525 | 0.03121259 | 6.96734596 | 17.0484791 | 11.50454 | 14.0078987 |
| -2.3660005 | 0.01798142 | 0.03122162 | 4.0984388 | 16.7778683 | 25.3099879 | 15.6888466 |
| -2.3659707 | 0.01798287 | 0.03122246 | 0 | 32.4732935 | 179.470823 | 45.3855918 |
| 2.36565067 | 0.01799842 | 0.03124442 | 139.346919 | 125.834012 | 378.499364 | 294.726189 |
| -2.3656657 | 0.01799769 | 0.03124442 | 7.37718984 | 14.8835929 | 17.2568099 | 10.0856871 |
| -2.365682 | 0.0179969 | 0.03124442 | 38.9351686 | 8.38893415 | 12.6549939 | 7.28410733 |
| -2.3654507 | 0.01800814 | 0.03125794 | 24.1807889 | 50.8748265 | 47.1686138 | 48.7474875 |
| 2.36545398 | 0.01800798 | 0.03125794 | 0 | 0.54122156 | 0 | 0 |
| -2.3650793 | 0.01802621 | 0.03128762 | 17.213443 | 4.60038325 | 9.20363196 | 44.2649599 |
| -2.3649401 | 0.01803299 | 0.03129769 | 8.60672148 | 17.3190899 | 13.8054479 | 8.96505518 |
| -2.3648699 | 0.0180364 | 0.03130194 | 7.78703372 | 15.4248144 | 9.20363196 | 17.3697944 |
| -2.3648224 | 0.01803872 | 0.03130428 | 33.6071982 | 53.851545 | 0 | 95.2537113 |
| -2.3642506 | 0.01806659 | 0.03135096 | 5.32797044 | 70.3588026 | 12.6549939 | 6.72379138 |
| -2.3641288 | 0.01807253 | 0.03135789 | 5.73781432 | 12.4480958 | 19.5577179 | 15.6888466 |
| -2.3641309 | 0.01807243 | 0.03135789 | 13.524848 | 7.8477126 | 12.6549939 | 12.3269509 |
| 2.36390882 | 0.01808326 | 0.03137483 | 0.40984388 | 1.3530539 | 2.30090799 | 0 |
| -2.3636282 | 0.01809697 | 0.03139692 | 2.45906328 | 14.6129821 | 39.1154358 | 22.9729539 |
| -2.3632651 | 0.0181147 | 0.03142601 | 10.6559409 | 17.8603114 | 5.75226998 | 20.1713742 |
| -2.3631411 | 0.01812077 | 0.03143484 | 35.6564176 | 103.914539 | 0 | 45.3855918 |
| -2.3630926 | 0.01812314 | 0.03143727 | 0 | 71.9824673 | 74.7795097 | 69.4791776 |
| -2.3627821 | 0.01813833 | 0.03146193 | 22.5414134 | 3.24732935 | 16.1063559 | 25.2142177 |
| -2.3626579 | 0.01814441 | 0.03147078 | 13.1150042 | 18.9427545 | 6.90272397 | 11.206319 |
| -2.3626268 | 0.01814593 | 0.03147173 | 3.68859492 | 23.8137486 | 13.8054479 | 23.5332698 |
| -2.3626034 | 0.01814708 | 0.03147203 | 0 | 70.3588026 | 69.0272397 | 76.763285 |
| -2.3625584 | 0.01814928 | 0.03147415 | 3.27875104 | 33.2851258 | 10.354086 | 28.0157974 |

| | | | | | | |
|---|---|---|---|---|---|---|
| -2.3620313 | 0.01817511 | 0.03151725 | 49.5911095 | 65.7584193 | 0 | 55.4712789 |
| -2.3615868 | 0.01819691 | 0.03155337 | 22.9512573 | 14.3423713 | 5.75226998 | 11.7666349 |
| -2.3612262 | 0.01821462 | 0.03158237 | 28.6890716 | 1.89427545 | 37.9649818 | 15.6888466 |
| -2.3611093 | 0.01822036 | 0.03159063 | 37.2957931 | 10.0125988 | 11.50454 | 6.72379138 |
| 2.36087695 | 0.01823178 | 0.03160873 | 2.86890716 | 0.54122156 | 4.60181598 | 1.68094785 |
| 2.36082548 | 0.01823431 | 0.03161142 | 42.6237635 | 44.9213893 | 70.1776937 | 66.6775979 |
| 2.36064376 | 0.01824325 | 0.03162522 | 120.494101 | 126.645845 | 51.7704298 | 85.7283401 |
| -2.3601742 | 0.01826636 | 0.03166358 | 178.282088 | 5.41221558 | 0 | 84.0473923 |
| -2.3600672 | 0.01827163 | 0.03167101 | 2.45906328 | 15.1542036 | 43.7172518 | 19.6110582 |
| -2.3599765 | 0.01827609 | 0.03167706 | 20.9020379 | 113.656527 | 0 | 63.3157022 |
| -2.3598704 | 0.01828132 | 0.03168441 | 9.42640924 | 14.3423713 | 17.2568099 | 7.84442328 |
| 2.35938004 | 0.0183055 | 0.03171781 | 0 | 0 | 0 | 0 |
| 2.35938004 | 0.0183055 | 0.03171781 | 0 | 0 | 0 | 0 |
| 2.35938004 | 0.0183055 | 0.03171781 | 0 | 0 | 0 | 0 |
| 2.35938004 | 0.0183055 | 0.03171781 | 0 | 0 | 0 | 0 |
| 2.35938004 | 0.0183055 | 0.03171781 | 0 | 0 | 0 | 0 |
| -2.3592915 | 0.01830987 | 0.03172367 | 11.4756286 | 11.3656527 | 10.354086 | 10.646003 |
| -2.3590452 | 0.01832202 | 0.03174303 | 457.795614 | 683.292217 | 571.775636 | 627.553862 |
| -2.3589099 | 0.0183287 | 0.0317529 | 30.3284471 | 80.6420122 | 0 | 68.9188617 |
| -2.358529 | 0.01834753 | 0.0317821 | 85.247527 | 43.0271139 | 0 | 47.0665397 |
| -2.358539 | 0.01834703 | 0.0317821 | 19.2626624 | 10.8244312 | 12.6549939 | 7.28410733 |
| -2.3581359 | 0.01836697 | 0.03181407 | 17.213443 | 11.6362635 | 8.05317797 | 10.646003 |
| -2.3579823 | 0.01837457 | 0.03182553 | 18.0331307 | 49.7923834 | 0 | 145.682147 |
| -2.3576392 | 0.01839156 | 0.03185325 | 13.1150042 | 12.9893174 | 24.1595339 | 5.60315949 |
| 2.35727406 | 0.01840965 | 0.03188288 | 0 | 0 | 0 | 0 |
| -2.3572446 | 0.01841112 | 0.0318837 | 8.60672148 | 11.9068743 | 11.50454 | 12.8872668 |
| -2.3569179 | 0.01842732 | 0.03191005 | 5.73781432 | 107.973701 | 0 | 142.320251 |
| -2.356833 | 0.01843153 | 0.03191564 | 6.96734596 | 7.30649104 | 12.6549939 | 48.7474875 |
| 2.35674299 | 0.018436 | 0.03192166 | 208.200691 | 130.163785 | 118.496762 | 107.580662 |
| -2.3566474 | 0.01844075 | 0.03192817 | 25.8201644 | 52.4984912 | 44.8677058 | 44.2649599 |
| -2.3556687 | 0.0184894 | 0.03201069 | 9.01656536 | 11.3656527 | 10.354086 | 14.5682147 |
| -2.3556101 | 0.01849231 | 0.03201384 | 224.184602 | 1019.39081 | 1073.37358 | 1016.97345 |
| -2.3555922 | 0.01849321 | 0.03201384 | 114.346443 | 51.6866588 | 0 | 25.7745336 |
| -2.3555199 | 0.01849681 | 0.03201732 | 222.545227 | 37.8855091 | 26.4604419 | 49.3078035 |
| -2.355512 | 0.0184972 | 0.03201732 | 39.3450125 | 15.966036 | 4.60181598 | 10.0856871 |
| -2.3551377 | 0.01851584 | 0.03204787 | 16.3937552 | 11.0950419 | 6.90272397 | 14.0078987 |
| 2.35473976 | 0.01853568 | 0.03208049 | 0 | 0 | 0 | 0 |
| -2.3547024 | 0.01853754 | 0.03208199 | 12.7051603 | 16.2366467 | 14.9559019 | 6.16347543 |
| -2.3543458 | 0.01855534 | 0.03211107 | 3.27875104 | 26.7904671 | 12.6549939 | 26.3348496 |
| 2.35430868 | 0.01855719 | 0.03211256 | 63.1159575 | 168.319905 | 0 | 189.947107 |
| -2.3542473 | 0.01856026 | 0.03211614 | 67.6242402 | 11.6362635 | 4.60181598 | 8.40473923 |
| -2.3540845 | 0.01856839 | 0.03212849 | 0 | 73.0649104 | 78.2308717 | 62.1950703 |
| -2.353934 | 0.01857591 | 0.03213978 | 7.78703372 | 18.1309222 | 10.354086 | 12.3269509 |

| | | | | | | |
|---|---|---|---|---|---|---|
| -2.3537869 | 0.01858326 | 0.03215078 | 6.55750208 | 9.74198805 | 24.1595339 | 14.5682147 |
| -2.3528817 | 0.01862856 | 0.03222742 | 3.68859492 | 18.401533 | 17.2568099 | 22.4126379 |
| -2.3526667 | 0.01863933 | 0.03224432 | 4.91812656 | 15.966036 | 14.9559019 | 19.0507423 |
| -2.3525993 | 0.01864271 | 0.03224844 | 7.78703372 | 13.2599282 | 10.354086 | 15.6888466 |
| 2.35257754 | 0.0186438 | 0.0322486 | 0 | 0.27061078 | 0 | 1.1206319 |
| -2.3525485 | 0.01864526 | 0.0322494 | 0 | 86.5954493 | 101.239952 | 36.4205367 |
| -2.3522795 | 0.01865875 | 0.03227049 | 19.2626624 | 17.5897006 | 4.60181598 | 14.0078987 |
| -2.3522654 | 0.01865945 | 0.03227049 | 259.84102 | 296.318803 | 178.320369 | 193.869318 |
| -2.3519645 | 0.01867456 | 0.03229488 | 7.78703372 | 9.47137727 | 17.2568099 | 14.0078987 |
| -2.3517447 | 0.01868559 | 0.03231224 | 30.3284471 | 76.8534613 | 0 | 70.0394936 |
| -2.3514518 | 0.01870031 | 0.03233423 | 0 | 126.645845 | 40.2658898 | 60.5141225 |
| -2.3514675 | 0.01869952 | 0.03233423 | 11.4756286 | 33.8263474 | 21.8586259 | 29.6967453 |
| -2.3513098 | 0.01870745 | 0.03234484 | 28.2792277 | 108.514922 | 0 | 49.8681194 |
| 2.35128049 | 0.01870892 | 0.03234565 | 0.81968776 | 0.81183234 | 1.150454 | 0 |
| 2.35126022 | 0.01870994 | 0.03234568 | 4.91812656 | 10.5538204 | 39.1154358 | 10.0856871 |
| -2.3511046 | 0.01871777 | 0.03235749 | 125.412227 | 189.968767 | 243.896247 | 214.601008 |
| -2.3508418 | 0.018731 | 0.03237861 | 23.3611012 | 13.2599282 | 8.05317797 | 8.40473923 |
| -2.3507055 | 0.01873786 | 0.032387 | 55.3289238 | 103.373318 | 47.1686138 | 0 |
| 2.35072544 | 0.01873685 | 0.032387 | 7.78703372 | 3.51794013 | 1.150454 | 5.60315949 |
| -2.3505468 | 0.01874585 | 0.03239909 | 11.0657848 | 28.9553534 | 28.7613499 | 25.7745336 |
| 2.35029974 | 0.0187583 | 0.03241886 | 0 | 0 | 0 | 0 |
| 2.3502016 | 0.01876325 | 0.03242568 | 8.1968776 | 8.65954493 | 11.50454 | 6.16347543 |
| -2.3498402 | 0.01878147 | 0.03245544 | 6.55750208 | 17.0484791 | 14.9559019 | 11.206319 |
| -2.349794 | 0.0187838 | 0.03245773 | 150.822548 | 262.221845 | 157.612197 | 360.843471 |
| -2.3496593 | 0.0187906 | 0.03246774 | 78.6900249 | 128.810731 | 86.2840497 | 79.5648647 |
| -2.3493751 | 0.01880495 | 0.03249079 | 40.1647002 | 54.9339882 | 0 | 75.6426531 |
| 2.34926863 | 0.01881033 | 0.03249835 | 77.4604933 | 59.8049822 | 0 | 38.1014845 |
| -2.3492061 | 0.01881349 | 0.03250207 | 6.55750208 | 10.0125988 | 21.8586259 | 14.5682147 |
| -2.3490536 | 0.0188212 | 0.03251364 | 27.0496961 | 17.5897006 | 32.2127119 | 14.0078987 |
| -2.3487908 | 0.01883449 | 0.03253486 | 0 | 25.708024 | 88.5849576 | 121.588561 |
| 2.34868281 | 0.01883995 | 0.03254255 | 29.9186032 | 22.1900839 | 29.9118039 | 21.292006 |
| -2.3482272 | 0.01886301 | 0.03258064 | 63.9356453 | 46.2744432 | 92.0363196 | 0 |
| 2.34804207 | 0.01887239 | 0.0325951 | 0.40984388 | 0 | 0 | 0 |
| -2.3480039 | 0.01887432 | 0.0325967 | 35.2465737 | 108.244312 | 66.7263317 | 0 |
| -2.3477437 | 0.01888751 | 0.03261621 | 109.018472 | 17.5897006 | 11.50454 | 24.0935858 |
| -2.3477411 | 0.01888764 | 0.03261621 | 36.4761053 | 79.2889583 | 0 | 56.5919108 |
| -2.3475747 | 0.01889608 | 0.03262905 | 11.0657848 | 10.2832096 | 10.354086 | 11.7666349 |
| -2.3473019 | 0.01890992 | 0.0326512 | 45.0828268 | 69.0057487 | 87.4345036 | 0 |
| -2.3469551 | 0.01892753 | 0.03267985 | 77.4604933 | 0 | 97.7885896 | 56.5919108 |
| -2.3468997 | 0.01893035 | 0.03268297 | 26.2300083 | 145.317988 | 0 | 33.6189569 |
| -2.3467736 | 0.01893676 | 0.03269228 | 12.2953164 | 11.0950419 | 9.20363196 | 11.206319 |
| -2.3466571 | 0.01894268 | 0.03270076 | 6.96734596 | 14.0717605 | 11.50454 | 15.1285306 |
| -2.3466262 | 0.01894425 | 0.03270172 | 25.4103206 | 24.8961917 | 25.3099879 | 12.3269509 |

| | | | | | | |
|---|---|---|---|---|---|---|
| -2.3465496 | 0.01894814 | 0.03270669 | 11.0657848 | 7.03588026 | 40.2658898 | 8.96505518 |
| -2.3464155 | 0.01895496 | 0.03271671 | 4.0984388 | 19.7545869 | 23.0090799 | 13.4475828 |
| 2.34618977 | 0.01896645 | 0.03273479 | 8.1968776 | 1.89427545 | 4.60181598 | 3.92221164 |
| -2.3460505 | 0.01897354 | 0.03274528 | 61.8864259 | 11.6362635 | 4.60181598 | 8.96505518 |
| -2.3457033 | 0.01899122 | 0.03277404 | 22.1315695 | 71.4412457 | 0 | 93.0124475 |
| -2.3455422 | 0.01899943 | 0.03278479 | 119.674413 | 3.78855091 | 20.7081719 | 13.4475828 |
| -2.3455412 | 0.01899948 | 0.03278479 | 22.1315695 | 8.93015571 | 16.1063559 | 6.72379138 |
| 2.34548299 | 0.01900245 | 0.03278816 | 43.0336074 | 33.8263474 | 34.5136199 | 31.3776931 |
| -2.3454187 | 0.01900572 | 0.03279205 | 5.32797044 | 15.4248144 | 31.0622579 | 10.0856871 |
| 2.34502758 | 0.01902567 | 0.03282472 | 0 | 1.3530539 | 0 | 1.1206319 |
| -2.3447576 | 0.01903945 | 0.03284674 | 21.3118818 | 9.20076649 | 12.6549939 | 7.84442328 |
| -2.3445887 | 0.01904808 | 0.03285987 | 10.6559409 | 11.9068743 | 11.50454 | 9.52537113 |
| -2.3441564 | 0.01907017 | 0.03289622 | 6.55750208 | 31.3908504 | 21.8586259 | 25.7745336 |
| -2.3440882 | 0.01907366 | 0.03290049 | 31.5579788 | 9.74198805 | 14.9559019 | 5.60315949 |
| -2.3437997 | 0.01908842 | 0.03292418 | 9.01656536 | 81.7244553 | 181.771731 | 0 |
| -2.3432999 | 0.01911401 | 0.03296397 | 2.45906328 | 17.8603114 | 27.6108959 | 23.5332698 |
| -2.3433004 | 0.01911398 | 0.03296397 | 78.2801811 | 25.4374132 | 23.0090799 | 28.5761134 |
| -2.3432894 | 0.01911455 | 0.03296397 | 31.5579788 | 15.966036 | 6.90272397 | 7.28410733 |
| -2.3432596 | 0.01911608 | 0.03296485 | 38.5253247 | 125.563402 | 0 | 28.5761134 |
| 2.34317714 | 0.0191203 | 0.03297037 | 0 | 0 | 19.5577179 | 0 |
| -2.3428747 | 0.01913581 | 0.03299535 | 11.0657848 | 25.708024 | 11.50454 | 6.72379138 |
| -2.3425332 | 0.01915333 | 0.03302379 | 20.0823501 | 27.3316887 | 18.4072639 | 17.3697944 |
| -2.3424921 | 0.01915544 | 0.03302567 | 3.68859492 | 28.6847426 | 21.8586259 | 11.7666349 |
| -2.3424624 | 0.01915696 | 0.03302653 | 401.237158 | 359.641725 | 248.498063 | 336.189569 |
| -2.3423797 | 0.01916121 | 0.03303209 | 8.60672148 | 11.6362635 | 14.9559019 | 10.0856871 |
| -2.3422729 | 0.01916669 | 0.03303978 | 270.087117 | 377.772648 | 276.108959 | 332.827673 |
| -2.3419385 | 0.01918388 | 0.03306763 | 3.68859492 | 10.0125988 | 33.3631659 | 22.9729539 |
| -2.3418636 | 0.01918773 | 0.0330725 | 11.4756286 | 12.7187066 | 8.05317797 | 12.3269509 |
| -2.3415923 | 0.01920168 | 0.03309478 | 6.55750208 | 20.0251977 | 16.1063559 | 9.52537113 |
| -2.3415282 | 0.01920498 | 0.0330987 | 11.4756286 | 217.029845 | 49.4695218 | 0 |
| -2.3414397 | 0.01920953 | 0.03310478 | 0 | 46.545054 | 57.5226998 | 114.304454 |
| -2.341314 | 0.019216 | 0.03311416 | 7.37718984 | 9.74198805 | 18.4072639 | 13.4475828 |
| -2.3406532 | 0.01925004 | 0.03317106 | 11.8854725 | 16.7778683 | 11.50454 | 7.28410733 |
| -2.3405708 | 0.01925429 | 0.0331766 | 8.1968776 | 11.9068743 | 9.20363196 | 20.1713742 |
| -2.3401593 | 0.01927552 | 0.03321141 | 197.134906 | 344.487522 | 491.243856 | 423.038541 |
| -2.3399827 | 0.01928463 | 0.03322534 | 17.213443 | 114.73897 | 0 | 65.556966 |
| -2.3399578 | 0.01928592 | 0.03322579 | 4.50828268 | 16.5072575 | 21.8586259 | 14.0078987 |
| -2.3396923 | 0.01929963 | 0.03324764 | 13.9346919 | 5.68282636 | 19.5577179 | 12.8872668 |
| -2.3396541 | 0.01930161 | 0.03324926 | 2.45906328 | 17.8603114 | 17.2568099 | 38.1014845 |
| 2.33960751 | 0.01930401 | 0.03325164 | 0 | 1.3530539 | 0 | 0 |
| 2.33946126 | 0.01931157 | 0.03326289 | 16.8035991 | 30.579018 | 25.3099879 | 27.4554815 |
| 2.33934385 | 0.01931765 | 0.03326979 | 0 | 0 | 0 | 0 |
| 2.33934385 | 0.01931765 | 0.03326979 | 0 | 0 | 0 | 0 |

| | | | | | | |
|---|---|---|---|---|---|---|
| 2.33883733 | 0.01934385 | 0.03331315 | 3.68859492 | 2.70610779 | 2.30090799 | 4.48252759 |
| 2.3384646 | 0.01936316 | 0.03334285 | 23.3611012 | 50.0629941 | 20.7081719 | 45.9459078 |
| -2.3384701 | 0.01936287 | 0.03334285 | 29.9186032 | 64.1347547 | 0 | 79.5648647 |
| 2.33843313 | 0.01936479 | 0.03334388 | 0 | 3.24732935 | 0 | 1.68094785 |
| -2.3383189 | 0.01937071 | 0.03335229 | 2.86890716 | 39.238563 | 24.1595339 | 10.646003 |
| 2.33806045 | 0.01938411 | 0.03337358 | 11.8854725 | 0 | 12.6549939 | 16.2491625 |
| -2.338012 | 0.01938663 | 0.03337614 | 18.8528185 | 51.9572696 | 0 | 128.312352 |
| -2.3378867 | 0.01939313 | 0.03338554 | 56.5584554 | 67.1114732 | 0 | 42.0236961 |
| -2.3378399 | 0.01939556 | 0.03338575 | 59.8372065 | 51.1454373 | 0 | 52.6696992 |
| -2.3378246 | 0.01939635 | 0.03338575 | 33.6071982 | 31.3908504 | 0 | 126.631404 |
| 2.33784963 | 0.01939505 | 0.03338575 | 42.6237635 | 26.2492456 | 29.9118039 | 25.7745336 |
| -2.3376538 | 0.01940522 | 0.03339923 | 0 | 38.1561199 | 121.948124 | 61.6347543 |
| 2.33721103 | 0.01942821 | 0.03343703 | 46.3123584 | 22.7313054 | 24.1595339 | 30.2570612 |
| 2.33675534 | 0.01945191 | 0.03347603 | 31.9678226 | 21.3782516 | 36.8145279 | 0 |
| -2.3364944 | 0.01946549 | 0.03349716 | 3.68859492 | 27.6022995 | 12.6549939 | 20.1713742 |
| -2.3364796 | 0.01946626 | 0.03349716 | 42.6237635 | 77.6652936 | 52.9208838 | 58.8331746 |
| -2.3361899 | 0.01948134 | 0.03352029 | 277.874151 | 7.30649104 | 35.6640739 | 0 |
| 2.33618157 | 0.01948178 | 0.03352029 | 3.27875104 | 2.16488623 | 1.150454 | 1.68094785 |
| -2.3360781 | 0.01948717 | 0.03352778 | 0 | 84.9717847 | 85.1335957 | 42.5840121 |
| -2.3359926 | 0.01949163 | 0.03353366 | 49.1812656 | 38.9679522 | 0 | 80.6854966 |
| -2.3357626 | 0.01950362 | 0.03355071 | 412.712787 | 707.647187 | 586.731538 | 708.799675 |
| -2.3357637 | 0.01950356 | 0.03355071 | 5.73781432 | 12.4480958 | 24.1595339 | 12.3269509 |
| -2.3355231 | 0.01951611 | 0.03357041 | 0 | 84.7011739 | 43.7172518 | 83.4870763 |
| -2.3354061 | 0.01952221 | 0.03357912 | 3.68859492 | 61.1580361 | 21.8586259 | 5.60315949 |
| -2.3352442 | 0.01953067 | 0.03359187 | 40.5745441 | 69.0057487 | 0 | 56.0315949 |
| -2.3351216 | 0.01953707 | 0.03360109 | 4.91812656 | 15.1542036 | 14.9559019 | 18.4904263 |
| -2.3349653 | 0.01954523 | 0.03361335 | 27.0496961 | 91.4664434 | 92.0363196 | 0 |
| 2.3348089 | 0.01955341 | 0.03362561 | 11.0657848 | 24.6255809 | 25.3099879 | 17.9301104 |
| -2.3345233 | 0.01956834 | 0.0336495 | 0 | 69.817581 | 60.9740618 | 75.6426531 |
| -2.3344303 | 0.0195732 | 0.03365607 | 40.5745441 | 75.7710182 | 0 | 50.4284354 |
| 2.33394558 | 0.01959857 | 0.0336979 | 0.40984388 | 1.08244312 | 0 | 0 |
| -2.3334875 | 0.01962258 | 0.03373737 | 37.705637 | 48.1687187 | 0 | 83.4870763 |
| -2.3333545 | 0.01962955 | 0.03374756 | 9.01656536 | 19.7545869 | 6.90272397 | 15.1285306 |
| -2.3328997 | 0.01965341 | 0.03378679 | 0 | 101.479042 | 56.3722458 | 53.2300151 |
| -2.3322686 | 0.01968657 | 0.03384199 | 13.524848 | 18.9427545 | 5.75226998 | 12.3269509 |
| -2.3322153 | 0.01968937 | 0.033845 | 69.6734596 | 50.3336049 | 73.6290557 | 0 |
| -2.3316893 | 0.01971704 | 0.03389077 | 13.1150042 | 14.3423713 | 6.90272397 | 11.7666349 |
| -2.3315958 | 0.01972197 | 0.03389382 | 40.1647002 | 53.3103235 | 0 | 71.7204414 |
| -2.3316141 | 0.019721 | 0.03389382 | 5.73781432 | 16.2366467 | 18.4072639 | 11.206319 |
| -2.3316144 | 0.01972098 | 0.03389382 | 32.3776665 | 94.1725511 | 0 | 47.6268556 |
| -2.3313295 | 0.01973599 | 0.03391611 | 4.0984388 | 20.83703 | 12.6549939 | 21.852322 |
| -2.3311209 | 0.01974698 | 0.0339332 | 42.2139196 | 41.9446708 | 0 | 84.6077082 |
| -2.3310946 | 0.01974837 | 0.03393378 | 48.3615778 | 87.9485032 | 0 | 34.7395888 |

| | | | | | | |
|---|---|---|---|---|---|---|
| -2.3309043 | 0.0197584 | 0.03394921 | 9.42640924 | 11.9068743 | 8.05317797 | 17.9301104 |
| 2.33048208 | 0.01978069 | 0.03398569 | 0 | 6.76526948 | 4.60181598 | 0 |
| -2.3304205 | 0.01978394 | 0.03398946 | 3.27875104 | 57.3694852 | 11.50454 | 12.8872668 |
| -2.3302623 | 0.01979229 | 0.03400201 | 0 | 68.7351379 | 52.9208838 | 85.1680242 |
| -2.3301237 | 0.01979962 | 0.03401278 | 24.5906328 | 8.65954493 | 6.90272397 | 14.0078987 |
| -2.3297429 | 0.01981974 | 0.03404515 | 6.96734596 | 27.0610779 | 8.05317797 | 14.5682147 |
| -2.3297273 | 0.01982057 | 0.03404515 | 86.0672148 | 0 | 60.9740618 | 76.202969 |
| -2.3294358 | 0.01983599 | 0.03406983 | 72.1325229 | 71.7118565 | 75.9299637 | 80.6854966 |
| 2.32936736 | 0.01983961 | 0.03407424 | 162.298176 | 102.832096 | 132.302209 | 109.821926 |
| -2.3292262 | 0.01984709 | 0.03408526 | 12.2953164 | 112.844695 | 0 | 80.1251807 |
| -2.3290175 | 0.01985814 | 0.03410243 | 48.7714217 | 45.4626109 | 0 | 68.9188617 |
| -2.3289302 | 0.01986277 | 0.03410856 | 34.8367298 | 8.11832337 | 10.354086 | 8.40473923 |
| -2.3288361 | 0.01986775 | 0.03411531 | 16.8035991 | 4.05916169 | 11.50454 | 33.058641 |
| 2.32844164 | 0.01988866 | 0.03414697 | 0.81968776 | 1.3530539 | 2.30090799 | 1.1206319 |
| -2.3284285 | 0.01988936 | 0.03414697 | 0 | 40.0503953 | 70.1776937 | 102.537819 |
| -2.3284371 | 0.0198889 | 0.03414697 | 7.78703372 | 10.5538204 | 13.8054479 | 12.8872668 |
| -2.3282978 | 0.0198963 | 0.03415706 | 5.73781432 | 43.5683354 | 9.20363196 | 11.7666349 |
| -2.3282436 | 0.01989917 | 0.03415836 | 10.6559409 | 10.0125988 | 10.354086 | 11.7666349 |
| 2.32825446 | 0.0198986 | 0.03415836 | 0 | 0 | 0 | 0 |
| -2.3280936 | 0.01990713 | 0.03417021 | 14.7543797 | 17.0484791 | 9.20363196 | 7.28410733 |
| -2.3280702 | 0.01990837 | 0.03417052 | 0 | 116.362635 | 82.8326877 | 26.3348496 |
| 2.32802188 | 0.01991094 | 0.03417311 | 3.27875104 | 1.62366467 | 5.75226998 | 2.80157974 |
| -2.3279248 | 0.0199161 | 0.03417833 | 0 | 133.411114 | 37.9649818 | 51.5490673 |
| -2.3279315 | 0.01991574 | 0.03417833 | 60.2470503 | 344.758133 | 303.719855 | 301.44998 |
| -2.32789 | 0.01991795 | 0.03417968 | 11.8854725 | 10.2832096 | 8.05317797 | 14.5682147 |
| -2.3277787 | 0.01992386 | 0.03418801 | 120.084257 | 215.676791 | 199.028541 | 247.099333 |
| -2.3277458 | 0.01992561 | 0.03418919 | 0 | 64.946587 | 33.3631659 | 122.709193 |
| -2.3276221 | 0.01993218 | 0.03419684 | 10.246097 | 25.9786348 | 8.05317797 | 9.52537113 |
| -2.3276289 | 0.01993182 | 0.03419684 | 8.60672148 | 15.966036 | 11.50454 | 9.52537113 |
| -2.3275847 | 0.01993417 | 0.03419843 | 60.2470503 | 74.1473535 | 0 | 33.058641 |
| -2.3275315 | 0.01993699 | 0.03420146 | 60.6568942 | 77.1240721 | 54.0713378 | 0 |
| -2.3274315 | 0.01994231 | 0.03420876 | 24.5906328 | 79.8301798 | 0 | 71.1601255 |
| -2.3273415 | 0.01994709 | 0.03421515 | 50.4107972 | 7.57710182 | 8.05317797 | 8.96505518 |
| -2.3271074 | 0.01995955 | 0.0342347 | 36.4761053 | 63.0523115 | 101.239952 | 0 |
| -2.3270767 | 0.01996118 | 0.03423568 | 81.1490882 | 36.803066 | 0 | 49.3078035 |
| -2.326992 | 0.01996569 | 0.03424159 | 12.7051603 | 9.74198805 | 8.05317797 | 14.5682147 |
| -2.3269198 | 0.01996953 | 0.03424637 | 5.73781432 | 9.47137727 | 18.4072639 | 20.7316901 |
| -2.3265195 | 0.01999085 | 0.0342811 | 13.1150042 | 10.8244312 | 11.50454 | 8.40473923 |
| -2.3263297 | 0.02000097 | 0.03429664 | 93.4444046 | 64.1347547 | 0 | 22.4126379 |
| -2.3259673 | 0.02002029 | 0.03432795 | 81.5589321 | 0 | 58.6731538 | 180.982051 |
| -2.3258013 | 0.02002915 | 0.03434131 | 9.01656536 | 18.401533 | 21.8586259 | 22.9729539 |
| -2.3255452 | 0.02004283 | 0.03436293 | 60.2470503 | 0 | 138.054479 | 39.7824324 |
| -2.325448 | 0.02004802 | 0.03437001 | 0 | 55.2045989 | 96.6381356 | 55.4712789 |

| | | | | | | |
|---|---|---|---|---|---|---|
| -2.3253114 | 0.02005532 | 0.03438069 | 12.7051603 | 8.38893415 | 27.6108959 | 7.28410733 |
| -2.3252902 | 0.02005645 | 0.03438081 | 0 | 83.6187308 | 50.6199758 | 71.1601255 |
| -2.3250758 | 0.02006791 | 0.03439863 | 94.2640924 | 103.102707 | 109.29313 | 103.65845 |
| -2.3246798 | 0.02008909 | 0.03443311 | 13.524848 | 10.5538204 | 5.75226998 | 24.6539017 |
| -2.3246155 | 0.02009253 | 0.03443717 | 12.2953164 | 8.38893415 | 10.354086 | 12.8872668 |
| -2.3244877 | 0.02009937 | 0.03444707 | 4.0984388 | 25.708024 | 17.2568099 | 12.8872668 |
| 2.32446432 | 0.02010062 | 0.03444739 | 22.9512573 | 11.6362635 | 9.20363196 | 15.6888466 |
| -2.3240432 | 0.02012318 | 0.03448238 | 4.0984388 | 43.0271139 | 27.6108959 | 5.60315949 |
| -2.3240582 | 0.02012237 | 0.03448238 | 1.22953164 | 21.6488623 | 36.8145279 | 26.8951655 |
| -2.3240087 | 0.02012503 | 0.03448372 | 5.73781432 | 12.7187066 | 18.4072639 | 13.4475828 |
| -2.3236332 | 0.02014516 | 0.03451639 | 6.55750208 | 14.3423713 | 20.7081719 | 9.52537113 |
| -2.3234174 | 0.02015674 | 0.0345344 | 4.50828268 | 15.1542036 | 25.3099879 | 12.8872668 |
| -2.3233482 | 0.02016045 | 0.03453892 | 32.3776665 | 76.3122397 | 0 | 58.2728587 |
| -2.3233095 | 0.02016253 | 0.03454065 | 52.4600166 | 66.5702517 | 0 | 42.5840121 |
| -2.3232701 | 0.02016465 | 0.03454129 | 0 | 47.3568864 | 105.841768 | 56.5919108 |
| -2.3232627 | 0.02016504 | 0.03454129 | 0 | 77.1240721 | 62.1245158 | 63.3157022 |
| -2.3232329 | 0.02016664 | 0.0345422 | 7.37718984 | 15.1542036 | 9.20363196 | 15.6888466 |
| -2.3231925 | 0.02016881 | 0.03454408 | 66.3947085 | 0 | 366.994825 | 182.662999 |
| -2.3226595 | 0.02019745 | 0.03459129 | 45.9025146 | 89.0309463 | 58.6731538 | 0 |
| -2.3223421 | 0.02021452 | 0.0346187 | 138.117388 | 349.358516 | 307.171217 | 341.232413 |
| -2.32231 | 0.02021625 | 0.03461982 | 190.16756 | 437.57763 | 370.446187 | 391.100532 |
| -2.3214358 | 0.02026333 | 0.03469861 | 5.32797044 | 21.3782516 | 10.354086 | 17.3697944 |
| -2.3214062 | 0.02026493 | 0.0346995 | 10.6559409 | 11.6362635 | 9.20363196 | 11.206319 |
| -2.3213312 | 0.02026898 | 0.03470459 | 6.1476582 | 13.8011497 | 14.9559019 | 12.8872668 |
| -2.321029 | 0.02028528 | 0.03473066 | 1.63937552 | 17.5897006 | 35.6640739 | 25.7745336 |
| 2.32093763 | 0.02029021 | 0.03473542 | 0 | 0 | 0 | 0 |
| 2.32093763 | 0.02029021 | 0.03473542 | 0 | 0 | 0 | 0 |
| -2.3209065 | 0.02029189 | 0.03473645 | 14.7543797 | 11.9068743 | 11.50454 | 7.28410733 |
| -2.3208012 | 0.02029758 | 0.03474435 | 11.0657848 | 9.74198805 | 8.05317797 | 17.9301104 |
| -2.3205321 | 0.02031211 | 0.03476738 | 39.3450125 | 90.3840002 | 0 | 39.2221164 |
| -2.3203557 | 0.02032164 | 0.03478185 | 0 | 44.3801678 | 79.3813257 | 81.8061285 |
| -2.3200603 | 0.02033762 | 0.03480735 | 9.01656536 | 11.3656527 | 12.6549939 | 10.0856871 |
| -2.3198718 | 0.02034781 | 0.03482295 | 13.524848 | 23.5431378 | 5.75226998 | 10.646003 |
| 2.31982708 | 0.02035023 | 0.03482525 | 1.22953164 | 0 | 0 | 0 |
| -2.3196443 | 0.02036013 | 0.03483956 | 0 | 58.1813175 | 32.2127119 | 131.113932 |
| -2.3196328 | 0.02036075 | 0.03483956 | 11.4756286 | 12.9893174 | 5.75226998 | 21.292006 |
| -2.3194961 | 0.02036815 | 0.03485037 | 0 | 84.7011739 | 111.594038 | 25.7745336 |
| -2.319469 | 0.02036962 | 0.03485104 | 9.83625312 | 11.6362635 | 10.354086 | 10.646003 |
| -2.3189882 | 0.02039568 | 0.03489377 | 62.2962697 | 20.2958084 | 0 | 99.1759229 |
| -2.318828 | 0.02040437 | 0.03490679 | 9.01656536 | 12.1774851 | 12.6549939 | 9.52537113 |
| -2.3186062 | 0.0204164 | 0.03492367 | 6.1476582 | 21.3782516 | 11.50454 | 12.3269509 |
| -2.3186141 | 0.02041597 | 0.03492367 | 29.5087594 | 75.5004074 | 96.6381356 | 0 |
| -2.3184828 | 0.0204231 | 0.03493328 | 38.9351686 | 73.0649104 | 0 | 50.4284354 |

| | | | | | | |
|---|---|---|---|---|---|---|
| -2.3182558 | 0.02043542 | 0.03495251 | 11.4756286 | 13.530539 | 21.8586259 | 5.60315949 |
| -2.3179759 | 0.02045063 | 0.03497667 | 1.22953164 | 12.9893174 | 52.9208838 | 28.5761134 |
| -2.3178791 | 0.02045589 | 0.03498381 | 25.0004767 | 7.03588026 | 14.9559019 | 7.84442328 |
| -2.3177366 | 0.02046364 | 0.03499521 | 6.55750208 | 15.6954252 | 13.8054479 | 11.206319 |
| -2.3177082 | 0.02046519 | 0.034996 | 17.6232868 | 53.0397127 | 0 | 121.588561 |
| -2.3175425 | 0.0204742 | 0.03500956 | 9.42640924 | 129.351952 | 0 | 75.6426531 |
| -2.3172798 | 0.02048849 | 0.03503214 | 0 | 33.2851258 | 123.098578 | 61.0744384 |
| -2.3172095 | 0.02049232 | 0.03503683 | 35.6564176 | 13.8011497 | 16.1063559 | 11.7666349 |
| -2.317056 | 0.02050068 | 0.03504927 | 8.1968776 | 16.2366467 | 12.6549939 | 8.96505518 |
| -2.3169717 | 0.02050528 | 0.0350497 | 27.45954 | 36.803066 | 0 | 119.907613 |
| 2.31698903 | 0.02050433 | 0.0350497 | 0 | 0 | 0 | 0 |
| 2.31698903 | 0.02050433 | 0.0350497 | 0 | 0 | 0 | 0 |
| 2.31698903 | 0.02050433 | 0.0350497 | 0 | 0 | 0 | 0 |
| 2.31684985 | 0.02051191 | 0.03505919 | 35.6564176 | 22.4606947 | 19.5577179 | 11.7666349 |
| -2.3164991 | 0.02053103 | 0.03509001 | 0 | 30.8496288 | 104.691314 | 79.0045488 |
| -2.3160871 | 0.02055352 | 0.03512658 | 13.524848 | 15.1542036 | 5.75226998 | 13.4475828 |
| -2.3159434 | 0.02056136 | 0.03513812 | 15.1642236 | 50.8748265 | 0 | 134.475828 |
| -2.315589 | 0.02058072 | 0.03516935 | 49.1812656 | 12.4480958 | 6.90272397 | 6.16347543 |
| -2.315368 | 0.0205928 | 0.03518813 | 6.55750208 | 11.0950419 | 18.4072639 | 12.3269509 |
| -2.3151109 | 0.02060686 | 0.03520988 | 0 | 60.6168145 | 58.6731538 | 81.2458125 |
| -2.3150955 | 0.02060771 | 0.03520988 | 63.5258014 | 61.1580361 | 0 | 36.4205367 |
| 2.31505172 | 0.0206101 | 0.0352121 | 16.8035991 | 7.57710182 | 3.45136199 | 3.92221164 |
| -2.3150089 | 0.02061244 | 0.03521424 | 80.3294005 | 197.545869 | 130.001301 | 158.009098 |
| -2.3147871 | 0.02062458 | 0.03523312 | 73.3620545 | 148.565318 | 134.603117 | 146.242463 |
| -2.3146098 | 0.0206343 | 0.03524785 | 13.524848 | 21.1076408 | 24.1595339 | 3.36189569 |
| 2.31418522 | 0.02065756 | 0.03528573 | 5.32797044 | 11.6362635 | 11.50454 | 11.7666349 |
| -2.3140568 | 0.02066461 | 0.03529589 | 224.184602 | 195.380983 | 251.949425 | 234.772382 |
| -2.3136658 | 0.02068606 | 0.03533066 | 10.6559409 | 9.47137727 | 8.05317797 | 20.1713742 |
| -2.3133843 | 0.02070152 | 0.03535519 | 3.27875104 | 4.87099402 | 40.2658898 | 39.2221164 |
| -2.3133158 | 0.02070528 | 0.03535788 | 9.01656536 | 11.3656527 | 11.50454 | 10.646003 |
| -2.3133279 | 0.02070462 | 0.03535788 | 3.27875104 | 18.6721438 | 18.4072639 | 20.1713742 |
| -2.3131622 | 0.02071372 | 0.03537042 | 59.0175187 | 217.300456 | 232.391707 | 0 |
| 2.31303142 | 0.02072091 | 0.03538083 | 24.1807889 | 15.6954252 | 21.8586259 | 26.3348496 |
| -2.3129635 | 0.02072465 | 0.03538533 | 31.9678226 | 7.03588026 | 4.60181598 | 23.5332698 |
| -2.3126028 | 0.02074449 | 0.03541734 | 86.4770587 | 125.292791 | 110.443584 | 131.113932 |
| -2.312548 | 0.0207475 | 0.03541874 | 29.9186032 | 26.7904671 | 179.470823 | 0 |
| -2.3125557 | 0.02074708 | 0.03541874 | 28.6890716 | 8.93015571 | 10.354086 | 7.84442328 |
| 2.31251209 | 0.02074948 | 0.03542024 | 29.5087594 | 7.57710182 | 5.75226998 | 5.60315949 |
| -2.3121674 | 0.02076846 | 0.03545076 | 14.7543797 | 12.9893174 | 10.354086 | 7.28410733 |
| -2.3118269 | 0.02078723 | 0.03548093 | 106.149565 | 462.203211 | 444.075242 | 327.224514 |
| -2.311576 | 0.02080106 | 0.03550266 | 25.8201644 | 73.8767427 | 0 | 68.9188617 |
| -2.3115151 | 0.02080442 | 0.03550652 | 3.27875104 | 27.0610779 | 36.8145279 | 7.84442328 |
| -2.3112841 | 0.02081717 | 0.03552452 | 4.91812656 | 9.20076649 | 21.8586259 | 21.852322 |

| | | | | | | |
|---|---|---|---|---|---|---|
| -2.3112938 | 0.02081663 | 0.03552452 | 28.2792277 | 10.8244312 | 5.75226998 | 11.7666349 |
| -2.3111779 | 0.02082303 | 0.03553265 | 9.01656536 | 12.4480958 | 6.90272397 | 24.0935858 |
| -2.3109139 | 0.02083761 | 0.03555564 | 57.3781432 | 19.7545869 | 144.957203 | 0 |
| -2.3107287 | 0.02084784 | 0.03557123 | 22.1315695 | 113.927138 | 0 | 46.5062237 |
| -2.310341 | 0.02086928 | 0.03560592 | 5.73781432 | 20.5664192 | 13.8054479 | 11.206319 |
| -2.3103181 | 0.02087055 | 0.03560621 | 22.1315695 | 48.4393295 | 0 | 110.382242 |
| -2.3101939 | 0.02087742 | 0.03561417 | 3.27875104 | 27.3316887 | 9.20363196 | 30.2570612 |
| 2.31020858 | 0.02087661 | 0.03561417 | 13.1150042 | 6.4946587 | 8.05317797 | 5.04284354 |
| -2.3100621 | 0.02088472 | 0.03562285 | 20.0823501 | 9.74198805 | 9.20363196 | 8.96505518 |
| -2.3100792 | 0.02088377 | 0.03562285 | 8.1968776 | 18.1309222 | 12.6549939 | 8.40473923 |
| -2.3098609 | 0.02089586 | 0.03563997 | 8.1968776 | 16.5072575 | 5.75226998 | 26.8951655 |
| -2.3095153 | 0.020915 | 0.03567074 | 13.524848 | 11.0950419 | 12.6549939 | 7.28410733 |
| 2.3092987 | 0.02092701 | 0.03568933 | 116.805506 | 79.2889583 | 72.4786017 | 81.8061285 |
| -2.3092329 | 0.02093066 | 0.03569367 | 6.96734596 | 12.9893174 | 12.6549939 | 12.3269509 |
| -2.3092074 | 0.02093208 | 0.0356942 | 27.0496961 | 40.0503953 | 0 | 110.942558 |
| -2.3091581 | 0.02093481 | 0.03569697 | 228.692885 | 6.22404792 | 0 | 33.058641 |
| 2.30908752 | 0.02093873 | 0.03570177 | 58.6076748 | 25.1668025 | 32.2127119 | 31.3776931 |
| -2.3090379 | 0.02094148 | 0.03570457 | 7.78703372 | 66.5702517 | 196.727633 | 81.8061285 |
| -2.3086542 | 0.02096278 | 0.035739 | 0 | 61.6992576 | 74.7795097 | 61.0744384 |
| 2.30853365 | 0.02096947 | 0.03574853 | 1.22953164 | 1.62366467 | 0 | 0 |
| -2.3081728 | 0.02098953 | 0.03578082 | 22.1315695 | 24.6255809 | 11.50454 | 29.6967453 |
| -2.307995 | 0.02099941 | 0.03579579 | 52.0501728 | 77.6652936 | 0 | 33.058641 |
| -2.3076977 | 0.02101596 | 0.0358221 | 11.4756286 | 21.9194731 | 5.75226998 | 12.3269509 |
| -2.3071666 | 0.02104553 | 0.03587062 | 0 | 116.362635 | 60.9740618 | 34.1792729 |
| -2.307007 | 0.02105443 | 0.03588389 | 49.1812656 | 52.4984912 | 0 | 53.7903311 |
| -2.3069369 | 0.02105834 | 0.03588866 | 64.3454891 | 43.0271139 | 12.6549939 | 84.0473923 |
| -2.3068646 | 0.02106237 | 0.03589363 | 117.625194 | 23.0019162 | 0 | 42.0236961 |
| 2.30670067 | 0.02107151 | 0.03590732 | 29.9186032 | 10.2832096 | 10.354086 | 9.52537113 |
| -2.3065897 | 0.02107771 | 0.03591597 | 0.81968776 | 12.7187066 | 41.4163438 | 48.1871716 |
| -2.3063036 | 0.02109368 | 0.03594129 | 4.0984388 | 3.24732935 | 28.7613499 | 59.3934906 |
| -2.3060923 | 0.02110547 | 0.03595949 | 121.313788 | 379.666923 | 322.127119 | 320.500723 |
| -2.3059098 | 0.02111567 | 0.03597497 | 0 | 68.4645271 | 83.9831417 | 47.0665397 |
| -2.3058564 | 0.02111866 | 0.03597816 | 7.37718984 | 11.9068743 | 13.8054479 | 11.206319 |
| -2.3056076 | 0.02113257 | 0.03599996 | 8.60672148 | 14.8835929 | 10.354086 | 10.0856871 |
| 2.30555525 | 0.0211355 | 0.03600115 | 0 | 0 | 0 | 0 |
| 2.30555525 | 0.0211355 | 0.03600115 | 0 | 0 | 0 | 0 |
| -2.3048836 | 0.0211731 | 0.03606329 | 29.5087594 | 50.6042157 | 0 | 85.7283401 |
| -2.3047369 | 0.02118131 | 0.03607539 | 172.954117 | 499.818109 | 476.287954 | 462.820974 |
| -2.3046795 | 0.02118453 | 0.03607896 | 8.1968776 | 8.11832337 | 12.6549939 | 19.0507423 |
| -2.3045856 | 0.02118979 | 0.03608602 | 24.1807889 | 5.41221558 | 9.20363196 | 16.8094785 |
| -2.3040978 | 0.02121715 | 0.03613071 | 37.705637 | 27.6022995 | 44.8677058 | 42.5840121 |
| -2.3038922 | 0.0212287 | 0.03614846 | 20.9020379 | 20.0251977 | 17.2568099 | 17.3697944 |
| -2.3037508 | 0.02123664 | 0.03616007 | 7.37718984 | 9.47137727 | 57.5226998 | 6.16347543 |

| | | | | | | |
|---|---|---|---|---|---|---|
| 2.30337357 | 0.02125783 | 0.03619426 | 0 | 1.89427545 | 0 | 0 |
| -2.3031563 | 0.02127005 | 0.03621315 | 0 | 30.0377965 | 139.204933 | 50.9887513 |
| -2.302993 | 0.02127924 | 0.03622687 | 0 | 90.1133895 | 47.1686138 | 62.1950703 |
| -2.3029221 | 0.02128322 | 0.03623176 | 43.0336074 | 107.70309 | 0 | 25.2142177 |
| -2.3028501 | 0.02128728 | 0.03623675 | 13.1150042 | 6.22404792 | 24.1595339 | 9.52537113 |
| -2.3027666 | 0.02129198 | 0.03624284 | 111.067691 | 28.4141318 | 12.6549939 | 19.0507423 |
| -2.3025997 | 0.02130137 | 0.03625692 | 0 | 46.0038325 | 41.4163438 | 123.269509 |
| -2.3025655 | 0.0213033 | 0.03625828 | 0 | 28.6847426 | 71.3281477 | 112.623506 |
| -2.3024714 | 0.0213086 | 0.0362654 | 6.1476582 | 11.0950419 | 17.2568099 | 13.4475828 |
| 2.30152172 | 0.02136216 | 0.03634314 | 0 | 0 | 0 | 0 |
| 2.30152172 | 0.02136216 | 0.03634314 | 0 | 0 | 0 | 0 |
| 2.30152172 | 0.02136216 | 0.03634314 | 0 | 0 | 0 | 0 |
| 2.30152172 | 0.02136216 | 0.03634314 | 0 | 0 | 0 | 0 |
| 2.30152172 | 0.02136216 | 0.03634314 | 0 | 0 | 0 | 0 |
| 2.30152172 | 0.02136216 | 0.03634314 | 0 | 0 | 0 | 0 |
| 2.30152172 | 0.02136216 | 0.03634314 | 0 | 0 | 0 | 0 |
| -2.3013932 | 0.02136942 | 0.03635357 | 16.3937552 | 10.8244312 | 20.7081719 | 5.04284354 |
| -2.3012712 | 0.02137631 | 0.03636337 | 12.7051603 | 14.8835929 | 23.0090799 | 4.48252759 |
| -2.3010456 | 0.02138906 | 0.03638314 | 38.5253247 | 10.2832096 | 6.90272397 | 8.40473923 |
| -2.3009926 | 0.02139205 | 0.03638631 | 25.8201644 | 73.3355211 | 0 | 66.1172819 |
| -2.300902 | 0.02139717 | 0.03639311 | 54.0993922 | 82.8068984 | 0 | 28.0157974 |
| -2.300166 | 0.02143882 | 0.03646202 | 16.8035991 | 24.3549701 | 29.9118039 | 14.0078987 |
| -2.2998564 | 0.02145636 | 0.03648993 | 0 | 85.2423954 | 66.7263317 | 45.9459078 |
| 2.29977282 | 0.02146109 | 0.03649607 | 0 | 0 | 0 | 0 |
| -2.2997024 | 0.02146509 | 0.03650094 | 15.5740674 | 3.24732935 | 12.6549939 | 37.5411686 |
| -2.2995983 | 0.02147099 | 0.03650905 | 0 | 61.4286469 | 49.4695218 | 86.2886561 |
| -2.2994854 | 0.02147739 | 0.03651801 | 65.9848647 | 37.8855091 | 83.9831417 | 0 |
| -2.2993098 | 0.02148736 | 0.03653303 | 18.4429746 | 4.60038325 | 26.4604419 | 9.52537113 |
| -2.2989679 | 0.02150676 | 0.03656255 | 9.01656536 | 20.83703 | 8.05317797 | 10.646003 |
| -2.298964 | 0.02150698 | 0.03656255 | 17.6232868 | 66.8408624 | 0 | 94.1330794 |
| -2.298887 | 0.02151136 | 0.03656806 | 0 | 67.9233056 | 85.1335957 | 44.8252759 |
| -2.2986579 | 0.02152437 | 0.03658826 | 80.7392443 | 145.047378 | 111.594038 | 117.666349 |
| -2.2986168 | 0.02152671 | 0.0365903 | 70.4931473 | 0 | 46.0181598 | 99.7362389 |
| -2.2984412 | 0.02153669 | 0.03660534 | 8.60672148 | 6.4946587 | 29.9118039 | 12.3269509 |
| -2.2981063 | 0.02155574 | 0.03663579 | 4.50828268 | 18.6721438 | 20.7081719 | 11.206319 |
| -2.2980301 | 0.02156008 | 0.03664124 | 67.2143963 | 41.4034492 | 0 | 47.0665397 |
| -2.2979761 | 0.02156315 | 0.03664453 | 35.2465737 | 285.764983 | 0 | 147.363094 |
| 2.29792379 | 0.02156613 | 0.03664766 | 1.63937552 | 4.60038325 | 12.6549939 | 7.28410733 |
| -2.2978826 | 0.02156847 | 0.03664971 | 35.6564176 | 78.7477367 | 0 | 45.3855918 |
| -2.2977545 | 0.02157576 | 0.03666018 | 795.506971 | 1179.05116 | 918.062288 | 1036.5845 |
| -2.2976204 | 0.0215834 | 0.03667123 | 32.7875104 | 60.3462037 | 0 | 64.99665 |
| -2.2971585 | 0.02160973 | 0.03671403 | 8.60672148 | 13.2599282 | 11.50454 | 9.52537113 |
| -2.2970559 | 0.02161558 | 0.03672204 | 9.83625312 | 39.5091738 | 33.3631659 | 34.1792729 |

| | | | | | | |
|---|---|---|---|---|---|---|
| -2.2970329 | 0.02161689 | 0.03672233 | 26.2300083 | 62.51109 | 0 | 75.0823371 |
| -2.2969428 | 0.02162203 | 0.03672913 | 184.429746 | 399.962732 | 406.11026 | 449.373391 |
| -2.2965418 | 0.02164492 | 0.03676456 | 7.78703372 | 7.8477126 | 28.7613499 | 11.206319 |
| -2.2965375 | 0.02164517 | 0.03676456 | 12.7051603 | 9.47137727 | 31.0622579 | 5.60315949 |
| -2.2964892 | 0.02164793 | 0.03676731 | 8.60672148 | 14.3423713 | 5.75226998 | 28.0157974 |
| -2.2962298 | 0.02166275 | 0.03679055 | 256.562269 | 4.87099402 | 0 | 25.7745336 |
| -2.295758 | 0.02168972 | 0.03683442 | 3.68859492 | 16.5072575 | 17.2568099 | 19.0507423 |
| 2.29570785 | 0.02169259 | 0.03683736 | 0.40984388 | 0.54122156 | 0 | 0 |
| -2.2956619 | 0.02169522 | 0.03683795 | 27.8693838 | 59.5343714 | 0 | 74.5220212 |
| 2.29567618 | 0.0216944 | 0.03683795 | 0.81968776 | 0 | 0 | 0 |
| -2.2954528 | 0.02170719 | 0.03685633 | 11.8854725 | 13.530539 | 43.7172518 | 3.36189569 |
| -2.2954199 | 0.02170907 | 0.03685759 | 11.4756286 | 1.89427545 | 36.8145279 | 29.6967453 |
| -2.2951559 | 0.02172419 | 0.03688133 | 30.738291 | 15.966036 | 204.780811 | 0 |
| -2.2950904 | 0.02172795 | 0.03688576 | 43.8532952 | 10.2832096 | 4.60181598 | 11.206319 |
| -2.2950425 | 0.02173069 | 0.03688847 | 20.9020379 | 74.4179643 | 0 | 74.5220212 |
| 2.29454588 | 0.02175916 | 0.03693487 | 36.8859492 | 20.2958084 | 20.7081719 | 17.3697944 |
| 2.29394889 | 0.02179343 | 0.03698785 | 0 | 0 | 0 | 0 |
| -2.2939423 | 0.02179381 | 0.03698785 | 306.973066 | 616.451355 | 601.68744 | 693.110828 |
| 2.29394889 | 0.02179343 | 0.03698785 | 0 | 0 | 0 | 0 |
| -2.2939113 | 0.0217956 | 0.03698893 | 2.0492194 | 6.4946587 | 65.5758777 | 23.5332698 |
| -2.2935994 | 0.02181352 | 0.03701741 | 138.527231 | 211.617629 | 179.470823 | 182.102683 |
| -2.29331 | 0.02183016 | 0.0370437 | 65.9848647 | 26.2492456 | 0 | 70.0394936 |
| -2.2931433 | 0.02183976 | 0.03705803 | 45.9025146 | 82.2656769 | 0 | 32.498325 |
| 2.29305375 | 0.02184491 | 0.03706288 | 0 | 0 | 1.150454 | 0 |
| 2.29305375 | 0.02184491 | 0.03706288 | 0 | 0 | 1.150454 | 0 |
| -2.292999 | 0.02184806 | 0.03706628 | 57.3781432 | 46.2744432 | 0 | 48.7474875 |
| -2.292768 | 0.02186136 | 0.0370869 | 24.1807889 | 96.6080482 | 77.0804177 | 0 |
| 2.29248181 | 0.02187786 | 0.03711264 | 29.0989155 | 37.0736767 | 47.1686138 | 40.3427483 |
| 2.29246481 | 0.02187884 | 0.03711264 | 3.68859492 | 2.70610779 | 2.30090799 | 3.36189569 |
| -2.2922908 | 0.02188887 | 0.03712771 | 0 | 56.8282636 | 86.2840497 | 51.5490673 |
| -2.2921695 | 0.02189587 | 0.03713763 | 6.1476582 | 11.3656527 | 14.9559019 | 14.0078987 |
| -2.2915925 | 0.02192917 | 0.03719216 | 52.8698605 | 54.9339882 | 0 | 44.2649599 |
| -2.2915233 | 0.02193317 | 0.03719699 | 2.45906328 | 8.65954493 | 39.1154358 | 28.0157974 |
| 2.2914058 | 0.02193995 | 0.03720654 | 4.0984388 | 3.51794013 | 4.60181598 | 4.48252759 |
| -2.291272 | 0.02194768 | 0.03721769 | 0.40984388 | 17.3190899 | 73.6290557 | 28.5761134 |
| -2.2911687 | 0.02195366 | 0.03722587 | 44.263139 | 32.2026827 | 0 | 84.6077082 |
| -2.2911455 | 0.021955 | 0.03722618 | 0 | 98.7729344 | 35.6640739 | 66.6775979 |
| -2.2911214 | 0.0219564 | 0.0372266 | 0 | 60.3462037 | 51.7704298 | 81.2458125 |
| -2.2906558 | 0.02198333 | 0.03727031 | 21.3118818 | 54.6633774 | 0 | 95.2537113 |
| 2.29040554 | 0.02199782 | 0.03729292 | 3.27875104 | 1.08244312 | 0 | 1.68094785 |
| -2.2903325 | 0.02200205 | 0.03729813 | 0 | 41.67406 | 97.7885896 | 58.8331746 |
| -2.2897068 | 0.02203832 | 0.03735765 | 111.477535 | 9.74198805 | 110.443584 | 0 |
| 2.28964846 | 0.0220417 | 0.03736143 | 2.45906328 | 5.41221558 | 2.30090799 | 2.80157974 |

| | | | | | | |
|---|---|---|---|---|---|---|
| -2.2892545 | 0.02206457 | 0.03739822 | 7.37718984 | 10.5538204 | 25.3099879 | 8.96505518 |
| -2.2891595 | 0.02207009 | 0.03740561 | 3.68859492 | 17.5897006 | 27.6108959 | 11.7666349 |
| -2.2890284 | 0.0220777 | 0.03741656 | 0 | 41.67406 | 94.3372276 | 61.0744384 |
| -2.2889213 | 0.02208393 | 0.03742514 | 14.3445358 | 97.6904913 | 0 | 70.5998095 |
| -2.2888177 | 0.02208995 | 0.03743338 | 0 | 62.51109 | 51.7704298 | 77.8839169 |
| -2.2886654 | 0.0220988 | 0.03744642 | 0 | 98.5023236 | 29.9118039 | 75.6426531 |
| -2.2885456 | 0.02210577 | 0.03745625 | 49.1812656 | 31.3908504 | 115.0454 | 0 |
| 2.28819319 | 0.02212627 | 0.03748903 | 26.2300083 | 6.4946587 | 2.30090799 | 7.28410733 |
| -2.288064 | 0.02213379 | 0.03749981 | 90.9853413 | 13.2599282 | 3.45136199 | 3.92221164 |
| -2.2880085 | 0.02213703 | 0.03750332 | 65.9848647 | 41.1328384 | 37.9649818 | 34.1792729 |
| -2.2879331 | 0.02214142 | 0.03750878 | 10.246097 | 12.7187066 | 5.75226998 | 22.4126379 |
| -2.287841 | 0.02214678 | 0.03751591 | 20.9020379 | 0 | 62.1245158 | 161.370993 |
| -2.2877384 | 0.02215276 | 0.03752407 | 4.0984388 | 14.3423713 | 16.1063559 | 19.6110582 |
| -2.2876794 | 0.0221562 | 0.03752792 | 4.50828268 | 3.24732935 | 59.8236078 | 24.0935858 |
| -2.2876315 | 0.02215899 | 0.03753067 | 24.5906328 | 17.0484791 | 10.354086 | 15.1285306 |
| -2.287241 | 0.02218176 | 0.03756728 | 15.1642236 | 107.70309 | 89.7354116 | 0 |
| -2.2871449 | 0.02218737 | 0.03757479 | 2.45906328 | 22.7313054 | 21.8586259 | 17.9301104 |
| -2.286878 | 0.02220294 | 0.03759921 | 2.45906328 | 16.5072575 | 23.0090799 | 23.5332698 |
| -2.2868452 | 0.02220486 | 0.03760048 | 18.8528185 | 68.1939163 | 0 | 84.0473923 |
| -2.2867163 | 0.02221239 | 0.03761125 | 38.5253247 | 70.3588026 | 73.6290557 | 0 |
| -2.2866314 | 0.02221735 | 0.03761768 | 0 | 74.9591858 | 51.7704298 | 64.4363341 |
| -2.286546 | 0.02222234 | 0.03762415 | 30.738291 | 63.3229223 | 0 | 62.1950703 |
| -2.2864552 | 0.02222764 | 0.03763116 | 3.27875104 | 15.4248144 | 17.2568099 | 23.5332698 |
| -2.2862937 | 0.02223708 | 0.03764517 | 7.37718984 | 15.1542036 | 8.05317797 | 16.2491625 |
| -2.2861767 | 0.02224392 | 0.03765477 | 63.5258014 | 93.3607188 | 87.4345036 | 90.7711837 |
| -2.2860896 | 0.02224902 | 0.03766143 | 4.50828268 | 17.8603114 | 12.6549939 | 17.3697944 |
| 2.28593887 | 0.02225783 | 0.03767438 | 1.22953164 | 1.08244312 | 0 | 1.68094785 |
| -2.2858172 | 0.02226495 | 0.0376811 | 2.0492194 | 40.5916169 | 6.90272397 | 37.5411686 |
| -2.2858111 | 0.02226531 | 0.0376811 | 232.38148 | 153.165701 | 224.338529 | 220.764484 |
| -2.2858147 | 0.0222651 | 0.0376811 | 4.50828268 | 16.7778683 | 12.6549939 | 18.4904263 |
| 2.2855924 | 0.02227812 | 0.0377008 | 0 | 2.16488623 | 2.30090799 | 2.24126379 |
| -2.285139 | 0.02230468 | 0.0377408 | 10.6559409 | 12.4480958 | 11.50454 | 7.84442328 |
| -2.2851291 | 0.02230526 | 0.0377408 | 0 | 68.4645271 | 31.0622579 | 103.098135 |
| -2.2851493 | 0.02230407 | 0.0377408 | 0 | 54.9339882 | 73.6290557 | 61.6347543 |
| -2.2849674 | 0.02231474 | 0.03775486 | 4.91812656 | 27.0610779 | 29.9118039 | 5.60315949 |
| -2.2848681 | 0.02232056 | 0.03776075 | 13.1150042 | 11.6362635 | 12.6549939 | 6.72379138 |
| -2.2848866 | 0.02231948 | 0.03776075 | 0 | 103.373318 | 80.5317797 | 24.6539017 |
| -2.2847965 | 0.02232476 | 0.03776588 | 2.0492194 | 15.4248144 | 25.3099879 | 28.0157974 |
| -2.2847305 | 0.02232863 | 0.03777045 | 12.7051603 | 96.3374374 | 0 | 76.202969 |
| -2.2845703 | 0.02233803 | 0.03778437 | 0 | 51.9572696 | 77.0804177 | 61.6347543 |
| -2.2845073 | 0.02234174 | 0.03778865 | 14.7543797 | 12.7187066 | 6.90272397 | 10.0856871 |
| -2.2844382 | 0.02234579 | 0.03779353 | 29.9186032 | 50.0629941 | 0 | 78.4442328 |
| -2.2843075 | 0.02235347 | 0.03780454 | 12.2953164 | 6.22404792 | 8.05317797 | 33.6189569 |

| | | | | | | |
|---|---|---|---|---|---|---|
| -2.284183 | 0.02236078 | 0.03781492 | 22.5414134 | 8.65954493 | 9.20363196 | 8.96505518 |
| -2.2840857 | 0.02236649 | 0.0378226 | 176.232868 | 213.241294 | 322.127119 | 356.921259 |
| -2.2840494 | 0.02236863 | 0.03782424 | 38.1154808 | 107.432479 | 155.311289 | 169.215416 |
| -2.2835 | 0.02240093 | 0.03787688 | 9.42640924 | 11.0950419 | 9.20363196 | 11.7666349 |
| -2.2834044 | 0.02240656 | 0.0378844 | 8.1968776 | 19.2133653 | 10.354086 | 8.96505518 |
| -2.2832568 | 0.02241525 | 0.03789711 | 0 | 22.7313054 | 157.612197 | 45.9459078 |
| -2.2830028 | 0.0224302 | 0.03792041 | 0 | 104.726372 | 33.3631659 | 62.7553862 |
| 2.28281803 | 0.02244109 | 0.03793683 | 11.0657848 | 15.6954252 | 10.354086 | 11.206319 |
| -2.282361 | 0.02246804 | 0.03798039 | 12.7051603 | 6.76526948 | 32.2127119 | 7.28410733 |
| 2.2823104 | 0.02247102 | 0.03798345 | 0.40984388 | 0.54122156 | 2.30090799 | 0 |
| 2.28216551 | 0.02247957 | 0.03798796 | 0 | 0 | 0 | 0 |
| 2.28216551 | 0.02247957 | 0.03798796 | 0 | 0 | 0 | 0 |
| 2.28223464 | 0.02247549 | 0.03798796 | 31.9678226 | 21.1076408 | 33.3631659 | 22.4126379 |
| 2.28222318 | 0.02247617 | 0.03798796 | 2.45906328 | 2.97671857 | 2.30090799 | 4.48252759 |
| 2.28216551 | 0.02247957 | 0.03798796 | 0 | 0 | 0 | 0 |
| -2.2820208 | 0.02248811 | 0.03800041 | 18.8528185 | 34.9087905 | 26.4604419 | 20.1713742 |
| -2.2816001 | 0.02251296 | 0.0380404 | 3.68859492 | 14.3423713 | 12.6549939 | 31.3776931 |
| -2.2815764 | 0.02251436 | 0.03804077 | 3.27875104 | 31.6614612 | 10.354086 | 20.1713742 |
| -2.2815436 | 0.0225163 | 0.03804206 | 5.32797044 | 11.9068743 | 16.1063559 | 15.1285306 |
| -2.2814927 | 0.02251931 | 0.03804515 | 6.1476582 | 11.0950419 | 17.2568099 | 12.3269509 |
| -2.2814659 | 0.02252089 | 0.03804584 | 21.3118818 | 51.416048 | 0 | 96.3743432 |
| 2.2813212 | 0.02252945 | 0.0380583 | 18.0331307 | 8.38893415 | 20.7081719 | 8.96505518 |
| 2.28098069 | 0.02254959 | 0.03808236 | 0 | 0 | 0 | 0 |
| 2.28098069 | 0.02254959 | 0.03808236 | 0 | 0 | 0 | 0 |
| 2.28098069 | 0.02254959 | 0.03808236 | 0 | 0 | 0 | 0 |
| 2.28098069 | 0.02254959 | 0.03808236 | 0 | 0 | 0 | 0 |
| 2.28098069 | 0.02254959 | 0.03808236 | 0 | 0 | 0 | 0 |
| -2.2804472 | 0.02258118 | 0.03813372 | 4.0984388 | 14.6129821 | 14.9559019 | 20.1713742 |
| -2.280393 | 0.02258439 | 0.03813714 | 95.0837801 | 0 | 40.2658898 | 75.6426531 |
| -2.2803523 | 0.0225868 | 0.03813923 | 9.01656536 | 12.1774851 | 17.2568099 | 7.28410733 |
| -2.2801918 | 0.02259631 | 0.03815329 | 72.5423667 | 26.7904671 | 0 | 58.8331746 |
| 2.28014865 | 0.02259887 | 0.03815562 | 0 | 1.3530539 | 0 | 0 |
| -2.2801219 | 0.02260046 | 0.0381563 | 4.50828268 | 13.530539 | 18.4072639 | 15.1285306 |
| -2.2799483 | 0.02261075 | 0.03817168 | 0 | 79.5595691 | 48.3190678 | 62.1950703 |
| -2.2797997 | 0.02261957 | 0.03818258 | 4.0984388 | 14.3423713 | 31.0622579 | 11.206319 |
| -2.2798129 | 0.02261879 | 0.03818258 | 6.96734596 | 12.4480958 | 14.9559019 | 10.0856871 |
| -2.2794853 | 0.02263824 | 0.03821208 | 13.1150042 | 16.5072575 | 8.05317797 | 7.84442328 |
| 2.27918803 | 0.02265589 | 0.03823988 | 29.0989155 | 45.7332217 | 18.4072639 | 49.8681194 |
| -2.2787622 | 0.0226812 | 0.03828061 | 13.524848 | 11.0950419 | 5.75226998 | 16.2491625 |
| -2.2787145 | 0.02268404 | 0.03828339 | 13.1150042 | 23.0019162 | 17.2568099 | 37.5411686 |
| -2.2786383 | 0.02268858 | 0.03828905 | 25.8201644 | 57.0988744 | 0 | 76.202969 |
| -2.278248 | 0.02271181 | 0.03832625 | 4.0984388 | 17.5897006 | 21.8586259 | 11.7666349 |
| -2.2777073 | 0.02274402 | 0.03837861 | 73.3620545 | 44.9213893 | 0 | 35.2999048 |

| | | | | | | |
|---|---|---|---|---|---|---|
| -2.2776863 | 0.02274527 | 0.03837871 | 66.8045524 | 37.0736767 | 0 | 47.6268556 |
| -2.2776633 | 0.02274665 | 0.03837902 | 50.4107972 | 81.1832337 | 47.1686138 | 0 |
| 2.27742846 | 0.02276065 | 0.03840065 | 16.8035991 | 7.8477126 | 2.30090799 | 12.3269509 |
| -2.2771599 | 0.02277668 | 0.03842568 | 29.5087594 | 63.3229223 | 0 | 61.6347543 |
| 2.27700214 | 0.0227861 | 0.03843956 | 15.1642236 | 10.8244312 | 13.8054479 | 7.28410733 |
| -2.2769669 | 0.0227882 | 0.0384411 | 30.3284471 | 64.1347547 | 0 | 59.3934906 |
| -2.2768649 | 0.02279429 | 0.03844937 | 31.9678226 | 29.4965749 | 10.354086 | 20.1713742 |
| -2.2767436 | 0.02280154 | 0.03845958 | 121.313788 | 11.9068743 | 0 | 57.1522268 |
| 2.27650534 | 0.02281578 | 0.03848159 | 35.6564176 | 26.7904671 | 23.0090799 | 21.292006 |
| 2.27615506 | 0.02283673 | 0.03851491 | 3.68859492 | 4.32977247 | 1.150454 | 2.24126379 |
| -2.2760875 | 0.02284077 | 0.03851972 | 5.32797044 | 15.1542036 | 18.4072639 | 10.646003 |
| -2.2758825 | 0.02285304 | 0.0385384 | 89.7558097 | 278.729103 | 253.099879 | 252.142177 |
| -2.2755655 | 0.02287203 | 0.0385684 | 1.63937552 | 23.5431378 | 20.7081719 | 26.8951655 |
| -2.2755359 | 0.0228738 | 0.03856938 | 4.91812656 | 16.2366467 | 16.1063559 | 12.3269509 |
| -2.2754354 | 0.02287982 | 0.03857571 | 91.8050291 | 30.0377965 | 0 | 39.2221164 |
| -2.275425 | 0.02288045 | 0.03857571 | 113.526755 | 0 | 43.7172518 | 54.910963 |
| -2.2754134 | 0.02288114 | 0.03857571 | 89.7558097 | 36.2618444 | 0 | 33.6189569 |
| 2.27535372 | 0.02288472 | 0.03857973 | 44.6729829 | 51.6866588 | 49.4695218 | 43.704644 |
| -2.2751639 | 0.0228961 | 0.0385969 | 7.37718984 | 14.3423713 | 8.05317797 | 16.2491625 |
| -2.2751185 | 0.02289882 | 0.03859947 | 18.4429746 | 10.2832096 | 5.75226998 | 13.4475828 |
| -2.2750077 | 0.02290547 | 0.03860664 | 15.1642236 | 7.8477126 | 10.354086 | 10.0856871 |
| -2.2750212 | 0.02290466 | 0.03860664 | 4.0984388 | 19.4839761 | 10.354086 | 22.9729539 |
| -2.2749326 | 0.02290998 | 0.03861222 | 1.22953164 | 25.4374132 | 20.7081719 | 32.498325 |
| -2.2745925 | 0.02293039 | 0.03864461 | 6.1476582 | 37.6148983 | 11.50454 | 7.84442328 |
| -2.2743038 | 0.02294773 | 0.03867181 | 7.78703372 | 16.7778683 | 5.75226998 | 22.9729539 |
| -2.2742759 | 0.0229494 | 0.03867262 | 11.4756286 | 46.2744432 | 179.470823 | 0 |
| -2.2742521 | 0.02295083 | 0.03867301 | 12.7051603 | 11.3656527 | 18.4072639 | 5.60315949 |
| -2.2740374 | 0.02296374 | 0.03869274 | 0 | 60.6168145 | 69.0272397 | 56.5919108 |
| -2.2739932 | 0.02296639 | 0.03869519 | 3.68859492 | 17.8603114 | 14.9559019 | 18.4904263 |
| -2.2738541 | 0.02297476 | 0.03870691 | 9.83625312 | 23.8137486 | 5.75226998 | 12.3269509 |
| -2.2738377 | 0.02297574 | 0.03870691 | 0 | 67.382084 | 58.6731538 | 59.9538065 |
| -2.2735623 | 0.02299232 | 0.03873281 | 35.6564176 | 58.7225391 | 0 | 55.4712789 |
| -2.27354 | 0.02299366 | 0.03873304 | 18.8528185 | 10.5538204 | 19.5577179 | 4.48252759 |
| -2.2733469 | 0.02300529 | 0.03875061 | 35.2465737 | 39.238563 | 117.346308 | 0 |
| -2.2732657 | 0.02301017 | 0.03875681 | 148.773328 | 656.50175 | 400.35799 | 470.105081 |
| -2.2732062 | 0.02301376 | 0.03876083 | 85.6573709 | 45.1920001 | 0 | 28.0157974 |
| -2.2730135 | 0.02302536 | 0.03877836 | 2.45906328 | 18.401533 | 40.2658898 | 11.7666349 |
| -2.2725312 | 0.02305444 | 0.03882531 | 66.3947085 | 62.7817008 | 46.0181598 | 0 |
| -2.2724934 | 0.02305672 | 0.03882712 | 2.45906328 | 37.8855091 | 24.1595339 | 9.52537113 |
| -2.2722885 | 0.02306909 | 0.03884593 | 8.60672148 | 6.76526948 | 16.1063559 | 15.1285306 |
| -2.2720886 | 0.02308116 | 0.03886422 | 8.60672148 | 18.1309222 | 10.354086 | 8.40473923 |
| -2.2720194 | 0.02308534 | 0.03886923 | 6.1476582 | 15.4248144 | 9.20363196 | 16.8094785 |
| -2.2718779 | 0.02309389 | 0.03888159 | 4.91812656 | 17.0484791 | 33.3631659 | 7.28410733 |

| | | | | | | |
|---|---|---|---|---|---|---|
| -2.2716068 | 0.02311027 | 0.03890715 | 7.78703372 | 14.0717605 | 11.50454 | 9.52537113 |
| 2.27145112 | 0.02311968 | 0.03892097 | 0 | 0 | 0 | 0.56031595 |
| -2.2711072 | 0.02314049 | 0.03895396 | 6.55750208 | 16.5072575 | 16.1063559 | 8.40473923 |
| -2.2709234 | 0.02315162 | 0.03897066 | 6.96734596 | 8.38893415 | 11.50454 | 25.2142177 |
| -2.2706457 | 0.02316843 | 0.03899693 | 168.035991 | 204.581749 | 234.692615 | 223.005748 |
| 2.27020045 | 0.02319542 | 0.03903829 | 0 | 0 | 0 | 0 |
| 2.27020045 | 0.02319542 | 0.03903829 | 0 | 0 | 0 | 0 |
| -2.2701164 | 0.02320052 | 0.03904346 | 3.27875104 | 14.8835929 | 24.1595339 | 16.2491625 |
| -2.27009 | 0.02320212 | 0.03904346 | 10.6559409 | 4.87099402 | 17.2568099 | 18.4904263 |
| -2.2701072 | 0.02320108 | 0.03904346 | 410.663568 | 1427.20125 | 992.841798 | 1139.68264 |
| 2.2698388 | 0.02321737 | 0.03906504 | 0 | 0.27061078 | 0 | 0 |
| 2.26984352 | 0.02321708 | 0.03906504 | 0 | 0 | 1.150454 | 0.56031595 |
| 2.26959165 | 0.02323237 | 0.03908825 | 0 | 0 | 8.05317797 | 5.60315949 |
| 2.26902344 | 0.0232669 | 0.03914431 | 4.50828268 | 2.97671857 | 1.150454 | 3.92221164 |
| 2.26876909 | 0.02328237 | 0.0391683 | 7.37718984 | 24.0843593 | 23.0090799 | 21.852322 |
| -2.2686981 | 0.02328669 | 0.03917352 | 18.0331307 | 24.8961917 | 8.05317797 | 5.04284354 |
| -2.2685392 | 0.02329636 | 0.0391875 | 27.45954 | 15.6954252 | 21.8586259 | 15.1285306 |
| -2.2685217 | 0.02329743 | 0.0391875 | 176.232868 | 409.97533 | 376.198456 | 398.944955 |
| -2.2681986 | 0.02331711 | 0.03921856 | 10.6559409 | 6.22404792 | 14.9559019 | 13.4475828 |
| -2.2680656 | 0.02332521 | 0.03923014 | 24.1807889 | 100.125988 | 0 | 40.9030642 |
| -2.2678511 | 0.02333828 | 0.03925009 | 6.96734596 | 90.9252218 | 0 | 105.339398 |
| -2.267747 | 0.02334463 | 0.03925872 | 24.5906328 | 0 | 106.992222 | 87.9696039 |
| -2.2674878 | 0.02336044 | 0.03928326 | 0 | 47.0862756 | 103.54086 | 42.5840121 |
| -2.2674489 | 0.02336282 | 0.03928521 | 11.0657848 | 10.2832096 | 21.8586259 | 6.16347543 |
| -2.2673807 | 0.02336698 | 0.03929016 | 4.91812656 | 15.4248144 | 29.9118039 | 8.40473923 |
| -2.2673044 | 0.02337163 | 0.03929594 | 46.3123584 | 27.3316887 | 119.647216 | 0 |
| -2.2672519 | 0.02337484 | 0.03929929 | 35.6564176 | 45.1920001 | 103.54086 | 0 |
| -2.2670151 | 0.02338931 | 0.03932156 | 70.0833035 | 41.67406 | 0 | 38.1014845 |
| -2.2668632 | 0.02339858 | 0.0393351 | 0 | 62.2404792 | 54.0713378 | 67.7982298 |
| -2.266711 | 0.02340789 | 0.0393487 | 174.183649 | 22.4606947 | 0 | 15.1285306 |
| -2.2666628 | 0.02341084 | 0.03935081 | 8.1968776 | 9.47137727 | 13.8054479 | 10.646003 |
| -2.2666506 | 0.02341158 | 0.03935081 | 63.1159575 | 70.0881918 | 32.2127119 | 49.8681194 |
| 2.26553118 | 0.0234801 | 0.03946393 | 0 | 8.38893415 | 17.2568099 | 0 |
| -2.2651951 | 0.02350071 | 0.03949651 | 55.3289238 | 47.3568864 | 0 | 43.144328 |
| 2.26509034 | 0.02350714 | 0.0395032 | 0.40984388 | 0 | 0 | 0.56031595 |
| 2.26509034 | 0.02350714 | 0.0395032 | 0.40984388 | 0 | 0 | 0.56031595 |
| 2.26496782 | 0.02351465 | 0.03951377 | 4.0984388 | 11.9068743 | 4.60181598 | 8.96505518 |
| -2.2649344 | 0.02351671 | 0.03951516 | 17.6232868 | 3.51794013 | 49.4695218 | 6.72379138 |
| -2.2647845 | 0.02352591 | 0.03952857 | 45.4926707 | 56.0164313 | 0 | 44.2649599 |
| -2.2647353 | 0.02352893 | 0.03952964 | 46.3123584 | 44.3801678 | 0 | 54.910963 |
| -2.2647343 | 0.02352899 | 0.03952964 | 4.50828268 | 20.5664192 | 13.8054479 | 12.8872668 |
| -2.2646014 | 0.02353715 | 0.03954129 | 4.91812656 | 11.3656527 | 23.0090799 | 12.8872668 |
| -2.2645459 | 0.02354056 | 0.03954495 | 58.6076748 | 47.6274971 | 52.9208838 | 47.0665397 |

| | | | | | | |
|---|---|---|---|---|---|---|
| -2.264361 | 0.02355192 | 0.03956198 | 2.0492194 | 27.6022995 | 20.7081719 | 17.3697944 |
| -2.2642089 | 0.02356127 | 0.03957563 | 36.4761053 | 6.22404792 | 10.354086 | 8.40473923 |
| -2.2641535 | 0.02356467 | 0.03957929 | 78.6900249 | 158.577917 | 130.001301 | 129.432984 |
| -2.2639404 | 0.02357778 | 0.03959925 | 14.7543797 | 21.3782516 | 19.5577179 | 19.0507423 |
| -2.2638456 | 0.02358361 | 0.03960698 | 1.63937552 | 30.8496288 | 13.8054479 | 29.1364293 |
| -2.263001 | 0.02363563 | 0.03969227 | 16.3937552 | 66.0290301 | 0 | 85.7283401 |
| -2.2628548 | 0.02364464 | 0.03970534 | 3.27875104 | 14.0717605 | 33.3631659 | 12.8872668 |
| -2.2626759 | 0.02365567 | 0.0397218 | 13.524848 | 7.8477126 | 9.20363196 | 11.7666349 |
| -2.2620852 | 0.02369214 | 0.03978096 | 0 | 16.2366467 | 142.656295 | 61.6347543 |
| -2.2618026 | 0.0237096 | 0.03980614 | 30.738291 | 0 | 62.1245158 | 121.588561 |
| -2.2618184 | 0.02370863 | 0.03980614 | 29.0989155 | 56.0164313 | 0 | 65.556966 |
| 2.26162516 | 0.02372057 | 0.03982249 | 81.5589321 | 59.2637606 | 75.9299637 | 62.1950703 |
| -2.2615748 | 0.02372369 | 0.03982565 | 6.55750208 | 95.525605 | 0 | 100.856871 |
| -2.2614788 | 0.02372963 | 0.03983293 | 4.0984388 | 49.5217726 | 14.9559019 | 6.72379138 |
| -2.2614644 | 0.02373052 | 0.03983293 | 39.7548564 | 0 | 113.894946 | 54.910963 |
| -2.2614448 | 0.02373173 | 0.03983293 | 86.4770587 | 17.5897006 | 0 | 61.6347543 |
| -2.2613438 | 0.02373798 | 0.0398382 | 29.9186032 | 57.640096 | 0 | 62.1950703 |
| -2.2613143 | 0.0237398 | 0.0398382 | 22.5414134 | 58.7225391 | 0 | 76.202969 |
| 2.26131734 | 0.02373961 | 0.0398382 | 11.4756286 | 0 | 0 | 0 |
| -2.2613533 | 0.02373739 | 0.0398382 | 4.0984388 | 33.2851258 | 16.1063559 | 8.96505518 |
| 2.2612319 | 0.0237449 | 0.03984469 | 1.63937552 | 2.43549701 | 0 | 1.1206319 |
| -2.2602034 | 0.02380863 | 0.03994955 | 0 | 50.6042157 | 40.2658898 | 99.1759229 |
| -2.2600633 | 0.02381733 | 0.03996206 | 26.6398522 | 6.76526948 | 20.7081719 | 5.04284354 |
| -2.2599375 | 0.02382513 | 0.03997308 | 2.0492194 | 21.1076408 | 26.4604419 | 17.3697944 |
| 2.25972618 | 0.02383825 | 0.03999302 | 3.27875104 | 2.43549701 | 1.150454 | 0 |
| -2.2592347 | 0.02386879 | 0.04003168 | 36.4761053 | 55.7458205 | 64.4254237 | 62.7553862 |
| 2.25917571 | 0.02387246 | 0.04003168 | 0 | 0 | 0 | 0 |
| 2.25917571 | 0.02387246 | 0.04003168 | 0 | 0 | 0 | 0 |
| 2.25917571 | 0.02387246 | 0.04003168 | 0 | 0 | 0 | 0 |
| 2.25917571 | 0.02387246 | 0.04003168 | 0 | 0 | 0 | 0 |
| 2.25917571 | 0.02387246 | 0.04003168 | 0 | 0 | 0 | 0 |
| 2.25917571 | 0.02387246 | 0.04003168 | 0 | 0 | 0 | 0 |
| -2.2591815 | 0.0238721 | 0.04003168 | 47.5418901 | 0 | 67.8767857 | 84.0473923 |
| 2.25917571 | 0.02387246 | 0.04003168 | 0 | 0 | 0 | 0 |
| -2.2590956 | 0.02387744 | 0.04003795 | 0 | 48.980551 | 65.5758777 | 67.7982298 |
| -2.2590528 | 0.0238801 | 0.04004034 | 18.4429746 | 73.6061319 | 102.390406 | 0 |
| -2.2590156 | 0.02388242 | 0.04004214 | 4.0984388 | 17.0484791 | 17.2568099 | 13.4475828 |
| -2.2588141 | 0.02389495 | 0.04006108 | 20.492194 | 104.996982 | 0 | 42.0236961 |
| 2.2587449 | 0.02389926 | 0.04006622 | 0.40984388 | 1.3530539 | 0 | 1.68094785 |
| -2.2583238 | 0.02392548 | 0.0401081 | 3.68859492 | 23.8137486 | 18.4072639 | 11.206319 |
| -2.2582103 | 0.02393255 | 0.04011787 | 46.3123584 | 82.5362876 | 0 | 26.3348496 |
| 2.25804884 | 0.02394261 | 0.04012016 | 0 | 0 | 0 | 0 |
| 2.25804884 | 0.02394261 | 0.04012016 | 0 | 0 | 0 | 0 |

| | | | | | | |
|---|---|---|---|---|---|---|
| 2.25804884 | 0.02394261 | 0.04012016 | 0 | 0 | 0 | 0 |
| 2.25804884 | 0.02394261 | 0.04012016 | 0 | 0 | 0 | 0 |
| 2.25804884 | 0.02394261 | 0.04012016 | 0 | 0 | 0 | 0 |
| 2.25804884 | 0.02394261 | 0.04012016 | 0 | 0 | 0 | 0 |
| 2.25804884 | 0.02394261 | 0.04012016 | 0 | 0 | 0 | 0 |
| 2.25797783 | 0.02394704 | 0.04012549 | 7.78703372 | 11.3656527 | 2.30090799 | 10.0856871 |
| 2.25791987 | 0.02395066 | 0.04012947 | 2.45906328 | 4.60038325 | 2.30090799 | 12.3269509 |
| -2.2577974 | 0.0239583 | 0.04014018 | 6.1476582 | 14.6129821 | 17.2568099 | 8.96505518 |
| 2.25763034 | 0.02396872 | 0.04015556 | 10.246097 | 5.41221558 | 3.45136199 | 2.80157974 |
| -2.2575922 | 0.02397109 | 0.04015746 | 283.611965 | 515.784145 | 434.87161 | 476.828872 |
| -2.2573594 | 0.02398563 | 0.04017972 | 8.60672148 | 16.2366467 | 5.75226998 | 17.9301104 |
| -2.2569408 | 0.02401177 | 0.04022143 | 72.9522106 | 0 | 65.5758777 | 57.7125427 |
| -2.256847 | 0.02401764 | 0.04022917 | 43.4434513 | 27.6022995 | 0 | 83.4870763 |
| 2.25654462 | 0.02403654 | 0.04025875 | 26.2300083 | 11.0950419 | 6.90272397 | 17.9301104 |
| -2.2564903 | 0.02403994 | 0.04026235 | 0 | 98.7729344 | 34.5136199 | 57.1522268 |
| -2.2563553 | 0.02404839 | 0.04027441 | 124.592539 | 28.4141318 | 0 | 22.9729539 |
| -2.2561472 | 0.02406142 | 0.04029174 | 8.1968776 | 10.8244312 | 12.6549939 | 9.52537113 |
| -2.2561302 | 0.02406248 | 0.04029174 | 22.1315695 | 56.2870421 | 0 | 78.4442328 |
| 2.25616821 | 0.0240601 | 0.04029174 | 0.40984388 | 0.54122156 | 0 | 0 |
| -2.2555249 | 0.0241004 | 0.04035315 | 6.55750208 | 6.76526948 | 16.1063559 | 22.9729539 |
| -2.2552107 | 0.02412011 | 0.04038405 | 67.2143963 | 45.4626109 | 57.5226998 | 0 |
| -2.2551161 | 0.02412604 | 0.04039189 | 6.1476582 | 30.579018 | 8.05317797 | 11.7666349 |
| -2.254756 | 0.02414865 | 0.04042764 | 44.263139 | 34.9087905 | 0 | 67.7982298 |
| 2.25456753 | 0.02416049 | 0.04044536 | 5.32797044 | 7.57710182 | 3.45136199 | 9.52537113 |
| 2.25426325 | 0.02417961 | 0.04047528 | 0 | 1.62366467 | 3.45136199 | 0.56031595 |
| -2.2538875 | 0.02420325 | 0.04051274 | 36.4761053 | 83.6187308 | 70.1776937 | 84.0473923 |
| -2.253814 | 0.02420787 | 0.04051838 | 22.5414134 | 12.9893174 | 5.75226998 | 8.96505518 |
| -2.253278 | 0.02424162 | 0.0405723 | 0 | 46.545054 | 65.5758777 | 68.9188617 |
| -2.2532625 | 0.0242426 | 0.0405723 | 6.55750208 | 8.11832337 | 14.9559019 | 17.3697944 |
| -2.2532391 | 0.02424408 | 0.04057267 | 7.78703372 | 9.74198805 | 12.6549939 | 11.206319 |
| 2.25315636 | 0.02424929 | 0.04057929 | 8.1968776 | 2.70610779 | 3.45136199 | 1.68094785 |
| -2.2530942 | 0.02425321 | 0.04058375 | 32.7875104 | 97.6904913 | 0 | 29.1364293 |
| -2.253014 | 0.02425826 | 0.0405901 | 5.73781432 | 17.0484791 | 9.20363196 | 15.6888466 |
| -2.2529334 | 0.02426335 | 0.04059651 | 4.50828268 | 21.1076408 | 11.50454 | 14.5682147 |
| -2.2526314 | 0.0242824 | 0.04062628 | 76.2309617 | 218.92412 | 177.169915 | 263.348496 |
| -2.2523398 | 0.02430081 | 0.04065496 | 4.50828268 | 16.7778683 | 12.6549939 | 15.6888466 |
| -2.2519745 | 0.02432388 | 0.04069146 | 22.5414134 | 8.65954493 | 19.5577179 | 4.48252759 |
| -2.2518005 | 0.02433488 | 0.04070609 | 6.55750208 | 18.9427545 | 8.05317797 | 14.0078987 |
| -2.2517963 | 0.02433515 | 0.04070609 | 33.1973543 | 61.9698684 | 79.3813257 | 0 |
| -2.2516834 | 0.02434229 | 0.04071553 | 0 | 20.2958084 | 77.0804177 | 104.218766 |
| -2.2516672 | 0.02434331 | 0.04071553 | 14.3445358 | 6.4946587 | 23.0090799 | 7.28410733 |
| -2.2516424 | 0.02434488 | 0.04071604 | 6.55750208 | 13.2599282 | 12.6549939 | 10.646003 |
| -2.2513877 | 0.02436099 | 0.04074087 | 21.7217256 | 4.87099402 | 11.50454 | 12.8872668 |

| | | | | | | |
|---|---|---|---|---|---|---|
| -2.2513303 | 0.02436463 | 0.04074484 | 0 | 94.4431619 | 64.4254237 | 30.8173772 |
| -2.2512011 | 0.0243728 | 0.04075641 | 0 | 115.280192 | 26.4604419 | 56.0315949 |
| -2.2506645 | 0.02440679 | 0.04081113 | 6.1476582 | 9.47137727 | 13.8054479 | 16.2491625 |
| -2.2505888 | 0.0244116 | 0.04081705 | 57.3781432 | 49.2511618 | 0 | 36.9808526 |
| -2.2505651 | 0.0244131 | 0.04081745 | 4.0984388 | 19.2133653 | 23.0090799 | 9.52537113 |
| -2.2505036 | 0.02441699 | 0.04082185 | 3.68859492 | 18.401533 | 17.2568099 | 14.0078987 |
| -2.2503661 | 0.02442571 | 0.04083431 | 27.8693838 | 26.2492456 | 31.0622579 | 39.2221164 |
| -2.2502709 | 0.02443175 | 0.04084229 | 6.96734596 | 26.2492456 | 6.90272397 | 12.8872668 |
| -2.2502372 | 0.02443389 | 0.04084375 | 2.86890716 | 4.87099402 | 29.9118039 | 44.2649599 |
| -2.250145 | 0.02443975 | 0.04085142 | 17.213443 | 9.74198805 | 8.05317797 | 8.96505518 |
| -2.2496174 | 0.02447324 | 0.0409053 | 182.79037 | 235.701989 | 169.116737 | 203.394689 |
| -2.2492176 | 0.02449865 | 0.04094565 | 9.83625312 | 12.1774851 | 6.90272397 | 13.4475828 |
| -2.2489922 | 0.024513 | 0.0409675 | 356.974019 | 973.386973 | 975.584988 | 940.770478 |
| -2.2488028 | 0.02452504 | 0.04098551 | 67.2143963 | 20.5664192 | 0 | 67.7982298 |
| 2.24872471 | 0.02453002 | 0.0409917 | 0.81968776 | 0 | 0 | 0 |
| -2.2483547 | 0.02455358 | 0.04102684 | 72.5423667 | 164.531354 | 161.063559 | 138.958355 |
| -2.2483647 | 0.02455294 | 0.04102684 | 5.73781432 | 24.8961917 | 5.75226998 | 21.292006 |
| -2.2481432 | 0.02456706 | 0.04104723 | 14.7543797 | 9.20076649 | 4.60181598 | 26.3348496 |
| -2.2477833 | 0.02459001 | 0.04108346 | 4.91812656 | 12.1774851 | 11.50454 | 22.4126379 |
| -2.2476508 | 0.02459847 | 0.04109545 | 122.953164 | 21.6488623 | 0 | 29.1364293 |
| -2.247267 | 0.02462297 | 0.04113426 | 15.1642236 | 12.9893174 | 11.50454 | 5.60315949 |
| -2.2472052 | 0.02462691 | 0.04113872 | 10.6559409 | 11.9068743 | 10.354086 | 7.84442328 |
| -2.2471773 | 0.0246287 | 0.04113957 | 0 | 71.1706349 | 46.0181598 | 62.1950703 |
| 2.24694889 | 0.02464329 | 0.04116182 | 0 | 2.43549701 | 0 | 2.24126379 |
| -2.2468021 | 0.02465268 | 0.04117536 | 9.83625312 | 17.5897006 | 4.60181598 | 18.4904263 |
| 2.24661141 | 0.02466487 | 0.0411936 | 0 | 1.62366467 | 0 | 0.56031595 |
| -2.2465106 | 0.02467132 | 0.04120011 | 10.246097 | 15.1542036 | 6.90272397 | 10.646003 |
| -2.24653 | 0.02467008 | 0.04120011 | 12.7051603 | 59.2637606 | 0 | 101.977503 |
| -2.2464366 | 0.02467605 | 0.04120588 | 7.37718984 | 11.0950419 | 8.05317797 | 20.7316901 |
| -2.2463556 | 0.02468124 | 0.04121241 | 2.45906328 | 33.2851258 | 23.0090799 | 10.0856871 |
| -2.2459109 | 0.02470972 | 0.04125783 | 7.78703372 | 8.65954493 | 9.20363196 | 23.5332698 |
| -2.2458383 | 0.02471437 | 0.04126346 | 8.60672148 | 0 | 143.806749 | 98.055291 |
| 2.24580746 | 0.02471634 | 0.04126462 | 9.01656536 | 13.2599282 | 9.20363196 | 44.8252759 |
| 2.2457043 | 0.02472295 | 0.04126713 | 0 | 0 | 0 | 0 |
| 2.2457043 | 0.02472295 | 0.04126713 | 0 | 0 | 0 | 0 |
| 2.2457043 | 0.02472295 | 0.04126713 | 0 | 0 | 0 | 0 |
| 2.2457043 | 0.02472295 | 0.04126713 | 0 | 0 | 0 | 0 |
| -2.2452991 | 0.02474894 | 0.04130836 | 13.1150042 | 11.3656527 | 12.6549939 | 6.16347543 |
| -2.2451591 | 0.02475792 | 0.04132122 | 111.477535 | 182.662276 | 180.621277 | 214.601008 |
| -2.2449555 | 0.02477099 | 0.0413409 | 7.78703372 | 19.4839761 | 6.90272397 | 12.8872668 |
| -2.2448813 | 0.02477575 | 0.04134671 | 9.01656536 | 110.138587 | 0 | 66.6775979 |
| -2.2446888 | 0.02478811 | 0.0413652 | 0 | 73.3355211 | 37.9649818 | 70.0394936 |
| -2.2442785 | 0.02481449 | 0.04140707 | 19.6725062 | 88.7603356 | 0 | 50.4284354 |

| | | | | | | |
|---|---|---|---|---|---|---|
| -2.2440985 | 0.02482606 | 0.04142424 | 26.6398522 | 7.30649104 | 10.354086 | 7.84442328 |
| -2.2438806 | 0.02484008 | 0.0414455 | 28.6890716 | 61.1580361 | 0 | 56.0315949 |
| -2.2438332 | 0.02484314 | 0.04144631 | 106.149565 | 397.256624 | 427.968886 | 333.387989 |
| 2.24383494 | 0.02484302 | 0.04144631 | 2.45906328 | 6.76526948 | 1.150454 | 3.92221164 |
| -2.2438027 | 0.0248451 | 0.04144744 | 0 | 51.6866588 | 60.9740618 | 64.4363341 |
| -2.2435726 | 0.02485991 | 0.04146786 | 40.1647002 | 35.7206228 | 0 | 68.9188617 |
| -2.2435766 | 0.02485966 | 0.04146786 | 67.6242402 | 27.8729103 | 0 | 51.5490673 |
| 2.24282363 | 0.02490819 | 0.04154625 | 9.83625312 | 18.1309222 | 21.8586259 | 12.8872668 |
| -2.242782 | 0.02491087 | 0.04154643 | 21.7217256 | 53.5809343 | 0 | 78.4442328 |
| -2.2428001 | 0.02490971 | 0.04154643 | 51.230485 | 77.9359044 | 40.2658898 | 0 |
| -2.2422452 | 0.02494553 | 0.04160209 | 7.78703372 | 27.3316887 | 3.45136199 | 24.0935858 |
| -2.2420472 | 0.02495832 | 0.04162127 | 1007.8061 | 1565.48336 | 1388.59797 | 1503.88801 |
| -2.2417896 | 0.02497497 | 0.04164688 | 87.7065903 | 0 | 106.992222 | 20.1713742 |
| 2.24161355 | 0.02498636 | 0.04166372 | 3.27875104 | 1.62366467 | 1.150454 | 0.56031595 |
| -2.2412917 | 0.02500718 | 0.04169629 | 131.559885 | 19.7545869 | 0 | 26.8951655 |
| -2.2411879 | 0.02501391 | 0.04170534 | 129.92051 | 299.566133 | 292.215315 | 316.578511 |
| -2.2410733 | 0.02502132 | 0.04171556 | 6.1476582 | 7.8477126 | 18.4072639 | 15.6888466 |
| -2.2406651 | 0.02504778 | 0.04175751 | 27.8693838 | 2.70610779 | 42.5667978 | 5.60315949 |
| -2.2405451 | 0.02505555 | 0.04176831 | 51.230485 | 47.8981079 | 66.7263317 | 0 |
| -2.2405112 | 0.02505775 | 0.04176982 | 4.91812656 | 22.4606947 | 8.05317797 | 17.9301104 |
| -2.2404721 | 0.02506029 | 0.0417719 | 41.8040758 | 73.0649104 | 0 | 31.3776931 |
| -2.2403781 | 0.02506638 | 0.0417799 | 11.0657848 | 8.93015571 | 6.90272397 | 17.9301104 |
| -2.2399393 | 0.02509486 | 0.04182304 | 2.86890716 | 9.20076649 | 28.7613499 | 24.0935858 |
| -2.2399418 | 0.0250947 | 0.04182304 | 62.2962697 | 41.9446708 | 27.6108959 | 30.8173772 |
| -2.2398685 | 0.02509946 | 0.04182855 | 82.3786199 | 24.6255809 | 0 | 44.8252759 |
| -2.2396043 | 0.02511662 | 0.04185283 | 257.7918 | 464.368097 | 394.60572 | 452.735286 |
| -2.2396154 | 0.0251159 | 0.04185283 | 4.0984388 | 16.2366467 | 11.50454 | 20.1713742 |
| -2.2394944 | 0.02512376 | 0.04186257 | 8.60672148 | 15.966036 | 21.8586259 | 5.04284354 |
| -2.2393097 | 0.02513577 | 0.04188041 | 7.37718984 | 13.2599282 | 8.05317797 | 14.5682147 |
| -2.2390859 | 0.02515033 | 0.04190251 | 7.78703372 | 9.20076649 | 10.354086 | 14.5682147 |
| -2.2385444 | 0.02518558 | 0.04195907 | 19.2626624 | 64.1347547 | 0 | 71.1601255 |
| -2.2381082 | 0.025214 | 0.04200425 | 0 | 40.0503953 | 44.8677058 | 99.1759229 |
| -2.2378185 | 0.02523289 | 0.04203356 | 211.479442 | 623.757846 | 646.555145 | 650.526816 |
| -2.2375979 | 0.02524729 | 0.04205537 | 13.9346919 | 15.4248144 | 9.20363196 | 6.16347543 |
| -2.2373698 | 0.02526218 | 0.04207801 | 2.0492194 | 19.2133653 | 21.8586259 | 20.7316901 |
| -2.2372376 | 0.02527081 | 0.04209021 | 35.6564176 | 75.5004074 | 57.5226998 | 0 |
| 2.23612319 | 0.0253437 | 0.04220944 | 271.726492 | 243.00848 | 326.728935 | 338.430833 |
| 2.23594126 | 0.02535562 | 0.04221404 | 0 | 0 | 0 | 0 |
| 2.23594126 | 0.02535562 | 0.04221404 | 0 | 0 | 0 | 0 |
| 2.23594126 | 0.02535562 | 0.04221404 | 0 | 0 | 0 | 0 |
| 2.23594126 | 0.02535562 | 0.04221404 | 0 | 0 | 0 | 0 |
| 2.23594126 | 0.02535562 | 0.04221404 | 0 | 0 | 0 | 0 |
| 2.23594126 | 0.02535562 | 0.04221404 | 0 | 0 | 0 | 0 |

| | | | | | | |
|---|---|---|---|---|---|---|
| 2.23594126 | 0.02535562 | 0.04221404 | 0 | 0 | 0 | 0 |
| -2.2358948 | 0.02535866 | 0.04221693 | 11.4756286 | 10.8244312 | 58.6731538 | 38.6618005 |
| -2.2353776 | 0.02539257 | 0.04227119 | 0 | 19.7545869 | 143.806749 | 44.8252759 |
| -2.235077 | 0.02541229 | 0.04230185 | 46.7222023 | 46.2744432 | 0 | 45.3855918 |
| -2.2349561 | 0.02542023 | 0.04230888 | 6.96734596 | 10.5538204 | 11.50454 | 12.3269509 |
| -2.2349878 | 0.02541815 | 0.04230888 | 10.6559409 | 5.1416048 | 13.8054479 | 17.9301104 |
| -2.2349528 | 0.02542045 | 0.04230888 | 6.1476582 | 15.1542036 | 18.4072639 | 7.84442328 |
| -2.234895 | 0.02542424 | 0.04231302 | 40.1647002 | 7.03588026 | 8.05317797 | 7.84442328 |
| -2.2343209 | 0.02546197 | 0.04237362 | 5.32797044 | 14.8835929 | 10.354086 | 15.6888466 |
| -2.2342586 | 0.02546606 | 0.04237825 | 219.266476 | 245.714587 | 349.738015 | 294.726189 |
| -2.2340441 | 0.02548017 | 0.04239954 | 6.1476582 | 13.530539 | 11.50454 | 11.7666349 |
| 2.23402311 | 0.02548155 | 0.04239965 | 114.346443 | 69.817581 | 43.7172518 | 57.1522268 |
| 2.23376563 | 0.0254985 | 0.04242566 | 13.1150042 | 4.32977247 | 6.90272397 | 5.04284354 |
| -2.2336364 | 0.02550701 | 0.04243764 | 37.2957931 | 10.0125988 | 4.60181598 | 10.0856871 |
| -2.2335615 | 0.02551194 | 0.04244365 | 25.0004767 | 13.8011497 | 4.60181598 | 9.52537113 |
| -2.2335147 | 0.02551502 | 0.04244659 | 245.08664 | 493.594061 | 436.022064 | 457.77813 |
| 2.23316487 | 0.02553807 | 0.04248057 | 0 | 0 | 0 | 0 |
| 2.23316487 | 0.02553807 | 0.04248057 | 0 | 0 | 0 | 0 |
| 2.23297772 | 0.02555041 | 0.04249671 | 1.63937552 | 0 | 4.60181598 | 0 |
| -2.2329861 | 0.02554986 | 0.04249671 | 7.37718984 | 12.4480958 | 10.354086 | 10.646003 |
| -2.2326474 | 0.02557221 | 0.04253077 | 4.91812656 | 15.966036 | 11.50454 | 14.5682147 |
| -2.2325939 | 0.02557574 | 0.04253445 | 0 | 54.6633774 | 54.0713378 | 64.99665 |
| -2.232517 | 0.02558081 | 0.0425407 | 32.3776665 | 7.57710182 | 5.75226998 | 11.7666349 |
| -2.2323667 | 0.02559074 | 0.04255501 | 41.3942319 | 47.3568864 | 0 | 49.3078035 |
| -2.2321274 | 0.02560655 | 0.04257911 | 65.5750208 | 57.3694852 | 41.4163438 | 0 |
| -2.2319269 | 0.02561979 | 0.04259456 | 27.45954 | 5.95343714 | 5.75226998 | 17.3697944 |
| -2.2319392 | 0.02561898 | 0.04259456 | 7.37718984 | 14.0717605 | 6.90272397 | 17.3697944 |
| -2.2319423 | 0.02561878 | 0.04259456 | 33.6071982 | 7.03588026 | 4.60181598 | 15.6888466 |
| -2.2318943 | 0.02562195 | 0.04259595 | 106.149565 | 172.108456 | 149.559019 | 180.982051 |
| -2.2315602 | 0.02564405 | 0.04263049 | 3.27875104 | 23.0019162 | 19.5577179 | 11.206319 |
| 2.23124212 | 0.0256651 | 0.04266109 | 0 | 0.54122156 | 0 | 0 |
| 2.23124212 | 0.0256651 | 0.04266109 | 0 | 0.54122156 | 0 | 0 |
| -2.230881 | 0.02568901 | 0.04269689 | 3.27875104 | 13.2599282 | 17.2568099 | 21.292006 |
| -2.2308769 | 0.02568928 | 0.04269689 | 257.381957 | 654.066253 | 531.509746 | 531.179519 |
| -2.2307943 | 0.02569476 | 0.04270379 | 0 | 60.8874253 | 72.4786017 | 42.0236961 |
| -2.2307551 | 0.02569735 | 0.04270591 | 51.230485 | 34.367569 | 0 | 53.7903311 |
| -2.2301623 | 0.02573667 | 0.04276905 | 63.5258014 | 5.68282636 | 5.75226998 | 7.84442328 |
| -2.2300871 | 0.02574166 | 0.04277361 | 72.9522106 | 156.954252 | 0 | 119.347297 |
| -2.230081 | 0.02574206 | 0.04277361 | 0 | 46.545054 | 54.0713378 | 73.9617052 |
| 2.2297735 | 0.02576248 | 0.04280534 | 0.40984388 | 0 | 0 | 0 |
| -2.2296503 | 0.02577067 | 0.04281674 | 167.626147 | 362.618444 | 392.304812 | 400.065587 |
| -2.2294031 | 0.0257871 | 0.04284183 | 4.0984388 | 25.9786348 | 12.6549939 | 11.7666349 |
| -2.2293482 | 0.02579074 | 0.04284568 | 18.0331307 | 25.708024 | 39.1154358 | 29.6967453 |

| | | | | | | |
|---|---|---|---|---|---|---|
| 2.22924101 | 0.02579787 | 0.04285532 | 2.86890716 | 2.43549701 | 5.75226998 | 1.68094785 |
| -2.2291805 | 0.0258019 | 0.0428598 | 6.1476582 | 12.4480958 | 16.1063559 | 9.52537113 |
| 2.2289443 | 0.02581761 | 0.04288369 | 0 | 1.89427545 | 0 | 0 |
| -2.2286894 | 0.02583458 | 0.04290967 | 5.73781432 | 10.2832096 | 33.3631659 | 8.40473923 |
| 2.22854132 | 0.02584444 | 0.04292384 | 1.22953164 | 0.27061078 | 1.150454 | 1.1206319 |
| -2.2283612 | 0.02585644 | 0.04294155 | 6.96734596 | 15.1542036 | 8.05317797 | 13.4475828 |
| -2.2282427 | 0.02586434 | 0.04295247 | 40.1647002 | 7.8477126 | 5.75226998 | 9.52537113 |
| 2.22808756 | 0.02587468 | 0.04296628 | 5.73781432 | 1.62366467 | 4.60181598 | 2.24126379 |
| -2.2280779 | 0.02587532 | 0.04296628 | 26.2300083 | 76.5828505 | 0 | 43.704644 |
| -2.2280563 | 0.02587676 | 0.04296647 | 41.8040758 | 60.3462037 | 0 | 36.9808526 |
| 2.22780263 | 0.02589368 | 0.04299235 | 0 | 0 | 0 | 0.56031595 |
| -2.2275052 | 0.02591353 | 0.04302309 | 0 | 61.6992576 | 50.6199758 | 59.9538065 |
| -2.2270265 | 0.0259455 | 0.04307396 | 4.0984388 | 26.2492456 | 28.7613499 | 5.60315949 |
| -2.2270009 | 0.02594721 | 0.04307459 | 11.8854725 | 77.3946828 | 0 | 77.3236009 |
| -2.2267036 | 0.02596709 | 0.04310537 | 0 | 58.7225391 | 75.9299637 | 40.3427483 |
| -2.2260447 | 0.02601118 | 0.04317635 | 7.37718984 | 126.375234 | 0 | 56.5919108 |
| -2.2258602 | 0.02602355 | 0.04319465 | 68.4439279 | 33.0145151 | 64.4254237 | 0 |
| -2.2258032 | 0.02602737 | 0.04319877 | 179.511619 | 115.821413 | 154.160835 | 113.183822 |
| -2.22557 | 0.02604299 | 0.04322248 | 40.1647002 | 6.4946587 | 6.90272397 | 9.52537113 |
| -2.2254486 | 0.02605114 | 0.04323377 | 52.4600166 | 60.3462037 | 48.3190678 | 0 |
| 2.22511584 | 0.02607346 | 0.0432686 | 37.2957931 | 17.5897006 | 20.7081719 | 22.4126379 |
| -2.22509 | 0.0260752 | 0.04326926 | 225.414134 | 234.078324 | 187.524001 | 231.970803 |
| -2.2250033 | 0.02608102 | 0.04327669 | 18.8528185 | 6.4946587 | 6.90272397 | 15.6888466 |
| -2.2249685 | 0.02608335 | 0.04327834 | 52.4600166 | 256.809629 | 98.9390436 | 170.336048 |
| 2.22483521 | 0.0260923 | 0.04329097 | 0 | 1.08244312 | 3.45136199 | 5.04284354 |
| 2.22480517 | 0.02609432 | 0.04329209 | 11.0657848 | 7.57710182 | 9.20363196 | 3.36189569 |
| -2.2246637 | 0.02610382 | 0.04330562 | 421.729352 | 769.617056 | 759.299637 | 757.547163 |
| 2.22461394 | 0.02610717 | 0.04330895 | 14.7543797 | 13.8011497 | 11.50454 | 8.96505518 |
| 2.22451493 | 0.02611382 | 0.04331776 | 4.91812656 | 2.70610779 | 6.90272397 | 6.16347543 |
| -2.2243242 | 0.02612664 | 0.0433368 | 204.102252 | 18.1309222 | 14.9559019 | 0 |
| -2.2242892 | 0.02612899 | 0.04333848 | 0 | 58.9931499 | 102.390406 | 25.2142177 |
| -2.2242142 | 0.02613404 | 0.04334461 | 18.4429746 | 7.30649104 | 9.20363196 | 9.52537113 |
| -2.2241269 | 0.02613991 | 0.04335213 | 31.5579788 | 9.47137727 | 24.1595339 | 26.3348496 |
| -2.2240566 | 0.02614463 | 0.04335774 | 12.7051603 | 7.57710182 | 8.05317797 | 13.4475828 |
| -2.223633 | 0.02617314 | 0.04340279 | 0 | 57.3694852 | 66.7263317 | 47.6268556 |
| -2.2235192 | 0.02618081 | 0.04341328 | 42.6237635 | 56.5576528 | 0 | 38.1014845 |
| -2.2234661 | 0.02618438 | 0.04341697 | 0 | 54.1221558 | 48.3190678 | 69.4791776 |
| 2.22316419 | 0.02620473 | 0.04344847 | 26.2300083 | 13.2599282 | 13.8054479 | 19.6110582 |
| -2.2230624 | 0.02621159 | 0.04345762 | 36.8859492 | 18.1309222 | 2.30090799 | 10.646003 |
| -2.2225833 | 0.02624391 | 0.04350257 | 67.6242402 | 583.978061 | 1190.71989 | 1046.67019 |
| -2.2226053 | 0.02624243 | 0.04350257 | 51.230485 | 36.2618444 | 0 | 49.3078035 |
| -2.2226227 | 0.02624125 | 0.04350257 | 5.73781432 | 13.2599282 | 10.354086 | 14.5682147 |
| -2.2225806 | 0.02624409 | 0.04350257 | 14.3445358 | 8.11832337 | 12.6549939 | 7.28410733 |

| | | | | | | |
|---|---|---|---|---|---|---|
| 2.22239654 | 0.02625652 | 0.04352093 | 0 | 0.54122156 | 0 | 0.56031595 |
| -2.2222574 | 0.02626591 | 0.04353427 | 1.22953164 | 33.8263474 | 6.90272397 | 48.7474875 |
| -2.221984 | 0.02628439 | 0.04356265 | 16.8035991 | 6.22404792 | 31.0622579 | 5.04284354 |
| -2.221726 | 0.02630183 | 0.04358932 | 67.2143963 | 41.9446708 | 0 | 31.3776931 |
| 2.22142582 | 0.02632213 | 0.04361849 | 0 | 0 | 0 | 0 |
| 2.22142582 | 0.02632213 | 0.04361849 | 0 | 0 | 0 | 0 |
| -2.2210837 | 0.02634529 | 0.04365239 | 50.0009534 | 43.5683354 | 0 | 42.0236961 |
| 2.22110154 | 0.02634409 | 0.04365239 | 52.0501728 | 55.4752097 | 54.0713378 | 51.5490673 |
| -2.2210326 | 0.02634876 | 0.04365589 | 2.86890716 | 21.9194731 | 17.2568099 | 14.5682147 |
| -2.2208402 | 0.02636178 | 0.04367523 | 0 | 74.688575 | 75.9299637 | 28.5761134 |
| -2.2205807 | 0.02637937 | 0.04370213 | 83.1983076 | 139.635162 | 153.010381 | 118.226665 |
| -2.2202805 | 0.02639973 | 0.04373361 | 0 | 60.3462037 | 56.3722458 | 53.2300151 |
| -2.2202189 | 0.02640391 | 0.0437383 | 29.0989155 | 11.9068743 | 4.60181598 | 9.52537113 |
| -2.2201359 | 0.02640954 | 0.04374538 | 32.7875104 | 4.60038325 | 4.60181598 | 24.0935858 |
| -2.2198718 | 0.02642747 | 0.04377283 | 3.27875104 | 16.5072575 | 21.8586259 | 12.8872668 |
| 2.21975202 | 0.02643561 | 0.04378406 | 5.73781432 | 5.95343714 | 11.50454 | 5.60315949 |
| -2.2195979 | 0.02644608 | 0.04379915 | 3.68859492 | 13.2599282 | 11.50454 | 28.0157974 |
| -2.2193603 | 0.02646222 | 0.04382364 | 27.0496961 | 34.367569 | 5.75226998 | 45.3855918 |
| -2.2192924 | 0.02646684 | 0.04382904 | 144.265046 | 263.033677 | 185.223093 | 218.52322 |
| -2.2192335 | 0.02647085 | 0.04383343 | 8.1968776 | 39.5091738 | 9.20363196 | 5.60315949 |
| -2.2192024 | 0.02647296 | 0.04383468 | 11.0657848 | 11.3656527 | 10.354086 | 7.28410733 |
| 2.21914276 | 0.02647701 | 0.04383915 | 6.55750208 | 3.24732935 | 4.60181598 | 3.36189569 |
| -2.2187877 | 0.02650117 | 0.0438769 | 9.42640924 | 21.6488623 | 10.354086 | 6.16347543 |
| -2.2187551 | 0.02650339 | 0.04387832 | 45.9025146 | 63.3229223 | 50.6199758 | 0 |
| -2.218734 | 0.02650483 | 0.04387845 | 12.2953164 | 25.9786348 | 26.4604419 | 16.2491625 |
| -2.2181827 | 0.02654238 | 0.04393836 | 132.379573 | 32.2026827 | 0 | 13.4475828 |
| -2.2179806 | 0.02655616 | 0.04395891 | 8.60672148 | 12.1774851 | 5.75226998 | 21.292006 |
| -2.2176829 | 0.02657646 | 0.04399027 | 88.1164342 | 206.746635 | 195.577179 | 210.678797 |
| -2.2176056 | 0.02658174 | 0.04399675 | 11.0657848 | 15.6954252 | 4.60181598 | 15.1285306 |
| -2.2174575 | 0.02659185 | 0.04401122 | 131.969729 | 6.76526948 | 0 | 53.2300151 |
| -2.2170828 | 0.02661744 | 0.04405132 | 0 | 41.9446708 | 89.7354116 | 43.704644 |
| 2.21703328 | 0.02662082 | 0.04405466 | 47.5418901 | 21.6488623 | 18.4072639 | 17.9301104 |
| -2.216966 | 0.02662541 | 0.04405598 | 0 | 89.0309463 | 32.2127119 | 56.0315949 |
| -2.2169617 | 0.02662571 | 0.04405598 | 15.9839113 | 10.8244312 | 10.354086 | 6.16347543 |
| -2.2169672 | 0.02662533 | 0.04405598 | 12.2953164 | 88.219114 | 0 | 62.7553862 |
| -2.2169274 | 0.02662805 | 0.04405759 | 0 | 29.7671857 | 127.700393 | 34.7395888 |
| -2.2168719 | 0.02663185 | 0.04406162 | 24.1807889 | 43.8389462 | 0 | 77.3236009 |
| 2.21660267 | 0.02665025 | 0.04408981 | 22.1315695 | 14.3423713 | 12.6549939 | 17.9301104 |
| -2.2164041 | 0.02666384 | 0.04411002 | 0 | 89.5721679 | 71.3281477 | 22.9729539 |
| 2.21635434 | 0.02666724 | 0.04411114 | 5.73781432 | 10.8244312 | 6.90272397 | 3.92221164 |
| -2.216367 | 0.02666638 | 0.04411114 | 0 | 82.2656769 | 79.3813257 | 22.4126379 |
| -2.2162191 | 0.0266765 | 0.04412419 | 0 | 46.0038325 | 28.7613499 | 109.821926 |
| -2.2161298 | 0.02668262 | 0.04413204 | 10.246097 | 9.74198805 | 9.20363196 | 9.52537113 |

| | | | | | | |
|---|---|---|---|---|---|---|
| -2.2161015 | 0.02668455 | 0.04413298 | 0 | 47.0862756 | 52.9208838 | 70.0394936 |
| -2.2160475 | 0.02668825 | 0.04413684 | 8.60672148 | 21.3782516 | 8.05317797 | 8.40473923 |
| -2.2157334 | 0.02670976 | 0.04417016 | 7.78703372 | 13.2599282 | 9.20363196 | 10.0856871 |
| -2.2155723 | 0.02672081 | 0.04418616 | 17.213443 | 7.30649104 | 9.20363196 | 9.52537113 |
| 2.21543097 | 0.0267305 | 0.04419765 | 0 | 0 | 0 | 0 |
| 2.21543097 | 0.0267305 | 0.04419765 | 0 | 0 | 0 | 0 |
| -2.2152531 | 0.0267427 | 0.04421556 | 4.0984388 | 23.5431378 | 8.05317797 | 19.6110582 |
| -2.2152287 | 0.02674437 | 0.04421606 | 6.55750208 | 10.5538204 | 21.8586259 | 8.40473923 |
| 2.21469308 | 0.02678114 | 0.04427458 | 0.40984388 | 0.54122156 | 0 | 0 |
| 2.21443189 | 0.02679908 | 0.04430198 | 8.1968776 | 8.11832337 | 2.30090799 | 9.52537113 |
| -2.2143745 | 0.02680303 | 0.04430624 | 313.940412 | 443.801678 | 449.827512 | 453.295602 |
| -2.2141595 | 0.02681781 | 0.0443284 | 14.3445358 | 7.8477126 | 17.2568099 | 6.16347543 |
| -2.2141176 | 0.02682069 | 0.04433089 | 14.3445358 | 18.9427545 | 14.9559019 | 11.206319 |
| -2.2138577 | 0.02683857 | 0.04435751 | 0 | 46.2744432 | 59.8236078 | 62.7553862 |
| -2.2138436 | 0.02683954 | 0.04435751 | 117.21535 | 34.367569 | 0 | 15.6888466 |
| -2.2137975 | 0.02684271 | 0.04436047 | 0 | 75.2297966 | 39.1154358 | 57.1522268 |
| -2.2137709 | 0.02684454 | 0.04436087 | 28.2792277 | 49.7923834 | 0 | 60.5141225 |
| -2.2137541 | 0.0268457 | 0.04436087 | 54.0993922 | 37.6148983 | 0 | 43.144328 |
| -2.2136401 | 0.02685355 | 0.04437157 | 40.984388 | 17.8603114 | 10.354086 | 24.0935858 |
| 2.21358966 | 0.02685702 | 0.04437504 | 0 | 13.2599282 | 0 | 0 |
| -2.2133718 | 0.02687202 | 0.04439756 | 10.6559409 | 79.0183475 | 0 | 76.202969 |
| -2.2128504 | 0.02690796 | 0.04445466 | 12.7051603 | 14.3423713 | 5.75226998 | 10.0856871 |
| -2.2118123 | 0.02697964 | 0.0445708 | 91.8050291 | 57.0988744 | 23.0090799 | 0 |
| -2.2117011 | 0.02698732 | 0.04458121 | 52.8698605 | 28.9553534 | 0 | 55.4712789 |
| -2.211579 | 0.02699577 | 0.04459288 | 0 | 65.4878086 | 34.5136199 | 72.2807574 |
| 2.21107715 | 0.02703049 | 0.04463654 | 0 | 0 | 0 | 0 |
| 2.21107715 | 0.02703049 | 0.04463654 | 0 | 0 | 0 | 0 |
| 2.21107715 | 0.02703049 | 0.04463654 | 0 | 0 | 0 | 0 |
| 2.21107715 | 0.02703049 | 0.04463654 | 0 | 0 | 0 | 0 |
| 2.21107715 | 0.02703049 | 0.04463654 | 0 | 0 | 0 | 0 |
| 2.21107715 | 0.02703049 | 0.04463654 | 0 | 0 | 0 | 0 |
| 2.21087087 | 0.02704478 | 0.04465784 | 143.035514 | 122.857294 | 153.010381 | 118.786981 |
| 2.21074781 | 0.02705331 | 0.04466964 | 1.22953164 | 0 | 2.30090799 | 0 |
| -2.2106824 | 0.02705784 | 0.04467484 | 20.9020379 | 2.70610779 | 12.6549939 | 21.852322 |
| -2.2106326 | 0.02706129 | 0.04467825 | 0 | 28.6847426 | 89.7354116 | 58.8331746 |
| -2.2103251 | 0.02708261 | 0.04471116 | 7.37718984 | 19.4839761 | 8.05317797 | 10.0856871 |
| -2.2102937 | 0.02708479 | 0.04471247 | 31.5579788 | 48.7099402 | 48.3190678 | 40.3427483 |
| -2.2101409 | 0.02709538 | 0.04472768 | 17.6232868 | 2.70610779 | 18.4072639 | 17.3697944 |
| -2.209902 | 0.02711197 | 0.04475276 | 19.2626624 | 100.937821 | 0 | 35.8602207 |
| -2.2098619 | 0.02711474 | 0.04475277 | 7.37718984 | 12.9893174 | 10.354086 | 9.52537113 |
| -2.2098785 | 0.02711359 | 0.04475277 | 38.9351686 | 48.4393295 | 0 | 45.9459078 |
| -2.2098091 | 0.02711841 | 0.04475654 | 43.0336074 | 55.7458205 | 58.6731538 | 0 |
| -2.209789 | 0.02711981 | 0.04475656 | 136.068168 | 12.4480958 | 0 | 30.8173772 |

| | | | | | | |
|---|---|---|---|---|---|---|
| -2.209617 | 0.02713176 | 0.0447722 | 0 | 45.4626109 | 56.3722458 | 66.1172819 |
| -2.2096125 | 0.02713206 | 0.0447722 | 7.78703372 | 14.6129821 | 10.354086 | 8.40473923 |
| 2.20933291 | 0.02715149 | 0.0447965 | 0 | 0 | 0 | 0 |
| -2.2093208 | 0.02715234 | 0.0447965 | 25.4103206 | 61.9698684 | 0 | 52.1093832 |
| 2.20933291 | 0.02715149 | 0.0447965 | 0 | 0 | 0 | 0 |
| 2.20933291 | 0.02715149 | 0.0447965 | 0 | 0 | 0 | 0 |
| 2.20923978 | 0.02715797 | 0.04480121 | 11.4756286 | 4.05916169 | 8.05317797 | 5.60315949 |
| -2.2092566 | 0.0271568 | 0.04480121 | 18.0331307 | 63.0523115 | 0 | 66.6775979 |
| -2.2091825 | 0.02716195 | 0.04480549 | 38.9351686 | 70.9000241 | 0 | 29.6967453 |
| -2.2090558 | 0.02717076 | 0.04481773 | 7.37718984 | 92.5488865 | 0 | 77.8839169 |
| -2.2089668 | 0.02717695 | 0.04482566 | 52.4600166 | 84.4305631 | 28.7613499 | 0 |
| -2.2086173 | 0.02720127 | 0.04486348 | 9.42640924 | 6.76526948 | 14.9559019 | 10.646003 |
| 2.20841576 | 0.0272153 | 0.04488423 | 0.81968776 | 0 | 0 | 1.1206319 |
| -2.2083967 | 0.02721663 | 0.04488423 | 7.37718984 | 89.0309463 | 110.443584 | 0 |
| -2.2081813 | 0.02723164 | 0.04490668 | 8.1968776 | 17.8603114 | 11.50454 | 6.72379138 |
| -2.2081112 | 0.02723652 | 0.04491244 | 78.2801811 | 14.0717605 | 90.8858656 | 0 |
| -2.2080106 | 0.02724354 | 0.04492171 | 4.0984388 | 21.6488623 | 11.50454 | 13.4475828 |
| 2.20787329 | 0.02725311 | 0.0449352 | 26.6398522 | 15.6954252 | 0 | 36.4205367 |
| -2.2077826 | 0.02725943 | 0.04494333 | 49.1812656 | 23.8137486 | 98.9390436 | 0 |
| -2.2074543 | 0.02728233 | 0.0449765 | 2.0492194 | 28.143521 | 48.3190678 | 5.04284354 |
| -2.2074705 | 0.0272812 | 0.0449765 | 42.2139196 | 41.67406 | 0 | 48.7474875 |
| -2.2074341 | 0.02728374 | 0.04497653 | 43.8532952 | 51.1454373 | 0 | 38.1014845 |
| -2.207265 | 0.02729555 | 0.04499139 | 43.4434513 | 41.9446708 | 0 | 47.0665397 |
| -2.2072847 | 0.02729418 | 0.04499139 | 4.50828268 | 10.8244312 | 19.5577179 | 13.4475828 |
| -2.2066432 | 0.027339 | 0.04506071 | 113.116911 | 25.9786348 | 0 | 21.852322 |
| -2.2066227 | 0.02734043 | 0.04506076 | 6.96734596 | 15.4248144 | 11.50454 | 8.40473923 |
| -2.2064832 | 0.02735019 | 0.04507225 | 7.37718984 | 11.3656527 | 18.4072639 | 7.28410733 |
| 2.20650027 | 0.02734899 | 0.04507225 | 0.40984388 | 0.27061078 | 0 | 1.68094785 |
| -2.2064304 | 0.02735388 | 0.04507603 | 43.8532952 | 35.4500121 | 0 | 54.350647 |
| -2.206394 | 0.02735643 | 0.04507793 | 29.0989155 | 57.0988744 | 0 | 49.8681194 |
| -2.2062627 | 0.02736561 | 0.04509076 | 5.73781432 | 11.0950419 | 12.6549939 | 12.8872668 |
| 2.20588824 | 0.02739183 | 0.04513165 | 1.63937552 | 0 | 0 | 1.1206319 |
| 2.20565617 | 0.02740808 | 0.04515613 | 31.1481349 | 0 | 39.1154358 | 17.3697944 |
| 2.20532336 | 0.02743141 | 0.04519226 | 0 | 0 | 0 | 13.4475828 |
| -2.2050425 | 0.02745111 | 0.0452201 | 4.91812656 | 24.6255809 | 9.20363196 | 12.3269509 |
| -2.2050555 | 0.0274502 | 0.0452201 | 0 | 70.0881918 | 34.5136199 | 65.556966 |
| -2.2046751 | 0.0274769 | 0.04526028 | 3.27875104 | 16.2366467 | 13.8054479 | 19.0507423 |
| -2.204439 | 0.02749349 | 0.04528529 | 4.50828268 | 15.966036 | 10.354086 | 16.8094785 |
| -2.2042016 | 0.02751017 | 0.04531001 | 9.83625312 | 9.20076649 | 9.20363196 | 10.0856871 |
| -2.2041855 | 0.0275113 | 0.04531001 | 7.78703372 | 20.0251977 | 35.6640739 | 2.80157974 |
| -2.2040997 | 0.02751733 | 0.04531763 | 64.755333 | 17.8603114 | 0 | 63.8760181 |
| -2.2037478 | 0.02754209 | 0.04535608 | 0 | 22.4606947 | 73.6290557 | 83.4870763 |
| -2.2036443 | 0.02754937 | 0.04536576 | 4.50828268 | 11.6362635 | 17.2568099 | 13.4475828 |

| | | | | | | |
|---|---|---|---|---|---|---|
| -2.2031911 | 0.02758128 | 0.045416 | 334.84245 | 524.173079 | 441.774334 | 499.801826 |
| 2.20312811 | 0.02758572 | 0.04541636 | 0 | 0 | 0 | 0 |
| 2.20312811 | 0.02758572 | 0.04541636 | 0 | 0 | 0 | 0 |
| 2.20312811 | 0.02758572 | 0.04541636 | 0 | 0 | 0 | 0 |
| -2.2030826 | 0.02758892 | 0.04541931 | 6.96734596 | 14.0717605 | 10.354086 | 9.52537113 |
| -2.2029248 | 0.02760005 | 0.04543531 | 0 | 34.0969582 | 59.8236078 | 76.202969 |
| -2.2028938 | 0.02760224 | 0.04543659 | 186.888809 | 252.479857 | 285.312591 | 279.597658 |
| -2.2027803 | 0.02761024 | 0.04544745 | 193.446311 | 202.146252 | 154.160835 | 192.18837 |
| 2.20247552 | 0.02763174 | 0.04548052 | 3.27875104 | 6.76526948 | 9.20363196 | 2.24126379 |
| -2.202375 | 0.02763883 | 0.04548987 | 93.8542485 | 21.9194731 | 0 | 34.1792729 |
| -2.2021273 | 0.02765632 | 0.04551633 | 0 | 28.143521 | 117.346308 | 38.6618005 |
| -2.2020284 | 0.0276633 | 0.04552551 | 25.0004767 | 12.9893174 | 4.60181598 | 8.96505518 |
| -2.2018015 | 0.02767933 | 0.04554745 | 42.6237635 | 0 | 94.3372276 | 47.6268556 |
| 2.20179978 | 0.02767945 | 0.04554745 | 3.68859492 | 1.08244312 | 0 | 1.1206319 |
| -2.201234 | 0.02771947 | 0.04561097 | 25.0004767 | 0 | 85.1335957 | 81.2458125 |
| 2.20106966 | 0.02773109 | 0.04562777 | 61.0667381 | 21.1076408 | 27.6108959 | 26.8951655 |
| -2.2008563 | 0.0277462 | 0.0456503 | 51.6403289 | 69.0057487 | 36.8145279 | 0 |
| -2.2006912 | 0.02775789 | 0.04566722 | 0 | 67.382084 | 101.239952 | 17.9301104 |
| -2.2001595 | 0.02779558 | 0.04572689 | 5.73781432 | 4.05916169 | 28.7613499 | 22.9729539 |
| -2.1999976 | 0.02780707 | 0.04574345 | 49.1812656 | 54.3927666 | 0 | 30.2570612 |
| 2.19991266 | 0.02781309 | 0.04575104 | 38.5253247 | 23.8137486 | 29.9118039 | 24.6539017 |
| -2.1997165 | 0.02782702 | 0.04576758 | 0 | 32.7439043 | 52.9208838 | 85.7283401 |
| -2.1996951 | 0.02782853 | 0.04576758 | 2.45906328 | 25.9786348 | 16.1063559 | 14.5682147 |
| -2.199691 | 0.02782882 | 0.04576758 | 4.0984388 | 17.0484791 | 17.2568099 | 10.646003 |
| -2.1997509 | 0.02782457 | 0.04576758 | 312.71088 | 513.619259 | 475.1375 | 503.724038 |
| -2.1996271 | 0.02783336 | 0.04577272 | 85.6573709 | 26.2492456 | 0 | 32.498325 |
| 2.19953294 | 0.02784005 | 0.04578138 | 10.246097 | 6.22404792 | 4.60181598 | 6.72379138 |
| 2.19941153 | 0.02784867 | 0.04579323 | 58.197831 | 76.3122397 | 67.8767857 | 58.2728587 |
| -2.198731 | 0.02789706 | 0.04587046 | 0 | 83.6187308 | 27.6108959 | 62.1950703 |
| -2.1985127 | 0.02791259 | 0.04589366 | 40.5745441 | 44.109557 | 0 | 45.9459078 |
| 2.19811809 | 0.02794069 | 0.04593753 | 17.213443 | 4.32977247 | 12.6549939 | 8.40473923 |
| 2.19801458 | 0.02794807 | 0.04594731 | 0 | 0 | 0 | 5.60315949 |
| -2.1979642 | 0.02795165 | 0.04595087 | 0 | 36.803066 | 55.2217918 | 75.6426531 |
| -2.1974482 | 0.02798846 | 0.04600902 | 31.9678226 | 58.7225391 | 0 | 42.5840121 |
| -2.1974027 | 0.0279917 | 0.04601201 | 18.4429746 | 36.2618444 | 0 | 98.055291 |
| 2.1972813 | 0.02800036 | 0.04602391 | 23.770945 | 19.4839761 | 18.4072639 | 15.1285306 |
| -2.1971303 | 0.02801114 | 0.04603929 | 3.68859492 | 13.8011497 | 17.2568099 | 14.5682147 |
| 2.19681279 | 0.02803382 | 0.04606249 | 0 | 0 | 0 | 0 |
| 2.19681279 | 0.02803382 | 0.04606249 | 0 | 0 | 0 | 0 |
| 2.19681279 | 0.02803382 | 0.04606249 | 0 | 0 | 0 | 0 |
| 2.19681279 | 0.02803382 | 0.04606249 | 0 | 0 | 0 | 0 |
| -2.1969026 | 0.02802741 | 0.04606249 | 64.755333 | 88.7603356 | 71.3281477 | 123.269509 |
| 2.19681279 | 0.02803382 | 0.04606249 | 0 | 0 | 0 | 0 |

| | | | | | | |
|---|---|---|---|---|---|---|
| -2.1962632 | 0.02807311 | 0.0461247 | 8.1968776 | 11.0950419 | 10.354086 | 8.96505518 |
| -2.1959427 | 0.02809605 | 0.04615875 | 10.246097 | 18.1309222 | 4.60181598 | 13.4475828 |
| -2.1959336 | 0.02809669 | 0.04615875 | 88.9361219 | 164.531354 | 161.063559 | 140.078987 |
| 2.19584542 | 0.02810301 | 0.04616678 | 0.81968776 | 1.08244312 | 0 | 0 |
| -2.1953993 | 0.02813497 | 0.04621693 | 35.2465737 | 59.5343714 | 60.9740618 | 0 |
| -2.1951689 | 0.02815148 | 0.04624077 | 11.8854725 | 77.9359044 | 0 | 66.6775979 |
| -2.1951569 | 0.02815235 | 0.04624077 | 264.349303 | 959.585823 | 707.529207 | 765.951902 |
| -2.1948996 | 0.0281708 | 0.04626873 | 4.0984388 | 10.8244312 | 19.5577179 | 14.5682147 |
| 2.19482934 | 0.02817584 | 0.04627465 | 0 | 1.89427545 | 0 | 0 |
| 2.19442166 | 0.02820511 | 0.04632037 | 16.8035991 | 10.0125988 | 10.354086 | 11.206319 |
| -2.1939892 | 0.02823618 | 0.04636904 | 23.770945 | 10.2832096 | 10.354086 | 5.04284354 |
| 2.19383445 | 0.02824731 | 0.0463826 | 4.0984388 | 4.32977247 | 0 | 1.68094785 |
| 2.19385064 | 0.02824615 | 0.0463826 | 0.81968776 | 7.8477126 | 4.60181598 | 3.36189569 |
| -2.1937938 | 0.02825023 | 0.04638503 | 40.984388 | 58.9931499 | 0 | 32.498325 |
| -2.1934449 | 0.02827534 | 0.04642389 | 23.3611012 | 70.6294134 | 0 | 44.8252759 |
| -2.1932205 | 0.0282915 | 0.04644806 | 100.411751 | 32.7439043 | 33.3631659 | 0 |
| -2.1931729 | 0.02829493 | 0.04645133 | 11.4756286 | 18.6721438 | 3.45136199 | 16.8094785 |
| 2.19300059 | 0.02830734 | 0.04646934 | 8.60672148 | 14.0717605 | 21.8586259 | 14.5682147 |
| -2.1928755 | 0.02831635 | 0.04648177 | 34.8367298 | 45.1920001 | 0 | 50.4284354 |
| -2.192827 | 0.02831984 | 0.04648515 | 6.1476582 | 9.20076649 | 14.9559019 | 11.7666349 |
| -2.1927071 | 0.02832849 | 0.04649697 | 3.68859492 | 15.6954252 | 9.20363196 | 26.3348496 |
| -2.1924063 | 0.02835018 | 0.04653021 | 70.0833035 | 25.4374132 | 65.5758777 | 0 |
| 2.19237523 | 0.02835242 | 0.04653153 | 7.37718984 | 4.60038325 | 3.45136199 | 6.72379138 |
| -2.1923379 | 0.02835512 | 0.04653359 | 16.8035991 | 0 | 163.364467 | 38.1014845 |
| -2.1922604 | 0.02836071 | 0.04654039 | 13.9346919 | 9.74198805 | 11.50454 | 6.16347543 |
| -2.1922074 | 0.02836454 | 0.04654431 | 4.0984388 | 26.2492456 | 12.6549939 | 10.0856871 |
| -2.1921775 | 0.02836669 | 0.04654548 | 31.1481349 | 20.5664192 | 9.20363196 | 25.2142177 |
| -2.1919006 | 0.02838669 | 0.0465757 | 9.83625312 | 22.1900839 | 2.30090799 | 28.0157974 |
| -2.1918824 | 0.028388 | 0.0465757 | 1.22953164 | 14.6129821 | 51.7704298 | 14.0078987 |
| -2.1917797 | 0.02839542 | 0.04658551 | 5.73781432 | 18.6721438 | 12.6549939 | 8.40473923 |
| 2.1913842 | 0.028424 | 0.04662926 | 0 | 0 | 11.50454 | 0 |
| -2.1913708 | 0.02842497 | 0.04662926 | 7.78703372 | 7.57710182 | 10.354086 | 16.2491625 |
| -2.1913394 | 0.02842724 | 0.04663061 | 0 | 33.2851258 | 62.1245158 | 71.1601255 |
| -2.1911568 | 0.02844045 | 0.04664634 | 13.524848 | 10.8244312 | 5.75226998 | 11.206319 |
| -2.1911469 | 0.02844116 | 0.04664634 | 55.7387677 | 8.65954493 | 5.75226998 | 5.04284354 |
| -2.1911754 | 0.0284391 | 0.04664634 | 211.069598 | 25.708024 | 450.977966 | 123.269509 |
| -2.1909814 | 0.02845314 | 0.04666361 | 4.50828268 | 107.161869 | 0 | 78.4442328 |
| 2.19088136 | 0.02846038 | 0.04667311 | 68.8537718 | 26.2492456 | 0 | 0 |
| -2.1908611 | 0.02846184 | 0.04667314 | 10.6559409 | 8.65954493 | 8.05317797 | 11.206319 |
| -2.1905752 | 0.02848255 | 0.04670472 | 4.91812656 | 17.8603114 | 14.9559019 | 8.96505518 |
| 2.1903812 | 0.0284966 | 0.04672065 | 0 | 0 | 0 | 0 |
| 2.1903812 | 0.0284966 | 0.04672065 | 0 | 0 | 0 | 0 |
| 2.1903812 | 0.0284966 | 0.04672065 | 0 | 0 | 0 | 0 |

| | | | | | | |
|---|---|---|---|---|---|---|
| 2.19011187 | 0.02851612 | 0.04675029 | 0 | 0.81183234 | 0 | 1.1206319 |
| -2.1900695 | 0.0285192 | 0.04675295 | 7.78703372 | 10.5538204 | 8.05317797 | 13.4475828 |
| -2.1895304 | 0.02855831 | 0.04681469 | 0 | 48.4393295 | 71.3281477 | 43.704644 |
| -2.1894329 | 0.02856539 | 0.04682392 | 46.7222023 | 25.4374132 | 0 | 62.7553862 |
| 2.18930845 | 0.02857443 | 0.04683636 | 23.770945 | 10.5538204 | 18.4072639 | 0 |
| -2.1890544 | 0.02859288 | 0.04686423 | 5.73781432 | 13.2599282 | 13.8054479 | 9.52537113 |
| -2.1886968 | 0.02861889 | 0.04690447 | 36.0662614 | 48.4393295 | 34.5136199 | 38.6618005 |
| -2.1886307 | 0.02862369 | 0.04690996 | 5.73781432 | 18.6721438 | 9.20363196 | 11.206319 |
| -2.1884755 | 0.02863498 | 0.0469237 | 2.0492194 | 26.2492456 | 21.8586259 | 12.3269509 |
| -2.1884909 | 0.02863386 | 0.0469237 | 0 | 28.143521 | 82.8326877 | 58.8331746 |
| -2.1884529 | 0.02863663 | 0.04692401 | 7.78703372 | 11.3656527 | 17.2568099 | 6.72379138 |
| -2.1882075 | 0.02865449 | 0.04694974 | 55.3289238 | 49.7923834 | 0 | 27.4554815 |
| -2.1881973 | 0.02865523 | 0.04694974 | 18.4429746 | 57.9107067 | 0 | 64.99665 |
| -2.1873429 | 0.0287175 | 0.04704839 | 9.01656536 | 77.6652936 | 0 | 76.202969 |
| -2.1873145 | 0.02871957 | 0.04704839 | 0 | 34.6381797 | 93.1867736 | 42.0236961 |
| -2.1873112 | 0.02871981 | 0.04704839 | 63.1159575 | 28.143521 | 0 | 42.0236961 |
| -2.1871336 | 0.02873277 | 0.04706722 | 29.5087594 | 35.7206228 | 100.089498 | 0 |
| -2.1868519 | 0.02875334 | 0.04709853 | 26.2300083 | 36.2618444 | 0 | 75.0823371 |
| 2.18680791 | 0.02875655 | 0.0471014 | 293.858062 | 270.340168 | 292.215315 | 298.088085 |
| -2.1867586 | 0.02876015 | 0.0471049 | 129.100822 | 20.0251977 | 0 | 19.0507423 |
| -2.1866685 | 0.02876673 | 0.04711329 | 33.1973543 | 36.2618444 | 92.0363196 | 0 |
| 2.18658256 | 0.02877301 | 0.04712119 | 36.8859492 | 33.0145151 | 19.5577179 | 36.9808526 |
| -2.1864545 | 0.02878237 | 0.04713413 | 36.4761053 | 11.0950419 | 3.45136199 | 10.0856871 |
| -2.1863343 | 0.02879116 | 0.04714612 | 9.42640924 | 8.93015571 | 6.90272397 | 16.8094785 |
| -2.1861427 | 0.02880516 | 0.04716667 | 0 | 51.9572696 | 71.3281477 | 39.7824324 |
| -2.1858744 | 0.0288248 | 0.04719642 | 57.3781432 | 29.2259641 | 0 | 44.8252759 |
| -2.1858421 | 0.02882716 | 0.0471979 | 20.0823501 | 18.1309222 | 9.20363196 | 13.4475828 |
| 2.18537248 | 0.02886155 | 0.0472518 | 0.81968776 | 0.81183234 | 0 | 0 |
| 2.18517333 | 0.02887614 | 0.0472733 | 9.42640924 | 17.8603114 | 17.2568099 | 14.5682147 |
| -2.1851145 | 0.02888045 | 0.04727795 | 11.4756286 | 11.9068743 | 4.60181598 | 16.8094785 |
| -2.1850289 | 0.02888673 | 0.04728583 | 4.50828268 | 14.8835929 | 18.4072639 | 9.52537113 |
| -2.1849624 | 0.02889161 | 0.04729142 | 20.0823501 | 56.5576528 | 0 | 61.6347543 |
| 2.18433807 | 0.02893742 | 0.0473568 | 0 | 0 | 0 | 0 |
| 2.18433807 | 0.02893742 | 0.0473568 | 0 | 0 | 0 | 0 |
| 2.18433807 | 0.02893742 | 0.0473568 | 0 | 0 | 0 | 0 |
| 2.18433807 | 0.02893742 | 0.0473568 | 0 | 0 | 0 | 0 |
| -2.1842584 | 0.02894327 | 0.04736398 | 11.8854725 | 99.0435452 | 0 | 47.0665397 |
| 2.18383368 | 0.02897447 | 0.04741264 | 0 | 0 | 0 | 0 |
| -2.1838089 | 0.02897629 | 0.04741321 | 20.0823501 | 8.65954493 | 6.90272397 | 8.96505518 |
| -2.1834562 | 0.02900223 | 0.04745325 | 13.524848 | 29.2259641 | 10.354086 | 3.36189569 |
| -2.1822463 | 0.02909136 | 0.04759667 | 9.42640924 | 7.03588026 | 9.20363196 | 15.1285306 |
| 2.18199559 | 0.02910986 | 0.04762453 | 0 | 0.81183234 | 2.30090799 | 0 |
| -2.1818185 | 0.02912293 | 0.04764349 | 5.32797044 | 18.9427545 | 11.50454 | 9.52537113 |

| | | | | | | |
|---|---|---|---|---|---|---|
| 2.18144273 | 0.02915069 | 0.04768649 | 0.81968776 | 0 | 0 | 0 |
| 2.18132999 | 0.02915902 | 0.0476977 | 0.40984388 | 0.81183234 | 0 | 0 |
| 2.18091987 | 0.02918934 | 0.04774489 | 34.8367298 | 22.4606947 | 4.60181598 | 0 |
| -2.1807982 | 0.02919835 | 0.0477451 | 8.60672148 | 9.74198805 | 9.20363196 | 10.0856871 |
| -2.1808077 | 0.02919765 | 0.0477451 | 0 | 110.409198 | 16.1063559 | 58.2728587 |
| -2.1808455 | 0.02919485 | 0.0477451 | 0 | 31.3908504 | 56.3722458 | 77.3236009 |
| -2.180891 | 0.02919148 | 0.0477451 | 142.62567 | 133.140503 | 125.399485 | 128.312352 |
| -2.1808146 | 0.02919714 | 0.0477451 | 26.6398522 | 9.47137727 | 8.05317797 | 6.16347543 |
| -2.1808406 | 0.02919521 | 0.0477451 | 5.73781432 | 3.24732935 | 16.1063559 | 45.9459078 |
| -2.1804802 | 0.02922188 | 0.04778117 | 13.524848 | 10.8244312 | 4.60181598 | 15.1285306 |
| -2.1803224 | 0.02923357 | 0.04779785 | 43.8532952 | 86.5954493 | 0 | 16.2491625 |
| -2.1802163 | 0.02924143 | 0.04780829 | 9.42640924 | 13.8011497 | 9.20363196 | 7.28410733 |
| -2.1801366 | 0.02924734 | 0.04781552 | 50.0009534 | 43.0271139 | 0 | 34.7395888 |
| -2.1800035 | 0.0292572 | 0.04782923 | 4.0984388 | 12.4480958 | 18.4072639 | 12.3269509 |
| -2.1797523 | 0.02927583 | 0.04785725 | 9.83625312 | 11.3656527 | 6.90272397 | 10.646003 |
| -2.1794688 | 0.02929686 | 0.04788921 | 127.051603 | 17.5897006 | 0 | 21.292006 |
| -2.1787657 | 0.02934908 | 0.04797214 | 3.27875104 | 18.1309222 | 13.8054479 | 15.1285306 |
| 2.17872775 | 0.0293519 | 0.04797286 | 20.9020379 | 26.5198564 | 13.8054479 | 35.8602207 |
| 2.17871982 | 0.02935249 | 0.04797286 | 90.5754975 | 106.079425 | 72.4786017 | 90.7711837 |
| -2.1785937 | 0.02936186 | 0.04798575 | 34.8367298 | 70.3588026 | 0 | 28.5761134 |
| -2.1785579 | 0.02936453 | 0.04798767 | 0 | 97.4198805 | 26.4604419 | 47.6268556 |
| 2.17851496 | 0.02936772 | 0.04799046 | 1.22953164 | 1.08244312 | 0 | 0 |
| -2.1783312 | 0.02938139 | 0.04800925 | 52.0501728 | 38.4267306 | 28.7613499 | 52.1093832 |
| -2.1783204 | 0.02938219 | 0.04800925 | 20.492194 | 41.9446708 | 106.992222 | 0 |
| -2.1780918 | 0.02939921 | 0.04802976 | 2.45906328 | 12.9893174 | 32.2127119 | 13.4475828 |
| -2.1781145 | 0.02939751 | 0.04802976 | 0 | 77.3946828 | 51.7704298 | 34.1792729 |
| -2.1780993 | 0.02939865 | 0.04802976 | 14.7543797 | 13.8011497 | 23.0090799 | 21.852322 |
| 2.17801731 | 0.02940475 | 0.04803638 | 0 | 1.3530539 | 0 | 0 |
| 2.17783019 | 0.02941868 | 0.04805631 | 0 | 0 | 5.75226998 | 0 |
| -2.1778135 | 0.02941992 | 0.04805631 | 9.01656536 | 9.20076649 | 10.354086 | 8.96505518 |
| -2.1776011 | 0.02943574 | 0.04807972 | 0.81968776 | 11.6362635 | 40.2658898 | 30.2570612 |
| -2.1775164 | 0.02944206 | 0.04808761 | 6.96734596 | 7.57710182 | 8.05317797 | 30.8173772 |
| -2.1773741 | 0.02945266 | 0.04810249 | 1.63937552 | 12.4480958 | 29.9118039 | 22.4126379 |
| -2.1772747 | 0.02946008 | 0.04811216 | 63.5258014 | 8.93015571 | 12.6549939 | 10.0856871 |
| -2.1772179 | 0.02946431 | 0.04811422 | 7.78703372 | 46.545054 | 19.5577179 | 28.0157974 |
| 2.17721918 | 0.02946422 | 0.04811422 | 6.55750208 | 0 | 0 | 0 |
| 2.17716247 | 0.02946845 | 0.0481161 | 0 | 0 | 0 | 0 |
| 2.17716247 | 0.02946845 | 0.0481161 | 0 | 0 | 0 | 0 |
| -2.1770513 | 0.02947674 | 0.04812721 | 12.7051603 | 5.95343714 | 6.90272397 | 22.4126379 |
| -2.1768135 | 0.02949448 | 0.04815374 | 7.37718984 | 12.1774851 | 16.1063559 | 6.72379138 |
| -2.1766859 | 0.02950401 | 0.04816686 | 88.1164342 | 236.24321 | 192.125817 | 191.628054 |
| -2.1766429 | 0.02950722 | 0.04816967 | 6.96734596 | 10.2832096 | 14.9559019 | 8.40473923 |
| 2.17649245 | 0.02951846 | 0.04818557 | 0.40984388 | 0.54122156 | 0 | 0 |

| | | | | | | |
|---|---|---|---|---|---|---|
| -2.1764314 | 0.02952302 | 0.04818922 | 9.83625312 | 13.530539 | 5.75226998 | 11.7666349 |
| -2.1764226 | 0.02952368 | 0.04818922 | 0 | 82.8068984 | 35.6640739 | 45.3855918 |
| 2.17621892 | 0.02953889 | 0.04820918 | 9.42640924 | 3.24732935 | 8.05317797 | 5.04284354 |
| -2.1762331 | 0.02953783 | 0.04820918 | 0 | 28.4141318 | 82.8326877 | 54.910963 |
| -2.1761514 | 0.02954394 | 0.04821498 | 14.3445358 | 5.41221558 | 16.1063559 | 8.40473923 |
| -2.1758328 | 0.02956777 | 0.04825143 | 11.0657848 | 11.6362635 | 12.6549939 | 5.60315949 |
| -2.1753868 | 0.02960114 | 0.04830345 | 64.755333 | 211.347019 | 134.603117 | 145.121831 |
| -2.175259 | 0.02961071 | 0.04831662 | 8.1968776 | 23.5431378 | 4.60181598 | 13.4475828 |
| -2.1750136 | 0.0296291 | 0.04834418 | 4.91812656 | 23.8137486 | 8.05317797 | 12.8872668 |
| -2.1746484 | 0.02965647 | 0.0483864 | 0 | 90.1133895 | 49.4695218 | 28.0157974 |
| 2.17446573 | 0.02967018 | 0.04840632 | 2.0492194 | 0.81183234 | 0 | 0 |
| 2.17440517 | 0.02967472 | 0.04841035 | 36.8859492 | 4.32977247 | 0 | 12.8872668 |
| -2.1743928 | 0.02967565 | 0.04841035 | 53.6895483 | 41.67406 | 52.9208838 | 0 |
| -2.1739454 | 0.02970924 | 0.04846269 | 0 | 51.416048 | 85.1335957 | 29.1364293 |
| -2.173854 | 0.0297161 | 0.04847144 | 208.200691 | 331.498204 | 247.347609 | 285.200818 |
| -2.1738335 | 0.02971764 | 0.04847151 | 7.78703372 | 13.530539 | 21.8586259 | 5.04284354 |
| -2.1736047 | 0.02973483 | 0.04849465 | 33.1973543 | 7.03588026 | 4.60181598 | 12.3269509 |
| -2.1736075 | 0.02973462 | 0.04849465 | 20.0823501 | 82.2656769 | 0 | 38.1014845 |
| -2.1735402 | 0.02973969 | 0.04850011 | 7.78703372 | 7.8477126 | 13.8054479 | 10.0856871 |
| 2.1732312 | 0.02976292 | 0.04853555 | 4.50828268 | 2.16488623 | 0 | 1.68094785 |
| 2.17315718 | 0.02976849 | 0.04854218 | 17.6232868 | 3.24732935 | 8.05317797 | 3.36189569 |
| 2.17307957 | 0.02977433 | 0.04854925 | 22.5414134 | 33.2851258 | 29.9118039 | 31.3776931 |
| -2.1730245 | 0.02977848 | 0.04855356 | 622.14301 | 1362.79588 | 896.203662 | 1078.04789 |
| -2.1729728 | 0.02978236 | 0.04855744 | 140.986295 | 287.118037 | 239.294431 | 281.838922 |
| -2.1729286 | 0.02978569 | 0.04856043 | 10.246097 | 77.3946828 | 0 | 66.6775979 |
| -2.1728783 | 0.02978948 | 0.04856415 | 15.1642236 | 9.47137727 | 18.4072639 | 16.2491625 |
| -2.1727479 | 0.02979929 | 0.04857769 | 46.3123584 | 34.367569 | 0 | 45.3855918 |
| -2.1726988 | 0.02980299 | 0.04858127 | 36.8859492 | 55.7458205 | 0 | 34.7395888 |
| -2.1726317 | 0.02980805 | 0.04858706 | 7.78703372 | 14.0717605 | 5.75226998 | 15.6888466 |
| -2.1724635 | 0.02982072 | 0.04860383 | 8.1968776 | 13.530539 | 35.6640739 | 57.7125427 |
| -2.1724551 | 0.02982135 | 0.04860383 | 6.96734596 | 0 | 126.549939 | 88.5299199 |
| -2.1722192 | 0.02983913 | 0.04863036 | 3.27875104 | 12.1774851 | 33.3631659 | 10.0856871 |
| -2.1721795 | 0.02984212 | 0.04863278 | 108.608628 | 276.564216 | 207.081719 | 251.581861 |
| 2.171933 | 0.02986072 | 0.04866063 | 0 | 1.62366467 | 0 | 0 |
| -2.1718743 | 0.02986515 | 0.04866539 | 8.1968776 | 18.6721438 | 0 | 172.577312 |
| -2.1716041 | 0.02988554 | 0.04869616 | 20.0823501 | 4.87099402 | 8.05317797 | 14.5682147 |
| -2.1710365 | 0.02992841 | 0.04876355 | 0 | 48.7099402 | 57.5226998 | 50.4284354 |
| -2.1706941 | 0.02995431 | 0.04880328 | 10.6559409 | 81.1832337 | 0 | 61.0744384 |
| 2.17036305 | 0.02997935 | 0.04882931 | 0 | 0 | 0 | 0 |
| 2.17036305 | 0.02997935 | 0.04882931 | 0 | 0 | 0 | 0 |
| 2.17036305 | 0.02997935 | 0.04882931 | 0 | 0 | 0 | 0 |
| 2.17036305 | 0.02997935 | 0.04882931 | 0 | 0 | 0 | 0 |
| 2.17036305 | 0.02997935 | 0.04882931 | 0 | 0 | 0 | 0 |

| | | | | | | |
|---|---|---|---|---|---|---|
| 2.17036305 | 0.02997935 | 0.04882931 | 0 | 0 | 0 | 0 |
| 2.17013516 | 0.02999661 | 0.04885495 | 15.9839113 | 10.2832096 | 17.2568099 | 7.28410733 |
| -2.170011 | 0.03000601 | 0.0488678 | 9.01656536 | 17.8603114 | 29.9118039 | 32.498325 |
| -2.1699649 | 0.03000951 | 0.04887102 | 7.78703372 | 12.4480958 | 6.90272397 | 12.8872668 |
| -2.1696803 | 0.03003108 | 0.04890369 | 0 | 60.075593 | 48.3190678 | 48.1871716 |
| -2.1692828 | 0.03006122 | 0.04895031 | 9.42640924 | 10.0125988 | 6.90272397 | 12.3269509 |
| 2.16916749 | 0.03006997 | 0.04896208 | 33.1973543 | 32.2026827 | 36.8145279 | 38.6618005 |
| 2.16903581 | 0.03007997 | 0.04897589 | 41.3942319 | 20.2958084 | 8.05317797 | 15.1285306 |
| 2.16860597 | 0.03011261 | 0.04902657 | 5.32797044 | 5.68282636 | 8.05317797 | 6.72379138 |
| -2.1685704 | 0.03011532 | 0.0490285 | 38.5253247 | 45.1920001 | 31.0622579 | 26.3348496 |
| -2.168196 | 0.03014378 | 0.04907236 | 13.1150042 | 14.6129821 | 5.75226998 | 8.40473923 |
| -2.1681482 | 0.03014741 | 0.0490758 | 9.01656536 | 2.43549701 | 31.0622579 | 19.6110582 |
| -2.167804 | 0.0301736 | 0.04911596 | 54.9190799 | 73.3355211 | 0 | 14.5682147 |
| -2.1677598 | 0.03017696 | 0.04911896 | 0 | 32.4732935 | 147.258111 | 59.3934906 |
| -2.167647 | 0.03018556 | 0.04913047 | 25.8201644 | 41.9446708 | 0 | 62.1950703 |
| 2.16752629 | 0.03019475 | 0.04914239 | 65.9848647 | 23.272527 | 34.5136199 | 25.7745336 |
| -2.1675108 | 0.03019593 | 0.04914239 | 0 | 81.4538445 | 49.4695218 | 31.3776931 |
| -2.1674359 | 0.03020163 | 0.04914919 | 75.00143 | 20.2958084 | 11.50454 | 28.0157974 |
| -2.1673169 | 0.0302107 | 0.04915817 | 8.60672148 | 11.9068743 | 26.4604419 | 4.48252759 |
| -2.1673395 | 0.03020898 | 0.04915817 | 1.63937552 | 21.1076408 | 16.1063559 | 23.5332698 |
| -2.1673036 | 0.03021171 | 0.04915817 | 38.5253247 | 4.05916169 | 6.90272397 | 12.3269509 |
| -2.1671912 | 0.03022028 | 0.04916057 | 22.9512573 | 13.8011497 | 3.45136199 | 10.646003 |
| 2.16716435 | 0.03022232 | 0.04916057 | 2.0492194 | 0 | 1.150454 | 2.24126379 |
| 2.16722939 | 0.03021737 | 0.04916057 | 2.0492194 | 0.54122156 | 4.60181598 | 2.80157974 |
| -2.1672407 | 0.03021651 | 0.04916057 | 15.5740674 | 12.7187066 | 17.2568099 | 3.36189569 |
| -2.1672107 | 0.0302188 | 0.04916057 | 44.263139 | 12.1774851 | 0 | 95.8140272 |
| -2.1671722 | 0.03022173 | 0.04916057 | 8.1968776 | 21.1076408 | 4.60181598 | 14.0078987 |
| -2.167091 | 0.03022792 | 0.0491672 | 9.42640924 | 13.530539 | 16.1063559 | 21.852322 |
| -2.1669234 | 0.0302407 | 0.0491855 | 3.68859492 | 8.11832337 | 24.1595339 | 17.3697944 |
| -2.1668188 | 0.03024867 | 0.049196 | 20.0823501 | 54.1221558 | 0 | 59.3934906 |
| -2.1666345 | 0.03026273 | 0.04921638 | 0 | 44.3801678 | 80.5317797 | 35.8602207 |
| 2.16654233 | 0.03026977 | 0.04922535 | 0 | 0.27061078 | 0 | 0 |
| -2.1662205 | 0.03029434 | 0.04926282 | 3.68859492 | 10.0125988 | 19.5577179 | 16.2491625 |
| 2.16619765 | 0.03029609 | 0.04926318 | 2.0492194 | 1.08244312 | 1.150454 | 1.68094785 |
| -2.1654159 | 0.03035585 | 0.04935539 | 3.68859492 | 17.0484791 | 16.1063559 | 11.206319 |
| -2.1654249 | 0.03035516 | 0.04935539 | 40.5745441 | 32.7439043 | 0 | 52.1093832 |
| -2.1652332 | 0.03036983 | 0.04937564 | 34.017042 | 51.9572696 | 0 | 39.2221164 |
| -2.1651981 | 0.03037252 | 0.04937752 | 68.8537718 | 13.530539 | 0 | 61.0744384 |
| -2.164807 | 0.03040247 | 0.04942372 | 58.197831 | 173.461509 | 179.470823 | 159.129729 |
| -2.1644185 | 0.03043224 | 0.04946963 | 0 | 44.9213893 | 49.4695218 | 61.0744384 |
| -2.1641785 | 0.03045065 | 0.04949706 | 8.1968776 | 12.9893174 | 6.90272397 | 11.206319 |
| -2.1640883 | 0.03045757 | 0.04950582 | 35.2465737 | 50.0629941 | 0 | 39.2221164 |
| -2.1640365 | 0.03046155 | 0.04950979 | 14.3445358 | 18.1309222 | 18.4072639 | 20.7316901 |

| | | | | | | |
|---|---|---|---|---|---|---|
| -2.1639179 | 0.03047065 | 0.0495221 | 17.6232868 | 54.1221558 | 0 | 64.4363341 |
| -2.1638211 | 0.03047808 | 0.04953167 | 13.1150042 | 10.0125988 | 5.75226998 | 11.206319 |
| -2.1637778 | 0.0304814 | 0.04953458 | 19.6725062 | 0 | 94.3372276 | 70.5998095 |
| -2.1636671 | 0.0304899 | 0.0495459 | 5.32797044 | 86.3248385 | 0 | 81.2458125 |
| -2.1636321 | 0.03049259 | 0.04954778 | 2.45906328 | 16.5072575 | 16.1063559 | 19.0507423 |
| -2.1635467 | 0.03049916 | 0.04955595 | 195.085687 | 351.523402 | 291.064861 | 290.243661 |
| 2.1634306 | 0.03050808 | 0.04955882 | 0 | 0 | 0 | 0 |
| 2.1634306 | 0.03050808 | 0.04955882 | 0 | 0 | 0 | 0 |
| 2.1634306 | 0.03050808 | 0.04955882 | 0 | 0 | 0 | 0 |
| -2.1634969 | 0.03050298 | 0.04955882 | 6.1476582 | 13.8011497 | 11.50454 | 8.96505518 |
| -2.1634238 | 0.0305086 | 0.04955882 | 266.808366 | 471.403977 | 415.313892 | 456.657498 |
| 2.16332419 | 0.03051625 | 0.04956876 | 19.6725062 | 11.9068743 | 19.5577179 | 9.52537113 |
| -2.1632151 | 0.03052464 | 0.04957988 | 48.7714217 | 48.7099402 | 0 | 28.5761134 |
| -2.1630188 | 0.03053974 | 0.04960192 | 3.27875104 | 12.9893174 | 19.5577179 | 14.0078987 |
| -2.1629722 | 0.03054332 | 0.04960524 | 12.7051603 | 2.43549701 | 28.7613499 | 14.5682147 |
| -2.1629146 | 0.03054775 | 0.04960993 | 0 | 58.4519283 | 41.4163438 | 55.4712789 |
| -2.1627458 | 0.03056074 | 0.04962854 | 15.1642236 | 5.95343714 | 8.05317797 | 12.8872668 |
| -2.1626587 | 0.03056744 | 0.04963692 | 40.5745441 | 63.8641439 | 0 | 25.2142177 |
| 2.16236645 | 0.03058994 | 0.04967096 | 2.0492194 | 0 | 0 | 0.56031595 |
| -2.1623183 | 0.03059365 | 0.04967448 | 15.9839113 | 11.6362635 | 8.05317797 | 6.16347543 |
| -2.1619481 | 0.03062218 | 0.0497183 | 10.6559409 | 9.74198805 | 4.60181598 | 22.9729539 |
| 2.16180095 | 0.03063352 | 0.04973422 | 0 | 1.3530539 | 1.150454 | 0 |
| 2.16100603 | 0.03069487 | 0.04982713 | 14.7543797 | 12.4480958 | 13.8054479 | 14.0078987 |
| -2.1610062 | 0.03069486 | 0.04982713 | 0 | 43.8389462 | 43.7172518 | 68.3585457 |
| -2.1609995 | 0.03069538 | 0.04982713 | 211.069598 | 412.410827 | 338.233475 | 373.730738 |
| -2.1607371 | 0.03071565 | 0.04985754 | 32.3776665 | 92.2782757 | 0 | 18.4904263 |
| -2.1604408 | 0.03073856 | 0.04989221 | 4.50828268 | 16.2366467 | 10.354086 | 13.4475828 |
| -2.1602962 | 0.03074975 | 0.04990786 | 0 | 49.2511618 | 71.3281477 | 36.4205367 |
| -2.1602526 | 0.03075312 | 0.04991083 | 9.01656536 | 12.7187066 | 8.05317797 | 8.40473923 |
| 2.15989306 | 0.03078095 | 0.04995348 | 8.1968776 | 6.4946587 | 4.60181598 | 6.16347543 |
| -2.159727 | 0.03079381 | 0.04997184 | 4.0984388 | 12.9893174 | 14.9559019 | 12.8872668 |
| -2.1596751 | 0.03079783 | 0.04997585 | 15.9839113 | 80.912623 | 0 | 43.704644 |

| GFP (+) 1 | GFP (+) 2 |
|---|---|
| 9875.70329 | 8406.5272 |
| 12970.6628 | 10271.4318 |
| 21966.9992 | 17572.0272 |
| 4017.26435 | 2602.19243 |
| 565.06198 | 542.123422 |
| 419.073322 | 274.675867 |
| 6964.51771 | 5153.78666 |
| 34.3502723 | 93.9680598 |
| 25.7627042 | 14.4566246 |
| 2753.17433 | 1575.77208 |
| 30.9152451 | 0 |
| 554.756898 | 520.438485 |
| 125.378494 | 93.9680598 |
| 66.983031 | 65.0548106 |
| 5461.6933 | 3903.28864 |
| 5362.07751 | 4018.94163 |
| 120.225953 | 130.109621 |
| 36.0677859 | 0 |
| 12142.8213 | 7907.77365 |
| 51.5254085 | 28.9132492 |
| 1076.88104 | 1149.30165 |
| 37.7852996 | 7.22831229 |
| 297.129856 | 281.904179 |
| 137.401089 | 108.424684 |
| 276.519692 | 267.447555 |
| 1337.94311 | 599.94992 |
| 3769.94239 | 3498.50315 |
| 826.12405 | 614.406545 |
| 14545.6228 | 7437.93335 |
| 5.15254085 | 7.22831229 |
| 601.129766 | 628.863169 |
| 1080.31606 | 621.634857 |
| 20.6101634 | 0 |
| 1.71751362 | 36.1415615 |
| 965.242652 | 571.036671 |
| 12.0225953 | 14.4566246 |
| 1470.19166 | 1301.09621 |
| 15.4576225 | 7.22831229 |
| 302.282396 | 202.392744 |
| 3.43502723 | 7.22831229 |
| 453.423595 | 477.068611 |
| 82.4406536 | 79.5114352 |

| | |
|---|---|
| 2394.21398 | 1474.57571 |
| 0 | 0 |
| 0 | 0 |
| 3443.6148 | 2060.069 |
| 8.58756808 | 0 |
| 0 | 0 |
| 41.2203268 | 14.4566246 |
| 1449.58149 | 19661.0094 |
| 431.095918 | 505.98186 |
| 924.022326 | 621.634857 |
| 748.835937 | 419.242113 |
| 34.3502723 | 36.1415615 |
| 388.158077 | 339.730678 |
| 0 | 0 |
| 235.299365 | 513.210173 |
| 0 | 0 |
| 25.7627042 | 7.22831229 |
| 0 | 21.6849369 |
| 0 | 0 |
| 0 | 0 |
| 0 | 0 |
| 36.0677859 | 137.337934 |
| 2612.33821 | 1040.87697 |
| 32.6327587 | 36.1415615 |
| 0 | 7.22831229 |
| 3.43502723 | 0 |
| 1238.32732 | 469.840299 |
| 22.327677 | 21.6849369 |
| 0 | 0 |
| 0 | 0 |
| 0 | 0 |
| 25.7627042 | 0 |
| 226.711797 | 274.675867 |
| 13.7401089 | 36.1415615 |
| 278.237206 | 419.242113 |
| 8.58756808 | 0 |
| 0 | 14.4566246 |
| 0 | 7.22831229 |
| 0 | 0 |
| 0 | 14.4566246 |
| 18.8926498 | 7.22831229 |
| 218.124229 | 195.164432 |
| 0 | 0 |

| | |
|---|---|
| 0 | 0 |
| 0 | 7.22831229 |
| 121.943467 | 166.251183 |
| 84.1581672 | 21.6849369 |
| 255.909529 | 318.045741 |
| 422.50835 | 332.502365 |
| 0 | 0 |
| 226.711797 | 159.02287 |
| 0 | 0 |
| 5.15254085 | 21.6849369 |
| 0 | 7.22831229 |
| 0 | 0 |
| 0 | 0 |
| 6.87005447 | 14.4566246 |
| 0 | 0 |
| 0 | 0 |
| 0 | 7.22831229 |
| 984.135302 | 484.296924 |
| 0 | 0 |
| 659.525229 | 383.100551 |
| 194.079039 | 166.251183 |
| 0 | 0 |
| 0 | 0 |
| 0 | 0 |
| 49.8078949 | 50.598186 |
| 0 | 0 |
| 0 | 0 |
| 0 | 0 |
| 15.4576225 | 14.4566246 |
| 10.3050817 | 0 |
| 1.71751362 | 14.4566246 |
| 0 | 7.22831229 |
| 408.768241 | 224.077681 |
| 15.4576225 | 0 |
| 0 | 0 |
| 231.864338 | 260.219243 |
| 0 | 0 |
| 0 | 7.22831229 |
| 0 | 0 |
| 0 | 0 |
| 0 | 0 |
| 0 | 0 |
| 37.7852996 | 0 |

| | |
|---|---|
| 0 | 7.22831229 |
| 0 | 0 |
| 5.15254085 | 0 |
| 274.802179 | 216.849369 |
| 0 | 0 |
| 0 | 0 |
| 0 | 0 |
| 6.87005447 | 0 |
| 0 | 0 |
| 12.0225953 | 0 |
| 106.485844 | 144.566246 |
| 0 | 0 |
| 0 | 0 |
| 0 | 0 |
| 5.15254085 | 14.4566246 |
| 0 | 0 |
| 0 | 0 |
| 0 | 0 |
| 0 | 0 |
| 237.016879 | 173.479495 |
| 0 | 0 |
| 0 | 0 |
| 219.841743 | 636.091482 |
| 0 | 7.22831229 |
| 0 | 0 |
| 0 | 0 |
| 0 | 0 |
| 3878.14575 | 1792.62145 |
| 0 | 0 |
| 0 | 0 |
| 0 | 0 |
| 0 | 0 |
| 1.71751362 | 0 |
| 0 | 0 |
| 0 | 0 |
| 305.717424 | 498.753548 |
| 0 | 0 |
| 99.6157898 | 187.93612 |
| 178.621416 | 303.589116 |
| 0 | 0 |
| 188.926498 | 303.589116 |
| 0 | 0 |
| 0 | 0 |

| | |
|---|---|
| 0 | 0 |
| 0 | 0 |
| 0 | 0 |
| 1.71751362 | 0 |
| 0 | 0 |
| 17.1751362 | 0 |
| 0 | 0 |
| 135.683576 | 245.762618 |
| 0 | 0 |
| 1.71751362 | 0 |
| 0 | 0 |
| 0 | 0 |
| 0 | 0 |
| 0 | 0 |
| 0 | 0 |
| 637.197552 | 881.8541 |
| 0 | 0 |
| 0 | 0 |
| 762.576046 | 209.621056 |
| 0 | 0 |
| 0 | 0 |
| 0 | 0 |
| 0 | 0 |
| 0 | 0 |
| 0 | 0 |
| 0 | 0 |
| 1.71751362 | 7.22831229 |
| 0 | 0 |
| 0 | 0 |
| 0 | 0 |
| 0 | 0 |
| 0 | 0 |
| 0 | 0 |
| 0 | 0 |
| 0 | 0 |
| 202.666607 | 202.392744 |
| 109.920871 | 101.196372 |
| 1.71751362 | 7.22831229 |
| 0 | 0 |
| 0 | 0 |
| 216.406716 | 231.305993 |
| 3.43502723 | 7.22831229 |
| 0 | 0 |

| | |
|---|---|
| 0 | 0 |
| 0 | 0 |
| 0 | 0 |
| 176.903902 | 252.99093 |
| 1035.66071 | 1091.47516 |
| 0 | 0 |
| 3.43502723 | 0 |
| 0 | 0 |
| 1.71751362 | 7.22831229 |
| 0 | 0 |
| 5.15254085 | 14.4566246 |
| 27.4802179 | 7.22831229 |
| 0 | 0 |
| 12.0225953 | 7.22831229 |
| 0 | 0 |
| 0 | 0 |
| 0 | 0 |
| 1.71751362 | 0 |
| 0 | 7.22831229 |
| 0 | 0 |
| 0 | 0 |
| 15.4576225 | 0 |
| 0 | 0 |
| 278.237206 | 101.196372 |
| 0 | 0 |
| 10.3050817 | 0 |
| 0 | 0 |
| 0 | 0 |
| 0 | 7.22831229 |
| 199.23158 | 187.93612 |
| 364.112887 | 440.92705 |
| 303.99991 | 166.251183 |
| 0 | 0 |
| 0 | 0 |
| 0 | 7.22831229 |
| 6.87005447 | 7.22831229 |
| 0 | 0 |
| 0 | 0 |
| 242.16942 | 252.99093 |
| 0 | 0 |
| 0 | 21.6849369 |
| 0 | 0 |
| 0 | 0 |

| | |
|---|---|
| 36.0677859 | 0 |
| 166.598821 | 238.534306 |
| 0 | 0 |
| 0 | 0 |
| 0 | 0 |
| 0 | 0 |
| 0 | 0 |
| 0 | 0 |
| 24.0451906 | 7.22831229 |
| 0 | 0 |
| 0 | 7.22831229 |
| 0 | 0 |
| 0 | 0 |
| 0 | 0 |
| 0 | 0 |
| 0 | 7.22831229 |
| 592.542198 | 289.132492 |
| 20.6101634 | 0 |
| 0 | 0 |
| 0 | 0 |
| 0 | 0 |
| 0 | 0 |
| 0 | 0 |
| 0 | 0 |
| 5.15254085 | 0 |
| 0 | 0 |
| 0 | 0 |
| 0 | 0 |
| 0 | 0 |
| 257.627042 | 571.036671 |
| 0 | 0 |
| 0 | 0 |
| 0 | 0 |
| 0 | 0 |
| 0 | 0 |
| 0 | 7.22831229 |
| 0 | 0 |
| 27.4802179 | 14.4566246 |
| 0 | 0 |
| 0 | 0 |
| 170.033848 | 643.319794 |
| 5.15254085 | 0 |
| 386.440564 | 159.02287 |

| | |
|---|---|
| 0 | 0 |
| 0 | 0 |
| 0 | 0 |
| 0 | 7.22831229 |
| 0 | 0 |
| 0 | 0 |
| 0 | 0 |
| 97.8982761 | 166.251183 |
| 0 | 0 |
| 61.8304902 | 7.22831229 |
| 0 | 0 |
| 0 | 0 |
| 0 | 0 |
| 0 | 0 |
| 0 | 0 |
| 0 | 0 |
| 0 | 0 |
| 0 | 0 |
| 56.6779493 | 7.22831229 |
| 0 | 0 |
| 32.6327587 | 28.9132492 |
| 0 | 0 |
| 0 | 0 |
| 0 | 0 |
| 0 | 0 |
| 0 | 0 |
| 1300.15781 | 650.548106 |
| 36.0677859 | 0 |
| 0 | 0 |
| 0 | 7.22831229 |
| 0 | 0 |
| 0 | 0 |
| 27.4802179 | 7.22831229 |
| 0 | 0 |
| 0 | 7.22831229 |
| 22.327677 | 21.6849369 |
| 0 | 0 |
| 1.71751362 | 0 |
| 0 | 0 |
| 810.666427 | 166.251183 |
| 206.101634 | 159.02287 |
| 0 | 0 |
| 0 | 7.22831229 |

| | |
|---|---|
| 202.666607 | 180.707807 |
| 164.881307 | 7.22831229 |
| 1.71751362 | 0 |
| 87.5931944 | 28.9132492 |
| 0 | 0 |
| 0 | 0 |
| 0 | 0 |
| 10.3050817 | 0 |
| 170.033848 | 245.762618 |
| 0 | 0 |
| 0 | 0 |
| 6.87005447 | 7.22831229 |
| 0 | 0 |
| 18.8926498 | 21.6849369 |
| 0 | 0 |
| 0 | 0 |
| 0 | 0 |
| 0 | 0 |
| 0 | 0 |
| 3.43502723 | 7.22831229 |
| 338.350182 | 245.762618 |
| 0 | 7.22831229 |
| 0 | 0 |
| 0 | 0 |
| 0 | 0 |
| 0 | 0 |
| 163.163794 | 397.557176 |
| 0 | 0 |
| 0 | 0 |
| 149.423685 | 491.525236 |
| 0 | 0 |
| 106.485844 | 72.2831229 |
| 0 | 0 |
| 0 | 0 |
| 0 | 0 |
| 0 | 0 |
| 0 | 0 |
| 0 | 0 |
| 0 | 7.22831229 |
| 0 | 0 |
| 0 | 0 |
| 0 | 0 |
| 0 | 0 |

| | |
|---|---|
| 0 | 0 |
| 0 | 0 |
| 0 | 0 |
| 0 | 0 |
| 0 | 0 |
| 0 | 0 |
| 0 | 0 |
| 0 | 0 |
| 24.0451906 | 0 |
| 20.6101634 | 21.6849369 |
| 0 | 0 |
| 0 | 0 |
| 0 | 0 |
| 3605.06108 | 1279.41128 |
| 0 | 0 |
| 18.8926498 | 0 |
| 0 | 0 |
| 0 | 0 |
| 384.72305 | 361.415615 |
| 652.655174 | 563.808359 |
| 0 | 7.22831229 |
| 0 | 0 |
| 0 | 0 |
| 0 | 7.22831229 |
| 0 | 0 |
| 0 | 0 |
| 0 | 0 |
| 13.7401089 | 0 |
| 42.9378404 | 0 |
| 0 | 0 |
| 0 | 0 |
| 15.4576225 | 7.22831229 |
| 0 | 0 |
| 0 | 0 |
| 3.43502723 | 0 |
| 0 | 0 |
| 0 | 0 |
| 0 | 0 |
| 0 | 0 |
| 0 | 0 |
| 0 | 0 |
| 0 | 0 |
| 0 | 0 |

| | |
|---|---|
| 0 | 0 |
| 53.2429221 | 50.598186 |
| 0 | 0 |
| 0 | 0 |
| 0 | 0 |
| 0 | 0 |
| 0 | 0 |
| 0 | 0 |
| 0 | 0 |
| 0 | 0 |
| 0 | 0 |
| 0 | 0 |
| 0 | 0 |
| 0 | 0 |
| 0 | 0 |
| 0 | 0 |
| 147.706171 | 187.93612 |
| 0 | 0 |
| 0 | 0 |
| 0 | 0 |
| 0 | 0 |
| 0 | 0 |
| 0 | 7.22831229 |
| 0 | 0 |
| 0 | 0 |
| 0 | 0 |
| 0 | 0 |
| 0 | 0 |
| 0 | 0 |
| 61.8304902 | 0 |
| 0 | 0 |
| 176.903902 | 151.794558 |
| 0 | 0 |
| 0 | 0 |
| 0 | 7.22831229 |
| 0 | 0 |
| 0 | 0 |
| 166.598821 | 173.479495 |
| 0 | 7.22831229 |
| 0 | 0 |
| 0 | 0 |
| 0 | 14.4566246 |
| 0 | 0 |

| | |
|---|---|
| 0 | 0 |
| 0 | 0 |
| 0 | 0 |
| 5.15254085 | 0 |
| 0 | 0 |
| 0 | 0 |
| 0 | 0 |
| 0 | 0 |
| 0 | 0 |
| 0 | 7.22831229 |
| 15.4576225 | 28.9132492 |
| 0 | 0 |
| 0 | 0 |
| 0 | 0 |
| 0 | 0 |
| 0 | 0 |
| 0 | 0 |
| 140.836117 | 101.196372 |
| 0 | 0 |
| 0 | 0 |
| 0 | 0 |
| 77.2881127 | 101.196372 |
| 0 | 0 |
| 0 | 0 |
| 0 | 0 |
| 0 | 701.146292 |
| 0 | 7.22831229 |
| 0 | 0 |
| 0 | 0 |
| 1.71751362 | 0 |
| 0 | 0 |
| 0 | 0 |
| 0 | 0 |
| 0 | 0 |
| 0 | 7.22831229 |
| 0 | 0 |
| 372.700455 | 0 |
| 5.15254085 | 7.22831229 |
| 1.71751362 | 7.22831229 |
| 0 | 0 |
| 0 | 0 |
| 355.525319 | 354.187302 |
| 1.71751362 | 7.22831229 |

| | |
|---|---|
| 0 | 0 |
| 0 | 0 |
| 0 | 7.22831229 |
| 0 | 0 |
| 1.71751362 | 0 |
| 0 | 7.22831229 |
| 3.43502723 | 0 |
| 132.248548 | 224.077681 |
| 322.89256 | 130.109621 |
| 0 | 0 |
| 0 | 0 |
| 15.4576225 | 57.8264983 |
| 0 | 0 |
| 3.43502723 | 0 |
| 0 | 0 |
| 30.9152451 | 7.22831229 |
| 0 | 0 |
| 0 | 0 |
| 0 | 0 |
| 0 | 0 |
| 5.15254085 | 7.22831229 |
| 0 | 0 |
| 0 | 0 |
| 0 | 0 |
| 0 | 0 |
| 0 | 0 |
| 0 | 0 |
| 0 | 0 |
| 0 | 0 |
| 0 | 0 |
| 0 | 0 |
| 357.242832 | 0 |
| 0 | 0 |
| 0 | 0 |
| 142.55363 | 173.479495 |
| 0 | 0 |
| 0 | 0 |
| 0 | 0 |
| 0 | 0 |
| 0 | 0 |
| 0 | 0 |
| 0 | 0 |
| 0 | 0 |

| | |
|---|---|
| 0 | 0 |
| 0 | 0 |
| 0 | 0 |
| 0 | 0 |
| 0 | 7.22831229 |
| 17.1751362 | 7.22831229 |
| 0 | 0 |
| 249.039474 | 224.077681 |
| 0 | 0 |
| 18.8926498 | 0 |
| 0 | 0 |
| 0 | 7.22831229 |
| 0 | 0 |
| 39.5028132 | 21.6849369 |
| 0 | 0 |
| 0 | 0 |
| 0 | 0 |
| 0 | 0 |
| 5.15254085 | 0 |
| 0 | 0 |
| 0 | 0 |
| 0 | 7.22831229 |
| 233.581852 | 195.164432 |
| 0 | 0 |
| 0 | 0 |
| 0 | 0 |
| 0 | 0 |
| 0 | 0 |
| 0 | 0 |
| 0 | 0 |
| 0 | 0 |
| 0 | 0 |
| 0 | 0 |
| 17.1751362 | 0 |
| 0 | 0 |
| 0 | 0 |
| 13.7401089 | 0 |
| 0 | 0 |
| 6.87005447 | 7.22831229 |
| 0 | 0 |
| 0 | 0 |
| 334.915155 | 0 |
| 0 | 0 |

| | |
|---|---|
| 314.304992 | 238.534306 |
| 0 | 0 |
| 0 | 0 |
| 0 | 0 |
| 0 | 0 |
| 200.949093 | 130.109621 |
| 0 | 0 |
| 0 | 0 |
| 0 | 0 |
| 0 | 0 |
| 12.0225953 | 7.22831229 |
| 0 | 0 |
| 0 | 0 |
| 0 | 0 |
| 0 | 0 |
| 417.355809 | 122.881309 |
| 0 | 0 |
| 0 | 0 |
| 0 | 0 |
| 1.71751362 | 14.4566246 |
| 0 | 0 |
| 0 | 0 |
| 326.327587 | 0 |
| 0 | 0 |
| 0 | 0 |
| 0 | 0 |
| 0 | 0 |
| 0 | 0 |
| 0 | 0 |
| 0 | 0 |
| 0 | 0 |
| 56.6779493 | 65.0548106 |
| 0 | 0 |
| 5.15254085 | 14.4566246 |
| 0 | 0 |
| 0 | 0 |
| 0 | 0 |
| 3.43502723 | 0 |
| 322.89256 | 0 |
| 3.43502723 | 0 |
| 5.15254085 | 7.22831229 |
| 0 | 0 |
| 0 | 0 |

| | |
|---|---|
| 121.943467 | 245.762618 |
| 444.836027 | 173.479495 |
| 0 | 0 |
| 0 | 0 |
| 0 | 0 |
| 0 | 0 |
| 0 | 0 |
| 1.71751362 | 0 |
| 17.1751362 | 0 |
| 0 | 0 |
| 0 | 0 |
| 0 | 0 |
| 0 | 0 |
| 10.3050817 | 7.22831229 |
| 0 | 0 |
| 123.66098 | 209.621056 |
| 0 | 14.4566246 |
| 0 | 0 |
| 0 | 0 |
| 0 | 0 |
| 135.683576 | 224.077681 |
| 224.994284 | 122.881309 |
| 0 | 0 |
| 1.71751362 | 0 |
| 1.71751362 | 0 |
| 0 | 0 |
| 0 | 0 |
| 0 | 0 |
| 0 | 0 |
| 0 | 0 |
| 3.43502723 | 0 |
| 0 | 0 |
| 17.1751362 | 14.4566246 |
| 0 | 0 |
| 0 | 0 |
| 0 | 7.22831229 |
| 1.71751362 | 7.22831229 |
| 0 | 0 |
| 0 | 0 |
| 0 | 0 |
| 0 | 0 |
| 0 | 0 |
| 1.71751362 | 0 |

| | |
|---|---|
| 6.87005447 | 7.22831229 |
| 0 | 0 |
| 0 | 0 |
| 0 | 0 |
| 0 | 0 |
| 0 | 0 |
| 0 | 0 |
| 0 | 0 |
| 0 | 0 |
| 0 | 0 |
| 0 | 7.22831229 |
| 0 | 0 |
| 0 | 0 |
| 0 | 0 |
| 1823.99946 | 19089.9728 |
| 0 | 0 |
| 17.1751362 | 0 |
| 1.71751362 | 0 |
| 0 | 0 |
| 0 | 0 |
| 0 | 0 |
| 0 | 0 |
| 0 | 0 |
| 0 | 0 |
| 0 | 0 |
| 0 | 0 |
| 0 | 0 |
| 0 | 0 |
| 0 | 0 |
| 0 | 0 |
| 25.7627042 | 0 |
| 0 | 0 |
| 427.66089 | 224.077681 |
| 0 | 0 |
| 0 | 0 |
| 0 | 0 |
| 0 | 7.22831229 |
| 0 | 0 |
| 34.3502723 | 0 |
| 1.71751362 | 21.6849369 |
| 0 | 0 |
| 0 | 0 |
| 0 | 0 |

| | |
|---|---|
| 0 | 0 |
| 144.271144 | 144.566246 |
| 0 | 0 |
| 0 | 7.22831229 |
| 0 | 0 |
| 0 | 0 |
| 0 | 0 |
| 0 | 0 |
| 0 | 0 |
| 0 | 0 |
| 0 | 0 |
| 0 | 0 |
| 0 | 0 |
| 0 | 0 |
| 0 | 0 |
| 0 | 0 |
| 0 | 0 |
| 0 | 0 |
| 0 | 0 |
| 44.655354 | 0 |
| 3.43502723 | 7.22831229 |
| 0 | 0 |
| 0 | 0 |
| 0 | 0 |
| 0 | 0 |
| 15.4576225 | 0 |
| 0 | 0 |
| 0 | 0 |
| 0 | 0 |
| 0 | 0 |
| 0 | 0 |
| 8.58756808 | 14.4566246 |
| 525.559167 | 440.92705 |
| 0 | 0 |
| 0 | 0 |
| 0 | 0 |
| 0 | 0 |
| 0 | 0 |
| 0 | 0 |
| 0 | 0 |
| 0 | 0 |
| 0 | 7.22831229 |
| 0 | 0 |

| | |
|---|---|
| 0 | 0 |
| 0 | 0 |
| 115.073412 | 231.305993 |
| 0 | 0 |
| 0 | 0 |
| 0 | 0 |
| 0 | 0 |
| 24.0451906 | 7.22831229 |
| 0 | 0 |
| 0 | 0 |
| 0 | 0 |
| 0 | 0 |
| 0 | 0 |
| 0 | 0 |
| 0 | 0 |
| 0 | 0 |
| 0 | 0 |
| 5.15254085 | 21.6849369 |
| 0 | 0 |
| 0 | 0 |
| 0 | 0 |
| 0 | 0 |
| 0 | 0 |
| 5.15254085 | 7.22831229 |
| 0 | 0 |
| 0 | 0 |
| 0 | 0 |
| 0 | 0 |
| 0 | 0 |
| 0 | 0 |
| 0 | 0 |
| 0 | 28.9132492 |
| 0 | 0 |
| 0 | 0 |
| 0 | 0 |
| 0 | 0 |
| 0 | 0 |
| 0 | 0 |
| 276.519692 | 0 |
| 0 | 0 |
| 3.43502723 | 0 |
| 0 | 0 |
| 0 | 0 |

| | |
|---|---|
| 0 | 0 |
| 0 | 0 |
| 0 | 0 |
| 0 | 0 |
| 1.71751362 | 0 |
| 0 | 0 |
| 0 | 0 |
| 0 | 0 |
| 0 | 0 |
| 0 | 0 |
| 0 | 0 |
| 0 | 0 |
| 0 | 0 |
| 0 | 0 |
| 0 | 0 |
| 0 | 0 |
| 0 | 0 |
| 0 | 0 |
| 0 | 0 |
| 0 | 0 |
| 0 | 0 |
| 0 | 0 |
| 0 | 0 |
| 140.836117 | 137.337934 |
| 0 | 0 |
| 0 | 0 |
| 0 | 0 |
| 0 | 0 |
| 0 | 0 |
| 0 | 0 |
| 0 | 0 |
| 0 | 0 |
| 0 | 0 |
| 0 | 0 |
| 0 | 0 |
| 0 | 0 |
| 0 | 0 |
| 0 | 0 |
| 0 | 0 |
| 0 | 14.4566246 |
| 0 | 0 |
| 0 | 0 |
| 1186.80191 | 1163.75828 |

| | |
|---|---|
| 0 | 0 |
| 0 | 0 |
| 0 | 0 |
| 1.71751362 | 7.22831229 |
| 0 | 0 |
| 0 | 0 |
| 0 | 0 |
| 36.0677859 | 0 |
| 0 | 0 |
| 0 | 0 |
| 0 | 0 |
| 0 | 0 |
| 0 | 0 |
| 0 | 0 |
| 0 | 0 |
| 0 | 0 |
| 0 | 0 |
| 0 | 0 |
| 0 | 0 |
| 0 | 0 |
| 0 | 0 |
| 25.7627042 | 0 |
| 0 | 0 |
| 0 | 0 |
| 0 | 0 |
| 0 | 0 |
| 0 | 0 |
| 0 | 0 |
| 0 | 0 |
| 0 | 0 |
| 0 | 0 |
| 0 | 0 |
| 30.9152451 | 0 |
| 0 | 0 |
| 0 | 0 |
| 0 | 0 |
| 360.677859 | 108.424684 |
| 1717.51362 | 693.91798 |
| 0 | 0 |
| 0 | 0 |
| 0 | 0 |
| 0 | 0 |
| 0 | 0 |

| | |
|---|---|
| 0 | 0 |
| 0 | 0 |
| 0 | 0 |
| 0 | 0 |
| 0 | 0 |
| 18.8926498 | 21.6849369 |
| 0 | 0 |
| 0 | 0 |
| 0 | 0 |
| 0 | 0 |
| 0 | 0 |
| 0 | 0 |
| 0 | 0 |
| 0 | 0 |
| 113.355899 | 108.424684 |
| 0 | 0 |
| 0 | 0 |
| 0 | 0 |
| 168.316334 | 245.762618 |
| 0 | 0 |
| 0 | 0 |
| 0 | 0 |
| 0 | 0 |
| 0 | 0 |
| 0 | 0 |
| 5.15254085 | 0 |
| 0 | 0 |
| 0 | 7.22831229 |
| 0 | 0 |
| 0 | 0 |
| 0 | 0 |
| 0 | 0 |
| 1.71751362 | 0 |
| 0 | 0 |
| 0 | 0 |
| 0 | 0 |
| 0 | 0 |
| 103.050817 | 14.4566246 |
| 0 | 0 |
| 0 | 0 |
| 0 | 0 |
| 0 | 0 |
| 0 | 0 |

| | |
|---|---|
| 0 | 0 |
| 0 | 0 |
| 0 | 0 |
| 3.43502723 | 0 |
| 0 | 0 |
| 0 | 0 |
| 0 | 0 |
| 0 | 0 |
| 0 | 0 |
| 0 | 0 |
| 0 | 0 |
| 0 | 0 |
| 678.417879 | 534.89511 |
| 0 | 0 |
| 0 | 0 |
| 0 | 7.22831229 |
| 0 | 0 |
| 0 | 0 |
| 17.1751362 | 0 |
| 0 | 0 |
| 0 | 0 |
| 0 | 0 |
| 0 | 0 |
| 0 | 7.22831229 |
| 22.327677 | 0 |
| 0 | 7.22831229 |
| 1.71751362 | 0 |
| 0 | 0 |
| 0 | 0 |
| 0 | 0 |
| 0 | 0 |
| 0 | 0 |
| 0 | 0 |
| 0 | 0 |
| 0 | 0 |
| 0 | 0 |
| 0 | 0 |
| 0 | 0 |
| 353.807805 | 187.93612 |
| 27.4802179 | 0 |
| 0 | 0 |
| 0 | 0 |
| 0 | 0 |

| | |
|---|---|
| 0 | 0 |
| 0 | 7.22831229 |
| 0 | 0 |
| 1.71751362 | 0 |
| 0 | 0 |
| 0 | 7.22831229 |
| 0 | 0 |
| 0 | 0 |
| 0 | 0 |
| 0 | 0 |
| 0 | 0 |
| 0 | 0 |
| 0 | 0 |
| 0 | 0 |
| 0 | 0 |
| 0 | 0 |
| 0 | 0 |
| 0 | 0 |
| 0 | 7.22831229 |
| 0 | 0 |
| 0 | 0 |
| 0 | 0 |
| 127.096008 | 137.337934 |
| 0 | 0 |
| 0 | 0 |
| 0 | 0 |
| 0 | 0 |
| 0 | 375.872239 |
| 0 | 0 |
| 0 | 0 |
| 1.71751362 | 0 |
| 0 | 0 |
| 0 | 0 |
| 0 | 0 |
| 0 | 7.22831229 |
| 0 | 0 |
| 0 | 0 |
| 0 | 0 |
| 0 | 0 |
| 0 | 0 |
| 0 | 0 |
| 0 | 0 |
| 0 | 0 |

| | |
|---|---|
| 0 | 0 |
| 0 | 0 |
| 0 | 0 |
| 0 | 0 |
| 0 | 0 |
| 0 | 0 |
| 0 | 0 |
| 0 | 7.22831229 |
| 0 | 0 |
| 0 | 0 |
| 0 | 0 |
| 0 | 0 |
| 0 | 0 |
| 0 | 0 |
| 0 | 0 |
| 0 | 0 |
| 0 | 0 |
| 0 | 0 |
| 0 | 0 |
| 0 | 0 |
| 0 | 0 |
| 0 | 0 |
| 0 | 0 |
| 0 | 0 |
| 0 | 0 |
| 0 | 0 |
| 34.3502723 | 0 |
| 0 | 0 |
| 25.7627042 | 0 |
| 0 | 0 |
| 1.71751362 | 0 |
| 22.327677 | 0 |
| 3.43502723 | 0 |
| 5.15254085 | 0 |
| 0 | 0 |
| 0 | 0 |
| 0 | 0 |
| 0 | 0 |
| 1.71751362 | 0 |
| 0 | 7.22831229 |
| 0 | 0 |
| 0 | 0 |
| 171.751362 | 180.707807 |

| | |
|---|---|
| 0 | 0 |
| 0 | 0 |
| 0 | 0 |
| 0 | 0 |
| 0 | 0 |
| 0 | 0 |
| 0 | 0 |
| 0 | 0 |
| 0 | 0 |
| 0 | 0 |
| 0 | 0 |
| 0 | 0 |
| 0 | 0 |
| 0 | 0 |
| 0 | 0 |
| 0 | 0 |
| 0 | 0 |
| 0 | 0 |
| 0 | 0 |
| 0 | 0 |
| 1.71751362 | 0 |
| 0 | 0 |
| 0 | 0 |
| 0 | 0 |
| 12.0225953 | 7.22831229 |
| 0 | 0 |
| 0 | 0 |
| 209.536661 | 173.479495 |
| 0 | 0 |
| 0 | 0 |
| 3.43502723 | 0 |
| 0 | 0 |
| 0 | 0 |
| 0 | 0 |
| 231.864338 | 101.196372 |
| 0 | 0 |
| 291.977315 | 86.7397475 |
| 0 | 0 |
| 0 | 0 |
| 0 | 0 |
| 0 | 0 |
| 0 | 0 |
| 0 | 0 |

| | |
|---|---|
| 0 | 0 |
| 1.71751362 | 7.22831229 |
| 0 | 0 |
| 0 | 0 |
| 0 | 0 |
| 0 | 0 |
| 0 | 0 |
| 0 | 0 |
| 0 | 0 |
| 0 | 0 |
| 0 | 0 |
| 0 | 0 |
| 0 | 0 |
| 0 | 0 |
| 1.71751362 | 0 |
| 0 | 0 |
| 0 | 0 |
| 0 | 7.22831229 |
| 0 | 0 |
| 0 | 0 |
| 0 | 0 |
| 0 | 0 |
| 0 | 0 |
| 0 | 0 |
| 1.71751362 | 0 |
| 0 | 0 |
| 115.073412 | 144.566246 |
| 0 | 0 |
| 0 | 0 |
| 0 | 0 |
| 0 | 0 |
| 0 | 0 |
| 0 | 7.22831229 |
| 0 | 0 |
| 0 | 0 |
| 0 | 0 |
| 0 | 0 |
| 0 | 0 |
| 0 | 0 |
| 0 | 0 |
| 0 | 0 |
| 0 | 0 |
| 0 | 0 |

| | |
|---|---|
| 0 | 0 |
| 0 | 0 |
| 0 | 0 |
| 0 | 0 |
| 0 | 0 |
| 0 | 0 |
| 0 | 0 |
| 0 | 0 |
| 0 | 0 |
| 0 | 0 |
| 0 | 0 |
| 0 | 0 |
| 0 | 0 |
| 0 | 0 |
| 1.71751362 | 7.22831229 |
| 6.87005447 | 28.9132492 |
| 0 | 0 |
| 0 | 0 |
| 1915.02768 | 1792.62145 |
| 0 | 0 |
| 0 | 0 |
| 0 | 0 |
| 0 | 0 |
| 0 | 0 |
| 0 | 0 |
| 0 | 0 |
| 0 | 0 |
| 0 | 0 |
| 508.38403 | 440.92705 |
| 0 | 0 |
| 0 | 0 |
| 0 | 0 |
| 1.71751362 | 79.5114352 |
| 0 | 0 |
| 0 | 0 |
| 0 | 0 |
| 0 | 0 |
| 0 | 0 |
| 0 | 0 |
| 0 | 0 |
| 0 | 0 |
| 22.327677 | 0 |
| 0 | 0 |

| | |
|---|---|
| 0 | 0 |
| 0 | 0 |
| 1.71751362 | 0 |
| 0 | 0 |
| 0 | 0 |
| 0 | 0 |
| 0 | 0 |
| 0 | 0 |
| 0 | 0 |
| 0 | 0 |
| 0 | 0 |
| 0 | 0 |
| 0 | 0 |
| 0 | 0 |
| 0 | 0 |
| 0 | 0 |
| 0 | 0 |
| 0 | 0 |
| 0 | 0 |
| 0 | 0 |
| 0 | 0 |
| 0 | 0 |
| 0 | 0 |
| 0 | 0 |
| 0 | 0 |
| 0 | 0 |
| 0 | 0 |
| 0 | 0 |
| 0 | 0 |
| 0 | 0 |
| 0 | 0 |
| 0 | 0 |
| 0 | 0 |
| 0 | 0 |
| 0 | 0 |
| 0 | 0 |
| 0 | 0 |
| 0 | 0 |
| 0 | 0 |
| 0 | 0 |
| 0 | 0 |
| 0 | 0 |
| 0 | 0 |

| | |
|---|---|
| 0 | 0 |
| 0 | 0 |
| 0 | 0 |
| 0 | 0 |
| 0 | 0 |
| 0 | 0 |
| 0 | 0 |
| 0 | 0 |
| 0 | 0 |
| 0 | 0 |
| 5.15254085 | 0 |
| 0 | 0 |
| 0 | 0 |
| 0 | 0 |
| 0 | 0 |
| 0 | 0 |
| 0 | 0 |
| 0 | 0 |
| 0 | 0 |
| 0 | 0 |
| 3.43502723 | 0 |
| 0 | 0 |
| 0 | 0 |
| 0 | 0 |
| 0 | 0 |
| 0 | 0 |
| 0 | 0 |
| 0 | 0 |
| 0 | 0 |
| 0 | 0 |
| 0 | 0 |
| 0 | 7.22831229 |
| 0 | 0 |
| 0 | 0 |
| 0 | 0 |
| 0 | 0 |
| 0 | 7.22831229 |
| 0 | 0 |
| 0 | 0 |
| 0 | 0 |
| 0 | 0 |
| 0 | 0 |
| 0 | 0 |

| | |
|---|---|
| 0 | 7.22831229 |
| 0 | 0 |
| 0 | 0 |
| 0 | 0 |
| 44.655354 | 0 |
| 0 | 0 |
| 0 | 0 |
| 0 | 0 |
| 0 | 0 |
| 0 | 0 |
| 0 | 0 |
| 0 | 0 |
| 0 | 0 |
| 0 | 0 |
| 0 | 0 |
| 0 | 0 |
| 5.15254085 | 7.22831229 |
| 0 | 0 |
| 0 | 0 |
| 0 | 7.22831229 |
| 0 | 0 |
| 0 | 0 |
| 0 | 0 |
| 0 | 0 |
| 0 | 0 |
| 0 | 0 |
| 314.304992 | 252.99093 |
| 0 | 0 |
| 0 | 0 |
| 0 | 0 |
| 0 | 0 |
| 0 | 0 |
| 0 | 0 |
| 0 | 0 |
| 0 | 0 |
| 0 | 0 |
| 0 | 0 |
| 0 | 0 |
| 0 | 0 |
| 0 | 0 |
| 0 | 0 |
| 0 | 0 |
| 0 | 0 |

| | |
|---|---|
| 0 | 0 |
| 0 | 0 |
| 0 | 0 |
| 0 | 0 |
| 0 | 0 |
| 0 | 0 |
| 0 | 0 |
| 0 | 0 |
| 0 | 0 |
| 0 | 0 |
| 0 | 7.22831229 |
| 0 | 0 |
| 0 | 0 |
| 0 | 0 |
| 0 | 0 |
| 0 | 0 |
| 0 | 0 |
| 0 | 0 |
| 0 | 0 |
| 0 | 0 |
| 0 | 0 |
| 0 | 0 |
| 0 | 0 |
| 0 | 0 |
| 0 | 0 |
| 0 | 0 |
| 0 | 0 |
| 1.71751362 | 0 |
| 0 | 0 |
| 0 | 0 |
| 0 | 0 |
| 0 | 0 |
| 0 | 0 |
| 0 | 0 |
| 0 | 0 |
| 0 | 0 |
| 0 | 0 |
| 0 | 0 |
| 0 | 0 |
| 0 | 0 |
| 0 | 0 |
| 0 | 0 |
| 0 | 0 |

| | |
|---|---|
| 0 | 0 |
| 0 | 0 |
| 0 | 0 |
| 0 | 0 |
| 0 | 0 |
| 0 | 0 |
| 0 | 0 |
| 0 | 0 |
| 0 | 0 |
| 0 | 0 |
| 1.71751362 | 0 |
| 0 | 0 |
| 0 | 0 |
| 24.0451906 | 0 |
| 0 | 0 |
| 0 | 0 |
| 0 | 0 |
| 0 | 0 |
| 0 | 0 |
| 0 | 0 |
| 63.5480038 | 0 |
| 0 | 0 |
| 0 | 0 |
| 0 | 0 |
| 96.1807625 | 180.707807 |
| 0 | 0 |
| 0 | 0 |
| 0 | 0 |
| 0 | 0 |
| 0 | 0 |
| 0 | 0 |
| 0 | 0 |
| 0 | 0 |
| 0 | 0 |
| 0 | 7.22831229 |
| 0 | 0 |
| 0 | 0 |
| 0 | 0 |
| 85.8756808 | 65.0548106 |
| 0 | 0 |
| 0 | 0 |
| 15.4576225 | 14.4566246 |
| 0 | 0 |

| | |
|---|---|
| 0 | 0 |
| 0 | 0 |
| 0 | 0 |
| 0 | 0 |
| 5.15254085 | 0 |
| 0 | 0 |
| 0 | 0 |
| 0 | 0 |
| 0 | 0 |
| 0 | 0 |
| 0 | 0 |
| 0 | 0 |
| 0 | 0 |
| 0 | 0 |
| 0 | 0 |
| 0 | 0 |
| 0 | 0 |
| 8.58756808 | 0 |
| 0 | 0 |
| 0 | 14.4566246 |
| 0 | 0 |
| 0 | 0 |
| 0 | 0 |
| 0 | 0 |
| 0 | 0 |
| 0 | 0 |
| 0 | 0 |
| 0 | 0 |
| 0 | 14.4566246 |
| 0 | 0 |
| 0 | 0 |
| 0 | 0 |
| 0 | 0 |
| 0 | 0 |
| 0 | 0 |
| 0 | 0 |
| 0 | 0 |
| 0 | 0 |
| 0 | 0 |
| 8.58756808 | 7.22831229 |
| 616.587388 | 412.013801 |
| 15.4576225 | 14.4566246 |
| 0 | 0 |

| | |
|---|---|
| 0 | 0 |
| 0 | 0 |
| 0 | 0 |
| 91.0282217 | 187.93612 |
| 0 | 0 |
| 0 | 0 |
| 108.203358 | 36.1415615 |
| 0 | 0 |
| 0 | 0 |
| 0 | 0 |
| 0 | 0 |
| 0 | 0 |
| 0 | 0 |
| 0 | 0 |
| 0 | 0 |
| 0 | 0 |
| 0 | 0 |
| 0 | 0 |
| 3.43502723 | 0 |
| 0 | 0 |
| 0 | 0 |
| 0 | 0 |
| 0 | 0 |
| 1.71751362 | 0 |
| 0 | 0 |
| 0 | 0 |
| 0 | 0 |
| 0 | 0 |
| 0 | 0 |
| 0 | 0 |
| 0 | 0 |
| 0 | 0 |
| 0 | 0 |
| 0 | 0 |
| 0 | 0 |
| 0 | 0 |
| 0 | 0 |
| 0 | 0 |
| 0 | 0 |
| 0 | 0 |
| 0 | 0 |
| 0 | 0 |
| 0 | 0 |

| | |
|---|---|
| 15.4576225 | 0 |
| 0 | 0 |
| 0 | 0 |
| 0 | 0 |
| 0 | 0 |
| 0 | 0 |
| 0 | 0 |
| 0 | 0 |
| 0 | 0 |
| 0 | 0 |
| 0 | 0 |
| 0 | 0 |
| 0 | 0 |
| 0 | 0 |
| 0 | 0 |
| 0 | 0 |
| 0 | 0 |
| 0 | 0 |
| 0 | 0 |
| 0 | 0 |
| 0 | 0 |
| 0 | 0 |
| 0 | 0 |
| 0 | 0 |
| 0 | 0 |
| 0 | 0 |
| 0 | 0 |
| 0 | 0 |
| 0 | 0 |
| 0 | 0 |
| 0 | 0 |
| 0 | 7.22831229 |
| 0 | 0 |
| 0 | 0 |
| 0 | 0 |
| 0 | 0 |
| 0 | 0 |
| 1.71751362 | 0 |
| 3.43502723 | 0 |
| 1.71751362 | 0 |
| 0 | 0 |
| 0 | 0 |
| 24.0451906 | 72.2831229 |

| | |
|---|---|
| 0 | 0 |
| 0 | 0 |
| 0 | 0 |
| 30.9152451 | 0 |
| 0 | 0 |
| 0 | 0 |
| 0 | 0 |
| 0 | 0 |
| 0 | 0 |
| 0 | 0 |
| 0 | 0 |
| 0 | 0 |
| 0 | 0 |
| 0 | 0 |
| 0 | 0 |
| 8.58756808 | 7.22831229 |
| 0 | 0 |
| 0 | 0 |
| 0 | 0 |
| 0 | 0 |
| 0 | 0 |
| 145.988657 | 101.196372 |
| 0 | 0 |
| 0 | 0 |
| 0 | 0 |
| 0 | 0 |
| 8.58756808 | 21.6849369 |
| 0 | 7.22831229 |
| 0 | 0 |
| 0 | 0 |
| 0 | 0 |
| 0 | 0 |
| 0 | 0 |
| 231.864338 | 137.337934 |
| 0 | 0 |
| 0 | 0 |
| 0 | 0 |
| 0 | 0 |
| 0 | 0 |
| 188.926498 | 209.621056 |
| 92.7457353 | 0 |
| 5.15254085 | 0 |
| 0 | 0 |

| | |
|---|---|
| 0 | 0 |
| 0 | 7.22831229 |
| 0 | 0 |
| 0 | 0 |
| 0 | 0 |
| 0 | 0 |
| 0 | 0 |
| 0 | 0 |
| 0 | 0 |
| 0 | 0 |
| 0 | 0 |
| 29.1977315 | 0 |
| 0 | 0 |
| 0 | 0 |
| 0 | 0 |
| 0 | 0 |
| 0 | 0 |
| 0 | 0 |
| 0 | 0 |
| 0 | 0 |
| 0 | 0 |
| 0 | 0 |
| 0 | 0 |
| 170.033848 | 93.9680598 |
| 0 | 0 |
| 0 | 0 |
| 0 | 0 |
| 1.71751362 | 0 |
| 0 | 0 |
| 0 | 0 |
| 0 | 0 |
| 0 | 0 |
| 0 | 0 |
| 0 | 0 |
| 0 | 0 |
| 0 | 0 |
| 0 | 0 |
| 0 | 0 |
| 0 | 0 |
| 0 | 0 |
| 0 | 0 |
| 0 | 0 |
| 0 | 0 |

| | |
|---|---|
| 0 | 0 |
| 1.71751362 | 0 |
| 0 | 0 |
| 0 | 0 |
| 1.71751362 | 0 |
| 0 | 0 |
| 0 | 0 |
| 0 | 0 |
| 113.355899 | 281.904179 |
| 30.9152451 | 57.8264983 |
| 3.43502723 | 0 |
| 0 | 0 |
| 0 | 0 |
| 0 | 0 |
| 0 | 0 |
| 0 | 0 |
| 0 | 0 |
| 0 | 0 |
| 0 | 0 |
| 0 | 0 |
| 0 | 0 |
| 0 | 0 |
| 0 | 0 |
| 0 | 0 |
| 0 | 0 |
| 0 | 0 |
| 0 | 0 |
| 1.71751362 | 7.22831229 |
| 0 | 0 |
| 0 | 0 |
| 0 | 0 |
| 0 | 0 |
| 0 | 0 |
| 0 | 0 |
| 0 | 0 |
| 0 | 0 |
| 0 | 0 |
| 0 | 0 |
| 0 | 0 |
| 0 | 0 |
| 75.5705991 | 101.196372 |
| 0 | 0 |
| 0 | 0 |

| | |
|---|---|
| 0 | 0 |
| 0 | 0 |
| 0 | 0 |
| 0 | 0 |
| 1.71751362 | 21.6849369 |
| 0 | 0 |
| 0 | 0 |
| 0 | 0 |
| 0 | 0 |
| 0 | 0 |
| 0 | 0 |
| 0 | 0 |
| 0 | 0 |
| 0 | 0 |
| 0 | 0 |
| 0 | 0 |
| 0 | 0 |
| 0 | 0 |
| 1.71751362 | 0 |
| 0 | 0 |
| 0 | 0 |
| 0 | 0 |
| 0 | 0 |
| 0 | 0 |
| 0 | 0 |
| 0 | 0 |
| 0 | 0 |
| 0 | 0 |
| 0 | 0 |
| 0 | 0 |
| 0 | 0 |
| 1.71751362 | 14.4566246 |
| 0 | 0 |
| 0 | 7.22831229 |
| 0 | 0 |
| 0 | 0 |
| 0 | 0 |
| 0 | 0 |
| 0 | 0 |
| 0 | 0 |
| 0 | 0 |
| 0 | 0 |
| 0 | 0 |
| 0 | 0 |

| | |
|---|---|
| 0 | 0 |
| 0 | 0 |
| 0 | 0 |
| 0 | 0 |
| 0 | 0 |
| 0 | 0 |
| 0 | 0 |
| 0 | 7.22831229 |
| 0 | 7.22831229 |
| 0 | 0 |
| 0 | 0 |
| 0 | 0 |
| 0 | 0 |
| 0 | 0 |
| 0 | 0 |
| 13.7401089 | 7.22831229 |
| 0 | 0 |
| 0 | 0 |
| 0 | 0 |
| 0 | 0 |
| 18.8926498 | 65.0548106 |
| 0 | 0 |
| 0 | 0 |
| 0 | 0 |
| 12.0225953 | 7.22831229 |
| 0 | 0 |
| 0 | 0 |
| 0 | 0 |
| 0 | 0 |
| 0 | 0 |
| 0 | 0 |
| 0 | 0 |
| 0 | 0 |
| 0 | 0 |
| 0 | 0 |
| 0 | 0 |
| 0 | 0 |
| 0 | 0 |
| 0 | 0 |
| 0 | 0 |
| 0 | 0 |
| 0 | 0 |
| 0 | 0 |

| | |
|---|---|
| 0 | 0 |
| 0 | 0 |
| 0 | 0 |
| 0 | 0 |
| 0 | 0 |
| 0 | 0 |
| 0 | 0 |
| 0 | 0 |
| 0 | 0 |
| 0 | 0 |
| 0 | 7.22831229 |
| 0 | 0 |
| 0 | 0 |
| 0 | 0 |
| 0 | 0 |
| 0 | 0 |
| 5.15254085 | 14.4566246 |
| 3.43502723 | 0 |
| 0 | 0 |
| 37.7852996 | 0 |
| 1.71751362 | 0 |
| 0 | 0 |
| 66.983031 | 7.22831229 |
| 0 | 0 |
| 204.38412 | 130.109621 |
| 0 | 0 |
| 0 | 0 |
| 0 | 0 |
| 0 | 0 |
| 0 | 0 |
| 0 | 0 |
| 0 | 0 |
| 0 | 0 |
| 0 | 0 |
| 0 | 0 |
| 0 | 0 |
| 0 | 0 |
| 0 | 0 |
| 0 | 0 |
| 0 | 0 |
| 0 | 0 |
| 0 | 0 |
| 0 | 0 |

| | |
|---|---|
| 0 | 0 |
| 0 | 0 |
| 0 | 0 |
| 56.6779493 | 0 |
| 0 | 0 |
| 0 | 0 |
| 0 | 0 |
| 0 | 0 |
| 0 | 0 |
| 0 | 0 |
| 0 | 0 |
| 3.43502723 | 0 |
| 0 | 0 |
| 0 | 14.4566246 |
| 0 | 0 |
| 0 | 0 |
| 0 | 0 |
| 0 | 0 |
| 0 | 0 |
| 0 | 0 |
| 0 | 0 |
| 0 | 0 |
| 0 | 0 |
| 0 | 0 |
| 0 | 0 |
| 0 | 0 |
| 0 | 0 |
| 0 | 0 |
| 0 | 0 |
| 0 | 0 |
| 0 | 0 |
| 0 | 0 |
| 0 | 0 |
| 0 | 0 |
| 0 | 0 |
| 0 | 0 |
| 0 | 0 |
| 0 | 0 |
| 0 | 0 |
| 0 | 0 |
| 0 | 7.22831229 |
| 0 | 0 |
| 0 | 0 |

| | |
|---|---|
| 0 | 0 |
| 77.2881127 | 318.045741 |
| 0 | 0 |
| 0 | 0 |
| 0 | 0 |
| 0 | 0 |
| 0 | 0 |
| 0 | 0 |
| 0 | 0 |
| 0 | 0 |
| 0 | 0 |
| 0 | 0 |
| 0 | 0 |
| 0 | 0 |
| 0 | 0 |
| 0 | 0 |
| 0 | 0 |
| 0 | 0 |
| 0 | 0 |
| 0 | 0 |
| 5.15254085 | 7.22831229 |
| 0 | 0 |
| 0 | 0 |
| 0 | 0 |
| 0 | 0 |
| 0 | 0 |
| 0 | 0 |
| 0 | 0 |
| 0 | 0 |
| 0 | 0 |
| 0 | 0 |
| 0 | 0 |
| 0 | 0 |
| 3.43502723 | 0 |
| 0 | 7.22831229 |
| 0 | 0 |
| 0 | 0 |
| 0 | 0 |
| 0 | 0 |
| 0 | 0 |
| 0 | 0 |
| 0 | 0 |
| 97.8982761 | 130.109621 |

| | |
|---|---|
| 1.71751362 | 0 |
| 0 | 0 |
| 0 | 0 |
| 0 | 0 |
| 0 | 0 |
| 0 | 0 |
| 0 | 0 |
| 0 | 0 |
| 0 | 0 |
| 0 | 0 |
| 0 | 0 |
| 1.71751362 | 0 |
| 0 | 0 |
| 0 | 0 |
| 0 | 0 |
| 0 | 0 |
| 0 | 0 |
| 0 | 0 |
| 0 | 0 |
| 0 | 0 |
| 0 | 0 |
| 0 | 0 |
| 0 | 14.4566246 |
| 0 | 0 |
| 0 | 0 |
| 0 | 0 |
| 0 | 0 |
| 0 | 0 |
| 0 | 7.22831229 |
| 0 | 0 |
| 0 | 0 |
| 0 | 21.6849369 |
| 0 | 0 |
| 0 | 0 |
| 0 | 0 |
| 0 | 0 |
| 0 | 0 |
| 0 | 0 |
| 0 | 0 |
| 0 | 0 |
| 0 | 7.22831229 |
| 0 | 0 |
| 0 | 0 |

| | |
|---|---|
| 0 | 0 |
| 0 | 0 |
| 0 | 0 |
| 0 | 0 |
| 0 | 0 |
| 0 | 0 |
| 0 | 0 |
| 0 | 0 |
| 0 | 0 |
| 0 | 0 |
| 0 | 0 |
| 0 | 0 |
| 0 | 0 |
| 407.050727 | 296.360804 |
| 0 | 0 |
| 0 | 0 |
| 0 | 0 |
| 0 | 0 |
| 0 | 0 |
| 0 | 0 |
| 0 | 0 |
| 0 | 0 |
| 0 | 0 |
| 20.6101634 | 0 |
| 0 | 0 |
| 0 | 0 |
| 0 | 0 |
| 0 | 0 |
| 0 | 0 |
| 0 | 0 |
| 0 | 0 |
| 0 | 0 |
| 0 | 0 |
| 0 | 0 |
| 0 | 0 |
| 0 | 0 |
| 0 | 0 |
| 0 | 0 |
| 0 | 0 |
| 0 | 0 |
| 0 | 0 |
| 0 | 0 |
| 0 | 0 |

| | |
|---|---|
| 0 | 0 |
| 0 | 0 |
| 0 | 0 |
| 0 | 0 |
| 0 | 0 |
| 0 | 7.22831229 |
| 0 | 0 |
| 0 | 0 |
| 0 | 0 |
| 0 | 0 |
| 0 | 0 |
| 0 | 0 |
| 111.638385 | 224.077681 |
| 3.43502723 | 0 |
| 0 | 0 |
| 187.208984 | 166.251183 |
| 0 | 0 |
| 0 | 0 |
| 0 | 0 |
| 0 | 0 |
| 0 | 0 |
| 0 | 0 |
| 247.321961 | 433.698738 |
| 0 | 0 |
| 0 | 0 |
| 0 | 0 |
| 0 | 0 |
| 0 | 0 |
| 0 | 0 |
| 0 | 0 |
| 0 | 0 |
| 0 | 0 |
| 6.87005447 | 0 |
| 0 | 0 |
| 0 | 0 |
| 0 | 0 |
| 0 | 0 |
| 0 | 0 |
| 0 | 0 |
| 0 | 0 |
| 0 | 7.22831229 |
| 0 | 0 |
| 0 | 0 |

| | |
|---|---|
| 0 | 0 |
| 15.4576225 | 14.4566246 |
| 0 | 7.22831229 |
| 0 | 0 |
| 0 | 0 |
| 0 | 0 |
| 0 | 0 |
| 0 | 0 |
| 0 | 0 |
| 0 | 0 |
| 0 | 0 |
| 0 | 0 |
| 0 | 7.22831229 |
| 68.7005447 | 115.652997 |
| 0 | 0 |
| 0 | 0 |
| 0 | 0 |
| 0 | 0 |
| 0 | 14.4566246 |
| 0 | 0 |
| 0 | 0 |
| 0 | 0 |
| 0 | 0 |
| 0 | 0 |
| 0 | 0 |
| 0 | 0 |
| 0 | 0 |
| 1.71751362 | 14.4566246 |
| 0 | 0 |
| 0 | 0 |
| 0 | 0 |
| 10.3050817 | 0 |
| 0 | 0 |
| 0 | 0 |
| 0 | 0 |
| 0 | 0 |
| 0 | 0 |
| 0 | 0 |
| 0 | 0 |
| 0 | 0 |
| 0 | 0 |
| 0 | 0 |
| 0 | 0 |

| | |
|---|---|
| 0 | 0 |
| 0 | 0 |
| 0 | 0 |
| 0 | 0 |
| 0 | 0 |
| 0 | 0 |
| 0 | 0 |
| 32.6327587 | 0 |
| 0 | 0 |
| 46.3728676 | 7.22831229 |
| 0 | 0 |
| 0 | 0 |
| 0 | 0 |
| 0 | 0 |
| 0 | 0 |
| 0 | 0 |
| 0 | 0 |
| 0 | 0 |
| 0 | 0 |
| 0 | 0 |
| 0 | 0 |
| 0 | 0 |
| 0 | 0 |
| 0 | 0 |
| 25.7627042 | 14.4566246 |
| 0 | 0 |
| 3.43502723 | 7.22831229 |
| 0 | 0 |
| 0 | 0 |
| 0 | 0 |
| 0 | 0 |
| 0 | 0 |
| 0 | 0 |
| 0 | 0 |
| 0 | 0 |
| 0 | 0 |
| 51.5254085 | 209.621056 |
| 0 | 0 |
| 0 | 0 |
| 0 | 0 |
| 1.71751362 | 0 |
| 0 | 0 |
| 0 | 0 |

| | |
|---|---|
| 0 | 0 |
| 0 | 0 |
| 0 | 0 |
| 0 | 0 |
| 0 | 0 |
| 0 | 0 |
| 0 | 0 |
| 0 | 7.22831229 |
| 15.4576225 | 0 |
| 0 | 0 |
| 0 | 0 |
| 0 | 0 |
| 0 | 0 |
| 0 | 0 |
| 12.0225953 | 0 |
| 0 | 0 |
| 1.71751362 | 0 |
| 0 | 0 |
| 0 | 0 |
| 0 | 0 |
| 0 | 0 |
| 0 | 0 |
| 0 | 0 |
| 0 | 0 |
| 0 | 0 |
| 0 | 14.4566246 |
| 0 | 0 |
| 0 | 0 |
| 0 | 0 |
| 10.3050817 | 7.22831229 |
| 0 | 7.22831229 |
| 0 | 0 |
| 0 | 0 |
| 0 | 0 |
| 0 | 0 |
| 0 | 0 |
| 0 | 0 |
| 0 | 0 |
| 0 | 0 |
| 0 | 0 |
| 0 | 0 |
| 1.71751362 | 7.22831229 |
| 0 | 0 |

| | |
|---|---|
| 0 | 0 |
| 0 | 0 |
| 149.423685 | 86.7397475 |
| 1.71751362 | 43.3698738 |
| 0 | 0 |
| 0 | 0 |
| 0 | 0 |
| 0 | 0 |
| 75.5705991 | 231.305993 |
| 0 | 0 |
| 0 | 0 |
| 0 | 0 |
| 0 | 0 |
| 0 | 0 |
| 0 | 0 |
| 0 | 0 |
| 0 | 0 |
| 0 | 0 |
| 0 | 0 |
| 0 | 0 |
| 0 | 0 |
| 0 | 0 |
| 0 | 0 |
| 0 | 0 |
| 0 | 0 |
| 0 | 0 |
| 0 | 0 |
| 0 | 0 |
| 0 | 0 |
| 1.71751362 | 0 |
| 0 | 7.22831229 |
| 0 | 0 |
| 0 | 0 |
| 0 | 0 |
| 0 | 0 |
| 0 | 0 |
| 0 | 0 |
| 0 | 0 |
| 0 | 0 |
| 0 | 0 |
| 0 | 0 |
| 0 | 0 |
| 0 | 0 |

| | |
|---|---|
| 0 | 0 |
| 0 | 0 |
| 465.44619 | 267.447555 |
| 0 | 0 |
| 0 | 0 |
| 0 | 14.4566246 |
| 0 | 0 |
| 0 | 0 |
| 0 | 0 |
| 0 | 0 |
| 0 | 0 |
| 0 | 0 |
| 0 | 0 |
| 0 | 0 |
| 0 | 7.22831229 |
| 0 | 0 |
| 0 | 0 |
| 0 | 0 |
| 0 | 0 |
| 0 | 0 |
| 0 | 0 |
| 0 | 0 |
| 0 | 0 |
| 0 | 0 |
| 0 | 0 |
| 0 | 0 |
| 0 | 0 |
| 0 | 0 |
| 0 | 0 |
| 22.327677 | 7.22831229 |
| 0 | 0 |
| 0 | 0 |
| 0 | 0 |
| 0 | 0 |
| 0 | 0 |
| 199.23158 | 130.109621 |
| 0 | 0 |
| 0 | 0 |
| 0 | 0 |
| 0 | 0 |
| 0 | 0 |
| 0 | 0 |
| 0 | 0 |

| | |
|---|---|
| 0 | 0 |
| 0 | 0 |
| 1.71751362 | 14.4566246 |
| 0 | 0 |
| 0 | 0 |
| 0 | 0 |
| 0 | 0 |
| 0 | 0 |
| 0 | 0 |
| 0 | 0 |
| 0 | 0 |
| 0 | 0 |
| 0 | 0 |
| 0 | 0 |
| 0 | 0 |
| 0 | 0 |
| 0 | 0 |
| 0 | 0 |
| 0 | 0 |
| 0 | 0 |
| 0 | 0 |
| 0 | 0 |
| 0 | 0 |
| 0 | 0 |
| 0 | 0 |
| 0 | 0 |
| 0 | 0 |
| 0 | 0 |
| 0 | 0 |
| 0 | 0 |
| 0 | 0 |
| 0 | 0 |
| 1.71751362 | 0 |
| 0 | 0 |
| 0 | 7.22831229 |
| 0 | 0 |
| 0 | 0 |
| 0 | 0 |
| 0 | 0 |
| 0 | 14.4566246 |
| 0 | 0 |
| 0 | 0 |
| 0 | 0 |

| | |
|---|---|
| 0 | 0 |
| 0 | 0 |
| 0 | 0 |
| 0 | 0 |
| 0 | 0 |
| 0 | 0 |
| 0 | 0 |
| 0 | 0 |
| 0 | 0 |
| 12.0225953 | 0 |
| 0 | 0 |
| 0 | 0 |
| 0 | 0 |
| 0 | 0 |
| 0 | 0 |
| 0 | 0 |
| 0 | 0 |
| 0 | 7.22831229 |
| 0 | 0 |
| 0 | 0 |
| 0 | 0 |
| 0 | 0 |
| 0 | 0 |
| 0 | 0 |
| 0 | 0 |
| 0 | 0 |
| 1.71751362 | 0 |
| 0 | 0 |
| 0 | 0 |
| 0 | 0 |
| 0 | 0 |
| 0 | 7.22831229 |
| 0 | 0 |
| 0 | 0 |
| 0 | 0 |
| 0 | 0 |
| 0 | 0 |
| 0 | 0 |
| 0 | 0 |
| 0 | 14.4566246 |
| 0 | 0 |
| 0 | 0 |
| 0 | 0 |

| | |
|---|---|
| 0 | 0 |
| 0 | 0 |
| 0 | 0 |
| 0 | 0 |
| 0 | 0 |
| 0 | 0 |
| 0 | 0 |
| 0 | 0 |
| 0 | 0 |
| 0 | 0 |
| 0 | 0 |
| 0 | 0 |
| 0 | 0 |
| 0 | 0 |
| 0 | 0 |
| 0 | 0 |
| 171.751362 | 79.5114352 |
| 0 | 7.22831229 |
| 0 | 0 |
| 0 | 0 |
| 0 | 0 |
| 0 | 0 |
| 0 | 0 |
| 0 | 0 |
| 0 | 0 |
| 0 | 0 |
| 0 | 0 |
| 0 | 0 |
| 0 | 0 |
| 0 | 0 |
| 0 | 0 |
| 0 | 0 |
| 0 | 0 |
| 1.71751362 | 7.22831229 |
| 0 | 0 |
| 0 | 0 |
| 0 | 0 |
| 0 | 0 |
| 0 | 7.22831229 |
| 37.7852996 | 0 |
| 0 | 0 |
| 0 | 0 |
| 3.43502723 | 0 |
| 0 | 0 |

| | |
|---|---|
| 262.779583 | 130.109621 |
| 0 | 0 |
| 0 | 0 |
| 0 | 0 |
| 0 | 0 |
| 0 | 0 |
| 0 | 0 |
| 0 | 0 |
| 0 | 0 |
| 0 | 0 |
| 0 | 0 |
| 0 | 0 |
| 0 | 0 |
| 0 | 0 |
| 0 | 0 |
| 0 | 7.22831229 |
| 0 | 0 |
| 0 | 0 |
| 0 | 0 |
| 0 | 0 |
| 0 | 0 |
| 0 | 0 |
| 0 | 0 |
| 135.683576 | 86.7397475 |
| 0 | 0 |
| 1.71751362 | 7.22831229 |
| 0 | 0 |
| 0 | 0 |
| 0 | 0 |
| 0 | 0 |
| 0 | 7.22831229 |
| 0 | 0 |
| 0 | 0 |
| 0 | 0 |
| 0 | 0 |
| 5.15254085 | 7.22831229 |
| 0 | 7.22831229 |
| 0 | 0 |
| 0 | 0 |
| 0 | 0 |
| 0 | 0 |
| 0 | 0 |
| 0 | 0 |

| | |
|---|---|
| 0 | 0 |
| 0 | 0 |
| 0 | 0 |
| 0 | 0 |
| 0 | 0 |
| 0 | 0 |
| 0 | 0 |
| 0 | 0 |
| 0 | 0 |
| 0 | 0 |
| 0 | 0 |
| 0 | 0 |
| 0 | 0 |
| 0 | 0 |
| 0 | 0 |
| 0 | 0 |
| 0 | 0 |
| 0 | 0 |
| 0 | 0 |
| 0 | 0 |
| 0 | 0 |
| 0 | 0 |
| 0 | 0 |
| 0 | 0 |
| 0 | 0 |
| 12.0225953 | 0 |
| 0 | 0 |
| 0 | 0 |
| 0 | 0 |
| 0 | 0 |
| 0 | 7.22831229 |
| 0 | 0 |
| 0 | 0 |
| 0 | 0 |
| 0 | 0 |
| 0 | 0 |
| 0 | 0 |
| 0 | 0 |
| 41.2203268 | 57.8264983 |
| 0 | 0 |
| 0 | 0 |
| 0 | 7.22831229 |
| 0 | 0 |

| | |
|---|---|
| 0 | 0 |
| 0 | 0 |
| 0 | 0 |
| 0 | 0 |
| 0 | 0 |
| 0 | 0 |
| 0 | 0 |
| 2526.46253 | 1077.01853 |
| 0 | 0 |
| 1.71751362 | 0 |
| 0 | 0 |
| 0 | 0 |
| 0 | 0 |
| 0 | 0 |
| 0 | 0 |
| 0 | 0 |
| 0 | 0 |
| 0 | 0 |
| 0 | 0 |
| 0 | 0 |
| 0 | 0 |
| 0 | 0 |
| 0 | 0 |
| 0 | 0 |
| 1.71751362 | 7.22831229 |
| 1.71751362 | 0 |
| 0 | 0 |
| 0 | 0 |
| 0 | 0 |
| 0 | 0 |
| 0 | 0 |
| 0 | 0 |
| 0 | 0 |
| 0 | 0 |
| 0 | 0 |
| 0 | 0 |
| 0 | 0 |
| 0 | 7.22831229 |
| 0 | 0 |
| 0 | 0 |
| 0 | 0 |
| 0 | 0 |
| 0 | 0 |

| | |
|---|---|
| 5.15254085 | 0 |
| 0 | 0 |
| 0 | 0 |
| 0 | 0 |
| 0 | 0 |
| 0 | 0 |
| 0 | 14.4566246 |
| 0 | 0 |
| 0 | 0 |
| 0 | 0 |
| 0 | 0 |
| 0 | 0 |
| 0 | 0 |
| 0 | 0 |
| 0 | 0 |
| 0 | 0 |
| 0 | 0 |
| 0 | 28.9132492 |
| 0 | 0 |
| 0 | 0 |
| 0 | 0 |
| 0 | 0 |
| 0 | 0 |
| 0 | 0 |
| 0 | 0 |
| 0 | 0 |
| 0 | 0 |
| 0 | 0 |
| 48.0903813 | 65.0548106 |
| 0 | 0 |
| 0 | 0 |
| 0 | 0 |
| 0 | 0 |
| 0 | 0 |
| 0 | 0 |
| 0 | 0 |
| 0 | 0 |
| 0 | 0 |
| 0 | 0 |
| 0 | 0 |
| 15.4576225 | 7.22831229 |
| 0 | 0 |
| 0 | 0 |

| | |
|---|---|
| 0 | 0 |
| 0 | 7.22831229 |
| 18.8926498 | 0 |
| 0 | 0 |
| 0 | 0 |
| 0 | 0 |
| 0 | 0 |
| 0 | 0 |
| 0 | 0 |
| 0 | 0 |
| 0 | 0 |
| 0 | 0 |
| 0 | 0 |
| 0 | 0 |
| 0 | 0 |
| 0 | 0 |
| 0 | 14.4566246 |
| 0 | 0 |
| 0 | 0 |
| 0 | 0 |
| 0 | 0 |
| 0 | 0 |
| 0 | 0 |
| 0 | 0 |
| 0 | 0 |
| 285.10726 | 86.7397475 |
| 0 | 0 |
| 0 | 0 |
| 0 | 0 |
| 0 | 0 |
| 0 | 0 |
| 0 | 7.22831229 |
| 0 | 0 |
| 0 | 0 |
| 0 | 0 |
| 0 | 0 |
| 0 | 0 |
| 0 | 0 |
| 0 | 0 |
| 0 | 0 |
| 0 | 0 |
| 0 | 0 |
| 0 | 0 |

| | |
|---|---|
| 341.78521 | 224.077681 |
| 0 | 0 |
| 0 | 0 |
| 0 | 0 |
| 0 | 0 |
| 0 | 0 |
| 0 | 0 |
| 0 | 0 |
| 75.5705991 | 166.251183 |
| 0 | 0 |
| 0 | 0 |
| 0 | 0 |
| 0 | 0 |
| 0 | 0 |
| 0 | 0 |
| 0 | 0 |
| 0 | 0 |
| 0 | 0 |
| 0 | 0 |
| 0 | 0 |
| 0 | 0 |
| 0 | 0 |
| 0 | 0 |
| 0 | 0 |
| 0 | 0 |
| 0 | 14.4566246 |
| 1.71751362 | 0 |
| 46.3728676 | 0 |
| 0 | 0 |
| 0 | 0 |
| 0 | 14.4566246 |
| 0 | 0 |
| 0 | 0 |
| 0 | 0 |
| 0 | 0 |
| 0 | 7.22831229 |
| 0 | 0 |
| 0 | 0 |
| 0 | 0 |
| 0 | 0 |
| 158.011253 | 187.93612 |
| 0 | 0 |
| 0 | 0 |

| | |
|---|---|
| 0 | 0 |
| 0 | 0 |
| 0 | 0 |
| 0 | 0 |
| 30.9152451 | 7.22831229 |
| 0 | 0 |
| 0 | 0 |
| 0 | 0 |
| 0 | 0 |
| 0 | 0 |
| 0 | 0 |
| 0 | 0 |
| 0 | 0 |
| 0 | 0 |
| 0 | 0 |
| 0 | 0 |
| 0 | 0 |
| 0 | 7.22831229 |
| 0 | 0 |
| 0 | 0 |
| 0 | 0 |
| 0 | 0 |
| 0 | 0 |
| 0 | 0 |
| 0 | 0 |
| 0 | 0 |
| 0 | 0 |
| 0 | 0 |
| 0 | 0 |
| 0 | 0 |
| 0 | 0 |
| 0 | 0 |
| 0 | 0 |
| 70.4180583 | 202.392744 |
| 0 | 0 |
| 0 | 0 |
| 0 | 0 |
| 1.71751362 | 7.22831229 |
| 0 | 0 |
| 0 | 0 |
| 0 | 0 |
| 0 | 0 |
| 54.9604357 | 0 |

| | |
|---|---|
| 0 | 0 |
| 108.203358 | 86.7397475 |
| 0 | 0 |
| 0 | 0 |
| 0 | 0 |
| 0 | 0 |
| 0 | 0 |
| 0 | 0 |
| 0 | 0 |
| 0 | 0 |
| 0 | 0 |
| 0 | 7.22831229 |
| 0 | 0 |
| 0 | 0 |
| 0 | 0 |
| 0 | 0 |
| 0 | 0 |
| 0 | 0 |
| 0 | 0 |
| 0 | 0 |
| 0 | 0 |
| 0 | 0 |
| 0 | 0 |
| 0 | 0 |
| 0 | 0 |
| 0 | 0 |
| 0 | 0 |
| 0 | 0 |
| 0 | 0 |
| 0 | 0 |
| 0 | 0 |
| 0 | 0 |
| 0 | 0 |
| 0 | 0 |
| 0 | 0 |
| 0 | 7.22831229 |
| 0 | 0 |
| 0 | 0 |
| 0 | 0 |
| 0 | 0 |
| 0 | 0 |
| 0 | 0 |
| 0 | 0 |

| | |
|---|---|
| 0 | 0 |
| 0 | 0 |
| 273.084665 | 187.93612 |
| 0 | 0 |
| 0 | 0 |
| 0 | 0 |
| 0 | 0 |
| 0 | 0 |
| 0 | 0 |
| 0 | 0 |
| 0 | 0 |
| 0 | 0 |
| 0 | 0 |
| 0 | 0 |
| 0 | 0 |
| 0 | 0 |
| 0 | 0 |
| 0 | 0 |
| 0 | 0 |
| 0 | 0 |
| 0 | 0 |
| 681.852906 | 455.383674 |
| 0 | 0 |
| 1.71751362 | 0 |
| 0 | 0 |
| 0 | 0 |
| 0 | 0 |
| 0 | 0 |
| 0 | 0 |
| 1.71751362 | 0 |
| 0 | 0 |
| 0 | 0 |
| 0 | 0 |
| 0 | 0 |
| 0 | 0 |
| 0 | 0 |
| 0 | 0 |
| 0 | 0 |
| 0 | 7.22831229 |
| 0 | 0 |
| 0 | 0 |
| 0 | 0 |
| 0 | 0 |

| | |
|---|---|
| 0 | 0 |
| 0 | 0 |
| 0 | 0 |
| 0 | 0 |
| 0 | 0 |
| 0 | 0 |
| 0 | 0 |
| 0 | 93.9680598 |
| 0 | 7.22831229 |
| 1.71751362 | 0 |
| 0 | 0 |
| 20.6101634 | 28.9132492 |
| 0 | 0 |
| 0 | 0 |
| 0 | 0 |
| 0 | 0 |
| 0 | 0 |
| 0 | 0 |
| 0 | 0 |
| 0 | 0 |
| 0 | 0 |
| 0 | 0 |
| 0 | 0 |
| 0 | 0 |
| 0 | 0 |
| 0 | 7.22831229 |
| 0 | 0 |
| 0 | 0 |
| 0 | 0 |
| 0 | 0 |
| 0 | 0 |
| 0 | 0 |
| 0 | 0 |
| 0 | 14.4566246 |
| 0 | 0 |
| 0 | 0 |
| 0 | 0 |
| 0 | 0 |
| 0 | 0 |
| 0 | 0 |
| 0 | 0 |
| 0 | 0 |
| 0 | 0 |

| | |
|---|---|
| 0 | 0 |
| 0 | 0 |
| 0 | 7.22831229 |
| 0 | 0 |
| 0 | 0 |
| 0 | 0 |
| 0 | 0 |
| 0 | 0 |
| 10.3050817 | 0 |
| 0 | 0 |
| 0 | 0 |
| 0 | 0 |
| 0 | 0 |
| 0 | 0 |
| 0 | 0 |
| 0 | 0 |
| 0 | 0 |
| 0 | 0 |
| 0 | 0 |
| 0 | 0 |
| 0 | 0 |
| 0 | 0 |
| 0 | 0 |
| 0 | 0 |
| 0 | 0 |
| 0 | 0 |
| 0 | 0 |
| 173.468875 | 72.2831229 |
| 0 | 0 |
| 0 | 0 |
| 0 | 0 |
| 0 | 0 |
| 0 | 0 |
| 0 | 0 |
| 0 | 0 |
| 0 | 0 |
| 0 | 0 |
| 0 | 0 |
| 0 | 0 |
| 0 | 0 |
| 0 | 0 |
| 0 | 0 |
| 0 | 7.22831229 |

| | |
|---|---|
| 0 | 0 |
| 0 | 0 |
| 0 | 0 |
| 0 | 0 |
| 0 | 0 |
| 0 | 0 |
| 0 | 0 |
| 0 | 0 |
| 0 | 0 |
| 151.141198 | 93.9680598 |
| 0 | 0 |
| 0 | 0 |
| 0 | 0 |
| 0 | 0 |
| 0 | 0 |
| 0 | 0 |
| 0 | 0 |
| 0 | 0 |
| 0 | 0 |
| 0 | 0 |
| 0 | 7.22831229 |
| 0 | 0 |
| 0 | 0 |
| 0 | 0 |
| 0 | 0 |
| 0 | 0 |
| 0 | 0 |
| 0 | 0 |
| 0 | 0 |
| 0 | 0 |
| 0 | 0 |
| 0 | 0 |
| 0 | 0 |
| 3.43502723 | 7.22831229 |
| 0 | 0 |
| 413.920782 | 274.675867 |
| 0 | 0 |
| 0 | 0 |
| 0 | 0 |
| 0 | 0 |
| 0 | 0 |
| 0 | 0 |
| 0 | 0 |

| | |
|---|---|
| 5.15254085 | 0 |
| 0 | 0 |
| 0 | 0 |
| 0 | 0 |
| 0 | 0 |
| 0 | 0 |
| 0 | 0 |
| 0 | 0 |
| 0 | 0 |
| 0 | 0 |
| 0 | 21.6849369 |
| 0 | 0 |
| 0 | 0 |
| 0 | 0 |
| 0 | 0 |
| 0 | 0 |
| 0 | 0 |
| 0 | 0 |
| 0 | 0 |
| 66.983031 | 346.95899 |
| 0 | 0 |
| 0 | 0 |
| 0 | 0 |
| 0 | 0 |
| 0 | 0 |
| 0 | 0 |
| 0 | 0 |
| 0 | 0 |
| 0 | 0 |
| 0 | 0 |
| 0 | 0 |
| 0 | 0 |
| 0 | 0 |
| 0 | 0 |
| 0 | 0 |
| 0 | 7.22831229 |
| 0 | 0 |
| 0 | 0 |
| 0 | 0 |
| 0 | 0 |
| 0 | 0 |
| 0 | 0 |
| 0 | 0 |

| | |
|---|---|
| 0 | 0 |
| 0 | 0 |
| 0 | 0 |
| 0 | 0 |
| 0 | 0 |
| 346.937751 | 383.100551 |
| 0 | 0 |
| 0 | 0 |
| 0 | 0 |
| 0 | 0 |
| 0 | 0 |
| 0 | 0 |
| 0 | 0 |
| 0 | 0 |
| 0 | 0 |
| 0 | 0 |
| 0 | 0 |
| 0 | 0 |
| 0 | 0 |
| 0 | 0 |
| 0 | 0 |
| 0 | 0 |
| 0 | 0 |
| 0 | 0 |
| 6.87005447 | 0 |
| 0 | 0 |
| 0 | 0 |
| 0 | 0 |
| 0 | 0 |
| 0 | 0 |
| 0 | 0 |
| 0 | 0 |
| 0 | 0 |
| 0 | 0 |
| 0 | 0 |
| 0 | 0 |
| 0 | 0 |
| 25.7627042 | 7.22831229 |
| 0 | 0 |
| 0 | 0 |
| 0 | 0 |
| 0 | 0 |
| 0 | 0 |

| | |
|---|---|
| 0 | 0 |
| 0 | 0 |
| 0 | 0 |
| 0 | 0 |
| 0 | 0 |
| 0 | 0 |
| 0 | 0 |
| 0 | 0 |
| 0 | 0 |
| 0 | 0 |
| 0 | 0 |
| 0 | 0 |
| 0 | 0 |
| 0 | 0 |
| 77.2881127 | 166.251183 |
| 0 | 0 |
| 0 | 0 |
| 0 | 0 |
| 0 | 0 |
| 0 | 0 |
| 0 | 0 |
| 0 | 0 |
| 1.71751362 | 0 |
| 0 | 0 |
| 0 | 0 |
| 0 | 0 |
| 0 | 0 |
| 0 | 0 |
| 0 | 0 |
| 0 | 0 |
| 0 | 0 |
| 0 | 0 |
| 0 | 0 |
| 0 | 0 |
| 0 | 0 |
| 0 | 0 |
| 0 | 7.22831229 |
| 0 | 0 |
| 3.43502723 | 7.22831229 |
| 0 | 0 |
| 0 | 0 |
| 0 | 0 |
| 0 | 0 |

| | |
|---|---|
| 0 | 7.22831229 |
| 0 | 0 |
| 0 | 0 |
| 0 | 0 |
| 0 | 0 |
| 0 | 0 |
| 0 | 7.22831229 |
| 0 | 0 |
| 0 | 0 |
| 0 | 0 |
| 0 | 0 |
| 0 | 0 |
| 0 | 0 |
| 3.43502723 | 14.4566246 |
| 0 | 0 |
| 207.819148 | 542.123422 |
| 58.395463 | 0 |
| 0 | 0 |
| 0 | 0 |
| 0 | 0 |
| 0 | 0 |
| 0 | 0 |
| 0 | 0 |
| 0 | 0 |
| 0 | 0 |
| 0 | 0 |
| 0 | 0 |
| 0 | 0 |
| 0 | 0 |
| 1.71751362 | 0 |
| 0 | 0 |
| 0 | 0 |
| 0 | 0 |
| 0 | 0 |
| 0 | 0 |
| 0 | 0 |
| 0 | 0 |
| 0 | 0 |
| 0 | 0 |
| 0 | 0 |
| 0 | 7.22831229 |
| 0 | 0 |
| 0 | 0 |

| | |
|---|---|
| 0 | 0 |
| 0 | 0 |
| 0 | 0 |
| 0 | 0 |
| 0 | 0 |
| 0 | 0 |
| 0 | 0 |
| 0 | 0 |
| 0 | 0 |
| 0 | 0 |
| 0 | 0 |
| 0 | 0 |
| 0 | 0 |
| 0 | 0 |
| 0 | 0 |
| 223.27677 | 238.534306 |
| 0 | 0 |
| 82.4406536 | 115.652997 |
| 0 | 0 |
| 0 | 0 |
| 0 | 0 |
| 0 | 0 |
| 0 | 0 |
| 0 | 0 |
| 0 | 0 |
| 0 | 0 |
| 0 | 0 |
| 0 | 0 |
| 0 | 0 |
| 0 | 0 |
| 0 | 0 |
| 0 | 0 |
| 12.0225953 | 21.6849369 |
| 0 | 0 |
| 0 | 0 |
| 0 | 0 |
| 0 | 0 |
| 0 | 0 |
| 0 | 0 |
| 0 | 0 |
| 5.15254085 | 0 |
| 0 | 0 |
| 0 | 0 |

| | |
|---|---|
| 0 | 0 |
| 0 | 0 |
| 0 | 0 |
| 0 | 0 |
| 0 | 0 |
| 0 | 0 |
| 0 | 0 |
| 0 | 0 |
| 0 | 0 |
| 0 | 0 |
| 1083.75109 | 795.114352 |
| 0 | 0 |
| 0 | 0 |
| 0 | 0 |
| 22.327677 | 86.7397475 |
| 0 | 0 |
| 0 | 0 |
| 0 | 0 |
| 0 | 0 |
| 0 | 0 |
| 0 | 0 |
| 8.58756808 | 14.4566246 |
| 0 | 0 |
| 0 | 0 |
| 0 | 0 |
| 0 | 0 |
| 0 | 0 |
| 0 | 0 |
| 0 | 0 |
| 0 | 0 |
| 1.71751362 | 0 |
| 0 | 0 |
| 0 | 0 |
| 0 | 0 |
| 0 | 0 |
| 0 | 0 |
| 0 | 0 |
| 0 | 0 |
| 0 | 0 |
| 0 | 0 |
| 0 | 0 |
| 24.0451906 | 0 |
| 0 | 0 |

| | |
|---|---|
| 0 | 7.22831229 |
| 0 | 0 |
| 0 | 0 |
| 0 | 0 |
| 0 | 0 |
| 5.15254085 | 115.652997 |
| 0 | 0 |
| 0 | 0 |
| 0 | 0 |
| 0 | 0 |
| 0 | 0 |
| 0 | 0 |
| 0 | 0 |
| 0 | 0 |
| 0 | 0 |
| 0 | 0 |
| 0 | 0 |
| 0 | 0 |
| 0 | 0 |
| 0 | 0 |
| 0 | 0 |
| 0 | 0 |
| 0 | 0 |
| 0 | 0 |
| 17.1751362 | 0 |
| 0 | 0 |
| 3.43502723 | 7.22831229 |
| 0 | 0 |
| 0 | 0 |
| 0 | 0 |
| 0 | 0 |
| 0 | 0 |
| 0 | 0 |
| 0 | 0 |
| 0 | 0 |
| 0 | 0 |
| 0 | 0 |
| 6.87005447 | 0 |
| 0 | 0 |
| 0 | 0 |
| 0 | 0 |
| 0 | 0 |
| 1.71751362 | 7.22831229 |

| | |
|---|---|
| 0 | 0 |
| 0 | 0 |
| 0 | 0 |
| 0 | 0 |
| 0 | 0 |
| 0 | 0 |
| 0 | 0 |
| 0 | 0 |
| 0 | 0 |
| 0 | 0 |
| 0 | 0 |
| 3.43502723 | 0 |
| 0 | 0 |
| 0 | 0 |
| 0 | 0 |
| 0 | 0 |
| 0 | 0 |
| 0 | 0 |
| 0 | 0 |
| 12.0225953 | 50.598186 |
| 0 | 0 |
| 0 | 0 |
| 0 | 0 |
| 0 | 0 |
| 0 | 0 |
| 173.468875 | 93.9680598 |
| 0 | 0 |
| 0 | 0 |
| 0 | 0 |
| 0 | 0 |
| 0 | 0 |
| 104.768331 | 86.7397475 |
| 0 | 0 |
| 0 | 0 |
| 0 | 14.4566246 |
| 0 | 0 |
| 0 | 0 |
| 0 | 0 |
| 0 | 0 |
| 0 | 0 |
| 0 | 0 |
| 0 | 0 |
| 0 | 0 |

| | |
|---|---|
| 0 | 0 |
| 0 | 0 |
| 0 | 0 |
| 0 | 0 |
| 0 | 0 |
| 0 | 7.22831229 |
| 0 | 0 |
| 0 | 0 |
| 1.71751362 | 14.4566246 |
| 0 | 0 |
| 0 | 0 |
| 0 | 0 |
| 0 | 0 |
| 0 | 0 |
| 0 | 0 |
| 0 | 0 |
| 0 | 0 |
| 0 | 14.4566246 |
| 0 | 0 |
| 0 | 0 |
| 0 | 0 |
| 0 | 0 |
| 290.259801 | 166.251183 |
| 0 | 0 |
| 0 | 0 |
| 0 | 0 |
| 0 | 0 |
| 0 | 0 |
| 0 | 7.22831229 |
| 130.531035 | 361.415615 |
| 0 | 0 |
| 0 | 0 |
| 0 | 0 |
| 3.43502723 | 43.3698738 |
| 0 | 0 |
| 0 | 0 |
| 0 | 0 |
| 0 | 0 |
| 0 | 0 |
| 0 | 0 |
| 0 | 0 |
| 0 | 0 |
| 0 | 0 |

| | |
|---|---|
| 0 | 0 |
| 0 | 0 |
| 0 | 0 |
| 0 | 0 |
| 0 | 0 |
| 0 | 0 |
| 0 | 0 |
| 0 | 0 |
| 0 | 0 |
| 0 | 0 |
| 0 | 0 |
| 0 | 0 |
| 0 | 0 |
| 0 | 0 |
| 113.355899 | 43.3698738 |
| 0 | 0 |
| 0 | 0 |
| 0 | 0 |
| 1.71751362 | 0 |
| 0 | 0 |
| 0 | 0 |
| 0 | 0 |
| 0 | 0 |
| 0 | 0 |
| 0 | 0 |
| 0 | 0 |
| 0 | 0 |
| 0 | 0 |
| 0 | 0 |
| 0 | 0 |
| 0 | 0 |
| 0 | 0 |
| 0 | 0 |
| 0 | 0 |
| 0 | 0 |
| 0 | 0 |
| 0 | 0 |
| 0 | 0 |
| 0 | 0 |
| 0 | 0 |
| 0 | 0 |
| 0 | 0 |
| 0 | 0 |

| | |
|---|---|
| 0 | 0 |
| 0 | 0 |
| 0 | 0 |
| 0 | 0 |
| 0 | 0 |
| 0 | 0 |
| 0 | 0 |
| 0 | 0 |
| 1.71751362 | 0 |
| 0 | 0 |
| 0 | 0 |
| 0 | 0 |
| 0 | 0 |
| 374.417968 | 281.904179 |
| 0 | 0 |
| 0 | 0 |
| 0 | 0 |
| 0 | 0 |
| 0 | 0 |
| 0 | 0 |
| 0 | 0 |
| 0 | 0 |
| 0 | 0 |
| 0 | 0 |
| 0 | 0 |
| 0 | 0 |
| 0 | 0 |
| 0 | 0 |
| 0 | 0 |
| 152.858712 | 231.305993 |
| 0 | 0 |
| 0 | 0 |
| 0 | 0 |
| 0 | 0 |
| 0 | 0 |
| 0 | 0 |
| 0 | 0 |
| 0 | 0 |
| 0 | 0 |
| 0 | 0 |
| 0 | 0 |
| 0 | 0 |
| 0 | 0 |

| | |
|---|---|
| 0 | 0 |
| 20.6101634 | 0 |
| 0 | 0 |
| 0 | 0 |
| 0 | 0 |
| 0 | 0 |
| 0 | 0 |
| 0 | 0 |
| 0 | 0 |
| 0 | 0 |
| 0 | 0 |
| 0 | 0 |
| 0 | 0 |
| 0 | 0 |
| 0 | 0 |
| 30.9152451 | 0 |
| 0 | 0 |
| 0 | 0 |
| 0 | 0 |
| 0 | 0 |
| 0 | 0 |
| 0 | 0 |
| 0 | 0 |
| 0 | 0 |
| 0 | 0 |
| 0 | 0 |
| 0 | 0 |
| 0 | 0 |
| 0 | 0 |
| 0 | 0 |
| 0 | 0 |
| 0 | 0 |
| 0 | 0 |
| 0 | 0 |
| 0 | 0 |
| 0 | 0 |
| 0 | 0 |
| 0 | 0 |
| 0 | 0 |
| 0 | 0 |
| 0 | 0 |
| 0 | 0 |
| 0 | 0 |

| | |
|---|---|
| 24.0451906 | 57.8264983 |
| 70.4180583 | 21.6849369 |
| 0 | 0 |
| 0 | 0 |
| 0 | 0 |
| 0 | 7.22831229 |
| 0 | 0 |
| 0 | 0 |
| 0 | 0 |
| 1.71751362 | 0 |
| 0 | 0 |
| 0 | 0 |
| 0 | 0 |
| 0 | 0 |
| 0 | 0 |
| 34.3502723 | 0 |
| 0 | 0 |
| 0 | 0 |
| 0 | 0 |
| 0 | 0 |
| 0 | 0 |
| 0 | 0 |
| 0 | 0 |
| 0 | 0 |
| 0 | 0 |
| 0 | 0 |
| 0 | 0 |
| 0 | 0 |
| 6.87005447 | 36.1415615 |
| 0 | 0 |
| 0 | 0 |
| 0 | 0 |
| 0 | 0 |
| 0 | 0 |
| 0 | 0 |
| 46.3728676 | 0 |
| 0 | 0 |
| 271.367151 | 412.013801 |
| 0 | 0 |
| 0 | 7.22831229 |
| 0 | 0 |
| 0 | 0 |
| 0 | 0 |

| | |
|---|---|
| 0 | 0 |
| 0 | 0 |
| 0 | 0 |
| 0 | 0 |
| 0 | 0 |
| 0 | 14.4566246 |
| 3.43502723 | 0 |
| 0 | 0 |
| 0 | 0 |
| 0 | 0 |
| 0 | 0 |
| 0 | 0 |
| 0 | 0 |
| 0 | 0 |
| 0 | 0 |
| 0 | 0 |
| 0 | 0 |
| 0 | 0 |
| 0 | 0 |
| 0 | 0 |
| 0 | 0 |
| 0 | 0 |
| 0 | 0 |
| 10.3050817 | 21.6849369 |
| 0 | 0 |
| 103.050817 | 57.8264983 |
| 0 | 0 |
| 0 | 0 |
| 0 | 0 |
| 0 | 7.22831229 |
| 0 | 0 |
| 0 | 0 |
| 0 | 0 |
| 0 | 0 |
| 0 | 0 |
| 0 | 0 |
| 46.3728676 | 0 |
| 0 | 0 |
| 0 | 0 |
| 0 | 0 |
| 0 | 0 |
| 0 | 0 |
| 0 | 0 |

| | |
|---|---|
| 0 | 0 |
| 0 | 0 |
| 0 | 0 |
| 0 | 0 |
| 0 | 0 |
| 0 | 0 |
| 0 | 0 |
| 0 | 0 |
| 273.084665 | 281.904179 |
| 0 | 0 |
| 0 | 0 |
| 0 | 0 |
| 0 | 0 |
| 0 | 0 |
| 63.5480038 | 173.479495 |
| 0 | 0 |
| 0 | 0 |
| 0 | 0 |
| 0 | 0 |
| 0 | 0 |
| 0 | 0 |
| 0 | 0 |
| 0 | 0 |
| 0 | 0 |
| 0 | 0 |
| 0 | 0 |
| 0 | 0 |
| 0 | 0 |
| 0 | 0 |
| 0 | 0 |
| 0 | 0 |
| 0 | 0 |
| 0 | 0 |
| 0 | 0 |
| 0 | 0 |
| 0 | 0 |
| 0 | 0 |
| 0 | 0 |
| 0 | 0 |
| 0 | 0 |
| 0 | 0 |
| 0 | 0 |
| 0 | 0 |

| | |
|---|---|
| 0 | 0 |
| 0 | 0 |
| 0 | 0 |
| 0 | 0 |
| 0 | 0 |
| 0 | 7.22831229 |
| 0 | 0 |
| 0 | 0 |
| 0 | 0 |
| 0 | 0 |
| 0 | 0 |
| 0 | 0 |
| 0 | 0 |
| 0 | 0 |
| 0 | 0 |
| 0 | 0 |
| 0 | 0 |
| 0 | 14.4566246 |
| 0 | 0 |
| 61.8304902 | 65.0548106 |
| 0 | 0 |
| 0 | 0 |
| 0 | 0 |
| 0 | 0 |
| 0 | 0 |
| 0 | 0 |
| 0 | 0 |
| 0 | 0 |
| 0 | 0 |
| 0 | 0 |
| 0 | 0 |
| 0 | 0 |
| 0 | 7.22831229 |
| 0 | 0 |
| 0 | 0 |
| 0 | 0 |
| 0 | 0 |
| 0 | 0 |
| 75.5705991 | 115.652997 |
| 0 | 0 |
| 0 | 0 |
| 0 | 0 |
| 0 | 7.22831229 |

| | |
|---|---|
| 0 | 0 |
| 0 | 0 |
| 0 | 0 |
| 0 | 0 |
| 0 | 0 |
| 0 | 0 |
| 0 | 0 |
| 0 | 0 |
| 0 | 0 |
| 0 | 0 |
| 0 | 0 |
| 0 | 0 |
| 0 | 0 |
| 0 | 0 |
| 0 | 0 |
| 0 | 0 |
| 0 | 0 |
| 0 | 0 |
| 0 | 0 |
| 0 | 0 |
| 0 | 0 |
| 0 | 0 |
| 66.983031 | 144.566246 |
| 0 | 0 |
| 0 | 0 |
| 0 | 0 |
| 0 | 0 |
| 0 | 0 |
| 0 | 0 |
| 0 | 0 |
| 0 | 0 |
| 0 | 0 |
| 0 | 0 |
| 0 | 0 |
| 0 | 0 |
| 0 | 0 |
| 0 | 0 |
| 0 | 0 |
| 0 | 0 |
| 0 | 0 |
| 0 | 0 |
| 0 | 0 |
| 0 | 0 |

| | |
|---|---|
| 5.15254085 | 7.22831229 |
| 17.1751362 | 0 |
| 0 | 0 |
| 0 | 0 |
| 0 | 0 |
| 0 | 0 |
| 288.542288 | 137.337934 |
| 0 | 0 |
| 0 | 0 |
| 0 | 0 |
| 0 | 0 |
| 0 | 0 |
| 0 | 0 |
| 0 | 0 |
| 0 | 0 |
| 0 | 0 |
| 0 | 0 |
| 0 | 0 |
| 0 | 0 |
| 0 | 0 |
| 0 | 0 |
| 0 | 0 |
| 0 | 0 |
| 0 | 0 |
| 0 | 0 |
| 0 | 0 |
| 0 | 0 |
| 0 | 0 |
| 0 | 0 |
| 0 | 0 |
| 3.43502723 | 0 |
| 22.327677 | 0 |
| 0 | 0 |
| 0 | 0 |
| 0 | 0 |
| 0 | 0 |
| 0 | 0 |
| 0 | 0 |
| 0 | 0 |
| 0 | 0 |
| 0 | 0 |
| 0 | 0 |
| 0 | 7.22831229 |

| | |
|---|---|
| 0 | 0 |
| 0 | 0 |
| 0 | 0 |
| 0 | 0 |
| 1.71751362 | 0 |
| 6.87005447 | 0 |
| 24.0451906 | 0 |
| 0 | 0 |
| 0 | 0 |
| 0 | 0 |
| 0 | 0 |
| 0 | 0 |
| 0 | 0 |
| 0 | 0 |
| 0 | 0 |
| 0 | 0 |
| 0 | 0 |
| 0 | 0 |
| 0 | 0 |
| 0 | 0 |
| 0 | 0 |
| 1.71751362 | 0 |
| 372.700455 | 744.516166 |
| 0 | 0 |
| 0 | 0 |
| 0 | 14.4566246 |
| 0 | 0 |
| 0 | 0 |
| 0 | 0 |
| 0 | 0 |
| 0 | 0 |
| 0 | 0 |
| 0 | 0 |
| 0 | 0 |
| 0 | 0 |
| 0 | 0 |
| 0 | 0 |
| 0 | 0 |
| 0 | 0 |
| 0 | 0 |
| 0 | 0 |
| 0 | 0 |
| 0 | 0 |

| | |
|---|---|
| 0 | 0 |
| 63.5480038 | 65.0548106 |
| 0 | 0 |
| 0 | 0 |
| 0 | 7.22831229 |
| 0 | 0 |
| 0 | 0 |
| 0 | 0 |
| 0 | 0 |
| 0 | 0 |
| 0 | 0 |
| 0 | 0 |
| 0 | 0 |
| 0 | 0 |
| 0 | 0 |
| 0 | 0 |
| 0 | 0 |
| 0 | 0 |
| 0 | 0 |
| 0 | 0 |
| 17.1751362 | 43.3698738 |
| 1.71751362 | 0 |
| 0 | 0 |
| 0 | 0 |
| 0 | 0 |
| 0 | 0 |
| 0 | 0 |
| 0 | 0 |
| 0 | 0 |
| 0 | 0 |
| 0 | 0 |
| 0 | 0 |
| 0 | 0 |
| 0 | 0 |
| 0 | 0 |
| 2043.8412 | 650.548106 |
| 0 | 0 |
| 0 | 0 |
| 0 | 0 |
| 0 | 0 |
| 0 | 0 |
| 39.5028132 | 0 |
| 0 | 0 |

| | |
|---|---|
| 0 | 0 |
| 0 | 0 |
| 0 | 0 |
| 0 | 0 |
| 0 | 0 |
| 0 | 0 |
| 0 | 0 |
| 0 | 0 |
| 0 | 0 |
| 0 | 0 |
| 0 | 0 |
| 0 | 0 |
| 0 | 0 |
| 10.3050817 | 0 |
| 0 | 0 |
| 3.43502723 | 7.22831229 |
| 0 | 0 |
| 1.71751362 | 0 |
| 0 | 7.22831229 |
| 0 | 7.22831229 |
| 0 | 0 |
| 0 | 0 |
| 0 | 0 |
| 0 | 0 |
| 0 | 0 |
| 0 | 0 |
| 0 | 0 |
| 0 | 0 |
| 0 | 7.22831229 |
| 0 | 0 |
| 0 | 0 |
| 0 | 0 |
| 0 | 7.22831229 |
| 0 | 0 |
| 0 | 0 |
| 0 | 0 |
| 0 | 0 |
| 0 | 0 |
| 0 | 0 |
| 0 | 0 |
| 0 | 0 |
| 0 | 0 |
| 0 | 0 |

| | |
|---|---|
| 0 | 0 |
| 0 | 0 |
| 0 | 0 |
| 0 | 7.22831229 |
| 0 | 0 |
| 0 | 0 |
| 5.15254085 | 0 |
| 0 | 0 |
| 0 | 0 |
| 0 | 0 |
| 0 | 7.22831229 |
| 0 | 0 |
| 0 | 0 |
| 0 | 0 |
| 0 | 0 |
| 0 | 0 |
| 0 | 0 |
| 104.768331 | 79.5114352 |
| 0 | 0 |
| 0 | 0 |
| 0 | 0 |
| 0 | 0 |
| 0 | 0 |
| 0 | 0 |
| 0 | 0 |
| 0 | 0 |
| 0 | 0 |
| 0 | 0 |
| 0 | 0 |
| 0 | 0 |
| 0 | 0 |
| 0 | 0 |
| 0 | 0 |
| 0 | 7.22831229 |
| 0 | 0 |
| 130.531035 | 1105.93178 |
| 0 | 0 |
| 0 | 0 |
| 0 | 0 |
| 0 | 0 |
| 0 | 0 |
| 1.71751362 | 14.4566246 |
| 0 | 0 |

| | |
|---|---|
| 0 | 0 |
| 1.71751362 | 0 |
| 0 | 0 |
| 0 | 0 |
| 221.559257 | 130.109621 |
| 0 | 0 |
| 551.321871 | 4149.05126 |
| 0 | 0 |
| 0 | 0 |
| 0 | 0 |
| 1.71751362 | 0 |
| 0 | 0 |
| 0 | 0 |
| 0 | 0 |
| 0 | 0 |
| 0 | 0 |
| 0 | 0 |
| 44.655354 | 0 |
| 0 | 0 |
| 0 | 0 |
| 0 | 0 |
| 0 | 0 |
| 1.71751362 | 0 |
| 0 | 0 |
| 0 | 0 |
| 0 | 0 |
| 1.71751362 | 0 |
| 0 | 0 |
| 0 | 0 |
| 0 | 0 |
| 0 | 0 |
| 0 | 0 |
| 0 | 0 |
| 0 | 0 |
| 0 | 0 |
| 0 | 0 |
| 0 | 0 |
| 37.7852996 | 0 |
| 0 | 0 |
| 0 | 0 |
| 0 | 0 |
| 0 | 0 |
| 32.6327587 | 0 |

| | |
|---|---|
| 0 | 0 |
| 0 | 0 |
| 0 | 0 |
| 0 | 14.4566246 |
| 0 | 0 |
| 0 | 0 |
| 0 | 0 |
| 0 | 0 |
| 0 | 0 |
| 0 | 7.22831229 |
| 0 | 0 |
| 0 | 0 |
| 0 | 0 |
| 0 | 0 |
| 0 | 0 |
| 0 | 0 |
| 0 | 0 |
| 0 | 0 |
| 0 | 0 |
| 0 | 0 |
| 1.71751362 | 0 |
| 0 | 0 |
| 0 | 0 |
| 0 | 7.22831229 |
| 0 | 0 |
| 77.2881127 | 101.196372 |
| 0 | 0 |
| 0 | 0 |
| 0 | 0 |
| 0 | 0 |
| 0 | 0 |
| 0 | 0 |
| 0 | 0 |
| 0 | 0 |
| 0 | 0 |
| 0 | 0 |
| 0 | 0 |
| 0 | 0 |
| 0 | 0 |
| 0 | 0 |
| 0 | 0 |
| 1.71751362 | 0 |
| 0 | 0 |

| | |
|---|---|
| 3.43502723 | 21.6849369 |
| 0 | 0 |
| 0 | 0 |
| 0 | 0 |
| 3.43502723 | 0 |
| 168.316334 | 166.251183 |
| 0 | 43.3698738 |
| 0 | 0 |
| 0 | 0 |
| 266.214611 | 50.598186 |
| 0 | 0 |
| 0 | 0 |
| 0 | 0 |
| 0 | 0 |
| 0 | 0 |
| 0 | 0 |
| 0 | 0 |
| 0 | 0 |
| 0 | 0 |
| 0 | 0 |
| 0 | 0 |
| 0 | 0 |
| 0 | 0 |
| 0 | 0 |
| 0 | 0 |
| 209.536661 | 202.392744 |
| 0 | 0 |
| 0 | 0 |
| 0 | 0 |
| 0 | 0 |
| 0 | 7.22831229 |
| 0 | 0 |
| 0 | 0 |
| 0 | 0 |
| 0 | 0 |
| 0 | 0 |
| 0 | 0 |
| 0 | 0 |
| 0 | 7.22831229 |
| 0 | 0 |
| 0 | 0 |
| 0 | 0 |
| 0 | 0 |

| | |
|---|---|
| 0 | 0 |
| 0 | 0 |
| 0 | 0 |
| 1.71751362 | 0 |
| 0 | 0 |
| 0 | 0 |
| 0 | 0 |
| 0 | 0 |
| 0 | 0 |
| 0 | 0 |
| 0 | 0 |
| 8.58756808 | 0 |
| 0 | 0 |
| 0 | 0 |
| 0 | 0 |
| 0 | 0 |
| 0 | 0 |
| 0 | 0 |
| 0 | 0 |
| 0 | 0 |
| 0 | 0 |
| 0 | 0 |
| 0 | 0 |
| 0 | 0 |
| 0 | 0 |
| 0 | 0 |
| 0 | 0 |
| 0 | 0 |
| 0 | 0 |
| 0 | 0 |
| 0 | 0 |
| 0 | 0 |
| 0 | 0 |
| 0 | 0 |
| 0 | 0 |
| 0 | 0 |
| 0 | 0 |
| 0 | 0 |
| 0 | 0 |
| 0 | 0 |
| 0 | 0 |
| 0 | 0 |
| 111.638385 | 72.2831229 |

| | |
|---|---|
| 0 | 0 |
| 0 | 0 |
| 0 | 0 |
| 0 | 0 |
| 0 | 0 |
| 0 | 0 |
| 0 | 7.22831229 |
| 0 | 0 |
| 0 | 0 |
| 0 | 0 |
| 0 | 0 |
| 0 | 0 |
| 0 | 0 |
| 0 | 0 |
| 0 | 0 |
| 5.15254085 | 0 |
| 0 | 14.4566246 |
| 0 | 0 |
| 0 | 0 |
| 0 | 0 |
| 0 | 0 |
| 0 | 0 |
| 0 | 0 |
| 0 | 0 |
| 0 | 0 |
| 0 | 7.22831229 |
| 0 | 0 |
| 0 | 0 |
| 0 | 0 |
| 0 | 0 |
| 0 | 0 |
| 0 | 0 |
| 120.225953 | 137.337934 |
| 0 | 0 |
| 0 | 0 |
| 0 | 0 |
| 0 | 0 |
| 0 | 0 |
| 0 | 0 |
| 0 | 0 |
| 0 | 0 |
| 0 | 0 |
| 0 | 0 |

| | |
|---|---|
| 0 | 0 |
| 0 | 0 |
| 0 | 0 |
| 0 | 0 |
| 0 | 0 |
| 0 | 0 |
| 0 | 0 |
| 0 | 7.22831229 |
| 224.994284 | 318.045741 |
| 0 | 0 |
| 0 | 0 |
| 0 | 0 |
| 0 | 0 |
| 0 | 0 |
| 0 | 0 |
| 0 | 0 |
| 0 | 0 |
| 0 | 0 |
| 0 | 0 |
| 0 | 0 |
| 0 | 0 |
| 0 | 0 |
| 0 | 0 |
| 0 | 0 |
| 0 | 0 |
| 0 | 0 |
| 0 | 0 |
| 0 | 0 |
| 0 | 7.22831229 |
| 0 | 0 |
| 0 | 0 |
| 0 | 0 |
| 0 | 0 |
| 0 | 0 |
| 0 | 0 |
| 0 | 0 |
| 0 | 0 |
| 0 | 0 |
| 731.660801 | 187.93612 |
| 0 | 0 |
| 0 | 0 |
| 0 | 0 |
| 0 | 0 |

| | |
|---|---|
| 0 | 0 |
| 0 | 0 |
| 0 | 0 |
| 0 | 0 |
| 0 | 0 |
| 0 | 0 |
| 0 | 0 |
| 0 | 0 |
| 0 | 14.4566246 |
| 0 | 0 |
| 0 | 0 |
| 0 | 0 |
| 0 | 0 |
| 0 | 0 |
| 0 | 0 |
| 0 | 0 |
| 0 | 0 |
| 0 | 0 |
| 0 | 0 |
| 0 | 0 |
| 0 | 0 |
| 0 | 0 |
| 0 | 0 |
| 0 | 0 |
| 0 | 0 |
| 0 | 0 |
| 32.6327587 | 0 |
| 0 | 0 |
| 0 | 0 |
| 0 | 0 |
| 0 | 0 |
| 0 | 0 |
| 0 | 0 |
| 0 | 0 |
| 0 | 0 |
| 0 | 0 |
| 94.4632489 | 79.5114352 |
| 0 | 0 |
| 0 | 0 |
| 0 | 7.22831229 |
| 0 | 0 |
| 0 | 0 |
| 0 | 0 |

| | |
|---|---|
| 0 | 0 |
| 0 | 0 |
| 0 | 0 |
| 0 | 0 |
| 0 | 0 |
| 0 | 0 |
| 0 | 14.4566246 |
| 0 | 0 |
| 0 | 0 |
| 0 | 0 |
| 0 | 0 |
| 0 | 0 |
| 0 | 0 |
| 1.71751362 | 0 |
| 0 | 0 |
| 0 | 0 |
| 0 | 0 |
| 0 | 0 |
| 0 | 0 |
| 0 | 0 |
| 216.406716 | 260.219243 |
| 0 | 0 |
| 0 | 0 |
| 1.71751362 | 7.22831229 |
| 0 | 0 |
| 0 | 0 |
| 0 | 0 |
| 0 | 0 |
| 0 | 0 |
| 0 | 0 |
| 0 | 0 |
| 0 | 0 |
| 0 | 0 |
| 0 | 0 |
| 34.3502723 | 14.4566246 |
| 0 | 0 |
| 0 | 0 |
| 0 | 0 |
| 0 | 0 |
| 0 | 0 |
| 0 | 0 |
| 108.203358 | 166.251183 |
| 0 | 0 |

| | |
|---|---|
| 0 | 7.22831229 |
| 46.3728676 | 21.6849369 |
| 0 | 0 |
| 0 | 0 |
| 192.361525 | 159.02287 |
| 0 | 0 |
| 0 | 0 |
| 0 | 0 |
| 0 | 0 |
| 0 | 7.22831229 |
| 0 | 0 |
| 0 | 0 |
| 15.4576225 | 0 |
| 0 | 0 |
| 0 | 0 |
| 0 | 0 |
| 0 | 0 |
| 0 | 0 |
| 0 | 0 |
| 29.1977315 | 57.8264983 |
| 0 | 0 |
| 0 | 0 |
| 0 | 0 |
| 0 | 0 |
| 0 | 0 |
| 0 | 0 |
| 0 | 0 |
| 0 | 0 |
| 0 | 0 |
| 0 | 0 |
| 0 | 0 |
| 0 | 0 |
| 0 | 0 |
| 0 | 0 |
| 1.71751362 | 0 |
| 0 | 0 |
| 0 | 0 |
| 0 | 0 |
| 0 | 0 |
| 0 | 0 |
| 0 | 0 |
| 0 | 0 |
| 0 | 0 |

| | |
|---|---|
| 0 | 0 |
| 0 | 0 |
| 0 | 0 |
| 0 | 0 |
| 0 | 0 |
| 0 | 0 |
| 0 | 0 |
| 0 | 0 |
| 1.71751362 | 0 |
| 0 | 0 |
| 0 | 0 |
| 0 | 0 |
| 0 | 0 |
| 0 | 0 |
| 0 | 0 |
| 0 | 0 |
| 53.2429221 | 0 |
| 0 | 0 |
| 0 | 0 |
| 3.43502723 | 0 |
| 0 | 0 |
| 0 | 0 |
| 0 | 0 |
| 0 | 0 |
| 0 | 0 |
| 0 | 0 |
| 0 | 0 |
| 0 | 0 |
| 0 | 0 |
| 0 | 0 |
| 0 | 0 |
| 0 | 0 |
| 0 | 0 |
| 0 | 0 |
| 0 | 0 |
| 0 | 0 |
| 0 | 0 |
| 0 | 0 |
| 0 | 0 |
| 0 | 7.22831229 |
| 1.71751362 | 0 |
| 0 | 0 |
| 0 | 14.4566246 |

| | |
|---|---|
| 0 | 0 |
| 0 | 0 |
| 0 | 0 |
| 0 | 0 |
| 0 | 0 |
| 0 | 0 |
| 0 | 0 |
| 0 | 0 |
| 0 | 0 |
| 0 | 0 |
| 0 | 0 |
| 0 | 0 |
| 221.559257 | 50.598186 |
| 75.5705991 | 224.077681 |
| 0 | 0 |
| 27.4802179 | 0 |
| 0 | 0 |
| 0 | 0 |
| 54.9604357 | 173.479495 |
| 0 | 0 |
| 0 | 0 |
| 0 | 0 |
| 0 | 0 |
| 0 | 0 |
| 0 | 0 |
| 0 | 0 |
| 0 | 0 |
| 0 | 14.4566246 |
| 0 | 0 |
| 0 | 0 |
| 68.7005447 | 108.424684 |
| 0 | 0 |
| 0 | 0 |
| 0 | 0 |
| 0 | 0 |
| 0 | 0 |
| 0 | 0 |
| 0 | 0 |
| 0 | 0 |
| 0 | 0 |
| 0 | 0 |
| 0 | 0 |
| 0 | 0 |

| | |
|---|---|
| 1.71751362 | 0 |
| 0 | 0 |
| 0 | 0 |
| 63.5480038 | 36.1415615 |
| 0 | 0 |
| 0 | 0 |
| 0 | 0 |
| 0 | 0 |
| 0 | 0 |
| 0 | 0 |
| 0 | 0 |
| 0 | 0 |
| 8.58756808 | 0 |
| 0 | 0 |
| 0 | 0 |
| 0 | 0 |
| 0 | 0 |
| 0 | 0 |
| 0 | 0 |
| 1.71751362 | 0 |
| 0 | 7.22831229 |
| 0 | 0 |
| 0 | 0 |
| 0 | 0 |
| 0 | 0 |
| 0 | 0 |
| 0 | 0 |
| 0 | 0 |
| 0 | 0 |
| 0 | 0 |
| 0 | 0 |
| 0 | 0 |
| 0 | 0 |
| 0 | 0 |
| 305.717424 | 433.698738 |
| 0 | 0 |
| 0 | 0 |
| 0 | 0 |
| 0 | 0 |
| 0 | 7.22831229 |
| 0 | 0 |
| 0 | 7.22831229 |
| 0 | 14.4566246 |

| | |
|---|---|
| 0 | 0 |
| 0 | 0 |
| 0 | 0 |
| 0 | 0 |
| 0 | 0 |
| 0 | 0 |
| 0 | 0 |
| 0 | 0 |
| 44.655354 | 28.9132492 |
| 0 | 0 |
| 0 | 0 |
| 0 | 0 |
| 0 | 0 |
| 0 | 0 |
| 0 | 0 |
| 0 | 0 |
| 0 | 0 |
| 0 | 0 |
| 0 | 0 |
| 0 | 0 |
| 0 | 0 |
| 0 | 0 |
| 0 | 0 |
| 0 | 0 |
| 27.4802179 | 57.8264983 |
| 0 | 0 |
| 0 | 0 |
| 0 | 0 |
| 0 | 0 |
| 0 | 0 |
| 0 | 0 |
| 0 | 0 |
| 0 | 0 |
| 0 | 0 |
| 0 | 0 |
| 0 | 0 |
| 0 | 0 |
| 0 | 0 |
| 192.361525 | 187.93612 |
| 0 | 0 |
| 0 | 0 |
| 0 | 0 |
| 0 | 0 |

| | |
|---|---|
| 0 | 0 |
| 0 | 0 |
| 6.87005447 | 0 |
| 0 | 0 |
| 0 | 0 |
| 0 | 0 |
| 0 | 0 |
| 0 | 0 |
| 0 | 0 |
| 32.6327587 | 21.6849369 |
| 0 | 0 |
| 0 | 7.22831229 |
| 0 | 0 |
| 0 | 0 |
| 0 | 0 |
| 0 | 0 |
| 0 | 0 |
| 0 | 0 |
| 0 | 0 |
| 0 | 0 |
| 212.971688 | 202.392744 |
| 0 | 0 |
| 0 | 0 |
| 0 | 0 |
| 0 | 0 |
| 0 | 0 |
| 0 | 0 |
| 0 | 0 |
| 0 | 0 |
| 0 | 0 |
| 0 | 0 |
| 0 | 0 |
| 0 | 0 |
| 0 | 0 |
| 0 | 0 |
| 0 | 0 |
| 0 | 0 |
| 0 | 0 |
| 0 | 0 |
| 0 | 0 |
| 0 | 0 |
| 0 | 0 |

| | |
|---|---|
| 0 | 0 |
| 0 | 0 |
| 0 | 0 |
| 0 | 0 |
| 0 | 14.4566246 |
| 0 | 0 |
| 0 | 0 |
| 0 | 14.4566246 |
| 0 | 0 |
| 0 | 0 |
| 0 | 0 |
| 0 | 0 |
| 0 | 0 |
| 0 | 0 |
| 0 | 0 |
| 0 | 0 |
| 0 | 0 |
| 0 | 0 |
| 0 | 0 |
| 0 | 0 |
| 0 | 0 |
| 0 | 0 |
| 0 | 0 |
| 0 | 0 |
| 0 | 0 |
| 0 | 0 |
| 0 | 0 |
| 0 | 0 |
| 0 | 0 |
| 0 | 0 |
| 0 | 0 |
| 0 | 0 |
| 0 | 0 |
| 594.259711 | 628.863169 |
| 3.43502723 | 0 |
| 0 | 0 |
| 0 | 0 |
| 0 | 0 |
| 0 | 0 |
| 911.99973 | 599.94992 |
| 0 | 0 |
| 103.050817 | 7.22831229 |

| | |
|---|---|
| 13.7401089 | 0 |
| 0 | 0 |
| 0 | 0 |
| 29.1977315 | 7.22831229 |
| 0 | 0 |
| 0 | 0 |
| 0 | 0 |
| 0 | 0 |
| 0 | 0 |
| 0 | 0 |
| 0 | 0 |
| 0 | 0 |
| 34.3502723 | 7.22831229 |
| 0 | 0 |
| 0 | 0 |
| 0 | 0 |
| 0 | 0 |
| 0 | 0 |
| 0 | 0 |
| 0 | 0 |
| 0 | 0 |
| 0 | 0 |
| 0 | 0 |
| 0 | 0 |
| 0 | 0 |
| 0 | 0 |
| 0 | 0 |
| 0 | 0 |
| 0 | 0 |
| 0 | 0 |
| 0 | 0 |
| 0 | 0 |
| 0 | 0 |
| 309.152451 | 108.424684 |
| 0 | 0 |
| 0 | 0 |
| 0 | 0 |
| 0 | 0 |
| 0 | 0 |
| 0 | 0 |
| 0 | 0 |
| 0 | 0 |
| 0 | 0 |

| | |
|---|---|
| 0 | 0 |
| 0 | 0 |
| 0 | 0 |
| 0 | 0 |
| 0 | 0 |
| 0 | 0 |
| 0 | 0 |
| 0 | 0 |
| 0 | 0 |
| 0 | 0 |
| 6.87005447 | 0 |
| 0 | 0 |
| 0 | 0 |
| 0 | 0 |
| 0 | 0 |
| 0 | 0 |
| 0 | 0 |
| 0 | 0 |
| 0 | 0 |
| 0 | 0 |
| 0 | 0 |
| 0 | 0 |
| 0 | 0 |
| 0 | 0 |
| 0 | 0 |
| 0 | 0 |
| 0 | 0 |
| 115.073412 | 57.8264983 |
| 0 | 0 |
| 0 | 0 |
| 0 | 0 |
| 0 | 0 |
| 0 | 0 |
| 0 | 0 |
| 0 | 0 |
| 0 | 0 |
| 0 | 0 |
| 0 | 0 |
| 97.8982761 | 0 |
| 0 | 7.22831229 |
| 0 | 0 |
| 0 | 0 |
| 0 | 0 |

| | |
|---|---|
| 0 | 0 |
| 0 | 0 |
| 0 | 0 |
| 0 | 0 |
| 0 | 0 |
| 0 | 0 |
| 0 | 0 |
| 0 | 0 |
| 0 | 0 |
| 0 | 0 |
| 168.316334 | 50.598186 |
| 0 | 0 |
| 0 | 0 |
| 0 | 0 |
| 0 | 0 |
| 0 | 0 |
| 0 | 0 |
| 0 | 0 |
| 0 | 0 |
| 0 | 0 |
| 0 | 0 |
| 73.8530855 | 43.3698738 |
| 73.8530855 | 43.3698738 |
| 0 | 0 |
| 0 | 0 |
| 0 | 0 |
| 0 | 0 |
| 1.71751362 | 0 |
| 0 | 0 |
| 0 | 0 |
| 3.43502723 | 0 |
| 0 | 0 |
| 8.58756808 | 7.22831229 |
| 0 | 0 |
| 0 | 0 |
| 0 | 0 |
| 0 | 0 |
| 0 | 0 |
| 0 | 0 |
| 0 | 14.4566246 |
| 0 | 0 |
| 0 | 0 |
| 0 | 0 |

| | |
|---|---|
| 0 | 0 |
| 0 | 0 |
| 0 | 7.22831229 |
| 1.71751362 | 0 |
| 0 | 0 |
| 0 | 0 |
| 0 | 0 |
| 0 | 0 |
| 0 | 7.22831229 |
| 0 | 0 |
| 0 | 0 |
| 0 | 0 |
| 0 | 0 |
| 0 | 0 |
| 0 | 0 |
| 0 | 0 |
| 0 | 0 |
| 0 | 0 |
| 0 | 0 |
| 0 | 0 |
| 0 | 0 |
| 0 | 0 |
| 0 | 0 |
| 0 | 0 |
| 0 | 0 |
| 0 | 0 |
| 0 | 0 |
| 0 | 7.22831229 |
| 0 | 0 |
| 350.372778 | 151.794558 |
| 0 | 0 |
| 0 | 0 |
| 0 | 0 |
| 0 | 0 |
| 0 | 0 |
| 0 | 0 |
| 0 | 0 |
| 0 | 7.22831229 |
| 0 | 0 |
| 0 | 0 |
| 0 | 0 |
| 0 | 0 |
| 0 | 0 |

| | |
|---|---|
| 0 | 0 |
| 0 | 0 |
| 0 | 0 |
| 0 | 0 |
| 0 | 0 |
| 0 | 0 |
| 0 | 0 |
| 0 | 0 |
| 0 | 7.22831229 |
| 0 | 0 |
| 0 | 0 |
| 376.135482 | 289.132492 |
| 0 | 0 |
| 0 | 0 |
| 0 | 0 |
| 0 | 0 |
| 0 | 0 |
| 0 | 0 |
| 0 | 0 |
| 0 | 0 |
| 0 | 0 |
| 0 | 0 |
| 0 | 0 |
| 0 | 0 |
| 0 | 0 |
| 0 | 0 |
| 468.881217 | 195.164432 |
| 0 | 7.22831229 |
| 0 | 0 |
| 0 | 0 |
| 0 | 0 |
| 0 | 0 |
| 3.43502723 | 0 |
| 0 | 0 |
| 1.71751362 | 7.22831229 |
| 0 | 0 |
| 0 | 0 |
| 0 | 0 |
| 0 | 0 |
| 0 | 0 |
| 0 | 0 |
| 0 | 0 |
| 0 | 0 |

| | |
|---|---|
| 0 | 0 |
| 0 | 0 |
| 0 | 0 |
| 3.43502723 | 21.6849369 |
| 0 | 7.22831229 |
| 0 | 0 |
| 0 | 0 |
| 0 | 0 |
| 0 | 0 |
| 0 | 0 |
| 0 | 0 |
| 0 | 0 |
| 0 | 0 |
| 0 | 0 |
| 170.033848 | 187.93612 |
| 183.773957 | 50.598186 |
| 0 | 0 |
| 0 | 0 |
| 0 | 0 |
| 0 | 0 |
| 1.71751362 | 0 |
| 0 | 0 |
| 0 | 0 |
| 0 | 0 |
| 0 | 0 |
| 0 | 0 |
| 0 | 0 |
| 0 | 0 |
| 372.700455 | 36.1415615 |
| 0 | 0 |
| 0 | 0 |
| 0 | 0 |
| 0 | 0 |
| 0 | 0 |
| 0 | 0 |
| 49.8078949 | 0 |
| 0 | 0 |
| 0 | 0 |
| 0 | 0 |
| 0 | 0 |
| 120.225953 | 86.7397475 |
| 0 | 0 |
| 0 | 21.6849369 |

| | |
|---|---|
| 0 | 0 |
| 0 | 0 |
| 0 | 0 |
| 0 | 0 |
| 0 | 7.22831229 |
| 0 | 0 |
| 0 | 0 |
| 0 | 0 |
| 0 | 0 |
| 0 | 0 |
| 1.71751362 | 0 |
| 0 | 7.22831229 |
| 0 | 0 |
| 0 | 0 |
| 0 | 0 |
| 0 | 0 |
| 0 | 0 |
| 0 | 0 |
| 0 | 0 |
| 0 | 0 |
| 0 | 0 |
| 0 | 0 |
| 0 | 0 |
| 0 | 0 |
| 0 | 0 |
| 204.38412 | 238.534306 |
| 0 | 0 |
| 0 | 0 |
| 0 | 0 |
| 0 | 0 |
| 0 | 0 |
| 0 | 0 |
| 0 | 0 |
| 0 | 0 |
| 0 | 0 |
| 0 | 0 |
| 0 | 0 |
| 0 | 0 |
| 0 | 0 |
| 0 | 0 |
| 1.71751362 | 0 |
| 0 | 0 |
| 0 | 7.22831229 |

| | |
|---|---|
| 0 | 0 |
| 0 | 0 |
| 0 | 0 |
| 0 | 0 |
| 0 | 0 |
| 0 | 0 |
| 0 | 7.22831229 |
| 0 | 0 |
| 0 | 0 |
| 0 | 0 |
| 0 | 0 |
| 0 | 0 |
| 0 | 0 |
| 0 | 0 |
| 0 | 0 |
| 0 | 0 |
| 0 | 7.22831229 |
| 0 | 0 |
| 0 | 0 |
| 0 | 0 |
| 0 | 0 |
| 0 | 0 |
| 8.58756808 | 0 |
| 0 | 0 |
| 0 | 0 |
| 0 | 0 |
| 0 | 0 |
| 0 | 0 |
| 0 | 0 |
| 0 | 0 |
| 0 | 0 |
| 0 | 0 |
| 0 | 0 |
| 0 | 0 |
| 0 | 0 |
| 0 | 0 |
| 139.118603 | 151.794558 |
| 0 | 0 |
| 0 | 0 |
| 0 | 0 |
| 0 | 0 |
| 0 | 0 |
| 10.3050817 | 7.22831229 |

| | |
|---|---|
| 0 | 0 |
| 0 | 0 |
| 0 | 0 |
| 0 | 0 |
| 0 | 50.598186 |
| 0 | 0 |
| 0 | 0 |
| 0 | 0 |
| 0 | 0 |
| 0 | 0 |
| 0 | 0 |
| 0 | 0 |
| 0 | 0 |
| 0 | 0 |
| 0 | 0 |
| 0 | 0 |
| 0 | 0 |
| 0 | 0 |
| 0 | 0 |
| 0 | 0 |
| 0 | 0 |
| 0 | 0 |
| 0 | 0 |
| 0 | 0 |
| 0 | 0 |
| 0 | 0 |
| 0 | 0 |
| 0 | 0 |
| 0 | 0 |
| 0 | 0 |
| 0 | 0 |
| 12.0225953 | 7.22831229 |
| 0 | 0 |
| 0 | 7.22831229 |
| 0 | 0 |
| 0 | 0 |
| 0 | 0 |
| 32.6327587 | 0 |
| 0 | 0 |
| 0 | 0 |
| 0 | 0 |
| 0 | 0 |
| 0 | 0 |

| | |
|---|---|
| 0 | 0 |
| 0 | 0 |
| 0 | 0 |
| 0 | 0 |
| 0 | 0 |
| 3026.25899 | 1207.12815 |
| 0 | 0 |
| 0 | 0 |
| 0 | 0 |
| 0 | 7.22831229 |
| 0 | 0 |
| 3.43502723 | 0 |
| 0 | 0 |
| 0 | 0 |
| 0 | 0 |
| 221.559257 | 180.707807 |
| 0 | 7.22831229 |
| 0 | 0 |
| 0 | 0 |
| 0 | 0 |
| 0 | 0 |
| 1.71751362 | 14.4566246 |
| 528.994194 | 180.707807 |
| 0 | 0 |
| 0 | 0 |
| 0 | 0 |
| 0 | 0 |
| 0 | 0 |
| 0 | 0 |
| 0 | 0 |
| 0 | 0 |
| 0 | 0 |
| 0 | 0 |
| 0 | 0 |
| 0 | 0 |
| 0 | 0 |
| 0 | 0 |
| 13.7401089 | 0 |
| 0 | 0 |
| 0 | 0 |
| 0 | 0 |
| 0 | 0 |
| 0 | 0 |

| | |
|---|---|
| 0 | 0 |
| 0 | 0 |
| 0 | 0 |
| 0 | 0 |
| 0 | 0 |
| 0 | 0 |
| 0 | 0 |
| 0 | 0 |
| 0 | 0 |
| 0 | 0 |
| 0 | 0 |
| 0 | 0 |
| 0 | 0 |
| 0 | 0 |
| 1.71751362 | 7.22831229 |
| 0 | 0 |
| 0 | 0 |
| 0 | 0 |
| 0 | 0 |
| 0 | 0 |
| 0 | 0 |
| 0 | 0 |
| 0 | 0 |
| 0 | 0 |
| 0 | 0 |
| 0 | 0 |
| 0 | 0 |
| 0 | 0 |
| 0 | 0 |
| 0 | 0 |
| 5.15254085 | 0 |
| 0 | 0 |
| 0 | 0 |
| 0 | 0 |
| 0 | 0 |
| 0 | 0 |
| 0 | 0 |
| 27.4802179 | 7.22831229 |
| 0 | 0 |
| 1.71751362 | 7.22831229 |
| 0 | 0 |
| 0 | 0 |
| 0 | 0 |

| | |
|---|---|
| 0 | 0 |
| 0 | 0 |
| 0 | 0 |
| 0 | 0 |
| 0 | 0 |
| 0 | 0 |
| 0 | 0 |
| 0 | 0 |
| 0 | 0 |
| 0 | 0 |
| 0 | 0 |
| 0 | 0 |
| 0 | 0 |
| 0 | 0 |
| 322.89256 | 159.02287 |
| 0 | 0 |
| 0 | 0 |
| 0 | 0 |
| 44.655354 | 0 |
| 0 | 0 |
| 0 | 0 |
| 0 | 0 |
| 0 | 0 |
| 0 | 0 |
| 6.87005447 | 0 |
| 166.598821 | 144.566246 |
| 0 | 0 |
| 0 | 0 |
| 0 | 0 |
| 0 | 0 |
| 0 | 0 |
| 0 | 14.4566246 |
| 42.9378404 | 7.22831229 |
| 0 | 0 |
| 0 | 0 |
| 0 | 0 |
| 0 | 0 |
| 0 | 0 |
| 0 | 0 |
| 0 | 0 |
| 0 | 7.22831229 |
| 0 | 0 |
| 0 | 0 |

| | |
|---|---|
| 0 | 0 |
| 0 | 0 |
| 0 | 0 |
| 0 | 0 |
| 0 | 0 |
| 261.06207 | 332.502365 |
| 0 | 0 |
| 0 | 0 |
| 0 | 0 |
| 0 | 0 |
| 0 | 7.22831229 |
| 0 | 0 |
| 0 | 0 |
| 0 | 0 |
| 0 | 0 |
| 0 | 0 |
| 0 | 0 |
| 0 | 0 |
| 0 | 0 |
| 0 | 0 |
| 0 | 0 |
| 0 | 21.6849369 |
| 0 | 0 |
| 0 | 0 |
| 3.43502723 | 21.6849369 |
| 0 | 7.22831229 |
| 0 | 0 |
| 0 | 0 |
| 0 | 0 |
| 0 | 0 |
| 0 | 0 |
| 0 | 0 |
| 0 | 0 |
| 0 | 0 |
| 0 | 0 |
| 1.71751362 | 0 |
| 0 | 0 |
| 0 | 0 |
| 1.71751362 | 0 |
| 0 | 0 |
| 0 | 0 |
| 0 | 0 |
| 0 | 0 |

| | |
|---|---|
| 0 | 0 |
| 0 | 0 |
| 0 | 0 |
| 0 | 0 |
| 0 | 0 |
| 0 | 0 |
| 0 | 0 |
| 0 | 0 |
| 0 | 0 |
| 0 | 0 |
| 0 | 0 |
| 0 | 0 |
| 0 | 0 |
| 0 | 0 |
| 0 | 7.22831229 |
| 1.71751362 | 36.1415615 |
| 0 | 0 |
| 0 | 0 |
| 13.7401089 | 7.22831229 |
| 0 | 0 |
| 0 | 0 |
| 0 | 0 |
| 0 | 0 |
| 0 | 0 |
| 0 | 0 |
| 0 | 0 |
| 0 | 0 |
| 0 | 0 |
| 0 | 0 |
| 0 | 0 |
| 0 | 0 |
| 0 | 0 |
| 0 | 0 |
| 0 | 0 |
| 0 | 0 |
| 0 | 0 |
| 0 | 0 |
| 0 | 0 |
| 0 | 0 |
| 0 | 0 |
| 0 | 0 |
| 0 | 0 |
| 0 | 36.1415615 |

| | |
|---|---|
| 0 | 7.22831229 |
| 0 | 0 |
| 1.71751362 | 0 |
| 0 | 0 |
| 0 | 0 |
| 0 | 0 |
| 0 | 0 |
| 0 | 0 |
| 0 | 0 |
| 0 | 7.22831229 |
| 0 | 0 |
| 0 | 0 |
| 0 | 0 |
| 0 | 0 |
| 0 | 0 |
| 0 | 0 |
| 0 | 0 |
| 0 | 0 |
| 0 | 0 |
| 0 | 7.22831229 |
| 0 | 0 |
| 0 | 0 |
| 0 | 0 |
| 0 | 0 |
| 0 | 0 |
| 0 | 0 |
| 0 | 0 |
| 0 | 0 |
| 0 | 0 |
| 0 | 0 |
| 0 | 0 |
| 3.43502723 | 14.4566246 |
| 0 | 0 |
| 0 | 0 |
| 0 | 0 |
| 0 | 0 |
| 0 | 0 |
| 0 | 0 |
| 0 | 0 |
| 333.197642 | 180.707807 |
| 0 | 0 |
| 0 | 0 |
| 0 | 0 |

| | |
|---|---|
| 0 | 0 |
| 0 | 0 |
| 0 | 0 |
| 0 | 0 |
| 0 | 0 |
| 0 | 0 |
| 0 | 7.22831229 |
| 0 | 0 |
| 0 | 0 |
| 0 | 0 |
| 0 | 0 |
| 0 | 0 |
| 0 | 0 |
| 0 | 0 |
| 0 | 0 |
| 0 | 0 |
| 0 | 0 |
| 5.15254085 | 0 |
| 0 | 0 |
| 0 | 50.598186 |
| 0 | 7.22831229 |
| 0 | 0 |
| 77.2881127 | 101.196372 |
| 0 | 0 |
| 0 | 0 |
| 0 | 0 |
| 0 | 0 |
| 0 | 0 |
| 0 | 0 |
| 0 | 0 |
| 0 | 0 |
| 0 | 0 |
| 118.50844 | 93.9680598 |
| 0 | 0 |
| 0 | 0 |
| 0 | 0 |
| 0 | 0 |
| 0 | 0 |
| 0 | 0 |
| 0 | 0 |
| 0 | 0 |
| 0 | 0 |
| 0 | 0 |

| | |
|---|---|
| 0 | 0 |
| 0 | 0 |
| 0 | 0 |
| 1.71751362 | 0 |
| 0 | 0 |
| 1.71751362 | 0 |
| 0 | 0 |
| 214.689202 | 43.3698738 |
| 3.43502723 | 0 |
| 0 | 0 |
| 0 | 0 |
| 0 | 0 |
| 0 | 0 |
| 0 | 0 |
| 0 | 7.22831229 |
| 0 | 0 |
| 0 | 0 |
| 0 | 0 |
| 0 | 0 |
| 0 | 0 |
| 0 | 0 |
| 0 | 0 |
| 0 | 0 |
| 0 | 0 |
| 0 | 0 |
| 0 | 7.22831229 |
| 0 | 0 |
| 0 | 0 |
| 0 | 0 |
| 0 | 0 |
| 0 | 0 |
| 0 | 0 |
| 0 | 0 |
| 0 | 0 |
| 0 | 0 |
| 0 | 0 |
| 0 | 0 |
| 0 | 0 |
| 0 | 0 |
| 0 | 0 |
| 0 | 0 |
| 0 | 0 |
| 0 | 0 |

| | |
|---|---|
| 0 | 0 |
| 0 | 0 |
| 0 | 0 |
| 0 | 0 |
| 0 | 0 |
| 0 | 0 |
| 0 | 0 |
| 0 | 0 |
| 0 | 0 |
| 0 | 0 |
| 0 | 0 |
| 0 | 0 |
| 0 | 0 |
| 0 | 0 |
| 0 | 0 |
| 0 | 0 |
| 0 | 0 |
| 0 | 0 |
| 0 | 0 |
| 0 | 0 |
| 0 | 0 |
| 0 | 0 |
| 0 | 0 |
| 0 | 0 |
| 0 | 0 |
| 0 | 0 |
| 0 | 0 |
| 0 | 0 |
| 0 | 0 |
| 0 | 0 |
| 0 | 0 |
| 0 | 0 |
| 0 | 0 |
| 0 | 0 |
| 0 | 7.22831229 |
| 0 | 0 |
| 0 | 0 |
| 0 | 0 |
| 0 | 0 |
| 0 | 0 |
| 0 | 0 |
| 0 | 0 |
| 250.756988 | 101.196372 |

| | |
|---|---|
| 0 | 0 |
| 0 | 0 |
| 0 | 0 |
| 0 | 0 |
| 0 | 0 |
| 1.71751362 | 7.22831229 |
| 0 | 0 |
| 2198.41743 | 1091.47516 |
| 0 | 0 |
| 0 | 0 |
| 0 | 0 |
| 0 | 0 |
| 46.3728676 | 101.196372 |
| 0 | 0 |
| 0 | 0 |
| 0 | 0 |
| 0 | 0 |
| 1.71751362 | 0 |
| 0 | 0 |
| 0 | 0 |
| 0 | 0 |
| 0 | 0 |
| 0 | 0 |
| 0 | 0 |
| 0 | 0 |
| 324.610074 | 36.1415615 |
| 0 | 0 |
| 0 | 0 |
| 6.87005447 | 0 |
| 0 | 0 |
| 0 | 0 |
| 0 | 0 |
| 0 | 0 |
| 0 | 0 |
| 0 | 7.22831229 |
| 0 | 0 |
| 0 | 0 |
| 0 | 0 |
| 0 | 0 |
| 0 | 0 |
| 0 | 0 |
| 0 | 0 |
| 0 | 0 |

| | |
|---|---|
| 0 | 0 |
| 0 | 0 |
| 0 | 0 |
| 0 | 0 |
| 0 | 0 |
| 0 | 0 |
| 0 | 0 |
| 0 | 0 |
| 0 | 0 |
| 0 | 0 |
| 0 | 0 |
| 0 | 0 |
| 0 | 0 |
| 0 | 0 |
| 0 | 0 |
| 0 | 0 |
| 0 | 0 |
| 549.604357 | 448.155362 |
| 0 | 0 |
| 0 | 0 |
| 0 | 0 |
| 0 | 0 |
| 0 | 0 |
| 0 | 0 |
| 0 | 0 |
| 0 | 0 |
| 0 | 0 |
| 0 | 0 |
| 0 | 0 |
| 0 | 0 |
| 3.43502723 | 0 |
| 27.4802179 | 0 |
| 0 | 0 |
| 0 | 0 |
| 0 | 0 |
| 0 | 0 |
| 0 | 0 |
| 0 | 0 |
| 0 | 0 |
| 0 | 0 |
| 0 | 0 |
| 0 | 0 |
| 0 | 0 |

| | |
|---|---|
| 0 | 0 |
| 0 | 0 |
| 0 | 0 |
| 0 | 0 |
| 0 | 0 |
| 0 | 0 |
| 0 | 0 |
| 0 | 0 |
| 0 | 0 |
| 0 | 0 |
| 0 | 0 |
| 0 | 0 |
| 0 | 0 |
| 0 | 0 |
| 0 | 0 |
| 0 | 0 |
| 0 | 0 |
| 0 | 0 |
| 0 | 0 |
| 0 | 0 |
| 0 | 0 |
| 0 | 0 |
| 0 | 0 |
| 0 | 0 |
| 0 | 0 |
| 0 | 0 |
| 0 | 0 |
| 0 | 0 |
| 0 | 0 |
| 0 | 0 |
| 0 | 0 |
| 0 | 0 |
| 0 | 0 |
| 0 | 0 |
| 49.8078949 | 144.566246 |
| 0 | 0 |
| 0 | 0 |
| 0 | 0 |
| 0 | 0 |
| 0 | 0 |
| 0 | 0 |
| 0 | 0 |
| 0 | 0 |

| | |
|---|---|
| 0 | 0 |
| 0 | 0 |
| 0 | 0 |
| 0 | 0 |
| 0 | 0 |
| 0 | 0 |
| 0 | 0 |
| 0 | 0 |
| 0 | 0 |
| 0 | 0 |
| 0 | 0 |
| 245.604447 | 130.109621 |
| 0 | 0 |
| 0 | 0 |
| 0 | 0 |
| 1.71751362 | 0 |
| 0 | 0 |
| 0 | 0 |
| 0 | 0 |
| 133.966062 | 0 |
| 0 | 0 |
| 0 | 0 |
| 0 | 0 |
| 0 | 0 |
| 0 | 0 |
| 0 | 0 |
| 0 | 0 |
| 0 | 0 |
| 0 | 0 |
| 0 | 0 |
| 0 | 0 |
| 0 | 0 |
| 0 | 0 |
| 0 | 0 |
| 0 | 0 |
| 0 | 0 |
| 0 | 0 |
| 0 | 0 |
| 142.55363 | 50.598186 |
| 0 | 0 |
| 0 | 0 |
| 0 | 0 |
| 0 | 0 |

| | |
|---|---|
| 0 | 0 |
| 0 | 0 |
| 0 | 0 |
| 0 | 0 |
| 0 | 0 |
| 0 | 0 |
| 0 | 0 |
| 0 | 0 |
| 0 | 0 |
| 145.988657 | 65.0548106 |
| 1.71751362 | 0 |
| 0 | 0 |
| 0 | 0 |
| 0 | 0 |
| 0 | 0 |
| 0 | 7.22831229 |
| 0 | 0 |
| 103.050817 | 115.652997 |
| 0 | 0 |
| 0 | 0 |
| 0 | 0 |
| 0 | 0 |
| 0 | 0 |
| 0 | 0 |
| 0 | 0 |
| 3.43502723 | 0 |
| 0 | 0 |
| 0 | 0 |
| 0 | 0 |
| 0 | 0 |
| 0 | 0 |
| 0 | 0 |
| 0 | 0 |
| 0 | 0 |
| 0 | 0 |
| 0 | 0 |
| 0 | 0 |
| 0 | 0 |
| 0 | 0 |
| 0 | 14.4566246 |
| 0 | 0 |
| 0 | 0 |
| 0 | 0 |

| | |
|---|---|
| 0 | 0 |
| 0 | 0 |
| 0 | 0 |
| 0 | 0 |
| 0 | 0 |
| 0 | 0 |
| 0 | 0 |
| 0 | 0 |
| 0 | 0 |
| 0 | 0 |
| 0 | 0 |
| 0 | 0 |
| 0 | 0 |
| 0 | 0 |
| 0 | 0 |
| 0 | 0 |
| 0 | 0 |
| 0 | 0 |
| 0 | 0 |
| 0 | 0 |
| 0 | 0 |
| 0 | 0 |
| 0 | 0 |
| 0 | 0 |
| 0 | 0 |
| 0 | 0 |
| 0 | 0 |
| 0 | 0 |
| 0 | 7.22831229 |
| 0 | 0 |
| 0 | 0 |
| 0 | 0 |
| 0 | 0 |
| 0 | 0 |
| 175.186389 | 252.99093 |
| 0 | 0 |
| 0 | 0 |
| 0 | 0 |
| 0 | 0 |
| 0 | 0 |
| 0 | 0 |
| 0 | 0 |
| 0 | 0 |

| | |
|---|---|
| 0 | 7.22831229 |
| 0 | 0 |
| 0 | 0 |
| 0 | 0 |
| 0 | 0 |
| 0 | 0 |
| 0 | 0 |
| 0 | 0 |
| 0 | 0 |
| 0 | 0 |
| 0 | 0 |
| 0 | 0 |
| 0 | 0 |
| 0 | 0 |
| 0 | 0 |
| 17.1751362 | 93.9680598 |
| 0 | 0 |
| 0 | 0 |
| 0 | 0 |
| 0 | 0 |
| 0 | 0 |
| 0 | 0 |
| 0 | 0 |
| 0 | 0 |
| 0 | 0 |
| 0 | 0 |
| 0 | 0 |
| 0 | 0 |
| 0 | 7.22831229 |
| 0 | 0 |
| 0 | 0 |
| 0 | 0 |
| 48.0903813 | 14.4566246 |
| 0 | 0 |
| 0 | 0 |
| 0 | 0 |
| 0 | 7.22831229 |
| 0 | 0 |
| 0 | 0 |
| 0 | 0 |
| 0 | 0 |
| 0 | 21.6849369 |
| 0 | 0 |

| | |
|---|---|
| 0 | 0 |
| 0 | 0 |
| 0 | 0 |
| 0 | 0 |
| 0 | 0 |
| 0 | 0 |
| 0 | 0 |
| 0 | 0 |
| 0 | 7.22831229 |
| 0 | 0 |
| 0 | 0 |
| 0 | 0 |
| 0 | 0 |
| 0 | 0 |
| 0 | 0 |
| 0 | 0 |
| 0 | 0 |
| 0 | 0 |
| 0 | 0 |
| 0 | 0 |
| 0 | 0 |
| 0 | 0 |
| 0 | 0 |
| 0 | 0 |
| 0 | 0 |
| 0 | 0 |
| 0 | 0 |
| 0 | 0 |
| 0 | 0 |
| 0 | 0 |
| 0 | 0 |
| 0 | 0 |
| 0 | 0 |
| 0 | 0 |
| 1.71751362 | 0 |
| 1.71751362 | 0 |
| 0 | 0 |
| 0 | 0 |
| 0 | 0 |
| 0 | 0 |
| 0 | 0 |
| 0 | 0 |
| 0 | 0 |

| | |
|---|---|
| 0 | 0 |
| 0 | 0 |
| 0 | 0 |
| 0 | 0 |
| 0 | 0 |
| 0 | 0 |
| 0 | 0 |
| 0 | 0 |
| 0 | 36.1415615 |
| 0 | 36.1415615 |
| 0 | 0 |
| 285.10726 | 527.666797 |
| 0 | 7.22831229 |
| 0 | 0 |
| 0 | 0 |
| 0 | 0 |
| 0 | 0 |
| 0 | 0 |
| 0 | 0 |
| 1.71751362 | 0 |
| 0 | 0 |
| 0 | 0 |
| 66.983031 | 0 |
| 0 | 7.22831229 |
| 0 | 0 |
| 0 | 0 |
| 0 | 0 |
| 0 | 7.22831229 |
| 0 | 0 |
| 463.728676 | 534.89511 |
| 0 | 0 |
| 0 | 0 |
| 0 | 0 |
| 0 | 0 |
| 0 | 0 |
| 0 | 0 |
| 0 | 0 |
| 0 | 0 |
| 0 | 0 |
| 0 | 0 |
| 0 | 0 |
| 0 | 0 |
| 0 | 0 |

| | |
|---|---|
| 0 | 0 |
| 0 | 0 |
| 0 | 0 |
| 0 | 0 |
| 0 | 0 |
| 0 | 0 |
| 0 | 0 |
| 0 | 0 |
| 661.242742 | 209.621056 |
| 0 | 0 |
| 0 | 0 |
| 173.468875 | 108.424684 |
| 0 | 0 |
| 0 | 0 |
| 0 | 0 |
| 0 | 0 |
| 0 | 0 |
| 0 | 0 |
| 0 | 0 |
| 0 | 0 |
| 0 | 0 |
| 0 | 0 |
| 0 | 0 |
| 0 | 0 |
| 0 | 0 |
| 0 | 0 |
| 0 | 0 |
| 0 | 0 |
| 0 | 7.22831229 |
| 0 | 0 |
| 0 | 0 |
| 0 | 0 |
| 0 | 7.22831229 |
| 0 | 0 |
| 0 | 0 |
| 0 | 0 |
| 0 | 0 |
| 0 | 0 |
| 0 | 0 |
| 0 | 0 |
| 0 | 0 |
| 0 | 0 |
| 0 | 0 |

| | |
|---|---|
| 0 | 0 |
| 0 | 0 |
| 1.71751362 | 0 |
| 0 | 0 |
| 0 | 0 |
| 0 | 0 |
| 0 | 0 |
| 0 | 0 |
| 15.4576225 | 14.4566246 |
| 0 | 0 |
| 0 | 0 |
| 0 | 0 |
| 0 | 0 |
| 0 | 0 |
| 0 | 0 |
| 0 | 0 |
| 0 | 0 |
| 0 | 0 |
| 0 | 0 |
| 0 | 0 |
| 0 | 0 |
| 711.050637 | 917.995661 |
| 0 | 0 |
| 0 | 0 |
| 0 | 0 |
| 0 | 0 |
| 0 | 0 |
| 486.056353 | 209.621056 |
| 0 | 0 |
| 0 | 0 |
| 0 | 0 |
| 0 | 0 |
| 0 | 0 |
| 0 | 0 |
| 0 | 0 |
| 0 | 0 |
| 0 | 0 |
| 0 | 0 |
| 0 | 7.22831229 |
| 0 | 0 |
| 0 | 0 |
| 0 | 0 |
| 0 | 0 |

| | |
|---|---|
| 0 | 0 |
| 0 | 0 |
| 0 | 0 |
| 0 | 0 |
| 0 | 0 |
| 0 | 0 |
| 0 | 14.4566246 |
| 0 | 0 |
| 0 | 0 |
| 0 | 0 |
| 0 | 0 |
| 0 | 0 |
| 0 | 0 |
| 0 | 0 |
| 0 | 0 |
| 0 | 0 |
| 0 | 0 |
| 0 | 0 |
| 0 | 0 |
| 0 | 0 |
| 0 | 0 |
| 3.43502723 | 0 |
| 0 | 0 |
| 0 | 0 |
| 0 | 0 |
| 0 | 0 |
| 0 | 0 |
| 0 | 0 |
| 0 | 0 |
| 0 | 0 |
| 0 | 0 |
| 0 | 0 |
| 0 | 0 |
| 0 | 0 |
| 0 | 0 |
| 0 | 0 |
| 0 | 0 |
| 0 | 0 |
| 0 | 7.22831229 |
| 0 | 0 |
| 0 | 0 |
| 0 | 0 |
| 0 | 0 |

| | |
|---|---|
| 0 | 0 |
| 0 | 0 |
| 0 | 0 |
| 0 | 0 |
| 0 | 0 |
| 0 | 0 |
| 0 | 0 |
| 0 | 0 |
| 0 | 0 |
| 0 | 0 |
| 0 | 0 |
| 0 | 0 |
| 0 | 0 |
| 0 | 0 |
| 0 | 0 |
| 0 | 0 |
| 145.988657 | 354.187302 |
| 0 | 0 |
| 0 | 0 |
| 0 | 0 |
| 0 | 0 |
| 0 | 0 |
| 0 | 0 |
| 0 | 0 |
| 0 | 0 |
| 0 | 0 |
| 0 | 0 |
| 0 | 0 |
| 0 | 0 |
| 0 | 0 |
| 0 | 0 |
| 0 | 0 |
| 0 | 7.22831229 |
| 0 | 0 |
| 0 | 0 |
| 0 | 0 |
| 0 | 0 |
| 0 | 0 |
| 0 | 0 |
| 0 | 0 |
| 0 | 21.6849369 |
| 3.43502723 | 28.9132492 |
| 3.43502723 | 7.22831229 |

| | |
|---|---|
| 0 | 0 |
| 0 | 0 |
| 0 | 0 |
| 1.71751362 | 0 |
| 0 | 0 |
| 0 | 0 |
| 0 | 0 |
| 0 | 7.22831229 |
| 0 | 0 |
| 0 | 0 |
| 0 | 0 |
| 1.71751362 | 0 |
| 0 | 0 |
| 0 | 0 |
| 0 | 0 |
| 0 | 0 |
| 0 | 0 |
| 0 | 7.22831229 |
| 0 | 0 |
| 0 | 0 |
| 0 | 0 |
| 0 | 0 |
| 0 | 0 |
| 0 | 0 |
| 0 | 0 |
| 0 | 0 |
| 5.15254085 | 14.4566246 |
| 118.50844 | 57.8264983 |
| 358.960346 | 115.652997 |
| 0 | 0 |
| 0 | 0 |
| 0 | 7.22831229 |
| 0 | 0 |
| 54.9604357 | 57.8264983 |
| 18.8926498 | 0 |
| 0 | 0 |
| 0 | 0 |
| 0 | 0 |
| 0 | 0 |
| 0 | 7.22831229 |
| 0 | 0 |
| 0 | 0 |
| 0 | 0 |

| | |
|---|---|
| 0 | 0 |
| 0 | 0 |
| 0 | 0 |
| 0 | 0 |
| 0 | 0 |
| 0 | 0 |
| 0 | 0 |
| 0 | 0 |
| 0 | 0 |
| 0 | 0 |
| 70.4180583 | 122.881309 |
| 0 | 0 |
| 0 | 0 |
| 0 | 0 |
| 0 | 0 |
| 0 | 0 |
| 0 | 0 |
| 0 | 0 |
| 0 | 0 |
| 0 | 0 |
| 0 | 0 |
| 0 | 0 |
| 0 | 43.3698738 |
| 0 | 0 |
| 0 | 0 |
| 0 | 0 |
| 0 | 0 |
| 0 | 7.22831229 |
| 0 | 0 |
| 0 | 0 |
| 0 | 0 |
| 0 | 0 |
| 0 | 7.22831229 |
| 0 | 0 |
| 0 | 0 |
| 0 | 0 |
| 0 | 0 |
| 0 | 0 |
| 0 | 0 |
| 0 | 0 |
| 0 | 0 |
| 0 | 0 |
| 0 | 0 |

| | |
|---|---|
| 0 | 0 |
| 283.389747 | 151.794558 |
| 0 | 0 |
| 0 | 0 |
| 0 | 0 |
| 0 | 0 |
| 1.71751362 | 0 |
| 0 | 0 |
| 6.87005447 | 43.3698738 |
| 0 | 0 |
| 0 | 0 |
| 0 | 0 |
| 0 | 0 |
| 0 | 0 |
| 0 | 0 |
| 0 | 0 |
| 0 | 0 |
| 0 | 0 |
| 0 | 0 |
| 0 | 0 |
| 0 | 0 |
| 0 | 0 |
| 0 | 0 |
| 0 | 0 |
| 0 | 0 |
| 0 | 0 |
| 0 | 0 |
| 0 | 0 |
| 0 | 0 |
| 0 | 0 |
| 0 | 0 |
| 0 | 0 |
| 0 | 0 |
| 0 | 0 |
| 0 | 0 |
| 0 | 0 |
| 0 | 0 |
| 0 | 0 |
| 0 | 0 |
| 0 | 0 |
| 0 | 0 |
| 0 | 0 |
| 0 | 0 |

| | |
|---|---|
| 0 | 0 |
| 0 | 0 |
| 0 | 0 |
| 0 | 0 |
| 0 | 0 |
| 0 | 0 |
| 0 | 0 |
| 0 | 0 |
| 0 | 0 |
| 0 | 0 |
| 0 | 0 |
| 0 | 0 |
| 0 | 0 |
| 0 | 0 |
| 0 | 0 |
| 0 | 0 |
| 0 | 0 |
| 874.214431 | 578.264983 |
| 0 | 0 |
| 0 | 0 |
| 0 | 0 |
| 0 | 0 |
| 0 | 0 |
| 0 | 0 |
| 0 | 0 |
| 0 | 0 |
| 0 | 0 |
| 0 | 0 |
| 0 | 0 |
| 0 | 0 |
| 0 | 0 |
| 0 | 0 |
| 0 | 0 |
| 0 | 0 |
| 0 | 0 |
| 0 | 0 |
| 0 | 0 |
| 0 | 0 |
| 0 | 7.22831229 |
| 0 | 0 |
| 0 | 0 |
| 0 | 0 |
| 0 | 0 |

| | |
|---|---|
| 0 | 0 |
| 0 | 0 |
| 0 | 0 |
| 0 | 0 |
| 0 | 0 |
| 0 | 0 |
| 0 | 0 |
| 0 | 0 |
| 0 | 0 |
| 0 | 0 |
| 0 | 0 |
| 0 | 0 |
| 0 | 0 |
| 0 | 0 |
| 61.8304902 | 86.7397475 |
| 0 | 0 |
| 0 | 0 |
| 0 | 0 |
| 0 | 0 |
| 496.361435 | 346.95899 |
| 37.7852996 | 0 |
| 0 | 0 |
| 0 | 0 |
| 0 | 0 |
| 0 | 0 |
| 0 | 0 |
| 0 | 0 |
| 1.71751362 | 0 |
| 0 | 0 |
| 0 | 0 |
| 0 | 0 |
| 0 | 0 |
| 0 | 0 |
| 0 | 0 |
| 0 | 0 |
| 0 | 0 |
| 0 | 0 |
| 0 | 0 |
| 0 | 0 |
| 0 | 0 |
| 0 | 0 |
| 0 | 0 |
| 0 | 0 |

| | |
|---|---|
| 1.71751362 | 0 |
| 0 | 0 |
| 0 | 0 |
| 0 | 0 |
| 0 | 0 |
| 0 | 0 |
| 0 | 0 |
| 0 | 0 |
| 0 | 0 |
| 0 | 0 |
| 0 | 0 |
| 0 | 0 |
| 3.43502723 | 0 |
| 0 | 0 |
| 0 | 0 |
| 0 | 0 |
| 0 | 0 |
| 0 | 0 |
| 0 | 0 |
| 0 | 0 |
| 0 | 0 |
| 0 | 0 |
| 0 | 0 |
| 0 | 0 |
| 0 | 0 |
| 0 | 0 |
| 0 | 0 |
| 0 | 0 |
| 0 | 0 |
| 0 | 0 |
| 0 | 0 |
| 0 | 0 |
| 0 | 0 |
| 0 | 0 |
| 0 | 0 |
| 0 | 0 |
| 0 | 0 |
| 0 | 0 |
| 0 | 0 |
| 0 | 0 |
| 0 | 0 |
| 0 | 0 |
| 355.525319 | 397.557176 |

| | |
|---|---|
| 5.15254085 | 0 |
| 0 | 0 |
| 0 | 0 |
| 0 | 0 |
| 0 | 0 |
| 0 | 0 |
| 0 | 0 |
| 0 | 0 |
| 0 | 0 |
| 0 | 0 |
| 0 | 0 |
| 0 | 0 |
| 1.71751362 | 65.0548106 |
| 0 | 0 |
| 0 | 0 |
| 0 | 0 |
| 0 | 0 |
| 0 | 0 |
| 0 | 0 |
| 0 | 0 |
| 0 | 0 |
| 0 | 0 |
| 0 | 0 |
| 0 | 0 |
| 0 | 0 |
| 0 | 0 |
| 0 | 0 |
| 0 | 0 |
| 0 | 0 |
| 0 | 0 |
| 0 | 0 |
| 0 | 0 |
| 0 | 0 |
| 85.8756808 | 65.0548106 |
| 0 | 0 |
| 0 | 0 |
| 0 | 0 |
| 0 | 0 |
| 0 | 0 |
| 0 | 0 |
| 0 | 0 |
| 0 | 0 |
| 0 | 0 |

| | |
|---|---|
| 0 | 0 |
| 0 | 0 |
| 0 | 0 |
| 0 | 0 |
| 0 | 0 |
| 0 | 0 |
| 0 | 0 |
| 0 | 0 |
| 1.71751362 | 0 |
| 0 | 0 |
| 0 | 0 |
| 0 | 0 |
| 0 | 0 |
| 0 | 0 |
| 0 | 0 |
| 0 | 0 |
| 0 | 0 |
| 0 | 0 |
| 0 | 0 |
| 0 | 0 |
| 0 | 0 |
| 0 | 0 |
| 0 | 0 |
| 0 | 0 |
| 1.71751362 | 0 |
| 0 | 7.22831229 |
| 0 | 0 |
| 0 | 0 |
| 0 | 0 |
| 0 | 0 |
| 0 | 0 |
| 0 | 0 |
| 0 | 7.22831229 |
| 0 | 0 |
| 0 | 0 |
| 0 | 0 |
| 0 | 0 |
| 0 | 21.6849369 |
| 0 | 0 |
| 0 | 0 |
| 0 | 0 |
| 0 | 0 |
| 0 | 0 |

| | |
|---|---|
| 0 | 0 |
| 0 | 0 |
| 0 | 0 |
| 0 | 0 |
| 0 | 0 |
| 0 | 0 |
| 0 | 0 |
| 0 | 0 |
| 0 | 0 |
| 0 | 0 |
| 0 | 7.22831229 |
| 0 | 0 |
| 0 | 0 |
| 0 | 0 |
| 0 | 0 |
| 0 | 0 |
| 0 | 0 |
| 0 | 0 |
| 0 | 0 |
| 0 | 0 |
| 0 | 0 |
| 0 | 0 |
| 0 | 0 |
| 1.71751362 | 0 |
| 0 | 0 |
| 0 | 0 |
| 0 | 0 |
| 0 | 0 |
| 0 | 0 |
| 0 | 0 |
| 0 | 0 |
| 92.7457353 | 57.8264983 |
| 0 | 7.22831229 |
| 0 | 0 |
| 0 | 0 |
| 0 | 0 |
| 0 | 0 |
| 0 | 0 |
| 0 | 0 |
| 0 | 0 |
| 0 | 7.22831229 |
| 0 | 0 |
| 0 | 0 |

| | |
|---|---|
| 0 | 0 |
| 0 | 0 |
| 0 | 0 |
| 0 | 0 |
| 0 | 0 |
| 0 | 0 |
| 0 | 0 |
| 0 | 0 |
| 0 | 7.22831229 |
| 0 | 0 |
| 0 | 0 |
| 0 | 0 |
| 0 | 0 |
| 0 | 7.22831229 |
| 0 | 0 |
| 0 | 0 |
| 0 | 0 |
| 0 | 0 |
| 0 | 0 |
| 0 | 0 |
| 0 | 0 |
| 0 | 0 |
| 0 | 14.4566246 |
| 0 | 0 |
| 0 | 0 |
| 0 | 0 |
| 0 | 0 |
| 1.71751362 | 14.4566246 |
| 0 | 0 |
| 0 | 0 |
| 0 | 0 |
| 0 | 0 |
| 0 | 0 |
| 0 | 0 |
| 0 | 0 |
| 0 | 0 |
| 0 | 0 |
| 0 | 0 |
| 0 | 0 |
| 0 | 0 |
| 0 | 0 |
| 0 | 0 |
| 0 | 0 |

| | |
|---|---|
| 0 | 0 |
| 0 | 0 |
| 0 | 0 |
| 0 | 0 |
| 0 | 0 |
| 0 | 0 |
| 1.71751362 | 0 |
| 0 | 0 |
| 0 | 0 |
| 0 | 0 |
| 0 | 0 |
| 0 | 0 |
| 44.655354 | 7.22831229 |
| 0 | 0 |
| 0 | 0 |
| 0 | 0 |
| 0 | 0 |
| 0 | 0 |
| 0 | 0 |
| 0 | 0 |
| 0 | 0 |
| 0 | 7.22831229 |
| 0 | 0 |
| 0 | 0 |
| 0 | 0 |
| 0 | 0 |
| 0 | 0 |
| 46.3728676 | 0 |
| 0 | 0 |
| 0 | 0 |
| 0 | 0 |
| 0 | 0 |
| 0 | 0 |
| 0 | 0 |
| 0 | 0 |
| 0 | 0 |
| 0 | 0 |
| 42.9378404 | 0 |
| 0 | 0 |
| 0 | 0 |
| 0 | 0 |
| 0 | 0 |
| 0 | 0 |

| | |
|---|---|
| 0 | 0 |
| 0 | 0 |
| 0 | 0 |
| 0 | 0 |
| 0 | 0 |
| 0 | 0 |
| 0 | 0 |
| 0 | 0 |
| 0 | 0 |
| 0 | 0 |
| 0 | 0 |
| 0 | 0 |
| 0 | 0 |
| 0 | 0 |
| 0 | 0 |
| 0 | 0 |
| 0 | 0 |
| 0 | 0 |
| 0 | 43.3698738 |
| 0 | 36.1415615 |
| 0 | 0 |
| 0 | 0 |
| 0 | 0 |
| 0 | 0 |
| 0 | 0 |
| 0 | 0 |
| 0 | 14.4566246 |
| 0 | 0 |
| 0 | 0 |
| 0 | 0 |
| 0 | 0 |
| 0 | 0 |
| 0 | 0 |
| 0 | 0 |
| 0 | 0 |
| 0 | 0 |
| 0 | 0 |
| 0 | 0 |
| 0 | 0 |
| 0 | 0 |
| 58.395463 | 0 |
| 0 | 0 |
| 0 | 0 |

| | |
|---|---|
| 0 | 0 |
| 0 | 0 |
| 0 | 7.22831229 |
| 0 | 0 |
| 0 | 0 |
| 0 | 0 |
| 0 | 0 |
| 0 | 0 |
| 0 | 0 |
| 0 | 0 |
| 0 | 0 |
| 0 | 0 |
| 0 | 0 |
| 0 | 0 |
| 0 | 0 |
| 0 | 0 |
| 0 | 7.22831229 |
| 0 | 0 |
| 0 | 0 |
| 0 | 0 |
| 0 | 0 |
| 0 | 0 |
| 0 | 0 |
| 0 | 0 |
| 0 | 0 |
| 0 | 0 |
| 0 | 0 |
| 0 | 0 |
| 0 | 0 |
| 1.71751362 | 0 |
| 0 | 0 |
| 0 | 0 |
| 0 | 0 |
| 0 | 0 |
| 84.1581672 | 122.881309 |
| 0 | 0 |
| 0 | 0 |
| 0 | 0 |
| 0 | 0 |
| 0 | 0 |
| 0 | 0 |
| 303.99991 | 93.9680598 |
| 0 | 0 |

| | |
|---|---|
| 0 | 0 |
| 0 | 0 |
| 0 | 0 |
| 0 | 0 |
| 0 | 0 |
| 0 | 0 |
| 0 | 0 |
| 0 | 0 |
| 0 | 0 |
| 0 | 0 |
| 0 | 0 |
| 6.87005447 | 21.6849369 |
| 0 | 0 |
| 0 | 0 |
| 0 | 0 |
| 0 | 0 |
| 0 | 0 |
| 0 | 0 |
| 0 | 0 |
| 85.8756808 | 7.22831229 |
| 0 | 0 |
| 0 | 0 |
| 0 | 0 |
| 0 | 0 |
| 66.983031 | 14.4566246 |
| 0 | 0 |
| 0 | 0 |
| 0 | 0 |
| 0 | 0 |
| 0 | 0 |
| 673.265338 | 542.123422 |
| 6.87005447 | 7.22831229 |
| 0 | 0 |
| 0 | 0 |
| 0 | 0 |
| 0 | 0 |
| 0 | 0 |
| 0 | 0 |
| 0 | 0 |
| 0 | 0 |
| 0 | 7.22831229 |
| 0 | 0 |
| 0 | 0 |

| | |
|---|---|
| 0 | 0 |
| 0 | 0 |
| 0 | 0 |
| 1.71751362 | 0 |
| 0 | 0 |
| 0 | 0 |
| 0 | 0 |
| 0 | 0 |
| 0 | 0 |
| 0 | 0 |
| 0 | 0 |
| 0 | 0 |
| 36.0677859 | 57.8264983 |
| 0 | 0 |
| 0 | 0 |
| 0 | 0 |
| 0 | 0 |
| 0 | 0 |
| 0 | 0 |
| 0 | 0 |
| 0 | 0 |
| 0 | 0 |
| 0 | 0 |
| 0 | 0 |
| 0 | 0 |
| 0 | 0 |
| 0 | 0 |
| 0 | 0 |
| 0 | 7.22831229 |
| 0 | 0 |
| 583.95463 | 325.274053 |
| 0 | 0 |
| 0 | 0 |
| 1.71751362 | 0 |
| 128.813521 | 21.6849369 |
| 27.4802179 | 0 |
| 0 | 0 |
| 0 | 0 |
| 0 | 0 |
| 0 | 0 |
| 0 | 0 |
| 0 | 0 |
| 0 | 0 |

| | |
|---|---|
| 0 | 0 |
| 410.485754 | 571.036671 |
| 0 | 0 |
| 0 | 0 |
| 0 | 0 |
| 0 | 0 |
| 0 | 0 |
| 0 | 0 |
| 0 | 0 |
| 0 | 0 |
| 0 | 0 |
| 0 | 0 |
| 0 | 0 |
| 0 | 0 |
| 0 | 0 |
| 0 | 0 |
| 0 | 0 |
| 6.87005447 | 0 |
| 0 | 0 |
| 0 | 0 |
| 0 | 0 |
| 0 | 0 |
| 0 | 0 |
| 0 | 0 |
| 0 | 0 |
| 0 | 0 |
| 0 | 0 |
| 0 | 0 |
| 0 | 0 |
| 0 | 0 |
| 0 | 0 |
| 30.9152451 | 21.6849369 |
| 0 | 0 |
| 0 | 7.22831229 |
| 0 | 0 |
| 0 | 0 |
| 0 | 0 |
| 0 | 0 |
| 0 | 0 |
| 0 | 0 |
| 0 | 0 |
| 0 | 0 |
| 0 | 0 |

| | |
|---|---|
| 0 | 0 |
| 0 | 0 |
| 8.58756808 | 159.02287 |
| 211.254175 | 281.904179 |
| 0 | 7.22831229 |
| 0 | 0 |
| 46.3728676 | 115.652997 |
| 0 | 0 |
| 0 | 0 |
| 0 | 0 |
| 0 | 0 |
| 0 | 0 |
| 0 | 0 |
| 0 | 0 |
| 0 | 0 |
| 0 | 0 |
| 0 | 0 |
| 0 | 0 |
| 6.87005447 | 28.9132492 |
| 0 | 14.4566246 |
| 0 | 0 |
| 0 | 0 |
| 0 | 0 |
| 0 | 0 |
| 0 | 0 |
| 0 | 0 |
| 0 | 0 |
| 0 | 0 |
| 0 | 0 |
| 0 | 0 |
| 0 | 0 |
| 0 | 0 |
| 0 | 0 |
| 41.2203268 | 28.9132492 |
| 0 | 0 |
| 0 | 0 |
| 0 | 0 |
| 0 | 0 |
| 0 | 0 |
| 0 | 0 |
| 0 | 0 |
| 0 | 0 |
| 0 | 0 |

| | |
|---|---|
| 0 | 0 |
| 0 | 7.22831229 |
| 0 | 0 |
| 0 | 0 |
| 0 | 0 |
| 0 | 0 |
| 0 | 0 |
| 0 | 0 |
| 0 | 0 |
| 0 | 0 |
| 0 | 0 |
| 0 | 0 |
| 0 | 0 |
| 0 | 0 |
| 0 | 0 |
| 0 | 0 |
| 0 | 0 |
| 0 | 0 |
| 0 | 0 |
| 0 | 0 |
| 32.6327587 | 202.392744 |
| 0 | 0 |
| 0 | 0 |
| 0 | 0 |
| 0 | 0 |
| 0 | 0 |
| 190.644011 | 14.4566246 |
| 0 | 0 |
| 5.15254085 | 36.1415615 |
| 0 | 0 |
| 0 | 0 |
| 0 | 0 |
| 0 | 0 |
| 0 | 0 |
| 0 | 0 |
| 0 | 0 |
| 0 | 0 |
| 0 | 0 |
| 0 | 0 |
| 0 | 0 |
| 0 | 0 |
| 0 | 0 |
| 0 | 0 |

| | |
|---|---|
| 954.937571 | 231.305993 |
| 0 | 0 |
| 0 | 0 |
| 0 | 0 |
| 0 | 0 |
| 73.8530855 | 0 |
| 0 | 0 |
| 0 | 14.4566246 |
| 0 | 0 |
| 0 | 0 |
| 0 | 0 |
| 0 | 0 |
| 0 | 0 |
| 97.8982761 | 202.392744 |
| 3.43502723 | 0 |
| 0 | 0 |
| 0 | 0 |
| 0 | 0 |
| 0 | 0 |
| 0 | 0 |
| 0 | 0 |
| 0 | 0 |
| 0 | 0 |
| 80.72314 | 310.817429 |
| 0 | 0 |
| 0 | 0 |
| 0 | 0 |
| 0 | 0 |
| 0 | 0 |
| 0 | 0 |
| 0 | 0 |
| 0 | 0 |
| 1.71751362 | 7.22831229 |
| 0 | 0 |
| 0 | 0 |
| 0 | 0 |
| 0 | 0 |
| 85.8756808 | 115.652997 |
| 0 | 0 |
| 0 | 0 |
| 0 | 0 |
| 0 | 0 |
| 0 | 0 |

| | |
|---|---|
| 0 | 0 |
| 0 | 7.22831229 |
| 0 | 0 |
| 0 | 0 |
| 0 | 0 |
| 0 | 0 |
| 0 | 0 |
| 0 | 0 |
| 0 | 0 |
| 0 | 0 |
| 0 | 0 |
| 24.0451906 | 7.22831229 |
| 0 | 0 |
| 0 | 0 |
| 0 | 0 |
| 0 | 0 |
| 0 | 0 |
| 0 | 0 |
| 0 | 0 |
| 0 | 0 |
| 3.43502723 | 0 |
| 0 | 0 |
| 0 | 0 |
| 0 | 0 |
| 0 | 0 |
| 0 | 0 |
| 0 | 0 |
| 0 | 0 |
| 0 | 0 |
| 0 | 0 |
| 29.1977315 | 57.8264983 |
| 0 | 0 |
| 0 | 0 |
| 0 | 0 |
| 135.683576 | 14.4566246 |
| 0 | 0 |
| 0 | 0 |
| 0 | 0 |
| 0 | 0 |
| 0 | 0 |
| 0 | 0 |
| 0 | 0 |
| 0 | 0 |

| | |
|---|---|
| 8.58756808 | 7.22831229 |
| 0 | 0 |
| 0 | 0 |
| 0 | 0 |
| 0 | 0 |
| 0 | 0 |
| 0 | 0 |
| 0 | 0 |
| 0 | 0 |
| 0 | 0 |
| 0 | 0 |
| 0 | 0 |
| 0 | 0 |
| 0 | 0 |
| 20.6101634 | 14.4566246 |
| 0 | 0 |
| 0 | 0 |
| 0 | 0 |
| 0 | 0 |
| 0 | 7.22831229 |
| 0 | 0 |
| 24.0451906 | 65.0548106 |
| 0 | 0 |
| 0 | 0 |
| 0 | 0 |
| 0 | 0 |
| 0 | 0 |
| 30.9152451 | 7.22831229 |
| 0 | 0 |
| 0 | 0 |
| 82.4406536 | 57.8264983 |
| 0 | 0 |
| 0 | 0 |
| 0 | 7.22831229 |
| 0 | 0 |
| 0 | 0 |
| 12.0225953 | 0 |
| 0 | 0 |
| 0 | 0 |
| 0 | 0 |
| 432.813431 | 375.872239 |
| 1.71751362 | 0 |
| 0 | 0 |

| | |
|---|---|
| 0 | 0 |
| 0 | 0 |
| 0 | 0 |
| 0 | 0 |
| 0 | 0 |
| 0 | 0 |
| 0 | 0 |
| 0 | 0 |
| 0 | 0 |
| 0 | 0 |
| 0 | 0 |
| 0 | 0 |
| 30.9152451 | 0 |
| 0 | 0 |
| 0 | 0 |
| 0 | 0 |
| 0 | 0 |
| 0 | 0 |
| 0 | 0 |
| 0 | 0 |
| 0 | 0 |
| 5.15254085 | 0 |
| 0 | 0 |
| 0 | 0 |
| 0 | 0 |
| 0 | 0 |
| 0 | 0 |
| 0 | 0 |
| 1.71751362 | 7.22831229 |
| 0 | 0 |
| 1.71751362 | 0 |
| 0 | 0 |
| 0 | 0 |
| 0 | 0 |
| 13.7401089 | 0 |
| 0 | 0 |
| 0 | 0 |
| 0 | 0 |
| 0 | 0 |
| 0 | 0 |
| 0 | 0 |
| 0 | 0 |
| 0 | 0 |

| | |
|---|---|
| 0 | 0 |
| 0 | 0 |
| 0 | 0 |
| 0 | 0 |
| 5.15254085 | 0 |
| 0 | 0 |
| 0 | 0 |
| 0 | 0 |
| 0 | 0 |
| 0 | 0 |
| 0 | 0 |
| 0 | 0 |
| 0 | 0 |
| 0 | 0 |
| 101.333303 | 137.337934 |
| 0 | 0 |
| 0 | 0 |
| 0 | 0 |
| 0 | 14.4566246 |
| 0 | 0 |
| 224.994284 | 93.9680598 |
| 0 | 0 |
| 0 | 0 |
| 0 | 0 |
| 1.71751362 | 0 |
| 0 | 0 |
| 0 | 0 |
| 0 | 0 |
| 0 | 0 |
| 0 | 0 |
| 0 | 0 |
| 0 | 0 |
| 1.71751362 | 0 |
| 0 | 0 |
| 0 | 0 |
| 0 | 21.6849369 |
| 334.915155 | 93.9680598 |
| 0 | 0 |
| 0 | 0 |
| 0 | 0 |
| 0 | 0 |
| 0 | 0 |
| 0 | 0 |

| | |
|---|---|
| 0 | 0 |
| 0 | 0 |
| 0 | 0 |
| 178.621416 | 180.707807 |
| 0 | 0 |
| 0 | 0 |
| 0 | 0 |
| 0 | 0 |
| 0 | 0 |
| 0 | 0 |
| 0 | 0 |
| 0 | 0 |
| 0 | 0 |
| 53.2429221 | 14.4566246 |
| 3.43502723 | 0 |
| 0 | 0 |
| 0 | 0 |
| 0 | 0 |
| 0 | 0 |
| 0 | 0 |
| 255.909529 | 187.93612 |
| 0 | 0 |
| 3.43502723 | 0 |
| 0 | 0 |
| 0 | 0 |
| 0 | 0 |
| 0 | 0 |
| 0 | 0 |
| 0 | 0 |
| 0 | 0 |
| 0 | 0 |
| 0 | 0 |
| 0 | 0 |
| 0 | 0 |
| 0 | 0 |
| 0 | 0 |
| 0 | 0 |
| 0 | 0 |
| 0 | 0 |
| 0 | 0 |
| 0 | 0 |
| 0 | 0 |
| 0 | 0 |

| | |
|---|---|
| 0 | 0 |
| 0 | 0 |
| 0 | 0 |
| 0 | 0 |
| 0 | 0 |
| 0 | 0 |
| 0 | 0 |
| 0 | 0 |
| 0 | 0 |
| 0 | 0 |
| 0 | 0 |
| 0 | 0 |
| 1183.36688 | 318.045741 |
| 0 | 0 |
| 0 | 0 |
| 128.813521 | 65.0548106 |
| 0 | 0 |
| 0 | 0 |
| 0 | 0 |
| 0 | 0 |
| 0 | 0 |
| 0 | 0 |
| 0 | 0 |
| 152.858712 | 144.566246 |
| 0 | 0 |
| 0 | 0 |
| 0 | 0 |
| 0 | 0 |
| 0 | 0 |
| 0 | 0 |
| 6.87005447 | 0 |
| 0 | 0 |
| 0 | 0 |
| 0 | 0 |
| 0 | 0 |
| 0 | 0 |
| 262.779583 | 166.251183 |
| 0 | 0 |
| 0 | 0 |
| 0 | 0 |
| 0 | 0 |
| 0 | 0 |
| 0 | 0 |

| | |
|---|---|
| 0 | 0 |
| 53.2429221 | 86.7397475 |
| 0 | 0 |
| 0 | 0 |
| 0 | 0 |
| 0 | 0 |
| 0 | 0 |
| 0 | 0 |
| 0 | 0 |
| 0 | 0 |
| 0 | 7.22831229 |
| 0 | 0 |
| 0 | 0 |
| 0 | 0 |
| 0 | 0 |
| 0 | 0 |
| 0 | 0 |
| 0 | 0 |
| 0 | 0 |
| 61.8304902 | 101.196372 |
| 0 | 0 |
| 0 | 0 |
| 0 | 0 |
| 0 | 0 |
| 0 | 0 |
| 0 | 0 |
| 0 | 0 |
| 0 | 0 |
| 0 | 0 |
| 0 | 0 |
| 0 | 7.22831229 |
| 0 | 0 |
| 0 | 0 |
| 1.71751362 | 0 |
| 0 | 0 |
| 0 | 0 |
| 0 | 0 |
| 0 | 0 |
| 0 | 0 |
| 0 | 0 |
| 0 | 0 |
| 0 | 0 |
| 1.71751362 | 0 |

| | |
|---|---|
| 0 | 0 |
| 0 | 0 |
| 0 | 0 |
| 0 | 0 |
| 0 | 0 |
| 0 | 0 |
| 0 | 0 |
| 0 | 0 |
| 0 | 0 |
| 0 | 0 |
| 0 | 0 |
| 0 | 0 |
| 0 | 0 |
| 0 | 0 |
| 0 | 0 |
| 0 | 0 |
| 0 | 0 |
| 0 | 0 |
| 0 | 0 |
| 0 | 0 |
| 0 | 0 |
| 0 | 0 |
| 0 | 0 |
| 0 | 0 |
| 0 | 0 |
| 0 | 0 |
| 0 | 7.22831229 |
| 0 | 0 |
| 0 | 0 |
| 0 | 0 |
| 0 | 0 |
| 0 | 0 |
| 0 | 0 |
| 0 | 0 |
| 0 | 0 |
| 3.43502723 | 7.22831229 |
| 358.960346 | 115.652997 |
| 0 | 0 |
| 0 | 0 |
| 0 | 0 |
| 0 | 0 |
| 0 | 0 |
| 0 | 0 |

| | |
|---|---|
| 0 | 0 |
| 0 | 0 |
| 0 | 0 |
| 0 | 0 |
| 0 | 0 |
| 0 | 0 |
| 0 | 0 |
| 82.4406536 | 79.5114352 |
| 0 | 0 |
| 5.15254085 | 0 |
| 1.71751362 | 0 |
| 0 | 0 |
| 1.71751362 | 0 |
| 0 | 0 |
| 0 | 0 |
| 0 | 0 |
| 0 | 0 |
| 0 | 0 |
| 0 | 0 |
| 0 | 0 |
| 0 | 0 |
| 961.807625 | 281.904179 |
| 0 | 0 |
| 0 | 0 |
| 0 | 0 |
| 0 | 0 |
| 0 | 0 |
| 0 | 0 |
| 0 | 0 |
| 170.033848 | 57.8264983 |
| 0 | 0 |
| 0 | 0 |
| 0 | 0 |
| 0 | 0 |
| 0 | 0 |
| 0 | 0 |
| 0 | 7.22831229 |
| 0 | 0 |
| 0 | 0 |
| 0 | 0 |
| 0 | 0 |
| 0 | 0 |
| 0 | 0 |

| | |
|---|---|
| 0 | 0 |
| 0 | 0 |
| 0 | 0 |
| 542.734303 | 209.621056 |
| 0 | 0 |
| 0 | 0 |
| 0 | 0 |
| 0 | 0 |
| 0 | 0 |
| 0 | 0 |
| 0 | 7.22831229 |
| 0 | 0 |
| 0 | 0 |
| 1.71751362 | 14.4566246 |
| 0 | 0 |
| 0 | 0 |
| 0 | 0 |
| 0 | 0 |
| 0 | 0 |
| 32.6327587 | 115.652997 |
| 202.666607 | 72.2831229 |
| 0 | 0 |
| 0 | 0 |
| 0 | 0 |
| 0 | 0 |
| 0 | 0 |
| 0 | 0 |
| 0 | 0 |
| 0 | 0 |
| 226.711797 | 0 |
| 0 | 0 |
| 0 | 0 |
| 0 | 0 |
| 0 | 0 |
| 0 | 0 |
| 0 | 0 |
| 0 | 0 |
| 0 | 0 |
| 0 | 0 |
| 0 | 0 |
| 0 | 0 |
| 0 | 0 |
| 0 | 0 |

| | |
|---|---|
| 0 | 0 |
| 0 | 7.22831229 |
| 0 | 0 |
| 0 | 0 |
| 0 | 0 |
| 0 | 0 |
| 0 | 0 |
| 0 | 0 |
| 0 | 0 |
| 0 | 0 |
| 0 | 0 |
| 0 | 0 |
| 0 | 0 |
| 0 | 0 |
| 0 | 0 |
| 0 | 0 |
| 44.655354 | 50.598186 |
| 0 | 0 |
| 0 | 0 |
| 0 | 0 |
| 0 | 0 |
| 0 | 0 |
| 0 | 0 |
| 0 | 0 |
| 0 | 0 |
| 0 | 0 |
| 0 | 0 |
| 0 | 0 |
| 0 | 0 |
| 0 | 0 |
| 0 | 0 |
| 0 | 0 |
| 0 | 0 |
| 0 | 0 |
| 0 | 0 |
| 0 | 0 |
| 0 | 0 |
| 0 | 0 |
| 0 | 0 |
| 0 | 0 |
| 0 | 0 |
| 0 | 0 |
| 0 | 0 |

| | |
|---|---|
| 0 | 0 |
| 0 | 0 |
| 0 | 0 |
| 0 | 0 |
| 0 | 0 |
| 0 | 0 |
| 0 | 0 |
| 0 | 0 |
| 0 | 0 |
| 0 | 0 |
| 49.8078949 | 346.95899 |
| 0 | 14.4566246 |
| 0 | 0 |
| 0 | 0 |
| 0 | 0 |
| 0 | 0 |
| 0 | 0 |
| 0 | 0 |
| 6.87005447 | 36.1415615 |
| 0 | 0 |
| 0 | 0 |
| 0 | 0 |
| 0 | 0 |
| 0 | 0 |
| 0 | 0 |
| 27.4802179 | 0 |
| 0 | 0 |
| 207.819148 | 173.479495 |
| 0 | 14.4566246 |
| 0 | 0 |
| 2306.62079 | 722.831229 |
| 0 | 0 |
| 132.248548 | 187.93612 |
| 0 | 0 |
| 0 | 0 |
| 0 | 0 |
| 0 | 0 |
| 0 | 0 |
| 0 | 0 |
| 0 | 0 |
| 0 | 0 |
| 0 | 0 |
| 0 | 0 |

| | |
|---|---|
| 0 | 0 |
| 0 | 7.22831229 |
| 0 | 0 |
| 0 | 0 |
| 0 | 0 |
| 0 | 0 |
| 0 | 0 |
| 0 | 0 |
| 0 | 0 |
| 0 | 0 |
| 0 | 0 |
| 0 | 0 |
| 0 | 0 |
| 0 | 0 |
| 0 | 0 |
| 0 | 0 |
| 0 | 36.1415615 |
| 63.5480038 | 0 |
| 0 | 0 |
| 0 | 0 |
| 0 | 0 |
| 0 | 0 |
| 0 | 0 |
| 0 | 0 |
| 0 | 0 |
| 745.40091 | 477.068611 |
| 0 | 0 |
| 0 | 0 |
| 0 | 0 |
| 0 | 0 |
| 0 | 0 |
| 0 | 0 |
| 139.118603 | 0 |
| 6.87005447 | 7.22831229 |
| 0 | 0 |
| 0 | 0 |
| 0 | 0 |
| 0 | 0 |
| 0 | 36.1415615 |
| 0 | 0 |
| 0 | 0 |
| 0 | 0 |
| 0 | 0 |

| | |
|---|---|
| 0 | 0 |
| 0 | 0 |
| 0 | 0 |
| 0 | 0 |
| 49.8078949 | 115.652997 |
| 0 | 0 |
| 0 | 0 |
| 0 | 0 |
| 0 | 0 |
| 0 | 0 |
| 0 | 0 |
| 0 | 0 |
| 0 | 0 |
| 197.514066 | 173.479495 |
| 0 | 0 |
| 0 | 0 |
| 0 | 0 |
| 0 | 0 |
| 0 | 0 |
| 0 | 0 |
| 0 | 0 |
| 0 | 0 |
| 0 | 0 |
| 0 | 0 |
| 0 | 0 |
| 0 | 0 |
| 0 | 21.6849369 |
| 0 | 0 |
| 0 | 0 |
| 0 | 0 |
| 0 | 0 |
| 0 | 0 |
| 0 | 0 |
| 58.395463 | 72.2831229 |
| 0 | 0 |
| 0 | 0 |
| 0 | 0 |
| 0 | 0 |
| 0 | 0 |
| 0 | 0 |
| 0 | 0 |
| 0 | 0 |
| 0 | 0 |

| | |
|---|---|
| 0 | 0 |
| 0 | 0 |
| 0 | 0 |
| 0 | 0 |
| 0 | 0 |
| 0 | 0 |
| 0 | 0 |
| 0 | 0 |
| 0 | 0 |
| 0 | 0 |
| 0 | 7.22831229 |
| 0 | 0 |
| 1.71751362 | 0 |
| 0 | 0 |
| 0 | 7.22831229 |
| 0 | 0 |
| 0 | 0 |
| 0 | 0 |
| 0 | 0 |
| 0 | 0 |
| 0 | 0 |
| 0 | 0 |
| 0 | 0 |
| 0 | 0 |
| 0 | 0 |
| 0 | 0 |
| 0 | 0 |
| 0 | 0 |
| 0 | 0 |
| 0 | 0 |
| 0 | 0 |
| 1.71751362 | 0 |
| 91.0282217 | 108.424684 |
| 0 | 0 |
| 0 | 0 |
| 0 | 0 |
| 0 | 0 |
| 0 | 0 |
| 0 | 0 |
| 0 | 0 |
| 0 | 0 |
| 0 | 0 |
| 0 | 0 |

| | |
|---|---|
| 3.43502723 | 14.4566246 |
| 0 | 0 |
| 0 | 0 |
| 0 | 0 |
| 0 | 0 |
| 0 | 0 |
| 0 | 0 |
| 0 | 0 |
| 0 | 0 |
| 0 | 0 |
| 0 | 7.22831229 |
| 0 | 0 |
| 0 | 0 |
| 0 | 0 |
| 0 | 0 |
| 3.43502723 | 0 |
| 0 | 0 |
| 87.5931944 | 267.447555 |
| 0 | 0 |
| 0 | 0 |
| 0 | 0 |
| 0 | 0 |
| 0 | 0 |
| 0 | 0 |
| 0 | 0 |
| 0 | 0 |
| 0 | 0 |
| 0 | 0 |
| 0 | 0 |
| 0 | 0 |
| 0 | 0 |
| 0 | 0 |
| 0 | 0 |
| 0 | 0 |
| 0 | 14.4566246 |
| 0 | 0 |
| 0 | 0 |
| 0 | 0 |
| 0 | 0 |
| 0 | 0 |
| 0 | 0 |
| 0 | 0 |
| 0 | 0 |

| | |
|---|---|
| 0 | 0 |
| 0 | 0 |
| 0 | 0 |
| 30.9152451 | 0 |
| 0 | 0 |
| 0 | 0 |
| 0 | 0 |
| 0 | 0 |
| 384.72305 | 209.621056 |
| 0 | 0 |
| 0 | 0 |
| 0 | 0 |
| 0 | 0 |
| 0 | 0 |
| 0 | 93.9680598 |
| 0 | 0 |
| 0 | 0 |
| 0 | 0 |
| 0 | 0 |
| 0 | 0 |
| 0 | 0 |
| 0 | 0 |
| 0 | 0 |
| 0 | 0 |
| 0 | 0 |
| 0 | 0 |
| 0 | 0 |
| 29.1977315 | 21.6849369 |
| 508.38403 | 180.707807 |
| 0 | 0 |
| 0 | 0 |
| 0 | 0 |
| 0 | 0 |
| 0 | 0 |
| 0 | 7.22831229 |
| 0 | 0 |
| 0 | 0 |
| 0 | 0 |
| 0 | 7.22831229 |
| 0 | 0 |
| 0 | 0 |
| 0 | 0 |
| 0 | 0 |

| | |
|---|---|
| 0 | 0 |
| 0 | 7.22831229 |
| 0 | 0 |
| 0 | 0 |
| 0 | 0 |
| 0 | 0 |
| 0 | 0 |
| 0 | 0 |
| 0 | 0 |
| 0 | 0 |
| 0 | 0 |
| 0 | 0 |
| 0 | 0 |
| 170.033848 | 7.22831229 |
| 0 | 0 |
| 0 | 0 |
| 0 | 14.4566246 |
| 0 | 0 |
| 0 | 0 |
| 0 | 0 |
| 0 | 0 |
| 0 | 0 |
| 0 | 0 |
| 0 | 0 |
| 0 | 0 |
| 0 | 0 |
| 0 | 0 |
| 0 | 0 |
| 0 | 0 |
| 386.440564 | 310.817429 |
| 0 | 43.3698738 |
| 0 | 0 |
| 0 | 0 |
| 0 | 0 |
| 0 | 0 |
| 0 | 0 |
| 0 | 0 |
| 0 | 0 |
| 0 | 0 |
| 0 | 137.337934 |
| 0 | 0 |
| 0 | 0 |
| 0 | 0 |

| | |
|---|---|
| 0 | 0 |
| 0 | 0 |
| 3.43502723 | 7.22831229 |
| 0 | 7.22831229 |
| 0 | 0 |
| 0 | 0 |
| 0 | 0 |
| 0 | 0 |
| 0 | 14.4566246 |
| 0 | 0 |
| 0 | 0 |
| 0 | 0 |
| 0 | 0 |
| 0 | 0 |
| 0 | 0 |
| 0 | 0 |
| 0 | 0 |
| 0 | 0 |
| 0 | 0 |
| 0 | 0 |
| 139.118603 | 7.22831229 |
| 0 | 0 |
| 0 | 0 |
| 0 | 0 |
| 0 | 0 |
| 0 | 0 |
| 0 | 0 |
| 0 | 0 |
| 0 | 0 |
| 0 | 0 |
| 0 | 0 |
| 0 | 0 |
| 0 | 0 |
| 0 | 0 |
| 0 | 0 |
| 0 | 86.7397475 |
| 0 | 0 |
| 0 | 0 |
| 0 | 0 |
| 13.7401089 | 0 |
| 0 | 0 |
| 0 | 0 |
| 0 | 0 |

| | |
|---|---|
| 185.491471 | 28.9132492 |
| 0 | 0 |
| 0 | 0 |
| 0 | 0 |
| 0 | 0 |
| 0 | 0 |
| 8.58756808 | 0 |
| 0 | 0 |
| 0 | 0 |
| 0 | 0 |
| 0 | 0 |
| 0 | 0 |
| 0 | 0 |
| 0 | 0 |
| 0 | 0 |
| 0 | 0 |
| 0 | 0 |
| 0 | 0 |
| 0 | 0 |
| 0 | 0 |
| 0 | 0 |
| 0 | 0 |
| 0 | 0 |
| 0 | 0 |
| 0 | 0 |
| 0 | 0 |
| 0 | 0 |
| 0 | 0 |
| 1374.01089 | 1366.15102 |
| 0 | 7.22831229 |
| 0 | 0 |
| 0 | 0 |
| 0 | 0 |
| 0 | 0 |
| 0 | 0 |
| 0 | 0 |
| 0 | 0 |
| 0 | 0 |
| 0 | 0 |
| 0 | 0 |
| 0 | 0 |
| 0 | 0 |
| 0 | 0 |

| | |
|---|---|
| 0 | 0 |
| 0 | 0 |
| 0 | 0 |
| 0 | 0 |
| 0 | 0 |
| 0 | 0 |
| 0 | 0 |
| 0 | 0 |
| 0 | 0 |
| 0 | 14.4566246 |
| 0 | 0 |
| 0 | 0 |
| 20.6101634 | 0 |
| 0 | 0 |
| 0 | 0 |
| 0 | 0 |
| 0 | 0 |
| 0 | 0 |
| 0 | 0 |
| 0 | 0 |
| 0 | 0 |
| 0 | 0 |
| 0 | 0 |
| 0 | 0 |
| 0 | 0 |
| 0 | 0 |
| 0 | 0 |
| 0 | 0 |
| 285.10726 | 195.164432 |
| 0 | 0 |
| 0 | 0 |
| 0 | 0 |
| 0 | 0 |
| 0 | 0 |
| 0 | 0 |
| 0 | 0 |
| 0 | 0 |
| 0 | 0 |
| 0 | 0 |
| 0 | 0 |
| 0 | 0 |
| 0 | 0 |
| 0 | 0 |

| | |
|---|---|
| 0 | 0 |
| 0 | 0 |
| 0 | 0 |
| 0 | 0 |
| 0 | 0 |
| 0 | 0 |
| 0 | 0 |
| 0 | 0 |
| 0 | 0 |
| 0 | 0 |
| 0 | 0 |
| 0 | 0 |
| 0 | 0 |
| 0 | 0 |
| 30.9152451 | 86.7397475 |
| 0 | 7.22831229 |
| 0 | 0 |
| 0 | 0 |
| 0 | 0 |
| 0 | 0 |
| 0 | 0 |
| 0 | 0 |
| 0 | 0 |
| 0 | 0 |
| 0 | 0 |
| 0 | 0 |
| 0 | 0 |
| 0 | 0 |
| 910.282217 | 361.415615 |
| 0 | 0 |
| 108.203358 | 72.2831229 |
| 0 | 0 |
| 63.5480038 | 209.621056 |
| 0 | 0 |
| 0 | 0 |
| 0 | 0 |
| 0 | 0 |
| 0 | 0 |
| 0 | 0 |
| 0 | 0 |
| 0 | 0 |
| 0 | 0 |
| 0 | 0 |

| | |
|---|---|
| 0 | 0 |
| 0 | 0 |
| 0 | 0 |
| 0 | 0 |
| 0 | 0 |
| 0 | 0 |
| 0 | 0 |
| 0 | 0 |
| 0 | 0 |
| 0 | 0 |
| 0 | 0 |
| 0 | 0 |
| 0 | 0 |
| 0 | 0 |
| 0 | 0 |
| 0 | 0 |
| 0 | 0 |
| 0 | 0 |
| 0 | 0 |
| 0 | 0 |
| 0 | 0 |
| 0 | 0 |
| 0 | 0 |
| 15.4576225 | 0 |
| 0 | 0 |
| 0 | 0 |
| 0 | 0 |
| 0 | 0 |
| 0 | 0 |
| 0 | 0 |
| 0 | 0 |
| 0 | 0 |
| 8.58756808 | 0 |
| 0 | 0 |
| 0 | 0 |
| 0 | 0 |
| 0 | 0 |
| 0 | 0 |
| 0 | 0 |
| 0 | 0 |
| 0 | 0 |
| 0 | 0 |
| 0 | 0 |

| | |
|---|---|
| 0 | 0 |
| 0 | 0 |
| 99.6157898 | 7.22831229 |
| 0 | 0 |
| 0 | 0 |
| 0 | 0 |
| 0 | 0 |
| 0 | 0 |
| 0 | 0 |
| 192.361525 | 130.109621 |
| 0 | 28.9132492 |
| 0 | 0 |
| 0 | 0 |
| 0 | 0 |
| 0 | 0 |
| 0 | 0 |
| 0 | 0 |
| 0 | 0 |
| 0 | 0 |
| 0 | 0 |
| 0 | 0 |
| 0 | 0 |
| 0 | 0 |
| 0 | 0 |
| 0 | 0 |
| 0 | 0 |
| 0 | 0 |
| 0 | 0 |
| 0 | 0 |
| 0 | 0 |
| 0 | 0 |
| 0 | 0 |
| 0 | 0 |
| 0 | 7.22831229 |
| 0 | 0 |
| 0 | 0 |
| 0 | 0 |
| 0 | 0 |
| 0 | 0 |
| 0 | 0 |
| 0 | 0 |
| 0 | 0 |
| 0 | 0 |

| | |
|---|---|
| 0 | 0 |
| 0 | 36.1415615 |
| 0 | 0 |
| 0 | 0 |
| 0 | 0 |
| 0 | 0 |
| 0 | 0 |
| 0 | 0 |
| 0 | 0 |
| 0 | 0 |
| 0 | 0 |
| 0 | 0 |
| 3.43502723 | 21.6849369 |
| 0 | 0 |
| 0 | 0 |
| 0 | 0 |
| 118.50844 | 173.479495 |
| 0 | 0 |
| 0 | 0 |
| 0 | 0 |
| 0 | 0 |
| 0 | 0 |
| 0 | 0 |
| 0 | 0 |
| 0 | 0 |
| 0 | 0 |
| 0 | 0 |
| 0 | 0 |
| 0 | 0 |
| 0 | 0 |
| 0 | 0 |
| 0 | 0 |
| 0 | 0 |
| 0 | 0 |
| 0 | 0 |
| 0 | 0 |
| 41.2203268 | 0 |
| 0 | 0 |
| 0 | 0 |
| 0 | 0 |
| 0 | 0 |
| 0 | 0 |
| 0 | 0 |

| | |
|---|---|
| 0 | 0 |
| 0 | 7.22831229 |
| 0 | 0 |
| 0 | 0 |
| 0 | 0 |
| 0 | 0 |
| 0 | 0 |
| 0 | 7.22831229 |
| 0 | 0 |
| 0 | 0 |
| 0 | 0 |
| 0 | 0 |
| 5.15254085 | 0 |
| 0 | 0 |
| 0 | 0 |
| 0 | 0 |
| 0 | 0 |
| 0 | 0 |
| 0 | 0 |
| 321.175046 | 2421.48462 |
| 0 | 0 |
| 310.869965 | 224.077681 |
| 0 | 0 |
| 0 | 7.22831229 |
| 0 | 0 |
| 34.3502723 | 130.109621 |
| 0 | 0 |
| 0 | 0 |
| 0 | 0 |
| 0 | 0 |
| 0 | 0 |
| 0 | 0 |
| 0 | 0 |
| 0 | 0 |
| 0 | 0 |
| 0 | 0 |
| 0 | 0 |
| 0 | 0 |
| 0 | 0 |
| 0 | 0 |
| 0 | 0 |
| 0 | 0 |
| 30.9152451 | 0 |

| | |
|---|---|
| 0 | 0 |
| 0 | 0 |
| 0 | 0 |
| 0 | 0 |
| 0 | 0 |
| 0 | 0 |
| 0 | 0 |
| 0 | 0 |
| 0 | 0 |
| 0 | 0 |
| 0 | 0 |
| 0 | 0 |
| 0 | 0 |
| 0 | 0 |
| 0 | 0 |
| 0 | 0 |
| 0 | 0 |
| 0 | 0 |
| 8.58756808 | 28.9132492 |
| 0 | 0 |
| 0 | 0 |
| 0 | 0 |
| 0 | 0 |
| 0 | 0 |
| 0 | 0 |
| 0 | 0 |
| 0 | 0 |
| 0 | 0 |
| 0 | 0 |
| 0 | 0 |
| 0 | 0 |
| 0 | 0 |
| 0 | 7.22831229 |
| 0 | 0 |
| 0 | 0 |
| 0 | 0 |
| 0 | 0 |
| 61.8304902 | 36.1415615 |
| 0 | 0 |
| 0 | 0 |
| 0 | 0 |
| 0 | 0 |
| 0 | 7.22831229 |

| | |
|---|---|
| 0 | 0 |
| 0 | 0 |
| 0 | 0 |
| 0 | 0 |
| 0 | 0 |
| 0 | 0 |
| 0 | 0 |
| 0 | 7.22831229 |
| 0 | 0 |
| 0 | 0 |
| 0 | 0 |
| 0 | 0 |
| 0 | 0 |
| 0 | 0 |
| 0 | 0 |
| 0 | 0 |
| 0 | 0 |
| 1.71751362 | 0 |
| 0 | 0 |
| 0 | 0 |
| 0 | 0 |
| 0 | 0 |
| 0 | 0 |
| 0 | 7.22831229 |
| 0 | 7.22831229 |
| 0 | 0 |
| 0 | 0 |
| 231.864338 | 72.2831229 |
| 0 | 0 |
| 0 | 0 |
| 0 | 0 |
| 0 | 0 |
| 0 | 7.22831229 |
| 0 | 0 |
| 0 | 0 |
| 0 | 0 |
| 0 | 0 |
| 0 | 0 |
| 0 | 0 |
| 0 | 0 |
| 1.71751362 | 0 |
| 0 | 0 |
| 0 | 21.6849369 |

| | |
|---|---|
| 0 | 0 |
| 0 | 0 |
| 0 | 0 |
| 0 | 0 |
| 0 | 0 |
| 0 | 0 |
| 0 | 0 |
| 0 | 0 |
| 0 | 0 |
| 0 | 28.9132492 |
| 0 | 0 |
| 0 | 0 |
| 0 | 0 |
| 0 | 0 |
| 0 | 0 |
| 0 | 0 |
| 0 | 0 |
| 0 | 0 |
| 0 | 0 |
| 0 | 0 |
| 0 | 0 |
| 0 | 0 |
| 0 | 0 |
| 0 | 57.8264983 |
| 0 | 0 |
| 0 | 0 |
| 0 | 0 |
| 0 | 0 |
| 0 | 14.4566246 |
| 0 | 0 |
| 0 | 0 |
| 0 | 0 |
| 0 | 0 |
| 0 | 0 |
| 0 | 0 |
| 0 | 0 |
| 0 | 0 |
| 0 | 0 |
| 0 | 0 |
| 0 | 0 |
| 0 | 0 |
| 0 | 0 |
| 0 | 0 |

| | |
|---|---|
| 0 | 0 |
| 0 | 21.6849369 |
| 0 | 0 |
| 116.790926 | 267.447555 |
| 0 | 0 |
| 168.316334 | 267.447555 |
| 0 | 0 |
| 0 | 0 |
| 25.7627042 | 0 |
| 0 | 0 |
| 0 | 0 |
| 1.71751362 | 21.6849369 |
| 3.43502723 | 7.22831229 |
| 0 | 0 |
| 0 | 0 |
| 0 | 0 |
| 0 | 0 |
| 0 | 0 |
| 0 | 0 |
| 0 | 0 |
| 0 | 0 |
| 0 | 0 |
| 0 | 0 |
| 0 | 0 |
| 0 | 0 |
| 0 | 0 |
| 0 | 0 |
| 0 | 0 |
| 0 | 0 |
| 0 | 0 |
| 0 | 0 |
| 0 | 0 |
| 0 | 0 |
| 1.71751362 | 0 |
| 0 | 7.22831229 |
| 0 | 0 |
| 0 | 0 |
| 0 | 0 |
| 0 | 0 |
| 0 | 0 |
| 0 | 0 |
| 0 | 0 |
| 0 | 0 |

| | |
|---|---|
| 0 | 0 |
| 0 | 0 |
| 0 | 0 |
| 0 | 0 |
| 3.43502723 | 7.22831229 |
| 0 | 0 |
| 0 | 0 |
| 0 | 0 |
| 0 | 0 |
| 0 | 0 |
| 0 | 14.4566246 |
| 214.689202 | 14.4566246 |
| 0 | 0 |
| 0 | 0 |
| 0 | 0 |
| 0 | 0 |
| 0 | 0 |
| 0 | 7.22831229 |
| 0 | 0 |
| 0 | 0 |
| 0 | 0 |
| 0 | 0 |
| 0 | 0 |
| 0 | 0 |
| 0 | 0 |
| 0 | 0 |
| 0 | 0 |
| 0 | 0 |
| 0 | 0 |
| 0 | 0 |
| 0 | 0 |
| 0 | 0 |
| 0 | 0 |
| 0 | 0 |
| 0 | 0 |
| 0 | 0 |
| 0 | 0 |
| 3.43502723 | 0 |
| 346.937751 | 50.598186 |
| 1.71751362 | 0 |
| 0 | 0 |
| 0 | 0 |

| | |
|---|---|
| 0 | 0 |
| 0 | 0 |
| 0 | 0 |
| 0 | 0 |
| 0 | 0 |
| 0 | 0 |
| 0 | 0 |
| 0 | 0 |
| 0 | 0 |
| 0 | 0 |
| 72.1355719 | 108.424684 |
| 0 | 0 |
| 60.1129766 | 7.22831229 |
| 0 | 0 |
| 0 | 0 |
| 0 | 0 |
| 0 | 0 |
| 139.118603 | 36.1415615 |
| 0 | 0 |
| 0 | 0 |
| 159.728766 | 238.534306 |
| 0 | 0 |
| 0 | 0 |
| 0 | 0 |
| 0 | 0 |
| 0 | 7.22831229 |
| 0 | 0 |
| 0 | 0 |
| 0 | 0 |
| 0 | 0 |
| 0 | 0 |
| 60.1129766 | 7.22831229 |
| 0 | 0 |
| 0 | 0 |
| 0 | 0 |
| 0 | 0 |
| 0 | 0 |
| 0 | 0 |
| 0 | 0 |
| 0 | 0 |
| 0 | 0 |
| 0 | 0 |
| 0 | 0 |

| | |
|---|---|
| 0 | 0 |
| 0 | 0 |
| 0 | 0 |
| 0 | 0 |
| 0 | 0 |
| 0 | 0 |
| 340.067696 | 209.621056 |
| 1.71751362 | 0 |
| 0 | 0 |
| 0 | 0 |
| 0 | 0 |
| 0 | 0 |
| 0 | 0 |
| 0 | 0 |
| 56.6779493 | 101.196372 |
| 0 | 0 |
| 0 | 0 |
| 0 | 0 |
| 0 | 0 |
| 0 | 0 |
| 0 | 0 |
| 0 | 0 |
| 0 | 0 |
| 0 | 7.22831229 |
| 3.43502723 | 0 |
| 0 | 0 |
| 0 | 0 |
| 0 | 0 |
| 0 | 0 |
| 0 | 0 |
| 0 | 0 |
| 0 | 0 |
| 0 | 0 |
| 0 | 0 |
| 0 | 0 |
| 0 | 0 |
| 0 | 0 |
| 0 | 0 |
| 0 | 0 |
| 0 | 0 |
| 0 | 28.9132492 |
| 0 | 0 |
| 6.87005447 | 0 |

| | |
|---|---|
| 0 | 0 |
| 0 | 0 |
| 0 | 0 |
| 0 | 0 |
| 0 | 0 |
| 1142.14655 | 1669.74014 |
| 0 | 0 |
| 0 | 0 |
| 0 | 0 |
| 0 | 0 |
| 0 | 0 |
| 0 | 0 |
| 0 | 0 |
| 0 | 0 |
| 0 | 0 |
| 0 | 0 |
| 0 | 0 |
| 0 | 0 |
| 0 | 0 |
| 0 | 0 |
| 0 | 0 |
| 0 | 0 |
| 0 | 0 |
| 0 | 0 |
| 355.525319 | 14.4566246 |
| 0 | 0 |
| 0 | 0 |
| 0 | 0 |
| 0 | 28.9132492 |
| 0 | 0 |
| 0 | 0 |
| 154.576225 | 28.9132492 |
| 0 | 0 |
| 0 | 0 |
| 0 | 0 |
| 0 | 0 |
| 0 | 0 |
| 0 | 0 |
| 0 | 0 |
| 0 | 0 |
| 0 | 0 |
| 0 | 0 |
| 0 | 0 |

| | |
|---|---|
| 0 | 0 |
| 0 | 0 |
| 0 | 0 |
| 0 | 0 |
| 0 | 0 |
| 0 | 0 |
| 0 | 0 |
| 0 | 0 |
| 0 | 0 |
| 0 | 0 |
| 0 | 0 |
| 0 | 0 |
| 0 | 0 |
| 0 | 0 |
| 0 | 0 |
| 0 | 0 |
| 0 | 0 |
| 53.2429221 | 14.4566246 |
| 0 | 0 |
| 0 | 0 |
| 0 | 0 |
| 0 | 0 |
| 5.15254085 | 0 |
| 0 | 0 |
| 0 | 0 |
| 0 | 0 |
| 29.1977315 | 14.4566246 |
| 0 | 0 |
| 1.71751362 | 0 |
| 0 | 0 |
| 0 | 0 |
| 0 | 0 |
| 0 | 0 |
| 0 | 7.22831229 |
| 0 | 0 |
| 29.1977315 | 0 |
| 0 | 0 |
| 0 | 0 |
| 0 | 0 |
| 1.71751362 | 7.22831229 |
| 1.71751362 | 14.4566246 |
| 0 | 0 |
| 0 | 0 |

| | |
|---|---|
| 0 | 0 |
| 80.72314 | 7.22831229 |
| 51.5254085 | 7.22831229 |
| 0 | 0 |
| 0 | 0 |
| 0 | 0 |
| 0 | 0 |
| 0 | 0 |
| 0 | 0 |
| 0 | 0 |
| 0 | 0 |
| 0 | 0 |
| 0 | 0 |
| 0 | 0 |
| 209.536661 | 36.1415615 |
| 1.71751362 | 0 |
| 0 | 0 |
| 0 | 0 |
| 0 | 0 |
| 0 | 0 |
| 0 | 0 |
| 0 | 0 |
| 0 | 0 |
| 3.43502723 | 0 |
| 0 | 0 |
| 0 | 0 |
| 0 | 0 |
| 0 | 0 |
| 1.71751362 | 43.3698738 |
| 54.9604357 | 7.22831229 |
| 0 | 0 |
| 0 | 0 |
| 0 | 0 |
| 49.8078949 | 72.2831229 |
| 231.864338 | 166.251183 |
| 0 | 0 |
| 0 | 0 |
| 0 | 0 |
| 168.316334 | 122.881309 |
| 0 | 0 |
| 0 | 0 |
| 0 | 0 |
| 0 | 0 |

| | |
|---|---|
| 0 | 0 |
| 0 | 0 |
| 0 | 0 |
| 0 | 0 |
| 0 | 0 |
| 0 | 0 |
| 0 | 0 |
| 0 | 0 |
| 0 | 0 |
| 0 | 0 |
| 0 | 0 |
| 0 | 0 |
| 0 | 0 |
| 0 | 0 |
| 0 | 0 |
| 0 | 0 |
| 0 | 0 |
| 149.423685 | 28.9132492 |
| 0 | 0 |
| 0 | 0 |
| 1.71751362 | 14.4566246 |
| 0 | 7.22831229 |
| 0 | 0 |
| 0 | 0 |
| 0 | 0 |
| 0 | 0 |
| 0 | 0 |
| 0 | 0 |
| 0 | 0 |
| 0 | 0 |
| 139.118603 | 14.4566246 |
| 0 | 0 |
| 0 | 0 |
| 0 | 0 |
| 0 | 0 |
| 0 | 0 |
| 0 | 0 |
| 0 | 0 |
| 53.2429221 | 7.22831229 |
| 0 | 0 |
| 0 | 0 |
| 0 | 0 |
| 0 | 0 |

| | |
|---|---|
| 1.71751362 | 0 |
| 0 | 0 |
| 0 | 7.22831229 |
| 0 | 0 |
| 0 | 0 |
| 0 | 0 |
| 0 | 0 |
| 0 | 0 |
| 0 | 0 |
| 0 | 0 |
| 0 | 0 |
| 0 | 0 |
| 0 | 0 |
| 0 | 0 |
| 0 | 0 |
| 0 | 0 |
| 0 | 0 |
| 0 | 0 |
| 0 | 0 |
| 240.451906 | 93.9680598 |
| 0 | 0 |
| 0 | 0 |
| 0 | 0 |
| 0 | 0 |
| 82.4406536 | 281.904179 |
| 0 | 0 |
| 0 | 0 |
| 0 | 0 |
| 0 | 0 |
| 0 | 0 |
| 0 | 0 |
| 0 | 0 |
| 0 | 0 |
| 0 | 0 |
| 0 | 0 |
| 0 | 0 |
| 0 | 0 |
| 0 | 0 |
| 178.621416 | 144.566246 |
| 0 | 0 |
| 0 | 0 |
| 20.6101634 | 36.1415615 |
| 32.6327587 | 0 |

| | |
|---|---|
| 0 | 0 |
| 1.71751362 | 7.22831229 |
| 0 | 0 |
| 0 | 0 |
| 0 | 0 |
| 0 | 0 |
| 0 | 0 |
| 0 | 0 |
| 158.011253 | 101.196372 |
| 0 | 0 |
| 0 | 0 |
| 0 | 0 |
| 0 | 0 |
| 0 | 0 |
| 0 | 0 |
| 0 | 93.9680598 |
| 0 | 0 |
| 0 | 0 |
| 0 | 0 |
| 0 | 0 |
| 0 | 0 |
| 0 | 0 |
| 0 | 0 |
| 0 | 0 |
| 0 | 0 |
| 0 | 21.6849369 |
| 0 | 0 |
| 0 | 0 |
| 0 | 0 |
| 0 | 0 |
| 0 | 0 |
| 0 | 0 |
| 0 | 0 |
| 10.3050817 | 14.4566246 |
| 0 | 0 |
| 0 | 0 |
| 0 | 0 |
| 0 | 0 |
| 0 | 7.22831229 |
| 0 | 0 |
| 0 | 0 |
| 0 | 0 |
| 0 | 0 |

| | |
|---|---|
| 0 | 0 |
| 0 | 0 |
| 0 | 0 |
| 0 | 0 |
| 0 | 0 |
| 360.677859 | 238.534306 |
| 0 | 0 |
| 305.717424 | 14.4566246 |
| 1.71751362 | 28.9132492 |
| 0 | 0 |
| 0 | 0 |
| 0 | 0 |
| 206.101634 | 151.794558 |
| 0 | 0 |
| 0 | 0 |
| 0 | 0 |
| 0 | 14.4566246 |
| 0 | 0 |
| 0 | 7.22831229 |
| 0 | 0 |
| 0 | 0 |
| 0 | 0 |
| 51.5254085 | 86.7397475 |
| 133.966062 | 36.1415615 |
| 0 | 0 |
| 0 | 0 |
| 0 | 0 |
| 283.389747 | 115.652997 |
| 0 | 0 |
| 0 | 0 |
| 0 | 0 |
| 0 | 0 |
| 0 | 0 |
| 0 | 0 |
| 0 | 0 |
| 0 | 0 |
| 53.2429221 | 65.0548106 |
| 13.7401089 | 303.589116 |
| 0 | 0 |
| 0 | 0 |
| 0 | 0 |
| 0 | 14.4566246 |
| 0 | 0 |

| | |
|---|---|
| 0 | 0 |
| 0 | 0 |
| 0 | 0 |
| 0 | 0 |
| 0 | 0 |
| 0 | 0 |
| 0 | 0 |
| 0 | 7.22831229 |
| 0 | 0 |
| 0 | 0 |
| 0 | 0 |
| 0 | 0 |
| 1.71751362 | 0 |
| 0 | 0 |
| 0 | 0 |
| 0 | 0 |
| 0 | 0 |
| 0 | 0 |
| 0 | 0 |
| 0 | 0 |
| 42.9378404 | 115.652997 |
| 0 | 0 |
| 0 | 0 |
| 0 | 0 |
| 0 | 0 |
| 0 | 0 |
| 8.58756808 | 0 |
| 0 | 0 |
| 0 | 0 |
| 154.576225 | 101.196372 |
| 48.0903813 | 72.2831229 |
| 0 | 0 |
| 0 | 0 |
| 0 | 0 |
| 5.15254085 | 0 |
| 434.530945 | 72.2831229 |
| 0 | 0 |
| 0 | 0 |
| 0 | 0 |
| 0 | 0 |
| 0 | 0 |
| 36.0677859 | 0 |
| 0 | 0 |

| | |
|---|---|
| 0 | 0 |
| 12.0225953 | 7.22831229 |
| 0 | 0 |
| 0 | 0 |
| 0 | 0 |
| 0 | 0 |
| 0 | 50.598186 |
| 0 | 0 |
| 0 | 0 |
| 0 | 0 |
| 0 | 0 |
| 0 | 0 |
| 0 | 7.22831229 |
| 0 | 0 |
| 0 | 0 |
| 0 | 0 |
| 0 | 0 |
| 10.3050817 | 7.22831229 |
| 0 | 0 |
| 0 | 0 |
| 0 | 0 |
| 0 | 0 |
| 0 | 0 |
| 0 | 0 |
| 0 | 0 |
| 0 | 0 |
| 8.58756808 | 0 |
| 68.7005447 | 101.196372 |
| 0 | 0 |
| 0 | 0 |
| 0 | 0 |
| 0 | 0 |
| 0 | 0 |
| 0 | 0 |
| 0 | 0 |
| 0 | 0 |
| 0 | 0 |
| 0 | 0 |
| 0 | 0 |
| 0 | 0 |
| 20.6101634 | 108.424684 |
| 0 | 0 |
| 0 | 0 |

| | |
|---|---|
| 259.344556 | 101.196372 |
| 1.71751362 | 7.22831229 |
| 0 | 0 |
| 0 | 0 |
| 104.768331 | 36.1415615 |
| 195.796552 | 21.6849369 |
| 0 | 0 |
| 0 | 0 |
| 0 | 0 |
| 0 | 0 |
| 0 | 0 |
| 0 | 0 |
| 0 | 0 |
| 0 | 0 |
| 0 | 0 |
| 0 | 0 |
| 0 | 0 |
| 0 | 0 |
| 0 | 0 |
| 0 | 0 |
| 0 | 0 |
| 13.7401089 | 14.4566246 |
| 0 | 0 |
| 0 | 0 |
| 41.2203268 | 14.4566246 |
| 0 | 0 |
| 0 | 0 |
| 0 | 0 |
| 0 | 0 |
| 0 | 0 |
| 0 | 0 |
| 0 | 0 |
| 0 | 0 |
| 0 | 0 |
| 0 | 0 |
| 0 | 0 |
| 0 | 0 |
| 0 | 0 |
| 0 | 0 |
| 0 | 0 |
| 0 | 0 |
| 0 | 0 |
| 37.7852996 | 28.9132492 |

| | |
|---|---|
| 0 | 0 |
| 0 | 0 |
| 79.0056264 | 14.4566246 |
| 0 | 0 |
| 0 | 7.22831229 |
| 0 | 0 |
| 0 | 0 |
| 0 | 0 |
| 0 | 0 |
| 0 | 0 |
| 0 | 0 |
| 0 | 0 |
| 0 | 0 |
| 0 | 0 |
| 0 | 0 |
| 0 | 0 |
| 25.7627042 | 72.2831229 |
| 0 | 0 |
| 159.728766 | 151.794558 |
| 68.7005447 | 57.8264983 |
| 0 | 0 |
| 0 | 0 |
| 0 | 0 |
| 0 | 0 |
| 0 | 0 |
| 0 | 0 |
| 0 | 0 |
| 0 | 0 |
| 0 | 0 |
| 0 | 0 |
| 0 | 0 |
| 0 | 0 |
| 0 | 0 |
| 0 | 0 |
| 0 | 0 |
| 0 | 0 |
| 0 | 0 |
| 0 | 0 |
| 0 | 0 |
| 0 | 0 |
| 0 | 0 |
| 0 | 0 |
| 0 | 0 |

| | |
|---|---|
| 0 | 0 |
| 0 | 0 |
| 0 | 0 |
| 0 | 0 |
| 0 | 0 |
| 0 | 0 |
| 0 | 0 |
| 0 | 7.22831229 |
| 0 | 0 |
| 0 | 0 |
| 1.71751362 | 0 |
| 0 | 0 |
| 0 | 0 |
| 0 | 0 |
| 0 | 0 |
| 0 | 0 |
| 0 | 0 |
| 0 | 0 |
| 0 | 0 |
| 0 | 0 |
| 0 | 0 |
| 0 | 0 |
| 0 | 0 |
| 0 | 0 |
| 0 | 0 |
| 0 | 21.6849369 |
| 39.5028132 | 0 |
| 0 | 7.22831229 |
| 0 | 0 |
| 0 | 0 |
| 0 | 0 |
| 0 | 0 |
| 1.71751362 | 0 |
| 0 | 0 |
| 0 | 0 |
| 0 | 0 |
| 0 | 0 |
| 0 | 0 |
| 0 | 0 |
| 0 | 0 |
| 0 | 0 |
| 0 | 0 |
| 0 | 0 |

| | |
|---|---|
| 427.66089 | 354.187302 |
| 0 | 0 |
| 17.1751362 | 0 |
| 0 | 0 |
| 0 | 0 |
| 0 | 0 |
| 0 | 0 |
| 0 | 0 |
| 0 | 0 |
| 0 | 0 |
| 0 | 21.6849369 |
| 0 | 0 |
| 0 | 0 |
| 0 | 0 |
| 0 | 0 |
| 0 | 0 |
| 0 | 0 |
| 0 | 0 |
| 0 | 0 |
| 0 | 0 |
| 0 | 0 |
| 0 | 0 |
| 214.689202 | 238.534306 |
| 0 | 0 |
| 0 | 0 |
| 0 | 0 |
| 0 | 0 |
| 0 | 0 |
| 0 | 0 |
| 149.423685 | 36.1415615 |
| 0 | 0 |
| 0 | 0 |
| 0 | 0 |
| 303.99991 | 144.566246 |
| 0 | 0 |
| 0 | 0 |
| 70.4180583 | 0 |
| 0 | 0 |
| 0 | 0 |
| 0 | 0 |
| 0 | 0 |
| 3.43502723 | 0 |
| 0 | 0 |

| | |
|---|---|
| 0 | 0 |
| 0 | 0 |
| 0 | 0 |
| 0 | 0 |
| 0 | 0 |
| 3.43502723 | 7.22831229 |
| 0 | 0 |
| 0 | 0 |
| 0 | 0 |
| 0 | 28.9132492 |
| 0 | 0 |
| 0 | 0 |
| 0 | 0 |
| 0 | 0 |
| 0 | 0 |
| 0 | 21.6849369 |
| 12.0225953 | 72.2831229 |
| 0 | 0 |
| 0 | 0 |
| 0 | 0 |
| 0 | 0 |
| 0 | 0 |
| 0 | 0 |
| 0 | 0 |
| 0 | 0 |
| 0 | 0 |
| 0 | 0 |
| 1.71751362 | 0 |
| 0 | 0 |
| 0 | 0 |
| 0 | 0 |
| 0 | 0 |
| 0 | 0 |
| 0 | 0 |
| 0 | 0 |
| 0 | 0 |
| 0 | 0 |
| 0 | 0 |
| 0 | 0 |
| 0 | 0 |
| 0 | 0 |
| 0 | 0 |
| 6.87005447 | 0 |

| | |
|---|---|
| 0 | 0 |
| 197.514066 | 296.360804 |
| 0 | 0 |
| 0 | 0 |
| 0 | 0 |
| 0 | 0 |
| 0 | 0 |
| 0 | 0 |
| 0 | 0 |
| 0 | 0 |
| 158.011253 | 101.196372 |
| 0 | 0 |
| 0 | 0 |
| 0 | 0 |
| 0 | 0 |
| 0 | 0 |
| 0 | 0 |
| 0 | 7.22831229 |
| 0 | 0 |
| 0 | 0 |
| 0 | 0 |
| 0 | 0 |
| 0 | 0 |
| 0 | 0 |
| 0 | 0 |
| 0 | 0 |
| 0 | 0 |
| 0 | 0 |
| 0 | 0 |
| 0 | 0 |
| 0 | 0 |
| 0 | 0 |
| 0 | 0 |
| 0 | 0 |
| 0 | 0 |
| 0 | 0 |
| 0 | 0 |
| 0 | 0 |
| 0 | 0 |
| 0 | 0 |
| 0 | 0 |
| 0 | 0 |
| 0 | 0 |
| 0 | 0 |

| | |
|---|---|
| 0 | 0 |
| 127.096008 | 14.4566246 |
| 0 | 0 |
| 0 | 0 |
| 0 | 0 |
| 0 | 0 |
| 1.71751362 | 0 |
| 0 | 0 |
| 0 | 0 |
| 0 | 0 |
| 185.491471 | 224.077681 |
| 0 | 86.7397475 |
| 0 | 0 |
| 0 | 0 |
| 0 | 0 |
| 0 | 0 |
| 0 | 0 |
| 77.2881127 | 43.3698738 |
| 77.2881127 | 43.3698738 |
| 5.15254085 | 0 |
| 0 | 0 |
| 0 | 0 |
| 0 | 0 |
| 1.71751362 | 0 |
| 91.0282217 | 274.675867 |
| 0 | 0 |
| 128.813521 | 28.9132492 |
| 0 | 0 |
| 128.813521 | 93.9680598 |
| 0 | 0 |
| 0 | 0 |
| 0 | 0 |
| 0 | 0 |
| 0 | 0 |
| 0 | 0 |
| 0 | 0 |
| 10.3050817 | 0 |
| 0 | 0 |
| 1.71751362 | 14.4566246 |
| 0 | 0 |
| 0 | 0 |
| 0 | 0 |
| 0 | 0 |

| | |
|---|---|
| 0 | 0 |
| 0 | 0 |
| 0 | 0 |
| 0 | 0 |
| 0 | 0 |
| 0 | 0 |
| 0 | 0 |
| 0 | 0 |
| 0 | 0 |
| 0 | 0 |
| 286.824774 | 50.598186 |
| 0 | 0 |
| 298.847369 | 173.479495 |
| 0 | 0 |
| 0 | 0 |
| 0 | 0 |
| 0 | 0 |
| 0 | 0 |
| 0 | 0 |
| 0 | 0 |
| 41.2203268 | 50.598186 |
| 0 | 0 |
| 0 | 0 |
| 0 | 0 |
| 0 | 0 |
| 0 | 50.598186 |
| 0 | 0 |
| 0 | 0 |
| 0 | 0 |
| 0 | 0 |
| 176.903902 | 21.6849369 |
| 0 | 0 |
| 0 | 0 |
| 1.71751362 | 0 |
| 0 | 0 |
| 0 | 14.4566246 |
| 0 | 0 |
| 0 | 0 |
| 0 | 0 |
| 0 | 0 |
| 0 | 0 |
| 0 | 0 |
| 0 | 0 |

| | |
|---|---|
| 0 | 0 |
| 0 | 0 |
| 0 | 0 |
| 0 | 0 |
| 0 | 0 |
| 0 | 14.4566246 |
| 0 | 0 |
| 0 | 0 |
| 0 | 0 |
| 0 | 0 |
| 0 | 65.0548106 |
| 0 | 7.22831229 |
| 0 | 0 |
| 0 | 0 |
| 0 | 0 |
| 0 | 0 |
| 0 | 0 |
| 0 | 0 |
| 1.71751362 | 0 |
| 0 | 0 |
| 0 | 21.6849369 |
| 0 | 0 |
| 259.344556 | 43.3698738 |
| 0 | 0 |
| 0 | 0 |
| 0 | 0 |
| 0 | 0 |
| 0 | 0 |
| 0 | 0 |
| 0 | 0 |
| 0 | 0 |
| 0 | 0 |
| 0 | 0 |
| 0 | 0 |
| 0 | 0 |
| 0 | 0 |
| 0 | 0 |
| 0 | 93.9680598 |
| 0 | 0 |
| 0 | 0 |
| 0 | 0 |
| 0 | 0 |
| 262.779583 | 166.251183 |

| | |
|---|---|
| 0 | 0 |
| 0 | 0 |
| 0 | 0 |
| 0 | 0 |
| 322.89256 | 621.634857 |
| 0 | 0 |
| 121.943467 | 72.2831229 |
| 0 | 0 |
| 0 | 0 |
| 0 | 0 |
| 0 | 0 |
| 419.073322 | 216.849369 |
| 0 | 0 |
| 0 | 0 |
| 0 | 0 |
| 0 | 0 |
| 0 | 0 |
| 0 | 0 |
| 0 | 0 |
| 0 | 7.22831229 |
| 1.71751362 | 0 |
| 0 | 0 |
| 0 | 0 |
| 0 | 0 |
| 0 | 0 |
| 0 | 0 |
| 0 | 0 |
| 0 | 0 |
| 0 | 7.22831229 |
| 0 | 0 |
| 582.237116 | 166.251183 |
| 0 | 0 |
| 0 | 0 |
| 235.299365 | 14.4566246 |
| 0 | 7.22831229 |
| 56.6779493 | 122.881309 |
| 0 | 0 |
| 0 | 0 |
| 0 | 0 |
| 0 | 0 |
| 0 | 0 |
| 0 | 0 |
| 0 | 0 |

| | |
|---|---|
| 0 | 0 |
| 0 | 0 |
| 0 | 0 |
| 12.0225953 | 0 |
| 0 | 0 |
| 0 | 0 |
| 0 | 0 |
| 0 | 0 |
| 1.71751362 | 21.6849369 |
| 113.355899 | 122.881309 |
| 0 | 0 |
| 0 | 0 |
| 0 | 0 |
| 0 | 0 |
| 101.333303 | 43.3698738 |
| 0 | 0 |
| 1.71751362 | 50.598186 |
| 0 | 0 |
| 0 | 0 |
| 0 | 0 |
| 0 | 0 |
| 0 | 0 |
| 0 | 0 |
| 0 | 0 |
| 1.71751362 | 7.22831229 |
| 0 | 0 |
| 0 | 0 |
| 0 | 0 |
| 0 | 0 |
| 0 | 0 |
| 61.8304902 | 50.598186 |
| 0 | 0 |
| 0 | 0 |
| 0 | 0 |
| 0 | 0 |
| 15.4576225 | 65.0548106 |
| 0 | 0 |
| 0 | 0 |
| 0 | 0 |
| 0 | 7.22831229 |
| 0 | 0 |
| 0 | 0 |
| 0 | 0 |

| | |
|---|---|
| 0 | 0 |
| 0 | 0 |
| 0 | 0 |
| 0 | 0 |
| 0 | 0 |
| 0 | 0 |
| 0 | 0 |
| 0 | 0 |
| 0 | 0 |
| 0 | 0 |
| 168.316334 | 108.424684 |
| 0 | 0 |
| 0 | 0 |
| 0 | 0 |
| 0 | 0 |
| 0 | 0 |
| 0 | 0 |
| 0 | 0 |
| 0 | 7.22831229 |
| 0 | 0 |
| 0 | 0 |
| 171.751362 | 122.881309 |
| 0 | 0 |
| 0 | 0 |
| 0 | 0 |
| 127.096008 | 93.9680598 |
| 0 | 0 |
| 0 | 0 |
| 0 | 0 |
| 0 | 0 |
| 0 | 0 |
| 0 | 0 |
| 0 | 0 |
| 0 | 0 |
| 0 | 0 |
| 0 | 7.22831229 |
| 0 | 0 |
| 0 | 7.22831229 |
| 0 | 0 |
| 0 | 0 |
| 0 | 0 |
| 0 | 0 |
| 0 | 0 |

| | |
|---|---|
| 60.1129766 | 50.598186 |
| 0 | 0 |
| 0 | 0 |
| 0 | 0 |
| 0 | 0 |
| 0 | 0 |
| 0 | 0 |
| 0 | 0 |
| 0 | 0 |
| 0 | 0 |
| 0 | 0 |
| 34.3502723 | 93.9680598 |
| 0 | 0 |
| 0 | 0 |
| 0 | 0 |
| 0 | 0 |
| 1.71751362 | 0 |
| 0 | 0 |
| 0 | 0 |
| 0 | 0 |
| 0 | 0 |
| 0 | 0 |
| 0 | 0 |
| 0 | 0 |
| 0 | 0 |
| 0 | 0 |
| 0 | 0 |
| 0 | 0 |
| 0 | 0 |
| 0 | 7.22831229 |
| 0 | 0 |
| 0 | 0 |
| 0 | 0 |
| 111.638385 | 303.589116 |
| 0 | 0 |
| 0 | 0 |
| 0 | 0 |
| 0 | 0 |
| 0 | 0 |
| 0 | 0 |
| 0 | 0 |
| 0 | 0 |
| 0 | 28.9132492 |

| | |
|---|---|
| 0 | 7.22831229 |
| 0 | 0 |
| 0 | 0 |
| 0 | 0 |
| 0 | 0 |
| 0 | 0 |
| 0 | 0 |
| 0 | 0 |
| 0 | 0 |
| 0 | 0 |
| 0 | 0 |
| 5.15254085 | 0 |
| 0 | 0 |
| 0 | 0 |
| 72.1355719 | 43.3698738 |
| 0 | 0 |
| 29.1977315 | 0 |
| 0 | 0 |
| 0 | 0 |
| 0 | 0 |
| 0 | 0 |
| 0 | 0 |
| 0 | 0 |
| 0 | 0 |
| 0 | 0 |
| 0 | 0 |
| 0 | 0 |
| 1.71751362 | 0 |
| 0 | 0 |
| 0 | 0 |
| 0 | 0 |
| 51.5254085 | 0 |
| 109.920871 | 173.479495 |
| 0 | 0 |
| 0 | 0 |
| 152.858712 | 0 |
| 0 | 0 |
| 0 | 0 |
| 0 | 0 |
| 0 | 0 |
| 0 | 0 |
| 0 | 0 |
| 0 | 0 |

| | |
|---|---|
| 0 | 0 |
| 0 | 0 |
| 0 | 7.22831229 |
| 0 | 0 |
| 0 | 0 |
| 0 | 0 |
| 0 | 0 |
| 0 | 0 |
| 0 | 0 |
| 0 | 0 |
| 0 | 0 |
| 0 | 0 |
| 0 | 0 |
| 0 | 0 |
| 0 | 0 |
| 0 | 0 |
| 0 | 0 |
| 0 | 0 |
| 0 | 14.4566246 |
| 0 | 0 |
| 0 | 0 |
| 0 | 0 |
| 0 | 0 |
| 0 | 0 |
| 0 | 0 |
| 0 | 0 |
| 0 | 0 |
| 0 | 0 |
| 0 | 0 |
| 0 | 0 |
| 0 | 0 |
| 0 | 0 |
| 0 | 7.22831229 |
| 0 | 0 |
| 1214.28213 | 643.319794 |
| 0 | 0 |
| 0 | 0 |
| 0 | 0 |
| 72.1355719 | 180.707807 |
| 0 | 0 |
| 0 | 0 |
| 0 | 0 |
| 0 | 0 |

| | |
|---|---|
| 0 | 0 |
| 0 | 0 |
| 0 | 0 |
| 0 | 0 |
| 0 | 0 |
| 34.3502723 | 0 |
| 0 | 0 |
| 0 | 7.22831229 |
| 0 | 0 |
| 417.355809 | 325.274053 |
| 0 | 0 |
| 144.271144 | 108.424684 |
| 0 | 0 |
| 0 | 0 |
| 0 | 0 |
| 0 | 0 |
| 0 | 0 |
| 0 | 0 |
| 0 | 72.2831229 |
| 0 | 0 |
| 0 | 0 |
| 0 | 0 |
| 0 | 0 |
| 0 | 7.22831229 |
| 0 | 0 |
| 0 | 0 |
| 0 | 0 |
| 51.5254085 | 57.8264983 |
| 0 | 0 |
| 0 | 0 |
| 0 | 0 |
| 0 | 0 |
| 0 | 0 |
| 0 | 0 |
| 3.43502723 | 0 |
| 0 | 0 |
| 0 | 0 |
| 0 | 0 |
| 0 | 0 |
| 0 | 0 |
| 0 | 0 |
| 0 | 0 |
| 0 | 0 |

| | |
|---|---|
| 0 | 0 |
| 0 | 0 |
| 178.621416 | 137.337934 |
| 0 | 0 |
| 0 | 0 |
| 0 | 0 |
| 0 | 0 |
| 0 | 0 |
| 0 | 14.4566246 |
| 0 | 0 |
| 0 | 0 |
| 0 | 50.598186 |
| 0 | 0 |
| 0 | 0 |
| 0 | 0 |
| 0 | 0 |
| 0 | 0 |
| 0 | 0 |
| 0 | 0 |
| 0 | 0 |
| 0 | 0 |
| 22.327677 | 0 |
| 0 | 0 |
| 0 | 0 |
| 0 | 0 |
| 0 | 0 |
| 0 | 0 |
| 0 | 0 |
| 0 | 0 |
| 0 | 0 |
| 1.71751362 | 0 |
| 0 | 0 |
| 0 | 0 |
| 0 | 0 |
| 0 | 0 |
| 170.033848 | 216.849369 |
| 1.71751362 | 0 |
| 0 | 0 |
| 132.248548 | 7.22831229 |
| 0 | 0 |
| 0 | 0 |
| 0 | 0 |
| 0 | 0 |

| | |
|---|---|
| 0 | 0 |
| 42.9378404 | 21.6849369 |
| 49.8078949 | 0 |
| 0 | 0 |
| 0 | 0 |
| 0 | 0 |
| 0 | 28.9132492 |
| 0 | 7.22831229 |
| 0 | 0 |
| 0 | 0 |
| 0 | 0 |
| 0 | 0 |
| 0 | 0 |
| 0 | 0 |
| 0 | 0 |
| 0 | 0 |
| 0 | 0 |
| 0 | 0 |
| 0 | 0 |
| 0 | 0 |
| 0 | 0 |
| 0 | 0 |
| 0 | 0 |
| 0 | 0 |
| 0 | 0 |
| 0 | 0 |
| 0 | 0 |
| 0 | 0 |
| 0 | 0 |
| 0 | 0 |
| 0 | 14.4566246 |
| 0 | 0 |
| 0 | 0 |
| 0 | 0 |
| 0 | 0 |
| 0 | 0 |
| 0 | 0 |
| 0 | 0 |
| 0 | 0 |
| 0 | 0 |
| 0 | 0 |
| 0 | 0 |
| 0 | 0 |

| | |
|---|---|
| 0 | 0 |
| 0 | 0 |
| 0 | 0 |
| 0 | 0 |
| 0 | 0 |
| 0 | 0 |
| 0 | 0 |
| 154.576225 | 21.6849369 |
| 0 | 0 |
| 0 | 0 |
| 68.7005447 | 202.392744 |
| 0 | 0 |
| 0 | 0 |
| 3.43502723 | 36.1415615 |
| 0 | 0 |
| 30.9152451 | 151.794558 |
| 0 | 0 |
| 0 | 0 |
| 0 | 0 |
| 0 | 0 |
| 0 | 0 |
| 0 | 0 |
| 0 | 0 |
| 0 | 0 |
| 0 | 0 |
| 0 | 0 |
| 0 | 0 |
| 18.8926498 | 7.22831229 |
| 0 | 0 |
| 0 | 0 |
| 0 | 0 |
| 63.5480038 | 101.196372 |
| 0 | 0 |
| 0 | 0 |
| 0 | 0 |
| 0 | 0 |
| 0 | 0 |
| 0 | 0 |
| 0 | 0 |
| 0 | 0 |
| 0 | 0 |
| 0 | 0 |
| 0 | 0 |

| | |
|---|---|
| 0 | 0 |
| 70.4180583 | 36.1415615 |
| 0 | 0 |
| 0 | 0 |
| 0 | 0 |
| 0 | 0 |
| 0 | 0 |
| 0 | 0 |
| 0 | 0 |
| 0 | 0 |
| 0 | 0 |
| 0 | 0 |
| 0 | 21.6849369 |
| 0 | 0 |
| 0 | 0 |
| 0 | 7.22831229 |
| 0 | 0 |
| 0 | 0 |
| 0 | 0 |
| 0 | 0 |
| 0 | 0 |
| 0 | 0 |
| 0 | 14.4566246 |
| 0 | 0 |
| 0 | 0 |
| 6.87005447 | 368.643927 |
| 0 | 0 |
| 3.43502723 | 0 |
| 0 | 0 |
| 0 | 0 |
| 0 | 0 |
| 0 | 0 |
| 0 | 0 |
| 0 | 14.4566246 |
| 0 | 0 |
| 0 | 0 |
| 0 | 0 |
| 0 | 0 |
| 0 | 14.4566246 |
| 0 | 0 |
| 0 | 0 |
| 82.4406536 | 43.3698738 |
| 0 | 0 |

| | |
|---|---|
| 0 | 0 |
| 0 | 0 |
| 0 | 0 |
| 0 | 0 |
| 0 | 0 |
| 0 | 0 |
| 0 | 0 |
| 0 | 0 |
| 0 | 0 |
| 152.858712 | 115.652997 |
| 0 | 0 |
| 0 | 0 |
| 0 | 0 |
| 0 | 0 |
| 0 | 0 |
| 0 | 0 |
| 0 | 0 |
| 1.71751362 | 36.1415615 |
| 0 | 0 |
| 0 | 0 |
| 649.220147 | 216.849369 |
| 0 | 0 |
| 1.71751362 | 0 |
| 0 | 0 |
| 0 | 0 |
| 0 | 0 |
| 0 | 0 |
| 0 | 0 |
| 0 | 0 |
| 0 | 0 |
| 0 | 0 |
| 0 | 7.22831229 |
| 0 | 0 |
| 0 | 0 |
| 0 | 0 |
| 0 | 0 |
| 0 | 0 |
| 0 | 0 |
| 590.824684 | 368.643927 |
| 0 | 0 |
| 0 | 0 |
| 0 | 0 |
| 0 | 0 |

| | |
|---|---|
| 0 | 0 |
| 0 | 0 |
| 0 | 0 |
| 0 | 0 |
| 30.9152451 | 43.3698738 |
| 0 | 0 |
| 0 | 0 |
| 0 | 0 |
| 0 | 0 |
| 0 | 0 |
| 0 | 0 |
| 0 | 0 |
| 1.71751362 | 0 |
| 137.401089 | 14.4566246 |
| 0 | 0 |
| 0 | 0 |
| 0 | 0 |
| 0 | 0 |
| 0 | 0 |
| 0 | 0 |
| 0 | 0 |
| 0 | 0 |
| 0 | 0 |
| 3.43502723 | 7.22831229 |
| 0 | 0 |
| 211.254175 | 303.589116 |
| 0 | 0 |
| 89.3107081 | 0 |
| 0 | 0 |
| 0 | 0 |
| 0 | 0 |
| 0 | 0 |
| 0 | 0 |
| 0 | 0 |
| 0 | 0 |
| 0 | 0 |
| 0 | 0 |
| 0 | 0 |
| 0 | 0 |
| 0 | 0 |
| 0 | 0 |
| 0 | 0 |
| 0 | 0 |

| | |
|---|---|
| 29.1977315 | 0 |
| 0 | 0 |
| 0 | 0 |
| 0 | 0 |
| 0 | 0 |
| 0 | 0 |
| 0 | 7.22831229 |
| 0 | 0 |
| 0 | 0 |
| 0 | 0 |
| 0 | 0 |
| 0 | 0 |
| 0 | 0 |
| 0 | 0 |
| 0 | 0 |
| 0 | 0 |
| 0 | 0 |
| 0 | 0 |
| 0 | 0 |
| 0 | 0 |
| 0 | 0 |
| 0 | 0 |
| 0 | 0 |
| 0 | 0 |
| 0 | 0 |
| 0 | 0 |
| 0 | 0 |
| 1.71751362 | 0 |
| 39.5028132 | 36.1415615 |
| 0 | 0 |
| 0 | 0 |
| 1.71751362 | 0 |
| 0 | 0 |
| 0 | 0 |
| 1.71751362 | 0 |
| 0 | 0 |
| 0 | 0 |
| 0 | 0 |
| 0 | 0 |
| 0 | 0 |
| 0 | 0 |
| 235.299365 | 621.634857 |
| 0 | 0 |

| | |
|---|---|
| 0 | 0 |
| 0 | 0 |
| 0 | 0 |
| 0 | 0 |
| 0 | 0 |
| 0 | 0 |
| 97.8982761 | 209.621056 |
| 0 | 0 |
| 0 | 0 |
| 0 | 0 |
| 0 | 0 |
| 0 | 0 |
| 0 | 0 |
| 0 | 0 |
| 0 | 0 |
| 0 | 0 |
| 0 | 0 |
| 0 | 0 |
| 0 | 0 |
| 0 | 0 |
| 0 | 0 |
| 0 | 7.22831229 |
| 0 | 7.22831229 |
| 0 | 0 |
| 0 | 0 |
| 0 | 0 |
| 0 | 0 |
| 0 | 0 |
| 0 | 0 |
| 0 | 0 |
| 0 | 0 |
| 0 | 0 |
| 0 | 0 |
| 0 | 0 |
| 0 | 0 |
| 0 | 0 |
| 0 | 0 |
| 0 | 0 |
| 0 | 0 |
| 0 | 0 |
| 0 | 0 |
| 0 | 0 |
| 0 | 0 |

| | |
|---|---|
| 0 | 0 |
| 0 | 7.22831229 |
| 0 | 0 |
| 5.15254085 | 0 |
| 0 | 0 |
| 0 | 0 |
| 27.4802179 | 0 |
| 0 | 0 |
| 0 | 0 |
| 0 | 0 |
| 0 | 0 |
| 0 | 0 |
| 0 | 0 |
| 0 | 0 |
| 0 | 0 |
| 0 | 0 |
| 0 | 0 |
| 0 | 0 |
| 0 | 0 |
| 0 | 0 |
| 0 | 0 |
| 0 | 0 |
| 0 | 0 |
| 0 | 0 |
| 0 | 0 |
| 0 | 21.6849369 |
| 0 | 0 |
| 0 | 0 |
| 0 | 0 |
| 0 | 0 |
| 206.101634 | 137.337934 |
| 0 | 0 |
| 0 | 0 |
| 0 | 0 |
| 0 | 0 |
| 0 | 0 |
| 0 | 0 |
| 0 | 0 |
| 0 | 0 |
| 0 | 0 |
| 0 | 0 |
| 39.5028132 | 93.9680598 |
| 0 | 0 |

| | |
|---|---|
| 0 | 0 |
| 0 | 0 |
| 164.881307 | 21.6849369 |
| 0 | 0 |
| 0 | 0 |
| 0 | 0 |
| 0 | 0 |
| 0 | 0 |
| 0 | 0 |
| 0 | 0 |
| 0 | 0 |
| 0 | 0 |
| 0 | 0 |
| 42.9378404 | 57.8264983 |
| 0 | 0 |
| 0 | 0 |
| 0 | 0 |
| 0 | 0 |
| 0 | 7.22831229 |
| 0 | 0 |
| 0 | 0 |
| 0 | 0 |
| 0 | 0 |
| 0 | 0 |
| 1.71751362 | 0 |
| 0 | 0 |
| 0 | 0 |
| 0 | 0 |
| 41.2203268 | 0 |
| 0 | 0 |
| 0 | 0 |
| 0 | 0 |
| 0 | 0 |
| 0 | 0 |
| 0 | 0 |
| 0 | 0 |
| 0 | 7.22831229 |
| 0 | 0 |
| 0 | 0 |
| 0 | 0 |
| 32.6327587 | 14.4566246 |
| 0 | 0 |
| 0 | 0 |

| | |
|---|---|
| 291.977315 | 122.881309 |
| 42.9378404 | 7.22831229 |
| 0 | 0 |
| 0 | 0 |
| 0 | 0 |
| 0 | 0 |
| 0 | 0 |
| 0 | 7.22831229 |
| 1.71751362 | 0 |
| 0 | 0 |
| 109.920871 | 274.675867 |
| 0 | 0 |
| 30.9152451 | 0 |
| 0 | 0 |
| 0 | 0 |
| 0 | 0 |
| 1.71751362 | 0 |
| 0 | 0 |
| 0 | 0 |
| 0 | 14.4566246 |
| 0 | 0 |
| 0 | 0 |
| 0 | 0 |
| 0 | 0 |
| 0 | 0 |
| 0 | 0 |
| 0 | 0 |
| 0 | 0 |
| 0 | 0 |
| 0 | 0 |
| 0 | 0 |
| 0 | 0 |
| 0 | 0 |
| 0 | 0 |
| 0 | 0 |
| 0 | 0 |
| 0 | 0 |
| 15.4576225 | 0 |
| 0 | 0 |
| 58.395463 | 0 |
| 22.327677 | 0 |
| 0 | 0 |
| 54.9604357 | 115.652997 |

| | |
|---|---|
| 0 | 0 |
| 0 | 101.196372 |
| 0 | 0 |
| 0 | 0 |
| 0 | 0 |
| 0 | 0 |
| 0 | 0 |
| 0 | 0 |
| 0 | 0 |
| 0 | 0 |
| 0 | 0 |
| 0 | 0 |
| 0 | 0 |
| 0 | 0 |
| 0 | 0 |
| 0 | 0 |
| 0 | 0 |
| 0 | 0 |
| 0 | 0 |
| 204.38412 | 115.652997 |
| 0 | 0 |
| 760.858532 | 874.625787 |
| 0 | 0 |
| 0 | 0 |
| 0 | 0 |
| 0 | 7.22831229 |
| 250.756988 | 166.251183 |
| 0 | 0 |
| 0 | 0 |
| 0 | 0 |
| 1167.90926 | 361.415615 |
| 0 | 0 |
| 0 | 0 |
| 0 | 0 |
| 0 | 0 |
| 0 | 0 |
| 0 | 0 |
| 70.4180583 | 86.7397475 |
| 0 | 0 |
| 0 | 0 |
| 0 | 0 |
| 0 | 0 |
| 0 | 0 |

| | |
|---|---|
| 1983.72823 | 20181.4479 |
| 0 | 0 |
| 0 | 0 |
| 1.71751362 | 0 |
| 0 | 0 |
| 0 | 0 |
| 1.71751362 | 0 |
| 0 | 0 |
| 0 | 0 |
| 135.683576 | 21.6849369 |
| 0 | 0 |
| 0 | 0 |
| 0 | 0 |
| 0 | 0 |
| 0 | 0 |
| 0 | 0 |
| 0 | 0 |
| 0 | 0 |
| 39.5028132 | 0 |
| 0 | 0 |
| 0 | 0 |
| 0 | 7.22831229 |
| 0 | 0 |
| 0 | 0 |
| 0 | 0 |
| 0 | 0 |
| 0 | 0 |
| 0 | 0 |
| 0 | 0 |
| 0 | 0 |
| 0 | 0 |
| 0 | 0 |
| 0 | 0 |
| 0 | 0 |
| 0 | 0 |
| 0 | 0 |
| 0 | 0 |
| 0 | 0 |
| 0 | 0 |
| 0 | 0 |
| 3.43502723 | 0 |
| 8.58756808 | 7.22831229 |
| 0 | 0 |

| | |
|---|---|
| 0 | 7.22831229 |
| 0 | 0 |
| 0 | 0 |
| 0 | 0 |
| 0 | 0 |
| 0 | 0 |
| 0 | 0 |
| 0 | 0 |
| 0 | 0 |
| 0 | 0 |
| 0 | 0 |
| 0 | 0 |
| 0 | 0 |
| 0 | 0 |
| 0 | 0 |
| 0 | 0 |
| 0 | 7.22831229 |
| 0 | 0 |
| 0 | 0 |
| 0 | 0 |
| 0 | 0 |
| 0 | 0 |
| 0 | 0 |
| 207.819148 | 130.109621 |
| 0 | 0 |
| 0 | 0 |
| 0 | 0 |
| 0 | 0 |
| 0 | 0 |
| 0 | 7.22831229 |
| 0 | 0 |
| 0 | 0 |
| 0 | 0 |
| 0 | 0 |
| 0 | 0 |
| 0 | 0 |
| 0 | 0 |
| 0 | 0 |
| 0 | 0 |
| 0 | 0 |
| 0 | 0 |
| 0 | 0 |
| 0 | 0 |

| | |
|---|---|
| 0 | 0 |
| 0 | 0 |
| 0 | 0 |
| 0 | 0 |
| 0 | 0 |
| 0 | 0 |
| 0 | 0 |
| 0 | 0 |
| 0 | 0 |
| 0 | 0 |
| 0 | 0 |
| 0 | 0 |
| 0 | 0 |
| 0 | 0 |
| 0 | 7.22831229 |
| 0 | 0 |
| 0 | 0 |
| 0 | 0 |
| 0 | 0 |
| 0 | 0 |
| 0 | 0 |
| 0 | 0 |
| 0 | 0 |
| 0 | 0 |
| 0 | 0 |
| 0 | 0 |
| 0 | 0 |
| 0 | 0 |
| 0 | 0 |
| 91.0282217 | 151.794558 |
| 0 | 0 |
| 0 | 0 |
| 0 | 0 |
| 0 | 0 |
| 0 | 0 |
| 0 | 0 |
| 0 | 7.22831229 |
| 0 | 7.22831229 |
| 0 | 0 |
| 0 | 0 |
| 0 | 0 |
| 0 | 0 |
| 0 | 0 |

| | |
|---|---|
| 0 | 0 |
| 0 | 0 |
| 0 | 0 |
| 0 | 0 |
| 0 | 0 |
| 207.819148 | 7.22831229 |
| 0 | 0 |
| 1.71751362 | 14.4566246 |
| 0 | 0 |
| 0 | 0 |
| 0 | 0 |
| 0 | 0 |
| 0 | 0 |
| 0 | 0 |
| 0 | 0 |
| 0 | 0 |
| 30.9152451 | 108.424684 |
| 0 | 0 |
| 0 | 0 |
| 0 | 0 |
| 0 | 0 |
| 0 | 0 |
| 0 | 0 |
| 0 | 0 |
| 0 | 0 |
| 0 | 0 |
| 0 | 0 |
| 0 | 0 |
| 0 | 0 |
| 163.163794 | 0 |
| 44.655354 | 57.8264983 |
| 0 | 0 |
| 41.2203268 | 72.2831229 |
| 0 | 7.22831229 |
| 0 | 0 |
| 687.005447 | 368.643927 |
| 3.43502723 | 28.9132492 |
| 0 | 0 |
| 0 | 0 |
| 0 | 0 |
| 0 | 0 |
| 0 | 0 |
| 0 | 0 |

| | |
|---|---|
| 0 | 0 |
| 0 | 0 |
| 0 | 0 |
| 0 | 0 |
| 0 | 0 |
| 0 | 0 |
| 0 | 0 |
| 0 | 0 |
| 0 | 0 |
| 0 | 0 |
| 0 | 0 |
| 0 | 0 |
| 147.706171 | 151.794558 |
| 97.8982761 | 195.164432 |
| 0 | 0 |
| 0 | 0 |
| 6.87005447 | 0 |
| 0 | 0 |
| 0 | 0 |
| 0 | 0 |
| 0 | 0 |
| 0 | 0 |
| 0 | 0 |
| 0 | 0 |
| 606.282307 | 325.274053 |
| 152.858712 | 7.22831229 |
| 0 | 0 |
| 0 | 0 |
| 0 | 0 |
| 0 | 0 |
| 0 | 0 |
| 0 | 0 |
| 0 | 0 |
| 104.768331 | 72.2831229 |
| 0 | 0 |
| 0 | 0 |
| 0 | 0 |
| 0 | 0 |
| 0 | 0 |
| 0 | 0 |
| 0 | 0 |
| 0 | 0 |
| 0 | 0 |

| | |
|---|---|
| 0 | 0 |
| 0 | 0 |
| 0 | 14.4566246 |
| 0 | 0 |
| 0 | 0 |
| 0 | 0 |
| 0 | 0 |
| 0 | 0 |
| 0 | 0 |
| 0 | 0 |
| 0 | 0 |
| 0 | 0 |
| 84.1581672 | 7.22831229 |
| 0 | 0 |
| 0 | 0 |
| 0 | 0 |
| 0 | 0 |
| 0 | 0 |
| 20.6101634 | 0 |
| 0 | 0 |
| 0 | 0 |
| 6.87005447 | 14.4566246 |
| 0 | 0 |
| 1.71751362 | 36.1415615 |
| 0 | 0 |
| 0 | 0 |
| 0 | 0 |
| 0 | 0 |
| 0 | 0 |
| 0 | 0 |
| 0 | 0 |
| 0 | 0 |
| 0 | 0 |
| 0 | 0 |
| 0 | 21.6849369 |
| 0 | 0 |
| 0 | 0 |
| 0 | 0 |
| 0 | 0 |
| 0 | 0 |
| 0 | 0 |
| 0 | 0 |
| 0 | 0 |

| | |
|---|---|
| 0 | 7.22831229 |
| 0 | 0 |
| 0 | 0 |
| 0 | 0 |
| 0 | 0 |
| 0 | 0 |
| 0 | 0 |
| 0 | 0 |
| 0 | 0 |
| 1.71751362 | 0 |
| 0 | 0 |
| 0 | 0 |
| 0 | 0 |
| 0 | 0 |
| 0 | 28.9132492 |
| 0 | 0 |
| 0 | 0 |
| 0 | 0 |
| 1.71751362 | 0 |
| 24.0451906 | 101.196372 |
| 0 | 7.22831229 |
| 135.683576 | 0 |
| 0 | 0 |
| 0 | 0 |
| 0 | 0 |
| 0 | 0 |
| 0 | 0 |
| 0 | 0 |
| 0 | 0 |
| 0 | 0 |
| 0 | 0 |
| 0 | 0 |
| 0 | 0 |
| 0 | 0 |
| 0 | 0 |
| 0 | 0 |
| 0 | 0 |
| 0 | 0 |
| 0 | 0 |
| 0 | 0 |
| 0 | 0 |
| 0 | 0 |
| 0 | 0 |

| | |
|---|---|
| 0 | 0 |
| 18.8926498 | 137.337934 |
| 0 | 0 |
| 0 | 0 |
| 0 | 0 |
| 0 | 0 |
| 0 | 0 |
| 0 | 0 |
| 1.71751362 | 65.0548106 |
| 0 | 0 |
| 0 | 0 |
| 0 | 0 |
| 0 | 7.22831229 |
| 32.6327587 | 43.3698738 |
| 0 | 0 |
| 0 | 0 |
| 0 | 0 |
| 0 | 0 |
| 13.7401089 | 79.5114352 |
| 0 | 0 |
| 0 | 0 |
| 91.0282217 | 281.904179 |
| 0 | 0 |
| 0 | 7.22831229 |
| 0 | 0 |
| 0 | 0 |
| 0 | 0 |
| 0 | 0 |
| 0 | 0 |
| 0 | 0 |
| 0 | 0 |
| 0 | 0 |
| 0 | 43.3698738 |
| 32.6327587 | 7.22831229 |
| 0 | 0 |
| 0 | 0 |
| 0 | 0 |
| 0 | 0 |
| 0 | 0 |
| 0 | 0 |
| 0 | 0 |
| 1.71751362 | 0 |
| 0 | 0 |

| | |
|---|---|
| 0 | 0 |
| 0 | 0 |
| 0 | 0 |
| 0 | 0 |
| 0 | 0 |
| 0 | 0 |
| 0 | 0 |
| 0 | 0 |
| 0 | 0 |
| 0 | 0 |
| 0 | 7.22831229 |
| 5.15254085 | 28.9132492 |
| 0 | 0 |
| 0 | 0 |
| 0 | 0 |
| 0 | 0 |
| 0 | 0 |
| 0 | 0 |
| 0 | 0 |
| 554.756898 | 375.872239 |
| 0 | 0 |
| 0 | 0 |
| 0 | 0 |
| 0 | 0 |
| 0 | 0 |
| 0 | 0 |
| 0 | 0 |
| 0 | 0 |
| 0 | 0 |
| 0 | 0 |
| 0 | 0 |
| 0 | 0 |
| 49.8078949 | 252.99093 |
| 0 | 7.22831229 |
| 0 | 0 |
| 340.067696 | 144.566246 |
| 0 | 0 |
| 0 | 0 |
| 0 | 0 |
| 0 | 7.22831229 |
| 0 | 0 |
| 0 | 0 |
| 0 | 0 |

| | |
|---|---|
| 185.491471 | 231.305993 |
| 0 | 0 |
| 0 | 0 |
| 3.43502723 | 21.6849369 |
| 0 | 7.22831229 |
| 0 | 0 |
| 0 | 0 |
| 0 | 0 |
| 0 | 0 |
| 0 | 0 |
| 0 | 0 |
| 0 | 0 |
| 1099.20871 | 1799.84976 |
| 0 | 0 |
| 0 | 0 |
| 0 | 0 |
| 0 | 0 |
| 24.0451906 | 7.22831229 |
| 0 | 0 |
| 0 | 0 |
| 99.6157898 | 368.643927 |
| 0 | 0 |
| 0 | 0 |
| 0 | 0 |
| 0 | 0 |
| 0 | 0 |
| 0 | 0 |
| 0 | 0 |
| 0 | 14.4566246 |
| 0 | 0 |
| 0 | 0 |
| 3.43502723 | 7.22831229 |
| 0 | 0 |
| 0 | 0 |
| 0 | 0 |
| 0 | 0 |
| 273.084665 | 1272.18296 |
| 0 | 0 |
| 0 | 0 |
| 0 | 0 |
| 0 | 0 |
| 0 | 0 |
| 0 | 0 |

| | |
|---|---|
| 0 | 0 |
| 0 | 0 |
| 0 | 0 |
| 0 | 0 |
| 0 | 0 |
| 0 | 28.9132492 |
| 0 | 0 |
| 0 | 0 |
| 0 | 0 |
| 0 | 0 |
| 0 | 0 |
| 0 | 0 |
| 0 | 0 |
| 0 | 0 |
| 0 | 0 |
| 0 | 0 |
| 0 | 0 |
| 192.361525 | 209.621056 |
| 0 | 0 |
| 13.7401089 | 0 |
| 0 | 0 |
| 0 | 0 |
| 0 | 0 |
| 0 | 0 |
| 0 | 0 |
| 0 | 0 |
| 0 | 0 |
| 0 | 0 |
| 0 | 0 |
| 0 | 0 |
| 0 | 0 |
| 0 | 0 |
| 0 | 0 |
| 1.71751362 | 0 |
| 0 | 0 |
| 0 | 0 |
| 0 | 0 |
| 0 | 0 |
| 0 | 0 |
| 0 | 0 |
| 178.621416 | 130.109621 |
| 0 | 0 |
| 61.8304902 | 36.1415615 |

| | |
|---|---|
| 0 | 0 |
| 0 | 0 |
| 70.4180583 | 0 |
| 0 | 0 |
| 0 | 0 |
| 87.5931944 | 28.9132492 |
| 87.5931944 | 28.9132492 |
| 0 | 0 |
| 0 | 0 |
| 0 | 0 |
| 0 | 0 |
| 54.9604357 | 36.1415615 |
| 0 | 0 |
| 0 | 0 |
| 0 | 0 |
| 0 | 0 |
| 0 | 0 |
| 0 | 0 |
| 0 | 0 |
| 20.6101634 | 0 |
| 0 | 0 |
| 0 | 0 |
| 0 | 0 |
| 0 | 0 |
| 0 | 0 |
| 79.0056264 | 0 |
| 0 | 0 |
| 0 | 0 |
| 0 | 0 |
| 0 | 0 |
| 281.672233 | 7.22831229 |
| 0 | 0 |
| 6.87005447 | 50.598186 |
| 0 | 0 |
| 0 | 0 |
| 0 | 0 |
| 0 | 0 |
| 300.564883 | 469.840299 |
| 0 | 0 |
| 0 | 0 |
| 1.71751362 | 0 |
| 0 | 0 |
| 0 | 0 |

| | |
|---|---|
| 0 | 7.22831229 |
| 0 | 0 |
| 0 | 0 |
| 22.327677 | 79.5114352 |
| 0 | 0 |
| 0 | 0 |
| 604.564793 | 72.2831229 |
| 0 | 0 |
| 1.71751362 | 0 |
| 6.87005447 | 0 |
| 0 | 7.22831229 |
| 0 | 0 |
| 0 | 0 |
| 0 | 0 |
| 0 | 0 |
| 0 | 0 |
| 0 | 0 |
| 0 | 0 |
| 0 | 0 |
| 0 | 0 |
| 0 | 0 |
| 0 | 0 |
| 0 | 0 |
| 0 | 0 |
| 0 | 0 |
| 0 | 0 |
| 0 | 0 |
| 0 | 0 |
| 0 | 0 |
| 0 | 0 |
| 0 | 0 |
| 0 | 0 |
| 0 | 0 |
| 0 | 0 |
| 0 | 0 |
| 0 | 0 |
| 0 | 0 |
| 0 | 0 |
| 0 | 0 |
| 0 | 0 |
| 0 | 0 |
| 0 | 0 |
| 0 | 0 |

| | |
|---|---|
| 0 | 0 |
| 0 | 0 |
| 0 | 0 |
| 0 | 0 |
| 0 | 0 |
| 0 | 0 |
| 1.71751362 | 0 |
| 0 | 0 |
| 0 | 0 |
| 0 | 7.22831229 |
| 0 | 0 |
| 0 | 0 |
| 13.7401089 | 202.392744 |
| 0 | 0 |
| 0 | 0 |
| 0 | 0 |
| 0 | 0 |
| 0 | 0 |
| 0 | 0 |
| 0 | 0 |
| 0 | 0 |
| 0 | 0 |
| 0 | 0 |
| 0 | 0 |
| 0 | 0 |
| 46.3728676 | 0 |
| 0 | 0 |
| 0 | 0 |
| 0 | 0 |
| 0 | 0 |
| 0 | 0 |
| 0 | 0 |
| 0 | 0 |
| 0 | 0 |
| 115.073412 | 21.6849369 |
| 0 | 0 |
| 139.118603 | 1452.89077 |
| 0 | 28.9132492 |
| 0 | 0 |
| 0 | 0 |
| 0 | 0 |
| 638.915065 | 296.360804 |
| 0 | 0 |

| | |
|---|---|
| 0 | 0 |
| 0 | 0 |
| 0 | 0 |
| 0 | 0 |
| 0 | 0 |
| 0 | 0 |
| 70.4180583 | 7.22831229 |
| 0 | 0 |
| 0 | 7.22831229 |
| 0 | 0 |
| 0 | 0 |
| 0 | 0 |
| 0 | 0 |
| 0 | 0 |
| 0 | 0 |
| 0 | 0 |
| 0 | 72.2831229 |
| 0 | 0 |
| 0 | 0 |
| 0 | 0 |
| 0 | 0 |
| 27.4802179 | 0 |
| 8.58756808 | 0 |
| 0 | 0 |
| 0 | 0 |
| 0 | 0 |
| 0 | 0 |
| 0 | 0 |
| 0 | 0 |
| 0 | 7.22831229 |
| 0 | 0 |
| 0 | 0 |
| 726.50826 | 173.479495 |
| 0 | 0 |
| 0 | 0 |
| 0 | 0 |
| 0 | 0 |
| 0 | 14.4566246 |
| 0 | 0 |
| 0 | 0 |
| 0 | 0 |
| 0 | 7.22831229 |
| 75.5705991 | 101.196372 |

| | |
|---|---|
| 0 | 0 |
| 0 | 0 |
| 0 | 0 |
| 0 | 0 |
| 0 | 0 |
| 0 | 0 |
| 0 | 0 |
| 0 | 0 |
| 0 | 0 |
| 0 | 0 |
| 0 | 0 |
| 0 | 0 |
| 712.768151 | 390.328864 |
| 0 | 0 |
| 0 | 0 |
| 0 | 0 |
| 0 | 0 |
| 0 | 0 |
| 0 | 0 |
| 0 | 0 |
| 0 | 0 |
| 0 | 0 |
| 0 | 0 |
| 0 | 0 |
| 3.43502723 | 0 |
| 0 | 0 |
| 0 | 0 |
| 0 | 0 |
| 0 | 0 |
| 0 | 0 |
| 10.3050817 | 86.7397475 |
| 0 | 0 |
| 0 | 0 |
| 0 | 0 |
| 0 | 0 |
| 0 | 0 |
| 0 | 0 |
| 0 | 0 |
| 0 | 0 |
| 0 | 0 |
| 0 | 0 |
| 0 | 0 |
| 0 | 0 |

| | |
|---|---|
| 113.355899 | 7.22831229 |
| 0 | 0 |
| 0 | 0 |
| 0 | 0 |
| 0 | 0 |
| 0 | 0 |
| 0 | 0 |
| 0 | 0 |
| 0 | 7.22831229 |
| 202.666607 | 122.881309 |
| 1.71751362 | 0 |
| 0 | 0 |
| 0 | 0 |
| 0 | 0 |
| 0 | 0 |
| 0 | 0 |
| 0 | 0 |
| 0 | 0 |
| 0 | 0 |
| 0 | 0 |
| 0 | 0 |
| 0 | 7.22831229 |
| 0 | 0 |
| 70.4180583 | 0 |
| 0 | 0 |
| 0 | 0 |
| 0 | 0 |
| 0 | 0 |
| 0 | 0 |
| 149.423685 | 173.479495 |
| 51.5254085 | 296.360804 |
| 1.71751362 | 0 |
| 0 | 0 |
| 0 | 0 |
| 0 | 0 |
| 0 | 0 |
| 0 | 0 |
| 0 | 0 |
| 0 | 0 |
| 0 | 28.9132492 |
| 340.067696 | 202.392744 |
| 1.71751362 | 0 |
| 104.768331 | 36.1415615 |

| | |
|---|---|
| 0 | 0 |
| 0 | 0 |
| 0 | 0 |
| 0 | 0 |
| 0 | 0 |
| 0 | 0 |
| 0 | 0 |
| 0 | 0 |
| 0 | 0 |
| 0 | 0 |
| 0 | 0 |
| 0 | 0 |
| 1.71751362 | 0 |
| 0 | 0 |
| 0 | 0 |
| 0 | 0 |
| 0 | 0 |
| 0 | 0 |
| 0 | 0 |
| 0 | 0 |
| 0 | 0 |
| 0 | 0 |
| 0 | 0 |
| 12.0225953 | 0 |
| 0 | 0 |
| 0 | 0 |
| 73.8530855 | 7.22831229 |
| 0 | 0 |
| 0 | 0 |
| 0 | 0 |
| 0 | 0 |
| 0 | 0 |
| 0 | 0 |
| 0 | 0 |
| 0 | 0 |
| 0 | 0 |
| 0 | 0 |
| 0 | 0 |
| 0 | 36.1415615 |
| 0 | 0 |
| 0 | 0 |
| 0 | 0 |
| 0 | 0 |

| | |
|---|---|
| 0 | 0 |
| 0 | 0 |
| 0 | 0 |
| 0 | 0 |
| 0 | 0 |
| 0 | 0 |
| 0 | 0 |
| 0 | 0 |
| 0 | 0 |
| 6.87005447 | 0 |
| 0 | 0 |
| 1.71751362 | 7.22831229 |
| 0 | 0 |
| 0 | 0 |
| 0 | 0 |
| 0 | 0 |
| 0 | 0 |
| 0 | 0 |
| 3.43502723 | 14.4566246 |
| 0 | 0 |
| 0 | 14.4566246 |
| 0 | 0 |
| 0 | 0 |
| 109.920871 | 151.794558 |
| 0 | 0 |
| 0 | 0 |
| 0 | 0 |
| 0 | 0 |
| 0 | 0 |
| 0 | 0 |
| 0 | 0 |
| 0 | 0 |
| 0 | 0 |
| 0 | 0 |
| 0 | 0 |
| 0 | 0 |
| 0 | 0 |
| 0 | 0 |
| 0 | 0 |
| 0 | 0 |
| 0 | 0 |
| 0 | 0 |
| 0 | 0 |

| | |
|---|---|
| 0 | 0 |
| 0 | 0 |
| 0 | 7.22831229 |
| 0 | 0 |
| 0 | 0 |
| 0 | 0 |
| 0 | 0 |
| 0 | 0 |
| 30.9152451 | 0 |
| 0 | 0 |
| 0 | 0 |
| 0 | 7.22831229 |
| 0 | 0 |
| 0 | 0 |
| 0 | 0 |
| 0 | 0 |
| 0 | 0 |
| 0 | 0 |
| 0 | 0 |
| 0 | 0 |
| 0 | 0 |
| 0 | 0 |
| 0 | 0 |
| 0 | 0 |
| 0 | 0 |
| 0 | 0 |
| 0 | 0 |
| 0 | 0 |
| 1.71751362 | 0 |
| 0 | 0 |
| 1.71751362 | 0 |
| 0 | 28.9132492 |
| 0 | 0 |
| 0 | 0 |
| 0 | 0 |
| 0 | 0 |
| 34.3502723 | 65.0548106 |
| 0 | 0 |
| 0 | 0 |
| 0 | 0 |
| 0 | 0 |
| 0 | 0 |
| 0 | 0 |

| | |
|---|---|
| 0 | 0 |
| 0 | 0 |
| 0 | 0 |
| 0 | 0 |
| 0 | 0 |
| 0 | 0 |
| 0 | 0 |
| 22.327677 | 50.598186 |
| 0 | 0 |
| 0 | 0 |
| 63.5480038 | 339.730678 |
| 0 | 0 |
| 0 | 0 |
| 0 | 0 |
| 0 | 0 |
| 182.056443 | 122.881309 |
| 0 | 0 |
| 0 | 0 |
| 0 | 0 |
| 0 | 0 |
| 0 | 0 |
| 0 | 0 |
| 0 | 0 |
| 283.389747 | 159.02287 |
| 0 | 0 |
| 63.5480038 | 115.652997 |
| 0 | 0 |
| 82.4406536 | 43.3698738 |
| 0 | 0 |
| 37.7852996 | 0 |
| 0 | 0 |
| 0 | 0 |
| 0 | 0 |
| 0 | 0 |
| 0 | 0 |
| 0 | 0 |
| 0 | 0 |
| 0 | 0 |
| 0 | 0 |
| 0 | 0 |
| 22.327677 | 14.4566246 |
| 5.15254085 | 7.22831229 |
| 0 | 0 |

| | |
|---|---|
| 0 | 0 |
| 0 | 0 |
| 0 | 0 |
| 0 | 0 |
| 0 | 0 |
| 0 | 0 |
| 0 | 0 |
| 0 | 0 |
| 0 | 0 |
| 17.1751362 | 0 |
| 0 | 0 |
| 0 | 0 |
| 0 | 0 |
| 0 | 0 |
| 0 | 0 |
| 0 | 0 |
| 0 | 0 |
| 0 | 0 |
| 0 | 0 |
| 0 | 0 |
| 34.3502723 | 7.22831229 |
| 5.15254085 | 7.22831229 |
| 0 | 0 |
| 0 | 0 |
| 0 | 0 |
| 123.66098 | 21.6849369 |
| 3.43502723 | 0 |
| 0 | 0 |
| 0 | 0 |
| 0 | 0 |
| 183.773957 | 419.242113 |
| 0 | 0 |
| 0 | 0 |
| 0 | 0 |
| 0 | 0 |
| 0 | 0 |
| 0 | 0 |
| 0 | 0 |
| 0 | 0 |
| 3.43502723 | 7.22831229 |
| 113.355899 | 86.7397475 |
| 0 | 7.22831229 |
| 0 | 0 |

| | |
|---|---|
| 446.55354 | 224.077681 |
| 0 | 0 |
| 108.203358 | 65.0548106 |
| 0 | 0 |
| 187.208984 | 809.570977 |
| 0 | 0 |
| 0 | 0 |
| 0 | 0 |
| 0 | 0 |
| 0 | 0 |
| 0 | 0 |
| 0 | 0 |
| 0 | 0 |
| 0 | 0 |
| 0 | 0 |
| 0 | 0 |
| 0 | 0 |
| 0 | 0 |
| 0 | 0 |
| 0 | 0 |
| 0 | 43.3698738 |
| 101.333303 | 50.598186 |
| 0 | 0 |
| 0 | 0 |
| 0 | 7.22831229 |
| 0 | 0 |
| 0 | 0 |
| 0 | 0 |
| 0 | 0 |
| 0 | 0 |
| 0 | 50.598186 |
| 0 | 0 |
| 0 | 0 |
| 0 | 0 |
| 0 | 0 |
| 0 | 0 |
| 0 | 0 |
| 0 | 0 |
| 0 | 0 |
| 0 | 0 |
| 0 | 0 |
| 0 | 0 |
| 0 | 0 |

| | |
|---|---|
| 0 | 0 |
| 0 | 0 |
| 0 | 0 |
| 0 | 0 |
| 0 | 0 |
| 3.43502723 | 0 |
| 80.72314 | 7.22831229 |
| 65.2655174 | 144.566246 |
| 0 | 0 |
| 0 | 0 |
| 123.66098 | 245.762618 |
| 0 | 0 |
| 0 | 0 |
| 0 | 0 |
| 0 | 0 |
| 0 | 0 |
| 0 | 0 |
| 17.1751362 | 375.872239 |
| 0 | 0 |
| 0 | 0 |
| 0 | 0 |
| 0 | 0 |
| 0 | 0 |
| 0 | 0 |
| 0 | 0 |
| 0 | 0 |
| 0 | 0 |
| 0 | 0 |
| 0 | 0 |
| 0 | 0 |
| 0 | 0 |
| 0 | 0 |
| 0 | 0 |
| 103.050817 | 21.6849369 |
| 0 | 0 |
| 0 | 0 |
| 0 | 0 |
| 0 | 0 |
| 0 | 0 |
| 0 | 0 |
| 94.4632489 | 7.22831229 |
| 0 | 0 |
| 0 | 0 |

| | |
|---|---|
| 0 | 0 |
| 0 | 0 |
| 0 | 0 |
| 0 | 0 |
| 305.717424 | 231.305993 |
| 1.71751362 | 0 |
| 0 | 0 |
| 0 | 0 |
| 0 | 0 |
| 0 | 0 |
| 0 | 0 |
| 0 | 0 |
| 0 | 0 |
| 0 | 0 |
| 0 | 0 |
| 0 | 0 |
| 0 | 0 |
| 0 | 0 |
| 109.920871 | 7.22831229 |
| 0 | 0 |
| 12.0225953 | 0 |
| 0 | 0 |
| 0 | 0 |
| 0 | 0 |
| 0 | 0 |
| 0 | 0 |
| 0 | 0 |
| 0 | 0 |
| 0 | 0 |
| 0 | 0 |
| 0 | 0 |
| 0 | 0 |
| 0 | 0 |
| 0 | 0 |
| 0 | 0 |
| 0 | 0 |
| 0 | 0 |
| 0 | 0 |
| 0 | 0 |
| 36.0677859 | 65.0548106 |
| 1.71751362 | 0 |
| 0 | 0 |
| 39.5028132 | 0 |

| | |
|---|---|
| 0 | 0 |
| 0 | 0 |
| 0 | 0 |
| 0 | 0 |
| 0 | 0 |
| 0 | 0 |
| 0 | 0 |
| 180.33893 | 187.93612 |
| 79.0056264 | 216.849369 |
| 0 | 0 |
| 106.485844 | 166.251183 |
| 0 | 0 |
| 0 | 0 |
| 0 | 0 |
| 0 | 0 |
| 0 | 0 |
| 0 | 0 |
| 0 | 0 |
| 0 | 0 |
| 0 | 0 |
| 94.4632489 | 151.794558 |
| 0 | 0 |
| 0 | 0 |
| 0 | 0 |
| 0 | 0 |
| 0 | 0 |
| 0 | 0 |
| 0 | 0 |
| 0 | 0 |
| 0 | 0 |
| 0 | 0 |
| 0 | 0 |
| 0 | 0 |
| 0 | 0 |
| 0 | 0 |
| 0 | 0 |
| 0 | 36.1415615 |
| 0 | 0 |
| 0 | 0 |
| 0 | 0 |
| 0 | 0 |
| 5.15254085 | 43.3698738 |
| 0 | 0 |

| | |
|---|---|
| 0 | 0 |
| 0 | 0 |
| 0 | 0 |
| 0 | 0 |
| 0 | 0 |
| 0 | 0 |
| 0 | 0 |
| 27.4802179 | 93.9680598 |
| 0 | 0 |
| 0 | 0 |
| 0 | 7.22831229 |
| 0 | 0 |
| 0 | 0 |
| 0 | 0 |
| 0 | 50.598186 |
| 0 | 0 |
| 0 | 0 |
| 0 | 0 |
| 0 | 0 |
| 0 | 0 |
| 0 | 0 |
| 0 | 7.22831229 |
| 0 | 0 |
| 0 | 0 |
| 0 | 0 |
| 0 | 0 |
| 0 | 0 |
| 6.87005447 | 224.077681 |
| 142.55363 | 122.881309 |
| 0 | 0 |
| 0 | 0 |
| 0 | 0 |
| 0 | 0 |
| 8.58756808 | 195.164432 |
| 0 | 0 |
| 0 | 0 |
| 0 | 0 |
| 0 | 0 |
| 0 | 0 |
| 0 | 0 |
| 0 | 0 |
| 0 | 0 |
| 0 | 0 |

| | |
|---|---|
| 0 | 0 |
| 0 | 0 |
| 0 | 0 |
| 0 | 0 |
| 0 | 0 |
| 1.71751362 | 0 |
| 0 | 0 |
| 0 | 0 |
| 8.58756808 | 0 |
| 63.5480038 | 50.598186 |
| 0 | 0 |
| 0 | 0 |
| 0 | 0 |
| 0 | 0 |
| 0 | 0 |
| 0 | 0 |
| 65.2655174 | 36.1415615 |
| 0 | 0 |
| 0 | 0 |
| 0 | 0 |
| 0 | 0 |
| 204.38412 | 151.794558 |
| 0 | 0 |
| 0 | 0 |
| 0 | 0 |
| 0 | 0 |
| 75.5705991 | 7.22831229 |
| 106.485844 | 21.6849369 |
| 0 | 0 |
| 1.71751362 | 7.22831229 |
| 0 | 0 |
| 0 | 0 |
| 0 | 0 |
| 0 | 0 |
| 0 | 0 |
| 0 | 0 |
| 63.5480038 | 14.4566246 |
| 0 | 0 |
| 0 | 0 |
| 0 | 7.22831229 |
| 0 | 0 |
| 103.050817 | 101.196372 |
| 0 | 0 |

| | |
|---|---|
| 0 | 0 |
| 0 | 0 |
| 0 | 0 |
| 0 | 0 |
| 0 | 0 |
| 0 | 0 |
| 0 | 0 |
| 0 | 0 |
| 0 | 0 |
| 0 | 0 |
| 0 | 0 |
| 0 | 0 |
| 0 | 0 |
| 0 | 0 |
| 0 | 0 |
| 0 | 0 |
| 0 | 0 |
| 0 | 0 |
| 0 | 0 |
| 18.8926498 | 43.3698738 |
| 0 | 0 |
| 0 | 0 |
| 0 | 0 |
| 3.43502723 | 28.9132492 |
| 0 | 0 |
| 0 | 0 |
| 0 | 0 |
| 144.271144 | 50.598186 |
| 1.71751362 | 7.22831229 |
| 0 | 7.22831229 |
| 0 | 0 |
| 0 | 0 |
| 0 | 0 |
| 0 | 0 |
| 0 | 0 |
| 0 | 0 |
| 0 | 0 |
| 0 | 0 |
| 0 | 0 |
| 0 | 0 |
| 0 | 0 |
| 0 | 0 |
| 0 | 0 |

| | |
|---|---|
| 0 | 0 |
| 0 | 0 |
| 0 | 0 |
| 867.344376 | 831.255914 |
| 0 | 0 |
| 242.16942 | 65.0548106 |
| 0 | 0 |
| 0 | 0 |
| 0 | 0 |
| 0 | 0 |
| 121.943467 | 0 |
| 0 | 0 |
| 0 | 0 |
| 0 | 0 |
| 0 | 0 |
| 0 | 0 |
| 0 | 0 |
| 0 | 0 |
| 0 | 0 |
| 0 | 0 |
| 0 | 0 |
| 254.192015 | 72.2831229 |
| 0 | 0 |
| 0 | 0 |
| 1.71751362 | 0 |
| 0 | 0 |
| 0 | 0 |
| 0 | 0 |
| 0 | 0 |
| 0 | 0 |
| 0 | 0 |
| 0 | 0 |
| 0 | 0 |
| 0 | 0 |
| 0 | 0 |
| 163.163794 | 115.652997 |
| 0 | 0 |
| 0 | 7.22831229 |
| 0 | 0 |
| 0 | 0 |
| 0 | 0 |
| 0 | 0 |
| 115.073412 | 332.502365 |

| | |
|---|---|
| 0 | 0 |
| 0 | 0 |
| 0 | 0 |
| 0 | 0 |
| 0 | 0 |
| 0 | 0 |
| 0 | 0 |
| 0 | 0 |
| 0 | 0 |
| 0 | 0 |
| 0 | 0 |
| 0 | 0 |
| 0 | 0 |
| 0 | 0 |
| 0 | 0 |
| 0 | 0 |
| 0 | 0 |
| 0 | 0 |
| 0 | 0 |
| 0 | 0 |
| 0 | 0 |
| 0 | 0 |
| 147.706171 | 21.6849369 |
| 0 | 0 |
| 0 | 0 |
| 0 | 0 |
| 0 | 0 |
| 0 | 0 |
| 0 | 0 |
| 0 | 0 |
| 0 | 0 |
| 0 | 0 |
| 0 | 0 |
| 0 | 0 |
| 0 | 0 |
| 0 | 0 |
| 73.8530855 | 14.4566246 |
| 0 | 0 |
| 0 | 0 |
| 0 | 0 |
| 0 | 0 |
| 0 | 0 |
| 68.7005447 | 7.22831229 |

| | |
|---|---|
| 133.966062 | 28.9132492 |
| 0 | 0 |
| 218.124229 | 7.22831229 |
| 0 | 0 |
| 0 | 0 |
| 0 | 0 |
| 0 | 0 |
| 0 | 0 |
| 0 | 0 |
| 0 | 0 |
| 0 | 0 |
| 0 | 0 |
| 0 | 0 |
| 0 | 0 |
| 0 | 0 |
| 0 | 0 |
| 0 | 0 |
| 0 | 0 |
| 0 | 0 |
| 0 | 0 |
| 0 | 0 |
| 0 | 0 |
| 0 | 0 |
| 61.8304902 | 21.6849369 |
| 79.0056264 | 130.109621 |
| 0 | 0 |
| 0 | 0 |
| 0 | 0 |
| 0 | 0 |
| 0 | 0 |
| 0 | 0 |
| 0 | 7.22831229 |
| 0 | 0 |
| 0 | 0 |
| 82.4406536 | 7.22831229 |
| 0 | 0 |
| 0 | 0 |
| 0 | 0 |
| 0 | 0 |
| 0 | 0 |
| 0 | 0 |
| 0 | 0 |
| 0 | 0 |

| | |
|---|---|
| 0 | 0 |
| 0 | 0 |
| 0 | 0 |
| 0 | 0 |
| 0 | 7.22831229 |
| 0 | 0 |
| 0 | 0 |
| 0 | 0 |
| 0 | 0 |
| 0 | 0 |
| 0 | 0 |
| 0 | 0 |
| 0 | 0 |
| 0 | 0 |
| 0 | 0 |
| 946.350003 | 303.589116 |
| 0 | 14.4566246 |
| 0 | 0 |
| 0 | 0 |
| 0 | 0 |
| 0 | 0 |
| 0 | 0 |
| 0 | 0 |
| 0 | 0 |
| 0 | 0 |
| 1.71751362 | 0 |
| 0 | 0 |
| 206.101634 | 14.4566246 |
| 0 | 0 |
| 0 | 0 |
| 0 | 0 |
| 0 | 0 |
| 0 | 0 |
| 0 | 0 |
| 0 | 0 |
| 0 | 0 |
| 0 | 0 |
| 65.2655174 | 14.4566246 |
| 0 | 0 |
| 0 | 0 |
| 0 | 0 |
| 0 | 0 |
| 0 | 0 |

| | |
|---|---|
| 0 | 0 |
| 0 | 0 |
| 0 | 86.7397475 |
| 0 | 0 |
| 0 | 0 |
| 0 | 21.6849369 |
| 1.71751362 | 0 |
| 0 | 0 |
| 0 | 0 |
| 0 | 0 |
| 0 | 0 |
| 0 | 0 |
| 0 | 0 |
| 0 | 0 |
| 0 | 0 |
| 0 | 0 |
| 0 | 0 |
| 0 | 0 |
| 0 | 0 |
| 0 | 0 |
| 0 | 0 |
| 0 | 0 |
| 0 | 0 |
| 0 | 0 |
| 0 | 0 |
| 0 | 0 |
| 0 | 0 |
| 0 | 0 |
| 0 | 0 |
| 15.4576225 | 14.4566246 |
| 0 | 0 |
| 0 | 0 |
| 0 | 0 |
| 0 | 0 |
| 0 | 0 |
| 0 | 0 |
| 0 | 0 |
| 0 | 0 |
| 73.8530855 | 28.9132492 |
| 0 | 0 |
| 142.55363 | 0 |
| 0 | 101.196372 |
| 0 | 0 |

| | |
|---|---|
| 0 | 0 |
| 0 | 0 |
| 0 | 0 |
| 0 | 0 |
| 0 | 0 |
| 0 | 0 |
| 46.3728676 | 0 |
| 5.15254085 | 0 |
| 0 | 0 |
| 94.4632489 | 7.22831229 |
| 0 | 0 |
| 0 | 0 |
| 0 | 0 |
| 127.096008 | 0 |
| 0 | 0 |
| 0 | 0 |
| 0 | 0 |
| 0 | 0 |
| 0 | 0 |
| 0 | 0 |
| 0 | 0 |
| 0 | 0 |
| 0 | 0 |
| 0 | 0 |
| 0 | 0 |
| 0 | 0 |
| 0 | 0 |
| 0 | 0 |
| 0 | 0 |
| 0 | 0 |
| 75.5705991 | 0 |
| 0 | 0 |
| 0 | 0 |
| 0 | 0 |
| 0 | 0 |
| 0 | 0 |
| 0 | 0 |
| 0 | 0 |
| 0 | 0 |
| 0 | 0 |
| 0 | 0 |
| 0 | 0 |
| 0 | 0 |

| | |
|---|---|
| 0 | 0 |
| 0 | 0 |
| 0 | 0 |
| 0 | 0 |
| 0 | 0 |
| 89.3107081 | 7.22831229 |
| 0 | 0 |
| 0 | 0 |
| 0 | 0 |
| 0 | 0 |
| 0 | 0 |
| 0 | 0 |
| 0 | 0 |
| 477.468785 | 679.461355 |
| 0 | 0 |
| 0 | 0 |
| 0 | 0 |
| 0 | 0 |
| 0 | 0 |
| 0 | 0 |
| 0 | 0 |
| 0 | 0 |
| 0 | 0 |
| 49.8078949 | 57.8264983 |
| 0 | 0 |
| 1.71751362 | 50.598186 |
| 0 | 0 |
| 0 | 0 |
| 0 | 0 |
| 0 | 0 |
| 0 | 0 |
| 1.71751362 | 0 |
| 0 | 0 |
| 0 | 0 |
| 0 | 7.22831229 |
| 0 | 0 |
| 0 | 0 |
| 0 | 0 |
| 0 | 0 |
| 0 | 0 |
| 0 | 0 |
| 0 | 0 |
| 0 | 14.4566246 |

| | |
|---|---|
| 0 | 0 |
| 0 | 0 |
| 0 | 0 |
| 180.33893 | 57.8264983 |
| 0 | 0 |
| 0 | 0 |
| 0 | 0 |
| 0 | 0 |
| 0 | 0 |
| 0 | 0 |
| 0 | 0 |
| 0 | 0 |
| 0 | 0 |
| 0 | 0 |
| 0 | 0 |
| 0 | 0 |
| 0 | 0 |
| 0 | 0 |
| 133.966062 | 14.4566246 |
| 0 | 0 |
| 0 | 0 |
| 0 | 0 |
| 0 | 0 |
| 3.43502723 | 0 |
| 0 | 0 |
| 0 | 0 |
| 53.2429221 | 36.1415615 |
| 0 | 0 |
| 0 | 0 |
| 0 | 0 |
| 0 | 0 |
| 0 | 7.22831229 |
| 0 | 0 |
| 0 | 0 |
| 0 | 0 |
| 0 | 0 |
| 0 | 0 |
| 0 | 0 |
| 0 | 0 |
| 0 | 0 |
| 489.491381 | 159.02287 |
| 0 | 0 |
| 0 | 0 |

| | |
|---|---|
| 0 | 0 |
| 0 | 0 |
| 0 | 0 |
| 0 | 0 |
| 6.87005447 | 93.9680598 |
| 0 | 0 |
| 0 | 0 |
| 0 | 0 |
| 0 | 0 |
| 0 | 0 |
| 0 | 0 |
| 0 | 0 |
| 0 | 0 |
| 0 | 0 |
| 0 | 0 |
| 0 | 0 |
| 130.531035 | 0 |
| 0 | 0 |
| 0 | 0 |
| 15.4576225 | 187.93612 |
| 0 | 0 |
| 0 | 0 |
| 0 | 0 |
| 0 | 0 |
| 0 | 0 |
| 0 | 0 |
| 0 | 0 |
| 65.2655174 | 14.4566246 |
| 0 | 0 |
| 0 | 28.9132492 |
| 0 | 0 |
| 0 | 0 |
| 0 | 0 |
| 0 | 0 |
| 0 | 0 |
| 0 | 0 |
| 0 | 0 |
| 20.6101634 | 0 |
| 0 | 0 |
| 0 | 0 |
| 0 | 0 |
| 0 | 0 |
| 5.15254085 | 36.1415615 |

| | |
|---|---|
| 0 | 0 |
| 0 | 0 |
| 0 | 0 |
| 0 | 0 |
| 0 | 0 |
| 0 | 0 |
| 39.5028132 | 0 |
| 0 | 0 |
| 0 | 0 |
| 0 | 0 |
| 0 | 0 |
| 0 | 0 |
| 0 | 0 |
| 0 | 0 |
| 29.1977315 | 101.196372 |
| 0 | 0 |
| 0 | 0 |
| 0 | 0 |
| 0 | 0 |
| 156.293739 | 0 |
| 0 | 0 |
| 0 | 0 |
| 0 | 0 |
| 30.9152451 | 0 |
| 0 | 0 |
| 0 | 0 |
| 0 | 0 |
| 0 | 0 |
| 0 | 0 |
| 0 | 0 |
| 0 | 0 |
| 0 | 0 |
| 219.841743 | 28.9132492 |
| 0 | 0 |
| 0 | 0 |
| 0 | 0 |
| 0 | 0 |
| 0 | 0 |
| 0 | 0 |
| 0 | 0 |
| 0 | 0 |
| 0 | 0 |
| 0 | 0 |

| | |
|---|---|
| 0 | 0 |
| 0 | 0 |
| 0 | 0 |
| 0 | 0 |
| 0 | 0 |
| 0 | 0 |
| 0 | 0 |
| 0 | 0 |
| 0 | 0 |
| 0 | 0 |
| 0 | 0 |
| 0 | 0 |
| 0 | 0 |
| 0 | 0 |
| 0 | 0 |
| 0 | 0 |
| 77.2881127 | 0 |
| 0 | 0 |
| 0 | 0 |
| 0 | 0 |
| 0 | 0 |
| 0 | 0 |
| 0 | 0 |
| 0 | 0 |
| 0 | 0 |
| 0 | 0 |
| 0 | 0 |
| 0 | 7.22831229 |
| 0 | 0 |
| 0 | 0 |
| 0 | 0 |
| 0 | 0 |
| 0 | 0 |
| 0 | 0 |
| 0 | 0 |
| 0 | 0 |
| 0 | 0 |
| 0 | 21.6849369 |
| 0 | 0 |
| 0 | 0 |
| 0 | 0 |
| 48.0903813 | 0 |
| 0 | 0 |

| | |
|---|---|
| 0 | 0 |
| 70.4180583 | 14.4566246 |
| 0 | 0 |
| 0 | 0 |
| 0 | 0 |
| 0 | 0 |
| 0 | 7.22831229 |
| 0 | 0 |
| 0 | 0 |
| 0 | 0 |
| 6.87005447 | 43.3698738 |
| 0 | 0 |
| 0 | 0 |
| 27.4802179 | 108.424684 |
| 0 | 0 |
| 0 | 0 |
| 0 | 0 |
| 0 | 0 |
| 206.101634 | 144.566246 |
| 0 | 0 |
| 0 | 0 |
| 0 | 0 |
| 0 | 0 |
| 0 | 0 |
| 0 | 0 |
| 0 | 0 |
| 0 | 0 |
| 0 | 0 |
| 20.6101634 | 14.4566246 |
| 0 | 0 |
| 0 | 0 |
| 0 | 0 |
| 0 | 0 |
| 0 | 0 |
| 0 | 0 |
| 0 | 57.8264983 |
| 0 | 0 |
| 0 | 0 |
| 36.0677859 | 14.4566246 |
| 0 | 0 |
| 68.7005447 | 209.621056 |
| 0 | 0 |
| 0 | 7.22831229 |

| | |
|---|---|
| 0 | 0 |
| 0 | 0 |
| 0 | 0 |
| 0 | 0 |
| 0 | 0 |
| 0 | 0 |
| 0 | 0 |
| 0 | 0 |
| 0 | 0 |
| 0 | 0 |
| 1.71751362 | 0 |
| 0 | 0 |
| 0 | 0 |
| 0 | 0 |
| 0 | 0 |
| 0 | 0 |
| 92.7457353 | 7.22831229 |
| 0 | 0 |
| 0 | 0 |
| 0 | 0 |
| 0 | 0 |
| 0 | 0 |
| 0 | 0 |
| 0 | 0 |
| 0 | 0 |
| 0 | 0 |
| 0 | 0 |
| 0 | 0 |
| 0 | 0 |
| 0 | 0 |
| 0 | 0 |
| 15.4576225 | 7.22831229 |
| 0 | 0 |
| 0 | 0 |
| 0 | 0 |
| 0 | 0 |
| 0 | 0 |
| 0 | 7.22831229 |
| 0 | 0 |
| 0 | 0 |
| 0 | 0 |
| 0 | 0 |
| 0 | 36.1415615 |

| | |
|---|---|
| 0 | 0 |
| 0 | 0 |
| 0 | 0 |
| 288.542288 | 65.0548106 |
| 0 | 0 |
| 0 | 0 |
| 0 | 0 |
| 0 | 0 |
| 0 | 0 |
| 0 | 0 |
| 544.451816 | 383.100551 |
| 36.0677859 | 216.849369 |
| 0 | 0 |
| 0 | 0 |
| 0 | 0 |
| 0 | 0 |
| 0 | 0 |
| 91.0282217 | 202.392744 |
| 0 | 0 |
| 0 | 0 |
| 6.87005447 | 180.707807 |
| 0 | 0 |
| 60.1129766 | 14.4566246 |
| 0 | 0 |
| 0 | 0 |
| 0 | 0 |
| 0 | 43.3698738 |
| 0 | 0 |
| 0 | 0 |
| 0 | 0 |
| 10.3050817 | 28.9132492 |
| 0 | 0 |
| 0 | 0 |
| 0 | 0 |
| 0 | 0 |
| 0 | 0 |
| 0 | 0 |
| 0 | 0 |
| 0 | 0 |
| 0 | 0 |
| 0 | 0 |
| 0 | 0 |
| 0 | 0 |

| | |
|---|---|
| 0 | 0 |
| 0 | 0 |
| 0 | 0 |
| 0 | 0 |
| 0 | 0 |
| 0 | 0 |
| 0 | 0 |
| 0 | 0 |
| 0 | 0 |
| 0 | 0 |
| 0 | 14.4566246 |
| 0 | 0 |
| 0 | 0 |
| 0 | 0 |
| 0 | 0 |
| 194.079039 | 216.849369 |
| 0 | 0 |
| 0 | 0 |
| 0 | 0 |
| 0 | 0 |
| 0 | 0 |
| 85.8756808 | 0 |
| 0 | 0 |
| 0 | 0 |
| 0 | 0 |
| 0 | 0 |
| 0 | 0 |
| 1.71751362 | 7.22831229 |
| 15.4576225 | 0 |
| 0 | 0 |
| 0 | 0 |
| 0 | 0 |
| 0 | 0 |
| 115.073412 | 14.4566246 |
| 8.58756808 | 122.881309 |
| 0 | 0 |
| 0 | 0 |
| 0 | 0 |
| 0 | 0 |
| 0 | 0 |
| 0 | 0 |
| 0 | 7.22831229 |
| 0 | 0 |

| | |
|---|---|
| 0 | 0 |
| 0 | 0 |
| 0 | 0 |
| 0 | 0 |
| 0 | 0 |
| 0 | 0 |
| 0 | 0 |
| 0 | 0 |
| 0 | 0 |
| 0 | 21.6849369 |
| 0 | 0 |
| 0 | 0 |
| 0 | 0 |
| 0 | 0 |
| 113.355899 | 7.22831229 |
| 0 | 0 |
| 1.71751362 | 0 |
| 0 | 0 |
| 1.71751362 | 14.4566246 |
| 0 | 0 |
| 0 | 7.22831229 |
| 0 | 0 |
| 0 | 0 |
| 0 | 0 |
| 166.598821 | 101.196372 |
| 104.768331 | 36.1415615 |
| 0 | 0 |
| 0 | 0 |
| 0 | 0 |
| 0 | 43.3698738 |
| 0 | 0 |
| 0 | 0 |
| 0 | 0 |
| 0 | 0 |
| 0 | 0 |
| 58.395463 | 0 |
| 0 | 0 |
| 0 | 0 |
| 0 | 0 |
| 0 | 0 |
| 0 | 0 |
| 0 | 0 |
| 0 | 0 |

| | |
|---|---|
| 58.395463 | 86.7397475 |
| 0 | 0 |
| 0 | 0 |
| 0 | 0 |
| 10.3050817 | 137.337934 |
| 0 | 0 |
| 0 | 0 |
| 85.8756808 | 101.196372 |
| 0 | 0 |
| 0 | 0 |
| 0 | 7.22831229 |
| 0 | 0 |
| 0 | 0 |
| 0 | 0 |
| 0 | 0 |
| 0 | 0 |
| 0 | 0 |
| 0 | 0 |
| 42.9378404 | 28.9132492 |
| 0 | 0 |
| 0 | 0 |
| 0 | 0 |
| 0 | 0 |
| 0 | 0 |
| 0 | 0 |
| 0 | 0 |
| 0 | 0 |
| 0 | 0 |
| 0 | 0 |
| 0 | 0 |
| 0 | 0 |
| 130.531035 | 397.557176 |
| 0 | 0 |
| 0 | 0 |
| 0 | 0 |
| 0 | 0 |
| 0 | 0 |
| 0 | 0 |
| 0 | 0 |
| 0 | 0 |
| 0 | 0 |
| 432.813431 | 57.8264983 |
| 0 | 0 |

| | |
|---|---|
| 0 | 0 |
| 0 | 0 |
| 0 | 0 |
| 0 | 0 |
| 0 | 0 |
| 0 | 0 |
| 0 | 0 |
| 0 | 0 |
| 0 | 0 |
| 219.841743 | 159.02287 |
| 0 | 0 |
| 0 | 0 |
| 3.43502723 | 0 |
| 0 | 0 |
| 0 | 0 |
| 0 | 0 |
| 0 | 0 |
| 3.43502723 | 0 |
| 0 | 0 |
| 0 | 0 |
| 0 | 14.4566246 |
| 0 | 0 |
| 0 | 0 |
| 0 | 14.4566246 |
| 0 | 0 |
| 0 | 0 |
| 0 | 0 |
| 0 | 0 |
| 0 | 0 |
| 0 | 0 |
| 0 | 0 |
| 0 | 0 |
| 0 | 0 |
| 0 | 0 |
| 0 | 0 |
| 91.0282217 | 122.881309 |
| 0 | 0 |
| 0 | 0 |
| 0 | 0 |
| 0 | 0 |
| 0 | 0 |
| 0 | 0 |
| 0 | 0 |

| | |
|---|---|
| 0 | 0 |
| 0 | 0 |
| 0 | 0 |
| 0 | 0 |
| 0 | 0 |
| 0 | 0 |
| 0 | 0 |
| 0 | 0 |
| 0 | 0 |
| 0 | 0 |
| 0 | 0 |
| 0 | 0 |
| 0 | 0 |
| 0 | 0 |
| 0 | 0 |
| 176.903902 | 7.22831229 |
| 0 | 0 |
| 0 | 0 |
| 0 | 0 |
| 0 | 0 |
| 0 | 0 |
| 0 | 0 |
| 0 | 0 |
| 0 | 0 |
| 0 | 0 |
| 0 | 0 |
| 0 | 0 |
| 46.3728676 | 0 |
| 44.655354 | 932.452286 |
| 0 | 0 |
| 0 | 0 |
| 0 | 0 |
| 0 | 0 |
| 0 | 0 |
| 0 | 0 |
| 182.056443 | 7.22831229 |
| 6.87005447 | 0 |
| 0 | 0 |
| 0 | 0 |
| 0 | 0 |
| 0 | 0 |
| 0 | 0 |
| 0 | 0 |

| | |
|---|---|
| 0 | 14.4566246 |
| 0 | 0 |
| 0 | 0 |
| 1.71751362 | 14.4566246 |
| 0 | 0 |
| 0 | 0 |
| 0 | 0 |
| 0 | 0 |
| 0 | 0 |
| 0 | 0 |
| 0 | 0 |
| 0 | 0 |
| 0 | 0 |
| 0 | 0 |
| 0 | 0 |
| 0 | 0 |
| 0 | 0 |
| 0 | 0 |
| 0 | 0 |
| 5.15254085 | 195.164432 |
| 0 | 0 |
| 0 | 0 |
| 0 | 0 |
| 0 | 0 |
| 419.073322 | 72.2831229 |
| 0 | 0 |
| 0 | 0 |
| 25.7627042 | 0 |
| 0 | 0 |
| 0 | 0 |
| 0 | 0 |
| 0 | 0 |
| 3.43502723 | 0 |
| 0 | 0 |
| 0 | 0 |
| 0 | 0 |
| 0 | 0 |
| 0 | 0 |
| 0 | 0 |
| 0 | 0 |
| 0 | 0 |
| 12.0225953 | 7.22831229 |
| 0 | 0 |

| | |
|---|---|
| 0 | 0 |
| 0 | 0 |
| 3.43502723 | 7.22831229 |
| 0 | 0 |
| 0 | 0 |
| 0 | 0 |
| 0 | 0 |
| 0 | 0 |
| 0 | 0 |
| 56.6779493 | 93.9680598 |
| 0 | 0 |
| 0 | 0 |
| 0 | 0 |
| 0 | 0 |
| 0 | 0 |
| 0 | 0 |
| 0 | 0 |
| 0 | 0 |
| 0 | 0 |
| 0 | 0 |
| 0 | 0 |
| 140.836117 | 137.337934 |
| 0 | 0 |
| 0 | 0 |
| 0 | 7.22831229 |
| 37.7852996 | 390.328864 |
| 0 | 0 |
| 75.5705991 | 65.0548106 |
| 0 | 7.22831229 |
| 0 | 0 |
| 0 | 0 |
| 0 | 7.22831229 |
| 0 | 21.6849369 |
| 0 | 0 |
| 0 | 0 |
| 0 | 0 |
| 18.8926498 | 7.22831229 |
| 0 | 72.2831229 |
| 0 | 0 |
| 0 | 0 |
| 0 | 0 |
| 0 | 0 |
| 0 | 0 |

| | |
|---|---|
| 0 | 0 |
| 0 | 0 |
| 22.327677 | 72.2831229 |
| 0 | 0 |
| 0 | 0 |
| 0 | 0 |
| 0 | 0 |
| 0 | 0 |
| 0 | 0 |
| 0 | 0 |
| 0 | 0 |
| 1.71751362 | 0 |
| 0 | 0 |
| 0 | 0 |
| 8.58756808 | 144.566246 |
| 0 | 0 |
| 0 | 0 |
| 0 | 0 |
| 0 | 0 |
| 0 | 0 |
| 0 | 0 |
| 0 | 0 |
| 0 | 0 |
| 51.5254085 | 130.109621 |
| 1.71751362 | 0 |
| 0 | 0 |
| 0 | 0 |
| 0 | 0 |
| 0 | 0 |
| 0 | 0 |
| 0 | 0 |
| 0 | 0 |
| 0 | 0 |
| 0 | 0 |
| 0 | 0 |
| 0 | 0 |
| 0 | 0 |
| 0 | 751.744478 |
| 0 | 0 |
| 0 | 14.4566246 |
| 0 | 0 |
| 0 | 0 |
| 0 | 0 |

| | |
|---|---|
| 0 | 14.4566246 |
| 0 | 0 |
| 0 | 65.0548106 |
| 0 | 0 |
| 0 | 0 |
| 0 | 0 |
| 0 | 0 |
| 0 | 0 |
| 194.079039 | 50.598186 |
| 13.7401089 | 231.305993 |
| 0 | 0 |
| 0 | 0 |
| 224.994284 | 57.8264983 |
| 61.8304902 | 28.9132492 |
| 0 | 7.22831229 |
| 0 | 0 |
| 0 | 0 |
| 0 | 0 |
| 0 | 0 |
| 819.253995 | 455.383674 |
| 13.7401089 | 79.5114352 |
| 0 | 0 |
| 0 | 0 |
| 20.6101634 | 0 |
| 0 | 0 |
| 0 | 0 |
| 0 | 0 |
| 0 | 0 |
| 0 | 0 |
| 0 | 0 |
| 0 | 0 |
| 0 | 0 |
| 0 | 0 |
| 142.55363 | 7.22831229 |
| 1.71751362 | 144.566246 |
| 0 | 0 |
| 0 | 0 |
| 0 | 0 |
| 0 | 0 |
| 0 | 0 |
| 0 | 0 |
| 0 | 0 |
| 224.994284 | 101.196372 |

| | |
|---|---|
| 0 | 0 |
| 8.58756808 | 50.598186 |
| 0 | 0 |
| 0 | 0 |
| 0 | 0 |
| 0 | 0 |
| 0 | 0 |
| 0 | 0 |
| 1.71751362 | 0 |
| 0 | 0 |
| 0 | 0 |
| 0 | 0 |
| 0 | 0 |
| 0 | 0 |
| 6.87005447 | 72.2831229 |
| 0 | 0 |
| 0 | 0 |
| 0 | 0 |
| 0 | 7.22831229 |
| 0 | 0 |
| 413.920782 | 28.9132492 |
| 115.073412 | 0 |
| 0 | 0 |
| 0 | 0 |
| 0 | 0 |
| 0 | 0 |
| 1.71751362 | 0 |
| 0 | 0 |
| 0 | 0 |
| 0 | 0 |
| 0 | 0 |
| 0 | 0 |
| 0 | 0 |
| 0 | 0 |
| 56.6779493 | 28.9132492 |
| 0 | 0 |
| 0 | 0 |
| 0 | 0 |
| 0 | 0 |
| 0 | 0 |
| 0 | 0 |
| 0 | 0 |
| 12.0225953 | 21.6849369 |

| | |
|---|---|
| 0 | 0 |
| 0 | 0 |
| 0 | 0 |
| 0 | 0 |
| 1.71751362 | 0 |
| 0 | 0 |
| 0 | 0 |
| 0 | 0 |
| 0 | 0 |
| 0 | 0 |
| 72.1355719 | 0 |
| 0 | 0 |
| 152.858712 | 252.99093 |
| 0 | 0 |
| 0 | 0 |
| 27.4802179 | 0 |
| 0 | 0 |
| 0 | 0 |
| 0 | 0 |
| 0 | 0 |
| 0 | 0 |
| 0 | 0 |
| 0 | 0 |
| 0 | 0 |
| 0 | 0 |
| 0 | 0 |
| 0 | 0 |
| 0 | 0 |
| 0 | 0 |
| 0 | 0 |
| 0 | 0 |
| 0 | 0 |
| 0 | 0 |
| 0 | 0 |
| 0 | 0 |
| 0 | 0 |
| 156.293739 | 0 |
| 449.988568 | 267.447555 |
| 0 | 0 |
| 12.0225953 | 0 |
| 0 | 0 |
| 0 | 0 |
| 0 | 0 |

| | |
|---|---|
| 0 | 0 |
| 0 | 0 |
| 518.689112 | 115.652997 |
| 0 | 0 |
| 0 | 0 |
| 94.4632489 | 7.22831229 |
| 0 | 0 |
| 0 | 0 |
| 0 | 0 |
| 0 | 0 |
| 0 | 14.4566246 |
| 0 | 0 |
| 0 | 0 |
| 0 | 0 |
| 0 | 0 |
| 0 | 0 |
| 0 | 0 |
| 0 | 0 |
| 0 | 0 |
| 0 | 0 |
| 0 | 0 |
| 0 | 0 |
| 0 | 0 |
| 161.44628 | 7.22831229 |
| 0 | 0 |
| 1399.7736 | 860.169163 |
| 0 | 0 |
| 0 | 0 |
| 0 | 0 |
| 0 | 0 |
| 0 | 0 |
| 0 | 0 |
| 0 | 0 |
| 0 | 0 |
| 0 | 0 |
| 314.304992 | 108.424684 |
| 0 | 0 |
| 0 | 0 |
| 0 | 0 |
| 0 | 0 |
| 0 | 0 |
| 0 | 0 |
| 0 | 0 |

| | |
|---|---|
| 0 | 0 |
| 0 | 0 |
| 0 | 0 |
| 0 | 0 |
| 0 | 0 |
| 0 | 0 |
| 0 | 0 |
| 0 | 0 |
| 0 | 0 |
| 0 | 0 |
| 61.8304902 | 0 |
| 0 | 0 |
| 0 | 0 |
| 0 | 7.22831229 |
| 0 | 0 |
| 0 | 0 |
| 0 | 0 |
| 60.1129766 | 0 |
| 0 | 0 |
| 0 | 0 |
| 0 | 7.22831229 |
| 0 | 0 |
| 6.87005447 | 0 |
| 0 | 0 |
| 0 | 0 |
| 0 | 0 |
| 0 | 0 |
| 0 | 0 |
| 0 | 0 |
| 420.790836 | 7.22831229 |
| 0 | 0 |
| 0 | 0 |
| 3.43502723 | 7.22831229 |
| 77.2881127 | 36.1415615 |
| 0 | 0 |
| 0 | 0 |
| 0 | 0 |
| 0 | 0 |
| 0 | 0 |
| 0 | 0 |
| 0 | 0 |
| 0 | 0 |
| 0 | 0 |

| | |
|---|---|
| 261.06207 | 252.99093 |
| 0 | 0 |
| 1.71751362 | 0 |
| 0 | 0 |
| 0 | 0 |
| 0 | 0 |
| 0 | 0 |
| 58.395463 | 7.22831229 |
| 0 | 0 |
| 0 | 0 |
| 0 | 0 |
| 0 | 7.22831229 |
| 0 | 0 |
| 0 | 0 |
| 0 | 0 |
| 0 | 0 |
| 0 | 0 |
| 0 | 0 |
| 0 | 0 |
| 0 | 0 |
| 3.43502723 | 0 |
| 0 | 0 |
| 0 | 0 |
| 0 | 0 |
| 271.367151 | 28.9132492 |
| 0 | 0 |
| 0 | 0 |
| 0 | 0 |
| 0 | 0 |
| 247.321961 | 50.598186 |
| 0 | 0 |
| 0 | 0 |
| 0 | 0 |
| 0 | 0 |
| 0 | 0 |
| 0 | 0 |
| 70.4180583 | 21.6849369 |
| 0 | 0 |
| 0 | 0 |
| 0 | 0 |
| 0 | 0 |
| 0 | 0 |
| 0 | 0 |

| | |
|---|---|
| 0 | 0 |
| 0 | 0 |
| 0 | 0 |
| 0 | 0 |
| 0 | 0 |
| 0 | 7.22831229 |
| 0 | 0 |
| 0 | 0 |
| 139.118603 | 101.196372 |
| 0 | 0 |
| 0 | 0 |
| 0 | 0 |
| 0 | 0 |
| 407.050727 | 469.840299 |
| 0 | 0 |
| 0 | 115.652997 |
| 0 | 0 |
| 0 | 0 |
| 0 | 0 |
| 1.71751362 | 14.4566246 |
| 0 | 0 |
| 0 | 0 |
| 0 | 0 |
| 0 | 7.22831229 |
| 0 | 0 |
| 5.15254085 | 0 |
| 0 | 0 |
| 111.638385 | 7.22831229 |
| 0 | 0 |
| 0 | 0 |
| 0 | 0 |
| 70.4180583 | 21.6849369 |
| 0 | 0 |
| 0 | 0 |
| 0 | 0 |
| 154.576225 | 7.22831229 |
| 163.163794 | 252.99093 |
| 0 | 0 |
| 0 | 7.22831229 |
| 0 | 0 |
| 0 | 0 |
| 0 | 0 |
| 0 | 0 |

| | |
|---|---|
| 3.43502723 | 0 |
| 0 | 0 |
| 0 | 0 |
| 42.9378404 | 43.3698738 |
| 0 | 14.4566246 |
| 0 | 7.22831229 |
| 0 | 0 |
| 0 | 0 |
| 46.3728676 | 122.881309 |
| 0 | 7.22831229 |
| 0 | 0 |
| 0 | 0 |
| 0 | 0 |
| 0 | 0 |
| 0 | 0 |
| 30.9152451 | 0 |
| 0 | 0 |
| 0 | 0 |
| 0 | 0 |
| 29.1977315 | 0 |
| 0 | 0 |
| 0 | 0 |
| 0 | 0 |
| 0 | 0 |
| 8.58756808 | 0 |
| 0 | 0 |
| 0 | 0 |
| 0 | 0 |
| 0 | 0 |
| 77.2881127 | 14.4566246 |
| 0 | 0 |
| 750.55345 | 693.91798 |
| 0 | 0 |
| 66.983031 | 0 |
| 0 | 14.4566246 |
| 0 | 0 |
| 0 | 0 |
| 0 | 0 |
| 51.5254085 | 122.881309 |
| 70.4180583 | 289.132492 |
| 10.3050817 | 108.424684 |
| 0 | 0 |
| 0 | 0 |

| | |
|---|---|
| 0 | 0 |
| 0 | 0 |
| 0 | 0 |
| 0 | 0 |
| 1016.76806 | 318.045741 |
| 0 | 0 |
| 0 | 0 |
| 0 | 0 |
| 0 | 0 |
| 1.71751362 | 0 |
| 0 | 0 |
| 0 | 0 |
| 0 | 0 |
| 0 | 0 |
| 0 | 0 |
| 704.180583 | 383.100551 |
| 8.58756808 | 0 |
| 0 | 0 |
| 0 | 0 |
| 0 | 0 |
| 0 | 0 |
| 0 | 0 |
| 22.327677 | 36.1415615 |
| 0 | 0 |
| 0 | 0 |
| 151.141198 | 7.22831229 |
| 0 | 0 |
| 0 | 0 |
| 0 | 0 |
| 0 | 0 |
| 0 | 0 |
| 836.429131 | 397.557176 |
| 0 | 0 |
| 3.43502723 | 7.22831229 |
| 0 | 0 |
| 0 | 0 |
| 0 | 0 |
| 0 | 0 |
| 0 | 0 |
| 0 | 0 |
| 0 | 0 |
| 0 | 0 |
| 0 | 0 |

| | |
|---|---|
| 0 | 0 |
| 0 | 0 |
| 0 | 0 |
| 0 | 0 |
| 0 | 0 |
| 0 | 0 |
| 0 | 0 |
| 0 | 7.22831229 |
| 0 | 0 |
| 0 | 0 |
| 0 | 0 |
| 0 | 0 |
| 0 | 0 |
| 0 | 0 |
| 0 | 0 |
| 68.7005447 | 79.5114352 |
| 0 | 0 |
| 0 | 0 |
| 0 | 0 |
| 0 | 0 |
| 0 | 0 |
| 24.0451906 | 0 |
| 0 | 0 |
| 1.71751362 | 14.4566246 |
| 0 | 0 |
| 0 | 0 |
| 0 | 0 |
| 0 | 0 |
| 0 | 0 |
| 0 | 0 |
| 0 | 0 |
| 0 | 0 |
| 0 | 0 |
| 77.2881127 | 72.2831229 |
| 0 | 0 |
| 0 | 0 |
| 0 | 0 |
| 0 | 0 |
| 0 | 14.4566246 |
| 0 | 0 |
| 0 | 0 |
| 0 | 7.22831229 |
| 0 | 0 |

| | |
|---|---|
| 216.406716 | 108.424684 |
| 0 | 0 |
| 0 | 0 |
| 0 | 0 |
| 0 | 0 |
| 106.485844 | 14.4566246 |
| 0 | 0 |
| 1.71751362 | 0 |
| 0 | 0 |
| 0 | 0 |
| 0 | 0 |
| 0 | 0 |
| 0 | 0 |
| 0 | 0 |
| 0 | 0 |
| 132.248548 | 7.22831229 |
| 0 | 0 |
| 0 | 0 |
| 0 | 0 |
| 0 | 0 |
| 0 | 0 |
| 3.43502723 | 57.8264983 |
| 0 | 0 |
| 0 | 0 |
| 30.9152451 | 86.7397475 |
| 0 | 0 |
| 0 | 0 |
| 0 | 0 |
| 0 | 0 |
| 0 | 0 |
| 0 | 0 |
| 0 | 0 |
| 0 | 0 |
| 0 | 0 |
| 0 | 0 |
| 0 | 0 |
| 0 | 0 |
| 0 | 0 |
| 0 | 0 |
| 0 | 0 |
| 0 | 0 |
| 0 | 0 |
| 58.395463 | 43.3698738 |

| | |
|---|---|
| 0 | 0 |
| 34.3502723 | 28.9132492 |
| 0 | 0 |
| 183.773957 | 144.566246 |
| 120.225953 | 0 |
| 0 | 0 |
| 0 | 0 |
| 0 | 0 |
| 0 | 0 |
| 0 | 0 |
| 116.790926 | 122.881309 |
| 0 | 0 |
| 132.248548 | 7.22831229 |
| 0 | 0 |
| 0 | 0 |
| 0 | 0 |
| 0 | 0 |
| 0 | 0 |
| 0 | 0 |
| 0 | 0 |
| 0 | 0 |
| 0 | 0 |
| 0 | 0 |
| 0 | 0 |
| 0 | 0 |
| 0 | 0 |
| 0 | 0 |
| 0 | 0 |
| 0 | 0 |
| 0 | 0 |
| 30.9152451 | 0 |
| 0 | 0 |
| 0 | 0 |
| 82.4406536 | 0 |
| 0 | 0 |
| 0 | 0 |
| 0 | 0 |
| 0 | 0 |
| 0 | 0 |
| 0 | 0 |
| 0 | 0 |
| 30.9152451 | 7.22831229 |
| 0 | 0 |

| | |
|---|---|
| 0 | 0 |
| 0 | 0 |
| 0 | 0 |
| 0 | 0 |
| 61.8304902 | 173.479495 |
| 0 | 0 |
| 0 | 0 |
| 79.0056264 | 65.0548106 |
| 0 | 0 |
| 0 | 0 |
| 0 | 0 |
| 0 | 0 |
| 0 | 0 |
| 27.4802179 | 108.424684 |
| 0 | 0 |
| 32.6327587 | 43.3698738 |
| 0 | 0 |
| 0 | 0 |
| 0 | 0 |
| 0 | 0 |
| 0 | 0 |
| 0 | 0 |
| 0 | 0 |
| 0 | 0 |
| 0 | 0 |
| 0 | 0 |
| 0 | 0 |
| 681.852906 | 115.652997 |
| 0 | 0 |
| 0 | 0 |
| 0 | 0 |
| 0 | 0 |
| 24.0451906 | 57.8264983 |
| 0 | 0 |
| 0 | 0 |
| 0 | 0 |
| 0 | 0 |
| 0 | 0 |
| 8.58756808 | 7.22831229 |
| 278.237206 | 130.109621 |
| 0 | 0 |
| 0 | 0 |
| 0 | 0 |

| | |
|---|---|
| 0 | 0 |
| 0 | 0 |
| 27.4802179 | 0 |
| 0 | 0 |
| 0 | 0 |
| 0 | 0 |
| 0 | 0 |
| 0 | 0 |
| 0 | 0 |
| 0 | 0 |
| 243.886934 | 28.9132492 |
| 0 | 0 |
| 0 | 0 |
| 345.220237 | 448.155362 |
| 0 | 0 |
| 0 | 0 |
| 0 | 0 |
| 0 | 0 |
| 0 | 0 |
| 273.084665 | 412.013801 |
| 0 | 0 |
| 0 | 0 |
| 0 | 0 |
| 0 | 0 |
| 0 | 0 |
| 0 | 0 |
| 0 | 0 |
| 0 | 0 |
| 0 | 0 |
| 0 | 0 |
| 30.9152451 | 0 |
| 0 | 0 |
| 0 | 0 |
| 0 | 0 |
| 140.836117 | 7.22831229 |
| 0 | 0 |
| 0 | 0 |
| 79.0056264 | 0 |
| 0 | 0 |
| 0 | 0 |
| 0 | 0 |
| 101.333303 | 173.479495 |
| 0 | 0 |

| | |
|---|---|
| 0 | 0 |
| 0 | 0 |
| 0 | 0 |
| 133.966062 | 14.4566246 |
| 1.71751362 | 0 |
| 0 | 0 |
| 17.1751362 | 130.109621 |
| 27.4802179 | 72.2831229 |
| 0 | 0 |
| 0 | 0 |
| 80.72314 | 0 |
| 0 | 0 |
| 0 | 0 |
| 0 | 0 |
| 0 | 0 |
| 0 | 0 |
| 0 | 0 |
| 1.71751362 | 0 |
| 0 | 0 |
| 0 | 0 |
| 0 | 0 |
| 0 | 0 |
| 0 | 0 |
| 0 | 0 |
| 5.15254085 | 0 |
| 0 | 0 |
| 0 | 57.8264983 |
| 0 | 0 |
| 0 | 0 |
| 0 | 0 |
| 0 | 0 |
| 0 | 36.1415615 |
| 0 | 0 |
| 0 | 0 |
| 0 | 0 |
| 0 | 0 |
| 0 | 0 |
| 0 | 0 |
| 29.1977315 | 130.109621 |
| 3.43502723 | 7.22831229 |
| 0 | 0 |
| 0 | 0 |
| 0 | 0 |

| | |
|---|---|
| 0 | 0 |
| 0 | 0 |
| 0 | 0 |
| 1.71751362 | 0 |
| 0 | 0 |
| 0 | 0 |
| 0 | 0 |
| 0 | 0 |
| 0 | 0 |
| 1.71751362 | 0 |
| 0 | 0 |
| 0 | 0 |
| 0 | 0 |
| 0 | 0 |
| 0 | 0 |
| 0 | 0 |
| 0 | 0 |
| 0 | 0 |
| 386.440564 | 831.255914 |
| 0 | 0 |
| 0 | 0 |
| 0 | 0 |
| 0 | 0 |
| 0 | 0 |
| 0 | 0 |
| 0 | 0 |
| 0 | 0 |
| 0 | 0 |
| 0 | 0 |
| 0 | 0 |
| 0 | 0 |
| 1.71751362 | 21.6849369 |
| 0 | 0 |
| 0 | 0 |
| 0 | 0 |
| 0 | 0 |
| 0 | 0 |
| 0 | 0 |
| 0 | 0 |
| 0 | 0 |
| 34.3502723 | 0 |
| 0 | 0 |
| 0 | 0 |

| | |
|---|---|
| 0 | 0 |
| 0 | 0 |
| 0 | 14.4566246 |
| 0 | 0 |
| 0 | 21.6849369 |
| 0 | 0 |
| 0 | 7.22831229 |
| 0 | 0 |
| 77.2881127 | 0 |
| 17.1751362 | 86.7397475 |
| 13.7401089 | 86.7397475 |
| 1149.01661 | 527.666797 |
| 0 | 0 |
| 0 | 0 |
| 0 | 0 |
| 0 | 0 |
| 216.406716 | 65.0548106 |
| 0 | 0 |
| 0 | 0 |
| 0 | 7.22831229 |
| 0 | 0 |
| 0 | 0 |
| 0 | 0 |
| 12.0225953 | 0 |
| 0 | 0 |
| 0 | 0 |
| 1.71751362 | 0 |
| 0 | 0 |
| 0 | 0 |
| 0 | 0 |
| 3.43502723 | 0 |
| 0 | 0 |
| 0 | 0 |
| 0 | 0 |
| 135.683576 | 7.22831229 |
| 0 | 50.598186 |
| 0 | 0 |
| 0 | 0 |
| 36.0677859 | 14.4566246 |
| 0 | 0 |
| 0 | 0 |
| 0 | 0 |
| 0 | 0 |

| | |
|---|---|
| 0 | 0 |
| 0 | 0 |
| 0 | 0 |
| 0 | 0 |
| 0 | 0 |
| 6.87005447 | 130.109621 |
| 1.71751362 | 7.22831229 |
| 0 | 0 |
| 0 | 0 |
| 0 | 0 |
| 0 | 0 |
| 0 | 0 |
| 0 | 0 |
| 0 | 0 |
| 0 | 0 |
| 0 | 0 |
| 0 | 0 |
| 0 | 0 |
| 0 | 21.6849369 |
| 0 | 43.3698738 |
| 0 | 0 |
| 0 | 0 |
| 437.965972 | 224.077681 |
| 0 | 0 |
| 0 | 0 |
| 0 | 0 |
| 0 | 0 |
| 0 | 0 |
| 0 | 0 |
| 0 | 0 |
| 285.10726 | 3115.4026 |
| 113.355899 | 7.22831229 |
| 0 | 0 |
| 0 | 0 |
| 0 | 0 |
| 0 | 0 |
| 0 | 0 |
| 0 | 0 |
| 267.932124 | 57.8264983 |
| 0 | 0 |
| 0 | 0 |
| 0 | 0 |
| 0 | 0 |

| | |
|---|---|
| 65.2655174 | 505.98186 |
| 0 | 14.4566246 |
| 0 | 0 |
| 0 | 0 |
| 0 | 0 |
| 0 | 14.4566246 |
| 0 | 0 |
| 0 | 0 |
| 0 | 0 |
| 0 | 0 |
| 0 | 0 |
| 0 | 0 |
| 0 | 0 |
| 0 | 14.4566246 |
| 0 | 0 |
| 79.0056264 | 440.92705 |
| 0 | 0 |
| 0 | 0 |
| 0 | 0 |
| 0 | 0 |
| 618.304902 | 260.219243 |
| 0 | 0 |
| 0 | 0 |
| 0 | 0 |
| 5.15254085 | 0 |
| 0 | 0 |
| 0 | 0 |
| 0 | 0 |
| 0 | 0 |
| 0 | 0 |
| 0 | 0 |
| 0 | 0 |
| 0 | 0 |
| 0 | 0 |
| 214.689202 | 0 |
| 97.8982761 | 303.589116 |
| 0 | 0 |
| 0 | 7.22831229 |
| 0 | 0 |
| 0 | 0 |
| 0 | 0 |
| 0 | 0 |
| 0 | 0 |

| | |
|---|---|
| 0 | 0 |
| 12.0225953 | 28.9132492 |
| 0 | 0 |
| 0 | 0 |
| 61.8304902 | 137.337934 |
| 27.4802179 | 28.9132492 |
| 0 | 0 |
| 8.58756808 | 0 |
| 0 | 0 |
| 17.1751362 | 0 |
| 0 | 0 |
| 235.299365 | 231.305993 |
| 5.15254085 | 0 |
| 0 | 0 |
| 0 | 0 |
| 197.514066 | 0 |
| 61.8304902 | 166.251183 |
| 199.23158 | 0 |
| 0 | 0 |
| 0 | 0 |
| 65.2655174 | 7.22831229 |
| 0 | 0 |
| 0 | 0 |
| 1.71751362 | 0 |
| 0 | 0 |
| 0 | 0 |
| 0 | 0 |
| 128.813521 | 419.242113 |
| 0 | 0 |
| 0 | 0 |
| 0 | 0 |
| 82.4406536 | 144.566246 |
| 0 | 7.22831229 |
| 0 | 0 |
| 0 | 0 |
| 0 | 0 |
| 63.5480038 | 0 |
| 0 | 0 |
| 0 | 0 |
| 0 | 0 |
| 233.581852 | 122.881309 |
| 0 | 0 |
| 0 | 0 |

| | |
|---|---|
| 0 | 0 |
| 0 | 0 |
| 121.943467 | 14.4566246 |
| 0 | 0 |
| 0 | 0 |
| 0 | 0 |
| 0 | 0 |
| 65.2655174 | 0 |
| 0 | 0 |
| 0 | 0 |
| 0 | 0 |
| 0 | 0 |
| 0 | 0 |
| 0 | 0 |
| 44.655354 | 0 |
| 0 | 0 |
| 0 | 0 |
| 0 | 0 |
| 120.225953 | 93.9680598 |
| 0 | 0 |
| 0 | 0 |
| 0 | 0 |
| 0 | 0 |
| 63.5480038 | 7.22831229 |
| 0 | 0 |
| 54.9604357 | 187.93612 |
| 0 | 0 |
| 0 | 0 |
| 0 | 0 |
| 0 | 0 |
| 123.66098 | 93.9680598 |
| 0 | 0 |
| 12.0225953 | 36.1415615 |
| 360.677859 | 252.99093 |
| 0 | 0 |
| 0 | 0 |
| 0 | 0 |
| 0 | 0 |
| 0 | 0 |
| 0 | 0 |
| 0 | 0 |
| 0 | 0 |
| 0 | 0 |

| | |
|---|---|
| 0 | 0 |
| 0 | 0 |
| 0 | 0 |
| 0 | 0 |
| 0 | 0 |
| 0 | 0 |
| 0 | 7.22831229 |
| 0 | 0 |
| 0 | 0 |
| 0 | 0 |
| 0 | 0 |
| 0 | 0 |
| 1.71751362 | 144.566246 |
| 0 | 0 |
| 0 | 0 |
| 0 | 0 |
| 0 | 0 |
| 0 | 0 |
| 0 | 0 |
| 12.0225953 | 0 |
| 0 | 0 |
| 0 | 0 |
| 0 | 0 |
| 0 | 0 |
| 0 | 0 |
| 0 | 0 |
| 0 | 0 |
| 0 | 0 |
| 0 | 0 |
| 0 | 0 |
| 0 | 0 |
| 0 | 0 |
| 0 | 0 |
| 0 | 0 |
| 341.78521 | 93.9680598 |
| 34.3502723 | 0 |
| 0 | 0 |
| 0 | 0 |
| 0 | 0 |
| 0 | 0 |
| 0 | 0 |
| 0 | 0 |
| 0 | 0 |

| | |
|---|---|
| 0 | 0 |
| 147.706171 | 36.1415615 |
| 163.163794 | 7.22831229 |
| 0 | 0 |
| 0 | 0 |
| 0 | 0 |
| 0 | 0 |
| 42.9378404 | 180.707807 |
| 0 | 0 |
| 0 | 0 |
| 0 | 0 |
| 0 | 0 |
| 0 | 0 |
| 22.327677 | 0 |
| 286.824774 | 36.1415615 |
| 0 | 0 |
| 0 | 0 |
| 192.361525 | 79.5114352 |
| 0 | 0 |
| 0 | 0 |
| 0 | 14.4566246 |
| 0 | 0 |
| 0 | 7.22831229 |
| 73.8530855 | 14.4566246 |
| 168.316334 | 65.0548106 |
| 0 | 0 |
| 24.0451906 | 0 |
| 104.768331 | 28.9132492 |
| 0 | 0 |
| 0 | 0 |
| 0 | 0 |
| 0 | 0 |
| 0 | 0 |
| 0 | 0 |
| 0 | 0 |
| 0 | 0 |
| 0 | 0 |
| 0 | 0 |
| 22.327677 | 14.4566246 |
| 0 | 0 |
| 0 | 0 |
| 6.87005447 | 57.8264983 |
| 0 | 0 |

| | |
|---|---|
| 0 | 0 |
| 0 | 0 |
| 0 | 0 |
| 0 | 0 |
| 0 | 0 |
| 125.378494 | 7.22831229 |
| 0 | 0 |
| 0 | 0 |
| 0 | 0 |
| 0 | 0 |
| 58.395463 | 7.22831229 |
| 0 | 0 |
| 0 | 0 |
| 0 | 0 |
| 0 | 0 |
| 0 | 0 |
| 1.71751362 | 7.22831229 |
| 0 | 0 |
| 0 | 0 |
| 0 | 0 |
| 0 | 0 |
| 0 | 0 |
| 0 | 0 |
| 0 | 0 |
| 0 | 0 |
| 0 | 0 |
| 0 | 0 |
| 0 | 0 |
| 0 | 0 |
| 0 | 0 |
| 0 | 0 |
| 0 | 0 |
| 0 | 0 |
| 0 | 0 |
| 0 | 0 |
| 0 | 0 |
| 0 | 0 |
| 0 | 0 |
| 0 | 0 |
| 0 | 0 |
| 3.43502723 | 14.4566246 |
| 0 | 0 |
| 0 | 0 |

| | |
|---|---|
| 0 | 7.22831229 |
| 0 | 14.4566246 |
| 0 | 0 |
| 0 | 0 |
| 0 | 0 |
| 0 | 0 |
| 0 | 0 |
| 0 | 0 |
| 0 | 0 |
| 0 | 0 |
| 0 | 0 |
| 0 | 0 |
| 0 | 0 |
| 0 | 0 |
| 0 | 0 |
| 0 | 0 |
| 0 | 0 |
| 0 | 0 |
| 516.971599 | 159.02287 |
| 0 | 57.8264983 |
| 0 | 0 |
| 0 | 0 |
| 0 | 0 |
| 408.768241 | 50.598186 |
| 0 | 0 |
| 0 | 28.9132492 |
| 0 | 0 |
| 0 | 0 |
| 0 | 0 |
| 0 | 0 |
| 0 | 0 |
| 0 | 0 |
| 0 | 0 |
| 0 | 0 |
| 0 | 0 |
| 0 | 0 |
| 1.71751362 | 0 |
| 0 | 0 |
| 0 | 7.22831229 |
| 0 | 0 |
| 5.15254085 | 0 |
| 161.44628 | 65.0548106 |
| 135.683576 | 115.652997 |

| | |
|---|---|
| 1.71751362 | 0 |
| 80.72314 | 50.598186 |
| 0 | 0 |
| 0 | 0 |
| 0 | 14.4566246 |
| 0 | 0 |
| 0 | 0 |
| 0 | 7.22831229 |
| 183.773957 | 50.598186 |
| 0 | 0 |
| 0 | 0 |
| 0 | 0 |
| 0 | 0 |
| 0 | 0 |
| 0 | 0 |
| 0 | 0 |
| 0 | 0 |
| 0 | 0 |
| 0 | 0 |
| 396.745645 | 86.7397475 |
| 0 | 0 |
| 0 | 0 |
| 0 | 14.4566246 |
| 115.073412 | 0 |
| 0 | 0 |
| 0 | 0 |
| 6.87005447 | 7.22831229 |
| 0 | 0 |
| 0 | 0 |
| 0 | 0 |
| 0 | 0 |
| 0 | 0 |
| 121.943467 | 7.22831229 |
| 0 | 0 |
| 0 | 7.22831229 |
| 0 | 0 |
| 0 | 0 |
| 0 | 0 |
| 0 | 0 |
| 175.186389 | 115.652997 |
| 0 | 0 |
| 0 | 0 |
| 0 | 0 |

| | |
|---|---|
| 0 | 0 |
| 0 | 0 |
| 0 | 0 |
| 0 | 0 |
| 61.8304902 | 14.4566246 |
| 0 | 0 |
| 0 | 0 |
| 0 | 0 |
| 0 | 0 |
| 0 | 0 |
| 32.6327587 | 28.9132492 |
| 0 | 0 |
| 0 | 0 |
| 0 | 0 |
| 0 | 0 |
| 0 | 0 |
| 0 | 0 |
| 0 | 0 |
| 0 | 0 |
| 0 | 0 |
| 116.790926 | 0 |
| 0 | 0 |
| 0 | 0 |
| 0 | 0 |
| 0 | 0 |
| 0 | 0 |
| 200.949093 | 14.4566246 |
| 0 | 0 |
| 0 | 0 |
| 0 | 0 |
| 42.9378404 | 166.251183 |
| 125.378494 | 448.155362 |
| 103.050817 | 50.598186 |
| 0 | 0 |
| 0 | 0 |
| 0 | 0 |
| 0 | 0 |
| 0 | 0 |
| 0 | 0 |
| 0 | 0 |
| 0 | 7.22831229 |
| 0 | 0 |
| 0 | 0 |

| | |
|---|---|
| 0 | 0 |
| 0 | 0 |
| 0 | 0 |
| 0 | 0 |
| 0 | 0 |
| 29.1977315 | 0 |
| 285.10726 | 751.744478 |
| 0 | 0 |
| 0 | 0 |
| 0 | 0 |
| 183.773957 | 245.762618 |
| 0 | 0 |
| 1688.31588 | 787.88604 |
| 0 | 0 |
| 0 | 0 |
| 0 | 0 |
| 0 | 0 |
| 0 | 0 |
| 0 | 0 |
| 0 | 0 |
| 486.056353 | 469.840299 |
| 77.2881127 | 14.4566246 |
| 0 | 0 |
| 0 | 0 |
| 0 | 0 |
| 1.71751362 | 0 |
| 44.655354 | 0 |
| 0 | 0 |
| 0 | 0 |
| 89.3107081 | 14.4566246 |
| 15.4576225 | 72.2831229 |
| 0 | 0 |
| 0 | 0 |
| 0 | 0 |
| 0 | 0 |
| 0 | 0 |
| 0 | 0 |
| 0 | 0 |
| 0 | 0 |
| 0 | 0 |
| 0 | 7.22831229 |
| 0 | 0 |
| 0 | 0 |

| | |
|---|---|
| 0 | 7.22831229 |
| 0 | 0 |
| 0 | 0 |
| 0 | 0 |
| 187.208984 | 0 |
| 0 | 0 |
| 0 | 0 |
| 0 | 0 |
| 0 | 0 |
| 0 | 0 |
| 0 | 0 |
| 0 | 0 |
| 0 | 0 |
| 0 | 7.22831229 |
| 54.9604357 | 7.22831229 |
| 0 | 0 |
| 0 | 0 |
| 0 | 0 |
| 0 | 0 |
| 44.655354 | 21.6849369 |
| 0 | 0 |
| 0 | 0 |
| 0 | 0 |
| 0 | 0 |
| 0 | 0 |
| 0 | 0 |
| 0 | 0 |
| 0 | 0 |
| 3.43502723 | 166.251183 |
| 0 | 0 |
| 0 | 7.22831229 |
| 0 | 0 |
| 0 | 0 |
| 0 | 0 |
| 0 | 0 |
| 0 | 0 |
| 32.6327587 | 7.22831229 |
| 0 | 0 |
| 0 | 0 |
| 0 | 0 |
| 0 | 0 |
| 0 | 0 |
| 0 | 0 |

| | |
|---|---|
| 0 | 0 |
| 0 | 0 |
| 0 | 0 |
| 0 | 0 |
| 113.355899 | 86.7397475 |
| 0 | 7.22831229 |
| 0 | 0 |
| 0 | 0 |
| 0 | 0 |
| 0 | 0 |
| 0 | 0 |
| 0 | 0 |
| 0 | 0 |
| 0 | 0 |
| 0 | 0 |
| 0 | 0 |
| 36.0677859 | 802.342664 |
| 0 | 28.9132492 |
| 0 | 0 |
| 0 | 0 |
| 0 | 0 |
| 0 | 0 |
| 70.4180583 | 21.6849369 |
| 0 | 0 |
| 0 | 7.22831229 |
| 0 | 0 |
| 0 | 0 |
| 0 | 0 |
| 420.790836 | 274.675867 |
| 0 | 7.22831229 |
| 0 | 0 |
| 0 | 0 |
| 0 | 0 |
| 0 | 0 |
| 0 | 0 |
| 13.7401089 | 180.707807 |
| 24.0451906 | 28.9132492 |
| 103.050817 | 0 |
| 0 | 0 |
| 0 | 0 |
| 0 | 0 |
| 0 | 0 |
| 0 | 0 |

| | |
|---|---|
| 0 | 0 |
| 0 | 0 |
| 0 | 0 |
| 0 | 0 |
| 42.9378404 | 21.6849369 |
| 0 | 0 |
| 1.71751362 | 21.6849369 |
| 0 | 0 |
| 0 | 0 |
| 0 | 0 |
| 0 | 0 |
| 0 | 0 |
| 0 | 0 |
| 15.4576225 | 0 |
| 206.101634 | 50.598186 |
| 0 | 0 |
| 0 | 0 |
| 0 | 0 |
| 0 | 0 |
| 0 | 0 |
| 1.71751362 | 0 |
| 0 | 0 |
| 41.2203268 | 14.4566246 |
| 0 | 0 |
| 0 | 0 |
| 0 | 0 |
| 0 | 0 |
| 0 | 0 |
| 0 | 0 |
| 0 | 0 |
| 0 | 0 |
| 0 | 0 |
| 0 | 0 |
| 0 | 0 |
| 0 | 0 |
| 3.43502723 | 151.794558 |
| 0 | 0 |
| 553.039384 | 159.02287 |
| 0 | 0 |
| 0 | 0 |
| 0 | 7.22831229 |
| 0 | 0 |
| 0 | 0 |

| | |
|---|---|
| 0 | 0 |
| 0 | 0 |
| 0 | 0 |
| 156.293739 | 0 |
| 0 | 0 |
| 0 | 0 |
| 0 | 0 |
| 0 | 0 |
| 0 | 0 |
| 0 | 0 |
| 0 | 0 |
| 72.1355719 | 202.392744 |
| 255.909529 | 86.7397475 |
| 0 | 0 |
| 1.71751362 | 57.8264983 |
| 0 | 0 |
| 0 | 0 |
| 0 | 0 |
| 0 | 0 |
| 34.3502723 | 0 |
| 0 | 0 |
| 0 | 0 |
| 0 | 0 |
| 0 | 0 |
| 1.71751362 | 0 |
| 54.9604357 | 7.22831229 |
| 0 | 7.22831229 |
| 0 | 0 |
| 0 | 0 |
| 0 | 0 |
| 0 | 0 |
| 0 | 14.4566246 |
| 0 | 0 |
| 0 | 0 |
| 0 | 0 |
| 0 | 0 |
| 0 | 0 |
| 0 | 0 |
| 0 | 0 |
| 0 | 0 |
| 0 | 7.22831229 |
| 0 | 0 |
| 0 | 0 |

| | |
|---|---|
| 0 | 0 |
| 0 | 65.0548106 |
| 0 | 0 |
| 72.1355719 | 7.22831229 |
| 0 | 28.9132492 |
| 0 | 0 |
| 0 | 0 |
| 0 | 0 |
| 0 | 0 |
| 0 | 0 |
| 0 | 0 |
| 0 | 0 |
| 0 | 0 |
| 0 | 0 |
| 0 | 0 |
| 0 | 0 |
| 0 | 0 |
| 0 | 0 |
| 0 | 0 |
| 0 | 0 |
| 0 | 0 |
| 0 | 0 |
| 0 | 0 |
| 235.299365 | 159.02287 |
| 0 | 0 |
| 0 | 0 |
| 0 | 0 |
| 0 | 0 |
| 0 | 0 |
| 0 | 0 |
| 0 | 0 |
| 0 | 0 |
| 0 | 0 |
| 0 | 0 |
| 0 | 0 |
| 0 | 0 |
| 0 | 0 |
| 0 | 0 |
| 0 | 0 |
| 0 | 0 |
| 0 | 0 |
| 0 | 0 |
| 0 | 0 |

| | |
|---|---|
| 237.016879 | 43.3698738 |
| 0 | 0 |
| 0 | 0 |
| 338.350182 | 187.93612 |
| 0 | 0 |
| 0 | 0 |
| 106.485844 | 137.337934 |
| 0 | 0 |
| 0 | 0 |
| 0 | 0 |
| 0 | 0 |
| 0 | 0 |
| 0 | 0 |
| 1.71751362 | 0 |
| 65.2655174 | 21.6849369 |
| 0 | 0 |
| 0 | 36.1415615 |
| 0 | 0 |
| 0 | 0 |
| 0 | 0 |
| 1.71751362 | 0 |
| 0 | 0 |
| 0 | 0 |
| 0 | 0 |
| 0 | 0 |
| 0 | 0 |
| 0 | 0 |
| 173.468875 | 0 |
| 0 | 0 |
| 0 | 0 |
| 63.5480038 | 7.22831229 |
| 0 | 0 |
| 0 | 0 |
| 0 | 0 |
| 0 | 0 |
| 0 | 0 |
| 0 | 0 |
| 0 | 0 |
| 0 | 0 |
| 0 | 0 |
| 0 | 0 |
| 1.71751362 | 0 |
| 0 | 0 |

| | |
|---|---|
| 0 | 0 |
| 245.604447 | 144.566246 |
| 0 | 0 |
| 185.491471 | 65.0548106 |
| 15.4576225 | 65.0548106 |
| 0 | 0 |
| 0 | 0 |
| 0 | 0 |
| 0 | 0 |
| 0 | 0 |
| 0 | 0 |
| 0 | 0 |
| 0 | 0 |
| 18.8926498 | 0 |
| 0 | 0 |
| 0 | 0 |
| 0 | 0 |
| 0 | 0 |
| 0 | 0 |
| 0 | 0 |
| 0 | 0 |
| 0 | 7.22831229 |
| 0 | 0 |
| 0 | 0 |
| 0 | 0 |
| 0 | 0 |
| 0 | 0 |
| 116.790926 | 151.794558 |
| 0 | 0 |
| 0 | 0 |
| 0 | 0 |
| 96.1807625 | 173.479495 |
| 0 | 0 |
| 44.655354 | 137.337934 |
| 0 | 0 |
| 0 | 0 |
| 0 | 0 |
| 0 | 0 |
| 18.8926498 | 318.045741 |
| 0 | 0 |
| 0 | 0 |
| 108.203358 | 7.22831229 |
| 0 | 0 |

| | |
|---|---|
| 111.638385 | 0 |
| 0 | 0 |
| 0 | 0 |
| 0 | 0 |
| 0 | 0 |
| 0 | 0 |
| 0 | 0 |
| 0 | 0 |
| 0 | 0 |
| 0 | 0 |
| 0 | 7.22831229 |
| 0 | 0 |
| 0 | 0 |
| 0 | 0 |
| 0 | 0 |
| 0 | 0 |
| 0 | 0 |
| 0 | 0 |
| 0 | 0 |
| 154.576225 | 50.598186 |
| 0 | 0 |
| 0 | 0 |
| 0 | 0 |
| 0 | 0 |
| 0 | 0 |
| 0 | 0 |
| 0 | 0 |
| 0 | 0 |
| 0 | 0 |
| 0 | 0 |
| 0 | 0 |
| 0 | 0 |
| 0 | 0 |
| 0 | 0 |
| 0 | 14.4566246 |
| 264.497097 | 28.9132492 |
| 0 | 0 |
| 0 | 0 |
| 0 | 0 |
| 0 | 0 |
| 0 | 0 |
| 0 | 0 |
| 13.7401089 | 0 |

| | |
|---|---|
| 0 | 0 |
| 0 | 0 |
| 0 | 0 |
| 0 | 0 |
| 0 | 0 |
| 0 | 0 |
| 0 | 0 |
| 0 | 0 |
| 0 | 0 |
| 0 | 0 |
| 0 | 0 |
| 0 | 0 |
| 0 | 0 |
| 104.768331 | 50.598186 |
| 0 | 0 |
| 0 | 21.6849369 |
| 0 | 0 |
| 0 | 0 |
| 0 | 0 |
| 0 | 0 |
| 0 | 0 |
| 0 | 0 |
| 164.881307 | 0 |
| 0 | 7.22831229 |
| 0 | 0 |
| 42.9378404 | 0 |
| 0 | 0 |
| 0 | 0 |
| 530.711707 | 274.675867 |
| 0 | 0 |
| 0 | 0 |
| 0 | 0 |
| 0 | 0 |
| 0 | 0 |
| 65.2655174 | 130.109621 |
| 0 | 0 |
| 0 | 7.22831229 |
| 161.44628 | 130.109621 |
| 0 | 0 |
| 0 | 0 |
| 92.7457353 | 7.22831229 |
| 0 | 0 |
| 0 | 0 |

| | |
|---|---|
| 0 | 0 |
| 0 | 0 |
| 0 | 0 |
| 0 | 0 |
| 0 | 0 |
| 0 | 0 |
| 0 | 0 |
| 0 | 0 |
| 0 | 0 |
| 0 | 0 |
| 0 | 0 |
| 0 | 0 |
| 0 | 0 |
| 0 | 0 |
| 0 | 0 |
| 63.5480038 | 108.424684 |
| 0 | 0 |
| 0 | 0 |
| 0 | 0 |
| 0 | 0 |
| 0 | 0 |
| 0 | 0 |
| 278.237206 | 7.22831229 |
| 264.497097 | 7.22831229 |
| 0 | 0 |
| 0 | 0 |
| 0 | 0 |
| 0 | 0 |
| 0 | 0 |
| 0 | 0 |
| 22.327677 | 0 |
| 0 | 0 |
| 0 | 0 |
| 0 | 0 |
| 0 | 0 |
| 20.6101634 | 202.392744 |
| 0 | 0 |
| 257.627042 | 36.1415615 |
| 0 | 0 |
| 0 | 0 |
| 0 | 0 |
| 0 | 0 |
| 1.71751362 | 14.4566246 |

| | |
|---|---|
| 0 | 0 |
| 0 | 0 |
| 0 | 0 |
| 0 | 0 |
| 0 | 0 |
| 0 | 7.22831229 |
| 0 | 0 |
| 0 | 0 |
| 0 | 0 |
| 0 | 0 |
| 867.344376 | 173.479495 |
| 0 | 0 |
| 0 | 0 |
| 0 | 0 |
| 0 | 0 |
| 0 | 0 |
| 0 | 0 |
| 0 | 0 |
| 0 | 0 |
| 0 | 0 |
| 0 | 0 |
| 0 | 0 |
| 0 | 0 |
| 154.576225 | 7.22831229 |
| 0 | 0 |
| 0 | 14.4566246 |
| 230.146825 | 346.95899 |
| 5.15254085 | 0 |
| 0 | 0 |
| 0 | 0 |
| 75.5705991 | 0 |
| 0 | 0 |
| 0 | 0 |
| 0 | 0 |
| 0 | 0 |
| 0 | 0 |
| 0 | 0 |
| 233.581852 | 79.5114352 |
| 0 | 0 |
| 0 | 14.4566246 |
| 0 | 0 |
| 331.480128 | 7.22831229 |
| 0 | 0 |

| | |
|---|---|
| 0 | 0 |
| 0 | 0 |
| 0 | 0 |
| 41.2203268 | 0 |
| 0 | 0 |
| 103.050817 | 7.22831229 |
| 0 | 0 |
| 0 | 0 |
| 135.683576 | 0 |
| 0 | 0 |
| 0 | 0 |
| 0 | 0 |
| 0 | 0 |
| 0 | 0 |
| 0 | 0 |
| 0 | 0 |
| 0 | 0 |
| 66.983031 | 14.4566246 |
| 0 | 0 |
| 0 | 0 |
| 0 | 0 |
| 0 | 0 |
| 0 | 0 |
| 0 | 0 |
| 0 | 0 |
| 8.58756808 | 14.4566246 |
| 0 | 65.0548106 |
| 0 | 0 |
| 0 | 0 |
| 0 | 0 |
| 0 | 0 |
| 32.6327587 | 86.7397475 |
| 0 | 0 |
| 54.9604357 | 7.22831229 |
| 523.841653 | 144.566246 |
| 0 | 0 |
| 0 | 0 |
| 0 | 0 |
| 0 | 0 |
| 0 | 0 |
| 41.2203268 | 0 |
| 0 | 0 |
| 0 | 0 |

| | |
|---|---|
| 145.988657 | 187.93612 |
| 0 | 0 |
| 0 | 0 |
| 0 | 0 |
| 0 | 0 |
| 0 | 0 |
| 0 | 0 |
| 0 | 0 |
| 0 | 0 |
| 0 | 43.3698738 |
| 0 | 0 |
| 0 | 0 |
| 0 | 7.22831229 |
| 0 | 0 |
| 0 | 0 |
| 0 | 7.22831229 |
| 319.457533 | 187.93612 |
| 0 | 0 |
| 0 | 0 |
| 80.72314 | 0 |
| 0 | 0 |
| 0 | 0 |
| 0 | 0 |
| 0 | 0 |
| 0 | 0 |
| 0 | 0 |
| 0 | 21.6849369 |
| 420.790836 | 72.2831229 |
| 5.15254085 | 0 |
| 41.2203268 | 79.5114352 |
| 0 | 0 |
| 303.99991 | 390.328864 |
| 0 | 7.22831229 |
| 0 | 0 |
| 0 | 0 |
| 0 | 0 |
| 0 | 0 |
| 0 | 0 |
| 0 | 0 |
| 0 | 0 |
| 0 | 0 |
| 0 | 0 |
| 0 | 0 |

| | |
|---|---|
| 0 | 0 |
| 0 | 0 |
| 0 | 0 |
| 0 | 0 |
| 0 | 0 |
| 0 | 0 |
| 0 | 0 |
| 0 | 7.22831229 |
| 0 | 0 |
| 0 | 0 |
| 0 | 0 |
| 0 | 0 |
| 0 | 0 |
| 0 | 0 |
| 1.71751362 | 173.479495 |
| 0 | 0 |
| 0 | 0 |
| 1.71751362 | 0 |
| 0 | 0 |
| 786.621236 | 7286.13879 |
| 0 | 0 |
| 0 | 7.22831229 |
| 0 | 0 |
| 376.135482 | 101.196372 |
| 44.655354 | 36.1415615 |
| 154.576225 | 21.6849369 |
| 0 | 0 |
| 0 | 0 |
| 714.485664 | 383.100551 |
| 0 | 0 |
| 0 | 0 |
| 0 | 0 |
| 0 | 0 |
| 0 | 0 |
| 0 | 0 |
| 0 | 0 |
| 5.15254085 | 115.652997 |
| 41.2203268 | 21.6849369 |
| 0 | 0 |
| 0 | 7.22831229 |
| 0 | 0 |
| 0 | 0 |
| 0 | 0 |

| | |
|---|---|
| 0 | 0 |
| 0 | 0 |
| 1.71751362 | 7.22831229 |
| 0 | 0 |
| 0 | 0 |
| 0 | 0 |
| 0 | 0 |
| 0 | 0 |
| 0 | 0 |
| 0 | 0 |
| 0 | 0 |
| 34.3502723 | 202.392744 |
| 0 | 0 |
| 1.71751362 | 0 |
| 0 | 0 |
| 0 | 0 |
| 0 | 0 |
| 39.5028132 | 86.7397475 |
| 0 | 0 |
| 0 | 0 |
| 1.71751362 | 0 |
| 72.1355719 | 21.6849369 |
| 0 | 0 |
| 0 | 0 |
| 0 | 0 |
| 0 | 0 |
| 0 | 0 |
| 0 | 0 |
| 0 | 0 |
| 0 | 0 |
| 243.886934 | 79.5114352 |
| 207.819148 | 65.0548106 |
| 0 | 0 |
| 0 | 0 |
| 0 | 14.4566246 |
| 0 | 0 |
| 0 | 0 |
| 0 | 0 |
| 0 | 0 |
| 0 | 0 |
| 0 | 0 |
| 0 | 0 |
| 0 | 0 |

| | |
|---|---|
| 0 | 0 |
| 0 | 0 |
| 0 | 0 |
| 0 | 0 |
| 0 | 0 |
| 0 | 0 |
| 0 | 0 |
| 0 | 0 |
| 0 | 0 |
| 6.87005447 | 187.93612 |
| 56.6779493 | 0 |
| 0 | 0 |
| 0 | 0 |
| 0 | 0 |
| 0 | 0 |
| 94.4632489 | 0 |
| 0 | 0 |
| 0 | 0 |
| 0 | 0 |
| 0 | 0 |
| 1.71751362 | 0 |
| 0 | 0 |
| 0 | 0 |
| 97.8982761 | 7.22831229 |
| 0 | 0 |
| 0 | 0 |
| 0 | 0 |
| 0 | 0 |
| 0 | 0 |
| 0 | 0 |
| 0 | 0 |
| 56.6779493 | 0 |
| 0 | 0 |
| 0 | 0 |
| 3.43502723 | 0 |
| 79.0056264 | 0 |
| 0 | 0 |
| 0 | 50.598186 |
| 0 | 7.22831229 |
| 101.333303 | 0 |
| 0 | 0 |
| 1.71751362 | 0 |
| 0 | 0 |

| | |
|---|---|
| 0 | 0 |
| 0 | 0 |
| 0 | 0 |
| 0 | 0 |
| 0 | 0 |
| 104.768331 | 144.566246 |
| 0 | 0 |
| 0 | 0 |
| 0 | 0 |
| 0 | 0 |
| 0 | 0 |
| 58.395463 | 43.3698738 |
| 0 | 0 |
| 0 | 0 |
| 0 | 0 |
| 0 | 0 |
| 1.71751362 | 0 |
| 0 | 0 |
| 0 | 0 |
| 89.3107081 | 0 |
| 0 | 0 |
| 0 | 0 |
| 0 | 0 |
| 503.23149 | 7.22831229 |
| 0 | 0 |
| 0 | 0 |
| 0 | 0 |
| 8.58756808 | 0 |
| 0 | 0 |
| 0 | 0 |
| 133.966062 | 79.5114352 |
| 0 | 0 |
| 0 | 0 |
| 0 | 0 |
| 0 | 0 |
| 206.101634 | 0 |
| 0 | 0 |
| 0 | 0 |
| 267.932124 | 180.707807 |
| 0 | 0 |
| 0 | 0 |
| 0 | 0 |
| 0 | 0 |

| | |
|---|---|
| 0 | 0 |
| 0 | 0 |
| 0 | 0 |
| 0 | 0 |
| 0 | 0 |
| 0 | 7.22831229 |
| 0 | 0 |
| 0 | 0 |
| 0 | 0 |
| 0 | 50.598186 |
| 0 | 0 |
| 75.5705991 | 7.22831229 |
| 0 | 0 |
| 0 | 0 |
| 0 | 0 |
| 0 | 0 |
| 87.5931944 | 0 |
| 0 | 0 |
| 0 | 0 |
| 0 | 0 |
| 0 | 0 |
| 0 | 0 |
| 3.43502723 | 7.22831229 |
| 0 | 0 |
| 8.58756808 | 0 |
| 0 | 0 |
| 0 | 0 |
| 0 | 0 |
| 0 | 0 |
| 0 | 0 |
| 0 | 0 |
| 0 | 0 |
| 0 | 0 |
| 0 | 0 |
| 0 | 0 |
| 0 | 0 |
| 930.89238 | 889.082412 |
| 0 | 0 |
| 0 | 0 |
| 266.214611 | 404.785488 |
| 0 | 0 |
| 130.531035 | 289.132492 |
| 0 | 0 |

| | |
|---|---|
| 0 | 0 |
| 0 | 0 |
| 0 | 0 |
| 0 | 0 |
| 56.6779493 | 137.337934 |
| 0 | 0 |
| 0 | 0 |
| 0 | 0 |
| 1.71751362 | 151.794558 |
| 0 | 0 |
| 0 | 0 |
| 0 | 0 |
| 0 | 0 |
| 0 | 0 |
| 121.943467 | 7.22831229 |
| 293.694828 | 65.0548106 |
| 0 | 0 |
| 0 | 0 |
| 0 | 0 |
| 0 | 0 |
| 0 | 0 |
| 0 | 0 |
| 0 | 0 |
| 176.903902 | 79.5114352 |
| 0 | 0 |
| 10.3050817 | 79.5114352 |
| 0 | 0 |
| 0 | 0 |
| 0 | 0 |
| 29.1977315 | 108.424684 |
| 0 | 0 |
| 0 | 0 |
| 0 | 0 |
| 326.327587 | 0 |
| 133.966062 | 79.5114352 |
| 0 | 0 |
| 0 | 0 |
| 0 | 0 |
| 0 | 0 |
| 0 | 0 |
| 5.15254085 | 36.1415615 |
| 61.8304902 | 79.5114352 |
| 0 | 0 |

| | |
|---|---|
| 0 | 21.6849369 |
| 10.3050817 | 72.2831229 |
| 142.55363 | 0 |
| 0 | 0 |
| 0 | 0 |
| 0 | 0 |
| 0 | 0 |
| 0 | 0 |
| 20.6101634 | 0 |
| 0 | 0 |
| 15.4576225 | 36.1415615 |
| 152.858712 | 36.1415615 |
| 0 | 0 |
| 0 | 0 |
| 0 | 0 |
| 0 | 0 |
| 0 | 0 |
| 0 | 0 |
| 0 | 0 |
| 154.576225 | 0 |
| 84.1581672 | 14.4566246 |
| 0 | 0 |
| 0 | 0 |
| 92.7457353 | 7.22831229 |
| 0 | 0 |
| 24.0451906 | 0 |
| 145.988657 | 50.598186 |
| 0 | 0 |
| 0 | 0 |
| 0 | 0 |
| 0 | 0 |
| 202.666607 | 28.9132492 |
| 0 | 0 |
| 130.531035 | 0 |
| 8.58756808 | 79.5114352 |
| 0 | 0 |
| 0 | 0 |
| 0 | 0 |
| 0 | 0 |
| 0 | 0 |
| 0 | 0 |
| 0 | 7.22831229 |
| 0 | 0 |

| | |
|---|---|
| 1.71751362 | 14.4566246 |
| 221.559257 | 36.1415615 |
| 0 | 0 |
| 0 | 0 |
| 10.3050817 | 57.8264983 |
| 34.3502723 | 0 |
| 0 | 0 |
| 0 | 0 |
| 0 | 0 |
| 0 | 0 |
| 0 | 0 |
| 41.2203268 | 108.424684 |
| 0 | 0 |
| 0 | 0 |
| 0 | 0 |
| 0 | 0 |
| 0 | 0 |
| 0 | 0 |
| 0 | 0 |
| 0 | 0 |
| 0 | 36.1415615 |
| 0 | 0 |
| 0 | 0 |
| 0 | 0 |
| 0 | 0 |
| 0 | 0 |
| 0 | 0 |
| 0 | 0 |
| 0 | 0 |
| 0 | 0 |
| 0 | 0 |
| 0 | 0 |
| 0 | 0 |
| 0 | 7.22831229 |
| 0 | 0 |
| 0 | 0 |
| 0 | 0 |
| 0 | 0 |
| 0 | 0 |
| 0 | 0 |
| 0 | 0 |
| 0 | 0 |
| 0 | 0 |

| | |
|---|---|
| 0 | 0 |
| 0 | 0 |
| 0 | 0 |
| 0 | 0 |
| 0 | 0 |
| 0 | 0 |
| 0 | 0 |
| 0 | 0 |
| 0 | 0 |
| 0 | 0 |
| 0 | 0 |
| 0 | 0 |
| 0 | 0 |
| 0 | 0 |
| 0 | 0 |
| 0 | 0 |
| 0 | 0 |
| 0 | 0 |
| 0 | 0 |
| 0 | 0 |
| 0 | 0 |
| 37.7852996 | 144.566246 |
| 0 | 0 |
| 0 | 0 |
| 0 | 0 |
| 0 | 0 |
| 0 | 0 |
| 0 | 0 |
| 0 | 0 |
| 0 | 0 |
| 0 | 0 |
| 0 | 0 |
| 0 | 0 |
| 0 | 0 |
| 0 | 14.4566246 |
| 0 | 0 |
| 0 | 0 |
| 0 | 0 |
| 0 | 0 |
| 0 | 0 |
| 0 | 0 |
| 0 | 0 |
| 0 | 0 |

| | |
|---|---|
| 0 | 0 |
| 0 | 0 |
| 0 | 0 |
| 0 | 0 |
| 0 | 7.22831229 |
| 173.468875 | 166.251183 |
| 249.039474 | 440.92705 |
| 0 | 0 |
| 27.4802179 | 43.3698738 |
| 66.983031 | 339.730678 |
| 0 | 0 |
| 48.0903813 | 50.598186 |
| 0 | 0 |
| 0 | 0 |
| 1.71751362 | 0 |
| 0 | 14.4566246 |
| 0 | 0 |
| 0 | 0 |
| 0 | 0 |
| 0 | 0 |
| 0 | 0 |
| 115.073412 | 65.0548106 |
| 0 | 0 |
| 0 | 0 |
| 0 | 0 |
| 0 | 0 |
| 0 | 0 |
| 0 | 0 |
| 0 | 0 |
| 0 | 0 |
| 0 | 0 |
| 0 | 0 |
| 0 | 0 |
| 0 | 0 |
| 0 | 0 |
| 0 | 0 |
| 0 | 0 |
| 0 | 0 |
| 109.920871 | 231.305993 |
| 0 | 0 |
| 0 | 0 |
| 231.864338 | 122.881309 |
| 0 | 0 |

| | |
|---|---|
| 89.3107081 | 14.4566246 |
| 0 | 0 |
| 44.655354 | 0 |
| 202.666607 | 0 |
| 0 | 0 |
| 0 | 0 |
| 0 | 0 |
| 0 | 0 |
| 0 | 0 |
| 0 | 0 |
| 0 | 0 |
| 0 | 0 |
| 3.43502723 | 36.1415615 |
| 0 | 0 |
| 0 | 0 |
| 0 | 0 |
| 6.87005447 | 0 |
| 182.056443 | 36.1415615 |
| 75.5705991 | 79.5114352 |
| 0 | 0 |
| 0 | 0 |
| 0 | 0 |
| 0 | 0 |
| 87.5931944 | 21.6849369 |
| 0 | 0 |
| 0 | 0 |
| 0 | 0 |
| 0 | 0 |
| 0 | 0 |
| 0 | 0 |
| 0 | 0 |
| 0 | 7.22831229 |
| 0 | 0 |
| 65.2655174 | 0 |
| 0 | 14.4566246 |
| 0 | 0 |
| 0 | 0 |
| 0 | 0 |
| 0 | 0 |
| 0 | 0 |
| 0 | 0 |
| 0 | 0 |
| 0 | 0 |

| | |
|---|---|
| 135.683576 | 36.1415615 |
| 15.4576225 | 0 |
| 0 | 0 |
| 0 | 0 |
| 0 | 0 |
| 0 | 0 |
| 0 | 0 |
| 0 | 0 |
| 0 | 0 |
| 0 | 0 |
| 0 | 0 |
| 0 | 0 |
| 27.4802179 | 7.22831229 |
| 0 | 0 |
| 0 | 0 |
| 0 | 0 |
| 0 | 0 |
| 0 | 0 |
| 0 | 0 |
| 0 | 0 |
| 0 | 0 |
| 0 | 0 |
| 0 | 0 |
| 0 | 0 |
| 0 | 0 |
| 0 | 0 |
| 0 | 28.9132492 |
| 0 | 0 |
| 144.271144 | 274.675867 |
| 0 | 7.22831229 |
| 0 | 0 |
| 0 | 0 |
| 123.66098 | 0 |
| 0 | 0 |
| 0 | 0 |
| 0 | 0 |
| 0 | 0 |
| 0 | 0 |
| 518.689112 | 101.196372 |
| 0 | 0 |
| 0 | 0 |
| 77.2881127 | 122.881309 |
| 0 | 0 |

| | |
|---|---|
| 0 | 0 |
| 0 | 0 |
| 0 | 0 |
| 0 | 0 |
| 0 | 0 |
| 0 | 0 |
| 0 | 0 |
| 0 | 0 |
| 0 | 7.22831229 |
| 0 | 0 |
| 0 | 0 |
| 0 | 0 |
| 0 | 0 |
| 0 | 7.22831229 |
| 0 | 0 |
| 0 | 0 |
| 0 | 0 |
| 0 | 0 |
| 0 | 0 |
| 0 | 0 |
| 0 | 0 |
| 0 | 0 |
| 1.71751362 | 65.0548106 |
| 0 | 0 |
| 0 | 0 |
| 0 | 0 |
| 0 | 0 |
| 70.4180583 | 0 |
| 0 | 0 |
| 120.225953 | 0 |
| 154.576225 | 0 |
| 0 | 7.22831229 |
| 0 | 0 |
| 0 | 0 |
| 0 | 0 |
| 0 | 0 |
| 0 | 57.8264983 |
| 96.1807625 | 28.9132492 |
| 37.7852996 | 7.22831229 |
| 0 | 7.22831229 |
| 0 | 0 |
| 0 | 0 |
| 0 | 0 |

| | |
|---|---|
| 1.71751362 | 144.566246 |
| 0 | 0 |
| 0 | 0 |
| 0 | 0 |
| 0 | 0 |
| 0 | 0 |
| 60.1129766 | 0 |
| 0 | 0 |
| 154.576225 | 0 |
| 0 | 0 |
| 0 | 0 |
| 0 | 0 |
| 154.576225 | 28.9132492 |
| 0 | 0 |
| 20.6101634 | 144.566246 |
| 84.1581672 | 43.3698738 |
| 60.1129766 | 0 |
| 0 | 0 |
| 0 | 0 |
| 0 | 0 |
| 0 | 0 |
| 0 | 0 |
| 0 | 0 |
| 0 | 7.22831229 |
| 0 | 0 |
| 0 | 50.598186 |
| 0 | 0 |
| 0 | 0 |
| 0 | 0 |
| 106.485844 | 43.3698738 |
| 0 | 0 |
| 0 | 0 |
| 0 | 0 |
| 0 | 0 |
| 0 | 0 |
| 8.58756808 | 0 |
| 0 | 0 |
| 70.4180583 | 93.9680598 |
| 439.683486 | 187.93612 |
| 3.43502723 | 0 |
| 0 | 0 |
| 75.5705991 | 65.0548106 |
| 0 | 0 |

| | |
|---|---|
| 0 | 0 |
| 0 | 0 |
| 20.6101634 | 65.0548106 |
| 0 | 0 |
| 869.06189 | 0 |
| 0 | 7.22831229 |
| 0 | 0 |
| 0 | 0 |
| 0 | 0 |
| 0 | 0 |
| 0 | 0 |
| 252.474502 | 477.068611 |
| 0 | 0 |
| 0 | 0 |
| 164.881307 | 101.196372 |
| 168.316334 | 0 |
| 5.15254085 | 0 |
| 0 | 0 |
| 0 | 0 |
| 0 | 7.22831229 |
| 0 | 0 |
| 0 | 0 |
| 0 | 0 |
| 0 | 0 |
| 0 | 0 |
| 0 | 0 |
| 0 | 0 |
| 8.58756808 | 72.2831229 |
| 0 | 0 |
| 56.6779493 | 0 |
| 0 | 0 |
| 0 | 0 |
| 0 | 0 |
| 0 | 0 |
| 0 | 0 |
| 0 | 0 |
| 0 | 0 |
| 0 | 0 |
| 171.751362 | 0 |
| 0 | 0 |
| 0 | 0 |
| 0 | 0 |
| 0 | 0 |

| | |
|---|---|
| 0 | 0 |
| 0 | 0 |
| 0 | 0 |
| 0 | 0 |
| 0 | 0 |
| 0 | 0 |
| 0 | 0 |
| 0 | 0 |
| 0 | 0 |
| 0 | 0 |
| 0 | 0 |
| 127.096008 | 0 |
| 0 | 0 |
| 0 | 0 |
| 0 | 0 |
| 175.186389 | 0 |
| 0 | 0 |
| 147.706171 | 0 |
| 0 | 0 |
| 133.966062 | 159.02287 |
| 0 | 0 |
| 0 | 0 |
| 29.1977315 | 93.9680598 |
| 0 | 0 |
| 0 | 0 |
| 0 | 0 |
| 108.203358 | 7546.35803 |
| 278.237206 | 0 |
| 0 | 0 |
| 0 | 0 |
| 0 | 0 |
| 441.400999 | 390.328864 |
| 0 | 0 |
| 0 | 0 |
| 29.1977315 | 7.22831229 |
| 0 | 0 |
| 0 | 0 |
| 0 | 0 |
| 158.011253 | 137.337934 |
| 0 | 0 |
| 0 | 0 |
| 0 | 0 |
| 0 | 0 |

| | |
|---|---|
| 0 | 0 |
| 0 | 0 |
| 0 | 28.9132492 |
| 0 | 0 |
| 0 | 0 |
| 0 | 0 |
| 68.7005447 | 7.22831229 |
| 79.0056264 | 0 |
| 247.321961 | 7.22831229 |
| 0 | 0 |
| 0 | 0 |
| 0 | 0 |
| 0 | 0 |
| 0 | 0 |
| 106.485844 | 0 |
| 0 | 0 |
| 0 | 0 |
| 0 | 0 |
| 1.71751362 | 28.9132492 |
| 0 | 0 |
| 0 | 0 |
| 0 | 0 |
| 0 | 0 |
| 0 | 0 |
| 0 | 0 |
| 0 | 7.22831229 |
| 127.096008 | 7.22831229 |
| 39.5028132 | 14.4566246 |
| 0 | 0 |
| 0 | 0 |
| 154.576225 | 14.4566246 |
| 0 | 0 |
| 0 | 0 |
| 0 | 0 |
| 0 | 0 |
| 145.988657 | 0 |
| 13.7401089 | 79.5114352 |
| 0 | 0 |
| 0 | 0 |
| 0 | 0 |
| 0 | 0 |
| 0 | 0 |
| 0 | 0 |

| | |
|---|---|
| 0 | 0 |
| 0 | 0 |
| 0 | 0 |
| 0 | 0 |
| 0 | 0 |
| 85.8756808 | 7.22831229 |
| 0 | 0 |
| 0 | 0 |
| 0 | 0 |
| 0 | 0 |
| 0 | 0 |
| 1.71751362 | 0 |
| 5.15254085 | 79.5114352 |
| 0 | 0 |
| 0 | 0 |
| 0 | 0 |
| 73.8530855 | 0 |
| 0 | 0 |
| 0 | 0 |
| 44.655354 | 0 |
| 0 | 0 |
| 0 | 0 |
| 0 | 0 |
| 104.768331 | 7.22831229 |
| 0 | 0 |
| 0 | 0 |
| 0 | 0 |
| 235.299365 | 86.7397475 |
| 0 | 0 |
| 0 | 0 |
| 0 | 0 |
| 0 | 65.0548106 |
| 0 | 0 |
| 0 | 0 |
| 0 | 0 |
| 0 | 0 |
| 1.71751362 | 0 |
| 1.71751362 | 50.598186 |
| 0 | 0 |
| 0 | 0 |
| 0 | 0 |
| 0 | 0 |
| 0 | 0 |

| | |
|---|---|
| 0 | 0 |
| 0 | 0 |
| 0 | 0 |
| 0 | 14.4566246 |
| 0 | 0 |
| 269.649638 | 173.479495 |
| 0 | 0 |
| 0 | 0 |
| 0 | 0 |
| 0 | 0 |
| 0 | 0 |
| 0 | 0 |
| 0 | 7.22831229 |
| 0 | 0 |
| 0 | 0 |
| 0 | 0 |
| 0 | 7.22831229 |
| 0 | 0 |
| 0 | 0 |
| 0 | 36.1415615 |
| 0 | 0 |
| 0 | 0 |
| 0 | 0 |
| 0 | 0 |
| 0 | 0 |
| 0 | 0 |
| 17.1751362 | 36.1415615 |
| 0 | 0 |
| 0 | 0 |
| 0 | 0 |
| 530.711707 | 852.94085 |
| 151.141198 | 0 |
| 0 | 0 |
| 0 | 0 |
| 0 | 0 |
| 0 | 0 |
| 0 | 0 |
| 97.8982761 | 0 |
| 0 | 0 |
| 0 | 0 |
| 0 | 0 |
| 0 | 0 |
| 185.491471 | 0 |

| | |
|---|---|
| 0 | 0 |
| 0 | 0 |
| 0 | 0 |
| 0 | 0 |
| 0 | 0 |
| 0 | 0 |
| 0 | 0 |
| 0 | 0 |
| 0 | 0 |
| 0 | 0 |
| 53.2429221 | 180.707807 |
| 0 | 0 |
| 0 | 166.251183 |
| 130.531035 | 14.4566246 |
| 0 | 0 |
| 214.689202 | 0 |
| 0 | 0 |
| 0 | 0 |
| 0 | 0 |
| 0 | 0 |
| 0 | 0 |
| 0 | 0 |
| 0 | 0 |
| 0 | 0 |
| 0 | 0 |
| 0 | 0 |
| 30.9152451 | 455.383674 |
| 0 | 0 |
| 0 | 0 |
| 0 | 0 |
| 118.50844 | 0 |
| 0 | 0 |
| 0 | 0 |
| 0 | 0 |
| 0 | 0 |
| 0 | 0 |
| 0 | 0 |
| 0 | 0 |
| 0 | 7.22831229 |
| 0 | 0 |
| 0 | 0 |
| 58.395463 | 21.6849369 |
| 0 | 0 |

| | |
|---|---|
| 0 | 0 |
| 0 | 0 |
| 0 | 0 |
| 0 | 0 |
| 0 | 0 |
| 0 | 0 |
| 10.3050817 | 21.6849369 |
| 42.9378404 | 195.164432 |
| 0 | 0 |
| 0 | 0 |
| 0 | 0 |
| 0 | 0 |
| 0 | 0 |
| 0 | 0 |
| 0 | 0 |
| 0 | 0 |
| 0 | 0 |
| 0 | 0 |
| 176.903902 | 0 |
| 0 | 0 |
| 0 | 0 |
| 51.5254085 | 14.4566246 |
| 0 | 0 |
| 1.71751362 | 101.196372 |
| 0 | 0 |
| 0 | 0 |
| 70.4180583 | 36.1415615 |
| 108.203358 | 0 |
| 1.71751362 | 0 |
| 333.197642 | 115.652997 |
| 0 | 0 |
| 0 | 0 |
| 0 | 7.22831229 |
| 0 | 0 |
| 0 | 0 |
| 0 | 0 |
| 0 | 0 |
| 41.2203268 | 130.109621 |
| 0 | 0 |
| 0 | 0 |
| 1.71751362 | 0 |
| 0 | 43.3698738 |
| 0 | 0 |

| | |
|---|---|
| 0 | 0 |
| 180.33893 | 0 |
| 145.988657 | 0 |
| 87.5931944 | 0 |
| 0 | 0 |
| 0 | 137.337934 |
| 0 | 0 |
| 0 | 0 |
| 0 | 0 |
| 0 | 0 |
| 0 | 0 |
| 0 | 0 |
| 0 | 0 |
| 0 | 0 |
| 0 | 0 |
| 0 | 0 |
| 0 | 0 |
| 0 | 0 |
| 0 | 0 |
| 0 | 0 |
| 0 | 0 |
| 135.683576 | 108.424684 |
| 0 | 0 |
| 0 | 0 |
| 5.15254085 | 0 |
| 0 | 0 |
| 0 | 0 |
| 0 | 0 |
| 66.983031 | 130.109621 |
| 0 | 0 |
| 0 | 0 |
| 0 | 0 |
| 0 | 0 |
| 0 | 0 |
| 0 | 0 |
| 13.7401089 | 50.598186 |
| 0 | 14.4566246 |
| 0 | 0 |
| 0 | 0 |
| 0 | 0 |
| 0 | 0 |
| 0 | 0 |
| 0 | 0 |

| | |
|---|---|
| 0 | 0 |
| 556.474412 | 231.305993 |
| 0 | 7.22831229 |
| 0 | 0 |
| 0 | 0 |
| 0 | 0 |
| 0 | 0 |
| 0 | 0 |
| 0 | 0 |
| 0 | 0 |
| 0 | 65.0548106 |
| 0 | 0 |
| 0 | 0 |
| 85.8756808 | 0 |
| 0 | 0 |
| 0 | 0 |
| 12.0225953 | 108.424684 |
| 6.87005447 | 28.9132492 |
| 0 | 0 |
| 0 | 0 |
| 0 | 0 |
| 0 | 0 |
| 0 | 0 |
| 0 | 0 |
| 0 | 0 |
| 0 | 0 |
| 0 | 0 |
| 0 | 0 |
| 0 | 0 |
| 0 | 0 |
| 0 | 0 |
| 0 | 0 |
| 0 | 0 |
| 0 | 0 |
| 0 | 0 |
| 0 | 0 |
| 0 | 0 |
| 0 | 0 |
| 0 | 0 |
| 0 | 0 |
| 0 | 0 |
| 0 | 0 |
| 0 | 0 |
| 0 | 0 |

| | |
|---|---|
| 0 | 0 |
| 0 | 0 |
| 0 | 0 |
| 0 | 0 |
| 0 | 0 |
| 0 | 36.1415615 |
| 0 | 0 |
| 0 | 28.9132492 |
| 0 | 0 |
| 125.378494 | 21.6849369 |
| 0 | 0 |
| 0 | 0 |
| 1.71751362 | 0 |
| 48.0903813 | 0 |
| 0 | 0 |
| 0 | 0 |
| 0 | 0 |
| 0 | 0 |
| 75.5705991 | 7.22831229 |
| 0 | 0 |
| 39.5028132 | 43.3698738 |
| 0 | 0 |
| 0 | 0 |
| 0 | 28.9132492 |
| 0 | 0 |
| 0 | 0 |
| 0 | 7.22831229 |
| 0 | 0 |
| 0 | 0 |
| 0 | 0 |
| 0 | 0 |
| 0 | 0 |
| 0 | 0 |
| 13.7401089 | 7.22831229 |
| 0 | 0 |
| 8.58756808 | 202.392744 |
| 0 | 0 |
| 0 | 0 |
| 0 | 0 |
| 0 | 36.1415615 |
| 0 | 7.22831229 |
| 0 | 0 |
| 0 | 0 |

| | |
|---|---|
| 0 | 0 |
| 0 | 0 |
| 0 | 0 |
| 0 | 0 |
| 0 | 0 |
| 0 | 0 |
| 0 | 0 |
| 29.1977315 | 0 |
| 0 | 0 |
| 18.8926498 | 0 |
| 0 | 0 |
| 0 | 0 |
| 13.7401089 | 0 |
| 0 | 0 |
| 0 | 0 |
| 0 | 0 |
| 173.468875 | 0 |
| 0 | 0 |
| 0 | 0 |
| 0 | 0 |
| 0 | 0 |
| 0 | 0 |
| 0 | 0 |
| 0 | 0 |
| 0 | 0 |
| 0 | 0 |
| 0 | 0 |
| 73.8530855 | 0 |
| 0 | 0 |
| 0 | 0 |
| 0 | 7.22831229 |
| 0 | 0 |
| 27.4802179 | 101.196372 |
| 0 | 0 |
| 0 | 0 |
| 142.55363 | 0 |
| 91.0282217 | 50.598186 |
| 0 | 0 |
| 0 | 0 |
| 0 | 7.22831229 |
| 0 | 0 |
| 0 | 0 |
| 0 | 0 |

| | |
|---|---|
| 0 | 0 |
| 0 | 0 |
| 0 | 0 |
| 0 | 0 |
| 0 | 7.22831229 |
| 0 | 0 |
| 0 | 0 |
| 79.0056264 | 7.22831229 |
| 49.8078949 | 115.652997 |
| 0 | 0 |
| 0 | 0 |
| 0 | 0 |
| 0 | 0 |
| 0 | 0 |
| 0 | 0 |
| 0 | 0 |
| 0 | 0 |
| 145.988657 | 57.8264983 |
| 0 | 0 |
| 0 | 0 |
| 97.8982761 | 28.9132492 |
| 0 | 0 |
| 0 | 0 |
| 0 | 0 |
| 0 | 0 |
| 41.2203268 | 0 |
| 0 | 0 |
| 0 | 0 |
| 0 | 0 |
| 0 | 0 |
| 0 | 0 |
| 3.43502723 | 93.9680598 |
| 0 | 0 |
| 0 | 0 |
| 0 | 0 |
| 0 | 0 |
| 0 | 0 |
| 0 | 0 |
| 0 | 7.22831229 |
| 0 | 115.652997 |
| 0 | 0 |
| 0 | 0 |
| 0 | 0 |

| | |
|---|---|
| 0 | 0 |
| 0 | 0 |
| 0 | 0 |
| 73.8530855 | 0 |
| 0 | 0 |
| 0 | 0 |
| 0 | 0 |
| 0 | 0 |
| 0 | 0 |
| 0 | 108.424684 |
| 82.4406536 | 0 |
| 477.468785 | 14.4566246 |
| 18.8926498 | 14.4566246 |
| 0 | 0 |
| 0 | 0 |
| 0 | 0 |
| 0 | 0 |
| 80.72314 | 151.794558 |
| 0 | 0 |
| 0 | 0 |
| 0 | 0 |
| 66.983031 | 144.566246 |
| 0 | 0 |
| 72.1355719 | 7.22831229 |
| 73.8530855 | 0 |
| 0 | 0 |
| 0 | 7.22831229 |
| 0 | 0 |
| 0 | 0 |
| 0 | 0 |
| 0 | 0 |
| 0 | 0 |
| 0 | 0 |
| 70.4180583 | 0 |
| 0 | 0 |
| 1.71751362 | 0 |
| 0 | 0 |
| 0 | 0 |
| 0 | 0 |
| 0 | 0 |
| 0 | 0 |
| 1.71751362 | 0 |
| 0 | 0 |

| | |
|---|---|
| 0 | 7.22831229 |
| 0 | 0 |
| 0 | 0 |
| 132.248548 | 0 |
| 0 | 0 |
| 0 | 0 |
| 0 | 0 |
| 0 | 0 |
| 242.16942 | 310.817429 |
| 0 | 0 |
| 0 | 0 |
| 0 | 0 |
| 0 | 0 |
| 0 | 0 |
| 259.344556 | 14.4566246 |
| 0 | 0 |
| 190.644011 | 7.22831229 |
| 0 | 0 |
| 0 | 0 |
| 0 | 0 |
| 0 | 0 |
| 121.943467 | 21.6849369 |
| 0 | 0 |
| 0 | 0 |
| 0 | 0 |
| 0 | 0 |
| 0 | 0 |
| 0 | 0 |
| 60.1129766 | 7.22831229 |
| 0 | 0 |
| 0 | 0 |
| 0 | 0 |
| 0 | 0 |
| 0 | 0 |
| 0 | 0 |
| 0 | 0 |
| 0 | 0 |
| 0 | 0 |
| 0 | 0 |
| 0 | 0 |
| 89.3107081 | 0 |
| 0 | 0 |
| 0 | 0 |

| | |
|---|---|
| 0 | 0 |
| 0 | 0 |
| 0 | 0 |
| 0 | 0 |
| 0 | 0 |
| 0 | 0 |
| 0 | 0 |
| 0 | 0 |
| 0 | 0 |
| 53.2429221 | 0 |
| 0 | 0 |
| 0 | 0 |
| 168.316334 | 824.027601 |
| 0 | 0 |
| 0 | 0 |
| 0 | 0 |
| 0 | 7.22831229 |
| 0 | 0 |
| 0 | 0 |
| 0 | 0 |
| 0 | 0 |
| 0 | 0 |
| 0 | 0 |
| 0 | 0 |
| 424.225863 | 303.589116 |
| 0 | 0 |
| 0 | 0 |
| 0 | 14.4566246 |
| 0 | 14.4566246 |
| 27.4802179 | 86.7397475 |
| 0 | 7.22831229 |
| 0 | 0 |
| 0 | 0 |
| 224.994284 | 0 |
| 0 | 0 |
| 0 | 0 |
| 0 | 0 |
| 0 | 0 |
| 0 | 0 |
| 0 | 36.1415615 |
| 0 | 0 |
| 29.1977315 | 0 |
| 92.7457353 | 0 |

| | |
|---|---|
| 0 | 0 |
| 87.5931944 | 0 |
| 0 | 0 |
| 121.943467 | 0 |
| 0 | 0 |
| 8.58756808 | 7.22831229 |
| 0 | 0 |
| 0 | 151.794558 |
| 51.5254085 | 7.22831229 |
| 0 | 0 |
| 0 | 7.22831229 |
| 0 | 0 |
| 113.355899 | 14.4566246 |
| 0 | 0 |
| 0 | 0 |
| 0 | 28.9132492 |
| 0 | 0 |
| 434.530945 | 14.4566246 |
| 58.395463 | 0 |
| 0 | 0 |
| 0 | 0 |
| 0 | 7.22831229 |
| 152.858712 | 122.881309 |
| 0 | 0 |
| 0 | 7.22831229 |
| 0 | 0 |
| 6.87005447 | 14.4566246 |
| 0 | 0 |
| 0 | 115.652997 |
| 0 | 0 |
| 0 | 0 |
| 0 | 0 |
| 0 | 0 |
| 0 | 0 |
| 0 | 0 |
| 0 | 0 |
| 0 | 0 |
| 0 | 0 |
| 0 | 0 |
| 0 | 0 |
| 0 | 0 |
| 18.8926498 | 122.881309 |
| 0 | 0 |

| | |
|---|---|
| 0 | 0 |
| 48.0903813 | 0 |
| 0 | 0 |
| 0 | 0 |
| 0 | 0 |
| 0 | 0 |
| 0 | 0 |
| 0 | 0 |
| 0 | 0 |
| 0 | 0 |
| 0 | 0 |
| 0 | 0 |
| 0 | 0 |
| 0 | 0 |
| 0 | 0 |
| 25.7627042 | 0 |
| 0 | 14.4566246 |
| 0 | 0 |
| 79.0056264 | 14.4566246 |
| 0 | 0 |
| 0 | 0 |
| 0 | 0 |
| 0 | 0 |
| 42.9378404 | 72.2831229 |
| 0 | 0 |
| 0 | 0 |
| 0 | 0 |
| 0 | 0 |
| 0 | 0 |
| 0 | 0 |
| 0 | 0 |
| 0 | 0 |
| 0 | 0 |
| 0 | 0 |
| 17.1751362 | 0 |
| 5.15254085 | 14.4566246 |
| 0 | 0 |
| 0 | 0 |
| 0 | 0 |
| 0 | 0 |
| 0 | 0 |
| 0 | 0 |
| 0 | 0 |

| | |
|---|---|
| 0 | 0 |
| 0 | 0 |
| 0 | 0 |
| 0 | 14.4566246 |
| 0 | 0 |
| 0 | 0 |
| 0 | 0 |
| 1.71751362 | 7.22831229 |
| 0 | 0 |
| 0 | 0 |
| 0 | 0 |
| 0 | 0 |
| 0 | 7.22831229 |
| 0 | 0 |
| 0 | 0 |
| 42.9378404 | 28.9132492 |
| 0 | 0 |
| 0 | 0 |
| 0 | 0 |
| 0 | 0 |
| 0 | 0 |
| 158.011253 | 7.22831229 |
| 267.932124 | 14.4566246 |
| 0 | 0 |
| 0 | 0 |
| 0 | 0 |
| 0 | 0 |
| 541.016789 | 303.589116 |
| 0 | 0 |
| 0 | 0 |
| 0 | 0 |
| 0 | 0 |
| 0 | 0 |
| 30.9152451 | 0 |
| 44.655354 | 43.3698738 |
| 0 | 0 |
| 0 | 0 |
| 0 | 0 |
| 0 | 0 |
| 0 | 0 |
| 0 | 0 |
| 0 | 0 |
| 159.728766 | 79.5114352 |

| | |
|---|---|
| 0 | 0 |
| 0 | 0 |
| 0 | 0 |
| 0 | 0 |
| 0 | 0 |
| 0 | 0 |
| 0 | 0 |
| 0 | 0 |
| 0 | 0 |
| 0 | 0 |
| 116.790926 | 93.9680598 |
| 0 | 0 |
| 3.43502723 | 0 |
| 3.43502723 | 86.7397475 |
| 0 | 0 |
| 44.655354 | 14.4566246 |
| 0 | 0 |
| 0 | 0 |
| 0 | 14.4566246 |
| 0 | 0 |
| 0 | 0 |
| 192.361525 | 14.4566246 |
| 0 | 0 |
| 6.87005447 | 65.0548106 |
| 0 | 0 |
| 0 | 0 |
| 0 | 0 |
| 0 | 0 |
| 1.71751362 | 0 |
| 0 | 0 |
| 0 | 0 |
| 0 | 0 |
| 0 | 0 |
| 1.71751362 | 0 |
| 0 | 43.3698738 |
| 1.71751362 | 151.794558 |
| 197.514066 | 79.5114352 |
| 30.9152451 | 0 |
| 0 | 0 |
| 106.485844 | 0 |
| 0 | 0 |
| 27.4802179 | 79.5114352 |
| 66.983031 | 0 |

| | |
|---|---|
| 0 | 0 |
| 0 | 0 |
| 0 | 0 |
| 25.7627042 | 93.9680598 |
| 0 | 0 |
| 0 | 0 |
| 0 | 0 |
| 0 | 0 |
| 0 | 7.22831229 |
| 0 | 0 |
| 0 | 0 |
| 0 | 0 |
| 116.790926 | 14.4566246 |
| 487.773867 | 404.785488 |
| 0 | 0 |
| 0 | 0 |
| 0 | 0 |
| 0 | 0 |
| 0 | 0 |
| 0 | 0 |
| 0 | 0 |
| 0 | 0 |
| 0 | 0 |
| 0 | 0 |
| 0 | 0 |
| 0 | 0 |
| 195.796552 | 65.0548106 |
| 0 | 0 |
| 18.8926498 | 72.2831229 |
| 109.920871 | 0 |
| 0 | 0 |
| 0 | 0 |
| 0 | 0 |
| 309.152451 | 57.8264983 |
| 10.3050817 | 21.6849369 |
| 0 | 0 |
| 0 | 0 |
| 0 | 130.109621 |
| 0 | 0 |
| 0 | 0 |
| 0 | 0 |
| 0 | 7.22831229 |
| 0 | 0 |

| | |
|---|---|
| 0 | 0 |
| 0 | 0 |
| 0 | 0 |
| 0 | 0 |
| 0 | 0 |
| 0 | 0 |
| 0 | 0 |
| 0 | 0 |
| 0 | 0 |
| 0 | 86.7397475 |
| 0 | 0 |
| 0 | 202.392744 |
| 0 | 0 |
| 0 | 0 |
| 0 | 0 |
| 51.5254085 | 50.598186 |
| 0 | 0 |
| 0 | 0 |
| 0 | 0 |
| 0 | 0 |
| 0 | 0 |
| 0 | 0 |
| 56.6779493 | 0 |
| 0 | 0 |
| 139.118603 | 0 |
| 0 | 0 |
| 0 | 0 |
| 0 | 7.22831229 |
| 115.073412 | 0 |
| 0 | 0 |
| 46.3728676 | 0 |
| 0 | 0 |
| 0 | 0 |
| 0 | 7.22831229 |
| 0 | 43.3698738 |
| 34.3502723 | 0 |
| 0 | 0 |
| 0 | 0 |
| 0 | 0 |
| 0 | 0 |
| 89.3107081 | 0 |
| 0 | 0 |
| 0 | 0 |

| | |
|---|---|
| 0 | 0 |
| 0 | 0 |
| 0 | 0 |
| 0 | 0 |
| 0 | 0 |
| 0 | 0 |
| 0 | 0 |
| 0 | 0 |
| 0 | 0 |
| 0 | 0 |
| 84.1581672 | 0 |
| 0 | 0 |
| 0 | 0 |
| 0 | 0 |
| 0 | 0 |
| 0 | 0 |
| 0 | 0 |
| 0 | 0 |
| 0 | 0 |
| 0 | 0 |
| 0 | 0 |
| 1.71751362 | 0 |
| 0 | 0 |
| 77.2881127 | 43.3698738 |
| 0 | 0 |
| 0 | 0 |
| 558.191925 | 346.95899 |
| 65.2655174 | 7.22831229 |
| 0 | 0 |
| 0 | 0 |
| 0 | 0 |
| 0 | 0 |
| 0 | 0 |
| 0 | 0 |
| 0 | 0 |
| 0 | 0 |
| 0 | 0 |
| 0 | 0 |
| 0 | 0 |
| 0 | 0 |
| 0 | 0 |
| 352.090291 | 43.3698738 |
| 66.983031 | 93.9680598 |

| | |
|---|---|
| 0 | 0 |
| 0 | 0 |
| 0 | 0 |
| 17.1751362 | 130.109621 |
| 0 | 57.8264983 |
| 0 | 0 |
| 0 | 0 |
| 0 | 0 |
| 0 | 0 |
| 0 | 0 |
| 0 | 0 |
| 0 | 0 |
| 0 | 0 |
| 0 | 0 |
| 0 | 0 |
| 0 | 0 |
| 121.943467 | 7.22831229 |
| 0 | 14.4566246 |
| 0 | 0 |
| 0 | 0 |
| 0 | 0 |
| 0 | 0 |
| 0 | 0 |
| 0 | 0 |
| 0 | 0 |
| 84.1581672 | 14.4566246 |
| 70.4180583 | 0 |
| 0 | 72.2831229 |
| 0 | 0 |
| 0 | 0 |
| 0 | 0 |
| 0 | 0 |
| 25.7627042 | 14.4566246 |
| 0 | 0 |
| 0 | 0 |
| 0 | 0 |
| 0 | 0 |
| 0 | 28.9132492 |
| 0 | 0 |
| 0 | 0 |
| 0 | 0 |
| 0 | 0 |
| 15.4576225 | 7.22831229 |

| | |
|---|---|
| 163.163794 | 187.93612 |
| 0 | 0 |
| 158.011253 | 0 |
| 0 | 0 |
| 0 | 0 |
| 0 | 0 |
| 0 | 0 |
| 0 | 0 |
| 0 | 0 |
| 106.485844 | 115.652997 |
| 0 | 0 |
| 0 | 0 |
| 0 | 7.22831229 |
| 0 | 0 |
| 0 | 0 |
| 0 | 0 |
| 386.440564 | 101.196372 |
| 0 | 0 |
| 41.2203268 | 0 |
| 0 | 0 |
| 0 | 0 |
| 0 | 0 |
| 0 | 0 |
| 0 | 0 |
| 0 | 0 |
| 0 | 0 |
| 1.71751362 | 7.22831229 |
| 0 | 0 |
| 0 | 0 |
| 0 | 187.93612 |
| 0 | 0 |
| 63.5480038 | 7.22831229 |
| 13.7401089 | 245.762618 |
| 58.395463 | 21.6849369 |
| 0 | 0 |
| 0 | 122.881309 |
| 0 | 0 |
| 204.38412 | 0 |
| 1.71751362 | 0 |
| 0 | 0 |
| 898.259621 | 166.251183 |
| 0 | 0 |
| 458.576136 | 137.337934 |

| | |
|---|---|
| 0 | 0 |
| 0 | 0 |
| 0 | 0 |
| 12.0225953 | 202.392744 |
| 0 | 0 |
| 0 | 0 |
| 0 | 0 |
| 0 | 0 |
| 132.248548 | 14.4566246 |
| 0 | 0 |
| 0 | 7.22831229 |
| 17.1751362 | 173.479495 |
| 0 | 0 |
| 0 | 0 |
| 0 | 0 |
| 0 | 0 |
| 0 | 0 |
| 0 | 0 |
| 0 | 0 |
| 187.208984 | 7.22831229 |
| 0 | 0 |
| 0 | 0 |
| 0 | 0 |
| 0 | 0 |
| 0 | 0 |
| 0 | 0 |
| 79.0056264 | 0 |
| 0 | 0 |
| 0 | 0 |
| 0 | 0 |
| 0 | 0 |
| 0 | 0 |
| 231.864338 | 693.91798 |
| 0 | 0 |
| 0 | 0 |
| 0 | 0 |
| 0 | 0 |
| 0 | 0 |
| 0 | 28.9132492 |
| 0 | 0 |
| 0 | 0 |
| 393.310618 | 751.744478 |
| 188.926498 | 86.7397475 |

| | |
|---|---|
| 0 | 0 |
| 0 | 0 |
| 0 | 0 |
| 0 | 0 |
| 89.3107081 | 0 |
| 0 | 0 |
| 0 | 0 |
| 0 | 0 |
| 0 | 0 |
| 0 | 0 |
| 0 | 0 |
| 0 | 0 |
| 0 | 0 |
| 0 | 0 |
| 0 | 0 |
| 0 | 0 |
| 0 | 0 |
| 0 | 0 |
| 0 | 0 |
| 0 | 0 |
| 0 | 0 |
| 195.796552 | 28.9132492 |
| 0 | 0 |
| 0 | 0 |
| 0 | 0 |
| 0 | 0 |
| 0 | 0 |
| 0 | 7.22831229 |
| 0 | 0 |
| 0 | 0 |
| 0 | 0 |
| 0 | 0 |
| 72.1355719 | 7.22831229 |
| 0 | 0 |
| 0 | 0 |
| 249.039474 | 79.5114352 |
| 0 | 0 |
| 0 | 0 |
| 0 | 0 |
| 465.44619 | 224.077681 |
| 0 | 0 |
| 0 | 0 |
| 0 | 0 |

| | |
|---|---|
| 0 | 0 |
| 0 | 0 |
| 0 | 0 |
| 0 | 0 |
| 0 | 0 |
| 0 | 0 |
| 1.71751362 | 7.22831229 |
| 0 | 7.22831229 |
| 0 | 0 |
| 0 | 0 |
| 0 | 0 |
| 0 | 0 |
| 0 | 0 |
| 0 | 0 |
| 0 | 0 |
| 0 | 0 |
| 0 | 0 |
| 0 | 0 |
| 0 | 0 |
| 1.71751362 | 0 |
| 0 | 0 |
| 0 | 0 |
| 0 | 0 |
| 0 | 0 |
| 0 | 0 |
| 0 | 0 |
| 0 | 0 |
| 0 | 0 |
| 0 | 0 |
| 0 | 0 |
| 0 | 0 |
| 0 | 0 |
| 0 | 0 |
| 0 | 0 |
| 0 | 0 |
| 216.406716 | 0 |
| 0 | 0 |
| 89.3107081 | 0 |
| 0 | 0 |
| 0 | 0 |
| 0 | 0 |
| 0 | 0 |
| 0 | 0 |

| | |
|---|---|
| 0 | 0 |
| 0 | 0 |
| 0 | 0 |
| 0 | 0 |
| 0 | 7.22831229 |
| 0 | 0 |
| 0 | 0 |
| 0 | 0 |
| 0 | 0 |
| 0 | 0 |
| 89.3107081 | 7.22831229 |
| 113.355899 | 0 |
| 0 | 0 |
| 0 | 0 |
| 0 | 0 |
| 0 | 0 |
| 0 | 0 |
| 17.1751362 | 0 |
| 20.6101634 | 0 |
| 0 | 0 |
| 0 | 0 |
| 0 | 0 |
| 10.3050817 | 0 |
| 0 | 0 |
| 0 | 0 |
| 91.0282217 | 0 |
| 0 | 0 |
| 85.8756808 | 0 |
| 0 | 0 |
| 0 | 0 |
| 0 | 0 |
| 0 | 0 |
| 0 | 0 |
| 0 | 7.22831229 |
| 0 | 0 |
| 0 | 0 |
| 0 | 0 |
| 0 | 0 |
| 0 | 144.566246 |
| 0 | 0 |
| 0 | 0 |
| 0 | 0 |
| 0 | 0 |

| | |
|---|---|
| 0 | 0 |
| 94.4632489 | 0 |
| 0 | 0 |
| 5.15254085 | 7.22831229 |
| 0 | 0 |
| 0 | 0 |
| 0 | 0 |
| 0 | 0 |
| 0 | 0 |
| 0 | 0 |
| 0 | 0 |
| 87.5931944 | 0 |
| 0 | 0 |
| 0 | 0 |
| 3.43502723 | 7.22831229 |
| 0 | 7.22831229 |
| 12.0225953 | 43.3698738 |
| 25.7627042 | 0 |
| 0 | 0 |
| 0 | 0 |
| 137.401089 | 122.881309 |
| 0 | 0 |
| 0 | 0 |
| 0 | 0 |
| 0 | 0 |
| 32.6327587 | 122.881309 |
| 0 | 0 |
| 3.43502723 | 0 |
| 0 | 0 |
| 0 | 0 |
| 212.971688 | 296.360804 |
| 0 | 0 |
| 0 | 0 |
| 0 | 0 |
| 0 | 0 |
| 0 | 0 |
| 0 | 0 |
| 0 | 0 |
| 0 | 0 |
| 0 | 0 |
| 0 | 0 |
| 151.141198 | 0 |
| 0 | 0 |

| | |
|---|---|
| 0 | 0 |
| 0 | 0 |
| 936.044921 | 527.666797 |
| 0 | 0 |
| 39.5028132 | 14.4566246 |
| 0 | 0 |
| 0 | 0 |
| 0 | 0 |
| 0 | 0 |
| 0 | 0 |
| 0 | 101.196372 |
| 34.3502723 | 7.22831229 |
| 0 | 0 |
| 0 | 0 |
| 197.514066 | 7.22831229 |
| 0 | 0 |
| 0 | 0 |
| 115.073412 | 7.22831229 |
| 0 | 0 |
| 0 | 0 |
| 0 | 0 |
| 0 | 0 |
| 0 | 7.22831229 |
| 0 | 0 |
| 0 | 0 |
| 0 | 0 |
| 0 | 0 |
| 0 | 0 |
| 0 | 0 |
| 0 | 0 |
| 0 | 0 |
| 245.604447 | 0 |
| 0 | 0 |
| 0 | 0 |
| 0 | 0 |
| 108.203358 | 0 |
| 0 | 0 |
| 0 | 0 |
| 15.4576225 | 50.598186 |
| 18.8926498 | 36.1415615 |
| 0 | 0 |
| 0 | 0 |
| 0 | 0 |

| | |
|---|---|
| 0 | 0 |
| 0 | 0 |
| 92.7457353 | 57.8264983 |
| 463.728676 | 455.383674 |
| 0 | 0 |
| 0 | 0 |
| 0 | 0 |
| 0 | 0 |
| 15.4576225 | 36.1415615 |
| 0 | 0 |
| 0 | 0 |
| 0 | 0 |
| 89.3107081 | 0 |
| 0 | 0 |
| 0 | 0 |
| 0 | 0 |
| 207.819148 | 0 |
| 24.0451906 | 21.6849369 |
| 164.881307 | 28.9132492 |
| 298.847369 | 7.22831229 |
| 0 | 0 |
| 0 | 0 |
| 0 | 0 |
| 0 | 0 |
| 0 | 0 |
| 0 | 0 |
| 0 | 0 |
| 49.8078949 | 455.383674 |
| 0 | 0 |
| 0 | 36.1415615 |
| 0 | 0 |
| 1.71751362 | 57.8264983 |
| 0 | 0 |
| 0 | 0 |
| 1.71751362 | 209.621056 |
| 0 | 0 |
| 0 | 0 |
| 0 | 0 |
| 0 | 0 |
| 0 | 0 |
| 0 | 0 |
| 104.768331 | 151.794558 |
| 0 | 0 |

| | |
|---|---|
| 0 | 0 |
| 0 | 0 |
| 0 | 0 |
| 154.576225 | 0 |
| 0 | 0 |
| 0 | 0 |
| 0 | 0 |
| 0 | 0 |
| 0 | 0 |
| 0 | 0 |
| 0 | 0 |
| 53.2429221 | 7.22831229 |
| 0 | 0 |
| 0 | 180.707807 |
| 115.073412 | 7.22831229 |
| 0 | 0 |
| 0 | 0 |
| 1.71751362 | 21.6849369 |
| 0 | 0 |
| 8.58756808 | 50.598186 |
| 0 | 0 |
| 0 | 0 |
| 0 | 0 |
| 0 | 0 |
| 0 | 0 |
| 63.5480038 | 108.424684 |
| 0 | 0 |
| 3.43502723 | 0 |
| 0 | 0 |
| 85.8756808 | 0 |
| 0 | 0 |
| 214.689202 | 79.5114352 |
| 29.1977315 | 202.392744 |
| 0 | 0 |
| 0 | 0 |
| 171.751362 | 0 |
| 0 | 0 |
| 214.689202 | 79.5114352 |
| 0 | 0 |
| 0 | 0 |
| 0 | 7.22831229 |
| 0 | 0 |
| 0 | 0 |

| | |
|---|---|
| 0 | 151.794558 |
| 0 | 0 |
| 0 | 0 |
| 0 | 0 |
| 0 | 0 |
| 84.1581672 | 0 |
| 0 | 0 |
| 0 | 0 |
| 0 | 0 |
| 0 | 0 |
| 0 | 0 |
| 84.1581672 | 0 |
| 0 | 0 |
| 0 | 0 |
| 0 | 0 |
| 0 | 0 |
| 0 | 0 |
| 0 | 0 |
| 0 | 0 |
| 0 | 0 |
| 0 | 0 |
| 0 | 0 |
| 0 | 0 |
| 0 | 0 |
| 0 | 0 |
| 0 | 0 |
| 0 | 0 |
| 0 | 0 |
| 0 | 0 |
| 0 | 0 |
| 0 | 0 |
| 0 | 0 |
| 0 | 0 |
| 0 | 0 |
| 0 | 0 |
| 415.638295 | 715.602917 |
| 0 | 0 |
| 0 | 0 |
| 0 | 0 |
| 0 | 0 |
| 92.7457353 | 36.1415615 |
| 0 | 0 |
| 0 | 0 |

| | |
|---|---|
| 70.4180583 | 7.22831229 |
| 0 | 0 |
| 0 | 0 |
| 0 | 0 |
| 0 | 0 |
| 0 | 0 |
| 0 | 0 |
| 8.58756808 | 0 |
| 0 | 0 |
| 0 | 0 |
| 0 | 0 |
| 0 | 0 |
| 0 | 0 |
| 0 | 0 |
| 0 | 72.2831229 |
| 0 | 0 |
| 0 | 0 |
| 0 | 7.22831229 |
| 0 | 0 |
| 1.71751362 | 57.8264983 |
| 0 | 0 |
| 223.27677 | 28.9132492 |
| 1.71751362 | 115.652997 |
| 219.841743 | 7.22831229 |
| 3.43502723 | 72.2831229 |
| 0 | 0 |
| 0 | 0 |
| 0 | 0 |
| 0 | 0 |
| 0 | 0 |
| 0 | 0 |
| 171.751362 | 21.6849369 |
| 0 | 0 |
| 54.9604357 | 65.0548106 |
| 0 | 0 |
| 0 | 0 |
| 0 | 21.6849369 |
| 0 | 0 |
| 0 | 0 |
| 0 | 0 |
| 0 | 0 |
| 0 | 0 |
| 125.378494 | 0 |

| | |
|---|---|
| 0 | 0 |
| 0 | 0 |
| 0 | 0 |
| 0 | 0 |
| 0 | 43.3698738 |
| 0 | 0 |
| 0 | 0 |
| 51.5254085 | 36.1415615 |
| 0 | 0 |
| 0 | 0 |
| 0 | 43.3698738 |
| 0 | 0 |
| 0 | 0 |
| 22.327677 | 166.251183 |
| 0 | 0 |
| 6.87005447 | 0 |
| 0 | 0 |
| 0 | 0 |
| 0 | 0 |
| 80.72314 | 28.9132492 |
| 0 | 0 |
| 94.4632489 | 260.219243 |
| 0 | 0 |
| 365.8304 | 542.123422 |
| 0 | 0 |
| 0 | 0 |
| 0 | 122.881309 |
| 0 | 0 |
| 0 | 0 |
| 79.0056264 | 57.8264983 |
| 0 | 0 |
| 0 | 0 |
| 0 | 0 |
| 0 | 0 |
| 123.66098 | 0 |
| 0 | 0 |
| 0 | 0 |
| 6.87005447 | 21.6849369 |
| 0 | 0 |
| 0 | 0 |
| 0 | 0 |
| 0 | 0 |
| 0 | 93.9680598 |

| | |
|---|---|
| 808.948913 | 231.305993 |
| 0 | 0 |
| 0 | 0 |
| 151.141198 | 0 |
| 132.248548 | 14.4566246 |
| 0 | 0 |
| 0 | 0 |
| 0 | 0 |
| 0 | 0 |
| 68.7005447 | 0 |
| 0 | 0 |
| 84.1581672 | 7.22831229 |
| 0 | 0 |
| 0 | 0 |
| 0 | 0 |
| 120.225953 | 0 |
| 226.711797 | 65.0548106 |
| 0 | 0 |
| 0 | 0 |
| 0 | 0 |
| 0 | 0 |
| 0 | 0 |
| 0 | 0 |
| 139.118603 | 130.109621 |
| 0 | 0 |
| 0 | 0 |
| 245.604447 | 86.7397475 |
| 87.5931944 | 14.4566246 |
| 0 | 86.7397475 |
| 0 | 0 |
| 0 | 0 |
| 0 | 0 |
| 0 | 72.2831229 |
| 0 | 0 |
| 0 | 7.22831229 |
| 0 | 0 |
| 3.43502723 | 0 |
| 0 | 0 |
| 0 | 0 |
| 0 | 0 |
| 0 | 0 |
| 0 | 0 |
| 0 | 0 |

| | |
|---|---|
| 0 | 0 |
| 1.71751362 | 0 |
| 0 | 0 |
| 0 | 0 |
| 0 | 0 |
| 247.321961 | 245.762618 |
| 352.090291 | 289.132492 |
| 0 | 0 |
| 0 | 0 |
| 0 | 0 |
| 0 | 0 |
| 0 | 0 |
| 0 | 0 |
| 0 | 0 |
| 0 | 0 |
| 0 | 0 |
| 0 | 0 |
| 0 | 0 |
| 0 | 0 |
| 0 | 0 |
| 0 | 0 |
| 164.881307 | 137.337934 |
| 0 | 0 |
| 0 | 0 |
| 0 | 0 |
| 0 | 0 |
| 0 | 0 |
| 0 | 7.22831229 |
| 0 | 0 |
| 0 | 0 |
| 0 | 0 |
| 0 | 0 |
| 0 | 0 |
| 0 | 0 |
| 84.1581672 | 7.22831229 |
| 0 | 0 |
| 0 | 0 |
| 0 | 0 |
| 0 | 0 |
| 3.43502723 | 0 |
| 0 | 0 |
| 0 | 0 |
| 0 | 0 |

| | |
|---|---|
| 0 | 0 |
| 0 | 0 |
| 0 | 0 |
| 0 | 0 |
| 0 | 0 |
| 0 | 0 |
| 0 | 0 |
| 118.50844 | 7.22831229 |
| 0 | 0 |
| 386.440564 | 245.762618 |
| 0 | 0 |
| 0 | 0 |
| 0 | 0 |
| 0 | 0 |
| 0 | 0 |
| 0 | 0 |
| 0 | 0 |
| 0 | 0 |
| 0 | 0 |
| 0 | 0 |
| 0 | 0 |
| 87.5931944 | 0 |
| 0 | 0 |
| 24.0451906 | 0 |
| 0 | 0 |
| 0 | 0 |
| 0 | 0 |
| 0 | 0 |
| 53.2429221 | 43.3698738 |
| 190.644011 | 0 |
| 3.43502723 | 0 |
| 0 | 115.652997 |
| 680.135392 | 426.470425 |
| 0 | 0 |
| 1.71751362 | 65.0548106 |
| 0 | 0 |
| 0 | 0 |
| 226.711797 | 260.219243 |
| 0 | 0 |
| 0 | 0 |
| 1.71751362 | 36.1415615 |
| 0 | 0 |
| 0 | 0 |

| | |
|---|---|
| 0 | 86.7397475 |
| 0 | 0 |
| 0 | 0 |
| 109.920871 | 43.3698738 |
| 0 | 0 |
| 0 | 0 |
| 0 | 0 |
| 66.983031 | 0 |
| 32.6327587 | 0 |
| 0 | 0 |
| 0 | 0 |
| 0 | 0 |
| 118.50844 | 7.22831229 |
| 18.8926498 | 0 |
| 0 | 0 |
| 0 | 0 |
| 0 | 0 |
| 1.71751362 | 7.22831229 |
| 0 | 0 |
| 0 | 0 |
| 250.756988 | 224.077681 |
| 0 | 0 |
| 0 | 0 |
| 0 | 0 |
| 0 | 0 |
| 0 | 0 |
| 77.2881127 | 0 |
| 0 | 0 |
| 0 | 0 |
| 0 | 0 |
| 0 | 0 |
| 0 | 0 |
| 0 | 7.22831229 |
| 0 | 0 |
| 0 | 7.22831229 |
| 0 | 7.22831229 |
| 257.627042 | 0 |
| 0 | 0 |
| 0 | 0 |
| 0 | 0 |
| 0 | 0 |
| 274.802179 | 0 |
| 0 | 0 |

| | |
|---|---|
| 0 | 0 |
| 0 | 0 |
| 0 | 0 |
| 231.864338 | 339.730678 |
| 0 | 0 |
| 0 | 0 |
| 0 | 0 |
| 1.71751362 | 0 |
| 0 | 0 |
| 0 | 0 |
| 0 | 0 |
| 17.1751362 | 0 |
| 0 | 0 |
| 82.4406536 | 0 |
| 0 | 0 |
| 0 | 0 |
| 0 | 0 |
| 0 | 0 |
| 0 | 0 |
| 0 | 0 |
| 0 | 0 |
| 0 | 0 |
| 128.813521 | 0 |
| 0 | 0 |
| 0 | 0 |
| 503.23149 | 72.2831229 |
| 0 | 0 |
| 6.87005447 | 108.424684 |
| 121.943467 | 86.7397475 |
| 1.71751362 | 7.22831229 |
| 51.5254085 | 0 |
| 37.7852996 | 0 |
| 0 | 0 |
| 936.044921 | 997.507096 |
| 13.7401089 | 0 |
| 0 | 0 |
| 0 | 0 |
| 53.2429221 | 0 |
| 218.124229 | 245.762618 |
| 130.531035 | 0 |
| 0 | 0 |
| 0 | 0 |
| 0 | 0 |

| | |
|---|---|
| 0 | 0 |
| 0 | 0 |
| 0 | 0 |
| 65.2655174 | 7.22831229 |
| 0 | 0 |
| 0 | 0 |
| 0 | 0 |
| 147.706171 | 28.9132492 |
| 0 | 0 |
| 0 | 0 |
| 0 | 0 |
| 75.5705991 | 36.1415615 |
| 0 | 0 |
| 0 | 0 |
| 0 | 0 |
| 0 | 0 |
| 0 | 7.22831229 |
| 0 | 0 |
| 0 | 0 |
| 0 | 0 |
| 34.3502723 | 187.93612 |
| 0 | 0 |
| 0 | 0 |
| 0 | 0 |
| 0 | 0 |
| 0 | 0 |
| 0 | 0 |
| 0 | 0 |
| 0 | 0 |
| 0 | 0 |
| 0 | 0 |
| 0 | 0 |
| 0 | 0 |
| 0 | 0 |
| 0 | 0 |
| 0 | 0 |
| 0 | 0 |
| 0 | 137.337934 |
| 0 | 0 |
| 0 | 0 |
| 0 | 36.1415615 |
| 68.7005447 | 50.598186 |
| 1.71751362 | 0 |

| | |
|---|---|
| 0 | 0 |
| 0 | 0 |
| 0 | 0 |
| 0 | 0 |
| 0 | 0 |
| 0 | 0 |
| 0 | 0 |
| 0 | 0 |
| 0 | 0 |
| 0 | 0 |
| 0 | 0 |
| 0 | 0 |
| 36.0677859 | 28.9132492 |
| 0 | 0 |
| 0 | 0 |
| 0 | 7.22831229 |
| 104.768331 | 7.22831229 |
| 0 | 0 |
| 249.039474 | 0 |
| 51.5254085 | 65.0548106 |
| 34.3502723 | 14.4566246 |
| 0 | 0 |
| 0 | 0 |
| 0 | 0 |
| 0 | 0 |
| 8.58756808 | 0 |
| 0 | 0 |
| 72.1355719 | 159.02287 |
| 0 | 0 |
| 0 | 0 |
| 0 | 0 |
| 0 | 0 |
| 91.0282217 | 0 |
| 0 | 0 |
| 17.1751362 | 79.5114352 |
| 0 | 0 |
| 0 | 0 |
| 0 | 0 |
| 185.491471 | 0 |
| 0 | 0 |
| 0 | 0 |
| 0 | 0 |
| 0 | 0 |

| | |
|---|---|
| 0 | 0 |
| 70.4180583 | 137.337934 |
| 0 | 0 |
| 0 | 0 |
| 0 | 0 |
| 0 | 0 |
| 113.355899 | 0 |
| 0 | 0 |
| 0 | 0 |
| 338.350182 | 419.242113 |
| 103.050817 | 72.2831229 |
| 0 | 86.7397475 |
| 0 | 0 |
| 0 | 0 |
| 338.350182 | 0 |
| 41.2203268 | 0 |
| 0 | 0 |
| 0 | 0 |
| 53.2429221 | 224.077681 |
| 0 | 0 |
| 0 | 0 |
| 0 | 0 |
| 0 | 0 |
| 195.796552 | 57.8264983 |
| 0 | 0 |
| 51.5254085 | 7.22831229 |
| 60.1129766 | 173.479495 |
| 0 | 0 |
| 0 | 0 |
| 0 | 0 |
| 0 | 0 |
| 94.4632489 | 0 |
| 269.649638 | 672.233043 |
| 0 | 0 |
| 0 | 0 |
| 0 | 0 |
| 0 | 0 |
| 3.43502723 | 0 |
| 0 | 0 |
| 0 | 0 |
| 36.0677859 | 0 |
| 0 | 0 |
| 44.655354 | 108.424684 |

| | |
|---|---|
| 0 | 0 |
| 0 | 0 |
| 0 | 0 |
| 0 | 281.904179 |
| 0 | 0 |
| 0 | 0 |
| 0 | 0 |
| 5.15254085 | 50.598186 |
| 0 | 0 |
| 0 | 0 |
| 0 | 0 |
| 3.43502723 | 65.0548106 |
| 0 | 0 |
| 204.38412 | 14.4566246 |
| 0 | 0 |
| 0 | 0 |
| 422.50835 | 151.794558 |
| 34.3502723 | 0 |
| 0 | 0 |
| 0 | 0 |
| 0 | 0 |
| 3.43502723 | 0 |
| 0 | 0 |
| 111.638385 | 0 |
| 56.6779493 | 57.8264983 |
| 0 | 0 |
| 0 | 0 |
| 0 | 0 |
| 0 | 0 |
| 0 | 0 |
| 0 | 0 |
| 0 | 7.22831229 |
| 0 | 0 |
| 0 | 0 |
| 0 | 0 |
| 80.72314 | 93.9680598 |
| 0 | 0 |
| 0 | 0 |
| 0 | 0 |
| 0 | 0 |
| 0 | 0 |
| 0 | 0 |
| 0 | 0 |

| | |
|---|---|
| 0 | 0 |
| 0 | 0 |
| 0 | 0 |
| 0 | 0 |
| 0 | 0 |
| 0 | 0 |
| 139.118603 | 28.9132492 |
| 32.6327587 | 137.337934 |
| 0 | 0 |
| 504.949003 | 281.904179 |
| 0 | 0 |
| 0 | 0 |
| 0 | 0 |
| 0 | 0 |
| 0 | 0 |
| 0 | 0 |
| 0 | 0 |
| 0 | 0 |
| 0 | 0 |
| 0 | 0 |
| 0 | 101.196372 |
| 0 | 0 |
| 0 | 0 |
| 0 | 0 |
| 0 | 0 |
| 0 | 7.22831229 |
| 0 | 0 |
| 48.0903813 | 390.328864 |
| 0 | 0 |
| 0 | 0 |
| 8.58756808 | 0 |
| 133.966062 | 0 |
| 0 | 0 |
| 0 | 0 |
| 0 | 0 |
| 0 | 14.4566246 |
| 0 | 0 |
| 0 | 0 |
| 111.638385 | 14.4566246 |
| 0 | 0 |
| 0 | 0 |
| 0 | 0 |
| 0 | 0 |

| | |
|---|---|
| 395.028132 | 209.621056 |
| 0 | 0 |
| 106.485844 | 0 |
| 0 | 0 |
| 99.6157898 | 21.6849369 |
| 89.3107081 | 7.22831229 |
| 152.858712 | 28.9132492 |
| 274.802179 | 79.5114352 |
| 0 | 0 |
| 85.8756808 | 0 |
| 0 | 0 |
| 249.039474 | 130.109621 |
| 0 | 0 |
| 0 | 0 |
| 0 | 0 |
| 0 | 0 |
| 142.55363 | 7.22831229 |
| 0 | 0 |
| 0 | 0 |
| 82.4406536 | 93.9680598 |
| 0 | 0 |
| 0 | 0 |
| 0 | 7.22831229 |
| 1.71751362 | 0 |
| 0 | 0 |
| 0 | 0 |
| 0 | 101.196372 |
| 0 | 7.22831229 |
| 218.124229 | 0 |
| 0 | 0 |
| 0 | 0 |
| 0 | 0 |
| 65.2655174 | 0 |
| 0 | 0 |
| 0 | 0 |
| 0 | 0 |
| 0 | 0 |
| 0 | 0 |
| 32.6327587 | 14.4566246 |
| 0 | 0 |
| 0 | 0 |
| 0 | 0 |
| 0 | 0 |

| | |
|---|---|
| 0 | 0 |
| 0 | 0 |
| 0 | 93.9680598 |
| 0 | 0 |
| 128.813521 | 50.598186 |
| 0 | 0 |
| 0 | 0 |
| 0 | 0 |
| 0 | 0 |
| 1.71751362 | 0 |
| 0 | 0 |
| 0 | 0 |
| 0 | 0 |
| 0 | 0 |
| 104.768331 | 0 |
| 0 | 0 |
| 0 | 0 |
| 0 | 0 |
| 84.1581672 | 0 |
| 0 | 0 |
| 0 | 0 |
| 176.903902 | 21.6849369 |
| 0 | 0 |
| 0 | 0 |
| 0 | 0 |
| 0 | 0 |
| 0 | 0 |
| 151.141198 | 0 |
| 70.4180583 | 0 |
| 0 | 0 |
| 0 | 0 |
| 63.5480038 | 21.6849369 |
| 0 | 0 |
| 0 | 0 |
| 0 | 0 |
| 103.050817 | 0 |
| 0 | 0 |
| 0 | 0 |
| 0 | 0 |
| 0 | 0 |
| 94.4632489 | 0 |
| 0 | 0 |
| 0 | 0 |

| | |
|---|---|
| 0 | 65.0548106 |
| 0 | 0 |
| 0 | 0 |
| 0 | 0 |
| 0 | 0 |
| 0 | 0 |
| 17.1751362 | 0 |
| 285.10726 | 14.4566246 |
| 0 | 0 |
| 0 | 0 |
| 0 | 0 |
| 0 | 0 |
| 0 | 0 |
| 118.50844 | 0 |
| 0 | 0 |
| 0 | 0 |
| 0 | 0 |
| 0 | 0 |
| 0 | 0 |
| 0 | 0 |
| 0 | 0 |
| 0 | 0 |
| 149.423685 | 101.196372 |
| 0 | 0 |
| 0 | 0 |
| 0 | 0 |
| 0 | 0 |
| 0 | 0 |
| 121.943467 | 0 |
| 0 | 0 |
| 103.050817 | 0 |
| 103.050817 | 0 |
| 0 | 0 |
| 0 | 0 |
| 0 | 14.4566246 |
| 132.248548 | 7.22831229 |
| 0 | 0 |
| 0 | 303.589116 |
| 127.096008 | 7.22831229 |
| 0 | 0 |
| 30.9152451 | 0 |
| 0 | 0 |
| 0 | 0 |

| | |
|---|---|
| 0 | 50.598186 |
| 0 | 0 |
| 0 | 0 |
| 56.6779493 | 14.4566246 |
| 0 | 43.3698738 |
| 116.790926 | 180.707807 |
| 0 | 0 |
| 0 | 0 |
| 0 | 0 |
| 0 | 0 |
| 0 | 0 |
| 0 | 7.22831229 |
| 139.118603 | 7.22831229 |
| 6.87005447 | 0 |
| 0 | 0 |
| 0 | 0 |
| 0 | 0 |
| 0 | 0 |
| 0 | 0 |
| 0 | 0 |
| 0 | 0 |
| 228.429311 | 122.881309 |
| 0 | 0 |
| 0 | 0 |
| 0 | 0 |
| 0 | 0 |
| 0 | 0 |
| 1.71751362 | 144.566246 |
| 0 | 0 |
| 0 | 0 |
| 68.7005447 | 0 |
| 0 | 0 |
| 152.858712 | 0 |
| 906.847189 | 144.566246 |
| 0 | 0 |
| 0 | 0 |
| 0 | 7.22831229 |
| 0 | 0 |
| 0 | 0 |
| 0 | 0 |
| 0 | 0 |
| 0 | 0 |
| 0 | 0 |

| | |
|---|---|
| 0 | 0 |
| 101.333303 | 0 |
| 0 | 7.22831229 |
| 104.768331 | 0 |
| 0 | 0 |
| 5.15254085 | 0 |
| 0 | 0 |
| 0 | 0 |
| 0 | 0 |
| 705.898096 | 361.415615 |
| 204.38412 | 79.5114352 |
| 0 | 0 |
| 25.7627042 | 7.22831229 |
| 0 | 0 |
| 0 | 0 |
| 0 | 0 |
| 94.4632489 | 7.22831229 |
| 120.225953 | 0 |
| 0 | 0 |
| 0 | 0 |
| 140.836117 | 0 |
| 0 | 0 |
| 0 | 0 |
| 0 | 0 |
| 0 | 0 |
| 0 | 0 |
| 0 | 72.2831229 |
| 0 | 93.9680598 |
| 212.971688 | 28.9132492 |
| 156.293739 | 0 |
| 0 | 0 |
| 0 | 0 |
| 0 | 0 |
| 0 | 0 |
| 0 | 0 |
| 0 | 0 |
| 0 | 0 |
| 0 | 72.2831229 |
| 0 | 0 |
| 0 | 0 |
| 0 | 0 |
| 0 | 0 |
| 0 | 0 |

| | |
|---|---|
| 0 | 0 |
| 0 | 0 |
| 0 | 101.196372 |
| 0 | 0 |
| 0 | 0 |
| 32.6327587 | 50.598186 |
| 0 | 0 |
| 221.559257 | 0 |
| 101.333303 | 0 |
| 0 | 0 |
| 0 | 0 |
| 0 | 0 |
| 0 | 0 |
| 0 | 0 |
| 0 | 0 |
| 0 | 0 |
| 0 | 0 |
| 5.15254085 | 0 |
| 0 | 0 |
| 0 | 0 |
| 0 | 0 |
| 0 | 0 |
| 0 | 0 |
| 94.4632489 | 0 |
| 103.050817 | 0 |
| 132.248548 | 0 |
| 0 | 0 |
| 0 | 0 |
| 0 | 0 |
| 0 | 0 |
| 0 | 0 |
| 0 | 0 |
| 163.163794 | 137.337934 |
| 0 | 0 |
| 334.915155 | 50.598186 |
| 484.33884 | 607.178233 |
| 22.327677 | 28.9132492 |
| 0 | 0 |
| 0 | 0 |
| 0 | 0 |
| 70.4180583 | 0 |
| 0 | 0 |
| 0 | 0 |

| | |
|---|---|
| 0 | 0 |
| 106.485844 | 7.22831229 |
| 0 | 0 |
| 0 | 0 |
| 94.4632489 | 7.22831229 |
| 255.909529 | 36.1415615 |
| 0 | 0 |
| 0 | 0 |
| 0 | 0 |
| 25.7627042 | 28.9132492 |
| 0 | 0 |
| 0 | 0 |
| 0 | 0 |
| 0 | 0 |
| 34.3502723 | 0 |
| 0 | 0 |
| 0 | 0 |
| 0 | 0 |
| 0 | 0 |
| 0 | 0 |
| 219.841743 | 65.0548106 |
| 0 | 0 |
| 0 | 0 |
| 0 | 0 |
| 0 | 0 |
| 0 | 0 |
| 0 | 0 |
| 0 | 0 |
| 63.5480038 | 0 |
| 24.0451906 | 180.707807 |
| 654.372688 | 578.264983 |
| 39.5028132 | 0 |
| 0 | 0 |
| 0 | 0 |
| 0 | 0 |
| 0 | 0 |
| 0 | 0 |
| 20.6101634 | 0 |
| 97.8982761 | 0 |
| 0 | 0 |
| 0 | 0 |
| 5.15254085 | 21.6849369 |
| 1.71751362 | 7.22831229 |

| | |
|---|---|
| 0 | 0 |
| 0 | 0 |
| 0 | 0 |
| 0 | 0 |
| 0 | 0 |
| 0 | 0 |
| 5.15254085 | 0 |
| 0 | 0 |
| 63.5480038 | 14.4566246 |
| 0 | 0 |
| 0 | 93.9680598 |
| 269.649638 | 0 |
| 37.7852996 | 108.424684 |
| 0 | 0 |
| 29.1977315 | 130.109621 |
| 0 | 0 |
| 0 | 0 |
| 0 | 0 |
| 0 | 0 |
| 168.316334 | 36.1415615 |
| 0 | 0 |
| 0 | 0 |
| 0 | 0 |
| 0 | 0 |
| 0 | 0 |
| 0 | 0 |
| 0 | 0 |
| 3.43502723 | 0 |
| 0 | 0 |
| 0 | 0 |
| 0 | 0 |
| 0 | 0 |
| 20.6101634 | 14.4566246 |
| 0 | 0 |
| 5.15254085 | 0 |
| 1.71751362 | 0 |
| 0 | 0 |
| 0 | 0 |
| 388.158077 | 0 |
| 0 | 0 |
| 0 | 0 |
| 0 | 0 |
| 657.807715 | 151.794558 |

| | |
|---|---|
| 0 | 0 |
| 0 | 0 |
| 0 | 0 |
| 1.71751362 | 0 |
| 0 | 7.22831229 |
| 158.011253 | 93.9680598 |
| 0 | 0 |
| 142.55363 | 14.4566246 |
| 0 | 0 |
| 202.666607 | 0 |
| 0 | 0 |
| 0 | 0 |
| 0 | 0 |
| 0 | 0 |
| 0 | 0 |
| 0 | 0 |
| 0 | 0 |
| 0 | 0 |
| 0 | 0 |
| 0 | 0 |
| 556.474412 | 1344.46609 |
| 0 | 0 |
| 125.378494 | 0 |
| 0 | 0 |
| 0 | 0 |
| 839.864158 | 325.274053 |
| 0 | 0 |
| 0 | 0 |
| 0 | 0 |
| 187.208984 | 50.598186 |
| 0 | 0 |
| 0 | 0 |
| 29.1977315 | 0 |
| 0 | 0 |
| 0 | 0 |
| 0 | 0 |
| 0 | 0 |
| 0 | 0 |
| 0 | 0 |
| 0 | 7.22831229 |
| 0 | 0 |
| 0 | 0 |
| 0 | 0 |

| | |
|---|---|
| 0 | 0 |
| 0 | 0 |
| 0 | 0 |
| 0 | 0 |
| 0 | 79.5114352 |
| 0 | 0 |
| 0 | 0 |
| 0 | 0 |
| 0 | 0 |
| 0 | 0 |
| 0 | 0 |
| 0 | 50.598186 |
| 0 | 0 |
| 0 | 0 |
| 0 | 0 |
| 63.5480038 | 0 |
| 0 | 0 |
| 0 | 7.22831229 |
| 0 | 0 |
| 13.7401089 | 65.0548106 |
| 0 | 0 |
| 0 | 0 |
| 0 | 0 |
| 0 | 0 |
| 0 | 0 |
| 243.886934 | 354.187302 |
| 0 | 0 |
| 0 | 0 |
| 0 | 0 |
| 0 | 0 |
| 484.33884 | 672.233043 |
| 92.7457353 | 0 |
| 0 | 0 |
| 0 | 0 |
| 0 | 0 |
| 0 | 0 |
| 752.270964 | 65.0548106 |
| 0 | 0 |
| 0 | 0 |
| 0 | 0 |
| 0 | 0 |
| 0 | 0 |
| 0 | 0 |

| | |
|---|---|
| 0 | 0 |
| 303.99991 | 173.479495 |
| 0 | 0 |
| 0 | 0 |
| 0 | 0 |
| 97.8982761 | 0 |
| 0 | 0 |
| 0 | 0 |
| 0 | 0 |
| 0 | 0 |
| 0 | 0 |
| 0 | 0 |
| 0 | 0 |
| 0 | 0 |
| 0 | 0 |
| 0 | 0 |
| 54.9604357 | 0 |
| 0 | 0 |
| 0 | 0 |
| 0 | 0 |
| 0 | 0 |
| 0 | 0 |
| 0 | 0 |
| 161.44628 | 36.1415615 |
| 0 | 0 |
| 0 | 0 |
| 0 | 50.598186 |
| 0 | 0 |
| 0 | 0 |
| 0 | 0 |
| 0 | 0 |
| 0 | 0 |
| 0 | 0 |
| 0 | 0 |
| 0 | 0 |
| 0 | 0 |
| 0 | 0 |
| 0 | 0 |
| 0 | 0 |
| 0 | 0 |
| 161.44628 | 0 |
| 0 | 0 |
| 130.531035 | 0 |

| | |
|---|---|
| 0 | 0 |
| 0 | 0 |
| 0 | 0 |
| 0 | 0 |
| 0 | 0 |
| 0 | 0 |
| 0 | 0 |
| 0 | 0 |
| 77.2881127 | 7.22831229 |
| 0 | 0 |
| 206.101634 | 0 |
| 0 | 0 |
| 0 | 0 |
| 0 | 0 |
| 400.180673 | 216.849369 |
| 283.389747 | 65.0548106 |
| 0 | 0 |
| 0 | 0 |
| 0 | 0 |
| 0 | 0 |
| 0 | 0 |
| 467.163704 | 130.109621 |
| 27.4802179 | 0 |
| 0 | 0 |
| 0 | 0 |
| 0 | 7.22831229 |
| 0 | 0 |
| 0 | 0 |
| 0 | 267.447555 |
| 3.43502723 | 332.502365 |
| 0 | 57.8264983 |
| 204.38412 | 0 |
| 0 | 0 |
| 0 | 14.4566246 |
| 0 | 0 |
| 152.858712 | 130.109621 |
| 63.5480038 | 0 |
| 0 | 79.5114352 |
| 25.7627042 | 0 |
| 0 | 0 |
| 0 | 0 |
| 0 | 0 |
| 0 | 0 |

| | |
|---|---|
| 3.43502723 | 0 |
| 0 | 0 |
| 221.559257 | 0 |
| 0 | 0 |
| 0 | 0 |
| 0 | 0 |
| 0 | 0 |
| 0 | 0 |
| 0 | 0 |
| 0 | 0 |
| 0 | 0 |
| 61.8304902 | 0 |
| 0 | 0 |
| 0 | 0 |
| 0 | 0 |
| 0 | 0 |
| 0 | 0 |
| 37.7852996 | 159.02287 |
| 0 | 0 |
| 13.7401089 | 187.93612 |
| 24.0451906 | 0 |
| 5.15254085 | 14.4566246 |
| 0 | 0 |
| 0 | 28.9132492 |
| 5.15254085 | 0 |
| 0 | 0 |
| 18.8926498 | 173.479495 |
| 0 | 0 |
| 0 | 0 |
| 0 | 0 |
| 0 | 101.196372 |
| 0 | 0 |
| 1.71751362 | 0 |
| 0 | 0 |
| 0 | 0 |
| 0 | 0 |
| 0 | 0 |
| 0 | 130.109621 |
| 0 | 0 |
| 0 | 0 |
| 0 | 0 |
| 0 | 0 |
| 358.960346 | 36.1415615 |

| | |
|---|---|
| 0 | 36.1415615 |
| 0 | 0 |
| 0 | 0 |
| 0 | 0 |
| 0 | 0 |
| 0 | 0 |
| 0 | 0 |
| 0 | 0 |
| 0 | 0 |
| 0 | 0 |
| 0 | 0 |
| 200.949093 | 0 |
| 200.949093 | 0 |
| 0 | 0 |
| 0 | 0 |
| 0 | 0 |
| 0 | 0 |
| 0 | 0 |
| 0 | 0 |
| 0 | 0 |
| 0 | 0 |
| 0 | 0 |
| 0 | 0 |
| 0 | 0 |
| 0 | 7.22831229 |
| 0 | 0 |
| 0 | 0 |
| 0 | 0 |
| 0 | 0 |
| 0 | 0 |
| 15.4576225 | 159.02287 |
| 0 | 0 |
| 101.333303 | 0 |
| 0 | 0 |
| 0 | 0 |
| 0 | 0 |
| 0 | 0 |
| 0 | 0 |
| 0 | 0 |
| 0 | 260.219243 |
| 0 | 0 |
| 0 | 0 |
| 547.886844 | 72.2831229 |

| | |
|---|---|
| 0 | 0 |
| 0 | 0 |
| 0 | 0 |
| 0 | 0 |
| 0 | 0 |
| 199.23158 | 0 |
| 0 | 0 |
| 17.1751362 | 21.6849369 |
| 0 | 0 |
| 0 | 0 |
| 0 | 0 |
| 0 | 0 |
| 0 | 0 |
| 170.033848 | 28.9132492 |
| 0 | 0 |
| 1.71751362 | 0 |
| 15.4576225 | 130.109621 |
| 0 | 0 |
| 0 | 0 |
| 0 | 0 |
| 0 | 14.4566246 |
| 0 | 0 |
| 0 | 0 |
| 0 | 0 |
| 42.9378404 | 7.22831229 |
| 20.6101634 | 28.9132492 |
| 0 | 0 |
| 0 | 0 |
| 0 | 0 |
| 0 | 7.22831229 |
| 0 | 0 |
| 197.514066 | 0 |
| 0 | 0 |
| 0 | 0 |
| 0 | 0 |
| 0 | 0 |
| 0 | 0 |
| 0 | 0 |
| 0 | 0 |
| 0 | 0 |
| 12.0225953 | 43.3698738 |
| 0 | 0 |
| 0 | 0 |

| | |
|---|---|
| 0 | 0 |
| 0 | 0 |
| 0 | 0 |
| 0 | 0 |
| 0 | 0 |
| 5.15254085 | 93.9680598 |
| 0 | 0 |
| 0 | 0 |
| 27.4802179 | 93.9680598 |
| 0 | 0 |
| 6.87005447 | 0 |
| 0 | 0 |
| 0 | 0 |
| 0 | 0 |
| 0 | 7.22831229 |
| 0 | 0 |
| 0 | 0 |
| 0 | 0 |
| 0 | 0 |
| 0 | 0 |
| 0 | 0 |
| 149.423685 | 0 |
| 0 | 0 |
| 0 | 0 |
| 0 | 0 |
| 0 | 0 |
| 0 | 0 |
| 482.621326 | 187.93612 |
| 0 | 0 |
| 0 | 0 |
| 0 | 0 |
| 0 | 0 |
| 0 | 0 |
| 0 | 0 |
| 0 | 0 |
| 137.401089 | 7.22831229 |
| 0 | 0 |
| 0 | 0 |
| 142.55363 | 0 |
| 0 | 0 |
| 0 | 0 |
| 0 | 0 |
| 0 | 0 |

| | |
|---|---|
| 34.3502723 | 7.22831229 |
| 0 | 0 |
| 0 | 0 |
| 20.6101634 | 93.9680598 |
| 0 | 0 |
| 75.5705991 | 0 |
| 1.71751362 | 57.8264983 |
| 0 | 0 |
| 3.43502723 | 14.4566246 |
| 0 | 0 |
| 0 | 0 |
| 0 | 0 |
| 0 | 0 |
| 0 | 0 |
| 194.079039 | 0 |
| 0 | 0 |
| 0 | 0 |
| 0 | 0 |
| 85.8756808 | 0 |
| 0 | 0 |
| 0 | 0 |
| 17.1751362 | 0 |
| 0 | 0 |
| 0 | 0 |
| 0 | 0 |
| 0 | 0 |
| 0 | 0 |
| 0 | 0 |
| 0 | 0 |
| 0 | 0 |
| 101.333303 | 0 |
| 0 | 0 |
| 0 | 0 |
| 0 | 0 |
| 0 | 0 |
| 0 | 0 |
| 0 | 0 |
| 0 | 0 |
| 0 | 0 |
| 3.43502723 | 0 |
| 56.6779493 | 28.9132492 |
| 49.8078949 | 122.881309 |
| 0 | 57.8264983 |

| | |
|---|---|
| 0 | 0 |
| 0 | 0 |
| 0 | 0 |
| 109.920871 | 0 |
| 0 | 0 |
| 192.361525 | 0 |
| 0 | 7.22831229 |
| 0 | 0 |
| 0 | 0 |
| 72.1355719 | 65.0548106 |
| 133.966062 | 72.2831229 |
| 0 | 0 |
| 127.096008 | 115.652997 |
| 0 | 0 |
| 0 | 0 |
| 0 | 14.4566246 |
| 0 | 72.2831229 |
| 0 | 0 |
| 0 | 0 |
| 0 | 0 |
| 0 | 0 |
| 0 | 0 |
| 0 | 0 |
| 65.2655174 | 50.598186 |
| 65.2655174 | 7.22831229 |
| 0 | 0 |
| 0 | 0 |
| 0 | 0 |
| 0 | 0 |
| 151.141198 | 173.479495 |
| 0 | 180.707807 |
| 89.3107081 | 7.22831229 |
| 0 | 0 |
| 0 | 0 |
| 0 | 0 |
| 0 | 0 |
| 56.6779493 | 0 |
| 0 | 0 |
| 0 | 0 |
| 0 | 65.0548106 |
| 0 | 0 |
| 0 | 151.794558 |
| 0 | 0 |

| | |
|---|---|
| 94.4632489 | 7.22831229 |
| 0 | 0 |
| 0 | 0 |
| 0 | 0 |
| 278.237206 | 108.424684 |
| 0 | 0 |
| 0 | 0 |
| 0 | 0 |
| 0 | 245.762618 |
| 0 | 245.762618 |
| 0 | 0 |
| 89.3107081 | 130.109621 |
| 0 | 0 |
| 0 | 0 |
| 0 | 0 |
| 0 | 0 |
| 1.71751362 | 79.5114352 |
| 0 | 0 |
| 0 | 0 |
| 60.1129766 | 0 |
| 12.0225953 | 14.4566246 |
| 0 | 0 |
| 0 | 0 |
| 0 | 0 |
| 0 | 0 |
| 1.71751362 | 65.0548106 |
| 1.71751362 | 7.22831229 |
| 0 | 0 |
| 0 | 0 |
| 116.790926 | 21.6849369 |
| 0 | 115.652997 |
| 54.9604357 | 28.9132492 |
| 0 | 0 |
| 0 | 0 |
| 0 | 0 |
| 0 | 0 |
| 0 | 0 |
| 0 | 0 |
| 0 | 0 |
| 242.16942 | 563.808359 |
| 376.135482 | 260.219243 |
| 80.72314 | 0 |
| 89.3107081 | 50.598186 |

| | |
|---|---|
| 0 | 0 |
| 0 | 0 |
| 0 | 0 |
| 0 | 0 |
| 145.988657 | 57.8264983 |
| 0 | 0 |
| 1.71751362 | 0 |
| 0 | 0 |
| 0 | 0 |
| 0 | 0 |
| 0 | 0 |
| 0 | 0 |
| 0 | 0 |
| 0 | 0 |
| 161.44628 | 1395.06427 |
| 254.192015 | 216.849369 |
| 0 | 0 |
| 0 | 0 |
| 0 | 0 |
| 0 | 130.109621 |
| 0 | 0 |
| 0 | 0 |
| 0 | 0 |
| 79.0056264 | 130.109621 |
| 0 | 0 |
| 0 | 0 |
| 0 | 0 |
| 0 | 0 |
| 0 | 0 |
| 41.2203268 | 0 |
| 0 | 0 |
| 583.95463 | 4199.64944 |
| 53.2429221 | 122.881309 |
| 0 | 0 |
| 252.474502 | 50.598186 |
| 36.0677859 | 0 |
| 0 | 0 |
| 0 | 0 |
| 0 | 0 |
| 0 | 0 |
| 158.011253 | 14.4566246 |
| 152.858712 | 144.566246 |
| 0 | 0 |

| | |
|---|---|
| 0 | 0 |
| 0 | 0 |
| 0 | 0 |
| 0 | 0 |
| 0 | 0 |
| 0 | 0 |
| 77.2881127 | 14.4566246 |
| 0 | 0 |
| 94.4632489 | 0 |
| 0 | 0 |
| 185.491471 | 0 |
| 0 | 0 |
| 0 | 115.652997 |
| 0 | 115.652997 |
| 0 | 21.6849369 |
| 0 | 0 |
| 0 | 0 |
| 0 | 0 |
| 0 | 187.93612 |
| 0 | 0 |
| 0 | 0 |
| 0 | 0 |
| 0 | 7.22831229 |
| 0 | 0 |
| 0 | 0 |
| 0 | 0 |
| 0 | 0 |
| 0 | 0 |
| 3.43502723 | 86.7397475 |
| 0 | 238.534306 |
| 0 | 238.534306 |
| 0 | 0 |
| 15.4576225 | 0 |
| 130.531035 | 36.1415615 |
| 0 | 0 |
| 0 | 0 |
| 0 | 0 |
| 0 | 0 |
| 0 | 0 |
| 0 | 0 |
| 37.7852996 | 122.881309 |
| 142.55363 | 57.8264983 |
| 0 | 0 |

| | |
|---|---|
| 10.3050817 | 14.4566246 |
| 0 | 0 |
| 5.15254085 | 79.5114352 |
| 0 | 0 |
| 0 | 0 |
| 91.0282217 | 0 |
| 0 | 0 |
| 0 | 0 |
| 183.773957 | 0 |
| 0 | 0 |
| 77.2881127 | 50.598186 |
| 0 | 0 |
| 0 | 0 |
| 0 | 0 |
| 61.8304902 | 50.598186 |
| 0 | 0 |
| 1.71751362 | 0 |
| 0 | 0 |
| 0 | 0 |
| 0 | 0 |
| 0 | 0 |
| 0 | 0 |
| 0 | 0 |
| 0 | 0 |
| 0 | 122.881309 |
| 0 | 0 |
| 0 | 0 |
| 60.1129766 | 0 |
| 0 | 0 |
| 0 | 0 |
| 0 | 0 |
| 0 | 21.6849369 |
| 0 | 0 |
| 0 | 0 |
| 0 | 0 |
| 0 | 0 |
| 85.8756808 | 0 |
| 0 | 0 |
| 250.756988 | 36.1415615 |
| 1.71751362 | 0 |
| 0 | 0 |
| 3.43502723 | 50.598186 |
| 5.15254085 | 14.4566246 |

| | |
|---|---|
| 0 | 14.4566246 |
| 0 | 0 |
| 0 | 0 |
| 63.5480038 | 14.4566246 |
| 285.10726 | 491.525236 |
| 0 | 0 |
| 0 | 0 |
| 0 | 86.7397475 |
| 0 | 57.8264983 |
| 194.079039 | 7.22831229 |
| 0 | 0 |
| 5.15254085 | 57.8264983 |
| 49.8078949 | 115.652997 |
| 0 | 7.22831229 |
| 0 | 0 |
| 212.971688 | 50.598186 |
| 0 | 0 |
| 120.225953 | 7.22831229 |
| 0 | 0 |
| 0 | 0 |
| 54.9604357 | 0 |
| 5.15254085 | 0 |
| 0 | 0 |
| 0 | 0 |
| 0 | 0 |
| 44.655354 | 0 |
| 0 | 0 |
| 0 | 0 |
| 0 | 0 |
| 0 | 0 |
| 0 | 0 |
| 0 | 0 |
| 0 | 0 |
| 121.943467 | 0 |
| 170.033848 | 57.8264983 |
| 94.4632489 | 166.251183 |
| 0 | 0 |
| 6.87005447 | 101.196372 |
| 0 | 0 |
| 0 | 0 |
| 0 | 0 |
| 12.0225953 | 0 |
| 0 | 231.305993 |

| | |
|---|---|
| 1.71751362 | 375.872239 |
| 0 | 0 |
| 0 | 0 |
| 0 | 0 |
| 3.43502723 | 0 |
| 0 | 0 |
| 0 | 0 |
| 0 | 0 |
| 13.7401089 | 0 |
| 0 | 0 |
| 0 | 0 |
| 0 | 0 |
| 0 | 0 |
| 0 | 0 |
| 0 | 0 |
| 0 | 0 |
| 0 | 0 |
| 0 | 115.652997 |
| 0 | 159.02287 |
| 0 | 0 |
| 0 | 0 |
| 0 | 0 |
| 460.293649 | 187.93612 |
| 0 | 0 |
| 0 | 0 |
| 0 | 0 |
| 0 | 0 |
| 0 | 0 |
| 0 | 0 |
| 0 | 0 |
| 0 | 0 |
| 0 | 0 |
| 0 | 0 |
| 0 | 0 |
| 0 | 0 |
| 0 | 0 |
| 68.7005447 | 7.22831229 |
| 1.71751362 | 0 |
| 72.1355719 | 0 |
| 0 | 0 |
| 0 | 0 |
| 60.1129766 | 245.762618 |
| 104.768331 | 7.22831229 |

| | |
|---|---|
| 238.734393 | 173.479495 |
| 0 | 0 |
| 0 | 0 |
| 27.4802179 | 43.3698738 |
| 3.43502723 | 0 |
| 75.5705991 | 0 |
| 0 | 0 |
| 0 | 0 |
| 5.15254085 | 0 |
| 8.58756808 | 0 |
| 17.1751362 | 0 |
| 0 | 0 |
| 0 | 0 |
| 0 | 0 |
| 34.3502723 | 151.794558 |
| 0 | 0 |
| 0 | 0 |
| 0 | 0 |
| 176.903902 | 0 |
| 0 | 0 |
| 111.638385 | 0 |
| 0 | 0 |
| 0 | 0 |
| 0 | 0 |
| 0 | 0 |
| 0 | 0 |
| 0 | 0 |
| 77.2881127 | 36.1415615 |
| 0 | 0 |
| 192.361525 | 542.123422 |
| 0 | 0 |
| 0 | 0 |
| 0 | 0 |
| 0 | 0 |
| 0 | 0 |
| 0 | 0 |
| 161.44628 | 14.4566246 |
| 242.16942 | 101.196372 |
| 0 | 0 |
| 0 | 0 |
| 0 | 0 |
| 0 | 0 |
| 0 | 0 |

| | |
|---|---|
| 0 | 0 |
| 0 | 0 |
| 0 | 0 |
| 0 | 0 |
| 0 | 0 |
| 1.71751362 | 43.3698738 |
| 0 | 0 |
| 0 | 0 |
| 175.186389 | 0 |
| 0 | 0 |
| 0 | 0 |
| 0 | 0 |
| 0 | 0 |
| 0 | 0 |
| 297.129856 | 14.4566246 |
| 0 | 0 |
| 0 | 0 |
| 0 | 0 |
| 0 | 0 |
| 0 | 0 |
| 0 | 224.077681 |
| 0 | 224.077681 |
| 1.71751362 | 0 |
| 0 | 0 |
| 0 | 0 |
| 0 | 0 |
| 0 | 0 |
| 0 | 0 |
| 0 | 0 |
| 0 | 0 |
| 0 | 0 |
| 0 | 0 |
| 0 | 0 |
| 0 | 0 |
| 0 | 0 |
| 0 | 0 |
| 0 | 0 |
| 142.55363 | 7.22831229 |
| 0 | 0 |
| 0 | 0 |
| 0 | 0 |
| 408.768241 | 115.652997 |
| 0 | 0 |

| | |
|---|---|
| 0 | 0 |
| 0 | 0 |
| 0 | 0 |
| 0 | 0 |
| 0 | 0 |
| 0 | 0 |
| 0 | 130.109621 |
| 0 | 0 |
| 0 | 0 |
| 63.5480038 | 0 |
| 0 | 7.22831229 |
| 82.4406536 | 0 |
| 0 | 0 |
| 3.43502723 | 0 |
| 132.248548 | 216.849369 |
| 0 | 0 |
| 0 | 0 |
| 0 | 0 |
| 0 | 0 |
| 0 | 0 |
| 183.773957 | 0 |
| 219.841743 | 0 |
| 0 | 0 |
| 85.8756808 | 7.22831229 |
| 0 | 0 |
| 0 | 0 |
| 0 | 0 |
| 37.7852996 | 151.794558 |
| 0 | 0 |
| 0 | 0 |
| 0 | 0 |
| 1.71751362 | 7.22831229 |
| 0 | 0 |
| 0 | 0 |
| 0 | 0 |
| 0 | 0 |
| 3.43502723 | 0 |
| 27.4802179 | 79.5114352 |
| 0 | 0 |
| 0 | 0 |
| 0 | 0 |
| 171.751362 | 0 |
| 171.751362 | 0 |

| | |
|---|---|
| 171.751362 | 0 |
| 0 | 0 |
| 0 | 0 |
| 182.056443 | 101.196372 |
| 0 | 0 |
| 0 | 0 |
| 0 | 0 |
| 109.920871 | 7.22831229 |
| 0 | 0 |
| 0 | 0 |
| 0 | 0 |
| 147.706171 | 36.1415615 |
| 5.15254085 | 7.22831229 |
| 0 | 0 |
| 44.655354 | 7.22831229 |
| 0 | 0 |
| 87.5931944 | 224.077681 |
| 0 | 93.9680598 |
| 0 | 0 |
| 159.728766 | 36.1415615 |
| 0 | 0 |
| 0 | 0 |
| 3.43502723 | 0 |
| 0 | 0 |
| 0 | 0 |
| 0 | 0 |
| 0 | 0 |
| 6.87005447 | 72.2831229 |
| 0 | 0 |
| 58.395463 | 0 |
| 0 | 0 |
| 0 | 0 |
| 115.073412 | 0 |
| 0 | 0 |
| 80.72314 | 0 |
| 0 | 0 |
| 46.3728676 | 21.6849369 |
| 0 | 0 |
| 0 | 0 |
| 170.033848 | 0 |
| 0 | 0 |
| 0 | 0 |
| 0 | 0 |

| | |
|---|---|
| 44.655354 | 86.7397475 |
| 0 | 0 |
| 0 | 0 |
| 273.084665 | 7.22831229 |
| 0 | 0 |
| 0 | 0 |
| 0 | 0 |
| 12.0225953 | 7.22831229 |
| 305.717424 | 14.4566246 |
| 0 | 0 |
| 168.316334 | 14.4566246 |
| 0 | 0 |
| 0 | 0 |
| 0 | 0 |
| 1.71751362 | 0 |
| 0 | 0 |
| 0 | 0 |
| 0 | 0 |
| 0 | 0 |
| 0 | 0 |
| 154.576225 | 21.6849369 |
| 0 | 0 |
| 0 | 0 |
| 0 | 0 |
| 657.807715 | 274.675867 |
| 0 | 0 |
| 0 | 0 |
| 0 | 0 |
| 168.316334 | 0 |
| 0 | 0 |
| 0 | 0 |
| 0 | 0 |
| 0 | 79.5114352 |
| 0 | 0 |
| 0 | 79.5114352 |
| 0 | 0 |
| 200.949093 | 0 |
| 0 | 0 |
| 0 | 0 |
| 1.71751362 | 0 |
| 0 | 0 |
| 0 | 0 |
| 133.966062 | 50.598186 |

| | |
|---|---|
| 0 | 0 |
| 0 | 0 |
| 5.15254085 | 0 |
| 0 | 0 |
| 642.350093 | 1272.18296 |
| 0 | 0 |
| 219.841743 | 72.2831229 |
| 0 | 0 |
| 0 | 0 |
| 0 | 0 |
| 80.72314 | 0 |
| 12.0225953 | 0 |
| 0 | 0 |
| 0 | 0 |
| 197.514066 | 21.6849369 |
| 0 | 0 |
| 0 | 0 |
| 0 | 0 |
| 0 | 0 |
| 10.3050817 | 21.6849369 |
| 0 | 0 |
| 0 | 0 |
| 0 | 0 |
| 103.050817 | 79.5114352 |
| 111.638385 | 0 |
| 0 | 0 |
| 0 | 0 |
| 0 | 0 |
| 0 | 0 |
| 0 | 0 |
| 0 | 0 |
| 0 | 0 |
| 0 | 0 |
| 0 | 0 |
| 0 | 0 |
| 0 | 0 |
| 0 | 0 |
| 0 | 0 |
| 1.71751362 | 57.8264983 |
| 166.598821 | 0 |
| 166.598821 | 0 |
| 0 | 0 |
| 0 | 0 |

| | |
|---|---|
| 0 | 0 |
| 0 | 0 |
| 0 | 0 |
| 6.87005447 | 144.566246 |
| 0 | 0 |
| 0 | 0 |
| 75.5705991 | 0 |
| 97.8982761 | 79.5114352 |
| 0 | 0 |
| 0 | 36.1415615 |
| 30.9152451 | 7.22831229 |
| 0 | 0 |
| 0 | 0 |
| 0 | 0 |
| 0 | 0 |
| 0 | 0 |
| 0 | 0 |
| 0 | 0 |
| 0 | 0 |
| 118.50844 | 209.621056 |
| 0 | 0 |
| 0 | 0 |
| 0 | 0 |
| 0 | 0 |
| 0 | 7.22831229 |
| 0 | 0 |
| 0 | 0 |
| 0 | 0 |
| 0 | 0 |
| 125.378494 | 0 |
| 0 | 0 |
| 0 | 0 |
| 0 | 202.392744 |
| 0 | 0 |
| 0 | 0 |
| 0 | 0 |
| 1.71751362 | 130.109621 |
| 0 | 0 |
| 0 | 0 |
| 0 | 0 |
| 295.412342 | 101.196372 |
| 0 | 0 |
| 0 | 0 |

| | |
|---|---|
| 0 | 0 |
| 116.790926 | 0 |
| 0 | 0 |
| 0 | 0 |
| 0 | 0 |
| 0 | 0 |
| 75.5705991 | 0 |
| 0 | 0 |
| 89.3107081 | 7.22831229 |
| 22.327677 | 43.3698738 |
| 154.576225 | 0 |
| 0 | 7.22831229 |
| 0 | 0 |
| 192.361525 | 361.415615 |
| 0 | 0 |
| 0 | 0 |
| 0 | 0 |
| 41.2203268 | 36.1415615 |
| 0 | 0 |
| 0 | 0 |
| 0 | 0 |
| 0 | 0 |
| 0 | 0 |
| 0 | 0 |
| 262.779583 | 173.479495 |
| 0 | 0 |
| 0 | 0 |
| 0 | 0 |
| 0 | 0 |
| 0 | 0 |
| 0 | 101.196372 |
| 0 | 101.196372 |
| 0 | 101.196372 |
| 0 | 0 |
| 0 | 0 |
| 163.163794 | 0 |
| 0 | 0 |
| 0 | 0 |
| 0 | 36.1415615 |
| 0 | 0 |
| 41.2203268 | 79.5114352 |
| 0 | 0 |
| 0 | 0 |

| | |
|---|---|
| 0 | 0 |
| 0 | 0 |
| 0 | 0 |
| 721.355719 | 281.904179 |
| 0 | 0 |
| 0 | 0 |
| 0 | 0 |
| 0 | 0 |
| 0 | 0 |
| 91.0282217 | 0 |
| 0 | 0 |
| 0 | 0 |
| 0 | 0 |
| 54.9604357 | 0 |
| 0 | 0 |
| 3.43502723 | 43.3698738 |
| 1.71751362 | 137.337934 |
| 0 | 0 |
| 0 | 0 |
| 0 | 0 |
| 0 | 28.9132492 |
| 0 | 0 |
| 0 | 0 |
| 75.5705991 | 21.6849369 |
| 0 | 0 |
| 0 | 0 |
| 0 | 0 |
| 128.813521 | 0 |
| 20.6101634 | 0 |
| 58.395463 | 289.132492 |
| 0 | 0 |
| 0 | 0 |
| 0 | 0 |
| 0 | 0 |
| 0 | 0 |
| 0 | 0 |
| 68.7005447 | 0 |
| 0 | 0 |
| 0 | 0 |
| 0 | 0 |
| 0 | 0 |
| 0 | 0 |
| 0 | 0 |

| | |
|---|---|
| 0 | 0 |
| 0 | 0 |
| 0 | 0 |
| 0 | 0 |
| 0 | 0 |
| 1.71751362 | 36.1415615 |
| 63.5480038 | 0 |
| 18.8926498 | 159.02287 |
| 0 | 0 |
| 0 | 0 |
| 20.6101634 | 14.4566246 |
| 0 | 0 |
| 80.72314 | 14.4566246 |
| 0 | 0 |
| 0 | 0 |
| 0 | 0 |
| 0 | 0 |
| 0 | 0 |
| 0 | 0 |
| 0 | 0 |
| 0 | 0 |
| 0 | 0 |
| 0 | 0 |
| 0 | 0 |
| 0 | 173.479495 |
| 0 | 0 |
| 0 | 7.22831229 |
| 159.728766 | 0 |
| 159.728766 | 0 |
| 0 | 0 |
| 0 | 202.392744 |
| 0 | 0 |
| 58.395463 | 0 |
| 0 | 0 |
| 0 | 0 |
| 0 | 0 |
| 0 | 0 |
| 41.2203268 | 0 |
| 0 | 0 |
| 0 | 0 |
| 0 | 0 |
| 0 | 0 |
| 1.71751362 | 0 |

| | |
|---|---|
| 92.7457353 | 28.9132492 |
| 0 | 0 |
| 264.497097 | 108.424684 |
| 87.5931944 | 7.22831229 |
| 0 | 0 |
| 0 | 0 |
| 0 | 0 |
| 257.627042 | 0 |
| 0 | 0 |
| 0 | 0 |
| 0 | 7.22831229 |
| 0 | 72.2831229 |
| 0 | 0 |
| 0 | 0 |
| 0 | 0 |
| 140.836117 | 0 |
| 0 | 0 |
| 0 | 166.251183 |
| 0 | 0 |
| 0 | 0 |
| 0 | 101.196372 |
| 0 | 0 |
| 0 | 0 |
| 0 | 0 |
| 27.4802179 | 224.077681 |
| 0 | 0 |
| 0 | 0 |
| 0 | 0 |
| 0 | 0 |
| 3.43502723 | 0 |
| 0 | 130.109621 |
| 158.011253 | 0 |
| 0 | 0 |
| 0 | 0 |
| 0 | 0 |
| 94.4632489 | 209.621056 |
| 0 | 0 |
| 65.2655174 | 0 |
| 27.4802179 | 325.274053 |
| 0 | 0 |
| 0 | 0 |
| 0 | 0 |
| 283.389747 | 0 |

| | |
|---|---|
| 37.7852996 | 0 |
| 0 | 0 |
| 152.858712 | 0 |
| 0 | 0 |
| 0 | 7.22831229 |
| 103.050817 | 115.652997 |
| 0 | 0 |
| 82.4406536 | 7.22831229 |
| 0 | 0 |
| 0 | 0 |
| 0 | 0 |
| 0 | 0 |
| 0 | 0 |
| 0 | 0 |
| 0 | 0 |
| 0 | 0 |
| 3.43502723 | 79.5114352 |
| 0 | 0 |
| 0 | 0 |
| 0 | 0 |
| 0 | 0 |
| 0 | 0 |
| 1.71751362 | 0 |
| 0 | 28.9132492 |
| 224.994284 | 28.9132492 |
| 0 | 0 |
| 0 | 0 |
| 0 | 0 |
| 0 | 0 |
| 0 | 50.598186 |
| 451.706081 | 43.3698738 |
| 0 | 0 |
| 0 | 0 |
| 0 | 0 |
| 0 | 0 |
| 0 | 0 |
| 0 | 0 |
| 156.293739 | 0 |
| 25.7627042 | 0 |
| 0 | 0 |
| 173.468875 | 0 |
| 0 | 0 |
| 46.3728676 | 0 |

| | |
|---|---|
| 48.0903813 | 166.251183 |
| 0 | 0 |
| 0 | 0 |
| 0 | 0 |
| 128.813521 | 0 |
| 0 | 0 |
| 6.87005447 | 72.2831229 |
| 206.101634 | 21.6849369 |
| 0 | 0 |
| 0 | 0 |
| 0 | 0 |
| 0 | 0 |
| 0 | 0 |
| 1.71751362 | 0 |
| 3.43502723 | 7.22831229 |
| 161.44628 | 21.6849369 |
| 73.8530855 | 0 |
| 0 | 0 |
| 0 | 0 |
| 0 | 0 |
| 0 | 202.392744 |
| 0 | 0 |
| 0 | 0 |
| 0 | 0 |
| 0 | 0 |
| 0 | 0 |
| 0 | 0 |
| 0 | 0 |
| 0 | 0 |
| 0 | 0 |
| 0 | 0 |
| 0 | 0 |
| 0 | 0 |
| 0 | 0 |
| 0 | 0 |
| 168.316334 | 115.652997 |
| 0 | 0 |
| 0 | 0 |
| 298.847369 | 7.22831229 |
| 0 | 0 |
| 0 | 0 |
| 0 | 0 |
| 0 | 93.9680598 |

| | |
|---|---|
| 0 | 0 |
| 0 | 93.9680598 |
| 0 | 195.164432 |
| 0 | 195.164432 |
| 0 | 0 |
| 0 | 0 |
| 1.71751362 | 0 |
| 0 | 0 |
| 188.926498 | 21.6849369 |
| 0 | 0 |
| 0 | 0 |
| 0 | 202.392744 |
| 0 | 0 |
| 0 | 0 |
| 0 | 0 |
| 238.734393 | 0 |
| 0 | 0 |
| 0 | 0 |
| 0 | 0 |
| 0 | 0 |
| 48.0903813 | 0 |
| 0 | 0 |
| 0 | 0 |
| 0 | 0 |
| 0 | 0 |
| 0 | 0 |
| 0 | 0 |
| 0 | 0 |
| 0 | 0 |
| 175.186389 | 14.4566246 |
| 0 | 0 |
| 0 | 0 |
| 0 | 0 |
| 0 | 0 |
| 0 | 0 |
| 0 | 0 |
| 0 | 0 |
| 0 | 0 |
| 0 | 0 |
| 0 | 0 |
| 72.1355719 | 0 |
| 0 | 0 |
| 0 | 0 |

| | |
|---|---|
| 0 | 0 |
| 0 | 0 |
| 0 | 0 |
| 0 | 0 |
| 0 | 0 |
| 0 | 0 |
| 27.4802179 | 101.196372 |
| 180.33893 | 72.2831229 |
| 0 | 0 |
| 0 | 14.4566246 |
| 290.259801 | 202.392744 |
| 0 | 0 |
| 0 | 0 |
| 0 | 7.22831229 |
| 0 | 0 |
| 0 | 0 |
| 0 | 0 |
| 0 | 0 |
| 0 | 0 |
| 0 | 0 |
| 0 | 0 |
| 192.361525 | 21.6849369 |
| 0 | 0 |
| 0 | 0 |
| 0 | 0 |
| 0 | 0 |
| 0 | 0 |
| 0 | 0 |
| 53.2429221 | 0 |
| 0 | 0 |
| 0 | 0 |
| 0 | 0 |
| 0 | 0 |
| 0 | 0 |
| 72.1355719 | 0 |
| 72.1355719 | 0 |
| 0 | 0 |
| 0 | 0 |
| 0 | 0 |
| 0 | 0 |
| 0 | 0 |
| 0 | 0 |
| 0 | 86.7397475 |

| | |
|---|---|
| 39.5028132 | 7.22831229 |
| 0 | 0 |
| 60.1129766 | 21.6849369 |
| 0 | 0 |
| 151.141198 | 0 |
| 151.141198 | 0 |
| 0 | 0 |
| 0 | 0 |
| 0 | 0 |
| 0 | 0 |
| 0 | 0 |
| 0 | 0 |
| 0 | 0 |
| 0 | 0 |
| 0 | 0 |
| 80.72314 | 0 |
| 0 | 0 |
| 1028.79066 | 498.753548 |
| 290.259801 | 108.424684 |
| 1.71751362 | 0 |
| 0 | 0 |
| 0 | 0 |
| 0 | 65.0548106 |
| 0 | 0 |
| 0 | 0 |
| 137.401089 | 101.196372 |
| 0 | 0 |
| 0 | 0 |
| 0 | 0 |
| 137.401089 | 21.6849369 |
| 0 | 0 |
| 1.71751362 | 0 |
| 0 | 0 |
| 0 | 0 |
| 0 | 0 |
| 0 | 187.93612 |
| 0 | 187.93612 |
| 383.005536 | 0 |
| 0 | 0 |
| 0 | 0 |
| 0 | 0 |
| 0 | 0 |
| 0 | 0 |

| | |
|---|---|
| 0 | 0 |
| 0 | 0 |
| 0 | 0 |
| 0 | 0 |
| 75.5705991 | 267.447555 |
| 0 | 0 |
| 0 | 0 |
| 0 | 115.652997 |
| 84.1581672 | 0 |
| 0 | 0 |
| 400.180673 | 137.337934 |
| 0 | 122.881309 |
| 0 | 0 |
| 0 | 0 |
| 65.2655174 | 0 |
| 0 | 0 |
| 0 | 0 |
| 0 | 0 |
| 0 | 0 |
| 0 | 0 |
| 15.4576225 | 43.3698738 |
| 0 | 0 |
| 0 | 0 |
| 46.3728676 | 0 |
| 0 | 0 |
| 0 | 0 |
| 34.3502723 | 202.392744 |
| 0 | 0 |
| 0 | 0 |
| 0 | 0 |
| 0 | 0 |
| 0 | 0 |
| 25.7627042 | 159.02287 |
| 0 | 0 |
| 0 | 0 |
| 147.706171 | 0 |
| 147.706171 | 0 |
| 0 | 0 |
| 75.5705991 | 108.424684 |
| 0 | 0 |
| 0 | 144.566246 |
| 0 | 7.22831229 |
| 0 | 0 |

| | |
|---|---|
| 0 | 0 |
| 17.1751362 | 195.164432 |
| 0 | 86.7397475 |
| 0 | 0 |
| 216.406716 | 0 |
| 305.717424 | 14.4566246 |
| 0 | 0 |
| 0 | 0 |
| 0 | 7.22831229 |
| 0 | 7.22831229 |
| 0 | 7.22831229 |
| 0 | 0 |
| 0 | 57.8264983 |
| 128.813521 | 14.4566246 |
| 163.163794 | 412.013801 |
| 0 | 0 |
| 0 | 0 |
| 0 | 0 |
| 0 | 0 |
| 0 | 0 |
| 128.813521 | 144.566246 |
| 0 | 0 |
| 0 | 7.22831229 |
| 0 | 0 |
| 0 | 86.7397475 |
| 0 | 0 |
| 63.5480038 | 14.4566246 |
| 0 | 0 |
| 0 | 0 |
| 41.2203268 | 86.7397475 |
| 0 | 0 |
| 149.423685 | 72.2831229 |
| 104.768331 | 36.1415615 |
| 0 | 0 |
| 158.011253 | 72.2831229 |
| 0 | 0 |
| 0 | 0 |
| 0 | 0 |
| 12.0225953 | 7.22831229 |
| 0 | 0 |
| 0 | 0 |
| 0 | 101.196372 |
| 61.8304902 | 0 |

| | |
|---|---|
| 218.124229 | 0 |
| 152.858712 | 7.22831229 |
| 101.333303 | 0 |
| 48.0903813 | 0 |
| 0 | 0 |
| 0 | 0 |
| 259.344556 | 7.22831229 |
| 218.124229 | 7.22831229 |
| 0 | 0 |
| 0 | 0 |
| 0 | 0 |
| 0 | 0 |
| 48.0903813 | 0 |
| 0 | 0 |
| 0 | 0 |
| 0 | 0 |
| 0 | 0 |
| 0 | 0 |
| 0 | 0 |
| 0 | 0 |
| 53.2429221 | 28.9132492 |
| 0 | 0 |
| 0 | 0 |
| 0 | 0 |
| 0 | 0 |
| 0 | 0 |
| 0 | 0 |
| 0 | 0 |
| 0 | 0 |
| 0 | 0 |
| 182.056443 | 7.22831229 |
| 85.8756808 | 21.6849369 |
| 0 | 224.077681 |
| 37.7852996 | 0 |
| 0 | 43.3698738 |
| 0 | 0 |
| 0 | 0 |
| 0 | 0 |
| 0 | 0 |
| 1.71751362 | 50.598186 |
| 1.71751362 | 50.598186 |
| 1.71751362 | 50.598186 |
| 1.71751362 | 50.598186 |

| | |
|---|---|
| 1.71751362 | 50.598186 |
| 0 | 0 |
| 1.71751362 | 7.22831229 |
| 0 | 0 |
| 0 | 0 |
| 0 | 65.0548106 |
| 182.056443 | 7.22831229 |
| 0 | 180.707807 |
| 0 | 0 |
| 144.271144 | 0 |
| 63.5480038 | 202.392744 |
| 0 | 0 |
| 0 | 0 |
| 0 | 0 |
| 0 | 0 |
| 0 | 0 |
| 0 | 0 |
| 61.8304902 | 0 |
| 0 | 0 |
| 0 | 0 |
| 0 | 0 |
| 0 | 0 |
| 0 | 0 |
| 17.1751362 | 0 |
| 211.254175 | 0 |
| 61.8304902 | 14.4566246 |
| 3.43502723 | 0 |
| 0 | 173.479495 |
| 0 | 0 |
| 0 | 0 |
| 0 | 0 |
| 41.2203268 | 7.22831229 |
| 116.790926 | 101.196372 |
| 1.71751362 | 180.707807 |
| 39.5028132 | 101.196372 |
| 0 | 93.9680598 |
| 1.71751362 | 0 |
| 1.71751362 | 0 |
| 0 | 72.2831229 |
| 0 | 0 |
| 0 | 0 |
| 420.790836 | 0 |
| 5.15254085 | 115.652997 |

| | |
|---|---|
| 0 | 0 |
| 0 | 0 |
| 0 | 0 |
| 97.8982761 | 0 |
| 0 | 0 |
| 0 | 0 |
| 0 | 0 |
| 0 | 0 |
| 0 | 7.22831229 |
| 0 | 0 |
| 1.71751362 | 72.2831229 |
| 0 | 0 |
| 0 | 0 |
| 0 | 0 |
| 0 | 0 |
| 0 | 0 |
| 0 | 0 |
| 0 | 0 |
| 161.44628 | 86.7397475 |
| 0 | 0 |
| 42.9378404 | 14.4566246 |
| 0 | 0 |
| 0 | 0 |
| 0 | 0 |
| 0 | 50.598186 |
| 1.71751362 | 0 |
| 0 | 0 |
| 0 | 144.566246 |
| 0 | 0 |
| 0 | 0 |
| 0 | 0 |
| 0 | 0 |
| 0 | 0 |
| 0 | 0 |
| 1.71751362 | 7.22831229 |
| 0 | 0 |
| 0 | 0 |
| 0 | 0 |
| 0 | 57.8264983 |
| 0 | 0 |
| 0 | 0 |
| 0 | 0 |
| 0 | 0 |

| | |
|---|---|
| 0 | 0 |
| 0 | 0 |
| 0 | 0 |
| 82.4406536 | 231.305993 |
| 0 | 122.881309 |
| 0 | 0 |
| 0 | 0 |
| 0 | 0 |
| 0 | 0 |
| 0 | 0 |
| 0 | 0 |
| 0 | 0 |
| 0 | 65.0548106 |
| 0 | 0 |
| 0 | 0 |
| 0 | 86.7397475 |
| 0 | 0 |
| 142.55363 | 79.5114352 |
| 219.841743 | 21.6849369 |
| 0 | 0 |
| 0 | 7.22831229 |
| 0 | 0 |
| 0 | 0 |
| 0 | 14.4566246 |
| 5.15254085 | 21.6849369 |
| 0 | 0 |
| 0 | 0 |
| 140.836117 | 0 |
| 0 | 0 |
| 0 | 0 |
| 0 | 0 |
| 82.4406536 | 14.4566246 |
| 0 | 0 |
| 0 | 0 |
| 0 | 0 |
| 0 | 0 |
| 0 | 7.22831229 |
| 0 | 0 |
| 0 | 0 |
| 106.485844 | 0 |
| 22.327677 | 7.22831229 |
| 66.983031 | 0 |
| 0 | 0 |

| | |
|---|---|
| 37.7852996 | 21.6849369 |
| 0 | 0 |
| 0 | 0 |
| 0 | 0 |
| 0 | 0 |
| 0 | 0 |
| 0 | 0 |
| 54.9604357 | 122.881309 |
| 0 | 0 |
| 0 | 0 |
| 1.71751362 | 0 |
| 0 | 0 |
| 0 | 0 |
| 0 | 0 |
| 0 | 0 |
| 66.983031 | 0 |
| 56.6779493 | 86.7397475 |
| 0 | 43.3698738 |
| 0 | 0 |
| 87.5931944 | 0 |
| 22.327677 | 7.22831229 |
| 0 | 0 |
| 0 | 0 |
| 0 | 0 |
| 0 | 0 |
| 0 | 0 |
| 6.87005447 | 7.22831229 |
| 0 | 0 |
| 135.683576 | 137.337934 |
| 0 | 0 |
| 0 | 0 |
| 0 | 0 |
| 0 | 0 |
| 0 | 0 |
| 0 | 0 |
| 0 | 0 |
| 0 | 173.479495 |
| 0 | 173.479495 |
| 0 | 173.479495 |
| 0 | 0 |
| 0 | 122.881309 |
| 0 | 0 |
| 0 | 0 |

| | |
|---|---|
| 0 | 0 |
| 0 | 0 |
| 73.8530855 | 332.502365 |
| 0 | 0 |
| 58.395463 | 0 |
| 0 | 0 |
| 0 | 0 |
| 0 | 0 |
| 0 | 0 |
| 0 | 0 |
| 0 | 0 |
| 137.401089 | 0 |
| 0 | 0 |
| 0 | 0 |
| 0 | 0 |
| 147.706171 | 115.652997 |
| 0 | 0 |
| 0 | 0 |
| 0 | 101.196372 |
| 0 | 0 |
| 30.9152451 | 7.22831229 |
| 0 | 0 |
| 0 | 0 |
| 0 | 0 |
| 0 | 93.9680598 |
| 0 | 0 |
| 0 | 0 |
| 0 | 0 |
| 139.118603 | 0 |
| 1.71751362 | 0 |
| 0 | 0 |
| 171.751362 | 14.4566246 |
| 0 | 0 |
| 159.728766 | 159.02287 |
| 0 | 0 |
| 0 | 0 |
| 0 | 0 |
| 0 | 0 |
| 0 | 0 |
| 0 | 0 |
| 0 | 0 |
| 0 | 0 |
| 0 | 0 |

| | |
|---|---|
| 0 | 0 |
| 0 | 7.22831229 |
| 0 | 0 |
| 137.401089 | 0 |
| 48.0903813 | 0 |
| 0 | 0 |
| 501.513976 | 968.593847 |
| 0 | 0 |
| 0 | 0 |
| 0 | 0 |
| 252.474502 | 14.4566246 |
| 534.146735 | 426.470425 |
| 1.71751362 | 0 |
| 0 | 0 |
| 0 | 0 |
| 0 | 0 |
| 0 | 0 |
| 0 | 86.7397475 |
| 0 | 0 |
| 0 | 86.7397475 |
| 0 | 0 |
| 0 | 0 |
| 0 | 0 |
| 0 | 0 |
| 63.5480038 | 0 |
| 0 | 397.557176 |
| 0 | 0 |
| 65.2655174 | 0 |
| 0 | 0 |
| 0 | 0 |
| 0 | 0 |
| 65.2655174 | 0 |
| 0 | 0 |
| 0 | 0 |
| 0 | 0 |
| 0 | 0 |
| 0 | 0 |
| 0 | 0 |
| 0 | 0 |
| 0 | 0 |
| 0 | 0 |
| 63.5480038 | 137.337934 |
| 0 | 0 |

| | |
|---|---|
| 0 | 0 |
| 0 | 0 |
| 0 | 0 |
| 0 | 0 |
| 0 | 0 |
| 0 | 0 |
| 1.71751362 | 144.566246 |
| 30.9152451 | 7.22831229 |
| 12.0225953 | 28.9132492 |
| 32.6327587 | 0 |
| 12.0225953 | 21.6849369 |
| 94.4632489 | 43.3698738 |
| 0 | 151.794558 |
| 0 | 0 |
| 0 | 7.22831229 |
| 0 | 0 |
| 0 | 187.93612 |
| 0 | 0 |
| 0 | 0 |
| 0 | 7.22831229 |
| 0 | 0 |
| 135.683576 | 0 |
| 0 | 0 |
| 0 | 0 |
| 0 | 0 |
| 0 | 0 |
| 0 | 0 |
| 0 | 0 |
| 5.15254085 | 86.7397475 |
| 0 | 0 |
| 115.073412 | 0 |
| 0 | 0 |
| 0 | 0 |
| 0 | 0 |
| 0 | 0 |
| 0 | 0 |
| 659.525229 | 159.02287 |
| 66.983031 | 0 |
| 190.644011 | 0 |
| 0 | 7.22831229 |
| 0 | 0 |
| 18.8926498 | 14.4566246 |
| 0 | 0 |

| | |
|---|---|
| 0 | 115.652997 |
| 0 | 0 |
| 48.0903813 | 28.9132492 |
| 0 | 0 |
| 0 | 0 |
| 0 | 0 |
| 0 | 0 |
| 68.7005447 | 252.99093 |
| 0 | 0 |
| 0 | 0 |
| 0 | 0 |
| 0 | 57.8264983 |
| 0 | 0 |
| 0 | 0 |
| 0 | 0 |
| 0 | 0 |
| 0 | 0 |
| 212.971688 | 14.4566246 |
| 0 | 0 |
| 0 | 0 |
| 1.71751362 | 0 |
| 0 | 0 |
| 0 | 0 |
| 0 | 0 |
| 0 | 0 |
| 53.2429221 | 0 |
| 0 | 0 |
| 0 | 0 |
| 133.966062 | 0 |
| 324.610074 | 0 |
| 0 | 0 |
| 0 | 7.22831229 |
| 0 | 0 |
| 0 | 14.4566246 |
| 0 | 0 |
| 63.5480038 | 0 |
| 0 | 0 |
| 0 | 0 |
| 106.485844 | 281.904179 |
| 0 | 0 |
| 0 | 0 |
| 0 | 137.337934 |
| 0 | 0 |

| | |
|---|---|
| 42.9378404 | 0 |
| 0 | 0 |
| 87.5931944 | 0 |
| 0 | 0 |
| 0 | 0 |
| 0 | 122.881309 |
| 0 | 0 |
| 30.9152451 | 14.4566246 |
| 0 | 0 |
| 0 | 0 |
| 0 | 0 |
| 0 | 86.7397475 |
| 44.655354 | 0 |
| 0 | 0 |
| 345.220237 | 65.0548106 |
| 0 | 0 |
| 0 | 50.598186 |
| 0 | 0 |
| 0 | 0 |
| 0 | 57.8264983 |
| 0 | 0 |
| 94.4632489 | 816.799289 |
| 0 | 137.337934 |
| 0 | 0 |
| 0 | 0 |
| 353.807805 | 14.4566246 |
| 0 | 0 |
| 54.9604357 | 108.424684 |
| 0 | 0 |
| 0 | 0 |
| 0 | 0 |
| 0 | 0 |
| 0 | 0 |
| 0 | 0 |
| 0 | 0 |
| 214.689202 | 14.4566246 |
| 0 | 0 |
| 185.491471 | 7.22831229 |
| 97.8982761 | 0 |
| 1.71751362 | 0 |
| 0 | 0 |
| 0 | 0 |
| 0 | 0 |

| | |
|---|---|
| 132.248548 | 0 |
| 0 | 0 |
| 132.248548 | 0 |
| 0 | 0 |
| 0 | 0 |
| 0 | 0 |
| 0 | 0 |
| 0 | 0 |
| 87.5931944 | 43.3698738 |
| 0 | 0 |
| 0 | 57.8264983 |
| 0 | 0 |
| 0 | 0 |
| 0 | 0 |
| 0 | 0 |
| 0 | 0 |
| 0 | 0 |
| 0 | 0 |
| 0 | 0 |
| 0 | 0 |
| 0 | 0 |
| 0 | 0 |
| 0 | 0 |
| 1.71751362 | 0 |
| 39.5028132 | 0 |
| 0 | 137.337934 |
| 170.033848 | 231.305993 |
| 0 | 0 |
| 0 | 0 |
| 60.1129766 | 7.22831229 |
| 0 | 0 |
| 61.8304902 | 0 |
| 0 | 0 |
| 0 | 0 |
| 0 | 7.22831229 |
| 0 | 0 |
| 0 | 0 |
| 0 | 0 |
| 0 | 0 |
| 0 | 0 |
| 0 | 0 |
| 0 | 108.424684 |
| 0 | 0 |

| | |
|---|---|
| 0 | 0 |
| 118.50844 | 0 |
| 0 | 0 |
| 0 | 0 |
| 0 | 0 |
| 0 | 0 |
| 0 | 0 |
| 0 | 0 |
| 0 | 0 |
| 0 | 0 |
| 0 | 0 |
| 58.395463 | 50.598186 |
| 255.909529 | 21.6849369 |
| 0 | 0 |
| 0 | 0 |
| 53.2429221 | 43.3698738 |
| 130.531035 | 0 |
| 0 | 0 |
| 75.5705991 | 14.4566246 |
| 15.4576225 | 14.4566246 |
| 0 | 0 |
| 0 | 0 |
| 0 | 0 |
| 0 | 28.9132492 |
| 0 | 0 |
| 8.58756808 | 86.7397475 |
| 1.71751362 | 0 |
| 135.683576 | 0 |
| 0 | 0 |
| 0 | 0 |
| 20.6101634 | 0 |
| 0 | 43.3698738 |
| 0 | 0 |
| 0 | 0 |
| 0 | 0 |
| 0 | 0 |
| 0 | 0 |
| 0 | 0 |
| 243.886934 | 0 |
| 121.943467 | 195.164432 |
| 0 | 0 |
| 0 | 0 |
| 0 | 0 |

| | |
|---|---|
| 290.259801 | 7.22831229 |
| 0 | 0 |
| 0 | 0 |
| 0 | 0 |
| 0 | 0 |
| 0 | 0 |
| 388.158077 | 0 |
| 0 | 0 |
| 0 | 0 |
| 0 | 0 |
| 0 | 0 |
| 0 | 0 |
| 0 | 93.9680598 |
| 0 | 0 |
| 0 | 0 |
| 0 | 0 |
| 0 | 0 |
| 0 | 0 |
| 0 | 0 |
| 0 | 0 |
| 60.1129766 | 0 |
| 0 | 0 |
| 0 | 0 |
| 128.813521 | 0 |
| 0 | 0 |
| 128.813521 | 0 |
| 0 | 0 |
| 0 | 0 |
| 0 | 0 |
| 0 | 159.02287 |
| 0 | 159.02287 |
| 0 | 159.02287 |
| 0 | 0 |
| 278.237206 | 7.22831229 |
| 73.8530855 | 0 |
| 0 | 0 |
| 0 | 0 |
| 0 | 0 |
| 0 | 0 |
| 0 | 0 |
| 5.15254085 | 50.598186 |
| 27.4802179 | 318.045741 |
| 0 | 0 |

| | |
|---|---|
| 0 | 0 |
| 0 | 0 |
| 8.58756808 | 72.2831229 |
| 53.2429221 | 0 |
| 0 | 36.1415615 |
| 0 | 0 |
| 0 | 0 |
| 0 | 0 |
| 1.71751362 | 65.0548106 |
| 0 | 0 |
| 0 | 0 |
| 0 | 0 |
| 0 | 0 |
| 0 | 0 |
| 1.71751362 | 14.4566246 |
| 0 | 0 |
| 0 | 0 |
| 0 | 0 |
| 0 | 0 |
| 56.6779493 | 289.132492 |
| 72.1355719 | 0 |
| 0 | 0 |
| 0 | 0 |
| 0 | 0 |
| 0 | 0 |
| 0 | 0 |
| 0 | 21.6849369 |
| 0 | 0 |
| 49.8078949 | 0 |
| 0 | 0 |
| 0 | 0 |
| 0 | 0 |
| 0 | 0 |
| 0 | 0 |
| 0 | 0 |
| 0 | 0 |
| 0 | 180.707807 |
| 127.096008 | 0 |
| 127.096008 | 0 |
| 127.096008 | 0 |
| 0 | 0 |
| 0 | 0 |
| 39.5028132 | 28.9132492 |

| | |
|---|---|
| 0 | 0 |
| 0 | 0 |
| 0 | 0 |
| 0 | 0 |
| 0 | 0 |
| 68.7005447 | 0 |
| 0 | 0 |
| 0 | 0 |
| 96.1807625 | 50.598186 |
| 0 | 0 |
| 0 | 0 |
| 34.3502723 | 115.652997 |
| 156.293739 | 0 |
| 0 | 0 |
| 0 | 0 |
| 92.7457353 | 0 |
| 0 | 0 |
| 0 | 0 |
| 0 | 0 |
| 0 | 0 |
| 0 | 0 |
| 0 | 0 |
| 30.9152451 | 0 |
| 1719.23113 | 665.004731 |
| 0 | 0 |
| 908.564703 | 592.721608 |
| 49.8078949 | 14.4566246 |
| 0 | 0 |
| 6.87005447 | 28.9132492 |
| 0 | 0 |
| 242.16942 | 36.1415615 |
| 273.084665 | 354.187302 |
| 0 | 0 |
| 295.412342 | 108.424684 |
| 0 | 0 |
| 125.378494 | 0 |
| 125.378494 | 0 |
| 125.378494 | 0 |
| 125.378494 | 0 |
| 0 | 0 |
| 0 | 0 |
| 0 | 0 |
| 0 | 0 |

| | |
|---|---|
| 0 | 0 |
| 10.3050817 | 151.794558 |
| 0 | 0 |
| 0 | 0 |
| 0 | 57.8264983 |
| 0 | 0 |
| 0 | 0 |
| 0 | 0 |
| 237.016879 | 0 |
| 77.2881127 | 115.652997 |
| 92.7457353 | 0 |
| 6.87005447 | 202.392744 |
| 0 | 0 |
| 0 | 0 |
| 0 | 0 |
| 0 | 0 |
| 48.0903813 | 0 |
| 0 | 0 |
| 0 | 0 |
| 0 | 0 |
| 0 | 0 |
| 657.807715 | 672.233043 |
| 0 | 0 |
| 0 | 0 |
| 12.0225953 | 0 |
| 0 | 101.196372 |
| 0 | 0 |
| 0 | 0 |
| 0 | 0 |
| 0 | 0 |
| 0 | 0 |
| 0 | 0 |
| 0 | 0 |
| 60.1129766 | 0 |
| 0 | 0 |
| 0 | 0 |
| 0 | 0 |
| 0 | 0 |
| 0 | 0 |
| 0 | 0 |
| 0 | 0 |
| 0 | 0 |
| 0 | 0 |

| | |
|---|---|
| 0 | 0 |
| 0 | 0 |
| 0 | 0 |
| 0 | 0 |
| 68.7005447 | 0 |
| 264.497097 | 72.2831229 |
| 212.971688 | 267.447555 |
| 0 | 0 |
| 0 | 0 |
| 0 | 0 |
| 0 | 0 |
| 0 | 151.794558 |
| 0 | 151.794558 |
| 0 | 151.794558 |
| 0 | 151.794558 |
| 0 | 151.794558 |
| 0 | 0 |
| 168.316334 | 0 |
| 0 | 0 |
| 0 | 0 |
| 0 | 0 |
| 0 | 0 |
| 0 | 0 |
| 0 | 0 |
| 1.71751362 | 43.3698738 |
| 0 | 0 |
| 0 | 0 |
| 0 | 0 |
| 283.389747 | 325.274053 |
| 0 | 14.4566246 |
| 0 | 0 |
| 0 | 137.337934 |
| 0 | 0 |
| 13.7401089 | 0 |
| 0 | 0 |
| 0 | 0 |
| 27.4802179 | 7.22831229 |
| 0 | 0 |
| 0 | 0 |
| 463.728676 | 2811.81348 |
| 0 | 0 |
| 0 | 0 |
| 0 | 0 |

| | |
|---|---|
| 0 | 0 |
| 0 | 0 |
| 0 | 0 |
| 0 | 0 |
| 51.5254085 | 0 |
| 0 | 0 |
| 0 | 0 |
| 0 | 57.8264983 |
| 0 | 0 |
| 0 | 0 |
| 0 | 0 |
| 0 | 7.22831229 |
| 0 | 0 |
| 0 | 57.8264983 |
| 127.096008 | 50.598186 |
| 80.72314 | 93.9680598 |
| 0 | 0 |
| 0 | 0 |
| 61.8304902 | 7.22831229 |
| 0 | 7.22831229 |
| 121.943467 | 0 |
| 29.1977315 | 28.9132492 |
| 0 | 0 |
| 73.8530855 | 0 |
| 0 | 28.9132492 |
| 0 | 0 |
| 595.977225 | 339.730678 |
| 0 | 0 |
| 0 | 7.22831229 |
| 0 | 0 |
| 37.7852996 | 144.566246 |
| 0 | 0 |
| 58.395463 | 0 |
| 0 | 0 |
| 0 | 7.22831229 |
| 0 | 0 |
| 0 | 0 |
| 0 | 0 |
| 0 | 0 |
| 0 | 0 |
| 0 | 0 |
| 0 | 0 |
| 0 | 7.22831229 |

| | |
|---|---|
| 0 | 0 |
| 0 | 0 |
| 106.485844 | 0 |
| 0 | 0 |
| 0 | 0 |
| 1.71751362 | 0 |
| 0 | 0 |
| 269.649638 | 28.9132492 |
| 0 | 0 |
| 0 | 101.196372 |
| 0 | 0 |
| 0 | 0 |
| 1.71751362 | 0 |
| 0 | 0 |
| 0 | 0 |
| 0 | 0 |
| 5.15254085 | 7.22831229 |
| 0 | 0 |
| 0 | 0 |
| 504.949003 | 267.447555 |
| 0 | 0 |
| 3.43502723 | 0 |
| 0 | 0 |
| 1.71751362 | 101.196372 |
| 0 | 0 |
| 144.271144 | 14.4566246 |
| 0 | 0 |
| 0 | 0 |
| 0 | 0 |
| 0 | 0 |
| 0 | 0 |
| 0 | 0 |
| 0 | 0 |
| 0 | 0 |
| 0 | 79.5114352 |
| 0 | 0 |
| 0 | 0 |
| 0 | 0 |
| 0 | 0 |
| 0 | 159.02287 |
| 48.0903813 | 101.196372 |
| 0 | 144.566246 |
| 0 | 144.566246 |

| | |
|---|---|
| 0 | 115.652997 |
| 286.824774 | 36.1415615 |
| 0 | 0 |
| 149.423685 | 0 |
| 0 | 0 |
| 29.1977315 | 296.360804 |
| 0 | 0 |
| 0 | 0 |
| 0 | 0 |
| 0 | 0 |
| 453.423595 | 7.22831229 |
| 0 | 0 |
| 61.8304902 | 122.881309 |
| 159.728766 | 361.415615 |
| 0 | 0 |
| 22.327677 | 7.22831229 |
| 0 | 0 |
| 48.0903813 | 0 |
| 0 | 0 |
| 0 | 0 |
| 171.751362 | 0 |
| 0 | 0 |
| 0 | 0 |
| 0 | 0 |
| 0 | 0 |
| 0 | 0 |
| 0 | 0 |
| 65.2655174 | 50.598186 |
| 0 | 0 |
| 0 | 0 |
| 0 | 72.2831229 |
| 0 | 0 |
| 0 | 0 |
| 0 | 0 |
| 0 | 0 |
| 0 | 0 |
| 0 | 0 |
| 0 | 0 |
| 0 | 0 |
| 0 | 0 |
| 0 | 0 |
| 0 | 0 |
| 0 | 0 |

| | |
|---|---|
| 0 | 0 |
| 80.72314 | 57.8264983 |
| 0 | 0 |
| 0 | 0 |
| 0 | 0 |
| 0 | 0 |
| 0 | 0 |
| 37.7852996 | 14.4566246 |
| 563.344466 | 151.794558 |
| 0 | 0 |
| 0 | 0 |
| 0 | 0 |
| 0 | 0 |
| 0 | 57.8264983 |
| 0 | 0 |
| 0 | 0 |
| 0 | 0 |
| 0 | 0 |
| 118.50844 | 0 |
| 0 | 0 |
| 0 | 0 |
| 68.7005447 | 0 |
| 0 | 0 |
| 65.2655174 | 0 |
| 0 | 0 |
| 0 | 50.598186 |
| 0 | 0 |
| 0 | 0 |
| 0 | 0 |
| 0 | 0 |
| 0 | 0 |
| 0 | 0 |
| 0 | 0 |
| 0 | 0 |
| 0 | 0 |
| 0 | 0 |
| 0 | 0 |
| 0 | 0 |
| 0 | 0 |
| 1.71751362 | 0 |
| 1.71751362 | 0 |
| 0 | 0 |
| 0 | 0 |

| | |
|---|---|
| 0 | 0 |
| 0 | 0 |
| 42.9378404 | 0 |
| 0 | 0 |
| 0 | 0 |
| 0 | 0 |
| 61.8304902 | 36.1415615 |
| 0 | 0 |
| 0 | 0 |
| 0 | 0 |
| 0 | 0 |
| 0 | 0 |
| 0 | 0 |
| 0 | 0 |
| 0 | 0 |
| 0 | 0 |
| 0 | 0 |
| 0 | 7.22831229 |
| 0 | 0 |
| 5.15254085 | 101.196372 |
| 108.203358 | 0 |
| 0 | 0 |
| 0 | 0 |
| 0 | 0 |
| 0 | 0 |
| 0 | 137.337934 |
| 0 | 137.337934 |
| 0 | 0 |
| 0 | 0 |
| 0 | 0 |
| 0 | 0 |
| 0 | 0 |
| 0 | 0 |
| 113.355899 | 0 |
| 0 | 0 |
| 0 | 0 |
| 0 | 0 |
| 0 | 0 |
| 0 | 0 |
| 0 | 0 |
| 0 | 0 |
| 0 | 0 |
| 0 | 0 |

| | |
|---|---|
| 0 | 0 |
| 0 | 0 |
| 0 | 0 |
| 0 | 0 |
| 0 | 0 |
| 0 | 0 |
| 0 | 0 |
| 1.71751362 | 0 |
| 0 | 0 |
| 0 | 0 |
| 116.790926 | 0 |
| 116.790926 | 0 |
| 116.790926 | 0 |
| 527.27668 | 0 |
| 0 | 0 |
| 0 | 0 |
| 0 | 0 |
| 0 | 0 |
| 0 | 0 |
| 0 | 0 |
| 0 | 0 |
| 202.666607 | 0 |
| 58.395463 | 7.22831229 |
| 46.3728676 | 0 |
| 0 | 0 |
| 199.23158 | 0 |
| 84.1581672 | 0 |
| 0 | 0 |
| 0 | 0 |
| 0 | 0 |
| 0 | 0 |
| 0 | 7.22831229 |
| 200.949093 | 14.4566246 |
| 0 | 0 |
| 53.2429221 | 7.22831229 |
| 0 | 0 |
| 0 | 0 |
| 139.118603 | 21.6849369 |
| 0 | 0 |
| 91.0282217 | 0 |
| 0 | 0 |
| 0 | 0 |
| 0 | 0 |

| | |
|---|---|
| 0 | 0 |
| 0 | 0 |
| 0 | 0 |
| 0 | 0 |
| 0 | 0 |
| 0 | 0 |
| 0 | 0 |
| 1.71751362 | 173.479495 |
| 0 | 0 |
| 0 | 0 |
| 0 | 0 |
| 0 | 0 |
| 163.163794 | 187.93612 |
| 0 | 0 |
| 0 | 0 |
| 0 | 0 |
| 54.9604357 | 216.849369 |
| 0 | 14.4566246 |
| 0 | 0 |
| 75.5705991 | 0 |
| 0 | 7.22831229 |
| 0 | 0 |
| 0 | 0 |
| 0 | 0 |
| 0 | 0 |
| 8.58756808 | 7.22831229 |
| 0 | 0 |
| 66.983031 | 50.598186 |
| 0 | 0 |
| 0 | 0 |
| 0 | 65.0548106 |
| 0 | 0 |
| 0 | 0 |
| 0 | 0 |
| 115.073412 | 0 |
| 115.073412 | 0 |
| 0 | 0 |
| 116.790926 | 0 |
| 0 | 0 |
| 0 | 0 |
| 0 | 14.4566246 |
| 0 | 7.22831229 |
| 0 | 0 |

| | |
|---|---|
| 37.7852996 | 14.4566246 |
| 0 | 0 |
| 0 | 0 |
| 0 | 0 |
| 0 | 0 |
| 1.71751362 | 7.22831229 |
| 0 | 0 |
| 0 | 0 |
| 0 | 0 |
| 0 | 130.109621 |
| 0 | 130.109621 |
| 0 | 130.109621 |
| 0 | 130.109621 |
| 0 | 130.109621 |
| 0 | 130.109621 |
| 0 | 130.109621 |
| 0 | 0 |
| 0 | 0 |
| 0 | 0 |
| 0 | 0 |
| 0 | 0 |
| 0 | 7.22831229 |
| 0 | 0 |
| 15.4576225 | 14.4566246 |
| 0 | 0 |
| 0 | 0 |
| 0 | 0 |
| 0 | 0 |
| 0 | 0 |
| 0 | 0 |
| 0 | 0 |
| 54.9604357 | 14.4566246 |
| 0 | 0 |
| 0 | 0 |
| 0 | 0 |
| 0 | 0 |
| 0 | 7.22831229 |
| 161.44628 | 0 |
| 0 | 0 |
| 365.8304 | 14.4566246 |
| 0 | 0 |
| 0 | 0 |
| 1.71751362 | 7.22831229 |

| | |
|---|---|
| 0 | 0 |
| 152.858712 | 151.794558 |
| 0 | 0 |
| 0 | 0 |
| 0 | 0 |
| 0 | 0 |
| 0 | 0 |
| 36.0677859 | 0 |
| 0 | 0 |
| 0 | 108.424684 |
| 0 | 0 |
| 0 | 0 |
| 0 | 0 |
| 0 | 0 |
| 0 | 0 |
| 51.5254085 | 93.9680598 |
| 113.355899 | 0 |
| 159.728766 | 0 |
| 113.355899 | 0 |
| 0 | 0 |
| 68.7005447 | 0 |
| 0 | 0 |
| 0 | 0 |
| 63.5480038 | 0 |
| 63.5480038 | 0 |
| 0 | 0 |
| 0 | 0 |
| 61.8304902 | 144.566246 |
| 3.43502723 | 50.598186 |
| 0 | 0 |
| 0 | 0 |
| 0 | 0 |
| 0 | 0 |
| 68.7005447 | 0 |
| 0 | 0 |
| 0 | 0 |
| 0 | 0 |
| 0 | 0 |
| 0 | 0 |
| 65.2655174 | 0 |
| 0 | 0 |
| 0 | 0 |
| 65.2655174 | 0 |

| | |
|---|---|
| 0 | 0 |
| 0 | 0 |
| 0 | 0 |
| 0 | 0 |
| 0 | 0 |
| 0 | 0 |
| 0 | 0 |
| 132.248548 | 21.6849369 |
| 0 | 0 |
| 12.0225953 | 14.4566246 |
| 0 | 0 |
| 0 | 0 |
| 0 | 0 |
| 0 | 0 |
| 1.71751362 | 0 |
| 0 | 0 |
| 0 | 0 |
| 0 | 0 |
| 0 | 0 |
| 0 | 0 |
| 0 | 0 |
| 0 | 0 |
| 0 | 0 |
| 0 | 0 |
| 34.3502723 | 0 |
| 0 | 0 |
| 0 | 65.0548106 |
| 0 | 0 |
| 108.203358 | 108.424684 |
| 0 | 0 |
| 82.4406536 | 0 |
| 0 | 0 |
| 0 | 0 |
| 0 | 0 |
| 0 | 0 |
| 0 | 0 |
| 0 | 0 |
| 0 | 0 |
| 0 | 0 |
| 0 | 0 |
| 0 | 0 |
| 0 | 0 |
| 0 | 0 |

| | |
|---|---|
| 0 | 0 |
| 109.920871 | 7.22831229 |
| 5.15254085 | 43.3698738 |
| 0 | 0 |
| 0 | 0 |
| 0 | 0 |
| 0 | 0 |
| 6.87005447 | 159.02287 |
| 0 | 0 |
| 44.655354 | 0 |
| 111.638385 | 0 |
| 111.638385 | 0 |
| 34.3502723 | 173.479495 |
| 0 | 101.196372 |
| 111.638385 | 0 |
| 5.15254085 | 0 |
| 0 | 0 |
| 0 | 0 |
| 0 | 0 |
| 0 | 0 |
| 0 | 0 |
| 30.9152451 | 79.5114352 |
| 0 | 122.881309 |
| 0 | 122.881309 |
| 0 | 122.881309 |
| 0 | 122.881309 |
| 0 | 122.881309 |
| 0 | 0 |
| 0 | 0 |
| 0 | 0 |
| 0 | 0 |
| 113.355899 | 0 |
| 0 | 0 |
| 0 | 0 |
| 0 | 0 |
| 0 | 0 |
| 0 | 0 |
| 274.802179 | 36.1415615 |
| 0 | 0 |
| 0 | 7.22831229 |
| 0 | 0 |
| 0 | 0 |
| 0 | 0 |

| | |
|---|---|
| 0 | 0 |
| 0 | 0 |
| 144.271144 | 7.22831229 |
| 0 | 0 |
| 120.225953 | 7.22831229 |
| 0 | 0 |
| 0 | 7.22831229 |
| 0 | 0 |
| 56.6779493 | 79.5114352 |
| 63.5480038 | 0 |
| 0 | 0 |
| 68.7005447 | 86.7397475 |
| 0 | 0 |
| 0 | 0 |
| 0 | 0 |
| 0 | 0 |
| 0 | 0 |
| 65.2655174 | 195.164432 |
| 0 | 0 |
| 0 | 0 |
| 0 | 0 |
| 0 | 0 |
| 0 | 0 |
| 0 | 0 |
| 0 | 0 |
| 0 | 0 |
| 0 | 0 |
| 0 | 0 |
| 0 | 0 |
| 0 | 0 |
| 0 | 0 |
| 0 | 0 |
| 0 | 0 |
| 0 | 0 |
| 161.44628 | 21.6849369 |
| 0 | 0 |
| 0 | 0 |
| 0 | 0 |
| 0 | 0 |
| 0 | 0 |
| 0 | 0 |
| 0 | 0 |
| 0 | 0 |

| | |
|---|---|
| 0 | 0 |
| 53.2429221 | 0 |
| 0 | 0 |
| 0 | 0 |
| 96.1807625 | 7.22831229 |
| 109.920871 | 0 |
| 109.920871 | 0 |
| 0 | 0 |
| 0 | 0 |
| 0 | 180.707807 |
| 51.5254085 | 0 |
| 48.0903813 | 0 |
| 96.1807625 | 79.5114352 |
| 20.6101634 | 14.4566246 |
| 156.293739 | 21.6849369 |
| 0 | 0 |
| 0 | 7.22831229 |
| 0 | 79.5114352 |
| 0 | 0 |
| 0 | 0 |
| 0 | 0 |
| 0 | 0 |
| 0 | 0 |
| 0 | 0 |
| 0 | 0 |
| 0 | 0 |
| 0 | 0 |
| 0 | 0 |
| 0 | 0 |
| 0 | 0 |
| 0 | 0 |
| 13.7401089 | 21.6849369 |
| 99.6157898 | 231.305993 |
| 0 | 0 |
| 0 | 50.598186 |
| 0 | 50.598186 |
| 149.423685 | 0 |
| 0 | 0 |
| 0 | 0 |
| 0 | 0 |
| 0 | 0 |
| 0 | 0 |
| 20.6101634 | 0 |

| | |
|---|---|
| 0 | 0 |
| 0 | 0 |
| 48.0903813 | 0 |
| 0 | 7.22831229 |
| 0 | 0 |
| 0 | 0 |
| 0 | 0 |
| 0 | 0 |
| 0 | 0 |
| 0 | 0 |
| 0 | 0 |
| 106.485844 | 202.392744 |
| 0 | 0 |
| 0 | 0 |
| 0 | 0 |
| 0 | 0 |
| 0 | 0 |
| 0 | 0 |
| 336.632669 | 36.1415615 |
| 0 | 0 |
| 53.2429221 | 0 |
| 0 | 0 |
| 0 | 0 |
| 0 | 0 |
| 0 | 122.881309 |
| 6.87005447 | 28.9132492 |
| 0 | 115.652997 |
| 0 | 115.652997 |
| 0 | 115.652997 |
| 0 | 115.652997 |
| 0 | 115.652997 |
| 0 | 115.652997 |
| 0 | 0 |
| 0 | 115.652997 |
| 0 | 0 |
| 0 | 0 |
| 0 | 0 |
| 0 | 0 |
| 41.2203268 | 0 |
| 0 | 0 |
| 0 | 0 |
| 108.203358 | 0 |
| 108.203358 | 0 |

| | |
|---|---|
| 108.203358 | 0 |
| 108.203358 | 0 |
| 108.203358 | 0 |
| 108.203358 | 0 |
| 108.203358 | 0 |
| 173.468875 | 0 |
| 133.966062 | 0 |
| 0 | 0 |
| 115.073412 | 0 |
| 137.401089 | 0 |
| 0 | 0 |
| 0 | 0 |
| 0 | 0 |
| 206.101634 | 7.22831229 |
| 0 | 0 |
| 0 | 0 |
| 0 | 0 |
| 0 | 0 |
| 34.3502723 | 0 |
| 0 | 0 |
| 0 | 0 |
| 0 | 0 |
| 0 | 0 |
| 61.8304902 | 7.22831229 |
| 0 | 108.424684 |
| 24.0451906 | 0 |
| 0 | 0 |
| 0 | 0 |
| 0 | 0 |
| 0 | 0 |
| 15.4576225 | 36.1415615 |
| 0 | 0 |
| 0 | 0 |
| 0 | 0 |
| 60.1129766 | 0 |
| 0 | 0 |
| 0 | 0 |
| 0 | 0 |
| 0 | 0 |
| 0 | 0 |
| 0 | 0 |
| 0 | 0 |
| 0 | 0 |

| | |
|---|---|
| 0 | 0 |
| 0 | 0 |
| 0 | 0 |
| 0 | 0 |
| 0 | 0 |
| 0 | 0 |
| 8.58756808 | 7.22831229 |
| 0 | 0 |
| 0 | 0 |
| 0 | 0 |
| 77.2881127 | 0 |
| 0 | 0 |
| 391.593105 | 238.534306 |
| 0 | 0 |
| 79.0056264 | 0 |
| 49.8078949 | 0 |
| 0 | 0 |
| 0 | 0 |
| 0 | 0 |
| 0 | 0 |
| 0 | 0 |
| 0 | 0 |
| 0 | 0 |
| 3.43502723 | 65.0548106 |
| 0 | 0 |
| 0 | 86.7397475 |
| 0 | 0 |
| 0 | 0 |
| 0 | 0 |
| 0 | 0 |
| 0 | 0 |
| 0 | 0 |
| 120.225953 | 57.8264983 |
| 106.485844 | 0 |
| 106.485844 | 0 |
| 106.485844 | 0 |
| 106.485844 | 0 |
| 0 | 0 |
| 18.8926498 | 93.9680598 |
| 0 | 0 |
| 0 | 0 |
| 0 | 0 |
| 0 | 0 |

| | |
|---|---|
| 0 | 0 |
| 0 | 0 |
| 92.7457353 | 0 |
| 84.1581672 | 0 |
| 0 | 0 |
| 0 | 0 |
| 0 | 0 |
| 300.564883 | 0 |
| 0 | 0 |
| 0 | 0 |
| 0 | 0 |
| 346.937751 | 0 |
| 0 | 0 |
| 44.655354 | 0 |
| 0 | 0 |
| 0 | 65.0548106 |
| 0 | 0 |
| 0 | 0 |
| 0 | 0 |
| 0 | 0 |
| 0 | 0 |
| 0 | 0 |
| 0 | 0 |
| 0 | 14.4566246 |
| 0 | 0 |
| 223.27677 | 159.02287 |
| 0 | 0 |
| 0 | 0 |
| 0 | 0 |
| 0 | 0 |
| 0 | 0 |
| 0 | 0 |
| 243.886934 | 57.8264983 |
| 0 | 0 |
| 0 | 0 |
| 0 | 0 |
| 695.593015 | 397.557176 |
| 0 | 108.424684 |
| 0 | 108.424684 |
| 0 | 108.424684 |
| 0 | 108.424684 |
| 0 | 108.424684 |
| 0 | 108.424684 |

| | |
|---|---|
| 0 | 108.424684 |
| 3.43502723 | 0 |
| 0 | 0 |
| 0 | 0 |
| 0 | 0 |
| 0 | 0 |
| 0 | 0 |
| 0 | 0 |
| 0 | 0 |
| 0 | 72.2831229 |
| 0 | 0 |
| 118.50844 | 303.589116 |
| 140.836117 | 0 |
| 0 | 0 |
| 0 | 0 |
| 132.248548 | 0 |
| 104.768331 | 0 |
| 104.768331 | 0 |
| 188.926498 | 0 |
| 0 | 0 |
| 0 | 0 |
| 0 | 0 |
| 0 | 0 |
| 0 | 0 |
| 0 | 0 |
| 0 | 0 |
| 0 | 0 |
| 0 | 0 |
| 1.71751362 | 57.8264983 |
| 0 | 0 |
| 65.2655174 | 0 |
| 65.2655174 | 0 |
| 0 | 0 |
| 149.423685 | 0 |
| 0 | 0 |
| 0 | 0 |
| 0 | 0 |
| 0 | 7.22831229 |
| 0 | 0 |
| 0 | 65.0548106 |
| 108.203358 | 0 |
| 0 | 0 |
| 6.87005447 | 0 |

| | |
|---|---|
| 0 | 101.196372 |
| 0 | 0 |
| 137.401089 | 0 |
| 0 | 0 |
| 0 | 43.3698738 |
| 0 | 0 |
| 0 | 0 |
| 0 | 115.652997 |
| 0 | 0 |
| 0 | 0 |
| 0 | 65.0548106 |
| 0 | 0 |
| 0 | 0 |
| 0 | 0 |
| 0 | 0 |
| 0 | 0 |
| 0 | 0 |
| 63.5480038 | 72.2831229 |
| 0 | 0 |
| 0 | 0 |
| 56.6779493 | 79.5114352 |
| 25.7627042 | 130.109621 |
| 0 | 0 |
| 0 | 36.1415615 |
| 61.8304902 | 7.22831229 |
| 53.2429221 | 14.4566246 |
| 1.71751362 | 187.93612 |
| 73.8530855 | 14.4566246 |
| 42.9378404 | 7.22831229 |
| 0 | 0 |
| 0 | 0 |
| 0 | 0 |
| 3.43502723 | 0 |
| 0 | 0 |
| 0 | 0 |
| 0 | 0 |
| 0 | 0 |
| 29.1977315 | 108.424684 |
| 0 | 0 |
| 139.118603 | 0 |
| 0 | 0 |
| 0 | 0 |
| 0 | 0 |

| | |
|---|---|
| 10.3050817 | 7.22831229 |
| 0 | 0 |
| 0 | 0 |
| 0 | 0 |
| 103.050817 | 0 |
| 103.050817 | 0 |
| 0 | 0 |
| 170.033848 | 65.0548106 |
| 0 | 0 |
| 0 | 0 |
| 49.8078949 | 0 |
| 0 | 0 |
| 0 | 0 |
| 0 | 0 |
| 0 | 0 |
| 120.225953 | 0 |
| 0 | 0 |
| 0 | 7.22831229 |
| 92.7457353 | 7.22831229 |
| 0 | 0 |
| 0 | 0 |
| 0 | 130.109621 |
| 0 | 0 |
| 0 | 0 |
| 0 | 7.22831229 |
| 0 | 0 |
| 0 | 0 |
| 63.5480038 | 0 |
| 0 | 0 |
| 0 | 0 |
| 0 | 0 |
| 44.655354 | 159.02287 |
| 0 | 0 |
| 0 | 0 |
| 0 | 0 |
| 0 | 0 |
| 0 | 0 |
| 30.9152451 | 79.5114352 |
| 0 | 0 |
| 32.6327587 | 21.6849369 |
| 0 | 0 |
| 0 | 0 |
| 0 | 0 |

| | |
|---|---|
| 0 | 0 |
| 0 | 0 |
| 0 | 0 |
| 0 | 0 |
| 101.333303 | 0 |
| 101.333303 | 0 |
| 0 | 0 |
| 0 | 0 |
| 32.6327587 | 0 |
| 1.71751362 | 144.566246 |
| 0 | 101.196372 |
| 0 | 0 |
| 1.71751362 | 0 |
| 0 | 0 |
| 0 | 0 |
| 0 | 0 |
| 0 | 0 |
| 0 | 0 |
| 0 | 7.22831229 |
| 358.960346 | 36.1415615 |
| 0 | 0 |
| 0 | 0 |
| 0 | 0 |
| 0 | 0 |
| 0 | 0 |
| 0 | 101.196372 |
| 0 | 101.196372 |
| 0 | 101.196372 |
| 0 | 101.196372 |
| 0 | 101.196372 |
| 0 | 101.196372 |
| 170.033848 | 462.611987 |
| 96.1807625 | 0 |
| 0 | 0 |
| 0 | 0 |
| 0 | 0 |
| 8.58756808 | 21.6849369 |
| 0 | 0 |
| 0 | 0 |
| 0 | 0 |
| 0 | 0 |
| 0 | 0 |
| 0 | 0 |

| | |
|---|---|
| 0 | 0 |
| 0 | 0 |
| 99.6157898 | 0 |
| 0 | 0 |
| 99.6157898 | 0 |
| 99.6157898 | 0 |
| 68.7005447 | 7.22831229 |
| 0 | 0 |
| 0 | 0 |
| 0 | 0 |
| 0 | 0 |
| 0 | 0 |
| 0 | 72.2831229 |
| 0 | 0 |
| 0 | 0 |
| 0 | 0 |
| 0 | 0 |
| 82.4406536 | 274.675867 |
| 0 | 0 |
| 0 | 0 |
| 0 | 0 |
| 0 | 0 |
| 0 | 0 |
| 0 | 0 |
| 0 | 0 |
| 0 | 0 |
| 0 | 0 |
| 32.6327587 | 0 |
| 0 | 0 |
| 0 | 0 |
| 0 | 0 |
| 72.1355719 | 0 |
| 279.954719 | 238.534306 |
| 257.627042 | 115.652997 |
| 0 | 0 |
| 0 | 0 |
| 0 | 0 |
| 0 | 0 |
| 0 | 0 |
| 0 | 0 |
| 0 | 0 |
| 0 | 0 |
| 0 | 0 |

| | |
|---|---|
| 0 | 108.424684 |
| 97.8982761 | 0 |
| 97.8982761 | 0 |
| 97.8982761 | 0 |
| 0 | 0 |
| 0 | 0 |
| 12.0225953 | 122.881309 |
| 75.5705991 | 0 |
| 1.71751362 | 108.424684 |
| 0 | 0 |
| 0 | 0 |
| 0 | 0 |
| 0 | 0 |
| 61.8304902 | 0 |
| 0 | 0 |
| 97.8982761 | 101.196372 |
| 0 | 0 |
| 0 | 0 |
| 0 | 0 |
| 0 | 0 |
| 68.7005447 | 65.0548106 |
| 0 | 0 |
| 0 | 0 |
| 0 | 0 |
| 0 | 108.424684 |
| 0 | 0 |
| 0 | 195.164432 |
| 266.214611 | 72.2831229 |
| 0 | 0 |
| 0 | 0 |
| 39.5028132 | 43.3698738 |
| 75.5705991 | 57.8264983 |
| 0 | 0 |
| 0 | 0 |
| 0 | 0 |
| 58.395463 | 36.1415615 |
| 0 | 0 |
| 96.1807625 | 0 |
| 96.1807625 | 0 |
| 96.1807625 | 0 |
| 96.1807625 | 0 |
| 39.5028132 | 21.6849369 |
| 96.1807625 | 0 |

| | |
|---|---|
| 0 | 0 |
| 0 | 0 |
| 72.1355719 | 28.9132492 |
| 15.4576225 | 7.22831229 |
| 0 | 0 |
| 0 | 0 |
| 185.491471 | 0 |
| 0 | 0 |
| 130.531035 | 0 |
| 30.9152451 | 43.3698738 |
| 0 | 0 |
| 44.655354 | 7.22831229 |
| 99.6157898 | 0 |
| 0 | 0 |
| 0 | 0 |
| 0 | 0 |
| 0 | 0 |
| 49.8078949 | 36.1415615 |
| 0 | 0 |
| 0 | 0 |
| 0 | 0 |
| 0 | 0 |
| 91.0282217 | 0 |
| 0 | 0 |
| 0 | 0 |
| 0 | 0 |
| 0 | 7.22831229 |
| 0 | 0 |
| 0 | 0 |
| 0 | 0 |
| 176.903902 | 137.337934 |
| 0 | 0 |
| 0 | 0 |
| 0 | 0 |
| 0 | 0 |
| 22.327677 | 50.598186 |
| 0 | 0 |
| 568.497007 | 657.776419 |
| 0 | 0 |
| 0 | 0 |
| 94.4632489 | 0 |
| 94.4632489 | 0 |
| 94.4632489 | 0 |

| | |
|---|---|
| 48.0903813 | 0 |
| 0 | 0 |
| 0 | 0 |
| 0 | 0 |
| 82.4406536 | 173.479495 |
| 0 | 0 |
| 6.87005447 | 21.6849369 |
| 0 | 0 |
| 0 | 0 |
| 0 | 0 |
| 0 | 0 |
| 0 | 0 |
| 0 | 0 |
| 0 | 0 |
| 0 | 0 |
| 0 | 0 |
| 0 | 0 |
| 0 | 0 |
| 759.141018 | 346.95899 |
| 0 | 0 |
| 0 | 0 |
| 58.395463 | 108.424684 |
| 0 | 0 |
| 0 | 0 |
| 0 | 0 |
| 0 | 0 |
| 1.71751362 | 0 |
| 0 | 57.8264983 |
| 70.4180583 | 21.6849369 |
| 0 | 0 |
| 0 | 0 |
| 0 | 0 |
| 0 | 93.9680598 |
| 0 | 93.9680598 |
| 0 | 93.9680598 |
| 0 | 93.9680598 |
| 0 | 0 |
| 92.7457353 | 0 |
| 0 | 0 |
| 0 | 0 |
| 0 | 0 |
| 0 | 101.196372 |
| 0 | 0 |

| | |
|---|---|
| 72.1355719 | 0 |
| 0 | 50.598186 |
| 305.717424 | 79.5114352 |
| 0 | 0 |
| 0 | 0 |
| 0 | 0 |
| 68.7005447 | 86.7397475 |
| 0 | 0 |
| 0 | 0 |
| 0 | 0 |
| 0 | 0 |
| 0 | 0 |
| 0 | 0 |
| 0 | 0 |
| 0 | 0 |
| 0 | 0 |
| 0 | 0 |
| 73.8530855 | 57.8264983 |
| 204.38412 | 130.109621 |
| 0 | 0 |
| 0 | 0 |
| 0 | 72.2831229 |
| 15.4576225 | 7.22831229 |
| 0 | 0 |
| 0 | 0 |
| 0 | 0 |
| 0 | 7.22831229 |
| 97.8982761 | 0 |
| 3.43502723 | 202.392744 |
| 0 | 0 |
| 0 | 0 |
| 0 | 0 |
| 0 | 0 |
| 1.71751362 | 0 |
| 0 | 7.22831229 |
| 46.3728676 | 108.424684 |
| 91.0282217 | 0 |
| 91.0282217 | 0 |
| 0 | 0 |
| 0 | 0 |
| 27.4802179 | 93.9680598 |
| 0 | 0 |
| 0 | 43.3698738 |

| | |
|---|---|
| 0 | 0 |
| 0 | 0 |
| 5.15254085 | 86.7397475 |
| 0 | 0 |
| 0 | 0 |
| 0 | 0 |
| 49.8078949 | 0 |
| 0 | 0 |
| 0 | 0 |
| 0 | 0 |
| 65.2655174 | 0 |
| 8.58756808 | 440.92705 |
| 0 | 0 |
| 0 | 0 |
| 101.333303 | 0 |
| 0 | 0 |
| 0 | 0 |
| 0 | 0 |
| 0 | 0 |
| 77.2881127 | 0 |
| 54.9604357 | 28.9132492 |
| 116.790926 | 36.1415615 |
| 274.802179 | 0 |
| 0 | 65.0548106 |
| 0 | 0 |
| 1.71751362 | 0 |
| 0 | 0 |
| 0 | 0 |
| 0 | 0 |
| 0 | 7.22831229 |
| 0 | 0 |
| 0 | 0 |
| 0 | 57.8264983 |
| 111.638385 | 0 |
| 0 | 0 |
| 0 | 0 |
| 0 | 0 |
| 0 | 0 |
| 89.3107081 | 0 |
| 89.3107081 | 0 |
| 89.3107081 | 0 |
| 89.3107081 | 0 |
| 89.3107081 | 0 |

| | |
|---|---|
| 89.3107081 | 0 |
| 219.841743 | 0 |
| 0 | 7.22831229 |
| 0 | 0 |
| 0 | 0 |
| 0 | 0 |
| 207.819148 | 28.9132492 |
| 113.355899 | 50.598186 |
| 0 | 166.251183 |
| 0 | 14.4566246 |
| 0 | 0 |
| 0 | 0 |
| 0 | 0 |
| 1.71751362 | 0 |
| 0 | 0 |
| 242.16942 | 43.3698738 |
| 0 | 0 |
| 5.15254085 | 0 |
| 0 | 0 |
| 0 | 0 |
| 0 | 0 |
| 0 | 0 |
| 0 | 93.9680598 |
| 27.4802179 | 7.22831229 |
| 0 | 0 |
| 0 | 0 |
| 0 | 0 |
| 1.71751362 | 0 |
| 0 | 0 |
| 0 | 0 |
| 0 | 0 |
| 0 | 57.8264983 |
| 0 | 0 |
| 29.1977315 | 0 |
| 0 | 0 |
| 0 | 0 |
| 0 | 0 |
| 0 | 0 |
| 58.395463 | 7.22831229 |
| 0 | 0 |
| 0 | 0 |
| 0 | 0 |
| 3.43502723 | 0 |

| | |
|---|---|
| 0 | 0 |
| 0 | 0 |
| 0 | 0 |
| 0 | 0 |
| 0 | 0 |
| 104.768331 | 0 |
| 87.5931944 | 0 |
| 87.5931944 | 0 |
| 87.5931944 | 0 |
| 0 | 0 |
| 0 | 101.196372 |
| 25.7627042 | 86.7397475 |
| 0 | 0 |
| 0 | 0 |
| 0 | 0 |
| 0 | 0 |
| 0 | 0 |
| 0 | 0 |
| 0 | 93.9680598 |
| 0 | 0 |
| 0 | 0 |
| 22.327677 | 7.22831229 |
| 8.58756808 | 144.566246 |
| 0 | 0 |
| 0 | 86.7397475 |
| 0 | 0 |
| 0 | 0 |
| 0 | 0 |
| 0 | 0 |
| 96.1807625 | 0 |
| 0 | 0 |
| 0 | 0 |
