## Supplemental Table 2 for "Downregulation of Neurodevelopmental Gene Expression in iPSC-Derived Cerebral Organoids Upon Infection by Human Cytomegalovirus"

| | Gene | baseMean | log2FoldChai | lfcSE | stat |
|---|---|---|---|---|---|
| 1 | NR_125368.1 | 2403.23548 | 4.41802698 | 0.23357971 | 18.9144297 |
| 2 | NM_001184735.1 | 617.987346 | 3.15989648 | 0.17258761 | 18.3089413 |
| 3 | NR_110665.1 | 290.573365 | 4.90520568 | 0.29602023 | 16.570508 |
| 4 | NM_001402.5 | 23469.2183 | -8.5222554 | 0.57816985 | -14.740055 |
| 5 | NM_152733.2 | 5318.08291 | 17.2971121 | 1.18796849 | 14.5602449 |
| 6 | NR_027451.1 | 1466.98746 | -6.1465951 | 0.42695454 | -14.396369 |
| 7 | NR_126435.1 | 5891.57004 | 3.86549111 | 0.27364106 | 14.1261372 |
| 8 | NR_047679.1 | 3993.24428 | 4.29058982 | 0.30583915 | 14.0289097 |
| 9 | NM_001190881.2 | 1508.56242 | 15.4795171 | 1.11042466 | 13.9401777 |
| 10 | NR_002728.3 | 3375.89376 | 3.24843477 | 0.23342677 | 13.9162905 |
| 11 | NR_047469.1 | 3350.31522 | 3.81738633 | 0.2758185 | 13.8402111 |
| 12 | NM_001039211.2 | 549.034786 | 3.99549078 | 0.29785642 | 13.4141501 |
| 13 | NR_034022.1 | 1477.7492 | 3.29552851 | 0.24683219 | 13.3512915 |
| 14 | NM_005348.3 | 4200.02503 | -10.588131 | 0.82206145 | -12.879975 |
| 15 | NM_006445.3 | 1023.18041 | -8.0380599 | 0.63206553 | -12.717131 |
| 16 | NM_001164436.1 | 708.306758 | 4.3115642 | 0.33997854 | 12.6818716 |
| 17 | NM_003360.4 | 188.251534 | 3.28335025 | 0.26658483 | 12.316343 |
| 18 | NR_125837.1 | 805.580412 | 4.03725098 | 0.34180071 | 11.8117105 |
| 19 | NM_001278593.1 | 243.182348 | 12.8441942 | 1.15145759 | 11.1547262 |
| 20 | NR_125813.1 | 288.73059 | 4.0750126 | 0.36851952 | 11.0577931 |
| 21 | NM_002046.5 | 5084.47208 | -13.934117 | 1.2709102 | -10.963888 |
| 22 | NM_012401.3 | 991.751465 | -8.5007821 | 0.78395516 | -10.843454 |
| 23 | NM_005346.4 | 510.350886 | -6.3839815 | 0.58907005 | -10.837389 |
| 24 | NM_080686.2 | 1074.26236 | -8.9229735 | 0.82696642 | -10.790007 |
| 25 | NM_007355.3 | 3664.28265 | -13.46156 | 1.24809757 | -10.785663 |
| 26 | NM_015276.1 | 1706.63899 | -9.5906075 | 0.89314122 | -10.738064 |
| 27 | NM_001961.3 | 7627.3252 | -13.557341 | 1.28219897 | -10.573508 |
| 28 | NM_178014.3 | 5310.05963 | -13.996751 | 1.32713413 | -10.546599 |
| 29 | NM_003118.3 | 5725.7326 | -14.105499 | 1.33944736 | -10.530835 |
| 30 | NM_003380.3 | 5226.45137 | -10.628101 | 1.01171544 | -10.50503 |
| 31 | NM_080817.4 | 1490.57948 | 5.75499144 | 0.5477422 | 10.5067521 |
| 32 | NM_001101.3 | 11227.7755 | -11.940934 | 1.14156959 | -10.460102 |
| 33 | NM_001418.3 | 3641.88219 | -13.452703 | 1.29162705 | -10.415315 |
| 34 | NM_004321.7 | 2320.60704 | -9.3273035 | 0.89601765 | -10.409732 |
| 35 | NM_001291969.1 | 316.830248 | 13.2284243 | 1.27121017 | 10.4061662 |
| 36 | NM_001145204.2 | 685.066967 | 2.75795675 | 0.26516766 | 10.4008037 |
| 37 | NM_002823.4 | 2304.9495 | -9.7079072 | 0.93409208 | -10.39288 |
| 38 | NM_000967.3 | 3566.37131 | -13.422469 | 1.2937522 | -10.374838 |
| 39 | NM_001428.3 | 2727.77685 | -13.035704 | 1.26469037 | -10.307427 |
| 40 | NM_006597.5 | 3483.70388 | -13.388648 | 1.29913251 | -10.305837 |
| 41 | NR_144568.1 | 5521.31141 | -14.053004 | 1.36299027 | -10.310422 |
| 42 | NM_001110556.1 | 2721.14242 | -13.032281 | 1.27335825 | -10.234575 |

| | | | | | |
|---|---|---|---|---|---|
| 43 | NM_002654.5 | 1648.57286 | -12.309356 | 1.20327784 | -10.229854 |
| 44 | NM_001025.4 | 1491.93423 | -9.1124241 | 0.89206951 | -10.214926 |
| 45 | NM_017758.3 | 319.20004 | -6.1470644 | 0.601653 | -10.21696 |
| 46 | NM_021109.3 | 5176.1158 | -13.959878 | 1.36695223 | -10.212411 |
| 47 | NM_022731.4 | 1082.60585 | -7.6859136 | 0.75483759 | -10.182208 |
| 48 | NM_016333.3 | 1632.10406 | -12.294937 | 1.20808516 | -10.177211 |
| 49 | NM_184041.2 | 1852.59883 | -12.477576 | 1.22601585 | -10.177336 |
| 50 | NM_005909.4 | 3667.50445 | -13.462802 | 1.33119446 | -10.113325 |
| 51 | NR_027701.1 | 102.767601 | 3.99064189 | 0.39496471 | 10.1037936 |
| 52 | NM_002228.3 | 514.231367 | -5.9612016 | 0.59027971 | -10.098944 |
| 53 | NM_001243960.1 | 282.727544 | 13.0645118 | 1.29414774 | 10.0950699 |
| 54 | NM_001404.4 | 4412.67035 | -11.960417 | 1.18838535 | -10.064427 |
| 55 | NR_130931.1 | 1149.7693 | 3.27636012 | 0.32732472 | 10.0095102 |
| 56 | NR_144567.1 | 1633.85242 | -12.29637 | 1.23387306 | -9.965669 |
| 57 | NM_000599.3 | 2689.91786 | -12.05364 | 1.21012249 | -9.9606775 |
| 58 | NR_033770.1 | 219.101263 | 5.42444435 | 0.54470239 | 9.95854705 |
| 59 | NR_110133.1 | 1768.26524 | 4.01704186 | 0.40348711 | 9.95581223 |
| 60 | NM_001206491.1 | 139.522184 | 12.0452631 | 1.21246443 | 9.93452905 |
| 61 | NR_110867.1 | 307.727949 | 3.37461028 | 0.33982686 | 9.93038127 |
| 62 | NM_014014.4 | 917.88286 | -7.9522955 | 0.80164016 | -9.9200313 |
| 63 | NM_002032.2 | 4325.77102 | -12.739084 | 1.28630421 | -9.9036325 |
| 64 | NM_003334.3 | 1664.14764 | -12.322877 | 1.24840483 | -9.8708985 |
| 65 | NM_002778.3 | 1446.63821 | -12.120947 | 1.23170474 | -9.8407892 |
| 66 | NR_046479.1 | 111.69815 | 11.7278842 | 1.19461395 | 9.81730056 |
| 67 | NM_001114173.2 | 197.194608 | 12.5444215 | 1.2807778 | 9.79437771 |
| 68 | NM_005381.2 | 1511.58138 | -12.184087 | 1.24387505 | -9.7952663 |
| 69 | NM_000597.2 | 2026.4961 | -12.606944 | 1.28738577 | -9.7926703 |
| 70 | NM_004343.3 | 2015.76047 | -12.599397 | 1.2879236 | -9.7827209 |
| 71 | NM_002568.3 | 1885.97746 | -10.716624 | 1.09847228 | -9.7559351 |
| 72 | NM_006158.4 | 4578.24775 | -9.0216531 | 0.92588524 | -9.7438136 |
| 73 | NR_027053.2 | 768.825588 | 3.41392278 | 0.35065076 | 9.73596299 |
| 74 | NM_001746.3 | 1711.05924 | -12.36292 | 1.27285158 | -9.7127743 |
| 75 | NM_005382.2 | 4399.08203 | -13.725231 | 1.41430445 | -9.7045804 |
| 76 | NM_006986.3 | 1294.82341 | -11.960869 | 1.23255238 | -9.7041468 |
| 77 | NM_000972.2 | 2107.59762 | -9.6515902 | 0.99529371 | -9.6972281 |
| 78 | NM_001975.2 | 1561.08983 | -12.23052 | 1.26221207 | -9.6897503 |
| 79 | NM_000175.4 | 1169.1764 | -11.813465 | 1.2221564 | -9.666083 |
| 80 | NM_004559.4 | 3640.99852 | -13.452338 | 1.3954664 | -9.6400301 |
| 81 | NM_001831.3 | 2409.1344 | -12.856567 | 1.33951583 | -9.5979207 |
| 82 | NM_198589.2 | 1248.89133 | -11.908708 | 1.241409 | -9.5928967 |
| 83 | NM_005347.4 | 1783.63002 | -10.638374 | 1.10941284 | -9.5891931 |
| 84 | NM_021104.1 | 2029.97737 | -12.609462 | 1.31640846 | -9.5786851 |
| 85 | NM_001002.3 | 2988.46631 | -13.167407 | 1.3755035 | -9.572791 |

| | | | | | |
|---|---|---|---|---|---|
| 86 | NM_001202515.1 | 585.097246 | 2.76948848 | 0.28927664 | 9.57384082 |
| 87 | NM_201413.2 | 1031.728 | -11.633384 | 1.21704948 | -9.5586781 |
| 88 | NM_014423.3 | 414.283956 | -7.1989855 | 0.75334752 | -9.5559956 |
| 89 | NM_012423.3 | 2282.77254 | -12.778782 | 1.33900826 | -9.5434677 |
| 90 | NM_000970.4 | 2077.88703 | -12.643112 | 1.32578284 | -9.5363367 |
| 91 | NR_106781.1 | 3438.54009 | -13.369853 | 1.40270883 | -9.5314525 |
| 92 | NM_002332.2 | 1416.40896 | -7.914752 | 0.83173467 | -9.5159578 |
| 93 | NM_003295.3 | 2732.90539 | -13.038447 | 1.37289282 | -9.4970614 |
| 94 | NM_006761.4 | 2182.21168 | -12.713797 | 1.33885028 | -9.4960559 |
| 95 | NM_002473.5 | 1325.54404 | -11.994762 | 1.2650071 | -9.4819718 |
| 96 | NM_001686.3 | 1658.63612 | -12.317992 | 1.29942551 | -9.4795676 |
| 97 | NM_000093.4 | 1444.96212 | -12.119184 | 1.27905042 | -9.4751417 |
| 98 | NM_003128.2 | 1197.9321 | -8.7469046 | 0.92383508 | -9.4680369 |
| 99 | NM_000202.7 | 1158.73138 | -11.800538 | 1.2469951 | -9.4631792 |
| 100 | NM_002074.4 | 1684.90124 | -12.340654 | 1.30405858 | -9.4632662 |
| 101 | NM_000291.3 | 1147.19673 | -11.785978 | 1.24893224 | -9.4368437 |
| 102 | NM_000615.6 | 1794.15502 | -9.2609149 | 0.98203079 | -9.4303712 |
| 103 | NM_000977.3 | 2563.35382 | -12.94603 | 1.37270321 | -9.4310484 |
| 104 | NM_001387.2 | 3082.90555 | -13.21231 | 1.40113268 | -9.4297351 |
| 105 | NM_032591.2 | 113.654952 | 11.7458057 | 1.2452346 | 9.43260469 |
| 106 | NM_001873.3 | 1105.04396 | -8.6357106 | 0.91605855 | -9.427029 |
| 107 | NM_021939.3 | 671.494273 | -8.2159836 | 0.87176552 | -9.4245338 |
| 108 | NM_002156.4 | 987.604572 | -11.570131 | 1.22815481 | -9.4207432 |
| 109 | NM_005184.3 | 1998.76783 | -12.587122 | 1.33695318 | -9.4147817 |
| 110 | NM_000516.5 | 2660.00012 | -12.999437 | 1.38158227 | -9.4090937 |
| 111 | NM_003217.2 | 1351.36473 | -12.022432 | 1.28152215 | -9.3813687 |
| 112 | NM_004651.3 | 1089.41576 | -11.711651 | 1.24924945 | -9.3749499 |
| 113 | NM_006082.2 | 3519.22653 | -12.441314 | 1.32793596 | -9.3689111 |
| 114 | NM_000138.4 | 449.889213 | -7.285165 | 0.77823133 | -9.3611818 |
| 115 | NM_001164782.1 | 118.00034 | 11.8029392 | 1.26472125 | 9.33244321 |
| 116 | NR_135276.1 | 528.399669 | 3.76587843 | 0.40388345 | 9.32417123 |
| 117 | NM_001010.2 | 1811.85105 | -12.445474 | 1.33523565 | -9.3208073 |
| 118 | NM_007158.5 | 1214.68824 | -11.868577 | 1.27362407 | -9.3187443 |
| 119 | NM_002520.6 | 1394.82672 | -12.068099 | 1.29555141 | -9.3150294 |
| 120 | NM_025081.2 | 341.512791 | -6.9448293 | 0.74559788 | -9.3144435 |
| 121 | NM_001493.2 | 1763.67939 | -12.406598 | 1.33331776 | -9.3050569 |
| 122 | NM_002136.3 | 2978.99563 | -8.5618858 | 0.92043099 | -9.3020399 |
| 123 | NM_001291311.1 | 1460.5651 | -12.134631 | 1.30607769 | -9.2908953 |
| 124 | NM_001417.6 | 1102.39457 | -11.728571 | 1.26374554 | -9.2808013 |
| 125 | NM_005507.2 | 2132.60467 | -12.680632 | 1.36625231 | -9.281325 |
| 126 | NM_007126.3 | 1082.82538 | -11.702879 | 1.26117305 | -9.2793604 |
| 127 | NM_003752.4 | 1241.30946 | -11.899887 | 1.28294211 | -9.2754668 |
| 128 | NM_001130438.2 | 1854.3832 | -12.478922 | 1.34587023 | -9.2720097 |

| | | | | | |
|---|---|---|---|---|---|
| 129 | NM_000088.3 | 9841.2894 | -10.076464 | 1.08690661 | -9.2707726 |
| 130 | NM_003299.2 | 1989.6724 | -11.618561 | 1.25333647 | -9.2701049 |
| 131 | NM_001306129.1 | 984.592589 | -11.56594 | 1.2479956 | -9.2676129 |
| 132 | NM_001923.4 | 894.539552 | -11.427352 | 1.23356225 | -9.263701 |
| 133 | NM_004046.5 | 1226.08044 | -11.882091 | 1.28332858 | -9.2588065 |
| 134 | NM_007029.3 | 3310.0458 | -13.314867 | 1.43796613 | -9.2595134 |
| 135 | NM_001940.3 | 952.94252 | -11.518404 | 1.24643446 | -9.2410828 |
| 136 | NM_015015.2 | 438.710782 | -6.5813601 | 0.71245793 | -9.2375421 |
| 137 | NM_001079524.1 | 413.547153 | -6.5264769 | 0.70688928 | -9.2326721 |
| 138 | NM_212482.2 | 1036.43736 | -11.639836 | 1.26144809 | -9.2273602 |
| 139 | NM_002824.5 | 2233.77393 | -12.747476 | 1.38161317 | -9.2265161 |
| 140 | NM_000612.5 | 1621.43641 | -12.285385 | 1.33259351 | -9.2191544 |
| 141 | NM_001042465.2 | 1377.88676 | -12.050514 | 1.3083374 | -9.2105557 |
| 142 | NM_007363.4 | 1094.43255 | -11.718162 | 1.27454206 | -9.1940173 |
| 143 | NR_109904.1 | 101.875378 | 3.03120941 | 0.33010148 | 9.1826593 |
| 144 | NM_012433.3 | 777.246552 | -11.224553 | 1.22255208 | -9.1812473 |
| 145 | NM_001256802.1 | 1404.65098 | -9.5798387 | 1.04390456 | -9.1769296 |
| 146 | NM_018834.5 | 1402.43361 | -12.075893 | 1.31717771 | -9.1680056 |
| 147 | NM_000971.3 | 1813.53342 | -9.1957752 | 1.00548291 | -9.1456305 |
| 148 | NM_006088.5 | 690.846852 | -11.05484 | 1.20887326 | -9.1447471 |
| 149 | NM_004134.6 | 793.874029 | -11.254989 | 1.23140549 | -9.1399535 |
| 150 | NM_001009944.2 | 937.770732 | -11.49538 | 1.25984249 | -9.1244579 |
| 151 | NM_001823.4 | 1680.21205 | -12.336589 | 1.35281916 | -9.1191709 |
| 152 | NM_001164319.1 | 407.210413 | -6.6325004 | 0.72738139 | -9.1183257 |
| 153 | NM_000998.4 | 1611.63428 | -12.276523 | 1.34675755 | -9.1156149 |
| 154 | NM_000365.5 | 1202.12297 | -11.853483 | 1.30078691 | -9.1125476 |
| 155 | NM_000980.3 | 1746.0414 | -12.392081 | 1.36038664 | -9.1092344 |
| 156 | NM_003290.2 | 899.195622 | -8.7046696 | 0.95580927 | -9.1071199 |
| 157 | NM_005801.3 | 1337.40386 | -12.007429 | 1.31936352 | -9.1009255 |
| 158 | NR_119374.1 | 208.032527 | 7.1078819 | 0.78100668 | 9.10092327 |
| 159 | NM_006039.4 | 1108.72883 | -9.340282 | 1.02637924 | -9.100225 |
| 160 | NM_004599.3 | 1560.26254 | -12.229803 | 1.34424385 | -9.0979049 |
| 161 | NM_012479.3 | 1758.20569 | -12.40211 | 1.36310129 | -9.0984511 |
| 162 | NM_001743.5 | 1767.90134 | -12.410047 | 1.36452596 | -9.0947682 |
| 163 | NM_003011.3 | 1320.00451 | -11.988517 | 1.31882899 | -9.090274 |
| 164 | NM_023009.6 | 2127.47734 | -12.677138 | 1.39646908 | -9.0779942 |
| 165 | NM_006888.4 | 1805.80739 | -10.610382 | 1.16995434 | -9.0690562 |
| 166 | NM_004932.3 | 295.589844 | -5.2480158 | 0.57910301 | -9.0623183 |
| 167 | NM_015382.3 | 342.956233 | -6.8932754 | 0.7606708 | -9.0621006 |
| 168 | NM_005313.4 | 1082.54231 | -11.702522 | 1.29147501 | -9.0613613 |
| 169 | NM_004396.4 | 1394.23705 | -10.184915 | 1.12416512 | -9.0599815 |
| 170 | NM_001040152.1 | 1478.00859 | -8.1834497 | 0.90342125 | -9.0582879 |
| 171 | NM_000852.3 | 903.319394 | -11.441402 | 1.26324129 | -9.0571786 |

| | | | | | |
|---|---|---|---|---|---|
| 172 | NM_001416.3 | 1326.89691 | -11.996069 | 1.32540621 | -9.0508623 |
| 173 | NM_005911.5 | 661.250252 | -10.991278 | 1.21709859 | -9.0307212 |
| 174 | NM_002375.4 | 738.832215 | -11.151734 | 1.23509632 | -9.0290398 |
| 175 | NM_005324.4 | 1528.0535 | -11.237517 | 1.24475063 | -9.0279267 |
| 176 | NM_001315537.1 | 1397.30589 | -12.070647 | 1.33861155 | -9.0172885 |
| 177 | NM_015626.8 | 777.254824 | -11.224316 | 1.24490738 | -9.0161854 |
| 178 | NM_001854.3 | 843.340982 | -11.342405 | 1.25910814 | -9.0082852 |
| 179 | NM_001903.4 | 616.0167 | -10.889473 | 1.20871854 | -9.009106 |
| 180 | NM_013336.3 | 987.188933 | -11.569481 | 1.28428427 | -9.0085048 |
| 181 | NM_021999.4 | 725.970706 | -11.126161 | 1.23495875 | -9.0093382 |
| 182 | NM_001261826.1 | 767.298876 | -11.205941 | 1.24406164 | -9.0075445 |
| 183 | NM_005885.3 | 796.447802 | -11.259612 | 1.25009428 | -9.0070104 |
| 184 | NM_201414.2 | 2114.49525 | -12.668329 | 1.4078651 | -8.9982549 |
| 185 | NM_005100.3 | 778.366549 | -11.226696 | 1.24837616 | -8.9930397 |
| 186 | NM_004068.3 | 1074.68375 | -11.691931 | 1.30025763 | -8.9920111 |
| 187 | NM_000398.6 | 791.216349 | -11.250225 | 1.25178675 | -8.9873335 |
| 188 | NM_003750.2 | 718.21075 | -11.110612 | 1.23678825 | -8.9834391 |
| 189 | NM_006805.3 | 1018.27613 | -11.614096 | 1.29299029 | -8.9823533 |
| 190 | NM_001330681.1 | 1453.61614 | -12.127644 | 1.35128343 | -8.9749076 |
| 191 | NM_001190702.1 | 12243.273 | -7.3613625 | 0.82064172 | -8.9702513 |
| 192 | NM_002128.5 | 1207.94192 | -11.860496 | 1.32228281 | -8.9697118 |
| 193 | NM_001005335.1 | 202.404484 | -5.6427637 | 0.62936633 | -8.9657857 |
| 194 | NM_002295.5 | 1647.67991 | -12.308438 | 1.3731207 | -8.9638425 |
| 195 | NM_001005271.2 | 1300.06616 | -11.966503 | 1.33511348 | -8.9629106 |
| 196 | NM_012215.3 | 693.535304 | -11.059928 | 1.23421761 | -8.9610846 |
| 197 | NR_002824.3 | 470.338808 | -7.7408936 | 0.8641811 | -8.9574901 |
| 198 | NM_031844.2 | 725.566635 | -11.124984 | 1.24207553 | -8.9567694 |
| 199 | NM_002211.3 | 1116.10749 | -11.746574 | 1.31196632 | -8.9534111 |
| 200 | NM_003932.4 | 868.237203 | -11.384165 | 1.27309743 | -8.9421002 |
| 201 | NM_003107.2 | 2252.89926 | -10.222548 | 1.14349446 | -8.9397443 |
| 202 | NM_032727.3 | 2064.29548 | -12.633675 | 1.4131629 | -8.9399991 |
| 203 | NM_004039.2 | 1097.01852 | -11.721696 | 1.31130141 | -8.9389792 |
| 204 | NR_039980.1 | 274.539095 | 2.50118738 | 0.27998833 | 8.93318456 |
| 205 | NM_001658.3 | 937.585075 | -11.495058 | 1.28694556 | -8.9320471 |
| 206 | NM_001539.3 | 964.30386 | -11.535565 | 1.29239545 | -8.9257242 |
| 207 | NM_016609.4 | 610.03662 | -10.875196 | 1.21857427 | -8.9245242 |
| 208 | NM_000975.3 | 1222.90425 | -11.878294 | 1.33195204 | -8.917959 |
| 209 | NM_000979.3 | 1314.34705 | -11.982337 | 1.34459805 | -8.9114641 |
| 210 | NM_001030006.1 | 931.399776 | -11.485468 | 1.28926255 | -8.9085563 |
| 211 | NM_001069.2 | 2090.08254 | -12.651573 | 1.42037399 | -8.9072127 |
| 212 | NM_001839.4 | 748.122033 | -11.169538 | 1.25576031 | -8.8946418 |
| 213 | NM_006317.4 | 1938.51013 | -12.542941 | 1.41028518 | -8.893904 |
| 214 | NM_004798.3 | 425.776212 | -4.3339002 | 0.48805345 | -8.8799704 |

| | | | | | |
|---|---|---|---|---|---|
| 215 | NM_002265.5 | 634.597469 | -10.932166 | 1.23162127 | -8.8762398 |
| 216 | NM_001303624.1 | 1268.79647 | -11.931497 | 1.34464983 | -8.8733115 |
| 217 | NM_001322257.1 | 63.9081747 | 10.9237398 | 1.23124347 | 8.87211997 |
| 218 | NM_001235.3 | 1065.06524 | -11.679068 | 1.31755484 | -8.8641989 |
| 219 | NM_005506.3 | 766.590703 | -11.204538 | 1.26408255 | -8.8637707 |
| 220 | NM_001273.3 | 570.429424 | -10.7784 | 1.21690268 | -8.857241 |
| 221 | NM_005566.3 | 908.073694 | -11.448685 | 1.29325515 | -8.8526112 |
| 222 | NM_004140.3 | 400.782808 | -6.3508388 | 0.71763089 | -8.8497289 |
| 223 | NM_005724.5 | 820.272307 | -11.302202 | 1.27920635 | -8.8353238 |
| 224 | NM_012117.2 | 643.019421 | -10.950675 | 1.23984549 | -8.83229 |
| 225 | NM_004181.4 | 1740.62005 | -12.387612 | 1.40316786 | -8.8283176 |
| 226 | NM_003255.4 | 1088.63804 | -11.710552 | 1.32656626 | -8.8277172 |
| 227 | NM_001003.2 | 1623.45638 | -12.287066 | 1.39200097 | -8.8269091 |
| 228 | NM_001664.3 | 659.210623 | -10.986875 | 1.24521907 | -8.8232468 |
| 229 | NM_001021.4 | 1472.72054 | -12.146474 | 1.3769665 | -8.8211832 |
| 230 | NM_005563.3 | 1890.28388 | -12.506604 | 1.41790359 | -8.8204898 |
| 231 | NM_002687.3 | 519.893153 | -10.644766 | 1.20704598 | -8.8188572 |
| 232 | NM_004522.2 | 1835.03819 | -12.463811 | 1.41404654 | -8.8142862 |
| 233 | NM_006098.4 | 2017.61142 | -11.638622 | 1.32054415 | -8.8135048 |
| 234 | NR_135520.1 | 591.264894 | 4.99853465 | 0.56731315 | 8.81089159 |
| 235 | NM_020638.2 | 383.947441 | 5.81413861 | 0.660021 | 8.80902067 |
| 236 | NM_002137.3 | 1823.20023 | -8.6572519 | 0.98292184 | -8.8076707 |
| 237 | NM_000188.2 | 743.351553 | -11.160102 | 1.26802265 | -8.8011847 |
| 238 | NM_052925.3 | 695.05385 | -11.063579 | 1.25740201 | -8.7987602 |
| 239 | NM_000993.4 | 1019.73258 | -8.1257508 | 0.92397586 | -8.7943324 |
| 240 | NM_001636.3 | 1024.37497 | -11.622776 | 1.3235616 | -8.7814393 |
| 241 | NM_000978.3 | 1105.46347 | -11.73264 | 1.33620579 | -8.7805636 |
| 242 | NM_153828.2 | 970.089751 | -11.544225 | 1.31536726 | -8.7764274 |
| 243 | NM_002808.4 | 710.22372 | -11.094393 | 1.26486342 | -8.7712181 |
| 244 | NM_014762.3 | 1316.20864 | -11.984435 | 1.3668745 | -8.7677653 |
| 245 | NM_001113202.1 | 1065.50886 | -11.679497 | 1.33316537 | -8.7607263 |
| 246 | NM_002972.3 | 983.01126 | -8.2965563 | 0.94708155 | -8.7601287 |
| 247 | NM_014819.4 | 949.877141 | -11.513761 | 1.31430555 | -8.7603377 |
| 248 | NM_001020.5 | 1217.6887 | -11.872116 | 1.35581831 | -8.7564209 |
| 249 | NM_001969.4 | 644.258275 | -10.953695 | 1.2509041 | -8.7566222 |
| 250 | NR_110744.1 | 130.20248 | 4.51824843 | 0.51601558 | 8.75603103 |
| 251 | NM_001013.3 | 988.030012 | -11.570587 | 1.32169518 | -8.7543539 |
| 252 | NM_006035.3 | 636.826252 | -10.93702 | 1.24939316 | -8.7538659 |
| 253 | NM_000701.7 | 672.050344 | -11.014718 | 1.25859096 | -8.7516268 |
| 254 | NM_001033026.1 | 695.387651 | -11.063793 | 1.26589415 | -8.7399035 |
| 255 | NM_001206799.1 | 860.112218 | -11.370555 | 1.30105598 | -8.7394818 |
| 256 | NM_002291.2 | 649.012269 | -10.964821 | 1.25461254 | -8.7396074 |
| 257 | NM_021960.4 | 577.543336 | -10.796167 | 1.2357068 | -8.7368355 |

| | | | | | |
|---|---|---|---|---|---|
| 258 | NR_046392.1 | 83.0167502 | 4.05131422 | 0.46372739 | 8.73641351 |
| 259 | NM_004265.3 | 1337.55755 | -12.007613 | 1.3748258 | -8.7339163 |
| 260 | NM_001012.1 | 2141.46388 | -11.724564 | 1.34249005 | -8.7334455 |
| 261 | NM_004481.4 | 586.144157 | -10.81761 | 1.23902066 | -8.7307745 |
| 262 | NM_000968.3 | 3226.0488 | -9.501575 | 1.08881235 | -8.7265496 |
| 263 | NM_015110.3 | 152.740196 | -4.4044573 | 0.50493695 | -8.7227866 |
| 264 | NM_004395.3 | 1486.47604 | -9.3847466 | 1.07595745 | -8.7222284 |
| 265 | NM_182810.2 | 823.533303 | -11.307938 | 1.29685079 | -8.7195369 |
| 266 | NM_001297654.1 | 514.701154 | -10.630299 | 1.21981428 | -8.7146867 |
| 267 | NM_006148.3 | 685.443055 | -11.043161 | 1.26748269 | -8.712672 |
| 268 | NM_002626.5 | 515.840184 | -10.632967 | 1.22130709 | -8.7062188 |
| 269 | NM_001320965.1 | 529.751097 | -10.671103 | 1.22662357 | -8.6995742 |
| 270 | NM_006086.3 | 2893.0439 | -10.452569 | 1.20152164 | -8.6994433 |
| 271 | NM_014944.4 | 884.930876 | -11.411709 | 1.31179479 | -8.6993097 |
| 272 | NM_006196.3 | 775.520964 | -11.221216 | 1.29045392 | -8.6955571 |
| 273 | NM_005610.2 | 669.66956 | -11.009389 | 1.26617846 | -8.6949739 |
| 274 | NM_002825.6 | 538.898262 | -10.696138 | 1.23027685 | -8.6940904 |
| 275 | NM_001015.4 | 2041.95164 | -11.655911 | 1.34076905 | -8.6934519 |
| 276 | NR_036647.1 | 148.440412 | 3.28339577 | 0.37798265 | 8.6866308 |
| 277 | NM_020134.3 | 1291.53644 | -11.957092 | 1.37659556 | -8.6859873 |
| 278 | NM_005722.3 | 799.038113 | -11.264331 | 1.29713797 | -8.6839884 |
| 279 | NM_001035006.2 | 1155.08702 | -11.795986 | 1.35913591 | -8.6790331 |
| 280 | NM_004356.3 | 817.482545 | -11.297313 | 1.30208556 | -8.6763214 |
| 281 | NM_033022.3 | 1417.41467 | -12.091235 | 1.39393776 | -8.6741568 |
| 282 | NM_001321678.1 | 66.6454159 | 10.9820705 | 1.26658467 | 8.67061693 |
| 283 | NM_206917.2 | 777.149257 | -11.22424 | 1.29480635 | -8.668663 |
| 284 | NM_000165.4 | 574.323799 | -7.4968266 | 0.86495834 | -8.667269 |
| 285 | NM_000484.3 | 772.601596 | -11.216097 | 1.29422111 | -8.666291 |
| 286 | NM_006429.3 | 783.474215 | -11.2359 | 1.297365 | -8.6605541 |
| 287 | NM_000969.3 | 1815.23077 | -11.486088 | 1.32646665 | -8.6591611 |
| 288 | NR_015362.1 | 89.2389975 | 3.45558263 | 0.3995688 | 8.6482795 |
| 289 | NM_007184.3 | 698.755883 | -11.070953 | 1.28023375 | -8.6476025 |
| 290 | NM_005022.3 | 770.106449 | -11.211214 | 1.29717267 | -8.6428079 |
| 291 | NM_016395.2 | 615.699763 | -7.9447885 | 0.91939255 | -8.6413453 |
| 292 | NM_001013251.2 | 642.595738 | -10.950141 | 1.26781505 | -8.6370177 |
| 293 | NM_001193416.2 | 541.945952 | -10.704123 | 1.23996284 | -8.6326161 |
| 294 | NM_016128.3 | 515.811833 | -10.633223 | 1.23297432 | -8.6240427 |
| 295 | NM_004500.3 | 763.759188 | -11.198999 | 1.29876366 | -8.6228148 |
| 296 | NM_002734.4 | 483.022539 | -10.538312 | 1.22243914 | -8.6207255 |
| 297 | NM_001128204.1 | 698.438304 | -11.070632 | 1.28451953 | -8.6185007 |
| 298 | NM_001023.3 | 1036.24919 | -11.639363 | 1.35092418 | -8.6158524 |
| 299 | NM_001145667.1 | 632.044725 | -10.926115 | 1.26820198 | -8.6154375 |
| 300 | NM_003633.3 | 1184.29603 | -11.83201 | 1.37329729 | -8.6157676 |

| | | | | | |
|---|---|---|---|---|---|
| 301 | NM_020755.3 | 1362.18053 | -7.5575013 | 0.87718213 | -8.6156581 |
| 302 | NM_030926.5 | 555.530855 | -10.740076 | 1.24639399 | -8.6169186 |
| 303 | NM_001155.4 | 504.860447 | -10.602622 | 1.23075956 | -8.6146984 |
| 304 | NM_005737.3 | 557.924568 | -10.746451 | 1.24782211 | -8.6121661 |
| 305 | NM_004995.3 | 600.005168 | -10.851504 | 1.26054632 | -8.6085722 |
| 306 | NM_001904.3 | 474.252462 | -10.511625 | 1.22122349 | -8.6074541 |
| 307 | NM_005560.4 | 683.110255 | -8.5048643 | 0.98803928 | -8.60782 |
| 308 | NM_002056.3 | 385.661267 | -4.9671084 | 0.57714837 | -8.6062938 |
| 309 | NM_005127.2 | 66.92639 | 4.6736366 | 0.54311666 | 8.60521678 |
| 310 | NM_002045.3 | 1557.2019 | -12.226976 | 1.42117842 | -8.6034066 |
| 311 | NM_004521.2 | 614.58513 | -10.885557 | 1.26557509 | -8.6012733 |
| 312 | NM_001271938.1 | 637.306656 | -10.93806 | 1.27182105 | -8.600314 |
| 313 | NM_000990.4 | 1398.83395 | -4.9881235 | 0.58012407 | -8.5983737 |
| 314 | NM_004924.5 | 611.571258 | -10.878927 | 1.26519487 | -8.5986176 |
| 315 | NM_004152.3 | 951.916736 | -11.516877 | 1.33955898 | -8.5975141 |
| 316 | NM_024536.5 | 495.653235 | -10.575774 | 1.2301231 | -8.59733 |
| 317 | NM_005066.2 | 669.619541 | -11.009307 | 1.28065636 | -8.5966133 |
| 318 | NM_006559.2 | 775.733467 | -11.221532 | 1.30561991 | -8.5947926 |
| 319 | NM_002901.2 | 559.0623 | -10.749269 | 1.25074441 | -8.594297 |
| 320 | NM_004484.3 | 665.191363 | -11.00018 | 1.28011155 | -8.5931414 |
| 321 | NM_031263.3 | 451.655256 | -10.440873 | 1.21543369 | -8.5902446 |
| 322 | NM_005617.3 | 1224.29395 | -11.879934 | 1.38316972 | -8.5889201 |
| 323 | NM_004637.5 | 636.247222 | -10.935735 | 1.27364778 | -8.5861535 |
| 324 | NM_002292.3 | 607.87337 | -10.870176 | 1.26645444 | -8.5831561 |
| 325 | NM_005742.3 | 670.675948 | -11.011791 | 1.28303021 | -8.5826435 |
| 326 | NM_020414.3 | 560.653666 | -10.753256 | 1.25287068 | -8.5828939 |
| 327 | NM_005275.3 | 312.681835 | -4.9584543 | 0.57789972 | -8.5801293 |
| 328 | NM_152296.4 | 1337.73853 | -12.007802 | 1.4005049 | -8.5739096 |
| 329 | NM_004772.2 | 1257.43072 | -11.918447 | 1.39029811 | -8.5725841 |
| 330 | NM_001130.5 | 987.078596 | -11.569178 | 1.35015945 | -8.5687492 |
| 331 | NM_006835.2 | 887.323261 | -11.415442 | 1.33226579 | -8.568442 |
| 332 | NM_152221.2 | 790.398625 | -11.248577 | 1.31299485 | -8.5671141 |
| 333 | NM_001064.3 | 748.183255 | -11.169511 | 1.30441161 | -8.5628733 |
| 334 | NM_014766.4 | 802.266502 | -11.270052 | 1.31639825 | -8.5612787 |
| 335 | NM_006029.4 | 722.962553 | -11.119972 | 1.29909789 | -8.5597643 |
| 336 | NM_000661.4 | 1543.1171 | -12.213834 | 1.42706182 | -8.5587277 |
| 337 | NM_001286968.1 | 714.940244 | -11.103799 | 1.29747729 | -8.5579913 |
| 338 | NM_006265.2 | 583.332994 | -10.81026 | 1.26340432 | -8.5564531 |
| 339 | NM_006617.1 | 1022.95291 | -11.620809 | 1.35816955 | -8.5562284 |
| 340 | NM_006943.3 | 699.172323 | -11.071659 | 1.29397151 | -8.5563392 |
| 341 | NM_006015.4 | 499.683358 | -10.58703 | 1.23745705 | -8.555473 |
| 342 | NM_020338.3 | 1087.77865 | -9.9345797 | 1.16154429 | -8.5529064 |
| 343 | NM_022551.2 | 2249.92567 | -10.956571 | 1.28105317 | -8.5527838 |

| | | | | | |
|---|---|---|---|---|---|
| 344 | NM_000973.4 | 2443.15431 | -9.0957238 | 1.06368115 | -8.5511751 |
| 345 | NM_001007.4 | 1301.3638 | -10.199378 | 1.19476083 | -8.536753 |
| 346 | NM_001970.4 | 808.113841 | -11.280603 | 1.32181312 | -8.5341889 |
| 347 | NM_006054.3 | 1158.463 | -11.800204 | 1.38291874 | -8.5328256 |
| 348 | NM_001330262.1 | 818.195181 | -9.5146458 | 1.1153365 | -8.5307401 |
| 349 | NM_033296.2 | 692.370633 | -11.057591 | 1.29639415 | -8.5294978 |
| 350 | NM_001028.2 | 868.274629 | -11.384213 | 1.33538583 | -8.5250365 |
| 351 | NM_001033044.3 | 396.857402 | -10.254674 | 1.20309141 | -8.5236036 |
| 352 | NM_002743.3 | 579.979148 | -10.802223 | 1.26840822 | -8.5163612 |
| 353 | NM_144665.3 | 587.492653 | -10.820742 | 1.2708858 | -8.5143311 |
| 354 | NM_001605.2 | 519.860266 | -10.644172 | 1.25051981 | -8.5117982 |
| 355 | NM_001655.4 | 464.366988 | -10.481525 | 1.23163833 | -8.5102296 |
| 356 | NR_121644.1 | 76.0245932 | 7.87520122 | 0.92537042 | 8.5103231 |
| 357 | NM_020728.2 | 229.519518 | -5.1310433 | 0.60303989 | -8.50863 |
| 358 | NM_001018006.1 | 891.886789 | -11.423107 | 1.34262465 | -8.5080423 |
| 359 | NM_002948.3 | 1113.30542 | -11.742824 | 1.38026101 | -8.5076835 |
| 360 | NM_005762.2 | 1295.11921 | -10.99889 | 1.29316876 | -8.5053785 |
| 361 | NM_001780.5 | 620.375918 | -10.899293 | 1.2816987 | -8.503787 |
| 362 | NR_026712.1 | 1035.90698 | -11.638868 | 1.36883042 | -8.5027832 |
| 363 | NM_017563.4 | 295.836676 | -5.647263 | 0.66439249 | -8.4998899 |
| 364 | NM_024911.6 | 570.066894 | -10.777622 | 1.26794273 | -8.500086 |
| 365 | NM_005273.3 | 750.723749 | -11.174336 | 1.31492763 | -8.4980616 |
| 366 | NM_001014809.2 | 1506.44964 | -12.179184 | 1.43334696 | -8.4970243 |
| 367 | NR_103849.2 | 61.599924 | 3.50411309 | 0.41244772 | 8.49589633 |
| 368 | NM_014742.3 | 287.145937 | -6.6621797 | 0.78430941 | -8.4943258 |
| 369 | NM_001286561.1 | 793.484402 | -10.290951 | 1.2115767 | -8.4938505 |
| 370 | NM_000382.2 | 199.65445 | -5.5749585 | 0.65638928 | -8.4933723 |
| 371 | NM_005567.3 | 519.963896 | -10.644956 | 1.25341645 | -8.4927529 |
| 372 | NM_004736.3 | 1143.76602 | -11.781782 | 1.38758851 | -8.4908331 |
| 373 | NM_016548.3 | 640.760717 | -10.945867 | 1.28935811 | -8.4893926 |
| 374 | NM_002567.3 | 958.008917 | -11.52609 | 1.35828627 | -8.4857593 |
| 375 | NM_001257293.1 | 531.902395 | -10.67694 | 1.25832517 | -8.4850408 |
| 376 | NM_032242.3 | 927.655421 | -11.479678 | 1.35290905 | -8.4851809 |
| 377 | NM_177433.2 | 804.572975 | -11.274406 | 1.33012949 | -8.476172 |
| 378 | NM_152436.2 | 74.342562 | 3.77131719 | 0.44495121 | 8.47579933 |
| 379 | NM_001280.2 | 580.995799 | -10.804465 | 1.27490724 | -8.4747068 |
| 380 | NM_001101417.3 | 46.2339386 | 10.4534812 | 1.2338521 | 8.47223197 |
| 381 | NM_005318.3 | 772.097814 | -11.214646 | 1.32403723 | -8.4700382 |
| 382 | NM_001037333.2 | 669.363655 | -11.008849 | 1.29986189 | -8.4692454 |
| 383 | NM_003344.3 | 546.47921 | -10.716189 | 1.26539967 | -8.4686198 |
| 384 | NM_000989.3 | 888.048932 | -11.416652 | 1.34821998 | -8.4679447 |
| 385 | NM_001009.3 | 1203.95751 | -11.855757 | 1.40022065 | -8.4670632 |
| 386 | NM_031372.3 | 670.494176 | -11.011169 | 1.30063242 | -8.4660112 |

| | | | | | |
|---|---|---|---|---|---|
| 387 | NM_001017392.4 | 393.910856 | -10.244273 | 1.21009426 | -8.4656818 |
| 388 | NM_016424.4 | 430.320332 | -10.371466 | 1.22575645 | -8.4612783 |
| 389 | NM_014739.2 | 453.643613 | -10.447535 | 1.23506276 | -8.4591129 |
| 390 | NM_004640.6 | 409.945633 | -10.301006 | 1.21788709 | -8.4580959 |
| 391 | NM_005520.2 | 549.058561 | -10.722982 | 1.2677742 | -8.4581165 |
| 392 | NM_001008938.3 | 591.811035 | -10.831224 | 1.2808168 | -8.4564973 |
| 393 | NM_001012614.1 | 525.439372 | -10.659556 | 1.26115626 | -8.4522086 |
| 394 | NM_024830.4 | 391.529819 | -10.235448 | 1.21120854 | -8.4506077 |
| 395 | NM_000598.4 | 1378.58673 | -12.051247 | 1.42700462 | -8.445135 |
| 396 | NM_005514.7 | 573.873019 | -10.78701 | 1.27759404 | -8.4432217 |
| 397 | NM_004161.4 | 576.182671 | -10.792597 | 1.27854066 | -8.4413403 |
| 398 | NM_021643.3 | 468.701937 | -10.494934 | 1.24329751 | -8.4412088 |
| 399 | NM_004911.4 | 520.441378 | -10.646063 | 1.26169649 | -8.4378955 |
| 400 | NM_002635.3 | 730.886363 | -11.135717 | 1.32009351 | -8.4355516 |
| 401 | NR_002315.1 | 834.954064 | -11.32775 | 1.34344094 | -8.4318929 |
| 402 | NM_015161.2 | 749.43014 | -11.171784 | 1.32523813 | -8.4300199 |
| 403 | NM_002415.1 | 899.063821 | -11.434424 | 1.35649894 | -8.4293642 |
| 404 | NM_004530.5 | 524.441209 | -10.6573 | 1.26456951 | -8.4276113 |
| 405 | NM_001011.3 | 957.415292 | -11.525179 | 1.36780653 | -8.4260301 |
| 406 | NM_006201.4 | 545.664233 | -10.714201 | 1.27168739 | -8.4251845 |
| 407 | NM_001967.3 | 1280.58881 | -10.982254 | 1.30376821 | -8.4234715 |
| 408 | NM_001219.4 | 601.390733 | -10.854582 | 1.28872783 | -8.4227108 |
| 409 | NM_004766.2 | 403.067236 | -10.277519 | 1.22028109 | -8.4222557 |
| 410 | NM_001142277.1 | 780.919337 | -11.2315 | 1.33380123 | -8.4206702 |
| 411 | NM_001204.6 | 400.268597 | -4.8220088 | 0.57262545 | -8.4208775 |
| 412 | NM_001901.2 | 1119.98266 | -11.751563 | 1.3954817 | -8.4211513 |
| 413 | NM_006819.2 | 554.704656 | -10.737871 | 1.27514565 | -8.4208973 |
| 414 | NM_001128619.2 | 548.935258 | -7.6226978 | 0.90537398 | -8.4193912 |
| 415 | NM_030571.3 | 980.631076 | -11.559766 | 1.37352639 | -8.4161223 |
| 416 | NM_000986.3 | 1208.66523 | -10.027895 | 1.19178169 | -8.4142043 |
| 417 | NM_003045.4 | 244.161206 | -6.1743312 | 0.73387647 | -8.4133113 |
| 418 | NM_001330700.1 | 566.7206 | -10.768557 | 1.27998673 | -8.4130227 |
| 419 | NM_005898.4 | 579.798901 | -10.801556 | 1.2840209 | -8.4122896 |
| 420 | NM_007209.3 | 966.080286 | -11.538156 | 1.37155774 | -8.4124466 |
| 421 | NM_001321483.1 | 981.282804 | -11.560705 | 1.37492445 | -8.4082471 |
| 422 | NM_006825.3 | 645.990571 | -10.957706 | 1.3031956 | -8.4083357 |
| 423 | NM_001012478.1 | 498.108071 | -8.2347684 | 0.97959598 | -8.4062906 |
| 424 | NM_001195532.1 | 488.763575 | -10.555775 | 1.25572322 | -8.4061317 |
| 425 | NM_006026.3 | 1088.52693 | -11.710291 | 1.39306039 | -8.4061619 |
| 426 | NM_007114.2 | 171.047415 | -5.0923932 | 0.60600688 | -8.4031937 |
| 427 | NM_001690.3 | 480.737082 | -4.6457854 | 0.55291527 | -8.4023459 |
| 428 | NM_002130.7 | 975.803819 | -11.552711 | 1.37502452 | -8.401822 |
| 429 | NM_002271.4 | 566.903653 | -10.769097 | 1.28201449 | -8.4001371 |

| | | | | | |
|---|---|---|---|---|---|
| 430 | NM_018984.3 | 169.588628 | -5.5841211 | 0.66505321 | -8.3965028 |
| 431 | NM_021103.3 | 2432.18937 | -11.908265 | 1.41824414 | -8.3964844 |
| 432 | NM_198576.3 | 879.557654 | -6.9749834 | 0.83073341 | -8.3961754 |
| 433 | NM_006601.6 | 738.983128 | -11.151507 | 1.32839299 | -8.394735 |
| 434 | NM_017952.5 | 177.423262 | -5.067463 | 0.60375008 | -8.3933124 |
| 435 | NM_002603.3 | 49.6754924 | 10.5575643 | 1.25792632 | 8.39283197 |
| 436 | NM_030801.3 | 652.611288 | -10.972176 | 1.30743384 | -8.3921464 |
| 437 | NM_018247.3 | 563.828503 | -10.761409 | 1.28239108 | -8.391675 |
| 438 | NM_004747.3 | 486.378635 | -10.548234 | 1.2571136 | -8.3908359 |
| 439 | NM_005561.3 | 707.462171 | -11.088841 | 1.32163504 | -8.3902441 |
| 440 | NM_001297698.1 | 61.9628445 | 10.8741152 | 1.29631629 | 8.38847376 |
| 441 | NM_020927.2 | 502.490713 | -10.595749 | 1.26338314 | -8.3868061 |
| 442 | NM_031157.3 | 608.416885 | -10.871064 | 1.29625869 | -8.3864926 |
| 443 | NM_001018108.3 | 491.401171 | -10.563036 | 1.25980878 | -8.3846343 |
| 444 | NM_021019.4 | 824.768995 | -11.310097 | 1.34912791 | -8.3832652 |
| 445 | NM_021149.3 | 673.881865 | -11.018644 | 1.31471023 | -8.3810436 |
| 446 | NM_014225.5 | 1321.01946 | -11.027493 | 1.31612142 | -8.3787807 |
| 447 | NM_004643.3 | 419.782522 | -10.335911 | 1.23362994 | -8.3784537 |
| 448 | NM_000982.3 | 993.64827 | -11.578774 | 1.38205815 | -8.3779212 |
| 449 | NM_001199780.1 | 60.0332917 | 8.93180027 | 1.06610903 | 8.37794263 |
| 450 | NM_015074.3 | 1007.72441 | -4.6949843 | 0.5604496 | -8.3771748 |
| 451 | NM_001537.3 | 554.589816 | -10.737505 | 1.28224946 | -8.3739598 |
| 452 | NM_001105079.2 | 282.608844 | -6.4381599 | 0.76941122 | -8.367645 |
| 453 | NM_001001701.3 | 240.753303 | -6.4128538 | 0.76649451 | -8.3664707 |
| 454 | NM_000389.4 | 536.693635 | -10.690312 | 1.27811447 | -8.3641275 |
| 455 | NM_001190487.2 | 955.118438 | -11.521928 | 1.37750104 | -8.3643696 |
| 456 | NM_005918.3 | 469.555499 | -10.49732 | 1.2554827 | -8.3611826 |
| 457 | NM_001318188.1 | 432.600392 | -10.378606 | 1.24170001 | -8.3583843 |
| 458 | NM_006621.5 | 375.003176 | -10.173172 | 1.21768241 | -8.3545361 |
| 459 | NM_006908.4 | 712.990466 | -11.099944 | 1.32913373 | -8.3512618 |
| 460 | NM_003681.4 | 388.780068 | -10.225162 | 1.2245874 | -8.3498838 |
| 461 | NM_004368.3 | 522.663899 | -10.652098 | 1.27698602 | -8.3415936 |
| 462 | NM_000942.4 | 630.701723 | -10.923167 | 1.30958952 | -8.3409093 |
| 463 | NM_004309.5 | 733.072278 | -11.140077 | 1.33556279 | -8.3411108 |
| 464 | NM_001004.3 | 1054.43165 | -11.664435 | 1.3986783 | -8.3396123 |
| 465 | NM_020123.3 | 502.824752 | -10.596123 | 1.27076322 | -8.3383929 |
| 466 | NM_004800.2 | 447.847634 | -10.429092 | 1.2509528 | -8.336919 |
| 467 | NM_080840.2 | 413.844861 | -10.314973 | 1.23739019 | -8.3360711 |
| 468 | NM_181689.2 | 635.119417 | -10.93297 | 1.31170165 | -8.3349517 |
| 469 | NM_001527.3 | 723.142493 | -11.120283 | 1.33440297 | -8.333527 |
| 470 | NM_006644.3 | 432.932905 | -10.38006 | 1.24582051 | -8.3319063 |
| 471 | NR_033320.2 | 576.973345 | -10.794496 | 1.29621622 | -8.3276973 |
| 472 | NM_001030.4 | 844.660814 | -11.344391 | 1.36315131 | -8.3221805 |

| | | | | | |
|---|---|---|---|---|---|
| 473 | NM_001126340.2 | 368.243816 | -10.14714 | 1.21955872 | -8.3203373 |
| 474 | NM_013436.4 | 420.684946 | -10.338288 | 1.24269763 | -8.3192308 |
| 475 | NM_170679.2 | 714.149541 | -11.10228 | 1.33450003 | -8.3194306 |
| 476 | NM_002951.4 | 522.224552 | -10.650931 | 1.28081582 | -8.31574 |
| 477 | NM_003900.4 | 513.383742 | -10.626426 | 1.27794244 | -8.3152616 |
| 478 | NM_001154.3 | 516.129921 | -10.634099 | 1.27909714 | -8.3137539 |
| 479 | NM_139215.2 | 457.967733 | -10.461384 | 1.25847291 | -8.3127606 |
| 480 | NM_001270366.1 | 886.364961 | -11.413966 | 1.37327981 | -8.311464 |
| 481 | NM_001142423.1 | 471.11157 | -10.50215 | 1.26425923 | -8.3069597 |
| 482 | NM_004339.3 | 682.853014 | -11.037703 | 1.32912466 | -8.3044902 |
| 483 | NM_001694.3 | 774.138433 | -11.218697 | 1.35147828 | -8.3010564 |
| 484 | NM_004446.2 | 453.94557 | -10.448611 | 1.25891022 | -8.2997271 |
| 485 | NM_012218.3 | 456.407088 | -8.0852181 | 0.97458887 | -8.2960295 |
| 486 | NM_006013.4 | 848.005766 | -11.350035 | 1.36831197 | -8.2949177 |
| 487 | NM_001278559.1 | 404.684592 | -4.8353978 | 0.58307907 | -8.292868 |
| 488 | NM_032737.3 | 477.039904 | -10.520131 | 1.26855526 | -8.2930015 |
| 489 | NM_001908.4 | 583.956857 | -10.812087 | 1.30385112 | -8.2924242 |
| 490 | NM_017590.5 | 583.287628 | -10.810296 | 1.30380539 | -8.2913419 |
| 491 | NM_000302.3 | 446.153199 | -10.423943 | 1.25726277 | -8.2909822 |
| 492 | NM_001016.3 | 1007.64739 | -11.598988 | 1.39912467 | -8.2901747 |
| 493 | NM_003129.3 | 713.243776 | -11.100484 | 1.33897167 | -8.290305 |
| 494 | NM_015338.5 | 390.965794 | -10.232947 | 1.23439386 | -8.2898554 |
| 495 | NM_002402.3 | 713.968205 | -11.102228 | 1.33933367 | -8.2893665 |
| 496 | NM_000859.2 | 919.331029 | -11.46668 | 1.38364356 | -8.2873074 |
| 497 | NM_003292.2 | 358.573473 | -10.109033 | 1.21980728 | -8.2874013 |
| 498 | NM_145290.3 | 200.305191 | -6.1693451 | 0.74444257 | -8.2872008 |
| 499 | NM_001079559.2 | 385.962145 | -7.2904823 | 0.88004807 | -8.2841865 |
| 500 | NM_006667.4 | 640.620092 | -10.945522 | 1.32134501 | -8.2836217 |
| 501 | NM_004539.3 | 461.806881 | -10.473328 | 1.26468406 | -8.2813791 |
| 502 | NM_003186.3 | 1263.95849 | -11.926006 | 1.44034973 | -8.2799377 |
| 503 | NM_014335.2 | 639.663347 | -10.943334 | 1.32223517 | -8.2763903 |
| 504 | NM_018206.5 | 454.134247 | -10.449289 | 1.26294668 | -8.2737375 |
| 505 | NM_001797.3 | 621.71978 | -10.902571 | 1.31784726 | -8.2730153 |
| 506 | NM_015710.4 | 567.612148 | -10.770894 | 1.30200096 | -8.2725702 |
| 507 | NM_014607.3 | 355.677096 | -10.096755 | 1.22091444 | -8.2698299 |
| 508 | NM_001320967.1 | 368.596424 | -6.8429888 | 0.82778296 | -8.266646 |
| 509 | NM_001127350.1 | 53.8673628 | 10.6740485 | 1.29143873 | 8.26523798 |
| 510 | NM_002821.4 | 440.492677 | -10.405235 | 1.2590072 | -8.2646352 |
| 511 | NM_003089.5 | 429.421335 | -10.368096 | 1.25449466 | -8.2647588 |
| 512 | NM_170782.2 | 124.404444 | 11.8800804 | 1.43783756 | 8.26246352 |
| 513 | NM_002868.3 | 440.54516 | -10.405268 | 1.25939634 | -8.2621074 |
| 514 | NM_001533.2 | 580.019538 | -10.802191 | 1.30747925 | -8.2618452 |
| 515 | NM_005721.4 | 504.135669 | -10.599854 | 1.2830789 | -8.2612644 |

| | | | | | |
|---|---|---|---|---|---|
| 516 | NM_001606.4 | 400.868118 | -10.269243 | 1.24330093 | -8.2596599 |
| 517 | NM_053056.2 | 437.412332 | -10.39568 | 1.25911638 | -8.2563294 |
| 518 | NM_003286.3 | 392.736114 | -10.239578 | 1.24042772 | -8.2548765 |
| 519 | NM_004494.2 | 413.964592 | -10.31528 | 1.24967133 | -8.2543948 |
| 520 | NM_006842.2 | 698.696312 | -10.107668 | 1.22453533 | -8.2542889 |
| 521 | NM_002139.3 | 467.511339 | -10.490933 | 1.27111143 | -8.253354 |
| 522 | NM_000018.3 | 407.109158 | -10.291976 | 1.24725692 | -8.251689 |
| 523 | NM_014730.3 | 480.350906 | -10.530246 | 1.27618263 | -8.2513628 |
| 524 | NM_004218.3 | 503.589389 | -7.8657613 | 0.95336533 | -8.2505217 |
| 525 | NM_001009184.1 | 493.566381 | -10.569371 | 1.28138279 | -8.2484107 |
| 526 | NM_194247.3 | 530.099702 | -10.672315 | 1.29386564 | -8.2483949 |
| 527 | NM_002631.3 | 641.157799 | -10.946779 | 1.3276105 | -8.2454748 |
| 528 | NM_006818.3 | 1047.83744 | -11.655405 | 1.41384363 | -8.243772 |
| 529 | NM_019020.3 | 525.276077 | -10.659175 | 1.29311076 | -8.2430487 |
| 530 | NM_001101662.1 | 421.075455 | -10.340143 | 1.25486675 | -8.2400324 |
| 531 | NM_001494.3 | 633.340918 | -10.929047 | 1.32674969 | -8.2374596 |
| 532 | NM_006431.2 | 686.871337 | -11.046053 | 1.34093933 | -8.2375487 |
| 533 | NM_003072.3 | 377.740293 | -10.183471 | 1.23635509 | -8.2366881 |
| 534 | NM_033161.3 | 501.770384 | -10.593196 | 1.28641922 | -8.2346377 |
| 535 | NM_006083.3 | 332.776042 | -10.000913 | 1.21489419 | -8.2319209 |
| 536 | NM_018926.2 | 467.530299 | -10.491264 | 1.27520399 | -8.2271258 |
| 537 | NM_012094.4 | 571.719774 | -10.781325 | 1.31061442 | -8.2261606 |
| 538 | NM_001006.4 | 2486.4006 | -9.5689765 | 1.16333477 | -8.2254711 |
| 539 | NM_000430.3 | 949.406236 | -10.550773 | 1.28273193 | -8.2252359 |
| 540 | NM_006717.2 | 512.087943 | -10.622346 | 1.2916935 | -8.223581 |
| 541 | NR_029405.1 | 516.071042 | 3.3680218 | 0.40965244 | 8.22165684 |
| 542 | NM_005642.2 | 431.398833 | -10.37493 | 1.26210021 | -8.2203692 |
| 543 | NM_183422.3 | 329.457526 | -9.9869669 | 1.21553214 | -8.2161274 |
| 544 | NM_002391.4 | 537.191742 | -10.691632 | 1.30150688 | -8.2148102 |
| 545 | NM_001025368.2 | 550.287473 | -10.726241 | 1.30638314 | -8.2106396 |
| 546 | NM_003768.4 | 1029.49134 | -11.629926 | 1.41640939 | -8.2108507 |
| 547 | NM_002270.3 | 504.596422 | -10.601023 | 1.29142046 | -8.2088084 |
| 548 | NM_021130.4 | 1742.4862 | -8.0889618 | 0.98546044 | -8.208307 |
| 549 | NR_110292.1 | 116.461993 | 3.4945213 | 0.42584036 | 8.20617677 |
| 550 | NM_001278116.1 | 1100.19446 | -11.725764 | 1.42900733 | -8.2055309 |
| 551 | NM_001304275.1 | 430.121509 | -10.370754 | 1.26404699 | -8.2044054 |
| 552 | NM_004168.3 | 302.328307 | -9.8621275 | 1.20212573 | -8.2039068 |
| 553 | NM_001127228.1 | 528.562287 | -10.668097 | 1.30043994 | -8.2034526 |
| 554 | NM_002741.3 | 510.301136 | -10.617293 | 1.29427648 | -8.2032649 |
| 555 | NM_003769.2 | 411.498214 | -10.306802 | 1.25663211 | -8.2019247 |
| 556 | NM_001111020.2 | 346.411962 | -10.058397 | 1.22715203 | -8.1965368 |
| 557 | NM_000551.3 | 444.716879 | -10.418639 | 1.27136153 | -8.1948676 |
| 558 | NM_005998.4 | 612.9142 | -10.881674 | 1.32871998 | -8.189592 |

| | | | | | |
|---|---|---|---|---|---|
| 559 | NR_103797.1 | 344.641251 | -10.051162 | 1.22743783 | -8.1887341 |
| 560 | NM_012293.2 | 807.317546 | -10.316905 | 1.26005152 | -8.1876851 |
| 561 | NM_001320547.1 | 367.618369 | -10.144715 | 1.23914472 | -8.1868686 |
| 562 | NM_007040.5 | 355.716473 | -10.096552 | 1.23329457 | -8.1866505 |
| 563 | NM_002337.3 | 367.965276 | -10.145837 | 1.23941724 | -8.1859739 |
| 564 | NM_006430.3 | 550.241169 | -10.726033 | 1.31028639 | -8.1860218 |
| 565 | NM_018031.4 | 362.687984 | -10.125416 | 1.23690601 | -8.1860836 |
| 566 | NM_002081.2 | 778.388922 | -8.5182611 | 1.04090682 | -8.1835001 |
| 567 | NM_002819.4 | 390.918015 | -10.233087 | 1.2510685 | -8.1794779 |
| 568 | NR_023312.2 | 352.358987 | -10.082371 | 1.23282791 | -8.1782471 |
| 569 | NM_002998.3 | 572.516931 | -10.783576 | 1.31881261 | -8.1767308 |
| 570 | NM_015578.2 | 239.863065 | -5.757382 | 0.70412751 | -8.1766184 |
| 571 | NM_001305105.1 | 625.993064 | -10.912179 | 1.33529333 | -8.1721214 |
| 572 | NM_002764.3 | 297.308125 | -9.8380565 | 1.20392143 | -8.1716766 |
| 573 | NM_014972.2 | 406.456166 | -10.289276 | 1.25953586 | -8.1691009 |
| 574 | NM_006854.3 | 455.840445 | -10.454681 | 1.27982126 | -8.1688602 |
| 575 | NM_030752.2 | 666.275657 | -11.002146 | 1.34693878 | -8.1682602 |
| 576 | NM_003134.5 | 534.086597 | -10.683141 | 1.30812684 | -8.1667474 |
| 577 | NM_004462.4 | 900.16539 | -11.436255 | 1.40038689 | -8.1664969 |
| 578 | NM_004910.2 | 306.995011 | -9.8851056 | 1.21046673 | -8.1663587 |
| 579 | NM_203504.2 | 475.33 | -10.514868 | 1.28766416 | -8.1658463 |
| 580 | NM_005051.2 | 343.84562 | -10.048172 | 1.23074459 | -8.1643029 |
| 581 | NM_001284306.1 | 68.1445699 | 11.0054488 | 1.34858437 | 8.1607418 |
| 582 | NM_001844.4 | 655.061491 | -10.977873 | 1.34519801 | -8.1607863 |
| 583 | NM_002862.3 | 552.69029 | -10.732527 | 1.31525888 | -8.1600109 |
| 584 | NM_000435.2 | 460.549653 | -10.469641 | 1.28335844 | -8.1580023 |
| 585 | NM_001080424.1 | 451.794156 | -10.44149 | 1.28001254 | -8.1573344 |
| 586 | NM_000371.3 | 2768.77005 | -13.057296 | 1.60086392 | -8.1564062 |
| 587 | NM_030649.2 | 666.634746 | -11.002918 | 1.34904189 | -8.1560978 |
| 588 | NM_000445.4 | 451.45019 | -10.440741 | 1.28024165 | -8.155289 |
| 589 | NR_130727.1 | 31.4492374 | 7.01502347 | 0.86020437 | 8.15506607 |
| 590 | NM_006227.3 | 383.896864 | -10.207604 | 1.251839 | -8.1540865 |
| 591 | NR_027001.1 | 345.819862 | -10.055898 | 1.23326068 | -8.153911 |
| 592 | NM_003170.4 | 332.729134 | -10.001012 | 1.22679034 | -8.1521767 |
| 593 | NM_001909.4 | 684.970468 | -10.079244 | 1.23644562 | -8.1517889 |
| 594 | NM_006904.6 | 396.598503 | -10.253669 | 1.25786738 | -8.1516293 |
| 595 | NM_020532.4 | 988.19234 | -11.570853 | 1.41955482 | -8.1510437 |
| 596 | NM_001677.3 | 412.303505 | -5.2469499 | 0.64393411 | -8.1482715 |
| 597 | NR_003249.2 | 384.273764 | -10.207859 | 1.25276102 | -8.1482888 |
| 598 | NM_022574.4 | 333.852035 | -10.005319 | 1.22872653 | -8.1428364 |
| 599 | NM_006325.4 | 785.580265 | -7.572316 | 0.92998627 | -8.1423955 |
| 600 | NM_003079.4 | 391.60326 | -10.235534 | 1.25716641 | -8.1417499 |
| 601 | NM_006141.2 | 458.22267 | -10.462136 | 1.28495512 | -8.1420248 |

| | | | | | |
|---|---|---|---|---|---|
| 602 | NM_032632.4 | 382.72479 | -10.202179 | 1.25308222 | -8.141668 |
| 603 | NM_024718.4 | 472.282017 | -10.505545 | 1.29040866 | -8.141254 |
| 604 | NM_002809.3 | 472.724397 | -10.507068 | 1.29069177 | -8.1406485 |
| 605 | NM_001077443.1 | 429.228608 | -10.367893 | 1.27364428 | -8.1403362 |
| 606 | NR_037946.1 | 355.078691 | -10.09439 | 1.24008275 | -8.1400942 |
| 607 | NM_003816.2 | 448.460492 | -10.431321 | 1.28196431 | -8.1369821 |
| 608 | NM_031992.1 | 760.564326 | -10.230702 | 1.25735742 | -8.13667 |
| 609 | NM_006310.3 | 402.638042 | -10.275471 | 1.26317448 | -8.1346414 |
| 610 | NM_016368.4 | 312.432891 | -9.9100276 | 1.21848184 | -8.1330942 |
| 611 | NM_017620.2 | 341.507964 | -10.037926 | 1.23428384 | -8.1325916 |
| 612 | NM_004427.3 | 338.870397 | -10.026697 | 1.23324098 | -8.130363 |
| 613 | NM_007359.4 | 394.891363 | -10.247378 | 1.26049485 | -8.1296471 |
| 614 | NM_001958.3 | 1078.78048 | -11.697378 | 1.43897466 | -8.1289671 |
| 615 | NM_000984.5 | 739.14797 | -8.8329745 | 1.08674331 | -8.1279309 |
| 616 | NM_001183.5 | 604.129163 | -10.86099 | 1.33662448 | -8.1256854 |
| 617 | NM_006371.4 | 278.007613 | -5.5909818 | 0.68822879 | -8.1237255 |
| 618 | NM_181697.2 | 497.630948 | -10.581151 | 1.30247938 | -8.1238531 |
| 619 | NM_004990.3 | 566.707889 | -10.76865 | 1.32563567 | -8.1233856 |
| 620 | NM_001037675.3 | 306.50563 | -9.882279 | 1.2169657 | -8.1204252 |
| 621 | NM_002715.2 | 553.399737 | -10.734341 | 1.32187357 | -8.1205501 |
| 622 | NM_001996.3 | 407.283783 | -10.292663 | 1.26765495 | -8.119452 |
| 623 | NM_000992.2 | 1216.7067 | -10.908763 | 1.34361543 | -8.1189622 |
| 624 | NM_006924.4 | 501.574246 | -10.592443 | 1.30487527 | -8.1175907 |
| 625 | NM_153649.3 | 379.722432 | -10.190718 | 1.25574203 | -8.1152956 |
| 626 | NM_021648.4 | 498.889993 | -10.58479 | 1.30502074 | -8.1108211 |
| 627 | NM_004793.3 | 334.92454 | -10.009527 | 1.23413666 | -8.1105503 |
| 628 | NM_001357.4 | 584.291188 | -8.9889729 | 1.10861316 | -8.1083044 |
| 629 | NM_001659.2 | 584.544037 | -10.813329 | 1.3336879 | -8.1078405 |
| 630 | NM_003142.4 | 366.598236 | -10.140199 | 1.25102446 | -8.105516 |
| 631 | NM_013974.2 | 559.764296 | -10.750804 | 1.32669691 | -8.1034363 |
| 632 | NM_148174.3 | 575.808053 | -10.79159 | 1.33180412 | -8.1029858 |
| 633 | NM_020117.10 | 368.310308 | -10.146653 | 1.25235544 | -8.1020554 |
| 634 | NM_001098272.2 | 763.806335 | -11.199168 | 1.38241106 | -8.1011849 |
| 635 | NM_001195553.1 | 1055.21892 | -11.665544 | 1.4402602 | -8.0996087 |
| 636 | NM_152713.4 | 335.520947 | -10.012703 | 1.23635373 | -8.0985747 |
| 637 | NM_000988.3 | 967.64117 | -11.540515 | 1.42525883 | -8.0971362 |
| 638 | NM_198475.2 | 555.671589 | -10.74017 | 1.3265321 | -8.096427 |
| 639 | NM_001145543.1 | 299.094174 | -9.8460265 | 1.21621496 | -8.0956301 |
| 640 | NM_182557.2 | 301.095563 | -7.4910212 | 0.92535929 | -8.0952569 |
| 641 | NM_005968.4 | 383.596976 | -10.205735 | 1.26115236 | -8.092389 |
| 642 | NM_005877.5 | 337.038042 | -10.019025 | 1.23830837 | -8.0908969 |
| 643 | NM_001102420.2 | 429.851266 | -10.369742 | 1.28188955 | -8.0894195 |
| 644 | NM_031407.6 | 652.011107 | -10.007821 | 1.23713615 | -8.0895068 |

| | | | | | |
|---|---|---|---|---|---|
| 645 | NM_006364.3 | 303.902315 | -9.8705774 | 1.22031378 | -8.0885568 |
| 646 | NM_002070.3 | 559.73646 | -10.750839 | 1.32922911 | -8.0880253 |
| 647 | NM_002586.4 | 328.014345 | -9.9803306 | 1.23402008 | -8.0876566 |
| 648 | NM_015060.2 | 220.481325 | -4.0189656 | 0.49697758 | -8.0868147 |
| 649 | NM_007245.3 | 345.828472 | -10.05668 | 1.24381579 | -8.0853452 |
| 650 | NM_030662.3 | 440.126723 | -10.404022 | 1.2869504 | -8.0842451 |
| 651 | NM_000938.2 | 413.967195 | -10.315509 | 1.27615509 | -8.0832723 |
| 652 | NM_001568.2 | 532.289035 | -10.678179 | 1.32104368 | -8.0831385 |
| 653 | NM_004930.4 | 503.212288 | -10.597241 | 1.3110062 | -8.083288 |
| 654 | NM_031370.2 | 306.719219 | -9.8835017 | 1.22275819 | -8.0829569 |
| 655 | NM_080426.3 | 405.34379 | -10.285395 | 1.27243548 | -8.0832346 |
| 656 | NM_003074.3 | 402.653505 | -10.275497 | 1.27138909 | -8.0821029 |
| 657 | NM_021029.5 | 770.810564 | -11.212382 | 1.38734269 | -8.0819122 |
| 658 | NM_014639.3 | 261.876483 | -6.0753049 | 0.75180787 | -8.0809275 |
| 659 | NM_001206654.1 | 364.148892 | -10.130685 | 1.253798 | -8.0799977 |
| 660 | NM_080391.3 | 389.08965 | -10.226039 | 1.26578507 | -8.0788118 |
| 661 | NM_130445.3 | 1197.62311 | -8.0526405 | 0.99693466 | -8.0774005 |
| 662 | NM_016176.3 | 412.325227 | -10.309955 | 1.27655509 | -8.0763883 |
| 663 | NM_001089.2 | 553.705598 | -10.735212 | 1.32935415 | -8.0755093 |
| 664 | NM_080677.2 | 537.212098 | -10.691616 | 1.32402244 | -8.075102 |
| 665 | NM_152879.2 | 317.587974 | -9.9331901 | 1.23029462 | -8.0738304 |
| 666 | NM_000999.3 | 702.913974 | -11.079368 | 1.37243696 | -8.0727701 |
| 667 | NM_005552.4 | 970.990197 | -11.545513 | 1.43019151 | -8.0727042 |
| 668 | NM_130853.2 | 679.048059 | -6.4909048 | 0.80417208 | -8.0715372 |
| 669 | NM_002660.2 | 466.284386 | -10.487341 | 1.29939865 | -8.0709191 |
| 670 | NM_021034.2 | 365.773546 | -10.137876 | 1.25641838 | -8.0688695 |
| 671 | NM_002086.4 | 546.380951 | -10.715929 | 1.32845778 | -8.0664433 |
| 672 | NM_001893.4 | 369.173615 | -10.150199 | 1.25904926 | -8.0617962 |
| 673 | NM_018448.4 | 509.245215 | -10.614326 | 1.3166043 | -8.061895 |
| 674 | NM_004331.2 | 510.910894 | -10.618835 | 1.31723464 | -8.0614606 |
| 675 | NM_013402.4 | 573.265071 | -6.9574549 | 0.86341771 | -8.0580405 |
| 676 | NM_006389.4 | 376.059606 | -10.176963 | 1.26327525 | -8.0560141 |
| 677 | NM_012181.4 | 490.417191 | -10.560086 | 1.31087622 | -8.0557465 |
| 678 | NM_012268.3 | 429.803629 | -10.369627 | 1.28722995 | -8.0557686 |
| 679 | NM_006009.3 | 11685.5929 | -11.020597 | 1.36843725 | -8.0534181 |
| 680 | NM_013236.3 | 527.95227 | -10.66642 | 1.32458724 | -8.0526369 |
| 681 | NM_021943.2 | 362.054617 | -10.122292 | 1.25731772 | -8.0507037 |
| 682 | NM_006472.5 | 301.046494 | -9.8560446 | 1.22427546 | -8.0505122 |
| 683 | NM_002076.3 | 512.303878 | -2.6121857 | 0.32455713 | -8.0484622 |
| 684 | NM_006870.3 | 672.36379 | -11.01528 | 1.36863513 | -8.0483687 |
| 685 | NM_016185.3 | 980.780481 | -11.559967 | 1.43660522 | -8.0467251 |
| 686 | NM_003973.4 | 1383.76746 | -11.094386 | 1.37879997 | -8.0464075 |
| 687 | NM_004048.2 | 404.954221 | -10.284169 | 1.2781529 | -8.0461176 |

| | | | | | |
|---|---|---|---|---|---|
| 688 | NM_007278.1 | 562.751066 | -10.758627 | 1.33825722 | -8.0392818 |
| 689 | NM_018215.3 | 347.174947 | -10.061637 | 1.25186115 | -8.0373427 |
| 690 | NM_019058.3 | 293.707584 | -9.8204162 | 1.22196843 | -8.0365548 |
| 691 | NM_014263.3 | 320.708574 | -9.9467929 | 1.23786127 | -8.0354666 |
| 692 | NR_110493.1 | 399.785998 | -10.265282 | 1.2777329 | -8.0339809 |
| 693 | NM_001018069.1 | 350.161235 | -10.073566 | 1.25471054 | -8.0285973 |
| 694 | NM_001305275.1 | 591.25939 | -10.829748 | 1.34890823 | -8.0285284 |
| 695 | NM_014286.3 | 786.614982 | -11.241721 | 1.40025511 | -8.0283378 |
| 696 | NM_001458.4 | 342.797976 | -10.043815 | 1.25139937 | -8.0260665 |
| 697 | NM_002628.4 | 359.128879 | -10.110699 | 1.25976739 | -8.0258457 |
| 698 | NM_001030009.1 | 878.452552 | -11.401003 | 1.42108591 | -8.0227401 |
| 699 | NM_001136135.1 | 533.602686 | -10.681827 | 1.33154573 | -8.0221259 |
| 700 | NM_024595.2 | 334.676267 | -10.00876 | 1.24770808 | -8.0217163 |
| 701 | NM_001163547.2 | 37.4387611 | 10.1505873 | 1.26546328 | 8.02124211 |
| 702 | NM_001141980.1 | 326.856926 | -9.9749962 | 1.24417418 | -8.0173631 |
| 703 | NM_001012279.2 | 693.379034 | -11.059647 | 1.37989501 | -8.0148466 |
| 704 | NM_017896.2 | 263.357479 | -6.8153777 | 0.85036699 | -8.0146311 |
| 705 | NM_002037.5 | 488.118069 | -10.553228 | 1.31678162 | -8.0144103 |
| 706 | NM_006460.2 | 162.972888 | -5.582258 | 0.69698034 | -8.0092044 |
| 707 | NM_001693.3 | 506.783279 | -10.607425 | 1.32448154 | -8.0087372 |
| 708 | NM_021239.2 | 442.578197 | -8.5922291 | 1.07284551 | -8.0088224 |
| 709 | NM_000454.4 | 383.983661 | -10.207141 | 1.27472076 | -8.0073547 |
| 710 | NM_178862.2 | 464.915928 | -10.482946 | 1.3092445 | -8.0068664 |
| 711 | NM_006585.3 | 472.53772 | -10.50648 | 1.31224167 | -8.0065134 |
| 712 | NM_003906.4 | 283.15801 | -7.3697192 | 0.92062567 | -8.0051202 |
| 713 | NM_014866.1 | 267.989073 | -9.6883398 | 1.21043845 | -8.0039921 |
| 714 | NM_013438.4 | 464.642329 | -10.481873 | 1.31020151 | -8.000199 |
| 715 | NM_013271.4 | 686.844779 | -11.046069 | 1.38081201 | -7.9996909 |
| 716 | NM_025181.4 | 255.302984 | -9.6184865 | 1.20247396 | -7.9989146 |
| 717 | NM_130854.2 | 270.249139 | -9.7009644 | 1.21290637 | -7.9981149 |
| 718 | NM_015526.2 | 613.919619 | -10.884069 | 1.36100856 | -7.9970613 |
| 719 | NM_002539.2 | 813.581037 | -11.290226 | 1.41196864 | -7.9960884 |
| 720 | NM_004040.3 | 575.45815 | -10.790787 | 1.34961415 | -7.9954606 |
| 721 | NM_003836.6 | 759.518292 | -11.191388 | 1.3998045 | -7.9949647 |
| 722 | NM_000289.5 | 495.072494 | -10.57362 | 1.32273235 | -7.9937717 |
| 723 | NM_001553.2 | 444.380298 | -10.418401 | 1.30337127 | -7.9934257 |
| 724 | NM_017514.4 | 459.874307 | -10.467366 | 1.30956135 | -7.9930324 |
| 725 | NM_001102.3 | 645.776174 | -10.957247 | 1.37092784 | -7.9925773 |
| 726 | NM_001286366.1 | 633.255114 | -10.928913 | 1.36746599 | -7.9920913 |
| 727 | NM_016201.3 | 343.324139 | -10.045801 | 1.25732307 | -7.9898324 |
| 728 | NM_058243.2 | 259.878008 | -9.6439905 | 1.20790176 | -7.9840851 |
| 729 | NM_012478.3 | 369.766408 | -10.15277 | 1.27167236 | -7.9837939 |
| 730 | NM_003366.3 | 434.146949 | -10.384112 | 1.3007085 | -7.9834278 |

| | | | | | |
|---|---|---|---|---|---|
| 731 | NM_001244701.1 | 276.653263 | -9.7352147 | 1.21983341 | -7.9807739 |
| 732 | NM_014275.4 | 322.065368 | -7.1571104 | 0.89682995 | -7.9804543 |
| 733 | NM_003754.2 | 507.408595 | -10.609171 | 1.32985057 | -7.9777169 |
| 734 | NM_006513.3 | 447.881964 | -10.42923 | 1.30764637 | -7.9755738 |
| 735 | NM_002373.5 | 616.21359 | -9.0622821 | 1.1363557 | -7.9748639 |
| 736 | NM_001190836.1 | 327.304953 | -9.9771931 | 1.25173373 | -7.9706992 |
| 737 | NM_001338.4 | 398.785285 | -10.261468 | 1.28754355 | -7.9698024 |
| 738 | NM_001286572.2 | 357.341958 | -10.103452 | 1.26779347 | -7.9693201 |
| 739 | NM_001078171.1 | 486.091473 | -10.547304 | 1.32382027 | -7.9673234 |
| 740 | NM_001085460.1 | 348.070195 | -10.065876 | 1.26334608 | -7.9676313 |
| 741 | NM_004630.3 | 304.767584 | -9.8730771 | 1.23917514 | -7.967459 |
| 742 | NM_004939.2 | 371.621359 | -10.159837 | 1.27520229 | -7.9672353 |
| 743 | NM_170665.3 | 772.271521 | -10.252196 | 1.2867604 | -7.9674472 |
| 744 | NM_005027.3 | 593.694634 | -9.0089887 | 1.1308699 | -7.9664236 |
| 745 | NM_005896.3 | 495.657451 | -10.575436 | 1.32786096 | -7.9642647 |
| 746 | NM_001748.4 | 316.182852 | -9.9275986 | 1.24665879 | -7.9633648 |
| 747 | NM_014413.3 | 318.75722 | -9.9383396 | 1.24805543 | -7.9630594 |
| 748 | NM_004157.3 | 302.062513 | -9.8609338 | 1.23850529 | -7.9619634 |
| 749 | NM_004663.4 | 431.354909 | -10.374953 | 1.30366504 | -7.9582968 |
| 750 | NM_031989.4 | 453.266956 | -10.446523 | 1.31265787 | -7.9582986 |
| 751 | NM_145296.1 | 583.295636 | -10.810194 | 1.3583925 | -7.9580782 |
| 752 | NM_019613.3 | 322.385982 | -9.9548006 | 1.25128159 | -7.9556837 |
| 753 | NM_004859.3 | 1370.37759 | -6.8716183 | 0.86377338 | -7.9553485 |
| 754 | NM_001031.4 | 626.913801 | -10.914317 | 1.37251722 | -7.9520435 |
| 755 | NM_139276.2 | 338.504555 | -10.025851 | 1.26090066 | -7.9513408 |
| 756 | NM_001297553.1 | 387.072312 | -10.218773 | 1.28529789 | -7.9505097 |
| 757 | NM_018240.6 | 328.112463 | -9.9806568 | 1.25544939 | -7.949868 |
| 758 | NM_005338.6 | 302.547058 | -9.8633782 | 1.24072827 | -7.9496683 |
| 759 | NM_000994.3 | 1125.49249 | -7.5593355 | 0.95093906 | -7.9493375 |
| 760 | NM_002340.5 | 576.835826 | -10.79422 | 1.35800304 | -7.9485976 |
| 761 | NM_015541.2 | 211.324656 | -6.503744 | 0.81822224 | -7.9486278 |
| 762 | NM_004187.3 | 327.283715 | -9.9760778 | 1.25521961 | -7.9476753 |
| 763 | NM_003144.4 | 350.174329 | -10.074231 | 1.26770142 | -7.9468483 |
| 764 | NM_001304.4 | 324.111459 | -9.9630112 | 1.25378858 | -7.9463247 |
| 765 | NM_001042681.1 | 320.605249 | -9.9466086 | 1.25194088 | -7.9449507 |
| 766 | NM_001394.6 | 691.204634 | -11.055249 | 1.39156488 | -7.9444725 |
| 767 | NM_184043.2 | 369.584483 | -10.151592 | 1.27819417 | -7.9421358 |
| 768 | NM_004689.3 | 336.337505 | -10.015782 | 1.26131106 | -7.9407707 |
| 769 | NM_002695.3 | 367.201541 | -10.142592 | 1.27737289 | -7.9401968 |
| 770 | NR_109890.1 | 105.548819 | 3.32000489 | 0.41828299 | 7.93722189 |
| 771 | NM_001039580.1 | 204.200643 | -6.517059 | 0.82112046 | -7.9367879 |
| 772 | NM_001546.3 | 541.380387 | -10.702688 | 1.34850479 | -7.9367071 |
| 773 | NM_053064.4 | 711.776271 | -11.09738 | 1.39850236 | -7.9351886 |

| | | | | | |
|---|---|---|---|---|---|
| 774 | NR_028272.1 | 400.777061 | -10.269419 | 1.29423177 | -7.9347605 |
| 775 | NM_003463.4 | 521.925312 | -10.649771 | 1.34276748 | -7.9312098 |
| 776 | NM_002673.5 | 304.119657 | -9.869727 | 1.24449428 | -7.930713 |
| 777 | NM_002092.3 | 300.524645 | -9.8535535 | 1.24274068 | -7.9288895 |
| 778 | NM_015172.3 | 637.402241 | -7.2798871 | 0.91820675 | -7.9283746 |
| 779 | NM_006311.3 | 331.078381 | -9.993243 | 1.26070078 | -7.9267366 |
| 780 | NM_001895.3 | 315.984117 | -9.9257811 | 1.25291242 | -7.9221668 |
| 781 | NM_031462.3 | 425.104955 | -10.354081 | 1.30707001 | -7.921596 |
| 782 | NM_001618.3 | 534.230661 | -7.4112818 | 0.93566568 | -7.9208653 |
| 783 | NM_001134231.1 | 558.780064 | -10.748386 | 1.35718131 | -7.9196388 |
| 784 | NM_000786.3 | 496.165377 | -10.576869 | 1.33575135 | -7.9182921 |
| 785 | NM_001067.3 | 305.39189 | -9.8761399 | 1.24721685 | -7.9185427 |
| 786 | NM_003405.3 | 687.807164 | -11.048054 | 1.39526728 | -7.9182349 |
| 787 | NM_005245.3 | 559.062661 | -9.7860894 | 1.23594686 | -7.9178885 |
| 788 | NM_032408.3 | 330.338028 | -9.9899423 | 1.2622237 | -7.9145577 |
| 789 | NM_015894.3 | 670.390292 | -11.01108 | 1.39136501 | -7.9138689 |
| 790 | NM_001093725.1 | 517.020248 | -10.63622 | 1.34478522 | -7.909233 |
| 791 | NM_004595.4 | 470.500922 | -10.50019 | 1.32762305 | -7.9090146 |
| 792 | NM_006016.5 | 296.199264 | -9.8334489 | 1.24365452 | -7.9068976 |
| 793 | NM_006623.3 | 387.274298 | -10.219308 | 1.29243081 | -7.9070446 |
| 794 | NM_000099.3 | 314.481893 | -9.9192634 | 1.25471757 | -7.9055746 |
| 795 | NM_018837.3 | 392.039488 | -10.237359 | 1.29524309 | -7.9038131 |
| 796 | NM_012096.2 | 175.524574 | -3.8479434 | 0.48689308 | -7.9030564 |
| 797 | NM_001080425.3 | 426.713399 | -10.359095 | 1.31107342 | -7.9012321 |
| 798 | NM_003564.2 | 278.250737 | -9.7428704 | 1.2330755 | -7.9012764 |
| 799 | NM_030810.4 | 369.941471 | -10.153555 | 1.2850596 | -7.9012328 |
| 800 | NM_181449.2 | 74.729041 | 6.44779627 | 0.81603773 | 7.90134578 |
| 801 | NM_001791.3 | 445.34955 | -10.420981 | 1.31898124 | -7.9007799 |
| 802 | NM_025235.3 | 277.847983 | -9.7405018 | 1.23311287 | -7.8991162 |
| 803 | NR_034009.1 | 294.979461 | -9.8266264 | 1.24400616 | -7.8991782 |
| 804 | NM_003343.5 | 283.291447 | -9.7679833 | 1.23662992 | -7.8988735 |
| 805 | NM_006812.3 | 380.58394 | -10.194071 | 1.2907567 | -7.897748 |
| 806 | NM_002911.3 | 292.901826 | -9.8162048 | 1.24301435 | -7.897097 |
| 807 | NM_004826.3 | 284.942206 | -9.7778578 | 1.23842306 | -7.89541 |
| 808 | NM_016463.8 | 325.806637 | -9.9706265 | 1.26292991 | -7.8948376 |
| 809 | NM_013291.2 | 488.790309 | -8.7695985 | 1.11083377 | -7.8946092 |
| 810 | NM_002254.6 | 809.374934 | -11.282847 | 1.4292371 | -7.8943142 |
| 811 | NM_012073.4 | 475.318261 | -10.514782 | 1.332072 | -7.8935536 |
| 812 | NM_145690.2 | 1765.19314 | -8.7939669 | 1.11429051 | -7.8919876 |
| 813 | NM_003753.3 | 336.814666 | -10.017949 | 1.26966749 | -7.8902149 |
| 814 | NM_014292.4 | 276.955411 | -9.7354371 | 1.23393287 | -7.8897624 |
| 815 | NR_037688.1 | 1089.5431 | -10.74916 | 1.3624132 | -7.8897944 |
| 816 | NM_001199456.1 | 323.694862 | -9.960476 | 1.26250782 | -7.8894371 |

| | | | | | |
|---|---|---|---|---|---|
| 817 | NM_005626.4 | 287.840858 | -9.7910049 | 1.24136194 | -7.8873088 |
| 818 | NM_139207.4 | 328.23016 | -9.9800699 | 1.26550689 | -7.8862233 |
| 819 | NM_007103.3 | 379.58659 | -10.190463 | 1.29223745 | -7.8859057 |
| 820 | NM_015043.3 | 285.222066 | -9.7781698 | 1.24001658 | -7.8855154 |
| 821 | NM_024293.5 | 373.709095 | -10.168048 | 1.28946674 | -7.8854673 |
| 822 | NM_002117.5 | 391.761011 | -10.236319 | 1.29832441 | -7.8842535 |
| 823 | NM_002094.3 | 332.289127 | -9.9985183 | 1.2681878 | -7.8840991 |
| 824 | NM_173595.3 | 341.671763 | -10.03893 | 1.2733942 | -7.8835994 |
| 825 | NM_000489.4 | 314.349977 | -9.9184764 | 1.258224 | -7.8829179 |
| 826 | NM_002140.4 | 266.861096 | -9.6831498 | 1.22847247 | -7.8822684 |
| 827 | NM_007118.3 | 391.946997 | -10.236622 | 1.29866827 | -7.8823991 |
| 828 | NM_015980.4 | 1039.90531 | -11.644419 | 1.47754288 | -7.8809347 |
| 829 | NM_001197293.2 | 670.080538 | -11.010392 | 1.39713238 | -7.8807076 |
| 830 | NM_014709.3 | 353.764367 | -8.290046 | 1.05195616 | -7.8806003 |
| 831 | NM_020151.3 | 389.023751 | -10.225812 | 1.29765711 | -7.8802114 |
| 832 | NM_002161.5 | 288.298452 | -9.7937013 | 1.24304553 | -7.8787953 |
| 833 | NM_004327.3 | 410.121273 | -10.302142 | 1.30764293 | -7.8784063 |
| 834 | NM_005570.3 | 268.950902 | -9.6945755 | 1.23051601 | -7.8784635 |
| 835 | NM_206833.3 | 492.69867 | -10.566752 | 1.34126578 | -7.8781941 |
| 836 | NM_033643.2 | 734.371488 | -8.6216162 | 1.09473305 | -7.8755421 |
| 837 | NM_020746.4 | 245.945858 | -9.5642181 | 1.21448619 | -7.8751148 |
| 838 | NM_024299.2 | 454.006146 | -10.448804 | 1.32706966 | -7.8735915 |
| 839 | NM_005137.2 | 400.070718 | -10.266256 | 1.30424158 | -7.8714371 |
| 840 | NM_001014.4 | 1126.71265 | -9.9578628 | 1.26521512 | -7.8704899 |
| 841 | NM_002807.3 | 395.212339 | -10.248732 | 1.30226046 | -7.8699559 |
| 842 | NM_198334.2 | 568.917888 | -10.774361 | 1.36906734 | -7.8698548 |
| 843 | NM_130440.3 | 302.658403 | -9.8647863 | 1.25354228 | -7.8695282 |
| 844 | NM_014599.5 | 413.67765 | -10.314207 | 1.31068812 | -7.869307 |
| 845 | NM_003021.3 | 419.282713 | -10.333926 | 1.31324572 | -7.8689962 |
| 846 | NM_203499.2 | 327.047462 | -5.9581699 | 0.75719086 | -7.8687821 |
| 847 | NM_005386.3 | 311.414794 | -9.9039956 | 1.25890093 | -7.8671763 |
| 848 | NM_015171.3 | 369.448687 | -10.151304 | 1.29031931 | -7.8672804 |
| 849 | NM_000182.4 | 382.783121 | -6.9127383 | 0.87876448 | -7.8664289 |
| 850 | NM_198318.4 | 428.305293 | -10.364683 | 1.31764951 | -7.8660395 |
| 851 | NM_002085.4 | 356.325568 | -10.099318 | 1.2840954 | -7.8649281 |
| 852 | NM_021102.3 | 268.681517 | -9.6915517 | 1.23235824 | -7.8642325 |
| 853 | NM_003062.3 | 349.322063 | -10.071181 | 1.280785 | -7.8632879 |
| 854 | NM_002792.3 | 478.210016 | -10.523678 | 1.33838867 | -7.8629462 |
| 855 | NM_005445.3 | 291.35844 | -9.8086584 | 1.24744536 | -7.8629964 |
| 856 | NR_026811.3 | 271.817743 | -9.707797 | 1.2348394 | -7.8615867 |
| 857 | NM_030915.3 | 415.865047 | -10.322199 | 1.3131784 | -7.86047 |
| 858 | NM_002107.4 | 710.91939 | -8.7839601 | 1.1176705 | -7.8591679 |
| 859 | NM_000820.3 | 271.664774 | -9.7080867 | 1.23530192 | -7.8588777 |

| | | | | | |
|---|---|---|---|---|---|
| 860 | NM_006044.3 | 302.137745 | -9.8616994 | 1.25495276 | -7.8582236 |
| 861 | NM_005271.4 | 327.064192 | -9.975597 | 1.26948255 | -7.8580025 |
| 862 | NM_014232.2 | 642.616183 | -10.950016 | 1.39364893 | -7.8570838 |
| 863 | NM_002047.3 | 347.011469 | -10.061032 | 1.28078786 | -7.8553462 |
| 864 | NM_002507.3 | 614.539199 | -10.885708 | 1.38610725 | -7.8534384 |
| 865 | NM_052932.2 | 230.346214 | -9.4698383 | 1.2058135 | -7.8534851 |
| 866 | NM_005751.4 | 288.753866 | -9.7954303 | 1.24732704 | -7.8531371 |
| 867 | NM_014330.3 | 261.073511 | -9.6510495 | 1.2291172 | -7.8520172 |
| 868 | NM_001317897.1 | 94.6327289 | 11.4883906 | 1.46326664 | 7.85119424 |
| 869 | NM_018243.3 | 837.259853 | -4.8640098 | 0.61955047 | -7.8508693 |
| 870 | NM_006305.3 | 508.322576 | -10.611628 | 1.3521567 | -7.8479279 |
| 871 | NM_006618.4 | 425.674415 | -10.355752 | 1.31961711 | -7.8475427 |
| 872 | NM_001009552.1 | 368.735496 | -10.148493 | 1.29324217 | -7.8473261 |
| 873 | NM_001099415.2 | 273.436203 | -9.717616 | 1.23858573 | -7.8457355 |
| 874 | NM_003791.3 | 311.271275 | -9.9044836 | 1.26272193 | -7.8437567 |
| 875 | NM_005443.4 | 346.404595 | -10.058444 | 1.28232561 | -7.8439078 |
| 876 | NM_015013.3 | 404.349332 | -7.942027 | 1.01254352 | -7.8436401 |
| 877 | NM_002880.3 | 369.886393 | -10.152981 | 1.29480174 | -7.8413406 |
| 878 | NM_001190274.1 | 322.889898 | -9.9568974 | 1.27008751 | -7.8395365 |
| 879 | NM_003590.4 | 295.064265 | -9.8270862 | 1.25354295 | -7.8394491 |
| 880 | NM_002444.2 | 651.895673 | -10.007674 | 1.27673233 | -7.8385063 |
| 881 | NM_017921.3 | 274.963962 | -9.7253684 | 1.24088192 | -7.8374648 |
| 882 | NM_001193417.2 | 256.191207 | -9.6235895 | 1.22792922 | -7.837251 |
| 883 | NM_007235.4 | 349.468179 | -6.4357955 | 0.82129411 | -7.8361642 |
| 884 | NM_005744.3 | 250.852824 | -9.5928797 | 1.22430369 | -7.835376 |
| 885 | NM_201599.2 | 257.037939 | -9.6290671 | 1.22893205 | -7.8353129 |
| 886 | NM_001001563.3 | 283.121021 | -9.7672864 | 1.24674031 | -7.8342589 |
| 887 | NM_001018.4 | 992.660883 | -9.6267175 | 1.22881209 | -7.8341657 |
| 888 | NM_002710.3 | 306.63066 | -9.8823132 | 1.2615768 | -7.8333029 |
| 889 | NR_027166.1 | 587.797582 | -10.821322 | 1.381703 | -7.8318728 |
| 890 | NM_001415.3 | 656.498878 | -4.6777678 | 0.59729406 | -7.8315995 |
| 891 | NM_031243.2 | 307.475665 | -9.8866208 | 1.26241714 | -7.8315008 |
| 892 | NM_002740.5 | 169.589365 | -5.3431558 | 0.6823019 | -7.8310727 |
| 893 | NM_004638.3 | 316.884871 | -9.9294688 | 1.26809567 | -7.8302206 |
| 894 | NM_006555.3 | 441.859046 | -10.40965 | 1.32957606 | -7.8293004 |
| 895 | NM_000358.2 | 860.616974 | -10.409171 | 1.32959169 | -7.828848 |
| 896 | NM_001300905.1 | 255.647055 | -9.6192619 | 1.2287378 | -7.8285716 |
| 897 | NM_013390.2 | 317.225642 | -9.9319093 | 1.26866068 | -7.8286569 |
| 898 | NM_058004.3 | 349.009212 | -10.069197 | 1.28623923 | -7.8284012 |
| 899 | NM_002950.3 | 591.758639 | -9.0546557 | 1.15676256 | -7.8275837 |
| 900 | NM_019116.2 | 344.026116 | -10.048668 | 1.28378656 | -7.8273668 |
| 901 | NM_014649.2 | 261.173262 | -9.6508858 | 1.23312004 | -7.8263961 |
| 902 | NM_001144831.1 | 409.402915 | -10.299492 | 1.31626084 | -7.824811 |

| | | | | | |
|---|---|---|---|---|---|
| 903 | NM_001330682.1 | 396.914486 | -10.254845 | 1.31063853 | -7.8243121 |
| 904 | NM_022766.5 | 415.272364 | -6.2263292 | 0.79576285 | -7.8243527 |
| 905 | NM_003746.2 | 409.052458 | -10.298219 | 1.3163407 | -7.8233687 |
| 906 | NM_001000.3 | 727.948529 | -11.129848 | 1.42268891 | -7.8231074 |
| 907 | NM_004787.3 | 286.86773 | -9.7874526 | 1.25113926 | -7.8228323 |
| 908 | NM_002714.3 | 254.563032 | -9.6137085 | 1.22900306 | -7.8223634 |
| 909 | NM_016107.4 | 332.892316 | -10.000993 | 1.27893027 | -7.8198108 |
| 910 | NM_001010858.2 | 748.271127 | -8.1673687 | 1.0448881 | -7.8165008 |
| 911 | NM_018060.3 | 245.259897 | -9.5601496 | 1.22308537 | -7.8164205 |
| 912 | NM_014686.3 | 138.766083 | -4.8817086 | 0.62457027 | -7.8161079 |
| 913 | NM_001252507.2 | 691.168237 | -6.205092 | 0.79419022 | -7.8131055 |
| 914 | NM_002266.3 | 426.280936 | -10.357914 | 1.32579076 | -7.8126306 |
| 915 | NM_006401.2 | 311.118309 | -9.9035397 | 1.26769461 | -7.8122441 |
| 916 | NM_003100.3 | 172.187231 | -4.1689991 | 0.53371123 | -7.8113387 |
| 917 | NM_005803.3 | 452.935214 | -10.445386 | 1.33728548 | -7.810887 |
| 918 | NM_001319236.1 | 682.78891 | -11.037461 | 1.41336656 | -7.8093406 |
| 919 | NM_020713.2 | 337.732898 | -10.021972 | 1.28337237 | -7.8090911 |
| 920 | NM_003378.3 | 784.688303 | -11.238233 | 1.43920574 | -7.8086355 |
| 921 | NM_016626.4 | 211.380057 | -5.9869472 | 0.76678069 | -7.8079002 |
| 922 | NM_001111077.1 | 306.876748 | -9.8827999 | 1.26584985 | -7.807245 |
| 923 | NM_016337.2 | 383.697861 | -10.206159 | 1.30738916 | -7.8065195 |
| 924 | NM_207577.1 | 423.226734 | -10.347527 | 1.3258495 | -7.8044507 |
| 925 | NM_033426.2 | 156.413972 | -3.0026321 | 0.38481442 | -7.8028055 |
| 926 | NM_139072.3 | 503.142187 | -10.597031 | 1.35811537 | -7.8027474 |
| 927 | NM_002414.4 | 446.695403 | -10.425533 | 1.33633776 | -7.8015703 |
| 928 | NM_007006.2 | 411.389388 | -10.306332 | 1.32131087 | -7.8000814 |
| 929 | NM_004731.4 | 35.9308404 | 10.0804202 | 1.29261017 | 7.79849987 |
| 930 | NM_002293.3 | 431.779288 | -8.511652 | 1.09164076 | -7.7971182 |
| 931 | NM_001017.2 | 761.431012 | -11.194752 | 1.43582511 | -7.7967381 |
| 932 | NM_079423.3 | 514.260871 | -10.628545 | 1.36341645 | -7.7955238 |
| 933 | NM_001190470.1 | 10547.132 | -7.5439814 | 0.9677907 | -7.7950546 |
| 934 | NM_002537.3 | 326.599628 | -9.9735563 | 1.27947172 | -7.795058 |
| 935 | NM_017794.4 | 148.845094 | -5.3841037 | 0.6907153 | -7.794968 |
| 936 | NM_152862.2 | 556.100252 | -7.7250544 | 0.9910604 | -7.7947363 |
| 937 | NM_004593.2 | 381.076813 | -10.195938 | 1.3082338 | -7.7936662 |
| 938 | NM_004819.2 | 301.080229 | -9.8561302 | 1.26471656 | -7.7931535 |
| 939 | NM_003565.2 | 337.177885 | -10.019487 | 1.28579694 | -7.7924335 |
| 940 | NM_003333.4 | 454.223422 | -10.44942 | 1.34107869 | -7.791802 |
| 941 | NM_003119.3 | 275.749634 | -9.7294497 | 1.24875202 | -7.7913385 |
| 942 | NM_003179.2 | 671.477242 | -11.013418 | 1.41363679 | -7.7908397 |
| 943 | NM_006876.2 | 329.308646 | -9.9855604 | 1.28178112 | -7.7903787 |
| 944 | NM_001040715.1 | 410.187821 | -10.302277 | 1.32248967 | -7.7900624 |
| 945 | NM_014396.3 | 246.678892 | -9.5686165 | 1.22859114 | -7.7882838 |

| | | | | | |
|---|---|---|---|---|---|
| 946 | NM_005766.3 | 424.223733 | -10.350887 | 1.3291727 | -7.7874659 |
| 947 | NM_015904.3 | 394.103557 | -10.24449 | 1.3155201 | -7.7874069 |
| 948 | NM_001291324.1 | 233.564364 | -5.899618 | 0.7577256 | -7.7859557 |
| 949 | NM_003365.2 | 381.35386 | -10.197212 | 1.30981237 | -7.7852465 |
| 950 | NM_005231.3 | 312.373662 | -9.9098995 | 1.2730137 | -7.7845976 |
| 951 | NM_015130.2 | 247.438009 | -9.5742781 | 1.23002781 | -7.78379 |
| 952 | NM_012319.3 | 490.204254 | -10.559353 | 1.3567098 | -7.7830593 |
| 953 | NM_001032296.3 | 227.340558 | -9.4518042 | 1.21454274 | -7.7821916 |
| 954 | NM_001145263.1 | 348.208374 | -10.066116 | 1.293754 | -7.7805485 |
| 955 | NM_004984.2 | 617.466347 | -10.892444 | 1.40012046 | -7.779648 |
| 956 | NM_003009.2 | 367.329701 | -10.143049 | 1.30388853 | -7.7790766 |
| 957 | NM_004738.4 | 269.958998 | -9.6990692 | 1.24682979 | -7.7789841 |
| 958 | NM_014294.5 | 252.207959 | -9.6015961 | 1.23457396 | -7.7772547 |
| 959 | NM_004518.5 | 616.408188 | -10.889897 | 1.40035885 | -7.7765042 |
| 960 | NM_014287.3 | 376.37941 | -10.178343 | 1.3088875 | -7.7763316 |
| 961 | NM_004958.3 | 248.939495 | -6.8231692 | 0.87766762 | -7.7742064 |
| 962 | NM_001530.3 | 328.271302 | -9.9808902 | 1.28388489 | -7.7739759 |
| 963 | NM_024757.4 | 277.133948 | -9.7364284 | 1.25247739 | -7.7737358 |
| 964 | NM_003737.3 | 737.585854 | -10.186414 | 1.31041499 | -7.7734261 |
| 965 | NM_004703.5 | 322.402231 | -9.9547356 | 1.28071246 | -7.7728107 |
| 966 | NM_001204054.1 | 30.5283349 | 9.84700184 | 1.26697084 | 7.77208249 |
| 967 | NM_023015.4 | 262.104892 | -9.6563933 | 1.24245622 | -7.772019 |
| 968 | NM_001641.3 | 336.742022 | -10.01751 | 1.28922413 | -7.7701853 |
| 969 | NM_001005332.1 | 443.06329 | -8.5439918 | 1.09964744 | -7.7697555 |
| 970 | NM_003640.4 | 233.86917 | -9.4905684 | 1.22146573 | -7.7698197 |
| 971 | NM_005964.3 | 893.922323 | -10.463911 | 1.34678294 | -7.76956 |
| 972 | NM_001142276.1 | 345.207795 | -10.053559 | 1.29447948 | -7.7664875 |
| 973 | NM_001040456.2 | 301.529399 | -9.8586645 | 1.26948502 | -7.7658769 |
| 974 | NR_003491.3 | 312.561652 | -9.909317 | 1.27601632 | -7.7658231 |
| 975 | NM_003061.2 | 478.762147 | -10.525258 | 1.35541418 | -7.7653447 |
| 976 | NM_001040134.1 | 558.64998 | -10.747922 | 1.38433363 | -7.763968 |
| 977 | NM_002317.6 | 326.744139 | -9.9746709 | 1.28479987 | -7.7635989 |
| 978 | NM_004905.2 | 307.896874 | -9.8885218 | 1.27378833 | -7.7630809 |
| 979 | NM_001678.4 | 227.488744 | -9.4510157 | 1.21760697 | -7.7619593 |
| 980 | NM_001106.3 | 219.731014 | -6.2539692 | 0.80574124 | -7.7617588 |
| 981 | NM_032208.2 | 269.644645 | -9.6975685 | 1.2495743 | -7.7606978 |
| 982 | NM_033446.2 | 311.20002 | -7.1040035 | 0.91540958 | -7.7604645 |
| 983 | NM_001129.4 | 391.097085 | -8.3763168 | 1.07942302 | -7.7599946 |
| 984 | NM_003498.5 | 435.497145 | -10.388675 | 1.3392089 | -7.7573223 |
| 985 | NM_006708.2 | 316.126687 | -9.9265215 | 1.27961921 | -7.7574027 |
| 986 | NM_001543.4 | 379.482313 | -10.190047 | 1.31372749 | -7.7565908 |
| 987 | NM_022748.11 | 341.22863 | -10.037198 | 1.29415537 | -7.7557903 |
| 988 | NM_001012761.2 | 408.334815 | -10.295868 | 1.3276764 | -7.7548024 |

| | | | | | |
|---|---|---|---|---|---|
| 989 | NM_198968.3 | 248.308348 | -9.5784915 | 1.23515117 | -7.7549143 |
| 990 | NM_006411.3 | 406.007138 | -10.287511 | 1.32675822 | -7.7538704 |
| 991 | NM_017798.3 | 250.308216 | -9.589516 | 1.23683879 | -7.7532465 |
| 992 | NM_006295.2 | 389.789706 | -10.228769 | 1.3194186 | -7.7524818 |
| 993 | NM_001042544.1 | 266.630178 | -9.681524 | 1.24916664 | -7.7503863 |
| 994 | NM_018509.3 | 331.06459 | -9.9931568 | 1.28939224 | -7.7502846 |
| 995 | NM_002147.3 | 504.976119 | -10.602387 | 1.36809546 | -7.7497425 |
| 996 | NM_017747.2 | 206.396554 | -6.5412283 | 0.84407286 | -7.7496015 |
| 997 | NM_002428.3 | 571.817776 | -10.781675 | 1.39131306 | -7.7492807 |
| 998 | NM_001199864.1 | 237.731193 | -9.5150167 | 1.22817914 | -7.7472547 |
| 999 | NM_002585.3 | 616.8307 | -10.89088 | 1.4057622 | -7.7473134 |
| 1000 | NM_003908.4 | 298.992234 | -9.8460766 | 1.2709363 | -7.7471047 |
| 1001 | NM_001867.2 | 436.997654 | -10.393651 | 1.34172557 | -7.7464802 |
| 1002 | NM_002184.3 | 383.524245 | -10.205623 | 1.31745216 | -7.7464849 |
| 1003 | NM_001792.4 | 746.760039 | -7.705635 | 0.99500639 | -7.7443071 |
| 1004 | NM_014615.3 | 352.407398 | -10.083508 | 1.30208608 | -7.744118 |
| 1005 | NM_001143998.1 | 275.502379 | -9.7281083 | 1.25623707 | -7.7438475 |
| 1006 | NM_014786.3 | 271.290284 | -9.7065527 | 1.2536385 | -7.7427047 |
| 1007 | NM_001100399.1 | 265.707015 | -9.6755312 | 1.24976065 | -7.7419074 |
| 1008 | NM_033008.2 | 339.846006 | -10.031009 | 1.2957159 | -7.7416733 |
| 1009 | NM_002225.3 | 212.928109 | -9.357383 | 1.20876311 | -7.7412877 |
| 1010 | NM_001619.3 | 327.351628 | -9.9766473 | 1.28882955 | -7.7408586 |
| 1011 | NM_001287.5 | 306.709834 | -9.8830072 | 1.27679809 | -7.740462 |
| 1012 | NM_014246.1 | 260.576221 | -6.8446757 | 0.88444735 | -7.7389295 |
| 1013 | NM_004136.3 | 253.706233 | -9.6088814 | 1.24167422 | -7.7386494 |
| 1014 | NM_057161.3 | 378.991159 | -10.188088 | 1.31669233 | -7.7376376 |
| 1015 | NM_015902.5 | 286.238747 | -9.7830663 | 1.26444346 | -7.7370532 |
| 1016 | NM_001626.5 | 214.549249 | -9.3673398 | 1.21078016 | -7.7366149 |
| 1017 | NM_001130144.2 | 221.417156 | -9.4123816 | 1.2167929 | -7.7354015 |
| 1018 | NM_003040.3 | 275.756943 | -9.730343 | 1.25794047 | -7.7351378 |
| 1019 | NM_001079864.2 | 263.997503 | -9.6667907 | 1.24993256 | -7.7338498 |
| 1020 | NM_015270.4 | 301.890923 | -9.8601362 | 1.27500868 | -7.7333875 |
| 1021 | NM_014631.2 | 238.078375 | -9.5176127 | 1.23089107 | -7.7322948 |
| 1022 | NM_001288732.1 | 224.202915 | -9.4321512 | 1.22016897 | -7.7302008 |
| 1023 | NM_031449.3 | 267.51667 | -9.6857198 | 1.25298671 | -7.7301058 |
| 1024 | NM_001039550.1 | 243.146186 | -9.547589 | 1.23517826 | -7.7297256 |
| 1025 | NM_031266.2 | 378.174775 | -10.184973 | 1.31779046 | -7.7288256 |
| 1026 | NM_002214.2 | 154.167422 | -3.7629159 | 0.48692291 | -7.72795 |
| 1027 | NM_032039.3 | 272.186829 | -9.7111098 | 1.25661886 | -7.7279676 |
| 1028 | NM_001252102.1 | 614.05834 | -10.884301 | 1.40853113 | -7.7274126 |
| 1029 | NM_015327.2 | 281.874952 | -9.7608489 | 1.26343792 | -7.7256261 |
| 1030 | NM_182931.2 | 257.684236 | -9.6313475 | 1.24669144 | -7.7255263 |
| 1031 | NM_033000.3 | 277.752759 | -9.7387642 | 1.26062656 | -7.7253363 |

| | | | | | |
|---|---|---|---|---|---|
| 1032 | NM_001322410.1 | 255.116946 | -9.6166617 | 1.24485188 | -7.7251453 |
| 1033 | NM_000801.4 | 378.260159 | -10.185365 | 1.31859824 | -7.7243886 |
| 1034 | NM_014389.2 | 321.498029 | -9.9508064 | 1.2882816 | -7.7240927 |
| 1035 | NM_001145638.2 | 411.148 | -10.30559 | 1.33431962 | -7.7234794 |
| 1036 | NM_005077.4 | 273.392206 | -9.7169125 | 1.25814988 | -7.7231757 |
| 1037 | NM_001332.3 | 532.710165 | -10.679206 | 1.38283374 | -7.7226973 |
| 1038 | NM_001171613.1 | 418.326323 | -10.330595 | 1.33774636 | -7.7223872 |
| 1039 | NM_000636.3 | 241.099097 | -9.5355505 | 1.23514245 | -7.7202031 |
| 1040 | NM_005499.2 | 344.332874 | -10.049581 | 1.30203438 | -7.7183685 |
| 1041 | NM_013403.2 | 313.710506 | -9.9149524 | 1.28483446 | -7.7169104 |
| 1042 | NM_020791.2 | 335.398943 | -10.011572 | 1.29745684 | -7.7163048 |
| 1043 | NM_181672.2 | 261.867539 | -9.6544697 | 1.25119969 | -7.7161702 |
| 1044 | NM_000391.3 | 246.020852 | -9.5658092 | 1.23979704 | -7.7156251 |
| 1045 | NM_014849.4 | 464.792162 | -10.482774 | 1.35871654 | -7.7152032 |
| 1046 | NM_004175.4 | 313.123738 | -9.9126873 | 1.28492926 | -7.7145783 |
| 1047 | NM_005937.3 | 325.028266 | -3.6362667 | 0.47134998 | -7.7145791 |
| 1048 | NM_004538.5 | 469.506705 | -10.49718 | 1.36083574 | -7.7137742 |
| 1049 | NM_000414.3 | 222.477959 | -9.419929 | 1.22126478 | -7.7132569 |
| 1050 | NM_001788.5 | 408.021506 | -10.294743 | 1.33478055 | -7.7126858 |
| 1051 | NM_014003.3 | 245.722411 | -6.4992288 | 0.84267887 | -7.7125808 |
| 1052 | NM_001856.3 | 190.200462 | -6.1110962 | 0.79239648 | -7.7121697 |
| 1053 | NM_020719.2 | 258.55708 | -9.6363651 | 1.24956266 | -7.7117903 |
| 1054 | NM_138370.2 | 304.158014 | -9.871378 | 1.28013739 | -7.7111863 |
| 1055 | NM_004656.3 | 358.056387 | -10.106146 | 1.31070635 | -7.7104582 |
| 1056 | NM_014901.4 | 317.170223 | -9.9313101 | 1.28810936 | -7.7099899 |
| 1057 | NM_030981.2 | 307.977514 | -9.8890398 | 1.28261451 | -7.7100639 |
| 1058 | NM_001098209.1 | 236.478583 | -9.5089289 | 1.23337359 | -7.7096908 |
| 1059 | NM_001252512.1 | 37.994524 | 10.1698798 | 1.3193922 | 7.70800358 |
| 1060 | NM_032800.2 | 382.220239 | -10.200626 | 1.3235301 | -7.7071358 |
| 1061 | NM_015177.1 | 275.914888 | -9.7301678 | 1.26314245 | -7.7031437 |
| 1062 | NM_177438.2 | 243.940647 | -9.5534776 | 1.24020998 | -7.703113 |
| 1063 | NM_005629.3 | 410.137552 | -10.302433 | 1.33764002 | -7.7019471 |
| 1064 | NM_002613.4 | 247.966311 | -9.5757832 | 1.24355494 | -7.7003299 |
| 1065 | NM_001001520.2 | 284.52434 | -9.7742737 | 1.269375 | -7.7000679 |
| 1066 | NM_001954.4 | 312.282579 | -9.9094687 | 1.28693428 | -7.7000581 |
| 1067 | NM_002796.2 | 352.089328 | -10.081952 | 1.30943045 | -7.6994939 |
| 1068 | NM_014680.3 | 365.91785 | -10.138183 | 1.31683322 | -7.6989117 |
| 1069 | NM_017871.5 | 253.955438 | -9.6103585 | 1.24829738 | -7.6987732 |
| 1070 | NM_019106.5 | 1149.32022 | -10.826532 | 1.4064855 | -7.6975781 |
| 1071 | NM_001100623.1 | 279.425429 | -9.7475643 | 1.26639741 | -7.6970817 |
| 1072 | NM_001300995.1 | 26.7292157 | 9.66708952 | 1.25596671 | 7.69693134 |
| 1073 | NM_030939.4 | 320.911932 | -9.9479768 | 1.29293375 | -7.6941118 |
| 1074 | NM_001308373.1 | 319.327161 | -9.9409562 | 1.29222726 | -7.6928854 |

| | | | | | |
|---|---|---|---|---|---|
| 1075 | NM_001288976.1 | 381.838162 | -10.198742 | 1.32577109 | -7.6926868 |
| 1076 | NM_001419.2 | 308.4173 | -9.8907691 | 1.28577859 | -7.6924357 |
| 1077 | NM_021259.2 | 235.510956 | -9.5027325 | 1.23545139 | -7.6917089 |
| 1078 | NM_001172437.2 | 439.510947 | -10.401944 | 1.35241131 | -7.6914054 |
| 1079 | NM_003025.3 | 293.305572 | -9.8183903 | 1.27679062 | -7.6898985 |
| 1080 | NM_001145056.1 | 349.105997 | -10.069923 | 1.30959575 | -7.6893373 |
| 1081 | NM_003870.3 | 212.906014 | -9.3576092 | 1.21719217 | -7.6878651 |
| 1082 | NM_001320214.1 | 380.729737 | -9.2314955 | 1.20093235 | -7.6869405 |
| 1083 | NM_005735.3 | 258.380335 | -9.635221 | 1.25350741 | -7.6866087 |
| 1084 | NM_015850.3 | 201.671503 | -9.2784725 | 1.20738324 | -7.6847783 |
| 1085 | NM_006047.5 | 257.854567 | -9.6319221 | 1.25356741 | -7.6836092 |
| 1086 | NM_001198802.1 | 279.825623 | -9.751035 | 1.26915073 | -7.6831181 |
| 1087 | NM_006947.3 | 306.200339 | -9.8803926 | 1.2859952 | -7.6830711 |
| 1088 | NM_002087.3 | 390.919895 | -10.233199 | 1.33231913 | -7.6807415 |
| 1089 | NM_016732.2 | 335.007674 | -10.010223 | 1.30335468 | -7.6803526 |
| 1090 | NM_001110.3 | 241.20504 | -9.5362287 | 1.24168522 | -7.6800694 |
| 1091 | NM_022818.4 | 428.448667 | -10.365211 | 1.3496384 | -7.6799915 |
| 1092 | NM_015331.2 | 272.352021 | -9.7118108 | 1.26464544 | -7.6794732 |
| 1093 | NM_004380.2 | 222.358774 | -9.4199234 | 1.22671095 | -7.6790082 |
| 1094 | NM_024038.3 | 394.28247 | -10.245182 | 1.33451229 | -7.6770985 |
| 1095 | NM_005065.5 | 329.276642 | -9.9853093 | 1.30073902 | -7.6766431 |
| 1096 | NM_007107.4 | 311.718369 | -9.9062935 | 1.29048676 | -7.6764008 |
| 1097 | NM_005777.2 | 225.684236 | -9.4398906 | 1.22994505 | -7.6750507 |
| 1098 | NM_014764.3 | 261.605566 | -9.6527907 | 1.25790052 | -7.6737314 |
| 1099 | NM_016292.2 | 287.664908 | -9.7903994 | 1.27583568 | -7.673715 |
| 1100 | NM_033133.4 | 294.073412 | -9.8224004 | 1.28022992 | -7.6723722 |
| 1101 | NM_001080533.2 | 199.132722 | -9.2606191 | 1.20710459 | -7.671762 |
| 1102 | NR_003580.2 | 440.104617 | -10.403871 | 1.35617169 | -7.6715002 |
| 1103 | NM_001098504.1 | 287.439208 | -9.7888827 | 1.27616459 | -7.6705487 |
| 1104 | NM_014812.2 | 387.72763 | -10.22098 | 1.33252713 | -7.6703731 |
| 1105 | NM_014268.3 | 508.226944 | -10.611503 | 1.3835637 | -7.6696889 |
| 1106 | NM_007353.2 | 318.666082 | -9.938052 | 1.29580143 | -7.669425 |
| 1107 | NM_014823.2 | 242.620606 | -7.1820419 | 0.93645139 | -7.6694231 |
| 1108 | NM_012249.3 | 223.871888 | -9.4294897 | 1.22958466 | -7.6688414 |
| 1109 | NM_006107.3 | 300.80799 | -9.8545857 | 1.28503924 | -7.6687041 |
| 1110 | NM_012111.2 | 274.488163 | -9.7228596 | 1.26793795 | -7.6682456 |
| 1111 | NM_001170634.1 | 394.150122 | -10.245014 | 1.33609539 | -7.6678758 |
| 1112 | NM_004247.3 | 301.827731 | -9.8598455 | 1.28590047 | -7.6676584 |
| 1113 | NM_016312.2 | 257.751544 | -9.6317244 | 1.25618467 | -7.667443 |
| 1114 | NM_032777.9 | 381.559617 | -7.1650962 | 0.93450536 | -7.6672607 |
| 1115 | NM_032808.6 | 361.505722 | -10.119964 | 1.32021235 | -7.665406 |
| 1116 | NM_001294.3 | 388.146678 | -10.222745 | 1.33384087 | -7.6641414 |
| 1117 | NM_014719.2 | 385.981807 | -10.214699 | 1.33298806 | -7.6630089 |

| | | | | | |
|---|---|---|---|---|---|
| 1118 | NM_024516.3 | 216.035859 | -9.3783901 | 1.22391213 | -7.6626335 |
| 1119 | NM_013253.4 | 487.075077 | -10.550186 | 1.37692431 | -7.6621395 |
| 1120 | NM_000884.2 | 339.065376 | -10.027549 | 1.30877939 | -7.6617566 |
| 1121 | NM_014298.4 | 282.534832 | -9.7649288 | 1.27451497 | -7.6616823 |
| 1122 | NM_001271641.1 | 450.923418 | -10.438866 | 1.3626462 | -7.660731 |
| 1123 | NM_006801.2 | 394.369741 | -8.4583146 | 1.10413526 | -7.6605782 |
| 1124 | NM_024612.4 | 200.081331 | -9.2657382 | 1.20959224 | -7.6602163 |
| 1125 | NM_003969.3 | 362.606694 | -10.124428 | 1.32174666 | -7.6598853 |
| 1126 | NM_001096.2 | 424.039436 | -10.350135 | 1.35131204 | -7.6593231 |
| 1127 | NM_001920.4 | 299.38397 | -9.8485577 | 1.28582028 | -7.6593579 |
| 1128 | NM_138769.2 | 293.740751 | -7.309041 | 0.95433265 | -7.658798 |
| 1129 | NM_004768.3 | 257.537499 | -9.6309191 | 1.25799919 | -7.6557435 |
| 1130 | NM_007270.4 | 248.520652 | -9.5797193 | 1.25134746 | -7.655523 |
| 1131 | NM_014646.2 | 212.17172 | -6.0268256 | 0.78729055 | -7.655148 |
| 1132 | NM_001131016.1 | 248.333724 | -9.5779598 | 1.25138911 | -7.6538622 |
| 1133 | NM_001670.2 | 303.295044 | -6.5563443 | 0.85671447 | -7.6528931 |
| 1134 | NM_032430.1 | 540.243743 | -10.699598 | 1.39845132 | -7.6510335 |
| 1135 | NM_022754.6 | 300.331069 | -9.8522992 | 1.28797489 | -7.6494498 |
| 1136 | NM_001195427.1 | 317.548917 | -9.9328259 | 1.29852225 | -7.6493306 |
| 1137 | NM_001152.4 | 381.659716 | -7.7283677 | 1.01038741 | -7.6489153 |
| 1138 | NM_031412.2 | 348.619789 | -10.067704 | 1.3162753 | -7.6486307 |
| 1139 | NM_001007553.2 | 238.897799 | -9.5221524 | 1.24499687 | -7.6483344 |
| 1140 | NM_016322.3 | 345.272057 | -10.053748 | 1.31480644 | -7.6465615 |
| 1141 | NM_139062.2 | 228.839471 | -9.4607755 | 1.2373384 | -7.6460696 |
| 1142 | NM_001033930.2 | 356.431802 | -10.099631 | 1.32095038 | -7.6457311 |
| 1143 | NM_002688.5 | 438.275982 | -10.397713 | 1.35994501 | -7.6456865 |
| 1144 | NM_013245.2 | 358.683864 | -10.108673 | 1.32210924 | -7.6458685 |
| 1145 | NM_000179.2 | 266.789031 | -9.6814723 | 1.26632756 | -7.6453144 |
| 1146 | NM_002812.4 | 385.61157 | -10.213277 | 1.33597593 | -7.644806 |
| 1147 | NM_015151.3 | 193.15075 | -9.2158541 | 1.20551732 | -7.6447297 |
| 1148 | NM_003133.5 | 688.687844 | -9.1216944 | 1.19329027 | -7.6441539 |
| 1149 | NM_001448.2 | 495.428944 | -10.574831 | 1.38348002 | -7.6436459 |
| 1150 | NM_001025091.1 | 278.10057 | -9.7417279 | 1.27465523 | -7.6426375 |
| 1151 | NM_001077488.3 | 545.916362 | -10.714799 | 1.40197667 | -7.6426373 |
| 1152 | NM_001286134.1 | 310.335552 | -9.8999689 | 1.2953607 | -7.6426349 |
| 1153 | NM_014971.1 | 455.144957 | -10.45243 | 1.36771609 | -7.6422512 |
| 1154 | NM_006224.3 | 387.192335 | -8.4314777 | 1.10331839 | -7.6419262 |
| 1155 | NM_021807.3 | 242.86546 | -9.5457965 | 1.24916461 | -7.6417442 |
| 1156 | NM_019609.4 | 231.162635 | -9.4762881 | 1.24011049 | -7.6414869 |
| 1157 | NM_005032.6 | 196.837118 | -9.2440607 | 1.20976369 | -7.6412119 |
| 1158 | NM_145716.3 | 421.725606 | -10.342018 | 1.35366454 | -7.6400153 |
| 1159 | NM_198830.1 | 462.967379 | -6.9800968 | 0.91370207 | -7.6393576 |
| 1160 | NM_004459.6 | 223.248784 | -9.4235614 | 1.23360785 | -7.6390251 |

| | | | | | |
|---|---|---|---|---|---|
| 1161 | NM_003339.2 | 366.581262 | -10.14014 | 1.3274463 | -7.6388322 |
| 1162 | NM_030912.2 | 220.468492 | -9.4064639 | 1.23144741 | -7.6385429 |
| 1163 | NM_005631.4 | 217.005475 | -9.3848903 | 1.22864183 | -7.6384265 |
| 1164 | NM_001270975.1 | 264.332282 | -9.6683513 | 1.26590291 | -7.6375141 |
| 1165 | NM_002794.4 | 337.493578 | -10.020857 | 1.31207338 | -7.6374212 |
| 1166 | NM_003200.3 | 296.977126 | -9.8359896 | 1.28804258 | -7.6363855 |
| 1167 | NM_144703.2 | 306.770787 | -9.8832106 | 1.29425198 | -7.6362337 |
| 1168 | NM_001135586.1 | 248.907798 | -9.5813092 | 1.25485717 | -7.6353782 |
| 1169 | NM_014713.4 | 453.515322 | -8.5636114 | 1.12174047 | -7.6342181 |
| 1170 | NM_014856.2 | 259.25414 | -9.640734 | 1.26298791 | -7.6332749 |
| 1171 | NM_017457.5 | 425.085457 | -10.353963 | 1.35668485 | -7.6318114 |
| 1172 | NM_002954.5 | 576.51041 | -10.793337 | 1.41435055 | -7.6313025 |
| 1173 | NR_126012.1 | 100.367263 | 4.16639758 | 0.5461678 | 7.62842033 |
| 1174 | NM_013446.3 | 294.895967 | -9.826162 | 1.28815179 | -7.6281088 |
| 1175 | NM_199141.1 | 318.638138 | -7.2006019 | 0.9441724 | -7.6263635 |
| 1176 | NM_001009998.3 | 229.017173 | -9.4614171 | 1.24067823 | -7.626004 |
| 1177 | NM_001199383.1 | 287.145528 | -9.7879064 | 1.28350088 | -7.6259445 |
| 1178 | NM_004516.3 | 357.363887 | -10.103207 | 1.32486516 | -7.6258379 |
| 1179 | NM_015197.3 | 413.145448 | -10.312597 | 1.35262633 | -7.6241287 |
| 1180 | NM_031262.3 | 256.923538 | -9.6271854 | 1.26272848 | -7.6241136 |
| 1181 | NM_005392.3 | 277.121163 | -9.7364052 | 1.27727053 | -7.6228215 |
| 1182 | NM_006315.5 | 268.375606 | -6.5809292 | 0.86333199 | -7.6227098 |
| 1183 | NM_002709.2 | 341.303362 | -10.037114 | 1.31681395 | -7.6222715 |
| 1184 | NM_003756.2 | 435.241224 | -10.387781 | 1.3629111 | -7.6217598 |
| 1185 | NR_037901.1 | 198.5769 | 3.46985527 | 0.45530221 | 7.62099369 |
| 1186 | NM_005216.4 | 607.858667 | -6.2921154 | 0.82570585 | -7.6202868 |
| 1187 | NM_004341.4 | 203.507281 | -9.2918004 | 1.21954796 | -7.6190528 |
| 1188 | NM_006426.2 | 464.218979 | -10.480815 | 1.37589481 | -7.6174535 |
| 1189 | NM_005255.3 | 217.290512 | -9.3856695 | 1.23217422 | -7.6171611 |
| 1190 | NM_006546.3 | 286.042832 | -9.7825962 | 1.28426881 | -7.6172497 |
| 1191 | NM_017780.3 | 218.947847 | -9.3958149 | 1.23356636 | -7.6167892 |
| 1192 | NM_174921.2 | 221.484904 | -9.4129845 | 1.23582573 | -7.6167572 |
| 1193 | NR_046252.1 | 299.994049 | -9.8507918 | 1.29328366 | -7.6168841 |
| 1194 | NM_006795.3 | 279.870398 | -9.7510346 | 1.28035727 | -7.6158701 |
| 1195 | NM_000455.4 | 246.946646 | -9.5700584 | 1.25710324 | -7.6127863 |
| 1196 | NM_004583.3 | 329.942617 | -9.9881347 | 1.31210127 | -7.61232 |
| 1197 | NM_017905.4 | 207.661122 | -9.3215939 | 1.22453515 | -7.6123531 |
| 1198 | NM_000014.5 | 558.502389 | -10.747746 | 1.41200736 | -7.6116783 |
| 1199 | NM_000527.4 | 388.417607 | -10.223923 | 1.34345829 | -7.6101525 |
| 1200 | NM_002210.4 | 302.041966 | -9.8606722 | 1.29571049 | -7.6102434 |
| 1201 | NM_002273.3 | 324.173235 | -9.9632593 | 1.30948345 | -7.6085415 |
| 1202 | NM_178537.4 | 776.029507 | -7.7668638 | 1.02081846 | -7.6084673 |
| 1203 | NM_006502.2 | 198.906597 | -9.2585094 | 1.21691371 | -7.6081889 |

| | | | | | |
|---|---|---|---|---|---|
| 1204 | NM_018649.2 | 359.355052 | -10.111402 | 1.32908708 | -7.6077797 |
| 1205 | NM_053275.3 | 486.337631 | -10.547925 | 1.38645065 | -7.6078622 |
| 1206 | NM_001009993.3 | 412.874255 | -10.311528 | 1.35543441 | -7.607545 |
| 1207 | NM_018026.3 | 324.003174 | -9.962156 | 1.30952437 | -7.6074613 |
| 1208 | NR_027409.1 | 690.945256 | -7.2469012 | 0.95265218 | -7.6070799 |
| 1209 | NM_001024736.1 | 228.164355 | -9.4568024 | 1.24332483 | -7.6060594 |
| 1210 | NM_006698.3 | 349.150985 | -10.069968 | 1.3239847 | -7.6058042 |
| 1211 | NM_016131.4 | 449.897514 | -10.435601 | 1.37206038 | -7.6057885 |
| 1212 | NM_015097.2 | 323.41824 | -9.9593888 | 1.30949499 | -7.6055188 |
| 1213 | NM_003190.4 | 228.527863 | -9.4595833 | 1.24381888 | -7.605274 |
| 1214 | NM_005340.6 | 404.691737 | -10.282857 | 1.35225307 | -7.6042398 |
| 1215 | NM_130839.2 | 280.290453 | -9.7529323 | 1.28289553 | -7.602281 |
| 1216 | NM_006925.4 | 187.852102 | -9.1766148 | 1.20730067 | -7.6009358 |
| 1217 | NM_014738.5 | 195.235111 | -9.2300757 | 1.21441665 | -7.6004192 |
| 1218 | NM_014970.3 | 378.961425 | -10.188119 | 1.34055527 | -7.599925 |
| 1219 | NM_002014.3 | 481.629583 | -10.533888 | 1.38611641 | -7.5995693 |
| 1220 | NM_015720.3 | 634.3488 | -10.931284 | 1.43845247 | -7.5993364 |
| 1221 | NM_018998.3 | 291.161784 | -9.8077275 | 1.29058559 | -7.5994398 |
| 1222 | NM_001042424.2 | 303.125351 | -9.8657731 | 1.29827366 | -7.5991475 |
| 1223 | NM_015229.3 | 228.316395 | -4.8943704 | 0.64410401 | -7.5987268 |
| 1224 | NM_018639.4 | 536.107412 | -10.688493 | 1.4066541 | -7.5985226 |
| 1225 | NM_015874.4 | 310.907655 | -9.9019576 | 1.30318366 | -7.5982825 |
| 1226 | NM_020898.2 | 213.803234 | -9.3623007 | 1.2321536 | -7.5983227 |
| 1227 | NM_014548.3 | 562.276255 | -10.757318 | 1.41586169 | -7.5977181 |
| 1228 | NM_015466.3 | 231.378178 | -9.4762604 | 1.24731815 | -7.5973082 |
| 1229 | NM_004342.6 | 660.948586 | -10.990721 | 1.44680336 | -7.5965547 |
| 1230 | NR_047532.1 | 97.7398881 | 3.5383959 | 0.46579887 | 7.59640292 |
| 1231 | NM_004943.1 | 234.953138 | -9.4987183 | 1.25048212 | -7.5960449 |
| 1232 | NM_013325.4 | 245.208974 | -9.5596986 | 1.25858905 | -7.5955679 |
| 1233 | NM_001197294.1 | 348.225258 | -10.066233 | 1.3253341 | -7.5952419 |
| 1234 | NM_006464.3 | 285.093995 | -9.7774652 | 1.28756516 | -7.5937634 |
| 1235 | NM_001552.2 | 223.224996 | -9.4259324 | 1.24136279 | -7.5932133 |
| 1236 | NM_000687.3 | 300.347607 | -9.8529663 | 1.2976602 | -7.5928708 |
| 1237 | NM_022075.4 | 256.614682 | -9.6260907 | 1.2679472 | -7.5918703 |
| 1238 | NM_001037283.1 | 229.751247 | -9.4659877 | 1.24699604 | -7.5910328 |
| 1239 | NM_020474.3 | 263.584022 | -9.664197 | 1.27314554 | -7.590803 |
| 1240 | NM_012305.3 | 286.206555 | -9.7829356 | 1.28886306 | -7.5903607 |
| 1241 | NR_002196.2 | 546.538604 | -10.716512 | 1.41191958 | -7.5900299 |
| 1242 | NM_006572.5 | 252.862989 | -9.6040523 | 1.26543026 | -7.5895548 |
| 1243 | NM_014502.4 | 309.438881 | -9.8956883 | 1.30409987 | -7.5881369 |
| 1244 | NM_004960.3 | 577.435856 | -10.795613 | 1.42287382 | -7.587189 |
| 1245 | NM_006793.4 | 246.833588 | -9.5696244 | 1.26125064 | -7.5874089 |
| 1246 | NM_016057.2 | 309.902238 | -9.8976857 | 1.3045313 | -7.5871585 |

| | | | | | |
|---|---|---|---|---|---|
| 1247 | NM_031991.3 | 188.934678 | -9.1838447 | 1.21043065 | -7.5872539 |
| 1248 | NM_024042.3 | 223.39878 | -7.0607129 | 0.93076384 | -7.5859339 |
| 1249 | NM_004285.3 | 187.628949 | -9.1750175 | 1.20958697 | -7.5852483 |
| 1250 | NM_001130965.2 | 206.345388 | -9.3107228 | 1.22751657 | -7.5850078 |
| 1251 | NM_032493.3 | 289.912963 | -9.8017018 | 1.29229612 | -7.5847181 |
| 1252 | NM_007165.4 | 303.074269 | -9.8656496 | 1.30091758 | -7.5836085 |
| 1253 | NM_022735.3 | 168.018963 | -5.9263001 | 0.78176889 | -7.5806292 |
| 1254 | NM_014501.2 | 405.552116 | -10.285823 | 1.35690931 | -7.5803321 |
| 1255 | NM_001033853.1 | 373.077606 | -10.165703 | 1.34110893 | -7.5800723 |
| 1256 | NM_001348.2 | 258.828658 | -9.6380297 | 1.27165828 | -7.5791035 |
| 1257 | NM_003051.3 | 267.839965 | -9.6871225 | 1.27824936 | -7.5784294 |
| 1258 | NM_004691.4 | 320.039584 | -9.9444769 | 1.31220673 | -7.5784376 |
| 1259 | NM_002080.3 | 298.627609 | -9.8444455 | 1.29916071 | -7.5775425 |
| 1260 | NM_006312.5 | 294.984953 | -9.8265697 | 1.29684924 | -7.5772645 |
| 1261 | NM_015658.3 | 314.518719 | -9.9191474 | 1.30905265 | -7.577348 |
| 1262 | NM_003076.4 | 301.549475 | -9.8581162 | 1.30109037 | -7.5768112 |
| 1263 | NM_014740.3 | 243.072699 | -9.5472878 | 1.26014169 | -7.5763605 |
| 1264 | NM_001316.3 | 323.479222 | -9.9595286 | 1.31458383 | -7.5761837 |
| 1265 | NM_199054.2 | 195.406626 | -9.2335109 | 1.21879373 | -7.5759422 |
| 1266 | NM_002804.4 | 311.009086 | -9.9030115 | 1.30724163 | -7.5755019 |
| 1267 | NM_001276252.1 | 211.855756 | -9.3491243 | 1.23441655 | -7.5737192 |
| 1268 | NM_001020658.1 | 215.830861 | -9.3750784 | 1.23795077 | -7.5730624 |
| 1269 | NM_003755.4 | 317.457706 | -9.9325643 | 1.31167218 | -7.5724441 |
| 1270 | NM_001284214.1 | 224.91299 | -9.4366168 | 1.24623659 | -7.572091 |
| 1271 | NM_183376.2 | 192.334639 | -9.2105007 | 1.21650504 | -7.5712803 |
| 1272 | NM_002810.2 | 343.987321 | -10.048449 | 1.32740848 | -7.5699751 |
| 1273 | NM_001302883.1 | 282.360842 | -7.020907 | 0.92749585 | -7.5697449 |
| 1274 | NM_006367.3 | 341.841894 | -10.039193 | 1.3262791 | -7.5694425 |
| 1275 | NM_012286.2 | 272.2102 | -9.7104977 | 1.28302662 | -7.5684304 |
| 1276 | NM_000987.3 | 1177.50608 | -8.3874299 | 1.10823765 | -7.5682593 |
| 1277 | NM_003139.3 | 238.389772 | -9.5191323 | 1.25783426 | -7.5678749 |
| 1278 | NM_019884.2 | 322.467471 | -9.9551852 | 1.31578014 | -7.5659944 |
| 1279 | NM_002442.3 | 248.330644 | -4.773063 | 0.63090323 | -7.5654439 |
| 1280 | NM_001144.5 | 227.887633 | -9.4545437 | 1.24981111 | -7.5647781 |
| 1281 | NM_012477.3 | 245.351944 | -9.5608262 | 1.26387388 | -7.5646996 |
| 1282 | NM_003146.2 | 512.434854 | -9.6606164 | 1.27722259 | -7.5637688 |
| 1283 | NM_002482.3 | 317.311502 | -9.9318837 | 1.31342153 | -7.5618402 |
| 1284 | NM_152742.2 | 514.563501 | -10.629398 | 1.40595869 | -7.5602492 |
| 1285 | NM_001037663.1 | 423.119793 | -10.347008 | 1.36870185 | -7.5597241 |
| 1286 | NM_001048183.1 | 208.530085 | -9.3257094 | 1.23364505 | -7.5594754 |
| 1287 | NM_003472.3 | 274.793539 | -9.723767 | 1.28629326 | -7.5595257 |
| 1288 | NM_006496.3 | 335.740951 | -6.97606 | 0.92283059 | -7.5594156 |
| 1289 | NM_014700.3 | 240.948431 | -9.5345592 | 1.26126018 | -7.5595499 |

| | | | | | |
|---|---|---|---|---|---|
| 1290 | NM_022780.3 | 213.376824 | -9.3588751 | 1.23820775 | -7.5584046 |
| 1291 | NM_003792.3 | 361.56438 | -10.120201 | 1.33931003 | -7.5562797 |
| 1292 | NM_012207.2 | 263.636039 | -9.6643089 | 1.2794737 | -7.553347 |
| 1293 | NM_001271803.1 | 368.448703 | -10.147449 | 1.34355065 | -7.5527105 |
| 1294 | NM_001330071.1 | 479.725659 | -10.528184 | 1.393951 | -7.5527649 |
| 1295 | NM_003345.4 | 316.495928 | -9.9280536 | 1.31456848 | -7.5523289 |
| 1296 | NM_007144.2 | 241.436883 | -9.5377541 | 1.26313616 | -7.5508519 |
| 1297 | NM_001098833.1 | 316.638159 | -9.9289606 | 1.31518726 | -7.5494653 |
| 1298 | NM_013233.2 | 185.730139 | -9.160541 | 1.2136389 | -7.5479955 |
| 1299 | NM_030920.4 | 200.335651 | -9.2671425 | 1.22792604 | -7.5469875 |
| 1300 | NM_003885.2 | 613.047918 | -10.881967 | 1.4420016 | -7.546432 |
| 1301 | NM_012235.3 | 251.567982 | -9.5969415 | 1.2719124 | -7.545285 |
| 1302 | NM_004761.4 | 209.259424 | -9.332833 | 1.23701802 | -7.5446217 |
| 1303 | NM_001037984.2 | 263.084678 | -9.6620197 | 1.28072651 | -7.5441709 |
| 1304 | NM_001282611.1 | 592.67294 | -10.833313 | 1.43602473 | -7.5439599 |
| 1305 | NM_181891.2 | 319.499766 | -9.9415614 | 1.31813185 | -7.5421601 |
| 1306 | NM_006826.3 | 1659.54616 | -8.5486114 | 1.13347828 | -7.5419278 |
| 1307 | NM_020461.3 | 248.046831 | -9.5767677 | 1.26982477 | -7.5418026 |
| 1308 | NM_025241.2 | 296.641562 | -9.8345577 | 1.30418927 | -7.5407442 |
| 1309 | NM_015024.4 | 271.635517 | -9.707632 | 1.28745039 | -7.5401989 |
| 1310 | NM_006628.5 | 361.851121 | -10.121263 | 1.34234796 | -7.5399701 |
| 1311 | NR_110005.1 | 91.7935708 | 7.14885191 | 0.948137 | 7.53989341 |
| 1312 | NM_005625.3 | 219.934848 | -9.4034009 | 1.24739752 | -7.5384156 |
| 1313 | NM_152758.5 | 204.610461 | -9.2982995 | 1.2337 | -7.536921 |
| 1314 | NM_006791.3 | 942.948346 | -7.7118579 | 1.02324683 | -7.5366546 |
| 1315 | NM_003145.3 | 343.228123 | -10.045134 | 1.33292634 | -7.5361508 |
| 1316 | NM_138423.3 | 248.493178 | -9.5793765 | 1.27147265 | -7.5340799 |
| 1317 | NM_014921.4 | 439.301915 | -3.8786609 | 0.51497549 | -7.5317388 |
| 1318 | NM_032552.3 | 300.274148 | -9.8522042 | 1.30821736 | -7.5310147 |
| 1319 | NM_001012426.1 | 201.551863 | -9.2761005 | 1.23178171 | -7.5306367 |
| 1320 | NM_198241.2 | 321.59645 | -9.9520392 | 1.32164211 | -7.5300561 |
| 1321 | NM_198896.1 | 452.480822 | -10.443862 | 1.38707875 | -7.529394 |
| 1322 | NM_001282659.1 | 242.534496 | -9.5444779 | 1.26766678 | -7.5291694 |
| 1323 | NM_002730.3 | 353.540688 | -10.088059 | 1.34000342 | -7.5283831 |
| 1324 | NM_020410.2 | 250.097923 | -6.8395668 | 0.90853574 | -7.5281208 |
| 1325 | NM_020729.2 | 584.275665 | 2.59497357 | 0.34472333 | 7.52769942 |
| 1326 | NM_001042539.2 | 343.485985 | -10.046061 | 1.33475139 | -7.5265409 |
| 1327 | NM_003481.2 | 271.672985 | -9.707754 | 1.28983395 | -7.5263595 |
| 1328 | NM_000941.2 | 262.638668 | -9.6591246 | 1.28353601 | -7.5254021 |
| 1329 | NM_001322033.1 | 184.328278 | -9.1496433 | 1.21583462 | -7.5254013 |
| 1330 | NM_015965.6 | 352.637821 | -10.084215 | 1.34026802 | -7.5240289 |
| 1331 | NM_002392.5 | 191.643751 | -9.2045765 | 1.22342136 | -7.5236355 |
| 1332 | NM_144498.2 | 186.46316 | -9.1648656 | 1.21824245 | -7.5230227 |

| | | | | | |
|---|---|---|---|---|---|
| 1333 | NM_138501.5 | 250.952198 | -9.5936764 | 1.27527163 | -7.5228493 |
| 1334 | NM_014608.4 | 180.828227 | -9.1211216 | 1.21260505 | -7.5219228 |
| 1335 | NM_001104647.1 | 251.186016 | -9.594419 | 1.27566577 | -7.5211072 |
| 1336 | NM_006936.2 | 289.448297 | -9.7992584 | 1.30330922 | -7.5187517 |
| 1337 | NM_004888.3 | 343.726411 | -10.047338 | 1.3364679 | -7.5178295 |
| 1338 | NM_024835.4 | 200.459219 | -9.2697644 | 1.23303625 | -7.5178361 |
| 1339 | NM_004986.3 | 225.427735 | -9.4393779 | 1.25563998 | -7.5175831 |
| 1340 | NM_001146688.1 | 235.839931 | -9.5032289 | 1.26434266 | -7.5163397 |
| 1341 | NM_021814.4 | 293.449087 | -9.819165 | 1.30638785 | -7.5162709 |
| 1342 | NM_003418.4 | 271.842941 | -9.709423 | 1.29195603 | -7.5152891 |
| 1343 | NM_006990.4 | 239.759747 | -7.0308381 | 0.935553 | -7.5151682 |
| 1344 | NM_006022.3 | 437.971658 | -10.396809 | 1.38353865 | -7.5146502 |
| 1345 | NM_139067.3 | 394.385376 | -10.245664 | 1.36354161 | -7.5140094 |
| 1346 | NM_017748.4 | 97.7501617 | -4.3604162 | 0.5803338 | -7.5136347 |
| 1347 | NM_001039705.2 | 301.265432 | -9.8573502 | 1.3123128 | -7.5114334 |
| 1348 | NM_001039960.2 | 320.518969 | -9.9461947 | 1.32418814 | -7.5111644 |
| 1349 | NM_001469.4 | 274.479532 | -9.7227738 | 1.29446813 | -7.5110183 |
| 1350 | NM_004052.3 | 381.61327 | -10.197739 | 1.35780482 | -7.5104599 |
| 1351 | NM_002383.3 | 598.760025 | -10.847932 | 1.44508311 | -7.5067875 |
| 1352 | NM_079834.3 | 187.79173 | -9.1760566 | 1.22249843 | -7.5059864 |
| 1353 | NM_206852.2 | 1018.85312 | -10.652216 | 1.41922298 | -7.5056675 |
| 1354 | NM_015213.3 | 276.088455 | -9.7310311 | 1.2965443 | -7.5053595 |
| 1355 | NM_005153.2 | 240.832509 | -9.533904 | 1.27042116 | -7.5045223 |
| 1356 | NM_002834.4 | 356.150484 | -8.2617389 | 1.10099999 | -7.5038501 |
| 1357 | NM_014262.4 | 224.127429 | -9.4309017 | 1.25679991 | -7.5039007 |
| 1358 | NM_006486.2 | 194.273509 | -9.2256937 | 1.22949966 | -7.5036163 |
| 1359 | NM_005016.5 | 298.32997 | -9.8420787 | 1.31173482 | -7.5031009 |
| 1360 | NM_177401.4 | 530.667369 | -9.7110653 | 1.29441596 | -7.5022756 |
| 1361 | NM_004385.4 | 260.294563 | -9.6462439 | 1.2858361 | -7.5019234 |
| 1362 | NM_005339.4 | 313.578167 | -9.9147342 | 1.3218438 | -7.5006852 |
| 1363 | NM_001938.2 | 189.194726 | -9.1861154 | 1.22474369 | -7.5004391 |
| 1364 | NM_004359.1 | 267.008806 | -9.6827676 | 1.29128821 | -7.4985332 |
| 1365 | NM_001127173.2 | 538.2712 | -10.694374 | 1.42627298 | -7.4981254 |
| 1366 | NM_015584.4 | 274.789714 | -9.7243273 | 1.29690946 | -7.4980772 |
| 1367 | NM_001360.2 | 432.073693 | -10.37742 | 1.3840794 | -7.497706 |
| 1368 | NM_031463.4 | 325.870962 | -9.9701612 | 1.32980181 | -7.497479 |
| 1369 | NM_001315507.1 | 217.223722 | -9.3849337 | 1.25177999 | -7.4972709 |
| 1370 | NM_006816.2 | 251.053409 | -9.5941129 | 1.27970369 | -7.4971362 |
| 1371 | NM_020297.3 | 752.277708 | 3.24469953 | 0.43279345 | 7.49710861 |
| 1372 | NM_005500.2 | 343.938304 | -10.048126 | 1.34065125 | -7.4949587 |
| 1373 | NM_005087.3 | 196.426572 | -9.239874 | 1.2328649 | -7.4946363 |
| 1374 | NM_003934.1 | 199.622848 | -9.2632358 | 1.23603173 | -7.4943349 |
| 1375 | NM_001288.4 | 221.255747 | -9.4122707 | 1.2559991 | -7.4938515 |

| | | | | | |
|---|---|---|---|---|---|
| 1376 | NM_014714.3 | 217.265449 | -9.3861377 | 1.25264445 | -7.4930582 |
| 1377 | NM_001130991.2 | 392.716855 | -10.239829 | 1.36659524 | -7.4929497 |
| 1378 | NM_020738.2 | 1281.45373 | -8.7925926 | 1.17348905 | -7.4926925 |
| 1379 | NM_005176.5 | 364.525058 | -10.131984 | 1.35228222 | -7.4925068 |
| 1380 | NM_015047.2 | 209.150261 | -9.3297554 | 1.24525381 | -7.492252 |
| 1381 | NM_018675.2 | 6108.05633 | 7.11365158 | 0.94950229 | 7.49197939 |
| 1382 | NM_001008895.2 | 172.770685 | -9.0555636 | 1.2088263 | -7.4912033 |
| 1383 | NM_014828.3 | 233.464235 | -9.4890791 | 1.26672831 | -7.4910137 |
| 1384 | NM_001164.4 | 407.18548 | -10.291594 | 1.37406493 | -7.4898895 |
| 1385 | NM_018426.2 | 265.785091 | -6.3839169 | 0.85234839 | -7.4897976 |
| 1386 | NM_003347.3 | 276.891964 | -9.7352182 | 1.29983128 | -7.4896014 |
| 1387 | NM_006666.2 | 221.043195 | -9.4104006 | 1.25648118 | -7.4894879 |
| 1388 | NM_004860.3 | 257.734786 | -9.6318794 | 1.28623269 | -7.4884424 |
| 1389 | NM_014500.4 | 269.91803 | -9.6982169 | 1.29513865 | -7.4881689 |
| 1390 | NM_001711.5 | 261.232954 | -7.2273658 | 0.96518696 | -7.4880475 |
| 1391 | NM_001135653.1 | 188.865336 | -9.1840126 | 1.22654401 | -7.4877155 |
| 1392 | NM_015208.4 | 385.023676 | -10.210792 | 1.36383111 | -7.4868446 |
| 1393 | NM_022765.3 | 351.75533 | -10.080623 | 1.34655027 | -7.4862579 |
| 1394 | NM_001306131.1 | 322.895411 | -9.9575306 | 1.33012631 | -7.4861542 |
| 1395 | NM_001042646.2 | 242.502348 | -9.5439575 | 1.2754175 | -7.4830065 |
| 1396 | NM_003373.3 | 481.572436 | -8.6304235 | 1.15336455 | -7.4828236 |
| 1397 | NM_001087.4 | 253.318528 | -9.6068704 | 1.28396904 | -7.4821667 |
| 1398 | NM_005103.4 | 501.211003 | -10.591382 | 1.41557506 | -7.4820349 |
| 1399 | NM_004344.1 | 179.511868 | -9.1116135 | 1.21843432 | -7.4781327 |
| 1400 | NM_001320153.1 | 253.204284 | -9.6062624 | 1.28461585 | -7.4779261 |
| 1401 | NM_006329.3 | 444.179807 | -10.417381 | 1.39312917 | -7.4776852 |
| 1402 | NM_000089.3 | 5665.0585 | -9.3827352 | 1.2549253 | -7.476728 |
| 1403 | NM_015039.3 | 375.405741 | -10.174427 | 1.360847 | -7.4765397 |
| 1404 | NM_004783.3 | 247.37008 | -9.5728561 | 1.28043568 | -7.4762491 |
| 1405 | NR_135154.1 | 367.627606 | -9.1809926 | 1.22801229 | -7.4763036 |
| 1406 | NM_002624.3 | 379.50989 | -10.189991 | 1.36301709 | -7.4760551 |
| 1407 | NM_002768.4 | 267.989703 | -9.6879949 | 1.29613718 | -7.4745135 |
| 1408 | NM_002902.2 | 337.070887 | -10.019052 | 1.34041607 | -7.4745835 |
| 1409 | NM_003909.4 | 223.027491 | -9.4227376 | 1.26069622 | -7.4742332 |
| 1410 | NM_001006935.2 | 310.171603 | -9.8989279 | 1.32446972 | -7.4738801 |
| 1411 | NM_001833.3 | 268.097864 | -9.6888215 | 1.29663726 | -7.4722683 |
| 1412 | NM_004985.4 | 200.874284 | -5.6790586 | 0.76001862 | -7.4722624 |
| 1413 | NM_006476.4 | 328.485651 | -9.9818092 | 1.33590042 | -7.471971 |
| 1414 | NM_001316374.1 | 299.986412 | -9.85085 | 1.31874874 | -7.469846 |
| 1415 | NM_003439.3 | 241.307135 | -9.5371189 | 1.27678031 | -7.4696632 |
| 1416 | NM_019094.5 | 217.750059 | -9.3880917 | 1.25683464 | -7.4696316 |
| 1417 | NM_006392.3 | 247.107844 | -9.5712187 | 1.28137653 | -7.4694818 |
| 1418 | NM_000875.4 | 240.104111 | -9.528712 | 1.27578811 | -7.468883 |

| | | | | | |
|---|---|---|---|---|---|
| 1419 | NM_001520.3 | 218.117719 | -9.3914024 | 1.25743897 | -7.4686745 |
| 1420 | NM_003795.5 | 290.480504 | -9.8044691 | 1.31283194 | -7.468183 |
| 1421 | NM_001098534.1 | 432.988279 | -8.6025899 | 1.15196291 | -7.4677664 |
| 1422 | NM_015705.5 | 226.007227 | -9.442141 | 1.26437981 | -7.4678043 |
| 1423 | NM_001134422.1 | 235.537021 | -9.5018687 | 1.27252923 | -7.4669158 |
| 1424 | NM_003663.3 | 274.073441 | -9.7203219 | 1.30206559 | -7.465309 |
| 1425 | NM_005481.2 | 216.788315 | -9.3823009 | 1.25679107 | -7.4652829 |
| 1426 | NM_002883.3 | 323.729496 | -9.9607687 | 1.3344179 | -7.4645047 |
| 1427 | NM_003692.4 | 451.53323 | -10.440853 | 1.39874864 | -7.4644237 |
| 1428 | NM_001099784.2 | 338.203516 | -10.024002 | 1.34300716 | -7.4638483 |
| 1429 | NM_001136025.4 | 274.947141 | -9.7260924 | 1.30313412 | -7.4636158 |
| 1430 | NM_001303275.1 | 226.68565 | -9.4477439 | 1.265923 | -7.4631268 |
| 1431 | NM_001386.5 | 724.978263 | -10.161415 | 1.36158514 | -7.4629301 |
| 1432 | NR_026656.1 | 493.634431 | -10.569635 | 1.4162698 | -7.4630092 |
| 1433 | NM_004373.3 | 486.496663 | -10.548448 | 1.41349707 | -7.4626601 |
| 1434 | NM_000284.3 | 230.942662 | -9.4734986 | 1.26947264 | -7.4625465 |
| 1435 | NM_004393.5 | 256.635186 | -9.6262806 | 1.29013609 | -7.4614459 |
| 1436 | NM_001242758.1 | 220.888992 | -9.4106921 | 1.26134185 | -7.4608577 |
| 1437 | NM_004866.5 | 265.960212 | -9.6769613 | 1.29704185 | -7.4607935 |
| 1438 | NM_033546.3 | 365.742606 | -10.136925 | 1.3587722 | -7.4603562 |
| 1439 | NM_006247.3 | 263.102782 | -9.6616447 | 1.29509616 | -7.4601755 |
| 1440 | NM_001024807.2 | 362.186713 | -10.122805 | 1.35696392 | -7.4598926 |
| 1441 | NM_003367.3 | 298.492539 | -9.8435533 | 1.31954125 | -7.4598299 |
| 1442 | NM_002627.4 | 305.949729 | -9.8796964 | 1.32444897 | -7.4594768 |
| 1443 | NM_020444.4 | 231.007318 | -9.4735411 | 1.27006485 | -7.4591003 |
| 1444 | NM_018071.4 | 231.897624 | -9.4801009 | 1.27106209 | -7.4584089 |
| 1445 | NM_018011.3 | 355.102 | -9.1296948 | 1.22455845 | -7.4554994 |
| 1446 | NM_001662.3 | 348.137336 | -10.065643 | 1.35023509 | -7.4547338 |
| 1447 | NM_002793.3 | 268.325806 | -9.6900104 | 1.30002166 | -7.4537299 |
| 1448 | NM_007260.2 | 215.549307 | -6.8860867 | 0.92391694 | -7.4531447 |
| 1449 | NM_017892.3 | 303.835319 | -8.076594 | 1.08383965 | -7.4518348 |
| 1450 | NM_016480.4 | 263.255643 | -9.6624632 | 1.29682129 | -7.4508826 |
| 1451 | NM_002959.6 | 407.355513 | -10.292391 | 1.38144516 | -7.4504521 |
| 1452 | NM_032801.4 | 223.195208 | -9.4245445 | 1.26514998 | -7.4493496 |
| 1453 | NM_005151.3 | 221.908718 | -9.4157979 | 1.26399891 | -7.4492136 |
| 1454 | NM_001518.4 | 222.396939 | -9.4199766 | 1.26463758 | -7.4487559 |
| 1455 | NM_000451.3 | 484.183708 | 6.32189247 | 0.84873051 | 7.44864523 |
| 1456 | NM_000637.3 | 185.123615 | -9.1545949 | 1.22913609 | -7.4479913 |
| 1457 | NM_001005.4 | 341.887155 | -10.039576 | 1.3479593 | -7.4479815 |
| 1458 | NM_001281720.1 | 454.313624 | -10.44983 | 1.40303281 | -7.4480299 |
| 1459 | NM_182729.2 | 247.580549 | -9.5736011 | 1.28538519 | -7.4480406 |
| 1460 | NM_014889.3 | 188.273997 | -9.1788883 | 1.23247589 | -7.4475195 |
| 1461 | NR_036592.1 | 196.964808 | -9.2435437 | 1.24124958 | -7.4469663 |

| | | | | | |
|---|---|---|---|---|---|
| 1462 | NM_017797.3 | 444.612894 | -9.455435 | 1.26995155 | -7.4455085 |
| 1463 | NM_001313.4 | 1460.26531 | -8.7093408 | 1.16982781 | -7.4449767 |
| 1464 | NM_032409.2 | 294.774094 | -9.825685 | 1.31979764 | -7.4448421 |
| 1465 | NM_006409.3 | 284.442573 | -9.7741915 | 1.31293195 | -7.444553 |
| 1466 | NM_003861.2 | 229.109842 | -9.4621435 | 1.27113055 | -7.4438802 |
| 1467 | NM_015114.2 | 212.797012 | -9.3552987 | 1.25694789 | -7.4428692 |
| 1468 | NM_006802.3 | 232.850295 | -9.4855216 | 1.2745291 | -7.4423735 |
| 1469 | NM_015909.3 | 188.013139 | -9.1769907 | 1.23310722 | -7.4421677 |
| 1470 | NM_175852.3 | 251.634769 | -7.79262 | 1.04717329 | -7.4415763 |
| 1471 | NM_006937.3 | 607.609423 | -9.9064205 | 1.33125493 | -7.441415 |
| 1472 | NM_032221.4 | 202.687418 | -5.9648726 | 0.80161665 | -7.4410538 |
| 1473 | NM_002860.3 | 218.871591 | -9.3955697 | 1.26289345 | -7.4397169 |
| 1474 | NM_033089.6 | 177.894178 | -9.0968753 | 1.2227357 | -7.4397724 |
| 1475 | NM_004739.3 | 301.796013 | -9.8594201 | 1.32536129 | -7.4390433 |
| 1476 | NM_001297595.1 | 254.317591 | -9.6126267 | 1.29242224 | -7.4376828 |
| 1477 | NM_025029.4 | 342.491503 | -10.041976 | 1.35019931 | -7.4374025 |
| 1478 | NM_002572.3 | 351.095176 | -10.077695 | 1.3550582 | -7.4370937 |
| 1479 | NM_001322434.1 | 227.589089 | -9.4521143 | 1.27105909 | -7.4364082 |
| 1480 | NM_015329.3 | 205.470919 | -9.3052065 | 1.2513779 | -7.4359684 |
| 1481 | NM_002609.3 | 213.703568 | -9.3629837 | 1.25916572 | -7.435863 |
| 1482 | NM_014344.3 | 422.359474 | -10.344487 | 1.39129112 | -7.4351709 |
| 1483 | NM_005973.4 | 231.596556 | -6.5611259 | 0.88247287 | -7.4349322 |
| 1484 | NM_030790.4 | 305.973262 | -9.8793702 | 1.32881472 | -7.4347237 |
| 1485 | NM_014613.2 | 205.646787 | -9.3062763 | 1.25182554 | -7.4341639 |
| 1486 | NM_014874.3 | 268.014442 | -9.6876918 | 1.30311647 | -7.4342486 |
| 1487 | NM_018959.3 | 241.879617 | -9.5400305 | 1.28345937 | -7.43306 |
| 1488 | NM_001098509.1 | 305.257658 | -9.8761314 | 1.32870529 | -7.4328984 |
| 1489 | NM_005015.3 | 190.481153 | -9.1958993 | 1.23723489 | -7.432622 |
| 1490 | NM_001030055.1 | 201.47586 | -9.278059 | 1.24841464 | -7.431873 |
| 1491 | NM_015544.2 | 184.06453 | -9.1469378 | 1.23078565 | -7.4317878 |
| 1492 | NM_005345.5 | 270.648944 | -9.7024386 | 1.30564366 | -7.4311536 |
| 1493 | NM_006391.2 | 474.34145 | -9.548906 | 1.28503176 | -7.4308716 |
| 1494 | NM_016404.2 | 241.814903 | -9.5396914 | 1.28381802 | -7.4307193 |
| 1495 | NM_002096.2 | 341.789087 | -8.2526263 | 1.11089892 | -7.4287824 |
| 1496 | NM_003099.4 | 198.052975 | -9.2519795 | 1.2454698 | -7.4285057 |
| 1497 | NM_001040158.1 | 289.464012 | -9.7988791 | 1.31925029 | -7.4276119 |
| 1498 | NM_001017405.2 | 207.595614 | -9.3200874 | 1.25483013 | -7.4273698 |
| 1499 | NM_001159673.1 | 227.46054 | -9.4511895 | 1.27251163 | -7.4271931 |
| 1500 | NM_001284280.1 | 192.80274 | -9.2126494 | 1.2405858 | -7.4260478 |
| 1501 | NM_018976.4 | 446.079675 | -10.423275 | 1.40374692 | -7.4253237 |
| 1502 | NM_000997.4 | 737.88326 | -10.186369 | 1.37193911 | -7.4247967 |
| 1503 | NM_001289123.1 | 416.312002 | -10.323741 | 1.39062068 | -7.4238371 |
| 1504 | NM_001032396.2 | 237.839443 | -9.5159379 | 1.28188388 | -7.4234009 |

| | | | | | |
|---|---|---|---|---|---|
| 1505 | NM_016263.3 | 258.833332 | -9.6379306 | 1.29831209 | -7.4234313 |
| 1506 | NM_054016.3 | 202.472379 | -9.2832027 | 1.2505111 | -7.4235268 |
| 1507 | NM_006294.4 | 354.265494 | -3.656621 | 0.49260672 | -7.4230028 |
| 1508 | NM_022003.3 | 595.933072 | -10.841148 | 1.46049426 | -7.4229312 |
| 1509 | NM_018847.3 | 249.23208 | -9.5836488 | 1.29131543 | -7.4216172 |
| 1510 | NM_145012.5 | 216.7492 | -9.3818979 | 1.26432554 | -7.4204765 |
| 1511 | NM_014747.2 | 372.14241 | -10.16203 | 1.36964063 | -7.4194861 |
| 1512 | NM_006145.2 | 253.648249 | -9.6091414 | 1.2955535 | -7.4170163 |
| 1513 | NM_001134382.2 | 195.41879 | -9.233133 | 1.24515623 | -7.4152405 |
| 1514 | NM_004699.3 | 252.102057 | -9.6000545 | 1.29464006 | -7.4152305 |
| 1515 | NM_006772.2 | 179.006366 | -9.1052729 | 1.22791885 | -7.4152073 |
| 1516 | NM_020531.2 | 389.511171 | -8.3994872 | 1.13285716 | -7.4144275 |
| 1517 | NM_145899.2 | 359.109085 | -10.110311 | 1.36374324 | -7.4136468 |
| 1518 | NM_002896.3 | 288.431529 | -9.7942718 | 1.32122546 | -7.4130208 |
| 1519 | NM_002931.3 | 202.77097 | -9.286198 | 1.25271399 | -7.4128637 |
| 1520 | NM_003971.5 | 205.689236 | -9.3058941 | 1.25539596 | -7.4127163 |
| 1521 | NM_001330063.1 | 208.021261 | -9.3235871 | 1.25782626 | -7.4124602 |
| 1522 | NM_022912.2 | 462.733662 | -10.476129 | 1.41337576 | -7.4121329 |
| 1523 | NM_173562.4 | 222.017828 | -9.4165279 | 1.27044304 | -7.4120032 |
| 1524 | NM_016080.3 | 222.887429 | -9.4223342 | 1.27130125 | -7.4115668 |
| 1525 | NM_001278912.1 | 280.203508 | -9.752461 | 1.31612066 | -7.4100052 |
| 1526 | NM_003331.4 | 178.66545 | -9.1039468 | 1.22861351 | -7.4099355 |
| 1527 | NM_001128217.2 | 182.503494 | -9.1324198 | 1.23250848 | -7.4096203 |
| 1528 | NM_020135.2 | 194.642467 | -9.2266024 | 1.24533255 | -7.4089467 |
| 1529 | NM_017455.3 | 267.75218 | -9.6870065 | 1.30753304 | -7.4086131 |
| 1530 | NM_021224.5 | 212.204065 | -9.3511585 | 1.26226022 | -7.4082652 |
| 1531 | NM_001101372.1 | 363.436533 | -10.127822 | 1.36712839 | -7.4080987 |
| 1532 | NM_002018.3 | 213.957881 | -9.3627285 | 1.26393452 | -7.4076056 |
| 1533 | NM_002305.3 | 339.874278 | -10.031366 | 1.35419919 | -7.4076002 |
| 1534 | NM_017811.3 | 403.141234 | -9.3142207 | 1.25758617 | -7.4064274 |
| 1535 | NM_006544.3 | 266.283724 | -9.6787077 | 1.30687009 | -7.4060213 |
| 1536 | NM_001640.3 | 244.281312 | -9.5548632 | 1.29024521 | -7.405463 |
| 1537 | NM_153273.3 | 212.446103 | -9.353362 | 1.26309599 | -7.4051078 |
| 1538 | NM_001177563.1 | 32.9291803 | 9.96216853 | 1.3453422 | 7.40493274 |
| 1539 | NM_006756.3 | 281.699394 | -9.7599844 | 1.31803966 | -7.4049246 |
| 1540 | NM_002827.3 | 225.549244 | -9.4397658 | 1.27489129 | -7.4043692 |
| 1541 | NM_007065.3 | 442.7813 | -9.4487503 | 1.2761575 | -7.4040628 |
| 1542 | NM_014392.4 | 456.430072 | -10.45649 | 1.4123473 | -7.4036252 |
| 1543 | NM_032333.4 | 193.39518 | -9.2169328 | 1.24495875 | -7.4034042 |
| 1544 | NM_005407.2 | 211.887501 | -9.3499578 | 1.26305418 | -7.4026577 |
| 1545 | NM_002088.4 | 292.739656 | -9.8153663 | 1.32596229 | -7.4024476 |
| 1546 | NM_015482.1 | 322.996352 | -9.9573796 | 1.34518854 | -7.4022186 |
| 1547 | NM_001349.3 | 193.321596 | -9.2160584 | 1.24522194 | -7.4011372 |

| | | | | | |
|---|---|---|---|---|---|
| 1548 | NM_181442.3 | 265.43846 | -9.6738144 | 1.30708209 | -7.4010765 |
| 1549 | NM_032772.5 | 185.149219 | -9.1563813 | 1.23721417 | -7.4008054 |
| 1550 | NM_016605.2 | 214.325747 | -9.3656416 | 1.26550611 | -7.4007083 |
| 1551 | NM_018235.2 | 236.546652 | -9.5083024 | 1.28482505 | -7.4004647 |
| 1552 | NM_016308.2 | 266.990902 | -9.6824657 | 1.3084131 | -7.4001595 |
| 1553 | NM_020524.3 | 215.474759 | -9.3748619 | 1.26719281 | -7.3981338 |
| 1554 | NM_006660.4 | 168.910359 | -9.0222045 | 1.21959788 | -7.3976879 |
| 1555 | NM_032476.3 | 164.859594 | -8.9887626 | 1.21508304 | -7.3976529 |
| 1556 | NM_001261.3 | 181.489316 | -9.1254025 | 1.23365722 | -7.3970325 |
| 1557 | NM_002227.3 | 185.143618 | -9.1541731 | 1.23753514 | -7.3971016 |
| 1558 | NM_006380.3 | 186.437615 | -9.1652697 | 1.23904029 | -7.3970716 |
| 1559 | NM_003680.3 | 369.436088 | -7.8232035 | 1.05770768 | -7.3963759 |
| 1560 | NM_005419.3 | 187.834924 | -9.1750541 | 1.24052105 | -7.3961293 |
| 1561 | NM_015933.5 | 269.990961 | -9.6988227 | 1.31139921 | -7.395782 |
| 1562 | NM_006838.3 | 229.268473 | -9.4630723 | 1.27953861 | -7.3956911 |
| 1563 | NM_001080457.1 | 446.118746 | -7.4086011 | 1.00185905 | -7.3948537 |
| 1564 | NM_004077.2 | 503.619206 | -9.6356063 | 1.30301749 | -7.3948404 |
| 1565 | NM_033200.2 | 230.433032 | -9.4704445 | 1.2807139 | -7.3946605 |
| 1566 | NM_016320.4 | 171.681453 | -9.0461697 | 1.22338075 | -7.3944025 |
| 1567 | NM_058164.3 | 390.484861 | -10.231165 | 1.38365537 | -7.3943018 |
| 1568 | NM_001999.3 | 451.904063 | -9.4786378 | 1.28202524 | -7.3934876 |
| 1569 | NM_006796.2 | 196.941804 | -9.2441524 | 1.25032014 | -7.3934284 |
| 1570 | NM_016289.3 | 266.513733 | -9.6800205 | 1.30932922 | -7.3931142 |
| 1571 | NM_004598.3 | 400.811292 | -10.269069 | 1.38917596 | -7.3922017 |
| 1572 | NM_014371.3 | 197.943935 | -9.2522007 | 1.25162143 | -7.3921718 |
| 1573 | NM_022130.3 | 219.112821 | -9.3975955 | 1.27135808 | -7.3917771 |
| 1574 | NM_005669.4 | 291.758884 | -9.8107813 | 1.32739605 | -7.3909978 |
| 1575 | NM_007263.3 | 276.099277 | -9.7311189 | 1.3166449 | -7.3908454 |
| 1576 | NM_003352.4 | 268.883969 | -9.6928587 | 1.3115267 | -7.3905157 |
| 1577 | NM_005124.3 | 194.417901 | -9.2255049 | 1.24830428 | -7.3904295 |
| 1578 | NM_138967.3 | 391.536014 | -10.235002 | 1.3849778 | -7.3900116 |
| 1579 | NM_001136157.1 | 199.796808 | -9.2658761 | 1.25393649 | -7.3894302 |
| 1580 | NM_006517.4 | 192.114078 | -9.2081488 | 1.24611703 | -7.3894735 |
| 1581 | NM_005243.3 | 257.11137 | -9.6283075 | 1.30303162 | -7.3891588 |
| 1582 | NM_004730.3 | 222.49046 | -9.4195549 | 1.27495704 | -7.3881351 |
| 1583 | NM_004429.4 | 228.096555 | -9.4550199 | 1.27980122 | -7.3878816 |
| 1584 | NM_015423.2 | 349.364784 | -10.07076 | 1.36315111 | -7.3878531 |
| 1585 | NM_002409.4 | 288.171682 | -9.7928747 | 1.32576928 | -7.3865603 |
| 1586 | NM_002855.4 | 207.96458 | -9.3222149 | 1.26220811 | -7.3856402 |
| 1587 | NM_001113239.2 | 231.003729 | -9.4737949 | 1.28276588 | -7.3854435 |
| 1588 | NM_003842.4 | 182.909422 | -9.1365105 | 1.23713424 | -7.3852216 |
| 1589 | NM_032389.4 | 208.413023 | -9.3256627 | 1.26283338 | -7.3847135 |
| 1590 | NM_198390.2 | 478.748002 | -9.562257 | 1.29495994 | -7.3842107 |

| | | | | | |
|---|---|---|---|---|---|
| 1591 | NM_022826.3 | 186.310881 | -9.1625899 | 1.24101421 | -7.3831466 |
| 1592 | NM_016255.2 | 201.576998 | -9.2771519 | 1.25654925 | -7.3830388 |
| 1593 | NM_199511.2 | 309.098731 | -9.8944586 | 1.34017304 | -7.3829709 |
| 1594 | NM_014172.5 | 253.485764 | -9.6078974 | 1.30141084 | -7.3826781 |
| 1595 | NM_001182.4 | 193.887549 | -9.2215668 | 1.24921291 | -7.3819016 |
| 1596 | NM_003801.3 | 269.188647 | -9.6947482 | 1.31332802 | -7.3818179 |
| 1597 | NM_004302.4 | 319.201953 | -9.9403878 | 1.34663658 | -7.381641 |
| 1598 | NM_007178.3 | 619.201864 | -9.1342586 | 1.23747392 | -7.3813746 |
| 1599 | NM_198291.2 | 265.362428 | -9.6738581 | 1.31072353 | -7.3805481 |
| 1600 | NM_016142.2 | 201.039077 | -9.2733291 | 1.25653421 | -7.3800849 |
| 1601 | NM_001271741.1 | 192.905982 | -9.2148379 | 1.24869035 | -7.3796021 |
| 1602 | NM_004355.3 | 547.256177 | -10.718408 | 1.45244606 | -7.3795566 |
| 1603 | NM_139015.4 | 276.663705 | -7.376936 | 0.99965771 | -7.3794619 |
| 1604 | NM_002817.3 | 287.561668 | -9.7897743 | 1.32680199 | -7.3784742 |
| 1605 | NM_006927.3 | 207.562172 | -4.0518289 | 0.54916117 | -7.3782145 |
| 1606 | NM_003469.4 | 487.552701 | -10.551687 | 1.43013296 | -7.3781163 |
| 1607 | NM_032960.3 | 194.741704 | -9.2277124 | 1.25074758 | -7.3777576 |
| 1608 | NM_015388.3 | 305.155942 | -7.5372166 | 1.02164451 | -7.3775335 |
| 1609 | NM_005891.2 | 434.484637 | -10.385382 | 1.40776612 | -7.3772068 |
| 1610 | NM_018128.4 | 166.066969 | -5.2109122 | 0.70644477 | -7.3762485 |
| 1611 | NM_001193646.1 | 250.334024 | -9.5897685 | 1.30010599 | -7.3761436 |
| 1612 | NR_123722.1 | 213.499624 | -6.8652898 | 0.93075249 | -7.3760639 |
| 1613 | NM_001145108.1 | 529.348752 | -10.670086 | 1.4467234 | -7.3753464 |
| 1614 | NM_000690.3 | 229.346379 | -9.463531 | 1.28325869 | -7.3746089 |
| 1615 | NM_022089.3 | 317.283694 | -9.931615 | 1.34689026 | -7.3737373 |
| 1616 | NM_001130138.1 | 562.732543 | -6.8566465 | 0.92989932 | -7.3735365 |
| 1617 | NM_017761.3 | 197.00984 | -9.2445511 | 1.25383005 | -7.3730496 |
| 1618 | NM_001102397.2 | 216.500591 | -9.3802136 | 1.27224967 | -7.3729346 |
| 1619 | NM_001312.3 | 290.829513 | -9.8060159 | 1.33032551 | -7.3711402 |
| 1620 | NM_006817.3 | 256.443512 | -9.6246731 | 1.30573668 | -7.3710674 |
| 1621 | NM_002078.4 | 175.927106 | -9.0809115 | 1.23198338 | -7.3709692 |
| 1622 | NM_001001998.2 | 179.566904 | -9.1107606 | 1.23627629 | -7.3695182 |
| 1623 | NM_145754.3 | 355.865919 | -10.097261 | 1.37026709 | -7.3688272 |
| 1624 | NM_007262.4 | 287.560996 | -9.7898562 | 1.3285716 | -7.368708 |
| 1625 | NM_005785.3 | 217.585872 | -9.3868334 | 1.27405136 | -7.3677041 |
| 1626 | NM_006989.5 | 266.178228 | -9.6778092 | 1.31377684 | -7.3664027 |
| 1627 | NM_001128914.1 | 315.387637 | -9.9232002 | 1.34712151 | -7.366225 |
| 1628 | NM_016073.3 | 377.75735 | -10.183351 | 1.38266792 | -7.3650012 |
| 1629 | NM_001679.3 | 272.427654 | -9.7117833 | 1.31870471 | -7.3646384 |
| 1630 | NM_022173.2 | 284.046169 | -9.7724079 | 1.32693929 | -7.3646232 |
| 1631 | NM_017870.3 | 416.180271 | -10.323204 | 1.40179707 | -7.3642644 |
| 1632 | NM_003959.2 | 364.215747 | -10.130663 | 1.37579551 | -7.363495 |
| 1633 | NM_002869.4 | 177.293944 | -9.0916624 | 1.23478003 | -7.3629814 |

| | | | | | |
|---|---|---|---|---|---|
| 1634 | NM_015571.3 | 188.207758 | -9.1784493 | 1.24666789 | -7.3623852 |
| 1635 | NM_020959.2 | 275.612027 | -9.7286218 | 1.32141348 | -7.3622843 |
| 1636 | NM_002708.3 | 234.800909 | -9.4975258 | 1.29018064 | -7.3613923 |
| 1637 | NM_173614.2 | 323.980607 | -9.9621875 | 1.35346967 | -7.3604808 |
| 1638 | NM_031286.3 | 215.412994 | -9.3733136 | 1.2737727 | -7.3587019 |
| 1639 | NM_001323619.1 | 182.160313 | -9.1312592 | 1.24093843 | -7.3583499 |
| 1640 | NM_003860.3 | 280.988069 | -9.7564343 | 1.32593526 | -7.3581529 |
| 1641 | NM_006324.2 | 246.228494 | -9.5658849 | 1.30013739 | -7.3575954 |
| 1642 | NM_152346.2 | 180.605107 | -9.119508 | 1.23949078 | -7.3574634 |
| 1643 | NM_006948.4 | 296.238995 | -9.8327271 | 1.33673687 | -7.3557686 |
| 1644 | NM_004060.3 | 220.654378 | -9.40684 | 1.27894086 | -7.3551798 |
| 1645 | NM_001278141.1 | 150.331406 | -6.0740809 | 0.82588158 | -7.3546633 |
| 1646 | NM_032853.3 | 204.608787 | -9.2990343 | 1.26448851 | -7.3539888 |
| 1647 | NM_003273.3 | 222.220579 | -9.4181079 | 1.28070494 | -7.3538467 |
| 1648 | NM_018179.4 | 260.689366 | -9.6474868 | 1.31201942 | -7.3531586 |
| 1649 | NM_001863.4 | 287.794238 | -6.9233935 | 0.94158238 | -7.3529344 |
| 1650 | NM_003751.3 | 195.577026 | -9.2339962 | 1.2559021 | -7.3524809 |
| 1651 | NM_004326.3 | 201.03836 | -9.2732088 | 1.2612782 | -7.3522311 |
| 1652 | NM_015339.4 | 169.682505 | -9.0299706 | 1.22825404 | -7.3518753 |
| 1653 | NM_014306.4 | 225.97181 | -9.4422166 | 1.28442305 | -7.3513291 |
| 1654 | NM_021070.4 | 218.617513 | -9.3955732 | 1.278095 | -7.3512323 |
| 1655 | NM_001199161.1 | 162.649938 | -8.9670193 | 1.21983195 | -7.3510284 |
| 1656 | NM_001292034.2 | 173.386821 | -9.0607446 | 1.23256876 | -7.3511068 |
| 1657 | NM_006510.4 | 221.362745 | -9.412342 | 1.28049571 | -7.3505455 |
| 1658 | NM_173468.3 | 124.141501 | -4.9617213 | 0.67501402 | -7.3505455 |
| 1659 | NM_004426.2 | 455.966201 | -6.7956374 | 0.92454749 | -7.3502307 |
| 1660 | NM_005342.3 | 273.107182 | -9.7154303 | 1.3218727 | -7.3497473 |
| 1661 | NM_021078.2 | 202.609285 | -6.9218206 | 0.94185097 | -7.3491675 |
| 1662 | NM_003799.2 | 183.00374 | -9.1373155 | 1.24340559 | -7.3486202 |
| 1663 | NM_006709.4 | 273.57949 | -9.7178241 | 1.32242423 | -7.3484922 |
| 1664 | NM_020830.4 | 174.005672 | -9.0647386 | 1.23367843 | -7.3477321 |
| 1665 | NM_001193483.2 | 180.38552 | -9.1177639 | 1.24099004 | -7.3471693 |
| 1666 | NM_012308.2 | 151.435377 | -8.8643772 | 1.20649098 | -7.3472387 |
| 1667 | NM_003714.2 | 171.674637 | -9.0457426 | 1.23122991 | -7.3469159 |
| 1668 | NM_001082968.1 | 207.522135 | -9.3189351 | 1.26843917 | -7.3467733 |
| 1669 | NM_020944.2 | 194.57679 | -9.2262969 | 1.25595762 | -7.3460256 |
| 1670 | NM_014325.3 | 617.884944 | -9.9306153 | 1.35185864 | -7.3458977 |
| 1671 | NM_014183.3 | 250.547263 | -9.5911764 | 1.30572327 | -7.3454894 |
| 1672 | NM_022149.4 | 229.146612 | -9.4622317 | 1.28847531 | -7.3437432 |
| 1673 | NM_004549.5 | 309.895555 | -9.8976727 | 1.3478965 | -7.343051 |
| 1674 | NM_032812.8 | 161.574357 | -8.958991 | 1.22005779 | -7.3430874 |
| 1675 | NM_001344.3 | 244.368309 | -9.5552446 | 1.30133132 | -7.3426686 |
| 1676 | NM_207122.1 | 238.100174 | -9.5178092 | 1.29628406 | -7.3423792 |

| | | | | | |
|---|---|---|---|---|---|
| 1677 | NM_024671.3 | 200.301136 | -9.2681686 | 1.26233387 | -7.3420897 |
| 1678 | NM_003581.4 | 225.561617 | -9.439305 | 1.28567473 | -7.3419076 |
| 1679 | NM_001845.5 | 1500.51246 | -7.8284138 | 1.06634528 | -7.3413499 |
| 1680 | NM_006178.3 | 335.486842 | -10.012255 | 1.36412848 | -7.3396714 |
| 1681 | NM_003348.3 | 340.604765 | -10.034062 | 1.36712161 | -7.3395533 |
| 1682 | NM_001144995.1 | 168.058255 | -9.0153587 | 1.22835866 | -7.3393538 |
| 1683 | NM_003017.4 | 493.088633 | -9.6049571 | 1.30874011 | -7.3390867 |
| 1684 | NM_004074.2 | 270.573254 | -9.7020917 | 1.32204226 | -7.3387153 |
| 1685 | NM_018084.4 | 244.431012 | -9.5555341 | 1.3020727 | -7.3387101 |
| 1686 | NM_001320.6 | 240.438015 | -9.5318731 | 1.29893375 | -7.3382288 |
| 1687 | NM_000108.4 | 191.713926 | -9.2051092 | 1.254423 | -7.3381222 |
| 1688 | NM_005984.4 | 228.233275 | -9.4561571 | 1.28877879 | -7.3373004 |
| 1689 | NM_005455.4 | 212.389945 | -9.3522627 | 1.27463384 | -7.3372151 |
| 1690 | NM_016628.4 | 185.395796 | -9.1573912 | 1.24809332 | -7.3371045 |
| 1691 | NM_000090.3 | 5846.35914 | -7.1745718 | 0.97790037 | -7.3367104 |
| 1692 | NM_005109.2 | 194.231796 | -9.223552 | 1.2571975 | -7.3365974 |
| 1693 | NM_001007237.2 | 388.596495 | -10.224053 | 1.3936043 | -7.3364104 |
| 1694 | NM_001330249.1 | 173.630162 | -9.0637985 | 1.23578701 | -7.3344341 |
| 1695 | NM_004776.3 | 269.163085 | -9.6945918 | 1.3219606 | -7.3334953 |
| 1696 | NM_007002.3 | 233.345254 | -9.4882481 | 1.29398066 | -7.3326043 |
| 1697 | NM_005186.3 | 252.800546 | -9.6036801 | 1.3097475 | -7.3324668 |
| 1698 | NM_006230.3 | 219.748627 | -9.4018773 | 1.28237482 | -7.3316141 |
| 1699 | NM_002815.3 | 214.963657 | -9.3699342 | 1.27804178 | -7.3314772 |
| 1700 | NM_002396.4 | 181.08811 | -9.1222881 | 1.24432823 | -7.3310947 |
| 1701 | NM_001204467.1 | 197.679179 | -9.2493068 | 1.26176073 | -7.330476 |
| 1702 | NM_014685.3 | 199.513254 | -9.2623607 | 1.26366226 | -7.3297755 |
| 1703 | NM_003676.3 | 199.444254 | -9.2624475 | 1.26379219 | -7.3290906 |
| 1704 | NM_000346.3 | 166.736955 | -9.0033375 | 1.22866504 | -7.3277396 |
| 1705 | NM_007046.3 | 217.272876 | -9.3865588 | 1.28099777 | -7.3275372 |
| 1706 | NM_020992.3 | 310.193951 | -9.8995856 | 1.35107834 | -7.3271736 |
| 1707 | NM_001304815.1 | 163.479595 | -5.8942482 | 0.80452912 | -7.3263329 |
| 1708 | NM_016272.3 | 173.494069 | -6.5918733 | 0.89986342 | -7.3254154 |
| 1709 | NM_001173431.1 | 346.354642 | -9.0942844 | 1.24155519 | -7.3249136 |
| 1710 | NM_001627.3 | 174.378837 | -9.0685384 | 1.23805279 | -7.3248398 |
| 1711 | NM_003106.3 | 258.71648 | -7.8154442 | 1.06699369 | -7.3247333 |
| 1712 | NM_014938.5 | 315.335504 | -3.6497324 | 0.49829806 | -7.3243962 |
| 1713 | NM_001080453.2 | 556.305362 | -7.0945603 | 0.96873933 | -7.3234978 |
| 1714 | NM_002775.4 | 290.512037 | -9.8050592 | 1.33890884 | -7.3231716 |
| 1715 | NM_015570.3 | 478.379333 | -10.524075 | 1.43711574 | -7.3230533 |
| 1716 | NM_006548.5 | 186.904622 | -9.1679199 | 1.25196436 | -7.3228281 |
| 1717 | NM_001137673.1 | 167.246204 | -9.0078203 | 1.2302511 | -7.3219364 |
| 1718 | NM_001318728.1 | 205.813265 | -9.3065007 | 1.27104225 | -7.3219444 |
| 1719 | NM_001242559.1 | 325.933383 | -9.9702838 | 1.36176223 | -7.321604 |

| | | | | | |
|---|---|---|---|---|---|
| 1720 | NM_017994.4 | 235.388657 | -7.1494607 | 0.97650025 | -7.3215145 |
| 1721 | NM_003131.3 | 196.364265 | -9.2394774 | 1.26220586 | -7.3201034 |
| 1722 | NM_020865.2 | 225.523121 | -9.4389323 | 1.28953055 | -7.3196655 |
| 1723 | NM_004397.5 | 259.479985 | -9.6409172 | 1.31719006 | -7.3193061 |
| 1724 | NM_001042368.2 | 310.939756 | -9.9026765 | 1.35310541 | -7.3184812 |
| 1725 | NM_001130072.1 | 230.115132 | -9.4682658 | 1.29378934 | -7.3182438 |
| 1726 | NM_004434.2 | 150.032392 | -8.8522255 | 1.20960704 | -7.3182655 |
| 1727 | NM_001018057.1 | 390.344259 | -10.231026 | 1.39804182 | -7.3181119 |
| 1728 | NM_015537.4 | 166.991282 | -9.0069697 | 1.2307935 | -7.3180186 |
| 1729 | NM_001163285.1 | 160.226361 | -8.9458528 | 1.22248456 | -7.3177634 |
| 1730 | NM_001916.4 | 247.812557 | -9.5751402 | 1.30857818 | -7.3172091 |
| 1731 | NM_006833.4 | 276.206504 | -9.7316596 | 1.33002867 | -7.3168796 |
| 1732 | NM_001330395.1 | 262.658937 | -9.659059 | 1.32021281 | -7.3162894 |
| 1733 | NM_001420.3 | 596.92585 | -10.843529 | 1.48219641 | -7.3158516 |
| 1734 | NM_003036.3 | 318.308858 | -5.7105036 | 0.78057968 | -7.3157216 |
| 1735 | NM_005880.3 | 236.43925 | -7.0518793 | 0.96394414 | -7.3156514 |
| 1736 | NM_052916.2 | 320.88013 | -9.9479018 | 1.35978496 | -7.3157905 |
| 1737 | NM_001076786.2 | 149.865208 | -5.4674988 | 0.74762225 | -7.3131837 |
| 1738 | NM_001318734.1 | 292.169844 | -9.8125388 | 1.34175626 | -7.3132052 |
| 1739 | NM_003321.4 | 469.545113 | -9.534602 | 1.30380365 | -7.3129125 |
| 1740 | NM_001166110.1 | 427.704854 | -7.098096 | 0.97075052 | -7.3119673 |
| 1741 | NM_015046.5 | 172.27995 | -6.6490863 | 0.90934535 | -7.3119484 |
| 1742 | NM_001122955.3 | 342.049579 | -10.040258 | 1.37315261 | -7.3118297 |
| 1743 | NM_000274.3 | 188.383797 | -9.1798747 | 1.25554903 | -7.3114426 |
| 1744 | NM_004628.4 | 151.933311 | -8.8708155 | 1.21328665 | -7.311393 |
| 1745 | NM_030927.2 | 294.53193 | -9.8243033 | 1.34390241 | -7.3102803 |
| 1746 | NM_006345.3 | 276.210624 | -6.9969298 | 0.95719905 | -7.3097959 |
| 1747 | NM_004965.6 | 659.598927 | -7.3492905 | 1.00541372 | -7.3097177 |
| 1748 | NM_018715.3 | 178.197858 | -9.0990011 | 1.24496024 | -7.308668 |
| 1749 | NM_013265.3 | 275.171835 | -9.7260548 | 1.33085705 | -7.3081138 |
| 1750 | NM_015277.5 | 421.792533 | -10.342412 | 1.41537739 | -7.3071765 |
| 1751 | NM_006096.3 | 275.052595 | -9.7256355 | 1.33100497 | -7.3069866 |
| 1752 | NM_007315.3 | 189.392967 | -9.1887105 | 1.25753334 | -7.3069319 |
| 1753 | NM_001270520.1 | 226.928916 | -9.4478427 | 1.29303118 | -7.30674 |
| 1754 | NM_032236.6 | 187.968284 | -9.1763556 | 1.25596628 | -7.3062117 |
| 1755 | NM_001271875.1 | 181.285239 | -9.1246909 | 1.24892521 | -7.3060347 |
| 1756 | NM_002116.7 | 233.557392 | -9.4902341 | 1.29901411 | -7.3057205 |
| 1757 | NM_016222.3 | 187.448652 | -9.1725529 | 1.25551414 | -7.3058141 |
| 1758 | NM_024294.3 | 192.412452 | -9.2107133 | 1.26075054 | -7.3057381 |
| 1759 | NM_170707.3 | 164.695422 | -8.9872695 | 1.23013973 | -7.3058932 |
| 1760 | NM_005006.6 | 175.964576 | -9.0809661 | 1.24311259 | -7.3050231 |
| 1761 | NM_002314.3 | 307.320559 | -9.8858828 | 1.35335473 | -7.3047241 |
| 1762 | NM_001212.3 | 250.6211 | -9.5914599 | 1.31334488 | -7.3030778 |

| | | | | | |
|---|---|---|---|---|---|
| 1763 | NM_006245.3 | 237.42085 | -9.5133579 | 1.30273286 | -7.302616 |
| 1764 | NM_133642.3 | 229.656066 | -9.4656914 | 1.29620877 | -7.3025978 |
| 1765 | NM_031490.4 | 147.678123 | -8.82778 | 1.20887502 | -7.3024753 |
| 1766 | NM_015125.4 | 396.703781 | -7.6710598 | 1.05050023 | -7.3022924 |
| 1767 | NM_002755.3 | 177.598314 | -9.0946261 | 1.2455593 | -7.3016403 |
| 1768 | NM_012324.4 | 405.357817 | -10.285193 | 1.40874996 | -7.3009357 |
| 1769 | NM_012137.3 | 202.002461 | -9.2804671 | 1.27118985 | -7.3006145 |
| 1770 | NM_138689.2 | 389.545788 | -10.227713 | 1.40093069 | -7.3006562 |
| 1771 | NM_014671.2 | 196.40595 | -9.2399102 | 1.26574303 | -7.299989 |
| 1772 | NM_005494.2 | 305.231367 | -9.8757443 | 1.35295599 | -7.2993832 |
| 1773 | NM_015170.2 | 209.444826 | -9.3338032 | 1.27870337 | -7.299428 |
| 1774 | NM_139323.3 | 906.772972 | -8.0075331 | 1.09702654 | -7.2993066 |
| 1775 | NM_005861.3 | 233.114491 | -9.4870893 | 1.29978208 | -7.2989845 |
| 1776 | NM_014077.3 | 252.696768 | -9.6034162 | 1.31574234 | -7.2988578 |
| 1777 | NM_016407.4 | 236.329872 | -9.5066293 | 1.30251453 | -7.2986743 |
| 1778 | NM_139279.5 | 190.281349 | -9.1954596 | 1.25987093 | -7.2987315 |
| 1779 | NM_005789.3 | 238.628213 | -9.5208937 | 1.30455796 | -7.2981761 |
| 1780 | NM_017745.5 | 180.54272 | -9.118367 | 1.24940515 | -7.2981667 |
| 1781 | NM_000362.4 | 490.171692 | -7.4790982 | 1.02524571 | -7.2949325 |
| 1782 | NM_001031684.2 | 233.653972 | -9.4903586 | 1.30095755 | -7.2949025 |
| 1783 | NM_004515.3 | 511.761917 | -7.3674045 | 1.01005183 | -7.2940856 |
| 1784 | NM_001031835.2 | 195.062568 | -9.2302647 | 1.26548848 | -7.2938354 |
| 1785 | NM_012247.4 | 202.486925 | -9.2836786 | 1.27282596 | -7.2937533 |
| 1786 | NM_001004431.2 | 164.429059 | -8.9847054 | 1.23187619 | -7.2935133 |
| 1787 | NM_021252.4 | 255.573852 | -9.6196289 | 1.31909766 | -7.2925828 |
| 1788 | NM_006823.3 | 445.029452 | -10.419836 | 1.4288566 | -7.2924294 |
| 1789 | NM_001277.2 | 166.459316 | -9.0010402 | 1.23437773 | -7.2919658 |
| 1790 | NM_032311.4 | 234.827191 | -9.4971394 | 1.30241841 | -7.2919266 |
| 1791 | NM_020337.2 | 187.485314 | -9.1722415 | 1.25789258 | -7.2917526 |
| 1792 | NM_022356.3 | 214.132138 | -9.365055 | 1.28434917 | -7.2916736 |
| 1793 | NM_152793.2 | 332.16364 | -9.9976496 | 1.37132899 | -7.2904822 |
| 1794 | NM_001673.4 | 292.318317 | -9.8134594 | 1.34610993 | -7.2902363 |
| 1795 | NM_005120.2 | 155.734101 | -8.9068004 | 1.22177165 | -7.2900696 |
| 1796 | NM_002805.5 | 233.346972 | -9.4884628 | 1.30157808 | -7.2899681 |
| 1797 | NM_002105.2 | 265.395903 | -9.6738969 | 1.32703031 | -7.2898839 |
| 1798 | NM_012109.2 | 482.39729 | -10.536243 | 1.44537221 | -7.2896398 |
| 1799 | NM_133443.3 | 229.460371 | -9.4637919 | 1.29828937 | -7.2894319 |
| 1800 | NM_018259.5 | 178.553727 | -9.1023566 | 1.24876321 | -7.2890973 |
| 1801 | NM_001878.3 | 197.00035 | -9.2457179 | 1.26848382 | -7.2887945 |
| 1802 | NM_001613.2 | 434.623511 | -10.385984 | 1.4249952 | -7.2884346 |
| 1803 | NM_030789.3 | 241.746886 | -9.5396824 | 1.30902036 | -7.2876501 |
| 1804 | NM_015178.2 | 157.324348 | -8.9193457 | 1.2239249 | -7.2874943 |
| 1805 | NM_020131.4 | 187.495317 | -9.1723779 | 1.25870145 | -7.2871751 |

| | | | | | |
|---|---|---|---|---|---|
| 1806 | NM_021095.2 | 176.303675 | -9.0857549 | 1.24680553 | -7.287227 |
| 1807 | NM_001032221.3 | 667.599013 | -10.041785 | 1.37810996 | -7.2866355 |
| 1808 | NM_020182.4 | 270.211115 | -9.7003067 | 1.33128079 | -7.2864468 |
| 1809 | NM_003634.3 | 268.368227 | -9.6901893 | 1.33018183 | -7.2848607 |
| 1810 | NM_004718.3 | 282.0694 | -9.7619305 | 1.34003904 | -7.2848105 |
| 1811 | NM_020335.2 | 431.515191 | -9.4124658 | 1.29216131 | -7.2842808 |
| 1812 | NM_004092.3 | 285.724396 | -9.7808502 | 1.34296458 | -7.2830292 |
| 1813 | NM_181523.2 | 157.564802 | -8.9227912 | 1.22514726 | -7.2830357 |
| 1814 | NM_005104.3 | 205.395571 | -9.305139 | 1.277711 | -7.2826633 |
| 1815 | NM_002887.3 | 181.715581 | -9.1280529 | 1.25350645 | -7.2820151 |
| 1816 | NM_014633.4 | 174.88588 | -9.0725319 | 1.24587398 | -7.2820622 |
| 1817 | NM_000528.3 | 175.181331 | -9.0755176 | 1.24646903 | -7.2809813 |
| 1818 | NM_015375.2 | 152.527635 | -8.8738414 | 1.21883836 | -7.2805728 |
| 1819 | NM_002319.4 | 155.341344 | -8.9030478 | 1.22287831 | -7.2804037 |
| 1820 | NM_003736.2 | 255.968822 | -9.62221 | 1.32217346 | -7.2775701 |
| 1821 | NM_024656.3 | 271.65523 | -7.1704538 | 0.98531377 | -7.2773303 |
| 1822 | NM_007362.3 | 237.572186 | -9.5140928 | 1.30747968 | -7.2766659 |
| 1823 | NM_016456.4 | 226.86944 | -9.4477171 | 1.29844481 | -7.2761792 |
| 1824 | NM_130809.4 | 155.027863 | -8.8981008 | 1.22294232 | -7.2759775 |
| 1825 | NM_001554.4 | 437.157698 | -8.0798046 | 1.11071491 | -7.274418 |
| 1826 | NM_007146.2 | 181.721702 | -9.1269439 | 1.25472555 | -7.2740559 |
| 1827 | NM_006814.4 | 224.300652 | -9.4313234 | 1.29679963 | -7.2727684 |
| 1828 | NM_024298.4 | 184.568522 | -9.1500649 | 1.2582045 | -7.2723194 |
| 1829 | NM_031484.3 | 160.243 | -8.9479877 | 1.23040549 | -7.2723893 |
| 1830 | NM_177924.4 | 278.754798 | -9.7452268 | 1.34011667 | -7.2719241 |
| 1831 | NM_032833.4 | 205.830285 | -4.4135194 | 0.60699716 | -7.271071 |
| 1832 | NM_001113363.1 | 305.228854 | -9.8755168 | 1.35825655 | -7.2707301 |
| 1833 | NM_020132.4 | 195.288236 | -9.2319065 | 1.26978335 | -7.270458 |
| 1834 | NM_012321.4 | 281.249408 | -9.7577045 | 1.3423792 | -7.2689628 |
| 1835 | NM_015185.2 | 194.476993 | -9.2251881 | 1.26927059 | -7.268102 |
| 1836 | NM_001759.3 | 822.185876 | -7.6785811 | 1.05662243 | -7.2671002 |
| 1837 | NM_021008.3 | 260.664288 | -9.6481293 | 1.32764826 | -7.2670824 |
| 1838 | NM_019044.4 | 178.945128 | -9.1051312 | 1.25295133 | -7.2669472 |
| 1839 | NM_015016.1 | 116.337548 | -2.8692278 | 0.39484901 | -7.2666456 |
| 1840 | NR_037612.1 | 347.186171 | -10.061639 | 1.38463322 | -7.2666454 |
| 1841 | NM_022051.2 | 224.548444 | -4.4424908 | 0.61148874 | -7.2650411 |
| 1842 | NR_026581.1 | 451.550584 | -9.4779463 | 1.30461648 | -7.2649291 |
| 1843 | NM_002957.5 | 164.732642 | -8.9865308 | 1.23701616 | -7.2646835 |
| 1844 | NM_001243332.1 | 235.536882 | -9.5015348 | 1.30810254 | -7.2636009 |
| 1845 | NM_006109.4 | 196.656201 | -9.2416207 | 1.27241721 | -7.2630428 |
| 1846 | NM_014614.2 | 187.133909 | -9.1700653 | 1.26270494 | -7.2622392 |
| 1847 | NM_001128846.1 | 196.486985 | -9.2392815 | 1.27233234 | -7.2616888 |
| 1848 | NM_005436.4 | 167.060648 | -9.0064785 | 1.24028398 | -7.2616261 |

| | | | | | |
|---|---|---|---|---|---|
| 1849 | NM_001032283.2 | 177.646365 | -9.0937942 | 1.25237222 | -7.2612551 |
| 1850 | NM_001170629.1 | 183.644422 | -9.1420552 | 1.25901295 | -7.2612877 |
| 1851 | NM_012121.4 | 166.92475 | -9.0047685 | 1.24010584 | -7.2612903 |
| 1852 | NM_001170937.1 | 276.9722 | -9.7362537 | 1.34122547 | -7.2592222 |
| 1853 | NM_014061.4 | 219.389749 | -9.3992939 | 1.29511698 | -7.2574864 |
| 1854 | NM_015908.5 | 166.677499 | -9.002244 | 1.24043946 | -7.2573022 |
| 1855 | NM_014737.2 | 195.225602 | -9.2310272 | 1.27201502 | -7.2570111 |
| 1856 | NM_001077516.1 | 193.036942 | -9.215148 | 1.26990806 | -7.2565474 |
| 1857 | NM_002593.3 | 232.939163 | -9.4867843 | 1.3073556 | -7.2564682 |
| 1858 | NM_015374.2 | 368.353736 | -10.146987 | 1.39854379 | -7.2553946 |
| 1859 | NM_001042486.3 | 289.566116 | -9.7997721 | 1.35093628 | -7.2540594 |
| 1860 | NM_005526.3 | 233.669194 | -9.490432 | 1.30828215 | -7.2541171 |
| 1861 | NM_032704.4 | 211.282541 | -9.3454033 | 1.28834067 | -7.2538293 |
| 1862 | NM_014931.3 | 322.639473 | -6.331147 | 0.87285519 | -7.2533761 |
| 1863 | NM_032233.2 | 149.406572 | -8.8459002 | 1.21955115 | -7.2534064 |
| 1864 | NM_001024.3 | 466.936783 | -10.489218 | 1.44627631 | -7.2525682 |
| 1865 | NM_006570.4 | 245.941478 | -9.5643021 | 1.31874012 | -7.2526057 |
| 1866 | NM_007313.2 | 156.738073 | -8.9132793 | 1.22910194 | -7.2518633 |
| 1867 | NM_002055.4 | 464.75054 | -10.482642 | 1.44557393 | -7.2515435 |
| 1868 | NM_002541.3 | 161.342698 | -8.9577339 | 1.23529656 | -7.2514845 |
| 1869 | NM_014862.3 | 411.814977 | -9.3440456 | 1.28857082 | -7.25148 |
| 1870 | NM_017873.3 | 145.286854 | -8.8047431 | 1.21425922 | -7.2511232 |
| 1871 | NM_003247.3 | 170.756436 | -9.0395872 | 1.24673556 | -7.2506051 |
| 1872 | NM_004711.4 | 233.984436 | -9.4921933 | 1.30919916 | -7.2503814 |
| 1873 | NM_001079514.2 | 154.476882 | -8.8943681 | 1.22701239 | -7.2488006 |
| 1874 | NM_003136.3 | 174.799879 | -9.0713035 | 1.25141407 | -7.2488425 |
| 1875 | NM_015160.2 | 172.071986 | -9.0490701 | 1.24848289 | -7.248053 |
| 1876 | NM_015457.2 | 178.428963 | -9.1021021 | 1.25597567 | -7.247037 |
| 1877 | NM_004615.3 | 275.671662 | -9.7290285 | 1.3425201 | -7.2468401 |
| 1878 | NM_002480.2 | 159.788449 | -8.942305 | 1.23399017 | -7.2466582 |
| 1879 | NM_178126.3 | 166.287505 | -8.9999239 | 1.24197485 | -7.2464623 |
| 1880 | NM_004448.3 | 187.747615 | -9.1763689 | 1.26645422 | -7.2457171 |
| 1881 | NM_006469.4 | 445.285611 | -9.4565995 | 1.30523398 | -7.2451374 |
| 1882 | NM_015214.2 | 163.699039 | -8.9767544 | 1.23903931 | -7.2449311 |
| 1883 | NM_024345.4 | 145.694511 | -8.8084542 | 1.2158507 | -7.2446841 |
| 1884 | NM_001242318.2 | 156.561751 | -8.9121342 | 1.23043645 | -7.2430674 |
| 1885 | NM_007007.2 | 308.548451 | -8.082635 | 1.11594455 | -7.2428643 |
| 1886 | NM_138346.2 | 176.516783 | -9.0861218 | 1.25453366 | -7.2426289 |
| 1887 | NM_001113407.2 | 304.255754 | -9.8713195 | 1.36298361 | -7.2424345 |
| 1888 | NM_015057.4 | 325.637615 | -6.9259688 | 0.95632662 | -7.2422629 |
| 1889 | NM_178502.3 | 205.606303 | -9.3058395 | 1.28510631 | -7.2412994 |
| 1890 | NM_017583.5 | 353.089366 | -9.1217456 | 1.25985802 | -7.2402965 |
| 1891 | NM_024551.2 | 211.448787 | -9.3460544 | 1.29085562 | -7.2402012 |

| | | | | | |
|---|---|---|---|---|---|
| 1892 | NM_001261420.1 | 219.253828 | -9.3993181 | 1.29824002 | -7.2400465 |
| 1893 | NM_005076.4 | 173.49762 | -9.0593813 | 1.25127542 | -7.2401177 |
| 1894 | NM_001142481.1 | 229.950701 | -9.4673937 | 1.30767742 | -7.2398541 |
| 1895 | NM_006503.3 | 253.398135 | -9.6075548 | 1.32703332 | -7.2398745 |
| 1896 | NM_006471.3 | 186.532995 | -9.1660279 | 1.26619371 | -7.2390408 |
| 1897 | NM_018230.2 | 176.761403 | -9.0876861 | 1.25541653 | -7.2387816 |
| 1898 | NM_001128148.2 | 212.469865 | -9.3534876 | 1.29215248 | -7.2386872 |
| 1899 | NM_019121.1 | 188.104128 | -9.1774802 | 1.2678998 | -7.2383324 |
| 1900 | NM_015999.5 | 154.577957 | -8.8953254 | 1.22896739 | -7.2380484 |
| 1901 | NM_004850.4 | 154.802087 | -8.8958762 | 1.22918059 | -7.237241 |
| 1902 | NM_012326.3 | 276.799034 | -9.7348365 | 1.34519118 | -7.236768 |
| 1903 | NM_182972.2 | 207.266407 | -9.3168657 | 1.28751046 | -7.2363417 |
| 1904 | NR_024077.2 | 150.711865 | -8.8595362 | 1.22434324 | -7.236154 |
| 1905 | NM_001031714.3 | 169.391013 | -9.0264764 | 1.24758004 | -7.2351882 |
| 1906 | NM_001068.3 | 343.118677 | -9.0809071 | 1.25519308 | -7.2346695 |
| 1907 | NM_001321473.1 | 176.932993 | -9.0901022 | 1.25646254 | -7.2346783 |
| 1908 | NM_006423.2 | 288.042396 | -9.7922519 | 1.35369441 | -7.2337241 |
| 1909 | NM_020150.4 | 229.732357 | -9.465993 | 1.30860447 | -7.2336548 |
| 1910 | NM_001242836.1 | 399.621478 | -7.3641656 | 1.01814231 | -7.2329434 |
| 1911 | NM_001689.4 | 340.486906 | -7.3146935 | 1.01138266 | -7.2323698 |
| 1912 | NM_003461.4 | 227.01586 | -9.4490878 | 1.30658148 | -7.2319162 |
| 1913 | NM_001284215.1 | 192.649995 | -9.2112181 | 1.27388887 | -7.2307863 |
| 1914 | NM_016026.3 | 314.599591 | -9.9195899 | 1.37195465 | -7.2302608 |
| 1915 | NM_018490.3 | 239.00361 | -9.5234309 | 1.31721465 | -7.2299765 |
| 1916 | NM_173701.1 | 217.086945 | -9.3842776 | 1.29795555 | -7.2300454 |
| 1917 | NM_001278556.1 | 304.629377 | -9.8729699 | 1.36569354 | -7.2292719 |
| 1918 | NM_001291964.1 | 158.635122 | -8.9310277 | 1.23555274 | -7.2283662 |
| 1919 | NM_001001391.1 | 472.27733 | -10.505803 | 1.45361797 | -7.2273479 |
| 1920 | NM_138473.2 | 138.216292 | -8.7335015 | 1.20844093 | -7.2270819 |
| 1921 | NM_012197.3 | 258.918977 | -7.2959467 | 1.00955018 | -7.2269282 |
| 1922 | NM_005004.3 | 252.567562 | -9.6027258 | 1.3288453 | -7.2263685 |
| 1923 | NR_138470.1 | 265.880887 | -9.6765336 | 1.3392586 | -7.2252914 |
| 1924 | NM_001199281.1 | 144.281515 | -8.7957522 | 1.21738124 | -7.2251418 |
| 1925 | NM_012398.2 | 269.546174 | -9.696324 | 1.34206597 | -7.2249236 |
| 1926 | NM_017789.4 | 270.293633 | -9.7005386 | 1.34270282 | -7.2246356 |
| 1927 | NM_022760.4 | 153.890484 | -8.8887347 | 1.23037618 | -7.2244041 |
| 1928 | NM_018650.4 | 151.843854 | -8.8671031 | 1.22741729 | -7.224196 |
| 1929 | NM_002356.6 | 1762.21342 | -8.3656632 | 1.15819296 | -7.2230306 |
| 1930 | NM_006565.3 | 171.888786 | -9.0470947 | 1.25270009 | -7.2220755 |
| 1931 | NM_012233.2 | 152.261394 | -8.8722481 | 1.22854339 | -7.2217621 |
| 1932 | NM_005234.3 | 176.099985 | -9.0833926 | 1.25780405 | -7.2216278 |
| 1933 | NM_053023.4 | 220.019876 | -9.4037281 | 1.30233375 | -7.2206745 |
| 1934 | NM_001308176.1 | 198.08663 | -9.2518017 | 1.28138452 | -7.2201603 |

| | | | | | |
|---|---|---|---|---|---|
| 1935 | NM_032905.4 | 173.543114 | -9.0613214 | 1.25503051 | -7.2200009 |
| 1936 | NM_001083913.1 | 171.728831 | -9.0462569 | 1.25297763 | -7.2198072 |
| 1937 | NM_002512.3 | 345.857255 | -10.056138 | 1.39288367 | -7.2196537 |
| 1938 | NM_006328.3 | 192.989102 | -9.2146002 | 1.276336 | -7.2195725 |
| 1939 | NM_001318187.1 | 174.239064 | -9.0680885 | 1.2562117 | -7.218599 |
| 1940 | NM_017595.5 | 197.769841 | -9.2502953 | 1.28155098 | -7.2180471 |
| 1941 | NM_001697.2 | 298.607219 | -9.8443018 | 1.36396089 | -7.217437 |
| 1942 | NM_004627.4 | 185.886636 | -9.1600858 | 1.26916068 | -7.2174358 |
| 1943 | NM_001170687.1 | 149.351753 | -8.8449499 | 1.22552657 | -7.2172649 |
| 1944 | NR_026943.1 | 227.913595 | -6.6518594 | 0.92171844 | -7.2168019 |
| 1945 | NM_145686.3 | 257.628807 | -9.6311962 | 1.3346907 | -7.2160511 |
| 1946 | NM_001128847.1 | 204.419245 | -9.2975888 | 1.28866792 | -7.2148835 |
| 1947 | NM_080645.2 | 544.283346 | -10.710503 | 1.48458187 | -7.2144913 |
| 1948 | NM_004172.4 | 158.420087 | -8.9284298 | 1.2375891 | -7.2143733 |
| 1949 | NM_004879.4 | 232.860945 | -9.4853798 | 1.31495956 | -7.2134385 |
| 1950 | NM_001286050.1 | 362.974182 | -10.125682 | 1.40374603 | -7.2133289 |
| 1951 | NM_001492.5 | 261.982842 | -9.6554882 | 1.33862684 | -7.2129797 |
| 1952 | NM_022720.6 | 180.790979 | -9.1204212 | 1.26456709 | -7.2122874 |
| 1953 | NM_002841.3 | 184.568597 | -6.7059907 | 0.92981601 | -7.2121696 |
| 1954 | NM_002893.3 | 350.629528 | -9.1116276 | 1.26341479 | -7.2119051 |
| 1955 | NM_001144952.1 | 147.226548 | -8.8259326 | 1.2238673 | -7.211511 |
| 1956 | NM_032595.4 | 447.840859 | -9.4650637 | 1.31249814 | -7.2114873 |
| 1957 | NM_017542.4 | 210.796149 | -9.3417167 | 1.29541936 | -7.2113456 |
| 1958 | NM_001164803.1 | 209.92895 | -9.3355696 | 1.29466985 | -7.2107724 |
| 1959 | NM_004047.4 | 333.70611 | -10.004526 | 1.38744722 | -7.2107433 |
| 1960 | NM_024731.3 | 151.830963 | -8.8687913 | 1.23004506 | -7.2101353 |
| 1961 | NM_001135699.1 | 280.298927 | -9.7530688 | 1.35284512 | -7.2093019 |
| 1962 | NM_006197.3 | 187.738146 | -9.1747829 | 1.27269831 | -7.208922 |
| 1963 | NM_017670.2 | 214.456352 | -9.3666776 | 1.29952149 | -7.2077897 |
| 1964 | NM_002938.4 | 181.808885 | -9.1281471 | 1.26655166 | -7.2070863 |
| 1965 | NM_053024.3 | 593.77151 | -9.8732085 | 1.36993912 | -7.2070418 |
| 1966 | NM_004378.2 | 265.274141 | -9.6738299 | 1.34241271 | -7.2063009 |
| 1967 | NM_007185.6 | 349.337254 | -10.070524 | 1.39752296 | -7.2059808 |
| 1968 | NM_005529.6 | 290.537346 | -5.3207323 | 0.73839584 | -7.2057993 |
| 1969 | NM_001080517.2 | 243.274347 | -9.5496 | 1.32542538 | -7.2049322 |
| 1970 | NM_001286.3 | 228.553494 | -9.4586744 | 1.31285391 | -7.2046663 |
| 1971 | NM_021098.2 | 260.181417 | -9.6451534 | 1.33877578 | -7.2044576 |
| 1972 | NM_002952.3 | 3913.22168 | -7.8789237 | 1.09366611 | -7.2041399 |
| 1973 | NM_016462.3 | 186.413604 | -9.1645783 | 1.27213473 | -7.2040941 |
| 1974 | NM_012104.4 | 195.42717 | -9.2331508 | 1.28178131 | -7.2033745 |
| 1975 | NM_021942.5 | 152.426873 | -8.8732639 | 1.23188951 | -7.2029705 |
| 1976 | NM_174907.3 | 181.21765 | -9.1232138 | 1.26666798 | -7.2025297 |
| 1977 | NM_005072.4 | 147.412651 | -8.8274609 | 1.22582059 | -7.2012666 |

| | | | | | |
|---|---|---|---|---|---|
| 1978 | NM_001146334.1 | 311.66749 | -9.9060754 | 1.37566068 | -7.2009584 |
| 1979 | NM_001321075.1 | 232.20572 | -9.4808188 | 1.31665866 | -7.2006656 |
| 1980 | NM_003150.3 | 179.852421 | -9.1121964 | 1.26547944 | -7.2005883 |
| 1981 | NM_012245.2 | 155.921009 | -8.9068986 | 1.23704192 | -7.200159 |
| 1982 | NM_006622.3 | 297.405247 | -9.8383734 | 1.36647362 | -7.1998268 |
| 1983 | NM_014811.3 | 157.748273 | -8.9238998 | 1.23946514 | -7.199799 |
| 1984 | NM_001136528.1 | 264.946746 | -9.6718636 | 1.34341627 | -7.1994539 |
| 1985 | NM_003091.3 | 226.619528 | -9.4460004 | 1.31205966 | -7.1993681 |
| 1986 | NM_006595.3 | 146.195979 | -8.812799 | 1.2240891 | -7.1994751 |
| 1987 | NM_014864.3 | 194.631664 | -9.2258375 | 1.28148825 | -7.199315 |
| 1988 | NM_016025.4 | 210.653865 | -9.3409789 | 1.29744209 | -7.1995343 |
| 1989 | NM_016211.4 | 135.038223 | -8.6986992 | 1.20824404 | -7.1994555 |
| 1990 | NM_024086.3 | 121.722939 | -3.4523843 | 0.47953668 | -7.1994164 |
| 1991 | NM_206962.3 | 186.597717 | -9.1654701 | 1.27316381 | -7.1989716 |
| 1992 | NM_001077198.2 | 195.320047 | -9.2320471 | 1.28267429 | -7.1974991 |
| 1993 | NM_003651.4 | 164.578806 | -8.9864256 | 1.2485369 | -7.1975651 |
| 1994 | NM_001128912.1 | 209.118574 | -9.329265 | 1.29628694 | -7.1969136 |
| 1995 | NM_004774.3 | 184.923833 | -9.1528253 | 1.27196864 | -7.1957948 |
| 1996 | NM_006360.5 | 240.324454 | -9.5308481 | 1.32457625 | -7.1953941 |
| 1997 | NM_001100423.1 | 149.114284 | -8.8438108 | 1.22911514 | -7.1952663 |
| 1998 | NM_006827.5 | 614.484617 | -4.7923306 | 0.66612487 | -7.1943427 |
| 1999 | NM_004756.4 | 196.414405 | -9.2393088 | 1.28427376 | -7.1941895 |
| 2000 | NM_198976.2 | 194.70901 | -9.2273065 | 1.28273117 | -7.1934842 |
| 2001 | NM_001304717.2 | 313.570286 | -8.1063138 | 1.12706287 | -7.1924238 |
| 2002 | NM_001456.3 | 831.183073 | -7.406023 | 1.02987579 | -7.1911808 |
| 2003 | NM_145725.2 | 154.59267 | -8.8959325 | 1.23705175 | -7.1912371 |
| 2004 | NM_001199698.1 | 208.214472 | -9.3249044 | 1.29683497 | -7.1905097 |
| 2005 | NM_001291411.1 | 181.931146 | -9.1300731 | 1.26972862 | -7.1905704 |
| 2006 | NM_001207044.1 | 412.639299 | -10.310856 | 1.43411141 | -7.1897177 |
| 2007 | NM_004844.4 | 231.813039 | -9.4789333 | 1.31860005 | -7.1886341 |
| 2008 | NM_001318100.1 | 187.55371 | -9.1729939 | 1.27623854 | -7.187523 |
| 2009 | NM_001010972.1 | 221.48491 | -9.4135912 | 1.30990929 | -7.1864451 |
| 2010 | NM_003325.3 | 197.129718 | -9.2451034 | 1.28645269 | -7.1865087 |
| 2011 | NM_001242560.1 | 383.084767 | -10.203583 | 1.41993662 | -7.1859427 |
| 2012 | NM_052876.3 | 216.21165 | -9.3785718 | 1.30514505 | -7.1858463 |
| 2013 | NM_025137.3 | 142.751579 | -8.7803511 | 1.2220208 | -7.1851077 |
| 2014 | NM_014752.2 | 228.385835 | -9.4575556 | 1.31631715 | -7.1848609 |
| 2015 | NM_001307924.2 | 371.62364 | -9.195288 | 1.279891 | -7.1844306 |
| 2016 | NM_015153.3 | 142.484554 | -8.7768191 | 1.22165654 | -7.184359 |
| 2017 | NM_001321208.1 | 218.684196 | -9.3948241 | 1.30771192 | -7.1841695 |
| 2018 | NM_004461.2 | 223.954501 | -9.4291669 | 1.31249573 | -7.1841505 |
| 2019 | NM_004386.2 | 560.250513 | -10.751981 | 1.49665437 | -7.1840109 |
| 2020 | NM_014827.5 | 194.59187 | -9.2259168 | 1.28427455 | -7.1837574 |

| | | | | | |
|---|---|---|---|---|---|
| 2021 | NM_001082579.1 | 357.614345 | -10.104902 | 1.40686485 | -7.1825675 |
| 2022 | NM_006302.2 | 194.324653 | -6.1036079 | 0.84977761 | -7.1825943 |
| 2023 | NM_001271223.2 | 138.270761 | -8.7330835 | 1.21599176 | -7.1818608 |
| 2024 | NM_002508.2 | 167.676057 | -9.0132965 | 1.25500527 | -7.1818794 |
| 2025 | NM_002814.3 | 187.736374 | -9.1738953 | 1.27740945 | -7.1816404 |
| 2026 | NM_023018.4 | 182.286995 | -9.132458 | 1.27165893 | -7.181531 |
| 2027 | NM_001032288.2 | 276.287809 | -9.7322654 | 1.35522462 | -7.1812932 |
| 2028 | NM_004607.2 | 245.11808 | -9.5593995 | 1.33116026 | -7.1812537 |
| 2029 | NM_012446.4 | 211.791773 | -9.3479634 | 1.30173135 | -7.1811772 |
| 2030 | NM_015695.2 | 153.139444 | -8.8816707 | 1.23685874 | -7.1808287 |
| 2031 | NM_001082959.1 | 139.564379 | -8.7462396 | 1.21805 | -7.1805259 |
| 2032 | NM_001830.3 | 294.588682 | -9.8244965 | 1.36821485 | -7.1805218 |
| 2033 | NM_001001973.2 | 235.883726 | -9.5037808 | 1.32362898 | -7.1800942 |
| 2034 | NM_013374.5 | 210.296295 | -5.5795014 | 0.77709304 | -7.1799657 |
| 2035 | NM_015354.2 | 189.824229 | -9.1905958 | 1.28004413 | -7.1799054 |
| 2036 | NM_001204077.1 | 188.036946 | -9.1768312 | 1.27816587 | -7.1796873 |
| 2037 | NM_014412.2 | 207.742587 | -9.3206537 | 1.29827752 | -7.179246 |
| 2038 | NM_007190.3 | 142.58902 | -8.7787493 | 1.22296502 | -7.1782505 |
| 2039 | NM_032876.5 | 160.159496 | -8.9472567 | 1.24644738 | -7.1782065 |
| 2040 | NM_020062.3 | 165.584222 | -8.9935635 | 1.25291852 | -7.1780913 |
| 2041 | NM_006646.5 | 245.010327 | -9.5585933 | 1.33177826 | -7.1773159 |
| 2042 | NM_001412.3 | 398.591023 | -9.2967035 | 1.29531054 | -7.1772005 |
| 2043 | NM_015251.2 | 241.651349 | -9.538859 | 1.32916063 | -7.1766036 |
| 2044 | NM_018979.3 | 178.396875 | -9.1011849 | 1.26816954 | -7.176631 |
| 2045 | NM_006079.4 | 198.991919 | -9.2582298 | 1.29007211 | -7.1765212 |
| 2046 | NM_002797.4 | 247.231261 | -9.5717287 | 1.33384052 | -7.1760668 |
| 2047 | NM_004381.4 | 208.104334 | -9.3233491 | 1.29930087 | -7.175666 |
| 2048 | NM_001202502.1 | 228.604923 | -9.459168 | 1.31826799 | -7.1754515 |
| 2049 | NM_206915.2 | 251.901814 | -9.598613 | 1.33772502 | -7.1753259 |
| 2050 | NM_003827.3 | 211.255853 | -9.3449652 | 1.30241708 | -7.1750942 |
| 2051 | NM_005548.2 | 239.567181 | -9.5263733 | 1.32781142 | -7.1744927 |
| 2052 | NR_130745.1 | 144.091735 | -8.792952 | 1.22564124 | -7.1741645 |
| 2053 | NM_001022.3 | 693.137267 | -10.096684 | 1.40751792 | -7.1733963 |
| 2054 | NM_001288738.1 | 215.189703 | -9.371739 | 1.30648513 | -7.1732458 |
| 2055 | NM_001142289.2 | 191.463946 | -9.2029256 | 1.28303693 | -7.1727675 |
| 2056 | NM_004162.4 | 177.251676 | -9.0916558 | 1.26752234 | -7.1727776 |
| 2057 | NM_001436.3 | 262.715486 | -9.6592475 | 1.34667242 | -7.1726779 |
| 2058 | NM_013237.3 | 252.821592 | -9.6040586 | 1.3389869 | -7.1726308 |
| 2059 | NM_004597.6 | 282.689447 | -9.7651498 | 1.36151799 | -7.1722517 |
| 2060 | NM_014726.2 | 225.52404 | -9.4392468 | 1.31617493 | -7.1717266 |
| 2061 | NM_183001.4 | 192.644163 | -9.2119277 | 1.28453094 | -7.1714331 |
| 2062 | NM_017588.2 | 176.953552 | -9.0889365 | 1.26741771 | -7.1712241 |
| 2063 | NM_007145.2 | 193.457714 | -9.2168516 | 1.28526692 | -7.1711576 |

| | | | | | |
|---|---|---|---|---|---|
| 2064 | NM_020764.3 | 245.823641 | -9.5637029 | 1.33380892 | -7.1702197 |
| 2065 | NM_014516.3 | 160.198323 | -8.9460532 | 1.2478521 | -7.1691614 |
| 2066 | NM_001329.3 | 180.264806 | -9.1165086 | 1.27166278 | -7.1689671 |
| 2067 | NM_001806.3 | 235.94954 | -9.5040822 | 1.32583195 | -7.1683912 |
| 2068 | NM_001009881.2 | 145.37325 | -8.8071808 | 1.22872838 | -7.1677198 |
| 2069 | NM_007111.4 | 157.072375 | -8.9168938 | 1.24408896 | -7.1674085 |
| 2070 | NM_004592.3 | 186.499779 | -9.1648379 | 1.27871913 | -7.1672017 |
| 2071 | NM_001291986.1 | 160.204064 | -8.9473407 | 1.2484194 | -7.166935 |
| 2072 | NM_001634.5 | 159.23559 | -8.9359354 | 1.24703306 | -7.1657566 |
| 2073 | NM_007371.3 | 379.177045 | -9.2245222 | 1.28739488 | -7.1652625 |
| 2074 | NM_001292043.1 | 226.806281 | -9.4464107 | 1.31837773 | -7.1651777 |
| 2075 | NM_080732.3 | 235.631876 | -6.0843816 | 0.84924856 | -7.1644297 |
| 2076 | NM_015629.3 | 183.087137 | -9.1387081 | 1.27572762 | -7.1635261 |
| 2077 | NM_002786.3 | 222.275507 | -9.4183626 | 1.31486451 | -7.1629909 |
| 2078 | NM_032512.3 | 335.143151 | -10.010875 | 1.39770106 | -7.1623863 |
| 2079 | NM_012203.1 | 206.810323 | -9.3143993 | 1.30051714 | -7.1620734 |
| 2080 | NM_005412.5 | 327.252523 | -9.9761201 | 1.39307092 | -7.1612435 |
| 2081 | NM_016343.3 | 166.472271 | -9.0006748 | 1.25693996 | -7.1607834 |
| 2082 | NM_052844.3 | 145.180828 | -8.8047949 | 1.22961899 | -7.160588 |
| 2083 | NM_001318509.1 | 305.320708 | -9.8763955 | 1.37929371 | -7.1604731 |
| 2084 | NM_032888.3 | 159.480051 | -6.569216 | 0.91746278 | -7.160199 |
| 2085 | NM_005708.4 | 180.674795 | -9.120175 | 1.27403347 | -7.158505 |
| 2086 | NM_194434.2 | 370.399417 | -5.2589607 | 0.73469621 | -7.1580072 |
| 2087 | NM_022737.2 | 318.693956 | -9.9382162 | 1.38855076 | -7.1572581 |
| 2088 | NM_001025370.2 | 161.516655 | -8.9584228 | 1.25201323 | -7.1552142 |
| 2089 | NM_032442.2 | 204.669537 | -9.2990072 | 1.29962662 | -7.1551375 |
| 2090 | NM_198880.2 | 166.589721 | -9.0029645 | 1.25826249 | -7.1550766 |
| 2091 | NM_001142495.1 | 641.29145 | -6.0230603 | 0.84179775 | -7.1549969 |
| 2092 | NM_002103.4 | 148.523098 | -8.837596 | 1.23528883 | -7.154275 |
| 2093 | NM_015164.2 | 277.105422 | -6.4500105 | 0.90164746 | -7.1535837 |
| 2094 | NM_014712.2 | 140.710857 | -8.7586934 | 1.22473193 | -7.1515188 |
| 2095 | NM_138962.3 | 177.47437 | -9.0945588 | 1.2717544 | -7.1511911 |
| 2096 | NM_004064.4 | 276.952287 | -9.7352427 | 1.36147737 | -7.1504991 |
| 2097 | NM_014216.5 | 222.711669 | -9.4208278 | 1.3174966 | -7.1505519 |
| 2098 | NM_004475.2 | 284.131712 | -7.9868732 | 1.11698099 | -7.150411 |
| 2099 | NM_014341.2 | 175.799374 | -9.0812123 | 1.27010074 | -7.1499937 |
| 2100 | NM_004631.4 | 151.157853 | -8.8637961 | 1.23970804 | -7.1499061 |
| 2101 | NM_080598.5 | 315.028577 | -9.9214073 | 1.38789579 | -7.1485247 |
| 2102 | NM_001282539.1 | 449.053767 | -10.432916 | 1.45946702 | -7.1484424 |
| 2103 | NM_005121.2 | 179.304804 | -6.3493756 | 0.88825794 | -7.1481214 |
| 2104 | NM_001039355.2 | 185.712163 | -9.1585819 | 1.28134358 | -7.1476394 |
| 2105 | NM_012336.3 | 207.739772 | -9.3199813 | 1.30399728 | -7.1472398 |
| 2106 | NM_000183.2 | 140.725138 | -8.7604467 | 1.22576818 | -7.1469034 |

| | | | | | |
|---|---|---|---|---|---|
| 2107 | NM_001014431.1 | 379.381887 | -8.2817309 | 1.15890176 | -7.146189 |
| 2108 | NM_021254.2 | 138.969286 | -8.7427232 | 1.22353646 | -7.1454538 |
| 2109 | NM_001345868.1 | 81.3999793 | 11.2691804 | 1.57717386 | 7.14517325 |
| 2110 | NM_001320596.1 | 267.872251 | -9.6877742 | 1.35590108 | -7.1448975 |
| 2111 | NM_001136195.1 | 215.179978 | -9.3711422 | 1.31159962 | -7.1448192 |
| 2112 | NM_022473.2 | 136.649444 | -8.7170695 | 1.22013401 | -7.1443541 |
| 2113 | NM_058163.2 | 148.303764 | -8.8343622 | 1.23663979 | -7.1438444 |
| 2114 | NM_014918.4 | 132.475691 | -8.6723342 | 1.2139805 | -7.1437179 |
| 2115 | NM_173587.3 | 323.765911 | -9.9605873 | 1.39436174 | -7.1434743 |
| 2116 | NM_001286077.1 | 240.471608 | -9.5317766 | 1.33448248 | -7.1426765 |
| 2117 | NM_002634.3 | 228.986474 | -9.4611703 | 1.32458855 | -7.1427239 |
| 2118 | NM_020307.3 | 140.880589 | -8.7603391 | 1.22654709 | -7.1422771 |
| 2119 | NM_002489.3 | 390.005088 | -7.8268909 | 1.09592946 | -7.1417835 |
| 2120 | NM_015050.2 | 170.737611 | -9.0380532 | 1.26564927 | -7.1410409 |
| 2121 | NM_015379.4 | 227.262478 | -9.4504995 | 1.32339516 | -7.1411018 |
| 2122 | NM_002746.2 | 187.869465 | -9.175712 | 1.28502152 | -7.1405123 |
| 2123 | NM_138635.3 | 313.143331 | -6.9544447 | 0.97394141 | -7.1405165 |
| 2124 | NM_001190987.1 | 205.446615 | -9.3042913 | 1.30316807 | -7.1397477 |
| 2125 | NM_019015.2 | 167.208251 | -9.0082559 | 1.26173227 | -7.1395938 |
| 2126 | NM_005516.5 | 185.506291 | -3.9090701 | 0.54753981 | -7.1393349 |
| 2127 | NM_014847.3 | 300.732649 | -8.8913822 | 1.24543724 | -7.1391652 |
| 2128 | NM_016410.5 | 199.464481 | -9.2619396 | 1.29751176 | -7.1382317 |
| 2129 | NM_004295.3 | 298.535781 | -8.8791393 | 1.24398015 | -7.1376856 |
| 2130 | NM_001168370.1 | 128.382364 | -8.6264883 | 1.2088623 | -7.1360388 |
| 2131 | NM_016306.5 | 209.084521 | -9.3303767 | 1.3074952 | -7.1360696 |
| 2132 | NM_020198.2 | 233.080526 | -7.6640908 | 1.07399099 | -7.1360848 |
| 2133 | NM_152727.5 | 130.174264 | -8.6476105 | 1.21181407 | -7.1360868 |
| 2134 | NM_004687.4 | 339.43659 | -8.1813148 | 1.14655265 | -7.135577 |
| 2135 | NM_015289.3 | 311.673722 | -8.9411708 | 1.25306226 | -7.1354561 |
| 2136 | NM_001252100.1 | 406.69128 | -10.28993 | 1.44226893 | -7.1345429 |
| 2137 | NM_005655.3 | 126.974914 | -8.610605 | 1.20691258 | -7.1344065 |
| 2138 | NM_004587.2 | 278.533006 | -8.7815338 | 1.23089848 | -7.134247 |
| 2139 | NM_003010.3 | 206.614942 | -9.312886 | 1.30545352 | -7.1338319 |
| 2140 | NM_001009999.2 | 222.399028 | -9.4190449 | 1.32045483 | -7.1331822 |
| 2141 | NM_000251.2 | 156.471424 | -8.9114686 | 1.24932398 | -7.1330326 |
| 2142 | NM_001011553.3 | 269.549254 | -9.6961919 | 1.35939167 | -7.1327434 |
| 2143 | NM_138387.3 | 185.031532 | -9.1543595 | 1.28348426 | -7.1324283 |
| 2144 | NM_016032.3 | 207.946047 | -9.3225532 | 1.30711471 | -7.1321615 |
| 2145 | NM_182649.1 | 181.397314 | -9.1244298 | 1.27938895 | -7.1318654 |
| 2146 | NM_183079.2 | 144.300483 | -8.7944966 | 1.23336881 | -7.1304678 |
| 2147 | NM_017526.4 | 136.948377 | -8.721168 | 1.22311489 | -7.1302934 |
| 2148 | NM_001142551.1 | 291.32714 | -9.8085234 | 1.37563996 | -7.130153 |
| 2149 | NM_001829.3 | 174.884743 | -9.071892 | 1.27243381 | -7.1295591 |

| | | | | | |
|---|---|---|---|---|---|
| 2150 | NM_022080.2 | 300.904465 | -9.8554685 | 1.38236655 | -7.1294177 |
| 2151 | NM_144599.4 | 203.345449 | -9.2898797 | 1.303039 | -7.129395 |
| 2152 | NM_014366.4 | 168.589573 | -9.0194961 | 1.26527692 | -7.128476 |
| 2153 | NM_001284389.1 | 175.641774 | -9.0771069 | 1.27337895 | -7.1283626 |
| 2154 | NM_015215.3 | 260.015603 | -9.6446169 | 1.35299521 | -7.1283452 |
| 2155 | NM_003028.2 | 176.016365 | -9.0814014 | 1.2740368 | -7.1280527 |
| 2156 | NM_015100.3 | 141.199894 | -8.7631866 | 1.22939708 | -7.128036 |
| 2157 | NM_016617.3 | 175.83038 | -9.0790354 | 1.27371325 | -7.1280058 |
| 2158 | NR_028052.1 | 797.956888 | -7.6381598 | 1.07169128 | -7.1272017 |
| 2159 | NM_001099285.1 | 287.754036 | -9.7906978 | 1.37382772 | -7.1265834 |
| 2160 | NM_015037.3 | 199.97501 | -9.2670105 | 1.30039418 | -7.1263089 |
| 2161 | NM_012106.3 | 189.879391 | -9.1911064 | 1.28998423 | -7.1249758 |
| 2162 | NM_001024401.2 | 1043.21663 | -5.4970644 | 0.77162508 | -7.1240096 |
| 2163 | NM_016303.2 | 151.527993 | -8.8662841 | 1.24462173 | -7.1236777 |
| 2164 | NM_014153.3 | 147.055998 | -8.8223637 | 1.23847425 | -7.1235746 |
| 2165 | NM_019023.2 | 127.317371 | -8.6146671 | 1.20973308 | -7.1211305 |
| 2166 | NM_001300792.1 | 324.839993 | -9.9655698 | 1.39948301 | -7.1208937 |
| 2167 | NM_021975.3 | 167.913854 | -9.0137781 | 1.26592017 | -7.1203369 |
| 2168 | NM_198252.2 | 136.477386 | -8.7161474 | 1.22420822 | -7.1198243 |
| 2169 | NM_001113493.1 | 187.503417 | -9.1728239 | 1.28840902 | -7.1194968 |
| 2170 | NM_004872.4 | 176.061305 | -9.0833498 | 1.27585166 | -7.1194404 |
| 2171 | NM_015670.5 | 203.784654 | -9.2931006 | 1.30531298 | -7.1194424 |
| 2172 | NM_006339.2 | 152.684727 | -8.8767028 | 1.24687815 | -7.1191421 |
| 2173 | NM_005080.3 | 185.108341 | -9.1551227 | 1.28607931 | -7.1186299 |
| 2174 | NM_014975.2 | 407.602015 | -6.954252 | 0.97696419 | -7.1182261 |
| 2175 | NM_003557.2 | 154.505316 | -8.8921155 | 1.24931747 | -7.1175788 |
| 2176 | NM_019102.3 | 314.181732 | -9.9177889 | 1.39343605 | -7.1175056 |
| 2177 | NM_001144062.1 | 168.800567 | -9.0226756 | 1.26787454 | -7.1163789 |
| 2178 | NM_003075.4 | 147.654337 | -8.8274804 | 1.24056316 | -7.1157041 |
| 2179 | NR_034144.1 | 79.6160688 | 3.7403903 | 0.52565973 | 7.11561122 |
| 2180 | NM_000700.2 | 206.790964 | -9.3149012 | 1.30918471 | -7.1150398 |
| 2181 | NM_001170718.1 | 176.386718 | -9.0844841 | 1.27715216 | -7.1130789 |
| 2182 | NM_000041.3 | 196.663095 | -9.2430159 | 1.29966669 | -7.1118356 |
| 2183 | NR_024019.3 | 138.999425 | -8.7414618 | 1.22915006 | -7.1117938 |
| 2184 | NM_002899.3 | 519.908158 | -7.1846825 | 1.01033076 | -7.1112182 |
| 2185 | NM_005336.5 | 173.570214 | -9.0628381 | 1.27447152 | -7.1110558 |
| 2186 | NM_001289823.1 | 159.814455 | -8.9430167 | 1.25767187 | -7.110771 |
| 2187 | NM_005899.4 | 169.609418 | -9.026581 | 1.26946027 | -7.110566 |
| 2188 | NM_138795.3 | 317.483148 | -9.9326431 | 1.39690721 | -7.110453 |
| 2189 | NM_014313.3 | 207.11946 | -9.3161835 | 1.31036175 | -7.1096272 |
| 2190 | NM_198935.2 | 218.836046 | -9.395366 | 1.32165347 | -7.1087968 |
| 2191 | NM_003375.4 | 276.454287 | -9.7329504 | 1.36923 | -7.1083386 |
| 2192 | NM_152493.2 | 242.798438 | -9.5457412 | 1.34289072 | -7.1083529 |

| | | | | | |
|---|---|---|---|---|---|
| 2193 | NM_002267.3 | 231.19613 | -9.4750036 | 1.33300436 | -7.1080064 |
| 2194 | NM_032272.4 | 211.299345 | -9.3450137 | 1.3148921 | -7.1070575 |
| 2195 | NM_177535.2 | 258.379203 | -9.6363442 | 1.35594151 | -7.106755 |
| 2196 | NM_031866.2 | 139.85164 | -8.7482539 | 1.23103013 | -7.1064499 |
| 2197 | NM_012179.3 | 153.141756 | -8.8807556 | 1.24968653 | -7.1063866 |
| 2198 | NM_016132.4 | 141.412986 | -8.7672766 | 1.23373537 | -7.1062861 |
| 2199 | NM_001001.4 | 189.08215 | -9.1851386 | 1.29255494 | -7.1061881 |
| 2200 | NM_001681.3 | 385.609093 | -9.2500977 | 1.30171411 | -7.10609 |
| 2201 | NM_005497.3 | 150.741366 | -8.8574213 | 1.24656959 | -7.1054367 |
| 2202 | NM_005628.2 | 198.092997 | -9.2521252 | 1.30213032 | -7.1053757 |
| 2203 | NM_018936.3 | 204.546416 | -9.2986541 | 1.30877103 | -7.1048746 |
| 2204 | NM_001198665.1 | 176.450234 | -9.0866483 | 1.27909445 | -7.1039697 |
| 2205 | NM_005439.2 | 204.562029 | -9.2987013 | 1.3089868 | -7.1037395 |
| 2206 | NM_018476.3 | 425.549663 | -10.355322 | 1.45795301 | -7.1026447 |
| 2207 | NM_033081.2 | 129.548837 | -8.6378652 | 1.21615461 | -7.1026045 |
| 2208 | NM_002856.2 | 220.504155 | -9.4071048 | 1.32450524 | -7.1023538 |
| 2209 | NM_003872.2 | 245.666245 | -9.5628779 | 1.34651313 | -7.1019567 |
| 2210 | NM_016039.2 | 213.685822 | -9.3614109 | 1.31835765 | -7.1008128 |
| 2211 | NM_001321092.1 | 216.308896 | -9.3789584 | 1.32090593 | -7.1003984 |
| 2212 | NM_012112.4 | 164.728782 | -8.9856519 | 1.265514 | -7.1003971 |
| 2213 | NM_014909.4 | 374.514945 | -8.3813553 | 1.18039022 | -7.1004954 |
| 2214 | NM_001105203.1 | 266.128829 | -9.6780304 | 1.36321046 | -7.0994397 |
| 2215 | NM_001170755.2 | 220.333835 | -6.1044708 | 0.85984654 | -7.0994887 |
| 2216 | NM_001079872.1 | 178.800276 | -9.104982 | 1.28267652 | -7.0984241 |
| 2217 | NM_001278216.1 | 137.12313 | -8.7225809 | 1.22884191 | -7.0982124 |
| 2218 | NM_003400.3 | 644.828919 | -7.1584827 | 1.00847673 | -7.0983122 |
| 2219 | NM_019845.2 | 366.616379 | -10.14046 | 1.42859763 | -7.0981921 |
| 2220 | NM_005088.2 | 151.795753 | -8.8688224 | 1.24947521 | -7.0980379 |
| 2221 | NM_001512.3 | 245.446302 | -9.5612246 | 1.34720113 | -7.0971026 |
| 2222 | NM_018013.3 | 220.762414 | -9.4086203 | 1.32572457 | -7.0969646 |
| 2223 | NM_005415.4 | 221.755993 | -7.5626699 | 1.06563911 | -7.0968397 |
| 2224 | NM_173689.6 | 145.592048 | -8.8100043 | 1.24146598 | -7.0964525 |
| 2225 | NM_002167.4 | 153.441459 | -8.8852821 | 1.25211667 | -7.0962094 |
| 2226 | NM_014949.3 | 139.556719 | -8.7466085 | 1.23271875 | -7.0953804 |
| 2227 | NM_015527.3 | 247.192759 | -6.8377988 | 0.96371862 | -7.0952233 |
| 2228 | NM_001099402.1 | 154.315437 | -8.8926688 | 1.25336127 | -7.0950563 |
| 2229 | NM_012295.3 | 156.835364 | -8.9167329 | 1.25677017 | -7.094959 |
| 2230 | NM_003632.2 | 334.365347 | -6.5226194 | 0.91939118 | -7.0944986 |
| 2231 | NM_017825.2 | 128.516046 | -8.627858 | 1.21614635 | -7.0944241 |
| 2232 | NM_181741.3 | 226.255996 | -9.4431058 | 1.33123469 | -7.0934943 |
| 2233 | NM_001284307.1 | 142.534368 | -8.7780067 | 1.2375212 | -7.0932172 |
| 2234 | NM_012470.3 | 187.005458 | -9.1689177 | 1.2927957 | -7.0923176 |
| 2235 | NM_020196.2 | 191.940932 | -9.2063418 | 1.29817192 | -7.0917739 |

| | | | | | |
|---|---|---|---|---|---|
| 2236 | NM_000143.3 | 180.280652 | -9.1162494 | 1.28553059 | -7.0914294 |
| 2237 | NM_015560.2 | 181.107909 | -9.1223003 | 1.28652322 | -7.0906612 |
| 2238 | NM_006136.2 | 222.971418 | -9.4225847 | 1.32890542 | -7.0904856 |
| 2239 | NM_017969.2 | 173.622789 | -9.0619727 | 1.27805581 | -7.0904358 |
| 2240 | NM_001688.4 | 316.181686 | -8.1333082 | 1.14718743 | -7.0897815 |
| 2241 | NM_031954.4 | 212.219536 | -9.3515608 | 1.31911141 | -7.0892881 |
| 2242 | NM_001270362.1 | 146.058292 | -8.8128333 | 1.24315225 | -7.0891022 |
| 2243 | NM_006886.3 | 288.938748 | -9.7968173 | 1.38198565 | -7.0889428 |
| 2244 | NM_015027.3 | 158.149773 | -8.926955 | 1.25941611 | -7.0881696 |
| 2245 | NM_014935.4 | 270.44983 | -4.6333628 | 0.65368246 | -7.0880941 |
| 2246 | NM_014647.3 | 148.560976 | -8.8359569 | 1.24660359 | -7.0880246 |
| 2247 | NM_001135999.1 | 270.418432 | -9.7011813 | 1.36872413 | -7.087755 |
| 2248 | NM_015048.1 | 142.17333 | -8.7729803 | 1.23779517 | -7.0875864 |
| 2249 | NM_181042.4 | 214.372971 | -9.366366 | 1.32162008 | -7.0870337 |
| 2250 | NM_001130691.1 | 229.555562 | -9.4644338 | 1.33552327 | -7.0866858 |
| 2251 | NM_015905.2 | 156.287772 | -8.9108289 | 1.25741378 | -7.0866321 |
| 2252 | NM_031459.4 | 157.492679 | -8.9213471 | 1.25890378 | -7.0865997 |
| 2253 | NM_001330332.1 | 171.836916 | 2.90582377 | 0.41007638 | 7.08605489 |
| 2254 | NM_001039538.1 | 268.86269 | -9.6925478 | 1.36792121 | -7.0856039 |
| 2255 | NM_021033.6 | 335.144363 | -7.6885647 | 1.08511158 | -7.0855061 |
| 2256 | NM_021135.5 | 244.321469 | -9.5548823 | 1.34854655 | -7.0853189 |
| 2257 | NM_005589.3 | 135.400469 | -8.7040672 | 1.22858957 | -7.0846012 |
| 2258 | NM_006861.6 | 202.940831 | -9.2867381 | 1.31085209 | -7.084505 |
| 2259 | NM_138373.4 | 154.973853 | -8.8989364 | 1.25619531 | -7.0840389 |
| 2260 | NM_020825.3 | 158.59901 | -5.5087304 | 0.77764518 | -7.083861 |
| 2261 | NM_207368.4 | 269.607104 | -9.6968807 | 1.36896725 | -7.0833548 |
| 2262 | NM_001004333.4 | 218.908813 | -9.3964032 | 1.32663675 | -7.0828757 |
| 2263 | NM_002576.4 | 225.372606 | -9.4380406 | 1.33252179 | -7.0828415 |
| 2264 | NM_014748.3 | 187.976684 | -9.1767008 | 1.29565255 | -7.0826864 |
| 2265 | NM_021210.4 | 194.301332 | -9.2242444 | 1.30246896 | -7.0821222 |
| 2266 | NM_013943.2 | 412.065356 | -6.9555273 | 0.98215358 | -7.0819141 |
| 2267 | NM_016016.3 | 169.690171 | -9.0283819 | 1.27484219 | -7.0819604 |
| 2268 | NM_020443.4 | 234.9289 | -9.4982129 | 1.34152358 | -7.0801684 |
| 2269 | NM_030630.2 | 169.462589 | -9.0270374 | 1.2749994 | -7.0800327 |
| 2270 | NM_002858.3 | 173.527496 | -9.0607778 | 1.27986564 | -7.0794758 |
| 2271 | NM_145230.3 | 330.11222 | -9.9888952 | 1.41101636 | -7.07922 |
| 2272 | NM_182917.4 | 176.061361 | -9.0830829 | 1.2830701 | -7.079179 |
| 2273 | NM_016219.4 | 210.631039 | -9.3406859 | 1.3198734 | -7.0769559 |
| 2274 | NM_001318797.1 | 155.968448 | -8.9080964 | 1.25894844 | -7.075823 |
| 2275 | NM_001843.3 | 314.430967 | -9.9185076 | 1.40190436 | -7.0750245 |
| 2276 | NM_002067.4 | 324.069466 | -6.6890572 | 0.94553235 | -7.0743823 |
| 2277 | NM_015246.3 | 142.717326 | -8.7793414 | 1.24104876 | -7.074131 |
| 2278 | NM_004431.4 | 132.847743 | -8.6773457 | 1.22666964 | -7.073906 |

| | | | | | |
|---|---|---|---|---|---|
| 2279 | NM_001409.3 | 141.884677 | -8.7712451 | 1.23996436 | -7.073788 |
| 2280 | NM_182976.3 | 137.896886 | -8.7296401 | 1.23410105 | -7.0736834 |
| 2281 | NM_017739.3 | 195.134743 | -9.23079 | 1.30500716 | -7.0733635 |
| 2282 | NM_006221.3 | 221.994619 | -9.4165565 | 1.33137448 | -7.0728083 |
| 2283 | NM_001379.3 | 161.623225 | -8.9583637 | 1.26663339 | -7.0725782 |
| 2284 | NM_002940.2 | 195.170766 | -9.2303569 | 1.30518725 | -7.0720556 |
| 2285 | NM_014952.4 | 134.046703 | -8.6870569 | 1.22840399 | -7.0718241 |
| 2286 | NM_002835.3 | 181.419376 | -9.124959 | 1.29035668 | -7.0716564 |
| 2287 | NM_207013.2 | 263.278152 | -9.6624347 | 1.36641765 | -7.0713626 |
| 2288 | NM_152988.2 | 205.530301 | -7.5080743 | 1.06177167 | -7.0712702 |
| 2289 | NM_001301059.1 | 257.552296 | -9.6308192 | 1.36207189 | -7.0707128 |
| 2290 | NM_000268.3 | 139.508714 | -8.7477889 | 1.23720458 | -7.0706083 |
| 2291 | NM_001159390.1 | 263.393233 | -9.6631337 | 1.36666622 | -7.070588 |
| 2292 | NM_000791.3 | 175.576439 | -9.0775019 | 1.28388507 | -7.0703384 |
| 2293 | NM_020225.2 | 143.906331 | -8.7901014 | 1.24324318 | -7.0702994 |
| 2294 | NM_033088.3 | 229.859139 | -9.4667576 | 1.33896226 | -7.0702199 |
| 2295 | NM_152743.3 | 163.005302 | -8.9709989 | 1.26885375 | -7.0701599 |
| 2296 | NM_018471.2 | 213.151711 | -9.3577412 | 1.32357607 | -7.0700441 |
| 2297 | NM_001142478.1 | 320.826689 | -9.9477369 | 1.4072327 | -7.0690064 |
| 2298 | NM_013341.4 | 283.671711 | -9.7701156 | 1.38209767 | -7.0690486 |
| 2299 | NM_001172630.1 | 280.685303 | -9.7548671 | 1.38008611 | -7.0683032 |
| 2300 | NM_018083.4 | 231.638476 | -9.4778149 | 1.34089784 | -7.0682602 |
| 2301 | NM_033201.2 | 292.977641 | -9.8167202 | 1.38896595 | -7.0676464 |
| 2302 | NM_015179.3 | 121.313002 | -8.5456882 | 1.20913819 | -7.067586 |
| 2303 | NM_001032.4 | 400.880467 | -10.269196 | 1.45318046 | -7.0667042 |
| 2304 | NM_006519.3 | 210.563268 | -9.3401128 | 1.32173056 | -7.0665785 |
| 2305 | NM_018332.4 | 158.778853 | -8.9333257 | 1.26417777 | -7.0665106 |
| 2306 | NM_006421.4 | 189.772971 | -9.1902462 | 1.30057309 | -7.066305 |
| 2307 | NM_003812.3 | 163.712599 | -8.9767685 | 1.27038155 | -7.0661988 |
| 2308 | NM_001135652.2 | 80.8321259 | 11.2577077 | 1.59326272 | 7.06582005 |
| 2309 | NM_001289135.2 | 178.428036 | -9.1022145 | 1.28821312 | -7.0657676 |
| 2310 | NM_012234.6 | 187.670115 | -9.1737776 | 1.29834439 | -7.0657506 |
| 2311 | NM_017582.6 | 216.95701 | -7.0363873 | 0.99589646 | -7.0653804 |
| 2312 | NM_000671.4 | 377.873934 | -9.2196394 | 1.30499391 | -7.0648907 |
| 2313 | NM_001283054.1 | 387.056553 | -10.218533 | 1.44640682 | -7.064771 |
| 2314 | NM_006432.3 | 182.063402 | -9.1312227 | 1.29254985 | -7.0645033 |
| 2315 | NM_016040.4 | 128.857226 | -8.6313443 | 1.22182131 | -7.0643262 |
| 2316 | NM_001190467.1 | 241.715961 | -9.5390337 | 1.35042804 | -7.0637112 |
| 2317 | NM_000925.3 | 153.321238 | -8.8819256 | 1.2574201 | -7.0636104 |
| 2318 | NM_005875.2 | 209.114422 | -9.3301894 | 1.32096975 | -7.0631363 |
| 2319 | NM_001098507.1 | 161.912925 | -8.959391 | 1.26863621 | -7.0622224 |
| 2320 | NM_019026.4 | 169.23803 | -9.0249447 | 1.27791442 | -7.062245 |
| 2321 | NM_006289.3 | 533.45984 | -6.8765465 | 0.97373901 | -7.0620017 |

| | | | | | |
|---|---|---|---|---|---|
| 2322 | NM_000100.3 | 138.35702 | -8.7349622 | 1.23691497 | -7.0618938 |
| 2323 | NM_001321291.1 | 132.809778 | -8.6750227 | 1.22852046 | -7.061358 |
| 2324 | NM_018045.7 | 206.998119 | -9.3154549 | 1.31939506 | -7.0603985 |
| 2325 | NM_002866.4 | 460.581099 | -10.469476 | 1.48297362 | -7.0597857 |
| 2326 | NM_003655.2 | 184.072059 | -9.1462013 | 1.29553156 | -7.0598059 |
| 2327 | NM_004229.3 | 129.797437 | -8.6438311 | 1.22442349 | -7.059511 |
| 2328 | NM_002717.3 | 166.223963 | -8.9988787 | 1.27484238 | -7.0588168 |
| 2329 | NM_017727.4 | 148.031527 | -8.8326335 | 1.25134735 | -7.0584986 |
| 2330 | NM_152295.4 | 208.285837 | -9.3249418 | 1.32117684 | -7.0580573 |
| 2331 | NM_002816.4 | 198.727427 | -9.2567667 | 1.31168263 | -7.0571696 |
| 2332 | NM_177983.2 | 388.721831 | -9.2617628 | 1.31244738 | -7.0568641 |
| 2333 | NM_001184823.1 | 197.143071 | -9.2452705 | 1.31014026 | -7.0567028 |
| 2334 | NM_198268.2 | 177.94873 | -9.0975087 | 1.2891953 | -7.0567343 |
| 2335 | NM_004261.4 | 178.119228 | -9.0985149 | 1.28936169 | -7.056604 |
| 2336 | NM_025238.3 | 220.897356 | -9.408949 | 1.3333785 | -7.0564727 |
| 2337 | NM_004255.3 | 272.152837 | -9.7103483 | 1.37617278 | -7.0560532 |
| 2338 | NM_005572.3 | 168.537215 | -9.0207732 | 1.27847962 | -7.05586 |
| 2339 | NM_015045.3 | 140.942687 | -8.7604053 | 1.24159369 | -7.0557746 |
| 2340 | NM_000487.5 | 129.963376 | -8.6430808 | 1.22500353 | -7.0555558 |
| 2341 | NM_021815.4 | 321.744171 | -9.9520739 | 1.4106233 | -7.0550897 |
| 2342 | NM_004612.3 | 252.668231 | -9.6036947 | 1.3612732 | -7.0549356 |
| 2343 | NM_001076785.2 | 235.11914 | -5.629292 | 0.79796674 | -7.0545447 |
| 2344 | NM_002129.3 | 209.589996 | -9.3334685 | 1.32323921 | -7.0535005 |
| 2345 | NM_022156.4 | 141.337032 | -8.7654885 | 1.24273134 | -7.0534059 |
| 2346 | NM_015336.2 | 208.219178 | -9.3237034 | 1.32191732 | -7.0531668 |
| 2347 | NM_015525.3 | 139.838082 | -8.7496086 | 1.24052562 | -7.0531462 |
| 2348 | NM_152265.4 | 243.651994 | -9.5506083 | 1.35423415 | -7.0524054 |
| 2349 | NM_001318119.1 | 136.877307 | -8.7173405 | 1.23613277 | -7.0521069 |
| 2350 | NM_022081.5 | 123.904939 | -8.5759346 | 1.21613687 | -7.0517841 |
| 2351 | NM_001819.2 | 321.964372 | -9.9530112 | 1.4115515 | -7.0511145 |
| 2352 | NM_206943.2 | 224.480545 | -9.432771 | 1.33785014 | -7.0506933 |
| 2353 | NM_001193489.1 | 163.026866 | -8.9699111 | 1.27225645 | -7.0503955 |
| 2354 | NM_015386.2 | 178.863763 | -9.1047615 | 1.291441 | -7.0500793 |
| 2355 | NM_002403.3 | 153.119705 | -8.8813693 | 1.25981063 | -7.0497654 |
| 2356 | NM_030913.4 | 215.020221 | -9.370298 | 1.32920208 | -7.0495661 |
| 2357 | NM_005040.3 | 153.152948 | -8.8815991 | 1.26001992 | -7.0487767 |
| 2358 | NM_019088.3 | 172.691679 | -9.0545381 | 1.28460202 | -7.0485161 |
| 2359 | NM_003819.3 | 162.972044 | -8.9707391 | 1.27280452 | -7.0480101 |
| 2360 | NM_005732.3 | 151.183152 | -8.8613372 | 1.25731277 | -7.0478384 |
| 2361 | NM_000404.3 | 143.124528 | -8.7849936 | 1.24650063 | -7.0477249 |
| 2362 | NM_015084.2 | 153.433212 | -8.8834039 | 1.2605232 | -7.0473943 |
| 2363 | NM_152421.3 | 197.550768 | -9.2481343 | 1.31228206 | -7.0473678 |
| 2364 | NM_033386.3 | 210.883564 | -9.3425875 | 1.32579861 | -7.0467622 |

| | | | | | |
|---|---|---|---|---|---|
| 2365 | NM_201630.1 | 150.965154 | -8.8616098 | 1.25761553 | -7.0463584 |
| 2366 | NM_002206.2 | 169.338947 | -9.0253605 | 1.28094483 | -7.045862 |
| 2367 | NM_014303.3 | 173.835249 | -9.0639454 | 1.28642816 | -7.0458232 |
| 2368 | NM_080703.2 | 166.045147 | -8.9988804 | 1.27718828 | -7.0458527 |
| 2369 | NM_002221.3 | 182.589128 | -9.1358095 | 1.2966389 | -7.0457623 |
| 2370 | NM_015322.4 | 339.459684 | -4.5243103 | 0.64217068 | -7.0453392 |
| 2371 | NM_014089.3 | 175.759407 | -9.0787578 | 1.28871323 | -7.0448239 |
| 2372 | NM_013366.3 | 141.64489 | -8.7684457 | 1.24472035 | -7.0445106 |
| 2373 | NM_001080434.1 | 310.121177 | -9.8987955 | 1.40533872 | -7.043708 |
| 2374 | NM_002806.3 | 176.638292 | -9.0865715 | 1.29008645 | -7.0433819 |
| 2375 | NM_033222.4 | 211.90016 | -9.3489177 | 1.32749497 | -7.0425259 |
| 2376 | NM_018198.3 | 175.175309 | -9.074781 | 1.28861501 | -7.0422748 |
| 2377 | NM_005689.2 | 150.355473 | -8.8537854 | 1.25726931 | -7.0420755 |
| 2378 | NM_006055.2 | 185.678074 | -9.1590181 | 1.30069769 | -7.0416194 |
| 2379 | NM_007285.6 | 274.335517 | -9.7218886 | 1.38063714 | -7.0415957 |
| 2380 | NM_014570.4 | 142.463234 | -8.7757796 | 1.24628129 | -7.0415722 |
| 2381 | NM_033063.1 | 140.758083 | -8.7604307 | 1.24410527 | -7.041551 |
| 2382 | NM_003038.4 | 233.264544 | -9.4879959 | 1.34747386 | -7.0413209 |
| 2383 | NM_207012.3 | 145.343785 | -8.8061915 | 1.25077228 | -7.0406034 |
| 2384 | NM_032575.2 | 126.207484 | -8.6008592 | 1.22195024 | -7.038633 |
| 2385 | NM_001282850.1 | 167.94694 | -9.0123388 | 1.28052836 | -7.0379845 |
| 2386 | NM_002791.2 | 260.863753 | -9.6492535 | 1.37103565 | -7.0379304 |
| 2387 | NM_005414.4 | 146.493967 | -8.8169255 | 1.25294958 | -7.0369356 |
| 2388 | NM_007224.3 | 205.785424 | -9.3069948 | 1.32258783 | -7.0369578 |
| 2389 | NM_019082.3 | 170.884044 | -9.0389908 | 1.28450298 | -7.0369559 |
| 2390 | NM_003195.4 | 178.556121 | -9.1023759 | 1.29359318 | -7.0365058 |
| 2391 | NM_000520.5 | 178.971861 | -9.1063282 | 1.29427796 | -7.0358365 |
| 2392 | NM_003607.3 | 185.127424 | -9.1541687 | 1.30108476 | -7.0357973 |
| 2393 | NM_001184970.1 | 129.456673 | -8.6399835 | 1.22803894 | -7.0355941 |
| 2394 | NM_032288.6 | 242.74934 | -9.5453119 | 1.35671102 | -7.0356264 |
| 2395 | NM_001081563.2 | 144.522454 | -8.7987275 | 1.2507483 | -7.0347707 |
| 2396 | NM_001145358.1 | 151.577715 | -8.8652971 | 1.26023857 | -7.0346182 |
| 2397 | NM_000919.3 | 144.188412 | -8.7943542 | 1.25020366 | -7.0343372 |
| 2398 | NM_001304809.1 | 156.398064 | -8.9109727 | 1.26690533 | -7.0336532 |
| 2399 | NM_014071.4 | 211.224919 | -9.3443225 | 1.32851912 | -7.0336379 |
| 2400 | NM_032590.4 | 195.117612 | -9.229589 | 1.31221644 | -7.0335873 |
| 2401 | NM_183352.2 | 208.33923 | -9.3251062 | 1.32593314 | -7.0328631 |
| 2402 | NM_000287.3 | 130.791349 | -8.6527848 | 1.23036368 | -7.032705 |
| 2403 | NM_018127.6 | 172.800796 | -9.0548825 | 1.28762103 | -7.0322574 |
| 2404 | NM_006710.4 | 193.620685 | -9.2189992 | 1.31104665 | -7.0317857 |
| 2405 | NM_017902.2 | 150.699663 | -8.8581028 | 1.25978803 | -7.0314232 |
| 2406 | NM_148415.2 | 156.514922 | -8.9113752 | 1.26748372 | -7.0307611 |
| 2407 | NM_003341.4 | 193.158305 | -9.2158451 | 1.31081252 | -7.0306355 |

| | | | | | |
|---|---|---|---|---|---|
| 2408 | NM_025082.3 | 174.982561 | -9.073068 | 1.29055063 | -7.0303851 |
| 2409 | NR_037611.1 | 138.112984 | -8.7322279 | 1.24211404 | -7.0301338 |
| 2410 | NM_001330247.1 | 132.980888 | -8.6778777 | 1.23443622 | -7.0298308 |
| 2411 | NM_198551.3 | 132.592185 | -8.6716842 | 1.23369587 | -7.0290292 |
| 2412 | NM_182977.2 | 158.619497 | -8.9315573 | 1.27068745 | -7.0289176 |
| 2413 | NM_004227.3 | 150.160566 | -8.8526976 | 1.25953834 | -7.0285257 |
| 2414 | NM_030811.3 | 141.803922 | -6.3876891 | 0.90882523 | -7.028512 |
| 2415 | NM_006004.3 | 252.435603 | -9.6018717 | 1.36619753 | -7.0281724 |
| 2416 | NM_014888.2 | 201.860178 | -9.2782385 | 1.32015788 | -7.0281279 |
| 2417 | NM_004124.2 | 235.327743 | -9.5003888 | 1.3517819 | -7.0280485 |
| 2418 | NM_017707.3 | 154.250682 | -8.8919302 | 1.26525962 | -7.0277515 |
| 2419 | NM_001271958.1 | 155.348919 | -8.9020584 | 1.26675184 | -7.0274683 |
| 2420 | NM_002487.2 | 453.439245 | -9.4839358 | 1.34956132 | -7.0274211 |
| 2421 | NM_001861.4 | 346.758824 | -10.059882 | 1.43158679 | -7.0270847 |
| 2422 | NM_001309840.1 | 1563.44827 | -8.19372 | 1.16618389 | -7.026096 |
| 2423 | NM_020928.1 | 159.552888 | -8.9400663 | 1.27244171 | -7.0259142 |
| 2424 | NM_015937.5 | 202.963028 | -9.2873035 | 1.32191664 | -7.0256348 |
| 2425 | NM_030796.4 | 308.660916 | -9.8917545 | 1.40798518 | -7.0254678 |
| 2426 | NM_052899.2 | 333.77195 | -10.004857 | 1.42410849 | -7.0253477 |
| 2427 | NM_016441.2 | 244.146028 | -6.4716226 | 0.92125264 | -7.0248077 |
| 2428 | NM_002038.3 | 178.447529 | -9.1027255 | 1.29583493 | -7.0246026 |
| 2429 | NM_018209.3 | 133.778395 | -8.6861681 | 1.23652964 | -7.0246339 |
| 2430 | NM_152277.2 | 192.999405 | -6.3505593 | 0.90403663 | -7.0246704 |
| 2431 | NM_001347423.1 | 1909.37366 | -7.992098 | 1.13778039 | -7.0242888 |
| 2432 | NR_110001.1 | 68.0870842 | 2.61874085 | 0.37281442 | 7.02424783 |
| 2433 | NM_015320.3 | 130.405241 | -8.6511417 | 1.23164554 | -7.0240515 |
| 2434 | NM_017607.3 | 133.522571 | -8.6827909 | 1.23656553 | -7.0216989 |
| 2435 | NM_018257.2 | 161.962847 | -8.96106 | 1.27618583 | -7.0217517 |
| 2436 | NM_001345840.1 | 275.279848 | -9.7269348 | 1.38529142 | -7.0215802 |
| 2437 | NM_032268.4 | 237.247278 | -9.5122514 | 1.3547356 | -7.0214818 |
| 2438 | NM_004463.2 | 169.72743 | -9.0291014 | 1.28598514 | -7.0211553 |
| 2439 | NM_001077180.2 | 221.585709 | -9.4134823 | 1.34080005 | -7.020795 |
| 2440 | NM_184234.2 | 151.937724 | -8.8712595 | 1.26362002 | -7.020512 |
| 2441 | NM_001300946.1 | 204.20891 | -6.5524221 | 0.93340226 | -7.0199338 |
| 2442 | NM_003250.5 | 292.619789 | -9.8146368 | 1.3981231 | -7.019866 |
| 2443 | NM_003780.4 | 277.090203 | -9.7361816 | 1.38701011 | -7.0195462 |
| 2444 | NM_015299.2 | 132.725754 | -8.6756817 | 1.23596364 | -7.0193665 |
| 2445 | NM_000310.3 | 201.830864 | -9.2791606 | 1.32204146 | -7.0188121 |
| 2446 | NM_006244.3 | 250.276557 | -9.589357 | 1.36628848 | -7.0185448 |
| 2447 | NM_005916.4 | 237.593504 | -9.5141174 | 1.3556674 | -7.0180322 |
| 2448 | NM_001165036.1 | 118.328798 | -8.5096794 | 1.21256085 | -7.0179401 |
| 2449 | NM_002789.5 | 218.984582 | -9.3966331 | 1.33908893 | -7.0171838 |
| 2450 | NM_001291813.1 | 121.998633 | -8.5523882 | 1.21888726 | -7.016554 |

| | | | | | |
|---|---|---|---|---|---|
| 2451 | NM_017721.4 | 146.808456 | -8.8199688 | 1.25710623 | -7.0160887 |
| 2452 | NM_004713.5 | 167.913167 | -9.013387 | 1.28468678 | -7.0160191 |
| 2453 | NM_080670.3 | 166.070266 | -8.9979285 | 1.28267098 | -7.0149934 |
| 2454 | NM_006428.4 | 164.654042 | -8.98513 | 1.28088005 | -7.0148099 |
| 2455 | NM_007208.3 | 190.176828 | -9.1930251 | 1.31053683 | -7.0147019 |
| 2456 | NM_022566.2 | 155.293921 | -8.9011002 | 1.26897663 | -7.0143925 |
| 2457 | NM_001190708.1 | 867.654821 | -8.1630746 | 1.16377645 | -7.0142978 |
| 2458 | NM_016019.4 | 226.703027 | -7.62174 | 1.08661051 | -7.0142337 |
| 2459 | NM_032287.2 | 320.863275 | -8.984291 | 1.28089659 | -7.0140643 |
| 2460 | NM_006319.4 | 174.654445 | -9.0702988 | 1.29321033 | -7.0137847 |
| 2461 | NM_020223.3 | 238.950378 | -9.5227837 | 1.35772684 | -7.01377 |
| 2462 | NM_001865.3 | 315.783432 | -9.9247945 | 1.41516722 | -7.0131602 |
| 2463 | NM_004436.2 | 133.248491 | -8.6806445 | 1.23778187 | -7.0130648 |
| 2464 | NM_012156.2 | 300.125568 | -9.851571 | 1.4050288 | -7.0116506 |
| 2465 | NM_015461.2 | 134.361857 | -8.6940747 | 1.23994791 | -7.0116451 |
| 2466 | NM_032102.3 | 169.197357 | -9.0246169 | 1.28707491 | -7.0117262 |
| 2467 | NM_014233.3 | 163.785967 | -8.9770608 | 1.28053858 | -7.010379 |
| 2468 | NM_002970.3 | 123.904784 | -8.5748064 | 1.22317772 | -7.0102703 |
| 2469 | NM_020747.2 | 147.234777 | -5.4002515 | 0.77034182 | -7.0102016 |
| 2470 | NM_006373.3 | 917.142785 | -7.6848675 | 1.09628777 | -7.0098999 |
| 2471 | NM_000581.3 | 335.020518 | -9.0469455 | 1.2908782 | -7.0083649 |
| 2472 | NM_015678.4 | 208.615364 | -9.3261411 | 1.33082456 | -7.0077916 |
| 2473 | NM_199368.1 | 151.624521 | -6.4983089 | 0.9274285 | -7.0068031 |
| 2474 | NM_021003.4 | 194.798528 | -9.2275428 | 1.31698939 | -7.006543 |
| 2475 | NM_005054.2 | 201.596125 | -7.4791444 | 1.06747154 | -7.006411 |
| 2476 | NM_052875.4 | 412.769869 | -9.347197 | 1.33413615 | -7.0061792 |
| 2477 | NM_005886.2 | 147.700707 | -8.8285083 | 1.26018716 | -7.005712 |
| 2478 | NM_153377.4 | 181.889404 | -9.129547 | 1.30326779 | -7.0051198 |
| 2479 | NM_001321983.1 | 52.9961875 | 10.6504544 | 1.52044781 | 7.00481417 |
| 2480 | NM_005611.3 | 137.322877 | -8.7231997 | 1.24530957 | -7.0048443 |
| 2481 | NM_014300.3 | 168.269735 | -9.0169089 | 1.28723241 | -7.0048803 |
| 2482 | NM_022832.3 | 117.947149 | -8.5044495 | 1.21406648 | -7.004929 |
| 2483 | NM_031892.2 | 172.400973 | -9.0522125 | 1.29225771 | -7.0049592 |
| 2484 | NM_015404.3 | 142.759413 | -8.7811095 | 1.25368235 | -7.0042539 |
| 2485 | NM_006694.3 | 198.165972 | -9.2527008 | 1.3210385 | -7.0041114 |
| 2486 | NM_003270.3 | 117.449926 | -8.4990892 | 1.21347247 | -7.0039407 |
| 2487 | NM_001130021.1 | 375.075608 | -10.17317 | 1.45264184 | -7.0032195 |
| 2488 | NM_001010942.2 | 197.07673 | -9.2443559 | 1.32005408 | -7.003013 |
| 2489 | NM_020919.3 | 132.918648 | -8.6767232 | 1.23910628 | -7.0024043 |
| 2490 | NM_001346581.1 | 171.308231 | -9.0419392 | 1.29140807 | -7.0016128 |
| 2491 | NM_001142565.1 | 177.994284 | -9.098962 | 1.29962314 | -7.0012311 |
| 2492 | NM_015662.2 | 146.897779 | -8.8220025 | 1.26014048 | -7.0008088 |
| 2493 | NM_015159.2 | 243.337414 | -9.5495325 | 1.36415168 | -7.0003451 |

| | | | | | |
|---|---|---|---|---|---|
| 2494 | NM_003689.3 | 117.410877 | -8.4967841 | 1.21388016 | -6.9996895 |
| 2495 | NM_018184.2 | 224.668718 | -9.4337987 | 1.34773965 | -6.9997189 |
| 2496 | NM_001126055.2 | 133.902903 | -8.6875348 | 1.24121574 | -6.9992141 |
| 2497 | NM_133171.4 | 187.334734 | -9.1716882 | 1.31045472 | -6.9988593 |
| 2498 | NM_015397.3 | 140.021743 | -5.4051702 | 0.77233766 | -6.9984549 |
| 2499 | NM_001329443.1 | 161.327493 | -8.9553414 | 1.27969188 | -6.998045 |
| 2500 | NM_003238.4 | 240.25278 | -9.5308274 | 1.36195892 | -6.9978817 |
| 2501 | NM_002109.5 | 203.717643 | -9.2924818 | 1.32792002 | -6.9977722 |
| 2502 | NM_001128916.1 | 169.018058 | -9.0226752 | 1.28940157 | -6.9975681 |
| 2503 | NM_198335.3 | 132.601579 | -8.6736337 | 1.23956035 | -6.9973469 |
| 2504 | NM_001347721.1 | 152.458311 | -8.8735884 | 1.2683051 | -6.9964146 |
| 2505 | NM_003198.2 | 153.267249 | -8.8815975 | 1.26948003 | -6.9962483 |
| 2506 | NM_020310.2 | 144.22682 | -8.7943348 | 1.25699799 | -6.9962998 |
| 2507 | NM_015949.2 | 180.17784 | -9.1151248 | 1.30318699 | -6.9944872 |
| 2508 | NM_153831.3 | 140.784456 | -8.7588242 | 1.25227812 | -6.9943123 |
| 2509 | NM_018064.3 | 205.456061 | -9.3044985 | 1.33044383 | -6.9935297 |
| 2510 | NM_006268.4 | 165.263611 | -8.990986 | 1.28574447 | -6.9928249 |
| 2511 | NM_003941.3 | 137.655637 | -8.726993 | 1.24810016 | -6.9922216 |
| 2512 | NM_004438.4 | 223.759363 | -9.4275389 | 1.34838781 | -6.9917118 |
| 2513 | NM_002865.2 | 432.702147 | -7.7579747 | 1.10961038 | -6.9916205 |
| 2514 | NM_016002.2 | 163.314249 | -8.9741494 | 1.28374426 | -6.9906053 |
| 2515 | NM_145173.3 | 314.320837 | -7.1221093 | 1.01883361 | -6.9904538 |
| 2516 | NM_001924.3 | 130.580986 | -8.6505864 | 1.23759026 | -6.989863 |
| 2517 | NM_002669.3 | 165.513263 | -8.9924933 | 1.28655138 | -6.9896107 |
| 2518 | NM_014520.3 | 156.201958 | -8.9092801 | 1.27471026 | -6.989259 |
| 2519 | NM_001136139.2 | 134.563982 | -8.6952435 | 1.24413212 | -6.9890033 |
| 2520 | NM_002213.4 | 298.755432 | -6.3564773 | 0.90949403 | -6.9890258 |
| 2521 | NM_014781.4 | 173.197488 | -9.0574883 | 1.29595875 | -6.9890251 |
| 2522 | NM_020199.2 | 139.583706 | -8.748432 | 1.25180401 | -6.9886595 |
| 2523 | NM_020851.2 | 510.41837 | -10.617637 | 1.51926762 | -6.9886548 |
| 2524 | NM_031885.3 | 119.769429 | -8.5261755 | 1.22001106 | -6.9886051 |
| 2525 | NM_014974.2 | 257.931652 | -7.8236095 | 1.11949314 | -6.9885283 |
| 2526 | NM_022462.4 | 32.0668453 | 9.93296195 | 1.42140868 | 6.98811123 |
| 2527 | NM_080861.3 | 192.401656 | -9.2096285 | 1.31790716 | -6.9880708 |
| 2528 | NM_005451.4 | 146.681344 | -8.8198423 | 1.26226369 | -6.9873216 |
| 2529 | NM_016208.3 | 201.973539 | -9.2801887 | 1.32819157 | -6.987086 |
| 2530 | NM_001308301.1 | 165.168659 | 2.40324046 | 0.3439645 | 6.98688508 |
| 2531 | NM_019858.1 | 149.880102 | -8.8489321 | 1.26650902 | -6.9868686 |
| 2532 | NM_001142397.1 | 197.642467 | -9.2486701 | 1.32380001 | -6.9864557 |
| 2533 | NM_003156.3 | 148.543042 | -8.8377062 | 1.26511457 | -6.9856964 |
| 2534 | NM_021136.2 | 157.474081 | -8.9214073 | 1.27719554 | -6.9851538 |
| 2535 | NM_016030.5 | 135.992218 | -8.7083361 | 1.24673184 | -6.9849311 |
| 2536 | NM_006034.4 | 258.811886 | -9.6376465 | 1.37993193 | -6.9841463 |

| | | | | | |
|---|---|---|---|---|---|
| 2537 | NM_015089.3 | 121.674071 | -8.548901 | 1.22407615 | -6.9839618 |
| 2538 | NM_014758.2 | 133.524711 | -8.6822726 | 1.24323299 | -6.9836247 |
| 2539 | NM_006097.4 | 201.103661 | -9.2746928 | 1.32830972 | -6.982327 |
| 2540 | NM_001153484.1 | 229.624524 | -9.4650751 | 1.35563351 | -6.9820309 |
| 2541 | NM_016111.3 | 155.268032 | -8.900885 | 1.27489061 | -6.9816853 |
| 2542 | NM_181336.3 | 173.493359 | -9.0608446 | 1.29786893 | -6.9813249 |
| 2543 | NM_001112732.2 | 251.486843 | -9.5962721 | 1.37469376 | -6.9806617 |
| 2544 | NM_176801.2 | 187.666922 | -9.1729718 | 1.31407878 | -6.9805342 |
| 2545 | NM_001330330.1 | 168.84219 | -9.0211083 | 1.29241074 | -6.9800629 |
| 2546 | NM_017449.4 | 264.499726 | -9.6691521 | 1.38526147 | -6.9800195 |
| 2547 | NM_020761.2 | 140.48031 | -8.7564182 | 1.25465068 | -6.9791682 |
| 2548 | NM_005829.4 | 132.623158 | -8.6726544 | 1.24272447 | -6.9787427 |
| 2549 | NM_130783.4 | 146.850893 | -8.8187511 | 1.26366365 | -6.9787171 |
| 2550 | NM_018061.2 | 152.6415 | -8.8756723 | 1.2718332 | -6.9786449 |
| 2551 | NM_001170535.1 | 153.860509 | -8.8872744 | 1.27366945 | -6.977693 |
| 2552 | NM_015033.2 | 236.754569 | -4.9629735 | 0.71126689 | -6.9776529 |
| 2553 | NM_144723.2 | 208.163379 | -9.3237177 | 1.33622583 | -6.9776512 |
| 2554 | NM_001256378.1 | 122.067789 | -8.5533827 | 1.22587208 | -6.977386 |
| 2555 | NM_024818.4 | 126.030794 | -8.5982536 | 1.23241609 | -6.9767456 |
| 2556 | NM_005928.3 | 235.891515 | -9.5046183 | 1.36236114 | -6.9765777 |
| 2557 | NR_104023.1 | 119.410564 | -8.5230146 | 1.22186846 | -6.9753945 |
| 2558 | NM_018358.2 | 137.902152 | -8.7304331 | 1.25164302 | -6.9751782 |
| 2559 | NM_032840.2 | 151.706084 | -8.8675169 | 1.27135272 | -6.9748676 |
| 2560 | NM_003016.4 | 127.276678 | -8.6150147 | 1.23522036 | -6.9744759 |
| 2561 | NM_130386.2 | 255.101163 | -7.8331391 | 1.12320169 | -6.9739382 |
| 2562 | NM_002890.2 | 140.842352 | -8.7597667 | 1.25620206 | -6.9732147 |
| 2563 | NM_015599.2 | 118.656045 | -8.513331 | 1.22093943 | -6.972771 |
| 2564 | NM_006913.3 | 226.303882 | -9.4441489 | 1.35463925 | -6.9717077 |
| 2565 | NM_014245.4 | 153.647883 | -8.8857903 | 1.27462011 | -6.9713244 |
| 2566 | NM_015332.3 | 360.159805 | -9.151546 | 1.31301237 | -6.9698856 |
| 2567 | NM_018174.5 | 204.190224 | -6.9487108 | 0.99701879 | -6.9694883 |
| 2568 | NM_032875.2 | 125.479722 | -8.5917923 | 1.23286085 | -6.9689878 |
| 2569 | NM_138352.2 | 262.375051 | -9.6573501 | 1.38581933 | -6.9686935 |
| 2570 | NM_000308.3 | 169.2611 | -9.025881 | 1.29521848 | -6.9686166 |
| 2571 | NM_001300.5 | 175.592948 | -9.0784392 | 1.30286218 | -6.9680733 |
| 2572 | NM_002839.3 | 130.008941 | -8.6459467 | 1.24097553 | -6.9670565 |
| 2573 | NM_001159702.2 | 258.437527 | -9.6359966 | 1.3831105 | -6.9669029 |
| 2574 | NM_006449.4 | 152.979602 | -8.8790027 | 1.27454885 | -6.9663887 |
| 2575 | NM_058167.2 | 150.522161 | -8.8556312 | 1.27118896 | -6.9664161 |
| 2576 | NM_015994.3 | 180.20645 | -9.1155325 | 1.30851622 | -6.9663122 |
| 2577 | NM_001103146.1 | 150.985002 | -8.8613123 | 1.27208012 | -6.9660017 |
| 2578 | NM_001128219.2 | 135.761282 | -8.7076217 | 1.25028543 | -6.9645071 |
| 2579 | NM_018683.3 | 146.547507 | -8.8169058 | 1.26607419 | -6.9639725 |

| | | | | | |
|---|---|---|---|---|---|
| 2580 | NM_031845.2 | 682.441273 | -10.074084 | 1.4467323 | -6.9633367 |
| 2581 | NM_003715.3 | 210.475207 | -5.8258534 | 0.83668088 | -6.9630531 |
| 2582 | NR_135167.1 | 178.198139 | -9.1003689 | 1.30695365 | -6.9630388 |
| 2583 | NM_020211.2 | 383.884203 | -10.206686 | 1.46607708 | -6.9619025 |
| 2584 | NM_203304.3 | 175.066171 | -9.0740837 | 1.30344236 | -6.9616302 |
| 2585 | NM_004186.4 | 111.090064 | -8.417611 | 1.20916756 | -6.9614926 |
| 2586 | NM_007005.4 | 218.700493 | -9.3953703 | 1.34963299 | -6.9614261 |
| 2587 | NM_177949.3 | 175.625827 | -9.0786685 | 1.3041428 | -6.9614068 |
| 2588 | NM_153029.3 | 139.380987 | -6.3715516 | 0.91532104 | -6.961002 |
| 2589 | NR_136326.1 | 122.751065 | -8.5635894 | 1.23022004 | -6.9610225 |
| 2590 | NM_001031732.3 | 125.712943 | -8.596779 | 1.23503611 | -6.9607512 |
| 2591 | NM_006406.1 | 208.518442 | -9.326037 | 1.33983326 | -6.9605953 |
| 2592 | NM_178152.2 | 484.715532 | -10.54305 | 1.51475525 | -6.9602336 |
| 2593 | NM_080607.2 | 290.773951 | -9.80587 | 1.40887519 | -6.96007 |
| 2594 | NM_020315.4 | 270.304617 | -9.7003639 | 1.39376456 | -6.9598297 |
| 2595 | NM_003073.4 | 233.72598 | -9.4905684 | 1.36366956 | -6.9595807 |
| 2596 | NM_020801.3 | 194.233697 | -9.2236819 | 1.32536292 | -6.9593632 |
| 2597 | NM_024649.4 | 124.737311 | -8.5871093 | 1.2338882 | -6.9593901 |
| 2598 | NM_002318.2 | 290.581567 | -6.4582003 | 0.92802055 | -6.9591135 |
| 2599 | NM_004192.3 | 131.814355 | -8.66435 | 1.24506277 | -6.9589664 |
| 2600 | NM_004566.3 | 197.559259 | -9.2482846 | 1.32906411 | -6.958494 |
| 2601 | NM_002826.4 | 195.728323 | -9.2354002 | 1.32729625 | -6.9580549 |
| 2602 | NM_001127231.2 | 228.331868 | -9.4563288 | 1.35913817 | -6.957592 |
| 2603 | NM_001282582.1 | 177.712132 | -9.0963609 | 1.30742937 | -6.9574396 |
| 2604 | NM_001256095.1 | 243.87341 | -9.552691 | 1.37311469 | -6.9569506 |
| 2605 | NM_201274.3 | 278.177461 | -9.7426182 | 1.40054506 | -6.9563047 |
| 2606 | NM_001137675.3 | 118.713421 | -8.5144858 | 1.2243417 | -6.9543378 |
| 2607 | NM_030782.4 | 234.380555 | -7.1571629 | 1.02917843 | -6.9542488 |
| 2608 | NM_003688.3 | 183.097506 | -9.1385373 | 1.31419832 | -6.9536973 |
| 2609 | NM_182978.3 | 241.020923 | -9.5352343 | 1.37131564 | -6.9533476 |
| 2610 | NM_031314.2 | 221.998305 | -9.416215 | 1.354531 | -6.9516423 |
| 2611 | NM_001018003.2 | 151.326422 | -8.8641992 | 1.27523642 | -6.9510242 |
| 2612 | NM_006575.4 | 139.050213 | -8.7412619 | 1.25755921 | -6.9509744 |
| 2613 | NM_001014794.2 | 140.717073 | -8.7591859 | 1.26026614 | -6.9502668 |
| 2614 | NM_017580.2 | 210.424072 | -7.5370232 | 1.08460058 | -6.9491233 |
| 2615 | NM_017999.4 | 132.460492 | -8.6716474 | 1.24791988 | -6.9488815 |
| 2616 | NM_012448.3 | 141.407778 | -8.7663258 | 1.26155588 | -6.9488208 |
| 2617 | NM_003359.3 | 116.295619 | -8.4860142 | 1.22139892 | -6.9477827 |
| 2618 | NM_153718.3 | 163.296752 | -8.9723876 | 1.29149025 | -6.9473134 |
| 2619 | NM_002355.3 | 191.949758 | -9.2068827 | 1.32526595 | -6.9471963 |
| 2620 | NM_015419.3 | 246.084499 | -7.7800975 | 1.1199192 | -6.9470168 |
| 2621 | NM_019004.1 | 198.836339 | -6.854071 | 0.98677383 | -6.9459392 |
| 2622 | NM_004114.3 | 288.213121 | -9.7931677 | 1.409942 | -6.9457947 |

| | | | | | |
|---|---|---|---|---|---|
| 2623 | NM_007055.3 | 209.024793 | -4.2807964 | 0.61641316 | -6.9446869 |
| 2624 | NM_005685.3 | 127.181565 | -8.6109691 | 1.24003202 | -6.9441506 |
| 2625 | NM_012317.3 | 207.38741 | -9.3183833 | 1.34195962 | -6.9438627 |
| 2626 | NM_005165.2 | 177.184195 | -9.0907245 | 1.3092543 | -6.9434368 |
| 2627 | NM_001040445.2 | 138.719556 | -8.7376323 | 1.25851345 | -6.9428199 |
| 2628 | NM_001242898.1 | 169.17224 | -9.0241148 | 1.29984633 | -6.9424474 |
| 2629 | NM_004544.3 | 186.174446 | -9.1622838 | 1.31972112 | -6.9425909 |
| 2630 | NM_015638.2 | 211.020468 | -9.3433904 | 1.34582333 | -6.9425088 |
| 2631 | NM_016641.3 | 179.896719 | -9.1128973 | 1.31262626 | -6.942492 |
| 2632 | NM_004231.3 | 241.652115 | -9.5389095 | 1.3740269 | -6.9423019 |
| 2633 | NM_170746.3 | 175.225486 | -9.0748792 | 1.30722313 | -6.9421043 |
| 2634 | NM_016252.3 | 216.160149 | -7.552576 | 1.08795605 | -6.9419863 |
| 2635 | NM_172020.4 | 255.471443 | -8.6536482 | 1.24659815 | -6.9418106 |
| 2636 | NM_183419.3 | 127.099653 | -8.6130627 | 1.24085379 | -6.941239 |
| 2637 | NM_006482.2 | 145.809065 | -8.8094258 | 1.26916874 | -6.9410989 |
| 2638 | NM_002787.4 | 203.460064 | -9.2906614 | 1.33861209 | -6.9405181 |
| 2639 | NM_015496.4 | 132.102343 | -8.6679232 | 1.24924775 | -6.9385141 |
| 2640 | NM_021079.4 | 285.717582 | -8.8173881 | 1.2710252 | -6.9372253 |
| 2641 | NM_000895.2 | 135.774265 | -8.7075023 | 1.25523976 | -6.9369236 |
| 2642 | NM_004499.3 | 202.634981 | -9.2847305 | 1.33872928 | -6.9354803 |
| 2643 | NM_006753.5 | 140.488212 | -8.7561204 | 1.26250468 | -6.9355151 |
| 2644 | NM_005864.3 | 135.432386 | -8.7022051 | 1.25497957 | -6.9341409 |
| 2645 | NM_001082576.1 | 249.115096 | -9.5821544 | 1.38193677 | -6.9338587 |
| 2646 | NM_017437.2 | 127.486887 | -8.6151317 | 1.24253913 | -6.9334893 |
| 2647 | NM_013382.5 | 137.73171 | -8.7285131 | 1.25893992 | -6.9332245 |
| 2648 | NM_174909.4 | 403.670844 | -7.0242662 | 1.01312905 | -6.9332394 |
| 2649 | NM_001161726.1 | 193.265836 | -9.2161713 | 1.32929507 | -6.9331268 |
| 2650 | NM_032246.4 | 233.230331 | -9.4872998 | 1.36842434 | -6.9330101 |
| 2651 | NM_001164665.1 | 131.373756 | -8.659659 | 1.24910438 | -6.9326945 |
| 2652 | NM_003022.2 | 162.752967 | -8.9688576 | 1.29397516 | -6.931244 |
| 2653 | NM_003972.2 | 210.454021 | -8.3724197 | 1.20810322 | -6.9302188 |
| 2654 | NM_138471.2 | 141.669706 | -8.7687525 | 1.26537831 | -6.9297477 |
| 2655 | NR_036447.1 | 1186.96341 | -6.3451833 | 0.91565563 | -6.9296612 |
| 2656 | NM_002226.4 | 159.89927 | -8.9428966 | 1.2905392 | -6.9295815 |
| 2657 | NM_001317122.1 | 149.316594 | -8.8434111 | 1.27620035 | -6.9294849 |
| 2658 | NM_005782.3 | 207.215892 | -9.3168381 | 1.34455154 | -6.9293276 |
| 2659 | NM_015140.3 | 162.992195 | -8.9702306 | 1.29456968 | -6.9291215 |
| 2660 | NM_004818.2 | 283.66207 | -8.804777 | 1.27083356 | -6.9283479 |
| 2661 | NM_152699.4 | 155.857491 | -8.9059095 | 1.285557 | -6.927666 |
| 2662 | NM_014041.3 | 158.217123 | -8.9280608 | 1.28878576 | -6.927498 |
| 2663 | NM_001012329.1 | 187.899255 | -9.175938 | 1.32459374 | -6.9273602 |
| 2664 | NM_001321688.1 | 139.828945 | -8.7506126 | 1.26343726 | -6.9260365 |
| 2665 | NM_017813.4 | 298.583576 | -8.8803296 | 1.28223362 | -6.9256721 |

| | | | | | |
|---|---|---|---|---|---|
| 2666 | NM_001173408.1 | 180.134749 | -9.1156354 | 1.31629758 | -6.9252086 |
| 2667 | NM_001330123.1 | 170.365846 | -9.0350291 | 1.30467191 | -6.925135 |
| 2668 | NM_002388.4 | 149.843321 | -8.8488628 | 1.27780738 | -6.9250365 |
| 2669 | NM_020239.3 | 158.523764 | -6.0860318 | 0.87883955 | -6.9250773 |
| 2670 | NM_024103.2 | 298.593026 | -8.880969 | 1.28245836 | -6.924957 |
| 2671 | NR_037600.1 | 210.656233 | -9.3405663 | 1.34892203 | -6.9244672 |
| 2672 | NM_004504.4 | 121.36099 | -8.5452915 | 1.23410802 | -6.9242654 |
| 2673 | NM_006372.4 | 134.85467 | -8.6969834 | 1.25612074 | -6.9236843 |
| 2674 | NM_006521.5 | 147.126312 | -8.8238253 | 1.27443454 | -6.923718 |
| 2675 | NM_000254.2 | 128.67258 | -8.6306187 | 1.24657859 | -6.9234454 |
| 2676 | NM_001286665.1 | 256.847285 | -9.6272839 | 1.39054276 | -6.9234001 |
| 2677 | NM_005160.3 | 167.638676 | -5.6418163 | 0.81496021 | -6.922812 |
| 2678 | NM_021962.4 | 224.60968 | -9.4329909 | 1.36258703 | -6.9228539 |
| 2679 | NM_032228.5 | 190.682763 | -9.1969052 | 1.32849505 | -6.9227997 |
| 2680 | NM_012416.3 | 130.725691 | -8.6526225 | 1.24990147 | -6.9226436 |
| 2681 | NM_001291345.1 | 163.391544 | -8.9735693 | 1.29631169 | -6.9223855 |
| 2682 | NM_006736.5 | 226.510872 | -9.4454627 | 1.36468753 | -6.9213373 |
| 2683 | NM_015631.5 | 121.083944 | -8.5428905 | 1.23433386 | -6.9210534 |
| 2684 | NM_001042466.2 | 197.159935 | -9.2454156 | 1.33586214 | -6.9209354 |
| 2685 | NM_012458.3 | 167.941791 | -9.0139414 | 1.30244407 | -6.9207896 |
| 2686 | NM_182547.3 | 127.045624 | -8.6117545 | 1.2443331 | -6.9207791 |
| 2687 | NM_001984.1 | 174.550196 | -9.0694098 | 1.31055348 | -6.9202898 |
| 2688 | NM_003629.3 | 222.352138 | -9.4191165 | 1.36110355 | -6.9202057 |
| 2689 | NM_002296.3 | 174.401245 | -9.0683408 | 1.31047462 | -6.9198905 |
| 2690 | NM_174913.2 | 164.209194 | -8.9813476 | 1.29790181 | -6.9198976 |
| 2691 | NM_001085454.1 | 361.525505 | -10.120001 | 1.46270802 | -6.918675 |
| 2692 | NM_002157.2 | 213.077786 | -6.6203112 | 0.95694319 | -6.9181862 |
| 2693 | NM_001145415.1 | 125.09773 | -8.5885555 | 1.24169958 | -6.9167741 |
| 2694 | NM_000611.5 | 167.260601 | -9.0079129 | 1.30237706 | -6.9165168 |
| 2695 | NM_031442.3 | 150.488747 | -8.8560352 | 1.28041534 | -6.9165332 |
| 2696 | NM_030957.3 | 198.586856 | -9.2556166 | 1.33830844 | -6.9159069 |
| 2697 | NM_001005388.2 | 320.578768 | -9.9466928 | 1.43825003 | -6.91583 |
| 2698 | NM_001077397.1 | 136.102629 | -8.7117925 | 1.2597637 | -6.915418 |
| 2699 | NM_003660.3 | 280.647661 | -9.7546875 | 1.41060072 | -6.9152719 |
| 2700 | NM_006356.2 | 198.643111 | -9.2561126 | 1.33850371 | -6.9152685 |
| 2701 | NM_020401.3 | 143.693285 | -3.8950794 | 0.56325115 | -6.9153509 |
| 2702 | NM_005650.3 | 196.891866 | -9.2430219 | 1.33668957 | -6.9148605 |
| 2703 | NM_001297624.1 | 47.3322354 | 10.4777771 | 1.51529143 | 6.91469434 |
| 2704 | NM_016529.5 | 233.081001 | -9.4868704 | 1.37198289 | -6.9147148 |
| 2705 | NM_024735.4 | 150.423846 | -8.8550741 | 1.28066072 | -6.9144574 |
| 2706 | NM_006110.2 | 250.210236 | -7.7461953 | 1.12044497 | -6.9134992 |
| 2707 | NM_001136018.3 | 148.783745 | -8.8400211 | 1.27874753 | -6.9130308 |
| 2708 | NM_003875.2 | 137.691494 | -8.7266834 | 1.26239177 | -6.9128171 |

| | | | | | |
|---|---|---|---|---|---|
| 2709 | NM_019009.3 | 197.21287 | -9.2459589 | 1.33752649 | -6.9127295 |
| 2710 | NM_001127.3 | 193.860452 | -9.2213752 | 1.33403341 | -6.912402 |
| 2711 | NM_006369.4 | 250.151276 | -8.6254066 | 1.24785557 | -6.9121834 |
| 2712 | NM_005000.4 | 225.669785 | -9.4399024 | 1.36589795 | -6.911133 |
| 2713 | NM_203462.2 | 276.962192 | -7.9561468 | 1.15128791 | -6.9106492 |
| 2714 | NM_002588.3 | 126.067572 | -8.602137 | 1.24480813 | -6.9104119 |
| 2715 | NM_016630.6 | 145.617938 | -8.8075952 | 1.27458113 | -6.910188 |
| 2716 | NM_001257181.1 | 132.976685 | -8.6792418 | 1.25607257 | -6.9098251 |
| 2717 | NM_016038.2 | 173.119256 | -9.057619 | 1.31099015 | -6.9089909 |
| 2718 | NM_020820.3 | 179.08426 | -5.9325028 | 0.85873969 | -6.9083832 |
| 2719 | NM_001204961.1 | 213.94017 | -9.363801 | 1.35546631 | -6.9081768 |
| 2720 | NM_001321022.1 | 27.4472244 | 9.68836007 | 1.40247259 | 6.9080566 |
| 2721 | NM_004444.4 | 138.666795 | -8.7390192 | 1.26511089 | -6.9077101 |
| 2722 | NM_005639.2 | 338.734045 | -10.025954 | 1.45154593 | -6.9070869 |
| 2723 | NM_016121.4 | 117.692985 | -8.500967 | 1.23076831 | -6.9070408 |
| 2724 | NM_006767.3 | 290.632025 | -8.8401888 | 1.27993907 | -6.9067263 |
| 2725 | NM_014236.3 | 160.633068 | -8.9495077 | 1.29579196 | -6.906593 |
| 2726 | NM_020156.4 | 87.1229843 | -3.9930968 | 0.57818821 | -6.9062232 |
| 2727 | NM_015476.3 | 214.831159 | -9.3687667 | 1.35663603 | -6.9058808 |
| 2728 | NM_030938.4 | 198.619543 | -9.2560113 | 1.34030991 | -6.9058739 |
| 2729 | NM_147187.2 | 156.098722 | -8.9097378 | 1.29021139 | -6.9056419 |
| 2730 | NM_001120.4 | 124.457769 | -8.581958 | 1.24275524 | -6.9055899 |
| 2731 | NM_018403.6 | 120.599363 | -8.5346512 | 1.23599119 | -6.9051068 |
| 2732 | NM_201269.2 | 129.989308 | -8.6438927 | 1.25190602 | -6.904586 |
| 2733 | NR_130152.1 | 118.751593 | -8.5133106 | 1.2330501 | -6.9042698 |
| 2734 | NM_014597.4 | 134.81988 | -8.6963948 | 1.25961289 | -6.9040218 |
| 2735 | NM_014815.3 | 140.438068 | -8.7572097 | 1.26843447 | -6.9039512 |
| 2736 | NM_004354.2 | 209.830364 | -9.3349352 | 1.35216025 | -6.9037196 |
| 2737 | NM_139030.3 | 197.845145 | -9.2506974 | 1.34026517 | -6.9021397 |
| 2738 | NM_001136203.1 | 208.065704 | -9.3228685 | 1.35077371 | -6.9018729 |
| 2739 | NM_015104.2 | 122.45235 | -8.5580967 | 1.23996292 | -6.9018973 |
| 2740 | NM_130847.2 | 138.837667 | -8.7407724 | 1.26643483 | -6.901873 |
| 2741 | NM_001033.4 | 188.101031 | -9.1771089 | 1.32983014 | -6.9009632 |
| 2742 | NM_144582.2 | 168.744025 | -9.0212109 | 1.30724818 | -6.9009168 |
| 2743 | NM_015283.1 | 169.195594 | -9.0237235 | 1.30764868 | -6.9007247 |
| 2744 | NM_019597.4 | 192.350335 | -9.209582 | 1.33461337 | -6.9005617 |
| 2745 | NM_017740.2 | 128.598777 | -8.6291582 | 1.25051909 | -6.900461 |
| 2746 | NM_004112.3 | 128.371438 | -8.6260795 | 1.25015125 | -6.9000286 |
| 2747 | NM_005146.4 | 249.901406 | -8.6236214 | 1.24982245 | -6.8998772 |
| 2748 | NM_005278.4 | 239.108378 | -9.5232773 | 1.38021968 | -6.8998272 |
| 2749 | NM_001199973.1 | 110.214291 | -8.4056812 | 1.21828109 | -6.8996238 |
| 2750 | NM_007213.2 | 199.296733 | -9.2611565 | 1.34240271 | -6.8989405 |
| 2751 | NM_014060.2 | 136.804012 | -8.7177688 | 1.26372587 | -6.8984651 |

| | | | | | |
|---|---|---|---|---|---|
| 2752 | NM_152753.3 | 119.532185 | -8.5249582 | 1.2357848 | -6.8984165 |
| 2753 | NM_019892.5 | 128.74949 | -8.630752 | 1.25113137 | -6.8983579 |
| 2754 | NM_015209.2 | 115.214853 | -8.4721783 | 1.22817804 | -6.8981679 |
| 2755 | NM_000269.2 | 266.235732 | -9.6787047 | 1.40310222 | -6.8980752 |
| 2756 | NM_003758.3 | 222.469849 | -9.4193378 | 1.36558396 | -6.8976629 |
| 2757 | NM_001282925.1 | 301.03882 | -9.8557793 | 1.42888479 | -6.8975325 |
| 2758 | NM_022917.4 | 156.123907 | -8.9087538 | 1.29167787 | -6.8970399 |
| 2759 | NM_018233.3 | 131.307518 | -8.659794 | 1.25560122 | -6.8969302 |
| 2760 | NM_016311.4 | 179.363806 | -9.1086376 | 1.32069221 | -6.8968663 |
| 2761 | NM_001519.3 | 147.645368 | -8.8270529 | 1.27987421 | -6.8968129 |
| 2762 | NM_006830.3 | 235.740235 | -9.5031271 | 1.37817756 | -6.8954302 |
| 2763 | NM_002556.2 | 225.827924 | -8.4769271 | 1.22945865 | -6.8948453 |
| 2764 | NM_138565.2 | 159.767972 | -7.1214497 | 1.0330026 | -6.8939321 |
| 2765 | NM_033505.3 | 202.599745 | -9.2844809 | 1.34687754 | -6.8933371 |
| 2766 | NM_152299.3 | 187.678922 | -9.1742065 | 1.33092093 | -6.8931266 |
| 2767 | NM_001008701.2 | 401.624473 | -10.271767 | 1.49020169 | -6.89287 |
| 2768 | NM_001287190.1 | 176.171827 | -9.0842486 | 1.31791304 | -6.8929044 |
| 2769 | NM_133635.4 | 137.008022 | -8.7221702 | 1.26539567 | -6.8928403 |
| 2770 | NM_004252.4 | 116.748687 | -8.4918139 | 1.23216727 | -6.8917704 |
| 2771 | NM_001033561.1 | 132.936187 | -8.6771165 | 1.25908637 | -6.8915975 |
| 2772 | NM_013234.3 | 210.606169 | -9.3402774 | 1.35533813 | -6.8914739 |
| 2773 | NM_016139.3 | 144.054422 | -8.7940074 | 1.27608339 | -6.891405 |
| 2774 | NM_004082.4 | 243.246801 | -9.5485037 | 1.38561006 | -6.8911911 |
| 2775 | NM_000311.3 | 172.847927 | -9.056686 | 1.31425397 | -6.8911231 |
| 2776 | NM_001004067.3 | 173.873745 | -9.0642338 | 1.31537231 | -6.8910025 |
| 2777 | NM_016530.2 | 150.099786 | -8.8509299 | 1.28445239 | -6.8908197 |
| 2778 | NM_133474.3 | 106.168171 | -3.4901791 | 0.50649742 | -6.8908132 |
| 2779 | NM_016374.5 | 135.959333 | -8.7088269 | 1.26388471 | -6.8905232 |
| 2780 | NM_001081491.1 | 120.07438 | -8.5319186 | 1.23822829 | -6.8904246 |
| 2781 | NM_001330564.1 | 154.618187 | -8.8958148 | 1.29108672 | -6.890176 |
| 2782 | NM_014283.4 | 175.36834 | -9.0762646 | 1.31730273 | -6.890037 |
| 2783 | NM_176812.4 | 281.832016 | -8.7972377 | 1.27684018 | -6.8898503 |
| 2784 | NR_104162.1 | 61.5330667 | 6.07417714 | 0.88162266 | 6.88976977 |
| 2785 | NM_018010.3 | 121.932948 | -8.5531989 | 1.24149871 | -6.8894142 |
| 2786 | NM_016141.3 | 201.822301 | -9.2789636 | 1.34687761 | -6.8892404 |
| 2787 | NM_018368.3 | 118.832057 | -8.5152013 | 1.23603185 | -6.8891439 |
| 2788 | NM_015294.4 | 167.736483 | -9.0114328 | 1.30813122 | -6.8887835 |
| 2789 | NM_002398.2 | 262.548005 | -9.658787 | 1.40212622 | -6.8886715 |
| 2790 | NM_152395.2 | 117.517798 | -8.4993213 | 1.23381254 | -6.888665 |
| 2791 | NM_007277.4 | 135.511412 | -8.7049354 | 1.26367515 | -6.8885864 |
| 2792 | NM_007173.5 | 256.138218 | -9.6231527 | 1.39701826 | -6.8883514 |
| 2793 | NM_006620.3 | 141.152725 | -8.7632061 | 1.27233376 | -6.8875058 |
| 2794 | NM_005461.4 | 249.021081 | -9.5825015 | 1.39135724 | -6.8871611 |

| | | | | | |
|---|---|---|---|---|---|
| 2795 | NM_013986.3 | 224.322779 | -9.4316643 | 1.36959796 | -6.8864474 |
| 2796 | NM_014117.2 | 144.249866 | -8.7946823 | 1.277144 | -6.8862104 |
| 2797 | NM_003575.3 | 151.166127 | -8.861843 | 1.28690726 | -6.8861551 |
| 2798 | NM_015906.3 | 128.947655 | -8.6322619 | 1.25362331 | -6.8858499 |
| 2799 | NM_015577.2 | 171.404618 | -9.0424674 | 1.31323449 | -6.8856457 |
| 2800 | NM_001037131.2 | 150.230067 | -8.8533227 | 1.28580349 | -6.88544 |
| 2801 | NM_001040260.3 | 209.884588 | -9.3352453 | 1.3559212 | -6.8847993 |
| 2802 | NM_001040006.1 | 173.340571 | -9.0595689 | 1.3159156 | -6.8846124 |
| 2803 | NM_001271977.1 | 193.682014 | -9.2195781 | 1.33919745 | -6.8844053 |
| 2804 | NM_001122659.2 | 156.540071 | -8.9129789 | 1.29467463 | -6.8843389 |
| 2805 | NM_005967.3 | 195.076074 | -9.2296323 | 1.34071777 | -6.8840979 |
| 2806 | NM_001302633.1 | 186.061777 | -9.1623315 | 1.33105166 | -6.8835281 |
| 2807 | NM_006765.3 | 143.744357 | -8.7903558 | 1.27708941 | -6.8831169 |
| 2808 | NM_152588.2 | 182.35931 | -9.1329494 | 1.32695443 | -6.8826398 |
| 2809 | NM_152419.2 | 180.667376 | -9.1197737 | 1.32506491 | -6.8825109 |
| 2810 | NM_003315.3 | 113.50415 | -8.4476704 | 1.22744536 | -6.8823189 |
| 2811 | NM_016061.2 | 191.648824 | -9.2042778 | 1.33737381 | -6.8823524 |
| 2812 | NM_001862.2 | 239.230839 | -9.5243129 | 1.38396954 | -6.8818805 |
| 2813 | NM_020920.3 | 168.423061 | -6.2663934 | 0.91060181 | -6.8815955 |
| 2814 | NM_001031689.2 | 141.206668 | -8.7629877 | 1.27344468 | -6.8813258 |
| 2815 | NM_002013.3 | 225.489794 | -9.4389875 | 1.37169189 | -6.8812738 |
| 2816 | NM_018089.2 | 123.156913 | -8.5647899 | 1.24467255 | -6.8811591 |
| 2817 | NM_001048241.2 | 210.81585 | -9.3417703 | 1.35774053 | -6.8803796 |
| 2818 | NM_006755.1 | 239.431858 | -7.1222427 | 1.03516793 | -6.8802776 |
| 2819 | NM_138773.3 | 192.22823 | -9.2084038 | 1.33853342 | -6.8794724 |
| 2820 | NM_022744.3 | 256.99183 | -8.6637461 | 1.25939497 | -6.8792923 |
| 2821 | NM_001398.2 | 200.063559 | -7.4734538 | 1.08644175 | -6.8788352 |
| 2822 | NM_177424.2 | 250.130231 | -9.5886251 | 1.39404581 | -6.8782712 |
| 2823 | NM_001281.2 | 231.06663 | -9.4742065 | 1.37756193 | -6.8775176 |
| 2824 | NM_000434.3 | 143.130701 | -8.783824 | 1.27720123 | -6.8774002 |
| 2825 | NM_001071.2 | 131.468516 | -8.6598555 | 1.25917053 | -6.8774287 |
| 2826 | NM_020680.3 | 127.180252 | -8.6132166 | 1.25247296 | -6.876968 |
| 2827 | NM_080820.5 | 228.971326 | -9.4610823 | 1.37576521 | -6.87696 |
| 2828 | NM_032940.2 | 155.456194 | -8.9024892 | 1.29465318 | -6.8763506 |
| 2829 | NM_001483.2 | 117.691021 | -8.5018037 | 1.23640968 | -6.8762028 |
| 2830 | NM_001081755.2 | 142.058247 | -8.7716952 | 1.27568289 | -6.8760781 |
| 2831 | NM_001346792.1 | 66.4327354 | 10.9776481 | 1.59659066 | 6.87568104 |
| 2832 | NM_002069.5 | 240.924176 | -9.534392 | 1.38670786 | -6.8755592 |
| 2833 | NM_013442.2 | 156.344624 | -8.9097996 | 1.29586875 | -6.8755417 |
| 2834 | NM_000381.3 | 121.223982 | -8.5412527 | 1.24230105 | -6.8753485 |
| 2835 | NM_001009984.2 | 220.149462 | -6.6713206 | 0.97036754 | -6.8750451 |
| 2836 | NM_005456.3 | 482.973993 | -7.0819873 | 1.03010154 | -6.8750381 |
| 2837 | NM_017824.4 | 165.750261 | -8.9947935 | 1.30843316 | -6.8744769 |

| | | | | | |
|---|---|---|---|---|---|
| 2838 | NM_004526.3 | 169.680653 | -9.0283508 | 1.31342035 | -6.8739233 |
| 2839 | NM_013285.2 | 144.259715 | -8.794921 | 1.27948913 | -6.8737755 |
| 2840 | NM_016076.4 | 110.528785 | -8.4106877 | 1.22362265 | -6.8735959 |
| 2841 | NM_020378.3 | 145.089135 | -8.8032191 | 1.28080682 | -6.8731826 |
| 2842 | NM_001277325.1 | 124.943491 | -8.5871761 | 1.249426 | -6.8728969 |
| 2843 | NM_017592.2 | 144.650923 | -8.7998447 | 1.28038061 | -6.8728351 |
| 2844 | NM_001024455.3 | 160.34675 | -8.9473059 | 1.30186335 | -6.8726921 |
| 2845 | NM_023019.3 | 337.736599 | -10.021916 | 1.45824397 | -6.872592 |
| 2846 | NM_001318852.1 | 347.420343 | -10.062784 | 1.4643534 | -6.8718274 |
| 2847 | NM_181892.3 | 121.829931 | -8.55257 | 1.24457281 | -6.8718921 |
| 2848 | NM_182895.4 | 126.198953 | -8.60199 | 1.25177376 | -6.8718408 |
| 2849 | NM_001293.2 | 149.871976 | -8.8494618 | 1.28788323 | -6.8713231 |
| 2850 | NM_001293696.1 | 196.669689 | -9.2414925 | 1.34492584 | -6.871377 |
| 2851 | NM_001308208.1 | 122.401098 | -8.5579354 | 1.24545731 | -6.8713198 |
| 2852 | NM_152925.2 | 109.266526 | -8.3931769 | 1.22151968 | -6.8710943 |
| 2853 | NM_018972.2 | 215.039006 | -9.3705196 | 1.36377902 | -6.8709956 |
| 2854 | NM_007030.2 | 134.740225 | -8.6961589 | 1.2656583 | -6.8708583 |
| 2855 | NM_024658.3 | 126.392469 | -8.6061605 | 1.25264833 | -6.8703724 |
| 2856 | NM_001127266.1 | 137.666903 | -8.7269283 | 1.27038782 | -6.8694994 |
| 2857 | NM_001696.3 | 185.793234 | -9.1594287 | 1.33336851 | -6.8693903 |
| 2858 | NM_022131.2 | 207.949151 | -9.3225684 | 1.35718206 | -6.8690625 |
| 2859 | NM_019107.3 | 194.538764 | -9.2263147 | 1.34333972 | -6.8681917 |
| 2860 | NM_030581.3 | 122.873656 | -8.5633861 | 1.24682689 | -6.8681436 |
| 2861 | NM_014820.4 | 272.165891 | -8.7463905 | 1.27349638 | -6.8680136 |
| 2862 | NM_002657.3 | 109.19999 | -8.3911913 | 1.22178976 | -6.8679502 |
| 2863 | NM_014679.4 | 124.237949 | -8.577632 | 1.24895441 | -6.8678504 |
| 2864 | NM_001127211.2 | 112.864128 | -8.4399155 | 1.22895066 | -6.8675788 |
| 2865 | NM_020314.5 | 135.735971 | -8.7079213 | 1.26811748 | -6.8668096 |
| 2866 | NM_199166.2 | 107.202115 | -8.3677303 | 1.2186096 | -6.8666211 |
| 2867 | NM_012289.3 | 155.03636 | -8.8983213 | 1.29597282 | -6.8661327 |
| 2868 | NM_003137.4 | 171.091385 | -9.0404678 | 1.31670928 | -6.8659559 |
| 2869 | NM_003001.3 | 153.771218 | -8.8868559 | 1.29435349 | -6.8658647 |
| 2870 | NM_003132.2 | 218.089483 | -9.3908513 | 1.36776704 | -6.8658266 |
| 2871 | NM_022064.4 | 150.908489 | -8.8593322 | 1.29052754 | -6.864892 |
| 2872 | NM_178031.2 | 618.975381 | -6.5798329 | 0.95851784 | -6.864591 |
| 2873 | NM_019096.4 | 132.916347 | -8.6755796 | 1.26385846 | -6.8643601 |
| 2874 | NM_005119.3 | 235.384004 | -8.5363541 | 1.24369401 | -6.8637092 |
| 2875 | NM_002720.2 | 167.241539 | -9.007526 | 1.31243403 | -6.8632219 |
| 2876 | NM_020839.3 | 144.041982 | -8.7921197 | 1.28105218 | -6.8632018 |
| 2877 | NM_001098787.1 | 123.967622 | -8.5777973 | 1.24983797 | -6.8631274 |
| 2878 | NM_001008529.2 | 287.413421 | -9.7893495 | 1.42641141 | -6.8629214 |
| 2879 | NM_003274.4 | 138.322867 | -8.7335992 | 1.27258065 | -6.8629043 |
| 2880 | NM_001135822.1 | 200.633927 | -9.2707712 | 1.35102223 | -6.8620419 |

| | | | | | |
|---|---|---|---|---|---|
| 2881 | NM_015173.3 | 105.395469 | -8.3426343 | 1.21583834 | -6.8616312 |
| 2882 | NM_001070.4 | 128.299482 | -8.625722 | 1.25711294 | -6.8615331 |
| 2883 | NM_001330351.1 | 155.652165 | -8.9047302 | 1.2978069 | -6.8613676 |
| 2884 | NM_201632.4 | 24.9765138 | 9.55125118 | 1.3921504 | 6.86078974 |
| 2885 | NM_006832.2 | 145.149933 | -8.8035533 | 1.28325302 | -6.860341 |
| 2886 | NM_016147.2 | 222.9019 | -7.5949047 | 1.10712084 | -6.8600504 |
| 2887 | NM_003254.2 | 189.900814 | -9.1919086 | 1.33993622 | -6.8599598 |
| 2888 | NM_015026.2 | 136.508058 | -8.7138183 | 1.27038152 | -6.8592137 |
| 2889 | NM_004417.3 | 128.899901 | -8.63214 | 1.25853745 | -6.8588663 |
| 2890 | NM_025161.5 | 140.255938 | -8.7540861 | 1.27636934 | -6.8585839 |
| 2891 | NM_001024936.1 | 247.995825 | -9.576139 | 1.39624061 | -6.8585163 |
| 2892 | NM_004618.4 | 111.8928 | -8.4287435 | 1.22901187 | -6.8581466 |
| 2893 | NM_014316.3 | 230.537469 | -9.4707568 | 1.38104324 | -6.857683 |
| 2894 | NM_003644.2 | 35.8165032 | 10.087091 | 1.47093376 | 6.85761062 |
| 2895 | NM_031946.6 | 175.825746 | -9.0796408 | 1.32410859 | -6.8571723 |
| 2896 | NM_018303.5 | 149.038706 | -8.8414083 | 1.28945572 | -6.8566979 |
| 2897 | NM_032932.5 | 240.733729 | -9.5332836 | 1.39038776 | -6.8565647 |
| 2898 | NM_002475.4 | 255.523667 | -9.619409 | 1.4029663 | -6.856479 |
| 2899 | NM_015343.4 | 117.88786 | -8.5038732 | 1.24027963 | -6.8564161 |
| 2900 | NM_001303256.2 | 163.676785 | -8.9765971 | 1.30929426 | -6.8560578 |
| 2901 | NM_148921.3 | 144.461746 | -8.7961597 | 1.2830699 | -6.8555577 |
| 2902 | NM_018425.3 | 142.875102 | -8.7811051 | 1.28100788 | -6.8548408 |
| 2903 | NM_015317.2 | 169.404335 | -9.026189 | 1.3167926 | -6.8546778 |
| 2904 | NM_006844.4 | 176.243287 | -9.0835643 | 1.32549063 | -6.8529826 |
| 2905 | NM_001964.2 | 250.003989 | -9.5884672 | 1.39923195 | -6.8526646 |
| 2906 | NM_006336.3 | 226.291792 | -8.4782383 | 1.23723134 | -6.8525894 |
| 2907 | NM_016274.5 | 209.358912 | -9.3318589 | 1.36181244 | -6.8525287 |
| 2908 | NM_001308068.1 | 160.47691 | -8.9491525 | 1.30599898 | -6.8523427 |
| 2909 | NM_018375.4 | 146.393457 | -8.8146178 | 1.28636995 | -6.8523194 |
| 2910 | NM_017519.2 | 171.790968 | -7.2165762 | 1.05318206 | -6.852164 |
| 2911 | NM_001303524.1 | 173.116123 | -9.0579231 | 1.32193866 | -6.8519995 |
| 2912 | NM_003562.4 | 177.878491 | -9.0967207 | 1.32760835 | -6.851961 |
| 2913 | NM_177947.2 | 201.406759 | -9.2769316 | 1.35429921 | -6.8499867 |
| 2914 | NM_001098416.3 | 127.709453 | -8.6189159 | 1.25834679 | -6.8493964 |
| 2915 | NM_020831.4 | 154.503359 | -8.8933533 | 1.29848973 | -6.8489978 |
| 2916 | NM_000415.2 | 44.3058746 | 10.395865 | 1.51791996 | 6.84875705 |
| 2917 | NM_004204.3 | 136.573583 | -8.7164592 | 1.2727286 | -6.8486393 |
| 2918 | NM_003249.4 | 149.875894 | -8.8496521 | 1.2922134 | -6.8484448 |
| 2919 | NM_001079803.2 | 162.844135 | -8.9688055 | 1.30967098 | -6.8481364 |
| 2920 | NM_006700.2 | 160.684876 | -8.9499469 | 1.30704274 | -6.8474784 |
| 2921 | NM_001040450.2 | 140.535915 | -8.755955 | 1.27873192 | -6.8473735 |
| 2922 | NM_133457.4 | 165.071927 | -8.9894498 | 1.31284605 | -6.8472993 |
| 2923 | NM_006758.2 | 168.520599 | -9.0188975 | 1.31719944 | -6.847025 |

| | | | | | |
|---|---|---|---|---|---|
| 2924 | NM_003560.2 | 112.888079 | -8.4432606 | 1.23317526 | -6.8467645 |
| 2925 | NM_006494.3 | 111.713385 | -8.4276633 | 1.23093992 | -6.8465269 |
| 2926 | NM_015649.2 | 154.943795 | -8.8975699 | 1.2996495 | -6.8461304 |
| 2927 | NM_025146.3 | 167.042888 | -9.0059181 | 1.31547712 | -6.846123 |
| 2928 | NM_001303241.1 | 139.940019 | -8.7507083 | 1.27829698 | -6.8455988 |
| 2929 | NM_020376.3 | 117.874996 | -8.5048567 | 1.24245199 | -6.8452196 |
| 2930 | NM_001039141.2 | 134.706025 | -8.6955736 | 1.27044584 | -6.8445055 |
| 2931 | NM_001113348.1 | 138.634894 | -8.7377122 | 1.27662114 | -6.8444051 |
| 2932 | NM_001300860.1 | 178.752348 | -9.1033041 | 1.33007381 | -6.8442097 |
| 2933 | NM_198597.2 | 339.790549 | -6.36991 | 0.93070016 | -6.8442129 |
| 2934 | NM_182553.2 | 238.867287 | -9.5221839 | 1.3913926 | -6.8436356 |
| 2935 | NM_017755.5 | 127.61972 | -8.6173052 | 1.25926534 | -6.8431211 |
| 2936 | NM_001242866.2 | 122.927808 | -8.562358 | 1.25127253 | -6.8429202 |
| 2937 | NM_013373.3 | 143.077887 | -8.7828201 | 1.28359121 | -6.842381 |
| 2938 | NM_001204063.1 | 109.618918 | -8.3996595 | 1.22763931 | -6.8421233 |
| 2939 | NM_025138.4 | 138.3775 | -8.7340112 | 1.27655652 | -6.8418524 |
| 2940 | NM_002446.3 | 247.565604 | -9.5737122 | 1.39937627 | -6.8414138 |
| 2941 | NM_006010.5 | 145.738956 | -8.8096714 | 1.28778539 | -6.8409469 |
| 2942 | NM_014669.4 | 148.74477 | -8.8387503 | 1.29203225 | -6.8409672 |
| 2943 | NM_022841.5 | 150.538507 | -8.8555141 | 1.29456343 | -6.8405409 |
| 2944 | NM_006742.2 | 109.699193 | -8.4007988 | 1.22812476 | -6.8403464 |
| 2945 | NM_004684.5 | 115.984302 | -8.4819206 | 1.24002001 | -6.8401482 |
| 2946 | NM_016282.3 | 134.905134 | -8.6976551 | 1.27159406 | -6.839962 |
| 2947 | NM_001003800.1 | 145.829443 | -8.8100128 | 1.28811324 | -6.8394708 |
| 2948 | NM_144586.6 | 170.135414 | -9.0324086 | 1.32063349 | -6.8394514 |
| 2949 | NM_004907.2 | 207.407538 | -8.3544856 | 1.22160751 | -6.8389278 |
| 2950 | NM_201575.3 | 267.989889 | -9.6882946 | 1.41663784 | -6.8389354 |
| 2951 | NM_006011.3 | 260.848109 | -9.649233 | 1.41104834 | -6.8383433 |
| 2952 | NM_033254.3 | 105.177947 | -8.3369064 | 1.21917133 | -6.8381746 |
| 2953 | NM_024006.5 | 128.493638 | -8.6279089 | 1.26180354 | -6.8377594 |
| 2954 | NM_001122630.1 | 144.52639 | -8.7990545 | 1.2868978 | -6.8374151 |
| 2955 | NM_130465.4 | 200.81333 | -9.2718806 | 1.35605658 | -6.8373848 |
| 2956 | NM_000402.4 | 161.383642 | -8.9561807 | 1.30989471 | -6.8373287 |
| 2957 | NM_001130066.1 | 174.779565 | -5.1564379 | 0.75426539 | -6.8363708 |
| 2958 | NM_024569.4 | 221.222402 | -9.4114031 | 1.37665861 | -6.8364103 |
| 2959 | NM_001303448.1 | 147.079886 | -8.8208084 | 1.29041334 | -6.8356457 |
| 2960 | NM_100264.2 | 166.567145 | -9.0015598 | 1.31687006 | -6.8355718 |
| 2961 | NM_178000.2 | 165.121049 | -8.9906678 | 1.31529581 | -6.8354721 |
| 2962 | NM_006804.3 | 171.632191 | -5.734059 | 0.83909043 | -6.8336603 |
| 2963 | NM_006280.2 | 210.492982 | -9.3396986 | 1.36675368 | -6.8334908 |
| 2964 | NM_014612.4 | 146.876819 | -8.8223361 | 1.29105257 | -6.8334445 |
| 2965 | NM_024036.4 | 262.998334 | -9.6609499 | 1.41414431 | -6.8316577 |
| 2966 | NM_001033057.1 | 108.979059 | -8.3882654 | 1.22787072 | -6.8315542 |

| | | | | | |
|---|---|---|---|---|---|
| 2967 | NM_001470.3 | 216.88733 | -9.3833559 | 1.37366193 | -6.8309063 |
| 2968 | NM_020962.2 | 129.841799 | -8.6443364 | 1.2655276 | -6.8306186 |
| 2969 | NM_001777.3 | 260.696408 | -9.6481884 | 1.4125177 | -6.8304903 |
| 2970 | NM_015005.2 | 181.58598 | -9.1270077 | 1.33624171 | -6.8303568 |
| 2971 | NM_001032363.3 | 148.203152 | -8.8337539 | 1.29332739 | -6.8302535 |
| 2972 | NM_005781.4 | 174.353746 | -9.0674318 | 1.32756768 | -6.8301089 |
| 2973 | NM_024790.6 | 104.872504 | -8.3327666 | 1.22009702 | -6.8295934 |
| 2974 | NM_015203.4 | 144.317962 | -8.7966791 | 1.28807955 | -6.829298 |
| 2975 | NM_004674.4 | 123.575255 | -8.5717182 | 1.25516154 | -6.8291753 |
| 2976 | NM_014282.3 | 172.431662 | -9.0520744 | 1.32551955 | -6.8290765 |
| 2977 | NM_002850.3 | 197.409582 | -9.2470723 | 1.35412917 | -6.8287963 |
| 2978 | NM_152903.4 | 202.839183 | -3.7915383 | 0.55529302 | -6.8279956 |
| 2979 | NM_018142.2 | 116.725881 | -8.4889975 | 1.24340469 | -6.8272201 |
| 2980 | NM_198963.2 | 136.224957 | -8.7113803 | 1.27610545 | -6.8265364 |
| 2981 | NM_004716.3 | 201.186078 | -9.2744869 | 1.35865044 | -6.8262495 |
| 2982 | NM_003749.2 | 243.316904 | -7.7187658 | 1.13075865 | -6.8261833 |
| 2983 | NM_001040438.2 | 150.640568 | -8.857142 | 1.29764788 | -6.8255358 |
| 2984 | NM_078480.2 | 184.128917 | -9.1468472 | 1.34020005 | -6.8249865 |
| 2985 | NM_020783.3 | 567.84382 | -8.7811023 | 1.28664347 | -6.824814 |
| 2986 | NM_181453.3 | 126.742567 | -8.6091583 | 1.261663 | -6.8236592 |
| 2987 | NM_015656.1 | 394.93992 | -7.9448307 | 1.16433884 | -6.8234696 |
| 2988 | NM_018051.4 | 121.527279 | -8.5483101 | 1.25280037 | -6.8233617 |
| 2989 | NM_012248.3 | 158.435384 | -8.9296968 | 1.30872781 | -6.8231887 |
| 2990 | NM_183006.3 | 132.629186 | -8.6733017 | 1.27119538 | -6.8229494 |
| 2991 | NM_005132.2 | 108.404729 | -8.3839437 | 1.22885947 | -6.8225407 |
| 2992 | NM_001017995.2 | 250.436977 | -8.6266764 | 1.26461673 | -6.8215739 |
| 2993 | NM_015509.3 | 298.937084 | -9.8457161 | 1.44335053 | -6.8214311 |
| 2994 | NM_015548.4 | 292.263965 | -8.848465 | 1.29719019 | -6.8212549 |
| 2995 | NM_001009925.1 | 142.034631 | -8.7735349 | 1.28621952 | -6.82118 |
| 2996 | NM_018907.3 | 203.49798 | -9.2913373 | 1.36221717 | -6.820746 |
| 2997 | NM_013400.3 | 127.102032 | -8.6125504 | 1.26279317 | -6.8202384 |
| 2998 | NM_016237.4 | 199.560008 | -9.2624967 | 1.35835642 | -6.8189001 |
| 2999 | NM_032991.2 | 117.620183 | -8.4991325 | 1.2464331 | -6.8187635 |
| 3000 | NM_023011.3 | 158.545068 | -8.9301562 | 1.30967214 | -6.8186197 |
| 3001 | NM_005632.2 | 128.088724 | -8.623045 | 1.26468917 | -6.8183117 |
| 3002 | NM_005219.4 | 151.360277 | -8.8639997 | 1.30010962 | -6.8178864 |
| 3003 | NM_005009.2 | 146.45416 | -8.8166605 | 1.29338968 | -6.8167086 |
| 3004 | NM_007346.3 | 199.474251 | -8.2953593 | 1.21696366 | -6.8164396 |
| 3005 | NM_016022.3 | 128.508986 | -8.6285408 | 1.2658358 | -6.8164771 |
| 3006 | NM_001321096.2 | 148.109124 | -8.8319109 | 1.29572831 | -6.816175 |
| 3007 | NM_013263.4 | 101.327833 | -8.2866296 | 1.21586 | -6.8154471 |
| 3008 | NM_052897.3 | 144.910709 | -6.3051956 | 0.92516199 | -6.8152341 |
| 3009 | NM_016485.4 | 232.583014 | -9.4833479 | 1.39156308 | -6.814889 |

| | | | | | |
|---|---|---|---|---|---|
| 3010 | NM_002496.3 | 175.776939 | -9.0796602 | 1.33248501 | -6.8140806 |
| 3011 | NM_000917.3 | 164.605715 | -8.9858988 | 1.3187472 | -6.8139661 |
| 3012 | NM_006111.2 | 105.65619 | -8.3475226 | 1.22507171 | -6.8139053 |
| 3013 | NM_012402.3 | 120.487781 | -8.5350597 | 1.25263719 | -6.8136726 |
| 3014 | NM_004690.3 | 116.374508 | -8.4832598 | 1.24506387 | -6.8135137 |
| 3015 | NM_001143667.1 | 127.091633 | -8.6118177 | 1.26412426 | -6.8124772 |
| 3016 | NM_018645.5 | 342.14508 | -10.040269 | 1.47388234 | -6.8121236 |
| 3017 | NM_030640.2 | 105.878157 | -8.3482261 | 1.22552923 | -6.8119355 |
| 3018 | NM_032019.5 | 138.280236 | -8.7340639 | 1.28220394 | -6.8117588 |
| 3019 | NM_001145161.2 | 253.062072 | -9.6053793 | 1.41022272 | -6.8112499 |
| 3020 | NM_002197.2 | 123.81822 | -8.5742247 | 1.25887079 | -6.8110443 |
| 3021 | NM_006887.4 | 175.433115 | -9.0778856 | 1.33282695 | -6.8110009 |
| 3022 | NM_024953.3 | 111.745841 | -8.4249563 | 1.23701776 | -6.8106995 |
| 3023 | NM_001283.3 | 249.307162 | -9.5838135 | 1.40719989 | -6.8105559 |
| 3024 | NM_018841.5 | 148.917723 | -8.8410823 | 1.29832657 | -6.8095983 |
| 3025 | NR_046251.1 | 147.763707 | -8.8282001 | 1.29648564 | -6.8093311 |
| 3026 | NM_004075.4 | 113.889991 | -8.4531422 | 1.24147583 | -6.8089462 |
| 3027 | NM_001095.3 | 213.910448 | -9.3628612 | 1.37514723 | -6.8086246 |
| 3028 | NM_024682.2 | 141.821715 | -8.7697578 | 1.28816149 | -6.8079646 |
| 3029 | NM_016265.3 | 101.425482 | -8.2874804 | 1.21735272 | -6.8077889 |
| 3030 | NM_016075.3 | 146.696315 | -8.818533 | 1.29538719 | -6.8076426 |
| 3031 | NM_032514.3 | 171.985457 | -9.0480536 | 1.32911642 | -6.8075704 |
| 3032 | NM_001195555.1 | 118.038122 | -8.5029449 | 1.24923065 | -6.8065452 |
| 3033 | NM_001092.4 | 163.510206 | -8.9768083 | 1.31893046 | -6.8061271 |
| 3034 | NM_130439.3 | 139.584905 | -8.7460409 | 1.28508943 | -6.8057839 |
| 3035 | NM_014824.2 | 141.509547 | -8.766336 | 1.2881226 | -6.8055137 |
| 3036 | NM_015456.4 | 287.636286 | -8.8250338 | 1.29674065 | -6.8055503 |
| 3037 | NR_110130.1 | 104.324731 | 3.39222936 | 0.49846523 | 6.80534797 |
| 3038 | NM_003707.2 | 150.133965 | -8.8516249 | 1.30089482 | -6.8042587 |
| 3039 | NM_002798.2 | 204.611417 | -9.2989501 | 1.36676185 | -6.803636 |
| 3040 | NR_134873.1 | 231.726495 | 3.3939751 | 0.49886814 | 6.80335109 |
| 3041 | NM_001301044.1 | 188.147071 | -9.1766018 | 1.34897051 | -6.8026704 |
| 3042 | NM_018028.3 | 112.26976 | -8.4347987 | 1.23995198 | -6.8025205 |
| 3043 | NM_018170.4 | 184.590796 | -9.1500246 | 1.34513049 | -6.8023323 |
| 3044 | NM_001300828.1 | 143.796067 | -8.7898379 | 1.29222448 | -6.8020983 |
| 3045 | NM_145245.4 | 191.141564 | -9.2004389 | 1.35259572 | -6.8020613 |
| 3046 | NM_018981.2 | 197.184033 | -9.2457818 | 1.35937071 | -6.8015161 |
| 3047 | NM_004927.3 | 119.869201 | -8.5274571 | 1.25398034 | -6.8003117 |
| 3048 | NM_001286771.2 | 151.611787 | -8.8662405 | 1.3040332 | -6.7990911 |
| 3049 | NM_033517.1 | 195.703338 | -8.269944 | 1.21635742 | -6.7989424 |
| 3050 | NM_001794.4 | 172.662566 | -9.0537201 | 1.33179915 | -6.7981122 |
| 3051 | NM_018091.5 | 110.522662 | -8.4094961 | 1.23703913 | -6.7980842 |
| 3052 | NM_001001410.2 | 169.942132 | -9.0308782 | 1.32850407 | -6.7977799 |

| | | | | | |
|---|---|---|---|---|---|
| 3053 | NM_012264.4 | 216.584035 | -8.4148938 | 1.23789921 | -6.7977213 |
| 3054 | NM_004548.2 | 166.838981 | -9.0043062 | 1.3246599 | -6.7974476 |
| 3055 | NM_000368.4 | 128.18182 | -8.6246233 | 1.26896247 | -6.7965944 |
| 3056 | NM_001134647.1 | 264.163438 | -9.6671135 | 1.42242921 | -6.7962001 |
| 3057 | NM_001134673.3 | 167.889256 | -9.0126233 | 1.32612807 | -6.7961937 |
| 3058 | NM_001284245.1 | 102.978188 | -8.3060193 | 1.22244847 | -6.7945762 |
| 3059 | NM_007085.4 | 2588.36156 | -8.5044304 | 1.25173024 | -6.79414 |
| 3060 | NM_001323342.1 | 119.994825 | -8.5310853 | 1.2556771 | -6.794012 |
| 3061 | NM_005013.2 | 114.82517 | -8.4652637 | 1.24609169 | -6.7934517 |
| 3062 | NM_001129995.1 | 137.937476 | -8.7285287 | 1.2850039 | -6.7926088 |
| 3063 | NM_013433.4 | 233.166531 | -9.4873858 | 1.39677527 | -6.7923495 |
| 3064 | NM_022913.3 | 138.51033 | -8.733934 | 1.28588894 | -6.7921371 |
| 3065 | NM_017623.4 | 109.560425 | -8.3965106 | 1.23641103 | -6.791035 |
| 3066 | NM_001008273.1 | 226.011963 | -9.4422975 | 1.39042073 | -6.7909642 |
| 3067 | NM_002945.3 | 226.417103 | -8.4809481 | 1.24889575 | -6.7907574 |
| 3068 | NM_012161.3 | 161.004328 | -8.9527819 | 1.31852872 | -6.7899787 |
| 3069 | NM_000214.2 | 136.726238 | -5.9525749 | 0.87684449 | -6.7886324 |
| 3070 | NM_000849.4 | 161.696706 | -8.9587397 | 1.31981581 | -6.7878711 |
| 3071 | NM_018677.3 | 169.790937 | -9.0298144 | 1.33028061 | -6.787902 |
| 3072 | NM_015680.5 | 163.683894 | -8.9768381 | 1.32249276 | -6.7878165 |
| 3073 | NM_001449.4 | 209.216252 | -9.3308912 | 1.37496638 | -6.7862686 |
| 3074 | NM_015062.4 | 111.884726 | -8.4272176 | 1.24182272 | -6.786168 |
| 3075 | NM_005197.3 | 197.620366 | -9.2481663 | 1.36283558 | -6.7859736 |
| 3076 | NM_004432.3 | 241.841191 | -9.5398899 | 1.40588614 | -6.7856775 |
| 3077 | NM_001276275.1 | 154.386462 | -8.8937396 | 1.31071773 | -6.7853966 |
| 3078 | NM_005576.3 | 123.635846 | -8.5742114 | 1.26363856 | -6.7853354 |
| 3079 | NM_015475.4 | 121.948981 | -8.5520441 | 1.2604145 | -6.7851045 |
| 3080 | NM_003897.3 | 114.991188 | -8.469864 | 1.24844021 | -6.7843569 |
| 3081 | NM_015051.2 | 129.205721 | -8.6359853 | 1.27292961 | -6.7843384 |
| 3082 | NM_015907.2 | 105.722119 | -8.3482877 | 1.23058762 | -6.7839848 |
| 3083 | NM_020896.3 | 154.73244 | -8.8954707 | 1.31131988 | -6.7836009 |
| 3084 | NM_014754.2 | 188.635416 | -9.1814529 | 1.35349639 | -6.7835074 |
| 3085 | NM_145185.3 | 114.251675 | -8.4571862 | 1.24682157 | -6.7829964 |
| 3086 | NM_013243.3 | 320.976907 | -9.9484155 | 1.46670855 | -6.7828169 |
| 3087 | NM_001320883.1 | 229.00279 | -9.4608525 | 1.39488798 | -6.7825177 |
| 3088 | NM_175614.4 | 228.238302 | -9.4564274 | 1.39423747 | -6.7825084 |
| 3089 | NM_005741.4 | 126.980955 | -8.60976 | 1.26952238 | -6.7818891 |
| 3090 | NM_006405.6 | 108.335198 | -8.3834417 | 1.23622276 | -6.7814976 |
| 3091 | NM_001330311.1 | 193.700032 | -9.2192795 | 1.35950739 | -6.7813383 |
| 3092 | NM_015959.3 | 160.072298 | -8.9445573 | 1.31912512 | -6.7806739 |
| 3093 | NM_014767.2 | 302.52606 | -8.8997634 | 1.31284838 | -6.7789728 |
| 3094 | NM_003940.2 | 85.4756655 | -4.6294787 | 0.68306957 | -6.777463 |
| 3095 | NM_001048200.2 | 102.869685 | -8.3089327 | 1.2260504 | -6.7769911 |

| | | | | | |
|---|---|---|---|---|---|
| 3096 | NM_003406.3 | 173.712155 | -9.0627505 | 1.33740973 | -6.7763455 |
| 3097 | NM_012399.4 | 158.620671 | -8.9306613 | 1.31792978 | -6.77628 |
| 3098 | NM_198076.5 | 198.95902 | -9.2581156 | 1.36625197 | -6.7762871 |
| 3099 | NM_182620.3 | 165.693152 | -8.9943147 | 1.32742674 | -6.7757523 |
| 3100 | NM_001317231.1 | 190.049506 | -9.1920987 | 1.35666905 | -6.7754908 |
| 3101 | NM_032025.4 | 153.441436 | -8.8830869 | 1.31124359 | -6.7745512 |
| 3102 | NM_145733.2 | 348.904193 | -10.068785 | 1.48630188 | -6.7743873 |
| 3103 | NM_005397.3 | 228.763168 | -9.4603435 | 1.39658399 | -6.7739166 |
| 3104 | NM_175907.5 | 103.360812 | -8.3143918 | 1.22741835 | -6.7738859 |
| 3105 | NM_001130067.1 | 229.610093 | -9.4652304 | 1.39744348 | -6.7732474 |
| 3106 | NM_001099625.1 | 136.739295 | -8.7173954 | 1.28708968 | -6.7729511 |
| 3107 | NM_030648.3 | 154.345549 | -8.8917385 | 1.31297376 | -6.7722134 |
| 3108 | NM_001282858.1 | 126.868415 | -8.6095936 | 1.2714911 | -6.7712575 |
| 3109 | NM_001085365.1 | 175.159469 | -9.0745458 | 1.34020709 | -6.7710027 |
| 3110 | NM_005805.5 | 172.548292 | -9.0529379 | 1.33704812 | -6.7708393 |
| 3111 | NM_017946.3 | 107.73786 | -8.3755811 | 1.23705813 | -6.7705639 |
| 3112 | NM_203364.2 | 348.704353 | -9.1034993 | 1.34483876 | -6.7692125 |
| 3113 | NM_007166.3 | 240.840845 | -9.5339087 | 1.40844433 | -6.7691058 |
| 3114 | NM_017838.3 | 187.549369 | -9.1732046 | 1.35516951 | -6.7690459 |
| 3115 | NM_003873.5 | 108.800074 | -8.3900129 | 1.23948122 | -6.7689714 |
| 3116 | NM_003478.3 | 120.479967 | -8.5347829 | 1.26095423 | -6.7685113 |
| 3117 | NM_001001794.3 | 106.387899 | -4.7733093 | 0.70528494 | -6.7679162 |
| 3118 | NM_007192.3 | 477.503215 | -6.7195521 | 0.99295649 | -6.767217 |
| 3119 | NM_002721.4 | 140.093791 | -8.7523318 | 1.2937217 | -6.7652354 |
| 3120 | NM_006763.2 | 99.1498644 | -8.2555079 | 1.22032555 | -6.7650046 |
| 3121 | NM_152902.4 | 211.730466 | -9.3478539 | 1.38208931 | -6.7635672 |
| 3122 | NM_005399.4 | 138.930733 | -8.7400806 | 1.29235309 | -6.76292 |
| 3123 | NM_213636.2 | 141.908013 | -8.7718135 | 1.29729221 | -6.7616328 |
| 3124 | NM_001202464.1 | 144.048399 | -8.7922634 | 1.30042246 | -6.7610824 |
| 3125 | NM_006808.2 | 173.159417 | -9.0580203 | 1.33985127 | -6.760467 |
| 3126 | NM_021158.4 | 117.897501 | -8.5031515 | 1.25785073 | -6.760064 |
| 3127 | NM_017836.3 | 150.090253 | -8.8515223 | 1.30942868 | -6.7598353 |
| 3128 | NM_016406.3 | 128.795684 | -8.6313595 | 1.27690536 | -6.7595922 |
| 3129 | NM_015878.5 | 155.627124 | -8.9032974 | 1.31730275 | -6.7587328 |
| 3130 | NM_004370.5 | 268.056808 | -8.7251863 | 1.29102564 | -6.7583369 |
| 3131 | NM_004450.2 | 332.363778 | -9.0354623 | 1.33693316 | -6.7583501 |
| 3132 | NM_018202.5 | 175.073643 | -9.0736433 | 1.34258376 | -6.7583443 |
| 3133 | NM_003653.3 | 170.573698 | -9.0358038 | 1.33707154 | -6.757906 |
| 3134 | NM_013354.6 | 149.776092 | -8.8482514 | 1.30942077 | -6.7573782 |
| 3135 | NM_001134693.1 | 163.568476 | -8.9760934 | 1.32838759 | -6.7571344 |
| 3136 | NM_001098635.1 | 315.564014 | -9.9238881 | 1.468678 | -6.757021 |
| 3137 | NM_001174116.1 | 145.667158 | -8.8069909 | 1.30347653 | -6.7565397 |
| 3138 | NM_002633.2 | 100.555896 | -8.2726205 | 1.22438971 | -6.7565256 |

| | | | | | |
|---|---|---|---|---|---|
| 3139 | NM_023924.4 | 130.278405 | -8.6473318 | 1.27983395 | -6.7566045 |
| 3140 | NM_003953.5 | 153.664387 | -8.8864971 | 1.3153315 | -6.7560893 |
| 3141 | NM_004424.4 | 138.853375 | -8.7390783 | 1.29352461 | -6.7560201 |
| 3142 | NM_145730.2 | 107.597965 | -8.3706091 | 1.23898676 | -6.7560118 |
| 3143 | NM_182641.3 | 124.480828 | -8.5831208 | 1.27046806 | -6.755873 |
| 3144 | NM_000546.5 | 142.176494 | -8.7737101 | 1.29874974 | -6.7555049 |
| 3145 | NM_004207.3 | 145.228663 | -8.8031488 | 1.30311858 | -6.7554472 |
| 3146 | NM_001014436.2 | 169.161123 | -9.0239649 | 1.3358933 | -6.7550043 |
| 3147 | NM_033656.3 | 114.731007 | -8.4629042 | 1.25291025 | -6.7545973 |
| 3148 | NM_018992.3 | 145.940021 | -8.8111163 | 1.30449935 | -6.7544046 |
| 3149 | NM_003930.4 | 202.429688 | -9.2835636 | 1.37448971 | -6.7541892 |
| 3150 | NM_024295.5 | 157.4357 | -8.9202907 | 1.32071925 | -6.7541157 |
| 3151 | NM_001933.4 | 164.080231 | -8.9807886 | 1.32983931 | -6.7532886 |
| 3152 | NM_006241.7 | 139.410158 | -8.7454387 | 1.29502342 | -6.7531124 |
| 3153 | NM_002790.3 | 170.025539 | -9.0315352 | 1.33740604 | -6.7530241 |
| 3154 | NM_002870.3 | 125.033248 | -8.5888975 | 1.27188365 | -6.7528956 |
| 3155 | NR_003260.1 | 55.2873475 | 2.4061127 | 0.35631588 | 6.75275175 |
| 3156 | NM_014238.1 | 100.99666 | -8.2798138 | 1.22630726 | -6.7518265 |
| 3157 | NR_036610.1 | 151.762071 | -8.8674813 | 1.31347843 | -6.7511435 |
| 3158 | NM_007024.4 | 129.421732 | -8.6383692 | 1.27959247 | -6.7508753 |
| 3159 | NM_033644.2 | 128.824832 | -8.6290984 | 1.27825402 | -6.7506913 |
| 3160 | NM_001031804.2 | 135.78628 | -8.7091804 | 1.29014579 | -6.7505398 |
| 3161 | NM_001320219.1 | 246.258992 | -9.565928 | 1.41711825 | -6.750268 |
| 3162 | NM_005573.3 | 181.639106 | -9.1267939 | 1.35206852 | -6.7502451 |
| 3163 | NM_002857.3 | 116.865122 | -8.4909697 | 1.2579612 | -6.7497866 |
| 3164 | NM_014884.4 | 188.426992 | -9.1800548 | 1.36018975 | -6.7490987 |
| 3165 | NM_016608.1 | 164.726199 | -8.9859496 | 1.33145271 | -6.7489814 |
| 3166 | NM_001112726.2 | 119.46515 | -8.5221778 | 1.26276278 | -6.7488351 |
| 3167 | NM_018047.2 | 140.73335 | -8.7588863 | 1.29793662 | -6.7483159 |
| 3168 | NR_003563.1 | 131.445012 | -8.6597149 | 1.28323322 | -6.7483562 |
| 3169 | NM_139315.2 | 133.473675 | -8.6817673 | 1.28652608 | -6.7482249 |
| 3170 | NR_027692.1 | 158.909351 | -8.9328893 | 1.32377873 | -6.748023 |
| 3171 | NM_001321.2 | 301.454477 | -9.8576056 | 1.46083016 | -6.7479477 |
| 3172 | NM_001201339.1 | 237.642241 | -9.5139398 | 1.40991522 | -6.7478807 |
| 3173 | NR_033711.1 | 96.5740664 | -8.2168294 | 1.21778661 | -6.7473475 |
| 3174 | NM_001080485.3 | 99.7888378 | -8.2655017 | 1.22502807 | -6.7471937 |
| 3175 | NM_006659.3 | 214.864766 | -9.3692939 | 1.38863617 | -6.7471193 |
| 3176 | NM_004319.2 | 189.083244 | -9.1848739 | 1.36131786 | -6.7470458 |
| 3177 | NM_016172.2 | 142.747944 | -8.7792389 | 1.30122925 | -6.746881 |
| 3178 | NM_007027.3 | 152.121359 | -8.8709193 | 1.31517685 | -6.7450391 |
| 3179 | NM_022777.3 | 100.430738 | -8.2720881 | 1.2263994 | -6.7450197 |
| 3180 | NM_001199179.1 | 164.306009 | -8.9818929 | 1.33168283 | -6.7447689 |
| 3181 | NM_015199.3 | 140.404227 | -8.754754 | 1.29806519 | -6.744464 |

| | | | | | |
|---|---|---|---|---|---|
| 3182 | NM_176096.2 | 153.201888 | -8.8808763 | 1.31678335 | -6.7443716 |
| 3183 | NM_020148.2 | 220.955949 | -9.4091567 | 1.39524004 | -6.7437548 |
| 3184 | NM_002408.3 | 112.06309 | -8.4311566 | 1.25025795 | -6.7435337 |
| 3185 | NM_001030007.1 | 215.619719 | -9.3747552 | 1.39021664 | -6.7433772 |
| 3186 | NM_014230.3 | 178.862699 | -8.1391535 | 1.20700463 | -6.7432662 |
| 3187 | NM_173854.5 | 134.152634 | -8.6912169 | 1.28889296 | -6.7431642 |
| 3188 | NM_022763.3 | 126.813124 | -8.6082209 | 1.27664071 | -6.7428689 |
| 3189 | NM_005698.3 | 132.713634 | -8.6744672 | 1.28657103 | -6.742315 |
| 3190 | NM_005716.3 | 118.626558 | -8.5126179 | 1.26257375 | -6.7422738 |
| 3191 | NM_006048.4 | 199.01053 | -8.2946444 | 1.23037921 | -6.7415349 |
| 3192 | NM_024407.4 | 177.594727 | -9.0943812 | 1.34905096 | -6.7413178 |
| 3193 | NM_207354.2 | 123.72752 | -8.5718848 | 1.27156934 | -6.7411855 |
| 3194 | NM_020857.2 | 134.622243 | -8.695423 | 1.28994271 | -6.7409373 |
| 3195 | NM_002138.3 | 304.957775 | -9.8744368 | 1.46508157 | -6.7398546 |
| 3196 | NM_003169.3 | 109.264568 | -8.3952511 | 1.24563864 | -6.7397163 |
| 3197 | NM_006292.3 | 175.105929 | -9.0740322 | 1.34661724 | -6.7383901 |
| 3198 | NM_032015.4 | 128.155119 | -8.6245946 | 1.28017376 | -6.73705 |
| 3199 | NM_001682.2 | 219.02157 | -8.4331752 | 1.2518394 | -6.7366271 |
| 3200 | NM_173354.4 | 100.413118 | -4.6830547 | 0.69516388 | -6.7366197 |
| 3201 | NM_044472.2 | 282.68159 | -9.7651676 | 1.44962572 | -6.7363372 |
| 3202 | NM_004786.2 | 183.252626 | -9.1397118 | 1.35681954 | -6.7361293 |
| 3203 | NM_006164.4 | 107.573498 | -8.3710518 | 1.24278626 | -6.7357132 |
| 3204 | NM_022733.2 | 187.133386 | -9.1698256 | 1.36147352 | -6.7352214 |
| 3205 | NM_013260.7 | 149.327274 | -8.8443849 | 1.31321725 | -6.7348985 |
| 3206 | NM_016417.2 | 131.280057 | -8.6585145 | 1.28561119 | -6.7349402 |
| 3207 | NM_006474.4 | 167.956886 | -9.0143516 | 1.33848316 | -6.7347516 |
| 3208 | NM_007234.4 | 193.157799 | -9.2156029 | 1.36857648 | -6.7337142 |
| 3209 | NM_006297.2 | 126.823932 | -8.6090329 | 1.2787127 | -6.732578 |
| 3210 | NM_178557.3 | 333.159888 | -9.0374372 | 1.34234021 | -6.7325981 |
| 3211 | NM_013242.2 | 141.426063 | -8.7656773 | 1.30206685 | -6.7321254 |
| 3212 | NM_004896.4 | 134.26803 | -8.6914817 | 1.29147555 | -6.7298848 |
| 3213 | NM_147164.2 | 200.177917 | -9.2670438 | 1.37708114 | -6.7294828 |
| 3214 | NM_012087.3 | 154.023259 | -8.889347 | 1.32096776 | -6.7294201 |
| 3215 | NM_001172640.1 | 172.062635 | -9.0476318 | 1.34450425 | -6.7293441 |
| 3216 | NM_001098616.2 | 202.94842 | -9.2867407 | 1.3800836 | -6.7291146 |
| 3217 | NM_005817.4 | 131.591012 | -8.6625999 | 1.28757212 | -6.727856 |
| 3218 | NM_005517.3 | 934.876764 | -7.5740742 | 1.12587135 | -6.7272998 |
| 3219 | NM_020428.3 | 141.935391 | -8.7713798 | 1.30384025 | -6.7273424 |
| 3220 | NM_015347.4 | 117.774349 | -8.5026043 | 1.26395283 | -6.726995 |
| 3221 | NM_006304.1 | 123.22312 | -8.5668531 | 1.27363059 | -6.7263249 |
| 3222 | NR_109994.1 | 557.704334 | 4.08665968 | 0.60759075 | 6.72600713 |
| 3223 | NM_001801.2 | 121.755201 | -8.5512114 | 1.27137518 | -6.7259543 |
| 3224 | NM_024045.1 | 122.86269 | -8.5630614 | 1.27328989 | -6.7251468 |

| | | | | | |
|---|---|---|---|---|---|
| 3225 | NM_194291.2 | 122.677909 | -8.5608913 | 1.27299951 | -6.7249761 |
| 3226 | NM_003592.2 | 227.050759 | -8.4848603 | 1.26177057 | -6.7245667 |
| 3227 | NM_003388.4 | 287.152793 | -9.7876599 | 1.45553675 | -6.7244334 |
| 3228 | NM_017801.2 | 110.190454 | -8.4065509 | 1.25014602 | -6.7244552 |
| 3229 | NM_001035254.2 | 100.955041 | -8.2799073 | 1.23134061 | -6.7243029 |
| 3230 | NM_001261406.1 | 124.605465 | -8.5812272 | 1.27615467 | -6.7242846 |
| 3231 | NM_004078.2 | 202.847127 | -9.2866993 | 1.38131806 | -6.723071 |
| 3232 | NM_024665.5 | 162.226203 | -8.9642054 | 1.33369038 | -6.7213542 |
| 3233 | NM_001301045.1 | 228.281669 | -9.4566272 | 1.40717679 | -6.7202837 |
| 3234 | NM_001242792.1 | 256.882706 | -9.6269609 | 1.43254094 | -6.7201995 |
| 3235 | NM_001145856.1 | 198.589072 | -9.2552969 | 1.37732266 | -6.7197739 |
| 3236 | NM_013379.2 | 163.384616 | -7.1753778 | 1.06783014 | -6.7195872 |
| 3237 | NM_022479.2 | 291.161339 | -8.8429519 | 1.31604475 | -6.7193398 |
| 3238 | NM_005503.3 | 288.805171 | -9.7960578 | 1.45812188 | -6.7182709 |
| 3239 | NM_014038.2 | 172.191689 | -9.0498702 | 1.34714598 | -6.7178096 |
| 3240 | NM_001320595.1 | 120.902645 | -8.5395667 | 1.27122974 | -6.7175636 |
| 3241 | NM_001846.3 | 1888.08003 | -7.1785083 | 1.06870196 | -6.7170348 |
| 3242 | NM_014967.4 | 115.736073 | -8.4771181 | 1.26204111 | -6.7169905 |
| 3243 | NM_005233.5 | 133.451676 | -8.6816431 | 1.29250118 | -6.7169325 |
| 3244 | NM_001915.3 | 99.2887347 | -8.2582388 | 1.22952595 | -6.7166039 |
| 3245 | NM_015665.5 | 115.740034 | -8.4766942 | 1.26211328 | -6.7162705 |
| 3246 | NM_021205.5 | 97.1186665 | -8.2259196 | 1.22477758 | -6.7162559 |
| 3247 | NM_004146.5 | 171.957633 | -9.0480202 | 1.34726507 | -6.7158427 |
| 3248 | NM_014255.5 | 129.456106 | -8.6385878 | 1.28634182 | -6.7156239 |
| 3249 | NM_015311.2 | 149.62816 | -8.8482407 | 1.3177249 | -6.714786 |
| 3250 | NM_001099695.1 | 129.619852 | -8.6398313 | 1.28672925 | -6.7145682 |
| 3251 | NM_002501.3 | 198.282949 | -9.2528597 | 1.37807505 | -6.7143366 |
| 3252 | NM_152531.4 | 142.304722 | -8.7749529 | 1.30696976 | -6.7139679 |
| 3253 | NM_001667.3 | 130.013426 | -8.6452411 | 1.28767025 | -6.7138625 |
| 3254 | NM_004069.4 | 176.203767 | -9.0831353 | 1.35290062 | -6.713823 |
| 3255 | NM_001031698.2 | 123.362868 | -8.5684192 | 1.27628586 | -6.713558 |
| 3256 | NM_030650.2 | 162.281776 | -8.9640386 | 1.33521338 | -6.7135625 |
| 3257 | NM_018638.4 | 271.985504 | -7.9016493 | 1.17699311 | -6.7134202 |
| 3258 | NM_001040446.2 | 95.568272 | -8.1983674 | 1.22122585 | -6.7132278 |
| 3259 | NM_006023.2 | 161.63462 | -8.9587811 | 1.33468673 | -6.7122726 |
| 3260 | NM_015321.2 | 194.710021 | -7.4376233 | 1.10807969 | -6.7121737 |
| 3261 | NM_001323036.2 | 93.6261521 | 11.4696459 | 1.70881585 | 6.71204328 |
| 3262 | NM_021627.2 | 108.417484 | -8.3831854 | 1.24903943 | -6.711706 |
| 3263 | NM_016275.3 | 218.869484 | -9.3960948 | 1.40001609 | -6.7114192 |
| 3264 | NM_018072.5 | 109.490737 | -8.3968487 | 1.25120996 | -6.7109829 |
| 3265 | NM_013255.4 | 128.283883 | -3.4226543 | 0.51001132 | -6.7109379 |
| 3266 | NM_015444.2 | 106.137245 | -8.3501208 | 1.24435898 | -6.7103793 |
| 3267 | NM_021242.5 | 128.995287 | -4.4166942 | 0.65821473 | -6.7101114 |

| | | | | | |
|---|---|---|---|---|---|
| 3268 | NR_027873.1 | 98.4641504 | -8.2448 | 1.22871219 | -6.7101149 |
| 3269 | NM_020722.1 | 308.780653 | -8.9276827 | 1.33053814 | -6.7098284 |
| 3270 | NM_000821.6 | 100.391093 | -8.2692706 | 1.23255788 | -6.7090323 |
| 3271 | NM_003796.3 | 112.292072 | -8.4319567 | 1.25682059 | -6.7089581 |
| 3272 | NM_001025107.2 | 248.501183 | -9.579652 | 1.42790181 | -6.7089011 |
| 3273 | NM_001170543.1 | 117.53122 | -8.4980647 | 1.26682769 | -6.7081457 |
| 3274 | NM_001076788.1 | 156.260916 | -8.9094688 | 1.32820842 | -6.7078846 |
| 3275 | NM_014372.4 | 340.317031 | -8.0932634 | 1.20657025 | -6.7076604 |
| 3276 | NM_001414.3 | 151.901547 | -8.8685688 | 1.32219907 | -6.7074384 |
| 3277 | NM_001015053.1 | 180.393006 | -9.1179294 | 1.35942098 | -6.7072154 |
| 3278 | NM_054012.3 | 168.737066 | -9.0214353 | 1.3450957 | -6.7069096 |
| 3279 | NM_003435.4 | 106.737987 | -8.3599909 | 1.24658374 | -6.7063211 |
| 3280 | NM_001115113.2 | 244.310016 | -8.5886855 | 1.28074382 | -6.7060136 |
| 3281 | NM_015859.3 | 111.978806 | -8.4287159 | 1.25702378 | -6.7052955 |
| 3282 | NM_003368.4 | 115.843685 | -8.4766933 | 1.26419166 | -6.705228 |
| 3283 | NM_003569.2 | 231.504899 | -9.4768996 | 1.41343882 | -6.7048531 |
| 3284 | NM_138736.2 | 221.103015 | -9.4103323 | 1.40373407 | -6.7037856 |
| 3285 | NM_002731.3 | 188.164565 | -9.1778445 | 1.36906788 | -6.7037176 |
| 3286 | NM_002573.3 | 285.555836 | -9.7796759 | 1.45887004 | -6.7035963 |
| 3287 | NM_001321358.1 | 154.426803 | -8.8923941 | 1.3266332 | -6.7029787 |
| 3288 | NM_022140.4 | 99.1716858 | -8.2543112 | 1.231434 | -6.7030074 |
| 3289 | NM_181701.3 | 141.923482 | -8.7705179 | 1.30844319 | -6.7030178 |
| 3290 | NM_003350.2 | 179.479345 | -9.1095526 | 1.35940871 | -6.7011139 |
| 3291 | NM_032558.2 | 108.321437 | -8.3821697 | 1.25087719 | -6.7010333 |
| 3292 | NM_000285.3 | 138.78809 | -8.7390728 | 1.30429498 | -6.7002273 |
| 3293 | NM_005997.2 | 192.534857 | -9.2109454 | 1.37479038 | -6.6998908 |
| 3294 | NM_002168.3 | 260.306469 | -8.6824297 | 1.29600761 | -6.6993663 |
| 3295 | NM_145056.2 | 194.881471 | -9.2285489 | 1.37769567 | -6.6985396 |
| 3296 | NM_006403.3 | 121.315415 | -8.5443183 | 1.27572681 | -6.6976082 |
| 3297 | NM_017893.3 | 116.32166 | -8.4841717 | 1.26682177 | -6.6972102 |
| 3298 | NM_001199867.1 | 114.251463 | -8.4604543 | 1.26329791 | -6.6971173 |
| 3299 | NM_001304504.1 | 131.580651 | -8.6615782 | 1.29367033 | -6.695352 |
| 3300 | NM_003559.4 | 863.363818 | -6.2593571 | 0.934895 | -6.6952515 |
| 3301 | NM_006285.2 | 145.476497 | -8.8067944 | 1.31540628 | -6.6951135 |
| 3302 | NM_022745.4 | 110.842082 | -8.4135201 | 1.25673093 | -6.6947665 |
| 3303 | NM_001014840.1 | 129.288393 | -8.6365998 | 1.29008328 | -6.6946064 |
| 3304 | NM_017830.3 | 120.261245 | -8.5320697 | 1.27446932 | -6.6946058 |
| 3305 | NM_001040000.2 | 270.481186 | -9.701214 | 1.44914086 | -6.6944589 |
| 3306 | NM_001199802.1 | 157.890642 | -8.9250493 | 1.33335025 | -6.6937021 |
| 3307 | NM_001278389.1 | 458.640716 | -9.4994717 | 1.41916896 | -6.6936862 |
| 3308 | NM_213595.3 | 124.076443 | -8.5776011 | 1.28148293 | -6.6934962 |
| 3309 | NM_001330177.1 | 96.4955445 | -8.2152638 | 1.22749157 | -6.6927253 |
| 3310 | NM_032436.3 | 111.566769 | -8.4251092 | 1.25886741 | -6.6926104 |

| | | | | | |
|---|---|---|---|---|---|
| 3311 | NM_001042545.1 | 104.197731 | -8.3278588 | 1.24435615 | -6.6925043 |
| 3312 | NM_002345.3 | 168.969878 | -9.0232462 | 1.34833984 | -6.6921156 |
| 3313 | NM_001017962.2 | 110.528887 | -8.4096152 | 1.25672053 | -6.6917146 |
| 3314 | NM_032016.3 | 211.989033 | -9.3500299 | 1.39729988 | -6.6914984 |
| 3315 | NM_021117.3 | 120.068149 | -8.5304366 | 1.27484048 | -6.6913757 |
| 3316 | NM_015701.4 | 121.411016 | -8.546861 | 1.27740889 | -6.6907793 |
| 3317 | NM_207370.2 | 186.264511 | -9.1633883 | 1.36956714 | -6.6907186 |
| 3318 | NM_006713.3 | 368.846019 | -9.1857527 | 1.37296069 | -6.6904703 |
| 3319 | NM_001191061.1 | 154.544783 | -8.8935156 | 1.3293771 | -6.689987 |
| 3320 | NM_001303512.1 | 190.110357 | -9.192651 | 1.37410643 | -6.6899119 |
| 3321 | NM_017452.3 | 141.225424 | -8.7637624 | 1.31002158 | -6.6897848 |
| 3322 | NM_001080429.2 | 195.382525 | -9.2323791 | 1.38011573 | -6.6895688 |
| 3323 | NM_018255.3 | 116.446237 | -8.4832328 | 1.26814 | -6.6895081 |
| 3324 | NM_005804.3 | 168.803761 | -9.021119 | 1.34858543 | -6.6893197 |
| 3325 | NM_014660.3 | 204.59235 | -9.298445 | 1.39010859 | -6.6890062 |
| 3326 | NM_024535.4 | 166.598099 | -9.0017551 | 1.34575803 | -6.6889849 |
| 3327 | NM_005956.3 | 145.986507 | -8.8117332 | 1.31737929 | -6.6888354 |
| 3328 | NM_016561.2 | 122.591074 | -8.5601644 | 1.27980253 | -6.6886603 |
| 3329 | NM_004374.3 | 198.408607 | -9.2544837 | 1.38365904 | -6.6884135 |
| 3330 | NM_001080543.1 | 115.639042 | -8.4752418 | 1.26719981 | -6.6881653 |
| 3331 | NM_016291.3 | 149.698916 | -8.8471714 | 1.32287109 | -6.687856 |
| 3332 | NM_006415.3 | 127.051002 | -8.6117224 | 1.28770027 | -6.6876762 |
| 3333 | NM_001177.5 | 155.857034 | -7.1063622 | 1.06261917 | -6.6875908 |
| 3334 | NM_001330701.1 | 138.275365 | -8.732058 | 1.3057148 | -6.6875692 |
| 3335 | NM_006648.3 | 178.399863 | -8.1338965 | 1.21631861 | -6.6873075 |
| 3336 | NM_031902.4 | 118.94136 | -8.5159211 | 1.27349844 | -6.6870291 |
| 3337 | NM_000665.4 | 186.301422 | -9.163786 | 1.37040918 | -6.6868977 |
| 3338 | NM_002917.1 | 137.292365 | -8.7234508 | 1.30456961 | -6.6868419 |
| 3339 | NM_004383.2 | 168.502891 | -9.0184628 | 1.34871256 | -6.6867197 |
| 3340 | NM_015532.4 | 129.447256 | -8.6383114 | 1.29189385 | -6.6865489 |
| 3341 | NM_001034996.2 | 208.930998 | -9.3287742 | 1.39529566 | -6.6858763 |
| 3342 | NM_002847.4 | 95.7315869 | -8.2040268 | 1.22706293 | -6.6859055 |
| 3343 | NM_198530.3 | 108.298849 | -8.3824579 | 1.25374891 | -6.6859144 |
| 3344 | NM_015565.2 | 116.681176 | -8.4879374 | 1.26955734 | -6.6857456 |
| 3345 | NM_017617.4 | 339.84857 | -7.0257853 | 1.05087063 | -6.6856805 |
| 3346 | NM_003372.6 | 190.429602 | -9.1950025 | 1.37541997 | -6.6852327 |
| 3347 | NM_000814.5 | 238.201819 | -9.5180758 | 1.42380788 | -6.6849439 |
| 3348 | NM_016068.2 | 158.697519 | -8.9321372 | 1.33626229 | -6.6844191 |
| 3349 | NM_133266.4 | 114.84876 | -8.4636848 | 1.26618781 | -6.6843834 |
| 3350 | NM_001031690.2 | 283.463345 | -9.7692047 | 1.46157905 | -6.6840071 |
| 3351 | NM_001166007.1 | 97.8502444 | -8.2341353 | 1.23196982 | -6.6837151 |
| 3352 | NM_015634.3 | 131.135416 | -8.6569761 | 1.29528329 | -6.6834616 |
| 3353 | NM_006712.4 | 135.955716 | -8.7084226 | 1.30306235 | -6.6830437 |

| | | | | | |
|---|---|---|---|---|---|
| 3354 | NM_172070.3 | 193.142255 | -8.2505724 | 1.23455201 | -6.6830497 |
| 3355 | NM_020442.5 | 107.98907 | -8.3790601 | 1.25382045 | -6.682823 |
| 3356 | NM_001960.5 | 207.413773 | -9.3184055 | 1.39464081 | -6.681581 |
| 3357 | NM_024865.3 | 122.643703 | -8.562503 | 1.28152858 | -6.6814764 |
| 3358 | NM_014947.4 | 100.290645 | -8.2717828 | 1.23806348 | -6.6812267 |
| 3359 | NM_031229.3 | 126.120513 | -8.6018276 | 1.28751751 | -6.6809403 |
| 3360 | NM_001257413.1 | 299.262303 | 2.63220025 | 0.39403885 | 6.68005265 |
| 3361 | NM_002811.4 | 227.933362 | -8.4906174 | 1.27115745 | -6.6794381 |
| 3362 | NM_152558.4 | 139.182371 | -8.7414115 | 1.3087751 | -6.6790784 |
| 3363 | NM_001141973.2 | 192.989625 | -9.2148286 | 1.37970978 | -6.6788166 |
| 3364 | NM_001330520.1 | 197.342098 | -8.2823967 | 1.24014071 | -6.6785944 |
| 3365 | NM_001535.4 | 147.156273 | -8.8237855 | 1.32121495 | -6.6785389 |
| 3366 | NM_174916.2 | 111.69424 | -8.425787 | 1.26161794 | -6.6785567 |
| 3367 | NM_004822.2 | 421.56635 | -9.3790223 | 1.40443151 | -6.6781628 |
| 3368 | NM_016479.4 | 163.707731 | -8.978506 | 1.3444732 | -6.6780847 |
| 3369 | NM_000642.2 | 88.8990475 | -8.0965125 | 1.21241408 | -6.6780093 |
| 3370 | NM_152726.2 | 116.685498 | -8.4881567 | 1.27110227 | -6.6777921 |
| 3371 | NM_006933.6 | 264.983201 | -7.8179141 | 1.17075207 | -6.6776855 |
| 3372 | NM_024927.4 | 101.35767 | -8.2833126 | 1.24047744 | -6.6775197 |
| 3373 | NM_080491.2 | 160.764431 | -8.9507767 | 1.34045607 | -6.6774114 |
| 3374 | NM_004318.3 | 112.07922 | -8.4322735 | 1.26303735 | -6.676187 |
| 3375 | NM_018241.2 | 117.653106 | -4.8998739 | 0.73396318 | -6.6759125 |
| 3376 | NM_030579.2 | 240.022477 | -7.7357957 | 1.15890315 | -6.6751011 |
| 3377 | NM_147223.2 | 122.07486 | -8.5513644 | 1.28111928 | -6.6749166 |
| 3378 | NM_014188.2 | 245.448609 | -8.5976709 | 1.28814418 | -6.6744632 |
| 3379 | NM_020313.3 | 145.184192 | -8.8038748 | 1.31912599 | -6.6740212 |
| 3380 | NM_001321967.1 | 150.903547 | -8.8594147 | 1.32753232 | -6.6735962 |
| 3381 | NM_001198832.2 | 211.902882 | -7.0166606 | 1.05141874 | -6.6735167 |
| 3382 | NM_153816.5 | 118.389265 | -8.5092374 | 1.27512299 | -6.6732679 |
| 3383 | NM_001001521.1 | 152.791436 | -8.8773825 | 1.33030878 | -6.6731744 |
| 3384 | NM_021090.3 | 116.143636 | -8.4842764 | 1.27139982 | -6.6731772 |
| 3385 | NM_016118.4 | 98.5461865 | -8.2452336 | 1.23560092 | -6.6730555 |
| 3386 | NM_006337.4 | 155.244646 | -8.8999753 | 1.33373281 | -6.6729822 |
| 3387 | NM_144973.3 | 172.384513 | -9.0511149 | 1.35639531 | -6.6729181 |
| 3388 | NM_001174063.1 | 107.565433 | -8.368984 | 1.25426692 | -6.6724107 |
| 3389 | NM_005471.4 | 123.74002 | -8.5729725 | 1.28494897 | -6.6718389 |
| 3390 | NM_001002800.2 | 105.701843 | -8.3461792 | 1.25099211 | -6.6716482 |
| 3391 | NM_001136035.2 | 128.346824 | -8.6259205 | 1.29292992 | -6.6716072 |
| 3392 | NM_002212.3 | 149.182884 | -8.8429757 | 1.32553404 | -6.6712551 |
| 3393 | NM_019059.3 | 195.641954 | -9.233932 | 1.38424022 | -6.6707583 |
| 3394 | NM_004208.3 | 89.3732629 | -8.1056999 | 1.21514037 | -6.6705872 |
| 3395 | NM_004901.4 | 151.522737 | -8.8658254 | 1.3292673 | -6.6697085 |
| 3396 | NM_018201.4 | 136.907749 | -8.7197793 | 1.30742772 | -6.669416 |

| | | | | | |
|---|---|---|---|---|---|
| 3397 | NR_038967.1 | 106.455052 | -8.3556909 | 1.25290779 | -6.669039 |
| 3398 | NM_078485.3 | 104.110237 | -8.325965 | 1.24850655 | -6.6687396 |
| 3399 | NM_182970.3 | 200.374903 | -9.2683182 | 1.38983664 | -6.6686386 |
| 3400 | NM_015201.4 | 207.265923 | -9.3174305 | 1.39722716 | -6.6685151 |
| 3401 | NM_001142640.1 | 119.131577 | -8.5180649 | 1.27740712 | -6.668246 |
| 3402 | NM_138440.2 | 164.53918 | -8.985095 | 1.34750954 | -6.6679268 |
| 3403 | NM_017548.4 | 122.9912 | -8.5637041 | 1.28440882 | -6.6674286 |
| 3404 | NM_139156.3 | 173.001605 | -8.0937121 | 1.21392743 | -6.6673773 |
| 3405 | NM_003379.4 | 168.995394 | -9.0230544 | 1.35336534 | -6.6671239 |
| 3406 | NM_015387.4 | 175.803581 | -9.0793885 | 1.36182723 | -6.6670634 |
| 3407 | NM_172231.3 | 130.545599 | -8.6499823 | 1.29758747 | -6.6662037 |
| 3408 | NM_152344.3 | 139.341988 | -8.7445382 | 1.3118665 | -6.6657226 |
| 3409 | NM_001201428.1 | 136.124941 | -8.7113921 | 1.30696785 | -6.6653453 |
| 3410 | NM_004882.3 | 96.0603132 | -8.207368 | 1.23144658 | -6.6648185 |
| 3411 | NM_016390.3 | 113.537895 | -8.4488355 | 1.26772765 | -6.6645509 |
| 3412 | NM_001014839.1 | 90.7995994 | -8.1248757 | 1.21920472 | -6.6640783 |
| 3413 | NM_032050.1 | 142.233406 | -8.7738324 | 1.3166038 | -6.6639884 |
| 3414 | NM_001079817.2 | 161.319034 | -7.0992565 | 1.06534369 | -6.663818 |
| 3415 | NM_018903.3 | 155.962902 | -8.9076411 | 1.33676513 | -6.6635798 |
| 3416 | NM_032581.3 | 154.129532 | -8.8890955 | 1.33398077 | -6.6635859 |
| 3417 | NM_020655.3 | 255.3854 | -9.6186894 | 1.44354623 | -6.6632362 |
| 3418 | NM_145239.2 | 253.197047 | -5.2710458 | 0.79107861 | -6.6631126 |
| 3419 | NM_152892.2 | 118.708737 | -8.5129342 | 1.27777938 | -6.662288 |
| 3420 | NM_001289405.1 | 139.97866 | -8.7512983 | 1.31374025 | -6.6613612 |
| 3421 | NM_001001323.1 | 259.120884 | -9.6396895 | 1.44721663 | -6.6608476 |
| 3422 | NM_001206652.1 | 208.135484 | -9.3231626 | 1.39973385 | -6.6606681 |
| 3423 | NM_015049.2 | 114.433748 | -8.4604291 | 1.27019999 | -6.6607063 |
| 3424 | NM_017520.3 | 293.127503 | -2.7254173 | 0.40922097 | -6.660014 |
| 3425 | NM_005134.3 | 138.483388 | -8.7345866 | 1.31156244 | -6.6596803 |
| 3426 | NM_014141.5 | 496.026718 | -5.2198187 | 0.78386764 | -6.6590563 |
| 3427 | NM_003340.6 | 105.444098 | -8.3415963 | 1.25269041 | -6.6589448 |
| 3428 | NM_006349.2 | 158.016604 | -8.9258372 | 1.34055142 | -6.6583326 |
| 3429 | NM_201568.2 | 108.161595 | -8.3785376 | 1.25835986 | -6.6583001 |
| 3430 | NM_007367.3 | 125.859837 | -8.5981902 | 1.29144696 | -6.6577958 |
| 3431 | NM_017432.4 | 100.170293 | -8.265752 | 1.24151184 | -6.6578117 |
| 3432 | NM_006547.2 | 141.207367 | -6.4041034 | 0.96191787 | -6.6576406 |
| 3433 | NM_001319239.1 | 107.655714 | -8.3728326 | 1.2576593 | -6.6574728 |
| 3434 | NM_001550.3 | 181.548056 | -9.1261567 | 1.3708388 | -6.6573522 |
| 3435 | NM_001042472.2 | 158.459642 | -8.9302051 | 1.3414194 | -6.6572804 |
| 3436 | NM_024589.2 | 171.523514 | -9.0440874 | 1.35854386 | -6.6571921 |
| 3437 | NM_005953.3 | 179.821169 | -9.1123201 | 1.36883099 | -6.6570089 |
| 3438 | NM_003483.4 | 276.505422 | -7.8967622 | 1.18629373 | -6.6566669 |
| 3439 | NM_013975.3 | 101.167799 | -8.2829603 | 1.24452282 | -6.6555311 |

| | | | | | |
|---|---|---|---|---|---|
| 3440 | NM_001321691.1 | 151.287156 | -8.8641105 | 1.33188946 | -6.65529 |
| 3441 | NM_001719.2 | 352.736993 | -2.8908279 | 0.43436737 | -6.6552602 |
| 3442 | NM_024755.3 | 130.299717 | -8.6475445 | 1.29934613 | -6.6553048 |
| 3443 | NR_036569.1 | 127.025332 | -8.6113638 | 1.29398943 | -6.654895 |
| 3444 | NM_001431.3 | 133.169035 | -8.680401 | 1.30450035 | -6.654196 |
| 3445 | NM_004938.3 | 117.666657 | -8.5010719 | 1.27755473 | -6.6541743 |
| 3446 | NM_001042616.2 | 156.928279 | -8.9158284 | 1.33995585 | -6.6538225 |
| 3447 | NM_001256447.1 | 123.724487 | -8.5738645 | 1.2886125 | -6.653563 |
| 3448 | NM_005834.4 | 131.437475 | -8.6607308 | 1.30176916 | -6.6530465 |
| 3449 | NM_173834.3 | 115.78526 | -8.4767183 | 1.27414138 | -6.6528867 |
| 3450 | NM_021244.4 | 119.662056 | -8.524464 | 1.28133039 | -6.6528228 |
| 3451 | NM_000292.2 | 106.246736 | -8.3553402 | 1.25592995 | -6.652712 |
| 3452 | NM_003412.3 | 193.182074 | -9.2168366 | 1.38580968 | -6.6508675 |
| 3453 | NM_198402.4 | 118.885622 | -8.5146718 | 1.2802601 | -6.6507359 |
| 3454 | NM_002882.3 | 170.370066 | -9.0342172 | 1.35839475 | -6.6506567 |
| 3455 | NM_197968.3 | 139.683217 | -8.7466558 | 1.315258 | -6.6501445 |
| 3456 | NM_001195001.1 | 108.511397 | -8.3859806 | 1.26112483 | -6.6496039 |
| 3457 | NM_004781.3 | 106.375862 | -8.3567758 | 1.25687646 | -6.6488442 |
| 3458 | NM_014870.3 | 104.852377 | -8.3348334 | 1.25361447 | -6.6486417 |
| 3459 | NM_003883.3 | 145.641189 | -5.6888633 | 0.85570109 | -6.6481898 |
| 3460 | NM_007364.3 | 161.430503 | -8.9567871 | 1.34726774 | -6.6481122 |
| 3461 | NM_001135998.2 | 181.08314 | -9.1226105 | 1.37226067 | -6.64787 |
| 3462 | NM_130811.3 | 279.593627 | -9.7494955 | 1.46660355 | -6.6476694 |
| 3463 | NM_003920.4 | 110.481887 | -8.4086688 | 1.2649561 | -6.6473997 |
| 3464 | NM_001242818.1 | 97.4145957 | -8.2263544 | 1.23758571 | -6.6470987 |
| 3465 | NM_001166347.1 | 93.4338664 | -8.1691393 | 1.22898918 | -6.6470393 |
| 3466 | NM_005717.3 | 338.685394 | -9.0613795 | 1.36330794 | -6.6466124 |
| 3467 | NM_001617.3 | 152.663355 | -8.8757902 | 1.33541204 | -6.6464806 |
| 3468 | NM_004553.4 | 158.142434 | -8.9271165 | 1.3431332 | -6.6464864 |
| 3469 | NM_001242362.1 | 103.401098 | -8.3140122 | 1.25090788 | -6.6463825 |
| 3470 | NM_021129.3 | 256.373302 | -9.6240852 | 1.44805935 | -6.6461953 |
| 3471 | NM_001243127.2 | 159.590123 | -2.8765713 | 0.4328176 | -6.6461514 |
| 3472 | NM_015516.3 | 94.1555073 | -8.1804684 | 1.23093608 | -6.6457296 |
| 3473 | NM_022459.4 | 94.3794482 | -8.1826547 | 1.23142429 | -6.6448703 |
| 3474 | NM_033121.1 | 99.5710444 | -8.2601582 | 1.2432977 | -6.6437493 |
| 3475 | NM_020447.4 | 105.503161 | -8.3445124 | 1.25603581 | -6.6435307 |
| 3476 | NM_006609.4 | 99.8801778 | -8.2641202 | 1.2439462 | -6.6434707 |
| 3477 | NM_014678.4 | 113.36189 | -8.446327 | 1.27148698 | -6.6428734 |
| 3478 | NM_019022.3 | 125.043371 | -8.5880528 | 1.29282046 | -6.6428813 |
| 3479 | NM_001272060.1 | 112.75892 | -8.4388366 | 1.27041445 | -6.6425855 |
| 3480 | NM_030809.2 | 220.102069 | -7.5560491 | 1.13757571 | -6.6422384 |
| 3481 | NM_002048.2 | 93.6993205 | -8.1717785 | 1.23038261 | -6.6416564 |
| 3482 | NM_015380.4 | 140.56124 | -8.7571415 | 1.31864431 | -6.6410187 |

| | | | | | |
|---|---|---|---|---|---|
| 3483 | NM_021991.2 | 141.113935 | -8.7631831 | 1.3195797 | -6.6408896 |
| 3484 | NM_033116.5 | 121.136497 | -8.5408567 | 1.28616632 | -6.6405539 |
| 3485 | NM_033001.3 | 178.601082 | -9.1025697 | 1.37076834 | -6.6404872 |
| 3486 | NM_015913.3 | 132.493368 | -8.6716874 | 1.30605108 | -6.6396235 |
| 3487 | NM_006045.2 | 802.874043 | -6.9624093 | 1.04871683 | -6.6389792 |
| 3488 | NM_001144774.2 | 227.325376 | -9.4505475 | 1.42394114 | -6.6368948 |
| 3489 | NM_001018050.3 | 140.880894 | -8.7602877 | 1.32004793 | -6.6363406 |
| 3490 | NM_005819.5 | 131.751826 | -8.6634167 | 1.30546877 | -6.6362497 |
| 3491 | NM_004890.2 | 133.775607 | -8.6862458 | 1.30898508 | -6.6358631 |
| 3492 | NM_022106.2 | 93.8969172 | -8.1726349 | 1.23160814 | -6.6357428 |
| 3493 | NM_021219.3 | 113.283475 | -8.4454197 | 1.27276046 | -6.6355138 |
| 3494 | NM_023935.2 | 142.288677 | -8.7746589 | 1.32239996 | -6.6354047 |
| 3495 | NM_138342.3 | 97.9782599 | -8.2353373 | 1.2414487 | -6.6336509 |
| 3496 | NM_170606.2 | 220.622526 | -8.442741 | 1.27277915 | -6.6333118 |
| 3497 | NM_006091.4 | 297.570299 | -9.8389783 | 1.48340378 | -6.6327041 |
| 3498 | NM_022101.3 | 98.728945 | -8.2476353 | 1.24356577 | -6.632247 |
| 3499 | NM_032975.3 | 90.6162812 | -8.1244244 | 1.22501703 | -6.632091 |
| 3500 | NM_004596.4 | 135.950292 | -8.7089949 | 1.31323117 | -6.6317303 |
| 3501 | NM_001406.3 | 176.587031 | -7.2746255 | 1.09696655 | -6.6315837 |
| 3502 | NM_004267.4 | 131.754012 | -8.6638674 | 1.30671405 | -6.6302703 |
| 3503 | NM_001346813.1 | 143.001593 | -8.7823441 | 1.32462862 | -6.6300425 |
| 3504 | NM_006003.2 | 213.639793 | -6.3282721 | 0.95448995 | -6.630004 |
| 3505 | NM_001244008.1 | 161.280721 | -8.9560429 | 1.35089555 | -6.6297079 |
| 3506 | NM_015235.2 | 121.916098 | -8.5515801 | 1.28992717 | -6.6295061 |
| 3507 | NM_006979.2 | 297.656245 | -6.2599144 | 0.94430717 | -6.6291082 |
| 3508 | NM_021215.3 | 93.6123497 | -8.1711416 | 1.2326171 | -6.6290997 |
| 3509 | NM_014982.2 | 158.090142 | -8.9265048 | 1.34657429 | -6.6290474 |
| 3510 | NM_001130040.1 | 170.666213 | -9.0384498 | 1.3634861 | -6.628927 |
| 3511 | NM_006368.4 | 149.610678 | -8.8469703 | 1.33464509 | -6.6287063 |
| 3512 | NM_001110514.1 | 117.12822 | -8.4962295 | 1.2818773 | -6.6279585 |
| 3513 | NM_005866.3 | 138.909973 | -8.7407937 | 1.31893241 | -6.6271734 |
| 3514 | NM_001913.4 | 94.3597159 | -8.1844807 | 1.23500822 | -6.6270658 |
| 3515 | NM_001282697.1 | 132.794182 | -8.6746356 | 1.3090775 | -6.6265256 |
| 3516 | NM_175075.4 | 90.0518055 | -8.1165514 | 1.22510252 | -6.6252018 |
| 3517 | NM_018686.5 | 135.800582 | -8.7068173 | 1.31430933 | -6.6246332 |
| 3518 | NM_007358.3 | 125.830792 | -8.5965219 | 1.29775218 | -6.6241629 |
| 3519 | NM_000071.2 | 195.896714 | -9.2356847 | 1.39427806 | -6.6239906 |
| 3520 | NM_004420.2 | 347.11413 | -8.2185429 | 1.24079284 | -6.6236221 |
| 3521 | NM_145728.2 | 126.410756 | -8.604537 | 1.29906433 | -6.623642 |
| 3522 | NM_032434.3 | 130.241627 | -8.646821 | 1.30552289 | -6.6232627 |
| 3523 | NM_005697.4 | 101.854449 | -8.294027 | 1.25233368 | -6.6228571 |
| 3524 | NM_003430.3 | 145.755416 | -8.8077273 | 1.32996232 | -6.622539 |
| 3525 | NM_005085.3 | 114.053869 | -8.4545146 | 1.27669422 | -6.6221923 |

| | | | | | |
|---|---|---|---|---|---|
| 3526 | NM_004239.4 | 98.8803079 | -8.2504467 | 1.24590082 | -6.6220735 |
| 3527 | NM_005035.3 | 224.955608 | -6.8576631 | 1.03559925 | -6.6219275 |
| 3528 | NM_024034.4 | 241.898614 | -9.5403552 | 1.44081968 | -6.6214775 |
| 3529 | NM_001301226.1 | 149.032347 | -8.8427028 | 1.33552546 | -6.6211413 |
| 3530 | NM_016491.3 | 135.949702 | -8.7091068 | 1.31536823 | -6.6210409 |
| 3531 | NM_020240.2 | 94.0152081 | -8.1742038 | 1.23458429 | -6.6210172 |
| 3532 | NM_014351.3 | 216.678033 | -9.3813401 | 1.41694306 | -6.6208307 |
| 3533 | NM_022095.3 | 96.2980836 | -8.2116467 | 1.2402835 | -6.620782 |
| 3534 | NM_032611.2 | 87.6285451 | -8.0756583 | 1.21973851 | -6.6208111 |
| 3535 | NM_018428.2 | 109.642989 | -8.3980055 | 1.26849861 | -6.6204294 |
| 3536 | NM_003893.4 | 99.8962339 | -8.2650995 | 1.24845233 | -6.6202764 |
| 3537 | NM_138638.4 | 140.329294 | -8.7549215 | 1.32254755 | -6.6197404 |
| 3538 | NM_002696.2 | 160.583213 | -8.9490144 | 1.3519138 | -6.6195156 |
| 3539 | NM_001018004.1 | 133.475022 | -8.6825767 | 1.3116917 | -6.619373 |
| 3540 | NM_001077484.1 | 180.237325 | -9.1153657 | 1.37709279 | -6.6192821 |
| 3541 | NM_013388.5 | 124.267733 | -8.5790454 | 1.2960997 | -6.6191246 |
| 3542 | NM_006390.3 | 99.4271241 | -8.2583919 | 1.24774113 | -6.6186741 |
| 3543 | NM_032223.3 | 258.316477 | -5.9894815 | 0.9049687 | -6.6184405 |
| 3544 | NM_001330221.1 | 393.793474 | -9.2791976 | 1.40213774 | -6.6178931 |
| 3545 | NM_020357.2 | 169.516385 | -9.0271418 | 1.36408117 | -6.6177453 |
| 3546 | NM_020651.3 | 139.331131 | -8.7442357 | 1.32143436 | -6.6172305 |
| 3547 | NM_004273.4 | 181.293004 | -8.1610606 | 1.23336535 | -6.6169044 |
| 3548 | NM_152787.4 | 142.1274 | -6.9262866 | 1.04676655 | -6.6168399 |
| 3549 | NM_003794.3 | 128.518866 | -8.6273824 | 1.30393287 | -6.6164314 |
| 3550 | NM_025083.3 | 126.537857 | -8.6053697 | 1.30063653 | -6.6162756 |
| 3551 | NM_002229.2 | 113.673458 | -8.4525179 | 1.27767478 | -6.6155473 |
| 3552 | NM_014363.5 | 153.700473 | -8.886178 | 1.34326448 | -6.6153599 |
| 3553 | NM_016062.3 | 166.356235 | -9.0002236 | 1.36050225 | -6.6153684 |
| 3554 | NM_001102575.1 | 91.267655 | -8.1332793 | 1.22959475 | -6.6146016 |
| 3555 | NM_006999.4 | 101.657045 | -8.2885485 | 1.25308966 | -6.6144896 |
| 3556 | NM_198526.3 | 115.309848 | -8.4714084 | 1.28074359 | -6.6144453 |
| 3557 | NM_002859.3 | 93.2290129 | -8.1674508 | 1.23484528 | -6.6141491 |
| 3558 | NM_005983.3 | 89.8726574 | -8.1125856 | 1.22661291 | -6.6138107 |
| 3559 | NM_006837.2 | 150.387781 | -8.8545857 | 1.33881571 | -6.613745 |
| 3560 | NM_001317812.1 | 93.1926172 | -8.1634509 | 1.23432741 | -6.6136836 |
| 3561 | NM_020372.3 | 90.4866268 | -8.1229049 | 1.22820492 | -6.6136398 |
| 3562 | NM_003646.3 | 146.548583 | -8.81717 | 1.33328527 | -6.6131159 |
| 3563 | NM_021637.2 | 264.97116 | -9.6717601 | 1.46256935 | -6.6128557 |
| 3564 | NM_018268.3 | 150.173145 | -8.8528851 | 1.33888554 | -6.6121298 |
| 3565 | NM_015284.3 | 149.92001 | -4.8812829 | 0.73823853 | -6.6120673 |
| 3566 | NM_002788.3 | 130.949013 | -8.6544591 | 1.30900739 | -6.611467 |
| 3567 | NM_207356.2 | 85.8915754 | -3.5574894 | 0.5380897 | -6.6113315 |
| 3568 | NM_023083.3 | 128.888094 | -8.6315971 | 1.30571575 | -6.6106249 |

| | | | | | |
|---|---|---|---|---|---|
| 3569 | NM_138394.3 | 112.921665 | -8.4404207 | 1.27682467 | -6.6104774 |
| 3570 | NM_005796.2 | 256.386803 | -7.7882474 | 1.17831348 | -6.6096566 |
| 3571 | NM_017864.3 | 100.384744 | -8.2702893 | 1.25134343 | -6.6091283 |
| 3572 | NM_181443.3 | 89.3492962 | -8.1014318 | 1.22589387 | -6.6085915 |
| 3573 | NM_000848.3 | 108.763848 | -8.3884701 | 1.26941529 | -6.608137 |
| 3574 | NM_018997.3 | 126.011288 | -8.5999248 | 1.30144657 | -6.6079737 |
| 3575 | NM_001271776.1 | 122.497595 | -8.558785 | 1.29529854 | -6.6075771 |
| 3576 | NM_001363.4 | 104.127735 | -8.3229817 | 1.25967506 | -6.607245 |
| 3577 | NM_032303.4 | 84.5737109 | -8.0258712 | 1.21473395 | -6.6071021 |
| 3578 | NM_001284239.1 | 120.528213 | -8.5350018 | 1.29181447 | -6.6069873 |
| 3579 | NM_005653.4 | 127.914584 | -8.6216181 | 1.30491888 | -6.6070146 |
| 3580 | NM_080546.4 | 86.8396426 | -8.0632827 | 1.22050931 | -6.6064901 |
| 3581 | NM_001042663.1 | 110.990718 | -8.4158268 | 1.2740716 | -6.6054583 |
| 3582 | NM_194356.2 | 93.7560389 | -8.1730965 | 1.2373629 | -6.6052542 |
| 3583 | NR_003594.1 | 54.7958064 | 7.86322851 | 1.19052829 | 6.60482291 |
| 3584 | NM_001105568.2 | 87.6001382 | -8.0773741 | 1.22314562 | -6.6037714 |
| 3585 | NM_001985.2 | 112.740417 | -8.4388028 | 1.277904 | -6.6036281 |
| 3586 | NM_001164273.1 | 100.897377 | -8.2808668 | 1.25419335 | -6.602544 |
| 3587 | NM_025135.4 | 155.467466 | -8.9018374 | 1.34845805 | -6.6014938 |
| 3588 | NM_018121.3 | 113.156682 | -8.4437075 | 1.27915373 | -6.6010107 |
| 3589 | NM_004550.4 | 216.961682 | -7.60321 | 1.15186371 | -6.6007896 |
| 3590 | NM_014278.3 | 88.1792813 | -8.0832496 | 1.22471173 | -6.6001243 |
| 3591 | NM_033257.3 | 131.902893 | -8.6651534 | 1.31288704 | -6.6000754 |
| 3592 | NM_001276380.1 | 98.065149 | -8.2363042 | 1.2479355 | -6.5999439 |
| 3593 | NM_033070.2 | 163.172262 | -8.9717541 | 1.35946472 | -6.5994755 |
| 3594 | NM_014497.4 | 212.847387 | -9.3548214 | 1.41753916 | -6.5993389 |
| 3595 | NM_021202.2 | 116.889624 | -8.4929308 | 1.28695065 | -6.5992668 |
| 3596 | NM_031431.3 | 113.505812 | -8.4484031 | 1.28034676 | -6.5985274 |
| 3597 | NM_014873.2 | 196.809236 | -4.3012707 | 0.65186066 | -6.598451 |
| 3598 | NM_001303513.1 | 178.812742 | -9.1039532 | 1.37984636 | -6.5978021 |
| 3599 | NM_013365.4 | 112.685534 | -8.4374074 | 1.27883575 | -6.5977257 |
| 3600 | NM_022749.5 | 183.844151 | -8.1801218 | 1.23991948 | -6.5973008 |
| 3601 | NM_000709.3 | 113.625067 | -8.4490338 | 1.28080002 | -6.5966846 |
| 3602 | NM_001143775.1 | 227.932774 | -6.7350885 | 1.02099734 | -6.5965779 |
| 3603 | NM_020933.4 | 99.2944558 | -8.2537333 | 1.25121008 | -6.5966007 |
| 3604 | NM_001001486.1 | 87.9941697 | -8.0816799 | 1.22516011 | -6.5964275 |
| 3605 | NM_006012.2 | 150.036814 | -8.8509991 | 1.34183678 | -6.5961816 |
| 3606 | NM_022805.3 | 184.596473 | -9.1503695 | 1.38737166 | -6.595471 |
| 3607 | NM_001995.3 | 103.283241 | -8.3126064 | 1.26052501 | -6.5945589 |
| 3608 | NM_003905.3 | 134.511172 | -8.692954 | 1.31819332 | -6.5945972 |
| 3609 | NM_020145.3 | 104.7919 | -8.3358703 | 1.26403758 | -6.5946381 |
| 3610 | NM_006195.5 | 243.655066 | -9.5510324 | 1.44835265 | -6.5944108 |
| 3611 | NM_152624.5 | 89.8936549 | -8.1139254 | 1.23042608 | -6.5944029 |

| | | | | | |
|---|---|---|---|---|---|
| 3612 | NM_005476.5 | 117.453008 | -8.4992431 | 1.2888963 | -6.5942024 |
| 3613 | NM_014814.2 | 132.150725 | -8.6673803 | 1.31444854 | -6.5939289 |
| 3614 | NM_019885.3 | 210.222501 | -9.3381959 | 1.41621067 | -6.5937901 |
| 3615 | NM_004551.2 | 141.042248 | -8.7621073 | 1.32886267 | -6.5936891 |
| 3616 | NM_017859.3 | 114.375341 | -8.4593185 | 1.28294975 | -6.5936476 |
| 3617 | NM_018008.3 | 218.940955 | -9.3960224 | 1.42501513 | -6.59363 |
| 3618 | NM_014601.3 | 128.899501 | -3.8087402 | 0.57765643 | -6.5934352 |
| 3619 | NM_178832.3 | 118.791786 | -8.5143201 | 1.29133669 | -6.5934161 |
| 3620 | NM_022060.2 | 109.596042 | -8.3994086 | 1.27392211 | -6.5933455 |
| 3621 | NM_012194.2 | 170.165941 | -9.0328122 | 1.37005864 | -6.5930114 |
| 3622 | NM_005749.3 | 87.549439 | -8.0763696 | 1.2250056 | -6.5929246 |
| 3623 | NM_194460.2 | 147.508709 | -8.8262504 | 1.33880725 | -6.5926222 |
| 3624 | NM_001134493.1 | 156.638983 | -8.9132499 | 1.35230504 | -6.5911534 |
| 3625 | NM_020659.3 | 193.239248 | -9.2157752 | 1.39822334 | -6.5910609 |
| 3626 | NM_015189.2 | 122.1241 | -8.5536965 | 1.29780853 | -6.5908771 |
| 3627 | NR_028325.1 | 124.707217 | -6.0875628 | 0.92363465 | -6.5908774 |
| 3628 | NM_017916.2 | 116.351292 | -8.4843292 | 1.28733943 | -6.5905922 |
| 3629 | NM_001098794.1 | 84.6348613 | -8.0244313 | 1.21756724 | -6.5905447 |
| 3630 | NM_002419.3 | 92.2269873 | -8.150372 | 1.236717 | -6.5903291 |
| 3631 | NM_001184772.2 | 84.9779786 | -8.0329886 | 1.21894022 | -6.5901415 |
| 3632 | NM_001983.3 | 91.7560134 | -8.1408421 | 1.23536796 | -6.5898116 |
| 3633 | NM_138391.4 | 122.389256 | -8.5572957 | 1.2985743 | -6.5897621 |
| 3634 | NM_032569.3 | 128.985023 | -8.6349327 | 1.31038247 | -6.5896277 |
| 3635 | NM_000297.3 | 121.07615 | -8.5426575 | 1.29649347 | -6.5890478 |
| 3636 | NM_001291303.1 | 181.761907 | -9.1273666 | 1.38523871 | -6.5890207 |
| 3637 | NR_002599.1 | 190.913465 | -9.1985368 | 1.39602791 | -6.589078 |
| 3638 | NM_007249.4 | 118.05955 | -8.5045917 | 1.29075846 | -6.5888329 |
| 3639 | NM_145693.2 | 150.2533 | -6.3759536 | 0.96773968 | -6.5885007 |
| 3640 | NM_012328.2 | 102.33914 | -8.2988439 | 1.25974375 | -6.5877238 |
| 3641 | NM_000255.3 | 110.33533 | -8.4076235 | 1.27631962 | -6.5873966 |
| 3642 | NM_001278563.2 | 149.326312 | -8.8455151 | 1.34285431 | -6.5870996 |
| 3643 | NM_014956.4 | 94.8346829 | -8.189327 | 1.24340779 | -6.5861956 |
| 3644 | NM_001853.3 | 144.190604 | -8.7949082 | 1.33540219 | -6.5859621 |
| 3645 | NM_139321.2 | 262.239993 | -7.8856948 | 1.19740716 | -6.585642 |
| 3646 | NM_001320845.1 | 103.3544 | -8.3124943 | 1.26223325 | -6.5855453 |
| 3647 | NM_014616.2 | 90.6931465 | -8.1258905 | 1.23390883 | -6.5854869 |
| 3648 | NM_001003713.2 | 149.787157 | -8.8489169 | 1.34373463 | -6.5853158 |
| 3649 | NM_015107.2 | 86.8805839 | -8.0612875 | 1.22413612 | -6.5852869 |
| 3650 | NM_012086.4 | 102.72125 | -8.3041981 | 1.26110211 | -6.5848737 |
| 3651 | NM_001834.3 | 116.099392 | -8.4824628 | 1.28820929 | -6.5846931 |
| 3652 | NM_006402.2 | 148.647454 | -8.8375741 | 1.34213964 | -6.5846904 |
| 3653 | NM_002904.5 | 141.838681 | -8.7699723 | 1.3318938 | -6.5845883 |
| 3654 | NM_004854.4 | 163.461192 | -8.974438 | 1.36318179 | -6.5834491 |

| | | | | | |
|---|---|---|---|---|---|
| 3655 | NM_004578.3 | 166.528928 | -9.0017979 | 1.36735081 | -6.5833858 |
| 3656 | NM_001892.5 | 202.100392 | -9.2808915 | 1.40991342 | -6.5825967 |
| 3657 | NM_001039618.2 | 90.9740977 | -8.1307295 | 1.23519573 | -6.5825434 |
| 3658 | NM_001329431.1 | 92.1137386 | -8.1452907 | 1.23743513 | -6.5823981 |
| 3659 | NM_020745.3 | 132.249657 | -3.0930415 | 0.46989393 | -6.5824249 |
| 3660 | NM_015080.3 | 296.056378 | -9.8317597 | 1.49366207 | -6.5823186 |
| 3661 | NM_175748.3 | 109.580784 | -8.3973563 | 1.27574878 | -6.5822962 |
| 3662 | NM_014851.3 | 83.9245291 | -8.015115 | 1.21779146 | -6.581681 |
| 3663 | NM_015323.4 | 85.2073936 | -8.035814 | 1.220931 | -6.5817102 |
| 3664 | NM_001291459.1 | 111.249128 | -8.419836 | 1.27930345 | -6.5815784 |
| 3665 | NM_003087.2 | 219.503023 | -9.400575 | 1.42830813 | -6.5816155 |
| 3666 | NM_001031710.2 | 184.470067 | -6.8112606 | 1.0350287 | -6.5807456 |
| 3667 | NM_030973.3 | 160.419584 | -7.9817572 | 1.21293367 | -6.5805389 |
| 3668 | NM_030940.3 | 135.574622 | -8.7043796 | 1.32275692 | -6.5804831 |
| 3669 | NM_004089.3 | 101.30997 | -8.2863067 | 1.25930219 | -6.580078 |
| 3670 | NM_001033519.1 | 107.772779 | -8.373284 | 1.27256415 | -6.5798521 |
| 3671 | NM_001048218.1 | 113.448387 | -8.4492618 | 1.28412289 | -6.5797922 |
| 3672 | NM_005493.2 | 214.917447 | -8.4056477 | 1.27759748 | -6.5792613 |
| 3673 | NM_014203.2 | 262.272294 | -9.6570664 | 1.4678053 | -6.5792557 |
| 3674 | NM_199002.1 | 97.8283988 | -8.2344859 | 1.25157266 | -6.5793111 |
| 3675 | NM_005663.4 | 140.629361 | -6.0034126 | 0.91249946 | -6.5790862 |
| 3676 | NM_031287.2 | 154.814195 | -8.8965992 | 1.35237063 | -6.5785215 |
| 3677 | NM_017489.2 | 104.077471 | -8.3235246 | 1.26532174 | -6.5781883 |
| 3678 | NM_000723.4 | 175.451087 | -9.0769291 | 1.37997722 | -6.5775934 |
| 3679 | NM_014771.3 | 88.2453157 | -8.0866392 | 1.22946658 | -6.5773558 |
| 3680 | NM_005253.3 | 129.429042 | -8.6391369 | 1.31349383 | -6.5772192 |
| 3681 | NM_001198774.1 | 130.808514 | -8.6534516 | 1.31579327 | -6.5766042 |
| 3682 | NM_006813.2 | 124.735456 | -8.5842976 | 1.30533697 | -6.5763077 |
| 3683 | NM_052948.3 | 159.559525 | -8.939751 | 1.3593889 | -6.5763013 |
| 3684 | NM_001085411.2 | 119.70747 | -8.5247097 | 1.29634949 | -6.5759348 |
| 3685 | NM_001289972.1 | 91.4369473 | -8.1340554 | 1.23694451 | -6.5759259 |
| 3686 | NM_173809.4 | 130.066464 | -8.645404 | 1.31473841 | -6.5757599 |
| 3687 | NM_206853.2 | 295.392631 | -7.8920159 | 1.20022199 | -6.5754635 |
| 3688 | NM_001144061.1 | 140.535102 | -8.7571727 | 1.33185549 | -6.5751673 |
| 3689 | NM_002795.3 | 148.738332 | -8.8386796 | 1.34425151 | -6.5751681 |
| 3690 | NM_001930.3 | 157.261436 | -8.9187627 | 1.35660123 | -6.5743437 |
| 3691 | NM_012406.3 | 176.382168 | -8.1201429 | 1.23512233 | -6.5743633 |
| 3692 | NM_144572.1 | 81.8078127 | -7.9759841 | 1.21321443 | -6.5742576 |
| 3693 | NM_003713.4 | 134.013132 | -6.8841034 | 1.04720079 | -6.5738141 |
| 3694 | NM_002735.2 | 180.826772 | -9.120483 | 1.387414 | -6.5737285 |
| 3695 | NM_175866.4 | 106.941785 | -8.3651017 | 1.27253128 | -6.5735922 |
| 3696 | NM_018046.4 | 117.926665 | -6.0854214 | 0.92574828 | -6.5735163 |
| 3697 | NM_007121.5 | 92.4574033 | -8.153872 | 1.2404428 | -6.573356 |

| | | | | | |
|---|---|---|---|---|---|
| 3698 | NM_152833.2 | 111.287747 | -8.4200347 | 1.28100012 | -6.5730163 |
| 3699 | NM_003849.3 | 143.185674 | -8.7837776 | 1.33639324 | -6.5727492 |
| 3700 | NM_014047.2 | 138.095804 | -8.7315044 | 1.32847168 | -6.5725935 |
| 3701 | NM_001300821.2 | 115.70641 | -8.4766837 | 1.28974252 | -6.5723845 |
| 3702 | NM_002645.3 | 101.36877 | -8.2878475 | 1.26104012 | -6.5722314 |
| 3703 | NM_004547.5 | 161.06568 | -8.9533702 | 1.36235998 | -6.5719562 |
| 3704 | NM_016100.4 | 94.2608352 | -8.1820308 | 1.24504513 | -6.5716741 |
| 3705 | NM_030645.1 | 154.200571 | -6.4245645 | 0.97769211 | -6.571153 |
| 3706 | NM_003336.3 | 150.96766 | -8.8597477 | 1.3483432 | -6.5708402 |
| 3707 | NM_021633.3 | 122.551283 | -8.5592174 | 1.30266826 | -6.5705273 |
| 3708 | NM_181716.2 | 211.555312 | -9.3466681 | 1.42252447 | -6.5704797 |
| 3709 | NM_016199.2 | 169.61619 | -9.0280081 | 1.37419109 | -6.569689 |
| 3710 | NM_001343.3 | 120.302058 | -8.5347747 | 1.29922609 | -6.569122 |
| 3711 | NM_024548.3 | 99.6486582 | -8.2587001 | 1.25723406 | -6.568944 |
| 3712 | NM_019843.3 | 94.3890217 | -8.1814622 | 1.24549092 | -6.5688653 |
| 3713 | NM_033028.4 | 91.3314137 | -8.1348391 | 1.23845204 | -6.568554 |
| 3714 | NM_001128827.2 | 148.046638 | -8.8315063 | 1.34457524 | -6.56825 |
| 3715 | NR_026550.2 | 140.282827 | -8.7541958 | 1.33283688 | -6.5680924 |
| 3716 | NM_007100.3 | 131.925725 | -8.665827 | 1.31943927 | -6.5678104 |
| 3717 | NM_005180.8 | 148.918166 | -8.8398475 | 1.34604786 | -6.5672609 |
| 3718 | NM_016079.3 | 130.854952 | -8.6529448 | 1.31766028 | -6.5669011 |
| 3719 | NM_032846.3 | 126.31646 | -8.6044198 | 1.31026192 | -6.5669464 |
| 3720 | NM_203434.2 | 147.457493 | -8.8264126 | 1.3440729 | -6.5669151 |
| 3721 | NM_175610.3 | 96.0631182 | -8.2108552 | 1.25042897 | -6.5664307 |
| 3722 | NM_015559.2 | 182.988207 | -6.3882261 | 0.97307777 | -6.5649697 |
| 3723 | NM_144711.5 | 150.771174 | -8.8584325 | 1.34936296 | -6.5648997 |
| 3724 | NM_013346.3 | 144.761385 | -8.7997618 | 1.34058498 | -6.5641209 |
| 3725 | NM_021204.4 | 165.104525 | -8.9884859 | 1.3693281 | -6.5641579 |
| 3726 | NM_006362.4 | 113.494183 | -8.4483234 | 1.28705708 | -6.5640627 |
| 3727 | NM_003587.4 | 108.666547 | -8.3874366 | 1.277796 | -6.5639872 |
| 3728 | NM_001042590.2 | 112.127853 | -8.4311508 | 1.28448048 | -6.5638606 |
| 3729 | NR_134537.1 | 158.37066 | -8.9290586 | 1.36052268 | -6.562962 |
| 3730 | NM_001145522.1 | 219.784672 | -9.4026358 | 1.43272024 | -6.5627857 |
| 3731 | NM_002799.3 | 289.8327 | -8.8361203 | 1.34650831 | -6.5622471 |
| 3732 | NM_001039877.1 | 163.169993 | -8.9729188 | 1.36741171 | -6.561973 |
| 3733 | NM_001164814.1 | 96.1716165 | -8.208803 | 1.2510678 | -6.5614374 |
| 3734 | NM_213674.1 | 178.714233 | -9.10401 | 1.3875152 | -6.5613767 |
| 3735 | NM_016151.3 | 98.1043649 | -8.2387838 | 1.25582702 | -6.5604448 |
| 3736 | NM_002892.3 | 134.941203 | -8.6971079 | 1.32573828 | -6.5601997 |
| 3737 | NM_002881.2 | 107.869722 | -8.375077 | 1.27666409 | -6.5601258 |
| 3738 | NM_024527.4 | 117.483427 | -8.4980442 | 1.29546008 | -6.5598658 |
| 3739 | NM_001330653.1 | 160.508585 | -8.9489308 | 1.36423576 | -6.5596659 |
| 3740 | NM_024678.5 | 96.2291535 | -8.2092465 | 1.25153039 | -6.5593664 |

| | | | | | |
|---|---|---|---|---|---|
| 3741 | NM_020782.1 | 214.184734 | -7.5313481 | 1.14828465 | -6.5587814 |
| 3742 | NM_020470.2 | 131.642051 | -8.6625578 | 1.32080695 | -6.5585344 |
| 3743 | NM_175885.3 | 93.4276286 | -8.165205 | 1.24498821 | -6.5584597 |
| 3744 | NM_001162383.1 | 190.49979 | -9.1956026 | 1.40213507 | -6.5582858 |
| 3745 | NM_001082575.2 | 182.6417 | -9.135064 | 1.39297212 | -6.5579661 |
| 3746 | NM_032752.2 | 166.615499 | -7.1739697 | 1.09398117 | -6.557672 |
| 3747 | NM_020319.2 | 105.298867 | -8.3402042 | 1.27183557 | -6.5576121 |
| 3748 | NM_001244926.1 | 104.43642 | -8.3283133 | 1.27004483 | -6.5574955 |
| 3749 | NM_021005.3 | 213.510154 | -7.4624188 | 1.13801145 | -6.5574198 |
| 3750 | NM_033630.2 | 148.748022 | -8.8387558 | 1.34792144 | -6.5573227 |
| 3751 | NM_018644.3 | 436.008925 | -10.390269 | 1.58463125 | -6.5569002 |
| 3752 | NM_207113.2 | 146.181855 | -8.8134147 | 1.34414661 | -6.5568849 |
| 3753 | NM_016146.5 | 134.177173 | -8.6900465 | 1.32555786 | -6.5557655 |
| 3754 | NM_025179.3 | 409.705157 | -7.3110035 | 1.11522407 | -6.5556364 |
| 3755 | NM_021976.4 | 115.597206 | -8.4764996 | 1.29305773 | -6.5553915 |
| 3756 | NM_153712.4 | 259.527052 | -5.2221581 | 0.79664734 | -6.5551692 |
| 3757 | NM_000466.2 | 90.4991836 | -8.1209441 | 1.23888451 | -6.5550453 |
| 3758 | NM_021227.3 | 152.04691 | -8.8707101 | 1.35333357 | -6.5547108 |
| 3759 | NM_002766.2 | 102.692753 | -8.3032403 | 1.26683005 | -6.5543442 |
| 3760 | NM_015368.3 | 134.073872 | -8.6887267 | 1.32567671 | -6.5541822 |
| 3761 | NM_001278628.1 | 86.168112 | -8.050542 | 1.22835391 | -6.5539271 |
| 3762 | NM_014773.4 | 105.375482 | -8.3425607 | 1.27292617 | -6.5538449 |
| 3763 | NM_025240.2 | 165.699025 | -8.9950307 | 1.37247945 | -6.5538546 |
| 3764 | NM_002335.3 | 194.160963 | -8.2570148 | 1.2599145 | -6.553631 |
| 3765 | NM_001135575.1 | 203.08422 | -9.2878275 | 1.41722177 | -6.5535456 |
| 3766 | NM_198236.2 | 135.548367 | -8.7056973 | 1.32844948 | -6.5532769 |
| 3767 | NM_007320.2 | 101.186343 | -8.2818164 | 1.26381038 | -6.553053 |
| 3768 | NM_004552.2 | 167.601426 | -9.0110122 | 1.37513838 | -6.552804 |
| 3769 | NM_022457.6 | 103.356222 | -8.3131342 | 1.26881931 | -6.5518661 |
| 3770 | NM_006442.3 | 303.382442 | -7.9249061 | 1.2096194 | -6.55157 |
| 3771 | NM_181784.2 | 109.833864 | -8.4019971 | 1.28258999 | -6.5508051 |
| 3772 | NM_139245.3 | 184.338426 | -9.1479033 | 1.39649575 | -6.5506131 |
| 3773 | NM_015897.3 | 132.001425 | -8.6661884 | 1.32296651 | -6.5505728 |
| 3774 | NM_005704.4 | 252.252673 | -9.6008337 | 1.46570609 | -6.550313 |
| 3775 | NM_017528.4 | 108.738999 | -8.3878146 | 1.28052655 | -6.5502856 |
| 3776 | NM_006899.4 | 107.040284 | -8.3647536 | 1.27703946 | -6.5501137 |
| 3777 | NM_001001852.3 | 137.797839 | -8.7283795 | 1.33269009 | -6.5494443 |
| 3778 | NM_032486.3 | 232.437324 | -9.4833732 | 1.44797021 | -6.5494257 |
| 3779 | NM_015076.4 | 115.88807 | -8.4776126 | 1.29453466 | -6.5487722 |
| 3780 | NM_025073.2 | 115.4592 | -8.4739151 | 1.29401774 | -6.5485309 |
| 3781 | NM_001300971.1 | 168.514012 | -9.018697 | 1.37735463 | -6.5478395 |
| 3782 | NM_001303614.1 | 87.5091397 | -8.0740484 | 1.23308943 | -6.5478206 |
| 3783 | NM_022092.2 | 104.912971 | -8.334515 | 1.27285351 | -6.547898 |

| | | | | | |
|---|---|---|---|---|---|
| 3784 | NM_032775.3 | 127.604033 | -8.6175552 | 1.31611041 | -6.5477449 |
| 3785 | NM_004281.3 | 100.271014 | -8.2710993 | 1.26323747 | -6.5475412 |
| 3786 | NM_001300801.1 | 130.11784 | -8.6457591 | 1.3206678 | -6.5465056 |
| 3787 | NM_001077446.3 | 89.9188155 | -8.1126774 | 1.23927625 | -6.5463027 |
| 3788 | NM_012453.2 | 102.505076 | -8.3017692 | 1.26824452 | -6.5458743 |
| 3789 | NM_002818.2 | 108.372844 | -8.3823456 | 1.28060065 | -6.5456359 |
| 3790 | NM_004330.3 | 96.4442441 | -8.2137118 | 1.25492927 | -6.5451591 |
| 3791 | NM_002165.3 | 89.1269248 | -8.1019601 | 1.23787611 | -6.5450493 |
| 3792 | NM_001146116.1 | 17.3026547 | 9.03795365 | 1.38090966 | 6.5449275 |
| 3793 | NM_004171.3 | 139.538025 | -8.7454469 | 1.33620456 | -6.5449911 |
| 3794 | NM_012088.2 | 130.804829 | -8.6533132 | 1.32214039 | -6.5449277 |
| 3795 | NM_000755.4 | 104.817783 | -8.3353259 | 1.27369761 | -6.5441953 |
| 3796 | NM_001085487.2 | 89.3879534 | -8.1045992 | 1.23852218 | -6.5437659 |
| 3797 | NM_000405.4 | 96.5154464 | -8.2151874 | 1.25557317 | -6.5429778 |
| 3798 | NM_020342.2 | 189.802803 | -7.3448519 | 1.12255003 | -6.5430063 |
| 3799 | NM_020404.2 | 125.438189 | -8.5941572 | 1.3135895 | -6.5424984 |
| 3800 | NM_002926.3 | 135.323074 | -8.7018773 | 1.3300968 | -6.5422887 |
| 3801 | NM_020382.4 | 105.822029 | -8.3478433 | 1.27596787 | -6.5423617 |
| 3802 | NM_020909.3 | 96.068478 | -8.2087837 | 1.25472317 | -6.5423066 |
| 3803 | NM_001002836.3 | 111.103951 | -8.4178019 | 1.28673349 | -6.5419933 |
| 3804 | NM_004147.3 | 133.39803 | -8.6816479 | 1.32713576 | -6.5416427 |
| 3805 | NM_003324.4 | 87.3872741 | -8.0690977 | 1.23351391 | -6.5415539 |
| 3806 | NM_001446.4 | 412.841477 | -10.311521 | 1.57634943 | -6.5413928 |
| 3807 | NM_018161.4 | 96.4610805 | -8.2147958 | 1.25585641 | -6.5411903 |
| 3808 | NM_001291412.1 | 111.359674 | -8.4206357 | 1.28735291 | -6.5410468 |
| 3809 | NM_001270546.1 | 125.523272 | -6.2299365 | 0.95244993 | -6.5409597 |
| 3810 | NM_003157.4 | 88.2287369 | -8.0846341 | 1.23601763 | -6.5408728 |
| 3811 | NM_001024660.4 | 128.588486 | -5.450973 | 0.83338968 | -6.5407254 |
| 3812 | NM_206926.1 | 426.451393 | -5.3488024 | 0.81782931 | -6.540243 |
| 3813 | NM_001322374.1 | 126.35318 | -8.602467 | 1.31532229 | -6.5401971 |
| 3814 | NM_001959.3 | 143.623259 | -8.7885552 | 1.34384168 | -6.5398739 |
| 3815 | NM_001614.3 | 10142.5826 | -9.5415778 | 1.45917922 | -6.5390034 |
| 3816 | NM_032621.3 | 195.648311 | -9.2340342 | 1.41217443 | -6.5388765 |
| 3817 | NM_001321178.1 | 81.8908093 | -7.9802978 | 1.22047403 | -6.5386871 |
| 3818 | NM_033129.3 | 161.897795 | -8.9605863 | 1.37040411 | -6.5386452 |
| 3819 | NR_036627.2 | 143.853458 | -8.7916846 | 1.34461282 | -6.5384506 |
| 3820 | NM_001610.3 | 122.632201 | -8.5605736 | 1.30934555 | -6.5380553 |
| 3821 | NM_014937.3 | 140.623888 | -8.7577296 | 1.33950131 | -6.5380523 |
| 3822 | NM_053045.1 | 114.571545 | -8.4618655 | 1.29424159 | -6.5380881 |
| 3823 | NM_001098536.1 | 235.410999 | -9.5012859 | 1.45330868 | -6.537693 |
| 3824 | NM_020814.2 | 185.005091 | -9.1532874 | 1.40008843 | -6.5376495 |
| 3825 | NM_031203.3 | 379.942264 | -6.7337517 | 1.03004587 | -6.5373319 |
| 3826 | NM_033140.3 | 180.903416 | -8.1531776 | 1.24720625 | -6.5371526 |

| | | | | | |
|---|---|---|---|---|---|
| 3827 | NM_007186.5 | 94.9280469 | -8.188435 | 1.25261613 | -6.5370665 |
| 3828 | NM_003869.5 | 99.6562845 | -8.2631922 | 1.26424699 | -6.5360584 |
| 3829 | NM_007275.2 | 122.238291 | -8.555934 | 1.30907983 | -6.5358382 |
| 3830 | NM_005594.5 | 132.979223 | -8.6769277 | 1.32780601 | -6.5347856 |
| 3831 | NM_019063.4 | 86.3752326 | -8.0568605 | 1.23302109 | -6.5342439 |
| 3832 | NM_001081550.1 | 182.661795 | -8.1673267 | 1.24994507 | -6.5341485 |
| 3833 | NM_002843.3 | 94.47608 | -8.1843508 | 1.25258711 | -6.5339574 |
| 3834 | NM_020940.3 | 107.808089 | -8.374225 | 1.28165539 | -6.5339131 |
| 3835 | NM_015190.4 | 134.524767 | -8.6936075 | 1.33062531 | -6.5334752 |
| 3836 | NM_003377.4 | 103.484307 | -8.3158749 | 1.27285909 | -6.5332251 |
| 3837 | NM_015470.2 | 102.120209 | -8.2971921 | 1.27000017 | -6.5332212 |
| 3838 | NM_032141.3 | 88.4544005 | -8.0888106 | 1.23810236 | -6.5332325 |
| 3839 | NM_001042678.1 | 147.060455 | -8.8234032 | 1.35056215 | -6.5331338 |
| 3840 | NM_004649.7 | 130.89014 | -8.6535509 | 1.32457361 | -6.5330842 |
| 3841 | NM_030877.4 | 120.844821 | -8.5386116 | 1.30706511 | -6.5326597 |
| 3842 | NM_001321381.1 | 112.344903 | -8.4324968 | 1.29098295 | -6.5318421 |
| 3843 | NM_003130.3 | 124.076379 | -8.5768846 | 1.31309139 | -6.531826 |
| 3844 | NM_014023.3 | 163.928119 | -8.0116526 | 1.22689779 | -6.5300082 |
| 3845 | NM_006636.3 | 148.053849 | -8.8319923 | 1.35274485 | -6.5289417 |
| 3846 | NM_001177701.2 | 98.6713746 | -8.2471906 | 1.26325825 | -6.5285073 |
| 3847 | NM_002616.2 | 162.924636 | -8.0040131 | 1.2260506 | -6.5282894 |
| 3848 | NM_004834.4 | 105.509803 | -8.344875 | 1.27849416 | -6.5271123 |
| 3849 | NM_032478.3 | 128.233695 | -8.6248365 | 1.32141706 | -6.5269601 |
| 3850 | NM_003626.3 | 108.112109 | -8.3795332 | 1.28386503 | -6.5268023 |
| 3851 | NM_001142299.1 | 88.4008919 | -8.0889085 | 1.23941492 | -6.5263928 |
| 3852 | NM_004389.3 | 126.165352 | -8.6022324 | 1.31814077 | -6.5260347 |
| 3853 | NM_001001487.1 | 102.906297 | -5.9300097 | 0.90878182 | -6.5252291 |
| 3854 | NM_001291694.1 | 192.866221 | -8.2460651 | 1.26378939 | -6.5248729 |
| 3855 | NM_024067.3 | 121.234356 | -8.5435203 | 1.30940914 | -6.5247141 |
| 3856 | NM_001024948.2 | 83.4180794 | -8.0017619 | 1.22642849 | -6.5244423 |
| 3857 | NM_024843.3 | 108.765924 | -8.3888347 | 1.28577204 | -6.5243561 |
| 3858 | NM_177972.2 | 336.53203 | -6.7656536 | 1.03711246 | -6.5235486 |
| 3859 | NM_181050.2 | 93.0620024 | -8.1611641 | 1.25108781 | -6.5232544 |
| 3860 | NM_001330619.1 | 95.9917221 | -8.2071234 | 1.25814077 | -6.5232155 |
| 3861 | NM_006052.1 | 117.721527 | -2.893129 | 0.44352049 | -6.5231011 |
| 3862 | NM_016029.3 | 119.564752 | -8.5241171 | 1.30700177 | -6.5218864 |
| 3863 | NM_005920.3 | 122.040989 | -8.5524674 | 1.31138708 | -6.5216956 |
| 3864 | NM_014703.2 | 84.8990697 | -8.029479 | 1.23119865 | -6.5216763 |
| 3865 | NM_015411.2 | 142.143069 | -8.773544 | 1.34535581 | -6.5213558 |
| 3866 | NM_015073.2 | 122.151499 | -3.1469279 | 0.48262672 | -6.5204179 |
| 3867 | NM_002706.5 | 87.4397016 | -8.0725765 | 1.23808771 | -6.5201976 |
| 3868 | NM_031469.3 | 102.050357 | -5.8877203 | 0.90301757 | -6.5200506 |
| 3869 | NM_001098511.2 | 192.96149 | -9.2139341 | 1.41323897 | -6.5197283 |

| | | | | | |
|---|---|---|---|---|---|
| 3870 | NM_005453.4 | 110.215972 | -8.4056058 | 1.28927878 | -6.5196185 |
| 3871 | NM_006929.4 | 178.389811 | -8.1363932 | 1.24813935 | -6.518818 |
| 3872 | NM_000019.3 | 126.224708 | -8.601715 | 1.31956119 | -6.5186177 |
| 3873 | NM_003093.2 | 148.514515 | -8.8365914 | 1.35566508 | -6.5182702 |
| 3874 | NM_019030.3 | 102.49073 | -8.2995599 | 1.27333852 | -6.5179524 |
| 3875 | NM_001849.3 | 628.032644 | -7.3994832 | 1.13530656 | -6.5176081 |
| 3876 | NM_005786.5 | 84.6476822 | -8.0274877 | 1.23171195 | -6.5173417 |
| 3877 | NM_015258.1 | 91.3964876 | -8.1369252 | 1.24855487 | -6.5170746 |
| 3878 | NR_104307.2 | 102.310572 | -8.2975615 | 1.27356796 | -6.515209 |
| 3879 | NM_138425.3 | 248.841379 | -9.5809776 | 1.47057869 | -6.5151071 |
| 3880 | NM_001194955.1 | 115.457582 | -8.4723375 | 1.30045838 | -6.5148855 |
| 3881 | NM_000725.3 | 150.663909 | -8.8572405 | 1.35956901 | -6.5147414 |
| 3882 | NM_002097.2 | 196.018509 | -9.2368911 | 1.41795541 | -6.5142324 |
| 3883 | NM_006116.2 | 109.766685 | -8.4001017 | 1.28958004 | -6.5138274 |
| 3884 | NM_016355.3 | 118.964759 | -8.5158136 | 1.30736676 | -6.5137143 |
| 3885 | NM_001243186.1 | 92.0185241 | -8.1466857 | 1.25072551 | -6.5135681 |
| 3886 | NM_015131.2 | 102.81807 | -8.3056228 | 1.2751587 | -6.5134032 |
| 3887 | NM_032635.3 | 132.262891 | -8.6697213 | 1.33105665 | -6.5134127 |
| 3888 | NM_033082.3 | 136.250326 | -8.7117004 | 1.3375142 | -6.5133517 |
| 3889 | NM_016169.3 | 86.4837654 | -8.0557087 | 1.23687341 | -6.5129613 |
| 3890 | NM_032204.4 | 105.034238 | -8.3374274 | 1.28015294 | -6.512837 |
| 3891 | NM_001145418.1 | 361.293861 | -2.3794307 | 0.365393 | -6.5119767 |
| 3892 | NM_002615.6 | 267.67695 | -9.687043 | 1.48758303 | -6.5119343 |
| 3893 | NM_006031.5 | 111.915017 | -8.4279644 | 1.29433981 | -6.5114001 |
| 3894 | NM_001303264.1 | 117.18726 | -8.495547 | 1.30476392 | -6.5111756 |
| 3895 | NM_032299.3 | 138.737394 | -8.7381611 | 1.34204004 | -6.5111031 |
| 3896 | NM_198317.2 | 116.720067 | -8.4891956 | 1.30404434 | -6.5098979 |
| 3897 | NM_015326.4 | 184.13104 | -9.1471628 | 1.40521053 | -6.5094607 |
| 3898 | NM_000754.3 | 129.118799 | -8.6350498 | 1.3265474 | -6.5094167 |
| 3899 | NM_001330420.1 | 107.36274 | -8.3683589 | 1.2855931 | -6.5093371 |
| 3900 | NM_005927.4 | 118.978702 | -8.5165596 | 1.30836946 | -6.5092926 |
| 3901 | NM_014248.3 | 161.018867 | -8.9530849 | 1.3755294 | -6.5088284 |
| 3902 | NM_004708.3 | 135.385136 | -8.7026908 | 1.33711509 | -6.5085577 |
| 3903 | NM_001029863.2 | 104.693474 | -8.3315695 | 1.2802375 | -6.5078312 |
| 3904 | NM_001284.3 | 117.98471 | -8.5039826 | 1.30678502 | -6.5075605 |
| 3905 | NM_006527.3 | 125.699279 | -8.5954363 | 1.32090467 | -6.5072343 |
| 3906 | NM_003164.4 | 103.352423 | -8.3137583 | 1.27763265 | -6.5071586 |
| 3907 | NM_144982.4 | 120.549666 | -5.4609367 | 0.83922913 | -6.5070867 |
| 3908 | NM_018946.3 | 100.00208 | -8.2664284 | 1.27064504 | -6.5056944 |
| 3909 | NM_001130674.2 | 91.8594691 | -8.14532 | 1.25207746 | -6.5054442 |
| 3910 | NM_001123329.1 | 80.0826696 | -7.9448724 | 1.22129597 | -6.5052801 |
| 3911 | NM_003094.3 | 119.704974 | -8.52502 | 1.31050825 | -6.5051251 |
| 3912 | NM_015540.3 | 122.91728 | -6.1995156 | 0.95311216 | -6.5044974 |

| | | | | | |
|---|---|---|---|---|---|
| 3913 | NM_000937.4 | 749.054498 | -6.9962711 | 1.07565025 | -6.5042249 |
| 3914 | NM_032360.3 | 142.906327 | -8.7810743 | 1.35005294 | -6.5042444 |
| 3915 | NM_004300.3 | 195.219079 | -7.3376595 | 1.12816386 | -6.5040725 |
| 3916 | NM_018434.5 | 97.8323363 | -8.2349011 | 1.26621062 | -6.5035792 |
| 3917 | NR_046331.1 | 139.964705 | -8.75065 | 1.34552646 | -6.5035139 |
| 3918 | NM_015694.2 | 98.5465125 | -8.2445024 | 1.26772052 | -6.5034069 |
| 3919 | NM_004546.2 | 167.451848 | -9.0097168 | 1.38540868 | -6.5032917 |
| 3920 | NM_001143835.1 | 82.2001244 | -7.9808224 | 1.22722392 | -6.5031509 |
| 3921 | NM_001166.4 | 110.998304 | -8.415743 | 1.29430881 | -6.5021136 |
| 3922 | NM_001569.3 | 128.315189 | -8.6253371 | 1.32672623 | -6.5012185 |
| 3923 | NM_001039802.1 | 81.2949205 | -4.3187278 | 0.66436237 | -6.5005605 |
| 3924 | NM_005831.4 | 81.0241654 | -7.9650343 | 1.22530764 | -6.5004363 |
| 3925 | NM_017653.3 | 108.524033 | -8.383936 | 1.28974655 | -6.5004523 |
| 3926 | NM_153812.2 | 86.3592045 | -8.0562345 | 1.23937114 | -6.5002599 |
| 3927 | NM_173798.3 | 145.301407 | -8.8052378 | 1.35472224 | -6.4996628 |
| 3928 | NM_015971.3 | 111.87851 | -8.4275889 | 1.29663748 | -6.4995722 |
| 3929 | NM_006363.4 | 78.3042388 | -7.9144692 | 1.21773677 | -6.4993268 |
| 3930 | NR_040722.1 | 132.978007 | -8.6753639 | 1.33480571 | -6.4993458 |
| 3931 | NM_014487.5 | 87.3298282 | -8.0700342 | 1.24168492 | -6.4992609 |
| 3932 | NM_020856.3 | 162.594508 | -8.967262 | 1.37981229 | -6.4989 |
| 3933 | NM_001668.3 | 83.404959 | -8.0064843 | 1.23200232 | -6.4987574 |
| 3934 | NM_001042537.1 | 210.219805 | -9.3380047 | 1.43717246 | -6.4974837 |
| 3935 | NM_001131024.1 | 120.221612 | -8.5308549 | 1.31298566 | -6.4972948 |
| 3936 | NM_174983.4 | 117.364077 | -8.4967982 | 1.30776012 | -6.4972147 |
| 3937 | NM_213619.2 | 147.991581 | -8.831455 | 1.35939393 | -6.496612 |
| 3938 | NM_005524.3 | 91.6604046 | -8.1407102 | 1.25308284 | -6.4965459 |
| 3939 | NM_001135643.1 | 204.754229 | -9.2994641 | 1.43160747 | -6.4958198 |
| 3940 | NM_001031723.3 | 100.979202 | -8.2792836 | 1.27456769 | -6.4957582 |
| 3941 | NM_000985.4 | 88.460918 | -8.090418 | 1.24552598 | -6.4955835 |
| 3942 | NM_002736.2 | 361.459676 | -9.1565488 | 1.40966328 | -6.4955575 |
| 3943 | NM_022575.3 | 189.153551 | -6.1490772 | 0.94677036 | -6.4947926 |
| 3944 | NM_138820.3 | 98.337799 | -8.2420127 | 1.26904406 | -6.4946624 |
| 3945 | NM_001172773.1 | 86.1252305 | -8.0480615 | 1.23930017 | -6.4940373 |
| 3946 | NM_021928.3 | 264.513077 | -3.6034935 | 0.55490484 | -6.4938945 |
| 3947 | NM_006848.2 | 143.472861 | -8.7861476 | 1.353019 | -6.4937356 |
| 3948 | NR_033850.1 | 49.3293628 | 2.58753944 | 0.39846948 | 6.49369546 |
| 3949 | NM_001035005.3 | 168.933492 | -9.0224878 | 1.38949061 | -6.493378 |
| 3950 | NM_001083956.1 | 86.5804177 | -8.0611574 | 1.24146312 | -6.4932717 |
| 3951 | NM_005542.5 | 215.279499 | -9.3721489 | 1.44336174 | -6.4932779 |
| 3952 | NM_058246.3 | 107.476421 | -8.3699497 | 1.28906008 | -6.493064 |
| 3953 | NM_014734.3 | 102.63433 | -5.925701 | 0.91262775 | -6.4930099 |
| 3954 | NM_005052.2 | 157.812819 | -8.9239618 | 1.37452055 | -6.4924179 |
| 3955 | NM_001037954.3 | 103.311845 | -8.3117035 | 1.28028611 | -6.4920672 |

| | | | | | |
|---|---|---|---|---|---|
| 3956 | NM_001080412.2 | 84.7991964 | -8.0271152 | 1.23652159 | -6.4916903 |
| 3957 | NM_016129.2 | 120.630907 | -8.5364385 | 1.31497867 | -6.4916935 |
| 3958 | NM_138453.3 | 367.998225 | -9.1814284 | 1.41449404 | -6.490963 |
| 3959 | NM_138389.3 | 82.8434247 | -7.9958294 | 1.23201281 | -6.4900537 |
| 3960 | NM_000903.2 | 99.4728778 | -8.2610125 | 1.27295535 | -6.4896326 |
| 3961 | NM_002966.2 | 169.328964 | -9.0261369 | 1.39093474 | -6.4892598 |
| 3962 | NM_021959.2 | 148.159811 | -8.8333158 | 1.36128359 | -6.4889608 |
| 3963 | NM_001007246.2 | 170.055721 | -9.0316775 | 1.39191161 | -6.4886861 |
| 3964 | NM_004567.3 | 102.768121 | -8.3040648 | 1.27980822 | -6.4885228 |
| 3965 | NM_003302.2 | 97.393719 | -8.229636 | 1.26847374 | -6.4878253 |
| 3966 | NM_001287491.1 | 197.394219 | -7.4656704 | 1.15076186 | -6.4875893 |
| 3967 | NM_002336.2 | 161.231271 | -7.125722 | 1.09840353 | -6.4873444 |
| 3968 | NM_031417.3 | 173.997744 | -9.0646701 | 1.39740254 | -6.4867995 |
| 3969 | NM_004529.3 | 148.342249 | -8.8349567 | 1.3619993 | -6.4867556 |
| 3970 | NM_001319113.1 | 129.543924 | -8.6397013 | 1.33196446 | -6.4864353 |
| 3971 | NM_006246.3 | 133.326621 | -8.6797494 | 1.33817321 | -6.4862675 |
| 3972 | NM_021160.2 | 141.441689 | -8.7656565 | 1.3515309 | -6.4857241 |
| 3973 | NM_004894.2 | 154.502989 | -8.8934363 | 1.37127404 | -6.4855281 |
| 3974 | NM_015077.3 | 235.7111 | -8.5392759 | 1.31672226 | -6.4852522 |
| 3975 | NR_125375.1 | 96.7086704 | -8.219967 | 1.26749323 | -6.4852157 |
| 3976 | NM_001136108.2 | 138.231747 | -8.7325303 | 1.34662864 | -6.4847353 |
| 3977 | NM_001006634.2 | 88.9317846 | -8.0968501 | 1.24865938 | -6.4844346 |
| 3978 | NM_017550.2 | 104.204593 | -8.3251572 | 1.2838962 | -6.4842915 |
| 3979 | NM_003313.3 | 120.342583 | -8.5329848 | 1.31614252 | -6.4833288 |
| 3980 | NM_004223.4 | 123.882503 | -8.5752118 | 1.32266796 | -6.4832687 |
| 3981 | NM_014110.4 | 129.55456 | -8.6393067 | 1.33255788 | -6.4832507 |
| 3982 | NM_025159.2 | 46.7735057 | 4.44847373 | 0.68614163 | 6.48331708 |
| 3983 | NM_139246.4 | 111.346606 | -8.4204403 | 1.29890944 | -6.4827001 |
| 3984 | NM_001321439.1 | 105.990398 | -8.3490251 | 1.2879117 | -6.4826068 |
| 3985 | NM_030627.3 | 115.547762 | -8.4735764 | 1.30728323 | -6.4818214 |
| 3986 | NM_001317955.1 | 95.7393366 | -8.2018829 | 1.26541639 | -6.4815684 |
| 3987 | NM_145886.3 | 81.051933 | -7.9629376 | 1.22867935 | -6.4808916 |
| 3988 | NM_006586.4 | 96.1738377 | -8.2102911 | 1.2668601 | -6.4808191 |
| 3989 | NM_004685.3 | 141.345994 | -8.764999 | 1.35249421 | -6.4806185 |
| 3990 | NM_182947.3 | 179.337244 | -9.10826 | 1.40548254 | -6.4805216 |
| 3991 | NM_199069.1 | 99.2878385 | -8.255752 | 1.27392643 | -6.4805563 |
| 3992 | NM_173611.3 | 126.709785 | -8.6064591 | 1.32808716 | -6.4803421 |
| 3993 | NM_001244950.1 | 142.653476 | -8.7799083 | 1.35489799 | -6.480125 |
| 3994 | NM_012394.3 | 183.231387 | -9.1393303 | 1.4103979 | -6.4799659 |
| 3995 | NM_153702.3 | 101.663294 | -8.2902293 | 1.27942214 | -6.4796669 |
| 3996 | NM_005189.2 | 120.774353 | -8.5376399 | 1.31767002 | -6.4793459 |
| 3997 | NM_021203.3 | 142.577728 | -8.7779639 | 1.35480469 | -6.479136 |
| 3998 | NM_016155.5 | 151.058727 | -8.8609632 | 1.36762696 | -6.479079 |

| | | | | | |
|---|---|---|---|---|---|
| 3999 | NM_020379.3 | 80.3391921 | -7.9531 | 1.22751614 | -6.4790187 |
| 4000 | NM_017586.3 | 143.209052 | -8.7837079 | 1.35579721 | -6.4786295 |
| 4001 | NM_021821.3 | 119.590254 | -8.5237079 | 1.31566803 | -6.4786159 |
| 4002 | NM_001166286.1 | 128.485337 | -8.6277948 | 1.33174643 | -6.4785567 |
| 4003 | NM_018052.3 | 122.486287 | -8.5587987 | 1.32117734 | -6.4781604 |
| 4004 | NM_001144072.1 | 108.388345 | -8.3819238 | 1.29390739 | -6.4779936 |
| 4005 | NM_003610.3 | 120.859068 | -8.5389209 | 1.31817538 | -6.4778338 |
| 4006 | NM_001516.4 | 86.4404468 | -8.0569756 | 1.24381258 | -6.4776444 |
| 4007 | NM_181575.4 | 141.831617 | -8.7706948 | 1.35409538 | -6.4771617 |
| 4008 | NM_015462.4 | 117.273536 | -8.4950393 | 1.31157524 | -6.4769744 |
| 4009 | NM_001135597.1 | 197.8626 | -9.2498275 | 1.42817238 | -6.4766884 |
| 4010 | NM_001142573.1 | 100.931567 | -8.2799672 | 1.27843413 | -6.4766475 |
| 4011 | NM_015000.3 | 92.6530834 | -8.1556914 | 1.25924715 | -6.4766407 |
| 4012 | NM_024333.2 | 183.957324 | -9.1452165 | 1.41201611 | -6.4767084 |
| 4013 | NM_003201.2 | 110.641765 | -8.4106543 | 1.29862766 | -6.476571 |
| 4014 | NM_030808.4 | 111.083427 | -8.4173488 | 1.29971929 | -6.4762821 |
| 4015 | NM_014302.3 | 139.730617 | -8.7485274 | 1.35094146 | -6.4758745 |
| 4016 | NM_003952.2 | 99.9183399 | -8.2656044 | 1.27643815 | -6.4755229 |
| 4017 | NM_016607.3 | 137.941834 | -8.7290023 | 1.34799322 | -6.4755535 |
| 4018 | NM_020967.2 | 93.2524802 | -8.1625581 | 1.26063555 | -6.4749547 |
| 4019 | NM_003968.3 | 111.934689 | -8.4283766 | 1.3016976 | -6.4749113 |
| 4020 | NM_138730.2 | 82.2342841 | -7.982794 | 1.23294023 | -6.4745994 |
| 4021 | NM_016454.3 | 116.943304 | -8.4919506 | 1.31160678 | -6.4744638 |
| 4022 | NM_001663.3 | 196.880282 | -8.2781564 | 1.27868265 | -6.4739726 |
| 4023 | NM_005581.4 | 101.884633 | -8.2954582 | 1.28135461 | -6.4739754 |
| 4024 | NM_020829.3 | 85.8012811 | -8.0435627 | 1.2424428 | -6.4739904 |
| 4025 | NM_194332.2 | 98.2727485 | -8.2394588 | 1.27279862 | -6.4734976 |
| 4026 | NM_001265605.1 | 215.695749 | -9.3747293 | 1.44818553 | -6.4734311 |
| 4027 | NM_001284230.1 | 123.116443 | -8.5657215 | 1.3232607 | -6.4731927 |
| 4028 | NM_004644.4 | 139.081003 | -8.7418892 | 1.35069606 | -6.4721365 |
| 4029 | NM_019594.3 | 147.827971 | -8.8306129 | 1.36440714 | -6.4721245 |
| 4030 | NM_016823.3 | 116.270803 | -8.4835274 | 1.31080245 | -6.4720106 |
| 4031 | NM_018226.4 | 104.668478 | -8.3321352 | 1.28740478 | -6.47204 |
| 4032 | NM_023936.1 | 151.012078 | -8.8604757 | 1.36922497 | -6.4711613 |
| 4033 | NM_001185076.1 | 110.048514 | -8.4039468 | 1.29869902 | -6.4710504 |
| 4034 | NM_001035521.2 | 105.099412 | -8.3378063 | 1.28851095 | -6.4708851 |
| 4035 | NM_004098.3 | 119.377514 | -8.5204489 | 1.31674166 | -6.4708584 |
| 4036 | NM_001102421.2 | 125.203763 | -8.5896991 | 1.32757231 | -6.4702306 |
| 4037 | NM_004214.4 | 165.558432 | -8.9932297 | 1.38994711 | -6.4701957 |
| 4038 | NM_020850.2 | 75.5239052 | -7.8618667 | 1.21508411 | -6.4702243 |
| 4039 | NM_002973.3 | 87.646214 | -8.0748524 | 1.24806887 | -6.4698773 |
| 4040 | NM_024663.3 | 76.0065793 | -7.8711049 | 1.21660918 | -6.469707 |
| 4041 | NM_000532.4 | 89.6347106 | -8.1077945 | 1.25331544 | -6.4690774 |

| | | | | | |
|---|---|---|---|---|---|
| 4042 | NM_001037442.3 | 79.4096626 | -7.9361722 | 1.2268107 | -6.468946 |
| 4043 | NM_016400.3 | 120.277012 | -8.5322033 | 1.31894176 | -6.4689765 |
| 4044 | NM_024552.2 | 115.622443 | -8.4750197 | 1.31025554 | -6.468219 |
| 4045 | NM_003188.3 | 152.330312 | -8.872946 | 1.37178713 | -6.4681653 |
| 4046 | NM_032041.2 | 213.349367 | -9.3594996 | 1.44716112 | -6.4674897 |
| 4047 | NM_001173464.1 | 258.548864 | -9.636423 | 1.49005595 | -6.4671551 |
| 4048 | NM_005005.2 | 203.067202 | -9.2879271 | 1.4362347 | -6.4668589 |
| 4049 | NM_001302632.1 | 89.6788648 | -8.1117848 | 1.2543906 | -6.4667137 |
| 4050 | NM_001042682.1 | 167.431465 | -6.6254269 | 1.02455327 | -6.4666495 |
| 4051 | NM_002691.3 | 98.4542395 | -8.2423828 | 1.27461368 | -6.4665733 |
| 4052 | NM_007373.3 | 132.825516 | -8.6754597 | 1.34170025 | -6.4660193 |
| 4053 | NM_020216.3 | 92.9769972 | -8.1613265 | 1.26219341 | -6.4659872 |
| 4054 | NM_153333.2 | 114.419094 | -8.4599491 | 1.30837628 | -6.4659908 |
| 4055 | NM_001078172.1 | 137.622426 | -8.7262848 | 1.34971302 | -6.4652891 |
| 4056 | NM_001128913.1 | 148.685716 | -8.8385445 | 1.36707726 | -6.4652853 |
| 4057 | NM_006701.4 | 110.656422 | -8.4117193 | 1.30106042 | -6.4652795 |
| 4058 | NM_138367.1 | 91.3205527 | -8.1345335 | 1.25822176 | -6.4651032 |
| 4059 | NM_024725.3 | 102.328405 | -8.2980652 | 1.2836118 | -6.4646221 |
| 4060 | NR_136702.1 | 263.053305 | -6.6491186 | 1.02856969 | -6.4644318 |
| 4061 | NM_003945.3 | 110.576366 | -8.4115096 | 1.30126525 | -6.4641007 |
| 4062 | NM_001164748.1 | 97.8550538 | -8.2369833 | 1.27429094 | -6.4639738 |
| 4063 | NR_024359.2 | 89.8702605 | -8.1122541 | 1.25509531 | -6.4634566 |
| 4064 | NM_001316342.1 | 114.811457 | -8.4651679 | 1.30972162 | -6.4633337 |
| 4065 | NM_014016.4 | 90.3500587 | -8.11728 | 1.25588528 | -6.4633929 |
| 4066 | NM_032013.3 | 130.382416 | -8.6479506 | 1.33800265 | -6.4633284 |
| 4067 | NM_012384.4 | 78.7803865 | -7.921362 | 1.22562334 | -6.4631292 |
| 4068 | NM_015889.4 | 126.914181 | -8.6090494 | 1.33203258 | -6.4630922 |
| 4069 | NM_199168.3 | 181.531949 | -9.1266037 | 1.41209882 | -6.463148 |
| 4070 | NM_000216.3 | 262.572482 | -8.6953784 | 1.3454133 | -6.4629794 |
| 4071 | NM_001330597.1 | 174.544579 | -9.0697089 | 1.40343057 | -6.4625277 |
| 4072 | NM_001321856.1 | 90.4922202 | -8.1238275 | 1.25709489 | -6.4623821 |
| 4073 | NM_014272.4 | 226.213247 | -8.4781049 | 1.31211287 | -6.4614143 |
| 4074 | NM_020895.4 | 248.50361 | -9.5791356 | 1.48277246 | -6.4602869 |
| 4075 | NM_001199172.1 | 157.962453 | -8.9254278 | 1.38165726 | -6.4599435 |
| 4076 | NM_004830.3 | 94.9953167 | -8.190405 | 1.26792974 | -6.4596679 |
| 4077 | NM_022157.3 | 112.581817 | -8.4373035 | 1.30618356 | -6.4595083 |
| 4078 | NM_000718.3 | 143.378935 | -8.7861222 | 1.36036896 | -6.4586318 |
| 4079 | NM_006991.4 | 82.9549376 | -7.9975593 | 1.23827217 | -6.4586441 |
| 4080 | NM_001346143.1 | 94.8664618 | -8.1918219 | 1.26837324 | -6.4585263 |
| 4081 | NM_004838.3 | 101.382282 | -8.286346 | 1.28310181 | -6.4580581 |
| 4082 | NM_020170.3 | 107.617023 | -8.3733275 | 1.29665413 | -6.4576415 |
| 4083 | NM_015268.3 | 83.4719047 | -8.0044255 | 1.2396573 | -6.4569663 |
| 4084 | NM_006589.2 | 112.649349 | -8.439049 | 1.30706188 | -6.4565031 |

| | | | | | |
|---|---|---|---|---|---|
| 4085 | NM_005274.2 | 105.53863 | -8.3439983 | 1.29244318 | -6.4559885 |
| 4086 | NM_020751.2 | 78.5494323 | -7.9192709 | 1.22668887 | -6.4558105 |
| 4087 | NM_015274.2 | 117.401187 | -8.4974543 | 1.31628362 | -6.4556408 |
| 4088 | NM_000305.2 | 116.026619 | -8.481157 | 1.31392991 | -6.4548017 |
| 4089 | NM_001282142.1 | 91.8273494 | -8.144375 | 1.26175417 | -6.4548033 |
| 4090 | NM_004236.3 | 93.7794865 | -8.1739867 | 1.26633043 | -6.4548609 |
| 4091 | NM_032023.3 | 95.5147853 | -8.1995337 | 1.27033193 | -6.4546388 |
| 4092 | NM_014804.2 | 76.0670638 | -7.8714036 | 1.21961707 | -6.4539959 |
| 4093 | NM_001199753.1 | 120.714773 | -8.5368893 | 1.32275923 | -6.4538497 |
| 4094 | NM_001145678.1 | 416.054708 | 4.48295344 | 0.69463831 | 6.45365126 |
| 4095 | NM_001164831.2 | 337.188426 | -9.0562787 | 1.40332318 | -6.4534519 |
| 4096 | NM_004036.4 | 120.332811 | -8.5327126 | 1.32229227 | -6.4529702 |
| 4097 | NM_021970.3 | 135.755048 | -8.7063688 | 1.34929592 | -6.4525273 |
| 4098 | NR_122104.1 | 70.3959379 | 4.08951999 | 0.6338001 | 6.45238146 |
| 4099 | NM_207044.1 | 160.638375 | -8.9503573 | 1.38716201 | -6.4522798 |
| 4100 | NM_018133.3 | 94.4925298 | -8.1831954 | 1.26827544 | -6.4522225 |
| 4101 | NM_015497.4 | 108.40195 | -8.3817288 | 1.2990687 | -6.4521059 |
| 4102 | NM_015530.4 | 109.874002 | -8.4012713 | 1.30209181 | -6.4521343 |
| 4103 | NM_006560.3 | 131.834814 | -8.6655198 | 1.34311251 | -6.4518197 |
| 4104 | NM_001997.4 | 689.984225 | -7.4418322 | 1.15357457 | -6.4511063 |
| 4105 | NM_004749.3 | 98.4294686 | -8.2432514 | 1.27783856 | -6.4509334 |
| 4106 | NM_138392.3 | 98.917964 | -8.2503934 | 1.27903423 | -6.4504868 |
| 4107 | NM_001142930.1 | 125.95673 | -8.5988678 | 1.33306983 | -6.4504257 |
| 4108 | NM_013328.3 | 111.380643 | -8.4214226 | 1.30561513 | -6.450157 |
| 4109 | NM_002146.4 | 151.522154 | -8.8661692 | 1.37463547 | -6.449833 |
| 4110 | NM_001174096.1 | 83.0329511 | -7.999468 | 1.24034268 | -6.4494015 |
| 4111 | NM_006353.2 | 127.754517 | -8.6189946 | 1.33644438 | -6.4491981 |
| 4112 | NM_001288730.1 | 109.14937 | -8.3921933 | 1.30131602 | -6.4490049 |
| 4113 | NM_005488.2 | 95.4610143 | -8.1989538 | 1.27134611 | -6.4490336 |
| 4114 | NM_030775.2 | 78.4664185 | -7.914763 | 1.22730129 | -6.4489161 |
| 4115 | NM_016145.3 | 145.970295 | -8.8116322 | 1.36641453 | -6.4487255 |
| 4116 | NM_175875.4 | 84.0859289 | -8.0185025 | 1.24344523 | -6.4486174 |
| 4117 | NM_012342.2 | 99.5634894 | -8.2619847 | 1.28123448 | -6.4484564 |
| 4118 | NM_020189.5 | 118.81128 | -8.5141166 | 1.3208018 | -6.4461728 |
| 4119 | NM_003101.5 | 85.2396034 | -8.0354192 | 1.24657399 | -6.4460026 |
| 4120 | NM_022048.4 | 91.4703524 | -8.1367515 | 1.26238836 | -6.4455216 |
| 4121 | NM_014755.2 | 86.4108365 | -8.0550343 | 1.24999368 | -6.44406 |
| 4122 | NM_001127358.1 | 110.08987 | -8.4043277 | 1.30430007 | -6.4435538 |
| 4123 | NM_012105.4 | 73.5840724 | -7.8227935 | 1.21409696 | -6.4433021 |
| 4124 | NM_030621.4 | 84.9425871 | -8.0276877 | 1.24592698 | -6.4431446 |
| 4125 | NM_020529.2 | 75.9609035 | -7.8706816 | 1.22157798 | -6.4430447 |
| 4126 | NM_001080855.2 | 115.915069 | -8.479102 | 1.316046 | -6.4428614 |
| 4127 | NM_138797.2 | 82.692215 | -7.9900575 | 1.24017154 | -6.4427035 |

| | | | | | |
|---|---|---|---|---|---|
| 4128 | NM_004090.3 | 206.737226 | -6.7286728 | 1.04441827 | -6.4425078 |
| 4129 | NM_024541.2 | 91.7558002 | -8.1426869 | 1.26407286 | -6.4416279 |
| 4130 | NM_003473.3 | 94.4140318 | -8.1822026 | 1.27022201 | -6.4415531 |
| 4131 | NM_001046.2 | 102.500936 | -8.3025345 | 1.28896914 | -6.4412205 |
| 4132 | NM_017778.2 | 103.81417 | -8.3190056 | 1.29160724 | -6.4408168 |
| 4133 | NM_001079518.1 | 83.6314866 | -8.0084049 | 1.24343238 | -6.4405632 |
| 4134 | NM_001048249.3 | 106.872978 | -8.3617602 | 1.29832319 | -6.4404304 |
| 4135 | NM_144604.3 | 182.656523 | -8.1705312 | 1.26870218 | -6.4400702 |
| 4136 | NM_139045.3 | 119.575115 | -8.5223615 | 1.32340112 | -6.4397418 |
| 4137 | NM_033274.4 | 335.15304 | -9.0464423 | 1.4048107 | -6.4396166 |
| 4138 | NM_006988.4 | 176.629164 | -9.0876568 | 1.41129939 | -6.4392126 |
| 4139 | NM_019008.5 | 123.981515 | -8.57415 | 1.33155554 | -6.4391982 |
| 4140 | NM_014618.2 | 171.487236 | -9.0442555 | 1.40465104 | -6.4387917 |
| 4141 | NM_001156.4 | 93.1488733 | -8.1652633 | 1.26816957 | -6.4386211 |
| 4142 | NM_031966.3 | 104.266456 | -8.3266958 | 1.29326251 | -6.4385195 |
| 4143 | NM_004357.4 | 116.434145 | -8.4858685 | 1.31803317 | -6.4382814 |
| 4144 | NM_005662.6 | 101.034855 | -8.283418 | 1.28661638 | -6.4381412 |
| 4145 | NM_024041.3 | 89.8062198 | -8.1097736 | 1.25970809 | -6.4378197 |
| 4146 | NM_032999.3 | 205.559693 | -9.3058418 | 1.44548774 | -6.4378559 |
| 4147 | NM_005902.3 | 124.108169 | -8.5792112 | 1.33266968 | -6.4376127 |
| 4148 | NM_201442.3 | 89.3965863 | -8.1068492 | 1.2592953 | -6.4376078 |
| 4149 | NM_001017974.1 | 95.6207399 | -8.2031033 | 1.27454079 | -6.4361246 |
| 4150 | NM_018840.4 | 120.71883 | -8.537114 | 1.32645741 | -6.4360257 |
| 4151 | NM_002492.3 | 101.506563 | -8.2861806 | 1.28760318 | -6.4353527 |
| 4152 | NM_001195626.1 | 85.2077213 | -8.033686 | 1.24851041 | -6.4346167 |
| 4153 | NM_013392.3 | 90.7593546 | -8.1248351 | 1.26276277 | -6.4341738 |
| 4154 | NM_181521.2 | 124.340667 | -6.7760966 | 1.05324401 | -6.4335487 |
| 4155 | NM_005094.3 | 152.546843 | -8.8753996 | 1.37958071 | -6.4334037 |
| 4156 | NM_024666.4 | 92.7567242 | -8.1600624 | 1.26839375 | -6.4333827 |
| 4157 | NM_016205.2 | 131.098866 | -8.656228 | 1.34558364 | -6.4330658 |
| 4158 | NM_001256494.1 | 98.8173186 | -8.2489144 | 1.28238961 | -6.4324557 |
| 4159 | NM_003329.3 | 270.651971 | -6.6934073 | 1.04057121 | -6.4324356 |
| 4160 | NM_024529.4 | 84.6590456 | -8.0247285 | 1.24776584 | -6.4312776 |
| 4161 | NM_001145774.2 | 144.929424 | -8.7996882 | 1.36841588 | -6.4305657 |
| 4162 | NR_028103.1 | 99.2842007 | -8.2581617 | 1.28431992 | -6.429988 |
| 4163 | NM_006366.2 | 145.209112 | -8.8037479 | 1.3691921 | -6.4298851 |
| 4164 | NM_012229.4 | 102.049959 | -8.2973685 | 1.29058154 | -6.4291703 |
| 4165 | NM_022491.2 | 175.340576 | -8.1088496 | 1.2612669 | -6.4291306 |
| 4166 | NM_014498.4 | 121.962011 | -8.5539839 | 1.33067413 | -6.4283085 |
| 4167 | NM_017736.4 | 91.4582824 | -8.1366657 | 1.26582231 | -6.4279683 |
| 4168 | NM_004508.3 | 176.076015 | -9.0822045 | 1.41296729 | -6.4277528 |
| 4169 | NM_015069.4 | 141.152974 | -8.7627068 | 1.36336318 | -6.4272726 |
| 4170 | NM_024523.5 | 82.0764649 | -7.9818719 | 1.2419119 | -6.4270838 |

| | | | | | |
|---|---|---|---|---|---|
| 4171 | NM_032989.2 | 131.167044 | -8.6569491 | 1.34696072 | -6.4270242 |
| 4172 | NM_016240.2 | 108.327725 | -8.3815094 | 1.30417675 | -6.4266668 |
| 4173 | NM_013232.3 | 94.7928636 | -8.1863062 | 1.27381593 | -6.4266006 |
| 4174 | NM_001135821.1 | 132.248998 | -8.6693949 | 1.34902793 | -6.4264014 |
| 4175 | NM_001128431.2 | 96.9196682 | -8.2223985 | 1.27948968 | -6.4263109 |
| 4176 | NM_012304.4 | 72.2938627 | -7.7981714 | 1.21363509 | -6.4254664 |
| 4177 | NR_034076.1 | 88.215817 | -8.0848889 | 1.2583434 | -6.4250259 |
| 4178 | NM_007051.2 | 120.43188 | -8.534114 | 1.32829768 | -6.4248505 |
| 4179 | NM_001079673.1 | 117.113661 | -5.9875356 | 0.93200206 | -6.4243802 |
| 4180 | NM_001256798.1 | 99.5983481 | -8.2589671 | 1.28556741 | -6.424375 |
| 4181 | NM_001981.2 | 82.1358565 | -7.9844705 | 1.24285278 | -6.4243091 |
| 4182 | NM_000858.5 | 113.967496 | -8.4538182 | 1.31594013 | -6.4241663 |
| 4183 | NM_012079.5 | 135.735385 | -8.7065183 | 1.35528016 | -6.4241465 |
| 4184 | NM_032926.2 | 119.984104 | -8.5284798 | 1.327588 | -6.424041 |
| 4185 | NM_001733.4 | 92.3809119 | -8.1541796 | 1.26940197 | -6.4236387 |
| 4186 | NR_024281.1 | 149.276646 | -8.8437113 | 1.37682919 | -6.4232451 |
| 4187 | NM_022776.4 | 85.1610141 | -8.0349433 | 1.25094148 | -6.4231168 |
| 4188 | NM_001142864.3 | 190.07931 | -8.2267088 | 1.2809226 | -6.422487 |
| 4189 | NR_073503.1 | 119.110809 | -8.5174755 | 1.32620238 | -6.4224553 |
| 4190 | NM_022834.4 | 97.9673406 | -8.2363025 | 1.28245858 | -6.4222757 |
| 4191 | NM_001330574.1 | 131.319467 | -8.6584648 | 1.34824369 | -6.4220325 |
| 4192 | NM_014456.4 | 147.330061 | -8.8264832 | 1.37442346 | -6.4219532 |
| 4193 | NM_005780.2 | 162.626448 | -8.0035447 | 1.24634623 | -6.4216062 |
| 4194 | NM_000945.3 | 343.589953 | -6.8006132 | 1.05905977 | -6.4213686 |
| 4195 | NM_006263.3 | 97.7409902 | -8.2330774 | 1.28215333 | -6.4212893 |
| 4196 | NM_014453.3 | 108.336015 | -8.3811725 | 1.30524155 | -6.4211658 |
| 4197 | NM_001444.2 | 184.24972 | -9.1475388 | 1.42461316 | -6.4210686 |
| 4198 | NM_001423.2 | 99.3258939 | -8.2581442 | 1.28650578 | -6.4190494 |
| 4199 | NM_006348.3 | 106.759175 | -8.3596744 | 1.30233584 | -6.4189851 |
| 4200 | NM_001348129.1 | 114.524989 | -8.4614102 | 1.3182275 | -6.4187784 |
| 4201 | NM_006910.4 | 168.343434 | -7.2294038 | 1.12628804 | -6.4187877 |
| 4202 | NM_024537.3 | 109.665163 | -8.3989149 | 1.3085214 | -6.4186302 |
| 4203 | NM_024876.3 | 77.0663812 | -7.8931473 | 1.22972504 | -6.4186278 |
| 4204 | NM_001142674.1 | 88.000669 | -8.0810516 | 1.25902554 | -6.418497 |
| 4205 | NM_020750.2 | 222.415903 | -5.6783378 | 0.88468674 | -6.4184728 |
| 4206 | NM_004027.2 | 160.03155 | -8.9439758 | 1.39355545 | -6.4180982 |
| 4207 | NM_001242352.1 | 104.202373 | -8.3252245 | 1.29717713 | -6.417955 |
| 4208 | NM_007188.4 | 112.592239 | -8.4367169 | 1.31460032 | -6.4177049 |
| 4209 | NM_022757.4 | 82.8528079 | -7.9918235 | 1.24533613 | -6.4174027 |
| 4210 | NM_005166.4 | 1075.39861 | -8.1332827 | 1.26752944 | -6.416642 |
| 4211 | NR_001564.2 | 174.429911 | -9.0696294 | 1.41346099 | -6.416611 |
| 4212 | NM_015202.3 | 88.9735681 | -8.0991336 | 1.26222177 | -6.4165695 |
| 4213 | NM_152289.2 | 79.9242693 | -7.9429864 | 1.23789688 | -6.416517 |

| | | | | | |
|---|---|---|---|---|---|
| 4214 | NM_001321882.1 | 88.8497999 | -8.093539 | 1.2614436 | -6.4160926 |
| 4215 | NM_001285878.1 | 82.3864135 | -7.9842046 | 1.2444844 | -6.4156727 |
| 4216 | NR_034031.1 | 123.164664 | -8.5657234 | 1.33522004 | -6.4152149 |
| 4217 | NM_178167.3 | 176.939597 | -8.1246218 | 1.26651493 | -6.4149435 |
| 4218 | NM_001145437.1 | 155.674562 | -8.9047808 | 1.38822852 | -6.414492 |
| 4219 | NM_000178.3 | 107.254099 | -8.366898 | 1.30442393 | -6.4142475 |
| 4220 | NM_006453.2 | 121.911999 | -8.5517433 | 1.33337379 | -6.4136128 |
| 4221 | NM_003330.3 | 111.299244 | -8.4200582 | 1.31291606 | -6.4132494 |
| 4222 | NM_003078.3 | 141.076112 | -8.7625426 | 1.36634608 | -6.4131209 |
| 4223 | NM_138408.3 | 97.9671381 | -8.2363384 | 1.2842965 | -6.4131128 |
| 4224 | NM_013396.4 | 78.9314871 | -7.925557 | 1.23585171 | -6.4130324 |
| 4225 | NM_003363.3 | 100.497053 | -8.2732265 | 1.29009808 | -6.4128663 |
| 4226 | NM_006185.3 | 110.88067 | -8.4166731 | 1.31265399 | -6.411951 |
| 4227 | NM_015014.3 | 98.0988007 | -8.2376669 | 1.28483918 | -6.4114381 |
| 4228 | NM_005539.4 | 109.877025 | -8.4014524 | 1.31040109 | -6.4113594 |
| 4229 | NM_001128852.1 | 134.878205 | -8.6987572 | 1.35692834 | -6.4106239 |
| 4230 | NM_005687.4 | 158.997921 | -8.9346727 | 1.39377361 | -6.4104189 |
| 4231 | NM_001163474.1 | 85.2058838 | -8.0323239 | 1.25308901 | -6.4100186 |
| 4232 | NM_001170689.1 | 85.831767 | -8.0460535 | 1.25528917 | -6.4097212 |
| 4233 | NM_004719.2 | 189.479776 | -5.5409351 | 0.86451441 | -6.4093033 |
| 4234 | NM_015881.5 | 119.307746 | -8.5216503 | 1.32964509 | -6.4089661 |
| 4235 | NM_001199174.1 | 108.456426 | -8.3830625 | 1.30805718 | -6.408789 |
| 4236 | NM_015231.2 | 149.664814 | -4.9190549 | 0.76766904 | -6.4077807 |
| 4237 | NM_031492.3 | 93.2255053 | -8.1654878 | 1.27440275 | -6.4073056 |
| 4238 | NM_024117.3 | 107.845742 | -8.3754759 | 1.30722817 | -6.4070497 |
| 4239 | NM_148977.2 | 75.0714721 | -7.8494731 | 1.22531241 | -6.4060994 |
| 4240 | NM_004055.4 | 124.43125 | -8.5812045 | 1.33962332 | -6.4056846 |
| 4241 | NM_003165.3 | 185.267795 | -9.1555463 | 1.42944555 | -6.4049633 |
| 4242 | NM_032520.4 | 109.10342 | -8.3916702 | 1.31021679 | -6.4047952 |
| 4243 | NM_139048.2 | 74.9754929 | -7.8511244 | 1.22592196 | -6.4042612 |
| 4244 | NM_014397.5 | 101.564432 | -8.2889488 | 1.29429545 | -6.4042169 |
| 4245 | NM_002360.3 | 86.2663262 | -8.0541559 | 1.25768406 | -6.403958 |
| 4246 | NM_001291310.1 | 116.008716 | -8.4797717 | 1.32419008 | -6.4037421 |
| 4247 | NM_006451.4 | 253.251135 | -8.6423451 | 1.3495743 | -6.4037564 |
| 4248 | NM_001198934.1 | 99.7117417 | -8.262601 | 1.29043012 | -6.4029822 |
| 4249 | NM_001142806.1 | 140.977828 | -8.7628469 | 1.36868691 | -6.402375 |
| 4250 | NM_014267.5 | 566.339814 | -2.6190797 | 0.40907959 | -6.4023719 |
| 4251 | NM_004725.3 | 148.66948 | -8.8379408 | 1.38047126 | -6.4021187 |
| 4252 | NM_001145337.2 | 98.5915428 | -8.2427241 | 1.28752124 | -6.4020102 |
| 4253 | NM_001080394.3 | 92.3069813 | -8.1503959 | 1.27320559 | -6.4014767 |
| 4254 | NM_030969.4 | 104.911736 | -8.334501 | 1.30203799 | -6.4011196 |
| 4255 | NM_005839.3 | 111.750496 | -8.4282248 | 1.3169643 | -6.3997367 |
| 4256 | NM_014795.3 | 216.760809 | -9.3815738 | 1.46601081 | -6.3993892 |

| | | | | | |
|---|---|---|---|---|---|
| 4257 | NM_016258.2 | 112.938692 | -8.439577 | 1.31899936 | -6.3984694 |
| 4258 | NM_001320851.1 | 76.1755678 | -7.8724434 | 1.23054775 | -6.3975115 |
| 4259 | NM_015353.2 | 144.128432 | -8.7929717 | 1.37454196 | -6.3970195 |
| 4260 | NM_005872.2 | 120.838843 | -8.538693 | 1.33480102 | -6.3969782 |
| 4261 | NM_001281534.1 | 163.099072 | -8.9710235 | 1.40242294 | -6.3968032 |
| 4262 | NM_198563.2 | 98.887963 | -8.2501543 | 1.28972991 | -6.3968078 |
| 4263 | NM_004184.3 | 147.341826 | -8.8254368 | 1.37971582 | -6.3965613 |
| 4264 | NM_002747.3 | 84.2176408 | -8.0201723 | 1.25385415 | -6.3964156 |
| 4265 | NM_006624.5 | 115.297097 | -8.4705051 | 1.32426335 | -6.3963902 |
| 4266 | NM_022830.2 | 88.6373949 | -8.0914223 | 1.26502863 | -6.3962365 |
| 4267 | NM_001164161.1 | 109.087787 | -8.389799 | 1.31175227 | -6.3958716 |
| 4268 | NM_004414.6 | 184.300116 | -9.147666 | 1.43024698 | -6.3958645 |
| 4269 | NM_080725.2 | 87.347424 | -8.0724621 | 1.26212933 | -6.3959073 |
| 4270 | NM_002485.4 | 82.6346994 | -7.9899927 | 1.24925506 | -6.3958057 |
| 4271 | NM_032227.3 | 84.2356505 | -8.0194989 | 1.25397918 | -6.3952409 |
| 4272 | NM_001042498.2 | 75.3498718 | -7.8589879 | 1.22890306 | -6.3951243 |
| 4273 | NM_003670.2 | 110.047477 | -6.0381427 | 0.94418334 | -6.3950956 |
| 4274 | NM_006094.4 | 91.7662049 | -8.1414164 | 1.27310896 | -6.3949094 |
| 4275 | NM_001321166.1 | 71.700776 | -7.7863754 | 1.21762054 | -6.3947471 |
| 4276 | NM_004493.2 | 119.547203 | -8.5234735 | 1.33288446 | -6.3947579 |
| 4277 | NM_001134398.1 | 146.409272 | -8.816194 | 1.37871406 | -6.394505 |
| 4278 | NM_078471.3 | 95.685009 | -8.2038046 | 1.28300199 | -6.394226 |
| 4279 | NM_024324.4 | 104.715315 | -8.3318729 | 1.3030676 | -6.394045 |
| 4280 | NR_040016.1 | 76.03034 | -7.8703632 | 1.23098798 | -6.3935337 |
| 4281 | NM_030914.3 | 131.172961 | -8.6570748 | 1.35412583 | -6.3931096 |
| 4282 | NM_014603.2 | 143.345283 | -7.82189 | 1.22359248 | -6.3925613 |
| 4283 | NM_016231.4 | 120.3347 | -8.5329153 | 1.33497194 | -6.3918312 |
| 4284 | NM_016423.2 | 122.250458 | -8.555576 | 1.33862726 | -6.3913056 |
| 4285 | NM_014319.4 | 120.758936 | -8.5376217 | 1.33594409 | -6.3907028 |
| 4286 | NM_020457.2 | 101.495112 | -8.286774 | 1.2967654 | -6.3903417 |
| 4287 | NM_001321305.1 | 81.6927307 | -7.9775442 | 1.24839658 | -6.3902323 |
| 4288 | NM_001001555.2 | 132.962366 | -7.7114842 | 1.20698747 | -6.3890342 |
| 4289 | NM_206876.1 | 332.945757 | -4.6353933 | 0.72554924 | -6.388806 |
| 4290 | NM_003047.4 | 89.5624905 | -8.1075917 | 1.26908321 | -6.3885423 |
| 4291 | NM_001287135.1 | 82.6880458 | -7.9920502 | 1.25105478 | -6.3882496 |
| 4292 | NM_005047.3 | 81.5783705 | -7.9726597 | 1.24804183 | -6.388135 |
| 4293 | NR_003700.1 | 151.915397 | -5.565184 | 0.87117976 | -6.3881007 |
| 4294 | NM_001005336.2 | 126.586467 | -8.606058 | 1.34724862 | -6.3878766 |
| 4295 | NM_004870.3 | 120.398838 | -8.5339694 | 1.33603334 | -6.3875423 |
| 4296 | NM_001193511.1 | 100.404156 | -8.2736142 | 1.29533882 | -6.3872201 |
| 4297 | NM_004913.2 | 123.441386 | -8.569787 | 1.34174368 | -6.3870523 |
| 4298 | NM_001240.3 | 73.0615333 | -7.811228 | 1.22299816 | -6.3869499 |
| 4299 | NM_001040441.2 | 103.468629 | -8.3147083 | 1.30183564 | -6.3869109 |

| | | | | | |
|---|---|---|---|---|---|
| 4300 | NM_001184979.1 | 174.783573 | -9.071413 | 1.4203334 | -6.3868195 |
| 4301 | NR_038834.1 | 139.544315 | 3.30814847 | 0.51796763 | 6.3867861 |
| 4302 | NM_002752.4 | 104.51509 | -8.3292834 | 1.30425425 | -6.3862421 |
| 4303 | NM_152795.3 | 86.8713255 | -8.0624126 | 1.2625122 | -6.3860076 |
| 4304 | NM_006868.3 | 136.054144 | -8.7105241 | 1.36402824 | -6.3858825 |
| 4305 | NM_213569.2 | 77.2851416 | -7.8953488 | 1.23647364 | -6.3853758 |
| 4306 | NM_024072.3 | 101.538525 | -4.6803178 | 0.73300634 | -6.3850987 |
| 4307 | NM_020808.4 | 182.720496 | -7.3015295 | 1.14356989 | -6.3848564 |
| 4308 | NM_182487.3 | 82.4624499 | -7.9905129 | 1.25158243 | -6.3843282 |
| 4309 | NM_001330231.1 | 114.320542 | -8.4583239 | 1.32499579 | -6.383661 |
| 4310 | NM_007241.3 | 132.746854 | -8.6742268 | 1.35894258 | -6.3830709 |
| 4311 | NM_022340.3 | 81.8939936 | -7.9771967 | 1.24980584 | -6.3827488 |
| 4312 | NM_001145135.1 | 77.9323039 | -7.9054973 | 1.23859787 | -6.3826182 |
| 4313 | NM_001310.3 | 86.6157626 | -8.0576464 | 1.26244764 | -6.3825589 |
| 4314 | NM_016143.4 | 103.060391 | -8.3113728 | 1.30222015 | -6.3824637 |
| 4315 | NM_015963.5 | 106.673112 | -8.3586581 | 1.3096477 | -6.3823714 |
| 4316 | NM_001114183.1 | 353.535982 | -10.087753 | 1.58061546 | -6.3821677 |
| 4317 | NM_001100412.1 | 166.701429 | -8.0346371 | 1.2589273 | -6.3821295 |
| 4318 | NM_005884.3 | 74.4725385 | -7.8400154 | 1.22844115 | -6.3820847 |
| 4319 | NM_020385.3 | 103.738097 | -8.3181714 | 1.30351285 | -6.3813497 |
| 4320 | NM_001286577.1 | 76.7916067 | -7.8866992 | 1.23594557 | -6.3811056 |
| 4321 | NM_182767.5 | 118.651948 | -8.5124195 | 1.33402584 | -6.3810005 |
| 4322 | NM_000476.2 | 168.204502 | -9.016457 | 1.41305766 | -6.3808132 |
| 4323 | NM_020142.3 | 117.583675 | -8.4983237 | 1.33189629 | -6.3806196 |
| 4324 | NM_022553.5 | 84.3837632 | -8.0195699 | 1.25686545 | -6.3806113 |
| 4325 | NM_003034.3 | 115.617302 | -8.4751601 | 1.32864505 | -6.3787993 |
| 4326 | NM_001142416.1 | 89.5004752 | -8.1041383 | 1.27053325 | -6.3785331 |
| 4327 | NM_001323964.1 | 69.7375436 | -7.7456259 | 1.2144172 | -6.3780602 |
| 4328 | NM_003489.3 | 146.441788 | -7.8512946 | 1.23103176 | -6.3778165 |
| 4329 | NM_001330782.1 | 175.709126 | -9.0797999 | 1.42366964 | -6.3777436 |
| 4330 | NM_005530.2 | 94.2547323 | -8.1799302 | 1.28258609 | -6.3776851 |
| 4331 | NM_022767.3 | 115.57886 | -8.4744466 | 1.32876362 | -6.3776931 |
| 4332 | NM_199040.3 | 128.358862 | -8.6250615 | 1.35252843 | -6.3769909 |
| 4333 | NM_001048210.2 | 82.5352962 | -7.9875718 | 1.25259419 | -6.3768233 |
| 4334 | NM_012208.3 | 88.9507286 | -8.0975527 | 1.26995398 | -6.3762568 |
| 4335 | NM_182706.4 | 166.800112 | -9.0035383 | 1.41203581 | -6.3762818 |
| 4336 | NM_015622.5 | 124.655116 | -8.5842573 | 1.34644666 | -6.3754901 |
| 4337 | NM_001099666.1 | 131.851039 | -6.8368282 | 1.07241524 | -6.3751688 |
| 4338 | NM_176869.2 | 85.2057216 | -8.0342974 | 1.26032647 | -6.3747748 |
| 4339 | NM_194278.3 | 79.8153098 | -7.9411152 | 1.24575025 | -6.3745644 |
| 4340 | NM_032142.3 | 76.6243434 | -7.8811422 | 1.23641811 | -6.3741724 |
| 4341 | NM_006275.5 | 6314.243 | 4.52890648 | 0.71058731 | 6.37346939 |
| 4342 | NM_014374.3 | 96.8108084 | -8.2208558 | 1.28989278 | -6.3732862 |

| | | | | | |
|---|---|---|---|---|---|
| 4343 | NM_173510.3 | 81.9751361 | -7.9792432 | 1.25197575 | -6.3733209 |
| 4344 | NM_001015881.1 | 121.177005 | -8.5427716 | 1.34051175 | -6.3727689 |
| 4345 | NM_018394.3 | 96.2267032 | -8.2095981 | 1.28823439 | -6.3727518 |
| 4346 | NM_203390.2 | 82.6573137 | -7.9897977 | 1.25373391 | -6.3728018 |
| 4347 | NM_001206736.1 | 178.460974 | -9.1014679 | 1.42843328 | -6.3716437 |
| 4348 | NM_003168.2 | 113.267216 | -8.4459037 | 1.32562903 | -6.3712423 |
| 4349 | NM_018110.3 | 142.890206 | -8.781013 | 1.37843138 | -6.3702938 |
| 4350 | NR_036443.1 | 97.5382247 | -8.2303309 | 1.29200151 | -6.3702177 |
| 4351 | NM_133369.2 | 156.050312 | -8.9072424 | 1.39833284 | -6.3699014 |
| 4352 | NM_018171.3 | 92.6333638 | -8.1532212 | 1.28010716 | -6.3691708 |
| 4353 | NM_001024921.3 | 150.898819 | -8.8591748 | 1.39102518 | -6.3688098 |
| 4354 | NM_015922.2 | 115.354883 | -8.4725535 | 1.3304627 | -6.3681255 |
| 4355 | NM_018137.2 | 78.7845198 | -7.9231779 | 1.2441898 | -6.3681425 |
| 4356 | NM_001271618.1 | 134.053159 | -8.6884818 | 1.36437964 | -6.3680823 |
| 4357 | NM_016938.4 | 107.961041 | -8.3759987 | 1.31532347 | -6.3680144 |
| 4358 | NM_052850.3 | 133.154253 | -8.6789884 | 1.36303348 | -6.3674066 |
| 4359 | NM_198836.2 | 260.919433 | -9.6496828 | 1.51554978 | -6.367117 |
| 4360 | NM_032382.4 | 82.1488114 | -7.9835177 | 1.25388295 | -6.3670359 |
| 4361 | NM_201412.2 | 138.399858 | -8.734409 | 1.371811 | -6.3670645 |
| 4362 | NM_001271184.1 | 126.778672 | -8.6077808 | 1.35198582 | -6.3667685 |
| 4363 | NM_001008530.2 | 87.2396536 | -8.0718717 | 1.26790184 | -6.3663223 |
| 4364 | NM_138499.3 | 72.5602805 | -7.8047551 | 1.22593822 | -6.3663527 |
| 4365 | NM_032431.2 | 86.5678941 | -8.0582776 | 1.26582127 | -6.3660469 |
| 4366 | NM_018433.5 | 79.2508662 | -7.9274988 | 1.24529676 | -6.3659515 |
| 4367 | NM_001199839.1 | 92.4886665 | -8.1533173 | 1.28079095 | -6.3658455 |
| 4368 | NM_014235.4 | 117.252685 | -8.4952703 | 1.33454164 | -6.365684 |
| 4369 | NM_005667.3 | 96.0203072 | -8.207037 | 1.28929971 | -6.3654997 |
| 4370 | NM_001281454.1 | 338.513791 | -6.566454 | 1.03159931 | -6.3653144 |
| 4371 | NM_003690.4 | 121.3573 | -8.5454643 | 1.34258091 | -6.3649529 |
| 4372 | NR_026771.1 | 82.8924369 | 3.65262857 | 0.57386266 | 6.36498736 |
| 4373 | NM_001031623.2 | 75.2317494 | -7.8557088 | 1.23424278 | -6.3648003 |
| 4374 | NM_006703.3 | 215.64185 | -7.6018548 | 1.19436382 | -6.3647731 |
| 4375 | NM_015641.3 | 115.89513 | -8.4794691 | 1.33243704 | -6.3638798 |
| 4376 | NM_016207.3 | 100.032745 | -8.2652355 | 1.29876918 | -6.3638987 |
| 4377 | NM_016045.2 | 99.234654 | -8.2542948 | 1.2970882 | -6.3637112 |
| 4378 | NM_001282918.1 | 112.536061 | -8.4361134 | 1.32571163 | -6.3634604 |
| 4379 | NM_052937.3 | 76.1031197 | -7.8725126 | 1.2371465 | -6.3634441 |
| 4380 | NM_014806.4 | 129.965216 | -8.6442652 | 1.35849402 | -6.3631235 |
| 4381 | NM_199359.2 | 97.1597057 | -8.2243879 | 1.2925799 | -6.3627695 |
| 4382 | NM_006037.3 | 90.5525035 | -5.7397112 | 0.90210996 | -6.3625405 |
| 4383 | NM_000026.3 | 106.9759 | -8.3636205 | 1.3145283 | -6.3624499 |
| 4384 | NM_178815.3 | 80.7786186 | -7.9563702 | 1.25053391 | -6.3623786 |
| 4385 | NM_005102.2 | 83.0863022 | -7.9998155 | 1.25740654 | -6.3621551 |

| | | | | | |
|---|---|---|---|---|---|
| 4386 | NM_001146069.1 | 130.317063 | -8.6486379 | 1.35940359 | -6.3620826 |
| 4387 | NM_001755.2 | 87.0170717 | -8.065216 | 1.26781245 | -6.3615213 |
| 4388 | NM_001802.1 | 98.9146735 | -5.1625879 | 0.81160878 | -6.3609316 |
| 4389 | NM_005163.2 | 96.5637936 | -8.213763 | 1.29129511 | -6.3608721 |
| 4390 | NM_022835.2 | 422.041399 | -2.3943796 | 0.37642763 | -6.3607967 |
| 4391 | NM_004952.4 | 164.319285 | -8.9823789 | 1.41217362 | -6.360676 |
| 4392 | NM_001199455.1 | 85.065113 | -8.0323591 | 1.26286259 | -6.3604379 |
| 4393 | NM_001031702.3 | 122.101264 | -8.5538584 | 1.34491543 | -6.3601459 |
| 4394 | NM_015886.4 | 166.595543 | -9.0026552 | 1.41554637 | -6.3598448 |
| 4395 | NM_015534.5 | 91.5280292 | -8.1379414 | 1.27970033 | -6.3592555 |
| 4396 | NM_001130921.1 | 98.5486431 | -8.2465062 | 1.2967924 | -6.3591568 |
| 4397 | NM_018067.4 | 124.562188 | -8.58279 | 1.34967198 | -6.3591674 |
| 4398 | NM_024047.4 | 83.0865586 | -7.9989369 | 1.2578679 | -6.3591232 |
| 4399 | NM_006985.2 | 99.7747725 | -8.2625815 | 1.29949801 | -6.3582871 |
| 4400 | NM_024077.4 | 101.152631 | -8.2808974 | 1.30239341 | -6.3582151 |
| 4401 | NM_025015.2 | 209.891037 | -9.3358644 | 1.46833223 | -6.3581417 |
| 4402 | NM_001919.3 | 88.3275409 | -8.0864939 | 1.27184134 | -6.3580996 |
| 4403 | NM_014367.3 | 99.6062925 | -8.2589037 | 1.29900093 | -6.357889 |
| 4404 | NM_013272.3 | 82.8646963 | -7.9961343 | 1.25778339 | -6.3573222 |
| 4405 | NM_006734.3 | 169.514163 | -9.026722 | 1.41994056 | -6.3571126 |
| 4406 | NM_017865.3 | 108.082393 | -8.3771228 | 1.31775127 | -6.3571351 |
| 4407 | NM_213611.2 | 109.632023 | -8.3990428 | 1.32120553 | -6.3571054 |
| 4408 | NM_052834.2 | 100.367281 | -8.2719882 | 1.30127363 | -6.35684 |
| 4409 | NM_004664.3 | 117.167121 | -8.4929402 | 1.33605837 | -6.3567134 |
| 4410 | NM_172353.2 | 80.2012698 | -7.946463 | 1.25011466 | -6.3565873 |
| 4411 | NM_139126.3 | 79.8869676 | -7.9417706 | 1.24938825 | -6.3565274 |
| 4412 | NM_001079520.1 | 216.43512 | -9.379456 | 1.47562908 | -6.3562423 |
| 4413 | NM_005452.5 | 115.911577 | -8.4791859 | 1.33401583 | -6.3561359 |
| 4414 | NM_014963.2 | 79.3170245 | -7.9304699 | 1.24776681 | -6.3557308 |
| 4415 | NM_153827.4 | 78.1094398 | -7.9127857 | 1.24497835 | -6.3557617 |
| 4416 | NM_032353.3 | 109.1952 | -8.3929972 | 1.32057828 | -6.3555469 |
| 4417 | NM_033055.2 | 154.928542 | -6.3424288 | 0.9979652 | -6.3553607 |
| 4418 | NM_207311.2 | 168.392137 | -6.2867268 | 0.98924382 | -6.3550832 |
| 4419 | NM_004044.6 | 115.081514 | -8.4686126 | 1.33267183 | -6.3546121 |
| 4420 | NM_007056.2 | 72.2241889 | -7.7939095 | 1.22655644 | -6.3543015 |
| 4421 | NM_007348.3 | 193.170345 | -4.8761131 | 0.76745629 | -6.3536037 |
| 4422 | NM_012097.3 | 91.9424069 | -8.1433113 | 1.28172814 | -6.3533842 |
| 4423 | NM_001275.3 | 201.161982 | -9.2743392 | 1.45976769 | -6.353298 |
| 4424 | NR_033886.1 | 33.0773407 | 4.21203229 | 0.66297945 | 6.35318685 |
| 4425 | NM_024598.3 | 146.150286 | -8.8132038 | 1.38730179 | -6.3527661 |
| 4426 | NM_001304808.1 | 77.3811967 | -7.8954438 | 1.24286778 | -6.3526016 |
| 4427 | NM_001003712.1 | 99.0644747 | -8.2537914 | 1.29953717 | -6.3513316 |
| 4428 | NM_001278196.1 | 270.345279 | -9.7004892 | 1.52730953 | -6.3513578 |

| | | | | | |
|---|---|---|---|---|---|
| 4429 | NM_003374.2 | 107.658708 | -8.3727672 | 1.31832408 | -6.351069 |
| 4430 | NM_005010.4 | 151.976503 | -8.869769 | 1.3965836 | -6.3510476 |
| 4431 | NM_015103.2 | 305.55163 | -8.1200703 | 1.27852722 | -6.3511126 |
| 4432 | NM_183415.2 | 76.3827518 | -7.8749717 | 1.23994924 | -6.3510436 |
| 4433 | NM_152267.3 | 84.7508257 | -8.0276141 | 1.26399279 | -6.3509968 |
| 4434 | NM_004814.2 | 101.261309 | -8.2836815 | 1.30439049 | -6.3506147 |
| 4435 | NM_147189.2 | 122.060658 | -8.5529855 | 1.34680487 | -6.3505751 |
| 4436 | NM_005776.2 | 138.055418 | -8.7311244 | 1.37488836 | -6.3504243 |
| 4437 | NM_001113482.1 | 154.259798 | -8.8911897 | 1.40011615 | -6.3503229 |
| 4438 | NM_001079839.2 | 106.323054 | -8.354968 | 1.3158951 | -6.349266 |
| 4439 | NM_001121.3 | 80.6486043 | -7.9590475 | 1.25360446 | -6.3489304 |
| 4440 | NR_038170.1 | 120.132932 | -8.5306491 | 1.34363417 | -6.3489373 |
| 4441 | NM_203388.2 | 106.559684 | -8.3576991 | 1.31649121 | -6.3484655 |
| 4442 | NM_023080.2 | 195.808705 | -8.2685551 | 1.30246606 | -6.3483843 |
| 4443 | NM_002235.3 | 135.692689 | -8.705654 | 1.37138486 | -6.3480751 |
| 4444 | NM_015040.3 | 169.436263 | -4.2162154 | 0.66417045 | -6.3480925 |
| 4445 | NM_015833.3 | 95.9900078 | -8.2088983 | 1.29315286 | -6.3479721 |
| 4446 | NM_001025105.2 | 79.1847097 | -7.9264695 | 1.24870423 | -6.3477558 |
| 4447 | NM_006590.3 | 114.162405 | -8.4561518 | 1.33215212 | -6.3477374 |
| 4448 | NM_018416.2 | 100.392146 | -8.2706013 | 1.30293647 | -6.3476627 |
| 4449 | NM_015713.4 | 114.366527 | -8.4591565 | 1.33267861 | -6.3474843 |
| 4450 | NM_021727.4 | 104.850376 | -8.3346669 | 1.31306819 | -6.3474745 |
| 4451 | NM_000271.4 | 96.8461363 | -8.2198438 | 1.29502469 | -6.3472487 |
| 4452 | NM_001018111.2 | 116.096236 | -6.1190227 | 0.96407243 | -6.3470571 |
| 4453 | NM_000757.5 | 76.9932819 | -7.8917415 | 1.2433901 | -6.3469554 |
| 4454 | NM_001099272.1 | 74.1820172 | -7.8338696 | 1.23429506 | -6.346837 |
| 4455 | NM_015187.4 | 170.329149 | -9.0339854 | 1.42338341 | -6.346839 |
| 4456 | NM_001282412.1 | 108.678696 | -8.3857691 | 1.32130427 | -6.3465844 |
| 4457 | NM_032701.3 | 80.8328246 | -7.957782 | 1.25387665 | -6.3465429 |
| 4458 | NM_181491.2 | 82.9954239 | -7.9980291 | 1.26023056 | -6.3464808 |
| 4459 | NM_019112.3 | 162.3276 | -7.1792977 | 1.13124982 | -6.3463414 |
| 4460 | NM_001195478.1 | 94.4365287 | -8.1846613 | 1.28967389 | -6.3463031 |
| 4461 | NM_001080509.2 | 162.029641 | -5.450175 | 0.85884032 | -6.3459702 |
| 4462 | NM_001024844.1 | 14.3945347 | 8.77485921 | 1.38286161 | 6.34543556 |
| 4463 | NM_007108.3 | 176.847198 | -9.0884851 | 1.43233087 | -6.3452414 |
| 4464 | NM_015569.4 | 160.1004 | -8.9448182 | 1.40973858 | -6.3450191 |
| 4465 | NM_004871.2 | 114.054449 | -8.4546049 | 1.33249091 | -6.3449625 |
| 4466 | NM_006540.3 | 86.6572838 | -8.0574537 | 1.26993326 | -6.3447851 |
| 4467 | NM_001306144.1 | 90.864685 | -8.1273143 | 1.28095619 | -6.3447246 |
| 4468 | NM_024792.2 | 76.5020464 | -7.879101 | 1.24187645 | -6.3445128 |
| 4469 | NM_145257.4 | 200.149103 | -9.2665658 | 1.46083754 | -6.3433239 |
| 4470 | NM_182789.3 | 126.024796 | -8.59903 | 1.35576051 | -6.3425877 |
| 4471 | NM_001035507.2 | 86.250178 | -8.0543272 | 1.26993148 | -6.3423321 |

| | | | | | |
|---|---|---|---|---|---|
| 4472 | NM_002027.2 | 97.3912565 | -8.2268821 | 1.29736092 | -6.3412439 |
| 4473 | NM_032428.1 | 71.7047069 | -7.7861367 | 1.2279116 | -6.3409587 |
| 4474 | NM_001308096.1 | 104.636352 | -8.32986 | 1.31369588 | -6.3407826 |
| 4475 | NM_001130831.1 | 135.835532 | -8.7066638 | 1.37317194 | -6.3405488 |
| 4476 | NM_016072.4 | 118.519617 | -8.5107015 | 1.34232408 | -6.3402733 |
| 4477 | NM_001134479.1 | 111.444955 | -8.4220111 | 1.32848708 | -6.3395506 |
| 4478 | NM_173050.3 | 112.049759 | -8.4294595 | 1.32966783 | -6.3395228 |
| 4479 | NM_015310.3 | 120.785866 | -8.5372792 | 1.34685288 | -6.3386873 |
| 4480 | NM_001129994.1 | 100.245587 | -8.2705555 | 1.30483972 | -6.3383689 |
| 4481 | NM_212471.2 | 107.670953 | -8.3744331 | 1.32122253 | -6.3383972 |
| 4482 | NM_001749.3 | 120.465976 | -8.5342616 | 1.34646456 | -6.3382742 |
| 4483 | NM_030819.3 | 109.614524 | -8.398296 | 1.32505113 | -6.338092 |
| 4484 | NM_000113.2 | 114.061325 | -8.4560215 | 1.33416941 | -6.3380419 |
| 4485 | NM_018244.4 | 88.4838214 | -8.089487 | 1.27644594 | -6.3375085 |
| 4486 | NM_130434.4 | 86.2710155 | -8.0536965 | 1.27081558 | -6.3374235 |
| 4487 | NR_038109.1 | 78.6583152 | -7.9197108 | 1.24976936 | -6.3369379 |
| 4488 | NM_004996.3 | 177.036084 | -8.1258465 | 1.28264287 | -6.3352369 |
| 4489 | NM_001040661.1 | 87.1187004 | -8.0666624 | 1.27347344 | -6.3343782 |
| 4490 | NM_016238.2 | 123.121684 | -8.5655114 | 1.35222069 | -6.3344035 |
| 4491 | NM_205842.2 | 201.991255 | -9.2803904 | 1.46514214 | -6.3341229 |
| 4492 | NM_022158.3 | 90.6638218 | -8.1245937 | 1.28271335 | -6.3339122 |
| 4493 | NM_001002029.3 | 111.38718 | -8.4236503 | 1.32997012 | -6.333714 |
| 4494 | NM_025010.4 | 99.3517505 | -8.2570344 | 1.30366205 | -6.3337231 |
| 4495 | NM_001033581.1 | 131.820722 | -8.6637874 | 1.3679073 | -6.3336071 |
| 4496 | NM_021260.3 | 92.9996503 | -8.1615305 | 1.28874719 | -6.3329182 |
| 4497 | NM_024069.3 | 119.404678 | -8.5216784 | 1.34562406 | -6.332882 |
| 4498 | NM_002513.2 | 89.7503584 | -8.1088173 | 1.28044755 | -6.3327993 |
| 4499 | NM_015484.4 | 88.0301021 | -8.0813459 | 1.27623132 | -6.3321952 |
| 4500 | NM_001256126.1 | 277.135491 | -9.7364354 | 1.53762769 | -6.332115 |
| 4501 | NM_001079862.2 | 127.058839 | -8.6115123 | 1.36003926 | -6.3318115 |
| 4502 | NM_033414.2 | 108.325472 | -8.3815206 | 1.32374736 | -6.3316618 |
| 4503 | NM_001324100.1 | 19.7040798 | 9.22957213 | 1.45771439 | 6.33153669 |
| 4504 | NM_003111.4 | 84.0889578 | -8.0135247 | 1.26566711 | -6.3314632 |
| 4505 | NM_001330618.1 | 103.472143 | -8.3139934 | 1.3131346 | -6.3314099 |
| 4506 | NM_007254.3 | 84.1645753 | -8.0177085 | 1.26634972 | -6.3313541 |
| 4507 | NM_018120.5 | 126.064127 | -8.5998436 | 1.35837938 | -6.3309586 |
| 4508 | NM_032477.2 | 124.982653 | -8.5874635 | 1.35670522 | -6.3296458 |
| 4509 | NM_014940.3 | 84.123416 | -8.0171 | 1.26663772 | -6.3294341 |
| 4510 | NM_015490.3 | 84.5925744 | -8.024275 | 1.26781533 | -6.3292143 |
| 4511 | NM_032286.2 | 112.861516 | -8.4402011 | 1.33358794 | -6.3289423 |
| 4512 | NR_023915.1 | 121.508604 | -4.9955361 | 0.78939965 | -6.3282725 |
| 4513 | NM_032116.4 | 98.1563574 | -8.2406924 | 1.30229501 | -6.327823 |
| 4514 | NM_004964.2 | 139.793158 | -5.4411109 | 0.85994232 | -6.3272975 |

| | | | | | |
|---|---|---|---|---|---|
| 4515 | NM_005620.1 | 122.389006 | -8.5582641 | 1.35263176 | -6.3271205 |
| 4516 | NM_001080395.2 | 89.3127149 | -8.1020875 | 1.28066506 | -6.3264688 |
| 4517 | NM_001171617.1 | 83.2345709 | -8.0040254 | 1.26518964 | -6.3263444 |
| 4518 | NM_144985.3 | 117.432791 | -8.497359 | 1.34329393 | -6.325763 |
| 4519 | NR_046262.1 | 134.783371 | -3.779156 | 0.59745225 | -6.3254527 |
| 4520 | NM_018695.3 | 87.729699 | -8.0746673 | 1.27661269 | -6.3250721 |
| 4521 | NR_103804.1 | 87.1419754 | -8.0703862 | 1.27601499 | -6.3246798 |
| 4522 | NM_080594.3 | 228.200964 | -8.490185 | 1.34252067 | -6.3240628 |
| 4523 | NM_002968.2 | 131.462205 | -8.6601051 | 1.36954723 | -6.3233344 |
| 4524 | NM_003470.2 | 377.236749 | -6.3802748 | 1.00903031 | -6.3231745 |
| 4525 | NM_024040.2 | 207.150491 | -8.3523071 | 1.3209124 | -6.3231347 |
| 4526 | NM_005760.2 | 153.323585 | -7.9141377 | 1.25184445 | -6.3219817 |
| 4527 | NM_006384.3 | 76.6744664 | -7.8849499 | 1.24726473 | -6.3217934 |
| 4528 | NM_007284.3 | 106.748343 | -8.3598773 | 1.32265948 | -6.3205061 |
| 4529 | NM_016603.3 | 101.204283 | -8.2833144 | 1.31065387 | -6.3199862 |
| 4530 | NM_032012.3 | 343.851134 | -5.1613661 | 0.81671064 | -6.3196998 |
| 4531 | NM_003090.3 | 94.7906878 | -8.1881856 | 1.29584401 | -6.318805 |
| 4532 | NM_001079521.1 | 76.8710966 | -7.883791 | 1.24772486 | -6.3185333 |
| 4533 | NM_005198.4 | 76.588844 | -7.8816858 | 1.24741477 | -6.3184163 |
| 4534 | NM_001042601.2 | 69.0612439 | -7.7347845 | 1.22422929 | -6.3180848 |
| 4535 | NM_016494.3 | 109.299473 | -8.3941359 | 1.32862883 | -6.3178938 |
| 4536 | NM_001265612.1 | 96.8713558 | -22.335347 | 3.53533464 | -6.3177461 |
| 4537 | NM_014062.2 | 75.7303352 | -7.8636715 | 1.24469163 | -6.3177668 |
| 4538 | NM_053041.2 | 113.876823 | -8.4536396 | 1.33808497 | -6.317715 |
| 4539 | NM_153232.3 | 110.670348 | -8.4118273 | 1.33149859 | -6.3175639 |
| 4540 | NM_014408.4 | 111.211489 | -8.4191841 | 1.33274616 | -6.3171701 |
| 4541 | NM_013386.4 | 84.1148001 | -8.0161953 | 1.26905299 | -6.316675 |
| 4542 | NM_002488.4 | 116.468391 | -8.4855989 | 1.34337785 | -6.3166137 |
| 4543 | NM_005659.6 | 106.009968 | -8.3504 | 1.32199495 | -6.3165143 |
| 4544 | NM_023928.4 | 74.710418 | -7.8438158 | 1.24181397 | -6.3164178 |
| 4545 | NM_016468.6 | 115.00722 | -8.4672701 | 1.34054636 | -6.3162829 |
| 4546 | NM_014010.4 | 172.039538 | -9.0489732 | 1.43266139 | -6.3161982 |
| 4547 | NM_006233.4 | 101.954975 | -8.2940338 | 1.31341565 | -6.3148584 |
| 4548 | NM_022756.5 | 198.509752 | -9.2548162 | 1.46561148 | -6.314645 |
| 4549 | NM_000345.3 | 158.112019 | -8.9267098 | 1.41369849 | -6.3144368 |
| 4550 | NM_001040202.1 | 134.52031 | -8.6934884 | 1.37676931 | -6.3144118 |
| 4551 | NM_016545.4 | 170.068783 | -7.2492238 | 1.14803512 | -6.3144617 |
| 4552 | NM_001261434.1 | 112.516602 | -8.4359904 | 1.33601094 | -6.3143123 |
| 4553 | NM_002149.3 | 148.629789 | -8.8377131 | 1.39963131 | -6.3143151 |
| 4554 | NM_003977.3 | 124.086273 | -8.5772339 | 1.35855587 | -6.3134937 |
| 4555 | NM_001135944.1 | 124.032117 | -8.5765299 | 1.35860939 | -6.3127268 |
| 4556 | NM_004311.3 | 246.906595 | -6.3070338 | 0.99916144 | -6.3123271 |
| 4557 | NM_001085466.1 | 91.2949045 | -8.1347664 | 1.28873514 | -6.3122097 |

| | | | | | |
|---|---|---|---|---|---|
| 4558 | NM_020780.1 | 168.281468 | -9.016292 | 1.42840125 | -6.3121564 |
| 4559 | NM_020960.4 | 378.439923 | -6.3854603 | 1.01161315 | -6.3121562 |
| 4560 | NM_002647.3 | 73.4270245 | -7.8199759 | 1.23891274 | -6.3119666 |
| 4561 | NM_201998.2 | 73.0693149 | -7.8140871 | 1.23807666 | -6.3114728 |
| 4562 | NM_178234.2 | 156.052802 | -8.9077441 | 1.41139695 | -6.3112961 |
| 4563 | NM_003904.4 | 119.931508 | -8.5279588 | 1.35124139 | -6.311203 |
| 4564 | NM_020840.2 | 117.517171 | -8.4987116 | 1.34660915 | -6.311194 |
| 4565 | NM_006218.3 | 98.5251505 | -8.2437289 | 1.3062547 | -6.310966 |
| 4566 | NM_014251.2 | 75.6273896 | -7.8659027 | 1.24638421 | -6.3109775 |
| 4567 | NM_032289.2 | 131.472967 | -8.6605699 | 1.37243963 | -6.3103467 |
| 4568 | NM_012202.4 | 173.583778 | -9.0617435 | 1.43605623 | -6.3101592 |
| 4569 | NR_027410.1 | 258.148878 | -5.4902045 | 0.87007962 | -6.3100025 |
| 4570 | NM_000421.3 | 97.8639296 | -8.2350418 | 1.30514203 | -6.3096902 |
| 4571 | NM_001130056.3 | 104.881566 | -8.3347957 | 1.32096257 | -6.3096381 |
| 4572 | NM_006033.3 | 92.6940999 | -8.1563208 | 1.29268246 | -6.3096089 |
| 4573 | NM_007013.3 | 82.3409749 | -7.9861759 | 1.26573742 | -6.3095045 |
| 4574 | NM_005327.4 | 92.7005684 | -8.1566133 | 1.29276563 | -6.3094293 |
| 4575 | NM_001142421.1 | 92.8519177 | -8.1604439 | 1.29339289 | -6.309331 |
| 4576 | NM_001110781.2 | 518.272278 | -6.7087763 | 1.06338753 | -6.3088724 |
| 4577 | NM_001271978.1 | 101.920796 | -8.2943278 | 1.31476349 | -6.3086083 |
| 4578 | NM_021818.3 | 73.9601574 | -4.9636541 | 0.78680817 | -6.3085951 |
| 4579 | NM_014547.4 | 119.833395 | -6.6920254 | 1.06092277 | -6.3077404 |
| 4580 | NM_022121.4 | 77.8101747 | -7.9073184 | 1.25379649 | -6.3067001 |
| 4581 | NM_001173523.1 | 134.195391 | -8.690949 | 1.37807033 | -6.3066077 |
| 4582 | NM_014222.2 | 114.50392 | -8.461327 | 1.34167066 | -6.3065604 |
| 4583 | NM_007266.3 | 105.04717 | -8.3370094 | 1.3219872 | -6.3064221 |
| 4584 | NM_001199417.1 | 190.953273 | -7.2877323 | 1.15570375 | -6.3058828 |
| 4585 | NM_004209.5 | 154.442694 | -8.8929262 | 1.41032096 | -6.3056045 |
| 4586 | NR_039666.1 | 149.640912 | -8.8484311 | 1.4033368 | -6.3052798 |
| 4587 | NM_025224.3 | 89.9237507 | -8.1139606 | 1.28700554 | -6.3045265 |
| 4588 | NM_001098816.2 | 201.283387 | -7.4370531 | 1.17965682 | -6.3044209 |
| 4589 | NM_018427.4 | 131.601841 | -8.6619396 | 1.37414994 | -6.3034894 |
| 4590 | NM_020988.2 | 374.536685 | -9.2067578 | 1.46059194 | -6.3034429 |
| 4591 | NM_025233.6 | 90.9752281 | -8.1283715 | 1.28952573 | -6.3033806 |
| 4592 | NM_016010.2 | 127.355535 | -8.614544 | 1.36689621 | -6.3022664 |
| 4593 | NR_038241.1 | 82.6882696 | -7.9886213 | 1.2676008 | -6.3021586 |
| 4594 | NM_003337.3 | 126.815676 | -8.6081914 | 1.36606521 | -6.3014499 |
| 4595 | NM_001258437.1 | 92.1474728 | -8.1461873 | 1.292851 | -6.3009483 |
| 4596 | NM_001751.5 | 116.488977 | -8.4863744 | 1.34685026 | -6.3009042 |
| 4597 | NM_194326.3 | 112.107082 | -8.4309803 | 1.33805262 | -6.3009333 |
| 4598 | NM_014142.3 | 84.4098783 | -8.0205392 | 1.27299448 | -6.3005294 |
| 4599 | NM_018927.3 | 156.467721 | -8.9115721 | 1.41455676 | -6.2999042 |
| 4600 | NM_001198999.1 | 170.075046 | -9.0321675 | 1.43370688 | -6.2998704 |

| | | | | | |
|---|---|---|---|---|---|
| 4601 | NM_001453.2 | 98.5642532 | -8.2474162 | 1.30930131 | -6.2990972 |
| 4602 | NM_032508.3 | 94.7906375 | -8.1874511 | 1.29983579 | -6.298835 |
| 4603 | NM_000474.3 | 135.793908 | -8.7082235 | 1.38258173 | -6.2985235 |
| 4604 | NM_022456.4 | 77.4327118 | -7.8961473 | 1.25370819 | -6.2982339 |
| 4605 | NM_021115.4 | 160.849305 | -8.9514972 | 1.42129358 | -6.2981339 |
| 4606 | NM_173470.2 | 119.74562 | -8.5258186 | 1.35381601 | -6.2976199 |
| 4607 | NM_024015.4 | 127.419617 | -8.6160925 | 1.36820123 | -6.2973869 |
| 4608 | NM_201521.2 | 114.929975 | -8.4662771 | 1.34447395 | -6.2970927 |
| 4609 | NM_012143.3 | 100.69116 | -8.2745159 | 1.31403208 | -6.2970426 |
| 4610 | NM_000543.4 | 132.414111 | -8.6709722 | 1.37707093 | -6.2966779 |
| 4611 | NM_000122.1 | 150.191087 | -7.8846075 | 1.25235094 | -6.2958451 |
| 4612 | NM_002196.2 | 101.820775 | -8.2897668 | 1.31681603 | -6.2953113 |
| 4613 | NM_004483.4 | 127.947639 | -8.6213326 | 1.36948386 | -6.2953152 |
| 4614 | NM_022752.5 | 89.972997 | -8.1136772 | 1.28886164 | -6.2952275 |
| 4615 | NM_139202.2 | 199.45735 | -9.2622956 | 1.47131682 | -6.2952421 |
| 4616 | NM_006579.2 | 133.725932 | -8.6851957 | 1.379826 | -6.2944137 |
| 4617 | NM_005406.2 | 135.32951 | -7.7327342 | 1.22852957 | -6.2943004 |
| 4618 | NM_033624.2 | 78.6066853 | -7.9204183 | 1.25843522 | -6.2938625 |
| 4619 | NM_138396.5 | 136.579607 | -8.7155874 | 1.38483605 | -6.2935879 |
| 4620 | NM_001013649.3 | 91.3046387 | -5.7483693 | 0.91340592 | -6.2933349 |
| 4621 | NM_152341.4 | 90.251803 | -8.1172931 | 1.28992025 | -6.2928643 |
| 4622 | NM_001024924.1 | 107.204753 | -8.366231 | 1.32949647 | -6.2927816 |
| 4623 | NM_024544.2 | 96.4153202 | -8.2142245 | 1.30543414 | -6.2923316 |
| 4624 | NM_002956.2 | 151.88727 | -8.8681163 | 1.40948446 | -6.2917446 |
| 4625 | NM_003462.3 | 142.636034 | -8.7794266 | 1.39544054 | -6.2915089 |
| 4626 | NM_001272004.1 | 70.2520599 | -7.7568923 | 1.23292515 | -6.2914543 |
| 4627 | NM_001135564.1 | 81.7940484 | -7.9750658 | 1.2676225 | -6.2913571 |
| 4628 | NM_001142685.1 | 115.235401 | -8.4715679 | 1.34653413 | -6.2913874 |
| 4629 | NM_005534.3 | 117.360348 | -6.1263898 | 0.97387928 | -6.2907076 |
| 4630 | NM_032306.3 | 121.607293 | -8.5477615 | 1.35885723 | -6.2904044 |
| 4631 | NM_001042462.1 | 81.6934717 | -7.9735269 | 1.26758865 | -6.290311 |
| 4632 | NM_001286646.1 | 70.0539485 | -7.7557635 | 1.23297077 | -6.290306 |
| 4633 | NM_138799.3 | 110.958075 | -8.4154841 | 1.33806738 | -6.2892828 |
| 4634 | NM_001077183.2 | 107.35238 | -8.3671881 | 1.33043991 | -6.2890387 |
| 4635 | NM_025176.5 | 161.858597 | -6.3924113 | 1.01646406 | -6.2888709 |
| 4636 | NM_080687.2 | 111.806423 | -8.4268244 | 1.339985 | -6.2887453 |
| 4637 | NM_182811.1 | 134.264976 | -7.7230419 | 1.22809266 | -6.2886475 |
| 4638 | NR_027700.2 | 82.3861409 | -7.9877981 | 1.2701866 | -6.2886808 |
| 4639 | NM_031215.2 | 89.5610272 | -8.1061267 | 1.28916725 | -6.2878782 |
| 4640 | NM_001321324.1 | 85.5449993 | -8.0435989 | 1.27923323 | -6.2878283 |
| 4641 | NM_003917.4 | 70.9408345 | -7.7694929 | 1.23564726 | -6.287792 |
| 4642 | NM_002949.3 | 112.387141 | -8.4341498 | 1.3414113 | -6.2875196 |
| 4643 | NM_003942.2 | 68.5169584 | -7.7189435 | 1.22773391 | -6.2871469 |

| | | | | | |
|---|---|---|---|---|---|
| 4644 | NM_001144382.1 | 86.2495757 | -8.0507735 | 1.28055971 | -6.2869177 |
| 4645 | NM_001145845.1 | 80.7473844 | -7.9544522 | 1.26523961 | -6.2869137 |
| 4646 | NM_001319943.1 | 79.4895753 | -7.9352379 | 1.26219659 | -6.2868478 |
| 4647 | NM_030974.3 | 96.8353814 | -8.2191508 | 1.30745561 | -6.2863708 |
| 4648 | NM_005903.6 | 103.422359 | -8.3151447 | 1.32275949 | -6.2862106 |
| 4649 | NM_014904.2 | 76.5895582 | -7.8802574 | 1.25359173 | -6.2861435 |
| 4650 | NR_015353.1 | 72.9705619 | -7.8114255 | 1.24264107 | -6.2861479 |
| 4651 | NM_152926.2 | 126.963479 | -8.6119929 | 1.37000385 | -6.2861085 |
| 4652 | NM_001321694.1 | 132.777484 | -8.6743787 | 1.38000815 | -6.2857446 |
| 4653 | NM_001100427.1 | 98.6860792 | -8.2454465 | 1.31206795 | -6.2843136 |
| 4654 | NM_002751.6 | 160.446626 | -8.9478428 | 1.42404754 | -6.2833877 |
| 4655 | NM_031209.2 | 106.272225 | -8.3533896 | 1.32962709 | -6.2825056 |
| 4656 | NM_001998.2 | 89.2370951 | -8.1044606 | 1.29010307 | -6.2820256 |
| 4657 | NM_181866.2 | 161.405794 | -8.9569636 | 1.42580472 | -6.2820409 |
| 4658 | NR_033465.1 | 124.032622 | -8.5764671 | 1.36530252 | -6.2817339 |
| 4659 | NM_018357.3 | 92.5051752 | -8.1536812 | 1.29802066 | -6.2816267 |
| 4660 | NM_015934.4 | 187.26915 | -8.2062747 | 1.30648869 | -6.2811678 |
| 4661 | NM_022726.3 | 135.298701 | -8.7019433 | 1.3854027 | -6.2811652 |
| 4662 | NM_023034.1 | 140.342123 | -5.7030347 | 0.90800014 | -6.2808742 |
| 4663 | NM_001297736.1 | 30.7088201 | 9.86585126 | 1.5708045 | 6.28076331 |
| 4664 | NM_021228.2 | 476.249645 | -6.0837991 | 0.96879033 | -6.2797893 |
| 4665 | NM_006377.3 | 140.780079 | -7.7924703 | 1.2408912 | -6.279737 |
| 4666 | NM_001135812.1 | 140.580206 | -5.2044481 | 0.82878966 | -6.2795765 |
| 4667 | NM_001003801.1 | 109.765146 | -8.3990132 | 1.33756104 | -6.2793495 |
| 4668 | NM_006234.4 | 100.510572 | -8.2728168 | 1.31751278 | -6.2791169 |
| 4669 | NM_001124758.2 | 105.550882 | -8.3434788 | 1.32880726 | -6.2789233 |
| 4670 | NM_001294338.1 | 77.0237654 | -7.8892409 | 1.25648173 | -6.2788346 |
| 4671 | NM_001306215.1 | 101.021909 | -8.2799473 | 1.31872506 | -6.2787517 |
| 4672 | NM_001166109.1 | 143.834156 | -8.7910307 | 1.40034735 | -6.2777501 |
| 4673 | NM_001077415.2 | 109.198191 | -8.3926213 | 1.33692485 | -6.2775565 |
| 4674 | NM_017907.2 | 142.137928 | -8.7732598 | 1.39762194 | -6.2772768 |
| 4675 | NM_017802.3 | 84.7281662 | -8.0274156 | 1.27891913 | -6.2767187 |
| 4676 | NM_020418.3 | 113.285507 | -8.4462226 | 1.34570779 | -6.2764165 |
| 4677 | NM_001011713.2 | 95.042756 | -8.1917525 | 1.30517283 | -6.2763738 |
| 4678 | NM_000318.2 | 66.5308166 | -7.6764383 | 1.22307891 | -6.276323 |
| 4679 | NM_001018068.1 | 157.747738 | -8.9224786 | 1.42181 | -6.2754366 |
| 4680 | NM_024831.7 | 77.2325832 | -7.8929155 | 1.25778387 | -6.2752558 |
| 4681 | NM_001145076.2 | 118.993213 | -8.5169238 | 1.35735143 | -6.2746637 |
| 4682 | NM_022464.4 | 75.4264975 | -7.8603555 | 1.2527139 | -6.2746613 |
| 4683 | NM_020320.4 | 68.0656345 | -7.7078528 | 1.22842959 | -6.2745581 |
| 4684 | NM_177441.2 | 88.7806077 | -8.0944924 | 1.29022092 | -6.2737259 |
| 4685 | NM_001039703.5 | 100.716218 | -8.2746409 | 1.31899667 | -6.2734358 |
| 4686 | NM_005262.2 | 98.6056832 | -8.2456322 | 1.31439669 | -6.2733208 |

| | | | | | |
|---|---|---|---|---|---|
| 4687 | NM_014434.3 | 95.0133045 | -8.1942975 | 1.30625611 | -6.273117 |
| 4688 | NM_006836.1 | 415.358431 | -5.848696 | 0.93235916 | -6.2730075 |
| 4689 | NM_001114617.1 | 94.1703215 | -8.1806059 | 1.30421707 | -6.2724267 |
| 4690 | NM_022730.3 | 108.964776 | -8.3895215 | 1.33751886 | -6.272451 |
| 4691 | NM_001308142.1 | 71.8610504 | -7.787357 | 1.24161881 | -6.2719387 |
| 4692 | NM_001040402.2 | 97.7572669 | -8.2311778 | 1.31240017 | -6.2718506 |
| 4693 | NM_016033.2 | 80.6831748 | -7.9560635 | 1.26858197 | -6.2716196 |
| 4694 | NM_017660.3 | 92.241754 | -8.1475536 | 1.29913819 | -6.2715065 |
| 4695 | NM_138821.2 | 77.8997317 | -7.9090151 | 1.26113967 | -6.2713237 |
| 4696 | NM_003683.5 | 92.9388916 | -8.1604363 | 1.30126456 | -6.2711585 |
| 4697 | NM_004541.3 | 108.721634 | -8.3866925 | 1.33735334 | -6.2711119 |
| 4698 | NM_030593.2 | 105.343637 | -8.3415221 | 1.33015633 | -6.271084 |
| 4699 | NM_018144.3 | 124.761661 | -8.5849934 | 1.36906271 | -6.2707086 |
| 4700 | NM_194071.3 | 121.197651 | -6.1905518 | 0.98727405 | -6.2703479 |
| 4701 | NM_014772.2 | 249.832074 | -8.6233036 | 1.37527059 | -6.2702596 |
| 4702 | NM_014905.4 | 143.187146 | -8.7840273 | 1.40093388 | -6.2701227 |
| 4703 | NM_002982.3 | 122.884929 | -8.5639954 | 1.36595041 | -6.2696239 |
| 4704 | NM_052848.2 | 85.108202 | -8.0336983 | 1.28143135 | -6.2693162 |
| 4705 | NM_170692.2 | 78.6930405 | -7.9224497 | 1.26373491 | -6.2690756 |
| 4706 | NR_033227.1 | 100.222092 | -8.2682662 | 1.318896 | -6.2690813 |
| 4707 | NM_016083.4 | 203.78554 | -9.2928167 | 1.48242624 | -6.2686536 |
| 4708 | NM_015528.2 | 85.3598479 | -8.0354416 | 1.28186719 | -6.2685445 |
| 4709 | NM_001127229.1 | 85.9029206 | -8.0464503 | 1.28366666 | -6.2683331 |
| 4710 | NM_001320546.1 | 84.8987055 | -8.0288836 | 1.28089727 | -6.2681714 |
| 4711 | NM_005586.3 | 104.720326 | -8.3321262 | 1.32939169 | -6.2676232 |
| 4712 | NM_000016.5 | 81.8230551 | -7.9789552 | 1.27313022 | -6.2671949 |
| 4713 | NM_030792.6 | 132.646653 | -8.6737457 | 1.38401965 | -6.2670683 |
| 4714 | NM_033554.3 | 186.704079 | -9.167065 | 1.46280202 | -6.2667845 |
| 4715 | NM_015884.3 | 104.765257 | -8.3333167 | 1.32982713 | -6.2664662 |
| 4716 | NM_018138.4 | 64.2616282 | -7.6281513 | 1.21733865 | -6.2662524 |
| 4717 | NM_177968.3 | 86.289993 | -4.2370965 | 0.67618204 | -6.2662067 |
| 4718 | NM_020897.2 | 141.234356 | -8.7640881 | 1.39866176 | -6.2660526 |
| 4719 | NM_001033564.2 | 85.5937647 | -4.1456771 | 0.66163269 | -6.2658288 |
| 4720 | NM_022068.3 | 82.7293236 | -7.9948681 | 1.27596098 | -6.2657622 |
| 4721 | NM_020141.3 | 87.6545273 | -8.074581 | 1.28872456 | -6.26556 |
| 4722 | NM_006690.3 | 225.651332 | -6.7336114 | 1.07471055 | -6.2655116 |
| 4723 | NM_014281.4 | 130.951689 | -8.6532824 | 1.38126825 | -6.2647371 |
| 4724 | NM_021639.4 | 131.555536 | -7.6962272 | 1.22850458 | -6.2647118 |
| 4725 | NM_001282730.1 | 77.9262522 | -7.9055876 | 1.26201079 | -6.2642789 |
| 4726 | NM_020824.3 | 533.521115 | -7.0647932 | 1.12780184 | -6.264215 |
| 4727 | NM_003731.2 | 116.581066 | -8.4872216 | 1.35491719 | -6.264015 |
| 4728 | NM_001322119.1 | 173.549143 | -2.0718284 | 0.33077243 | -6.2636067 |
| 4729 | NM_138793.3 | 93.0986644 | -8.1619635 | 1.30311314 | -6.2634343 |

| | | | | | |
|---|---|---|---|---|---|
| 4730 | NM_001012732.1 | 78.0388305 | -7.9105071 | 1.26303486 | -6.2630948 |
| 4731 | NM_020718.3 | 88.4524858 | -8.0878918 | 1.29155323 | -6.2621436 |
| 4732 | NM_001135655.1 | 199.591092 | -9.2628448 | 1.47919122 | -6.262101 |
| 4733 | NM_001320044.1 | 120.664449 | -8.5365663 | 1.36326666 | -6.2618463 |
| 4734 | NM_001311162.1 | 168.228756 | -9.0165373 | 1.43998111 | -6.2615664 |
| 4735 | NM_001204298.1 | 189.189192 | -7.2701954 | 1.16115843 | -6.2611571 |
| 4736 | NM_013364.5 | 68.4473144 | -7.7220728 | 1.23337289 | -6.2609393 |
| 4737 | NM_001316772.1 | 153.759848 | -8.8869198 | 1.41946999 | -6.260731 |
| 4738 | NM_015099.3 | 109.017709 | -8.3905406 | 1.34018165 | -6.2607488 |
| 4739 | NM_016930.3 | 75.666347 | -7.8656703 | 1.25641983 | -6.2603837 |
| 4740 | NM_001206426.1 | 149.655084 | -8.8474066 | 1.41325185 | -6.2603184 |
| 4741 | NM_203331.2 | 132.424245 | -8.6719115 | 1.38535826 | -6.2596888 |
| 4742 | NM_005711.4 | 201.337935 | -9.2761189 | 1.48190513 | -6.2595902 |
| 4743 | NM_003309.3 | 572.54475 | -3.6454124 | 0.58240004 | -6.2592928 |
| 4744 | NM_005359.5 | 229.54791 | -5.9838777 | 0.95598992 | -6.2593523 |
| 4745 | NM_007202.3 | 67.105488 | -7.6909635 | 1.22871855 | -6.259337 |
| 4746 | NR_024384.1 | 117.251923 | -8.4948469 | 1.35716075 | -6.2592784 |
| 4747 | NM_001282507.1 | 112.92212 | -8.4399182 | 1.3484019 | -6.2592008 |
| 4748 | NM_153326.2 | 82.5535299 | -7.9915906 | 1.27685627 | -6.258802 |
| 4749 | NM_001281482.1 | 95.4383217 | -8.200085 | 1.31020225 | -6.2586407 |
| 4750 | NM_004603.3 | 146.050677 | -8.8123632 | 1.40804886 | -6.2585635 |
| 4751 | NM_005900.2 | 117.497512 | -8.4982865 | 1.35786432 | -6.2585682 |
| 4752 | NM_031454.1 | 97.2534629 | -8.2251478 | 1.31421085 | -6.2586212 |
| 4753 | NR_003615.2 | 68.5253301 | -7.7231607 | 1.234044 | -6.258416 |
| 4754 | NM_024017.4 | 173.846234 | -9.0642083 | 1.44838679 | -6.2581407 |
| 4755 | NM_001039690.3 | 85.7774015 | -8.0430839 | 1.28528552 | -6.2578188 |
| 4756 | NM_016613.6 | 130.080363 | -8.6458932 | 1.38161756 | -6.257805 |
| 4757 | NM_139280.3 | 85.9802875 | -8.0483356 | 1.28618015 | -6.2575492 |
| 4758 | NM_018969.5 | 190.618429 | -8.2297618 | 1.31523868 | -6.2572383 |
| 4759 | NM_000609.6 | 143.34906 | -8.7861871 | 1.40440902 | -6.2561454 |
| 4760 | NR_002578.2 | 375.446684 | -6.9396201 | 1.10932808 | -6.2556967 |
| 4761 | NM_014377.1 | 82.6142022 | -7.9901087 | 1.27727183 | -6.2556055 |
| 4762 | NM_006378.3 | 118.355642 | -8.5087376 | 1.36027254 | -6.2551712 |
| 4763 | NM_000098.2 | 69.4970303 | -7.7445466 | 1.2381159 | -6.2551063 |
| 4764 | NM_000416.2 | 84.7608875 | -8.0283316 | 1.28350142 | -6.2550236 |
| 4765 | NM_030576.3 | 136.593734 | -8.7155222 | 1.39340209 | -6.2548508 |
| 4766 | NM_018238.3 | 89.6653644 | -8.1083177 | 1.29637168 | -6.2546242 |
| 4767 | NM_178150.2 | 139.059186 | -7.776548 | 1.2433662 | -6.254431 |
| 4768 | NM_001277062.1 | 104.257963 | -8.3248795 | 1.33108461 | -6.2542076 |
| 4769 | NM_020690.5 | 78.5068762 | -7.9160849 | 1.26572532 | -6.2541886 |
| 4770 | NM_015156.3 | 106.339778 | -5.9435173 | 0.9503563 | -6.2539884 |
| 4771 | NM_017613.3 | 78.2003145 | -7.9098499 | 1.26477505 | -6.2539579 |
| 4772 | NM_015143.2 | 107.712565 | -8.3726103 | 1.33890293 | -6.2533363 |

| | | | | | |
|---|---|---|---|---|---|
| 4773 | NM_001131025.1 | 76.662087 | -7.8794978 | 1.26010141 | -6.2530664 |
| 4774 | NM_005002.4 | 116.894711 | -8.4914081 | 1.35795329 | -6.2530929 |
| 4775 | NR_026908.1 | 65.5877508 | 2.61619508 | 0.41844154 | 6.25223561 |
| 4776 | NM_004193.2 | 87.5393281 | -8.0769726 | 1.29189214 | -6.2520487 |
| 4777 | NM_001126.3 | 132.795734 | -4.9468517 | 0.79124608 | -6.2519762 |
| 4778 | NM_022061.3 | 80.8747037 | -7.9589598 | 1.27303266 | -6.2519683 |
| 4779 | NM_004041.4 | 126.035353 | -8.598783 | 1.37551567 | -6.2513159 |
| 4780 | NM_001003703.1 | 94.8503291 | -8.189531 | 1.31007523 | -6.2511914 |
| 4781 | NM_001318942.1 | 99.3372454 | -8.2563348 | 1.32080115 | -6.2510051 |
| 4782 | NM_031903.2 | 90.1010575 | -8.1150357 | 1.29822253 | -6.2508819 |
| 4783 | NM_017953.3 | 80.4606681 | -7.9526954 | 1.27230138 | -6.250638 |
| 4784 | NM_012425.3 | 139.098255 | -8.7418589 | 1.39857774 | -6.2505348 |
| 4785 | NM_012474.4 | 128.135305 | -8.6231202 | 1.37961288 | -6.2503912 |
| 4786 | NM_052920.1 | 151.688771 | -7.8986855 | 1.26370705 | -6.2504087 |
| 4787 | NM_134442.4 | 92.3814285 | -8.149928 | 1.30389421 | -6.2504519 |
| 4788 | NM_138761.3 | 100.09845 | -8.2675244 | 1.32283446 | -6.2498556 |
| 4789 | NM_005190.3 | 92.5025225 | -8.1524433 | 1.30445678 | -6.2496845 |
| 4790 | NM_015722.3 | 74.8690319 | -7.8503229 | 1.25611372 | -6.2496912 |
| 4791 | NR_036605.1 | 88.5394141 | -8.090632 | 1.29455878 | -6.2497216 |
| 4792 | NM_017482.3 | 168.234204 | -9.0161264 | 1.44267444 | -6.2495918 |
| 4793 | NM_001321365.1 | 104.205574 | -8.3241712 | 1.33199373 | -6.2494072 |
| 4794 | NM_003716.3 | 165.318011 | -8.9912748 | 1.43879906 | -6.2491525 |
| 4795 | NM_005049.2 | 81.5346618 | -7.9716186 | 1.27566302 | -6.2490003 |
| 4796 | NM_001301073.1 | 93.1780237 | -8.1640667 | 1.30649859 | -6.248814 |
| 4797 | NM_152527.4 | 118.628119 | -8.5117382 | 1.36216235 | -6.2486958 |
| 4798 | NM_152857.2 | 92.7589176 | -8.1572442 | 1.30546822 | -6.2485199 |
| 4799 | NM_004582.3 | 99.2526172 | -8.255247 | 1.32123711 | -6.2481192 |
| 4800 | NM_018077.2 | 72.2327804 | -7.7965775 | 1.24782597 | -6.2481289 |
| 4801 | NM_001037163.1 | 112.948285 | -8.4412236 | 1.35109525 | -6.2476895 |
| 4802 | NM_017750.3 | 75.5155619 | -7.8610941 | 1.25823635 | -6.2477087 |
| 4803 | NM_001080470.1 | 113.859042 | -8.4530227 | 1.35307996 | -6.2472455 |
| 4804 | NM_001136265.1 | 103.633877 | -8.3173292 | 1.33135738 | -6.2472551 |
| 4805 | NM_080862.2 | 132.218207 | -8.6691905 | 1.38768038 | -6.2472531 |
| 4806 | NM_016063.2 | 78.4994092 | -7.9175819 | 1.26745523 | -6.2468336 |
| 4807 | NM_007122.4 | 153.937517 | -8.8879023 | 1.42286501 | -6.2464831 |
| 4808 | NM_025069.2 | 267.773671 | -7.0500828 | 1.12866154 | -6.2464101 |
| 4809 | NM_001178030.1 | 75.5150521 | -7.8622824 | 1.25870828 | -6.2463102 |
| 4810 | NM_018405.3 | 81.8004228 | -7.9767714 | 1.27706783 | -6.2461611 |
| 4811 | NM_014850.3 | 96.7232361 | -8.217223 | 1.31559342 | -6.2460202 |
| 4812 | NM_001198916.1 | 77.3756013 | -7.8988271 | 1.26476726 | -6.2452811 |
| 4813 | NM_022488.4 | 104.780924 | -8.3332242 | 1.33432562 | -6.2452704 |
| 4814 | NM_138462.2 | 119.114882 | -8.5179304 | 1.36392392 | -6.2451653 |
| 4815 | NM_000238.3 | 127.373384 | -8.6151299 | 1.37951469 | -6.245044 |

| | | | | | |
|---|---|---|---|---|---|
| 4816 | NM_024102.3 | 105.245535 | -8.3399598 | 1.33547368 | -6.2449451 |
| 4817 | NM_145166.3 | 101.556905 | -5.5198432 | 0.88388971 | -6.2449457 |
| 4818 | NM_001042414.2 | 110.53773 | -8.4097513 | 1.34692601 | -6.2436624 |
| 4819 | NM_033138.3 | 90.6954218 | -8.1269301 | 1.30168376 | -6.2433983 |
| 4820 | NM_003491.3 | 119.400343 | -8.5220068 | 1.36513433 | -6.2426141 |
| 4821 | NM_015117.2 | 89.25744 | -8.1015525 | 1.29787797 | -6.2421527 |
| 4822 | NM_001159773.1 | 90.5515474 | -8.1248882 | 1.30168226 | -6.2418368 |
| 4823 | NM_001282533.1 | 217.081497 | -8.42014 | 1.34902607 | -6.2416436 |
| 4824 | NM_001002909.3 | 126.927043 | -8.6096789 | 1.37950793 | -6.2411232 |
| 4825 | NM_017945.3 | 108.551409 | -8.3841946 | 1.34338345 | -6.2411031 |
| 4826 | NM_031918.3 | 69.702638 | -7.7453739 | 1.24114167 | -6.2405236 |
| 4827 | NM_001005920.2 | 85.0725145 | -8.0304391 | 1.2868887 | -6.2401971 |
| 4828 | NM_002230.2 | 113.119514 | -8.4441047 | 1.3532303 | -6.2399613 |
| 4829 | NM_016097.4 | 104.261481 | -8.3254546 | 1.33428098 | -6.2396562 |
| 4830 | NM_003896.3 | 110.397611 | -8.4092176 | 1.34773057 | -6.2395392 |
| 4831 | NM_003214.3 | 85.435376 | -8.0404789 | 1.2886449 | -6.2394838 |
| 4832 | NM_003314.2 | 97.4498682 | -8.2290949 | 1.31893162 | -6.2392127 |
| 4833 | NM_001003796.1 | 200.144834 | -8.3005022 | 1.33039271 | -6.2391368 |
| 4834 | NM_032342.2 | 129.81856 | -8.6424386 | 1.38524536 | -6.2389226 |
| 4835 | NM_000876.3 | 519.896892 | -3.3688252 | 0.54004237 | -6.2380757 |
| 4836 | NM_000921.4 | 132.423019 | -8.6715132 | 1.3901484 | -6.2378327 |
| 4837 | NM_001289022.1 | 109.300108 | -8.3942264 | 1.34569582 | -6.2378335 |
| 4838 | NM_001243028.2 | 74.9855027 | -7.8506315 | 1.25863295 | -6.2374273 |
| 4839 | NM_015483.2 | 153.701458 | -7.9202772 | 1.2698116 | -6.237364 |
| 4840 | NM_016573.3 | 94.7242709 | -8.1875176 | 1.31267982 | -6.2372541 |
| 4841 | NM_001039697.1 | 151.532441 | -6.3004597 | 1.01015211 | -6.2371396 |
| 4842 | NM_014853.2 | 73.2850409 | -7.8165327 | 1.25322272 | -6.2371457 |
| 4843 | NM_004492.2 | 109.673599 | -8.3988132 | 1.3466037 | -6.2370341 |
| 4844 | NM_153223.3 | 74.0724864 | -7.8307964 | 1.25553301 | -6.2370295 |
| 4845 | NM_006806.4 | 74.7451896 | -7.8443159 | 1.25777461 | -6.2366626 |
| 4846 | NM_000390.3 | 69.7153573 | -7.7471108 | 1.24224469 | -6.2363807 |
| 4847 | NM_001258401.2 | 62.5727385 | 10.8847267 | 1.74542006 | 6.23616454 |
| 4848 | NM_007372.3 | 69.1123962 | -7.7317798 | 1.23985209 | -6.2360501 |
| 4849 | NM_198560.2 | 155.485802 | -8.9029511 | 1.42766784 | -6.2360101 |
| 4850 | NM_014384.2 | 82.5265359 | -7.9884254 | 1.28107252 | -6.2357324 |
| 4851 | NM_024113.4 | 97.4729413 | -8.2289955 | 1.31968109 | -6.235594 |
| 4852 | NM_012141.2 | 73.9640706 | -7.8291403 | 1.25568147 | -6.2349732 |
| 4853 | NM_000374.4 | 102.955493 | -8.3080847 | 1.33251591 | -6.2348859 |
| 4854 | NM_014426.3 | 70.6874018 | -7.7656042 | 1.24550667 | -6.2348957 |
| 4855 | NM_170674.4 | 167.529029 | -9.0109621 | 1.44540594 | -6.2342085 |
| 4856 | NM_001628.3 | 190.188701 | -8.2289613 | 1.32002879 | -6.2339256 |
| 4857 | NM_153757.3 | 135.589126 | -8.7050078 | 1.39655812 | -6.2331869 |
| 4858 | NM_152371.3 | 84.676655 | -8.0272454 | 1.28784897 | -6.2330643 |

| | | | | | |
|---|---|---|---|---|---|
| 4859 | NM_006074.4 | 74.4436538 | -7.8435344 | 1.25841551 | -6.2328653 |
| 4860 | NM_000819.4 | 112.336319 | -8.4341799 | 1.35323666 | -6.2325978 |
| 4861 | NM_001002251.2 | 82.0141424 | -7.9789937 | 1.28020661 | -6.2325828 |
| 4862 | NM_033100.3 | 84.9971558 | -8.0338025 | 1.2890044 | -6.232564 |
| 4863 | NR_109842.1 | 111.201477 | 3.09661779 | 0.49684066 | 6.23261753 |
| 4864 | NR_125376.1 | 97.8623487 | -8.2351763 | 1.32133469 | -6.2324681 |
| 4865 | NR_027039.1 | 30.1858856 | 4.91450497 | 0.78864998 | 6.23154134 |
| 4866 | NM_001171135.1 | 82.7554051 | -7.9918032 | 1.28255357 | -6.2311652 |
| 4867 | NM_001127511.2 | 87.0959168 | -8.0656792 | 1.29449651 | -6.2307461 |
| 4868 | NM_001322393.1 | 84.0209934 | -8.0168821 | 1.28666157 | -6.2307621 |
| 4869 | NM_014916.3 | 149.169257 | -7.0305681 | 1.12843132 | -6.2303908 |
| 4870 | NM_017645.4 | 86.3628815 | -8.0540785 | 1.29270903 | -6.2303878 |
| 4871 | NM_014699.3 | 78.2270845 | -7.9125641 | 1.27007761 | -6.2299847 |
| 4872 | NM_001042479.1 | 70.5210589 | -7.7591057 | 1.24557347 | -6.2293441 |
| 4873 | NM_001281501.1 | 136.522001 | -8.7158312 | 1.39927091 | -6.2288375 |
| 4874 | NM_001164094.1 | 91.6472014 | -8.1398845 | 1.30681859 | -6.2287792 |
| 4875 | NM_006118.3 | 110.133575 | -8.4046978 | 1.3493355 | -6.228768 |
| 4876 | NM_138300.3 | 139.409559 | -6.9558162 | 1.11679691 | -6.2283627 |
| 4877 | NM_001164817.1 | 109.280636 | -8.3920751 | 1.34744833 | -6.2281239 |
| 4878 | NM_005647.3 | 101.607865 | -8.2896087 | 1.33122639 | -6.2270466 |
| 4879 | NM_001001485.2 | 69.5212065 | -7.7404154 | 1.24312839 | -6.2265615 |
| 4880 | NM_002437.4 | 81.1750894 | -7.9645233 | 1.2791242 | -6.2265441 |
| 4881 | NM_152415.2 | 85.8650935 | -8.0446025 | 1.29203813 | -6.2262888 |
| 4882 | NM_005862.2 | 116.602548 | -6.1319708 | 0.98492923 | -6.2257984 |
| 4883 | NM_020245.4 | 110.04473 | -8.404414 | 1.34995661 | -6.2256919 |
| 4884 | NM_001145468.3 | 168.043017 | -9.015183 | 1.44809546 | -6.2255447 |
| 4885 | NM_025195.3 | 65.9281194 | -7.6683601 | 1.23182014 | -6.2252271 |
| 4886 | NM_001328682.1 | 210.550385 | -9.3399533 | 1.50036009 | -6.2251411 |
| 4887 | NM_001330211.1 | 89.9329255 | -8.1124229 | 1.30318898 | -6.2250549 |
| 4888 | NM_174938.5 | 124.366709 | -8.5806142 | 1.37843119 | -6.224913 |
| 4889 | NM_198398.1 | 179.952933 | -9.1132656 | 1.46401844 | -6.2248298 |
| 4890 | NM_024812.2 | 112.464767 | -8.4351412 | 1.35521086 | -6.2242279 |
| 4891 | NM_022762.4 | 92.2849127 | -8.1499808 | 1.30942567 | -6.2240881 |
| 4892 | NM_002499.3 | 115.073999 | -22.569775 | 3.62625026 | -6.2239981 |
| 4893 | NM_000153.3 | 121.930209 | -7.5873122 | 1.21916004 | -6.2233931 |
| 4894 | NM_001866.2 | 132.301784 | -8.6700132 | 1.39317431 | -6.2232078 |
| 4895 | NM_004655.3 | 88.3052308 | -8.0885282 | 1.29980891 | -6.2228595 |
| 4896 | NM_001184985.1 | 65.1524168 | -7.6485289 | 1.22913442 | -6.2226952 |
| 4897 | NM_001271287.1 | 90.2020559 | -8.1160276 | 1.30441498 | -6.2219675 |
| 4898 | NM_024604.2 | 81.0986687 | -7.9616004 | 1.27970419 | -6.2214381 |
| 4899 | NM_001040431.2 | 113.290021 | -8.4458922 | 1.35756559 | -6.2213511 |
| 4900 | NM_001303457.1 | 74.3431216 | -7.8376676 | 1.26017456 | -6.2195095 |
| 4901 | NM_014802.2 | 70.8424474 | -7.767196 | 1.24891013 | -6.2191793 |

| | | | | | |
|---|---|---|---|---|---|
| 4902 | NM_178511.5 | 70.5744956 | -7.7636175 | 1.24833716 | -6.2191672 |
| 4903 | NM_007357.2 | 66.1674498 | -7.6702248 | 1.23336159 | -6.2189587 |
| 4904 | NM_012080.4 | 72.5588896 | -7.8039777 | 1.2548959 | -6.2188248 |
| 4905 | NM_001172673.1 | 274.660942 | -8.759855 | 1.40864028 | -6.21866 |
| 4906 | NM_001018020.1 | 74.9874626 | -7.8504156 | 1.26240914 | -6.2185985 |
| 4907 | NM_015330.4 | 92.9168079 | -8.157298 | 1.31183498 | -6.2182348 |
| 4908 | NM_016505.3 | 103.652306 | -8.3172926 | 1.33757381 | -6.2181934 |
| 4909 | NM_017971.3 | 111.046309 | -8.4168914 | 1.35376852 | -6.2173786 |
| 4910 | NM_001303274.1 | 200.635179 | -9.2701554 | 1.49135545 | -6.2159262 |
| 4911 | NM_000393.4 | 955.168177 | -7.3633776 | 1.18460751 | -6.2158795 |
| 4912 | NM_001128226.2 | 75.362692 | -7.8567461 | 1.26407179 | -6.2154271 |
| 4913 | NM_012227.3 | 112.094172 | -8.4302262 | 1.3564042 | -6.2151283 |
| 4914 | NM_020416.3 | 139.658409 | -8.7474361 | 1.40749496 | -6.214897 |
| 4915 | NM_001144825.1 | 146.559341 | -8.8173447 | 1.41882063 | -6.2145591 |
| 4916 | NM_001278203.1 | 29.5974805 | 9.81662092 | 1.57969717 | 6.21424225 |
| 4917 | NM_014673.4 | 77.0704563 | -7.8925038 | 1.27012782 | -6.2139445 |
| 4918 | NM_138792.3 | 82.895211 | -7.9952555 | 1.2867468 | -6.2135422 |
| 4919 | NM_004301.4 | 84.6218896 | -8.0230021 | 1.29128528 | -6.213191 |
| 4920 | NM_152278.3 | 166.747945 | -9.0034814 | 1.44909125 | -6.2131915 |
| 4921 | NM_015989.4 | 87.7015625 | -8.0743645 | 1.29955892 | -6.2131577 |
| 4922 | NM_001040181.1 | 86.493168 | -8.0564637 | 1.29669571 | -6.2130719 |
| 4923 | NM_012142.4 | 81.0535072 | -7.965515 | 1.28217108 | -6.2125212 |
| 4924 | NM_001025159.2 | 184.836545 | -9.1525743 | 1.47331544 | -6.21223 |
| 4925 | NM_012200.3 | 107.33445 | -8.3675443 | 1.3469662 | -6.2121412 |
| 4926 | NM_012129.4 | 82.0524196 | -7.9778724 | 1.28426682 | -6.2120054 |
| 4927 | NM_138785.3 | 74.1314836 | -7.8352509 | 1.26131361 | -6.2119768 |
| 4928 | NM_018222.4 | 189.056054 | -3.167599 | 0.50992818 | -6.2118532 |
| 4929 | NR_026920.1 | 119.00168 | 6.45088117 | 1.0385331 | 6.21153162 |
| 4930 | NM_153633.2 | 134.467918 | -8.6935779 | 1.39964287 | -6.211283 |
| 4931 | NM_017814.2 | 78.3863864 | -7.9153246 | 1.27439862 | -6.2110273 |
| 4932 | NM_001282182.1 | 21.6128786 | 9.35391314 | 1.50602679 | 6.21098724 |
| 4933 | NM_014840.2 | 223.867693 | -8.4645462 | 1.36289394 | -6.2107153 |
| 4934 | NM_001304438.1 | 159.397762 | -8.9386252 | 1.43949525 | -6.2095551 |
| 4935 | NM_003239.4 | 77.0427353 | -7.893008 | 1.2711555 | -6.2093174 |
| 4936 | NM_017575.4 | 129.929204 | -7.6790298 | 1.23669969 | -6.2092923 |
| 4937 | NM_152359.2 | 70.6472049 | -7.7676878 | 1.25131247 | -6.2076324 |
| 4938 | NM_172351.2 | 83.6213426 | -8.005746 | 1.28973194 | -6.2072945 |
| 4939 | NM_022450.3 | 75.7493921 | -7.8678193 | 1.26757717 | -6.2069746 |
| 4940 | NM_001217.4 | 98.0531758 | -8.2367648 | 1.32713781 | -6.2064126 |
| 4941 | NR_028327.1 | 62.9671655 | -7.5988851 | 1.22436949 | -6.206366 |
| 4942 | NM_032110.2 | 98.3877084 | -8.2410061 | 1.32785106 | -6.2062729 |
| 4943 | NM_130475.2 | 142.133114 | -8.7731351 | 1.41375754 | -6.2055443 |
| 4944 | NM_080797.3 | 82.3557839 | -7.9872478 | 1.28717009 | -6.2052777 |

| | | | | | |
|---|---|---|---|---|---|
| 4945 | NM_001135694.2 | 237.504531 | -9.5138444 | 1.53328087 | -6.2048934 |
| 4946 | NM_014043.3 | 85.2479168 | -8.0348906 | 1.29502016 | -6.2044522 |
| 4947 | NM_182540.4 | 80.1800746 | -7.947061 | 1.28095721 | -6.2040019 |
| 4948 | NM_005618.3 | 96.2533943 | -8.2099386 | 1.32334236 | -6.2039415 |
| 4949 | NM_001007099.2 | 90.9208979 | -8.128286 | 1.31021818 | -6.2037652 |
| 4950 | NM_138333.4 | 60.8326399 | -7.5492941 | 1.21688322 | -6.2037951 |
| 4951 | NM_004645.2 | 79.9378995 | -7.9413328 | 1.28010253 | -6.2036694 |
| 4952 | NM_001286060.1 | 98.1636144 | -8.2376006 | 1.32797121 | -6.203147 |
| 4953 | NM_007269.3 | 64.5679076 | -7.6376072 | 1.23163561 | -6.2011907 |
| 4954 | NM_015472.4 | 92.015441 | -8.1477083 | 1.3139368 | -6.2009895 |
| 4955 | NM_001009996.2 | 72.345105 | -7.800772 | 1.25803632 | -6.2007526 |
| 4956 | NM_003642.3 | 76.0188472 | -7.8685377 | 1.26896508 | -6.200752 |
| 4957 | NM_133372.2 | 84.1677268 | -8.0162162 | 1.29277153 | -6.2007988 |
| 4958 | NM_006675.4 | 131.084279 | -8.6578722 | 1.39631494 | -6.2005153 |
| 4959 | NM_020197.2 | 66.4192526 | -7.677102 | 1.23815384 | -6.2004428 |
| 4960 | NM_173542.3 | 120.018592 | -8.5293972 | 1.37567507 | -6.200154 |
| 4961 | NM_022373.4 | 82.2154681 | -5.1304688 | 0.82753781 | -6.199679 |
| 4962 | NM_024092.2 | 102.826194 | -5.5312093 | 0.89218771 | -6.1996027 |
| 4963 | NR_110514.1 | 226.106205 | 2.39983251 | 0.38710798 | 6.19938781 |
| 4964 | NM_001142334.1 | 164.792047 | -8.9870427 | 1.44970434 | -6.1992246 |
| 4965 | NM_001244014.1 | 107.135484 | -8.3664683 | 1.34974688 | -6.1985461 |
| 4966 | NM_001101802.1 | 64.8814943 | -7.6437797 | 1.23321209 | -6.1982686 |
| 4967 | NM_001130158.2 | 111.277068 | -8.4197937 | 1.35843039 | -6.1981783 |
| 4968 | NM_001347217.1 | 116.912079 | -8.4910666 | 1.36996001 | -6.1980397 |
| 4969 | NM_025099.5 | 82.6046212 | -7.9888908 | 1.28894294 | -6.1980175 |
| 4970 | NM_001287060.1 | 79.6171331 | -7.9379378 | 1.28074979 | -6.1978834 |
| 4971 | NM_020943.2 | 62.0323467 | -7.5767582 | 1.22251425 | -6.1976849 |
| 4972 | NM_003664.4 | 87.685733 | -8.0791977 | 1.30368165 | -6.1972167 |
| 4973 | NM_016096.3 | 107.509902 | -8.3700211 | 1.35060719 | -6.1972283 |
| 4974 | NM_019095.5 | 86.3139908 | -8.0536275 | 1.29958479 | -6.1970774 |
| 4975 | NM_203284.2 | 76.8360885 | -7.8853393 | 1.27243471 | -6.1970483 |
| 4976 | NM_001322815.1 | 26.1757507 | 9.63922947 | 1.5555712 | 6.19658521 |
| 4977 | NM_005063.4 | 2707.33577 | -8.5188363 | 1.37478701 | -6.1964772 |
| 4978 | NM_153260.2 | 269.525934 | 1.82230903 | 0.29409726 | 6.19628012 |
| 4979 | NM_015885.3 | 79.5711488 | -7.9343458 | 1.28053111 | -6.1961367 |
| 4980 | NM_183394.2 | 100.057355 | -8.2668223 | 1.33421101 | -6.1960381 |
| 4981 | NM_001172665.1 | 85.7621724 | -8.0438129 | 1.29825922 | -6.195845 |
| 4982 | NM_013301.2 | 89.8572867 | -8.1107911 | 1.30916223 | -6.1954057 |
| 4983 | NM_001757.3 | 76.301453 | -7.8751974 | 1.27136107 | -6.1943043 |
| 4984 | NM_014173.3 | 108.415147 | -8.3832756 | 1.35341225 | -6.1941775 |
| 4985 | NM_025078.4 | 79.8029908 | -7.9399189 | 1.28187149 | -6.1940054 |
| 4986 | NM_144999.3 | 91.4210481 | -8.1357191 | 1.31352339 | -6.1938137 |
| 4987 | NM_024958.2 | 97.0165509 | -8.2206645 | 1.32733431 | -6.193364 |

| | | | | | |
|---|---|---|---|---|---|
| 4988 | NM_001031685.2 | 104.705029 | -8.3308653 | 1.34516678 | -6.1931839 |
| 4989 | NM_001025603.1 | 97.7784557 | -8.2333485 | 1.32950283 | -6.1928026 |
| 4990 | NM_001320704.1 | 78.4833624 | -7.9185386 | 1.27870592 | -6.192619 |
| 4991 | NM_001722.2 | 80.0129133 | -7.9450771 | 1.28298493 | -6.1926504 |
| 4992 | NM_014801.3 | 91.8426567 | -8.1431182 | 1.31503802 | -6.1923063 |
| 4993 | NM_001084.4 | 173.066184 | -7.1466974 | 1.15413131 | -6.1922741 |
| 4994 | NR_027856.1 | 71.258412 | -7.7777011 | 1.25607507 | -6.1920671 |
| 4995 | NM_014783.5 | 71.700375 | -7.7853481 | 1.25732164 | -6.19201 |
| 4996 | NM_001142353.2 | 154.986064 | -8.8976415 | 1.43710109 | -6.1913818 |
| 4997 | NM_001255.2 | 79.261024 | -7.9295093 | 1.2808379 | -6.1908765 |
| 4998 | NM_006129.4 | 109.680123 | -8.399783 | 1.35682738 | -6.1907529 |
| 4999 | NR_109905.1 | 20.7953882 | 9.31009817 | 1.5038657 | 6.19077764 |
| 5000 | NM_018288.3 | 63.5269711 | -7.6146129 | 1.23004377 | -6.1905219 |
| 5001 | NM_020921.3 | 102.31457 | -8.2984013 | 1.34070494 | -6.1895806 |
| 5002 | NM_001287520.1 | 69.2040266 | -7.732463 | 1.2492975 | -6.1894489 |
| 5003 | NM_032281.2 | 399.059258 | -7.5770278 | 1.22420679 | -6.1893365 |
| 5004 | NM_033087.3 | 66.3544709 | -7.6742242 | 1.23990359 | -6.1893717 |
| 5005 | NR_073148.1 | 71.4820522 | -7.782095 | 1.25733644 | -6.1893498 |
| 5006 | NM_003591.3 | 98.8492963 | -8.2493937 | 1.33291516 | -6.1889863 |
| 5007 | NM_003185.3 | 79.1905301 | -7.9284748 | 1.28107536 | -6.1889215 |
| 5008 | NM_001142749.2 | 66.7467604 | -7.6831045 | 1.24143639 | -6.1888829 |
| 5009 | NM_004099.5 | 62.9586984 | -7.5978554 | 1.22768346 | -6.1887739 |
| 5010 | NM_016041.4 | 79.3374033 | -7.9308417 | 1.28149404 | -6.1887464 |
| 5011 | NM_018346.2 | 94.8311466 | -8.1885452 | 1.32317184 | -6.1885728 |
| 5012 | NM_015946.4 | 83.2831973 | -8.0033558 | 1.29331245 | -6.1882616 |
| 5013 | NM_004554.4 | 68.177847 | -7.7155127 | 1.24682537 | -6.1881262 |
| 5014 | NM_001635.3 | 104.479924 | -8.3289261 | 1.34598819 | -6.1879638 |
| 5015 | NM_006856.2 | 80.258629 | -7.9486238 | 1.28460403 | -6.1876061 |
| 5016 | NM_004906.4 | 113.68383 | -8.4499592 | 1.36571148 | -6.1872213 |
| 5017 | NM_175921.5 | 94.553394 | -8.1848859 | 1.32290625 | -6.1870491 |
| 5018 | NM_001190.3 | 74.5869202 | -7.8446516 | 1.26793182 | -6.1869665 |
| 5019 | NM_014452.4 | 162.299669 | -8.0007925 | 1.29321921 | -6.1867257 |
| 5020 | NM_001173463.1 | 126.776889 | -8.6068006 | 1.39118653 | -6.1866619 |
| 5021 | NM_145295.3 | 93.3185358 | -8.1657095 | 1.31990346 | -6.186596 |
| 5022 | NM_006526.2 | 75.0689897 | -7.8532825 | 1.26945948 | -6.1863199 |
| 5023 | NM_175859.2 | 82.0042724 | -7.9812647 | 1.29014424 | -6.1863352 |
| 5024 | NM_033278.3 | 110.464217 | -8.4091986 | 1.35933326 | -6.1862671 |
| 5025 | NM_014886.4 | 129.752668 | -8.6415564 | 1.39696934 | -6.1859313 |
| 5026 | NM_018364.4 | 105.532545 | -8.3428508 | 1.34876699 | -6.185539 |
| 5027 | NM_001320852.1 | 88.9314762 | -8.0955001 | 1.30882255 | -6.1853305 |
| 5028 | NM_021178.4 | 107.031305 | -8.3636601 | 1.35219132 | -6.1852639 |
| 5029 | NM_004073.3 | 80.5587982 | -7.9548565 | 1.2862591 | -6.18449 |
| 5030 | NM_173546.2 | 153.664564 | -7.9219442 | 1.28093667 | -6.1844933 |

| | | | | | |
|---|---|---|---|---|---|
| 5031 | NM_024624.5 | 80.3546886 | -7.9481156 | 1.28519012 | -6.184389 |
| 5032 | NM_001114382.2 | 187.069556 | -9.1693388 | 1.4827175 | -6.1841441 |
| 5033 | NM_014206.3 | 113.558725 | -8.449566 | 1.36636322 | -6.1839823 |
| 5034 | NM_032283.2 | 102.525784 | -8.3012852 | 1.34241086 | -6.1838632 |
| 5035 | NM_020451.2 | 70.1889929 | -7.7529706 | 1.25377956 | -6.1836792 |
| 5036 | NM_001113490.1 | 75.4471149 | -7.8614712 | 1.27147933 | -6.1829328 |
| 5037 | NM_001122772.2 | 218.119215 | -9.3908846 | 1.51887052 | -6.1828079 |
| 5038 | NM_006335.2 | 162.301288 | -7.9970149 | 1.29343586 | -6.1827688 |
| 5039 | NM_001168241.1 | 153.046411 | -8.8799303 | 1.4362643 | -6.1826576 |
| 5040 | NM_144568.3 | 96.3541831 | -8.2123884 | 1.32841606 | -6.1820906 |
| 5041 | NM_018229.3 | 81.7351798 | -7.9744747 | 1.2899846 | -6.1818371 |
| 5042 | NM_005827.2 | 85.1890827 | -8.0347583 | 1.29981609 | -6.1814578 |
| 5043 | NM_001852.3 | 83.1619802 | -8.0009462 | 1.29443591 | -6.1810292 |
| 5044 | NM_001077195.1 | 80.5979533 | -7.9559305 | 1.28722507 | -6.1806833 |
| 5045 | NM_005702.3 | 109.431408 | -8.3958082 | 1.35839053 | -6.1807028 |
| 5046 | NM_014681.5 | 78.7775797 | -7.9215836 | 1.28165729 | -6.1807346 |
| 5047 | NM_021009.6 | 3906.42585 | -7.6434571 | 1.23673196 | -6.1803668 |
| 5048 | NM_173552.3 | 63.215743 | -7.6055607 | 1.2306474 | -6.1801298 |
| 5049 | NM_001127649.2 | 106.635577 | -8.3593638 | 1.3526738 | -6.1798815 |
| 5050 | NM_012412.4 | 92.8256477 | -8.1574783 | 1.32005707 | -6.1796407 |
| 5051 | NM_006938.3 | 131.158313 | -8.6567619 | 1.40092938 | -6.1792993 |
| 5052 | NM_199129.2 | 82.4258956 | -7.9878481 | 1.29275047 | -6.1789559 |
| 5053 | NM_025128.4 | 152.868534 | -6.5447042 | 1.05921405 | -6.1788306 |
| 5054 | NM_145021.5 | 67.7313112 | -7.7016752 | 1.24647871 | -6.1787459 |
| 5055 | NM_001320714.1 | 154.122846 | -8.889384 | 1.43874764 | -6.1785568 |
| 5056 | NM_001615.3 | 153.412468 | -8.8837002 | 1.43782375 | -6.1785738 |
| 5057 | NM_024948.3 | 88.5095787 | -8.0893195 | 1.30925106 | -6.1785854 |
| 5058 | NM_007039.3 | 79.1576647 | -7.9287271 | 1.283316 | -6.1783123 |
| 5059 | NM_198859.3 | 88.5585909 | -8.090635 | 1.30952706 | -6.1782877 |
| 5060 | NM_175061.3 | 87.2254589 | -8.0691939 | 1.30620198 | -6.1776004 |
| 5061 | NM_006782.3 | 85.196785 | -8.0345735 | 1.30062066 | -6.1774918 |
| 5062 | NM_022458.3 | 143.735186 | -6.9903298 | 1.13163422 | -6.177199 |
| 5063 | NM_014333.3 | 193.199693 | -9.2168784 | 1.49210245 | -6.1771083 |
| 5064 | NM_001111322.1 | 89.991084 | -8.1133065 | 1.3134697 | -6.1770031 |
| 5065 | NM_021189.4 | 182.771363 | -9.1365026 | 1.4791167 | -6.1769991 |
| 5066 | NM_001301268.1 | 71.105127 | -7.7731335 | 1.2584116 | -6.1769404 |
| 5067 | NM_014296.2 | 115.680288 | -7.5068055 | 1.21538764 | -6.1764702 |
| 5068 | NM_001080123.1 | 98.8749882 | -8.2505205 | 1.33591089 | -6.1759512 |
| 5069 | NM_183357.2 | 138.09869 | -8.7308896 | 1.41373808 | -6.1757476 |
| 5070 | NM_001256736.2 | 55.9417521 | 10.7197762 | 1.73589857 | 6.17534712 |
| 5071 | NM_006461.3 | 81.4979753 | -7.9702301 | 1.29068204 | -6.1752081 |
| 5072 | NM_201381.2 | 32.8067897 | 9.95649767 | 1.61238926 | 6.17499629 |
| 5073 | NM_001242809.1 | 73.8813003 | -7.8306245 | 1.26823553 | -6.1744245 |

| | | | | | |
|---|---|---|---|---|---|
| 5074 | NM_016437.2 | 77.3349898 | -7.8953069 | 1.27870946 | -6.1744338 |
| 5075 | NM_019070.4 | 117.275146 | -8.4961062 | 1.37601973 | -6.1744073 |
| 5076 | NM_032301.2 | 61.7689529 | -7.5730447 | 1.22659177 | -6.1740547 |
| 5077 | NM_006693.3 | 88.0563185 | -8.0810904 | 1.3089172 | -6.1738744 |
| 5078 | NM_022831.2 | 173.601588 | -8.0965497 | 1.3114443 | -6.1737656 |
| 5079 | NM_004998.3 | 67.6768087 | -7.7053047 | 1.2482602 | -6.1728354 |
| 5080 | NM_016733.2 | 121.640263 | -8.5493357 | 1.38501032 | -6.1727596 |
| 5081 | NM_023934.3 | 122.947956 | -8.5636532 | 1.38744909 | -6.1722288 |
| 5082 | NM_201379.2 | 189.453642 | -9.1888716 | 1.48883499 | -6.1718536 |
| 5083 | NM_133471.3 | 94.5578591 | -8.1845852 | 1.3261503 | -6.1716875 |
| 5084 | NM_014868.4 | 478.400288 | -6.0120458 | 0.97415812 | -6.1715298 |
| 5085 | NM_015935.4 | 84.257039 | -8.0183947 | 1.2993123 | -6.1712605 |
| 5086 | NM_001013845.1 | 80.438381 | -7.9517337 | 1.28857755 | -6.1709391 |
| 5087 | NM_005643.3 | 95.6744122 | -8.2022617 | 1.32917721 | -6.1709317 |
| 5088 | NM_138369.2 | 174.532725 | -8.1048499 | 1.31346483 | -6.1705877 |
| 5089 | NM_016286.3 | 112.255336 | -8.4325043 | 1.36658153 | -6.1705095 |
| 5090 | NM_001098785.1 | 177.758143 | -9.0956053 | 1.47408195 | -6.1703525 |
| 5091 | NM_001245005.1 | 90.1817829 | -8.1151112 | 1.3151833 | -6.1703272 |
| 5092 | NM_001080497.2 | 240.866025 | -4.0747589 | 0.66053098 | -6.1689142 |
| 5093 | NM_001256864.1 | 131.259732 | -8.659704 | 1.40384789 | -6.1685487 |
| 5094 | NM_002359.3 | 167.722432 | -8.0447718 | 1.30417185 | -6.1684906 |
| 5095 | NM_002803.3 | 210.154661 | -5.8537815 | 0.94900038 | -6.1683658 |
| 5096 | NM_032014.2 | 112.379965 | -8.4341117 | 1.36731703 | -6.1683659 |
| 5097 | NM_198426.2 | 88.1190993 | -8.0841046 | 1.31057848 | -6.1683483 |
| 5098 | NM_001142782.1 | 83.3096393 | -8.0016427 | 1.29723913 | -6.1682095 |
| 5099 | NM_001134381.1 | 107.904847 | -8.3758094 | 1.35794616 | -6.1679981 |
| 5100 | NM_207333.2 | 127.974933 | -3.0778559 | 0.49904868 | -6.1674463 |
| 5101 | NM_001135651.2 | 65.4132448 | -7.654142 | 1.24112458 | -6.1671021 |
| 5102 | NM_012250.5 | 82.7834527 | -7.9925599 | 1.2962265 | -6.1660211 |
| 5103 | NM_001142610.1 | 77.6629048 | -7.9010039 | 1.28138507 | -6.1659872 |
| 5104 | NM_005688.3 | 116.995143 | -7.5239599 | 1.22029317 | -6.1656986 |
| 5105 | NM_012151.3 | 67.3661798 | -7.6956509 | 1.24813915 | -6.1656994 |
| 5106 | NM_018145.2 | 81.1698721 | -7.9653005 | 1.29190157 | -6.1655629 |
| 5107 | NM_020841.4 | 166.808753 | -9.0035117 | 1.46034291 | -6.1653408 |
| 5108 | NM_005125.1 | 87.0713766 | -8.0660873 | 1.30849863 | -6.1643834 |
| 5109 | NM_016047.3 | 111.44615 | -8.421919 | 1.36624192 | -6.1642955 |
| 5110 | NM_005067.5 | 99.9931488 | -8.2656181 | 1.34092131 | -6.1641336 |
| 5111 | NM_024946.2 | 228.609609 | -6.1964175 | 1.00529563 | -6.1637764 |
| 5112 | NM_001278579.1 | 119.083185 | -8.5183449 | 1.38205633 | -6.1635295 |
| 5113 | NM_001284254.1 | 68.2738941 | -7.7188109 | 1.25234599 | -6.1634812 |
| 5114 | NM_001241.3 | 88.9635097 | -8.0961786 | 1.31361872 | -6.1632638 |
| 5115 | NM_012429.4 | 65.9427381 | -7.6689866 | 1.24430378 | -6.1632752 |
| 5116 | NM_022875.2 | 68.023749 | -7.7119761 | 1.25128952 | -6.1632228 |

| | | | | | |
|---|---|---|---|---|---|
| 5117 | NR_024542.1 | 97.9399014 | -8.2356185 | 1.33630053 | -6.1629987 |
| 5118 | NM_001042576.1 | 90.3532989 | -8.1220767 | 1.31790082 | -6.1628892 |
| 5119 | NM_024619.3 | 85.1901678 | -8.0340948 | 1.30364178 | -6.1628086 |
| 5120 | NR_003038.2 | 130.301069 | -8.6477694 | 1.40330897 | -6.162413 |
| 5121 | NM_002374.3 | 1008.33973 | -7.5667602 | 1.22812071 | -6.1612512 |
| 5122 | NM_005402.3 | 188.361573 | -8.2124531 | 1.33292873 | -6.1612095 |
| 5123 | NM_001498.3 | 103.175579 | -8.3121925 | 1.34925297 | -6.1605886 |
| 5124 | NM_015252.3 | 110.959307 | -8.4153961 | 1.36603631 | -6.1604484 |
| 5125 | NM_006706.3 | 97.8843428 | -8.2332094 | 1.33648123 | -6.1603629 |
| 5126 | NM_030780.4 | 74.2582736 | -7.8368451 | 1.2721485 | -6.1603226 |
| 5127 | NM_018714.2 | 105.992165 | -6.5224149 | 1.05879636 | -6.1602166 |
| 5128 | NM_023010.3 | 95.3199349 | -8.1959907 | 1.33051561 | -6.1600109 |
| 5129 | NM_018119.3 | 87.6877867 | -8.0762838 | 1.31114482 | -6.1597191 |
| 5130 | NM_001199378.1 | 76.5951418 | -22.009472 | 3.573361 | -6.1593194 |
| 5131 | NM_001130440.1 | 111.069523 | -8.4163716 | 1.36648204 | -6.1591527 |
| 5132 | NM_015448.2 | 59.617058 | -7.5187958 | 1.22082265 | -6.1587944 |
| 5133 | NM_017412.3 | 170.093243 | -9.0315466 | 1.46654192 | -6.1583964 |
| 5134 | NM_001009813.2 | 145.685644 | -8.8085598 | 1.4304704 | -6.1578064 |
| 5135 | NM_004145.3 | 201.510385 | -3.9509157 | 0.6416175 | -6.1577431 |
| 5136 | NM_001176.3 | 98.6532457 | -8.246508 | 1.33934963 | -6.1570988 |
| 5137 | NM_000386.3 | 158.354582 | -7.964154 | 1.29350405 | -6.1570383 |
| 5138 | NM_001005373.3 | 90.8569713 | -8.126514 | 1.3199352 | -6.1567522 |
| 5139 | NM_144607.5 | 68.3026299 | -7.7182425 | 1.25368596 | -6.1564401 |
| 5140 | NM_182705.2 | 84.3492739 | -4.5518462 | 0.73935993 | -6.1564686 |
| 5141 | NM_001675.4 | 68.2211657 | -7.7107729 | 1.25249918 | -6.1563097 |
| 5142 | NR_036570.1 | 74.5238954 | -7.8433156 | 1.27409038 | -6.156012 |
| 5143 | NM_024909.3 | 187.008633 | -9.16885 | 1.48959626 | -6.1552585 |
| 5144 | NM_001880.3 | 131.539858 | -8.6606213 | 1.40705616 | -6.1551355 |
| 5145 | NM_006699.3 | 161.455388 | -7.989019 | 1.29801439 | -6.1548 |
| 5146 | NM_007279.2 | 70.4662453 | -7.7598747 | 1.26095336 | -6.1539744 |
| 5147 | NM_001079818.2 | 83.8784446 | -8.0152886 | 1.30246862 | -6.1539206 |
| 5148 | NM_019027.3 | 174.804483 | 3.06951745 | 0.49884594 | 6.15323729 |
| 5149 | NM_001300847.1 | 88.9657647 | -8.0949895 | 1.31558687 | -6.1531395 |
| 5150 | NM_015061.3 | 62.0772788 | -7.5770106 | 1.23140304 | -6.1531524 |
| 5151 | NM_152735.3 | 69.6986617 | -7.7472222 | 1.25905667 | -6.1531958 |
| 5152 | NM_022662.3 | 127.589146 | -7.6523347 | 1.24368988 | -6.1529283 |
| 5153 | NM_133646.2 | 100.198205 | -5.7961638 | 0.94203499 | -6.1528115 |
| 5154 | NM_020946.1 | 114.394985 | -6.5958735 | 1.07201758 | -6.1527662 |
| 5155 | NM_001320803.1 | 64.0379454 | -7.6219065 | 1.23893964 | -6.1519595 |
| 5156 | NM_001331037.1 | 79.8696167 | -7.9429556 | 1.29127598 | -6.1512456 |
| 5157 | NM_024632.5 | 76.0616749 | -7.8694986 | 1.27937031 | -6.1510718 |
| 5158 | NM_018718.2 | 72.4089532 | -7.8001397 | 1.2681656 | -6.1507265 |
| 5159 | NM_003031.3 | 110.926621 | -8.4148809 | 1.36817496 | -6.1504421 |

| | | | | | |
|---|---|---|---|---|---|
| 5160 | NM_002249.5 | 64.8707454 | -7.6391094 | 1.24208475 | -6.1502321 |
| 5161 | NM_006122.3 | 95.6469688 | -8.2021455 | 1.33373545 | -6.1497545 |
| 5162 | NM_001321304.1 | 77.7494159 | -7.90538 | 1.28551491 | -6.1495825 |
| 5163 | NM_005131.2 | 70.1158612 | -7.7531488 | 1.26093675 | -6.1487214 |
| 5164 | NM_001127493.1 | 156.594098 | -8.9128287 | 1.44966354 | -6.1482051 |
| 5165 | NM_078469.2 | 88.8684902 | -8.0954495 | 1.31671179 | -6.1482319 |
| 5166 | NM_017822.3 | 96.7456082 | -8.2173051 | 1.33658383 | -6.1479908 |
| 5167 | NM_005504.6 | 133.255716 | -8.6796302 | 1.41181284 | -6.1478617 |
| 5168 | NM_001007225.2 | 82.2217276 | -7.9858819 | 1.29908171 | -6.1473284 |
| 5169 | NM_001256405.1 | 105.739252 | -5.5878027 | 0.90898294 | -6.1473131 |
| 5170 | NM_018262.3 | 88.5942132 | -8.0930237 | 1.3166147 | -6.1468429 |
| 5171 | NM_001017977.2 | 84.9935823 | -8.0304308 | 1.30649569 | -6.1465421 |
| 5172 | NM_015355.3 | 106.436028 | -8.3555603 | 1.35941315 | -6.1464465 |
| 5173 | NM_001172173.1 | 99.1996777 | -8.2540302 | 1.34292552 | -6.1463053 |
| 5174 | NM_031452.3 | 87.2994461 | -8.0702436 | 1.31306021 | -6.1461337 |
| 5175 | NM_004284.4 | 69.2288699 | -7.7379036 | 1.25908047 | -6.1456784 |
| 5176 | NM_002466.3 | 96.2757054 | -8.2109478 | 1.33622624 | -6.1448784 |
| 5177 | NM_015607.3 | 86.4452549 | -8.0578711 | 1.31133718 | -6.1447744 |
| 5178 | NM_139281.2 | 95.8255927 | -5.10959 | 0.83154469 | -6.1446967 |
| 5179 | NM_001330658.1 | 77.3780597 | -7.8964051 | 1.28509477 | -6.1446092 |
| 5180 | NM_000930.4 | 92.5308438 | -8.1543541 | 1.32716025 | -6.1442121 |
| 5181 | NM_003878.2 | 104.008863 | -8.3223431 | 1.35452105 | -6.1441224 |
| 5182 | NM_024836.2 | 134.652288 | -7.729519 | 1.25820829 | -6.1432746 |
| 5183 | NM_001161841.1 | 80.0090051 | -7.9472057 | 1.29368899 | -6.1430574 |
| 5184 | NM_005460.3 | 73.4406084 | -7.819255 | 1.27286854 | -6.1430185 |
| 5185 | NM_024329.5 | 126.936503 | -4.8551792 | 0.7903993 | -6.1426918 |
| 5186 | NM_001142424.1 | 86.081124 | -8.0495794 | 1.31050975 | -6.1423269 |
| 5187 | NM_006323.4 | 80.5271353 | -7.9506115 | 1.29446655 | -6.1419984 |
| 5188 | NM_144772.2 | 107.183865 | -8.3662537 | 1.3621583 | -6.14191 |
| 5189 | NM_001321149.1 | 85.8728099 | -8.0489213 | 1.31052317 | -6.1417619 |
| 5190 | NM_001661.3 | 115.36005 | -8.4712562 | 1.37929721 | -6.1417193 |
| 5191 | NM_003992.4 | 69.0104234 | -7.7312606 | 1.25881383 | -6.1417029 |
| 5192 | NM_017698.2 | 67.7833526 | -7.7075768 | 1.25494334 | -6.1417727 |
| 5193 | NM_198389.2 | 130.578292 | -8.6516723 | 1.40888097 | -6.1408113 |
| 5194 | NM_033318.4 | 118.296312 | -8.5081909 | 1.38566616 | -6.1401448 |
| 5195 | NM_032448.2 | 60.2919182 | -7.5383058 | 1.22777534 | -6.1398088 |
| 5196 | NM_004775.3 | 60.7152255 | -7.5479993 | 1.22938941 | -6.1396326 |
| 5197 | NM_002084.4 | 116.353409 | -8.485064 | 1.3820548 | -6.1394555 |
| 5198 | NM_000479.4 | 94.888176 | -8.1905853 | 1.33421749 | -6.1388682 |
| 5199 | NM_001324331.1 | 108.047658 | -8.3769727 | 1.36464154 | -6.1385884 |
| 5200 | NM_014175.3 | 96.0720916 | -8.2082889 | 1.33720097 | -6.1384109 |
| 5201 | NM_007204.4 | 73.3692834 | -7.8175786 | 1.27366049 | -6.1378826 |
| 5202 | NM_001130969.1 | 401.745123 | -10.272207 | 1.67359519 | -6.1378085 |

| | | | | | |
|---|---|---|---|---|---|
| 5203 | NM_001163.3 | 258.522094 | -8.6725405 | 1.41298644 | -6.137738 |
| 5204 | NM_004232.3 | 76.7790503 | -7.8851694 | 1.28483068 | -6.1371273 |
| 5205 | NM_001077665.2 | 74.0443189 | -7.8308311 | 1.27603133 | -6.1368643 |
| 5206 | NM_003887.2 | 115.492647 | -8.4752028 | 1.38107237 | -6.1366826 |
| 5207 | NM_001076683.1 | 155.612527 | -7.9372457 | 1.29362867 | -6.1356445 |
| 5208 | NM_000170.2 | 86.0004017 | -8.0476093 | 1.31173162 | -6.1351035 |
| 5209 | NM_020383.3 | 127.709473 | -8.6181679 | 1.40473277 | -6.1350942 |
| 5210 | NM_001018139.2 | 94.1962837 | -8.1804385 | 1.3334426 | -6.1348262 |
| 5211 | NM_000788.2 | 113.125124 | -8.44332 | 1.37634966 | -6.1345748 |
| 5212 | NM_018335.4 | 63.9315658 | -7.6181396 | 1.24187713 | -6.1343746 |
| 5213 | NM_032603.4 | 64.9809289 | -7.6460594 | 1.24642365 | -6.1343986 |
| 5214 | NM_003825.3 | 63.9630213 | -7.6244489 | 1.24306038 | -6.1336111 |
| 5215 | NM_144631.5 | 68.9484407 | -7.729396 | 1.26020502 | -6.1334433 |
| 5216 | NM_000367.4 | 62.3146253 | -7.5829618 | 1.2363591 | -6.1333005 |
| 5217 | NM_014953.4 | 104.3767 | -8.3278524 | 1.35788972 | -6.1329372 |
| 5218 | NM_032383.4 | 63.8655535 | -7.6208855 | 1.24273364 | -6.1323564 |
| 5219 | NM_024509.1 | 90.7190664 | -8.1253102 | 1.32501224 | -6.1322529 |
| 5220 | NM_031896.4 | 319.029138 | -7.2596655 | 1.18394528 | -6.1317576 |
| 5221 | NR_015389.1 | 147.285775 | -7.8560512 | 1.28124713 | -6.1315659 |
| 5222 | NM_001136263.1 | 81.7020727 | -7.9743194 | 1.30059465 | -6.1312872 |
| 5223 | NR_130156.1 | 64.95207 | -7.6426975 | 1.24661195 | -6.1307751 |
| 5224 | NM_013283.4 | 66.5151572 | -7.6758475 | 1.25209241 | -6.1304162 |
| 5225 | NM_001278640.1 | 119.340924 | -8.5208318 | 1.3900208 | -6.1300031 |
| 5226 | NM_005053.3 | 127.610468 | -8.6171916 | 1.40578863 | -6.1297918 |
| 5227 | NM_032172.2 | 109.654861 | -6.0416553 | 0.98562915 | -6.1297449 |
| 5228 | NM_001145306.1 | 74.764772 | -7.8448313 | 1.2798203 | -6.129635 |
| 5229 | NM_014688.3 | 71.2444047 | -7.7764214 | 1.26879234 | -6.1289946 |
| 5230 | NM_001323685.1 | 91.265387 | -8.1343343 | 1.32727642 | -6.128591 |
| 5231 | NM_013375.3 | 74.5995432 | -7.8418482 | 1.27957366 | -6.1284851 |
| 5232 | NM_006272.2 | 136.247194 | -8.7125822 | 1.42171596 | -6.1282158 |
| 5233 | NM_017638.2 | 59.2256772 | -7.5115128 | 1.22576348 | -6.1280279 |
| 5234 | NM_001012994.1 | 159.471644 | -5.4209859 | 0.88463814 | -6.1279134 |
| 5235 | NM_001042522.2 | 94.7906005 | -8.1891268 | 1.33650491 | -6.1272703 |
| 5236 | NM_173852.3 | 92.3849957 | -8.152166 | 1.33052783 | -6.1270165 |
| 5237 | NM_000270.3 | 92.9483753 | -8.1610105 | 1.3320013 | -6.1268788 |
| 5238 | NM_001286611.1 | 119.451345 | -8.5226842 | 1.39105807 | -6.1267638 |
| 5239 | NM_006400.4 | 147.208916 | -8.8235168 | 1.44015683 | -6.126775 |
| 5240 | NM_001127320.2 | 161.212566 | -7.9874051 | 1.30370282 | -6.1267069 |
| 5241 | NM_015921.2 | 85.9192519 | -8.0477784 | 1.31359236 | -6.1265417 |
| 5242 | NM_002063.3 | 152.741471 | -8.877158 | 1.44903502 | -6.126255 |
| 5243 | NM_014655.3 | 111.571968 | -8.4240533 | 1.37507528 | -6.1262488 |
| 5244 | NM_001282757.1 | 66.0290476 | -7.6662577 | 1.25143713 | -6.1259631 |
| 5245 | NM_020961.3 | 68.0228951 | -7.7086221 | 1.25840265 | -6.1257199 |

| | | | | | |
|---|---|---|---|---|---|
| 5246 | NM_003798.3 | 61.9372714 | -7.5778701 | 1.23718676 | -6.1250817 |
| 5247 | NM_004287.4 | 87.8162926 | -8.0796319 | 1.31915101 | -6.1248726 |
| 5248 | NM_002532.5 | 86.3373944 | -8.0530702 | 1.31492526 | -6.1243558 |
| 5249 | NM_018049.2 | 181.375272 | -6.1005916 | 0.99612462 | -6.1243257 |
| 5250 | NM_005385.3 | 319.958418 | -5.2277804 | 0.85365398 | -6.124004 |
| 5251 | NM_015434.3 | 76.302826 | -7.8739285 | 1.28580302 | -6.123744 |
| 5252 | NM_001130142.1 | 65.1468761 | -7.6475064 | 1.24884297 | -6.1236733 |
| 5253 | NM_032822.2 | 71.9139567 | -7.790178 | 1.2721486 | -6.1236384 |
| 5254 | NM_001037172.2 | 116.429984 | -8.4852008 | 1.38565823 | -6.1235885 |
| 5255 | NM_153321.2 | 138.763983 | -8.7389092 | 1.42709261 | -6.1235754 |
| 5256 | NM_002813.6 | 89.1961365 | -8.1015623 | 1.32309487 | -6.1231908 |
| 5257 | NM_001270454.1 | 68.9086628 | -7.7299152 | 1.2624301 | -6.1230441 |
| 5258 | NM_030806.3 | 172.742836 | -7.2793447 | 1.18884529 | -6.1230378 |
| 5259 | NM_024686.4 | 58.6143209 | -7.4940994 | 1.22403551 | -6.1224526 |
| 5260 | NM_001305626.1 | 63.5487732 | -7.6155976 | 1.24389904 | -6.1223599 |
| 5261 | NM_003153.4 | 84.616234 | -8.0267763 | 1.31106889 | -6.1223147 |
| 5262 | NM_022494.2 | 76.8348542 | -7.8857085 | 1.28821492 | -6.1214231 |
| 5263 | NM_001160103.1 | 60.6334623 | -7.5470968 | 1.23299881 | -6.1209279 |
| 5264 | NM_001300846.1 | 82.9345417 | -7.9961535 | 1.30636119 | -6.1209362 |
| 5265 | NM_018113.3 | 102.259906 | -8.2979084 | 1.35571308 | -6.1206966 |
| 5266 | NM_173566.2 | 118.06852 | -7.5357507 | 1.2312237 | -6.1205374 |
| 5267 | NM_032169.4 | 77.6416539 | -7.9018681 | 1.29106621 | -6.1204206 |
| 5268 | NM_001308319.1 | 103.813969 | -8.3210351 | 1.35959255 | -6.1202418 |
| 5269 | NM_000495.4 | 122.39643 | -8.5580991 | 1.39833835 | -6.1201919 |
| 5270 | NM_001307943.1 | 63.4607724 | -7.608944 | 1.24326127 | -6.1201488 |
| 5271 | NM_001270965.1 | 133.094641 | -8.6769174 | 1.41782321 | -6.1198867 |
| 5272 | NM_001301191.1 | 140.72131 | -8.7583078 | 1.43112804 | -6.1198632 |
| 5273 | NM_024095.4 | 81.817394 | -7.9755227 | 1.30325454 | -6.1196968 |
| 5274 | NM_033112.3 | 93.7885418 | -8.1736337 | 1.33562187 | -6.1197214 |
| 5275 | NM_001127671.1 | 164.216206 | -8.0161503 | 1.30990137 | -6.1196594 |
| 5276 | NM_001042573.2 | 125.153401 | -7.6212329 | 1.24537672 | -6.1196205 |
| 5277 | NM_001134225.1 | 67.9046837 | -7.708952 | 1.2597434 | -6.1194621 |
| 5278 | NM_152334.2 | 71.9372037 | -7.7890305 | 1.27294751 | -6.1188937 |
| 5279 | NM_025158.4 | 72.5794147 | -7.8007263 | 1.27489833 | -6.1187047 |
| 5280 | NM_001330542.1 | 77.8239043 | -7.90277 | 1.29177577 | -6.1177568 |
| 5281 | NM_006554.4 | 110.552851 | -8.4102592 | 1.37471644 | -6.1178138 |
| 5282 | NM_018192.3 | 84.193552 | -8.0197499 | 1.31089318 | -6.1177753 |
| 5283 | NM_018151.4 | 86.5569821 | -8.0550459 | 1.31671262 | -6.1175429 |
| 5284 | NM_001100588.1 | 77.7794296 | -7.9031122 | 1.29193209 | -6.1172814 |
| 5285 | NM_181349.2 | 66.5440097 | -7.6755189 | 1.25474777 | -6.1171807 |
| 5286 | NM_021132.3 | 136.346454 | -8.7131038 | 1.424395 | -6.1170559 |
| 5287 | NM_003916.4 | 127.453728 | -6.8269491 | 1.11606091 | -6.1170041 |
| 5288 | NM_001199097.1 | 172.745084 | -9.0553696 | 1.48037524 | -6.1169421 |

| | | | | | |
|---|---|---|---|---|---|
| 5289 | NM_021934.4 | 88.7614411 | -8.0935715 | 1.32317986 | -6.1167584 |
| 5290 | NM_001271789.1 | 127.981886 | -8.6228643 | 1.40982072 | -6.1162842 |
| 5291 | NM_031432.3 | 95.1147272 | -8.1926297 | 1.33952597 | -6.1160663 |
| 5292 | NR_045058.1 | 83.9406405 | -8.0116416 | 1.31005615 | -6.1154948 |
| 5293 | NM_018838.4 | 104.622758 | -8.3307332 | 1.36230884 | -6.1151575 |
| 5294 | NM_152367.2 | 422.495899 | 2.94546789 | 0.48166706 | 6.11515325 |
| 5295 | NM_203372.1 | 212.85898 | -8.3915206 | 1.37226387 | -6.1150926 |
| 5296 | NM_182640.2 | 68.8270349 | -7.7279332 | 1.26377113 | -6.1149784 |
| 5297 | NM_001271044.2 | 185.000693 | -9.1535316 | 1.49693039 | -6.1148679 |
| 5298 | NM_014049.4 | 69.1394398 | -7.7343849 | 1.26486758 | -6.1147783 |
| 5299 | NM_018400.3 | 136.566381 | -8.715345 | 1.42534204 | -6.1145639 |
| 5300 | NM_133476.4 | 77.1907097 | -7.8903289 | 1.29044716 | -6.1144145 |
| 5301 | NM_012090.5 | 659.009295 | -5.1249687 | 0.83819832 | -6.1142674 |
| 5302 | NM_001134375.1 | 70.7548299 | -7.7649785 | 1.27001987 | -6.1140607 |
| 5303 | NM_153033.4 | 178.470396 | -8.1369754 | 1.33090989 | -6.113844 |
| 5304 | NM_030793.4 | 84.1149861 | -8.0164455 | 1.31122239 | -6.1137192 |
| 5305 | NM_001010000.2 | 89.7795908 | -8.1105028 | 1.32680974 | -6.1127851 |
| 5306 | NM_001204450.1 | 78.1757449 | -7.9129176 | 1.29458379 | -6.1123256 |
| 5307 | NM_005646.3 | 107.078613 | -6.5632796 | 1.07377797 | -6.1123246 |
| 5308 | NM_022338.3 | 85.7838483 | -8.0456156 | 1.31644352 | -6.1116299 |
| 5309 | NM_001040455.1 | 117.54134 | -7.5308029 | 1.23223536 | -6.1114971 |
| 5310 | NM_004992.3 | 243.380467 | -5.4069728 | 0.88474606 | -6.1113273 |
| 5311 | NM_005627.3 | 94.5587686 | -8.1861379 | 1.33957114 | -6.1110139 |
| 5312 | NM_012473.3 | 148.025932 | -6.4990261 | 1.06353253 | -6.110792 |
| 5313 | NM_003907.2 | 145.933266 | -5.5153248 | 0.90258559 | -6.1105837 |
| 5314 | NR_033302.1 | 61.5199591 | -7.564767 | 1.23800237 | -6.1104624 |
| 5315 | NM_014976.1 | 202.381662 | -5.7987523 | 0.9490173 | -6.1102704 |
| 5316 | NM_001300759.1 | 91.9862586 | -8.1446377 | 1.33305639 | -6.1097473 |
| 5317 | NM_003184.3 | 94.1400821 | -5.4085103 | 0.88524625 | -6.109611 |
| 5318 | NM_020195.2 | 79.3596778 | -7.9318833 | 1.29829234 | -6.109474 |
| 5319 | NM_012124.2 | 70.4907637 | -7.7613328 | 1.27058274 | -6.1084828 |
| 5320 | NM_001946.3 | 77.3699186 | -7.8967446 | 1.29283169 | -6.1080995 |
| 5321 | NM_001170880.1 | 104.773561 | -8.3326405 | 1.36424508 | -6.1078765 |
| 5322 | NM_001896.2 | 73.1762275 | -7.8147399 | 1.27950195 | -6.107642 |
| 5323 | NM_014458.3 | 100.860009 | -8.2782564 | 1.35539621 | -6.1076284 |
| 5324 | NM_207111.3 | 72.6885517 | -7.8024458 | 1.27749417 | -6.1076175 |
| 5325 | NM_005981.4 | 67.7928237 | -7.7028804 | 1.26123194 | -6.1074258 |
| 5326 | NM_001134367.3 | 60.7181939 | -7.5439788 | 1.23530303 | -6.1069864 |
| 5327 | NM_020246.3 | 107.43164 | -8.369402 | 1.37047692 | -6.1069267 |
| 5328 | NM_023071.3 | 132.056266 | -8.6664587 | 1.41913809 | -6.1068466 |
| 5329 | NR_036684.1 | 74.6568656 | -7.8422343 | 1.28421266 | -6.1066477 |
| 5330 | NM_001040284.2 | 59.0536122 | -7.5068158 | 1.22934201 | -6.1063688 |
| 5331 | NM_020998.3 | 57.9313424 | -7.479102 | 1.22480647 | -6.1063541 |

| | | | | | |
|---|---|---|---|---|---|
| 5332 | NM_024321.4 | 71.3973011 | -7.7783627 | 1.27387693 | -6.106055 |
| 5333 | NM_001199260.1 | 64.1030194 | -7.6251448 | 1.24893704 | -6.1053076 |
| 5334 | NM_000147.4 | 64.449303 | -7.6345574 | 1.25051168 | -6.1051468 |
| 5335 | NM_001282149.1 | 191.158268 | -9.199973 | 1.50703954 | -6.104666 |
| 5336 | NM_016200.4 | 135.909092 | -5.6610578 | 0.9273768 | -6.1043773 |
| 5337 | NM_005148.3 | 102.487758 | -6.4982916 | 1.06470694 | -6.1033617 |
| 5338 | NM_183381.2 | 81.7244529 | -7.9748005 | 1.30671788 | -6.1029245 |
| 5339 | NM_032125.2 | 98.7844772 | -8.2484766 | 1.35160395 | -6.1027319 |
| 5340 | NM_001007559.2 | 66.8927732 | -7.6893734 | 1.26008373 | -6.1022718 |
| 5341 | NM_178151.2 | 101.70718 | -8.290204 | 1.35853726 | -6.1023016 |
| 5342 | NM_181659.2 | 63.5293836 | -7.6141075 | 1.24775854 | -6.1022283 |
| 5343 | NM_001609.3 | 66.2908894 | -7.6711813 | 1.25712197 | -6.1021774 |
| 5344 | NM_001136193.1 | 93.8481993 | -8.1720107 | 1.33922327 | -6.1020525 |
| 5345 | NM_018285.3 | 98.4495845 | -8.2427973 | 1.35084406 | -6.1019607 |
| 5346 | NM_139157.2 | 77.8792349 | -7.9056105 | 1.29561177 | -6.1018359 |
| 5347 | NM_001252499.2 | 78.2848337 | -7.911137 | 1.29655399 | -6.1016642 |
| 5348 | NM_000400.3 | 139.544964 | -6.9621835 | 1.14110942 | -6.1012409 |
| 5349 | NM_138619.3 | 85.3111409 | -8.0356812 | 1.31708853 | -6.1010942 |
| 5350 | NM_003110.5 | 59.7407864 | -7.5245991 | 1.23334211 | -6.1009829 |
| 5351 | NM_001077619.1 | 193.268518 | -5.4477118 | 0.89297125 | -6.1006576 |
| 5352 | NM_003443.2 | 97.8222296 | -8.2332226 | 1.34956272 | -6.1006595 |
| 5353 | NM_032145.4 | 67.1179043 | -7.6899469 | 1.26058068 | -6.1003211 |
| 5354 | NM_001199058.1 | 90.1508547 | -8.1163329 | 1.33049738 | -6.1002247 |
| 5355 | NM_003404.4 | 85.7677831 | -8.0429571 | 1.31850733 | -6.1000474 |
| 5356 | NM_015898.3 | 69.9606876 | -7.7468728 | 1.2701532 | -6.0991641 |
| 5357 | NM_006420.2 | 131.393265 | -5.137921 | 0.84241231 | -6.0990574 |
| 5358 | NM_144949.2 | 66.2580795 | -7.6717823 | 1.25796765 | -6.0985529 |
| 5359 | NM_001039708.1 | 72.6632336 | -7.803513 | 1.27960264 | -6.0983877 |
| 5360 | NM_015471.3 | 76.660975 | -7.8804883 | 1.29240183 | -6.0975528 |
| 5361 | NM_000837.1 | 81.9709455 | -7.9793451 | 1.30874734 | -6.0969332 |
| 5362 | NM_013448.2 | 70.9296368 | -7.7707567 | 1.27459768 | -6.096635 |
| 5363 | NM_019012.5 | 66.4727163 | -7.6741054 | 1.25875711 | -6.0965736 |
| 5364 | NM_001271836.1 | 61.5198702 | -7.5633023 | 1.24059253 | -6.0965241 |
| 5365 | NM_170744.4 | 105.815602 | -8.3490818 | 1.36961306 | -6.095942 |
| 5366 | NM_207299.1 | 158.546469 | -8.9306985 | 1.46502142 | -6.0959508 |
| 5367 | NM_005380.7 | 69.8848758 | -7.7522094 | 1.27175122 | -6.0956965 |
| 5368 | NM_174902.3 | 96.4224239 | -8.2135279 | 1.34747422 | -6.0954991 |
| 5369 | NM_014860.2 | 81.5274654 | -7.9703669 | 1.30764094 | -6.0952259 |
| 5370 | NM_001278.4 | 68.675404 | -7.7239048 | 1.26723164 | -6.095101 |
| 5371 | NM_001168243.1 | 171.167674 | -9.0411952 | 1.4833716 | -6.0950305 |
| 5372 | NR_037771.1 | 54.268474 | 4.36520013 | 0.7161947 | 6.09499086 |
| 5373 | NM_030885.3 | 272.15402 | -7.9059365 | 1.29721251 | -6.0945577 |
| 5374 | NM_130468.3 | 69.3121646 | -7.7400108 | 1.27005469 | -6.0942342 |

| | | | | | |
|---|---|---|---|---|---|
| 5375 | NM_001039465.1 | 66.8276419 | -7.6841402 | 1.26094583 | -6.0939495 |
| 5376 | NM_006571.3 | 92.8188061 | -8.158588 | 1.33888489 | -6.0935694 |
| 5377 | NM_015344.2 | 158.625175 | -7.9635177 | 1.30708843 | -6.0925623 |
| 5378 | NM_020456.3 | 66.6946329 | -7.6791386 | 1.26054516 | -6.0919186 |
| 5379 | NM_001077352.1 | 60.429959 | -7.5393056 | 1.23764983 | -6.0916306 |
| 5380 | NM_001033025.2 | 86.9830857 | -8.0652103 | 1.32417165 | -6.0907589 |
| 5381 | NM_012306.3 | 259.708064 | -6.5489825 | 1.07525093 | -6.090655 |
| 5382 | NM_021982.2 | 127.288933 | -3.7812894 | 0.62084912 | -6.0905127 |
| 5383 | NM_004962.4 | 262.379788 | -7.0077998 | 1.15061543 | -6.0904796 |
| 5384 | NM_182905.4 | 70.0753932 | -7.7525378 | 1.27301098 | -6.0899222 |
| 5385 | NM_033415.3 | 101.771534 | -8.2913908 | 1.36162966 | -6.0893142 |
| 5386 | NM_080430.3 | 79.0143048 | -7.9264708 | 1.30178344 | -6.0889319 |
| 5387 | NR_028104.1 | 73.255662 | -7.8146537 | 1.2834499 | -6.0887875 |
| 5388 | NM_020652.2 | 84.2365998 | -8.0179341 | 1.31689733 | -6.0885036 |
| 5389 | NM_013399.2 | 119.017384 | -8.5164085 | 1.39894439 | -6.0877392 |
| 5390 | NM_001289971.1 | 134.085842 | -8.6880891 | 1.42724957 | -6.0872949 |
| 5391 | NM_005871.3 | 83.5673879 | -8.0059177 | 1.31523507 | -6.0870622 |
| 5392 | NM_007043.6 | 77.5765798 | -7.8984535 | 1.29761366 | -6.0869069 |
| 5393 | NM_005569.3 | 59.2297522 | -7.5067007 | 1.23326433 | -6.0868546 |
| 5394 | NM_016056.3 | 65.569863 | -7.6573964 | 1.25804061 | -6.0867641 |
| 5395 | NM_021137.4 | 221.808858 | -6.3179915 | 1.03802228 | -6.0865663 |
| 5396 | NM_020662.3 | 68.4358421 | -7.7178113 | 1.26813011 | -6.0859775 |
| 5397 | NM_001798.4 | 68.7927358 | -7.7240704 | 1.2691819 | -6.0858656 |
| 5398 | NM_003850.2 | 136.916549 | -6.3817945 | 1.04873801 | -6.0852133 |
| 5399 | NM_016166.2 | 130.852391 | -7.6881405 | 1.26342174 | -6.0851735 |
| 5400 | NR_026922.1 | 85.4982595 | -8.038812 | 1.32110535 | -6.0849137 |
| 5401 | NM_001007468.2 | 94.2358539 | -8.1820425 | 1.34468976 | -6.0847065 |
| 5402 | NM_177996.2 | 114.336211 | -8.46008 | 1.39038037 | -6.0847234 |
| 5403 | NM_001206665.2 | 117.633035 | -8.4996915 | 1.39694258 | -6.084496 |
| 5404 | NM_152769.2 | 103.487884 | -8.3151752 | 1.36672566 | -6.084012 |
| 5405 | NM_018208.3 | 75.7584637 | -7.8654199 | 1.29282777 | -6.0838884 |
| 5406 | NM_032313.3 | 69.5500552 | -7.7416206 | 1.27247425 | -6.0839114 |
| 5407 | NM_001017408.2 | 79.9500598 | -7.939978 | 1.30515346 | -6.0835589 |
| 5408 | NM_006505.4 | 99.0961728 | -8.253434 | 1.35669208 | -6.0834984 |
| 5409 | NM_002455.4 | 72.1972831 | -7.797153 | 1.28176024 | -6.0831603 |
| 5410 | NR_135599.1 | 80.3698581 | -7.9495941 | 1.30687736 | -6.0828922 |
| 5411 | NM_181671.2 | 84.9415992 | -8.030086 | 1.32013112 | -6.0827943 |
| 5412 | NM_001400.4 | 61.0544272 | -7.5537093 | 1.2418627 | -6.0825639 |
| 5413 | NM_031862.3 | 100.926292 | -8.2813367 | 1.36151525 | -6.0824414 |
| 5414 | NM_001034841.3 | 69.4762994 | -7.7439495 | 1.27317278 | -6.0824027 |
| 5415 | NM_003171.4 | 63.7874797 | -7.6162441 | 1.25223495 | -6.0821207 |
| 5416 | NM_001080419.2 | 116.977116 | -7.526492 | 1.2375074 | -6.0819774 |
| 5417 | NM_005089.3 | 75.2299652 | -7.8539148 | 1.29144781 | -6.0814806 |

| | | | | | |
|---|---|---|---|---|---|
| 5418 | NM_001145670.1 | 65.3643681 | -7.6528788 | 1.25843578 | -6.0812628 |
| 5419 | NM_018092.4 | 161.889573 | -8.9608856 | 1.47361633 | -6.080881 |
| 5420 | NM_007198.3 | 96.2510312 | -5.1316804 | 0.8439261 | -6.0807225 |
| 5421 | NM_001163735.1 | 70.763863 | -7.7660604 | 1.27720862 | -6.0804948 |
| 5422 | NM_001305043.1 | 117.492136 | -8.4971588 | 1.39751008 | -6.0802129 |
| 5423 | NM_001317724.1 | 90.5313394 | -8.1216004 | 1.33577858 | -6.0800499 |
| 5424 | NM_004799.3 | 84.1319874 | -8.0181312 | 1.31879539 | -6.0798903 |
| 5425 | NM_001253389.1 | 59.0638658 | -7.5093597 | 1.23517161 | -6.0796084 |
| 5426 | NM_147782.3 | 95.753081 | -8.2007903 | 1.34890019 | -6.0796124 |
| 5427 | NM_001008215.2 | 73.6413138 | -7.8239003 | 1.2869587 | -6.0793716 |
| 5428 | NM_001905.3 | 113.111597 | -8.4437601 | 1.38902375 | -6.078917 |
| 5429 | NM_000744.6 | 157.936381 | -8.925054 | 1.46826551 | -6.0786376 |
| 5430 | NM_032882.5 | 74.8951189 | -7.8488727 | 1.29149642 | -6.0773477 |
| 5431 | NM_033027.3 | 57.8007122 | -7.4747538 | 1.23002742 | -6.0769001 |
| 5432 | NM_032320.6 | 94.1305603 | -8.1781862 | 1.34586167 | -6.0765429 |
| 5433 | NM_001166266.1 | 87.72743 | -8.0778152 | 1.32939178 | -6.076324 |
| 5434 | NM_001077628.2 | 119.991761 | -5.7832871 | 0.95185747 | -6.0757911 |
| 5435 | NM_031206.4 | 80.5623488 | -7.953782 | 1.30911071 | -6.0757138 |
| 5436 | NM_001142479.1 | 138.313465 | -8.7335522 | 1.43746623 | -6.0756573 |
| 5437 | NM_138439.2 | 80.5262543 | -7.9542363 | 1.30923278 | -6.0754943 |
| 5438 | NM_007344.3 | 60.9516811 | -7.5499676 | 1.2426987 | -6.0754611 |
| 5439 | NM_181675.3 | 129.238076 | -8.6364489 | 1.42161681 | -6.0750892 |
| 5440 | NM_001957.3 | 72.9436697 | -7.8129266 | 1.28607191 | -6.0750309 |
| 5441 | NM_170784.2 | 73.4451945 | -7.820564 | 1.28734621 | -6.0749501 |
| 5442 | NM_004225.2 | 98.8202711 | -8.2479484 | 1.35780343 | -6.074479 |
| 5443 | NM_001128228.2 | 71.4919483 | -7.78189 | 1.28109346 | -6.0744124 |
| 5444 | NM_003187.4 | 113.821398 | -8.4524266 | 1.39152247 | -6.0742293 |
| 5445 | NM_024083.3 | 80.1644593 | -7.9458424 | 1.30814319 | -6.0741381 |
| 5446 | NM_203418.2 | 137.238933 | -8.7233622 | 1.43636853 | -6.0732061 |
| 5447 | NM_000431.3 | 97.0807343 | -8.2231853 | 1.35404559 | -6.0730491 |
| 5448 | NM_001136053.2 | 79.2325796 | -7.930299 | 1.30607038 | -6.0718772 |
| 5449 | NM_015503.2 | 68.7909237 | -7.7270346 | 1.27265089 | -6.0716058 |
| 5450 | NM_015542.3 | 117.927927 | -7.5343828 | 1.24112929 | -6.0705865 |
| 5451 | NR_036659.1 | 115.229648 | -8.4700317 | 1.39526605 | -6.0705495 |
| 5452 | NM_001099409.1 | 60.4658452 | -7.5397908 | 1.24205755 | -6.0704037 |
| 5453 | NM_153280.2 | 92.0024505 | -8.1455064 | 1.34191496 | -6.0700615 |
| 5454 | NM_016309.2 | 107.954483 | -8.3764138 | 1.38000261 | -6.0698536 |
| 5455 | NM_004884.3 | 162.995692 | -8.003532 | 1.31859581 | -6.0697387 |
| 5456 | NM_033549.4 | 133.916738 | -7.7180029 | 1.27160596 | -6.0694926 |
| 5457 | NM_001002860.3 | 64.4167179 | -7.6349504 | 1.25794187 | -6.0693985 |
| 5458 | NM_001195518.1 | 156.554405 | -7.944475 | 1.3089827 | -6.0691979 |
| 5459 | NM_006489.2 | 109.488232 | -8.3971919 | 1.38391586 | -6.0677041 |
| 5460 | NM_006095.2 | 118.852451 | -8.5148022 | 1.40337042 | -6.0673947 |

| | | | | | |
|---|---|---|---|---|---|
| 5461 | NM_020779.3 | 67.0298804 | -7.6907742 | 1.26754567 | -6.0674533 |
| 5462 | NR_045209.1 | 81.498804 | -7.9689435 | 1.31339911 | -6.0674196 |
| 5463 | NM_014961.4 | 187.74819 | -9.1744816 | 1.51214544 | -6.0671952 |
| 5464 | NM_183377.1 | 135.490489 | -8.7042987 | 1.43472099 | -6.0668929 |
| 5465 | NM_014051.3 | 103.669196 | -8.317876 | 1.37103644 | -6.0668526 |
| 5466 | NM_020862.1 | 124.234294 | -8.5789228 | 1.41409062 | -6.0667419 |
| 5467 | NM_003635.3 | 63.2264434 | -7.605979 | 1.25384127 | -6.0661419 |
| 5468 | NM_003477.2 | 77.2918959 | -7.8933873 | 1.30128118 | -6.0658583 |
| 5469 | NM_001013438.2 | 153.913914 | -7.9206842 | 1.30583126 | -6.0656262 |
| 5470 | NM_024031.3 | 98.5370191 | -8.2432726 | 1.35914827 | -6.0650282 |
| 5471 | NM_001013843.2 | 79.0300199 | -7.9257339 | 1.30682002 | -6.0649009 |
| 5472 | NM_020923.2 | 100.012192 | -8.2680096 | 1.363317 | -6.0646274 |
| 5473 | NM_001001790.2 | 115.473019 | -8.4732532 | 1.39719838 | -6.0644596 |
| 5474 | NM_022753.3 | 70.7060398 | -7.7675537 | 1.28084749 | -6.0643861 |
| 5475 | NM_015698.5 | 70.3289557 | -7.7589598 | 1.27944143 | -6.0643338 |
| 5476 | NM_001254952.2 | 132.709897 | -8.6743821 | 1.43043673 | -6.0641494 |
| 5477 | NM_005881.3 | 68.3056831 | -7.7186464 | 1.27288099 | -6.0639183 |
| 5478 | NM_012279.3 | 63.855354 | -7.617055 | 1.25613639 | -6.0638757 |
| 5479 | NM_004095.3 | 128.388991 | -8.6262913 | 1.42263044 | -6.0636206 |
| 5480 | NM_001190946.2 | 89.6913435 | -8.1085797 | 1.33728569 | -6.0634611 |
| 5481 | NM_001174100.1 | 104.40202 | -8.3277607 | 1.37345369 | -6.063372 |
| 5482 | NM_014166.3 | 82.4108069 | -7.9865012 | 1.31717463 | -6.0633579 |
| 5483 | NM_001015877.1 | 110.891182 | -7.4491956 | 1.2286809 | -6.0627585 |
| 5484 | NR_038419.1 | 46.8737593 | 5.36937673 | 0.88564667 | 6.06266239 |
| 5485 | NM_004922.3 | 68.7021295 | -7.7250951 | 1.27422004 | -6.0626068 |
| 5486 | NM_001174061.1 | 104.961905 | -8.3340094 | 1.37485915 | -6.0617187 |
| 5487 | NM_001288655.1 | 59.9797246 | -7.5292716 | 1.24218201 | -6.0613272 |
| 5488 | NM_014239.3 | 83.1657352 | -7.9999055 | 1.31986143 | -6.0611707 |
| 5489 | NM_033394.2 | 99.5553656 | -8.2611085 | 1.36295708 | -6.0611655 |
| 5490 | NM_024561.4 | 62.8423921 | -7.593922 | 1.25289417 | -6.0611041 |
| 5491 | NM_001065.3 | 98.8315298 | -8.2504001 | 1.36122326 | -6.0610191 |
| 5492 | NM_006357.3 | 176.825153 | -8.1236514 | 1.34036328 | -6.0607833 |
| 5493 | NR_045600.1 | 92.2666024 | -8.1503128 | 1.34479481 | -6.0606367 |
| 5494 | NM_012323.3 | 60.8454062 | -7.5494672 | 1.24569659 | -6.0604382 |
| 5495 | NM_003104.5 | 84.9324154 | -8.0301843 | 1.32505454 | -6.060267 |
| 5496 | NM_017495.5 | 60.7932157 | -7.5508662 | 1.24596713 | -6.0602451 |
| 5497 | NR_003089.1 | 62.6698975 | -7.5923889 | 1.25280845 | -6.0602951 |
| 5498 | NM_002505.4 | 126.515058 | -8.6048644 | 1.41992319 | -6.0600914 |
| 5499 | NM_001204386.1 | 65.839064 | -7.6589733 | 1.26385537 | -6.0600077 |
| 5500 | NM_001294335.1 | 176.559958 | -9.0856339 | 1.4992979 | -6.0599257 |
| 5501 | NM_014817.3 | 71.4606 | -7.7844944 | 1.28461473 | -6.0597892 |
| 5502 | NM_022365.3 | 83.7005295 | -8.0092037 | 1.32174916 | -6.0595489 |
| 5503 | NM_198242.2 | 138.544291 | -8.7355799 | 1.44169482 | -6.0592435 |

| | | | | | |
|---|---|---|---|---|---|
| 5504 | NM_014906.4 | 93.5997968 | -8.171713 | 1.34884088 | -6.0583224 |
| 5505 | NM_018297.3 | 59.9025729 | -7.5302645 | 1.24301006 | -6.0580881 |
| 5506 | NM_001317781.1 | 54.6853343 | -7.3936284 | 1.22046376 | -6.0580483 |
| 5507 | NM_015391.3 | 100.138913 | -8.2679934 | 1.36485311 | -6.0577899 |
| 5508 | NM_033280.3 | 140.00021 | -8.7510991 | 1.44460484 | -6.0577805 |
| 5509 | NM_032390.4 | 86.8232523 | -8.0613654 | 1.33090215 | -6.0570684 |
| 5510 | NM_024100.3 | 148.336882 | -6.928513 | 1.1439041 | -6.0569002 |
| 5511 | NM_017644.3 | 155.348473 | -5.3969016 | 0.89108904 | -6.0565233 |
| 5512 | NM_001242860.2 | 132.447686 | -8.6713087 | 1.43188052 | -6.0558885 |
| 5513 | NM_025132.3 | 83.0025827 | -7.998562 | 1.32079173 | -6.0558843 |
| 5514 | NM_023926.4 | 57.6662216 | -7.4682783 | 1.23324047 | -6.0558168 |
| 5515 | NM_001251974.1 | 108.016257 | -8.3779321 | 1.38347345 | -6.0557231 |
| 5516 | NM_032880.4 | 90.99903 | -8.1299276 | 1.34256193 | -6.0555326 |
| 5517 | NM_004263.4 | 123.127499 | -8.566245 | 1.41465153 | -6.0553746 |
| 5518 | NR_028024.1 | 76.0798646 | -7.8696864 | 1.29967907 | -6.0550997 |
| 5519 | NM_001143852.1 | 68.5001201 | -7.7183961 | 1.27472109 | -6.0549686 |
| 5520 | NM_199341.3 | 67.8640564 | -7.7065961 | 1.27278578 | -6.0549043 |
| 5521 | NM_001031727.3 | 57.025045 | -7.4532824 | 1.23099346 | -6.0546888 |
| 5522 | NM_183050.3 | 56.2802258 | -7.4379696 | 1.22847423 | -6.0546403 |
| 5523 | NM_001008272.1 | 115.686411 | -8.4760515 | 1.39994251 | -6.0545711 |
| 5524 | NM_052874.4 | 236.351327 | -5.803669 | 0.95862315 | -6.0541716 |
| 5525 | NM_001136472.1 | 116.981434 | -7.5282111 | 1.24354926 | -6.0538101 |
| 5526 | NM_001321209.1 | 75.2187314 | -7.8545304 | 1.29745485 | -6.0537986 |
| 5527 | NM_006702.4 | 98.7492717 | -8.2476411 | 1.36243849 | -6.0535879 |
| 5528 | NM_002676.2 | 91.6630377 | -8.1401847 | 1.34478521 | -6.0531486 |
| 5529 | NM_001037637.1 | 92.6226616 | -8.1552455 | 1.34737067 | -6.0527112 |
| 5530 | NM_003936.4 | 202.659566 | -7.4991812 | 1.2391797 | -6.0517302 |
| 5531 | NM_006084.4 | 71.7970345 | -7.7911105 | 1.28743885 | -6.0516354 |
| 5532 | NM_002490.4 | 177.390472 | -8.1280265 | 1.34317253 | -6.0513645 |
| 5533 | NM_001076677.2 | 138.853639 | -8.7396733 | 1.44427394 | -6.0512574 |
| 5534 | NM_001023567.4 | 166.144983 | -8.9982408 | 1.48704201 | -6.0511006 |
| 5535 | NM_020441.2 | 114.559201 | -6.6199301 | 1.09400981 | -6.0510701 |
| 5536 | NM_016353.4 | 71.0319253 | -7.7708834 | 1.28431818 | -6.0505905 |
| 5537 | NM_001142730.2 | 62.217912 | -7.5821502 | 1.25320716 | -6.050197 |
| 5538 | NM_014602.2 | 63.9224216 | -7.6195867 | 1.25952365 | -6.0495781 |
| 5539 | NM_198868.2 | 140.596538 | -8.7575629 | 1.44765318 | -6.0494896 |
| 5540 | NM_015373.3 | 59.8367809 | -7.5274215 | 1.24439985 | -6.0490376 |
| 5541 | NR_033230.2 | 77.6061216 | -7.8993153 | 1.30587752 | -6.0490476 |
| 5542 | NM_001242824.1 | 110.380176 | -8.4087424 | 1.39022968 | -6.0484556 |
| 5543 | NM_016451.4 | 100.635472 | -8.2740056 | 1.36794792 | -6.04848 |
| 5544 | NM_001160226.1 | 132.061137 | -8.6671303 | 1.4329701 | -6.0483678 |
| 5545 | NM_001048.3 | 115.757853 | -8.4777529 | 1.40168817 | -6.0482446 |
| 5546 | NM_003687.3 | 62.433211 | -7.5878185 | 1.25456328 | -6.0481752 |

| | | | | | |
|---|---|---|---|---|---|
| 5547 | NM_001318785.1 | 86.3463373 | -8.0535817 | 1.33160989 | -6.0480038 |
| 5548 | NM_024907.6 | 61.486358 | -7.5641564 | 1.25069259 | -6.0479741 |
| 5549 | NM_001270448.1 | 67.635028 | -7.7054045 | 1.27408236 | -6.0478072 |
| 5550 | NM_022482.4 | 68.4906603 | -7.7187942 | 1.27638449 | -6.0473895 |
| 5551 | NM_001029.3 | 492.750424 | -7.1391642 | 1.18067117 | -6.0466998 |
| 5552 | NM_001278380.1 | 66.3836504 | -7.6747797 | 1.26926589 | -6.0466288 |
| 5553 | NM_003765.2 | 69.7089835 | -7.7459202 | 1.28105343 | -6.0465239 |
| 5554 | NM_032656.3 | 76.9086817 | -7.8872097 | 1.30441795 | -6.0465357 |
| 5555 | NM_032380.4 | 80.8531297 | -7.9598371 | 1.31644247 | -6.0464755 |
| 5556 | NM_001382.3 | 71.2720672 | -7.7794342 | 1.28669119 | -6.0460771 |
| 5557 | NM_001278480.1 | 72.2214557 | -7.7950871 | 1.28928637 | -6.0460479 |
| 5558 | NM_144564.4 | 90.2706341 | -8.1180927 | 1.34282111 | -6.0455504 |
| 5559 | NM_032038.2 | 107.206298 | -8.3651728 | 1.38376539 | -6.0452248 |
| 5560 | NM_001407.2 | 395.788257 | -6.8120726 | 1.12689086 | -6.0450154 |
| 5561 | NM_006327.3 | 91.9624282 | -8.1442815 | 1.34732413 | -6.0447826 |
| 5562 | NM_020187.2 | 68.8387101 | -7.7251691 | 1.27809215 | -6.0442974 |
| 5563 | NM_001308321.1 | 905.056836 | 3.85429377 | 0.63769983 | 6.0440564 |
| 5564 | NM_015453.2 | 92.8585089 | -8.1582252 | 1.34980273 | -6.044013 |
| 5565 | NM_001142641.1 | 339.170149 | -4.915677 | 0.81332522 | -6.0439254 |
| 5566 | NM_030667.2 | 119.659247 | -8.5242115 | 1.41048178 | -6.0434751 |
| 5567 | NM_005700.4 | 92.3035966 | -8.1507246 | 1.34869268 | -6.0434261 |
| 5568 | NM_001270377.1 | 188.659344 | -8.2147702 | 1.35934903 | -6.0431647 |
| 5569 | NM_015449.3 | 97.9592822 | -8.2378701 | 1.36322444 | -6.0429301 |
| 5570 | NM_001136204.2 | 175.657215 | -6.2776077 | 1.03888383 | -6.0426464 |
| 5571 | NM_014858.3 | 131.55002 | -8.6612229 | 1.43336109 | -6.0425967 |
| 5572 | NM_024096.1 | 110.270164 | -8.4066933 | 1.39143091 | -6.0417612 |
| 5573 | NM_025140.2 | 78.5788787 | -7.9175474 | 1.31076529 | -6.0404005 |
| 5574 | NM_001190965.2 | 149.753788 | -8.8484697 | 1.46488871 | -6.0403699 |
| 5575 | NR_073080.1 | 108.463391 | -8.3834801 | 1.38810369 | -6.0395201 |
| 5576 | NM_006858.3 | 67.7454211 | -7.7058323 | 1.27599573 | -6.0390737 |
| 5577 | NM_001003940.1 | 75.4318637 | -7.8619932 | 1.30190581 | -6.0388341 |
| 5578 | NM_012239.5 | 61.5892267 | -7.5675597 | 1.25323078 | -6.0384407 |
| 5579 | NM_001244808.1 | 82.5716655 | -7.9903757 | 1.32333084 | -6.0380786 |
| 5580 | NM_152298.3 | 63.2405827 | -7.6008689 | 1.25883625 | -6.0380124 |
| 5581 | NM_032124.4 | 82.7936639 | -7.9925558 | 1.32375209 | -6.0378041 |
| 5582 | NM_173680.3 | 72.8363791 | -7.8074004 | 1.29309638 | -6.0377559 |
| 5583 | NM_024639.4 | 98.2482939 | -8.2414914 | 1.36504765 | -6.0375119 |
| 5584 | NM_014598.3 | 261.641046 | -3.1905818 | 0.52846777 | -6.0374198 |
| 5585 | NM_198244.2 | 82.9658627 | -7.997735 | 1.32475837 | -6.0371273 |
| 5586 | NM_198273.2 | 56.7677277 | -7.4528038 | 1.23452798 | -6.0369663 |
| 5587 | NM_052854.3 | 66.3396119 | -7.675954 | 1.27151456 | -6.0368589 |
| 5588 | NR_135468.1 | 75.3113702 | -7.8560489 | 1.30139772 | -6.0366241 |
| 5589 | NM_057175.3 | 163.35861 | -4.7291018 | 0.7834113 | -6.0365504 |

| | | | | | |
|---|---|---|---|---|---|
| 5590 | NM_000320.2 | 118.581182 | -8.5115692 | 1.41012734 | -6.0360287 |
| 5591 | NM_001146054.1 | 105.9447 | -8.3497669 | 1.38332023 | -6.0360333 |
| 5592 | NM_032439.3 | 127.215687 | -8.6132525 | 1.42696053 | -6.0360832 |
| 5593 | NM_001297707.1 | 180.497669 | -9.1179142 | 1.51059846 | -6.0359615 |
| 5594 | NM_001310137.1 | 102.320172 | -8.2981 | 1.37479061 | -6.035901 |
| 5595 | NM_030583.3 | 98.9826027 | -8.2519242 | 1.36721724 | -6.0355618 |
| 5596 | NM_015137.5 | 60.705728 | -7.5495005 | 1.25092941 | -6.0351131 |
| 5597 | NM_002946.4 | 90.7814909 | -8.125627 | 1.34646202 | -6.0347985 |
| 5598 | NM_001253383.1 | 1216.4127 | -7.7668709 | 1.28706979 | -6.0345374 |
| 5599 | NM_005869.3 | 73.2556733 | -7.8157907 | 1.29527029 | -6.0341002 |
| 5600 | NM_198467.2 | 80.9682007 | -7.9604897 | 1.31925758 | -6.0340678 |
| 5601 | NM_012133.5 | 91.8446366 | -5.7757349 | 0.95719914 | -6.0339951 |
| 5602 | NM_025114.3 | 141.337838 | -6.8616222 | 1.13719066 | -6.0338362 |
| 5603 | NM_033301.2 | 71.5439761 | -7.7833197 | 1.28994882 | -6.0338206 |
| 5604 | NM_015124.4 | 152.055062 | -7.0590514 | 1.16994179 | -6.0336774 |
| 5605 | NM_001206998.1 | 65.6583751 | -7.6585564 | 1.26936442 | -6.0333788 |
| 5606 | NM_015675.3 | 86.0692364 | -8.05071 | 1.3343574 | -6.0333986 |
| 5607 | NM_138384.2 | 81.4342688 | -7.970001 | 1.32098284 | -6.0333872 |
| 5608 | NM_002145.3 | 93.6859706 | -8.1729524 | 1.35473932 | -6.0328598 |
| 5609 | NM_014727.2 | 268.640824 | -5.8786537 | 0.97453169 | -6.0322858 |
| 5610 | NM_001039469.2 | 61.8440882 | -7.5718694 | 1.25524121 | -6.0322027 |
| 5611 | NM_001002027.1 | 112.339917 | -8.4337403 | 1.39823756 | -6.0316934 |
| 5612 | NM_001347.3 | 131.454107 | -5.2933741 | 0.87762463 | -6.0314785 |
| 5613 | NM_001043351.1 | 102.104262 | -8.2972736 | 1.37587525 | -6.0305421 |
| 5614 | NM_001142467.1 | 136.291924 | -8.7123704 | 1.44480171 | -6.0301495 |
| 5615 | NM_024779.4 | 70.7228483 | -7.7656751 | 1.28782349 | -6.0300772 |
| 5616 | NM_004542.3 | 98.2485436 | -8.2404701 | 1.36657026 | -6.0300376 |
| 5617 | NM_024066.2 | 78.672943 | -7.9181439 | 1.31315452 | -6.0298645 |
| 5618 | NM_003071.3 | 66.1217987 | -7.6686654 | 1.27182166 | -6.0296704 |
| 5619 | NM_003598.1 | 70.7030794 | -7.76465 | 1.287746 | -6.029644 |
| 5620 | NM_002106.3 | 471.525008 | -6.7235158 | 1.11522213 | -6.0288579 |
| 5621 | NM_001286379.1 | 66.2406702 | -7.6707069 | 1.27236185 | -6.028715 |
| 5622 | NM_017560.1 | 142.821652 | -7.8142822 | 1.29617682 | -6.0287162 |
| 5623 | NM_005648.3 | 108.206988 | -8.3792437 | 1.38997067 | -6.0283601 |
| 5624 | NM_001455.3 | 78.4883977 | -7.914744 | 1.31295139 | -6.0282079 |
| 5625 | NM_001242783.1 | 74.3186042 | -7.8375489 | 1.3001956 | -6.0279768 |
| 5626 | NM_003095.2 | 109.706209 | -8.399277 | 1.39354265 | -6.0272838 |
| 5627 | NM_006852.3 | 80.3178465 | -7.9477313 | 1.31863485 | -6.0272419 |
| 5628 | NM_080748.2 | 101.061219 | -8.2810105 | 1.37393667 | -6.0272142 |
| 5629 | NM_001322246.1 | 127.688582 | -8.6184684 | 1.43004382 | -6.0267163 |
| 5630 | NR_024565.1 | 79.0073915 | -7.9256154 | 1.31541565 | -6.0251795 |
| 5631 | NM_001256574.1 | 72.617422 | -7.8077469 | 1.29588426 | -6.0250342 |
| 5632 | NM_001198972.1 | 121.581249 | -8.5487634 | 1.41893608 | -6.0247699 |

| | | | | | |
|---|---|---|---|---|---|
| 5633 | NM_001170464.2 | 107.494815 | -8.3693062 | 1.38949653 | -6.0232653 |
| 5634 | NM_002611.4 | 74.3039962 | -7.8387928 | 1.30152712 | -6.0227656 |
| 5635 | NM_001098205.1 | 123.106166 | -8.5668909 | 1.42247277 | -6.0225342 |
| 5636 | NM_000199.3 | 87.194604 | -4.9326918 | 0.81910641 | -6.0220402 |
| 5637 | NM_001130445.2 | 63.5795546 | -7.6084022 | 1.26344042 | -6.0219715 |
| 5638 | NM_031420.3 | 81.8936039 | -7.9775056 | 1.32479751 | -6.0216792 |
| 5639 | NM_004517.3 | 76.5266516 | -7.8797687 | 1.30868562 | -6.0211319 |
| 5640 | NM_022455.4 | 78.2440658 | -7.9116062 | 1.3140139 | -6.0209456 |
| 5641 | NM_152390.2 | 171.940685 | -8.0837187 | 1.342645 | -6.0207416 |
| 5642 | NM_001168303.1 | 74.3898832 | -7.8403475 | 1.30224338 | -6.0206468 |
| 5643 | NM_003685.2 | 665.238208 | -7.1630885 | 1.18976259 | -6.0206032 |
| 5644 | NM_014774.2 | 121.122124 | -5.0153666 | 0.83313001 | -6.0199087 |
| 5645 | NM_018360.2 | 123.053525 | -7.5981188 | 1.26243349 | -6.018629 |
| 5646 | NM_017926.3 | 124.171128 | -7.61015 | 1.26450191 | -6.0182985 |
| 5647 | NM_016174.4 | 82.9272202 | -7.9962427 | 1.3286758 | -6.0182045 |
| 5648 | NM_001004356.2 | 79.2484165 | -7.9309359 | 1.31784406 | -6.0181141 |
| 5649 | NM_053042.2 | 99.3097153 | -4.4659162 | 0.74208525 | -6.0180636 |
| 5650 | NM_005113.3 | 118.923311 | -7.5504751 | 1.25466717 | -6.0179108 |
| 5651 | NM_146387.1 | 89.6858416 | -8.1088246 | 1.3474656 | -6.0178342 |
| 5652 | NM_001160372.3 | 97.0119306 | -8.2228706 | 1.36642983 | -6.0177775 |
| 5653 | NM_024761.4 | 54.3324254 | -7.3845904 | 1.22721709 | -6.0173465 |
| 5654 | NM_032193.3 | 74.18455 | -7.8356592 | 1.30219638 | -6.0172638 |
| 5655 | NM_001031812.3 | 65.4525863 | -7.6545432 | 1.27217703 | -6.0168852 |
| 5656 | NM_031450.3 | 72.8520618 | -7.8090492 | 1.29814042 | -6.0155658 |
| 5657 | NM_018116.3 | 75.2169083 | -7.8555733 | 1.3060796 | -6.0146206 |
| 5658 | NM_007067.4 | 79.3853156 | -7.9352749 | 1.31937684 | -6.0144112 |
| 5659 | NM_000836.2 | 81.0355101 | -7.962651 | 1.32395659 | -6.014284 |
| 5660 | NM_006688.4 | 175.201669 | -8.1110425 | 1.34865562 | -6.0141687 |
| 5661 | NM_032982.3 | 76.3761915 | -7.8792042 | 1.31025219 | -6.013502 |
| 5662 | NM_001324188.1 | 62.2095709 | -7.581514 | 1.26084546 | -6.0130399 |
| 5663 | NM_001135734.2 | 73.4418432 | -7.820209 | 1.30058844 | -6.0128238 |
| 5664 | NM_014160.4 | 112.492661 | -7.4692961 | 1.24223246 | -6.0128006 |
| 5665 | NM_002495.3 | 84.4567729 | -8.022431 | 1.33423891 | -6.0127395 |
| 5666 | NM_013306.4 | 88.9454157 | -8.0967328 | 1.34660098 | -6.0127187 |
| 5667 | NM_020873.6 | 156.992751 | -5.415329 | 0.90067107 | -6.012549 |
| 5668 | NM_001032295.1 | 62.3388991 | -7.587367 | 1.26202653 | -6.0120503 |
| 5669 | NM_001252078.1 | 85.3169563 | -8.0370544 | 1.33683467 | -6.0120033 |
| 5670 | NM_024814.3 | 78.1153143 | -7.9088624 | 1.31555563 | -6.0118039 |
| 5671 | NM_001134338.2 | 89.2282062 | -8.1020031 | 1.34774427 | -6.0115285 |
| 5672 | NM_020912.1 | 57.4839407 | -7.4680794 | 1.2423256 | -6.0113705 |
| 5673 | NM_206920.2 | 68.9190205 | -7.7287082 | 1.28572809 | -6.011153 |
| 5674 | NM_020871.3 | 79.3345052 | -7.9306428 | 1.31936439 | -6.0109572 |
| 5675 | NM_177939.2 | 111.639097 | -7.460243 | 1.24115997 | -6.0107022 |

| | | | | | |
|---|---|---|---|---|---|
| 5676 | NM_001001795.1 | 90.3056921 | -8.118111 | 1.35062637 | -6.0106268 |
| 5677 | NM_020812.3 | 55.1929577 | -7.4114175 | 1.23326422 | -6.0095942 |
| 5678 | NM_007247.5 | 73.0567465 | -7.815428 | 1.30061126 | -6.0090423 |
| 5679 | NM_005384.2 | 108.335245 | -8.3805525 | 1.39467252 | -6.0089751 |
| 5680 | NM_138709.2 | 64.1924794 | -7.6268683 | 1.26924959 | -6.0089586 |
| 5681 | NM_033380.2 | 238.572569 | -9.5207655 | 1.5845352 | -6.0085541 |
| 5682 | NM_001015052.2 | 82.9447097 | -7.9959135 | 1.33097835 | -6.0075459 |
| 5683 | NM_006099.3 | 113.689621 | -7.487606 | 1.24636433 | -6.007558 |
| 5684 | NM_001078645.2 | 106.194374 | -8.3516177 | 1.39026377 | -6.0072181 |
| 5685 | NM_024419.4 | 61.7124383 | -7.5670364 | 1.25966453 | -6.0071838 |
| 5686 | NM_025196.3 | 74.2287306 | -7.8358535 | 1.30445953 | -6.0069732 |
| 5687 | NM_053051.3 | 70.3076806 | -7.7558145 | 1.29132934 | -6.0060701 |
| 5688 | NM_001047.3 | 112.441072 | -8.4350399 | 1.40456773 | -6.0054348 |
| 5689 | NM_021174.5 | 61.2532434 | -7.5555343 | 1.25830079 | -6.0045534 |
| 5690 | NM_130463.3 | 153.552772 | -8.8844237 | 1.47967605 | -6.0043032 |
| 5691 | NM_001242868.1 | 106.697334 | -8.3585248 | 1.3921294 | -6.0041292 |
| 5692 | NM_022552.4 | 62.5493607 | -7.5848714 | 1.26339267 | -6.003574 |
| 5693 | NM_001242911.1 | 68.9515322 | -7.7295325 | 1.28752438 | -6.0034067 |
| 5694 | NM_016579.3 | 102.530195 | -8.3019284 | 1.38289659 | -6.0032894 |
| 5695 | NM_020784.2 | 89.7330042 | -8.109202 | 1.35081948 | -6.0031722 |
| 5696 | NM_014034.2 | 90.6186236 | -8.1232197 | 1.35327673 | -6.0026301 |
| 5697 | NM_018321.3 | 68.3597677 | -7.7152379 | 1.28542521 | -6.0020901 |
| 5698 | NM_001001872.3 | 67.4502948 | -7.6969046 | 1.28241144 | -6.0018995 |
| 5699 | NM_006457.4 | 143.36147 | -8.7867434 | 1.46400186 | -6.0018663 |
| 5700 | NM_130799.2 | 73.2578741 | -7.8160816 | 1.30228914 | -6.001802 |
| 5701 | NM_001001975.1 | 118.917151 | -8.5153881 | 1.41882421 | -6.0017218 |
| 5702 | NM_025154.5 | 67.1298664 | -7.6922857 | 1.28170777 | -6.0015909 |
| 5703 | NM_003705.4 | 92.9414164 | -8.1599169 | 1.35981234 | -6.0007669 |
| 5704 | NM_133502.2 | 72.5110877 | -7.8008418 | 1.30018811 | -5.9997794 |
| 5705 | NM_001032382.1 | 79.4320006 | -7.932077 | 1.32215244 | -5.9993665 |
| 5706 | NM_001004125.2 | 75.7702978 | -7.8639502 | 1.31087941 | -5.9989882 |
| 5707 | NM_018999.3 | 108.271826 | -21.206809 | 3.53515552 | -5.9988335 |
| 5708 | NM_002595.4 | 64.270188 | -7.6301947 | 1.27197963 | -5.9986768 |
| 5709 | NM_001258429.1 | 99.4380063 | -8.2575303 | 1.37656531 | -5.9986477 |
| 5710 | NM_001019.4 | 101.101701 | -8.2813589 | 1.38056996 | -5.9985073 |
| 5711 | NM_004447.5 | 70.3553912 | -7.758424 | 1.29338679 | -5.9985335 |
| 5712 | NM_024809.4 | 66.9817293 | -7.6872058 | 1.28154488 | -5.9983899 |
| 5713 | NM_005030.5 | 80.8642496 | -7.9589909 | 1.32689245 | -5.9982186 |
| 5714 | NR_132964.1 | 606.954235 | -6.5722742 | 1.09596907 | -5.9967697 |
| 5715 | NM_024692.5 | 52.4652022 | -7.3364508 | 1.2234976 | -5.9962936 |
| 5716 | NM_003704.3 | 85.4353593 | -8.0377584 | 1.34051391 | -5.9960275 |
| 5717 | NM_001128919.2 | 133.904077 | -7.7180827 | 1.28725784 | -5.995755 |
| 5718 | NM_002276.4 | 60.938446 | -7.553822 | 1.2599719 | -5.9952306 |

| | | | | | |
|---|---|---|---|---|---|
| 5719 | NM_001293233.1 | 114.019646 | -8.4542953 | 1.4102859 | -5.9947385 |
| 5720 | NM_033023.4 | 902.798318 | 4.97742093 | 0.83039548 | 5.99403667 |
| 5721 | NR_144402.1 | 61.6476385 | -7.5683026 | 1.26289373 | -5.9928262 |
| 5722 | NM_145212.3 | 156.0677 | -3.3807104 | 0.56417186 | -5.9923414 |
| 5723 | NM_001540.3 | 418.753655 | -6.8104311 | 1.13655555 | -5.9921674 |
| 5724 | NM_001318323.1 | 54.0927878 | 2.04356313 | 0.34104904 | 5.99199199 |
| 5725 | NM_000154.1 | 57.2085619 | -7.4591976 | 1.24491767 | -5.9917196 |
| 5726 | NM_001308824.1 | 56.818838 | -7.4524527 | 1.24379476 | -5.9917061 |
| 5727 | NM_018135.3 | 52.5088378 | -7.3376677 | 1.22485868 | -5.9906239 |
| 5728 | NM_014604.3 | 96.3984969 | -8.2128528 | 1.37095788 | -5.9905946 |
| 5729 | NM_005933.3 | 249.657462 | -5.4748198 | 0.91393942 | -5.9903531 |
| 5730 | NM_001143981.1 | 132.010087 | -8.6664091 | 1.44675237 | -5.9902505 |
| 5731 | NM_001316990.1 | 60.7058431 | -7.5432962 | 1.25927171 | -5.9902054 |
| 5732 | NM_001242882.1 | 82.7170365 | -7.9908886 | 1.33401037 | -5.9901248 |
| 5733 | NM_003383.4 | 67.4793877 | -7.6988624 | 1.2852618 | -5.9901122 |
| 5734 | NM_206831.2 | 95.691653 | -8.2020962 | 1.3695128 | -5.9890613 |
| 5735 | NM_004789.3 | 232.379736 | -9.4820882 | 1.58327243 | -5.9889177 |
| 5736 | NM_015915.4 | 119.110837 | -8.5179366 | 1.42227966 | -5.9889323 |
| 5737 | NM_000021.3 | 103.724857 | -8.3184225 | 1.38900457 | -5.9887654 |
| 5738 | NM_013254.3 | 56.9277262 | -7.4527136 | 1.24457404 | -5.9881641 |
| 5739 | NM_175884.5 | 80.3041876 | -7.9484522 | 1.32735924 | -5.9881696 |
| 5740 | NM_001171093.1 | 76.7985006 | -7.8847817 | 1.31673651 | -5.9881241 |
| 5741 | NM_018246.2 | 71.1581079 | -7.77194 | 1.29791707 | -5.9880097 |
| 5742 | NM_001256763.1 | 114.77033 | -8.4644147 | 1.41374913 | -5.9872113 |
| 5743 | NM_018648.3 | 102.573489 | -8.3030651 | 1.38687975 | -5.9868673 |
| 5744 | NM_003597.4 | 62.7754348 | -7.591972 | 1.26815257 | -5.9866393 |
| 5745 | NM_022461.4 | 102.85591 | -4.9246637 | 0.82264063 | -5.9864095 |
| 5746 | NM_001142426.1 | 156.044986 | -8.9080494 | 1.48814119 | -5.9860243 |
| 5747 | NM_030955.3 | 51.5283283 | -7.3107049 | 1.22131802 | -5.9859142 |
| 5748 | NM_019005.3 | 66.9360671 | -7.6863752 | 1.28417962 | -5.9854362 |
| 5749 | NM_001135050.1 | 108.563826 | -8.3840514 | 1.40090808 | -5.9847263 |
| 5750 | NM_145858.2 | 61.9912169 | -7.5759912 | 1.26590688 | -5.9846355 |
| 5751 | NM_152783.4 | 61.3487857 | -7.5651428 | 1.26409338 | -5.9846392 |
| 5752 | NM_016338.4 | 62.5596935 | -7.588518 | 1.26800676 | -5.9846037 |
| 5753 | NM_000868.3 | 110.90088 | -8.4165429 | 1.40641097 | -5.9844122 |
| 5754 | NM_001322040.1 | 68.0397475 | -7.7110904 | 1.28855356 | -5.9842995 |
| 5755 | NM_020162.3 | 58.6840427 | -7.4981421 | 1.25307069 | -5.9838141 |
| 5756 | NM_001199344.1 | 77.7157687 | -7.9039982 | 1.32096134 | -5.9835197 |
| 5757 | NM_014169.3 | 70.300823 | -7.7574627 | 1.29665177 | -5.9826877 |
| 5758 | NM_030818.3 | 54.3674955 | -7.3849151 | 1.23442729 | -5.9824627 |
| 5759 | NM_173602.2 | 236.826357 | -6.3954365 | 1.06902669 | -5.9824853 |
| 5760 | NR_049735.2 | 64.5889271 | 2.93923919 | 0.49130965 | 5.98245764 |
| 5761 | NM_001322271.1 | 43.1004888 | 10.34885 | 1.72989917 | 5.98234288 |

| | | | | | |
|---|---|---|---|---|---|
| 5762 | NM_013432.4 | 76.291067 | -4.2753312 | 0.71469371 | -5.9820468 |
| 5763 | NM_016156.5 | 99.3532657 | -8.2569572 | 1.38035698 | -5.981755 |
| 5764 | NM_001025203.1 | 75.7121875 | -7.8630025 | 1.31453763 | -5.9815728 |
| 5765 | NM_006500.2 | 134.058757 | -7.7206832 | 1.29075532 | -5.9815234 |
| 5766 | NM_032118.3 | 68.4167976 | -7.7183467 | 1.29051917 | -5.9808075 |
| 5767 | NM_001128160.1 | 91.121733 | -8.1326555 | 1.35994118 | -5.9801524 |
| 5768 | NM_001172415.1 | 150.817312 | -7.8932229 | 1.31995015 | -5.9799401 |
| 5769 | NM_001136049.2 | 51.3565722 | -7.3057395 | 1.22173002 | -5.9798314 |
| 5770 | NM_002093.3 | 84.0213312 | -8.0155157 | 1.34042383 | -5.9798367 |
| 5771 | NM_002435.2 | 54.9476292 | -7.4025286 | 1.23806001 | -5.9791355 |
| 5772 | NM_001013836.1 | 84.4701391 | -8.0220607 | 1.34168286 | -5.9791035 |
| 5773 | NM_004176.4 | 70.2566282 | -7.7562718 | 1.29728686 | -5.978841 |
| 5774 | NM_017867.2 | 83.8894593 | -8.0115412 | 1.34014439 | -5.978118 |
| 5775 | NM_005718.4 | 97.6367459 | -8.2317068 | 1.37716595 | -5.9772802 |
| 5776 | NM_001786.4 | 72.4334831 | -7.7995409 | 1.30487176 | -5.977247 |
| 5777 | NM_133173.2 | 121.824904 | -8.5503081 | 1.43048386 | -5.977214 |
| 5778 | NM_000366.5 | 71.9190336 | -7.7891481 | 1.30317224 | -5.9770673 |
| 5779 | NM_001004342.3 | 140.437571 | -8.7554933 | 1.46498199 | -5.9765194 |
| 5780 | NM_020546.2 | 127.390867 | -6.2618059 | 1.04774568 | -5.976456 |
| 5781 | NM_005993.4 | 420.74904 | -6.2733157 | 1.0496849 | -5.9763798 |
| 5782 | NM_018708.2 | 120.428818 | -6.7455768 | 1.12870152 | -5.9764044 |
| 5783 | NM_016525.4 | 79.00576 | -7.9252581 | 1.32611079 | -5.9763167 |
| 5784 | NM_016475.4 | 56.0742568 | -7.433141 | 1.24382042 | -5.9760564 |
| 5785 | NM_004441.4 | 137.640751 | -8.7263405 | 1.46032577 | -5.9756122 |
| 5786 | NM_181553.2 | 71.4407594 | -7.7817793 | 1.30225571 | -5.9756154 |
| 5787 | NM_013247.4 | 70.9973179 | -7.769723 | 1.30029978 | -5.9753321 |
| 5788 | NM_025234.2 | 83.2125635 | -8.0005782 | 1.33896534 | -5.9751944 |
| 5789 | NM_003002.3 | 65.8523754 | -7.6624921 | 1.28246221 | -5.9748287 |
| 5790 | NM_058229.3 | 107.291771 | -8.3678225 | 1.40065663 | -5.9742141 |
| 5791 | NM_014299.2 | 75.43827 | -7.8587934 | 1.31547388 | -5.9741159 |
| 5792 | NM_001171944.1 | 99.8919266 | -7.2967544 | 1.22151146 | -5.9735456 |
| 5793 | NM_022104.3 | 133.468803 | -7.7132481 | 1.29124663 | -5.9734894 |
| 5794 | NM_018115.3 | 68.8335789 | -7.7256466 | 1.29337046 | -5.9732667 |
| 5795 | NM_019045.4 | 63.440543 | -7.6106136 | 1.27428629 | -5.9724519 |
| 5796 | NM_005550.3 | 149.575204 | -8.846606 | 1.48132098 | -5.9721061 |
| 5797 | NM_004705.3 | 146.412335 | -5.4745987 | 0.9167078 | -5.9720215 |
| 5798 | NM_001330358.1 | 83.7899479 | -8.0108078 | 1.34145427 | -5.9717338 |
| 5799 | NM_017943.3 | 73.5668667 | -7.824069 | 1.31022088 | -5.9715649 |
| 5800 | NR_102706.1 | 86.2124187 | -8.0524717 | 1.34850716 | -5.9713971 |
| 5801 | NM_003115.4 | 71.0032108 | -7.7716443 | 1.30154287 | -5.9711013 |
| 5802 | NM_018325.4 | 89.0009042 | -8.0967005 | 1.35599163 | -5.9710549 |
| 5803 | NM_001656.3 | 72.3306196 | -7.7975239 | 1.30618862 | -5.9696768 |
| 5804 | NM_021183.4 | 143.571693 | -6.2002543 | 1.03862532 | -5.9696737 |

| | | | | | |
|---|---|---|---|---|---|
| 5805 | NM_016497.3 | 153.366042 | -7.9177867 | 1.32634539 | -5.9696266 |
| 5806 | NM_012280.3 | 93.4377929 | -8.1677357 | 1.36831448 | -5.9691948 |
| 5807 | NM_005536.3 | 64.8960344 | -7.6424086 | 1.28037324 | -5.9688912 |
| 5808 | NM_014223.4 | 95.8087591 | -8.2038018 | 1.37446075 | -5.9687422 |
| 5809 | NM_032667.6 | 67.2160652 | -7.693435 | 1.28899706 | -5.9685435 |
| 5810 | NM_005154.4 | 51.547839 | -7.3081336 | 1.22454262 | -5.9680517 |
| 5811 | NR_073024.1 | 79.4081368 | -7.9337173 | 1.32945027 | -5.9676676 |
| 5812 | NM_013235.4 | 76.5933467 | -7.880753 | 1.32066219 | -5.9672739 |
| 5813 | NR_104034.1 | 321.291146 | 4.53207717 | 0.75950554 | 5.96714167 |
| 5814 | NM_005944.6 | 133.24922 | -8.6802195 | 1.45468032 | -5.9670976 |
| 5815 | NM_004565.2 | 74.0509088 | -7.8326286 | 1.31274648 | -5.9665965 |
| 5816 | NR_024113.1 | 93.5602598 | -8.1686385 | 1.36916226 | -5.9661581 |
| 5817 | NM_015529.3 | 87.6628731 | -8.0765383 | 1.35375234 | -5.9660383 |
| 5818 | NR_073599.1 | 78.216257 | -7.9136281 | 1.32644515 | -5.9660425 |
| 5819 | NM_001321179.1 | 56.4804133 | -7.4449062 | 1.24788833 | -5.9660036 |
| 5820 | NM_018566.3 | 54.7626467 | -7.396491 | 1.23979669 | -5.9658902 |
| 5821 | NM_001012991.2 | 70.9163444 | -7.7697785 | 1.30246842 | -5.9654256 |
| 5822 | NM_014640.4 | 152.327049 | -8.8740478 | 1.4876338 | -5.9652098 |
| 5823 | NM_001146070.1 | 54.1803198 | -7.3831201 | 1.23771988 | -5.9650978 |
| 5824 | NM_005740.2 | 61.638461 | -7.5677834 | 1.26876386 | -5.9646902 |
| 5825 | NM_001308210.1 | 61.6460379 | -7.5701635 | 1.26917373 | -5.9646393 |
| 5826 | NM_003467.2 | 56.5718014 | -7.4452076 | 1.24826839 | -5.9644285 |
| 5827 | NM_005675.4 | 71.683028 | -7.7883722 | 1.305844 | -5.964244 |
| 5828 | NM_015239.2 | 75.2046241 | -7.8556328 | 1.31716155 | -5.9640618 |
| 5829 | NM_005202.3 | 61.7227396 | -7.5731364 | 1.26994894 | -5.9633393 |
| 5830 | NM_012098.2 | 72.2748405 | -7.7986561 | 1.30776185 | -5.9633611 |
| 5831 | NM_024116.3 | 56.1558781 | -7.4339141 | 1.24659776 | -5.9633623 |
| 5832 | NM_001135245.1 | 82.7061298 | -7.9899112 | 1.33991707 | -5.9629894 |
| 5833 | NM_007176.3 | 150.25853 | -7.0777878 | 1.18694785 | -5.9630149 |
| 5834 | NM_182828.3 | 91.3125669 | -8.1361843 | 1.36452288 | -5.9626588 |
| 5835 | NM_017849.3 | 103.871511 | -8.3190106 | 1.3952676 | -5.9623047 |
| 5836 | NR_003662.3 | 78.6114761 | -7.9217313 | 1.32877535 | -5.9616784 |
| 5837 | NM_016218.3 | 57.8582275 | -7.4745219 | 1.25378589 | -5.9615617 |
| 5838 | NM_024105.3 | 63.5379504 | -7.6129696 | 1.27706741 | -5.9612903 |
| 5839 | NM_178860.4 | 179.714772 | -9.1113329 | 1.52843681 | -5.9612101 |
| 5840 | NM_005725.5 | 59.2095656 | -7.5067056 | 1.25928879 | -5.9610675 |
| 5841 | NM_001128920.2 | 80.8060651 | -7.9610487 | 1.33581612 | -5.959689 |
| 5842 | NM_002263.3 | 69.1646592 | -7.7329769 | 1.29759688 | -5.9594601 |
| 5843 | NM_052940.4 | 56.0039143 | -7.4309223 | 1.24693808 | -5.9593354 |
| 5844 | NM_003291.3 | 116.535641 | -6.6511374 | 1.11615291 | -5.9589841 |
| 5845 | NM_000808.3 | 155.613736 | -6.0632513 | 1.01751093 | -5.9589053 |
| 5846 | NM_002749.3 | 76.2841121 | -7.8740572 | 1.32142389 | -5.9587671 |
| 5847 | NM_002853.3 | 65.3636274 | -7.6512788 | 1.28407366 | -5.9585981 |

| | | | | | |
|---|---|---|---|---|---|
| 5848 | NM_032349.3 | 90.2339792 | -8.1173386 | 1.36232716 | -5.9584356 |
| 5849 | NM_138349.3 | 64.7665446 | -7.6402309 | 1.28225551 | -5.9584309 |
| 5850 | NM_030768.2 | 79.6520566 | -7.9368536 | 1.33204546 | -5.9583955 |
| 5851 | NM_001130053.3 | 136.473742 | -8.7143649 | 1.46264808 | -5.9579369 |
| 5852 | NM_000142.4 | 210.293769 | -9.3379024 | 1.56734538 | -5.9577822 |
| 5853 | NR_136661.1 | 57.2779988 | -7.4612607 | 1.25239158 | -5.9576101 |
| 5854 | NM_016070.3 | 84.4602298 | -8.0221251 | 1.3465413 | -5.9575782 |
| 5855 | NM_178191.2 | 63.2555803 | -7.6050844 | 1.27661202 | -5.9572402 |
| 5856 | NM_017728.3 | 76.6826697 | -7.8834919 | 1.32342831 | -5.9568711 |
| 5857 | NM_005606.6 | 105.680463 | -8.3455921 | 1.40118906 | -5.9560785 |
| 5858 | NM_017415.2 | 50.2571654 | -7.2726235 | 1.22113548 | -5.9556238 |
| 5859 | NM_000107.2 | 57.6866534 | -7.4689065 | 1.2542217 | -5.955013 |
| 5860 | NM_003143.2 | 84.3644372 | -8.0201217 | 1.34701565 | -5.9539929 |
| 5861 | NM_015950.4 | 66.731016 | -7.6809712 | 1.29013487 | -5.9536188 |
| 5862 | NM_004714.2 | 66.3682024 | -7.6759974 | 1.2893394 | -5.9534343 |
| 5863 | NM_005851.4 | 67.4221901 | -7.6972292 | 1.29290831 | -5.9534224 |
| 5864 | NM_001040440.2 | 129.676469 | -8.6403641 | 1.45143274 | -5.9529896 |
| 5865 | NM_001286626.1 | 61.3129703 | -7.5576791 | 1.26957784 | -5.9529072 |
| 5866 | NM_198285.2 | 59.8178448 | -7.5255062 | 1.26418309 | -5.952861 |
| 5867 | NM_022453.2 | 72.1073699 | -7.7937167 | 1.30924814 | -5.9528186 |
| 5868 | NR_110492.1 | 106.881263 | -8.3623335 | 1.40477927 | -5.952774 |
| 5869 | NM_016099.2 | 151.768626 | -7.8996335 | 1.32711333 | -5.9524935 |
| 5870 | NM_001014433.2 | 80.5819353 | -7.9547252 | 1.33660296 | -5.9514497 |
| 5871 | NM_052911.2 | 56.6582291 | -7.444811 | 1.25097337 | -5.9512146 |
| 5872 | NM_006785.3 | 64.9619518 | -7.644625 | 1.2846229 | -5.9508709 |
| 5873 | NM_175839.2 | 61.5408031 | -7.565044 | 1.2713352 | -5.9504716 |
| 5874 | NM_016581.4 | 94.6647909 | -8.1860292 | 1.37573016 | -5.950316 |
| 5875 | NM_003617.3 | 111.821577 | -8.4278015 | 1.41665553 | -5.9490831 |
| 5876 | NM_001278356.1 | 64.4021296 | -7.6296034 | 1.28250152 | -5.9490014 |
| 5877 | NM_001312673.1 | 54.9132688 | -7.4038419 | 1.24458367 | -5.9488503 |
| 5878 | NM_001042412.1 | 73.6261788 | -7.8231745 | 1.31513603 | -5.9485668 |
| 5879 | NM_001163280.1 | 72.241244 | -7.7971834 | 1.31082722 | -5.9482922 |
| 5880 | NM_014665.3 | 117.176776 | -7.5269107 | 1.26542031 | -5.9481507 |
| 5881 | NM_024896.2 | 54.9381949 | -7.4046719 | 1.24486629 | -5.9481665 |
| 5882 | NM_016946.4 | 50.2263506 | -7.2739847 | 1.22290529 | -5.9481178 |
| 5883 | NM_015443.3 | 117.965947 | -7.5330045 | 1.26646688 | -5.948047 |
| 5884 | NM_022343.3 | 111.357066 | -8.4216271 | 1.4159971 | -5.9474889 |
| 5885 | NM_001015048.2 | 108.976557 | -7.4170194 | 1.24709743 | -5.9474258 |
| 5886 | NM_020536.4 | 62.0316331 | -7.5772654 | 1.27408353 | -5.9472281 |
| 5887 | NM_001077199.2 | 118.29146 | -8.5079679 | 1.43072157 | -5.9466273 |
| 5888 | NM_001042610.2 | 115.171336 | -8.4698795 | 1.42447065 | -5.9459839 |
| 5889 | NM_022143.4 | 68.7521752 | -7.7257257 | 1.29935068 | -5.9458357 |
| 5890 | NM_000034.3 | 55.0143931 | -7.4008022 | 1.24471482 | -5.9457814 |

| | | | | | |
|---|---|---|---|---|---|
| 5891 | NM_139273.3 | 60.3100131 | -7.5375163 | 1.26771355 | -5.9457567 |
| 5892 | NM_016458.3 | 82.4866414 | -7.9881885 | 1.34357445 | -5.9454752 |
| 5893 | NM_018431.4 | 103.030589 | -8.3081109 | 1.39738833 | -5.9454561 |
| 5894 | NM_183075.2 | 55.0322739 | -7.4054149 | 1.24556936 | -5.9454055 |
| 5895 | NM_001760.4 | 82.4571909 | -7.987269 | 1.34344694 | -5.945355 |
| 5896 | NM_003718.4 | 66.0266776 | -7.66635 | 1.28949545 | -5.9452323 |
| 5897 | NM_018275.4 | 90.0792581 | -8.1153519 | 1.36501879 | -5.9452309 |
| 5898 | NM_000920.3 | 69.4255915 | -7.7380325 | 1.30156585 | -5.9451717 |
| 5899 | NM_018030.4 | 79.9920656 | -7.9456031 | 1.33652817 | -5.9449575 |
| 5900 | NM_032830.2 | 84.5579033 | -8.0227635 | 1.34963082 | -5.9444134 |
| 5901 | NM_001315.2 | 65.8270975 | -7.6600322 | 1.28865155 | -5.944223 |
| 5902 | NM_003423.3 | 62.8821236 | -7.5929353 | 1.27743981 | -5.9438693 |
| 5903 | NR_045634.1 | 72.3391231 | -7.7970024 | 1.31199833 | -5.9428448 |
| 5904 | NM_006545.4 | 77.6455836 | -7.89981 | 1.32932464 | -5.9427244 |
| 5905 | NM_001331012.1 | 90.7024129 | -8.1259147 | 1.36739405 | -5.9426283 |
| 5906 | NM_015602.3 | 107.696271 | -2.7000576 | 0.45443346 | -5.9415907 |
| 5907 | NM_015205.2 | 73.4466842 | -7.8190929 | 1.31603876 | -5.941385 |
| 5908 | NM_198682.2 | 21.6317909 | 6.23985723 | 1.05065852 | 5.93899645 |
| 5909 | NM_016106.3 | 66.9362045 | -7.6835746 | 1.29377238 | -5.9388921 |
| 5910 | NM_153329.3 | 68.4613967 | -7.7190848 | 1.29987829 | -5.9383135 |
| 5911 | NM_206854.2 | 90.024886 | -8.1148331 | 1.36663945 | -5.937801 |
| 5912 | NM_002718.4 | 92.973943 | -8.1600831 | 1.3743067 | -5.9375997 |
| 5913 | NM_001042405.1 | 217.269404 | -8.4205162 | 1.41830304 | -5.937036 |
| 5914 | NM_153207.4 | 63.5182123 | -7.6108717 | 1.28198454 | -5.936789 |
| 5915 | NM_001206612.1 | 120.097256 | -8.5292 | 1.43669396 | -5.9366854 |
| 5916 | NM_030641.3 | 191.052062 | 2.75739947 | 0.46447368 | 5.93661076 |
| 5917 | NM_021170.3 | 85.0289071 | -8.0305944 | 1.35275447 | -5.936476 |
| 5918 | NM_001114394.1 | 55.6170382 | -7.4169544 | 1.24939412 | -5.9364409 |
| 5919 | NM_001256442.1 | 102.661417 | -8.3053947 | 1.39911178 | -5.936191 |
| 5920 | NM_001127183.2 | 66.1224586 | -7.6716956 | 1.29249734 | -5.9355601 |
| 5921 | NM_172170.4 | 71.6118807 | -7.7814413 | 1.31104435 | -5.9352998 |
| 5922 | NM_152704.3 | 96.3774783 | -8.211888 | 1.38363201 | -5.9350231 |
| 5923 | NM_001687.4 | 83.821564 | -8.0113973 | 1.34989829 | -5.9348155 |
| 5924 | NM_001304997.1 | 53.5919708 | -7.3646504 | 1.24100598 | -5.9344197 |
| 5925 | NM_001139500.1 | 91.4483579 | -8.1374502 | 1.37125914 | -5.9342906 |
| 5926 | NM_024417.4 | 55.2406797 | -7.40872 | 1.24846493 | -5.9342636 |
| 5927 | NM_022077.3 | 68.7803878 | -7.7254838 | 1.30189667 | -5.9340222 |
| 5928 | NM_015523.3 | 68.2463367 | -7.7144358 | 1.30008887 | -5.9337757 |
| 5929 | NM_001607.3 | 79.8100649 | -7.9406412 | 1.33824502 | -5.9336229 |
| 5930 | NM_001001433.2 | 62.5747493 | -7.5904212 | 1.27927517 | -5.9333765 |
| 5931 | NM_004961.3 | 92.1067554 | -8.1485171 | 1.37334741 | -5.9333254 |
| 5932 | NM_005370.4 | 112.086532 | -6.639862 | 1.11914767 | -5.9329633 |
| 5933 | NM_021237.4 | 90.2856317 | -8.1178438 | 1.36830398 | -5.9327781 |

| | | | | | |
|---|---|---|---|---|---|
| 5934 | NM_145251.3 | 63.7521566 | -7.6156955 | 1.28371281 | -5.932554 |
| 5935 | NM_152243.2 | 88.2269694 | -8.0861645 | 1.36321877 | -5.9316705 |
| 5936 | NR_027293.2 | 57.1665448 | -7.4589841 | 1.25752918 | -5.9314601 |
| 5937 | NR_109792.1 | 120.121589 | -8.5301996 | 1.43815593 | -5.9313454 |
| 5938 | NM_001282790.1 | 127.936239 | -7.6541717 | 1.29050194 | -5.9311586 |
| 5939 | NM_018994.2 | 145.143397 | -6.2104783 | 1.04710831 | -5.9310754 |
| 5940 | NM_015603.2 | 72.2919982 | -5.4155029 | 0.91310092 | -5.930892 |
| 5941 | NM_130443.3 | 74.0949071 | -7.8333254 | 1.32076822 | -5.9308857 |
| 5942 | NM_001319944.1 | 68.0890909 | -7.7103686 | 1.30009859 | -5.930603 |
| 5943 | NM_016133.3 | 53.6931292 | -7.3728326 | 1.24325801 | -5.9302515 |
| 5944 | NM_144571.2 | 83.5449735 | -5.2089925 | 0.87838971 | -5.9301611 |
| 5945 | NM_004423.3 | 536.839503 | -4.1671492 | 0.70273632 | -5.9298902 |
| 5946 | NM_032848.2 | 57.6181201 | -7.4715179 | 1.26000967 | -5.9297306 |
| 5947 | NM_175931.2 | 83.2480416 | -8.0021273 | 1.34948683 | -5.9297557 |
| 5948 | NM_001079841.2 | 114.17262 | -8.4571555 | 1.42630085 | -5.9294331 |
| 5949 | NM_001319661.1 | 69.0975988 | -7.735021 | 1.30454171 | -5.9293014 |
| 5950 | NM_005842.3 | 75.9489776 | -7.8687937 | 1.32717323 | -5.9289877 |
| 5951 | NM_078470.5 | 75.7830109 | -7.8648726 | 1.32656596 | -5.928746 |
| 5952 | NM_019052.3 | 54.468564 | -7.3894818 | 1.24641158 | -5.9286049 |
| 5953 | NM_005729.3 | 102.643144 | -7.3339764 | 1.23711186 | -5.928305 |
| 5954 | NM_001136044.1 | 84.5047752 | -8.0224865 | 1.35339604 | -5.9276711 |
| 5955 | NM_024297.2 | 120.0488 | -7.5601006 | 1.27541668 | -5.9275535 |
| 5956 | NM_017769.4 | 75.5131497 | -7.8590497 | 1.3258834 | -5.9274063 |
| 5957 | NM_018125.3 | 88.8116535 | -8.0953071 | 1.36573662 | -5.9274292 |
| 5958 | NM_017553.2 | 74.3554527 | -7.837326 | 1.32229542 | -5.9270613 |
| 5959 | NM_181481.4 | 82.3668502 | -7.9883215 | 1.34800938 | -5.9260133 |
| 5960 | NM_024701.3 | 71.9618031 | -7.7933602 | 1.31516107 | -5.9257838 |
| 5961 | NR_110730.1 | 58.6909122 | -7.4987407 | 1.26552823 | -5.925384 |
| 5962 | NM_007212.3 | 92.8965695 | -8.1591186 | 1.3770105 | -5.9252406 |
| 5963 | NR_104013.1 | 54.2837238 | -7.3878882 | 1.24685083 | -5.9252382 |
| 5964 | NM_001201338.1 | 125.256426 | -8.5917988 | 1.4500574 | -5.9251439 |
| 5965 | NR_033266.1 | 80.3265353 | -5.5372697 | 0.93462597 | -5.9245836 |
| 5966 | NM_007066.4 | 61.3483342 | -7.5607835 | 1.27624828 | -5.9242262 |
| 5967 | NM_017746.3 | 74.2997366 | -7.836952 | 1.32286976 | -5.9242052 |
| 5968 | NM_178439.4 | 65.1969897 | -4.0368507 | 0.68149283 | -5.923541 |
| 5969 | NM_001281533.1 | 120.353693 | -8.5325742 | 1.44046779 | -5.9234745 |
| 5970 | NM_001141969.1 | 72.0217374 | -7.790806 | 1.3152949 | -5.923239 |
| 5971 | NM_152316.2 | 75.3367336 | -7.856871 | 1.32656336 | -5.9227258 |
| 5972 | NM_003824.3 | 68.9386984 | -7.7293893 | 1.30508454 | -5.92252 |
| 5973 | NM_018407.4 | 669.772887 | -6.3524108 | 1.07260931 | -5.9223902 |
| 5974 | NM_003081.4 | 194.110415 | -9.2225263 | 1.55726891 | -5.9222439 |
| 5975 | NM_006766.4 | 250.912489 | -5.7792328 | 0.97586892 | -5.9221404 |
| 5976 | NM_004928.2 | 54.476247 | -7.3867394 | 1.24731681 | -5.9221036 |

| | | | | | |
|---|---|---|---|---|---|
| 5977 | NM_001198869.1 | 94.8378529 | -8.1913825 | 1.38336061 | -5.9213646 |
| 5978 | NM_001587.3 | 146.68368 | -8.8198432 | 1.48951997 | -5.9212655 |
| 5979 | NM_005400.2 | 120.955747 | -8.5401257 | 1.44230009 | -5.921185 |
| 5980 | NM_001330396.1 | 96.8698816 | -6.3534536 | 1.07308205 | -5.9207529 |
| 5981 | NM_005034.3 | 99.7518549 | -8.2619543 | 1.39543749 | -5.920691 |
| 5982 | NM_003818.3 | 341.010754 | -4.0112605 | 0.67754974 | -5.9202451 |
| 5983 | NM_018989.1 | 111.394315 | -7.4502638 | 1.25850402 | -5.9199364 |
| 5984 | NM_033416.2 | 78.3604955 | -7.9151116 | 1.33708873 | -5.9196607 |
| 5985 | NM_033025.5 | 73.1547049 | -7.816274 | 1.32044564 | -5.9194213 |
| 5986 | NR_109843.1 | 55.3950925 | -7.4135042 | 1.25242173 | -5.9193353 |
| 5987 | NM_001130709.1 | 66.4069102 | -7.6753225 | 1.29696603 | -5.9179056 |
| 5988 | NM_000084.4 | 96.213517 | -5.4270959 | 0.91713632 | -5.9174364 |
| 5989 | NM_198055.1 | 51.7151828 | -7.3175986 | 1.23670476 | -5.9170133 |
| 5990 | NM_016091.3 | 808.958295 | -7.4287734 | 1.25552312 | -5.916875 |
| 5991 | NM_138732.2 | 175.790795 | -7.3133485 | 1.23614629 | -5.9162484 |
| 5992 | NR_028025.1 | 62.5867461 | -7.5903049 | 1.28295487 | -5.9162681 |
| 5993 | NM_001136223.1 | 91.7092825 | -8.140682 | 1.37599397 | -5.9162192 |
| 5994 | NM_016271.4 | 75.3582549 | -7.8567902 | 1.32809825 | -5.9158201 |
| 5995 | NM_015342.3 | 56.6039178 | -7.4429157 | 1.25815098 | -5.9157572 |
| 5996 | NM_014157.3 | 80.451025 | -7.95353 | 1.34456843 | -5.9153033 |
| 5997 | NM_005431.1 | 107.79226 | 1.67325218 | 0.28287461 | 5.91517273 |
| 5998 | NM_000249.3 | 72.4532466 | -7.7998155 | 1.3186466 | -5.9150158 |
| 5999 | NM_005800.4 | 65.6359339 | -7.6582392 | 1.29473896 | -5.9148905 |
| 6000 | NM_001198801.1 | 71.4295248 | -7.7828574 | 1.31591554 | -5.914405 |
| 6001 | NM_003748.3 | 112.221283 | -8.4330066 | 1.42588546 | -5.9142244 |
| 6002 | NM_144664.4 | 61.3312751 | -7.5612822 | 1.27849799 | -5.9141917 |
| 6003 | NM_006645.2 | 57.0231172 | -7.4555814 | 1.26066289 | -5.9140167 |
| 6004 | NM_001242833.1 | 111.479838 | -8.4232425 | 1.42433129 | -5.9138225 |
| 6005 | NM_001359.1 | 52.1971104 | -7.3316092 | 1.23974147 | -5.913821 |
| 6006 | NM_002095.4 | 60.9319244 | -7.5507453 | 1.27679552 | -5.913825 |
| 6007 | NM_004435.2 | 82.3929841 | -7.9862874 | 1.35048381 | -5.9136491 |
| 6008 | NM_001040025.2 | 75.0651012 | -7.851609 | 1.32773407 | -5.9135403 |
| 6009 | NM_002515.2 | 93.2024993 | -8.1639614 | 1.38055923 | -5.9135177 |
| 6010 | NM_012265.2 | 80.6303159 | -7.9551521 | 1.34523698 | -5.9135693 |
| 6011 | NM_019109.4 | 77.4066188 | -7.8960357 | 1.33526227 | -5.9134717 |
| 6012 | NM_001077207.3 | 105.780262 | -8.3471652 | 1.41158228 | -5.9133395 |
| 6013 | NM_001032381.1 | 72.975139 | -7.8104335 | 1.32088679 | -5.9130226 |
| 6014 | NM_001991.4 | 75.4287761 | -7.8611397 | 1.32951937 | -5.912768 |
| 6015 | NM_032314.3 | 78.3076459 | -7.9123579 | 1.338338 | -5.9120774 |
| 6016 | NM_000156.5 | 91.6546852 | -8.1402876 | 1.37692646 | -5.9119262 |
| 6017 | NM_000038.5 | 273.806939 | -8.7552398 | 1.48097565 | -5.9118054 |
| 6018 | NM_018697.3 | 159.304281 | -6.2224258 | 1.05260604 | -5.9114479 |
| 6019 | NM_015726.3 | 120.804444 | -5.2333744 | 0.88535375 | -5.9110547 |

| | | | | | |
|---|---|---|---|---|---|
| 6020 | NM_023039.4 | 59.4645572 | -7.5140564 | 1.27158921 | -5.9091854 |
| 6021 | NM_005050.3 | 61.3916158 | -7.5602309 | 1.27944991 | -5.9089699 |
| 6022 | NM_006556.3 | 91.9634037 | -8.1445638 | 1.37837018 | -5.9088363 |
| 6023 | NM_145201.5 | 63.218077 | -7.6076394 | 1.28754488 | -5.9086402 |
| 6024 | NM_173808.2 | 173.114086 | -9.0574796 | 1.53294065 | -5.9085651 |
| 6025 | NM_001278530.1 | 69.112173 | -7.7341599 | 1.30929903 | -5.9070997 |
| 6026 | NM_016587.3 | 430.189096 | -4.8813419 | 0.82635377 | -5.907085 |
| 6027 | NM_178148.3 | 72.1124321 | -7.794126 | 1.31951341 | -5.9068183 |
| 6028 | NM_001114618.1 | 69.4848919 | -7.7372521 | 1.30990366 | -5.9067337 |
| 6029 | NM_003358.2 | 140.159773 | -7.788081 | 1.31853362 | -5.906623 |
| 6030 | NM_153613.2 | 148.378895 | -6.1151092 | 1.03534428 | -5.9063533 |
| 6031 | NM_001135178.2 | 58.6409206 | -7.4934139 | 1.26876952 | -5.9060482 |
| 6032 | NM_004107.4 | 64.8985824 | -7.6448405 | 1.29448632 | -5.9056943 |
| 6033 | NM_001297600.1 | 62.1157667 | -7.5821668 | 1.28403219 | -5.9049663 |
| 6034 | NM_052950.3 | 49.6540805 | -7.2548271 | 1.22863979 | -5.9047633 |
| 6035 | NM_021184.3 | 57.6266651 | -7.4715583 | 1.26537769 | -5.9046073 |
| 6036 | NM_017931.2 | 53.318906 | -7.3612957 | 1.24675566 | -5.9043612 |
| 6037 | NM_012175.3 | 78.2196939 | -7.9115698 | 1.34000644 | -5.9041282 |
| 6038 | NM_014285.6 | 60.2492187 | -7.5325839 | 1.27582286 | -5.9040986 |
| 6039 | NM_015002.2 | 255.736282 | -5.2316809 | 0.88610812 | -5.9041112 |
| 6040 | NM_006070.5 | 78.3738521 | -7.9123134 | 1.34020593 | -5.9038042 |
| 6041 | NM_001033859.2 | 65.0335007 | -7.6488631 | 1.29566977 | -5.9034047 |
| 6042 | NM_003958.3 | 65.4147262 | -7.6531714 | 1.2964063 | -5.9033741 |
| 6043 | NM_015442.2 | 66.710062 | -7.6801791 | 1.30114052 | -5.9026516 |
| 6044 | NM_007124.2 | 97.7919063 | -7.2692167 | 1.23154244 | -5.9025304 |
| 6045 | NM_018389.4 | 74.6168847 | -7.8444146 | 1.32900022 | -5.9024931 |
| 6046 | NM_030954.3 | 57.3410604 | -7.4665604 | 1.26505914 | -5.9021433 |
| 6047 | NM_001281775.2 | 65.4957873 | -7.6586474 | 1.29767089 | -5.9018411 |
| 6048 | NM_001318005.1 | 59.9185184 | -7.5266828 | 1.27536562 | -5.9015883 |
| 6049 | NM_016539.3 | 65.5417392 | -7.6543211 | 1.29718372 | -5.9007225 |
| 6050 | NM_001318798.1 | 56.5895885 | -7.4462133 | 1.26197966 | -5.9004225 |
| 6051 | NM_006537.3 | 75.2847109 | -7.8562693 | 1.33147658 | -5.9004187 |
| 6052 | NM_033182.5 | 62.0116351 | -7.5725114 | 1.28338941 | -5.9004004 |
| 6053 | NM_001277074.1 | 73.1482962 | -7.8142792 | 1.32438317 | -5.900316 |
| 6054 | NM_003427.4 | 54.7536478 | -7.3969005 | 1.25367147 | -5.9001905 |
| 6055 | NR_015404.1 | 70.7944326 | -7.7662093 | 1.31628038 | -5.9001178 |
| 6056 | NM_001011649.2 | 69.7161573 | -7.7437829 | 1.31251348 | -5.8999644 |
| 6057 | NM_001206528.2 | 121.61535 | -8.5487036 | 1.44904215 | -5.8995548 |
| 6058 | NM_030937.4 | 76.1150943 | -7.871876 | 1.33433307 | -5.8994836 |
| 6059 | NM_017411.3 | 56.4340603 | -7.4438585 | 1.26182119 | -5.8992974 |
| 6060 | NM_153485.2 | 58.9458763 | -7.5016244 | 1.27165279 | -5.8991138 |
| 6061 | NR_033781.1 | 92.9659311 | 3.14899381 | 0.53382333 | 5.89894382 |
| 6062 | NM_002370.3 | 60.8495487 | -7.5479005 | 1.27966395 | -5.8983458 |

| | | | | | |
|---|---|---|---|---|---|
| 6063 | NM_015267.3 | 158.549206 | -7.1599113 | 1.21392289 | -5.8981599 |
| 6064 | NM_001278710.1 | 85.7197682 | -8.0424604 | 1.36367486 | -5.8976378 |
| 6065 | NM_001017440.2 | 50.3399217 | -7.2755513 | 1.2336866 | -5.8974065 |
| 6066 | NM_205861.2 | 62.8760708 | -7.5957165 | 1.28798044 | -5.897385 |
| 6067 | NM_181485.2 | 67.8050772 | -7.7048043 | 1.30653256 | -5.8971391 |
| 6068 | NM_000157.3 | 79.3112506 | -7.9313928 | 1.34501207 | -5.8968934 |
| 6069 | NM_001318729.1 | 173.647582 | -8.0942699 | 1.37277975 | -5.8962626 |
| 6070 | NM_013330.4 | 50.8575322 | -7.2884576 | 1.23616605 | -5.8960183 |
| 6071 | NM_014887.2 | 101.251162 | -7.316907 | 1.24103518 | -5.8958095 |
| 6072 | NM_018571.5 | 96.8522422 | -7.2490828 | 1.22968256 | -5.8950847 |
| 6073 | NM_203350.2 | 113.245746 | -7.4756097 | 1.2682032 | -5.8946466 |
| 6074 | NM_017966.4 | 78.9139019 | -4.5306237 | 0.76863938 | -5.8943424 |
| 6075 | NM_006633.3 | 55.6855244 | -7.4223499 | 1.25930069 | -5.8940251 |
| 6076 | NM_002406.3 | 74.7525533 | -7.848693 | 1.33172241 | -5.8936404 |
| 6077 | NM_001260.2 | 64.7586171 | -7.6366143 | 1.29579877 | -5.8933643 |
| 6078 | NM_016538.2 | 52.1402724 | -7.3267187 | 1.24324446 | -5.8932245 |
| 6079 | NM_001206999.1 | 57.738328 | -7.4706348 | 1.2676896 | -5.8931104 |
| 6080 | NM_001322898.1 | 90.8769233 | -8.128197 | 1.37926785 | -5.8931243 |
| 6081 | NM_002061.3 | 84.6760143 | -8.0269425 | 1.3620789 | -5.8931553 |
| 6082 | NM_005876.4 | 155.607807 | -7.0394746 | 1.19454855 | -5.893 |
| 6083 | NR_026659.1 | 73.4904503 | 1.7566836 | 0.29809984 | 5.89293701 |
| 6084 | NM_024583.4 | 66.3211717 | -7.6728244 | 1.30209153 | -5.8926921 |
| 6085 | NM_001004439.1 | 237.114499 | -8.5480045 | 1.45078182 | -5.8919986 |
| 6086 | NM_021214.1 | 96.7616652 | -8.2182646 | 1.39487544 | -5.8917552 |
| 6087 | NM_006869.3 | 110.939051 | -8.4157794 | 1.42852891 | -5.8912209 |
| 6088 | NM_001167734.1 | 101.185229 | -8.2819126 | 1.40582578 | -5.8911372 |
| 6089 | NM_001254738.1 | 76.1700312 | -7.86981 | 1.33599544 | -5.8905965 |
| 6090 | NM_001008493.2 | 97.9923534 | -8.2351072 | 1.39805997 | -5.890382 |
| 6091 | NM_007317.2 | 120.286652 | -7.5653327 | 1.28435727 | -5.8903647 |
| 6092 | NM_032312.3 | 69.3302302 | -7.7374355 | 1.31369669 | -5.8898188 |
| 6093 | NM_138357.2 | 62.459153 | -7.5874038 | 1.28823437 | -5.8897698 |
| 6094 | NM_182923.3 | 73.4102313 | -7.8184171 | 1.32753752 | -5.8894132 |
| 6095 | NM_032412.3 | 56.854761 | -7.4553109 | 1.26609101 | -5.8884478 |
| 6096 | NM_004765.3 | 79.5472359 | -7.9359386 | 1.34775374 | -5.8882705 |
| 6097 | NM_025191.3 | 77.0067035 | -7.892055 | 1.34031241 | -5.8882205 |
| 6098 | NM_020453.3 | 47.8216707 | -7.2020578 | 1.22315554 | -5.8880965 |
| 6099 | NM_024546.3 | 75.7875857 | -7.8654762 | 1.33585089 | -5.8879896 |
| 6100 | NM_001257361.1 | 55.3014551 | -7.4120158 | 1.2588587 | -5.8878854 |
| 6101 | NM_001282112.1 | 63.9913108 | -7.6225955 | 1.29470904 | -5.887497 |
| 6102 | NM_130466.3 | 62.6761835 | -7.5932289 | 1.2897307 | -5.8874531 |
| 6103 | NM_015399.3 | 77.3135282 | -7.8941631 | 1.34087342 | -5.8873291 |
| 6104 | NM_017832.3 | 48.5828636 | -7.2243531 | 1.22710107 | -5.8873334 |
| 6105 | NM_024591.4 | 70.8393921 | -7.7690715 | 1.31965691 | -5.8871904 |

| | | | | | |
|---|---|---|---|---|---|
| 6106 | NM_206836.2 | 50.5296065 | -7.2842143 | 1.23732802 | -5.887052 |
| 6107 | NM_007240.2 | 72.7064419 | -7.8053205 | 1.32587089 | -5.8869386 |
| 6108 | NM_001042618.1 | 78.0762745 | -7.9093408 | 1.34361167 | -5.886627 |
| 6109 | NM_024571.3 | 82.8133487 | -7.9938441 | 1.35810339 | -5.886035 |
| 6110 | NM_022039.3 | 50.6818148 | -7.2836928 | 1.23745961 | -5.8860045 |
| 6111 | NM_001130967.2 | 49.6433286 | -7.2547784 | 1.23256345 | -5.885927 |
| 6112 | NM_024721.4 | 97.5905957 | -7.2646717 | 1.23424735 | -5.8859124 |
| 6113 | NM_001079537.1 | 80.9351791 | -7.9596768 | 1.35234839 | -5.8858183 |
| 6114 | NM_001305424.1 | 86.5321751 | -8.0559427 | 1.36891885 | -5.8848943 |
| 6115 | NM_138426.3 | 108.163844 | -7.4072542 | 1.25879135 | -5.8844178 |
| 6116 | NM_145698.4 | 73.4252652 | -7.81967 | 1.32893068 | -5.884182 |
| 6117 | NM_017807.3 | 66.1399643 | -7.6695366 | 1.30347871 | -5.8838986 |
| 6118 | NM_025205.4 | 118.519245 | -6.7251666 | 1.14323061 | -5.8825985 |
| 6119 | NM_199342.3 | 93.6427307 | -8.1705732 | 1.38893548 | -5.8826153 |
| 6120 | NM_032663.4 | 74.3557473 | -7.8363096 | 1.33213196 | -5.8825325 |
| 6121 | NM_016098.3 | 85.4117268 | -8.0382073 | 1.36656779 | -5.8820407 |
| 6122 | NM_152227.2 | 60.5377554 | -7.5418027 | 1.28220336 | -5.8819084 |
| 6123 | NR_135223.1 | 61.5056178 | -7.5647726 | 1.28612325 | -5.8818411 |
| 6124 | NM_001171941.2 | 55.2265726 | -7.4133721 | 1.26040856 | -5.8817215 |
| 6125 | NM_001317919.1 | 76.3384083 | -7.8788904 | 1.33971558 | -5.8810172 |
| 6126 | NM_004220.2 | 75.6765268 | -7.8653291 | 1.33742929 | -5.8809308 |
| 6127 | NM_003180.2 | 90.8828836 | -8.1275684 | 1.38215889 | -5.880343 |
| 6128 | NM_007246.3 | 73.0541636 | -7.8117751 | 1.32851133 | -5.8800968 |
| 6129 | NM_017549.4 | 69.5027301 | -7.7436433 | 1.31716451 | -5.8790252 |
| 6130 | NM_024296.4 | 87.8779959 | -8.0793046 | 1.37428584 | -5.8789114 |
| 6131 | NM_015188.1 | 75.2933106 | -7.8556035 | 1.33637802 | -5.8782795 |
| 6132 | NM_025106.3 | 81.7787459 | -7.976654 | 1.35701547 | -5.8780863 |
| 6133 | NM_001762.3 | 641.518599 | -6.7822005 | 1.15392131 | -5.8775243 |
| 6134 | NM_001199873.1 | 80.682493 | -7.9551707 | 1.35358591 | -5.8771081 |
| 6135 | NM_032985.4 | 64.09327 | -7.6199955 | 1.29660584 | -5.8768789 |
| 6136 | NM_001142419.1 | 63.2076523 | -7.603426 | 1.2938098 | -5.8767726 |
| 6137 | NM_004635.4 | 53.107906 | -7.3547695 | 1.25149529 | -5.8767856 |
| 6138 | NM_001301085.1 | 64.5967412 | -7.637688 | 1.2996503 | -5.8767255 |
| 6139 | NM_173348.1 | 56.5473241 | -7.4438433 | 1.26666933 | -5.876706 |
| 6140 | NM_032194.2 | 60.0378514 | -7.5278207 | 1.28096572 | -5.8766761 |
| 6141 | NM_025065.6 | 62.7986572 | -7.5939865 | 1.29224072 | -5.8766036 |
| 6142 | NM_052951.2 | 71.2570092 | -7.7774262 | 1.32354136 | -5.8762245 |
| 6143 | NM_004535.2 | 136.650281 | -6.3882191 | 1.08716591 | -5.8760296 |
| 6144 | NM_006157.4 | 88.5142501 | -8.089643 | 1.37671391 | -5.8760524 |
| 6145 | NM_001199135.1 | 62.6417613 | -7.5903695 | 1.29179092 | -5.8758499 |
| 6146 | NM_016101.4 | 49.6113122 | -7.2527876 | 1.23434483 | -5.8758196 |
| 6147 | NM_000521.3 | 105.551427 | -7.3754171 | 1.25528764 | -5.8754798 |
| 6148 | NM_032511.3 | 90.9943756 | -8.130618 | 1.38392722 | -5.8750329 |

| | | | | | |
|---|---|---|---|---|---|
| 6149 | NM_032182.3 | 63.4455888 | -7.6083629 | 1.29506377 | -5.8748944 |
| 6150 | NM_024319.3 | 71.5375835 | -7.7818575 | 1.32469524 | -5.8744511 |
| 6151 | NM_145246.4 | 63.6326872 | -7.6126929 | 1.29597287 | -5.8741144 |
| 6152 | NM_030665.3 | 358.735224 | -6.5911466 | 1.12212298 | -5.8738184 |
| 6153 | NM_001083614.1 | 56.546618 | -7.4429407 | 1.26722438 | -5.8734197 |
| 6154 | NM_001145659.1 | 25.9581018 | 9.62658187 | 1.63906739 | 5.8732069 |
| 6155 | NM_001282146.1 | 62.4686967 | -7.5854509 | 1.29171328 | -5.8723953 |
| 6156 | NM_004807.2 | 602.441963 | -6.201214 | 1.0560421 | -5.8721277 |
| 6157 | NM_001102564.1 | 56.2703358 | -7.4349689 | 1.26648797 | -5.8705405 |
| 6158 | NM_001114172.1 | 99.6498903 | -8.2613305 | 1.40730629 | -5.8703145 |
| 6159 | NM_001317899.1 | 85.42986 | -8.0381135 | 1.36927738 | -5.8703325 |
| 6160 | NM_025164.4 | 62.1712217 | -7.5801508 | 1.29140484 | -5.8696937 |
| 6161 | NM_030803.6 | 74.1931238 | -7.8346321 | 1.33478317 | -5.8695916 |
| 6162 | NM_004707.3 | 88.0998319 | -8.0836152 | 1.37724898 | -5.8693927 |
| 6163 | NM_001145199.1 | 59.3902132 | -7.5143903 | 1.28027645 | -5.8693498 |
| 6164 | NM_032351.5 | 74.3250296 | -7.8382742 | 1.33549175 | -5.8692045 |
| 6165 | NM_018453.3 | 65.5422731 | -7.6566033 | 1.30484694 | -5.8678172 |
| 6166 | NM_001282281.1 | 87.2138411 | -8.0680995 | 1.37503188 | -5.8675727 |
| 6167 | NM_015305.3 | 58.2461642 | -7.4872841 | 1.27620782 | -5.866822 |
| 6168 | NM_000748.2 | 116.690539 | -8.4884785 | 1.44698257 | -5.8663308 |
| 6169 | NM_005224.2 | 53.7685994 | -7.3746547 | 1.25715825 | -5.8661308 |
| 6170 | NM_000155.3 | 53.9659798 | -7.3770602 | 1.25763747 | -5.8658082 |
| 6171 | NM_019606.5 | 96.6244429 | -7.2447017 | 1.23512939 | -5.8655407 |
| 6172 | NM_001243042.1 | 69.5606574 | -7.7443993 | 1.32042639 | -5.8650746 |
| 6173 | NR_003265.3 | 47.7475005 | -7.1999816 | 1.22760063 | -5.8650846 |
| 6174 | NM_022063.2 | 65.3820046 | -7.6505916 | 1.30447774 | -5.8648694 |
| 6175 | NM_004443.3 | 108.262007 | -5.0674766 | 0.86406696 | -5.8646805 |
| 6176 | NM_001136499.1 | 59.6843617 | -7.5181172 | 1.28194709 | -5.864608 |
| 6177 | NM_016524.3 | 54.6207956 | -7.3959588 | 1.26114664 | -5.8644717 |
| 6178 | NM_001284364.2 | 88.0692769 | -8.0825466 | 1.37825361 | -5.8643391 |
| 6179 | NM_020813.3 | 64.4686919 | -7.630958 | 1.30130611 | -5.8640761 |
| 6180 | NM_032813.3 | 111.681363 | -7.4580148 | 1.2718374 | -5.8639687 |
| 6181 | NM_007187.4 | 72.6984042 | -7.8043203 | 1.33094953 | -5.8637237 |
| 6182 | NM_006058.4 | 108.989346 | -8.388798 | 1.43064834 | -5.8636338 |
| 6183 | NM_012310.4 | 54.1660682 | -7.3809255 | 1.2587724 | -5.8635902 |
| 6184 | NM_001034850.2 | 76.4755505 | -7.8797187 | 1.34388907 | -5.8633699 |
| 6185 | NM_006443.2 | 52.1367415 | -7.3271147 | 1.24964411 | -5.8633611 |
| 6186 | NM_018207.2 | 66.8980047 | -7.6866525 | 1.31108234 | -5.8628298 |
| 6187 | NM_032339.4 | 93.0047592 | -8.1612042 | 1.39209725 | -5.8625245 |
| 6188 | NM_182943.2 | 100.992201 | -8.2810899 | 1.4126455 | -5.8621147 |
| 6189 | NM_020689.3 | 191.704394 | -5.8589796 | 0.99947062 | -5.8620828 |
| 6190 | NM_018337.3 | 65.8201821 | -7.6619064 | 1.30705769 | -5.8619496 |
| 6191 | NM_006067.4 | 79.8570311 | -7.9402495 | 1.35459967 | -5.8616945 |

| | | | | | |
|---|---|---|---|---|---|
| 6192 | NM_001571.5 | 51.8837857 | -7.316626 | 1.24831708 | -5.8611919 |
| 6193 | NM_033428.2 | 87.9567043 | -8.0805724 | 1.37867084 | -5.8611325 |
| 6194 | NM_018685.4 | 64.8114456 | -7.6400166 | 1.3036606 | -5.8604338 |
| 6195 | NM_001080414.3 | 108.451497 | -4.0208188 | 0.68613804 | -5.8600728 |
| 6196 | NM_033510.2 | 119.134603 | -8.5184555 | 1.45366818 | -5.8599725 |
| 6197 | NM_005509.5 | 67.8793664 | -7.7062411 | 1.31508749 | -5.8598696 |
| 6198 | NM_020644.2 | 75.478845 | -7.8599709 | 1.3413543 | -5.8597276 |
| 6199 | NM_017637.5 | 53.5344752 | -7.3668053 | 1.25729037 | -5.8592713 |
| 6200 | NM_005859.4 | 52.5207732 | -7.3373155 | 1.25254379 | -5.8579313 |
| 6201 | NM_001135190.1 | 73.4383416 | -4.3730008 | 0.74651852 | -5.8578598 |
| 6202 | NM_001017395.3 | 85.81823 | -8.0453722 | 1.37351966 | -5.857486 |
| 6203 | NM_007265.2 | 60.8870292 | -7.5494017 | 1.28886499 | -5.8574031 |
| 6204 | NM_000194.2 | 58.3243325 | -7.4859153 | 1.27815079 | -5.8568327 |
| 6205 | NM_033449.2 | 59.1230137 | -7.5087867 | 1.28218223 | -5.8562555 |
| 6206 | NM_001098426.1 | 54.2663428 | -7.3858473 | 1.26123926 | -5.856024 |
| 6207 | NM_004313.3 | 107.155678 | -8.3656128 | 1.42855975 | -5.8559769 |
| 6208 | NM_020932.2 | 154.824463 | -7.9319026 | 1.35459804 | -5.8555397 |
| 6209 | NM_021809.6 | 54.2837195 | -7.3805714 | 1.26043824 | -5.8555597 |
| 6210 | NM_000146.3 | 2656.01625 | -6.9857488 | 1.1930218 | -5.8555081 |
| 6211 | NM_004897.4 | 70.5217334 | -7.7626806 | 1.32587077 | -5.8547792 |
| 6212 | NM_018584.5 | 453.07348 | -7.3696908 | 1.2587477 | -5.8547799 |
| 6213 | NM_198925.2 | 58.0074849 | -7.4800544 | 1.27767384 | -5.8544318 |
| 6214 | NM_022443.4 | 57.7089064 | -7.471852 | 1.27633749 | -5.854135 |
| 6215 | NM_001346231.1 | 76.6535494 | -7.882388 | 1.3464782 | -5.8540777 |
| 6216 | NM_021218.2 | 68.8812496 | -7.7264801 | 1.31989069 | -5.8538787 |
| 6217 | NM_020827.2 | 110.992003 | -6.6271588 | 1.13214527 | -5.8536294 |
| 6218 | NM_001113434.4 | 61.9984649 | -4.4442142 | 0.75925209 | -5.8534105 |
| 6219 | NM_030949.2 | 60.6453058 | -7.5438142 | 1.28880183 | -5.8533547 |
| 6220 | NM_012329.2 | 183.034596 | -8.1708776 | 1.39594214 | -5.8533068 |
| 6221 | NM_001034023.1 | 64.1636507 | -7.6274804 | 1.30315202 | -5.853101 |
| 6222 | NM_003276.2 | 79.0306459 | -7.9256889 | 1.35430739 | -5.8522082 |
| 6223 | NM_001014286.2 | 68.5803701 | -7.7217154 | 1.31960511 | -5.8515349 |
| 6224 | NM_174893.3 | 63.9766991 | -7.6193449 | 1.30217894 | -5.8512272 |
| 6225 | NM_015436.3 | 77.1085325 | -7.8912075 | 1.34867944 | -5.8510624 |
| 6226 | NM_019080.2 | 102.942986 | -8.3072803 | 1.41979707 | -5.8510336 |
| 6227 | NM_001287000.1 | 62.2999212 | -7.5809735 | 1.29569061 | -5.8509134 |
| 6228 | NM_002719.3 | 67.8617059 | -7.7077212 | 1.31736059 | -5.8508819 |
| 6229 | NM_001172688.1 | 48.6617961 | -7.2282344 | 1.23542788 | -5.8507943 |
| 6230 | NM_001172624.1 | 194.739525 | -9.2282427 | 1.57727865 | -5.8507371 |
| 6231 | NM_002075.3 | 51.7374125 | -7.3190412 | 1.251107 | -5.8500522 |
| 6232 | NM_004604.4 | 57.6349394 | -7.4714596 | 1.27716793 | -5.8500213 |
| 6233 | NM_003092.4 | 135.707199 | -7.7400585 | 1.32311322 | -5.8498837 |
| 6234 | NM_014836.4 | 57.2115951 | -7.4642896 | 1.2759745 | -5.8498736 |

| | | | | | |
|---|---|---|---|---|---|
| 6235 | NM_139265.3 | 55.3783462 | -7.4161634 | 1.2677871 | -5.8496915 |
| 6236 | NM_031480.2 | 83.0450366 | -7.9965423 | 1.36705429 | -5.8494695 |
| 6237 | NM_012071.3 | 86.8759972 | -8.0622603 | 1.37830685 | -5.8493943 |
| 6238 | NM_004053.3 | 69.7549513 | -7.7465018 | 1.32435082 | -5.8492823 |
| 6239 | NM_080592.3 | 84.2033835 | -8.0174483 | 1.37068942 | -5.8492085 |
| 6240 | NM_005604.3 | 76.790637 | -7.8834392 | 1.34782209 | -5.8490206 |
| 6241 | NM_012138.3 | 104.123125 | -5.2638012 | 0.90002011 | -5.8485373 |
| 6242 | NR_003264.2 | 48.1007089 | -7.2092211 | 1.2326504 | -5.8485529 |
| 6243 | NM_152441.2 | 63.4978931 | -7.6094645 | 1.30110262 | -5.8484737 |
| 6244 | NR_003098.1 | 125.267032 | -7.6242312 | 1.30366902 | -5.8482874 |
| 6245 | NM_015463.2 | 103.671522 | -8.3176535 | 1.42237802 | -5.8477095 |
| 6246 | NM_001002292.3 | 85.6155783 | -8.0427708 | 1.37537982 | -5.8476726 |
| 6247 | NM_002121.5 | 115.942673 | -8.4797936 | 1.45026267 | -5.8470743 |
| 6248 | NM_201542.4 | 74.4187696 | -7.8397299 | 1.34080053 | -5.8470516 |
| 6249 | NM_015412.3 | 53.2755137 | -7.3580072 | 1.25853698 | -5.8464768 |
| 6250 | NM_001388.4 | 69.9977624 | -7.7515095 | 1.32588594 | -5.8462868 |
| 6251 | NM_006468.7 | 70.4042802 | -7.7601333 | 1.32742564 | -5.8460022 |
| 6252 | NM_001113378.1 | 62.715485 | -7.5901629 | 1.29843328 | -5.8456318 |
| 6253 | NM_080796.3 | 51.8995798 | -7.3219179 | 1.25255161 | -5.8456018 |
| 6254 | NM_000976.3 | 925.309846 | -6.8964801 | 1.17979908 | -5.84547 |
| 6255 | NM_001308317.1 | 93.6568341 | -8.1707214 | 1.39784285 | -5.845236 |
| 6256 | NM_001098790.1 | 56.8760611 | -7.4500157 | 1.27462069 | -5.8448884 |
| 6257 | NM_003020.4 | 116.684937 | -8.4880083 | 1.45222517 | -5.8448294 |
| 6258 | NM_153631.2 | 92.9851165 | -8.1619979 | 1.39646253 | -5.8447668 |
| 6259 | NR_125420.1 | 36.5620087 | 2.90919344 | 0.49775272 | 5.84465606 |
| 6260 | NM_001198905.1 | 65.8919405 | -7.6622912 | 1.31104259 | -5.8444259 |
| 6261 | NM_001243781.1 | 80.1351913 | -7.9487039 | 1.36004545 | -5.8444399 |
| 6262 | NM_178833.5 | 54.1672665 | -7.3813642 | 1.26306094 | -5.8440285 |
| 6263 | NM_015955.3 | 61.6828006 | -7.5684083 | 1.29514852 | -5.8436605 |
| 6264 | NM_001320977.1 | 76.2971317 | -7.8752127 | 1.34766558 | -5.8435956 |
| 6265 | NM_006225.3 | 64.9951493 | -7.6454475 | 1.30835126 | -5.8435741 |
| 6266 | NM_004272.4 | 78.6145325 | -7.9177864 | 1.35497378 | -5.8434979 |
| 6267 | NM_001164315.1 | 86.7842994 | -8.0603749 | 1.37938283 | -5.8434647 |
| 6268 | NM_018674.5 | 70.4732276 | -7.7615999 | 1.32838605 | -5.8428797 |
| 6269 | NM_145253.2 | 62.6344344 | -7.5911809 | 1.29926184 | -5.8426875 |
| 6270 | NM_001320663.1 | 86.2329676 | -8.0511016 | 1.37800475 | -5.842579 |
| 6271 | NM_198098.3 | 497.203488 | -10.58007 | 1.81094158 | -5.8423033 |
| 6272 | NM_001166278.1 | 67.2021182 | -7.6921326 | 1.3166783 | -5.8420744 |
| 6273 | NM_001282874.1 | 123.707562 | -8.5730986 | 1.46749081 | -5.8420118 |
| 6274 | NM_001346494.1 | 82.3500248 | -7.9854536 | 1.36692097 | -5.8419278 |
| 6275 | NM_014463.2 | 92.9684803 | -8.1605068 | 1.3968907 | -5.8419079 |
| 6276 | NM_001454.3 | 189.739228 | -8.2265544 | 1.4082355 | -5.8417462 |
| 6277 | NM_020773.2 | 47.3908812 | -7.1894079 | 1.23077074 | -5.8413868 |

| | | | | | |
|---|---|---|---|---|---|
| 6278 | NM_001319956.1 | 89.7319763 | -8.1108557 | 1.38853411 | -5.8413082 |
| 6279 | NM_007326.4 | 41.4871448 | 10.287339 | 1.76121499 | 5.84104666 |
| 6280 | NM_018356.2 | 69.1163056 | -7.7350758 | 1.32428528 | -5.8409438 |
| 6281 | NM_001199637.1 | 120.177242 | -8.5302875 | 1.46059512 | -5.8402821 |
| 6282 | NR_123740.1 | 73.317852 | -7.8168784 | 1.33852837 | -5.8399049 |
| 6283 | NM_024923.3 | 106.021338 | -7.3849471 | 1.26458658 | -5.8398114 |
| 6284 | NM_003730.4 | 50.1122813 | -7.2701256 | 1.24498479 | -5.8395297 |
| 6285 | NM_018590.4 | 84.1486696 | -8.0171822 | 1.37301602 | -5.8391032 |
| 6286 | NM_183238.3 | 65.1469938 | -7.6482154 | 1.30997906 | -5.8384257 |
| 6287 | NM_153365.2 | 68.3771435 | -7.7179217 | 1.32202243 | -5.8379658 |
| 6288 | NM_001163424.1 | 109.724055 | -8.3998662 | 1.43892299 | -5.8376065 |
| 6289 | NM_001099857.2 | 62.1427271 | -7.58162 | 1.29880644 | -5.8373748 |
| 6290 | NM_170713.2 | 61.964248 | -7.575715 | 1.29782764 | -5.8372273 |
| 6291 | NM_014966.3 | 75.5282099 | -7.8581769 | 1.34641898 | -5.8363534 |
| 6292 | NM_012158.2 | 87.6525242 | -5.7138999 | 0.97906861 | -5.8360567 |
| 6293 | NM_001301056.1 | 93.3549996 | -8.1672056 | 1.39949512 | -5.8358228 |
| 6294 | NM_024090.2 | 77.4018876 | -7.8976568 | 1.35345106 | -5.8351994 |
| 6295 | NM_023007.2 | 59.7956316 | -7.5241113 | 1.28976868 | -5.8336905 |
| 6296 | NM_017615.2 | 60.8120103 | -7.5484987 | 1.29405844 | -5.8331977 |
| 6297 | NM_153000.4 | 55.5574272 | -7.4189214 | 1.27184635 | -5.83319 |
| 6298 | NM_005730.3 | 432.579117 | -4.450237 | 0.7629236 | -5.8331359 |
| 6299 | NM_006711.4 | 80.5569266 | -7.9534666 | 1.36357094 | -5.832822 |
| 6300 | NM_018971.2 | 149.928395 | -7.884476 | 1.35174194 | -5.8328264 |
| 6301 | NM_001313998.1 | 71.0343227 | -7.770758 | 1.33225545 | -5.8327838 |
| 6302 | NM_173551.4 | 83.944318 | -6.1527846 | 1.05504485 | -5.8317754 |
| 6303 | NM_004366.5 | 51.4165347 | -7.3048278 | 1.25264096 | -5.8315415 |
| 6304 | NM_006108.3 | 62.5513897 | -7.5921204 | 1.30192005 | -5.8314797 |
| 6305 | NM_002231.3 | 73.5408041 | -7.8250174 | 1.34200475 | -5.8308418 |
| 6306 | NM_032305.2 | 70.0672294 | -7.7510916 | 1.32938033 | -5.830605 |
| 6307 | NR_033936.3 | 59.4124028 | -7.5149949 | 1.28893724 | -5.8303808 |
| 6308 | NM_001142295.1 | 47.2371628 | -7.1868876 | 1.23272049 | -5.8301031 |
| 6309 | NM_001170738.1 | 67.3655997 | -7.6963445 | 1.32013663 | -5.8299606 |
| 6310 | NM_004864.3 | 59.7707219 | -7.5275715 | 1.29123293 | -5.8297549 |
| 6311 | NM_007129.3 | 146.704821 | -8.8196517 | 1.51286981 | -5.8297493 |
| 6312 | NM_001110303.3 | 132.248667 | -6.077282 | 1.04266471 | -5.8286062 |
| 6313 | NM_001100913.2 | 62.1546989 | -7.5779673 | 1.30016168 | -5.8284807 |
| 6314 | NM_001007228.1 | 100.145479 | -8.2677806 | 1.41852188 | -5.8284477 |
| 6315 | NM_002493.4 | 89.4386695 | -8.1040472 | 1.39063659 | -5.8275808 |
| 6316 | NM_015361.3 | 92.770155 | -8.1571308 | 1.3997539 | -5.8275464 |
| 6317 | NM_032112.2 | 86.1156014 | -8.0497438 | 1.38132205 | -5.8275648 |
| 6318 | NM_001282307.1 | 96.5924734 | -8.2149578 | 1.40973665 | -5.8272996 |
| 6319 | NM_001006617.1 | 70.5704798 | -7.7602099 | 1.3317159 | -5.8272263 |
| 6320 | NM_004710.4 | 54.8948795 | -7.4002452 | 1.26995767 | -5.827159 |

| | | | | | |
|---|---|---|---|---|---|
| 6321 | NR_109988.1 | 62.7655507 | -7.5919602 | 1.30285542 | -5.8271701 |
| 6322 | NM_000158.3 | 52.4309 | -7.3380781 | 1.25934044 | -5.8269216 |
| 6323 | NM_138376.2 | 59.1385232 | -7.5092111 | 1.2887117 | -5.8269132 |
| 6324 | NM_024326.3 | 82.6167258 | -7.9900777 | 1.37142668 | -5.8261064 |
| 6325 | NM_001282772.1 | 52.9967596 | -7.3498371 | 1.26158897 | -5.8258571 |
| 6326 | NM_001134773.2 | 60.9475193 | -7.5508385 | 1.29611831 | -5.8257325 |
| 6327 | NM_052831.2 | 56.0769148 | -7.4326268 | 1.27588868 | -5.8254509 |
| 6328 | NM_001040627.1 | 59.0764836 | -7.5080818 | 1.28889412 | -5.8252122 |
| 6329 | NM_001128309.1 | 12.8644711 | 8.5924161 | 1.47506459 | 5.82511175 |
| 6330 | NM_172081.2 | 90.3879806 | -8.1180146 | 1.39364431 | -5.8250262 |
| 6331 | NM_001020820.2 | 57.4463231 | -7.4680923 | 1.28209812 | -5.8248992 |
| 6332 | NM_013337.3 | 71.546666 | -7.7832619 | 1.33621945 | -5.8248381 |
| 6333 | NM_005915.5 | 122.503731 | -7.5884869 | 1.30294604 | -5.8240991 |
| 6334 | NM_001143887.1 | 85.5150945 | -8.0394287 | 1.38040018 | -5.8239841 |
| 6335 | NM_001039848.3 | 65.4471093 | -7.6549067 | 1.31446532 | -5.8235897 |
| 6336 | NM_005510.3 | 54.8145964 | -7.3988141 | 1.27054398 | -5.8233436 |
| 6337 | NM_016504.2 | 70.2633418 | -7.7563479 | 1.33197113 | -5.8232102 |
| 6338 | NM_172316.2 | 130.305342 | -8.6479422 | 1.48511609 | -5.8230749 |
| 6339 | NR_073110.1 | 110.400393 | -8.4083677 | 1.44399942 | -5.8229717 |
| 6340 | NM_030634.2 | 65.1293806 | -7.6479314 | 1.31341776 | -5.8229237 |
| 6341 | NM_001198838.1 | 56.7859582 | -7.4532891 | 1.28003282 | -5.8227328 |
| 6342 | NM_001130860.3 | 74.7293486 | -7.8474184 | 1.34781338 | -5.822333 |
| 6343 | NM_001193388.3 | 45.9531009 | -7.1426146 | 1.22677904 | -5.8222503 |
| 6344 | NM_001044305.2 | 184.196082 | -9.1468364 | 1.57104909 | -5.8221201 |
| 6345 | NM_181836.5 | 214.458624 | -6.2483428 | 1.07329053 | -5.8216696 |
| 6346 | NM_015939.4 | 59.8510944 | -7.5236276 | 1.29240819 | -5.821402 |
| 6347 | NM_001317934.1 | 53.482739 | -7.3652173 | 1.26520992 | -5.8213401 |
| 6348 | NM_001048205.1 | 54.2190104 | -7.3859812 | 1.26880809 | -5.8211965 |
| 6349 | NM_000126.3 | 120.282272 | -5.9531501 | 1.0228188 | -5.820337 |
| 6350 | NM_033453.3 | 59.8642229 | -7.5262735 | 1.293123 | -5.8202302 |
| 6351 | NM_002622.4 | 162.482163 | -7.1995586 | 1.2370495 | -5.8199439 |
| 6352 | NM_001330437.1 | 52.6072758 | -7.337923 | 1.26089213 | -5.8196279 |
| 6353 | NM_153042.3 | 90.996357 | -7.1557404 | 1.22965569 | -5.8193041 |
| 6354 | NM_001128206.1 | 77.6123718 | -7.9020191 | 1.35792246 | -5.8191976 |
| 6355 | NM_001001330.2 | 94.4695264 | -3.5797438 | 0.61516631 | -5.819148 |
| 6356 | NM_025147.4 | 53.5936251 | -7.3641615 | 1.26551058 | -5.8191228 |
| 6357 | NM_001724.4 | 74.2984861 | -7.837109 | 1.34688312 | -5.8187001 |
| 6358 | NM_021110.3 | 66.6525568 | -7.6833138 | 1.32046464 | -5.8186441 |
| 6359 | NM_001142420.1 | 82.4794312 | -7.9880039 | 1.3728371 | -5.8186101 |
| 6360 | NM_003863.3 | 66.8577479 | -7.6850187 | 1.32108533 | -5.8172008 |
| 6361 | NM_001010906.1 | 94.2794099 | 2.12543246 | 0.36537228 | 5.81716943 |
| 6362 | NM_016224.4 | 97.7026892 | -7.26208 | 1.24845853 | -5.8168372 |
| 6363 | NM_001427.3 | 122.465943 | -8.5586676 | 1.47137897 | -5.8167663 |

| | | | | | |
|---|---|---|---|---|---|
| 6364 | NM_032375.5 | 65.8341763 | -7.6630874 | 1.31742793 | -5.8167033 |
| 6365 | NM_014362.3 | 46.5198579 | -7.1628635 | 1.23151814 | -5.8162874 |
| 6366 | NM_001163435.2 | 64.9730693 | -7.6421756 | 1.31393857 | -5.8162351 |
| 6367 | NM_148957.3 | 102.06142 | -8.2953689 | 1.426255 | -5.8161891 |
| 6368 | NM_017572.3 | 53.2014634 | -7.3561101 | 1.2647994 | -5.8160291 |
| 6369 | NR_027928.2 | 48.6483402 | -7.2228612 | 1.24190911 | -5.8159339 |
| 6370 | NM_001042683.2 | 57.4226792 | -7.4632353 | 1.28328851 | -5.8157112 |
| 6371 | NM_001161346.1 | 75.6578437 | -7.8620778 | 1.3519008 | -5.815573 |
| 6372 | NM_001163809.1 | 49.4955669 | -7.2494304 | 1.24658992 | -5.8154091 |
| 6373 | NM_003033.3 | 60.1248534 | -7.534114 | 1.29555316 | -5.8153646 |
| 6374 | NM_019592.6 | 191.577571 | -5.513716 | 0.94820239 | -5.8149147 |
| 6375 | NM_181837.2 | 51.9664797 | -7.3210579 | 1.2591343 | -5.8143583 |
| 6376 | NM_002346.2 | 85.8086566 | -8.0476473 | 1.38428194 | -5.8135898 |
| 6377 | NM_015972.3 | 77.592179 | -7.8994518 | 1.35887301 | -5.8132377 |
| 6378 | NM_138431.2 | 62.8024763 | -7.5927898 | 1.30622458 | -5.8127751 |
| 6379 | NM_015964.3 | 125.273599 | -8.5921322 | 1.47827433 | -5.8122718 |
| 6380 | NM_001145658.1 | 62.8129607 | -7.5908913 | 1.30622379 | -5.8113252 |
| 6381 | NM_020971.2 | 107.24651 | -8.3662043 | 1.43965626 | -5.8112513 |
| 6382 | NM_001290261.1 | 90.4501707 | -8.1189954 | 1.39721059 | -5.8108602 |
| 6383 | NM_021925.3 | 60.4320532 | -7.5389282 | 1.29751075 | -5.8103011 |
| 6384 | NR_104473.1 | 72.8174399 | -7.8077276 | 1.34378668 | -5.8102433 |
| 6385 | NM_205768.2 | 211.362598 | -9.3453449 | 1.60845728 | -5.8101294 |
| 6386 | NM_001008495.3 | 50.0392898 | -7.270798 | 1.25148363 | -5.8097428 |
| 6387 | NM_000918.3 | 1370.75185 | -7.3312841 | 1.2619531 | -5.8094743 |
| 6388 | NM_001300908.1 | 100.425969 | -8.2714339 | 1.42377982 | -5.8094895 |
| 6389 | NM_194255.2 | 56.5309255 | -7.44284 | 1.28118547 | -5.8093384 |
| 6390 | NM_014050.3 | 75.6356024 | -7.8617048 | 1.35330382 | -5.8092681 |
| 6391 | NM_024713.2 | 68.4240949 | -7.7180504 | 1.32875049 | -5.8085024 |
| 6392 | NM_001105549.1 | 51.5007942 | -7.3130387 | 1.25912308 | -5.8080412 |
| 6393 | NM_001193315.1 | 65.1244537 | -7.6471554 | 1.31668926 | -5.8078665 |
| 6394 | NM_004240.3 | 53.1177498 | -7.356616 | 1.26674377 | -5.8075012 |
| 6395 | NM_014962.3 | 61.6328107 | -7.5694489 | 1.30340653 | -5.8074351 |
| 6396 | NM_013239.4 | 101.978495 | -7.3269557 | 1.26181294 | -5.8066893 |
| 6397 | NM_003859.2 | 58.4966724 | -7.4905822 | 1.2900169 | -5.8065768 |
| 6398 | NM_014190.3 | 304.437326 | -9.8722141 | 1.70021041 | -5.8064661 |
| 6399 | NM_001330218.1 | 53.4337314 | -7.3586094 | 1.26743617 | -5.8059014 |
| 6400 | NM_001347681.1 | 91.2668498 | -8.1333341 | 1.40093198 | -5.8056595 |
| 6401 | NM_004846.3 | 63.7688651 | -7.6155811 | 1.31177485 | -5.8055551 |
| 6402 | NM_001077351.1 | 55.9717846 | -7.4261569 | 1.27916393 | -5.8054771 |
| 6403 | NM_153487.3 | 96.8325641 | -7.2540162 | 1.24952596 | -5.8054146 |
| 6404 | NR_027383.1 | 64.4111958 | -7.6288551 | 1.31410694 | -5.8053534 |
| 6405 | NM_001324304.1 | 105.875959 | -8.3479876 | 1.43800391 | -5.8052607 |
| 6406 | NM_001198557.1 | 69.505796 | -7.7386074 | 1.33311624 | -5.8049007 |

| | | | | | |
|---|---|---|---|---|---|
| 6407 | NM_001286834.1 | 59.6850096 | -7.5203798 | 1.29553387 | -5.80485 |
| 6408 | NR_033787.2 | 83.0629835 | -6.1928845 | 1.06695777 | -5.8042452 |
| 6409 | NM_001145524.1 | 68.7075419 | -7.7234665 | 1.33080636 | -5.803599 |
| 6410 | NM_001322118.1 | 86.6205788 | -8.059067 | 1.38866197 | -5.8034764 |
| 6411 | NM_020230.6 | 51.1776421 | -7.2963911 | 1.25725915 | -5.8034106 |
| 6412 | NM_152641.3 | 111.516208 | -7.4552027 | 1.2846314 | -5.8033789 |
| 6413 | NM_014186.3 | 81.1856377 | -7.9645054 | 1.37248392 | -5.8029863 |
| 6414 | NM_001105539.2 | 86.700263 | -8.057915 | 1.38868059 | -5.8025691 |
| 6415 | NM_001029991.1 | 51.2845423 | -7.3007453 | 1.25823033 | -5.8023918 |
| 6416 | NM_021145.3 | 61.5987652 | -7.5625966 | 1.30339522 | -5.8022282 |
| 6417 | NM_002977.3 | 95.5289951 | -8.2003124 | 1.41333 | -5.8021215 |
| 6418 | NM_001174128.1 | 52.0575912 | -7.3261296 | 1.26269015 | -5.802001 |
| 6419 | NM_021998.4 | 88.0265498 | -5.2627437 | 0.90707767 | -5.8018667 |
| 6420 | NM_001297705.1 | 58.804303 | -7.5019811 | 1.29306093 | -5.8017228 |
| 6421 | NM_006238.4 | 66.3539992 | -7.673688 | 1.32267741 | -5.8016323 |
| 6422 | NM_138578.2 | 76.1327569 | -7.8710239 | 1.356706 | -5.8015693 |
| 6423 | NM_032816.4 | 45.5906067 | -7.1312641 | 1.22920854 | -5.8015087 |
| 6424 | NM_130787.2 | 282.166823 | -6.2599014 | 1.07904371 | -5.8013419 |
| 6425 | NM_001193582.1 | 191.642546 | -9.2040976 | 1.58677433 | -5.8005082 |
| 6426 | NR_016021.1 | 148.593426 | -3.4596587 | 0.59645226 | -5.800395 |
| 6427 | NM_002668.2 | 46.4716371 | -7.1633487 | 1.23499751 | -5.800294 |
| 6428 | NM_183353.2 | 68.4145457 | -7.714669 | 1.33012493 | -5.7999582 |
| 6429 | NM_001278554.1 | 79.582152 | -7.9355774 | 1.36833625 | -5.7994351 |
| 6430 | NM_006775.2 | 47.2005569 | -7.1831673 | 1.23863515 | -5.79926 |
| 6431 | NM_001300865.1 | 77.0848159 | -7.8909803 | 1.36070149 | -5.7992002 |
| 6432 | NM_001172.3 | 106.428509 | -8.3556586 | 1.44085043 | -5.7991159 |
| 6433 | NM_015210.3 | 99.176962 | -7.2836678 | 1.25606625 | -5.7987927 |
| 6434 | NM_001024809.3 | 64.5137638 | -7.6357996 | 1.31704943 | -5.797656 |
| 6435 | NM_001199054.1 | 76.6051877 | -7.8835958 | 1.35986546 | -5.7973351 |
| 6436 | NM_017819.3 | 73.0657205 | -7.812635 | 1.34771166 | -5.7969633 |
| 6437 | NM_004228.6 | 94.8028927 | -5.0340114 | 0.86849117 | -5.7962725 |
| 6438 | NR_027768.1 | 55.484192 | -7.4180068 | 1.27979367 | -5.7962522 |
| 6439 | NM_015646.5 | 72.6425942 | -7.805423 | 1.34671941 | -5.7958792 |
| 6440 | NM_178026.2 | 272.717368 | -6.0752014 | 1.04820757 | -5.7958 |
| 6441 | NM_001024662.2 | 73.7330701 | -7.8279511 | 1.35069166 | -5.7955131 |
| 6442 | NM_000281.3 | 66.3914849 | -7.6760065 | 1.32449387 | -5.7954262 |
| 6443 | NM_016396.2 | 122.238079 | -7.5882098 | 1.30971922 | -5.7937684 |
| 6444 | NR_110439.1 | 77.6151699 | -7.8991832 | 1.3634439 | -5.793552 |
| 6445 | NM_022346.4 | 77.9739634 | -7.9052041 | 1.36461232 | -5.7930036 |
| 6446 | NM_001136156.1 | 61.2945485 | -7.5588483 | 1.30484145 | -5.7929248 |
| 6447 | NM_001329857.1 | 74.0722807 | -7.8309012 | 1.35182093 | -5.7928539 |
| 6448 | NM_003055.2 | 275.004827 | -7.0413155 | 1.21552353 | -5.7928254 |
| 6449 | NM_022167.3 | 68.4799442 | -7.7225131 | 1.33315381 | -5.7926648 |

| | | | | | |
|---|---|---|---|---|---|
| 6450 | NM_001912.4 | 52.2677378 | -7.3295184 | 1.26538879 | -5.7923054 |
| 6451 | NM_031438.3 | 44.956592 | -7.1108218 | 1.22766812 | -5.7921369 |
| 6452 | NM_021134.3 | 81.9395974 | -7.9782835 | 1.37744768 | -5.7920773 |
| 6453 | NM_016474.4 | 55.6091985 | -7.4181456 | 1.28089165 | -5.7913919 |
| 6454 | NM_015055.3 | 60.7754326 | -7.5465715 | 1.30314482 | -5.7910459 |
| 6455 | NR_104318.1 | 93.1875468 | -8.1637224 | 1.4097602 | -5.7908589 |
| 6456 | NM_145868.1 | 47.5683694 | -7.1948369 | 1.24247148 | -5.7907461 |
| 6457 | NM_020312.3 | 94.5883383 | -7.2147339 | 1.24595844 | -5.7905093 |
| 6458 | NM_001348098.1 | 74.3066048 | -7.8366696 | 1.35353131 | -5.7897956 |
| 6459 | NM_001018072.1 | 50.8296958 | -7.2935147 | 1.25976519 | -5.7895826 |
| 6460 | NM_020191.2 | 57.2723229 | -7.4617418 | 1.28888259 | -5.7893108 |
| 6461 | NM_031937.2 | 52.648232 | -7.3405454 | 1.26800894 | -5.7890328 |
| 6462 | NM_001330167.1 | 66.2279012 | -7.6728507 | 1.32560759 | -5.7881765 |
| 6463 | NM_001035235.3 | 88.9153757 | -5.7377419 | 0.99139667 | -5.7875339 |
| 6464 | NM_006182.2 | 49.256219 | -7.2490468 | 1.2525463 | -5.7874482 |
| 6465 | NM_001321645.1 | 46.0516329 | -7.1444103 | 1.23456707 | -5.7869763 |
| 6466 | NM_007223.2 | 66.1377739 | -7.6702304 | 1.32559692 | -5.7862464 |
| 6467 | NM_014159.6 | 258.070119 | -5.2480479 | 0.90709267 | -5.7855698 |
| 6468 | NM_001012508.3 | 63.5390719 | -7.6111924 | 1.31557065 | -5.7854684 |
| 6469 | NM_033637.3 | 88.6424361 | -8.0925558 | 1.39883374 | -5.7852163 |
| 6470 | NM_001204478.1 | 44.9638321 | -7.1113941 | 1.22924752 | -5.7851604 |
| 6471 | NM_001282597.2 | 165.683689 | -8.9941968 | 1.55474873 | -5.7849842 |
| 6472 | NM_001161662.1 | 60.1627017 | -7.5364288 | 1.30278979 | -5.7848387 |
| 6473 | NM_001284498.1 | 60.3286174 | -7.5391941 | 1.30372069 | -5.7828292 |
| 6474 | NM_031267.3 | 88.7288379 | -7.1214275 | 1.23150962 | -5.7826811 |
| 6475 | NM_018316.2 | 55.0524728 | -7.4041223 | 1.28041442 | -5.7825983 |
| 6476 | NM_032251.5 | 47.2709066 | -7.184186 | 1.24249746 | -5.7820529 |
| 6477 | NM_024844.4 | 69.7878288 | -7.7453442 | 1.33966908 | -5.7815354 |
| 6478 | NM_134323.1 | 60.5513857 | -7.5417177 | 1.30451873 | -5.7812261 |
| 6479 | NM_001167582.1 | 104.679943 | -7.3667627 | 1.27426611 | -5.7811807 |
| 6480 | NR_002809.2 | 54.9296778 | -7.4011044 | 1.28023344 | -5.7810585 |
| 6481 | NM_001330729.1 | 86.4796857 | -8.0564757 | 1.3936242 | -5.7809528 |
| 6482 | NM_002828.3 | 44.1931466 | -7.0859598 | 1.22573688 | -5.7809795 |
| 6483 | NM_017762.2 | 47.6900532 | -7.1991771 | 1.2453325 | -5.7809276 |
| 6484 | NM_015367.3 | 60.4005524 | -7.538108 | 1.30397913 | -5.7808502 |
| 6485 | NM_001042454.2 | 69.9131649 | -7.750652 | 1.3409572 | -5.7799399 |
| 6486 | NM_001826.2 | 59.4085249 | -7.5132124 | 1.29987822 | -5.7799356 |
| 6487 | NM_001304963.1 | 69.6823171 | -7.7437418 | 1.33978167 | -5.7798535 |
| 6488 | NM_001330290.1 | 53.1139881 | -7.3511062 | 1.27189605 | -5.7796439 |
| 6489 | NM_001286389.1 | 91.5072632 | -8.1375914 | 1.40799632 | -5.7795545 |
| 6490 | NM_032051.1 | 46.7859341 | -7.1711798 | 1.24090132 | -5.7790089 |
| 6491 | NR_130914.1 | 199.374335 | 2.99591678 | 0.51841888 | 5.77895003 |
| 6492 | NM_003804.4 | 55.5637079 | -7.4218075 | 1.2843142 | -5.7788099 |

| | | | | | |
|---|---|---|---|---|---|
| 6493 | NM_001186.3 | 90.4042232 | -8.1198835 | 1.40516395 | -5.7786022 |
| 6494 | NM_198545.3 | 85.5327606 | -8.0393002 | 1.39150091 | -5.7774308 |
| 6495 | NM_006253.4 | 122.564605 | -7.5894025 | 1.31363535 | -5.7774043 |
| 6496 | NM_024617.3 | 56.9144681 | -7.4507258 | 1.28964505 | -5.7773461 |
| 6497 | NM_001146345.1 | 51.8224938 | -7.3191642 | 1.2669112 | -5.7771722 |
| 6498 | NM_003069.4 | 85.7240923 | -8.0460271 | 1.3928689 | -5.776586 |
| 6499 | NM_015176.3 | 66.4346242 | -7.6748913 | 1.32863707 | -5.7765145 |
| 6500 | NM_015917.2 | 51.7355624 | -7.3170755 | 1.2669367 | -5.7754073 |
| 6501 | NM_052879.4 | 61.193748 | -7.5548363 | 1.30812893 | -5.7752994 |
| 6502 | NM_000900.4 | 118.158274 | -8.5078416 | 1.47329254 | -5.774713 |
| 6503 | NM_006243.3 | 43.8190946 | -7.0756997 | 1.22532853 | -5.7745327 |
| 6504 | NM_152834.2 | 54.5040876 | -7.390633 | 1.27990285 | -5.7743704 |
| 6505 | NM_032368.4 | 70.0781697 | -7.7558294 | 1.34318467 | -5.7742093 |
| 6506 | NM_058169.5 | 50.081062 | -7.2669836 | 1.25855082 | -5.7740884 |
| 6507 | NM_021974.4 | 76.7623863 | -7.8837022 | 1.36540665 | -5.7738859 |
| 6508 | NM_032357.3 | 81.1544131 | -7.9636405 | 1.37936088 | -5.7734278 |
| 6509 | NM_014109.3 | 70.0268478 | -7.7497474 | 1.34237871 | -5.7731454 |
| 6510 | NM_001037335.2 | 74.0288513 | -7.8338289 | 1.35706648 | -5.7726198 |
| 6511 | NM_015568.3 | 104.001825 | -8.3225106 | 1.44173193 | -5.7725784 |
| 6512 | NM_021873.3 | 49.1701444 | -7.2377849 | 1.25384854 | -5.7724555 |
| 6513 | NM_006893.2 | 52.550326 | -7.3395339 | 1.27163492 | -5.7717304 |
| 6514 | NM_018034.3 | 69.2097467 | -7.7349115 | 1.34016329 | -5.7716187 |
| 6515 | NM_001193655.1 | 54.23905 | -7.3821207 | 1.27904507 | -5.7715876 |
| 6516 | NM_001042679.1 | 60.1857801 | -7.5341379 | 1.30541153 | -5.7714657 |
| 6517 | NM_003338.4 | 84.9728162 | -8.0307432 | 1.39150005 | -5.7712849 |
| 6518 | NM_001007563.2 | 733.168211 | -4.435128 | 0.76851753 | -5.7710173 |
| 6519 | NM_144736.4 | 47.1770302 | -7.1835506 | 1.24476412 | -5.7710136 |
| 6520 | NM_014380.2 | 44.9209304 | -7.1118065 | 1.23243562 | -5.7705298 |
| 6521 | NM_032627.4 | 49.9528331 | -7.2610563 | 1.25830764 | -5.7704937 |
| 6522 | NM_006923.3 | 65.6151724 | -7.6584097 | 1.3271968 | -5.7703648 |
| 6523 | NM_001174101.1 | 62.8397936 | -7.5947425 | 1.31630376 | -5.7697491 |
| 6524 | NM_031426.3 | 62.2203516 | -7.5796465 | 1.31389736 | -5.7688269 |
| 6525 | NM_023937.3 | 67.0174735 | -7.6891032 | 1.33288612 | -5.7687623 |
| 6526 | NM_001135733.1 | 87.7004862 | -8.0747301 | 1.39974002 | -5.7687356 |
| 6527 | NM_173633.2 | 95.327863 | -8.1965091 | 1.42086056 | -5.7686936 |
| 6528 | NM_006346.2 | 49.0651758 | -7.2375615 | 1.25483894 | -5.7677215 |
| 6529 | NM_001206916.1 | 95.3854283 | -8.1967922 | 1.42122039 | -5.7674322 |
| 6530 | NM_018264.3 | 56.0712832 | -7.4304435 | 1.28836427 | -5.7673468 |
| 6531 | NM_022497.4 | 57.601291 | -7.4714962 | 1.29554972 | -5.7670471 |
| 6532 | NR_036624.1 | 71.1379305 | -7.7775494 | 1.348613 | -5.7670728 |
| 6533 | NM_001291107.1 | 145.832496 | -8.8097603 | 1.52794046 | -5.7657746 |
| 6534 | NM_032350.5 | 72.3085394 | -7.7979711 | 1.35245239 | -5.7658008 |
| 6535 | NM_145109.2 | 49.1231179 | -7.2399902 | 1.25568978 | -5.7657475 |

| | | | | | |
|---|---|---|---|---|---|
| 6536 | NM_152240.2 | 49.0653586 | -7.2357272 | 1.25494862 | -5.7657557 |
| 6537 | NM_018214.4 | 45.2709649 | -7.1238799 | 1.23556538 | -5.7656843 |
| 6538 | NM_001017969.2 | 151.425437 | -5.5864129 | 0.96891665 | -5.765628 |
| 6539 | NM_001100418.1 | 69.0865737 | -7.7313504 | 1.34097639 | -5.7654635 |
| 6540 | NM_145030.3 | 46.6203622 | -7.16485 | 1.24272157 | -5.7654508 |
| 6541 | NM_015092.4 | 330.864498 | -3.4224133 | 0.59364192 | -5.7651139 |
| 6542 | NM_014663.2 | 270.273185 | -5.8972473 | 1.02297779 | -5.7647852 |
| 6543 | NM_003211.4 | 159.908669 | -5.6699866 | 0.98358609 | -5.7646064 |
| 6544 | NM_000713.2 | 58.4212178 | -7.4900929 | 1.29936416 | -5.7644293 |
| 6545 | NM_006950.3 | 111.481734 | -8.4221684 | 1.46122085 | -5.7637888 |
| 6546 | NM_017866.5 | 61.3876992 | -7.5613274 | 1.31187152 | -5.7637713 |
| 6547 | NM_052853.3 | 76.6777329 | -7.8817832 | 1.36762255 | -5.7631275 |
| 6548 | NM_006334.3 | 126.740267 | -8.6077578 | 1.49361226 | -5.7630471 |
| 6549 | NM_003193.4 | 72.1410973 | -7.7937762 | 1.35240169 | -5.7629152 |
| 6550 | NM_001293697.1 | 103.196892 | -8.3104383 | 1.44213409 | -5.7625975 |
| 6551 | NM_001164766.1 | 63.4077328 | -4.397383 | 0.76314286 | -5.7622016 |
| 6552 | NM_152388.3 | 73.4582713 | -7.8184693 | 1.35693271 | -5.7618696 |
| 6553 | NM_201433.1 | 125.316404 | -8.5925232 | 1.49128186 | -5.7618371 |
| 6554 | NM_001080522.2 | 59.875554 | -7.5271792 | 1.30641066 | -5.7617252 |
| 6555 | NR_029426.1 | 53.6165864 | -7.3650903 | 1.2782871 | -5.7616871 |
| 6556 | NM_002410.4 | 330.219205 | -4.8066096 | 0.83427296 | -5.7614352 |
| 6557 | NM_175744.4 | 84.115116 | -8.0159278 | 1.39135283 | -5.7612473 |
| 6558 | NM_001300909.1 | 54.4577079 | -7.3924339 | 1.28316634 | -5.7610877 |
| 6559 | NM_001242854.1 | 44.0288931 | -7.0799424 | 1.22893842 | -5.7610229 |
| 6560 | NM_004242.3 | 46.1024849 | -7.146464 | 1.2405196 | -5.7608634 |
| 6561 | NM_015058.1 | 51.9428002 | -7.3240534 | 1.27152159 | -5.76007 |
| 6562 | NM_001024649.1 | 163.573927 | -8.9768864 | 1.55852523 | -5.7598595 |
| 6563 | NM_001033910.2 | 43.3100097 | -7.0607512 | 1.22585021 | -5.7598809 |
| 6564 | NM_032795.2 | 66.0770335 | -7.6671643 | 1.33114608 | -5.7598219 |
| 6565 | NM_001018056.2 | 87.0694756 | -8.0660234 | 1.40057331 | -5.7590869 |
| 6566 | NM_024653.3 | 66.397507 | -7.6762474 | 1.33306734 | -5.7583343 |
| 6567 | NM_006043.1 | 99.3078571 | -8.2562323 | 1.43383184 | -5.7581594 |
| 6568 | NM_024309.3 | 53.8961123 | -7.3722271 | 1.28033255 | -5.7580565 |
| 6569 | NM_001098577.2 | 45.8009971 | -7.1428812 | 1.2405679 | -5.7577511 |
| 6570 | NM_001136191.2 | 59.8059001 | -7.5280065 | 1.30747549 | -5.7576655 |
| 6571 | NM_020205.3 | 46.6015392 | -7.1687778 | 1.24510298 | -5.7575782 |
| 6572 | NM_021241.2 | 70.2064009 | -7.7565537 | 1.34720364 | -5.7575213 |
| 6573 | NM_032444.3 | 88.1726416 | -4.1926128 | 0.72820228 | -5.7574838 |
| 6574 | NM_022769.4 | 73.8076759 | -7.8267156 | 1.35942517 | -5.7573714 |
| 6575 | NM_148170.4 | 45.2069768 | -7.1207954 | 1.23684478 | -5.7572264 |
| 6576 | NM_006996.2 | 54.7918853 | -7.396982 | 1.28506746 | -5.756104 |
| 6577 | NM_001135629.2 | 65.9173615 | -7.6636162 | 1.33145046 | -5.7558403 |
| 6578 | NM_018952.4 | 158.855787 | -7.969326 | 1.38466527 | -5.755417 |

| | | | | | |
|---|---|---|---|---|---|
| 6579 | NM_001193360.1 | 71.444031 | -7.7781449 | 1.35155081 | -5.7549778 |
| 6580 | NM_005001.4 | 62.7257994 | -7.592873 | 1.31937963 | -5.7548812 |
| 6581 | NM_177402.4 | 164.946385 | -6.2827639 | 1.09176866 | -5.754666 |
| 6582 | NM_001134937.1 | 54.9841711 | -7.399084 | 1.28582093 | -5.7543658 |
| 6583 | NM_145306.2 | 86.3454618 | -8.0538701 | 1.39967566 | -5.7540974 |
| 6584 | NM_001038707.1 | 54.2390779 | -7.3843545 | 1.28335721 | -5.7539354 |
| 6585 | NM_017643.2 | 110.284079 | -5.3562111 | 0.93087296 | -5.7539657 |
| 6586 | NM_018211.3 | 54.9849322 | -7.4014336 | 1.28632924 | -5.7539185 |
| 6587 | NM_031216.3 | 77.6633108 | -7.9003428 | 1.37304227 | -5.7538962 |
| 6588 | NM_015111.1 | 84.5121255 | -7.0504141 | 1.22534149 | -5.7538361 |
| 6589 | NM_006922.3 | 121.736678 | -8.5497223 | 1.48606211 | -5.7532738 |
| 6590 | NM_012120.2 | 50.3142774 | -7.2777037 | 1.26504747 | -5.7529096 |
| 6591 | NM_145214.2 | 89.1166155 | -7.1274067 | 1.23898173 | -5.7526326 |
| 6592 | NM_021120.3 | 105.838604 | -8.3472966 | 1.45116417 | -5.752138 |
| 6593 | NM_020159.4 | 91.7339082 | -8.1418862 | 1.41547858 | -5.7520377 |
| 6594 | NM_080390.3 | 126.123038 | -8.6004668 | 1.49523246 | -5.7519262 |
| 6595 | NM_153611.4 | 59.337646 | -7.5156423 | 1.30666189 | -5.7517881 |
| 6596 | NM_016154.4 | 77.1794991 | -7.891775 | 1.37209092 | -5.7516415 |
| 6597 | NM_138610.2 | 74.4600595 | -7.8426652 | 1.3635602 | -5.751609 |
| 6598 | NR_103529.1 | 89.1768749 | -8.1007857 | 1.40851531 | -5.751294 |
| 6599 | NM_053053.3 | 66.8438835 | -7.684555 | 1.33623492 | -5.7509012 |
| 6600 | NM_152653.3 | 62.4201116 | -7.586147 | 1.31912523 | -5.7508922 |
| 6601 | NM_005028.4 | 70.5604501 | -7.7623715 | 1.34985671 | -5.7505152 |
| 6602 | NM_139012.2 | 55.4970379 | -7.4192517 | 1.29025788 | -5.7502084 |
| 6603 | NM_020175.2 | 62.274526 | -7.5819484 | 1.31870301 | -5.7495496 |
| 6604 | NM_198243.2 | 52.0119874 | -7.3204714 | 1.27323543 | -5.7495034 |
| 6605 | NM_170686.2 | 54.0931853 | -7.3770981 | 1.28311511 | -5.7493658 |
| 6606 | NM_002039.3 | 48.540654 | -7.2240758 | 1.25669524 | -5.7484707 |
| 6607 | NM_004753.6 | 105.568501 | -7.379588 | 1.28378428 | -5.7483085 |
| 6608 | NM_006014.4 | 71.7443799 | -7.7861697 | 1.35453643 | -5.7482173 |
| 6609 | NM_025092.4 | 83.7621248 | -5.6517823 | 0.98324801 | -5.748074 |
| 6610 | NM_199044.3 | 51.9544807 | -7.3191858 | 1.27339725 | -5.7477631 |
| 6611 | NM_001042635.1 | 66.3930655 | -7.6746219 | 1.33528501 | -5.7475534 |
| 6612 | NM_003096.3 | 83.7526346 | -8.0090864 | 1.39356344 | -5.747199 |
| 6613 | NR_103464.1 | 65.1167255 | -7.6471113 | 1.33063312 | -5.746972 |
| 6614 | NM_001300815.1 | 53.1885761 | -7.3536295 | 1.27958606 | -5.7468815 |
| 6615 | NM_019007.3 | 63.3474454 | -7.6082735 | 1.32390815 | -5.7468288 |
| 6616 | NM_006715.3 | 61.6327949 | -7.5669401 | 1.31691022 | -5.7459802 |
| 6617 | NM_025150.4 | 62.5761022 | -7.5857706 | 1.32018536 | -5.745989 |
| 6618 | NM_015635.3 | 43.2390686 | -7.0570213 | 1.22834767 | -5.7451335 |
| 6619 | NM_001145073.2 | 48.3594336 | -7.2204737 | 1.25681297 | -5.7450662 |
| 6620 | NM_001002762.2 | 52.9119144 | -7.3483387 | 1.27912305 | -5.7448255 |
| 6621 | NM_178566.4 | 91.6457452 | -5.6598734 | 0.98527802 | -5.744443 |

| | | | | | |
|---|---|---|---|---|---|
| 6622 | NR_028028.1 | 12.7468836 | 8.59701624 | 1.49677994 | 5.74367414 |
| 6623 | NM_005326.4 | 76.0417289 | -7.8704883 | 1.37031052 | -5.7435802 |
| 6624 | NR_015433.1 | 63.3323123 | -7.6056454 | 1.32420692 | -5.7435475 |
| 6625 | NM_016048.2 | 67.5856342 | -7.7005202 | 1.34075784 | -5.7434087 |
| 6626 | NM_020803.4 | 65.0442524 | -7.6467784 | 1.33146476 | -5.7431324 |
| 6627 | NM_181514.1 | 65.9656104 | -7.665659 | 1.33477879 | -5.7430183 |
| 6628 | NM_006788.3 | 184.981415 | -5.488899 | 0.95581009 | -5.7426669 |
| 6629 | NM_006358.3 | 65.8036122 | -7.6618364 | 1.33425794 | -5.7423952 |
| 6630 | NM_152405.4 | 91.7184089 | -7.1668834 | 1.24807375 | -5.7423556 |
| 6631 | NM_001270401.1 | 67.5999698 | -7.6976154 | 1.34058461 | -5.741984 |
| 6632 | NM_001347701.1 | 45.3009293 | -7.125546 | 1.24096391 | -5.7419446 |
| 6633 | NM_016188.4 | 133.885771 | -8.6867965 | 1.51302684 | -5.7413366 |
| 6634 | NR_033688.1 | 54.8238956 | -7.401507 | 1.28920218 | -5.741153 |
| 6635 | NM_018197.2 | 55.0712805 | -7.4030367 | 1.28948998 | -5.7410579 |
| 6636 | NM_033215.4 | 64.833712 | -7.6401515 | 1.33081799 | -5.7409439 |
| 6637 | NM_016564.3 | 98.6077245 | -8.2458962 | 1.43634417 | -5.7408916 |
| 6638 | NM_013300.2 | 68.7739915 | -7.725639 | 1.3458898 | -5.740172 |
| 6639 | NM_014460.3 | 110.466392 | -8.4095157 | 1.46508424 | -5.7399537 |
| 6640 | NM_001308120.1 | 46.3293475 | -7.1582455 | 1.24712228 | -5.7398105 |
| 6641 | NM_006824.2 | 144.589045 | -6.0059608 | 1.04641089 | -5.7395817 |
| 6642 | NM_001318934.1 | 51.7534036 | -7.3106485 | 1.27376973 | -5.73938 |
| 6643 | NM_001286045.1 | 42.6104921 | -7.0339332 | 1.22568428 | -5.7387806 |
| 6644 | NM_030935.4 | 60.5675202 | -7.5450864 | 1.31486938 | -5.7382783 |
| 6645 | NM_005308.2 | 79.5739015 | -7.9363071 | 1.38305914 | -5.7382268 |
| 6646 | NM_015004.3 | 54.4056721 | -7.3876676 | 1.28750723 | -5.737962 |
| 6647 | NM_021992.2 | 124.847051 | -8.5858056 | 1.49643623 | -5.7375018 |
| 6648 | NM_024827.3 | 50.3261156 | -7.2771543 | 1.2684298 | -5.737136 |
| 6649 | NM_025222.3 | 796.061907 | -5.8075626 | 1.01233468 | -5.736801 |
| 6650 | NM_005649.2 | 57.0322259 | -7.4541373 | 1.29936493 | -5.7367543 |
| 6651 | NM_030918.5 | 67.0008964 | -7.6873812 | 1.34008591 | -5.7364838 |
| 6652 | NM_002536.2 | 72.8090905 | -7.8077649 | 1.3610966 | -5.7363782 |
| 6653 | NM_016446.3 | 56.5275654 | -7.442647 | 1.29762228 | -5.7356036 |
| 6654 | NM_002430.2 | 430.05654 | -7.1798728 | 1.25184528 | -5.7354315 |
| 6655 | NM_032450.2 | 69.4687463 | -7.7425961 | 1.34997774 | -5.7353509 |
| 6656 | NM_004824.3 | 53.0018387 | -7.3522724 | 1.28193587 | -5.7352888 |
| 6657 | NM_007032.5 | 133.492131 | -2.1532071 | 0.37544742 | -5.7350429 |
| 6658 | NM_198516.2 | 58.9173443 | -7.5054074 | 1.30876851 | -5.7347096 |
| 6659 | NM_078483.3 | 75.228721 | -7.8557462 | 1.36990228 | -5.7345304 |
| 6660 | NM_007318.2 | 96.3818439 | -7.2480823 | 1.26397112 | -5.7343734 |
| 6661 | NM_002969.5 | 93.1858654 | -8.1644613 | 1.42382437 | -5.7341773 |
| 6662 | NM_001199763.1 | 90.9207772 | -8.1297036 | 1.41782245 | -5.7339363 |
| 6663 | NM_001199818.1 | 72.1281781 | -7.7938422 | 1.35929231 | -5.73375 |
| 6664 | NM_001167.3 | 78.0127993 | -7.9059367 | 1.37886366 | -5.7336609 |

| | | | | | |
|---|---|---|---|---|---|
| 6665 | NM_001256155.2 | 53.2913676 | -7.3584826 | 1.28338352 | -5.7336583 |
| 6666 | NM_014606.2 | 71.1597006 | -7.772758 | 1.35569309 | -5.7334201 |
| 6667 | NR_024271.1 | 49.6043267 | -7.2522073 | 1.26494977 | -5.7331979 |
| 6668 | NM_001323934.1 | 63.719441 | -7.6167989 | 1.32858332 | -5.7330231 |
| 6669 | NR_135475.1 | 43.156926 | -7.0556757 | 1.23071636 | -5.7329828 |
| 6670 | NM_001161576.2 | 48.6544918 | -7.22934 | 1.26110159 | -5.7325596 |
| 6671 | NM_015617.2 | 49.0224503 | -7.2397791 | 1.26316987 | -5.7314375 |
| 6672 | NR_135125.1 | 72.2713424 | -7.7983876 | 1.36070885 | -5.7311214 |
| 6673 | NM_006649.3 | 48.263481 | -7.2151535 | 1.25895983 | -5.7310435 |
| 6674 | NM_001114093.1 | 86.6945603 | -8.0602364 | 1.40644787 | -5.7309173 |
| 6675 | NM_001323068.1 | 108.243252 | -8.3791592 | 1.46228975 | -5.7301634 |
| 6676 | NM_001197109.1 | 97.9191113 | -8.2350962 | 1.43716623 | -5.730093 |
| 6677 | NM_020870.3 | 60.6954453 | -7.5449288 | 1.31699287 | -5.7289063 |
| 6678 | NM_014946.3 | 118.444657 | -8.5092369 | 1.48538881 | -5.7286259 |
| 6679 | NM_144695.3 | 51.3697194 | -7.3043741 | 1.27506106 | -5.7286465 |
| 6680 | NM_001166005.1 | 101.109326 | -8.28141 | 1.44565452 | -5.7284848 |
| 6681 | NM_001330660.1 | 62.3402015 | -7.5870865 | 1.32449808 | -5.7282729 |
| 6682 | NM_012417.3 | 122.006489 | -8.5530588 | 1.49317861 | -5.7280882 |
| 6683 | NM_001284351.2 | 47.3057814 | -7.1890727 | 1.25517936 | -5.7275263 |
| 6684 | NM_001263.3 | 44.3064999 | -7.0935376 | 1.23855812 | -5.7272545 |
| 6685 | NM_015974.2 | 46.0557352 | -7.1492346 | 1.24828221 | -5.7272583 |
| 6686 | NM_001128631.2 | 52.0596755 | -7.3247154 | 1.27894621 | -5.7271489 |
| 6687 | NM_001126111.2 | 53.5108299 | -7.3633057 | 1.28576572 | -5.7267865 |
| 6688 | NM_018953.3 | 84.2013374 | -8.0187424 | 1.40023475 | -5.7267129 |
| 6689 | NM_178565.4 | 51.846307 | -7.3164699 | 1.27763674 | -5.726565 |
| 6690 | NM_203352.2 | 63.2702799 | -7.6073528 | 1.32853942 | -5.7261024 |
| 6691 | NM_001256828.1 | 52.2501593 | -7.3323746 | 1.28056539 | -5.7258885 |
| 6692 | NR_038849.1 | 456.360499 | 4.64484881 | 0.81121242 | 5.72581075 |
| 6693 | NM_001319837.1 | 59.1734808 | -7.509594 | 1.31155004 | -5.7257396 |
| 6694 | NM_014674.2 | 95.9741502 | -7.2388137 | 1.26433674 | -5.7253842 |
| 6695 | NM_024640.3 | 71.3876262 | -7.7789975 | 1.35869166 | -5.7253589 |
| 6696 | NM_175736.4 | 73.710074 | -7.8261356 | 1.3669861 | -5.7251026 |
| 6697 | NM_002577.4 | 399.834962 | -4.7280957 | 0.82586558 | -5.7250184 |
| 6698 | NM_017906.2 | 59.0651965 | -7.505462 | 1.31100609 | -5.7249635 |
| 6699 | NM_152773.4 | 62.4289026 | -7.5844541 | 1.32482385 | -5.7248774 |
| 6700 | NM_001007069.1 | 61.0631272 | -7.5508281 | 1.31899525 | -5.7246818 |
| 6701 | NM_003271.4 | 54.980141 | -7.3992074 | 1.29261155 | -5.7242312 |
| 6702 | NM_015492.4 | 45.417477 | -7.127864 | 1.24527118 | -5.7239452 |
| 6703 | NM_014320.2 | 65.7794544 | -7.6614941 | 1.33851886 | -5.7238596 |
| 6704 | NM_001111067.2 | 70.7588464 | -7.7673827 | 1.35709648 | -5.7235302 |
| 6705 | NM_004317.3 | 164.300512 | -5.9678613 | 1.0427767 | -5.7230482 |
| 6706 | NM_032646.5 | 43.4918068 | -7.063671 | 1.23429297 | -5.722848 |
| 6707 | NM_001842.4 | 75.1406453 | -7.8528956 | 1.37221821 | -5.7227747 |

| | | | | | |
|---|---|---|---|---|---|
| 6708 | NM_003426.3 | 96.5032508 | -8.2130643 | 1.43517533 | -5.7226905 |
| 6709 | NM_003611.2 | 59.071868 | -7.503022 | 1.31110597 | -5.7226664 |
| 6710 | NM_001330178.1 | 46.3937454 | -7.1530762 | 1.24998254 | -5.7225409 |
| 6711 | NM_152864.3 | 51.1841044 | -7.2996393 | 1.27566577 | -5.7222193 |
| 6712 | NM_032837.2 | 60.7978095 | -7.5502046 | 1.31948574 | -5.7220813 |
| 6713 | NM_024112.3 | 146.841879 | -7.8543675 | 1.37265083 | -5.7220433 |
| 6714 | NM_001204142.1 | 53.7440139 | -7.3695902 | 1.28793656 | -5.7220134 |
| 6715 | NM_004666.2 | 34.7449049 | 6.48725245 | 1.13379025 | 5.72173948 |
| 6716 | NM_015513.4 | 44.897609 | -7.1108287 | 1.24285647 | -5.7213595 |
| 6717 | NM_033288.3 | 63.8634796 | -7.6170589 | 1.33133624 | -5.7213637 |
| 6718 | NM_145269.4 | 63.0118917 | -7.5984776 | 1.3281889 | -5.7209314 |
| 6719 | NM_003609.4 | 68.3572467 | -7.7155314 | 1.34871642 | -5.7206476 |
| 6720 | NM_001303503.1 | 54.1154716 | -7.3786522 | 1.28994578 | -5.7201259 |
| 6721 | NM_018136.4 | 59.1959788 | -7.5082965 | 1.31261677 | -5.720098 |
| 6722 | NM_019857.4 | 60.1243937 | -7.5287294 | 1.31621449 | -5.7199867 |
| 6723 | NM_002617.3 | 58.9582615 | -7.5033968 | 1.31180914 | -5.7198845 |
| 6724 | NM_001177413.1 | 281.760382 | -6.3141057 | 1.10389523 | -5.7198415 |
| 6725 | NM_020347.3 | 54.4590612 | -7.3894668 | 1.29191471 | -5.7197791 |
| 6726 | NM_002748.3 | 464.484982 | -5.1565898 | 0.90157681 | -5.7195235 |
| 6727 | NM_032259.3 | 91.2995084 | -7.1674246 | 1.25322853 | -5.7191681 |
| 6728 | NR_028408.1 | 69.2521992 | -7.735291 | 1.35257893 | -5.7189202 |
| 6729 | NM_001256265.1 | 75.7637282 | -7.8663745 | 1.37565322 | -5.7182831 |
| 6730 | NM_005955.2 | 61.4801328 | -3.9518452 | 0.6911005 | -5.7181918 |
| 6731 | NM_144679.2 | 58.8403838 | -7.5002802 | 1.31165991 | -5.7181592 |
| 6732 | NM_006159.2 | 120.541809 | -8.5338769 | 1.49242475 | -5.7181288 |
| 6733 | NM_001242837.1 | 82.5037877 | -7.9885954 | 1.39720451 | -5.7175563 |
| 6734 | NM_014017.3 | 64.8297083 | -7.6402113 | 1.33633662 | -5.7172805 |
| 6735 | NM_014808.3 | 49.0270917 | -7.235725 | 1.26559365 | -5.7172577 |
| 6736 | NM_194280.3 | 61.0719339 | -7.5543359 | 1.32138398 | -5.7169876 |
| 6737 | NM_001024845.2 | 74.2959608 | -7.8363835 | 1.37074894 | -5.7168627 |
| 6738 | NM_018107.4 | 65.7447923 | -7.6633504 | 1.3404924 | -5.7168175 |
| 6739 | NM_016103.3 | 79.3079575 | -7.9316979 | 1.38744106 | -5.7167819 |
| 6740 | NM_173537.3 | 44.9089704 | -7.1092657 | 1.24360084 | -5.7166782 |
| 6741 | NM_024046.4 | 262.091455 | -9.6558367 | 1.68915325 | -5.716377 |
| 6742 | NM_001008239.2 | 49.5775546 | -7.2551234 | 1.26920866 | -5.7162574 |
| 6743 | NM_002180.2 | 69.2043764 | -7.7346569 | 1.35319993 | -5.7158271 |
| 6744 | NM_021248.2 | 62.3702613 | -7.5856308 | 1.32727182 | -5.7152052 |
| 6745 | NM_032862.4 | 58.5921088 | -7.4935336 | 1.31118733 | -5.7150747 |
| 6746 | NM_017426.3 | 49.5593484 | -7.2565426 | 1.26973204 | -5.7150189 |
| 6747 | NM_001323572.1 | 93.2657487 | -8.1647966 | 1.42866493 | -5.7149836 |
| 6748 | NM_178314.4 | 98.6325828 | -7.2742524 | 1.27287457 | -5.7148226 |
| 6749 | NM_153747.1 | 61.9299473 | -7.5757931 | 1.32576166 | -5.7142949 |
| 6750 | NM_001008395.3 | 67.6949915 | -7.7041191 | 1.34823713 | -5.7142167 |

| | | | | | |
|---|---|---|---|---|---|
| 6751 | NR_126511.1 | 61.9293508 | -7.5730526 | 1.32529487 | -5.7142398 |
| 6752 | NM_024098.3 | 58.2627697 | -7.486852 | 1.3102785 | -5.7139394 |
| 6753 | NM_001330988.1 | 81.099421 | -7.9628536 | 1.39376707 | -5.7131882 |
| 6754 | NM_005926.2 | 124.762918 | -5.0746094 | 0.88830707 | -5.7126748 |
| 6755 | NM_007097.3 | 147.78898 | -8.8291759 | 1.5455331 | -5.7127058 |
| 6756 | NM_017900.2 | 72.3352743 | -7.7978038 | 1.36500373 | -5.7126612 |
| 6757 | NM_001441.2 | 67.458677 | -7.6967218 | 1.34741844 | -5.7121987 |
| 6758 | NM_001185077.2 | 54.4858608 | -7.3923213 | 1.29414246 | -5.7121388 |
| 6759 | NM_030815.2 | 115.360082 | -6.6125288 | 1.1576389 | -5.7120825 |
| 6760 | NM_004929.3 | 61.1734476 | -7.5600384 | 1.32356154 | -5.7118904 |
| 6761 | NM_001145301.2 | 49.7275554 | -7.2555972 | 1.27029526 | -5.7117408 |
| 6762 | NM_001146213.1 | 114.021214 | -7.487108 | 1.31084404 | -5.7116696 |
| 6763 | NM_001301228.1 | 71.9452167 | -7.7905107 | 1.36412335 | -5.7110016 |
| 6764 | NM_198793.2 | 2077.49568 | 3.50492382 | 0.61371756 | 5.71097205 |
| 6765 | NM_174931.3 | 65.5253768 | -7.6547584 | 1.34040586 | -5.7107766 |
| 6766 | NM_012459.2 | 84.6890075 | -8.0261814 | 1.40551039 | -5.7105102 |
| 6767 | NM_001199022.2 | 48.1861088 | -7.2134719 | 1.26321919 | -5.7103882 |
| 6768 | NM_001128610.2 | 43.6247378 | -7.0733325 | 1.2387199 | -5.7101952 |
| 6769 | NM_001009880.1 | 452.861263 | -3.5286493 | 0.61797214 | -5.7100459 |
| 6770 | NM_022827.3 | 60.6227871 | -7.5455137 | 1.32143973 | -5.7100703 |
| 6771 | NM_001077621.1 | 82.6615358 | -7.9905242 | 1.39940756 | -5.7099336 |
| 6772 | NM_014667.3 | 62.7748787 | -7.5968468 | 1.33060379 | -5.7093229 |
| 6773 | NM_001130720.2 | 104.183824 | -8.3247846 | 1.45811266 | -5.7092876 |
| 6774 | NM_006301.3 | 79.2854772 | -5.5645623 | 0.97468074 | -5.7091128 |
| 6775 | NM_004329.2 | 106.237072 | -7.38751 | 1.29413173 | -5.7084684 |
| 6776 | NM_014008.4 | 53.859596 | -7.3741355 | 1.29191422 | -5.7079142 |
| 6777 | NM_003108.3 | 1705.47069 | -3.6515973 | 0.63975163 | -5.7078359 |
| 6778 | NM_000375.2 | 71.9873724 | -7.7916543 | 1.36519006 | -5.707377 |
| 6779 | NM_005174.3 | 67.9303166 | -7.7100603 | 1.35090663 | -5.7073228 |
| 6780 | NM_004045.3 | 82.4235041 | -7.9874305 | 1.39956209 | -5.7070927 |
| 6781 | NM_001002262.3 | 62.201323 | -7.581126 | 1.32840618 | -5.7069337 |
| 6782 | NM_001319.6 | 315.610963 | -6.4045052 | 1.12226394 | -5.7067727 |
| 6783 | NM_001007257.2 | 147.535996 | -5.3328038 | 0.93451547 | -5.7064906 |
| 6784 | NM_012219.4 | 52.7221428 | -7.3377223 | 1.28599678 | -5.7058636 |
| 6785 | NM_001303103.1 | 83.767023 | -8.008038 | 1.40366479 | -5.7050929 |
| 6786 | NM_015469.2 | 80.8480776 | -7.9588765 | 1.39505808 | -5.7050503 |
| 6787 | NM_001139441.1 | 135.255352 | -7.7331556 | 1.35552002 | -5.7049365 |
| 6788 | NM_032521.2 | 275.857954 | 2.58968889 | 0.45398177 | 5.70438961 |
| 6789 | NM_004148.3 | 51.3798018 | -7.3074683 | 1.28105512 | -5.7042575 |
| 6790 | NM_001319050.1 | 50.8719271 | -7.2917043 | 1.27830164 | -5.7042126 |
| 6791 | NM_017854.1 | 84.038441 | -8.0143512 | 1.40500654 | -5.704138 |
| 6792 | NM_024165.2 | 66.7673147 | -7.6862461 | 1.34751119 | -5.7040313 |
| 6793 | NM_021126.5 | 79.4656062 | -7.9336939 | 1.3909263 | -5.7038923 |

| | | | | | |
|---|---|---|---|---|---|
| 6794 | NM_001256891.1 | 47.5388034 | -7.1960787 | 1.26164397 | -5.7037317 |
| 6795 | NM_006085.5 | 69.0761443 | -7.7302511 | 1.35530191 | -5.7037115 |
| 6796 | NM_001214909.1 | 79.594001 | -7.9349695 | 1.39126322 | -5.703428 |
| 6797 | NM_018385.2 | 120.772144 | -5.2425229 | 0.91934649 | -5.7024451 |
| 6798 | NM_145345.2 | 49.4852616 | -7.2531356 | 1.2719618 | -5.7023219 |
| 6799 | NM_032998.2 | 75.9396063 | -7.8681404 | 1.37984523 | -5.7021905 |
| 6800 | NR_102326.1 | 46.4383118 | -7.1600251 | 1.25567879 | -5.7021152 |
| 6801 | NM_001144757.2 | 74.1355639 | -7.8349455 | 1.37413946 | -5.7017106 |
| 6802 | NM_001288767.1 | 59.5506884 | -7.5161427 | 1.31834062 | -5.7012145 |
| 6803 | NM_020368.2 | 90.2257317 | -6.3008051 | 1.10517683 | -5.7011737 |
| 6804 | NM_001082967.2 | 97.5472321 | -8.2295756 | 1.44360321 | -5.7007186 |
| 6805 | NM_033114.3 | 149.667584 | -7.8821727 | 1.38278024 | -5.7002353 |
| 6806 | NM_014177.2 | 65.0546319 | -7.6450442 | 1.34120495 | -5.7001312 |
| 6807 | NM_001282431.1 | 91.4935253 | -6.2889533 | 1.10353124 | -5.6989355 |
| 6808 | NM_025126.3 | 63.7398438 | -7.6149191 | 1.33619523 | -5.6989569 |
| 6809 | NM_000373.3 | 63.3736792 | -7.6069989 | 1.33486018 | -5.6987234 |
| 6810 | NM_171982.4 | 59.8432183 | -7.5231053 | 1.32021583 | -5.6983905 |
| 6811 | NM_148414.2 | 66.3512678 | -7.6751901 | 1.34704403 | -5.697802 |
| 6812 | NM_001322477.1 | 55.4451034 | -7.4162835 | 1.30161675 | -5.6977474 |
| 6813 | NM_001007156.2 | 79.842439 | -7.9410626 | 1.39379632 | -5.697434 |
| 6814 | NM_007041.3 | 56.8502428 | -7.4474819 | 1.30720893 | -5.6972392 |
| 6815 | NM_024954.4 | 54.0846194 | -7.3799241 | 1.29537967 | -5.6971128 |
| 6816 | NM_001330289.1 | 171.492357 | -9.0434963 | 1.58740505 | -5.6970313 |
| 6817 | NM_054035.2 | 42.6410075 | -7.0388757 | 1.23555405 | -5.6969387 |
| 6818 | NM_001010924.1 | 372.327362 | -6.3810859 | 1.12012511 | -5.6967618 |
| 6819 | NM_004279.2 | 140.681425 | -5.2601252 | 0.92339058 | -5.6965333 |
| 6820 | NM_001330074.1 | 93.1958736 | -8.1662638 | 1.43357887 | -5.6964175 |
| 6821 | NM_144710.4 | 46.0245886 | -7.1423937 | 1.25383436 | -5.6964412 |
| 6822 | NM_012225.3 | 86.2840197 | -8.0528634 | 1.41382405 | -5.6958031 |
| 6823 | NM_001551.2 | 124.421544 | -7.6153372 | 1.33703436 | -5.695693 |
| 6824 | NM_002582.3 | 57.560529 | -7.4670916 | 1.31100878 | -5.6956839 |
| 6825 | NM_198395.1 | 78.8991052 | -7.9237143 | 1.39119488 | -5.6956178 |
| 6826 | NM_017599.3 | 126.00647 | -8.5991533 | 1.51008739 | -5.6944739 |
| 6827 | NM_001278198.1 | 56.5103075 | -7.4406352 | 1.30669445 | -5.6942426 |
| 6828 | NM_032403.2 | 111.810757 | -8.4277435 | 1.48004656 | -5.6942421 |
| 6829 | NM_001301072.1 | 42.8267591 | -7.0438983 | 1.23709998 | -5.6938796 |
| 6830 | NM_001242374.1 | 59.0612043 | -7.5054665 | 1.31819486 | -5.6937458 |
| 6831 | NM_006395.2 | 48.1448373 | -7.2124906 | 1.26673912 | -5.6937459 |
| 6832 | NM_018158.2 | 95.8407416 | -7.2381208 | 1.27129692 | -5.6934935 |
| 6833 | NM_017423.2 | 66.5228078 | -7.6781335 | 1.34859374 | -5.693437 |
| 6834 | NM_001100165.1 | 53.5974728 | -7.3693807 | 1.29438146 | -5.6933608 |
| 6835 | NM_000593.5 | 53.9824484 | -7.3753737 | 1.29567819 | -5.6922882 |
| 6836 | NM_003502.3 | 43.2199655 | -7.0597442 | 1.24027579 | -5.6920761 |

| | | | | | |
|---|---|---|---|---|---|
| 6837 | NM_001282129.1 | 63.2865789 | -7.6018592 | 1.33554442 | -5.6919554 |
| 6838 | NM_052907.3 | 71.7324527 | -7.7852051 | 1.36778303 | -5.6918422 |
| 6839 | NM_002131.3 | 83.2347171 | -8.0018818 | 1.40588018 | -5.6917239 |
| 6840 | NM_001130475.2 | 47.7279647 | -7.1999673 | 1.26500743 | -5.6916403 |
| 6841 | NM_004852.2 | 287.329733 | -7.090176 | 1.24572379 | -5.6916116 |
| 6842 | NM_139056.3 | 114.155278 | -8.4578307 | 1.48611909 | -5.69122 |
| 6843 | NM_012230.3 | 76.5599911 | -7.8800807 | 1.38466361 | -5.6909712 |
| 6844 | NM_001198777.1 | 58.2953393 | -7.4858961 | 1.31542764 | -5.690846 |
| 6845 | NM_148956.3 | 67.940375 | -7.7078957 | 1.3545017 | -5.6905766 |
| 6846 | NM_001135702.1 | 56.3635937 | -7.4409016 | 1.30774175 | -5.6898861 |
| 6847 | NM_018318.4 | 55.5408937 | -7.4168562 | 1.30353365 | -5.689808 |
| 6848 | NM_001286245.1 | 65.0486051 | -7.6439799 | 1.34346825 | -5.6897361 |
| 6849 | NR_073527.1 | 76.9726594 | -7.8861618 | 1.38605466 | -5.689647 |
| 6850 | NM_003310.3 | 70.665953 | -7.764101 | 1.36461494 | -5.6895911 |
| 6851 | NM_015313.2 | 61.4836226 | -7.5612109 | 1.32895801 | -5.6895784 |
| 6852 | NM_017840.3 | 56.2421064 | -7.4329803 | 1.30645782 | -5.6894147 |
| 6853 | NM_001164463.1 | 52.8993736 | -7.3517048 | 1.29223154 | -5.6891544 |
| 6854 | NR_024467.1 | 71.5500494 | 4.8596785 | 0.85433356 | 5.68826832 |
| 6855 | NM_004760.2 | 68.3255003 | -7.7152501 | 1.3564711 | -5.6877364 |
| 6856 | NR_024031.2 | 71.4116014 | -7.7794971 | 1.36792187 | -5.6870917 |
| 6857 | NM_014902.5 | 57.2994014 | -7.4625649 | 1.31223464 | -5.6869135 |
| 6858 | NM_022899.4 | 87.6300838 | -7.1085544 | 1.25006963 | -5.6865268 |
| 6859 | NM_016226.4 | 78.5586332 | -7.9177352 | 1.39238455 | -5.6864572 |
| 6860 | NM_018090.4 | 74.5221673 | -5.069329 | 0.89148587 | -5.6863818 |
| 6861 | NM_022170.1 | 59.2320043 | -7.5137947 | 1.32137569 | -5.6863425 |
| 6862 | NM_006696.3 | 63.948715 | -7.622469 | 1.34049731 | -5.6862993 |
| 6863 | NM_021729.5 | 93.8874019 | -7.2030048 | 1.26674138 | -5.6862474 |
| 6864 | NM_001321646.1 | 71.6015484 | -7.7826078 | 1.36868938 | -5.6861754 |
| 6865 | NM_017636.3 | 59.3387764 | -7.5128332 | 1.3213547 | -5.6857051 |
| 6866 | NM_017853.2 | 49.2937216 | -7.2422993 | 1.27381761 | -5.6855073 |
| 6867 | NM_203495.3 | 98.5955751 | -8.2453371 | 1.4503175 | -5.6851945 |
| 6868 | NM_022787.3 | 396.331209 | 3.3274619 | 0.585389 | 5.68418929 |
| 6869 | NM_021626.2 | 54.9715682 | -7.4050242 | 1.30278516 | -5.6839949 |
| 6870 | NR_027447.1 | 141.991832 | 2.63175282 | 0.46301618 | 5.68393268 |
| 6871 | NM_152740.3 | 45.7034525 | -7.1410345 | 1.25636561 | -5.6838825 |
| 6872 | NM_031909.2 | 64.8506823 | -7.6408095 | 1.34435888 | -5.6836084 |
| 6873 | NR_110909.1 | 22.6551189 | 9.42311537 | 1.65810657 | 5.68305774 |
| 6874 | NM_001171624.1 | 94.9069876 | -8.1906945 | 1.44129502 | -5.6828716 |
| 6875 | NM_007010.4 | 78.4860941 | -7.917223 | 1.39341762 | -5.6818737 |
| 6876 | NM_001084392.1 | 80.8285924 | -7.9586258 | 1.40077906 | -5.6815711 |
| 6877 | NM_004470.3 | 55.1008841 | -7.406549 | 1.30360609 | -5.6815851 |
| 6878 | NR_076392.1 | 92.8674862 | -8.1601475 | 1.43624913 | -5.6815683 |
| 6879 | NM_152913.2 | 102.505819 | -8.3020447 | 1.46131601 | -5.6812111 |

| | | | | | |
|---|---|---|---|---|---|
| 6880 | NM_017742.5 | 42.4228547 | -7.0277717 | 1.23714803 | -5.6806231 |
| 6881 | NM_001012968.2 | 48.2052652 | -7.2134111 | 1.26991694 | -5.6802227 |
| 6882 | NM_003425.3 | 44.1672079 | -7.0882061 | 1.24797017 | -5.6797881 |
| 6883 | NM_003194.4 | 52.5586657 | -7.3399568 | 1.29234596 | -5.6795603 |
| 6884 | NM_015340.3 | 92.3194315 | -7.1799837 | 1.26419559 | -5.679488 |
| 6885 | NM_001322384.1 | 61.7102034 | -7.5692039 | 1.33276401 | -5.679328 |
| 6886 | NM_000191.2 | 44.2332094 | -7.0940084 | 1.24910667 | -5.6792655 |
| 6887 | NM_013241.2 | 45.7319403 | -7.1387901 | 1.25711929 | -5.6786895 |
| 6888 | NM_021163.3 | 64.0370531 | -7.6255456 | 1.34283595 | -5.6786874 |
| 6889 | NM_147188.2 | 58.2764841 | -7.4845679 | 1.31802828 | -5.6786095 |
| 6890 | NM_005857.4 | 121.077359 | -6.7291535 | 1.18517813 | -5.6777571 |
| 6891 | NM_001330645.1 | 118.589028 | -8.5128985 | 1.49950446 | -5.6771412 |
| 6892 | NM_001308152.1 | 46.8535901 | -7.1769429 | 1.26419023 | -5.6771067 |
| 6893 | NM_004457.3 | 107.746819 | -7.4029421 | 1.30405243 | -5.6768746 |
| 6894 | NM_001300826.1 | 66.9642314 | -7.6852391 | 1.35384109 | -5.6766183 |
| 6895 | NM_001348015.1 | 74.849575 | -7.8482558 | 1.38276367 | -5.6757752 |
| 6896 | NM_001248008.1 | 71.5781252 | -7.7825965 | 1.37122189 | -5.6756653 |
| 6897 | NM_001301084.1 | 123.100922 | -8.5652094 | 1.50914958 | -5.6755205 |
| 6898 | NM_000553.4 | 43.6259073 | -7.0695549 | 1.24565087 | -5.6753903 |
| 6899 | NM_004815.3 | 51.4448667 | -7.3095296 | 1.28807774 | -5.6747581 |
| 6900 | NM_012168.5 | 44.4444765 | -7.0989663 | 1.2510077 | -5.6745984 |
| 6901 | NM_001010862.2 | 69.6877892 | -7.7442195 | 1.36479642 | -5.6742671 |
| 6902 | NM_139025.4 | 47.6261707 | -7.1961946 | 1.26831047 | -5.6738431 |
| 6903 | NM_003647.2 | 79.6303117 | -3.1453753 | 0.55437532 | -5.6737291 |
| 6904 | NM_001042535.3 | 94.3201802 | -8.1810994 | 1.44204859 | -5.6732481 |
| 6905 | NM_080655.2 | 67.8110418 | -7.7048863 | 1.35820322 | -5.6728523 |
| 6906 | NM_198216.1 | 64.5772069 | -7.6351414 | 1.34594072 | -5.6727174 |
| 6907 | NM_001171137.1 | 71.4250534 | -7.7790082 | 1.37131573 | -5.6726602 |
| 6908 | NM_001300947.1 | 45.4167568 | -7.1260385 | 1.25626124 | -5.6724178 |
| 6909 | NM_001303142.1 | 41.9906558 | -7.0152685 | 1.23676593 | -5.6722686 |
| 6910 | NM_007062.2 | 169.635935 | -5.7638819 | 1.01615021 | -5.6722735 |
| 6911 | NM_001025248.1 | 101.529781 | -7.3156667 | 1.28977727 | -5.6720388 |
| 6912 | NM_001271592.1 | 127.926063 | -7.6544585 | 1.34950704 | -5.6720404 |
| 6913 | NM_001079670.2 | 61.1675672 | -7.5567344 | 1.33230499 | -5.6719253 |
| 6914 | NM_012162.3 | 61.9058841 | -7.5741999 | 1.33549877 | -5.671439 |
| 6915 | NM_022893.3 | 243.386233 | -9.5490044 | 1.68379302 | -5.6711272 |
| 6916 | NM_025203.2 | 56.0211371 | -7.4287113 | 1.31001948 | -5.6706876 |
| 6917 | NM_024573.2 | 55.5491849 | -7.418075 | 1.30817789 | -5.6705399 |
| 6918 | NM_001320493.1 | 36.4597066 | 3.74539644 | 0.66053307 | 5.6702633 |
| 6919 | NM_022037.2 | 102.327364 | -8.2973869 | 1.46348591 | -5.6696049 |
| 6920 | NM_025075.3 | 68.2835201 | -7.7150731 | 1.36082115 | -5.6694247 |
| 6921 | NM_003927.4 | 48.6455107 | -7.2274544 | 1.2748209 | -5.6693881 |
| 6922 | NM_025212.3 | 66.9625945 | -7.6894295 | 1.35631875 | -5.6693381 |

| | | | | | |
|---|---|---|---|---|---|
| 6923 | NM_016277.4 | 67.3113536 | -7.6940489 | 1.35728378 | -5.6687106 |
| 6924 | NM_152737.3 | 49.3548862 | -7.2466339 | 1.27863953 | -5.6674565 |
| 6925 | NM_017908.3 | 46.7855435 | -7.1694234 | 1.26524551 | -5.6664287 |
| 6926 | NM_032047.4 | 53.5630201 | -7.3646115 | 1.29974527 | -5.6661961 |
| 6927 | NM_017872.4 | 44.5235059 | -7.0987043 | 1.25282949 | -5.6661376 |
| 6928 | NM_002533.3 | 45.7770139 | -7.1336316 | 1.25900921 | -5.6660678 |
| 6929 | NM_024493.3 | 43.5006893 | -7.0648282 | 1.24686595 | -5.6660687 |
| 6930 | NM_021045.2 | 90.7970998 | -8.1257159 | 1.43421324 | -5.6656261 |
| 6931 | NM_001348022.1 | 47.2980009 | -7.181218 | 1.267579 | -5.6653021 |
| 6932 | NM_000235.3 | 54.56124 | -7.3877119 | 1.30408595 | -5.6650498 |
| 6933 | NM_001330125.1 | 116.482009 | -8.4849852 | 1.49777717 | -5.6650518 |
| 6934 | NM_006236.2 | 415.276336 | -6.4600249 | 1.14036308 | -5.6648843 |
| 6935 | NM_152755.1 | 47.5613605 | -7.1967431 | 1.27045972 | -5.6646764 |
| 6936 | NM_001135700.1 | 51.5180254 | -7.305736 | 1.28972892 | -5.6645515 |
| 6937 | NM_031925.2 | 76.8707444 | -7.8860934 | 1.39223191 | -5.6643533 |
| 6938 | NM_014923.4 | 94.7363578 | -7.2222262 | 1.27507239 | -5.6641696 |
| 6939 | NR_047665.1 | 59.6958873 | -7.5188929 | 1.32748938 | -5.6639948 |
| 6940 | NM_001040458.1 | 44.159176 | -7.0915514 | 1.25204749 | -5.6639636 |
| 6941 | NM_015920.3 | 135.556809 | -5.1208929 | 0.90413313 | -5.6638704 |
| 6942 | NM_007112.4 | 76.9381333 | -7.89016 | 1.39316098 | -5.6634949 |
| 6943 | NM_001276293.1 | 46.2549347 | -7.1551256 | 1.26349025 | -5.6629844 |
| 6944 | NM_003392.4 | 68.1944442 | -7.7159554 | 1.36260388 | -5.6626548 |
| 6945 | NM_015608.2 | 57.713784 | -7.4722156 | 1.31974641 | -5.6618571 |
| 6946 | NM_001286635.1 | 108.672563 | -8.3864231 | 1.48128477 | -5.6615873 |
| 6947 | NM_015879.2 | 79.2916757 | -7.9316338 | 1.40113782 | -5.6608519 |
| 6948 | NM_001800.3 | 99.5529933 | -8.2589987 | 1.45898088 | -5.6607999 |
| 6949 | NM_012263.4 | 57.5714732 | -7.4679451 | 1.31924108 | -5.6607888 |
| 6950 | NM_006822.2 | 45.2854277 | -7.1222966 | 1.25819017 | -5.6607473 |
| 6951 | NM_004661.3 | 95.1075685 | -5.6295821 | 0.99452307 | -5.6605847 |
| 6952 | NM_014473.2 | 65.2380151 | -7.6481685 | 1.35114705 | -5.6605005 |
| 6953 | NM_001253792.1 | 44.447269 | -7.0959109 | 1.25359089 | -5.6604679 |
| 6954 | NM_032340.3 | 86.1271356 | -8.0500868 | 1.42216508 | -5.6604447 |
| 6955 | NM_002124.3 | 103.605174 | -8.3176916 | 1.46978352 | -5.659127 |
| 6956 | NM_001316973.1 | 110.552305 | -8.4093447 | 1.48603025 | -5.6589324 |
| 6957 | NM_003998.3 | 43.6853589 | -7.0733252 | 1.24999316 | -5.6586911 |
| 6958 | NM_134269.2 | 63.6651834 | -7.616503 | 1.34602417 | -5.6585188 |
| 6959 | NM_001077269.1 | 50.3308271 | -7.280174 | 1.28663542 | -5.6583038 |
| 6960 | NM_032188.2 | 101.727768 | -7.3185602 | 1.29342676 | -5.6582718 |
| 6961 | NM_053050.4 | 62.9423423 | -7.5984113 | 1.34304944 | -5.6575812 |
| 6962 | NM_001023563.3 | 51.1800261 | -7.2982539 | 1.29008243 | -5.6571996 |
| 6963 | NM_001143993.1 | 46.1874232 | -7.1482389 | 1.26356093 | -5.6572174 |
| 6964 | NM_001286657.1 | 44.2267419 | -7.0907509 | 1.25340416 | -5.6571943 |
| 6965 | NM_201997.2 | 60.0603562 | -7.526677 | 1.33046721 | -5.6571683 |

| | | | | | |
|---|---|---|---|---|---|
| 6966 | NM_001083600.2 | 66.7222164 | -7.6815235 | 1.3578547 | -5.6571027 |
| 6967 | NM_018367.6 | 90.8334409 | -8.1279849 | 1.43684972 | -5.6568093 |
| 6968 | NM_152705.2 | 62.8797763 | -7.5965631 | 1.34294049 | -5.656664 |
| 6969 | NM_001321077.1 | 75.1083959 | -7.8523503 | 1.38840451 | -5.6556646 |
| 6970 | NM_024644.4 | 55.6664702 | -7.4210281 | 1.31217324 | -5.6555246 |
| 6971 | NM_030628.1 | 86.4141745 | -7.0883284 | 1.25334315 | -5.6555369 |
| 6972 | NM_006845.3 | 45.0286244 | -7.1168986 | 1.25844115 | -5.6553289 |
| 6973 | NM_033632.3 | 55.2869721 | -7.4099094 | 1.31027001 | -5.6552538 |
| 6974 | NM_014055.3 | 42.289342 | -7.0279876 | 1.24276704 | -5.6551126 |
| 6975 | NM_001347219.1 | 51.0869647 | -7.30068 | 1.29106346 | -5.6547802 |
| 6976 | NM_001322885.1 | 63.4689949 | -7.6107148 | 1.34591366 | -5.6546828 |
| 6977 | NM_012258.3 | 93.0237426 | -8.1603089 | 1.44310967 | -5.65467 |
| 6978 | NM_022916.4 | 111.35307 | -3.1895824 | 0.56406235 | -5.6546628 |
| 6979 | NM_016484.4 | 73.0566472 | -7.8133698 | 1.38184713 | -5.6542939 |
| 6980 | NM_030755.4 | 114.385715 | -4.5266163 | 0.80057155 | -5.6542307 |
| 6981 | NM_006397.2 | 59.4009733 | -7.513022 | 1.3287699 | -5.6541181 |
| 6982 | NM_005388.4 | 78.0418812 | -5.48968 | 0.97093034 | -5.6540411 |
| 6983 | NM_006541.4 | 92.3876829 | -8.1513487 | 1.4416814 | -5.6540569 |
| 6984 | NM_058174.2 | 46.1358364 | -7.1546904 | 1.26549635 | -5.6536634 |
| 6985 | NR_003285.2 | 121.508065 | -8.5462523 | 1.51163138 | -5.6536616 |
| 6986 | NM_001297569.1 | 30.7563561 | 9.86583352 | 1.74508166 | 5.65350824 |
| 6987 | NM_001304762.1 | 42.7947598 | -7.0442421 | 1.2459923 | -5.6535197 |
| 6988 | NM_139283.1 | 53.5143464 | -7.3624596 | 1.30230591 | -5.6534026 |
| 6989 | NR_003256.2 | 59.6118208 | -7.521095 | 1.33039797 | -5.6532671 |
| 6990 | NM_001130107.1 | 61.4114513 | -7.5647148 | 1.3381742 | -5.653012 |
| 6991 | NM_001114632.1 | 42.1112499 | -7.0199051 | 1.24184176 | -5.6528177 |
| 6992 | NM_198219.2 | 59.839159 | -7.5217475 | 1.33082446 | -5.6519456 |
| 6993 | NM_001321701.1 | 55.540974 | -7.4212094 | 1.31310367 | -5.6516554 |
| 6994 | NM_207115.1 | 80.1544477 | -7.9468027 | 1.40612554 | -5.6515599 |
| 6995 | NM_014621.2 | 86.8375743 | -8.0629053 | 1.4267282 | -5.6513254 |
| 6996 | NM_013437.4 | 90.1893425 | -7.1494218 | 1.26519633 | -5.6508398 |
| 6997 | NM_015500.1 | 46.5451971 | -7.1675675 | 1.26841272 | -5.6508165 |
| 6998 | NM_032784.4 | 47.5993232 | -7.1913856 | 1.27272661 | -5.6503773 |
| 6999 | NM_001205293.1 | 120.667406 | -8.5365647 | 1.51103081 | -5.6494974 |
| 7000 | NM_182691.2 | 65.2894862 | -7.6482252 | 1.35396243 | -5.6487721 |
| 7001 | NM_001318477.1 | 103.880735 | 1.78688931 | 0.31634125 | 5.64861307 |
| 7002 | NM_032482.2 | 394.848733 | -6.0650722 | 1.07374411 | -5.6485266 |
| 7003 | NM_138766.2 | 73.5791348 | -7.8237757 | 1.38512665 | -5.648419 |
| 7004 | NM_018844.3 | 78.7577508 | -7.9209931 | 1.40242576 | -5.6480659 |
| 7005 | NM_019002.3 | 43.7919493 | -7.0723325 | 1.25216737 | -5.6480728 |
| 7006 | NM_016440.3 | 44.6040297 | -7.0977788 | 1.2566848 | -5.6480183 |
| 7007 | NM_006591.2 | 43.4648941 | -7.0657862 | 1.25102887 | -5.6479801 |
| 7008 | NM_001308248.1 | 54.9567617 | -7.4021671 | 1.31068323 | -5.6475638 |

| | | | | | |
|---|---|---|---|---|---|
| 7009 | NR_036599.1 | 39.9435161 | -6.9432786 | 1.2294834 | -5.6473138 |
| 7010 | NM_001037738.2 | 95.4133235 | -7.2266989 | 1.2797441 | -5.6469875 |
| 7011 | NM_002845.3 | 121.636107 | -7.584279 | 1.34310536 | -5.6468236 |
| 7012 | NM_005225.2 | 66.2831103 | -7.6709876 | 1.35845041 | -5.6468661 |
| 7013 | NM_016542.3 | 41.9144877 | -7.0147102 | 1.24223528 | -5.6468451 |
| 7014 | NM_005904.3 | 41.0729661 | -6.9806188 | 1.23621977 | -5.6467458 |
| 7015 | NM_032900.5 | 48.079492 | -7.2072387 | 1.2764137 | -5.6464755 |
| 7016 | NM_001271822.1 | 44.4281628 | -7.0989617 | 1.2574597 | -5.6454785 |
| 7017 | NM_001013839.3 | 92.0489453 | -8.1443162 | 1.44265786 | -5.6453553 |
| 7018 | NM_005763.3 | 114.795255 | -3.6550098 | 0.64743857 | -5.6453385 |
| 7019 | NM_023948.4 | 58.726774 | -7.4981606 | 1.32821378 | -5.6452965 |
| 7020 | NM_005003.2 | 139.883358 | -7.7840148 | 1.37887652 | -5.6451862 |
| 7021 | NM_018445.5 | 101.469292 | -7.3160228 | 1.29598684 | -5.6451366 |
| 7022 | NM_001258277.1 | 66.6026831 | -7.6806249 | 1.36058186 | -5.6451031 |
| 7023 | NM_001761.2 | 63.8092823 | -7.6199017 | 1.34986512 | -5.6449357 |
| 7024 | NM_001008220.1 | 139.728112 | -8.7479398 | 1.54994461 | -5.6440338 |
| 7025 | NM_003203.4 | 46.2021095 | -7.1512596 | 1.26708691 | -5.6438588 |
| 7026 | NM_000633.2 | 86.2466618 | -6.2571562 | 1.10871119 | -5.6436304 |
| 7027 | NM_018270.5 | 57.8651654 | -7.4742622 | 1.3243923 | -5.6435409 |
| 7028 | NM_019091.3 | 55.661661 | -7.4203252 | 1.31490172 | -5.6432546 |
| 7029 | NR_028502.1 | 47.2268522 | -7.1883207 | 1.27385848 | -5.6429508 |
| 7030 | NM_019056.6 | 66.2526816 | -7.6715435 | 1.35967933 | -5.6421711 |
| 7031 | NM_000097.5 | 54.1838487 | -7.3824162 | 1.30851552 | -5.6418255 |
| 7032 | NM_001270764.1 | 68.4163941 | -7.719138 | 1.36821446 | -5.6417603 |
| 7033 | NM_018256.3 | 64.4081783 | -7.631046 | 1.35261091 | -5.6417156 |
| 7034 | NM_001162499.1 | 60.8804163 | -7.5496716 | 1.33827231 | -5.6413568 |
| 7035 | NM_001025780.2 | 87.7115838 | -7.1022976 | 1.25909099 | -5.6408136 |
| 7036 | NM_001184727.1 | 65.9046815 | -7.6611268 | 1.3581695 | -5.6407737 |
| 7037 | NM_001286086.1 | 63.3248934 | -7.6072802 | 1.34863152 | -5.6407403 |
| 7038 | NM_198256.3 | 41.7806582 | -7.0068798 | 1.24219207 | -5.6407379 |
| 7039 | NM_001674.3 | 53.615842 | -7.3683252 | 1.30634488 | -5.6404135 |
| 7040 | NM_001318040.1 | 112.98874 | -8.4417412 | 1.49666059 | -5.6403845 |
| 7041 | NM_015072.4 | 72.9903243 | -6.8428856 | 1.21327139 | -5.6400288 |
| 7042 | NM_018387.4 | 97.6833323 | -7.2622435 | 1.2876354 | -5.6399844 |
| 7043 | NM_001308232.1 | 59.0673976 | 5.25399608 | 0.93159185 | 5.6398047 |
| 7044 | NM_020753.4 | 87.2234298 | -7.1020879 | 1.25927516 | -5.6398221 |
| 7045 | NM_024513.3 | 43.2543149 | -7.060846 | 1.25200712 | -5.6396213 |
| 7046 | NM_001031703.2 | 59.7125687 | -7.5218751 | 1.33382442 | -5.6393292 |
| 7047 | NM_006600.3 | 401.070234 | -5.3216542 | 0.94368681 | -5.6392165 |
| 7048 | NM_048368.3 | 49.8159812 | -7.2578102 | 1.28704672 | -5.6391195 |
| 7049 | NM_001008392.1 | 45.2125087 | -7.1174231 | 1.26218296 | -5.6389789 |
| 7050 | NR_034035.2 | 50.5206005 | -7.2812367 | 1.2914227 | -5.6381514 |
| 7051 | NM_006809.4 | 80.5453416 | -6.9878611 | 1.23941547 | -5.6380296 |

| | | | | | |
|---|---|---|---|---|---|
| 7052 | NM_007110.4 | 51.0480417 | -7.2930112 | 1.29354537 | -5.6380019 |
| 7053 | NM_023072.2 | 86.6749813 | -7.0917487 | 1.25788899 | -5.6378176 |
| 7054 | NM_001017528.2 | 41.2535922 | -6.9874449 | 1.23949475 | -5.6373332 |
| 7055 | NM_014187.3 | 57.3085953 | -7.4635254 | 1.32394771 | -5.6373264 |
| 7056 | NM_018710.2 | 76.6280553 | -7.8817171 | 1.39828501 | -5.6367028 |
| 7057 | NM_152283.4 | 85.8001856 | -8.0446714 | 1.42728663 | -5.6363391 |
| 7058 | NM_032111.3 | 59.8198684 | -7.5239046 | 1.33494232 | -5.6361271 |
| 7059 | NM_001006613.1 | 52.0880796 | -7.3214156 | 1.29902184 | -5.6360989 |
| 7060 | NM_139021.2 | 134.320224 | -8.6925426 | 1.54231548 | -5.6360341 |
| 7061 | NM_002990.4 | 86.0075812 | 2.93969686 | 0.52160843 | 5.63583083 |
| 7062 | NM_016024.2 | 57.2367779 | -7.4607353 | 1.32381244 | -5.6357948 |
| 7063 | NR_037646.1 | 64.0735611 | -7.620469 | 1.3521602 | -5.6357737 |
| 7064 | NM_205767.2 | 70.2300383 | -7.7549662 | 1.376042 | -5.6357046 |
| 7065 | NM_006777.3 | 94.9381095 | -7.2183261 | 1.28085213 | -5.6355655 |
| 7066 | NM_015508.4 | 43.2250672 | -7.0597875 | 1.25276937 | -5.6353449 |
| 7067 | NR_024110.1 | 47.8100475 | -7.197113 | 1.27714647 | -5.6353074 |
| 7068 | NM_001319139.1 | 64.1816239 | -7.6227942 | 1.35281028 | -5.6347844 |
| 7069 | NM_147150.2 | 61.6430301 | -7.5681366 | 1.34312468 | -5.6347238 |
| 7070 | NM_004623.4 | 60.822981 | -7.5478695 | 1.33962706 | -5.6343066 |
| 7071 | NM_001011880.2 | 71.5253264 | -7.7832247 | 1.38142478 | -5.6342008 |
| 7072 | NM_020706.2 | 67.8396457 | -7.7036634 | 1.36730426 | -5.6341984 |
| 7073 | NM_017659.3 | 56.5662927 | -7.4433041 | 1.32112762 | -5.6340538 |
| 7074 | NM_004629.1 | 59.1097013 | -7.506861 | 1.33243877 | -5.6339256 |
| 7075 | NM_000485.2 | 53.970332 | -7.3764889 | 1.30946094 | -5.6332256 |
| 7076 | NM_183398.2 | 50.7173126 | -7.2842614 | 1.29327412 | -5.6324187 |
| 7077 | NR_051963.1 | 44.1755024 | -7.0859425 | 1.25809319 | -5.6322875 |
| 7078 | NR_001434.4 | 51.4758364 | -7.3104322 | 1.29797381 | -5.6321878 |
| 7079 | NM_020709.2 | 61.4533489 | -7.5628519 | 1.34289623 | -5.6317471 |
| 7080 | NM_032492.3 | 63.1216504 | -7.6020604 | 1.34986452 | -5.631721 |
| 7081 | NM_021947.2 | 65.6485702 | -7.6576121 | 1.35978378 | -5.6314925 |
| 7082 | NM_021194.2 | 48.7023519 | -7.2310609 | 1.2840783 | -5.631324 |
| 7083 | NR_111952.1 | 54.2801945 | -7.3842626 | 1.31140318 | -5.6308104 |
| 7084 | NM_001329451.1 | 45.4523978 | -7.1291427 | 1.26626877 | -5.6300391 |
| 7085 | NM_001005210.2 | 97.6266893 | -8.2307306 | 1.4620185 | -5.6297035 |
| 7086 | NM_001257.4 | 97.9161741 | -8.2354936 | 1.46291031 | -5.6295273 |
| 7087 | NM_032164.3 | 40.6576437 | -6.9659059 | 1.23750596 | -5.6289877 |
| 7088 | NM_001008537.2 | 85.7962681 | -8.0443498 | 1.42913632 | -5.6288191 |
| 7089 | NM_014587.4 | 55.993934 | -7.4259674 | 1.31928057 | -5.6288007 |
| 7090 | NM_002393.4 | 66.7560127 | -7.6803821 | 1.36453237 | -5.6285818 |
| 7091 | NM_138288.3 | 104.519553 | -2.899488 | 0.51513617 | -5.6285856 |
| 7092 | NM_001308036.1 | 58.9878827 | -7.5041063 | 1.33343006 | -5.6276714 |
| 7093 | NR_003579.1 | 48.7360841 | -7.2299522 | 1.28484273 | -5.6271106 |
| 7094 | NM_001258311.1 | 124.341998 | -7.6106952 | 1.35256908 | -5.626844 |

| | | | | | |
|---|---|---|---|---|---|
| 7095 | NM_001584.2 | 104.814721 | -8.3335579 | 1.48103603 | -5.6268435 |
| 7096 | NM_001258372.1 | 49.5032235 | -7.2549918 | 1.28938456 | -5.6267091 |
| 7097 | NM_013318.3 | 1738.09311 | -6.1172086 | 1.08718667 | -5.6266405 |
| 7098 | NM_001130848.1 | 70.5434474 | -7.7655033 | 1.38022214 | -5.6262707 |
| 7099 | NM_152271.4 | 100.451377 | -8.2714542 | 1.47030661 | -5.6256662 |
| 7100 | NM_017887.2 | 61.5834929 | -7.5663178 | 1.3451429 | -5.6249175 |
| 7101 | NM_001256575.1 | 95.0556834 | -8.1922803 | 1.45646034 | -5.6247877 |
| 7102 | NM_002861.4 | 98.6585973 | -8.2461671 | 1.46609197 | -5.6245906 |
| 7103 | NM_001114600.2 | 68.2149615 | -7.7143004 | 1.37154619 | -5.6245284 |
| 7104 | NM_145685.2 | 50.1070298 | -7.2680031 | 1.29221937 | -5.6244344 |
| 7105 | NM_018704.2 | 125.320454 | -6.6575606 | 1.18373924 | -5.6241784 |
| 7106 | NR_024057.2 | 67.3984606 | -7.6974294 | 1.36874196 | -5.6237257 |
| 7107 | NM_138401.3 | 60.9861211 | -7.5518234 | 1.34289671 | -5.6235326 |
| 7108 | NM_018343.2 | 51.5493198 | -7.3111072 | 1.30014383 | -5.6233065 |
| 7109 | NM_030934.4 | 48.2528086 | -7.2129868 | 1.28289265 | -5.6224399 |
| 7110 | NM_001300862.1 | 57.1464809 | -7.4574263 | 1.32643362 | -5.6221632 |
| 7111 | NM_006857.2 | 74.577431 | -7.8419568 | 1.39488899 | -5.6219218 |
| 7112 | NM_006963.4 | 101.157438 | -7.3145064 | 1.30107043 | -5.6219142 |
| 7113 | NM_016162.3 | 58.1739983 | -7.4845659 | 1.33142854 | -5.6214552 |
| 7114 | NM_020243.4 | 128.044148 | -7.6531751 | 1.36142507 | -5.6214442 |
| 7115 | NM_015381.6 | 61.3703237 | -7.5620476 | 1.34526494 | -5.6212329 |
| 7116 | NM_001130924.2 | 42.9929394 | -7.0480363 | 1.2538657 | -5.6210456 |
| 7117 | NM_138701.3 | 58.7272355 | -7.4973972 | 1.33385695 | -5.6208405 |
| 7118 | NM_003595.3 | 43.2215508 | -7.0603204 | 1.25611284 | -5.6207692 |
| 7119 | NM_017777.3 | 49.4555422 | -7.2505071 | 1.2899648 | -5.6207015 |
| 7120 | NM_012212.3 | 45.481293 | -7.1326092 | 1.26904489 | -5.6204546 |
| 7121 | NM_014989.5 | 90.2669185 | -8.1185311 | 1.44449551 | -5.6203228 |
| 7122 | NM_020693.3 | 46.4451139 | -7.1558403 | 1.2733111 | -5.619868 |
| 7123 | NM_017955.3 | 47.9637038 | -7.2063601 | 1.28236511 | -5.6195853 |
| 7124 | NM_001304420.2 | 48.6825092 | -7.2230184 | 1.28533618 | -5.6195558 |
| 7125 | NM_024122.4 | 62.1849124 | -7.5786953 | 1.34873553 | -5.6191115 |
| 7126 | NM_016467.4 | 42.567301 | -7.0344462 | 1.25188595 | -5.6190791 |
| 7127 | NM_145804.2 | 87.970661 | -7.1076732 | 1.26493975 | -5.6189816 |
| 7128 | NR_104586.1 | 47.3827072 | -7.1905227 | 1.27971751 | -5.6188359 |
| 7129 | NM_022828.4 | 49.2412938 | -7.2430795 | 1.28908583 | -5.6187721 |
| 7130 | NM_001105251.2 | 44.9092823 | -7.1063242 | 1.26483339 | -5.6183876 |
| 7131 | NM_005822.3 | 99.9433827 | -8.2650834 | 1.47108806 | -5.6183472 |
| 7132 | NM_002504.5 | 53.7455209 | -7.3699073 | 1.31186815 | -5.6178719 |
| 7133 | NM_001264573.1 | 48.7103946 | -7.2285697 | 1.28680309 | -5.6174637 |
| 7134 | NM_144609.2 | 59.5706762 | -7.5178654 | 1.33829995 | -5.6174742 |
| 7135 | NM_020701.3 | 61.1075589 | -7.5584473 | 1.34554581 | -5.6173839 |
| 7136 | NM_001329545.1 | 41.4254482 | -6.9942878 | 1.24516316 | -5.6171657 |
| 7137 | NM_138360.3 | 196.707543 | -7.4910335 | 1.33362337 | -5.6170532 |

| | | | | | |
|---|---|---|---|---|---|
| 7138 | NM_032291.3 | 97.7593989 | -8.2329222 | 1.46573693 | -5.6169167 |
| 7139 | NM_017640.5 | 45.7077995 | -7.1313485 | 1.26968506 | -5.6166279 |
| 7140 | NM_001243159.1 | 46.2201931 | -7.1540294 | 1.2738933 | -5.615878 |
| 7141 | NM_006532.3 | 41.9136413 | -7.0109576 | 1.24843523 | -5.615796 |
| 7142 | NM_019556.2 | 39.0968443 | -6.9125555 | 1.23096429 | -5.6155614 |
| 7143 | NM_001326.2 | 95.0623456 | -7.2250801 | 1.28669476 | -5.6152246 |
| 7144 | NM_001071775.2 | 64.5557608 | -7.6334345 | 1.3597498 | -5.6138522 |
| 7145 | NM_001271999.1 | 56.7781153 | -7.4458231 | 1.32646295 | -5.6132914 |
| 7146 | NM_000181.3 | 57.586054 | -7.470117 | 1.33080197 | -5.6132446 |
| 7147 | NM_001130106.1 | 68.5569079 | -7.7197868 | 1.3752904 | -5.6132049 |
| 7148 | NM_030805.3 | 48.3792028 | -7.2178314 | 1.28586875 | -5.6131945 |
| 7149 | NM_032781.3 | 57.8636003 | -7.4787809 | 1.33234697 | -5.6132382 |
| 7150 | NM_173469.3 | 66.9250033 | -7.6864419 | 1.36936525 | -5.6131423 |
| 7151 | NM_005044.4 | 275.015971 | -1.8383522 | 0.32752188 | -5.6129143 |
| 7152 | NM_032831.3 | 48.6436405 | -7.2300103 | 1.28811099 | -5.6128783 |
| 7153 | NM_012420.2 | 71.8537866 | -5.4201452 | 0.96573102 | -5.6124792 |
| 7154 | NM_018307.4 | 57.5684994 | -7.4695385 | 1.33104283 | -5.6117943 |
| 7155 | NM_015420.6 | 107.756326 | -7.402581 | 1.31913154 | -5.611708 |
| 7156 | NM_001135191.1 | 79.2164403 | -7.927013 | 1.41260616 | -5.6116229 |
| 7157 | NM_203505.2 | 73.7245556 | -7.8269687 | 1.39482188 | -5.6114468 |
| 7158 | NR_130944.1 | 48.2505625 | -7.2113663 | 1.28513017 | -5.6113897 |
| 7159 | NR_036521.1 | 62.4062617 | -7.5849563 | 1.3517205 | -5.6113348 |
| 7160 | NM_005883.2 | 1283.04898 | -6.5595886 | 1.16913515 | -5.6106333 |
| 7161 | NM_032376.3 | 47.9712352 | -7.2065441 | 1.28450196 | -5.6103799 |
| 7162 | NM_032377.3 | 94.4281902 | -7.2156973 | 1.28617489 | -5.6101992 |
| 7163 | NM_004545.3 | 75.442531 | -7.8592574 | 1.40097527 | -5.6098473 |
| 7164 | NM_001478.4 | 68.9948796 | -7.7304091 | 1.37805658 | -5.6096456 |
| 7165 | NM_018446.3 | 45.445571 | -7.1297782 | 1.27106264 | -5.6093052 |
| 7166 | NM_001142418.1 | 54.6523352 | -7.3919261 | 1.317839 | -5.6091268 |
| 7167 | NM_016167.4 | 49.4165068 | -7.2493529 | 1.29243308 | -5.6090741 |
| 7168 | NM_018410.4 | 49.9463728 | -7.2627264 | 1.29484137 | -5.60897 |
| 7169 | NM_005726.5 | 53.1061534 | -7.353431 | 1.31116379 | -5.6083237 |
| 7170 | NM_003612.4 | 86.2524567 | -8.0519021 | 1.43575218 | -5.608142 |
| 7171 | NM_016059.4 | 122.85821 | -7.596468 | 1.35454411 | -5.6081363 |
| 7172 | NM_016500.3 | 51.5660228 | -7.3110413 | 1.30365718 | -5.6081011 |
| 7173 | NM_006811.3 | 304.485255 | -4.055582 | 0.72317263 | -5.6080414 |
| 7174 | NM_001195305.1 | 59.1170732 | -7.5068253 | 1.33859443 | -5.6079909 |
| 7175 | NM_014939.3 | 124.356799 | -5.073742 | 0.9049404 | -5.6067139 |
| 7176 | NM_018074.5 | 53.1844413 | -7.3535971 | 1.31156524 | -5.6067338 |
| 7177 | NM_172226.2 | 204.460016 | -9.2977133 | 1.65837693 | -5.6065139 |
| 7178 | NM_176806.3 | 68.2556769 | -7.7155711 | 1.37622397 | -5.6063339 |
| 7179 | NM_001172681.1 | 69.0201284 | -7.7300956 | 1.37891224 | -5.6059373 |
| 7180 | NM_000714.5 | 88.1450329 | -8.0831919 | 1.4419526 | -5.6057265 |

| | | | | | |
|---|---|---|---|---|---|
| 7181 | NM_153832.2 | 95.4707763 | -6.4095662 | 1.14340441 | -5.6056861 |
| 7182 | NM_001408.2 | 556.290488 | -5.0677508 | 0.90408944 | -5.6053644 |
| 7183 | NM_018684.3 | 70.9450849 | -7.7703763 | 1.38625808 | -5.6052884 |
| 7184 | NM_020815.2 | 64.7525992 | -7.6396885 | 1.3629374 | -5.6053113 |
| 7185 | NM_001198690.1 | 58.1451164 | -7.4846267 | 1.33531462 | -5.6051409 |
| 7186 | NM_001134771.1 | 66.3963686 | -7.6741692 | 1.36917848 | -5.6049444 |
| 7187 | NM_004519.3 | 137.653854 | -8.726352 | 1.55692505 | -5.6048632 |
| 7188 | NM_002237.3 | 102.18459 | -8.2964643 | 1.48027914 | -5.6046621 |
| 7189 | NM_004813.2 | 64.7668598 | -7.6385991 | 1.36294471 | -5.604482 |
| 7190 | NM_207009.3 | 56.2152738 | -7.4345405 | 1.32656279 | -5.6043638 |
| 7191 | NM_018444.3 | 67.7783472 | -7.7055131 | 1.37493928 | -5.604257 |
| 7192 | NM_031284.4 | 64.9845074 | -7.6451339 | 1.36417828 | -5.6042044 |
| 7193 | NM_001160259.1 | 83.9578033 | -8.0141442 | 1.43003893 | -5.6041441 |
| 7194 | NM_007084.3 | 77.6073828 | -7.8985502 | 1.40944092 | -5.6040307 |
| 7195 | NM_032036.2 | 60.1528913 | -7.5323188 | 1.34410641 | -5.6039602 |
| 7196 | NM_032421.2 | 234.44005 | -5.6882883 | 1.01504493 | -5.6039768 |
| 7197 | NM_182543.3 | 40.108655 | -6.9458037 | 1.23958561 | -5.6033271 |
| 7198 | NM_001199198.2 | 78.8160602 | -7.9219747 | 1.41393392 | -5.6027899 |
| 7199 | NR_037608.1 | 114.45236 | -6.6832045 | 1.19284313 | -5.6027522 |
| 7200 | NM_012228.3 | 61.6260277 | -7.5673394 | 1.35068574 | -5.6025907 |
| 7201 | NM_001163484.1 | 64.2699854 | -7.6311721 | 1.36209311 | -5.6025334 |
| 7202 | NM_007222.4 | 83.4906788 | -8.0052515 | 1.42886425 | -5.6025277 |
| 7203 | NM_001077497.2 | 62.0239498 | -7.5757919 | 1.35227172 | -5.6022705 |
| 7204 | NM_004394.2 | 261.518422 | -6.1412331 | 1.096306 | -5.6017509 |
| 7205 | NM_017775.3 | 169.5476 | -5.1672761 | 0.92251049 | -5.6013196 |
| 7206 | NM_015711.3 | 91.3363429 | -7.1670581 | 1.27963458 | -5.6008631 |
| 7207 | NM_007342.2 | 53.3378002 | -7.3589056 | 1.3139211 | -5.600721 |
| 7208 | NM_001769.3 | 41.9916558 | -7.017306 | 1.25299372 | -5.6004319 |
| 7209 | NM_001122679.1 | 78.7821033 | -7.9237542 | 1.4148648 | -5.6003614 |
| 7210 | NM_182533.2 | 55.5602982 | -7.4180289 | 1.32459145 | -5.6002391 |
| 7211 | NM_012424.4 | 58.6248429 | -7.4936814 | 1.33817444 | -5.5999287 |
| 7212 | NM_018224.3 | 67.9793733 | -7.7080956 | 1.37646598 | -5.5999173 |
| 7213 | NM_001514.5 | 59.0807058 | -7.5062896 | 1.34045116 | -5.5998232 |
| 7214 | NM_152374.1 | 158.997773 | -7.9698706 | 1.4232418 | -5.5998008 |
| 7215 | NM_007221.3 | 50.293274 | -7.2732668 | 1.29889733 | -5.5995702 |
| 7216 | NM_002953.3 | 41.9486333 | -7.0164049 | 1.25310155 | -5.5992309 |
| 7217 | NM_052868.5 | 62.7293325 | -7.595366 | 1.35654107 | -5.5990682 |
| 7218 | NM_000355.3 | 40.4070901 | -6.9598236 | 1.24304474 | -5.599013 |
| 7219 | NM_001330327.1 | 40.9205668 | -6.9816291 | 1.24696305 | -5.5989061 |
| 7220 | NM_001270941.1 | 143.862299 | -8.7903613 | 1.57005955 | -5.5987439 |
| 7221 | NM_001303514.1 | 171.505563 | -9.0438074 | 1.61532489 | -5.5987544 |
| 7222 | NM_002894.2 | 40.259151 | -6.9560801 | 1.24243086 | -5.5987664 |
| 7223 | NR_024493.2 | 53.0811475 | -7.3525985 | 1.31325006 | -5.5987802 |

| | | | | | |
|---|---|---|---|---|---|
| 7224 | NR_038366.1 | 85.8049421 | -8.0458305 | 1.43712625 | -5.5985551 |
| 7225 | NM_002357.3 | 56.9087118 | -7.4536128 | 1.33139289 | -5.5983571 |
| 7226 | NM_006653.4 | 62.5864292 | -7.5892301 | 1.35562621 | -5.5983206 |
| 7227 | NM_024408.3 | 363.639554 | -4.7956248 | 0.85667435 | -5.5979554 |
| 7228 | NM_001142568.2 | 81.2744623 | -7.968866 | 1.42354975 | -5.5978837 |
| 7229 | NM_003939.4 | 60.025613 | -7.5272215 | 1.34466832 | -5.5978277 |
| 7230 | NM_005952.3 | 89.8482572 | -8.1113868 | 1.44914382 | -5.5973649 |
| 7231 | NR_073396.1 | 202.845229 | 2.64735016 | 0.47299744 | 5.59696511 |
| 7232 | NM_003957.3 | 86.5261817 | -8.056649 | 1.4395237 | -5.5967463 |
| 7233 | NM_005223.3 | 197.156531 | 1.51952642 | 0.27151223 | 5.59653029 |
| 7234 | NM_001323523.1 | 101.075516 | -8.2815579 | 1.47991966 | -5.595951 |
| 7235 | NM_015087.4 | 71.6308441 | -7.7874148 | 1.39161214 | -5.5959664 |
| 7236 | NM_004579.4 | 44.9258434 | -7.111923 | 1.27094209 | -5.5957884 |
| 7237 | NM_003449.4 | 44.0933455 | -7.0868909 | 1.26648159 | -5.5957315 |
| 7238 | NM_001122.3 | 39.8447102 | -6.9378733 | 1.23994797 | -5.5952938 |
| 7239 | NM_002141.4 | 72.1065479 | -7.7946113 | 1.39314195 | -5.5949871 |
| 7240 | NM_152437.2 | 313.788528 | -5.9796811 | 1.06875824 | -5.5949801 |
| 7241 | NM_003888.3 | 78.0242459 | -7.9088592 | 1.41371687 | -5.5943728 |
| 7242 | NM_020844.2 | 49.3581128 | -7.2447588 | 1.29506932 | -5.5941089 |
| 7243 | NM_213725.1 | 112.277504 | -8.4333477 | 1.50753873 | -5.5941168 |
| 7244 | NM_006416.4 | 64.9000705 | -7.6417582 | 1.3660775 | -5.593942 |
| 7245 | NM_001177984.2 | 74.711805 | -7.8436661 | 1.40219736 | -5.5938389 |
| 7246 | NM_004487.4 | 115.418044 | -8.4722399 | 1.51459235 | -5.5937427 |
| 7247 | NM_006821.5 | 39.2044847 | -6.9123744 | 1.23582084 | -5.5933466 |
| 7248 | NR_033183.1 | 66.5903314 | -7.6795301 | 1.37305563 | -5.5930218 |
| 7249 | NM_006577.5 | 44.8517342 | -7.1060123 | 1.27055275 | -5.5928511 |
| 7250 | NM_015655.3 | 53.1726295 | -7.3580369 | 1.31586137 | -5.5918025 |
| 7251 | NM_001111.4 | 96.9328306 | -8.2191642 | 1.47009182 | -5.5909189 |
| 7252 | NM_001080482.2 | 58.2715336 | -7.4869178 | 1.33944093 | -5.5895841 |
| 7253 | NM_016339.4 | 44.6982133 | -7.1001065 | 1.27024453 | -5.5895588 |
| 7254 | NM_001330698.1 | 80.0730031 | -7.9449981 | 1.42144916 | -5.5893649 |
| 7255 | NM_020828.1 | 51.3621193 | -7.3034999 | 1.30678479 | -5.588908 |
| 7256 | NM_153265.2 | 50.5778179 | -7.286317 | 1.30371101 | -5.588905 |
| 7257 | NM_032731.3 | 70.9256767 | -7.7700385 | 1.39034075 | -5.5885858 |
| 7258 | NM_001281956.1 | 76.6882434 | -7.8832416 | 1.41061289 | -5.5885223 |
| 7259 | NM_001018138.1 | 75.5081016 | -7.8603328 | 1.40661487 | -5.58812 |
| 7260 | NM_182715.2 | 82.8354969 | -7.0222759 | 1.25668355 | -5.5879429 |
| 7261 | NM_001278270.1 | 92.0912538 | -7.1806236 | 1.28507113 | -5.5877247 |
| 7262 | NM_020533.2 | 69.130576 | -7.7336683 | 1.38404883 | -5.5877135 |
| 7263 | NM_001002030.1 | 59.8440206 | -7.5252519 | 1.34678272 | -5.5875768 |
| 7264 | NM_007242.5 | 58.6562528 | -7.4950564 | 1.34138608 | -5.587546 |
| 7265 | NM_001145928.1 | 45.3546396 | -7.1279419 | 1.27576736 | -5.58718 |
| 7266 | NM_005830.3 | 51.8880328 | -7.3174014 | 1.30970143 | -5.587076 |

| | | | | | |
|---|---|---|---|---|---|
| 7267 | NM_001300883.2 | 45.8393394 | -7.1395388 | 1.277983 | -5.5865679 |
| 7268 | NM_020905.3 | 46.1547025 | -7.1497097 | 1.27986857 | -5.5862843 |
| 7269 | NM_007259.4 | 51.3939995 | -7.304092 | 1.30755815 | -5.5860552 |
| 7270 | NM_016651.5 | 105.653305 | -8.3447929 | 1.49387014 | -5.586023 |
| 7271 | NM_001198843.1 | 53.9380367 | -7.3748752 | 1.32025894 | -5.5859309 |
| 7272 | NM_170663.4 | 152.818415 | -8.8773314 | 1.58928438 | -5.5857413 |
| 7273 | NM_001100420.1 | 40.9388604 | -6.976575 | 1.24908628 | -5.5853427 |
| 7274 | NM_174936.3 | 49.3941973 | -7.2485642 | 1.29799623 | -5.5844263 |
| 7275 | NM_032315.2 | 52.4387608 | -7.3336406 | 1.31325472 | -5.5843246 |
| 7276 | NM_001142333.1 | 74.7655491 | -7.8452016 | 1.40494353 | -5.5839978 |
| 7277 | NM_004111.5 | 89.4152929 | -6.313184 | 1.130612 | -5.5838643 |
| 7278 | NM_001149.3 | 39.9439769 | -6.9426122 | 1.24342542 | -5.5834569 |
| 7279 | NM_015254.3 | 40.2588454 | -6.9581735 | 1.24633776 | -5.5828956 |
| 7280 | NM_148893.2 | 50.9559432 | -7.2915622 | 1.30607039 | -5.5828248 |
| 7281 | NM_014462.2 | 60.9119797 | -7.5502281 | 1.35241764 | -5.5827637 |
| 7282 | NM_004264.4 | 61.2697162 | -7.5578734 | 1.35385406 | -5.5824875 |
| 7283 | NM_007044.3 | 38.8678082 | -6.9003214 | 1.23605959 | -5.5825152 |
| 7284 | NR_024058.1 | 59.2479921 | -7.509752 | 1.34523488 | -5.5824839 |
| 7285 | NM_000031.5 | 89.6619544 | -6.2895925 | 1.1266897 | -5.5823645 |
| 7286 | NM_001080432.2 | 257.916454 | -4.3504441 | 0.77931573 | -5.5823897 |
| 7287 | NM_001254757.1 | 40.3442478 | -6.9545317 | 1.24579933 | -5.5823851 |
| 7288 | NM_006286.4 | 73.3307511 | -7.8199973 | 1.40082677 | -5.5824156 |
| 7289 | NM_138782.2 | 43.6041242 | -7.0681917 | 1.2661511 | -5.5824235 |
| 7290 | NM_032352.3 | 56.7543171 | -7.4469261 | 1.33406966 | -5.5821119 |
| 7291 | NM_001136050.2 | 45.0496465 | -7.1182304 | 1.2752031 | -5.5820366 |
| 7292 | NM_014943.3 | 37.7775478 | -6.8636564 | 1.2299028 | -5.5806495 |
| 7293 | NM_005941.4 | 45.3739164 | -7.1261399 | 1.27697423 | -5.5804884 |
| 7294 | NM_016004.4 | 65.1665579 | -7.6453133 | 1.37005006 | -5.5803168 |
| 7295 | NM_001313959.1 | 75.8497286 | -7.8679229 | 1.41003291 | -5.5799569 |
| 7296 | NM_002733.4 | 100.236229 | -7.2980362 | 1.30795591 | -5.5797265 |
| 7297 | NM_022742.4 | 62.5740734 | -7.5901397 | 1.36033815 | -5.5795978 |
| 7298 | NM_016399.2 | 70.3914786 | -7.7582584 | 1.39048422 | -5.5795372 |
| 7299 | NM_173547.3 | 69.0574047 | -7.7303343 | 1.38550764 | -5.5794238 |
| 7300 | NM_017456.3 | 94.4358551 | -8.18339 | 1.46690048 | -5.5786947 |
| 7301 | NM_014284.2 | 58.424762 | -7.4904566 | 1.34273667 | -5.5785001 |
| 7302 | NM_016044.2 | 50.2615705 | -7.2727114 | 1.30378526 | -5.5781513 |
| 7303 | NM_005837.2 | 91.1390052 | -8.1318106 | 1.45794173 | -5.5775965 |
| 7304 | NM_139070.2 | 72.4841538 | -7.8010134 | 1.39865933 | -5.5774936 |
| 7305 | NM_001001992.1 | 50.7098945 | -7.2857418 | 1.30640459 | -5.5769413 |
| 7306 | NM_015316.2 | 105.161003 | -7.3716717 | 1.32184405 | -5.5768089 |
| 7307 | NM_023075.5 | 44.3085063 | -7.0883751 | 1.27116483 | -5.5762832 |
| 7308 | NM_001168401.1 | 92.0923972 | -8.1465645 | 1.46119116 | -5.5752901 |
| 7309 | NM_016138.4 | 41.795952 | -7.0057499 | 1.25665591 | -5.574915 |

| | | | | | |
|---|---|---|---|---|---|
| 7310 | NM_003641.3 | 83.1033833 | -8.0007158 | 1.43516011 | -5.5747897 |
| 7311 | NM_001321770.1 | 54.8027446 | -7.4011815 | 1.32766491 | -5.5745855 |
| 7312 | NM_001330431.1 | 58.8686471 | -7.5008431 | 1.34560521 | -5.5743267 |
| 7313 | NM_006520.2 | 50.0488963 | -7.2678705 | 1.30388118 | -5.5740283 |
| 7314 | NM_001077399.2 | 52.5613532 | -7.3385081 | 1.31664828 | -5.5736283 |
| 7315 | NM_005605.4 | 54.195401 | -7.3810286 | 1.32434121 | -5.5733587 |
| 7316 | NM_178841.3 | 61.9542171 | -7.5732629 | 1.3588281 | -5.5733782 |
| 7317 | NM_018216.2 | 110.117145 | -5.8092411 | 1.04234114 | -5.5732628 |
| 7318 | NM_006810.3 | 57.3819902 | -7.465378 | 1.33957425 | -5.5729483 |
| 7319 | NM_022065.4 | 48.5296186 | -7.22293 | 1.29617375 | -5.5725013 |
| 7320 | NM_018122.4 | 48.0614755 | -7.2093498 | 1.29377445 | -5.572339 |
| 7321 | NM_001099922.2 | 50.7596426 | -7.2845192 | 1.30730344 | -5.5721717 |
| 7322 | NR_122070.1 | 19.4914747 | 3.08531713 | 0.55373154 | 5.57186452 |
| 7323 | NM_032042.5 | 57.2054186 | -7.4610991 | 1.33907347 | -5.571837 |
| 7324 | NM_001145365.1 | 74.1989177 | -7.8332926 | 1.4060046 | -5.5713136 |
| 7325 | NM_052840.4 | 72.817344 | -7.8063381 | 1.40125369 | -5.570967 |
| 7326 | NM_194318.3 | 42.7140685 | -7.042823 | 1.26423715 | -5.5708084 |
| 7327 | NM_144684.3 | 46.7580551 | -7.1655252 | 1.28631821 | -5.5705697 |
| 7328 | NM_004336.4 | 53.0156885 | -7.3493401 | 1.31934171 | -5.5704599 |
| 7329 | NM_001113361.1 | 39.9639929 | -6.9398827 | 1.24593326 | -5.5700277 |
| 7330 | NM_019037.2 | 57.9570109 | -7.4777861 | 1.34264053 | -5.5694625 |
| 7331 | NM_001010983.2 | 43.4076123 | -7.066128 | 1.26874717 | -5.5693744 |
| 7332 | NM_001198953.1 | 49.8280351 | -7.2591552 | 1.30343622 | -5.5692446 |
| 7333 | NM_001300952.1 | 49.2605191 | -7.245546 | 1.30101781 | -5.5691367 |
| 7334 | NM_001167740.1 | 66.3436904 | -7.6737962 | 1.377977 | -5.5688855 |
| 7335 | NM_198507.2 | 42.6870378 | -7.0379102 | 1.26381446 | -5.5687843 |
| 7336 | NR_024408.1 | 59.5248796 | -5.1335305 | 0.92195314 | -5.5681035 |
| 7337 | NM_001145297.3 | 47.1702228 | -7.1860901 | 1.29073301 | -5.5674489 |
| 7338 | NM_015384.4 | 45.3589452 | -7.1300497 | 1.28073842 | -5.5671397 |
| 7339 | NM_001195479.1 | 67.235668 | -7.6942506 | 1.38224717 | -5.5664795 |
| 7340 | NM_004935.3 | 85.5097087 | -8.0396895 | 1.44440637 | -5.5660856 |
| 7341 | NM_007167.3 | 70.4986926 | -5.340508 | 0.95949359 | -5.5659652 |
| 7342 | NM_001324276.1 | 49.3678149 | -7.2474593 | 1.30223946 | -5.5653815 |
| 7343 | NM_016115.4 | 52.862164 | -7.3452014 | 1.31991043 | -5.5649241 |
| 7344 | NR_033322.3 | 38.4864762 | -6.8851769 | 1.23761624 | -5.5632568 |
| 7345 | NM_202468.2 | 89.2527622 | -8.1013932 | 1.45629302 | -5.5630241 |
| 7346 | NR_028482.1 | 46.9271798 | -7.1738391 | 1.28967546 | -5.562515 |
| 7347 | NM_000965.4 | 38.580193 | -6.8956181 | 1.23972125 | -5.5622327 |
| 7348 | NM_024805.2 | 63.3998778 | -7.6077683 | 1.36779084 | -5.5620845 |
| 7349 | NM_006885.3 | 49.8790941 | -7.2658 | 1.30633756 | -5.5619621 |
| 7350 | NM_007048.5 | 37.4633884 | -6.8496008 | 1.23155181 | -5.5617642 |
| 7351 | NM_030965.2 | 57.9607347 | -7.4790142 | 1.34484177 | -5.5612596 |
| 7352 | NM_018164.2 | 60.8789979 | -7.5494648 | 1.35757362 | -5.5609984 |

| | | | | | |
|---|---|---|---|---|---|
| 7353 | NM_005229.4 | 56.282198 | -7.4360802 | 1.33719637 | -5.5609486 |
| 7354 | NM_018668.4 | 62.6309887 | -7.593236 | 1.36552646 | -5.5606656 |
| 7355 | NM_001320485.1 | 61.7875984 | -7.5697015 | 1.36138042 | -5.5603132 |
| 7356 | NM_002765.4 | 43.8599906 | -7.076612 | 1.27277854 | -5.5599712 |
| 7357 | NM_001078173.1 | 60.5639478 | -7.5419097 | 1.35658436 | -5.5594845 |
| 7358 | NM_020218.1 | 67.4819215 | -7.6979433 | 1.38466877 | -5.5594113 |
| 7359 | NM_002387.2 | 46.2881466 | -7.1575719 | 1.28762922 | -5.5587212 |
| 7360 | NM_031443.3 | 65.7304673 | -7.659994 | 1.37804998 | -5.5585749 |
| 7361 | NM_181705.3 | 76.9094414 | -7.8884166 | 1.41915185 | -5.558543 |
| 7362 | NM_001330124.1 | 49.389106 | -7.2484774 | 1.30411227 | -5.5581698 |
| 7363 | NM_022839.4 | 56.4167057 | -7.4402011 | 1.33865134 | -5.5579828 |
| 7364 | NM_001012979.2 | 104.312231 | -8.3263525 | 1.49811305 | -5.5578933 |
| 7365 | NM_020154.2 | 131.232342 | -5.6328199 | 1.01351033 | -5.5577331 |
| 7366 | NM_001040101.1 | 120.54495 | -8.5354353 | 1.5358025 | -5.5576386 |
| 7367 | NM_020232.4 | 56.1547487 | -7.4331616 | 1.33756231 | -5.5572451 |
| 7368 | NM_201592.2 | 89.8920058 | -8.1107393 | 1.45962059 | -5.5567449 |
| 7369 | NM_005082.4 | 86.5881713 | -6.2688061 | 1.12816172 | -5.5566556 |
| 7370 | NM_020360.3 | 70.4199311 | -7.7605297 | 1.39664671 | -5.5565446 |
| 7371 | NR_015432.1 | 48.0787967 | -7.2087742 | 1.29743534 | -5.5561722 |
| 7372 | NM_005682.6 | 124.704692 | -8.5844038 | 1.54514604 | -5.5557232 |
| 7373 | NM_002333.3 | 436.090215 | -5.7029383 | 1.02660795 | -5.5551278 |
| 7374 | NM_003162.3 | 87.200225 | -7.0949431 | 1.27725552 | -5.5548346 |
| 7375 | NM_014878.4 | 53.3881078 | -7.3608959 | 1.32520456 | -5.5545356 |
| 7376 | NR_135550.1 | 46.8162893 | 3.00011124 | 0.54012723 | 5.55445285 |
| 7377 | NM_001163546.2 | 65.8661097 | -7.6630196 | 1.3796837 | -5.5541858 |
| 7378 | NM_001173462.1 | 61.9177491 | -7.5737176 | 1.36361115 | -5.5541622 |
| 7379 | NM_003174.3 | 74.8424797 | -6.8757186 | 1.23793324 | -5.5541917 |
| 7380 | NM_000189.4 | 324.257674 | -6.1785996 | 1.11245331 | -5.5540306 |
| 7381 | NM_203298.2 | 58.8311608 | -7.499758 | 1.35038653 | -5.5537862 |
| 7382 | NM_004257.5 | 80.4132464 | -7.9519019 | 1.43182922 | -5.5536665 |
| 7383 | NM_001098204.1 | 112.240018 | -7.4660327 | 1.34435468 | -5.5536182 |
| 7384 | NM_021075.3 | 40.4046041 | -6.9602269 | 1.25345354 | -5.5528399 |
| 7385 | NM_016246.2 | 44.4214141 | -7.0989794 | 1.27864771 | -5.5519431 |
| 7386 | NM_016371.3 | 48.4421016 | -7.2217908 | 1.30078971 | -5.5518511 |
| 7387 | NM_001253.3 | 196.847934 | -3.6849113 | 0.66376953 | -5.5514921 |
| 7388 | NR_135260.1 | 103.97817 | -8.3214424 | 1.49896414 | -5.551462 |
| 7389 | NM_145057.3 | 43.3606186 | -7.0652276 | 1.27269717 | -5.5513816 |
| 7390 | NM_016640.3 | 48.8008885 | -7.2302602 | 1.30248642 | -5.5511213 |
| 7391 | NM_017818.3 | 52.3747454 | -7.3308139 | 1.32062351 | -5.5510248 |
| 7392 | NM_005907.3 | 44.8519104 | -7.1139307 | 1.28160379 | -5.5508034 |
| 7393 | NM_015679.2 | 62.1881887 | -7.5797836 | 1.36560379 | -5.5504998 |
| 7394 | NM_001271314.1 | 56.9875606 | -7.4535446 | 1.3428689 | -5.5504634 |
| 7395 | NM_001114090.1 | 52.7487858 | -7.3408086 | 1.32256429 | -5.5504362 |

| | | | | | |
|---|---|---|---|---|---|
| 7396 | NM_001008892.2 | 62.0411428 | -7.5760305 | 1.36507262 | -5.5499102 |
| 7397 | NM_006166.3 | 108.41514 | -3.6200739 | 0.65228993 | -5.5497928 |
| 7398 | NM_001040092.2 | 55.4122612 | -7.4172623 | 1.33652922 | -5.5496447 |
| 7399 | NM_054025.2 | 111.501437 | -8.4221723 | 1.51759416 | -5.5496868 |
| 7400 | NM_182961.3 | 62.044054 | -7.5749741 | 1.36494523 | -5.5496543 |
| 7401 | NM_014846.3 | 99.094634 | -6.4696551 | 1.16594022 | -5.5488738 |
| 7402 | NM_017909.3 | 51.4601242 | -7.3077103 | 1.31709889 | -5.5483384 |
| 7403 | NR_134455.1 | 45.617687 | -7.1335526 | 1.28580086 | -5.5479451 |
| 7404 | NM_052917.3 | 84.2972568 | -8.018966 | 1.44556788 | -5.5472774 |
| 7405 | NM_001321431.1 | 114.577718 | -6.687044 | 1.20566998 | -5.5463303 |
| 7406 | NM_001517.4 | 92.6785902 | -7.1893524 | 1.29626023 | -5.5462261 |
| 7407 | NM_001300867.1 | 49.8509098 | -7.2602715 | 1.30923396 | -5.5454348 |
| 7408 | NM_004832.2 | 52.849222 | -7.3460285 | 1.32469822 | -5.5454354 |
| 7409 | NM_018073.7 | 57.4219263 | -7.4641531 | 1.34609486 | -5.5450424 |
| 7410 | NM_139162.3 | 61.2502775 | -7.5566814 | 1.36278369 | -5.5450336 |
| 7411 | NM_001137667.1 | 64.0217481 | -7.6200839 | 1.37425678 | -5.5448764 |
| 7412 | NM_001172646.1 | 41.1324673 | -6.9874285 | 1.26039763 | -5.5438287 |
| 7413 | NM_015282.2 | 77.9300255 | -7.905802 | 1.4261165 | -5.5435878 |
| 7414 | NM_021904.3 | 102.362674 | -8.2991718 | 1.49719056 | -5.5431633 |
| 7415 | NR_024356.2 | 92.5160663 | -8.1536591 | 1.47103364 | -5.5428094 |
| 7416 | NM_023004.5 | 63.7117077 | -7.614506 | 1.37385615 | -5.5424333 |
| 7417 | NM_003766.4 | 45.2129747 | -7.1226229 | 1.28520978 | -5.5419925 |
| 7418 | NM_001322503.1 | 78.4429625 | 3.09603943 | 0.55865867 | 5.54191601 |
| 7419 | NM_002979.4 | 39.049782 | -6.9136532 | 1.2476096 | -5.5415197 |
| 7420 | NM_001247.4 | 41.1589351 | -6.9903144 | 1.2614794 | -5.5413624 |
| 7421 | NM_001204375.1 | 83.8085685 | -8.0093132 | 1.44538188 | -5.5413129 |
| 7422 | NM_013293.4 | 108.153238 | -7.4113101 | 1.33752333 | -5.5410698 |
| 7423 | NM_017446.3 | 47.6637964 | -7.1953192 | 1.29863102 | -5.5406956 |
| 7424 | NM_005860.2 | 95.0361059 | -6.4079954 | 1.15661476 | -5.5403023 |
| 7425 | NM_024075.4 | 59.5680738 | -7.5171791 | 1.35685584 | -5.5401457 |
| 7426 | NM_015028.3 | 150.529627 | -8.8559482 | 1.59852674 | -5.5400688 |
| 7427 | NM_001077418.2 | 88.2051936 | -7.1195051 | 1.28523949 | -5.5394385 |
| 7428 | NR_046084.1 | 38.6731752 | -6.8949037 | 1.24470658 | -5.5393808 |
| 7429 | NM_001248004.1 | 42.1984211 | -7.0175017 | 1.26688474 | -5.5391792 |
| 7430 | NM_006427.3 | 59.7284028 | -7.5222769 | 1.3580869 | -5.5388774 |
| 7431 | NM_012297.4 | 66.6025951 | -7.679854 | 1.38658786 | -5.538671 |
| 7432 | NM_198513.1 | 92.6107008 | -8.1539932 | 1.47245271 | -5.5376945 |
| 7433 | NM_014161.4 | 69.718776 | -7.7461765 | 1.39883431 | -5.537594 |
| 7434 | NM_182765.4 | 55.1282677 | -7.4060228 | 1.33744957 | -5.5374221 |
| 7435 | NM_017765.2 | 42.4744123 | -7.0331283 | 1.27013937 | -5.5372886 |
| 7436 | NM_024572.3 | 66.2168797 | -7.6721789 | 1.38566374 | -5.5368259 |
| 7437 | NM_032304.3 | 48.9582397 | -7.2354551 | 1.30690757 | -5.5363174 |
| 7438 | NM_001318875.1 | 54.2669425 | -7.3861764 | 1.33416654 | -5.5361727 |

| | | | | | |
|---|---|---|---|---|---|
| 7439 | NM_005513.2 | 60.2763675 | -3.0762726 | 0.55567976 | -5.5360529 |
| 7440 | NM_015464.2 | 70.4442504 | -7.7620662 | 1.40213866 | -5.5358763 |
| 7441 | NM_006574.3 | 67.2993303 | -7.6948875 | 1.39007654 | -5.5355854 |
| 7442 | NM_001362.3 | 47.7732962 | -7.2018589 | 1.30104467 | -5.5354433 |
| 7443 | NM_144693.2 | 56.6952608 | -7.4463952 | 1.3453143 | -5.5350599 |
| 7444 | NM_153756.2 | 42.5219954 | -7.0327885 | 1.2705907 | -5.5350543 |
| 7445 | NM_032140.2 | 36.3932819 | -6.8090794 | 1.2301945 | -5.5349617 |
| 7446 | NM_173575.3 | 79.1757298 | -7.9284233 | 1.43244099 | -5.5349039 |
| 7447 | NM_001145416.1 | 43.4022502 | -7.0629507 | 1.27608952 | -5.5348395 |
| 7448 | NM_152652.2 | 57.9462127 | -7.4786176 | 1.35120982 | -5.5347567 |
| 7449 | NM_032869.3 | 50.9714381 | -7.2938424 | 1.31796376 | -5.5341752 |
| 7450 | NM_145648.3 | 57.955257 | -7.4773001 | 1.35111385 | -5.5341747 |
| 7451 | NR_036608.1 | 50.2339899 | -7.2731488 | 1.31423349 | -5.5341375 |
| 7452 | NM_177938.2 | 43.741896 | -7.0714814 | 1.27787203 | -5.5337946 |
| 7453 | NM_019886.3 | 60.1308201 | -7.5332871 | 1.36139583 | -5.5335024 |
| 7454 | NM_001005367.2 | 66.8089589 | -7.6840542 | 1.38872004 | -5.5331917 |
| 7455 | NM_153348.2 | 40.0057684 | -6.9444232 | 1.2550665 | -5.5331118 |
| 7456 | NM_032466.3 | 53.4011392 | -7.3578675 | 1.32988003 | -5.5327303 |
| 7457 | NM_002693.2 | 64.2583294 | -7.6281855 | 1.37881449 | -5.5324234 |
| 7458 | NM_001082969.1 | 63.4148391 | -7.6070311 | 1.37505206 | -5.5321768 |
| 7459 | NM_001487.3 | 74.1970428 | -7.8350119 | 1.41632213 | -5.531942 |
| 7460 | NM_012174.1 | 50.1987105 | -7.2725673 | 1.31473897 | -5.5315675 |
| 7461 | NM_001305066.1 | 52.7291835 | -7.341653 | 1.32723956 | -5.5315206 |
| 7462 | NR_026903.1 | 50.6502949 | -7.2835822 | 1.31675815 | -5.5314503 |
| 7463 | NM_002923.3 | 58.5026868 | -7.4934185 | 1.354727 | -5.5313126 |
| 7464 | NR_033464.1 | 74.0825752 | -7.8339915 | 1.41641588 | -5.5308554 |
| 7465 | NM_001114.4 | 60.2455799 | -7.5351888 | 1.36239715 | -5.5308313 |
| 7466 | NM_201267.1 | 52.0755605 | -7.3275715 | 1.32486659 | -5.5307995 |
| 7467 | NM_001290039.1 | 75.1173733 | -7.8508493 | 1.41954065 | -5.5305562 |
| 7468 | NM_002852.3 | 82.3991803 | -7.9872041 | 1.44420937 | -5.5305029 |
| 7469 | NM_025040.3 | 47.151155 | -7.1798565 | 1.29829912 | -5.5302021 |
| 7470 | NM_001025088.1 | 103.426149 | -8.3127831 | 1.50318731 | -5.5301046 |
| 7471 | NM_006909.2 | 98.4924787 | -7.2735289 | 1.31531119 | -5.529892 |
| 7472 | NM_001171155.1 | 67.3972476 | -7.6967422 | 1.3920155 | -5.5292073 |
| 7473 | NM_001199954.1 | 133.832581 | -7.7207395 | 1.39639053 | -5.5290689 |
| 7474 | NM_031921.5 | 60.25909 | -7.5333942 | 1.36252897 | -5.5289791 |
| 7475 | NM_017975.4 | 64.124912 | -7.6236861 | 1.37888006 | -5.5288972 |
| 7476 | NM_006918.4 | 393.213501 | -4.5069915 | 0.81522967 | -5.528493 |
| 7477 | NM_018928.2 | 67.6256846 | -7.7020277 | 1.39343103 | -5.5273835 |
| 7478 | NM_152715.3 | 41.2062529 | -6.9872262 | 1.26413738 | -5.5272681 |
| 7479 | NM_016192.3 | 78.381349 | -7.9146862 | 1.43200118 | -5.527011 |
| 7480 | NM_022755.5 | 45.1424834 | -7.1186712 | 1.28799781 | -5.526928 |
| 7481 | NM_080668.3 | 50.1050642 | -7.2705375 | 1.31549579 | -5.5268421 |

| | | | | | |
|---|---|---|---|---|---|
| 7482 | NM_198525.2 | 83.1150366 | -7.0257051 | 1.27142598 | -5.5258467 |
| 7483 | NM_023078.4 | 41.0554677 | -6.9818149 | 1.26357109 | -5.5254627 |
| 7484 | NM_001033045.3 | 45.3801964 | -7.1251157 | 1.2895864 | -5.5251169 |
| 7485 | NM_001321706.1 | 93.197553 | -8.1645463 | 1.47772579 | -5.5250753 |
| 7486 | NM_001110199.2 | 87.4537129 | -8.0718509 | 1.46097359 | -5.5249807 |
| 7487 | NM_001005862.2 | 57.228625 | -7.4613297 | 1.35061595 | -5.5243903 |
| 7488 | NM_022739.3 | 82.8695383 | -7.0258136 | 1.27194345 | -5.5236839 |
| 7489 | NM_014067.3 | 54.4057147 | -7.3861243 | 1.33722216 | -5.5234833 |
| 7490 | NM_016478.4 | 52.1964562 | -7.3282151 | 1.32679422 | -5.5232492 |
| 7491 | NM_032842.3 | 57.1204959 | -7.4611193 | 1.35101125 | -5.5226182 |
| 7492 | NM_016360.3 | 46.1251344 | -7.1498048 | 1.29465124 | -5.5225721 |
| 7493 | NM_139179.3 | 65.1266188 | -7.6481445 | 1.38489566 | -5.5225421 |
| 7494 | NM_001099289.2 | 75.0281804 | -7.8505722 | 1.42170006 | -5.5219609 |
| 7495 | NR_029407.1 | 55.7712007 | -7.4235933 | 1.34447022 | -5.5215751 |
| 7496 | NM_001799.3 | 37.9981855 | -6.8662668 | 1.24359734 | -5.5212942 |
| 7497 | NM_001297704.1 | 86.0859483 | -8.048171 | 1.45769727 | -5.5211539 |
| 7498 | NM_138358.2 | 46.346681 | -7.1560306 | 1.2961161 | -5.521134 |
| 7499 | NM_005389.2 | 113.074069 | -8.4433226 | 1.52950872 | -5.5202841 |
| 7500 | NM_001083619.1 | 90.8132659 | -8.1264835 | 1.47214627 | -5.5201604 |
| 7501 | NM_001003722.1 | 82.9108085 | -7.0268796 | 1.27304099 | -5.5197591 |
| 7502 | NM_016262.4 | 46.6148803 | -7.1628399 | 1.29770137 | -5.5196365 |
| 7503 | NM_024580.5 | 39.2686826 | -6.9142121 | 1.25267675 | -5.5195502 |
| 7504 | NM_212492.2 | 53.825581 | -7.3707993 | 1.33542665 | -5.519434 |
| 7505 | NM_005908.3 | 76.9046824 | -6.0897111 | 1.10339886 | -5.5190479 |
| 7506 | NM_017650.2 | 56.4383351 | -7.4404945 | 1.34815841 | -5.5190061 |
| 7507 | NM_001308147.1 | 87.2467906 | -7.0979196 | 1.28617199 | -5.5186395 |
| 7508 | NM_178568.3 | 47.7334104 | -7.1963825 | 1.30402092 | -5.5186097 |
| 7509 | NM_078468.2 | 41.7823237 | -7.0051137 | 1.26940171 | -5.5184373 |
| 7510 | NM_031231.3 | 43.5061269 | -7.0686757 | 1.28110189 | -5.517653 |
| 7511 | NR_038269.1 | 41.6662803 | -7.0000499 | 1.26879114 | -5.5171018 |
| 7512 | NM_020664.3 | 51.8067834 | -7.3177738 | 1.32649225 | -5.5166352 |
| 7513 | NM_201280.2 | 43.081261 | -7.0479035 | 1.27758286 | -5.5165921 |
| 7514 | NM_004477.2 | 59.831226 | -7.5247551 | 1.36404177 | -5.5165137 |
| 7515 | NM_001195044.1 | 55.699747 | -7.4174703 | 1.34465989 | -5.5162427 |
| 7516 | NM_031297.4 | 68.8566825 | -7.7274432 | 1.40091964 | -5.5159789 |
| 7517 | NM_001097615.2 | 96.4579566 | -7.2472814 | 1.31397242 | -5.5155507 |
| 7518 | NM_001130963.1 | 48.7597308 | -7.225336 | 1.3101752 | -5.5147861 |
| 7519 | NM_025201.4 | 40.4128558 | -6.96055 | 1.26216154 | -5.5147853 |
| 7520 | NM_001289103.1 | 58.4040611 | -7.4911813 | 1.35838913 | -5.5147536 |
| 7521 | NM_001014763.1 | 71.006065 | -7.7716405 | 1.4093669 | -5.5142777 |
| 7522 | NM_001039803.2 | 51.790759 | -7.3172897 | 1.32700496 | -5.5141389 |
| 7523 | NM_020686.5 | 150.636531 | -6.8830396 | 1.2482615 | -5.5141007 |
| 7524 | NM_001318819.1 | 48.8446436 | -7.2278206 | 1.31091983 | -5.5135489 |

| | | | | | |
|---|---|---|---|---|---|
| 7525 | NM_001014447.2 | 47.4155037 | -7.1911394 | 1.30427999 | -5.5134936 |
| 7526 | NM_030767.5 | 71.2163108 | -7.7755136 | 1.41027255 | -5.5134829 |
| 7527 | NM_001242865.2 | 47.7952242 | -7.2015414 | 1.30624712 | -5.5131539 |
| 7528 | NM_032175.3 | 42.9137934 | -7.040733 | 1.27714419 | -5.5128725 |
| 7529 | NM_005469.3 | 52.9767727 | -7.3478779 | 1.33296151 | -5.5124457 |
| 7530 | NM_001130971.1 | 43.8925424 | -7.0732027 | 1.28317307 | -5.5122749 |
| 7531 | NM_032458.2 | 45.6302183 | -7.134061 | 1.29428706 | -5.511962 |
| 7532 | NM_001532.2 | 49.6695602 | -7.2553468 | 1.31635699 | -5.5116863 |
| 7533 | NM_006024.6 | 55.970381 | -7.4279265 | 1.3476985 | -5.511564 |
| 7534 | NM_002015.3 | 53.607596 | -7.3696245 | 1.3371414 | -5.5114773 |
| 7535 | NM_024089.2 | 47.4614749 | -7.192611 | 1.30503985 | -5.511411 |
| 7536 | NM_018180.2 | 70.1793166 | -6.783358 | 1.23080197 | -5.5113317 |
| 7537 | NM_133496.4 | 46.0374568 | -7.1463989 | 1.29703476 | -5.5097975 |
| 7538 | NM_016639.2 | 126.858803 | -7.64423 | 1.38742656 | -5.5096466 |
| 7539 | NM_001081562.2 | 45.4255594 | -7.1245479 | 1.29339655 | -5.5084018 |
| 7540 | NM_004957.5 | 78.01538 | -7.90674 | 1.4353948 | -5.5084078 |
| 7541 | NM_002601.3 | 65.6452605 | -7.6582858 | 1.39036781 | -5.5081006 |
| 7542 | NM_001135244.1 | 55.9807256 | -7.4285591 | 1.34869204 | -5.5079729 |
| 7543 | NM_001321387.2 | 52.472431 | -7.3375864 | 1.33218617 | -5.5079287 |
| 7544 | NM_001014765.1 | 36.1252254 | -6.7957931 | 1.23386117 | -5.5077453 |
| 7545 | NM_001203258.1 | 62.4861199 | -7.5885581 | 1.37791324 | -5.507283 |
| 7546 | NM_001160329.1 | 79.3843059 | -7.9323308 | 1.44040246 | -5.5070239 |
| 7547 | NM_001142572.1 | 62.1674361 | -4.4631319 | 0.810549 | -5.5063073 |
| 7548 | NM_017619.3 | 37.8034612 | -6.8634561 | 1.24669387 | -5.5053259 |
| 7549 | NM_001143826.2 | 92.8875454 | -8.1601348 | 1.48228438 | -5.5051075 |
| 7550 | NM_001145176.1 | 80.2365173 | -7.9474069 | 1.44366317 | -5.5050285 |
| 7551 | NM_001322956.1 | 84.778813 | -8.0271841 | 1.45819664 | -5.5048708 |
| 7552 | NM_003000.2 | 101.630963 | -7.3220269 | 1.33041751 | -5.5035557 |
| 7553 | NM_001330518.1 | 59.9896622 | -7.5291216 | 1.36811624 | -5.5032762 |
| 7554 | NM_033547.3 | 83.771634 | -7.0407651 | 1.27940452 | -5.5031579 |
| 7555 | NM_020465.3 | 94.8924855 | -8.1900277 | 1.48826048 | -5.5030875 |
| 7556 | NM_173832.5 | 52.8343642 | -7.3447001 | 1.33477012 | -5.5025956 |
| 7557 | NM_006386.4 | 1109.63555 | -6.6116158 | 1.20155394 | -5.5025543 |
| 7558 | NM_001083601.2 | 48.2271053 | -7.2108225 | 1.31047308 | -5.5024575 |
| 7559 | NM_001256627.1 | 63.7691711 | -7.6163868 | 1.38422816 | -5.5022626 |
| 7560 | NM_172230.2 | 69.9854718 | -7.7521371 | 1.40890875 | -5.502228 |
| 7561 | NM_002510.2 | 62.2090456 | -7.5834233 | 1.37825208 | -5.5022034 |
| 7562 | NM_001318730.1 | 70.2885224 | -7.7554002 | 1.40954697 | -5.5020516 |
| 7563 | NM_017719.4 | 61.612835 | -7.5671332 | 1.37539795 | -5.5017773 |
| 7564 | NM_004182.3 | 58.1272802 | -7.4830181 | 1.36030918 | -5.5009686 |
| 7565 | NM_015926.5 | 45.9239071 | -7.1451761 | 1.29894973 | -5.5007334 |
| 7566 | NM_005483.2 | 74.8054013 | -5.4871729 | 0.99754765 | -5.5006624 |
| 7567 | NM_001001132.1 | 107.875147 | -8.3744596 | 1.52247419 | -5.5005593 |

| | | | | | |
|---|---|---|---|---|---|
| 7568 | NM_004365.3 | 42.1879329 | -7.0205403 | 1.27633712 | -5.5005375 |
| 7569 | NM_006473.3 | 83.1035533 | -5.6478635 | 1.02687807 | -5.5000332 |
| 7570 | NM_001039613.2 | 48.010797 | -7.2066821 | 1.31031172 | -5.4999753 |
| 7571 | NM_001199163.1 | 44.0072475 | -7.0818533 | 1.28761088 | -5.4999949 |
| 7572 | NM_020431.3 | 109.379851 | -5.8875518 | 1.07048199 | -5.4999074 |
| 7573 | NM_006705.3 | 99.8245027 | -8.2623686 | 1.50239463 | -5.4994663 |
| 7574 | NM_017919.2 | 71.2325131 | -6.804762 | 1.2373744 | -5.4993557 |
| 7575 | NM_001099269.2 | 59.8235169 | -7.5225305 | 1.36815023 | -5.498322 |
| 7576 | NM_033259.2 | 85.2867263 | -8.0363014 | 1.46162187 | -5.4982082 |
| 7577 | NM_014643.3 | 117.023108 | -5.4140888 | 0.98474318 | -5.4979703 |
| 7578 | NR_045262.1 | 41.2810457 | 2.28472119 | 0.41557756 | 5.49770111 |
| 7579 | NM_025129.4 | 47.6372981 | -7.1944425 | 1.3087302 | -5.4972694 |
| 7580 | NM_004723.3 | 130.227973 | -7.6775322 | 1.39668473 | -5.4969686 |
| 7581 | NM_001323631.1 | 40.249957 | -6.9543684 | 1.26513676 | -5.4969301 |
| 7582 | NM_001282689.1 | 65.1044222 | -5.2753557 | 0.95975983 | -5.4965373 |
| 7583 | NM_012166.2 | 94.7132521 | -7.2209154 | 1.31374964 | -5.4964166 |
| 7584 | NM_001319228.1 | 53.5469236 | -7.3601486 | 1.33911634 | -5.4962727 |
| 7585 | NM_001242659.1 | 84.2436057 | -5.6704598 | 1.03178153 | -5.495795 |
| 7586 | NM_024111.4 | 55.0804107 | -7.4047887 | 1.34743618 | -5.4954652 |
| 7587 | NR_038861.1 | 44.8103742 | -7.1113377 | 1.29407261 | -5.4953158 |
| 7588 | NM_148894.2 | 320.654854 | -5.4552354 | 0.99277962 | -5.4949108 |
| 7589 | NM_015958.2 | 52.9842495 | -7.349153 | 1.33756757 | -5.4944163 |
| 7590 | NM_153759.3 | 87.2912859 | -5.275246 | 0.96010664 | -5.4944376 |
| 7591 | NM_001099658.1 | 100.938491 | -8.2791571 | 1.50686529 | -5.4942915 |
| 7592 | NM_021805.2 | 39.7822403 | -6.9338501 | 1.26201734 | -5.4942589 |
| 7593 | NM_001033504.1 | 46.5316037 | -7.1627877 | 1.30369285 | -5.4942295 |
| 7594 | NM_001159944.2 | 74.4179966 | -7.8395332 | 1.42689115 | -5.4941354 |
| 7595 | NM_144567.4 | 46.040872 | -7.1472917 | 1.30089419 | -5.4941376 |
| 7596 | NM_001130111.1 | 255.125824 | -4.8322044 | 0.87956331 | -5.4938676 |
| 7597 | NM_005690.4 | 63.6189397 | -7.6116843 | 1.38549313 | -5.4938448 |
| 7598 | NM_004724.3 | 44.632396 | -7.1029481 | 1.29303702 | -5.4932287 |
| 7599 | NM_005126.4 | 50.8901223 | -7.2890907 | 1.32697762 | -5.493002 |
| 7600 | NM_001143944.1 | 56.468495 | -7.4407472 | 1.35460763 | -5.492917 |
| 7601 | NR_109840.1 | 50.9870284 | 10.5885157 | 1.92769355 | 5.4928418 |
| 7602 | NM_024051.3 | 44.3850526 | -7.0914436 | 1.2911074 | -5.492528 |
| 7603 | NM_001135993.1 | 44.2152779 | -7.0859057 | 1.29012063 | -5.4924366 |
| 7604 | NM_014517.4 | 67.9772534 | -7.7071065 | 1.40328594 | -5.4921854 |
| 7605 | NM_001135740.1 | 47.3199511 | -7.1842002 | 1.30808853 | -5.4921361 |
| 7606 | NM_001276270.1 | 44.9454313 | -7.1090576 | 1.29449322 | -5.4917689 |
| 7607 | NM_001131017.1 | 59.8605921 | -7.5243428 | 1.37016425 | -5.4915627 |
| 7608 | NM_018102.4 | 37.6689304 | -6.8525843 | 1.24790093 | -5.4912887 |
| 7609 | NM_003246.3 | 684.74146 | -3.7547126 | 0.68379922 | -5.4909576 |
| 7610 | NM_014798.2 | 43.6486175 | -7.0704487 | 1.28767938 | -5.4908457 |

| | | | | | |
|---|---|---|---|---|---|
| 7611 | NM_021971.2 | 38.7503712 | -6.8998679 | 1.25670956 | -5.4904236 |
| 7612 | NM_001006684.1 | 52.3368253 | -7.3316967 | 1.33550288 | -5.4898397 |
| 7613 | NM_006064.4 | 51.6403597 | -7.3144943 | 1.33237346 | -5.4898229 |
| 7614 | NM_181482.4 | 54.0647885 | -7.3759903 | 1.3436598 | -5.4894775 |
| 7615 | NM_016535.3 | 49.8360739 | -7.2616323 | 1.32310676 | -5.4883192 |
| 7616 | NM_198204.1 | 57.8366081 | -7.4752221 | 1.36207662 | -5.4881069 |
| 7617 | NM_001199933.1 | 36.5837686 | -6.8193346 | 1.24258732 | -5.4880124 |
| 7618 | NM_014934.4 | 24.0041386 | 9.49492324 | 1.73019729 | 5.48776912 |
| 7619 | NM_014154.4 | 53.4923498 | -7.3616903 | 1.34150928 | -5.4876178 |
| 7620 | NM_016656.3 | 54.4999445 | -7.3885625 | 1.34640479 | -5.4876235 |
| 7621 | NM_020650.2 | 46.5322314 | -7.1661617 | 1.30597167 | -5.487226 |
| 7622 | NM_005033.2 | 44.661053 | -7.1015873 | 1.29421492 | -5.4871778 |
| 7623 | NM_001144888.1 | 48.6074887 | -7.2288787 | 1.31756095 | -5.4865611 |
| 7624 | NM_016228.3 | 54.5878291 | -7.3927314 | 1.34748209 | -5.48633 |
| 7625 | NM_152261.3 | 77.171079 | -7.8916009 | 1.43844752 | -5.4861931 |
| 7626 | NM_001142964.1 | 48.2551824 | -7.2155027 | 1.31526841 | -5.4859545 |
| 7627 | NM_005466.3 | 72.1448648 | -6.8204535 | 1.24327803 | -5.4858635 |
| 7628 | NM_001112734.3 | 44.187231 | -7.084678 | 1.29149504 | -5.4856409 |
| 7629 | NM_001282799.1 | 81.0405082 | -5.6093192 | 1.02257864 | -5.4854648 |
| 7630 | NM_003013.2 | 96.3476207 | -4.8721911 | 0.88825841 | -5.4851055 |
| 7631 | NM_001201380.2 | 36.9078617 | -6.8305589 | 1.24533756 | -5.4849056 |
| 7632 | NM_032324.2 | 50.3371126 | -7.2743655 | 1.32629007 | -5.4847471 |
| 7633 | NR_027155.2 | 42.7149053 | -7.0382578 | 1.28326089 | -5.4846664 |
| 7634 | NM_001024226.1 | 75.7324662 | -7.8632794 | 1.43372181 | -5.4845224 |
| 7635 | NM_012062.4 | 135.506405 | -8.7034299 | 1.58704127 | -5.4840602 |
| 7636 | NM_004874.3 | 52.4214788 | -7.3304156 | 1.3367498 | -5.4837604 |
| 7637 | NM_014015.3 | 80.0056583 | -5.1887645 | 0.94621201 | -5.4837229 |
| 7638 | NM_001144758.2 | 96.7346218 | -8.2179515 | 1.49867701 | -5.4834707 |
| 7639 | NM_001304996.1 | 43.3216791 | -7.0538362 | 1.28642089 | -5.4833035 |
| 7640 | NM_152686.3 | 186.13161 | -3.2556635 | 0.59376804 | -5.4830561 |
| 7641 | NM_001047160.2 | 42.4156475 | -7.0252526 | 1.28127193 | -5.4830301 |
| 7642 | NM_021177.4 | 72.8354795 | -7.8082753 | 1.42417767 | -5.4826553 |
| 7643 | NM_024635.3 | 46.1795997 | -7.153368 | 1.30477285 | -5.4824623 |
| 7644 | NM_001346592.1 | 52.8935589 | -7.3469027 | 1.3401346 | -5.4822126 |
| 7645 | NM_017503.4 | 51.3256632 | -7.302934 | 1.33215876 | -5.4820298 |
| 7646 | NM_004564.2 | 51.989115 | -7.3227915 | 1.33585361 | -5.481732 |
| 7647 | NM_022078.2 | 45.4971356 | -7.12888 | 1.30062683 | -5.481111 |
| 7648 | NM_016836.3 | 76.1378161 | -7.8734359 | 1.43651059 | -5.4809453 |
| 7649 | NM_001098638.1 | 66.4714142 | -4.7341572 | 0.8637839 | -5.4807194 |
| 7650 | NM_022821.3 | 56.3411031 | -7.4398427 | 1.35750272 | -5.4805361 |
| 7651 | NM_001827.2 | 69.1356994 | -7.7324793 | 1.41091297 | -5.4804793 |
| 7652 | NM_003482.3 | 240.834165 | -5.5841945 | 1.01893331 | -5.4804318 |
| 7653 | NM_001257363.2 | 47.0058483 | -7.1745028 | 1.30912873 | -5.4803646 |

| | | | | | |
|---|---|---|---|---|---|
| 7654 | NM_138615.2 | 327.986922 | -6.0498679 | 1.10394921 | -5.480205 |
| 7655 | NM_021255.2 | 70.5628748 | -7.7608363 | 1.41634482 | -5.4794822 |
| 7656 | NM_001282736.1 | 19.9457761 | 9.24635042 | 1.68746544 | 5.47943099 |
| 7657 | NM_022804.2 | 55.6553251 | -7.4198852 | 1.35430124 | -5.4787554 |
| 7658 | NM_006594.4 | 56.075681 | -7.4279696 | 1.3557937 | -5.4786872 |
| 7659 | NM_001287387.1 | 71.8093902 | -7.7887263 | 1.42180813 | -5.4780431 |
| 7660 | NM_203394.2 | 37.0977777 | -6.836878 | 1.24810289 | -5.477816 |
| 7661 | NM_001040457.2 | 38.3851043 | -6.8813762 | 1.25627542 | -5.4776016 |
| 7662 | NM_145918.2 | 55.4659168 | -7.4194554 | 1.3545027 | -5.4776232 |
| 7663 | NM_006815.3 | 466.65027 | -6.6413322 | 1.21266291 | -5.4766515 |
| 7664 | NM_016546.3 | 42.0261875 | -7.0195533 | 1.28179847 | -5.4763314 |
| 7665 | NM_001278433.1 | 59.8223174 | -5.1301978 | 0.93685776 | -5.4759623 |
| 7666 | NM_198446.2 | 67.2097689 | -7.69148 | 1.40464616 | -5.475742 |
| 7667 | NM_139076.2 | 38.2869527 | -6.8809374 | 1.25664619 | -5.4756363 |
| 7668 | NM_001199803.2 | 39.0488893 | -6.9084201 | 1.26173261 | -5.4753441 |
| 7669 | NM_138609.2 | 156.730543 | -5.4351384 | 0.99266191 | -5.4753167 |
| 7670 | NM_018087.4 | 53.1110988 | -7.3526654 | 1.3430108 | -5.4747627 |
| 7671 | NM_000262.2 | 79.2904545 | -6.1148404 | 1.11699085 | -5.4743872 |
| 7672 | NM_017657.4 | 56.2712717 | -7.434682 | 1.35813999 | -5.4741647 |
| 7673 | NM_001229.4 | 38.9911699 | -6.9025903 | 1.26096642 | -5.4740477 |
| 7674 | NM_001145408.1 | 49.6581214 | -7.2536685 | 1.32514422 | -5.473871 |
| 7675 | NM_003488.3 | 129.116739 | -7.6654338 | 1.40037707 | -5.4738356 |
| 7676 | NM_006270.4 | 42.8044651 | -7.0442923 | 1.2869876 | -5.4734734 |
| 7677 | NM_197964.4 | 39.3796515 | -6.9209537 | 1.26448126 | -5.4733541 |
| 7678 | NM_005484.3 | 43.9664254 | -7.075869 | 1.29284568 | -5.4730963 |
| 7679 | NM_053279.2 | 39.2603315 | -6.9226019 | 1.26486298 | -5.4730054 |
| 7680 | NM_001282684.1 | 51.3748113 | -7.3015184 | 1.33416117 | -5.4727409 |
| 7681 | NM_001042497.1 | 53.7722826 | -7.3666176 | 1.34607923 | -5.4726478 |
| 7682 | NM_133631.3 | 73.0732439 | -7.8138221 | 1.42781019 | -5.4725916 |
| 7683 | NM_024104.3 | 97.4135066 | -7.2604775 | 1.32679787 | -5.4721806 |
| 7684 | NM_001345882.1 | 46.299885 | -7.1508883 | 1.30678285 | -5.472132 |
| 7685 | NM_003085.4 | 105.214548 | -8.3386949 | 1.52384179 | -5.4721526 |
| 7686 | NR_036442.1 | 52.5298664 | -7.3367384 | 1.34078811 | -5.4719596 |
| 7687 | NM_015478.6 | 34.8832964 | -6.7485818 | 1.23331734 | -5.471894 |
| 7688 | NM_003135.2 | 58.7657092 | -7.4988498 | 1.37048657 | -5.4716697 |
| 7689 | NM_001330272.1 | 41.5425225 | -7.0011908 | 1.27954213 | -5.4716376 |
| 7690 | NR_135601.1 | 21.3871763 | 9.32545387 | 1.70441555 | 5.47134991 |
| 7691 | NM_032828.3 | 98.5162507 | -8.2433381 | 1.50674305 | -5.4709647 |
| 7692 | NM_002863.4 | 43.9227573 | -7.084163 | 1.29488872 | -5.4708663 |
| 7693 | NM_138980.3 | 70.6173223 | -7.7628716 | 1.41908068 | -5.4703526 |
| 7694 | NM_001113547.1 | 62.489018 | -7.5876732 | 1.38712 | -5.4700914 |
| 7695 | NM_001289950.2 | 53.7189676 | -7.3660666 | 1.34661503 | -5.4700612 |
| 7696 | NM_020649.2 | 41.5985082 | -6.9981816 | 1.2795542 | -5.4692342 |

| | | | | | |
|---|---|---|---|---|---|
| 7697 | NM_001293215.1 | 55.8858375 | -7.4271271 | 1.35799824 | -5.4691728 |
| 7698 | NM_001114403.2 | 68.2635819 | -4.942344 | 0.90385729 | -5.4680579 |
| 7699 | NM_001194946.1 | 270.386656 | -9.7009553 | 1.77429365 | -5.4675027 |
| 7700 | NM_001122821.1 | 41.7059961 | -7.0092792 | 1.28199579 | -5.4674744 |
| 7701 | NM_002996.4 | 52.7654802 | -7.3474511 | 1.34386662 | -5.4673961 |
| 7702 | NM_001258023.1 | 117.434601 | -8.4972088 | 1.55417065 | -5.4673589 |
| 7703 | NM_003921.4 | 44.2703498 | -7.0876463 | 1.29644406 | -5.4669897 |
| 7704 | NM_001164326.1 | 41.5069312 | -6.9999382 | 1.28046148 | -5.4667308 |
| 7705 | NM_014573.2 | 211.427125 | -4.9801824 | 0.91099465 | -5.4667526 |
| 7706 | NM_178587.2 | 49.5040018 | -7.2525625 | 1.32683366 | -5.4660676 |
| 7707 | NM_001297429.1 | 28.9469054 | 9.76844956 | 1.78716954 | 5.46587738 |
| 7708 | NM_001291931.1 | 43.5115319 | -7.062262 | 1.29209988 | -5.4657245 |
| 7709 | NM_001079855.1 | 80.0325648 | -7.9442804 | 1.45349825 | -5.4656278 |
| 7710 | NM_016108.3 | 52.171014 | -7.3282916 | 1.34089733 | -5.4652146 |
| 7711 | NR_003697.1 | 55.5927929 | -7.4183986 | 1.35754557 | -5.4645669 |
| 7712 | NM_000317.2 | 58.7699171 | -7.4998244 | 1.37261755 | -5.463885 |
| 7713 | NM_001017919.1 | 44.6890492 | -7.1015492 | 1.29986341 | -5.4633042 |
| 7714 | NM_018622.6 | 51.3176989 | -7.3027334 | 1.33670245 | -5.4632453 |
| 7715 | NM_001258329.1 | 76.633285 | -7.8830177 | 1.44308494 | -5.4626152 |
| 7716 | NM_006383.3 | 63.3857802 | -7.6071453 | 1.39257836 | -5.4626336 |
| 7717 | NM_017594.4 | 125.822889 | -6.8310786 | 1.25050794 | -5.4626431 |
| 7718 | NM_014409.3 | 58.0534411 | -7.4804575 | 1.36940977 | -5.4625413 |
| 7719 | NM_001199155.1 | 48.6771962 | -7.2225421 | 1.32223277 | -5.4623832 |
| 7720 | NM_001297551.1 | 45.9205984 | -7.1423976 | 1.30760198 | -5.4622107 |
| 7721 | NM_001305077.1 | 56.0991492 | -7.4340689 | 1.3610191 | -5.4621342 |
| 7722 | NM_004058.4 | 67.0854069 | -7.6917328 | 1.40820622 | -5.4620784 |
| 7723 | NM_001135156.2 | 156.632283 | -8.9131219 | 1.63184583 | -5.4619877 |
| 7724 | NM_002491.2 | 69.5450096 | -7.7419377 | 1.4174155 | -5.4620101 |
| 7725 | NM_032765.3 | 42.1334276 | -7.0173089 | 1.28474715 | -5.4620155 |
| 7726 | NM_032631.3 | 73.6265667 | -7.8248103 | 1.43263996 | -5.4618121 |
| 7727 | NM_017668.2 | 36.8846151 | -6.8263924 | 1.24992266 | -5.4614518 |
| 7728 | NM_001130137.1 | 34.6671487 | -6.7378725 | 1.23375418 | -5.4612763 |
| 7729 | NM_017661.3 | 37.9079933 | -6.8705894 | 1.2581058 | -5.4610585 |
| 7730 | NM_006627.2 | 51.3394701 | -7.3040154 | 1.33756227 | -5.4606919 |
| 7731 | NM_018269.3 | 72.1857777 | -7.7952287 | 1.42762133 | -5.4602915 |
| 7732 | NM_001198633.1 | 74.9882205 | -7.8493319 | 1.43759428 | -5.4600467 |
| 7733 | NM_021644.3 | 57.8052076 | -7.4724329 | 1.36878886 | -5.4591568 |
| 7734 | NM_000152.4 | 41.6031717 | -6.9984541 | 1.28208889 | -5.4586341 |
| 7735 | NM_014057.4 | 52.0617302 | -7.3275761 | 1.34245158 | -5.4583541 |
| 7736 | NM_147184.3 | 38.5844947 | -6.8950186 | 1.26326187 | -5.4581071 |
| 7737 | NM_006463.4 | 48.9110489 | -7.2347187 | 1.32553541 | -5.4579596 |
| 7738 | NM_003584.2 | 37.6642961 | -6.8575605 | 1.25652301 | -5.4575685 |
| 7739 | NM_145914.2 | 40.9120174 | -6.9747967 | 1.27800203 | -5.4575787 |

| | | | | | |
|---|---|---|---|---|---|
| 7740 | NM_001017981.1 | 85.1754175 | -5.5832567 | 1.02306672 | -5.457373 |
| 7741 | NM_032459.2 | 49.813409 | -7.259052 | 1.33027672 | -5.4567985 |
| 7742 | NM_000932.2 | 58.686252 | -7.4963379 | 1.37381717 | -5.4565761 |
| 7743 | NM_001291867.1 | 50.6501409 | -7.2870536 | 1.33549133 | -5.4564589 |
| 7744 | NM_001080396.2 | 71.7921242 | -7.7876036 | 1.42752079 | -5.4553346 |
| 7745 | NM_032509.3 | 45.8558509 | -7.1401495 | 1.30897514 | -5.4547633 |
| 7746 | NM_032634.3 | 39.2778844 | -6.9185107 | 1.268339 | -5.4547804 |
| 7747 | NM_006191.2 | 384.371259 | -6.2960924 | 1.15425032 | -5.4547028 |
| 7748 | NM_145261.3 | 39.8641092 | -6.9386214 | 1.2720929 | -5.4544926 |
| 7749 | NM_001300989.1 | 87.4927136 | -8.0725329 | 1.48001472 | -5.4543599 |
| 7750 | NM_017585.3 | 65.8122662 | -7.6634145 | 1.40508044 | -5.4540753 |
| 7751 | NM_001080430.3 | 42.398307 | -7.0235125 | 1.28789143 | -5.4534974 |
| 7752 | NM_002467.4 | 113.061643 | -7.4735037 | 1.37042002 | -5.4534403 |
| 7753 | NM_001135005.2 | 72.8500832 | -7.8091858 | 1.43203264 | -5.4532178 |
| 7754 | NR_110257.1 | 379.957392 | 3.4386556 | 0.63058699 | 5.45310266 |
| 7755 | NM_012388.3 | 64.4854914 | -7.6300725 | 1.39924959 | -5.4529746 |
| 7756 | NR_121647.1 | 47.3432639 | 4.3394936 | 0.79584252 | 5.4527039 |
| 7757 | NM_015987.4 | 43.1304948 | -7.0527158 | 1.29358693 | -5.4520617 |
| 7758 | NM_003437.4 | 39.499898 | -6.9261191 | 1.2703932 | -5.4519491 |
| 7759 | NM_174891.3 | 35.3194477 | -6.7687974 | 1.24154221 | -5.4519269 |
| 7760 | NM_001278074.1 | 59.0870346 | -7.5090483 | 1.37735915 | -5.4517722 |
| 7761 | NM_001321155.1 | 45.9590309 | -7.1445204 | 1.31078829 | -5.4505526 |
| 7762 | NM_203464.2 | 50.3729732 | -7.2782696 | 1.33534718 | -5.4504699 |
| 7763 | NM_001025077.2 | 186.537741 | -9.1650347 | 1.68157608 | -5.4502647 |
| 7764 | NM_003851.2 | 88.7608346 | -7.1253325 | 1.3073348 | -5.4502737 |
| 7765 | NM_001031740.2 | 52.2431105 | -7.331078 | 1.34509446 | -5.4502328 |
| 7766 | NM_033641.3 | 150.0088 | -8.8515313 | 1.62426145 | -5.449573 |
| 7767 | NM_020774.3 | 379.465489 | -6.0113922 | 1.10314916 | -5.4493014 |
| 7768 | NM_001932.4 | 74.0925584 | -7.8337474 | 1.43773482 | -5.4486733 |
| 7769 | NM_001137550.1 | 55.6163174 | -7.4204928 | 1.36192059 | -5.4485503 |
| 7770 | NM_014478.4 | 88.9699096 | -8.0963179 | 1.48597912 | -5.4484736 |
| 7771 | NR_134562.1 | 37.166075 | -6.8363943 | 1.25476264 | -5.4483566 |
| 7772 | NM_001317920.1 | 77.0538346 | -7.8885854 | 1.44792377 | -5.4482049 |
| 7773 | NM_021927.2 | 68.8294205 | -7.7274663 | 1.4184216 | -5.4479333 |
| 7774 | NR_003604.2 | 62.6707636 | -7.5912971 | 1.39345451 | -5.4478255 |
| 7775 | NR_003272.2 | 54.0591678 | -7.3793274 | 1.35455927 | -5.44777 |
| 7776 | NM_002436.3 | 43.3697269 | -7.059464 | 1.2959012 | -5.4475326 |
| 7777 | NM_018290.3 | 36.8951876 | -6.8258516 | 1.25302711 | -5.4474892 |
| 7778 | NM_018474.4 | 34.2865852 | -6.7193829 | 1.23358465 | -5.4470383 |
| 7779 | NM_001282313.1 | 100.374698 | -8.271212 | 1.51851054 | -5.4469243 |
| 7780 | NM_020190.3 | 54.2878995 | -7.3877847 | 1.3563724 | -5.446723 |
| 7781 | NM_001330091.1 | 75.0855958 | -7.8518015 | 1.44161574 | -5.4465287 |
| 7782 | NM_015351.1 | 114.848617 | -5.4749796 | 1.00523146 | -5.4464865 |

| | | | | | |
|---|---|---|---|---|---|
| 7783 | NM_000203.4 | 46.8313074 | -7.1761991 | 1.31760218 | -5.446408 |
| 7784 | NM_198573.2 | 108.952356 | -8.3892203 | 1.54038931 | -5.4461688 |
| 7785 | NM_005811.4 | 141.669821 | -5.6605644 | 1.03938924 | -5.4460487 |
| 7786 | NM_005679.3 | 77.5885613 | -7.9019764 | 1.45105019 | -5.4456948 |
| 7787 | NM_033267.4 | 43.4990308 | -7.0668277 | 1.29770957 | -5.4456157 |
| 7788 | NM_003580.3 | 40.3221906 | -6.9576014 | 1.27765758 | -5.4455916 |
| 7789 | NM_207336.2 | 58.1662431 | -7.4816851 | 1.37392027 | -5.4455016 |
| 7790 | NR_003584.3 | 58.7275145 | -7.4967612 | 1.37668862 | -5.4455024 |
| 7791 | NM_001127190.1 | 50.6976982 | -7.2878789 | 1.33842158 | -5.4451295 |
| 7792 | NM_017759.4 | 65.4164715 | -5.8075849 | 1.06660105 | -5.4449459 |
| 7793 | NM_178570.2 | 69.7273425 | -7.744719 | 1.42239407 | -5.4448476 |
| 7794 | NM_013326.4 | 50.5210124 | -7.2822752 | 1.33754583 | -5.4445052 |
| 7795 | NM_001203247.1 | 55.0688355 | -7.4016403 | 1.3595487 | -5.4441891 |
| 7796 | NM_057169.4 | 39.868195 | -6.9396563 | 1.27468688 | -5.4442047 |
| 7797 | NM_018263.4 | 37.4725346 | -6.8549984 | 1.25918932 | -5.4439776 |
| 7798 | NM_001145077.1 | 55.542074 | -7.4171459 | 1.36252837 | -5.4436635 |
| 7799 | NM_018203.2 | 44.7936404 | -7.1059471 | 1.30540754 | -5.4434703 |
| 7800 | NM_152434.2 | 48.4780309 | -7.2191161 | 1.3262206 | -5.443375 |
| 7801 | NR_027788.1 | 42.2769486 | -7.0252895 | 1.29066532 | -5.4431536 |
| 7802 | NM_001128150.1 | 52.1354243 | -7.3256831 | 1.34586719 | -5.4430951 |
| 7803 | NM_177556.2 | 63.674431 | -7.6150991 | 1.39903805 | -5.4430965 |
| 7804 | NM_001184971.1 | 50.7794644 | -7.2890014 | 1.3391918 | -5.442836 |
| 7805 | NM_001086521.1 | 66.1284736 | -7.6689212 | 1.40900237 | -5.4428022 |
| 7806 | NM_018130.2 | 35.5548667 | -6.7762587 | 1.24512747 | -5.4422208 |
| 7807 | NM_001261843.1 | 50.5211263 | -7.2831019 | 1.33828609 | -5.4421114 |
| 7808 | NM_152288.2 | 38.2297642 | -6.8835757 | 1.26491216 | -5.4419397 |
| 7809 | NM_001317898.1 | 71.5030744 | -7.781351 | 1.429925 | -5.4417896 |
| 7810 | NM_023947.3 | 75.6774564 | -7.8640483 | 1.44530845 | -5.4410865 |
| 7811 | NM_001290118.1 | 49.63543 | 10.5564051 | 1.94018007 | 5.44094088 |
| 7812 | NM_003216.3 | 41.126867 | -6.9808688 | 1.28302891 | -5.4409287 |
| 7813 | NM_025182.3 | 49.4439811 | -7.2500832 | 1.33254599 | -5.4407752 |
| 7814 | NM_053002.5 | 44.9581181 | -7.1134116 | 1.30742847 | -5.4407654 |
| 7815 | NM_001040437.2 | 102.104584 | -6.5172589 | 1.19793865 | -5.4403945 |
| 7816 | NM_022773.2 | 42.5109469 | -7.034723 | 1.2930504 | -5.440409 |
| 7817 | NM_001135110.2 | 87.4426265 | -7.1032222 | 1.3057338 | -5.4400232 |
| 7818 | NM_015669.4 | 76.7702905 | -7.8835282 | 1.44925013 | -5.4397292 |
| 7819 | NM_015158.3 | 66.9741826 | -7.689798 | 1.41367364 | -5.439585 |
| 7820 | NM_021121.3 | 58.9121996 | -7.5046756 | 1.37971635 | -5.4392887 |
| 7821 | NM_016003.3 | 80.2953072 | -7.9488809 | 1.46139197 | -5.4392532 |
| 7822 | NM_201398.2 | 61.6933178 | -7.56838 | 1.39146589 | -5.4391416 |
| 7823 | NR_003255.2 | 331.564204 | 1.32635897 | 0.24385391 | 5.43915409 |
| 7824 | NR_037601.1 | 53.4227221 | -7.3593052 | 1.35310105 | -5.4388438 |
| 7825 | NM_006186.3 | 74.032257 | -7.8315179 | 1.43996265 | -5.4386951 |

| | | | | | |
|---|---|---|---|---|---|
| 7826 | NM_001306177.1 | 46.3182538 | -7.1591242 | 1.31638821 | -5.4384597 |
| 7827 | NM_001042413.1 | 36.0235649 | -6.7922432 | 1.24894483 | -5.4383853 |
| 7828 | NM_032711.3 | 42.9583997 | -7.0459106 | 1.29564253 | -5.4381594 |
| 7829 | NM_175918.3 | 71.0577281 | -3.4952291 | 0.64275168 | -5.4379151 |
| 7830 | NM_017647.3 | 144.364382 | -5.6849069 | 1.04562247 | -5.4368638 |
| 7831 | NM_013349.4 | 46.4897382 | -7.1600477 | 1.31696363 | -5.4367847 |
| 7832 | NM_021100.4 | 43.7339717 | -7.0726235 | 1.30096766 | -5.436433 |
| 7833 | NM_001321018.1 | 55.4920912 | -7.4149437 | 1.36394217 | -5.4364062 |
| 7834 | NM_014417.4 | 96.1136984 | -7.240599 | 1.33190475 | -5.4362739 |
| 7835 | NM_033286.3 | 53.579315 | -7.3651763 | 1.35495284 | -5.4357437 |
| 7836 | NM_001319674.1 | 37.9757454 | -6.8632686 | 1.26263072 | -5.4356895 |
| 7837 | NM_000996.2 | 758.872582 | -7.0797605 | 1.30262189 | -5.4350081 |
| 7838 | NM_024093.2 | 53.3281568 | -7.3561261 | 1.35350099 | -5.4348879 |
| 7839 | NM_001320815.1 | 69.3666266 | -7.7370491 | 1.42360789 | -5.4348175 |
| 7840 | NM_016302.3 | 91.2978825 | -7.1634428 | 1.31821034 | -5.4342183 |
| 7841 | NM_207363.2 | 36.3722271 | -6.8060166 | 1.25250237 | -5.4339351 |
| 7842 | NM_138570.3 | 40.1686864 | -6.9509658 | 1.27934625 | -5.433217 |
| 7843 | NM_001017963.2 | 60.7816757 | -7.5469305 | 1.3890967 | -5.4329771 |
| 7844 | NM_022903.3 | 43.7963843 | -7.0753232 | 1.30234318 | -5.432764 |
| 7845 | NM_001329984.1 | 168.503965 | -9.0179655 | 1.65995204 | -5.4326663 |
| 7846 | NM_001306130.1 | 73.6334587 | -7.8265497 | 1.44077101 | -5.4321954 |
| 7847 | NM_000745.3 | 50.8375638 | -7.2896207 | 1.34202571 | -5.4318041 |
| 7848 | NM_001173476.1 | 70.126505 | -7.7534393 | 1.4274193 | -5.4317882 |
| 7849 | NM_002380.4 | 61.0697083 | -7.5563581 | 1.39112734 | -5.4318234 |
| 7850 | NM_172080.2 | 133.827901 | -8.6862246 | 1.59932298 | -5.4311885 |
| 7851 | NM_012291.4 | 42.4605732 | -7.0308278 | 1.29466384 | -5.4306204 |
| 7852 | NR_037713.1 | 64.5958832 | -7.6345598 | 1.40605389 | -5.4297775 |
| 7853 | NM_022727.5 | 51.2114409 | -7.301928 | 1.34503218 | -5.4288129 |
| 7854 | NM_015204.2 | 126.203088 | -7.6350621 | 1.40652084 | -5.428332 |
| 7855 | NM_001099270.1 | 74.9392933 | -6.8742896 | 1.26638096 | -5.4282951 |
| 7856 | NM_001300981.1 | 38.8350419 | -6.904429 | 1.27197704 | -5.4281082 |
| 7857 | NM_005528.3 | 41.5976107 | -7.0013822 | 1.28993877 | -5.4276857 |
| 7858 | NM_015242.4 | 101.596896 | -8.2894512 | 1.52725708 | -5.4276725 |
| 7859 | NR_026052.1 | 195.721851 | -1.9655274 | 0.36213676 | -5.427583 |
| 7860 | NM_004763.4 | 50.9670077 | -7.292546 | 1.34365331 | -5.4274015 |
| 7861 | NM_182913.2 | 65.7342167 | -7.6580812 | 1.41102879 | -5.4273033 |
| 7862 | NM_001029896.1 | 56.4453759 | -7.4413134 | 1.37114978 | -5.4270609 |
| 7863 | NM_014746.4 | 178.775064 | -5.0066831 | 0.92253708 | -5.4270806 |
| 7864 | NM_024339.4 | 61.7120465 | -7.5677125 | 1.39452924 | -5.4267148 |
| 7865 | NM_001303451.1 | 18.6641374 | 9.15403422 | 1.68692921 | 5.42644833 |
| 7866 | NM_001243965.1 | 85.0154769 | -8.0322933 | 1.48026864 | -5.4262403 |
| 7867 | NM_017827.3 | 41.6684776 | -7.0015348 | 1.29038433 | -5.4259298 |
| 7868 | NM_052860.2 | 89.0423875 | -8.0974357 | 1.49239519 | -5.4257986 |

| | | | | | |
|---|---|---|---|---|---|
| 7869 | NM_013401.3 | 38.7673999 | -6.9044514 | 1.27260097 | -5.4254645 |
| 7870 | NM_138704.3 | 83.230271 | -7.0278242 | 1.29544467 | -5.4250285 |
| 7871 | NM_172315.2 | 146.232465 | -6.832506 | 1.25946912 | -5.4249095 |
| 7872 | NM_006807.4 | 49.5928558 | -7.2538409 | 1.33714656 | -5.424866 |
| 7873 | NM_021647.7 | 44.3188215 | -7.089771 | 1.30690587 | -5.424852 |
| 7874 | NM_001013642.2 | 77.6368482 | -7.9003955 | 1.45642095 | -5.4245275 |
| 7875 | NM_002417.4 | 56.0320281 | -7.4338511 | 1.37042978 | -5.424467 |
| 7876 | NM_182527.2 | 67.4562786 | -7.6952858 | 1.4186298 | -5.4244496 |
| 7877 | NM_001290187.1 | 44.3118904 | -7.0906512 | 1.30717786 | -5.4243967 |
| 7878 | NM_001168368.1 | 47.982337 | -7.209612 | 1.32941791 | -5.4231344 |
| 7879 | NM_001003787.2 | 44.8763203 | -7.1099642 | 1.3112411 | -5.4223165 |
| 7880 | NM_025232.3 | 38.6611868 | -6.8973822 | 1.27204159 | -5.422293 |
| 7881 | NM_001040167.1 | 48.9358017 | -7.2344394 | 1.33428577 | -5.4219565 |
| 7882 | NM_203374.1 | 36.3255809 | -6.8029609 | 1.25476264 | -5.4217114 |
| 7883 | NM_015693.3 | 36.1171626 | -6.7985346 | 1.25414805 | -5.4208389 |
| 7884 | NM_001242488.1 | 140.185734 | -4.1777734 | 0.77073857 | -5.4204805 |
| 7885 | NM_014585.5 | 49.5441558 | -7.2544917 | 1.3384397 | -5.420111 |
| 7886 | NM_001318484.1 | 30.5818631 | 4.79762168 | 0.88516636 | 5.42002258 |
| 7887 | NM_003619.3 | 63.4818967 | -6.636341 | 1.22454073 | -5.419453 |
| 7888 | NM_002716.4 | 47.8924013 | -7.2020533 | 1.32898171 | -5.4192268 |
| 7889 | NM_031220.3 | 43.1475237 | -7.0553563 | 1.30194123 | -5.4191051 |
| 7890 | NM_152269.4 | 56.3143571 | -7.4393133 | 1.37281342 | -5.4190272 |
| 7891 | NM_020832.2 | 45.8963585 | -7.1465915 | 1.31881646 | -5.4189432 |
| 7892 | NM_001080398.1 | 320.089413 | -4.8907688 | 0.90254742 | -5.4188497 |
| 7893 | NM_001322402.1 | 63.604125 | -7.6132554 | 1.40525264 | -5.4177129 |
| 7894 | NM_001184720.1 | 52.1633769 | -7.3280814 | 1.35275083 | -5.4171701 |
| 7895 | NM_002610.4 | 91.3643528 | -7.1661671 | 1.32288672 | -5.4170678 |
| 7896 | NM_001166242.1 | 49.3678904 | -7.2449023 | 1.33746811 | -5.4168785 |
| 7897 | NM_001330089.1 | 25.7320061 | 9.60654022 | 1.77377554 | 5.41587139 |
| 7898 | NM_152499.2 | 33.1907091 | -6.6751815 | 1.23269836 | -5.4150973 |
| 7899 | NM_001190438.1 | 55.8281899 | -7.4230618 | 1.37098762 | -5.4143901 |
| 7900 | NM_024854.3 | 48.5907883 | -7.2234729 | 1.33414461 | -5.4143103 |
| 7901 | NM_004269.3 | 58.1196409 | -7.4819241 | 1.38190354 | -5.4142159 |
| 7902 | NM_025189.3 | 41.1276161 | -6.9879167 | 1.29067619 | -5.4141517 |
| 7903 | NM_004849.3 | 53.1991707 | -7.3533683 | 1.35818162 | -5.4141274 |
| 7904 | NM_013387.3 | 126.548403 | -5.5840687 | 1.03140178 | -5.4140576 |
| 7905 | NM_003098.2 | 65.9596305 | -5.8642144 | 1.0833785 | -5.4128953 |
| 7906 | NM_001134389.1 | 42.97463 | -7.0456929 | 1.30173209 | -5.4125522 |
| 7907 | NM_001085375.1 | 36.1960683 | -6.8028141 | 1.2570666 | -5.4116576 |
| 7908 | NM_007019.3 | 47.9177291 | -7.2032881 | 1.33117563 | -5.411223 |
| 7909 | NM_032359.3 | 69.618852 | -7.7430699 | 1.4310906 | -5.4106078 |
| 7910 | NM_013231.5 | 48.7132517 | -7.2244483 | 1.33527241 | -5.4104677 |
| 7911 | NM_000190.3 | 50.1764034 | -7.2710258 | 1.34393861 | -5.4102366 |

| | | | | | |
|---|---|---|---|---|---|
| 7912 | NM_017847.5 | 46.930838 | -7.1768586 | 1.32661302 | -5.4099112 |
| 7913 | NM_001033505.1 | 39.1069195 | -6.9108282 | 1.27747019 | -5.4097765 |
| 7914 | NM_014180.3 | 41.0686906 | -6.9822934 | 1.29070453 | -5.4096761 |
| 7915 | NM_001166426.2 | 34.5056348 | -6.7343111 | 1.2448824 | -5.4095962 |
| 7916 | NM_016175.3 | 34.7630978 | -6.7406322 | 1.24608815 | -5.4094345 |
| 7917 | NM_015261.2 | 80.4504083 | -4.8848919 | 0.90305444 | -5.4092995 |
| 7918 | NM_001142784.2 | 45.6249335 | -7.134392 | 1.31905778 | -5.4087032 |
| 7919 | NM_006978.2 | 47.392041 | -7.1867644 | 1.32885076 | -5.4082555 |
| 7920 | NR_024448.2 | 67.9669613 | -5.3434384 | 0.98811525 | -5.4077077 |
| 7921 | NM_001190452.1 | 3583.28735 | -7.5706234 | 1.40000881 | -5.4075541 |
| 7922 | NM_001321146.1 | 37.8711957 | -6.8616086 | 1.26896048 | -5.4072674 |
| 7923 | NM_001014432.1 | 59.1562006 | -7.5045106 | 1.38795313 | -5.4068905 |
| 7924 | NR_033327.1 | 96.0434059 | -8.2089283 | 1.51851491 | -5.4058925 |
| 7925 | NM_025152.2 | 36.6358273 | -6.8225036 | 1.26205624 | -5.4058633 |
| 7926 | NM_173555.3 | 35.349462 | -6.7610914 | 1.25080481 | -5.4053928 |
| 7927 | NM_004563.3 | 56.1778806 | -7.4339934 | 1.37535641 | -5.4051396 |
| 7928 | NM_001289175.1 | 49.6197692 | -7.2551972 | 1.34228593 | -5.4051056 |
| 7929 | NM_005270.4 | 69.2902005 | -4.5941879 | 0.8500008 | -5.4049218 |
| 7930 | NM_025103.2 | 38.7369313 | -6.8990202 | 1.27644391 | -5.4048754 |
| 7931 | NM_033419.4 | 44.1359495 | -7.0877213 | 1.31141676 | -5.4046292 |
| 7932 | NR_132749.1 | 59.5862662 | -7.5172477 | 1.39108998 | -5.4038544 |
| 7933 | NM_003876.2 | 45.7677994 | -7.1399311 | 1.32145001 | -5.4031034 |
| 7934 | NM_013323.2 | 49.2086327 | -7.2442728 | 1.34076572 | -5.4030862 |
| 7935 | NM_004845.4 | 75.3364422 | -7.8564631 | 1.45415154 | -5.4027816 |
| 7936 | NM_001029885.1 | 88.2934814 | -6.302098 | 1.16653019 | -5.4024302 |
| 7937 | NM_001330665.1 | 45.5587389 | -7.1338449 | 1.32065044 | -5.4017662 |
| 7938 | NM_014765.2 | 688.794244 | -6.6813585 | 1.23696354 | -5.4014191 |
| 7939 | NM_001195104.1 | 18.1551445 | 9.11167445 | 1.68698977 | 5.40114385 |
| 7940 | NM_024336.2 | 61.6054198 | -7.5677426 | 1.40118622 | -5.4009542 |
| 7941 | NM_017556.2 | 86.0941493 | -8.0500354 | 1.49051292 | -5.4008491 |
| 7942 | NR_033677.1 | 55.0995341 | -7.4046315 | 1.37121104 | -5.4000671 |
| 7943 | NM_001272071.1 | 125.219467 | -7.6208166 | 1.41142482 | -5.3993784 |
| 7944 | NM_181507.1 | 50.1689408 | -4.4639145 | 0.82676539 | -5.3992518 |
| 7945 | NM_199337.2 | 42.385102 | -7.0294446 | 1.30198135 | -5.3990363 |
| 7946 | NR_110834.1 | 59.7716039 | -7.525645 | 1.39394024 | -5.398829 |
| 7947 | NM_015102.4 | 42.3775126 | -7.0287544 | 1.30197799 | -5.3985202 |
| 7948 | NM_001171887.1 | 53.939788 | -7.3737827 | 1.36602173 | -5.3979981 |
| 7949 | NM_054013.3 | 37.2162485 | -6.8383165 | 1.26684341 | -5.3979177 |
| 7950 | NM_001328675.1 | 78.0047558 | -7.9068215 | 1.46481438 | -5.3978317 |
| 7951 | NM_145112.2 | 50.8215582 | -7.2893415 | 1.35042253 | -5.3978228 |
| 7952 | NM_001304482.1 | 60.3648208 | -7.5385137 | 1.39660468 | -5.3977434 |
| 7953 | NM_012244.3 | 165.065686 | -8.9896337 | 1.66554246 | -5.3974209 |
| 7954 | NM_014957.2 | 47.8997843 | -7.2034776 | 1.33470959 | -5.3970374 |

| | | | | | |
|---|---|---|---|---|---|
| 7955 | NM_003566.3 | 100.829109 | -6.5017111 | 1.20476193 | -5.3966771 |
| 7956 | NM_177537.2 | 77.1307062 | -6.9221822 | 1.28275668 | -5.396333 |
| 7957 | NM_003507.1 | 157.626209 | -5.2582745 | 0.97464133 | -5.3950868 |
| 7958 | NM_024028.3 | 76.8851507 | -7.8868885 | 1.4619368 | -5.3948218 |
| 7959 | NM_017634.3 | 39.3324208 | -6.9206415 | 1.2828359 | -5.3947987 |
| 7960 | NM_018066.3 | 42.6314724 | -7.0365134 | 1.30437336 | -5.3945547 |
| 7961 | NM_020676.6 | 65.9221421 | -7.6631566 | 1.42069855 | -5.3939357 |
| 7962 | NM_006080.2 | 129.142824 | -7.6692652 | 1.42185818 | -5.3938328 |
| 7963 | NM_001968.4 | 79.3099769 | -7.9325527 | 1.47071363 | -5.3936759 |
| 7964 | NM_006917.4 | 61.0144371 | -7.5540152 | 1.4005569 | -5.3935797 |
| 7965 | NM_199478.2 | 52.3496473 | -7.3313473 | 1.35935635 | -5.393249 |
| 7966 | NM_001145436.1 | 87.3639566 | -8.0726596 | 1.49693679 | -5.3927859 |
| 7967 | NM_001034173.3 | 86.1001554 | -8.0497425 | 1.49275434 | -5.3925434 |
| 7968 | NM_145691.3 | 40.3219659 | -6.9540463 | 1.28961001 | -5.3923638 |
| 7969 | NM_031298.3 | 53.6728153 | -7.3684347 | 1.36648353 | -5.3922602 |
| 7970 | NM_194314.2 | 89.9883395 | -7.1411272 | 1.32436732 | -5.3921047 |
| 7971 | NM_152932.2 | 44.3929136 | -7.0983426 | 1.31650989 | -5.3917883 |
| 7972 | NM_001330170.1 | 47.1221987 | -7.180599 | 1.33180424 | -5.3916325 |
| 7973 | NM_001039590.2 | 41.4516217 | -6.9947376 | 1.29754778 | -5.3907361 |
| 7974 | NM_203290.3 | 42.1232705 | -7.0172861 | 1.3017844 | -5.3905133 |
| 7975 | NM_001145426.1 | 39.8408933 | -6.9430983 | 1.2880658 | -5.3903288 |
| 7976 | NM_148172.2 | 51.2576687 | -7.3039424 | 1.35504783 | -5.3901731 |
| 7977 | NM_014018.2 | 41.5826361 | -6.9991632 | 1.29858748 | -5.389828 |
| 7978 | NM_181471.2 | 55.122073 | -7.4037443 | 1.37371851 | -5.3895643 |
| 7979 | NM_017734.4 | 54.1746481 | -7.3828569 | 1.36987059 | -5.3894557 |
| 7980 | NM_001301210.1 | 157.871423 | -3.9308833 | 0.72938853 | -5.3892858 |
| 7981 | NM_194279.3 | 41.1436661 | -6.984027 | 1.29591756 | -5.3892525 |
| 7982 | NM_005843.5 | 94.6995054 | -4.5180979 | 0.83847052 | -5.3884995 |
| 7983 | NM_000232.4 | 286.24467 | -3.2573318 | 0.60454728 | -5.3880514 |
| 7984 | NM_012154.3 | 33.1781316 | -6.6720919 | 1.23846236 | -5.3873998 |
| 7985 | NM_001164381.1 | 62.8324485 | -7.5950197 | 1.40978175 | -5.3873727 |
| 7986 | NM_001304562.1 | 70.1209971 | -7.753913 | 1.4392831 | -5.3873439 |
| 7987 | NM_001161009.2 | 54.9774339 | -7.4014749 | 1.37400707 | -5.3867808 |
| 7988 | NM_001320752.1 | 54.0564468 | -7.3772521 | 1.36954826 | -5.3866317 |
| 7989 | NM_001185082.1 | 34.6318346 | -6.7329044 | 1.25000283 | -5.3863113 |
| 7990 | NR_024550.1 | 69.9057435 | -7.7464044 | 1.43816582 | -5.3863082 |
| 7991 | NM_145059.2 | 53.1929966 | -4.5553261 | 0.84578385 | -5.3859223 |
| 7992 | NM_001321312.1 | 46.186017 | -7.1474232 | 1.32717427 | -5.3854444 |
| 7993 | NM_001331078.1 | 33.8967185 | -6.703797 | 1.24488044 | -5.385093 |
| 7994 | NM_012437.5 | 50.123749 | -7.2693132 | 1.34991019 | -5.3850347 |
| 7995 | NM_003440.3 | 42.1761879 | -7.0168835 | 1.30306044 | -5.3849256 |
| 7996 | NM_001286566.1 | 123.562235 | -8.5708428 | 1.59165241 | -5.3848709 |
| 7997 | NM_001271507.1 | 37.7891565 | -6.8562054 | 1.27328142 | -5.384674 |

| | | | | | |
|---|---|---|---|---|---|
| 7998 | NM_001002914.2 | 64.087323 | -5.8224781 | 1.08138671 | -5.3842701 |
| 7999 | NM_001004106.2 | 70.6320656 | -7.7655059 | 1.44236691 | -5.3838631 |
| 8000 | NM_014595.2 | 41.829647 | -7.0087873 | 1.30181432 | -5.3838609 |
| 8001 | NM_020408.5 | 60.8219008 | -7.5483374 | 1.40206602 | -5.3837246 |
| 8002 | NM_015480.2 | 48.1419974 | -7.2112398 | 1.33967382 | -5.3828325 |
| 8003 | NM_001001396.2 | 65.4462881 | -7.6538463 | 1.4219606 | -5.3826008 |
| 8004 | NR_121625.1 | 44.5170436 | 3.77982308 | 0.70224902 | 5.38245405 |
| 8005 | NM_001330556.1 | 85.5375005 | -8.039816 | 1.49389109 | -5.3817953 |
| 8006 | NM_024942.3 | 45.0888379 | -7.1156071 | 1.32216266 | -5.381794 |
| 8007 | NM_001042548.1 | 66.8672236 | -7.684365 | 1.42791125 | -5.3815425 |
| 8008 | NM_024928.4 | 44.6965992 | -7.107092 | 1.3206423 | -5.381542 |
| 8009 | NM_002040.3 | 34.6640713 | -6.7326282 | 1.25112831 | -5.3812452 |
| 8010 | NM_004973.3 | 85.1136865 | -7.0571292 | 1.31145412 | -5.3811484 |
| 8011 | NM_001297659.1 | 50.3068183 | -7.2743253 | 1.3518971 | -5.3808277 |
| 8012 | NM_198207.2 | 44.8235943 | -7.1100224 | 1.32140021 | -5.380673 |
| 8013 | NM_001322250.1 | 98.4019514 | -7.2763588 | 1.35241778 | -5.3802596 |
| 8014 | NM_001100429.1 | 90.8567659 | -8.1271211 | 1.51067521 | -5.3797938 |
| 8015 | NR_027164.1 | 81.7620149 | -7.9758707 | 1.48256537 | -5.3797767 |
| 8016 | NM_031210.5 | 64.2052447 | -7.6264897 | 1.41825765 | -5.3773654 |
| 8017 | NM_001330637.1 | 67.0879102 | -7.6922077 | 1.43051033 | -5.3772472 |
| 8018 | NM_001206941.1 | 126.924119 | -8.6096732 | 1.60114787 | -5.3771881 |
| 8019 | NM_144969.2 | 56.9787096 | -7.4540383 | 1.3862788 | -5.3770124 |
| 8020 | NM_033395.1 | 70.6235975 | -6.7950039 | 1.263873 | -5.3763345 |
| 8021 | NM_001301169.1 | 88.5580911 | -8.089986 | 1.50479144 | -5.376151 |
| 8022 | NM_173540.2 | 37.6998059 | -6.8581238 | 1.27573711 | -5.3758127 |
| 8023 | NM_014339.6 | 44.8787422 | -7.1084815 | 1.3223782 | -5.3755284 |
| 8024 | NM_002412.4 | 37.7075508 | -6.8595807 | 1.27630445 | -5.3745646 |
| 8025 | NM_003277.3 | 79.8201577 | -7.9421961 | 1.47786781 | -5.374091 |
| 8026 | NM_004128.2 | 87.4854526 | -7.1052604 | 1.32215653 | -5.3739933 |
| 8027 | NM_014167.4 | 44.7899559 | -7.1042884 | 1.32200072 | -5.3738915 |
| 8028 | NM_001024678.3 | 52.3258427 | -7.3309375 | 1.36422594 | -5.3736975 |
| 8029 | NM_020640.3 | 83.2156328 | -7.0266661 | 1.3076127 | -5.37366 |
| 8030 | NM_001002857.1 | 56.0289298 | -7.4333641 | 1.38334913 | -5.3734548 |
| 8031 | NM_006341.3 | 74.2907264 | -4.6996863 | 0.87464466 | -5.3732521 |
| 8032 | NM_003848.3 | 37.7190087 | -6.8632579 | 1.27731494 | -5.3731916 |
| 8033 | NM_003457.3 | 83.1629003 | -8.0011912 | 1.48914525 | -5.3730092 |
| 8034 | NM_001145023.1 | 37.0428858 | -6.8371985 | 1.2725565 | -5.3728054 |
| 8035 | NM_018944.2 | 42.9339232 | -7.0440448 | 1.31105685 | -5.3727989 |
| 8036 | NM_003304.4 | 36.8607419 | -6.8280146 | 1.27094044 | -5.3724112 |
| 8037 | NM_001082537.2 | 40.1423082 | -6.9469899 | 1.2930986 | -5.372359 |
| 8038 | NM_014289.3 | 70.2923171 | -7.7585245 | 1.44425074 | -5.3720066 |
| 8039 | NM_001031801.1 | 45.2225013 | -7.1206092 | 1.32553207 | -5.3718875 |
| 8040 | NM_183413.2 | 86.1191004 | -8.0508043 | 1.49870002 | -5.3718584 |

| | | | | | |
|---|---|---|---|---|---|
| 8041 | NM_002623.3 | 72.2479261 | -7.7959796 | 1.45129924 | -5.3717244 |
| 8042 | NM_006642.3 | 43.6877124 | -7.0701147 | 1.3161838 | -5.3716773 |
| 8043 | NM_006388.3 | 40.3993272 | -6.958393 | 1.2954708 | -5.3713237 |
| 8044 | NM_018075.4 | 64.4896245 | -7.6335134 | 1.42116678 | -5.3713002 |
| 8045 | NM_022465.3 | 76.4906934 | -6.9050116 | 1.28557032 | -5.371166 |
| 8046 | NM_003026.3 | 136.43381 | -8.7138249 | 1.62239289 | -5.3709709 |
| 8047 | NM_001166373.1 | 72.9902935 | -7.8114272 | 1.45469183 | -5.3698158 |
| 8048 | NM_021222.2 | 44.7990902 | -7.1108345 | 1.32421827 | -5.3698357 |
| 8049 | NM_001282625.1 | 38.5536285 | -6.8890994 | 1.28294403 | -5.3697584 |
| 8050 | NM_031288.3 | 62.3531723 | -6.6092872 | 1.23083339 | -5.3697659 |
| 8051 | NM_014243.2 | 52.5244054 | -7.3353384 | 1.3660882 | -5.3695936 |
| 8052 | NM_016283.4 | 42.9692478 | -7.0477855 | 1.31260582 | -5.3693084 |
| 8053 | NM_018317.3 | 40.2081061 | -6.9491977 | 1.29426787 | -5.3692113 |
| 8054 | NM_183065.2 | 34.268119 | -6.7178361 | 1.25122408 | -5.3690112 |
| 8055 | NM_014790.4 | 52.8881465 | -7.3501824 | 1.36901019 | -5.3689757 |
| 8056 | NM_001193465.1 | 52.5288433 | -7.339194 | 1.36701974 | -5.3687549 |
| 8057 | NM_018082.5 | 48.4228814 | -7.2189927 | 1.34471 | -5.3684383 |
| 8058 | NM_001098.2 | 423.577813 | -6.0647443 | 1.1297292 | -5.3683168 |
| 8059 | NM_145255.3 | 54.6221148 | -7.3929684 | 1.37723702 | -5.367971 |
| 8060 | NM_173467.4 | 39.3327399 | -6.918685 | 1.28890913 | -5.367861 |
| 8061 | NM_021190.3 | 87.1163041 | -8.0664558 | 1.50281435 | -5.3675664 |
| 8062 | NM_005940.4 | 61.8722116 | -7.5738449 | 1.41108378 | -5.3673956 |
| 8063 | NM_030674.3 | 76.5469058 | -7.8804281 | 1.46830871 | -5.3670104 |
| 8064 | NM_001201483.1 | 41.3091553 | -6.9913865 | 1.3028732 | -5.3661296 |
| 8065 | NM_014914.4 | 349.166861 | -4.5827576 | 0.85410966 | -5.3655377 |
| 8066 | NM_024418.2 | 91.8593403 | -8.1449833 | 1.51807194 | -5.3653474 |
| 8067 | NM_032689.4 | 36.8956656 | -6.8266765 | 1.27236014 | -5.3653649 |
| 8068 | NM_201430.2 | 66.4764949 | -7.6755924 | 1.43074658 | -5.3647463 |
| 8069 | NM_006375.3 | 40.5598397 | -6.9632144 | 1.29805937 | -5.3643266 |
| 8070 | NM_004889.3 | 50.7041874 | -7.2855031 | 1.35816581 | -5.3642221 |
| 8071 | NM_025251.2 | 45.5111379 | -7.1336821 | 1.32987725 | -5.3641658 |
| 8072 | NM_001242795.1 | 47.7453486 | -7.1996479 | 1.34220837 | -5.3640315 |
| 8073 | NM_015219.4 | 243.39221 | -5.5630246 | 1.03712239 | -5.3639037 |
| 8074 | NM_032294.2 | 50.7214243 | -7.2837104 | 1.35791159 | -5.3639062 |
| 8075 | NM_018057.6 | 55.7428213 | -7.4207945 | 1.38365749 | -5.363173 |
| 8076 | NM_001004105.2 | 44.2469404 | -7.0885625 | 1.32172156 | -5.3631285 |
| 8077 | NM_018843.3 | 41.756158 | -7.0048072 | 1.30612952 | -5.3630265 |
| 8078 | NM_017812.3 | 64.7984989 | -6.6614596 | 1.24228854 | -5.3622483 |
| 8079 | NM_014890.3 | 58.805202 | -7.5010877 | 1.39890875 | -5.3620994 |
| 8080 | NM_001283044.2 | 50.1560155 | -7.2662565 | 1.3551359 | -5.3620131 |
| 8081 | NM_172109.2 | 63.0331562 | -7.5983061 | 1.41711361 | -5.3618186 |
| 8082 | NM_020821.2 | 72.4576447 | -7.7999375 | 1.45485917 | -5.3613007 |
| 8083 | NM_001320360.1 | 63.6727762 | -7.6148386 | 1.4203847 | -5.36111 |

| | | | | | |
|---|---|---|---|---|---|
| 8084 | NM_133328.3 | 40.282227 | -6.9531527 | 1.29695722 | -5.3611272 |
| 8085 | NM_145265.2 | 42.3812552 | -7.0272405 | 1.31080124 | -5.3610268 |
| 8086 | NM_016841.4 | 1634.45863 | -6.6510213 | 1.24070588 | -5.3606752 |
| 8087 | NM_057159.2 | 37.7123601 | -6.8597282 | 1.27994626 | -5.3593877 |
| 8088 | NM_145080.3 | 67.9225975 | -7.7070684 | 1.4380718 | -5.3593071 |
| 8089 | NM_001122606.1 | 78.6539546 | -7.9197606 | 1.47805699 | -5.3582241 |
| 8090 | NM_003086.3 | 121.394888 | -4.8578462 | 0.90663151 | -5.3581264 |
| 8091 | NM_017664.3 | 94.0400696 | -7.2053823 | 1.34476989 | -5.3580783 |
| 8092 | NM_005147.5 | 51.8802061 | -7.3204273 | 1.36628462 | -5.3579081 |
| 8093 | NM_001346092.1 | 65.2599741 | -6.6788922 | 1.24667678 | -5.3573567 |
| 8094 | NM_016498.4 | 35.6679997 | -6.7759049 | 1.26479054 | -5.3573336 |
| 8095 | NM_001317078.1 | 57.4324322 | -7.4652075 | 1.39355389 | -5.3569565 |
| 8096 | NM_020236.3 | 63.2298548 | -7.6039733 | 1.41947498 | -5.3568914 |
| 8097 | NM_018463.3 | 43.7199937 | -7.071311 | 1.32007862 | -5.3567347 |
| 8098 | NM_001244262.1 | 55.8271894 | -4.3127131 | 0.80513846 | -5.3564862 |
| 8099 | NM_001128126.2 | 56.4654474 | -7.4411133 | 1.38928787 | -5.356063 |
| 8100 | NM_031464.4 | 148.423763 | -7.0816758 | 1.32218667 | -5.3560332 |
| 8101 | NM_024664.3 | 50.4614857 | -7.2763339 | 1.35855745 | -5.3559266 |
| 8102 | NM_001278442.1 | 39.0837069 | -6.9116139 | 1.29055015 | -5.3555562 |
| 8103 | NM_001297696.1 | 147.597385 | 3.29635279 | 0.61550421 | 5.35553251 |
| 8104 | NM_001168618.1 | 59.6582757 | -6.5537016 | 1.22374206 | -5.35546 |
| 8105 | NM_024562.1 | 44.7834461 | -7.1057041 | 1.32686626 | -5.3552527 |
| 8106 | NM_001207051.1 | 51.221766 | -7.3030515 | 1.36376128 | -5.3550805 |
| 8107 | NM_021198.2 | 37.6450122 | -6.8532761 | 1.27979551 | -5.3549775 |
| 8108 | NM_198183.2 | 59.1564869 | -7.5075775 | 1.4020407 | -5.35475 |
| 8109 | NM_001098844.2 | 75.3960527 | 2.76177739 | 0.51576574 | 5.35471279 |
| 8110 | NM_203371.1 | 76.5942711 | -7.8821593 | 1.4721374 | -5.354228 |
| 8111 | NM_139169.4 | 81.992341 | -5.1588876 | 0.96355377 | -5.3540215 |
| 8112 | NM_001206559.1 | 44.8558939 | -7.105792 | 1.32726934 | -5.3536926 |
| 8113 | NM_024065.4 | 45.2308441 | -7.1208481 | 1.33020593 | -5.3531923 |
| 8114 | NM_031890.3 | 72.3662235 | -7.7992943 | 1.45712305 | -5.3525297 |
| 8115 | NM_022478.3 | 49.9558228 | -7.2628486 | 1.35703285 | -5.3520065 |
| 8116 | NM_005012.3 | 39.157401 | -6.9164546 | 1.29231824 | -5.3519747 |
| 8117 | NM_012475.4 | 54.3940618 | -7.3871017 | 1.38028811 | -5.3518549 |
| 8118 | NM_001278728.1 | 41.0354069 | -6.9809121 | 1.30468245 | -5.3506599 |
| 8119 | NM_002836.3 | 49.9698289 | -7.2641815 | 1.35785666 | -5.3497411 |
| 8120 | NM_001329429.1 | 57.8109432 | -7.4747575 | 1.39739572 | -5.3490628 |
| 8121 | NM_003864.3 | 51.9138154 | -7.3185261 | 1.36838821 | -5.348282 |
| 8122 | NM_016341.3 | 38.6571836 | -6.8907191 | 1.28865378 | -5.347223 |
| 8123 | NM_001130089.1 | 88.7163946 | -7.1248168 | 1.33245971 | -5.3471161 |
| 8124 | NM_019065.2 | 44.4958958 | -7.0986093 | 1.32757163 | -5.3470631 |
| 8125 | NM_017626.4 | 44.7986339 | -7.1071373 | 1.32922693 | -5.3468202 |
| 8126 | NM_002830.3 | 85.4442299 | -5.6970633 | 1.06555579 | -5.346565 |

| | | | | | |
|---|---|---|---|---|---|
| 8127 | NM_016408.3 | 48.197894 | -7.2092823 | 1.34839836 | -5.3465523 |
| 8128 | NM_001276405.1 | 38.6108123 | -6.8895446 | 1.28867156 | -5.3462378 |
| 8129 | NM_001127325.1 | 66.2397511 | -7.6708849 | 1.43499672 | -5.3455766 |
| 8130 | NM_001099403.1 | 147.919427 | -8.8303419 | 1.65199053 | -5.345274 |
| 8131 | NM_177538.2 | 559.027036 | 1.17369401 | 0.21959075 | 5.34491542 |
| 8132 | NM_001005861.2 | 61.1457177 | -7.5578393 | 1.41413013 | -5.3445147 |
| 8133 | NM_020806.4 | 38.5622374 | -6.8900957 | 1.28920862 | -5.3444381 |
| 8134 | NM_017831.3 | 272.434064 | 2.69864884 | 0.50495773 | 5.34430646 |
| 8135 | NM_199287.2 | 71.938477 | -6.8155382 | 1.27529919 | -5.3442661 |
| 8136 | NM_001284513.1 | 43.5110675 | -7.0632966 | 1.32173538 | -5.3439567 |
| 8137 | NM_001199340.1 | 38.0044835 | -6.8660769 | 1.28492443 | -5.3435647 |
| 8138 | NM_001356.4 | 33.3883221 | -6.6830809 | 1.25080307 | -5.343032 |
| 8139 | NM_005360.4 | 42.5837787 | -7.0388152 | 1.31742895 | -5.3428424 |
| 8140 | NM_014628.2 | 47.6205409 | -7.1940774 | 1.34651731 | -5.3427292 |
| 8141 | NM_174889.4 | 54.9391389 | -7.4003001 | 1.38515767 | -5.3425687 |
| 8142 | NR_073007.1 | 58.2208128 | -7.4847848 | 1.40111719 | -5.342012 |
| 8143 | NM_004642.3 | 178.809065 | -6.4233264 | 1.2025198 | -5.3415556 |
| 8144 | NM_016507.3 | 85.64407 | -8.0417917 | 1.50559055 | -5.3412873 |
| 8145 | NM_201446.2 | 36.1941434 | -6.7981737 | 1.27297021 | -5.3404028 |
| 8146 | NM_152540.3 | 49.1813575 | -7.2414946 | 1.35610718 | -5.3399132 |
| 8147 | NM_006153.5 | 47.9556644 | -7.2061447 | 1.34962307 | -5.3393758 |
| 8148 | NM_138328.2 | 47.2747319 | -7.1854249 | 1.3457647 | -5.3392877 |
| 8149 | NM_005007.3 | 46.9109717 | -7.1721829 | 1.34334757 | -5.3390374 |
| 8150 | NM_002928.3 | 73.860514 | -6.8577644 | 1.28452396 | -5.3387594 |
| 8151 | NM_025155.2 | 43.6813202 | -7.069171 | 1.32421443 | -5.3383884 |
| 8152 | NM_018412.3 | 58.0681417 | -7.4813039 | 1.4014772 | -5.338156 |
| 8153 | NM_001822.5 | 51.45893 | -7.3087744 | 1.36917211 | -5.3380976 |
| 8154 | NM_032178.2 | 38.1521559 | -6.8747544 | 1.28796656 | -5.3376808 |
| 8155 | NM_203414.2 | 51.2568672 | -7.2994624 | 1.36755833 | -5.3375877 |
| 8156 | NM_001277324.1 | 43.1733234 | -7.0508064 | 1.32109778 | -5.3370814 |
| 8157 | NM_001077525.2 | 41.8885632 | -7.0086751 | 1.31325383 | -5.3368778 |
| 8158 | NM_030816.4 | 127.401844 | -4.7516975 | 0.89044795 | -5.3363001 |
| 8159 | NM_014346.3 | 41.3109804 | -6.9901769 | 1.31000047 | -5.336011 |
| 8160 | NM_016470.7 | 86.70629 | -7.086718 | 1.32825971 | -5.3353406 |
| 8161 | NR_132355.1 | 47.0685842 | -7.1795986 | 1.3456742 | -5.3353171 |
| 8162 | NM_016565.2 | 48.5468434 | -7.2236175 | 1.35415132 | -5.3344242 |
| 8163 | NM_181783.3 | 74.0862388 | -6.8645649 | 1.2870618 | -5.3335162 |
| 8164 | NM_001130996.1 | 92.9347506 | -8.1602569 | 1.530013 | -5.333456 |
| 8165 | NM_020210.3 | 32.4941211 | -6.6406156 | 1.24509256 | -5.3334313 |
| 8166 | NM_021032.4 | 56.2745334 | -7.4360805 | 1.39426439 | -5.3333361 |
| 8167 | NM_203318.1 | 47.4552248 | -7.1908486 | 1.34831471 | -5.3332123 |
| 8168 | NM_018320.4 | 52.7162523 | -7.3423721 | 1.37673959 | -5.3331597 |
| 8169 | NM_001256443.1 | 35.4923059 | -6.7774391 | 1.27082708 | -5.333093 |

| | | | | | |
|---|---|---|---|---|---|
| 8170 | NM_203389.2 | 59.7328074 | -7.5219552 | 1.41059471 | -5.3324709 |
| 8171 | NM_012241.4 | 36.1615875 | -6.7987189 | 1.27497618 | -5.3324281 |
| 8172 | NM_052886.2 | 60.0297207 | -7.5278951 | 1.41174792 | -5.3323225 |
| 8173 | NM_181718.3 | 66.8137711 | -7.6837343 | 1.4409992 | -5.3322266 |
| 8174 | NM_138477.2 | 90.9085985 | -3.1932451 | 0.59889001 | -5.3319392 |
| 8175 | NM_000052.6 | 42.42831 | -7.033605 | 1.31918515 | -5.33178 |
| 8176 | NM_053005.5 | 46.7088387 | -7.1660992 | 1.34413162 | -5.3313969 |
| 8177 | NM_006299.4 | 35.5521235 | -6.7752939 | 1.27087005 | -5.3312248 |
| 8178 | NM_019053.5 | 41.7230479 | -7.0042435 | 1.3139288 | -5.3307634 |
| 8179 | NM_153498.3 | 248.440688 | -3.4679505 | 0.65057787 | -5.3305695 |
| 8180 | NM_006089.2 | 37.5839993 | -6.8498202 | 1.28506358 | -5.3303357 |
| 8181 | NM_003390.3 | 87.1100225 | -6.1454478 | 1.15299604 | -5.3299818 |
| 8182 | NM_018093.3 | 73.3672564 | -6.849989 | 1.28518129 | -5.3299788 |
| 8183 | NR_130945.1 | 36.9246511 | -6.8232911 | 1.28058083 | -5.3282783 |
| 8184 | NM_001258330.1 | 35.168007 | -6.7640233 | 1.26950195 | -5.3280921 |
| 8185 | NM_007324.3 | 50.5647246 | -7.2797162 | 1.36633283 | -5.3279231 |
| 8186 | NM_012392.3 | 53.1934982 | -7.3537728 | 1.38024998 | -5.3278557 |
| 8187 | NM_018413.5 | 76.1275823 | -7.871396 | 1.47744887 | -5.3276943 |
| 8188 | NM_014056.3 | 294.158439 | -2.7240011 | 0.51129677 | -5.3276322 |
| 8189 | NR_038335.1 | 76.6975237 | -7.8845717 | 1.48011914 | -5.3269845 |
| 8190 | NM_016050.4 | 42.0582013 | -7.0145627 | 1.31682798 | -5.3268633 |
| 8191 | NM_032890.3 | 33.4215749 | -6.690306 | 1.25606891 | -5.3263845 |
| 8192 | NM_145119.3 | 45.1005542 | -7.1182509 | 1.33649906 | -5.3260426 |
| 8193 | NM_205845.2 | 68.6526907 | -7.7224848 | 1.44998167 | -5.3259189 |
| 8194 | NM_032199.2 | 38.1813135 | -6.8811311 | 1.29209756 | -5.3255507 |
| 8195 | NM_001127496.1 | 84.2060946 | -6.1731058 | 1.15932246 | -5.3247531 |
| 8196 | NM_020904.2 | 50.2794412 | -7.2767813 | 1.366653 | -5.3245274 |
| 8197 | NM_017983.6 | 47.5026249 | -7.1957159 | 1.35147539 | -5.3243411 |
| 8198 | NR_040039.1 | 48.8960721 | 2.87111411 | 0.53926718 | 5.32410318 |
| 8199 | NM_144563.2 | 41.0408453 | -6.9814247 | 1.3114019 | -5.3236347 |
| 8200 | NM_174905.3 | 49.8302058 | -7.2592959 | 1.36360751 | -5.3235963 |
| 8201 | NM_001276414.1 | 35.9494559 | -6.7947624 | 1.27643785 | -5.3232223 |
| 8202 | NM_022073.3 | 71.6560985 | -6.8082312 | 1.27910491 | -5.3226527 |
| 8203 | NM_001076778.2 | 67.8932976 | -7.7051376 | 1.44813117 | -5.3207456 |
| 8204 | NM_078474.2 | 60.015336 | -7.5271176 | 1.41469185 | -5.3206764 |
| 8205 | NM_003656.4 | 63.3084148 | -7.6061507 | 1.42961851 | -5.3204058 |
| 8206 | NM_001127896.1 | 73.3564856 | -7.8191343 | 1.46970528 | -5.3202056 |
| 8207 | NM_015952.3 | 48.8508488 | -7.2315135 | 1.35928484 | -5.3200869 |
| 8208 | NM_001330323.1 | 52.2125399 | -7.3303175 | 1.37791745 | -5.3198524 |
| 8209 | NM_003335.2 | 37.1413735 | -6.8428975 | 1.28638348 | -5.3194849 |
| 8210 | NM_001286247.1 | 83.4445397 | -7.0306375 | 1.32172558 | -5.3192868 |
| 8211 | NM_181351.4 | 148.642073 | -8.8379823 | 1.66157801 | -5.3190294 |
| 8212 | NM_002916.3 | 51.9211871 | -7.3192028 | 1.37617659 | -5.3185055 |

| | | | | | |
|---|---|---|---|---|---|
| 8213 | NM_017910.3 | 51.174285 | -7.2992731 | 1.37250251 | -5.318222 |
| 8214 | NM_032968.4 | 99.9220458 | -8.2646975 | 1.55407752 | -5.3180728 |
| 8215 | NM_002915.3 | 41.4486452 | -6.9926034 | 1.31488219 | -5.3180455 |
| 8216 | NM_004440.3 | 55.9290238 | -7.429868 | 1.3972199 | -5.3176082 |
| 8217 | NM_023940.2 | 56.2508722 | -7.434306 | 1.39823072 | -5.3169379 |
| 8218 | NM_001321693.1 | 48.1819152 | -7.21341 | 1.35674133 | -5.3167172 |
| 8219 | NM_001316311.1 | 49.7974341 | -7.2608605 | 1.36567399 | -5.3166865 |
| 8220 | NM_001322215.1 | 40.3752916 | -6.9590784 | 1.30907193 | -5.3160398 |
| 8221 | NM_032630.2 | 42.4560253 | -7.0308098 | 1.32257656 | -5.3159946 |
| 8222 | NM_001320464.1 | 49.8674318 | -7.2633031 | 1.36637166 | -5.3157595 |
| 8223 | NR_002909.2 | 44.0983585 | -7.0871347 | 1.33336182 | -5.3152374 |
| 8224 | NR_110049.2 | 89.9218293 | 1.71692852 | 0.32302289 | 5.31519153 |
| 8225 | NM_003422.2 | 35.9543418 | -6.7943762 | 1.27843576 | -5.3146012 |
| 8226 | NM_001322324.1 | 64.9962377 | -7.645278 | 1.43855348 | -5.3145594 |
| 8227 | NM_012330.3 | 42.5328337 | -7.0288503 | 1.32257149 | -5.3145334 |
| 8228 | NM_001300799.1 | 83.1651822 | -7.9989518 | 1.50513433 | -5.3144438 |
| 8229 | NM_033375.4 | 86.6311142 | -8.0601851 | 1.51667425 | -5.3143812 |
| 8230 | NM_006232.4 | 41.1912947 | -6.9882652 | 1.3150196 | -5.314191 |
| 8231 | NM_001039614.2 | 63.3359244 | -7.6061568 | 1.43138 | -5.3138627 |
| 8232 | NM_003721.3 | 45.2863234 | -7.1218779 | 1.34032917 | -5.313529 |
| 8233 | NM_001135746.1 | 41.6081813 | -6.9982713 | 1.31711574 | -5.3133306 |
| 8234 | NM_025141.3 | 63.4085721 | -7.6091304 | 1.43210023 | -5.3132667 |
| 8235 | NM_001495.4 | 44.5713328 | -7.0991225 | 1.33612786 | -5.313206 |
| 8236 | NM_001128220.2 | 37.1026386 | -6.8362657 | 1.2866978 | -5.3130313 |
| 8237 | NM_004083.5 | 61.7047615 | -7.5684835 | 1.42451135 | -5.3130384 |
| 8238 | NM_001040439.1 | 179.847066 | -8.1465026 | 1.53333113 | -5.3129441 |
| 8239 | NR_144509.1 | 52.9747636 | -7.350378 | 1.38363463 | -5.3123692 |
| 8240 | NM_002202.2 | 98.2312861 | -7.2693596 | 1.36855155 | -5.3117178 |
| 8241 | NM_001031711.2 | 369.811443 | -6.1002791 | 1.1484696 | -5.3116592 |
| 8242 | NR_125956.1 | 41.3149004 | 4.96382982 | 0.93453481 | 5.31155157 |
| 8243 | NM_020975.4 | 114.654518 | -7.4975991 | 1.41158257 | -5.3114846 |
| 8244 | NM_006480.4 | 31.5506852 | -6.6022487 | 1.24313525 | -5.3109657 |
| 8245 | NM_207332.2 | 35.4094789 | -6.7713431 | 1.27502663 | -5.3107464 |
| 8246 | NM_001130058.1 | 58.1224393 | -7.4825153 | 1.40902295 | -5.3104283 |
| 8247 | NR_109832.1 | 85.5725713 | 4.44475149 | 0.83698486 | 5.31043239 |
| 8248 | NM_206925.2 | 57.4355447 | -7.4647727 | 1.40575878 | -5.3101377 |
| 8249 | NM_020365.4 | 59.6457023 | -7.5196052 | 1.41617525 | -5.3097985 |
| 8250 | NM_177533.4 | 53.6184163 | -7.3651052 | 1.3871315 | -5.3095941 |
| 8251 | NR_040104.1 | 41.4595991 | -6.9939017 | 1.31725852 | -5.3094374 |
| 8252 | NM_001079539.1 | 51.2487289 | -7.2996943 | 1.37505166 | -5.3086691 |
| 8253 | NM_001290117.1 | 33.2950612 | -6.6843469 | 1.25922917 | -5.3082847 |
| 8254 | NR_026874.2 | 55.0182985 | -7.4050426 | 1.39500192 | -5.308267 |
| 8255 | NM_016221.3 | 111.786461 | -6.6160439 | 1.24649486 | -5.3077185 |

| | | | | | |
|---|---|---|---|---|---|
| 8256 | NM_006123.4 | 37.429836 | -6.8476604 | 1.29014529 | -5.3076661 |
| 8257 | NM_033542.3 | 62.0084484 | -6.6083669 | 1.2450574 | -5.3076805 |
| 8258 | NM_145806.3 | 38.8671291 | -6.9019649 | 1.3003948 | -5.3075919 |
| 8259 | NM_181861.1 | 55.9083507 | -7.4239442 | 1.39886188 | -5.3071317 |
| 8260 | NM_018050.2 | 38.8394807 | -6.9007597 | 1.30028924 | -5.3070959 |
| 8261 | NM_000663.4 | 99.1344904 | -6.4789028 | 1.22084846 | -5.3068853 |
| 8262 | NM_001319121.1 | 46.5228186 | -7.1631269 | 1.34983936 | -5.3066514 |
| 8263 | NM_018569.5 | 65.8465538 | -7.6603129 | 1.44356556 | -5.3065223 |
| 8264 | NM_001006605.4 | 48.7211401 | -7.2281468 | 1.36230811 | -5.3058091 |
| 8265 | NM_015458.3 | 241.988204 | -3.7910717 | 0.71452089 | -5.3057535 |
| 8266 | NM_016627.4 | 63.2465266 | -7.6040092 | 1.43317876 | -5.3056949 |
| 8267 | NM_018691.3 | 35.867467 | -6.7858688 | 1.279083 | -5.3052607 |
| 8268 | NM_013312.2 | 54.7827313 | -7.3955124 | 1.39404206 | -5.3050856 |
| 8269 | NM_001282311.1 | 32.8358456 | -6.6632571 | 1.2562384 | -5.3041342 |
| 8270 | NM_000313.3 | 71.3857321 | -6.8136991 | 1.28478172 | -5.3033904 |
| 8271 | NM_006676.7 | 48.7117117 | -7.2293198 | 1.36332203 | -5.3027236 |
| 8272 | NM_018217.2 | 38.9592831 | -6.9097583 | 1.30305961 | -5.3027185 |
| 8273 | NM_001271.3 | 394.053045 | -4.67383 | 0.88146782 | -5.3023263 |
| 8274 | NM_000683.3 | 37.7006861 | -6.8613987 | 1.294062 | -5.3022179 |
| 8275 | NM_001291285.1 | 46.1142422 | -7.1503046 | 1.3485649 | -5.3021583 |
| 8276 | NM_001308325.1 | 49.0173403 | -7.2364661 | 1.36482309 | -5.3021275 |
| 8277 | NM_001321076.1 | 100.172926 | -8.2678681 | 1.55934525 | -5.3021408 |
| 8278 | NM_201222.2 | 50.0444067 | -7.2713122 | 1.37145716 | -5.301888 |
| 8279 | NR_028040.1 | 57.1572335 | -7.4565632 | 1.4064292 | -5.3017693 |
| 8280 | NM_015271.4 | 333.951082 | -6.2311472 | 1.17532015 | -5.3016594 |
| 8281 | NR_026865.1 | 44.4323131 | -7.0944216 | 1.33815371 | -5.3016493 |
| 8282 | NM_001130970.1 | 238.894751 | -9.5222385 | 1.79612608 | -5.3015424 |
| 8283 | NM_001005333.1 | 32.3433213 | -6.6336664 | 1.25131579 | -5.3013527 |
| 8284 | NM_175575.5 | 194.433035 | -9.2257656 | 1.74027413 | -5.3013289 |
| 8285 | NM_020888.2 | 83.5463039 | -8.0064214 | 1.51035206 | -5.3010299 |
| 8286 | NM_001789.2 | 42.5859196 | -7.0326106 | 1.3268209 | -5.3003465 |
| 8287 | NM_145887.3 | 43.3371233 | -7.0587001 | 1.33180464 | -5.3001018 |
| 8288 | NM_001145432.1 | 40.6414947 | -6.9655075 | 1.31422998 | -5.3000675 |
| 8289 | NM_001323600.1 | 34.2653143 | -6.7220889 | 1.26831579 | -5.300012 |
| 8290 | NM_017454.3 | 42.0910598 | -7.0196216 | 1.32444925 | -5.3000306 |
| 8291 | NM_032052.1 | 42.7063234 | -7.0350243 | 1.32750391 | -5.2994377 |
| 8292 | NM_005845.4 | 38.4262856 | -6.8852847 | 1.29925568 | -5.299407 |
| 8293 | NM_001010875.3 | 39.430967 | -6.9233496 | 1.30645044 | -5.2993588 |
| 8294 | NM_020630.4 | 67.5857224 | -7.7021515 | 1.45341394 | -5.2993516 |
| 8295 | NM_022892.1 | 125.718603 | -4.2434345 | 0.80082325 | -5.2988403 |
| 8296 | NR_036434.1 | 42.3515739 | -7.0244298 | 1.32576124 | -5.2984124 |
| 8297 | NR_110837.1 | 186.9075 | 2.01602821 | 0.38051844 | 5.29810912 |
| 8298 | NM_001258392.2 | 45.1543207 | -7.1174453 | 1.34341766 | -5.2980138 |

| | | | | | |
|---|---|---|---|---|---|
| 8299 | NM_001145191.1 | 88.2026662 | -4.6226528 | 0.87255168 | -5.2978556 |
| 8300 | NM_017754.3 | 123.346816 | -3.2199387 | 0.60780248 | -5.2976729 |
| 8301 | NM_018372.3 | 35.4336508 | -6.7677286 | 1.27751464 | -5.2975742 |
| 8302 | NM_178496.3 | 53.6931233 | -7.3683059 | 1.3909887 | -5.2971717 |
| 8303 | NM_001008910.3 | 34.9008616 | -6.7435422 | 1.27312334 | -5.2968491 |
| 8304 | NM_007276.4 | 39.9238589 | -6.939814 | 1.31027098 | -5.2964724 |
| 8305 | NM_014978.2 | 57.0432008 | -7.4554737 | 1.40763027 | -5.2964716 |
| 8306 | NM_001324116.1 | 35.0375407 | -6.750387 | 1.27476919 | -5.2953798 |
| 8307 | NM_007274.3 | 223.787071 | -6.1993104 | 1.1707145 | -5.2953221 |
| 8308 | NM_019896.2 | 58.7990437 | -7.4991732 | 1.41663656 | -5.2936465 |
| 8309 | NR_033338.1 | 64.1986152 | -7.6266743 | 1.44081313 | -5.2933126 |
| 8310 | NM_206949.2 | 33.1914625 | -6.6779647 | 1.2616783 | -5.2929219 |
| 8311 | NM_003244.3 | 32.2334361 | -6.6328545 | 1.25317916 | -5.2928222 |
| 8312 | NM_138465.3 | 42.9534819 | -7.0449813 | 1.33112097 | -5.2925177 |
| 8313 | NM_001301699.1 | 50.5083836 | -7.2814317 | 1.37588562 | -5.292178 |
| 8314 | NM_173548.2 | 34.6971346 | -6.7381394 | 1.27329529 | -5.2918907 |
| 8315 | NM_002599.4 | 86.5657579 | -8.0564013 | 1.52242537 | -5.2918202 |
| 8316 | NM_001282602.1 | 69.8626793 | -7.7465576 | 1.46398227 | -5.2914285 |
| 8317 | NM_020238.2 | 77.9085269 | -6.9372277 | 1.31104111 | -5.2913884 |
| 8318 | NM_022006.1 | 61.4075718 | -7.56314 | 1.42943114 | -5.2910138 |
| 8319 | NR_003945.1 | 37.3113538 | 2.51563535 | 0.47546291 | 5.29091814 |
| 8320 | NM_152789.3 | 56.3655028 | -7.4355576 | 1.40546262 | -5.2904698 |
| 8321 | NM_001111125.2 | 50.5436559 | -7.2795533 | 1.37607184 | -5.2900968 |
| 8322 | NM_138493.2 | 53.6641511 | -7.3668798 | 1.3925994 | -5.2900208 |
| 8323 | NM_001321039.1 | 31.4889869 | -6.6023819 | 1.24818818 | -5.2895726 |
| 8324 | NM_001314063.1 | 126.28829 | -8.602152 | 1.62628177 | -5.2894598 |
| 8325 | NM_001897.4 | 41.5661636 | -7.0042868 | 1.32422569 | -5.2893452 |
| 8326 | NM_017941.5 | 50.7177915 | -7.2828303 | 1.37697576 | -5.289004 |
| 8327 | NM_033117.3 | 94.3587217 | -7.2148744 | 1.36414967 | -5.288917 |
| 8328 | NM_001265609.1 | 32.7316529 | -6.6496258 | 1.25731755 | -5.2887401 |
| 8329 | NM_001080512.2 | 57.2633252 | -4.549562 | 0.8602916 | -5.2883953 |
| 8330 | NR_073502.1 | 58.1314464 | -7.4855066 | 1.41550185 | -5.2882352 |
| 8331 | NM_004280.4 | 49.0881098 | -7.2391632 | 1.36901236 | -5.2878728 |
| 8332 | NM_018899.5 | 62.4859063 | -7.5864158 | 1.43472789 | -5.2877035 |
| 8333 | NM_001130136.1 | 107.733902 | -8.3732469 | 1.58356337 | -5.2875983 |
| 8334 | NR_126523.1 | 96.5703023 | 4.73969221 | 0.89637923 | 5.28759711 |
| 8335 | NM_001252053.1 | 46.204696 | -7.1548902 | 1.35363324 | -5.2856933 |
| 8336 | NM_023933.2 | 48.8949099 | -7.2323603 | 1.36832695 | -5.2855499 |
| 8337 | NM_032829.2 | 43.2311419 | -7.0593727 | 1.3356399 | -5.2853862 |
| 8338 | NM_001253380.1 | 60.3180509 | -7.5392729 | 1.42652497 | -5.285062 |
| 8339 | NM_139078.2 | 62.1918379 | -7.5805655 | 1.43433837 | -5.2850608 |
| 8340 | NM_003720.3 | 40.1538789 | -6.9499102 | 1.31517624 | -5.2843945 |
| 8341 | NM_006769.3 | 316.799017 | -6.3784291 | 1.20715183 | -5.2838665 |

| | | | | | |
|---|---|---|---|---|---|
| 8342 | NM_015691.3 | 115.423756 | -4.3125055 | 0.81616265 | -5.28388 |
| 8343 | NM_001242916.1 | 56.9983782 | -7.4529031 | 1.41063856 | -5.2833541 |
| 8344 | NM_004701.3 | 48.1735096 | -7.2114153 | 1.36505332 | -5.2828818 |
| 8345 | NM_201378.3 | 97.3032868 | -8.2245229 | 1.55692441 | -5.2825448 |
| 8346 | NM_022719.2 | 92.2031802 | -5.4173907 | 1.02556172 | -5.2823644 |
| 8347 | NM_018663.2 | 45.7185452 | -7.1354042 | 1.35088842 | -5.2820086 |
| 8348 | NM_001288814.1 | 49.3766917 | -7.2458198 | 1.37196078 | -5.2813607 |
| 8349 | NM_001017926.2 | 44.3300766 | -7.0915121 | 1.34275226 | -5.2813258 |
| 8350 | NM_003582.2 | 31.5644725 | -6.605677 | 1.25081572 | -5.2810953 |
| 8351 | NM_001317238.1 | 52.3000724 | -7.332333 | 1.38843778 | -5.280995 |
| 8352 | NM_000824.4 | 48.1691847 | -7.2106497 | 1.36544376 | -5.2808105 |
| 8353 | NM_015957.3 | 41.445684 | -6.9921312 | 1.32418035 | -5.2803466 |
| 8354 | NM_005680.2 | 30.1476998 | -6.5390034 | 1.23840937 | -5.280163 |
| 8355 | NM_198897.1 | 35.3035071 | -6.7623293 | 1.28082125 | -5.2796823 |
| 8356 | NM_001308115.1 | 33.5176575 | -6.6897088 | 1.2671522 | -5.2793254 |
| 8357 | NM_001136159.1 | 64.3310573 | -4.5423652 | 0.86044456 | -5.2790911 |
| 8358 | NM_001199861.1 | 31.6866769 | -6.6096086 | 1.2520354 | -5.2790908 |
| 8359 | NM_013393.1 | 70.3012542 | -6.7889731 | 1.28611027 | -5.2786866 |
| 8360 | NM_012267.4 | 40.7100287 | -6.9704779 | 1.32052965 | -5.278547 |
| 8361 | NM_001130825.1 | 68.1055001 | -7.7119951 | 1.4610271 | -5.278475 |
| 8362 | NR_144445.1 | 100.041535 | -4.8969508 | 0.92772902 | -5.2784279 |
| 8363 | NM_176814.4 | 30.9280867 | -6.5734802 | 1.2454432 | -5.2780248 |
| 8364 | NM_024337.3 | 48.8486667 | -7.2343077 | 1.37065836 | -5.2779802 |
| 8365 | NM_130843.3 | 68.1072163 | -7.712926 | 1.46134174 | -5.2779755 |
| 8366 | NM_144489.3 | 30.4627213 | -6.5536402 | 1.24172755 | -5.2778407 |
| 8367 | NM_019848.4 | 34.4392373 | -6.7336577 | 1.27589195 | -5.2776081 |
| 8368 | NM_014044.6 | 50.2088866 | -7.2715405 | 1.37789947 | -5.2772649 |
| 8369 | NM_003981.3 | 37.2965881 | -6.8391751 | 1.29599059 | -5.2771796 |
| 8370 | NM_015945.11 | 57.9420693 | -7.4787288 | 1.41720532 | -5.2770962 |
| 8371 | NM_001278719.1 | 51.5186546 | -7.3106041 | 1.38537042 | -5.2770032 |
| 8372 | NM_022451.10 | 37.7688929 | -6.8587773 | 1.29983079 | -5.2766694 |
| 8373 | NM_006466.3 | 34.5154226 | -6.7315476 | 1.27590384 | -5.2759051 |
| 8374 | NM_014393.2 | 111.476717 | -6.4594285 | 1.2244314 | -5.2754515 |
| 8375 | NM_025202.3 | 32.7418574 | -6.658352 | 1.26220179 | -5.2751882 |
| 8376 | NM_017913.3 | 65.6130955 | -5.8144875 | 1.10231131 | -5.2748143 |
| 8377 | NR_047519.1 | 78.0421509 | -6.9399585 | 1.31572122 | -5.274642 |
| 8378 | NM_001321368.1 | 42.476125 | -7.032085 | 1.33321237 | -5.2745422 |
| 8379 | NM_052910.2 | 63.1134243 | -7.6008027 | 1.4411295 | -5.2741983 |
| 8380 | NM_022736.2 | 46.9625416 | -7.1729876 | 1.36003353 | -5.2741256 |
| 8381 | NM_001011655.2 | 40.958478 | -6.9776025 | 1.32314467 | -5.2734993 |
| 8382 | NM_003501.2 | 36.1667626 | -6.7951384 | 1.28857873 | -5.2733591 |
| 8383 | NM_017566.3 | 42.5964818 | -7.0306689 | 1.3332669 | -5.2732644 |
| 8384 | NM_001083914.2 | 54.1518772 | -7.380179 | 1.39958562 | -5.2731172 |

| | | | | | |
|---|---|---|---|---|---|
| 8385 | NM_182916.2 | 43.5213002 | -7.0649009 | 1.33983553 | -5.2729613 |
| 8386 | NM_052963.2 | 46.8042525 | -7.1697691 | 1.35980664 | -5.2726387 |
| 8387 | NM_001112802.1 | 37.8729513 | -6.8600887 | 1.30127316 | -5.2718283 |
| 8388 | NM_020188.3 | 60.7467322 | -7.5457921 | 1.43135691 | -5.2717754 |
| 8389 | NM_182894.2 | 63.4847517 | -7.6112245 | 1.44390642 | -5.2712727 |
| 8390 | NM_018486.2 | 37.6473703 | -6.8569516 | 1.30088215 | -5.2710014 |
| 8391 | NM_001177387.1 | 34.7939906 | -6.7364433 | 1.27805931 | -5.2708378 |
| 8392 | NM_178865.4 | 100.788532 | -7.3054132 | 1.38607931 | -5.2705593 |
| 8393 | NM_001256523.1 | 55.526788 | -7.4187167 | 1.40758325 | -5.2705349 |
| 8394 | NM_002484.3 | 35.7077406 | -6.7808144 | 1.28659981 | -5.2703369 |
| 8395 | NR_027512.1 | 37.7309533 | -6.8611387 | 1.30184663 | -5.2703126 |
| 8396 | NM_005238.3 | 37.1420468 | -6.8410356 | 1.29811152 | -5.2699907 |
| 8397 | NM_207318.3 | 251.796172 | -5.1196368 | 0.97147242 | -5.2699765 |
| 8398 | NM_032366.4 | 40.7502294 | -6.9683227 | 1.32237175 | -5.2695641 |
| 8399 | NM_153208.2 | 55.7531008 | -7.4207737 | 1.40824671 | -5.2695125 |
| 8400 | NM_001260492.1 | 53.9083171 | -7.3744274 | 1.39949077 | -5.2693648 |
| 8401 | NM_001320836.1 | 34.9144518 | -6.7466146 | 1.2803822 | -5.2692193 |
| 8402 | NM_030653.3 | 40.3169906 | -6.9572283 | 1.32040307 | -5.2690186 |
| 8403 | NM_001168300.1 | 62.9718716 | -7.5994303 | 1.44242557 | -5.2685078 |
| 8404 | NM_001500.3 | 49.8369674 | -7.2610482 | 1.37829364 | -5.2681432 |
| 8405 | NM_001318338.1 | 63.0973025 | -7.6034225 | 1.44341662 | -5.2676562 |
| 8406 | NM_001654.4 | 94.6579822 | -4.6669044 | 0.88595291 | -5.2676665 |
| 8407 | NM_001098811.1 | 75.4353922 | -7.8587156 | 1.4919221 | -5.2675107 |
| 8408 | NM_014155.4 | 101.117499 | -4.7687299 | 0.90530852 | -5.2675191 |
| 8409 | NM_001289912.1 | 32.47746 | -6.6449539 | 1.26159559 | -5.267103 |
| 8410 | NM_001348078.1 | 31.7045593 | -6.6143959 | 1.25578947 | -5.2671216 |
| 8411 | NM_024641.3 | 34.0524857 | -6.7137051 | 1.27467711 | -5.2669849 |
| 8412 | NM_001144950.1 | 44.0962684 | -7.0881088 | 1.34579502 | -5.2668561 |
| 8413 | NM_020692.2 | 80.4855751 | -6.9846573 | 1.32616098 | -5.2668246 |
| 8414 | NM_001256007.2 | 38.0772823 | -6.8757704 | 1.30554499 | -5.2665902 |
| 8415 | NM_006875.3 | 40.8105704 | -6.9755043 | 1.32448513 | -5.2665781 |
| 8416 | NM_004589.3 | 64.040534 | -6.6463728 | 1.26199699 | -5.266552 |
| 8417 | NM_006288.4 | 75.5666409 | -7.8615051 | 1.49276974 | -5.2663883 |
| 8418 | NM_001317956.1 | 35.2528257 | -6.7608107 | 1.28378205 | -5.2663228 |
| 8419 | NM_001100814.2 | 44.98272 | -7.1115346 | 1.35040263 | -5.2662328 |
| 8420 | NR_073365.1 | 43.1046013 | -7.0543825 | 1.3395726 | -5.2661442 |
| 8421 | NM_014650.3 | 48.2898275 | -7.2181866 | 1.37072517 | -5.265962 |
| 8422 | NM_014078.5 | 48.9731743 | -7.234674 | 1.37406514 | -5.2651609 |
| 8423 | NM_001003891.2 | 41.42728 | -6.9995179 | 1.32949328 | -5.2648013 |
| 8424 | NM_013335.3 | 35.6930454 | -6.7787839 | 1.28762244 | -5.2645743 |
| 8425 | NM_001283053.1 | 51.0131497 | -7.2938598 | 1.38550325 | -5.2644119 |
| 8426 | NM_001292033.1 | 68.3377391 | 2.44692753 | 0.46481974 | 5.26425046 |
| 8427 | NM_004853.2 | 46.5813959 | -7.1639447 | 1.3608626 | -5.2642675 |

| | | | | | |
|---|---|---|---|---|---|
| 8428 | NM_001040427.1 | 47.578042 | -7.1942271 | 1.36663232 | -5.2642009 |
| 8429 | NM_014936.4 | 39.7037157 | -6.9333634 | 1.31717409 | -5.263817 |
| 8430 | NM_015167.2 | 34.5018147 | -6.7298968 | 1.27855813 | -5.2636612 |
| 8431 | NM_012309.4 | 37.7746026 | -6.8589142 | 1.30307712 | -5.2636288 |
| 8432 | NR_045106.2 | 30.4643219 | -6.5471321 | 1.24389153 | -5.2634269 |
| 8433 | NM_017828.4 | 30.1253305 | -6.5320177 | 1.24105429 | -5.2632812 |
| 8434 | NM_018315.4 | 43.4813945 | -7.0613924 | 1.34163519 | -5.2632731 |
| 8435 | NM_007361.3 | 115.507941 | -8.4749094 | 1.61025624 | -5.2630812 |
| 8436 | NM_001278463.1 | 78.1603906 | -7.9120762 | 1.50343617 | -5.2626619 |
| 8437 | NM_001308191.1 | 47.9128076 | -7.2029078 | 1.36871202 | -5.2625444 |
| 8438 | NM_003711.3 | 42.2865639 | -7.0241413 | 1.33476093 | -5.2624714 |
| 8439 | NM_004071.3 | 56.3745068 | -7.4384994 | 1.41350183 | -5.2624618 |
| 8440 | NM_007238.4 | 45.3066521 | -7.1232338 | 1.35362789 | -5.2623279 |
| 8441 | NM_019051.2 | 44.263421 | -7.0890328 | 1.34718405 | -5.2621116 |
| 8442 | NM_173794.3 | 42.2819567 | -7.0239249 | 1.33480753 | -5.2621256 |
| 8443 | NM_001323543.1 | 34.2235384 | -6.7215855 | 1.27766197 | -5.2608481 |
| 8444 | NM_004979.5 | 32.2234218 | -6.6314408 | 1.26066542 | -5.2602703 |
| 8445 | NR_037941.1 | 53.9930384 | -7.3798806 | 1.4029645 | -5.2602048 |
| 8446 | NM_001252065.1 | 61.8303002 | -7.5727228 | 1.43964223 | -5.2601422 |
| 8447 | NM_005788.3 | 36.030801 | -6.7901878 | 1.29095239 | -5.2598282 |
| 8448 | NM_032122.4 | 45.116316 | -7.1151986 | 1.35281865 | -5.2595362 |
| 8449 | NM_004794.2 | 62.5105271 | -7.5877459 | 1.44272783 | -5.2593052 |
| 8450 | NM_003528.2 | 77.3997977 | -6.9277826 | 1.31725523 | -5.2592561 |
| 8451 | NR_026956.1 | 123.162131 | 1.15355663 | 0.21933804 | 5.25926377 |
| 8452 | NM_002579.2 | 48.1019512 | -7.2140174 | 1.37177723 | -5.2588841 |
| 8453 | NM_018059.4 | 35.7398226 | -6.7813111 | 1.28951536 | -5.258806 |
| 8454 | NM_004362.2 | 61.5945216 | -6.5959844 | 1.25443782 | -5.2581198 |
| 8455 | NM_001321690.1 | 69.6021904 | -7.7420053 | 1.47240665 | -5.2580619 |
| 8456 | NM_001174118.1 | 48.7072611 | -7.224347 | 1.37397773 | -5.2579797 |
| 8457 | NM_057167.3 | 69.1668786 | -7.7360331 | 1.47131724 | -5.257896 |
| 8458 | NM_021949.3 | 102.526122 | -8.3026469 | 1.57910891 | -5.2578051 |
| 8459 | NM_176824.2 | 32.7667097 | -6.656227 | 1.26604131 | -5.2575117 |
| 8460 | NM_014165.3 | 59.3931785 | -7.5135035 | 1.42933897 | -5.2566282 |
| 8461 | NM_003012.4 | 465.042237 | -6.320428 | 1.20243432 | -5.2563603 |
| 8462 | NM_001109662.3 | 467.079499 | -4.9341614 | 0.93877947 | -5.2559324 |
| 8463 | NM_181723.2 | 52.2004844 | -7.3279608 | 1.39423698 | -5.2558933 |
| 8464 | NM_015041.2 | 51.5017378 | -7.3121362 | 1.39142125 | -5.2551564 |
| 8465 | NM_004468.4 | 39.5721476 | -6.9292783 | 1.31867733 | -5.2547186 |
| 8466 | NM_001100619.2 | 34.3423769 | -6.729676 | 1.28072063 | -5.2546011 |
| 8467 | NM_002913.4 | 134.099464 | -5.0158246 | 0.95455858 | -5.2546011 |
| 8468 | NM_001039503.2 | 33.7460493 | -6.6960962 | 1.27438108 | -5.2543908 |
| 8469 | NM_001161780.1 | 74.7294762 | -7.8454824 | 1.49314287 | -5.2543414 |
| 8470 | NM_178454.4 | 55.7167312 | -6.4491759 | 1.22745434 | -5.2541066 |

| | | | | | |
|---|---|---|---|---|---|
| 8471 | NM_001297741.1 | 59.3031787 | -7.5128712 | 1.42995459 | -5.2539229 |
| 8472 | NM_020964.2 | 96.0116595 | -3.397724 | 0.64672428 | -5.2537443 |
| 8473 | NM_024900.4 | 37.2898516 | -6.8434393 | 1.30266321 | -5.2534218 |
| 8474 | NM_015135.2 | 178.521089 | -5.2871102 | 1.00658495 | -5.2525226 |
| 8475 | NM_021101.4 | 72.855018 | -7.8108121 | 1.48719239 | -5.2520522 |
| 8476 | NM_080669.5 | 34.6862606 | -6.7391319 | 1.28329363 | -5.2514341 |
| 8477 | NM_017956.3 | 36.7557629 | -6.8202186 | 1.29883656 | -5.2510214 |
| 8478 | NR_023384.1 | 44.7308999 | -7.1028106 | 1.35266414 | -5.2509787 |
| 8479 | NM_003797.4 | 47.4673182 | -7.1886613 | 1.36908483 | -5.2507056 |
| 8480 | NM_013338.4 | 49.7187169 | -7.2575003 | 1.38219131 | -5.2507205 |
| 8481 | NM_001745.3 | 228.035045 | -5.1240123 | 0.97592266 | -5.2504287 |
| 8482 | NM_018905.2 | 70.0539968 | -7.752487 | 1.47662417 | -5.2501423 |
| 8483 | NM_018555.5 | 29.9638702 | -6.5273715 | 1.2434125 | -5.2495624 |
| 8484 | NM_004805.3 | 90.4491164 | -4.3905893 | 0.83637843 | -5.2495247 |
| 8485 | NM_006412.3 | 30.32092 | -6.5483203 | 1.24747435 | -5.2492625 |
| 8486 | NM_017998.2 | 51.4342263 | -7.3060509 | 1.39182402 | -5.2492634 |
| 8487 | NM_001286835.1 | 38.2826785 | -6.8798433 | 1.3107143 | -5.2489267 |
| 8488 | NM_001146175.1 | 54.0320546 | -7.3772413 | 1.40549232 | -5.2488664 |
| 8489 | NM_020166.4 | 40.3767107 | -6.9550563 | 1.32507282 | -5.2488107 |
| 8490 | NM_001166159.1 | 33.2397805 | -6.6783765 | 1.27237499 | -5.2487486 |
| 8491 | NM_153035.2 | 57.1820975 | -2.7721892 | 0.52818655 | -5.2485039 |
| 8492 | NM_001018136.2 | 57.8747813 | -7.476818 | 1.42464583 | -5.2481942 |
| 8493 | NR_037709.1 | 30.5001162 | -6.5539616 | 1.24890592 | -5.2477624 |
| 8494 | NM_152305.2 | 45.1535815 | -7.1195805 | 1.35679406 | -5.2473553 |
| 8495 | NM_022782.3 | 33.6685928 | -6.6954692 | 1.27599182 | -5.2472666 |
| 8496 | NM_001199563.1 | 31.6882208 | -6.604666 | 1.25874305 | -5.2470327 |
| 8497 | NM_001320633.1 | 38.4358986 | -6.8818103 | 1.31164182 | -5.2467146 |
| 8498 | NM_031300.3 | 38.8900143 | -6.9003342 | 1.31523441 | -5.2464672 |
| 8499 | NM_001122842.2 | 52.7455616 | -7.3415753 | 1.39945591 | -5.2460212 |
| 8500 | NM_213720.2 | 32.2216024 | -6.6281575 | 1.26353794 | -5.2457131 |
| 8501 | NM_001042631.2 | 41.7516075 | -7.0048174 | 1.3353972 | -5.2454936 |
| 8502 | NM_021221.2 | 30.4193065 | -6.5504181 | 1.24878414 | -5.2454366 |
| 8503 | NM_004569.3 | 46.3404017 | -7.155503 | 1.3642129 | -5.2451512 |
| 8504 | NM_024525.4 | 46.9228226 | -7.1722139 | 1.36744941 | -5.2449574 |
| 8505 | NM_024581.5 | 29.6954269 | -6.5176691 | 1.24271305 | -5.2447096 |
| 8506 | NM_173601.1 | 47.7116681 | -7.1986949 | 1.37259196 | -5.2445993 |
| 8507 | NM_001206856.2 | 35.6427093 | -6.7721963 | 1.2913021 | -5.2444709 |
| 8508 | NM_001318860.1 | 52.8404171 | -7.3449887 | 1.40060154 | -5.2441672 |
| 8509 | NM_203459.2 | 104.454923 | -8.3279916 | 1.58808177 | -5.2440572 |
| 8510 | NM_001014446.2 | 61.7786817 | -7.571784 | 1.44395401 | -5.2437847 |
| 8511 | NM_001099696.2 | 59.8590358 | -7.5280813 | 1.43564588 | -5.2436895 |
| 8512 | NM_016586.2 | 40.0176443 | -6.9432012 | 1.32422183 | -5.2432312 |
| 8513 | NM_031212.3 | 66.5954605 | -6.7086417 | 1.27948748 | -5.2432257 |

| | | | | | |
|---|---|---|---|---|---|
| 8514 | NM_004886.3 | 38.3263354 | -6.8805182 | 1.31233147 | -5.2429728 |
| 8515 | NR_038222.1 | 16.5700992 | 3.77636118 | 0.72028803 | 5.24284872 |
| 8516 | NM_001278651.1 | 42.1960218 | -7.0199497 | 1.3389923 | -5.2427111 |
| 8517 | NM_002971.5 | 131.46994 | -4.9856466 | 0.9510718 | -5.2421349 |
| 8518 | NM_032507.3 | 50.7099957 | -7.2857081 | 1.38990247 | -5.2418844 |
| 8519 | NM_015986.3 | 34.0583292 | -6.7089419 | 1.28009273 | -5.2409812 |
| 8520 | NM_001039577.3 | 39.584339 | -6.9336627 | 1.32302738 | -5.2407553 |
| 8521 | NM_006038.3 | 58.1696294 | -6.5165824 | 1.2434998 | -5.2405174 |
| 8522 | NM_006214.3 | 38.8001887 | -6.9005391 | 1.31685143 | -5.2401804 |
| 8523 | NM_004898.3 | 60.4277814 | -6.5699082 | 1.25385768 | -5.239756 |
| 8524 | NM_024955.5 | 78.706741 | -7.9223292 | 1.51205296 | -5.2394522 |
| 8525 | NM_001007097.2 | 207.917887 | -5.3702825 | 1.02499197 | -5.239341 |
| 8526 | NM_016148.3 | 325.31363 | -5.4278209 | 1.03605308 | -5.2389409 |
| 8527 | NM_016464.4 | 32.0932715 | -6.6265901 | 1.26491816 | -5.2387501 |
| 8528 | NM_022114.3 | 99.225925 | -8.2555413 | 1.57592838 | -5.2385257 |
| 8529 | NM_020242.2 | 40.9460921 | -6.9752119 | 1.33161224 | -5.2381704 |
| 8530 | NM_005966.3 | 55.76069 | -7.4202046 | 1.41663349 | -5.2379141 |
| 8531 | NM_020346.2 | 44.5283794 | -7.0971982 | 1.35506087 | -5.2375493 |
| 8532 | NM_025125.3 | 44.8141187 | -7.1055061 | 1.35664236 | -5.2375676 |
| 8533 | NM_020437.4 | 50.5414426 | -7.2812968 | 1.39022372 | -5.2375001 |
| 8534 | NM_178140.3 | 35.0191289 | -6.7493252 | 1.28864883 | -5.2375209 |
| 8535 | NM_001242840.1 | 55.2349959 | -7.409933 | 1.41479711 | -5.2374527 |
| 8536 | NM_018341.2 | 35.0327396 | -6.7496608 | 1.28874337 | -5.2373971 |
| 8537 | NM_006759.3 | 62.0251831 | -7.5777675 | 1.44707133 | -5.2366233 |
| 8538 | NM_018977.3 | 43.6060845 | -7.0705913 | 1.35036148 | -5.236073 |
| 8539 | NM_080476.4 | 72.7238454 | -6.0167017 | 1.14910052 | -5.2360099 |
| 8540 | NR_047657.1 | 53.4353739 | -7.3586131 | 1.40538216 | -5.2360229 |
| 8541 | NM_033212.3 | 31.8690691 | -6.6221519 | 1.26492426 | -5.2352161 |
| 8542 | NM_001083613.1 | 43.0321015 | -7.049186 | 1.3465083 | -5.2351597 |
| 8543 | NM_001122636.1 | 60.6474587 | -7.5443034 | 1.44108711 | -5.2351474 |
| 8544 | NM_001080488.1 | 66.904685 | -7.6866746 | 1.46835834 | -5.2348765 |
| 8545 | NM_145115.2 | 60.1376832 | -6.5628404 | 1.25376307 | -5.2345141 |
| 8546 | NM_001319193.1 | 53.3129662 | -6.3853724 | 1.21994311 | -5.2341559 |
| 8547 | NM_020931.3 | 44.7115718 | -7.1017117 | 1.35693969 | -5.2336237 |
| 8548 | NM_138356.2 | 57.6080404 | -7.4703483 | 1.42753523 | -5.2330395 |
| 8549 | NM_020798.2 | 73.0268006 | -6.8379116 | 1.3067751 | -5.2326614 |
| 8550 | NM_001134337.2 | 75.5914741 | -6.8913943 | 1.31700429 | -5.2326286 |
| 8551 | NM_005692.4 | 41.6521213 | -7.0015447 | 1.33813773 | -5.232305 |
| 8552 | NM_001035258.1 | 62.3056061 | -7.5836366 | 1.44943943 | -5.2321169 |
| 8553 | NM_001304749.1 | 31.5639985 | -6.6010689 | 1.26164094 | -5.2321296 |
| 8554 | NM_018491.3 | 47.0200137 | -7.1789411 | 1.37231592 | -5.2312598 |
| 8555 | NM_130836.2 | 44.341659 | -7.0889045 | 1.35511993 | -5.2312008 |
| 8556 | NM_001079669.2 | 70.9164948 | -7.770699 | 1.48563851 | -5.2305449 |

| | | | | | |
|---|---|---|---|---|---|
| 8557 | NM_003983.5 | 72.0130308 | -7.7939804 | 1.49023761 | -5.2300253 |
| 8558 | NM_017914.3 | 48.0648 | -7.2093083 | 1.37849975 | -5.229822 |
| 8559 | NM_001001567.1 | 47.0325275 | -7.1767248 | 1.37228218 | -5.2297734 |
| 8560 | NM_002516.3 | 364.974158 | -5.9049626 | 1.12917901 | -5.22943 |
| 8561 | NM_032832.5 | 158.533514 | -5.1064432 | 0.9766966 | -5.2282799 |
| 8562 | NR_003263.2 | 29.0004333 | -6.4816974 | 1.23979959 | -5.2280203 |
| 8563 | NM_001204160.2 | 53.4010407 | -7.3599232 | 1.40780227 | -5.2279523 |
| 8564 | NM_002053.2 | 34.4699964 | -6.7350736 | 1.28829496 | -5.2278972 |
| 8565 | NM_001010867.3 | 48.5422896 | -7.2195671 | 1.38109225 | -5.2274329 |
| 8566 | NM_198066.3 | 75.4520272 | -3.3463739 | 0.64019384 | -5.2271261 |
| 8567 | NM_001352.4 | 35.2662624 | -6.7601682 | 1.29331826 | -5.226995 |
| 8568 | NM_001329631.1 | 44.9985679 | -7.1170096 | 1.36161706 | -5.2268804 |
| 8569 | NR_024403.1 | 33.2704932 | -6.6759284 | 1.27726403 | -5.2267411 |
| 8570 | NM_020362.4 | 217.316124 | -5.7535131 | 1.10080051 | -5.2266629 |
| 8571 | NM_005694.1 | 43.7256793 | -7.0727437 | 1.3532575 | -5.2264582 |
| 8572 | NM_005733.2 | 67.5659264 | -6.7258226 | 1.2868872 | -5.2264274 |
| 8573 | NM_173659.3 | 47.7139127 | -7.1977688 | 1.37736777 | -5.2257421 |
| 8574 | NM_001203245.2 | 44.4015914 | -7.0970748 | 1.35834214 | -5.2248065 |
| 8575 | NM_005480.3 | 40.1953916 | -6.9501143 | 1.33029289 | -5.2244993 |
| 8576 | NM_001017964.1 | 46.7592432 | -7.1684756 | 1.37214826 | -5.2242719 |
| 8577 | NM_001145797.1 | 51.4885981 | -7.3069719 | 1.39867599 | -5.2242063 |
| 8578 | NM_024833.2 | 33.9892819 | -6.7052248 | 1.28349324 | -5.2241995 |
| 8579 | NM_001142272.1 | 52.0181579 | -7.3233963 | 1.4019141 | -5.2238552 |
| 8580 | NM_015094.2 | 160.930965 | -5.2975054 | 1.01411976 | -5.2237473 |
| 8581 | NM_020695.3 | 193.947143 | -4.9407235 | 0.94583423 | -5.2236675 |
| 8582 | NM_001102368.2 | 41.8798975 | -7.0117984 | 1.34233787 | -5.2235719 |
| 8583 | NM_138443.3 | 45.1904334 | -7.1188366 | 1.36284675 | -5.2235049 |
| 8584 | NM_006851.2 | 51.7587806 | -7.3168359 | 1.4008483 | -5.2231465 |
| 8585 | NM_001142533.1 | 47.5243442 | -7.1899938 | 1.37669302 | -5.2226558 |
| 8586 | NM_014425.4 | 33.9898423 | -6.7094073 | 1.28485597 | -5.2219139 |
| 8587 | NM_021930.5 | 50.9627366 | -7.2960719 | 1.39723993 | -5.2217746 |
| 8588 | NM_014637.3 | 34.1381446 | -6.7151003 | 1.28622504 | -5.2207818 |
| 8589 | NM_001346413.1 | 38.8434127 | -6.9011936 | 1.32200151 | -5.2202615 |
| 8590 | NR_034181.1 | 40.4013257 | -6.9571163 | 1.33292936 | -5.2194186 |
| 8591 | NM_020309.3 | 316.967896 | -8.0508789 | 1.54275211 | -5.2185175 |
| 8592 | NM_001077366.1 | 56.1602224 | -7.4349932 | 1.4247824 | -5.2183359 |
| 8593 | NR_144491.1 | 13.4691157 | 8.65702655 | 1.65920841 | 5.2175643 |
| 8594 | NM_001319990.1 | 40.5357419 | -6.9678661 | 1.33558975 | -5.2170707 |
| 8595 | NM_002553.3 | 45.8752445 | -7.1397745 | 1.36856416 | -5.2169819 |
| 8596 | NM_004219.3 | 39.7641408 | -6.9354715 | 1.3294225 | -5.2169055 |
| 8597 | NM_014000.2 | 33.1880949 | -6.6808915 | 1.28063103 | -5.2168746 |
| 8598 | NM_015166.3 | 69.0445241 | -7.7310213 | 1.48210315 | -5.2162505 |
| 8599 | NR_026580.1 | 42.8940866 | -7.0457956 | 1.35077716 | -5.2161051 |

| | | | | | |
|---|---|---|---|---|---|
| 8600 | NM_032836.2 | 75.5157288 | -6.07335 | 1.16435529 | -5.2160625 |
| 8601 | NM_003054.4 | 32.6532198 | -6.6519168 | 1.27544289 | -5.2153781 |
| 8602 | NM_001083908.1 | 32.5372514 | -6.6399711 | 1.2732637 | -5.2149221 |
| 8603 | NM_001012241.1 | 413.511431 | -6.0329806 | 1.15689944 | -5.2147839 |
| 8604 | NM_002399.3 | 55.1626267 | -7.4070148 | 1.42043192 | -5.2146215 |
| 8605 | NM_001042469.1 | 34.5064671 | -6.734215 | 1.29143694 | -5.2145132 |
| 8606 | NM_032687.3 | 36.9555604 | -6.8298681 | 1.30978394 | -5.2144998 |
| 8607 | NM_014079.3 | 94.8483947 | 3.16264433 | 0.60655081 | 5.21414571 |
| 8608 | NM_016008.3 | 32.0720219 | -6.6314053 | 1.27190602 | -5.2137542 |
| 8609 | NM_018231.2 | 97.6751352 | -3.876513 | 0.74353231 | -5.2136443 |
| 8610 | NR_073000.1 | 41.0025498 | -6.9821699 | 1.33928442 | -5.2133586 |
| 8611 | NM_002840.4 | 70.3061104 | -7.7552266 | 1.48758609 | -5.213296 |
| 8612 | NM_019110.4 | 32.2953046 | -6.63445 | 1.27288206 | -5.2121483 |
| 8613 | NM_138353.3 | 152.010145 | -5.7304837 | 1.09951189 | -5.2118434 |
| 8614 | NM_001002755.2 | 47.8297371 | -7.2016943 | 1.3818744 | -5.2115405 |
| 8615 | NM_006223.3 | 42.2574521 | -7.0215877 | 1.34738448 | -5.2112725 |
| 8616 | NM_001083893.1 | 42.4207401 | -7.0279064 | 1.34869869 | -5.2108795 |
| 8617 | NM_006531.4 | 32.7183763 | -6.6530818 | 1.27687889 | -5.2104251 |
| 8618 | NR_039988.1 | 94.7683334 | 3.0109471 | 0.57789355 | 5.21021057 |
| 8619 | NM_001744.5 | 96.9898886 | -8.2201508 | 1.57784715 | -5.2097257 |
| 8620 | NM_001300750.1 | 35.8360766 | -6.7898533 | 1.30339974 | -5.2093407 |
| 8621 | NM_001005619.1 | 60.8173095 | -7.5491022 | 1.44916855 | -5.2092645 |
| 8622 | NM_001162536.2 | 52.5893083 | -7.3356161 | 1.40819875 | -5.2092193 |
| 8623 | NM_001142444.1 | 35.8502651 | -6.7880361 | 1.3031005 | -5.2091424 |
| 8624 | NM_015997.3 | 31.3685457 | -6.592679 | 1.26566254 | -5.2088759 |
| 8625 | NM_001330740.1 | 109.742429 | -7.4295189 | 1.42644038 | -5.2084328 |
| 8626 | NM_172016.2 | 31.8721815 | -6.6185798 | 1.27085299 | -5.2079823 |
| 8627 | NM_001031725.5 | 31.738645 | -6.6132034 | 1.26990406 | -5.2076402 |
| 8628 | NM_007014.4 | 43.4868828 | -7.0652163 | 1.35671515 | -5.20759 |
| 8629 | NM_198458.2 | 36.3777404 | -6.8062002 | 1.30707879 | -5.2071843 |
| 8630 | NM_022486.4 | 43.2870061 | -7.0558456 | 1.35506035 | -5.2070342 |
| 8631 | NM_020724.1 | 59.733261 | -5.1546704 | 0.9900053 | -5.2067099 |
| 8632 | NM_001283020.1 | 43.9392209 | -7.0751946 | 1.35891163 | -5.2065156 |
| 8633 | NR_104184.1 | 54.8695251 | -2.8377562 | 0.54506379 | -5.2062828 |
| 8634 | NM_001139498.1 | 71.2230876 | -7.7755962 | 1.49353526 | -5.2061685 |
| 8635 | NM_152729.2 | 49.3069018 | -7.2482388 | 1.3922591 | -5.206099 |
| 8636 | NM_017629.3 | 147.041789 | -4.291785 | 0.82439561 | -5.2059775 |
| 8637 | NM_194298.2 | 73.2598184 | -7.8189625 | 1.50212798 | -5.2052572 |
| 8638 | NM_001083602.1 | 56.6176867 | -7.4456606 | 1.43042588 | -5.2052055 |
| 8639 | NM_001303.3 | 37.6752177 | -6.8589048 | 1.3178193 | -5.2047385 |
| 8640 | NM_001198569.1 | 38.9799041 | -6.9111843 | 1.32804922 | -5.2040122 |
| 8641 | NM_001267046.1 | 36.4605311 | -6.8074042 | 1.3084827 | -5.2025175 |
| 8642 | NM_005230.3 | 42.0957635 | -7.0210972 | 1.34955724 | -5.202519 |

| | | | | | |
|---|---|---|---|---|---|
| 8643 | NM_001282198.1 | 30.1722926 | -6.5343671 | 1.25607511 | -5.2022105 |
| 8644 | NM_003596.3 | 71.0100122 | -5.2565809 | 1.01048722 | -5.2020261 |
| 8645 | NM_014064.3 | 49.2804907 | -7.2435433 | 1.39245515 | -5.201994 |
| 8646 | NM_002910.5 | 33.7058688 | -6.699181 | 1.2878921 | -5.2016633 |
| 8647 | NM_001163213.1 | 41.9620836 | -7.0107648 | 1.34780483 | -5.2016172 |
| 8648 | NM_001291447.1 | 43.8851655 | -7.0787417 | 1.36091283 | -5.2014659 |
| 8649 | NM_006849.2 | 78.0259418 | -5.9868201 | 1.15112333 | -5.2008503 |
| 8650 | NM_001282680.1 | 45.5904761 | -7.1372145 | 1.37248658 | -5.2002071 |
| 8651 | NM_001330112.1 | 40.2270487 | -6.9531686 | 1.33709054 | -5.2002227 |
| 8652 | NM_001207008.1 | 131.308363 | -4.9856201 | 0.95874309 | -5.2001628 |
| 8653 | NR_045072.1 | 72.4840188 | -7.8011524 | 1.50023556 | -5.1999517 |
| 8654 | NM_001039673.2 | 83.1535614 | -8.0000073 | 1.53868053 | -5.1992647 |
| 8655 | NM_001294340.1 | 29.6754093 | -6.5095294 | 1.25202669 | -5.1991938 |
| 8656 | NM_001320846.1 | 57.0465911 | -7.4564461 | 1.43429311 | -5.1986905 |
| 8657 | NM_004454.2 | 75.3076684 | -6.8819816 | 1.32396901 | -5.197993 |
| 8658 | NM_014903.5 | 122.365918 | -7.5871491 | 1.45970354 | -5.1977329 |
| 8659 | NM_207644.2 | 39.2301634 | -6.9179428 | 1.33101231 | -5.1975048 |
| 8660 | NR_102336.1 | 71.7897932 | 9.43871808 | 1.81605585 | 5.19737215 |
| 8661 | NR_102759.1 | 34.7560552 | -6.740146 | 1.2968966 | -5.1971344 |
| 8662 | NM_017948.5 | 61.990699 | -7.5754559 | 1.457731 | -5.1967447 |
| 8663 | NM_004316.3 | 39.9936585 | -6.9393682 | 1.33552677 | -5.1959784 |
| 8664 | NM_022728.3 | 447.165095 | -10.426757 | 2.00670129 | -5.1959686 |
| 8665 | NM_015260.3 | 43.4992545 | -7.0680559 | 1.36030574 | -5.1959318 |
| 8666 | NM_203288.1 | 48.2352786 | -7.213006 | 1.38824035 | -5.1957905 |
| 8667 | NM_020733.1 | 252.51027 | -4.7065484 | 0.90592684 | -5.1952853 |
| 8668 | NM_001003897.1 | 39.0413485 | -6.9077298 | 1.32971248 | -5.1949049 |
| 8669 | NM_030797.3 | 223.96997 | -8.4649257 | 1.62972666 | -5.194077 |
| 8670 | NM_012232.5 | 439.473447 | -6.5416018 | 1.25972969 | -5.1928614 |
| 8671 | NM_012213.2 | 33.9654455 | -6.71166 | 1.29257986 | -5.1924528 |
| 8672 | NM_001258374.1 | 45.9303436 | -7.145626 | 1.37627494 | -5.1920048 |
| 8673 | NM_001173984.2 | 61.4379234 | -7.5625381 | 1.45658459 | -5.1919663 |
| 8674 | NM_003729.3 | 154.389129 | -2.2342376 | 0.43035381 | -5.1916297 |
| 8675 | NM_001006639.1 | 42.7381091 | -7.0362991 | 1.35555994 | -5.1906956 |
| 8676 | NR_027948.2 | 81.6834556 | -7.9716256 | 1.53576885 | -5.1906415 |
| 8677 | NM_001330474.1 | 49.6291114 | -7.2544414 | 1.39761703 | -5.1905788 |
| 8678 | NM_014751.5 | 43.2177616 | -7.0587138 | 1.35996901 | -5.190349 |
| 8679 | NM_032620.3 | 52.8684884 | -7.3442998 | 1.41526928 | -5.1893303 |
| 8680 | NM_001278677.1 | 41.8740203 | -7.0115265 | 1.3511505 | -5.1893009 |
| 8681 | NM_025194.2 | 63.9685444 | -6.6537203 | 1.28221586 | -5.1892356 |
| 8682 | NM_001198800.2 | 41.3374498 | -6.9904559 | 1.34711564 | -5.1892025 |
| 8683 | NM_002382.4 | 49.1842892 | -7.2434977 | 1.39596136 | -5.1888955 |
| 8684 | NR_125930.1 | 20.8415851 | 3.96904043 | 0.76493036 | 5.18876049 |
| 8685 | NM_003496.3 | 194.550318 | -5.7067213 | 1.0998666 | -5.1885577 |

| | | | | | |
|---|---|---|---|---|---|
| 8686 | NR_037861.1 | 32.4414662 | -6.6454809 | 1.28098292 | -5.1877982 |
| 8687 | NM_001100818.1 | 54.8350216 | -7.3989796 | 1.42623404 | -5.1877738 |
| 8688 | NM_001184700.1 | 15.8159037 | 8.88518045 | 1.71276811 | 5.18761436 |
| 8689 | NM_001201372.1 | 55.1989461 | -7.4072713 | 1.42791246 | -5.1874828 |
| 8690 | NM_002166.4 | 321.821782 | -6.3454815 | 1.22323043 | -5.1874785 |
| 8691 | NR_037718.1 | 56.4636235 | -7.4433053 | 1.43496456 | -5.1871004 |
| 8692 | NM_019028.2 | 39.0819818 | -6.907169 | 1.33165076 | -5.1869223 |
| 8693 | NM_001706.4 | 30.752561 | -6.5613656 | 1.26504259 | -5.1866757 |
| 8694 | NM_003675.3 | 43.6038131 | -7.0663058 | 1.36240308 | -5.1866484 |
| 8695 | NM_016065.3 | 214.330846 | -2.625837 | 0.5063027 | -5.1862985 |
| 8696 | NM_003668.3 | 33.5212842 | -6.6890371 | 1.2898368 | -5.1859561 |
| 8697 | NM_001166037.1 | 51.5369896 | -7.3089072 | 1.40945105 | -5.185641 |
| 8698 | NM_001242487.1 | 66.6845542 | -7.6780517 | 1.48068897 | -5.1854588 |
| 8699 | NM_001018109.2 | 31.0661922 | -6.5844265 | 1.26984022 | -5.1852402 |
| 8700 | NM_025045.5 | 29.1566373 | -6.493302 | 1.2522966 | -5.1851151 |
| 8701 | NM_014580.4 | 47.5116908 | -7.1933061 | 1.3873095 | -5.1850767 |
| 8702 | NM_005116.5 | 361.314984 | -5.367279 | 1.03519249 | -5.1848125 |
| 8703 | NM_145893.2 | 57.3939848 | -7.4629913 | 1.43941006 | -5.184757 |
| 8704 | NM_001321003.1 | 43.0086231 | -4.1812576 | 0.80648849 | -5.1845223 |
| 8705 | NM_001145939.1 | 72.4893478 | -7.8006726 | 1.50475857 | -5.1840028 |
| 8706 | NM_000229.1 | 30.2148447 | -6.5442914 | 1.26248002 | -5.1836792 |
| 8707 | NM_015416.4 | 46.9284153 | -7.1716413 | 1.38350341 | -5.1836817 |
| 8708 | NM_001199847.1 | 53.581554 | -4.9873207 | 0.96222063 | -5.1831363 |
| 8709 | NM_023943.3 | 62.4667031 | -7.584314 | 1.46328992 | -5.1830563 |
| 8710 | NM_001164160.1 | 57.3810248 | -7.4674255 | 1.44080391 | -5.1828187 |
| 8711 | NM_012407.3 | 49.4153575 | -7.2499372 | 1.39885648 | -5.1827599 |
| 8712 | NR_040072.1 | 31.0918643 | -6.5835298 | 1.270495 | -5.181862 |
| 8713 | NM_001199323.1 | 23.9661879 | 9.50691491 | 1.83486749 | 5.18125421 |
| 8714 | NM_003450.2 | 30.6996179 | -6.5593046 | 1.26600133 | -5.1811198 |
| 8715 | NM_152293.2 | 172.224167 | -3.2853908 | 0.63411734 | -5.1810455 |
| 8716 | NM_013248.2 | 42.830507 | -7.0411469 | 1.35916204 | -5.1805058 |
| 8717 | NM_007059.3 | 40.3605991 | -6.9551496 | 1.34262507 | -5.180262 |
| 8718 | NM_001098208.1 | 65.2552911 | -7.649613 | 1.47675938 | -5.1799996 |
| 8719 | NM_001564.3 | 38.9927037 | -6.9038503 | 1.33280712 | -5.179932 |
| 8720 | NM_001330580.1 | 36.45783 | -6.8097043 | 1.31478454 | -5.179331 |
| 8721 | NM_001164387.1 | 43.836357 | -7.0745899 | 1.36603963 | -5.1789053 |
| 8722 | NM_001277842.1 | 51.2453656 | -7.3000957 | 1.40976702 | -5.1782284 |
| 8723 | NM_001163788.2 | 67.0347996 | -7.6880671 | 1.48474003 | -5.178056 |
| 8724 | NM_058182.4 | 46.7411855 | -3.9987218 | 0.77229656 | -5.1777025 |
| 8725 | NM_001145373.2 | 38.1298943 | -6.873558 | 1.32761303 | -5.1773807 |
| 8726 | NM_000712.3 | 62.6494553 | -7.5910969 | 1.46640202 | -5.176682 |
| 8727 | NM_025072.6 | 80.961739 | -6.1806036 | 1.19398823 | -5.176436 |
| 8728 | NM_001009185.2 | 41.9312583 | -7.0091093 | 1.35413517 | -5.176078 |

| | | | | | |
|---|---|---|---|---|---|
| 8729 | NM_017816.2 | 36.9002522 | -6.8263887 | 1.31894949 | -5.1756256 |
| 8730 | NM_207360.2 | 104.3897 | 2.56948246 | 0.49653713 | 5.17480426 |
| 8731 | NM_017883.5 | 48.4298262 | -7.2226758 | 1.39580268 | -5.174568 |
| 8732 | NM_001330731.1 | 33.5920289 | -6.6916154 | 1.29323632 | -5.1743176 |
| 8733 | NM_148920.2 | 50.7561008 | -7.2873789 | 1.40861547 | -5.1734338 |
| 8734 | NM_022492.4 | 31.3921451 | -6.5920654 | 1.27431987 | -5.1730068 |
| 8735 | NM_003486.6 | 889.938112 | -5.9871856 | 1.15757647 | -5.1721728 |
| 8736 | NM_001034852.2 | 42.8537623 | -7.0417291 | 1.36148964 | -5.1720769 |
| 8737 | NM_003192.2 | 82.4929283 | -4.9844901 | 0.96375662 | -5.1719387 |
| 8738 | NM_001193495.1 | 265.700467 | -4.9752706 | 0.96208989 | -5.1713157 |
| 8739 | NM_001040118.2 | 31.9240583 | -6.6247313 | 1.28107834 | -5.1712148 |
| 8740 | NM_203437.3 | 47.5859328 | -7.1932879 | 1.39108062 | -5.1710072 |
| 8741 | NM_001007527.1 | 42.1459498 | -7.0178625 | 1.35716862 | -5.1709584 |
| 8742 | NM_153688.3 | 44.1153171 | -7.0871709 | 1.37073728 | -5.170335 |
| 8743 | NM_002738.6 | 80.9164672 | -6.9912361 | 1.35231863 | -5.1698142 |
| 8744 | NM_006499.4 | 37.3508641 | -6.8401127 | 1.32317335 | -5.1694759 |
| 8745 | NR_120685.1 | 43.5298109 | -7.0659607 | 1.36701514 | -5.1688972 |
| 8746 | NM_016373.3 | 32.993239 | -6.664156 | 1.2893348 | -5.1686777 |
| 8747 | NM_018935.3 | 42.75019 | -7.0380048 | 1.36166585 | -5.1686724 |
| 8748 | NM_001206747.1 | 33.0362919 | -6.6685996 | 1.29031252 | -5.1682049 |
| 8749 | NM_080416.3 | 39.186937 | -6.9133745 | 1.33791703 | -5.167267 |
| 8750 | NM_032928.3 | 44.1408103 | -7.0863486 | 1.3714895 | -5.1668997 |
| 8751 | NM_001160258.1 | 64.3357891 | -7.6300804 | 1.47691766 | -5.1662192 |
| 8752 | NM_001243143.1 | 43.3894385 | -7.0565298 | 1.36594793 | -5.1660313 |
| 8753 | NM_017691.4 | 50.1686658 | -7.2714175 | 1.40753661 | -5.1660592 |
| 8754 | NM_018340.2 | 30.755037 | -6.5679701 | 1.27137454 | -5.1660387 |
| 8755 | NM_005792.2 | 44.3077635 | -7.0909556 | 1.37266927 | -5.1658151 |
| 8756 | NM_199363.2 | 45.5308067 | -7.1307674 | 1.38042823 | -5.1656198 |
| 8757 | NM_001135556.1 | 77.3554076 | -7.8954637 | 1.5285536 | -5.1653169 |
| 8758 | NM_012063.3 | 40.7600922 | -6.9747557 | 1.35041454 | -5.1648997 |
| 8759 | NM_001286836.1 | 37.783565 | -6.8618059 | 1.32866282 | -5.1644449 |
| 8760 | NM_001024956.2 | 30.6773001 | -6.5566403 | 1.26958672 | -5.1643894 |
| 8761 | NM_017622.2 | 36.9446408 | -6.82706 | 1.32194501 | -5.1644054 |
| 8762 | NM_016532.3 | 73.1272521 | -6.8390239 | 1.32432232 | -5.1641687 |
| 8763 | NM_003599.3 | 31.3581722 | -6.5948258 | 1.27708733 | -5.1639584 |
| 8764 | NM_001130012.2 | 45.3610759 | -7.1263214 | 1.38017572 | -5.1633435 |
| 8765 | NR_015370.2 | 75.349187 | -6.8882404 | 1.33420202 | -5.1628166 |
| 8766 | NM_145689.2 | 48.225967 | -7.2173113 | 1.39804691 | -5.1624243 |
| 8767 | NM_198850.3 | 32.7137645 | -6.6524526 | 1.28869919 | -5.1621454 |
| 8768 | NM_016329.3 | 32.3636056 | -6.6364007 | 1.28559775 | -5.1621129 |
| 8769 | NM_014872.2 | 134.843815 | -5.028732 | 0.97420869 | -5.1618632 |
| 8770 | NM_001330418.1 | 80.8691338 | -5.6244256 | 1.08984185 | -5.1607723 |
| 8771 | NM_014782.6 | 42.0663436 | -7.0185998 | 1.35998559 | -5.1607898 |

| | | | | | |
|---|---|---|---|---|---|
| 8772 | NM_002559.3 | 92.1370332 | -7.1812387 | 1.39151329 | -5.1607403 |
| 8773 | NM_004307.1 | 48.657462 | -7.229552 | 1.40093313 | -5.1605261 |
| 8774 | NR_110761.1 | 48.5037304 | 3.36616092 | 0.65230665 | 5.16039642 |
| 8775 | NM_005644.3 | 38.6040327 | -6.8916531 | 1.33553718 | -5.1602105 |
| 8776 | NM_001425.2 | 51.80896 | -7.3187233 | 1.41835467 | -5.1600093 |
| 8777 | NM_001042463.2 | 32.4451533 | -6.6389112 | 1.28667668 | -5.1597354 |
| 8778 | NM_182565.3 | 108.189934 | -5.0981171 | 0.98806572 | -5.1596943 |
| 8779 | NM_006347.3 | 43.0555328 | -7.0498518 | 1.36635371 | -5.1596097 |
| 8780 | NM_003762.4 | 40.1474666 | -6.947763 | 1.3467454 | -5.1589284 |
| 8781 | NM_032271.2 | 341.696114 | -6.1908297 | 1.20006324 | -5.1587529 |
| 8782 | NM_012113.2 | 34.636497 | -6.7417264 | 1.30685888 | -5.1587257 |
| 8783 | NM_003516.2 | 28.9533934 | -6.4790781 | 1.25596377 | -5.1586505 |
| 8784 | NM_001253382.1 | 77.0853102 | -7.8903316 | 1.52957844 | -5.1585008 |
| 8785 | NM_152285.3 | 33.6333363 | -6.6978196 | 1.2984167 | -5.1584515 |
| 8786 | NM_001256335.1 | 39.7435429 | -6.9349522 | 1.34444722 | -5.1582182 |
| 8787 | NM_001320193.1 | 53.0037616 | -7.3501817 | 1.42507481 | -5.1577515 |
| 8788 | NM_001011658.3 | 37.9133654 | -6.8651103 | 1.3310465 | -5.1576788 |
| 8789 | NM_001122670.1 | 33.0978178 | -6.6709541 | 1.29339838 | -5.1576948 |
| 8790 | NM_152230.4 | 40.7168662 | -6.9665269 | 1.35074563 | -5.1575417 |
| 8791 | NM_002941.3 | 106.065298 | -6.0389971 | 1.17108119 | -5.1567707 |
| 8792 | NM_001193268.1 | 49.7648569 | -7.2578731 | 1.40746171 | -5.1567109 |
| 8793 | NM_145043.2 | 31.9578566 | -6.6175903 | 1.28337035 | -5.1564151 |
| 8794 | NM_015163.5 | 64.2337397 | -7.6259994 | 1.47894556 | -5.156376 |
| 8795 | NM_170750.2 | 47.3462929 | -7.1870261 | 1.39390245 | -5.1560467 |
| 8796 | NR_037882.1 | 32.7431725 | -6.6578486 | 1.29135321 | -5.1557146 |
| 8797 | NM_004503.3 | 58.865982 | -7.5028067 | 1.45525474 | -5.1556655 |
| 8798 | NM_024330.1 | 40.0729298 | -6.9449603 | 1.34708702 | -5.1555395 |
| 8799 | NM_014708.4 | 58.3569286 | -5.1203709 | 0.99328199 | -5.1550023 |
| 8800 | NM_152581.3 | 36.0080151 | -6.7919547 | 1.31761533 | -5.1547326 |
| 8801 | NM_005901.5 | 102.556314 | -3.5309879 | 0.6850441 | -5.1543951 |
| 8802 | NM_001145450.1 | 32.5021984 | -6.6504518 | 1.2903149 | -5.1541309 |
| 8803 | NR_144366.1 | 36.4086125 | -6.8034625 | 1.32001267 | -5.1540888 |
| 8804 | NM_005706.3 | 46.9031244 | -7.1712227 | 1.39146492 | -5.1537215 |
| 8805 | NM_199245.2 | 29.245744 | -6.4945731 | 1.26023176 | -5.1534752 |
| 8806 | NM_153271.1 | 28.9776651 | -6.4835308 | 1.25811814 | -5.1533562 |
| 8807 | NM_002521.2 | 63.1897726 | -7.6045062 | 1.47590515 | -5.1524355 |
| 8808 | NM_005868.5 | 37.4974559 | -6.8514071 | 1.32973979 | -5.152442 |
| 8809 | NM_001545.2 | 41.2168555 | -6.9868644 | 1.35614046 | -5.1520212 |
| 8810 | NM_001009921.2 | 91.4479268 | -8.1359398 | 1.57919653 | -5.1519489 |
| 8811 | NM_001329222.1 | 286.024507 | -3.8979208 | 0.75661307 | -5.1518021 |
| 8812 | NM_004206.3 | 34.9012679 | -6.7457029 | 1.30948119 | -5.1514317 |
| 8813 | NM_002846.3 | 78.5964776 | -7.9162452 | 1.53703236 | -5.1503439 |
| 8814 | NM_022872.2 | 37.2551497 | -6.8449017 | 1.32907036 | -5.1501425 |

| | | | | | |
|---|---|---|---|---|---|
| 8815 | NM_016301.3 | 38.9240204 | -6.9032831 | 1.3404124 | -5.1501188 |
| 8816 | NM_024828.3 | 33.1291577 | -6.6709947 | 1.29550001 | -5.149359 |
| 8817 | NM_001136201.1 | 41.4031847 | -6.9949378 | 1.35855089 | -5.1488228 |
| 8818 | NM_018204.4 | 77.9930031 | -7.9087139 | 1.53607472 | -5.1486518 |
| 8819 | NM_001039887.2 | 29.156077 | -6.4923578 | 1.26107046 | -5.1482911 |
| 8820 | NM_020438.4 | 49.5019032 | -7.2501961 | 1.40830753 | -5.1481626 |
| 8821 | NM_001099642.1 | 35.6112953 | -6.7752265 | 1.31607202 | -5.1480666 |
| 8822 | NM_012243.2 | 42.1884467 | -7.0257291 | 1.36476766 | -5.1479305 |
| 8823 | NM_001010935.2 | 32.0040194 | -6.6285367 | 1.2878736 | -5.1468845 |
| 8824 | NM_001037806.3 | 207.921887 | -5.7933282 | 1.12562112 | -5.1467835 |
| 8825 | NM_001321072.1 | 44.2636819 | -7.0927211 | 1.3781785 | -5.1464459 |
| 8826 | NM_022126.3 | 32.9901411 | -6.6630457 | 1.29470487 | -5.1463819 |
| 8827 | NM_001321328.1 | 29.6456032 | -6.5077559 | 1.26468289 | -5.145761 |
| 8828 | NM_001014834.2 | 29.0384776 | -6.4856788 | 1.26044516 | -5.1455461 |
| 8829 | NM_001167607.1 | 30.7841208 | 9.85711619 | 1.91572269 | 5.14537737 |
| 8830 | NM_001001331.2 | 108.037434 | -8.3783922 | 1.62837657 | -5.1452424 |
| 8831 | NM_001304347.1 | 57.760488 | -7.4751702 | 1.45285235 | -5.1451686 |
| 8832 | NM_001017979.2 | 39.7904396 | -6.9353094 | 1.3480169 | -5.1448238 |
| 8833 | NM_012381.3 | 33.0321751 | -6.6634848 | 1.29531125 | -5.1443117 |
| 8834 | NM_001012978.2 | 64.9919438 | -7.6442214 | 1.48602891 | -5.1440597 |
| 8835 | NM_020137.4 | 322.710338 | -6.1109503 | 1.18803166 | -5.1437605 |
| 8836 | NM_002395.5 | 44.966836 | -7.1134754 | 1.38305614 | -5.143302 |
| 8837 | NM_004792.2 | 202.41431 | -4.8959266 | 0.95195594 | -5.143018 |
| 8838 | NM_152551.3 | 57.5727658 | -4.3614858 | 0.84808651 | -5.1427369 |
| 8839 | NM_001145125.1 | 61.2583988 | -7.5584029 | 1.46975138 | -5.1426405 |
| 8840 | NM_173647.3 | 58.2932251 | -6.5104185 | 1.26606048 | -5.142265 |
| 8841 | NM_016352.3 | 36.5714773 | -6.8166519 | 1.32566791 | -5.142051 |
| 8842 | NM_001330566.1 | 58.4467273 | -7.4894484 | 1.45672765 | -5.1412825 |
| 8843 | NR_111971.1 | 149.203479 | -5.3737787 | 1.04523451 | -5.1412182 |
| 8844 | NM_004650.2 | 31.9348378 | -6.6178184 | 1.2873495 | -5.140654 |
| 8845 | NM_006834.4 | 41.1871283 | -6.9860899 | 1.35902228 | -5.1405264 |
| 8846 | NM_000538.3 | 33.5398442 | -6.6883796 | 1.30111743 | -5.1404888 |
| 8847 | NR_027237.1 | 51.1583346 | -7.2969686 | 1.41951811 | -5.1404548 |
| 8848 | NR_073415.2 | 57.0638679 | 3.53825405 | 0.68831612 | 5.14044921 |
| 8849 | NM_001127453.1 | 50.1913778 | -7.2704944 | 1.41455116 | -5.1397889 |
| 8850 | NM_145911.2 | 32.0797234 | -6.6232806 | 1.28875397 | -5.1392902 |
| 8851 | NM_001010911.2 | 38.0501409 | -6.8714177 | 1.33711213 | -5.1389989 |
| 8852 | NM_015394.4 | 59.4268476 | -5.0090443 | 0.97476481 | -5.1387209 |
| 8853 | NM_138284.1 | 40.0318953 | -6.9439762 | 1.35137609 | -5.1384483 |
| 8854 | NM_001284325.1 | 34.1308936 | -6.7212372 | 1.30809953 | -5.1381695 |
| 8855 | NM_001002002.2 | 36.067436 | -6.7929574 | 1.32212151 | -5.1379221 |
| 8856 | NM_001142282.1 | 47.0009316 | -7.1803224 | 1.39753318 | -5.1378547 |
| 8857 | NM_138740.3 | 44.0014384 | -7.0803064 | 1.37806461 | -5.1378624 |

| | | | | | |
|---|---|---|---|---|---|
| 8858 | NM_014733.4 | 110.692987 | -6.5282256 | 1.27064864 | -5.1377111 |
| 8859 | NM_018210.3 | 32.8273709 | -6.6630309 | 1.29693803 | -5.1375091 |
| 8860 | NM_001079843.2 | 47.9362316 | -3.8073464 | 0.74109423 | -5.137466 |
| 8861 | NM_001109903.1 | 59.7975751 | -7.5230827 | 1.46436058 | -5.1374523 |
| 8862 | NM_001283009.1 | 54.4289991 | -7.3879209 | 1.43815464 | -5.1370838 |
| 8863 | NM_032326.3 | 37.8080529 | -6.8616645 | 1.33570678 | -5.1371039 |
| 8864 | NM_001127213.1 | 60.9755197 | -7.5493392 | 1.46958512 | -5.1370548 |
| 8865 | NM_021983.4 | 69.2623072 | -7.7367988 | 1.50608738 | -5.1370185 |
| 8866 | NM_001002252.2 | 37.7344735 | -6.8559362 | 1.3347436 | -5.1365193 |
| 8867 | NM_024585.3 | 35.3964058 | -6.7680478 | 1.31772543 | -5.1361594 |
| 8868 | NM_139058.2 | 66.4676303 | -6.7035319 | 1.30526947 | -5.1357456 |
| 8869 | NM_001297573.1 | 42.8436098 | -7.0470135 | 1.37223532 | -5.1354264 |
| 8870 | NM_006907.3 | 43.795902 | -7.0752115 | 1.37774789 | -5.1353456 |
| 8871 | NM_001142273.1 | 109.053771 | -8.3914687 | 1.6342869 | -5.1346362 |
| 8872 | NM_007131.4 | 27.7456711 | -6.4205725 | 1.2504643 | -5.1345508 |
| 8873 | NM_014521.2 | 264.429098 | -5.6022446 | 1.09116777 | -5.1341735 |
| 8874 | NM_014805.3 | 274.507302 | -5.6585254 | 1.10219118 | -5.1338874 |
| 8875 | NR_045196.2 | 28.208565 | -6.4437942 | 1.25518802 | -5.1337283 |
| 8876 | NM_000899.4 | 60.4037759 | -7.5388024 | 1.46891438 | -5.1322273 |
| 8877 | NM_031940.3 | 40.1260919 | -6.944606 | 1.35316595 | -5.132117 |
| 8878 | NM_004540.3 | 90.0249325 | -7.1472044 | 1.39266345 | -5.1320399 |
| 8879 | NM_024522.2 | 285.264096 | -6.5728632 | 1.28080369 | -5.1318271 |
| 8880 | NM_080678.2 | 46.8044319 | -7.1691455 | 1.39701429 | -5.1317624 |
| 8881 | NM_001303433.1 | 47.0838677 | -7.1746421 | 1.39826933 | -5.1310873 |
| 8882 | NM_014930.2 | 45.8182181 | -7.136281 | 1.39092436 | -5.1306032 |
| 8883 | NM_172014.3 | 52.5169981 | 2.18182891 | 0.42528046 | 5.13032957 |
| 8884 | NM_139286.3 | 37.6650584 | -6.8553991 | 1.33630454 | -5.1301173 |
| 8885 | NM_018381.3 | 42.2968567 | -7.0279601 | 1.36999634 | -5.1299116 |
| 8886 | NM_006493.2 | 73.5487426 | -4.7607266 | 0.92807787 | -5.1296629 |
| 8887 | NM_001184777.1 | 54.7328211 | -7.3941506 | 1.44153677 | -5.1293527 |
| 8888 | NM_024913.4 | 37.3045572 | -6.8467936 | 1.33484066 | -5.1292966 |
| 8889 | NM_001308154.1 | 113.910891 | -8.4539383 | 1.64828353 | -5.1289345 |
| 8890 | NM_198449.2 | 32.4758307 | -6.6472926 | 1.29608209 | -5.128759 |
| 8891 | NM_003275.3 | 53.3313292 | -7.3590906 | 1.43488381 | -5.1287014 |
| 8892 | NM_005249.4 | 551.893084 | -10.73037 | 2.09249734 | -5.1280209 |
| 8893 | NM_130802.2 | 38.0162968 | -6.8677984 | 1.33931316 | -5.1278511 |
| 8894 | NM_004117.3 | 31.7443316 | -6.616821 | 1.29048304 | -5.1273987 |
| 8895 | NM_020374.3 | 53.3645894 | -7.3573245 | 1.43492331 | -5.1273294 |
| 8896 | NM_024690.2 | 33.5073327 | 6.13930443 | 1.19737958 | 5.12728337 |
| 8897 | NM_173686.2 | 30.4599388 | -6.5558957 | 1.27871606 | -5.1269362 |
| 8898 | NM_006439.4 | 38.5290966 | -6.8920725 | 1.34430261 | -5.1268758 |
| 8899 | NM_015127.4 | 34.42252 | -6.7301761 | 1.31272768 | -5.1268639 |
| 8900 | NM_052918.4 | 91.1939742 | -8.1318956 | 1.58633525 | -5.126215 |

| | | | | | |
|---|---|---|---|---|---|
| 8901 | NM_178564.3 | 295.265371 | -5.9871708 | 1.16799417 | -5.126028 |
| 8902 | NM_207303.4 | 91.0389575 | -7.1579325 | 1.39651412 | -5.1255712 |
| 8903 | NM_001083.3 | 54.5498704 | -7.3948008 | 1.44278744 | -5.125357 |
| 8904 | NM_014945.4 | 55.2021884 | -7.4080668 | 1.44545575 | -5.1250734 |
| 8905 | NM_001198941.1 | 40.754572 | -6.9752813 | 1.36102378 | -5.1250253 |
| 8906 | NM_021105.2 | 34.5620027 | -6.7387234 | 1.31490253 | -5.1248843 |
| 8907 | NM_032332.3 | 32.8119786 | -6.6532271 | 1.29858883 | -5.1234286 |
| 8908 | NM_001099271.1 | 33.5310313 | -6.6886385 | 1.30563267 | -5.1229099 |
| 8909 | NM_153456.3 | 66.2108745 | -6.7029196 | 1.30843318 | -5.1228597 |
| 8910 | NM_001301168.1 | 38.168129 | -6.8774964 | 1.34256355 | -5.1226599 |
| 8911 | NM_152272.4 | 73.1388664 | -6.8455939 | 1.33634854 | -5.1226112 |
| 8912 | NR_038862.1 | 55.9073234 | 3.37800467 | 0.659452 | 5.12244203 |
| 8913 | NM_032536.2 | 56.5722517 | -7.4427953 | 1.45299721 | -5.1223741 |
| 8914 | NM_152264.4 | 49.3768864 | -7.2474679 | 1.4148723 | -5.1223478 |
| 8915 | NM_006396.2 | 43.5181481 | -7.06394 | 1.37906221 | -5.122278 |
| 8916 | NM_004333.4 | 29.6325005 | -6.5132747 | 1.2715964 | -5.1221242 |
| 8917 | NM_001201461.1 | 50.9229589 | -7.2894026 | 1.42323934 | -5.1216984 |
| 8918 | NM_032343.2 | 90.8071915 | -8.1265464 | 1.5867185 | -5.1216057 |
| 8919 | NM_182776.2 | 115.075584 | -4.9762886 | 0.97168836 | -5.1212805 |
| 8920 | NM_005441.2 | 29.0253564 | -6.4788795 | 1.26516763 | -5.1209653 |
| 8921 | NM_198080.3 | 50.8955059 | -7.2905043 | 1.42367062 | -5.1209207 |
| 8922 | NM_017547.3 | 49.6270138 | -7.2536756 | 1.41668722 | -5.1201673 |
| 8923 | NR_052003.1 | 36.7632964 | -6.8253344 | 1.33303594 | -5.1201428 |
| 8924 | NM_001318069.1 | 41.7266634 | -7.0066701 | 1.36848212 | -5.1200304 |
| 8925 | NM_203500.1 | 34.4025102 | -6.7280372 | 1.31423019 | -5.119375 |
| 8926 | NM_001164277.1 | 26.9703468 | -6.3719173 | 1.24470076 | -5.1192363 |
| 8927 | NM_015960.2 | 30.6464248 | -6.5589885 | 1.28130937 | -5.1189734 |
| 8928 | NM_001160302.1 | 51.3974129 | -7.3047968 | 1.42706377 | -5.1187599 |
| 8929 | NM_213590.2 | 34.2121844 | -6.7129465 | 1.31146842 | -5.118649 |
| 8930 | NM_015686.2 | 48.3561883 | -7.217231 | 1.40999939 | -5.1186057 |
| 8931 | NM_013376.3 | 29.9575651 | -6.5275326 | 1.27528342 | -5.1184956 |
| 8932 | NM_001007793.2 | 68.6560662 | -4.9709529 | 0.97118423 | -5.1184448 |
| 8933 | NM_022163.3 | 38.7315912 | -6.8975036 | 1.34769035 | -5.1180181 |
| 8934 | NM_020809.3 | 31.1938559 | -6.587338 | 1.28716825 | -5.1176977 |
| 8935 | NM_001173982.1 | 45.2870909 | -7.1253538 | 1.39250255 | -5.1169413 |
| 8936 | NM_016173.4 | 45.7137975 | -7.1330187 | 1.3940464 | -5.1167728 |
| 8937 | NM_001323408.1 | 34.7509667 | -6.7449353 | 1.31822767 | -5.1166695 |
| 8938 | NM_004632.3 | 40.1827926 | -6.947447 | 1.3578485 | -5.1165112 |
| 8939 | NM_007157.3 | 60.5530999 | -4.4550889 | 0.87077348 | -5.1162432 |
| 8940 | NM_001288629.1 | 78.5400036 | -7.915474 | 1.5471693 | -5.1161007 |
| 8941 | NM_032501.3 | 36.3651156 | -6.803744 | 1.32988222 | -5.1160501 |
| 8942 | NM_024629.3 | 42.9992115 | -7.0457734 | 1.3772492 | -5.1158305 |
| 8943 | NM_020810.3 | 59.822773 | -5.7019339 | 1.11462132 | -5.1155794 |

| | | | | | |
|---|---|---|---|---|---|
| 8944 | NM_006993.2 | 48.5411196 | -7.2227938 | 1.41195001 | -5.1154742 |
| 8945 | NM_138335.2 | 31.7086817 | -6.6050555 | 1.2911877 | -5.1154883 |
| 8946 | NM_145798.2 | 59.6050407 | -6.5500625 | 1.28043664 | -5.1154913 |
| 8947 | NM_016223.4 | 33.3135938 | -6.6781008 | 1.30555422 | -5.1151462 |
| 8948 | NM_003659.3 | 189.329773 | -4.7364732 | 0.92602456 | -5.1148462 |
| 8949 | NM_003042.3 | 28.2684128 | -6.4387789 | 1.25887628 | -5.1147035 |
| 8950 | NM_021047.2 | 31.0241594 | -6.5728904 | 1.2851193 | -5.114615 |
| 8951 | NR_027946.2 | 77.4338496 | -7.8984099 | 1.54428469 | -5.1146074 |
| 8952 | NM_199413.2 | 56.1036236 | -7.4307766 | 1.45293868 | -5.1143085 |
| 8953 | NM_005614.3 | 279.74968 | -5.2340318 | 1.02348746 | -5.1139188 |
| 8954 | NR_132426.1 | 41.3854198 | -6.996292 | 1.36809733 | -5.1138847 |
| 8955 | NM_001321228.1 | 31.0957229 | -6.5768946 | 1.28612948 | -5.1137112 |
| 8956 | NM_001006946.1 | 45.2245995 | -7.1220845 | 1.39276571 | -5.1136271 |
| 8957 | NM_017940.4 | 273.184333 | -3.6716888 | 0.71803175 | -5.1135466 |
| 8958 | NM_005836.2 | 28.7298259 | -6.4613064 | 1.26362543 | -5.1133083 |
| 8959 | NM_001038618.2 | 27.1821318 | -6.3888356 | 1.24954882 | -5.112914 |
| 8960 | NM_001297774.1 | 42.8062586 | -7.0402052 | 1.37696413 | -5.1128458 |
| 8961 | NM_006198.2 | 36.24809 | -6.8044221 | 1.33084758 | -5.1128486 |
| 8962 | NM_014007.3 | 42.2949037 | -7.0207586 | 1.37322083 | -5.1126217 |
| 8963 | NM_001033566.2 | 37.2423754 | -6.8405604 | 1.33802114 | -5.1124457 |
| 8964 | NM_001042388.2 | 51.7299128 | -7.317595 | 1.43137078 | -5.1122987 |
| 8965 | NM_022474.3 | 44.5140122 | -7.1002174 | 1.38900543 | -5.1117276 |
| 8966 | NM_001271662.1 | 42.1656822 | -7.0208865 | 1.3735296 | -5.1115655 |
| 8967 | NM_016648.3 | 42.2836931 | -7.0246206 | 1.3743261 | -5.1113201 |
| 8968 | NM_016067.3 | 37.3388778 | -6.8428362 | 1.33878897 | -5.1112134 |
| 8969 | NM_019855.4 | 19.7140818 | 8.47086703 | 1.65730609 | 5.11122664 |
| 8970 | NM_021211.3 | 45.2055912 | -7.1189369 | 1.3928142 | -5.1111892 |
| 8971 | NM_001281731.1 | 33.6825895 | -6.6925916 | 1.30941942 | -5.1111137 |
| 8972 | NM_012198.4 | 55.5262457 | -7.4156478 | 1.45097593 | -5.1108 |
| 8973 | NM_001284499.1 | 81.4959329 | 7.1353386 | 1.39613823 | 5.11076802 |
| 8974 | NM_203453.3 | 45.1254038 | -7.1168525 | 1.39258617 | -5.1105293 |
| 8975 | NM_001291941.1 | 62.334309 | -7.5831454 | 1.48390822 | -5.1102523 |
| 8976 | NM_024613.3 | 29.4043343 | -6.4946366 | 1.27094979 | -5.1100654 |
| 8977 | NR_034041.1 | 92.7783667 | -4.773049 | 0.93423532 | -5.1090436 |
| 8978 | NM_001320672.1 | 15.5480787 | 3.90301584 | 0.76400846 | 5.10860294 |
| 8979 | NM_001207067.1 | 65.8378637 | -7.6615431 | 1.49989542 | -5.1080515 |
| 8980 | NM_016034.4 | 99.9669541 | -4.978648 | 0.97469627 | -5.1078968 |
| 8981 | NR_033243.2 | 35.1853476 | -6.7554734 | 1.32255056 | -5.1079131 |
| 8982 | NM_194449.3 | 118.04716 | -4.634888 | 0.90745551 | -5.1075649 |
| 8983 | NM_004954.4 | 51.79706 | -7.3147506 | 1.43215668 | -5.1075072 |
| 8984 | NM_001127195.1 | 74.9335382 | -7.8495669 | 1.53694358 | -5.1072577 |
| 8985 | NM_000136.2 | 41.9403992 | -7.0134237 | 1.37325606 | -5.1071493 |
| 8986 | NM_182491.2 | 44.4619226 | -7.0965872 | 1.38966332 | -5.1066953 |

| | | | | | |
|---|---|---|---|---|---|
| 8987 | NM_001301007.1 | 57.3477646 | -7.4614078 | 1.46115115 | -5.106527 |
| 8988 | NM_203463.2 | 55.1126876 | -7.4092787 | 1.45096624 | -5.1064446 |
| 8989 | NM_213622.2 | 38.5685144 | -6.8885112 | 1.34904735 | -5.1062042 |
| 8990 | NM_148923.3 | 42.9539451 | -7.0465368 | 1.38024076 | -5.1052954 |
| 8991 | NM_177477.3 | 39.6214359 | -6.9321739 | 1.35787546 | -5.1051618 |
| 8992 | NM_001303007.1 | 29.1411922 | -6.4896479 | 1.27130557 | -5.1047113 |
| 8993 | NM_015009.2 | 49.6129326 | -7.2581321 | 1.42204697 | -5.104003 |
| 8994 | NM_001345873.1 | 51.5097864 | -7.3077107 | 1.43178015 | -5.1039335 |
| 8995 | NM_018004.2 | 34.1288559 | -6.7191965 | 1.3165044 | -5.1038162 |
| 8996 | NM_144778.3 | 38.9826701 | -6.9050507 | 1.35293448 | -5.1037583 |
| 8997 | NM_001031827.2 | 47.9927541 | -7.2074642 | 1.41220099 | -5.1037099 |
| 8998 | NM_031280.3 | 85.2409135 | -6.08835 | 1.19297985 | -5.103481 |
| 8999 | NM_001024218.1 | 84.7772426 | -8.0274742 | 1.57294933 | -5.1034538 |
| 9000 | NM_005772.4 | 29.8706019 | -6.52278 | 1.27817037 | -5.1032164 |
| 9001 | NM_001271441.1 | 27.4052074 | -6.4006664 | 1.25433858 | -5.1028219 |
| 9002 | NM_000037.3 | 63.7316773 | -7.6163063 | 1.49258801 | -5.1027519 |
| 9003 | NM_000169.2 | 36.469172 | -6.811052 | 1.33483088 | -5.102558 |
| 9004 | NM_024638.3 | 30.3864896 | -6.5440871 | 1.28253368 | -5.1024681 |
| 9005 | NM_022717.3 | 35.6683119 | -6.7778613 | 1.32844571 | -5.1020988 |
| 9006 | NM_001256441.1 | 37.429309 | -6.8466027 | 1.34195221 | -5.1019721 |
| 9007 | NM_002101.4 | 40.9624008 | -6.9795577 | 1.36818074 | -5.1013419 |
| 9008 | NM_024855.3 | 35.7532536 | -6.7801888 | 1.32925293 | -5.1007514 |
| 9009 | NM_001303471.2 | 43.0022664 | -7.046715 | 1.3815111 | -5.1007299 |
| 9010 | NM_021127.2 | 27.883133 | -6.4283142 | 1.26028068 | -5.1007004 |
| 9011 | NM_003370.3 | 76.0221558 | -4.8137076 | 0.94378977 | -5.1004024 |
| 9012 | NM_001199562.1 | 31.0029886 | -6.5830385 | 1.29071525 | -5.1003028 |
| 9013 | NM_001130528.2 | 51.1310182 | -7.3016048 | 1.43176753 | -5.0997139 |
| 9014 | NM_018212.5 | 742.746304 | -3.243097 | 0.63593743 | -5.0997108 |
| 9015 | NM_032824.2 | 26.297156 | -6.3414662 | 1.24352415 | -5.0995923 |
| 9016 | NM_001100122.1 | 57.1216921 | -7.4587809 | 1.46266953 | -5.09943 |
| 9017 | NM_001289930.1 | 34.0146607 | -6.7064662 | 1.31514417 | -5.0994153 |
| 9018 | NM_178498.3 | 80.3270183 | 3.56003331 | 0.69815962 | 5.09916819 |
| 9019 | NM_001308263.1 | 65.8390764 | -6.6951588 | 1.31301021 | -5.0990912 |
| 9020 | NM_001113525.1 | 30.7587205 | -6.5644384 | 1.28742696 | -5.0988822 |
| 9021 | NM_021243.2 | 73.6270265 | -7.8232286 | 1.53435985 | -5.0986922 |
| 9022 | NM_001093767.2 | 69.2681695 | -7.7352992 | 1.51712411 | -5.0986595 |
| 9023 | NM_001127510.2 | 47.2947021 | -7.189213 | 1.41002986 | -5.0986246 |
| 9024 | NM_001142446.1 | 54.2290752 | -7.3836808 | 1.44825598 | -5.0983258 |
| 9025 | NM_004116.4 | 62.0256815 | -7.5770904 | 1.48618905 | -5.0983355 |
| 9026 | NM_001042551.1 | 63.9221009 | -7.6188619 | 1.49450438 | -5.0979188 |
| 9027 | NM_000033.3 | 76.1027035 | -6.0918624 | 1.19505087 | -5.0975758 |
| 9028 | NM_018719.4 | 40.6525686 | -6.9639679 | 1.36628921 | -5.096994 |
| 9029 | NM_032026.3 | 35.6396724 | -6.777433 | 1.32986699 | -5.096324 |

| | | | | | |
|---|---|---|---|---|---|
| 9030 | NR_023392.1 | 47.5905566 | -4.3889253 | 0.86120283 | -5.0962736 |
| 9031 | NM_032907.4 | 41.756043 | -7.0061188 | 1.37486113 | -5.0958738 |
| 9032 | NM_152829.2 | 31.8453456 | 9.91729732 | 1.9461474 | 5.09586135 |
| 9033 | NM_014895.3 | 28.325571 | -6.4433192 | 1.26470034 | -5.0947398 |
| 9034 | NM_145169.2 | 30.2545995 | -6.5420444 | 1.28423295 | -5.0941259 |
| 9035 | NM_003431.3 | 26.5525414 | -6.3523433 | 1.24702752 | -5.093988 |
| 9036 | NM_001289969.1 | 45.7940544 | -7.1361224 | 1.40094471 | -5.0937931 |
| 9037 | NM_003289.3 | 44.4078553 | -7.0976754 | 1.39350076 | -5.0934133 |
| 9038 | NM_181501.1 | 105.784109 | -5.0506558 | 0.99169087 | -5.092974 |
| 9039 | NM_024292.3 | 39.5502935 | -6.9270923 | 1.36025683 | -5.0924885 |
| 9040 | NR_040071.1 | 45.8317817 | 1.9733022 | 0.38749665 | 5.09243681 |
| 9041 | NM_001018005.1 | 60.5661089 | -7.5445134 | 1.48157776 | -5.0922156 |
| 9042 | NM_006184.5 | 495.446456 | -6.1020011 | 1.19839239 | -5.0918223 |
| 9043 | NM_001013703.3 | 192.862914 | -3.9303318 | 0.77190021 | -5.0917615 |
| 9044 | NM_022066.3 | 286.500474 | -5.1805352 | 1.01744675 | -5.0917016 |
| 9045 | NM_016069.9 | 46.6140102 | -7.1639608 | 1.40710477 | -5.0912774 |
| 9046 | NM_001174084.1 | 30.1262635 | -6.5343116 | 1.28345173 | -5.0912017 |
| 9047 | NM_004404.4 | 779.276111 | -6.7219848 | 1.32042119 | -5.0907884 |
| 9048 | NM_001008703.2 | 48.4544383 | -7.2207234 | 1.4184262 | -5.0906585 |
| 9049 | NM_014653.3 | 37.1363091 | -6.8358976 | 1.3428589 | -5.0905554 |
| 9050 | NM_001318525.1 | 51.2232028 | -7.3010988 | 1.4344532 | -5.0898132 |
| 9051 | NM_002134.3 | 61.4141583 | -7.5624456 | 1.48589236 | -5.0894976 |
| 9052 | NM_001040108.1 | 35.9513311 | -6.783806 | 1.33300483 | -5.0891084 |
| 9053 | NM_152791.4 | 38.9319004 | -6.9004028 | 1.35638679 | -5.0873415 |
| 9054 | NM_001330611.1 | 30.6799699 | -6.5637862 | 1.29031233 | -5.0869747 |
| 9055 | NM_003944.3 | 33.3194036 | -6.6866069 | 1.31464957 | -5.0862276 |
| 9056 | NM_017934.6 | 273.283635 | -5.1071745 | 1.00417691 | -5.085931 |
| 9057 | NM_018294.5 | 41.9864258 | -7.0120476 | 1.37873845 | -5.0858432 |
| 9058 | NM_032790.3 | 27.123563 | -6.3855004 | 1.25565157 | -5.0854079 |
| 9059 | NR_120580.1 | 25.7610662 | 5.00406986 | 0.98406045 | 5.08512446 |
| 9060 | NM_032593.2 | 43.0042962 | -7.0477946 | 1.38610199 | -5.0846147 |
| 9061 | NM_001034.3 | 109.650952 | -2.5271704 | 0.49705894 | -5.0842469 |
| 9062 | NM_172171.2 | 73.5725569 | -7.8228255 | 1.53872642 | -5.0839613 |
| 9063 | NM_001282749.1 | 72.543853 | -7.8017717 | 1.5346553 | -5.0837291 |
| 9064 | NM_001111019.2 | 75.6717101 | -6.8956195 | 1.35644931 | -5.0835807 |
| 9065 | NM_005645.3 | 34.9371905 | -6.7498543 | 1.32778378 | -5.0835493 |
| 9066 | NM_001286177.1 | 52.1675715 | -7.3257294 | 1.44109108 | -5.0834603 |
| 9067 | NM_001270998.1 | 40.5942272 | -6.9636781 | 1.36997724 | -5.0830611 |
| 9068 | NM_145160.2 | 36.4937313 | -6.8141926 | 1.34077833 | -5.0822664 |
| 9069 | NM_001256012.1 | 84.3485281 | -8.0216259 | 1.57841718 | -5.0820696 |
| 9070 | NM_001167671.2 | 67.4474527 | -7.695921 | 1.51442564 | -5.0817424 |
| 9071 | NM_052965.3 | 48.0486756 | -7.2073822 | 1.41848287 | -5.0810498 |
| 9072 | NM_182476.2 | 39.1153927 | -6.9100635 | 1.3601645 | -5.0803145 |

| | | | | | |
|---|---|---|---|---|---|
| 9073 | NM_001033088.1 | 73.4330372 | -7.8208768 | 1.5394719 | -5.0802335 |
| 9074 | NM_014698.2 | 30.5011053 | -6.5590291 | 1.29133115 | -5.0792774 |
| 9075 | NM_001267547.2 | 40.1176859 | -6.9461669 | 1.36769566 | -5.0787372 |
| 9076 | NM_198043.2 | 42.9941334 | -7.0489179 | 1.38801874 | -5.0784025 |
| 9077 | NM_001159676.1 | 36.2865759 | -6.801617 | 1.33934394 | -5.0783199 |
| 9078 | NM_000750.3 | 45.586731 | -7.1331105 | 1.40462891 | -5.0782882 |
| 9079 | NM_001316951.1 | 28.570509 | -6.4560839 | 1.27131654 | -5.0782662 |
| 9080 | NM_013443.4 | 36.4988934 | -6.8128561 | 1.34160038 | -5.078156 |
| 9081 | NM_018095.5 | 30.1428836 | -6.5374397 | 1.28739435 | -5.0780398 |
| 9082 | NM_052811.3 | 53.9442028 | -7.3747839 | 1.45235336 | -5.0778165 |
| 9083 | NR_130947.1 | 41.3000647 | 10.2856902 | 2.02573457 | 5.07751132 |
| 9084 | NM_006352.4 | 73.3558324 | -7.8168872 | 1.53960425 | -5.0772055 |
| 9085 | NR_126432.1 | 44.6103154 | -7.1030285 | 1.39900514 | -5.0771997 |
| 9086 | NM_001286631.1 | 51.6844806 | -7.312704 | 1.44033922 | -5.0770707 |
| 9087 | NM_006506.3 | 45.7429527 | -7.1360885 | 1.40555694 | -5.077054 |
| 9088 | NM_006747.3 | 51.851971 | -5.4718889 | 1.07788577 | -5.0765016 |
| 9089 | NM_183057.2 | 39.4342331 | -6.9211123 | 1.36338921 | -5.0764024 |
| 9090 | NM_002822.4 | 151.427708 | -5.2099928 | 1.0263795 | -5.0760882 |
| 9091 | NM_002833.3 | 196.644076 | -5.4412712 | 1.0719673 | -5.0759675 |
| 9092 | NM_002431.3 | 32.3326064 | -6.6327181 | 1.3066971 | -5.0759415 |
| 9093 | NM_012262.3 | 147.428859 | -5.0014886 | 0.98545333 | -5.0753176 |
| 9094 | NM_014066.3 | 38.8500811 | -6.9009003 | 1.35987413 | -5.074661 |
| 9095 | NM_001163417.1 | 56.7148648 | -7.444768 | 1.46706422 | -5.0746026 |
| 9096 | NM_030569.6 | 65.4257999 | -6.6867498 | 1.31770102 | -5.0745577 |
| 9097 | NM_001032291.2 | 53.9129455 | -7.3747849 | 1.45337003 | -5.0742652 |
| 9098 | NM_001128430.1 | 82.88772 | -6.178171 | 1.21765388 | -5.0738318 |
| 9099 | NM_133265.2 | 74.5775989 | -4.5245557 | 0.89176201 | -5.0737255 |
| 9100 | NM_001243246.1 | 29.3284245 | -6.5006453 | 1.28139589 | -5.0730967 |
| 9101 | NM_015896.3 | 75.947703 | -7.8704536 | 1.55157289 | -5.0725645 |
| 9102 | NM_017450.2 | 138.959981 | -5.3185055 | 1.04848927 | -5.0725416 |
| 9103 | NM_001130026.2 | 76.4059055 | 1.81926604 | 0.35866275 | 5.07235844 |
| 9104 | NM_032683.2 | 40.6750391 | -6.9686398 | 1.37391024 | -5.0721216 |
| 9105 | NR_130950.1 | 23.633643 | 9.49069503 | 1.87114569 | 5.07213044 |
| 9106 | NM_017841.2 | 74.0756625 | -6.8579219 | 1.35213781 | -5.0719104 |
| 9107 | NM_001145099.1 | 44.1791712 | -7.0864699 | 1.39728006 | -5.0716174 |
| 9108 | NM_130442.3 | 55.4713091 | -7.4156686 | 1.4624171 | -5.0708301 |
| 9109 | NM_024681.3 | 35.2793455 | -6.7676649 | 1.33471177 | -5.0705066 |
| 9110 | NM_206538.3 | 348.714897 | -6.1546426 | 1.21383279 | -5.0704205 |
| 9111 | NM_175609.2 | 65.8285435 | -7.6615537 | 1.51141176 | -5.0691373 |
| 9112 | NM_201543.2 | 31.9913646 | -6.6245813 | 1.30691297 | -5.0688772 |
| 9113 | NM_001008860.2 | 43.8465234 | -7.0765806 | 1.39612493 | -5.0687302 |
| 9114 | NM_013251.3 | 62.7406176 | -7.5937188 | 1.49823491 | -5.0684434 |
| 9115 | NM_024709.4 | 39.1243699 | -6.9112021 | 1.36365978 | -5.0681278 |

| | | | | | |
|---|---|---|---|---|---|
| 9116 | NM_006051.3 | 38.0298535 | -6.8734906 | 1.35623469 | -5.0680687 |
| 9117 | NM_017444.5 | 29.5717173 | -6.5019715 | 1.28293314 | -5.0680517 |
| 9118 | NM_032207.3 | 29.2874072 | -6.4931015 | 1.2812743 | -5.0676905 |
| 9119 | NM_058180.4 | 27.5831579 | -6.4125394 | 1.26544755 | -5.0674083 |
| 9120 | NM_025247.5 | 56.0590687 | -3.6968814 | 0.72958574 | -5.0670965 |
| 9121 | NM_033196.2 | 45.1616462 | -7.1170389 | 1.40456489 | -5.0670773 |
| 9122 | NM_001810.5 | 272.398417 | -5.6491366 | 1.1149588 | -5.0666774 |
| 9123 | NM_024308.3 | 27.7941508 | -6.4217797 | 1.26747809 | -5.0665804 |
| 9124 | NM_005783.3 | 49.2248907 | -7.2435433 | 1.42978323 | -5.0661829 |
| 9125 | NM_001279354.1 | 63.1951024 | -7.604689 | 1.50128894 | -5.06544 |
| 9126 | NR_036477.1 | 30.4971727 | -6.5507717 | 1.29322631 | -5.0654488 |
| 9127 | NM_000067.2 | 105.693151 | -8.3468004 | 1.64782679 | -5.0653385 |
| 9128 | NM_003312.5 | 26.329981 | -6.3396716 | 1.25162212 | -5.0651642 |
| 9129 | NM_013449.3 | 87.6340171 | -5.3519896 | 1.05666186 | -5.0649975 |
| 9130 | NM_003585.4 | 51.232877 | -6.3328889 | 1.25050218 | -5.0642766 |
| 9131 | NM_001256661.1 | 26.480598 | -6.3503338 | 1.25407426 | -5.0637622 |
| 9132 | NR_036462.1 | 26.3913456 | -6.3448239 | 1.25301205 | -5.0636575 |
| 9133 | NM_015566.2 | 57.759647 | -6.4943957 | 1.2825649 | -5.0636001 |
| 9134 | NM_001080469.1 | 34.8703701 | -6.7456061 | 1.33239956 | -5.0627501 |
| 9135 | NM_015088.2 | 99.6668969 | -4.8765201 | 0.96327891 | -5.0624176 |
| 9136 | NM_001142529.1 | 29.1113056 | -6.4845293 | 1.28093722 | -5.0623318 |
| 9137 | NM_173664.5 | 41.7920376 | -7.0062684 | 1.38413255 | -5.0618478 |
| 9138 | NM_080597.3 | 57.5572056 | -5.6265993 | 1.11158023 | -5.0618022 |
| 9139 | NM_152766.3 | 41.4551159 | -6.9958691 | 1.38210118 | -5.0617634 |
| 9140 | NM_001330373.1 | 54.8983283 | -7.3978442 | 1.46157742 | -5.061548 |
| 9141 | NM_152222.1 | 33.8042983 | -6.7008927 | 1.32402161 | -5.0610146 |
| 9142 | NM_177925.3 | 32.9513175 | -6.6646647 | 1.31689631 | -5.060888 |
| 9143 | NM_005101.3 | 32.443216 | -6.6472262 | 1.31351065 | -5.0606565 |
| 9144 | NM_013410.3 | 48.5358636 | -7.2226216 | 1.4273192 | -5.0602708 |
| 9145 | NM_001099638.1 | 52.6785581 | -7.3443853 | 1.45166272 | -5.0592918 |
| 9146 | NM_015545.3 | 113.441774 | -2.2112234 | 0.43709941 | -5.0588569 |
| 9147 | NM_001166271.2 | 42.9119069 | -7.0418646 | 1.39200886 | -5.0587786 |
| 9148 | NM_001165414.1 | 51.2308997 | -7.3010165 | 1.4434674 | -5.0579712 |
| 9149 | NR_046567.1 | 43.3541128 | 2.27131695 | 0.44908578 | 5.05764611 |
| 9150 | NM_020192.4 | 32.1415121 | -6.6273467 | 1.31046945 | -5.057231 |
| 9151 | NM_006165.3 | 35.6096209 | -6.7802854 | 1.34096704 | -5.0562655 |
| 9152 | NM_000239.2 | 225.7723 | 3.03547656 | 0.6003448 | 5.05622198 |
| 9153 | NM_006190.4 | 34.9511317 | -6.7482313 | 1.33472316 | -5.0559034 |
| 9154 | NM_152407.3 | 67.4849949 | -6.7275615 | 1.33063178 | -5.0559152 |
| 9155 | NM_182760.3 | 38.4346051 | -6.8858437 | 1.36197002 | -5.0557968 |
| 9156 | NR_126481.1 | 17.239479 | 6.4188152 | 1.26974802 | 5.05518821 |
| 9157 | NM_014184.3 | 36.3484783 | -6.8066918 | 1.34659803 | -5.0547318 |
| 9158 | NM_144643.3 | 32.0284867 | -6.6202491 | 1.30994341 | -5.0538436 |

| | | | | | |
|---|---|---|---|---|---|
| 9159 | NR_027107.2 | 26.8465779 | -6.3716417 | 1.26079038 | -5.0536884 |
| 9160 | NM_016472.4 | 27.9387858 | -6.4325237 | 1.27289498 | -5.0534598 |
| 9161 | NM_001319842.1 | 31.5331254 | -6.5985218 | 1.30579928 | -5.0532435 |
| 9162 | NM_015892.4 | 37.4981896 | -6.8463251 | 1.35489189 | -5.0530416 |
| 9163 | NM_005433.3 | 159.866324 | -4.0184092 | 0.79528663 | -5.0527811 |
| 9164 | NM_012325.2 | 507.547382 | -6.4984574 | 1.28628116 | -5.0521283 |
| 9165 | NM_001170649.1 | 26.7924827 | -6.3622392 | 1.25932948 | -5.0520847 |
| 9166 | NM_001129993.2 | 42.279735 | -7.0263452 | 1.39085513 | -5.0518167 |
| 9167 | NM_152424.3 | 66.7439715 | -6.7070582 | 1.32769124 | -5.0516702 |
| 9168 | NM_001170766.1 | 46.0430337 | -7.1460284 | 1.41459586 | -5.0516395 |
| 9169 | NM_001163287.1 | 30.8442368 | -6.5752406 | 1.30161658 | -5.0515956 |
| 9170 | NM_003083.3 | 44.5263399 | -7.098052 | 1.4051609 | -5.0514158 |
| 9171 | NM_052954.3 | 27.9441357 | -6.4290759 | 1.27295097 | -5.0505291 |
| 9172 | NM_001085049.2 | 98.9994304 | -8.2516615 | 1.63386453 | -5.0503952 |
| 9173 | NM_015428.3 | 35.0884909 | -6.7498042 | 1.33653532 | -5.050225 |
| 9174 | NM_017950.3 | 63.7225598 | -7.6175754 | 1.50854051 | -5.0496326 |
| 9175 | NM_016213.4 | 37.0500434 | -6.8319948 | 1.35306369 | -5.0492781 |
| 9176 | NR_024158.1 | 48.866117 | -6.2607374 | 1.24012502 | -5.0484728 |
| 9177 | NM_012343.3 | 40.026833 | -6.9423144 | 1.37519069 | -5.0482558 |
| 9178 | NM_033489.2 | 28.7471007 | -6.4692233 | 1.28154128 | -5.0480023 |
| 9179 | NM_001017998.3 | 51.3986294 | -7.3063346 | 1.44738982 | -5.0479384 |
| 9180 | NM_175617.3 | 56.5684434 | -7.4441601 | 1.47481373 | -5.0475256 |
| 9181 | NM_020143.3 | 43.2620445 | -7.0569208 | 1.39813705 | -5.0473741 |
| 9182 | NM_001172509.1 | 66.4106565 | -7.6741425 | 1.52044415 | -5.0473031 |
| 9183 | NM_001002248.2 | 46.0180565 | -7.1447586 | 1.41559871 | -5.0471638 |
| 9184 | NM_173829.3 | 113.13088 | -3.1031265 | 0.61482866 | -5.0471403 |
| 9185 | NM_016299.3 | 31.8040782 | -6.6127042 | 1.31035028 | -5.0465165 |
| 9186 | NM_001002009.2 | 40.246387 | -6.9493644 | 1.37712447 | -5.0462863 |
| 9187 | NM_001085372.2 | 54.9444486 | -4.0299852 | 0.79870957 | -5.0456202 |
| 9188 | NM_006593.3 | 288.6115 | -7.5521616 | 1.49677571 | -5.0456201 |
| 9189 | NM_001346363.1 | 30.7795192 | -6.5710283 | 1.30234654 | -5.0455298 |
| 9190 | NM_032758.3 | 67.7995981 | -4.6216906 | 0.91605787 | -5.045195 |
| 9191 | NM_001320942.1 | 27.9033156 | -6.419361 | 1.27256015 | -5.0444461 |
| 9192 | NM_199191.2 | 26.406674 | -6.3493436 | 1.25871609 | -5.0443016 |
| 9193 | NM_145062.2 | 42.938361 | -7.0457812 | 1.39691297 | -5.0438226 |
| 9194 | NM_001307931.1 | 83.4553782 | 2.66511488 | 0.52841714 | 5.04358144 |
| 9195 | NR_046330.1 | 30.3036067 | -6.548512 | 1.29853212 | -5.0430112 |
| 9196 | NM_001243349.1 | 37.2251963 | -6.8417137 | 1.35670006 | -5.042908 |
| 9197 | NM_018025.2 | 38.6503752 | -6.8914059 | 1.36667011 | -5.0424794 |
| 9198 | NR_039969.1 | 27.8494619 | -6.4257029 | 1.27437047 | -5.0422566 |
| 9199 | NM_018947.5 | 619.856044 | -2.9357034 | 0.58224228 | -5.0420649 |
| 9200 | NM_198686.2 | 44.8626893 | -7.1082743 | 1.41001306 | -5.0412826 |
| 9201 | NM_015690.4 | 82.0407923 | -7.9821512 | 1.58342989 | -5.0410513 |

| | | | | | |
|---|---|---|---|---|---|
| 9202 | NM_003954.4 | 47.2756582 | -3.7927637 | 0.75238835 | -5.0409655 |
| 9203 | NM_001153.4 | 69.8275849 | -7.7494621 | 1.53736242 | -5.0407516 |
| 9204 | NM_001278604.1 | 37.4023127 | -6.8440945 | 1.35782767 | -5.0404736 |
| 9205 | NR_104457.1 | 26.9953418 | -6.3717325 | 1.26411559 | -5.0404667 |
| 9206 | NM_016457.4 | 38.9394786 | -6.9092301 | 1.37077631 | -5.0403776 |
| 9207 | NM_019099.4 | 44.765057 | -7.1047392 | 1.40958983 | -5.0402883 |
| 9208 | NM_001145445.1 | 54.2522325 | -4.610109 | 0.91471196 | -5.039957 |
| 9209 | NM_001042697.1 | 31.9840386 | -6.6176103 | 1.31313535 | -5.0395493 |
| 9210 | NM_145175.3 | 59.2212363 | -6.5330987 | 1.29646462 | -5.0391647 |
| 9211 | NR_004858.1 | 28.6367213 | -6.4610917 | 1.28228277 | -5.0387417 |
| 9212 | NM_001331036.1 | 36.0412614 | -6.792896 | 1.34823875 | -5.038348 |
| 9213 | NM_004975.3 | 63.6585613 | -5.8232226 | 1.15585701 | -5.0380129 |
| 9214 | NR_026717.1 | 41.7099341 | -7.0009453 | 1.38961876 | -5.0380331 |
| 9215 | NM_144579.2 | 25.9975276 | -6.3174068 | 1.25397408 | -5.0379086 |
| 9216 | NM_033427.2 | 101.51491 | -7.3155147 | 1.45222505 | -5.0374525 |
| 9217 | NM_014723.3 | 80.1080472 | -6.9782479 | 1.38550896 | -5.0365953 |
| 9218 | NM_032793.4 | 54.6824203 | -7.3942018 | 1.4681281 | -5.0364827 |
| 9219 | NM_007057.3 | 42.4486986 | -7.0289799 | 1.39564874 | -5.0363531 |
| 9220 | NM_001025235.1 | 34.7989737 | -6.7472669 | 1.33975493 | -5.0361948 |
| 9221 | NM_001310159.1 | 25.8782753 | -6.3160914 | 1.25413518 | -5.0362126 |
| 9222 | NM_001348149.1 | 37.8920236 | -6.8659704 | 1.36332603 | -5.0361911 |
| 9223 | NM_003763.5 | 63.5620127 | -7.6146605 | 1.51198328 | -5.0362068 |
| 9224 | NM_138361.5 | 37.161273 | -6.8343803 | 1.35704637 | -5.0362172 |
| 9225 | NM_001009894.2 | 43.5075671 | -7.0632236 | 1.4026067 | -5.0357835 |
| 9226 | NM_001299.5 | 50.3077589 | -7.276423 | 1.44503163 | -5.0354766 |
| 9227 | NM_198264.1 | 34.6163866 | -6.7341403 | 1.33733867 | -5.0354787 |
| 9228 | NM_021937.4 | 65.6322699 | -5.7911217 | 1.15010825 | -5.035284 |
| 9229 | NM_001004128.2 | 46.6928634 | -7.1670246 | 1.423403 | -5.0351338 |
| 9230 | NM_001698.2 | 32.8686103 | -6.6667061 | 1.32405952 | -5.0350502 |
| 9231 | NM_001948.3 | 34.9936746 | -6.7530945 | 1.34126759 | -5.03486 |
| 9232 | NM_005938.3 | 25.5438562 | -6.300154 | 1.25132995 | -5.0347664 |
| 9233 | NM_032420.3 | 80.3786242 | -6.9827285 | 1.38695049 | -5.034591 |
| 9234 | NM_001430.4 | 103.485861 | -8.3163191 | 1.65192498 | -5.0343201 |
| 9235 | NM_153617.1 | 37.9795867 | -6.87051 | 1.3647536 | -5.0342494 |
| 9236 | NM_000086.2 | 40.7533015 | -6.9700243 | 1.3845355 | -5.0341969 |
| 9237 | NR_033669.1 | 35.0894054 | -6.7542627 | 1.34176408 | -5.0338676 |
| 9238 | NM_024574.3 | 38.9143636 | -6.9078984 | 1.37235186 | -5.0336204 |
| 9239 | NM_031435.3 | 38.2475583 | -6.8772704 | 1.36626578 | -5.0336256 |
| 9240 | NM_004715.4 | 41.1310566 | -6.9815008 | 1.38704139 | -5.033376 |
| 9241 | NM_001142725.1 | 26.7290101 | -6.3584665 | 1.26331722 | -5.0331512 |
| 9242 | NM_133259.3 | 479.363448 | -5.9621737 | 1.18465343 | -5.0328422 |
| 9243 | NM_007327.3 | 81.2564558 | -7.965368 | 1.58273437 | -5.0326626 |
| 9244 | NM_001201362.1 | 37.6358286 | -6.8589126 | 1.3629537 | -5.0323885 |

| | | | | | |
|---|---|---|---|---|---|
| 9245 | NM_001005415.1 | 44.297393 | -7.090417 | 1.40906988 | -5.0319839 |
| 9246 | NM_032881.2 | 73.0579189 | -6.8446101 | 1.36026428 | -5.0318238 |
| 9247 | NM_006670.4 | 77.4595624 | -7.8987846 | 1.56986838 | -5.0314948 |
| 9248 | NM_018983.3 | 39.7860177 | -6.9348587 | 1.37850913 | -5.0306948 |
| 9249 | NM_001199580.1 | 44.9412485 | -7.1131189 | 1.41397922 | -5.0305682 |
| 9250 | NR_038889.1 | 26.732974 | -6.3573301 | 1.26378397 | -5.0303931 |
| 9251 | NM_001168.2 | 45.319949 | -7.1221381 | 1.41596286 | -5.0298905 |
| 9252 | NM_183393.2 | 118.988712 | -5.8210195 | 1.15745982 | -5.0291331 |
| 9253 | NM_001277285.1 | 33.3428735 | -6.6805221 | 1.32846952 | -5.0287357 |
| 9254 | NM_014676.2 | 120.516708 | -5.8637964 | 1.16606286 | -5.0287138 |
| 9255 | NM_005531.2 | 31.6262081 | -6.6067016 | 1.31382356 | -5.0286064 |
| 9256 | NM_015101.3 | 43.4986419 | -7.0675003 | 1.40545774 | -5.028611 |
| 9257 | NM_012260.3 | 27.5658563 | -6.412768 | 1.27527167 | -5.0285505 |
| 9258 | NM_001184897.1 | 25.3099464 | -6.2878496 | 1.25045585 | -5.0284459 |
| 9259 | NM_005600.2 | 37.6038977 | -6.8573452 | 1.36376416 | -5.0282485 |
| 9260 | NM_005613.5 | 34.1815814 | -6.7188662 | 1.33629017 | -5.0279994 |
| 9261 | NR_133003.1 | 54.6803134 | -7.3915781 | 1.47018062 | -5.0276667 |
| 9262 | NM_181504.3 | 78.1781023 | -7.9104893 | 1.57349393 | -5.0273402 |
| 9263 | NM_006931.2 | 257.292917 | -3.8097877 | 0.75781974 | -5.0273007 |
| 9264 | NM_001184819.1 | 45.0553433 | -7.1198483 | 1.41634852 | -5.0269042 |
| 9265 | NM_015346.3 | 82.1479225 | -4.6780771 | 0.93064446 | -5.0267071 |
| 9266 | NM_138364.3 | 25.4203097 | -6.2886564 | 1.25107294 | -5.0266105 |
| 9267 | NM_080621.4 | 70.0447052 | -4.9900508 | 0.99274387 | -5.0265239 |
| 9268 | NM_001308230.1 | 32.8174004 | -6.653362 | 1.32380845 | -5.025925 |
| 9269 | NM_014394.2 | 420.719077 | -5.5217941 | 1.09868011 | -5.0258433 |
| 9270 | NM_001167604.1 | 34.4670017 | -6.7306525 | 1.33927616 | -5.0255897 |
| 9271 | NM_001008695.1 | 30.4162537 | -6.5447977 | 1.30230858 | -5.0255353 |
| 9272 | NM_001122823.1 | 26.8594879 | -6.3684991 | 1.26726308 | -5.0253962 |
| 9273 | NM_133264.4 | 174.583015 | -2.9956376 | 0.59610166 | -5.0253804 |
| 9274 | NM_018086.3 | 29.6788541 | -6.5201359 | 1.29746542 | -5.0252868 |
| 9275 | NM_012435.2 | 387.371211 | -6.3564213 | 1.26492146 | -5.025151 |
| 9276 | NR_027435.1 | 59.4892142 | 2.34153404 | 0.46599464 | 5.02480899 |
| 9277 | NM_001144060.1 | 49.6939222 | -5.4018704 | 1.07510394 | -5.0245099 |
| 9278 | NM_001128161.1 | 40.8855729 | -6.9780813 | 1.38886119 | -5.0243187 |
| 9279 | NM_147175.3 | 97.6900518 | -6.253277 | 1.24468504 | -5.0239834 |
| 9280 | NM_015589.5 | 30.3023095 | -6.5498351 | 1.30384505 | -5.0234766 |
| 9281 | NM_004127.5 | 49.8585383 | -7.2630083 | 1.4458761 | -5.0232577 |
| 9282 | NM_004748.4 | 33.1309056 | -6.6751245 | 1.32887625 | -5.0231348 |
| 9283 | NM_015941.3 | 39.8737065 | -6.9398892 | 1.38169699 | -5.0227288 |
| 9284 | NM_003149.2 | 50.0516568 | -7.2674332 | 1.44698143 | -5.0224786 |
| 9285 | NR_033426.1 | 29.1825365 | -6.4941474 | 1.29302374 | -5.0224502 |
| 9286 | NM_022481.5 | 35.2346913 | -6.7545348 | 1.34498393 | -5.0220189 |
| 9287 | NM_001012759.2 | 44.7525932 | -7.1052434 | 1.41486051 | -5.0218685 |

| | | | | | |
|---|---|---|---|---|---|
| 9288 | NM_001005910.2 | 32.020945 | -6.6161513 | 1.31748206 | -5.0218151 |
| 9289 | NM_003815.4 | 35.9218746 | -6.7902908 | 1.35221309 | -5.0216129 |
| 9290 | NM_020200.6 | 38.3684347 | -6.8803585 | 1.37023611 | -5.0212941 |
| 9291 | NM_032192.3 | 30.8163458 | -6.5619593 | 1.30684546 | -5.0212205 |
| 9292 | NM_001252049.1 | 55.0248499 | -7.4002867 | 1.47381815 | -5.0211667 |
| 9293 | NM_145177.2 | 49.0236303 | -6.2677392 | 1.24827921 | -5.0211036 |
| 9294 | NM_001145443.2 | 56.3424179 | -7.436728 | 1.4811491 | -5.0209179 |
| 9295 | NM_182914.2 | 113.715017 | -8.4504248 | 1.68316315 | -5.0205619 |
| 9296 | NM_001270393.1 | 73.5982513 | -5.4007796 | 1.07576135 | -5.0204254 |
| 9297 | NM_004568.5 | 37.1057384 | -6.8365838 | 1.36181763 | -5.0201904 |
| 9298 | NM_198149.2 | 40.0915282 | -6.9472438 | 1.38386597 | -5.020171 |
| 9299 | NR_135199.1 | 48.6416337 | -7.2259304 | 1.43943219 | -5.0199867 |
| 9300 | NM_001146343.1 | 33.7414926 | -6.6973774 | 1.33416875 | -5.0198878 |
| 9301 | NM_030758.3 | 33.4267902 | -6.6819603 | 1.33117438 | -5.019598 |
| 9302 | NM_004581.5 | 35.4342338 | -6.7680794 | 1.34855732 | -5.0187555 |
| 9303 | NM_019593.3 | 145.234377 | -4.4396028 | 0.88467964 | -5.0183169 |
| 9304 | NM_001319066.1 | 39.1126079 | -6.9096838 | 1.37692367 | -5.0182039 |
| 9305 | NM_022132.4 | 103.185636 | -4.6211866 | 0.92089935 | -5.0181233 |
| 9306 | NM_001281300.1 | 154.805885 | -5.2463816 | 1.04556146 | -5.0177649 |
| 9307 | NM_005853.5 | 40.8912208 | -6.9768386 | 1.39048199 | -5.0175685 |
| 9308 | NM_016332.3 | 35.5299554 | -6.772825 | 1.34991672 | -5.0172169 |
| 9309 | NM_025080.3 | 83.2145208 | -7.032277 | 1.40162448 | -5.0172333 |
| 9310 | NM_001318219.1 | 87.1742181 | -4.5505632 | 0.90706661 | -5.0167905 |
| 9311 | NM_001318975.1 | 19.6710058 | 9.21058344 | 1.8360498 | 5.01652158 |
| 9312 | NM_001077440.1 | 42.9709938 | -7.046412 | 1.40466765 | -5.0164265 |
| 9313 | NM_001322206.1 | 47.0377027 | -7.1759375 | 1.43048994 | -5.0164194 |
| 9314 | NM_001127218.2 | 36.8958217 | -6.8245358 | 1.3605607 | -5.0159731 |
| 9315 | NM_001321512.1 | 30.303555 | -6.5446941 | 1.30477074 | -5.0159725 |
| 9316 | NM_001127660.1 | 99.8562981 | -8.2649957 | 1.64799488 | -5.0151829 |
| 9317 | NM_015129.5 | 31.6868559 | -6.6136422 | 1.31874153 | -5.0151163 |
| 9318 | NM_181889.2 | 47.1623837 | -7.1811584 | 1.43200559 | -5.0147559 |
| 9319 | NM_207291.2 | 49.5320415 | -7.2539972 | 1.44708232 | -5.0128435 |
| 9320 | NM_194249.2 | 30.7128749 | -6.5617368 | 1.30899184 | -5.0128172 |
| 9321 | NM_017633.2 | 29.6677061 | -6.5189211 | 1.30047097 | -5.0127387 |
| 9322 | NM_020978.4 | 30.5304292 | -6.5523958 | 1.30715441 | -5.0127175 |
| 9323 | NM_004275.4 | 34.4135465 | -6.7278982 | 1.34219864 | -5.012595 |
| 9324 | NM_012301.3 | 66.2866581 | -7.6725645 | 1.53066109 | -5.0125822 |
| 9325 | NM_032174.5 | 48.1829219 | -7.2129001 | 1.43895339 | -5.0126016 |
| 9326 | NM_012256.3 | 47.6827452 | -6.2170163 | 1.24033743 | -5.0123589 |
| 9327 | NM_012145.3 | 38.0442007 | -6.8708616 | 1.37080906 | -5.0122674 |
| 9328 | NM_001031737.2 | 30.7232217 | -6.5690788 | 1.31061105 | -5.0122261 |
| 9329 | NM_024603.3 | 43.6295708 | -7.0673159 | 1.41001585 | -5.0122244 |
| 9330 | NM_012110.3 | 27.6451365 | -6.4156564 | 1.28005945 | -5.0119988 |

| | | | | | |
|---|---|---|---|---|---|
| 9331 | NR_131336.1 | 42.1690439 | -7.0221356 | 1.40109398 | -5.0118948 |
| 9332 | NR_130704.1 | 34.0955182 | -6.7142501 | 1.33970875 | -5.0117237 |
| 9333 | NM_005105.4 | 494.920884 | -4.3018566 | 0.85841363 | -5.0114029 |
| 9334 | NM_138555.3 | 30.3819254 | -6.5429833 | 1.30567555 | -5.0111862 |
| 9335 | NM_001271148.1 | 33.1317874 | 9.97456042 | 1.99054668 | 5.01096534 |
| 9336 | NM_030763.2 | 31.0316164 | -6.5715841 | 1.31152837 | -5.0106305 |
| 9337 | NM_004104.4 | 1978.77213 | -7.1997318 | 1.43704038 | -5.010111 |
| 9338 | NM_001328650.1 | 53.1309048 | -7.3505015 | 1.46717979 | -5.0099528 |
| 9339 | NR_040244.1 | 79.8514193 | 2.65794847 | 0.53063455 | 5.00900004 |
| 9340 | NR_028027.1 | 31.7922503 | -6.6164454 | 1.32093612 | -5.0089065 |
| 9341 | NM_018913.2 | 44.4794286 | -7.0968006 | 1.41708921 | -5.0080126 |
| 9342 | NM_001099679.1 | 47.474542 | -7.1934414 | 1.43644512 | -5.007808 |
| 9343 | NM_001346218.1 | 51.7012752 | -7.3122428 | 1.46020778 | -5.0076728 |
| 9344 | NM_001270771.1 | 28.5126808 | -6.4558562 | 1.28955953 | -5.0062491 |
| 9345 | NM_004065.2 | 47.2998961 | -7.182589 | 1.43476667 | -5.0061025 |
| 9346 | NM_020244.2 | 61.8369295 | -6.594992 | 1.31747079 | -5.0057975 |
| 9347 | NM_017875.2 | 90.5777416 | -4.8308671 | 0.96509464 | -5.005589 |
| 9348 | NM_018017.3 | 45.6434411 | -7.1360701 | 1.42568056 | -5.005378 |
| 9349 | NM_001325.2 | 30.3866655 | -6.5517974 | 1.30899777 | -5.0052014 |
| 9350 | NM_172056.2 | 37.9955724 | -6.8689501 | 1.37249142 | -5.0047308 |
| 9351 | NM_001206.2 | 54.7624265 | -6.4230956 | 1.28348157 | -5.0044315 |
| 9352 | NR_040289.1 | 32.4198885 | 3.60039395 | 0.71945894 | 5.00430775 |
| 9353 | NM_002560.2 | 26.5660887 | -6.3562529 | 1.27018104 | -5.0042102 |
| 9354 | NM_001011663.1 | 28.83457 | -6.4714256 | 1.29325989 | -5.0039638 |
| 9355 | NM_001270.2 | 124.564253 | -4.392588 | 0.87782143 | -5.0039654 |
| 9356 | NR_024209.1 | 35.8902892 | -6.783285 | 1.35564789 | -5.0037219 |
| 9357 | NM_201589.3 | 107.859667 | -6.6192827 | 1.32295956 | -5.0033901 |
| 9358 | NM_032490.4 | 24.6427867 | -6.2420367 | 1.24759557 | -5.0032533 |
| 9359 | NM_001318082.1 | 36.8003681 | -6.819907 | 1.36314546 | -5.0030662 |
| 9360 | NM_022898.2 | 98.2074651 | -8.2394509 | 1.6469576 | -5.0028312 |
| 9361 | NR_026963.1 | 28.1774945 | -6.4313432 | 1.28567221 | -5.0023195 |
| 9362 | NM_001142616.1 | 53.8005465 | -7.3699847 | 1.47335854 | -5.0021665 |
| 9363 | NM_152409.3 | 12.0952827 | 8.53012909 | 1.70528203 | 5.00218083 |
| 9364 | NM_181652.2 | 45.433007 | -7.1281869 | 1.42506649 | -5.002003 |
| 9365 | NM_020409.2 | 73.837521 | -5.4938656 | 1.09841807 | -5.0016162 |
| 9366 | NM_001812.2 | 25.7842438 | -6.3086976 | 1.26145513 | -5.0011272 |
| 9367 | NM_001278231.1 | 39.8377402 | -6.9385459 | 1.38741953 | -5.0010439 |
| 9368 | NM_001195144.1 | 55.4215175 | -7.4128913 | 1.4823676 | -5.0007105 |
| 9369 | NM_001167675.1 | 66.6077777 | -7.6784858 | 1.53565 | -5.0001535 |
| 9370 | NM_213618.1 | 56.4301352 | -7.4432616 | 1.4887971 | -4.9995138 |
| 9371 | NM_001304779.1 | 31.5807404 | 9.90564972 | 1.98134301 | 4.99946233 |
| 9372 | NM_014624.3 | 34.2124615 | -6.7199835 | 1.344324 | -4.9987826 |
| 9373 | NM_018380.3 | 32.5567142 | -6.6475835 | 1.32990456 | -4.9985417 |

| | | | | | |
|---|---|---|---|---|---|
| 9374 | NM_080593.2 | 25.798857 | -6.317509 | 1.26389981 | -4.9984255 |
| 9375 | NM_001013625.3 | 43.9830266 | -7.0840543 | 1.41736111 | -4.9980589 |
| 9376 | NM_138421.2 | 37.0768576 | -6.8328041 | 1.36716671 | -4.9977842 |
| 9377 | NM_001145860.1 | 37.0840586 | -6.8335801 | 1.36733135 | -4.9977499 |
| 9378 | NM_001286440.1 | 36.1764707 | -6.8018928 | 1.36101945 | -4.9976456 |
| 9379 | NR_003239.1 | 32.6883789 | -6.6538687 | 1.33151733 | -4.9972077 |
| 9380 | NM_001014443.2 | 29.7940683 | -6.5207092 | 1.3048981 | -4.9971022 |
| 9381 | NM_001288634.1 | 44.9369436 | -7.1153034 | 1.42391711 | -4.9969927 |
| 9382 | NM_018139.2 | 33.1050711 | -6.6764691 | 1.33614884 | -4.9968005 |
| 9383 | NM_025215.5 | 37.3147402 | -6.8406613 | 1.36905233 | -4.9966397 |
| 9384 | NM_002907.3 | 41.3870317 | -6.9916975 | 1.39939558 | -4.9962267 |
| 9385 | NM_001278431.1 | 44.2132029 | -7.0919432 | 1.41947917 | -4.9961587 |
| 9386 | NM_001130415.1 | 30.4206241 | -6.5478476 | 1.31072053 | -4.9956093 |
| 9387 | NM_024528.3 | 54.1647672 | -5.520723 | 1.10512574 | -4.995561 |
| 9388 | NM_139033.2 | 30.0317218 | -6.5252415 | 1.30620289 | -4.9955803 |
| 9389 | NM_001278511.1 | 52.8203165 | -7.3445982 | 1.47027986 | -4.9953743 |
| 9390 | NM_001130519.2 | 65.6414504 | -7.6564838 | 1.53276175 | -4.995221 |
| 9391 | NM_004742.2 | 47.6827262 | -7.1992292 | 1.44123298 | -4.9951877 |
| 9392 | NM_001265596.1 | 39.6097812 | -6.9314098 | 1.38768434 | -4.9949471 |
| 9393 | NM_002897.4 | 87.6941752 | -5.0423083 | 1.00956675 | -4.994527 |
| 9394 | NM_001616.4 | 85.5248762 | -7.0657351 | 1.41470468 | -4.9944947 |
| 9395 | NM_172232.3 | 30.8605516 | -6.5698643 | 1.31550505 | -4.9941764 |
| 9396 | NM_014026.4 | 25.5536111 | -6.2960344 | 1.26068116 | -4.9941528 |
| 9397 | NM_018328.4 | 54.1452724 | -6.407707 | 1.28305124 | -4.9941162 |
| 9398 | NM_001260508.1 | 304.245033 | 4.42599433 | 0.88625779 | 4.99402587 |
| 9399 | NM_005133.2 | 38.265443 | -6.8782014 | 1.37740581 | -4.9935911 |
| 9400 | NM_024819.6 | 35.2047686 | -6.7580461 | 1.35340463 | -4.9933671 |
| 9401 | NM_022555.3 | 58.6619181 | -7.4975077 | 1.50156885 | -4.9931161 |
| 9402 | NM_001196.3 | 38.0258917 | -6.8703754 | 1.37604203 | -4.9928529 |
| 9403 | NM_001077181.1 | 25.8570594 | -6.3139136 | 1.26467285 | -4.9925272 |
| 9404 | NM_015194.2 | 57.9368271 | -6.5031519 | 1.30259428 | -4.9924616 |
| 9405 | NM_016950.2 | 48.5653985 | -7.2266616 | 1.44754874 | -4.9923442 |
| 9406 | NM_000017.3 | 26.6544133 | -6.3536056 | 1.27268571 | -4.9922817 |
| 9407 | NR_145420.1 | 184.007887 | 5.75830504 | 1.15348669 | 4.99208623 |
| 9408 | NM_018451.4 | 38.4953609 | -6.8864935 | 1.37950213 | -4.9920137 |
| 9409 | NM_020235.6 | 48.6588201 | -7.2221183 | 1.44678836 | -4.9918278 |
| 9410 | NM_018019.2 | 58.4923788 | -4.726015 | 0.94689661 | -4.9910571 |
| 9411 | NM_033113.2 | 47.1385944 | -4.7970962 | 0.96113096 | -4.9910953 |
| 9412 | NR_135639.1 | 136.483226 | 3.33233712 | 0.66765993 | 4.99106948 |
| 9413 | NM_003442.5 | 27.4925265 | -6.3972318 | 1.28176223 | -4.990966 |
| 9414 | NM_052839.3 | 26.5540418 | -6.3510717 | 1.27262277 | -4.9905376 |
| 9415 | NR_015442.1 | 58.1042505 | -6.5022955 | 1.3030301 | -4.9901346 |
| 9416 | NM_015052.4 | 52.0321361 | -7.3228384 | 1.46760964 | -4.9896364 |

| | | | | | |
|---|---|---|---|---|---|
| 9417 | NM_001326295.1 | 29.7137627 | -6.5151275 | 1.30575194 | -4.9895599 |
| 9418 | NM_203413.2 | 32.9289861 | -6.6647406 | 1.33573963 | -4.9895507 |
| 9419 | NM_001288580.1 | 38.7783303 | -6.8947876 | 1.38186264 | -4.9894884 |
| 9420 | NM_000280.4 | 45.812433 | -7.1353403 | 1.43009291 | -4.9894243 |
| 9421 | NM_016287.3 | 621.970838 | -2.7239409 | 0.54596089 | -4.9892601 |
| 9422 | NM_001289746.1 | 70.0655161 | -6.7824568 | 1.35949506 | -4.9889529 |
| 9423 | NM_001282391.1 | 47.2289957 | -7.18387 | 1.44007606 | -4.9885351 |
| 9424 | NR_027671.2 | 39.5092887 | -6.9282569 | 1.38885207 | -4.9884772 |
| 9425 | NM_001023560.3 | 38.5769316 | -6.8887661 | 1.38126293 | -4.9872953 |
| 9426 | NM_032898.4 | 55.0694516 | -3.2668583 | 0.65506682 | -4.9870612 |
| 9427 | NM_001001994.2 | 111.157627 | -7.4513796 | 1.49425358 | -4.9866901 |
| 9428 | NM_080656.2 | 80.6658325 | -6.000247 | 1.20341844 | -4.9860022 |
| 9429 | NM_001161358.1 | 42.5636241 | -7.0317022 | 1.41030777 | -4.9859345 |
| 9430 | NM_003630.2 | 34.9987635 | -6.7488426 | 1.35357102 | -4.9859538 |
| 9431 | NM_031478.5 | 299.620682 | -6.2456641 | 1.25273409 | -4.9856264 |
| 9432 | NM_001322412.1 | 46.2714732 | -7.1542644 | 1.43499661 | -4.9855619 |
| 9433 | NM_001197319.2 | 22.8618368 | 9.4232154 | 1.89031303 | 4.98500263 |
| 9434 | NM_000408.4 | 42.2287213 | -7.0203895 | 1.40835098 | -4.9848295 |
| 9435 | NM_001330495.1 | 46.9323013 | -7.1726387 | 1.43892283 | -4.9847279 |
| 9436 | NM_022165.2 | 52.1432874 | -7.326474 | 1.46982788 | -4.9845796 |
| 9437 | NM_213594.2 | 45.4589308 | -7.1270282 | 1.42985674 | -4.9844351 |
| 9438 | NM_002023.4 | 33.8154627 | -6.7054431 | 1.34545515 | -4.9837731 |
| 9439 | NM_001206840.1 | 30.3260987 | -6.550482 | 1.31438685 | -4.9836789 |
| 9440 | NM_001284388.1 | 39.4374722 | -6.9184576 | 1.38835472 | -4.9832061 |
| 9441 | NM_153047.3 | 29.4060852 | -6.4982789 | 1.30403415 | -4.9832122 |
| 9442 | NM_138632.2 | 28.144278 | -6.4383428 | 1.29205129 | -4.9830396 |
| 9443 | NR_144477.1 | 38.2128216 | -6.8815007 | 1.38110569 | -4.9826025 |
| 9444 | NM_152654.2 | 231.007853 | 2.71032062 | 0.54398227 | 4.98236939 |
| 9445 | NR_026718.1 | 88.8989037 | -4.8034791 | 0.96410539 | -4.9823175 |
| 9446 | NM_001024858.2 | 37.9691483 | -6.8680578 | 1.37854382 | -4.9821106 |
| 9447 | NM_012139.3 | 28.6045558 | -6.4550455 | 1.29565746 | -4.9820618 |
| 9448 | NM_016567.3 | 38.6491844 | -6.8918025 | 1.38341392 | -4.9817357 |
| 9449 | NM_018112.2 | 34.293346 | -6.7195227 | 1.34888663 | -4.9815326 |
| 9450 | NM_032319.2 | 36.5083181 | -6.8117442 | 1.36746586 | -4.9812901 |
| 9451 | NM_006716.3 | 34.5188223 | -6.7285298 | 1.35079114 | -4.981177 |
| 9452 | NM_001114748.1 | 32.7513616 | -6.6525729 | 1.33562431 | -4.9808714 |
| 9453 | NM_001042724.1 | 126.507279 | -5.4624653 | 1.09671314 | -4.9807603 |
| 9454 | NM_018452.5 | 79.3539585 | -6.1588268 | 1.23656751 | -4.9805828 |
| 9455 | NM_006530.3 | 41.8489039 | -7.0069199 | 1.40689385 | -4.9804183 |
| 9456 | NM_004416.2 | 138.790149 | -6.0570549 | 1.21620433 | -4.9802938 |
| 9457 | NR_024122.1 | 44.4951127 | -7.0981222 | 1.42524706 | -4.9802749 |
| 9458 | NM_152526.5 | 25.9931475 | -6.3229547 | 1.26980116 | -4.9794841 |
| 9459 | NM_001135610.1 | 34.812846 | -6.7454282 | 1.3547203 | -4.9792036 |

| | | | | | |
|---|---|---|---|---|---|
| 9460 | NM_024078.2 | 63.596648 | -6.6424146 | 1.33421037 | -4.9785362 |
| 9461 | NM_017735.4 | 37.3684417 | -6.8455496 | 1.37505584 | -4.9783794 |
| 9462 | NM_001161345.1 | 50.5116057 | -7.2817067 | 1.46276074 | -4.9780573 |
| 9463 | NM_016085.4 | 31.3371378 | -6.5915526 | 1.32417791 | -4.9778452 |
| 9464 | NM_001143779.1 | 36.8716217 | -6.8206081 | 1.37021993 | -4.9777469 |
| 9465 | NM_001160246.1 | 49.7248842 | -7.2569607 | 1.4579378 | -4.9775516 |
| 9466 | NM_031434.3 | 26.8131538 | -6.3662838 | 1.27900384 | -4.977533 |
| 9467 | NM_001242858.2 | 57.2657091 | -7.4610011 | 1.49913381 | -4.9768747 |
| 9468 | NM_016618.2 | 33.1984233 | -6.6761749 | 1.34145957 | -4.9767992 |
| 9469 | NM_080649.2 | 39.1450408 | -6.9092883 | 1.38839722 | -4.9764492 |
| 9470 | NM_024936.2 | 28.114315 | -6.4367711 | 1.2934997 | -4.9762447 |
| 9471 | NM_017440.4 | 29.7226768 | -6.5089766 | 1.30806347 | -4.9760404 |
| 9472 | NM_014264.4 | 28.100311 | -6.4337924 | 1.29310578 | -4.9754571 |
| 9473 | NM_199076.2 | 29.276432 | -6.4960707 | 1.30569011 | -4.975201 |
| 9474 | NM_017510.5 | 409.901685 | -5.920947 | 1.19012787 | -4.9750512 |
| 9475 | NM_001303257.2 | 31.7960936 | -6.6159153 | 1.32983345 | -4.9749954 |
| 9476 | NM_003178.5 | 55.6542191 | -7.4199698 | 1.49178682 | -4.9738808 |
| 9477 | NM_052936.4 | 30.8389342 | -6.5640048 | 1.31977389 | -4.9735829 |
| 9478 | NM_001467.5 | 37.0281101 | -6.8294838 | 1.37321816 | -4.9733422 |
| 9479 | NM_001200003.1 | 38.6886269 | -6.8937772 | 1.3861674 | -4.9732645 |
| 9480 | NM_018218.2 | 99.8443198 | -4.5720868 | 0.91934265 | -4.973213 |
| 9481 | NM_004486.4 | 408.5003 | -3.2937329 | 0.66233321 | -4.9729243 |
| 9482 | NM_016046.4 | 32.9707634 | -6.6668398 | 1.34066796 | -4.9727748 |
| 9483 | NM_001293186.1 | 33.262146 | -6.67703 | 1.3427612 | -4.9726117 |
| 9484 | NM_012257.3 | 38.8146441 | -6.9030361 | 1.38821937 | -4.972583 |
| 9485 | NR_036640.1 | 39.202555 | -6.9134158 | 1.39036522 | -4.9723739 |
| 9486 | NR_110091.1 | 41.7669196 | 2.50944211 | 0.50467739 | 4.9723688 |
| 9487 | NM_032861.3 | 62.2947887 | -6.6140745 | 1.33022075 | -4.9721631 |
| 9488 | NM_001002010.2 | 37.2670448 | -6.8426673 | 1.37620478 | -4.9721287 |
| 9489 | NM_198799.3 | 26.3460462 | -6.3408531 | 1.27532096 | -4.9719665 |
| 9490 | NM_001256875.1 | 34.6121117 | -6.7344245 | 1.3545165 | -4.9718291 |
| 9491 | NM_001281789.1 | 43.3427671 | -7.0574748 | 1.4195008 | -4.9718005 |
| 9492 | NM_018079.4 | 46.3829284 | -6.173933 | 1.24179305 | -4.971789 |
| 9493 | NM_001206602.1 | 37.9019744 | -6.8682214 | 1.38164289 | -4.9710539 |
| 9494 | NM_032730.5 | 26.4665972 | -6.3438546 | 1.27616364 | -4.9710354 |
| 9495 | NM_172251.2 | 41.4392529 | -6.9946434 | 1.40726714 | -4.9703736 |
| 9496 | NM_001282474.1 | 32.5200853 | -6.648173 | 1.33779193 | -4.969512 |
| 9497 | NM_007349.3 | 55.7496708 | -5.5587367 | 1.11859522 | -4.9693907 |
| 9498 | NM_152943.2 | 40.3183067 | -6.9524343 | 1.39918086 | -4.9689318 |
| 9499 | NM_000094.3 | 192.178206 | -5.2606246 | 1.05878322 | -4.9685569 |
| 9500 | NM_001324.2 | 27.2809404 | -6.3934136 | 1.28676986 | -4.9685758 |
| 9501 | NM_181643.5 | 66.8175174 | -7.6859255 | 1.54690242 | -4.968591 |
| 9502 | NM_006928.4 | 36.8048679 | -6.8272297 | 1.37419646 | -4.9681613 |

| | | | | | |
|---|---|---|---|---|---|
| 9503 | NM_001303426.1 | 36.9892139 | -6.8291882 | 1.37460113 | -4.9681235 |
| 9504 | NM_022836.3 | 25.9768575 | -6.3238175 | 1.27293692 | -4.9678954 |
| 9505 | NM_001319115.1 | 33.9788435 | -6.7087892 | 1.3505671 | -4.9673868 |
| 9506 | NM_001135774.1 | 40.1877963 | -6.9531376 | 1.39977164 | -4.9673371 |
| 9507 | NM_001318784.1 | 68.9583731 | -7.728802 | 1.55592188 | -4.9673458 |
| 9508 | NM_001220473.2 | 46.7363816 | -7.1644565 | 1.44240941 | -4.9670062 |
| 9509 | NM_022822.2 | 66.5841801 | -7.6810296 | 1.54654316 | -4.9665796 |
| 9510 | NR_110851.1 | 52.6358054 | 3.07132134 | 0.61840812 | 4.96649578 |
| 9511 | NM_001105244.1 | 63.6638047 | -7.6121451 | 1.53273233 | -4.9663891 |
| 9512 | NM_003800.4 | 54.2116285 | -7.3846942 | 1.48694139 | -4.9663653 |
| 9513 | NM_145729.2 | 37.4736965 | -6.8500843 | 1.37928966 | -4.9663856 |
| 9514 | NM_015976.4 | 33.3235795 | -6.6779218 | 1.34467866 | -4.9661841 |
| 9515 | NR_132313.1 | 36.4567309 | 3.32101871 | 0.6688277 | 4.96543238 |
| 9516 | NM_001002258.4 | 26.4390322 | -6.3497091 | 1.27884234 | -4.9652009 |
| 9517 | NM_032346.1 | 40.6817308 | -6.9671142 | 1.4032848 | -4.9648612 |
| 9518 | NM_000688.5 | 38.8802231 | -6.9042829 | 1.39078285 | -4.9643141 |
| 9519 | NM_015139.2 | 65.1284207 | -3.1043155 | 0.62534747 | -4.9641449 |
| 9520 | NM_001025201.3 | 34.8182258 | -6.7417451 | 1.3581525 | -4.9639088 |
| 9521 | NM_001197113.1 | 40.8907335 | -6.9785097 | 1.40588359 | -4.9637891 |
| 9522 | NM_001199282.2 | 42.8880003 | -7.0439965 | 1.4191425 | -4.9635582 |
| 9523 | NM_004335.3 | 40.9142374 | -6.9794693 | 1.40617797 | -4.9634324 |
| 9524 | NM_001287045.1 | 67.7411963 | -7.7032235 | 1.55206081 | -4.9632227 |
| 9525 | NM_001007075.2 | 40.5462753 | -6.9649758 | 1.40339985 | -4.9629304 |
| 9526 | NM_001301098.1 | 28.6238754 | -6.459471 | 1.30159957 | -4.9627175 |
| 9527 | NM_032027.2 | 43.5940892 | -7.0656242 | 1.42379047 | -4.9625449 |
| 9528 | NR_033436.1 | 83.4473925 | -4.7053238 | 0.94819963 | -4.9623767 |
| 9529 | NM_001029884.2 | 58.8431376 | -7.5025108 | 1.5119811 | -4.9620401 |
| 9530 | NM_001224.4 | 38.3057119 | -6.8783952 | 1.38639005 | -4.9613709 |
| 9531 | NM_001170794.1 | 283.297866 | -4.6482173 | 0.93689772 | -4.9612857 |
| 9532 | NM_001002246.2 | 35.1006299 | -6.7538121 | 1.36132927 | -4.9611892 |
| 9533 | NM_001318249.1 | 54.2739307 | 6.9302186 | 1.39696867 | 4.96089766 |
| 9534 | NM_018353.4 | 30.0387338 | -6.5266238 | 1.31564346 | -4.9607846 |
| 9535 | NM_025077.3 | 34.8332963 | -6.7450875 | 1.35970924 | -4.9606837 |
| 9536 | NM_001122633.2 | 27.3821284 | -6.4010401 | 1.29036688 | -4.9606357 |
| 9537 | NM_001201366.1 | 56.8535374 | -7.4523302 | 1.50235679 | -4.9604264 |
| 9538 | NM_007214.4 | 369.096605 | -4.5866144 | 0.92467694 | -4.9602345 |
| 9539 | NM_144721.5 | 26.9172442 | -6.3767222 | 1.2856204 | -4.960035 |
| 9540 | NM_001330232.1 | 64.0787339 | -7.6249464 | 1.53739693 | -4.9596472 |
| 9541 | NR_038301.1 | 31.8925559 | -6.6166373 | 1.33412635 | -4.9595282 |
| 9542 | NM_003783.3 | 58.579544 | -7.4937671 | 1.51114278 | -4.9590066 |
| 9543 | NM_001270399.1 | 40.2757614 | -6.9506088 | 1.40161981 | -4.958983 |
| 9544 | NM_021076.3 | 104.988177 | -5.3203007 | 1.07289893 | -4.9588088 |
| 9545 | NR_003561.2 | 132.83243 | 1.81341695 | 0.36574245 | 4.95818016 |

| | | | | | |
|---|---|---|---|---|---|
| 9546 | NM_001002245.2 | 37.2631438 | -6.8424507 | 1.3800639 | -4.958068 |
| 9547 | NM_152640.4 | 28.3173442 | -6.4494436 | 1.30079247 | -4.9580881 |
| 9548 | NM_002975.2 | 24.4016637 | -6.23508 | 1.25763488 | -4.9577824 |
| 9549 | NM_032138.5 | 52.1450913 | -6.3441723 | 1.27976125 | -4.9573092 |
| 9550 | NM_014324.5 | 33.9034507 | -6.7077158 | 1.35310814 | -4.9572652 |
| 9551 | NM_001037495.1 | 40.0326269 | -6.9436168 | 1.40076874 | -4.9570044 |
| 9552 | NM_175063.5 | 35.8174909 | -6.7856984 | 1.36890959 | -4.9570099 |
| 9553 | NM_020760.2 | 33.0387144 | -6.6681183 | 1.34529818 | -4.9566099 |
| 9554 | NM_003112.4 | 40.2649463 | -6.9492628 | 1.40206707 | -4.956441 |
| 9555 | NM_001039840.2 | 31.6266751 | -6.6056288 | 1.33277931 | -4.956281 |
| 9556 | NM_020440.3 | 340.408318 | -5.8634436 | 1.18306669 | -4.9561396 |
| 9557 | NR_045026.1 | 27.4057935 | -6.3972605 | 1.29088726 | -4.9557082 |
| 9558 | NM_001013663.1 | 39.5418593 | -6.9264114 | 1.39767397 | -4.9556703 |
| 9559 | NM_178826.3 | 26.3675041 | -6.3492018 | 1.28134491 | -4.9551076 |
| 9560 | NM_001098484.2 | 26.0885374 | -6.3206854 | 1.2756826 | -4.9547477 |
| 9561 | NR_109807.1 | 25.4266276 | -6.2935059 | 1.27021759 | -4.9546676 |
| 9562 | NM_007215.3 | 23.8324101 | -6.1949766 | 1.25052799 | -4.9538888 |
| 9563 | NR_037188.1 | 27.0908183 | -6.3882008 | 1.28953518 | -4.9538786 |
| 9564 | NM_022768.4 | 25.75155 | -6.3054331 | 1.27286509 | -4.9537324 |
| 9565 | NM_172359.2 | 40.0054388 | -6.9460286 | 1.40228935 | -4.9533491 |
| 9566 | NM_007080.2 | 33.5886368 | -6.6932393 | 1.35142495 | -4.9527273 |
| 9567 | NM_001330785.1 | 63.3758033 | -7.6063011 | 1.53580598 | -4.9526445 |
| 9568 | NM_004277.4 | 27.4072402 | -6.4052756 | 1.29339022 | -4.9523148 |
| 9569 | NM_001294348.1 | 56.7758445 | -7.4465632 | 1.50384474 | -4.9516835 |
| 9570 | NM_001319143.1 | 34.3958613 | -6.7252759 | 1.35823438 | -4.9514841 |
| 9571 | NM_001288615.1 | 24.3926773 | -6.2278396 | 1.25780128 | -4.9513701 |
| 9572 | NM_001001991.2 | 27.628029 | -6.4139195 | 1.29550195 | -4.9509146 |
| 9573 | NM_001144879.1 | 25.3499998 | 9.58904305 | 1.93695237 | 4.95058278 |
| 9574 | NM_032355.3 | 44.0739812 | -7.0836093 | 1.43086527 | -4.9505775 |
| 9575 | NM_003242.5 | 29.4835799 | -6.5080249 | 1.31468311 | -4.9502613 |
| 9576 | NM_000964.3 | 24.9851013 | -6.2678761 | 1.26642355 | -4.9492732 |
| 9577 | NM_014481.3 | 26.0911047 | -6.3249523 | 1.27805022 | -4.9489075 |
| 9578 | NM_001201556.1 | 35.13983 | -6.7561987 | 1.36522137 | -4.9487935 |
| 9579 | NM_022366.2 | 28.3701216 | -6.4441733 | 1.30224643 | -4.9485053 |
| 9580 | NM_181882.2 | 27.2134033 | -6.3925499 | 1.29190382 | -4.9481624 |
| 9581 | NM_152760.2 | 46.9008055 | -7.1723187 | 1.44962794 | -4.9476962 |
| 9582 | NM_001174120.1 | 39.7994788 | -6.9319831 | 1.40109775 | -4.9475371 |
| 9583 | NM_018479.3 | 50.6992507 | -7.2872092 | 1.47291106 | -4.9474876 |
| 9584 | NM_001004127.2 | 27.3027736 | -6.3953199 | 1.29275014 | -4.9470657 |
| 9585 | NM_032740.3 | 52.9426565 | -6.367943 | 1.28729114 | -4.9467776 |
| 9586 | NM_001191057.3 | 36.5360234 | -6.8138163 | 1.37743923 | -4.9467273 |
| 9587 | NM_001206646.1 | 33.7820261 | -6.7047727 | 1.35543831 | -4.9465717 |
| 9588 | NM_001001976.2 | 33.1816331 | -6.6768072 | 1.34982575 | -4.9464216 |

| | | | | | |
|---|---|---|---|---|---|
| 9589 | NM_001099678.1 | 252.876473 | -1.5554969 | 0.31452907 | -4.945479 |
| 9590 | NM_173824.3 | 54.590645 | -6.4120792 | 1.29671616 | -4.9448595 |
| 9591 | NM_001142302.1 | 27.6996654 | 9.71587816 | 1.96495661 | 4.94457645 |
| 9592 | NM_001135654.1 | 54.4214053 | -7.3872026 | 1.49405528 | -4.9443971 |
| 9593 | NM_024520.2 | 28.814487 | -6.4709426 | 1.30874986 | -4.9443693 |
| 9594 | NM_001033719.2 | 28.6521815 | -6.4602679 | 1.30664725 | -4.944156 |
| 9595 | NM_030761.4 | 67.5309411 | -6.724957 | 1.36018021 | -4.9441662 |
| 9596 | NM_003678.4 | 44.0691888 | -7.0832255 | 1.43282972 | -4.9435222 |
| 9597 | NM_007069.3 | 31.5514943 | -6.6044983 | 1.33600919 | -4.9434527 |
| 9598 | NM_001098206.1 | 31.3680813 | -6.5936283 | 1.33383869 | -4.9433476 |
| 9599 | NM_001303037.1 | 69.0795557 | -5.8570587 | 1.18485397 | -4.9432747 |
| 9600 | NM_016401.3 | 51.9381918 | -7.3191731 | 1.48074698 | -4.9428925 |
| 9601 | NM_001077427.3 | 48.0898363 | -7.2106582 | 1.45906094 | -4.9419857 |
| 9602 | NM_134446.3 | 43.3967679 | -7.0601305 | 1.42864378 | -4.941841 |
| 9603 | NM_003155.2 | 44.5878112 | -5.2542239 | 1.06322564 | -4.9417769 |
| 9604 | NM_018339.5 | 176.562254 | -3.6880279 | 0.74631913 | -4.9416231 |
| 9605 | NR_002217.1 | 30.4145558 | -6.5493207 | 1.32534138 | -4.9416104 |
| 9606 | NM_033411.4 | 29.3785186 | -6.4986883 | 1.31512068 | -4.9415148 |
| 9607 | NM_014932.3 | 37.2191694 | -6.8366518 | 1.3836411 | -4.9410586 |
| 9608 | NM_144638.2 | 28.942866 | -6.4751098 | 1.31057034 | -4.9406808 |
| 9609 | NM_001329943.1 | 33.457477 | -6.6829615 | 1.35266438 | -4.940591 |
| 9610 | NM_173564.3 | 243.425239 | -5.5528519 | 1.12395359 | -4.9404637 |
| 9611 | NM_023106.2 | 58.2985872 | -7.4887638 | 1.51583102 | -4.9403685 |
| 9612 | NM_174976.2 | 46.2060135 | -7.1527242 | 1.44789583 | -4.940082 |
| 9613 | NM_152307.2 | 56.2894306 | -6.4582236 | 1.30741813 | -4.9396772 |
| 9614 | NM_178326.2 | 37.7297865 | -6.8638285 | 1.38970967 | -4.9390377 |
| 9615 | NM_001317961.1 | 33.2784848 | -6.6759138 | 1.35172866 | -4.9387973 |
| 9616 | NR_133948.1 | 51.6791379 | -7.3110643 | 1.48047011 | -4.9383397 |
| 9617 | NM_080867.2 | 90.0756214 | -3.8493222 | 0.77950122 | -4.9381863 |
| 9618 | NM_024630.2 | 38.4741209 | -6.8889841 | 1.39523594 | -4.9375047 |
| 9619 | NM_001048209.1 | 71.4172547 | -7.7797426 | 1.57592086 | -4.9366328 |
| 9620 | NM_174858.2 | 42.6330088 | -7.0362556 | 1.42550709 | -4.9359667 |
| 9621 | NR_038281.1 | 48.6109734 | 2.41766462 | 0.48980679 | 4.9359557 |
| 9622 | NR_033399.1 | 30.2228683 | -6.5342536 | 1.32396323 | -4.935374 |
| 9623 | NM_022734.2 | 38.6834148 | -6.8977143 | 1.39762204 | -4.9353217 |
| 9624 | NM_001317990.1 | 45.5876107 | -7.1360843 | 1.44606352 | -4.9348346 |
| 9625 | NM_007171.3 | 48.0860812 | -7.2053025 | 1.46008449 | -4.9348531 |
| 9626 | NM_001243571.1 | 38.4634133 | -6.8910053 | 1.39652664 | -4.9343887 |
| 9627 | NM_001048212.3 | 31.9358907 | -6.6189924 | 1.3417716 | -4.9330247 |
| 9628 | NM_001198536.1 | 158.184027 | 3.25298703 | 0.65944289 | 4.93293213 |
| 9629 | NM_000353.2 | 50.7248172 | 2.93197081 | 0.59437084 | 4.93289813 |
| 9630 | NM_001330488.1 | 43.8294816 | -7.074979 | 1.4342991 | -4.9327082 |
| 9631 | NR_130898.1 | 33.5412332 | -6.6896063 | 1.35648663 | -4.9315682 |

| | | | | | |
|---|---|---|---|---|---|
| 9632 | NR_024241.1 | 25.6211488 | -6.3036246 | 1.27825955 | -4.9314121 |
| 9633 | NM_001330760.1 | 53.9172711 | -7.3724767 | 1.49509372 | -4.9311134 |
| 9634 | NM_001257096.1 | 51.9043596 | -7.3207244 | 1.48472661 | -4.9306885 |
| 9635 | NM_005334.2 | 358.650663 | -4.4538381 | 0.90331415 | -4.9305528 |
| 9636 | NM_021948.4 | 49.0133898 | -7.2368844 | 1.46790024 | -4.9300928 |
| 9637 | NM_153335.5 | 30.8703827 | -6.5700185 | 1.33268184 | -4.9299227 |
| 9638 | NR_034123.1 | 25.3605906 | -6.2898355 | 1.27585899 | -4.929883 |
| 9639 | NM_001199863.1 | 34.8227102 | -6.7412803 | 1.36744697 | -4.9298295 |
| 9640 | NM_000512.4 | 37.5937667 | -6.8542213 | 1.39049502 | -4.9293389 |
| 9641 | NM_001142599.1 | 25.542771 | -6.3014421 | 1.27842871 | -4.9290524 |
| 9642 | NM_001303062.1 | 27.150135 | -6.3908496 | 1.29670397 | -4.928534 |
| 9643 | NM_001289413.1 | 54.1484565 | -7.3804264 | 1.49757282 | -4.9282588 |
| 9644 | NM_001323960.1 | 27.6640549 | -6.4078139 | 1.30021303 | -4.9282801 |
| 9645 | NR_110555.1 | 41.9108501 | -6.0393722 | 1.22545864 | -4.9282546 |
| 9646 | NM_005071.2 | 45.0460882 | -7.115011 | 1.44377507 | -4.9280606 |
| 9647 | NM_001160418.1 | 23.9787954 | 3.23316477 | 0.65617564 | 4.92728557 |
| 9648 | NM_001317968.1 | 25.3851374 | -6.2867863 | 1.27594128 | -4.9271752 |
| 9649 | NM_033393.2 | 26.7287704 | -6.3622281 | 1.29133326 | -4.9268677 |
| 9650 | NR_132419.1 | 43.5989222 | -7.0692707 | 1.43489866 | -4.926669 |
| 9651 | NM_001291604.1 | 19.318616 | 9.20121538 | 1.86765066 | 4.92662552 |
| 9652 | NM_001280787.1 | 38.2992458 | -6.8780723 | 1.39625121 | -4.9260995 |
| 9653 | NM_001278158.1 | 25.7964333 | -6.3093104 | 1.2808303 | -4.9259534 |
| 9654 | NR_045861.1 | 24.3863959 | -6.2285581 | 1.26451675 | -4.925643 |
| 9655 | NM_000696.3 | 159.939693 | -4.9819697 | 1.01150859 | -4.9252867 |
| 9656 | NM_001330164.1 | 40.048279 | -6.9490813 | 1.41095529 | -4.9250897 |
| 9657 | NM_022079.2 | 48.4313736 | -7.2167529 | 1.46536145 | -4.9248961 |
| 9658 | NM_052924.2 | 125.931229 | -3.7395848 | 0.75932337 | -4.924891 |
| 9659 | NM_001011551.2 | 28.5206302 | -6.4549924 | 1.31069288 | -4.9248703 |
| 9660 | NM_015575.3 | 47.1866979 | -7.1781596 | 1.45762817 | -4.9245478 |
| 9661 | NM_198316.1 | 49.6348456 | -5.4505387 | 1.10681179 | -4.9245398 |
| 9662 | NM_003183.5 | 149.091421 | -3.9923102 | 0.81079556 | -4.9239419 |
| 9663 | NM_138395.3 | 27.4648659 | -6.4008935 | 1.29996614 | -4.9238925 |
| 9664 | NM_016125.3 | 25.1789897 | -6.2778079 | 1.27505359 | -4.923564 |
| 9665 | NM_001031734.3 | 39.894717 | -6.9406619 | 1.409943 | -4.9226543 |
| 9666 | NR_003613.1 | 30.6839248 | -6.5628727 | 1.33319354 | -4.9226706 |
| 9667 | NM_003622.3 | 33.3121073 | -6.6775457 | 1.3565284 | -4.9225256 |
| 9668 | NM_001256166.1 | 35.5131694 | -6.7744185 | 1.37633748 | -4.9220621 |
| 9669 | NM_153354.4 | 37.3724001 | -6.8437289 | 1.3905011 | -4.9217717 |
| 9670 | NM_001322006.1 | 48.8638872 | -7.2305333 | 1.4691229 | -4.9216668 |
| 9671 | NM_021021.3 | 41.762197 | -7.0094102 | 1.42422598 | -4.9215576 |
| 9672 | NM_001740.4 | 42.1406695 | -7.0191855 | 1.42622768 | -4.9215042 |
| 9673 | NM_017680.4 | 25.64021 | -6.3094897 | 1.28206347 | -4.9213552 |
| 9674 | NM_001143828.1 | 66.4939158 | 2.98906307 | 0.60741786 | 4.92093381 |

| | | | | | |
|---|---|---|---|---|---|
| 9675 | NM_145810.2 | 35.4424708 | -6.7695914 | 1.375759 | -4.920623 |
| 9676 | NM_015269.2 | 40.2694373 | -6.9505424 | 1.41262467 | -4.9203037 |
| 9677 | NM_006949.3 | 29.2285931 | -6.4958888 | 1.32027095 | -4.920118 |
| 9678 | NR_074083.1 | 29.1276701 | -6.4824012 | 1.31759569 | -4.9198712 |
| 9679 | NM_021080.3 | 60.1109224 | -7.530479 | 1.53081153 | -4.9192724 |
| 9680 | NM_030917.3 | 30.5391276 | -6.5577261 | 1.33308179 | -4.9192226 |
| 9681 | NM_017425.3 | 29.7775478 | -6.5221209 | 1.32586432 | -4.9191466 |
| 9682 | NM_001113496.1 | 39.0127707 | -6.9099985 | 1.40477479 | -4.9189369 |
| 9683 | NM_018480.4 | 36.5604343 | -6.8141548 | 1.38529713 | -4.918912 |
| 9684 | NM_152414.4 | 117.858592 | -8.5025901 | 1.7285804 | -4.9188283 |
| 9685 | NM_001077238.1 | 33.7266545 | -6.6964721 | 1.36146329 | -4.9185844 |
| 9686 | NR_003287.2 | 22646.9502 | -6.8941166 | 1.40169754 | -4.9184053 |
| 9687 | NM_001329632.1 | 30.235978 | -6.5352213 | 1.3287854 | -4.9181917 |
| 9688 | NM_001304833.1 | 26.5352196 | -6.356828 | 1.29254795 | -4.9180597 |
| 9689 | NM_001143912.1 | 29.2740039 | -6.4932341 | 1.32054638 | -4.9170815 |
| 9690 | NM_001267065.1 | 28.9921875 | -6.4733786 | 1.31662649 | -4.9166401 |
| 9691 | NM_001003697.1 | 35.3392092 | -6.7656385 | 1.37615626 | -4.9163301 |
| 9692 | NM_018398.2 | 28.4811278 | -6.4542858 | 1.31304172 | -4.9155223 |
| 9693 | NM_000983.3 | 1040.8391 | -6.9678729 | 1.41755709 | -4.915409 |
| 9694 | NM_001005909.2 | 91.2927651 | -5.1080229 | 1.03919073 | -4.9153853 |
| 9695 | NM_201526.1 | 46.5947019 | -7.1661671 | 1.45793724 | -4.9152782 |
| 9696 | NM_206866.2 | 36.5525873 | -6.817724 | 1.38708492 | -4.9151453 |
| 9697 | NM_007370.6 | 31.3878418 | -6.5929183 | 1.34135655 | -4.9151124 |
| 9698 | NM_017990.4 | 38.413852 | -6.8904482 | 1.40189392 | -4.9150995 |
| 9699 | NM_031904.4 | 36.0807911 | -6.8004849 | 1.38365974 | -4.9148535 |
| 9700 | NM_033529.3 | 36.8877766 | -6.8308201 | 1.38983169 | -4.9148542 |
| 9701 | NM_012237.3 | 49.9446995 | -7.263436 | 1.4779018 | -4.9146946 |
| 9702 | NM_019887.5 | 77.7612135 | -5.1743903 | 1.05286277 | -4.9145914 |
| 9703 | NR_073499.1 | 41.6679951 | -7.0026835 | 1.42487768 | -4.9145857 |
| 9704 | NM_006303.3 | 39.9416151 | -6.939727 | 1.41210286 | -4.9144628 |
| 9705 | NM_001172705.1 | 30.8300516 | -6.568404 | 1.33673956 | -4.91375 |
| 9706 | NM_002390.5 | 49.9672808 | -7.2641082 | 1.47838968 | -4.9135274 |
| 9707 | NM_001143688.2 | 29.7676859 | -6.5236438 | 1.32775802 | -4.9132776 |
| 9708 | NM_001167934.1 | 36.2495843 | -6.8012372 | 1.38429212 | -4.9131517 |
| 9709 | NM_001345869.1 | 36.7026479 | -6.8158748 | 1.38730991 | -4.9130153 |
| 9710 | NR_102705.1 | 30.720628 | -6.565679 | 1.33645546 | -4.9127556 |
| 9711 | NM_138355.3 | 30.2061681 | -6.5337572 | 1.33017262 | -4.9119618 |
| 9712 | NM_005809.5 | 771.432252 | -6.5013798 | 1.32359287 | -4.9119181 |
| 9713 | NM_138774.3 | 203.945871 | -5.5027023 | 1.12029973 | -4.9118126 |
| 9714 | NM_018352.2 | 32.8520923 | -6.6559117 | 1.35512578 | -4.911656 |
| 9715 | NM_001962.2 | 100.973348 | -4.0599754 | 0.82667386 | -4.9112179 |
| 9716 | NM_005921.1 | 46.0559562 | -5.3412901 | 1.08769366 | -4.9106566 |
| 9717 | NM_182498.3 | 448.17741 | -6.1611616 | 1.25470971 | -4.910428 |

| | | | | | |
|---|---|---|---|---|---|
| 9718 | NM_174890.3 | 23.6416184 | -6.1856584 | 1.25978685 | -4.9100833 |
| 9719 | NM_015510.4 | 31.2059854 | -6.5859204 | 1.34142103 | -4.9096594 |
| 9720 | NM_080759.5 | 89.8684191 | -4.8226617 | 0.98229021 | -4.9096098 |
| 9721 | NM_001243351.1 | 34.4676614 | -6.7309613 | 1.37105098 | -4.9093443 |
| 9722 | NM_173573.2 | 56.364595 | -7.4402263 | 1.5155241 | -4.9093421 |
| 9723 | NM_015393.3 | 115.416147 | -7.5078296 | 1.52976684 | -4.9078261 |
| 9724 | NM_001199388.2 | 26.1985134 | -6.3370167 | 1.29162433 | -4.9062383 |
| 9725 | NM_019589.2 | 342.228614 | -5.1317364 | 1.04596438 | -4.9062248 |
| 9726 | NM_015954.3 | 29.1591504 | -6.4870117 | 1.32223616 | -4.9060916 |
| 9727 | NM_001316920.1 | 30.1822833 | -6.5351962 | 1.33207126 | -4.906041 |
| 9728 | NM_004977.2 | 26.9546391 | -6.3773462 | 1.29989262 | -4.9060562 |
| 9729 | NM_014472.4 | 23.6463909 | -6.1874362 | 1.2611995 | -4.9059932 |
| 9730 | NM_145271.3 | 26.4325874 | -6.3427278 | 1.2928807 | -4.9058879 |
| 9731 | NM_005522.4 | 51.2714717 | -7.3029651 | 1.48862615 | -4.9058423 |
| 9732 | NM_023112.3 | 23.9143671 | -6.2025458 | 1.26432267 | -4.905825 |
| 9733 | NM_001127322.1 | 30.9112544 | -6.5721145 | 1.33972766 | -4.90556 |
| 9734 | NM_024615.3 | 29.2023673 | -6.4892599 | 1.32288249 | -4.9053941 |
| 9735 | NM_001199267.1 | 37.3078907 | -6.8451383 | 1.39544676 | -4.9053382 |
| 9736 | NM_001375.2 | 71.1018667 | -3.8554309 | 0.78599059 | -4.9051871 |
| 9737 | NM_001013694.2 | 31.9715691 | -6.6193393 | 1.34961151 | -4.9046257 |
| 9738 | NM_001322437.1 | 25.0667363 | -6.2667461 | 1.27774587 | -4.9045325 |
| 9739 | NM_001193523.1 | 25.6448633 | -6.3052069 | 1.2856789 | -4.9041848 |
| 9740 | NM_001284295.1 | 28.976756 | -6.4768321 | 1.32100658 | -4.9029522 |
| 9741 | NM_006120.3 | 50.0833049 | -7.2692822 | 1.48280633 | -4.9023814 |
| 9742 | NM_001142798.2 | 44.8191912 | -7.1077152 | 1.44990862 | -4.9021815 |
| 9743 | NM_001317395.1 | 38.8221656 | -6.8993896 | 1.40741355 | -4.9021765 |
| 9744 | NM_001198938.1 | 23.4575527 | -6.1796791 | 1.26062395 | -4.9020797 |
| 9745 | NM_021195.4 | 47.5710561 | -7.1946428 | 1.46769212 | -4.9020109 |
| 9746 | NM_001184749.2 | 26.825523 | -6.3648538 | 1.29846488 | -4.9018298 |
| 9747 | NM_001271562.1 | 29.5198817 | -6.5044375 | 1.32693416 | -4.901854 |
| 9748 | NM_080764.3 | 27.3864027 | -6.3928373 | 1.30417198 | -4.9018361 |
| 9749 | NR_027906.1 | 30.2581701 | -6.5380256 | 1.33385285 | -4.9016093 |
| 9750 | NM_001329639.1 | 27.7301943 | -6.419915 | 1.30986254 | -4.9012128 |
| 9751 | NM_020177.2 | 103.09257 | -4.1153433 | 0.83970088 | -4.9009634 |
| 9752 | NM_001083965.1 | 38.0911296 | -6.8705448 | 1.40191416 | -4.9008313 |
| 9753 | NM_001143978.2 | 34.6596092 | -6.7370261 | 1.37471022 | -4.9006881 |
| 9754 | NM_001277145.1 | 25.4591197 | -6.2982723 | 1.28523201 | -4.9004944 |
| 9755 | NM_145231.3 | 27.8410132 | -6.4268315 | 1.31153455 | -4.9002381 |
| 9756 | NM_001198793.1 | 38.6410494 | -6.8918514 | 1.4064439 | -4.9001964 |
| 9757 | NM_005333.4 | 37.7032098 | -6.8619595 | 1.40047227 | -4.8997468 |
| 9758 | NM_020979.4 | 68.679478 | -6.7542713 | 1.37850521 | -4.8997067 |
| 9759 | NM_002192.3 | 101.633121 | -4.7474575 | 0.96916563 | -4.8984997 |
| 9760 | NR_034105.3 | 31.9541139 | -6.6226323 | 1.35199095 | -4.8984294 |

| | | | | | |
|---|---|---|---|---|---|
| 9761 | NM_005008.3 | 74.5095627 | -6.872985 | 1.40313121 | -4.8983195 |
| 9762 | NM_144617.2 | 26.7773615 | -6.3659744 | 1.29969274 | -4.8980611 |
| 9763 | NM_004978.4 | 42.7584266 | -7.0396912 | 1.43728513 | -4.8979086 |
| 9764 | NM_001033714.2 | 28.1144206 | -6.4319461 | 1.31326003 | -4.8976943 |
| 9765 | NM_001144759.2 | 284.746091 | -5.5973983 | 1.14286606 | -4.8976853 |
| 9766 | NM_001271634.1 | 51.6810446 | -4.1599116 | 0.84943166 | -4.8972882 |
| 9767 | NM_001099733.1 | 45.4118018 | -7.1283114 | 1.45561604 | -4.8971097 |
| 9768 | NM_018035.2 | 42.0042561 | -7.0126337 | 1.4320064 | -4.8970687 |
| 9769 | NM_000263.3 | 96.4026461 | -3.833644 | 0.78287882 | -4.8968549 |
| 9770 | NM_001195753.1 | 24.3992183 | -6.2249369 | 1.27120912 | -4.896863 |
| 9771 | NM_019625.3 | 41.63381 | -6.9988068 | 1.42928707 | -4.8967117 |
| 9772 | NM_152396.3 | 31.5666512 | -6.6023145 | 1.34837131 | -4.8965106 |
| 9773 | NM_000124.3 | 34.8687074 | -6.7456746 | 1.37771306 | -4.8962841 |
| 9774 | NM_000853.3 | 31.077229 | -6.5740337 | 1.34265767 | -4.8962843 |
| 9775 | NR_034126.1 | 134.028839 | 2.57858298 | 0.52665535 | 4.89614883 |
| 9776 | NM_005482.2 | 69.151034 | -6.7576358 | 1.3803359 | -4.8956459 |
| 9777 | NM_001008390.1 | 34.782641 | -6.742064 | 1.37727468 | -4.8952211 |
| 9778 | NM_006831.2 | 32.1494652 | -6.6256526 | 1.35362114 | -4.8947614 |
| 9779 | NM_001330360.1 | 31.5979024 | -6.6062519 | 1.34972766 | -4.8945073 |
| 9780 | NM_006113.4 | 27.0360696 | -6.3809849 | 1.30371488 | -4.8944635 |
| 9781 | NM_019111.4 | 435.867645 | -6.3575957 | 1.29911937 | -4.8937733 |
| 9782 | NM_015279.1 | 47.2778413 | -7.1837701 | 1.46795453 | -4.8937279 |
| 9783 | NM_001145165.1 | 43.4886465 | -7.0616961 | 1.44301978 | -4.8936932 |
| 9784 | NM_003738.4 | 33.9280438 | -6.7068347 | 1.370545 | -4.8935531 |
| 9785 | NR_003246.4 | 27.0927134 | -6.3862783 | 1.30510766 | -4.8932962 |
| 9786 | NM_000795.3 | 42.1766723 | -7.0203395 | 1.43472981 | -4.8931439 |
| 9787 | NM_015082.1 | 32.2784838 | -6.6361415 | 1.35631493 | -4.8927733 |
| 9788 | NM_001172225.2 | 30.7322256 | -6.5635165 | 1.34178435 | -4.8916329 |
| 9789 | NM_001195563.1 | 25.4837411 | -6.2883205 | 1.28570714 | -4.8909431 |
| 9790 | NM_001330333.1 | 46.4616856 | -7.1574782 | 1.4634245 | -4.8909104 |
| 9791 | NM_012382.2 | 77.0268379 | -4.0905635 | 0.83645535 | -4.8903549 |
| 9792 | NM_001039672.2 | 34.8102477 | -6.7390693 | 1.37820046 | -4.8897599 |
| 9793 | NM_001376.4 | 1635.48849 | -5.5800961 | 1.14123385 | -4.8895291 |
| 9794 | NM_004251.4 | 36.5926695 | -6.8182507 | 1.39469794 | -4.8886934 |
| 9795 | NM_001145033.1 | 53.9613656 | -7.3738893 | 1.50837591 | -4.8886284 |
| 9796 | NM_001083924.1 | 19.2879796 | 6.03963063 | 1.23545924 | 4.88857133 |
| 9797 | NM_001281453.1 | 40.1840192 | -6.9476108 | 1.42137374 | -4.8879549 |
| 9798 | NM_001242793.1 | 37.1958944 | -6.8405448 | 1.39968593 | -4.8871998 |
| 9799 | NM_012300.2 | 42.3530832 | -7.0249076 | 1.4374314 | -4.8871255 |
| 9800 | NM_133467.2 | 25.5553134 | -6.3013997 | 1.28953954 | -4.8865502 |
| 9801 | NR_026982.1 | 40.6977499 | -6.9686084 | 1.42608749 | -4.8865224 |
| 9802 | NM_144635.4 | 45.8942757 | -7.1410472 | 1.46155312 | -4.8859306 |
| 9803 | NM_024698.5 | 38.2391527 | -6.8825909 | 1.40866658 | -4.8858907 |

| | | | | | |
|---|---|---|---|---|---|
| 9804 | NM_001042475.2 | 59.3395354 | -7.5121174 | 1.53755473 | -4.8857561 |
| 9805 | NM_001288786.1 | 45.4501106 | -7.1308142 | 1.45960302 | -4.8854477 |
| 9806 | NM_019083.2 | 27.1487336 | -6.3860356 | 1.30723222 | -4.8851577 |
| 9807 | NM_001308195.1 | 35.289171 | -6.7651766 | 1.38494441 | -4.8848001 |
| 9808 | NM_001348096.1 | 41.0787916 | -6.9858437 | 1.4302197 | -4.8844549 |
| 9809 | NM_144580.2 | 35.748537 | -6.7843068 | 1.38898234 | -4.8843723 |
| 9810 | NM_001039583.1 | 42.8341535 | -7.0383727 | 1.44101729 | -4.8843083 |
| 9811 | NM_005475.2 | 32.4900104 | -6.6453247 | 1.36059761 | -4.884122 |
| 9812 | NM_001253387.1 | 62.5685234 | -7.5887442 | 1.55390258 | -4.883668 |
| 9813 | NM_001198812.1 | 56.432131 | -7.4406217 | 1.52375995 | -4.8830669 |
| 9814 | NM_018196.3 | 26.3238215 | -6.3372022 | 1.29793819 | -4.8825146 |
| 9815 | NM_021269.2 | 29.3188799 | -6.4922682 | 1.32977163 | -4.882243 |
| 9816 | NM_001323393.1 | 53.5339349 | -7.3638149 | 1.50842176 | -4.881801 |
| 9817 | NM_001300984.1 | 23.3197656 | -6.1632814 | 1.26257567 | -4.8815145 |
| 9818 | NM_001164097.1 | 789.52644 | -6.5362282 | 1.33905334 | -4.8812307 |
| 9819 | NM_139178.3 | 27.0699625 | -6.3808953 | 1.30741104 | -4.8805579 |
| 9820 | NR_027158.1 | 54.5349214 | -7.392239 | 1.51489893 | -4.8796912 |
| 9821 | NM_001048230.1 | 45.5104798 | -7.129718 | 1.46116684 | -4.8794688 |
| 9822 | NM_001144037.1 | 29.0907693 | -6.4847149 | 1.32931057 | -4.8782542 |
| 9823 | NM_014799.3 | 54.9924233 | -7.4034238 | 1.51768223 | -4.8781119 |
| 9824 | NM_001330213.1 | 31.1311349 | -6.5852578 | 1.35002838 | -4.8778662 |
| 9825 | NM_032901.3 | 30.5800539 | -6.5544859 | 1.34372434 | -4.8778501 |
| 9826 | NM_001098497.2 | 29.6253899 | -6.5101434 | 1.33469003 | -4.8776444 |
| 9827 | NM_001329139.1 | 89.2665024 | -8.1023253 | 1.66118049 | -4.8774504 |
| 9828 | NM_001134369.2 | 38.2931492 | -6.8768343 | 1.41001404 | -4.8771389 |
| 9829 | NM_017772.3 | 58.0112487 | -5.6268801 | 1.15384003 | -4.8766553 |
| 9830 | NM_144720.3 | 60.4231652 | -7.5378554 | 1.54569615 | -4.8766735 |
| 9831 | NM_001017958.2 | 57.9227728 | -7.4805681 | 1.53398268 | -4.8765662 |
| 9832 | NM_024121.2 | 107.455706 | -3.6763456 | 0.75389107 | -4.8764944 |
| 9833 | NM_012450.3 | 66.2105057 | -7.6727739 | 1.57345922 | -4.8763729 |
| 9834 | NM_178336.2 | 34.3852152 | -6.7239267 | 1.37886998 | -4.8764037 |
| 9835 | NR_137176.1 | 67.9766223 | -7.7090164 | 1.58088687 | -4.8763872 |
| 9836 | NM_017988.5 | 34.3597146 | -6.7234135 | 1.37897347 | -4.8756656 |
| 9837 | NM_020122.4 | 252.574306 | -2.3168948 | 0.47520983 | -4.8755195 |
| 9838 | NM_017666.4 | 37.6368562 | -3.9970306 | 0.81996874 | -4.8746134 |
| 9839 | NM_000027.3 | 24.580448 | -6.2345194 | 1.27899196 | -4.8745571 |
| 9840 | NM_032424.2 | 36.9701855 | -6.8259859 | 1.4005372 | -4.8738341 |
| 9841 | NM_016370.2 | 74.0728374 | -6.8575521 | 1.40708551 | -4.8735859 |
| 9842 | NM_138444.3 | 295.302775 | -5.5384189 | 1.13644415 | -4.8734634 |
| 9843 | NM_001164317.1 | 37.2125228 | -6.8415871 | 1.40401062 | -4.8728884 |
| 9844 | NM_001286117.1 | 99.4531347 | -5.1425519 | 1.05535176 | -4.8728321 |
| 9845 | NR_028324.1 | 28.4198511 | -6.4514325 | 1.32407832 | -4.8723949 |
| 9846 | NM_001042492.2 | 67.1580961 | -7.6909794 | 1.57858115 | -4.8720837 |

| | | | | | |
|---|---|---|---|---|---|
| 9847 | NM_001256577.2 | 62.3442724 | -7.5849573 | 1.55684183 | -4.8720153 |
| 9848 | NM_032936.3 | 30.418334 | -6.5520452 | 1.34493286 | -4.8716522 |
| 9849 | NR_135469.1 | 17.8863159 | 9.08433625 | 1.86495565 | 4.87107361 |
| 9850 | NM_001122957.2 | 37.9319952 | -6.8655789 | 1.40950104 | -4.8709286 |
| 9851 | NM_002524.4 | 231.141299 | -4.3205282 | 0.88705186 | -4.8706602 |
| 9852 | NM_006283.2 | 48.6412726 | -6.2591296 | 1.28533962 | -4.869631 |
| 9853 | NR_104025.1 | 33.9037105 | -6.7025816 | 1.37641565 | -4.8695912 |
| 9854 | NR_024173.1 | 115.578931 | 3.04999144 | 0.62636342 | 4.86936395 |
| 9855 | NM_032827.6 | 24.2524699 | -6.2267448 | 1.27888534 | -4.8688844 |
| 9856 | NM_005038.2 | 153.291745 | -2.7750188 | 0.57001779 | -4.868302 |
| 9857 | NR_027783.2 | 30.5743937 | -6.5519125 | 1.34586019 | -4.8681969 |
| 9858 | NM_001077700.2 | 30.5502502 | -6.5562963 | 1.34679288 | -4.8680806 |
| 9859 | NM_173804.4 | 23.5018552 | -6.1795176 | 1.26943517 | -4.8679269 |
| 9860 | NM_006117.2 | 36.0865265 | -6.7945267 | 1.3958029 | -4.8678267 |
| 9861 | NM_177937.2 | 26.3501041 | -6.3350715 | 1.30152934 | -4.8674058 |
| 9862 | NM_001025071.1 | 31.4919178 | -6.5984557 | 1.35566738 | -4.8673117 |
| 9863 | NM_012456.2 | 38.5045714 | -6.8888001 | 1.41539898 | -4.8670376 |
| 9864 | NM_001199111.1 | 64.1448696 | -7.6250803 | 1.56682903 | -4.8665682 |
| 9865 | NM_001008212.1 | 40.0406329 | -6.9492535 | 1.42800672 | -4.8664011 |
| 9866 | NM_144574.3 | 24.4951031 | -6.2315431 | 1.28052736 | -4.8663881 |
| 9867 | NM_000204.4 | 43.5976741 | -7.0723099 | 1.45338036 | -4.8661108 |
| 9868 | NM_004614.4 | 33.9064135 | -6.7008462 | 1.37713439 | -4.8657896 |
| 9869 | NR_037148.1 | 30.6771944 | -6.5559612 | 1.34765091 | -4.8647325 |
| 9870 | NM_020861.2 | 46.8814203 | -6.1991125 | 1.27430645 | -4.8646952 |
| 9871 | NM_001127691.2 | 69.6165623 | 2.44009629 | 0.50160171 | 4.86460921 |
| 9872 | NM_001142395.1 | 31.2991275 | -6.5894075 | 1.35468373 | -4.8641667 |
| 9873 | NM_001142546.1 | 103.742556 | -5.6464301 | 1.16087465 | -4.8639447 |
| 9874 | NM_019032.5 | 40.2727779 | -6.9573697 | 1.43050811 | -4.8635654 |
| 9875 | NM_001330640.1 | 39.1293781 | -6.9116617 | 1.42128394 | -4.8629704 |
| 9876 | NM_020165.3 | 32.2304738 | -6.6301736 | 1.36357458 | -4.8623476 |
| 9877 | NM_001271781.1 | 34.6719951 | -6.7395719 | 1.38611948 | -4.8621869 |
| 9878 | NR_038361.1 | 23.5000587 | -6.1765586 | 1.27034836 | -4.8620983 |
| 9879 | NM_001127448.1 | 112.055892 | -8.4310764 | 1.73405646 | -4.8620542 |
| 9880 | NM_052857.3 | 52.4625011 | -5.5224732 | 1.13607872 | -4.8609952 |
| 9881 | NM_001114100.2 | 89.3440974 | -8.103173 | 1.66718841 | -4.8603823 |
| 9882 | NM_001166112.1 | 49.1138282 | -7.2430597 | 1.49029895 | -4.8601388 |
| 9883 | NM_006478.4 | 43.2831322 | -7.058626 | 1.45237222 | -4.8600668 |
| 9884 | NM_001318122.1 | 30.0950501 | -6.5368088 | 1.34507861 | -4.8597968 |
| 9885 | NR_033416.1 | 30.187346 | -6.5351897 | 1.34481978 | -4.8595282 |
| 9886 | NM_004233.3 | 29.3495521 | -6.494536 | 1.33647235 | -4.8594615 |
| 9887 | NM_018465.3 | 34.9794631 | -6.7494199 | 1.38899249 | -4.8592199 |
| 9888 | NM_000168.5 | 142.590703 | -4.9591064 | 1.02070567 | -4.8585077 |
| 9889 | NM_003846.2 | 48.7354182 | -7.2290983 | 1.48808669 | -4.857982 |

| | | | | | |
|---|---|---|---|---|---|
| 9890 | NM_014583.3 | 32.9978906 | -6.6673203 | 1.37245059 | -4.8579674 |
| 9891 | NM_017632.3 | 38.8554622 | -6.901704 | 1.42073566 | -4.8578382 |
| 9892 | NM_153701.2 | 108.138627 | 2.02061644 | 0.41597666 | 4.85752356 |
| 9893 | NM_199442.1 | 41.3529987 | -6.9912646 | 1.43928936 | -4.857442 |
| 9894 | NM_001044370.1 | 155.836594 | -8.9058095 | 1.83346854 | -4.8573561 |
| 9895 | NR_024024.1 | 27.0063167 | -6.3802721 | 1.31359138 | -4.8571209 |
| 9896 | NM_017820.4 | 43.7171575 | -5.1820698 | 1.06697791 | -4.8567733 |
| 9897 | NM_000439.4 | 41.2630114 | -6.9892026 | 1.43912326 | -4.8565699 |
| 9898 | NM_022152.5 | 25.5351741 | -6.3006447 | 1.29743616 | -4.8562271 |
| 9899 | NM_002528.6 | 36.5759278 | -6.8125259 | 1.40287541 | -4.8561162 |
| 9900 | NM_032309.3 | 35.9478807 | -6.7868083 | 1.39759278 | -4.8560699 |
| 9901 | NM_001330104.1 | 26.5838741 | -6.3497026 | 1.30758738 | -4.8560446 |
| 9902 | NM_001142351.1 | 52.0913549 | -7.326265 | 1.50876486 | -4.855803 |
| 9903 | NM_003993.3 | 33.7929369 | -6.7021993 | 1.38029385 | -4.8556322 |
| 9904 | NM_001031696.3 | 31.1067904 | -6.5868646 | 1.35657411 | -4.855514 |
| 9905 | NM_022838.3 | 37.4816839 | -6.8483129 | 1.4108466 | -4.854045 |
| 9906 | NM_001170820.3 | 39.5152906 | -6.9292099 | 1.42757094 | -4.8538463 |
| 9907 | NM_002583.2 | 26.7323607 | -6.3671905 | 1.31180907 | -4.8537479 |
| 9908 | NM_030636.2 | 32.9893174 | -6.6640876 | 1.37297571 | -4.8537549 |
| 9909 | NM_001300731.1 | 45.128469 | -7.1187088 | 1.46680425 | -4.8532098 |
| 9910 | NM_001244889.1 | 68.9909502 | -7.729638 | 1.59270764 | -4.853143 |
| 9911 | NR_134482.1 | 25.6717828 | -6.29599 | 1.29734979 | -4.8529626 |
| 9912 | NM_004914.3 | 92.2360171 | -3.8387485 | 0.79101888 | -4.8529164 |
| 9913 | NM_000164.3 | 39.6615744 | -6.9338646 | 1.42885773 | -4.8527327 |
| 9914 | NM_018366.2 | 50.302347 | -6.3023166 | 1.29873738 | -4.852649 |
| 9915 | NM_001114107.4 | 29.1231649 | -6.4897736 | 1.33744108 | -4.852381 |
| 9916 | NR_026589.1 | 74.9208915 | 2.11859743 | 0.4366192 | 4.85227724 |
| 9917 | NM_020341.3 | 45.2070935 | -7.1187009 | 1.46714424 | -4.8520798 |
| 9918 | NM_017576.2 | 43.5861466 | -7.0690197 | 1.45695608 | -4.8519099 |
| 9919 | NM_001100878.1 | 22.0737167 | -6.0898237 | 1.25532859 | -4.851179 |
| 9920 | NM_014379.3 | 113.659596 | 2.80753806 | 0.5787519 | 4.85102176 |
| 9921 | NM_181332.2 | 81.7070135 | -7.9744414 | 1.6439966 | -4.8506435 |
| 9922 | NM_001286485.1 | 30.8656583 | -6.5736133 | 1.35526139 | -4.8504395 |
| 9923 | NM_001128917.1 | 35.9387933 | -6.7909323 | 1.40030206 | -4.8496196 |
| 9924 | NM_024852.3 | 26.0007217 | -6.3184311 | 1.30291955 | -4.8494407 |
| 9925 | NM_005713.2 | 60.6966548 | -7.5442463 | 1.55573703 | -4.8493069 |
| 9926 | NM_001330308.1 | 28.7282071 | -6.4652241 | 1.33331261 | -4.8489934 |
| 9927 | NM_001320036.1 | 24.600516 | -6.2477951 | 1.28850126 | -4.8488855 |
| 9928 | NM_138414.2 | 34.5329815 | -6.7312826 | 1.38821923 | -4.8488613 |
| 9929 | NM_001321953.1 | 25.5050139 | -6.2997314 | 1.29923371 | -4.8488054 |
| 9930 | NM_001142298.1 | 53.179224 | -6.3805014 | 1.31592272 | -4.84869 |
| 9931 | NM_024741.2 | 34.7468769 | -6.7441157 | 1.39108086 | -4.8481119 |
| 9932 | NM_006704.4 | 42.1981672 | -7.0215203 | 1.44832992 | -4.8480116 |

| | | | | | |
|---|---|---|---|---|---|
| 9933 | NM_001318750.1 | 58.0917574 | -6.5115038 | 1.34318545 | -4.847807 |
| 9934 | NM_138717.2 | 23.1925573 | -6.1632106 | 1.27134555 | -4.8477856 |
| 9935 | NM_206893.3 | 143.420528 | 3.77840023 | 0.77942273 | 4.84769059 |
| 9936 | NR_046088.1 | 147.303879 | 4.96617967 | 1.02456271 | 4.84712124 |
| 9937 | NM_014550.3 | 25.7793343 | -6.3154747 | 1.30300564 | -4.8468514 |
| 9938 | NM_014322.2 | 31.8795199 | -6.6178399 | 1.36550787 | -4.8464312 |
| 9939 | NM_001244896.1 | 45.6304496 | -7.1345422 | 1.47213427 | -4.8463936 |
| 9940 | NM_001130864.1 | 41.1885815 | -5.9973513 | 1.23755606 | -4.8461249 |
| 9941 | NM_152361.2 | 29.495761 | -6.5036616 | 1.34205042 | -4.8460636 |
| 9942 | NR_026812.1 | 15.3091992 | 6.95833366 | 1.43591416 | 4.84592593 |
| 9943 | NM_001288765.1 | 45.65595 | -7.1328693 | 1.4719749 | -4.8457819 |
| 9944 | NM_001139490.1 | 100.590846 | -2.0158575 | 0.4160562 | -4.8451568 |
| 9945 | NR_033710.1 | 27.8926943 | -6.4237047 | 1.32596969 | -4.8445336 |
| 9946 | NR_037802.1 | 25.7964168 | -6.3038937 | 1.30140978 | -4.8438961 |
| 9947 | NM_024597.3 | 25.4363424 | -6.2962758 | 1.30010986 | -4.8428798 |
| 9948 | NM_016202.2 | 22.88384 | -6.1364917 | 1.2671675 | -4.842684 |
| 9949 | NM_001024934.1 | 39.8736203 | -6.9400432 | 1.43326463 | -4.8421227 |
| 9950 | NM_001286414.2 | 44.9866837 | -7.1156449 | 1.46966494 | -4.8416783 |
| 9951 | NM_005748.5 | 42.1605425 | -7.0181174 | 1.44960956 | -4.8413846 |
| 9952 | NM_003172.3 | 41.1769851 | -6.985438 | 1.44296657 | -4.8410255 |
| 9953 | NR_109841.1 | 22.8768329 | 2.91841624 | 0.60286323 | 4.84092596 |
| 9954 | NM_001080501.2 | 25.9231433 | -6.3178914 | 1.30511375 | -4.8408741 |
| 9955 | NM_014563.5 | 317.058681 | 1.94365203 | 0.40151237 | 4.84082726 |
| 9956 | NM_001288661.1 | 36.3584064 | -6.807483 | 1.40628304 | -4.8407631 |
| 9957 | NM_001284201.1 | 29.5194941 | -6.5064124 | 1.34418778 | -4.8404044 |
| 9958 | NM_005598.3 | 81.6106915 | -6.2051527 | 1.282018 | -4.8401447 |
| 9959 | NM_001303421.1 | 31.9039781 | -6.6131103 | 1.36643179 | -4.8396929 |
| 9960 | NM_000635.3 | 38.4475572 | -6.889968 | 1.42402696 | -4.8383691 |
| 9961 | NR_134528.1 | 39.9373956 | -6.9385267 | 1.43415536 | -4.838058 |
| 9962 | NM_000137.2 | 28.5816187 | -6.4625339 | 1.33590026 | -4.8375871 |
| 9963 | NM_194455.1 | 26.2227266 | -6.3328809 | 1.30910103 | -4.83758 |
| 9964 | NM_003779.3 | 33.7785311 | -6.7040804 | 1.38591757 | -4.8372865 |
| 9965 | NM_018154.2 | 26.8661696 | -6.368997 | 1.3167291 | -4.8369835 |
| 9966 | NM_013322.2 | 55.9383723 | -7.4268708 | 1.53546013 | -4.8369024 |
| 9967 | NR_135527.1 | 23.4201413 | -6.169389 | 1.27549848 | -4.8368454 |
| 9968 | NM_016434.3 | 28.9202377 | -6.4782387 | 1.33937903 | -4.8367479 |
| 9969 | NM_001297442.1 | 55.4686192 | -7.4169946 | 1.53365161 | -4.8361665 |
| 9970 | NM_001105520.1 | 25.2994271 | -6.2799141 | 1.2985461 | -4.8361118 |
| 9971 | NM_001008216.1 | 43.6040483 | -7.0675392 | 1.46149283 | -4.8358357 |
| 9972 | NM_005634.2 | 47.9391165 | -7.2023333 | 1.48940603 | -4.8357084 |
| 9973 | NM_201522.2 | 27.8812732 | -6.4246707 | 1.32861148 | -4.8356278 |
| 9974 | NM_015090.3 | 61.5794992 | -7.5659214 | 1.56464475 | -4.8355523 |
| 9975 | NM_199444.1 | 34.7171325 | -6.7427297 | 1.39444261 | -4.8354301 |

| | | | | | |
|---|---|---|---|---|---|
| 9976 | NM_002386.3 | 45.3160043 | -6.1425257 | 1.27044198 | -4.8349517 |
| 9977 | NM_002999.3 | 72.9850417 | -3.2291688 | 0.66790088 | -4.8348025 |
| 9978 | NM_001143780.2 | 30.3516512 | -6.5470407 | 1.35420552 | -4.834599 |
| 9979 | NM_001199856.1 | 39.8962509 | -6.9409259 | 1.43569635 | -4.8345361 |
| 9980 | NM_001001974.2 | 31.4336199 | -6.5953357 | 1.3643584 | -4.83402 |
| 9981 | NM_144963.3 | 102.543656 | -4.6996462 | 0.97229329 | -4.8335685 |
| 9982 | NM_001290293.2 | 24.3491485 | -6.2235855 | 1.2876384 | -4.8333333 |
| 9983 | NM_001318508.1 | 26.7328708 | -6.3687245 | 1.31774968 | -4.8330306 |
| 9984 | NR_029408.1 | 36.4726772 | -6.8093593 | 1.40901802 | -4.8326985 |
| 9985 | NM_016334.3 | 29.6008252 | -6.509486 | 1.34708821 | -4.8322641 |
| 9986 | NM_016955.3 | 22.5746065 | -6.117254 | 1.26599325 | -4.8319799 |
| 9987 | NM_078629.3 | 37.9183099 | -6.8666738 | 1.42138285 | -4.8309812 |
| 9988 | NM_001127716.1 | 40.8611569 | -6.9762743 | 1.444087 | -4.8309238 |
| 9989 | NR_037947.1 | 30.3595986 | -6.5408616 | 1.35396547 | -4.8308925 |
| 9990 | NM_001171509.2 | 64.3583131 | -7.6295989 | 1.57954112 | -4.8302629 |
| 9991 | NM_001195738.1 | 49.4028307 | -7.2463064 | 1.50029027 | -4.8299363 |
| 9992 | NM_003088.3 | 1480.59918 | -7.0770144 | 1.46525484 | -4.8298864 |
| 9993 | NM_004773.3 | 30.2465108 | -6.5342948 | 1.35320237 | -4.8287639 |
| 9994 | NM_001199.3 | 35.6779379 | -6.781844 | 1.40457438 | -4.8283979 |
| 9995 | NM_015918.3 | 33.5489413 | -6.6879154 | 1.38512876 | -4.8283709 |
| 9996 | NR_136703.1 | 26.0433971 | -6.3204552 | 1.30918307 | -4.8277856 |
| 9997 | NM_001301237.1 | 34.9734273 | -6.7477572 | 1.39777783 | -4.8274891 |
| 9998 | NM_001145847.1 | 35.829803 | -6.7792808 | 1.40441386 | -4.8271247 |
| 9999 | NM_001136232.2 | 26.9625614 | -6.3820886 | 1.32224236 | -4.8267162 |
| 10000 | NM_001973.3 | 89.1321922 | -1.7856466 | 0.36995043 | -4.8267185 |
| 10001 | NM_022359.5 | 42.4822443 | -7.0289121 | 1.45631847 | -4.8264938 |
| 10002 | NM_012460.3 | 35.9785927 | -6.7884823 | 1.40652435 | -4.8264236 |
| 10003 | NM_001018161.1 | 30.7985966 | -6.5640698 | 1.36005312 | -4.8263334 |
| 10004 | NM_194282.3 | 30.181346 | -6.5389825 | 1.35498756 | -4.8258616 |
| 10005 | NM_206824.2 | 32.508319 | -6.6452101 | 1.37718795 | -4.825202 |
| 10006 | NM_001159291.1 | 37.9508936 | -6.866447 | 1.42304765 | -4.8251701 |
| 10007 | NM_001276290.1 | 28.1920607 | -6.4455537 | 1.33585062 | -4.8250557 |
| 10008 | NM_001102426.2 | 26.0370753 | -6.3311731 | 1.31218151 | -4.8249217 |
| 10009 | NM_001199693.1 | 47.4283196 | -7.1926183 | 1.49079919 | -4.8246728 |
| 10010 | NM_001270966.1 | 47.3560214 | -7.1879864 | 1.48983504 | -4.8246861 |
| 10011 | NM_152515.4 | 26.5763535 | -6.3601049 | 1.31830929 | -4.8244406 |
| 10012 | NM_152544.2 | 27.8523799 | -6.4209966 | 1.33096004 | -4.8243346 |
| 10013 | NR_030727.1 | 38.0451415 | 4.10852483 | 0.85167714 | 4.8240403 |
| 10014 | NM_006658.4 | 23.6902579 | -6.1840767 | 1.28196935 | -4.8238881 |
| 10015 | NM_145063.3 | 24.921435 | -6.25533 | 1.29697849 | -4.8230021 |
| 10016 | NR_037672.1 | 26.6274291 | -6.3631546 | 1.3193603 | -4.8229089 |
| 10017 | NM_178834.4 | 38.4706757 | -6.8912253 | 1.42887389 | -4.8228366 |
| 10018 | NM_178526.4 | 46.9801574 | -6.2018276 | 1.28627086 | -4.8215565 |

| | | | | | |
|---|---|---|---|---|---|
| 10019 | NM_001085426.2 | 34.181717 | -6.722623 | 1.39435716 | -4.8213063 |
| 10020 | NM_001143787.1 | 38.4935416 | -6.8855246 | 1.42823219 | -4.821012 |
| 10021 | NM_001297723.1 | 22.5747381 | -6.1213933 | 1.26974171 | -4.8209752 |
| 10022 | NM_001330322.1 | 25.6728017 | -6.2967954 | 1.30629789 | -4.8203365 |
| 10023 | NM_001166105.2 | 32.19918 | -6.631673 | 1.37586046 | -4.8200186 |
| 10024 | NM_001143984.1 | 29.0280406 | -6.48001 | 1.34442273 | -4.81992 |
| 10025 | NM_001289789.1 | 54.0646514 | -7.3781553 | 1.53084427 | -4.8196642 |
| 10026 | NM_001202413.1 | 95.2153961 | -7.2278479 | 1.49969361 | -4.8195497 |
| 10027 | NM_007109.2 | 28.0331527 | -6.4235095 | 1.33288388 | -4.8192567 |
| 10028 | NM_031446.4 | 26.9052505 | -6.3715007 | 1.32229793 | -4.8185061 |
| 10029 | NM_153334.6 | 42.9295188 | -7.0498461 | 1.46334053 | -4.8176388 |
| 10030 | NM_002207.2 | 26.7234627 | -6.3620926 | 1.32068874 | -4.8172536 |
| 10031 | NM_139177.3 | 27.5061192 | -6.4039768 | 1.32940342 | -4.8171809 |
| 10032 | NM_003258.4 | 38.3331963 | -6.8828155 | 1.42881185 | -4.8171601 |
| 10033 | NM_012339.3 | 23.660811 | -6.1916472 | 1.28540476 | -4.8168852 |
| 10034 | NM_001080792.2 | 37.6430495 | -6.856411 | 1.42342769 | -4.8168313 |
| 10035 | NM_014182.4 | 33.1810337 | -6.6751874 | 1.38583056 | -4.8167414 |
| 10036 | NM_001310315.1 | 37.1369204 | -6.8362678 | 1.41939302 | -4.8163319 |
| 10037 | NM_001256495.1 | 29.8060167 | -6.5168925 | 1.35333764 | -4.8154225 |
| 10038 | NM_017679.3 | 25.9979533 | -6.319599 | 1.31236647 | -4.8154225 |
| 10039 | NR_037676.1 | 190.976448 | 3.62440033 | 0.75268104 | 4.81532034 |
| 10040 | NM_006602.3 | 28.1810662 | -6.4397726 | 1.33741928 | -4.8150738 |
| 10041 | NM_018640.4 | 97.2395764 | -8.224985 | 1.70830883 | -4.8146944 |
| 10042 | NM_001282974.1 | 225.451022 | -5.4268281 | 1.1273262 | -4.8138933 |
| 10043 | NM_013367.2 | 41.8808143 | -7.0085344 | 1.45621989 | -4.812827 |
| 10044 | NM_078476.3 | 41.0466085 | -6.978794 | 1.45013902 | -4.8125 |
| 10045 | NM_138317.2 | 27.4281572 | -6.3951542 | 1.32888219 | -4.8124313 |
| 10046 | NM_001451.2 | 71.3497632 | -6.8046761 | 1.41401325 | -4.8123143 |
| 10047 | NM_152516.3 | 33.9200507 | -6.7072317 | 1.39395118 | -4.811669 |
| 10048 | NM_000456.2 | 23.0129566 | -6.1426099 | 1.27663921 | -4.8115473 |
| 10049 | NM_024578.2 | 22.4385679 | -6.1114412 | 1.27017233 | -4.8115055 |
| 10050 | NM_030802.3 | 28.1402083 | -6.4422336 | 1.33898996 | -4.8112636 |
| 10051 | NM_001665.3 | 29.11221 | -6.4886207 | 1.34863943 | -4.8112346 |
| 10052 | NM_001039885.2 | 33.4908388 | -6.6903422 | 1.39058552 | -4.8111692 |
| 10053 | NM_003455.3 | 34.9848584 | -6.7453527 | 1.40201581 | -4.8111816 |
| 10054 | NM_024940.7 | 23.6834779 | -6.1925173 | 1.28717973 | -4.8109189 |
| 10055 | NM_130906.2 | 38.6119509 | -6.8914621 | 1.43262426 | -4.8103765 |
| 10056 | NR_002186.1 | 25.7636423 | -6.3055695 | 1.31089027 | -4.8101429 |
| 10057 | NM_033657.2 | 47.0342458 | -7.17774 | 1.49232469 | -4.809771 |
| 10058 | NM_201286.3 | 25.7540372 | -6.3121335 | 1.31246526 | -4.8093719 |
| 10059 | NM_014337.3 | 42.2727087 | -7.0271302 | 1.46119865 | -4.8091546 |
| 10060 | NM_001077188.1 | 61.3383214 | -7.5610281 | 1.57240398 | -4.8085786 |
| 10061 | NM_006630.2 | 31.8225222 | -6.6111709 | 1.37486687 | -4.8085899 |

| | | | | | |
|---|---|---|---|---|---|
| 10062 | NM_018930.3 | 66.4504715 | -3.9387734 | 0.81913761 | -4.8084392 |
| 10063 | NM_199351.2 | 70.9474437 | -3.4108169 | 0.70934493 | -4.8084039 |
| 10064 | NM_006200.5 | 48.0197439 | -7.208988 | 1.49929576 | -4.8082495 |
| 10065 | NM_001145312.1 | 22.5693704 | -6.1198675 | 1.2728367 | -4.8080539 |
| 10066 | NM_020703.2 | 27.6511102 | -6.4083853 | 1.33285357 | -4.808019 |
| 10067 | NM_001100119.1 | 31.3097129 | -6.5839766 | 1.36939319 | -4.8079519 |
| 10068 | NM_025185.3 | 275.850561 | -5.8719378 | 1.22147969 | -4.8072332 |
| 10069 | NM_001199346.1 | 56.6180534 | -7.4433463 | 1.5483868 | -4.807162 |
| 10070 | NM_001276318.1 | 57.6991485 | -5.5408693 | 1.15264586 | -4.8070873 |
| 10071 | NM_147686.3 | 28.7374094 | -6.4699019 | 1.3461201 | -4.8063333 |
| 10072 | NM_014181.2 | 90.7584868 | -4.2356889 | 0.8813445 | -4.8059401 |
| 10073 | NM_003741.3 | 25.6560095 | -6.3036624 | 1.31169287 | -4.8057457 |
| 10074 | NM_080685.2 | 28.8299156 | -6.4642286 | 1.34518962 | -4.8054404 |
| 10075 | NR_037630.1 | 25.0540049 | -6.2663164 | 1.30405614 | -4.8052505 |
| 10076 | NR_033268.1 | 23.9193191 | -6.2049131 | 1.29128533 | -4.8052223 |
| 10077 | NM_198082.2 | 42.2877417 | -5.2091291 | 1.08416009 | -4.8047601 |
| 10078 | NM_001202546.2 | 34.3541153 | -6.7235387 | 1.39948628 | -4.8042905 |
| 10079 | NM_016937.3 | 34.2429008 | -6.7175897 | 1.39836222 | -4.8038981 |
| 10080 | NM_133499.2 | 140.970088 | -5.7776897 | 1.20272653 | -4.8038266 |
| 10081 | NM_017918.4 | 24.7625091 | -6.2459029 | 1.3003701 | -4.8031733 |
| 10082 | NR_027303.1 | 80.5911892 | -6.1891825 | 1.28861954 | -4.8029557 |
| 10083 | NM_002312.3 | 42.2645895 | -7.0264545 | 1.46301334 | -4.8027276 |
| 10084 | NM_001101426.3 | 22.2982197 | -6.1020969 | 1.27062612 | -4.802433 |
| 10085 | NM_031923.3 | 54.053391 | -6.4048751 | 1.33382501 | -4.8018856 |
| 10086 | NM_001170670.1 | 29.7426464 | 9.80838221 | 2.04321488 | 4.80046534 |
| 10087 | NM_001330490.1 | 22.5692938 | -6.1211563 | 1.27511384 | -4.8004783 |
| 10088 | NM_005011.4 | 30.918519 | -6.5700537 | 1.36868405 | -4.8002705 |
| 10089 | NM_145859.1 | 28.179227 | -6.4396616 | 1.34153631 | -4.8002142 |
| 10090 | NM_001329615.1 | 25.9591496 | -6.3138562 | 1.31535418 | -4.8001187 |
| 10091 | NM_001113498.2 | 32.6556106 | -6.6508063 | 1.38561257 | -4.7999033 |
| 10092 | NM_213649.1 | 33.777054 | -6.6977191 | 1.39538205 | -4.7999178 |
| 10093 | NM_001144000.1 | 21.8955312 | -6.074077 | 1.26569516 | -4.7990047 |
| 10094 | NM_014672.3 | 34.9427111 | -6.7464717 | 1.40580811 | -4.798999 |
| 10095 | NM_198309.3 | 23.5893117 | -6.1881002 | 1.28948108 | -4.7989073 |
| 10096 | NM_183380.3 | 68.6324159 | -7.7239212 | 1.60954429 | -4.7988249 |
| 10097 | NM_003224.5 | 28.6766228 | -6.4670028 | 1.34765959 | -4.7986916 |
| 10098 | NM_001319206.1 | 46.3646125 | -7.1551569 | 1.49108308 | -4.7986306 |
| 10099 | NR_015453.2 | 32.3281401 | -6.6360427 | 1.38290723 | -4.7986174 |
| 10100 | NR_038854.1 | 21.2292541 | -6.0335966 | 1.25740128 | -4.7984654 |
| 10101 | NM_001286760.1 | 66.8338919 | -6.7097789 | 1.39836127 | -4.7983157 |
| 10102 | NM_005828.4 | 803.562561 | -3.0318078 | 0.63186274 | -4.7982062 |
| 10103 | NM_001198951.1 | 25.2724285 | -6.2782545 | 1.30846797 | -4.7981721 |
| 10104 | NM_178562.4 | 29.5780855 | -6.5086441 | 1.35652871 | -4.7980143 |

| | | | | | |
|---|---|---|---|---|---|
| 10105 | NM_032826.4 | 156.827331 | -4.0926869 | 0.85301951 | -4.7978819 |
| 10106 | NM_005252.3 | 118.160058 | -5.0331539 | 1.04904924 | -4.7978243 |
| 10107 | NM_001135635.1 | 30.6396839 | -6.5614848 | 1.36766544 | -4.7975803 |
| 10108 | NM_001134450.1 | 28.5286353 | -6.4494081 | 1.34434108 | -4.7974493 |
| 10109 | NR_002211.1 | 45.023284 | -6.145772 | 1.28108793 | -4.7973069 |
| 10110 | NM_001083591.1 | 92.2230014 | -8.1482465 | 1.69852917 | -4.7972367 |
| 10111 | NM_025188.3 | 27.1450736 | -6.3823133 | 1.33048202 | -4.7969933 |
| 10112 | NR_047594.1 | 27.7011278 | -6.4121889 | 1.33673109 | -4.7969176 |
| 10113 | NM_015149.4 | 26.0507828 | -6.3237665 | 1.31838684 | -4.7965941 |
| 10114 | NM_006090.4 | 27.5835669 | -6.4026898 | 1.33500882 | -4.7959906 |
| 10115 | NM_005410.2 | 27.4034664 | -6.4014052 | 1.33486027 | -4.7955621 |
| 10116 | NM_001348136.1 | 68.9426091 | -7.7297138 | 1.61189565 | -4.7954182 |
| 10117 | NM_001127202.3 | 34.6248474 | -6.7367907 | 1.40489162 | -4.7952387 |
| 10118 | NM_001145044.1 | 43.5116611 | -6.0924656 | 1.27055325 | -4.7951281 |
| 10119 | NM_004215.4 | 22.223431 | -6.097842 | 1.27167732 | -4.7951174 |
| 10120 | NM_144660.2 | 34.4233172 | -6.7228015 | 1.40208488 | -4.7948606 |
| 10121 | NM_002726.4 | 159.164015 | -4.953408 | 1.03309959 | -4.7947052 |
| 10122 | NM_019013.2 | 42.3562145 | -7.0252125 | 1.46531743 | -4.7943281 |
| 10123 | NM_001031720.3 | 24.8501434 | -6.2615573 | 1.30610166 | -4.7940811 |
| 10124 | NM_016540.3 | 22.8373929 | -6.1296034 | 1.27858036 | -4.7940697 |
| 10125 | NR_104481.1 | 28.1209682 | 9.72750975 | 2.02918286 | 4.79380637 |
| 10126 | NR_135198.1 | 179.885539 | 2.81469725 | 0.58718444 | 4.79354879 |
| 10127 | NM_203456.2 | 29.799464 | -6.5190296 | 1.36004002 | -4.793263 |
| 10128 | NM_004839.3 | 25.8250187 | -6.3197377 | 1.31852211 | -4.7930465 |
| 10129 | NM_001031738.2 | 31.2579763 | -6.5933924 | 1.375656 | -4.7929078 |
| 10130 | NM_006440.4 | 25.2115439 | -6.2796656 | 1.31023418 | -4.7927811 |
| 10131 | NM_001333.3 | 23.9071445 | -6.2065136 | 1.29498259 | -4.7927391 |
| 10132 | NR_033908.1 | 27.1060017 | -6.3881246 | 1.33290386 | -4.7926372 |
| 10133 | NM_207520.1 | 47.7878893 | -7.2020175 | 1.5028861 | -4.7921247 |
| 10134 | NM_207346.2 | 47.260404 | -6.2121728 | 1.29641041 | -4.7918258 |
| 10135 | NM_001130845.1 | 29.0170863 | -6.4830919 | 1.35311131 | -4.791248 |
| 10136 | NM_001172574.1 | 33.2186909 | -6.6761644 | 1.39350552 | -4.7909135 |
| 10137 | NM_153719.3 | 35.5037777 | -6.7752441 | 1.41422622 | -4.7907782 |
| 10138 | NM_001319846.1 | 45.2453322 | -7.1193501 | 1.48605922 | -4.790758 |
| 10139 | NM_022485.4 | 23.3485736 | -6.1731313 | 1.28864084 | -4.7904203 |
| 10140 | NM_006260.4 | 172.610722 | -1.8045214 | 0.37670142 | -4.7903229 |
| 10141 | NM_182764.2 | 59.0820151 | -5.7228751 | 1.19473423 | -4.7900822 |
| 10142 | NR_028301.1 | 35.9003165 | -6.7868204 | 1.4169425 | -4.7897642 |
| 10143 | NM_145060.3 | 219.468012 | 2.75137394 | 0.57444378 | 4.78963137 |
| 10144 | NM_003503.3 | 42.3736793 | -7.0233086 | 1.46640906 | -4.7894607 |
| 10145 | NM_032134.2 | 28.6928849 | -6.4623173 | 1.34954805 | -4.7885048 |
| 10146 | NM_001671.4 | 34.9638764 | -6.7482045 | 1.40930315 | -4.7883271 |
| 10147 | NM_000784.3 | 90.2364413 | -8.1188066 | 1.69566307 | -4.7879834 |

| | | | | | |
|---|---|---|---|---|---|
| 10148 | NM_002003.4 | 143.800648 | 2.33461851 | 0.4875981 | 4.78799756 |
| 10149 | NM_015421.3 | 26.3537303 | -6.3414985 | 1.32446679 | -4.7879634 |
| 10150 | NM_001281460.1 | 24.3566725 | -6.2208751 | 1.29928295 | -4.7879294 |
| 10151 | NM_002997.4 | 53.7889435 | -6.3921971 | 1.33507196 | -4.7879045 |
| 10152 | NM_001166279.1 | 39.5237389 | -6.9242752 | 1.44632016 | -4.787512 |
| 10153 | NM_001317786.1 | 22.47128 | -6.1101148 | 1.27627192 | -4.787471 |
| 10154 | NR_036435.1 | 27.0044509 | -6.3695542 | 1.33059888 | -4.786983 |
| 10155 | NM_001242819.1 | 41.470078 | -6.997874 | 1.46207916 | -4.7862484 |
| 10156 | NR_034170.1 | 25.2447134 | -6.2837387 | 1.31287448 | -4.7862448 |
| 10157 | NM_182501.3 | 26.4861162 | -6.3439308 | 1.32546452 | -4.7861944 |
| 10158 | NM_022897.4 | 27.814626 | -6.4185005 | 1.34106964 | -4.7861053 |
| 10159 | NM_012322.2 | 45.6789758 | -7.1349915 | 1.49085826 | -4.7858282 |
| 10160 | NM_004785.5 | 23.3649369 | -6.1661481 | 1.28860523 | -4.7851335 |
| 10161 | NM_019591.3 | 28.9967484 | -6.4741293 | 1.35298013 | -4.7850883 |
| 10162 | NM_002801.3 | 34.466266 | -6.7296967 | 1.40639598 | -4.7850653 |
| 10163 | NM_016013.3 | 32.591142 | -6.6483749 | 1.38958619 | -4.7844279 |
| 10164 | NM_181655.3 | 28.7568013 | -6.4639037 | 1.35107278 | -4.784275 |
| 10165 | NM_004453.3 | 25.0493734 | -6.2702233 | 1.31083431 | -4.7833836 |
| 10166 | NM_001226.3 | 27.4334076 | -6.3986291 | 1.33777607 | -4.7830345 |
| 10167 | NM_201541.1 | 26.4858022 | -6.3520711 | 1.32814299 | -4.7826711 |
| 10168 | NR_040252.1 | 27.2561984 | -6.3873231 | 1.33550946 | -4.7826865 |
| 10169 | NM_004059.4 | 31.3142254 | -6.5856897 | 1.37724465 | -4.7817864 |
| 10170 | NM_001177711.1 | 34.7004323 | -6.7393517 | 1.40949757 | -4.7813858 |
| 10171 | NM_004976.4 | 67.4204168 | -6.7267104 | 1.40685269 | -4.7813893 |
| 10172 | NM_138734.2 | 342.250112 | -6.1696222 | 1.29037942 | -4.7812466 |
| 10173 | NM_006281.3 | 22.08329 | -6.0937957 | 1.27458793 | -4.7809928 |
| 10174 | NM_016310.4 | 32.3288151 | -6.6349946 | 1.38778987 | -4.7809793 |
| 10175 | NM_002594.4 | 30.9731577 | -6.5717261 | 1.37461358 | -4.7807807 |
| 10176 | NM_001902.5 | 20.6060871 | -5.9866316 | 1.25246889 | -4.7798645 |
| 10177 | NM_003068.4 | 29.4864324 | -6.5076487 | 1.36146649 | -4.7798816 |
| 10178 | NM_133455.3 | 42.4681758 | -7.027504 | 1.47032087 | -4.7795717 |
| 10179 | NM_001289745.1 | 21.1196027 | -6.01809 | 1.25946311 | -4.7782979 |
| 10180 | NM_133277.3 | 133.861312 | 2.90365282 | 0.60768496 | 4.77822062 |
| 10181 | NR_038406.1 | 78.5854615 | 4.61953668 | 0.96682089 | 4.77806877 |
| 10182 | NM_001199570.1 | 54.8789097 | -7.4016296 | 1.54910896 | -4.7779916 |
| 10183 | NM_000282.3 | 40.1131846 | -6.9498044 | 1.45464717 | -4.7776564 |
| 10184 | NM_004419.3 | 40.0800279 | -6.9449093 | 1.45376314 | -4.7771945 |
| 10185 | NM_018163.2 | 27.2995814 | -6.3911995 | 1.33802102 | -4.7766062 |
| 10186 | NM_001243963.1 | 83.5063595 | -5.5808891 | 1.1684509 | -4.7763146 |
| 10187 | NM_024055.4 | 21.7655431 | -6.0668542 | 1.2702473 | -4.7761205 |
| 10188 | NM_001317951.1 | 27.5280696 | -6.4061082 | 1.34131097 | -4.7760052 |
| 10189 | NM_001201477.1 | 49.9195386 | -6.2782839 | 1.31456484 | -4.7759408 |
| 10190 | NM_001011725.1 | 45.4705682 | -6.1536972 | 1.2885859 | -4.7755429 |

| | | | | | |
|---|---|---|---|---|---|
| 10191 | NM_006677.2 | 52.7400028 | -7.3447144 | 1.53806621 | -4.7752914 |
| 10192 | NM_001278225.1 | 27.7445477 | -6.4172321 | 1.34385696 | -4.7752345 |
| 10193 | NM_182679.2 | 44.5998912 | -7.1011696 | 1.48719046 | -4.7748892 |
| 10194 | NM_003772.3 | 41.4630977 | -6.0242427 | 1.26174955 | -4.7745154 |
| 10195 | NM_001199461.1 | 33.7155419 | -6.6949092 | 1.4025044 | -4.7735388 |
| 10196 | NM_001198719.1 | 38.6636483 | -6.8943304 | 1.44435987 | -4.7732775 |
| 10197 | NM_130797.3 | 53.6892 | -7.3669423 | 1.54345949 | -4.7730066 |
| 10198 | NM_014466.2 | 56.8406675 | -7.4525651 | 1.56141756 | -4.7729482 |
| 10199 | NR_027504.1 | 22.7471075 | -6.1313905 | 1.28499704 | -4.7715211 |
| 10200 | NM_001160037.1 | 39.975899 | -6.9452476 | 1.45564055 | -4.7712655 |
| 10201 | NM_080611.4 | 27.0398138 | -6.3793943 | 1.33712677 | -4.770972 |
| 10202 | NM_003487.3 | 51.8661437 | -6.3422555 | 1.32940691 | -4.7707406 |
| 10203 | NM_001317343.1 | 45.3880367 | -6.1491833 | 1.28896864 | -4.7706229 |
| 10204 | NM_012333.4 | 21.4407886 | -6.0463325 | 1.26742501 | -4.7705643 |
| 10205 | NM_019555.2 | 38.2606754 | -6.8791415 | 1.44199808 | -4.7705622 |
| 10206 | NR_022008.1 | 23.0310536 | -6.1501645 | 1.28919807 | -4.770535 |
| 10207 | NM_001286439.1 | 24.0704026 | -6.2104644 | 1.30193484 | -4.7701806 |
| 10208 | NM_006005.3 | 138.864126 | -4.9222391 | 1.03201771 | -4.7695297 |
| 10209 | NM_012180.2 | 26.8732823 | -6.3695975 | 1.33557603 | -4.7691762 |
| 10210 | NM_024057.3 | 24.9370867 | -6.2605949 | 1.31271786 | -4.7691855 |
| 10211 | NM_016647.2 | 36.7622683 | -6.8207369 | 1.43022833 | -4.7689846 |
| 10212 | NM_133489.2 | 80.5829665 | -5.9670819 | 1.25126473 | -4.7688405 |
| 10213 | NM_148965.1 | 29.41045 | -6.4964254 | 1.36235714 | -4.7685186 |
| 10214 | NM_015148.3 | 36.2076815 | -6.796802 | 1.42544156 | -4.7682081 |
| 10215 | NM_001112736.1 | 177.033112 | -5.0064284 | 1.04998518 | -4.7680943 |
| 10216 | NM_020710.2 | 268.924686 | -4.1156209 | 0.86323079 | -4.7676947 |
| 10217 | NM_001143853.1 | 36.8814211 | -6.8264723 | 1.43185493 | -4.7675726 |
| 10218 | NM_001258218.1 | 26.3096382 | -6.3366442 | 1.32932457 | -4.7668149 |
| 10219 | NM_001287439.1 | 37.9454799 | -6.863335 | 1.43984641 | -4.7667133 |
| 10220 | NR_036573.1 | 46.6063347 | -3.2667215 | 0.68535135 | -4.7664916 |
| 10221 | NM_001127715.2 | 40.5814197 | -6.9598964 | 1.46019249 | -4.7664239 |
| 10222 | NM_173582.4 | 619.05219 | -3.1974554 | 0.67083943 | -4.7663499 |
| 10223 | NM_001042399.1 | 26.7035902 | -6.3633428 | 1.33521138 | -4.7657943 |
| 10224 | NM_001098525.2 | 40.7174478 | -6.9650393 | 1.46154062 | -4.7655462 |
| 10225 | NM_175876.4 | 81.1558196 | -2.6321345 | 0.55237271 | -4.7651423 |
| 10226 | NM_001282764.1 | 32.415998 | -6.6384038 | 1.3931346 | -4.7650843 |
| 10227 | NM_018682.3 | 84.5708223 | -6.0286329 | 1.26516947 | -4.7650794 |
| 10228 | NM_013284.3 | 23.1357051 | -6.1608906 | 1.29299155 | -4.7648344 |
| 10229 | NM_004390.4 | 24.55969 | -6.2475277 | 1.31119444 | -4.7647607 |
| 10230 | NR_045286.1 | 51.1812448 | -7.2985065 | 1.53193202 | -4.7642496 |
| 10231 | NM_016510.5 | 25.3416888 | -6.2803348 | 1.31833253 | -4.7638473 |
| 10232 | NM_001134405.1 | 39.6245569 | -6.9287996 | 1.45446827 | -4.7638025 |
| 10233 | NM_001017390.2 | 23.4243998 | -6.1693665 | 1.29508386 | -4.7636811 |

| | | | | | |
|---|---|---|---|---|---|
| 10234 | NM_003173.3 | 26.481221 | -6.3469193 | 1.33235309 | -4.7636917 |
| 10235 | NM_145052.3 | 26.983569 | -6.3738473 | 1.33803907 | -4.7635734 |
| 10236 | NR_037193.1 | 31.9585309 | -6.6174022 | 1.38929311 | -4.7631433 |
| 10237 | NM_001143668.1 | 33.8849961 | -6.704884 | 1.40782838 | -4.762572 |
| 10238 | NM_173514.3 | 30.9117378 | -6.5750828 | 1.38061265 | -4.7624385 |
| 10239 | NM_032947.4 | 21.6540377 | -6.0631602 | 1.2731337 | -4.7623908 |
| 10240 | NM_001025253.2 | 56.3574622 | -7.4372618 | 1.56179117 | -4.7620079 |
| 10241 | NR_073008.1 | 25.6811952 | -6.3013712 | 1.32340301 | -4.7614907 |
| 10242 | NM_024908.3 | 36.0396496 | -6.7916055 | 1.4266422 | -4.7605528 |
| 10243 | NM_016095.2 | 35.2199852 | -6.7579355 | 1.41959361 | -4.7604719 |
| 10244 | NM_001184774.1 | 30.1570723 | -6.5339552 | 1.37258803 | -4.7603178 |
| 10245 | NM_001278222.1 | 27.0217468 | -6.3787653 | 1.34007574 | -4.7600035 |
| 10246 | NM_138775.2 | 27.3272041 | -6.3931725 | 1.34312474 | -4.7599246 |
| 10247 | NM_004857.3 | 51.4443132 | -2.7190166 | 0.57124034 | -4.759847 |
| 10248 | NM_080916.2 | 24.8476295 | -6.2542844 | 1.31400325 | -4.7597176 |
| 10249 | NM_152511.4 | 24.1013979 | -6.204827 | 1.30378213 | -4.7590981 |
| 10250 | NM_001037281.1 | 37.8874197 | -6.8620039 | 1.44207081 | -4.7584376 |
| 10251 | NM_001322929.1 | 37.6248891 | -6.8509418 | 1.43983637 | -4.7581392 |
| 10252 | NM_017443.4 | 73.2026541 | -7.8167029 | 1.64287437 | -4.7579432 |
| 10253 | NM_001128918.2 | 35.686547 | -6.7803115 | 1.42525324 | -4.7572679 |
| 10254 | NM_198471.2 | 21.912797 | -6.0765478 | 1.27740044 | -4.7569639 |
| 10255 | NM_144576.3 | 29.4061226 | -6.4933495 | 1.36503226 | -4.7569202 |
| 10256 | NM_000885.5 | 25.9723269 | -6.32757 | 1.33025252 | -4.7566683 |
| 10257 | NM_005239.5 | 107.642945 | -4.3102733 | 0.90615566 | -4.7566588 |
| 10258 | NM_172373.3 | 29.166706 | -6.4942033 | 1.36532119 | -4.7565389 |
| 10259 | NM_020459.1 | 59.6560256 | -5.6961078 | 1.19765977 | -4.7560317 |
| 10260 | NM_145313.3 | 22.5600055 | -6.1136098 | 1.28557711 | -4.7555372 |
| 10261 | NM_001271006.1 | 31.2581562 | -6.5848382 | 1.38473694 | -4.755299 |
| 10262 | NM_001002913.2 | 22.0860921 | -6.0934898 | 1.28149335 | -4.7549913 |
| 10263 | NM_001113402.1 | 37.7928395 | -6.8590811 | 1.44261724 | -4.7546091 |
| 10264 | NM_001216.2 | 74.7244207 | -7.8444549 | 1.64998343 | -4.7542628 |
| 10265 | NM_001134938.1 | 32.5536814 | -6.6461551 | 1.39841003 | -4.7526512 |
| 10266 | NM_006092.2 | 22.5926233 | -6.1248268 | 1.28876073 | -4.7524934 |
| 10267 | NM_002514.3 | 39.3195924 | -5.9358987 | 1.2490279 | -4.7524148 |
| 10268 | NM_033396.2 | 327.989938 | -5.8723955 | 1.23568745 | -4.7523308 |
| 10269 | NM_000309.3 | 23.3548541 | -6.1743637 | 1.2992803 | -4.7521414 |
| 10270 | NM_016042.3 | 25.9271921 | -6.3178771 | 1.32957204 | -4.7518125 |
| 10271 | NM_014291.3 | 34.7714668 | -6.7424398 | 1.41911786 | -4.7511486 |
| 10272 | NM_001304748.1 | 28.0606917 | -6.4385961 | 1.35530289 | -4.7506695 |
| 10273 | NM_001951.3 | 36.3012073 | -6.8045429 | 1.43241141 | -4.750411 |
| 10274 | NM_007034.4 | 22.1642671 | -6.094037 | 1.28286977 | -4.7503162 |
| 10275 | NR_002319.2 | 81.346576 | 3.02207156 | 0.63619795 | 4.75020638 |
| 10276 | NM_016429.3 | 22.9469636 | -6.148087 | 1.29440642 | -4.7497346 |

| | | | | | |
|---|---|---|---|---|---|
| 10277 | NM_014830.2 | 83.9495995 | -4.0736185 | 0.85775409 | -4.7491683 |
| 10278 | NM_003259.3 | 55.2960659 | -4.8104814 | 1.01308065 | -4.7483696 |
| 10279 | NM_194324.2 | 40.6905775 | -6.9673193 | 1.46730622 | -4.7483744 |
| 10280 | NM_015562.1 | 108.182389 | -3.1512384 | 0.66365411 | -4.7483145 |
| 10281 | NM_080752.3 | 25.5038461 | -6.2932296 | 1.32537327 | -4.7482696 |
| 10282 | NR_027122.1 | 177.836268 | 2.04403853 | 0.43048206 | 4.74825487 |
| 10283 | NM_002745.4 | 425.441407 | -5.7912736 | 1.21972389 | -4.7480202 |
| 10284 | NM_001346768.1 | 166.323243 | -5.3672111 | 1.13045413 | -4.7478362 |
| 10285 | NM_004966.3 | 34.6947604 | -6.7384696 | 1.41930094 | -4.7477384 |
| 10286 | NM_003618.3 | 72.5094517 | -7.8009043 | 1.64340761 | -4.7467861 |
| 10287 | NM_005505.4 | 32.9473788 | -6.6684904 | 1.40486364 | -4.7467172 |
| 10288 | NM_152385.2 | 29.0736314 | -6.4797862 | 1.36512534 | -4.7466603 |
| 10289 | NM_203475.2 | 53.3050875 | -7.3572025 | 1.55031006 | -4.7456329 |
| 10290 | NM_001303525.1 | 26.4763177 | -6.3547742 | 1.33916556 | -4.7453238 |
| 10291 | NM_006156.2 | 221.992725 | -5.850566 | 1.23301928 | -4.7449104 |
| 10292 | NM_001267783.1 | 36.514072 | -6.8135831 | 1.43599439 | -4.7448536 |
| 10293 | NM_001002247.2 | 45.886548 | -7.1407433 | 1.50495152 | -4.7448328 |
| 10294 | NM_202469.2 | 35.9533206 | -6.7919629 | 1.43152467 | -4.7445657 |
| 10295 | NM_001023587.2 | 29.640294 | -6.5100431 | 1.3723164 | -4.7438354 |
| 10296 | NM_001161376.1 | 21.7061711 | -6.0604586 | 1.27757638 | -4.7437152 |
| 10297 | NR_110051.1 | 79.4035321 | 2.48132452 | 0.52307663 | 4.74371129 |
| 10298 | NM_001322030.1 | 25.5136469 | -6.292918 | 1.32659437 | -4.7436641 |
| 10299 | NR_024278.1 | 57.0121731 | -4.6327563 | 0.97666491 | -4.7434451 |
| 10300 | NM_001114121.2 | 21.4732246 | -6.048289 | 1.27521458 | -4.7429578 |
| 10301 | NM_182526.2 | 43.311268 | -7.0574882 | 1.48804867 | -4.7427805 |
| 10302 | NM_001114123.2 | 39.4594464 | -6.9234282 | 1.45984155 | -4.7425888 |
| 10303 | NM_001033050.2 | 30.8509937 | -6.5655025 | 1.38449215 | -4.7421739 |
| 10304 | NM_021138.3 | 47.2850941 | -6.2110516 | 1.30995247 | -4.7414328 |
| 10305 | NM_152945.3 | 24.928513 | -6.2600322 | 1.32042598 | -4.7409187 |
| 10306 | NM_015910.5 | 20.6230972 | -5.9921112 | 1.26406491 | -4.7403509 |
| 10307 | NM_003141.3 | 21.9126718 | -6.0802212 | 1.28267038 | -4.7402835 |
| 10308 | NM_152484.2 | 40.4348096 | -5.9800399 | 1.26156658 | -4.7401699 |
| 10309 | NM_001319999.1 | 58.8180832 | -7.4988144 | 1.58202613 | -4.7400067 |
| 10310 | NM_001270551.1 | 37.6751911 | -6.8575395 | 1.44681538 | -4.7397474 |
| 10311 | NM_014176.3 | 43.0386105 | -7.0478619 | 1.48698985 | -4.7396839 |
| 10312 | NM_001297658.1 | 24.9258679 | -6.2591563 | 1.32060449 | -4.7396146 |
| 10313 | NM_001297419.1 | 25.6066042 | -6.2955769 | 1.32835078 | -4.7393934 |
| 10314 | NM_001207055.1 | 22.8811743 | -6.1329298 | 1.29406659 | -4.7392691 |
| 10315 | NM_174923.2 | 46.8031349 | -7.1702921 | 1.51294984 | -4.7392795 |
| 10316 | NR_103772.1 | 22.4687054 | -6.1052233 | 1.28826975 | -4.7390876 |
| 10317 | NM_017917.3 | 30.7253546 | -6.5619979 | 1.38466195 | -4.7390614 |
| 10318 | NM_001033678.3 | 22.71835 | -6.1232223 | 1.29216078 | -4.7387465 |
| 10319 | NM_001039366.1 | 42.5914239 | -7.0365628 | 1.4851149 | -4.7380595 |

| | | | | | |
|---|---|---|---|---|---|
| 10320 | NM_194330.2 | 47.72117 | -7.1955563 | 1.51868438 | -4.7380196 |
| 10321 | NM_001317915.1 | 40.3454376 | -6.9572968 | 1.46845831 | -4.7378238 |
| 10322 | NR_103498.1 | 27.8956207 | -6.4252088 | 1.35617432 | -4.7377455 |
| 10323 | NM_001320973.1 | 29.9967964 | -6.5256573 | 1.37746674 | -4.7374337 |
| 10324 | NM_001012974.2 | 27.0399816 | -6.3842088 | 1.34765296 | -4.7372795 |
| 10325 | NM_004299.4 | 24.6926781 | -6.251908 | 1.32009416 | -4.7359561 |
| 10326 | NM_145309.5 | 31.5418834 | -6.6016984 | 1.39397811 | -4.7358695 |
| 10327 | NM_001083909.2 | 24.3419839 | -6.2289898 | 1.31546805 | -4.735189 |
| 10328 | NM_016257.3 | 52.0816778 | -7.3241119 | 1.54673547 | -4.7352065 |
| 10329 | NM_020795.3 | 754.820775 | -4.6246459 | 0.97674562 | -4.7347496 |
| 10330 | NM_003420.3 | 29.2497579 | -6.4907274 | 1.37095514 | -4.7344565 |
| 10331 | NM_001310121.1 | 22.2353762 | -6.0899466 | 1.28652655 | -4.7336346 |
| 10332 | NM_031905.4 | 29.3146067 | -6.4979582 | 1.3727455 | -4.7335491 |
| 10333 | NM_198220.2 | 31.2021868 | -6.5841423 | 1.3909566 | -4.7335354 |
| 10334 | NM_198155.4 | 46.3082151 | -7.1566577 | 1.51197228 | -4.733326 |
| 10335 | NM_001098207.1 | 36.1967419 | -6.7988198 | 1.4366262 | -4.7324905 |
| 10336 | NM_024650.3 | 27.8782093 | -6.4206057 | 1.35676432 | -4.7322926 |
| 10337 | NM_032892.4 | 44.349462 | -7.0946873 | 1.49938529 | -4.7317306 |
| 10338 | NM_001288645.1 | 67.2950439 | -7.6941304 | 1.62637279 | -4.7308529 |
| 10339 | NM_052935.4 | 29.0538251 | -6.4810318 | 1.36996484 | -4.7308015 |
| 10340 | NM_001146314.1 | 21.1883592 | -6.0237782 | 1.27333698 | -4.7307023 |
| 10341 | NR_131180.1 | 16.9763113 | 8.9893535 | 1.900322 | 4.73043699 |
| 10342 | NM_001100117.2 | 38.406164 | -6.8859982 | 1.45588727 | -4.7297606 |
| 10343 | NM_001256277.1 | 33.4463477 | -6.6875119 | 1.41394757 | -4.7296746 |
| 10344 | NR_145128.1 | 29.7939034 | -6.5158916 | 1.37765844 | -4.7296859 |
| 10345 | NM_001080849.2 | 35.4782375 | -6.7692751 | 1.43131359 | -4.7294144 |
| 10346 | NM_080822.2 | 34.7257834 | -6.7385735 | 1.4251207 | -4.728423 |
| 10347 | NM_153685.3 | 49.0579717 | -7.2354973 | 1.53041418 | -4.7278034 |
| 10348 | NM_138783.3 | 42.7100928 | -6.0621951 | 1.2822747 | -4.7276883 |
| 10349 | NM_153741.1 | 31.9242245 | -6.6180796 | 1.39992235 | -4.7274619 |
| 10350 | NM_001142448.1 | 35.4606212 | -6.774759 | 1.43308599 | -4.7273919 |
| 10351 | NM_001301771.1 | 28.2531456 | -6.4434827 | 1.36301765 | -4.7273656 |
| 10352 | NM_052880.4 | 35.6154512 | -6.7797157 | 1.43424665 | -4.7270221 |
| 10353 | NM_001185011.1 | 31.2252604 | -6.5819191 | 1.39246346 | -4.7268164 |
| 10354 | NM_198212.2 | 49.8673037 | 7.12400777 | 1.50717955 | 4.72671473 |
| 10355 | NM_001309842.1 | 45.5963484 | -7.1308193 | 1.50881758 | -4.7260977 |
| 10356 | NM_004096.4 | 484.533037 | -4.9584546 | 1.04917908 | -4.7260326 |
| 10357 | NM_001307936.1 | 32.1913803 | -6.6244147 | 1.40206837 | -4.7247444 |
| 10358 | NM_001257389.1 | 38.5237118 | -6.8890122 | 1.45822341 | -4.7242502 |
| 10359 | NR_036576.1 | 27.3164805 | -6.3933143 | 1.35331873 | -4.7241748 |
| 10360 | NR_046341.1 | 56.5368852 | -5.5018924 | 1.16469692 | -4.7238833 |
| 10361 | NM_015409.4 | 312.473675 | -4.3182027 | 0.9141942 | -4.7235069 |
| 10362 | NR_110281.1 | 37.9046745 | 3.74621664 | 0.79313684 | 4.72329171 |

| | | | | | |
|---|---|---|---|---|---|
| 10363 | NM_013339.3 | 26.2292809 | -6.3333114 | 1.34096959 | -4.7229344 |
| 10364 | NM_030751.5 | 46.5482554 | -7.1593267 | 1.5158615 | -4.7229425 |
| 10365 | NM_001130035.1 | 21.3662327 | -6.0416331 | 1.27926669 | -4.7227315 |
| 10366 | NM_016245.4 | 29.6373715 | -6.5113439 | 1.37877005 | -4.7225742 |
| 10367 | NM_001323607.1 | 34.6901647 | -6.7380543 | 1.42681674 | -4.7224385 |
| 10368 | NM_006020.2 | 25.5911109 | -6.2992561 | 1.33402837 | -4.7219806 |
| 10369 | NR_131187.1 | 65.5121288 | 4.06376477 | 0.86069157 | 4.72151106 |
| 10370 | NM_003862.2 | 37.19837 | -6.8406397 | 1.44905013 | -4.7207751 |
| 10371 | NM_003948.4 | 27.1707999 | -6.3790668 | 1.35133009 | -4.7205837 |
| 10372 | NM_001303255.1 | 23.1962923 | -6.1599853 | 1.30513556 | -4.7198049 |
| 10373 | NM_032565.4 | 31.0733349 | -6.5804136 | 1.39450763 | -4.7188079 |
| 10374 | NM_002125.3 | 49.505255 | -7.2524539 | 1.53700458 | -4.7185636 |
| 10375 | NM_001145639.1 | 34.7055403 | -6.7402124 | 1.42845393 | -4.7185368 |
| 10376 | NM_001330523.1 | 28.5947567 | -6.4622857 | 1.36958256 | -4.7184346 |
| 10377 | NM_004667.5 | 357.001859 | -4.6821077 | 0.99229985 | -4.7184403 |
| 10378 | NM_001017928.3 | 25.4693949 | -6.2925851 | 1.33368725 | -4.7181864 |
| 10379 | NM_006002.4 | 32.9560022 | -6.6632386 | 1.41224787 | -4.7181793 |
| 10380 | NM_016592.3 | 39.4038405 | -6.9236412 | 1.46743317 | -4.7181986 |
| 10381 | NM_177454.3 | 187.033715 | -4.3679166 | 0.92578112 | -4.7180878 |
| 10382 | NM_002072.4 | 409.598962 | -4.798623 | 1.01711175 | -4.7178916 |
| 10383 | NR_135227.1 | 21.2923211 | -6.0314687 | 1.27845351 | -4.717785 |
| 10384 | NM_001001660.2 | 34.6373252 | -6.7352381 | 1.42773994 | -4.7174124 |
| 10385 | NM_001172418.1 | 27.1356475 | -6.3867578 | 1.35386585 | -4.7174229 |
| 10386 | NM_001042734.3 | 25.7548832 | -6.3145956 | 1.33862586 | -4.7172222 |
| 10387 | NM_001143819.2 | 39.1144484 | -6.9119906 | 1.46532628 | -4.7170318 |
| 10388 | NM_001134388.1 | 25.5193221 | -6.2911489 | 1.33371867 | -4.7169985 |
| 10389 | NM_001265589.1 | 26.4844005 | -6.3420831 | 1.34454202 | -4.7169095 |
| 10390 | NM_005745.7 | 25.8648398 | -6.3102484 | 1.33792154 | -4.7164562 |
| 10391 | NR_038190.1 | 22.6416588 | -6.116839 | 1.29693571 | -4.7163779 |
| 10392 | NM_033452.2 | 20.42736 | -5.9807054 | 1.2681845 | -4.7159584 |
| 10393 | NM_172341.3 | 94.4267128 | -4.4978058 | 0.9537886 | -4.7157261 |
| 10394 | NM_001303143.1 | 34.322408 | -6.7176275 | 1.42461535 | -4.7153974 |
| 10395 | NM_001173516.1 | 18.8369416 | 9.16565044 | 1.94421733 | 4.71431373 |
| 10396 | NM_021825.4 | 30.2458654 | -6.541219 | 1.38765983 | -4.7138491 |
| 10397 | NM_015358.2 | 47.7996462 | -7.2026741 | 1.52806338 | -4.7135965 |
| 10398 | NM_005393.2 | 34.6486123 | -6.7365136 | 1.42918351 | -4.7135399 |
| 10399 | NR_024127.1 | 28.1966104 | -6.4379662 | 1.36587062 | -4.7134524 |
| 10400 | NM_053035.2 | 29.0957148 | -6.4849938 | 1.37588178 | -4.7133365 |
| 10401 | NM_032494.2 | 27.1357065 | -6.3819664 | 1.35403682 | -4.7132887 |
| 10402 | NM_001040151.1 | 333.982436 | -3.9826443 | 0.84500936 | -4.7131363 |
| 10403 | NM_001160243.1 | 23.5923786 | -6.1888371 | 1.31311678 | -4.7130897 |
| 10404 | NM_001324266.1 | 37.3854538 | -6.8424022 | 1.4518062 | -4.7130272 |
| 10405 | NM_005491.4 | 22.8657729 | -6.1290579 | 1.30048742 | -4.7128929 |

| | | | | | |
|---|---|---|---|---|---|
| 10406 | NM_001271783.1 | 46.1405373 | -7.1496745 | 1.51722405 | -4.7123393 |
| 10407 | NM_001986.2 | 32.0612281 | -6.6256016 | 1.40601006 | -4.712343 |
| 10408 | NM_020158.3 | 39.8705855 | -6.9376888 | 1.4724253 | -4.7117424 |
| 10409 | NR_134858.1 | 141.943754 | 2.86057449 | 0.60712106 | 4.71170363 |
| 10410 | NM_021168.4 | 48.6316403 | -7.2265464 | 1.53380422 | -4.7115181 |
| 10411 | NM_015325.2 | 161.688962 | -5.0069698 | 1.06279689 | -4.7111258 |
| 10412 | NM_178003.2 | 62.4308438 | -7.5846995 | 1.61000934 | -4.7109661 |
| 10413 | NM_153339.2 | 29.6768029 | -6.5111022 | 1.38218721 | -4.7107238 |
| 10414 | NM_001950.3 | 138.197015 | -4.9174941 | 1.04391337 | -4.7106343 |
| 10415 | NM_001286069.1 | 31.7925823 | -6.6098488 | 1.40320837 | -4.7105255 |
| 10416 | NM_005612.4 | 44.4785252 | -7.099318 | 1.50714159 | -4.7104519 |
| 10417 | NM_006331.7 | 32.5053066 | -6.643703 | 1.41055027 | -4.710008 |
| 10418 | NM_007045.3 | 45.5906158 | -7.1352654 | 1.51507252 | -4.7095207 |
| 10419 | NM_001168647.2 | 31.3201982 | -6.5892953 | 1.39929311 | -4.7090172 |
| 10420 | NM_001949.4 | 36.5091813 | -6.8098415 | 1.44614362 | -4.7089662 |
| 10421 | NM_017793.2 | 33.8197597 | -6.698947 | 1.42273058 | -4.7085141 |
| 10422 | NM_001206641.2 | 28.8051241 | -6.4698962 | 1.37420363 | -4.7081059 |
| 10423 | NM_006042.2 | 29.9454538 | -6.5282716 | 1.38666239 | -4.7079027 |
| 10424 | NM_020744.3 | 22.4474473 | -6.1155892 | 1.29902525 | -4.7078294 |
| 10425 | NM_030795.3 | 46.4769627 | -7.1607676 | 1.52103931 | -4.7078123 |
| 10426 | NM_020458.3 | 31.432363 | -6.6020148 | 1.4024295 | -4.7075556 |
| 10427 | NM_001099858.1 | 24.25854 | -6.2257181 | 1.32265457 | -4.7069871 |
| 10428 | NR_028039.1 | 34.1739181 | -6.7209384 | 1.42787707 | -4.7069447 |
| 10429 | NM_016144.3 | 28.4138486 | -6.4492525 | 1.37018967 | -4.706832 |
| 10430 | NM_001017922.1 | 22.8159935 | -6.1391272 | 1.3043521 | -4.7066488 |
| 10431 | NM_005854.2 | 21.7769665 | -6.0634699 | 1.28836682 | -4.7063226 |
| 10432 | NM_020399.3 | 94.25336 | -8.1808581 | 1.73850893 | -4.705675 |
| 10433 | NM_016235.2 | 158.339847 | -5.3856081 | 1.1445658 | -4.7053722 |
| 10434 | NM_005086.4 | 21.1717639 | -6.0305506 | 1.28177212 | -4.7048539 |
| 10435 | NM_001282671.1 | 25.747947 | -6.307501 | 1.34067922 | -4.7047056 |
| 10436 | NM_080605.3 | 203.470467 | -5.727877 | 1.21751202 | -4.7045753 |
| 10437 | NM_005833.3 | 26.9047036 | -6.370758 | 1.35419736 | -4.7044531 |
| 10438 | NM_001267774.1 | 32.5075361 | -6.6434805 | 1.41221327 | -4.704304 |
| 10439 | NM_001291994.1 | 59.583311 | -7.5182895 | 1.59823472 | -4.704121 |
| 10440 | NM_001300907.1 | 28.9120658 | -6.4716414 | 1.37579693 | -4.703922 |
| 10441 | NM_018958.2 | 31.7103232 | 2.29442073 | 0.4877683 | 4.70391518 |
| 10442 | NR_134855.1 | 23.7819809 | -6.1945928 | 1.31689862 | -4.7039254 |
| 10443 | NM_005269.2 | 23.1563998 | -6.1603553 | 1.30988286 | -4.7029819 |
| 10444 | NM_001101801.2 | 45.9483515 | -6.1592169 | 1.3096894 | -4.7028073 |
| 10445 | NM_001244580.1 | 46.0896207 | -7.1469629 | 1.51971805 | -4.7028216 |
| 10446 | NM_001331007.1 | 48.382946 | -7.2209762 | 1.53553182 | -4.7025898 |
| 10447 | NM_014636.2 | 52.0750816 | -7.3215802 | 1.55700061 | -4.7023618 |
| 10448 | NM_001012754.3 | 28.4766619 | -6.4586515 | 1.37352264 | -4.7022534 |

| | | | | | |
|---|---|---|---|---|---|
| 10449 | NM_001134707.1 | 20.9549616 | -6.0172458 | 1.27991103 | -4.7013001 |
| 10450 | NM_001080397.2 | 38.7971294 | -6.9000041 | 1.46777646 | -4.7009911 |
| 10451 | NM_138457.2 | 228.190059 | -2.6630906 | 0.56656341 | -4.7004281 |
| 10452 | NM_058230.2 | 24.495446 | -6.2322301 | 1.32602689 | -4.6999274 |
| 10453 | NM_002802.2 | 302.725499 | -5.5810039 | 1.18751512 | -4.6997329 |
| 10454 | NR_038970.1 | 213.992214 | -5.4333161 | 1.15615531 | -4.6994691 |
| 10455 | NM_016104.3 | 34.5304047 | -6.7297916 | 1.43226307 | -4.698712 |
| 10456 | NM_001358.2 | 331.504701 | -5.8841949 | 1.25230741 | -4.6986824 |
| 10457 | NM_007200.4 | 50.6921083 | -6.3076744 | 1.34245056 | -4.6986269 |
| 10458 | NR_052010.1 | 27.5204721 | -6.4091396 | 1.36405564 | -4.698591 |
| 10459 | NM_005079.3 | 32.7462719 | -6.6593267 | 1.4175629 | -4.6977293 |
| 10460 | NM_152683.3 | 20.067127 | -5.9434144 | 1.26519076 | -4.6976429 |
| 10461 | NM_001707.3 | 32.4751861 | -6.6436465 | 1.41428802 | -4.6975202 |
| 10462 | NM_178448.3 | 54.4463502 | -6.4140314 | 1.36564523 | -4.696704 |
| 10463 | NM_002246.2 | 164.090016 | -3.8671667 | 0.82346942 | -4.6961874 |
| 10464 | NM_004170.5 | 61.9133352 | -5.7967159 | 1.23434877 | -4.6961734 |
| 10465 | NM_001278789.1 | 26.3743481 | -6.3466247 | 1.35153986 | -4.6958472 |
| 10466 | NM_138285.4 | 25.1267848 | -6.2710812 | 1.33554796 | -4.6955118 |
| 10467 | NM_001330374.1 | 12.0902933 | 8.50960926 | 1.81235599 | 4.6953299 |
| 10468 | NM_001170416.1 | 20.5456562 | -5.9851417 | 1.27470799 | -4.6953042 |
| 10469 | NM_176867.3 | 26.0372174 | 6.79497439 | 1.44723076 | 4.69515613 |
| 10470 | NM_014562.3 | 22.5044183 | -6.1117067 | 1.30173947 | -4.6950306 |
| 10471 | NM_016551.2 | 186.363802 | -4.0471074 | 0.86207619 | -4.6946052 |
| 10472 | NM_016126.3 | 21.8832321 | -6.0754215 | 1.29417472 | -4.6944369 |
| 10473 | NM_001128833.1 | 176.474608 | -5.0044511 | 1.06608759 | -4.6942213 |
| 10474 | NM_015365.2 | 27.4170802 | -6.3968754 | 1.36272036 | -4.6941952 |
| 10475 | NM_001037340.2 | 36.9789535 | -6.8306472 | 1.4551494 | -4.6941209 |
| 10476 | NR_034075.1 | 27.2771994 | -6.3838686 | 1.35999053 | -4.6940537 |
| 10477 | NR_135262.1 | 67.4695613 | -7.6971772 | 1.63979812 | -4.6939785 |
| 10478 | NM_004451.4 | 38.1983746 | -6.8759232 | 1.46489126 | -4.6938113 |
| 10479 | NM_144651.4 | 41.3451161 | -6.9925881 | 1.48983233 | -4.6935403 |
| 10480 | NM_172177.3 | 21.1371506 | -6.0262005 | 1.28396456 | -4.6934321 |
| 10481 | NM_152600.2 | 141.397864 | -4.9535063 | 1.05547118 | -4.6931705 |
| 10482 | NM_001145648.1 | 44.4077249 | -7.0932803 | 1.51151927 | -4.692815 |
| 10483 | NM_177948.2 | 31.9324427 | -6.6226279 | 1.4113124 | -4.6925315 |
| 10484 | NM_001042432.1 | 24.8643889 | -6.2588359 | 1.33403348 | -4.6916633 |
| 10485 | NM_001142315.1 | 23.2675416 | -6.1571701 | 1.31246234 | -4.6913118 |
| 10486 | NM_175039.3 | 25.5245438 | -6.2946113 | 1.34179919 | -4.6911723 |
| 10487 | NM_001288997.1 | 13.6953977 | 8.70857214 | 1.85670341 | 4.69034099 |
| 10488 | NM_001276312.1 | 31.5290184 | -6.6033549 | 1.40787495 | -4.6902993 |
| 10489 | NM_021961.5 | 210.20012 | -5.1459644 | 1.09770377 | -4.6879354 |
| 10490 | NM_005962.4 | 24.7214869 | -6.2565555 | 1.33464088 | -4.6878195 |
| 10491 | NM_001099693.1 | 24.8716473 | -6.2542465 | 1.33417698 | -4.6877188 |

| | | | | | |
|---|---|---|---|---|---|
| 10492 | NM_007281.3 | 45.8073137 | -7.1406668 | 1.5232966 | -4.6876405 |
| 10493 | NM_020350.4 | 29.3063287 | -6.5001278 | 1.38665635 | -4.6876271 |
| 10494 | NM_180982.2 | 21.704363 | -6.0588242 | 1.29272108 | -4.6868766 |
| 10495 | NR_110533.1 | 21.7863919 | -6.0599301 | 1.29307082 | -4.6864642 |
| 10496 | NM_022341.1 | 23.3655406 | -6.164764 | 1.31553914 | -4.6861122 |
| 10497 | NM_001278228.1 | 32.800291 | -6.6566182 | 1.42050569 | -4.6860905 |
| 10498 | NM_001163636.2 | 49.0137981 | -7.2384359 | 1.54476002 | -4.6857996 |
| 10499 | NM_001278182.1 | 78.6206625 | -7.9173102 | 1.68966345 | -4.6857321 |
| 10500 | NM_001323956.1 | 49.5415665 | -7.2522455 | 1.54780755 | -4.6854957 |
| 10501 | NM_001272082.1 | 311.330013 | -6.0320653 | 1.28741946 | -4.6853924 |
| 10502 | NM_001288656.1 | 35.7549577 | -6.7837702 | 1.44787146 | -4.68534 |
| 10503 | NM_001258420.1 | 28.3329249 | -6.44469 | 1.3755601 | -4.6851388 |
| 10504 | NM_052885.3 | 25.8353482 | -6.3154194 | 1.34817428 | -4.6844236 |
| 10505 | NM_001160147.1 | 41.2286007 | -6.9862624 | 1.49145165 | -4.6842031 |
| 10506 | NM_005357.3 | 32.8981711 | -6.660356 | 1.42188041 | -4.6841886 |
| 10507 | NM_015221.3 | 24.7931021 | -6.2606891 | 1.33662563 | -4.6839511 |
| 10508 | NM_032889.4 | 61.5908371 | -4.8253641 | 1.03025589 | -4.683656 |
| 10509 | NM_012385.2 | 30.0624086 | -6.5343238 | 1.39517006 | -4.6835321 |
| 10510 | NM_001348124.1 | 37.0260884 | -6.8292319 | 1.45832383 | -4.6829324 |
| 10511 | NM_001330612.1 | 32.490339 | -6.6390272 | 1.41773734 | -4.682833 |
| 10512 | NM_198205.1 | 31.7808366 | -6.6129003 | 1.41222306 | -4.6826175 |
| 10513 | NM_001085347.2 | 23.8341634 | -6.1970691 | 1.32346457 | -4.6824593 |
| 10514 | NM_001164473.2 | 80.8001887 | -7.9572876 | 1.69946771 | -4.6822235 |
| 10515 | NM_001171136.1 | 52.0088096 | -7.3195464 | 1.56329823 | -4.6821177 |
| 10516 | NM_017711.3 | 27.584568 | -6.4088187 | 1.36912418 | -4.6809623 |
| 10517 | NM_203458.4 | 58.6986261 | -4.8799173 | 1.04259343 | -4.6805563 |
| 10518 | NM_001098576.1 | 33.4824198 | -6.6852821 | 1.42835671 | -4.6804009 |
| 10519 | NM_001291379.1 | 35.69026 | -6.7762713 | 1.44796379 | -4.6798624 |
| 10520 | NR_038270.1 | 21.6899606 | -6.0640758 | 1.29592138 | -4.6793547 |
| 10521 | NM_183235.2 | 28.1494914 | -6.4391971 | 1.37609299 | -4.6793328 |
| 10522 | NM_024540.3 | 29.9194715 | -6.5234501 | 1.39412796 | -4.6792334 |
| 10523 | NM_001144058.1 | 29.4964017 | -6.5103108 | 1.39144326 | -4.6788188 |
| 10524 | NR_027078.1 | 25.0901568 | 2.84234882 | 0.60750752 | 4.67870557 |
| 10525 | NM_175854.7 | 111.835399 | -2.0693693 | 0.44232153 | -4.6784276 |
| 10526 | NR_135104.1 | 25.0577394 | -6.2610154 | 1.33830107 | -4.678331 |
| 10527 | NM_147128.3 | 38.0245621 | -5.8929087 | 1.25965042 | -4.6782096 |
| 10528 | NR_120613.1 | 30.8754815 | -6.5714212 | 1.40483189 | -4.6777278 |
| 10529 | NM_001243756.1 | 24.1455491 | -6.2220163 | 1.33041527 | -4.6767475 |
| 10530 | NM_014911.3 | 332.238142 | -2.2531517 | 0.48178059 | -4.6767174 |
| 10531 | NM_003838.4 | 20.8057419 | -6.0039391 | 1.28384079 | -4.6765449 |
| 10532 | NM_032844.4 | 25.364137 | -6.286845 | 1.34433505 | -4.6765462 |
| 10533 | NM_001100912.1 | 25.9920846 | -6.323004 | 1.35209234 | -4.6764587 |
| 10534 | NM_020209.3 | 22.972166 | -6.147127 | 1.31449276 | -4.6764251 |

| | | | | | |
|---|---|---|---|---|---|
| 10535 | NM_001271895.1 | 27.2182324 | -6.3924876 | 1.36698379 | -4.6763449 |
| 10536 | NM_001283049.1 | 26.0756051 | -6.3297948 | 1.35372717 | -4.6758275 |
| 10537 | NM_203411.1 | 21.0317963 | -6.0242832 | 1.28843638 | -4.6756544 |
| 10538 | NM_024769.3 | 47.265177 | -6.2082537 | 1.32796645 | -4.675008 |
| 10539 | NM_207327.4 | 24.5923094 | -6.238594 | 1.33448875 | -4.6748944 |
| 10540 | NM_001318234.1 | 59.0323161 | -7.5045888 | 1.60546928 | -4.6743895 |
| 10541 | NM_001297599.1 | 55.3592473 | -7.4112207 | 1.58574636 | -4.6736483 |
| 10542 | NM_134324.2 | 25.3223967 | -6.2833447 | 1.34456846 | -4.6731311 |
| 10543 | NM_173630.3 | 32.3318728 | -6.6317813 | 1.41934718 | -4.6724166 |
| 10544 | NM_001012643.3 | 29.3456734 | -6.4945826 | 1.39016834 | -4.6717958 |
| 10545 | NM_021723.4 | 29.2785042 | -6.488034 | 1.38883592 | -4.6715626 |
| 10546 | NM_001171740.2 | 23.6707619 | -6.1801683 | 1.32307682 | -4.6710578 |
| 10547 | NM_001243203.1 | 39.7529187 | -6.9334426 | 1.48442514 | -4.6707931 |
| 10548 | NM_013397.5 | 23.3274275 | -6.1657893 | 1.32007392 | -4.6707909 |
| 10549 | NM_017879.1 | 22.5894754 | -6.1170655 | 1.30973625 | -4.670456 |
| 10550 | NM_015042.1 | 259.577676 | -5.3506266 | 1.14564642 | -4.6703996 |
| 10551 | NM_014174.2 | 26.5811196 | -6.3529551 | 1.36042061 | -4.6698463 |
| 10552 | NM_017518.7 | 28.8121373 | -6.4699218 | 1.38549782 | -4.6697452 |
| 10553 | NM_152998.2 | 29.1467923 | -6.4849686 | 1.38873186 | -4.6697054 |
| 10554 | NM_000797.3 | 19.4519613 | -5.8987523 | 1.26327854 | -4.6693996 |
| 10555 | NM_020423.6 | 24.4743417 | -6.2268461 | 1.33358859 | -4.6692407 |
| 10556 | NM_001083588.1 | 20.4492449 | -5.9738128 | 1.2795695 | -4.6686115 |
| 10557 | NM_001024939.3 | 19.4615027 | -5.9086052 | 1.26596981 | -4.667256 |
| 10558 | NM_031218.3 | 26.8871094 | -6.3685325 | 1.3645178 | -4.6672403 |
| 10559 | NM_001251971.1 | 290.758344 | -4.2470476 | 0.91001085 | -4.6670296 |
| 10560 | NM_001318134.1 | 30.0993254 | -6.5365438 | 1.40062561 | -4.6668744 |
| 10561 | NM_000904.4 | 31.8320961 | -6.6134532 | 1.4173634 | -4.6660251 |
| 10562 | NM_030922.6 | 38.9580921 | -6.9043505 | 1.47986682 | -4.6655215 |
| 10563 | NM_001142651.2 | 54.6821714 | -7.3943797 | 1.584999 | -4.6652267 |
| 10564 | NM_002269.2 | 41.3517402 | -6.0166543 | 1.28967986 | -4.6652309 |
| 10565 | NR_110864.1 | 29.6355077 | -6.5162279 | 1.39677479 | -4.6651958 |
| 10566 | NM_001174051.2 | 52.2738624 | -7.3288403 | 1.57119531 | -4.6644999 |
| 10567 | NM_001032387.1 | 27.7519763 | -6.4195883 | 1.37633353 | -4.6642679 |
| 10568 | NM_020116.4 | 30.6618962 | -6.5639943 | 1.40729219 | -4.6642725 |
| 10569 | NM_133462.3 | 29.3417805 | -6.4962519 | 1.39281136 | -4.664129 |
| 10570 | NR_046357.1 | 49.8481832 | -7.26036 | 1.55674907 | -4.6637959 |
| 10571 | NM_001282551.1 | 31.6482547 | -6.6075191 | 1.41689223 | -4.6633887 |
| 10572 | NM_001039349.2 | 57.8065583 | -4.7287514 | 1.01402493 | -4.6633483 |
| 10573 | NM_001320848.1 | 23.1927357 | -6.1610689 | 1.32131205 | -4.6628416 |
| 10574 | NM_005934.3 | 514.360618 | -5.8291482 | 1.25013886 | -4.6628006 |
| 10575 | NM_001001438.2 | 37.6091432 | -6.8529292 | 1.46973131 | -4.6627089 |
| 10576 | NM_012341.2 | 149.598421 | -4.4478704 | 0.95401496 | -4.6622648 |
| 10577 | NM_004217.3 | 27.8852048 | -6.4218175 | 1.37767752 | -4.6613358 |

| | | | | | |
|---|---|---|---|---|---|
| 10578 | NM_001010866.3 | 61.8800744 | -5.6114741 | 1.20384856 | -4.6612791 |
| 10579 | NM_016093.3 | 44.9321719 | -7.1100057 | 1.52537365 | -4.6611568 |
| 10580 | NM_016466.5 | 29.5070507 | -6.5037564 | 1.39541276 | -4.6608119 |
| 10581 | NR_132421.1 | 24.724259 | 9.53777915 | 2.04649486 | 4.66054391 |
| 10582 | NM_203349.3 | 32.9573599 | -6.6650593 | 1.4305826 | -4.6589825 |
| 10583 | NM_001318089.1 | 27.8353636 | -6.418389 | 1.37769721 | -4.6587806 |
| 10584 | NM_018916.3 | 53.3207235 | -7.3570918 | 1.57923994 | -4.6586282 |
| 10585 | NR_030737.1 | 25.4354679 | -6.2893797 | 1.35008562 | -4.6585043 |
| 10586 | NM_012463.3 | 123.523836 | -2.1003391 | 0.45089997 | -4.6581043 |
| 10587 | NM_001142677.1 | 32.9211492 | -6.6639685 | 1.43092926 | -4.6570915 |
| 10588 | NM_000044.3 | 29.9004508 | -6.5243749 | 1.40114365 | -4.656464 |
| 10589 | NM_001271629.1 | 20.7708925 | -5.9980301 | 1.28842082 | -4.6553347 |
| 10590 | NM_173666.3 | 27.1342566 | -6.3783321 | 1.37013496 | -4.6552583 |
| 10591 | NM_001142522.2 | 26.050136 | -6.3188567 | 1.35754932 | -4.6546056 |
| 10592 | NM_019114.4 | 24.5196029 | -6.2415051 | 1.34093531 | -4.6545907 |
| 10593 | NM_148908.3 | 39.806289 | -6.9350405 | 1.4899695 | -4.6544848 |
| 10594 | NM_173473.3 | 46.3654961 | -7.1552137 | 1.53727837 | -4.6544685 |
| 10595 | NM_000290.3 | 20.4958713 | -5.9785439 | 1.28456036 | -4.6541557 |
| 10596 | NM_016576.4 | 21.3299411 | -6.0437048 | 1.29856239 | -4.6541505 |
| 10597 | NM_001297567.1 | 86.5423814 | 4.31490082 | 0.92728922 | 4.65324167 |
| 10598 | NM_152658.2 | 22.9785983 | -6.1440116 | 1.32048645 | -4.6528395 |
| 10599 | NM_001118890.1 | 44.0549229 | -7.0827526 | 1.52231067 | -4.6526328 |
| 10600 | NM_001170418.1 | 63.2732036 | 3.71961212 | 0.79946899 | 4.65260339 |
| 10601 | NM_020421.3 | 22.9920783 | -6.1401428 | 1.31974805 | -4.6525114 |
| 10602 | NM_001807.4 | 19.1047411 | -5.8784788 | 1.26359613 | -4.6521817 |
| 10603 | NR_138044.1 | 19.8018155 | -5.9231334 | 1.27328254 | -4.651861 |
| 10604 | NM_014449.1 | 41.35022 | -6.990146 | 1.5026963 | -4.6517357 |
| 10605 | NM_021111.2 | 51.5227336 | -4.5000037 | 0.96738007 | -4.6517433 |
| 10606 | NM_002208.4 | 32.2337873 | -6.6328968 | 1.42604454 | -4.651255 |
| 10607 | NM_001145374.1 | 34.0269254 | -6.7117377 | 1.44307484 | -4.6509977 |
| 10608 | NM_001320542.1 | 27.5946479 | -6.4127834 | 1.3788051 | -4.6509716 |
| 10609 | NM_014682.2 | 42.1332301 | -7.0167838 | 1.50875622 | -4.6507075 |
| 10610 | NM_001164811.1 | 26.3543592 | -6.3459316 | 1.36460135 | -4.6503923 |
| 10611 | NM_005641.3 | 49.906312 | -6.2853394 | 1.35160402 | -4.6502817 |
| 10612 | NR_002570.3 | 26.5180893 | -6.3474627 | 1.36499834 | -4.6501615 |
| 10613 | NM_033148.3 | 24.1651191 | -6.2172176 | 1.33703617 | -4.6499995 |
| 10614 | NR_049762.1 | 39.1130538 | -6.9069378 | 1.48544049 | -4.6497573 |
| 10615 | NM_001077350.2 | 47.4181176 | -7.1871928 | 1.54584579 | -4.6493595 |
| 10616 | NM_021229.3 | 26.5094041 | -6.3530159 | 1.36647993 | -4.6491835 |
| 10617 | NM_001304998.1 | 141.695441 | 12.0675002 | 2.59582571 | 4.64880986 |
| 10618 | NM_001003945.2 | 29.5191404 | -6.5092699 | 1.40030734 | -4.648458 |
| 10619 | NM_002498.2 | 24.1561384 | -6.2149407 | 1.33705484 | -4.6482317 |
| 10620 | NR_138095.1 | 24.1665914 | -6.2142879 | 1.33694011 | -4.6481423 |

| | | | | | |
|---|---|---|---|---|---|
| 10621 | NM_138386.2 | 19.4967224 | -5.9118025 | 1.27204588 | -4.6474759 |
| 10622 | NM_017703.2 | 32.9663074 | -6.6623759 | 1.43362346 | -4.6472285 |
| 10623 | NM_024316.2 | 28.0829665 | -6.4335732 | 1.38440502 | -4.6471756 |
| 10624 | NR_033403.1 | 19.8945699 | -5.9394296 | 1.27807846 | -4.6471557 |
| 10625 | NM_001666.4 | 20.3270395 | -5.9600994 | 1.28261562 | -4.646832 |
| 10626 | NM_005787.5 | 32.5867222 | -6.6510014 | 1.43132616 | -4.6467406 |
| 10627 | NM_001018000.3 | 29.3350356 | -6.4941908 | 1.39766038 | -4.6464727 |
| 10628 | NM_003456.2 | 24.4990771 | -6.2288591 | 1.34059906 | -4.6463251 |
| 10629 | NR_003226.1 | 24.3692493 | -6.2363271 | 1.34224339 | -4.6461969 |
| 10630 | NM_080911.2 | 29.6477774 | -6.5057909 | 1.40030754 | -4.6459729 |
| 10631 | NM_001122769.2 | 38.8617467 | -5.9175564 | 1.27371872 | -4.6458895 |
| 10632 | NM_001190991.1 | 22.2329467 | -6.095834 | 1.312131 | -4.6457511 |
| 10633 | NM_007175.6 | 178.878688 | -3.3701205 | 0.72548647 | -4.6453251 |
| 10634 | NM_001013436.2 | 32.5140311 | -6.6434536 | 1.43021093 | -4.6450866 |
| 10635 | NM_173651.3 | 21.2342849 | -6.0249916 | 1.2972712 | -4.6443578 |
| 10636 | NM_001311178.1 | 21.3298673 | -6.0418885 | 1.3009381 | -4.6442552 |
| 10637 | NM_001174164.1 | 35.4710503 | -6.7698634 | 1.45771026 | -4.6441763 |
| 10638 | NM_001322224.1 | 27.9619575 | -6.42374 | 1.38321475 | -4.6440655 |
| 10639 | NM_145865.2 | 44.9768601 | 2.43134579 | 0.52355824 | 4.64388789 |
| 10640 | NM_012338.3 | 27.9869855 | -6.4299131 | 1.3846784 | -4.6436148 |
| 10641 | NM_001218.4 | 65.3903021 | -7.6539625 | 1.6482939 | -4.6435666 |
| 10642 | NM_007086.3 | 32.3491121 | -6.6357309 | 1.42902319 | -4.6435432 |
| 10643 | NM_004332.3 | 30.6854931 | -6.5613332 | 1.41301373 | -4.6435028 |
| 10644 | NM_014353.4 | 27.0965609 | -6.3757455 | 1.37309957 | -4.6433235 |
| 10645 | NM_018048.4 | 35.5088931 | -6.7696391 | 1.45794318 | -4.6432805 |
| 10646 | NM_145870.2 | 25.521161 | -6.2948715 | 1.35573501 | -4.643143 |
| 10647 | NM_001006622.2 | 27.469666 | -6.4016115 | 1.37878061 | -4.6429515 |
| 10648 | NM_138340.4 | 19.9455672 | -5.9375231 | 1.27885695 | -4.642836 |
| 10649 | NM_001006109.1 | 29.0413059 | -6.484707 | 1.39673184 | -4.6427717 |
| 10650 | NM_000817.2 | 27.636947 | -6.4152185 | 1.3817945 | -4.6426719 |
| 10651 | NM_001321099.1 | 62.3103948 | -7.5806829 | 1.63284205 | -4.6426309 |
| 10652 | NM_198097.3 | 116.883805 | -4.8313467 | 1.04067138 | -4.6425286 |
| 10653 | NM_001006943.2 | 18.9441773 | -5.865616 | 1.26348295 | -4.642418 |
| 10654 | NM_178558.4 | 24.154934 | -6.2123318 | 1.33818912 | -4.6423422 |
| 10655 | NM_001481.2 | 25.4642127 | -6.2896723 | 1.35488397 | -4.6422221 |
| 10656 | NM_014421.2 | 39.5180191 | -5.9583495 | 1.28354988 | -4.6420865 |
| 10657 | NM_001185176.1 | 12.9198742 | 8.62526402 | 1.85841547 | 4.64119255 |
| 10658 | NM_007331.1 | 52.5159346 | -5.3948282 | 1.16242435 | -4.6410144 |
| 10659 | NM_015237.3 | 28.791152 | -6.4674834 | 1.39361189 | -4.6408067 |
| 10660 | NM_014687.3 | 143.742612 | -2.1624198 | 0.46598785 | -4.6405067 |
| 10661 | NM_001256067.1 | 19.8501074 | -5.9389397 | 1.27987214 | -4.6402602 |
| 10662 | NM_002205.4 | 123.305939 | -5.1057348 | 1.10037431 | -4.6399982 |
| 10663 | NM_080571.1 | 51.6878741 | -7.3127718 | 1.57607427 | -4.6398649 |

| | | | | | |
|---|---|---|---|---|---|
| 10664 | NM_032345.2 | 19.7194452 | -5.9219912 | 1.27636255 | -4.6397406 |
| 10665 | NM_130384.2 | 24.0307097 | -6.2068464 | 1.33776354 | -4.6397186 |
| 10666 | NM_173828.4 | 34.2612893 | -6.7190509 | 1.44822678 | -4.6395019 |
| 10667 | NM_001330724.1 | 24.3695497 | -6.2311217 | 1.34320455 | -4.6389969 |
| 10668 | NM_001303099.1 | 27.6821064 | -6.4125564 | 1.38236779 | -4.6388208 |
| 10669 | NM_001177651.1 | 28.0285566 | -6.4264547 | 1.38545643 | -4.6385109 |
| 10670 | NM_014297.4 | 30.883731 | -6.5689317 | 1.41627407 | -4.6381783 |
| 10671 | NM_002268.4 | 382.12462 | -1.8586655 | 0.40074566 | -4.6380178 |
| 10672 | NR_120424.1 | 97.2496752 | 2.05438326 | 0.44296276 | 4.6378239 |
| 10673 | NM_017880.1 | 22.9719123 | -6.1447797 | 1.32523416 | -4.6367501 |
| 10674 | NM_001190849.1 | 24.2626717 | -6.2212004 | 1.34186217 | -4.636244 |
| 10675 | NM_001308028.1 | 40.5422465 | -6.9613462 | 1.50154131 | -4.6361336 |
| 10676 | NM_003408.2 | 28.1618643 | -6.4312114 | 1.3873078 | -4.6357495 |
| 10677 | NM_001252092.1 | 27.3397186 | -6.3981298 | 1.38020436 | -4.6356394 |
| 10678 | NM_003955.4 | 95.9600575 | -5.0574369 | 1.09098867 | -4.6356456 |
| 10679 | NM_001320616.1 | 36.7262194 | -6.8198771 | 1.47129326 | -4.6352942 |
| 10680 | NM_001115.2 | 21.1867639 | -6.0323616 | 1.30146633 | -4.6350501 |
| 10681 | NM_021824.3 | 22.7228597 | -6.1297721 | 1.32254988 | -4.6348135 |
| 10682 | NM_001271969.1 | 42.7943775 | -7.042802 | 1.5196341 | -4.634538 |
| 10683 | NM_001195215.1 | 28.6639834 | -6.4664579 | 1.39529435 | -4.6344758 |
| 10684 | NM_014584.2 | 214.469156 | -2.7989494 | 0.60399236 | -4.6340809 |
| 10685 | NM_145330.2 | 31.0970082 | -6.5811635 | 1.42017554 | -4.6340493 |
| 10686 | NM_025268.3 | 21.4550623 | -6.045135 | 1.30470561 | -4.6333326 |
| 10687 | NM_020777.2 | 171.477663 | -4.9631955 | 1.07130429 | -4.6328532 |
| 10688 | NM_033046.2 | 22.4283885 | -6.1051154 | 1.31782428 | -4.6327234 |
| 10689 | NM_001144775.2 | 51.2762725 | -7.3029267 | 1.57639827 | -4.632666 |
| 10690 | NR_125755.1 | 20.6798436 | -5.9975075 | 1.29467049 | -4.6324586 |
| 10691 | NM_033445.2 | 35.1022923 | -6.7535237 | 1.45820611 | -4.6313917 |
| 10692 | NR_034053.2 | 30.8350316 | -6.5665341 | 1.41786773 | -4.6312741 |
| 10693 | NM_003272.3 | 42.0687236 | -6.0424882 | 1.30479937 | -4.6309711 |
| 10694 | NM_007137.3 | 31.1405765 | -6.5829178 | 1.4215114 | -4.6309286 |
| 10695 | NM_021967.2 | 33.7053602 | -6.6992137 | 1.44664161 | -4.6308731 |
| 10696 | NM_001206670.1 | 65.4765619 | 4.58089709 | 0.98921903 | 4.63082183 |
| 10697 | NM_001142296.1 | 39.8338595 | -6.9338957 | 1.49741317 | -4.6305828 |
| 10698 | NM_001143942.1 | 18.9158261 | -5.8643481 | 1.2664753 | -4.630448 |
| 10699 | NM_182665.3 | 24.3481453 | -6.2309731 | 1.345843 | -4.629792 |
| 10700 | NM_002452.3 | 38.2577637 | -6.8767526 | 1.48533909 | -4.6297527 |
| 10701 | NM_004891.3 | 24.2854854 | -6.2207094 | 1.34372569 | -4.6294488 |
| 10702 | NM_012140.4 | 27.0573881 | -6.3858855 | 1.37965879 | -4.6285977 |
| 10703 | NM_018401.2 | 28.764047 | -6.4693684 | 1.39776287 | -4.6283733 |
| 10704 | NM_001040058.1 | 32.2363418 | -6.631205 | 1.43285427 | -4.6279689 |
| 10705 | NM_031448.4 | 31.4157285 | -6.5897232 | 1.42392328 | -4.6278639 |
| 10706 | NM_001200016.1 | 20.2016585 | -5.9584716 | 1.28758614 | -4.6276295 |

| | | | | | |
|---|---|---|---|---|---|
| 10707 | NM_001304331.1 | 69.4462547 | -7.7394577 | 1.67277927 | -4.6267059 |
| 10708 | NM_005746.2 | 158.51377 | -4.9800428 | 1.07646591 | -4.6262894 |
| 10709 | NR_036542.1 | 27.0595309 | -6.3733064 | 1.3777724 | -4.625805 |
| 10710 | NM_001135058.1 | 33.9001282 | -6.7022686 | 1.44894286 | -4.6256266 |
| 10711 | NM_024874.4 | 277.985425 | -3.0403727 | 0.65736595 | -4.6250839 |
| 10712 | NM_016305.3 | 37.0845365 | -6.8331713 | 1.47758166 | -4.6245643 |
| 10713 | NM_153182.3 | 23.6303151 | -6.1766299 | 1.33562365 | -4.6245287 |
| 10714 | NM_000380.3 | 44.1992792 | -6.1124771 | 1.32176085 | -4.6244955 |
| 10715 | NM_152260.2 | 40.8923352 | -5.9992527 | 1.29728461 | -4.6244692 |
| 10716 | NM_001099673.2 | 27.7520389 | -6.4174088 | 1.38772647 | -4.6244047 |
| 10717 | NM_152750.4 | 29.0677584 | -6.476092 | 1.40046787 | -4.6242346 |
| 10718 | NM_001303253.1 | 41.1371294 | -6.9841721 | 1.51036489 | -4.6241621 |
| 10719 | NM_001001890.2 | 28.8274845 | -6.4741862 | 1.40023902 | -4.6236293 |
| 10720 | NM_001107.4 | 24.9235724 | -6.2605443 | 1.35404676 | -4.6235806 |
| 10721 | NM_203486.2 | 64.4555642 | -6.661196 | 1.44079619 | -4.6232743 |
| 10722 | NM_000425.4 | 40.1154255 | -6.9511226 | 1.5035333 | -4.6231916 |
| 10723 | NM_001042552.2 | 35.2168691 | -6.7556522 | 1.46129734 | -4.6230511 |
| 10724 | NR_132353.1 | 25.4827042 | -6.2906558 | 1.36073765 | -4.6229747 |
| 10725 | NM_001321356.1 | 18.9962799 | -5.8675999 | 1.26924367 | -4.6229105 |
| 10726 | NM_001025930.3 | 24.5496168 | -6.2449148 | 1.35088368 | -4.6228368 |
| 10727 | NM_014494.2 | 64.4378834 | -7.6320363 | 1.65105457 | -4.6225222 |
| 10728 | NM_015564.2 | 53.7166855 | -6.3872969 | 1.38195343 | -4.6219335 |
| 10729 | NM_001987.4 | 67.437636 | -5.7158665 | 1.23673059 | -4.6217555 |
| 10730 | NM_015356.4 | 263.101723 | -4.2661323 | 0.92307274 | -4.6216643 |
| 10731 | NM_001040454.1 | 23.6097609 | -6.1810011 | 1.33746994 | -4.6214131 |
| 10732 | NM_003454.3 | 21.7644033 | -6.0647336 | 1.31231401 | -4.6214043 |
| 10733 | NM_005638.5 | 36.1735504 | -6.8014594 | 1.47171832 | -4.621441 |
| 10734 | NM_020765.2 | 485.52921 | -4.5186833 | 0.97776828 | -4.6214256 |
| 10735 | NM_001130029.1 | 47.9443103 | -7.20516 | 1.55917072 | -4.6211488 |
| 10736 | NM_199334.3 | 65.0660615 | -5.8740345 | 1.27115161 | -4.6210337 |
| 10737 | NM_006176.2 | 29.7012647 | -6.5143974 | 1.40984722 | -4.6206407 |
| 10738 | NM_001178005.1 | 59.3272562 | 1.66926498 | 0.36126975 | 4.62055008 |
| 10739 | NM_024870.3 | 46.6921958 | -3.2975701 | 0.71371991 | -4.620258 |
| 10740 | NM_001127392.2 | 21.1244856 | -6.0175053 | 1.302526 | -4.6198734 |
| 10741 | NM_001202412.1 | 27.1418635 | -6.3779638 | 1.38056851 | -4.6198097 |
| 10742 | NM_024740.2 | 36.3597626 | -6.8014067 | 1.47227407 | -4.6196607 |
| 10743 | NM_003060.3 | 37.3575781 | -5.8625578 | 1.26905963 | -4.6196078 |
| 10744 | NM_147173.2 | 30.5838884 | -6.5571793 | 1.41988291 | -4.6181127 |
| 10745 | NM_020817.1 | 32.3200272 | -6.6394774 | 1.43797206 | -4.6172506 |
| 10746 | NM_018418.4 | 22.821681 | -6.1274234 | 1.32707954 | -4.6172239 |
| 10747 | NR_036614.1 | 44.5829246 | -7.1027002 | 1.53840225 | -4.6169331 |
| 10748 | NM_006321.3 | 61.3750011 | -7.5608269 | 1.63770673 | -4.616716 |
| 10749 | NR_134867.1 | 35.4063217 | -6.7701191 | 1.46643251 | -4.6167274 |

| | | | | | |
|---|---|---|---|---|---|
| 10750 | NR_104220.1 | 65.8385551 | 6.22987307 | 1.34946048 | 4.61656579 |
| 10751 | NR_024330.1 | 25.6797252 | -6.3097341 | 1.36681751 | -4.6163691 |
| 10752 | NM_001134406.1 | 30.2261725 | -6.5401357 | 1.41681016 | -4.6160988 |
| 10753 | NM_014593.3 | 159.919508 | -4.9939084 | 1.081853 | -4.6160693 |
| 10754 | NM_006951.4 | 28.0436978 | -6.4302332 | 1.3931112 | -4.6157358 |
| 10755 | NM_032231.5 | 26.2993414 | -6.3416288 | 1.37396666 | -4.6155624 |
| 10756 | NM_001316984.1 | 23.5418109 | -6.1705865 | 1.33692105 | -4.6155205 |
| 10757 | NM_152718.2 | 23.7261149 | -6.1877369 | 1.34064279 | -4.6155001 |
| 10758 | NM_207395.2 | 24.0198946 | -6.2076557 | 1.34502536 | -4.6152704 |
| 10759 | NM_001330617.1 | 22.4688358 | -6.1081445 | 1.32359731 | -4.6148058 |
| 10760 | NM_019071.2 | 26.4363285 | -6.3428262 | 1.37457419 | -4.6143935 |
| 10761 | NM_019590.4 | 41.9148609 | -7.0135698 | 1.51994692 | -4.6143518 |
| 10762 | NR_033797.2 | 27.749639 | -6.4143282 | 1.39007933 | -4.6143613 |
| 10763 | NM_001033602.2 | 19.3981217 | -5.901472 | 1.27899037 | -4.6141645 |
| 10764 | NM_015627.2 | 28.0453097 | -6.437258 | 1.39510329 | -4.6141802 |
| 10765 | NM_017787.4 | 26.0752065 | -6.3317824 | 1.37228318 | -4.6140494 |
| 10766 | NR_104223.1 | 23.393086 | -6.1685475 | 1.33715 | -4.6132053 |
| 10767 | NM_001242856.1 | 19.9117499 | 9.22828694 | 2.00051585 | 4.61295366 |
| 10768 | NM_013355.4 | 22.3832089 | -6.1076403 | 1.32402729 | -4.6129263 |
| 10769 | NM_138446.1 | 45.5553937 | -5.3342516 | 1.15641376 | -4.6127535 |
| 10770 | NM_004732.3 | 42.2763453 | -6.0444514 | 1.31041021 | -4.6126407 |
| 10771 | NM_001018055.2 | 60.2027292 | -5.7579013 | 1.24839789 | -4.6122325 |
| 10772 | NM_001193350.1 | 32.7018238 | -6.6546883 | 1.44289775 | -4.6120304 |
| 10773 | NM_001011671.2 | 21.6081093 | -6.0524045 | 1.31235577 | -4.6118626 |
| 10774 | NM_001170721.1 | 19.3635916 | -5.9044489 | 1.28034024 | -4.6116249 |
| 10775 | NM_130393.3 | 43.269968 | -7.0570347 | 1.53034937 | -4.611388 |
| 10776 | NM_001017980.3 | 184.523954 | -2.8446778 | 0.61690237 | -4.6112285 |
| 10777 | NM_001078.3 | 23.6324602 | -6.1859312 | 1.34155324 | -4.6110218 |
| 10778 | NM_001301267.1 | 36.6341762 | -6.818034 | 1.47868832 | -4.6108662 |
| 10779 | NM_001318727.1 | 267.901072 | -5.0063453 | 1.08580438 | -4.6107249 |
| 10780 | NM_001142471.1 | 44.0886365 | -6.0994521 | 1.32291002 | -4.6106326 |
| 10781 | NM_012383.4 | 29.4751425 | -6.5024232 | 1.4104528 | -4.6101671 |
| 10782 | NR_038279.1 | 26.9920083 | -6.374744 | 1.38278906 | -4.6100625 |
| 10783 | NM_032479.3 | 47.6499793 | -6.2224954 | 1.34977955 | -4.6100087 |
| 10784 | NM_001204051.1 | 20.9257933 | -6.0012096 | 1.30191378 | -4.6095292 |
| 10785 | NM_152379.3 | 19.9879712 | -5.9392602 | 1.28855166 | -4.6092527 |
| 10786 | NM_001033555.2 | 44.5276487 | -7.0987047 | 1.54017773 | -4.6090166 |
| 10787 | NM_001305115.1 | 46.7294371 | -7.1660555 | 1.55528255 | -4.6075586 |
| 10788 | NM_018482.3 | 212.42777 | -3.4231168 | 0.74294388 | -4.6075039 |
| 10789 | NM_024109.3 | 20.8192564 | -6.00808 | 1.30398541 | -4.6074749 |
| 10790 | NM_001204404.1 | 42.3305961 | -7.0238946 | 1.52460004 | -4.6070408 |
| 10791 | NM_001085384.2 | 22.2266519 | -6.1035742 | 1.32487687 | -4.6068992 |
| 10792 | NM_001113528.1 | 35.6010441 | -6.779765 | 1.47165642 | -4.6068939 |

| | | | | | |
|---|---|---|---|---|---|
| 10793 | NM_032949.2 | 29.1628347 | -6.4853183 | 1.40785339 | -4.6065296 |
| 10794 | NM_018056.3 | 31.4462901 | -6.5975323 | 1.4323265 | -4.6061651 |
| 10795 | NM_194281.3 | 20.1556469 | -5.9599436 | 1.29409872 | -4.6054784 |
| 10796 | NM_001320627.1 | 29.674114 | -6.5168278 | 1.41511216 | -4.605167 |
| 10797 | NM_001130020.1 | 62.777103 | -6.6251907 | 1.43870478 | -4.6049689 |
| 10798 | NM_015398.2 | 26.0950654 | -6.3217992 | 1.37286589 | -4.6048192 |
| 10799 | NM_020987.4 | 243.55394 | -4.396042 | 0.9546858 | -4.6047003 |
| 10800 | NM_001704.2 | 101.76363 | -6.243761 | 1.35603835 | -4.6044132 |
| 10801 | NM_001321958.1 | 26.8747754 | -6.365836 | 1.38256349 | -4.6043715 |
| 10802 | NM_171998.3 | 68.4235393 | -5.7313264 | 1.24481046 | -4.604176 |
| 10803 | NM_001318192.1 | 108.333255 | -8.3821894 | 1.82064555 | -4.6039656 |
| 10804 | NM_183043.2 | 30.5542717 | -6.5490263 | 1.42246982 | -4.6039826 |
| 10805 | NM_000286.2 | 21.3229873 | -6.0409041 | 1.31212185 | -4.60392 |
| 10806 | NM_020663.4 | 36.6361329 | -5.8449195 | 1.26963337 | -4.6036278 |
| 10807 | NM_181303.1 | 20.910097 | -6.0110662 | 1.30581216 | -4.6033161 |
| 10808 | NR_073090.1 | 27.4620758 | -6.3956007 | 1.38945505 | -4.6029562 |
| 10809 | NM_020202.4 | 43.0163909 | -6.0652374 | 1.31772703 | -4.6028026 |
| 10810 | NM_133177.3 | 48.2554676 | -7.2148985 | 1.56758413 | -4.602559 |
| 10811 | NM_020311.2 | 51.6426046 | -6.3325508 | 1.37597987 | -4.6022118 |
| 10812 | NM_001491.2 | 30.6968602 | -6.5609532 | 1.42566263 | -4.6020377 |
| 10813 | NM_020371.2 | 26.8020706 | -6.368049 | 1.38378955 | -4.6018913 |
| 10814 | NM_001198868.1 | 39.9493604 | -6.9448816 | 1.50919051 | -4.6017263 |
| 10815 | NM_001278689.1 | 22.0058736 | -6.0807 | 1.32151637 | -4.6013051 |
| 10816 | NM_172115.2 | 20.2389487 | -5.9667822 | 1.29686908 | -4.6009133 |
| 10817 | NM_001127230.1 | 47.1388426 | -7.1810933 | 1.5609591 | -4.6004366 |
| 10818 | NR_024048.2 | 19.6966888 | -5.9158834 | 1.28595335 | -4.6003872 |
| 10819 | NM_001313905.1 | 19.2058125 | -5.8834523 | 1.27905638 | -4.5998382 |
| 10820 | NM_021724.4 | 26.5000479 | -6.3468842 | 1.38001004 | -4.599158 |
| 10821 | NM_001291106.1 | 49.3895068 | -7.2499263 | 1.57658812 | -4.598491 |
| 10822 | NM_001253823.1 | 18.9973674 | -5.8629251 | 1.2751505 | -4.5978299 |
| 10823 | NR_045553.1 | 24.7082107 | -6.2512399 | 1.3596574 | -4.5976581 |
| 10824 | NM_001256449.1 | 26.187096 | -6.3342295 | 1.37777973 | -4.5974182 |
| 10825 | NM_001199140.1 | 93.3162564 | -4.677984 | 1.01758569 | -4.5971402 |
| 10826 | NM_001080114.1 | 40.9009526 | 2.53064093 | 0.55052627 | 4.59676684 |
| 10827 | NM_001320126.1 | 21.37644 | -6.0315201 | 1.3122577 | -4.5962924 |
| 10828 | NM_001008707.1 | 68.9116475 | -7.7275289 | 1.68140082 | -4.5958875 |
| 10829 | NM_001202414.1 | 24.3316571 | -6.2192357 | 1.35341892 | -4.5952038 |
| 10830 | NM_001308293.1 | 36.1510413 | -5.824536 | 1.26753547 | -4.5951661 |
| 10831 | NM_001267062.1 | 38.2624772 | -6.8797639 | 1.49741915 | -4.5944142 |
| 10832 | NM_001166660.1 | 57.2236085 | -7.4606225 | 1.62389178 | -4.5942855 |
| 10833 | NM_178580.2 | 34.9414367 | -6.745992 | 1.46853767 | -4.5936799 |
| 10834 | NM_001289153.1 | 21.3642359 | -6.0308562 | 1.31304257 | -4.5930393 |
| 10835 | NM_004101.3 | 19.66693 | -5.9236097 | 1.28985211 | -4.592472 |

| | | | | | |
|---|---|---|---|---|---|
| 10836 | NM_001098398.1 | 62.2568012 | -7.5837536 | 1.65143986 | -4.5922069 |
| 10837 | NM_001002877.1 | 26.7630023 | -6.3628266 | 1.38569475 | -4.5917953 |
| 10838 | NM_001093756.1 | 26.2170742 | -6.3324776 | 1.37926583 | -4.5911944 |
| 10839 | NM_001135599.3 | 34.6888995 | -6.7409442 | 1.4684535 | -4.5905057 |
| 10840 | NR_034108.1 | 20.7485206 | -5.9976519 | 1.30655294 | -4.5904393 |
| 10841 | NM_001321708.1 | 29.9645236 | -6.5240374 | 1.42125168 | -4.5903463 |
| 10842 | NR_073526.1 | 49.7035047 | -7.2548175 | 1.5805882 | -4.5899479 |
| 10843 | NM_148175.2 | 24.3224846 | -6.2260497 | 1.35646839 | -4.5898966 |
| 10844 | NR_138575.1 | 24.552926 | -6.2456674 | 1.360742 | -4.5898983 |
| 10845 | NM_001003694.1 | 25.2471813 | -6.2804418 | 1.36848057 | -4.589354 |
| 10846 | NM_001270805.1 | 34.7009476 | -6.7385909 | 1.4684393 | -4.5889476 |
| 10847 | NM_001124.2 | 19.9911657 | -5.9474964 | 1.29611541 | -4.5887089 |
| 10848 | NM_018105.2 | 25.82311 | -6.3121239 | 1.37567928 | -4.5883688 |
| 10849 | NM_020972.2 | 23.194167 | -6.1642174 | 1.3434765 | -4.5882585 |
| 10850 | NM_016230.3 | 29.4272034 | -6.4990049 | 1.4165417 | -4.5879376 |
| 10851 | NR_120551.1 | 22.0920092 | 4.10569057 | 0.89499029 | 4.58741353 |
| 10852 | NM_001276271.1 | 24.9543652 | -6.2629777 | 1.3653985 | -4.586923 |
| 10853 | NM_001033549.2 | 32.0143534 | -6.6163057 | 1.44250313 | -4.5866838 |
| 10854 | NM_002658.4 | 23.7381765 | -6.1894448 | 1.34949396 | -4.5864932 |
| 10855 | NM_001079910.1 | 49.4883352 | -6.2724397 | 1.36778702 | -4.5858307 |
| 10856 | NM_002404.2 | 307.441193 | -4.5182286 | 0.98527238 | -4.585766 |
| 10857 | NR_003110.2 | 21.3860975 | -6.0420263 | 1.31757663 | -4.5857115 |
| 10858 | NM_001291878.1 | 93.2926659 | -8.165588 | 1.78081718 | -4.5853039 |
| 10859 | NM_001083112.2 | 33.5211855 | -6.6849103 | 1.45797559 | -4.5850633 |
| 10860 | NM_001098673.1 | 21.5789998 | -6.050066 | 1.31951793 | -4.5850578 |
| 10861 | NM_001172896.1 | 35.7117105 | -6.7820803 | 1.47922432 | -4.5848896 |
| 10862 | NM_001625.3 | 50.7779742 | -2.9736082 | 0.64860942 | -4.5845899 |
| 10863 | NM_173872.3 | 27.2728696 | -6.3961142 | 1.39526654 | -4.5841522 |
| 10864 | NM_001300977.1 | 24.8476708 | -6.2570471 | 1.36497453 | -4.5840029 |
| 10865 | NM_006958.2 | 23.8548184 | -6.1951477 | 1.35146679 | -4.5840177 |
| 10866 | NR_027260.1 | 413.154016 | -4.0388669 | 0.88109198 | -4.5839333 |
| 10867 | NR_073403.1 | 18.1180607 | -5.8047974 | 1.26638984 | -4.5837365 |
| 10868 | NM_198705.2 | 27.1786143 | -6.3829843 | 1.39266727 | -4.5832802 |
| 10869 | NM_000264.3 | 155.615718 | -5.7011382 | 1.24398441 | -4.5829659 |
| 10870 | NM_001039374.4 | 22.1919572 | -6.0985222 | 1.33082129 | -4.5825253 |
| 10871 | NM_014735.4 | 25.3659394 | -6.2911874 | 1.37308105 | -4.5818034 |
| 10872 | NM_017846.4 | 42.5412561 | -6.0474349 | 1.32015131 | -4.580865 |
| 10873 | NM_020201.3 | 30.7887719 | -6.5658319 | 1.43333762 | -4.5807993 |
| 10874 | NM_201265.1 | 33.3376683 | -6.6808452 | 1.45845548 | -4.5807673 |
| 10875 | NM_016941.3 | 32.2575187 | -6.6303157 | 1.44758765 | -4.5802516 |
| 10876 | NM_018477.2 | 267.15464 | -3.0973857 | 0.67628922 | -4.579972 |
| 10877 | NM_054033.3 | 35.521916 | -6.7727934 | 1.47887188 | -4.5797026 |
| 10878 | NM_001134364.1 | 115.643823 | -5.2994835 | 1.15718201 | -4.5796456 |

| | | | | | |
|---|---|---|---|---|---|
| 10879 | NM_001006945.1 | 25.8782447 | -6.3141056 | 1.37876725 | -4.5795297 |
| 10880 | NM_001317123.1 | 14.2498149 | 8.76049219 | 1.91303357 | 4.5793719 |
| 10881 | NM_032811.2 | 32.0824867 | -6.6252979 | 1.44677552 | -4.5793544 |
| 10882 | NM_020160.2 | 32.63063 | -6.6488526 | 1.45198811 | -4.5791371 |
| 10883 | NM_001168399.1 | 28.773353 | -6.4655076 | 1.41202193 | -4.5789003 |
| 10884 | NM_201999.2 | 35.3458352 | -6.7636121 | 1.47732395 | -4.5782864 |
| 10885 | NM_001321266.1 | 30.6391842 | -6.5562018 | 1.43211472 | -4.5779865 |
| 10886 | NM_001001420.2 | 91.1355244 | -3.798228 | 0.82978085 | -4.577387 |
| 10887 | NR_045574.1 | 21.3041333 | -6.0417538 | 1.31992671 | -4.5773403 |
| 10888 | NM_018696.2 | 23.021463 | -6.1481851 | 1.34320988 | -4.5772334 |
| 10889 | NM_020422.5 | 20.7693463 | -5.9930034 | 1.3093154 | -4.5772038 |
| 10890 | NM_001329877.1 | 65.2577799 | 1.95957908 | 0.42813333 | 4.57702998 |
| 10891 | NR_120416.1 | 27.0897531 | 2.92129513 | 0.63828223 | 4.57680787 |
| 10892 | NM_033315.3 | 105.190245 | -4.8639269 | 1.06276534 | -4.5766706 |
| 10893 | NM_001300778.1 | 25.3871842 | -6.2858462 | 1.37364756 | -4.5760255 |
| 10894 | NM_020133.2 | 211.737505 | -5.2854543 | 1.15503689 | -4.5760047 |
| 10895 | NR_038911.1 | 22.9478233 | -6.1396236 | 1.34185374 | -4.575479 |
| 10896 | NM_153376.2 | 21.5739328 | -6.0552139 | 1.32345955 | -4.575292 |
| 10897 | NM_001003652.3 | 55.0845986 | -7.4047439 | 1.6184881 | -4.5750994 |
| 10898 | NM_182546.3 | 32.6518111 | -6.6540503 | 1.45450419 | -4.5747893 |
| 10899 | NR_110000.1 | 88.7767263 | 3.06572629 | 0.67014495 | 4.57472119 |
| 10900 | NM_020717.3 | 19.4798827 | -5.9136438 | 1.29271836 | -4.5745802 |
| 10901 | NM_021253.3 | 21.3514049 | -6.0398406 | 1.32033011 | -4.5744928 |
| 10902 | NM_005599.3 | 24.2453764 | -6.2249493 | 1.36081419 | -4.57443 |
| 10903 | NR_135786.1 | 24.5971992 | -6.2384322 | 1.36392504 | -4.5738821 |
| 10904 | NM_001303413.1 | 24.3461042 | -6.2232364 | 1.36063406 | -4.5737767 |
| 10905 | NM_001035.2 | 23.3326951 | -6.1635191 | 1.34764128 | -4.5735607 |
| 10906 | NM_015001.2 | 316.835634 | -5.4477181 | 1.19121128 | -4.5732593 |
| 10907 | NM_015895.4 | 25.2460945 | -6.2770017 | 1.37258609 | -4.5731206 |
| 10908 | NM_001308027.1 | 25.0631503 | -6.2677625 | 1.37063857 | -4.5728777 |
| 10909 | NM_178040.3 | 51.533329 | -7.3087827 | 1.59828727 | -4.5728843 |
| 10910 | NM_001330404.1 | 25.5165996 | -6.2921551 | 1.3761218 | -4.5723824 |
| 10911 | NR_135249.1 | 25.562105 | -6.3046461 | 1.37894716 | -4.5720723 |
| 10912 | NM_001002031.2 | 19.6651501 | -5.9250962 | 1.29607324 | -4.5715751 |
| 10913 | NR_109997.1 | 365.342018 | 2.41942686 | 0.52936374 | 4.57044309 |
| 10914 | NR_103492.1 | 32.2062559 | 9.92249804 | 2.17112313 | 4.57021432 |
| 10915 | NM_001011657.3 | 37.6098036 | -5.036895 | 1.10212345 | -4.5701732 |
| 10916 | NM_002183.3 | 19.4497408 | -5.9068175 | 1.29249014 | -4.5701065 |
| 10917 | NM_001202519.1 | 29.196678 | -6.4908715 | 1.42031684 | -4.5700166 |
| 10918 | NR_135778.1 | 25.4188728 | -6.290447 | 1.37649853 | -4.5698901 |
| 10919 | NM_001270468.1 | 32.1820814 | -6.6281957 | 1.45042692 | -4.569824 |
| 10920 | NM_016053.3 | 36.5451068 | -6.811774 | 1.49064482 | -4.5696828 |
| 10921 | NM_016487.4 | 23.7207303 | -6.188433 | 1.35425202 | -4.5696317 |

| | | | | | |
|---|---|---|---|---|---|
| 10922 | NM_001207062.1 | 55.7843766 | -6.4529973 | 1.41219299 | -4.5694869 |
| 10923 | NM_032725.3 | 49.7216876 | -5.3146473 | 1.16314258 | -4.569214 |
| 10924 | NM_001139442.1 | 26.3022313 | -6.3400125 | 1.38768161 | -4.5687804 |
| 10925 | NM_001292004.1 | 20.0072308 | -5.9438513 | 1.30097708 | -4.5687594 |
| 10926 | NM_001168280.1 | 51.2474139 | -5.4222677 | 1.18683162 | -4.5686916 |
| 10927 | NM_018361.3 | 217.345437 | -4.2491872 | 0.93012962 | -4.5683818 |
| 10928 | NM_019034.2 | 18.624794 | -5.8423159 | 1.27892835 | -4.5681338 |
| 10929 | NM_133466.3 | 101.885796 | -1.7116614 | 0.37470247 | -4.5680549 |
| 10930 | NR_045014.1 | 106.256445 | -8.3531444 | 1.82870013 | -4.5678043 |
| 10931 | NM_014279.4 | 43.3054304 | -7.0551168 | 1.54454429 | -4.567766 |
| 10932 | NM_001271043.2 | 59.1527005 | -7.5080455 | 1.64402968 | -4.5668552 |
| 10933 | NM_173576.2 | 32.7054139 | -6.6550527 | 1.45732651 | -4.5666175 |
| 10934 | NM_020911.1 | 249.532985 | -5.8194043 | 1.27436144 | -4.5665257 |
| 10935 | NM_144653.4 | 194.978296 | -4.6532679 | 1.01902055 | -4.5664123 |
| 10936 | NM_003616.2 | 26.1906987 | -6.3279704 | 1.38580271 | -4.5662852 |
| 10937 | NM_001142726.1 | 111.387651 | -4.9524043 | 1.08456842 | -4.5662443 |
| 10938 | NM_001307941.1 | 32.371847 | -6.6403394 | 1.45467497 | -4.5648268 |
| 10939 | NM_032437.3 | 38.9515715 | -5.9158681 | 1.29598681 | -4.5647595 |
| 10940 | NM_001330449.1 | 24.4324264 | -6.2318905 | 1.36528451 | -4.5645362 |
| 10941 | NM_001040272.5 | 48.3408814 | -6.2480283 | 1.36885779 | -4.5644101 |
| 10942 | NR_015430.1 | 21.8544831 | 2.59803769 | 0.56923772 | 4.56406456 |
| 10943 | NM_001184792.1 | 20.8391106 | -6.0071884 | 1.31621227 | -4.5639967 |
| 10944 | NM_198536.2 | 26.7106172 | -6.3606911 | 1.39370005 | -4.5638881 |
| 10945 | NM_138559.1 | 62.6723429 | -7.5911316 | 1.66331396 | -4.5638597 |
| 10946 | NM_018157.3 | 47.0373689 | -6.2075431 | 1.36048654 | -4.5627376 |
| 10947 | NM_001348081.1 | 30.5883503 | -6.558398 | 1.43748178 | -4.5624216 |
| 10948 | NM_001145434.1 | 25.1901113 | -6.2737705 | 1.37530249 | -4.5617387 |
| 10949 | NM_022044.2 | 41.1802534 | -6.0027951 | 1.31607302 | -4.5611413 |
| 10950 | NR_132114.1 | 23.8665496 | -6.1980873 | 1.35888906 | -4.561143 |
| 10951 | NM_025250.2 | 1666.89568 | -5.4709819 | 1.19951631 | -4.56099 |
| 10952 | NM_015651.2 | 29.4198228 | -6.4988967 | 1.42495308 | -4.5607794 |
| 10953 | NM_032387.4 | 21.0289143 | -6.0186591 | 1.31966695 | -4.5607409 |
| 10954 | NR_024079.1 | 51.2482645 | 2.40908754 | 0.52833346 | 4.55978608 |
| 10955 | NM_001122764.1 | 47.2727466 | -4.828716 | 1.05905978 | -4.5594367 |
| 10956 | NM_001321112.1 | 20.3563872 | -5.9694602 | 1.30934959 | -4.5591034 |
| 10957 | NM_001042371.2 | 115.647118 | -4.8195434 | 1.05720997 | -4.5587381 |
| 10958 | NM_014290.2 | 23.1750841 | -6.1511358 | 1.3493257 | -4.5586739 |
| 10959 | NM_152446.3 | 17.6059262 | -5.7605894 | 1.26367373 | -4.558605 |
| 10960 | NM_018085.4 | 595.2821 | -2.0366999 | 0.44685021 | -4.557903 |
| 10961 | NM_001300962.1 | 49.3348733 | -7.2485733 | 1.59059563 | -4.557144 |
| 10962 | NM_001318828.1 | 34.0121116 | -6.70701 | 1.47175949 | -4.5571373 |
| 10963 | NM_001320035.1 | 58.4468058 | -6.5228768 | 1.43142768 | -4.5569028 |
| 10964 | NM_017723.2 | 17.7153355 | -5.7722308 | 1.26690959 | -4.5561505 |

| | | | | | |
|---|---|---|---|---|---|
| 10965 | NM_173618.2 | 32.412546 | -6.6388775 | 1.45713905 | -4.5561043 |
| 10966 | NR_135077.1 | 26.5049335 | -6.3469561 | 1.3931143 | -4.5559479 |
| 10967 | NM_153322.2 | 36.019779 | -6.795489 | 1.49157403 | -4.555918 |
| 10968 | NM_001145862.1 | 19.7582561 | -5.9322634 | 1.30225824 | -4.5553664 |
| 10969 | NM_001714.2 | 22.5888179 | -6.1159934 | 1.34263071 | -4.5552313 |
| 10970 | NM_000186.3 | 19.8092585 | -5.9369878 | 1.30343379 | -4.5548825 |
| 10971 | NR_015449.1 | 26.799158 | -6.3705126 | 1.39871222 | -4.5545556 |
| 10972 | NM_016248.3 | 289.250816 | -5.2038964 | 1.14273851 | -4.5538821 |
| 10973 | NM_001323384.1 | 34.5039185 | -6.7280266 | 1.4774699 | -4.5537487 |
| 10974 | NM_133436.3 | 28.2407943 | -6.4459865 | 1.41583578 | -4.5527784 |
| 10975 | NM_181839.2 | 59.2566673 | -5.7147 | 1.25528524 | -4.5525112 |
| 10976 | NR_036623.1 | 23.0320213 | -6.1474104 | 1.35033879 | -4.5524948 |
| 10977 | NM_001114118.2 | 216.873701 | -2.4755587 | 0.54382204 | -4.5521485 |
| 10978 | NM_002204.3 | 200.872012 | -5.343136 | 1.17393816 | -4.5514629 |
| 10979 | NR_033734.1 | 22.1091245 | -6.0871469 | 1.33743869 | -4.5513465 |
| 10980 | NM_016315.3 | 26.9286228 | -6.3742614 | 1.40059903 | -4.5510966 |
| 10981 | NM_001018024.2 | 23.0560537 | -6.1472353 | 1.35079213 | -4.5508374 |
| 10982 | NM_016307.3 | 24.5833821 | -6.2437431 | 1.37204171 | -4.5506948 |
| 10983 | NM_032472.3 | 18.7736197 | -5.8603278 | 1.28795129 | -4.550116 |
| 10984 | NM_001256799.2 | 25.9984823 | -6.3232394 | 1.38971909 | -4.5500126 |
| 10985 | NM_001291814.1 | 30.8716875 | -6.565416 | 1.4429507 | -4.5499933 |
| 10986 | NM_001320717.1 | 25.1782036 | -6.2714226 | 1.3785306 | -4.5493532 |
| 10987 | NM_001267727.1 | 19.2331375 | -5.8899774 | 1.29469971 | -4.5493 |
| 10988 | NM_014881.4 | 20.9162503 | -6.0159183 | 1.32246704 | -4.5490119 |
| 10989 | NM_001039707.1 | 22.139755 | -6.0828697 | 1.33720894 | -4.5489299 |
| 10990 | NM_170601.4 | 89.6197397 | -3.3986935 | 0.74721925 | -4.5484554 |
| 10991 | NM_001300965.1 | 27.6283868 | -6.409819 | 1.40929122 | -4.5482572 |
| 10992 | NM_001243794.1 | 29.5878518 | -6.5088974 | 1.43115282 | -4.54801 |
| 10993 | NM_001195226.1 | 50.7323795 | -7.2887237 | 1.60270157 | -4.5477735 |
| 10994 | NM_177995.2 | 23.1397685 | -6.1511858 | 1.35258927 | -4.5477115 |
| 10995 | NR_036577.1 | 21.2956037 | -6.0398637 | 1.32817988 | -4.5474742 |
| 10996 | NM_001258026.1 | 24.7647261 | -6.2513647 | 1.374742 | -4.5473003 |
| 10997 | NM_001271953.1 | 24.6406126 | -6.2438664 | 1.37320802 | -4.5469195 |
| 10998 | NM_203451.2 | 20.6545976 | -5.9939188 | 1.3183712 | -4.5464576 |
| 10999 | NM_138415.4 | 45.7305728 | -7.1346561 | 1.56932683 | -4.5463162 |
| 11000 | NM_138972.3 | 61.1359713 | -7.5551112 | 1.66189359 | -4.546086 |
| 11001 | NR_110138.1 | 51.5924291 | 2.82065068 | 0.62046938 | 4.54599497 |
| 11002 | NM_001322342.1 | 22.8068874 | -6.1309046 | 1.34868835 | -4.5458275 |
| 11003 | NM_001136494.1 | 47.3582723 | -7.1892781 | 1.58198886 | -4.5444556 |
| 11004 | NM_001330635.1 | 41.9698419 | -7.0115008 | 1.54286385 | -4.5444715 |
| 11005 | NM_005739.3 | 30.1827561 | -6.5334776 | 1.43772165 | -4.5443272 |
| 11006 | NM_007156.4 | 21.4208563 | -6.0386019 | 1.32881699 | -4.5443442 |
| 11007 | NM_005319.3 | 24.1711127 | -6.217076 | 1.36811029 | -4.5442798 |

| | | | | | |
|---|---|---|---|---|---|
| 11008 | NM_001006623.2 | 23.8486365 | -6.1960843 | 1.36360094 | -4.5439132 |
| 11009 | NM_001142475.1 | 45.0310919 | -7.1161064 | 1.56616266 | -4.5436573 |
| 11010 | NM_153340.4 | 65.4781829 | -4.9288846 | 1.08480429 | -4.5435703 |
| 11011 | NM_024834.3 | 58.1879491 | -7.4840443 | 1.6472122 | -4.5434609 |
| 11012 | NM_001134651.1 | 38.810118 | -6.8976771 | 1.51825703 | -4.543155 |
| 11013 | NM_001005914.2 | 41.8466128 | -7.012411 | 1.54361734 | -4.5428428 |
| 11014 | NM_052990.2 | 39.0672057 | -5.9360158 | 1.30667604 | -4.5428366 |
| 11015 | NM_006491.2 | 21.308921 | -6.0343615 | 1.32837102 | -4.5426777 |
| 11016 | NM_001080450.2 | 40.8113365 | -5.9847848 | 1.31748122 | -4.5425959 |
| 11017 | NM_001346590.1 | 22.6973729 | -6.1298921 | 1.34951485 | -4.5422932 |
| 11018 | NM_001159643.1 | 8.47804692 | 7.98019741 | 1.75694075 | 4.5420982 |
| 11019 | NM_001308023.1 | 105.458689 | -3.6871198 | 0.81188239 | -4.5414457 |
| 11020 | NM_014442.2 | 35.1382504 | 2.39236429 | 0.52680538 | 4.54126773 |
| 11021 | NM_001172432.1 | 29.0573709 | -6.4841167 | 1.42783866 | -4.541211 |
| 11022 | NR_108087.1 | 53.942239 | 3.13163343 | 0.68960623 | 4.54119077 |
| 11023 | NM_003728.3 | 72.5814773 | -4.21052 | 0.92723718 | -4.540931 |
| 11024 | NM_001204859.1 | 31.2330382 | -6.5857021 | 1.45030908 | -4.5408956 |
| 11025 | NM_020300.4 | 23.3103425 | -6.1660263 | 1.35797596 | -4.5406005 |
| 11026 | NM_032449.2 | 35.306967 | -6.7639349 | 1.4897254 | -4.5403904 |
| 11027 | NM_001304536.1 | 46.7066749 | -7.1697336 | 1.57919661 | -4.5401146 |
| 11028 | NM_001316766.1 | 21.312166 | -6.0359552 | 1.3295614 | -4.5398093 |
| 11029 | NM_014718.3 | 636.385925 | -6.0733643 | 1.33782256 | -4.5397383 |
| 11030 | NM_001285829.1 | 20.2278775 | -5.9589701 | 1.31263497 | -4.5397008 |
| 11031 | NR_033194.1 | 24.5079403 | -6.2328751 | 1.37299165 | -4.5396307 |
| 11032 | NM_003776.3 | 33.2751493 | -6.6778667 | 1.47112677 | -4.5392871 |
| 11033 | NM_144976.3 | 20.4429686 | -5.9795268 | 1.31738204 | -4.5389467 |
| 11034 | NM_001320194.1 | 22.8504448 | -6.1307617 | 1.35070766 | -4.5389257 |
| 11035 | NM_001190201.1 | 19.8026804 | -5.9278881 | 1.30610415 | -4.5386029 |
| 11036 | NM_001128615.1 | 24.5523598 | -6.2417729 | 1.37531437 | -4.5384336 |
| 11037 | NM_001199723.1 | 28.22671 | -6.44591 | 1.42036877 | -4.5381946 |
| 11038 | NM_001282778.1 | 34.5510178 | -6.7296265 | 1.48289935 | -4.5381546 |
| 11039 | NM_017803.4 | 22.3022213 | -6.1013831 | 1.3444722 | -4.5381252 |
| 11040 | NM_020990.4 | 33.1439659 | -6.674011 | 1.47074511 | -4.5378434 |
| 11041 | NM_001272067.1 | 24.253671 | -6.2260579 | 1.37217281 | -4.5373716 |
| 11042 | NM_144998.3 | 30.5948224 | -6.5577047 | 1.44537138 | -4.5370379 |
| 11043 | NR_003654.2 | 35.8150563 | -4.3934134 | 0.9684476 | -4.5365525 |
| 11044 | NM_003080.2 | 17.6016608 | -5.7612726 | 1.27003606 | -4.5363063 |
| 11045 | NM_001282306.1 | 28.7563733 | -6.4710582 | 1.4265259 | -4.5362361 |
| 11046 | NM_001323302.1 | 37.1756021 | -6.8341999 | 1.50669577 | -4.5358858 |
| 11047 | NM_007255.2 | 76.8952281 | -5.2117171 | 1.14914245 | -4.5353098 |
| 11048 | NM_022166.3 | 155.637268 | -4.9558432 | 1.09274334 | -4.5352307 |
| 11049 | NM_016654.4 | 30.5043599 | -6.5522799 | 1.44498674 | -4.5344914 |
| 11050 | NM_022124.5 | 51.1311437 | -7.2999157 | 1.6100257 | -4.5340367 |

| | | | | | |
|---|---|---|---|---|---|
| 11051 | NR_045587.1 | 36.9244619 | -6.8293222 | 1.50631602 | -4.5337912 |
| 11052 | NM_001030010.2 | 18.8715062 | -5.8556342 | 1.29160004 | -4.5336281 |
| 11053 | NM_001323295.1 | 35.1419858 | -6.7520734 | 1.48936776 | -4.5335166 |
| 11054 | NM_002889.3 | 59.1578374 | -7.5101709 | 1.65665553 | -4.5333329 |
| 11055 | NM_001258022.1 | 53.4120123 | -7.3611931 | 1.62394639 | -4.532904 |
| 11056 | NM_004361.3 | 38.0029527 | -6.8682569 | 1.51540431 | -4.5322934 |
| 11057 | NM_004501.3 | 728.236376 | -6.071806 | 1.33973841 | -4.5320832 |
| 11058 | NM_001135213.2 | 34.7713412 | -6.7366885 | 1.48649515 | -4.5319276 |
| 11059 | NM_001135662.1 | 29.1193208 | -6.4811327 | 1.43013997 | -4.5318171 |
| 11060 | NM_001007278.2 | 121.567853 | 2.09431419 | 0.4621584 | 4.53159395 |
| 11061 | NM_001200.3 | 26.4747913 | -6.3537839 | 1.40227846 | -4.531043 |
| 11062 | NM_015175.2 | 129.849378 | -3.7729238 | 0.83280204 | -4.5303969 |
| 11063 | NM_001315530.1 | 25.440772 | -6.2936773 | 1.38926706 | -4.5302142 |
| 11064 | NM_022840.4 | 23.5763804 | -6.1777691 | 1.36369809 | -4.530159 |
| 11065 | NM_001270875.1 | 56.2471569 | -7.4378259 | 1.64201292 | -4.5297 |
| 11066 | NM_001993.4 | 58.859481 | -7.503105 | 1.65642637 | -4.5296942 |
| 11067 | NM_015155.2 | 184.707249 | -4.9548529 | 1.09386415 | -4.5296785 |
| 11068 | NM_153689.5 | 74.7917467 | -4.5569137 | 1.00600596 | -4.5297085 |
| 11069 | NM_203384.1 | 38.1755522 | -6.8746187 | 1.51773745 | -4.5295177 |
| 11070 | NM_001045.5 | 41.9422849 | 2.53769243 | 0.56025964 | 4.52949358 |
| 11071 | NM_015407.4 | 53.8818038 | -5.0344715 | 1.11156677 | -4.529167 |
| 11072 | NM_001256829.1 | 20.9177165 | -6.0093409 | 1.32697784 | -4.5285917 |
| 11073 | NM_004315.5 | 27.9836186 | -6.4223539 | 1.41822065 | -4.5284589 |
| 11074 | NM_022169.4 | 23.6436654 | -6.1789798 | 1.36453178 | -4.5282784 |
| 11075 | NM_001002018.2 | 26.6474165 | -6.3535765 | 1.40319722 | -4.5279284 |
| 11076 | NM_014891.6 | 440.269968 | -4.8296788 | 1.06671914 | -4.5276011 |
| 11077 | NM_014445.3 | 336.642475 | -5.5398289 | 1.22376132 | -4.5268868 |
| 11078 | NM_001323599.1 | 21.186976 | -6.032314 | 1.33263384 | -4.5266103 |
| 11079 | NM_007117.4 | 19.1092176 | -5.8823853 | 1.29954002 | -4.5265134 |
| 11080 | NM_000127.2 | 173.140638 | -4.9818304 | 1.10062813 | -4.5263521 |
| 11081 | NM_001300959.1 | 93.7975591 | -5.0136799 | 1.10772755 | -4.5260948 |
| 11082 | NM_003999.2 | 38.8369012 | -6.9024525 | 1.52526727 | -4.5254052 |
| 11083 | NM_000221.2 | 31.9242242 | -6.6174893 | 1.46239336 | -4.525109 |
| 11084 | NM_014345.2 | 140.856176 | -5.2102759 | 1.15143153 | -4.5250418 |
| 11085 | NM_001184876.2 | 50.7493052 | -7.2846996 | 1.60987166 | -4.5250188 |
| 11086 | NM_001305840.1 | 25.9752223 | -6.3189252 | 1.39675687 | -4.523998 |
| 11087 | NM_000981.3 | 1342.84919 | -6.820852 | 1.50775753 | -4.5238388 |
| 11088 | NM_001165972.1 | 37.2080866 | -6.8370576 | 1.51138416 | -4.523706 |
| 11089 | NM_007148.4 | 88.6436809 | -7.123722 | 1.57490757 | -4.5232635 |
| 11090 | NM_001128167.2 | 66.0997884 | -7.6662226 | 1.6950307 | -4.5227633 |
| 11091 | NR_045494.1 | 48.5196359 | -7.2247487 | 1.5975427 | -4.5224135 |
| 11092 | NM_144589.2 | 31.8080607 | -6.6128356 | 1.46240086 | -4.5219035 |
| 11093 | NM_201429.2 | 63.9899953 | -7.6205659 | 1.68540205 | -4.5215122 |

| | | | | | |
|---|---|---|---|---|---|
| 11094 | NM_003915.5 | 25.6100231 | -6.2991812 | 1.39354164 | -4.5202676 |
| 11095 | NM_001003684.1 | 22.6745588 | -6.1239224 | 1.35487642 | -4.5199122 |
| 11096 | NM_001076678.2 | 43.7699775 | -7.0712289 | 1.56445667 | -4.5199263 |
| 11097 | NM_003747.2 | 326.339638 | -4.4813847 | 0.99147219 | -4.5199298 |
| 11098 | NM_021165.3 | 23.7094486 | -6.1880928 | 1.36905777 | -4.5199647 |
| 11099 | NM_018942.2 | 40.5935366 | -6.9647409 | 1.54093675 | -4.5198097 |
| 11100 | NM_018078.3 | 22.4394473 | -6.1075115 | 1.35133325 | -4.5196191 |
| 11101 | NM_182984.4 | 21.0436022 | -6.0254791 | 1.3332396 | -4.519427 |
| 11102 | NR_144529.1 | 20.8522514 | -6.0047007 | 1.32871654 | -4.5191736 |
| 11103 | NM_024832.4 | 20.0622515 | -5.9560967 | 1.31807307 | -4.518791 |
| 11104 | NR_103803.1 | 30.4960065 | -6.5488933 | 1.4494948 | -4.5180523 |
| 11105 | NM_152274.4 | 29.2541967 | -6.4909916 | 1.43669948 | -4.5179884 |
| 11106 | NM_001287215.1 | 22.8097828 | -6.132977 | 1.35748248 | -4.5179051 |
| 11107 | NM_001242921.1 | 21.564574 | -6.0555366 | 1.34037034 | -4.5178086 |
| 11108 | NM_001286752.1 | 18.199246 | -5.8076762 | 1.28555165 | -4.517653 |
| 11109 | NM_001330111.1 | 45.8570118 | -5.3517715 | 1.18473295 | -4.5172809 |
| 11110 | NM_133640.4 | 40.2043564 | -6.951185 | 1.53890542 | -4.516967 |
| 11111 | NM_014948.3 | 25.5430698 | -6.303741 | 1.39582097 | -4.516153 |
| 11112 | NR_002940.2 | 134.737891 | -2.0016717 | 0.44322642 | -4.5161381 |
| 11113 | NM_016209.4 | 26.3190011 | -6.3342738 | 1.40265353 | -4.5159219 |
| 11114 | NM_001292018.1 | 34.1187897 | -6.7126174 | 1.48657688 | -4.5154862 |
| 11115 | NM_001322427.1 | 23.5470089 | -6.1798209 | 1.36896629 | -4.5142243 |
| 11116 | NM_003774.4 | 23.042318 | -6.1557404 | 1.36366913 | -4.5141012 |
| 11117 | NM_001267536.2 | 36.8168463 | -5.8430607 | 1.29449545 | -4.5137746 |
| 11118 | NM_174934.3 | 23.0540485 | -6.1540365 | 1.3634121 | -4.5137024 |
| 11119 | NM_201551.2 | 48.7993699 | -7.2319125 | 1.60230993 | -4.5134293 |
| 11120 | NR_047521.1 | 36.0536201 | -6.796155 | 1.50597152 | -4.5128045 |
| 11121 | NM_022658.3 | 40.701157 | -6.9699331 | 1.54462409 | -4.5123814 |
| 11122 | NM_182835.2 | 33.773528 | -6.6996785 | 1.48478425 | -4.5122236 |
| 11123 | NM_133444.2 | 20.0142214 | -5.9532482 | 1.31937341 | -4.5121784 |
| 11124 | NR_048569.1 | 44.303669 | -7.0895152 | 1.57138408 | -4.5116374 |
| 11125 | NM_080627.3 | 450.832511 | -4.0059659 | 0.88797748 | -4.5113372 |
| 11126 | NM_001164093.1 | 27.8830988 | -6.4173618 | 1.42252615 | -4.5112434 |
| 11127 | NM_002036.3 | 31.8020469 | -6.6125918 | 1.46581128 | -4.5112164 |
| 11128 | NM_177559.2 | 17.5931045 | -5.7556471 | 1.27591712 | -4.5109882 |
| 11129 | NM_015341.4 | 26.2843635 | -6.3400761 | 1.40552249 | -4.5108322 |
| 11130 | NM_001171132.2 | 24.6825382 | -6.2500455 | 1.38576958 | -4.5101621 |
| 11131 | NM_001257370.1 | 19.1387869 | -5.8770055 | 1.30309124 | -4.5100491 |
| 11132 | NM_025180.3 | 23.4731364 | -6.1674471 | 1.36758677 | -4.50973 |
| 11133 | NM_001301837.1 | 28.1510933 | -6.439771 | 1.4281211 | -4.5092612 |
| 11134 | NM_001122964.2 | 41.7060878 | -7.0068279 | 1.55394252 | -4.509065 |
| 11135 | NM_024570.3 | 36.0307312 | -6.7914965 | 1.50634245 | -4.5086007 |
| 11136 | NM_015557.2 | 146.680579 | -4.2805229 | 0.94976224 | -4.5069416 |

| | | | | | |
|---|---|---|---|---|---|
| 11137 | NM_152911.3 | 22.4386184 | -6.1117787 | 1.35607984 | -4.5069461 |
| 11138 | NM_001317824.1 | 17.3288138 | 9.04235035 | 2.00635109 | 4.50686344 |
| 11139 | NM_001018078.2 | 23.4549376 | -6.1728087 | 1.36973598 | -4.5065683 |
| 11140 | NM_001082480.2 | 19.3414898 | -5.8938283 | 1.30791252 | -4.5062863 |
| 11141 | NM_001032284.2 | 126.046073 | -4.4677081 | 0.99145205 | -4.5062271 |
| 11142 | NM_033036.2 | 52.609783 | -6.3670775 | 1.41307018 | -4.5058466 |
| 11143 | NM_006568.2 | 25.0471961 | -6.2671862 | 1.39091998 | -4.5057849 |
| 11144 | NM_001122898.2 | 30.3895647 | -6.5477508 | 1.45322578 | -4.5056666 |
| 11145 | NM_001205.2 | 25.0951522 | -6.27075 | 1.39174504 | -4.5056744 |
| 11146 | NM_173519.2 | 36.803755 | -6.8222484 | 1.51415151 | -4.5056577 |
| 11147 | NR_125921.1 | 62.1937065 | 2.43029686 | 0.53939627 | 4.50558705 |
| 11148 | NM_001178074.1 | 60.2441651 | -6.5620577 | 1.45663843 | -4.5049324 |
| 11149 | NM_001204890.1 | 42.3664596 | -7.0251475 | 1.5594303 | -4.5049448 |
| 11150 | NM_153026.2 | 46.9290694 | -7.175499 | 1.59287107 | -4.5047582 |
| 11151 | NR_077228.1 | 59.0333678 | 2.31879191 | 0.5147469 | 4.50472242 |
| 11152 | NM_001330235.1 | 20.0051919 | -5.9416115 | 1.31903948 | -4.5044986 |
| 11153 | NM_006855.3 | 31.7269354 | -5.6215204 | 1.24813076 | -4.5039515 |
| 11154 | NM_001128922.1 | 26.6738794 | -6.3634926 | 1.41318019 | -4.5029591 |
| 11155 | NM_001317232.1 | 34.2062756 | -6.721395 | 1.49269789 | -4.5028502 |
| 11156 | NM_032857.4 | 19.0269941 | -5.8716609 | 1.30416594 | -4.5022346 |
| 11157 | NM_001286129.1 | 18.6530622 | -5.8418961 | 1.29783685 | -4.5012562 |
| 11158 | NR_024084.1 | 23.2253618 | -6.1568516 | 1.36785988 | -4.5010836 |
| 11159 | NM_057180.2 | 28.690527 | -6.4618367 | 1.4356314 | -4.5010416 |
| 11160 | NM_015721.2 | 78.9729512 | -4.4218166 | 0.98244778 | -4.5008159 |
| 11161 | NM_031307.3 | 21.6588779 | -6.0558608 | 1.34550549 | -4.5008072 |
| 11162 | NM_022915.3 | 98.9093346 | -3.4225189 | 0.76046736 | -4.5005467 |
| 11163 | NM_001199708.1 | 22.6336596 | -6.1138427 | 1.35849036 | -4.5004682 |
| 11164 | NM_006668.1 | 24.7869983 | -6.2544374 | 1.38976871 | -4.5003441 |
| 11165 | NM_080648.2 | 42.3578177 | -7.0289778 | 1.56187766 | -4.5003383 |
| 11166 | NM_001280794.2 | 34.6721322 | -6.7356936 | 1.4967444 | -4.5002297 |
| 11167 | NM_020737.2 | 34.8750615 | -6.7460148 | 1.49909742 | -4.500051 |
| 11168 | NM_001185118.1 | 29.9106818 | -6.5179995 | 1.44849516 | -4.4998421 |
| 11169 | NM_145715.2 | 17.6105045 | -5.767176 | 1.28167597 | -4.4997145 |
| 11170 | NM_001270482.1 | 27.3759539 | -6.4006198 | 1.4224792 | -4.4996228 |
| 11171 | NR_104010.1 | 36.3112984 | -6.8038601 | 1.51261659 | -4.4980732 |
| 11172 | NM_199424.2 | 37.0744504 | -6.8335682 | 1.51923319 | -4.4980377 |
| 11173 | NM_006656.5 | 19.9770193 | -5.9437401 | 1.32143328 | -4.4979495 |
| 11174 | NM_033364.3 | 35.8529197 | -6.788329 | 1.50948773 | -4.4971077 |
| 11175 | NM_020993.4 | 55.4163675 | -7.4113341 | 1.64808798 | -4.4969286 |
| 11176 | NR_045839.1 | 22.2029231 | -6.0948599 | 1.35550268 | -4.4963835 |
| 11177 | NM_018348.5 | 18.249558 | -5.8099532 | 1.29229552 | -4.4958395 |
| 11178 | NR_110290.1 | 25.1700458 | -6.2823163 | 1.39751382 | -4.4953518 |
| 11179 | NR_047492.1 | 61.0116534 | 3.0641092 | 0.68166626 | 4.49502843 |

| | | | | | |
|---|---|---|---|---|---|
| 11180 | NM_005440.4 | 240.86344 | -2.9883557 | 0.66483362 | -4.4948926 |
| 11181 | NM_017573.4 | 35.7266597 | -4.3979159 | 0.97856122 | -4.4942675 |
| 11182 | NM_002663.4 | 27.6234544 | -6.4152283 | 1.42755091 | -4.49387 |
| 11183 | NM_001322949.1 | 26.9936313 | -6.3737181 | 1.41834904 | -4.4937585 |
| 11184 | NM_017784.4 | 24.9570565 | -6.2612539 | 1.39357111 | -4.4929561 |
| 11185 | NM_001252393.1 | 23.1325441 | -6.1610065 | 1.37135653 | -4.4926366 |
| 11186 | NM_198315.2 | 18.3174458 | -5.8251349 | 1.29665236 | -4.4924415 |
| 11187 | NM_181726.3 | 17.8008691 | -5.7768001 | 1.28591699 | -4.4923585 |
| 11188 | NM_001301069.1 | 368.955267 | -5.6132693 | 1.24957567 | -4.4921404 |
| 11189 | NM_203506.2 | 10.5925377 | 8.33668432 | 1.85588905 | 4.49201655 |
| 11190 | NM_001033002.3 | 31.7178714 | -6.6085579 | 1.4713321 | -4.4915474 |
| 11191 | NM_001164715.1 | 23.9156917 | -6.1941364 | 1.37906219 | -4.491557 |
| 11192 | NM_017868.3 | 24.4556846 | -6.2362302 | 1.38847564 | -4.4914221 |
| 11193 | NM_001303112.1 | 27.5668864 | -6.4017184 | 1.42542972 | -4.4910796 |
| 11194 | NM_001136258.1 | 31.0343364 | -6.5810771 | 1.46549523 | -4.4906848 |
| 11195 | NM_016354.3 | 19.623586 | -5.9190123 | 1.31823071 | -4.4901187 |
| 11196 | NM_153692.3 | 44.8451496 | 3.77025224 | 0.83975241 | 4.48971885 |
| 11197 | NM_198794.2 | 36.0583338 | -6.7934023 | 1.51325942 | -4.4892516 |
| 11198 | NM_181657.3 | 32.5141025 | -5.6603117 | 1.26089697 | -4.4891152 |
| 11199 | NM_003922.3 | 332.26031 | -4.4650783 | 0.99465419 | -4.4890761 |
| 11200 | NM_001268.3 | 25.5715998 | -6.3008851 | 1.40367319 | -4.4888548 |
| 11201 | NM_016049.3 | 21.4694613 | -6.0485208 | 1.34770922 | -4.4880013 |
| 11202 | NM_145034.4 | 149.111012 | 2.19245061 | 0.48854037 | 4.48775728 |
| 11203 | NR_110765.1 | 48.6941763 | 4.55212381 | 1.01434769 | 4.48773518 |
| 11204 | NM_001080514.2 | 21.9311935 | -6.0797764 | 1.35476418 | -4.4877009 |
| 11205 | NM_001123.3 | 21.2445808 | -6.0298036 | 1.34365833 | -4.4876018 |
| 11206 | NM_001253816.1 | 40.0480902 | -6.9452729 | 1.54767536 | -4.4875515 |
| 11207 | NM_006155.2 | 21.3353284 | -6.0329622 | 1.3444622 | -4.487268 |
| 11208 | NM_020445.5 | 60.3052087 | -7.5348573 | 1.67917402 | -4.4872403 |
| 11209 | NM_001270977.1 | 35.7585821 | -6.7858209 | 1.51231765 | -4.4870341 |
| 11210 | NM_001346858.1 | 18.7033396 | -5.8505237 | 1.30387079 | -4.4870426 |
| 11211 | NM_001171134.2 | 18.092083 | -5.8009525 | 1.29284886 | -4.4869533 |
| 11212 | NM_001085.4 | 43.7904444 | -7.0768674 | 1.5772313 | -4.4868926 |
| 11213 | NM_001010938.1 | 48.512873 | -6.2485922 | 1.39264556 | -4.4868503 |
| 11214 | NM_001320817.1 | 31.21166 | -5.5965978 | 1.24736893 | -4.4867221 |
| 11215 | NM_014382.3 | 43.4775681 | -7.065202 | 1.57468769 | -4.4867322 |
| 11216 | NM_001321035.1 | 32.6431771 | -6.6461146 | 1.48129763 | -4.4866842 |
| 11217 | NM_032317.2 | 40.5090079 | -3.5429911 | 0.78967818 | -4.4866266 |
| 11218 | NM_001042473.3 | 21.621957 | -6.0587062 | 1.35041471 | -4.4865523 |
| 11219 | NM_001130858.2 | 25.5048851 | -6.2919926 | 1.40243573 | -4.4864749 |
| 11220 | NM_000212.2 | 27.8536516 | -6.4253198 | 1.43220573 | -4.4863106 |
| 11221 | NM_001144765.1 | 220.453453 | 2.44432589 | 0.54491019 | 4.48574081 |
| 11222 | NM_001318054.1 | 17.1291078 | -5.719018 | 1.2749296 | -4.485752 |

| | | | | | |
|---|---|---|---|---|---|
| 11223 | NR_134302.1 | 20.6195574 | -5.9847902 | 1.33423728 | -4.4855517 |
| 11224 | NM_001164380.1 | 26.314493 | -6.3382467 | 1.41308038 | -4.4854113 |
| 11225 | NM_001322820.1 | 40.4947298 | -6.9641154 | 1.55264333 | -4.4853285 |
| 11226 | NM_001134830.1 | 42.0606097 | -6.038216 | 1.34645682 | -4.4845225 |
| 11227 | NM_024884.2 | 610.590573 | 2.14653913 | 0.47865608 | 4.48451241 |
| 11228 | NM_001322019.1 | 21.2703924 | -6.0393312 | 1.34672661 | -4.4844523 |
| 11229 | NR_134862.1 | 25.4551555 | -6.2898772 | 1.40262841 | -4.4843504 |
| 11230 | NM_003831.3 | 235.508879 | -2.4345392 | 0.54297599 | -4.4836959 |
| 11231 | NM_001921.2 | 31.7608743 | -6.6088645 | 1.47409373 | -4.4833407 |
| 11232 | NM_018845.3 | 20.597348 | -5.9782633 | 1.3334359 | -4.4833526 |
| 11233 | NM_152328.4 | 24.6187842 | -6.248683 | 1.3937709 | -4.4832927 |
| 11234 | NR_144404.1 | 21.019948 | -6.0092384 | 1.34041511 | -4.4831175 |
| 11235 | NR_045019.1 | 21.6449336 | -6.059126 | 1.35155568 | -4.4830754 |
| 11236 | NM_178006.3 | 20.2203486 | -5.9619159 | 1.32993868 | -4.4828502 |
| 11237 | NM_001297652.1 | 21.4115469 | -6.0481664 | 1.34919624 | -4.4827922 |
| 11238 | NR_144431.1 | 31.6547265 | -6.6071983 | 1.47404821 | -4.4823489 |
| 11239 | NM_019610.5 | 27.9879874 | -6.4310858 | 1.43479811 | -4.4822235 |
| 11240 | NM_032273.3 | 30.7958282 | -6.5645874 | 1.46466111 | -4.4819838 |
| 11241 | NM_001130677.1 | 32.0941075 | -6.6239037 | 1.47794529 | -4.4818328 |
| 11242 | NM_002632.5 | 25.2588536 | -6.2861501 | 1.40282474 | -4.4810659 |
| 11243 | NM_005138.2 | 17.8982029 | -5.790094 | 1.29215408 | -4.4809626 |
| 11244 | NM_003371.3 | 151.291825 | -4.7769214 | 1.06609374 | -4.4807705 |
| 11245 | NM_025108.2 | 31.0620962 | -6.577629 | 1.46810907 | -4.4803409 |
| 11246 | NM_138459.3 | 219.231193 | -5.5315568 | 1.23467948 | -4.4801561 |
| 11247 | NM_019854.4 | 32.4724212 | -6.6425284 | 1.48277173 | -4.4798051 |
| 11248 | NM_001004722.3 | 27.0723223 | -6.3798009 | 1.42437746 | -4.47901 |
| 11249 | NM_015278.4 | 27.2234502 | -6.3936862 | 1.42761801 | -4.4785693 |
| 11250 | NM_144775.2 | 109.222594 | -2.2238489 | 0.49659669 | -4.4781791 |
| 11251 | NM_001243784.1 | 22.3627404 | -6.1028319 | 1.36300005 | -4.4774994 |
| 11252 | NM_004782.3 | 142.694028 | -1.6801284 | 0.37524275 | -4.4774441 |
| 11253 | NM_001130912.2 | 51.5481957 | -3.2994485 | 0.73691231 | -4.4773964 |
| 11254 | NR_038285.1 | 21.2468983 | -6.0305554 | 1.34690103 | -4.477356 |
| 11255 | NM_170725.2 | 16.9884795 | -5.7143085 | 1.27637091 | -4.4769968 |
| 11256 | NM_001282686.1 | 26.6043509 | -6.3596583 | 1.42098028 | -4.4755429 |
| 11257 | NM_003910.3 | 170.90807 | -5.1007666 | 1.13972723 | -4.4754275 |
| 11258 | NM_001077489.3 | 244.434902 | -5.2659889 | 1.17668341 | -4.475281 |
| 11259 | NM_001195446.1 | 67.2405351 | -4.3922562 | 0.98152099 | -4.4749488 |
| 11260 | NM_022062.2 | 100.330018 | -4.4332613 | 0.99074186 | -4.4746886 |
| 11261 | NM_001034025.1 | 35.5670731 | -6.774507 | 1.51413343 | -4.474181 |
| 11262 | NM_007350.3 | 348.695989 | -3.7529319 | 0.83879773 | -4.4741799 |
| 11263 | NM_001130683.3 | 20.1902895 | -5.9615103 | 1.33256642 | -4.473706 |
| 11264 | NM_001330195.1 | 25.3898177 | -6.2878805 | 1.4055535 | -4.4735974 |
| 11265 | NM_006370.2 | 167.799897 | -4.0265725 | 0.90009687 | -4.473488 |

| | | | | | |
|---|---|---|---|---|---|
| 11266 | NR_033709.1 | 23.9550308 | -6.1990173 | 1.38585804 | -4.4730536 |
| 11267 | NM_001161352.1 | 26.9490288 | -6.3748864 | 1.42533032 | -4.4725677 |
| 11268 | NM_182572.3 | 16.7202466 | -5.6848023 | 1.27104079 | -4.4725569 |
| 11269 | NM_014666.3 | 62.3593715 | -5.7689346 | 1.28986908 | -4.4724962 |
| 11270 | NM_001136562.2 | 46.6365724 | -6.1936562 | 1.38489989 | -4.4722772 |
| 11271 | NM_018694.3 | 22.2121544 | -6.0944701 | 1.36280518 | -4.4720039 |
| 11272 | NR_045557.1 | 28.2122446 | -6.4446232 | 1.44113658 | -4.4719031 |
| 11273 | NM_022369.3 | 30.6650024 | -6.5570569 | 1.46636799 | -4.4716312 |
| 11274 | NM_001134445.1 | 37.4515513 | -6.8490444 | 1.53179773 | -4.4712459 |
| 11275 | NM_001003788.2 | 17.2341235 | -5.7275958 | 1.28100972 | -4.4711572 |
| 11276 | NM_001258038.1 | 24.4490635 | -6.2390839 | 1.39551718 | -4.4708041 |
| 11277 | NM_001130926.2 | 24.1391342 | -6.2104807 | 1.38922909 | -4.4704511 |
| 11278 | NM_001142541.2 | 20.2368988 | -5.9529302 | 1.33178021 | -4.4699044 |
| 11279 | NM_001145850.1 | 16.9760447 | -5.7124314 | 1.27802098 | -4.4697478 |
| 11280 | NM_001660.3 | 551.52522 | -6.2092645 | 1.38919183 | -4.4696955 |
| 11281 | NM_001310148.1 | 26.3740659 | -6.3429912 | 1.41933514 | -4.4689876 |
| 11282 | NM_022113.5 | 18.2726928 | -5.8216307 | 1.303083 | -4.4675824 |
| 11283 | NM_001282285.1 | 23.330519 | -6.1702618 | 1.38114889 | -4.4674849 |
| 11284 | NM_001321187.1 | 20.1183226 | -5.954092 | 1.33275034 | -4.4675224 |
| 11285 | NM_014431.2 | 48.7150581 | -4.1472855 | 0.92832553 | -4.4674905 |
| 11286 | NR_015380.2 | 17.9552274 | -5.7865124 | 1.2953367 | -4.4671879 |
| 11287 | NR_134983.1 | 21.2373295 | -6.0384909 | 1.35214323 | -4.4658663 |
| 11288 | NM_001103167.1 | 20.9389837 | -6.0108981 | 1.34605443 | -4.4655684 |
| 11289 | NM_002886.3 | 123.376662 | -2.922204 | 0.65448381 | -4.4648988 |
| 11290 | NM_024672.5 | 19.6451498 | -5.9124813 | 1.32427192 | -4.4647034 |
| 11291 | NR_027702.1 | 41.1890573 | -6.9862745 | 1.56480022 | -4.4646431 |
| 11292 | NM_001204872.1 | 22.5042857 | -6.1204474 | 1.3709563 | -4.4643636 |
| 11293 | NM_001040143.1 | 51.7585078 | -7.3145023 | 1.63845033 | -4.4642808 |
| 11294 | NM_001289045.1 | 18.5373059 | -5.8379389 | 1.30786364 | -4.4637214 |
| 11295 | NM_001289061.1 | 21.8754778 | -6.0780766 | 1.36165786 | -4.4637326 |
| 11296 | NM_001256455.1 | 17.8130704 | -5.7818116 | 1.29535212 | -4.4635057 |
| 11297 | NM_002154.3 | 477.91236 | -5.7241031 | 1.28243706 | -4.4634573 |
| 11298 | NM_031232.3 | 21.2742114 | -6.027947 | 1.35053161 | -4.4633883 |
| 11299 | NR_046418.2 | 17.7348912 | -5.7642705 | 1.29157498 | -4.4629778 |
| 11300 | NR_024397.1 | 23.9623593 | -6.2045821 | 1.39027146 | -4.4628565 |
| 11301 | NM_001007464.2 | 23.8442623 | -6.2017251 | 1.38964078 | -4.4628261 |
| 11302 | NM_145294.4 | 109.211064 | -4.5657896 | 1.0232039 | -4.4622481 |
| 11303 | NM_001047434.2 | 38.6360274 | -5.9109685 | 1.32468911 | -4.4621552 |
| 11304 | NM_001123378.2 | 24.9993218 | -6.2685854 | 1.40497452 | -4.4617075 |
| 11305 | NM_001048265.1 | 19.9069112 | -5.9446742 | 1.33246034 | -4.4614267 |
| 11306 | NM_001037494.1 | 28.6390281 | -6.4611156 | 1.44844795 | -4.4607165 |
| 11307 | NM_206825.1 | 25.789534 | -6.3051419 | 1.41357885 | -4.4604104 |
| 11308 | NM_006859.3 | 18.7686138 | -5.8549827 | 1.31275312 | -4.4600791 |

| | | | | | |
|---|---|---|---|---|---|
| 11309 | NM_030786.2 | 28.6134465 | -6.4620363 | 1.44892429 | -4.4598854 |
| 11310 | NM_015414.3 | 26.6253933 | -6.3588175 | 1.42586455 | -4.4596224 |
| 11311 | NM_001201459.1 | 26.3470558 | -6.3388946 | 1.42143655 | -4.4594988 |
| 11312 | NM_033407.3 | 18.785675 | -5.8537553 | 1.31269697 | -4.4593348 |
| 11313 | NM_001282755.1 | 18.1855296 | -5.8078777 | 1.30249371 | -4.4590447 |
| 11314 | NM_183372.5 | 19.6283389 | -5.9109708 | 1.32576813 | -4.4585253 |
| 11315 | NM_001077692.1 | 18.3655703 | -5.8173028 | 1.30495558 | -4.457855 |
| 11316 | NM_152447.4 | 28.5631168 | -6.4551708 | 1.44806474 | -4.4577916 |
| 11317 | NM_001308287.1 | 48.8843454 | 7.00956185 | 1.57259604 | 4.45731877 |
| 11318 | NM_001282963.1 | 23.7079915 | 9.47822254 | 2.12649446 | 4.45720537 |
| 11319 | NM_015245.2 | 295.567549 | -5.7100893 | 1.28108816 | -4.4572181 |
| 11320 | NM_014619.4 | 20.483301 | -5.9858209 | 1.34304375 | -4.4569069 |
| 11321 | NM_001278698.1 | 19.9690783 | -5.935564 | 1.33178518 | -4.4568479 |
| 11322 | NM_005584.4 | 18.4314372 | -5.8279065 | 1.30768157 | -4.456671 |
| 11323 | NM_182697.2 | 45.6531162 | -7.1363676 | 1.6013829 | -4.4563781 |
| 11324 | NM_012261.3 | 18.4999191 | -5.8341473 | 1.30925439 | -4.4560838 |
| 11325 | NM_001161364.1 | 40.9811154 | -6.9796343 | 1.56635951 | -4.4559594 |
| 11326 | NM_001303523.1 | 9.57415843 | 8.19084937 | 1.83840418 | 4.45541273 |
| 11327 | NR_033690.1 | 19.1600542 | -5.8792431 | 1.31958203 | -4.4553828 |
| 11328 | NM_014338.3 | 38.1838079 | -6.8740187 | 1.54296152 | -4.4550811 |
| 11329 | NM_001142778.1 | 10.1739999 | 8.25433443 | 1.85284449 | 4.45495262 |
| 11330 | NM_001267040.1 | 84.3535819 | -4.7504923 | 1.06652791 | -4.454166 |
| 11331 | NM_024514.4 | 22.1961777 | -6.0921456 | 1.36774758 | -4.4541447 |
| 11332 | NM_001025242.1 | 41.9977401 | -7.0171024 | 1.57551624 | -4.4538433 |
| 11333 | NM_001278784.1 | 37.2578523 | -6.8415732 | 1.53628582 | -4.4533205 |
| 11334 | NM_032385.4 | 16.8814877 | -5.7059193 | 1.28126767 | -4.453339 |
| 11335 | NM_001100169.1 | 37.7544962 | -6.8624136 | 1.54101056 | -4.4531905 |
| 11336 | NM_001199302.1 | 131.97025 | -5.0265028 | 1.12887488 | -4.452666 |
| 11337 | NM_007008.2 | 19.9134265 | -5.9422179 | 1.3346882 | -4.4521393 |
| 11338 | NM_024657.4 | 41.5089411 | -7.0005626 | 1.57246652 | -4.4519629 |
| 11339 | NM_000641.3 | 31.7115955 | -6.6092701 | 1.48474653 | -4.4514467 |
| 11340 | NM_001204376.1 | 24.5443239 | -6.240208 | 1.40186266 | -4.451369 |
| 11341 | NM_013441.3 | 18.4334841 | -5.8281773 | 1.30936927 | -4.4511334 |
| 11342 | NM_004562.2 | 22.5069864 | -6.109498 | 1.37265866 | -4.4508501 |
| 11343 | NR_002781.1 | 19.0592269 | -5.8725479 | 1.31959218 | -4.4502748 |
| 11344 | NM_001173524.1 | 16.4152766 | -5.6589286 | 1.27164451 | -4.4500869 |
| 11345 | NR_026846.1 | 41.2496917 | -4.1772894 | 0.93871321 | -4.4500166 |
| 11346 | NM_001308237.1 | 17.9438153 | -5.7838794 | 1.29982493 | -4.4497372 |
| 11347 | NM_194259.2 | 27.7723785 | -6.4147033 | 1.44164784 | -4.4495633 |
| 11348 | NM_001321744.1 | 48.3851123 | -7.2161194 | 1.62184613 | -4.4493243 |
| 11349 | NM_001322414.1 | 25.9326232 | -6.3130549 | 1.41891863 | -4.4492015 |
| 11350 | NM_000857.3 | 47.4426468 | -7.1888294 | 1.6157802 | -4.4491382 |
| 11351 | NM_001318239.1 | 20.9931192 | -6.0194242 | 1.35317642 | -4.4483662 |

| | | | | | |
|---|---|---|---|---|---|
| 11352 | NR_038885.1 | 47.2524079 | 4.43448827 | 0.9969687 | 4.44797141 |
| 11353 | NM_001242859.2 | 79.490443 | -7.935102 | 1.78399039 | -4.4479511 |
| 11354 | NM_017696.2 | 17.136867 | -5.7182767 | 1.2857245 | -4.4475132 |
| 11355 | NR_040000.1 | 37.9636447 | 4.69253769 | 1.05509367 | 4.44750813 |
| 11356 | NM_145111.3 | 34.7183285 | -5.7477469 | 1.29253235 | -4.4468882 |
| 11357 | NR_038118.1 | 17.136827 | -5.725432 | 1.28752027 | -4.4468675 |
| 11358 | NM_001330983.1 | 75.9501672 | -7.8697555 | 1.77000181 | -4.446185 |
| 11359 | NM_015888.4 | 49.1101241 | -5.4375316 | 1.22296708 | -4.4461798 |
| 11360 | NM_001320042.1 | 18.1936365 | -5.800827 | 1.30472552 | -4.4460133 |
| 11361 | NM_053013.3 | 16.7778048 | -5.6935126 | 1.28061295 | -4.4459277 |
| 11362 | NM_000841.3 | 30.7339444 | -6.5641618 | 1.47669456 | -4.4451723 |
| 11363 | NM_001256732.2 | 57.8533982 | -7.4752939 | 1.68170135 | -4.4450782 |
| 11364 | NR_110907.1 | 65.2599745 | 2.3549408 | 0.52980968 | 4.44488066 |
| 11365 | NM_001060.5 | 73.9757005 | 1.75115484 | 0.39408206 | 4.44362991 |
| 11366 | NM_004712.4 | 363.295811 | -5.795915 | 1.30440636 | -4.4433354 |
| 11367 | NM_001122838.1 | 23.0791984 | -6.1427819 | 1.38256246 | -4.4430411 |
| 11368 | NM_001330234.1 | 18.8005486 | -5.8486631 | 1.31641183 | -4.4428825 |
| 11369 | NR_026808.1 | 53.3182669 | 1.76659229 | 0.39763192 | 4.44278295 |
| 11370 | NM_001289399.1 | 22.7797994 | -6.138334 | 1.38185863 | -4.4420853 |
| 11371 | NM_001204468.1 | 47.1778246 | -7.1838403 | 1.6172723 | -4.4419486 |
| 11372 | NM_020939.1 | 27.4336187 | -6.3944198 | 1.43959152 | -4.4418293 |
| 11373 | NM_145165.3 | 23.2157546 | -6.1540505 | 1.38577998 | -4.4408568 |
| 11374 | NM_199250.2 | 30.811516 | -6.5622381 | 1.47770034 | -4.440845 |
| 11375 | NM_144570.2 | 265.757912 | -3.9452461 | 0.88849214 | -4.4403837 |
| 11376 | NR_109819.1 | 22.1693522 | -6.0894775 | 1.37140789 | -4.4403109 |
| 11377 | NM_001099650.1 | 57.6008313 | -4.2461812 | 0.95636139 | -4.4399337 |
| 11378 | NM_001135580.1 | 17.7701306 | -5.7761159 | 1.30099283 | -4.4397753 |
| 11379 | NM_001243749.1 | 18.2977865 | -5.8192752 | 1.31083401 | -4.4393685 |
| 11380 | NM_001307981.1 | 16.7466709 | -5.6940624 | 1.28264396 | -4.4393165 |
| 11381 | NR_024117.2 | 29.021813 | -5.504162 | 1.23987376 | -4.4392923 |
| 11382 | NR_027840.1 | 22.6694847 | -6.1293872 | 1.38075191 | -4.4391662 |
| 11383 | NM_207309.2 | 50.3875135 | -2.4437301 | 0.5505008 | -4.4391037 |
| 11384 | NM_015291.3 | 35.0597651 | -6.7545578 | 1.52163645 | -4.4390089 |
| 11385 | NM_001288764.1 | 30.2107398 | -6.5440456 | 1.47431586 | -4.4386998 |
| 11386 | NM_001008661.2 | 22.1408092 | -6.0939274 | 1.37295022 | -4.438564 |
| 11387 | NM_001286722.1 | 51.216194 | -6.3159574 | 1.42316255 | -4.4379733 |
| 11388 | NM_004491.4 | 576.980232 | -6.1138958 | 1.37765448 | -4.4379022 |
| 11389 | NM_001001436.2 | 51.0937495 | -5.4302151 | 1.22377166 | -4.4372781 |
| 11390 | NM_001203.2 | 24.3246177 | -6.222134 | 1.40228069 | -4.437153 |
| 11391 | NM_001330454.1 | 36.6544337 | -6.8156758 | 1.5360615 | -4.4371113 |
| 11392 | NM_020770.2 | 18.2779093 | -5.8221959 | 1.31218542 | -4.4370222 |
| 11393 | NM_173543.2 | 18.6322343 | -5.8433635 | 1.31697584 | -4.4369558 |
| 11394 | NM_001345880.1 | 28.3745528 | -6.4457962 | 1.45278692 | -4.436849 |

| | | | | | |
|---|---|---|---|---|---|
| 11395 | NR_033833.1 | 19.1337569 | 3.54281776 | 0.79849638 | 4.43686138 |
| 11396 | NM_173039.2 | 20.9048661 | -6.0061924 | 1.3537353 | -4.4367554 |
| 11397 | NM_020727.4 | 40.1808258 | -6.949609 | 1.56653665 | -4.4362888 |
| 11398 | NM_001321547.1 | 21.5648389 | -6.0484238 | 1.36341698 | -4.4362245 |
| 11399 | NM_005159.4 | 22.0511467 | -6.0890462 | 1.37257261 | -4.4362288 |
| 11400 | NM_015086.1 | 70.2746916 | -7.7558972 | 1.74834797 | -4.436129 |
| 11401 | NM_001316321.1 | 64.3951248 | -3.7451501 | 0.84439807 | -4.4352897 |
| 11402 | NM_001329846.1 | 32.3449492 | -6.6335318 | 1.49572819 | -4.4349848 |
| 11403 | NM_001297565.1 | 38.8762629 | -5.9259684 | 1.33636073 | -4.4344078 |
| 11404 | NM_018015.5 | 23.805902 | -6.2010078 | 1.39843062 | -4.434262 |
| 11405 | NM_138393.2 | 19.4986616 | -5.9150354 | 1.33393826 | -4.4342647 |
| 11406 | NM_001276469.1 | 40.1454673 | -6.9472815 | 1.5668754 | -4.4338443 |
| 11407 | NM_003745.1 | 20.6559129 | -5.9905167 | 1.35113996 | -4.4336759 |
| 11408 | NM_032538.2 | 151.74612 | -5.2408555 | 1.18215783 | -4.433296 |
| 11409 | NM_001085447.1 | 27.6375573 | -6.413566 | 1.44669221 | -4.4332623 |
| 11410 | NM_001080449.2 | 22.8034644 | -6.1315552 | 1.38313151 | -4.4330964 |
| 11411 | NM_021082.3 | 23.6339745 | -6.1890228 | 1.39609385 | -4.4330994 |
| 11412 | NM_001286373.1 | 21.9011254 | -6.0687563 | 1.36912122 | -4.4325924 |
| 11413 | NM_001134486.2 | 40.2053525 | 4.24303925 | 0.95724449 | 4.43255543 |
| 11414 | NM_057092.2 | 35.4623058 | -4.8925006 | 1.10395965 | -4.4317749 |
| 11415 | NM_001318083.1 | 17.2573925 | -5.7288862 | 1.29303547 | -4.4305716 |
| 11416 | NM_025139.5 | 26.4212764 | -6.3498607 | 1.43325869 | -4.4303661 |
| 11417 | NM_018717.4 | 105.471248 | -4.6832779 | 1.05720952 | -4.4298484 |
| 11418 | NM_001098510.1 | 20.0991968 | -5.9458899 | 1.34226741 | -4.4297358 |
| 11419 | NR_049753.2 | 19.8621039 | -5.9409126 | 1.3413583 | -4.4290274 |
| 11420 | NR_046176.2 | 20.9057599 | -6.011755 | 1.35745442 | -4.4286975 |
| 11421 | NM_001282791.1 | 47.369435 | -6.2065797 | 1.40157536 | -4.4282883 |
| 11422 | NM_001170640.1 | 17.2176941 | -5.7341972 | 1.29491905 | -4.4282283 |
| 11423 | NM_001075098.3 | 17.8298641 | -5.7807821 | 1.30545319 | -4.4281803 |
| 11424 | NM_001193538.2 | 20.6177576 | -5.9852766 | 1.35172348 | -4.4278853 |
| 11425 | NR_001568.1 | 748.059765 | -5.8993497 | 1.33233873 | -4.4278152 |
| 11426 | NM_002629.3 | 1014.05835 | -6.5264618 | 1.4741615 | -4.4272366 |
| 11427 | NM_152482.2 | 82.9234651 | -4.7236676 | 1.06699762 | -4.4270647 |
| 11428 | NM_001171797.1 | 40.4438713 | -6.9586134 | 1.57211633 | -4.4262713 |
| 11429 | NR_027149.1 | 20.2394833 | -5.959201 | 1.34633055 | -4.426254 |
| 11430 | NM_000478.5 | 17.6753519 | -5.7742558 | 1.30459453 | -4.4260923 |
| 11431 | NM_024817.2 | 18.1429953 | -5.7985549 | 1.31011355 | -4.4259941 |
| 11432 | NM_020818.4 | 35.7541718 | -6.7823311 | 1.53246273 | -4.4257723 |
| 11433 | NR_027487.2 | 20.8626159 | -6.0054368 | 1.35704689 | -4.4253717 |
| 11434 | NM_006980.4 | 18.4827263 | -5.8312666 | 1.31776483 | -4.4251193 |
| 11435 | NM_017621.3 | 57.9128544 | -4.1596562 | 0.94003755 | -4.4249894 |
| 11436 | NM_001012393.2 | 88.6178415 | -8.090618 | 1.82850106 | -4.424727 |
| 11437 | NM_014234.4 | 16.4948675 | -5.6726688 | 1.28209125 | -4.4245437 |

| | | | | | |
|---|---|---|---|---|---|
| 11438 | NM_001080417.2 | 163.131735 | -4.1214142 | 0.93153435 | -4.4243287 |
| 11439 | NM_020853.1 | 163.388963 | -2.9971318 | 0.6774191 | -4.4243391 |
| 11440 | NM_015450.2 | 23.8074464 | -6.1971039 | 1.4009405 | -4.4235311 |
| 11441 | NM_001317177.1 | 46.6379435 | -7.1658902 | 1.61997725 | -4.4234512 |
| 11442 | NM_003840.4 | 68.7344169 | -3.6704955 | 0.82982116 | -4.4232368 |
| 11443 | NM_001318226.1 | 21.6249859 | -6.0491363 | 1.36774776 | -4.4226987 |
| 11444 | NR_136651.1 | 20.5911688 | -5.9877597 | 1.35400025 | -4.4222738 |
| 11445 | NM_001281504.1 | 21.5019001 | -6.0404846 | 1.36605971 | -4.4218305 |
| 11446 | NM_001276282.3 | 23.2457892 | -6.1551841 | 1.39216709 | -4.421297 |
| 11447 | NM_001288766.1 | 36.5789437 | -4.9850186 | 1.12754262 | -4.4211354 |
| 11448 | NM_001128224.2 | 57.5525072 | -5.1436242 | 1.16344794 | -4.421018 |
| 11449 | NM_138467.2 | 18.2503552 | -5.8147265 | 1.31524047 | -4.4210368 |
| 11450 | NM_019104.2 | 42.4286054 | -6.0557243 | 1.36979807 | -4.4208883 |
| 11451 | NM_052943.3 | 18.3947157 | -5.8317445 | 1.3192074 | -4.4206427 |
| 11452 | NM_006258.3 | 17.1088317 | -5.7258069 | 1.29535142 | -4.420273 |
| 11453 | NM_001193389.1 | 15.1223359 | 8.84025396 | 2.00000864 | 4.42010788 |
| 11454 | NM_005773.2 | 19.3178269 | -5.8910145 | 1.33289564 | -4.4197118 |
| 11455 | NM_152262.2 | 26.4949823 | -6.3460274 | 1.43587009 | -4.419639 |
| 11456 | NM_005258.2 | 17.657291 | -5.7713951 | 1.30588867 | -4.4195154 |
| 11457 | NM_030651.3 | 31.5665881 | -6.6006134 | 1.49361161 | -4.4192301 |
| 11458 | NM_012345.2 | 39.014638 | -5.9207105 | 1.33978585 | -4.4191469 |
| 11459 | NM_001015049.2 | 31.2928845 | -6.5909006 | 1.4915266 | -4.4188958 |
| 11460 | NM_133460.2 | 19.067741 | -5.8679624 | 1.32795642 | -4.4187914 |
| 11461 | NM_152279.3 | 218.820159 | 1.81527292 | 0.41082723 | 4.41857988 |
| 11462 | NR_134535.1 | 23.8042966 | -6.1989079 | 1.40294154 | -4.4185076 |
| 11463 | NM_001284382.1 | 37.9949493 | -6.8681079 | 1.55441842 | -4.4184422 |
| 11464 | NM_033157.3 | 28.0900448 | -6.4337396 | 1.45614783 | -4.4183286 |
| 11465 | NM_001271878.1 | 31.2630781 | -6.5889057 | 1.49137543 | -4.418006 |
| 11466 | NM_016027.2 | 17.721024 | -5.7703282 | 1.30609265 | -4.4180084 |
| 11467 | NM_001123363.3 | 30.2117441 | -6.5398594 | 1.48029905 | -4.4179312 |
| 11468 | NM_012331.4 | 19.5577078 | -5.9145932 | 1.33898068 | -4.4172356 |
| 11469 | NM_001289808.1 | 27.5462559 | -6.404835 | 1.4500189 | -4.41707 |
| 11470 | NM_001282673.1 | 38.5941991 | -6.8918421 | 1.56029174 | -4.4170215 |
| 11471 | NM_016606.3 | 46.5634825 | -7.1657497 | 1.62238988 | -4.4167865 |
| 11472 | NM_152457.2 | 25.7403189 | -6.3033613 | 1.42714786 | -4.4167542 |
| 11473 | NR_134612.1 | 30.5576578 | -6.5547521 | 1.48411192 | -4.4166158 |
| 11474 | NM_001271442.1 | 15.9296258 | -5.6137492 | 1.27109491 | -4.4164674 |
| 11475 | NR_136405.1 | 22.1305794 | -6.097902 | 1.38096561 | -4.4156799 |
| 11476 | NM_145291.3 | 18.8529767 | -5.8574294 | 1.32656584 | -4.4154834 |
| 11477 | NM_152698.2 | 36.1903995 | -6.7951562 | 1.53930534 | -4.4144304 |
| 11478 | NM_014875.2 | 21.9048374 | -6.0687246 | 1.37476484 | -4.4143729 |
| 11479 | NM_001113567.2 | 28.7327129 | -6.4616595 | 1.46381626 | -4.4142558 |
| 11480 | NM_001324020.1 | 31.1139546 | -6.5767009 | 1.49016301 | -4.4134104 |

| | | | | | |
|---|---|---|---|---|---|
| 11481 | NM_001243247.1 | 28.627651 | -6.4669266 | 1.46535315 | -4.4132205 |
| 11482 | NM_001098633.3 | 39.8712288 | -5.9624692 | 1.35108584 | -4.4130943 |
| 11483 | NM_001282930.1 | 45.3993506 | -7.1256603 | 1.61466685 | -4.4130839 |
| 11484 | NM_152912.4 | 20.7585966 | -6.0026326 | 1.36024824 | -4.4128949 |
| 11485 | NM_001345876.1 | 24.2181351 | -6.222682 | 1.4102811 | -4.41237 |
| 11486 | NM_145065.2 | 20.9735717 | -6.0173622 | 1.3638354 | -4.4120883 |
| 11487 | NM_138794.4 | 29.0539639 | -6.4783522 | 1.46846526 | -4.4116483 |
| 11488 | NM_001135055.2 | 27.8973431 | -6.4219983 | 1.45571381 | -4.4115802 |
| 11489 | NM_001329990.1 | 39.6483895 | -6.9299782 | 1.57089232 | -4.4114915 |
| 11490 | NM_021931.3 | 37.1538278 | -6.8368069 | 1.54985347 | -4.4112602 |
| 11491 | NM_014997.3 | 227.423883 | -4.0471854 | 0.91777189 | -4.4097945 |
| 11492 | NM_001346810.1 | 33.1483136 | -6.6723922 | 1.5131028 | -4.4097415 |
| 11493 | NM_020317.4 | 19.8032153 | -5.9326323 | 1.34552263 | -4.4091657 |
| 11494 | NM_001008.3 | 30.3542718 | -6.5468912 | 1.4849249 | -4.4089039 |
| 11495 | NM_006019.3 | 21.9689729 | -6.0847223 | 1.38009597 | -4.4089124 |
| 11496 | NM_001080826.2 | 123.539161 | -4.9276714 | 1.11771618 | -4.4086965 |
| 11497 | NM_024608.3 | 21.1198152 | -6.0186893 | 1.36518513 | -4.4086983 |
| 11498 | NM_001177465.2 | 20.7479281 | -5.9896219 | 1.3587058 | -4.4083288 |
| 11499 | NM_001317991.1 | 25.8640184 | -6.3157309 | 1.4327456 | -4.4081315 |
| 11500 | NM_177417.2 | 20.6661009 | -5.9943114 | 1.36002208 | -4.4075104 |
| 11501 | NR_121583.1 | 18.3583274 | 3.02525482 | 0.68639039 | 4.40748419 |
| 11502 | NM_001127207.1 | 39.6347505 | -5.941467 | 1.34815142 | -4.4071214 |
| 11503 | NM_152402.2 | 21.9370654 | -6.072101 | 1.37784583 | -4.4069524 |
| 11504 | NM_001146082.2 | 17.7499509 | -5.7699878 | 1.3095327 | -4.4061426 |
| 11505 | NM_001286734.1 | 41.6146106 | -5.1168091 | 1.16135675 | -4.4058892 |
| 11506 | NM_018146.3 | 32.4894799 | -5.6532777 | 1.28337666 | -4.4050028 |
| 11507 | NR_108047.1 | 124.972699 | 3.34130691 | 0.7585464 | 4.40488136 |
| 11508 | NM_001271819.1 | 377.13517 | 6.80921939 | 1.54589133 | 4.40472059 |
| 11509 | NM_182836.2 | 36.0091527 | -5.8028279 | 1.31758363 | -4.4041439 |
| 11510 | NM_152478.2 | 29.7315897 | -6.5142058 | 1.47918535 | -4.4039145 |
| 11511 | NM_004915.3 | 20.0349463 | -5.945927 | 1.35020069 | -4.4037357 |
| 11512 | NM_001321532.1 | 37.0121537 | -6.8299231 | 1.55096449 | -4.4036618 |
| 11513 | NM_001405.3 | 1020.2651 | 4.11369497 | 0.9342579 | 4.40316851 |
| 11514 | NM_001290050.1 | 104.507192 | -2.6874658 | 0.61037453 | -4.4029783 |
| 11515 | NM_012471.2 | 28.548206 | -6.4591613 | 1.46700596 | -4.402955 |
| 11516 | NM_025160.6 | 122.848126 | -4.5979946 | 1.04448826 | -4.4021506 |
| 11517 | NM_001134870.1 | 162.178692 | -4.0231744 | 0.91403191 | -4.4015689 |
| 11518 | NM_001195263.1 | 24.3291897 | -6.2264763 | 1.41464766 | -4.4014326 |
| 11519 | NM_172037.4 | 88.9901361 | -4.3573933 | 0.99001932 | -4.4013215 |
| 11520 | NR_024159.1 | 19.0348528 | -5.872736 | 1.33436338 | -4.401152 |
| 11521 | NM_001262.2 | 22.0805237 | -6.0861187 | 1.38285723 | -4.4011186 |
| 11522 | NM_001324255.1 | 963.931994 | -11.534939 | 2.62142511 | -4.4002551 |
| 11523 | NM_012414.3 | 258.792081 | -5.1482931 | 1.17009182 | -4.3999052 |

| | | | | | |
|---|---|---|---|---|---|
| 11524 | NM_178169.3 | 27.4678279 | -6.4059122 | 1.45608449 | -4.3994097 |
| 11525 | NM_001077490.2 | 22.0024415 | -6.0861418 | 1.38343199 | -4.3993068 |
| 11526 | NM_015633.2 | 23.7456802 | -6.1870715 | 1.40644222 | -4.399094 |
| 11527 | NM_153015.2 | 26.5247834 | -6.3478765 | 1.44337046 | -4.3979537 |
| 11528 | NM_001008225.2 | 17.3134156 | -5.7341479 | 1.30385407 | -4.3978449 |
| 11529 | NM_003724.3 | 119.513929 | -3.7140017 | 0.84450391 | -4.3978503 |
| 11530 | NM_016077.4 | 24.8820707 | -6.260865 | 1.42363897 | -4.3977898 |
| 11531 | NM_007098.3 | 21.1693401 | -6.0256251 | 1.37026816 | -4.3974057 |
| 11532 | NM_138720.2 | 18.4889279 | -5.835264 | 1.32699341 | -4.3973572 |
| 11533 | NM_003070.4 | 26.7488405 | -6.3648076 | 1.44748313 | -4.3971549 |
| 11534 | NM_013377.3 | 17.887897 | -5.7820456 | 1.31528227 | -4.3960492 |
| 11535 | NM_145008.2 | 20.8683452 | -6.0006315 | 1.3651277 | -4.3956558 |
| 11536 | NM_000465.3 | 19.6271832 | -5.9077797 | 1.34404773 | -4.3955133 |
| 11537 | NM_001198805.1 | 19.5399118 | -5.9105081 | 1.34476686 | -4.3951917 |
| 11538 | NM_003743.4 | 45.3313369 | -7.125306 | 1.62127097 | -4.394889 |
| 11539 | NM_016196.3 | 28.7943381 | -6.4674124 | 1.47159159 | -4.3948419 |
| 11540 | NM_001321109.1 | 17.0801184 | -5.7241415 | 1.30249201 | -4.3947613 |
| 11541 | NM_001313893.1 | 34.8256878 | -6.7433709 | 1.53452992 | -4.3944213 |
| 11542 | NM_004102.4 | 37.7502517 | -6.8598686 | 1.5611509 | -4.3941099 |
| 11543 | NR_033334.1 | 15.8923016 | -5.6143941 | 1.27774986 | -4.3939696 |
| 11544 | NR_033998.1 | 19.3697779 | -5.903185 | 1.34352398 | -4.393807 |
| 11545 | NR_024072.2 | 31.9312244 | 1.94162987 | 0.44195984 | 4.39322691 |
| 11546 | NM_001320309.1 | 34.1134024 | -6.7126356 | 1.52824578 | -4.3923796 |
| 11547 | NM_014711.5 | 20.8791444 | -5.9983437 | 1.36586009 | -4.3916239 |
| 11548 | NR_109846.1 | 21.9877551 | -6.0760848 | 1.38361029 | -4.3914712 |
| 11549 | NM_080676.5 | 24.2955646 | -6.2237583 | 1.41724518 | -4.3914478 |
| 11550 | NM_007074.3 | 87.1613792 | -5.3787012 | 1.22482996 | -4.391386 |
| 11551 | NM_018252.2 | 75.2358221 | -5.8179476 | 1.32514809 | -4.3904131 |
| 11552 | NM_003407.3 | 28.887997 | -6.4787329 | 1.47582161 | -4.389916 |
| 11553 | NM_014507.3 | 17.6019276 | -5.7574697 | 1.31163296 | -4.3895433 |
| 11554 | NM_033412.3 | 19.7551451 | -5.9296295 | 1.35096369 | -4.389185 |
| 11555 | NM_153704.5 | 18.4534534 | -5.8312267 | 1.32858942 | -4.3890359 |
| 11556 | NM_001308295.1 | 30.3954146 | -6.5470809 | 1.49172381 | -4.3889364 |
| 11557 | NM_001538.3 | 37.1201773 | -5.8545518 | 1.3339297 | -4.3889508 |
| 11558 | NM_052841.3 | 17.4521706 | -5.755895 | 1.31144972 | -4.3889559 |
| 11559 | NM_005677.3 | 22.2184073 | -6.0984594 | 1.38962235 | -4.3885732 |
| 11560 | NM_000082.3 | 24.0341142 | -6.209493 | 1.41497937 | -4.3883983 |
| 11561 | NM_001318794.1 | 21.0047618 | -6.0181902 | 1.37145903 | -4.3881662 |
| 11562 | NM_015044.4 | 306.880121 | -1.8928444 | 0.4314487 | -4.387183 |
| 11563 | NM_032847.2 | 34.7934315 | -5.7670957 | 1.31454818 | -4.3871315 |
| 11564 | NM_178812.3 | 384.974444 | -3.9052909 | 0.89017449 | -4.3871071 |
| 11565 | NM_018295.4 | 18.0077137 | -5.7952271 | 1.3211351 | -4.3865514 |
| 11566 | NM_032108.3 | 327.442407 | -5.8496298 | 1.3335984 | -4.3863503 |

| | | | | | |
|---|---|---|---|---|---|
| 11567 | NM_152744.3 | 129.59514 | -4.2192103 | 0.9618989 | -4.3863345 |
| 11568 | NM_018334.4 | 38.302615 | -5.9072726 | 1.34678953 | -4.3861884 |
| 11569 | NM_001033117.2 | 254.685721 | -5.706488 | 1.30113762 | -4.3857682 |
| 11570 | NM_002122.3 | 34.755429 | -6.7433276 | 1.53754889 | -4.3857647 |
| 11571 | NM_004769.3 | 20.1731729 | -5.9500611 | 1.35689186 | -4.3850666 |
| 11572 | NM_030919.2 | 35.077251 | -5.7783851 | 1.31777398 | -4.3849591 |
| 11573 | NM_032789.4 | 34.2733548 | -6.7235339 | 1.53336504 | -4.3848227 |
| 11574 | NM_001199397.1 | 34.8282917 | -6.7460055 | 1.53851268 | -4.3847578 |
| 11575 | NM_000966.5 | 20.338972 | -5.9745355 | 1.36257734 | -4.3847313 |
| 11576 | NR_015381.1 | 19.2923764 | -5.8930277 | 1.34400132 | -4.384689 |
| 11577 | NM_182524.3 | 23.5402329 | -6.171982 | 1.40773884 | -4.3843231 |
| 11578 | NM_001163147.1 | 27.7527106 | -6.4180078 | 1.46388467 | -4.3842305 |
| 11579 | NM_001752.3 | 48.5441887 | -7.2224848 | 1.64748149 | -4.3839551 |
| 11580 | NM_014984.3 | 32.1966054 | -6.6354118 | 1.51375741 | -4.383405 |
| 11581 | NM_001003698.3 | 21.0813262 | -6.0228269 | 1.37411165 | -4.3830695 |
| 11582 | NM_014454.2 | 39.8751543 | -5.9531196 | 1.35828623 | -4.3828167 |
| 11583 | NM_003285.2 | 30.0495525 | -6.5318596 | 1.49036019 | -4.3827389 |
| 11584 | NM_001316943.1 | 31.6051311 | -5.6220116 | 1.28281319 | -4.3825645 |
| 11585 | NM_016134.3 | 17.8603432 | -5.7897852 | 1.32113541 | -4.3824313 |
| 11586 | NM_018036.6 | 114.6347 | -4.9169847 | 1.12207299 | -4.3820543 |
| 11587 | NM_032167.4 | 78.6223391 | -4.4211772 | 1.00898095 | -4.3818243 |
| 11588 | NM_001257971.1 | 21.2342869 | -6.032198 | 1.37672231 | -4.3815648 |
| 11589 | NM_001282732.1 | 21.6271972 | 5.39855715 | 1.23216572 | 4.38135639 |
| 11590 | NM_181539.4 | 16.3410487 | 3.36811755 | 0.76877555 | 4.38114552 |
| 11591 | NR_015384.2 | 33.6894008 | -6.6956798 | 1.52853521 | -4.3804551 |
| 11592 | NM_152835.4 | 38.0738607 | -6.8691076 | 1.56832143 | -4.3799106 |
| 11593 | NR_026551.1 | 23.8672472 | -6.1959084 | 1.41462653 | -4.3798899 |
| 11594 | NM_001134456.1 | 57.2454238 | -7.4605602 | 1.703596 | -4.3793013 |
| 11595 | NM_032626.5 | 15.7760188 | -5.6058888 | 1.28015637 | -4.3790656 |
| 11596 | NM_003182.2 | 110.049891 | -5.1631438 | 1.1790635 | -4.379021 |
| 11597 | NM_006965.3 | 276.732344 | -5.2498088 | 1.19895766 | -4.378644 |
| 11598 | NM_033629.4 | 15.4854476 | -5.5781873 | 1.27395234 | -4.3786467 |
| 11599 | NM_147168.1 | 30.4975882 | -6.555322 | 1.49710342 | -4.3786701 |
| 11600 | NM_198971.2 | 46.3017157 | -4.395121 | 1.00380511 | -4.3784605 |
| 11601 | NM_024771.3 | 150.684293 | -3.1824666 | 0.72687767 | -4.3782699 |
| 11602 | NM_000101.3 | 19.506596 | -5.9091064 | 1.34967615 | -4.3781661 |
| 11603 | NM_194283.3 | 20.5997438 | -5.9777752 | 1.36544545 | -4.3778938 |
| 11604 | NM_014648.3 | 215.81421 | -2.1434173 | 0.48962321 | -4.3776873 |
| 11605 | NM_018117.11 | 170.181933 | -5.1737356 | 1.18185569 | -4.3776374 |
| 11606 | NM_003924.3 | 130.976636 | -5.3018272 | 1.21122484 | -4.3772445 |
| 11607 | NM_004856.6 | 22.9074894 | -6.145385 | 1.40408288 | -4.3767965 |
| 11608 | NM_021004.3 | 16.3649482 | -5.6491834 | 1.29071058 | -4.376801 |
| 11609 | NR_103535.1 | 98.0162605 | 3.57026856 | 0.81579748 | 4.3764153 |

| | | | | | |
|---|---|---|---|---|---|
| 11610 | NM_178229.4 | 51.7928408 | -4.282024 | 0.97844135 | -4.3763727 |
| 11611 | NM_001171025.1 | 29.0455556 | -6.4870892 | 1.48234358 | -4.3762386 |
| 11612 | NM_032131.5 | 19.2631242 | -5.8916046 | 1.34641463 | -4.3757728 |
| 11613 | NM_207310.2 | 22.0694419 | -6.0856441 | 1.39083641 | -4.3755283 |
| 11614 | NM_001141945.2 | 24.1037535 | -6.2186386 | 1.42133355 | -4.3752141 |
| 11615 | NM_031301.3 | 41.2267916 | -6.9903825 | 1.59780758 | -4.3749839 |
| 11616 | NM_206839.2 | 34.289481 | -6.7203501 | 1.53613705 | -4.3748376 |
| 11617 | NR_126510.1 | 24.6332148 | -6.2432904 | 1.42717306 | -4.3745854 |
| 11618 | NM_019100.4 | 20.4159577 | -5.9785472 | 1.36683101 | -4.3740207 |
| 11619 | NM_006946.2 | 641.934614 | -4.9242941 | 1.12582393 | -4.3739469 |
| 11620 | NM_001081461.1 | 91.6174131 | -3.2781422 | 0.74950032 | -4.3737702 |
| 11621 | NM_005575.2 | 18.7137888 | -5.8475962 | 1.33697278 | -4.3737586 |
| 11622 | NM_016621.3 | 145.412366 | -5.009665 | 1.14541302 | -4.3736756 |
| 11623 | NM_001130997.2 | 21.720487 | -6.0634377 | 1.38639219 | -4.3735371 |
| 11624 | NM_001270976.1 | 19.3179808 | -5.9001197 | 1.34929029 | -4.3727578 |
| 11625 | NM_017546.4 | 97.0377123 | -4.3856088 | 1.00297154 | -4.3726154 |
| 11626 | NM_004425.3 | 19.8789881 | -5.9350007 | 1.35733016 | -4.3725549 |
| 11627 | NR_004053.3 | 15.9362121 | -5.6173691 | 1.28471484 | -4.3724638 |
| 11628 | NM_001290259.1 | 34.5958729 | -5.7408635 | 1.31307451 | -4.3720775 |
| 11629 | NM_003706.2 | 20.1639467 | -5.9545269 | 1.36194064 | -4.3720899 |
| 11630 | NM_000159.3 | 25.4468516 | -6.2906336 | 1.43898808 | -4.3715676 |
| 11631 | NM_004260.3 | 102.054883 | -4.559292 | 1.04297071 | -4.3714478 |
| 11632 | NM_001308117.1 | 20.9711624 | -6.0143765 | 1.37609223 | -4.3706202 |
| 11633 | NM_198282.3 | 34.6549543 | -6.7400135 | 1.54240362 | -4.3698117 |
| 11634 | NM_032373.4 | 39.1112585 | -6.9098503 | 1.58141789 | -4.3694019 |
| 11635 | NM_015604.3 | 25.2161282 | -6.2742822 | 1.43605112 | -4.3691218 |
| 11636 | NM_001007794.2 | 19.0377947 | -5.8815919 | 1.34663596 | -4.3676183 |
| 11637 | NM_001013672.4 | 18.7187878 | -5.8547725 | 1.34050167 | -4.3675981 |
| 11638 | NM_001272013.1 | 36.3389644 | -6.8088316 | 1.559021 | -4.3673764 |
| 11639 | NM_001536.5 | 24.3323688 | -6.2234589 | 1.42502963 | -4.3672488 |
| 11640 | NM_014229.2 | 22.0513577 | -6.0816962 | 1.39259008 | -4.3671834 |
| 11641 | NM_173810.3 | 20.950698 | -6.0088833 | 1.37592507 | -4.3671589 |
| 11642 | NR_134301.1 | 138.688439 | -5.5238936 | 1.26496939 | -4.3668199 |
| 11643 | NM_006497.3 | 25.8331532 | -6.3186029 | 1.44705929 | -4.3665128 |
| 11644 | NM_207446.2 | 36.8049524 | -5.8514671 | 1.34021131 | -4.366078 |
| 11645 | NR_024284.1 | 19.1404778 | -5.8807492 | 1.34694846 | -4.3659793 |
| 11646 | NM_052969.1 | 25.735398 | -6.3053415 | 1.44421546 | -4.3659285 |
| 11647 | NM_214462.4 | 19.9734525 | -5.9498426 | 1.36281779 | -4.3658387 |
| 11648 | NM_001331010.1 | 26.3058957 | -6.3403764 | 1.45228443 | -4.3657952 |
| 11649 | NR_023314.4 | 28.2427185 | -6.4390955 | 1.47497038 | -4.3655761 |
| 11650 | NM_030762.2 | 16.8321042 | -5.7002308 | 1.30583409 | -4.365203 |
| 11651 | NM_032534.3 | 20.8200811 | -6.0076905 | 1.37629998 | -4.3651025 |
| 11652 | NR_135534.1 | 17.7426074 | -5.7686662 | 1.32157327 | -4.3649991 |

| | | | | | |
|---|---|---|---|---|---|
| 11653 | NM_024039.2 | 20.5629733 | -5.9853642 | 1.3712615 | -4.3648598 |
| 11654 | NM_001162429.2 | 37.9629681 | -6.8678278 | 1.57352985 | -4.3645997 |
| 11655 | NM_019895.2 | 35.8736344 | -6.7886447 | 1.55561505 | -4.3639618 |
| 11656 | NR_126447.1 | 80.5745002 | 3.50088335 | 0.80225248 | 4.36381741 |
| 11657 | NM_001109938.2 | 20.9594138 | -6.0126683 | 1.37785198 | -4.3637984 |
| 11658 | NM_001330260.1 | 35.2249635 | -6.7617651 | 1.5496693 | -4.36336 |
| 11659 | NM_001144823.2 | 38.4170787 | -6.881652 | 1.57716056 | -4.3633173 |
| 11660 | NM_002884.3 | 91.4631096 | -4.353971 | 0.99787514 | -4.3632423 |
| 11661 | NM_032792.3 | 19.462584 | -5.9056703 | 1.3535329 | -4.3631523 |
| 11662 | NM_018502.4 | 21.3954716 | -6.0353269 | 1.38326003 | -4.3631181 |
| 11663 | NM_001122824.1 | 35.4001572 | -4.9260111 | 1.12907513 | -4.3628727 |
| 11664 | NM_018909.3 | 44.1617556 | -7.0857533 | 1.62411991 | -4.3628265 |
| 11665 | NM_015969.2 | 25.2050253 | -6.2778409 | 1.43908546 | -4.3623823 |
| 11666 | NM_014005.4 | 85.4126151 | 1.84304798 | 0.42250205 | 4.36222263 |
| 11667 | NM_001004354.2 | 36.2198802 | -5.8199602 | 1.33419699 | -4.3621446 |
| 11668 | NM_002555.5 | 22.3483462 | -6.1064169 | 1.40002663 | -4.3616434 |
| 11669 | NM_001194998.1 | 16.6832702 | -5.6769246 | 1.30172738 | -4.3610703 |
| 11670 | NM_006828.3 | 125.665855 | -4.6345902 | 1.06277998 | -4.3608181 |
| 11671 | NM_001205294.1 | 138.582299 | -5.5150571 | 1.26475615 | -4.3605695 |
| 11672 | NM_001171168.1 | 23.99086 | -6.2006781 | 1.42240052 | -4.3593053 |
| 11673 | NM_152896.2 | 90.5039731 | -4.9473999 | 1.13491708 | -4.3592611 |
| 11674 | NR_033248.1 | 20.8673766 | -6.0044537 | 1.37767561 | -4.3583944 |
| 11675 | NR_046329.1 | 24.9476636 | -6.265427 | 1.437559 | -4.358379 |
| 11676 | NM_001290295.1 | 25.0735118 | -6.2692935 | 1.43845629 | -4.3583483 |
| 11677 | NM_001270433.1 | 19.3731609 | -5.9058911 | 1.35508906 | -4.3583047 |
| 11678 | NM_001195608.1 | 23.6818867 | -6.182322 | 1.41855491 | -4.3581831 |
| 11679 | NR_027084.1 | 9.90470556 | 4.90686703 | 1.12595974 | 4.35794181 |
| 11680 | NM_052883.1 | 15.8736796 | -5.6143704 | 1.28840849 | -4.3576012 |
| 11681 | NM_001286208.1 | 18.5072319 | -5.8260241 | 1.33738846 | -4.3562691 |
| 11682 | NM_016084.4 | 29.6174821 | -6.508544 | 1.49415136 | -4.3560138 |
| 11683 | NM_001040667.2 | 28.9340126 | -6.4807109 | 1.48798214 | -4.3553687 |
| 11684 | NM_001199013.1 | 23.0348949 | -6.1505105 | 1.41239975 | -4.3546528 |
| 11685 | NM_033331.2 | 28.1409686 | -6.4373643 | 1.47858951 | -4.3537197 |
| 11686 | NM_003260.4 | 31.2440924 | -6.582026 | 1.51186205 | -4.353589 |
| 11687 | NR_123713.1 | 19.7366882 | -5.9229767 | 1.3608258 | -4.352487 |
| 11688 | NM_031207.5 | 23.9659988 | -6.2012901 | 1.42477927 | -4.3524567 |
| 11689 | NM_001301243.1 | 22.6576799 | -6.1174758 | 1.40560407 | -4.3522041 |
| 11690 | NM_001304466.1 | 18.4606433 | -5.8346038 | 1.34076883 | -4.3516852 |
| 11691 | NM_001135570.1 | 60.2651176 | -5.2184113 | 1.19918332 | -4.3516377 |
| 11692 | NM_001321702.1 | 85.3368176 | -8.0369066 | 1.84697888 | -4.3513798 |
| 11693 | NM_001287816.1 | 46.0313571 | -6.1762103 | 1.41942199 | -4.3512151 |
| 11694 | NR_037945.1 | 21.2041617 | -6.0227644 | 1.38425494 | -4.350907 |
| 11695 | NM_004460.4 | 20.2244457 | -5.9657089 | 1.37135808 | -4.3502196 |

| | | | | | |
|---|---|---|---|---|---|
| 11696 | NM_000064.3 | 27.5458682 | -6.4077003 | 1.47304634 | -4.3499652 |
| 11697 | NM_001318736.1 | 21.0912161 | -6.0285373 | 1.38589588 | -4.3499208 |
| 11698 | NM_207322.2 | 64.7777287 | 2.5463569 | 0.58540211 | 4.34975695 |
| 11699 | NM_030961.2 | 61.5192761 | -3.2570421 | 0.74890233 | -4.349088 |
| 11700 | NM_022845.2 | 51.4357013 | -5.5318234 | 1.27200452 | -4.3489023 |
| 11701 | NM_001257281.1 | 20.2977385 | -5.9652437 | 1.37170909 | -4.3487673 |
| 11702 | NM_001040199.1 | 18.0783666 | -5.8004566 | 1.3339452 | -4.348347 |
| 11703 | NM_001796.4 | 38.8968195 | -5.9194004 | 1.36136768 | -4.3481276 |
| 11704 | NM_003946.6 | 21.8181105 | -6.0614589 | 1.39409247 | -4.3479604 |
| 11705 | NM_001037317.1 | 32.7689031 | -6.6555482 | 1.53078416 | -4.3478032 |
| 11706 | NM_001168407.1 | 20.0706109 | -5.9513861 | 1.36889257 | -4.3475918 |
| 11707 | NM_001278438.1 | 24.6492239 | -6.2406248 | 1.43541782 | -4.3476016 |
| 11708 | NM_001308392.1 | 33.3929371 | -6.6813988 | 1.53681011 | -4.347576 |
| 11709 | NM_031217.3 | 19.8402972 | -5.9339729 | 1.36492456 | -4.3474731 |
| 11710 | NM_001166114.1 | 73.9849415 | -3.7712017 | 0.8674813 | -4.3473003 |
| 11711 | NR_134476.1 | 92.7425717 | 1.90102418 | 0.43731238 | 4.34706233 |
| 11712 | NM_001098671.1 | 21.510903 | -6.0452645 | 1.39066234 | -4.3470398 |
| 11713 | NM_001164278.1 | 22.2980322 | -6.1039351 | 1.40424043 | -4.3467877 |
| 11714 | NM_006343.2 | 24.3967229 | -6.2353314 | 1.4345238 | -4.3466211 |
| 11715 | NM_001318915.1 | 27.634096 | -6.4081774 | 1.47430668 | -4.3465701 |
| 11716 | NM_152245.2 | 25.0964329 | -6.2740756 | 1.44349606 | -4.3464446 |
| 11717 | NM_025211.3 | 24.0084858 | -6.2007246 | 1.4267591 | -4.3460207 |
| 11718 | NM_001297754.1 | 18.7721268 | -5.8517214 | 1.34648127 | -4.345936 |
| 11719 | NM_016558.3 | 18.3677302 | -5.8267807 | 1.34077205 | -4.3458399 |
| 11720 | NR_033519.1 | 27.7170137 | -6.4101379 | 1.47502629 | -4.3457787 |
| 11721 | NM_020528.2 | 26.7479036 | -6.3645318 | 1.46454527 | -4.3457392 |
| 11722 | NR_003290.2 | 16.8161161 | -5.6905847 | 1.30961176 | -4.3452456 |
| 11723 | NR_024417.1 | 28.0113984 | 2.01616378 | 0.46401906 | 4.3450021 |
| 11724 | NM_001301871.1 | 28.7733953 | -6.4703544 | 1.48921241 | -4.3448164 |
| 11725 | NM_014301.4 | 20.1188931 | -5.9430424 | 1.36788157 | -4.3447054 |
| 11726 | NM_001198535.1 | 29.435848 | -6.4993475 | 1.49600123 | -4.3444801 |
| 11727 | NM_016544.2 | 18.8190474 | -5.8608074 | 1.34905714 | -4.344373 |
| 11728 | NM_001286659.2 | 44.5205257 | -5.2317579 | 1.20456513 | -4.3432752 |
| 11729 | NM_024787.3 | 62.5288564 | -4.5493826 | 1.04752228 | -4.3429936 |
| 11730 | NM_021963.3 | 77.3998044 | -4.6214022 | 1.06413602 | -4.342868 |
| 11731 | NM_025112.4 | 28.9301353 | -5.4950028 | 1.26549638 | -4.3421719 |
| 11732 | NM_003202.4 | 25.4568613 | -6.2959278 | 1.45014074 | -4.3415978 |
| 11733 | NM_199298.1 | 26.0329762 | -6.3224254 | 1.45632572 | -4.3413539 |
| 11734 | NR_045620.1 | 16.2312898 | -5.6471498 | 1.3009611 | -4.3407522 |
| 11735 | NM_005670.3 | 23.301632 | -6.1634561 | 1.41999877 | -4.3404658 |
| 11736 | NM_001201480.1 | 22.9578028 | -6.1458536 | 1.41601338 | -4.3402511 |
| 11737 | NM_001172303.2 | 25.4841382 | -6.292742 | 1.45028057 | -4.3389825 |
| 11738 | NM_001271581.1 | 40.7006249 | -5.1513448 | 1.18723528 | -4.3389418 |

| | | | | | |
|---|---|---|---|---|---|
| 11739 | NM_016281.3 | 21.0669275 | -6.0121203 | 1.38567475 | -4.3387673 |
| 11740 | NM_001105207.2 | 29.7884464 | -6.5192661 | 1.502623 | -4.3385907 |
| 11741 | NM_000245.3 | 25.5104652 | -6.2960817 | 1.45126586 | -4.3383379 |
| 11742 | NM_032515.4 | 95.0246515 | -4.5259523 | 1.04331671 | -4.3380426 |
| 11743 | NM_001346445.1 | 290.379678 | -5.4227989 | 1.25006356 | -4.3380186 |
| 11744 | NM_001008223.1 | 25.622554 | -6.3035674 | 1.45313587 | -4.3379064 |
| 11745 | NM_002133.2 | 59.7402962 | -6.5552385 | 1.51121263 | -4.337734 |
| 11746 | NM_033363.1 | 22.0547201 | -6.0886629 | 1.40380571 | -4.3372547 |
| 11747 | NR_033255.1 | 19.418569 | -5.8941536 | 1.35907142 | -4.3368976 |
| 11748 | NM_001101672.1 | 20.200106 | -5.9590308 | 1.37431766 | -4.3359923 |
| 11749 | NM_005895.3 | 505.017991 | -2.6225801 | 0.60497845 | -4.3349976 |
| 11750 | NM_198321.3 | 222.115021 | -3.8764689 | 0.8943065 | -4.3346089 |
| 11751 | NM_176880.5 | 25.3057161 | -6.2819914 | 1.44954774 | -4.3337596 |
| 11752 | NM_004626.2 | 18.7342065 | -5.8565723 | 1.35150694 | -4.3333646 |
| 11753 | NM_058216.2 | 21.6562266 | -6.0567649 | 1.39774614 | -4.3332367 |
| 11754 | NR_038244.1 | 15.2617398 | -5.5505873 | 1.28098161 | -4.3330734 |
| 11755 | NM_001011724.1 | 22.1919637 | -6.0900294 | 1.40564594 | -4.3325486 |
| 11756 | NM_001303269.1 | 32.0780131 | -5.6383973 | 1.30140011 | -4.3325625 |
| 11757 | NM_001008727.3 | 16.2833493 | -5.6404424 | 1.30208489 | -4.3318546 |
| 11758 | NM_001083607.1 | 20.794686 | -5.9913097 | 1.38317348 | -4.3315678 |
| 11759 | NM_005073.3 | 71.4222309 | 2.6152142 | 0.60375781 | 4.3315617 |
| 11760 | NM_133503.3 | 34.0712191 | -6.7156547 | 1.55059447 | -4.3310194 |
| 11761 | NM_001193482.1 | 64.2920329 | 10.9284672 | 2.52346768 | 4.33073396 |
| 11762 | NM_005664.3 | 19.7936427 | -5.9242739 | 1.36804099 | -4.3304798 |
| 11763 | NM_001270523.1 | 48.8244558 | -7.2332808 | 1.67036841 | -4.3303505 |
| 11764 | NM_001320147.1 | 176.144309 | -5.6609859 | 1.30729195 | -4.330315 |
| 11765 | NM_032810.3 | 96.3764121 | -4.9614144 | 1.1457455 | -4.3302936 |
| 11766 | NM_001126108.1 | 39.4495624 | 4.67676009 | 1.08010853 | 4.32989828 |
| 11767 | NM_006087.3 | 80.8260611 | -5.6711191 | 1.30977473 | -4.3298432 |
| 11768 | NM_032941.2 | 33.4836604 | -4.8746549 | 1.12603477 | -4.3290447 |
| 11769 | NM_017737.4 | 495.799515 | -5.2663754 | 1.21668691 | -4.3284557 |
| 11770 | NM_175040.3 | 17.4695682 | -5.7438643 | 1.32728463 | -4.3275302 |
| 11771 | NM_020800.2 | 35.0677152 | -6.7578411 | 1.56161697 | -4.3274639 |
| 11772 | NM_001166357.1 | 34.2114756 | -6.7176091 | 1.55243241 | -4.3271508 |
| 11773 | NM_003269.4 | 22.7991439 | -6.1279451 | 1.41619004 | -4.3270641 |
| 11774 | NM_001287429.1 | 30.5046694 | -6.5547932 | 1.51506025 | -4.3264241 |
| 11775 | NM_014329.4 | 338.404864 | -5.3726559 | 1.24201708 | -4.3257504 |
| 11776 | NR_024247.1 | 25.4586282 | -6.2909005 | 1.45438221 | -4.3254796 |
| 11777 | NM_032017.2 | 20.2959758 | -5.9672911 | 1.37976474 | -4.3248613 |
| 11778 | NR_103729.1 | 33.3244768 | -6.6766006 | 1.54378038 | -4.3248384 |
| 11779 | NM_001330663.1 | 18.5954017 | -5.8375953 | 1.34998514 | -4.3241922 |
| 11780 | NM_004278.3 | 16.449579 | -5.6700437 | 1.31126841 | -4.3240908 |
| 11781 | NM_017837.3 | 15.2649032 | -5.5555741 | 1.28488494 | -4.3237911 |

| | | | | | |
|---|---|---|---|---|---|
| 11782 | NM_005139.2 | 17.2765042 | -5.7403241 | 1.32763016 | -4.3237374 |
| 11783 | NM_001304947.1 | 18.3100642 | -5.8170102 | 1.34552836 | -4.3232163 |
| 11784 | NM_002065.6 | 22.8641773 | -6.1420017 | 1.4208086 | -4.3228917 |
| 11785 | NM_001300797.1 | 21.4458446 | -6.0470188 | 1.39911545 | -4.3220299 |
| 11786 | NM_052865.3 | 25.4211406 | -6.2894511 | 1.4552414 | -4.3219298 |
| 11787 | NM_022342.4 | 16.7466023 | -5.6970896 | 1.31843744 | -4.3210921 |
| 11788 | NM_001330763.1 | 19.9399699 | -5.9295203 | 1.37232258 | -4.3207919 |
| 11789 | NM_006567.4 | 17.951463 | -5.7864778 | 1.33921705 | -4.3207917 |
| 11790 | NM_007243.2 | 15.891447 | -5.6119918 | 1.29894061 | -4.3204375 |
| 11791 | NM_201650.2 | 23.771754 | -6.1959317 | 1.43411092 | -4.3203992 |
| 11792 | NM_024969.3 | 59.3371113 | -6.5414928 | 1.51419364 | -4.3201165 |
| 11793 | NM_015012.3 | 174.450278 | -2.3790431 | 0.55071951 | -4.3198817 |
| 11794 | NM_000816.3 | 40.6043396 | -6.9648716 | 1.61229963 | -4.319837 |
| 11795 | NM_013280.4 | 32.1419074 | -4.2348133 | 0.98043073 | -4.3193397 |
| 11796 | NM_001276336.1 | 37.5900473 | -6.8495289 | 1.58598776 | -4.3187779 |
| 11797 | NM_006983.1 | 15.6469021 | -5.5895875 | 1.29432386 | -4.3185386 |
| 11798 | NM_000329.2 | 28.5939412 | -6.4636422 | 1.49684037 | -4.3181907 |
| 11799 | NM_001320925.2 | 23.4945341 | -6.1714923 | 1.42921828 | -4.3180894 |
| 11800 | NM_015193.4 | 39.7080203 | -6.9316197 | 1.6055616 | -4.3172556 |
| 11801 | NR_033328.1 | 28.6787836 | -6.4625934 | 1.4969155 | -4.3172733 |
| 11802 | NM_001278346.1 | 18.0039563 | -5.790131 | 1.34141892 | -4.3164226 |
| 11803 | NR_026680.3 | 23.0973438 | -6.1506013 | 1.42500492 | -4.3161965 |
| 11804 | NM_014720.3 | 16.8439335 | -5.6878811 | 1.31788257 | -4.3159241 |
| 11805 | NM_014369.3 | 46.0439837 | -6.1724868 | 1.43030785 | -4.3154953 |
| 11806 | NM_000411.6 | 23.5444345 | -6.1833301 | 1.43283021 | -4.315466 |
| 11807 | NM_181468.2 | 20.1947954 | -5.9634014 | 1.38190293 | -4.3153548 |
| 11808 | NM_003834.2 | 21.5736268 | -6.045782 | 1.40116508 | -4.3148249 |
| 11809 | NM_172229.2 | 18.9948054 | -5.8625355 | 1.35870268 | -4.3148038 |
| 11810 | NM_005371.5 | 20.3231202 | -5.9692835 | 1.38364064 | -4.3141863 |
| 11811 | NR_103446.1 | 17.5248078 | -5.7576605 | 1.33459071 | -4.3141769 |
| 11812 | NM_001006621.1 | 30.2523404 | -6.5397389 | 1.51595501 | -4.31394 |
| 11813 | NM_001129996.1 | 19.080178 | -5.8737025 | 1.36171089 | -4.3134725 |
| 11814 | NM_001429.3 | 277.014838 | -5.1600313 | 1.19642675 | -4.3128686 |
| 11815 | NM_001025593.2 | 22.1843367 | -6.0844803 | 1.41081045 | -4.3127554 |
| 11816 | NM_032367.3 | 15.1164275 | -5.5377397 | 1.28420985 | -4.3121766 |
| 11817 | NR_103833.1 | 18.0596163 | -5.8018471 | 1.34557584 | -4.3117949 |
| 11818 | NM_001276294.1 | 24.2321304 | -6.2273669 | 1.44433714 | -4.3115743 |
| 11819 | NM_015697.7 | 22.8886422 | -6.1361775 | 1.42344741 | -4.3107862 |
| 11820 | NM_001145642.4 | 55.3277322 | -7.4122913 | 1.71976764 | -4.3100539 |
| 11821 | NM_001261430.1 | 16.4686944 | -5.6560257 | 1.31238895 | -4.3097175 |
| 11822 | NM_005541.4 | 35.0224629 | -6.7550767 | 1.56747734 | -4.3095211 |
| 11823 | NM_001142425.1 | 32.8712143 | -6.6642172 | 1.54653405 | -4.3091306 |
| 11824 | NM_001943.4 | 112.599975 | -2.265824 | 0.52581993 | -4.3091255 |

| | | | | | |
|---|---|---|---|---|---|
| 11825 | NM_001278214.1 | 42.5659451 | -7.0310243 | 1.63167137 | -4.3090934 |
| 11826 | NM_003781.3 | 24.1275842 | -6.2105186 | 1.44128832 | -4.309005 |
| 11827 | NM_018393.3 | 29.3565105 | -6.4946461 | 1.50779265 | -4.3073868 |
| 11828 | NM_033211.3 | 79.7253744 | -5.2299553 | 1.21431113 | -4.3069319 |
| 11829 | NM_001184991.1 | 26.1471667 | -6.3272225 | 1.46914271 | -4.3067447 |
| 11830 | NR_136669.1 | 20.3602316 | -5.9618115 | 1.38443777 | -4.3063052 |
| 11831 | NM_001029882.3 | 232.377315 | -2.8804146 | 0.6688895 | -4.3062638 |
| 11832 | NM_201266.1 | 31.9867292 | -6.6232001 | 1.53827481 | -4.3056026 |
| 11833 | NM_144671.4 | 43.6013139 | -7.067861 | 1.64162223 | -4.3054126 |
| 11834 | NR_036641.1 | 24.785993 | -6.2590465 | 1.45377637 | -4.3053709 |
| 11835 | NM_014675.4 | 212.575895 | -4.9534148 | 1.15075392 | -4.3044953 |
| 11836 | NM_001253875.1 | 41.8252709 | -7.0039741 | 1.62733617 | -4.3039504 |
| 11837 | NM_033132.3 | 38.5074042 | -6.8913814 | 1.60140173 | -4.3033433 |
| 11838 | NM_001284281.1 | 32.0059108 | -5.6419239 | 1.31125467 | -4.3026911 |
| 11839 | NM_001301016.2 | 18.8636942 | -5.8657316 | 1.36327521 | -4.302676 |
| 11840 | NM_004983.2 | 36.1777126 | -6.7966986 | 1.57965209 | -4.3026554 |
| 11841 | NM_014883.3 | 39.0208912 | -5.1081978 | 1.18722865 | -4.3026234 |
| 11842 | NR_037868.1 | 150.22486 | 1.88232836 | 0.43752809 | 4.30218856 |
| 11843 | NM_001319106.1 | 31.1584068 | -6.5835427 | 1.53028512 | -4.3021674 |
| 11844 | NM_001146071.1 | 16.6582467 | -5.6717793 | 1.31872702 | -4.3009503 |
| 11845 | NM_032878.3 | 19.1891441 | -5.8749559 | 1.36628834 | -4.2999386 |
| 11846 | NR_038367.1 | 43.0882078 | -6.078156 | 1.41362007 | -4.2997097 |
| 11847 | NM_001242485.1 | 22.028627 | -6.0787124 | 1.41379748 | -4.2995638 |
| 11848 | NM_018347.2 | 19.7438697 | -5.9332561 | 1.37999087 | -4.2994894 |
| 11849 | NM_001031694.2 | 17.4993122 | -5.7590884 | 1.33951182 | -4.2993935 |
| 11850 | NM_001293212.1 | 22.504272 | -6.1085312 | 1.42093379 | -4.2989555 |
| 11851 | NM_002606.2 | 22.8845797 | -6.1314821 | 1.4263481 | -4.2987277 |
| 11852 | NM_004203.4 | 25.2188844 | -6.2752795 | 1.45994839 | -4.2982886 |
| 11853 | NM_019852.4 | 116.888837 | -4.5255554 | 1.05287869 | -4.2982686 |
| 11854 | NM_015676.1 | 21.1884549 | -6.024882 | 1.40176423 | -4.2980709 |
| 11855 | NM_202001.2 | 49.2825838 | -3.83423 | 0.89217107 | -4.2976399 |
| 11856 | NM_017570.4 | 31.7659296 | -4.2180672 | 0.98155514 | -4.2973308 |
| 11857 | NM_145061.5 | 17.8527 | -5.7770298 | 1.34432566 | -4.297344 |
| 11858 | NM_003355.2 | 80.1045193 | -4.2606598 | 0.9915011 | -4.2971811 |
| 11859 | NM_138447.2 | 60.9084708 | -4.8738913 | 1.13422921 | -4.2970956 |
| 11860 | NM_024079.4 | 40.0149135 | -5.9603942 | 1.3871432 | -4.2968846 |
| 11861 | NM_001348137.1 | 48.6663802 | -7.2281337 | 1.68242539 | -4.2962581 |
| 11862 | NM_032435.2 | 14.8425852 | -5.5116534 | 1.28289892 | -4.2962491 |
| 11863 | NM_021632.3 | 18.9347345 | -5.8636048 | 1.36495635 | -4.2958185 |
| 11864 | NM_020153.3 | 15.8329788 | -5.6117756 | 1.30649062 | -4.295305 |
| 11865 | NM_001040146.3 | 21.362842 | -6.0439083 | 1.40723479 | -4.2948827 |
| 11866 | NM_004282.3 | 18.3337492 | -5.8206693 | 1.35537282 | -4.2945153 |
| 11867 | NR_026678.1 | 18.6179473 | -5.8410569 | 1.36021089 | -4.2942289 |

| | | | | | |
|---|---|---|---|---|---|
| 11868 | NM_001709.4 | 22.5040948 | -6.1133263 | 1.42362241 | -4.2942048 |
| 11869 | NR_036556.1 | 21.4547888 | -6.0528488 | 1.40959198 | -4.2940432 |
| 11870 | NM_002123.4 | 26.9626048 | -6.3765479 | 1.48518813 | -4.2934277 |
| 11871 | NM_000073.2 | 26.9854133 | 2.33579618 | 0.5440755 | 4.29314714 |
| 11872 | NR_110204.1 | 17.5013621 | -5.7515299 | 1.33972409 | -4.2930704 |
| 11873 | NM_173174.2 | 18.0250561 | -5.7875551 | 1.34831391 | -4.2924389 |
| 11874 | NM_001321351.1 | 31.394388 | -6.5914105 | 1.53581312 | -4.2918051 |
| 11875 | NM_001100595.1 | 46.9922448 | -4.3482627 | 1.0131836 | -4.2916828 |
| 11876 | NM_138436.3 | 22.9300919 | -6.1391472 | 1.43047925 | -4.2916716 |
| 11877 | NR_038863.2 | 27.3058798 | 4.05691802 | 0.9453015 | 4.29166569 |
| 11878 | NM_005657.2 | 45.2258906 | -7.1205026 | 1.65928417 | -4.2913099 |
| 11879 | NM_001244606.1 | 18.7233304 | -5.8494277 | 1.36315341 | -4.2911001 |
| 11880 | NM_020730.2 | 17.0509435 | -5.7198142 | 1.33313854 | -4.2904875 |
| 11881 | NM_014822.3 | 93.0806871 | -4.6890336 | 1.09291121 | -4.2904068 |
| 11882 | NR_134281.1 | 67.5370503 | 2.76885423 | 0.64542722 | 4.28995576 |
| 11883 | NM_033495.3 | 27.0662251 | -6.3839019 | 1.48831897 | -4.2893372 |
| 11884 | NR_029377.1 | 26.4955665 | -6.3456612 | 1.4794244 | -4.2892771 |
| 11885 | NM_001330512.1 | 17.5555956 | -5.7627402 | 1.34369906 | -4.2887134 |
| 11886 | NM_002874.4 | 528.227834 | -5.9829843 | 1.39506174 | -4.2886878 |
| 11887 | NM_032470.3 | 15.6399801 | -5.5818317 | 1.30159431 | -4.2884574 |
| 11888 | NM_001037536.2 | 27.7500136 | -6.4181241 | 1.49662078 | -4.2884104 |
| 11889 | NM_001100594.1 | 45.6066175 | -4.7866591 | 1.11622201 | -4.2882679 |
| 11890 | NM_001271420.1 | 15.4431573 | -5.5713173 | 1.29921487 | -4.2882185 |
| 11891 | NM_013259.2 | 28.6803979 | -6.4635223 | 1.50745144 | -4.2877151 |
| 11892 | NM_001130085.1 | 22.1213838 | -6.0915671 | 1.42075206 | -4.2875652 |
| 11893 | NM_145860.1 | 20.7294786 | -5.9958536 | 1.39845227 | -4.2874925 |
| 11894 | NM_004270.4 | 21.9451087 | -6.0768176 | 1.41746209 | -4.2871112 |
| 11895 | NM_033020.2 | 109.031284 | -4.7408543 | 1.10587125 | -4.2869858 |
| 11896 | NM_001011546.1 | 26.3878476 | -6.3439885 | 1.47995881 | -4.286598 |
| 11897 | NM_001005242.2 | 24.1710382 | -6.2159406 | 1.45038543 | -4.2857164 |
| 11898 | NM_021965.3 | 25.9004525 | -6.3215881 | 1.47536809 | -4.2847532 |
| 11899 | NM_001271948.1 | 16.4684855 | -5.6685696 | 1.32304015 | -4.2845031 |
| 11900 | NM_006662.2 | 314.291035 | -4.0829178 | 0.95301186 | -4.2842255 |
| 11901 | NM_002306.3 | 15.1181441 | -5.5432531 | 1.29390053 | -4.2841416 |
| 11902 | NM_001199878.1 | 16.3454746 | -5.662176 | 1.32178329 | -4.2837401 |
| 11903 | NM_152259.3 | 23.5701936 | -6.1751062 | 1.44151947 | -4.283748 |
| 11904 | NM_000055.3 | 20.8333585 | -6.0011158 | 1.40101181 | -4.2834156 |
| 11905 | NM_001144035.1 | 20.3463896 | -5.970397 | 1.39394053 | -4.2831074 |
| 11906 | NR_119375.1 | 51.6537635 | 3.53494868 | 0.82533286 | 4.28305823 |
| 11907 | NM_001197122.1 | 21.134484 | -6.0304688 | 1.40809043 | -4.2827283 |
| 11908 | NM_001080383.1 | 174.053109 | 7.75642067 | 1.81132361 | 4.28218382 |
| 11909 | NR_134875.1 | 22.5896082 | -6.1224009 | 1.42975176 | -4.2821426 |
| 11910 | NM_001136039.2 | 19.5885285 | -5.9090346 | 1.37997525 | -4.281986 |

| | | | | | |
|---|---|---|---|---|---|
| 11911 | NM_001030059.2 | 16.4931385 | -5.6717097 | 1.32456221 | -4.2819504 |
| 11912 | NM_001127898.3 | 23.5980623 | -6.1789873 | 1.44314851 | -4.2816019 |
| 11913 | NM_001013635.3 | 229.048666 | -5.4143076 | 1.26463121 | -4.2813332 |
| 11914 | NM_032564.4 | 20.1103041 | -5.9500503 | 1.38976529 | -4.2813347 |
| 11915 | NM_001198670.1 | 20.7586752 | -5.9885488 | 1.39899292 | -4.2806141 |
| 11916 | NM_018420.2 | 19.6869263 | -5.9254118 | 1.38426593 | -4.2805444 |
| 11917 | NM_001318746.1 | 7.53317653 | 7.80933151 | 1.82474991 | 4.27967222 |
| 11918 | NM_001258373.1 | 24.7128607 | -6.2532149 | 1.46118455 | -4.2795518 |
| 11919 | NM_013372.6 | 26.5427439 | -6.3530671 | 1.48455802 | -4.2794333 |
| 11920 | NM_014030.3 | 71.6854505 | -4.5070486 | 1.05325536 | -4.2791603 |
| 11921 | NM_012421.3 | 134.590218 | -5.1259285 | 1.19793673 | -4.2789643 |
| 11922 | NM_001204159.2 | 32.2293193 | -6.6351324 | 1.55090783 | -4.2782249 |
| 11923 | NM_001146156.1 | 277.116177 | -5.354186 | 1.25174729 | -4.2773698 |
| 11924 | NM_001304476.1 | 19.6331736 | -5.9097978 | 1.38168399 | -4.2772427 |
| 11925 | NM_001276723.1 | 20.9014151 | -6.0026643 | 1.40353915 | -4.2768057 |
| 11926 | NM_005879.2 | 19.2994154 | -5.889088 | 1.3770063 | -4.2767328 |
| 11927 | NR_033902.1 | 435.323684 | -4.4816091 | 1.0479314 | -4.2766245 |
| 11928 | NM_023002.2 | 15.4882578 | -5.5737754 | 1.30346058 | -4.2761365 |
| 11929 | NM_001256053.1 | 50.3641875 | -5.2861771 | 1.23627976 | -4.2758745 |
| 11930 | NM_001201397.1 | 20.2214956 | -5.9629097 | 1.39476419 | -4.2752099 |
| 11931 | NM_001145065.1 | 20.4638889 | -5.9711231 | 1.39673831 | -4.2750478 |
| 11932 | NR_003286.2 | 7295.21794 | -6.0207334 | 1.40834247 | -4.2750493 |
| 11933 | NM_001164465.3 | 14.4074875 | -5.4686667 | 1.27939621 | -4.2744121 |
| 11934 | NM_005187.5 | 18.9330749 | -5.8630611 | 1.3716634 | -4.2744168 |
| 11935 | NM_153200.2 | 19.6404906 | -5.9187491 | 1.38468852 | -4.2744264 |
| 11936 | NM_001258276.1 | 21.3016321 | -6.0427347 | 1.41396025 | -4.2736242 |
| 11937 | NR_120526.1 | 322.510912 | 4.61218762 | 1.07923269 | 4.27358037 |
| 11938 | NM_001243795.1 | 19.0131381 | -5.87053 | 1.37380273 | -4.2731972 |
| 11939 | NM_001319957.1 | 157.434187 | -4.6878672 | 1.09704471 | -4.2731779 |
| 11940 | NM_001143960.2 | 17.5594755 | -5.7535251 | 1.34650535 | -4.2729314 |
| 11941 | NM_001099334.2 | 30.9464905 | -6.5726466 | 1.53825983 | -4.2727805 |
| 11942 | NM_001117.4 | 20.6086113 | -5.9867777 | 1.40118486 | -4.2726537 |
| 11943 | NR_003227.1 | 28.5121384 | -6.4535079 | 1.51044594 | -4.2725845 |
| 11944 | NM_130901.2 | 40.5559209 | -5.171893 | 1.21050288 | -4.2725161 |
| 11945 | NM_001164638.1 | 17.8636737 | -5.7826472 | 1.35355411 | -4.2721951 |
| 11946 | NR_002786.1 | 22.3833924 | -6.1055571 | 1.42923129 | -4.2719168 |
| 11947 | NM_145267.2 | 19.7670813 | -5.9275138 | 1.38757805 | -4.2718417 |
| 11948 | NM_175732.2 | 60.0702827 | -4.0026571 | 0.93724858 | -4.2706462 |
| 11949 | NM_001003676.2 | 17.5553384 | -5.751589 | 1.34687879 | -4.2703093 |
| 11950 | NM_001005368.2 | 15.3901015 | -5.5596157 | 1.30191852 | -4.2703254 |
| 11951 | NM_032440.3 | 45.524017 | -7.1318935 | 1.67010157 | -4.2703352 |
| 11952 | NM_001271886.1 | 22.8213264 | -6.1323992 | 1.43610364 | -4.2701647 |
| 11953 | NM_005307.2 | 20.4948771 | -5.9750311 | 1.39938576 | -4.2697527 |

| | | | | | |
|---|---|---|---|---|---|
| 11954 | NM_138476.3 | 19.9427928 | -5.9409253 | 1.39140739 | -4.2697239 |
| 11955 | NM_001289861.1 | 24.5019601 | -6.2346354 | 1.46038 | -4.2691871 |
| 11956 | NM_001193421.1 | 30.3657344 | -5.5507768 | 1.3003463 | -4.2686912 |
| 11957 | NM_004455.2 | 14.4772066 | -5.4769383 | 1.28327374 | -4.2679423 |
| 11958 | NM_021830.4 | 37.2877319 | -5.0476005 | 1.18272161 | -4.2677841 |
| 11959 | NM_030907.3 | 16.0577362 | -5.6304417 | 1.31932782 | -4.2676594 |
| 11960 | NM_138330.2 | 37.7157239 | -5.8813944 | 1.37821438 | -4.2674017 |
| 11961 | NM_001130102.2 | 16.964253 | -5.7064257 | 1.33744718 | -4.266655 |
| 11962 | NM_001278280.1 | 28.649154 | -6.4593148 | 1.51405111 | -4.2662462 |
| 11963 | NM_001288666.1 | 31.0676216 | -6.5810713 | 1.54261273 | -4.266185 |
| 11964 | NM_198391.2 | 17.3360651 | -5.743149 | 1.34622216 | -4.2661228 |
| 11965 | NM_014641.2 | 165.541645 | -5.1269029 | 1.2019462 | -4.2655011 |
| 11966 | NM_000081.3 | 66.6293504 | -3.89582 | 0.91335093 | -4.2654141 |
| 11967 | NM_181833.2 | 62.4377167 | 10.8859632 | 2.55215131 | 4.26540666 |
| 11968 | NM_022145.4 | 18.5108672 | -5.8384313 | 1.36880056 | -4.2653631 |
| 11969 | NM_138770.2 | 19.4039179 | -5.905672 | 1.38457002 | -4.2653473 |
| 11970 | NR_033935.1 | 18.8542374 | -5.8611188 | 1.37419287 | -4.2651355 |
| 11971 | NM_080422.2 | 19.2615616 | -5.8876549 | 1.38046675 | -4.2649741 |
| 11972 | NM_001134423.2 | 29.3548412 | -6.5011224 | 1.52434331 | -4.2648676 |
| 11973 | NM_001289095.1 | 14.5769831 | -5.4891178 | 1.28708792 | -4.2647575 |
| 11974 | NM_152832.2 | 30.2844019 | -6.542331 | 1.53409551 | -4.2646178 |
| 11975 | NR_135043.1 | 25.3616905 | -6.2861019 | 1.47404163 | -4.2645349 |
| 11976 | NM_052837.2 | 23.2811701 | -6.1659662 | 1.44597288 | -4.2642336 |
| 11977 | NM_005227.2 | 18.0277252 | -5.7933312 | 1.35861922 | -4.2641317 |
| 11978 | NM_001128205.1 | 42.2203891 | -5.2374028 | 1.22827887 | -4.2640177 |
| 11979 | NM_001301076.1 | 20.3093559 | -5.969848 | 1.40010473 | -4.2638581 |
| 11980 | NM_001166158.1 | 40.7798024 | 4.3679089 | 1.02456113 | 4.26319989 |
| 11981 | NR_109790.1 | 31.7812078 | -6.6084425 | 1.55012632 | -4.2631638 |
| 11982 | NM_006800.3 | 15.1696288 | -5.5494399 | 1.30172859 | -4.2631313 |
| 11983 | NR_040038.1 | 68.6179803 | 2.30807417 | 0.54152457 | 4.26217809 |
| 11984 | NM_001170881.1 | 25.6358173 | -6.3058982 | 1.4796193 | -4.2618383 |
| 11985 | NM_000689.4 | 29.8120708 | -6.5226231 | 1.53048163 | -4.2618108 |
| 11986 | NM_178011.4 | 28.1740576 | -6.438912 | 1.51087434 | -4.2617125 |
| 11987 | NM_001271849.1 | 15.0911822 | -5.5453895 | 1.30136344 | -4.2612151 |
| 11988 | NM_001242575.2 | 62.0022634 | 2.49356484 | 0.5852658 | 4.2605682 |
| 11989 | NM_001289978.1 | 31.219245 | -6.5851762 | 1.54581758 | -4.2599956 |
| 11990 | NM_031891.3 | 30.978179 | -5.5968198 | 1.31380851 | -4.2599966 |
| 11991 | NM_002558.3 | 16.5192469 | 7.45712329 | 1.75055217 | 4.25986921 |
| 11992 | NM_001277313.1 | 15.6325996 | -5.596635 | 1.31383149 | -4.2597815 |
| 11993 | NM_018846.4 | 22.9179763 | -6.1376441 | 1.44085253 | -4.259731 |
| 11994 | NM_153603.3 | 73.3846898 | -4.2032627 | 0.98689651 | -4.2590714 |
| 11995 | NM_001198534.1 | 31.3679606 | -6.5911133 | 1.54760647 | -4.2589078 |
| 11996 | NM_001286259.1 | 14.6402873 | -5.4917201 | 1.28954773 | -4.2586404 |

| | | | | | |
|---|---|---|---|---|---|
| 11997 | NM_014323.2 | 15.4401704 | -5.5673855 | 1.30735964 | -4.2584958 |
| 11998 | NM_004936.3 | 39.0184678 | -5.9358305 | 1.39401011 | -4.2580972 |
| 11999 | NM_199037.4 | 90.3106363 | 2.00266184 | 0.47032391 | 4.25804816 |
| 12000 | NM_152356.3 | 30.3783018 | -2.8350048 | 0.66582989 | -4.2578516 |
| 12001 | NM_001350.4 | 23.9358876 | -6.1973806 | 1.45565127 | -4.2574624 |
| 12002 | NM_002350.3 | 16.7141077 | -5.6940125 | 1.33778881 | -4.2562865 |
| 12003 | NR_103834.1 | 24.8872839 | -6.2543079 | 1.46952316 | -4.2560118 |
| 12004 | NM_001278913.1 | 29.2213105 | -6.4940728 | 1.525991 | -4.2556429 |
| 12005 | NM_006417.4 | 25.2607512 | -6.2834652 | 1.47659225 | -4.2553827 |
| 12006 | NM_001238.3 | 16.598744 | -5.6680874 | 1.33210676 | -4.2549799 |
| 12007 | NM_001306188.1 | 15.3069943 | -5.5645169 | 1.30778385 | -4.2549209 |
| 12008 | NM_020734.3 | 30.4920756 | -6.5529377 | 1.54009683 | -4.2548868 |
| 12009 | NM_022823.2 | 115.100008 | -4.8260056 | 1.13422163 | -4.2549053 |
| 12010 | NM_138690.2 | 27.8360201 | -6.4248782 | 1.51011674 | -4.2545573 |
| 12011 | NM_153018.2 | 21.1366004 | -6.021157 | 1.41542895 | -4.2539451 |
| 12012 | NM_152836.2 | 15.7803836 | -5.5928719 | 1.31514312 | -4.2526717 |
| 12013 | NM_001330354.1 | 16.7278034 | -5.6799819 | 1.33573118 | -4.2523391 |
| 12014 | NR_026641.1 | 19.2699505 | -5.8870297 | 1.38451388 | -4.2520554 |
| 12015 | NR_003551.1 | 15.4259081 | -5.5600816 | 1.3076353 | -4.2520125 |
| 12016 | NM_001284206.1 | 28.5299253 | -6.4531059 | 1.51774343 | -4.2517766 |
| 12017 | NM_013286.4 | 364.890877 | -4.0641702 | 0.95588209 | -4.2517485 |
| 12018 | NM_014999.3 | 243.024254 | -2.768567 | 0.65125033 | -4.2511565 |
| 12019 | NR_046347.1 | 18.805698 | -5.8618556 | 1.37908351 | -4.2505443 |
| 12020 | NM_001303506.1 | 15.3995238 | -5.5585653 | 1.30775811 | -4.2504537 |
| 12021 | NM_024949.5 | 88.970966 | -4.2563226 | 1.00138492 | -4.2504361 |
| 12022 | NM_003603.6 | 22.8244888 | -6.1285964 | 1.44210109 | -4.2497689 |
| 12023 | NM_003902.4 | 172.191481 | -4.8580844 | 1.14316742 | -4.2496701 |
| 12024 | NM_182513.3 | 180.446796 | 1.56926072 | 0.36928901 | 4.24941083 |
| 12025 | NM_001146218.1 | 20.4933932 | -5.9809428 | 1.40755687 | -4.249166 |
| 12026 | NM_001135674.1 | 19.9059109 | -5.9311823 | 1.39585371 | -4.2491432 |
| 12027 | NM_144964.3 | 25.0037295 | -6.266415 | 1.47476521 | -4.2490934 |
| 12028 | NM_031922.4 | 39.6319296 | -6.9260599 | 1.63002998 | -4.2490383 |
| 12029 | NM_001146698.1 | 23.3155013 | -6.1687067 | 1.45213875 | -4.2480147 |
| 12030 | NM_020805.2 | 32.5197149 | -6.6422286 | 1.56377488 | -4.2475606 |
| 12031 | NR_049755.1 | 20.0478921 | -5.9478516 | 1.40046713 | -4.2470483 |
| 12032 | NM_002338.4 | 230.04953 | -2.6807895 | 0.63123853 | -4.2468725 |
| 12033 | NM_207435.2 | 65.3098311 | -3.1662646 | 0.74555009 | -4.2468837 |
| 12034 | NM_153347.2 | 15.7766531 | -5.601657 | 1.31915382 | -4.2464017 |
| 12035 | NM_001287192.1 | 105.306554 | -5.2414637 | 1.23438586 | -4.2462117 |
| 12036 | NM_018273.3 | 17.1966857 | -5.7251919 | 1.3483143 | -4.246185 |
| 12037 | NM_001277200.1 | 18.1775193 | -5.8082725 | 1.36790051 | -4.2461221 |
| 12038 | NM_173593.3 | 15.3469155 | -5.5632597 | 1.31028852 | -4.245828 |
| 12039 | NM_018310.3 | 61.6885665 | -4.2707928 | 1.00588427 | -4.2458093 |

| | | | | | |
|---|---|---|---|---|---|
| 12040 | NM_001164095.2 | 17.2386906 | -5.7244303 | 1.34826775 | -4.2457667 |
| 12041 | NM_001258040.2 | 18.5646434 | -5.8422795 | 1.37617917 | -4.2452899 |
| 12042 | NM_002160.3 | 704.737391 | -4.8636266 | 1.14581124 | -4.2447015 |
| 12043 | NM_001290307.2 | 16.3163682 | -5.659808 | 1.33350595 | -4.2443066 |
| 12044 | NR_003370.2 | 16.9206104 | -5.702337 | 1.3436668 | -4.2438624 |
| 12045 | NM_001288984.1 | 30.2422143 | -6.5407968 | 1.54132786 | -4.2436116 |
| 12046 | NM_024899.3 | 17.4368423 | -5.7487704 | 1.35480406 | -4.2432486 |
| 12047 | NM_001329609.1 | 19.0546762 | -5.8757051 | 1.38475719 | -4.2431302 |
| 12048 | NM_001031717.3 | 19.885967 | -5.9392581 | 1.39980363 | -4.2429224 |
| 12049 | NM_001204080.1 | 31.0020929 | -6.5735445 | 1.54933409 | -4.2428193 |
| 12050 | NM_001271951.1 | 20.2170517 | -5.9529901 | 1.40312631 | -4.2426616 |
| 12051 | NM_001002880.1 | 15.1992708 | -5.5481604 | 1.30798331 | -4.2417669 |
| 12052 | NM_138383.2 | 575.059744 | -5.9595854 | 1.40500269 | -4.2416897 |
| 12053 | NM_001044264.2 | 17.4524434 | -5.745357 | 1.35462759 | -4.2412815 |
| 12054 | NM_001130028.1 | 23.6622781 | -6.1837646 | 1.45819401 | -4.2407008 |
| 12055 | NM_145243.4 | 14.7423787 | -5.5111517 | 1.29991274 | -4.239632 |
| 12056 | NM_016256.3 | 26.2027194 | -6.3325171 | 1.49368481 | -4.239527 |
| 12057 | NM_022895.2 | 37.6962293 | -6.8554172 | 1.61703106 | -4.2395087 |
| 12058 | NM_020799.3 | 21.3823554 | -6.0403489 | 1.42481289 | -4.239398 |
| 12059 | NM_016448.3 | 26.4567576 | -6.347297 | 1.49730302 | -4.2391533 |
| 12060 | NM_138625.3 | 28.7865055 | -4.0625906 | 0.95842083 | -4.2388379 |
| 12061 | NM_003213.3 | 18.8046643 | -5.8516976 | 1.38054996 | -4.2386714 |
| 12062 | NM_001270639.1 | 17.6175916 | -5.753037 | 1.3573643 | -4.2383883 |
| 12063 | NM_001330624.1 | 55.5022609 | -6.4343322 | 1.51813533 | -4.2383127 |
| 12064 | NM_001296.4 | 90.7876786 | 3.0648733 | 0.72314062 | 4.23828123 |
| 12065 | NM_001195278.1 | 221.596774 | -5.2408199 | 1.23673899 | -4.2376119 |
| 12066 | NM_001134363.2 | 36.1230992 | -5.0029219 | 1.18084485 | -4.2367309 |
| 12067 | NM_001270378.1 | 14.7830366 | -5.5104766 | 1.30091393 | -4.2358502 |
| 12068 | NM_032726.3 | 14.2096089 | -5.4515037 | 1.28702482 | -4.2357409 |
| 12069 | NM_006721.3 | 27.242532 | -6.3900012 | 1.5086417 | -4.2355989 |
| 12070 | NM_001145544.1 | 28.0072473 | -6.4344446 | 1.51916064 | -4.2355261 |
| 12071 | NM_001135146.1 | 19.191385 | -5.8923372 | 1.39129761 | -4.2351379 |
| 12072 | NM_054026.3 | 18.2586511 | -5.8105865 | 1.37204146 | -4.2349934 |
| 12073 | NM_198194.2 | 19.6613469 | 9.22391299 | 2.17802887 | 4.23498197 |
| 12074 | NM_001320893.1 | 26.1514921 | -6.3362078 | 1.49617615 | -4.2349344 |
| 12075 | NR_024080.1 | 20.6304409 | -5.9813227 | 1.41239663 | -4.2348746 |
| 12076 | NM_001097612.1 | 28.8058889 | -6.4716526 | 1.52822222 | -4.2347589 |
| 12077 | NM_001301700.1 | 21.1105859 | -6.0238788 | 1.42248085 | -4.2347697 |
| 12078 | NM_152586.3 | 22.6810763 | -6.1215705 | 1.44554529 | -4.234783 |
| 12079 | NM_001134232.1 | 158.343209 | -4.8563589 | 1.14684626 | -4.2345335 |
| 12080 | NM_016391.7 | 21.0682904 | -6.0200261 | 1.42186586 | -4.2338917 |
| 12081 | NM_001033583.2 | 20.9729762 | -6.0099358 | 1.41958909 | -4.2335742 |
| 12082 | NM_001293228.1 | 28.5401828 | -6.4546259 | 1.52470309 | -4.2333658 |

| | | | | | |
|---|---|---|---|---|---|
| 12083 | NM_022786.2 | 27.7454694 | -6.4164715 | 1.5156948 | -4.2333532 |
| 12084 | NM_000062.2 | 19.7689343 | -5.9343961 | 1.40185523 | -4.2332446 |
| 12085 | NM_000121.3 | 16.2423559 | -5.6490831 | 1.33452558 | -4.2330272 |
| 12086 | NM_001304961.1 | 34.8588604 | -6.7434607 | 1.59310849 | -4.2328948 |
| 12087 | NM_032865.5 | 18.6385936 | 3.30311012 | 0.78038793 | 4.23265149 |
| 12088 | NM_001003407.1 | 23.1255087 | -6.1524157 | 1.45384351 | -4.231828 |
| 12089 | NM_001278740.1 | 19.5651848 | -5.9139849 | 1.39774901 | -4.2310778 |
| 12090 | NM_003811.3 | 14.4208275 | -5.477214 | 1.29471569 | -4.2304376 |
| 12091 | NM_001301097.1 | 40.8973688 | -6.9726833 | 1.64828434 | -4.2302673 |
| 12092 | NM_001193464.1 | 21.9576447 | -6.0726974 | 1.43555454 | -4.2302101 |
| 12093 | NM_004037.7 | 35.179401 | -4.9631052 | 1.17328356 | -4.2300987 |
| 12094 | NM_004360.4 | 15.9687927 | -5.6229154 | 1.32933545 | -4.2298694 |
| 12095 | NM_004346.3 | 21.6597633 | -6.0654184 | 1.43406417 | -4.2295307 |
| 12096 | NM_001159969.1 | 158.503666 | -4.7187351 | 1.11572328 | -4.2293059 |
| 12097 | NM_025246.2 | 22.0369211 | -6.0814963 | 1.43798816 | -4.2291699 |
| 12098 | NR_135607.1 | 20.0576258 | -5.955405 | 1.40817048 | -4.229179 |
| 12099 | NR_132272.1 | 27.2295929 | -6.3922841 | 1.51166366 | -4.2286418 |
| 12100 | NM_021944.3 | 16.4107425 | -5.6580124 | 1.33803983 | -4.2285829 |
| 12101 | NM_000396.3 | 19.2672244 | -5.8962855 | 1.39444086 | -4.2284228 |
| 12102 | NM_182552.4 | 67.322061 | -2.9328112 | 0.69359256 | -4.2284352 |
| 12103 | NM_001142702.1 | 50.6259326 | -7.2819197 | 1.72226305 | -4.2281112 |
| 12104 | NM_058237.1 | 13.8737297 | -5.4171055 | 1.28131992 | -4.227754 |
| 12105 | NM_001301131.1 | 24.1624617 | -6.2163181 | 1.47049118 | -4.2273753 |
| 12106 | NM_005949.3 | 18.1209812 | -5.8001288 | 1.37204449 | -4.227362 |
| 12107 | NM_001007169.3 | 149.096118 | 2.84161116 | 0.67224066 | 4.2270742 |
| 12108 | NR_024052.2 | 30.6966182 | -6.5601815 | 1.55195878 | -4.2270333 |
| 12109 | NM_001305051.1 | 42.9516666 | -7.0458486 | 1.66704285 | -4.2265552 |
| 12110 | NM_006342.2 | 104.669687 | -4.3308083 | 1.02467863 | -4.226504 |
| 12111 | NM_001018070.2 | 39.133832 | -6.9107263 | 1.63513687 | -4.2263901 |
| 12112 | NM_001294333.1 | 16.6044217 | -5.683794 | 1.34490377 | -4.2261715 |
| 12113 | NM_001322248.1 | 23.0506471 | -6.1417072 | 1.45327282 | -4.2261213 |
| 12114 | NM_001142928.1 | 25.8223301 | -6.3086413 | 1.49280859 | -4.2260216 |
| 12115 | NM_001199165.2 | 19.7670749 | -5.9336812 | 1.40408379 | -4.2260164 |
| 12116 | NM_001142650.1 | 25.8177083 | -6.3108645 | 1.49336807 | -4.2259271 |
| 12117 | NM_001079811.2 | 21.5629906 | -6.0574216 | 1.43346174 | -4.2257295 |
| 12118 | NM_014280.2 | 252.893829 | -5.2182397 | 1.23490605 | -4.2256168 |
| 12119 | NM_001136103.2 | 30.354568 | -6.5454981 | 1.54904398 | -4.2255082 |
| 12120 | NM_001317026.1 | 41.0404528 | -5.9952641 | 1.41887719 | -4.225358 |
| 12121 | NM_018024.2 | 18.9686583 | -5.8704228 | 1.38943516 | -4.2250426 |
| 12122 | NM_018370.2 | 29.0961901 | -4.6602403 | 1.10309866 | -4.2246813 |
| 12123 | NM_001331022.1 | 20.5235513 | -5.9732707 | 1.41407178 | -4.2241637 |
| 12124 | NM_018131.4 | 18.6692863 | -5.8426336 | 1.38325583 | -4.2238272 |
| 12125 | NM_001143957.2 | 17.6803376 | -5.7600016 | 1.36377804 | -4.2235624 |

| | | | | | |
|---|---|---|---|---|---|
| 12126 | NM_001098797.1 | 28.7267687 | -6.4641063 | 1.53071117 | -4.2229432 |
| 12127 | NM_005418.3 | 23.9367504 | -6.1973974 | 1.46761967 | -4.2227544 |
| 12128 | NM_001085451.1 | 17.1814002 | -5.7214619 | 1.35500658 | -4.2224606 |
| 12129 | NM_014145.4 | 19.2087491 | -5.8762896 | 1.39176843 | -4.2221748 |
| 12130 | NR_028038.1 | 116.994687 | -3.8992072 | 0.92351336 | -4.2221449 |
| 12131 | NM_001185093.1 | 15.5650016 | -5.5787725 | 1.32136202 | -4.2219864 |
| 12132 | NM_001304509.1 | 18.9988026 | -5.8610698 | 1.38842221 | -4.2213887 |
| 12133 | NR_144640.1 | 18.093775 | -5.7935773 | 1.37243556 | -4.2213838 |
| 12134 | NM_001272088.1 | 27.5179034 | -6.4042518 | 1.51723775 | -4.2209942 |
| 12135 | NM_004514.3 | 337.764323 | -5.6822738 | 1.34621176 | -4.2209361 |
| 12136 | NR_038879.1 | 14.4764636 | -5.4793758 | 1.29822323 | -4.220673 |
| 12137 | NM_171997.2 | 15.7478863 | -5.5958611 | 1.32588108 | -4.2204849 |
| 12138 | NM_001101669.2 | 18.6340187 | -5.850575 | 1.38625393 | -4.2204208 |
| 12139 | NM_152354.5 | 21.1581676 | -6.0192714 | 1.42640993 | -4.2198749 |
| 12140 | NM_021938.3 | 25.9487692 | -6.3181915 | 1.49745679 | -4.2192814 |
| 12141 | NM_172349.2 | 176.850403 | -4.7838862 | 1.13383599 | -4.2192047 |
| 12142 | NM_001286119.1 | 16.1640907 | -5.6286572 | 1.33456718 | -4.21759 |
| 12143 | NM_001042406.1 | 25.1511435 | -6.2737078 | 1.48757757 | -4.2173987 |
| 12144 | NM_001286564.1 | 30.6741827 | -6.5551519 | 1.55437745 | -4.2172202 |
| 12145 | NM_002245.3 | 26.0895878 | -6.3303649 | 1.50108075 | -4.2172048 |
| 12146 | NM_005107.3 | 25.1002998 | -6.2712502 | 1.48708385 | -4.2171463 |
| 12147 | NM_001144994.1 | 39.2222724 | -5.9382572 | 1.40817491 | -4.2169883 |
| 12148 | NM_152219.3 | 15.4008701 | -5.5674477 | 1.32037336 | -4.2165707 |
| 12149 | NM_194429.2 | 32.74425 | -6.6523419 | 1.57767199 | -4.2165558 |
| 12150 | NM_004881.4 | 19.4991135 | -5.9150134 | 1.4029204 | -4.2162145 |
| 12151 | NM_005779.2 | 77.0159052 | -4.7163869 | 1.11864836 | -4.2161478 |
| 12152 | NR_138033.1 | 11.0488189 | 8.38153519 | 1.98832889 | 4.21536659 |
| 12153 | NM_032551.4 | 27.8450214 | -6.4242853 | 1.52434286 | -4.2144622 |
| 12154 | NM_015020.3 | 27.4125642 | -6.3948974 | 1.51740786 | -4.2143563 |
| 12155 | NM_080622.3 | 18.4826349 | -5.828297 | 1.38309355 | -4.2139572 |
| 12156 | NM_001320470.1 | 16.0581068 | -5.6268609 | 1.33533981 | -4.2138045 |
| 12157 | NM_015070.5 | 20.4153189 | -5.9792449 | 1.41896046 | -4.2138207 |
| 12158 | NM_001258282.1 | 14.8943882 | -5.5277149 | 1.31187997 | -4.2135828 |
| 12159 | NM_001243160.1 | 14.4301843 | -5.4761096 | 1.29997385 | -4.2124768 |
| 12160 | NM_016403.3 | 117.578113 | -4.6909625 | 1.1136527 | -4.212231 |
| 12161 | NM_004498.2 | 23.1430273 | -6.155862 | 1.46147572 | -4.2120864 |
| 12162 | NR_027387.1 | 38.5798531 | -5.9161168 | 1.40471613 | -4.2116102 |
| 12163 | NM_007022.4 | 23.4190688 | -6.1671366 | 1.46440601 | -4.2113571 |
| 12164 | NM_001195007.1 | 15.9345631 | -5.6061321 | 1.33127487 | -4.2111004 |
| 12165 | NR_015382.2 | 57.8040009 | -4.3174427 | 1.025482 | -4.2101594 |
| 12166 | NM_007018.4 | 29.7303392 | -5.5248125 | 1.31232072 | -4.209956 |
| 12167 | NM_001136020.2 | 25.7400427 | -6.3030962 | 1.49724814 | -4.2097873 |
| 12168 | NM_001286606.1 | 17.4827162 | -5.7547544 | 1.36702417 | -4.2096947 |

| | | | | | |
|---|---|---|---|---|---|
| 12169 | NM_001177519.2 | 21.9148404 | -6.0820507 | 1.44481442 | -4.2095723 |
| 12170 | NM_001001683.3 | 16.474653 | -5.6603235 | 1.34470157 | -4.209353 |
| 12171 | NM_173690.4 | 61.9352404 | -7.5733767 | 1.79937734 | -4.2088874 |
| 12172 | NM_002842.4 | 18.8889477 | -5.8676216 | 1.3941749 | -4.2086697 |
| 12173 | NM_001348090.1 | 20.1375851 | -5.952293 | 1.41452994 | -4.2079654 |
| 12174 | NM_018593.4 | 24.3286731 | -6.229864 | 1.48062494 | -4.2075909 |
| 12175 | NM_001286512.1 | 30.2434888 | -6.5379789 | 1.55391188 | -4.2074322 |
| 12176 | NR_028383.1 | 19.2878116 | -5.8946587 | 1.40103348 | -4.2073647 |
| 12177 | NM_001127394.3 | 25.0827366 | -6.2717303 | 1.49070004 | -4.2072383 |
| 12178 | NM_006168.2 | 23.4085154 | -6.1711607 | 1.46710935 | -4.20634 |
| 12179 | NR_132352.1 | 21.8857133 | -6.0719388 | 1.44351772 | -4.2063487 |
| 12180 | NM_020924.3 | 16.5941279 | -5.6732921 | 1.34878378 | -4.206228 |
| 12181 | NM_006320.4 | 248.945306 | -5.1951276 | 1.23512435 | -4.2061575 |
| 12182 | NR_000029.3 | 13.8597034 | -5.4103396 | 1.28638352 | -4.2058527 |
| 12183 | NM_153221.2 | 24.1663578 | -5.2343186 | 1.24458623 | -4.2056697 |
| 12184 | NM_004728.3 | 230.491816 | -5.0788819 | 1.20765515 | -4.205573 |
| 12185 | NM_001261413.1 | 193.702767 | -5.340334 | 1.26989212 | -4.2053446 |
| 12186 | NR_024553.1 | 19.2951868 | -5.8887658 | 1.4003227 | -4.205292 |
| 12187 | NM_001300849.1 | 14.0141959 | -5.4336478 | 1.29215325 | -4.205111 |
| 12188 | NM_001002811.2 | 173.7129 | -4.7570104 | 1.13127048 | -4.2050159 |
| 12189 | NM_001318339.1 | 61.9305961 | -7.5724183 | 1.80100331 | -4.2045555 |
| 12190 | NM_001288953.1 | 32.7233073 | -3.848756 | 0.91552207 | -4.2038921 |
| 12191 | NM_018901.3 | 35.9760999 | -6.7902985 | 1.61534221 | -4.2036285 |
| 12192 | NR_003659.2 | 149.666876 | -4.6451565 | 1.10504284 | -4.2035986 |
| 12193 | NM_000944.4 | 60.8888322 | -7.5493781 | 1.79630903 | -4.2027168 |
| 12194 | NM_001322131.1 | 21.6919406 | -6.0526005 | 1.44023013 | -4.2025231 |
| 12195 | NM_001098502.1 | 18.4551059 | -5.8339749 | 1.38829705 | -4.2022526 |
| 12196 | NR_015439.1 | 33.9753294 | -5.7322444 | 1.36414754 | -4.2020707 |
| 12197 | NM_013299.3 | 81.7359614 | -4.7133473 | 1.12173114 | -4.2018512 |
| 12198 | NM_198721.3 | 22.6622552 | -6.1199959 | 1.45674928 | -4.2011319 |
| 12199 | NM_001318810.1 | 22.6761041 | -6.1273518 | 1.45860797 | -4.2008215 |
| 12200 | NM_138779.4 | 17.1437118 | -5.7167241 | 1.36109241 | -4.2000999 |
| 12201 | NM_001080421.2 | 296.432163 | -5.4574912 | 1.29947633 | -4.1997619 |
| 12202 | NM_152772.2 | 18.0041563 | -5.7978025 | 1.38051357 | -4.1997432 |
| 12203 | NM_001143841.1 | 47.3268487 | -4.5298033 | 1.07866066 | -4.1994702 |
| 12204 | NM_001320990.1 | 15.299186 | -5.5514436 | 1.32195858 | -4.1994081 |
| 12205 | NM_001137560.1 | 375.74171 | -5.4110314 | 1.2885467 | -4.1993289 |
| 12206 | NM_001282982.1 | 25.2247565 | -6.2734955 | 1.49396096 | -4.1992365 |
| 12207 | NM_002079.2 | 179.707037 | -4.9249023 | 1.17284898 | -4.1990933 |
| 12208 | NM_016071.3 | 18.4766725 | -5.827313 | 1.38789357 | -4.1986743 |
| 12209 | NM_015426.4 | 15.0215144 | -5.5208084 | 1.31512532 | -4.1979333 |
| 12210 | NM_001102594.1 | 14.1174694 | -5.4388469 | 1.29560952 | -4.1979059 |
| 12211 | NM_006768.4 | 95.4910627 | -3.9472976 | 0.9403577 | -4.1976554 |

| | | | | | |
|---|---|---|---|---|---|
| 12212 | NM_001347869.1 | 13.4911984 | -5.3750195 | 1.28057822 | -4.1973379 |
| 12213 | NR_002188.3 | 18.7678699 | -5.8504101 | 1.39388466 | -4.1971981 |
| 12214 | NM_001170686.1 | 25.1805363 | -6.2713812 | 1.49420138 | -4.1971459 |
| 12215 | NM_006516.2 | 604.088219 | -6.1710015 | 1.47047643 | -4.1966001 |
| 12216 | NM_001319670.1 | 16.6277902 | -5.6861934 | 1.35513469 | -4.1960356 |
| 12217 | NM_014750.4 | 25.7096938 | -5.3116211 | 1.26588475 | -4.1959753 |
| 12218 | NM_024602.5 | 133.966604 | -4.7398985 | 1.12977399 | -4.1954396 |
| 12219 | NM_207511.2 | 24.0269998 | 2.20997926 | 0.52680779 | 4.19503908 |
| 12220 | NM_003551.2 | 17.9910236 | -5.7979896 | 1.38216976 | -4.1948462 |
| 12221 | NM_002758.3 | 16.6083934 | -5.6749908 | 1.35301144 | -4.1943406 |
| 12222 | NM_005637.3 | 133.958429 | -2.3657676 | 0.56422615 | -4.1929422 |
| 12223 | NM_001007227.1 | 18.1955349 | -5.813005 | 1.38643448 | -4.192773 |
| 12224 | NR_036541.1 | 15.6576653 | -5.5826046 | 1.3315394 | -4.1925944 |
| 12225 | NR_137177.1 | 34.4229149 | -6.7249759 | 1.60419731 | -4.1921127 |
| 12226 | NM_000946.2 | 22.7104166 | -6.1226095 | 1.46060585 | -4.1918287 |
| 12227 | NM_007023.3 | 17.0562647 | -5.713376 | 1.36300506 | -4.1917496 |
| 12228 | NM_001321091.1 | 17.8368625 | -5.7756377 | 1.37799348 | -4.1913389 |
| 12229 | NM_001172702.1 | 14.8628335 | -5.5215664 | 1.31739658 | -4.1912712 |
| 12230 | NM_001037305.1 | 17.8984665 | -5.7806485 | 1.37925435 | -4.1911403 |
| 12231 | NM_001323529.1 | 26.3000816 | -6.3414892 | 1.51321255 | -4.1907458 |
| 12232 | NM_001174153.1 | 25.2841818 | -6.2772231 | 1.49791074 | -4.1906523 |
| 12233 | NM_025221.5 | 26.2609488 | -6.3374287 | 1.51228799 | -4.1906229 |
| 12234 | NR_045680.1 | 36.1383667 | -4.9622467 | 1.18418495 | -4.1904321 |
| 12235 | NM_212552.2 | 16.6301049 | -5.6810184 | 1.35622721 | -4.1888397 |
| 12236 | NM_015063.2 | 127.316305 | -5.3901941 | 1.28690981 | -4.1884785 |
| 12237 | NM_005057.3 | 38.3334545 | -6.8829119 | 1.64343124 | -4.188135 |
| 12238 | NM_006219.2 | 70.8748727 | -4.6010011 | 1.09864223 | -4.1878975 |
| 12239 | NR_138463.1 | 16.5381602 | -5.6609327 | 1.35175595 | -4.1878363 |
| 12240 | NM_005178.4 | 14.824154 | -5.5203049 | 1.31820929 | -4.1877302 |
| 12241 | NM_005398.6 | 22.6988724 | -6.1257444 | 1.4628755 | -4.187468 |
| 12242 | NM_001077701.2 | 28.3380736 | -6.4422705 | 1.5385282 | -4.1872944 |
| 12243 | NM_001306206.1 | 25.5047631 | -6.2898153 | 1.50230935 | -4.1867644 |
| 12244 | NM_001139457.2 | 17.0424374 | -5.7127564 | 1.36453801 | -4.1865865 |
| 12245 | NM_015871.4 | 31.6674032 | -5.628058 | 1.34434818 | -4.1864586 |
| 12246 | NM_001206540.2 | 16.6296752 | -5.6774984 | 1.35618172 | -4.1863847 |
| 12247 | NM_018153.3 | 20.1922696 | -5.9629162 | 1.42489351 | -4.1848154 |
| 12248 | NM_002873.1 | 19.501076 | -5.899966 | 1.40986226 | -4.1847819 |
| 12249 | NM_001042588.1 | 28.788014 | -5.4887935 | 1.31168536 | -4.1845351 |
| 12250 | NR_111973.1 | 94.6819712 | 2.42847777 | 0.58037259 | 4.18434262 |
| 12251 | NM_001277066.1 | 28.1437929 | -6.4355684 | 1.53833423 | -4.1834656 |
| 12252 | NR_033342.1 | 24.0775802 | -6.209579 | 1.48431662 | -4.1834599 |
| 12253 | NM_001130442.2 | 20.3287749 | -5.9700825 | 1.42709125 | -4.1833923 |
| 12254 | NM_001163297.1 | 29.3741433 | -5.5247635 | 1.32081318 | -4.1828501 |

| | | | | | |
|---|---|---|---|---|---|
| 12255 | NR_026658.1 | 22.3924468 | -6.1096411 | 1.46071054 | -4.1826501 |
| 12256 | NM_001242761.1 | 24.8052137 | -6.2507228 | 1.49453031 | -4.1823995 |
| 12257 | NM_001167738.1 | 251.736739 | -3.9712467 | 0.94962856 | -4.1818947 |
| 12258 | NR_037653.1 | 14.7466893 | -5.5035794 | 1.31617495 | -4.1814953 |
| 12259 | NM_015122.2 | 22.2773802 | -6.0965249 | 1.45803233 | -4.1813372 |
| 12260 | NM_002026.3 | 591.423423 | -10.830224 | 2.59014984 | -4.1813119 |
| 12261 | NM_001012659.1 | 137.736675 | 1.86474068 | 0.44602431 | 4.18080506 |
| 12262 | NM_002885.2 | 37.6504651 | -6.8579911 | 1.64036188 | -4.1807794 |
| 12263 | NM_006143.2 | 25.6493339 | -6.302907 | 1.50766954 | -4.1805627 |
| 12264 | NM_004770.2 | 22.2043297 | -6.0914952 | 1.45717256 | -4.1803527 |
| 12265 | NM_015192.3 | 60.7167985 | -4.2491871 | 1.01646958 | -4.1803387 |
| 12266 | NM_001199263.1 | 20.4335862 | -5.9712305 | 1.42842816 | -4.1802806 |
| 12267 | NM_017974.3 | 30.6844448 | -5.5722582 | 1.33300381 | -4.1802268 |
| 12268 | NM_002646.3 | 190.523463 | -5.0142383 | 1.1997434 | -4.1794257 |
| 12269 | NM_032310.4 | 29.6469241 | -6.5106626 | 1.55782275 | -4.1793347 |
| 12270 | NR_002166.1 | 18.555543 | -5.8357106 | 1.39642876 | -4.1790249 |
| 12271 | NM_018530.2 | 14.8227841 | -5.5037722 | 1.31702826 | -4.1789325 |
| 12272 | NM_001287747.1 | 18.8165206 | -5.856253 | 1.40152372 | -4.1784901 |
| 12273 | NM_015704.2 | 223.77006 | -2.3625567 | 0.56545399 | -4.1781591 |
| 12274 | NM_001101667.1 | 17.6840214 | -5.7738748 | 1.38195371 | -4.1780522 |
| 12275 | NM_001347919.1 | 26.5475145 | -6.3507669 | 1.52010586 | -4.1778451 |
| 12276 | NM_001321433.1 | 16.097459 | -5.6343924 | 1.34871236 | -4.1776086 |
| 12277 | NM_001278647.1 | 15.91364 | -5.6162473 | 1.34437731 | -4.1775826 |
| 12278 | NR_051991.1 | 16.910765 | -5.6999926 | 1.36444925 | -4.1775043 |
| 12279 | NM_022103.3 | 17.9882048 | -5.7839899 | 1.38477559 | -4.1768428 |
| 12280 | NR_111980.1 | 473.562088 | 2.97465034 | 0.71217674 | 4.17684289 |
| 12281 | NM_001085457.1 | 22.5722959 | -6.1181521 | 1.46495916 | -4.1763295 |
| 12282 | NM_001567.3 | 351.269117 | -5.5575188 | 1.33080216 | -4.1760669 |
| 12283 | NM_003385.4 | 42.5646853 | -7.0321334 | 1.68393351 | -4.1760161 |
| 12284 | NM_001278379.1 | 97.1406836 | -4.7610452 | 1.14011085 | -4.1759494 |
| 12285 | NM_005465.4 | 509.538895 | -5.4634358 | 1.30836381 | -4.1757772 |
| 12286 | NR_003500.1 | 48.9438634 | -4.9077634 | 1.17531906 | -4.175686 |
| 12287 | NM_032447.4 | 27.2872227 | -6.3939466 | 1.53126539 | -4.1755966 |
| 12288 | NM_002744.4 | 27.1208362 | -5.4075492 | 1.29507917 | -4.1754584 |
| 12289 | NM_001242915.1 | 34.5469631 | -6.7341432 | 1.61295177 | -4.1750431 |
| 12290 | NM_178013.3 | 13.8426441 | -5.4130064 | 1.29663 | -4.1746731 |
| 12291 | NM_001195200.1 | 20.4466391 | -5.9734717 | 1.43102649 | -4.1742565 |
| 12292 | NR_046360.1 | 24.7154826 | -6.2468418 | 1.49674659 | -4.1736135 |
| 12293 | NM_001178034.1 | 40.5916226 | -5.9896839 | 1.43525709 | -4.1732481 |
| 12294 | NM_005374.4 | 34.083816 | -6.7136262 | 1.60876295 | -4.1731607 |
| 12295 | NM_001242823.2 | 64.3373814 | -7.6309753 | 1.82897879 | -4.1722601 |
| 12296 | NM_001135004.2 | 33.7850685 | -6.6991069 | 1.60566712 | -4.1721642 |
| 12297 | NM_001134321.1 | 19.8695513 | -5.9349132 | 1.42257885 | -4.1719397 |

| | | | | | |
|---|---|---|---|---|---|
| 12298 | NM_001030001.2 | 16.6292051 | -5.6787394 | 1.3611929 | -4.1718844 |
| 12299 | NM_030573.2 | 16.0382227 | -5.629429 | 1.34947387 | -4.1715732 |
| 12300 | NM_024610.5 | 16.6254979 | -5.67555 | 1.36056808 | -4.1714561 |
| 12301 | NM_001142799.2 | 20.4502746 | -5.9782875 | 1.43320766 | -4.171264 |
| 12302 | NR_110835.1 | 27.3141252 | -6.3961755 | 1.53356246 | -4.1707956 |
| 12303 | NM_005671.3 | 15.5572106 | -5.5826274 | 1.33857236 | -4.1705832 |
| 12304 | NM_001257180.1 | 38.8504401 | -6.9038105 | 1.65540407 | -4.1704685 |
| 12305 | NM_002381.4 | 15.5932421 | -5.5863689 | 1.33958994 | -4.1702082 |
| 12306 | NM_001197123.1 | 14.4037713 | -5.4725319 | 1.31231697 | -4.1701296 |
| 12307 | NM_015352.1 | 219.1771 | -2.868127 | 0.68781111 | -4.1699341 |
| 12308 | NM_006479.4 | 20.3098203 | -5.9618798 | 1.42976528 | -4.1698312 |
| 12309 | NM_018191.3 | 131.478035 | -4.1578285 | 0.99717668 | -4.1696006 |
| 12310 | NM_001206427.1 | 21.1178526 | -6.0231698 | 1.4446357 | -4.1693347 |
| 12311 | NM_001144955.1 | 30.1347412 | -5.5401334 | 1.32885066 | -4.1691167 |
| 12312 | NM_004328.4 | 19.7630951 | -5.9241186 | 1.4211026 | -4.1686776 |
| 12313 | NM_016594.2 | 15.0241362 | -5.5371727 | 1.32838007 | -4.1683647 |
| 12314 | NM_014638.3 | 31.7630001 | -6.6098648 | 1.58590726 | -4.167876 |
| 12315 | NM_001286582.1 | 47.9077353 | -7.201019 | 1.72778316 | -4.1677794 |
| 12316 | NM_001267776.1 | 16.1388137 | -5.6408456 | 1.35363656 | -4.1671788 |
| 12317 | NM_013381.2 | 18.9186377 | -5.8639495 | 1.40738193 | -4.1665658 |
| 12318 | NM_176793.1 | 15.3479345 | -5.557148 | 1.33380662 | -4.1663821 |
| 12319 | NM_023927.2 | 28.8589103 | -6.4773511 | 1.55472962 | -4.1662235 |
| 12320 | NM_024044.3 | 66.4723528 | -4.3924239 | 1.05435346 | -4.1659881 |
| 12321 | NM_015319.2 | 16.5039541 | -5.6570407 | 1.35794183 | -4.1658932 |
| 12322 | NM_021128.4 | 143.311925 | -3.7825246 | 0.9080791 | -4.1654131 |
| 12323 | NM_001328.2 | 150.306924 | -4.6159109 | 1.10832929 | -4.1647468 |
| 12324 | NM_001542.3 | 54.9827488 | -6.4262507 | 1.54301936 | -4.1647246 |
| 12325 | NM_001242846.1 | 20.6490564 | -5.9942615 | 1.43953715 | -4.16402 |
| 12326 | NM_001319839.1 | 39.9260272 | -5.9636258 | 1.43220398 | -4.1639501 |
| 12327 | NM_001330227.1 | 20.8232895 | -6.0039013 | 1.44190895 | -4.163856 |
| 12328 | NM_032554.3 | 105.335633 | 3.5972522 | 0.86400563 | 4.1634592 |
| 12329 | NM_021035.2 | 113.880347 | -2.6091872 | 0.62673025 | -4.1631743 |
| 12330 | NM_004126.3 | 17.2987368 | -5.7303477 | 1.37652176 | -4.1629184 |
| 12331 | NM_025027.4 | 19.1059867 | -5.8822077 | 1.41302716 | -4.1628413 |
| 12332 | NR_024249.1 | 14.5328367 | -5.4749743 | 1.31530971 | -4.1624982 |
| 12333 | NR_033349.1 | 19.4967797 | -5.8974455 | 1.41701171 | -4.161889 |
| 12334 | NM_001304793.1 | 31.0861565 | -5.583617 | 1.34163545 | -4.1617989 |
| 12335 | NM_001300733.1 | 14.9466693 | -5.5222225 | 1.32695541 | -4.1615735 |
| 12336 | NM_001257234.1 | 23.8618887 | -6.1935005 | 1.4885878 | -4.1606552 |
| 12337 | NM_006511.2 | 15.2874829 | -5.5561678 | 1.33551579 | -4.160316 |
| 12338 | NM_153213.4 | 15.226158 | -5.5479185 | 1.33355106 | -4.1602596 |
| 12339 | NM_001320608.1 | 15.5675154 | -5.5820832 | 1.34178344 | -4.1601968 |
| 12340 | NM_001080.3 | 123.731348 | -3.6037205 | 0.86632688 | -4.15977 |

| | | | | | |
|---|---|---|---|---|---|
| 12341 | NM_138417.2 | 30.5543657 | -5.5617116 | 1.33754886 | -4.1581372 |
| 12342 | NM_004507.3 | 15.6170321 | -5.5852652 | 1.34337525 | -4.157636 |
| 12343 | NM_199051.2 | 23.8082108 | -6.1958729 | 1.49029771 | -4.1574733 |
| 12344 | NM_001142284.1 | 46.3266498 | -7.1552748 | 1.72131361 | -4.1568688 |
| 12345 | NM_001031806.1 | 18.8416604 | -5.8605966 | 1.40988798 | -4.1567818 |
| 12346 | NM_001243227.1 | 47.3406786 | -7.1835085 | 1.72829234 | -4.1564198 |
| 12347 | NM_005815.4 | 14.7700579 | -5.5072129 | 1.3250434 | -4.1562509 |
| 12348 | NM_001002257.2 | 34.4481279 | -4.2359441 | 1.01919391 | -4.156171 |
| 12349 | NM_022490.2 | 68.045137 | -3.0452508 | 0.73273148 | -4.1560256 |
| 12350 | NM_001135934.1 | 28.1273859 | -6.4357517 | 1.54870259 | -4.1555763 |
| 12351 | NM_152695.5 | 26.0918175 | -5.3480854 | 1.2873452 | -4.1543523 |
| 12352 | NM_001134875.1 | 15.6274569 | -5.5791152 | 1.34335067 | -4.1531339 |
| 12353 | NM_130800.2 | 24.4306774 | -6.2310566 | 1.50044125 | -4.1528161 |
| 12354 | NR_047674.1 | 18.1638266 | -5.8012258 | 1.39693175 | -4.1528341 |
| 12355 | NM_020935.2 | 71.5057263 | -3.000133 | 0.72244885 | -4.1527272 |
| 12356 | NM_213568.1 | 22.5010905 | -6.1134558 | 1.47217689 | -4.1526639 |
| 12357 | NM_016453.3 | 34.8721563 | -6.7425365 | 1.62371867 | -4.1525276 |
| 12358 | NM_001267699.1 | 30.7575135 | -4.1003244 | 0.98754697 | -4.1520298 |
| 12359 | NM_001271869.1 | 35.3867562 | -6.7686339 | 1.63023631 | -4.1519342 |
| 12360 | NM_006992.3 | 15.3473341 | -5.5686191 | 1.34132915 | -4.1515679 |
| 12361 | NM_199190.2 | 28.0881527 | -6.4290275 | 1.54870775 | -4.1512206 |
| 12362 | NR_109820.1 | 26.2020935 | -6.3306886 | 1.52513686 | -4.1508987 |
| 12363 | NM_001159287.1 | 39.2903099 | -6.9159266 | 1.66615152 | -4.1508389 |
| 12364 | NM_001324366.1 | 23.9148817 | -6.1984427 | 1.49332589 | -4.1507635 |
| 12365 | NM_001321829.1 | 34.2721728 | -6.7197589 | 1.61910526 | -4.1502916 |
| 12366 | NM_173615.4 | 64.625765 | -7.6373435 | 1.84045288 | -4.1497088 |
| 12367 | NM_207340.1 | 27.265805 | -5.4091725 | 1.30354369 | -4.1495905 |
| 12368 | NR_103449.1 | 19.4654023 | 3.58843336 | 0.86478716 | 4.14949891 |
| 12369 | NM_001136113.1 | 16.4559795 | -5.6656794 | 1.36542072 | -4.1494019 |
| 12370 | NM_012393.2 | 174.760492 | -2.8172507 | 0.67897902 | -4.1492455 |
| 12371 | NM_212539.1 | 19.4434204 | -5.9087988 | 1.4242461 | -4.1487204 |
| 12372 | NM_021994.2 | 37.1096476 | -3.58612 | 0.86441328 | -4.1486174 |
| 12373 | NM_017458.3 | 33.6518897 | -6.6906681 | 1.61276757 | -4.1485632 |
| 12374 | NM_001171894.2 | 34.0617081 | -5.7400376 | 1.38366334 | -4.1484351 |
| 12375 | NM_001302348.1 | 15.976317 | -5.6175998 | 1.35428427 | -4.1480211 |
| 12376 | NM_001142305.1 | 36.9682455 | -5.8571734 | 1.41205673 | -4.1479732 |
| 12377 | NR_038995.1 | 54.2451993 | 3.80283316 | 0.91682428 | 4.14783208 |
| 12378 | NM_006580.3 | 45.783801 | 3.79978653 | 0.9161125 | 4.14772915 |
| 12379 | NR_015363.2 | 27.5800753 | -5.4227472 | 1.3075027 | -4.147408 |
| 12380 | NM_023070.2 | 20.6688229 | -5.9894898 | 1.44416234 | -4.1473799 |
| 12381 | NM_001304835.1 | 13.7995628 | -5.4034477 | 1.30296315 | -4.1470456 |
| 12382 | NM_000954.5 | 64.5384308 | -4.930308 | 1.18916617 | -4.1460211 |
| 12383 | NM_001243194.1 | 354.221217 | -4.3818045 | 1.05696442 | -4.1456499 |

| | | | | | |
|---|---|---|---|---|---|
| 12384 | NM_053043.2 | 228.729319 | -4.9692864 | 1.19867912 | -4.1456352 |
| 12385 | NM_001242935.2 | 22.24552 | -6.101903 | 1.47194609 | -4.1454664 |
| 12386 | NM_001310160.1 | 18.4227135 | -5.827389 | 1.40582894 | -4.1451622 |
| 12387 | NM_022151.4 | 325.678604 | -4.1881801 | 1.01038243 | -4.1451434 |
| 12388 | NM_005486.2 | 24.5340893 | -6.242057 | 1.50588135 | -4.1451187 |
| 12389 | NR_024181.1 | 13.1558292 | -5.3356604 | 1.28722138 | -4.1450993 |
| 12390 | NM_212543.1 | 14.3841784 | -5.4598729 | 1.31730588 | -4.1447268 |
| 12391 | NR_130139.1 | 34.7808115 | -6.7420486 | 1.62673903 | -4.1445176 |
| 12392 | NM_001198798.2 | 20.9262098 | -6.0099195 | 1.45046033 | -4.1434567 |
| 12393 | NM_001144776.2 | 54.9381073 | -7.4022148 | 1.78651051 | -4.1433928 |
| 12394 | NM_001289404.1 | 19.3333476 | -5.8968783 | 1.42321618 | -4.1433469 |
| 12395 | NM_001319119.1 | 18.4448991 | -5.8181816 | 1.40425002 | -4.1432661 |
| 12396 | NM_001144942.1 | 26.7377378 | -6.3673041 | 1.53691277 | -4.1429184 |
| 12397 | NM_001330596.1 | 18.30286 | -5.8168999 | 1.40406144 | -4.1429098 |
| 12398 | NM_001281767.2 | 37.6725781 | -6.855241 | 1.65476852 | -4.142719 |
| 12399 | NM_020657.3 | 47.5720593 | -4.1308487 | 0.99716547 | -4.142591 |
| 12400 | NM_024605.3 | 13.9479082 | -5.4282625 | 1.3105173 | -4.1420761 |
| 12401 | NM_032788.2 | 49.6519512 | -3.3012517 | 0.79700444 | -4.1420744 |
| 12402 | NM_080651.3 | 16.3495684 | -5.6514942 | 1.36451261 | -4.1417677 |
| 12403 | NM_001242929.1 | 24.7985638 | -6.2557006 | 1.51090791 | -4.1403586 |
| 12404 | NM_006333.3 | 14.7197244 | -5.5030535 | 1.32912064 | -4.1403717 |
| 12405 | NM_001319042.1 | 40.8936648 | -6.9736869 | 1.68442449 | -4.1401007 |
| 12406 | NM_001301849.1 | 29.9843396 | -6.5300629 | 1.57739935 | -4.1397652 |
| 12407 | NR_103551.2 | 27.1361391 | -6.3808561 | 1.54144478 | -4.1395295 |
| 12408 | NM_001123385.1 | 82.7793839 | -4.3203102 | 1.04371582 | -4.1393549 |
| 12409 | NM_002394.5 | 18.2096755 | -5.8101841 | 1.40372102 | -4.1391302 |
| 12410 | NM_003847.2 | 14.1909673 | -5.4490011 | 1.31647526 | -4.1390836 |
| 12411 | NM_001136005.1 | 13.5703111 | -5.3796789 | 1.29986704 | -4.1386378 |
| 12412 | NR_120634.1 | 75.0913244 | 2.07253802 | 0.5007856 | 4.13857352 |
| 12413 | NM_001172623.1 | 271.213354 | -4.7561508 | 1.14929867 | -4.1383071 |
| 12414 | NM_181353.2 | 14.895558 | -5.5210187 | 1.33412333 | -4.1383121 |
| 12415 | NM_001330241.1 | 18.7497913 | -5.8503045 | 1.41384967 | -4.1378547 |
| 12416 | NR_027749.1 | 17.1342238 | -5.7256453 | 1.3838028 | -4.1376165 |
| 12417 | NM_007232.2 | 25.5045822 | -6.2959063 | 1.52166016 | -4.1375246 |
| 12418 | NM_015681.4 | 14.9865217 | -5.5333727 | 1.33741594 | -4.1373611 |
| 12419 | NM_001320586.1 | 16.3134317 | -5.6573413 | 1.3674208 | -4.1372351 |
| 12420 | NM_001145536.1 | 13.7708463 | -5.4087329 | 1.30739525 | -4.1370297 |
| 12421 | NM_018475.4 | 112.039754 | -4.6530234 | 1.12487396 | -4.1364843 |
| 12422 | NM_001346167.1 | 22.3220469 | -6.0937557 | 1.47318712 | -4.1364438 |
| 12423 | NM_003821.5 | 25.6857011 | -5.3240315 | 1.28711557 | -4.1364052 |
| 12424 | NM_001205254.1 | 30.5195801 | -6.5568544 | 1.58524289 | -4.1361828 |
| 12425 | NM_001145313.2 | 17.768738 | -5.778992 | 1.39724418 | -4.1359929 |
| 12426 | NM_144606.5 | 13.9424606 | -5.4165233 | 1.30977675 | -4.1354554 |

| | | | | | |
|---|---|---|---|---|---|
| 12427 | NM_001204284.1 | 23.316512 | -6.1664247 | 1.49112459 | -4.1354188 |
| 12428 | NM_001242534.1 | 17.9953927 | -5.7975163 | 1.40204919 | -4.1350306 |
| 12429 | NM_001267580.1 | 32.455902 | -6.6437016 | 1.60670511 | -4.1349851 |
| 12430 | NM_001323682.1 | 20.2841245 | -5.9682249 | 1.44346594 | -4.1346489 |
| 12431 | NM_030978.2 | 109.070554 | -4.5783854 | 1.10766059 | -4.133383 |
| 12432 | NM_001271512.3 | 18.4814362 | -5.8274697 | 1.40986871 | -4.1333421 |
| 12433 | NR_033823.2 | 14.7998733 | -5.5179817 | 1.3350171 | -4.1332667 |
| 12434 | NM_005602.5 | 36.8319702 | -6.8273632 | 1.65227634 | -4.1320953 |
| 12435 | NR_002451.2 | 14.2666223 | -5.4641041 | 1.32236243 | -4.1320775 |
| 12436 | NM_006569.5 | 29.0974058 | -6.4835276 | 1.56921186 | -4.1317095 |
| 12437 | NM_032229.2 | 27.0961515 | -5.4073909 | 1.30874856 | -4.1317264 |
| 12438 | NM_016055.5 | 26.5642448 | -6.3528227 | 1.53769415 | -4.1313955 |
| 12439 | NM_001282759.1 | 25.967126 | -5.3422879 | 1.29312545 | -4.131299 |
| 12440 | NR_135105.1 | 32.1751915 | -5.6532531 | 1.36847493 | -4.1310608 |
| 12441 | NM_001271610.1 | 15.0629153 | -5.5339546 | 1.3397318 | -4.1306437 |
| 12442 | NR_026587.1 | 22.4463541 | -6.1156295 | 1.48070903 | -4.1302034 |
| 12443 | NM_001199635.1 | 27.8860575 | -6.4246191 | 1.55568127 | -4.1297785 |
| 12444 | NM_000352.4 | 24.8892345 | -6.2596154 | 1.51598586 | -4.1290724 |
| 12445 | NM_203386.2 | 25.1107686 | -6.2661122 | 1.51756431 | -4.1290588 |
| 12446 | NM_016127.5 | 1078.27116 | -6.3003146 | 1.52631334 | -4.127799 |
| 12447 | NM_004085.3 | 18.47311 | -5.829789 | 1.41244544 | -4.1274437 |
| 12448 | NM_001282394.1 | 128.505264 | -4.6786466 | 1.13366774 | -4.1269999 |
| 12449 | NM_001318093.1 | 26.741045 | -6.359744 | 1.54106959 | -4.1268377 |
| 12450 | NM_016578.3 | 223.031563 | -4.9316678 | 1.19507762 | -4.1266506 |
| 12451 | NM_002114.3 | 75.6109518 | -3.0843189 | 0.74764184 | -4.1253964 |
| 12452 | NM_003416.3 | 22.5019165 | -6.1114351 | 1.48143329 | -4.1253529 |
| 12453 | NM_001203263.1 | 17.5300824 | -5.7428548 | 1.39218613 | -4.1250625 |
| 12454 | NM_001130420.2 | 27.5647309 | -6.4048842 | 1.55268945 | -4.1250259 |
| 12455 | NM_001321697.1 | 16.2648319 | -5.6491525 | 1.36976994 | -4.1241616 |
| 12456 | NM_005512.2 | 19.1358362 | -5.8805791 | 1.42592619 | -4.1240417 |
| 12457 | NM_000224.2 | 137.765621 | -3.91501 | 0.94933726 | -4.1239401 |
| 12458 | NM_001267708.1 | 14.7866925 | -5.5156748 | 1.33760771 | -4.123537 |
| 12459 | NM_025057.2 | 31.2410451 | -5.5997743 | 1.35827503 | -4.1227102 |
| 12460 | NM_213603.2 | 15.5990022 | -5.5828921 | 1.35420097 | -4.1226467 |
| 12461 | NM_022372.4 | 23.0527991 | -6.1448208 | 1.49057304 | -4.1224553 |
| 12462 | NM_001004304.3 | 129.53766 | -4.5527431 | 1.10440777 | -4.1223389 |
| 12463 | NR_027241.2 | 31.3418747 | -5.6109755 | 1.36133814 | -4.1216619 |
| 12464 | NM_001161476.1 | 26.671225 | -5.3795191 | 1.30536275 | -4.1210913 |
| 12465 | NR_073449.1 | 23.0908227 | -6.1546245 | 1.49345577 | -4.1210625 |
| 12466 | NM_001111112.1 | 13.1046379 | -5.3364149 | 1.29496271 | -4.1209023 |
| 12467 | NR_104225.1 | 17.9376788 | -5.7808455 | 1.40285467 | -4.1207729 |
| 12468 | NM_001303507.1 | 27.3977541 | -6.4019309 | 1.55361078 | -4.1206788 |
| 12469 | NM_016020.3 | 21.4825876 | -6.0451554 | 1.46724522 | -4.1200716 |

| | | | | | |
|---|---|---|---|---|---|
| 12470 | NR_144476.1 | 19.1276244 | -5.8771965 | 1.4265299 | -4.1199252 |
| 12471 | NM_017857.3 | 20.5210733 | -5.9857935 | 1.45296128 | -4.1197199 |
| 12472 | NM_001346516.1 | 14.8674941 | -5.5259767 | 1.34138062 | -4.1196187 |
| 12473 | NM_005754.2 | 160.41249 | -5.0722415 | 1.23125592 | -4.1195672 |
| 12474 | NR_037642.1 | 16.4188064 | -5.6557912 | 1.37291342 | -4.1195541 |
| 12475 | NM_001039712.1 | 17.8381415 | -5.7866993 | 1.40489098 | -4.1189668 |
| 12476 | NM_001330419.1 | 22.8874941 | -6.1380911 | 1.49021243 | -4.118937 |
| 12477 | NR_034101.1 | 23.0494666 | -6.1494473 | 1.49315516 | -4.1184248 |
| 12478 | NM_004177.4 | 24.8075654 | -6.2607306 | 1.52107553 | -4.1159893 |
| 12479 | NM_004999.3 | 41.9576586 | -7.0150181 | 1.70440362 | -4.1158198 |
| 12480 | NR_003605.1 | 26.3848458 | -6.3433975 | 1.54125245 | -4.115742 |
| 12481 | NM_001130688.1 | 32.3913118 | -6.6356639 | 1.61229256 | -4.1156699 |
| 12482 | NM_001173479.1 | 19.8812126 | -5.9427825 | 1.44415083 | -4.1150705 |
| 12483 | NM_018140.3 | 16.3703463 | -5.6490349 | 1.37279019 | -4.1150024 |
| 12484 | NM_006154.3 | 29.0655564 | -5.5068242 | 1.33850172 | -4.1141704 |
| 12485 | NR_130107.1 | 52.601555 | 2.26687499 | 0.55101423 | 4.11400447 |
| 12486 | NM_021973.2 | 25.2939044 | -6.2832848 | 1.52730656 | -4.1139644 |
| 12487 | NM_001244584.2 | 34.8126717 | -6.7427431 | 1.63907658 | -4.113745 |
| 12488 | NR_104080.1 | 20.3315544 | -5.9647524 | 1.45006147 | -4.113448 |
| 12489 | NM_001105530.1 | 166.327686 | -4.9360873 | 1.2002431 | -4.1125729 |
| 12490 | NM_001202439.2 | 66.9964731 | -3.8513753 | 0.9365009 | -4.1125164 |
| 12491 | NR_131907.1 | 16.2790009 | -5.6417938 | 1.37220035 | -4.1114942 |
| 12492 | NM_016090.3 | 22.2582648 | -6.0992558 | 1.48350219 | -4.1113898 |
| 12493 | NM_198123.1 | 23.6910479 | -6.1899809 | 1.50590429 | -4.1104743 |
| 12494 | NM_001164637.2 | 19.5561219 | -5.915087 | 1.4390532 | -4.1104019 |
| 12495 | NM_001347902.1 | 20.48634 | -5.978715 | 1.45496595 | -4.1091786 |
| 12496 | NM_001318909.1 | 17.3480374 | -5.7477146 | 1.39898674 | -4.108484 |
| 12497 | NR_134893.1 | 18.4115634 | -5.8205162 | 1.41670655 | -4.108484 |
| 12498 | NM_001134831.1 | 30.2789398 | -6.5464543 | 1.5934562 | -4.1083365 |
| 12499 | NM_001199875.1 | 13.7523925 | -5.4129369 | 1.31758157 | -4.1082366 |
| 12500 | NM_001033578.2 | 17.6353723 | -5.7666115 | 1.40368122 | -4.1082059 |
| 12501 | NM_013417.3 | 168.476916 | -5.1393953 | 1.25103684 | -4.1081087 |
| 12502 | NM_178009.4 | 20.1592222 | -5.9488434 | 1.44818667 | -4.1077877 |
| 12503 | NM_018205.3 | 43.4126072 | -7.0612029 | 1.71919806 | -4.1072655 |
| 12504 | NM_007021.3 | 27.3048912 | -6.3908793 | 1.55602631 | -4.1071794 |
| 12505 | NM_017665.3 | 18.2832274 | -5.8083997 | 1.41426205 | -4.107018 |
| 12506 | NM_001284528.1 | 20.388557 | -5.968012 | 1.45335124 | -4.1063797 |
| 12507 | NM_144666.2 | 30.1548732 | -5.5612989 | 1.35432993 | -4.1063103 |
| 12508 | NM_001330466.1 | 15.2215479 | -5.5568689 | 1.35329473 | -4.1061779 |
| 12509 | NM_001331039.1 | 18.6842133 | -5.8370265 | 1.42158879 | -4.105988 |
| 12510 | NM_001164382.1 | 16.0899916 | -5.6290055 | 1.37117355 | -4.1052466 |
| 12511 | NM_001003398.1 | 57.4574053 | -3.9400623 | 0.95978544 | -4.105149 |
| 12512 | NM_001135865.1 | 199.027669 | -5.3716372 | 1.30851576 | -4.1051376 |

| | | | | | |
|---|---|---|---|---|---|
| 12513 | NR_028090.1 | 51.6248928 | -2.7227255 | 0.66327774 | -4.1049552 |
| 12514 | NM_001195328.1 | 16.5941625 | -5.6687404 | 1.38097666 | -4.1048778 |
| 12515 | NM_001317771.1 | 15.9091839 | -5.6203806 | 1.36921464 | -4.1048207 |
| 12516 | NM_002194.3 | 30.0397607 | -5.5487578 | 1.35184038 | -4.1045954 |
| 12517 | NM_152562.3 | 19.9101755 | -5.9379368 | 1.4467104 | -4.1044406 |
| 12518 | NM_002590.3 | 42.592271 | -5.2573428 | 1.28089946 | -4.1044149 |
| 12519 | NM_005261.3 | 24.0079594 | -6.2099154 | 1.51302579 | -4.1043024 |
| 12520 | NM_199122.2 | 19.3038426 | -5.8922398 | 1.4356593 | -4.1042048 |
| 12521 | NM_001243313.1 | 55.7825354 | -5.1132778 | 1.2460163 | -4.1037005 |
| 12522 | NM_178012.4 | 3793.40479 | -6.9439262 | 1.69218037 | -4.1035378 |
| 12523 | NM_000505.3 | 25.3265564 | -6.2822402 | 1.53103227 | -4.103271 |
| 12524 | NM_003097.4 | 664.284659 | -6.0316118 | 1.46998969 | -4.1031661 |
| 12525 | NM_020647.3 | 19.160367 | -5.8782479 | 1.43262179 | -4.10314 |
| 12526 | NM_001243788.1 | 36.0525609 | -4.3934053 | 1.07112775 | -4.1016632 |
| 12527 | NM_006133.2 | 101.542118 | -4.6400458 | 1.13133398 | -4.1013935 |
| 12528 | NM_001271907.1 | 15.9161741 | -5.6218016 | 1.37084056 | -4.1009887 |
| 12529 | NM_001160328.1 | 20.4779176 | -5.9761073 | 1.45729072 | -4.100834 |
| 12530 | NM_001304959.1 | 23.6612019 | -6.188975 | 1.50930284 | -4.1005521 |
| 12531 | NR_030717.1 | 14.6984458 | -5.4953564 | 1.34038469 | -4.0998353 |
| 12532 | NM_030799.8 | 186.949643 | -3.1455146 | 0.76728738 | -4.0995261 |
| 12533 | NM_001136019.2 | 15.1385629 | -5.5520387 | 1.35432277 | -4.0994944 |
| 12534 | NM_182743.2 | 26.2306722 | -6.3405218 | 1.54670919 | -4.0993626 |
| 12535 | NM_001128302.2 | 22.7258171 | -6.1302827 | 1.49565419 | -4.09873 |
| 12536 | NM_001482.2 | 64.2532371 | -4.1177397 | 1.00464117 | -4.0987169 |
| 12537 | NM_000161.2 | 21.6716904 | -6.0591684 | 1.4784406 | -4.0983509 |
| 12538 | NM_001190478.1 | 49.886678 | 3.64971465 | 0.89055947 | 4.09822677 |
| 12539 | NM_014569.3 | 19.223073 | -5.8934445 | 1.4381104 | -4.0980473 |
| 12540 | NM_005619.4 | 24.8377965 | -6.253384 | 1.52600702 | -4.0978737 |
| 12541 | NM_004234.4 | 14.6160323 | -5.4902628 | 1.33981107 | -4.0977888 |
| 12542 | NM_025248.2 | 228.36725 | -4.6291311 | 1.12970101 | -4.0976604 |
| 12543 | NM_001282866.1 | 17.6491884 | -5.7679962 | 1.40774724 | -4.0973237 |
| 12544 | NM_001258.2 | 17.1181075 | -5.7174166 | 1.39541362 | -4.0972917 |
| 12545 | NM_001042665.1 | 17.6662929 | -5.7687107 | 1.40801643 | -4.0970478 |
| 12546 | NM_006714.4 | 16.8349691 | -5.6891581 | 1.38861455 | -4.0970031 |
| 12547 | NM_017815.2 | 19.2463448 | -5.8886402 | 1.43753793 | -4.0963373 |
| 12548 | NM_058179.3 | 325.46468 | -5.3199854 | 1.29900886 | -4.0954189 |
| 12549 | NM_001195790.1 | 10.6337386 | 8.31624455 | 2.03063934 | 4.09538236 |
| 12550 | NM_014170.2 | 17.5744357 | -5.7565729 | 1.40563007 | -4.0953684 |
| 12551 | NM_017845.4 | 22.351493 | -6.1009367 | 1.48992413 | -4.0947969 |
| 12552 | NM_001242739.1 | 13.2467641 | -5.3498553 | 1.30654844 | -4.0946475 |
| 12553 | NR_134519.1 | 21.6637985 | -6.0633375 | 1.48079013 | -4.0946636 |
| 12554 | NM_001243101.1 | 19.3934767 | -5.8974416 | 1.44040462 | -4.0942951 |
| 12555 | NM_001039548.2 | 44.6572353 | -4.7684301 | 1.16486825 | -4.093536 |

| | | | | | |
|---|---|---|---|---|---|
| 12556 | NM_002077.3 | 92.1899467 | -3.8292211 | 0.93544959 | -4.0934553 |
| 12557 | NM_001166168.1 | 15.6168504 | -5.5945231 | 1.36671153 | -4.0934191 |
| 12558 | NM_032786.2 | 18.5124187 | -5.8327801 | 1.42515396 | -4.0927369 |
| 12559 | NR_027073.1 | 44.9676402 | -4.3671292 | 1.06705263 | -4.0927027 |
| 12560 | NM_001317163.1 | 14.0059608 | -5.4388071 | 1.32910293 | -4.0920887 |
| 12561 | NM_001199957.1 | 19.5304687 | -5.9134522 | 1.44510192 | -4.0920658 |
| 12562 | NR_135677.1 | 35.332572 | -6.7619311 | 1.65249876 | -4.0919433 |
| 12563 | NM_203397.2 | 15.3838653 | -5.5631605 | 1.35973391 | -4.0913597 |
| 12564 | NM_003505.1 | 165.295231 | -3.5928154 | 0.8782019 | -4.0911041 |
| 12565 | NM_001282864.1 | 15.4733485 | -5.5729773 | 1.3622601 | -4.0909789 |
| 12566 | NR_136746.1 | 27.9750064 | -6.4266189 | 1.57099286 | -4.0908008 |
| 12567 | NM_001077243.2 | 20.69169 | -5.9966311 | 1.46593952 | -4.0906402 |
| 12568 | NM_001305624.1 | 24.8999494 | -6.2594529 | 1.53018398 | -4.0906538 |
| 12569 | NM_001199355.1 | 26.2806448 | -6.336208 | 1.54901321 | -4.0904803 |
| 12570 | NR_036492.1 | 21.0172386 | -6.019609 | 1.47176983 | -4.0900478 |
| 12571 | NR_015360.1 | 14.0022135 | -5.4257716 | 1.32666363 | -4.089787 |
| 12572 | NM_004087.2 | 23.7184149 | -6.1959304 | 1.51506613 | -4.0895445 |
| 12573 | NM_006902.4 | 23.5796695 | -6.1859365 | 1.51276375 | -4.0891623 |
| 12574 | NM_022761.2 | 19.2564529 | -5.8893681 | 1.44025739 | -4.0891081 |
| 12575 | NM_201524.3 | 29.6738541 | -6.5138653 | 1.59309335 | -4.0888158 |
| 12576 | NM_001330993.1 | 14.0643614 | -5.4332796 | 1.32895862 | -4.0883738 |
| 12577 | NM_152308.2 | 16.8874749 | -5.6980685 | 1.39377941 | -4.088214 |
| 12578 | NM_015186.3 | 34.8536979 | -5.7643293 | 1.41003245 | -4.0880827 |
| 12579 | NM_025193.3 | 15.1291036 | -5.5375404 | 1.35466348 | -4.0877609 |
| 12580 | NM_173812.4 | 36.9995304 | -4.9946034 | 1.22188313 | -4.0876278 |
| 12581 | NM_001167676.1 | 12.8156303 | -5.298281 | 1.29640978 | -4.0868875 |
| 12582 | NR_027407.1 | 113.760722 | 7.18337621 | 1.75778373 | 4.08660981 |
| 12583 | NM_003458.3 | 211.316012 | -3.7340226 | 0.91378036 | -4.0863459 |
| 12584 | NR_033827.1 | 86.5736453 | 1.9081592 | 0.46698016 | 4.08616759 |
| 12585 | NM_003805.4 | 15.7560033 | -5.5974488 | 1.36990357 | -4.0860167 |
| 12586 | NM_001164760.1 | 20.7367128 | -5.9989326 | 1.46844978 | -4.0852147 |
| 12587 | NR_126046.1 | 15.1791673 | -5.5376997 | 1.35560355 | -4.0850437 |
| 12588 | NM_001143679.1 | 16.1310641 | -5.6310468 | 1.3784893 | -4.0849405 |
| 12589 | NM_003492.2 | 15.8346461 | -5.6088244 | 1.37342471 | -4.0838237 |
| 12590 | NM_001199251.2 | 14.4737324 | -5.4726436 | 1.34017149 | -4.0835398 |
| 12591 | NM_170783.3 | 18.6172158 | -5.8357808 | 1.42912365 | -4.083468 |
| 12592 | NM_001013690.4 | 47.3535488 | -4.6310354 | 1.1341298 | -4.0833381 |
| 12593 | NM_001127692.2 | 17.937305 | -5.7880147 | 1.4176492 | -4.0828258 |
| 12594 | NR_015366.4 | 13.7145185 | -5.4060004 | 1.32416522 | -4.0825724 |
| 12595 | NM_019042.4 | 20.4020774 | -5.9785242 | 1.46459044 | -4.0820451 |
| 12596 | NM_001080410.2 | 217.041003 | -5.2145591 | 1.27762716 | -4.0814404 |
| 12597 | NM_001323312.1 | 18.424294 | -5.823541 | 1.42703449 | -4.0808691 |
| 12598 | NM_001172668.1 | 18.4007858 | -5.8203511 | 1.42632782 | -4.0806545 |

| | | | | | |
|---|---|---|---|---|---|
| 12599 | NM_030671.2 | 30.2411303 | -6.5382035 | 1.60272675 | -4.0794249 |
| 12600 | NM_001286265.1 | 19.2174032 | -5.8884807 | 1.44351468 | -4.0792662 |
| 12601 | NR_040037.1 | 47.4525267 | -2.2581938 | 0.55361273 | -4.0790135 |
| 12602 | NR_046098.1 | 53.50241 | 3.80889022 | 0.93378728 | 4.0789699 |
| 12603 | NM_001290144.1 | 30.5445032 | -6.5523695 | 1.60641144 | -4.0788863 |
| 12604 | NR_033704.1 | 16.4702791 | -5.6643975 | 1.38872444 | -4.0788492 |
| 12605 | NM_033655.3 | 24.4171271 | -5.2387438 | 1.28442841 | -4.0786577 |
| 12606 | NM_152432.2 | 22.5985043 | -5.119888 | 1.25542328 | -4.0782165 |
| 12607 | NM_058190.3 | 16.2662017 | -5.641385 | 1.38332449 | -4.0781357 |
| 12608 | NM_001220778.1 | 19.7290166 | -5.9300412 | 1.4543216 | -4.0775308 |
| 12609 | NR_103844.1 | 16.5251518 | -5.6660501 | 1.38968408 | -4.0772217 |
| 12610 | NM_005870.4 | 293.52383 | -5.1656266 | 1.26705188 | -4.0768864 |
| 12611 | NM_001099338.1 | 29.2134442 | -5.5101899 | 1.35162433 | -4.076717 |
| 12612 | NM_001322194.1 | 21.6368009 | -6.0620555 | 1.48710871 | -4.0764037 |
| 12613 | NM_020860.3 | 72.7016559 | -5.0777348 | 1.24567483 | -4.0762923 |
| 12614 | NM_015312.3 | 294.194867 | -4.6587856 | 1.14294289 | -4.0761316 |
| 12615 | NM_001318873.1 | 17.7898403 | -5.7704653 | 1.41573836 | -4.0759405 |
| 12616 | NM_003123.4 | 196.458634 | 2.20387486 | 0.54085811 | 4.07477452 |
| 12617 | NM_001207014.1 | 14.8895419 | -5.5091726 | 1.35202688 | -4.0747508 |
| 12618 | NM_016210.4 | 14.9091572 | -5.5225818 | 1.35556598 | -4.0740044 |
| 12619 | NM_001142310.1 | 29.2592622 | -6.4917757 | 1.59347315 | -4.0739787 |
| 12620 | NM_024071.3 | 26.7100413 | -6.3646411 | 1.56249222 | -4.0733906 |
| 12621 | NM_021141.3 | 821.627164 | -5.8991749 | 1.44841064 | -4.0728608 |
| 12622 | NM_002665.4 | 12.5674633 | -5.2768433 | 1.29598904 | -4.0716728 |
| 12623 | NM_001031849.2 | 31.8272617 | -6.6112252 | 1.62379158 | -4.071474 |
| 12624 | NR_027231.1 | 27.6817358 | -5.412215 | 1.32932065 | -4.0714143 |
| 12625 | NM_001145145.1 | 16.383091 | -5.6539562 | 1.38953663 | -4.0689508 |
| 12626 | NR_125372.1 | 16.5278486 | -5.6780546 | 1.39571448 | -4.0682064 |
| 12627 | NM_001309474.1 | 14.959816 | -5.5281416 | 1.35913621 | -4.0673933 |
| 12628 | NM_023931.3 | 19.4645917 | -5.9044217 | 1.45169552 | -4.067259 |
| 12629 | NM_133458.3 | 73.4928915 | -4.816321 | 1.18421293 | -4.0671072 |
| 12630 | NM_022833.3 | 217.64981 | -4.9421296 | 1.21557615 | -4.0656685 |
| 12631 | NM_032344.3 | 18.8587375 | -5.8529934 | 1.43969293 | -4.0654457 |
| 12632 | NM_020804.4 | 15.2058254 | -5.5367875 | 1.36207294 | -4.0649714 |
| 12633 | NM_015324.3 | 34.0837103 | -4.7482604 | 1.16812438 | -4.0648586 |
| 12634 | NM_018354.2 | 15.2429414 | -5.54609 | 1.36477667 | -4.0637345 |
| 12635 | NM_013363.3 | 12.7034732 | -5.2832465 | 1.3001568 | -4.0635456 |
| 12636 | NM_001314025.1 | 16.8028099 | -5.6859106 | 1.39938731 | -4.0631429 |
| 12637 | NM_001015509.2 | 16.2988788 | -5.6421971 | 1.38872644 | -4.0628571 |
| 12638 | NM_148912.3 | 21.7058593 | -6.0683505 | 1.49368423 | -4.0626729 |
| 12639 | NM_080647.1 | 19.2675861 | -5.8938857 | 1.45077646 | -4.0625733 |
| 12640 | NM_001285439.1 | 24.5191335 | -6.2330139 | 1.53431134 | -4.0624179 |
| 12641 | NM_001308420.1 | 14.9422613 | -5.5306097 | 1.36145781 | -4.0622703 |

| | | | | | |
|---|---|---|---|---|---|
| 12642 | NM_138418.3 | 18.4660721 | -5.8206961 | 1.4329262 | -4.0621046 |
| 12643 | NM_012426.4 | 488.80881 | -2.8673888 | 0.7059529 | -4.0617283 |
| 12644 | NM_018557.2 | 81.615134 | -4.1304095 | 1.01691892 | -4.0616901 |
| 12645 | NM_000440.2 | 79.9411647 | 1.99773322 | 0.49187238 | 4.06148688 |
| 12646 | NM_001195150.2 | 19.451504 | -5.9056901 | 1.45410422 | -4.061394 |
| 12647 | NM_000363.4 | 14.3903377 | -5.4613679 | 1.34481731 | -4.0610482 |
| 12648 | NM_053052.3 | 24.7195885 | -6.246752 | 1.53838814 | -4.0605825 |
| 12649 | NM_001312920.1 | 12.7034076 | -5.2955551 | 1.30415198 | -4.0605352 |
| 12650 | NM_024996.5 | 134.173295 | -4.6092857 | 1.13524193 | -4.0601792 |
| 12651 | NM_001302082.1 | 21.2152742 | -6.0287441 | 1.48505049 | -4.0596223 |
| 12652 | NR_036522.1 | 20.9842855 | -6.0142183 | 1.48165443 | -4.0591235 |
| 12653 | NM_022351.4 | 32.6667179 | -5.6721992 | 1.39752873 | -4.0587353 |
| 12654 | NM_001267616.1 | 41.8998978 | -7.0092838 | 1.72700972 | -4.0586244 |
| 12655 | NM_001304421.2 | 20.2911925 | -5.9671349 | 1.47026933 | -4.0585318 |
| 12656 | NM_001204088.1 | 15.0033707 | -5.5241199 | 1.36113941 | -4.0584527 |
| 12657 | NM_001321133.1 | 17.0054208 | -5.7072352 | 1.40625945 | -4.0584511 |
| 12658 | NR_046473.1 | 36.1203986 | 2.84438682 | 0.70087406 | 4.05834224 |
| 12659 | NM_144657.4 | 19.4750158 | -5.9014911 | 1.45422584 | -4.0581668 |
| 12660 | NM_001685.4 | 31.7765455 | -6.6101342 | 1.62886344 | -4.0581267 |
| 12661 | NM_020425.5 | 16.4428697 | -5.6677225 | 1.39675193 | -4.0577875 |
| 12662 | NM_173474.3 | 18.5477133 | -5.8301981 | 1.43694981 | -4.0573429 |
| 12663 | NM_001320210.1 | 19.1851519 | -5.877927 | 1.44873624 | -4.0572789 |
| 12664 | NM_152268.3 | 13.838108 | -5.4201434 | 1.33594374 | -4.0571644 |
| 12665 | NM_014839.4 | 51.4914924 | -7.3077135 | 1.80169879 | -4.0560129 |
| 12666 | NM_001127398.2 | 21.2460075 | -6.0251486 | 1.48551794 | -4.0559245 |
| 12667 | NM_001167575.1 | 14.6963874 | -5.5084399 | 1.35824918 | -4.0555444 |
| 12668 | NM_212535.2 | 18.4139547 | -5.8290144 | 1.43737289 | -4.0553251 |
| 12669 | NM_001300755.1 | 14.0028149 | -5.4227332 | 1.33720483 | -4.0552749 |
| 12670 | NM_001300756.1 | 19.252156 | -5.890357 | 1.45251877 | -4.0552708 |
| 12671 | NM_152517.2 | 17.595605 | -5.7663522 | 1.42194687 | -4.0552515 |
| 12672 | NM_006242.3 | 34.0802186 | -4.9206197 | 1.21348587 | -4.054946 |
| 12673 | NM_032127.3 | 19.9001835 | -5.9397521 | 1.46490294 | -4.0547069 |
| 12674 | NM_004675.3 | 15.7113328 | -5.6023433 | 1.38173562 | -4.0545696 |
| 12675 | NR_015427.2 | 193.267629 | -3.6278201 | 0.89478314 | -4.0544127 |
| 12676 | NM_014405.3 | 469.376007 | -5.8062727 | 1.43215334 | -4.0542255 |
| 12677 | NM_001319218.1 | 17.9933203 | -5.7854823 | 1.42705062 | -4.0541535 |
| 12678 | NM_019605.3 | 18.0877512 | -5.8019227 | 1.4313623 | -4.053427 |
| 12679 | NR_036511.1 | 15.3413503 | -5.5603301 | 1.37175663 | -4.0534377 |
| 12680 | NM_015873.3 | 14.1100027 | -5.435328 | 1.34119739 | -4.0525936 |
| 12681 | NM_001256167.1 | 27.2166047 | -6.3849454 | 1.57582335 | -4.0518155 |
| 12682 | NM_152643.7 | 135.791455 | -4.8359873 | 1.19366278 | -4.0513848 |
| 12683 | NM_021912.4 | 18.2840915 | -5.8067037 | 1.43338492 | -4.0510428 |
| 12684 | NM_014231.4 | 53.499837 | -5.3228629 | 1.31397345 | -4.0509669 |

| | | | | | |
|---|---|---|---|---|---|
| 12685 | NM_024334.2 | 242.752782 | -5.06199 | 1.24957906 | -4.0509562 |
| 12686 | NM_001318391.1 | 18.7354689 | -5.8575442 | 1.44619348 | -4.0503185 |
| 12687 | NM_001267574.2 | 21.236983 | -6.0310531 | 1.48922304 | -4.0497984 |
| 12688 | NM_003579.3 | 16.5288938 | -5.660486 | 1.39782957 | -4.0494822 |
| 12689 | NM_001085428.2 | 829.146971 | 7.2051075 | 1.77929519 | 4.04941662 |
| 12690 | NM_024328.5 | 15.0767166 | -5.5354606 | 1.36723922 | -4.0486409 |
| 12691 | NM_130470.2 | 20.8927086 | -5.9995168 | 1.48185876 | -4.0486428 |
| 12692 | NM_001174124.1 | 25.9726129 | -6.3146276 | 1.55977704 | -4.0484168 |
| 12693 | NM_001330585.1 | 98.6047996 | -1.7872631 | 0.44149216 | -4.0482329 |
| 12694 | NM_001318733.1 | 13.8665167 | -5.4177713 | 1.33833561 | -4.048141 |
| 12695 | NM_001753.4 | 22.537161 | -6.1193493 | 1.51165202 | -4.0481204 |
| 12696 | NM_000123.3 | 140.51019 | -4.5541713 | 1.12508038 | -4.047863 |
| 12697 | NM_001242313.1 | 16.2142419 | -5.6364753 | 1.39247017 | -4.0478248 |
| 12698 | NR_135106.1 | 15.0113861 | -5.5183932 | 1.3634158 | -4.0474763 |
| 12699 | NM_020845.2 | 31.8857623 | -6.6191942 | 1.6355107 | -4.0471727 |
| 12700 | NM_001017371.4 | 100.87577 | -4.4613194 | 1.10234355 | -4.0471225 |
| 12701 | NM_152450.2 | 14.805964 | -5.5041351 | 1.36029375 | -4.0462842 |
| 12702 | NM_145071.2 | 12.8503317 | -5.3015363 | 1.31051554 | -4.0453823 |
| 12703 | NR_109913.1 | 13.7377762 | -5.4009286 | 1.33515889 | -4.0451579 |
| 12704 | NM_001318129.1 | 23.3709242 | -6.1653573 | 1.52418998 | -4.0450058 |
| 12705 | NM_001178137.1 | 30.3009084 | -5.5612281 | 1.37497599 | -4.0446002 |
| 12706 | NR_144398.1 | 39.8345911 | 4.62570925 | 1.14386296 | 4.04393658 |
| 12707 | NM_024667.2 | 184.667121 | -4.8554904 | 1.20069211 | -4.0439096 |
| 12708 | NM_001322152.1 | 43.9251273 | -7.0757973 | 1.74975128 | -4.0438875 |
| 12709 | NM_001142648.1 | 26.7375556 | -6.3656328 | 1.57418738 | -4.043758 |
| 12710 | NM_001848.2 | 1259.57439 | -5.3134656 | 1.3140144 | -4.043689 |
| 12711 | NM_016589.3 | 120.455557 | -4.5828501 | 1.13344703 | -4.0432856 |
| 12712 | NM_201628.2 | 145.052899 | -5.5546524 | 1.37383919 | -4.0431605 |
| 12713 | NM_001323981.1 | 25.0118849 | -6.2610686 | 1.5486033 | -4.0430422 |
| 12714 | NM_001316952.1 | 16.2048626 | -5.6340643 | 1.39366284 | -4.0426308 |
| 12715 | NM_006040.2 | 53.9829423 | -5.5882421 | 1.38234265 | -4.0425882 |
| 12716 | NM_001303412.1 | 18.9866994 | -5.871108 | 1.4529422 | -4.0408408 |
| 12717 | NM_024717.5 | 14.2498172 | -5.4450242 | 1.34759019 | -4.0405638 |
| 12718 | NM_001278230.1 | 24.7551351 | -5.2676634 | 1.30374984 | -4.0403943 |
| 12719 | NM_181503.2 | 62.4016578 | -4.3004522 | 1.06440511 | -4.0402401 |
| 12720 | NR_036467.1 | 16.9466671 | -5.7040161 | 1.41182445 | -4.0401737 |
| 12721 | NM_005954.2 | 24.0028784 | -6.2034849 | 1.53548649 | -4.0400778 |
| 12722 | NM_017882.2 | 52.8336587 | -4.3047548 | 1.0655721 | -4.0398531 |
| 12723 | NR_024593.1 | 72.5778072 | 1.56159609 | 0.3865699 | 4.03962157 |
| 12724 | NR_135192.1 | 20.5757035 | -5.9835671 | 1.48121706 | -4.0396288 |
| 12725 | NM_178232.3 | 20.3534447 | -5.9716239 | 1.47832734 | -4.0394463 |
| 12726 | NR_073520.1 | 22.2906315 | -5.1180913 | 1.26718716 | -4.0389387 |
| 12727 | NM_001319184.1 | 18.2960896 | -5.8139035 | 1.43952154 | -4.0387749 |

| | | | | | |
|---|---|---|---|---|---|
| 12728 | NM_001080506.1 | 17.9894326 | -5.7926174 | 1.43439319 | -4.0383748 |
| 12729 | NM_000674.2 | 27.7488547 | -6.4170863 | 1.58917333 | -4.0380028 |
| 12730 | NM_001282179.1 | 31.5888544 | -6.6022434 | 1.6352516 | -4.037448 |
| 12731 | NM_001303534.1 | 15.2546664 | -5.5541371 | 1.37569316 | -4.0373372 |
| 12732 | NR_046292.1 | 15.9051576 | -5.6224034 | 1.39260922 | -4.037316 |
| 12733 | NM_145897.2 | 38.4722504 | -6.8894216 | 1.70655375 | -4.0370376 |
| 12734 | NM_001005386.2 | 20.0102308 | -5.9487567 | 1.47365905 | -4.0367252 |
| 12735 | NM_001329138.1 | 38.5714412 | -6.8882679 | 1.70646752 | -4.0365655 |
| 12736 | NM_001270427.1 | 7.68096023 | 7.86359258 | 1.94814631 | 4.03644867 |
| 12737 | NM_001199662.1 | 17.1045114 | -5.717176 | 1.41639438 | -4.0364294 |
| 12738 | NR_036478.1 | 13.8843165 | -5.4228938 | 1.34369844 | -4.0357967 |
| 12739 | NM_001105573.1 | 146.444794 | -3.4082831 | 0.84454117 | -4.0356625 |
| 12740 | NM_031455.3 | 21.4894077 | -6.0530102 | 1.49991064 | -4.0355806 |
| 12741 | NR_048551.1 | 31.0297399 | -6.5765027 | 1.62985518 | -4.0350227 |
| 12742 | NM_001130929.1 | 215.717584 | -3.6908477 | 0.91473851 | -4.0348664 |
| 12743 | NM_001345944.1 | 20.8458821 | -6.0043259 | 1.48834902 | -4.034219 |
| 12744 | NM_173620.2 | 17.1849248 | -5.7259789 | 1.41936028 | -4.034197 |
| 12745 | NM_001128203.1 | 15.8103603 | -5.610519 | 1.39083205 | -4.0339299 |
| 12746 | NM_015395.2 | 266.637256 | -2.6624237 | 0.66002436 | -4.0338265 |
| 12747 | NM_003995.3 | 92.9779535 | -3.7296437 | 0.92468483 | -4.0334215 |
| 12748 | NM_024097.3 | 18.4427139 | -5.8247871 | 1.44438828 | -4.0327017 |
| 12749 | NM_002155.4 | 22.2013648 | -6.0955164 | 1.51155028 | -4.0326257 |
| 12750 | NR_144421.1 | 15.1521427 | -5.5512123 | 1.37660709 | -4.0325322 |
| 12751 | NR_027040.1 | 21.8232904 | -6.0668719 | 1.50450457 | -4.0324716 |
| 12752 | NM_016426.6 | 55.0169732 | -3.8756762 | 0.96114444 | -4.0323557 |
| 12753 | NM_001317998.1 | 13.8773342 | -5.410156 | 1.34175734 | -4.0321419 |
| 12754 | NM_002098.5 | 28.2601738 | -3.1883153 | 0.79074078 | -4.0320612 |
| 12755 | NM_153341.3 | 18.7213481 | -5.8549128 | 1.4521092 | -4.0320058 |
| 12756 | NM_001330189.1 | 13.5523498 | -5.3711567 | 1.33214017 | -4.0319757 |
| 12757 | NM_000746.5 | 26.303933 | -6.339724 | 1.57279941 | -4.0308535 |
| 12758 | NM_001282750.1 | 33.318812 | -5.6848238 | 1.41045097 | -4.0305008 |
| 12759 | NM_018032.4 | 18.0455213 | -5.7943211 | 1.4377604 | -4.030102 |
| 12760 | NM_198182.2 | 12.5985345 | -5.2687849 | 1.3076993 | -4.0290493 |
| 12761 | NM_001271696.1 | 15.8054917 | -5.6001063 | 1.3899729 | -4.028932 |
| 12762 | NM_148178.2 | 16.3796312 | -5.6555721 | 1.40421848 | -4.0275585 |
| 12763 | NM_001127362.1 | 13.2019541 | -5.3505395 | 1.32849884 | -4.0275078 |
| 12764 | NM_020883.1 | 118.637785 | -4.7129757 | 1.1701962 | -4.0275089 |
| 12765 | NM_178335.2 | 13.8457713 | -5.4078286 | 1.34282342 | -4.0272075 |
| 12766 | NM_001171020.1 | 14.6846678 | -5.5005524 | 1.3659127 | -4.027016 |
| 12767 | NM_001322207.1 | 23.1920424 | -6.1592587 | 1.52955373 | -4.0268338 |
| 12768 | NM_006407.3 | 290.682329 | -5.4604977 | 1.35610815 | -4.0265946 |
| 12769 | NM_005701.3 | 13.8571373 | -5.4210451 | 1.34642497 | -4.0262512 |
| 12770 | NM_001329717.1 | 21.0873735 | -6.0219952 | 1.49592752 | -4.0255929 |

| | | | | | |
|---|---|---|---|---|---|
| 12771 | NM_001123377.1 | 164.207205 | -4.91992 | 1.22224878 | -4.0253017 |
| 12772 | NR_037669.1 | 17.1335744 | -5.7214687 | 1.42140195 | -4.0252292 |
| 12773 | NM_019048.2 | 136.05096 | -4.6322579 | 1.15083783 | -4.0251179 |
| 12774 | NR_045018.1 | 17.117666 | -5.7224913 | 1.42171148 | -4.0250722 |
| 12775 | NR_073562.1 | 21.7799295 | -5.0638295 | 1.25815862 | -4.0247941 |
| 12776 | NM_203446.2 | 25.8837931 | -6.3174939 | 1.56968278 | -4.0246947 |
| 12777 | NR_015364.2 | 12.3801654 | -5.2586858 | 1.3066171 | -4.0246571 |
| 12778 | NM_014620.5 | 26.1016291 | -6.3313686 | 1.57329418 | -4.0242751 |
| 12779 | NM_001572.3 | 15.4451164 | -5.5782592 | 1.38624683 | -4.0240014 |
| 12780 | NM_174869.2 | 13.0655283 | -5.3281873 | 1.32415538 | -4.0238385 |
| 12781 | NM_001142287.1 | 35.513641 | -4.8690765 | 1.21020509 | -4.0233482 |
| 12782 | NM_001080950.1 | 117.152513 | -3.2028113 | 0.79607799 | -4.0232381 |
| 12783 | NM_144641.3 | 18.5794842 | -5.8436723 | 1.45260359 | -4.0228954 |
| 12784 | NR_038253.1 | 14.5642407 | -5.4818775 | 1.36274513 | -4.0226726 |
| 12785 | NM_020233.4 | 16.7764083 | -5.6865529 | 1.41378136 | -4.0222294 |
| 12786 | NM_001242850.1 | 26.9148953 | -6.3673554 | 1.58307117 | -4.0221536 |
| 12787 | NM_006515.3 | 15.0062088 | -5.5261407 | 1.37395337 | -4.022073 |
| 12788 | NM_080927.3 | 204.239693 | -4.9017925 | 1.21883068 | -4.0217174 |
| 12789 | NM_031488.4 | 120.865183 | -3.4743159 | 0.86393041 | -4.0215228 |
| 12790 | NR_134539.1 | 17.1214706 | -5.7132901 | 1.42074114 | -4.0213449 |
| 12791 | NM_017694.3 | 53.9081549 | -4.6577946 | 1.15829861 | -4.0212382 |
| 12792 | NM_032712.2 | 23.7319364 | -6.1929724 | 1.54022237 | -4.0208301 |
| 12793 | NM_198531.4 | 35.5265049 | -6.770382 | 1.684669 | -4.0188203 |
| 12794 | NM_001258315.1 | 14.1612107 | -5.4438402 | 1.3550506 | -4.0174442 |
| 12795 | NM_000850.4 | 15.212567 | -5.5541018 | 1.38250822 | -4.0174096 |
| 12796 | NM_021964.2 | 160.30352 | -4.7570754 | 1.18414666 | -4.0173026 |
| 12797 | NR_003698.1 | 14.0530284 | -5.4343879 | 1.35277608 | -4.0172117 |
| 12798 | NM_001317836.1 | 23.4960173 | -6.172834 | 1.53678114 | -4.0167294 |
| 12799 | NM_002126.4 | 13.828629 | -5.4067374 | 1.34614277 | -4.0164666 |
| 12800 | NM_001199942.1 | 29.6325176 | -6.5133634 | 1.62204979 | -4.0155139 |
| 12801 | NM_152709.4 | 18.5977704 | -5.843053 | 1.4552709 | -4.0150965 |
| 12802 | NM_145331.2 | 30.4298189 | -6.5525809 | 1.63243515 | -4.0139916 |
| 12803 | NM_000450.2 | 19.6312646 | 8.46433896 | 2.10883514 | 4.01375091 |
| 12804 | NM_001134373.2 | 213.45138 | -3.429438 | 0.85460956 | -4.0128711 |
| 12805 | NM_015435.4 | 24.1476055 | -6.2198558 | 1.55001782 | -4.012764 |
| 12806 | NM_001284418.1 | 20.8843696 | -6.006359 | 1.49694461 | -4.0124123 |
| 12807 | NM_017551.2 | 107.060896 | -4.321083 | 1.07693176 | -4.0124018 |
| 12808 | NR_027780.1 | 25.511955 | -6.2995317 | 1.57013388 | -4.0120985 |
| 12809 | NM_001304369.1 | 40.108251 | 2.45717148 | 0.61245743 | 4.01198739 |
| 12810 | NM_052923.1 | 29.2890441 | -4.6866531 | 1.16834904 | -4.0113467 |
| 12811 | NM_001172772.1 | 18.1827787 | -5.8015585 | 1.44646926 | -4.0108412 |
| 12812 | NM_018012.3 | 109.709635 | -4.1020622 | 1.02283783 | -4.0104718 |
| 12813 | NM_001320640.1 | 31.7980631 | -6.6139471 | 1.64920513 | -4.0103847 |

| | | | | | |
|---|---|---|---|---|---|
| 12814 | NM_001286696.1 | 17.7593515 | -5.7619873 | 1.43683368 | -4.0101978 |
| 12815 | NM_001127584.1 | 30.3156131 | -6.5381951 | 1.63042651 | -4.0101134 |
| 12816 | NM_001145440.2 | 13.5761166 | -5.3850432 | 1.34286631 | -4.0101112 |
| 12817 | NR_037719.1 | 15.5130814 | -5.5702411 | 1.38913864 | -4.0098525 |
| 12818 | NM_013238.2 | 47.1668055 | -4.1249984 | 1.02872374 | -4.0098213 |
| 12819 | NM_001206836.1 | 7.19371678 | 7.74274758 | 1.93105254 | 4.00959964 |
| 12820 | NM_025049.3 | 19.3604634 | -5.8982574 | 1.47105043 | -4.0095549 |
| 12821 | NM_182502.3 | 29.2879311 | 2.77912012 | 0.69317808 | 4.00924411 |
| 12822 | NM_001319235.1 | 28.5188091 | -6.455712 | 1.61048059 | -4.0085624 |
| 12823 | NM_144966.5 | 21.9510935 | -5.0777387 | 1.26685027 | -4.0081601 |
| 12824 | NM_004398.3 | 76.6070579 | -4.6831318 | 1.16863462 | -4.0073533 |
| 12825 | NM_016598.2 | 19.5122491 | -5.9039007 | 1.47332766 | -4.0071879 |
| 12826 | NM_001136158.1 | 19.5422235 | -5.9176955 | 1.47690024 | -4.006835 |
| 12827 | NM_001267723.1 | 13.3237068 | -5.360307 | 1.33787223 | -4.0065911 |
| 12828 | NR_024534.1 | 18.9789266 | -5.8695139 | 1.46522384 | -4.005882 |
| 12829 | NM_001098634.1 | 72.6824466 | -7.8065248 | 1.94895097 | -4.0055009 |
| 12830 | NM_001304782.1 | 96.9467593 | 3.48799565 | 0.87082099 | 4.00541063 |
| 12831 | NM_003023.4 | 27.8158002 | -6.4247064 | 1.60405845 | -4.005282 |
| 12832 | NM_002193.3 | 20.0496601 | -5.94121 | 1.48335735 | -4.0052452 |
| 12833 | NM_001433.3 | 11.8057835 | -5.1804098 | 1.29349117 | -4.0049827 |
| 12834 | NR_026800.2 | 24.8383644 | 8.85416223 | 2.21082937 | 4.00490528 |
| 12835 | NR_027295.2 | 14.0105564 | -5.4268526 | 1.35512713 | -4.0046816 |
| 12836 | NM_001308221.1 | 8.87214378 | 8.08278413 | 2.01856631 | 4.00422026 |
| 12837 | NM_001304434.1 | 26.2167076 | -6.329064 | 1.58064869 | -4.0040928 |
| 12838 | NM_001014812.2 | 20.4657937 | -5.9738188 | 1.49200194 | -4.0038948 |
| 12839 | NR_105019.1 | 15.0138056 | -5.5264396 | 1.38028528 | -4.0038387 |
| 12840 | NR_038956.1 | 18.9850478 | -5.8705191 | 1.46630472 | -4.0036147 |
| 12841 | NR_026694.2 | 16.4673752 | -5.6563103 | 1.41286559 | -4.0034313 |
| 12842 | NM_001189.3 | 58.2596665 | -4.7817538 | 1.19448523 | -4.0031921 |
| 12843 | NM_001242821.1 | 23.3323731 | -6.1629394 | 1.53961457 | -4.0029105 |
| 12844 | NM_001001556.2 | 27.13907 | -4.5751687 | 1.14297669 | -4.0028538 |
| 12845 | NM_001330185.1 | 27.3553917 | -6.3997185 | 1.59880247 | -4.00282 |
| 12846 | NM_001256630.1 | 18.0496592 | -5.7889248 | 1.44634258 | -4.0024575 |
| 12847 | NM_001242336.1 | 18.683672 | -5.8399068 | 1.45924581 | -4.0020034 |
| 12848 | NM_001018116.2 | 26.9307261 | -5.3873199 | 1.34628444 | -4.0016209 |
| 12849 | NM_052958.2 | 15.8353666 | -5.6051021 | 1.40075421 | -4.0014887 |
| 12850 | NM_001304493.1 | 22.3745196 | -6.0975079 | 1.52393155 | -4.0011692 |
| 12851 | NM_000813.2 | 39.5557545 | -6.926242 | 1.73106962 | -4.0011343 |
| 12852 | NM_015257.2 | 21.1626022 | -6.028637 | 1.50677688 | -4.0010151 |
| 12853 | NM_001001550.2 | 32.3764505 | -5.6640102 | 1.41569252 | -4.000876 |
| 12854 | NM_004722.3 | 60.8039736 | -4.1489535 | 1.03702034 | -4.000841 |
| 12855 | NM_205850.2 | 13.6349727 | -5.3921993 | 1.34778718 | -4.0007795 |
| 12856 | NM_001127244.1 | 13.8583911 | -5.40654 | 1.35145117 | -4.0005441 |

| | | | | | |
|---|---|---|---|---|---|
| 12857 | NM_080629.2 | 34.7368114 | -4.9509939 | 1.23758788 | -4.0005191 |
| 12858 | NM_001190214.1 | 12.064711 | -5.2219568 | 1.30532649 | -4.0004986 |
| 12859 | NM_001142560.1 | 16.8049791 | -5.6932067 | 1.42325151 | -4.000141 |
| 12860 | NM_001243288.1 | 12.6769465 | -5.2861503 | 1.32152715 | -4.0000316 |
| 12861 | NM_152420.2 | 22.5389508 | -6.1105543 | 1.52772323 | -3.9997783 |
| 12862 | NM_001018053.1 | 13.0988662 | -5.3430403 | 1.33588542 | -3.9996247 |
| 12863 | NM_001201536.1 | 16.4127216 | -5.6520851 | 1.41331705 | -3.9991629 |
| 12864 | NM_001014795.2 | 15.4622464 | -5.5693623 | 1.39276703 | -3.9987752 |
| 12865 | NM_001143761.1 | 19.6231408 | -5.9172288 | 1.4798304 | -3.9985858 |
| 12866 | NM_001033521.1 | 17.6808107 | -5.7620364 | 1.441055 | -3.9984847 |
| 12867 | NM_002753.4 | 23.6544755 | -6.1909334 | 1.54851067 | -3.9979921 |
| 12868 | NM_018957.3 | 24.6663689 | -4.4251112 | 1.10685975 | -3.9978969 |
| 12869 | NM_002939.3 | 15.7315821 | -5.5885546 | 1.39792815 | -3.9977409 |
| 12870 | NM_006981.3 | 49.9789974 | -6.2836283 | 1.57185029 | -3.9975997 |
| 12871 | NM_001267868.1 | 21.9690342 | -5.0917875 | 1.27390168 | -3.9970019 |
| 12872 | NR_132999.1 | 28.0282387 | -2.8792549 | 0.7204406 | -3.9965195 |
| 12873 | NM_145058.2 | 13.1825612 | -5.3457399 | 1.33774715 | -3.9960765 |
| 12874 | NM_001127895.1 | 15.7333603 | -5.6004765 | 1.40153751 | -3.9959519 |
| 12875 | NM_173177.2 | 17.9331566 | -5.7819976 | 1.44705685 | -3.9956949 |
| 12876 | NM_001242906.1 | 22.2918328 | -6.1025849 | 1.52745732 | -3.9952573 |
| 12877 | NM_173627.4 | 17.2929689 | -5.7360355 | 1.43572177 | -3.9952278 |
| 12878 | NM_032854.3 | 13.6091559 | -5.3985153 | 1.35126708 | -3.9951505 |
| 12879 | NM_015836.3 | 19.8133261 | -5.9364172 | 1.48602266 | -3.9948363 |
| 12880 | NM_003342.4 | 205.87648 | -5.1754432 | 1.29555788 | -3.9947603 |
| 12881 | NM_001197222.1 | 31.7694054 | -6.6081663 | 1.65426452 | -3.9946249 |
| 12882 | NM_001286272.1 | 15.0550135 | -5.5315236 | 1.38488879 | -3.9942006 |
| 12883 | NM_001322978.1 | 28.9472847 | -6.4753705 | 1.62122402 | -3.9941245 |
| 12884 | NM_175850.2 | 12.7322319 | -5.2971419 | 1.32624616 | -3.9940865 |
| 12885 | NM_001271918.1 | 16.8559079 | -5.7001634 | 1.42726392 | -3.9937697 |
| 12886 | NM_001288754.1 | 32.8499306 | -6.6570762 | 1.66712161 | -3.9931557 |
| 12887 | NM_001040023.1 | 22.9742501 | -6.1413708 | 1.53801027 | -3.9930623 |
| 12888 | NM_024848.2 | 17.2997443 | -5.7380853 | 1.43712809 | -3.9927445 |
| 12889 | NM_152905.3 | 30.1996831 | -6.5412225 | 1.63828341 | -3.9927295 |
| 12890 | NM_005226.3 | 147.904142 | -3.5278862 | 0.88358762 | -3.992684 |
| 12891 | NM_024945.2 | 52.4153171 | -3.1406177 | 0.78660126 | -3.9926426 |
| 12892 | NM_000177.4 | 35.170147 | -6.7612884 | 1.69364666 | -3.9921482 |
| 12893 | NM_014578.3 | 13.4239759 | -5.3751271 | 1.34657028 | -3.9917167 |
| 12894 | NM_001025236.1 | 21.9187182 | -6.0762166 | 1.52233858 | -3.99137 |
| 12895 | NR_046437.1 | 12.816445 | -5.3106087 | 1.33081984 | -3.9904791 |
| 12896 | NM_001271471.2 | 15.1400781 | -5.5320971 | 1.38658956 | -3.9897149 |
| 12897 | NM_018950.2 | 13.075198 | -5.3361164 | 1.33755405 | -3.9894585 |
| 12898 | NM_001171868.1 | 13.731319 | -5.3906276 | 1.35157996 | -3.9883897 |
| 12899 | NM_001293104.1 | 16.4544357 | -5.6670542 | 1.4210349 | -3.9879768 |

| | | | | | |
|---|---|---|---|---|---|
| 12900 | NM_001324309.1 | 16.7428974 | -5.6966546 | 1.42846083 | -3.9879669 |
| 12901 | NM_018044.4 | 20.4627096 | -5.9730522 | 1.49783683 | -3.9877857 |
| 12902 | NM_182981.2 | 12.5070691 | -5.2709905 | 1.32185316 | -3.9875764 |
| 12903 | NM_023037.2 | 163.499392 | -4.7239829 | 1.18472666 | -3.9874033 |
| 12904 | NM_006576.3 | 13.2042461 | -5.3443771 | 1.34040377 | -3.9871397 |
| 12905 | NM_001303031.1 | 17.9433503 | -5.7909913 | 1.45249488 | -3.9869271 |
| 12906 | NM_001308382.1 | 13.8935841 | -5.4224403 | 1.36021521 | -3.9864577 |
| 12907 | NR_033714.1 | 23.3835498 | -6.1655217 | 1.54698024 | -3.9855206 |
| 12908 | NM_033345.3 | 21.6331287 | -6.0517341 | 1.51849222 | -3.9853574 |
| 12909 | NM_033362.3 | 26.1562921 | -6.3297909 | 1.58833075 | -3.9851844 |
| 12910 | NM_145232.3 | 48.5577207 | -4.4960325 | 1.12818563 | -3.9851886 |
| 12911 | NR_110858.1 | 15.0329153 | -5.5422455 | 1.3908705 | -3.9847315 |
| 12912 | NM_170774.1 | 9.95481909 | 8.21829689 | 2.06247414 | 3.98467875 |
| 12913 | NM_001199697.1 | 26.1564067 | -6.3351636 | 1.59019441 | -3.9838925 |
| 12914 | NM_017421.3 | 21.7641694 | -6.0622505 | 1.52181153 | -3.9835751 |
| 12915 | NM_001286767.1 | 47.7515182 | 10.4974421 | 2.63527523 | 3.98343291 |
| 12916 | NM_001320247.1 | 35.1331982 | -6.7548341 | 1.69576751 | -3.9833492 |
| 12917 | NR_027624.1 | 20.1435647 | -5.9507794 | 1.49413568 | -3.982757 |
| 12918 | NM_001006640.1 | 18.9295746 | -5.863467 | 1.47228128 | -3.9825725 |
| 12919 | NM_001320310.1 | 35.2832837 | -6.760226 | 1.69746534 | -3.9825414 |
| 12920 | NM_152732.4 | 16.0578407 | -5.6310012 | 1.41405452 | -3.982167 |
| 12921 | NM_001282512.1 | 61.3860995 | 2.68326511 | 0.67386148 | 3.98192387 |
| 12922 | NM_001099646.1 | 17.0630778 | -5.7194327 | 1.43665403 | -3.9810787 |
| 12923 | NM_001113207.1 | 17.2576286 | -5.7253052 | 1.43815133 | -3.9810172 |
| 12924 | NM_001300914.1 | 17.728808 | -5.7764967 | 1.45101132 | -3.9810142 |
| 12925 | NR_144335.1 | 15.4122489 | -5.5611959 | 1.39708833 | -3.9805614 |
| 12926 | NM_001270550.1 | 13.2435547 | -5.3485632 | 1.34396626 | -3.9796856 |
| 12927 | NM_001014445.2 | 48.8337264 | -5.4708199 | 1.3747726 | -3.9794362 |
| 12928 | NM_004836.6 | 65.0408679 | -3.9479007 | 0.99208924 | -3.9793806 |
| 12929 | NR_132751.1 | 13.8377888 | -5.4150643 | 1.36078817 | -3.9793587 |
| 12930 | NM_001080461.1 | 13.9370504 | -5.4333234 | 1.36561677 | -3.978659 |
| 12931 | NM_001319652.1 | 23.2800053 | -6.1550383 | 1.54700642 | -3.9786766 |
| 12932 | NM_000048.3 | 18.4125736 | -5.8308218 | 1.4657664 | -3.9780021 |
| 12933 | NM_001160305.2 | 20.7810716 | -5.9961225 | 1.50731899 | -3.978005 |
| 12934 | NM_183323.2 | 16.3677318 | -5.6565547 | 1.42211014 | -3.9775785 |
| 12935 | NM_001710.5 | 23.0849669 | -6.1556531 | 1.547604 | -3.9775376 |
| 12936 | NM_014001.4 | 24.9554628 | -6.2678258 | 1.57583775 | -3.9774563 |
| 12937 | NM_015059.2 | 194.158169 | -4.1483629 | 1.04301381 | -3.9772847 |
| 12938 | NR_135229.1 | 30.4799733 | -6.5476489 | 1.64645938 | -3.9768056 |
| 12939 | NR_135781.1 | 16.4973417 | -5.6737756 | 1.42685334 | -3.9764252 |
| 12940 | NM_001127203.3 | 24.1394956 | -6.2094057 | 1.56178641 | -3.9758354 |
| 12941 | NM_001143980.1 | 11.9671097 | -5.2018515 | 1.30837388 | -3.9758142 |
| 12942 | NM_001080503.2 | 12.965009 | -5.3257903 | 1.33967081 | -3.975447 |

| | | | | | |
|---|---|---|---|---|---|
| 12943 | NM_001291428.1 | 14.5514044 | -5.4776963 | 1.37800073 | -3.975104 |
| 12944 | NM_032364.5 | 149.001089 | -2.4004694 | 0.60391221 | -3.9748648 |
| 12945 | NM_001277064.1 | 24.7712111 | -6.2542853 | 1.57364433 | -3.9743957 |
| 12946 | NM_001005290.3 | 19.9752414 | -5.9468314 | 1.49654449 | -3.9737084 |
| 12947 | NM_022475.2 | 34.8812046 | -5.7589635 | 1.44927835 | -3.9736766 |
| 12948 | NM_001242481.1 | 16.2406042 | -5.634779 | 1.41812049 | -3.9734134 |
| 12949 | NM_001014842.2 | 15.0963834 | -5.5407876 | 1.39457817 | -3.9730922 |
| 12950 | NM_001173989.2 | 13.2053482 | -5.3549692 | 1.34803986 | -3.9724116 |
| 12951 | NM_001039479.1 | 246.975014 | -4.9965502 | 1.25785975 | -3.9722634 |
| 12952 | NM_017715.3 | 15.6448058 | -5.5856675 | 1.40622062 | -3.9721132 |
| 12953 | NR_001445.2 | 25.0911691 | -6.2674388 | 1.57800065 | -3.9717593 |
| 12954 | NM_001278622.1 | 17.8789933 | -5.7906626 | 1.45801964 | -3.9715944 |
| 12955 | NM_001008222.2 | 15.0973922 | -5.5346517 | 1.39365747 | -3.9713142 |
| 12956 | NM_001322046.1 | 24.8800743 | -5.2549596 | 1.32338363 | -3.9708513 |
| 12957 | NM_001330237.1 | 18.1537773 | -5.8034635 | 1.46152603 | -3.9708246 |
| 12958 | NM_001171888.1 | 11.9133931 | -5.1989319 | 1.30928972 | -3.9708033 |
| 12959 | NR_110996.1 | 38.997888 | 3.17957437 | 0.80086168 | 3.97019166 |
| 12960 | NM_018323.3 | 21.2822116 | -6.0316664 | 1.51926485 | -3.9701217 |
| 12961 | NM_032347.2 | 15.4901653 | -5.5692087 | 1.40294497 | -3.9696559 |
| 12962 | NM_001309242.1 | 78.8816326 | -3.1315135 | 0.78888061 | -3.9695658 |
| 12963 | NR_073169.1 | 15.5333822 | -5.5860743 | 1.40724194 | -3.9695195 |
| 12964 | NM_001143937.1 | 12.7265998 | -5.2894842 | 1.33265101 | -3.9691443 |
| 12965 | NM_145731.3 | 12.2270622 | -5.238072 | 1.31974905 | -3.968991 |
| 12966 | NM_004506.3 | 34.4489412 | -5.7369384 | 1.44551472 | -3.9687858 |
| 12967 | NR_026545.2 | 17.6740778 | -5.7730636 | 1.4546817 | -3.9686095 |
| 12968 | NM_001270381.1 | 15.8730292 | -5.6144279 | 1.41474299 | -3.9685144 |
| 12969 | NM_001271285.1 | 17.8361557 | -5.7805052 | 1.45671953 | -3.9681662 |
| 12970 | NM_001018089.2 | 77.6984391 | -2.2220056 | 0.55996761 | -3.9680966 |
| 12971 | NM_001346141.1 | 17.359931 | -5.7339037 | 1.44505991 | -3.9679349 |
| 12972 | NM_018653.3 | 15.46274 | -5.5769932 | 1.40552278 | -3.9679138 |
| 12973 | NM_025263.3 | 22.3835778 | -6.1107761 | 1.54004254 | -3.9679269 |
| 12974 | NR_045513.1 | 13.5631154 | -5.377846 | 1.35535982 | -3.9678364 |
| 12975 | NM_001320179.1 | 18.0252369 | -5.7871215 | 1.45853464 | -3.9677642 |
| 12976 | NM_181640.2 | 14.8209837 | -5.5145115 | 1.39004168 | -3.9671555 |
| 12977 | NM_005591.3 | 21.6087684 | -5.056444 | 1.27464133 | -3.9669543 |
| 12978 | NM_001346393.1 | 18.7431045 | -5.8408567 | 1.47239775 | -3.9669014 |
| 12979 | NM_144689.4 | 24.0668899 | -6.2052539 | 1.56440824 | -3.9665183 |
| 12980 | NM_014706.3 | 225.576933 | -3.7421994 | 0.94354959 | -3.9660866 |
| 12981 | NM_212476.2 | 352.361989 | -10.083094 | 2.54246049 | -3.9658804 |
| 12982 | NM_001244705.1 | 28.9419006 | -4.6697368 | 1.17756327 | -3.9655932 |
| 12983 | NM_001103161.1 | 18.6693686 | -5.8430626 | 1.47382406 | -3.9645591 |
| 12984 | NM_001010879.3 | 15.6910647 | -5.5946681 | 1.41125315 | -3.9643264 |
| 12985 | NM_207406.3 | 30.2609142 | -6.538922 | 1.64944288 | -3.9643216 |

| | | | | | |
|---|---|---|---|---|---|
| 12986 | NM_005458.7 | 243.385723 | -3.5252057 | 0.88925013 | -3.9642453 |
| 12987 | NM_000943.4 | 27.0662419 | -5.3898361 | 1.36003744 | -3.9630056 |
| 12988 | NM_133491.4 | 21.4334859 | -6.0483377 | 1.52627044 | -3.9628218 |
| 12989 | NM_001098210.1 | 335.732955 | -5.1340505 | 1.29563314 | -3.9625804 |
| 12990 | NM_001167608.1 | 18.884697 | -5.8650377 | 1.48013948 | -3.9624899 |
| 12991 | NM_170709.2 | 41.1550851 | 10.2851324 | 2.59578188 | 3.96224832 |
| 12992 | NM_020859.3 | 200.101933 | -4.2881947 | 1.08252231 | -3.9612992 |
| 12993 | NM_001156474.1 | 15.9516686 | -5.6234882 | 1.41979372 | -3.9607783 |
| 12994 | NM_001001740.3 | 19.6156581 | -5.9156722 | 1.49359238 | -3.9607006 |
| 12995 | NR_130154.1 | 25.3939717 | -6.2865806 | 1.58726559 | -3.9606356 |
| 12996 | NM_001040653.3 | 97.4948238 | -4.2585638 | 1.07522863 | -3.9606124 |
| 12997 | NM_000629.2 | 197.766614 | -3.1276461 | 0.78983204 | -3.9598876 |
| 12998 | NR_024130.2 | 60.9071379 | 1.87006938 | 0.47229321 | 3.95955173 |
| 12999 | NM_001165884.2 | 8.31503514 | 7.9932886 | 2.01895036 | 3.95913081 |
| 13000 | NM_005070.3 | 94.3603136 | -4.9599245 | 1.25277963 | -3.9591357 |
| 13001 | NM_001318833.1 | 27.3892215 | -4.5825927 | 1.15760043 | -3.9586999 |
| 13002 | NM_001330284.1 | 20.9998037 | -6.0160536 | 1.51984733 | -3.9583276 |
| 13003 | NM_004741.4 | 157.399728 | -5.3167445 | 1.34340927 | -3.9576506 |
| 13004 | NR_024466.1 | 11.794069 | -5.1799553 | 1.30894224 | -3.9573597 |
| 13005 | NR_024525.2 | 12.1928163 | -5.2311365 | 1.32191938 | -3.9572281 |
| 13006 | NM_001284352.1 | 21.7418746 | -6.0681374 | 1.53348612 | -3.9570866 |
| 13007 | NM_003490.3 | 37.4945267 | -6.8506643 | 1.73128788 | -3.9569758 |
| 13008 | NM_001301838.1 | 17.2225513 | -5.7211018 | 1.44588895 | -3.9568058 |
| 13009 | NM_001013257.2 | 16.4274324 | -5.6683211 | 1.43265922 | -3.9565034 |
| 13010 | NR_024248.1 | 16.7185772 | -5.6804442 | 1.43578125 | -3.9563438 |
| 13011 | NM_012415.3 | 14.4147879 | -5.4643391 | 1.38130026 | -3.9559386 |
| 13012 | NM_001272083.1 | 12.9097379 | -5.3079788 | 1.34189986 | -3.95557 |
| 13013 | NR_109973.1 | 12.9939582 | -5.3217795 | 1.34576147 | -3.9544746 |
| 13014 | NM_006435.2 | 130.894301 | -4.7773285 | 1.20816609 | -3.9541985 |
| 13015 | NM_032850.4 | 13.6943214 | -5.4013034 | 1.36611092 | -3.953781 |
| 13016 | NM_001159699.1 | 11.4389018 | -5.1383186 | 1.29961744 | -3.9537163 |
| 13017 | NR_110285.1 | 31.0573434 | 2.17633425 | 0.5504556 | 3.95369624 |
| 13018 | NM_001322089.1 | 19.4046527 | -5.8986698 | 1.49211343 | -3.9532315 |
| 13019 | NM_014305.3 | 12.4124894 | -5.2462134 | 1.32706501 | -3.9532452 |
| 13020 | NM_001145399.1 | 32.2680395 | -4.7545941 | 1.20285383 | -3.9527613 |
| 13021 | NM_001242397.1 | 34.7117923 | -6.739688 | 1.70506947 | -3.9527351 |
| 13022 | NM_024753.4 | 75.2454556 | -2.9186446 | 0.7385154 | -3.952043 |
| 13023 | NM_199185.3 | 30.5768378 | -6.5589359 | 1.6599962 | -3.9511753 |
| 13024 | NM_198310.3 | 16.045677 | -5.6334206 | 1.42600389 | -3.9504946 |
| 13025 | NR_135579.1 | 18.4039515 | -5.826495 | 1.47514439 | -3.9497794 |
| 13026 | NR_024322.1 | 13.4002173 | -5.3587538 | 1.35685602 | -3.9493902 |
| 13027 | NM_001323413.1 | 16.4219899 | -5.6544793 | 1.43187034 | -3.9490163 |
| 13028 | NM_001256725.1 | 14.1886967 | -5.4539746 | 1.38115154 | -3.9488604 |

| | | | | | |
|---|---|---|---|---|---|
| 13029 | NM_001278159.1 | 13.0973208 | -5.3362188 | 1.35135293 | -3.9487973 |
| 13030 | NM_016556.3 | 16.1846974 | -5.6307241 | 1.4261926 | -3.9480811 |
| 13031 | NR_027121.1 | 74.028402 | 2.32574614 | 0.58910266 | 3.94794708 |
| 13032 | NM_001204199.1 | 17.2993771 | -5.7320586 | 1.45195437 | -3.9478228 |
| 13033 | NR_028099.1 | 17.8025431 | -5.7845154 | 1.46523917 | -3.9478302 |
| 13034 | NM_000462.3 | 30.5259505 | -6.5505055 | 1.65957981 | -3.9470867 |
| 13035 | NM_152665.2 | 22.4714161 | -6.1164414 | 1.54980128 | -3.9465972 |
| 13036 | NR_133646.1 | 16.0733767 | -5.6286186 | 1.42625707 | -3.9464264 |
| 13037 | NM_001172477.1 | 20.0301486 | -5.9490893 | 1.50753399 | -3.9462389 |
| 13038 | NM_080283.3 | 56.8880465 | 3.21662552 | 0.81518709 | 3.94587395 |
| 13039 | NM_001287526.1 | 62.52122 | 4.0854148 | 1.03555249 | 3.94515474 |
| 13040 | NM_001048223.2 | 19.1640456 | -5.8801454 | 1.49067427 | -3.9446213 |
| 13041 | NR_047673.1 | 19.4222836 | -5.9028547 | 1.496512 | -3.9444085 |
| 13042 | NM_001013653.2 | 15.8613578 | -5.6071234 | 1.42168939 | -3.9439863 |
| 13043 | NM_001185181.2 | 17.8767399 | -5.7862382 | 1.46710606 | -3.9439808 |
| 13044 | NM_145274.3 | 15.4055622 | -5.569021 | 1.41207736 | -3.9438498 |
| 13045 | NR_037883.1 | 34.4093194 | 2.42188076 | 0.61409093 | 3.94384716 |
| 13046 | NM_001282.2 | 256.993967 | -5.0573184 | 1.28237089 | -3.9437252 |
| 13047 | NM_001308440.1 | 30.0475425 | -5.5488422 | 1.40719749 | -3.9431865 |
| 13048 | NM_001346492.1 | 15.4913324 | -5.5769636 | 1.4143489 | -3.9431314 |
| 13049 | NR_036538.1 | 13.7679671 | -5.4093655 | 1.3719413 | -3.9428549 |
| 13050 | NM_001201.3 | 11.4577547 | -5.1392855 | 1.30359733 | -3.9423872 |
| 13051 | NM_001145636.1 | 13.2713006 | -5.3627319 | 1.36039948 | -3.9420273 |
| 13052 | NM_001136498.1 | 26.3395867 | -5.3564948 | 1.35882791 | -3.9419964 |
| 13053 | NR_037665.1 | 350.362654 | -4.2479608 | 1.07764061 | -3.9419086 |
| 13054 | NM_001302688.1 | 13.9716532 | -5.4187224 | 1.37465662 | -3.9418734 |
| 13055 | NM_024524.3 | 218.069577 | -4.8092574 | 1.22013818 | -3.9415678 |
| 13056 | NM_001190992.1 | 21.6857992 | -5.0824509 | 1.28953786 | -3.9412964 |
| 13057 | NM_012164.3 | 222.020596 | -3.7873162 | 0.96094326 | -3.9412485 |
| 13058 | NM_172218.2 | 34.0754737 | -5.7400386 | 1.45651107 | -3.9409509 |
| 13059 | NM_001130487.1 | 18.3103683 | -5.8237414 | 1.47786109 | -3.9406554 |
| 13060 | NM_017898.4 | 16.9666784 | -5.7094566 | 1.44901367 | -3.9402365 |
| 13061 | NM_024501.2 | 18.761531 | -5.8521547 | 1.48524529 | -3.9401941 |
| 13062 | NM_018955.3 | 1290.63154 | -6.0492776 | 1.53543001 | -3.9397938 |
| 13063 | NM_006492.2 | 14.9550943 | -5.5273753 | 1.40306104 | -3.9395116 |
| 13064 | NM_001193524.1 | 18.2842403 | -5.81992 | 1.47732959 | -3.9394865 |
| 13065 | NM_015942.4 | 31.4354087 | -5.6033423 | 1.42236795 | -3.9394464 |
| 13066 | NM_001280792.1 | 17.7388005 | -5.7774366 | 1.46675781 | -3.9389166 |
| 13067 | NM_024531.4 | 16.4421476 | -5.6632328 | 1.43789908 | -3.9385468 |
| 13068 | NR_045033.1 | 19.4308317 | -5.9018927 | 1.49853325 | -3.9384463 |
| 13069 | NM_001100419.1 | 24.8852648 | -6.2600089 | 1.58947419 | -3.9384149 |
| 13070 | NM_001171161.1 | 105.344365 | 2.43827056 | 0.61916509 | 3.93799747 |
| 13071 | NM_015417.4 | 25.6242111 | -6.3050889 | 1.6013614 | -3.9373304 |

| | | | | | |
|---|---|---|---|---|---|
| 13072 | NR_136664.1 | 15.2585242 | -5.5533659 | 1.41045517 | -3.9372863 |
| 13073 | NM_001289401.1 | 21.3554252 | -6.0332231 | 1.53236537 | -3.9371962 |
| 13074 | NM_001033081.2 | 35.0697068 | -6.7516909 | 1.71487201 | -3.9371398 |
| 13075 | NM_173631.3 | 13.1986337 | -5.3476665 | 1.35840732 | -3.936718 |
| 13076 | NM_001330305.1 | 31.5384155 | -5.6198531 | 1.42755451 | -3.9366995 |
| 13077 | NM_006103.3 | 11.7824405 | -5.1865158 | 1.31775549 | -3.9358711 |
| 13078 | NR_038103.1 | 16.0077667 | -5.6172937 | 1.42739573 | -3.9353443 |
| 13079 | NR_103783.1 | 13.9990924 | -5.434492 | 1.38105424 | -3.9350315 |
| 13080 | NM_006036.4 | 16.9007407 | -5.7065226 | 1.45031658 | -3.9346738 |
| 13081 | NM_001267609.1 | 21.8640702 | -6.0742755 | 1.54398924 | -3.9341437 |
| 13082 | NM_016361.4 | 11.5761873 | -5.1592304 | 1.31141784 | -3.9340859 |
| 13083 | NM_182734.2 | 18.0859874 | -5.7974218 | 1.47367055 | -3.9340012 |
| 13084 | NM_001037165.1 | 338.286193 | -1.4385802 | 0.36568421 | -3.9339412 |
| 13085 | NM_006000.2 | 17.6583057 | -5.7658901 | 1.46573063 | -3.9337993 |
| 13086 | NM_001164213.1 | 59.5130743 | -4.7928399 | 1.21838605 | -3.9337613 |
| 13087 | NM_004352.3 | 87.8496672 | -4.0970073 | 1.04149476 | -3.9337762 |
| 13088 | NM_173216.2 | 136.409903 | -4.7784915 | 1.21499378 | -3.9329349 |
| 13089 | NM_001287492.2 | 18.6005261 | -5.8426497 | 1.48558101 | -3.9329055 |
| 13090 | NM_014402.4 | 220.182891 | -2.7519876 | 0.69982988 | -3.9323665 |
| 13091 | NM_001305163.1 | 25.4159964 | -6.283636 | 1.5979694 | -3.932263 |
| 13092 | NM_181526.2 | 15.5133631 | -5.5808801 | 1.41939366 | -3.9318762 |
| 13093 | NM_001302545.1 | 22.844059 | -6.1409756 | 1.5618612 | -3.9318318 |
| 13094 | NM_001194986.1 | 28.3789717 | -5.4595866 | 1.38859077 | -3.9317463 |
| 13095 | NM_080680.2 | 16.6832823 | -5.6838792 | 1.44587011 | -3.9311133 |
| 13096 | NR_144632.1 | 14.8163622 | -5.5176158 | 1.40368321 | -3.9308127 |
| 13097 | NM_001888.4 | 22.3715252 | -5.0997159 | 1.29743435 | -3.9306157 |
| 13098 | NR_002836.2 | 131.101967 | 2.62669485 | 0.6683368 | 3.93019632 |
| 13099 | NM_002526.3 | 17.4334646 | -5.7510176 | 1.46329888 | -3.9301729 |
| 13100 | NM_005479.3 | 16.6029686 | -5.6759848 | 1.44427134 | -3.9299989 |
| 13101 | NM_194313.2 | 16.042225 | -5.6327634 | 1.43346248 | -3.9294809 |
| 13102 | NM_001130057.3 | 21.9688157 | -6.0815206 | 1.54785136 | -3.9290081 |
| 13103 | NM_006748.3 | 64.8734807 | -7.6410464 | 1.94479735 | -3.9289679 |
| 13104 | NM_020147.3 | 14.3677483 | -5.4650768 | 1.39108555 | -3.9286418 |
| 13105 | NR_038397.2 | 58.0918188 | -4.7618066 | 1.21207811 | -3.9286302 |
| 13106 | NM_021732.2 | 13.0472962 | -5.3335145 | 1.3577322 | -3.9282522 |
| 13107 | NR_073091.1 | 15.9526228 | -5.6147094 | 1.42931776 | -3.9282444 |
| 13108 | NM_032151.4 | 154.520481 | 1.73772102 | 0.44241199 | 3.92783432 |
| 13109 | NR_038938.1 | 15.048897 | -5.5330806 | 1.40870191 | -3.9277867 |
| 13110 | NR_132442.1 | 75.3216907 | 4.15250343 | 1.05724939 | 3.92764799 |
| 13111 | NR_026752.1 | 113.585414 | -3.7128947 | 0.94547637 | -3.9270095 |
| 13112 | NR_110787.1 | 254.364057 | 3.55258053 | 0.90473201 | 3.92666614 |
| 13113 | NM_004093.3 | 502.777451 | -5.8998245 | 1.50254267 | -3.9265604 |
| 13114 | NM_001330602.1 | 13.9169701 | -5.4162274 | 1.37965995 | -3.9257698 |

| | | | | | |
|---|---|---|---|---|---|
| 13115 | NM_152730.5 | 29.9073411 | -3.1663474 | 0.80655109 | -3.9257865 |
| 13116 | NM_001026.4 | 19.4194702 | -5.9048251 | 1.50418502 | -3.9255976 |
| 13117 | NR_023351.2 | 46.4972939 | 6.5381748 | 1.66554883 | 3.92553775 |
| 13118 | NM_001319844.1 | 21.5613136 | -6.0519422 | 1.5420628 | -3.9245757 |
| 13119 | NM_145315.4 | 14.1469678 | -5.4399899 | 1.38648629 | -3.92358 |
| 13120 | NM_183425.2 | 27.4022922 | -6.3975855 | 1.63080825 | -3.9229539 |
| 13121 | NM_173485.5 | 46.4575275 | -4.4346223 | 1.1305488 | -3.9225394 |
| 13122 | NM_001288798.1 | 11.1781346 | -5.1088121 | 1.30244168 | -3.9224882 |
| 13123 | NM_001282175.1 | 36.8886953 | -6.8269544 | 1.74055816 | -3.9222788 |
| 13124 | NR_033297.1 | 13.3825773 | -5.3611237 | 1.36687532 | -3.9221746 |
| 13125 | NM_014892.4 | 173.857911 | -4.6600246 | 1.18814429 | -3.9221033 |
| 13126 | NM_001102657.1 | 22.8151405 | -5.1292208 | 1.30784135 | -3.9218983 |
| 13127 | NM_001306147.1 | 24.1735009 | -6.2162874 | 1.58520517 | -3.9214403 |
| 13128 | NM_021980.4 | 253.408007 | -3.3222621 | 0.84728902 | -3.9210494 |
| 13129 | NM_001167947.2 | 14.1664383 | -5.4444259 | 1.38852127 | -3.9210245 |
| 13130 | NM_031244.3 | 15.115658 | -5.5344414 | 1.41152582 | -3.9208928 |
| 13131 | NM_014979.3 | 205.73904 | -3.7908779 | 0.9669125 | -3.9206008 |
| 13132 | NM_001011554.2 | 18.0702009 | -5.7883496 | 1.47651652 | -3.9202742 |
| 13133 | NR_027345.1 | 11.9028523 | -5.1857365 | 1.3228074 | -3.9202506 |
| 13134 | NR_138039.1 | 60.5004315 | 2.13472785 | 0.54454003 | 3.92024046 |
| 13135 | NM_001287763.1 | 47.8702851 | -7.2016246 | 1.83743674 | -3.9193864 |
| 13136 | NM_001193452.1 | 12.2276746 | -5.2410543 | 1.33732554 | -3.9190565 |
| 13137 | NM_001136225.1 | 11.4083631 | -5.1390893 | 1.31131411 | -3.9190376 |
| 13138 | NM_001287487.1 | 14.9031807 | -5.515596 | 1.40744913 | -3.9188599 |
| 13139 | NM_001258304.1 | 14.4819766 | -5.4849733 | 1.39980335 | -3.9183884 |
| 13140 | NM_020685.4 | 27.6776558 | -6.4108809 | 1.63613515 | -3.9183077 |
| 13141 | NM_001301030.1 | 13.7359055 | -5.4070218 | 1.37998032 | -3.9181876 |
| 13142 | NM_001330676.1 | 12.4189784 | -5.2559779 | 1.34160743 | -3.917672 |
| 13143 | NR_072977.1 | 17.2231059 | -5.7282379 | 1.46216311 | -3.9176463 |
| 13144 | NM_057749.2 | 15.0156618 | -5.5311194 | 1.41186854 | -3.9175881 |
| 13145 | NM_005192.3 | 14.5992726 | -5.4828487 | 1.39964679 | -3.9173088 |
| 13146 | NM_001003679.3 | 17.204156 | -5.7323392 | 1.4634901 | -3.9168964 |
| 13147 | NM_033406.2 | 12.5369102 | -5.265958 | 1.34470478 | -3.91607 |
| 13148 | NR_024494.1 | 13.2115897 | -5.339335 | 1.36343647 | -3.9160864 |
| 13149 | NM_025207.4 | 12.1355034 | -5.2219604 | 1.33361446 | -3.9156447 |
| 13150 | NM_003221.3 | 13.6617705 | -5.4045655 | 1.38028688 | -3.9155378 |
| 13151 | NM_017897.2 | 11.1897692 | -5.1093341 | 1.30489502 | -3.9155136 |
| 13152 | NM_001142966.1 | 54.2900024 | -3.9941495 | 1.02011988 | -3.9153726 |
| 13153 | NR_037845.1 | 56.0698094 | 2.26623173 | 0.57880161 | 3.91538601 |
| 13154 | NR_104246.1 | 25.3051 | -6.2837478 | 1.60501852 | -3.9150625 |
| 13155 | NM_201995.2 | 19.5831656 | -5.9169031 | 1.51146202 | -3.9146886 |
| 13156 | NM_058175.2 | 17.5771254 | -5.7632464 | 1.47251187 | -3.9138879 |
| 13157 | NM_001288727.1 | 16.0849657 | -5.6289543 | 1.43824694 | -3.9137607 |

| | | | | | |
|---|---|---|---|---|---|
| 13158 | NM_001134340.1 | 19.8510854 | -5.9343621 | 1.51631835 | -3.913665 |
| 13159 | NR_038192.1 | 31.9991172 | -6.6200814 | 1.69154628 | -3.9136271 |
| 13160 | NR_131202.1 | 16.0313402 | 5.89627104 | 1.50666583 | 3.9134564 |
| 13161 | NM_004556.2 | 42.1510626 | -4.3374341 | 1.10835971 | -3.9133812 |
| 13162 | NM_001102560.1 | 11.5945303 | -5.163146 | 1.31942293 | -3.913185 |
| 13163 | NM_001145520.1 | 17.0163757 | -5.7066971 | 1.45838663 | -3.9130207 |
| 13164 | NM_017848.5 | 14.1726749 | -5.4428791 | 1.39096259 | -3.9130306 |
| 13165 | NM_001282997.1 | 15.9250923 | -5.6088621 | 1.43355982 | -3.9125413 |
| 13166 | NM_001256470.1 | 31.1284781 | -6.5801671 | 1.68195368 | -3.9122166 |
| 13167 | NR_024547.1 | 14.1774327 | -5.4390613 | 1.39040282 | -3.9118601 |
| 13168 | NM_005112.4 | 17.558433 | -5.7519769 | 1.47067909 | -3.9111027 |
| 13169 | NM_024674.5 | 133.969778 | -1.8851078 | 0.48209544 | -3.9102377 |
| 13170 | NM_001330200.1 | 13.316352 | -5.3526236 | 1.3689323 | -3.9100718 |
| 13171 | NM_001012507.3 | 14.7514554 | -5.502988 | 1.40743426 | -3.9099432 |
| 13172 | NR_036444.1 | 20.4126794 | -5.975545 | 1.52837664 | -3.9097333 |
| 13173 | NM_001321163.1 | 14.8794934 | -5.5118989 | 1.40980354 | -3.9096929 |
| 13174 | NM_001282989.1 | 12.9338863 | -5.3231791 | 1.36168479 | -3.9092594 |
| 13175 | NM_013275.5 | 251.128063 | -9.5945876 | 2.45452966 | -3.9089312 |
| 13176 | NR_038275.2 | 25.6771058 | 2.95127767 | 0.75506085 | 3.90866203 |
| 13177 | NM_001239.3 | 47.142343 | -4.1308693 | 1.05698276 | -3.9081709 |
| 13178 | NM_024587.3 | 16.9398656 | -5.6951815 | 1.45790054 | -3.9064266 |
| 13179 | NM_181463.2 | 13.8500842 | -5.4090391 | 1.38487966 | -3.9057827 |
| 13180 | NR_110438.1 | 12.0524239 | -5.2162928 | 1.33559291 | -3.9056008 |
| 13181 | NM_024307.2 | 11.8875753 | -5.1940134 | 1.32995065 | -3.9054181 |
| 13182 | NM_001146015.1 | 26.2243321 | -5.3486341 | 1.36955947 | -3.9053683 |
| 13183 | NM_022090.4 | 26.9550666 | -4.48872 | 1.14948166 | -3.9049949 |
| 13184 | NR_136715.1 | 63.6284635 | -3.9919234 | 1.02228746 | -3.9048933 |
| 13185 | NM_001346100.1 | 13.9458097 | -5.4329106 | 1.39154888 | -3.9042183 |
| 13186 | NM_001253852.2 | 21.7640629 | -6.0696375 | 1.5547965 | -3.9038147 |
| 13187 | NM_001291641.1 | 17.9451692 | -5.7890447 | 1.48298512 | -3.9036431 |
| 13188 | NM_203346.4 | 241.1248 | -9.5359509 | 2.442951 | -3.9034557 |
| 13189 | NM_004855.4 | 12.2770203 | -5.2366538 | 1.34162993 | -3.9032029 |
| 13190 | NM_203291.1 | 16.8807751 | -5.7001751 | 1.46041588 | -3.9031177 |
| 13191 | NR_134489.1 | 14.5053527 | -5.4786562 | 1.40365609 | -3.9031328 |
| 13192 | NM_001521.3 | 15.9548178 | -5.6245287 | 1.44108558 | -3.9029803 |
| 13193 | NM_001100873.1 | 12.2995011 | -5.2479403 | 1.34463382 | -3.9028769 |
| 13194 | NM_004189.3 | 18.9620405 | -5.8691839 | 1.50391933 | -3.9025922 |
| 13195 | NM_004617.3 | 19.54925 | -5.9129043 | 1.51515556 | -3.9025064 |
| 13196 | NM_017704.2 | 25.4710535 | -5.2910193 | 1.35587969 | -3.9022779 |
| 13197 | NM_001323635.1 | 14.839063 | -5.516246 | 1.4137013 | -3.9019884 |
| 13198 | NM_005472.4 | 36.8764815 | 1.51144573 | 0.38736028 | 3.90191201 |
| 13199 | NM_177439.2 | 15.5438443 | -5.5844201 | 1.43123739 | -3.9018126 |
| 13200 | NM_018183.4 | 177.120764 | -2.7398159 | 0.70234234 | -3.9009692 |

| | | | | | |
|---|---|---|---|---|---|
| 13201 | NM_182700.5 | 19.2540589 | -5.8829171 | 1.50819767 | -3.9006274 |
| 13202 | NM_194271.2 | 14.9554563 | -5.5184173 | 1.41475124 | -3.9006273 |
| 13203 | NM_018659.2 | 20.6172839 | -5.9889406 | 1.53560207 | -3.9000603 |
| 13204 | NM_033015.3 | 23.8658405 | -6.195531 | 1.58900791 | -3.8989932 |
| 13205 | NM_001286616.1 | 22.2824284 | -5.1147405 | 1.31184765 | -3.898883 |
| 13206 | NM_018014.3 | 23.571617 | -6.1753682 | 1.58394831 | -3.8987183 |
| 13207 | NR_103523.1 | 23.6027867 | -6.180125 | 1.58531603 | -3.8983552 |
| 13208 | NM_001018052.3 | 15.6277211 | -5.5899584 | 1.43403714 | -3.8980569 |
| 13209 | NM_001014841.1 | 139.233579 | -5.3242675 | 1.3658963 | -3.8980027 |
| 13210 | NR_133680.1 | 64.8046464 | 4.99664417 | 1.28190702 | 3.89782106 |
| 13211 | NM_152731.2 | 16.8120519 | -5.6938432 | 1.46086163 | -3.8975924 |
| 13212 | NM_052927.2 | 17.4083453 | -5.7508636 | 1.47556452 | -3.8973989 |
| 13213 | NR_036432.1 | 15.3967751 | -5.5651429 | 1.42794358 | -3.8973129 |
| 13214 | NM_001324167.1 | 12.5144302 | -5.2758001 | 1.35371755 | -3.897268 |
| 13215 | NM_032020.4 | 92.9427919 | -4.1871901 | 1.07444065 | -3.8970883 |
| 13216 | NM_001112800.1 | 39.4545516 | -6.9249406 | 1.77718713 | -3.8965737 |
| 13217 | NM_145055.3 | 15.3434872 | -5.5504866 | 1.42449096 | -3.8964702 |
| 13218 | NM_018176.3 | 66.4547538 | -3.4326292 | 0.88101096 | -3.8962389 |
| 13219 | NM_009590.3 | 15.0782055 | -5.5349261 | 1.42069956 | -3.895916 |
| 13220 | NM_030647.1 | 71.6634697 | -4.028945 | 1.0341418 | -3.8959309 |
| 13221 | NM_001136137.1 | 16.0713552 | -5.6247935 | 1.44410783 | -3.8949955 |
| 13222 | NM_004286.4 | 302.001318 | -3.7042353 | 0.95108654 | -3.8947405 |
| 13223 | NR_103707.1 | 13.2027511 | -5.3416566 | 1.37151468 | -3.8947134 |
| 13224 | NM_003262.3 | 497.499848 | -3.3322039 | 0.85580081 | -3.8936676 |
| 13225 | NM_001134433.1 | 15.7245873 | -5.597043 | 1.43767342 | -3.8931255 |
| 13226 | NM_001286124.1 | 25.7516267 | -6.3105662 | 1.62105677 | -3.8928718 |
| 13227 | NR_109833.1 | 75.4528795 | 2.86504939 | 0.73611222 | 3.89213671 |
| 13228 | NM_001184824.1 | 16.5909452 | -5.6829013 | 1.46013592 | -3.8920358 |
| 13229 | NM_182692.2 | 180.914609 | -4.8296399 | 1.24098361 | -3.8917838 |
| 13230 | NM_001324192.1 | 13.5353583 | -5.3855778 | 1.38388578 | -3.8916346 |
| 13231 | NM_001042728.2 | 14.9803826 | -5.5342236 | 1.42250335 | -3.890482 |
| 13232 | NM_016602.2 | 11.1904187 | -5.1099231 | 1.3134958 | -3.8903231 |
| 13233 | NM_201538.1 | 23.5463294 | -6.1732069 | 1.58706123 | -3.8897094 |
| 13234 | NM_004528.3 | 216.470535 | -5.2396442 | 1.34711591 | -3.8895274 |
| 13235 | NM_001135032.1 | 15.3998844 | -5.5705772 | 1.43229692 | -3.8892614 |
| 13236 | NM_001146068.1 | 54.6165881 | -3.6526365 | 0.93921401 | -3.8890354 |
| 13237 | NM_173685.2 | 31.3658597 | -4.6122588 | 1.18596102 | -3.8890476 |
| 13238 | NM_001303529.1 | 12.0378712 | -5.2218786 | 1.34282633 | -3.8887222 |
| 13239 | NR_003606.2 | 21.4252226 | -6.0450636 | 1.55456131 | -3.8885978 |
| 13240 | NR_027139.1 | 11.8072211 | -5.1811449 | 1.3324026 | -3.8885731 |
| 13241 | NM_001271099.1 | 24.8434994 | -6.2551322 | 1.60869364 | -3.8883303 |
| 13242 | NM_017752.2 | 19.2417297 | -5.8925215 | 1.51578464 | -3.8874398 |
| 13243 | NM_001257999.2 | 14.4256051 | -5.4820715 | 1.41038365 | -3.8869364 |

| | | | | | |
|---|---|---|---|---|---|
| 13244 | NM_001099339.1 | 12.7940921 | -5.2959428 | 1.36254448 | -3.8868036 |
| 13245 | NM_001485.3 | 29.1972133 | -5.5107366 | 1.41803191 | -3.8861866 |
| 13246 | NM_001164796.1 | 24.3570406 | -5.2408874 | 1.34864817 | -3.8860301 |
| 13247 | NM_033064.4 | 807.319962 | -2.2949625 | 0.59058752 | -3.8858975 |
| 13248 | NM_001079802.1 | 19.6991227 | -5.9260966 | 1.52505335 | -3.885829 |
| 13249 | NM_001040214.2 | 53.4145699 | -7.3608497 | 1.89429248 | -3.8858042 |
| 13250 | NM_001102566.1 | 12.128012 | -5.2119259 | 1.34162832 | -3.8847763 |
| 13251 | NR_003083.2 | 11.8595402 | -5.1950049 | 1.33727278 | -3.8847758 |
| 13252 | NR_028050.1 | 20.047038 | -5.9407702 | 1.52935655 | -3.88449 |
| 13253 | NM_001032279.1 | 12.6811264 | -5.2918666 | 1.36237162 | -3.8843048 |
| 13254 | NM_194292.2 | 23.0467652 | -5.1601174 | 1.32849796 | -3.8841741 |
| 13255 | NM_001164283.2 | 213.062584 | 2.05391191 | 0.52879988 | 3.88410056 |
| 13256 | NM_001005339.1 | 19.1670862 | -5.8844074 | 1.51514952 | -3.883714 |
| 13257 | NM_001320220.1 | 47.3769102 | -6.2116106 | 1.59952416 | -3.8834115 |
| 13258 | NM_001329494.1 | 12.803108 | -5.3044521 | 1.36603064 | -3.8831135 |
| 13259 | NM_001143990.1 | 17.833692 | -5.768285 | 1.48561385 | -3.882762 |
| 13260 | NM_001076680.1 | 25.078397 | -5.2666382 | 1.35665414 | -3.8820787 |
| 13261 | NM_003803.3 | 11.9107623 | -5.2063304 | 1.34129354 | -3.8815743 |
| 13262 | NM_001271838.1 | 18.5266754 | -5.8248914 | 1.50092115 | -3.8808777 |
| 13263 | NM_001206572.1 | 18.5033945 | -5.8389551 | 1.50459712 | -3.8807432 |
| 13264 | NM_016442.3 | 13.2183495 | -5.340665 | 1.37621441 | -3.8806925 |
| 13265 | NM_001330692.1 | 16.9760891 | -5.7085521 | 1.47115319 | -3.8803248 |
| 13266 | NM_004023.2 | 22.6134231 | -6.1146455 | 1.57582228 | -3.8802888 |
| 13267 | NM_001294339.1 | 28.7057374 | -4.0864727 | 1.05319172 | -3.8800843 |
| 13268 | NR_004859.1 | 11.9315053 | -5.2010807 | 1.34049826 | -3.8799608 |
| 13269 | NM_001293562.1 | 11.4738537 | -5.1344574 | 1.32336329 | -3.8798548 |
| 13270 | NM_033160.6 | 13.6052487 | -5.3981487 | 1.39134339 | -3.8798105 |
| 13271 | NM_001321847.1 | 24.728404 | -6.2482202 | 1.610454 | -3.8797881 |
| 13272 | NM_001270732.1 | 14.2702841 | -5.4491872 | 1.40452215 | -3.8797446 |
| 13273 | NM_198853.2 | 12.3619762 | -5.2570061 | 1.35505392 | -3.8795549 |
| 13274 | NR_138077.1 | 29.8162967 | -6.5197547 | 1.68108274 | -3.8783068 |
| 13275 | NM_003643.3 | 42.1186544 | 2.20220755 | 0.56784727 | 3.87816875 |
| 13276 | NR_026967.1 | 26.1309392 | -5.3421826 | 1.37749594 | -3.8781839 |
| 13277 | NM_001166348.1 | 17.0510217 | -5.7198923 | 1.47513022 | -3.8775508 |
| 13278 | NM_021922.2 | 31.6069674 | -4.7646946 | 1.2288082 | -3.8774925 |
| 13279 | NM_001308229.1 | 23.0875129 | -4.2894836 | 1.10628035 | -3.8773929 |
| 13280 | NM_001258213.2 | 17.9524591 | -5.7840363 | 1.49176975 | -3.8772983 |
| 13281 | NM_002542.5 | 14.8185178 | -5.5099788 | 1.42108841 | -3.8772949 |
| 13282 | NM_032453.1 | 216.492501 | 1.70626856 | 0.44006506 | 3.87730979 |
| 13283 | NM_014260.3 | 23.6283979 | -6.178837 | 1.59372852 | -3.8769696 |
| 13284 | NM_199421.1 | 17.7704427 | -5.7781677 | 1.49043375 | -3.8768363 |
| 13285 | NM_000958.2 | 12.528399 | -5.2775662 | 1.36151161 | -3.876255 |
| 13286 | NM_199336.2 | 15.4099129 | -5.5694543 | 1.43706205 | -3.8755837 |

| | | | | | |
|---|---|---|---|---|---|
| 13287 | NM_001242822.1 | 15.9827063 | -5.6177616 | 1.44960777 | -3.8753666 |
| 13288 | NR_134857.1 | 20.5710588 | -5.9902795 | 1.54586329 | -3.8750383 |
| 13289 | NR_122038.1 | 15.4929297 | -5.5734369 | 1.43830364 | -3.8750071 |
| 13290 | NM_024108.2 | 12.0211626 | -5.2155847 | 1.34602416 | -3.8748076 |
| 13291 | NM_001164751.1 | 13.2258671 | -5.3436928 | 1.37926727 | -3.8742983 |
| 13292 | NM_001306212.1 | 13.7505181 | -5.4107449 | 1.39658735 | -3.8742617 |
| 13293 | NM_000333.3 | 18.4730445 | -5.8358484 | 1.50634556 | -3.8741764 |
| 13294 | NM_001297566.1 | 17.7566557 | -5.7749858 | 1.49064399 | -3.874155 |
| 13295 | NM_020752.2 | 20.5586963 | -5.9792722 | 1.54345057 | -3.8739642 |
| 13296 | NM_001297739.1 | 15.571003 | -5.5837538 | 1.4414566 | -3.8736885 |
| 13297 | NM_001037144.5 | 16.0984312 | -5.6392406 | 1.45611967 | -3.8727865 |
| 13298 | NM_001256192.1 | 11.8897649 | -5.2004046 | 1.34304662 | -3.8720954 |
| 13299 | NR_002775.2 | 121.958544 | 1.98813723 | 0.51350215 | 3.87172131 |
| 13300 | NR_122072.1 | 69.9494709 | 2.96955928 | 0.76701462 | 3.87158104 |
| 13301 | NM_001007073.1 | 20.4918492 | -5.9797017 | 1.54477494 | -3.870921 |
| 13302 | NM_001006933.1 | 16.4031911 | -5.6584631 | 1.46184059 | -3.8707798 |
| 13303 | NM_152387.3 | 11.297667 | -5.1222365 | 1.32348348 | -3.8702686 |
| 13304 | NR_037717.1 | 17.5600899 | -5.753152 | 1.48663844 | -3.8699067 |
| 13305 | NM_025209.3 | 16.3928031 | -5.6603647 | 1.46270121 | -3.8698024 |
| 13306 | NM_152426.3 | 104.236017 | 2.18938312 | 0.56581851 | 3.86940879 |
| 13307 | NM_001323951.1 | 13.0431139 | -5.3150359 | 1.3737632 | -3.8689608 |
| 13308 | NM_032718.4 | 66.550647 | -2.6416994 | 0.68279457 | -3.868952 |
| 13309 | NR_024543.1 | 19.2664838 | -5.8924324 | 1.52305467 | -3.8688253 |
| 13310 | NM_001014764.2 | 14.5559686 | -5.4828766 | 1.41724103 | -3.8686973 |
| 13311 | NM_198833.1 | 17.4643003 | -5.7415009 | 1.4845216 | -3.8675765 |
| 13312 | NM_024091.3 | 22.1455736 | -5.0971842 | 1.31803957 | -3.8672467 |
| 13313 | NM_017806.3 | 24.706572 | -6.2421001 | 1.61413753 | -3.8671427 |
| 13314 | NM_001077268.1 | 13.6041151 | -5.3928153 | 1.39467194 | -3.8667268 |
| 13315 | NM_016078.5 | 13.9296269 | -5.4194687 | 1.4016568 | -3.8664734 |
| 13316 | NR_110395.1 | 29.3192832 | 2.38690961 | 0.61741974 | 3.86594314 |
| 13317 | NR_024529.1 | 14.2588745 | -5.4652088 | 1.41381385 | -3.8655787 |
| 13318 | NM_018289.3 | 19.6395326 | -4.9206381 | 1.27296161 | -3.8655039 |
| 13319 | NM_148918.2 | 15.1076395 | -5.5375408 | 1.43278106 | -3.8648897 |
| 13320 | NM_002912.4 | 161.122503 | -4.6539316 | 1.2042357 | -3.8646351 |
| 13321 | NM_001166054.1 | 6.83538528 | 7.70710338 | 1.99433038 | 3.86450684 |
| 13322 | NM_133452.2 | 161.453127 | -4.8256897 | 1.24884871 | -3.8641107 |
| 13323 | NR_103831.1 | 73.1038772 | 2.38276944 | 0.61670184 | 3.86373003 |
| 13324 | NM_000839.3 | 19.6942245 | -5.9176711 | 1.53169641 | -3.8634752 |
| 13325 | NR_133652.1 | 12.751695 | -5.2938323 | 1.37045881 | -3.8628175 |
| 13326 | NM_001287750.1 | 13.9208374 | -5.4308003 | 1.4060719 | -3.8623916 |
| 13327 | NM_001286451.1 | 14.885176 | -5.5175161 | 1.42855065 | -3.8623175 |
| 13328 | NR_037807.1 | 10.8404988 | -5.0558777 | 1.30915283 | -3.8619461 |
| 13329 | NM_006942.1 | 11.5423516 | -5.1562373 | 1.33522961 | -3.8616858 |

| | | | | | |
|---|---|---|---|---|---|
| 13330 | NR_027408.1 | 23.5718006 | -6.1840414 | 1.60162406 | -3.8611067 |
| 13331 | NM_001032730.1 | 12.2144877 | -5.2323812 | 1.35528578 | -3.8607217 |
| 13332 | NM_181712.4 | 13.7079918 | -5.4030251 | 1.39948239 | -3.860731 |
| 13333 | NM_001025237.1 | 13.0274295 | -5.3256467 | 1.37945585 | -3.8606866 |
| 13334 | NM_001166686.1 | 12.3838787 | -5.2427306 | 1.35800425 | -3.8606143 |
| 13335 | NM_177963.3 | 11.7989264 | -5.1858997 | 1.34364477 | -3.8595765 |
| 13336 | NM_024789.3 | 39.0940488 | -4.886895 | 1.26622097 | -3.859433 |
| 13337 | NR_023318.1 | 17.6617272 | -5.7600989 | 1.49257949 | -3.8591572 |
| 13338 | NM_001105529.1 | 97.4086022 | -4.9924265 | 1.29390239 | -3.8584259 |
| 13339 | NM_020922.4 | 16.286002 | -5.6545653 | 1.46564609 | -3.85807 |
| 13340 | NM_014518.3 | 17.3707104 | -5.7409587 | 1.48810215 | -3.8579063 |
| 13341 | NM_001318926.1 | 12.0821993 | -5.2211641 | 1.35347921 | -3.8575872 |
| 13342 | NM_001098533.2 | 15.860359 | -5.6129357 | 1.45527688 | -3.8569538 |
| 13343 | NM_001324087.1 | 12.4470196 | -5.2607555 | 1.36402033 | -3.8568014 |
| 13344 | NM_001256600.1 | 21.7845972 | -6.0621678 | 1.57185555 | -3.8566953 |
| 13345 | NM_001301820.1 | 20.6343571 | -5.9925381 | 1.55381883 | -3.8566517 |
| 13346 | NM_001242498.1 | 12.0000802 | -5.2168836 | 1.35278508 | -3.8564023 |
| 13347 | NM_016217.2 | 126.510095 | -3.6569205 | 0.94829311 | -3.8563187 |
| 13348 | NM_000640.2 | 14.9366207 | -5.5192082 | 1.43129647 | -3.8560901 |
| 13349 | NM_004625.3 | 24.0546926 | -6.2075098 | 1.61011949 | -3.85531 |
| 13350 | NM_001099400.1 | 24.4674772 | -6.2335487 | 1.61689806 | -3.8552515 |
| 13351 | NM_030577.2 | 15.9976663 | -5.6234522 | 1.45864789 | -3.8552499 |
| 13352 | NM_000288.3 | 12.1487294 | -5.2265627 | 1.35571827 | -3.8551983 |
| 13353 | NR_003189.2 | 18.816065 | -5.8531648 | 1.51841681 | -3.8547813 |
| 13354 | NM_021020.3 | 246.865558 | -4.270933 | 1.1079848 | -3.8546855 |
| 13355 | NM_018487.2 | 12.0581073 | -5.2215478 | 1.35460622 | -3.854661 |
| 13356 | NM_001136202.1 | 12.9539348 | -5.3242438 | 1.38137285 | -3.8543133 |
| 13357 | NM_001199379.1 | 195.847831 | -9.235942 | 2.39631863 | -3.8542212 |
| 13358 | NM_201532.2 | 16.5574581 | -5.6698447 | 1.47119861 | -3.8538948 |
| 13359 | NM_001322310.1 | 19.1145055 | -5.8822658 | 1.52649098 | -3.853456 |
| 13360 | NM_032601.3 | 12.836006 | -5.3025697 | 1.37608275 | -3.85338 |
| 13361 | NM_016284.4 | 190.704211 | -9.1975486 | 2.3869467 | -3.8532694 |
| 13362 | NM_020318.2 | 20.1359332 | -5.9526752 | 1.54541419 | -3.8518316 |
| 13363 | NM_030971.3 | 255.808961 | -4.2466598 | 1.10274274 | -3.8509978 |
| 13364 | NM_001300929.1 | 15.1584489 | -5.5396637 | 1.43912269 | -3.8493339 |
| 13365 | NM_004537.6 | 405.350152 | -5.3162776 | 1.38132054 | -3.8486922 |
| 13366 | NM_152345.4 | 295.276488 | -5.4739562 | 1.42250962 | -3.8480978 |
| 13367 | NM_024766.4 | 11.900474 | -5.1974871 | 1.35076662 | -3.8478054 |
| 13368 | NM_001271685.1 | 22.5706959 | -6.1109275 | 1.5884741 | -3.8470426 |
| 13369 | NM_001166113.1 | 21.8319614 | -6.0726166 | 1.57873935 | -3.8464973 |
| 13370 | NM_144988.3 | 13.8486601 | -5.4181477 | 1.40860642 | -3.8464596 |
| 13371 | NM_001014440.3 | 38.6552561 | -6.8966518 | 1.79315123 | -3.8461072 |
| 13372 | NM_001166600.2 | 19.4820001 | -4.9034097 | 1.27498366 | -3.8458608 |

| | | | | | |
|---|---|---|---|---|---|
| 13373 | NM_001098728.1 | 41.92552 | -3.7636903 | 0.978666 | -3.8457352 |
| 13374 | NM_173662.2 | 20.2396878 | -5.9586126 | 1.54954572 | -3.8453932 |
| 13375 | NM_001204143.1 | 16.4989648 | -5.6751483 | 1.4759871 | -3.844985 |
| 13376 | NM_001161546.1 | 10.6930564 | -5.0470516 | 1.31279018 | -3.8445227 |
| 13377 | NM_002143.2 | 53.3816109 | -6.3880051 | 1.66169604 | -3.8442681 |
| 13378 | NM_001013619.3 | 112.310886 | 2.21635021 | 0.57663715 | 3.84357859 |
| 13379 | NM_021216.4 | 86.6107343 | -4.2337866 | 1.10166273 | -3.8430878 |
| 13380 | NM_001136131.2 | 12.6421678 | -5.2778658 | 1.37338047 | -3.8429743 |
| 13381 | NM_001195639.1 | 15.3280041 | -5.5582176 | 1.44639978 | -3.8427949 |
| 13382 | NM_001311199.1 | 13.8473561 | -5.416488 | 1.40951608 | -3.8427997 |
| 13383 | NM_021191.2 | 21.0432057 | -6.0111327 | 1.56431409 | -3.8426636 |
| 13384 | NM_001042510.1 | 20.2771314 | -5.9608394 | 1.55128435 | -3.8425189 |
| 13385 | NM_001199341.1 | 19.4861483 | -5.9085571 | 1.53775425 | -3.8423286 |
| 13386 | NR_026849.1 | 13.5397407 | -5.3777351 | 1.39969882 | -3.8420659 |
| 13387 | NM_133325.2 | 13.1104646 | -5.3336847 | 1.38832548 | -3.8418114 |
| 13388 | NR_121676.1 | 30.4610044 | 2.67923675 | 0.6974111 | 3.84168926 |
| 13389 | NM_001288567.1 | 17.9886239 | -5.7971778 | 1.50906428 | -3.8415711 |
| 13390 | NM_198128.2 | 16.1813143 | -5.6322641 | 1.46624318 | -3.8412892 |
| 13391 | NM_001318841.1 | 18.6946817 | -5.8482436 | 1.52265613 | -3.840817 |
| 13392 | NM_018132.3 | 12.1908513 | -5.2296376 | 1.36163382 | -3.8407077 |
| 13393 | NR_026900.1 | 18.9979668 | -5.8675985 | 1.52782709 | -3.840486 |
| 13394 | NM_001256183.1 | 344.325372 | -4.1207827 | 1.07303754 | -3.8402969 |
| 13395 | NM_004130.3 | 14.4951075 | -5.4801979 | 1.42718317 | -3.83987 |
| 13396 | NM_001242699.1 | 22.6278171 | -6.1257093 | 1.59568695 | -3.8389167 |
| 13397 | NM_030923.4 | 48.2696124 | -4.1718888 | 1.08682004 | -3.8386197 |
| 13398 | NR_001545.2 | 40.4966632 | -3.2867756 | 0.85635451 | -3.8381016 |
| 13399 | NM_001270731.1 | 21.7866901 | -6.0648666 | 1.5802895 | -3.83782 |
| 13400 | NM_014399.3 | 400.596249 | -5.4747396 | 1.42655065 | -3.8377464 |
| 13401 | NM_005795.5 | 12.0730079 | -5.2234055 | 1.36108529 | -3.8376768 |
| 13402 | NM_175062.3 | 15.2200817 | -5.5426434 | 1.4444467 | -3.8372087 |
| 13403 | NM_001276373.1 | 7.7515914 | 7.86228189 | 2.04898735 | 3.83715492 |
| 13404 | NM_006284.3 | 147.096973 | -4.9301612 | 1.28540296 | -3.8354985 |
| 13405 | NM_001160148.1 | 29.6818579 | -6.5094909 | 1.69726935 | -3.8352727 |
| 13406 | NM_001202234.1 | 66.7628248 | -2.9810443 | 0.77728256 | -3.8352132 |
| 13407 | NM_001251989.1 | 13.8433327 | -5.411277 | 1.41109513 | -3.8348067 |
| 13408 | NM_001282728.1 | 16.8385613 | -5.6963071 | 1.48545975 | -3.8347098 |
| 13409 | NM_002234.3 | 15.2077194 | -5.5556373 | 1.44896639 | -3.8342072 |
| 13410 | NM_206937.1 | 30.1391508 | -6.532273 | 1.70428513 | -3.8328522 |
| 13411 | NM_001098621.1 | 12.7654527 | -5.2900409 | 1.38020727 | -3.8327873 |
| 13412 | NM_001127711.2 | 18.1536468 | -5.7987808 | 1.5129447 | -3.8327778 |
| 13413 | NR_111943.1 | 19.0201485 | -5.8685618 | 1.53127934 | -3.8324567 |
| 13414 | NM_032679.2 | 13.3916693 | -5.3583834 | 1.39832294 | -3.8320071 |
| 13415 | NM_003512.3 | 11.3045572 | -5.1274527 | 1.33825354 | -3.8314509 |

| | | | | | |
|---|---|---|---|---|---|
| 13416 | NM_153634.2 | 12.8328148 | -5.3136812 | 1.38685915 | -3.8314498 |
| 13417 | NM_001280559.1 | 15.5399327 | -5.5721081 | 1.45463311 | -3.8305934 |
| 13418 | NM_007191.4 | 18.7303029 | -5.8530072 | 1.52798773 | -3.8305328 |
| 13419 | NR_036471.1 | 13.5036182 | -5.3799409 | 1.40475537 | -3.8298063 |
| 13420 | NM_001137551.1 | 23.542575 | -6.177759 | 1.6132152 | -3.8294699 |
| 13421 | NM_001048201.2 | 26.1462856 | -6.330281 | 1.65312697 | -3.829277 |
| 13422 | NM_000418.3 | 21.7420052 | -5.0754629 | 1.32547045 | -3.8291785 |
| 13423 | NM_001003700.1 | 5.45022773 | 7.33940084 | 1.91698045 | 3.82862581 |
| 13424 | NM_002689.3 | 59.3321983 | -2.5026302 | 0.6536632 | -3.8286234 |
| 13425 | NM_001032386.1 | 20.9938932 | -6.0147979 | 1.57108663 | -3.8284318 |
| 13426 | NM_003441.2 | 11.3436758 | -5.1248328 | 1.33874243 | -3.8280947 |
| 13427 | NM_004980.4 | 14.3555526 | -5.4663944 | 1.42799376 | -3.828024 |
| 13428 | NM_017729.3 | 15.0490705 | -5.5394868 | 1.44711285 | -3.8279577 |
| 13429 | NM_032582.3 | 218.81334 | -4.2908692 | 1.12098623 | -3.8277626 |
| 13430 | NM_001270469.1 | 15.3332298 | -5.5517899 | 1.450421 | -3.8277093 |
| 13431 | NM_021159.4 | 24.3180578 | -6.2294368 | 1.62748383 | -3.827649 |
| 13432 | NR_110090.1 | 57.6930901 | 2.9428689 | 0.76887875 | 3.82748113 |
| 13433 | NM_001040182.1 | 18.5687814 | -5.8303105 | 1.52331281 | -3.8273889 |
| 13434 | NM_174908.3 | 286.327025 | -2.952346 | 0.77137644 | -3.8273738 |
| 13435 | NM_152900.2 | 18.4326989 | -5.8325196 | 1.52395892 | -3.8272158 |
| 13436 | NM_001319054.1 | 14.4991708 | -5.4803668 | 1.43232129 | -3.8262133 |
| 13437 | NM_001008237.2 | 11.1088168 | -5.0919946 | 1.33092377 | -3.82591 |
| 13438 | NM_138412.3 | 14.3165462 | -5.4566883 | 1.42648625 | -3.8252653 |
| 13439 | NM_002347.4 | 17.9551549 | -5.7940644 | 1.51498522 | -3.8245023 |
| 13440 | NM_145802.3 | 217.915428 | -5.2452296 | 1.3717063 | -3.8238722 |
| 13441 | NM_001291861.2 | 532.505384 | -5.5871622 | 1.46136943 | -3.8232374 |
| 13442 | NR_024611.2 | 18.7399848 | -5.8492198 | 1.5302333 | -3.8224366 |
| 13443 | NM_145298.5 | 103.963931 | 1.80764325 | 0.47303062 | 3.82140851 |
| 13444 | NM_000436.3 | 157.706659 | -4.7394707 | 1.2402962 | -3.821241 |
| 13445 | NR_033240.1 | 10.8597898 | -5.0604587 | 1.32439361 | -3.8209628 |
| 13446 | NM_005982.3 | 14.8617038 | -5.5219006 | 1.44522989 | -3.8207766 |
| 13447 | NR_132739.1 | 97.5885664 | -4.2663881 | 1.11665914 | -3.8206718 |
| 13448 | NM_001008213.1 | 16.3556664 | -5.6462978 | 1.47826362 | -3.8195473 |
| 13449 | NM_001128848.1 | 262.81321 | -9.6600905 | 2.52942614 | -3.8190839 |
| 13450 | NM_001146267.1 | 321.781601 | 2.9421415 | 0.77047273 | 3.8186186 |
| 13451 | NR_135779.1 | 7.92727339 | 7.92412753 | 2.07520396 | 3.81848131 |
| 13452 | NM_173557.2 | 19.8916296 | -4.9250406 | 1.2898731 | -3.8182366 |
| 13453 | NR_001591.1 | 33.6845634 | 1.82824648 | 0.47892321 | 3.81741049 |
| 13454 | NM_001324150.1 | 25.6529629 | -4.4897481 | 1.17652978 | -3.8160939 |
| 13455 | NM_001146105.1 | 14.4327226 | -5.4786267 | 1.43574419 | -3.8158794 |
| 13456 | NR_002453.5 | 11.591526 | -5.1626509 | 1.35294883 | -3.8158508 |
| 13457 | NR_046551.1 | 24.372554 | 2.41112751 | 0.63190229 | 3.81566507 |
| 13458 | NM_001142465.2 | 15.8905238 | -5.6106059 | 1.47052374 | -3.8153793 |

| | | | | | |
|---|---|---|---|---|---|
| 13459 | NR_104228.1 | 11.1527468 | -5.0999438 | 1.33672812 | -3.8152439 |
| 13460 | NM_130834.2 | 18.5663143 | -5.8415053 | 1.53115576 | -3.8150954 |
| 13461 | NM_173497.3 | 16.7643039 | -5.6848227 | 1.49044355 | -3.8141818 |
| 13462 | NR_026562.3 | 10.7421998 | -5.0532221 | 1.32491519 | -3.8139967 |
| 13463 | NR_110017.1 | 25.2105986 | 1.82611194 | 0.4788527 | 3.81351498 |
| 13464 | NM_001320847.1 | 15.7788025 | -5.5977904 | 1.4681167 | -3.8129056 |
| 13465 | NM_001163079.1 | 26.5789967 | -5.355932 | 1.405292 | -3.8112592 |
| 13466 | NM_001040125.1 | 15.2194732 | -5.5406178 | 1.45375832 | -3.8112372 |
| 13467 | NM_014247.2 | 195.530227 | -4.6474801 | 1.21943924 | -3.8111617 |
| 13468 | NM_001130698.1 | 13.8872627 | -5.4168241 | 1.42134276 | -3.8110611 |
| 13469 | NM_001278542.1 | 14.9125258 | -5.5134481 | 1.4469054 | -3.8105104 |
| 13470 | NM_024094.2 | 16.9379623 | -5.7013782 | 1.49621966 | -3.8105221 |
| 13471 | NM_145278.4 | 56.0053236 | 5.9515064 | 1.56212179 | 3.80988629 |
| 13472 | NM_174937.3 | 42.5435576 | -7.0315947 | 1.84594529 | -3.8092108 |
| 13473 | NM_152594.2 | 141.753036 | -3.6132051 | 0.94858959 | -3.8090288 |
| 13474 | NM_001286436.1 | 12.1139026 | -5.2293558 | 1.37294218 | -3.8088682 |
| 13475 | NM_001013838.1 | 18.7894356 | -5.8528543 | 1.53689631 | -3.8082298 |
| 13476 | NM_173658.2 | 10.3869413 | -4.9961949 | 1.31200904 | -3.8080491 |
| 13477 | NM_001008224.1 | 19.631684 | -5.923676 | 1.55560726 | -3.8079509 |
| 13478 | NM_001421.3 | 13.1060341 | -5.338677 | 1.40199024 | -3.8079274 |
| 13479 | NM_001001711.2 | 45.1352715 | 9.65153295 | 2.5346287 | 3.80786856 |
| 13480 | NM_198476.3 | 47.1578521 | -3.192566 | 0.83857229 | -3.8071447 |
| 13481 | NM_001330651.1 | 17.0359303 | -5.7153634 | 1.50126931 | -3.8070207 |
| 13482 | NM_001136116.1 | 15.8958201 | -5.6086002 | 1.47325034 | -3.8069566 |
| 13483 | NR_045147.1 | 17.7669926 | -5.7667117 | 1.51481615 | -3.8068724 |
| 13484 | NM_001142451.1 | 14.5199371 | -5.479296 | 1.43934298 | -3.8068036 |
| 13485 | NM_005596.3 | 520.909948 | -5.7906032 | 1.52132304 | -3.8062943 |
| 13486 | NR_040254.1 | 20.4478601 | 5.41329079 | 1.42220161 | 3.80627526 |
| 13487 | NM_001020819.2 | 21.0055753 | -6.018741 | 1.58132733 | -3.8061323 |
| 13488 | NM_005019.4 | 30.015344 | -6.5272255 | 1.71500149 | -3.8059591 |
| 13489 | NM_145719.2 | 15.2365396 | -5.5447847 | 1.45693936 | -3.8057759 |
| 13490 | NR_033184.1 | 22.6140499 | -6.1167335 | 1.60737417 | -3.8054198 |
| 13491 | NM_001141936.2 | 19.4439779 | -5.904984 | 1.55176275 | -3.8053395 |
| 13492 | NM_019050.2 | 36.5761405 | -3.4597262 | 0.90918162 | -3.8053191 |
| 13493 | NM_152449.3 | 18.9351198 | -5.8575615 | 1.53932748 | -3.8052732 |
| 13494 | NM_001321345.1 | 15.1109308 | -5.5381923 | 1.45551207 | -3.8049786 |
| 13495 | NM_001018066.2 | 12.7688181 | -5.2934781 | 1.39124341 | -3.804854 |
| 13496 | NM_001166304.1 | 13.2518093 | -5.3577127 | 1.4082238 | -3.8045889 |
| 13497 | NM_001286458.1 | 14.0051842 | -5.4345734 | 1.42864679 | -3.8040007 |
| 13498 | NM_033102.2 | 11.5920163 | -5.1667669 | 1.35829108 | -3.8038731 |
| 13499 | NM_001290040.1 | 23.7083933 | -6.1835355 | 1.62560129 | -3.8038451 |
| 13500 | NM_152547.4 | 12.1182736 | -5.2121637 | 1.37040378 | -3.8033781 |
| 13501 | NM_001171816.1 | 10.6934832 | -5.0388833 | 1.32504003 | -3.8028159 |

| | | | | | |
|---|---|---|---|---|---|
| 13502 | NR_026698.1 | 16.0987552 | -5.6379802 | 1.48263501 | -3.8026758 |
| 13503 | NM_006041.2 | 63.14289 | -2.7352717 | 0.7193533 | -3.8024037 |
| 13504 | NM_002547.2 | 495.3826 | 1.38252007 | 0.36364021 | 3.80188999 |
| 13505 | NR_038988.1 | 10.4911597 | -5.0122629 | 1.31858366 | -3.8012475 |
| 13506 | NM_001130714.2 | 35.124902 | -6.753471 | 1.77676448 | -3.8009939 |
| 13507 | NM_030639.2 | 27.9937089 | -6.4258025 | 1.69058539 | -3.8009334 |
| 13508 | NM_001037161.1 | 11.7022801 | -5.1675531 | 1.35958451 | -3.8008326 |
| 13509 | NM_003364.3 | 19.6277645 | -5.9205331 | 1.55776089 | -3.8006688 |
| 13510 | NR_135777.1 | 13.1781706 | -5.3322685 | 1.40309505 | -3.8003616 |
| 13511 | NM_001860.2 | 11.6956048 | -5.1734015 | 1.36136481 | -3.8001581 |
| 13512 | NR_134950.1 | 11.1582937 | -5.0957211 | 1.34107133 | -3.7997391 |
| 13513 | NM_032168.2 | 107.825671 | -4.2828271 | 1.12719 | -3.7995609 |
| 13514 | NM_032432.4 | 14.6277949 | -5.4858465 | 1.44382106 | -3.7995335 |
| 13515 | NM_001007022.2 | 21.6632963 | -6.0646194 | 1.59635252 | -3.7990477 |
| 13516 | NM_001318966.1 | 17.3667722 | -5.7412206 | 1.51149892 | -3.7983624 |
| 13517 | NM_025165.2 | 13.273018 | -5.360862 | 1.41136946 | -3.7983406 |
| 13518 | NM_001171109.1 | 15.0615938 | -5.5318326 | 1.45670101 | -3.7975072 |
| 13519 | NM_001286522.1 | 15.1620573 | -5.5359928 | 1.45788726 | -3.7972709 |
| 13520 | NM_031477.4 | 12.3418646 | -5.2464232 | 1.38163033 | -3.7972698 |
| 13521 | NR_073083.1 | 27.9092634 | -6.428075 | 1.69307988 | -3.7966755 |
| 13522 | NM_001136043.2 | 21.1878596 | -6.023779 | 1.58687075 | -3.7960111 |
| 13523 | NM_001083617.1 | 118.965305 | -4.2940439 | 1.13125278 | -3.7958306 |
| 13524 | NM_000303.2 | 63.1627773 | -3.9126862 | 1.03096006 | -3.7951869 |
| 13525 | NR_037576.1 | 13.4949849 | -5.3770836 | 1.41687585 | -3.7950281 |
| 13526 | NM_001258324.1 | 17.0461985 | -5.7104704 | 1.50473404 | -3.7950031 |
| 13527 | NM_003287.3 | 16.655085 | -5.6800183 | 1.49675883 | -3.7948788 |
| 13528 | NM_014241.3 | 16.7223268 | -5.6855749 | 1.49824826 | -3.7948149 |
| 13529 | NM_173194.2 | 17.7638091 | -5.7680964 | 1.52015041 | -3.7944248 |
| 13530 | NM_181699.2 | 11.4711799 | -5.1501593 | 1.3580488 | -3.7923227 |
| 13531 | NR_029383.1 | 10.3556309 | -4.9980629 | 1.31802629 | -3.7920814 |
| 13532 | NM_005373.2 | 118.275753 | 1.81786465 | 0.47939455 | 3.7920011 |
| 13533 | NM_002546.3 | 10.4650915 | -5.0149765 | 1.3228625 | -3.7910036 |
| 13534 | NM_016495.5 | 13.1342849 | -5.3309427 | 1.40648753 | -3.7902524 |
| 13535 | NM_001080493.3 | 11.9359471 | -5.200256 | 1.37210259 | -3.7899906 |
| 13536 | NM_001203261.1 | 13.2509497 | -5.343798 | 1.4100114 | -3.789897 |
| 13537 | NM_145812.2 | 16.5193054 | -5.6693517 | 1.49590894 | -3.7899043 |
| 13538 | NM_017676.2 | 13.4001067 | -5.3600176 | 1.41465557 | -3.7889206 |
| 13539 | NM_022746.3 | 21.3691598 | -5.0501274 | 1.33314623 | -3.7881271 |
| 13540 | NM_001198546.1 | 41.8356013 | -7.0085144 | 1.85018158 | -3.7880144 |
| 13541 | NM_001193535.1 | 12.5711169 | -5.2618242 | 1.38907887 | -3.7879953 |
| 13542 | NM_001282450.1 | 20.7783356 | -5.0099218 | 1.32265258 | -3.7877837 |
| 13543 | NM_145233.3 | 11.1364139 | -5.0996614 | 1.34634118 | -3.7877928 |
| 13544 | NM_001286095.1 | 26.2289574 | -6.3360481 | 1.67283046 | -3.7876212 |

| | | | | | |
|---|---|---|---|---|---|
| 13545 | NM_006786.3 | 16.9414757 | -5.7046016 | 1.50618916 | -3.7874403 |
| 13546 | NM_018124.3 | 97.0104357 | -3.0941805 | 0.8172191 | -3.7862313 |
| 13547 | NM_015093.5 | 12.2875446 | -5.2316623 | 1.38199882 | -3.7855765 |
| 13548 | NR_110602.1 | 17.5310439 | -5.7465246 | 1.51806497 | -3.7854273 |
| 13549 | NM_024645.2 | 13.3394273 | -5.3584888 | 1.41556859 | -3.7853968 |
| 13550 | NM_001164756.1 | 19.4276313 | -5.8999344 | 1.55876538 | -3.7850048 |
| 13551 | NM_012334.2 | 256.058386 | -1.4863418 | 0.39271054 | -3.7848279 |
| 13552 | NM_006650.3 | 194.984308 | -5.3213241 | 1.40607282 | -3.7845295 |
| 13553 | NM_001040451.2 | 15.4682764 | -5.5772971 | 1.47388522 | -3.7840783 |
| 13554 | NM_001037808.2 | 19.8344024 | -5.9365475 | 1.56887531 | -3.7839511 |
| 13555 | NM_001024732.2 | 14.8896384 | -5.509022 | 1.45611965 | -3.783358 |
| 13556 | NM_004422.2 | 163.525967 | -4.571366 | 1.20849381 | -3.7826971 |
| 13557 | NM_005537.5 | 11.2866245 | -5.127352 | 1.35547885 | -3.7826869 |
| 13558 | NM_001271954.1 | 14.2144656 | -5.4602786 | 1.44354079 | -3.7825592 |
| 13559 | NM_001128214.1 | 45.1650663 | -3.1205061 | 0.82510484 | -3.7819511 |
| 13560 | NR_125362.1 | 40.151948 | 1.60892124 | 0.42542567 | 3.78190917 |
| 13561 | NM_002178.2 | 15.7088946 | -5.60239 | 1.48151969 | -3.7815157 |
| 13562 | NR_037701.1 | 127.415467 | -4.1200969 | 1.08967839 | -3.781021 |
| 13563 | NR_027406.1 | 214.069563 | -4.7860508 | 1.26586138 | -3.7808649 |
| 13564 | NM_001012511.2 | 14.7827484 | -5.50705 | 1.45658911 | -3.7807848 |
| 13565 | NM_002309.4 | 44.950593 | -4.8025351 | 1.27025526 | -3.7807638 |
| 13566 | NM_001201543.1 | 21.4710863 | -6.0515518 | 1.60086824 | -3.7801686 |
| 13567 | NM_001128845.1 | 233.268569 | -9.4880046 | 2.51002422 | -3.780045 |
| 13568 | NM_005633.3 | 190.926544 | -4.1945049 | 1.10971205 | -3.7798137 |
| 13569 | NM_001324195.1 | 14.0249215 | -5.4267887 | 1.43576352 | -3.7797232 |
| 13570 | NM_022572.4 | 22.6377223 | -5.145744 | 1.3616799 | -3.7789674 |
| 13571 | NR_037702.1 | 15.08242 | -5.5356548 | 1.46491122 | -3.778833 |
| 13572 | NM_001319958.1 | 11.5384022 | -5.1405513 | 1.36037973 | -3.7787621 |
| 13573 | NM_001301083.1 | 15.8580804 | -5.6047462 | 1.48325275 | -3.7786859 |
| 13574 | NM_001004196.3 | 22.8535053 | -6.1316626 | 1.62275057 | -3.7785613 |
| 13575 | NM_001025100.1 | 11.4731334 | -5.1532025 | 1.36384785 | -3.7784291 |
| 13576 | NR_144326.1 | 10.8054218 | -5.0579055 | 1.33866814 | -3.7783117 |
| 13577 | NM_001008739.1 | 16.3712822 | -5.6561934 | 1.49714944 | -3.7779752 |
| 13578 | NM_001324047.1 | 19.9592273 | -5.9431296 | 1.57313485 | -3.7778895 |
| 13579 | NM_016304.2 | 268.104948 | -5.3334847 | 1.41180864 | -3.7777674 |
| 13580 | NR_130172.1 | 23.1866841 | -6.1556318 | 1.62947407 | -3.7776801 |
| 13581 | NM_080388.2 | 15.0688286 | -5.5393263 | 1.46634908 | -3.7776314 |
| 13582 | NR_145460.1 | 24.9141024 | 4.86574478 | 1.28806315 | 3.77756695 |
| 13583 | NM_020648.5 | 157.090711 | -4.5102953 | 1.19417024 | -3.7769283 |
| 13584 | NM_052846.1 | 28.5966353 | -4.0911846 | 1.08328834 | -3.7766349 |
| 13585 | NM_001330128.1 | 23.7703063 | -6.190252 | 1.63910666 | -3.7766011 |
| 13586 | NM_005206.4 | 151.215546 | -3.271095 | 0.86615524 | -3.776569 |
| 13587 | NR_046106.1 | 19.615293 | -4.9036003 | 1.29858803 | -3.7761015 |

| | | | | | |
|---|---|---|---|---|---|
| 13588 | NM_001347722.1 | 12.3930978 | -5.2549517 | 1.3916602 | -3.7760308 |
| 13589 | NM_001129979.1 | 11.8314639 | -5.182442 | 1.37293564 | -3.7747159 |
| 13590 | NM_001013439.2 | 13.9321161 | -5.4153894 | 1.43497992 | -3.7738433 |
| 13591 | NR_131235.1 | 41.6060701 | -6.9991024 | 1.85470962 | -3.7736917 |
| 13592 | NR_046308.1 | 12.9810905 | -5.3196392 | 1.40986371 | -3.7731585 |
| 13593 | NM_014845.5 | 119.268859 | -3.8756432 | 1.02729413 | -3.7726714 |
| 13594 | NM_005295.2 | 42.3139839 | -7.0240039 | 1.86190945 | -3.7724734 |
| 13595 | NM_032807.4 | 11.8634178 | -5.1946502 | 1.37714787 | -3.7720352 |
| 13596 | NR_002838.1 | 11.7439097 | -5.1766605 | 1.37247465 | -3.7717713 |
| 13597 | NR_109929.1 | 24.0124048 | -6.2120208 | 1.64730461 | -3.7710213 |
| 13598 | NM_005515.3 | 18.489747 | -5.8325796 | 1.5472364 | -3.7696758 |
| 13599 | NM_015488.4 | 18.968399 | -5.8669525 | 1.55644919 | -3.7694469 |
| 13600 | NM_001321926.1 | 103.281672 | -4.6821817 | 1.24223345 | -3.7691641 |
| 13601 | NM_001258024.1 | 10.4195475 | -5.0062324 | 1.32841269 | -3.7685822 |
| 13602 | NM_007076.2 | 12.0835433 | -5.2154369 | 1.38395121 | -3.7685121 |
| 13603 | NR_037178.1 | 114.986532 | -4.0665724 | 1.07917919 | -3.7682087 |
| 13604 | NM_001297697.1 | 17.5136458 | -5.7483806 | 1.5259141 | -3.7671718 |
| 13605 | NM_015315.4 | 775.950658 | -5.561188 | 1.47627747 | -3.7670344 |
| 13606 | NR_135548.1 | 12.6248101 | -5.2730828 | 1.39990017 | -3.7667563 |
| 13607 | NM_001261382.1 | 14.9630315 | -5.5194534 | 1.46537129 | -3.7665904 |
| 13608 | NM_017858.2 | 15.1350517 | -5.5365716 | 1.46993808 | -3.7665339 |
| 13609 | NM_033018.3 | 88.8613357 | -4.1231367 | 1.09467485 | -3.7665401 |
| 13610 | NM_001301132.1 | 12.4510179 | -5.2576844 | 1.39591747 | -3.7664723 |
| 13611 | NM_000831.3 | 341.930219 | -5.6817741 | 1.50879119 | -3.7657789 |
| 13612 | NM_001184766.1 | 18.7361486 | -4.8435829 | 1.28637011 | -3.7653105 |
| 13613 | NM_018106.3 | 21.1812666 | -6.029895 | 1.60145143 | -3.7652687 |
| 13614 | NM_018925.2 | 11.0844427 | -5.0940044 | 1.35308851 | -3.7647237 |
| 13615 | NM_001098783.2 | 12.0385117 | -5.2132812 | 1.3849611 | -3.7642076 |
| 13616 | NM_006150.4 | 15.8855777 | -5.6159781 | 1.49195391 | -3.7641767 |
| 13617 | NR_026880.1 | 14.3993029 | -5.474134 | 1.45438192 | -3.7638903 |
| 13618 | NM_153336.2 | 12.1850909 | -5.2271062 | 1.38882586 | -3.7636873 |
| 13619 | NM_001321367.1 | 99.8198951 | 2.11279903 | 0.56145549 | 3.76307483 |
| 13620 | NM_175573.2 | 18.285396 | -5.8207445 | 1.54681913 | -3.7630415 |
| 13621 | NM_000991.4 | 544.770978 | -3.7960311 | 1.00879964 | -3.7629188 |
| 13622 | NM_001143830.1 | 16.4243555 | -5.6589336 | 1.50386081 | -3.7629371 |
| 13623 | NM_018386.4 | 12.8546393 | -5.3141541 | 1.41225888 | -3.7628754 |
| 13624 | NM_145901.2 | 14.9489409 | -5.5252391 | 1.46844611 | -3.7626434 |
| 13625 | NM_004452.3 | 15.3372923 | -5.5611079 | 1.47810421 | -3.7623246 |
| 13626 | NM_001284501.1 | 11.9254442 | -5.1916148 | 1.3800111 | -3.7620094 |
| 13627 | NR_104230.1 | 12.1185581 | -5.2261611 | 1.38931137 | -3.7616917 |
| 13628 | NM_001205228.1 | 33.6748838 | -4.3506538 | 1.15667883 | -3.7613326 |
| 13629 | NR_024206.1 | 19.6321725 | -5.9185102 | 1.57356877 | -3.7612021 |
| 13630 | NM_000419.4 | 14.0594596 | -5.4415051 | 1.44675995 | -3.7611665 |

| | | | | | |
|---|---|---|---|---|---|
| 13631 | NM_153464.2 | 19.7392728 | -5.9293852 | 1.57648219 | -3.7611495 |
| 13632 | NM_001330306.1 | 14.1581951 | -5.4413756 | 1.4467891 | -3.7610013 |
| 13633 | NM_001306080.1 | 19.5187225 | -5.9135386 | 1.57310281 | -3.7591559 |
| 13634 | NM_020957.3 | 25.1773379 | -4.3850698 | 1.16663 | -3.7587494 |
| 13635 | NM_001063.3 | 10.7349702 | -5.0426213 | 1.34183187 | -3.7580128 |
| 13636 | NM_014844.4 | 135.516609 | -3.3444166 | 0.8899393 | -3.7580278 |
| 13637 | NM_005164.3 | 17.1144032 | -5.7128186 | 1.5203103 | -3.7576662 |
| 13638 | NM_183008.2 | 25.628338 | -6.3047078 | 1.67796093 | -3.7573627 |
| 13639 | NM_170710.4 | 12.8404223 | -5.3146918 | 1.41481223 | -3.7564644 |
| 13640 | NM_001324105.1 | 14.3908007 | -5.4673779 | 1.45577968 | -3.7556355 |
| 13641 | NM_004143.3 | 14.3205121 | -5.4596949 | 1.45381918 | -3.7554154 |
| 13642 | NR_073574.1 | 15.2992251 | -5.5629547 | 1.48131824 | -3.7554082 |
| 13643 | NM_001079846.1 | 115.614677 | -3.311159 | 0.88183973 | -3.7548308 |
| 13644 | NR_026856.1 | 13.354852 | -5.3545243 | 1.42622592 | -3.7543311 |
| 13645 | NR_033872.1 | 20.5651427 | -4.9928459 | 1.32992981 | -3.7542176 |
| 13646 | NM_014317.4 | 15.3982474 | -5.5618806 | 1.48179987 | -3.7534628 |
| 13647 | NM_001324320.1 | 323.739488 | -5.0841628 | 1.35474618 | -3.7528527 |
| 13648 | NM_001014972.2 | 156.301357 | -8.9106028 | 2.37458123 | -3.7524944 |
| 13649 | NM_182919.3 | 10.5828098 | -5.0162114 | 1.33693353 | -3.7520276 |
| 13650 | NM_006293.3 | 101.284655 | -4.1877563 | 1.11626328 | -3.7515847 |
| 13651 | NR_102752.1 | 10.5321284 | -5.0128307 | 1.33649149 | -3.7507389 |
| 13652 | NM_144732.4 | 114.603717 | -4.25186 | 1.13368119 | -3.75049 |
| 13653 | NM_001330240.1 | 11.5871693 | -5.1557296 | 1.37468871 | -3.7504706 |
| 13654 | NM_001307976.1 | 22.8666928 | -4.3215099 | 1.15228025 | -3.7503983 |
| 13655 | NM_001127695.2 | 19.6680918 | -5.9260865 | 1.58015724 | -3.7503144 |
| 13656 | NM_001143936.1 | 17.4546638 | -5.7461728 | 1.53252001 | -3.7494928 |
| 13657 | NM_001130690.2 | 100.706681 | -4.8372143 | 1.2902943 | -3.7489233 |
| 13658 | NR_109948.1 | 60.0943265 | 3.78253769 | 1.00898784 | 3.7488437 |
| 13659 | NM_018369.2 | 17.5484953 | -5.7507858 | 1.53412873 | -3.7485679 |
| 13660 | NM_017860.4 | 10.7964232 | -5.0611151 | 1.35020857 | -3.748395 |
| 13661 | NM_001136205.2 | 10.7527837 | -5.0579665 | 1.34954393 | -3.747908 |
| 13662 | NR_024407.1 | 48.0661535 | 1.80594035 | 0.48191123 | 3.74745435 |
| 13663 | NM_001197181.1 | 17.6308885 | -5.7572116 | 1.53639233 | -3.7472275 |
| 13664 | NM_001172774.1 | 141.597668 | -4.4604905 | 1.19040887 | -3.7470239 |
| 13665 | NM_001014985.2 | 14.3066142 | -5.4543528 | 1.45585394 | -3.7464972 |
| 13666 | NM_001278374.1 | 21.2063872 | -6.0271479 | 1.60878481 | -3.7463978 |
| 13667 | NM_001165037.1 | 18.9755235 | -5.8648964 | 1.56557814 | -3.7461537 |
| 13668 | NM_022110.3 | 14.354513 | -5.4671988 | 1.45962511 | -3.7456185 |
| 13669 | NM_001171743.2 | 18.85594 | -5.8577644 | 1.56396843 | -3.7454493 |
| 13670 | NM_001321196.1 | 11.8677457 | -5.1909538 | 1.38593097 | -3.7454635 |
| 13671 | NM_144987.3 | 11.9068151 | -5.1986055 | 1.38798687 | -3.7454285 |
| 13672 | NM_001287059.1 | 53.6529032 | -4.6678872 | 1.2463262 | -3.7453174 |
| 13673 | NM_018948.3 | 110.087308 | -4.5194053 | 1.20669464 | -3.7452767 |

| | | | | | |
|---|---|---|---|---|---|
| 13674 | NM_001131036.1 | 16.3058684 | -5.6498271 | 1.50856178 | -3.7451745 |
| 13675 | NM_001031722.3 | 10.6949495 | -5.0415282 | 1.34619188 | -3.7450294 |
| 13676 | NR_134951.1 | 11.722462 | -5.183552 | 1.38413736 | -3.7449694 |
| 13677 | NM_001204827.1 | 11.6325315 | -5.1612217 | 1.37827896 | -3.7446859 |
| 13678 | NM_001282931.1 | 23.8719729 | -6.2025075 | 1.65644261 | -3.7444747 |
| 13679 | NM_003124.4 | 11.9598627 | -5.2001652 | 1.38877709 | -3.7444203 |
| 13680 | NR_046079.1 | 14.4160071 | -5.4628969 | 1.45894869 | -3.7444065 |
| 13681 | NM_001290214.2 | 25.7686529 | -6.3072413 | 1.68445961 | -3.7443707 |
| 13682 | NM_016060.2 | 21.7952106 | -5.082419 | 1.35751333 | -3.7439183 |
| 13683 | NM_152788.4 | 16.0949546 | -5.6257971 | 1.50281061 | -3.743517 |
| 13684 | NM_024944.2 | 36.19304 | -4.7687823 | 1.27416528 | -3.7426716 |
| 13685 | NR_033440.1 | 15.1529813 | -5.5462753 | 1.48204839 | -3.7423038 |
| 13686 | NM_001322084.1 | 13.1544616 | -5.3361527 | 1.42592086 | -3.7422503 |
| 13687 | NR_040070.1 | 12.2026262 | -5.2358094 | 1.39911413 | -3.7422318 |
| 13688 | NM_002674.3 | 36.3222761 | -6.8055327 | 1.8189891 | -3.7413818 |
| 13689 | NM_017987.4 | 19.018411 | -5.8699125 | 1.5689642 | -3.741266 |
| 13690 | NR_027297.1 | 19.1754845 | -4.8944853 | 1.3084443 | -3.7406906 |
| 13691 | NM_002449.4 | 20.8369585 | -6.0073575 | 1.60624759 | -3.7399947 |
| 13692 | NM_001270574.1 | 15.6951799 | -5.5927162 | 1.49555011 | -3.7395713 |
| 13693 | NM_001308213.1 | 15.3314324 | -5.5587884 | 1.48649576 | -3.7395252 |
| 13694 | NR_130726.1 | 30.3612378 | 2.7903226 | 0.74625676 | 3.7390919 |
| 13695 | NM_004274.4 | 346.814869 | -2.8646778 | 0.76615341 | -3.7390394 |
| 13696 | NM_001161573.1 | 7.21968962 | 7.76757689 | 2.07766796 | 3.73860359 |
| 13697 | NM_001165937.1 | 26.5449422 | -6.352333 | 1.69920738 | -3.7384095 |
| 13698 | NM_001039380.3 | 11.6466264 | -5.1605781 | 1.38044318 | -3.7383488 |
| 13699 | NM_001276468.1 | 25.1184039 | -6.2708499 | 1.67743446 | -3.7383576 |
| 13700 | NM_001002000.2 | 14.1956771 | -5.4401255 | 1.455363 | -3.7379853 |
| 13701 | NM_145003.4 | 10.050114 | -4.949457 | 1.32412895 | -3.7378965 |
| 13702 | NM_178153.2 | 1064.9375 | -5.274535 | 1.41109801 | -3.7378941 |
| 13703 | NR_036484.1 | 28.156594 | -6.4393241 | 1.72281455 | -3.7376769 |
| 13704 | NM_203376.1 | 13.4008319 | -5.3623553 | 1.43488604 | -3.7371298 |
| 13705 | NM_001330422.1 | 20.2163931 | -5.961342 | 1.59524717 | -3.7369394 |
| 13706 | NM_001265582.1 | 12.8479421 | -5.3051593 | 1.41971341 | -3.7367819 |
| 13707 | NM_001197223.1 | 30.7783742 | -6.5627596 | 1.75632789 | -3.7366369 |
| 13708 | NM_001282658.1 | 12.8205698 | -5.3086416 | 1.42095422 | -3.7359695 |
| 13709 | NM_001284194.1 | 11.3686727 | -5.1215779 | 1.37097216 | -3.7357271 |
| 13710 | NM_019040.4 | 15.5946423 | -5.5838737 | 1.49472533 | -3.7357189 |
| 13711 | NM_003558.3 | 22.7346013 | -6.1310521 | 1.64155353 | -3.7349084 |
| 13712 | NM_139284.2 | 39.3965885 | -5.9503147 | 1.59316153 | -3.7349099 |
| 13713 | NM_175623.3 | 18.1600392 | -5.7997941 | 1.55300711 | -3.7345573 |
| 13714 | NM_001330336.1 | 17.182914 | -5.7281474 | 1.53411931 | -3.7338344 |
| 13715 | NM_001100170.1 | 167.121015 | -3.8789013 | 1.03891369 | -3.7336127 |
| 13716 | NR_038883.2 | 10.9473202 | -5.0764758 | 1.35979551 | -3.7332642 |

| | | | | | |
|---|---|---|---|---|---|
| 13717 | NM_001321705.1 | 36.6010153 | -6.8175642 | 1.8263224 | -3.7329467 |
| 13718 | NM_001002033.2 | 11.1333387 | -5.108794 | 1.36864638 | -3.7327348 |
| 13719 | NM_001297653.1 | 210.064512 | -9.3368081 | 2.50136774 | -3.7326811 |
| 13720 | NM_016381.5 | 11.3379749 | -5.1334846 | 1.3753842 | -3.7324005 |
| 13721 | NM_001321194.1 | 25.1090961 | -6.2690988 | 1.67971218 | -3.7322459 |
| 13722 | NM_001111033.1 | 91.6238152 | 7.29549458 | 1.95482692 | 3.73204118 |
| 13723 | NM_001731.2 | 178.880187 | -4.6089117 | 1.23509033 | -3.7316394 |
| 13724 | NM_001321484.1 | 15.8064624 | -5.6005418 | 1.50103225 | -3.7311269 |
| 13725 | NM_033418.3 | 10.8207461 | -5.0504466 | 1.35369428 | -3.730862 |
| 13726 | NR_110725.1 | 152.309029 | 3.48172008 | 0.93328777 | 3.73059647 |
| 13727 | NM_001136501.2 | 22.1516474 | -4.2654333 | 1.14354671 | -3.7300035 |
| 13728 | NM_016074.4 | 12.5228776 | -5.2712313 | 1.41338167 | -3.7295172 |
| 13729 | NR_138150.1 | 9.81349323 | -4.9149211 | 1.31793496 | -3.7292592 |
| 13730 | NM_002118.4 | 18.9720774 | -5.8694826 | 1.57398309 | -3.7290633 |
| 13731 | NM_001278237.1 | 12.1259301 | -5.2155552 | 1.39888914 | -3.7283549 |
| 13732 | NR_109978.1 | 16.9716818 | -5.701348 | 1.52923069 | -3.7282459 |
| 13733 | NR_046258.1 | 54.4398819 | -3.6779582 | 0.98651732 | -3.7282247 |
| 13734 | NM_001145268.1 | 11.3168269 | -5.1177152 | 1.3728218 | -3.7278802 |
| 13735 | NR_104041.1 | 19.3689512 | -5.8991602 | 1.58259512 | -3.7275233 |
| 13736 | NM_001318849.1 | 122.093522 | -4.6164728 | 1.23850449 | -3.7274575 |
| 13737 | NM_001284285.1 | 18.4709349 | -5.8252393 | 1.562866 | -3.7272801 |
| 13738 | NM_001288668.1 | 11.5433676 | -5.1446048 | 1.38041135 | -3.7268636 |
| 13739 | NR_024497.2 | 26.3338524 | -4.5446059 | 1.21951535 | -3.7265672 |
| 13740 | NM_138571.4 | 234.824177 | -2.0816113 | 0.55860269 | -3.7264612 |
| 13741 | NM_032457.3 | 90.6901773 | -4.1569933 | 1.11555142 | -3.7264022 |
| 13742 | NR_027154.1 | 24.9829185 | -4.41423 | 1.18461595 | -3.7262963 |
| 13743 | NM_194315.1 | 13.2004305 | -5.3547857 | 1.43708112 | -3.7261541 |
| 13744 | NR_045631.1 | 14.9815983 | -5.5223681 | 1.48208685 | -3.7260759 |
| 13745 | NM_001001329.2 | 63.7148104 | -3.4478671 | 0.92541168 | -3.7257657 |
| 13746 | NM_001100388.1 | 23.1803568 | -6.1603754 | 1.65366282 | -3.7252911 |
| 13747 | NM_001322362.1 | 29.9297179 | -6.5249434 | 1.75158969 | -3.7251552 |
| 13748 | NR_110010.1 | 15.782096 | -5.5925455 | 1.50139195 | -3.7249071 |
| 13749 | NM_001199653.1 | 13.0022981 | -5.320799 | 1.42870721 | -3.7242053 |
| 13750 | NM_012074.4 | 14.0176601 | -5.4335924 | 1.4591264 | -3.7238669 |
| 13751 | NM_182703.5 | 30.9861123 | -3.0261806 | 0.81268822 | -3.7236674 |
| 13752 | NM_012266.5 | 155.477244 | -4.8477126 | 1.30187345 | -3.7236435 |
| 13753 | NM_001255980.1 | 9.91299071 | -4.9388331 | 1.32654832 | -3.7230706 |
| 13754 | NM_018982.4 | 22.0164581 | -5.0777901 | 1.36402512 | -3.7226515 |
| 13755 | NR_046285.1 | 12.0723245 | -5.2119232 | 1.40017515 | -3.7223366 |
| 13756 | NM_001267612.1 | 23.7778249 | -6.1973109 | 1.6649559 | -3.7222072 |
| 13757 | NM_006441.3 | 10.772126 | -5.0591976 | 1.35937772 | -3.7217012 |
| 13758 | NR_144485.1 | 10.9496599 | -5.0798085 | 1.36540135 | -3.7203776 |
| 13759 | NM_001080436.1 | 74.1100753 | -3.9963659 | 1.07437137 | -3.7197248 |

| | | | | | |
|---|---|---|---|---|---|
| 13760 | NR_024468.1 | 21.2077213 | 5.2325715 | 1.40691532 | 3.71918012 |
| 13761 | NR_027418.1 | 269.150741 | -3.4164373 | 0.91862304 | -3.7190852 |
| 13762 | NM_001025247.1 | 18.3118256 | -4.8271981 | 1.29819818 | -3.7183831 |
| 13763 | NM_024301.4 | 99.3592803 | -1.8008054 | 0.48433332 | -3.7181119 |
| 13764 | NM_032834.3 | 13.3164658 | -5.3510902 | 1.43919756 | -3.7181068 |
| 13765 | NM_014002.3 | 10.9904008 | -5.0829504 | 1.36725744 | -3.717625 |
| 13766 | NM_001130687.2 | 11.3686562 | -5.1220093 | 1.37785559 | -3.7173774 |
| 13767 | NR_136256.1 | 24.9827771 | 2.77095527 | 0.74546118 | 3.71710203 |
| 13768 | NM_003914.3 | 13.5537822 | -5.3706287 | 1.44498744 | -3.7167304 |
| 13769 | NR_028407.2 | 9.90587803 | -4.9370568 | 1.32846696 | -3.7163565 |
| 13770 | NM_001033028.1 | 24.1941261 | -4.4050676 | 1.18538027 | -3.7161641 |
| 13771 | NR_046243.1 | 40.7685617 | -4.1269154 | 1.11063782 | -3.7158066 |
| 13772 | NM_001318063.1 | 11.0575402 | -5.0894244 | 1.3698633 | -3.715279 |
| 13773 | NM_139201.2 | 10.7077629 | -5.043413 | 1.35750592 | -3.7152052 |
| 13774 | NM_001282875.1 | 174.312687 | -4.0935147 | 1.10187459 | -3.7150459 |
| 13775 | NM_014391.2 | 75.7107774 | -3.0064476 | 0.80939329 | -3.7144459 |
| 13776 | NR_110723.1 | 11.324708 | -5.1249482 | 1.38033417 | -3.7128315 |
| 13777 | NR_073484.1 | 17.7517948 | -5.7727336 | 1.55519432 | -3.711905 |
| 13778 | NM_001161500.1 | 17.6397162 | -5.7547895 | 1.55038874 | -3.7118365 |
| 13779 | NM_001285450.1 | 10.24836 | -4.9889052 | 1.34419801 | -3.7114362 |
| 13780 | NM_007309.3 | 11.3804787 | -5.121282 | 1.3799397 | -3.7112361 |
| 13781 | NM_005775.4 | 12.080347 | -5.2120379 | 1.40440891 | -3.7111968 |
| 13782 | NM_080832.2 | 22.1899887 | -4.2623567 | 1.14856492 | -3.7110281 |
| 13783 | NR_120311.1 | 54.8662586 | 3.06366601 | 0.82573954 | 3.71020867 |
| 13784 | NM_032451.1 | 128.654418 | -3.0304127 | 0.81685009 | -3.7098762 |
| 13785 | NM_001347995.1 | 15.0958991 | -5.5435665 | 1.49432879 | -3.7097368 |
| 13786 | NM_152792.2 | 11.4317512 | -5.1358316 | 1.38443414 | -3.7096973 |
| 13787 | NM_212554.3 | 29.280025 | -4.1300511 | 1.11364865 | -3.7085764 |
| 13788 | NM_001206943.1 | 19.1795414 | -5.8803814 | 1.58598092 | -3.7077252 |
| 13789 | NR_034107.2 | 55.5472294 | 2.71097429 | 0.73129958 | 3.70706393 |
| 13790 | NM_001300970.1 | 15.4339555 | -5.5586738 | 1.4995677 | -3.7068509 |
| 13791 | NM_001270530.1 | 20.7608797 | -5.9967794 | 1.61797802 | -3.7063417 |
| 13792 | NM_002413.4 | 10.2462955 | -4.9831142 | 1.34464457 | -3.7058969 |
| 13793 | NM_001302084.1 | 19.1548479 | -5.8812702 | 1.58720733 | -3.7054203 |
| 13794 | NM_001304723.1 | 12.3036261 | -5.2413278 | 1.4145943 | -3.7051809 |
| 13795 | NM_014028.3 | 145.589244 | -3.2513712 | 0.87752512 | -3.7051603 |
| 13796 | NM_001283021.1 | 11.1367668 | -5.0998661 | 1.37651997 | -3.704898 |
| 13797 | NM_001002879.1 | 11.1193667 | -5.0844461 | 1.37247132 | -3.7045919 |
| 13798 | NM_032673.2 | 27.144162 | -4.0090597 | 1.08228261 | -3.7042633 |
| 13799 | NR_075083.2 | 114.109598 | 2.3023149 | 0.62155494 | 3.70412133 |
| 13800 | NM_001321301.1 | 16.4441655 | -5.6555034 | 1.52697171 | -3.7037382 |
| 13801 | NM_001079512.3 | 26.1626969 | -3.3694628 | 0.90984842 | -3.7033232 |
| 13802 | NM_001127323.1 | 23.1193912 | -6.1550668 | 1.66225179 | -3.7028487 |

| | | | | | |
|---|---|---|---|---|---|
| 13803 | NM_001143782.1 | 10.1672055 | -4.9718316 | 1.34270885 | -3.7028367 |
| 13804 | NM_001320610.1 | 15.2132263 | -5.5566201 | 1.50074183 | -3.7025823 |
| 13805 | NM_001304479.1 | 15.0582733 | -5.5344458 | 1.49476688 | -3.7025478 |
| 13806 | NM_024590.3 | 33.1680432 | -4.8979939 | 1.32306022 | -3.7020189 |
| 13807 | NM_001037666.2 | 16.9465099 | -4.7218876 | 1.27572057 | -3.7013494 |
| 13808 | NM_014205.3 | 41.5890116 | -3.6752081 | 0.99309567 | -3.7007594 |
| 13809 | NM_001031679.2 | 27.9470283 | -6.4288525 | 1.73726859 | -3.7005518 |
| 13810 | NR_036480.1 | 10.8104866 | -5.0641799 | 1.36855012 | -3.7003978 |
| 13811 | NR_047005.1 | 33.0187701 | 4.77135115 | 1.28952389 | 3.70008744 |
| 13812 | NR_130940.1 | 10.8633689 | -5.0627151 | 1.36841266 | -3.6996991 |
| 13813 | NM_152487.2 | 15.4177092 | -5.5768002 | 1.50762221 | -3.69907 |
| 13814 | NM_003384.2 | 68.3224983 | -4.4625202 | 1.20647558 | -3.6988069 |
| 13815 | NR_037185.1 | 10.8068479 | -5.0596631 | 1.36794142 | -3.6987425 |
| 13816 | NM_001253699.1 | 12.4752918 | -5.2713951 | 1.42525854 | -3.6985536 |
| 13817 | NM_024697.2 | 13.6203823 | -5.3888385 | 1.45706387 | -3.6984229 |
| 13818 | NM_015995.3 | 246.073714 | -5.0292837 | 1.35998182 | -3.6980522 |
| 13819 | NM_018221.4 | 135.21026 | -8.7015289 | 2.35386717 | -3.696695 |
| 13820 | NM_001005209.2 | 84.2340666 | -4.4262831 | 1.19755234 | -3.6961082 |
| 13821 | NM_001320038.1 | 14.1085304 | -5.4345627 | 1.47049248 | -3.6957432 |
| 13822 | NM_001347828.1 | 25.0759771 | -6.2649686 | 1.69517909 | -3.6957562 |
| 13823 | NM_014629.3 | 64.3682014 | -3.7729745 | 1.02107529 | -3.6950992 |
| 13824 | NM_001032282.3 | 11.4960147 | -5.1547225 | 1.39505117 | -3.6950061 |
| 13825 | NM_031219.3 | 12.6210167 | -5.283416 | 1.42988119 | -3.6950035 |
| 13826 | NM_012204.3 | 194.458608 | -2.991004 | 0.80948509 | -3.6949464 |
| 13827 | NM_001002296.1 | 18.0647635 | -5.8034667 | 1.57079403 | -3.694607 |
| 13828 | NM_012214.2 | 94.0170492 | -3.7027954 | 1.00229694 | -3.6943098 |
| 13829 | NM_144633.2 | 38.0632583 | -4.1317387 | 1.11846285 | -3.6941224 |
| 13830 | NM_018291.3 | 12.1637664 | -5.2345319 | 1.41716191 | -3.6936725 |
| 13831 | NM_001145304.1 | 9.49220153 | -4.86735 | 1.31787954 | -3.6933194 |
| 13832 | NM_080626.5 | 11.8980883 | -5.1828566 | 1.40362041 | -3.6924916 |
| 13833 | NM_001256644.1 | 17.7453296 | -5.7650369 | 1.56134261 | -3.6923587 |
| 13834 | NM_024099.3 | 10.2714496 | -4.9863803 | 1.35046546 | -3.6923419 |
| 13835 | NM_001318876.1 | 10.3778585 | -5.0031107 | 1.35520372 | -3.6917776 |
| 13836 | NM_024989.3 | 136.78602 | -8.7182074 | 2.36177672 | -3.6913767 |
| 13837 | NM_015600.4 | 13.2228848 | -5.3502501 | 1.44940148 | -3.6913513 |
| 13838 | NM_001256139.1 | 20.5469702 | -5.9796782 | 1.62033464 | -3.690397 |
| 13839 | NR_121638.1 | 21.3822957 | 6.71774548 | 1.82043938 | 3.69017806 |
| 13840 | NM_001561.5 | 177.257275 | 2.67651018 | 0.72535542 | 3.68992922 |
| 13841 | NM_018003.2 | 56.7961736 | -2.3487204 | 0.63656158 | -3.6896986 |
| 13842 | NM_001135769.2 | 32.6875977 | -4.3035877 | 1.16643045 | -3.6895365 |
| 13843 | NM_001244678.1 | 12.6769904 | -5.2903216 | 1.43389385 | -3.6894792 |
| 13844 | NR_138072.1 | 21.9396601 | -6.0706532 | 1.64557521 | -3.6890767 |
| 13845 | NM_001199298.1 | 23.9380815 | -6.2021956 | 1.68141458 | -3.6886771 |

| | | | | | |
|---|---|---|---|---|---|
| 13846 | NM_014623.3 | 16.1221499 | -5.6343798 | 1.52762422 | -3.6883284 |
| 13847 | NM_016094.3 | 173.911585 | -4.5674327 | 1.23843248 | -3.6880757 |
| 13848 | NM_024025.2 | 21.3217469 | -6.039494 | 1.6377246 | -3.6877348 |
| 13849 | NM_173811.3 | 12.0471274 | -5.2137463 | 1.41390725 | -3.6874741 |
| 13850 | NM_001173539.1 | 15.5625605 | -5.5794489 | 1.51314667 | -3.6873153 |
| 13851 | NM_001014838.1 | 34.6917083 | -3.9854219 | 1.08085551 | -3.6872846 |
| 13852 | NM_004887.4 | 1752.24391 | 3.63726818 | 0.98651181 | 3.68699911 |
| 13853 | NM_001322386.1 | 20.4963903 | -5.9841235 | 1.62307775 | -3.6868989 |
| 13854 | NM_001248005.1 | 27.2811486 | 9.68868844 | 2.62789544 | 3.68686223 |
| 13855 | NM_000259.3 | 16.6099135 | -5.6774858 | 1.54003752 | -3.6865893 |
| 13856 | NM_003786.3 | 13.8478341 | -5.4169056 | 1.46936191 | -3.6865701 |
| 13857 | NM_001314059.1 | 16.6330294 | -5.6858564 | 1.54234693 | -3.6864964 |
| 13858 | NM_181651.2 | 28.8719619 | -6.4762464 | 1.75674834 | -3.6864964 |
| 13859 | NM_001278648.1 | 17.1908613 | -5.7268839 | 1.55395849 | -3.6853519 |
| 13860 | NM_001174085.1 | 14.0812577 | -5.4335932 | 1.47438354 | -3.6853322 |
| 13861 | NM_001170414.2 | 10.8087789 | -5.0572413 | 1.37232725 | -3.685157 |
| 13862 | NM_001329640.1 | 27.63038 | 9.7110765 | 2.63600739 | 3.68400959 |
| 13863 | NR_134646.1 | 8.95716477 | 7.29307001 | 1.97985442 | 3.68363952 |
| 13864 | NM_001040165.2 | 10.8503305 | -5.0558842 | 1.3728742 | -3.6827003 |
| 13865 | NM_001199877.1 | 10.3411952 | -4.9894916 | 1.35488681 | -3.6825893 |
| 13866 | NM_033356.3 | 74.2717863 | 1.53648317 | 0.41725267 | 3.68238066 |
| 13867 | NM_032341.4 | 18.0558194 | -4.8109335 | 1.30648738 | -3.6823421 |
| 13868 | NR_015401.2 | 28.804649 | -4.6766614 | 1.27010267 | -3.6821129 |
| 13869 | NM_001684.4 | 232.595293 | -4.6630372 | 1.26648826 | -3.6818638 |
| 13870 | NM_001258306.1 | 18.2720542 | -4.7985362 | 1.30330064 | -3.6818337 |
| 13871 | NR_033773.1 | 9.6675058 | -4.8956416 | 1.32969615 | -3.6817747 |
| 13872 | NM_007099.3 | 30.5925311 | -4.5468036 | 1.23504494 | -3.6814884 |
| 13873 | NM_001128931.1 | 9.91645049 | -4.9321491 | 1.33972286 | -3.6814697 |
| 13874 | NM_001318327.1 | 27.0608788 | -3.5825337 | 0.97331807 | -3.680743 |
| 13875 | NM_199296.2 | 23.2102379 | -6.1621445 | 1.67419119 | -3.6806696 |
| 13876 | NM_020673.2 | 186.388592 | -1.8093453 | 0.49158719 | -3.6806193 |
| 13877 | NM_015454.2 | 71.4480332 | -3.9391224 | 1.07058267 | -3.6794192 |
| 13878 | NM_015021.1 | 252.541046 | -4.7873382 | 1.30121215 | -3.6791373 |
| 13879 | NM_005630.2 | 11.4565442 | -5.1503629 | 1.39992593 | -3.6790253 |
| 13880 | NM_024813.2 | 32.2308157 | -4.6069152 | 1.2522491 | -3.6789127 |
| 13881 | NM_001317227.1 | 13.8028339 | -5.4039864 | 1.46907527 | -3.6784952 |
| 13882 | NM_001330385.1 | 18.3536575 | -5.8189869 | 1.58193373 | -3.6784012 |
| 13883 | NM_001301075.1 | 14.862702 | -5.5061762 | 1.49718393 | -3.6776886 |
| 13884 | NR_038840.1 | 14.5930844 | -5.4874382 | 1.49211521 | -3.6776237 |
| 13885 | NR_138046.1 | 24.3335175 | -5.2206887 | 1.41964233 | -3.6774676 |
| 13886 | NM_001287421.1 | 11.7743532 | -5.1737958 | 1.40703393 | -3.6770938 |
| 13887 | NM_139343.2 | 188.002154 | -9.176742 | 2.49595499 | -3.6766456 |
| 13888 | NM_022072.3 | 11.4811112 | -5.1499644 | 1.40073126 | -3.6766256 |

| | | | | | |
|---|---|---|---|---|---|
| 13889 | NM_001206729.1 | 10.4624577 | -5.0191868 | 1.36519848 | -3.6765254 |
| 13890 | NM_001319220.1 | 45.253933 | 2.85245125 | 0.77592939 | 3.67617374 |
| 13891 | NM_001346145.1 | 26.601934 | -6.3529423 | 1.72822197 | -3.6759991 |
| 13892 | NM_001243532.1 | 30.8456612 | -6.5704338 | 1.78743314 | -3.6759047 |
| 13893 | NR_126168.1 | 65.9948245 | 2.15095769 | 0.58515186 | 3.67589652 |
| 13894 | NM_001320623.1 | 27.1262576 | -6.3826545 | 1.73665875 | -3.6752496 |
| 13895 | NM_005097.3 | 20.8533087 | -6.007416 | 1.63462531 | -3.6751027 |
| 13896 | NM_019117.4 | 15.5980133 | -5.588413 | 1.5212908 | -3.6734679 |
| 13897 | NM_198279.3 | 69.3160035 | -2.9330641 | 0.798481 | -3.6733049 |
| 13898 | NR_027775.2 | 12.5477309 | -5.2578709 | 1.43140531 | -3.673223 |
| 13899 | NM_016018.4 | 23.3938805 | -6.1662915 | 1.67905486 | -3.6724777 |
| 13900 | NM_017522.4 | 62.4178319 | -3.8992174 | 1.06183761 | -3.6721409 |
| 13901 | NR_051974.1 | 14.4975533 | -5.4793141 | 1.49225303 | -3.6718398 |
| 13902 | NM_001004308.2 | 48.0235542 | 4.69407517 | 1.27843647 | 3.67173129 |
| 13903 | NM_001274.5 | 13.9258916 | -5.4103815 | 1.47381626 | -3.6710014 |
| 13904 | NM_024773.2 | 12.1833396 | -5.2367943 | 1.42653464 | -3.6709899 |
| 13905 | NM_003943.4 | 16.5480449 | -4.6752126 | 1.27361671 | -3.670816 |
| 13906 | NM_053277.2 | 193.439843 | -5.8290182 | 1.58794407 | -3.6707956 |
| 13907 | NM_001242501.1 | 15.1196156 | -5.5367084 | 1.50856544 | -3.6701811 |
| 13908 | NR_046293.1 | 12.4048268 | -5.2446206 | 1.42906051 | -3.669978 |
| 13909 | NM_001316994.1 | 10.67485 | -5.0410962 | 1.37378653 | -3.6694902 |
| 13910 | NR_135263.1 | 36.8442082 | -6.8241118 | 1.85973531 | -3.6693995 |
| 13911 | NM_001100817.1 | 12.8325087 | 6.89731528 | 1.87984389 | 3.6690894 |
| 13912 | NM_001282506.1 | 9.85295871 | -4.9307991 | 1.34405365 | -3.6686029 |
| 13913 | NR_130112.1 | 12.6811718 | -5.2805237 | 1.43941471 | -3.6685214 |
| 13914 | NM_031948.4 | 13.7755693 | -5.4090237 | 1.47461063 | -3.668103 |
| 13915 | NR_134538.1 | 12.5527896 | -5.2753443 | 1.43829838 | -3.6677677 |
| 13916 | NM_001130156.2 | 18.9071096 | -5.8680514 | 1.59997218 | -3.6675959 |
| 13917 | NM_001099670.2 | 17.6903347 | -5.7605254 | 1.57070675 | -3.6674735 |
| 13918 | NM_001324066.1 | 24.480853 | -6.2334837 | 1.69969173 | -3.6674202 |
| 13919 | NM_001013628.2 | 10.4627223 | -5.0139635 | 1.36724849 | -3.6671926 |
| 13920 | NM_001145316.1 | 12.3405489 | -5.240228 | 1.42899787 | -3.6670649 |
| 13921 | NM_017770.3 | 42.2222698 | -4.2956903 | 1.17149254 | -3.6668525 |
| 13922 | NM_001130413.3 | 13.0532424 | -5.3376444 | 1.45572748 | -3.6666508 |
| 13923 | NM_004560.3 | 44.2732769 | -4.0506205 | 1.1047371 | -3.6665923 |
| 13924 | NM_017593.3 | 10.1790421 | -4.9713924 | 1.35596501 | -3.6663132 |
| 13925 | NR_138604.1 | 15.1218731 | -5.5440184 | 1.51226713 | -3.6660312 |
| 13926 | NM_003316.3 | 73.5369289 | -4.1597612 | 1.13475023 | -3.6657946 |
| 13927 | NM_138796.3 | 18.0068197 | -5.7971156 | 1.5814013 | -3.6658093 |
| 13928 | NM_001014979.2 | 13.6379872 | -5.3973145 | 1.47248687 | -3.6654415 |
| 13929 | NR_122105.1 | 9.71784545 | -4.8954804 | 1.3356221 | -3.6653185 |
| 13930 | NM_000347.5 | 18.9670978 | -5.8655712 | 1.60039107 | -3.6650862 |
| 13931 | NR_135293.1 | 75.9115744 | 3.19633661 | 0.87211922 | 3.66502257 |

| | | | | | |
|---|---|---|---|---|---|
| 13932 | NM_001267556.1 | 21.4004296 | -6.0445883 | 1.64937923 | -3.6647656 |
| 13933 | NR_002715.1 | 223.680516 | -4.3079728 | 1.17563141 | -3.6643907 |
| 13934 | NM_152479.5 | 105.308545 | -4.9906975 | 1.3622101 | -3.6636767 |
| 13935 | NM_013434.4 | 18.3300558 | -5.8136798 | 1.58725199 | -3.6627327 |
| 13936 | NM_001145453.2 | 149.186566 | -8.8434069 | 2.41475888 | -3.6622319 |
| 13937 | NM_173826.3 | 15.9372871 | -5.6195204 | 1.53461067 | -3.6618541 |
| 13938 | NR_073595.1 | 11.732573 | -5.1718417 | 1.41238735 | -3.6617728 |
| 13939 | NM_153700.2 | 17.9922536 | -4.8010552 | 1.31116056 | -3.6616836 |
| 13940 | NM_032429.3 | 18.1142776 | -5.8069944 | 1.58600137 | -3.6614056 |
| 13941 | NM_001252124.1 | 12.4591067 | -5.2688551 | 1.43927783 | -3.660763 |
| 13942 | NM_133450.3 | 13.5764326 | -5.3781032 | 1.4692936 | -3.6603325 |
| 13943 | NM_001256310.1 | 115.014468 | -4.3886465 | 1.1993942 | -3.6590527 |
| 13944 | NM_001134222.1 | 19.0327045 | -5.8751434 | 1.6058043 | -3.658692 |
| 13945 | NM_001330.3 | 10.6893145 | -5.0497151 | 1.3803377 | -3.6583186 |
| 13946 | NM_001172435.1 | 32.5379523 | -6.6472993 | 1.81704686 | -3.6582982 |
| 13947 | NM_001257191.2 | 208.346464 | -4.6599703 | 1.27387628 | -3.6581027 |
| 13948 | NM_001128849.1 | 17.8437027 | -5.7864472 | 1.58188046 | -3.6579548 |
| 13949 | NM_153769.2 | 11.9824512 | -5.2016396 | 1.4220699 | -3.6577946 |
| 13950 | NR_037776.2 | 13.0985585 | -5.3254431 | 1.45624974 | -3.6569573 |
| 13951 | NM_003744.5 | 14.5647114 | -5.4942942 | 1.50261637 | -3.656485 |
| 13952 | NM_024768.2 | 11.5716337 | -5.1539659 | 1.40974839 | -3.6559473 |
| 13953 | NM_001003692.1 | 12.133168 | -5.2309098 | 1.43120408 | -3.6549014 |
| 13954 | NM_018081.2 | 11.913778 | -5.1882197 | 1.41951714 | -3.6549187 |
| 13955 | NM_012419.4 | 15.759568 | -5.596499 | 1.53180061 | -3.6535427 |
| 13956 | NM_014045.4 | 305.861357 | -4.8621563 | 1.33083156 | -3.6534723 |
| 13957 | NR_104295.1 | 10.4264611 | -5.0112666 | 1.37172656 | -3.6532547 |
| 13958 | NM_001290263.1 | 24.4258689 | -4.4337352 | 1.21366351 | -3.6531832 |
| 13959 | NM_019118.4 | 15.0192236 | -5.5332809 | 1.51464873 | -3.6531777 |
| 13960 | NM_024707.2 | 10.218326 | -4.9711963 | 1.36085162 | -3.6530039 |
| 13961 | NM_152548.2 | 21.3409615 | -6.0414643 | 1.65406803 | -3.6524884 |
| 13962 | NM_001080495.2 | 719.851234 | -4.0806509 | 1.11724442 | -3.6524245 |
| 13963 | NM_006465.3 | 112.206477 | -4.2230855 | 1.15623699 | -3.6524394 |
| 13964 | NM_001288980.1 | 17.087335 | -5.7156892 | 1.56492017 | -3.6523839 |
| 13965 | NM_005306.2 | 46.5117773 | 2.9263381 | 0.80132874 | 3.65185716 |
| 13966 | NR_027706.1 | 9.6530608 | -4.8872266 | 1.33850472 | -3.6512584 |
| 13967 | NM_182642.2 | 49.5952336 | -3.9632974 | 1.08551761 | -3.6510669 |
| 13968 | NR_046412.1 | 13.6600444 | -5.3990157 | 1.47893665 | -3.6506065 |
| 13969 | NM_001172650.2 | 10.5608819 | -5.0161303 | 1.37422045 | -3.6501642 |
| 13970 | NM_032452.2 | 32.4083651 | -6.6393542 | 1.81897403 | -3.6500545 |
| 13971 | NM_018237.3 | 123.472768 | -8.5705463 | 2.34816707 | -3.6498878 |
| 13972 | NM_133644.3 | 16.3070854 | -5.6569357 | 1.55002534 | -3.6495763 |
| 13973 | NR_038934.1 | 10.9182869 | -5.0734685 | 1.39023518 | -3.6493599 |
| 13974 | NM_001330671.1 | 11.6648156 | -5.1615147 | 1.41443693 | -3.6491657 |

| | | | | | |
|---|---|---|---|---|---|
| 13975 | NR_045585.1 | 13.2720456 | -5.3528402 | 1.46703333 | -3.6487516 |
| 13976 | NM_001242532.1 | 17.1995494 | -5.7225832 | 1.5686052 | -3.6481985 |
| 13977 | NM_001025356.2 | 126.661543 | -8.6073359 | 2.35939706 | -3.6481083 |
| 13978 | NM_001287804.1 | 10.7109635 | -5.0455621 | 1.38374888 | -3.6462989 |
| 13979 | NM_022469.3 | 30.7024608 | -4.6705361 | 1.28091721 | -3.6462435 |
| 13980 | NM_020184.3 | 62.2096234 | -3.9633256 | 1.08712753 | -3.645686 |
| 13981 | NM_024742.2 | 23.5341176 | -3.6210206 | 0.99323122 | -3.6456975 |
| 13982 | NM_000141.4 | 138.433254 | -8.7354952 | 2.39621876 | -3.6455332 |
| 13983 | NM_018039.2 | 9.66521354 | -4.8916458 | 1.34181767 | -3.6455369 |
| 13984 | NM_175078.2 | 16.8542419 | 8.22470036 | 2.25641577 | 3.64502875 |
| 13985 | NM_014983.2 | 178.793148 | -4.7513567 | 1.30357286 | -3.6448724 |
| 13986 | NM_021903.2 | 145.848955 | -4.6264514 | 1.26935909 | -3.6447144 |
| 13987 | NM_001005362.2 | 125.329347 | -8.5921017 | 2.35746272 | -3.6446395 |
| 13988 | NR_038334.1 | 20.797516 | -4.1687885 | 1.14383301 | -3.6445779 |
| 13989 | NM_016724.2 | 15.9998104 | -5.6267113 | 1.54393269 | -3.644402 |
| 13990 | NR_024368.1 | 302.452729 | -1.3683666 | 0.37554883 | -3.6436451 |
| 13991 | NM_198195.1 | 33.5082598 | -4.9128226 | 1.34841074 | -3.643417 |
| 13992 | NM_005387.6 | 14.0853529 | -5.439464 | 1.49306939 | -3.6431421 |
| 13993 | NM_001319838.1 | 12.9403354 | -5.3157034 | 1.45911272 | -3.6431067 |
| 13994 | NR_027782.1 | 24.3784482 | -4.4191987 | 1.21308389 | -3.6429457 |
| 13995 | NR_037860.1 | 12.8845297 | -5.311356 | 1.45810153 | -3.6426516 |
| 13996 | NM_001002017.2 | 16.1904769 | -5.6354857 | 1.54712148 | -3.6425619 |
| 13997 | NM_001282568.1 | 12.4227194 | -5.256824 | 1.44326387 | -3.6423167 |
| 13998 | NM_003899.3 | 20.9317851 | -6.0144105 | 1.65131766 | -3.6421887 |
| 13999 | NM_001130478.1 | 15.4583219 | -5.5728394 | 1.53019216 | -3.6419213 |
| 14000 | NM_030625.2 | 64.160519 | -2.8576767 | 0.78482922 | -3.6411446 |
| 14001 | NM_138811.3 | 16.5027627 | -4.6725339 | 1.28333825 | -3.6409216 |
| 14002 | NM_001032410.1 | 27.1910396 | -6.3859152 | 1.75412406 | -3.6405151 |
| 14003 | NM_001256876.1 | 22.3315421 | -4.2716058 | 1.17348117 | -3.6401145 |
| 14004 | NM_001142370.1 | 34.018143 | -3.8725513 | 1.06386393 | -3.6400814 |
| 14005 | NM_001278716.1 | 11.4869717 | -5.1432648 | 1.41305427 | -3.6398212 |
| 14006 | NM_178858.4 | 20.1232681 | -3.528444 | 0.96944343 | -3.6396595 |
| 14007 | NM_001243998.1 | 70.1732405 | 1.32484638 | 0.36402512 | 3.63943669 |
| 14008 | NM_078628.1 | 16.2458956 | -4.6563381 | 1.27944978 | -3.6393285 |
| 14009 | NM_001025205.1 | 16.0896463 | -5.6364946 | 1.54882596 | -3.6392046 |
| 14010 | NM_001329953.1 | 10.5437969 | -5.0190514 | 1.37920836 | -3.6390814 |
| 14011 | NM_001098500.2 | 18.6984629 | -5.8482352 | 1.60709976 | -3.6389995 |
| 14012 | NM_001320966.1 | 163.344629 | -8.9738448 | 2.46607088 | -3.6389241 |
| 14013 | NM_001301196.1 | 18.6004835 | -5.8434355 | 1.6059952 | -3.6385137 |
| 14014 | NM_001227.4 | 9.8459281 | -4.9140402 | 1.35064691 | -3.6382864 |
| 14015 | NM_001306140.1 | 17.0291162 | -5.714023 | 1.5705978 | -3.6381198 |
| 14016 | NM_001099627.1 | 29.7661808 | -5.5269084 | 1.5191941 | -3.6380528 |
| 14017 | NM_001282696.1 | 12.138799 | -5.2213998 | 1.43527645 | -3.6379053 |

| | | | | | |
|---|---|---|---|---|---|
| 14018 | NM_018457.3 | 98.9952222 | -4.1569888 | 1.14300096 | -3.6369075 |
| 14019 | NR_121180.1 | 11.2435229 | 5.13306464 | 1.41153982 | 3.63650007 |
| 14020 | NM_001199917.1 | 14.0315417 | -5.4274961 | 1.49251809 | -3.6364692 |
| 14021 | NM_021249.4 | 19.1218382 | -5.8832144 | 1.61797863 | -3.6361508 |
| 14022 | NR_075079.1 | 16.8192338 | -4.6940922 | 1.29106056 | -3.635842 |
| 14023 | NM_001288770.1 | 9.60528417 | -4.8783397 | 1.34182298 | -3.635606 |
| 14024 | NM_016553.4 | 13.4629491 | -5.369726 | 1.47702946 | -3.6354901 |
| 14025 | NR_126327.1 | 11.5416017 | -5.1558679 | 1.41835202 | -3.6351117 |
| 14026 | NM_001271286.1 | 14.2787323 | -5.4671768 | 1.50407219 | -3.6349165 |
| 14027 | NM_015272.4 | 20.5062432 | -5.9790095 | 1.6452611 | -3.6340794 |
| 14028 | NR_024489.1 | 10.1397797 | -4.9729154 | 1.36842435 | -3.6340448 |
| 14029 | NM_015512.4 | 42.3873332 | -4.1874205 | 1.15233531 | -3.633856 |
| 14030 | NM_020204.2 | 9.37735101 | -4.8504015 | 1.33486675 | -3.6336222 |
| 14031 | NM_001288992.1 | 13.6256345 | -5.3988203 | 1.48588793 | -3.6333967 |
| 14032 | NR_026984.1 | 49.3835498 | -3.7265693 | 1.02567335 | -3.6332905 |
| 14033 | NM_001109.4 | 13.2062359 | -5.347509 | 1.47189318 | -3.6330823 |
| 14034 | NM_001145785.1 | 9.66798699 | -4.888947 | 1.34576414 | -3.6328408 |
| 14035 | NM_052978.4 | 13.9359791 | -5.4206969 | 1.49224772 | -3.6325717 |
| 14036 | NM_006068.4 | 23.0874947 | 2.18131197 | 0.60056167 | 3.63211988 |
| 14037 | NR_135622.1 | 11.6577725 | -5.1649299 | 1.42217588 | -3.6317097 |
| 14038 | NR_049780.1 | 11.6749077 | -5.1593499 | 1.42084837 | -3.6311756 |
| 14039 | NR_036533.1 | 14.0726725 | -5.4336162 | 1.49654959 | -3.6307626 |
| 14040 | NM_001160113.1 | 121.489145 | 2.59331125 | 0.71433284 | 3.63039623 |
| 14041 | NM_001314042.1 | 150.235788 | -3.5721423 | 0.98397408 | -3.6303216 |
| 14042 | NR_109909.1 | 9.55807088 | -4.8757081 | 1.34318075 | -3.6299717 |
| 14043 | NM_001884.3 | 41.4654762 | -4.5984826 | 1.26683344 | -3.6299031 |
| 14044 | NM_021140.3 | 18.0630775 | -5.8021134 | 1.59851974 | -3.6296789 |
| 14045 | NM_002643.3 | 28.7071006 | -4.1037075 | 1.13082698 | -3.6289437 |
| 14046 | NR_027052.1 | 16.9068875 | -4.6843776 | 1.29114627 | -3.6280766 |
| 14047 | NM_001243804.1 | 15.401282 | -5.5721011 | 1.53584744 | -3.6280304 |
| 14048 | NM_018833.2 | 13.197344 | -5.3540027 | 1.47645329 | -3.6262595 |
| 14049 | NM_212478.2 | 178.987461 | -9.1059628 | 2.51129098 | -3.6260087 |
| 14050 | NM_001281444.1 | 36.7616681 | 6.29300158 | 1.73558653 | 3.62586449 |
| 14051 | NM_206900.2 | 160.553945 | -5.0376296 | 1.38943155 | -3.6256767 |
| 14052 | NM_001321924.1 | 9.27222662 | -4.8311235 | 1.3325243 | -3.6255425 |
| 14053 | NM_001166693.2 | 19.2163635 | -5.8910673 | 1.62530049 | -3.6246019 |
| 14054 | NM_015295.2 | 245.416955 | -4.2502525 | 1.17261349 | -3.624598 |
| 14055 | NM_001190316.1 | 14.8189918 | -5.4997974 | 1.51753514 | -3.6241648 |
| 14056 | NM_001271635.1 | 11.5643558 | -5.1648694 | 1.42517007 | -3.6240372 |
| 14057 | NM_019108.3 | 133.770032 | -2.7200372 | 0.75058038 | -3.6239119 |
| 14058 | NR_125391.1 | 23.8123911 | 2.10623419 | 0.58126275 | 3.62354922 |
| 14059 | NM_018149.6 | 80.4072481 | -3.9730595 | 1.09648862 | -3.6234389 |
| 14060 | NM_001142554.1 | 13.4325405 | -5.3697007 | 1.48211969 | -3.6229873 |

| | | | | | |
|---|---|---|---|---|---|
| 14061 | NM_001319945.1 | 15.921244 | -5.6181602 | 1.5509025 | -3.6225103 |
| 14062 | NM_152309.2 | 35.1881278 | -4.0128604 | 1.10777367 | -3.6224551 |
| 14063 | NM_015474.3 | 74.683892 | -3.7957566 | 1.04799268 | -3.6219304 |
| 14064 | NM_016531.5 | 151.342341 | -4.3571092 | 1.20304073 | -3.6217471 |
| 14065 | NM_012089.2 | 70.8497017 | -2.7304207 | 0.75392335 | -3.6216157 |
| 14066 | NR_125408.1 | 10.1527727 | -4.967158 | 1.37175205 | -3.6210319 |
| 14067 | NM_001166059.1 | 12.2939646 | -5.2442715 | 1.44831156 | -3.6209554 |
| 14068 | NM_024959.3 | 11.8398921 | -5.1852867 | 1.43204155 | -3.6209052 |
| 14069 | NM_147202.1 | 120.220541 | -8.5320054 | 2.35660944 | -3.620458 |
| 14070 | NM_001080423.3 | 65.8132582 | -4.4092379 | 1.21806494 | -3.619871 |
| 14071 | NM_001199196.1 | 20.119025 | -5.9522861 | 1.64440436 | -3.6197216 |
| 14072 | NR_036625.1 | 382.751887 | -5.2256549 | 1.44365927 | -3.6197287 |
| 14073 | NM_001256552.1 | 9.77502193 | -4.9208213 | 1.35946569 | -3.619673 |
| 14074 | NM_032237.4 | 11.5939863 | -5.1459889 | 1.4219828 | -3.6188826 |
| 14075 | NM_001135953.1 | 14.9127128 | -5.524875 | 1.52675263 | -3.61871 |
| 14076 | NM_001143906.1 | 12.0028024 | -5.1955679 | 1.43578332 | -3.6186295 |
| 14077 | NM_001206844.1 | 46.8135113 | -3.5872341 | 0.9913785 | -3.6184303 |
| 14078 | NM_001184888.1 | 11.8631899 | -5.2009171 | 1.43769369 | -3.6175419 |
| 14079 | NM_016481.4 | 11.3362656 | -5.1279141 | 1.41772309 | -3.6170068 |
| 14080 | NM_001330130.1 | 10.9606766 | -5.0837582 | 1.40552983 | -3.6169693 |
| 14081 | NM_000621.4 | 11.284397 | -5.1192281 | 1.41537115 | -3.6168803 |
| 14082 | NM_001145971.1 | 11.2291888 | -5.1135837 | 1.41393453 | -3.6165633 |
| 14083 | NM_002286.5 | 9.63895756 | -4.8827677 | 1.35012922 | -3.6165188 |
| 14084 | NM_014770.3 | 18.612844 | -5.8402478 | 1.61489192 | -3.6164945 |
| 14085 | NM_001324122.1 | 19.4941563 | -5.902158 | 1.63202345 | -3.6164664 |
| 14086 | NM_001346048.1 | 14.6830984 | -5.4941817 | 1.51926816 | -3.6163344 |
| 14087 | NR_024116.2 | 14.4089559 | -5.4674639 | 1.51198973 | -3.6160721 |
| 14088 | NM_001003931.3 | 12.4716268 | -5.2609627 | 1.45520151 | -3.6152812 |
| 14089 | NR_027471.1 | 18.3739845 | -4.8321374 | 1.3366824 | -3.6150228 |
| 14090 | NM_001171948.1 | 11.510076 | -5.1505212 | 1.42480853 | -3.6148866 |
| 14091 | NM_016547.2 | 15.2573136 | -5.559704 | 1.53807642 | -3.6147125 |
| 14092 | NM_002024.5 | 13.5137413 | -5.3705779 | 1.48579612 | -3.614613 |
| 14093 | NM_001315508.1 | 46.5848319 | -2.4524406 | 0.67873733 | -3.6132396 |
| 14094 | NM_182688.2 | 20.3770577 | -5.9734573 | 1.65324936 | -3.6131617 |
| 14095 | NM_001017975.4 | 10.9302775 | -5.0687106 | 1.40296536 | -3.6128552 |
| 14096 | NM_001164211.1 | 18.4355499 | -5.8225102 | 1.61168296 | -3.6126896 |
| 14097 | NM_181785.3 | 12.6083798 | -5.2815143 | 1.46194607 | -3.6126601 |
| 14098 | NM_020226.3 | 20.4327095 | -5.9694592 | 1.65247678 | -3.6124315 |
| 14099 | NM_001142292.1 | 17.0991882 | -5.7237692 | 1.5845762 | -3.6121767 |
| 14100 | NM_007096.3 | 22.3894437 | -6.1024916 | 1.68955982 | -3.6118825 |
| 14101 | NM_005450.4 | 32.8939194 | -2.5391713 | 0.70314036 | -3.6111869 |
| 14102 | NM_001077526.2 | 13.0948646 | -5.3346793 | 1.4775512 | -3.610487 |
| 14103 | NM_024763.4 | 22.7137495 | -5.1493574 | 1.42626044 | -3.6103907 |

| | | | | | |
|---|---|---|---|---|---|
| 14104 | NR_122074.1 | 17.6744249 | 6.72386493 | 1.86264865 | 3.60984071 |
| 14105 | NM_001145783.1 | 35.6496606 | -5.0104638 | 1.38826757 | -3.6091485 |
| 14106 | NM_007257.5 | 355.998684 | -4.300077 | 1.19152277 | -3.608892 |
| 14107 | NR_038292.1 | 99.6099695 | 2.2404849 | 0.62095265 | 3.60814133 |
| 14108 | NM_174945.2 | 12.4616523 | -5.2581575 | 1.45737091 | -3.6079748 |
| 14109 | NM_001331097.1 | 96.2552866 | 2.35255191 | 0.65213149 | 3.6074809 |
| 14110 | NM_020787.3 | 18.7597448 | -4.8609935 | 1.34759578 | -3.60716 |
| 14111 | NM_145244.3 | 13.0806974 | -5.3364967 | 1.47942688 | -3.6071379 |
| 14112 | NM_001256534.1 | 15.3423096 | -5.5577953 | 1.54086249 | -3.6069379 |
| 14113 | NM_001024683.1 | 11.6528847 | -5.1670603 | 1.4327119 | -3.6064894 |
| 14114 | NR_028057.1 | 13.699664 | -5.406825 | 1.49929219 | -3.6062517 |
| 14115 | NR_120457.1 | 13.4462735 | -5.374118 | 1.49031566 | -3.6060266 |
| 14116 | NM_004097.2 | 65.0825016 | -7.645756 | 2.12035303 | -3.6058882 |
| 14117 | NR_026682.1 | 26.0262962 | 1.9156525 | 0.53130128 | 3.60558611 |
| 14118 | NM_145905.2 | 13.333129 | -5.3508452 | 1.4842457 | -3.6050939 |
| 14119 | NR_024388.1 | 83.2114582 | -3.5001813 | 0.97109657 | -3.6043597 |
| 14120 | NM_001813.2 | 13.9471384 | -5.4185308 | 1.50351715 | -3.6039036 |
| 14121 | NM_001195737.1 | 18.8352906 | -5.8556649 | 1.62489897 | -3.6037101 |
| 14122 | NM_001244644.1 | 24.1888262 | 9.51952711 | 2.64171876 | 3.60353541 |
| 14123 | NM_032873.4 | 62.7728007 | -3.9782961 | 1.10400228 | -3.6035216 |
| 14124 | NM_004163.4 | 23.6673644 | -5.2054841 | 1.444576 | -3.6034685 |
| 14125 | NM_001135256.1 | 66.5818916 | 7.0514478 | 1.95690074 | 3.6033753 |
| 14126 | NM_014574.3 | 84.8801015 | -4.4293815 | 1.22941402 | -3.6028396 |
| 14127 | NR_036750.2 | 52.4258924 | -3.6261376 | 1.00650573 | -3.6026994 |
| 14128 | NR_104414.1 | 10.2527988 | -4.9833128 | 1.38323464 | -3.6026518 |
| 14129 | NM_021817.2 | 15.8165905 | -5.6015947 | 1.55504335 | -3.6022113 |
| 14130 | NM_001167674.1 | 16.0856579 | -5.6267904 | 1.56208437 | -3.602104 |
| 14131 | NM_001162501.1 | 21.804298 | -6.0723037 | 1.68581753 | -3.6019934 |
| 14132 | NM_016089.2 | 50.0806695 | -3.3239053 | 0.92279219 | -3.6020085 |
| 14133 | NR_137175.1 | 28.5006922 | -6.45786 | 1.79300733 | -3.6016919 |
| 14134 | NM_001145064.2 | 255.261741 | -2.1816268 | 0.60575834 | -3.6014804 |
| 14135 | NR_024383.1 | 165.461137 | -8.9923817 | 2.49686063 | -3.6014752 |
| 14136 | NM_001319655.1 | 26.2330758 | -4.5426778 | 1.26145284 | -3.6011475 |
| 14137 | NR_109849.1 | 10.9538113 | -5.0750947 | 1.40962397 | -3.6003181 |
| 14138 | NR_024018.1 | 14.9834969 | -5.5336618 | 1.53703429 | -3.6002201 |
| 14139 | NM_176787.4 | 12.1326057 | -5.2190881 | 1.44991927 | -3.5995715 |
| 14140 | NM_001979.5 | 15.887873 | -5.6166403 | 1.56068996 | -3.5988188 |
| 14141 | NM_198120.2 | 15.5625238 | -5.5779147 | 1.55017845 | -3.5982404 |
| 14142 | NR_036682.1 | 13.7213438 | -5.3919265 | 1.49852146 | -3.5981643 |
| 14143 | NM_020981.3 | 9.70588885 | -4.9091909 | 1.36451696 | -3.59775 |
| 14144 | NR_038958.1 | 485.105407 | 2.4652556 | 0.68530824 | 3.59729457 |
| 14145 | NM_001320371.1 | 12.7321773 | -5.2897062 | 1.47049325 | -3.5972326 |
| 14146 | NM_198838.1 | 13.8684659 | -5.4224923 | 1.50757956 | -3.59682 |

| | | | | | |
|---|---|---|---|---|---|
| 14147 | NM_005025.4 | 14.8326098 | -5.5072573 | 1.53126051 | -3.5965515 |
| 14148 | NM_006598.2 | 184.901732 | -3.3977664 | 0.94476006 | -3.5964332 |
| 14149 | NM_003966.2 | 79.853997 | -2.7135983 | 0.75466542 | -3.5957635 |
| 14150 | NM_012464.4 | 10.6604398 | -5.0424448 | 1.40232786 | -3.5957674 |
| 14151 | NM_017884.5 | 20.2714597 | -5.9592338 | 1.65775399 | -3.5947636 |
| 14152 | NR_073424.1 | 18.9778479 | -5.8699807 | 1.63300789 | -3.594582 |
| 14153 | NM_001099432.1 | 24.2434982 | -6.2223271 | 1.73114307 | -3.5943459 |
| 14154 | NM_001145511.1 | 31.5428256 | -6.5976012 | 1.83573797 | -3.5939776 |
| 14155 | NM_002119.3 | 67.4295935 | -2.6866595 | 0.74761628 | -3.5936343 |
| 14156 | NR_026595.2 | 12.2667864 | -5.2333048 | 1.45628065 | -3.5936101 |
| 14157 | NM_001145673.1 | 12.8248209 | -5.3112809 | 1.47819547 | -3.5930843 |
| 14158 | NR_046325.1 | 11.5942306 | -5.1584265 | 1.43573813 | -3.5928742 |
| 14159 | NM_182958.2 | 11.4649178 | -5.143611 | 1.43163817 | -3.5928149 |
| 14160 | NM_001321047.1 | 16.9717525 | -5.7043989 | 1.58784234 | -3.5925474 |
| 14161 | NM_020178.4 | 19.567907 | -5.9096502 | 1.64522686 | -3.5919972 |
| 14162 | NM_001321970.1 | 19.2797081 | -5.8882708 | 1.63955325 | -3.5913873 |
| 14163 | NM_017724.2 | 43.847217 | -4.3425693 | 1.20927609 | -3.5910487 |
| 14164 | NM_030759.4 | 59.266259 | -3.8227284 | 1.06453106 | -3.5909976 |
| 14165 | NM_001293798.2 | 24.0629991 | 9.50880468 | 2.64812688 | 3.59076627 |
| 14166 | NR_036634.1 | 17.4523015 | -5.7514386 | 1.60185545 | -3.5904854 |
| 14167 | NM_033256.2 | 9.02946688 | -4.7894312 | 1.3341974 | -3.5897471 |
| 14168 | NR_131338.1 | 28.7613573 | 9.76688588 | 2.72116544 | 3.58922899 |
| 14169 | NM_001005417.2 | 15.8131567 | -5.613411 | 1.56400787 | -3.5891194 |
| 14170 | NM_001145812.1 | 21.2248242 | -6.0293456 | 1.6799213 | -3.5890643 |
| 14171 | NM_001204193.1 | 12.5937848 | -5.2646008 | 1.46684417 | -3.5890662 |
| 14172 | NM_001289133.1 | 29.1705413 | -5.5062196 | 1.53421624 | -3.5889462 |
| 14173 | NM_203370.1 | 14.785687 | -5.504722 | 1.53404332 | -3.5883746 |
| 14174 | NM_130838.1 | 42.6944578 | -5.287406 | 1.47354921 | -3.5882113 |
| 14175 | NR_033854.1 | 58.4243475 | 1.52103396 | 0.42391211 | 3.58808798 |
| 14176 | NM_013304.2 | 10.080542 | -4.9601396 | 1.38241401 | -3.5880275 |
| 14177 | NR_104317.1 | 9.20531805 | -4.8266333 | 1.34520605 | -3.5880253 |
| 14178 | NR_034137.1 | 8.96243001 | -4.7934882 | 1.33598407 | -3.587983 |
| 14179 | NM_015032.3 | 171.556286 | -4.4550502 | 1.24180775 | -3.5875523 |
| 14180 | NM_001821.3 | 169.489657 | -3.5184324 | 0.98082107 | -3.5872317 |
| 14181 | NR_045863.1 | 9.80554331 | -4.9272502 | 1.3735938 | -3.5871232 |
| 14182 | NR_104476.1 | 25.7702852 | -6.3132586 | 1.76013702 | -3.5867995 |
| 14183 | NM_001168254.1 | 17.7986385 | -5.7712283 | 1.60906611 | -3.5866944 |
| 14184 | NR_138074.1 | 13.1642348 | -5.3288049 | 1.48571348 | -3.5866975 |
| 14185 | NM_001286581.1 | 117.46358 | -8.4985674 | 2.36950874 | -3.5866369 |
| 14186 | NM_001320953.1 | 11.3135897 | -5.1329183 | 1.43113577 | -3.5866047 |
| 14187 | NR_033761.1 | 14.270165 | -5.4469031 | 1.51886118 | -3.5861757 |
| 14188 | NM_001270643.1 | 18.298138 | -5.8206072 | 1.623094 | -3.5861183 |
| 14189 | NR_136638.1 | 8.97262088 | -4.783695 | 1.33399286 | -3.5859975 |

| | | | | | |
|---|---|---|---|---|---|
| 14190 | NM_018058.6 | 17.7761404 | -5.7798484 | 1.61184405 | -3.5858608 |
| 14191 | NM_013959.3 | 14.7209168 | -5.4988211 | 1.53357565 | -3.5856211 |
| 14192 | NM_001104631.1 | 20.7954438 | -6.0018063 | 1.67417214 | -3.5849398 |
| 14193 | NM_001345907.1 | 9.00488233 | -4.7862828 | 1.33519689 | -3.5847018 |
| 14194 | NM_001131023.1 | 17.5110918 | -5.7507702 | 1.60454894 | -3.5840416 |
| 14195 | NM_001318821.1 | 109.026896 | -8.3911059 | 2.34139787 | -3.5838018 |
| 14196 | NR_135629.1 | 12.8338789 | -5.2983048 | 1.47841751 | -3.5837676 |
| 14197 | NM_052953.3 | 19.9286071 | -5.9348032 | 1.65615653 | -3.5834796 |
| 14198 | NM_001566.2 | 11.8075955 | -5.1833707 | 1.44653929 | -3.5832906 |
| 14199 | NM_001166621.1 | 14.720452 | -5.5086073 | 1.53754072 | -3.5827391 |
| 14200 | NM_031461.5 | 131.996212 | -4.3589672 | 1.21677255 | -3.582401 |
| 14201 | NR_105020.1 | 38.5849659 | -6.8906348 | 1.92349624 | -3.582349 |
| 14202 | NM_001114734.1 | 12.2088967 | -5.2406538 | 1.46320307 | -3.5816312 |
| 14203 | NM_003620.3 | 117.507856 | -1.9769834 | 0.55207139 | -3.5810284 |
| 14204 | NM_001130009.2 | 19.396345 | -5.8960641 | 1.64657377 | -3.5808078 |
| 14205 | NM_001303117.1 | 11.7601371 | -5.1839083 | 1.44770525 | -3.5807761 |
| 14206 | NM_001318943.1 | 9.09428552 | -4.8052421 | 1.34199305 | -3.5806759 |
| 14207 | NM_001318802.1 | 11.0010329 | -5.0827485 | 1.41953818 | -3.5805648 |
| 14208 | NM_001304960.1 | 11.3472426 | -5.1248831 | 1.43184408 | -3.5792187 |
| 14209 | NM_025004.2 | 8.95569287 | -4.7849281 | 1.33698989 | -3.5788813 |
| 14210 | NR_027855.1 | 28.1846005 | -5.4580347 | 1.5251727 | -3.5786339 |
| 14211 | NR_104345.1 | 18.1323959 | -5.8000461 | 1.62082754 | -3.5784474 |
| 14212 | NM_002937.4 | 10.2823185 | -4.9930413 | 1.39535535 | -3.5783295 |
| 14213 | NM_001304350.1 | 13.0781222 | -5.3250191 | 1.48821379 | -3.5781278 |
| 14214 | NR_028323.1 | 18.0755567 | -5.801816 | 1.62158998 | -3.5778563 |
| 14215 | NM_181742.3 | 26.1044016 | -6.3310737 | 1.76954264 | -3.5778023 |
| 14216 | NR_073135.1 | 21.520182 | -6.0479991 | 1.69042204 | -3.5778042 |
| 14217 | NR_027422.1 | 11.0579444 | -5.0881563 | 1.42217015 | -3.5777409 |
| 14218 | NM_004555.3 | 13.123454 | -5.3458912 | 1.49459033 | -3.5768271 |
| 14219 | NM_001267804.1 | 11.603619 | -5.1487184 | 1.43947724 | -3.5767973 |
| 14220 | NM_001308226.1 | 16.0519751 | -5.6263968 | 1.57306213 | -3.5767162 |
| 14221 | NR_038282.1 | 11.6487957 | -5.167377 | 1.44479358 | -3.5765504 |
| 14222 | NM_001319183.1 | 8.85612144 | -4.7686156 | 1.33342018 | -3.5762287 |
| 14223 | NM_001136180.1 | 9.06677974 | -4.8084348 | 1.34458535 | -3.576147 |
| 14224 | NM_138930.3 | 22.1351273 | -6.0910651 | 1.70365414 | -3.5752944 |
| 14225 | NM_020128.2 | 10.4661401 | -5.0011073 | 1.39899925 | -3.5747748 |
| 14226 | NM_001017923.1 | 12.2463614 | -5.2363618 | 1.46491325 | -3.5745201 |
| 14227 | NR_135268.1 | 22.685718 | -6.1217604 | 1.71264036 | -3.5744576 |
| 14228 | NM_001300773.1 | 24.1407429 | -6.2105952 | 1.73758918 | -3.5742598 |
| 14229 | NM_001317334.1 | 16.2628265 | -5.6479561 | 1.58026859 | -3.5740482 |
| 14230 | NM_001039999.2 | 9.59737938 | -4.8945083 | 1.36962792 | -3.5736044 |
| 14231 | NM_001330381.1 | 10.1529862 | -4.9742219 | 1.39195885 | -3.5735409 |
| 14232 | NR_022014.1 | 15.0056822 | 3.58821677 | 1.00424138 | 3.57306207 |

| | | | | | |
|---|---|---|---|---|---|
| 14233 | NM_001323032.2 | 17.9539426 | -5.7931708 | 1.62135691 | -3.5730386 |
| 14234 | NM_139034.2 | 20.6777883 | -5.9948977 | 1.67824754 | -3.5721176 |
| 14235 | NM_004585.4 | 10.7074922 | -5.0503893 | 1.41401629 | -3.5716628 |
| 14236 | NM_001278262.1 | 10.1959331 | -4.9640633 | 1.38985454 | -3.5716423 |
| 14237 | NM_001195076.1 | 11.7713662 | -5.1714445 | 1.44794883 | -3.5715658 |
| 14238 | NM_001321180.1 | 9.09555273 | -4.8146675 | 1.34806227 | -3.5715468 |
| 14239 | NM_133343.1 | 11.0847874 | -5.0825991 | 1.42311725 | -3.5714549 |
| 14240 | NR_037675.1 | 10.6956299 | -5.0507932 | 1.41460004 | -3.5704744 |
| 14241 | NR_110799.1 | 53.4393887 | 4.3747994 | 1.22528634 | 3.57043023 |
| 14242 | NM_001034194.1 | 14.4332357 | -5.4714344 | 1.53245722 | -3.5703668 |
| 14243 | NM_001809.3 | 12.3845325 | -5.2556683 | 1.47215908 | -3.570041 |
| 14244 | NM_006454.2 | 357.575955 | -3.9627323 | 1.11009302 | -3.5697299 |
| 14245 | NR_003505.3 | 8.73729294 | -4.7542205 | 1.33186046 | -3.5696086 |
| 14246 | NM_001077654.2 | 10.5309218 | -5.0245965 | 1.40769129 | -3.5693881 |
| 14247 | NR_038284.1 | 8.79391117 | -4.7632504 | 1.33452079 | -3.5692591 |
| 14248 | NM_006114.2 | 21.4568347 | -6.0456334 | 1.69399031 | -3.5688713 |
| 14249 | NM_152485.3 | 10.3074295 | -4.9836684 | 1.39655388 | -3.5685472 |
| 14250 | NR_026677.1 | 79.314806 | -3.954098 | 1.10806797 | -3.5684616 |
| 14251 | NM_001290222.1 | 9.2970503 | -4.8436812 | 1.35742704 | -3.568281 |
| 14252 | NM_001330103.1 | 73.6269977 | -3.2508306 | 0.91103291 | -3.5682911 |
| 14253 | NR_045022.1 | 12.6837945 | -5.2801475 | 1.47976148 | -3.5682423 |
| 14254 | NM_153214.2 | 9.65345374 | -4.9009321 | 1.3735317 | -3.5681245 |
| 14255 | NM_003548.2 | 10.7409951 | -5.0428355 | 1.41336434 | -3.5679657 |
| 14256 | NM_152927.2 | 11.0808803 | -5.0993432 | 1.42921565 | -3.5679313 |
| 14257 | NM_001334.2 | 22.1635789 | -5.0930645 | 1.42759877 | -3.5675742 |
| 14258 | NM_001282957.1 | 23.5624612 | -6.1838238 | 1.73352069 | -3.567205 |
| 14259 | NM_001320825.1 | 10.1024352 | -4.9533796 | 1.3887412 | -3.5668126 |
| 14260 | NM_014189.3 | 152.964438 | -8.8792509 | 2.48948514 | -3.5667017 |
| 14261 | NM_001282195.1 | 16.98065 | -5.7130247 | 1.60191925 | -3.5663624 |
| 14262 | NM_023923.3 | 150.368179 | 4.30533133 | 1.20726418 | 3.56618823 |
| 14263 | NM_144628.3 | 191.743257 | -2.999524 | 0.8411646 | -3.565918 |
| 14264 | NM_021217.2 | 21.1031326 | -4.1169543 | 1.15467529 | -3.565465 |
| 14265 | NR_038344.1 | 9.46877852 | -4.8735503 | 1.36689443 | -3.5654182 |
| 14266 | NM_001303531.1 | 57.3181616 | 2.04948278 | 0.57488623 | 3.56502323 |
| 14267 | NM_003395.2 | 10.1268894 | -4.9727497 | 1.39497941 | -3.5647478 |
| 14268 | NM_001163678.1 | 10.0536786 | -4.9506622 | 1.38919219 | -3.5636985 |
| 14269 | NR_132415.1 | 12.3574686 | -5.2397468 | 1.47045426 | -3.5633524 |
| 14270 | NR_028514.1 | 11.9972176 | -5.20883 | 1.4617915 | -3.5633194 |
| 14271 | NM_001310321.1 | 9.31325269 | -4.8383128 | 1.35784488 | -3.5632294 |
| 14272 | NM_001082578.1 | 181.788309 | -9.1280474 | 2.56203669 | -3.562809 |
| 14273 | NM_001301861.1 | 37.1587319 | -3.7782613 | 1.06050774 | -3.5626909 |
| 14274 | NM_017768.4 | 130.075348 | -4.4544047 | 1.25054942 | -3.5619582 |
| 14275 | NM_032361.3 | 159.316536 | -4.3426381 | 1.21917876 | -3.5619371 |

| | | | | | |
|---|---|---|---|---|---|
| 14276 | NM_153374.2 | 13.0412508 | -5.3240931 | 1.49483346 | -3.5616631 |
| 14277 | NM_001286479.2 | 20.0696986 | 9.25121347 | 2.5978744 | 3.56107033 |
| 14278 | NM_001282471.1 | 27.9805385 | 4.57656961 | 1.28517589 | 3.56104534 |
| 14279 | NM_001012662.2 | 27.8563602 | -4.4020047 | 1.2363507 | -3.5604823 |
| 14280 | NM_001276418.1 | 119.538863 | -2.9310599 | 0.82324151 | -3.5603888 |
| 14281 | NM_001190476.1 | 288.635703 | -4.1151887 | 1.1559684 | -3.5599492 |
| 14282 | NM_144997.5 | 198.421087 | -2.0313981 | 0.57063825 | -3.5598702 |
| 14283 | NM_006194.3 | 13.7530803 | -5.4081313 | 1.51951386 | -3.5591194 |
| 14284 | NR_110861.1 | 14.3024649 | -5.4597651 | 1.53402189 | -3.5591181 |
| 14285 | NM_001286625.1 | 83.1307553 | -4.1842305 | 1.17569045 | -3.5589559 |
| 14286 | NM_182796.2 | 13.1801308 | -5.3512171 | 1.50368209 | -3.5587424 |
| 14287 | NR_002330.1 | 9.50727548 | -4.8667493 | 1.36763702 | -3.5585094 |
| 14288 | NM_207325.2 | 18.3136747 | -5.8228411 | 1.63633633 | -3.5584622 |
| 14289 | NM_024565.6 | 11.1323598 | -5.0841802 | 1.42878018 | -3.5584062 |
| 14290 | NM_001346509.1 | 16.3211136 | -5.6482583 | 1.58745712 | -3.5580541 |
| 14291 | NM_001122961.1 | 13.9882773 | -5.4328465 | 1.52708015 | -3.5576695 |
| 14292 | NM_001292008.1 | 36.6089652 | 6.79187632 | 1.90912525 | 3.55758551 |
| 14293 | NR_110251.1 | 13.4154825 | 3.05963165 | 0.86019651 | 3.55689846 |
| 14294 | NM_030958.2 | 17.5452175 | -5.754088 | 1.61776102 | -3.556822 |
| 14295 | NM_178033.1 | 16.335177 | -5.6498337 | 1.58859709 | -3.5564925 |
| 14296 | NM_017970.3 | 44.2975146 | -3.8922328 | 1.0945119 | -3.5561356 |
| 14297 | NR_138423.1 | 10.8081656 | -5.0585897 | 1.42273185 | -3.5555468 |
| 14298 | NM_014168.3 | 13.5385729 | -5.377919 | 1.51266372 | -3.5552641 |
| 14299 | NM_001172697.1 | 14.3810458 | -5.460847 | 1.53599723 | -3.5552453 |
| 14300 | NM_001199876.1 | 11.4880789 | -5.1470171 | 1.44778103 | -3.5551074 |
| 14301 | NM_001136537.1 | 17.2367276 | -5.7355949 | 1.6133876 | -3.5550012 |
| 14302 | NM_005654.5 | 646.853679 | -4.4505278 | 1.25192378 | -3.5549511 |
| 14303 | NM_024782.2 | 37.6251182 | -3.5212958 | 0.99059446 | -3.5547299 |
| 14304 | NM_001282162.1 | 10.2556974 | -4.9833058 | 1.40200562 | -3.5544121 |
| 14305 | NM_172103.3 | 13.7552529 | -5.4072391 | 1.52135846 | -3.5542176 |
| 14306 | NM_001702.2 | 151.267157 | -4.7128323 | 1.32601992 | -3.5541188 |
| 14307 | NM_005906.5 | 14.2177855 | -5.4559153 | 1.5351468 | -3.5540023 |
| 14308 | NM_013451.3 | 165.104031 | -8.9895558 | 2.52993438 | -3.5532763 |
| 14309 | NM_139164.2 | 102.053992 | -8.2957676 | 2.33494624 | -3.5528731 |
| 14310 | NM_020468.5 | 22.4463454 | -5.1298629 | 1.44393449 | -3.5526978 |
| 14311 | NR_037144.1 | 9.38310217 | -4.8560568 | 1.3669327 | -3.5525207 |
| 14312 | NM_145260.2 | 16.1928606 | -5.644216 | 1.58888201 | -3.5523191 |
| 14313 | NM_001010897.2 | 20.4473642 | -5.9742805 | 1.68184011 | -3.5522286 |
| 14314 | NM_001346372.1 | 19.8451982 | -5.9309038 | 1.66964043 | -3.5522042 |
| 14315 | NM_001271779.1 | 9.39859789 | -4.8581492 | 1.36772159 | -3.5520015 |
| 14316 | NM_015982.3 | 12.0170699 | -5.2093743 | 1.46691827 | -3.5512369 |
| 14317 | NM_001256666.1 | 27.0914955 | -5.4061232 | 1.52235137 | -3.5511665 |
| 14318 | NM_015832.4 | 20.9506452 | -5.0002699 | 1.40829339 | -3.5505882 |

| | | | | | |
|---|---|---|---|---|---|
| 14319 | NM_021813.3 | 22.561349 | -6.1202016 | 1.72379034 | -3.5504327 |
| 14320 | NR_049763.1 | 146.979393 | -2.8298848 | 0.79710654 | -3.5501964 |
| 14321 | NM_002908.3 | 35.4466607 | -2.5532807 | 0.71920677 | -3.5501344 |
| 14322 | NM_006006.4 | 9.01408818 | -4.7992382 | 1.3518842 | -3.5500365 |
| 14323 | NR_136290.1 | 12.9080543 | -5.3073422 | 1.49512706 | -3.54976 |
| 14324 | NM_001025108.1 | 239.311133 | -3.5340296 | 0.99567953 | -3.5493645 |
| 14325 | NM_000790.3 | 12.0520618 | -5.2152187 | 1.46940076 | -3.5492146 |
| 14326 | NM_001006625.1 | 22.5791259 | 9.4148807 | 2.65302365 | 3.54873606 |
| 14327 | NM_020762.3 | 11.3006532 | -5.1219233 | 1.44333524 | -3.5486719 |
| 14328 | NM_001204103.1 | 21.6207911 | -6.050848 | 1.70532956 | -3.5481986 |
| 14329 | NR_033397.4 | 25.5470398 | -3.9266035 | 1.1067536 | -3.547857 |
| 14330 | NM_144495.2 | 12.5588289 | -5.2795394 | 1.48824433 | -3.547495 |
| 14331 | NM_024863.5 | 19.34927 | -5.8953385 | 1.66204753 | -3.5470336 |
| 14332 | NM_012428.3 | 14.1489809 | -5.4379262 | 1.53315287 | -3.546891 |
| 14333 | NM_018234.2 | 21.994906 | -4.1684075 | 1.17533571 | -3.5465676 |
| 14334 | NM_001008660.2 | 101.42534 | -8.2868703 | 2.3367646 | -3.5463009 |
| 14335 | NM_001320252.1 | 10.7846407 | -5.0519927 | 1.42461132 | -3.5462253 |
| 14336 | NM_001166139.2 | 41.3224746 | -4.1682248 | 1.17543274 | -3.5461194 |
| 14337 | NR_104006.1 | 12.2268464 | -5.2229016 | 1.47284755 | -3.5461251 |
| 14338 | NM_001243280.1 | 28.1812334 | -4.660163 | 1.31418506 | -3.5460478 |
| 14339 | NM_001134435.2 | 33.1531814 | -6.6751433 | 1.88251575 | -3.5458632 |
| 14340 | NM_001265611.1 | 30.2900751 | 9.84206393 | 2.77566561 | 3.5458392 |
| 14341 | NM_004034.3 | 21.8023771 | -6.0629464 | 1.70994669 | -3.5456932 |
| 14342 | NM_001317843.1 | 11.9240191 | -5.194921 | 1.46532297 | -3.5452396 |
| 14343 | NM_024887.3 | 21.1985603 | -6.0267662 | 1.70011937 | -3.5449077 |
| 14344 | NR_039973.1 | 87.5979205 | 3.02938944 | 0.85457439 | 3.54491015 |
| 14345 | NM_024997.3 | 11.0024321 | -5.0744394 | 1.43150529 | -3.5448276 |
| 14346 | NM_001256409.1 | 22.7845343 | -6.1297141 | 1.72922506 | -3.5447752 |
| 14347 | NR_130136.1 | 18.8334104 | -5.8596565 | 1.65336411 | -3.5440811 |
| 14348 | NM_198580.2 | 115.639992 | -3.6595638 | 1.03280347 | -3.5433303 |
| 14349 | NM_001243198.2 | 6.46649528 | 7.58787869 | 2.14151093 | 3.54323603 |
| 14350 | NM_001242597.1 | 11.5722657 | -5.152009 | 1.4541848 | -3.5428846 |
| 14351 | NR_002323.2 | 128.323989 | -8.6258123 | 2.43474687 | -3.5427963 |
| 14352 | NM_001178119.1 | 18.6416445 | -5.8370297 | 1.64777329 | -3.5423743 |
| 14353 | NM_001001936.2 | 13.2289062 | -5.3445742 | 1.50876881 | -3.5423414 |
| 14354 | NR_003502.2 | 18.4258711 | -3.9034327 | 1.10222962 | -3.541397 |
| 14355 | NM_145040.2 | 10.3217523 | -4.9995741 | 1.41194925 | -3.5409021 |
| 14356 | NM_001256469.1 | 120.464675 | -4.0727141 | 1.15025776 | -3.5406969 |
| 14357 | NM_006262.3 | 353.868255 | -5.2406527 | 1.48011378 | -3.5407094 |
| 14358 | NM_001317171.1 | 16.2265929 | -5.6447294 | 1.59432882 | -3.5405052 |
| 14359 | NM_024806.3 | 13.3698539 | -5.3698858 | 1.51697264 | -3.5398699 |
| 14360 | NM_003311.3 | 12.8390011 | -5.3035468 | 1.49824465 | -3.5398403 |
| 14361 | NM_024775.9 | 12.581705 | -5.2769358 | 1.49079084 | -3.5396889 |

| | | | | | |
|---|---|---|---|---|---|
| 14362 | NR_040023.1 | 88.2659132 | -3.1619146 | 0.89329949 | -3.5395908 |
| 14363 | NM_001178093.1 | 101.187711 | -8.2834632 | 2.34031515 | -3.5394648 |
| 14364 | NM_170733.3 | 13.0928372 | -5.3353517 | 1.50772371 | -3.53868 |
| 14365 | NM_153210.4 | 12.1113271 | -5.2279268 | 1.47740051 | -3.5385982 |
| 14366 | NM_014913.3 | 146.878579 | -3.8330765 | 1.08325622 | -3.5384763 |
| 14367 | NM_194252.2 | 9.78883701 | -4.9176672 | 1.3898413 | -3.538294 |
| 14368 | NM_015703.4 | 313.200911 | -1.7688064 | 0.49993196 | -3.5380943 |
| 14369 | NM_004956.4 | 17.9600891 | -5.7843331 | 1.63508154 | -3.5376419 |
| 14370 | NR_038131.1 | 12.1859108 | -5.2250367 | 1.47705872 | -3.5374604 |
| 14371 | NM_017413.4 | 9.3070738 | -4.8483062 | 1.37061462 | -3.5373227 |
| 14372 | NM_032316.3 | 176.541739 | -1.5876059 | 0.44883331 | -3.5371838 |
| 14373 | NM_016570.2 | 76.6874017 | -4.0585508 | 1.14740851 | -3.5371454 |
| 14374 | NM_006509.3 | 21.6107206 | -4.2507716 | 1.20190479 | -3.5366958 |
| 14375 | NM_001142936.1 | 20.4390813 | -4.979508 | 1.40819339 | -3.5360967 |
| 14376 | NM_001305654.1 | 14.4351404 | -5.4756128 | 1.54861873 | -3.5358043 |
| 14377 | NM_001286585.1 | 16.0767254 | -5.6238114 | 1.59055055 | -3.535764 |
| 14378 | NR_126059.1 | 26.0938837 | -2.4738679 | 0.69968396 | -3.5356933 |
| 14379 | NM_020439.2 | 12.2340149 | -5.2368713 | 1.48122915 | -3.5354903 |
| 14380 | NM_015134.3 | 446.167025 | -4.4280395 | 1.25271388 | -3.5347573 |
| 14381 | NM_001044723.2 | 18.7351666 | -5.8474054 | 1.65435347 | -3.5345563 |
| 14382 | NM_017823.4 | 11.9846832 | -5.21042 | 1.47416293 | -3.534494 |
| 14383 | NM_138622.3 | 10.5054879 | -5.0204169 | 1.42040299 | -3.5345018 |
| 14384 | NM_001282787.1 | 237.148883 | -5.7845691 | 1.63666776 | -3.5343576 |
| 14385 | NM_013995.2 | 232.910685 | -2.201468 | 0.6228965 | -3.5342437 |
| 14386 | NM_001286691.1 | 88.3976663 | -2.6968128 | 0.76314711 | -3.5338047 |
| 14387 | NM_014728.3 | 11.003516 | -5.0784541 | 1.43712685 | -3.533755 |
| 14388 | NM_001286094.1 | 13.3890265 | -5.3664649 | 1.51866172 | -3.5336802 |
| 14389 | NM_001303460.1 | 10.1858315 | -4.9688575 | 1.40618974 | -3.5335612 |
| 14390 | NM_001992.4 | 286.936275 | -5.0825781 | 1.4384653 | -3.5333338 |
| 14391 | NM_152355.2 | 48.9425377 | -3.7187685 | 1.05262289 | -3.5328593 |
| 14392 | NM_001015055.1 | 15.2240645 | -5.5551348 | 1.57244944 | -3.5327907 |
| 14393 | NM_001170635.1 | 104.421486 | -4.2450021 | 1.20196417 | -3.531721 |
| 14394 | NM_001330187.1 | 11.1294519 | -5.1058782 | 1.44573886 | -3.5316739 |
| 14395 | NM_001012301.3 | 9.06999443 | -4.8127337 | 1.36311159 | -3.5306968 |
| 14396 | NR_026991.1 | 21.1889645 | -3.5736404 | 1.01219307 | -3.5305917 |
| 14397 | NM_001286056.1 | 10.7397681 | -5.0382807 | 1.42714564 | -3.5303199 |
| 14398 | NM_001282768.1 | 11.5511634 | -5.1505652 | 1.45930251 | -3.5294705 |
| 14399 | NM_001197260.1 | 12.4101223 | -5.2554932 | 1.48907925 | -3.5293576 |
| 14400 | NM_001243611.1 | 12.6412393 | -5.2765885 | 1.49509545 | -3.5292653 |
| 14401 | NM_001199173.1 | 11.2309934 | -5.114302 | 1.44920217 | -3.5290466 |
| 14402 | NM_001293274.1 | 9.48617961 | -4.8733776 | 1.38093834 | -3.5290334 |
| 14403 | NM_024838.4 | 42.2419108 | -2.4523471 | 0.69492174 | -3.5289544 |
| 14404 | NR_135522.1 | 10.1326678 | 5.70805071 | 1.61752879 | 3.52887117 |

| | | | | | |
|---|---|---|---|---|---|
| 14405 | NM_001282418.1 | 37.5106113 | -6.8481548 | 1.94064619 | -3.5288013 |
| 14406 | NM_001201407.1 | 19.3382121 | -5.889982 | 1.66954988 | -3.5278862 |
| 14407 | NM_199242.2 | 9.33736733 | -4.8316959 | 1.36975719 | -3.5274105 |
| 14408 | NM_001285987.1 | 80.9527864 | 2.28180571 | 0.6470067 | 3.52671112 |
| 14409 | NM_014825.2 | 177.839634 | -2.9173913 | 0.82725335 | -3.5265995 |
| 14410 | NM_181711.3 | 20.2530687 | -4.1352881 | 1.1726461 | -3.5264587 |
| 14411 | NM_001193533.1 | 23.9942725 | 9.50766812 | 2.69626741 | 3.52623338 |
| 14412 | NM_206907.3 | 23.059203 | -6.1454051 | 1.74281689 | -3.5261335 |
| 14413 | NM_001302694.1 | 15.7884775 | -5.6021298 | 1.58875873 | -3.5261048 |
| 14414 | NM_001319041.1 | 17.2268047 | -5.7256293 | 1.62382291 | -3.5260183 |
| 14415 | NM_032532.2 | 55.2735157 | -3.626809 | 1.02866761 | -3.5257347 |
| 14416 | NM_001170797.1 | 10.036258 | -4.9472188 | 1.40326922 | -3.5254951 |
| 14417 | NM_001206839.1 | 137.694207 | -4.5429958 | 1.2886329 | -3.5254383 |
| 14418 | NR_073074.1 | 21.241264 | -5.0258848 | 1.42573339 | -3.5251225 |
| 14419 | NM_001102654.1 | 13.0358004 | -5.3331428 | 1.51290971 | -3.5250899 |
| 14420 | NM_001008710.2 | 13.1622919 | -5.3486705 | 1.51749142 | -3.5246793 |
| 14421 | NM_001286174.1 | 17.2582536 | -5.7355223 | 1.62770361 | -3.5236896 |
| 14422 | NM_001323424.1 | 10.9478386 | -5.0704017 | 1.43897358 | -3.5236239 |
| 14423 | NM_002517.3 | 10.5910667 | -5.0312962 | 1.42796293 | -3.5234081 |
| 14424 | NM_202494.2 | 18.7247104 | -4.840882 | 1.37401135 | -3.5231747 |
| 14425 | NM_004371.3 | 728.192649 | -4.2358574 | 1.20239247 | -3.5228576 |
| 14426 | NM_001271168.1 | 10.2972418 | -4.9821519 | 1.41428197 | -3.522743 |
| 14427 | NR_125949.1 | 133.571035 | 4.3173744 | 1.22557416 | 3.52273615 |
| 14428 | NM_152362.2 | 249.031934 | 1.29280154 | 0.36702082 | 3.52242017 |
| 14429 | NM_001316981.1 | 10.8964503 | -5.0621537 | 1.4371348 | -3.5223931 |
| 14430 | NM_015306.2 | 242.970295 | -4.8435498 | 1.37511106 | -3.5222972 |
| 14431 | NM_000117.2 | 195.395226 | -4.3084179 | 1.2232004 | -3.5222503 |
| 14432 | NR_027157.1 | 11.5300923 | -5.1552199 | 1.4636895 | -3.5220721 |
| 14433 | NM_001322796.1 | 10.1689825 | -4.9550248 | 1.40711002 | -3.5214196 |
| 14434 | NR_136596.1 | 13.5520864 | -5.3878128 | 1.53036457 | -3.5206074 |
| 14435 | NR_120449.1 | 9.23449524 | -4.8300672 | 1.37213627 | -3.5201075 |
| 14436 | NM_176792.2 | 11.6137708 | -5.1669558 | 1.46792164 | -3.5199125 |
| 14437 | NM_001185090.1 | 16.6572249 | -5.6715418 | 1.61158657 | -3.5192288 |
| 14438 | NM_031421.3 | 19.0096911 | -5.8744338 | 1.66924047 | -3.5192256 |
| 14439 | NM_001243942.1 | 15.9366354 | -5.6130221 | 1.59505153 | -3.5190224 |
| 14440 | NM_001166006.1 | 10.419763 | -5.0017179 | 1.42141563 | -3.5188285 |
| 14441 | NM_001288667.1 | 13.2275861 | -5.3434887 | 1.5185876 | -3.5187227 |
| 14442 | NM_001098623.2 | 26.9729264 | -4.3547127 | 1.23770687 | -3.5183716 |
| 14443 | NM_001128921.2 | 13.6540042 | -5.3964698 | 1.53381199 | -3.5183385 |
| 14444 | NR_002612.1 | 79.5592953 | 2.14381063 | 0.60935451 | 3.51816649 |
| 14445 | NM_001730.4 | 14.8755116 | -5.5226304 | 1.56979465 | -3.5180591 |
| 14446 | NM_152329.3 | 17.0897213 | -5.7134364 | 1.6241115 | -3.5178843 |
| 14447 | NM_152400.2 | 18.8472275 | -5.8537994 | 1.66413724 | -3.5176182 |

| | | | | | |
|---|---|---|---|---|---|
| 14448 | NM_181426.1 | 25.0867619 | -4.4798538 | 1.27356022 | -3.5175831 |
| 14449 | NM_001144936.1 | 597.571862 | -4.7431642 | 1.3485176 | -3.5173172 |
| 14450 | NM_001172428.1 | 12.8530273 | -5.3071684 | 1.50905319 | -3.5168862 |
| 14451 | NM_002918.4 | 73.5035919 | -3.3736815 | 0.95934309 | -3.5166579 |
| 14452 | NM_145755.2 | 15.9907918 | -5.6163603 | 1.59709107 | -3.5166187 |
| 14453 | NR_136586.1 | 25.7693351 | -6.3065892 | 1.79343378 | -3.5164885 |
| 14454 | NM_001278221.2 | 9.84711847 | -4.9145128 | 1.39770869 | -3.516121 |
| 14455 | NM_198044.2 | 11.4918089 | -5.1425852 | 1.4627196 | -3.5157697 |
| 14456 | NM_005791.2 | 117.951914 | -4.1875902 | 1.19131114 | -3.5151104 |
| 14457 | NM_022087.3 | 281.929587 | -4.6772902 | 1.33082168 | -3.5145882 |
| 14458 | NM_001267798.1 | 16.1209203 | 6.68606463 | 1.9026382 | 3.51410196 |
| 14459 | NM_198491.2 | 15.2564995 | -5.5592825 | 1.58217889 | -3.5136877 |
| 14460 | NR_144490.1 | 10.9453031 | -5.0730828 | 1.4439093 | -3.5134359 |
| 14461 | NM_001297721.1 | 11.8403365 | 8.46966714 | 2.41085251 | 3.51314198 |
| 14462 | NM_017414.3 | 17.0277732 | -4.700628 | 1.33806627 | -3.5130009 |
| 14463 | NR_046477.1 | 10.210492 | -4.9833661 | 1.41883381 | -3.5122973 |
| 14464 | NR_029434.1 | 35.6106841 | -4.0559462 | 1.15480128 | -3.512246 |
| 14465 | NM_001168331.1 | 9.01543745 | -4.7914158 | 1.36429487 | -3.5120089 |
| 14466 | NR_024014.1 | 13.1694246 | -5.3375612 | 1.51984177 | -3.511919 |
| 14467 | NM_001288570.1 | 12.5815942 | -5.2720767 | 1.50121529 | -3.5118725 |
| 14468 | NM_016035.4 | 85.8326969 | -3.9398434 | 1.12194281 | -3.5116259 |
| 14469 | NM_145010.3 | 12.0513094 | -5.2167395 | 1.48564465 | -3.5114316 |
| 14470 | NM_001172651.1 | 12.1406992 | -5.2254177 | 1.48817961 | -3.5112816 |
| 14471 | NM_000172.3 | 13.3846735 | 4.99861197 | 1.42361004 | 3.51122276 |
| 14472 | NM_001318186.1 | 165.854005 | -8.0283687 | 2.28660292 | -3.5110463 |
| 14473 | NM_002690.2 | 82.1268892 | -4.3403197 | 1.23633556 | -3.5106324 |
| 14474 | NM_001165924.1 | 21.5461223 | 6.0010136 | 1.70961928 | 3.51014619 |
| 14475 | NM_001243121.1 | 11.8364814 | -5.197805 | 1.48080164 | -3.5101291 |
| 14476 | NM_001324215.1 | 24.2292945 | -6.2232431 | 1.77322933 | -3.5095534 |
| 14477 | NM_020884.4 | 8.82887743 | -4.774677 | 1.36052983 | -3.5094247 |
| 14478 | NM_019090.2 | 16.0671859 | -4.5965536 | 1.30978572 | -3.5093936 |
| 14479 | NM_002692.3 | 9.64666542 | -4.8853549 | 1.39213405 | -3.5092561 |
| 14480 | NM_024693.4 | 17.7341306 | -5.7748741 | 1.64560566 | -3.5092697 |
| 14481 | NM_001348121.1 | 31.9910176 | 9.9216463 | 2.82760272 | 3.50885442 |
| 14482 | NM_031942.4 | 22.1755279 | -6.089752 | 1.7355691 | -3.5087926 |
| 14483 | NM_001620.2 | 103.991436 | -8.3227622 | 2.3722259 | -3.5084189 |
| 14484 | NM_001166269.1 | 13.3700966 | -5.3657559 | 1.529667 | -3.5077935 |
| 14485 | NM_005108.3 | 9.10691944 | -4.8032368 | 1.36931918 | -3.5077554 |
| 14486 | NM_001319086.1 | 10.5184919 | -5.0231625 | 1.43202368 | -3.507737 |
| 14487 | NM_001165931.1 | 22.6538523 | -6.121382 | 1.74545487 | -3.5070411 |
| 14488 | NM_001288566.1 | 12.1843759 | -5.2279105 | 1.49083163 | -3.5067075 |
| 14489 | NM_024704.4 | 11.5911841 | -5.1647885 | 1.47295474 | -3.5064136 |
| 14490 | NM_014364.4 | 10.0416798 | -4.9417208 | 1.40947219 | -3.506079 |

| | | | | | |
|---|---|---|---|---|---|
| 14491 | NM_001144766.2 | 83.4444362 | -4.3656023 | 1.24521328 | -3.5059073 |
| 14492 | NM_001207020.1 | 10.0050799 | -4.9394927 | 1.40891941 | -3.5058731 |
| 14493 | NM_006862.3 | 11.7537857 | -5.1801933 | 1.47768922 | -3.5056041 |
| 14494 | NM_004902.3 | 132.414066 | -8.6710781 | 2.47354382 | -3.5055284 |
| 14495 | NR_033819.1 | 11.7839982 | -5.1727182 | 1.47566124 | -3.5053561 |
| 14496 | NR_026918.1 | 20.4200812 | 7.03501808 | 2.0069922 | 3.50525431 |
| 14497 | NM_005738.4 | 13.2203388 | -5.3420885 | 1.52409645 | -3.5050856 |
| 14498 | NM_173600.2 | 20.2706527 | 4.18313838 | 1.19349944 | 3.50493537 |
| 14499 | NM_001244015.1 | 17.724397 | -5.7750748 | 1.64788874 | -3.5045296 |
| 14500 | NM_001206985.2 | 12.9834145 | -5.3113713 | 1.51559878 | -3.5044706 |
| 14501 | NM_001270491.1 | 141.232608 | -7.7992926 | 2.22567194 | -3.5042418 |
| 14502 | NM_001163692.1 | 9.03789505 | -4.7939237 | 1.3680873 | -3.5041066 |
| 14503 | NM_001318845.1 | 22.1228107 | -6.0863673 | 1.73696093 | -3.5040324 |
| 14504 | NM_001286242.1 | 201.839028 | -3.335417 | 0.95188661 | -3.5040066 |
| 14505 | NM_004237.3 | 30.8596298 | -3.4750468 | 0.99181293 | -3.5037321 |
| 14506 | NM_032177.3 | 289.282376 | -1.0836225 | 0.30928132 | -3.5036791 |
| 14507 | NM_001316935.1 | 11.6159001 | -5.1494766 | 1.46977163 | -3.5035896 |
| 14508 | NM_001206855.1 | 13.072957 | -5.3238093 | 1.51956581 | -3.5035069 |
| 14509 | NM_003413.3 | 23.9477947 | -6.206981 | 1.77175549 | -3.5032943 |
| 14510 | NM_001193343.1 | 13.8071813 | -5.411188 | 1.54497138 | -3.5024519 |
| 14511 | NM_001199707.1 | 9.54714214 | -4.8759682 | 1.39221511 | -3.5023095 |
| 14512 | NM_015335.4 | 311.353146 | -5.0550759 | 1.44341146 | -3.5021725 |
| 14513 | NM_001142676.1 | 11.2847707 | -5.1295073 | 1.46488341 | -3.5016488 |
| 14514 | NM_001256916.1 | 28.025773 | 2.04039463 | 0.58276997 | 3.50120068 |
| 14515 | NM_014760.3 | 218.475231 | -4.3429273 | 1.24043441 | -3.5011342 |
| 14516 | NM_001100159.2 | 10.9261444 | -5.0783315 | 1.45054062 | -3.5009923 |
| 14517 | NM_004759.4 | 10.8487043 | -5.0698723 | 1.44814057 | -3.5009532 |
| 14518 | NM_001032999.2 | 250.313749 | -1.5532302 | 0.44367447 | -3.5008329 |
| 14519 | NM_152774.3 | 22.6308829 | -5.1380596 | 1.46768116 | -3.5008009 |
| 14520 | NM_000361.2 | 18.2725413 | -3.9823931 | 1.13763332 | -3.5005946 |
| 14521 | NM_001206927.1 | 8.19673675 | 5.40501753 | 1.54408997 | 3.50045504 |
| 14522 | NM_018365.2 | 32.7077742 | -4.7230738 | 1.34930876 | -3.5003655 |
| 14523 | NM_001130867.1 | 10.520499 | -5.0233912 | 1.43532539 | -3.4998275 |
| 14524 | NM_001267594.1 | 22.9963614 | -4.2255238 | 1.20734732 | -3.4998411 |
| 14525 | NM_152357.2 | 66.79026 | -2.3967869 | 0.68492951 | -3.4993191 |
| 14526 | NM_001080532.1 | 13.4962059 | -5.376923 | 1.53676359 | -3.4988615 |
| 14527 | NR_046322.1 | 20.1438299 | -4.1335272 | 1.18145692 | -3.4986694 |
| 14528 | NM_001006681.1 | 16.7113625 | -5.6843518 | 1.62484544 | -3.4983954 |
| 14529 | NM_001319225.1 | 11.6956984 | -5.1756589 | 1.47955008 | -3.4981302 |
| 14530 | NM_000246.3 | 15.8206763 | -5.6036094 | 1.60194013 | -3.4980143 |
| 14531 | NM_001931.4 | 136.714365 | -3.8030916 | 1.08721048 | -3.498027 |
| 14532 | NM_031371.3 | 13.6728399 | -5.4029616 | 1.54479112 | -3.4975354 |
| 14533 | NM_000692.4 | 98.1707947 | -4.3384535 | 1.2405176 | -3.4972929 |

| | | | | | |
|---|---|---|---|---|---|
| 14534 | NM_012216.3 | 23.2716499 | -6.1638085 | 1.76283621 | -3.4965293 |
| 14535 | NR_026789.1 | 9.68119045 | -4.8895388 | 1.39847192 | -3.4963439 |
| 14536 | NM_001243374.1 | 165.824356 | -4.3487718 | 1.24381576 | -3.4963151 |
| 14537 | NM_001098173.1 | 18.4045845 | 3.87381133 | 1.10811615 | 3.49585316 |
| 14538 | NM_152398.3 | 22.4064299 | -6.1023766 | 1.74564286 | -3.4957761 |
| 14539 | NM_001317077.1 | 24.1938864 | -5.2391436 | 1.49877198 | -3.4956242 |
| 14540 | NM_175733.3 | 14.4146789 | -4.4802449 | 1.28177971 | -3.4953315 |
| 14541 | NM_001162422.1 | 24.8664 | 9.55894222 | 2.73529099 | 3.49467105 |
| 14542 | NM_001098531.2 | 12.2269903 | -5.233492 | 1.49787069 | -3.4939545 |
| 14543 | NR_023313.2 | 105.400548 | -8.3423325 | 2.38783185 | -3.4936851 |
| 14544 | NR_131157.1 | 12.7420945 | -5.2988372 | 1.51671567 | -3.493626 |
| 14545 | NR_024071.1 | 9.02096704 | -4.7944672 | 1.37249385 | -3.4932523 |
| 14546 | NM_003773.4 | 72.3457991 | -4.1445268 | 1.18648542 | -3.4931123 |
| 14547 | NM_001161586.1 | 13.8475671 | -5.4056455 | 1.54765575 | -3.4927958 |
| 14548 | NR_136240.1 | 15.614438 | -4.5874563 | 1.31382707 | -3.4916744 |
| 14549 | NM_001199800.1 | 17.6757468 | 4.41770552 | 1.26533573 | 3.49133072 |
| 14550 | NM_001253912.1 | 11.680146 | -5.1632703 | 1.47903991 | -3.4909608 |
| 14551 | NM_001330069.1 | 17.1015065 | -5.7201849 | 1.63859404 | -3.4909103 |
| 14552 | NM_001145710.1 | 8.53948822 | -4.7149551 | 1.35096575 | -3.4900626 |
| 14553 | NM_001025778.1 | 10.399914 | -5.0119978 | 1.43608773 | -3.4900359 |
| 14554 | NM_018249.5 | 94.8012057 | -8.1893827 | 2.34655729 | -3.4899564 |
| 14555 | NM_003673.3 | 11.3668783 | -5.1255689 | 1.46880945 | -3.4896078 |
| 14556 | NM_001281710.1 | 13.7183742 | -5.3893639 | 1.54462817 | -3.4891011 |
| 14557 | NM_005888.3 | 11.0179239 | -5.0787058 | 1.45560597 | -3.4890663 |
| 14558 | NM_001164384.1 | 12.4327595 | -5.2546765 | 1.50608139 | -3.4889725 |
| 14559 | NR_002822.3 | 9.66732316 | -4.8994644 | 1.40441784 | -3.4886087 |
| 14560 | NM_002592.2 | 12.4091372 | -5.252525 | 1.50571799 | -3.4883856 |
| 14561 | NM_001142316.1 | 12.2955478 | -5.2390872 | 1.50188868 | -3.4883326 |
| 14562 | NM_017712.3 | 91.8569687 | -8.143899 | 2.3348834 | -3.4879254 |
| 14563 | NM_001145054.1 | 8.52915769 | -4.7191477 | 1.35301092 | -3.4878859 |
| 14564 | NR_024629.1 | 8.3496703 | -4.6873433 | 1.34400693 | -3.4875886 |
| 14565 | NM_182924.3 | 75.7830723 | -3.886977 | 1.11455481 | -3.4874705 |
| 14566 | NM_012307.3 | 11.2813074 | -5.1181709 | 1.46769572 | -3.4872153 |
| 14567 | NM_001304834.1 | 9.57311092 | -4.8933096 | 1.40334806 | -3.4868824 |
| 14568 | NM_024952.7 | 11.0868999 | -5.0968658 | 1.4619287 | -3.4863983 |
| 14569 | NM_024811.3 | 11.5639555 | -5.1622576 | 1.48088236 | -3.4859336 |
| 14570 | NR_015395.2 | 12.4700121 | -5.267705 | 1.51131871 | -3.4855024 |
| 14571 | NM_001321964.1 | 12.0502975 | -5.2118766 | 1.49565274 | -3.4846836 |
| 14572 | NM_018442.3 | 13.9967682 | -5.4325695 | 1.55913028 | -3.4843589 |
| 14573 | NM_002739.4 | 13.5052956 | -5.3692614 | 1.54101267 | -3.4842422 |
| 14574 | NR_134469.1 | 170.600573 | -3.7864682 | 1.08677115 | -3.4841449 |
| 14575 | NM_006207.2 | 9.78331576 | -4.9198577 | 1.41210755 | -3.4840531 |
| 14576 | NM_001305182.1 | 9.16498966 | -4.8266246 | 1.38542005 | -3.4838709 |

| | | | | | |
|---|---|---|---|---|---|
| 14577 | NM_001301240.1 | 23.8863431 | -6.2006042 | 1.77981974 | -3.4838383 |
| 14578 | NM_020464.1 | 92.2692314 | -4.0537486 | 1.16363858 | -3.4836835 |
| 14579 | NR_104227.1 | 8.77253526 | -4.7549383 | 1.36500033 | -3.4834704 |
| 14580 | NM_015056.2 | 203.977994 | -4.7507914 | 1.36385399 | -3.4833578 |
| 14581 | NM_001304968.1 | 33.0477035 | 3.57878966 | 1.02743284 | 3.48323463 |
| 14582 | NM_001242704.1 | 8.88303822 | -4.7611354 | 1.36692235 | -3.483106 |
| 14583 | NR_038989.1 | 34.6765963 | 4.46697806 | 1.28246516 | 3.48311844 |
| 14584 | NM_001099294.1 | 158.445266 | -4.3648973 | 1.25332207 | -3.4826621 |
| 14585 | NM_001166588.2 | 16.6736662 | -4.6675811 | 1.3402928 | -3.4825086 |
| 14586 | NM_175605.4 | 11.0969577 | -5.1019237 | 1.46502922 | -3.4824723 |
| 14587 | NM_001146702.1 | 16.5792964 | -5.6801225 | 1.63140796 | -3.4817303 |
| 14588 | NM_001145364.2 | 11.5185634 | -5.1421923 | 1.47704158 | -3.4814134 |
| 14589 | NM_020677.4 | 11.8385601 | -5.1748928 | 1.48660267 | -3.4810194 |
| 14590 | NM_001124767.1 | 10.513332 | -5.021029 | 1.44262412 | -3.4804832 |
| 14591 | NM_024684.3 | 10.5363138 | -5.0147175 | 1.44085186 | -3.4803839 |
| 14592 | NM_001083953.1 | 12.2549258 | -5.2423904 | 1.50628922 | -3.4803345 |
| 14593 | NR_120332.1 | 18.7931157 | 3.20562907 | 0.92116711 | 3.47996473 |
| 14594 | NM_024070.3 | 9.19447462 | -4.8282426 | 1.387491 | -3.479837 |
| 14595 | NM_001101312.1 | 10.5224495 | -5.0236781 | 1.44370387 | -3.4797151 |
| 14596 | NM_003970.3 | 16.0298586 | -4.6266128 | 1.32962176 | -3.4796458 |
| 14597 | NM_016377.3 | 17.3996108 | -4.7280544 | 1.35902151 | -3.4790137 |
| 14598 | NM_018227.5 | 170.6646 | -4.4504495 | 1.27945157 | -3.478404 |
| 14599 | NM_001319240.1 | 21.3237959 | -4.2226408 | 1.21435653 | -3.4772661 |
| 14600 | NM_003205.3 | 89.3358258 | -8.1037974 | 2.33073119 | -3.4769335 |
| 14601 | NM_001303048.1 | 21.5035631 | -6.0503569 | 1.74025876 | -3.4766996 |
| 14602 | NR_107045.1 | 8.67680081 | 6.68753996 | 1.92383913 | 3.47614302 |
| 14603 | NM_002458.2 | 17.8996075 | 4.10311634 | 1.18067374 | 3.47523298 |
| 14604 | NM_001278173.1 | 9.81890374 | -4.9218574 | 1.41650172 | -3.4746568 |
| 14605 | NM_001012967.2 | 8.32069521 | -4.6817748 | 1.34741387 | -3.4746375 |
| 14606 | NM_001308340.1 | 22.9041944 | -4.285952 | 1.23367589 | -3.4741313 |
| 14607 | NM_001318872.1 | 9.91455647 | -4.9407615 | 1.4222567 | -3.4738887 |
| 14608 | NM_020056.4 | 17.9417032 | -5.789395 | 1.6667708 | -3.47342 |
| 14609 | NM_001330383.1 | 9.28645499 | -4.8392985 | 1.39350594 | -3.4727505 |
| 14610 | NM_001267608.1 | 14.4204006 | -5.4617807 | 1.57281424 | -3.4726165 |
| 14611 | NM_133645.2 | 13.7299465 | -5.4103108 | 1.55837326 | -3.4717682 |
| 14612 | NM_001198696.1 | 22.4519879 | -6.1120977 | 1.76071354 | -3.4713754 |
| 14613 | NM_032918.2 | 9.39372594 | -4.8500223 | 1.39714265 | -3.4713866 |
| 14614 | NM_080798.3 | 14.8045028 | -5.5051496 | 1.58595942 | -3.4711793 |
| 14615 | NM_194285.2 | 94.9718109 | -2.8817478 | 0.83025009 | -3.4709395 |
| 14616 | NM_001322240.1 | 18.1003835 | -5.7925924 | 1.66904548 | -3.4706019 |
| 14617 | NR_047510.2 | 9.9071473 | -4.9297112 | 1.4204164 | -3.4706099 |
| 14618 | NM_001145645.2 | 107.260674 | 4.86962886 | 1.40316142 | 3.47046947 |
| 14619 | NM_054027.4 | 145.35444 | -3.3386786 | 0.96203182 | -3.4704451 |

| | | | | | |
|---|---|---|---|---|---|
| 14620 | NM_001167935.1 | 15.8179907 | -5.6097576 | 1.61654179 | -3.4702212 |
| 14621 | NM_001082533.1 | 12.8557413 | -5.3015777 | 1.52797276 | -3.4696807 |
| 14622 | NM_176875.3 | 10.6502789 | -5.0349428 | 1.451255 | -3.4693715 |
| 14623 | NM_001002021.2 | 9.69526294 | -4.9033561 | 1.41333639 | -3.4693483 |
| 14624 | NM_001164883.1 | 13.4579993 | -5.3708179 | 1.54824781 | -3.4689653 |
| 14625 | NM_001025089.1 | 174.008443 | -2.6862297 | 0.77440946 | -3.4687459 |
| 14626 | NR_126419.1 | 91.8838656 | 3.24646029 | 0.93606359 | 3.46820484 |
| 14627 | NM_001242933.1 | 65.1169501 | 3.42037728 | 0.98632429 | 3.46780192 |
| 14628 | NR_109845.1 | 5.66787662 | 7.39752174 | 2.13324069 | 3.46773891 |
| 14629 | NM_001321746.1 | 108.236547 | -8.3805701 | 2.41698319 | -3.467368 |
| 14630 | NM_001320599.1 | 13.035044 | -5.3164931 | 1.5334542 | -3.4670048 |
| 14631 | NM_001282484.1 | 10.279224 | -4.9888033 | 1.43910709 | -3.4665963 |
| 14632 | NM_001301035.1 | 10.625564 | -5.0433115 | 1.45483937 | -3.4665762 |
| 14633 | NM_001039753.2 | 30.2049709 | -2.8485271 | 0.82172797 | -3.4665086 |
| 14634 | NM_022806.3 | 18.941488 | -5.8633782 | 1.69144189 | -3.466497 |
| 14635 | NR_103814.2 | 11.8111022 | 2.97628055 | 0.85859608 | 3.46645022 |
| 14636 | NM_001098832.1 | 16.8759159 | -5.6889236 | 1.64119343 | -3.4663334 |
| 14637 | NM_198947.3 | 12.524669 | -5.2764208 | 1.52249264 | -3.4656462 |
| 14638 | NR_003528.3 | 9.05412524 | -4.7999036 | 1.3850205 | -3.4655831 |
| 14639 | NM_001256617.1 | 64.1138173 | -3.9562349 | 1.14159938 | -3.4655195 |
| 14640 | NR_136234.1 | 19.9236237 | -4.9477639 | 1.42771015 | -3.4655241 |
| 14641 | NM_001199201.1 | 13.3603893 | -5.3663774 | 1.54858988 | -3.4653316 |
| 14642 | NM_001316307.1 | 25.5068543 | -6.2940827 | 1.81649241 | -3.464965 |
| 14643 | NM_139355.2 | 9.80852712 | -4.9079002 | 1.41650091 | -3.4648055 |
| 14644 | NM_013281.3 | 41.62078 | -3.462243 | 0.99930942 | -3.4646356 |
| 14645 | NM_153247.2 | 92.4546524 | -8.1532291 | 2.3535201 | -3.46427 |
| 14646 | NM_001018067.1 | 91.4483791 | -8.1374763 | 2.34932659 | -3.4637484 |
| 14647 | NM_001353.5 | 145.062443 | -4.6252275 | 1.33544413 | -3.4634377 |
| 14648 | NM_017799.3 | 58.3448585 | -3.6289239 | 1.04778937 | -3.4634097 |
| 14649 | NM_001258462.1 | 12.2715225 | -5.2259705 | 1.50934675 | -3.4624055 |
| 14650 | NM_016622.3 | 105.5785 | -4.1363348 | 1.19470696 | -3.462217 |
| 14651 | NM_015106.3 | 125.455902 | -8.5930926 | 2.48202512 | -3.4621296 |
| 14652 | NM_054110.4 | 12.7124301 | -5.29491 | 1.52978378 | -3.4612146 |
| 14653 | NM_001080480.2 | 19.5035529 | -5.9065879 | 1.70673232 | -3.4607582 |
| 14654 | NM_001198695.1 | 17.8819079 | -5.7870964 | 1.67234011 | -3.4604781 |
| 14655 | NM_014388.6 | 59.2082328 | -2.4249426 | 0.70075637 | -3.4604647 |
| 14656 | NM_002779.4 | 63.2711588 | -4.9433932 | 1.42856279 | -3.4603962 |
| 14657 | NM_012177.4 | 11.1681963 | -5.1107494 | 1.47708565 | -3.4600224 |
| 14658 | NM_018725.3 | 16.8972488 | -4.6794233 | 1.35271205 | -3.45929 |
| 14659 | NM_001303501.1 | 13.9621366 | -5.4337531 | 1.5708079 | -3.4592092 |
| 14660 | NM_001142683.2 | 9.99455774 | -4.9541964 | 1.43219738 | -3.4591575 |
| 14661 | NM_001256197.1 | 9.04023864 | -4.7953322 | 1.38629764 | -3.4590928 |
| 14662 | NR_120596.1 | 10.1846515 | -4.9646723 | 1.435249 | -3.4591017 |

| | | | | | |
|---|---|---|---|---|---|
| 14663 | NM_018672.4 | 23.1265897 | -3.7216183 | 1.07596337 | -3.4588708 |
| 14664 | NM_152373.3 | 9.23099011 | -4.8206842 | 1.3937292 | -3.4588385 |
| 14665 | NM_001286072.1 | 9.71568458 | -4.9130226 | 1.42046292 | -3.4587475 |
| 14666 | NR_102756.1 | 9.28847739 | -4.8458282 | 1.40104375 | -3.4587272 |
| 14667 | NM_001313990.1 | 16.4387226 | -5.6677663 | 1.6387969 | -3.4584922 |
| 14668 | NM_024875.4 | 97.3151777 | 3.4935531 | 1.01016425 | 3.45840106 |
| 14669 | NR_134559.1 | 93.8732286 | 2.11575981 | 0.61183649 | 3.45804776 |
| 14670 | NM_001130914.1 | 18.4884127 | -3.9836388 | 1.15210666 | -3.4576996 |
| 14671 | NM_005207.3 | 300.398812 | -3.1363526 | 0.90712685 | -3.4574576 |
| 14672 | NR_001281.2 | 23.5420482 | -6.1772541 | 1.78683107 | -3.4571003 |
| 14673 | NM_017751.4 | 88.7808703 | -8.0947418 | 2.34151218 | -3.4570573 |
| 14674 | NM_001142300.1 | 14.4744807 | -5.4755284 | 1.58392929 | -3.4569273 |
| 14675 | NM_001170717.1 | 18.975752 | -5.8655257 | 1.69673765 | -3.4569432 |
| 14676 | NM_001286084.1 | 11.5957994 | -5.1626594 | 1.49353859 | -3.4566629 |
| 14677 | NR_073411.1 | 9.96945807 | -4.9246602 | 1.42493038 | -3.4560707 |
| 14678 | NM_001243609.1 | 146.712882 | -8.8187158 | 2.55175373 | -3.4559432 |
| 14679 | NM_001286774.1 | 18.5369032 | -5.8330885 | 1.68785547 | -3.455917 |
| 14680 | NM_001078170.2 | 16.0224243 | -5.6244584 | 1.62751223 | -3.4558625 |
| 14681 | NM_001143757.1 | 21.7525738 | -6.0582579 | 1.75308345 | -3.4557727 |
| 14682 | NM_001321226.1 | 18.8867931 | -5.8673543 | 1.69783682 | -3.4557822 |
| 14683 | NM_013327.4 | 10.16152 | -4.9742169 | 1.43951275 | -3.4554865 |
| 14684 | NM_019024.2 | 131.270524 | -4.5058521 | 1.30419375 | -3.4548947 |
| 14685 | NM_213613.3 | 8.38630281 | -4.6870223 | 1.35695507 | -3.4540733 |
| 14686 | NM_152236.2 | 21.9924124 | -6.0804882 | 1.76069788 | -3.4534535 |
| 14687 | NM_001300789.1 | 12.1346683 | -5.2315336 | 1.51503103 | -3.4530868 |
| 14688 | NM_022650.2 | 40.3327015 | 7.10498553 | 2.0581257 | 3.45216307 |
| 14689 | NM_001320804.1 | 54.408107 | 1.96218431 | 0.56840205 | 3.45210631 |
| 14690 | NM_172209.2 | 181.60982 | 1.29048101 | 0.37389522 | 3.45145096 |
| 14691 | NM_001318939.1 | 9.16584107 | -4.8138002 | 1.39477254 | -3.4513156 |
| 14692 | NR_039979.1 | 19.4132518 | 4.27638179 | 1.23929626 | 3.45065334 |
| 14693 | NM_001001669.2 | 10.0849386 | -4.965522 | 1.43907769 | -3.4504892 |
| 14694 | NM_020473.3 | 31.818793 | 9.91338267 | 2.87382766 | 3.44953972 |
| 14695 | NM_001037330.1 | 47.1802408 | -3.2672023 | 0.94718869 | -3.449368 |
| 14696 | NM_001242618.1 | 11.1703752 | -5.1009903 | 1.47888485 | -3.449214 |
| 14697 | NM_001198852.1 | 97.0699521 | -2.3182528 | 0.67238246 | -3.4478187 |
| 14698 | NR_046455.1 | 56.5545562 | 4.25998457 | 1.2356274 | 3.44762877 |
| 14699 | NM_138927.2 | 85.6256538 | -8.0425559 | 2.3328268 | -3.4475581 |
| 14700 | NM_031916.4 | 13.8194056 | -5.4142813 | 1.57059489 | -3.4472806 |
| 14701 | NM_001317061.1 | 10.6000959 | -5.0298753 | 1.4591472 | -3.4471336 |
| 14702 | NR_027175.1 | 8.68803247 | -4.734232 | 1.37372011 | -3.4462857 |
| 14703 | NM_181673.2 | 125.430081 | -8.5930288 | 2.49343205 | -3.4462655 |
| 14704 | NR_024170.1 | 34.9467489 | -3.7581655 | 1.09058049 | -3.446023 |
| 14705 | NR_103478.1 | 14.6136194 | -5.4964113 | 1.59512034 | -3.4457659 |

| | | | | | |
|---|---|---|---|---|---|
| 14706 | NM_001243789.1 | 8.31018219 | -4.674859 | 1.35670831 | -3.4457363 |
| 14707 | NM_001199181.1 | 21.9085864 | -6.0784997 | 1.76411029 | -3.4456461 |
| 14708 | NM_001144997.1 | 14.2472473 | -5.4576354 | 1.58425667 | -3.4449187 |
| 14709 | NM_054014.3 | 8.85030975 | -4.7668265 | 1.38381441 | -3.4447007 |
| 14710 | NM_171999.3 | 259.601316 | -3.8064066 | 1.10500922 | -3.4446831 |
| 14711 | NM_001017420.2 | 9.64637498 | -4.8818298 | 1.4172631 | -3.4445473 |
| 14712 | NM_013409.2 | 72.1228559 | -4.1965372 | 1.21831166 | -3.4445515 |
| 14713 | NM_001038628.1 | 13.2753832 | -5.3452283 | 1.55183404 | -3.4444587 |
| 14714 | NM_015944.3 | 11.8356039 | -5.1928622 | 1.50766411 | -3.4443097 |
| 14715 | NM_001013620.3 | 29.7527088 | -3.4187175 | 0.99257554 | -3.4442895 |
| 14716 | NR_024254.1 | 7.97415527 | -4.6192967 | 1.34161274 | -3.4430924 |
| 14717 | NM_032307.4 | 8.76317847 | -4.7563229 | 1.38190386 | -3.4418623 |
| 14718 | NM_025151.4 | 98.4475399 | -3.2836084 | 0.95435082 | -3.4406723 |
| 14719 | NM_178238.3 | 18.6048483 | -4.8448307 | 1.4084248 | -3.4398931 |
| 14720 | NM_001279353.1 | 10.4775612 | -4.9987226 | 1.45359302 | -3.4388735 |
| 14721 | NM_033388.1 | 19.2234584 | -4.8763099 | 1.418019 | -3.4388185 |
| 14722 | NR_033385.1 | 10.38921 | -5.0069078 | 1.45601408 | -3.438777 |
| 14723 | NM_001243962.1 | 45.6724089 | -4.4385325 | 1.29074002 | -3.4387502 |
| 14724 | NM_001012758.2 | 9.03216033 | -4.7888335 | 1.39269219 | -3.4385441 |
| 14725 | NM_006634.2 | 16.7302557 | -4.6710173 | 1.35853578 | -3.4382733 |
| 14726 | NM_001258316.1 | 63.8992189 | -2.8514242 | 0.82935496 | -3.4381228 |
| 14727 | NM_004298.3 | 13.4238357 | -5.3722231 | 1.56255938 | -3.4380921 |
| 14728 | NM_001164758.1 | 14.5395978 | -5.4899755 | 1.59689017 | -3.4379168 |
| 14729 | NR_145426.1 | 10.8720473 | -5.0676086 | 1.47405805 | -3.4378623 |
| 14730 | NR_121578.1 | 32.2132809 | 1.87493519 | 0.54539287 | 3.43776991 |
| 14731 | NM_001204388.1 | 19.7508498 | 9.2244418 | 2.68338082 | 3.43761934 |
| 14732 | NM_001330081.1 | 19.3189169 | -5.8913941 | 1.71441289 | -3.4363916 |
| 14733 | NM_030929.4 | 9.04201255 | -4.8082798 | 1.39925402 | -3.4363166 |
| 14734 | NR_052022.1 | 13.1695694 | -5.3429429 | 1.55503009 | -3.4359096 |
| 14735 | NR_135091.1 | 37.7887364 | 3.03416376 | 0.88309954 | 3.43581174 |
| 14736 | NM_001935.3 | 9.10786752 | -4.8036366 | 1.39827535 | -3.4354011 |
| 14737 | NM_001003810.1 | 84.6256971 | -8.0256089 | 2.33649642 | -3.4348903 |
| 14738 | NM_005296.2 | 9.46520172 | -4.8688112 | 1.41757488 | -3.434606 |
| 14739 | NR_137424.1 | 8.13023093 | -4.6391637 | 1.35085961 | -3.4342308 |
| 14740 | NM_001308393.1 | 11.5560703 | -5.1635036 | 1.50362216 | -3.4340433 |
| 14741 | NM_181291.2 | 12.2259068 | -5.2223158 | 1.52080865 | -3.4339072 |
| 14742 | NM_001172677.1 | 10.8455387 | -5.0468423 | 1.46974345 | -3.4338253 |
| 14743 | NM_004791.2 | 15.2063132 | -5.5508244 | 1.61651776 | -3.4338159 |
| 14744 | NM_001144932.2 | 10.5551299 | -5.0328008 | 1.4656674 | -3.4337946 |
| 14745 | NM_001320458.1 | 62.2655268 | 3.11588239 | 0.90746514 | 3.43361111 |
| 14746 | NM_198503.3 | 21.8280494 | -6.0654462 | 1.76654064 | -3.4335164 |
| 14747 | NM_001042698.1 | 36.1096142 | -3.1109855 | 0.90621742 | -3.432935 |
| 14748 | NM_022908.2 | 10.9301378 | -5.0677864 | 1.47631413 | -3.432729 |

| | | | | | |
|---|---|---|---|---|---|
| 14749 | NM_001195156.1 | 9.82472871 | -4.9289864 | 1.43600972 | -3.4324186 |
| 14750 | NM_032021.2 | 8.31566348 | -4.6868705 | 1.36550406 | -3.4323373 |
| 14751 | NM_015138.4 | 484.08551 | -3.8059115 | 1.10892671 | -3.4320677 |
| 14752 | NM_001321210.1 | 9.07353031 | -4.8093292 | 1.40143588 | -3.4317155 |
| 14753 | NR_104194.1 | 13.1232734 | -5.3350631 | 1.55505221 | -3.4307936 |
| 14754 | NM_018923.2 | 11.8864756 | -5.1959347 | 1.51480692 | -3.430097 |
| 14755 | NM_001330208.1 | 17.7547517 | -5.7658721 | 1.68107782 | -3.4298663 |
| 14756 | NM_001267698.1 | 8.09694443 | -4.6426058 | 1.35367881 | -3.4296214 |
| 14757 | NM_145238.3 | 14.020672 | -4.4339034 | 1.29294753 | -3.4292988 |
| 14758 | NM_006332.4 | 18.5215026 | -4.8516142 | 1.4148209 | -3.4291367 |
| 14759 | NM_012099.1 | 142.108206 | 1.75039021 | 0.51048001 | 3.42891038 |
| 14760 | NM_145176.2 | 65.5137504 | -2.8364213 | 0.82720559 | -3.4289194 |
| 14761 | NR_047570.1 | 46.8600141 | 1.82405436 | 0.53196196 | 3.42891879 |
| 14762 | NM_001039763.3 | 9.01486284 | -4.8014572 | 1.40030604 | -3.4288627 |
| 14763 | NM_031886.2 | 100.59682 | 3.35837087 | 0.97954387 | 3.4285048 |
| 14764 | NM_002608.3 | 22.6738475 | -5.1211528 | 1.49385693 | -3.4281414 |
| 14765 | NM_033645.2 | 140.237124 | -7.7881799 | 2.27188731 | -3.4280661 |
| 14766 | NM_001271082.1 | 9.94973116 | -4.9360126 | 1.44005634 | -3.4276524 |
| 14767 | NM_001010898.3 | 93.0099904 | -3.9261068 | 1.14546956 | -3.4275087 |
| 14768 | NM_024409.3 | 8.41104951 | -4.7052473 | 1.37283913 | -3.4273843 |
| 14769 | NM_139354.2 | 10.679975 | -5.0295993 | 1.46750017 | -3.4273245 |
| 14770 | NM_001243879.1 | 24.2493204 | -5.2202737 | 1.52323406 | -3.4270989 |
| 14771 | NM_001305005.2 | 30.6906309 | -6.5633725 | 1.91514592 | -3.4270874 |
| 14772 | NM_003204.2 | 100.859426 | -8.2783708 | 2.41610817 | -3.4263246 |
| 14773 | NM_172159.3 | 11.2726606 | -5.1056799 | 1.49019967 | -3.4261717 |
| 14774 | NM_015198.4 | 9.99829683 | -4.9552443 | 1.44640597 | -3.4259015 |
| 14775 | NM_001277333.1 | 49.7511745 | 2.20253979 | 0.64309111 | 3.42492653 |
| 14776 | NM_173511.3 | 174.650807 | -3.6582526 | 1.06814922 | -3.4248516 |
| 14777 | NM_001280797.1 | 10.9906037 | -5.0656258 | 1.47919296 | -3.4245875 |
| 14778 | NR_024252.1 | 9.89975252 | -4.9289176 | 1.43931276 | -3.4244938 |
| 14779 | NM_001324302.1 | 15.1357422 | -5.5330987 | 1.61586384 | -3.4242357 |
| 14780 | NR_002776.4 | 9.41018707 | -4.8459002 | 1.41548876 | -3.4234819 |
| 14781 | NM_012170.3 | 10.1787487 | -4.9805225 | 1.45486198 | -3.4233643 |
| 14782 | NM_001003683.2 | 29.863693 | -6.5206449 | 1.90477532 | -3.4233145 |
| 14783 | NM_133368.2 | 84.5104957 | -8.0236796 | 2.34392135 | -3.4231863 |
| 14784 | NM_006561.3 | 151.709191 | -4.2720982 | 1.24803878 | -3.4230492 |
| 14785 | NM_199340.3 | 17.3050991 | -4.7253659 | 1.38048276 | -3.4229807 |
| 14786 | NM_014994.2 | 85.6770643 | -8.0434764 | 2.35000169 | -3.4227535 |
| 14787 | NM_194457.1 | 16.4283589 | -5.657481 | 1.65319668 | -3.4221464 |
| 14788 | NM_002742.2 | 19.4590937 | -4.9055553 | 1.43356717 | -3.4219222 |
| 14789 | NM_201613.2 | 9.16033132 | -4.8147921 | 1.40721149 | -3.4215128 |
| 14790 | NM_001003690.1 | 8.21028655 | -4.6606967 | 1.36223574 | -3.4213584 |
| 14791 | NR_024187.2 | 12.7614007 | -5.2883359 | 1.54595068 | -3.4207662 |

| | | | | | |
|---|---|---|---|---|---|
| 14792 | NM_001207011.1 | 13.1029624 | -5.3375361 | 1.56038388 | -3.4206558 |
| 14793 | NM_177400.2 | 13.7347856 | -5.4036141 | 1.57972434 | -3.4206057 |
| 14794 | NR_046085.1 | 9.92719169 | 5.36012049 | 1.56708912 | 3.42043119 |
| 14795 | NM_015983.3 | 78.0191935 | -1.7355717 | 0.50744196 | -3.4202369 |
| 14796 | NM_007147.3 | 31.4660276 | -3.2380385 | 0.94675645 | -3.4201389 |
| 14797 | NR_110124.1 | 39.6520401 | 4.61421996 | 1.34915573 | 3.42007959 |
| 14798 | NM_001007067.1 | 13.0163269 | -5.3276161 | 1.55779797 | -3.419966 |
| 14799 | NM_015191.2 | 90.8429031 | -3.9039705 | 1.14154289 | -3.419907 |
| 14800 | NR_024037.1 | 8.29226097 | -4.6731989 | 1.36657449 | -3.4196445 |
| 14801 | NR_040733.1 | 12.6523835 | -5.2879769 | 1.54653587 | -3.4192397 |
| 14802 | NM_001323027.1 | 21.9274794 | -2.5189938 | 0.73672411 | -3.4191819 |
| 14803 | NM_153695.3 | 18.6656289 | -4.8508281 | 1.41884647 | -3.4188535 |
| 14804 | NR_026647.1 | 16.3828367 | 5.94512392 | 1.73896624 | 3.41876902 |
| 14805 | NM_006603.4 | 79.3373691 | -7.9325851 | 2.32063066 | -3.4182885 |
| 14806 | NM_001314014.1 | 12.4039329 | -5.2444733 | 1.53430761 | -3.4181368 |
| 14807 | NM_001319658.1 | 10.7271883 | -5.0342394 | 1.4728441 | -3.4180396 |
| 14808 | NM_001171930.1 | 14.0744863 | -5.4373631 | 1.59096084 | -3.41766 |
| 14809 | NM_001323954.1 | 13.167231 | -5.3396785 | 1.56253483 | -3.417318 |
| 14810 | NM_001303111.1 | 10.5025923 | -5.0073987 | 1.46535474 | -3.4171921 |
| 14811 | NM_007375.3 | 423.073952 | -4.5529755 | 1.33261486 | -3.4165727 |
| 14812 | NR_104607.1 | 23.0488969 | 2.59832063 | 0.76055918 | 3.41632934 |
| 14813 | NR_038903.1 | 9.03614666 | -4.792905 | 1.40305646 | -3.4160458 |
| 14814 | NM_148919.3 | 7.79489866 | -4.587363 | 1.34311317 | -3.4154702 |
| 14815 | NM_152989.4 | 27.1710222 | -6.3810069 | 1.86828161 | -3.4154417 |
| 14816 | NM_001128303.1 | 8.23109825 | -4.6713814 | 1.36778928 | -3.4152786 |
| 14817 | NR_037715.1 | 8.63439814 | -4.7283236 | 1.38451121 | -3.4151573 |
| 14818 | NM_003414.5 | 14.9267054 | -4.5196515 | 1.32368602 | -3.4144438 |
| 14819 | NM_001297426.1 | 12.3486357 | -5.2429559 | 1.53569928 | -3.4140511 |
| 14820 | NR_126344.1 | 83.1952902 | 2.90264006 | 0.85031056 | 3.41362343 |
| 14821 | NM_022571.5 | 11.4371287 | -5.1451615 | 1.5073017 | -3.4134914 |
| 14822 | NR_003099.1 | 14.2078195 | -5.452044 | 1.59738673 | -3.4131021 |
| 14823 | NM_001202560.2 | 11.2802149 | -5.1098751 | 1.49718962 | -3.4129779 |
| 14824 | NM_183237.2 | 11.5896433 | -5.1585306 | 1.51144276 | -3.4129845 |
| 14825 | NM_001320676.1 | 12.9324476 | -5.3175438 | 1.55829863 | -3.4124036 |
| 14826 | NM_001136136.1 | 12.4298124 | -5.2529107 | 1.53944152 | -3.4122184 |
| 14827 | NM_001284236.1 | 101.078417 | -8.2818263 | 2.4274452 | -3.4117459 |
| 14828 | NM_000679.3 | 11.7620375 | -5.1813154 | 1.518965 | -3.4110828 |
| 14829 | NM_133433.3 | 186.489725 | -3.8800636 | 1.13755542 | -3.4108788 |
| 14830 | NM_016009.4 | 80.7355651 | -7.95786 | 2.33326243 | -3.4106151 |
| 14831 | NM_001190983.1 | 16.8481634 | -4.698136 | 1.37755663 | -3.4104848 |
| 14832 | NM_005723.3 | 213.20106 | -4.6175687 | 1.35405715 | -3.4101727 |
| 14833 | NM_001127617.2 | 9.01948128 | -4.7885642 | 1.40435074 | -3.4098064 |
| 14834 | NM_001178079.1 | 13.5874672 | -5.3836777 | 1.57916361 | -3.4091956 |

| | | | | | |
|---|---|---|---|---|---|
| 14835 | NM_001184940.1 | 109.992331 | -8.4032654 | 2.46519711 | -3.40876 |
| 14836 | NM_001034955.1 | 10.3636074 | -4.9844654 | 1.46256839 | -3.4080221 |
| 14837 | NM_000453.2 | 76.6015487 | 1.48557319 | 0.43598314 | 3.40740968 |
| 14838 | NM_025136.3 | 400.691471 | 1.19380023 | 0.35039006 | 3.40706074 |
| 14839 | NM_032280.2 | 8.74114277 | -4.7432492 | 1.39242796 | -3.4064593 |
| 14840 | NM_182746.2 | 223.400466 | -3.1447903 | 0.92323912 | -3.4062576 |
| 14841 | NM_175932.2 | 12.3559955 | -5.2564303 | 1.54333657 | -3.4058872 |
| 14842 | NM_001017971.2 | 33.5012484 | -2.02168 | 0.59362015 | -3.4056795 |
| 14843 | NM_001198780.1 | 103.114106 | -8.3102484 | 2.44021831 | -3.4055348 |
| 14844 | NM_003102.2 | 14.5231484 | -5.4879172 | 1.61152362 | -3.4054215 |
| 14845 | NM_001123384.1 | 11.9896226 | -5.2116974 | 1.53057081 | -3.4050678 |
| 14846 | NM_201520.2 | 9.13085543 | -4.8230972 | 1.41645463 | -3.4050489 |
| 14847 | NR_037631.1 | 13.1780919 | -5.3413829 | 1.56877388 | -3.4048138 |
| 14848 | NM_005568.4 | 17.6571229 | -3.9397998 | 1.15719377 | -3.4046155 |
| 14849 | NM_001271950.1 | 12.6482993 | -5.2777035 | 1.55036012 | -3.4041791 |
| 14850 | NM_145005.6 | 7.9489725 | -4.6106186 | 1.3544031 | -3.4041701 |
| 14851 | NM_152318.2 | 13.7934002 | -5.4031903 | 1.58739444 | -3.4038108 |
| 14852 | NM_001278562.1 | 9.17055907 | -4.8280024 | 1.418469 | -3.4036714 |
| 14853 | NM_024512.4 | 129.75719 | 2.39648558 | 0.70425217 | 3.40287995 |
| 14854 | NM_006867.3 | 8.24181035 | -4.6711045 | 1.37270981 | -3.4028347 |
| 14855 | NM_201566.2 | 10.0920759 | -4.9543778 | 1.45598423 | -3.4027689 |
| 14856 | NR_135006.1 | 96.6635438 | 3.70977204 | 1.09025731 | 3.40265733 |
| 14857 | NM_001142699.1 | 11.6549323 | -5.1594715 | 1.51648251 | -3.4022625 |
| 14858 | NR_104128.1 | 9.18356432 | -4.8252727 | 1.41858686 | -3.4014644 |
| 14859 | NM_001345993.1 | 8.41776091 | -4.6830006 | 1.37678618 | -3.4014001 |
| 14860 | NM_001457.3 | 126.456801 | -8.604838 | 2.52996842 | -3.4011642 |
| 14861 | NR_104189.1 | 14.5349226 | 4.29047673 | 1.26171504 | 3.40051168 |
| 14862 | NM_005923.3 | 8.11783613 | -4.6535333 | 1.36852474 | -3.4004013 |
| 14863 | NM_033518.3 | 10.5548537 | -5.0101887 | 1.47342651 | -3.4003655 |
| 14864 | NM_001291821.1 | 12.5078868 | -5.2552866 | 1.54593321 | -3.3994267 |
| 14865 | NM_001184742.1 | 10.0313463 | -4.9419833 | 1.45382521 | -3.3992968 |
| 14866 | NR_028137.1 | 31.4765868 | -6.5968284 | 1.94078571 | -3.3990504 |
| 14867 | NM_001284362.1 | 8.61963764 | 4.65608638 | 1.36999298 | 3.3986206 |
| 14868 | NM_005950.2 | 17.2710939 | -5.7280757 | 1.68542488 | -3.3985944 |
| 14869 | NM_031483.6 | 81.2557068 | -7.9670416 | 2.34424427 | -3.3985544 |
| 14870 | NM_001134316.1 | 8.24491635 | -4.6728364 | 1.37516831 | -3.3980106 |
| 14871 | NM_024016.3 | 192.953794 | -4.550899 | 1.33927907 | -3.3980214 |
| 14872 | NM_001206938.1 | 209.303191 | -3.5269239 | 1.0380727 | -3.3975693 |
| 14873 | NM_014309.2 | 226.6613 | -4.65511 | 1.37018167 | -3.39744 |
| 14874 | NM_001128636.2 | 109.751083 | -3.1416446 | 0.92492079 | -3.3966634 |
| 14875 | NR_033932.1 | 9.36926069 | -4.8530837 | 1.42889818 | -3.3963817 |
| 14876 | NM_001113756.2 | 49.4069797 | -7.2497701 | 2.13462431 | -3.3962745 |
| 14877 | NM_001206847.1 | 14.0461088 | -5.4312885 | 1.59940751 | -3.3958128 |

| | | | | | |
|---|---|---|---|---|---|
| 14878 | NM_001204173.1 | 8.35930919 | -4.6840586 | 1.37943975 | -3.3956239 |
| 14879 | NM_002563.4 | 8.5105469 | -4.700746 | 1.3844423 | -3.3954076 |
| 14880 | NR_040724.1 | 77.6528365 | -7.9016598 | 2.327359 | -3.3951186 |
| 14881 | NR_136271.1 | 8.79699151 | -4.7645822 | 1.40349167 | -3.3948062 |
| 14882 | NM_001330261.1 | 81.6936454 | -7.9746948 | 2.34911825 | -3.3947609 |
| 14883 | NM_001242312.1 | 72.628531 | -2.5033482 | 0.73745142 | -3.394594 |
| 14884 | NM_001279359.1 | 21.2363633 | -6.0255677 | 1.7750779 | -3.394537 |
| 14885 | NR_024199.2 | 12.0180178 | -5.2136015 | 1.53610708 | -3.3940352 |
| 14886 | NR_136334.1 | 43.1325734 | 2.3721671 | 0.69906888 | 3.39332386 |
| 14887 | NM_057166.4 | 227.056986 | -4.2784449 | 1.2609577 | -3.3930122 |
| 14888 | NM_001002269.2 | 9.19090388 | -4.8308552 | 1.42383716 | -3.3928425 |
| 14889 | NM_144722.3 | 12.9830232 | -5.325601 | 1.56979044 | -3.3925554 |
| 14890 | NM_001282951.2 | 10.7160115 | -5.0493273 | 1.48848327 | -3.3922634 |
| 14891 | NM_001330766.1 | 23.7440522 | -3.3050109 | 0.97442655 | -3.3917496 |
| 14892 | NM_001281425.1 | 30.9886926 | -5.5835741 | 1.64627341 | -3.3916445 |
| 14893 | NM_001127208.2 | 81.896055 | -3.1855464 | 0.93926817 | -3.3915196 |
| 14894 | NM_020937.3 | 14.0787494 | -5.4301361 | 1.6010872 | -3.3915305 |
| 14895 | NM_198969.1 | 17.254828 | -5.7343993 | 1.69099935 | -3.3911304 |
| 14896 | NR_138426.1 | 14.6798468 | -5.4902885 | 1.61904684 | -3.3910622 |
| 14897 | NM_032290.3 | 45.3025235 | -2.9119058 | 0.85872576 | -3.3909612 |
| 14898 | NM_001284367.1 | 9.20971692 | -4.8325073 | 1.4253314 | -3.3904447 |
| 14899 | NM_001286632.1 | 77.2052962 | -7.8932882 | 2.32834084 | -3.3900914 |
| 14900 | NM_153257.4 | 16.9157444 | -5.6965331 | 1.6805823 | -3.3896187 |
| 14901 | NM_006749.4 | 12.794043 | -5.298161 | 1.56336825 | -3.3889399 |
| 14902 | NM_023008.3 | 114.353446 | -2.3984923 | 0.70773829 | -3.3889536 |
| 14903 | NR_024246.2 | 42.7194335 | 4.02089544 | 1.18680809 | 3.38799126 |
| 14904 | NM_002557.3 | 10.426412 | -4.9954602 | 1.47451234 | -3.3878727 |
| 14905 | NM_018975.3 | 627.904771 | -5.3302342 | 1.5733489 | -3.3878272 |
| 14906 | NM_001194.3 | 123.802403 | -4.0004409 | 1.18089428 | -3.3876368 |
| 14907 | NR_002773.1 | 24.6088859 | 2.84205064 | 0.83902694 | 3.3873175 |
| 14908 | NM_001257197.1 | 19.2082559 | -5.8807216 | 1.73671518 | -3.3861175 |
| 14909 | NM_145799.3 | 11.0241284 | -5.0921114 | 1.50391183 | -3.3859109 |
| 14910 | NR_038942.1 | 12.0360622 | -5.2083633 | 1.53828423 | -3.3858264 |
| 14911 | NM_001286137.1 | 32.4377755 | 5.48985861 | 1.62151318 | 3.38563921 |
| 14912 | NM_001374.2 | 12.1903865 | -5.2315776 | 1.54587219 | -3.3842239 |
| 14913 | NM_001001395.2 | 107.804514 | -8.3741945 | 2.47451588 | -3.3841749 |
| 14914 | NM_001318230.1 | 29.6181378 | -4.7360234 | 1.39952162 | -3.3840302 |
| 14915 | NM_030821.4 | 133.007713 | -1.5618958 | 0.46158509 | -3.3837657 |
| 14916 | NR_033179.1 | 12.1845615 | -5.2196907 | 1.54294354 | -3.3829434 |
| 14917 | NM_003806.2 | 10.9116208 | -5.0753698 | 1.50030242 | -3.3828978 |
| 14918 | NM_018835.2 | 7.8018862 | -4.5852868 | 1.35547603 | -3.382787 |
| 14919 | NM_001300806.1 | 10.03193 | -4.9434625 | 1.46157112 | -3.3822935 |
| 14920 | NM_020161.3 | 45.6465827 | 3.45468715 | 1.021412 | 3.38226607 |

| | | | | | |
|---|---|---|---|---|---|
| 14921 | NR_132742.1 | 42.219141 | -2.3467139 | 0.69382958 | -3.3822627 |
| 14922 | NM_006296.6 | 10.1242177 | -4.9724663 | 1.47017625 | -3.3822246 |
| 14923 | NR_130910.1 | 9.24456918 | -4.8378632 | 1.43057395 | -3.3817638 |
| 14924 | NM_198480.3 | 12.2361887 | -5.2252239 | 1.54520025 | -3.3815836 |
| 14925 | NM_001281738.1 | 8.2295406 | -4.6543079 | 1.3764604 | -3.3813598 |
| 14926 | NM_005281.3 | 9.44935008 | -4.8631409 | 1.43823228 | -3.3813321 |
| 14927 | NM_001195056.1 | 10.9485405 | -5.0843418 | 1.50398158 | -3.3805878 |
| 14928 | NM_001328674.1 | 105.552146 | -8.3438753 | 2.46817822 | -3.3805806 |
| 14929 | NM_006475.2 | 94.1574074 | -8.1793104 | 2.41951944 | -3.3805516 |
| 14930 | NM_197962.2 | 10.1226087 | -4.9664519 | 1.46925218 | -3.3802584 |
| 14931 | NM_025213.2 | 9.53381887 | -4.8764029 | 1.44263041 | -3.3802163 |
| 14932 | NM_001243657.1 | 9.02648004 | -4.808376 | 1.42276862 | -3.379591 |
| 14933 | NM_138970.4 | 18.0535511 | 9.09433448 | 2.69099226 | 3.37954688 |
| 14934 | NR_130789.1 | 79.9186449 | -7.9431051 | 2.35044988 | -3.3793978 |
| 14935 | NM_024821.3 | 8.45593883 | -4.696481 | 1.38990281 | -3.3789996 |
| 14936 | NM_001308358.1 | 16.7127305 | -5.6844106 | 1.68281804 | -3.3779116 |
| 14937 | NM_021133.3 | 9.25348753 | -4.8246419 | 1.42834418 | -3.3777867 |
| 14938 | NM_022495.5 | 102.684993 | -2.9290551 | 0.86715031 | -3.3777941 |
| 14939 | NM_030773.3 | 11.2900948 | -5.1127279 | 1.51367126 | -3.3777003 |
| 14940 | NM_001278730.1 | 9.22343098 | -4.8336353 | 1.43106581 | -3.3776471 |
| 14941 | NM_012413.3 | 9.15817554 | -4.8226877 | 1.42800954 | -3.3772097 |
| 14942 | NR_103819.1 | 43.2350296 | 6.63662858 | 1.96558875 | 3.37640749 |
| 14943 | NM_001163558.2 | 17.589878 | 6.60962131 | 1.95761054 | 3.37637196 |
| 14944 | NM_001281715.1 | 9.5840929 | -4.8892185 | 1.4481671 | -3.3761425 |
| 14945 | NM_173471.3 | 10.0911619 | -4.9575848 | 1.46842471 | -3.3761246 |
| 14946 | NM_181435.5 | 15.0728042 | -5.5298734 | 1.63795074 | -3.3760926 |
| 14947 | NM_138378.2 | 14.5090892 | -5.4854355 | 1.62503629 | -3.3755773 |
| 14948 | NM_000110.3 | 9.16947147 | -4.8140849 | 1.42634645 | -3.3751161 |
| 14949 | NM_153810.4 | 149.029226 | -3.209725 | 0.95108135 | -3.3748165 |
| 14950 | NM_205835.3 | 11.8676641 | -5.1935779 | 1.53900524 | -3.374633 |
| 14951 | NR_046223.1 | 10.7299669 | -5.0413742 | 1.49390258 | -3.3746338 |
| 14952 | NM_006732.2 | 15.9906087 | -5.6246918 | 1.66709418 | -3.3739496 |
| 14953 | NM_001284303.1 | 10.0278209 | -4.9536383 | 1.46836273 | -3.3735794 |
| 14954 | NM_001267713.1 | 8.27359886 | -4.6572986 | 1.38058074 | -3.3734344 |
| 14955 | NM_020424.4 | 9.75269468 | -4.9013563 | 1.45298414 | -3.3733034 |
| 14956 | NM_033177.3 | 13.8276769 | -5.4133574 | 1.60476609 | -3.3733 |
| 14957 | NR_110128.1 | 12.6074997 | -5.2833575 | 1.56622768 | -3.3733011 |
| 14958 | NM_033504.3 | 10.4809996 | -5.0098172 | 1.48545356 | -3.3725842 |
| 14959 | NM_145025.4 | 16.3274255 | -4.6366486 | 1.37481151 | -3.3725704 |
| 14960 | NM_000029.3 | 8.9768697 | -4.7958396 | 1.42207299 | -3.3724286 |
| 14961 | NM_001135751.1 | 7.5423749 | -4.5374353 | 1.34549975 | -3.3723048 |
| 14962 | NM_001244973.1 | 107.063384 | -8.3646011 | 2.48042241 | -3.3722486 |
| 14963 | NM_001144872.1 | 27.0967168 | -4.5775031 | 1.35766613 | -3.371597 |

| | | | | | |
|---|---|---|---|---|---|
| 14964 | NR_135218.1 | 12.2686487 | -5.2485547 | 1.55669117 | -3.3716095 |
| 14965 | NM_004490.2 | 10.4746155 | -5.0127013 | 1.48720169 | -3.3705592 |
| 14966 | NM_001346870.1 | 8.5142377 | -4.7111656 | 1.39793788 | -3.3700822 |
| 14967 | NM_145045.4 | 10.8618807 | -5.0667751 | 1.50347701 | -3.3700383 |
| 14968 | NM_001304264.1 | 17.0257071 | -4.7220078 | 1.40118295 | -3.3700152 |
| 14969 | NM_001099455.1 | 43.4489396 | 6.41253592 | 1.90287399 | 3.36992148 |
| 14970 | NM_001145720.1 | 15.7490001 | -4.6014482 | 1.36550826 | -3.3697696 |
| 14971 | NM_001284301.2 | 11.863457 | -5.1993489 | 1.5429307 | -3.3697877 |
| 14972 | NM_006354.3 | 101.149718 | -8.2823753 | 2.45808561 | -3.3694414 |
| 14973 | NM_001040142.1 | 23.8229656 | -6.1961448 | 1.83914134 | -3.3690422 |
| 14974 | NR_109848.1 | 11.0202273 | -5.0802325 | 1.50795295 | -3.3689596 |
| 14975 | NM_001099850.2 | 9.74643918 | 4.9126277 | 1.45824278 | 3.36886818 |
| 14976 | NM_001955.4 | 9.94407836 | -4.931694 | 1.46389555 | -3.3688838 |
| 14977 | NM_000572.2 | 40.7877273 | 3.09960168 | 0.92019002 | 3.36843653 |
| 14978 | NM_138420.2 | 195.139892 | -3.263009 | 0.96876579 | -3.3682125 |
| 14979 | NM_001012727.1 | 7.93361341 | -4.608429 | 1.3682632 | -3.3680866 |
| 14980 | NM_001330342.1 | 9.80718382 | -4.9016818 | 1.45544939 | -3.3678133 |
| 14981 | NR_029193.1 | 18.7287761 | -4.856041 | 1.44224201 | -3.3670084 |
| 14982 | NM_001243228.1 | 16.2569696 | -5.6442896 | 1.676532 | -3.3666459 |
| 14983 | NM_006638.3 | 11.5804208 | -5.1495898 | 1.52966665 | -3.3664784 |
| 14984 | NM_174894.2 | 13.4763248 | -5.3780034 | 1.59755054 | -3.3664058 |
| 14985 | NM_001272053.1 | 332.888932 | -3.545807 | 1.05335739 | -3.3661956 |
| 14986 | NM_001288592.1 | 8.62174991 | -4.7419449 | 1.40882931 | -3.3658761 |
| 14987 | NM_022480.3 | 111.42407 | -4.504621 | 1.33832958 | -3.3658532 |
| 14988 | NM_138468.5 | 16.886193 | -5.6923582 | 1.69120374 | -3.3658619 |
| 14989 | NM_001163922.1 | 33.2940265 | -3.3352343 | 0.99106069 | -3.3653179 |
| 14990 | NM_003852.3 | 100.765396 | -8.2768534 | 2.45972292 | -3.3649535 |
| 14991 | NM_001204424.1 | 13.0230097 | -5.3291197 | 1.58375352 | -3.3648668 |
| 14992 | NM_001256172.1 | 9.80737074 | -4.9133653 | 1.46026046 | -3.3647185 |
| 14993 | NM_000949.6 | 9.22993734 | -4.8375082 | 1.43775555 | -3.3646249 |
| 14994 | NM_134261.2 | 15.3169101 | -5.5580365 | 1.65221649 | -3.363988 |
| 14995 | NM_001100166.1 | 5.08978453 | 7.27784103 | 2.16383225 | 3.36340353 |
| 14996 | NM_017672.5 | 74.3660379 | -7.8392027 | 2.33095418 | -3.3630874 |
| 14997 | NM_003621.3 | 23.462573 | -4.3860897 | 1.30432271 | -3.3627335 |
| 14998 | NM_004496.3 | 12.3147591 | -5.2459275 | 1.5600849 | -3.362591 |
| 14999 | NM_025115.3 | 18.7470232 | -4.8424552 | 1.44012431 | -3.3625258 |
| 15000 | NR_130972.1 | 12.2534343 | -5.2373023 | 1.5576382 | -3.3623356 |
| 15001 | NM_001330683.1 | 100.945337 | -8.2792848 | 2.46244115 | -3.3622265 |
| 15002 | NM_001198842.1 | 9.65417117 | -4.8886391 | 1.45435359 | -3.3613827 |
| 15003 | NM_014652.3 | 160.49294 | -4.2840566 | 1.27451065 | -3.3613345 |
| 15004 | NM_001323072.1 | 15.2485328 | -4.5492266 | 1.35344099 | -3.3612301 |
| 15005 | NM_198439.2 | 8.27935304 | -4.6689642 | 1.38917441 | -3.3609633 |
| 15006 | NM_001184743.1 | 10.0858038 | -4.9544088 | 1.47412833 | -3.3609074 |

| | | | | | |
|---|---|---|---|---|---|
| 15007 | NM_000359.2 | 9.49014234 | -4.8655222 | 1.44769706 | -3.3608704 |
| 15008 | NM_024700.3 | 90.8651093 | -1.7170362 | 0.51089744 | -3.3608236 |
| 15009 | NM_002641.3 | 12.3278575 | -5.2522383 | 1.56291296 | -3.3605443 |
| 15010 | NM_001320537.1 | 34.9965803 | -3.6927376 | 1.09902644 | -3.360008 |
| 15011 | NR_110644.1 | 8.66890975 | -4.7373189 | 1.41005821 | -3.3596619 |
| 15012 | NR_145328.1 | 9.97180998 | -4.9315305 | 1.4681079 | -3.3591063 |
| 15013 | NM_182612.3 | 102.619978 | -8.303085 | 2.47187812 | -3.3590188 |
| 15014 | NM_001242607.1 | 22.9200299 | -6.1376318 | 1.82759233 | -3.3583156 |
| 15015 | NM_001324086.1 | 16.4268343 | -5.6510851 | 1.68284668 | -3.3580511 |
| 15016 | NM_004094.4 | 189.287127 | -4.1286894 | 1.22962057 | -3.3576938 |
| 15017 | NM_003297.3 | 68.766756 | -2.9265874 | 0.87167042 | -3.3574472 |
| 15018 | NR_027263.1 | 9.84345039 | -4.9232811 | 1.46653227 | -3.3570902 |
| 15019 | NM_002264.3 | 265.090799 | -2.9565159 | 0.88070896 | -3.3569726 |
| 15020 | NR_104239.1 | 10.778496 | -5.0637827 | 1.50843802 | -3.356971 |
| 15021 | NM_001080391.1 | 66.5842505 | 1.81349487 | 0.54026687 | 3.35666495 |
| 15022 | NM_031293.2 | 20.9663424 | -5.004821 | 1.49108845 | -3.3564883 |
| 15023 | NM_001282144.1 | 15.2724356 | -5.5617691 | 1.65705676 | -3.3564144 |
| 15024 | NM_001322800.1 | 20.4922018 | -4.9885374 | 1.48627208 | -3.3564093 |
| 15025 | NM_024496.3 | 650.106719 | -3.6707232 | 1.09371793 | -3.3561881 |
| 15026 | NM_152613.2 | 20.4754939 | 8.51390767 | 2.53687662 | 3.35605902 |
| 15027 | NM_001128853.1 | 77.6420166 | -7.901417 | 2.35451593 | -3.3558562 |
| 15028 | NM_001284333.1 | 14.0131734 | -5.43934 | 1.6209305 | -3.3556898 |
| 15029 | NM_001289076.1 | 22.2580917 | 9.40059893 | 2.80161229 | 3.35542465 |
| 15030 | NR_037862.1 | 13.4855365 | -5.379892 | 1.60341979 | -3.3552611 |
| 15031 | NM_001303100.1 | 12.822261 | -5.3019445 | 1.58025189 | -3.3551262 |
| 15032 | NR_126343.1 | 7.82996755 | -4.5939179 | 1.36963406 | -3.3541207 |
| 15033 | NM_005969.3 | 477.405741 | -3.862091 | 1.15149107 | -3.3539913 |
| 15034 | NM_199282.2 | 7.94098254 | -4.6110325 | 1.37480887 | -3.3539444 |
| 15035 | NM_001243076.2 | 52.2811388 | -3.6365532 | 1.0843782 | -3.3535838 |
| 15036 | NM_005128.3 | 17.2738041 | 9.02803802 | 2.69248638 | 3.35304872 |
| 15037 | NM_031430.2 | 7.50071154 | -4.5330658 | 1.35241506 | -3.3518303 |
| 15038 | NM_001291018.1 | 10.9488964 | -5.0751271 | 1.51450861 | -3.3510058 |
| 15039 | NR_133635.1 | 8.47352242 | -4.7059002 | 1.40446141 | -3.3506796 |
| 15040 | NR_039992.2 | 11.8403047 | -5.1874854 | 1.54830281 | -3.3504334 |
| 15041 | NR_024451.1 | 122.663305 | 2.08376743 | 0.62206567 | 3.34975476 |
| 15042 | NM_001261412.1 | 99.6217333 | -8.2605851 | 2.46610064 | -3.3496545 |
| 15043 | NM_144989.2 | 10.5422822 | -5.0087969 | 1.49538717 | -3.3494984 |
| 15044 | NM_001113512.2 | 22.7288792 | -6.1259817 | 1.82917146 | -3.3490473 |
| 15045 | NM_198566.3 | 9.404512 | -4.8456325 | 1.44694581 | -3.3488694 |
| 15046 | NM_001030018.1 | 21.9401274 | -6.0778065 | 1.81491783 | -3.3488053 |
| 15047 | NR_046438.2 | 55.1144989 | -4.1478602 | 1.23868034 | -3.3486123 |
| 15048 | NM_001320724.1 | 12.4912967 | -5.2743187 | 1.57520969 | -3.348328 |
| 15049 | NM_018689.2 | 77.4511876 | -7.897895 | 2.3592163 | -3.3476774 |

| | | | | | |
|---|---|---|---|---|---|
| 15050 | NM_022750.3 | 11.7247387 | -5.1779123 | 1.54673946 | -3.3476306 |
| 15051 | NM_001195733.1 | 75.5593739 | -7.8621778 | 2.34874267 | -3.3473986 |
| 15052 | NR_126047.1 | 8.00502954 | -4.6286187 | 1.38301446 | -3.3467609 |
| 15053 | NM_001119.4 | 218.434673 | -4.6828925 | 1.39924759 | -3.3467219 |
| 15054 | NM_005929.5 | 15.6629678 | -4.5741176 | 1.36679588 | -3.3465989 |
| 15055 | NM_001005476.2 | 52.8146583 | -4.0825123 | 1.21997967 | -3.3463774 |
| 15056 | NR_015419.2 | 18.1731468 | -4.7913958 | 1.43193657 | -3.346095 |
| 15057 | NM_001143964.3 | 14.9550141 | -5.5239672 | 1.6509975 | -3.3458362 |
| 15058 | NM_001004051.3 | 73.6569508 | -7.8254563 | 2.3388994 | -3.3457858 |
| 15059 | NM_139211.4 | 11.7252037 | -5.1673799 | 1.5446456 | -3.3453498 |
| 15060 | NM_001199092.1 | 10.3374603 | 7.51402422 | 2.24617403 | 3.34525469 |
| 15061 | NM_001080379.1 | 9.52992782 | -4.8784215 | 1.45838706 | -3.3450801 |
| 15062 | NM_012242.2 | 14.3354377 | -5.4656608 | 1.63393731 | -3.345086 |
| 15063 | NM_001146264.2 | 9.7818606 | -4.9152075 | 1.46941602 | -3.3450074 |
| 15064 | NM_152416.3 | 11.2877123 | -5.1095111 | 1.52770356 | -3.3445698 |
| 15065 | NM_001135690.2 | 95.8015309 | -3.870837 | 1.15737603 | -3.3444939 |
| 15066 | NR_002944.2 | 7.57137867 | -4.5478873 | 1.35981728 | -3.3444841 |
| 15067 | NM_001166419.1 | 8.78919015 | -4.7428477 | 1.41814746 | -3.3443967 |
| 15068 | NM_133625.4 | 9.71906959 | -4.8929249 | 1.46303741 | -3.3443607 |
| 15069 | NM_001291398.1 | 76.556955 | -7.8811057 | 2.35672991 | -3.3440853 |
| 15070 | NM_006300.3 | 17.9687296 | -3.9626577 | 1.1850131 | -3.343978 |
| 15071 | NR_135008.1 | 9.42643186 | -4.8707964 | 1.45690348 | -3.3432527 |
| 15072 | NM_005358.5 | 9.80510119 | -4.9235792 | 1.47288156 | -3.3428208 |
| 15073 | NR_046226.1 | 8.8193781 | -4.7540972 | 1.42276227 | -3.3414558 |
| 15074 | NM_000266.3 | 15.7310538 | -5.5947222 | 1.67455984 | -3.3410106 |
| 15075 | NM_183009.2 | 9.44703629 | -4.8679007 | 1.45701189 | -3.3410164 |
| 15076 | NM_001317157.1 | 8.00877674 | -4.6232329 | 1.3838119 | -3.3409403 |
| 15077 | NM_006896.3 | 10.2551106 | -4.9860624 | 1.49244682 | -3.3408644 |
| 15078 | NM_001135241.2 | 12.022659 | -5.2156105 | 1.56129188 | -3.3405736 |
| 15079 | NM_024864.4 | 9.98942646 | -4.9374697 | 1.47833048 | -3.3398958 |
| 15080 | NM_015075.1 | 12.0550536 | -5.2215185 | 1.56375974 | -3.3390798 |
| 15081 | NM_001197323.1 | 14.1774973 | -5.4451344 | 1.63118548 | -3.3381455 |
| 15082 | NM_013246.2 | 8.86159669 | -4.7699508 | 1.42912468 | -3.337673 |
| 15083 | NM_002187.2 | 19.7025654 | 4.06964503 | 1.21962335 | 3.33680477 |
| 15084 | NM_001195605.1 | 164.759934 | -3.6089356 | 1.08157935 | -3.3367275 |
| 15085 | NM_001302621.1 | 13.3956146 | -5.3703247 | 1.60959596 | -3.3364427 |
| 15086 | NR_026921.1 | 17.0203565 | -3.8884377 | 1.16567804 | -3.3357733 |
| 15087 | NR_136736.1 | 8.87868685 | -4.7801633 | 1.43299507 | -3.3357849 |
| 15088 | NR_038456.1 | 25.4757493 | -4.5138642 | 1.35342987 | -3.3351297 |
| 15089 | NM_001282286.1 | 15.6133731 | -5.5808179 | 1.67371762 | -3.3343844 |
| 15090 | NM_001085481.1 | 8.27618967 | -4.6611473 | 1.39800485 | -3.3341425 |
| 15091 | NM_012476.2 | 12.1997833 | -5.2280391 | 1.56819944 | -3.3337846 |
| 15092 | NM_014722.4 | 23.2593746 | -6.1574747 | 1.84720576 | -3.3333995 |

| | | | | | |
|---|---|---|---|---|---|
| 15093 | NM_001193493.1 | 10.521279 | -5.0246571 | 1.50738965 | -3.3333499 |
| 15094 | NR_046361.1 | 9.61020111 | -4.8971791 | 1.46917246 | -3.3332909 |
| 15095 | NM_198333.2 | 75.7596848 | 4.40906625 | 1.32277849 | 3.33318562 |
| 15096 | NM_001271957.1 | 9.52202014 | -4.876893 | 1.46328641 | -3.3328355 |
| 15097 | NM_032356.5 | 13.2999164 | -5.3494157 | 1.60517006 | -3.3326162 |
| 15098 | NM_173487.2 | 19.6142856 | -4.921996 | 1.4769169 | -3.3326154 |
| 15099 | NM_016335.4 | 39.0846031 | -5.1691949 | 1.55132456 | -3.3321169 |
| 15100 | NM_017881.2 | 9.52953694 | -4.8594155 | 1.45835523 | -3.3321206 |
| 15101 | NM_001146028.1 | 397.315846 | -5.1506169 | 1.54615043 | -3.3312521 |
| 15102 | NR_003572.2 | 128.834079 | -2.2284736 | 0.66899381 | -3.3310825 |
| 15103 | NM_001286365.1 | 146.079048 | -4.4448782 | 1.33451131 | -3.330716 |
| 15104 | NM_013362.3 | 17.8127826 | -3.2423815 | 0.97360553 | -3.3302825 |
| 15105 | NM_023073.3 | 102.277006 | -3.0536986 | 0.91705375 | -3.3299014 |
| 15106 | NM_170677.4 | 94.4348226 | -8.1833412 | 2.45759769 | -3.3298132 |
| 15107 | NM_001163811.1 | 11.1143378 | -5.0881723 | 1.52816546 | -3.3295951 |
| 15108 | NM_006231.3 | 172.267872 | -3.5373961 | 1.06262946 | -3.3289084 |
| 15109 | NM_152335.2 | 9.47898067 | -4.856029 | 1.45880469 | -3.3287725 |
| 15110 | NM_032483.3 | 8.0541822 | -4.6360132 | 1.39272879 | -3.3287265 |
| 15111 | NM_001346576.1 | 14.3228678 | -5.4525187 | 1.63805527 | -3.3286536 |
| 15112 | NR_028064.1 | 10.6112769 | 5.2399732 | 1.57429956 | 3.32844736 |
| 15113 | NM_015716.4 | 71.4129748 | -7.7809155 | 2.33773975 | -3.3283925 |
| 15114 | NM_001305040.1 | 10.7410845 | -5.0380218 | 1.51366866 | -3.3283518 |
| 15115 | NM_005250.2 | 26.1147164 | -4.551852 | 1.36793263 | -3.3275411 |
| 15116 | NM_001308090.1 | 70.5627799 | -7.7635825 | 2.33352651 | -3.3269742 |
| 15117 | NM_152280.4 | 1019.01777 | -3.9168625 | 1.17738387 | -3.3267506 |
| 15118 | NM_001269.4 | 34.2778629 | -3.9730875 | 1.19434482 | -3.3265833 |
| 15119 | NM_022770.3 | 11.6392513 | -5.158315 | 1.55074135 | -3.3263542 |
| 15120 | NM_001042470.1 | 11.9122308 | -5.1928461 | 1.56124586 | -3.3260912 |
| 15121 | NR_133644.1 | 19.2283905 | -5.8817811 | 1.76839797 | -3.3260506 |
| 15122 | NM_033198.3 | 204.562584 | -3.3610163 | 1.01052801 | -3.3260001 |
| 15123 | NR_033896.1 | 8.58109723 | -4.7188727 | 1.41877548 | -3.326018 |
| 15124 | NM_001029840.2 | 19.8868349 | -4.9434839 | 1.48664023 | -3.3252725 |
| 15125 | NR_002934.3 | 13.5383795 | -4.3837214 | 1.31834998 | -3.3251576 |
| 15126 | NM_017658.4 | 121.263977 | -1.0429388 | 0.31366284 | -3.3250314 |
| 15127 | NM_001203251.1 | 16.6477246 | -5.6756645 | 1.70714674 | -3.3246495 |
| 15128 | NM_152721.5 | 188.446192 | -3.0999793 | 0.93256755 | -3.3241337 |
| 15129 | NM_022370.3 | 170.589577 | -4.4865942 | 1.34978133 | -3.3239415 |
| 15130 | NM_001287251.1 | 9.52197987 | -4.8596253 | 1.46207494 | -3.3237867 |
| 15131 | NM_001202.5 | 15.1106417 | -5.5464075 | 1.66885452 | -3.3234817 |
| 15132 | NM_001270767.1 | 10.8067188 | -5.0582852 | 1.52201884 | -3.3234051 |
| 15133 | NM_001184750.2 | 518.323785 | 6.64733063 | 2.00041848 | 3.32297002 |
| 15134 | NM_183421.1 | 16.3071832 | -5.652244 | 1.70097423 | -3.3229451 |
| 15135 | NM_001318760.1 | 10.3887614 | -5.0081914 | 1.50739484 | -3.3224151 |

| | | | | | |
|---|---|---|---|---|---|
| 15136 | NR_073154.1 | 8.97102121 | -4.7965915 | 1.4439004 | -3.3219684 |
| 15137 | NM_032217.4 | 91.4563076 | -8.1374079 | 2.44965799 | -3.3218547 |
| 15138 | NM_152605.3 | 16.775779 | -4.6880539 | 1.41133821 | -3.3217084 |
| 15139 | NM_001136026.2 | 10.4774923 | -4.9996141 | 1.50524488 | -3.3214623 |
| 15140 | NM_001118886.1 | 7.93812097 | -4.6221963 | 1.39163111 | -3.3214235 |
| 15141 | NM_002942.4 | 93.7139993 | -8.1723622 | 2.46054706 | -3.3213598 |
| 15142 | NM_199001.3 | 8.6649339 | -4.7496075 | 1.43011198 | -3.3211438 |
| 15143 | NM_030891.4 | 50.1777033 | -2.4145922 | 0.72705099 | -3.3210768 |
| 15144 | NM_020902.1 | 84.9636519 | -4.0734188 | 1.22727075 | -3.3190873 |
| 15145 | NM_003913.4 | 236.481131 | -4.4156885 | 1.3304694 | -3.3188952 |
| 15146 | NM_001331028.1 | 17.4052863 | -4.7536316 | 1.43243204 | -3.3185739 |
| 15147 | NM_052909.3 | 120.972812 | -2.0497477 | 0.61765786 | -3.3185811 |
| 15148 | NM_001271100.1 | 17.8953784 | -5.7853504 | 1.74347511 | -3.3182868 |
| 15149 | NM_001001342.2 | 15.218164 | -5.5516785 | 1.67318786 | -3.3180246 |
| 15150 | NM_015264.1 | 18.9482841 | -5.8600551 | 1.7661666 | -3.3179515 |
| 15151 | NR_034002.1 | 9.75607166 | -4.9039485 | 1.47799839 | -3.3179661 |
| 15152 | NM_002523.2 | 100.089163 | -4.3503696 | 1.31120772 | -3.317834 |
| 15153 | NM_153028.3 | 7.85537189 | -4.6050155 | 1.38825166 | -3.3171331 |
| 15154 | NM_001346030.1 | 7.96829844 | -4.6135787 | 1.39090246 | -3.3169678 |
| 15155 | NM_001322311.1 | 13.8917851 | -5.408837 | 1.63121628 | -3.3158307 |
| 15156 | NM_001198978.1 | 10.9697347 | -5.0867092 | 1.53410452 | -3.3157514 |
| 15157 | NM_001278471.1 | 16.8839307 | 8.99368724 | 2.71242377 | 3.31573825 |
| 15158 | NR_027413.2 | 40.4293597 | 1.98315885 | 0.59809919 | 3.31576917 |
| 15159 | NM_080744.1 | 7.43071039 | -4.5185312 | 1.36282929 | -3.3155519 |
| 15160 | NR_038429.1 | 10.008983 | -4.9338535 | 1.48824244 | -3.3152216 |
| 15161 | NM_001172713.1 | 83.1594265 | -3.7691297 | 1.13695365 | -3.3151129 |
| 15162 | NM_002354.2 | 13.7357025 | -5.4008083 | 1.62922882 | -3.3149477 |
| 15163 | NM_005215.3 | 249.999302 | -2.5446856 | 0.76764989 | -3.3149039 |
| 15164 | NR_036650.1 | 17.0141468 | -3.8043207 | 1.14767466 | -3.3148076 |
| 15165 | NM_145247.4 | 10.9583659 | -5.0697735 | 1.52957262 | -3.3145033 |
| 15166 | NM_001015882.2 | 13.098504 | -5.339939 | 1.61109831 | -3.3144712 |
| 15167 | NM_001277291.1 | 9.58584741 | -4.8700841 | 1.46945115 | -3.3142198 |
| 15168 | NR_144444.1 | 10.5334717 | -5.0213379 | 1.51508276 | -3.3142334 |
| 15169 | NM_198141.2 | 20.698343 | -3.1689988 | 0.95619026 | -3.3141927 |
| 15170 | NM_207514.2 | 20.2969335 | -5.9690324 | 1.80135036 | -3.3136432 |
| 15171 | NM_144967.3 | 12.748146 | -5.2860511 | 1.59589953 | -3.3122706 |
| 15172 | NM_152924.4 | 345.007404 | -3.5437041 | 1.07021682 | -3.311202 |
| 15173 | NM_018558.3 | 11.3770859 | -5.137993 | 1.55187833 | -3.310822 |
| 15174 | NM_001025436.2 | 68.5944323 | -2.5404906 | 0.76735142 | -3.3107264 |
| 15175 | NM_001256187.1 | 12.5829694 | -5.2742098 | 1.59322745 | -3.3103935 |
| 15176 | NM_001322383.1 | 9.49475341 | -4.864984 | 1.4696161 | -3.3103774 |
| 15177 | NM_014390.3 | 607.156726 | -3.9405341 | 1.19087856 | -3.3089303 |
| 15178 | NM_001127181.2 | 12.4224105 | -5.2489678 | 1.58689435 | -3.3076983 |

| | | | | | |
|---|---|---|---|---|---|
| 15179 | NM_058166.4 | 8.29517385 | -4.6610119 | 1.40942481 | -3.3070313 |
| 15180 | NM_003550.2 | 9.7992732 | -4.9145089 | 1.48617245 | -3.3068228 |
| 15181 | NM_001130182.1 | 26.5919568 | -3.240986 | 0.98014714 | -3.3066322 |
| 15182 | NM_001008393.3 | 42.4688411 | -3.5034216 | 1.05966196 | -3.3061691 |
| 15183 | NR_109972.1 | 13.7475786 | 8.00652892 | 2.42182487 | 3.30599004 |
| 15184 | NM_057162.2 | 8.17719611 | -4.6515037 | 1.40721428 | -3.3054694 |
| 15185 | NM_001131035.1 | 10.175213 | -4.9730265 | 1.50459452 | -3.305227 |
| 15186 | NM_013249.2 | 9.42399184 | -4.8693736 | 1.47338319 | -3.3048929 |
| 15187 | NM_001040629.1 | 15.048107 | -5.5241387 | 1.67152834 | -3.304843 |
| 15188 | NM_212551.4 | 48.7194562 | -3.7257073 | 1.1274318 | -3.3045966 |
| 15189 | NM_001039846.1 | 8.54543109 | -4.7269664 | 1.43059686 | -3.3041918 |
| 15190 | NM_001318360.1 | 64.9062271 | -7.6430691 | 2.3134192 | -3.3037977 |
| 15191 | NM_153362.2 | 72.1747448 | -4.1509917 | 1.25654117 | -3.3035063 |
| 15192 | NM_001098519.1 | 17.0613564 | -5.7189381 | 1.7313076 | -3.3032478 |
| 15193 | NM_001195192.1 | 50.8824661 | 3.85422779 | 1.16686617 | 3.30305897 |
| 15194 | NM_001289172.1 | 8.02646805 | -4.6143051 | 1.39697241 | -3.3030753 |
| 15195 | NM_003353.3 | 7.965288 | -4.6207723 | 1.39892213 | -3.3030947 |
| 15196 | NM_203401.1 | 13.4429078 | -5.3691822 | 1.62569186 | -3.302706 |
| 15197 | NM_019006.3 | 11.0283337 | -5.0923163 | 1.54193506 | -3.3025491 |
| 15198 | NR_045999.1 | 4.48437389 | 7.10549461 | 2.15167544 | 3.30230781 |
| 15199 | NM_001288985.1 | 7.3559543 | -4.5053635 | 1.36459369 | -3.3016154 |
| 15200 | NM_001318203.1 | 20.7835734 | -5.9989361 | 1.81707775 | -3.3014196 |
| 15201 | NM_000227.4 | 7.63829343 | -4.5459159 | 1.37731782 | -3.3005569 |
| 15202 | NM_001291163.1 | 12.1505139 | -5.2307446 | 1.58482559 | -3.3005175 |
| 15203 | NM_001317946.1 | 15.7579649 | -5.5958605 | 1.69559235 | -3.3002393 |
| 15204 | NM_015840.3 | 70.0527808 | -7.75316 | 2.34943894 | -3.3000049 |
| 15205 | NM_153604.3 | 16.0245356 | 3.3871677 | 1.02664697 | 3.29925263 |
| 15206 | NM_001135219.1 | 11.1933277 | -5.1049592 | 1.54734274 | -3.2991781 |
| 15207 | NM_001142777.1 | 9.0360163 | -4.8093741 | 1.45788904 | -3.2988616 |
| 15208 | NM_032804.5 | 122.663119 | -4.3986672 | 1.33347278 | -3.2986554 |
| 15209 | NM_018922.2 | 8.3739984 | -4.6895439 | 1.42168854 | -3.2985733 |
| 15210 | NM_138270.3 | 92.9432436 | -8.1602494 | 2.47391341 | -3.2985186 |
| 15211 | NR_024491.1 | 12.000293 | 3.95943694 | 1.20038194 | 3.29848092 |
| 15212 | NM_032561.4 | 10.4432638 | -5.0139345 | 1.52016283 | -3.2982878 |
| 15213 | NR_036461.1 | 7.31746559 | -4.4857533 | 1.36023933 | -3.2977677 |
| 15214 | NR_033373.1 | 16.083819 | -3.7081342 | 1.12456862 | -3.2973836 |
| 15215 | NM_025231.2 | 9.42950918 | -4.8693081 | 1.47679082 | -3.2972226 |
| 15216 | NM_001202425.1 | 10.8464954 | -5.0573002 | 1.53401051 | -3.2967833 |
| 15217 | NM_014217.3 | 10.9389366 | -5.0673724 | 1.53718769 | -3.2965215 |
| 15218 | NR_046687.1 | 30.4088464 | 3.44192049 | 1.04411915 | 3.29648249 |
| 15219 | NM_025230.4 | 88.9962519 | -8.0976124 | 2.45656983 | -3.2963086 |
| 15220 | NM_001287821.1 | 14.1182419 | -5.4443083 | 1.65171578 | -3.2961532 |
| 15221 | NM_144668.5 | 14.7923816 | -5.5134638 | 1.67271286 | -3.2961209 |

| | | | | | |
|---|---|---|---|---|---|
| 15222 | NM_207307.2 | 16.4628026 | -5.6671949 | 1.71988406 | -3.2951029 |
| 15223 | NM_032298.2 | 18.2519893 | -5.8161969 | 1.76549182 | -3.2943777 |
| 15224 | NM_005236.2 | 58.858711 | -3.0781685 | 0.93456777 | -3.2936814 |
| 15225 | NR_038420.1 | 8.63088704 | -4.7361548 | 1.43795622 | -3.2936711 |
| 15226 | NM_001161779.1 | 18.1313897 | -5.7975208 | 1.76026513 | -3.2935497 |
| 15227 | NM_175038.2 | 18.4299883 | 9.11848474 | 2.76936967 | 3.292621 |
| 15228 | NM_001319100.1 | 64.6500577 | -7.637344 | 2.3199496 | -3.2920301 |
| 15229 | NR_037803.2 | 7.3246666 | -4.4994159 | 1.3668188 | -3.2918891 |
| 15230 | NR_103782.1 | 11.0063948 | -5.0735053 | 1.54141755 | -3.2914542 |
| 15231 | NM_001278205.1 | 16.3863078 | -3.7177224 | 1.1295499 | -3.2913308 |
| 15232 | NM_001278244.1 | 14.1070206 | 8.74398675 | 2.65681582 | 3.29115278 |
| 15233 | NM_001256761.2 | 107.41728 | -8.3695478 | 2.54311709 | -3.2910588 |
| 15234 | NM_001321471.1 | 10.2476118 | -4.9654568 | 1.50881991 | -3.290954 |
| 15235 | NM_006483.2 | 41.5060203 | -3.6989577 | 1.12399511 | -3.290902 |
| 15236 | NM_001135862.2 | 87.4008242 | -8.0716834 | 2.45276071 | -3.2908564 |
| 15237 | NM_174958.2 | 22.7330375 | 9.42940602 | 2.86559663 | 3.29055594 |
| 15238 | NM_001103176.1 | 12.7539569 | -5.2951069 | 1.60937495 | -3.2901636 |
| 15239 | NM_145204.3 | 9.27947276 | -4.8406488 | 1.47125613 | -3.2901469 |
| 15240 | NM_018462.4 | 364.864186 | -4.7629707 | 1.44767318 | -3.290087 |
| 15241 | NM_000020.2 | 7.67804632 | -4.5685638 | 1.38861304 | -3.2900193 |
| 15242 | NM_018190.3 | 46.8548013 | -3.9295759 | 1.19440296 | -3.2899918 |
| 15243 | NM_014059.2 | 8.37151325 | -4.6860508 | 1.42440014 | -3.2898416 |
| 15244 | NM_001312911.1 | 9.64985725 | -4.8858732 | 1.48552884 | -3.2889791 |
| 15245 | NM_173074.2 | 8.03807095 | -4.6380748 | 1.41033581 | -3.2886315 |
| 15246 | NM_001193478.1 | 13.9824491 | -4.425027 | 1.34566912 | -3.288347 |
| 15247 | NM_005167.5 | 10.1068084 | -4.9603182 | 1.50848885 | -3.2882697 |
| 15248 | NM_001282960.1 | 95.5619669 | -8.2001723 | 2.49390916 | -3.2880798 |
| 15249 | NM_001784.4 | 10.0801376 | -4.9659288 | 1.51030958 | -3.2880204 |
| 15250 | NM_018424.3 | 24.5597271 | -5.2427308 | 1.59451319 | -3.2879821 |
| 15251 | NM_001039490.1 | 19.378185 | -5.8997408 | 1.79450053 | -3.2876785 |
| 15252 | NM_052845.3 | 116.954612 | -2.5399243 | 0.77259966 | -3.2875038 |
| 15253 | NM_018565.3 | 12.4279566 | -5.2502234 | 1.59705949 | -3.2874313 |
| 15254 | NM_001329922.1 | 16.7144954 | -5.6906993 | 1.73115671 | -3.2872237 |
| 15255 | NR_144937.1 | 52.8645567 | 5.01843298 | 1.52665524 | 3.28720777 |
| 15256 | NM_001085430.2 | 8.80908157 | -4.7611154 | 1.4484655 | -3.2870065 |
| 15257 | NM_001308429.1 | 10.873816 | -5.0562137 | 1.53833592 | -3.2868073 |
| 15258 | NM_001289181.1 | 10.0721907 | -4.9409899 | 1.50379648 | -3.2856772 |
| 15259 | NM_006314.2 | 10.9474957 | -5.0685306 | 1.54296417 | -3.2849308 |
| 15260 | NM_001080471.1 | 20.6727104 | -3.6100884 | 1.09900362 | -3.284874 |
| 15261 | NM_001012333.2 | 11.4213347 | -5.1425133 | 1.565775 | -3.2843246 |
| 15262 | NR_135685.1 | 11.5872678 | 6.73889867 | 2.05185752 | 3.28429173 |
| 15263 | NM_024754.4 | 10.3237449 | -4.9961509 | 1.52139429 | -3.283929 |
| 15264 | NM_017569.3 | 8.09309245 | -4.6394517 | 1.41289895 | -3.2836402 |

| | | | | | |
|---|---|---|---|---|---|
| 15265 | NR_120563.1 | 42.0997716 | 2.01070887 | 0.61241322 | 3.28325516 |
| 15266 | NM_001017989.2 | 7.62576428 | -4.5582184 | 1.3884932 | -3.2828526 |
| 15267 | NM_004899.4 | 9.85585419 | -4.9106881 | 1.49589522 | -3.2827754 |
| 15268 | NM_002397.4 | 90.0774308 | -8.1149484 | 2.47202817 | -3.2827087 |
| 15269 | NM_001163391.1 | 8.34227249 | -4.6813635 | 1.4261002 | -3.2826329 |
| 15270 | NR_104387.1 | 7.79574727 | -4.5811482 | 1.39584715 | -3.2819841 |
| 15271 | NM_001080515.2 | 17.6678788 | -5.7622622 | 1.75581977 | -3.2818073 |
| 15272 | NM_001346890.1 | 29.6589356 | -2.9756942 | 0.90675608 | -3.281692 |
| 15273 | NM_004843.3 | 30.9276322 | -3.3892857 | 1.03310123 | -3.2806908 |
| 15274 | NM_001242713.2 | 10.3774648 | -5.0090949 | 1.52696882 | -3.2804173 |
| 15275 | NM_001160042.1 | 16.2570633 | -5.6496073 | 1.72238996 | -3.2800977 |
| 15276 | NM_033223.4 | 10.4793992 | -5.0136509 | 1.52862628 | -3.2798408 |
| 15277 | NM_182832.2 | 9.7390964 | 4.20952715 | 1.28353167 | 3.27964417 |
| 15278 | NM_001145155.1 | 9.24561481 | -4.8400945 | 1.47595659 | -3.2792933 |
| 15279 | NM_001098622.2 | 9.54293337 | -4.8766611 | 1.48713588 | -3.2792303 |
| 15280 | NM_000413.3 | 8.88445003 | -4.7763025 | 1.4565556 | -3.2791763 |
| 15281 | NM_139159.4 | 155.099862 | -2.4015311 | 0.73236653 | -3.2791383 |
| 15282 | NM_018703.3 | 86.8440996 | -7.0903069 | 2.16258402 | -3.2786272 |
| 15283 | NM_181310.1 | 16.5192641 | 8.96132822 | 2.73362746 | 3.27818196 |
| 15284 | NM_024556.3 | 32.1597096 | -3.8962395 | 1.18887365 | -3.2772528 |
| 15285 | NR_029406.1 | 51.1941026 | 1.65814146 | 0.50599823 | 3.27697085 |
| 15286 | NM_001078651.2 | 13.6738278 | -5.3964399 | 1.64711547 | -3.2762973 |
| 15287 | NR_038924.1 | 8.20964233 | -4.662081 | 1.42298128 | -3.2762771 |
| 15288 | NM_173527.2 | 45.7355739 | -3.6265085 | 1.10718014 | -3.2754457 |
| 15289 | NM_001306.3 | 15.0620037 | -5.5397248 | 1.691357 | -3.2753137 |
| 15290 | NR_136513.1 | 14.6271893 | 6.03734401 | 1.84328351 | 3.27532035 |
| 15291 | NM_032276.3 | 21.4340708 | -4.2463972 | 1.2965703 | -3.2750999 |
| 15292 | NM_001010872.2 | 55.269626 | 2.61773242 | 0.7994028 | 3.27461003 |
| 15293 | NM_199420.3 | 12.5959556 | -4.2796581 | 1.30700415 | -3.2744028 |
| 15294 | NM_201598.3 | 17.9098784 | -5.7820358 | 1.76647256 | -3.2732101 |
| 15295 | NM_001286740.1 | 15.2015731 | -5.5424358 | 1.69330027 | -3.2731559 |
| 15296 | NM_001083926.1 | 10.9966712 | -5.0918582 | 1.55581311 | -3.2727955 |
| 15297 | NM_001114633.1 | 87.4381475 | -3.8497097 | 1.17635119 | -3.2725854 |
| 15298 | NM_145276.2 | 55.5254476 | 2.30969733 | 0.70583617 | 3.27228533 |
| 15299 | NM_005045.3 | 88.4513034 | -8.0892126 | 2.47261753 | -3.271518 |
| 15300 | NR_134283.1 | 7.69147135 | -4.5697285 | 1.39718637 | -3.270665 |
| 15301 | NR_130747.1 | 10.2378326 | -4.9741031 | 1.52134278 | -3.2695479 |
| 15302 | NM_015146.1 | 68.1955378 | -7.7142044 | 2.35945698 | -3.269483 |
| 15303 | NM_001018038.2 | 13.7014155 | -5.4050286 | 1.6532337 | -3.2693676 |
| 15304 | NM_032331.3 | 10.4612724 | -5.0104966 | 1.53312222 | -3.2681652 |
| 15305 | NM_001242852.1 | 9.37658455 | -4.8607661 | 1.48756685 | -3.2675951 |
| 15306 | NM_001256486.1 | 63.2260103 | -7.6052355 | 2.32774625 | -3.2672099 |
| 15307 | NR_110386.1 | 83.2010766 | 3.03089806 | 0.92773463 | 3.26698817 |

| | | | | | |
|---|---|---|---|---|---|
| 15308 | NR_125384.1 | 7.07938683 | -4.4448876 | 1.36093414 | -3.2660564 |
| 15309 | NR_120513.1 | 24.8511283 | 3.59592084 | 1.10109982 | 3.26575372 |
| 15310 | NR_134895.1 | 10.0651449 | -4.9617427 | 1.51949171 | -3.2653964 |
| 15311 | NM_000924.3 | 16.1096318 | -5.6351651 | 1.72624963 | -3.2643976 |
| 15312 | NR_038433.2 | 62.0574167 | 2.89926522 | 0.88826005 | 3.26398244 |
| 15313 | NM_001127893.1 | 15.3212092 | -4.5588079 | 1.39676537 | -3.2638323 |
| 15314 | NM_001320678.1 | 17.4309072 | -4.7504582 | 1.45563424 | -3.2634972 |
| 15315 | NM_024332.3 | 8.75370791 | -4.7621306 | 1.45925614 | -3.263396 |
| 15316 | NM_001271711.1 | 12.8091035 | -5.2943735 | 1.62241162 | -3.2632739 |
| 15317 | NM_018890.3 | 33.3246325 | 2.42078309 | 0.74184218 | 3.26320498 |
| 15318 | NM_181581.2 | 10.1242041 | -4.9476494 | 1.51651401 | -3.2625148 |
| 15319 | NM_080879.2 | 7.52117845 | -4.5342435 | 1.3900247 | -3.2619877 |
| 15320 | NM_001170637.3 | 96.3146228 | -8.2115706 | 2.51742901 | -3.2618876 |
| 15321 | NM_001282601.1 | 22.7804971 | -6.1286932 | 1.87901841 | -3.2616462 |
| 15322 | NM_018159.3 | 155.53862 | -4.3154354 | 1.32332859 | -3.261046 |
| 15323 | NM_001497.3 | 139.169889 | -1.8669375 | 0.57253428 | -3.260831 |
| 15324 | NM_005297.3 | 16.2707584 | -4.6564466 | 1.42847073 | -3.2597424 |
| 15325 | NM_152745.2 | 10.2283704 | -4.9804857 | 1.5279718 | -3.2595403 |
| 15326 | NM_001130055.3 | 8.85129573 | -4.7682461 | 1.46302964 | -3.2591589 |
| 15327 | NR_136158.1 | 57.5091592 | -2.8130104 | 0.86324019 | -3.2586648 |
| 15328 | NR_136399.1 | 12.2453776 | -5.2320358 | 1.60576946 | -3.2582733 |
| 15329 | NM_001199629.1 | 14.5455694 | -5.481757 | 1.68249054 | -3.2581206 |
| 15330 | NM_005994.3 | 133.111557 | -4.2744306 | 1.31203444 | -3.2578646 |
| 15331 | NM_033240.2 | 7.34136685 | -4.5016074 | 1.38194204 | -3.2574502 |
| 15332 | NM_005444.2 | 81.5082106 | -7.9708964 | 2.44726035 | -3.2570692 |
| 15333 | NM_052966.3 | 59.6699859 | -2.8190387 | 0.86556627 | -3.2568722 |
| 15334 | NM_020866.2 | 15.1621047 | -5.5404603 | 1.70147574 | -3.2562676 |
| 15335 | NM_022164.2 | 13.4069445 | -5.3696123 | 1.64906321 | -3.2561592 |
| 15336 | NR_033839.1 | 7.02849498 | -4.4314952 | 1.36096179 | -3.2561496 |
| 15337 | NM_022496.4 | 37.4784096 | -2.4771721 | 0.76079321 | -3.2560387 |
| 15338 | NR_002821.2 | 7.53563775 | -4.528222 | 1.39075131 | -3.2559538 |
| 15339 | NM_001256008.2 | 82.9196683 | -7.9955428 | 2.45587748 | -3.2556766 |
| 15340 | NM_173632.3 | 77.9725218 | -2.967519 | 0.91150032 | -3.2556423 |
| 15341 | NM_001323916.1 | 9.25467584 | -4.8345934 | 1.48499899 | -3.2556207 |
| 15342 | NM_001282907.1 | 8.96508667 | -4.7909593 | 1.47190115 | -3.2549464 |
| 15343 | NM_033276.3 | 9.40953157 | -4.8590924 | 1.49285309 | -3.2549033 |
| 15344 | NM_005309.2 | 7.61409274 | -4.5447667 | 1.39631631 | -3.254826 |
| 15345 | NM_080552.2 | 8.20492442 | -4.6713144 | 1.43550253 | -3.2541318 |
| 15346 | NM_001289165.1 | 10.7540589 | -5.0434476 | 1.54999039 | -3.2538573 |
| 15347 | NR_002305.1 | 10.082551 | -4.9579092 | 1.52375029 | -3.2537544 |
| 15348 | NM_001039842.2 | 7.38201944 | -4.4951451 | 1.38159228 | -3.2535974 |
| 15349 | NM_003549.3 | 8.12895775 | -4.6515368 | 1.4296646 | -3.2535861 |
| 15350 | NM_001320901.1 | 15.2622525 | 8.8485857 | 2.719962 | 3.25320195 |

| | | | | | |
|---|---|---|---|---|---|
| 15351 | NM_001105570.1 | 132.348604 | 1.28897375 | 0.39628694 | 3.25262736 |
| 15352 | NM_001316306.1 | 11.1525388 | -5.1028324 | 1.56890693 | -3.2524762 |
| 15353 | NM_001083537.1 | 11.4083455 | -5.1257226 | 1.57616476 | -3.2520221 |
| 15354 | NR_027356.2 | 10.5442494 | -5.0161161 | 1.54274045 | -3.2514323 |
| 15355 | NM_001286371.1 | 9.23171029 | -4.8329952 | 1.48677263 | -3.250662 |
| 15356 | NM_001005753.2 | 14.4082436 | -5.4783803 | 1.68536344 | -3.2505632 |
| 15357 | NM_007049.4 | 61.6910183 | -7.5698452 | 2.32878951 | -3.2505493 |
| 15358 | NR_038280.1 | 7.19652127 | -4.4626721 | 1.37323392 | -3.2497538 |
| 15359 | NM_005175.2 | 9.05344726 | -4.8065614 | 1.47919868 | -3.2494359 |
| 15360 | NM_001329914.1 | 17.1398499 | -5.7194907 | 1.76045392 | -3.2488727 |
| 15361 | NM_020769.2 | 9.00427512 | -4.7968974 | 1.47663286 | -3.2485376 |
| 15362 | NM_206923.3 | 8.93965859 | -4.7830402 | 1.47237154 | -3.248528 |
| 15363 | NM_001143854.1 | 66.0360689 | -7.6680285 | 2.36048817 | -3.2484927 |
| 15364 | NR_003494.1 | 18.7980519 | -2.8445567 | 0.87567209 | -3.2484268 |
| 15365 | NR_045639.1 | 8.43971683 | -4.7092446 | 1.4498014 | -3.2481998 |
| 15366 | NM_004602.3 | 59.1283676 | -7.5085928 | 2.31182264 | -3.2479104 |
| 15367 | NM_001136040.2 | 85.7991709 | -8.0447456 | 2.47701508 | -3.247758 |
| 15368 | NM_147127.4 | 7.3324288 | -4.5023722 | 1.38632794 | -3.2476963 |
| 15369 | NM_001329140.1 | 151.058978 | -4.4874297 | 1.38173841 | -3.2476695 |
| 15370 | NR_037918.2 | 9.80889123 | -4.9229029 | 1.51589373 | -3.247525 |
| 15371 | NM_001392.4 | 14.5848687 | -5.4916653 | 1.6912777 | -3.2470512 |
| 15372 | NM_001318739.1 | 10.5146843 | -5.0187144 | 1.54566282 | -3.2469659 |
| 15373 | NM_001123040.1 | 103.35948 | 2.34722081 | 0.72292512 | 3.2468381 |
| 15374 | NM_001317032.1 | 39.4221902 | 2.64090905 | 0.81356333 | 3.24610137 |
| 15375 | NM_001323987.1 | 20.2862316 | -5.9631513 | 1.83702905 | -3.2460844 |
| 15376 | NM_001882.3 | 13.054206 | -5.3298496 | 1.64193829 | -3.2460718 |
| 15377 | NM_001642.2 | 119.004548 | -8.5170355 | 2.62395884 | -3.2458724 |
| 15378 | NM_006172.3 | 8.57558497 | -4.710447 | 1.45129344 | -3.2456889 |
| 15379 | NM_001039111.2 | 33.0889642 | -2.9259088 | 0.90158735 | -3.245286 |
| 15380 | NM_001164815.1 | 62.7218588 | -7.5935964 | 2.33990074 | -3.2452643 |
| 15381 | NM_001039476.2 | 35.674344 | -4.6589092 | 1.43578152 | -3.2448594 |
| 15382 | NM_001163813.1 | 8.27595358 | -4.6587835 | 1.43579759 | -3.2447356 |
| 15383 | NM_001130141.1 | 10.5707228 | -5.0344506 | 1.55202544 | -3.2437938 |
| 15384 | NM_001254748.1 | 10.1083658 | -4.9605778 | 1.52931817 | -3.2436532 |
| 15385 | NR_045564.1 | 11.0233797 | -5.0797327 | 1.56610913 | -3.2435369 |
| 15386 | NM_016580.3 | 7.9624807 | -4.6028486 | 1.41912263 | -3.2434467 |
| 15387 | NM_001099645.1 | 8.32217286 | -4.6924597 | 1.44690312 | -3.2431057 |
| 15388 | NR_034006.2 | 18.5538878 | -3.4289428 | 1.05743371 | -3.2427024 |
| 15389 | NR_110849.1 | 18.4114521 | 4.8640401 | 1.49998865 | 3.24271794 |
| 15390 | NM_017791.2 | 8.15398745 | -4.6585221 | 1.43675063 | -3.2424013 |
| 15391 | NM_201574.2 | 50.4443757 | -3.5857204 | 1.10591587 | -3.2423085 |
| 15392 | NM_031369.2 | 83.7405699 | -8.0097041 | 2.47082179 | -3.2417166 |
| 15393 | NM_138813.3 | 7.15588297 | -4.4506037 | 1.37299212 | -3.2415362 |

| | | | | | |
|---|---|---|---|---|---|
| 15394 | NM_016422.3 | 149.156209 | -3.1222498 | 0.96327384 | -3.2412899 |
| 15395 | NM_001322272.1 | 11.3926985 | -5.1339143 | 1.58418379 | -3.2407315 |
| 15396 | NR_046342.1 | 16.6486898 | -5.6739159 | 1.75081254 | -3.2407329 |
| 15397 | NM_032868.5 | 14.9331862 | -5.5207165 | 1.70369031 | -3.240446 |
| 15398 | NM_001317079.1 | 7.18062967 | -4.474068 | 1.38082028 | -3.2401523 |
| 15399 | NM_001145409.1 | 59.6195904 | -7.5205828 | 2.32125695 | -3.2398752 |
| 15400 | NM_001012973.1 | 7.05084994 | -4.4434017 | 1.37202053 | -3.2385826 |
| 15401 | NM_138341.2 | 10.6928777 | -5.0281897 | 1.55274402 | -3.2382605 |
| 15402 | NM_144659.5 | 8.5544995 | -4.7085536 | 1.4541089 | -3.2381024 |
| 15403 | NM_001184943.1 | 89.6881656 | -8.1087667 | 2.50424969 | -3.2380025 |
| 15404 | NM_145020.4 | 16.1779268 | -4.6505285 | 1.43626771 | -3.237926 |
| 15405 | NM_177991.2 | 13.7660556 | -5.4079962 | 1.6704892 | -3.2373728 |
| 15406 | NR_104049.1 | 10.6322895 | -5.023862 | 1.55190708 | -3.2372183 |
| 15407 | NM_006558.2 | 149.077491 | -3.7746949 | 1.16613932 | -3.2369159 |
| 15408 | NM_001001894.2 | 63.1814047 | -7.6040596 | 2.34928088 | -3.2367605 |
| 15409 | NM_182925.4 | 8.61107474 | -4.725617 | 1.45999411 | -3.236737 |
| 15410 | NM_001304435.1 | 8.96757765 | -4.7901314 | 1.47996297 | -3.2366562 |
| 15411 | NR_015361.1 | 41.157305 | 2.7748193 | 0.85745132 | 3.23612461 |
| 15412 | NM_001278732.1 | 7.07139686 | -4.4453134 | 1.37374727 | -3.2359033 |
| 15413 | NM_001321463.1 | 104.711664 | -4.5560865 | 1.40798108 | -3.2359004 |
| 15414 | NR_137178.1 | 9.2636981 | -4.8329787 | 1.49369551 | -3.235585 |
| 15415 | NM_025256.6 | 75.1409058 | -7.8536325 | 2.42758564 | -3.2351619 |
| 15416 | NM_001035512.1 | 12.6303933 | -5.2829177 | 1.63312002 | -3.2348619 |
| 15417 | NM_000844.3 | 22.6416903 | -4.2714798 | 1.32047953 | -3.2347944 |
| 15418 | NM_024594.3 | 195.460214 | -4.3982107 | 1.35978602 | -3.2344874 |
| 15419 | NR_027055.1 | 7.50428531 | -4.5318034 | 1.40130543 | -3.2339869 |
| 15420 | NM_001285448.1 | 95.1862764 | -3.1455656 | 0.97267455 | -3.2339343 |
| 15421 | NM_025076.4 | 60.9733134 | -7.5529302 | 2.33565554 | -3.2337518 |
| 15422 | NR_027032.1 | 10.6526479 | -5.0300255 | 1.55559744 | -3.2335007 |
| 15423 | NM_021738.2 | 7.46644827 | -4.5312038 | 1.40134676 | -3.2334637 |
| 15424 | NM_021913.4 | 78.8410625 | -7.9230508 | 2.45061941 | -3.2330809 |
| 15425 | NR_135502.1 | 57.6124074 | -7.4710994 | 2.31094504 | -3.2329195 |
| 15426 | NM_001286815.1 | 7.76747797 | -4.5764034 | 1.41576347 | -3.2324633 |
| 15427 | NM_020336.3 | 73.1578958 | -7.8149839 | 2.41769253 | -3.2324143 |
| 15428 | NR_125906.1 | 54.2296284 | 1.50764866 | 0.46644531 | 3.23220885 |
| 15429 | NR_001459.2 | 9.11652354 | -4.8228973 | 1.49220207 | -3.2320671 |
| 15430 | NM_032427.3 | 46.3448258 | -3.4508644 | 1.0680477 | -3.2310022 |
| 15431 | NM_025218.3 | 9.9410085 | -4.9395916 | 1.52889045 | -3.2308342 |
| 15432 | NM_001282593.1 | 17.6374628 | 3.08286515 | 0.95421387 | 3.23079055 |
| 15433 | NM_032592.3 | 7.98911317 | -4.6275933 | 1.4323538 | -3.2307613 |
| 15434 | NM_001243750.1 | 8.99015219 | -4.7812608 | 1.48013567 | -3.2302855 |
| 15435 | NM_024719.3 | 7.58210507 | -4.5316176 | 1.40287281 | -3.2302412 |
| 15436 | NM_183387.2 | 23.3095885 | -3.3696571 | 1.04330929 | -3.2297777 |

| | | | | | |
|---|---|---|---|---|---|
| 15437 | NM_001287061.1 | 10.8193063 | -5.064761 | 1.56819655 | -3.2296724 |
| 15438 | NM_001031721.3 | 9.20996717 | -4.8342225 | 1.4968262 | -3.2296485 |
| 15439 | NM_030771.1 | 53.4261383 | -3.500551 | 1.08397431 | -3.2293672 |
| 15440 | NM_001145109.1 | 61.7548437 | -7.5714221 | 2.34458994 | -3.2293161 |
| 15441 | NM_001130110.1 | 9.06313725 | -4.7951816 | 1.48498784 | -3.229105 |
| 15442 | NM_016932.4 | 10.0895231 | -4.9663798 | 1.5380125 | -3.2290894 |
| 15443 | NM_001282585.1 | 9.72306204 | -4.9145896 | 1.52199542 | -3.2290436 |
| 15444 | NM_001330229.1 | 10.2724235 | -4.9779471 | 1.54162968 | -3.2290161 |
| 15445 | NM_012192.3 | 104.213237 | -1.7344736 | 0.53717615 | -3.2288731 |
| 15446 | NR_024580.1 | 12.0091564 | -4.2158648 | 1.30567476 | -3.2288782 |
| 15447 | NM_206933.2 | 12.8638448 | 5.51306323 | 1.70750534 | 3.22872386 |
| 15448 | NM_001102398.2 | 79.8763143 | -7.9414751 | 2.45974632 | -3.2285749 |
| 15449 | NM_012287.5 | 136.354532 | -2.6393467 | 0.81768383 | -3.2278328 |
| 15450 | NM_001136022.2 | 61.6596776 | -7.5689787 | 2.34492932 | -3.2278068 |
| 15451 | NM_003782.3 | 7.16904647 | -4.468969 | 1.38452768 | -3.2277932 |
| 15452 | NM_001166262.1 | 12.2221359 | -5.2259908 | 1.61918119 | -3.2275516 |
| 15453 | NM_000922.3 | 29.7431003 | -3.2292577 | 1.00059443 | -3.2273393 |
| 15454 | NM_001289003.1 | 50.372569 | -2.4176221 | 0.74918731 | -3.2269928 |
| 15455 | NM_201555.1 | 31.333793 | -2.4417775 | 0.75670813 | -3.2268418 |
| 15456 | NM_001130145.2 | 58.9403744 | -7.5040594 | 2.32558543 | -3.2267399 |
| 15457 | NM_005115.4 | 58.5694401 | -7.4948826 | 2.32282201 | -3.226628 |
| 15458 | NR_135107.1 | 20.2427628 | 9.25357878 | 2.86792667 | 3.2265744 |
| 15459 | NM_183078.2 | 45.5816569 | 2.51865623 | 0.78062712 | 3.22645237 |
| 15460 | NM_001253384.1 | 218.550439 | 2.37388008 | 0.73578221 | 3.22633525 |
| 15461 | NM_001190880.2 | 8.98623674 | -4.7971501 | 1.4870016 | -3.2260558 |
| 15462 | NM_001320329.1 | 8.66625006 | -4.7333551 | 1.46725492 | -3.2259937 |
| 15463 | NM_001146256.1 | 15.2118388 | 8.84239103 | 2.74123815 | 3.22569239 |
| 15464 | NM_015447.3 | 461.235945 | -4.9634993 | 1.53895765 | -3.2252345 |
| 15465 | NM_014661.3 | 205.552749 | -3.3740072 | 1.04619893 | -3.2250149 |
| 15466 | NM_152657.3 | 7.95210719 | -4.6158891 | 1.43132497 | -3.2249065 |
| 15467 | NM_001171162.1 | 32.1378394 | -2.5483175 | 0.79020585 | -3.224878 |
| 15468 | NM_001321703.1 | 22.2195693 | -5.1123839 | 1.58541307 | -3.2246384 |
| 15469 | NM_001308384.1 | 10.7974762 | -5.0548354 | 1.56760166 | -3.2245663 |
| 15470 | NM_001282622.1 | 61.5369709 | -7.5663421 | 2.34659271 | -3.2243951 |
| 15471 | NM_024010.2 | 13.2441297 | -5.3471124 | 1.65834307 | -3.2243704 |
| 15472 | NM_031901.5 | 67.7030217 | -3.2937897 | 1.02186558 | -3.2233102 |
| 15473 | NM_000144.4 | 164.374617 | 1.49043936 | 0.4624229 | 3.22310888 |
| 15474 | NR_104432.1 | 13.6101308 | -5.3904285 | 1.67270588 | -3.22258 |
| 15475 | NM_033158.4 | 77.3237447 | -3.7745993 | 1.17137571 | -3.2223643 |
| 15476 | NM_005095.2 | 260.499833 | -4.6562128 | 1.44508742 | -3.2220977 |
| 15477 | NM_152720.2 | 9.85384043 | -4.930382 | 1.53031643 | -3.2218055 |
| 15478 | NM_178310.3 | 7.27104959 | -4.4805884 | 1.39098305 | -3.2211668 |
| 15479 | NM_001291594.1 | 13.8623069 | -5.4195622 | 1.68254975 | -3.2210413 |

| | | | | | |
|---|---|---|---|---|---|
| 15480 | NM_001144951.1 | 7.47630874 | -4.5315295 | 1.40707484 | -3.220532 |
| 15481 | NM_015459.4 | 56.04782 | -7.4314173 | 2.30770146 | -3.220268 |
| 15482 | NM_001271520.1 | 8.35598265 | -4.6858475 | 1.45514757 | -3.2201872 |
| 15483 | NM_001271007.1 | 12.5244148 | -5.2679424 | 1.63594892 | -3.2201143 |
| 15484 | NR_024537.1 | 71.2691061 | 4.18826183 | 1.30066399 | 3.22009516 |
| 15485 | NR_026787.1 | 13.4764031 | 8.67789002 | 2.69493216 | 3.22007736 |
| 15486 | NM_005736.3 | 618.754657 | -4.0406349 | 1.25501409 | -3.2195933 |
| 15487 | NM_001256512.1 | 10.8259359 | -5.0639066 | 1.57293932 | -3.2193909 |
| 15488 | NM_207317.2 | 14.8260581 | -5.5167681 | 1.71379465 | -3.2190368 |
| 15489 | NM_001270896.1 | 11.1547714 | -5.0942826 | 1.58262154 | -3.2188887 |
| 15490 | NM_001276351.1 | 4.12838937 | 6.28268101 | 1.95181969 | 3.21888391 |
| 15491 | NM_001134745.2 | 13.4156202 | -4.3288299 | 1.34515053 | -3.2181007 |
| 15492 | NM_001201575.1 | 15.067885 | 8.83143294 | 2.74462811 | 3.21771569 |
| 15493 | NM_001243403.1 | 7.87709782 | -4.5929988 | 1.42749895 | -3.2175146 |
| 15494 | NM_006330.3 | 93.3971983 | -3.836309 | 1.19235519 | -3.2174213 |
| 15495 | NR_103810.1 | 23.7620818 | 2.64496251 | 0.8222402 | 3.21677597 |
| 15496 | NM_001330443.1 | 77.651464 | -7.9011327 | 2.45666688 | -3.2162003 |
| 15497 | NM_001348097.1 | 94.3273613 | -8.1814694 | 2.54416015 | -3.215784 |
| 15498 | NM_001164098.1 | 81.2838012 | -7.9671989 | 2.47770734 | -3.2155528 |
| 15499 | NM_024106.2 | 75.4025367 | -7.8586107 | 2.44423368 | -3.2151634 |
| 15500 | NR_125824.1 | 11.3653451 | 6.09252627 | 1.89497292 | 3.21509939 |
| 15501 | NM_001271788.1 | 72.9910999 | -7.8114366 | 2.42975738 | -3.2149039 |
| 15502 | NM_001284335.1 | 12.7680173 | -5.3045575 | 1.6500238 | -3.214837 |
| 15503 | NM_001100.3 | 10.2663378 | -4.9855796 | 1.55091175 | -3.214612 |
| 15504 | NM_001322886.1 | 83.099969 | -7.9991745 | 2.48853114 | -3.2144161 |
| 15505 | NM_001324058.1 | 12.4237541 | -4.2558525 | 1.32409723 | -3.214154 |
| 15506 | NM_004734.4 | 12.7989916 | -5.2999494 | 1.6495238 | -3.2130178 |
| 15507 | NM_001037132.2 | 30.2366579 | -5.5605737 | 1.73098481 | -3.2123758 |
| 15508 | NM_006387.5 | 302.482984 | -4.5436534 | 1.41444085 | -3.2123319 |
| 15509 | NM_001284260.1 | 9.38814238 | -4.8484475 | 1.50961089 | -3.21172 |
| 15510 | NM_006954.1 | 74.9142095 | -7.8492573 | 2.4442839 | -3.2112707 |
| 15511 | NM_001302371.1 | 75.0159134 | -6.8774374 | 2.14168428 | -3.2112284 |
| 15512 | NM_003757.3 | 387.382312 | -3.2121731 | 1.00037694 | -3.2109628 |
| 15513 | NM_004558.4 | 7.79449747 | -4.5932472 | 1.43051172 | -3.210912 |
| 15514 | NM_020526.4 | 17.1308603 | -3.8979205 | 1.21397884 | -3.2108636 |
| 15515 | NR_136700.1 | 10.3204398 | -4.9968314 | 1.55630957 | -3.2106925 |
| 15516 | NM_153618.1 | 9.04683318 | -4.8125016 | 1.49902592 | -3.2104192 |
| 15517 | NR_046722.1 | 24.6275937 | 5.40739859 | 1.68458781 | 3.20992385 |
| 15518 | NM_001199787.1 | 23.5805999 | -2.3817323 | 0.74202016 | -3.2097946 |
| 15519 | NM_001012456.1 | 9.98224198 | -4.9460269 | 1.54105405 | -3.209509 |
| 15520 | NR_144687.1 | 7.88528458 | -4.5894778 | 1.43005182 | -3.2093087 |
| 15521 | NR_144442.1 | 8.72692865 | -4.7590091 | 1.48290183 | -3.2092543 |
| 15522 | NM_001170460.1 | 14.040487 | -5.4407341 | 1.69560092 | -3.2087351 |

| | | | | | |
|---|---|---|---|---|---|
| 15523 | NM_001324216.1 | 12.2338579 | -5.2370842 | 1.6325338 | -3.2079484 |
| 15524 | NM_018933.3 | 8.99126242 | -4.7893655 | 1.49345883 | -3.2068949 |
| 15525 | NM_001248007.1 | 8.06183572 | -4.6348724 | 1.44565888 | -3.2060623 |
| 15526 | NM_001321949.1 | 9.08075991 | -4.7984225 | 1.49684072 | -3.2057002 |
| 15527 | NR_037888.1 | 7.02605193 | -4.4235322 | 1.37996799 | -3.2055325 |
| 15528 | NM_206808.3 | 9.30025459 | -4.8489463 | 1.51269332 | -3.2055052 |
| 15529 | NM_001322502.1 | 10.6166136 | -5.0282978 | 1.56875733 | -3.2052745 |
| 15530 | NR_036530.1 | 8.61879993 | -4.7209344 | 1.47315959 | -3.204632 |
| 15531 | NR_036510.1 | 10.9241611 | -5.0803391 | 1.58537674 | -3.2044996 |
| 15532 | NM_001130701.1 | 17.819415 | 6.16148168 | 1.92304957 | 3.20401604 |
| 15533 | NM_016203.3 | 9.28846809 | -4.8283096 | 1.50700896 | -3.2039024 |
| 15534 | NM_001145335.1 | 6.81949696 | -4.3940099 | 1.37149714 | -3.2038054 |
| 15535 | NM_016824.4 | 58.2915392 | -7.4880911 | 2.33741033 | -3.2035843 |
| 15536 | NM_152682.3 | 56.5543827 | -7.4444151 | 2.3237847 | -3.2035735 |
| 15537 | NM_018299.4 | 60.0758366 | -7.5314674 | 2.35101349 | -3.2034982 |
| 15538 | NM_018429.2 | 156.013184 | -4.1471663 | 1.29463975 | -3.2033361 |
| 15539 | NM_198524.2 | 11.3163587 | -5.1249771 | 1.59989373 | -3.2033235 |
| 15540 | NM_014865.3 | 218.922639 | -3.4453801 | 1.07558931 | -3.2032488 |
| 15541 | NM_024738.2 | 382.08353 | -2.1900296 | 0.68370065 | -3.2031995 |
| 15542 | NM_024798.2 | 144.47336 | 1.83291468 | 0.57231744 | 3.20261895 |
| 15543 | NM_001099335.1 | 8.59849199 | -4.7346421 | 1.47868258 | -3.2019327 |
| 15544 | NM_001134473.2 | 154.373676 | -7.2154913 | 2.25355068 | -3.2018323 |
| 15545 | NM_001079522.1 | 76.5533663 | -7.8803246 | 2.46135757 | -3.2016171 |
| 15546 | NM_001142482.1 | 62.4214313 | -7.5866453 | 2.37005777 | -3.2010382 |
| 15547 | NM_014941.3 | 12.3875278 | -4.2408384 | 1.32505421 | -3.2005018 |
| 15548 | NM_138422.3 | 8.06454333 | -4.6338016 | 1.44793252 | -3.2002884 |
| 15549 | NM_014809.3 | 11.7430198 | -5.1761049 | 1.61751077 | -3.2000435 |
| 15550 | NR_000011.1 | 8.71006234 | -4.7505337 | 1.4846322 | -3.1998051 |
| 15551 | NM_198484.4 | 73.2684544 | -7.817118 | 2.44316444 | -3.1995874 |
| 15552 | NM_199285.2 | 11.0083364 | -5.0796761 | 1.58759901 | -3.1995964 |
| 15553 | NM_032302.3 | 36.8296475 | -3.5120913 | 1.09768266 | -3.1995507 |
| 15554 | NR_073058.1 | 26.5443581 | 2.28481687 | 0.7142766 | 3.19878444 |
| 15555 | NM_001164496.1 | 52.3189439 | -5.0801134 | 1.58818752 | -3.1986861 |
| 15556 | NM_181838.1 | 8.18592357 | -4.6590439 | 1.45677646 | -3.1981873 |
| 15557 | NM_018374.3 | 15.7924451 | -5.5960342 | 1.74978332 | -3.1981298 |
| 15558 | NM_001142501.1 | 11.2595846 | -5.1085657 | 1.59784796 | -3.1971538 |
| 15559 | NM_170607.2 | 7.4633217 | -4.5168772 | 1.41290931 | -3.1968628 |
| 15560 | NM_032261.4 | 34.8374824 | -3.4229694 | 1.0707375 | -3.1968334 |
| 15561 | NM_001243770.1 | 7.45590126 | -4.5175632 | 1.41315305 | -3.1967969 |
| 15562 | NM_015232.1 | 9.62591854 | -4.8920902 | 1.5303138 | -3.1967889 |
| 15563 | NM_006841.5 | 16.0586635 | -3.1941451 | 0.9994143 | -3.1960171 |
| 15564 | NM_001171163.1 | 9.34620488 | -4.8476215 | 1.51714827 | -3.1952194 |
| 15565 | NM_177959.2 | 30.7261498 | -4.2478806 | 1.32957755 | -3.19491 |

| | | | | | |
|---|---|---|---|---|---|
| 15566 | NR_073609.1 | 17.5858819 | -3.3527255 | 1.04940321 | -3.1948878 |
| 15567 | NM_001101338.1 | 15.0739501 | -4.5376072 | 1.42046334 | -3.1944557 |
| 15568 | NM_001330131.1 | 7.16615901 | -4.4644856 | 1.39783927 | -3.1938476 |
| 15569 | NR_144394.1 | 13.0794512 | -5.3320881 | 1.66956693 | -3.1936953 |
| 15570 | NR_027508.1 | 20.9233023 | 4.79303042 | 1.50080796 | 3.19363339 |
| 15571 | NR_110331.1 | 9.96665829 | -4.9407005 | 1.54710742 | -3.1935084 |
| 15572 | NM_198240.1 | 70.6749229 | -6.7961893 | 2.12850639 | -3.1929381 |
| 15573 | NM_033290.3 | 53.2445187 | -7.3574496 | 2.30464529 | -3.1924434 |
| 15574 | NM_015465.4 | 53.7365892 | -7.3707267 | 2.3092437 | -3.1918358 |
| 15575 | NM_001305564.1 | 7.10122358 | -4.4510598 | 1.39461277 | -3.1916098 |
| 15576 | NR_034125.1 | 50.4592587 | 2.36073411 | 0.73967505 | 3.19158273 |
| 15577 | NM_014797.2 | 62.8221204 | -1.3762997 | 0.43124766 | -3.191437 |
| 15578 | NM_018166.2 | 9.55421065 | -4.8862492 | 1.53105734 | -3.1914214 |
| 15579 | NM_001177358.1 | 10.1026909 | -4.9647292 | 1.55568571 | -3.1913446 |
| 15580 | NR_038941.1 | 75.7185424 | 2.14830153 | 0.67324994 | 3.19094204 |
| 15581 | NM_016227.3 | 7.90494912 | -4.6051861 | 1.44327838 | -3.1907816 |
| 15582 | NM_020354.3 | 88.6819965 | -3.2882043 | 1.03062141 | -3.1905065 |
| 15583 | NM_001321518.1 | 11.3383127 | -5.1230096 | 1.60573168 | -3.1904518 |
| 15584 | NM_001146155.1 | 11.4184555 | -5.138673 | 1.6109436 | -3.1898528 |
| 15585 | NM_001307929.1 | 9.85653971 | -4.9268823 | 1.54475481 | -3.1894267 |
| 15586 | NM_001320572.1 | 74.1787363 | -7.8355409 | 2.45696179 | -3.1891179 |
| 15587 | NM_007280.1 | 9.57810173 | -4.8841662 | 1.53155072 | -3.1890332 |
| 15588 | NM_001136129.2 | 100.038254 | -8.2668702 | 2.59233998 | -3.1889607 |
| 15589 | NM_207380.2 | 23.9178522 | -3.4256089 | 1.07435468 | -3.188527 |
| 15590 | NM_020345.3 | 60.6139574 | -3.1944326 | 1.00189881 | -3.1883785 |
| 15591 | NM_130391.3 | 81.7520765 | -7.9749786 | 2.50132149 | -3.1883061 |
| 15592 | NM_001290051.1 | 11.4055527 | -5.1217252 | 1.60651477 | -3.1880972 |
| 15593 | NM_001282206.1 | 12.7746467 | -5.2960594 | 1.66129229 | -3.1879155 |
| 15594 | NR_037884.1 | 12.45606 | -4.2554308 | 1.33506554 | -3.1874321 |
| 15595 | NR_110650.1 | 22.6794744 | 4.44186373 | 1.3935728 | 3.18739269 |
| 15596 | NM_001278494.1 | 8.04362383 | -4.6146372 | 1.44804872 | -3.1867969 |
| 15597 | NM_001207012.1 | 9.63984083 | -4.8770073 | 1.53040972 | -3.1867331 |
| 15598 | NM_001320034.1 | 57.1470032 | -7.459402 | 2.3408466 | -3.1866257 |
| 15599 | NM_006264.2 | 57.408659 | -7.4660028 | 2.34299859 | -3.1865161 |
| 15600 | NR_028504.1 | 8.55951184 | -4.723673 | 1.48243397 | -3.1864306 |
| 15601 | NM_015250.3 | 54.3158798 | -3.709056 | 1.16407044 | -3.1862814 |
| 15602 | NM_000234.2 | 54.131227 | -7.3812523 | 2.31669893 | -3.1861077 |
| 15603 | NR_033339.1 | 62.2540218 | -3.1178182 | 0.97887687 | -3.1850974 |
| 15604 | NM_199246.1 | 75.0023013 | -6.8772982 | 2.15925677 | -3.1850303 |
| 15605 | NM_001190265.1 | 7.85430679 | -4.5939771 | 1.44250706 | -3.1847172 |
| 15606 | NR_110825.1 | 50.3937868 | 2.24305005 | 0.70438124 | 3.18442614 |
| 15607 | NM_001079535.1 | 9.85997397 | -4.9189292 | 1.5448206 | -3.1841427 |
| 15608 | NM_003280.2 | 7.86901657 | -4.6087765 | 1.44748759 | -3.1839834 |

| | | | | | |
|---|---|---|---|---|---|
| 15609 | NM_001330282.1 | 12.3263965 | -5.2448671 | 1.64732699 | -3.1838652 |
| 15610 | NM_001297671.1 | 53.8661206 | -7.3741758 | 2.31613195 | -3.1838323 |
| 15611 | NM_021952.4 | 15.9234597 | -5.6135978 | 1.76322681 | -3.1837072 |
| 15612 | NM_001278711.1 | 9.27134947 | -4.8265599 | 1.51623061 | -3.1832624 |
| 15613 | NM_004912.3 | 15.5392025 | -5.5810712 | 1.75330607 | -3.18317 |
| 15614 | NM_002111.8 | 300.433471 | -3.281454 | 1.0309187 | -3.1830386 |
| 15615 | NM_153681.2 | 7.47627173 | -4.5198275 | 1.42001477 | -3.182944 |
| 15616 | NM_181466.2 | 8.56296469 | -4.7191369 | 1.48278185 | -3.1826239 |
| 15617 | NM_013321.3 | 93.952035 | -3.4117178 | 1.07204757 | -3.1824313 |
| 15618 | NR_033870.2 | 19.2366759 | 2.6608154 | 0.83611322 | 3.18236257 |
| 15619 | NR_130740.1 | 12.5960552 | -4.2415895 | 1.33312577 | -3.1816874 |
| 15620 | NM_001143760.1 | 15.4816584 | -5.5774638 | 1.75304728 | -3.1815821 |
| 15621 | NM_006961.3 | 11.7465844 | -4.1271697 | 1.29721769 | -3.1815552 |
| 15622 | NM_005544.2 | 178.179219 | -3.5519348 | 1.11660057 | -3.1810255 |
| 15623 | NM_001942.3 | 14.0090709 | 8.73318246 | 2.74559189 | 3.18080138 |
| 15624 | NR_073409.1 | 7.85818236 | -4.6095496 | 1.44926612 | -3.1806095 |
| 15625 | NM_015292.2 | 244.790104 | -2.8148337 | 0.88506389 | -3.1803734 |
| 15626 | NR_023921.2 | 10.5048445 | -5.0060755 | 1.57428726 | -3.1798996 |
| 15627 | NM_001178087.1 | 56.9937613 | -7.4555714 | 2.34491457 | -3.179464 |
| 15628 | NM_005826.4 | 227.157366 | -4.5875712 | 1.4429683 | -3.1792599 |
| 15629 | NM_006393.2 | 16.0730382 | -3.8012274 | 1.19573719 | -3.1789823 |
| 15630 | NM_001321380.1 | 65.176223 | -1.9603886 | 0.61670024 | -3.1788356 |
| 15631 | NR_046757.1 | 9.20301973 | -4.8185757 | 1.51592242 | -3.1786427 |
| 15632 | NM_014214.2 | 22.8342481 | -3.4166819 | 1.0749608 | -3.1784246 |
| 15633 | NM_203406.1 | 43.4780112 | -3.8134742 | 1.19983448 | -3.1783336 |
| 15634 | NM_001320884.1 | 40.8037001 | -5.2399339 | 1.64884157 | -3.1779487 |
| 15635 | NR_036519.1 | 7.61852029 | -4.5578906 | 1.43428476 | -3.1778143 |
| 15636 | NM_003475.3 | 12.4076331 | -5.2522989 | 1.65303212 | -3.1773725 |
| 15637 | NM_152263.3 | 205.391595 | 2.0883717 | 0.65726375 | 3.1773724 |
| 15638 | NR_038341.1 | 18.0484744 | 2.87674653 | 0.90542774 | 3.17722377 |
| 15639 | NM_001105521.3 | 12.3848539 | -5.2558351 | 1.65441504 | -3.176854 |
| 15640 | NM_032369.3 | 10.7038721 | -5.0484374 | 1.58912292 | -3.1768703 |
| 15641 | NM_001306158.1 | 57.7083771 | -7.4735653 | 2.35260717 | -3.1767162 |
| 15642 | NM_001548.4 | 42.0970403 | -2.9393381 | 0.92529631 | -3.1766452 |
| 15643 | NM_152445.2 | 72.8740213 | -2.898862 | 0.91258297 | -3.1765462 |
| 15644 | NM_001330205.1 | 7.95384217 | -4.6214838 | 1.45496951 | -3.1763441 |
| 15645 | NM_015517.4 | 17.3924582 | -5.7445065 | 1.80873297 | -3.1759837 |
| 15646 | NM_018376.3 | 8.26716632 | -4.6614408 | 1.46778427 | -3.1758351 |
| 15647 | NM_001282383.1 | 11.2462478 | -5.1011558 | 1.60630835 | -3.1757015 |
| 15648 | NM_001164755.1 | 11.4174626 | -5.14522 | 1.62039578 | -3.175286 |
| 15649 | NM_015133.4 | 86.402382 | -8.054552 | 2.53672345 | -3.1751794 |
| 15650 | NR_034011.1 | 9.57758352 | -4.8825552 | 1.53775645 | -3.1751161 |
| 15651 | NM_001177802.1 | 14.1490777 | -5.4404076 | 1.71365509 | -3.174739 |

| | | | | | |
|---|---|---|---|---|---|
| 15652 | NM_001278342.1 | 16.4012118 | -5.6552459 | 1.78136038 | -3.1746781 |
| 15653 | NM_001079685.2 | 69.834255 | -7.7478998 | 2.44091876 | -3.1741736 |
| 15654 | NM_001558.3 | 11.60023 | -5.1547608 | 1.62396476 | -3.1741827 |
| 15655 | NM_001104546.1 | 26.2479132 | -3.128574 | 0.98572794 | -3.1738717 |
| 15656 | NM_052901.4 | 26.4236794 | -3.2461181 | 1.02281579 | -3.1737075 |
| 15657 | NM_001304724.1 | 9.72327851 | -4.888546 | 1.54039262 | -3.1735714 |
| 15658 | NM_001305142.1 | 13.695337 | -5.3875634 | 1.69762603 | -3.1735867 |
| 15659 | NR_037155.2 | 7.53236047 | -4.5454902 | 1.43241392 | -3.1733077 |
| 15660 | NM_015507.3 | 10.7261923 | -5.0489678 | 1.59111122 | -3.1732337 |
| 15661 | NM_001080477.3 | 321.034343 | -4.7257429 | 1.4894432 | -3.1728252 |
| 15662 | NM_001329686.1 | 9.66841067 | -4.893408 | 1.54262365 | -3.1721334 |
| 15663 | NM_001330245.1 | 63.617711 | -7.6135969 | 2.40018533 | -3.1720871 |
| 15664 | NM_001112741.1 | 9.98878829 | -4.9428341 | 1.55834791 | -3.1718425 |
| 15665 | NM_001779.2 | 8.75469293 | -4.7491056 | 1.49780442 | -3.1707114 |
| 15666 | NM_001279352.1 | 15.2875457 | 5.54420748 | 1.74871233 | 3.17045142 |
| 15667 | NM_001289068.1 | 11.9608477 | -5.1971534 | 1.63928794 | -3.1703725 |
| 15668 | NM_001258217.1 | 13.051943 | -5.3330685 | 1.68216821 | -3.1703539 |
| 15669 | NM_014508.2 | 8.85220008 | -4.769989 | 1.50469766 | -3.1700647 |
| 15670 | NM_203351.1 | 12.7663631 | -5.2961981 | 1.67080958 | -3.1698394 |
| 15671 | NM_138430.4 | 7.0007071 | -4.4372946 | 1.39991207 | -3.1696952 |
| 15672 | NR_102424.1 | 40.8330957 | 1.68702792 | 0.53224692 | 3.16963397 |
| 15673 | NM_181456.2 | 12.4417499 | -5.2514129 | 1.65686663 | -3.1694844 |
| 15674 | NM_001031617.2 | 97.9374011 | -2.5669891 | 0.8099552 | -3.1692976 |
| 15675 | NM_052813.4 | 9.24581064 | -4.8263506 | 1.52294644 | -3.1690876 |
| 15676 | NR_135120.1 | 8.67814863 | -4.735755 | 1.49438599 | -3.1690306 |
| 15677 | NM_001270428.1 | 10.2544445 | -4.974144 | 1.56979332 | -3.1686617 |
| 15678 | NM_001346818.1 | 72.1464337 | 4.92898385 | 1.55563275 | 3.16847523 |
| 15679 | NM_001317960.1 | 12.1076351 | -5.2151972 | 1.64609972 | -3.1682146 |
| 15680 | NR_038133.1 | 15.1401389 | -5.5434879 | 1.74985152 | -3.1679762 |
| 15681 | NM_017986.3 | 33.1422715 | 1.99849275 | 0.63087907 | 3.16779052 |
| 15682 | NM_001010908.1 | 8.68841634 | -4.7380575 | 1.4957612 | -3.1676564 |
| 15683 | NM_144578.3 | 332.529077 | -2.0221985 | 0.6384426 | -3.1673928 |
| 15684 | NM_001261395.1 | 15.7899309 | 8.89449299 | 2.80837925 | 3.16712673 |
| 15685 | NM_001322288.1 | 10.2853773 | -4.9943871 | 1.57712095 | -3.1667749 |
| 15686 | NM_000201.2 | 47.0146665 | -3.4785841 | 1.09855006 | -3.166523 |
| 15687 | NM_033119.4 | 51.1880484 | -2.3957718 | 0.75673293 | -3.165941 |
| 15688 | NM_001083620.1 | 94.607231 | -4.2485091 | 1.34210807 | -3.1655492 |
| 15689 | NR_027455.5 | 32.6997833 | -4.5329449 | 1.43215809 | -3.1651149 |
| 15690 | NM_006743.4 | 578.090504 | -3.065608 | 0.96863945 | -3.1648598 |
| 15691 | NM_001015051.3 | 9.20052028 | -4.8232042 | 1.52410036 | -3.1646237 |
| 15692 | NM_001319044.1 | 14.4455281 | 8.77479542 | 2.77321761 | 3.16412076 |
| 15693 | NM_001282167.1 | 8.17554718 | -4.6538619 | 1.47092062 | -3.163911 |
| 15694 | NM_002552.4 | 51.9498054 | -7.3219715 | 2.31420311 | -3.1639278 |

| | | | | | |
|---|---|---|---|---|---|
| 15695 | NM_001127364.2 | 10.9279052 | -5.0691724 | 1.60229724 | -3.1636904 |
| 15696 | NM_134268.4 | 35.2755225 | -3.5299416 | 1.11576728 | -3.1636898 |
| 15697 | NM_001346102.1 | 7.90441867 | -4.5992241 | 1.45390536 | -3.1633587 |
| 15698 | NM_024053.4 | 11.1839567 | -5.1044049 | 1.61365084 | -3.1632648 |
| 15699 | NM_004421.2 | 127.468907 | -4.0127064 | 1.26861322 | -3.1630652 |
| 15700 | NM_080672.4 | 17.569419 | -4.7753627 | 1.50975155 | -3.1630123 |
| 15701 | NM_001080392.1 | 129.36109 | -4.2603336 | 1.34719756 | -3.1623674 |
| 15702 | NM_145341.3 | 7.1937765 | -4.4711024 | 1.4138441 | -3.162373 |
| 15703 | NM_001101387.1 | 8.05167861 | -4.6335743 | 1.4653055 | -3.1621899 |
| 15704 | NM_032121.5 | 187.839334 | -1.5652344 | 0.49501483 | -3.161995 |
| 15705 | NM_080604.2 | 12.2436148 | -5.2286433 | 1.65358761 | -3.1619995 |
| 15706 | NM_001330475.1 | 26.0235201 | -3.5814435 | 1.1327818 | -3.1616358 |
| 15707 | NM_001301774.1 | 12.3131178 | 8.54816716 | 2.70381173 | 3.16152455 |
| 15708 | NR_024373.2 | 9.7421389 | -4.9101017 | 1.55308953 | -3.1615059 |
| 15709 | NM_001271959.1 | 10.9240751 | -5.077211 | 1.60595716 | -3.1614859 |
| 15710 | NR_028330.1 | 26.0240216 | -3.2237421 | 1.0197173 | -3.1614077 |
| 15711 | NM_001199699.1 | 45.3733681 | -2.8903154 | 0.91427595 | -3.1613162 |
| 15712 | NM_015150.1 | 151.166333 | -4.3040463 | 1.36155555 | -3.1611242 |
| 15713 | NM_080702.2 | 94.1722086 | -8.1788739 | 2.58733464 | -3.1611195 |
| 15714 | NM_001146016.1 | 17.3556048 | -5.7367665 | 1.81483314 | -3.1610435 |
| 15715 | NM_001198961.1 | 7.74413814 | -4.57288 | 1.44693145 | -3.1603985 |
| 15716 | NM_001320444.1 | 8.25952342 | -4.6655895 | 1.47631472 | -3.1602946 |
| 15717 | NM_015226.2 | 155.072369 | -3.2167384 | 1.01788781 | -3.1602091 |
| 15718 | NM_031861.2 | 18.970975 | -4.0495419 | 1.28145581 | -3.1601104 |
| 15719 | NM_001323329.1 | 71.1843346 | -7.7752418 | 2.46048898 | -3.1600393 |
| 15720 | NM_021902.3 | 13.1174419 | -5.3410915 | 1.69020086 | -3.1600336 |
| 15721 | NM_024642.4 | 26.7755493 | -4.6022137 | 1.45638546 | -3.1600245 |
| 15722 | NM_000104.3 | 80.3512754 | -3.5282145 | 1.11681847 | -3.1591656 |
| 15723 | NR_038915.1 | 8.2258049 | -4.6582947 | 1.47455095 | -3.1591277 |
| 15724 | NM_002503.4 | 10.6441851 | -5.0341788 | 1.59367784 | -3.1588434 |
| 15725 | NM_001287507.1 | 7.17600835 | -4.4685999 | 1.41479105 | -3.1584875 |
| 15726 | NM_005204.3 | 11.6434703 | -5.1708969 | 1.63714373 | -3.1584868 |
| 15727 | NM_001043229.1 | 7.77704218 | -4.5770295 | 1.44918526 | -3.1583467 |
| 15728 | NM_003222.3 | 26.7977609 | -4.4086761 | 1.39597181 | -3.1581412 |
| 15729 | NM_001204832.2 | 8.58405333 | -4.7146991 | 1.49302129 | -3.1578244 |
| 15730 | NM_001129840.1 | 68.6102863 | -7.7225395 | 2.44563581 | -3.1576817 |
| 15731 | NM_001258379.1 | 63.6586238 | -7.6145767 | 2.41148996 | -3.1576232 |
| 15732 | NM_178450.4 | 30.0670576 | -3.7728736 | 1.19484077 | -3.1576372 |
| 15733 | NM_001039457.2 | 17.3623092 | -5.7435402 | 1.81896304 | -3.1575904 |
| 15734 | NM_004056.5 | 8.7339779 | -4.7507132 | 1.50459769 | -3.1574641 |
| 15735 | NR_040085.1 | 27.0454236 | 1.55136281 | 0.49147022 | 3.15657539 |
| 15736 | NM_014567.3 | 52.1033953 | -7.3261347 | 2.32099297 | -3.1564657 |
| 15737 | NR_024371.1 | 14.5213928 | -5.4860455 | 1.73828565 | -3.1560092 |

| | | | | | |
|---|---|---|---|---|---|
| 15738 | NM_000076.2 | 13.6616076 | -5.4012972 | 1.71162253 | -3.1556591 |
| 15739 | NM_015511.4 | 17.2422089 | -5.7282173 | 1.81521076 | -3.1556762 |
| 15740 | NM_001317930.1 | 10.9035653 | -5.0735026 | 1.60827392 | -3.1546259 |
| 15741 | NM_004993.5 | 51.4684747 | -7.3084571 | 2.31694643 | -3.1543488 |
| 15742 | NM_001145196.1 | 6.78547905 | 6.93705872 | 2.19969676 | 3.15364319 |
| 15743 | NM_001145290.1 | 94.7687974 | 2.30656514 | 0.73149505 | 3.15322045 |
| 15744 | NM_144683.3 | 78.6777041 | -3.8197343 | 1.21137018 | -3.1532345 |
| 15745 | NM_001127598.2 | 11.591251 | -5.1560225 | 1.63523529 | -3.1530768 |
| 15746 | NM_001323387.1 | 17.5284862 | -5.7510659 | 1.82424102 | -3.1525801 |
| 15747 | NM_001330614.1 | 69.186319 | -7.7346709 | 2.4535157 | -3.1524848 |
| 15748 | NR_015410.1 | 8.67030112 | -4.7319875 | 1.50126233 | -3.1520058 |
| 15749 | NR_038926.1 | 11.2688471 | -5.1126837 | 1.62207495 | -3.1519405 |
| 15750 | NM_001330572.1 | 8.16556755 | -4.6348562 | 1.4705163 | -3.1518564 |
| 15751 | NM_001329917.1 | 456.949081 | 2.27375654 | 0.72146556 | 3.15158014 |
| 15752 | NM_001113561.2 | 18.7227371 | -3.450543 | 1.09514324 | -3.1507687 |
| 15753 | NM_205847.2 | 12.9978841 | -5.3284708 | 1.69120317 | -3.1506982 |
| 15754 | NM_020807.2 | 73.8681852 | -3.2356299 | 1.02702123 | -3.1504995 |
| 15755 | NR_122047.1 | 48.4138947 | 2.61469157 | 0.82993501 | 3.15047747 |
| 15756 | NM_001017415.1 | 9.51672554 | -4.8736449 | 1.54710739 | -3.1501659 |
| 15757 | NM_001278263.1 | 14.1934596 | 8.74134951 | 2.77509163 | 3.14993185 |
| 15758 | NM_001318104.1 | 7.49920521 | -4.5130895 | 1.4327524 | -3.1499438 |
| 15759 | NR_110751.1 | 32.9877297 | 2.24385628 | 0.71251698 | 3.1491969 |
| 15760 | NR_038458.1 | 10.744298 | -5.040051 | 1.60052309 | -3.1490024 |
| 15761 | NM_002867.3 | 426.131001 | -2.130915 | 0.67673414 | -3.1488215 |
| 15762 | NM_001130065.1 | 73.3958873 | -7.8204078 | 2.48375095 | -3.148628 |
| 15763 | NM_001281496.1 | 13.4317859 | -5.3730771 | 1.70651364 | -3.1485697 |
| 15764 | NR_033680.1 | 14.2700324 | 8.75452897 | 2.78055058 | 3.14848759 |
| 15765 | NM_001145848.1 | 49.5493896 | -7.2537911 | 2.30398255 | -3.1483707 |
| 15766 | NM_001329539.1 | 9.30183774 | -4.8265661 | 1.53327067 | -3.1478892 |
| 15767 | NM_001317782.1 | 10.2529947 | -4.983286 | 1.58320628 | -3.1475911 |
| 15768 | NM_001204857.1 | 8.35476699 | -4.6876122 | 1.4893032 | -3.1475204 |
| 15769 | NM_001321279.1 | 7.99463051 | -4.6265493 | 1.47006696 | -3.1471691 |
| 15770 | NM_001024960.2 | 78.9305749 | -7.9247524 | 2.51822125 | -3.1469643 |
| 15771 | NM_001130448.2 | 7.92351899 | -4.615039 | 1.46650004 | -3.146975 |
| 15772 | NM_001011645.2 | 13.6532629 | -4.3955413 | 1.39684725 | -3.1467588 |
| 15773 | NM_006901.3 | 90.9671492 | -3.1102334 | 0.98850838 | -3.1463905 |
| 15774 | NM_001320628.1 | 12.9908861 | -5.3125498 | 1.68848878 | -3.1463341 |
| 15775 | NM_005660.2 | 7.94818052 | -4.6153982 | 1.46694267 | -3.1462704 |
| 15776 | NM_001035267.1 | 177.227275 | -4.2638811 | 1.35524831 | -3.1461992 |
| 15777 | NM_020351.3 | 91.1998303 | -3.9218972 | 1.24661907 | -3.146027 |
| 15778 | NM_001329755.1 | 7.37222468 | -4.5110784 | 1.43397701 | -3.1458512 |
| 15779 | NR_030770.1 | 9.05503639 | -4.8099309 | 1.52902403 | -3.1457523 |
| 15780 | NR_103867.1 | 54.2237038 | -7.3836331 | 2.34727179 | -3.1456234 |

| | | | | | |
|---|---|---|---|---|---|
| 15781 | NM_025074.6 | 105.704079 | -3.9221833 | 1.24688721 | -3.1455799 |
| 15782 | NM_001033569.1 | 17.9374353 | -5.7859591 | 1.83991699 | -3.1446848 |
| 15783 | NR_045121.1 | 34.9467043 | -3.1977593 | 1.01709552 | -3.1440108 |
| 15784 | NM_006322.5 | 66.0997355 | -7.668611 | 2.43913562 | -3.1439871 |
| 15785 | NM_052903.4 | 74.9297478 | -2.6531146 | 0.84397162 | -3.1436064 |
| 15786 | NR_027420.1 | 43.83814 | 2.47328985 | 0.78678204 | 3.14355146 |
| 15787 | NM_030824.2 | 7.16998319 | -4.4533885 | 1.41676332 | -3.1433539 |
| 15788 | NM_001297669.1 | 9.17521835 | -4.8059366 | 1.52895054 | -3.1432911 |
| 15789 | NM_001272095.1 | 7.71388368 | -4.5824161 | 1.4579996 | -3.1429474 |
| 15790 | NR_024615.1 | 26.1853922 | 3.14730688 | 1.00157628 | 3.14235365 |
| 15791 | NM_001252079.1 | 8.07119539 | -4.6485039 | 1.4794415 | -3.1420667 |
| 15792 | NM_182975.3 | 6.81667219 | -4.4017521 | 1.40120202 | -3.1414115 |
| 15793 | NR_002318.2 | 6.66725661 | -4.3506173 | 1.38491673 | -3.1414288 |
| 15794 | NM_001083335.1 | 16.0429878 | -5.6286813 | 1.79179917 | -3.1413572 |
| 15795 | NM_001330539.1 | 8.2923114 | -4.677249 | 1.4890448 | -3.141107 |
| 15796 | NM_021116.2 | 311.656846 | -4.1378674 | 1.31732218 | -3.1411203 |
| 15797 | NR_033694.1 | 23.103412 | 3.41459782 | 1.08707972 | 3.14107397 |
| 15798 | NM_001281766.2 | 66.8144932 | -7.684019 | 2.44715662 | -3.1399784 |
| 15799 | NM_001301170.1 | 19.9215286 | -4.9319763 | 1.57070882 | -3.1399686 |
| 15800 | NR_134284.1 | 53.0764809 | -2.7850734 | 0.88699424 | -3.1399002 |
| 15801 | NM_001277961.1 | 9.47132701 | -4.8653087 | 1.54961055 | -3.1396977 |
| 15802 | NR_052852.1 | 101.229846 | -8.2839328 | 2.63888827 | -3.1391753 |
| 15803 | NM_001207065.1 | 16.438974 | 8.96456652 | 2.85579267 | 3.1390817 |
| 15804 | NM_152572.2 | 10.2881711 | -4.9920589 | 1.59035992 | -3.1389491 |
| 15805 | NM_001040712.2 | 55.1443375 | -7.40791 | 2.36034192 | -3.1384902 |
| 15806 | NR_015415.1 | 15.2929546 | -3.0308495 | 0.96605576 | -3.1373442 |
| 15807 | NM_003823.3 | 8.65852414 | -4.7340031 | 1.50920344 | -3.1367561 |
| 15808 | NM_001320698.1 | 7.45316482 | -4.5309026 | 1.44448732 | -3.1366856 |
| 15809 | NM_001313943.1 | 12.9468329 | -4.2838989 | 1.36582392 | -3.1364943 |
| 15810 | NM_005855.3 | 7.76347964 | -4.5828321 | 1.46122568 | -3.1362931 |
| 15811 | NM_001286042.1 | 15.1496024 | -5.5367951 | 1.76541118 | -3.1362638 |
| 15812 | NM_002851.2 | 308.278558 | -4.89707 | 1.56155501 | -3.1360215 |
| 15813 | NM_198474.3 | 13.7802914 | -3.5548603 | 1.13383376 | -3.1352571 |
| 15814 | NR_132119.1 | 42.0361041 | -2.1319588 | 0.68004447 | -3.1350285 |
| 15815 | NR_077247.1 | 6.75514219 | -4.3760588 | 1.39600137 | -3.1347096 |
| 15816 | NM_004969.3 | 47.8302537 | -7.2028367 | 2.29828198 | -3.1340091 |
| 15817 | NM_001288662.1 | 10.9310883 | -5.0768725 | 1.62009869 | -3.133681 |
| 15818 | NM_001265604.1 | 8.46115239 | -4.7162244 | 1.50505881 | -3.1335815 |
| 15819 | NR_125977.1 | 140.525581 | 2.47367194 | 0.78941372 | 3.13355582 |
| 15820 | NM_005609.3 | 15.460279 | -5.5773021 | 1.77994444 | -3.1334136 |
| 15821 | NR_003595.1 | 23.05448 | 3.92741695 | 1.25350089 | 3.13315848 |
| 15822 | NM_001324227.1 | 10.8448486 | -5.0623744 | 1.61618249 | -3.1323037 |
| 15823 | NM_001288965.1 | 12.2108795 | -5.2327694 | 1.67062607 | -3.1322206 |

| | | | | | |
|---|---|---|---|---|---|
| 15824 | NM_138463.3 | 6.53669094 | -4.339485 | 1.38560843 | -3.1318264 |
| 15825 | NR_002185.3 | 17.9727074 | 4.10260065 | 1.31006634 | 3.13159763 |
| 15826 | NM_015120.4 | 103.072614 | -3.3638656 | 1.07429815 | -3.1312216 |
| 15827 | NM_001077206.3 | 66.1807573 | -7.6702762 | 2.4497612 | -3.1310302 |
| 15828 | NM_013264.4 | 33.7812466 | -3.6511992 | 1.16616133 | -3.1309555 |
| 15829 | NM_001323511.1 | 90.908376 | -2.7898831 | 0.89116866 | -3.1305893 |
| 15830 | NM_006940.5 | 76.5564728 | -7.8803578 | 2.51722027 | -3.1305794 |
| 15831 | NM_024786.2 | 63.9061648 | -3.4906756 | 1.11504361 | -3.1305283 |
| 15832 | NM_001134233.1 | 11.5871513 | -4.0991936 | 1.30962555 | -3.1300502 |
| 15833 | NM_001293810.1 | 10.4398597 | -5.0073549 | 1.59995482 | -3.1296852 |
| 15834 | NM_020155.3 | 7.20059895 | -4.4825671 | 1.43244456 | -3.1293128 |
| 15835 | NM_213596.2 | 15.0946219 | -4.5405387 | 1.45127 | -3.1286657 |
| 15836 | NR_104101.1 | 6.36118154 | -4.2924702 | 1.37198154 | -3.1286647 |
| 15837 | NM_030962.3 | 191.32642 | -3.439303 | 1.0993096 | -3.1286028 |
| 15838 | NM_015485.4 | 12.0725129 | -5.2059806 | 1.66401109 | -3.1285733 |
| 15839 | NM_001243234.1 | 48.5346426 | -7.2238258 | 2.30901638 | -3.1285295 |
| 15840 | NM_001190329.1 | 7.83287145 | -4.5954378 | 1.46890687 | -3.1284746 |
| 15841 | NM_001323377.1 | 13.0372754 | 8.62458346 | 2.75697716 | 3.12827526 |
| 15842 | NM_018929.2 | 8.82430137 | -4.7607782 | 1.52202478 | -3.1279242 |
| 15843 | NM_002522.3 | 144.052973 | -4.5842666 | 1.46572565 | -3.1276431 |
| 15844 | NM_207315.3 | 9.08109733 | -4.8101034 | 1.53797324 | -3.1275599 |
| 15845 | NM_004758.3 | 72.2000892 | -7.7955108 | 2.49332839 | -3.126548 |
| 15846 | NM_001167959.1 | 10.6941064 | -5.0389607 | 1.61167939 | -3.126528 |
| 15847 | NR_110698.1 | 97.3067551 | 2.57520602 | 0.82370927 | 3.1263531 |
| 15848 | NM_032088.1 | 9.05797205 | -4.7928823 | 1.53308623 | -3.1262966 |
| 15849 | NM_001167890.1 | 12.5234374 | -5.2728625 | 1.68719906 | -3.1252166 |
| 15850 | NM_007271.3 | 77.2721257 | -3.7916279 | 1.2132831 | -3.1250974 |
| 15851 | NM_001001996.2 | 52.3609115 | -7.3333887 | 2.34666193 | -3.1250299 |
| 15852 | NM_004294.2 | 19.8113338 | -3.1102777 | 0.99529834 | -3.1249702 |
| 15853 | NM_001321230.1 | 7.81293085 | -4.5897897 | 1.46908909 | -3.1242419 |
| 15854 | NR_122073.1 | 9.61365052 | 2.45501575 | 0.7858126 | 3.12417457 |
| 15855 | NM_001320418.1 | 10.2978656 | -4.9749402 | 1.59272312 | -3.1235436 |
| 15856 | NM_207305.4 | 8.83443776 | -4.7644559 | 1.52541277 | -3.123388 |
| 15857 | NR_047662.2 | 108.962187 | 1.25926241 | 0.40330325 | 3.1223711 |
| 15858 | NR_111929.1 | 8.32963554 | -4.679354 | 1.49871862 | -3.1222365 |
| 15859 | NM_032370.2 | 7.98903825 | -4.6170342 | 1.478855 | -3.1220331 |
| 15860 | NR_024264.1 | 9.61741037 | -4.8897097 | 1.56632289 | -3.1217763 |
| 15861 | NM_181352.1 | 18.4996493 | -5.8352106 | 1.86927444 | -3.1216447 |
| 15862 | NM_198150.2 | 40.2337514 | -1.5821184 | 0.50683542 | -3.1215623 |
| 15863 | NM_002481.3 | 72.3131889 | -4.0095822 | 1.28463483 | -3.1211844 |
| 15864 | NM_001163315.2 | 92.3170242 | -2.6539478 | 0.85034867 | -3.1210113 |
| 15865 | NM_001025434.1 | 8.04911043 | -4.6267278 | 1.48270649 | -3.120461 |
| 15866 | NM_207510.3 | 11.4589037 | -5.1418347 | 1.64800778 | -3.1200306 |

| | | | | | |
|---|---|---|---|---|---|
| 15867 | NM_001082973.2 | 10.9655758 | -5.0821351 | 1.62903975 | -3.1197121 |
| 15868 | NM_001204453.1 | 6.9305438 | -4.4040755 | 1.41194885 | -3.1191466 |
| 15869 | NM_001303504.1 | 9.96991088 | -4.9432975 | 1.58481739 | -3.119159 |
| 15870 | NM_001146266.1 | 10.2237518 | -4.9749237 | 1.59509798 | -3.1188828 |
| 15871 | NM_000047.2 | 10.7699982 | -5.0563194 | 1.62127324 | -3.1187336 |
| 15872 | NM_001300949.1 | 260.959713 | 2.66957512 | 0.8559788 | 3.11873977 |
| 15873 | NM_001243425.1 | 7.80372802 | -4.5834296 | 1.46985914 | -3.1182781 |
| 15874 | NR_110748.1 | 16.2967616 | 8.17526909 | 2.62171455 | 3.11829108 |
| 15875 | NM_001135935.1 | 59.5729038 | -7.5190799 | 2.41161295 | -3.1178635 |
| 15876 | NM_001301189.1 | 12.8292912 | -5.3010447 | 1.70036915 | -3.1175846 |
| 15877 | NM_183243.2 | 12.6084446 | -5.2837969 | 1.69492729 | -3.1174181 |
| 15878 | NM_001142929.1 | 13.6004758 | -5.3856431 | 1.72804109 | -3.1166175 |
| 15879 | NM_001317112.1 | 3.70673868 | 6.79247639 | 2.17984217 | 3.11604045 |
| 15880 | NM_001282979.1 | 74.7766574 | -7.8469528 | 2.51896395 | -3.1151509 |
| 15881 | NM_145663.2 | 7.15150077 | -4.4592968 | 1.43153992 | -3.1150349 |
| 15882 | NM_001042595.2 | 6.32046626 | -4.2864373 | 1.37609772 | -3.1149222 |
| 15883 | NM_001312909.1 | 12.0930782 | -5.2225653 | 1.67694718 | -3.1143291 |
| 15884 | NM_001303098.1 | 11.0372295 | -5.0956672 | 1.63634096 | -3.114062 |
| 15885 | NM_015533.3 | 85.7572473 | -3.1988165 | 1.0272663 | -3.1139117 |
| 15886 | NM_001305102.1 | 52.138593 | -7.3272401 | 2.35343158 | -3.1134281 |
| 15887 | NM_001031709.2 | 6.58433411 | -4.3361654 | 1.39290128 | -3.1130458 |
| 15888 | NM_001010986.2 | 63.5736606 | -7.6125515 | 2.44595599 | -3.1123011 |
| 15889 | NM_001322376.1 | 61.6368969 | 5.27833906 | 1.69597016 | 3.11228297 |
| 15890 | NM_001004352.2 | 22.5075952 | 4.94615576 | 1.58950434 | 3.11175984 |
| 15891 | NM_001319120.1 | 47.8709517 | -7.2039156 | 2.31532973 | -3.1113994 |
| 15892 | NM_001300958.1 | 46.8498858 | -2.0963692 | 0.67381588 | -3.1111899 |
| 15893 | NM_001323324.1 | 47.6534698 | -7.1974248 | 2.31345021 | -3.1111216 |
| 15894 | NM_138322.3 | 48.7400218 | 6.97821123 | 2.2430015 | 3.11110413 |
| 15895 | NM_001010917.2 | 116.941696 | -4.3083959 | 1.38486644 | -3.1110552 |
| 15896 | NM_024620.3 | 115.788236 | -2.6633813 | 0.85615326 | -3.1108697 |
| 15897 | NR_110616.1 | 8.37802468 | -4.6827251 | 1.50530276 | -3.1108195 |
| 15898 | NM_012240.2 | 9.31013673 | -4.8366529 | 1.55489967 | -3.1105884 |
| 15899 | NM_001318521.1 | 8.87754477 | -4.7748527 | 1.53519288 | -3.1102624 |
| 15900 | NM_033408.3 | 7.88903392 | -4.5982632 | 1.47850885 | -3.1100681 |
| 15901 | NM_001035260.2 | 70.8815542 | 6.46411127 | 2.07865699 | 3.10975371 |
| 15902 | NM_001167820.1 | 8.49179365 | -4.6961533 | 1.51025929 | -3.1095013 |
| 15903 | NM_001198995.1 | 9.18893376 | -4.832167 | 1.55401116 | -3.1094802 |
| 15904 | NM_001345844.1 | 7.97665258 | -4.601256 | 1.48016983 | -3.1086001 |
| 15905 | NM_004428.2 | 10.8995275 | -5.0764968 | 1.63348964 | -3.1077618 |
| 15906 | NM_001170570.1 | 54.2389661 | 6.2551567 | 2.01276937 | 3.10773642 |
| 15907 | NM_001142588.1 | 9.18081239 | -4.8192356 | 1.55079765 | -3.1075851 |
| 15908 | NM_177990.2 | 72.0885939 | -3.4726723 | 1.11756976 | -3.1073427 |
| 15909 | NM_001329629.1 | 62.3343896 | -7.5840569 | 2.44078631 | -3.1072187 |

| | | | | | |
|---|---|---|---|---|---|
| 15910 | NM_001319960.1 | 64.6509237 | -7.6363488 | 2.45772239 | -3.1070835 |
| 15911 | NM_001009923.1 | 6.74086553 | -4.3752947 | 1.40834055 | -3.1067022 |
| 15912 | NM_001320824.1 | 11.9204356 | -5.1934055 | 1.67179559 | -3.1064835 |
| 15913 | NM_014400.2 | 7.34861895 | -4.4924569 | 1.44649796 | -3.1057471 |
| 15914 | NM_001130841.1 | 8.2407513 | -4.6680674 | 1.50306792 | -3.1056929 |
| 15915 | NR_028106.1 | 8.08352773 | -4.6445612 | 1.49564774 | -3.1053844 |
| 15916 | NM_001130414.1 | 101.499768 | -4.1155329 | 1.32535063 | -3.1052408 |
| 15917 | NM_001331020.1 | 21.4077195 | -3.5438626 | 1.14127707 | -3.1051729 |
| 15918 | NM_001718.5 | 6.58279918 | -4.349378 | 1.40087253 | -3.1047636 |
| 15919 | NM_001244949.1 | 58.5069852 | -7.492581 | 2.41335145 | -3.1046373 |
| 15920 | NM_001318485.1 | 7.0175891 | -4.4424575 | 1.43095755 | -3.1045348 |
| 15921 | NR_125352.1 | 7.98432034 | -4.625229 | 1.48992729 | -3.104332 |
| 15922 | NM_002871.4 | 64.3640826 | -2.6837113 | 0.86456139 | -3.1041305 |
| 15923 | NM_199248.2 | 9.36511819 | -4.834775 | 1.5575335 | -3.1041226 |
| 15924 | NM_002191.3 | 6.54706129 | -4.3334748 | 1.39629765 | -3.1035466 |
| 15925 | NM_015433.2 | 15.8539789 | -3.0501129 | 0.98283835 | -3.1033719 |
| 15926 | NM_022041.3 | 12.9993538 | -4.3105935 | 1.38909738 | -3.1031615 |
| 15927 | NM_001136196.1 | 192.431755 | -3.9163128 | 1.26209147 | -3.103034 |
| 15928 | NM_001128608.1 | 61.7567242 | -7.5706478 | 2.43989862 | -3.1028534 |
| 15929 | NM_001308422.1 | 9.67894111 | -4.9048296 | 1.58091328 | -3.1025292 |
| 15930 | NR_135852.1 | 12.8940876 | 8.61375672 | 2.77655321 | 3.10232006 |
| 15931 | NM_001184768.1 | 9.39237342 | -4.8467031 | 1.56242006 | -3.1020487 |
| 15932 | NM_005840.2 | 27.6373782 | -3.3247524 | 1.07184183 | -3.1019058 |
| 15933 | NM_001320264.1 | 12.9727329 | -5.3169907 | 1.7141512 | -3.1018213 |
| 15934 | NR_033992.2 | 7.02926359 | -4.4262726 | 1.42727216 | -3.1012113 |
| 15935 | NM_001282524.1 | 7.45157038 | -4.5202903 | 1.45767509 | -3.1010273 |
| 15936 | NM_001102668.2 | 9.09108736 | -4.817414 | 1.55352277 | -3.1009613 |
| 15937 | NM_006724.3 | 31.4175284 | -3.874206 | 1.2495467 | -3.1004891 |
| 15938 | NM_153605.3 | 26.274009 | -2.6864545 | 0.86649929 | -3.100354 |
| 15939 | NM_001450.3 | 12.6757975 | -5.2899866 | 1.70642618 | -3.1000383 |
| 15940 | NM_006729.4 | 7.96752364 | -4.6226513 | 1.49138284 | -3.0995739 |
| 15941 | NM_213566.1 | 269.775695 | 1.21764777 | 0.39287392 | 3.0993347 |
| 15942 | NR_103839.1 | 73.0025321 | 2.92029224 | 0.9423314 | 3.09900767 |
| 15943 | NM_145288.2 | 6.85496174 | -4.3883783 | 1.41607032 | -3.0989833 |
| 15944 | NM_033512.2 | 55.2394612 | -3.4024451 | 1.09814835 | -3.0983474 |
| 15945 | NM_021615.4 | 243.34663 | 1.80247373 | 0.58176743 | 3.098272 |
| 15946 | NM_001142291.2 | 46.8293314 | -7.1723966 | 2.31510836 | -3.0980824 |
| 15947 | NR_024004.1 | 6.62985866 | -4.3438081 | 1.40211043 | -3.0980499 |
| 15948 | NM_001193530.1 | 12.5969882 | 8.57065036 | 2.7667318 | 3.0977525 |
| 15949 | NM_001287188.1 | 69.2677465 | -7.7362079 | 2.49742922 | -3.0976685 |
| 15950 | NM_001256865.1 | 61.1438951 | -7.5560279 | 2.43931674 | -3.0976001 |
| 15951 | NR_023389.1 | 10.3332239 | -4.9899338 | 1.61128436 | -3.0968673 |
| 15952 | NM_133494.2 | 65.379432 | -3.5286736 | 1.13948989 | -3.0967134 |

| | | | | | |
|---|---|---|---|---|---|
| 15953 | NM_001033886.2 | 10.4312039 | -5.0018016 | 1.61542312 | -3.0962796 |
| 15954 | NR_104662.1 | 175.136468 | 2.68443339 | 0.86701365 | 3.09618353 |
| 15955 | NM_004268.4 | 118.112391 | -4.1257193 | 1.33264468 | -3.0958885 |
| 15956 | NM_001319300.1 | 11.6944483 | -5.1665023 | 1.66925969 | -3.095086 |
| 15957 | NM_024549.5 | 13.8380614 | -4.426664 | 1.43038391 | -3.0947384 |
| 15958 | NM_004863.3 | 253.741146 | -1.9633303 | 0.63449302 | -3.0943292 |
| 15959 | NM_031308.3 | 43.6936564 | -2.7903181 | 0.90178441 | -3.0942186 |
| 15960 | NM_001164386.1 | 51.1083215 | -7.298342 | 2.35891182 | -3.0939444 |
| 15961 | NM_001244766.1 | 12.587471 | -5.2743356 | 1.70472171 | -3.093957 |
| 15962 | NM_012482.4 | 53.6933395 | -7.3693423 | 2.38192364 | -3.0938617 |
| 15963 | NM_005288.3 | 122.856741 | -2.8073324 | 0.90747304 | -3.0935711 |
| 15964 | NM_030798.4 | 115.632536 | -3.863786 | 1.2491447 | -3.0931453 |
| 15965 | NM_000344.3 | 61.4901566 | -7.5640725 | 2.44545907 | -3.0931094 |
| 15966 | NM_005429.4 | 6.98716719 | -4.4373972 | 1.43507289 | -3.0921058 |
| 15967 | NM_001328638.1 | 6.80161784 | -4.3890088 | 1.41954075 | -3.0918512 |
| 15968 | NR_033344.1 | 6.5688016 | -4.3401637 | 1.4038267 | -3.0916663 |
| 15969 | NM_001281740.2 | 8.70890913 | -4.7480213 | 1.53576334 | -3.0916361 |
| 15970 | NM_001185012.1 | 7.13021599 | -4.4474726 | 1.43864356 | -3.0914347 |
| 15971 | NM_145113.2 | 10.999458 | -5.0804861 | 1.64361621 | -3.0910416 |
| 15972 | NM_001134848.1 | 21.7632567 | -2.502144 | 0.80951178 | -3.0909297 |
| 15973 | NM_153682.2 | 7.2664251 | -4.4880246 | 1.45244097 | -3.0899876 |
| 15974 | NM_031936.4 | 7.24088944 | -4.4796876 | 1.44979379 | -3.0898792 |
| 15975 | NM_174941.5 | 35.2538052 | 3.53694204 | 1.14474135 | 3.0897303 |
| 15976 | NM_001317102.1 | 12.5775394 | -5.271818 | 1.70681997 | -3.0886784 |
| 15977 | NM_002405.3 | 9.95133149 | -4.9224798 | 1.59374409 | -3.0886262 |
| 15978 | NR_036488.1 | 25.3051201 | -2.8731384 | 0.93030273 | -3.0883908 |
| 15979 | NM_004584.2 | 54.5476509 | -3.3832232 | 1.09568053 | -3.0877826 |
| 15980 | NM_001321612.1 | 248.445204 | -2.3140134 | 0.74942463 | -3.08772 |
| 15981 | NM_001243235.1 | 135.739001 | -6.9191152 | 2.24145791 | -3.0868816 |
| 15982 | NM_022160.2 | 8.79438411 | -4.768618 | 1.54481163 | -3.0868605 |
| 15983 | NM_001288747.1 | 7.22204263 | -4.4866578 | 1.45351417 | -3.0867658 |
| 15984 | NM_001278916.1 | 48.7691907 | -7.2309922 | 2.34315407 | -3.086008 |
| 15985 | NM_001300919.1 | 41.8019366 | 1.97419916 | 0.63973596 | 3.08595932 |
| 15986 | NR_015434.1 | 50.8414011 | -3.4301238 | 1.1115961 | -3.0857645 |
| 15987 | NR_038338.2 | 10.4330033 | -5.0019581 | 1.62122378 | -3.0852977 |
| 15988 | NM_001282116.1 | 112.802537 | -4.0592512 | 1.31571194 | -3.0852128 |
| 15989 | NM_012320.3 | 83.2540994 | -3.8926014 | 1.26191077 | -3.0846883 |
| 15990 | NM_153234.4 | 56.8793416 | -2.4983429 | 0.81002084 | -3.0842946 |
| 15991 | NM_001282958.1 | 69.1614474 | -7.7343918 | 2.50848867 | -3.0832875 |
| 15992 | NM_001322029.1 | 7.46470593 | -4.5353618 | 1.47117573 | -3.0828144 |
| 15993 | NM_145297.3 | 6.83654708 | -4.3909182 | 1.42431544 | -3.0828271 |
| 15994 | NM_001008800.2 | 63.8598477 | -7.6192559 | 2.47154298 | -3.0827932 |
| 15995 | NM_001282723.1 | 7.9371953 | -4.6035033 | 1.493499 | -3.0823612 |

| | | | | | |
|---|---|---|---|---|---|
| 15996 | NM_001293197.1 | 9.24610425 | -4.8250039 | 1.56557625 | -3.0819348 |
| 15997 | NM_001270891.1 | 6.45703143 | -4.3202054 | 1.40200523 | -3.0814474 |
| 15998 | NM_001282652.1 | 9.89205594 | -4.9218493 | 1.59737453 | -3.0812119 |
| 15999 | NR_003136.1 | 8.30176766 | -4.6794396 | 1.51873162 | -3.0811498 |
| 16000 | NM_173827.3 | 44.0145366 | -7.08302 | 2.29915187 | -3.0807099 |
| 16001 | NM_003452.3 | 11.8041411 | -5.1909338 | 1.68500829 | -3.0806577 |
| 16002 | NM_001190411.1 | 10.5197469 | -5.0220484 | 1.63038381 | -3.080286 |
| 16003 | NR_046870.1 | 14.1430458 | 8.73489823 | 2.83573882 | 3.0802901 |
| 16004 | NM_001289984.1 | 10.0545496 | -4.9576699 | 1.60976107 | -3.0797551 |
| 16005 | NM_017774.3 | 80.4980883 | -1.2701007 | 0.41240882 | -3.0797127 |
| 16006 | NM_145018.3 | 15.0736049 | -3.6918461 | 1.19905702 | -3.0789579 |
| 16007 | NR_034113.1 | 86.8786049 | 1.72686985 | 0.56093077 | 3.07857929 |
| 16008 | NM_001135217.1 | 11.9249638 | -5.2041347 | 1.6906068 | -3.0782644 |
| 16009 | NR_039674.1 | 9.48719852 | 4.35619539 | 1.41522396 | 3.07809614 |
| 16010 | NM_002477.1 | 24.836002 | -3.1477255 | 1.02282755 | -3.0774743 |
| 16011 | NM_005474.4 | 238.269743 | -4.6525367 | 1.51180449 | -3.0774724 |
| 16012 | NM_198488.3 | 58.3070366 | -2.778904 | 0.90299403 | -3.0774335 |
| 16013 | NM_181887.2 | 59.7750706 | -7.5233587 | 2.44479155 | -3.0773007 |
| 16014 | NM_001144869.2 | 7.89247954 | -4.5924707 | 1.4924537 | -3.0771278 |
| 16015 | NM_001184782.1 | 10.1212511 | -4.9500845 | 1.60892539 | -3.0766402 |
| 16016 | NM_001280801.1 | 17.657838 | -4.7437567 | 1.54201924 | -3.0763279 |
| 16017 | NM_025245.2 | 11.8584746 | -5.1837491 | 1.6852369 | -3.0759765 |
| 16018 | NM_001248.3 | 6.78746329 | -4.3860113 | 1.42626469 | -3.0751735 |
| 16019 | NM_001005855.1 | 6.25264338 | -4.2653726 | 1.38708094 | -3.0750712 |
| 16020 | NM_001145009.1 | 9.36916174 | -4.8467817 | 1.57614782 | -3.0750807 |
| 16021 | NR_024597.1 | 7.83286009 | -4.5861236 | 1.49160574 | -3.0746219 |
| 16022 | NM_207343.3 | 36.6208281 | -4.1135848 | 1.33801283 | -3.0743986 |
| 16023 | NM_001331009.1 | 54.5073449 | -7.3905188 | 2.40393327 | -3.0743444 |
| 16024 | NM_001159703.1 | 9.87656584 | -4.9358987 | 1.60566536 | -3.0740519 |
| 16025 | NM_003288.3 | 52.8591622 | -7.346252 | 2.38986084 | -3.0739246 |
| 16026 | NR_037583.1 | 7.79554494 | -4.5715955 | 1.48743659 | -3.0734725 |
| 16027 | NM_170676.4 | 63.262669 | -7.6049776 | 2.47440917 | -3.0734519 |
| 16028 | NM_001281293.1 | 57.7807521 | -7.4741756 | 2.43203385 | -3.0732202 |
| 16029 | NM_001017524.2 | 28.6849814 | -2.5049411 | 0.81520881 | -3.07276 |
| 16030 | NR_104169.2 | 6.39143813 | -4.2922468 | 1.39699382 | -3.072488 |
| 16031 | NM_006206.5 | 43.3259928 | -7.0602118 | 2.29801404 | -3.0723101 |
| 16032 | NM_031914.2 | 9.07684144 | -4.8094632 | 1.565418 | -3.0723188 |
| 16033 | NM_000447.2 | 48.9342468 | -7.2358675 | 2.35538697 | -3.0720504 |
| 16034 | NM_001301867.1 | 7.01424612 | -4.4442506 | 1.44686466 | -3.0716423 |
| 16035 | NM_001322884.1 | 59.7051341 | -7.5219022 | 2.44941278 | -3.0709002 |
| 16036 | NM_005434.4 | 17.2405507 | -2.4277176 | 0.79055462 | -3.0709044 |
| 16037 | NR_145416.1 | 8.03122298 | -4.6182723 | 1.50426642 | -3.070116 |
| 16038 | NM_001143970.1 | 16.3123528 | -4.6480696 | 1.51432693 | -3.0693964 |

| | | | | | |
|---|---|---|---|---|---|
| 16039 | NM_015392.3 | 453.57933 | -4.850119 | 1.58017541 | -3.0693548 |
| 16040 | NM_021131.4 | 82.027951 | -7.9798139 | 2.59988914 | -3.0692901 |
| 16041 | NM_198571.2 | 15.0174029 | -4.5042253 | 1.46752322 | -3.0692702 |
| 16042 | NM_015984.3 | 58.5574901 | -7.4936772 | 2.44163885 | -3.0691178 |
| 16043 | NM_001318154.1 | 57.9658971 | -7.4792401 | 2.43698945 | -3.069049 |
| 16044 | NM_207034.2 | 9.42018733 | -4.8471271 | 1.57950316 | -3.0687669 |
| 16045 | NM_001320598.1 | 10.226993 | -4.9643578 | 1.61784444 | -3.0685013 |
| 16046 | NM_001002812.2 | 13.5614963 | 8.67439272 | 2.82713526 | 3.06826236 |
| 16047 | NM_001286048.1 | 11.9684824 | 8.49816693 | 2.76987392 | 3.06806995 |
| 16048 | NM_015567.1 | 49.7632943 | -3.6331805 | 1.18431538 | -3.0677474 |
| 16049 | NM_054022.3 | 13.7662151 | 8.70750316 | 2.83841017 | 3.06773956 |
| 16050 | NR_120329.1 | 61.8549634 | 3.610434 | 1.17706622 | 3.06731595 |
| 16051 | NM_020775.4 | 58.5201417 | -7.49299 | 2.44319307 | -3.0668841 |
| 16052 | NM_004654.3 | 30.0050212 | -3.8005679 | 1.23931785 | -3.0666611 |
| 16053 | NR_046219.1 | 14.3791129 | 4.3774922 | 1.42752699 | 3.06648648 |
| 16054 | NM_001281778.2 | 58.118024 | -7.4829799 | 2.4409292 | -3.0656276 |
| 16055 | NR_126029.1 | 11.6895524 | 6.90325088 | 2.25181759 | 3.06563502 |
| 16056 | NM_001042461.2 | 12.9951589 | -5.3226235 | 1.7362863 | -3.0655218 |
| 16057 | NM_018287.6 | 43.7567032 | -7.0743564 | 2.30788896 | -3.0652932 |
| 16058 | NM_001320512.1 | 8.10962451 | -4.6264213 | 1.50995646 | -3.0639435 |
| 16059 | NM_020654.4 | 46.0560855 | -7.1481365 | 2.33303364 | -3.0638806 |
| 16060 | NM_001001670.2 | 4.43607189 | 7.08781646 | 2.31371136 | 3.06339701 |
| 16061 | NM_001033506.1 | 7.5675741 | -4.5443944 | 1.48353423 | -3.0632218 |
| 16062 | NM_001039582.3 | 76.3309194 | -7.8767834 | 2.57187198 | -3.0626654 |
| 16063 | NM_201436.2 | 7.28776097 | -4.4814371 | 1.46329304 | -3.0625698 |
| 16064 | NM_001045556.2 | 62.4281247 | -7.585715 | 2.47693575 | -3.0625401 |
| 16065 | NM_022758.5 | 40.0342927 | -3.6858925 | 1.20367431 | -3.0622009 |
| 16066 | NR_038947.1 | 7.29680758 | 5.07688671 | 1.65806539 | 3.06193395 |
| 16067 | NM_001112718.3 | 72.2078152 | -7.7956549 | 2.54612793 | -3.0617687 |
| 16068 | NM_001288616.1 | 7.71673942 | -4.5775031 | 1.49534063 | -3.0611775 |
| 16069 | NM_001159279.1 | 115.246704 | 2.60809647 | 0.85204639 | 3.06097943 |
| 16070 | NM_001199830.1 | 124.089661 | 7.01290178 | 2.29111748 | 3.06090886 |
| 16071 | NM_139319.2 | 43.3756095 | -7.0618129 | 2.3072022 | -3.060769 |
| 16072 | NM_001163545.2 | 43.6850827 | -7.0720942 | 2.31101831 | -3.0601636 |
| 16073 | NM_017491.3 | 661.136594 | -4.8147059 | 1.57340704 | -3.060051 |
| 16074 | NM_001009877.2 | 7.98777407 | -4.6142911 | 1.50794401 | -3.0599884 |
| 16075 | NM_001033925.1 | 45.9053861 | -7.1433824 | 2.33462581 | -3.0597547 |
| 16076 | NR_027028.2 | 6.18793873 | -4.2531577 | 1.39022868 | -3.0593223 |
| 16077 | NM_138802.2 | 10.8259902 | -5.0557974 | 1.65266102 | -3.0591859 |
| 16078 | NM_001001664.2 | 54.0562691 | -2.1004211 | 0.68662898 | -3.0590336 |
| 16079 | NM_022740.4 | 46.1905756 | -7.1522943 | 2.3383846 | -3.0586475 |
| 16080 | NM_031361.2 | 54.8869469 | -6.4295748 | 2.10220041 | -3.0584975 |
| 16081 | NM_002697.3 | 58.9122175 | -7.5024465 | 2.45303447 | -3.058435 |

| | | | | | |
|---|---|---|---|---|---|
| 16082 | NM_207040.1 | 56.7732153 | -7.4490967 | 2.43628734 | -3.0575608 |
| 16083 | NM_001318492.1 | 7.85827961 | -4.5790299 | 1.49777664 | -3.0572182 |
| 16084 | NM_001142301.1 | 14.308059 | -3.6278043 | 1.18680553 | -3.0567807 |
| 16085 | NM_001304794.1 | 9.5505992 | -4.889306 | 1.5997465 | -3.0563005 |
| 16086 | NM_138345.2 | 15.7926481 | -4.6054671 | 1.50697945 | -3.0560915 |
| 16087 | NM_000787.3 | 7.41124501 | -4.5232924 | 1.48025333 | -3.0557556 |
| 16088 | NM_001267066.1 | 45.6223568 | -7.1348242 | 2.33503162 | -3.0555579 |
| 16089 | NR_045789.1 | 7.40877402 | -4.5212468 | 1.48007178 | -3.0547484 |
| 16090 | NM_001305082.1 | 6.7184389 | -4.3735901 | 1.43201564 | -3.0541497 |
| 16091 | NR_046294.1 | 12.6934351 | -5.2931977 | 1.73312906 | -3.0541278 |
| 16092 | NR_024540.1 | 216.952307 | -4.3841241 | 1.43567359 | -3.0537053 |
| 16093 | NM_003199.2 | 63.7919138 | -7.6170563 | 2.49484169 | -3.0531221 |
| 16094 | NM_153612.3 | 22.1085359 | -4.1689389 | 1.36547412 | -3.0531072 |
| 16095 | NR_039996.1 | 24.0047562 | 2.46428063 | 0.80717296 | 3.05297717 |
| 16096 | NM_001166263.1 | 7.83965135 | -4.5826573 | 1.50128461 | -3.0524907 |
| 16097 | NM_001301063.1 | 54.9376297 | -7.4026532 | 2.42511805 | -3.0524919 |
| 16098 | NR_117086.1 | 12.0992474 | -5.2088234 | 1.70642502 | -3.0524772 |
| 16099 | NM_001032280.2 | 6.53966953 | -4.338789 | 1.42148774 | -3.0522873 |
| 16100 | NM_139312.2 | 10.1542634 | -4.9698993 | 1.6284127 | -3.05199 |
| 16101 | NM_014855.2 | 148.80341 | -3.311383 | 1.08501297 | -3.0519294 |
| 16102 | NM_003628.4 | 56.4972885 | -7.4419402 | 2.43873288 | -3.0515602 |
| 16103 | NM_001243270.1 | 9.09264787 | -4.7994105 | 1.57298181 | -3.0511545 |
| 16104 | NM_000602.4 | 198.744805 | -2.941649 | 0.96434715 | -3.0504046 |
| 16105 | NR_003545.1 | 10.1938913 | -4.9644253 | 1.62751171 | -3.0503162 |
| 16106 | NM_001321732.1 | 9.80562014 | -4.9243791 | 1.61440848 | -3.0502684 |
| 16107 | NM_001318136.1 | 8.79680887 | -4.7649859 | 1.5621742 | -3.050227 |
| 16108 | NM_013450.3 | 41.6064716 | -7.0018489 | 2.29563354 | -3.0500726 |
| 16109 | NM_004733.3 | 38.025574 | -3.6127934 | 1.18454335 | -3.0499461 |
| 16110 | NM_005892.3 | 36.3920583 | -3.5033775 | 1.14867772 | -3.049922 |
| 16111 | NM_001077365.1 | 43.4014449 | -7.0627471 | 2.31577532 | -3.0498413 |
| 16112 | NM_014702.4 | 7.33373117 | 5.60081005 | 1.83643537 | 3.04982693 |
| 16113 | NM_001001662.2 | 8.4787441 | -4.7171459 | 1.54673223 | -3.0497495 |
| 16114 | NR_037929.1 | 8.85001864 | -4.7753723 | 1.56589353 | -3.049615 |
| 16115 | NM_001142602.1 | 6.71880912 | -4.362806 | 1.43078064 | -3.0492487 |
| 16116 | NR_144634.1 | 8.6278836 | -4.7145601 | 1.5461594 | -3.049207 |
| 16117 | NM_001167856.2 | 78.8000117 | -7.922369 | 2.59931567 | -3.0478672 |
| 16118 | NM_001317721.1 | 8.51575825 | -4.69766 | 1.54156449 | -3.0473328 |
| 16119 | NR_126330.1 | 8.06781767 | -4.640042 | 1.5226987 | -3.0472489 |
| 16120 | NM_153027.1 | 8.88829683 | -4.7685506 | 1.56519471 | -3.0466182 |
| 16121 | NR_134285.1 | 6.27768649 | -4.2684681 | 1.40106932 | -3.0465788 |
| 16122 | NM_018167.4 | 7.81180316 | -4.5971385 | 1.50906038 | -3.0463583 |
| 16123 | NM_001282437.1 | 12.9819484 | -5.3127459 | 1.74406304 | -3.0461892 |
| 16124 | NM_021156.3 | 276.615996 | -2.5033995 | 0.82205282 | -3.0453025 |

| | | | | | |
|---|---|---|---|---|---|
| 16125 | NM_032119.3 | 41.6281631 | -7.0025617 | 2.29956258 | -3.0451712 |
| 16126 | NR_111941.1 | 10.2299164 | -4.9763139 | 1.63463251 | -3.0443013 |
| 16127 | NM_001143789.1 | 52.5463464 | -3.4851551 | 1.14487613 | -3.0441329 |
| 16128 | NM_004205.4 | 14.8256301 | -5.5161813 | 1.81207882 | -3.0441177 |
| 16129 | NR_026544.1 | 109.113169 | 3.03115775 | 0.99577233 | 3.0440269 |
| 16130 | NM_000898.4 | 9.41730093 | -4.8659549 | 1.5986583 | -3.0437742 |
| 16131 | NM_001169125.1 | 62.4672881 | -7.5867308 | 2.49272946 | -3.0435436 |
| 16132 | NM_017891.4 | 34.9589531 | -3.4400428 | 1.13044528 | -3.0430865 |
| 16133 | NM_020660.2 | 14.2089684 | -4.4540371 | 1.46370332 | -3.0429917 |
| 16134 | NM_032504.1 | 9.56327883 | -4.8710904 | 1.60077597 | -3.0429557 |
| 16135 | NM_148169.2 | 32.6671555 | 4.52769415 | 1.48797553 | 3.04285524 |
| 16136 | NM_001272000.1 | 55.2086076 | -7.4088683 | 2.43491152 | -3.0427669 |
| 16137 | NR_036481.2 | 43.1692897 | 3.44992808 | 1.13388594 | 3.04257064 |
| 16138 | NM_001320899.1 | 9.17466631 | -4.8291629 | 1.5879127 | -3.0412018 |
| 16139 | NM_006169.2 | 46.4426 | -3.8975807 | 1.28179944 | -3.0407102 |
| 16140 | NM_001076682.3 | 7.67872041 | -4.5612985 | 1.50017056 | -3.04052 |
| 16141 | NM_000783.3 | 9.84596423 | -4.92518 | 1.61986427 | -3.0404893 |
| 16142 | NM_001297549.1 | 40.8585095 | -6.9757763 | 2.29443173 | -3.0403068 |
| 16143 | NR_110291.1 | 27.1763261 | 3.76549069 | 1.23857473 | 3.04018045 |
| 16144 | NM_014470.3 | 25.9671299 | -4.2026162 | 1.38241546 | -3.0400529 |
| 16145 | NM_181304.2 | 7.98405267 | -4.6060546 | 1.51532495 | -3.0396481 |
| 16146 | NM_153051.2 | 68.9224165 | -7.7283825 | 2.54267694 | -3.0394669 |
| 16147 | NM_018934.3 | 61.5111781 | -3.5784469 | 1.17737474 | -3.039344 |
| 16148 | NM_138691.2 | 11.408881 | 3.11773431 | 1.02581802 | 3.03926646 |
| 16149 | NM_001287505.1 | 12.4883542 | -4.2182747 | 1.38797622 | -3.0391549 |
| 16150 | NM_003253.2 | 53.521882 | -3.7305153 | 1.22748934 | -3.0391427 |
| 16151 | NM_001257190.2 | 173.16332 | -2.9355254 | 0.96608364 | -3.0385831 |
| 16152 | NM_006839.2 | 43.5224078 | -7.066768 | 2.32567242 | -3.0385913 |
| 16153 | NM_178824.3 | 9.29756114 | -4.8479126 | 1.59556699 | -3.0383635 |
| 16154 | NM_001013627.2 | 6.21888999 | -4.2591139 | 1.40196357 | -3.0379633 |
| 16155 | NR_136740.1 | 51.691967 | -7.3136081 | 2.40749294 | -3.0378524 |
| 16156 | NM_001127196.1 | 305.912516 | -2.0241645 | 0.66637912 | -3.0375569 |
| 16157 | NM_001243224.1 | 9.89538544 | -4.9380545 | 1.62578129 | -3.0373424 |
| 16158 | NR_046095.1 | 15.1795708 | -5.5433804 | 1.82616069 | -3.0355382 |
| 16159 | NM_001323526.1 | 59.2416509 | -7.5101405 | 2.47417157 | -3.0354162 |
| 16160 | NM_152534.4 | 9.71492519 | -4.9094505 | 1.6175289 | -3.0351548 |
| 16161 | NM_018378.2 | 6.60525678 | -4.351627 | 1.43375919 | -3.035117 |
| 16162 | NR_027240.1 | 15.4723031 | 2.17846877 | 0.71782581 | 3.03481532 |
| 16163 | NM_001271893.3 | 7.21314985 | -4.4745575 | 1.47451448 | -3.0345972 |
| 16164 | NM_024081.5 | 95.8587693 | 2.22017043 | 0.73166051 | 3.03442705 |
| 16165 | NM_015891.2 | 103.405392 | -2.9189394 | 0.96196357 | -3.0343555 |
| 16166 | NM_152756.4 | 40.2562966 | -6.9543341 | 2.29193438 | -3.034264 |
| 16167 | NM_031910.3 | 101.151472 | -3.2310207 | 1.06488884 | -3.0341389 |

| | | | | | |
|---|---|---|---|---|---|
| 16168 | NR_040772.1 | 6.06200567 | -4.215557 | 1.38939472 | -3.034096 |
| 16169 | NM_001145873.1 | 6.45024297 | -4.3130954 | 1.42155768 | -3.0340629 |
| 16170 | NM_001199142.1 | 8.20829911 | -4.6710104 | 1.5396409 | -3.033831 |
| 16171 | NM_213654.3 | 10.6402213 | -5.0350759 | 1.65975333 | -3.033629 |
| 16172 | NM_024014.3 | 9.07540914 | -4.8102025 | 1.58572071 | -3.0334487 |
| 16173 | NM_001319062.1 | 10.4618737 | -5.0000225 | 1.64831574 | -3.033413 |
| 16174 | NM_018260.2 | 7.45314233 | -4.5217903 | 1.4908147 | -3.0331002 |
| 16175 | NM_001008781.2 | 213.412637 | -1.9720719 | 0.6502115 | -3.0329698 |
| 16176 | NM_001135163.1 | 29.0925832 | 3.02678349 | 0.99803015 | 3.03275757 |
| 16177 | NM_001311312.1 | 7.49582757 | -4.5324255 | 1.49449297 | -3.0327513 |
| 16178 | NM_001317813.1 | 8.68580916 | -4.751776 | 1.56686258 | -3.0326693 |
| 16179 | NM_001081677.1 | 92.6009109 | -3.6197463 | 1.19360004 | -3.0326291 |
| 16180 | NM_197977.2 | 9.09565728 | -4.8071579 | 1.58516348 | -3.0325944 |
| 16181 | NM_203383.1 | 42.8112111 | -7.0429226 | 2.32263919 | -3.032293 |
| 16182 | NM_153635.2 | 6.91757342 | -4.4074225 | 1.45362131 | -3.0320294 |
| 16183 | NM_183004.4 | 43.3736228 | -3.3525054 | 1.10584508 | -3.031623 |
| 16184 | NR_135657.1 | 9.92957622 | -4.9319721 | 1.62699357 | -3.0313409 |
| 16185 | NR_110804.1 | 26.5005342 | 3.02638752 | 0.9983724 | 3.03132131 |
| 16186 | NM_001324375.1 | 20.0418003 | 9.24723889 | 3.0506366 | 3.03124892 |
| 16187 | NM_014515.5 | 40.7039897 | -6.9701314 | 2.30048776 | -3.0298494 |
| 16188 | NM_001304952.1 | 7.639851 | -4.5379666 | 1.49780327 | -3.0297481 |
| 16189 | NM_001321590.1 | 64.5218127 | -7.6332375 | 2.5194265 | -3.029752 |
| 16190 | NR_121564.1 | 19.1378503 | 5.82206213 | 1.92173799 | 3.02958164 |
| 16191 | NM_001301701.1 | 9.31805791 | -4.8483344 | 1.60046556 | -3.0293276 |
| 16192 | NM_006731.2 | 53.907498 | -7.3744548 | 2.4343588 | -3.0293212 |
| 16193 | NM_017899.3 | 9.90736261 | -4.9377412 | 1.63006819 | -3.0291623 |
| 16194 | NM_001243782.1 | 3.58569381 | 6.7554152 | 2.2304929 | 3.02866474 |
| 16195 | NM_015378.3 | 51.1486589 | -7.2984915 | 2.41028913 | -3.0280564 |
| 16196 | NM_177405.2 | 118.732774 | -1.143994 | 0.37781935 | -3.0278862 |
| 16197 | NM_001206987.2 | 6.99484939 | -4.439029 | 1.46615907 | -3.0276585 |
| 16198 | NM_017782.4 | 54.4173068 | -7.3876286 | 2.44014942 | -3.0275312 |
| 16199 | NM_173465.3 | 7.79378018 | -4.592599 | 1.51707962 | -3.027263 |
| 16200 | NM_001793.5 | 42.4711361 | -7.0315706 | 2.32317547 | -3.0267066 |
| 16201 | NM_001171946.1 | 6.11759884 | -4.2376402 | 1.40010171 | -3.0266659 |
| 16202 | NM_178523.4 | 11.4733113 | -5.1472858 | 1.7007038 | -3.0265622 |
| 16203 | NR_111951.1 | 8.4314109 | -4.6950017 | 1.55126213 | -3.0265689 |
| 16204 | NM_001282234.1 | 7.12602462 | -4.4695195 | 1.47695542 | -3.0261709 |
| 16205 | NR_026788.1 | 16.4776904 | -3.7922851 | 1.25328201 | -3.0258833 |
| 16206 | NM_001244992.1 | 55.8556184 | -7.4252679 | 2.45407411 | -3.0256902 |
| 16207 | NM_012259.2 | 6.68977969 | -4.3541336 | 1.43915566 | -3.0254779 |
| 16208 | NM_014900.4 | 8.73456572 | -4.7566895 | 1.57262247 | -3.0246862 |
| 16209 | NM_001261408.1 | 11.1237598 | -5.0963007 | 1.68509607 | -3.0243383 |
| 16210 | NR_046228.1 | 21.3304849 | 2.63903734 | 0.8727239 | 3.02390865 |

| | | | | | |
|---|---|---|---|---|---|
| 16211 | NM_000294.2 | 116.096607 | -0.9518659 | 0.31480767 | -3.0236426 |
| 16212 | NM_001166108.1 | 6.80845143 | -4.3974745 | 1.45472373 | -3.0228932 |
| 16213 | NM_005761.2 | 52.9629636 | -3.7147009 | 1.22885943 | -3.0228852 |
| 16214 | NR_073470.1 | 10.1416156 | -4.9596144 | 1.64073452 | -3.0228013 |
| 16215 | NM_001025234.1 | 11.8603769 | -5.1980065 | 1.71970702 | -3.0226117 |
| 16216 | NM_002703.4 | 95.3910981 | -3.7661911 | 1.24608186 | -3.0224267 |
| 16217 | NM_001330441.1 | 11.8928619 | 8.48569651 | 2.8080433 | 3.02192509 |
| 16218 | NM_152595.4 | 6.22111957 | -4.2508509 | 1.40677972 | -3.0216891 |
| 16219 | NR_073468.1 | 53.7784895 | -7.3710996 | 2.43978253 | -3.0212117 |
| 16220 | NR_049784.1 | 7.80885325 | -4.5983579 | 1.52215338 | -3.0209557 |
| 16221 | NM_001323311.1 | 6.92143144 | -4.4201015 | 1.46319343 | -3.0208593 |
| 16222 | NM_018195.3 | 41.0470627 | -6.9823913 | 2.31142015 | -3.0208231 |
| 16223 | NM_003625.4 | 53.1006786 | -7.3526062 | 2.4340068 | -3.020783 |
| 16224 | NM_203497.3 | 21.7628085 | -6.0630729 | 2.00727534 | -3.0205487 |
| 16225 | NR_001587.1 | 7.64774047 | -4.5536052 | 1.50760042 | -3.0204324 |
| 16226 | NM_001134888.2 | 8.05047202 | -4.6433129 | 1.53733692 | -3.0203613 |
| 16227 | NR_144321.1 | 10.3964378 | -5.0078858 | 1.65821154 | -3.0200524 |
| 16228 | NR_104644.1 | 52.7202141 | 2.06422594 | 0.68352184 | 3.01998533 |
| 16229 | NM_001018064.2 | 97.3385324 | -3.9092633 | 1.2947207 | -3.0193874 |
| 16230 | NM_020448.4 | 39.4511444 | -6.9251141 | 2.2935977 | -3.0193238 |
| 16231 | NM_021179.2 | 13.6574967 | -5.3835451 | 1.78309594 | -3.0192123 |
| 16232 | NM_001286515.1 | 6.7322177 | -4.3653226 | 1.44586111 | -3.0191853 |
| 16233 | NM_006062.2 | 98.8013351 | -3.5837294 | 1.18711955 | -3.0188446 |
| 16234 | NM_001242768.1 | 40.9036097 | -6.9774601 | 2.31141728 | -3.0186934 |
| 16235 | NM_000826.3 | 64.5252448 | -7.6332091 | 2.52914283 | -3.0181012 |
| 16236 | NM_001291157.1 | 47.0372455 | -7.1789212 | 2.37869637 | -3.0180065 |
| 16237 | NM_016577.3 | 596.084926 | -4.8018212 | 1.59108058 | -3.0179623 |
| 16238 | NM_001322980.1 | 116.857247 | 3.78541215 | 1.25435333 | 3.01781966 |
| 16239 | NR_046359.1 | 9.82437271 | -4.9128342 | 1.62793753 | -3.0178272 |
| 16240 | NR_109763.1 | 6.71547763 | -4.3622109 | 1.44561739 | -3.0175418 |
| 16241 | NM_001168236.1 | 48.8167458 | -7.2316179 | 2.39658866 | -3.0174631 |
| 16242 | NM_001134296.1 | 9.9489776 | -4.9359759 | 1.6358468 | -3.0173828 |
| 16243 | NM_020395.3 | 9.86540092 | -4.923309 | 1.63170362 | -3.0172814 |
| 16244 | NM_001009936.2 | 7.63410412 | -4.5536328 | 1.50920762 | -3.0172342 |
| 16245 | NM_015036.2 | 102.748414 | -3.8827916 | 1.28688385 | -3.0172044 |
| 16246 | NM_001145299.3 | 53.1014845 | -7.352575 | 2.437015 | -3.0170413 |
| 16247 | NM_002878.3 | 12.1152152 | -5.2153658 | 1.72925449 | -3.015962 |
| 16248 | NM_138789.3 | 6.85479967 | -4.4080343 | 1.46155985 | -3.0159794 |
| 16249 | NR_023917.1 | 10.5274573 | -4.0006069 | 1.32646757 | -3.0159854 |
| 16250 | NM_001526.4 | 9.80729074 | -4.9143768 | 1.6294989 | -3.0158822 |
| 16251 | NM_014829.3 | 44.0242798 | -7.0829769 | 2.34857904 | -3.0158563 |
| 16252 | NM_001304342.1 | 56.0555114 | -7.4312111 | 2.46420306 | -3.0156651 |
| 16253 | NR_034082.1 | 6.13722031 | -4.2304407 | 1.40290143 | -3.0154939 |

| | | | | | |
|---|---|---|---|---|---|
| 16254 | NM_198532.2 | 36.9776285 | 2.14096292 | 0.71000511 | 3.01541904 |
| 16255 | NM_080413.2 | 12.1092013 | -5.2147108 | 1.7294347 | -3.015269 |
| 16256 | NM_001324094.1 | 66.5657674 | -3.862652 | 1.281082 | -3.0151481 |
| 16257 | NM_001286432.1 | 39.6623884 | -6.9328383 | 2.29941346 | -3.0150464 |
| 16258 | NR_133638.1 | 31.844661 | 3.35915797 | 1.11425858 | 3.01470237 |
| 16259 | NM_001194956.1 | 126.587944 | -2.5129201 | 0.83359103 | -3.0145719 |
| 16260 | NR_026675.1 | 23.6944116 | 2.33390212 | 0.77444761 | 3.01363463 |
| 16261 | NM_000387.5 | 38.2943256 | -3.3560519 | 1.1136853 | -3.0134652 |
| 16262 | NM_001305052.1 | 37.5034037 | 6.21296498 | 2.06180435 | 3.01336301 |
| 16263 | NM_032741.4 | 41.7088686 | -7.0052803 | 2.32474723 | -3.0133514 |
| 16264 | NM_001099401.1 | 7.09334533 | -4.4580743 | 1.47948515 | -3.0132606 |
| 16265 | NM_001164712.1 | 6.57687068 | -4.3507306 | 1.44398628 | -3.0129999 |
| 16266 | NM_203341.2 | 16.4490214 | -5.6652642 | 1.88027138 | -3.0130034 |
| 16267 | NM_053285.1 | 15.1954291 | -5.5520343 | 1.84287526 | -3.0127022 |
| 16268 | NM_001346007.1 | 8.3990598 | -4.6992191 | 1.55989234 | -3.0125279 |
| 16269 | NR_004401.2 | 27.9580462 | 3.0947877 | 1.02734679 | 3.01240801 |
| 16270 | NM_001321191.1 | 68.0367592 | -3.5966876 | 1.19402502 | -3.0122381 |
| 16271 | NM_133374.2 | 40.723564 | -6.9709863 | 2.31441053 | -3.0119921 |
| 16272 | NR_046028.1 | 7.08272362 | -4.4488005 | 1.47715338 | -3.0117391 |
| 16273 | NM_201517.2 | 12.3863599 | -5.2583681 | 1.74631816 | -3.0111169 |
| 16274 | NR_045215.1 | 6.21324663 | -4.2681773 | 1.4175903 | -3.010868 |
| 16275 | NR_026893.2 | 54.3339436 | -7.3855312 | 2.45316277 | -3.0106161 |
| 16276 | NM_001115016.2 | 52.2103005 | -7.3291201 | 2.43455656 | -3.0104538 |
| 16277 | NM_198181.3 | 39.8743951 | -6.9404571 | 2.30595971 | -3.0097912 |
| 16278 | NM_001220.4 | 11.3246622 | -5.1220363 | 1.70186861 | -3.0096544 |
| 16279 | NM_001330292.1 | 12.1774281 | -4.2305457 | 1.40578688 | -3.0093791 |
| 16280 | NM_001330239.1 | 45.0370289 | -7.1157997 | 2.36460309 | -3.0092998 |
| 16281 | NM_001166247.1 | 11.0533689 | -5.0830121 | 1.68915467 | -3.0092046 |
| 16282 | NM_001142279.2 | 6.72022725 | -4.3579836 | 1.44833075 | -3.0089698 |
| 16283 | NM_012418.3 | 6.35443009 | -4.3004976 | 1.42946816 | -3.0084599 |
| 16284 | NR_003109.1 | 8.91791032 | -4.7753955 | 1.58774637 | -3.0076564 |
| 16285 | NR_135065.1 | 38.8603515 | 3.74759492 | 1.24609442 | 3.00747267 |
| 16286 | NM_003015.3 | 7.89492268 | -4.6169404 | 1.53525617 | -3.0072769 |
| 16287 | NM_178516.3 | 7.73045253 | -4.5633643 | 1.51758037 | -3.0070001 |
| 16288 | NM_003394.3 | 17.9093087 | -3.3828479 | 1.12508451 | -3.00675 |
| 16289 | NM_001242895.1 | 166.982504 | 2.0725643 | 0.68931711 | 3.00669208 |
| 16290 | NR_027271.1 | 6.04559586 | -4.2238969 | 1.40514697 | -3.0060178 |
| 16291 | NM_001322189.1 | 9.28851042 | -4.8357307 | 1.60875052 | -3.0058922 |
| 16292 | NR_073472.1 | 10.0908867 | -4.9443731 | 1.64497477 | -3.005744 |
| 16293 | NM_001313903.1 | 11.3903903 | 8.43091779 | 2.80502951 | 3.00564317 |
| 16294 | NM_133639.3 | 11.4965185 | -5.1404018 | 1.71040907 | -3.005364 |
| 16295 | NM_207171.2 | 153.409481 | -4.1272489 | 1.37344786 | -3.0050277 |
| 16296 | NR_145459.1 | 85.0717735 | -3.347849 | 1.11414819 | -3.0048507 |

| | | | | | |
|---|---|---|---|---|---|
| 16297 | NM_001256404.1 | 6.98361996 | -4.4263615 | 1.47337352 | -3.0042358 |
| 16298 | NM_001184773.1 | 47.750464 | -7.2005544 | 2.39691654 | -3.0040906 |
| 16299 | NM_002218.4 | 10.4521334 | -4.9992995 | 1.66425684 | -3.0039231 |
| 16300 | NM_020954.3 | 17.9146034 | -2.4749128 | 0.82390923 | -3.0038659 |
| 16301 | NM_001318868.1 | 9.36189121 | -4.850445 | 1.61509449 | -3.0031958 |
| 16302 | NM_000231.2 | 8.56676004 | -4.7211163 | 1.57212153 | -3.0030225 |
| 16303 | NM_001159531.1 | 39.8838907 | -6.940855 | 2.31130542 | -3.0030021 |
| 16304 | NM_001317941.2 | 52.7498745 | -7.3428039 | 2.44520011 | -3.002946 |
| 16305 | NM_197941.3 | 28.2436089 | -2.5553231 | 0.85106051 | -3.0025163 |
| 16306 | NM_024311.3 | 12.9582319 | -5.3104615 | 1.76887126 | -3.0021752 |
| 16307 | NM_001243646.1 | 7.0000659 | -4.4415079 | 1.47946585 | -3.0021023 |
| 16308 | NM_024815.3 | 11.0616051 | -4.0343552 | 1.34396679 | -3.0018265 |
| 16309 | NR_027691.1 | 7.73004515 | -4.5727174 | 1.52343025 | -3.0015929 |
| 16310 | NM_001286686.1 | 11.4686525 | -5.1319637 | 1.71008468 | -3.0009997 |
| 16311 | NM_001330230.1 | 9.40627575 | -4.8644533 | 1.6209575 | -3.0009753 |
| 16312 | NM_001300861.1 | 11.4713523 | -5.148392 | 1.71566988 | -3.0008057 |
| 16313 | NM_001286823.1 | 7.31717212 | -4.4795811 | 1.49290488 | -3.0005804 |
| 16314 | NM_001100880.2 | 11.3974391 | 1.96414437 | 0.65466219 | 3.00024104 |
| 16315 | NM_018248.2 | 6.8591728 | -4.3870568 | 1.46226854 | -3.0001718 |
| 16316 | NR_110740.1 | 17.0388558 | 4.42875565 | 1.47620794 | 3.0000893 |
| 16317 | NM_014447.3 | 8.36218721 | -4.6793942 | 1.55979975 | -2.9999967 |
| 16318 | NM_207499.2 | 114.402043 | 2.45022989 | 0.81680866 | 2.99975994 |
| 16319 | NM_001130992.1 | 18.6764775 | -3.0232411 | 1.00806362 | -2.9990579 |
| 16320 | NM_001328685.1 | 52.4001454 | -7.3332275 | 2.44554421 | -2.9986076 |
| 16321 | NR_038886.1 | 29.5806495 | 3.63600224 | 1.21258905 | 2.9985445 |
| 16322 | NR_037575.1 | 10.036414 | -4.9541918 | 1.65228748 | -2.9983837 |
| 16323 | NM_130474.2 | 15.2668484 | 8.85483039 | 2.95393462 | 2.99763926 |
| 16324 | NM_139314.2 | 14.328573 | -5.465039 | 1.82324977 | -2.9974165 |
| 16325 | NM_178128.5 | 50.8307776 | 2.3099416 | 0.77069231 | 2.99722933 |
| 16326 | NM_001001522.1 | 30.7504634 | -3.2345563 | 1.07941275 | -2.996589 |
| 16327 | NM_001252007.1 | 6.43477291 | -4.3079567 | 1.43780916 | -2.996195 |
| 16328 | NM_001319134.1 | 52.8995871 | -7.3467928 | 2.45227275 | -2.9959118 |
| 16329 | NM_001276277.2 | 52.0144508 | -7.3230972 | 2.44441088 | -2.9958536 |
| 16330 | NR_135086.1 | 9.16142716 | -4.8148765 | 1.60718712 | -2.9958407 |
| 16331 | NM_001206769.1 | 9.57704002 | -4.8814273 | 1.62984882 | -2.9950185 |
| 16332 | NM_001291281.2 | 165.065068 | -4.1775788 | 1.39496303 | -2.9947595 |
| 16333 | NM_003014.3 | 113.544828 | -3.9359745 | 1.31430322 | -2.9947233 |
| 16334 | NM_001042493.2 | 14.4369377 | -5.4690229 | 1.8264202 | -2.9943946 |
| 16335 | NM_032531.3 | 46.1041331 | -7.1497131 | 2.38793333 | -2.9941008 |
| 16336 | NM_139266.2 | 17.8340199 | -5.7804848 | 1.93068031 | -2.9940145 |
| 16337 | NM_001122870.2 | 8.74543081 | -4.7434325 | 1.58445371 | -2.9937337 |
| 16338 | NM_013250.2 | 7.05042818 | -4.4387069 | 1.48267826 | -2.9937088 |
| 16339 | NM_018676.3 | 7.30579921 | -4.4938633 | 1.50137397 | -2.9931672 |

| | | | | | |
|---|---|---|---|---|---|
| 16340 | NM_016826.2 | 6.44718209 | -4.3145187 | 1.44168717 | -2.9926872 |
| 16341 | NM_000545.6 | 7.62876248 | -4.5489261 | 1.52048292 | -2.991764 |
| 16342 | NM_001243333.1 | 49.533562 | -7.2530793 | 2.42450114 | -2.9915759 |
| 16343 | NM_001042717.2 | 73.6455631 | 5.43519026 | 1.81694055 | 2.99139686 |
| 16344 | NR_138534.1 | 8.9961803 | -4.7791882 | 1.59764802 | -2.99139 |
| 16345 | NM_198332.1 | 67.7372977 | -2.7736227 | 0.92728452 | -2.9911236 |
| 16346 | NM_001170423.1 | 17.8606073 | -3.9121601 | 1.30798558 | -2.990981 |
| 16347 | NR_038457.1 | 9.66339627 | -4.8920649 | 1.63567251 | -2.9908584 |
| 16348 | NM_001318066.1 | 49.1225767 | -7.2410362 | 2.42147506 | -2.990341 |
| 16349 | NM_001347822.1 | 38.5039862 | -6.8901892 | 2.30415919 | -2.9903269 |
| 16350 | NM_001271522.1 | 7.35083121 | -4.4971112 | 1.50391401 | -2.9902715 |
| 16351 | NM_025198.4 | 13.9497228 | -5.4244202 | 1.81401153 | -2.9902898 |
| 16352 | NM_001169107.1 | 16.2554046 | -3.5981129 | 1.2033607 | -2.9900536 |
| 16353 | NR_073133.1 | 6.75075771 | -4.3845568 | 1.46640423 | -2.9900055 |
| 16354 | NM_138434.2 | 8.68221473 | -4.7322268 | 1.58287993 | -2.989631 |
| 16355 | NM_001004351.4 | 6.90800022 | -4.4236 | 1.48021784 | -2.9884791 |
| 16356 | NM_006181.2 | 6.73061729 | -4.376932 | 1.46474833 | -2.9881803 |
| 16357 | NM_001160233.1 | 12.7639003 | -5.2925203 | 1.77117699 | -2.9881375 |
| 16358 | NR_027626.1 | 18.4907533 | 9.13116541 | 3.05580677 | 2.98813573 |
| 16359 | NM_005477.2 | 62.844298 | -3.9488029 | 1.32151937 | -2.988078 |
| 16360 | NR_033915.1 | 6.85133156 | -4.4066425 | 1.47494615 | -2.9876633 |
| 16361 | NM_004430.2 | 5.99922285 | -4.1987617 | 1.40575919 | -2.9868286 |
| 16362 | NM_001134999.1 | 52.0513644 | -7.3241769 | 2.45235871 | -2.9865847 |
| 16363 | NM_001330509.1 | 9.83687317 | -4.9101764 | 1.64429589 | -2.9861879 |
| 16364 | NM_013352.3 | 37.9380741 | -6.8688381 | 2.30040376 | -2.9859272 |
| 16365 | NM_003105.5 | 139.561535 | -3.6710864 | 1.22951108 | -2.9858099 |
| 16366 | NM_013334.3 | 6.06851505 | -4.232912 | 1.41779177 | -2.9855668 |
| 16367 | NM_153251.3 | 62.0639851 | -7.5774024 | 2.53802735 | -2.985548 |
| 16368 | NR_026818.1 | 7.01006072 | -4.4438593 | 1.48863778 | -2.9851851 |
| 16369 | NM_001143823.2 | 9.04326928 | -4.7991083 | 1.60765854 | -2.985154 |
| 16370 | NM_001195243.1 | 6.53681305 | -4.3355231 | 1.45248598 | -2.9848984 |
| 16371 | NM_012176.2 | 6.50999057 | -4.3280301 | 1.45004744 | -2.9847507 |
| 16372 | NM_033430.2 | 9.94477501 | -4.934085 | 1.65311408 | -2.9847214 |
| 16373 | NM_032148.3 | 31.7859358 | -1.9895058 | 0.6666288 | -2.9844282 |
| 16374 | NM_002285.2 | 43.2389082 | -7.0575691 | 2.36536771 | -2.9837091 |
| 16375 | NM_001007471.2 | 9.61924393 | -4.8961098 | 1.64098363 | -2.9836433 |
| 16376 | NR_023353.1 | 13.0778131 | -4.3276845 | 1.45046266 | -2.983658 |
| 16377 | NM_144508.4 | 14.2572402 | -5.4499818 | 1.82664983 | -2.9835942 |
| 16378 | NM_015685.5 | 6.22661517 | -4.2469511 | 1.42354418 | -2.9833644 |
| 16379 | NM_001347890.1 | 7.52553802 | -4.5346923 | 1.52023491 | -2.9828892 |
| 16380 | NM_001172779.1 | 6.96009748 | -4.4274062 | 1.48441083 | -2.9826017 |
| 16381 | NM_018941.3 | 148.829697 | -1.2488514 | 0.41874328 | -2.9823795 |
| 16382 | NM_001135254.1 | 8.34750729 | -4.6903735 | 1.57290682 | -2.981978 |

| | | | | | |
|---|---|---|---|---|---|
| 16383 | NM_005446.3 | 7.4751646 | -4.5371754 | 1.52165514 | -2.981737 |
| 16384 | NM_201626.2 | 8.12579136 | -4.6418097 | 1.55692899 | -2.9813882 |
| 16385 | NM_001033030.1 | 8.85365281 | -4.7794522 | 1.60363065 | -2.9803947 |
| 16386 | NM_001271096.1 | 42.2214636 | -7.0231847 | 2.3567092 | -2.9800812 |
| 16387 | NR_136747.1 | 7.37584866 | -4.5019967 | 1.5107601 | -2.9799547 |
| 16388 | NM_001025161.2 | 11.0507942 | -5.0871202 | 1.70782875 | -2.9787062 |
| 16389 | NM_014506.2 | 39.638687 | -6.9319536 | 2.32723031 | -2.9786281 |
| 16390 | NM_001319993.1 | 13.8046542 | -3.4589783 | 1.16138647 | -2.9783181 |
| 16391 | NM_001135861.2 | 10.0090999 | -4.9325416 | 1.65618138 | -2.9782617 |
| 16392 | NM_001199868.1 | 78.8095221 | -2.7504418 | 0.92360759 | -2.9779334 |
| 16393 | NM_001330193.1 | 9.39932971 | -4.8556689 | 1.63056692 | -2.9779022 |
| 16394 | NR_138070.1 | 8.12734879 | -4.6422226 | 1.55906727 | -2.977564 |
| 16395 | NM_031485.3 | 98.7185883 | -2.7759073 | 0.93231006 | -2.9774507 |
| 16396 | NR_135066.1 | 13.699695 | 2.18811382 | 0.73497528 | 2.97712576 |
| 16397 | NR_126441.1 | 14.4976529 | 2.36751479 | 0.79525893 | 2.97703642 |
| 16398 | NM_001320782.1 | 9.71725851 | -4.8934286 | 1.64378286 | -2.9769313 |
| 16399 | NM_001008708.3 | 6.76918275 | -4.372918 | 1.46910457 | -2.9765873 |
| 16400 | NM_014929.3 | 729.756019 | 4.20047732 | 1.41199951 | 2.97484334 |
| 16401 | NM_001347821.1 | 8.87356678 | -4.7752538 | 1.60593845 | -2.9734974 |
| 16402 | NM_000773.3 | 7.04376898 | -4.4470268 | 1.49564672 | -2.9733136 |
| 16403 | NM_003627.5 | 6.23872794 | -4.2753871 | 1.43802573 | -2.973095 |
| 16404 | NR_004862.1 | 36.4331499 | -6.8104808 | 2.2908504 | -2.9729051 |
| 16405 | NM_003903.4 | 41.0122061 | -6.9813551 | 2.34904511 | -2.971997 |
| 16406 | NM_001316975.1 | 8.8040755 | -4.7620533 | 1.60234025 | -2.9719364 |
| 16407 | NM_020877.3 | 14.5377345 | -4.4578936 | 1.50011968 | -2.9716919 |
| 16408 | NM_001029858.3 | 309.039467 | -3.5950204 | 1.20978805 | -2.9716117 |
| 16409 | NM_184231.2 | 36.3006795 | -6.8051584 | 2.29024969 | -2.9713609 |
| 16410 | NR_134249.1 | 43.7760885 | 3.54100688 | 1.19179985 | 2.97114224 |
| 16411 | NM_033450.2 | 9.54090094 | -4.8766077 | 1.64150976 | -2.9708064 |
| 16412 | NM_014220.2 | 7.28847923 | -4.4978358 | 1.51434827 | -2.9701462 |
| 16413 | NM_148896.4 | 8.93591434 | -4.7739905 | 1.60779682 | -2.9692747 |
| 16414 | NM_005932.3 | 43.2458054 | -2.4445149 | 0.82332123 | -2.9690901 |
| 16415 | NR_046236.1 | 7.02983083 | -4.4307033 | 1.49232066 | -2.9690022 |
| 16416 | NM_024068.3 | 174.974121 | -2.4318637 | 0.81910568 | -2.9689254 |
| 16417 | NM_001184797.1 | 9.4754807 | -4.8625707 | 1.63790213 | -2.9687798 |
| 16418 | NM_001080496.2 | 36.108485 | -3.0566236 | 1.02959811 | -2.9687541 |
| 16419 | NM_001144996.1 | 37.4752987 | -6.851273 | 2.30786877 | -2.968658 |
| 16420 | NM_001080452.1 | 65.2788311 | -3.5175273 | 1.18491225 | -2.9685973 |
| 16421 | NM_000196.3 | 20.0978106 | -3.5880464 | 1.20872755 | -2.9684493 |
| 16422 | NM_001270497.1 | 8.36450196 | -4.6866956 | 1.57890822 | -2.9683142 |
| 16423 | NM_001318253.1 | 13.6175467 | -5.3824621 | 1.8134922 | -2.9680095 |
| 16424 | NM_001258308.1 | 45.6081973 | -7.1341334 | 2.40375588 | -2.9679109 |
| 16425 | NM_001136036.2 | 10.3419883 | -4.9882283 | 1.68073456 | -2.9678858 |

| | | | | | |
|---|---|---|---|---|---|
| 16426 | NM_005873.2 | 10.805267 | -5.0500707 | 1.70172835 | -2.9676127 |
| 16427 | NM_130852.2 | 14.4913249 | 8.77966913 | 2.95854266 | 2.96756551 |
| 16428 | NM_139285.3 | 10.654783 | -5.0408189 | 1.69866425 | -2.9675193 |
| 16429 | NM_198552.2 | 46.0010545 | -3.2752593 | 1.1037251 | -2.9674593 |
| 16430 | NM_014311.2 | 10.5672422 | -5.0192623 | 1.69153282 | -2.9672864 |
| 16431 | NR_037658.1 | 9.06431626 | -4.7920171 | 1.61508269 | -2.9670413 |
| 16432 | NM_019619.3 | 36.0378314 | -6.7946718 | 2.29011297 | -2.9669592 |
| 16433 | NM_005195.3 | 60.7447131 | -3.4187266 | 1.15233701 | -2.9667767 |
| 16434 | NM_001303458.2 | 40.1871912 | -6.9516848 | 2.34364267 | -2.966188 |
| 16435 | NM_006414.4 | 9.50598276 | -4.8615914 | 1.63913754 | -2.9659448 |
| 16436 | NR_144395.1 | 280.202438 | 1.24935662 | 0.42124077 | 2.9658967 |
| 16437 | NM_000332.3 | 50.1191807 | -7.2689972 | 2.45091384 | -2.9658314 |
| 16438 | NM_001300943.1 | 49.0909798 | -7.2390943 | 2.44096496 | -2.9656691 |
| 16439 | NM_203472.2 | 8.27018104 | -4.6743375 | 1.57704596 | -2.963983 |
| 16440 | NM_006212.2 | 43.728968 | -1.9427332 | 0.6554639 | -2.9639057 |
| 16441 | NM_201612.2 | 6.96092258 | -4.4359056 | 1.49679034 | -2.9636119 |
| 16442 | NM_024917.5 | 9.17670728 | -4.8081037 | 1.62252454 | -2.9633473 |
| 16443 | NM_001177306.1 | 51.0893172 | -7.2973716 | 2.46272817 | -2.9631251 |
| 16444 | NM_152545.2 | 50.9613101 | -7.2928874 | 2.46125883 | -2.9630721 |
| 16445 | NR_026598.1 | 11.1657216 | -5.1042497 | 1.72283476 | -2.9627042 |
| 16446 | NM_080631.3 | 10.7114708 | 8.34182226 | 2.81609124 | 2.96219886 |
| 16447 | NM_015266.2 | 36.2106946 | -6.8015165 | 2.29643412 | -2.961773 |
| 16448 | NM_153346.4 | 27.736188 | 1.88060403 | 0.63497673 | 2.96168967 |
| 16449 | NM_001135243.1 | 39.5558495 | -6.9290955 | 2.33959382 | -2.961666 |
| 16450 | NM_001005739.1 | 43.6953655 | -7.0713832 | 2.38807431 | -2.9611236 |
| 16451 | NM_152503.5 | 10.289706 | -3.9369923 | 1.32980237 | -2.9605845 |
| 16452 | NM_001330334.1 | 7.23298552 | -4.4866445 | 1.51548711 | -2.9605297 |
| 16453 | NR_039994.2 | 6.46400761 | -4.3022694 | 1.45343112 | -2.9600779 |
| 16454 | NM_001322331.1 | 9.43044051 | -4.8502061 | 1.63910336 | -2.9590605 |
| 16455 | NR_104612.1 | 9.71978688 | -4.8966145 | 1.65499538 | -2.9586877 |
| 16456 | NM_032180.2 | 35.9095737 | -6.7895873 | 2.2949057 | -2.9585474 |
| 16457 | NM_022918.3 | 35.4735832 | -6.7719513 | 2.28914383 | -2.95829 |
| 16458 | NR_037158.1 | 36.3290475 | 1.90705617 | 0.64465884 | 2.95824093 |
| 16459 | NM_012231.4 | 48.1164719 | -7.2102497 | 2.4375432 | -2.9579987 |
| 16460 | NM_001278429.3 | 14.8119501 | 4.48864747 | 1.51757087 | 2.95778443 |
| 16461 | NM_001308243.1 | 23.5589932 | 3.07755998 | 1.0405458 | 2.9576401 |
| 16462 | NM_005710.2 | 11.0949472 | -4.0762341 | 1.3786852 | -2.9566097 |
| 16463 | NM_001330233.1 | 15.8162695 | -5.5997157 | 1.89417757 | -2.9562781 |
| 16464 | NM_198401.3 | 14.0210145 | -5.4287985 | 1.83647226 | -2.9561015 |
| 16465 | NM_198867.1 | 8.8746032 | -4.7780904 | 1.61636136 | -2.9560781 |
| 16466 | NM_176794.1 | 6.8997485 | -4.4196799 | 1.49583793 | -2.9546516 |
| 16467 | NM_001723.5 | 11.1232173 | -4.0489725 | 1.37049355 | -2.95439 |
| 16468 | NM_002342.2 | 6.24322111 | -4.2695955 | 1.44520754 | -2.9543131 |

| | | | | | |
|---|---|---|---|---|---|
| 16469 | NM_006256.3 | 43.0692689 | -7.0506574 | 2.3866835 | -2.9541652 |
| 16470 | NM_001080466.1 | 35.5653826 | -3.0313623 | 1.02627999 | -2.9537381 |
| 16471 | NM_001256706.1 | 6.73945867 | -4.3605153 | 1.47626954 | -2.9537393 |
| 16472 | NM_181306.2 | 7.16014219 | -4.4592843 | 1.50970115 | -2.953753 |
| 16473 | NM_001318336.1 | 46.3522436 | -7.1575357 | 2.4234115 | -2.9534958 |
| 16474 | NM_006106.4 | 46.1746493 | -6.1797502 | 2.0923468 | -2.9535019 |
| 16475 | NM_153826.3 | 39.7086837 | -6.9343653 | 2.34784901 | -2.9534971 |
| 16476 | NM_012455.2 | 10.9582813 | -4.0337205 | 1.36576937 | -2.9534419 |
| 16477 | NR_027440.1 | 6.44183743 | -4.3010006 | 1.45632382 | -2.9533271 |
| 16478 | NM_080821.2 | 204.647744 | -2.1852427 | 0.73995401 | -2.9532142 |
| 16479 | NM_007314.3 | 40.8281856 | -6.9743796 | 2.36179126 | -2.9530042 |
| 16480 | NM_001286116.1 | 73.9310044 | 4.39295072 | 1.48763151 | 2.9529831 |
| 16481 | NM_021800.2 | 33.6303044 | -3.6580965 | 1.23885005 | -2.9528162 |
| 16482 | NM_001329805.1 | 47.6693646 | 7.53063564 | 2.55046495 | 2.95265208 |
| 16483 | NM_007365.2 | 30.3534515 | -3.8284781 | 1.29664701 | -2.9525986 |
| 16484 | NR_135259.1 | 8.46591623 | -4.6909925 | 1.58882328 | -2.9524948 |
| 16485 | NM_002334.3 | 111.363496 | -2.8935352 | 0.98006273 | -2.952398 |
| 16486 | NM_001257330.1 | 8.69041624 | -4.7388551 | 1.6053433 | -2.9519263 |
| 16487 | NM_001080400.1 | 7.13519132 | -4.4587239 | 1.51046575 | -2.9518868 |
| 16488 | NM_002767.3 | 140.818623 | -4.1897297 | 1.41938907 | -2.9517838 |
| 16489 | NM_201595.2 | 60.5877092 | -6.5719026 | 2.22647602 | -2.951706 |
| 16490 | NM_006309.3 | 14.720902 | -4.4998848 | 1.52464861 | -2.9514242 |
| 16491 | NM_001282758.1 | 6.98688825 | -4.4266271 | 1.49990606 | -2.9512695 |
| 16492 | NR_038965.1 | 6.98562395 | 2.97293646 | 1.00771265 | 2.95018273 |
| 16493 | NM_001174087.1 | 50.8136157 | -7.288838 | 2.47069813 | -2.9501127 |
| 16494 | NR_135591.1 | 36.024046 | -6.7939929 | 2.30297448 | -2.9500947 |
| 16495 | NM_017805.2 | 6.69738491 | -4.3632674 | 1.47922273 | -2.9497028 |
| 16496 | NM_001318827.1 | 35.8881236 | -6.7886423 | 2.3015554 | -2.9495889 |
| 16497 | NM_016183.3 | 154.1968 | -1.9739388 | 0.66922319 | -2.9495971 |
| 16498 | NM_198965.1 | 7.6900962 | -4.5641981 | 1.54745577 | -2.9494853 |
| 16499 | NM_001307980.1 | 8.54966973 | -4.7205608 | 1.60143167 | -2.9477129 |
| 16500 | NM_001171806.1 | 11.4222229 | -5.1367482 | 1.74264668 | -2.9476705 |
| 16501 | NM_001077663.2 | 10.2074878 | -4.9662706 | 1.68491096 | -2.9474974 |
| 16502 | NM_021831.5 | 29.9453002 | -3.1923609 | 1.08313145 | -2.9473439 |
| 16503 | NM_001100910.1 | 7.27086695 | -4.486493 | 1.52226491 | -2.9472485 |
| 16504 | NM_022128.2 | 6.60561889 | -4.3498746 | 1.47590478 | -2.9472597 |
| 16505 | NM_001242839.1 | 6.023477 | -4.2179224 | 1.43146281 | -2.9465819 |
| 16506 | NM_001048171.1 | 8.02956572 | -4.6171367 | 1.56702137 | -2.9464414 |
| 16507 | NM_173042.2 | 10.3669624 | -3.9284098 | 1.33328651 | -2.9464108 |
| 16508 | NR_135140.1 | 10.2245863 | -4.9733537 | 1.68796329 | -2.9463637 |
| 16509 | NR_103866.1 | 44.0948729 | -7.0841929 | 2.40441649 | -2.9463252 |
| 16510 | NM_001204458.1 | 39.5961124 | -6.9301459 | 2.35277459 | -2.9455205 |
| 16511 | NM_152228.2 | 6.5288744 | -4.3316749 | 1.47067341 | -2.9453683 |

| | | | | | |
|---|---|---|---|---|---|
| 16512 | NM_018225.2 | 278.590016 | -1.9606543 | 0.6656786 | -2.9453468 |
| 16513 | NM_005096.3 | 44.0640279 | -7.0833924 | 2.40498636 | -2.9452942 |
| 16514 | NM_001256702.1 | 17.0132152 | 9.01157484 | 3.05969211 | 2.94525544 |
| 16515 | NM_017951.4 | 50.7816435 | -7.2878227 | 2.47442149 | -2.9452633 |
| 16516 | NR_002912.1 | 6.47962012 | -4.3004067 | 1.46029456 | -2.9448899 |
| 16517 | NM_024841.4 | 8.6228459 | -4.7280865 | 1.60572159 | -2.9445245 |
| 16518 | NM_021709.2 | 7.27129166 | -4.468855 | 1.51771388 | -2.9444648 |
| 16519 | NM_001077624.1 | 15.2622151 | -3.5009444 | 1.18904995 | -2.9443207 |
| 16520 | NM_001300735.1 | 36.3216089 | -6.8060855 | 2.31168885 | -2.9442048 |
| 16521 | NM_016014.3 | 6.60972429 | -4.3480613 | 1.47683969 | -2.9441661 |
| 16522 | NM_000267.3 | 202.556919 | -2.5767341 | 0.87521858 | -2.9441035 |
| 16523 | NM_198468.2 | 53.4041389 | -1.2091111 | 0.41068975 | -2.9440986 |
| 16524 | NR_125386.1 | 43.3293811 | 1.37004316 | 0.46547419 | 2.94332784 |
| 16525 | NM_014683.3 | 193.875699 | -2.4109607 | 0.81921191 | -2.9430244 |
| 16526 | NM_139166.4 | 8.42085049 | 5.04210678 | 1.71341963 | 2.94271566 |
| 16527 | NM_001319224.1 | 5.96762509 | -4.1991111 | 1.42722405 | -2.9421527 |
| 16528 | NR_131977.1 | 15.7216335 | 3.37305409 | 1.1464781 | 2.94210075 |
| 16529 | NM_005574.3 | 6.92558057 | -4.4145283 | 1.50050307 | -2.9420322 |
| 16530 | NM_001286495.1 | 7.6174841 | -4.5608471 | 1.55026155 | -2.9419856 |
| 16531 | NR_029388.2 | 6.24565281 | -4.2610213 | 1.44841822 | -2.9418446 |
| 16532 | NM_177534.2 | 7.26344143 | -4.4957113 | 1.52825202 | -2.9417342 |
| 16533 | NM_153620.2 | 10.6649395 | -5.0436935 | 1.71475449 | -2.9413502 |
| 16534 | NM_024061.3 | 8.38227365 | -4.6839329 | 1.59250342 | -2.9412388 |
| 16535 | NM_001162426.1 | 49.6261043 | -7.2552016 | 2.46712046 | -2.9407569 |
| 16536 | NM_005977.3 | 34.5498056 | -6.7338715 | 2.28989348 | -2.940692 |
| 16537 | NM_148179.2 | 5.88103769 | -4.1881596 | 1.42424532 | -2.9406167 |
| 16538 | NM_001024629.2 | 4.64949737 | 7.10288492 | 2.41563192 | 2.94038378 |
| 16539 | NM_001321007.1 | 12.3551579 | -5.2402058 | 1.78219063 | -2.9403172 |
| 16540 | NM_001127890.4 | 8.42072446 | -4.6936453 | 1.59650959 | -2.9399418 |
| 16541 | NR_111930.1 | 6.22263718 | -4.2523341 | 1.4464898 | -2.9397609 |
| 16542 | NM_001199381.1 | 8.02298778 | -4.6342808 | 1.57645894 | -2.9396775 |
| 16543 | NM_133465.3 | 35.9905579 | -6.7929305 | 2.31080898 | -2.9396331 |
| 16544 | NM_001042603.2 | 180.862252 | -3.2276643 | 1.0980183 | -2.939536 |
| 16545 | NM_152504.3 | 7.76027328 | -4.5828586 | 1.55914105 | -2.9393483 |
| 16546 | NR_028414.1 | 21.0110796 | 4.71856738 | 1.60531297 | 2.93934421 |
| 16547 | NM_001297578.1 | 55.9079208 | -7.4275011 | 2.52714175 | -2.9390916 |
| 16548 | NR_077058.1 | 17.5420796 | 3.62511705 | 1.23348691 | 2.93891814 |
| 16549 | NM_152522.5 | 63.2320138 | -3.4832648 | 1.18527187 | -2.9387898 |
| 16550 | NR_037577.1 | 8.35067781 | -4.680018 | 1.59249264 | -2.9388004 |
| 16551 | NM_001329126.1 | 46.7207464 | -7.168184 | 2.43927732 | -2.9386507 |
| 16552 | NM_018559.3 | 5.8559319 | -4.1695538 | 1.4188992 | -2.9385835 |
| 16553 | NM_001033046.3 | 43.6418743 | -7.0696004 | 2.40585949 | -2.9384926 |
| 16554 | NR_125885.1 | 51.1195539 | 2.84968342 | 0.96990771 | 2.9380975 |

| | | | | | |
|---|---|---|---|---|---|
| 16555 | NM_001289910.1 | 43.4321219 | -7.0626262 | 2.40397157 | -2.9378992 |
| 16556 | NM_001004303.4 | 12.5446607 | -5.2750023 | 1.79568125 | -2.937605 |
| 16557 | NM_003568.2 | 13.0295005 | -3.3871846 | 1.1531375 | -2.937364 |
| 16558 | NM_001300932.1 | 8.52847646 | -4.7124671 | 1.60438048 | -2.9372503 |
| 16559 | NM_004458.2 | 71.6792299 | -6.8162137 | 2.32068571 | -2.937155 |
| 16560 | NM_001184987.1 | 45.8929045 | -7.1422269 | 2.43177528 | -2.9370423 |
| 16561 | NM_001130955.1 | 8.26451202 | -4.6772632 | 1.59263331 | -2.9368111 |
| 16562 | NR_033912.1 | 9.18613816 | 7.36159821 | 2.5068164 | 2.93663238 |
| 16563 | NM_178819.3 | 382.7404 | -2.9716574 | 1.01196545 | -2.9365206 |
| 16564 | NM_001164586.1 | 7.53404848 | -4.5362643 | 1.54481968 | -2.9364361 |
| 16565 | NM_173619.3 | 11.2893736 | -5.1253203 | 1.74551718 | -2.9362761 |
| 16566 | NM_145805.2 | 9.10222328 | -4.8114564 | 1.63874382 | -2.9360638 |
| 16567 | NM_001317101.1 | 10.9727833 | -5.0763152 | 1.72904733 | -2.935903 |
| 16568 | NR_121565.1 | 47.3278848 | 2.98396164 | 1.01649168 | 2.93554951 |
| 16569 | NM_001317138.1 | 9.04680464 | -4.8014651 | 1.63576127 | -2.9353092 |
| 16570 | NM_001308236.1 | 8.90899174 | -4.7768546 | 1.62744547 | -2.9351857 |
| 16571 | NM_024421.2 | 5.55383481 | -4.0975342 | 1.39614769 | -2.9348859 |
| 16572 | NM_032411.2 | 7.74928069 | -4.5815004 | 1.56109551 | -2.9347983 |
| 16573 | NM_001314033.2 | 8.31506822 | -4.6673145 | 1.59035786 | -2.9347574 |
| 16574 | NM_001271863.1 | 10.5439897 | -5.0236312 | 1.71186647 | -2.9345929 |
| 16575 | NM_153813.2 | 68.2099435 | -3.7749121 | 1.28642617 | -2.934418 |
| 16576 | NM_001317784.1 | 148.755866 | -2.5986499 | 0.88561738 | -2.9342806 |
| 16577 | NM_001257392.1 | 38.0632202 | -6.8737566 | 2.34278903 | -2.9340058 |
| 16578 | NM_006269.1 | 17.6515503 | 5.96063289 | 2.03184461 | 2.93360666 |
| 16579 | NM_003004.2 | 5.97353324 | -4.2021009 | 1.43243221 | -2.9335426 |
| 16580 | NM_004997.2 | 9.36298186 | 6.57931791 | 2.24299072 | 2.93327915 |
| 16581 | NM_003268.5 | 16.0690952 | 4.05567941 | 1.3827158 | 2.93312582 |
| 16582 | NM_001261830.1 | 8.68427024 | -4.7428533 | 1.61717936 | -2.9327936 |
| 16583 | NR_038392.1 | 9.62208287 | -4.8947021 | 1.66907576 | -2.9325823 |
| 16584 | NM_001165893.1 | 34.9017994 | -6.7484542 | 2.3012436 | -2.9325249 |
| 16585 | NR_051996.1 | 16.4114644 | 6.33461724 | 2.16023694 | 2.9323715 |
| 16586 | NR_033874.1 | 6.95856461 | -4.41953 | 1.50718411 | -2.9323093 |
| 16587 | NM_001195302.1 | 10.7978212 | -5.0596928 | 1.72553218 | -2.9322506 |
| 16588 | NM_002376.6 | 54.7272637 | -7.3956513 | 2.5221636 | -2.9322647 |
| 16589 | NM_015659.2 | 487.087963 | -1.6162767 | 0.55120015 | -2.9322864 |
| 16590 | NM_001167989.1 | 13.7695746 | -5.411178 | 1.84543034 | -2.9322039 |
| 16591 | NM_001105194.1 | 47.5228102 | -7.192945 | 2.45339569 | -2.9318324 |
| 16592 | NM_001320037.1 | 9.94694827 | -4.9332318 | 1.68290734 | -2.9313746 |
| 16593 | NR_136151.1 | 7.15880213 | -4.464548 | 1.523082 | -2.9312591 |
| 16594 | NM_021147.4 | 15.8852062 | -5.6156206 | 1.91579093 | -2.9312283 |
| 16595 | NM_001190317.1 | 34.7516804 | -6.7423613 | 2.3004624 | -2.9308722 |
| 16596 | NM_001321560.1 | 64.3529732 | -7.6291595 | 2.60302919 | -2.9308774 |
| 16597 | NM_004035.6 | 186.272476 | -1.7403402 | 0.59381155 | -2.9307954 |

| | | | | | |
|---|---|---|---|---|---|
| 16598 | NM_001177383.1 | 45.9673516 | -7.1442875 | 2.4376853 | -2.9307669 |
| 16599 | NM_006994.4 | 20.1750869 | -3.165353 | 1.08026106 | -2.9301741 |
| 16600 | NM_001292030.1 | 88.578425 | 4.65122662 | 1.58739086 | 2.93010797 |
| 16601 | NM_001278742.1 | 18.6917158 | 9.13853928 | 3.11886967 | 2.93008052 |
| 16602 | NM_001242415.1 | 17.2940545 | -3.3346748 | 1.13813046 | -2.9299583 |
| 16603 | NR_134658.1 | 8.7185049 | -4.7575795 | 1.62377939 | -2.9299421 |
| 16604 | NM_001167741.1 | 35.376163 | -6.7677899 | 2.30990254 | -2.9299028 |
| 16605 | NR_045682.1 | 62.7817293 | 2.24729428 | 0.76701794 | 2.92991095 |
| 16606 | NM_006340.2 | 6.96853391 | -4.4097009 | 1.50517403 | -2.929695 |
| 16607 | NM_001297763.1 | 74.3839574 | 1.02518102 | 0.34997006 | 2.92933923 |
| 16608 | NM_004197.1 | 35.0029199 | -6.7525607 | 2.30522675 | -2.9292393 |
| 16609 | NM_020936.2 | 45.4584083 | -7.128403 | 2.43366207 | -2.929085 |
| 16610 | NM_017686.3 | 63.4127109 | -1.2579823 | 0.42955909 | -2.9285431 |
| 16611 | NM_002494.3 | 8.27262446 | -4.6768498 | 1.59726941 | -2.9280282 |
| 16612 | NR_026899.1 | 28.3084233 | -3.0995825 | 1.0586907 | -2.9277508 |
| 16613 | NM_018988.3 | 31.4224214 | -2.9022018 | 0.99136047 | -2.927494 |
| 16614 | NM_032714.2 | 8.29110967 | -4.671285 | 1.59566366 | -2.9274872 |
| 16615 | NR_135074.1 | 11.6769068 | -5.1581327 | 1.76195518 | -2.9275051 |
| 16616 | NM_001301839.1 | 15.4172757 | -5.5701924 | 1.90282351 | -2.92733 |
| 16617 | NR_026887.1 | 7.99977299 | -4.615385 | 1.57684879 | -2.9269674 |
| 16618 | NM_020882.2 | 7.49042717 | -4.5182253 | 1.54371167 | -2.9268583 |
| 16619 | NM_001142579.1 | 55.3962863 | 1.16704336 | 0.39878529 | 2.92649548 |
| 16620 | NM_014838.2 | 172.823062 | -4.1797334 | 1.42830996 | -2.926349 |
| 16621 | NM_001012516.2 | 40.5577339 | -6.9648584 | 2.38068759 | -2.9255659 |
| 16622 | NM_001243013.1 | 9.37001183 | -4.8577535 | 1.66052075 | -2.9254398 |
| 16623 | NM_152372.3 | 8.43928543 | 2.46994663 | 0.84440573 | 2.92507092 |
| 16624 | NM_021069.4 | 45.5846503 | -7.132703 | 2.43851342 | -2.925021 |
| 16625 | NM_021188.2 | 45.0035215 | -7.1137862 | 2.43249076 | -2.9244864 |
| 16626 | NM_030930.3 | 47.7573759 | -3.5593401 | 1.2172361 | -2.9241164 |
| 16627 | NR_027982.1 | 47.0746893 | 3.13279094 | 1.07137666 | 2.92407987 |
| 16628 | NM_001127605.2 | 36.9969315 | -6.8327517 | 2.33684138 | -2.9239262 |
| 16629 | NM_014334.3 | 246.075828 | -1.5896063 | 0.54368273 | -2.9237756 |
| 16630 | NM_018467.3 | 45.9586193 | -3.4554947 | 1.18186491 | -2.9237646 |
| 16631 | NM_007026.3 | 53.561087 | -3.3625917 | 1.15014933 | -2.9236131 |
| 16632 | NM_152666.2 | 9.67102141 | -4.9051081 | 1.67780224 | -2.9235318 |
| 16633 | NM_001346228.1 | 9.84309233 | -4.9170885 | 1.68194313 | -2.923457 |
| 16634 | NM_000273.2 | 11.5502491 | -5.1576776 | 1.7643255 | -2.9233141 |
| 16635 | NM_001136469.2 | 32.6408445 | -3.6159414 | 1.23701985 | -2.923107 |
| 16636 | NM_001168379.1 | 20.7482783 | -4.9895127 | 1.70701968 | -2.922938 |
| 16637 | NM_152464.2 | 95.2943419 | -2.220262 | 0.75962051 | -2.9228568 |
| 16638 | NR_145419.1 | 35.173195 | -6.7596998 | 2.3130041 | -2.9224763 |
| 16639 | NM_001321383.1 | 5.81105688 | -4.1627945 | 1.42456055 | -2.9221604 |
| 16640 | NM_001145784.1 | 16.3215614 | -2.5873865 | 0.88549153 | -2.9219777 |

| | | | | | |
|---|---|---|---|---|---|
| 16641 | NM_152523.2 | 9.98213764 | 8.23638155 | 2.81900698 | 2.92173153 |
| 16642 | NM_000321.2 | 147.119069 | -4.0906652 | 1.40010096 | -2.9216931 |
| 16643 | NM_016297.3 | 382.967192 | -1.8072021 | 0.61855845 | -2.9216351 |
| 16644 | NM_003574.5 | 11.5099097 | -5.1440173 | 1.76086192 | -2.9213064 |
| 16645 | NM_001018054.2 | 47.3934555 | -7.1894704 | 2.46125035 | -2.9210643 |
| 16646 | NR_104143.1 | 8.61744977 | -4.730897 | 1.61974989 | -2.9207577 |
| 16647 | NM_015601.3 | 33.5673102 | -6.6924081 | 2.29181846 | -2.9201301 |
| 16648 | NM_139241.3 | 44.5083941 | -7.0979518 | 2.43088838 | -2.9199003 |
| 16649 | NM_001207000.1 | 9.93230152 | -4.9295546 | 1.68846766 | -2.9195434 |
| 16650 | NM_002581.4 | 71.3724525 | -3.6780988 | 1.259988 | -2.9191538 |
| 16651 | NM_001012729.1 | 56.681237 | 2.46321641 | 0.8440294 | 2.91840119 |
| 16652 | NM_006820.3 | 59.3009378 | -2.7900157 | 0.9560186 | -2.9183697 |
| 16653 | NR_073556.1 | 20.0298344 | -3.1461919 | 1.07831551 | -2.9176914 |
| 16654 | NM_001321363.1 | 38.2660409 | -6.8810047 | 2.35887945 | -2.917065 |
| 16655 | NM_001329474.1 | 6.94443571 | -4.4153319 | 1.51367255 | -2.9169664 |
| 16656 | NM_001281438.1 | 9.63139114 | -4.8929207 | 1.67744419 | -2.9168903 |
| 16657 | NM_001257359.1 | 220.092404 | -4.3996377 | 1.50859667 | -2.9163777 |
| 16658 | NR_026761.2 | 5.93232541 | -4.1871413 | 1.4357802 | -2.9162829 |
| 16659 | NM_001257320.2 | 17.1811443 | -5.7196418 | 1.96157819 | -2.9158368 |
| 16660 | NR_026742.1 | 5.6215777 | -4.1202751 | 1.41322384 | -2.9155149 |
| 16661 | NM_206855.2 | 85.3986481 | -3.596512 | 1.23382042 | -2.9149396 |
| 16662 | NM_004746.3 | 45.0623244 | -7.1155004 | 2.44113584 | -2.9148318 |
| 16663 | NM_152477.4 | 15.4505817 | -3.0959644 | 1.06234443 | -2.9142755 |
| 16664 | NR_029410.1 | 12.4744291 | -3.4103728 | 1.17028783 | -2.9141316 |
| 16665 | NM_182762.3 | 46.5026419 | 1.42008462 | 0.48731884 | 2.914077 |
| 16666 | NR_052019.1 | 9.12702182 | -4.8160324 | 1.65271421 | -2.914014 |
| 16667 | NR_135742.1 | 6.81070454 | -4.3775801 | 1.50227544 | -2.9139664 |
| 16668 | NM_002425.2 | 14.9486742 | -4.5024442 | 1.54550845 | -2.9132447 |
| 16669 | NR_110115.2 | 135.82157 | 1.48227451 | 0.50881129 | 2.91321073 |
| 16670 | NR_036512.1 | 6.12636878 | -4.2462836 | 1.45764598 | -2.9131104 |
| 16671 | NM_001282746.1 | 31.8504987 | -4.8798219 | 1.67520216 | -2.9129749 |
| 16672 | NM_207304.2 | 33.2127831 | -6.6770242 | 2.29224884 | -2.9128706 |
| 16673 | NM_001282973.1 | 10.1664319 | -4.9619399 | 1.70360046 | -2.9126195 |
| 16674 | NM_058172.5 | 49.3033397 | -7.245597 | 2.4880327 | -2.9121792 |
| 16675 | NM_000905.3 | 12.4319992 | -5.2540592 | 1.80441356 | -2.9117821 |
| 16676 | NM_145273.3 | 25.4965575 | 3.23115893 | 1.10982192 | 2.91142108 |
| 16677 | NM_006873.3 | 46.7465238 | -2.3611508 | 0.81103419 | -2.911284 |
| 16678 | NM_001037872.2 | 43.1462053 | -7.0526213 | 2.42257201 | -2.9112122 |
| 16679 | NM_001346446.1 | 33.4935072 | -6.6892562 | 2.29797488 | -2.9109353 |
| 16680 | NM_213599.2 | 40.4743055 | -6.9609601 | 2.39131274 | -2.9109367 |
| 16681 | NM_152486.2 | 353.597111 | -4.4494087 | 1.52878742 | -2.9104169 |
| 16682 | NM_001700.4 | 5.81865908 | -4.166311 | 1.43164483 | -2.9101569 |
| 16683 | NR_024210.1 | 8.30012787 | -4.6645186 | 1.60300995 | -2.90985 |

| | | | | | |
|---|---|---|---|---|---|
| 16684 | NM_001347923.1 | 11.9531913 | -5.1948274 | 1.78546494 | -2.9095096 |
| 16685 | NM_003826.2 | 122.205776 | -3.3972239 | 1.16764559 | -2.9094649 |
| 16686 | NM_001242769.1 | 8.22546853 | -4.6737362 | 1.6064564 | -2.9093452 |
| 16687 | NM_001294336.1 | 375.815532 | -3.5434692 | 1.21803168 | -2.9091766 |
| 16688 | NM_130807.2 | 181.849865 | -1.5097378 | 0.5189717 | -2.9090947 |
| 16689 | NM_001098424.1 | 33.5335231 | -6.6907002 | 2.30009204 | -2.9088837 |
| 16690 | NM_001323525.1 | 45.7214078 | -7.1365675 | 2.45338998 | -2.9088598 |
| 16691 | NM_018836.3 | 7.50142264 | -4.5333478 | 1.55880016 | -2.908229 |
| 16692 | NM_001320641.1 | 48.9789411 | -7.2360125 | 2.48838337 | -2.9079171 |
| 16693 | NM_001288973.1 | 158.231501 | -3.9473633 | 1.35748088 | -2.9078593 |
| 16694 | NM_001881.3 | 8.28965172 | -4.6732442 | 1.60724101 | -2.9076188 |
| 16695 | NM_033034.2 | 58.1910759 | -2.4722062 | 0.85031463 | -2.9074017 |
| 16696 | NM_006973.2 | 43.5570393 | -7.0667245 | 2.43102902 | -2.9068861 |
| 16697 | NR_109796.1 | 44.3546686 | -7.0926574 | 2.44000373 | -2.9068224 |
| 16698 | NM_019596.5 | 46.8065023 | -7.1710535 | 2.46780729 | -2.9058401 |
| 16699 | NM_005793.4 | 33.3596698 | -6.683282 | 2.30004302 | -2.9057204 |
| 16700 | NM_001079881.1 | 9.86812977 | -4.9199848 | 1.69329101 | -2.9055754 |
| 16701 | NM_000678.3 | 8.53668822 | -4.7210679 | 1.62501 | -2.9052546 |
| 16702 | NM_001330588.1 | 6.49358091 | -4.3148496 | 1.48536067 | -2.9049171 |
| 16703 | NM_001244735.1 | 8.28906789 | -4.6792714 | 1.6111811 | -2.9042492 |
| 16704 | NM_001012271.1 | 32.4306539 | -2.4704946 | 0.85078823 | -2.9037715 |
| 16705 | NM_001347729.1 | 6.29923583 | -4.2739274 | 1.47191036 | -2.9036601 |
| 16706 | NM_001321217.1 | 10.0040474 | -3.8921095 | 1.34047721 | -2.9035253 |
| 16707 | NM_001278284.1 | 35.8534327 | -6.7875344 | 2.3377724 | -2.9034197 |
| 16708 | NM_001290018.1 | 7.01011195 | -4.414424 | 1.52041716 | -2.9034295 |
| 16709 | NR_046661.1 | 17.9476578 | 2.90287672 | 0.9998291 | 2.90337289 |
| 16710 | NM_001286779.1 | 34.3673943 | -6.7262146 | 2.31672214 | -2.9033325 |
| 16711 | NR_135264.1 | 13.6093771 | -5.3833876 | 1.85422185 | -2.9033136 |
| 16712 | NM_001320437.1 | 32.6744868 | -6.6534043 | 2.29174652 | -2.9032026 |
| 16713 | NR_003266.2 | 43.6051937 | -2.9797576 | 1.02643667 | -2.9030116 |
| 16714 | NM_001319307.1 | 43.8782536 | -7.0775221 | 2.43805584 | -2.9029368 |
| 16715 | NM_001099692.1 | 7.39146951 | -4.5073738 | 1.55273074 | -2.9028689 |
| 16716 | NM_145301.2 | 7.00012016 | -4.4157934 | 1.52119108 | -2.9028525 |
| 16717 | NM_144598.4 | 47.0357791 | -7.1773883 | 2.47305892 | -2.902231 |
| 16718 | NM_001323516.1 | 33.4325465 | -6.6864195 | 2.3041681 | -2.9018801 |
| 16719 | NM_001033522.1 | 11.6576846 | -5.1690076 | 1.78134275 | -2.9017479 |
| 16720 | NM_032997.2 | 7.80102558 | -4.5949653 | 1.58369699 | -2.9014169 |
| 16721 | NM_015147.2 | 48.8899451 | -7.2328596 | 2.49314106 | -2.9011032 |
| 16722 | NM_001290043.1 | 33.4606303 | -6.6877639 | 2.30536833 | -2.9009524 |
| 16723 | NM_001347969.1 | 35.5513574 | -6.7751038 | 2.33563406 | -2.9007557 |
| 16724 | NR_024418.1 | 6.64519739 | -4.3533321 | 1.50102718 | -2.9002354 |
| 16725 | NM_002506.2 | 5.62507147 | -4.1187868 | 1.42016845 | -2.90021 |
| 16726 | NM_001261403.2 | 7.38151045 | -4.4969191 | 1.55072974 | -2.8998728 |

| | | | | | |
|---|---|---|---|---|---|
| 16727 | NM_001814.5 | 81.4334211 | -3.5224571 | 1.21473206 | -2.8997811 |
| 16728 | NM_201444.2 | 12.6268729 | -5.2853421 | 1.82267095 | -2.8997785 |
| 16729 | NM_001303052.1 | 38.961224 | -6.9070809 | 2.3820946 | -2.899583 |
| 16730 | NM_145011.3 | 43.2943673 | -7.0579916 | 2.43414973 | -2.8995717 |
| 16731 | NM_033387.3 | 7.02610539 | -4.4369789 | 1.53037695 | -2.8992719 |
| 16732 | NM_001135214.2 | 35.6913907 | -6.7807817 | 2.33900588 | -2.8990016 |
| 16733 | NM_020714.2 | 235.512886 | 1.4238424 | 0.49117019 | 2.89887788 |
| 16734 | NM_001005745.1 | 78.0056436 | -3.2362294 | 1.11646176 | -2.8986478 |
| 16735 | NM_001271754.1 | 15.1646996 | -4.5516701 | 1.57026788 | -2.8986583 |
| 16736 | NM_001290327.1 | 8.06530774 | -4.6330148 | 1.59834706 | -2.8986288 |
| 16737 | NM_007106.3 | 218.641468 | -4.2680527 | 1.47250918 | -2.8984897 |
| 16738 | NM_030783.2 | 38.7447716 | -6.8990535 | 2.38025416 | -2.8984525 |
| 16739 | NM_138448.3 | 9.91282465 | -4.9308255 | 1.70151445 | -2.897904 |
| 16740 | NM_001252052.1 | 38.2533551 | -6.8806093 | 2.37436515 | -2.8978733 |
| 16741 | NM_001166534.1 | 8.82837059 | 5.69471563 | 1.96529042 | 2.89764585 |
| 16742 | NM_002463.1 | 6.34265075 | -4.2701456 | 1.47386983 | -2.8972339 |
| 16743 | NR_134575.1 | 14.960263 | 4.01519614 | 1.38612274 | 2.89671038 |
| 16744 | NM_020700.1 | 89.7280129 | -2.7023352 | 0.93294665 | -2.8965592 |
| 16745 | NR_024490.1 | 98.1144846 | -1.2589723 | 0.4346676 | -2.8964025 |
| 16746 | NM_001130100.1 | 40.8545371 | -6.9749014 | 2.4085337 | -2.8959119 |
| 16747 | NM_001204423.1 | 13.6155151 | -4.3810988 | 1.51288728 | -2.8958528 |
| 16748 | NR_040584.2 | 55.0661063 | -1.7790332 | 0.61436053 | -2.8957479 |
| 16749 | NM_001013660.2 | 50.4092454 | 2.91738545 | 1.00750254 | 2.89566063 |
| 16750 | NM_024747.5 | 74.8561148 | -2.8190335 | 0.97367267 | -2.8952579 |
| 16751 | NM_001135669.1 | 46.7032899 | -7.1667594 | 2.47610552 | -2.8943675 |
| 16752 | NM_152349.2 | 14.589542 | -4.4843271 | 1.54946024 | -2.8941221 |
| 16753 | NM_004334.2 | 6.61898957 | -4.3497619 | 1.50307307 | -2.8939125 |
| 16754 | NM_001142450.1 | 34.3680547 | -6.726402 | 2.3243567 | -2.8938768 |
| 16755 | NM_180699.3 | 9.63850474 | -3.8810715 | 1.34122201 | -2.8936831 |
| 16756 | NM_001270400.1 | 5.75735987 | -4.1386856 | 1.43029637 | -2.8935861 |
| 16757 | NR_028077.1 | 9.64897515 | -4.8838246 | 1.68781252 | -2.8935824 |
| 16758 | NM_001177382.1 | 41.4118689 | -6.9946894 | 2.41738919 | -2.8934892 |
| 16759 | NM_018023.4 | 327.000452 | -2.8787816 | 0.99507613 | -2.8930264 |
| 16760 | NM_001166284.1 | 7.49390855 | -4.5264697 | 1.56485924 | -2.892573 |
| 16761 | NM_001001668.3 | 59.8412046 | -2.3448892 | 0.81071343 | -2.8923774 |
| 16762 | NM_178813.5 | 5.8607442 | -4.1782126 | 1.44467865 | -2.8921398 |
| 16763 | NM_001199694.2 | 9.37378794 | -4.8617825 | 1.68146268 | -2.8914008 |
| 16764 | NM_001291965.1 | 13.6118563 | -4.3922373 | 1.51911407 | -2.891315 |
| 16765 | NM_004945.3 | 44.5506375 | -7.0989345 | 2.45543461 | -2.8911112 |
| 16766 | NM_172069.3 | 42.8875915 | -3.1650716 | 1.09490747 | -2.8907206 |
| 16767 | NM_001008405.2 | 7.39821579 | -4.4911735 | 1.55375065 | -2.8905369 |
| 16768 | NM_016247.3 | 40.4584003 | 1.96907087 | 0.68123914 | 2.89042534 |
| 16769 | NM_000180.3 | 15.9429287 | 5.21439028 | 1.80414996 | 2.89021998 |

| | | | | | |
|---|---|---|---|---|---|
| 16770 | NR_136606.1 | 38.8786553 | -6.9037118 | 2.38884861 | -2.8899746 |
| 16771 | NM_001100626.1 | 40.6388049 | -6.967108 | 2.41105409 | -2.8896523 |
| 16772 | NM_001303012.1 | 21.2779268 | -3.9115398 | 1.35380732 | -2.8892884 |
| 16773 | NM_182985.4 | 7.44738712 | -4.5293225 | 1.56807119 | -2.8884674 |
| 16774 | NM_001242798.1 | 43.4338053 | -7.062173 | 2.44522644 | -2.8881468 |
| 16775 | NM_001143808.1 | 12.7852068 | 8.58785896 | 2.97413048 | 2.88751924 |
| 16776 | NM_001005782.1 | 11.7341239 | -5.1750382 | 1.79230211 | -2.8873694 |
| 16777 | NM_001695.4 | 240.494597 | -2.101396 | 0.7278904 | -2.8869676 |
| 16778 | NR_027141.2 | 6.51826493 | -4.3247543 | 1.49872669 | -2.885619 |
| 16779 | NM_004224.3 | 6.16780739 | -4.2614095 | 1.4769265 | -2.8853227 |
| 16780 | NM_197958.2 | 6.25829809 | -4.2733701 | 1.4810808 | -2.8853052 |
| 16781 | NM_032445.2 | 85.4804457 | -3.8529933 | 1.33562539 | -2.8847859 |
| 16782 | NR_104485.1 | 6.44415219 | -4.3164385 | 1.49634472 | -2.8846551 |
| 16783 | NM_024056.3 | 42.5583048 | -7.0335521 | 2.43836377 | -2.8845377 |
| 16784 | NR_138032.1 | 82.1046226 | 2.59212645 | 0.89862854 | 2.8845361 |
| 16785 | NM_001055.3 | 11.0582316 | -5.0948054 | 1.76635041 | -2.8843685 |
| 16786 | NM_183005.4 | 7.38403859 | -4.4929446 | 1.55768742 | -2.8843686 |
| 16787 | NM_001130041.1 | 46.9946229 | -7.176319 | 2.48811231 | -2.8842424 |
| 16788 | NM_001316978.1 | 5.56154267 | -4.0989403 | 1.42144366 | -2.883646 |
| 16789 | NM_001256015.1 | 5.4741753 | -4.0751213 | 1.41330429 | -2.8833998 |
| 16790 | NM_001286704.1 | 7.94496289 | -4.6247199 | 1.60401442 | -2.8832159 |
| 16791 | NM_001288624.1 | 19.8125394 | 1.80482398 | 0.62609289 | 2.88267764 |
| 16792 | NM_005640.2 | 7.78172692 | -4.5900309 | 1.5926324 | -2.8820404 |
| 16793 | NM_014315.2 | 175.249613 | -4.1980021 | 1.45666056 | -2.8819357 |
| 16794 | NM_001130916.2 | 52.953304 | -2.5268384 | 0.87682069 | -2.8818189 |
| 16795 | NM_017453.3 | 42.7818998 | -7.041826 | 2.44378534 | -2.8815239 |
| 16796 | NR_024603.1 | 6.18399754 | -4.250709 | 1.47541697 | -2.8810222 |
| 16797 | NR_110317.1 | 10.9592732 | -5.0681913 | 1.7595045 | -2.8804651 |
| 16798 | NM_001160367.1 | 43.5898429 | -7.0675393 | 2.45400306 | -2.8800043 |
| 16799 | NR_038242.1 | 35.4890072 | -6.7727201 | 2.35172916 | -2.8798895 |
| 16800 | NM_001256092.1 | 33.10112 | -6.6722117 | 2.31691016 | -2.8797887 |
| 16801 | NM_001130082.2 | 153.827932 | -5.2890769 | 1.83687942 | -2.8793816 |
| 16802 | NM_004820.4 | 33.0688073 | -1.4492023 | 0.50333266 | -2.8792138 |
| 16803 | NM_001003962.2 | 43.1653547 | -7.0541523 | 2.45030224 | -2.8788907 |
| 16804 | NM_001172220.1 | 41.7792768 | -7.0066526 | 2.43387832 | -2.8788015 |
| 16805 | NM_016612.3 | 293.758257 | 6.58779179 | 2.28884477 | 2.87821694 |
| 16806 | NM_001145310.3 | 6.26678363 | -4.2845948 | 1.48884288 | -2.8778019 |
| 16807 | NM_145014.2 | 21.9661083 | -2.2617259 | 0.78600096 | -2.8775104 |
| 16808 | NM_001114635.1 | 20.1191525 | -4.0123751 | 1.39440355 | -2.8774849 |
| 16809 | NM_001114089.3 | 46.0036943 | -7.1452708 | 2.4834043 | -2.877208 |
| 16810 | NM_004018.2 | 31.3862522 | -6.5954913 | 2.292534 | -2.8769437 |
| 16811 | NM_080474.2 | 43.2926389 | 3.68506656 | 1.28094462 | 2.8768352 |
| 16812 | NM_020463.3 | 42.6656597 | -6.0631716 | 2.10772952 | -2.8766365 |

| | | | | | |
|---|---|---|---|---|---|
| 16813 | NM_001199295.1 | 31.176176 | -6.5858074 | 2.28970976 | -2.876263 |
| 16814 | NM_001024216.1 | 162.649347 | 1.61532301 | 0.56167393 | 2.8759088 |
| 16815 | NM_001290072.1 | 6.56474958 | -4.3205069 | 1.50246318 | -2.8756158 |
| 16816 | NR_024344.1 | 134.693412 | -4.1211129 | 1.43315049 | -2.8755619 |
| 16817 | NM_001205028.1 | 17.2215255 | -3.8015837 | 1.32207034 | -2.8754776 |
| 16818 | NM_001145265.1 | 12.1011711 | -5.2171155 | 1.8144386 | -2.8753332 |
| 16819 | NM_001330301.1 | 43.4142959 | -7.0618893 | 2.45617479 | -2.8751575 |
| 16820 | NR_002223.3 | 11.4962816 | 5.01757604 | 1.74519566 | 2.87507937 |
| 16821 | NM_001317803.1 | 10.2038425 | -4.9825973 | 1.73341229 | -2.8744444 |
| 16822 | NM_006208.2 | 99.7654243 | 2.28859594 | 0.79639707 | 2.87368703 |
| 16823 | NM_001197295.1 | 6.02654392 | -4.227865 | 1.47146577 | -2.8732337 |
| 16824 | NM_021030.2 | 35.1176132 | -3.0155917 | 1.04955039 | -2.8732224 |
| 16825 | NM_001126051.1 | 5.98620418 | -4.2132157 | 1.46641513 | -2.8731398 |
| 16826 | NM_205548.2 | 6.05119093 | -4.2249864 | 1.47055718 | -2.8730515 |
| 16827 | NM_024503.4 | 41.6792243 | -7.0030433 | 2.43765844 | -2.8728567 |
| 16828 | NM_001142345.1 | 15.3838937 | 5.39038139 | 1.87646124 | 2.87263134 |
| 16829 | NR_015341.2 | 121.699077 | -2.9400678 | 1.02348777 | -2.8725969 |
| 16830 | NM_001317851.1 | 32.0113222 | -6.6236751 | 2.30585768 | -2.8725429 |
| 16831 | NM_006350.3 | 10.6252924 | -5.0318089 | 1.75190993 | -2.8721847 |
| 16832 | NM_001222.3 | 35.5150328 | -6.7733471 | 2.35838182 | -2.8720316 |
| 16833 | NM_001135914.1 | 6.09399334 | -4.2332009 | 1.47396896 | -2.8719743 |
| 16834 | NM_001329237.1 | 44.9825364 | -7.1145097 | 2.47774594 | -2.8713637 |
| 16835 | NM_022111.3 | 40.9589911 | -6.9781053 | 2.43086629 | -2.8706249 |
| 16836 | NM_001270470.1 | 9.5985772 | -4.8921476 | 1.70433448 | -2.8704152 |
| 16837 | NM_001137610.1 | 5.8243938 | -4.1756161 | 1.4547564 | -2.8703198 |
| 16838 | NM_001289931.1 | 12.7378815 | 7.41602054 | 2.5837117 | 2.87029723 |
| 16839 | NM_001130919.1 | 32.5758947 | -6.6488129 | 2.31647648 | -2.8702268 |
| 16840 | NM_001101391.1 | 5.71070777 | -4.1230037 | 1.43649025 | -2.8701926 |
| 16841 | NM_001322436.1 | 7.94247743 | -4.5987848 | 1.60234393 | -2.870036 |
| 16842 | NM_033244.3 | 9.80345712 | -4.9047172 | 1.70912676 | -2.8697211 |
| 16843 | NM_198441.2 | 3.41769264 | 6.71127366 | 2.33866331 | 2.86970494 |
| 16844 | NM_019035.4 | 35.4802941 | -6.7724233 | 2.36038453 | -2.8692034 |
| 16845 | NM_006434.2 | 34.4553139 | -6.7296392 | 2.34560242 | -2.8690451 |
| 16846 | NM_024654.4 | 527.684723 | 1.39330161 | 0.48581454 | 2.86797018 |
| 16847 | NM_001003794.2 | 41.6381911 | -7.0013808 | 2.44190814 | -2.8671762 |
| 16848 | NM_001271723.1 | 9.48751178 | -4.874674 | 1.70018238 | -2.8671477 |
| 16849 | NM_001134670.1 | 81.2693657 | 2.19944821 | 0.7671384 | 2.86708136 |
| 16850 | NM_001202498.1 | 24.1395959 | 6.19527766 | 2.16089323 | 2.86699851 |
| 16851 | NM_001023570.3 | 32.1190204 | -6.6289913 | 2.31256228 | -2.8665136 |
| 16852 | NM_020169.3 | 7.08969988 | -4.4612501 | 1.55636878 | -2.8664479 |
| 16853 | NM_053284.2 | 6.71798108 | -4.3793006 | 1.5278445 | -2.8663261 |
| 16854 | NM_024926.3 | 39.2360455 | -6.9163951 | 2.41317187 | -2.8661013 |
| 16855 | NM_001129731.2 | 11.6528872 | -5.1617759 | 1.80108599 | -2.8659242 |

| | | | | | |
|---|---|---|---|---|---|
| 16856 | NM_152679.3 | 18.6914862 | -3.474508 | 1.2125028 | -2.865567 |
| 16857 | NM_000574.4 | 8.80741326 | -4.7713176 | 1.66516659 | -2.8653695 |
| 16858 | NM_023012.5 | 148.983801 | -4.10605 | 1.43306276 | -2.8652269 |
| 16859 | NM_001134439.1 | 32.9168988 | -6.6643885 | 2.32602939 | -2.8651351 |
| 16860 | NM_001297734.1 | 33.0629588 | -6.6702777 | 2.32826322 | -2.8649156 |
| 16861 | NR_024061.1 | 85.777743 | 1.41841446 | 0.49510475 | 2.86487749 |
| 16862 | NM_015440.4 | 43.4864497 | -7.0648244 | 2.46608791 | -2.8647902 |
| 16863 | NM_001199052.1 | 7.34159992 | -4.4998094 | 1.57084723 | -2.8645748 |
| 16864 | NR_073585.1 | 9.38798209 | -4.8593738 | 1.69640602 | -2.8645111 |
| 16865 | NR_138153.1 | 7.16712456 | -4.4457143 | 1.55224793 | -2.8640491 |
| 16866 | NM_182920.1 | 30.9514511 | -6.575401 | 2.2959626 | -2.8638973 |
| 16867 | NM_000728.3 | 21.5691588 | -3.7124217 | 1.2963095 | -2.863839 |
| 16868 | NR_024420.1 | 6.23224423 | -4.2592687 | 1.48725494 | -2.8638457 |
| 16869 | NM_001242617.1 | 8.32526764 | -4.6756498 | 1.63279483 | -2.8635869 |
| 16870 | NM_001015072.3 | 5.39669405 | -4.0479035 | 1.41360649 | -2.8635293 |
| 16871 | NM_001313972.1 | 7.86144623 | -4.5989197 | 1.60602371 | -2.8635441 |
| 16872 | NM_024429.1 | 40.4618545 | -6.9603233 | 2.43096856 | -2.8631894 |
| 16873 | NM_001101376.2 | 9.82842644 | -4.9265283 | 1.72090982 | -2.8627463 |
| 16874 | NR_026962.1 | 10.4544005 | -3.995484 | 1.39569223 | -2.8627257 |
| 16875 | NM_001014980.2 | 7.37258679 | -4.5080818 | 1.57486429 | -2.8625208 |
| 16876 | NM_001004346.3 | 7.3920136 | -4.5182416 | 1.57846719 | -2.8624235 |
| 16877 | NM_005135.2 | 68.9247848 | 5.81083639 | 2.03005404 | 2.86240478 |
| 16878 | NM_001258320.1 | 7.89642885 | -4.592423 | 1.60457819 | -2.8620749 |
| 16879 | NM_001345871.1 | 33.2542026 | -6.6787695 | 2.3339552 | -2.8615671 |
| 16880 | NM_001142274.1 | 207.823234 | -2.8390228 | 0.99216495 | -2.8614423 |
| 16881 | NM_001252641.1 | 12.3623889 | -3.2934108 | 1.15115225 | -2.8609689 |
| 16882 | NM_001164404.1 | 5.49819335 | -4.0876422 | 1.42913812 | -2.8602149 |
| 16883 | NM_001323534.1 | 30.68837 | -6.56293 | 2.29474711 | -2.8599797 |
| 16884 | NM_013310.3 | 7.28792009 | -4.4894399 | 1.56981413 | -2.8598545 |
| 16885 | NM_080743.4 | 83.7230954 | -1.8353237 | 0.64177126 | -2.8597786 |
| 16886 | NR_040090.1 | 12.7782239 | -2.8214208 | 0.98670674 | -2.8594319 |
| 16887 | NM_001321123.1 | 42.2439869 | -7.0227789 | 2.4561889 | -2.8592177 |
| 16888 | NR_125872.1 | 8.91461246 | -4.7776293 | 1.67106263 | -2.8590366 |
| 16889 | NM_175888.3 | 7.0377203 | -4.4466599 | 1.55572953 | -2.8582474 |
| 16890 | NR_027058.1 | 178.061478 | 1.37184867 | 0.47996294 | 2.85823876 |
| 16891 | NM_001242861.2 | 43.158001 | -7.0528456 | 2.46766915 | -2.8581002 |
| 16892 | NM_015297.2 | 44.3347987 | -7.0923435 | 2.48162299 | -2.8579456 |
| 16893 | NM_173163.2 | 7.83717387 | -4.5993044 | 1.6101159 | -2.8565052 |
| 16894 | NM_001321980.1 | 6.15347933 | -4.22587 | 1.47950609 | -2.8562708 |
| 16895 | NM_001206947.1 | 50.6499502 | -7.2835978 | 2.55017482 | -2.8561171 |
| 16896 | NR_138051.1 | 32.3363374 | -6.6381524 | 2.32424145 | -2.8560511 |
| 16897 | NM_015715.4 | 6.63884183 | -4.3492354 | 1.52285913 | -2.855967 |
| 16898 | NM_001258312.1 | 40.9501914 | -6.9773153 | 2.4432548 | -2.8557461 |

| | | | | | |
|---|---|---|---|---|---|
| 16899 | NM_175856.4 | 16.5972826 | -3.5947146 | 1.25893313 | -2.8553658 |
| 16900 | NR_028045.1 | 6.8553456 | -4.3936912 | 1.5388508 | -2.8551768 |
| 16901 | NM_007181.5 | 10.0636206 | -3.8861508 | 1.36116959 | -2.8550086 |
| 16902 | NM_001556.2 | 32.9369594 | -6.6646157 | 2.33439498 | -2.8549649 |
| 16903 | NM_002217.3 | 13.2612151 | -4.3197141 | 1.51306245 | -2.8549477 |
| 16904 | NM_001278718.1 | 9.30614098 | -4.8458115 | 1.69739057 | -2.8548594 |
| 16905 | NM_001134387.1 | 9.76163804 | -4.9063582 | 1.71862778 | -2.8548114 |
| 16906 | NM_182639.3 | 6.96027319 | -4.4211776 | 1.54873568 | -2.8547012 |
| 16907 | NM_001277444.1 | 10.4476317 | 8.30954513 | 2.91098882 | 2.85454381 |
| 16908 | NM_001329848.1 | 7.80977029 | -4.5790276 | 1.60413673 | -2.854512 |
| 16909 | NM_133373.4 | 128.804496 | -3.3156396 | 1.16157202 | -2.8544417 |
| 16910 | NM_032986.3 | 42.4339779 | -6.0462054 | 2.11825882 | -2.854328 |
| 16911 | NM_152326.2 | 60.9768382 | -3.2968981 | 1.15546017 | -2.8533204 |
| 16912 | NM_000603.4 | 5.91632903 | -4.1879332 | 1.46786258 | -2.8530826 |
| 16913 | NR_033363.1 | 70.698048 | -3.8172713 | 1.33853423 | -2.8518294 |
| 16914 | NM_001128159.2 | 341.507253 | -1.8517413 | 0.64934107 | -2.8517236 |
| 16915 | NM_001297701.1 | 9.26691286 | -4.8443913 | 1.69921644 | -2.850956 |
| 16916 | NM_017895.7 | 127.563475 | -3.3198004 | 1.16453387 | -2.8507546 |
| 16917 | NR_134869.1 | 10.5056259 | -3.9518808 | 1.3863173 | -2.8506322 |
| 16918 | NM_004375.4 | 31.789196 | -6.6140347 | 2.32027313 | -2.8505414 |
| 16919 | NM_012082.3 | 65.6837386 | -4.0263394 | 1.41264766 | -2.8502078 |
| 16920 | NR_038440.1 | 8.04773854 | 7.93767911 | 2.7850441 | 2.85010895 |
| 16921 | NR_040059.1 | 11.2052946 | -5.1051966 | 1.79123777 | -2.8500943 |
| 16922 | NM_000550.2 | 23.0709841 | -3.0495854 | 1.07003624 | -2.8499833 |
| 16923 | NR_109956.1 | 30.6256744 | 4.43307913 | 1.55547689 | 2.84998071 |
| 16924 | NM_001256526.1 | 7.03105734 | -4.4445962 | 1.55962163 | -2.8497913 |
| 16925 | NM_014591.4 | 5.83590319 | -4.1722373 | 1.46415102 | -2.8495949 |
| 16926 | NM_018313.4 | 100.869021 | -3.6585077 | 1.28386585 | -2.8496027 |
| 16927 | NM_001288709.1 | 11.2835022 | -5.1231131 | 1.79811858 | -2.849152 |
| 16928 | NM_001306219.2 | 30.0209585 | -6.531475 | 2.29251473 | -2.8490439 |
| 16929 | NR_130897.1 | 5.63724986 | -4.1231409 | 1.44747751 | -2.8485009 |
| 16930 | NM_001080521.2 | 7.28161902 | -4.4769127 | 1.57187692 | -2.8481318 |
| 16931 | NR_033982.1 | 6.26018895 | -4.269012 | 1.49907039 | -2.8477729 |
| 16932 | NM_001261411.1 | 6.67931806 | -4.3750371 | 1.5364332 | -2.8475284 |
| 16933 | NM_001326580.1 | 11.8379338 | -4.1601986 | 1.46102322 | -2.8474555 |
| 16934 | NM_147148.2 | 7.21097246 | -4.4693574 | 1.5698933 | -2.8469179 |
| 16935 | NM_001077242.1 | 6.44929186 | -4.3083238 | 1.5133498 | -2.846879 |
| 16936 | NR_135542.1 | 51.1214281 | -7.2974847 | 2.56366825 | -2.8465012 |
| 16937 | NM_181578.4 | 7.33093677 | -4.4779121 | 1.57321823 | -2.8463388 |
| 16938 | NM_004842.3 | 14.8092925 | -5.5054828 | 1.93434264 | -2.8461776 |
| 16939 | NM_001329201.1 | 8.91916182 | -4.781834 | 1.68039727 | -2.8456569 |
| 16940 | NM_005608.2 | 5.26683734 | -4.0146343 | 1.41085085 | -2.8455412 |
| 16941 | NM_024119.2 | 11.3421395 | -5.1310267 | 1.80334529 | -2.8452824 |

| | | | | | |
|---|---|---|---|---|---|
| 16942 | NM_004196.4 | 6.06125667 | -4.2104004 | 1.47981724 | -2.8452165 |
| 16943 | NM_145284.5 | 29.5172457 | -6.5069721 | 2.28697717 | -2.8452283 |
| 16944 | NR_038302.1 | 7.87954906 | -4.6107649 | 1.62056798 | -2.8451537 |
| 16945 | NM_022817.2 | 91.5030926 | -0.8480978 | 0.2981945 | -2.8441094 |
| 16946 | NM_001100590.1 | 278.819081 | -3.8516457 | 1.35427388 | -2.844067 |
| 16947 | NM_001278194.1 | 51.7445757 | -7.3162857 | 2.57250435 | -2.8440324 |
| 16948 | NM_006252.3 | 60.7960899 | -1.8836998 | 0.66240376 | -2.8437335 |
| 16949 | NM_001034116.1 | 31.2488435 | -6.5892715 | 2.31731875 | -2.8434895 |
| 16950 | NM_001135208.1 | 8.86235933 | -4.7752351 | 1.67972746 | -2.842863 |
| 16951 | NR_003262.1 | 97.2908438 | 1.78640662 | 0.62839773 | 2.84279613 |
| 16952 | NM_006906.1 | 41.2598611 | -6.9887979 | 2.45846115 | -2.842753 |
| 16953 | NM_018661.4 | 14.225374 | 8.75329282 | 3.07946108 | 2.84247555 |
| 16954 | NR_110206.1 | 6.89264408 | -4.3975765 | 1.54718186 | -2.8423139 |
| 16955 | NM_022153.1 | 6.64696828 | -4.3634743 | 1.53549715 | -2.8417339 |
| 16956 | NM_182523.1 | 8.80946027 | -4.7552604 | 1.67340283 | -2.8416711 |
| 16957 | NM_033208.3 | 55.0143766 | -3.2640941 | 1.14866482 | -2.8416419 |
| 16958 | NM_001317902.1 | 6.52381239 | 5.00359888 | 1.76084595 | 2.84158809 |
| 16959 | NM_006028.4 | 5.2410226 | -4.0129532 | 1.41222426 | -2.8415835 |
| 16960 | NM_001320591.1 | 38.8213481 | -6.9001214 | 2.4285572 | -2.8412431 |
| 16961 | NM_001039664.1 | 9.31684469 | -3.780366 | 1.3306025 | -2.8410934 |
| 16962 | NM_001281749.1 | 54.2928254 | -3.2430868 | 1.14155361 | -2.8409413 |
| 16963 | NR_144356.1 | 7.76600261 | -4.5684755 | 1.60827722 | -2.840602 |
| 16964 | NM_001346877.1 | 7.68068249 | -4.5769267 | 1.61132671 | -2.8404709 |
| 16965 | NM_001077498.2 | 38.9569287 | -6.905842 | 2.43126752 | -2.8404287 |
| 16966 | NR_131192.1 | 7.58505595 | 3.83131403 | 1.34911682 | 2.83986825 |
| 16967 | NM_000722.3 | 129.957867 | -4.067019 | 1.43225579 | -2.8395899 |
| 16968 | NM_001256755.1 | 8.72800375 | 8.05519781 | 2.8368727 | 2.83946396 |
| 16969 | NM_001101357.1 | 8.52728513 | -4.7224263 | 1.66315318 | -2.8394416 |
| 16970 | NR_036505.1 | 5.80525049 | -4.1636065 | 1.4663916 | -2.8393551 |
| 16971 | NM_000247.2 | 7.73134332 | -4.5782721 | 1.61262112 | -2.8390253 |
| 16972 | NM_003506.3 | 29.3445643 | -6.4986154 | 2.28913625 | -2.8388941 |
| 16973 | NM_001257335.1 | 6.30878875 | -4.2938485 | 1.51253234 | -2.8388474 |
| 16974 | NM_000995.4 | 16.8693331 | -3.8300337 | 1.34920199 | -2.83874 |
| 16975 | NM_001305203.1 | 36.079274 | -6.7962734 | 2.39444801 | -2.8383466 |
| 16976 | NM_001508.2 | 9.24538371 | -4.8213148 | 1.69867524 | -2.8382793 |
| 16977 | NM_001190716.1 | 44.9531333 | -7.1128253 | 2.50666053 | -2.8375702 |
| 16978 | NM_199077.2 | 6.5087974 | -4.338593 | 1.52898865 | -2.8375573 |
| 16979 | NR_022007.2 | 6.20400375 | -4.2387394 | 1.49384548 | -2.8374685 |
| 16980 | NM_001079804.2 | 42.4338732 | -7.0301981 | 2.47765727 | -2.8374377 |
| 16981 | NR_037613.1 | 6.65133418 | -4.3452193 | 1.5315416 | -2.8371539 |
| 16982 | NM_001258299.1 | 7.50538927 | -4.5243945 | 1.59482624 | -2.83692 |
| 16983 | NM_173607.3 | 81.0387958 | -1.5673867 | 0.55253365 | -2.8367262 |
| 16984 | NM_001033551.2 | 39.2640838 | -6.9180458 | 2.43895333 | -2.8364814 |

| | | | | | |
|---|---|---|---|---|---|
| 16985 | NM_001257396.1 | 7.11934174 | -4.4395015 | 1.56518307 | -2.8364104 |
| 16986 | NM_017709.3 | 10.4353037 | -5.0118246 | 1.76699051 | -2.8363619 |
| 16987 | NR_024259.1 | 24.0817747 | 2.69797592 | 0.95134283 | 2.83596599 |
| 16988 | NM_001330489.1 | 29.1933849 | -6.4910868 | 2.28888856 | -2.8359121 |
| 16989 | NM_018293.2 | 48.6829747 | -3.2124302 | 1.13287937 | -2.835633 |
| 16990 | NM_001202404.1 | 3.99578473 | 6.88123386 | 2.42700151 | 2.83528207 |
| 16991 | NM_001130923.1 | 31.0307904 | -6.5790515 | 2.3205795 | -2.8350899 |
| 16992 | NM_199461.3 | 61.2763723 | -1.5040606 | 0.53069695 | -2.8341232 |
| 16993 | NM_001282763.1 | 36.8583665 | -6.8267158 | 2.40929643 | -2.8334894 |
| 16994 | NM_001007090.2 | 5.90268664 | -4.1765739 | 1.47427746 | -2.8329633 |
| 16995 | NM_004405.3 | 5.68553636 | -4.1318355 | 1.45858013 | -2.8327793 |
| 16996 | NM_001319167.1 | 8.98334449 | -4.7865396 | 1.68972072 | -2.83274 |
| 16997 | NM_005686.2 | 56.5517465 | -2.6029506 | 0.91892679 | -2.8325984 |
| 16998 | NM_020121.3 | 70.6973774 | -3.4129376 | 1.20491504 | -2.8325131 |
| 16999 | NM_001204140.1 | 38.5316101 | -6.889683 | 2.43240292 | -2.8324596 |
| 17000 | NM_001184702.1 | 29.901018 | -6.5253623 | 2.30431072 | -2.8318066 |
| 17001 | NM_173791.4 | 249.096278 | -2.1040931 | 0.74307488 | -2.831603 |
| 17002 | NM_001308050.1 | 31.5603427 | -6.6036497 | 2.3323073 | -2.8313806 |
| 17003 | NR_027412.1 | 33.1703054 | 2.18280224 | 0.77101776 | 2.83106612 |
| 17004 | NM_001031850.3 | 9.88131023 | 8.21580274 | 2.90247469 | 2.83061995 |
| 17005 | NM_001321350.1 | 61.973386 | -3.2558842 | 1.15027244 | -2.8305331 |
| 17006 | NM_001037164.2 | 29.9144547 | -6.5262879 | 2.30573009 | -2.8304648 |
| 17007 | NM_001220474.2 | 40.1736042 | -6.9499275 | 2.45605843 | -2.8297077 |
| 17008 | NM_015275.2 | 138.131572 | -3.0513401 | 1.07844012 | -2.8294015 |
| 17009 | NM_001306076.1 | 6.9813527 | -4.4391851 | 1.56899864 | -2.829311 |
| 17010 | NM_001300885.1 | 32.9835179 | -6.6667605 | 2.35652435 | -2.829065 |
| 17011 | NM_001300814.1 | 7.66520144 | 7.16415703 | 2.53289176 | 2.82844974 |
| 17012 | NM_002600.3 | 9.08034377 | -4.8150408 | 1.70257445 | -2.8280941 |
| 17013 | NM_005850.4 | 155.664968 | -3.8541644 | 1.3628881 | -2.827939 |
| 17014 | NR_024623.1 | 36.4203278 | 3.68700623 | 1.30382597 | 2.82783618 |
| 17015 | NM_001322963.1 | 8.52782995 | -4.7142498 | 1.667373 | -2.8273517 |
| 17016 | NM_001078650.2 | 6.89129695 | -4.4141343 | 1.56127099 | -2.8272698 |
| 17017 | NM_022664.2 | 13.8376123 | 8.71347281 | 3.08229835 | 2.82694011 |
| 17018 | NR_120488.1 | 13.8376123 | 8.71347281 | 3.08229835 | 2.82694011 |
| 17019 | NM_001134492.1 | 5.36890617 | -4.056465 | 1.43494195 | -2.8269192 |
| 17020 | NM_001318532.1 | 6.99636472 | -4.4378233 | 1.56990393 | -2.826812 |
| 17021 | NR_037599.1 | 9.81334609 | -3.9107607 | 1.38387945 | -2.8259403 |
| 17022 | NM_001198629.1 | 38.4848407 | -6.8880425 | 2.43750249 | -2.8258607 |
| 17023 | NM_001308357.1 | 8.51793247 | -4.7089162 | 1.66656982 | -2.8255139 |
| 17024 | NM_181868.1 | 31.9175898 | -6.6192506 | 2.34286116 | -2.825285 |
| 17025 | NR_033873.1 | 24.4430297 | -2.5164205 | 0.89076157 | -2.8250214 |
| 17026 | NM_001271186.1 | 30.0302182 | -6.5319435 | 2.31258612 | -2.824519 |
| 17027 | NM_001287511.1 | 5.75562586 | -4.1611463 | 1.47349365 | -2.8240001 |

| | | | | | |
|---|---|---|---|---|---|
| 17028 | NM_001142311.1 | 41.2640376 | -6.9891633 | 2.47569194 | -2.8231151 |
| 17029 | NR_120425.1 | 12.9193366 | -5.303667 | 1.87875689 | -2.8229661 |
| 17030 | NR_131972.1 | 5.80257967 | -4.1592306 | 1.47348819 | -2.8227105 |
| 17031 | NM_002073.2 | 312.940631 | -3.9590055 | 1.40270707 | -2.8224036 |
| 17032 | NM_001198784.1 | 5.63767973 | -4.116445 | 1.45872872 | -2.8219401 |
| 17033 | NM_001294337.1 | 7.93393166 | -4.6181497 | 1.63652101 | -2.8219312 |
| 17034 | NM_201440.1 | 8.6948005 | -4.7314582 | 1.67669568 | -2.8218944 |
| 17035 | NR_135583.1 | 13.8211061 | -5.405103 | 1.91549986 | -2.8217716 |
| 17036 | NM_016220.4 | 46.1504118 | 1.97873293 | 0.70143307 | 2.82098611 |
| 17037 | NM_014063.6 | 29.2019148 | -6.4915385 | 2.30133388 | -2.8207721 |
| 17038 | NM_004523.3 | 106.570542 | -2.1458049 | 0.76073062 | -2.8207158 |
| 17039 | NM_001276.2 | 6.8892385 | -4.4158209 | 1.56581841 | -2.820136 |
| 17040 | NM_001145795.1 | 38.6801313 | -6.8952559 | 2.44523786 | -2.8198712 |
| 17041 | NM_199184.1 | 6.15723169 | -4.2426765 | 1.50460869 | -2.8197873 |
| 17042 | NM_001012455.1 | 9.40373907 | -3.8342681 | 1.35991189 | -2.8194975 |
| 17043 | NM_001164832.2 | 7.88299999 | -4.6079415 | 1.63441888 | -2.8193149 |
| 17044 | NM_033083.6 | 142.163707 | -3.7840661 | 1.34218726 | -2.819328 |
| 17045 | NM_001303429.1 | 40.1888314 | -6.949957 | 2.46549074 | -2.818894 |
| 17046 | NM_004415.3 | 64.1359622 | -3.9609786 | 1.40520941 | -2.8187817 |
| 17047 | NM_152429.4 | 320.787973 | -3.9144099 | 1.38871978 | -2.8187183 |
| 17048 | NM_001010861.2 | 38.791148 | -6.8990499 | 2.44784778 | -2.8184146 |
| 17049 | NM_001040097.1 | 7.61377361 | -4.5425608 | 1.61175339 | -2.8183969 |
| 17050 | NM_001286210.1 | 34.6598485 | 4.45750221 | 1.58158617 | 2.81837456 |
| 17051 | NM_001289975.1 | 39.0952527 | -6.910275 | 2.45203661 | -2.8181777 |
| 17052 | NM_001270374.1 | 8.23609923 | -4.6707327 | 1.65740797 | -2.8180947 |
| 17053 | NM_001284350.1 | 8.77813085 | 3.78042844 | 1.34173183 | 2.81757378 |
| 17054 | NR_033201.2 | 7.61266561 | -4.5591202 | 1.61823835 | -2.8173354 |
| 17055 | NM_001042574.2 | 33.9571661 | -6.7084509 | 2.38144219 | -2.8169699 |
| 17056 | NM_001005361.2 | 55.4787558 | -7.4149356 | 2.63227211 | -2.8169335 |
| 17057 | NM_181573.2 | 5.26414086 | -4.0119534 | 1.42430802 | -2.8167737 |
| 17058 | NM_001272051.1 | 36.6219822 | -3.5273752 | 1.25238486 | -2.8165265 |
| 17059 | NM_024029.4 | 6.82914913 | -4.3996665 | 1.56225249 | -2.8162327 |
| 17060 | NM_001135189.1 | 50.9246951 | -6.3122605 | 2.24142869 | -2.8161772 |
| 17061 | NM_001258280.1 | 12.004792 | -5.2028737 | 1.84817582 | -2.81514 |
| 17062 | NM_014876.5 | 243.287002 | -2.1874101 | 0.77702445 | -2.815111 |
| 17063 | NM_001029859.2 | 47.5958082 | -2.7509366 | 0.97729326 | -2.8148527 |
| 17064 | NM_001193269.1 | 9.20499656 | -4.8136927 | 1.71016898 | -2.8147468 |
| 17065 | NR_073387.1 | 48.4559432 | 1.59076583 | 0.5652251 | 2.81439346 |
| 17066 | NR_138258.1 | 6.47513226 | -4.3104805 | 1.53167997 | -2.8142174 |
| 17067 | NM_144588.6 | 40.5699251 | -6.9644698 | 2.47481341 | -2.8141393 |
| 17068 | NM_001284261.1 | 10.7073225 | -4.0423062 | 1.43644863 | -2.8140973 |
| 17069 | NM_001242924.1 | 10.1310957 | -3.9627766 | 1.40836168 | -2.8137493 |
| 17070 | NM_014964.4 | 10.0103154 | -4.9539097 | 1.7611977 | -2.8128073 |

| | | | | | |
|---|---|---|---|---|---|
| 17071 | NM_199361.2 | 36.1783441 | -6.7988854 | 2.4171372 | -2.8127842 |
| 17072 | NM_001042633.2 | 29.0149013 | -6.4820344 | 2.30452801 | -2.8127384 |
| 17073 | NM_001099274.1 | 15.6218738 | -4.5893911 | 1.63167431 | -2.8126882 |
| 17074 | NM_001282685.1 | 6.7661047 | -4.3703695 | 1.5538777 | -2.8125569 |
| 17075 | NM_001330246.1 | 37.3359862 | -6.8441454 | 2.43392008 | -2.8119845 |
| 17076 | NM_001270876.1 | 55.8213614 | 7.02356049 | 2.49860636 | 2.81099119 |
| 17077 | NM_153050.2 | 42.8321391 | -7.0436187 | 2.50578433 | -2.8109437 |
| 17078 | NM_032855.3 | 42.0312906 | 1.80178972 | 0.64101687 | 2.81083043 |
| 17079 | NM_174887.3 | 6.38612732 | -4.3044582 | 1.5314172 | -2.8107679 |
| 17080 | NR_137639.1 | 10.2772174 | -4.9887039 | 1.7748727 | -2.810739 |
| 17081 | NR_026878.1 | 5.46149005 | -4.084456 | 1.45321136 | -2.8106414 |
| 17082 | NM_001346409.1 | 42.3383714 | -7.0263483 | 2.5004916 | -2.8099868 |
| 17083 | NM_001282956.1 | 6.54398309 | -4.3421779 | 1.5454154 | -2.8097157 |
| 17084 | NM_001199779.1 | 28.3100542 | -6.4468441 | 2.29452879 | -2.8096593 |
| 17085 | NM_178868.4 | 5.93270419 | -4.2053219 | 1.49680191 | -2.8095381 |
| 17086 | NM_024824.4 | 27.8684973 | -6.4241775 | 2.28662732 | -2.8094554 |
| 17087 | NM_001319078.1 | 39.2021479 | -6.9150002 | 2.46173452 | -2.8089951 |
| 17088 | NR_134657.1 | 8.176604 | 5.86809419 | 2.08951151 | 2.80835696 |
| 17089 | NM_001005463.2 | 93.6299283 | -3.7717538 | 1.34306247 | -2.8083234 |
| 17090 | NM_201428.2 | 36.8099188 | -6.8238407 | 2.43011244 | -2.808035 |
| 17091 | NR_144376.1 | 14.1778054 | -3.3820514 | 1.20464535 | -2.8075079 |
| 17092 | NM_005919.3 | 5.41324034 | -4.0678026 | 1.44891871 | -2.8074747 |
| 17093 | NM_001199636.1 | 11.22465 | -5.1019087 | 1.81738925 | -2.8072735 |
| 17094 | NM_001322315.1 | 9.01691015 | 4.35379487 | 1.55104011 | 2.8070163 |
| 17095 | NM_199202.2 | 4.36465148 | 5.66065115 | 2.01690376 | 2.8066045 |
| 17096 | NM_001271867.1 | 8.43369461 | -4.6888631 | 1.67069572 | -2.8065332 |
| 17097 | NM_001130523.2 | 45.7121788 | -7.1362452 | 2.54284114 | -2.8064062 |
| 17098 | NR_136155.1 | 27.8097036 | -6.4210334 | 2.28828693 | -2.8060438 |
| 17099 | NM_001127372.2 | 34.8295346 | -6.7455098 | 2.40436511 | -2.8055264 |
| 17100 | NR_034114.1 | 5.07319841 | -3.9682186 | 1.41443775 | -2.8055096 |
| 17101 | NM_198461.3 | 291.140891 | -3.9201898 | 1.39734988 | -2.8054461 |
| 17102 | NM_001025070.1 | 8.11370647 | -4.6509715 | 1.65787056 | -2.8053888 |
| 17103 | NM_001316898.1 | 36.8439563 | -6.8253546 | 2.43294611 | -2.8053867 |
| 17104 | NM_181829.2 | 34.1490097 | -6.7163459 | 2.39414715 | -2.8053188 |
| 17105 | NM_152778.2 | 59.8509908 | -2.2586367 | 0.805268 | -2.8048261 |
| 17106 | NM_015318.3 | 178.165113 | -3.8703661 | 1.37999775 | -2.8046177 |
| 17107 | NM_001145461.1 | 8.23678968 | -4.6548737 | 1.65976384 | -2.8045398 |
| 17108 | NR_039891.2 | 5.39074292 | -4.0638628 | 1.4491844 | -2.8042413 |
| 17109 | NM_015227.4 | 49.6954467 | -3.2495307 | 1.15887292 | -2.804044 |
| 17110 | NM_001308267.1 | 36.8556916 | -6.825475 | 2.43424815 | -2.8039356 |
| 17111 | NM_001282338.1 | 44.6646475 | -3.8507686 | 1.37339645 | -2.8038288 |
| 17112 | NM_001319170.1 | 8.70573706 | -4.7414257 | 1.69107096 | -2.8038005 |
| 17113 | NM_001164178.1 | 35.9221956 | -6.7901437 | 2.42182971 | -2.8037247 |

| | | | | | |
|---|---|---|---|---|---|
| 17114 | NM_001201552.1 | 36.5536932 | -6.813902 | 2.4304623 | -2.8035416 |
| 17115 | NM_001290223.1 | 29.6339397 | -6.5123822 | 2.32293644 | -2.8035129 |
| 17116 | NM_004212.3 | 27.5823333 | 1.4325249 | 0.51101074 | 2.80331659 |
| 17117 | NM_015296.2 | 39.3889211 | -6.9211686 | 2.46893581 | -2.8033003 |
| 17118 | NR_037805.1 | 51.2886539 | 1.78488179 | 0.63674626 | 2.80312882 |
| 17119 | NM_001288802.1 | 39.4931244 | -6.9252564 | 2.47065969 | -2.8029989 |
| 17120 | NM_153742.4 | 8.6495055 | 8.02915102 | 2.86453849 | 2.80294751 |
| 17121 | NM_001330563.1 | 6.52508502 | -4.3087387 | 1.53729082 | -2.802813 |
| 17122 | NM_003433.3 | 9.02959379 | -3.7337868 | 1.33216921 | -2.8027872 |
| 17123 | NM_001243144.1 | 7.34185615 | -4.4908674 | 1.60259543 | -2.8022465 |
| 17124 | NM_173589.3 | 6.60300131 | -4.3421729 | 1.54984474 | -2.8016825 |
| 17125 | NM_001258384.2 | 6.44429478 | -4.2955679 | 1.53386019 | -2.8004951 |
| 17126 | NM_000106.5 | 7.77936276 | -4.5831234 | 1.63700307 | -2.7997036 |
| 17127 | NM_001257207.1 | 9.18082626 | -4.8270194 | 1.72433938 | -2.7993442 |
| 17128 | NM_001300852.1 | 50.1088948 | -4.7715954 | 1.70455194 | -2.7993253 |
| 17129 | NM_013329.3 | 8.35154796 | -4.6916416 | 1.6761907 | -2.7989904 |
| 17130 | NM_080805.3 | 6.80247465 | -4.4019175 | 1.57273395 | -2.7988952 |
| 17131 | NM_138455.3 | 10.6026751 | -3.9865223 | 1.42433023 | -2.798875 |
| 17132 | NM_173855.4 | 5.35374085 | -4.036475 | 1.44231889 | -2.7986009 |
| 17133 | NM_001012984.2 | 5.74232293 | -4.1565626 | 1.48545493 | -2.7981748 |
| 17134 | NM_173078.4 | 36.8822298 | -6.8264442 | 2.4396912 | -2.7980772 |
| 17135 | NM_032864.3 | 132.834787 | -3.6791429 | 1.31492018 | -2.7979972 |
| 17136 | NM_001272020.1 | 27.5151 | -6.405692 | 2.28969492 | -2.7976181 |
| 17137 | NR_024480.1 | 31.9009563 | 2.05848605 | 0.73579815 | 2.79762332 |
| 17138 | NR_045385.1 | 33.0654878 | 2.23414225 | 0.79860362 | 2.79756089 |
| 17139 | NM_001267606.1 | 20.2159056 | -3.1819865 | 1.13747584 | -2.7974102 |
| 17140 | NM_001146197.1 | 6.20112524 | 5.54364933 | 1.98179163 | 2.79729172 |
| 17141 | NM_178507.2 | 61.5164341 | -3.6157891 | 1.29264187 | -2.7972087 |
| 17142 | NM_001145292.1 | 55.3181253 | -2.5283406 | 0.90396573 | -2.796943 |
| 17143 | NR_027673.1 | 6.94867546 | -4.4162841 | 1.5790326 | -2.796829 |
| 17144 | NM_001318085.1 | 27.7602454 | -6.4184795 | 2.29529211 | -2.7963672 |
| 17145 | NM_001303416.1 | 8.41822772 | -4.6815274 | 1.6744506 | -2.7958588 |
| 17146 | NR_026723.1 | 6.44610283 | -4.3020725 | 1.53880616 | -2.7957209 |
| 17147 | NR_136632.1 | 10.1324237 | -3.9546734 | 1.41455277 | -2.7957058 |
| 17148 | NM_001303487.1 | 33.8481765 | -6.7029046 | 2.39765763 | -2.7956054 |
| 17149 | NM_001256721.1 | 27.8596487 | -6.4237146 | 2.297809 | -2.7955825 |
| 17150 | NM_001297436.1 | 6.43615403 | -4.2935125 | 1.53584746 | -2.7955332 |
| 17151 | NM_001384.4 | 29.9996361 | -6.5302901 | 2.33615402 | -2.7953166 |
| 17152 | NM_001256737.1 | 7.11649448 | -4.4378441 | 1.58767898 | -2.7951772 |
| 17153 | NM_173822.4 | 41.2753987 | -6.9887213 | 2.50035402 | -2.7950927 |
| 17154 | NM_005996.3 | 28.7883063 | -6.471017 | 2.31515876 | -2.795064 |
| 17155 | NM_001310142.1 | 8.93332547 | 7.30478595 | 2.6135337 | 2.7949844 |
| 17156 | NM_002028.3 | 69.0488114 | -3.376505 | 1.20809355 | -2.7949036 |

| | | | | | |
|---|---|---|---|---|---|
| 17157 | NR_110812.1 | 19.7959436 | 4.46922327 | 1.59908182 | 2.79486841 |
| 17158 | NM_000856.5 | 8.33703327 | -4.6705094 | 1.67120847 | -2.7946899 |
| 17159 | NM_032557.6 | 36.5776848 | -6.8151413 | 2.43861693 | -2.7946748 |
| 17160 | NM_001331070.1 | 27.2880383 | -6.3939007 | 2.28835966 | -2.7940978 |
| 17161 | NM_015640.3 | 264.5452 | -3.8445075 | 1.37600198 | -2.7939694 |
| 17162 | NR_003142.2 | 7.2341523 | -4.4737247 | 1.60119743 | -2.7939869 |
| 17163 | NM_001304743.1 | 169.896392 | -3.7973614 | 1.3593878 | -2.7934349 |
| 17164 | NM_001346946.1 | 5.9200957 | -4.1873721 | 1.4990013 | -2.7934413 |
| 17165 | NR_027279.1 | 12.6083127 | 8.58244278 | 3.07238184 | 2.79341671 |
| 17166 | NM_001282397.1 | 30.1544651 | -3.092938 | 1.10726002 | -2.7933258 |
| 17167 | NR_037858.1 | 10.0749189 | -3.8996099 | 1.39611639 | -2.793184 |
| 17168 | NM_001024847.2 | 12.6734328 | -5.2872949 | 1.89304677 | -2.7930081 |
| 17169 | NM_201591.2 | 1004.98766 | -2.7079901 | 0.96960462 | -2.7928807 |
| 17170 | NM_001033723.2 | 173.634568 | -3.1475278 | 1.12701644 | -2.7927967 |
| 17171 | NM_001012514.2 | 33.4098985 | -6.6856363 | 2.39408624 | -2.7925629 |
| 17172 | NR_047653.1 | 41.6995487 | -7.0032457 | 2.50846197 | -2.7918485 |
| 17173 | NM_001286680.1 | 7.18925693 | -4.4689983 | 1.60087837 | -2.7915914 |
| 17174 | NM_001317921.1 | 27.4509785 | -6.4024822 | 2.29385537 | -2.7911447 |
| 17175 | NR_037566.1 | 11.0844381 | -4.0819467 | 1.46245413 | -2.7911622 |
| 17176 | NR_136559.1 | 30.4513518 | 3.75587283 | 1.34573981 | 2.79093536 |
| 17177 | NM_001206671.3 | 37.2080805 | -6.8389722 | 2.450553 | -2.7907873 |
| 17178 | NM_022902.4 | 114.340852 | -3.597053 | 1.28901685 | -2.7905399 |
| 17179 | NM_001330320.1 | 5.72622598 | -4.1414519 | 1.48412916 | -2.7904929 |
| 17180 | NM_001267561.1 | 18.646182 | -5.8457502 | 2.09493791 | -2.7904169 |
| 17181 | NM_201380.3 | 27.2134732 | -6.3899948 | 2.28996498 | -2.7904334 |
| 17182 | NM_001282802.1 | 36.7551236 | -6.8214258 | 2.4446493 | -2.7903495 |
| 17183 | NM_001178020.2 | 5.62545002 | -4.1162015 | 1.47525793 | -2.7901572 |
| 17184 | NM_001288999.1 | 30.5215968 | -6.5553326 | 2.34987072 | -2.7896567 |
| 17185 | NM_194296.1 | 5.03271091 | -3.954312 | 1.41772075 | -2.7892037 |
| 17186 | NR_026732.1 | 17.6835172 | -5.7638112 | 2.06647236 | -2.7892031 |
| 17187 | NM_001100621.2 | 27.5020725 | -6.4051608 | 2.29652947 | -2.789061 |
| 17188 | NR_027347.1 | 6.44558462 | -4.2987195 | 1.54136533 | -2.7889037 |
| 17189 | NR_028582.1 | 10.5167684 | -5.0171514 | 1.79905266 | -2.7887741 |
| 17190 | NM_001142483.1 | 7.19752598 | -4.4660717 | 1.60178432 | -2.7881854 |
| 17191 | NM_001319998.1 | 38.8163507 | -1.8446102 | 0.66158397 | -2.7881725 |
| 17192 | NM_001244883.1 | 5.06698251 | -3.9698597 | 1.42414076 | -2.7875473 |
| 17193 | NM_001258291.1 | 18.3209733 | 5.21453533 | 1.87073387 | 2.78742766 |
| 17194 | NM_001002249.2 | 8.44799104 | -4.7097791 | 1.6898542 | -2.787092 |
| 17195 | NM_001267611.1 | 27.5406463 | -6.4071244 | 2.29902997 | -2.7868816 |
| 17196 | NM_080660.3 | 54.7692842 | 1.38580764 | 0.49729504 | 2.78669102 |
| 17197 | NM_139055.3 | 77.6461819 | -3.4517112 | 1.23890038 | -2.7861087 |
| 17198 | NR_110278.1 | 12.5046125 | 3.83774285 | 1.37758691 | 2.78584445 |
| 17199 | NM_000481.3 | 28.1746364 | -6.4397026 | 2.31205988 | -2.7852664 |

| | | | | | |
|---|---|---|---|---|---|
| 17200 | NR_046681.1 | 5.48286294 | -4.088384 | 1.46787023 | -2.785249 |
| 17201 | NR_028326.1 | 42.3720104 | -3.3750548 | 1.21183502 | -2.7850778 |
| 17202 | NM_001184704.1 | 27.696326 | -6.4152825 | 2.30348691 | -2.785031 |
| 17203 | NM_001286789.1 | 6.9820646 | -4.4137511 | 1.58492748 | -2.7848284 |
| 17204 | NM_012210.3 | 35.8099846 | -6.7838338 | 2.43643677 | -2.7843258 |
| 17205 | NM_018349.3 | 5.04491496 | -3.9569484 | 1.42118582 | -2.7842583 |
| 17206 | NM_153839.6 | 11.751018 | 2.07112764 | 0.74388214 | 2.78421477 |
| 17207 | NR_120595.1 | 6.5739064 | -4.3350393 | 1.55702608 | -2.784179 |
| 17208 | NM_001134836.1 | 15.6731152 | 1.83655982 | 0.65965732 | 2.78411192 |
| 17209 | NM_000829.3 | 31.8994324 | -6.6189104 | 2.37753876 | -2.7839338 |
| 17210 | NM_001146191.1 | 13.8316921 | -4.4037918 | 1.581968 | -2.7837427 |
| 17211 | NM_052957.4 | 6.29893639 | -4.2843206 | 1.53908567 | -2.7836791 |
| 17212 | NM_001321234.1 | 28.788241 | -6.4711842 | 2.32481412 | -2.7835276 |
| 17213 | NM_144597.2 | 101.52622 | 1.8932176 | 0.68028976 | 2.78295765 |
| 17214 | NM_001127221.1 | 37.3861139 | -3.3238643 | 1.19449884 | -2.7826434 |
| 17215 | NM_001286526.1 | 14.4444309 | -3.0540236 | 1.09752885 | -2.7826363 |
| 17216 | NM_020843.2 | 122.706538 | -2.4641584 | 0.88555549 | -2.7826132 |
| 17217 | NM_001012267.2 | 7.74932205 | -4.5762426 | 1.64464179 | -2.7825163 |
| 17218 | NM_173484.3 | 21.1825762 | 1.88029087 | 0.67574995 | 2.7825246 |
| 17219 | NM_001271944.1 | 6.75204128 | -4.3656482 | 1.56912738 | -2.782214 |
| 17220 | NM_001320742.1 | 22.5895631 | -1.9817511 | 0.71240389 | -2.7817803 |
| 17221 | NM_001006115.2 | 51.1044555 | -3.6826553 | 1.323979 | -2.7815058 |
| 17222 | NR_145444.1 | 28.6319492 | 2.9670017 | 1.06669205 | 2.7814979 |
| 17223 | NM_000573.3 | 33.3507319 | 1.86410094 | 0.67018504 | 2.78147202 |
| 17224 | NM_003006.4 | 7.38038718 | -4.5111554 | 1.622002 | -2.7812267 |
| 17225 | NM_018097.2 | 39.2328604 | -6.9154284 | 2.48689634 | -2.7807466 |
| 17226 | NM_001128426.2 | 31.7178267 | -6.6105295 | 2.37740502 | -2.7805651 |
| 17227 | NM_030643.4 | 12.4765649 | -5.2661874 | 1.89396973 | -2.7805024 |
| 17228 | NR_038837.1 | 12.5376107 | 7.11409039 | 2.55856917 | 2.78049563 |
| 17229 | NM_001321221.1 | 137.052813 | 1.82807708 | 0.6575912 | 2.77995979 |
| 17230 | NM_004720.5 | 185.987778 | -4.1481982 | 1.492197 | -2.7799267 |
| 17231 | NM_002651.3 | 28.0577648 | -6.433877 | 2.31495948 | -2.7792612 |
| 17232 | NM_001166702.1 | 9.52602735 | -3.8447346 | 1.38343525 | -2.7791215 |
| 17233 | NR_109781.1 | 9.17863585 | -3.8158533 | 1.37307498 | -2.7790567 |
| 17234 | NM_001163436.2 | 7.44209465 | -4.5148636 | 1.62466911 | -2.7789434 |
| 17235 | NM_025003.3 | 5.26154385 | -4.0183651 | 1.44612319 | -2.7787156 |
| 17236 | NM_153268.3 | 16.3649843 | -3.8491182 | 1.3853677 | -2.7784091 |
| 17237 | NM_183399.2 | 21.7016623 | 5.52335881 | 1.987966 | 2.77839702 |
| 17238 | NR_126169.1 | 43.9018556 | 2.52055773 | 0.90742121 | 2.77771526 |
| 17239 | NM_016631.3 | 36.4830958 | -6.810571 | 2.45236756 | -2.7771412 |
| 17240 | NM_015702.2 | 161.622771 | -4.0750839 | 1.46745927 | -2.7769656 |
| 17241 | NM_001037293.2 | 133.872978 | 2.38656448 | 0.85945919 | 2.77682119 |
| 17242 | NM_001058.3 | 7.50423009 | -4.5344757 | 1.63304603 | -2.776698 |

| | | | | | |
|---|---|---|---|---|---|
| 17243 | NM_016328.2 | 26.7326603 | -6.3643306 | 2.29218879 | -2.7765299 |
| 17244 | NM_001271734.1 | 8.21251209 | -4.6660655 | 1.6805951 | -2.7764365 |
| 17245 | NM_003950.3 | 49.2933099 | 1.9647403 | 0.70765783 | 2.77639875 |
| 17246 | NM_001326358.1 | 31.1915123 | -6.5859655 | 2.3721451 | -2.7763755 |
| 17247 | NR_047576.2 | 5.74984601 | -4.1476494 | 1.49396572 | -2.7762681 |
| 17248 | NM_001018037.1 | 34.9919561 | -6.7509504 | 2.4317686 | -2.7761484 |
| 17249 | NM_001198910.1 | 7.48826748 | -4.5281822 | 1.63124013 | -2.7759139 |
| 17250 | NM_181333.3 | 8.6101903 | -4.7376909 | 1.70675407 | -2.7758486 |
| 17251 | NM_001126181.1 | 9.16287599 | -4.823655 | 1.73782294 | -2.7756884 |
| 17252 | NM_001308102.1 | 2.40588377 | 5.42552625 | 1.95469607 | 2.77563675 |
| 17253 | NM_001318324.1 | 27.3084645 | -6.3947332 | 2.30404207 | -2.7754412 |
| 17254 | NM_001004065.4 | 36.0217479 | -6.7933323 | 2.44860098 | -2.7743729 |
| 17255 | NR_036678.1 | 25.9350486 | 1.16740895 | 0.42083699 | 2.77401696 |
| 17256 | NM_004622.2 | 215.924282 | -3.9957002 | 1.440527 | -2.7737767 |
| 17257 | NM_001190482.1 | 27.4840682 | 1.46193955 | 0.52729499 | 2.77252694 |
| 17258 | NM_001286109.1 | 5.55606138 | -4.0818687 | 1.47226884 | -2.7725022 |
| 17259 | NM_001142327.1 | 27.265118 | -6.3925429 | 2.30573251 | -2.7724564 |
| 17260 | NM_001151.3 | 297.080227 | -2.5115547 | 0.9059253 | -2.772364 |
| 17261 | NR_131761.1 | 5.6571791 | -4.1135986 | 1.4839528 | -2.7720549 |
| 17262 | NR_004405.1 | 5.756181 | -4.138581 | 1.49318435 | -2.7716477 |
| 17263 | NR_026857.1 | 307.489239 | -3.2821754 | 1.18434008 | -2.7713116 |
| 17264 | NM_001103148.1 | 27.1857746 | -6.3883989 | 2.30523154 | -2.7712613 |
| 17265 | NM_213647.2 | 9.77625072 | -4.9175547 | 1.77455575 | -2.7711469 |
| 17266 | NR_136599.1 | 45.7875164 | -7.1384807 | 2.57626795 | -2.7708611 |
| 17267 | NR_133933.1 | 5.16575664 | -4.0016359 | 1.44497691 | -2.7693425 |
| 17268 | NM_006307.4 | 28.3303407 | -6.4478435 | 2.32857647 | -2.7690065 |
| 17269 | NM_001256020.1 | 9.63371405 | 6.72966331 | 2.43066992 | 2.76864549 |
| 17270 | NM_001301015.2 | 8.71751972 | -4.741923 | 1.71275034 | -2.7686014 |
| 17271 | NR_027248.3 | 8.27824527 | 7.17890273 | 2.59296798 | 2.76860446 |
| 17272 | NM_004941.2 | 26.5973527 | -6.356709 | 2.29605071 | -2.7685403 |
| 17273 | NM_080881.2 | 37.8121226 | -3.1483608 | 1.13739928 | -2.7680348 |
| 17274 | NM_001010852.3 | 54.2868379 | -3.5536742 | 1.28402205 | -2.7676115 |
| 17275 | NM_012396.4 | 77.2995306 | -2.6752605 | 0.96666939 | -2.7675031 |
| 17276 | NM_080630.3 | 37.3904085 | -6.8476875 | 2.47436808 | -2.767449 |
| 17277 | NM_001330317.1 | 8.22316166 | -4.6573274 | 1.68293348 | -2.7673865 |
| 17278 | NM_001889.3 | 26.2005746 | -6.3352029 | 2.28949357 | -2.7670761 |
| 17279 | NM_001130071.1 | 6.14849575 | -4.2415337 | 1.53335326 | -2.7661817 |
| 17280 | NM_053016.5 | 35.084135 | -6.7550624 | 2.44223853 | -2.7659307 |
| 17281 | NM_004249.3 | 5.81522798 | -4.177332 | 1.51031507 | -2.7658679 |
| 17282 | NM_001195308.1 | 8.90528133 | -4.7767842 | 1.72715246 | -2.7656992 |
| 17283 | NM_021722.5 | 30.2990818 | -6.5445497 | 2.36633396 | -2.7656915 |
| 17284 | NM_032776.2 | 36.0276552 | -6.792004 | 2.45587207 | -2.765618 |
| 17285 | NM_001286660.1 | 5.24749193 | -4.0033137 | 1.44754738 | -2.7655839 |

| | | | | | |
|---|---|---|---|---|---|
| 17286 | NM_001324220.1 | 34.903358 | -6.7474242 | 2.44002881 | -2.7653052 |
| 17287 | NR_073475.1 | 27.6173693 | -6.411085 | 2.31868372 | -2.7649674 |
| 17288 | NR_040535.1 | 76.09687 | 2.18532715 | 0.79040388 | 2.7648234 |
| 17289 | NM_001322242.1 | 27.0862913 | -6.3830153 | 2.30872754 | -2.764733 |
| 17290 | NR_026885.1 | 4.95500189 | -3.9309442 | 1.42191153 | -2.7645491 |
| 17291 | NR_033314.1 | 8.49680662 | -3.6985312 | 1.33826017 | -2.7636862 |
| 17292 | NM_153693.4 | 9.39046539 | -4.848216 | 1.75430687 | -2.7636077 |
| 17293 | NM_001018159.1 | 6.43722429 | -4.3196725 | 1.56320046 | -2.7633516 |
| 17294 | NR_024585.1 | 50.2160613 | 1.19394408 | 0.43213192 | 2.76291576 |
| 17295 | NM_001346875.1 | 6.80199956 | -4.3798299 | 1.5855379 | -2.7623622 |
| 17296 | NR_045669.1 | 5.79921103 | -4.1638311 | 1.50739734 | -2.7622651 |
| 17297 | NM_014149.3 | 89.6815994 | -3.7060026 | 1.3416875 | -2.762195 |
| 17298 | NR_134970.1 | 69.3712338 | 5.57719681 | 2.01912325 | 2.76218741 |
| 17299 | NM_001345970.1 | 27.1683063 | -6.3877328 | 2.31259895 | -2.7621446 |
| 17300 | NM_001113203.2 | 5.06084357 | -3.96195 | 1.43439516 | -2.762105 |
| 17301 | NM_001102592.1 | 5.7830865 | -4.153777 | 1.5041744 | -2.7614996 |
| 17302 | NR_029385.1 | 133.388948 | 2.92812965 | 1.06046286 | 2.76118077 |
| 17303 | NM_014140.3 | 26.1952538 | -6.3349825 | 2.29474438 | -2.7606485 |
| 17304 | NR_036504.1 | 95.0535307 | 1.22347001 | 0.44321452 | 2.76044659 |
| 17305 | NM_001243996.2 | 26.6759066 | -6.3611327 | 2.304931 | -2.7597931 |
| 17306 | NM_172071.3 | 42.2293086 | -7.021209 | 2.54427402 | -2.7596119 |
| 17307 | NM_022088.4 | 23.0703869 | 4.01133229 | 1.45363049 | 2.7595268 |
| 17308 | NM_030632.2 | 107.27837 | -3.7835206 | 1.37110118 | -2.7594758 |
| 17309 | NM_024019.3 | 79.0016432 | -4.2486492 | 1.53990395 | -2.7590352 |
| 17310 | NM_002502.5 | 26.7327487 | -6.3638925 | 2.30671591 | -2.758854 |
| 17311 | NM_001297771.1 | 37.8378717 | 1.3583714 | 0.49237964 | 2.75878874 |
| 17312 | NM_080681.2 | 31.0106189 | -6.5768543 | 2.38404402 | -2.7586967 |
| 17313 | NM_001297423.1 | 29.3144349 | -3.1694943 | 1.14899642 | -2.7584893 |
| 17314 | NM_001261436.1 | 13.2598976 | 5.21027199 | 1.88885977 | 2.75842181 |
| 17315 | NR_144951.1 | 26.2855482 | -6.3397672 | 2.2983952 | -2.7583451 |
| 17316 | NR_134934.1 | 5.63050439 | -4.1003308 | 1.48659134 | -2.7582098 |
| 17317 | NR_135825.1 | 7.2643031 | 2.30303536 | 0.83500586 | 2.75810682 |
| 17318 | NM_001159508.1 | 125.916762 | 3.56919215 | 1.29418608 | 2.75786628 |
| 17319 | NR_136547.1 | 7.88029812 | -4.5914578 | 1.66506249 | -2.7575288 |
| 17320 | NM_000069.2 | 19.0698751 | 7.5858953 | 2.75107739 | 2.75742708 |
| 17321 | NM_001303228.1 | 27.6927462 | 2.40294632 | 0.87151192 | 2.75721567 |
| 17322 | NM_172003.3 | 34.5580037 | -6.732638 | 2.44205524 | -2.7569556 |
| 17323 | NM_001289398.1 | 9.93941103 | -4.9340738 | 1.78971409 | -2.7569061 |
| 17324 | NM_012100.3 | 35.1918334 | -6.7582207 | 2.45145015 | -2.7568257 |
| 17325 | NR_144328.1 | 6.56913186 | -4.3445426 | 1.57600672 | -2.7566777 |
| 17326 | NM_199229.2 | 33.9216588 | -6.7060325 | 2.43289243 | -2.7564032 |
| 17327 | NM_001318766.1 | 8.41057923 | -4.6943819 | 1.70310685 | -2.7563637 |
| 17328 | NM_015132.4 | 106.234592 | -3.7692043 | 1.36747059 | -2.756333 |

| | | | | | |
|---|---|---|---|---|---|
| 17329 | NM_001320707.1 | 27.7906745 | -6.4200005 | 2.3293803 | -2.756098 |
| 17330 | NM_001300780.1 | 25.7075658 | -6.3077494 | 2.28894494 | -2.7557454 |
| 17331 | NM_147190.3 | 35.5123851 | -6.7723885 | 2.45759373 | -2.755699 |
| 17332 | NM_001321229.1 | 34.0261726 | -6.7097226 | 2.43506045 | -2.7554645 |
| 17333 | NR_048561.1 | 40.747226 | -6.9698226 | 2.52964583 | -2.7552563 |
| 17334 | NM_001146019.1 | 30.9663525 | -6.5750552 | 2.38666666 | -2.7549114 |
| 17335 | NR_144512.1 | 6.93768744 | -4.4120005 | 1.60155932 | -2.7548155 |
| 17336 | NR_144327.1 | 8.77801361 | -4.7670999 | 1.73057795 | -2.7546288 |
| 17337 | NM_001077239.1 | 27.6345337 | -6.4116917 | 2.327652 | -2.7545749 |
| 17338 | NM_001301047.2 | 10.3152623 | 8.27661746 | 3.00489187 | 2.75438112 |
| 17339 | NR_075071.1 | 10.5820914 | -5.0312082 | 1.82686702 | -2.754009 |
| 17340 | NM_001257134.1 | 7.19410602 | -4.4700778 | 1.62316973 | -2.7539189 |
| 17341 | NM_001320670.1 | 34.6889923 | -6.7388078 | 2.44710164 | -2.7537915 |
| 17342 | NM_198537.3 | 48.4929391 | -3.2149388 | 1.16752372 | -2.753639 |
| 17343 | NM_002135.4 | 6.31107571 | -4.2934667 | 1.55921487 | -2.7536081 |
| 17344 | NM_001286645.1 | 11.7378941 | -5.1737472 | 1.87892757 | -2.7535639 |
| 17345 | NM_004923.3 | 4.94680982 | -3.9347275 | 1.42905122 | -2.7533845 |
| 17346 | NM_003403.4 | 255.007222 | -3.7210536 | 1.3514815 | -2.7533145 |
| 17347 | NM_001291937.1 | 27.6308249 | -6.4119589 | 2.32883159 | -2.7532944 |
| 17348 | NM_004750.4 | 6.54249342 | -4.3383057 | 1.57583216 | -2.7530252 |
| 17349 | NM_001242476.1 | 5.08061885 | -3.9659988 | 1.44064628 | -2.7529303 |
| 17350 | NM_001282714.1 | 11.1855457 | -5.1110212 | 1.85667797 | -2.7527775 |
| 17351 | NM_001256006.1 | 8.75845848 | -4.7384283 | 1.72143643 | -2.7526014 |
| 17352 | NM_178548.3 | 12.7682971 | -3.4551787 | 1.2552703 | -2.7525376 |
| 17353 | NM_001166392.1 | 77.6654177 | -3.8196311 | 1.3877171 | -2.7524566 |
| 17354 | NM_001128626.1 | 34.4923811 | -6.7304732 | 2.44578886 | -2.7518619 |
| 17355 | NR_037159.1 | 6.18766776 | -4.2505766 | 1.54472841 | -2.751666 |
| 17356 | NR_135175.1 | 4.9212485 | -3.9231905 | 1.42589256 | -2.7513928 |
| 17357 | NM_015554.2 | 25.4697797 | -6.2945043 | 2.28782202 | -2.7513086 |
| 17358 | NM_001326421.1 | 8.19852565 | -4.6607085 | 1.69405661 | -2.7512118 |
| 17359 | NR_034146.1 | 33.4758495 | -2.9472975 | 1.07154693 | -2.7505072 |
| 17360 | NM_003833.4 | 6.21438243 | -4.2459816 | 1.54388299 | -2.7501965 |
| 17361 | NM_001282681.1 | 33.9639126 | -6.7073758 | 2.43905884 | -2.7499852 |
| 17362 | NM_001324305.1 | 10.414594 | -5.0076688 | 1.82125579 | -2.7495692 |
| 17363 | NM_001321918.1 | 7.96444248 | -4.6020995 | 1.6741265 | -2.7489557 |
| 17364 | NM_019041.6 | 33.5946872 | -6.6916484 | 2.43442886 | -2.748755 |
| 17365 | NM_139170.2 | 6.09739671 | -4.2416387 | 1.54317501 | -2.748644 |
| 17366 | NM_004322.3 | 5.54851264 | -4.0945425 | 1.49004225 | -2.7479372 |
| 17367 | NM_006049.3 | 5.96610144 | -4.1882617 | 1.52424113 | -2.7477685 |
| 17368 | NM_024007.4 | 25.5191228 | -6.2970675 | 2.29201821 | -2.7473898 |
| 17369 | NR_075090.1 | 26.3243749 | 4.61720206 | 1.68063597 | 2.74729457 |
| 17370 | NM_020741.2 | 5.19079975 | -4.0049612 | 1.45789498 | -2.7470849 |
| 17371 | NR_040046.1 | 62.9590347 | 1.6778984 | 0.61078835 | 2.74710284 |

| | | | | | |
|---|---|---|---|---|---|
| 17372 | NR_126431.1 | 9.49879216 | -4.8774079 | 1.77563495 | -2.7468529 |
| 17373 | NM_001300900.1 | 7.6441041 | -4.5649805 | 1.66209784 | -2.7465173 |
| 17374 | NM_014981.1 | 9.74097886 | -3.8916016 | 1.41692827 | -2.7465057 |
| 17375 | NM_001204747.1 | 25.4652064 | -6.29408 | 2.29186021 | -2.7462757 |
| 17376 | NM_001256932.1 | 5.34129171 | -4.0537422 | 1.47618319 | -2.746097 |
| 17377 | NM_001199280.1 | 30.4724922 | -6.5516492 | 2.38605044 | -2.7458134 |
| 17378 | NM_005169.3 | 8.45240738 | -4.7044764 | 1.71366668 | -2.7452692 |
| 17379 | NR_135845.1 | 33.7081182 | -6.6961504 | 2.43942121 | -2.7449751 |
| 17380 | NR_103527.2 | 5.58502519 | -4.1168752 | 1.49984011 | -2.744876 |
| 17381 | NR_027501.1 | 53.9409038 | 2.43127078 | 0.88582873 | 2.74462849 |
| 17382 | NM_001009955.3 | 240.644154 | -4.1866789 | 1.52562023 | -2.7442471 |
| 17383 | NM_001146171.1 | 8.8672427 | -4.7703766 | 1.73841443 | -2.7440963 |
| 17384 | NM_005562.2 | 5.23912644 | -4.0146113 | 1.46308935 | -2.7439276 |
| 17385 | NM_032378.5 | 53.5642674 | -2.5917061 | 0.94454265 | -2.7438741 |
| 17386 | NM_178237.2 | 33.164039 | -6.6735818 | 2.43235787 | -2.7436677 |
| 17387 | NR_136339.1 | 5.02554389 | -3.9472068 | 1.43872035 | -2.7435539 |
| 17388 | NM_001855.4 | 33.6568797 | -3.8575767 | 1.40625462 | -2.7431567 |
| 17389 | NM_001330210.1 | 6.44204573 | -4.2972129 | 1.56661514 | -2.7429921 |
| 17390 | NM_001330303.1 | 27.3863651 | -6.3985284 | 2.33269145 | -2.742981 |
| 17391 | NM_182932.2 | 33.1607107 | -6.6733432 | 2.43310158 | -2.742731 |
| 17392 | NM_020749.4 | 6.340375 | -4.2976075 | 1.56697765 | -2.7426093 |
| 17393 | NM_031471.5 | 5.22732357 | -3.9986307 | 1.45810875 | -2.7423405 |
| 17394 | NM_174855.3 | 9.14135231 | -4.8197359 | 1.75771653 | -2.7420439 |
| 17395 | NM_001001415.3 | 28.3661124 | -6.4495798 | 2.35330363 | -2.7406493 |
| 17396 | NM_001145.4 | 4.93018655 | -3.9211585 | 1.43077345 | -2.7405866 |
| 17397 | NM_018670.3 | 16.5782733 | -3.8715865 | 1.41279792 | -2.7403682 |
| 17398 | NM_001033677.1 | 10.7313615 | -5.0458089 | 1.84132269 | -2.7403176 |
| 17399 | NM_033429.2 | 28.677599 | -6.4653081 | 2.35952548 | -2.7400883 |
| 17400 | NM_153499.2 | 31.7433599 | -6.6107147 | 2.41258173 | -2.7400998 |
| 17401 | NR_104155.1 | 5.42687406 | 6.59739632 | 2.40774467 | 2.74007306 |
| 17402 | NM_002958.3 | 69.4617113 | -3.7816236 | 1.38017181 | -2.7399658 |
| 17403 | NM_001198850.1 | 6.18038786 | -4.246794 | 1.55000795 | -2.739853 |
| 17404 | NM_001242409.1 | 32.9991025 | -6.6662871 | 2.4331335 | -2.7397951 |
| 17405 | NM_022347.3 | 168.74577 | -1.1666197 | 0.42585748 | -2.7394603 |
| 17406 | NM_001007027.2 | 10.9038653 | -4.0581979 | 1.48191556 | -2.7384812 |
| 17407 | NM_016827.2 | 7.66421812 | -4.5676179 | 1.66832244 | -2.7378508 |
| 17408 | NM_001281484.1 | 26.6262815 | -6.358626 | 2.322838 | -2.7374384 |
| 17409 | NR_048568.1 | 9.89727259 | -4.9220171 | 1.79814709 | -2.7372717 |
| 17410 | NM_003573.2 | 40.8441022 | -6.9729025 | 2.54859239 | -2.7359819 |
| 17411 | NM_005999.2 | 137.083273 | -1.8833671 | 0.68842681 | -2.735755 |
| 17412 | NM_175902.4 | 25.6240434 | -6.3032658 | 2.30401785 | -2.7357712 |
| 17413 | NM_130810.3 | 8.92277475 | -4.7886947 | 1.75077512 | -2.7351855 |
| 17414 | NR_136637.1 | 8.78361312 | -3.7474621 | 1.37013309 | -2.7351081 |

| | | | | | |
|---|---|---|---|---|---|
| 17415 | NM_133180.2 | 10.4900969 | -5.0107142 | 1.83201915 | -2.7350774 |
| 17416 | NM_015029.2 | 8.57388494 | 8.01067905 | 2.92901035 | 2.73494392 |
| 17417 | NM_020715.2 | 55.9098176 | -3.1618287 | 1.1563275 | -2.7343713 |
| 17418 | NM_001257976.1 | 35.4602956 | -6.770792 | 2.47630121 | -2.734236 |
| 17419 | NM_001024808.2 | 263.439916 | -4.221791 | 1.54416293 | -2.7340321 |
| 17420 | NM_001282461.1 | 6.15285546 | -4.2268335 | 1.54603552 | -2.7339821 |
| 17421 | NM_198189.2 | 7.73951299 | -4.5872614 | 1.67830108 | -2.7332768 |
| 17422 | NM_032756.2 | 6.38623592 | -4.279746 | 1.56580565 | -2.7332549 |
| 17423 | NM_001162371.2 | 539.156626 | -4.4143316 | 1.61509688 | -2.7331683 |
| 17424 | NM_001287533.1 | 25.2095802 | -2.9160174 | 1.06704535 | -2.7327962 |
| 17425 | NM_001488.4 | 6.00720448 | -4.1939757 | 1.53469292 | -2.7327784 |
| 17426 | NR_033922.1 | 7.29441411 | 4.4083178 | 1.61346064 | 2.73221278 |
| 17427 | NM_001303542.2 | 44.9442992 | -7.1126266 | 2.60340969 | -2.7320427 |
| 17428 | NM_015914.6 | 25.0745237 | -6.2716887 | 2.29559228 | -2.7320569 |
| 17429 | NM_006477.4 | 5.76182738 | -4.1378509 | 1.51461882 | -2.7319421 |
| 17430 | NM_001099337.1 | 8.15838543 | -4.6610406 | 1.70615648 | -2.7318951 |
| 17431 | NM_152233.3 | 197.001459 | -4.1089322 | 1.50417122 | -2.7316918 |
| 17432 | NM_001286783.1 | 8.43858086 | -4.6889438 | 1.71668249 | -2.7313984 |
| 17433 | NM_001282908.1 | 7.55511957 | -4.5306069 | 1.65906337 | -2.7308221 |
| 17434 | NM_001170724.1 | 7.2076306 | 3.33071955 | 1.2200515 | 2.72998275 |
| 17435 | NM_001308240.1 | 6.93599855 | -4.4235549 | 1.62074381 | -2.7293363 |
| 17436 | NM_001199671.1 | 30.3331988 | -6.5446201 | 2.39793239 | -2.7292763 |
| 17437 | NM_000741.3 | 5.50353196 | -4.0883198 | 1.49801471 | -2.7291587 |
| 17438 | NM_001331071.1 | 32.4460078 | -6.6417721 | 2.43387903 | -2.7288834 |
| 17439 | NM_001271785.1 | 35.9584868 | -6.7897246 | 2.48815441 | -2.7288196 |
| 17440 | NM_024712.3 | 11.7506256 | -3.3353186 | 1.22235816 | -2.7285936 |
| 17441 | NM_001258031.1 | 7.59632025 | -4.5544804 | 1.66966895 | -2.7277745 |
| 17442 | NM_001322243.1 | 98.3673495 | -3.1345826 | 1.14914911 | -2.7277423 |
| 17443 | NR_033202.2 | 8.61501491 | -4.735781 | 1.73696824 | -2.7264638 |
| 17444 | NR_110172.1 | 7.20198516 | -4.4609713 | 1.63630735 | -2.7262429 |
| 17445 | NM_017922.3 | 114.81575 | -2.636466 | 0.96718759 | -2.7259096 |
| 17446 | NM_001321982.1 | 10.0336948 | -4.9472364 | 1.81494841 | -2.7258276 |
| 17447 | NM_001001931.2 | 15.2764236 | -4.535446 | 1.66397381 | -2.7256715 |
| 17448 | NM_015113.3 | 213.922051 | -3.9056249 | 1.43327162 | -2.7249719 |
| 17449 | NM_033103.4 | 78.9773269 | -3.3758418 | 1.23896781 | -2.7247212 |
| 17450 | NM_001330343.1 | 48.865967 | 3.5009603 | 1.28493611 | 2.72461818 |
| 17451 | NM_001907.2 | 7.49277798 | -4.52197 | 1.65980681 | -2.7243954 |
| 17452 | NM_004671.4 | 25.0747637 | -6.2717035 | 2.30217079 | -2.7242564 |
| 17453 | NM_004016.2 | 45.5250892 | -7.1307074 | 2.6176418 | -2.7240959 |
| 17454 | NM_181515.1 | 9.65326203 | -4.8935367 | 1.7965968 | -2.7237813 |
| 17455 | NR_033660.1 | 41.6310671 | -6.0293342 | 2.21372675 | -2.7236127 |
| 17456 | NM_001199189.1 | 6.06922931 | -4.2223121 | 1.55033446 | -2.7234846 |
| 17457 | NM_138348.5 | 68.3107005 | -2.6551057 | 0.97493837 | -2.7233575 |

| | | | | | |
|---|---|---|---|---|---|
| 17458 | NM_001243279.2 | 32.3338086 | -6.6365034 | 2.43756519 | -2.7225953 |
| 17459 | NM_014350.3 | 7.94104315 | -4.6221516 | 1.69771506 | -2.7225721 |
| 17460 | NM_001172517.1 | 34.2062292 | -6.7172969 | 2.46756292 | -2.7222393 |
| 17461 | NM_018712.3 | 31.8904587 | -6.6168831 | 2.43083272 | -2.7220644 |
| 17462 | NM_138957.3 | 7.47665744 | -4.529803 | 1.6641681 | -2.7219624 |
| 17463 | NM_001278212.1 | 25.329037 | -6.2867195 | 2.31009658 | -2.7214098 |
| 17464 | NM_031934.5 | 30.3063019 | -6.5432791 | 2.40460137 | -2.7211492 |
| 17465 | NR_109914.1 | 5.86113411 | -4.1889997 | 1.53961641 | -2.7208074 |
| 17466 | NM_001188.3 | 57.0871645 | -3.1971962 | 1.17514695 | -2.7206778 |
| 17467 | NM_001127258.2 | 18.7712561 | -2.3120739 | 0.84995053 | -2.7202453 |
| 17468 | NR_120317.1 | 32.1936142 | -2.9837815 | 1.09708418 | -2.719738 |
| 17469 | NM_198527.3 | 5.34406214 | -4.0413935 | 1.48601309 | -2.7196218 |
| 17470 | NM_000276.3 | 36.66834 | -6.8179067 | 2.50739399 | -2.7191206 |
| 17471 | NM_001270407.1 | 11.1769649 | 8.40832908 | 3.09228441 | 2.71913187 |
| 17472 | NM_024341.2 | 33.651224 | -6.6945661 | 2.46234278 | -2.7187791 |
| 17473 | NM_001314049.1 | 10.8010269 | -5.0513116 | 1.85800127 | -2.7186804 |
| 17474 | NM_001242849.1 | 25.1921897 | -6.2789049 | 2.30980855 | -2.718366 |
| 17475 | NM_006135.2 | 290.911664 | -4.0331627 | 1.48372283 | -2.7182723 |
| 17476 | NM_001079684.2 | 24.2626872 | -6.2244641 | 2.28990644 | -2.7182176 |
| 17477 | NM_016516.2 | 31.9024535 | -6.6184468 | 2.43545036 | -2.7175453 |
| 17478 | NM_000306.3 | 4.70859315 | 6.44776734 | 2.37286111 | 2.71729657 |
| 17479 | NM_001079823.1 | 66.8952223 | -2.8004666 | 1.03061952 | -2.7172653 |
| 17480 | NM_144629.2 | 14.4213307 | -2.5635697 | 0.94344972 | -2.7172298 |
| 17481 | NM_138639.1 | 24.3105994 | -6.2272212 | 2.29197177 | -2.7169712 |
| 17482 | NM_152995.5 | 26.170984 | -6.3333632 | 2.33110283 | -2.7168957 |
| 17483 | NM_001134396.1 | 6.38010238 | -4.3047397 | 1.58458608 | -2.7166336 |
| 17484 | NM_152440.4 | 6.55591631 | -4.3298657 | 1.59383343 | -2.7166363 |
| 17485 | NM_002468.4 | 32.0316915 | -6.6229923 | 2.43806474 | -2.7164957 |
| 17486 | NM_001297767.1 | 7.71651129 | -4.5760652 | 1.68463255 | -2.7163581 |
| 17487 | NM_005246.3 | 33.2621767 | -6.6787903 | 2.45886516 | -2.7162085 |
| 17488 | NM_020247.4 | 55.301397 | -3.1456651 | 1.15824197 | -2.7158963 |
| 17489 | NM_001322056.1 | 34.5817379 | -6.7335103 | 2.4794542 | -2.7157228 |
| 17490 | NR_047558.1 | 5.09850328 | -3.97759 | 1.46502196 | -2.7150378 |
| 17491 | NM_178120.4 | 5.91517796 | -4.1987743 | 1.54655351 | -2.7149234 |
| 17492 | NM_012135.2 | 20.7251186 | -2.872984 | 1.05823844 | -2.7148739 |
| 17493 | NM_001270552.1 | 26.4754074 | -6.3501047 | 2.33911669 | -2.7147447 |
| 17494 | NM_001206850.1 | 10.0768646 | -4.9534763 | 1.82475298 | -2.714601 |
| 17495 | NR_136893.1 | 7.74612543 | -4.5727707 | 1.68475143 | -2.7142109 |
| 17496 | NM_181846.2 | 25.5485807 | -6.2985599 | 2.32086221 | -2.7138879 |
| 17497 | NR_024421.1 | 5.55750201 | -4.0899244 | 1.50707157 | -2.7138223 |
| 17498 | NM_017877.3 | 200.685128 | -2.0235399 | 0.74566883 | -2.7137246 |
| 17499 | NM_001330269.1 | 9.44875965 | -3.8620788 | 1.42352068 | -2.7130472 |
| 17500 | NM_001318829.1 | 32.0554064 | -6.6247299 | 2.44187664 | -2.7129666 |

| | | | | | |
|---|---|---|---|---|---|
| 17501 | NR_130649.1 | 19.3633493 | -3.5490191 | 1.30819074 | -2.7129217 |
| 17502 | NR_135539.1 | 5.99647203 | -4.1983207 | 1.54764663 | -2.7127127 |
| 17503 | NM_001282530.1 | 5.54880611 | -4.1092124 | 1.51503197 | -2.7122942 |
| 17504 | NM_001321726.1 | 5.22259961 | -4.0031282 | 1.4760098 | -2.7121284 |
| 17505 | NM_001308257.1 | 4.97414217 | -3.9368816 | 1.45164863 | -2.7120072 |
| 17506 | NM_004235.5 | 11.4156547 | -4.0964908 | 1.51062307 | -2.7117889 |
| 17507 | NR_144372.1 | 5.49446576 | -4.0776592 | 1.50391228 | -2.7113677 |
| 17508 | NM_020826.2 | 262.356708 | -4.1902616 | 1.54549135 | -2.7112812 |
| 17509 | NM_022467.3 | 7.58434514 | -4.5470203 | 1.67706607 | -2.7112947 |
| 17510 | NM_012290.4 | 155.600279 | -3.8833889 | 1.43245647 | -2.7109996 |
| 17511 | NR_045418.1 | 31.4613042 | -6.5971737 | 2.43451987 | -2.7098459 |
| 17512 | NM_005036.4 | 31.7009295 | -6.6094385 | 2.43907144 | -2.7098175 |
| 17513 | NR_103768.1 | 7.95122743 | -3.5806209 | 1.3214873 | -2.7095387 |
| 17514 | NM_032144.2 | 6.34703825 | -4.2726769 | 1.57703882 | -2.7093036 |
| 17515 | NM_001024940.2 | 7.06701857 | -4.4451769 | 1.64079935 | -2.7091532 |
| 17516 | NM_001282583.1 | 7.44444024 | -4.530243 | 1.67274068 | -2.7082757 |
| 17517 | NM_001300969.1 | 24.9921066 | -6.2673103 | 2.31432671 | -2.7080491 |
| 17518 | NM_015852.3 | 61.347954 | -3.1906428 | 1.17827039 | -2.7079037 |
| 17519 | NM_001256435.2 | 5.84847444 | -4.1584585 | 1.53590029 | -2.7075055 |
| 17520 | NR_026845.1 | 5.93222292 | -4.1779913 | 1.5437016 | -2.706476 |
| 17521 | NM_001166305.1 | 6.24135872 | -4.2603325 | 1.57418492 | -2.7063736 |
| 17522 | NR_103738.1 | 5.61712146 | -4.0970768 | 1.51393191 | -2.7062491 |
| 17523 | NM_001304532.1 | 26.0828447 | -6.3288162 | 2.33866859 | -2.706162 |
| 17524 | NM_006398.3 | 7.06867266 | -4.454257 | 1.64596918 | -2.7061607 |
| 17525 | NR_103525.1 | 7.21398394 | -4.4702876 | 1.65187542 | -2.7061893 |
| 17526 | NM_005311.4 | 34.3928431 | -6.7256056 | 2.48584364 | -2.7055626 |
| 17527 | NM_173648.3 | 36.1519247 | 2.08674822 | 0.77140307 | 2.70513342 |
| 17528 | NM_017771.4 | 24.224359 | -6.2222877 | 2.30048234 | -2.7047753 |
| 17529 | NR_121608.1 | 4.90963963 | -3.9185661 | 1.44892178 | -2.7044704 |
| 17530 | NM_001128933.2 | 22.3049557 | -2.6963894 | 0.99710363 | -2.7042218 |
| 17531 | NM_002461.2 | 571.005903 | -4.4690456 | 1.65298144 | -2.7036272 |
| 17532 | NM_001007259.2 | 7.0212706 | -4.4270089 | 1.63746691 | -2.7035715 |
| 17533 | NM_001346471.1 | 32.0487436 | -6.6240161 | 2.4508638 | -2.7027272 |
| 17534 | NM_001002020.2 | 5.50022614 | -4.0847851 | 1.51172801 | -2.7020635 |
| 17535 | NM_001135703.2 | 47.1008206 | -6.1975676 | 2.293897 | -2.7017637 |
| 17536 | NM_001330082.1 | 29.7095792 | -6.5156466 | 2.41166352 | -2.7017229 |
| 17537 | NM_001305011.1 | 8.92255749 | 8.08626001 | 2.99326924 | 2.70148101 |
| 17538 | NR_045492.1 | 7.18102267 | -4.4523234 | 1.64815737 | -2.7013946 |
| 17539 | NM_001320703.1 | 31.2950319 | -6.5893857 | 2.43928155 | -2.7013633 |
| 17540 | NR_136749.1 | 5.62043864 | -4.1163881 | 1.52396 | -2.701113 |
| 17541 | NM_001099771.2 | 8.00760185 | -4.6332368 | 1.71545297 | -2.7008824 |
| 17542 | NM_001289807.1 | 25.2545441 | -6.2820366 | 2.32603804 | -2.7007454 |
| 17543 | NM_001174117.1 | 42.5954894 | -6.0619711 | 2.24463636 | -2.7006473 |

| | | | | | |
|---|---|---|---|---|---|
| 17544 | NM_018908.2 | 6.05671535 | -4.2089626 | 1.55883848 | -2.7000633 |
| 17545 | NM_022476.3 | 23.9349134 | -6.205059 | 2.2981555 | -2.700017 |
| 17546 | NR_110466.1 | 33.3801428 | -6.6835134 | 2.47538975 | -2.6999843 |
| 17547 | NM_001256513.1 | 35.4294935 | -3.2520664 | 1.20453395 | -2.6998545 |
| 17548 | NM_001127582.1 | 24.3866361 | -6.2317326 | 2.30845503 | -2.6995252 |
| 17549 | NM_001307978.1 | 18.027102 | 1.8372106 | 0.68062827 | 2.69928635 |
| 17550 | NM_001190882.2 | 37.555129 | -6.8533057 | 2.53916783 | -2.6990361 |
| 17551 | NM_001242796.2 | 9.39420331 | -4.8593202 | 1.80044664 | -2.6989526 |
| 17552 | NM_005734.4 | 23.5348791 | -6.1805167 | 2.29029085 | -2.6985729 |
| 17553 | NR_002834.1 | 18.540307 | -4.049635 | 1.5009604 | -2.6980292 |
| 17554 | NM_001288759.2 | 6.45246739 | -4.2936969 | 1.59147022 | -2.6979436 |
| 17555 | NM_001276289.1 | 8.76582946 | -4.7546885 | 1.76236209 | -2.6979067 |
| 17556 | NM_006226.3 | 56.4855098 | -3.3154303 | 1.22891692 | -2.6978474 |
| 17557 | NM_001002814.2 | 30.0105826 | -3.7050982 | 1.37344391 | -2.6976698 |
| 17558 | NR_135304.1 | 6.25366247 | -4.279839 | 1.58657821 | -2.6975279 |
| 17559 | NM_001321857.1 | 24.4900444 | -6.2381964 | 2.31279096 | -2.6972591 |
| 17560 | NR_136748.1 | 33.098397 | -6.6696566 | 2.4729471 | -2.6970478 |
| 17561 | NM_002773.4 | 6.2794441 | -4.271248 | 1.58391491 | -2.6966398 |
| 17562 | NM_015938.4 | 99.3454622 | -3.5486997 | 1.31602952 | -2.6965198 |
| 17563 | NR_132346.1 | 5.6497215 | -4.1210766 | 1.52839263 | -2.6963468 |
| 17564 | NM_001206846.1 | 19.4656713 | 6.51601732 | 2.41681142 | 2.69612154 |
| 17565 | NM_001202521.1 | 24.9510457 | -6.2643767 | 2.32371902 | -2.6958409 |
| 17566 | NR_002806.2 | 10.0127258 | -3.9194141 | 1.45400343 | -2.6956017 |
| 17567 | NM_001035505.1 | 10.7661417 | -3.9991314 | 1.48361179 | -2.6955376 |
| 17568 | NM_001291712.1 | 43.0899422 | 3.40929177 | 1.26494515 | 2.69520918 |
| 17569 | NR_024412.1 | 5.99280167 | -4.2142746 | 1.56366672 | -2.6951233 |
| 17570 | NM_001316309.1 | 14.4562996 | -3.66964 | 1.36159723 | -2.6950995 |
| 17571 | NM_001135248.1 | 24.6761631 | -6.2483864 | 2.3187499 | -2.694722 |
| 17572 | NM_138481.1 | 6.32874342 | -4.2734216 | 1.58591031 | -2.6946174 |
| 17573 | NM_145283.2 | 15.9661909 | 1.61827004 | 0.6007332 | 2.69382489 |
| 17574 | NM_001243089.1 | 26.6016335 | -6.356734 | 2.35986575 | -2.6936846 |
| 17575 | NM_001206480.2 | 7.76705119 | -4.5617064 | 1.69365912 | -2.6934029 |
| 17576 | NM_152342.3 | 43.1680283 | -3.3879068 | 1.25787909 | -2.6933485 |
| 17577 | NR_033890.1 | 39.8677932 | 2.53435448 | 0.94101043 | 2.69322677 |
| 17578 | NR_103730.1 | 25.5163337 | 3.38803173 | 1.25808063 | 2.69301636 |
| 17579 | NM_001081004.2 | 8.78309252 | -4.7614746 | 1.76812983 | -2.692944 |
| 17580 | NM_001329544.1 | 11.5330343 | -5.144265 | 1.91044476 | -2.6927054 |
| 17581 | NM_080603.4 | 142.895776 | 0.80695249 | 0.29968324 | 2.69268476 |
| 17582 | NM_001112706.2 | 11.1054103 | -5.1016382 | 1.89469949 | -2.6925844 |
| 17583 | NM_207418.2 | 6.83686613 | -4.379111 | 1.62655031 | -2.692269 |
| 17584 | NM_000075.3 | 295.760358 | -3.2002138 | 1.18873424 | -2.6921188 |
| 17585 | NM_001172557.1 | 6.3341251 | -4.2920075 | 1.59427823 | -2.692132 |
| 17586 | NM_032843.4 | 25.9708441 | -6.3223011 | 2.34845464 | -2.6921112 |

| | | | | | |
|---|---|---|---|---|---|
| 17587 | NM_001320137.1 | 8.08676697 | -4.6405985 | 1.72380751 | -2.692063 |
| 17588 | NM_024586.5 | 23.5579067 | -6.1817531 | 2.2964562 | -2.6918663 |
| 17589 | NM_024611.5 | 25.1945175 | -6.2786616 | 2.33253351 | -2.6917777 |
| 17590 | NM_001319138.1 | 12.305077 | -3.2908029 | 1.22269295 | -2.6914385 |
| 17591 | NM_001195203.2 | 31.6809603 | -6.6090654 | 2.45593655 | -2.6910571 |
| 17592 | NM_024906.2 | 128.502889 | 3.15337117 | 1.17180992 | 2.69102618 |
| 17593 | NR_028351.1 | 26.2570768 | 2.18985669 | 0.81379459 | 2.69092067 |
| 17594 | NM_001278277.1 | 9.21658205 | -4.8171652 | 1.79036003 | -2.6906126 |
| 17595 | NM_001199214.1 | 6.2025172 | -4.2703489 | 1.58716841 | -2.6905455 |
| 17596 | NM_001281805.1 | 62.5676417 | 1.57494018 | 0.58541209 | 2.69031031 |
| 17597 | NM_138812.3 | 5.91298922 | -4.1786431 | 1.55321395 | -2.6903203 |
| 17598 | NR_046593.1 | 104.864403 | 2.02894186 | 0.75425682 | 2.68998809 |
| 17599 | NM_173799.3 | 11.487156 | 6.0888831 | 2.26359669 | 2.68991518 |
| 17600 | NM_001300950.1 | 5.58372399 | -4.096538 | 1.52306047 | -2.6896752 |
| 17601 | NM_001329454.1 | 5.00720393 | -3.9466271 | 1.46740789 | -2.6895229 |
| 17602 | NM_001324244.1 | 23.4431376 | -6.1748487 | 2.29596471 | -2.6894353 |
| 17603 | NM_001258326.1 | 24.136365 | -6.2171515 | 2.31190712 | -2.6891874 |
| 17604 | NR_107053.1 | 18.3259607 | 3.78669505 | 1.4081159 | 2.68919274 |
| 17605 | NM_175634.2 | 31.9990571 | -6.6217041 | 2.46255036 | -2.6889619 |
| 17606 | NR_024460.1 | 23.3542986 | -6.1692779 | 2.29432574 | -2.6889285 |
| 17607 | NM_014757.4 | 148.001667 | -3.4079219 | 1.26747316 | -2.6887527 |
| 17608 | NM_001256105.1 | 45.601751 | -3.2780225 | 1.21925723 | -2.6885405 |
| 17609 | NM_001317744.1 | 7.04487707 | -4.4441739 | 1.65301783 | -2.6885215 |
| 17610 | NM_152565.1 | 15.7194985 | 1.92178052 | 0.71485971 | 2.6883324 |
| 17611 | NM_025262.3 | 6.60980643 | -4.3571469 | 1.62084765 | -2.6881903 |
| 17612 | NM_001318042.1 | 34.0472136 | -6.7123661 | 2.49704504 | -2.6881238 |
| 17613 | NR_136733.1 | 96.299061 | 4.62376236 | 1.72033647 | 2.68770816 |
| 17614 | NM_012095.5 | 29.6843444 | -6.5133573 | 2.423427 | -2.6876639 |
| 17615 | NM_012118.3 | 44.9243914 | -2.1638269 | 0.80531017 | -2.6869484 |
| 17616 | NR_001288.1 | 40.4129413 | 2.26331773 | 0.84245193 | 2.68658382 |
| 17617 | NM_001324319.1 | 93.4591598 | 3.75485719 | 1.39785419 | 2.68615799 |
| 17618 | NM_171846.3 | 22.9365736 | -6.1434785 | 2.28713123 | -2.6861067 |
| 17619 | NM_001145714.1 | 26.3588193 | -6.3439629 | 2.36191856 | -2.6859363 |
| 17620 | NM_006877.3 | 20.8679735 | -4.245929 | 1.58084215 | -2.6858653 |
| 17621 | NM_001141970.1 | 23.7130588 | -6.1916189 | 2.3053542 | -2.685756 |
| 17622 | NM_001307937.1 | 8.86325638 | -4.7687091 | 1.77564549 | -2.68562 |
| 17623 | NR_132660.1 | 39.3629447 | 1.87040392 | 0.69646862 | 2.68555376 |
| 17624 | NM_001278694.1 | 6.38495538 | -4.3025569 | 1.60216806 | -2.6854591 |
| 17625 | NM_001278387.1 | 10.2902589 | -4.9853389 | 1.85662928 | -2.6851558 |
| 17626 | NM_004437.3 | 7.69257869 | -4.5642532 | 1.70007244 | -2.6847404 |
| 17627 | NR_036520.1 | 12.391417 | -2.8047863 | 1.04478405 | -2.6845608 |
| 17628 | NM_138363.2 | 25.4939871 | -6.2960813 | 2.34548393 | -2.6843421 |
| 17629 | NM_006183.4 | 9.41279233 | -4.8468463 | 1.80590766 | -2.6838838 |

| | | | | | |
|---|---|---|---|---|---|
| 17630 | NM_001297609.1 | 29.9500657 | -6.5263576 | 2.43180466 | -2.6837508 |
| 17631 | NM_016001.2 | 90.9244129 | -3.5016599 | 1.30484512 | -2.6835828 |
| 17632 | NR_023390.1 | 137.638547 | 1.07916673 | 0.40224399 | 2.68286603 |
| 17633 | NM_001267571.1 | 9.66851205 | -4.8882731 | 1.82220516 | -2.682614 |
| 17634 | NR_104598.1 | 6.9493188 | -4.4208937 | 1.6480913 | -2.6824326 |
| 17635 | NM_001100176.1 | 23.639606 | -6.186727 | 2.30677084 | -2.681986 |
| 17636 | NM_005168.4 | 191.749947 | -4.0674765 | 1.51668769 | -2.6818155 |
| 17637 | NR_027285.1 | 5.97459229 | -4.2026853 | 1.56719473 | -2.6816612 |
| 17638 | NM_018911.2 | 6.72587061 | -4.3597772 | 1.62601706 | -2.6812616 |
| 17639 | NM_183373.3 | 99.8347257 | -2.386458 | 0.89017975 | -2.6808721 |
| 17640 | NM_001308170.1 | 29.8944981 | -6.5237683 | 2.43360876 | -2.6806973 |
| 17641 | NM_012155.2 | 23.4730917 | -6.1769525 | 2.30436432 | -2.6805451 |
| 17642 | NM_018043.5 | 34.6207344 | -3.2169958 | 1.20041131 | -2.6799113 |
| 17643 | NM_001145205.1 | 31.777285 | -3.0733206 | 1.14691552 | -2.6796399 |
| 17644 | NM_032037.3 | 5.40406015 | -4.0464706 | 1.51010051 | -2.6796035 |
| 17645 | NM_198970.1 | 24.9285486 | -6.2630742 | 2.33736723 | -2.6795422 |
| 17646 | NM_001123395.1 | 32.6742692 | -6.6531794 | 2.4831404 | -2.6793408 |
| 17647 | NM_001177665.1 | 23.6263237 | -6.185832 | 2.30902609 | -2.6789788 |
| 17648 | NM_001202470.2 | 7.49898792 | -4.5326514 | 1.69209473 | -2.678722 |
| 17649 | NR_046215.1 | 15.6633151 | 4.60794345 | 1.72025901 | 2.67863353 |
| 17650 | NR_104587.1 | 27.2267045 | -6.3890709 | 2.38527551 | -2.6785463 |
| 17651 | NM_000530.7 | 6.49508496 | -4.3321701 | 1.61746837 | -2.6783647 |
| 17652 | NR_109821.1 | 6.09126906 | -4.2192985 | 1.5753715 | -2.6782879 |
| 17653 | NM_001322219.1 | 5.64465298 | -4.1316334 | 1.5426773 | -2.6782227 |
| 17654 | NR_145450.1 | 79.8555669 | 2.38985276 | 0.89232799 | 2.67822235 |
| 17655 | NM_001258455.1 | 23.339702 | -6.1683732 | 2.3032238 | -2.6781476 |
| 17656 | NM_001270409.1 | 6.11155119 | -4.2151742 | 1.57406458 | -2.6778915 |
| 17657 | NM_001146285.1 | 8.15127031 | -4.6374824 | 1.73192062 | -2.677653 |
| 17658 | NM_001135644.1 | 6.11761831 | -4.2406433 | 1.58390145 | -2.6773404 |
| 17659 | NM_032728.3 | 9.54095871 | -3.8619573 | 1.44283238 | -2.67665 |
| 17660 | NM_001320849.1 | 5.15909648 | -3.9797011 | 1.48710178 | -2.6761458 |
| 17661 | NR_038272.1 | 5.2907529 | -4.013268 | 1.49969711 | -2.6760524 |
| 17662 | NM_017925.6 | 26.6617226 | -6.3598364 | 2.37660513 | -2.6760173 |
| 17663 | NM_001098784.1 | 100.143532 | -3.5620257 | 1.33113391 | -2.6759334 |
| 17664 | NM_175737.3 | 88.7221667 | 1.77803905 | 0.6645243 | 2.67565693 |
| 17665 | NM_001286730.1 | 84.0774377 | 5.88336043 | 2.19895376 | 2.67552712 |
| 17666 | NR_134854.1 | 4.88104008 | -3.8967536 | 1.45664031 | -2.6751653 |
| 17667 | NM_173206.3 | 22.6345103 | -6.1242616 | 2.28949897 | -2.6749353 |
| 17668 | NM_032489.2 | 4.88972772 | -3.9139295 | 1.46327996 | -2.6747646 |
| 17669 | NR_028443.1 | 5.62773926 | -4.1152728 | 1.53854619 | -2.6747801 |
| 17670 | NM_001306079.1 | 31.5898724 | -6.6035784 | 2.46889228 | -2.674713 |
| 17671 | NM_000865.2 | 5.26415222 | -4.0319242 | 1.50745727 | -2.6746524 |
| 17672 | NR_121570.1 | 9.48256814 | 2.00776136 | 0.75072285 | 2.67443754 |

| | | | | | |
|---|---|---|---|---|---|
| 17673 | NM_001277348.1 | 8.42348288 | -3.6188831 | 1.35323564 | -2.6742446 |
| 17674 | NM_021151.3 | 33.3783762 | -2.8137556 | 1.05217023 | -2.6742399 |
| 17675 | NM_001197221.1 | 22.7809156 | -6.1336342 | 2.29377804 | -2.6740313 |
| 17676 | NM_001005389.1 | 6.99119878 | -4.4280717 | 1.65602304 | -2.6739191 |
| 17677 | NM_025204.3 | 22.8167876 | -6.1357532 | 2.29506385 | -2.6734564 |
| 17678 | NR_103518.2 | 8.01388103 | -3.5470938 | 1.32688328 | -2.6732523 |
| 17679 | NR_003003.2 | 4.97761028 | -3.9383751 | 1.47369078 | -2.6724569 |
| 17680 | NM_001323627.1 | 34.079569 | -2.9833619 | 1.11638514 | -2.6723411 |
| 17681 | NM_001329921.1 | 11.1128788 | -5.1021693 | 1.90927368 | -2.672309 |
| 17682 | NM_005144.4 | 31.853102 | -6.6144409 | 2.47530888 | -2.6721679 |
| 17683 | NR_046417.1 | 13.5535308 | 4.63909118 | 1.73617365 | 2.67202026 |
| 17684 | NM_182678.2 | 23.799264 | -6.1964031 | 2.31926619 | -2.6717085 |
| 17685 | NR_024162.1 | 47.5080463 | 2.6804099 | 1.00330413 | 2.67158264 |
| 17686 | NM_001317950.1 | 23.7777185 | -6.1956864 | 2.31952832 | -2.6710975 |
| 17687 | NM_024820.2 | 6.39444843 | -4.3022881 | 1.61099391 | -2.67058 |
| 17688 | NR_045375.2 | 5.21553826 | -3.9954418 | 1.49617044 | -2.6704456 |
| 17689 | NM_000534.4 | 23.8168271 | -6.1973934 | 2.320972 | -2.6701715 |
| 17690 | NM_001301302.1 | 7.59596905 | -4.5464455 | 1.70278317 | -2.6700085 |
| 17691 | NM_001206833.1 | 29.6954686 | -6.51442 | 2.44007659 | -2.6697605 |
| 17692 | NM_001135673.3 | 23.504191 | -6.178368 | 2.31430398 | -2.6696441 |
| 17693 | NR_038841.1 | 114.707313 | 2.68228082 | 1.00476386 | 2.6695634 |
| 17694 | NR_073460.1 | 17.7199329 | 3.44106322 | 1.28911233 | 2.66932768 |
| 17695 | NM_001077664.2 | 87.7215436 | -4.0118753 | 1.50303863 | -2.6691765 |
| 17696 | NM_001306085.1 | 22.3434127 | -6.1056744 | 2.2878107 | -2.6687848 |
| 17697 | NR_125916.1 | 46.9378954 | 2.33978313 | 0.87678517 | 2.66859341 |
| 17698 | NM_130773.3 | 14.8208445 | -2.5559985 | 0.95842605 | -2.6668709 |
| 17699 | NM_001130713.2 | 22.9313546 | -6.1428416 | 2.30378313 | -2.6664149 |
| 17700 | NM_001042428.1 | 29.827005 | -6.5209182 | 2.44562752 | -2.6663579 |
| 17701 | NM_001145157.1 | 6.3534225 | -4.2803623 | 1.60535252 | -2.6663068 |
| 17702 | NM_001171133.2 | 4.77479418 | -3.8698956 | 1.45162442 | -2.6659069 |
| 17703 | NM_181465.2 | 5.45636999 | -4.0775436 | 1.52960757 | -2.6657449 |
| 17704 | NM_001042702.3 | 5.40353066 | -4.0670699 | 1.52576533 | -2.6655933 |
| 17705 | NM_001284408.1 | 15.8907466 | 3.65378123 | 1.37075041 | 2.66553357 |
| 17706 | NM_001323645.1 | 22.7647048 | -6.1326115 | 2.30081973 | -2.6654029 |
| 17707 | NM_001157.2 | 22.2722561 | -6.1010529 | 2.28924227 | -2.6650971 |
| 17708 | NM_178493.5 | 5.43861083 | -4.0516185 | 1.52025939 | -2.6650837 |
| 17709 | NM_173179.3 | 22.2880043 | -6.1021461 | 2.28979682 | -2.6649291 |
| 17710 | NM_080866.2 | 15.9390454 | 1.64965687 | 0.61918142 | 2.66425447 |
| 17711 | NM_032753.3 | 17.6032502 | 2.24631338 | 0.84315969 | 2.66416125 |
| 17712 | NM_001242848.1 | 24.0087599 | -6.2090707 | 2.33063195 | -2.6641147 |
| 17713 | NM_001206845.1 | 53.4051568 | 4.73932622 | 1.77905717 | 2.66395386 |
| 17714 | NM_001317188.1 | 6.73539235 | -4.3624222 | 1.63788185 | -2.6634535 |
| 17715 | NM_001025249.1 | 7.89288892 | -4.6034171 | 1.72841967 | -2.6633677 |

| | | | | | |
|---|---|---|---|---|---|
| 17716 | NM_198938.2 | 8.86837873 | -4.7695088 | 1.79087336 | -2.6632306 |
| 17717 | NM_001324132.1 | 24.0803734 | -6.2132599 | 2.33311198 | -2.6630783 |
| 17718 | NM_013342.3 | 43.2873466 | -3.1935701 | 1.19924108 | -2.6629926 |
| 17719 | NM_001193277.1 | 7.31687336 | 7.78707816 | 2.92467263 | 2.66254694 |
| 17720 | NM_001308056.1 | 33.5090978 | -6.6876612 | 2.51189243 | -2.6623995 |
| 17721 | NM_052859.3 | 84.7387501 | -2.0669198 | 0.77637104 | -2.6622835 |
| 17722 | NM_004841.3 | 44.3161325 | -2.9760789 | 1.11796951 | -2.6620394 |
| 17723 | NM_015308.3 | 118.889787 | -4.7339778 | 1.7784242 | -2.6618946 |
| 17724 | NM_001097599.1 | 7.17618796 | -4.4611811 | 1.67611339 | -2.6616225 |
| 17725 | NM_001347829.1 | 26.0174033 | -6.3250225 | 2.37644539 | -2.6615476 |
| 17726 | NM_032927.3 | 11.4330978 | -4.1163417 | 1.54679071 | -2.6612144 |
| 17727 | NM_001271824.1 | 8.9726315 | -4.7881462 | 1.79960186 | -2.6606697 |
| 17728 | NM_152559.2 | 5.52799744 | -4.1008129 | 1.54138448 | -2.6604737 |
| 17729 | NM_001080779.1 | 5.23926587 | -3.997647 | 1.50262802 | -2.6604369 |
| 17730 | NR_144474.1 | 103.908072 | -2.6943661 | 1.012831 | -2.6602327 |
| 17731 | NM_032484.4 | 29.1571395 | -3.1706223 | 1.19203734 | -2.6598347 |
| 17732 | NM_001190489.1 | 5.32290903 | 4.88940622 | 1.83832353 | 2.65970932 |
| 17733 | NM_001128225.2 | 30.8252369 | -6.5687366 | 2.46974433 | -2.6596828 |
| 17734 | NR_037648.1 | 23.1022415 | -6.1534349 | 2.31379932 | -2.6594506 |
| 17735 | NM_001199893.1 | 6.70121727 | -4.3734297 | 1.64452011 | -2.6593957 |
| 17736 | NM_012454.3 | 50.1096689 | -3.9414113 | 1.48206167 | -2.6594111 |
| 17737 | NM_001256835.1 | 22.9110624 | -6.141901 | 2.30962262 | -2.6592661 |
| 17738 | NM_000193.3 | 5.90256667 | -4.1931497 | 1.57714841 | -2.6586906 |
| 17739 | NM_001034853.1 | 9.89286121 | -4.9347917 | 1.85615137 | -2.6586149 |
| 17740 | NM_001146289.1 | 31.8487104 | -6.6162733 | 2.48881832 | -2.6583995 |
| 17741 | NR_034179.1 | 4.80058327 | -3.8757142 | 1.45802249 | -2.6581992 |
| 17742 | NM_207005.2 | 24.0621592 | -6.212864 | 2.33727984 | -2.6581601 |
| 17743 | NM_021048.4 | 67.5712092 | 3.60397766 | 1.35595906 | 2.65788087 |
| 17744 | NM_007334.2 | 26.8022184 | 9.66141342 | 3.63557213 | 2.65746713 |
| 17745 | NM_173653.3 | 5.28100023 | -4.0308141 | 1.5168141 | -2.6574213 |
| 17746 | NM_001306087.1 | 7.06730185 | -4.4398052 | 1.67075613 | -2.6573628 |
| 17747 | NM_018250.3 | 29.4149732 | -6.5000069 | 2.44614516 | -2.657245 |
| 17748 | NM_001282410.1 | 28.7407029 | -6.4669161 | 2.43375569 | -2.6571756 |
| 17749 | NM_001348120.1 | 29.1712133 | -6.4883982 | 2.44245344 | -2.6565084 |
| 17750 | NM_001319840.1 | 30.3688297 | -6.5465937 | 2.46478825 | -2.6560471 |
| 17751 | NM_001816.3 | 36.4181459 | 1.7784953 | 0.66960627 | 2.65603143 |
| 17752 | NM_014885.4 | 6.43424651 | -4.2874758 | 1.61439095 | -2.6557854 |
| 17753 | NR_136695.1 | 5.40457002 | -4.0413788 | 1.52183385 | -2.6555979 |
| 17754 | NM_001287244.1 | 14.5759287 | -3.6773753 | 1.38479317 | -2.6555412 |
| 17755 | NR_126050.1 | 4.87898694 | -3.9079592 | 1.4716356 | -2.655521 |
| 17756 | NM_178273.1 | 11.4525747 | 8.42855271 | 3.17417721 | 2.6553504 |
| 17757 | NM_001161630.1 | 8.52484133 | 6.83858559 | 2.57553072 | 2.65521413 |
| 17758 | NM_001039199.2 | 37.1162261 | -6.8365429 | 2.574864 | -2.6551083 |

| | | | | | |
|---|---|---|---|---|---|
| 17759 | NM_001193427.1 | 7.65575338 | 7.85523162 | 2.95870002 | 2.65496048 |
| 17760 | NM_001267828.1 | 7.65575338 | 7.85523162 | 2.95870002 | 2.65496048 |
| 17761 | NM_001317200.1 | 23.4596915 | -6.1759636 | 2.32629103 | -2.6548542 |
| 17762 | NM_001277372.2 | 8.52784935 | -4.7201366 | 1.77828699 | -2.6543165 |
| 17763 | NM_005765.2 | 390.407504 | -4.1706085 | 1.57124838 | -2.6543279 |
| 17764 | NM_001172745.1 | 5.36103831 | -4.0515652 | 1.52641772 | -2.6542965 |
| 17765 | NM_173176.2 | 29.3172791 | -6.4968561 | 2.44795325 | -2.6539952 |
| 17766 | NR_026815.1 | 6.40982941 | -4.2954734 | 1.61852023 | -2.653951 |
| 17767 | NM_001346031.1 | 21.8664301 | -6.0746058 | 2.28903187 | -2.6537882 |
| 17768 | NM_001136558.1 | 25.7163701 | -6.3077352 | 2.37693625 | -2.653725 |
| 17769 | NM_001261835.1 | 27.6961779 | -6.4146637 | 2.41745283 | -2.6534804 |
| 17770 | NM_178517.4 | 6.78315872 | -4.3705292 | 1.64709108 | -2.6534836 |
| 17771 | NR_073537.1 | 6.63235597 | -4.3318108 | 1.63259919 | -2.6533216 |
| 17772 | NM_203487.2 | 24.0909083 | -6.2137407 | 2.34193009 | -2.6532562 |
| 17773 | NM_001291416.1 | 22.0051328 | -6.0838634 | 2.29346189 | -2.6526987 |
| 17774 | NM_173480.2 | 11.5671406 | -3.1846195 | 1.20055898 | -2.652614 |
| 17775 | NR_037808.1 | 16.8871934 | 3.42976691 | 1.29315957 | 2.65223797 |
| 17776 | NM_001256847.2 | 31.3592941 | -5.6027626 | 2.11272337 | -2.6519149 |
| 17777 | NM_001243146.1 | 7.45901862 | -4.5259916 | 1.70681453 | -2.6517185 |
| 17778 | NR_135490.1 | 5.95578227 | -4.2042593 | 1.58556605 | -2.6515826 |
| 17779 | NR_103777.1 | 139.382281 | 2.06316508 | 0.77832723 | 2.65076822 |
| 17780 | NM_001202431.1 | 14.2011541 | -3.0024201 | 1.13267966 | -2.650723 |
| 17781 | NM_001311311.1 | 5.06159257 | -3.971067 | 1.49828033 | -2.6504165 |
| 17782 | NM_001100592.2 | 28.4717509 | -6.4547693 | 2.43605051 | -2.6496861 |
| 17783 | NM_022648.5 | 22.1918263 | -6.0959501 | 2.30073398 | -2.6495675 |
| 17784 | NM_001321759.1 | 30.0925004 | -6.532859 | 2.46568591 | -2.6495098 |
| 17785 | NM_001172828.1 | 178.875445 | -3.9637599 | 1.49629103 | -2.6490568 |
| 17786 | NM_001301020.1 | 6.77848306 | -4.3664161 | 1.6484538 | -2.648795 |
| 17787 | NM_021625.4 | 41.7913788 | -7.0079422 | 2.64601835 | -2.6484859 |
| 17788 | NM_001287436.2 | 7.72281945 | -4.577554 | 1.72842726 | -2.6483926 |
| 17789 | NM_022460.3 | 56.9238214 | -3.1962838 | 1.20704628 | -2.6480209 |
| 17790 | NM_001174127.1 | 35.6357185 | -5.7886139 | 2.18612266 | -2.6478907 |
| 17791 | NM_152882.3 | 27.0888646 | -6.3825126 | 2.4106951 | -2.6475818 |
| 17792 | NM_001040424.2 | 22.1903372 | -6.09562 | 2.30249588 | -2.6473967 |
| 17793 | NM_001145966.1 | 118.718355 | -3.582084 | 1.35326396 | -2.6469958 |
| 17794 | NM_007172.3 | 177.53169 | -2.8398353 | 1.07306196 | -2.6464784 |
| 17795 | NR_033990.1 | 27.1064643 | 2.85211664 | 1.07788112 | 2.64604007 |
| 17796 | NM_001139499.1 | 4.9918705 | -3.9401283 | 1.48924848 | -2.6457159 |
| 17797 | NM_001321877.1 | 23.764044 | -6.1948339 | 2.34162466 | -2.6455281 |
| 17798 | NM_145235.3 | 23.7436853 | -4.0132622 | 1.51707061 | -2.6454024 |
| 17799 | NR_046203.2 | 7.9266461 | 3.29774104 | 1.24687186 | 2.64481151 |
| 17800 | NM_001114396.1 | 178.0357 | 3.45052144 | 1.30466958 | 2.64474737 |
| 17801 | NR_024607.1 | 6.59388379 | -4.3388819 | 1.64065746 | -2.6445995 |

| | | | | | |
|---|---|---|---|---|---|
| 17802 | NM_001272103.1 | 10.8737091 | -4.0051307 | 1.51497042 | -2.6437022 |
| 17803 | NM_001324212.1 | 30.3626012 | -6.5473317 | 2.47658779 | -2.6436905 |
| 17804 | NM_133367.4 | 89.2396743 | -1.8875862 | 0.71397678 | -2.6437641 |
| 17805 | NM_198504.2 | 4.91844939 | -3.9309072 | 1.48687018 | -2.6437461 |
| 17806 | NR_026794.1 | 18.5124482 | 3.63955486 | 1.37667535 | 2.64372778 |
| 17807 | NM_001314051.1 | 12.8686977 | -3.2104341 | 1.21448662 | -2.6434496 |
| 17808 | NM_002439.4 | 66.5581387 | -3.2107734 | 1.21462297 | -2.6434322 |
| 17809 | NM_152490.4 | 78.9324432 | -3.2784315 | 1.24021017 | -2.6434483 |
| 17810 | NM_001304533.1 | 6.11548833 | -4.2352548 | 1.60259313 | -2.6427511 |
| 17811 | NM_020771.3 | 24.1514269 | -6.2180848 | 2.35303262 | -2.6425833 |
| 17812 | NM_001040159.1 | 27.9899034 | -6.4285304 | 2.43338614 | -2.6418045 |
| 17813 | NM_001257972.1 | 5.5492073 | -4.099091 | 1.55173416 | -2.6416193 |
| 17814 | NM_001171606.1 | 21.5707591 | -6.0548305 | 2.29260409 | -2.6410275 |
| 17815 | NR_125822.1 | 5.00961528 | -3.9429414 | 1.49315543 | -2.6406772 |
| 17816 | NM_000307.4 | 6.00484055 | -4.1988591 | 1.59014194 | -2.6405562 |
| 17817 | NM_001271560.1 | 6.3192955 | -4.2768497 | 1.61977506 | -2.6403973 |
| 17818 | NM_001134776.1 | 10.7768035 | -5.055262 | 1.91558209 | -2.6390213 |
| 17819 | NR_073064.1 | 9.35655524 | -4.8482676 | 1.83761442 | -2.6383487 |
| 17820 | NM_001330102.1 | 28.5755522 | -6.4582656 | 2.44794445 | -2.6382403 |
| 17821 | NM_001142477.1 | 26.7647587 | -6.3634243 | 2.41203683 | -2.6381954 |
| 17822 | NM_024050.5 | 194.912611 | -2.5589185 | 0.96997434 | -2.6381301 |
| 17823 | NM_001135676.1 | 6.1362271 | -4.232788 | 1.60456655 | -2.6379635 |
| 17824 | NM_001330203.1 | 30.8284196 | -6.5678925 | 2.48980943 | -2.6379097 |
| 17825 | NM_017890.4 | 35.6393051 | -6.7762254 | 2.569063 | -2.6376252 |
| 17826 | NM_178820.4 | 142.793656 | 1.66413107 | 0.63093496 | 2.63756358 |
| 17827 | NM_058192.2 | 23.5478673 | -6.181252 | 2.34363236 | -2.6374666 |
| 17828 | NM_001007074.1 | 4.98254771 | -3.9531245 | 1.49905507 | -2.6370776 |
| 17829 | NM_001286769.1 | 22.0083448 | -6.0835759 | 2.30695123 | -2.6370631 |
| 17830 | NM_001173456.1 | 9.13795234 | -3.7589261 | 1.42544781 | -2.6370142 |
| 17831 | NR_045791.1 | 41.6698065 | 4.31447452 | 1.63618332 | 2.63691388 |
| 17832 | NM_001320568.1 | 27.8680134 | -6.422563 | 2.43584883 | -2.6366837 |
| 17833 | NM_001347712.1 | 5.21333735 | -4.0084197 | 1.52030868 | -2.6365828 |
| 17834 | NM_005990.3 | 82.4743656 | -2.4414249 | 0.92598824 | -2.6365614 |
| 17835 | NM_024947.3 | 169.867507 | -1.4414266 | 0.54679097 | -2.6361566 |
| 17836 | NM_001166161.1 | 21.4099768 | -6.0440647 | 2.29310616 | -2.6357544 |
| 17837 | NM_201382.3 | 21.4598974 | -6.0477714 | 2.29557313 | -2.6345366 |
| 17838 | NM_001171091.1 | 21.4504264 | -6.0467547 | 2.29537123 | -2.6343254 |
| 17839 | NM_001320557.1 | 9.40791441 | -4.8494575 | 1.84120309 | -2.6338526 |
| 17840 | NM_000334.4 | 15.40388 | 3.55643341 | 1.35030873 | 2.63379281 |
| 17841 | NM_001098819.3 | 25.0382173 | -6.2696992 | 2.38064195 | -2.633617 |
| 17842 | NM_001320924.2 | 8.21652104 | -4.6648763 | 1.77135695 | -2.6335044 |
| 17843 | NR_046100.1 | 3.91913824 | 4.8701436 | 1.84934926 | 2.63343637 |
| 17844 | NM_031475.2 | 8.80937689 | -4.7677278 | 1.81050286 | -2.6333721 |

| | | | | | |
|---|---|---|---|---|---|
| 17845 | NM_001172685.1 | 11.9052102 | -4.1523527 | 1.57687458 | -2.6332803 |
| 17846 | NM_001256488.1 | 142.026114 | -2.8949483 | 1.09935739 | -2.6333095 |
| 17847 | NM_004485.3 | 113.612372 | 5.87098859 | 2.22952833 | 2.63328728 |
| 17848 | NM_173561.2 | 4.75242492 | -3.8819857 | 1.47433411 | -2.6330434 |
| 17849 | NM_001319847.1 | 5.94202676 | -4.1810895 | 1.58811 | -2.6327456 |
| 17850 | NM_032250.3 | 6.48215446 | -4.2987577 | 1.63290116 | -2.6325891 |
| 17851 | NM_001242832.1 | 37.1411604 | -5.8495617 | 2.22239415 | -2.6320991 |
| 17852 | NM_015853.4 | 10.1499783 | -3.9345434 | 1.49504256 | -2.6317267 |
| 17853 | NM_001301033.1 | 9.0205429 | -3.7862192 | 1.43875794 | -2.6315888 |
| 17854 | NR_136541.1 | 32.868391 | 2.72230278 | 1.03451318 | 2.63148197 |
| 17855 | NM_000369.2 | 5.45405523 | -4.0601615 | 1.54298418 | -2.6313695 |
| 17856 | NM_022783.3 | 6.28477047 | -4.2702673 | 1.62295651 | -2.6311656 |
| 17857 | NM_001171934.1 | 17.6871264 | -3.4052382 | 1.29428115 | -2.630988 |
| 17858 | NM_181093.3 | 6.01973814 | -4.2261152 | 1.60660282 | -2.6304667 |
| 17859 | NM_001098670.1 | 10.1037731 | -4.9539822 | 1.88334172 | -2.6304213 |
| 17860 | NM_014433.2 | 4.47960766 | -3.7880518 | 1.44011345 | -2.6303843 |
| 17861 | NM_001322165.1 | 8.71557238 | -4.7390958 | 1.80172789 | -2.6303061 |
| 17862 | NM_182648.1 | 37.2491304 | -5.8530542 | 2.22541457 | -2.6300961 |
| 17863 | NM_001321207.1 | 30.4051608 | -6.5483036 | 2.48979376 | -2.6300586 |
| 17864 | NR_036487.1 | 4.86953068 | -3.9023347 | 1.48375978 | -2.6300313 |
| 17865 | NM_001242857.2 | 30.9005155 | -6.5726274 | 2.49930463 | -2.6297824 |
| 17866 | NM_181674.2 | 6.7921194 | -4.3676687 | 1.66110005 | -2.6293833 |
| 17867 | NR_027246.1 | 15.7099322 | 3.9883804 | 1.51692866 | 2.62924718 |
| 17868 | NM_001113491.1 | 25.2166076 | -6.2796806 | 2.38850336 | -2.6291278 |
| 17869 | NM_001301726.1 | 59.5350648 | -3.1453282 | 1.19653599 | -2.6286951 |
| 17870 | NR_015375.2 | 4.74098947 | -3.8682099 | 1.47157556 | -2.6286179 |
| 17871 | NM_001040161.2 | 7.05260659 | -4.4299671 | 1.68555524 | -2.6281945 |
| 17872 | NM_001345975.1 | 30.0259859 | -6.5288729 | 2.48418583 | -2.6281741 |
| 17873 | NM_004510.3 | 7.70253389 | -3.5284592 | 1.34259488 | -2.6280893 |
| 17874 | NM_001160308.2 | 21.9037168 | -6.0766729 | 2.31244442 | -2.6278136 |
| 17875 | NM_182944.2 | 4.80402572 | -3.8809716 | 1.47688877 | -2.6278022 |
| 17876 | NR_046851.1 | 80.104616 | 2.58230621 | 0.98297386 | 2.62703446 |
| 17877 | NM_032153.5 | 24.9095133 | -6.262445 | 2.38388285 | -2.6269936 |
| 17878 | NM_001284242.1 | 24.8410805 | 9.55154531 | 3.63596159 | 2.6269654 |
| 17879 | NM_005681.3 | 6.43564415 | -4.2998339 | 1.63698211 | -2.6266835 |
| 17880 | NM_153435.1 | 25.8959258 | -6.3173306 | 2.40508818 | -2.6266524 |
| 17881 | NM_007152.4 | 24.1172546 | -6.215817 | 2.36656063 | -2.6265192 |
| 17882 | NM_001510.3 | 7.71702002 | -3.5472541 | 1.35071352 | -2.6262076 |
| 17883 | NM_001178032.2 | 12.3103971 | -3.3431176 | 1.27302146 | -2.6261282 |
| 17884 | NM_006794.3 | 4.62960707 | -3.8356261 | 1.46060378 | -2.6260551 |
| 17885 | NM_001198773.2 | 23.5617861 | -6.1816251 | 2.35424125 | -2.6257399 |
| 17886 | NR_038450.1 | 13.9105245 | 3.09672645 | 1.17966904 | 2.62508073 |
| 17887 | NM_033544.2 | 8.0256336 | -4.6252 | 1.76227489 | -2.6245622 |

| | | | | | |
|---|---|---|---|---|---|
| 17888 | NR_038391.1 | 6.07416954 | -4.2215607 | 1.60851943 | -2.6245009 |
| 17889 | NR_040092.1 | 140.484375 | -1.2134227 | 0.46234493 | -2.6244965 |
| 17890 | NM_144584.2 | 5.31414708 | -4.0226855 | 1.53286226 | -2.6242968 |
| 17891 | NR_027355.2 | 6.95105584 | -4.4153484 | 1.68253382 | -2.6242256 |
| 17892 | NM_001142286.1 | 20.8840975 | -6.0084878 | 2.29014573 | -2.6236268 |
| 17893 | NM_016179.2 | 6.22312656 | -4.2679102 | 1.62673497 | -2.6236051 |
| 17894 | NM_130808.2 | 39.0705928 | -3.5909856 | 1.3689185 | -2.6232282 |
| 17895 | NM_001330999.1 | 21.5018348 | -6.050641 | 2.30663041 | -2.6231515 |
| 17896 | NR_136649.1 | 41.0634031 | -3.6609386 | 1.39585048 | -2.6227297 |
| 17897 | NM_007037.5 | 10.3936349 | -3.9556523 | 1.50856816 | -2.6221237 |
| 17898 | NM_001347872.1 | 5.77487482 | -4.1653168 | 1.58869533 | -2.6218475 |
| 17899 | NM_001308177.1 | 29.608352 | -6.5093274 | 2.48280622 | -2.6217622 |
| 17900 | NM_001008391.3 | 5.51374849 | -4.0761554 | 1.55479001 | -2.6216758 |
| 17901 | NM_001161577.1 | 5.16265595 | -4.0006545 | 1.52604414 | -2.6215851 |
| 17902 | NM_007219.4 | 27.4781596 | -6.4015046 | 2.44192206 | -2.6215024 |
| 17903 | NR_027086.1 | 10.0604348 | 2.77848902 | 1.05990855 | 2.62144222 |
| 17904 | NM_001242877.2 | 23.2828206 | -6.1650131 | 2.35179826 | -2.6214039 |
| 17905 | NR_120450.1 | 8.93061122 | -3.7511551 | 1.43099537 | -2.6213607 |
| 17906 | NM_001307975.1 | 27.3579542 | -6.3957731 | 2.43995155 | -2.6212705 |
| 17907 | NM_001128620.1 | 7.01347729 | -4.4332887 | 1.69129921 | -2.6212327 |
| 17908 | NM_172108.4 | 75.1337197 | -3.3290882 | 1.27006565 | -2.6211938 |
| 17909 | NM_005914.3 | 4.88089231 | -3.9038802 | 1.48948215 | -2.6209648 |
| 17910 | NM_175922.3 | 7.9780829 | -4.6262349 | 1.76517322 | -2.620839 |
| 17911 | NM_001029997.3 | 21.5519905 | -6.0535004 | 2.30988067 | -2.6206983 |
| 17912 | NM_001161566.2 | 10.7988621 | 8.34329022 | 3.18363576 | 2.62067989 |
| 17913 | NM_170771.2 | 6.91717718 | 2.68710609 | 1.02533659 | 2.62070633 |
| 17914 | NM_001316893.1 | 6.06176352 | -4.206339 | 1.60548788 | -2.6199755 |
| 17915 | NM_001034024.1 | 28.6537415 | -6.4615438 | 2.46636881 | -2.6198611 |
| 17916 | NM_001143992.1 | 21.4849679 | -6.0490037 | 2.30890004 | -2.6198638 |
| 17917 | NM_032780.3 | 27.5728355 | -6.4066051 | 2.44542932 | -2.6198284 |
| 17918 | NM_001256675.1 | 6.42133564 | -4.3032473 | 1.64292547 | -2.619259 |
| 17919 | NM_001320868.1 | 5.03509962 | -3.960809 | 1.51228978 | -2.6190807 |
| 17920 | NM_001303473.1 | 4.83233786 | -3.8985366 | 1.48895103 | -2.6183108 |
| 17921 | NM_003638.2 | 20.7874556 | -6.0018277 | 2.29242062 | -2.618118 |
| 17922 | NM_138576.3 | 25.429728 | -6.291695 | 2.40326027 | -2.6179832 |
| 17923 | NM_001346766.1 | 29.203317 | -6.4895088 | 2.47891029 | -2.6178877 |
| 17924 | NM_019098.4 | 5.86375395 | 4.88760825 | 1.8670361 | 2.61784346 |
| 17925 | NM_001256496.1 | 21.1513414 | -6.0268871 | 2.302325 | -2.6177395 |
| 17926 | NM_022036.2 | 19.1255536 | -2.7497543 | 1.05048255 | -2.6176107 |
| 17927 | NM_006279.3 | 30.9814105 | -6.5763969 | 2.51252474 | -2.6174456 |
| 17928 | NM_001174102.1 | 7.05092087 | -4.4337353 | 1.6939799 | -2.6173482 |
| 17929 | NM_139278.2 | 6.03562671 | -4.2167013 | 1.61122139 | -2.6170837 |
| 17930 | NM_018355.3 | 10.2190655 | -4.9678672 | 1.89835446 | -2.6169334 |

| | | | | | |
|---|---|---|---|---|---|
| 17931 | NR_104219.1 | 21.9486233 | -2.5165173 | 0.96178971 | -2.6164943 |
| 17932 | NR_002830.1 | 4.68560143 | -3.8476477 | 1.47073641 | -2.6161368 |
| 17933 | NM_153636.2 | 68.5287616 | -3.4961203 | 1.3364607 | -2.6159545 |
| 17934 | NM_001278412.1 | 6.46904473 | -4.3276269 | 1.65449631 | -2.6156764 |
| 17935 | NM_001330166.1 | 5.88343856 | -4.1618953 | 1.59125887 | -2.6154734 |
| 17936 | NM_001145115.2 | 5.8565248 | -4.1718263 | 1.59507967 | -2.6154344 |
| 17937 | NM_001329456.1 | 8.35653882 | -4.6726543 | 1.78661328 | -2.6153698 |
| 17938 | NM_031208.3 | 109.308438 | -2.7218098 | 1.04069037 | -2.6153887 |
| 17939 | NM_001128428.2 | 27.999805 | -6.4284328 | 2.45804372 | -2.6152638 |
| 17940 | NM_001322012.1 | 21.3871674 | -6.0426542 | 2.31054012 | -2.6152561 |
| 17941 | NM_052817.2 | 24.728402 | -6.2504228 | 2.38995402 | -2.61529 |
| 17942 | NM_052941.4 | 62.3467402 | 1.96592862 | 0.75176296 | 2.61509108 |
| 17943 | NM_001303024.1 | 5.56138569 | -4.1032047 | 1.5691409 | -2.6149371 |
| 17944 | NM_001276501.1 | 4.99543506 | -3.9415737 | 1.50758006 | -2.6145037 |
| 17945 | NM_001321391.1 | 7.58472686 | -4.5419128 | 1.73733659 | -2.6142964 |
| 17946 | NM_021806.3 | 27.5619965 | -6.4057999 | 2.4503908 | -2.6141952 |
| 17947 | NM_001318891.1 | 25.9726964 | -6.3200597 | 2.41763618 | -2.6141484 |
| 17948 | NR_120591.1 | 6.65782815 | -4.3603379 | 1.66798237 | -2.6141391 |
| 17949 | NM_001001502.2 | 11.1418354 | -5.0927683 | 1.94831943 | -2.6139288 |
| 17950 | NM_005824.2 | 9.29695988 | -4.8247364 | 1.84578111 | -2.6139266 |
| 17951 | NM_001009555.3 | 20.5203338 | -5.9830502 | 2.28920426 | -2.6135939 |
| 17952 | NM_001308043.1 | 89.6593938 | 0.96501728 | 0.36927844 | 2.61325107 |
| 17953 | NM_000891.2 | 64.2815559 | 1.80854597 | 0.69208565 | 2.61318232 |
| 17954 | NM_001286216.1 | 38.8618218 | -4.957896 | 1.89726472 | -2.6131809 |
| 17955 | NM_024993.5 | 26.7259897 | -6.3618363 | 2.4346366 | -2.6130537 |
| 17956 | NM_001321026.1 | 7.6491954 | -4.5517514 | 1.74197642 | -2.6129811 |
| 17957 | NM_001178108.1 | 21.2449952 | -6.0332357 | 2.30918805 | -2.6127087 |
| 17958 | NM_001134403.1 | 25.3932981 | -6.2903899 | 2.40788836 | -2.6124093 |
| 17959 | NM_001010892.2 | 24.7955716 | -6.2557014 | 2.39466482 | -2.6123495 |
| 17960 | NM_178034.3 | 5.43960187 | 4.43508693 | 1.69790166 | 2.61209883 |
| 17961 | NM_001330727.1 | 7.26743563 | -4.4828315 | 1.71651077 | -2.6115953 |
| 17962 | NM_001286583.1 | 8.20921834 | 7.94556576 | 3.04250588 | 2.61152026 |
| 17963 | NM_032979.4 | 28.0320033 | -6.4301631 | 2.46223024 | -2.6115198 |
| 17964 | NR_003525.2 | 4.68362829 | -3.8591531 | 1.47798128 | -2.6110974 |
| 17965 | NR_103871.1 | 27.0224371 | -6.3780832 | 2.44325974 | -2.610481 |
| 17966 | NM_001305156.1 | 21.2971432 | -6.0367238 | 2.31253095 | -2.6104402 |
| 17967 | NM_001139443.1 | 6.0406041 | -4.2115565 | 1.6134365 | -2.610302 |
| 17968 | NM_001256668.1 | 26.7494007 | -6.3634793 | 2.43784514 | -2.6102886 |
| 17969 | NM_012486.2 | 30.4073584 | -6.5469982 | 2.50929726 | -2.6090963 |
| 17970 | NM_003790.2 | 21.2098614 | -6.0303701 | 2.31146062 | -2.6089002 |
| 17971 | NM_001195752.1 | 26.8016089 | -6.365849 | 2.44011254 | -2.6088342 |
| 17972 | NM_001323326.1 | 27.7534402 | -6.4156474 | 2.45924677 | -2.6087855 |
| 17973 | NM_001301224.1 | 7.83872008 | -4.5873794 | 1.75856064 | -2.6085989 |

| | | | | | |
|---|---|---|---|---|---|
| 17974 | NR_039987.1 | 11.2238586 | 3.82139437 | 1.46492329 | 2.60859691 |
| 17975 | NM_182965.2 | 5.50992432 | -4.0872744 | 1.56689722 | -2.6085147 |
| 17976 | NM_033480.2 | 146.387134 | -3.6489981 | 1.39912011 | -2.6080664 |
| 17977 | NM_138350.3 | 5.48049901 | -4.0887873 | 1.56779005 | -2.6079942 |
| 17978 | NM_004489.4 | 135.395092 | -3.8072433 | 1.45985871 | -2.6079533 |
| 17979 | NM_001166359.1 | 22.6465125 | -6.124515 | 2.34862241 | -2.6077053 |
| 17980 | NR_135823.1 | 42.3906222 | -2.1007596 | 0.80576586 | -2.6071589 |
| 17981 | NM_025217.3 | 5.78954964 | -4.1554135 | 1.59387578 | -2.6071125 |
| 17982 | NM_001105198.1 | 6.82137874 | -4.3824117 | 1.68099831 | -2.6070292 |
| 17983 | NR_122035.1 | 60.1487319 | -3.2871267 | 1.26124578 | -2.6062539 |
| 17984 | NM_001204083.1 | 4.98468085 | -3.9443965 | 1.51357671 | -2.6060103 |
| 17985 | NM_001130112.3 | 6.80368226 | -4.3746643 | 1.67889971 | -2.6056734 |
| 17986 | NM_001318763.1 | 24.6479836 | -6.2455373 | 2.3969197 | -2.6056515 |
| 17987 | NM_001042417.1 | 26.4688701 | -6.3479986 | 2.43636054 | -2.6055251 |
| 17988 | NM_000618.4 | 25.0356826 | -6.2692726 | 2.40636771 | -2.6052845 |
| 17989 | NM_022877.2 | 23.5336552 | 9.47335931 | 3.63625733 | 2.60524997 |
| 17990 | NM_001750.6 | 26.7456362 | -6.3634526 | 2.44259938 | -2.605197 |
| 17991 | NM_148887.2 | 33.8577356 | -2.1666244 | 0.83166817 | -2.6051549 |
| 17992 | NM_003567.3 | 32.2953064 | -2.8986155 | 1.11269619 | -2.6050376 |
| 17993 | NM_001242830.1 | 7.66455685 | -4.5658734 | 1.7528249 | -2.6048657 |
| 17994 | NM_002216.2 | 4.97445296 | -3.9403509 | 1.51283029 | -2.6046219 |
| 17995 | NM_001282401.1 | 6.1027834 | -4.2222717 | 1.62116818 | -2.6044624 |
| 17996 | NM_015962.4 | 126.617743 | -0.8730697 | 0.33522896 | -2.6043981 |
| 17997 | NR_120375.1 | 7.78677185 | 4.03921771 | 1.55127064 | 2.60381239 |
| 17998 | NM_206840.2 | 27.0031823 | -6.3774367 | 2.4494147 | -2.6036574 |
| 17999 | NM_001037339.2 | 9.26371306 | -4.8250795 | 1.85327493 | -2.6035422 |
| 18000 | NM_001244585.1 | 6.62226471 | -4.3304793 | 1.66334672 | -2.6034736 |
| 18001 | NR_073132.1 | 5.96463309 | -4.2029958 | 1.61437878 | -2.6034756 |
| 18002 | NM_001195125.1 | 5.5875225 | -4.0886232 | 1.5707632 | -2.6029533 |
| 18003 | NM_001297597.1 | 52.6328917 | -3.0705577 | 1.17963757 | -2.602967 |
| 18004 | NM_001256964.1 | 6.32717258 | -4.2781247 | 1.64385957 | -2.6024879 |
| 18005 | NM_001242786.1 | 7.25952898 | -4.4693557 | 1.71738019 | -2.6024265 |
| 18006 | NM_001289159.1 | 9.26904745 | 6.6706298 | 2.56337535 | 2.60228367 |
| 18007 | NM_001311179.1 | 6.00060914 | -4.205443 | 1.61633429 | -2.6018398 |
| 18008 | NR_030684.1 | 7.35967215 | -4.5096957 | 1.73352644 | -2.6014577 |
| 18009 | NM_016476.11 | 8.27152514 | -4.6647342 | 1.79321177 | -2.6013292 |
| 18010 | NM_001301410.1 | 4.70869404 | -3.8505551 | 1.48025777 | -2.6012733 |
| 18011 | NM_001142555.2 | 27.1588359 | -6.3843574 | 2.45443373 | -2.6011529 |
| 18012 | NM_001321618.1 | 6.934101 | 7.72470273 | 2.96973805 | 2.60113943 |
| 18013 | NM_024744.15 | 5.07905524 | -3.9748655 | 1.52817271 | -2.6010578 |
| 18014 | NM_001320134.1 | 26.3874369 | -6.3435224 | 2.43892248 | -2.6009529 |
| 18015 | NM_001282490.1 | 5.73382287 | -4.14126 | 1.5922598 | -2.6008695 |
| 18016 | NM_198700.2 | 11.1804346 | -5.1013842 | 1.96151081 | -2.6007423 |

| | | | | | |
|---|---|---|---|---|---|
| 18017 | NR_040001.2 | 6.03285325 | -4.2188859 | 1.62235946 | -2.6004631 |
| 18018 | NM_152868.2 | 82.9010204 | -3.5794123 | 1.37646368 | -2.6004408 |
| 18019 | NM_003867.3 | 9.11397238 | -4.8007119 | 1.84616713 | -2.6003669 |
| 18020 | NR_003187.3 | 11.2927797 | 5.06090913 | 1.94638391 | 2.60015976 |
| 18021 | NM_002252.4 | 7.85877603 | -3.5814155 | 1.37764694 | -2.5996613 |
| 18022 | NR_047470.1 | 4.67199074 | -3.8478983 | 1.48022157 | -2.5995421 |
| 18023 | NM_001321922.1 | 20.3568539 | -5.971348 | 2.29746825 | -2.5990993 |
| 18024 | NM_001301135.1 | 5.50979086 | -4.0748029 | 1.56782901 | -2.5990098 |
| 18025 | NM_001039584.1 | 20.8549217 | -6.0065918 | 2.3112138 | -2.5988906 |
| 18026 | NM_014959.3 | 22.343459 | -6.105455 | 2.34944219 | -2.5986828 |
| 18027 | NM_001292024.1 | 18.888516 | 2.88960797 | 1.11210887 | 2.59831392 |
| 18028 | NM_001302622.1 | 34.8939167 | -5.7716824 | 2.22152031 | -2.5980777 |
| 18029 | NM_144650.2 | 9.37008617 | -3.794792 | 1.46061607 | -2.5980763 |
| 18030 | NM_000789.3 | 28.5585499 | 3.49279044 | 1.34442105 | 2.59798851 |
| 18031 | NM_174911.4 | 282.300675 | -3.2468943 | 1.24977775 | -2.5979774 |
| 18032 | NM_001323615.1 | 20.4546389 | -5.9785026 | 2.30124415 | -2.5979436 |
| 18033 | NR_038230.1 | 11.375933 | 4.82950379 | 1.85900628 | 2.59789536 |
| 18034 | NM_003356.3 | 5.51763534 | -4.0764719 | 1.56922711 | -2.5977577 |
| 18035 | NR_073084.1 | 5.88532558 | -4.1911227 | 1.6134906 | -2.5975501 |
| 18036 | NM_175883.2 | 81.8228278 | 1.97191848 | 0.75924185 | 2.59722047 |
| 18037 | NM_001317911.1 | 9.14020637 | 8.1188946 | 3.12646893 | 2.59682561 |
| 18038 | NM_001198844.1 | 76.9768716 | -3.4715981 | 1.33690393 | -2.5967446 |
| 18039 | NM_001282279.1 | 29.0836278 | -6.4847711 | 2.49741802 | -2.5965902 |
| 18040 | NM_032995.2 | 49.9561431 | -3.1040465 | 1.19546281 | -2.5965229 |
| 18041 | NM_080865.3 | 4.833642 | -3.8781051 | 1.49366339 | -2.5963715 |
| 18042 | NM_001190919.1 | 33.3254202 | -5.7014031 | 2.1963681 | -2.5958322 |
| 18043 | NR_135668.1 | 4.47513713 | -3.7832721 | 1.45743032 | -2.5958511 |
| 18044 | NR_134473.1 | 5.95800883 | -4.1953433 | 1.6163216 | -2.5956117 |
| 18045 | NM_017541.3 | 5.39381493 | -4.0554197 | 1.56242917 | -2.5955863 |
| 18046 | NM_001207073.1 | 5.32923542 | -4.0440784 | 1.55834136 | -2.5951172 |
| 18047 | NR_110734.1 | 15.470766 | 5.81192038 | 2.23979966 | 2.59483939 |
| 18048 | NM_022487.3 | 20.9083202 | -6.0095901 | 2.31617541 | -2.5946179 |
| 18049 | NM_001007024.1 | 25.4611126 | -6.293284 | 2.42556277 | -2.5945666 |
| 18050 | NM_005282.2 | 14.1729495 | 2.75935815 | 1.06362961 | 2.59428481 |
| 18051 | NM_001294306.1 | 12.0791891 | -3.3170427 | 1.27864014 | -2.5941957 |
| 18052 | NM_173508.3 | 6.65657547 | -4.3553704 | 1.67893707 | -2.5941237 |
| 18053 | NM_001330426.1 | 69.9877678 | -3.4088168 | 1.31418334 | -2.593867 |
| 18054 | NM_052838.4 | 4.77603271 | -3.8822314 | 1.49676589 | -2.5937466 |
| 18055 | NR_038951.1 | 6.21983504 | -4.2550526 | 1.64057158 | -2.5936404 |
| 18056 | NM_023932.3 | 12.0580187 | -5.2100198 | 2.0088262 | -2.5935642 |
| 18057 | NM_001286279.1 | 4.9639513 | -3.9498207 | 1.52308108 | -2.5933096 |
| 18058 | NM_006458.3 | 6.18306613 | -4.262214 | 1.64384356 | -2.5928343 |
| 18059 | NM_014907.2 | 4.4793799 | -3.7945453 | 1.46348653 | -2.5928119 |

| | | | | | |
|---|---|---|---|---|---|
| 18060 | NM_024966.2 | 5.07962476 | -3.9756748 | 1.53335656 | -2.5927921 |
| 18061 | NM_144994.7 | 19.515602 | -2.780912 | 1.07265791 | -2.5925433 |
| 18062 | NR_144555.1 | 77.962495 | 2.47263656 | 0.95388014 | 2.59218791 |
| 18063 | NM_177542.3 | 5.95491131 | -4.1863385 | 1.61547256 | -2.5914018 |
| 18064 | NM_194328.2 | 29.1548104 | -6.4885244 | 2.50408673 | -2.591174 |
| 18065 | NM_001318955.1 | 5.11653017 | -3.9779764 | 1.53527631 | -2.5910492 |
| 18066 | NR_040052.1 | 21.0650558 | 3.13937636 | 1.21165688 | 2.59097804 |
| 18067 | NM_006745.4 | 463.198534 | -4.1455078 | 1.60014466 | -2.5907082 |
| 18068 | NM_001127380.2 | 15.9576629 | 3.40533947 | 1.31448396 | 2.59062839 |
| 18069 | NM_001301129.1 | 15.1075581 | 2.76601241 | 1.06779816 | 2.59038881 |
| 18070 | NM_198490.2 | 21.4054876 | -6.0434 | 2.33307573 | -2.5903145 |
| 18071 | NR_073546.1 | 27.5107135 | -6.4038516 | 2.47225244 | -2.5902903 |
| 18072 | NM_001302645.1 | 47.3803562 | 4.36546171 | 1.68541518 | 2.59014025 |
| 18073 | NM_207386.3 | 33.9510897 | -6.7075218 | 2.58975842 | -2.5900183 |
| 18074 | NM_001037171.1 | 6.59966385 | -4.3549287 | 1.68159177 | -2.5897657 |
| 18075 | NM_001049.2 | 17.1811478 | -3.3634222 | 1.29876522 | -2.5897076 |
| 18076 | NR_034072.1 | 77.1530561 | -1.5680073 | 0.60555727 | -2.5893625 |
| 18077 | NR_046706.1 | 5.24687728 | -4.0012166 | 1.54532813 | -2.5892343 |
| 18078 | NM_003793.3 | 66.5372724 | -3.2137093 | 1.24160938 | -2.5883417 |
| 18079 | NR_047666.1 | 21.3570955 | -6.0402066 | 2.3337058 | -2.5882468 |
| 18080 | NM_001160223.1 | 6.37928525 | -4.3019026 | 1.6622164 | -2.588052 |
| 18081 | NM_001195798.1 | 21.3911226 | -6.0426163 | 2.33481926 | -2.5880446 |
| 18082 | NM_031205.3 | 7.38901826 | -4.4945487 | 1.7368625 | -2.5877401 |
| 18083 | NM_001322799.1 | 15.288687 | 4.12766378 | 1.59537058 | 2.58727586 |
| 18084 | NM_001242354.1 | 19.840995 | -5.9346308 | 2.29382173 | -2.5872241 |
| 18085 | NM_001242883.1 | 26.4656499 | -6.3478214 | 2.45378711 | -2.5869487 |
| 18086 | NM_001137664.1 | 48.6220927 | -3.2255266 | 1.2469955 | -2.5866385 |
| 18087 | NM_199138.2 | 89.6357735 | -2.4942938 | 0.96433964 | -2.5865304 |
| 18088 | NR_134561.1 | 4.57891733 | -3.8124983 | 1.47409177 | -2.5863371 |
| 18089 | NM_001014990.2 | 8.51604161 | -4.710724 | 1.82155783 | -2.5860963 |
| 18090 | NM_001271762.1 | 7.28444616 | -4.4904837 | 1.7365961 | -2.5857963 |
| 18091 | NM_001330341.1 | 54.2142368 | -5.6925927 | 2.20216249 | -2.5850012 |
| 18092 | NM_001320970.1 | 34.0625084 | -6.7106878 | 2.59612453 | -2.5848867 |
| 18093 | NM_001323296.1 | 2.42240855 | 6.16855236 | 2.3873164 | 2.58388555 |
| 18094 | NM_001282534.1 | 6.69699197 | -4.3548471 | 1.68565565 | -2.5834737 |
| 18095 | NR_001575.1 | 35.1599252 | 1.48123997 | 0.57351598 | 2.58273531 |
| 18096 | NM_001190844.1 | 10.6887624 | -3.1676677 | 1.2265193 | -2.5826481 |
| 18097 | NM_001320453.1 | 11.3329235 | -5.1153003 | 1.98069233 | -2.582582 |
| 18098 | NM_015687.4 | 6.07949496 | -4.2392472 | 1.64151127 | -2.582527 |
| 18099 | NM_000337.5 | 21.0205868 | -6.0177338 | 2.33051454 | -2.5821481 |
| 18100 | NM_015348.1 | 232.798109 | -2.9379527 | 1.13793561 | -2.5818269 |
| 18101 | NM_005447.3 | 4.3445601 | -3.7373648 | 1.44776818 | -2.5814663 |
| 18102 | NM_032796.3 | 232.602971 | -3.9410644 | 1.52683709 | -2.5811951 |

| | | | | | |
|---|---|---|---|---|---|
| 18103 | NM_001258275.2 | 6.0776973 | -4.2263742 | 1.63740791 | -2.5811371 |
| 18104 | NM_001288757.1 | 8.63526039 | -4.7325133 | 1.83349107 | -2.5811488 |
| 18105 | NM_005157.5 | 271.865136 | -3.4781561 | 1.34767468 | -2.5808574 |
| 18106 | NM_001270661.1 | 102.36387 | 5.54966832 | 2.15046178 | 2.5806868 |
| 18107 | NM_007139.3 | 10.6305698 | -5.0220018 | 1.94607995 | -2.5805732 |
| 18108 | NR_136658.1 | 23.1782116 | -6.158649 | 2.38718999 | -2.5798738 |
| 18109 | NM_001039350.2 | 7.91192973 | -4.6051583 | 1.78505661 | -2.5798388 |
| 18110 | NM_001197114.1 | 25.288665 | -6.2821399 | 2.43537524 | -2.5795367 |
| 18111 | NM_001306210.1 | 28.2773625 | -6.4432867 | 2.49810137 | -2.5792735 |
| 18112 | NM_001347702.1 | 25.2967031 | -6.2826887 | 2.43595219 | -2.5791511 |
| 18113 | NM_033482.3 | 6.97209344 | 6.26849331 | 2.43054026 | 2.57905347 |
| 18114 | NM_181808.3 | 4.92864026 | -3.9162122 | 1.5185352 | -2.5789407 |
| 18115 | NM_001321006.1 | 57.4099998 | 1.43268446 | 0.55563705 | 2.57845382 |
| 18116 | NM_021958.3 | 6.2375942 | -4.2706724 | 1.65663998 | -2.5779122 |
| 18117 | NR_144447.1 | 6.74795468 | -4.3854326 | 1.70159235 | -2.5772522 |
| 18118 | NR_144550.1 | 7.75301533 | -3.5525456 | 1.3786538 | -2.5768221 |
| 18119 | NR_040013.1 | 11.3827361 | -3.2758786 | 1.27142285 | -2.5765453 |
| 18120 | NM_001329556.1 | 9.35531635 | -4.8470893 | 1.88152672 | -2.576147 |
| 18121 | NM_025236.3 | 5.40302079 | -4.0699032 | 1.57988828 | -2.5760703 |
| 18122 | NM_001081003.2 | 5.14427292 | -3.9694216 | 1.54091161 | -2.5760216 |
| 18123 | NM_001329787.1 | 92.5672974 | 5.75801036 | 2.23522047 | 2.57603688 |
| 18124 | NM_001039937.1 | 2.37410654 | 6.14106837 | 2.38404856 | 2.57589903 |
| 18125 | NM_001320709.1 | 25.0999226 | -6.2725957 | 2.43563138 | -2.5753469 |
| 18126 | NM_001039651.1 | 4.7752641 | -3.8915743 | 1.51144489 | -2.5747378 |
| 18127 | NM_176818.2 | 185.238986 | -2.7115926 | 1.05321371 | -2.5745891 |
| 18128 | NM_001278712.1 | 5.13704029 | -3.9879373 | 1.54904077 | -2.574456 |
| 18129 | NM_001308421.1 | 5.52086139 | -4.0961091 | 1.59144629 | -2.573828 |
| 18130 | NM_001136115.2 | 8.80074662 | 8.06444376 | 3.13387616 | 2.57331284 |
| 18131 | NM_031468.3 | 8.80074662 | 8.06444376 | 3.13387616 | 2.57331284 |
| 18132 | NM_001199850.1 | 19.8994322 | -5.9385377 | 2.30780751 | -2.5732379 |
| 18133 | NM_001242486.1 | 41.5470606 | 3.29473895 | 1.28050702 | 2.57299562 |
| 18134 | NM_000256.3 | 6.59372084 | 5.73525677 | 2.22910192 | 2.57290019 |
| 18135 | NM_030953.3 | 5.39716927 | -4.0668805 | 1.58064891 | -2.5729183 |
| 18136 | NM_001197297.1 | 71.5164443 | -3.222924 | 1.25272382 | -2.5727331 |
| 18137 | NM_005228.4 | 21.9331418 | -6.079102 | 2.36342842 | -2.572154 |
| 18138 | NM_001083606.1 | 21.8105782 | -6.071163 | 2.36046973 | -2.5720148 |
| 18139 | NM_004031.2 | 4.63070594 | -3.8376097 | 1.492153 | -2.5718607 |
| 18140 | NM_003898.3 | 20.0925924 | -5.9528231 | 2.31538409 | -2.5709873 |
| 18141 | NM_003059.2 | 5.2357631 | -4.0219043 | 1.56497145 | -2.5699538 |
| 18142 | NM_001330583.1 | 24.9519147 | -6.2627306 | 2.43706794 | -2.5697809 |
| 18143 | NM_000843.3 | 50.5542568 | 2.22363128 | 0.86534364 | 2.56965114 |
| 18144 | NM_138295.3 | 15.6917915 | 3.52800573 | 1.37309663 | 2.56937906 |
| 18145 | NM_144701.2 | 26.9926672 | -1.834898 | 0.71418168 | -2.5692314 |

| | | | | | |
|---|---|---|---|---|---|
| 18146 | NM_001621.4 | 79.9092972 | -2.108538 | 0.82069867 | -2.5691987 |
| 18147 | NM_001324214.1 | 8.88438018 | -4.7606869 | 1.85322985 | -2.5688594 |
| 18148 | NR_110258.2 | 18.5028095 | 7.29718385 | 2.84116358 | 2.56837864 |
| 18149 | NM_001286717.1 | 2.39931339 | 6.19469282 | 2.41199641 | 2.56828442 |
| 18150 | NM_001286707.1 | 62.981404 | 4.77162567 | 1.85812469 | 2.56797927 |
| 18151 | NM_001330673.1 | 5.57594749 | -4.1086605 | 1.60010961 | -2.5677369 |
| 18152 | NR_104359.1 | 35.2995584 | 2.30825101 | 0.89894449 | 2.56773475 |
| 18153 | NM_001284277.1 | 30.1564134 | -6.537414 | 2.54602642 | -2.5676929 |
| 18154 | NM_021979.3 | 6.57307608 | -4.3470836 | 1.69300976 | -2.567666 |
| 18155 | NM_001300734.1 | 8.84366006 | -4.7689563 | 1.85735186 | -2.5676106 |
| 18156 | NM_018292.4 | 76.8774902 | -1.7420303 | 0.67849566 | -2.5674892 |
| 18157 | NR_027456.1 | 56.1054986 | -3.4591715 | 1.34762692 | -2.5668614 |
| 18158 | NM_014310.3 | 8.34537469 | -3.657115 | 1.42484684 | -2.5666723 |
| 18159 | NM_019599.2 | 8.67402721 | 5.82301297 | 2.26871532 | 2.56665652 |
| 18160 | NM_001006630.1 | 21.8714189 | -6.0745882 | 2.36710319 | -2.5662541 |
| 18161 | NM_001267595.1 | 26.1436297 | -6.3298492 | 2.46708235 | -2.5657227 |
| 18162 | NM_001198900.1 | 26.6698741 | -6.3599382 | 2.47883606 | -2.5656954 |
| 18163 | NM_001282063.1 | 8.79339459 | -4.763924 | 1.85694119 | -2.5654684 |
| 18164 | NM_001195304.1 | 4.88473595 | -3.8957963 | 1.51883501 | -2.5649898 |
| 18165 | NM_001261443.1 | 23.7754246 | -6.1943235 | 2.41505577 | -2.564878 |
| 18166 | NM_001135179.1 | 195.724461 | -1.4916613 | 0.58161989 | -2.564667 |
| 18167 | NR_102368.1 | 4.59929613 | -3.8312936 | 1.49399462 | -2.5644628 |
| 18168 | NM_001308189.1 | 23.980601 | -6.2079588 | 2.42204098 | -2.5631105 |
| 18169 | NM_014920.3 | 89.654617 | -3.3909776 | 1.3230024 | -2.5630925 |
| 18170 | NM_001199729.2 | 13.9448267 | -5.4186346 | 2.11438578 | -2.5627464 |
| 18171 | NM_001316345.1 | 6.43173164 | -4.3014124 | 1.67860291 | -2.5624955 |
| 18172 | NM_001169118.1 | 30.5573601 | -6.556407 | 2.55869566 | -2.5624021 |
| 18173 | NM_001278210.1 | 23.205572 | -6.1577252 | 2.40376484 | -2.5617004 |
| 18174 | NM_001326411.1 | 23.2316816 | -6.1595994 | 2.40447485 | -2.5617233 |
| 18175 | NM_203415.1 | 24.3505975 | -6.2283123 | 2.43130454 | -2.5617162 |
| 18176 | NM_005293.2 | 6.36010744 | -4.2956408 | 1.67694182 | -2.561592 |
| 18177 | NM_148907.2 | 65.2877344 | -3.2983896 | 1.28764741 | -2.5615627 |
| 18178 | NM_001194937.1 | 22.4423276 | -6.1124963 | 2.3865269 | -2.5612518 |
| 18179 | NM_015514.1 | 4.87769709 | -3.9031586 | 1.52393654 | -2.5612343 |
| 18180 | NM_001199919.1 | 18.9419364 | -5.8677375 | 2.29100834 | -2.561203 |
| 18181 | NM_001289119.1 | 27.6094284 | -6.4083745 | 2.50248653 | -2.5608028 |
| 18182 | NM_001278293.2 | 20.2954994 | -5.9669275 | 2.33033643 | -2.5605434 |
| 18183 | NM_003666.3 | 26.8014629 | -2.1524954 | 0.84077884 | -2.5601208 |
| 18184 | NM_198489.2 | 6.64789085 | -4.3522189 | 1.70047581 | -2.5594124 |
| 18185 | NM_001336.3 | 28.5157127 | -3.6021665 | 1.40749458 | -2.5592756 |
| 18186 | NM_006306.3 | 328.424284 | -3.9476017 | 1.54259794 | -2.5590607 |
| 18187 | NR_138133.1 | 7.32983393 | -4.5050607 | 1.76057143 | -2.5588628 |
| 18188 | NM_031935.2 | 57.4123615 | -2.4488687 | 0.95708737 | -2.5586679 |

| | | | | | |
|---|---|---|---|---|---|
| 18189 | NM_001346201.1 | 132.905741 | -2.1084012 | 0.8241161 | -2.5583788 |
| 18190 | NM_001145412.3 | 19.278099 | -5.8931599 | 2.30350098 | -2.5583492 |
| 18191 | NM_001297778.1 | 19.6315644 | -5.9188141 | 2.31364844 | -2.5582167 |
| 18192 | NM_001145462.1 | 4.99134927 | -3.9272449 | 1.53524954 | -2.5580499 |
| 18193 | NM_001195131.1 | 60.6972355 | 1.13561924 | 0.44393874 | 2.55805391 |
| 18194 | NM_001195259.1 | 4.73273775 | -3.862275 | 1.5098335 | -2.5580801 |
| 18195 | NM_199512.2 | 23.1570408 | -6.1572829 | 2.40749377 | -2.5575488 |
| 18196 | NM_017436.5 | 6.31637222 | -4.2927212 | 1.67849387 | -2.557484 |
| 18197 | NM_001286586.1 | 19.4097638 | -5.9030707 | 2.30851616 | -2.5570844 |
| 18198 | NM_001321315.1 | 7.55550569 | 7.82527171 | 3.06046114 | 2.55689301 |
| 18199 | NM_053044.4 | 7.07884392 | -4.4547706 | 1.74226171 | -2.5568894 |
| 18200 | NM_001130721.1 | 47.7761179 | -4.8907613 | 1.91293901 | -2.556674 |
| 18201 | NM_182911.3 | 6.97867428 | -4.4328202 | 1.73391458 | -2.556539 |
| 18202 | NM_012163.2 | 4.91777736 | -3.9135093 | 1.53080736 | -2.5565002 |
| 18203 | NM_001256010.2 | 22.4310748 | -6.1105055 | 2.39038506 | -2.556285 |
| 18204 | NM_001083961.1 | 19.7572831 | -5.928046 | 2.31957065 | -2.5556652 |
| 18205 | NM_001184776.1 | 25.6150504 | -6.3007338 | 2.46547448 | -2.5555867 |
| 18206 | NM_001185056.1 | 21.7166864 | 5.26200752 | 2.05902687 | 2.55557981 |
| 18207 | NM_001143676.1 | 8.02874527 | -4.6267985 | 1.81049083 | -2.5555493 |
| 18208 | NM_005587.3 | 23.3914296 | -6.1710466 | 2.41490424 | -2.5554001 |
| 18209 | NM_001166695.2 | 26.6442451 | -6.3569544 | 2.48810485 | -2.5549383 |
| 18210 | NM_014634.3 | 144.260167 | -2.3916986 | 0.93612527 | -2.5548916 |
| 18211 | NM_014293.3 | 394.477449 | -1.6488352 | 0.645389 | -2.5547928 |
| 18212 | NM_001024212.1 | 5.62081734 | -4.0987905 | 1.60438561 | -2.5547415 |
| 18213 | NM_001345998.1 | 7.56631286 | -4.5312344 | 1.77372607 | -2.5546416 |
| 18214 | NM_001136052.2 | 19.2448327 | -5.8903595 | 2.30595266 | -2.5544148 |
| 18215 | NM_001012716.2 | 4.70604888 | -3.842415 | 1.5044728 | -2.5539943 |
| 18216 | NR_104183.1 | 4.48574659 | -3.7958536 | 1.48624107 | -2.5539959 |
| 18217 | NM_139320.1 | 4.88885043 | -3.9166849 | 1.53361889 | -2.5538841 |
| 18218 | NM_001321051.1 | 7.16500905 | -4.4711666 | 1.75133132 | -2.5530101 |
| 18219 | NM_001287336.1 | 19.6642097 | -5.9211795 | 2.31933223 | -2.5529674 |
| 18220 | NM_001271933.1 | 21.4547306 | -2.5544567 | 1.00064549 | -2.5528089 |
| 18221 | NR_046462.1 | 49.0642886 | -3.24236 | 1.27032707 | -2.552382 |
| 18222 | NM_001242450.1 | 20.3205966 | -5.9692181 | 2.33873331 | -2.5523295 |
| 18223 | NM_030567.4 | 7.45707408 | -4.5164045 | 1.76965612 | -2.5521368 |
| 18224 | NM_001329976.1 | 19.8566984 | -5.9360773 | 2.32624053 | -2.55179 |
| 18225 | NM_001242875.2 | 18.9564776 | -5.8685807 | 2.29986177 | -2.5517102 |
| 18226 | NR_038991.1 | 12.2874721 | -2.3159039 | 0.90760268 | -2.5516715 |
| 18227 | NR_033374.1 | 112.633728 | -1.0064199 | 0.39442611 | -2.5516055 |
| 18228 | NM_001136262.1 | 779.108079 | -4.3095401 | 1.68914609 | -2.5513128 |
| 18229 | NM_080654.2 | 6.34922486 | -4.2800996 | 1.67776286 | -2.5510755 |
| 18230 | NM_001316963.1 | 5.12342731 | -3.9893224 | 1.56383076 | -2.5509937 |
| 18231 | NM_000855.2 | 137.407259 | -2.9959347 | 1.17446282 | -2.5508979 |

| | | | | | |
|---|---|---|---|---|---|
| 18232 | NM_003901.3 | 275.060679 | -4.0201483 | 1.57608559 | -2.550717 |
| 18233 | NR_024111.1 | 24.8594807 | -6.2581387 | 2.45429806 | -2.5498691 |
| 18234 | NM_001261407.1 | 20.6586365 | -5.9927401 | 2.35029963 | -2.5497771 |
| 18235 | NM_000834.3 | 10.0823309 | -3.9320124 | 1.54216659 | -2.5496678 |
| 18236 | NM_001256672.1 | 24.1725436 | -6.2172187 | 2.43863009 | -2.5494718 |
| 18237 | NM_021569.3 | 24.8259051 | -6.2554187 | 2.45380609 | -2.5492718 |
| 18238 | NM_001289098.1 | 25.8243719 | -6.3113237 | 2.47577763 | -2.5492288 |
| 18239 | NR_024182.1 | 8.78631379 | 5.0892417 | 1.99658234 | 2.54897661 |
| 18240 | NR_026781.1 | 5.56161346 | -4.0998424 | 1.6086162 | -2.5486766 |
| 18241 | NM_022096.5 | 6.79019758 | -4.37093 | 1.71505774 | -2.5485614 |
| 18242 | NR_110139.1 | 4.7459269 | -3.8831926 | 1.52380306 | -2.548356 |
| 18243 | NM_001145343.1 | 19.3692517 | -5.9002263 | 2.31539016 | -2.5482644 |
| 18244 | NR_034021.1 | 53.798355 | -2.6197981 | 1.0281397 | -2.5480955 |
| 18245 | NM_001184.3 | 74.7837102 | -2.7872825 | 1.09391519 | -2.5479877 |
| 18246 | NM_198941.2 | 20.5473165 | -5.9848531 | 2.34908014 | -2.5477432 |
| 18247 | NM_198977.1 | 19.2485366 | -5.8907487 | 2.31274307 | -2.547083 |
| 18248 | NM_014468.3 | 45.8200018 | 1.74941792 | 0.68687786 | 2.54691268 |
| 18249 | NM_001308155.1 | 31.5546795 | -6.6007437 | 2.59196017 | -2.5466224 |
| 18250 | NM_001324330.1 | 32.4723168 | -6.6441059 | 2.60951943 | -2.5461033 |
| 18251 | NM_003063.2 | 6.60735202 | -4.3445986 | 1.70649332 | -2.5459218 |
| 18252 | NM_018327.2 | 11.6937574 | -4.1332016 | 1.62349256 | -2.5458704 |
| 18253 | NM_015595.3 | 22.2040139 | -6.0945285 | 2.39438053 | -2.5453467 |
| 18254 | NM_001318747.1 | 18.7334875 | -5.8520281 | 2.29912809 | -2.5453249 |
| 18255 | NM_001184999.1 | 20.265092 | 9.25706301 | 3.63716386 | 2.54513224 |
| 18256 | NR_045787.1 | 24.2440848 | -6.2216339 | 2.44482806 | -2.5448145 |
| 18257 | NM_001348122.1 | 19.4825549 | -5.9083476 | 2.32174809 | -2.5447841 |
| 18258 | NR_122112.1 | 42.4599196 | 3.09205819 | 1.2150925 | 2.54471012 |
| 18259 | NM_013389.2 | 4.90817032 | -3.9056607 | 1.53505547 | -2.5443124 |
| 18260 | NR_135752.1 | 24.1066102 | -6.215029 | 2.44269713 | -2.5443306 |
| 18261 | NM_001282208.1 | 9.57641342 | -4.8759069 | 1.91646698 | -2.5442165 |
| 18262 | NM_174882.2 | 17.5333948 | 3.2723987 | 1.28650303 | 2.54363856 |
| 18263 | NR_027426.2 | 46.1927134 | 2.55163447 | 1.00314755 | 2.54362827 |
| 18264 | NR_033907.2 | 41.2472781 | 3.28082018 | 1.28983919 | 2.54358855 |
| 18265 | NM_005556.3 | 5.91792274 | -4.1911977 | 1.64778681 | -2.5435315 |
| 18266 | NM_001728.3 | 5.66127713 | -4.1289129 | 1.62341774 | -2.5433459 |
| 18267 | NM_006952.3 | 18.7106976 | 3.29086354 | 1.29392276 | 2.54332302 |
| 18268 | NR_026879.1 | 6.61008832 | -4.3461191 | 1.70892851 | -2.5431837 |
| 18269 | NM_001166237.1 | 4.63400063 | -3.8474466 | 1.51296371 | -2.5429867 |
| 18270 | NM_001031836.2 | 10.7128742 | 5.66595446 | 2.22810446 | 2.54294831 |
| 18271 | NM_001317900.1 | 20.6547137 | -5.9927066 | 2.35734378 | -2.5421437 |
| 18272 | NM_001146096.1 | 6.78895501 | -4.3784293 | 1.72239387 | -2.5420604 |
| 18273 | NM_001290114.1 | 24.9189558 | -6.2606618 | 2.46352558 | -2.5413423 |
| 18274 | NM_001320666.1 | 22.0856198 | -6.0875915 | 2.3956068 | -2.541148 |

| | | | | | |
|---|---|---|---|---|---|
| 18275 | NM_001130524.1 | 4.65429109 | -3.8500994 | 1.5151504 | -2.5410675 |
| 18276 | NM_001040197.1 | 5.09340976 | -3.9575468 | 1.55748676 | -2.5409826 |
| 18277 | NM_001171795.1 | 25.985895 | -6.3220507 | 2.48825191 | -2.5407599 |
| 18278 | NM_001130005.1 | 19.6624069 | -5.9218746 | 2.33082325 | -2.5406794 |
| 18279 | NM_001324237.1 | 5.70922113 | -4.1296023 | 1.62552914 | -2.5404665 |
| 18280 | NM_198450.5 | 138.280186 | -1.1558014 | 0.45499551 | -2.540248 |
| 18281 | NM_001172219.1 | 18.9082644 | -5.8653384 | 2.30907446 | -2.5401253 |
| 18282 | NM_001319103.1 | 26.1387216 | -6.3310229 | 2.49257257 | -2.5399553 |
| 18283 | NM_020185.4 | 5.1066214 | -3.9869555 | 1.56975512 | -2.5398583 |
| 18284 | NM_001199039.1 | 6.47166932 | -4.3229608 | 1.70209997 | -2.5397808 |
| 18285 | NM_001323606.1 | 5.22183631 | -4.0142127 | 1.58064441 | -2.5396051 |
| 18286 | NR_038978.1 | 5.20290117 | -4.0131304 | 1.58025276 | -2.5395497 |
| 18287 | NM_000043.5 | 24.0697598 | -6.2114244 | 2.44593543 | -2.5394883 |
| 18288 | NM_033390.1 | 57.6695928 | -3.0980344 | 1.22010845 | -2.5391467 |
| 18289 | NM_001144826.1 | 114.622773 | -3.784195 | 1.49043553 | -2.538986 |
| 18290 | NM_001194991.1 | 23.1589613 | -6.1546511 | 2.42407896 | -2.5389648 |
| 18291 | NM_020796.4 | 204.718746 | -3.0434701 | 1.19870811 | -2.5389584 |
| 18292 | NM_021570.3 | 4.56494756 | -3.8238732 | 1.50615362 | -2.5388335 |
| 18293 | NR_024422.1 | 6.46867495 | -4.3078724 | 1.69681925 | -2.5387927 |
| 18294 | NM_001300888.1 | 7.51894598 | -4.5272847 | 1.78331209 | -2.5386945 |
| 18295 | NM_000743.4 | 86.8477923 | -3.4353573 | 1.3532773 | -2.5385464 |
| 18296 | NR_026757.1 | 155.266042 | -3.1536137 | 1.24234558 | -2.5384351 |
| 18297 | NM_001014797.2 | 26.3248846 | -6.3403938 | 2.49784641 | -2.5383441 |
| 18298 | NM_001168466.1 | 24.1805601 | -6.2168402 | 2.44928891 | -2.5382225 |
| 18299 | NM_001017398.1 | 5.96162493 | -4.2060542 | 1.65714791 | -2.5381284 |
| 18300 | NM_207401.2 | 101.638232 | 2.49736941 | 0.98393495 | 2.53814484 |
| 18301 | NM_001282547.1 | 232.906348 | -3.9417124 | 1.55319829 | -2.5378038 |
| 18302 | NM_001282280.1 | 25.6859703 | -6.3046908 | 2.48465333 | -2.5374529 |
| 18303 | NM_001039591.2 | 406.661935 | -3.976234 | 1.56703055 | -2.5374324 |
| 18304 | NM_001242547.1 | 23.7050225 | -6.1890842 | 2.43934273 | -2.5371934 |
| 18305 | NM_001291328.1 | 5.43242891 | -4.0611714 | 1.60084874 | -2.5368864 |
| 18306 | NR_027318.2 | 5.68793009 | 4.60655859 | 1.81598644 | 2.53667015 |
| 18307 | NR_132738.1 | 4.84283053 | -3.9145104 | 1.54355332 | -2.5360384 |
| 18308 | NM_138959.2 | 22.3605889 | -6.105367 | 2.40765657 | -2.5358131 |
| 18309 | NM_033631.3 | 18.6090587 | -5.8424454 | 2.30413984 | -2.5356297 |
| 18310 | NM_018917.3 | 5.74589877 | -4.136631 | 1.63144107 | -2.5355688 |
| 18311 | NM_001256394.1 | 9.00931687 | -4.7848208 | 1.88715348 | -2.5354699 |
| 18312 | NM_031419.3 | 18.27159 | -5.815928 | 2.29385505 | -2.5354383 |
| 18313 | NM_001321679.1 | 21.0958804 | -6.0221585 | 2.37522272 | -2.5354079 |
| 18314 | NM_001256718.1 | 24.4522045 | -6.2327383 | 2.4586007 | -2.5350755 |
| 18315 | NM_213597.3 | 5.00089069 | -3.9593168 | 1.56189271 | -2.534948 |
| 18316 | NM_018939.3 | 5.70743874 | -4.1218871 | 1.62609623 | -2.5348359 |
| 18317 | NR_044993.1 | 58.7490336 | 1.10945102 | 0.43771233 | 2.53465794 |

| | | | | | |
|---|---|---|---|---|---|
| 18318 | NM_001330762.1 | 38.8325008 | -3.0227622 | 1.19266401 | -2.5344625 |
| 18319 | NM_001203246.1 | 21.5823235 | -6.0549464 | 2.38906445 | -2.5344425 |
| 18320 | NR_137280.1 | 6.11843824 | -4.2335066 | 1.67047557 | -2.5343122 |
| 18321 | NM_005069.4 | 4.81485484 | -3.900089 | 1.53903804 | -2.5341083 |
| 18322 | NM_001284243.1 | 22.585424 | -6.1205228 | 2.4154177 | -2.5339397 |
| 18323 | NM_000046.3 | 140.348154 | -3.0366631 | 1.19842389 | -2.5338806 |
| 18324 | NM_001199514.1 | 6.70344774 | -4.3652194 | 1.72303129 | -2.5334534 |
| 18325 | NM_020348.2 | 24.0357882 | -6.2083934 | 2.45065768 | -2.5333581 |
| 18326 | NM_005098.3 | 4.33536863 | -3.7499512 | 1.48055393 | -2.5328029 |
| 18327 | NR_120628.1 | 20.9962695 | 1.75241357 | 0.69190398 | 2.53274099 |
| 18328 | NR_038864.1 | 32.86059 | 3.19988506 | 1.26391172 | 2.53173145 |
| 18329 | NM_001286580.1 | 26.0530344 | -6.3257309 | 2.49866285 | -2.5316464 |
| 18330 | NM_001291484.2 | 55.0889382 | 2.06474652 | 0.8155902 | 2.53159799 |
| 18331 | NM_033497.2 | 19.7209065 | -5.9256931 | 2.34090976 | -2.5313633 |
| 18332 | NM_018223.2 | 7.28150761 | -4.4959325 | 1.77612589 | -2.5313141 |
| 18333 | NM_001320296.1 | 4.69582932 | -3.8524904 | 1.5224094 | -2.5305219 |
| 18334 | NR_104487.1 | 23.5799771 | -6.1825524 | 2.44333445 | -2.530375 |
| 18335 | NM_022117.3 | 221.42974 | -3.3165426 | 1.310754 | -2.5302556 |
| 18336 | NM_001012751.2 | 27.367072 | -6.3950155 | 2.52757872 | -2.5300955 |
| 18337 | NM_001130859.2 | 25.1932859 | -6.2786072 | 2.48186363 | -2.5297954 |
| 18338 | NM_001330145.1 | 23.309558 | -6.1642286 | 2.43695581 | -2.529479 |
| 18339 | NM_021223.2 | 5.27400967 | -4.0247252 | 1.59131638 | -2.5291797 |
| 18340 | NM_172169.2 | 25.8551435 | -6.3136567 | 2.49656489 | -2.5289376 |
| 18341 | NM_001001560.2 | 23.4897261 | -6.175803 | 2.44236639 | -2.5286145 |
| 18342 | NM_001166208.1 | 18.0391493 | -5.797367 | 2.29270753 | -2.5286117 |
| 18343 | NR_125421.1 | 8.29177964 | 3.96961319 | 1.56991253 | 2.52855692 |
| 18344 | NM_001242539.2 | 7.59496456 | -4.5522961 | 1.8005115 | -2.5283349 |
| 18345 | NM_018271.4 | 7.78734786 | -4.5928662 | 1.81658214 | -2.5283009 |
| 18346 | NM_005519.1 | 5.50096076 | -4.0721142 | 1.61069351 | -2.5281745 |
| 18347 | NM_001317194.1 | 4.6195526 | -3.821907 | 1.51183938 | -2.5279848 |
| 18348 | NM_001282729.1 | 21.2249289 | -6.031014 | 2.38600486 | -2.5276621 |
| 18349 | NM_025149.5 | 18.86372 | -5.8621572 | 2.31940146 | -2.5274439 |
| 18350 | NM_001300988.1 | 25.3186841 | -6.2826788 | 2.48586604 | -2.5273602 |
| 18351 | NM_004636.3 | 25.0322785 | -6.2665213 | 2.47974951 | -2.5270784 |
| 18352 | NM_001318765.1 | 20.8572929 | -6.0057678 | 2.3768033 | -2.5268257 |
| 18353 | NR_003610.1 | 142.997923 | -2.9825207 | 1.18039124 | -2.5267222 |
| 18354 | NM_001330504.1 | 95.801497 | 3.88035073 | 1.53583902 | 2.5265348 |
| 18355 | NM_007104.4 | 1039.63113 | -4.4124622 | 1.74684186 | -2.5259655 |
| 18356 | NR_104165.1 | 4.4319788 | -3.7612727 | 1.48933604 | -2.5254695 |
| 18357 | NR_024380.1 | 67.5714051 | -1.4673542 | 0.5810304 | -2.5254344 |
| 18358 | NM_001329909.1 | 36.1096797 | -6.796969 | 2.69157338 | -2.5252772 |
| 18359 | NM_003809.2 | 5.04375858 | -3.9673578 | 1.57118413 | -2.525075 |
| 18360 | NM_001321524.1 | 19.3897488 | -5.9009357 | 2.33753696 | -2.5244246 |

| | | | | | |
|---|---|---|---|---|---|
| 18361 | NR_104118.1 | 10.8123122 | 3.82695617 | 1.51614227 | 2.52414054 |
| 18362 | NM_001271983.1 | 12.3643668 | 2.99213664 | 1.18621093 | 2.52243219 |
| 18363 | NM_178130.3 | 5.75486344 | -4.1552114 | 1.64774514 | -2.5217561 |
| 18364 | NR_024101.1 | 4.52179225 | -3.8123436 | 1.51200442 | -2.5213839 |
| 18365 | NM_018455.5 | 7.18703567 | -4.4736459 | 1.77455655 | -2.5209937 |
| 18366 | NM_198456.2 | 5.50041689 | -4.0699663 | 1.61443106 | -2.5209911 |
| 18367 | NM_001289797.1 | 4.28419468 | -3.7313735 | 1.48027903 | -2.5207231 |
| 18368 | NM_001267050.1 | 19.3259691 | -5.8965703 | 2.33975517 | -2.5201655 |
| 18369 | NM_199436.1 | 167.001757 | -2.7558247 | 1.09358514 | -2.519991 |
| 18370 | NM_000442.4 | 44.4189712 | 2.2971888 | 0.91167296 | 2.51975094 |
| 18371 | NM_001346550.1 | 23.0268122 | -6.1471444 | 2.43969379 | -2.5196377 |
| 18372 | NM_001012752.2 | 20.4526207 | -5.9787627 | 2.37373161 | -2.5187189 |
| 18373 | NM_021088.3 | 19.1006569 | -5.8801883 | 2.33467211 | -2.5186356 |
| 18374 | NM_001204183.1 | 149.756331 | 2.23507341 | 0.88746362 | 2.51849583 |
| 18375 | NM_181702.2 | 20.8542312 | -6.0060594 | 2.38490315 | -2.5183662 |
| 18376 | NM_001148.4 | 475.446808 | -3.6172056 | 1.43635107 | -2.5183297 |
| 18377 | NM_001307977.1 | 23.2851281 | -6.1627239 | 2.44755642 | -2.5179088 |
| 18378 | NM_001242834.1 | 18.4489493 | -5.8296078 | 2.31530101 | -2.5178617 |
| 18379 | NM_001272011.1 | 17.9619624 | -5.7911571 | 2.30029636 | -2.51757 |
| 18380 | NM_021173.4 | 24.7337588 | -6.2508111 | 2.48286009 | -2.5175849 |
| 18381 | NM_005155.6 | 18.6116425 | -5.8426398 | 2.32084109 | -2.5174665 |
| 18382 | NM_001010887.2 | 11.7160682 | -2.7140254 | 1.07812115 | -2.5173659 |
| 18383 | NM_001167931.1 | 6.19305489 | -4.257415 | 1.69176567 | -2.5165512 |
| 18384 | NM_001286798.1 | 5.4738909 | -4.0781171 | 1.6206846 | -2.5162929 |
| 18385 | NM_001037160.2 | 5.53664733 | -4.090421 | 1.62562603 | -2.5162128 |
| 18386 | NR_038383.1 | 15.6151864 | 3.16597769 | 1.25823836 | 2.51619867 |
| 18387 | NM_001320445.1 | 27.1812657 | -6.3846606 | 2.53747347 | -2.5161487 |
| 18388 | NM_013956.4 | 19.1612373 | -5.8840532 | 2.3385746 | -2.5160853 |
| 18389 | NR_110706.1 | 38.5386772 | 2.35947692 | 0.93781294 | 2.51593555 |
| 18390 | NM_001172766.2 | 17.833565 | -5.7809326 | 2.29802607 | -2.5156079 |
| 18391 | NR_026873.1 | 16.7424321 | -2.5244835 | 1.0035255 | -2.5156147 |
| 18392 | NM_001202559.1 | 23.3742512 | -6.1692661 | 2.45251825 | -2.5154822 |
| 18393 | NR_033489.1 | 20.9585096 | -6.0130466 | 2.39045918 | -2.5154358 |
| 18394 | NM_001256293.1 | 5.56952114 | -4.1022407 | 1.63084977 | -2.5154007 |
| 18395 | NM_001242314.1 | 25.455122 | -6.2902725 | 2.50075679 | -2.5153476 |
| 18396 | NM_017673.6 | 18.9291383 | -5.8664823 | 2.332662 | -2.5149303 |
| 18397 | NR_002814.2 | 5.57705269 | -4.0884188 | 1.62629076 | -2.5139532 |
| 18398 | NM_001135024.1 | 24.7558176 | -6.2517717 | 2.48700641 | -2.5137738 |
| 18399 | NM_024894.3 | 24.9098697 | -6.2600256 | 2.49093647 | -2.5131213 |
| 18400 | NM_006578.3 | 203.631846 | -3.0935805 | 1.23120056 | -2.5126536 |
| 18401 | NM_001290.4 | 19.8960455 | -5.9383357 | 2.3633939 | -2.5126305 |
| 18402 | NM_001290185.1 | 6.9054998 | -4.4106973 | 1.75558797 | -2.5123761 |
| 18403 | NR_073096.1 | 4.67167464 | -3.8387437 | 1.52803818 | -2.5122041 |

| | | | | | |
|---|---|---|---|---|---|
| 18404 | NR_111932.1 | 9.06803729 | -4.8000352 | 1.91069243 | -2.5121967 |
| 18405 | NM_001734.4 | 23.1209163 | -6.1531679 | 2.44950952 | -2.512 |
| 18406 | NM_001178110.1 | 17.6118283 | -5.762977 | 2.29438476 | -2.5117744 |
| 18407 | NM_001008388.4 | 144.581564 | -2.9765962 | 1.18513007 | -2.5116198 |
| 18408 | NM_001039920.2 | 18.8935119 | -5.8634444 | 2.33480417 | -2.5113217 |
| 18409 | NM_002519.2 | 39.3264075 | -2.5075973 | 0.99853657 | -2.5112724 |
| 18410 | NM_053046.3 | 7.15151485 | -4.4473474 | 1.77143976 | -2.5105835 |
| 18411 | NR_103448.1 | 4.43485172 | -3.7651212 | 1.49972185 | -2.5105464 |
| 18412 | NM_001267619.1 | 25.6004825 | -6.2984244 | 2.50881519 | -2.5105175 |
| 18413 | NM_001281458.1 | 18.7619826 | -5.8534812 | 2.33161391 | -2.5104848 |
| 18414 | NM_016270.3 | 15.3812543 | -2.7347875 | 1.08941038 | -2.5103372 |
| 18415 | NM_001077445.2 | 23.0890461 | -6.150479 | 2.45014198 | -2.5102541 |
| 18416 | NM_002985.2 | 439.639647 | 1.63943296 | 0.6531388 | 2.51008354 |
| 18417 | NR_109773.1 | 5.57493165 | -4.1114329 | 1.63808436 | -2.509903 |
| 18418 | NM_006883.2 | 59.3065021 | 1.69842544 | 0.67676494 | 2.50962386 |
| 18419 | NM_001318918.1 | 18.7498884 | -5.8524273 | 2.33218598 | -2.5094171 |
| 18420 | NM_138573.3 | 4.41849324 | -3.7684544 | 1.50179477 | -2.5093005 |
| 18421 | NM_014659.5 | 21.522085 | -6.0490281 | 2.41079626 | -2.5091412 |
| 18422 | NR_104676.1 | 4.15464271 | -3.6710901 | 1.46309395 | -2.5091281 |
| 18423 | NM_001164407.1 | 288.655766 | 1.46535964 | 0.58404013 | 2.50900505 |
| 18424 | NM_001330650.1 | 23.9855927 | 2.77588926 | 1.10645776 | 2.50880726 |
| 18425 | NM_013348.3 | 7.2246351 | -3.4401566 | 1.37135005 | -2.5085911 |
| 18426 | NM_213589.2 | 182.85738 | -2.7203363 | 1.08440457 | -2.5085991 |
| 18427 | NM_001300903.1 | 5.11649934 | -3.9556656 | 1.57690869 | -2.5084938 |
| 18428 | NM_003362.3 | 80.3411635 | -3.5269325 | 1.40616563 | -2.5081914 |
| 18429 | NM_198437.2 | 4.6811309 | -3.844441 | 1.53293943 | -2.5078884 |
| 18430 | NR_037705.1 | 18.3517446 | -5.8216033 | 2.32165313 | -2.507525 |
| 18431 | NR_136566.1 | 20.0744736 | -5.9508465 | 2.37320359 | -2.5075162 |
| 18432 | NM_001321183.1 | 7.66554592 | -4.5513719 | 1.8152704 | -2.5072694 |
| 18433 | NM_172215.2 | 7.04669906 | 7.72411526 | 3.08084467 | 2.50714206 |
| 18434 | NM_001346032.1 | 7.85587623 | 7.9027912 | 3.15254874 | 2.5067943 |
| 18435 | NM_201278.2 | 25.1898224 | -6.2754236 | 2.50355351 | -2.5066066 |
| 18436 | NM_018688.4 | 56.010137 | -3.0527437 | 1.21802661 | -2.506303 |
| 18437 | NM_001310327.1 | 212.679295 | 3.16987475 | 1.2648952 | 2.50603746 |
| 18438 | NM_003459.4 | 4.83971229 | -3.9059309 | 1.55860123 | -2.5060489 |
| 18439 | NM_001251888.1 | 5.62527977 | -4.1096796 | 1.63995013 | -2.5059784 |
| 18440 | NM_032338.3 | 77.196365 | -1.6529286 | 0.659603 | -2.5059446 |
| 18441 | NM_001305655.1 | 18.2374569 | -5.8130437 | 2.31980167 | -2.5058365 |
| 18442 | NM_001310123.1 | 77.308616 | -3.4709213 | 1.38526861 | -2.5055944 |
| 18443 | NM_001145258.1 | 111.881017 | 1.93084393 | 0.7706345 | 2.50552488 |
| 18444 | NM_001301409.1 | 4.35969143 | -3.7486567 | 1.4966671 | -2.5046697 |
| 18445 | NM_194260.2 | 4.49249804 | -3.7829335 | 1.51037072 | -2.5046391 |
| 18446 | NM_198428.2 | 25.0006544 | -6.2658355 | 2.50168651 | -2.5046446 |

| | | | | | |
|---|---|---|---|---|---|
| 18447 | NM_001012642.2 | 5.55348797 | -4.1050873 | 1.63905148 | -2.5045506 |
| 18448 | NR_134878.1 | 14.8129207 | 3.40258881 | 1.35873701 | 2.50422913 |
| 18449 | NM_022112.2 | 18.303954 | 9.10978526 | 3.63786341 | 2.50415814 |
| 18450 | NR_104233.1 | 18.303954 | 9.10978526 | 3.63786341 | 2.50415814 |
| 18451 | NM_004505.3 | 4.36100693 | -3.7519093 | 1.49849709 | -2.5037815 |
| 18452 | NM_001286565.1 | 172.148904 | -1.8202619 | 0.72701147 | -2.5037595 |
| 18453 | NM_001042351.2 | 21.4300423 | -6.0454573 | 2.41505446 | -2.5032385 |
| 18454 | NM_016623.4 | 20.0622124 | -5.9501611 | 2.37695502 | -2.5032704 |
| 18455 | NM_017785.4 | 22.7140777 | -6.1266639 | 2.44749226 | -2.5032414 |
| 18456 | NM_001042517.1 | 22.6624758 | -6.1241985 | 2.44657162 | -2.5031756 |
| 18457 | NM_020725.1 | 18.5105304 | -5.8341266 | 2.33082268 | -2.5030332 |
| 18458 | NM_001315531.1 | 5.799991 | -4.1673734 | 1.66545428 | -2.5022442 |
| 18459 | NM_001031726.3 | 23.5955468 | -6.1831803 | 2.4713397 | -2.5019548 |
| 18460 | NM_001171906.1 | 7.2613498 | -3.3859941 | 1.35343089 | -2.5017857 |
| 18461 | NM_001321788.1 | 22.9372282 | -6.1416217 | 2.45489437 | -2.5017866 |
| 18462 | NM_001323067.1 | 30.2711313 | -6.5429645 | 2.61553084 | -2.5015818 |
| 18463 | NM_015973.4 | 5.19573113 | -4.0021177 | 1.59982306 | -2.5016002 |
| 18464 | NM_001278304.1 | 70.3204363 | -2.3909806 | 0.95586608 | -2.5013761 |
| 18465 | NM_145660.2 | 121.353249 | 3.29781352 | 1.3184337 | 2.5013116 |
| 18466 | NM_001288799.1 | 10.6714855 | -5.0405856 | 2.01541352 | -2.5010181 |
| 18467 | NM_001039670.2 | 22.519034 | -6.1150596 | 2.44523472 | -2.5008068 |
| 18468 | NM_002686.4 | 4.69808457 | -3.841043 | 1.53597867 | -2.5007137 |
| 18469 | NM_001256356.1 | 10.2431423 | -3.9708258 | 1.58804029 | -2.5004566 |
| 18470 | NM_001278700.1 | 6.94806295 | -4.4217134 | 1.76838785 | -2.5004206 |
| 18471 | NR_024066.2 | 15.4255121 | 1.4676118 | 0.58695654 | 2.50037559 |
| 18472 | NM_007250.5 | 23.1219372 | -6.1519743 | 2.46048504 | -2.5003096 |
| 18473 | NM_000607.2 | 9.01657416 | 6.68304751 | 2.67293746 | 2.50026333 |
| 18474 | NR_037164.1 | 6.53855865 | -4.3399624 | 1.73646744 | -2.4993054 |
| 18475 | NM_197961.3 | 22.989307 | -6.143729 | 2.45824061 | -2.4992383 |
| 18476 | NM_003298.4 | 23.9613008 | -6.2054608 | 2.48313975 | -2.4990381 |
| 18477 | NM_001010860.1 | 11.9834438 | -3.2441726 | 1.29831718 | -2.498752 |
| 18478 | NM_001331035.1 | 23.0129948 | -6.1451665 | 2.45939432 | -2.4986504 |
| 18479 | NM_178505.7 | 5.58448965 | -4.1172882 | 1.64782585 | -2.4986186 |
| 18480 | NM_001329605.1 | 5.23795568 | -4.0026374 | 1.60198478 | -2.4985489 |
| 18481 | NM_001310319.1 | 22.3901483 | -6.1069285 | 2.44430183 | -2.4984347 |
| 18482 | NR_038261.1 | 4.55918718 | -3.8138576 | 1.52653507 | -2.4983754 |
| 18483 | NM_001171166.1 | 21.750195 | -6.0667508 | 2.42856794 | -2.4980774 |
| 18484 | NM_080662.3 | 4.89213074 | -3.8991142 | 1.56087171 | -2.4980363 |
| 18485 | NR_028093.1 | 41.2629515 | 2.72637139 | 1.09141572 | 2.49801367 |
| 18486 | NM_001242764.1 | 19.8257001 | -5.9328286 | 2.37538031 | -2.4976331 |
| 18487 | NR_125804.1 | 5.84981037 | -4.1840936 | 1.67523377 | -2.4976177 |
| 18488 | NM_001321993.1 | 21.0647131 | -6.0185205 | 2.40992588 | -2.4973882 |
| 18489 | NM_004947.4 | 183.957058 | -3.6504091 | 1.46180129 | -2.4971993 |

| | | | | | |
|---|---|---|---|---|---|
| 18490 | NR_109795.1 | 5.59698709 | -4.1021977 | 1.64274724 | -2.497157 |
| 18491 | NM_001142590.1 | 5.78026778 | -4.1660877 | 1.66850001 | -2.496906 |
| 18492 | NM_020367.5 | 22.2262485 | -6.0955585 | 2.44164075 | -2.496501 |
| 18493 | NM_001205019.1 | 18.6123141 | -5.8420985 | 2.34016887 | -2.4964431 |
| 18494 | NR_045562.1 | 18.2349138 | -5.8125483 | 2.32853781 | -2.4962224 |
| 18495 | NM_001172663.1 | 6.24386725 | -4.2648107 | 1.70852008 | -2.4962017 |
| 18496 | NM_004067.3 | 18.033385 | -5.7968295 | 2.32272997 | -2.4956967 |
| 18497 | NM_001287335.1 | 22.2744355 | -6.0985706 | 2.4436646 | -2.495666 |
| 18498 | NM_001647.3 | 6.98151042 | -4.4352819 | 1.77720225 | -2.495654 |
| 18499 | NM_032885.5 | 17.9554739 | -5.7906328 | 2.32034769 | -2.4955884 |
| 18500 | NM_199189.2 | 23.8141605 | -6.1962441 | 2.48292481 | -2.4955424 |
| 18501 | NM_153373.3 | 17.4738716 | -5.7514007 | 2.30473068 | -2.4954763 |
| 18502 | NM_183397.2 | 222.19693 | 3.35689629 | 1.34518461 | 2.49549115 |
| 18503 | NM_001039569.1 | 22.4558571 | -6.1095096 | 2.44836022 | -2.4953475 |
| 18504 | NR_037615.1 | 5.94229168 | -4.1774049 | 1.67442734 | -2.494826 |
| 18505 | NM_032328.3 | 7.28708284 | -4.4954811 | 1.80197268 | -2.4947554 |
| 18506 | NM_001195000.1 | 17.6722347 | -5.7678895 | 2.3122439 | -2.4944987 |
| 18507 | NM_017817.2 | 4.19636041 | -3.6837687 | 1.47678326 | -2.4944545 |
| 18508 | NM_001286678.1 | 5.76500505 | -4.136956 | 1.65855774 | -2.4943093 |
| 18509 | NM_153006.2 | 4.39763325 | -3.7643667 | 1.50931519 | -2.4940892 |
| 18510 | NM_001144770.1 | 28.9187748 | -6.4740377 | 2.59628053 | -2.4935817 |
| 18511 | NM_198855.3 | 24.6879625 | -6.2476166 | 2.50548806 | -2.4935727 |
| 18512 | NR_111944.1 | 4.27972414 | -3.7263898 | 1.49443726 | -2.493507 |
| 18513 | NR_027715.1 | 17.3384836 | 2.90661905 | 1.16586264 | 2.49310591 |
| 18514 | NM_001278696.1 | 6.00944807 | 7.50588131 | 3.01080649 | 2.49298032 |
| 18515 | NM_001042496.1 | 24.3355902 | -6.2269059 | 2.49782641 | -2.4929298 |
| 18516 | NM_001330643.1 | 23.565552 | -6.1814516 | 2.47971336 | -2.4928089 |
| 18517 | NM_001033559.2 | 5.5858623 | -4.1109125 | 1.64915091 | -2.4927449 |
| 18518 | NM_024577.3 | 34.8449494 | 2.16418099 | 0.86831836 | 2.49238193 |
| 18519 | NM_001330753.1 | 5.98424121 | 7.49602928 | 3.00772555 | 2.4922584 |
| 18520 | NM_001172780.1 | 5.2401832 | -4.0169524 | 1.61181429 | -2.4921931 |
| 18521 | NM_207168.1 | 7.2891331 | -4.4856172 | 1.79988522 | -2.4921685 |
| 18522 | NM_001184892.1 | 22.2658301 | -2.1911198 | 0.87921062 | -2.4921444 |
| 18523 | NM_001142769.1 | 5.624564 | 7.42444788 | 2.97921665 | 2.49208056 |
| 18524 | NR_073085.1 | 4.88533114 | -3.8963066 | 1.56382221 | -2.4915278 |
| 18525 | NR_144462.1 | 9.34049073 | 3.7950597 | 1.52322446 | 2.49146451 |
| 18526 | NM_001252607.1 | 28.5894375 | -6.4590665 | 2.59282789 | -2.4911281 |
| 18527 | NM_001271985.1 | 10.7625482 | -4.0386131 | 1.62119411 | -2.4911349 |
| 18528 | NM_001347953.1 | 22.2675182 | -6.0991412 | 2.44831312 | -2.4911606 |
| 18529 | NM_052952.2 | 9.1329411 | 3.96100802 | 1.59005638 | 2.49111168 |
| 18530 | NM_002144.3 | 6.80301246 | -4.3934976 | 1.76373891 | -2.4910136 |
| 18531 | NR_023924.2 | 46.6892957 | -3.0167477 | 1.21105885 | -2.4910001 |
| 18532 | NR_134920.1 | 10.3853419 | -2.9875611 | 1.19932619 | -2.491033 |

| | | | | | |
|---|---|---|---|---|---|
| 18533 | NM_001346350.1 | 29.0898851 | -6.4839102 | 2.60318533 | -2.4907601 |
| 18534 | NM_001287046.1 | 4.47929156 | -3.7774971 | 1.51681158 | -2.4904195 |
| 18535 | NM_001243650.1 | 394.584101 | 1.20645161 | 0.48450904 | 2.49004975 |
| 18536 | NM_001040662.1 | 9.87054834 | -3.8530135 | 1.54740574 | -2.4899827 |
| 18537 | NM_001280553.1 | 17.6502414 | 9.05714941 | 3.6381312 | 2.48950599 |
| 18538 | NM_001330120.1 | 5.16073921 | -3.9865757 | 1.60146243 | -2.4893345 |
| 18539 | NM_033124.4 | 18.293805 | -5.8175707 | 2.33702394 | -2.4893073 |
| 18540 | NM_001099225.1 | 7.59369304 | -3.5288834 | 1.41770907 | -2.489145 |
| 18541 | NM_005495.2 | 17.0424605 | 8.23116895 | 3.30688592 | 2.48909976 |
| 18542 | NM_001204137.1 | 18.4553536 | -5.8301059 | 2.34246751 | -2.4888737 |
| 18543 | NM_001006600.2 | 57.6267437 | -3.1011343 | 1.24604528 | -2.4887814 |
| 18544 | NM_001297761.1 | 5.8035164 | -4.148685 | 1.66695744 | -2.4887768 |
| 18545 | NM_001195381.1 | 4.71927785 | -3.8645773 | 1.55304505 | -2.4883871 |
| 18546 | NM_014791.3 | 9.69929142 | -4.8935177 | 1.96654186 | -2.4883873 |
| 18547 | NM_153638.3 | 26.3649949 | -6.3418323 | 2.54965877 | -2.4873259 |
| 18548 | NR_028492.1 | 4.28524539 | -3.7158072 | 1.49389938 | -2.4873209 |
| 18549 | NM_080625.3 | 4.36619778 | -3.76078 | 1.51203867 | -2.4872247 |
| 18550 | NR_033722.1 | 4.5659213 | -3.8193543 | 1.53560994 | -2.4871904 |
| 18551 | NM_000222.2 | 26.2672695 | -6.3353457 | 2.54741412 | -2.4869713 |
| 18552 | NM_001649.3 | 51.4200605 | -3.3240295 | 1.33661145 | -2.4869078 |
| 18553 | NM_001113206.1 | 4.77324789 | -3.8665981 | 1.55490242 | -2.4867143 |
| 18554 | NR_103726.1 | 64.415658 | -2.333314 | 0.93832695 | -2.4866749 |
| 18555 | NR_024337.1 | 8.89530557 | -3.7593193 | 1.51187459 | -2.4865285 |
| 18556 | NM_173549.2 | 4.73325293 | -3.8608477 | 1.55301507 | -2.4860336 |
| 18557 | NR_026802.1 | 22.1799207 | 2.30267649 | 0.9263374 | 2.48578595 |
| 18558 | NM_001294314.1 | 7.71108002 | -3.5515615 | 1.42883561 | -2.4856334 |
| 18559 | NM_001142468.2 | 6.20543303 | -4.2410901 | 1.70655266 | -2.4851797 |
| 18560 | NM_001301188.1 | 21.7656462 | -6.0658377 | 2.44105869 | -2.4849209 |
| 18561 | NM_015168.1 | 164.354636 | -3.7183776 | 1.49638302 | -2.4849103 |
| 18562 | NM_001330402.1 | 21.7667656 | -6.0655682 | 2.44102401 | -2.4848458 |
| 18563 | NM_001122752.1 | 53.8108875 | -3.3949 | 1.36631394 | -2.4847144 |
| 18564 | NM_001280790.1 | 5.7664291 | -4.1543672 | 1.67231311 | -2.4842041 |
| 18565 | NM_001024935.1 | 17.2548366 | -5.7333964 | 2.30859622 | -2.483499 |
| 18566 | NM_001304941.1 | 24.2437738 | -6.221706 | 2.5052818 | -2.4834356 |
| 18567 | NM_001099456.2 | 4.52520602 | -3.8099396 | 1.534197 | -2.4833444 |
| 18568 | NM_001256760.2 | 32.3740109 | -6.6397764 | 2.6737103 | -2.4833567 |
| 18569 | NM_001346873.1 | 16.8968631 | -5.7030025 | 2.29654437 | -2.4832973 |
| 18570 | NM_203330.2 | 16.8651035 | -5.700504 | 2.29552603 | -2.4833106 |
| 18571 | NM_018254.3 | 16.9121651 | -5.7045871 | 2.29722002 | -2.4832567 |
| 18572 | NM_015425.4 | 205.582921 | -1.6800887 | 0.67665112 | -2.4829468 |
| 18573 | NM_001258461.1 | 19.3057408 | -5.8954507 | 2.37455236 | -2.482763 |
| 18574 | NM_001193636.1 | 7.10374353 | -4.4381773 | 1.78768692 | -2.4826368 |
| 18575 | NM_001318792.1 | 20.1164198 | -5.9529683 | 2.39840253 | -2.4820556 |

| | | | | | |
|---|---|---|---|---|---|
| 18576 | NM_153266.3 | 55.0409038 | -3.2851876 | 1.32363894 | -2.4819364 |
| 18577 | NM_001282980.1 | 21.7542189 | -6.0666856 | 2.44452531 | -2.4817438 |
| 18578 | NM_001034172.2 | 16.0813444 | 5.74820628 | 2.31668927 | 2.48121591 |
| 18579 | NM_001198954.1 | 10.9103311 | -3.2367427 | 1.30453811 | -2.4811408 |
| 18580 | NR_040068.1 | 4.73095558 | -3.8740128 | 1.56145702 | -2.4810243 |
| 18581 | NM_001144899.1 | 40.9920786 | 1.78911472 | 0.7211717 | 2.48084432 |
| 18582 | NM_001012981.4 | 123.032628 | -2.6915641 | 1.08497667 | -2.4807576 |
| 18583 | NM_001318503.1 | 4.67852578 | -3.8623905 | 1.55701602 | -2.4806363 |
| 18584 | NM_004210.4 | 60.1653628 | -3.2942719 | 1.32818812 | -2.4802751 |
| 18585 | NM_004804.2 | 368.415209 | -2.8573756 | 1.15207019 | -2.4802096 |
| 18586 | NM_001145354.1 | 22.0808737 | -6.0851093 | 2.45415519 | -2.4795128 |
| 18587 | NM_001282342.1 | 17.5934042 | -5.7611145 | 2.3236516 | -2.4793366 |
| 18588 | NR_036476.1 | 17.4782716 | -5.7517221 | 2.32019282 | -2.4789845 |
| 18589 | NM_016591.2 | 5.47794196 | -4.0748758 | 1.64380496 | -2.478929 |
| 18590 | NM_001256080.1 | 7.51641648 | 7.83937537 | 3.16243183 | 2.4789073 |
| 18591 | NR_125838.1 | 27.4821259 | 2.34213731 | 0.94524165 | 2.47781856 |
| 18592 | NM_058195.3 | 5.95913056 | -4.2030246 | 1.69629731 | -2.4777641 |
| 18593 | NM_018842.4 | 16.857663 | -3.1158604 | 1.25758014 | -2.4776635 |
| 18594 | NM_001079866.1 | 21.5594358 | -6.0521428 | 2.44272074 | -2.4776237 |
| 18595 | NM_178863.4 | 107.817677 | -3.6115079 | 1.45773581 | -2.4774777 |
| 18596 | NM_001079675.2 | 18.4355232 | -5.8283305 | 2.35260843 | -2.4773908 |
| 18597 | NM_001323614.1 | 19.7312337 | -5.9261235 | 2.39228555 | -2.4771807 |
| 18598 | NM_006984.4 | 4.66381902 | -3.8424383 | 1.55114004 | -2.4771705 |
| 18599 | NR_036549.1 | 6.32350959 | -4.2795178 | 1.72790241 | -2.4767127 |
| 18600 | NM_006974.2 | 24.2562067 | -6.222484 | 2.51277141 | -2.476343 |
| 18601 | NM_152304.1 | 41.1585948 | 5.49775231 | 2.2204634 | 2.47594818 |
| 18602 | NR_121183.1 | 9.90828904 | 1.9730297 | 0.79710195 | 2.47525388 |
| 18603 | NR_126032.1 | 22.4983547 | -6.1123956 | 2.46944214 | -2.4752131 |
| 18604 | NR_033321.2 | 292.718657 | -3.0293889 | 1.22400237 | -2.4749861 |
| 18605 | NM_173674.2 | 13.8501508 | -2.7374058 | 1.10622895 | -2.4745382 |
| 18606 | NM_007293.2 | 218.069137 | -3.8154509 | 1.54206792 | -2.4742431 |
| 18607 | NM_030791.3 | 53.8723295 | -3.1159742 | 1.25934835 | -2.4742751 |
| 18608 | NM_182503.2 | 13.7417109 | -2.6970598 | 1.09004955 | -2.4742543 |
| 18609 | NM_018932.3 | 17.3590081 | -3.5555503 | 1.4370959 | -2.4741218 |
| 18610 | NM_153705.4 | 122.605578 | -2.1050123 | 0.85080771 | -2.474134 |
| 18611 | NM_001293307.2 | 9.65443898 | 6.28407269 | 2.53999339 | 2.4740508 |
| 18612 | NM_001288581.1 | 102.22803 | 2.8928559 | 1.16931195 | 2.47398129 |
| 18613 | NM_207348.2 | 85.4184872 | 1.32972079 | 0.53749558 | 2.47391951 |
| 18614 | NR_134623.1 | 6.98487247 | -3.3848629 | 1.36829478 | -2.4737819 |
| 18615 | NM_032961.2 | 120.846802 | -2.7922545 | 1.12896283 | -2.4732918 |
| 18616 | NM_001163438.1 | 4.18521844 | -3.6951223 | 1.49423038 | -2.4729268 |
| 18617 | NM_001252156.1 | 27.4022957 | -6.3965399 | 2.58682984 | -2.4727332 |
| 18618 | NM_006741.3 | 70.4020853 | -3.5392059 | 1.43135109 | -2.472633 |

| | | | | | |
|---|---|---|---|---|---|
| 18619 | NM_001258454.1 | 19.0475517 | -5.8752108 | 2.37668744 | -2.4720166 |
| 18620 | NM_181864.2 | 7.7435933 | -4.5665766 | 1.84730749 | -2.4720176 |
| 18621 | NR_046165.1 | 4.07143811 | -3.6513039 | 1.47705308 | -2.4720194 |
| 18622 | NM_001330324.1 | 19.596898 | -5.9169842 | 2.39361193 | -2.4719898 |
| 18623 | NM_001174150.1 | 4.91487878 | -3.9138896 | 1.58343751 | -2.4717676 |
| 18624 | NM_001320365.1 | 18.3687027 | -5.8230432 | 2.35587338 | -2.4717131 |
| 18625 | NR_134583.1 | 17.0793629 | -5.7181597 | 2.31347857 | -2.4716718 |
| 18626 | NR_138081.1 | 5.29244681 | -4.0340768 | 1.63219364 | -2.4715676 |
| 18627 | NM_001267035.1 | 34.6734803 | -3.0233857 | 1.22344287 | -2.4712112 |
| 18628 | NM_031496.1 | 9.4767285 | -4.8570471 | 1.96567298 | -2.4709334 |
| 18629 | NM_001321626.1 | 25.4911648 | -6.2926853 | 2.547049 | -2.4705788 |
| 18630 | NM_152476.2 | 9.66915959 | -4.890585 | 1.97960135 | -2.4704898 |
| 18631 | NR_046478.1 | 8.31515619 | -4.6745917 | 1.89218809 | -2.4704688 |
| 18632 | NM_032860.4 | 59.4682328 | -3.1015248 | 1.25551087 | -2.4703289 |
| 18633 | NM_198570.4 | 5.03307213 | -3.963819 | 1.6047552 | -2.4700459 |
| 18634 | NM_001346354.1 | 23.8820566 | -6.2012418 | 2.51087262 | -2.4697556 |
| 18635 | NM_004767.3 | 21.5201335 | 2.73626654 | 1.10798779 | 2.46958186 |
| 18636 | NM_022477.3 | 22.3135581 | -6.1019518 | 2.47085452 | -2.4695715 |
| 18637 | NM_001330600.1 | 7.04780434 | -4.4446599 | 1.80011285 | -2.4691007 |
| 18638 | NM_018914.2 | 21.3281401 | -6.0360986 | 2.4447765 | -2.4689777 |
| 18639 | NM_001281717.1 | 5.4367373 | -4.0499489 | 1.6404713 | -2.4687716 |
| 18640 | NR_037654.1 | 4.50681172 | -3.8049632 | 1.54123965 | -2.468768 |
| 18641 | NM_133338.2 | 7.1958039 | -4.4560539 | 1.80503539 | -2.4686795 |
| 18642 | NM_001145026.1 | 11.3323408 | 4.6721201 | 1.89317662 | 2.46787333 |
| 18643 | NM_024579.3 | 6.36236255 | -4.3005708 | 1.74263067 | -2.4678613 |
| 18644 | NM_001346749.1 | 7.44290763 | 7.82333171 | 3.17023693 | 2.46774354 |
| 18645 | NM_001289790.2 | 139.250881 | 5.68302503 | 2.30308571 | 2.46756993 |
| 18646 | NR_003010.1 | 4.01931607 | -3.6327353 | 1.47256156 | -2.4669497 |
| 18647 | NR_027404.1 | 5.10383673 | -3.9552222 | 1.60327591 | -2.4669629 |
| 18648 | NM_006803.3 | 112.926944 | -3.0352776 | 1.23055973 | -2.4665829 |
| 18649 | NM_001136134.1 | 5.16900826 | -3.9804279 | 1.6137691 | -2.4665412 |
| 18650 | NR_135689.1 | 5.96954485 | -4.2099062 | 1.70729218 | -2.4658381 |
| 18651 | NR_038431.1 | 6.66989572 | -4.3543282 | 1.7660605 | -2.46556 |
| 18652 | NM_001006641.3 | 24.9861622 | -6.264535 | 2.5408444 | -2.4655327 |
| 18653 | NM_012469.3 | 482.202325 | -4.0120193 | 1.62735103 | -2.4653681 |
| 18654 | NR_135148.1 | 4.87254023 | -3.9059572 | 1.58441224 | -2.4652405 |
| 18655 | NR_047655.1 | 22.3362659 | -6.1041267 | 2.47631035 | -2.4650088 |
| 18656 | NR_104268.1 | 4.67600272 | -3.8524199 | 1.56294701 | -2.4648435 |
| 18657 | NM_001164749.1 | 25.6368357 | -6.3003926 | 2.55642786 | -2.4645298 |
| 18658 | NM_001033580.2 | 5.42113298 | -4.0728229 | 1.6526754 | -2.4643816 |
| 18659 | NM_001242778.1 | 22.5098534 | -6.1136129 | 2.48126057 | -2.4639141 |
| 18660 | NM_018443.3 | 24.075795 | -6.2100001 | 2.52051364 | -2.4637836 |
| 18661 | NM_001276394.1 | 25.6665391 | 3.04882733 | 1.23749087 | 2.46371703 |

| | | | | | |
|---|---|---|---|---|---|
| 18662 | NM_001193532.1 | 6.28985449 | -4.2803242 | 1.7375368 | -2.4634437 |
| 18663 | NM_012144.3 | 23.9479866 | -6.2054144 | 2.51905086 | -2.4633939 |
| 18664 | NM_001301029.1 | 23.0917279 | -6.15059 | 2.49688179 | -2.4633084 |
| 18665 | NR_033412.1 | 4.36986497 | -3.7497674 | 1.52227505 | -2.4632653 |
| 18666 | NM_001195291.2 | 6.37096828 | -4.3070686 | 1.74861808 | -2.4631271 |
| 18667 | NM_017929.5 | 19.5003292 | -5.9074578 | 2.39839297 | -2.46309 |
| 18668 | NR_134527.1 | 5.64818354 | -4.1242052 | 1.67465243 | -2.4627231 |
| 18669 | NM_001330694.1 | 21.1746983 | -6.0256368 | 2.44676118 | -2.4626992 |
| 18670 | NM_001145264.1 | 6.30760369 | -4.2900791 | 1.74204107 | -2.4626739 |
| 18671 | NM_001144913.1 | 26.8470521 | -2.7806231 | 1.12967394 | -2.4614387 |
| 18672 | NM_001100829.2 | 247.980185 | -1.7573848 | 0.71397302 | -2.4614163 |
| 18673 | NM_001199184.1 | 17.9673498 | -5.7910262 | 2.35283619 | -2.461296 |
| 18674 | NM_052988.4 | 20.0172119 | -5.9471743 | 2.41640028 | -2.461171 |
| 18675 | NM_005699.3 | 6.41018239 | -4.3146178 | 1.75349299 | -2.4605846 |
| 18676 | NM_002198.2 | 223.287435 | 1.74307224 | 0.70840728 | 2.46055101 |
| 18677 | NM_001166049.1 | 43.5154019 | -1.8468351 | 0.75065063 | -2.4603125 |
| 18678 | NM_001098632.2 | 17.0245377 | -5.7139387 | 2.32253942 | -2.4602117 |
| 18679 | NM_005257.5 | 4.47285622 | -3.7981699 | 1.54404002 | -2.4598908 |
| 18680 | NM_022909.3 | 41.6792791 | -3.3084778 | 1.34498654 | -2.4598594 |
| 18681 | NM_001320776.1 | 4.36359774 | -3.7570533 | 1.52743236 | -2.4597183 |
| 18682 | NM_020340.4 | 196.334803 | -2.1120311 | 0.85864842 | -2.4597158 |
| 18683 | NM_003318.4 | 20.1454 | -5.9537488 | 2.42065441 | -2.4595616 |
| 18684 | NM_006626.5 | 64.3707874 | -3.165288 | 1.28693778 | -2.4595501 |
| 18685 | NM_173344.2 | 97.9545923 | -3.4684749 | 1.41018551 | -2.4595877 |
| 18686 | NM_001130726.3 | 22.559341 | -6.1170976 | 2.48722358 | -2.459408 |
| 18687 | NR_026541.1 | 4.45322037 | -3.7735756 | 1.53468433 | -2.4588611 |
| 18688 | NM_002737.2 | 178.837809 | -2.2028548 | 0.89597184 | -2.4586206 |
| 18689 | NM_001272049.1 | 51.8450188 | -2.6587211 | 1.08141536 | -2.4585568 |
| 18690 | NR_028350.1 | 4.3748167 | -3.7390717 | 1.52086188 | -2.4585215 |
| 18691 | NM_001288733.1 | 16.3428161 | 8.94574641 | 3.63873112 | 2.45847965 |
| 18692 | NR_110505.1 | 16.3428161 | 8.94574641 | 3.63873112 | 2.45847965 |
| 18693 | NM_020429.2 | 23.2140717 | -6.1600232 | 2.50564791 | -2.4584552 |
| 18694 | NM_001167869.1 | 22.1598803 | -6.0919606 | 2.47800179 | -2.4584166 |
| 18695 | NM_001286051.1 | 22.8857023 | -5.1567382 | 2.09805602 | -2.4578649 |
| 18696 | NM_001389.3 | 20.4454381 | -5.9771341 | 2.43199282 | -2.4577104 |
| 18697 | NM_003919.2 | 18.5925497 | -5.8410408 | 2.37664455 | -2.4576838 |
| 18698 | NM_032438.3 | 16.9492455 | -5.7072829 | 2.32227255 | -2.4576284 |
| 18699 | NM_001323031.2 | 23.593521 | -6.1814174 | 2.51521611 | -2.4576089 |
| 18700 | NM_000296.3 | 397.214049 | -3.1394516 | 1.27767721 | -2.4571555 |
| 18701 | NM_003608.3 | 11.9640916 | 3.42407766 | 1.39354451 | 2.45709961 |
| 18702 | NR_023363.1 | 26.8145542 | 3.11419663 | 1.26742686 | 2.45710165 |
| 18703 | NR_144323.1 | 4.97554578 | -3.9288505 | 1.59901127 | -2.4570499 |
| 18704 | NM_030587.2 | 21.0172665 | -6.0157137 | 2.44885614 | -2.4565403 |

| | | | | | |
|---|---|---|---|---|---|
| 18705 | NR_046901.1 | 12.7430108 | 5.63202332 | 2.29294252 | 2.4562427 |
| 18706 | NM_001291832.1 | 32.3553154 | 3.64483571 | 1.48432105 | 2.45555751 |
| 18707 | NM_001324062.1 | 16.9763827 | -5.7103257 | 2.32563969 | -2.4553785 |
| 18708 | NM_001317742.1 | 7.14902365 | -4.442212 | 1.80923077 | -2.4553042 |
| 18709 | NM_199344.2 | 4.60067409 | -3.8221688 | 1.55669699 | -2.4553069 |
| 18710 | NR_033242.1 | 4.32937746 | -3.7341799 | 1.52086894 | -2.4552937 |
| 18711 | NM_001440.3 | 340.590789 | -3.0343087 | 1.23593942 | -2.4550627 |
| 18712 | NM_006635.3 | 8.17711766 | -3.5632237 | 1.4516619 | -2.4545823 |
| 18713 | NM_017705.3 | 4.92457187 | -3.9338203 | 1.60264768 | -2.4545758 |
| 18714 | NM_006078.3 | 62.8854418 | -2.7085234 | 1.10355498 | -2.454362 |
| 18715 | NM_032043.2 | 153.706142 | 1.30810011 | 0.53305616 | 2.45396301 |
| 18716 | NM_030568.4 | 4.61161396 | -3.8176364 | 1.55571965 | -2.453936 |
| 18717 | NM_016518.2 | 65.7804632 | -2.7949051 | 1.1389757 | -2.453876 |
| 18718 | NM_001282387.1 | 5.85415187 | -4.1856999 | 1.70590459 | -2.4536542 |
| 18719 | NM_001303268.1 | 21.2713047 | -6.033282 | 2.45901694 | -2.4535341 |
| 18720 | NR_002771.1 | 4.33885105 | -3.7208939 | 1.51663812 | -2.4533828 |
| 18721 | NR_038271.1 | 4.96072223 | -3.9174841 | 1.59681693 | -2.4533082 |
| 18722 | NR_137287.1 | 21.8307454 | -6.0721428 | 2.47516559 | -2.4532269 |
| 18723 | NM_001309443.1 | 109.463511 | -3.6229211 | 1.47681475 | -2.4531995 |
| 18724 | NM_015630.3 | 129.479723 | -3.4451934 | 1.40447583 | -2.4530101 |
| 18725 | NR_047678.1 | 20.7179375 | -5.9943298 | 2.4437482 | -2.4529245 |
| 18726 | NM_001257994.1 | 7.40656712 | -4.498024 | 1.8338942 | -2.4527173 |
| 18727 | NR_073030.1 | 18.1071241 | -5.8025032 | 2.36598241 | -2.452471 |
| 18728 | NR_027148.1 | 6.69081005 | -4.3591351 | 1.77758818 | -2.452275 |
| 18729 | NM_000845.2 | 19.7680966 | -5.9290213 | 2.41817913 | -2.4518536 |
| 18730 | NM_001102653.1 | 121.249156 | -3.3419364 | 1.36302686 | -2.4518493 |
| 18731 | NR_135261.1 | 9.67747996 | -4.8902449 | 1.99448488 | -2.4518837 |
| 18732 | NM_016848.5 | 236.807264 | -3.4121871 | 1.39175019 | -2.4517238 |
| 18733 | NR_135282.1 | 16.0385334 | -5.628064 | 2.29576389 | -2.4514995 |
| 18734 | NR_102317.1 | 4.085567 | -3.6598126 | 1.49290129 | -2.4514766 |
| 18735 | NM_001011666.2 | 20.7797865 | -5.9984014 | 2.44703746 | -2.4512912 |
| 18736 | NM_000163.4 | 17.426144 | -5.7472877 | 2.34476783 | -2.4511116 |
| 18737 | NM_182500.2 | 6.48802663 | -4.3282021 | 1.76601832 | -2.4508251 |
| 18738 | NM_001330450.1 | 115.441504 | -2.2447579 | 0.91597671 | -2.4506714 |
| 18739 | NR_136745.1 | 33.8080378 | -4.9928526 | 2.0373502 | -2.45066 |
| 18740 | NR_103507.2 | 27.2283649 | -5.4068641 | 2.2064755 | -2.4504528 |
| 18741 | NM_001270436.1 | 19.5335742 | -5.9108556 | 2.4121762 | -2.4504245 |
| 18742 | NM_052847.2 | 91.7355657 | -2.2150602 | 0.90401219 | -2.4502548 |
| 18743 | NR_003135.2 | 4.81532984 | -3.8971549 | 1.59056425 | -2.4501713 |
| 18744 | NM_001099431.1 | 15.8982319 | 8.91299816 | 3.63891643 | 2.449355 |
| 18745 | NM_001286702.1 | 15.8982319 | 8.91299816 | 3.63891643 | 2.449355 |
| 18746 | NM_000554.5 | 123.05488 | 1.67507838 | 0.68394724 | 2.44913392 |
| 18747 | NM_001329641.1 | 16.0324653 | -5.6274592 | 2.29808394 | -2.4487614 |

| | | | | | |
|---|---|---|---|---|---|
| 18748 | NM_153809.2 | 10.4108018 | 1.9807333 | 0.80905395 | 2.44820916 |
| 18749 | NM_018309.4 | 19.0556154 | -5.8737351 | 2.39935539 | -2.4480471 |
| 18750 | NM_001184744.1 | 21.831073 | -2.995979 | 1.22388486 | -2.4479255 |
| 18751 | NM_145249.2 | 4.59868959 | -3.8105296 | 1.55708168 | -2.4472252 |
| 18752 | NM_001077394.1 | 6.82066159 | -4.3780937 | 1.78902472 | -2.4471958 |
| 18753 | NR_047025.1 | 6.09087141 | 3.60080594 | 1.47147851 | 2.44706662 |
| 18754 | NR_131917.1 | 48.062907 | 2.04482573 | 0.83566352 | 2.44694866 |
| 18755 | NM_001012334.2 | 6.10291686 | -4.2313387 | 1.7294333 | -2.446662 |
| 18756 | NM_001204836.1 | 16.139734 | -5.636828 | 2.30442202 | -2.4460919 |
| 18757 | NM_001256280.1 | 5.66998832 | 7.42101317 | 3.03383591 | 2.44608258 |
| 18758 | NM_001244871.1 | 21.6045616 | -6.0551077 | 2.47564145 | -2.4458743 |
| 18759 | NM_001099434.1 | 4.26814396 | -3.7275998 | 1.52432343 | -2.4454126 |
| 18760 | NM_001303114.1 | 5.66633355 | -4.1049109 | 1.67885956 | -2.4450591 |
| 18761 | NM_001346339.1 | 4.64070075 | -3.8410714 | 1.57099134 | -2.4449985 |
| 18762 | NM_001320298.1 | 19.490854 | -5.9069656 | 2.41612586 | -2.4448088 |
| 18763 | NR_110803.1 | 11.2902659 | 2.41607025 | 0.98848142 | 2.44422423 |
| 18764 | NR_033974.1 | 8.32980753 | -3.6708565 | 1.502134 | -2.443761 |
| 18765 | NR_104292.1 | 13.2119204 | -2.4794924 | 1.01461765 | -2.4437702 |
| 18766 | NR_036473.1 | 36.8704689 | 1.76583607 | 0.72269803 | 2.44339405 |
| 18767 | NM_020039.3 | 6.88780384 | -4.4005336 | 1.80128793 | -2.4429929 |
| 18768 | NM_145172.4 | 20.2451171 | -5.9633517 | 2.44107475 | -2.4429206 |
| 18769 | NR_132780.1 | 20.4802321 | -5.977363 | 2.44727936 | -2.4424523 |
| 18770 | NM_001199479.1 | 5.47977567 | -4.0796059 | 1.67034248 | -2.4423769 |
| 18771 | NR_108088.1 | 8.20494454 | 2.86312695 | 1.17233417 | 2.44224472 |
| 18772 | NM_004248.2 | 6.5186005 | -4.3306104 | 1.7732651 | -2.4421675 |
| 18773 | NR_134503.1 | 4.49791528 | 3.62152787 | 1.48293356 | 2.44213764 |
| 18774 | NM_018910.2 | 17.1021639 | -5.7207651 | 2.34257083 | -2.4420884 |
| 18775 | NM_025163.3 | 26.0223629 | -2.7564265 | 1.12875079 | -2.4420151 |
| 18776 | NM_001319107.1 | 29.1353269 | -6.4854556 | 2.65600739 | -2.4418063 |
| 18777 | NM_001199741.1 | 7.38254835 | -4.5170181 | 1.84995162 | -2.4416953 |
| 18778 | NM_001040100.1 | 4.56671859 | -3.8152517 | 1.56276612 | -2.4413453 |
| 18779 | NM_001282604.1 | 4.41801205 | -3.7811837 | 1.54887616 | -2.4412434 |
| 18780 | NM_001243737.1 | 20.3382905 | -5.9678569 | 2.44466755 | -2.4411732 |
| 18781 | NR_024321.1 | 6.84376812 | -3.3674018 | 1.37950978 | -2.4410133 |
| 18782 | NM_016952.4 | 182.142574 | -1.8677686 | 0.76520132 | -2.4408852 |
| 18783 | NR_110917.1 | 19.564093 | 1.51224636 | 0.61969409 | 2.4403111 |
| 18784 | NR_033263.1 | 26.6510815 | -1.9661479 | 0.80615407 | -2.4389233 |
| 18785 | NR_040087.2 | 15.9251545 | -5.618065 | 2.30368187 | -2.438733 |
| 18786 | NM_001177637.2 | 18.8586963 | -5.8599182 | 2.40310854 | -2.4384742 |
| 18787 | NR_037595.1 | 5.60903719 | -4.1129711 | 1.68671797 | -2.4384462 |
| 18788 | NM_001198899.1 | 7.46449252 | -4.5075384 | 1.84856733 | -2.4383956 |
| 18789 | NM_005170.2 | 4.98602179 | -3.9238313 | 1.60927188 | -2.438265 |
| 18790 | NM_002239.3 | 20.4453991 | -5.9753242 | 2.45067612 | -2.438235 |

| | | | | | |
|---|---|---|---|---|---|
| 18791 | NM_001143788.1 | 20.4730879 | -5.9769828 | 2.45150751 | -2.4380846 |
| 18792 | NM_001248001.1 | 31.5391055 | 4.47708296 | 1.83640893 | 2.43795535 |
| 18793 | NM_001282915.1 | 15.8319618 | -5.6091885 | 2.30116834 | -2.4375394 |
| 18794 | NM_001009932.2 | 27.6307374 | -2.6451978 | 1.08542278 | -2.4370207 |
| 18795 | NM_001267563.1 | 4.18871221 | -3.6930697 | 1.5155623 | -2.4367654 |
| 18796 | NM_001050.2 | 123.06305 | -1.8834149 | 0.7729854 | -2.4365465 |
| 18797 | NM_001014449.2 | 15.6905559 | -5.5964872 | 2.2972348 | -2.4361842 |
| 18798 | NM_001261415.2 | 24.1942207 | -6.2165895 | 2.55199241 | -2.4359749 |
| 18799 | NM_001318748.1 | 20.1702725 | -5.956085 | 2.4452482 | -2.4357793 |
| 18800 | NM_032750.2 | 6.21424006 | -4.2720383 | 1.75403664 | -2.4355468 |
| 18801 | NR_120684.1 | 52.2156384 | -1.4757522 | 0.60611095 | -2.4347888 |
| 18802 | NM_000496.2 | 4.70790508 | -3.8668054 | 1.58820472 | -2.4347021 |
| 18803 | NM_001098831.1 | 54.4106585 | 3.01377645 | 1.23800273 | 2.43438594 |
| 18804 | NM_005545.3 | 16.1697722 | -5.6399237 | 2.31677994 | -2.4343804 |
| 18805 | NM_001242537.1 | 5.86108464 | 7.46026326 | 3.06483662 | 2.43414713 |
| 18806 | NM_001244974.1 | 5.72129483 | -4.1522263 | 1.70586932 | -2.4340823 |
| 18807 | NR_110256.1 | 7.24566246 | 7.78115205 | 3.19678731 | 2.43405373 |
| 18808 | NM_003935.4 | 14.4015221 | -2.2263607 | 0.91476728 | -2.4338001 |
| 18809 | NR_120455.1 | 6.92087319 | -4.407839 | 1.8112204 | -2.4336293 |
| 18810 | NM_001281476.2 | 20.4177769 | -5.9729501 | 2.45453133 | -2.4334381 |
| 18811 | NM_005951.2 | 6.94938161 | -4.4175857 | 1.81553581 | -2.4332132 |
| 18812 | NM_003677.4 | 305.05094 | -2.457396 | 1.01022071 | -2.4325338 |
| 18813 | NM_138347.4 | 18.5339017 | -5.8345447 | 2.39884814 | -2.4322276 |
| 18814 | NM_016444.2 | 6.64296161 | -4.3589206 | 1.79261671 | -2.4315966 |
| 18815 | NM_014690.4 | 26.0094529 | -3.4500781 | 1.41891941 | -2.4314828 |
| 18816 | NM_001083608.1 | 20.1420545 | -5.9540156 | 2.44875113 | -2.4314498 |
| 18817 | NM_001081753.2 | 22.3606733 | -6.1067901 | 2.51168303 | -2.4313538 |
| 18818 | NM_005559.3 | 132.227562 | -2.8145734 | 1.15761237 | -2.4313609 |
| 18819 | NM_001201574.1 | 5.69519517 | 7.43146577 | 3.05658193 | 2.43129938 |
| 18820 | NR_135078.1 | 6.30091866 | -4.2597922 | 1.75206675 | -2.4312956 |
| 18821 | NR_030725.1 | 20.0598557 | -5.947631 | 2.44656867 | -2.4310092 |
| 18822 | NM_001005465.1 | 6.05986178 | 7.52487965 | 3.09549563 | 2.43091271 |
| 18823 | NM_001193630.1 | 6.05986178 | 7.52487965 | 3.09549563 | 2.43091271 |
| 18824 | NM_025219.2 | 544.023219 | -3.5957119 | 1.4791953 | -2.4308568 |
| 18825 | NM_001318052.1 | 95.1082025 | -3.423243 | 1.40846747 | -2.4304736 |
| 18826 | NM_015314.2 | 18.9718452 | -5.8673662 | 2.41425857 | -2.4302973 |
| 18827 | NM_207645.3 | 79.6459069 | -2.9126677 | 1.19848492 | -2.4302915 |
| 18828 | NR_027302.1 | 5.14548262 | -3.9932022 | 1.64332441 | -2.4299537 |
| 18829 | NM_033326.3 | 16.0227598 | -2.8071711 | 1.15525496 | -2.4299148 |
| 18830 | NR_110003.1 | 46.5430691 | 2.3218744 | 0.95566499 | 2.42959032 |
| 18831 | NR_045116.1 | 4.35827343 | -3.7263815 | 1.53415985 | -2.4289395 |
| 18832 | NM_001105565.2 | 4.35241875 | -3.7437805 | 1.54174075 | -2.4282815 |
| 18833 | NM_145655.3 | 21.021823 | 5.18818611 | 2.13664431 | 2.42819363 |

| | | | | | |
|---|---|---|---|---|---|
| 18834 | NM_001293236.1 | 16.1741751 | -5.6399197 | 2.3227618 | -2.4281094 |
| 18835 | NM_001330712.1 | 19.8789615 | -5.9348946 | 2.44442513 | -2.4279306 |
| 18836 | NM_005577.2 | 8.06624635 | 5.41962982 | 2.23219862 | 2.4279335 |
| 18837 | NR_026844.1 | 41.441112 | 3.12318461 | 1.28653386 | 2.42759613 |
| 18838 | NR_027163.1 | 21.7093995 | -6.0609952 | 2.4968412 | -2.4274652 |
| 18839 | NM_000825.3 | 4.35353517 | -3.7580642 | 1.54840933 | -2.4270483 |
| 18840 | NM_001321058.1 | 19.1603081 | -5.8841312 | 2.42496646 | -2.4264794 |
| 18841 | NM_001242817.1 | 20.9008248 | -6.0069756 | 2.47642977 | -2.4256596 |
| 18842 | NR_040028.1 | 5.33340732 | -4.0153974 | 1.65547828 | -2.4255211 |
| 18843 | NM_001242870.1 | 15.0353908 | 8.82501084 | 3.63943559 | 2.42482952 |
| 18844 | NM_176817.4 | 15.0353908 | 8.82501084 | 3.63943559 | 2.42482952 |
| 18845 | NM_001309434.1 | 4.91414416 | -3.9286332 | 1.62027411 | -2.4246719 |
| 18846 | NM_001024465.2 | 15.4863239 | -5.5775956 | 2.30039195 | -2.4246284 |
| 18847 | NM_001109977.1 | 6.96476504 | -4.4300403 | 1.82712189 | -2.4246003 |
| 18848 | NM_181459.3 | 6.30627483 | -4.265186 | 1.75936113 | -2.4242811 |
| 18849 | NM_001146337.2 | 51.963961 | 5.2304486 | 2.15791519 | 2.42384345 |
| 18850 | NM_031271.3 | 32.5520289 | 2.56879171 | 1.05982495 | 2.42378867 |
| 18851 | NR_040671.1 | 3.93539115 | -3.605664 | 1.4876022 | -2.4238093 |
| 18852 | NR_144688.1 | 55.4356268 | 6.41942615 | 2.64849454 | 2.42380192 |
| 18853 | NM_001162495.2 | 19.9124386 | -5.9372214 | 2.449905 | -2.4234496 |
| 18854 | NM_001286555.1 | 20.7945748 | -6.0009478 | 2.47657277 | -2.4230856 |
| 18855 | NM_001256468.1 | 16.2884789 | -5.6503083 | 2.33208165 | -2.4228604 |
| 18856 | NM_001308192.1 | 19.4867935 | -5.9082561 | 2.43921524 | -2.4221954 |
| 18857 | NM_001348094.1 | 20.4361945 | -5.9736454 | 2.46627846 | -2.4221293 |
| 18858 | NM_005201.3 | 14.4134554 | 8.06817406 | 3.33102432 | 2.42213004 |
| 18859 | NR_120420.1 | 7.94152999 | -3.5834294 | 1.47943156 | -2.4221664 |
| 18860 | NR_110767.1 | 7.53006827 | -4.521705 | 1.8669323 | -2.4219973 |
| 18861 | NM_024622.5 | 15.3907305 | -5.568544 | 2.29918517 | -2.4219641 |
| 18862 | NM_001195071.1 | 16.2018126 | -5.6430798 | 2.33016665 | -2.4217495 |
| 18863 | NM_001111045.1 | 15.3614733 | -5.5658391 | 2.29830392 | -2.4217159 |
| 18864 | NR_045104.1 | 15.4274053 | -5.5722779 | 2.301074 | -2.4215987 |
| 18865 | NM_002919.3 | 3.98065394 | -3.6083659 | 1.49009601 | -2.421566 |
| 18866 | NM_001198902.1 | 21.706812 | -6.0623803 | 2.50361425 | -2.4214514 |
| 18867 | NM_001282656.1 | 17.3120071 | -5.7378264 | 2.36994031 | -2.4210848 |
| 18868 | NM_033272.3 | 5.72456886 | -4.1246825 | 1.70371098 | -2.420999 |
| 18869 | NM_001079842.2 | 19.7509843 | -5.9252452 | 2.44750093 | -2.4209369 |
| 18870 | NM_144781.2 | 20.1019625 | -5.9517213 | 2.45866919 | -2.4207085 |
| 18871 | NM_001301089.1 | 15.5063473 | -5.5791346 | 2.30503573 | -2.4204113 |
| 18872 | NM_001321136.1 | 17.6665063 | 3.20524435 | 1.32427352 | 2.4203794 |
| 18873 | NM_020653.3 | 15.8647685 | -5.6123578 | 2.31885051 | -2.4203189 |
| 18874 | NM_001077349.1 | 2.05985365 | 5.96870392 | 2.46610187 | 2.42029901 |
| 18875 | NM_014435.3 | 5.80678528 | -4.1666594 | 1.72164211 | -2.4201658 |
| 18876 | NR_026730.1 | 93.4810797 | 1.23922201 | 0.5120628 | 2.42005865 |

| | | | | | |
|---|---|---|---|---|---|
| 18877 | NM_003841.4 | 4.98548117 | -3.9461782 | 1.63063517 | -2.4200252 |
| 18878 | NR_015399.1 | 7.50600709 | -4.5246213 | 1.8696683 | -2.4200128 |
| 18879 | NM_001032394.2 | 21.455491 | -6.0451183 | 2.49812112 | -2.419866 |
| 18880 | NM_012316.4 | 363.427873 | -2.1078297 | 0.87108796 | -2.4197668 |
| 18881 | NM_001231.4 | 4.90986438 | -3.9084583 | 1.61524929 | -2.4197245 |
| 18882 | NM_001305108.1 | 21.4592812 | -6.0450884 | 2.4984094 | -2.4195748 |
| 18883 | NR_138111.1 | 5.43844854 | -4.0578273 | 1.67708865 | -2.4195664 |
| 18884 | NM_001130725.1 | 18.2534017 | -5.8147234 | 2.40341093 | -2.419363 |
| 18885 | NR_135117.1 | 6.82869338 | -4.3930527 | 1.81578634 | -2.4193665 |
| 18886 | NM_001079530.1 | 4.79483122 | -3.8822272 | 1.60469288 | -2.4192961 |
| 18887 | NM_001270679.1 | 16.3866234 | 6.45512936 | 2.66840893 | 2.41909299 |
| 18888 | NM_001007253.3 | 83.2224466 | -3.3891801 | 1.40113148 | -2.418888 |
| 18889 | NR_044998.1 | 37.8340409 | -2.1744457 | 0.89908156 | -2.4185188 |
| 18890 | NM_018177.5 | 115.800718 | -2.1495102 | 0.88882992 | -2.4183594 |
| 18891 | NM_001144918.1 | 20.2171349 | -5.9599439 | 2.46452593 | -2.4182922 |
| 18892 | NM_001142405.1 | 22.3208956 | -6.1003068 | 2.52268421 | -2.4181809 |
| 18893 | NM_032239.3 | 6.55124655 | -4.3407472 | 1.79504026 | -2.4181893 |
| 18894 | NM_001318151.1 | 7.66499586 | -4.5536402 | 1.88358819 | -2.4175349 |
| 18895 | NM_001308160.1 | 32.396262 | -4.9297321 | 2.03980239 | -2.4167695 |
| 18896 | NM_153240.4 | 88.4572151 | -2.2298355 | 0.92269235 | -2.416662 |
| 18897 | NM_152292.4 | 7.70681103 | -4.5537564 | 1.88439398 | -2.4165628 |
| 18898 | NM_007287.2 | 19.6555756 | -5.918342 | 2.44916741 | -2.416471 |
| 18899 | NR_105018.1 | 22.8287363 | 1.92530146 | 0.7967563 | 2.4164245 |
| 18900 | NM_001286476.1 | 4.01492554 | -3.6280136 | 1.50149701 | -2.4162643 |
| 18901 | NM_001265579.1 | 20.2080949 | -5.9608616 | 2.46715595 | -2.4160863 |
| 18902 | NM_001301828.1 | 4.95087917 | -3.9435701 | 1.63247184 | -2.4157048 |
| 18903 | NM_001243765.1 | 127.091999 | -2.2295411 | 0.92295335 | -2.4156596 |
| 18904 | NM_001130006.1 | 52.4218848 | -4.681663 | 1.93807621 | -2.4156238 |
| 18905 | NM_001007088.1 | 18.1304674 | -5.8022206 | 2.4019772 | -2.4156019 |
| 18906 | NM_001329226.1 | 77.8112121 | 3.81763361 | 1.58044145 | 2.4155489 |
| 18907 | NM_080704.3 | 20.9070304 | -6.009634 | 2.48870571 | -2.4147628 |
| 18908 | NM_001286792.1 | 19.0279431 | -5.8742274 | 2.43270225 | -2.4146923 |
| 18909 | NM_001303229.1 | 20.6685783 | -5.9898345 | 2.48076034 | -2.4145156 |
| 18910 | NM_145867.1 | 4.11559583 | -3.6662223 | 1.51842612 | -2.4144884 |
| 18911 | NR_024413.1 | 10.465802 | 4.10109153 | 1.69853569 | 2.41448653 |
| 18912 | NM_001282857.1 | 100.034799 | -2.1616969 | 0.89532813 | -2.4144186 |
| 18913 | NM_197966.2 | 8.34338581 | -4.6756451 | 1.9365506 | -2.4144193 |
| 18914 | NM_014899.3 | 277.012299 | -1.9829637 | 0.82131177 | -2.4143861 |
| 18915 | NR_027133.1 | 31.189919 | 1.48318363 | 0.61443484 | 2.41389896 |
| 18916 | NM_001330661.1 | 6.56755799 | -4.3416299 | 1.79871129 | -2.4137447 |
| 18917 | NM_015328.3 | 21.8807418 | -6.0749574 | 2.51688383 | -2.4136821 |
| 18918 | NM_001161012.2 | 19.1082346 | -5.8795922 | 2.43660759 | -2.4130238 |
| 18919 | NR_036472.1 | 7.76121597 | -4.5706466 | 1.89428903 | -2.412856 |

| | | | | | |
|---|---|---|---|---|---|
| 18920 | NM_001330059.1 | 16.9100507 | -5.7035874 | 2.3639293 | -2.4127572 |
| 18921 | NM_001112812.1 | 21.0477186 | -5.034977 | 2.08689839 | -2.4126604 |
| 18922 | NM_052931.4 | 5.61957461 | 7.39991283 | 3.06712402 | 2.41265523 |
| 18923 | NR_024053.2 | 15.1616154 | -5.5470186 | 2.29935917 | -2.4124194 |
| 18924 | NM_133334.2 | 193.65422 | -3.7554763 | 1.55675798 | -2.41237 |
| 18925 | NR_131177.1 | 6.88894201 | -4.4050603 | 1.82613299 | -2.4122341 |
| 18926 | NM_001254732.2 | 19.6452918 | -5.9195856 | 2.45410561 | -2.4121152 |
| 18927 | NM_001001483.2 | 19.9620287 | -5.9416968 | 2.4640642 | -2.4113401 |
| 18928 | NR_040058.1 | 40.9800205 | 2.24475709 | 0.93102147 | 2.41106909 |
| 18929 | NR_046311.1 | 7.11985549 | -3.4255501 | 1.4209026 | -2.4108268 |
| 18930 | NM_001286517.1 | 15.5153587 | -5.5804366 | 2.31480689 | -2.4107569 |
| 18931 | NM_001290149.1 | 4.44507128 | -3.7568618 | 1.55864513 | -2.4103382 |
| 18932 | NR_046202.1 | 6.4863962 | -4.3202236 | 1.79273792 | -2.4098467 |
| 18933 | NM_152386.3 | 26.7266282 | -6.3632921 | 2.6409382 | -2.4094816 |
| 18934 | NR_024261.2 | 8.80576275 | -3.7439401 | 1.55392895 | -2.409338 |
| 18935 | NM_145037.2 | 146.899213 | -0.797505 | 0.33101492 | -2.4092721 |
| 18936 | NM_001172672.1 | 6.5623333 | 7.62018952 | 3.16288603 | 2.409252 |
| 18937 | NM_001126105.2 | 3.95900196 | -3.617804 | 1.50189328 | -2.4088289 |
| 18938 | NR_133664.1 | 27.6465096 | 2.69991822 | 1.12083925 | 2.40883625 |
| 18939 | NM_001164239.1 | 19.7099909 | -5.9211809 | 2.45853885 | -2.4084146 |
| 18940 | NM_001256484.1 | 18.3125737 | -5.8189248 | 2.41609805 | -2.4083976 |
| 18941 | NM_033224.4 | 239.119014 | -1.8458231 | 0.76642764 | -2.4083462 |
| 18942 | NM_001080475.2 | 100.241025 | -2.0724152 | 0.86057163 | -2.4081845 |
| 18943 | NM_001193451.1 | 70.440594 | -2.4997302 | 1.03805043 | -2.4081009 |
| 18944 | NM_198559.1 | 6.43135708 | -4.3168596 | 1.79264089 | -2.4081006 |
| 18945 | NR_136587.1 | 4.14650196 | -3.6687301 | 1.52366555 | -2.4078316 |
| 18946 | NM_001320597.1 | 17.1352064 | -5.723072 | 2.37699762 | -2.4076894 |
| 18947 | NM_001303625.1 | 20.9977683 | -6.0130378 | 2.49803704 | -2.4071052 |
| 18948 | NM_001172501.1 | 4.59879223 | -3.8014256 | 1.57931911 | -2.4070028 |
| 18949 | NM_000513.2 | 14.3816782 | 8.76063027 | 3.63983594 | 2.40687504 |
| 18950 | NM_001291278.1 | 14.3816782 | 8.76063027 | 3.63983594 | 2.40687504 |
| 18951 | NM_003561.2 | 14.3816782 | 8.76063027 | 3.63983594 | 2.40687504 |
| 18952 | NM_020525.4 | 14.3816782 | 8.76063027 | 3.63983594 | 2.40687504 |
| 18953 | NR_134924.1 | 14.3816782 | 8.76063027 | 3.63983594 | 2.40687504 |
| 18954 | NM_001006620.1 | 17.9234826 | -5.7871854 | 2.40456567 | -2.4067487 |
| 18955 | NR_031671.1 | 11.677981 | 4.07262427 | 1.69224083 | 2.40664579 |
| 18956 | NM_001099668.1 | 4.56995893 | -3.8210132 | 1.58773358 | -2.4065833 |
| 18957 | NM_001297608.1 | 20.3563178 | -5.9711011 | 2.48133814 | -2.4064036 |
| 18958 | NR_046058.1 | 15.1758921 | -5.5482872 | 2.3056848 | -2.4063511 |
| 18959 | NM_001161403.1 | 15.3646565 | -5.5665682 | 2.31336493 | -2.4062646 |
| 18960 | NM_020403.4 | 24.0090491 | -6.2068059 | 2.57974695 | -2.4059747 |
| 18961 | NM_001214907.1 | 16.2140956 | -5.6438818 | 2.34667544 | -2.4050543 |
| 18962 | NM_001346093.1 | 7.57860084 | 7.82919875 | 3.2555131 | 2.4049047 |

| | | | | | |
|---|---|---|---|---|---|
| 18963 | NM_001190918.1 | 20.0839901 | -5.9494095 | 2.47399064 | -2.4047825 |
| 18964 | NM_001204344.1 | 62.0880922 | 1.73530398 | 0.72170335 | 2.40445603 |
| 18965 | NM_020394.4 | 8.51486216 | -3.6155933 | 1.50376508 | -2.4043605 |
| 18966 | NM_207411.4 | 34.9742704 | 1.61344148 | 0.67117809 | 2.40389475 |
| 18967 | NM_152470.2 | 82.9028289 | -3.1841808 | 1.3246046 | -2.4038726 |
| 18968 | NM_001321127.1 | 4.86498619 | -3.912283 | 1.62752849 | -2.4038185 |
| 18969 | NM_001323294.1 | 20.3577574 | -5.9703819 | 2.48399995 | -2.4035354 |
| 18970 | NM_001006937.2 | 19.6328276 | -5.9165761 | 2.46164976 | -2.4035004 |
| 18971 | NM_001318853.1 | 18.3707779 | -5.8233842 | 2.42288877 | -2.4034881 |
| 18972 | NM_001174138.2 | 3.91874222 | -3.5928122 | 1.4949347 | -2.4033238 |
| 18973 | NM_024816.2 | 55.8160731 | -3.1954145 | 1.32958005 | -2.4033261 |
| 18974 | NM_213601.2 | 155.50367 | -1.561777 | 0.64987254 | -2.4032051 |
| 18975 | NM_053055.4 | 72.4014527 | -1.2913062 | 0.53733737 | -2.4031573 |
| 18976 | NM_001039395.3 | 9.22474291 | -3.8208595 | 1.59001437 | -2.4030346 |
| 18977 | NM_001080851.1 | 3.84555506 | -3.5593943 | 1.48125869 | -2.4029525 |
| 18978 | NM_001302652.1 | 21.4009588 | -6.0395878 | 2.51343205 | -2.4029246 |
| 18979 | NM_001168214.1 | 5.81955281 | -4.1629205 | 1.73290964 | -2.4022721 |
| 18980 | NM_001017370.2 | 20.6078034 | -2.6030102 | 1.08365711 | -2.4020608 |
| 18981 | NR_131782.1 | 24.48948 | -2.9000436 | 1.20733755 | -2.4020156 |
| 18982 | NM_000767.4 | 22.4595656 | 2.12141702 | 0.88325425 | 2.4018192 |
| 18983 | NM_001258325.1 | 21.700319 | -6.0605564 | 2.52376456 | -2.4013953 |
| 18984 | NM_001164392.1 | 15.6462613 | -5.5917599 | 2.32943291 | -2.4004812 |
| 18985 | NM_080722.4 | 22.2227794 | -6.0946999 | 2.53901772 | -2.4004165 |
| 18986 | NM_001199398.1 | 21.2818814 | -6.0320078 | 2.51298899 | -2.400332 |
| 18987 | NM_001555.4 | 17.9678024 | -5.7891161 | 2.41210846 | -2.4000232 |
| 18988 | NM_052818.2 | 5.14988539 | -3.9987453 | 1.66625968 | -2.3998332 |
| 18989 | NM_152313.3 | 90.9370706 | -3.2483759 | 1.35358873 | -2.3998249 |
| 18990 | NM_013279.3 | 16.3421877 | -5.655597 | 2.35678868 | -2.3997048 |
| 18991 | NM_001346500.1 | 5.58659706 | -4.088117 | 1.70392161 | -2.3992401 |
| 18992 | NM_001278642.1 | 138.990363 | -3.3012137 | 1.37603181 | -2.3990824 |
| 18993 | NR_134459.1 | 4.11438297 | -3.64872 | 1.52092722 | -2.3990103 |
| 18994 | NM_033213.4 | 16.5553418 | -5.6734061 | 2.36500681 | -2.3988963 |
| 18995 | NM_000283.3 | 12.5766779 | -5.2769869 | 2.20032297 | -2.3982783 |
| 18996 | NM_138706.4 | 25.5186642 | 1.74652131 | 0.72826237 | 2.39820341 |
| 18997 | NM_001130846.3 | 52.1060595 | 1.72853606 | 0.72080448 | 2.3980651 |
| 18998 | NM_032117.3 | 4.51482432 | -3.7939485 | 1.58209077 | -2.39806 |
| 18999 | NR_033234.1 | 16.0788462 | -5.6309644 | 2.34813388 | -2.3980593 |
| 19000 | NR_046719.1 | 4.34901331 | -3.7588487 | 1.56742609 | -2.3981027 |
| 19001 | NM_018489.2 | 192.159589 | -3.1351103 | 1.30737637 | -2.3980167 |
| 19002 | NM_001282434.1 | 7.88800362 | -4.5882396 | 1.91349425 | -2.3978329 |
| 19003 | NR_126448.1 | 7.06871482 | -4.4320816 | 1.8484975 | -2.3976671 |
| 19004 | NM_017646.5 | 15.3490711 | -5.5645328 | 2.32101565 | -2.397456 |
| 19005 | NR_106789.1 | 13.9594231 | 8.72563974 | 3.64006111 | 2.39711353 |

| | | | | | |
|---|---|---|---|---|---|
| 19006 | NM_153615.1 | 18.9709299 | -5.8673526 | 2.44779708 | -2.3969931 |
| 19007 | NR_134293.1 | 5.35163034 | -4.0383007 | 1.68479302 | -2.396912 |
| 19008 | NR_135840.1 | 4.7875789 | -3.8727717 | 1.6157698 | -2.3968586 |
| 19009 | NM_001164270.1 | 5.43765973 | -4.0526286 | 1.69084254 | -2.3968102 |
| 19010 | NM_133174.2 | 15.1645179 | -5.5470828 | 2.31439828 | -2.3967711 |
| 19011 | NM_001271521.1 | 4.79146575 | -3.8727096 | 1.6160639 | -2.396384 |
| 19012 | NM_033031.2 | 4.17716579 | -3.6742482 | 1.5333948 | -2.3961528 |
| 19013 | NM_001198827.1 | 19.4849839 | -5.9053959 | 2.46465537 | -2.3960331 |
| 19014 | NM_001320541.1 | 16.1825233 | -5.6406442 | 2.35424929 | -2.3959418 |
| 19015 | NM_021063.3 | 5.00626396 | -3.9380751 | 1.64366281 | -2.3959142 |
| 19016 | NM_004833.1 | 4.17930727 | -3.6798573 | 1.53626445 | -2.3953281 |
| 19017 | NM_001004340.3 | 8.73758167 | 5.42896184 | 2.26657389 | 2.39522826 |
| 19018 | NR_109867.1 | 4.82118983 | -3.8895859 | 1.62387714 | -2.3952464 |
| 19019 | NM_015645.4 | 11.5933684 | -5.1613516 | 2.15496439 | -2.3950983 |
| 19020 | NM_015672.1 | 5.22226618 | -4.0087261 | 1.67386102 | -2.3948978 |
| 19021 | NM_153477.2 | 6.14917072 | -4.2326957 | 1.76747844 | -2.3947651 |
| 19022 | NM_001195082.1 | 3.80833356 | -3.5472166 | 1.48139944 | -2.3945038 |
| 19023 | NM_001083113.1 | 4.82463 | -3.8955685 | 1.62690078 | -2.3944721 |
| 19024 | NM_001161425.1 | 19.0775864 | -5.8748107 | 2.45382569 | -2.3941434 |
| 19025 | NM_001039938.1 | 5.72157597 | -4.1348688 | 1.72713517 | -2.3940621 |
| 19026 | NR_036565.1 | 4.69059851 | -3.8672099 | 1.61539141 | -2.393977 |
| 19027 | NM_001034845.2 | 8.95383138 | -3.7185851 | 1.55341166 | -2.3938182 |
| 19028 | NM_001004317.3 | 41.7477038 | -3.1532707 | 1.31732436 | -2.3936935 |
| 19029 | NM_006235.2 | 53.959043 | 1.76102236 | 0.73575433 | 2.39349234 |
| 19030 | NM_016520.2 | 245.402605 | -1.359042 | 0.5678315 | -2.3933896 |
| 19031 | NM_001318058.1 | 6.87966265 | -4.4147615 | 1.84458684 | -2.3933606 |
| 19032 | NM_199367.2 | 16.1624861 | -2.8049796 | 1.17204033 | -2.3932449 |
| 19033 | NM_030882.3 | 88.0011486 | -1.7738529 | 0.74119988 | -2.393218 |
| 19034 | NR_026683.1 | 4.08389558 | -3.6439872 | 1.52271712 | -2.3930822 |
| 19035 | NM_001161563.2 | 18.3983888 | -5.8228879 | 2.43329187 | -2.3930084 |
| 19036 | NM_024793.2 | 30.3403176 | 3.24468976 | 1.35593088 | 2.39296104 |
| 19037 | NM_198834.2 | 22.3677547 | -6.1069474 | 2.55220095 | -2.3928161 |
| 19038 | NM_198038.2 | 25.2767224 | 5.34995529 | 2.236283 | 2.39234269 |
| 19039 | NM_001139488.1 | 14.7654057 | -5.5090665 | 2.30338548 | -2.3917258 |
| 19040 | NM_001301860.1 | 18.9431184 | -5.8665 | 2.45334126 | -2.3912287 |
| 19041 | NR_002815.2 | 15.040595 | 3.22793586 | 1.34995085 | 2.39115065 |
| 19042 | NM_001040708.1 | 18.5297208 | -5.8361284 | 2.44119668 | -2.3906834 |
| 19043 | NR_104247.1 | 28.562929 | -2.1671966 | 0.90662343 | -2.3904044 |
| 19044 | NM_002944.2 | 3.39011557 | 5.9014623 | 2.46886681 | 2.39035264 |
| 19045 | NM_183356.3 | 19.0267542 | -5.8710168 | 2.45612659 | -2.390356 |
| 19046 | NM_018902.4 | 21.1129276 | -6.0231056 | 2.51979873 | -2.3903122 |
| 19047 | NR_024561.1 | 28.6140524 | 1.8807398 | 0.7868845 | 2.3901091 |
| 19048 | NM_001169117.1 | 4.74874865 | -3.8657553 | 1.6175155 | -2.389934 |

| | | | | | |
|---|---|---|---|---|---|
| 19049 | NM_001012270.1 | 7.63263761 | -4.5486229 | 1.90330777 | -2.3898515 |
| 19050 | NM_032266.3 | 7.70681165 | -3.479991 | 1.45615295 | -2.3898527 |
| 19051 | NM_001288998.1 | 19.7771161 | -5.9260475 | 2.48001074 | -2.3895249 |
| 19052 | NM_000761.4 | 107.583897 | 2.42976165 | 1.01705455 | 2.38901803 |
| 19053 | NM_205858.1 | 6.21686861 | -4.239567 | 1.77471653 | -2.38887 |
| 19054 | NM_001003699.3 | 15.5463871 | -5.5836221 | 2.3374371 | -2.3887796 |
| 19055 | NR_024586.1 | 16.2685239 | -2.2543166 | 0.94376398 | -2.3886444 |
| 19056 | NR_110084.1 | 10.2069981 | -2.3547876 | 0.98586429 | -2.3885515 |
| 19057 | NM_014761.3 | 18.5849307 | -5.8363994 | 2.44381684 | -2.3882311 |
| 19058 | NM_001277092.1 | 6.23589475 | -4.2498408 | 1.7795789 | -2.388116 |
| 19059 | NM_001146276.1 | 13.7279655 | 8.6932431 | 3.64027449 | 2.38807352 |
| 19060 | NR_126340.1 | 13.7279655 | 8.6932431 | 3.64027449 | 2.38807352 |
| 19061 | NR_002803.2 | 7.54651887 | -3.4475502 | 1.44366613 | -2.3880523 |
| 19062 | NM_198514.3 | 41.6492197 | -1.2885153 | 0.53971171 | -2.3874141 |
| 19063 | NM_001313741.1 | 69.7090887 | -5.1566571 | 2.16028371 | -2.3870277 |
| 19064 | NR_038254.1 | 13.57839 | 8.6872955 | 3.64031419 | 2.38641366 |
| 19065 | NM_001283037.1 | 5.64652664 | -4.1195821 | 1.72632999 | -2.3863236 |
| 19066 | NM_017733.4 | 17.005435 | -5.7129164 | 2.39404308 | -2.3863048 |
| 19067 | NM_001316314.1 | 57.2852554 | -2.565828 | 1.07525769 | -2.3862447 |
| 19068 | NM_001244683.1 | 6.49803724 | 7.6304337 | 3.19807373 | 2.38594677 |
| 19069 | NM_001331030.1 | 6.49803724 | 7.6304337 | 3.19807373 | 2.38594677 |
| 19070 | NR_040020.1 | 13.5716614 | 8.68506022 | 3.64032915 | 2.38578982 |
| 19071 | NM_001320793.1 | 12.3424636 | -2.8356799 | 1.18858188 | -2.3857674 |
| 19072 | NM_024869.2 | 4.89231552 | -3.9111229 | 1.6395185 | -2.3855314 |
| 19073 | NM_001324081.1 | 5.49955095 | -4.0839814 | 1.7124735 | -2.3848436 |
| 19074 | NM_005930.3 | 19.3286916 | -5.8939087 | 2.47151475 | -2.3847354 |
| 19075 | NM_012347.4 | 5.35573542 | 7.34209178 | 3.07879188 | 2.38473143 |
| 19076 | NM_001082970.1 | 25.2162075 | -6.276229 | 2.63223071 | -2.3843765 |
| 19077 | NM_001204527.1 | 7.53305439 | -4.5376501 | 1.90306646 | -2.3843887 |
| 19078 | NM_198089.2 | 14.5944037 | -5.4919563 | 2.30355143 | -2.3841258 |
| 19079 | NM_017490.3 | 14.7891183 | -5.5115213 | 2.31185254 | -2.384028 |
| 19080 | NM_052906.4 | 158.08441 | -3.6563323 | 1.53379616 | -2.383845 |
| 19081 | NR_130138.1 | 7.20670832 | -4.4784336 | 1.87887792 | -2.3835681 |
| 19082 | NR_120363.1 | 51.1243357 | 2.02441364 | 0.84939709 | 2.38335363 |
| 19083 | NR_038932.1 | 7.34032024 | 4.19086918 | 1.75848276 | 2.38323018 |
| 19084 | NM_002441.4 | 6.46942394 | -4.3123741 | 1.80954072 | -2.3831319 |
| 19085 | NM_024680.3 | 4.52243042 | -3.7916656 | 1.5911608 | -2.3829556 |
| 19086 | NM_178837.4 | 19.0734313 | -5.8774662 | 2.46689594 | -2.3825351 |
| 19087 | NM_001311175.1 | 38.27486 | 1.56080832 | 0.65519305 | 2.38221135 |
| 19088 | NM_021785.4 | 16.417743 | -5.6619909 | 2.37678173 | -2.382209 |
| 19089 | NR_104051.1 | 5.90922684 | -4.1984693 | 1.76243057 | -2.3822041 |
| 19090 | NR_134656.1 | 5.93299972 | -4.1945408 | 1.76091724 | -2.3820204 |
| 19091 | NM_001256214.1 | 21.7139783 | -6.063892 | 2.5457348 | -2.381981 |

| | | | | | |
|---|---|---|---|---|---|
| 19092 | NM_001281773.2 | 19.0972073 | -5.8774014 | 2.46776909 | -2.3816659 |
| 19093 | NM_001313992.1 | 4.95802561 | -3.9354023 | 1.65237591 | -2.3816629 |
| 19094 | NM_001304464.1 | 5.93023539 | -4.1751611 | 1.75307574 | -2.3816205 |
| 19095 | NM_001284235.1 | 4.16722532 | -3.6732243 | 1.54240215 | -2.3814958 |
| 19096 | NM_021038.4 | 20.1226257 | -5.9538528 | 2.50030896 | -2.3812468 |
| 19097 | NM_001251874.1 | 3.93287112 | -3.6017788 | 1.51286083 | -2.3807734 |
| 19098 | NM_207352.3 | 164.055578 | 1.81656215 | 0.76301227 | 2.3807771 |
| 19099 | NM_182614.2 | 3.86115025 | -3.5777796 | 1.50279597 | -2.3807488 |
| 19100 | NM_005331.4 | 6.13507581 | -4.2321077 | 1.77783279 | -2.3804869 |
| 19101 | NR_110042.1 | 10.2230691 | 6.31710918 | 2.6537011 | 2.3804901 |
| 19102 | NM_001318831.1 | 19.3127909 | -5.8936781 | 2.47594675 | -2.3803735 |
| 19103 | NM_001167929.1 | 19.2021799 | -5.8858924 | 2.47285961 | -2.3801968 |
| 19104 | NM_000118.3 | 5.06010571 | -3.9524017 | 1.66069275 | -2.3799717 |
| 19105 | NM_172193.2 | 8.7870059 | -2.8696272 | 1.20595507 | -2.3795474 |
| 19106 | NM_001077637.1 | 6.80779227 | -3.3649679 | 1.41415336 | -2.3794929 |
| 19107 | NM_001281727.1 | 7.10559973 | -4.4455163 | 1.86834836 | -2.379383 |
| 19108 | NM_001127688.2 | 17.7615094 | -5.7743614 | 2.42698536 | -2.3792321 |
| 19109 | NM_001168467.1 | 18.1234963 | -5.8038183 | 2.4395669 | -2.3790363 |
| 19110 | NM_133342.2 | 15.1758746 | -5.5479609 | 2.33268657 | -2.3783568 |
| 19111 | NM_001293195.1 | 3.86736616 | -3.5756036 | 1.50341812 | -2.3783161 |
| 19112 | NR_119379.1 | 7.02674287 | 2.08653743 | 0.87739367 | 2.37810859 |
| 19113 | NR_028043.2 | 4.57873469 | -3.8226999 | 1.60760653 | -2.3778828 |
| 19114 | NM_001145414.3 | 6.88878159 | -3.3664451 | 1.41604743 | -2.3773533 |
| 19115 | NM_001258245.1 | 5.8779676 | -4.1835117 | 1.7600646 | -2.3769081 |
| 19116 | NM_175709.4 | 17.0093838 | -5.7101325 | 2.40237286 | -2.3768719 |
| 19117 | NM_000134.3 | 7.05992346 | 3.49446974 | 1.470252 | 2.37678285 |
| 19118 | NM_001318758.1 | 19.5965368 | -5.9149123 | 2.48869893 | -2.3767086 |
| 19119 | NM_181305.2 | 17.9515433 | -5.7901815 | 2.43623919 | -2.3766884 |
| 19120 | NR_033698.1 | 15.6314218 | -5.5907299 | 2.35234725 | -2.3766601 |
| 19121 | NM_001170938.1 | 14.5529274 | -5.4882072 | 2.30944909 | -2.376414 |
| 19122 | NM_001243197.1 | 4.11611404 | -3.6737591 | 1.54609947 | -2.3761466 |
| 19123 | NM_001278195.1 | 6.95436497 | -4.4074369 | 1.85503825 | -2.3759278 |
| 19124 | NM_001324164.1 | 5.03457307 | -3.9465694 | 1.66123464 | -2.3756845 |
| 19125 | NM_001297655.1 | 18.3137366 | -5.8162197 | 2.4485178 | -2.3754043 |
| 19126 | NM_014969.5 | 19.2948906 | -5.8908435 | 2.47993379 | -2.3754036 |
| 19127 | NR_110729.1 | 22.384989 | 1.41707303 | 0.59658948 | 2.37528998 |
| 19128 | NM_001102450.2 | 14.4682821 | -5.4797882 | 2.3070265 | -2.3752602 |
| 19129 | NM_001079882.1 | 14.9103773 | -5.523122 | 2.32532979 | -2.3751994 |
| 19130 | NM_004308.3 | 240.48197 | -1.3997275 | 0.58938003 | -2.3749151 |
| 19131 | NM_001039592.1 | 18.8228837 | -5.8583727 | 2.46683839 | -2.3748506 |
| 19132 | NR_135169.1 | 90.3204084 | 4.49043324 | 1.89086757 | 2.37480049 |
| 19133 | NR_135592.1 | 6.72530282 | -3.3281029 | 1.40174475 | -2.3742574 |
| 19134 | NR_027266.1 | 9.3802243 | 2.18827542 | 0.92169431 | 2.37418784 |

| | | | | | |
|---|---|---|---|---|---|
| 19135 | NM_001002758.1 | 13.1838996 | 8.64330663 | 3.64061289 | 2.37413504 |
| 19136 | NM_001034833.1 | 18.322643 | -5.8174486 | 2.45035495 | -2.3741248 |
| 19137 | NM_004676.2 | 13.1838996 | 8.64330663 | 3.64061289 | 2.37413504 |
| 19138 | NM_001330452.1 | 6.86539572 | -4.4085984 | 1.85736223 | -2.3735803 |
| 19139 | NM_001198645.1 | 42.073376 | 5.21003785 | 2.19513512 | 2.37344744 |
| 19140 | NR_120394.1 | 4.22580239 | -3.7093264 | 1.56294395 | -2.3732946 |
| 19141 | NM_017533.2 | 6.42452838 | 7.61158522 | 3.20767799 | 2.37292684 |
| 19142 | NM_001252151.1 | 14.5658896 | -5.4896434 | 2.31350953 | -2.372864 |
| 19143 | NM_183241.2 | 15.9523205 | -5.6198951 | 2.36841482 | -2.3728509 |
| 19144 | NM_001037582.2 | 486.265932 | -1.3843475 | 0.58347972 | -2.3725718 |
| 19145 | NM_003540.3 | 13.7942421 | 7.92153782 | 3.33903368 | 2.37240429 |
| 19146 | NR_028100.1 | 16.4896793 | -5.668497 | 2.38946234 | -2.3722897 |
| 19147 | NM_182507.2 | 4.79203211 | -3.8877293 | 1.63887863 | -2.3721887 |
| 19148 | NM_001100391.1 | 5.36197819 | -4.0281435 | 1.69822108 | -2.3719783 |
| 19149 | NM_001193529.2 | 4.24005125 | -3.6869855 | 1.55464449 | -2.371594 |
| 19150 | NM_001301778.1 | 15.0291191 | -5.5349056 | 2.33395896 | -2.3714665 |
| 19151 | NM_001080508.2 | 8.61824505 | -2.7364613 | 1.15401966 | -2.3712432 |
| 19152 | NM_001321032.1 | 17.7239054 | -5.7708266 | 2.43395174 | -2.37097 |
| 19153 | NM_005156.6 | 23.5244551 | -6.1786773 | 2.6067573 | -2.3702542 |
| 19154 | NM_001290355.2 | 14.8617262 | -5.5179843 | 2.32820523 | -2.3700593 |
| 19155 | NM_021074.4 | 183.354738 | -3.6724106 | 1.54952573 | -2.3700224 |
| 19156 | NM_001301009.1 | 18.977325 | -5.8684535 | 2.47620236 | -2.369941 |
| 19157 | NR_028136.2 | 6.4649495 | -4.3126359 | 1.81993528 | -2.3696644 |
| 19158 | NM_001130157.2 | 7.55582374 | -4.5486416 | 1.91984055 | -2.3692809 |
| 19159 | NM_001206648.1 | 53.9054346 | 3.12800663 | 1.32026739 | 2.36922206 |
| 19160 | NM_001178138.1 | 16.5295499 | -5.6707172 | 2.39396436 | -2.3687559 |
| 19161 | NM_001199480.1 | 18.7336864 | -5.8481702 | 2.46915196 | -2.3684934 |
| 19162 | NM_001308217.1 | 13.0742529 | 8.6225552 | 3.64075697 | 2.36834132 |
| 19163 | NM_001003828.2 | 4.46085427 | -3.7908738 | 1.60090718 | -2.3679535 |
| 19164 | NM_005468.2 | 12.7974695 | 3.16665598 | 1.33743274 | 2.36771233 |
| 19165 | NM_001005366.1 | 14.3570218 | -5.4686425 | 2.31003282 | -2.3673441 |
| 19166 | NM_203400.4 | 7.190803 | -4.4647432 | 1.88597918 | -2.3673343 |
| 19167 | NM_001303282.1 | 18.1981023 | -5.8078135 | 2.45360532 | -2.3670529 |
| 19168 | NM_020633.3 | 4.52781733 | -3.7818164 | 1.59790792 | -2.3667299 |
| 19169 | NR_046253.1 | 16.4623049 | -5.6657516 | 2.39393245 | -2.3667132 |
| 19170 | NM_001198593.1 | 5.39878635 | -4.0675607 | 1.71891739 | -2.3663503 |
| 19171 | NM_006404.4 | 5.12587037 | -3.992736 | 1.68753388 | -2.3660183 |
| 19172 | NR_027634.1 | 4.65280748 | -3.8553846 | 1.62964844 | -2.3657769 |
| 19173 | NR_109921.1 | 16.6517732 | -5.6826659 | 2.40202372 | -2.3657826 |
| 19174 | NM_001006932.2 | 16.9208255 | -5.7041904 | 2.41132729 | -2.3655811 |
| 19175 | NM_198156.2 | 192.340232 | 1.43522136 | 0.60680858 | 2.36519622 |
| 19176 | NR_029431.1 | 15.2057949 | -5.5511614 | 2.34709348 | -2.3651216 |
| 19177 | NM_001324025.1 | 18.2716513 | -5.8132889 | 2.45815524 | -2.364899 |

| | | | | | |
|---|---|---|---|---|---|
| 19178 | NM_015576.2 | 20.3441548 | -5.9701797 | 2.52459779 | -2.3648043 |
| 19179 | NR_002920.1 | 4.17719978 | -3.687345 | 1.55930758 | -2.3647323 |
| 19180 | NM_016357.4 | 42.2296604 | -2.8400264 | 1.20136204 | -2.3640054 |
| 19181 | NR_004380.1 | 4.06103075 | -3.6343446 | 1.53735517 | -2.3640241 |
| 19182 | NR_028344.1 | 38.2239707 | -1.6650751 | 0.70433427 | -2.3640411 |
| 19183 | NM_000360.3 | 10.8442999 | -4.0177561 | 1.69961515 | -2.3639211 |
| 19184 | NM_002699.3 | 29.8035917 | -2.9894277 | 1.26473771 | -2.3636741 |
| 19185 | NM_001286829.1 | 25.836215 | -6.3119295 | 2.6704822 | -2.3635917 |
| 19186 | NM_015286.5 | 46.3004145 | -3.4494966 | 1.45970069 | -2.3631533 |
| 19187 | NM_001278641.1 | 4.74260655 | -3.8742099 | 1.63960326 | -2.3628948 |
| 19188 | NM_021238.2 | 14.1127024 | -5.4439522 | 2.30409776 | -2.3627262 |
| 19189 | NM_080650.3 | 10.9306274 | -3.2442228 | 1.37319568 | -2.362535 |
| 19190 | NM_016821.2 | 18.3113224 | -5.8155803 | 2.46165083 | -2.3624716 |
| 19191 | NM_001005161.3 | 12.7961379 | 8.60030815 | 3.64091376 | 2.36212905 |
| 19192 | NM_001010876.1 | 12.7961379 | 8.60030815 | 3.64091376 | 2.36212905 |
| 19193 | NM_001323901.1 | 12.7961379 | 8.60030815 | 3.64091376 | 2.36212905 |
| 19194 | NR_037181.1 | 12.7961379 | 8.60030815 | 3.64091376 | 2.36212905 |
| 19195 | NM_006915.2 | 86.5858872 | -2.3076711 | 0.97697468 | -2.3620582 |
| 19196 | NR_102267.1 | 18.2107219 | -5.8090474 | 2.45948465 | -2.3618962 |
| 19197 | NM_001178009.2 | 17.5799009 | -5.7593421 | 2.43850468 | -2.3618335 |
| 19198 | NM_015831.2 | 5.39743383 | -4.0574752 | 1.71796438 | -2.3617924 |
| 19199 | NM_182486.1 | 5.04109609 | -3.9690313 | 1.68056639 | -2.3617224 |
| 19200 | NR_045214.1 | 12.6941538 | -2.4056459 | 1.01863382 | -2.3616395 |
| 19201 | NR_027514.2 | 18.1695548 | -5.8042894 | 2.45786975 | -2.3615122 |
| 19202 | NM_001318198.1 | 4.22272205 | -3.7030388 | 1.56812867 | -2.3614381 |
| 19203 | NR_104096.1 | 28.6684133 | 1.41038214 | 0.59736703 | 2.36099764 |
| 19204 | NR_047694.1 | 16.6831167 | -5.6844176 | 2.40784002 | -2.3607954 |
| 19205 | NM_006385.4 | 19.6808262 | -5.9204056 | 2.50796834 | -2.3606381 |
| 19206 | NR_003672.2 | 71.5788991 | 1.56569089 | 0.66332889 | 2.36035382 |
| 19207 | NR_073179.1 | 4.8309425 | -3.8757037 | 1.64201107 | -2.3603396 |
| 19208 | NM_001099639.1 | 17.1041878 | -5.7200436 | 2.42351276 | -2.3602284 |
| 19209 | NM_001322844.1 | 18.2519077 | -5.8111679 | 2.46217592 | -2.3601758 |
| 19210 | NR_136709.1 | 18.7198649 | -5.8470473 | 2.47799958 | -2.3595836 |
| 19211 | NM_003787.4 | 21.4270417 | -6.041764 | 2.56069914 | -2.3594197 |
| 19212 | NM_001010926.3 | 20.8469386 | -3.261774 | 1.38252359 | -2.3592899 |
| 19213 | NM_001287239.1 | 4.54195228 | -3.8039099 | 1.61238825 | -2.3591774 |
| 19214 | NM_001321968.1 | 20.1550293 | -5.9560233 | 2.52472312 | -2.3590798 |
| 19215 | NR_024362.1 | 3.80031793 | -3.5525895 | 1.50599861 | -2.3589594 |
| 19216 | NM_024312.4 | 273.396068 | -2.9173679 | 1.23687405 | -2.358662 |
| 19217 | NM_001102667.2 | 6.84166907 | -4.3953139 | 1.86352814 | -2.3585981 |
| 19218 | NM_001321173.1 | 3.87381286 | -3.5784191 | 1.51727782 | -2.3584469 |
| 19219 | NM_001323291.1 | 3.88704197 | -3.5685011 | 1.51313823 | -2.3583444 |
| 19220 | NR_138042.1 | 23.2987578 | 2.57354763 | 1.09147279 | 2.35786695 |

| | | | | | |
|---|---|---|---|---|---|
| 19221 | NM_001308375.1 | 4.67267404 | -3.8357236 | 1.62702007 | -2.3575146 |
| 19222 | NM_007042.4 | 16.4639421 | -5.6654008 | 2.40312604 | -2.357513 |
| 19223 | NM_001289156.1 | 3.82839928 | -3.5705572 | 1.5148263 | -2.3570737 |
| 19224 | NM_173489.4 | 9.92933046 | 6.43777765 | 2.73132661 | 2.35701495 |
| 19225 | NM_006539.3 | 6.24417214 | -4.2524374 | 1.80420282 | -2.3569619 |
| 19226 | NM_001127464.2 | 16.8486897 | -2.9163714 | 1.23746651 | -2.3567276 |
| 19227 | NR_038252.2 | 18.5790442 | 1.57208237 | 0.66723467 | 2.35611615 |
| 19228 | NM_001320952.1 | 17.343252 | -5.736956 | 2.43494414 | -2.3560935 |
| 19229 | NM_153836.3 | 25.3525475 | -2.9642993 | 1.25825795 | -2.3558756 |
| 19230 | NM_001161013.2 | 21.6889767 | 3.98079281 | 1.68989227 | 2.35564887 |
| 19231 | NR_135131.1 | 8.7277107 | -2.8141333 | 1.19485544 | -2.3552082 |
| 19232 | NM_004292.2 | 28.4718253 | -2.7034148 | 1.14789238 | -2.3551118 |
| 19233 | NM_001048195.2 | 14.951116 | -5.5263074 | 2.34654623 | -2.3550814 |
| 19234 | NR_027669.1 | 18.2286561 | -5.8087451 | 2.46656809 | -2.3549908 |
| 19235 | NM_001193272.1 | 63.6121074 | -1.4942162 | 0.6345082 | -2.3549203 |
| 19236 | NM_001270699.1 | 16.7757468 | -5.6892834 | 2.41627962 | -2.3545633 |
| 19237 | NM_001321784.1 | 16.8628347 | -5.6994587 | 2.42058409 | -2.3545799 |
| 19238 | NM_001330499.1 | 6.10864912 | -3.1783045 | 1.35011993 | -2.3540905 |
| 19239 | NM_001258032.1 | 4.1120764 | -3.6702998 | 1.55915583 | -2.3540301 |
| 19240 | NR_103720.1 | 4.52787684 | -3.8125914 | 1.61969376 | -2.3538964 |
| 19241 | NM_205843.2 | 14.6879641 | -5.5006264 | 2.33700914 | -2.3537034 |
| 19242 | NM_033127.2 | 4.10124219 | -3.6734009 | 1.56093488 | -2.3533339 |
| 19243 | NM_198590.2 | 5.48600876 | -4.084741 | 1.73575224 | -2.3532973 |
| 19244 | NM_001242462.1 | 5.57280458 | 7.39972815 | 3.14451903 | 2.35321462 |
| 19245 | NM_001324318.1 | 16.7500052 | -5.6880933 | 2.4174632 | -2.3529183 |
| 19246 | NM_001198806.1 | 4.14657577 | -3.6817078 | 1.56489833 | -2.3526818 |
| 19247 | NM_001320362.1 | 17.9789171 | -5.7889734 | 2.46072287 | -2.3525499 |
| 19248 | NM_001031743.2 | 16.4981635 | -2.8802833 | 1.22436105 | -2.3524787 |
| 19249 | NM_001302891.1 | 5.41249451 | -4.049314 | 1.72128544 | -2.3524942 |
| 19250 | NM_183420.1 | 19.5674647 | -5.911016 | 2.51271937 | -2.3524378 |
| 19251 | NR_033966.1 | 52.7192772 | 2.89634184 | 1.23125125 | 2.35235646 |
| 19252 | NM_001242897.1 | 17.7153366 | -5.7682342 | 2.4521405 | -2.3523261 |
| 19253 | NR_036662.1 | 17.9461559 | -5.790089 | 2.46168904 | -2.3520798 |
| 19254 | NM_001308396.1 | 4.71673216 | -3.8638461 | 1.64320991 | -2.3514014 |
| 19255 | NM_006898.4 | 157.975886 | -3.654338 | 1.55410103 | -2.351416 |
| 19256 | NM_001289097.1 | 4.74269489 | -3.8834721 | 1.65182876 | -2.3510137 |
| 19257 | NM_001282851.1 | 5.13735616 | -3.9764178 | 1.69175671 | -2.3504667 |
| 19258 | NM_001305134.1 | 7.35874354 | -4.5088694 | 1.9188769 | -2.3497439 |
| 19259 | NM_001320605.1 | 12.4083761 | 8.55598809 | 3.64123336 | 2.34975 |
| 19260 | NR_045566.1 | 5.30044806 | -4.0172779 | 1.7096858 | -2.3497171 |
| 19261 | NR_135179.1 | 11.3092483 | 6.50552889 | 2.76865349 | 2.34970859 |
| 19262 | NM_001163942.1 | 24.4014325 | 2.15037638 | 0.91517967 | 2.34967675 |
| 19263 | NR_111974.1 | 14.3964012 | -5.4725722 | 2.32920325 | -2.3495469 |

| | | | | | |
|---|---|---|---|---|---|
| 19264 | NM_173157.2 | 14.004913 | -5.4323722 | 2.31226803 | -2.3493696 |
| 19265 | NM_001282666.1 | 16.271431 | -5.6465778 | 2.40364763 | -2.3491704 |
| 19266 | NM_001301108.1 | 19.9279329 | -5.9367007 | 2.52724616 | -2.3490789 |
| 19267 | NM_001144891.1 | 5.54299736 | -4.080813 | 1.73745913 | -2.3487246 |
| 19268 | NM_001300758.1 | 18.1809368 | -5.8060262 | 2.47202658 | -2.3486908 |
| 19269 | NM_001271052.1 | 6.9248882 | 7.69814399 | 3.277749 | 2.34860692 |
| 19270 | NM_001079534.1 | 17.7551006 | -5.771745 | 2.45766129 | -2.3484705 |
| 19271 | NM_001318495.1 | 31.9868087 | 3.27691002 | 1.39535909 | 2.34843493 |
| 19272 | NM_001009186.1 | 19.0364412 | -4.884779 | 2.08077261 | -2.3475795 |
| 19273 | NM_001142293.1 | 12.4205402 | 8.54822086 | 3.64129038 | 2.34758011 |
| 19274 | NM_001204300.1 | 12.4205402 | 8.54822086 | 3.64129038 | 2.34758011 |
| 19275 | NM_001290058.1 | 12.4205402 | 8.54822086 | 3.64129038 | 2.34758011 |
| 19276 | NM_001328124.1 | 12.4205402 | 8.54822086 | 3.64129038 | 2.34758011 |
| 19277 | NR_047588.1 | 12.4205402 | 8.54822086 | 3.64129038 | 2.34758011 |
| 19278 | NM_001330759.1 | 8.47133765 | -4.6971766 | 2.00090913 | -2.3475212 |
| 19279 | NM_012269.2 | 21.086093 | 2.8484303 | 1.21345069 | 2.34738034 |
| 19280 | NM_001030005.2 | 3.9393948 | -3.5934341 | 1.53088995 | -2.3472844 |
| 19281 | NM_001135917.1 | 15.273421 | -5.5577584 | 2.3682124 | -2.3468159 |
| 19282 | NM_001278245.1 | 14.2771521 | -5.4609474 | 2.32710401 | -2.3466709 |
| 19283 | NM_145054.4 | 17.5769395 | -5.759644 | 2.45439535 | -2.3466651 |
| 19284 | NR_134665.1 | 45.0436544 | 3.48673689 | 1.48582584 | 2.34666594 |
| 19285 | NR_034104.1 | 5.05202461 | -3.9548223 | 1.68538278 | -2.3465425 |
| 19286 | NM_001168682.1 | 22.2012499 | -2.7638261 | 1.17785638 | -2.3464882 |
| 19287 | NM_001199984.1 | 19.7644516 | -5.9276854 | 2.52636289 | -2.3463317 |
| 19288 | NR_110586.1 | 16.0663629 | 3.01332141 | 1.28427749 | 2.34631646 |
| 19289 | NM_021988.5 | 4.99407657 | -3.9513643 | 1.68409381 | -2.3462852 |
| 19290 | NM_001146340.1 | 8.64046617 | 3.84704886 | 1.63972934 | 2.3461487 |
| 19291 | NM_001288704.1 | 32.2183238 | 2.09369393 | 0.89245647 | 2.34598998 |
| 19292 | NM_052972.2 | 42.3816217 | 1.75844234 | 0.74955165 | 2.34599223 |
| 19293 | NR_138421.1 | 43.8728531 | -2.4181788 | 1.03092335 | -2.3456437 |
| 19294 | NM_004445.5 | 15.3749429 | -5.5672865 | 2.37355389 | -2.3455488 |
| 19295 | NM_001347720.1 | 5.63517423 | -4.1165665 | 1.75522477 | -2.3453216 |
| 19296 | NM_001271905.1 | 17.5820489 | -5.757203 | 2.45491805 | -2.3451712 |
| 19297 | NM_001322802.1 | 4.90592621 | -3.9129621 | 1.66873627 | -2.3448655 |
| 19298 | NM_001318359.1 | 19.4140773 | -5.9022168 | 2.51736436 | -2.3446017 |
| 19299 | NR_110141.1 | 7.23044398 | 3.01829605 | 1.28735641 | 2.3445691 |
| 19300 | NR_134494.1 | 4.8569932 | -3.9054467 | 1.66577938 | -2.3445162 |
| 19301 | NR_027405.1 | 49.4522811 | -2.3466218 | 1.00093055 | -2.3444401 |
| 19302 | NM_174969.2 | 19.8570723 | -5.9328108 | 2.53089353 | -2.3441566 |
| 19303 | NM_004291.3 | 5.69200267 | -4.1247177 | 1.7596931 | -2.3439984 |
| 19304 | NM_138777.4 | 17.5991731 | -5.7590598 | 2.4570824 | -2.3438611 |
| 19305 | NR_033142.1 | 18.7440563 | -5.8492968 | 2.49586573 | -2.3435944 |
| 19306 | NM_000304.3 | 17.8057074 | -5.7766236 | 2.46489573 | -2.343557 |

| | | | | | |
|---|---|---|---|---|---|
| 19307 | NM_001308244.1 | 4.47074357 | -3.7729244 | 1.61030254 | -2.342991 |
| 19308 | NM_024519.3 | 266.083135 | -3.2872353 | 1.403106 | -2.3428275 |
| 19309 | NM_001280561.1 | 4.67717857 | -3.8472411 | 1.64234895 | -2.3425235 |
| 19310 | NM_001272087.1 | 6.65241729 | -3.2286309 | 1.37847583 | -2.3421745 |
| 19311 | NM_006216.3 | 14.4273265 | -5.4756676 | 2.33787377 | -2.3421571 |
| 19312 | NM_001308431.1 | 14.2774819 | -5.4598838 | 2.33119269 | -2.3420989 |
| 19313 | NR_135298.1 | 3.76260388 | -3.5254036 | 1.50538738 | -2.3418581 |
| 19314 | NM_013439.2 | 4.28844872 | -3.7040429 | 1.58210136 | -2.3412172 |
| 19315 | NM_001304508.1 | 4.8573236 | -3.8951747 | 1.66411199 | -2.3406926 |
| 19316 | NR_138466.1 | 14.2865732 | -5.4613723 | 2.33331475 | -2.3406068 |
| 19317 | NM_007345.3 | 13.9034886 | -5.4221996 | 2.31664928 | -2.3405354 |
| 19318 | NM_001330529.1 | 4.42030725 | -3.7433903 | 1.59951081 | -2.3403345 |
| 19319 | NM_001301078.1 | 4.17997324 | -3.6815329 | 1.57324769 | -2.3400847 |
| 19320 | NR_104400.2 | 7.2685481 | 6.44124487 | 2.75271483 | 2.33996083 |
| 19321 | NM_001331034.1 | 15.0213815 | -5.533684 | 2.36516751 | -2.3396584 |
| 19322 | NM_007283.6 | 17.9505452 | -5.7868898 | 2.47336135 | -2.3396864 |
| 19323 | NM_181441.2 | 18.5082117 | -5.8327155 | 2.49296752 | -2.3396677 |
| 19324 | NR_040027.1 | 4.06070034 | -3.6536512 | 1.56158521 | -2.3397066 |
| 19325 | NM_014924.4 | 102.513343 | -1.1223952 | 0.47978634 | -2.3393647 |
| 19326 | NM_199133.3 | 14.254796 | -5.4581615 | 2.33319155 | -2.3393542 |
| 19327 | NM_003409.4 | 18.6834362 | -5.8466721 | 2.49934963 | -2.3392774 |
| 19328 | NM_001290264.1 | 14.5547407 | -5.4886513 | 2.34677856 | -2.3388024 |
| 19329 | NM_001105574.1 | 5.58214607 | -4.1133115 | 1.75885045 | -2.3386363 |
| 19330 | NM_198510.2 | 8.08040091 | 5.35653074 | 2.29068123 | 2.33840077 |
| 19331 | NM_001318896.1 | 17.7301949 | -5.7686946 | 2.46697345 | -2.3383691 |
| 19332 | NM_198217.2 | 4.58022664 | -3.8246811 | 1.63582529 | -2.3380743 |
| 19333 | NM_001318162.1 | 6.3534646 | -4.2831008 | 1.83211611 | -2.3377889 |
| 19334 | NM_004165.2 | 26.0311577 | -3.4397756 | 1.47139752 | -2.3377609 |
| 19335 | NM_001322498.1 | 60.7616755 | 2.01125049 | 0.86039205 | 2.33759772 |
| 19336 | NM_001347865.1 | 4.7741507 | -3.8838373 | 1.66149356 | -2.3375578 |
| 19337 | NM_002829.3 | 14.5658272 | -5.4897784 | 2.34896591 | -2.3371043 |
| 19338 | NM_001033575.1 | 13.7327699 | -2.6742862 | 1.14458939 | -2.336459 |
| 19339 | NM_012445.3 | 18.4410131 | -5.8287158 | 2.49519554 | -2.3359756 |
| 19340 | NM_001320322.1 | 9.07445767 | -3.7369313 | 1.59974529 | -2.3359539 |
| 19341 | NM_001300897.1 | 4.74711705 | -3.8579314 | 1.6515907 | -2.3358883 |
| 19342 | NM_001193654.1 | 17.4929785 | -5.7503063 | 2.46174625 | -2.3358648 |
| 19343 | NR_106782.1 | 9.7123257 | 2.83872499 | 1.21531117 | 2.33580095 |
| 19344 | NM_001101340.1 | 10.3481014 | -2.5174466 | 1.07843899 | -2.3343431 |
| 19345 | NR_120567.1 | 25.413154 | 1.95574362 | 0.83780686 | 2.33436095 |
| 19346 | NM_001256281.2 | 16.3453877 | -5.6540821 | 2.42266614 | -2.3338264 |
| 19347 | NM_001006656.3 | 14.1910626 | -5.451083 | 2.33583798 | -2.3336734 |
| 19348 | NM_014594.2 | 69.8791981 | -3.0952663 | 1.32637379 | -2.3336305 |
| 19349 | NM_001316668.1 | 16.05474 | -5.630092 | 2.41274535 | -2.3334796 |

| | | | | | |
|---|---|---|---|---|---|
| 19350 | NM_001329916.1 | 24.9522402 | 4.57751696 | 1.96216488 | 2.33289108 |
| 19351 | NM_001267728.1 | 14.0004589 | -5.4317923 | 2.32846532 | -2.3327778 |
| 19352 | NM_016379.3 | 10.0777637 | 3.63915926 | 1.56002344 | 2.33275935 |
| 19353 | NM_001286745.1 | 8.03694648 | -4.6200194 | 1.98051453 | -2.3327369 |
| 19354 | NM_004686.4 | 84.7551575 | -3.2224538 | 1.38143246 | -2.3326901 |
| 19355 | NM_014271.3 | 3.72193992 | -3.5060276 | 1.50300265 | -2.3326823 |
| 19356 | NR_026870.1 | 13.066031 | 2.14011548 | 0.91749761 | 2.33255702 |
| 19357 | NR_145451.1 | 40.7040331 | 2.09675158 | 0.89896789 | 2.33239875 |
| 19358 | NM_001164144.2 | 19.3733396 | -5.8968014 | 2.52839727 | -2.332229 |
| 19359 | NM_001321309.1 | 13.553337 | -5.3854382 | 2.30967593 | -2.3316857 |
| 19360 | NM_001114753.2 | 13.6954061 | -5.4007076 | 2.31631332 | -2.3315963 |
| 19361 | NR_126521.1 | 5.61317747 | -4.1097701 | 1.76280015 | -2.3313874 |
| 19362 | NM_001297732.1 | 17.4331317 | -5.7445484 | 2.46410948 | -2.3312878 |
| 19363 | NM_001017363.1 | 5.31930925 | -4.0279956 | 1.72783291 | -2.3312414 |
| 19364 | NM_001321837.1 | 6.2035324 | -3.20966 | 1.37691914 | -2.3310446 |
| 19365 | NM_001197027.1 | 3.88272525 | -3.5802291 | 1.53615491 | -2.3306433 |
| 19366 | NM_000927.4 | 5.22247965 | -4.0181044 | 1.72413694 | -2.3305019 |
| 19367 | NM_031424.5 | 18.4256224 | -5.8258679 | 2.49989399 | -2.330446 |
| 19368 | NR_036551.1 | 45.6117907 | 2.88341083 | 1.23762989 | 2.32978441 |
| 19369 | NM_001032383.1 | 13.5261156 | -5.3827564 | 2.31049214 | -2.3297012 |
| 19370 | NM_001330432.1 | 13.4769557 | -5.3775455 | 2.30853084 | -2.3294233 |
| 19371 | NM_001320882.1 | 87.782896 | -8.0781363 | 3.46812468 | -2.329252 |
| 19372 | NR_073048.1 | 17.3960128 | -5.7413113 | 2.46502871 | -2.3291052 |
| 19373 | NR_037931.1 | 11.9592591 | 2.2094782 | 0.9488544 | 2.32857454 |
| 19374 | NM_001278512.1 | 4.74709139 | -3.8599925 | 1.65769935 | -2.3285239 |
| 19375 | NR_126534.1 | 5.34977725 | -4.0556032 | 1.74171177 | -2.3285157 |
| 19376 | NM_153188.2 | 98.629763 | -2.66527 | 1.14466064 | -2.3284369 |
| 19377 | NM_001330685.1 | 6.86194559 | -4.4037755 | 1.89140343 | -2.3283111 |
| 19378 | NM_001346473.1 | 19.5411079 | -5.9100549 | 2.53924872 | -2.3274817 |
| 19379 | NM_001193600.1 | 17.5529113 | -5.754465 | 2.47242062 | -2.327462 |
| 19380 | NM_174952.2 | 8.10074572 | 5.27299149 | 2.2656227 | 2.32739171 |
| 19381 | NR_120378.1 | 15.7559881 | -2.4352243 | 1.04645214 | -2.3271244 |
| 19382 | NM_001297733.1 | 23.3793376 | -6.1667844 | 2.65029753 | -2.3268272 |
| 19383 | NM_018711.4 | 227.752196 | -2.6250139 | 1.12819885 | -2.3267298 |
| 19384 | NM_001162995.2 | 4.40136614 | -3.7766076 | 1.62317155 | -2.3266842 |
| 19385 | NM_001128843.2 | 13.2343902 | 2.78065111 | 1.19525809 | 2.32640225 |
| 19386 | NM_004173.2 | 4.61663441 | -3.838417 | 1.64992564 | -2.3264182 |
| 19387 | NM_031951.4 | 4.46715441 | 3.92885125 | 1.68881997 | 2.32638844 |
| 19388 | NM_001279355.1 | 11.7668276 | 8.46985045 | 3.64188317 | 2.32567879 |
| 19389 | NM_001316312.1 | 11.7668276 | 8.46985045 | 3.64188317 | 2.32567879 |
| 19390 | NM_001319207.1 | 11.7668276 | 8.46985045 | 3.64188317 | 2.32567879 |
| 19391 | NM_001329522.1 | 11.7668276 | 8.46985045 | 3.64188317 | 2.32567879 |
| 19392 | NM_001716.4 | 11.7668276 | 8.46985045 | 3.64188317 | 2.32567879 |

| | | | | | |
|---|---|---|---|---|---|
| 19393 | NM_014438.4 | 11.7668276 | 8.46985045 | 3.64188317 | 2.32567879 |
| 19394 | NM_017602.3 | 20.967917 | -6.0105345 | 2.5842964 | -2.3257914 |
| 19395 | NM_025142.1 | 11.7668276 | 8.46985045 | 3.64188317 | 2.32567879 |
| 19396 | NR_046575.1 | 11.7668276 | 8.46985045 | 3.64188317 | 2.32567879 |
| 19397 | NR_132393.1 | 11.7668276 | 8.46985045 | 3.64188317 | 2.32567879 |
| 19398 | NR_135162.1 | 4.23432487 | -3.6903174 | 1.58664929 | -2.3258557 |
| 19399 | NM_001130148.1 | 13.4532446 | -5.3751366 | 2.31125396 | -2.3256365 |
| 19400 | NM_001308046.1 | 17.7925661 | -4.7888015 | 2.05916757 | -2.3256007 |
| 19401 | NM_001024594.2 | 4.2814182 | -3.725227 | 1.60196272 | -2.3254143 |
| 19402 | NM_001012977.2 | 4.32165228 | -3.7472166 | 1.61159778 | -2.3251562 |
| 19403 | NM_001010989.2 | 18.9531552 | -5.8677147 | 2.52381875 | -2.3249351 |
| 19404 | NR_046290.1 | 3.93980734 | -3.6052029 | 1.55095665 | -2.3245027 |
| 19405 | NR_046346.1 | 15.590914 | -5.5869889 | 2.40370237 | -2.3243264 |
| 19406 | NM_001324315.1 | 15.702203 | -5.5945016 | 2.40726978 | -2.3240028 |
| 19407 | NM_014869.6 | 43.3416494 | -2.7655731 | 1.19007293 | -2.3238686 |
| 19408 | NM_004809.4 | 15.0100943 | -5.5319144 | 2.38050169 | -2.3238439 |
| 19409 | NM_001040283.2 | 11.6328526 | 8.46304282 | 3.64193619 | 2.3237757 |
| 19410 | NM_001317929.1 | 11.6328526 | 8.46304282 | 3.64193619 | 2.3237757 |
| 19411 | NM_001146333.1 | 15.8234769 | -5.606679 | 2.41277611 | -2.3237461 |
| 19412 | NR_126364.1 | 16.0158267 | -1.727144 | 0.74329594 | -2.323629 |
| 19413 | NR_038875.2 | 41.3062457 | 3.02582538 | 1.30221216 | 2.32360399 |
| 19414 | NM_001320341.1 | 4.95124865 | -3.9258029 | 1.68983866 | -2.3231821 |
| 19415 | NM_004103.4 | 15.7353923 | -5.5987105 | 2.4100706 | -2.3230484 |
| 19416 | NM_005114.2 | 28.5799733 | -2.7111223 | 1.16704734 | -2.3230611 |
| 19417 | NR_138122.1 | 13.7354094 | -5.4050335 | 2.32688803 | -2.3228593 |
| 19418 | NM_014800.10 | 14.6516893 | -5.4977076 | 2.3668327 | -2.3228121 |
| 19419 | NR_144385.1 | 5.45547617 | -4.0519728 | 1.7444357 | -2.3227986 |
| 19420 | NM_001006627.1 | 17.871255 | -5.7822507 | 2.48942533 | -2.3227251 |
| 19421 | NM_001012664.2 | 17.132967 | -5.7209212 | 2.4631875 | -2.3225683 |
| 19422 | NM_018253.3 | 6.1973846 | -4.2600095 | 1.834273 | -2.3224512 |
| 19423 | NR_130151.1 | 6.64678515 | -3.2948734 | 1.41883122 | -2.3222448 |
| 19424 | NM_001261448.1 | 18.0418599 | -5.7933447 | 2.49531709 | -2.3216868 |
| 19425 | NM_001123376.2 | 13.7665817 | -5.4076574 | 2.32938459 | -2.3214962 |
| 19426 | NM_153500.1 | 11.5416315 | 8.4534915 | 3.642011 | 2.32110543 |
| 19427 | NM_001265577.1 | 46.7908234 | 1.7269883 | 0.74417426 | 2.32067729 |
| 19428 | NM_001287033.1 | 14.7495382 | -5.5077467 | 2.37378378 | -2.3202394 |
| 19429 | NR_144508.1 | 12.466472 | 7.7763593 | 3.35161509 | 2.32018268 |
| 19430 | NM_001286246.1 | 16.6754808 | -5.6827832 | 2.44969288 | -2.3197942 |
| 19431 | NM_002754.4 | 14.8262135 | -5.5151135 | 2.37749184 | -2.3197192 |
| 19432 | NM_003563.3 | 14.8853125 | -5.5205152 | 2.38002503 | -2.3195198 |
| 19433 | NR_121658.1 | 289.656226 | 4.31800126 | 1.86179291 | 2.31927044 |
| 19434 | NM_001145404.1 | 39.4813918 | 3.6685099 | 1.58178002 | 2.31922888 |
| 19435 | NM_001130090.1 | 13.9548822 | -5.4276994 | 2.34047455 | -2.3190594 |

| | | | | | |
|---|---|---|---|---|---|
| 19436 | NR_136663.1 | 13.4926471 | -5.3786158 | 2.31946758 | -2.3189011 |
| 19437 | NM_139029.1 | 14.6454422 | -5.4964745 | 2.37069744 | -2.3185053 |
| 19438 | NM_001320267.1 | 5.97750517 | -4.1946496 | 1.80922355 | -2.3184806 |
| 19439 | NM_001308273.1 | 3.67942493 | -3.4987163 | 1.50908487 | -2.3184357 |
| 19440 | NM_080836.3 | 140.443463 | -3.2848075 | 1.4168286 | -2.3184227 |
| 19441 | NM_022337.2 | 6.97303174 | -3.3324169 | 1.43751645 | -2.3181765 |
| 19442 | NM_018968.3 | 5.04310182 | -3.962574 | 1.70975378 | -2.3176285 |
| 19443 | NM_001193552.1 | 15.4281815 | -5.571042 | 2.40401833 | -2.3173875 |
| 19444 | NM_173462.3 | 65.2409736 | 1.37926517 | 0.59517642 | 2.31740559 |
| 19445 | NM_000213.4 | 14.1640974 | -5.4495763 | 2.35174797 | -2.317245 |
| 19446 | NM_001297752.1 | 4.64490879 | -3.8303855 | 1.65302067 | -2.3172036 |
| 19447 | NM_173683.3 | 4.09654087 | -3.6647571 | 1.58161642 | -2.317096 |
| 19448 | NR_040021.1 | 29.3862419 | 1.20548587 | 0.52037877 | 2.31655468 |
| 19449 | NM_001017416.1 | 18.7977485 | -5.8521218 | 2.52707118 | -2.3157725 |
| 19450 | NR_104636.1 | 56.0285574 | 2.74952724 | 1.18741422 | 2.31555862 |
| 19451 | NM_182623.2 | 13.069438 | -2.4653151 | 1.06469097 | -2.3155218 |
| 19452 | NM_198849.2 | 34.2410669 | -2.822436 | 1.21911255 | -2.3151563 |
| 19453 | NR_136618.1 | 7.70793963 | -4.5711519 | 1.97448851 | -2.3151069 |
| 19454 | NM_001287339.1 | 5.81911774 | 7.47211394 | 3.22864244 | 2.31432068 |
| 19455 | NM_138319.3 | 17.0176607 | -5.7109234 | 2.46779204 | -2.3141834 |
| 19456 | NR_126056.1 | 14.6807415 | -5.5009784 | 2.37725944 | -2.314 |
| 19457 | NM_003485.3 | 3.91479292 | -3.6070173 | 1.55911035 | -2.3135099 |
| 19458 | NM_024120.4 | 15.9322999 | -5.6178698 | 2.42833808 | -2.3134628 |
| 19459 | NM_001320679.1 | 6.07815284 | -4.2232456 | 1.8255865 | -2.3133637 |
| 19460 | NR_024334.1 | 4.57635129 | -3.8200683 | 1.65132114 | -2.3133406 |
| 19461 | NM_001322028.1 | 5.41251384 | -4.0655727 | 1.75759357 | -2.3131472 |
| 19462 | NR_138480.1 | 6.75882106 | 5.15014103 | 2.22719775 | 2.31238606 |
| 19463 | NM_001040022.1 | 18.1815223 | -5.8048262 | 2.5108681 | -2.3118802 |
| 19464 | NM_016178.2 | 4.25731483 | -3.7058307 | 1.6030494 | -2.3117383 |
| 19465 | NM_005727.3 | 4.16131415 | -3.6608434 | 1.58364708 | -2.3116536 |
| 19466 | NM_001319946.1 | 14.3869274 | -5.4711793 | 2.36707241 | -2.3113696 |
| 19467 | NR_037852.1 | 24.5717009 | -5.2419261 | 2.26814306 | -2.3111091 |
| 19468 | NM_001277061.1 | 17.8384083 | -5.7778432 | 2.50019244 | -2.3109594 |
| 19469 | NM_001007226.1 | 16.7620711 | -5.6880554 | 2.46142151 | -2.3108823 |
| 19470 | NM_001297550.1 | 16.4301697 | 3.48285281 | 1.50715961 | 2.31087191 |
| 19471 | NR_110067.1 | 4.99676494 | -3.9600378 | 1.71365411 | -2.3108734 |
| 19472 | NM_001282352.1 | 19.7857063 | -3.1899224 | 1.38046635 | -2.3107571 |
| 19473 | NM_001321848.1 | 5.09953667 | -3.977249 | 1.72119189 | -2.3107528 |
| 19474 | NM_001282282.1 | 33.721455 | 3.32460946 | 1.43889467 | 2.3105301 |
| 19475 | NM_001204864.1 | 5.63271536 | 6.20228857 | 2.68443247 | 2.31046548 |
| 19476 | NM_001330220.1 | 18.3400223 | -5.816858 | 2.51763054 | -2.3104494 |
| 19477 | NR_052002.1 | 4.11675221 | -3.6503581 | 1.5800115 | -2.3103364 |
| 19478 | NR_026543.1 | 4.12599184 | -3.656248 | 1.58318798 | -2.3094213 |

| | | | | | |
|---|---|---|---|---|---|
| 19479 | NM_001146686.2 | 7.02364066 | -4.4430734 | 1.92398775 | -2.3093044 |
| 19480 | NM_001130679.2 | 30.8762797 | -2.8620796 | 1.2394102 | -2.309227 |
| 19481 | NR_021493.1 | 12.4588128 | 5.26781705 | 2.28141787 | 2.30901017 |
| 19482 | NM_001198973.1 | 17.2690377 | -5.730989 | 2.4820551 | -2.3089693 |
| 19483 | NM_178001.2 | 13.6973619 | -5.4009901 | 2.33918734 | -2.3089173 |
| 19484 | NM_178857.5 | 11.2749924 | 3.55437573 | 1.53944572 | 2.3088672 |
| 19485 | NM_033382.2 | 4.2035901 | -3.7091 | 1.6067021 | -2.3085175 |
| 19486 | NM_001128172.2 | 2.03464679 | 5.904898 | 2.55814983 | 2.30826902 |
| 19487 | NR_130744.1 | 5.06172603 | -3.9769199 | 1.72350353 | -2.3074626 |
| 19488 | NM_032446.2 | 18.9790829 | -5.8657257 | 2.54222619 | -2.3073186 |
| 19489 | NM_001284244.1 | 10.6567626 | 6.05398435 | 2.62395309 | 2.30719992 |
| 19490 | NM_182542.2 | 6.82779899 | 5.14281027 | 2.22909533 | 2.3071289 |
| 19491 | NM_001305141.1 | 15.6790273 | -5.5950869 | 2.42543565 | -2.3068379 |
| 19492 | NM_001007214.1 | 11.3783607 | -4.0756724 | 1.76683857 | -2.3067599 |
| 19493 | NM_001039706.2 | 17.5390567 | -3.5409628 | 1.53512787 | -2.306624 |
| 19494 | NM_001013258.1 | 5.23394075 | -4.0237743 | 1.74447457 | -2.3065824 |
| 19495 | NM_017417.1 | 5.1622055 | -3.9903653 | 1.72999103 | -2.3065815 |
| 19496 | NR_037923.1 | 17.4850576 | -5.7483571 | 2.49252296 | -2.3062404 |
| 19497 | NM_001323550.1 | 8.56133281 | -3.6534154 | 1.58429189 | -2.3060242 |
| 19498 | NM_001289824.1 | 12.9408769 | -5.319063 | 2.30669874 | -2.3059201 |
| 19499 | NR_125854.1 | 41.5992116 | 3.29233012 | 1.42790549 | 2.3057059 |
| 19500 | NM_001134878.1 | 4.65276235 | -3.8503853 | 1.67001242 | -2.3056028 |
| 19501 | NR_144393.1 | 7.97634379 | -4.6190917 | 2.00356642 | -2.3054348 |
| 19502 | NM_001168648.2 | 19.7364927 | -5.9231816 | 2.56979699 | -2.304922 |
| 19503 | NM_001286842.1 | 13.2467747 | -5.3527111 | 2.32231523 | -2.3049029 |
| 19504 | NM_001001557.3 | 58.2039271 | -2.5543231 | 1.10849652 | -2.3043131 |
| 19505 | NR_026743.1 | 10.5435451 | 3.54544004 | 1.53896644 | 2.30377995 |
| 19506 | NM_007226.2 | 4.32198268 | -3.732855 | 1.62043227 | -2.3036168 |
| 19507 | NM_213662.1 | 77.8627497 | -2.0109385 | 0.87298793 | -2.3035123 |
| 19508 | NM_032574.3 | 4.04345939 | -3.6220018 | 1.57252932 | -2.3032969 |
| 19509 | NM_001278248.1 | 4.55283184 | -3.8268773 | 1.66162702 | -2.3030904 |
| 19510 | NM_001270507.1 | 16.5283369 | -5.6690668 | 2.46154498 | -2.3030523 |
| 19511 | NR_121600.1 | 4.08674285 | -3.6517241 | 1.58562992 | -2.3030116 |
| 19512 | NM_031486.2 | 13.3658568 | -5.3658968 | 2.33008905 | -2.302872 |
| 19513 | NM_001172703.2 | 16.7840278 | -5.6895436 | 2.47076303 | -2.3027476 |
| 19514 | NM_001009181.1 | 11.113115 | 8.38697605 | 3.64254587 | 2.30250389 |
| 19515 | NM_001191002.1 | 11.113115 | 8.38697605 | 3.64254587 | 2.30250389 |
| 19516 | NR_027500.1 | 11.113115 | 8.38697605 | 3.64254587 | 2.30250389 |
| 19517 | NR_047477.2 | 11.113115 | 8.38697605 | 3.64254587 | 2.30250389 |
| 19518 | NR_047672.1 | 11.113115 | 8.38697605 | 3.64254587 | 2.30250389 |
| 19519 | NM_181801.3 | 3.8857348 | -3.5843194 | 1.55684318 | -2.3022996 |
| 19520 | NM_001265601.1 | 4.7914656 | -3.8917961 | 1.69060233 | -2.3020175 |
| 19521 | NR_024872.2 | 13.2042471 | -5.3480631 | 2.32330722 | -2.3019182 |

| | | | | | |
|---|---|---|---|---|---|
| 19522 | NM_015054.1 | 99.3096962 | -3.1643351 | 1.37484185 | -2.3015993 |
| 19523 | NM_001323309.1 | 131.237065 | 5.38905476 | 2.34168896 | 2.30135379 |
| 19524 | NM_181646.3 | 9.04757247 | 3.57853142 | 1.55519159 | 2.30102287 |
| 19525 | NM_153350.3 | 194.540016 | -3.665792 | 1.59316548 | -2.3009487 |
| 19526 | NR_049757.1 | 15.8508235 | -5.6075189 | 2.43703297 | -2.3009615 |
| 19527 | NM_001256651.1 | 5.04148253 | 7.25867552 | 3.15480719 | 2.30083016 |
| 19528 | NR_038866.1 | 32.2599988 | 1.17077215 | 0.50887403 | 2.30071112 |
| 19529 | NM_001288649.1 | 4.55788275 | 7.12276036 | 3.0960832 | 2.30057137 |
| 19530 | NM_014775.3 | 16.7862721 | -5.6927947 | 2.47509535 | -2.3000304 |
| 19531 | NM_018319.3 | 35.3253789 | -2.1234734 | 0.92330998 | -2.2998488 |
| 19532 | NM_001321082.1 | 21.888671 | -5.0667287 | 2.20340774 | -2.2994967 |
| 19533 | NM_001144962.1 | 16.416698 | -5.6592936 | 2.46142516 | -2.2991938 |
| 19534 | NR_046072.1 | 19.6199621 | 4.15903598 | 1.80898024 | 2.29910526 |
| 19535 | NR_024236.1 | 30.8837546 | 2.26886362 | 0.98691805 | 2.29893821 |
| 19536 | NM_006651.3 | 142.143788 | -2.4944045 | 1.08506481 | -2.298853 |
| 19537 | NM_004780.2 | 14.1132532 | -5.4432559 | 2.36917778 | -2.2975295 |
| 19538 | NM_182898.3 | 17.0557387 | -5.715545 | 2.48768886 | -2.2975321 |
| 19539 | NM_005376.4 | 5.01849065 | -3.9360475 | 1.71321669 | -2.2974604 |
| 19540 | NM_001304480.1 | 17.3919401 | -5.7444 | 2.50094436 | -2.2968924 |
| 19541 | NM_001284431.1 | 10.862712 | 8.36630105 | 3.64271718 | 2.2967199 |
| 19542 | NM_134263.2 | 16.5710466 | -5.6742409 | 2.47067177 | -2.2966389 |
| 19543 | NM_014841.2 | 16.3140884 | -5.6501101 | 2.46022033 | -2.296587 |
| 19544 | NM_001320357.1 | 4.39973151 | -3.759612 | 1.63719188 | -2.2963784 |
| 19545 | NM_181489.5 | 326.018601 | -0.9391499 | 0.40899504 | -2.2962379 |
| 19546 | NM_001243740.2 | 45.6626043 | 1.19864238 | 0.52201155 | 2.29619895 |
| 19547 | NM_007197.3 | 6.71236722 | -3.2704245 | 1.42427748 | -2.296199 |
| 19548 | NM_174959.3 | 9.76963883 | 3.94278738 | 1.71738012 | 2.29581519 |
| 19549 | NR_028594.1 | 6.82667177 | -3.3536466 | 1.46077702 | -2.2957964 |
| 19550 | NM_001174129.1 | 4.816865 | -3.9024236 | 1.70011896 | -2.2953827 |
| 19551 | NM_001204387.1 | 6.89031796 | -4.3930876 | 1.91389326 | -2.2953671 |
| 19552 | NR_027312.1 | 7.65596889 | 4.18295198 | 1.82239366 | 2.29530649 |
| 19553 | NM_001345881.1 | 191.771342 | 1.31605835 | 0.57343058 | 2.29506135 |
| 19554 | NM_001257135.1 | 18.3927604 | -4.8330011 | 2.10592963 | -2.294949 |
| 19555 | NM_206967.2 | 4.91499255 | -3.9056783 | 1.70195906 | -2.2948133 |
| 19556 | NM_001303477.1 | 5.05951671 | -3.9485536 | 1.72087451 | -2.2945041 |
| 19557 | NM_152519.3 | 16.1755049 | -5.6368144 | 2.45755845 | -2.2936644 |
| 19558 | NM_001144956.1 | 3.99423594 | -3.6031782 | 1.57095365 | -2.2936247 |
| 19559 | NM_007335.3 | 17.9623161 | -5.7908951 | 2.52520149 | -2.2932408 |
| 19560 | NM_024963.5 | 954.298852 | 2.67880439 | 1.1682815 | 2.29294429 |
| 19561 | NM_001204456.1 | 17.2608875 | -5.7290697 | 2.49862079 | -2.2928929 |
| 19562 | NM_170695.3 | 4.8342546 | -3.9078682 | 1.70465779 | -2.2924649 |
| 19563 | NM_005676.4 | 87.8124928 | -8.0783456 | 3.52393365 | -2.2924227 |
| 19564 | NR_046298.2 | 17.0528231 | -5.7146453 | 2.49287794 | -2.2923887 |

| | | | | | |
|---|---|---|---|---|---|
| 19565 | NM_001329911.1 | 16.2001468 | -5.6381679 | 2.45960144 | -2.2923095 |
| 19566 | NR_102747.1 | 18.0981945 | -5.7981871 | 2.5295046 | -2.2922224 |
| 19567 | NM_001284237.1 | 9.07126046 | -4.8078793 | 2.09757532 | -2.2921128 |
| 19568 | NM_017567.4 | 102.013096 | -3.3709765 | 1.47074256 | -2.2920235 |
| 19569 | NR_110759.1 | 12.9294677 | 2.6805493 | 1.16958694 | 2.2918769 |
| 19570 | NM_001286167.1 | 19.7204652 | -5.9231874 | 2.58455502 | -2.2917629 |
| 19571 | NR_132737.1 | 79.0387905 | -2.5471851 | 1.11159533 | -2.291468 |
| 19572 | NM_138771.3 | 26.6152416 | -2.8065147 | 1.22479925 | -2.2914079 |
| 19573 | NR_002206.3 | 636.704615 | -3.1804129 | 1.38836211 | -2.2907662 |
| 19574 | NM_005554.3 | 5.84852475 | 6.16090456 | 2.6895784 | 2.2906581 |
| 19575 | NM_001318384.1 | 16.5273146 | -5.6688344 | 2.47479985 | -2.2906234 |
| 19576 | NM_001289010.1 | 16.4809029 | 4.84487052 | 2.11518845 | 2.29051483 |
| 19577 | NM_018436.3 | 11.8127594 | 7.69789892 | 3.36097315 | 2.29037799 |
| 19578 | NM_014705.3 | 105.726682 | -3.424431 | 1.49520801 | -2.2902706 |
| 19579 | NM_001256164.1 | 43.855275 | 6.02259527 | 2.63041627 | 2.28959779 |
| 19580 | NM_181709.4 | 14.9194469 | -5.5210421 | 2.41154193 | -2.2894241 |
| 19581 | NR_033441.1 | 16.102562 | -5.6311284 | 2.4598085 | -2.2892548 |
| 19582 | NR_033805.1 | 192.824451 | -1.6723669 | 0.73055086 | -2.2891861 |
| 19583 | NR_135680.1 | 5.67689173 | 4.85330174 | 2.12018497 | 2.28909355 |
| 19584 | NM_001323621.1 | 16.6252956 | -5.6755059 | 2.47959362 | -2.2888855 |
| 19585 | NR_003276.2 | 4.8318793 | 3.27456976 | 1.43081855 | 2.2885989 |
| 19586 | NM_000638.3 | 4.29801581 | -3.7388354 | 1.63385352 | -2.2883541 |
| 19587 | NM_001135154.1 | 10.9371032 | -3.1763677 | 1.38817086 | -2.2881677 |
| 19588 | NM_001306199.1 | 4.77696943 | -3.8660368 | 1.68956988 | -2.2881781 |
| 19589 | NM_001317095.1 | 13.0125206 | -5.3262944 | 2.32772652 | -2.288196 |
| 19590 | NM_001323564.1 | 3.71053316 | -3.4970786 | 1.52829864 | -2.2882168 |
| 19591 | NM_031302.4 | 12.8181218 | -5.3055968 | 2.31900656 | -2.2878749 |
| 19592 | NR_125369.1 | 4.93847823 | -3.9110053 | 1.70953933 | -2.2877539 |
| 19593 | NR_030761.1 | 12.7220862 | -5.2946288 | 2.31464303 | -2.2874494 |
| 19594 | NM_007163.3 | 45.6174234 | 2.19716217 | 0.96061766 | 2.287239 |
| 19595 | NM_203355.2 | 13.0161038 | -5.3274006 | 2.32937959 | -2.287047 |
| 19596 | NM_001033855.2 | 17.7471436 | -5.7691673 | 2.52303301 | -2.2866 |
| 19597 | NM_001204911.1 | 3.90607961 | -3.591287 | 1.5706099 | -2.2865557 |
| 19598 | NR_049739.1 | 3.8396439 | -3.5440475 | 1.55001453 | -2.2864608 |
| 19599 | NR_110933.1 | 4.7712266 | -3.866821 | 1.69122598 | -2.2864011 |
| 19600 | NM_001134778.1 | 3.77252702 | -3.5504655 | 1.55290915 | -2.2863318 |
| 19601 | NM_033503.3 | 15.1766149 | -5.5477571 | 2.42666994 | -2.2861606 |
| 19602 | NM_052905.3 | 301.337752 | -3.6005269 | 1.57495311 | -2.2861169 |
| 19603 | NM_001320818.1 | 19.033842 | 4.54643226 | 1.98889888 | 2.28590418 |
| 19604 | NM_001145315.1 | 15.9093818 | -5.6132612 | 2.45609234 | -2.2854439 |
| 19605 | NR_073065.1 | 13.180703 | -5.3448454 | 2.33921846 | -2.2848851 |
| 19606 | NM_001330451.1 | 4.78684745 | -3.8748989 | 1.69602313 | -2.284697 |
| 19607 | NR_034138.1 | 5.709651 | -4.128965 | 1.80736873 | -2.2845172 |

| | | | | | |
|---|---|---|---|---|---|
| 19608 | NM_198129.2 | 4.25624767 | -3.7172724 | 1.62719077 | -2.2844724 |
| 19609 | NM_001289397.1 | 13.7912072 | -5.4096991 | 2.36815215 | -2.2843545 |
| 19610 | NR_110995.1 | 11.9444482 | 2.81982583 | 1.23441912 | 2.28433422 |
| 19611 | NM_001130518.1 | 6.96544729 | -4.4076791 | 1.92965967 | -2.2841743 |
| 19612 | NM_001131019.2 | 6.26523115 | -4.2770152 | 1.87250604 | -2.2841129 |
| 19613 | NR_027183.2 | 17.2172194 | -5.7265358 | 2.50754032 | -2.2837263 |
| 19614 | NM_001164385.1 | 23.7542005 | -5.2107771 | 2.28175634 | -2.2836694 |
| 19615 | NM_019595.3 | 15.6946227 | -5.5960168 | 2.45086546 | -2.2832819 |
| 19616 | NM_001281719.1 | 13.3202521 | -5.3602289 | 2.34774151 | -2.2831427 |
| 19617 | NM_001318368.1 | 16.3277046 | -5.6517958 | 2.47545186 | -2.283137 |
| 19618 | NM_001008712.2 | 18.631001 | -2.738813 | 1.19966548 | -2.2829806 |
| 19619 | NM_001304461.1 | 12.0797356 | 5.10797265 | 2.23741743 | 2.28297705 |
| 19620 | NM_004570.5 | 6.61483546 | 6.30182647 | 2.76032818 | 2.282999 |
| 19621 | NM_001330427.1 | 56.4471489 | 3.00966245 | 1.31876149 | 2.28218861 |
| 19622 | NM_001010922.2 | 122.131044 | 2.5748033 | 1.1282971 | 2.28202598 |
| 19623 | NM_014361.3 | 14.3792221 | -5.4708906 | 2.39769155 | -2.2817325 |
| 19624 | NM_001322253.1 | 17.1074039 | -5.717687 | 2.5059407 | -2.281653 |
| 19625 | NR_110370.1 | 20.0575404 | 2.5683415 | 1.12571687 | 2.28151639 |
| 19626 | NM_024705.2 | 5.29043377 | -4.0111681 | 1.75813522 | -2.2814901 |
| 19627 | NM_001330734.1 | 12.5342874 | -5.2728288 | 2.31122947 | -2.2813956 |
| 19628 | NM_080600.2 | 10.4695674 | 8.31132946 | 3.64318474 | 2.28133627 |
| 19629 | NR_131982.1 | 4.44330556 | -3.7705197 | 1.65289476 | -2.2811614 |
| 19630 | NM_001304969.1 | 19.7273929 | -5.9247656 | 2.59758623 | -2.2808735 |
| 19631 | NM_015689.4 | 16.2868754 | -5.6504449 | 2.47791287 | -2.2803243 |
| 19632 | NR_029411.1 | 25.4920326 | -2.5191351 | 1.10472552 | -2.2803267 |
| 19633 | NM_004588.4 | 32.0104191 | -1.5194523 | 0.66637938 | -2.2801611 |
| 19634 | NM_175611.2 | 14.4023662 | -5.4736114 | 2.40053056 | -2.2801673 |
| 19635 | NR_110764.1 | 32.1580632 | 2.37081599 | 1.03979736 | 2.28007503 |
| 19636 | NM_001202468.1 | 4.36551123 | -3.7397026 | 1.64041392 | -2.279731 |
| 19637 | NM_207361.5 | 39.5190164 | -2.2719494 | 0.99660504 | -2.2796888 |
| 19638 | NM_016359.4 | 17.276131 | -5.7317472 | 2.51441343 | -2.2795564 |
| 19639 | NM_001256545.1 | 55.5109102 | -3.1759994 | 1.39327763 | -2.2795165 |
| 19640 | NM_172208.2 | 4.75261058 | -3.8759055 | 1.70047204 | -2.2793115 |
| 19641 | NM_001031677.3 | 50.8817338 | -2.9080672 | 1.27586901 | -2.2792835 |
| 19642 | NR_103456.1 | 19.0193916 | -2.475038 | 1.08590604 | -2.2792377 |
| 19643 | NM_001145080.2 | 6.16541511 | -3.1861015 | 1.39789681 | -2.2792108 |
| 19644 | NM_001322217.1 | 18.1966576 | -5.8054917 | 2.54719969 | -2.2791663 |
| 19645 | NM_001007470.1 | 20.2106135 | -5.9606958 | 2.61537375 | -2.279099 |
| 19646 | NR_135136.1 | 4.0217561 | -3.6339462 | 1.59449641 | -2.2790558 |
| 19647 | NM_014455.3 | 68.3814219 | -2.1153632 | 0.92820586 | -2.2789807 |
| 19648 | NM_133370.3 | 60.8859592 | -2.708278 | 1.18841639 | -2.2788966 |
| 19649 | NM_014697.2 | 18.276602 | -5.8152369 | 2.55190138 | -2.2787859 |
| 19650 | NR_136681.1 | 16.4066265 | -5.6589794 | 2.48361106 | -2.2785288 |

| | | | | | |
|---|---|---|---|---|---|
| 19651 | NM_001318126.1 | 10.001823 | 6.32320972 | 2.77523834 | 2.27843844 |
| 19652 | NM_001171.5 | 7.28888215 | -3.4729365 | 1.52438011 | -2.2782615 |
| 19653 | NR_136183.1 | 16.0174558 | -5.6225461 | 2.46805015 | -2.2781329 |
| 19654 | NM_001003795.2 | 116.775008 | -1.9546949 | 0.85805546 | -2.278052 |
| 19655 | NM_001044369.2 | 4.73843242 | -3.8535585 | 1.6916008 | -2.2780543 |
| 19656 | NM_001282168.1 | 10.4594023 | 8.29905073 | 3.64329161 | 2.27789911 |
| 19657 | NM_001347698.1 | 10.4594023 | 8.29905073 | 3.64329161 | 2.27789911 |
| 19658 | NR_002159.1 | 10.4594023 | 8.29905073 | 3.64329161 | 2.27789911 |
| 19659 | NR_027012.1 | 10.4594023 | 8.29905073 | 3.64329161 | 2.27789911 |
| 19660 | NR_036632.1 | 5.78042778 | -4.1629299 | 1.82764087 | -2.2777614 |
| 19661 | NM_001303494.1 | 59.4792066 | -3.9803523 | 1.74752714 | -2.2777056 |
| 19662 | NM_001308304.1 | 13.1246883 | -5.3390539 | 2.34413437 | -2.2776228 |
| 19663 | NM_152375.2 | 4.52780908 | -3.8153674 | 1.67526497 | -2.277471 |
| 19664 | NR_034150.1 | 17.8317825 | 1.62991936 | 0.71570511 | 2.27736164 |
| 19665 | NM_001042553.2 | 15.4279701 | -5.5712432 | 2.44644658 | -2.2772797 |
| 19666 | NM_001206952.1 | 35.4200873 | 5.29499959 | 2.3252563 | 2.27716815 |
| 19667 | NM_206929.3 | 17.2799871 | -5.7351198 | 2.51856372 | -2.277139 |
| 19668 | NM_001287485.1 | 16.0290666 | -5.6261085 | 2.47072802 | -2.2771055 |
| 19669 | NR_024399.1 | 4.66253146 | -3.8367131 | 1.68501664 | -2.2769586 |
| 19670 | NM_001031712.2 | 44.937532 | -2.8284736 | 1.24228817 | -2.2768257 |
| 19671 | NM_001145356.1 | 17.1047801 | -5.7174477 | 2.51160854 | -2.2764088 |
| 19672 | NM_024676.4 | 30.498565 | -2.8267407 | 1.24208995 | -2.2757939 |
| 19673 | NM_015932.5 | 179.992586 | -3.0638408 | 1.34635975 | -2.275648 |
| 19674 | NM_021953.3 | 15.8991251 | -5.6127435 | 2.46654861 | -2.2755455 |
| 19675 | NM_016319.3 | 16.5333807 | -4.6771322 | 2.05548251 | -2.2754424 |
| 19676 | NM_001324191.1 | 4.33364287 | -3.7329845 | 1.64058912 | -2.2753927 |
| 19677 | NM_001013437.1 | 25.4721424 | -2.9412945 | 1.29277869 | -2.2751725 |
| 19678 | NM_001199515.1 | 14.7280881 | -5.5051181 | 2.41997565 | -2.2748651 |
| 19679 | NM_001198533.1 | 15.1156631 | -5.5388851 | 2.43490668 | -2.2747833 |
| 19680 | NM_001282169.1 | 5.405976 | -4.0651051 | 1.78703086 | -2.2747817 |
| 19681 | NM_004653.4 | 15.9239891 | -5.6147384 | 2.46836461 | -2.2746795 |
| 19682 | NM_001257990.1 | 15.40417 | -5.5699785 | 2.4489826 | -2.2744051 |
| 19683 | NM_001018112.1 | 6.93080501 | -3.3576295 | 1.47630714 | -2.2743434 |
| 19684 | NR_026925.1 | 3.67148628 | -3.4913462 | 1.5351082 | -2.2743323 |
| 19685 | NM_001253901.1 | 15.6497453 | -5.5894131 | 2.45762878 | -2.2743114 |
| 19686 | NM_001292017.1 | 18.9559086 | -5.8681798 | 2.58037525 | -2.2741575 |
| 19687 | NM_199426.1 | 12.5072057 | -5.2702254 | 2.31748363 | -2.2741155 |
| 19688 | NR_049743.1 | 13.8277193 | -5.4142945 | 2.38086563 | -2.2740865 |
| 19689 | NM_012236.3 | 15.7331204 | -5.597703 | 2.46161613 | -2.2739951 |
| 19690 | NM_001652.3 | 26.1062098 | 2.27685138 | 1.00128793 | 2.27392274 |
| 19691 | NR_145132.1 | 5.27612526 | -4.0148594 | 1.76566586 | -2.27385 |
| 19692 | NM_001324024.1 | 16.2453996 | -5.6418993 | 2.48197111 | -2.2731527 |
| 19693 | NM_001159772.1 | 3.8019412 | -3.5492959 | 1.56160433 | -2.2728522 |

| | | | | | |
|---|---|---|---|---|---|
| 19694 | NM_002153.2 | 6.80957781 | -4.3969593 | 1.9346044 | -2.272795 |
| 19695 | NM_001320772.1 | 15.0950233 | -5.5377212 | 2.4366699 | -2.2726596 |
| 19696 | NM_001024943.1 | 16.2993826 | -5.6467957 | 2.48497541 | -2.2723749 |
| 19697 | NR_027270.1 | 8.14150288 | 5.00821639 | 2.2046613 | 2.27164888 |
| 19698 | NM_145263.3 | 17.4166273 | -5.7449905 | 2.52913046 | -2.2715279 |
| 19699 | NM_001330629.1 | 17.5357317 | -5.7544996 | 2.53342297 | -2.2714326 |
| 19700 | NM_181313.3 | 13.9029082 | -5.4220494 | 2.38715894 | -2.2713399 |
| 19701 | NM_003654.5 | 57.7910221 | -2.466202 | 1.08600989 | -2.2708835 |
| 19702 | NM_001133.2 | 10.6175425 | 6.29605067 | 2.77258375 | 2.27082434 |
| 19703 | NM_000947.4 | 25.4631819 | -2.3049971 | 1.01508599 | -2.2707407 |
| 19704 | NM_001134400.1 | 10.1837925 | 8.27350354 | 3.6435169 | 2.27074658 |
| 19705 | NM_001242402.2 | 10.1837925 | 8.27350354 | 3.6435169 | 2.27074658 |
| 19706 | NM_001306207.1 | 18.413304 | -5.823026 | 2.56431927 | -2.2707882 |
| 19707 | NM_001279360.1 | 4.54237013 | -3.8138115 | 1.67966993 | -2.270572 |
| 19708 | NM_001211.5 | 73.8500078 | -3.0015923 | 1.32212698 | -2.2702754 |
| 19709 | NM_001323083.1 | 4.6362415 | -3.8138889 | 1.6799998 | -2.2701722 |
| 19710 | NR_023350.1 | 12.7847508 | -5.3015572 | 2.33532442 | -2.2701588 |
| 19711 | NM_001099220.2 | 94.4393877 | -3.2544121 | 1.43363168 | -2.2700476 |
| 19712 | NM_001242926.1 | 12.7552782 | -5.2977848 | 2.33394207 | -2.269887 |
| 19713 | NM_000935.2 | 77.4595434 | -3.2721057 | 1.44186217 | -2.2693609 |
| 19714 | NM_173207.2 | 4.44099382 | -3.7579521 | 1.65632843 | -2.2688447 |
| 19715 | NM_001284518.1 | 13.6186508 | -5.3913963 | 2.37639288 | -2.2687311 |
| 19716 | NM_001278685.1 | 16.1933314 | -5.6414688 | 2.48682406 | -2.2685436 |
| 19717 | NR_034121.1 | 37.7806077 | 1.87101527 | 0.82477831 | 2.26850686 |
| 19718 | NR_027261.1 | 16.0659508 | -5.6261132 | 2.48031969 | -2.2683016 |
| 19719 | NR_045563.1 | 5.39110193 | -4.0298326 | 1.77703696 | -2.2677258 |
| 19720 | NM_001174097.2 | 4.22056324 | -3.712945 | 1.63732708 | -2.2676868 |
| 19721 | NM_001305208.1 | 14.2676663 | -5.4582673 | 2.40708774 | -2.2675814 |
| 19722 | NM_006177.3 | 4.56868339 | -3.7981406 | 1.67502885 | -2.2675076 |
| 19723 | NR_102432.2 | 10.5109864 | 2.86626963 | 1.26409933 | 2.2674402 |
| 19724 | NR_135558.1 | 3.79158199 | -3.5502924 | 1.5658023 | -2.2673951 |
| 19725 | NR_073156.1 | 3.62340491 | -3.4877465 | 1.53839365 | -2.2671353 |
| 19726 | NM_021061.4 | 15.0852264 | -5.5384244 | 2.44315526 | -2.2669146 |
| 19727 | NM_001085354.2 | 50.2877408 | -2.2551515 | 0.99504974 | -2.2663706 |
| 19728 | NR_038444.1 | 56.6139121 | 1.92648022 | 0.8500975 | 2.26618738 |
| 19729 | NM_001318139.1 | 16.0516473 | -5.6262385 | 2.48299604 | -2.2659071 |
| 19730 | NM_001282382.1 | 4.56409592 | -3.8202484 | 1.68614853 | -2.2656654 |
| 19731 | NM_145697.2 | 22.6056507 | -2.7785732 | 1.2265039 | -2.2654418 |
| 19732 | NR_002312.1 | 7.1512699 | -4.4498011 | 1.96419931 | -2.2654529 |
| 19733 | NM_198839.2 | 132.655793 | -2.4168259 | 1.06687609 | -2.2653295 |
| 19734 | NM_001006618.1 | 3.48059515 | -3.418127 | 1.50902344 | -2.2651252 |
| 19735 | NM_031938.5 | 12.2280072 | -5.2374384 | 2.31294967 | -2.2643979 |
| 19736 | NM_024808.3 | 15.674979 | -5.5917831 | 2.46968244 | -2.2641709 |

| | | | | | |
|---|---|---|---|---|---|
| 19737 | NR_135178.1 | 20.6894498 | 3.93264441 | 1.73689228 | 2.2641844 |
| 19738 | NM_001287501.1 | 48.7369588 | -4.621284 | 2.04190248 | -2.2632246 |
| 19739 | NM_001329712.1 | 15.7281581 | -5.5956943 | 2.4724908 | -2.263181 |
| 19740 | NM_001101676.1 | 15.1965852 | -5.5468601 | 2.45126388 | -2.2628572 |
| 19741 | NR_110942.1 | 4.49850565 | -3.7948508 | 1.67708777 | -2.2627622 |
| 19742 | NR_027340.1 | 10.5920412 | -2.3895177 | 1.05604141 | -2.2627121 |
| 19743 | NM_001252386.1 | 16.5556756 | -5.6700518 | 2.50622526 | -2.2623871 |
| 19744 | NR_125831.1 | 10.6582577 | 6.48291735 | 2.8654976 | 2.26240544 |
| 19745 | NM_004571.4 | 15.8691357 | -5.6108346 | 2.48027209 | -2.2621851 |
| 19746 | NM_001199881.1 | 5.25490801 | 7.29775229 | 3.22690805 | 2.26153091 |
| 19747 | NR_120683.1 | 4.66242882 | -3.8439801 | 1.69982992 | -2.261391 |
| 19748 | NM_003419.4 | 5.34150192 | -4.0394584 | 1.78634818 | -2.261294 |
| 19749 | NR_033262.1 | 4.26916068 | -3.7047346 | 1.638504 | -2.261047 |
| 19750 | NM_016835.4 | 5.16034613 | -3.987248 | 1.76347014 | -2.2610238 |
| 19751 | NM_001193466.1 | 6.10085273 | -4.2273182 | 1.86974377 | -2.2609078 |
| 19752 | NM_001287599.1 | 13.146064 | -5.3407689 | 2.36242612 | -2.2607136 |
| 19753 | NM_207196.2 | 16.0930439 | -5.6286941 | 2.49005468 | -2.2604701 |
| 19754 | NM_001304564.1 | 14.6655428 | 3.59835824 | 1.59192121 | 2.26038715 |
| 19755 | NM_002448.3 | 373.014288 | 2.19626143 | 0.9716872 | 2.26025559 |
| 19756 | NR_144629.1 | 12.0452686 | 3.22590424 | 1.42737543 | 2.26002507 |
| 19757 | NM_005597.3 | 35.0778709 | -4.5348763 | 2.0068514 | -2.2596971 |
| 19758 | NM_001170703.1 | 5.40614913 | 7.36311088 | 3.25866638 | 2.25954732 |
| 19759 | NM_001004330.2 | 7.42284185 | 5.09370324 | 2.25444479 | 2.25940473 |
| 19760 | NM_030972.3 | 4.47542454 | -3.785827 | 1.67569437 | -2.2592586 |
| 19761 | NM_018096.4 | 13.0293376 | -5.3278174 | 2.3584492 | -2.2590342 |
| 19762 | NM_001304351.1 | 14.5672564 | 4.88193434 | 2.16124474 | 2.25885308 |
| 19763 | NM_017730.3 | 12.329365 | -5.2494999 | 2.32400762 | -2.2588135 |
| 19764 | NM_001289087.1 | 11.1590467 | 7.61492959 | 3.37141942 | 2.25867169 |
| 19765 | NM_017416.1 | 3.57268649 | -3.4485565 | 1.52681397 | -2.2586619 |
| 19766 | NR_135640.1 | 3.87111335 | -3.564059 | 1.57792888 | -2.2586943 |
| 19767 | NM_000842.4 | 14.2106832 | -5.45297 | 2.41444453 | -2.2584781 |
| 19768 | NM_001242395.1 | 3.7277003 | -3.520558 | 1.55888267 | -2.2583855 |
| 19769 | NM_176895.2 | 5.43128389 | -4.0757584 | 1.80470668 | -2.2584049 |
| 19770 | NM_199247.2 | 4.17413589 | -3.6808355 | 1.63011387 | -2.2580236 |
| 19771 | NM_003586.2 | 20.0137434 | -5.9429963 | 2.63201283 | -2.2579663 |
| 19772 | NM_018464.4 | 72.7055122 | -3.0681209 | 1.3589295 | -2.2577484 |
| 19773 | NR_024617.1 | 3.70905257 | -3.5255651 | 1.56171797 | -2.2574915 |
| 19774 | NR_131234.1 | 6.29015193 | -4.2757658 | 1.89431903 | -2.2571519 |
| 19775 | NR_135121.1 | 9.84433272 | 8.22476499 | 3.64395786 | 2.25709663 |
| 19776 | NM_001105564.1 | 15.6323292 | -5.587053 | 2.47544851 | -2.2569862 |
| 19777 | NM_031895.5 | 219.589416 | -0.9284821 | 0.41139278 | -2.2569237 |
| 19778 | NM_145049.4 | 110.375578 | -3.3517695 | 1.48519397 | -2.2567891 |
| 19779 | NM_005349.3 | 17.2951503 | -5.7344359 | 2.54110926 | -2.2566664 |

| | | | | | |
|---|---|---|---|---|---|
| 19780 | NM_001395.3 | 15.4555428 | -5.5707539 | 2.46866363 | -2.2565869 |
| 19781 | NM_181804.2 | 6.18910336 | -4.2372176 | 1.87773103 | -2.2565626 |
| 19782 | NM_024043.3 | 15.5634164 | -5.5799519 | 2.47341432 | -2.2559714 |
| 19783 | NM_001346392.1 | 12.562579 | -5.275615 | 2.33864747 | -2.2558402 |
| 19784 | NM_001142618.1 | 12.1195694 | -5.224296 | 2.31592214 | -2.2558168 |
| 19785 | NR_130916.1 | 4.3418606 | -3.7294833 | 1.65336145 | -2.2556975 |
| 19786 | NM_001257145.1 | 15.6962068 | -5.5937102 | 2.4805469 | -2.255031 |
| 19787 | NM_001347885.1 | 16.737839 | -5.6851981 | 2.52114174 | -2.2550093 |
| 19788 | NM_001164276.1 | 7.98444353 | -4.605036 | 2.04232149 | -2.2548047 |
| 19789 | NM_005759.5 | 270.485126 | -3.631376 | 1.61072067 | -2.2545039 |
| 19790 | NR_034083.2 | 19.4748817 | 1.33632772 | 0.59281552 | 2.25420501 |
| 19791 | NM_014654.3 | 1031.19686 | -1.8417839 | 0.81712399 | -2.2539834 |
| 19792 | NR_037839.1 | 10.2050396 | 2.61711272 | 1.16130293 | 2.25360038 |
| 19793 | NM_001286198.1 | 29.1831182 | -1.8481294 | 0.82013902 | -2.2534343 |
| 19794 | NM_080616.4 | 263.959701 | -3.6598087 | 1.6241412 | -2.2533808 |
| 19795 | NR_102338.1 | 7.61394504 | -4.5383276 | 2.01404636 | -2.2533382 |
| 19796 | NR_136332.1 | 32.2678904 | 2.97924206 | 1.32213775 | 2.25335224 |
| 19797 | NM_001310339.1 | 4.53329546 | -3.8133987 | 1.69283184 | -2.2526742 |
| 19798 | NM_001281726.1 | 3.51764021 | -3.4313068 | 1.52332184 | -2.252516 |
| 19799 | NM_199121.2 | 3.67418276 | -3.4956201 | 1.55186148 | -2.2525336 |
| 19800 | NR_077234.1 | 3.77719967 | -3.5513658 | 1.5766705 | -2.2524464 |
| 19801 | NM_001318781.1 | 41.8567952 | -2.7114111 | 1.20378577 | -2.2524034 |
| 19802 | NR_033341.1 | 72.5025054 | -2.1939351 | 0.9740483 | -2.2523884 |
| 19803 | NM_017438.4 | 15.6674285 | -5.5898798 | 2.4821611 | -2.2520214 |
| 19804 | NM_001306077.1 | 125.763661 | 1.28952484 | 0.5726209 | 2.25196957 |
| 19805 | NM_000914.4 | 9.80568967 | 8.20541944 | 3.64413704 | 2.25167697 |
| 19806 | NM_001005213.1 | 9.80568967 | 8.20541944 | 3.64413704 | 2.25167697 |
| 19807 | NM_001005496.1 | 9.80568967 | 8.20541944 | 3.64413704 | 2.25167697 |
| 19808 | NM_001178002.2 | 9.80568967 | 8.20541944 | 3.64413704 | 2.25167697 |
| 19809 | NM_001252634.1 | 9.80568967 | 8.20541944 | 3.64413704 | 2.25167697 |
| 19810 | NM_001256478.1 | 9.80568967 | 8.20541944 | 3.64413704 | 2.25167697 |
| 19811 | NM_001286157.1 | 9.80568967 | 8.20541944 | 3.64413704 | 2.25167697 |
| 19812 | NM_001294347.1 | 9.80568967 | 8.20541944 | 3.64413704 | 2.25167697 |
| 19813 | NM_001330458.1 | 9.80568967 | 8.20541944 | 3.64413704 | 2.25167697 |
| 19814 | NM_001939.2 | 13.4180834 | -5.37093 | 2.38528907 | -2.2516893 |
| 19815 | NM_030968.4 | 12.0984162 | -5.2221287 | 2.31896611 | -2.2519211 |
| 19816 | NM_181486.2 | 9.80568967 | 8.20541944 | 3.64413704 | 2.25167697 |
| 19817 | NR_104448.1 | 9.80568967 | 8.20541944 | 3.64413704 | 2.25167697 |
| 19818 | NR_135481.1 | 9.80568967 | 8.20541944 | 3.64413704 | 2.25167697 |
| 19819 | NM_145720.3 | 6.59817435 | -3.2327666 | 1.43579786 | -2.2515471 |
| 19820 | NR_027716.2 | 17.2070196 | -5.7272675 | 2.54377865 | -2.2514803 |
| 19821 | NM_001247996.1 | 13.4605387 | -5.375836 | 2.38773673 | -2.2514358 |
| 19822 | NM_001110792.1 | 12.0657954 | -5.2177773 | 2.31762879 | -2.251343 |

| | | | | | |
|---|---|---|---|---|---|
| 19823 | NR_026651.1 | 4.12686611 | -3.6449974 | 1.61908198 | -2.2512741 |
| 19824 | NM_000449.3 | 11.9326359 | -5.2021115 | 2.31080351 | -2.2512132 |
| 19825 | NM_002540.4 | 12.1923401 | -5.2329443 | 2.32467798 | -2.2510405 |
| 19826 | NM_001001671.3 | 5.37503168 | -4.0578086 | 1.8027054 | -2.2509549 |
| 19827 | NM_001281535.1 | 39.075386 | -4.7019336 | 2.08887343 | -2.2509423 |
| 19828 | NM_145647.3 | 12.0887796 | -5.2211244 | 2.31968125 | -2.2507939 |
| 19829 | NR_103806.1 | 9.69404385 | 8.20052918 | 3.64418271 | 2.25030681 |
| 19830 | NM_030631.3 | 5.00972308 | -3.9557109 | 1.757967 | -2.2501622 |
| 19831 | NM_012294.3 | 41.8545828 | -2.8510092 | 1.26708178 | -2.2500593 |
| 19832 | NM_001317233.1 | 16.2138416 | -5.641742 | 2.50789858 | -2.2495894 |
| 19833 | NM_001190981.1 | 37.2221024 | -2.2967335 | 1.02100821 | -2.2494761 |
| 19834 | NM_153711.3 | 6.48462802 | -3.2740034 | 1.45544701 | -2.2494831 |
| 19835 | NM_003447.3 | 4.06491443 | -3.6553882 | 1.62504276 | -2.2494104 |
| 19836 | NM_001281445.1 | 12.1304173 | -5.225986 | 2.3233144 | -2.2493667 |
| 19837 | NM_001145901.1 | 3.74427225 | -3.5431772 | 1.57548473 | -2.2489441 |
| 19838 | NM_001318946.1 | 16.1054908 | -5.6301961 | 2.50346699 | -2.2489596 |
| 19839 | NM_001300836.1 | 3.57500441 | -3.4566938 | 1.53706799 | -2.248888 |
| 19840 | NM_181787.2 | 150.729511 | -0.8443016 | 0.37551687 | -2.2483719 |
| 19841 | NM_001135811.1 | 20.8822996 | -6.0035207 | 2.67029173 | -2.248264 |
| 19842 | NR_033191.2 | 19.7290265 | -2.8401114 | 1.26329215 | -2.2481826 |
| 19843 | NM_001164595.1 | 6.38077949 | -4.2994684 | 1.91307411 | -2.2474134 |
| 19844 | NM_001127889.4 | 16.1609932 | -5.6381055 | 2.50932088 | -2.2468651 |
| 19845 | NM_052893.1 | 12.803163 | -5.3038469 | 2.36079314 | -2.2466377 |
| 19846 | NM_006643.3 | 15.687313 | -5.5959474 | 2.49122271 | -2.2462654 |
| 19847 | NM_001290076.1 | 20.6857686 | -5.993423 | 2.66876903 | -2.2457631 |
| 19848 | NM_001317916.1 | 5.84825264 | -4.1540423 | 1.849877 | -2.2455776 |
| 19849 | NR_138473.1 | 7.22700968 | -4.4663476 | 1.98913896 | -2.2453673 |
| 19850 | NM_001008492.2 | 6.39677955 | -4.2938948 | 1.91236614 | -2.2453309 |
| 19851 | NM_018027.4 | 120.00201 | -4.974467 | 2.21567817 | -2.2451217 |
| 19852 | NR_104104.1 | 12.4208313 | -5.2593787 | 2.3428273 | -2.2448854 |
| 19853 | NM_024980.4 | 3.45331412 | -3.405498 | 1.5170269 | -2.2448501 |
| 19854 | NM_001136474.2 | 16.3030315 | -5.6465697 | 2.51558953 | -2.2446308 |
| 19855 | NM_001177381.1 | 16.3906863 | -5.6560826 | 2.51982492 | -2.2446332 |
| 19856 | NM_001300913.1 | 5.03675982 | -3.9389452 | 1.75486912 | -2.2445806 |
| 19857 | NM_001243777.1 | 15.3297343 | -5.5608448 | 2.4775509 | -2.2444926 |
| 19858 | NM_001278176.1 | 15.8657347 | -4.6164199 | 2.05716406 | -2.2440699 |
| 19859 | NR_033971.1 | 7.69378431 | 3.53815232 | 1.57685868 | 2.24379798 |
| 19860 | NM_018286.2 | 5.82543714 | -4.173083 | 1.85985504 | -2.2437679 |
| 19861 | NM_021213.3 | 11.9957658 | -5.2101268 | 2.32215363 | -2.2436615 |
| 19862 | NM_006553.3 | 12.513643 | -5.2700494 | 2.34921873 | -2.24332 |
| 19863 | NM_001320755.1 | 25.5631614 | -2.7027719 | 1.20482633 | -2.2432875 |
| 19864 | NM_001306196.1 | 12.0812602 | -5.2194927 | 2.3267517 | -2.243253 |
| 19865 | NR_110009.1 | 9.50487297 | 8.17432165 | 3.64443011 | 2.24296293 |

| | | | | | |
|---|---|---|---|---|---|
| 19866 | NM_001320108.1 | 26.0457242 | 4.98680935 | 2.22357562 | 2.24269835 |
| 19867 | NR_026825.2 | 10.7724936 | 4.41705759 | 1.96951752 | 2.24271048 |
| 19868 | NM_033410.3 | 12.4468008 | -5.2620968 | 2.34668069 | -2.2423574 |
| 19869 | NM_021614.3 | 5.2208142 | -3.9947507 | 1.78159432 | -2.2422336 |
| 19870 | NM_007218.3 | 118.241553 | -3.3298557 | 1.48511384 | -2.2421552 |
| 19871 | NR_036446.1 | 40.4757191 | 2.1167888 | 0.94421627 | 2.24184741 |
| 19872 | NR_135500.1 | 70.1062894 | -1.3115919 | 0.58505847 | -2.2418134 |
| 19873 | NM_018054.5 | 12.122853 | -5.2251329 | 2.33084915 | -2.2417293 |
| 19874 | NM_001318813.1 | 12.5210578 | -5.27199 | 2.35179834 | -2.2416846 |
| 19875 | NM_001031719.2 | 4.43613021 | -3.754653 | 1.67508891 | -2.2414649 |
| 19876 | NM_001319679.1 | 12.5739314 | -5.2768808 | 2.35424081 | -2.2414363 |
| 19877 | NM_001320384.1 | 4.46995461 | -3.7869893 | 1.6896537 | -2.2412813 |
| 19878 | NM_014915.2 | 15.7629621 | -5.5981568 | 2.49774857 | -2.2412811 |
| 19879 | NM_001142427.1 | 20.7489059 | -5.9942183 | 2.67522129 | -2.2406439 |
| 19880 | NM_024921.3 | 11.2915999 | 6.26319546 | 2.7955163 | 2.24044319 |
| 19881 | NM_203404.1 | 12.0485349 | -5.2156182 | 2.32794005 | -2.2404435 |
| 19882 | NM_001164462.1 | 14.8781834 | 2.98965997 | 1.33443333 | 2.24039665 |
| 19883 | NM_006237.3 | 13.0033749 | -4.3226072 | 1.92952848 | -2.2402401 |
| 19884 | NM_001164501.1 | 16.0440413 | -5.6266238 | 2.51173753 | -2.2401321 |
| 19885 | NM_138787.3 | 4.03270733 | -3.6509368 | 1.62998361 | -2.2398611 |
| 19886 | NR_126152.1 | 15.6170456 | 2.26346936 | 1.01066431 | 2.23958571 |
| 19887 | NR_002796.2 | 9.51493511 | 4.96602835 | 2.21747001 | 2.23950192 |
| 19888 | NM_016569.3 | 16.4149967 | -5.6568211 | 2.52606204 | -2.2393833 |
| 19889 | NM_012224.2 | 16.2348107 | -5.6429488 | 2.51989009 | -2.2393631 |
| 19890 | NM_001289968.1 | 19.48364 | -4.9214219 | 2.19776068 | -2.2392893 |
| 19891 | NM_015574.1 | 76.7399506 | -7.8838408 | 3.52065241 | -2.2393124 |
| 19892 | NM_018373.2 | 160.054605 | -1.6727558 | 0.74700227 | -2.2392914 |
| 19893 | NR_024532.1 | 11.9164849 | -5.2004722 | 2.32255874 | -2.2391133 |
| 19894 | NR_126358.1 | 3.74988995 | -3.5326208 | 1.57771848 | -2.2390692 |
| 19895 | NR_040089.1 | 3.54486992 | -3.4525872 | 1.5421207 | -2.2388567 |
| 19896 | NM_138424.1 | 26.3702494 | -3.3259858 | 1.48559557 | -2.2388232 |
| 19897 | NM_022100.2 | 67.8226682 | -2.3143072 | 1.03386545 | -2.2384994 |
| 19898 | NM_145326.2 | 49.9134193 | 0.73411601 | 0.32796613 | 2.23838972 |
| 19899 | NM_001242845.1 | 14.3084883 | -5.4607541 | 2.43992177 | -2.2380857 |
| 19900 | NM_001308164.1 | 13.0060672 | -5.3261786 | 2.38004615 | -2.2378468 |
| 19901 | NR_024555.1 | 17.6616636 | -5.7630166 | 2.57582125 | -2.2373511 |
| 19902 | NM_004063.3 | 16.0237628 | 2.07512289 | 0.92760444 | 2.23707737 |
| 19903 | NR_033947.1 | 3.98547434 | -3.6060602 | 1.6120253 | -2.2369749 |
| 19904 | NR_046254.1 | 12.7260196 | -5.2940696 | 2.36685283 | -2.2367549 |
| 19905 | NM_001145547.1 | 4.86874769 | -3.8926613 | 1.74035479 | -2.2367056 |
| 19906 | NM_001035003.1 | 6.00512797 | -4.2040301 | 1.87958204 | -2.2366835 |
| 19907 | NM_002564.3 | 18.2694996 | 2.58816212 | 1.15746958 | 2.23605196 |
| 19908 | NM_001242831.1 | 3.40265837 | -3.3937194 | 1.51783688 | -2.235892 |

| | | | | | |
|---|---|---|---|---|---|
| 19909 | NM_181786.3 | 4.15391937 | -3.661283 | 1.63759774 | -2.2357646 |
| 19910 | NM_001346098.1 | 3.91504332 | -3.5870353 | 1.60443774 | -2.2356962 |
| 19911 | NM_001142429.1 | 13.0519421 | -5.3304729 | 2.38438363 | -2.2355769 |
| 19912 | NM_001278487.1 | 15.711867 | -5.5958477 | 2.50313036 | -2.2355399 |
| 19913 | NM_018937.4 | 16.8742816 | -2.5653575 | 1.14759548 | -2.2354197 |
| 19914 | NM_032971.2 | 112.381654 | 0.7773942 | 0.34777659 | 2.23532643 |
| 19915 | NM_000666.2 | 11.9991539 | -5.2104977 | 2.33121975 | -2.235095 |
| 19916 | NM_033258.1 | 4.47335171 | -3.7665558 | 1.68520643 | -2.2350709 |
| 19917 | NM_001099952.2 | 13.2968202 | -5.3581405 | 2.397387 | -2.2349919 |
| 19918 | NM_175874.3 | 3.4895332 | -3.4155537 | 1.52828384 | -2.2348949 |
| 19919 | NR_002184.2 | 64.8486256 | -1.6315864 | 0.73022028 | -2.2343757 |
| 19920 | NM_001304273.1 | 3.41159642 | -3.3906088 | 1.51769337 | -2.2340539 |
| 19921 | NR_075080.1 | 11.6954969 | -5.1730155 | 2.3155314 | -2.2340511 |
| 19922 | NM_080742.2 | 20.714383 | -3.1028716 | 1.38898957 | -2.2339057 |
| 19923 | NM_001256827.1 | 9.3062821 | 8.1417645 | 3.64474374 | 2.2338373 |
| 19924 | NM_001286197.1 | 9.3062821 | 8.1417645 | 3.64474374 | 2.2338373 |
| 19925 | NM_001146106.1 | 12.7730161 | -2.3996103 | 1.07425291 | -2.233748 |
| 19926 | NR_038880.1 | 43.5544572 | 1.35381683 | 0.60609344 | 2.23367678 |
| 19927 | NM_018478.3 | 4.00242802 | -3.6025381 | 1.61297884 | -2.2334689 |
| 19928 | NM_001286189.1 | 15.2496151 | -5.5539245 | 2.48681171 | -2.2333514 |
| 19929 | NR_144545.1 | 4.7601779 | -3.8816295 | 1.73811946 | -2.2332352 |
| 19930 | NR_138537.1 | 19.4238039 | -4.9083148 | 2.19794739 | -2.2331357 |
| 19931 | NM_001330646.1 | 13.9261177 | -5.4205061 | 2.42756992 | -2.2328939 |
| 19932 | NM_016216.3 | 68.9298537 | -2.2298555 | 0.99864322 | -2.2328851 |
| 19933 | NM_001145674.1 | 16.7756345 | -5.6875204 | 2.54737373 | -2.2326996 |
| 19934 | NM_014779.3 | 18.8059834 | -5.853342 | 2.62180848 | -2.232559 |
| 19935 | NM_004982.3 | 10.307452 | -2.3803095 | 1.06628092 | -2.2323474 |
| 19936 | NM_001258390.1 | 5.8262901 | -4.1596235 | 1.86346563 | -2.2321976 |
| 19937 | NM_130801.2 | 14.9860667 | -5.5307072 | 2.47781254 | -2.2320927 |
| 19938 | NM_001190228.1 | 4.18594177 | -3.7032928 | 1.65927717 | -2.2318711 |
| 19939 | NM_000059.3 | 15.1214602 | -2.3661657 | 1.06021833 | -2.2317722 |
| 19940 | NM_001300894.1 | 16.3909138 | -5.6585212 | 2.53567564 | -2.2315635 |
| 19941 | NM_138992.2 | 5.30685369 | 7.33205529 | 3.28561108 | 2.23156518 |
| 19942 | NR_038975.1 | 4.50403524 | -3.7990217 | 1.70247574 | -2.2314689 |
| 19943 | NR_034091.1 | 14.660737 | 6.01334923 | 2.69489498 | 2.23138536 |
| 19944 | NR_038459.1 | 5.80778367 | 5.54733327 | 2.48605049 | 2.231384 |
| 19945 | NM_000327.3 | 7.06525654 | -3.3587675 | 1.50530934 | -2.2312806 |
| 19946 | NM_145206.3 | 20.9694547 | -2.6529886 | 1.18902472 | -2.2312308 |
| 19947 | NR_003574.1 | 4.30082922 | -3.7112733 | 1.66337314 | -2.231173 |
| 19948 | NR_135321.1 | 4.41987578 | 4.9713191 | 2.22823076 | 2.23106116 |
| 19949 | NM_032983.3 | 77.5767636 | 4.04049782 | 1.81132595 | 2.23068511 |
| 19950 | NM_001002243.2 | 15.6348137 | -4.5993068 | 2.06219602 | -2.2302957 |
| 19951 | NR_002726.2 | 7.30032533 | 6.09965971 | 2.73568244 | 2.22966658 |

| | | | | | |
|---|---|---|---|---|---|
| 19952 | NM_001271661.1 | 84.6357082 | 1.93306488 | 0.86710738 | 2.22932583 |
| 19953 | NM_001165967.1 | 14.6619805 | -5.4948778 | 2.46491234 | -2.2292386 |
| 19954 | NM_001243537.1 | 4.15179847 | -3.6757082 | 1.64921656 | -2.2287602 |
| 19955 | NR_027027.1 | 12.0759248 | -2.8108139 | 1.26122156 | -2.228644 |
| 19956 | NM_205853.3 | 3.69188241 | -3.4919185 | 1.56692153 | -2.2285216 |
| 19957 | NM_001199573.1 | 9.16541322 | 8.12204943 | 3.6449371 | 2.2283099 |
| 19958 | NM_001330191.1 | 9.16541322 | 8.12204943 | 3.6449371 | 2.2283099 |
| 19959 | NR_046227.1 | 9.16541322 | 8.12204943 | 3.6449371 | 2.2283099 |
| 19960 | NM_017442.3 | 3.57887674 | -3.4513589 | 1.54895748 | -2.2281818 |
| 19961 | NM_014908.3 | 73.0654652 | -3.1830296 | 1.42868562 | -2.2279426 |
| 19962 | NM_001317199.1 | 12.0467229 | -5.2152161 | 2.3409186 | -2.2278502 |
| 19963 | NM_001242801.1 | 4.29266281 | -3.7104045 | 1.66556229 | -2.2277188 |
| 19964 | NM_032107.4 | 17.74746 | -2.3582005 | 1.05868612 | -2.2274784 |
| 19965 | NM_001256412.1 | 1.7203939 | 5.69858007 | 2.55850574 | 2.22730791 |
| 19966 | NR_033996.1 | 1.7203939 | 5.69858007 | 2.55850574 | 2.22730791 |
| 19967 | NM_001135638.1 | 12.8776319 | -5.3107127 | 2.38466367 | -2.227028 |
| 19968 | NM_001290557.1 | 5.68442548 | -4.1361401 | 1.85730448 | -2.2269585 |
| 19969 | NM_001301647.1 | 11.6462163 | -5.1675112 | 2.32056536 | -2.2268329 |
| 19970 | NM_001003681.2 | 90.3100554 | -2.2654426 | 1.01744963 | -2.2265894 |
| 19971 | NM_001287242.1 | 5.42969941 | -4.0714243 | 1.82917537 | -2.225825 |
| 19972 | NM_001318812.1 | 7.36690257 | -4.4922233 | 2.01831374 | -2.2257309 |
| 19973 | NR_104191.1 | 4.98441567 | 2.72700102 | 1.22535166 | 2.22548441 |
| 19974 | NM_001204878.1 | 13.7206162 | -5.4023293 | 2.42761841 | -2.2253618 |
| 19975 | NM_000702.3 | 228.404105 | -2.7042715 | 1.2152683 | -2.2252465 |
| 19976 | NM_022361.4 | 3.67767653 | -3.4941248 | 1.57032738 | -2.2250932 |
| 19977 | NR_029449.1 | 10.5053341 | 7.52690562 | 3.38287487 | 2.22500267 |
| 19978 | NR_031680.1 | 10.5053341 | 7.52690562 | 3.38287487 | 2.22500267 |
| 19979 | NR_036539.1 | 3.99841007 | -3.6366913 | 1.63457166 | -2.2248589 |
| 19980 | NM_001166012.2 | 4.54791705 | -3.8068808 | 1.71117016 | -2.2247237 |
| 19981 | NM_001282806.1 | 8.86533048 | -3.708489 | 1.66693258 | -2.2247384 |
| 19982 | NM_001301182.1 | 16.8936804 | -5.701477 | 2.56288463 | -2.2246327 |
| 19983 | NM_001005407.1 | 194.208084 | -2.9987546 | 1.34814483 | -2.2243565 |
| 19984 | NR_033691.1 | 3.41978849 | -3.3848546 | 1.5217901 | -2.2242585 |
| 19985 | NM_001306141.1 | 3.93197133 | -3.5864437 | 1.61247132 | -2.2241907 |
| 19986 | NM_001347894.1 | 17.5928855 | -5.7564096 | 2.58815977 | -2.2241323 |
| 19987 | NM_001033873.1 | 9.15197702 | 8.10528326 | 3.64510362 | 2.22360846 |
| 19988 | NM_001162427.1 | 9.15197702 | 8.10528326 | 3.64510362 | 2.22360846 |
| 19989 | NM_001178075.1 | 16.1656346 | -5.6380136 | 2.53537907 | -2.2237359 |
| 19990 | NM_001242393.1 | 9.15197702 | 8.10528326 | 3.64510362 | 2.22360846 |
| 19991 | NM_001286838.1 | 9.15197702 | 8.10528326 | 3.64510362 | 2.22360846 |
| 19992 | NM_001300967.1 | 9.15197702 | 8.10528326 | 3.64510362 | 2.22360846 |
| 19993 | NM_001302504.1 | 9.15197702 | 8.10528326 | 3.64510362 | 2.22360846 |
| 19994 | NM_001320785.1 | 9.15197702 | 8.10528326 | 3.64510362 | 2.22360846 |

| | | | | | |
|---|---|---|---|---|---|
| 19995 | NM_017779.5 | 12.6842259 | -3.4971944 | 1.57269644 | -2.2236932 |
| 19996 | NR_026951.1 | 3.77077863 | -3.5446295 | 1.59410114 | -2.2235913 |
| 19997 | NM_001184891.1 | 4.06176242 | -3.6614474 | 1.64672494 | -2.2234723 |
| 19998 | NR_074080.1 | 13.763062 | -5.4060459 | 2.43192835 | -2.2229462 |
| 19999 | NM_001289125.1 | 14.6947744 | -5.4976379 | 2.47343503 | -2.2226733 |
| 20000 | NM_001272039.1 | 20.9240859 | -6.0063037 | 2.70241894 | -2.2225657 |
| 20001 | NM_022772.3 | 42.4959112 | -3.0231034 | 1.3602563 | -2.2224513 |
| 20002 | NM_015396.5 | 14.9515287 | -5.5216098 | 2.48513902 | -2.2218515 |
| 20003 | NM_001003680.3 | 14.4302284 | -5.4726597 | 2.46320731 | -2.2217617 |
| 20004 | NR_110048.1 | 8.63952785 | 6.05001823 | 2.72305497 | 2.22177602 |
| 20005 | NM_001277817.1 | 15.5103005 | -5.5784317 | 2.51119769 | -2.2214228 |
| 20006 | NM_006661.3 | 13.567689 | -5.3848862 | 2.42416726 | -2.2213344 |
| 20007 | NM_001164508.1 | 4.45823175 | -3.7745855 | 1.69934657 | -2.2211982 |
| 20008 | NM_153261.5 | 11.7802846 | -5.1830158 | 2.33349694 | -2.2211367 |
| 20009 | NM_001080448.2 | 5.59899488 | -4.0893729 | 1.84131413 | -2.2208991 |
| 20010 | NM_001322101.1 | 13.3146709 | -5.3587705 | 2.41292011 | -2.2208653 |
| 20011 | NM_013430.2 | 14.5185644 | -5.4806716 | 2.4678602 | -2.2208193 |
| 20012 | NR_134456.1 | 42.9698943 | 2.98183237 | 1.34268679 | 2.22079519 |
| 20013 | NM_178324.2 | 24.0035982 | -1.6285055 | 0.73342315 | -2.2204174 |
| 20014 | NM_020682.3 | 238.784228 | 1.23520845 | 0.55642122 | 2.21991615 |
| 20015 | NM_000959.3 | 4.09375605 | -3.6475988 | 1.6432893 | -2.2196936 |
| 20016 | NM_001312871.1 | 11.4218199 | -5.1392668 | 2.31534946 | -2.2196506 |
| 20017 | NM_001172292.1 | 13.8371144 | 4.53630197 | 2.04372509 | 2.21962435 |
| 20018 | NR_117084.1 | 4.0198999 | -3.6068871 | 1.62516557 | -2.2193967 |
| 20019 | NM_001287260.1 | 135.674666 | 3.94123361 | 1.77584572 | 2.21935586 |
| 20020 | NM_001297712.1 | 4.33735618 | 7.0343825 | 3.16954654 | 2.21936558 |
| 20021 | NM_001318804.1 | 4.95463978 | -3.9317547 | 1.77163831 | -2.2192762 |
| 20022 | NM_032133.4 | 4.36201238 | -3.7579384 | 1.69342155 | -2.2191394 |
| 20023 | NM_001320356.1 | 13.1875361 | -5.3461482 | 2.4091746 | -2.2190788 |
| 20024 | NM_170711.2 | 6.20216385 | -4.2493554 | 1.91506267 | -2.2189119 |
| 20025 | NM_003739.5 | 6.54736515 | -4.333782 | 1.95327008 | -2.2187316 |
| 20026 | NM_005585.4 | 45.8779079 | -2.7402887 | 1.23512098 | -2.2186399 |
| 20027 | NM_152435.2 | 9.12091145 | 3.94617757 | 1.7786604 | 2.21862339 |
| 20028 | NM_001289058.1 | 11.6010712 | -5.1613654 | 2.32652287 | -2.218489 |
| 20029 | NM_004079.4 | 12.6076561 | -5.2812806 | 2.38061592 | -2.2184514 |
| 20030 | NM_001347901.1 | 12.4849093 | -5.2670661 | 2.37472048 | -2.2179731 |
| 20031 | NM_001172218.1 | 11.5613297 | -5.1564169 | 2.32491232 | -2.2178974 |
| 20032 | NM_000869.5 | 4.34623683 | -3.7520198 | 1.69172828 | -2.2178619 |
| 20033 | NR_144521.1 | 18.1714809 | 2.14063109 | 0.96518024 | 2.21785632 |
| 20034 | NM_001291582.1 | 4.61143434 | -3.83285 | 1.72836518 | -2.2176159 |
| 20035 | NM_001043352.1 | 21.1249999 | -2.6698588 | 1.20398698 | -2.2175147 |
| 20036 | NM_021908.2 | 26.302943 | -2.9769695 | 1.34258378 | -2.2173435 |
| 20037 | NM_001142402.2 | 14.2612333 | -5.4579285 | 2.46167494 | -2.2171605 |

| | | | | | |
|---|---|---|---|---|---|
| 20038 | NM_001301074.1 | 14.4056266 | -5.4705907 | 2.46739762 | -2.21715 |
| 20039 | NM_001144919.1 | 14.5451799 | -5.4866723 | 2.47478516 | -2.2170297 |
| 20040 | NM_017984.4 | 5.12180913 | -3.9761244 | 1.79353753 | -2.2169173 |
| 20041 | NM_001127615.2 | 8.91852034 | 8.08050485 | 3.64535327 | 2.21665892 |
| 20042 | NM_001318721.1 | 8.91852034 | 8.08050485 | 3.64535327 | 2.21665892 |
| 20043 | NM_004981.1 | 8.91852034 | 8.08050485 | 3.64535327 | 2.21665892 |
| 20044 | NM_001242475.1 | 5.06668938 | 7.26970773 | 3.2798302 | 2.21648905 |
| 20045 | NR_134291.1 | 5.06668938 | 7.26970773 | 3.2798302 | 2.21648905 |
| 20046 | NM_001242482.1 | 14.6776629 | -5.4996495 | 2.48192858 | -2.2158774 |
| 20047 | NM_004688.2 | 6.19457791 | -3.2117611 | 1.44947388 | -2.2158116 |
| 20048 | NM_001099221.1 | 9.87252882 | 6.08521104 | 2.74661735 | 2.21552924 |
| 20049 | NM_001286209.1 | 14.4420102 | -5.4770915 | 2.47262812 | -2.2150891 |
| 20050 | NM_001320048.1 | 13.7780416 | -2.4265413 | 1.09546085 | -2.2150872 |
| 20051 | NM_003369.3 | 106.71485 | -2.7392445 | 1.23670981 | -2.2149452 |
| 20052 | NM_012327.5 | 35.0197777 | -3.0301934 | 1.36813742 | -2.2148311 |
| 20053 | NM_001007561.2 | 498.850215 | -1.0223763 | 0.46161783 | -2.2147679 |
| 20054 | NM_001171204.2 | 31.4167216 | -2.8728414 | 1.29731806 | -2.2144465 |
| 20055 | NM_017677.3 | 4.52124233 | -3.7931579 | 1.71313583 | -2.2141606 |
| 20056 | NM_001002911.3 | 4.85707851 | -3.9075638 | 1.76486703 | -2.214084 |
| 20057 | NM_001329411.1 | 30.7686761 | 1.40221036 | 0.63348186 | 2.21349726 |
| 20058 | NR_125363.1 | 3.53071536 | -3.4467451 | 1.55716984 | -2.2134676 |
| 20059 | NM_001323327.1 | 65.1551859 | -7.6481485 | 3.45538269 | -2.2134013 |
| 20060 | NM_007195.2 | 94.9835503 | -3.0263241 | 1.36740376 | -2.2131898 |
| 20061 | NM_001291744.1 | 3.4687072 | -3.4292677 | 1.54954023 | -2.2130873 |
| 20062 | NM_001323894.1 | 11.5759102 | -5.1581841 | 2.33088839 | -2.2129692 |
| 20063 | NM_001302769.1 | 15.2496081 | -5.5516843 | 2.50881143 | -2.2128743 |
| 20064 | NM_004466.5 | 3.97307652 | -3.619219 | 1.63554896 | -2.2128467 |
| 20065 | NR_104210.1 | 3.96983617 | -3.6123697 | 1.63275831 | -2.2124338 |
| 20066 | NM_173500.3 | 59.1066622 | -2.8553036 | 1.29059863 | -2.212387 |
| 20067 | NR_003614.3 | 4.34263209 | -3.7505625 | 1.69541294 | -2.2121823 |
| 20068 | NR_036540.1 | 4.2757348 | -3.7100692 | 1.67743966 | -2.2117453 |
| 20069 | NM_001130447.1 | 12.0336168 | -5.2133207 | 2.3577707 | -2.2111229 |
| 20070 | NM_053274.2 | 16.7093497 | -5.6854738 | 2.57139108 | -2.2110498 |
| 20071 | NM_001145396.1 | 5.5691609 | 7.38152322 | 3.33860016 | 2.21096354 |
| 20072 | NR_135654.1 | 4.2284137 | -3.6817436 | 1.66526032 | -2.2109117 |
| 20073 | NM_001040061.2 | 36.9557384 | 1.76179354 | 0.7969189 | 2.21075637 |
| 20074 | NR_120501.1 | 9.83746689 | 7.51298427 | 3.39866653 | 2.21056823 |
| 20075 | NM_020666.2 | 46.7076639 | -2.3184265 | 1.04880254 | -2.2105463 |
| 20076 | NM_001130520.2 | 11.9146336 | -5.1994395 | 2.35234422 | -2.2103226 |
| 20077 | NM_001330706.1 | 14.3231198 | -5.4621434 | 2.47134305 | -2.2101923 |
| 20078 | NM_001204180.1 | 3.75399623 | -3.5077898 | 1.58787957 | -2.2091032 |
| 20079 | NR_003138.3 | 15.1696336 | -1.968221 | 0.89104536 | -2.2088898 |
| 20080 | NR_126016.1 | 3.35073844 | -3.3667242 | 1.52427376 | -2.2087399 |

| | | | | | |
|---|---|---|---|---|---|
| 20081 | NM_014701.3 | 49.1423067 | -2.8554361 | 1.29300303 | -2.2083754 |
| 20082 | NM_001300833.2 | 11.460673 | -5.144473 | 2.32961463 | -2.2082936 |
| 20083 | NM_145866.1 | 134.357811 | -3.4206826 | 1.54909314 | -2.208184 |
| 20084 | NM_170589.4 | 11.5727445 | -5.1573925 | 2.33624745 | -2.207554 |
| 20085 | NR_024349.1 | 150.868378 | 1.18040221 | 0.53470802 | 2.20756407 |
| 20086 | NM_001525.2 | 5.20833939 | -4.0023364 | 1.81318934 | -2.2073461 |
| 20087 | NM_001166412.1 | 102.830439 | -3.0875709 | 1.39890493 | -2.2071342 |
| 20088 | NM_001166358.1 | 16.1630705 | -5.6345906 | 2.55313081 | -2.2069338 |
| 20089 | NM_153332.3 | 62.2791656 | -2.2443389 | 1.01699595 | -2.2068317 |
| 20090 | NR_002813.1 | 15.7066649 | 4.84851735 | 2.19742744 | 2.20645163 |
| 20091 | NM_001318088.1 | 13.5693526 | -5.3861562 | 2.44132635 | -2.2062418 |
| 20092 | NM_144978.2 | 11.3077154 | -5.1248991 | 2.32292158 | -2.2062299 |
| 20093 | NM_001318502.1 | 18.0479264 | -5.7953966 | 2.62697512 | -2.2061102 |
| 20094 | NM_201525.3 | 13.8209648 | -5.4134914 | 2.45400865 | -2.2059789 |
| 20095 | NM_001692.3 | 3.58555954 | -3.4677007 | 1.5720869 | -2.2057945 |
| 20096 | NM_001003792.2 | 28.8615701 | -2.7421407 | 1.24327858 | -2.2055722 |
| 20097 | NM_001322239.1 | 13.232463 | -5.3479152 | 2.42474491 | -2.2055579 |
| 20098 | NM_022764.2 | 11.3347943 | -5.128448 | 2.32521658 | -2.2055786 |
| 20099 | NM_001174156.1 | 165.35126 | -2.0547932 | 0.93166266 | -2.2055121 |
| 20100 | NM_001198621.2 | 4.60289413 | 4.85224791 | 2.20031685 | 2.20524963 |
| 20101 | NM_170754.2 | 14.2722355 | -5.4601056 | 2.47600599 | -2.2052069 |
| 20102 | NR_110566.1 | 3.89493763 | -3.5980359 | 1.63168574 | -2.2051035 |
| 20103 | NM_001297699.1 | 3.71226423 | -3.5254549 | 1.59893152 | -2.2048817 |
| 20104 | NM_001171083.2 | 20.2056048 | -4.9469449 | 2.2436736 | -2.2048416 |
| 20105 | NM_173528.2 | 5.29385456 | -4.0352121 | 1.83025243 | -2.2047298 |
| 20106 | NM_001326346.1 | 3.50862518 | -3.446267 | 1.56336093 | -2.2043963 |
| 20107 | NM_001135943.1 | 13.1124035 | -5.3354957 | 2.42051922 | -2.2042773 |
| 20108 | NM_001330142.1 | 4.02747946 | -3.6237702 | 1.64398804 | -2.2042558 |
| 20109 | NM_014991.4 | 252.530127 | -1.8992557 | 0.86162689 | -2.204267 |
| 20110 | NR_110622.1 | 4.32562724 | -3.7348136 | 1.69438692 | -2.2042271 |
| 20111 | NR_135594.1 | 13.4783975 | 3.55129748 | 1.61134251 | 2.20393706 |
| 20112 | NM_001080393.1 | 11.1935729 | -5.1102752 | 2.31885983 | -2.2037879 |
| 20113 | NM_017669.2 | 11.6193736 | -2.963242 | 1.34474821 | -2.2035665 |
| 20114 | NM_001201554.1 | 14.2042744 | -5.4490601 | 2.4728622 | -2.2035438 |
| 20115 | NM_001321124.1 | 4.30288237 | -3.7043331 | 1.68123776 | -2.2033369 |
| 20116 | NR_051984.1 | 19.1294989 | 4.0464588 | 1.83651389 | 2.20333689 |
| 20117 | NM_001037639.2 | 13.6471433 | -5.391744 | 2.44741053 | -2.2030403 |
| 20118 | NM_014877.3 | 98.2091738 | -3.536384 | 1.60523847 | -2.2030272 |
| 20119 | NM_001318395.1 | 3.98899377 | -3.6028612 | 1.63551161 | -2.2028955 |
| 20120 | NR_110718.1 | 3.64048069 | -3.4772491 | 1.57858914 | -2.2027575 |
| 20121 | NM_201437.2 | 12.4647392 | -5.2656618 | 2.39057194 | -2.2026786 |
| 20122 | NR_130733.1 | 3.4478955 | -3.4016173 | 1.54430934 | -2.2026787 |
| 20123 | NM_178818.2 | 14.3210038 | -5.4624408 | 2.48049287 | -2.2021594 |

| | | | | | |
|---|---|---|---|---|---|
| 20124 | NM_001163377.1 | 112.010287 | 3.22549035 | 1.46471656 | 2.20212595 |
| 20125 | NM_001278720.1 | 11.1260065 | -5.1013841 | 2.31686293 | -2.2018497 |
| 20126 | NR_125909.1 | 12.5566132 | 4.84338833 | 2.19988609 | 2.20165414 |
| 20127 | NM_001304732.1 | 3.70529402 | -3.5076979 | 1.59338277 | -2.2014157 |
| 20128 | NM_001324022.1 | 13.6280304 | -5.3910117 | 2.44897874 | -2.2013305 |
| 20129 | NR_138076.1 | 12.1793154 | -5.2313769 | 2.37690643 | -2.2009183 |
| 20130 | NM_001330355.1 | 13.3043354 | -5.3595358 | 2.43539153 | -2.2006875 |
| 20131 | NM_001330058.1 | 72.462615 | -3.7506717 | 1.70481669 | -2.200044 |
| 20132 | NM_152383.4 | 54.2663583 | -2.7907015 | 1.26851123 | -2.1999817 |
| 20133 | NM_001320934.1 | 14.2405103 | -5.453891 | 2.47912955 | -2.1999217 |
| 20134 | NM_001317158.1 | 14.7784775 | -5.5051785 | 2.50326485 | -2.1991994 |
| 20135 | NM_018129.3 | 180.961368 | 1.46411671 | 0.66586738 | 2.19881129 |
| 20136 | NM_001323304.1 | 8.53075859 | 8.01652727 | 3.64601789 | 2.19870761 |
| 20137 | NM_033126.2 | 8.53075859 | 8.01652727 | 3.64601789 | 2.19870761 |
| 20138 | NM_170706.3 | 8.53075859 | 8.01652727 | 3.64601789 | 2.19870761 |
| 20139 | NM_012238.4 | 91.6378714 | -3.0389928 | 1.38249486 | -2.1981946 |
| 20140 | NM_001159597.2 | 11.2880393 | -5.1216536 | 2.32995937 | -2.1981729 |
| 20141 | NM_133475.1 | 52.9706567 | -2.8603792 | 1.3013291 | -2.1980444 |
| 20142 | NM_001282552.2 | 13.9738951 | -5.4302028 | 2.47066054 | -2.1978749 |
| 20143 | NM_001204082.1 | 30.2554569 | 3.82990594 | 1.74281686 | 2.1975378 |
| 20144 | NM_001185119.1 | 8.48649373 | 8.01145651 | 3.64607183 | 2.19728433 |
| 20145 | NR_039999.1 | 34.5224546 | 1.40420801 | 0.639244 | 2.19666984 |
| 20146 | NR_027123.1 | 3.53024847 | -3.428105 | 1.56062396 | -2.1966246 |
| 20147 | NM_006608.2 | 13.3943934 | -5.3647216 | 2.44310756 | -2.1958598 |
| 20148 | NM_020139.3 | 72.2022067 | -2.8844811 | 1.31363326 | -2.195804 |
| 20149 | NM_001271518.1 | 4.36256303 | 7.048904 | 3.21023029 | 2.19576272 |
| 20150 | NR_038939.1 | 4.03567153 | -3.622046 | 1.65017054 | -2.1949525 |
| 20151 | NR_138043.1 | 3.67218094 | -3.5014522 | 1.5953245 | -2.1948213 |
| 20152 | NM_001271874.1 | 105.768954 | -2.2696357 | 1.03412934 | -2.1947309 |
| 20153 | NM_001142805.1 | 11.198773 | -5.1110758 | 2.32904066 | -2.1944983 |
| 20154 | NM_001042369.2 | 15.1688848 | -5.5427999 | 2.52619727 | -2.1941279 |
| 20155 | NM_001321080.1 | 15.0905281 | -5.5357322 | 2.5230395 | -2.1940727 |
| 20156 | NM_002636.4 | 16.7193187 | -5.6859902 | 2.59152753 | -2.194069 |
| 20157 | NM_024733.3 | 78.0603351 | 2.03718509 | 0.92855084 | 2.19394028 |
| 20158 | NR_015407.1 | 3.49220402 | -3.4248826 | 1.56113098 | -2.193847 |
| 20159 | NR_038386.1 | 21.9488344 | 2.12051583 | 0.96665584 | 2.19366163 |
| 20160 | NM_001017424.2 | 8.49826438 | 7.9976795 | 3.64621934 | 2.19341701 |
| 20161 | NM_001166420.1 | 8.49826438 | 7.9976795 | 3.64621934 | 2.19341701 |
| 20162 | NM_001207018.1 | 5.67202809 | -4.1161182 | 1.87651537 | -2.1934903 |
| 20163 | NM_001276480.1 | 8.49826438 | 7.9976795 | 3.64621934 | 2.19341701 |
| 20164 | NM_001278407.1 | 8.49826438 | 7.9976795 | 3.64621934 | 2.19341701 |
| 20165 | NM_001288636.1 | 8.49826438 | 7.9976795 | 3.64621934 | 2.19341701 |
| 20166 | NM_001291510.1 | 8.49826438 | 7.9976795 | 3.64621934 | 2.19341701 |

| | | | | | |
|---|---|---|---|---|---|
| 20167 | NM_001323557.1 | 8.49826438 | 7.9976795 | 3.64621934 | 2.19341701 |
| 20168 | NR_073482.1 | 8.49826438 | 7.9976795 | 3.64621934 | 2.19341701 |
| 20169 | NR_104617.1 | 8.49826438 | 7.9976795 | 3.64621934 | 2.19341701 |
| 20170 | NM_001284247.1 | 13.0927454 | -5.336277 | 2.43296556 | -2.193322 |
| 20171 | NM_001128423.1 | 11.8871042 | -5.1964864 | 2.36929401 | -2.1932636 |
| 20172 | NM_001330559.1 | 4.34082418 | -3.7203837 | 1.69627204 | -2.1932706 |
| 20173 | NM_001330725.1 | 4.12921486 | -3.6783244 | 1.67710834 | -2.1932539 |
| 20174 | NM_201515.1 | 11.3854272 | -5.1339555 | 2.34112432 | -2.1929444 |
| 20175 | NM_001282332.1 | 5.15567349 | -3.9861383 | 1.81789163 | -2.192726 |
| 20176 | NM_001198595.1 | 11.7004191 | -5.1734468 | 2.35958005 | -2.1925286 |
| 20177 | NM_001242901.1 | 3.45847629 | -3.4090993 | 1.5554052 | -2.1917757 |
| 20178 | NR_034157.1 | 17.2104781 | 2.83708048 | 1.29447952 | 2.19167661 |
| 20179 | NM_173588.3 | 6.14330668 | -3.1251836 | 1.42606326 | -2.1914761 |
| 20180 | NM_002891.4 | 15.4736435 | -5.5739007 | 2.54352547 | -2.1914074 |
| 20181 | NM_032548.3 | 5.80359426 | -4.1453337 | 1.89177408 | -2.1912414 |
| 20182 | NM_001290021.1 | 6.08173141 | -3.1151228 | 1.42227163 | -2.1902446 |
| 20183 | NR_038286.1 | 3.97866642 | -3.5890743 | 1.63876641 | -2.1901073 |
| 20184 | NM_001308258.1 | 7.96845706 | -3.5411719 | 1.61704811 | -2.1898989 |
| 20185 | NR_051965.1 | 5.16488531 | -3.9806964 | 1.81779644 | -2.1898472 |
| 20186 | NM_001159379.1 | 15.5785469 | -5.5822015 | 2.54931137 | -2.18969 |
| 20187 | NM_001173978.1 | 4.41868193 | -3.7718831 | 1.72260033 | -2.1896449 |
| 20188 | NM_001303030.1 | 27.7582503 | -1.8878012 | 0.86223597 | -2.1894252 |
| 20189 | NM_002677.3 | 12.3020613 | -2.6741206 | 1.2213718 | -2.1894403 |
| 20190 | NM_001329588.1 | 11.0008113 | -5.085423 | 2.32276317 | -2.1893851 |
| 20191 | NR_135265.1 | 6.3605946 | -4.282731 | 1.95615075 | -2.1893666 |
| 20192 | NM_207660.3 | 13.7466508 | -5.4022899 | 2.46765929 | -2.1892366 |
| 20193 | NM_001308139.1 | 5.04403464 | -3.9545916 | 1.80653737 | -2.189045 |
| 20194 | NR_002594.1 | 72.3248725 | 0.7571483 | 0.34592621 | 2.18875671 |
| 20195 | NM_012083.2 | 33.3274472 | -2.954516 | 1.35001093 | -2.1885127 |
| 20196 | NM_016614.2 | 91.8054988 | -2.2373234 | 1.0223085 | -2.1885013 |
| 20197 | NM_001005781.1 | 4.49353976 | -3.7974695 | 1.73527382 | -2.1883978 |
| 20198 | NM_032456.2 | 3.85216088 | -3.582131 | 1.63707653 | -2.1881268 |
| 20199 | NM_001142502.1 | 11.1041873 | -5.0988952 | 2.33034686 | -2.1880413 |
| 20200 | NM_198498.2 | 4.80073849 | 7.19624277 | 3.28900853 | 2.18796719 |
| 20201 | NM_001251.2 | 12.3800926 | -5.2554177 | 2.40200794 | -2.1879269 |
| 20202 | NM_001197247.2 | 4.7882139 | -3.8749039 | 1.77167617 | -2.1871401 |
| 20203 | NM_020344.3 | 14.3623289 | -5.4658859 | 2.49929101 | -2.1869746 |
| 20204 | NM_001205179.1 | 6.05679527 | -4.2008647 | 1.92105446 | -2.1867494 |
| 20205 | NR_037949.1 | 11.1362684 | -5.1031328 | 2.33376392 | -2.1866534 |
| 20206 | NM_016058.3 | 13.6703484 | -5.3941677 | 2.46697095 | -2.186555 |
| 20207 | NR_134464.1 | 13.8389524 | -5.4128174 | 2.47611268 | -2.1860142 |
| 20208 | NM_020180.3 | 60.8254302 | -7.5489749 | 3.45336701 | -2.1859753 |
| 20209 | NM_178581.2 | 14.2544164 | -5.4574969 | 2.49675713 | -2.1858341 |

| | | | | | |
|---|---|---|---|---|---|
| 20210 | NM_022470.3 | 183.109551 | -1.5273819 | 0.69884207 | -2.1855895 |
| 20211 | NM_001130822.2 | 11.0630556 | -5.0936366 | 2.33061415 | -2.1855341 |
| 20212 | NM_012430.4 | 60.5455023 | -1.4001528 | 0.64064267 | -2.1855441 |
| 20213 | NR_037716.1 | 11.3485664 | -5.1304542 | 2.34771059 | -2.1853009 |
| 20214 | NM_001304799.1 | 3.66267956 | -3.4872382 | 1.59604291 | -2.1849276 |
| 20215 | NR_126328.1 | 15.6168288 | 5.1387338 | 2.35193722 | 2.18489412 |
| 20216 | NR_109901.1 | 5.34042886 | -4.0266793 | 1.84351375 | -2.1842415 |
| 20217 | NR_003082.1 | 18.6098592 | 1.29895392 | 0.59471732 | 2.18415353 |
| 20218 | NM_001076781.2 | 11.307942 | -5.1241639 | 2.34714609 | -2.1831466 |
| 20219 | NR_037916.1 | 562.966169 | 1.27534727 | 0.58419389 | 2.18308902 |
| 20220 | NM_033026.5 | 69.3462325 | -3.0897859 | 1.41534297 | -2.1830652 |
| 20221 | NR_110745.1 | 27.4713889 | 1.8274541 | 0.83724279 | 2.18270509 |
| 20222 | NM_001291429.1 | 12.0098376 | -5.2121752 | 2.38801089 | -2.182643 |
| 20223 | NM_001289934.1 | 14.0078588 | -5.4297137 | 2.48807517 | -2.1822949 |
| 20224 | NR_029427.2 | 17.4796035 | -5.7469023 | 2.6335182 | -2.1822148 |
| 20225 | NM_001199866.1 | 6.12094869 | -3.101352 | 1.42126559 | -2.1821059 |
| 20226 | NM_002303.5 | 3.32841747 | -3.3613827 | 1.54048422 | -2.1820299 |
| 20227 | NM_003248.5 | 4.85863682 | -3.9002211 | 1.78771079 | -2.1816846 |
| 20228 | NM_001040274.2 | 13.0740819 | -3.110989 | 1.42600501 | -2.1816116 |
| 20229 | NM_198992.3 | 5.49254545 | -3.0239038 | 1.38622692 | -2.1813916 |
| 20230 | NM_030652.3 | 8.14703398 | 7.95280655 | 3.64670959 | 2.18081708 |
| 20231 | NM_021052.2 | 3.80550576 | -3.5509718 | 1.62838687 | -2.1806684 |
| 20232 | NM_020759.2 | 83.1652871 | -2.1130057 | 0.96919345 | -2.1801692 |
| 20233 | NM_015520.1 | 13.6794345 | -5.3955697 | 2.47491491 | -2.1801031 |
| 20234 | NM_024315.3 | 3.93475917 | -3.6125736 | 1.65717878 | -2.179954 |
| 20235 | NM_001346121.1 | 8.14299683 | 7.94957881 | 3.64674544 | 2.17991054 |
| 20236 | NR_136524.1 | 8.14299683 | 7.94957881 | 3.64674544 | 2.17991054 |
| 20237 | NM_001282180.1 | 12.9339218 | -5.3183758 | 2.44039054 | -2.1793134 |
| 20238 | NM_000022.3 | 11.2613239 | -5.1184761 | 2.34871759 | -2.1792642 |
| 20239 | NM_001040647.1 | 11.9794807 | -5.2066975 | 2.38924055 | -2.179227 |
| 20240 | NR_033995.2 | 19.0413741 | 2.68182891 | 1.2308836 | 2.17878353 |
| 20241 | NM_173517.5 | 208.42601 | -3.3342579 | 1.53047425 | -2.1785783 |
| 20242 | NM_020130.4 | 26.6185247 | -2.8155324 | 1.29244168 | -2.1784599 |
| 20243 | NM_001114091.2 | 17.6976165 | -5.7646551 | 2.64699084 | -2.1778145 |
| 20244 | NR_125963.1 | 10.1603675 | -2.2430295 | 1.02993872 | -2.1778282 |
| 20245 | NM_001256667.1 | 10.801218 | -5.0585792 | 2.3228581 | -2.1777392 |
| 20246 | NM_001319087.1 | 4.95479374 | -3.9240303 | 1.80255734 | -2.176924 |
| 20247 | NM_001330377.1 | 4.02536983 | -3.6063861 | 1.65668625 | -2.1768673 |
| 20248 | NM_002164.5 | 84.7169276 | 2.86865632 | 1.31798725 | 2.1765433 |
| 20249 | NR_003570.2 | 25.4851095 | 1.69671767 | 0.77973337 | 2.17602291 |
| 20250 | NM_022549.3 | 130.380304 | -1.6874902 | 0.77557657 | -2.1757879 |
| 20251 | NM_000173.6 | 3.67683411 | -3.4929753 | 1.6055773 | -2.1755261 |
| 20252 | NM_001281439.1 | 16.0474121 | -5.62804 | 2.58697086 | -2.1755328 |

| | | | | | |
|---|---|---|---|---|---|
| 20253 | NM_001130146.1 | 14.0357473 | -5.4307596 | 2.49667472 | -2.1751971 |
| 20254 | NM_001173452.1 | 16.0276289 | -5.6217572 | 2.58449906 | -2.1751825 |
| 20255 | NM_024939.2 | 4.2121156 | -3.699476 | 1.7008872 | -2.1750273 |
| 20256 | NR_024576.2 | 10.6890896 | -5.0436083 | 2.31889377 | -2.1750062 |
| 20257 | NM_016819.3 | 5.85598536 | -4.1764951 | 1.92045483 | -2.1747427 |
| 20258 | NM_006291.2 | 51.548338 | -3.062105 | 1.40810881 | -2.1746224 |
| 20259 | NR_109915.1 | 5.39808867 | -4.0589615 | 1.86714236 | -2.1738897 |
| 20260 | NM_152734.3 | 146.474846 | -3.1535461 | 1.45102916 | -2.1733168 |
| 20261 | NM_152475.2 | 117.112673 | 1.48299624 | 0.68238923 | 2.17324098 |
| 20262 | NM_001113182.2 | 71.5216042 | -7.7820893 | 3.58132534 | -2.1729635 |
| 20263 | NM_001031701.2 | 226.743467 | 1.83201445 | 0.84316902 | 2.17277248 |
| 20264 | NM_001291633.1 | 12.6647901 | -5.2863327 | 2.43321368 | -2.1725723 |
| 20265 | NM_001099756.1 | 6.17403421 | -4.2452684 | 1.95409901 | -2.172494 |
| 20266 | NM_001271844.1 | 10.8102279 | -5.0602262 | 2.32923011 | -2.1724887 |
| 20267 | NR_003148.3 | 37.340417 | -2.2515819 | 1.03653947 | -2.1722104 |
| 20268 | NM_004482.3 | 3.53401005 | -3.4598206 | 1.5927905 | -2.1721756 |
| 20269 | NM_001080511.2 | 5.3667843 | -4.0411089 | 1.86054892 | -2.1719982 |
| 20270 | NM_001136497.2 | 12.5230817 | -5.2706806 | 2.42706334 | -2.1716288 |
| 20271 | NM_001166579.1 | 3.61574837 | -3.4769206 | 1.60148943 | -2.1710544 |
| 20272 | NM_016300.4 | 14.0594608 | -5.4330202 | 2.50278483 | -2.17079 |
| 20273 | NM_018111.2 | 3.8737072 | -3.5801442 | 1.64940436 | -2.1705679 |
| 20274 | NM_000501.3 | 67.4971843 | -7.6990464 | 3.54721841 | -2.1704461 |
| 20275 | NR_110940.1 | 4.36785179 | -3.7367711 | 1.72179489 | -2.1702766 |
| 20276 | NM_001205251.1 | 4.72722963 | 7.1697897 | 3.30384191 | 2.17013704 |
| 20277 | NM_004200.3 | 76.4703269 | -2.415061 | 1.11295673 | -2.1699505 |
| 20278 | NM_017693.3 | 133.11642 | -3.1778988 | 1.46453448 | -2.1699037 |
| 20279 | NM_000830.4 | 4.21370931 | -3.7054727 | 1.70784304 | -2.1696799 |
| 20280 | NM_001079871.1 | 13.2918426 | -5.3570315 | 2.46913112 | -2.1696019 |
| 20281 | NM_001284511.1 | 14.4998791 | -5.4781114 | 2.52516134 | -2.1694105 |
| 20282 | NR_033984.1 | 47.5321144 | 1.3719211 | 0.63257541 | 2.16878664 |
| 20283 | NM_019069.3 | 46.6095531 | -2.2119058 | 1.02003462 | -2.1684615 |

| pvalue | padj | Mock 1 | Mock 2 | Mock 3 | Mock 4 | GFP (-) 1 |
|---|---|---|---|---|---|---|
| 8.68E-80 | 2.92E-75 | 236.319035 | 267.818473 | 225.516616 | 256.556692 | 4156.00571 |
| 7.02E-75 | 1.18E-70 | 135.682493 | 134.798999 | 138.279736 | 153.562866 | 1352.06818 |
| 1.14E-61 | 1.28E-57 | 20.5774398 | 22.0216187 | 27.8415575 | 17.6295737 | 624.945398 |
| 3.56E-49 | 3.00E-45 | 20555.5762 | 51488.7688 | 42451.8788 | 49454.6659 | 59.4054561 |
| 5.03E-48 | 3.38E-44 | 0 | 0 | 0 | 0 | 9804.27647 |
| 5.45E-47 | 3.06E-43 | 1754.22674 | 2962.90869 | 2547.50251 | 2895.88946 | 42.7719284 |
| 2.62E-45 | 1.26E-41 | 916.339114 | 771.423975 | 840.815037 | 927.872303 | 9545.26868 |
| 1.04E-44 | 4.36E-41 | 381.647203 | 458.22762 | 471.450374 | 472.287002 | 6838.7561 |
| 3.61E-44 | 1.35E-40 | 0 | 0 | 0 | 0 | 3276.80496 |
| 5.04E-44 | 1.70E-40 | 779.049008 | 677.109163 | 711.81582 | 739.978161 | 5976.18888 |
| 1.46E-43 | 4.46E-40 | 431.48319 | 501.603536 | 516.924918 | 577.136572 | 6004.7035 |
| 5.00E-41 | 1.40E-37 | 60.4462293 | 68.5117025 | 98.3735032 | 71.4461673 | 998.011662 |
| 1.16E-40 | 3.01E-37 | 353.674746 | 267.373592 | 301.616873 | 314.084774 | 2506.91025 |
| 5.84E-38 | 1.40E-34 | 4239.91716 | 9173.22759 | 7696.33455 | 8278.47668 | 9.50487297 |
| 4.75E-37 | 1.07E-33 | 1495.72265 | 1987.50669 | 1754.94617 | 1906.77758 | 11.8810912 |
| 7.45E-37 | 1.57E-33 | 74.5932191 | 75.4073608 | 86.3088283 | 76.549465 | 1304.54382 |
| 7.40E-35 | 1.46E-31 | 47.5853294 | 36.2578166 | 35.2659728 | 38.9706367 | 349.304082 |
| 3.40E-32 | 6.35E-29 | 99.9934963 | 110.552974 | 105.797919 | 107.633187 | 1613.45219 |
| 6.79E-29 | 1.20E-25 | 0 | 0 | 0 | 0 | 520.391795 |
| 2.01E-28 | 3.38E-25 | 38.5826995 | 35.8129354 | 42.6903882 | 31.0837221 | 411.085756 |
| 5.70E-28 | 9.13E-25 | 5205.77073 | 10396.4284 | 9342.69865 | 10646.4068 | 0 |
| 2.14E-27 | 3.28E-24 | 1451.99559 | 2086.7152 | 1524.7893 | 1866.87907 | 11.8810912 |
| 2.29E-27 | 3.35E-24 | 637.257587 | 1098.1892 | 835.246725 | 972.410173 | 16.6335277 |
| 3.84E-27 | 5.38E-24 | 1577.06784 | 2168.57334 | 1614.81034 | 2149.88013 | 9.50487297 |
| 4.02E-27 | 5.41E-24 | 4322.22692 | 7710.90314 | 6460.16939 | 7156.67907 | 0 |
| 6.74E-27 | 8.73E-24 | 1500.22397 | 3642.01982 | 3260.24638 | 3533.80166 | 4.75243649 |
| 3.95E-26 | 4.93E-23 | 7111.43457 | 16503.09 | 13769.5063 | 16004.8693 | 2.37621824 |
| 5.27E-26 | 6.33E-23 | 4076.26221 | 12321.4293 | 9509.74799 | 11262.9779 | 0 |
| 6.23E-26 | 7.23E-23 | 4708.37543 | 15855.1205 | 8380.30881 | 11136.3234 | 0 |
| 8.19E-26 | 8.89E-23 | 4509.03149 | 12669.3264 | 8763.59425 | 10626.9215 | 0 |
| 8.04E-26 | 8.89E-23 | 97.0997938 | 48.4920492 | 52.8989593 | 52.4247851 | 1620.58084 |
| 1.32E-25 | 1.38E-22 | 8179.21078 | 25956.5927 | 20939.6354 | 23500.6857 | 0 |
| 2.11E-25 | 2.15E-22 | 3335.47437 | 7940.46183 | 6756.21796 | 7461.02119 | 0 |
| 2.24E-25 | 2.22E-22 | 1549.09539 | 5359.48363 | 4295.95232 | 5018.86128 | 7.12865473 |
| 2.32E-25 | 2.23E-22 | 0 | 0 | 0 | 0 | 449.105248 |
| 2.46E-25 | 2.30E-22 | 207.060488 | 211.541003 | 169.833501 | 199.492545 | 1527.90833 |
| 2.67E-25 | 2.43E-22 | 1536.55601 | 4628.32141 | 4803.59672 | 5152.93883 | 2.37621824 |
| 3.23E-25 | 2.86E-22 | 3205.90081 | 7572.98997 | 6843.45484 | 7342.25353 | 0 |
| 6.52E-25 | 5.44E-22 | 2961.86523 | 4711.73663 | 5413.32683 | 6007.50922 | 0 |
| 6.63E-25 | 5.44E-22 | 3107.83645 | 7954.0307 | 6229.08447 | 7094.97556 | 0 |
| 6.32E-25 | 5.44E-22 | 3606.51784 | 9423.25082 | 14016.3681 | 11603.0431 | 0 |
| 1.39E-24 | 1.11E-21 | 3069.89679 | 6715.25905 | 4235.62895 | 5027.21214 | 0 |

| | | | | | | |
|---|---|---|---|---|---|---|
| 1.46E-24 | 1.14E-21 | 3104.94275 | 2849.68643 | 2673.71757 | 2911.66329 | 0 |
| 1.70E-24 | 1.27E-21 | 1159.0886 | 3284.11291 | 2888.09757 | 3100.02136 | 9.50487297 |
| 1.66E-24 | 1.27E-21 | 526.010804 | 519.398783 | 512.284658 | 656.005718 | 7.12865473 |
| 1.74E-24 | 1.28E-21 | 3132.59368 | 10816.6187 | 11040.1056 | 11243.4926 | 0 |
| 2.38E-24 | 1.70E-21 | 1040.4468 | 2166.79381 | 2108.53396 | 2238.02799 | 19.0097459 |
| 2.51E-24 | 1.72E-21 | 3017.81015 | 2996.9421 | 2483.46693 | 2926.50924 | 0 |
| 2.50E-24 | 1.72E-21 | 2685.03437 | 3176.45166 | 3078.34821 | 4028.3576 | 0 |
| 4.82E-24 | 3.24E-21 | 2688.89264 | 7833.69034 | 7776.14701 | 7373.80119 | 0 |
| 5.31E-24 | 3.51E-21 | 15.7546023 | 14.4586385 | 11.136623 | 13.9180845 | 197.226114 |
| 5.58E-24 | 3.61E-21 | 556.876963 | 1090.84867 | 900.210359 | 1006.74145 | 16.6335277 |
| 5.81E-24 | 3.69E-21 | 0 | 0 | 0 | 0 | 449.105248 |
| 7.93E-24 | 4.94E-21 | 3187.57403 | 9584.52025 | 8443.41634 | 9668.42939 | 4.75243649 |
| 1.38E-23 | 8.47E-21 | 167.834743 | 262.479899 | 241.293498 | 302.022435 | 2663.74065 |
| 2.15E-23 | 1.29E-20 | 3757.63341 | 2545.83258 | 2642.16381 | 2491.33713 | 0 |
| 2.27E-23 | 1.34E-20 | 4061.15065 | 5226.01928 | 4586.43257 | 4953.44629 | 2.37621824 |
| 2.31E-23 | 1.34E-20 | 7.07349491 | 13.123995 | 10.2085711 | 15.7738291 | 392.07601 |
| 2.38E-23 | 1.36E-20 | 299.658966 | 220.438627 | 215.308045 | 205.987651 | 2359.58472 |
| 2.95E-23 | 1.65E-20 | 0 | 0 | 0 | 0 | 223.364515 |
| 3.07E-23 | 1.69E-20 | 43.405537 | 63.1731283 | 56.6111669 | 82.5806349 | 658.212453 |
| 3.41E-23 | 1.85E-20 | 1354.8958 | 1761.72949 | 1520.14904 | 1771.77216 | 16.6335277 |
| 4.01E-23 | 2.14E-20 | 4100.05487 | 9679.50238 | 7017.00054 | 9481.46312 | 2.37621824 |
| 5.57E-23 | 2.93E-20 | 2090.53927 | 3781.04519 | 2666.29316 | 3111.15583 | 0 |
| 7.51E-23 | 3.89E-20 | 2563.17734 | 3188.90833 | 2024.08123 | 2350.30054 | 0 |
| 9.48E-23 | 4.84E-20 | 0 | 0 | 0 | 0 | 311.28459 |
| 1.19E-22 | 5.89E-20 | 0 | 0 | 0 | 0 | 266.136443 |
| 1.18E-22 | 5.89E-20 | 1830.42757 | 3161.54814 | 2682.07004 | 2907.02392 | 0 |
| 1.21E-22 | 5.90E-20 | 1945.53262 | 3481.86259 | 4230.06064 | 4528.01684 | 0 |
| 1.34E-22 | 6.42E-20 | 2079.6075 | 5025.6003 | 3138.67158 | 3866.44389 | 0 |
| 1.74E-22 | 8.24E-20 | 2131.0511 | 3827.53527 | 3283.44768 | 3955.05569 | 4.75243649 |
| 1.96E-22 | 9.16E-20 | 2311.42522 | 13068.3848 | 7444.83248 | 9178.97675 | 14.2573095 |
| 2.12E-22 | 9.76E-20 | 187.447615 | 126.568697 | 155.912722 | 128.510314 | 1102.56526 |
| 2.66E-22 | 1.21E-19 | 1766.12307 | 3792.6121 | 3031.94561 | 3386.7339 | 0 |
| 2.88E-22 | 1.28E-19 | 2305.63782 | 12631.0666 | 7171.05716 | 8685.81262 | 0 |
| 2.89E-22 | 1.28E-19 | 1758.40653 | 2755.59406 | 2094.61318 | 2455.15011 | 0 |
| 3.10E-22 | 1.35E-19 | 1579.96155 | 4623.20527 | 4042.59415 | 4495.54131 | 11.8810912 |
| 3.33E-22 | 1.44E-19 | 1670.63089 | 3028.52867 | 3050.50665 | 3177.96264 | 0 |
| 4.20E-22 | 1.79E-19 | 1843.93152 | 1819.78649 | 1999.02383 | 2521.49298 | 0 |
| 5.42E-22 | 2.28E-19 | 1910.80819 | 7381.02374 | 7576.61585 | 8618.54188 | 0 |
| 8.16E-22 | 3.39E-19 | 1898.59034 | 6501.49364 | 3790.16403 | 4673.69279 | 0 |
| 8.56E-22 | 3.51E-19 | 1559.06258 | 2607.89351 | 2112.24616 | 2463.03703 | 0 |
| 8.88E-22 | 3.60E-19 | 2061.60225 | 4199.45594 | 2766.52276 | 3453.07677 | 4.75243649 |
| 9.83E-22 | 3.94E-19 | 1615.65054 | 4330.25101 | 3995.2635 | 4268.67653 | 0 |
| 1.04E-21 | 4.07E-19 | 1737.82909 | 6212.76575 | 6148.34395 | 6820.32536 | 0 |

| | | | | | | |
|---|---|---|---|---|---|---|
| 1.03E-21 | 4.07E-19 | 203.202218 | 171.946578 | 169.833501 | 124.334889 | 957.615952 |
| 1.19E-21 | 4.61E-19 | 1819.49581 | 2117.63444 | 1538.71008 | 1746.25567 | 0 |
| 1.22E-21 | 4.68E-19 | 539.836271 | 730.494906 | 774.923351 | 840.652306 | 9.50487297 |
| 1.38E-21 | 5.22E-19 | 1614.04293 | 4639.44344 | 4607.77777 | 5118.14362 | 0 |
| 1.48E-21 | 5.53E-19 | 1576.10328 | 4253.73145 | 4122.40662 | 4592.9679 | 0 |
| 1.55E-21 | 5.73E-19 | 12472.8222 | 3970.11969 | 3391.1017 | 4235.73706 | 0 |
| 1.80E-21 | 6.58E-19 | 1616.29359 | 3256.75271 | 2236.60512 | 2770.6267 | 7.12865473 |
| 2.16E-21 | 7.80E-19 | 1630.7621 | 6004.56135 | 5146.97593 | 6348.03836 | 0 |
| 2.18E-21 | 7.80E-19 | 1546.52321 | 4840.97461 | 4252.33388 | 4635.65002 | 0 |
| 2.50E-21 | 8.84E-19 | 1741.04432 | 3296.56958 | 1829.19033 | 2412.00405 | 0 |
| 2.55E-21 | 8.95E-19 | 1449.10189 | 3491.64998 | 3197.13885 | 3472.56209 | 0 |
| 2.66E-21 | 9.24E-19 | 1766.12307 | 3700.5217 | 1951.69318 | 2696.39691 | 0 |
| 2.85E-21 | 9.79E-19 | 1218.89178 | 2725.78702 | 2043.57032 | 2383.70395 | 0 |
| 2.99E-21 | 1.01E-18 | 1366.79213 | 2292.91763 | 2100.18149 | 2351.22841 | 0 |
| 2.98E-21 | 1.01E-18 | 1432.38272 | 3559.04948 | 3330.77833 | 3472.09816 | 0 |
| 3.84E-21 | 1.28E-18 | 1419.52182 | 1819.34161 | 2299.71265 | 2491.80107 | 0 |
| 4.09E-21 | 1.32E-18 | 901.870602 | 4281.75896 | 3532.1656 | 3827.00931 | 7.12865473 |
| 4.06E-21 | 1.32E-18 | 1515.97857 | 5245.14917 | 5289.89593 | 5892.45306 | 0 |
| 4.11E-21 | 1.32E-18 | 1604.07573 | 7735.59404 | 5712.15955 | 6528.50952 | 0 |
| 4.00E-21 | 1.32E-18 | 0 | 0 | 0 | 0 | 275.641316 |
| 4.22E-21 | 1.34E-18 | 795.446655 | 2443.73235 | 2193.91473 | 2288.1331 | 9.50487297 |
| 4.32E-21 | 1.36E-18 | 788.37316 | 1665.41272 | 976.310617 | 1258.65878 | 7.12865473 |
| 4.48E-21 | 1.40E-18 | 1365.82756 | 2029.99285 | 1610.17008 | 1907.24152 | 0 |
| 4.74E-21 | 1.46E-18 | 1440.42078 | 4504.64444 | 3766.03468 | 4280.27493 | 0 |
| 5.00E-21 | 1.53E-18 | 1509.54812 | 6321.98408 | 5297.32034 | 5491.14829 | 0 |
| 6.51E-21 | 1.97E-18 | 1319.2068 | 2900.18044 | 2433.35213 | 2806.81372 | 0 |
| 6.92E-21 | 2.08E-18 | 1330.13857 | 2409.03162 | 1792.9963 | 2093.74385 | 0 |
| 7.33E-21 | 2.18E-18 | 2469.61429 | 7759.39519 | 7005.86392 | 7396.998 | 0 |
| 7.88E-21 | 2.33E-18 | 749.790461 | 1071.49633 | 566.111669 | 745.545395 | 7.12865473 |
| 1.03E-20 | 3.03E-18 | 0 | 0 | 0 | 0 | 154.454186 |
| 1.12E-20 | 3.24E-18 | 118.963324 | 71.1809896 | 74.2441534 | 65.4149973 | 855.438568 |
| 1.15E-20 | 3.32E-18 | 1317.92071 | 4082.89707 | 3418.94326 | 3863.19633 | 0 |
| 1.18E-20 | 3.36E-18 | 1226.92985 | 2556.28729 | 2283.93577 | 2435.66479 | 0 |
| 1.22E-20 | 3.44E-18 | 1260.68971 | 3096.81793 | 2559.56719 | 2846.71222 | 0 |
| 1.23E-20 | 3.44E-18 | 638.222155 | 635.290332 | 516.924918 | 588.27104 | 11.8810912 |
| 1.34E-20 | 3.72E-18 | 1296.05718 | 4014.38537 | 3354.90768 | 3680.40549 | 0 |
| 1.38E-20 | 3.80E-18 | 2543.88599 | 5986.98855 | 6191.03434 | 6091.01773 | 30.8908372 |
| 1.53E-20 | 4.18E-18 | 1441.70687 | 3839.99195 | 2113.17421 | 2829.08265 | 0 |
| 1.68E-20 | 4.53E-18 | 1172.27102 | 2117.18956 | 2183.70616 | 2243.59523 | 0 |
| 1.67E-20 | 4.53E-18 | 1323.70812 | 5091.44272 | 4037.02584 | 4476.05599 | 0 |
| 1.70E-20 | 4.55E-18 | 1228.21594 | 2438.83866 | 1805.98903 | 2106.73406 | 0 |
| 1.77E-20 | 4.68E-18 | 1213.74742 | 2815.20814 | 2252.382 | 2407.82863 | 0 |
| 1.83E-20 | 4.80E-18 | 1259.08209 | 3873.35804 | 3938.65234 | 3909.58995 | 0 |

| | | | | | | |
|---|---|---|---|---|---|---|
| 1.85E-20 | 4.81E-18 | 10769.396 | 25955.0356 | 13066.0429 | 19047.3626 | 0 |
| 1.86E-20 | 4.81E-18 | 2331.35962 | 4504.86688 | 3222.19626 | 3866.90782 | 2.37621824 |
| 1.90E-20 | 4.89E-18 | 1742.97345 | 2270.67357 | 1246.37372 | 1632.12738 | 0 |
| 1.97E-20 | 5.03E-18 | 1201.20805 | 1886.96355 | 1457.04151 | 1716.56376 | 0 |
| 2.07E-20 | 5.19E-18 | 1203.4587 | 2745.36179 | 2129.87915 | 2503.86341 | 0 |
| 2.05E-20 | 5.19E-18 | 1400.23047 | 8273.90028 | 6278.27122 | 7217.91864 | 0 |
| 2.44E-20 | 6.08E-18 | 1121.47047 | 1849.81597 | 1799.49267 | 1899.81854 | 0 |
| 2.52E-20 | 6.24E-18 | 389.04222 | 736.500802 | 967.958149 | 953.388791 | 9.50487297 |
| 2.64E-20 | 6.48E-18 | 372.644573 | 760.969267 | 825.038154 | 913.954218 | 9.50487297 |
| 2.77E-20 | 6.76E-18 | 1339.78424 | 2512.02161 | 1467.25008 | 1936.00556 | 0 |
| 2.80E-20 | 6.77E-18 | 1255.86687 | 4653.67963 | 4720.07205 | 5006.79894 | 0 |
| 2.99E-20 | 7.20E-18 | 1550.05995 | 4628.09897 | 2064.91552 | 3106.98041 | 0 |
| 3.24E-20 | 7.74E-18 | 1219.85635 | 3264.09325 | 2231.96486 | 2929.29286 | 0 |
| 3.78E-20 | 8.97E-18 | 1098.32085 | 2270.89601 | 2075.12409 | 2216.68693 | 0 |
| 4.21E-20 | 9.90E-18 | 29.5800696 | 22.4664998 | 27.8415575 | 19.9492545 | 166.335277 |
| 4.26E-20 | 9.96E-18 | 1097.6778 | 1578.66088 | 1320.61788 | 1443.7693 | 0 |
| 4.44E-20 | 1.03E-17 | 943.990049 | 3170.22332 | 2720.12017 | 2986.35701 | 0 |
| 4.82E-20 | 1.11E-17 | 1111.18175 | 2870.81829 | 2966.98198 | 2868.05329 | 0 |
| 5.93E-20 | 1.36E-17 | 1357.46798 | 3824.64355 | 3580.4243 | 3917.9408 | 14.2573095 |
| 5.98E-20 | 1.36E-17 | 1201.20805 | 1309.73021 | 1062.61944 | 1262.37027 | 0 |
| 6.25E-20 | 1.41E-17 | 1037.87462 | 1548.85385 | 1397.64619 | 1572.74355 | 0 |
| 7.21E-20 | 1.62E-17 | 1111.82479 | 1828.01679 | 1469.10618 | 2155.44736 | 0 |
| 7.57E-20 | 1.69E-17 | 1129.50853 | 3035.64676 | 3959.99753 | 3636.33155 | 0 |
| 7.63E-20 | 1.69E-17 | 960.066173 | 683.115059 | 484.443101 | 699.615716 | 9.50487297 |
| 7.82E-20 | 1.72E-17 | 1097.99932 | 3506.33106 | 3122.8947 | 3554.21486 | 0 |
| 8.05E-20 | 1.76E-17 | 1075.17123 | 2070.25459 | 2645.87602 | 2623.55894 | 0 |
| 8.30E-20 | 1.80E-17 | 1106.03739 | 3647.13595 | 3473.69833 | 3995.41814 | 0 |
| 8.46E-20 | 1.82E-17 | 895.440152 | 2256.65981 | 1382.79736 | 1749.96716 | 9.50487297 |
| 8.96E-20 | 1.91E-17 | 1057.16597 | 2828.33213 | 2534.50979 | 2941.81914 | 0 |
| 8.96E-20 | 1.91E-17 | 6.10892743 | 2.66928711 | 3.71220767 | 1.39180845 | 147.325531 |
| 9.01E-20 | 1.91E-17 | 918.589771 | 3016.51687 | 1694.6228 | 2122.04396 | 4.75243649 |
| 9.21E-20 | 1.92E-17 | 1089.63974 | 3603.76004 | 2889.02562 | 3339.41242 | 0 |
| 9.16E-20 | 1.92E-17 | 1101.85759 | 4074.22189 | 3418.94326 | 3712.41708 | 0 |
| 9.48E-20 | 1.97E-17 | 1103.14368 | 4155.19027 | 3404.09443 | 3712.88102 | 0 |
| 9.88E-20 | 2.04E-17 | 1039.16071 | 2711.9957 | 2637.52355 | 2851.35159 | 0 |
| 1.11E-19 | 2.27E-17 | 1103.14368 | 4638.10879 | 4527.03725 | 4624.05162 | 0 |
| 1.20E-19 | 2.45E-17 | 1248.47185 | 3651.13988 | 3803.15676 | 3930.93101 | 2.37621824 |
| 1.28E-19 | 2.58E-17 | 349.816476 | 680.890653 | 465.882062 | 535.382319 | 19.0097459 |
| 1.28E-19 | 2.58E-17 | 438.878207 | 733.164193 | 544.766475 | 667.604122 | 7.12865473 |
| 1.29E-19 | 2.58E-17 | 1081.60168 | 2657.7202 | 1707.61553 | 2130.85874 | 0 |
| 1.30E-19 | 2.60E-17 | 1348.78687 | 2686.41503 | 2789.72406 | 2925.58137 | 0 |
| 1.33E-19 | 2.62E-17 | 1179.02299 | 3366.41593 | 2665.36511 | 3111.15583 | 21.3859642 |
| 1.34E-19 | 2.63E-17 | 1018.90479 | 2050.45738 | 1495.09164 | 1758.78195 | 0 |

| | | | | | | |
|---|---|---|---|---|---|---|
| 1.42E-19 | 2.77E-17 | 1025.01372 | 2987.37716 | 2425.92771 | 2849.95978 | 0 |
| 1.71E-19 | 3.32E-17 | 950.420499 | 1277.69876 | 1172.12957 | 1228.50293 | 0 |
| 1.73E-19 | 3.35E-17 | 1205.38784 | 1628.93246 | 1006.93633 | 1330.56888 | 0 |
| 1.75E-19 | 3.36E-17 | 1785.41442 | 2784.51134 | 2992.96743 | 3131.10509 | 2.37621824 |
| 1.93E-19 | 3.69E-17 | 995.75517 | 3146.64462 | 2714.55186 | 2924.18956 | 0 |
| 1.95E-19 | 3.70E-17 | 934.022851 | 1396.70448 | 1510.86852 | 1599.18791 | 0 |
| 2.09E-19 | 3.89E-17 | 1085.13842 | 2068.25263 | 1272.35918 | 1477.63664 | 0 |
| 2.08E-19 | 3.89E-17 | 1065.20403 | 1205.40557 | 972.598409 | 1068.90889 | 0 |
| 2.09E-19 | 3.89E-17 | 997.362783 | 2301.37037 | 1603.67371 | 2007.91566 | 0 |
| 2.07E-19 | 3.89E-17 | 978.071433 | 1560.19832 | 1173.98567 | 1369.53952 | 0 |
| 2.11E-19 | 3.89E-17 | 947.526796 | 1610.46989 | 1294.63242 | 1518.46302 | 0 |
| 2.12E-19 | 3.89E-17 | 921.161951 | 1593.5644 | 1474.6745 | 1585.73377 | 0 |
| 2.29E-19 | 4.19E-17 | 1055.87988 | 5226.01928 | 3988.76714 | 4530.80045 | 0 |
| 2.40E-19 | 4.37E-17 | 988.038631 | 1770.84956 | 1238.94931 | 1450.72835 | 0 |
| 2.43E-19 | 4.39E-17 | 953.957246 | 2414.37019 | 1934.98825 | 2219.47055 | 0 |
| 2.53E-19 | 4.56E-17 | 937.238076 | 1713.015 | 1337.32281 | 1550.93855 | 0 |
| 2.62E-19 | 4.70E-17 | 939.810256 | 1518.15704 | 1186.05035 | 1383.4576 | 0 |
| 2.65E-19 | 4.72E-17 | 926.306311 | 2056.90816 | 1947.05292 | 2197.66555 | 0 |
| 2.84E-19 | 5.02E-17 | 976.785343 | 3054.55422 | 2743.32147 | 3400.65199 | 0 |
| 2.96E-19 | 5.21E-17 | 49103.2371 | 12268.4884 | 11052.1703 | 12891.3938 | 78.415202 |
| 2.97E-19 | 5.21E-17 | 934.665896 | 2382.33874 | 2562.35134 | 2576.23745 | 0 |
| 3.08E-19 | 5.37E-17 | 278.116959 | 400.615507 | 373.076871 | 343.312752 | 7.12865473 |
| 3.14E-19 | 5.44E-17 | 977.428388 | 3671.82686 | 3200.85106 | 3683.65304 | 0 |
| 3.16E-19 | 5.46E-17 | 962.638353 | 2317.16365 | 3007.81626 | 2812.84489 | 0 |
| 3.22E-19 | 5.52E-17 | 880.971639 | 1345.3207 | 1325.25814 | 1303.19665 | 0 |
| 3.32E-19 | 5.67E-17 | 605.105338 | 1072.3861 | 807.405168 | 797.97018 | 9.50487297 |
| 3.34E-19 | 5.68E-17 | 922.769564 | 1191.83669 | 1405.99865 | 1558.36153 | 0 |
| 3.45E-19 | 5.83E-17 | 1020.19088 | 2871.04073 | 1717.8241 | 2203.69672 | 0 |
| 3.82E-19 | 6.42E-17 | 889.331224 | 1829.12899 | 1562.83943 | 1796.36078 | 0 |
| 3.90E-19 | 6.50E-17 | 940.774824 | 4071.33016 | 5675.03747 | 5071.28607 | 0 |
| 3.89E-19 | 6.50E-17 | 1030.15808 | 5468.25708 | 3696.43079 | 4255.22238 | 0 |
| 3.93E-19 | 6.51E-17 | 1029.51503 | 2915.52885 | 1692.7667 | 2041.31907 | 0 |
| 4.14E-19 | 6.83E-17 | 105.137856 | 86.9742716 | 78.8844129 | 94.6429749 | 627.321616 |
| 4.18E-19 | 6.86E-17 | 906.050394 | 2042.67196 | 1604.60176 | 2009.77141 | 0 |
| 4.43E-19 | 7.23E-17 | 888.045134 | 2125.19742 | 1770.72306 | 1966.16141 | 0 |
| 4.48E-19 | 7.28E-17 | 906.693439 | 1227.87207 | 995.799707 | 1139.89112 | 0 |
| 4.75E-19 | 7.68E-17 | 899.941467 | 2602.55493 | 2404.58252 | 2653.25085 | 0 |
| 5.04E-19 | 8.11E-17 | 908.301052 | 2801.41682 | 2537.29394 | 2953.41754 | 0 |
| 5.17E-19 | 8.28E-17 | 871.004442 | 1970.37877 | 1702.04722 | 1976.368 | 0 |
| 5.23E-19 | 8.34E-17 | 972.284028 | 5162.84615 | 4012.89649 | 4482.55109 | 0 |
| 5.86E-19 | 9.30E-17 | 929.521536 | 1761.95193 | 1173.05762 | 1372.32314 | 0 |
| 5.90E-19 | 9.32E-17 | 934.022851 | 4226.37126 | 4123.33467 | 4285.84217 | 0 |
| 6.69E-19 | 1.05E-16 | 410.90575 | 831.482935 | 783.275818 | 849.931029 | 47.5243649 |

| | | | | | | |
|---|---|---|---|---|---|---|
| 6.92E-19 | 1.08E-16 | 892.224927 | 1376.68483 | 1010.64854 | 1162.624 | 0 |
| 7.10E-19 | 1.11E-16 | 909.265619 | 3107.0502 | 2195.77084 | 2669.48861 | 0 |
| 7.18E-19 | 1.11E-16 | 0 | 0 | 0 | 0 | 209.107205 |
| 7.71E-19 | 1.19E-16 | 993.504513 | 2828.10969 | 1522.00514 | 2111.83736 | 0 |
| 7.74E-19 | 1.19E-16 | 828.884995 | 1659.18438 | 1393.00593 | 1485.05962 | 0 |
| 8.20E-19 | 1.25E-16 | 981.608181 | 1174.48633 | 972.598409 | 864.31305 | 0 |
| 8.55E-19 | 1.30E-16 | 1012.15282 | 1172.26192 | 2000.87993 | 2171.22119 | 0 |
| 8.77E-19 | 1.33E-16 | 573.596133 | 735.166158 | 694.182834 | 778.948798 | 19.0097459 |
| 9.98E-19 | 1.51E-16 | 820.203888 | 1803.77076 | 1449.61709 | 1668.3144 | 0 |
| 1.03E-18 | 1.54E-16 | 798.340358 | 1183.82883 | 1329.8984 | 1189.06836 | 0 |
| 1.06E-18 | 1.59E-16 | 890.938837 | 4336.47935 | 3428.22378 | 3528.69837 | 0 |
| 1.07E-18 | 1.59E-16 | 848.81939 | 2596.77148 | 1952.62123 | 2222.25416 | 0 |
| 1.08E-18 | 1.59E-16 | 867.467695 | 3682.72645 | 3276.95132 | 3537.04922 | 0 |
| 1.11E-18 | 1.64E-16 | 805.09233 | 1407.82651 | 1147.07217 | 1254.48335 | 0 |
| 1.13E-18 | 1.66E-16 | 852.999182 | 3179.34339 | 2960.48562 | 3316.21561 | 0 |
| 1.14E-18 | 1.67E-16 | 879.364027 | 4451.03625 | 3896.89 | 4004.69686 | 0 |
| 1.16E-18 | 1.68E-16 | 977.428388 | 957.161869 | 828.750362 | 875.911454 | 0 |
| 1.20E-18 | 1.75E-16 | 872.290532 | 4329.36125 | 3779.02741 | 3864.58814 | 0 |
| 1.21E-18 | 1.75E-16 | 1497.33026 | 4482.62282 | 3755.82611 | 4385.1245 | 2.37621824 |
| 1.24E-18 | 1.79E-16 | 56.9094818 | 40.4841878 | 38.9781805 | 29.2279775 | 1119.19879 |
| 1.26E-18 | 1.81E-16 | 16.0761248 | 13.5688761 | 18.5610383 | 14.3820207 | 743.75631 |
| 1.28E-18 | 1.82E-16 | 2059.67311 | 3662.7068 | 3480.19469 | 3538.44103 | 21.3859642 |
| 1.35E-18 | 1.92E-16 | 781.621188 | 1537.28693 | 1352.17164 | 1532.38111 | 0 |
| 1.38E-18 | 1.96E-16 | 1671.27393 | 1251.89565 | 881.649321 | 1060.55804 | 0 |
| 1.44E-18 | 2.03E-16 | 647.224785 | 2193.26424 | 2027.79344 | 2252.87395 | 14.2573095 |
| 1.61E-18 | 2.26E-16 | 815.38105 | 2341.85456 | 1764.22669 | 2249.16246 | 0 |
| 1.63E-18 | 2.27E-16 | 799.304925 | 2396.13006 | 2120.59863 | 2422.21065 | 0 |
| 1.69E-18 | 2.35E-16 | 808.307555 | 2328.953 | 1728.03267 | 1925.33503 | 0 |
| 1.77E-18 | 2.45E-16 | 774.869216 | 1572.43255 | 1247.30178 | 1376.9625 | 0 |
| 1.82E-18 | 2.51E-16 | 861.358767 | 3463.17758 | 2216.18798 | 2672.73617 | 0 |
| 1.94E-18 | 2.66E-16 | 778.727486 | 2163.4572 | 2184.63421 | 2331.7431 | 0 |
| 1.95E-18 | 2.66E-16 | 879.364027 | 1961.03626 | 1980.46279 | 2045.95843 | 14.2573095 |
| 1.95E-18 | 2.66E-16 | 768.760288 | 1915.2135 | 1900.65033 | 2064.51587 | 0 |
| 2.02E-18 | 2.72E-16 | 787.408593 | 2561.18098 | 2518.7329 | 2656.4984 | 0 |
| 2.01E-18 | 2.72E-16 | 747.218281 | 1319.74003 | 1166.56126 | 1276.28835 | 0 |
| 2.02E-18 | 2.72E-16 | 17.6837373 | 11.3444702 | 8.35246725 | 12.0623399 | 168.711495 |
| 2.05E-18 | 2.75E-16 | 768.117243 | 2148.1088 | 2017.58487 | 1982.39917 | 0 |
| 2.06E-18 | 2.75E-16 | 756.542433 | 1321.96444 | 1096.95737 | 1282.31952 | 0 |
| 2.10E-18 | 2.80E-16 | 766.831153 | 1492.35393 | 1156.35269 | 1288.81463 | 0 |
| 2.33E-18 | 3.08E-16 | 732.428246 | 1398.26156 | 1366.09242 | 1370.93133 | 0 |
| 2.34E-18 | 3.08E-16 | 753.970253 | 1719.24334 | 1633.37137 | 1914.20056 | 0 |
| 2.34E-18 | 3.08E-16 | 1344.9286 | 1373.34822 | 765.642831 | 1059.16623 | 0 |
| 2.40E-18 | 3.14E-16 | 764.258973 | 1208.74218 | 945.684903 | 1124.11729 | 0 |

| | | | | | | |
|---|---|---|---|---|---|---|
| 2.41E-18 | 3.14E-16 | 15.4330798 | 10.4547078 | 10.2085711 | 6.95904227 | 159.206622 |
| 2.46E-18 | 3.20E-16 | 800.912538 | 3303.2428 | 2558.63914 | 2700.1084 | 0 |
| 2.47E-18 | 3.20E-16 | 1383.18978 | 4347.37894 | 4382.26115 | 4875.04108 | 2.37621824 |
| 2.53E-18 | 3.26E-16 | 803.80624 | 1316.62587 | 918.771398 | 1063.80559 | 0 |
| 2.63E-18 | 3.37E-16 | 2424.92267 | 6802.23332 | 6213.30758 | 7120.49205 | 21.3859642 |
| 2.71E-18 | 3.47E-16 | 281.975229 | 253.804716 | 245.933758 | 247.277969 | 11.8810912 |
| 2.73E-18 | 3.48E-16 | 910.230187 | 3381.98677 | 2943.78068 | 3158.47732 | 0 |
| 2.79E-18 | 3.55E-16 | 754.934821 | 1880.73521 | 1434.76826 | 1694.29482 | 0 |
| 2.92E-18 | 3.69E-16 | 927.592401 | 1069.93925 | 812.973479 | 792.402946 | 0 |
| 2.97E-18 | 3.74E-16 | 738.537174 | 1524.38538 | 1197.18697 | 1337.99186 | 0 |
| 3.14E-18 | 3.94E-16 | 720.210391 | 992.529923 | 920.627502 | 977.513471 | 0 |
| 3.33E-18 | 4.15E-16 | 713.779941 | 984.966943 | 1050.55477 | 958.956025 | 0 |
| 3.34E-18 | 4.15E-16 | 1481.57566 | 6811.79826 | 5668.54111 | 6275.66432 | 0 |
| 3.34E-18 | 4.15E-16 | 772.297036 | 2168.3509 | 1467.25008 | 1786.61812 | 0 |
| 3.45E-18 | 4.27E-16 | 719.567346 | 1628.26514 | 1437.55242 | 1643.26185 | 0 |
| 3.47E-18 | 4.28E-16 | 706.706446 | 1274.58459 | 1324.33009 | 1382.06579 | 0 |
| 3.50E-18 | 4.29E-16 | 716.995166 | 1097.29944 | 943.8288 | 1014.16443 | 0 |
| 3.52E-18 | 4.30E-16 | 1341.07033 | 4212.13506 | 3980.41467 | 4757.66523 | 2.37621824 |
| 3.73E-18 | 4.55E-16 | 44.048582 | 29.3621582 | 23.2012979 | 27.3722329 | 354.056518 |
| 3.75E-18 | 4.56E-16 | 756.863956 | 3042.31998 | 2502.95602 | 2738.6151 | 0 |
| 3.82E-18 | 4.62E-16 | 717.316689 | 1767.29051 | 1484.88307 | 1623.77653 | 0 |
| 3.99E-18 | 4.81E-16 | 738.215651 | 2484.43898 | 2271.87109 | 2591.08341 | 0 |
| 4.09E-18 | 4.91E-16 | 735.643471 | 1904.53635 | 1395.79008 | 1686.40791 | 0 |
| 4.17E-18 | 4.99E-16 | 751.076551 | 2868.59388 | 3035.65782 | 3266.57444 | 0 |
| 4.30E-18 | 5.13E-16 | 0 | 0 | 0 | 0 | 235.245606 |
| 4.37E-18 | 5.20E-16 | 702.848177 | 1681.42844 | 1470.96229 | 1584.80589 | 0 |
| 4.43E-18 | 5.24E-16 | 1188.66867 | 1246.77952 | 709.031665 | 861.529433 | 14.2573095 |
| 4.46E-18 | 5.27E-16 | 1005.72237 | 2128.08915 | 954.037371 | 1320.36229 | 0 |
| 4.69E-18 | 5.52E-16 | 695.131637 | 1670.30641 | 1563.76748 | 1555.11398 | 0 |
| 4.75E-18 | 5.57E-16 | 1292.84196 | 3926.96622 | 3497.82768 | 3986.60335 | 2.37621824 |
| 5.23E-18 | 6.11E-16 | 14.7900348 | 13.7913167 | 16.7049345 | 22.2689353 | 135.44444 |
| 5.26E-18 | 6.12E-16 | 718.281256 | 1632.26907 | 1170.27347 | 1370.46739 | 0 |
| 5.48E-18 | 6.36E-16 | 726.640841 | 1859.82579 | 1283.4958 | 1520.7827 | 0 |
| 5.56E-18 | 6.42E-16 | 600.2825 | 1482.78899 | 995.799707 | 1214.58484 | 0 |
| 5.77E-18 | 6.65E-16 | 725.676274 | 1499.47203 | 1026.42542 | 1246.59644 | 0 |
| 6.00E-18 | 6.89E-16 | 670.374404 | 1097.96676 | 1029.20958 | 996.070917 | 0 |
| 6.46E-18 | 7.40E-16 | 722.461049 | 1087.06718 | 798.124649 | 1003.02996 | 0 |
| 6.53E-18 | 7.45E-16 | 697.382294 | 1295.27157 | 1704.83137 | 1648.82908 | 0 |
| 6.65E-18 | 7.56E-16 | 687.093574 | 984.299622 | 825.966206 | 883.798368 | 0 |
| 6.78E-18 | 7.68E-16 | 1538.48514 | 1480.78702 | 688.614522 | 1181.18144 | 0 |
| 6.94E-18 | 7.76E-16 | 694.810114 | 2309.82311 | 1983.24695 | 2265.86416 | 0 |
| 6.97E-18 | 7.76E-16 | 670.695927 | 1372.01357 | 1133.15139 | 1248.45218 | 0 |
| 6.95E-18 | 7.76E-16 | 699.632952 | 2665.28318 | 2448.20096 | 2476.95511 | 0 |

| | | | | | | |
|---|---|---|---|---|---|---|
| 6.95E-18 | 7.76E-16 | 937.238076 | 3248.74485 | 2475.11446 | 2838.36137 | 28.5146189 |
| 6.88E-18 | 7.76E-16 | 700.597519 | 1256.56691 | 957.749578 | 973.801982 | 0 |
| 7.01E-18 | 7.79E-16 | 963.602921 | 1037.9078 | 678.405951 | 854.106455 | 0 |
| 7.17E-18 | 7.93E-16 | 743.360011 | 1294.60425 | 851.023608 | 1016.48411 | 0 |
| 7.40E-18 | 8.16E-16 | 908.622574 | 1448.97802 | 749.865949 | 1092.56964 | 0 |
| 7.47E-18 | 8.19E-16 | 663.622432 | 878.64034 | 852.879712 | 924.62475 | 0 |
| 7.45E-18 | 8.19E-16 | 1650.6965 | 1213.19099 | 838.958933 | 1065.1974 | 0 |
| 7.55E-18 | 8.24E-16 | 585.170943 | 679.333569 | 626.435044 | 746.937204 | 38.0194919 |
| 7.62E-18 | 8.29E-16 | 8.0380624 | 5.7834554 | 4.64025958 | 4.17542536 | 80.7914203 |
| 7.74E-18 | 8.40E-16 | 735.643471 | 4049.53099 | 2895.52198 | 3219.71689 | 0 |
| 7.88E-18 | 8.53E-16 | 654.619802 | 1158.69305 | 1186.05035 | 1302.73271 | 0 |
| 7.95E-18 | 8.57E-16 | 682.592259 | 1472.55672 | 1213.89191 | 1092.1057 | 0 |
| 8.09E-18 | 8.66E-16 | 900.584512 | 2915.08396 | 2682.99809 | 3065.22615 | 78.415202 |
| 8.07E-18 | 8.66E-16 | 800.269493 | 1524.60782 | 861.232179 | 1094.88932 | 0 |
| 8.15E-18 | 8.69E-16 | 673.589629 | 2067.80775 | 1898.79422 | 2023.22556 | 0 |
| 8.16E-18 | 8.69E-16 | 717.316689 | 1060.59674 | 764.71478 | 926.94443 | 0 |
| 8.21E-18 | 8.71E-16 | 651.7261 | 1309.73021 | 1320.61788 | 1405.2626 | 0 |
| 8.34E-18 | 8.83E-16 | 661.371775 | 1486.57048 | 1589.75293 | 1692.43908 | 0 |
| 8.38E-18 | 8.84E-16 | 691.916412 | 1267.24406 | 912.275034 | 1042.0006 | 0 |
| 8.46E-18 | 8.90E-16 | 868.753785 | 1731.25513 | 855.663867 | 1200.66676 | 0 |
| 8.68E-18 | 9.10E-16 | 673.589629 | 818.58138 | 896.498152 | 772.917628 | 0 |
| 8.78E-18 | 9.17E-16 | 686.129007 | 2702.20832 | 2487.17914 | 2694.54117 | 0 |
| 8.99E-18 | 9.37E-16 | 670.052882 | 1449.86778 | 1090.461 | 1243.34889 | 0 |
| 9.23E-18 | 9.57E-16 | 786.444026 | 1502.80864 | 844.527244 | 1121.33368 | 0 |
| 9.27E-18 | 9.57E-16 | 683.235304 | 1579.32821 | 1109.95009 | 1322.21803 | 0 |
| 9.25E-18 | 9.57E-16 | 658.15655 | 1183.16151 | 952.181267 | 1131.07634 | 0 |
| 9.48E-18 | 9.75E-16 | 294.193084 | 639.961584 | 578.176344 | 627.241677 | 23.7621824 |
| 1.00E-17 | 1.03E-15 | 691.916412 | 3231.17205 | 2593.90511 | 2847.17616 | 0 |
| 1.01E-17 | 1.03E-15 | 687.415097 | 2456.85634 | 2615.2503 | 3042.49328 | 0 |
| 1.05E-17 | 1.07E-15 | 663.943954 | 1929.67214 | 2066.77162 | 2249.16246 | 0 |
| 1.05E-17 | 1.07E-15 | 661.693297 | 1650.28676 | 1877.44903 | 2021.83375 | 0 |
| 1.06E-17 | 1.08E-15 | 649.475442 | 1536.17473 | 1574.9041 | 1772.2361 | 0 |
| 1.10E-17 | 1.11E-15 | 663.30091 | 1716.35161 | 1278.85554 | 1578.77472 | 0 |
| 1.12E-17 | 1.12E-15 | 642.72347 | 1638.4974 | 1733.60098 | 1601.04366 | 0 |
| 1.13E-17 | 1.14E-15 | 640.472812 | 1568.65106 | 1355.88385 | 1495.73015 | 0 |
| 1.14E-17 | 1.14E-15 | 684.842917 | 3217.15829 | 3224.05236 | 3675.76613 | 0 |
| 1.15E-17 | 1.15E-15 | 643.366515 | 1344.43094 | 1402.28645 | 1614.49781 | 0 |
| 1.16E-17 | 1.15E-15 | 625.361255 | 1131.11041 | 1134.07944 | 1192.77985 | 0 |
| 1.17E-17 | 1.15E-15 | 723.104094 | 2791.62944 | 1687.19839 | 1958.73843 | 0 |
| 1.17E-17 | 1.15E-15 | 635.971497 | 1403.3777 | 1332.68255 | 1522.17451 | 0 |
| 1.17E-17 | 1.16E-15 | 630.505615 | 964.502409 | 895.5701 | 1007.20538 | 0 |
| 1.20E-17 | 1.18E-15 | 912.802366 | 2500.67714 | 1963.75786 | 2232.46076 | 4.75243649 |
| 1.20E-17 | 1.18E-15 | 1002.50714 | 4868.55725 | 4594.78504 | 5278.20159 | 0 |

| | | | | | | |
|---|---|---|---|---|---|---|
| 1.22E-17 | 1.19E-15 | 2027.52086 | 5202.44058 | 4672.7414 | 5177.99138 | 21.3859642 |
| 1.38E-17 | 1.35E-15 | 923.091086 | 2936.43826 | 2411.07888 | 2834.18595 | 4.75243649 |
| 1.41E-17 | 1.37E-15 | 632.113227 | 1804.66053 | 1582.32852 | 1637.69461 | 0 |
| 1.43E-17 | 1.38E-15 | 653.655235 | 2688.19456 | 2310.84927 | 2456.54192 | 0 |
| 1.45E-17 | 1.41E-15 | 910.230187 | 1923.66624 | 1296.48853 | 1592.22887 | 4.75243649 |
| 1.47E-17 | 1.42E-15 | 623.110597 | 1526.83223 | 1315.97762 | 1380.67399 | 0 |
| 1.53E-17 | 1.47E-15 | 634.042362 | 1928.78238 | 1639.86774 | 1875.22992 | 0 |
| 1.55E-17 | 1.48E-15 | 670.374404 | 680.890653 | 687.68647 | 739.050289 | 0 |
| 1.65E-17 | 1.57E-15 | 646.58174 | 1339.75969 | 940.116592 | 1133.39602 | 0 |
| 1.68E-17 | 1.60E-15 | 634.042362 | 1351.99392 | 996.727759 | 1129.68453 | 0 |
| 1.71E-17 | 1.63E-15 | 606.71295 | 1054.36841 | 933.620228 | 1044.32028 | 0 |
| 1.74E-17 | 1.64E-15 | 630.184092 | 982.075216 | 780.491662 | 857.817944 | 0 |
| 1.73E-17 | 1.64E-15 | 1.60761248 | 0.22244059 | 0 | 0.9278723 | 130.692003 |
| 1.76E-17 | 1.66E-15 | 471.351979 | 381.485616 | 339.667002 | 384.603069 | 21.3859642 |
| 1.77E-17 | 1.66E-15 | 814.416483 | 2609.22815 | 1151.71243 | 1667.85046 | 0 |
| 1.77E-17 | 1.66E-15 | 638.543677 | 2300.92549 | 2235.67707 | 2617.9917 | 0 |
| 1.81E-17 | 1.69E-15 | 1122.75656 | 2665.72806 | 2498.31576 | 2776.65787 | 2.37621824 |
| 1.84E-17 | 1.71E-15 | 626.968867 | 1429.18081 | 1056.12308 | 1230.35867 | 0 |
| 1.85E-17 | 1.72E-15 | 629.541047 | 2190.81739 | 2067.69967 | 2363.29075 | 0 |
| 1.90E-17 | 1.75E-15 | 635.649975 | 497.599605 | 443.608816 | 467.183704 | 21.3859642 |
| 1.89E-17 | 1.75E-15 | 794.80361 | 1442.3048 | 749.865949 | 1003.4939 | 0 |
| 1.93E-17 | 1.78E-15 | 611.214265 | 1656.29265 | 1418.06333 | 1569.496 | 0 |
| 1.95E-17 | 1.79E-15 | 683.235304 | 3899.38359 | 2506.66823 | 3455.86039 | 0 |
| 1.96E-17 | 1.80E-15 | 8.0380624 | 11.7893514 | 12.0646749 | 13.4541484 | 130.692003 |
| 1.99E-17 | 1.82E-15 | 303.838759 | 608.37502 | 505.788295 | 577.600508 | 7.12865473 |
| 2.00E-17 | 1.82E-15 | 1238.18313 | 1289.71055 | 1575.83215 | 1446.08898 | 0 |
| 2.01E-17 | 1.83E-15 | 395.794193 | 326.320349 | 307.185185 | 342.848816 | 7.12865473 |
| 2.02E-17 | 1.83E-15 | 751.719596 | 1226.53743 | 696.96699 | 964.523259 | 0 |
| 2.05E-17 | 1.86E-15 | 630.184092 | 2689.97408 | 2319.20174 | 2367.00224 | 0 |
| 2.08E-17 | 1.87E-15 | 606.391428 | 1446.75361 | 1171.20152 | 1260.97846 | 0 |
| 2.14E-17 | 1.93E-15 | 615.394058 | 2134.31748 | 1897.86617 | 2058.4847 | 0 |
| 2.16E-17 | 1.93E-15 | 595.13814 | 950.711092 | 1111.8062 | 1065.66134 | 0 |
| 2.15E-17 | 1.93E-15 | 628.57648 | 2267.78184 | 1718.75215 | 1878.47748 | 0 |
| 2.33E-17 | 2.08E-15 | 669.088314 | 2113.18563 | 1245.44567 | 1604.29121 | 0 |
| 2.33E-17 | 2.08E-15 | 10.6102424 | 9.78738607 | 12.9927268 | 12.9902122 | 152.077968 |
| 2.36E-17 | 2.09E-15 | 584.206375 | 1131.77773 | 1147.07217 | 1203.91431 | 0 |
| 2.41E-17 | 2.13E-15 | 0 | 0 | 0 | 0 | 85.5438568 |
| 2.45E-17 | 2.17E-15 | 643.366515 | 1201.1792 | 1844.03916 | 1716.09982 | 0 |
| 2.47E-17 | 2.18E-15 | 593.85205 | 1440.52528 | 1225.02853 | 1426.13973 | 0 |
| 2.48E-17 | 2.18E-15 | 587.4216 | 1182.49419 | 1041.27425 | 1014.16443 | 0 |
| 2.50E-17 | 2.19E-15 | 599.960978 | 1823.34554 | 1849.60747 | 1943.42854 | 0 |
| 2.52E-17 | 2.20E-15 | 614.751013 | 2577.86403 | 2562.35134 | 2672.73617 | 0 |
| 2.54E-17 | 2.21E-15 | 586.13551 | 1298.38574 | 1387.43762 | 1421.50037 | 0 |

| | | | | | | |
|---|---|---|---|---|---|---|
| 2.55E-17 | 2.21E-15 | 774.869216 | 644.855277 | 638.499719 | 699.15178 | 0 |
| 2.64E-17 | 2.29E-15 | 587.100078 | 828.591207 | 758.218416 | 838.332625 | 0 |
| 2.69E-17 | 2.33E-15 | 583.56333 | 850.167944 | 814.829583 | 926.94443 | 0 |
| 2.72E-17 | 2.34E-15 | 625.361255 | 746.510628 | 835.246725 | 662.500824 | 0 |
| 2.72E-17 | 2.34E-15 | 580.348106 | 1180.71466 | 1042.2023 | 1040.14485 | 0 |
| 2.76E-17 | 2.36E-15 | 587.4216 | 1274.80704 | 1045.91451 | 1234.5341 | 0 |
| 2.86E-17 | 2.45E-15 | 575.203746 | 1085.95497 | 963.31789 | 1053.599 | 0 |
| 2.90E-17 | 2.47E-15 | 612.178833 | 734.276396 | 647.780238 | 746.473267 | 0 |
| 3.04E-17 | 2.59E-15 | 694.488592 | 4016.60978 | 2110.39006 | 2828.61871 | 0 |
| 3.09E-17 | 2.62E-15 | 632.756272 | 1364.67303 | 866.80049 | 1152.88134 | 0 |
| 3.14E-17 | 2.66E-15 | 576.168313 | 1253.67518 | 1045.91451 | 1157.5207 | 0 |
| 3.14E-17 | 2.66E-15 | 597.388798 | 1024.33893 | 775.851403 | 883.334432 | 0 |
| 3.23E-17 | 2.72E-15 | 626.647345 | 1212.96855 | 795.340493 | 1008.13326 | 0 |
| 3.30E-17 | 2.77E-15 | 582.920285 | 1664.96783 | 1359.59606 | 1508.72036 | 0 |
| 3.40E-17 | 2.85E-15 | 585.492465 | 1794.20582 | 1575.83215 | 1889.14801 | 0 |
| 3.46E-17 | 2.89E-15 | 571.988521 | 1569.9857 | 1570.26384 | 1533.77292 | 0 |
| 3.48E-17 | 2.90E-15 | 592.244438 | 1684.54261 | 1883.01734 | 2133.64236 | 0 |
| 3.53E-17 | 2.94E-15 | 735.321949 | 1319.74003 | 707.175561 | 908.85092 | 0 |
| 3.58E-17 | 2.97E-15 | 586.457033 | 1967.48704 | 1931.27604 | 2216.68693 | 0 |
| 3.60E-17 | 2.98E-15 | 588.064645 | 1243.44291 | 907.634775 | 1080.5073 | 0 |
| 3.65E-17 | 3.02E-15 | 1057.16597 | 2628.80292 | 2472.33031 | 2801.24648 | 0 |
| 3.68E-17 | 3.03E-15 | 636.614542 | 1485.23584 | 889.073736 | 1198.81101 | 0 |
| 3.69E-17 | 3.04E-15 | 610.892743 | 827.923885 | 642.211927 | 740.442097 | 0 |
| 3.74E-17 | 3.05E-15 | 872.612054 | 2128.31159 | 800.908804 | 1664.60291 | 0 |
| 3.74E-17 | 3.05E-15 | 317.985749 | 803.900301 | 796.268545 | 812.816137 | 30.8908372 |
| 3.73E-17 | 3.05E-15 | 693.202502 | 3230.28228 | 1595.32125 | 2321.07257 | 0 |
| 3.74E-17 | 3.05E-15 | 579.062016 | 1245.66732 | 942.900748 | 1115.30251 | 0 |
| 3.78E-17 | 3.08E-15 | 415.085542 | 1196.95283 | 1057.05113 | 1159.37644 | 9.50487297 |
| 3.89E-17 | 3.15E-15 | 581.634195 | 2236.41772 | 1955.40539 | 2090.96023 | 0 |
| 3.96E-17 | 3.20E-15 | 739.823264 | 2572.52545 | 2476.04251 | 2665.31319 | 2.37621824 |
| 3.99E-17 | 3.22E-15 | 335.026441 | 522.06807 | 377.71713 | 458.368917 | 2.37621824 |
| 4.00E-17 | 3.22E-15 | 553.661738 | 1110.64588 | 1144.28801 | 1158.44857 | 0 |
| 4.02E-17 | 3.22E-15 | 555.590873 | 1216.97248 | 1113.6623 | 1172.36665 | 0 |
| 4.02E-17 | 3.22E-15 | 577.454403 | 1950.35911 | 2083.47655 | 2151.27193 | 0 |
| 4.16E-17 | 3.32E-15 | 574.560701 | 2154.78202 | 2059.3472 | 2080.2897 | 0 |
| 4.16E-17 | 3.32E-15 | 603.176203 | 1588.67071 | 1031.99373 | 1298.09335 | 0 |
| 4.23E-17 | 3.35E-15 | 521.831011 | 1055.48061 | 919.69945 | 982.616768 | 7.12865473 |
| 4.24E-17 | 3.35E-15 | 756.220911 | 1155.35644 | 617.154525 | 892.613155 | 0 |
| 4.24E-17 | 3.35E-15 | 607.677518 | 1834.91245 | 2578.12823 | 2598.97032 | 0 |
| 4.34E-17 | 3.43E-15 | 318.307271 | 264.926746 | 274.703367 | 317.332327 | 16.6335277 |
| 4.38E-17 | 3.45E-15 | 414.76402 | 1049.25227 | 869.584646 | 932.511664 | 19.0097459 |
| 4.40E-17 | 3.46E-15 | 607.99904 | 2469.53546 | 1615.73839 | 2137.35385 | 0 |
| 4.46E-17 | 3.50E-15 | 549.160423 | 1192.05913 | 1097.88542 | 1129.22059 | 0 |

| | | | | | | |
|---|---|---|---|---|---|---|
| 4.60E-17 | 3.59E-15 | 245.321665 | 328.099874 | 297.904665 | 294.599456 | 4.75243649 |
| 4.60E-17 | 3.59E-15 | 989.003198 | 5761.43378 | 4991.06321 | 5281.44915 | 2.37621824 |
| 4.61E-17 | 3.59E-15 | 1050.73552 | 2122.30569 | 1331.7545 | 1616.81749 | 26.1384007 |
| 4.67E-17 | 3.63E-15 | 562.021323 | 1392.70055 | 1544.27839 | 1673.88163 | 0 |
| 4.73E-17 | 3.66E-15 | 218.313775 | 357.017151 | 300.688821 | 337.281582 | 9.50487297 |
| 4.75E-17 | 3.67E-15 | 0 | 0 | 0 | 0 | 87.920075 |
| 4.77E-17 | 3.68E-15 | 554.626306 | 1394.92496 | 1405.99865 | 1212.7291 | 0 |
| 4.79E-17 | 3.69E-15 | 568.451773 | 1256.12203 | 939.18854 | 1183.03719 | 0 |
| 4.83E-17 | 3.71E-15 | 558.806098 | 1053.03376 | 833.390621 | 959.419961 | 0 |
| 4.85E-17 | 3.72E-15 | 611.535788 | 1854.70966 | 1135.93555 | 1350.0542 | 0 |
| 4.92E-17 | 3.77E-15 | 0 | 0 | 0 | 0 | 206.730987 |
| 5.00E-17 | 3.81E-15 | 1153.30119 | 990.305518 | 585.60076 | 788.227521 | 0 |
| 5.01E-17 | 3.81E-15 | 547.874333 | 1228.76183 | 1155.42464 | 1326.85739 | 0 |
| 5.09E-17 | 3.86E-15 | 553.661738 | 1059.92942 | 849.167504 | 977.049535 | 0 |
| 5.15E-17 | 3.90E-15 | 578.418971 | 2052.90423 | 1468.17813 | 1673.88163 | 0 |
| 5.25E-17 | 3.97E-15 | 578.740493 | 1664.96783 | 1142.43191 | 1331.03282 | 0 |
| 5.35E-17 | 4.03E-15 | 1009.25912 | 2996.71966 | 2470.4742 | 2768.30701 | 2.37621824 |
| 5.36E-17 | 4.04E-15 | 572.953088 | 915.120597 | 721.096339 | 729.30763 | 0 |
| 5.39E-17 | 4.04E-15 | 562.021323 | 2099.39431 | 2031.50565 | 2262.61661 | 0 |
| 5.39E-17 | 4.04E-15 | 0.96456749 | 0 | 0 | 0 | 111.682257 |
| 5.42E-17 | 4.05E-15 | 736.286516 | 2039.78023 | 2021.29708 | 2061.26832 | 85.5438568 |
| 5.57E-17 | 4.16E-15 | 545.945198 | 1217.19492 | 993.015551 | 1125.97304 | 0 |
| 5.88E-17 | 4.38E-15 | 486.142014 | 548.983382 | 408.342843 | 520.536362 | 14.2573095 |
| 5.94E-17 | 4.41E-15 | 308.018551 | 502.493298 | 407.414792 | 453.26562 | 9.50487297 |
| 6.06E-17 | 4.48E-15 | 567.487206 | 1216.08272 | 852.879712 | 1120.40581 | 0 |
| 6.04E-17 | 4.48E-15 | 3253.80766 | 1297.27354 | 917.843346 | 1216.90452 | 0 |
| 6.21E-17 | 4.58E-15 | 533.405821 | 940.256384 | 846.383348 | 966.842939 | 0 |
| 6.36E-17 | 4.68E-15 | 544.016063 | 745.175985 | 919.69945 | 819.311243 | 0 |
| 6.57E-17 | 4.83E-15 | 566.522638 | 690.233158 | 613.442317 | 754.824118 | 0 |
| 6.75E-17 | 4.95E-15 | 541.122361 | 1577.32624 | 1322.47398 | 1550.01068 | 0 |
| 6.83E-17 | 5.00E-15 | 545.302153 | 759.634623 | 654.276601 | 762.247097 | 0 |
| 7.33E-17 | 5.35E-15 | 554.947828 | 1223.20082 | 854.735816 | 1025.76283 | 0 |
| 7.37E-17 | 5.36E-15 | 578.097448 | 1573.54475 | 978.16672 | 1285.10314 | 0 |
| 7.36E-17 | 5.36E-15 | 556.233918 | 1805.32785 | 1267.71892 | 1502.22526 | 0 |
| 7.45E-17 | 5.40E-15 | 545.302153 | 2228.40986 | 2188.34642 | 2418.96309 | 0 |
| 7.53E-17 | 5.45E-15 | 525.689281 | 1080.17152 | 908.562827 | 1005.34964 | 0 |
| 7.63E-17 | 5.51E-15 | 525.689281 | 922.683577 | 788.844129 | 897.716453 | 0 |
| 7.68E-17 | 5.53E-15 | 518.937309 | 792.555831 | 780.491662 | 804.929223 | 0 |
| 7.75E-17 | 5.57E-15 | 524.724714 | 1226.31499 | 1317.83372 | 1376.9625 | 0 |
| 7.85E-17 | 5.63E-15 | 526.332326 | 1550.41093 | 1493.23553 | 1492.01866 | 0 |
| 7.96E-17 | 5.69E-15 | 516.365129 | 835.486865 | 812.973479 | 865.704858 | 0 |
| 8.24E-17 | 5.89E-15 | 526.332326 | 1114.64981 | 1081.18048 | 1316.6508 | 0 |
| 8.64E-17 | 6.16E-15 | 527.296894 | 1785.30819 | 1756.80228 | 1843.21833 | 0 |

| | | | | | | |
|---|---|---|---|---|---|---|
| 8.77E-17 | 6.24E-15 | 556.555441 | 728.048059 | 584.672708 | 708.430503 | 0 |
| 8.85E-17 | 6.27E-15 | 532.441254 | 759.634623 | 920.627502 | 732.091247 | 0 |
| 8.84E-17 | 6.27E-15 | 525.046236 | 1608.46792 | 1358.66801 | 1506.86462 | 0 |
| 9.12E-17 | 6.44E-15 | 557.198486 | 1249.89369 | 816.685687 | 1031.794 | 0 |
| 9.16E-17 | 6.46E-15 | 603.819248 | 1286.15151 | 733.161014 | 970.554429 | 0 |
| 9.27E-17 | 6.53E-15 | 593.209005 | 1277.47632 | 743.369585 | 998.854534 | 0 |
| 9.35E-17 | 6.57E-15 | 520.544921 | 979.183488 | 787.916077 | 918.129643 | 0 |
| 9.45E-17 | 6.63E-15 | 530.190596 | 2055.79596 | 1705.75942 | 1912.80875 | 0 |
| 9.82E-17 | 6.87E-15 | 512.828381 | 990.750399 | 832.46257 | 961.739642 | 0 |
| 1.00E-16 | 7.00E-15 | 538.871704 | 1699.89101 | 1188.83451 | 1352.37388 | 0 |
| 1.03E-16 | 7.19E-15 | 534.691911 | 1864.71949 | 1349.38749 | 1670.17014 | 0 |
| 1.04E-16 | 7.25E-15 | 517.008174 | 903.998568 | 774.923351 | 981.688896 | 0 |
| 1.08E-16 | 7.47E-15 | 440.48582 | 982.520097 | 793.484389 | 970.554429 | 2.37621824 |
| 1.09E-16 | 7.52E-15 | 523.760146 | 1584.88922 | 1972.11032 | 1855.28067 | 0 |
| 1.11E-16 | 7.62E-15 | 476.174817 | 779.209395 | 721.096339 | 778.484862 | 16.6335277 |
| 1.10E-16 | 7.62E-15 | 501.896616 | 1012.54958 | 890.001788 | 934.831345 | 0 |
| 1.11E-16 | 7.63E-15 | 553.340216 | 1415.38949 | 888.145685 | 1230.82261 | 0 |
| 1.12E-16 | 7.69E-15 | 513.149904 | 1339.98213 | 1055.19503 | 1174.68634 | 0 |
| 1.12E-16 | 7.70E-15 | 558.806098 | 1029.89994 | 669.125432 | 865.240922 | 0 |
| 1.13E-16 | 7.72E-15 | 524.724714 | 2379.44702 | 1958.18954 | 2191.17044 | 0 |
| 1.13E-16 | 7.72E-15 | 520.544921 | 1699.89101 | 1302.05684 | 1470.21366 | 0 |
| 1.13E-16 | 7.72E-15 | 500.289004 | 754.51849 | 728.520755 | 753.43231 | 0 |
| 1.14E-16 | 7.74E-15 | 2137.16003 | 1205.62801 | 667.269328 | 987.720066 | 0 |
| 1.16E-16 | 7.83E-15 | 533.084299 | 2313.82704 | 1672.34955 | 1916.0563 | 0 |
| 1.16E-16 | 7.83E-15 | 597.71032 | 691.122921 | 522.493229 | 698.687844 | 0 |
| 1.16E-16 | 7.83E-15 | 333.740351 | 388.381274 | 338.73895 | 329.394667 | 11.8810912 |
| 1.19E-16 | 8.02E-15 | 406.082913 | 769.422009 | 721.096339 | 788.691457 | 0 |
| 1.19E-16 | 8.04E-15 | 506.719454 | 1455.4288 | 1216.67606 | 1305.51633 | 0 |
| 1.22E-16 | 8.18E-15 | 497.716824 | 967.171696 | 841.743089 | 926.016558 | 0 |
| 1.23E-16 | 8.26E-15 | 592.887483 | 3734.77755 | 1955.40539 | 2564.63904 | 0 |
| 1.27E-16 | 8.49E-15 | 500.932049 | 1341.31677 | 1225.95658 | 1409.43803 | 0 |
| 1.30E-16 | 8.67E-15 | 508.970111 | 1015.88619 | 780.491662 | 873.591773 | 0 |
| 1.31E-16 | 8.70E-15 | 611.214265 | 1674.75522 | 833.390621 | 1232.67835 | 0 |
| 1.31E-16 | 8.72E-15 | 493.215509 | 1178.26782 | 1108.09399 | 1193.70772 | 0 |
| 1.34E-16 | 8.90E-15 | 508.005544 | 700.687866 | 612.514265 | 668.531994 | 0 |
| 1.38E-16 | 9.13E-15 | 740.144786 | 728.937821 | 464.95401 | 631.881038 | 14.2573095 |
| 1.39E-16 | 9.22E-15 | 0 | 0 | 0 | 0 | 85.5438568 |
| 1.40E-16 | 9.23E-15 | 505.754886 | 993.197245 | 799.052701 | 785.443904 | 0 |
| 1.40E-16 | 9.23E-15 | 492.572464 | 811.0184 | 913.203086 | 789.155393 | 0 |
| 1.43E-16 | 9.38E-15 | 0 | 0 | 0 | 0 | 114.058476 |
| 1.43E-16 | 9.39E-15 | 486.463537 | 891.764335 | 821.325947 | 884.262304 | 0 |
| 1.43E-16 | 9.39E-15 | 500.932049 | 1290.82276 | 1023.64126 | 1244.74069 | 0 |
| 1.44E-16 | 9.42E-15 | 491.286374 | 1059.2621 | 916.915294 | 1061.48591 | 0 |

| | | | | | | |
|---|---|---|---|---|---|---|
| 1.46E-16 | 9.53E-15 | 494.501599 | 831.482935 | 702.535301 | 777.55699 | 0 |
| 1.50E-16 | 9.78E-15 | 1034.65939 | 813.465247 | 541.054268 | 672.707419 | 0 |
| 1.52E-16 | 9.88E-15 | 487.428104 | 787.662138 | 709.959716 | 764.102841 | 0 |
| 1.53E-16 | 9.89E-15 | 502.539661 | 698.908341 | 793.484389 | 902.81975 | 0 |
| 1.53E-16 | 9.89E-15 | 959.744651 | 1364.00571 | 1180.48204 | 1382.06579 | 0 |
| 1.54E-16 | 9.95E-15 | 481.640699 | 927.132389 | 902.994515 | 960.811769 | 0 |
| 1.56E-16 | 1.01E-14 | 972.605551 | 683.3375 | 585.60076 | 608.220294 | 0 |
| 1.57E-16 | 1.01E-14 | 499.645959 | 1079.28175 | 816.685687 | 966.842939 | 0 |
| 1.58E-16 | 1.01E-14 | 462.992394 | 1083.50813 | 905.778671 | 1063.34166 | 9.50487297 |
| 1.61E-16 | 1.03E-14 | 495.144644 | 1110.86832 | 858.448023 | 990.503683 | 0 |
| 1.61E-16 | 1.03E-14 | 508.327066 | 1015.66375 | 909.490879 | 1277.21622 | 0 |
| 1.65E-16 | 1.05E-14 | 502.539661 | 1562.20028 | 1168.41736 | 1254.94729 | 0 |
| 1.67E-16 | 1.06E-14 | 501.575094 | 2393.01589 | 2077.90824 | 2362.36288 | 0 |
| 1.68E-16 | 1.07E-14 | 490.964852 | 1165.81114 | 922.483605 | 1097.67293 | 0 |
| 1.72E-16 | 1.09E-14 | 482.283744 | 877.973018 | 753.578157 | 833.693264 | 0 |
| 1.76E-16 | 1.11E-14 | 489.357239 | 1470.55476 | 1192.54671 | 1280.92771 | 0 |
| 1.76E-16 | 1.11E-14 | 483.891357 | 1477.22797 | 1383.72541 | 1463.25462 | 0 |
| 1.77E-16 | 1.12E-14 | 491.286374 | 717.815792 | 647.780238 | 787.299649 | 0 |
| 1.80E-16 | 1.13E-14 | 499.002914 | 1181.60443 | 854.735816 | 977.049535 | 0 |
| 1.84E-16 | 1.16E-14 | 505.111841 | 649.52653 | 555.903098 | 618.890826 | 0 |
| 1.92E-16 | 1.20E-14 | 497.395302 | 1085.51009 | 781.419714 | 908.386984 | 0 |
| 1.93E-16 | 1.21E-14 | 475.853294 | 1244.33267 | 1108.09399 | 1173.75846 | 0 |
| 1.94E-16 | 1.22E-14 | 1184.48888 | 4998.46255 | 5398.478 | 5807.08881 | 0 |
| 1.95E-16 | 1.22E-14 | 881.936207 | 2068.25263 | 1792.06825 | 1901.21035 | 2.37621824 |
| 1.98E-16 | 1.23E-14 | 470.065889 | 1057.26014 | 1016.21685 | 1041.07272 | 0 |
| 2.01E-16 | 1.25E-14 | 132.145746 | 93.2026082 | 87.2368802 | 100.210209 | 722.370346 |
| 2.03E-16 | 1.26E-14 | 468.458277 | 839.713236 | 825.966206 | 885.654113 | 0 |
| 2.10E-16 | 1.30E-14 | 653.976757 | 599.922278 | 498.363879 | 553.939765 | 0 |
| 2.13E-16 | 1.31E-14 | 508.005544 | 1354.66321 | 902.066463 | 995.606981 | 0 |
| 2.20E-16 | 1.36E-14 | 1442.67144 | 521.400749 | 950.325163 | 937.614962 | 0 |
| 2.20E-16 | 1.36E-14 | 487.428104 | 2434.1674 | 2033.36175 | 2251.48214 | 0 |
| 2.23E-16 | 1.37E-14 | 463.956962 | 1007.211 | 1046.84256 | 1014.16443 | 0 |
| 2.24E-16 | 1.38E-14 | 1026.29981 | 4114.2612 | 3409.66274 | 3613.59868 | 0 |
| 2.28E-16 | 1.40E-14 | 26.6863672 | 16.6830444 | 20.4171422 | 22.7328714 | 190.097459 |
| 2.30E-16 | 1.40E-14 | 491.607897 | 2762.04484 | 2152.1524 | 2295.55608 | 0 |
| 2.32E-16 | 1.42E-14 | 466.529142 | 862.402177 | 778.635558 | 903.283687 | 0 |
| 2.33E-16 | 1.42E-14 | 517.329696 | 538.306234 | 538.270112 | 522.392106 | 0 |
| 2.34E-16 | 1.42E-14 | 466.207619 | 1134.22458 | 995.799707 | 1103.7041 | 0 |
| 2.34E-16 | 1.42E-14 | 466.207619 | 988.303552 | 993.015551 | 1124.58123 | 0 |
| 2.37E-16 | 1.43E-14 | 462.670872 | 793.890474 | 773.067247 | 850.858902 | 0 |
| 2.47E-16 | 1.50E-14 | 474.888727 | 623.945862 | 618.082577 | 707.966567 | 0 |
| 2.51E-16 | 1.52E-14 | 461.706304 | 818.58138 | 943.8288 | 888.901666 | 0 |
| 2.62E-16 | 1.58E-14 | 461.384782 | 1329.08254 | 1261.22256 | 1238.70952 | 0 |

| | | | | | | |
|---|---|---|---|---|---|---|
| 2.64E-16 | 1.59E-14 | 467.172187 | 673.550114 | 606.945954 | 664.820505 | 0 |
| 2.66E-16 | 1.60E-14 | 959.101606 | 1945.9103 | 1251.01398 | 1492.4826 | 0 |
| 2.68E-16 | 1.61E-14 | 505.754886 | 797.671964 | 560.543358 | 709.358375 | 0 |
| 2.69E-16 | 1.61E-14 | 482.926789 | 601.701803 | 641.283875 | 764.102841 | 0 |
| 2.70E-16 | 1.61E-14 | 476.174817 | 776.317668 | 612.514265 | 710.750184 | 0 |
| 2.70E-16 | 1.61E-14 | 456.883467 | 1138.00607 | 1101.59763 | 1155.20102 | 0 |
| 2.70E-16 | 1.61E-14 | 555.590873 | 833.4849 | 549.406735 | 600.33338 | 0 |
| 2.76E-16 | 1.64E-14 | 530.190596 | 1796.65267 | 1501.588 | 1610.78632 | 9.50487297 |
| 2.85E-16 | 1.69E-14 | 471.030457 | 847.943538 | 657.060757 | 760.391352 | 0 |
| 2.88E-16 | 1.71E-14 | 502.861184 | 619.941931 | 778.635558 | 565.074232 | 0 |
| 2.92E-16 | 1.72E-14 | 523.438624 | 1525.27514 | 859.376075 | 1099.52868 | 0 |
| 2.92E-16 | 1.72E-14 | 331.489694 | 425.751294 | 469.59427 | 430.996685 | 16.6335277 |
| 3.03E-16 | 1.79E-14 | 458.491079 | 1344.2085 | 1192.54671 | 1386.70516 | 0 |
| 3.04E-16 | 1.79E-14 | 526.332326 | 496.487402 | 510.428554 | 547.908595 | 0 |
| 3.11E-16 | 1.82E-14 | 463.635439 | 876.415934 | 686.758419 | 818.383371 | 0 |
| 3.11E-16 | 1.82E-14 | 464.600007 | 1022.5594 | 775.851403 | 927.872303 | 0 |
| 3.13E-16 | 1.83E-14 | 456.240422 | 1462.32445 | 1315.04957 | 1430.31515 | 0 |
| 3.17E-16 | 1.85E-14 | 456.883467 | 1210.5217 | 981.878928 | 1089.32208 | 0 |
| 3.17E-16 | 1.85E-14 | 466.207619 | 2141.21314 | 1763.29864 | 1930.43833 | 0 |
| 3.18E-16 | 1.85E-14 | 544.016063 | 571.672323 | 469.59427 | 563.682424 | 0 |
| 3.19E-16 | 1.85E-14 | 448.845405 | 1012.32714 | 962.389838 | 903.747623 | 0 |
| 3.23E-16 | 1.88E-14 | 480.997654 | 698.24102 | 545.694527 | 681.986142 | 0 |
| 3.33E-16 | 1.92E-14 | 0 | 0 | 0 | 0 | 68.9103291 |
| 3.33E-16 | 1.92E-14 | 539.193226 | 1828.90655 | 928.979969 | 1288.35069 | 0 |
| 3.35E-16 | 1.93E-14 | 453.668242 | 1258.12399 | 1036.63399 | 1120.40581 | 0 |
| 3.41E-16 | 1.96E-14 | 488.071149 | 1120.21082 | 722.952443 | 892.613155 | 0 |
| 3.42E-16 | 1.97E-14 | 461.063259 | 773.203499 | 897.426204 | 1030.86613 | 0 |
| 3.45E-16 | 1.98E-14 | 12467.9994 | 3301.90815 | 1053.33893 | 2558.14394 | 0 |
| 3.46E-16 | 1.98E-14 | 452.382152 | 1514.15311 | 1361.45216 | 1338.4558 | 0 |
| 3.48E-16 | 1.99E-14 | 486.463537 | 896.213147 | 698.823093 | 1078.65155 | 0 |
| 3.49E-16 | 1.99E-14 | 0.3215225 | 0.66732178 | 0.92805192 | 0.46393615 | 73.6627655 |
| 3.52E-16 | 2.01E-14 | 804.127763 | 797.449524 | 465.882062 | 619.818698 | 0 |
| 3.52E-16 | 2.01E-14 | 453.989765 | 628.172233 | 626.435044 | 712.141992 | 0 |
| 3.57E-16 | 2.03E-14 | 500.610526 | 692.680005 | 504.860243 | 630.953166 | 0 |
| 3.59E-16 | 2.03E-14 | 963.281398 | 1539.06646 | 1034.77789 | 1253.09154 | 0 |
| 3.59E-16 | 2.03E-14 | 443.701045 | 803.67786 | 728.520755 | 800.289861 | 0 |
| 3.61E-16 | 2.04E-14 | 459.455647 | 2269.56136 | 1951.69318 | 2236.63619 | 0 |
| 3.69E-16 | 2.08E-14 | 306.732461 | 1016.33107 | 722.952443 | 784.979968 | 14.2573095 |
| 3.69E-16 | 2.08E-14 | 441.77191 | 731.384668 | 787.916077 | 728.843694 | 0 |
| 3.86E-16 | 2.17E-14 | 453.34672 | 684.227262 | 595.809331 | 603.580933 | 0 |
| 3.88E-16 | 2.18E-14 | 500.932049 | 1744.37913 | 1570.26384 | 1664.13897 | 16.6335277 |
| 3.90E-16 | 2.18E-14 | 449.48845 | 828.813647 | 671.909588 | 791.011138 | 0 |
| 3.89E-16 | 2.18E-14 | 446.91627 | 1022.11452 | 822.253998 | 916.273899 | 0 |

| | | | | | | |
|---|---|---|---|---|---|---|
| 3.90E-16 | 2.18E-14 | 453.025197 | 678.221366 | 702.535301 | 845.291668 | 0 |
| 3.91E-16 | 2.18E-14 | 436.949072 | 940.256384 | 961.461786 | 967.306875 | 0 |
| 3.93E-16 | 2.19E-14 | 444.665612 | 1036.7956 | 852.879712 | 974.729854 | 0 |
| 3.94E-16 | 2.19E-14 | 451.739107 | 933.583166 | 727.592703 | 891.685283 | 0 |
| 3.95E-16 | 2.19E-14 | 456.561944 | 740.059851 | 591.169071 | 697.759972 | 0 |
| 4.05E-16 | 2.25E-14 | 491.286374 | 1101.97069 | 677.477899 | 868.488475 | 0 |
| 4.06E-16 | 2.25E-14 | 829.206517 | 1580.44041 | 1363.30827 | 1548.61887 | 2.37621824 |
| 4.13E-16 | 2.28E-14 | 437.270595 | 847.276217 | 757.290364 | 776.629117 | 0 |
| 4.18E-16 | 2.31E-14 | 470.708934 | 619.941931 | 507.644399 | 588.734976 | 0 |
| 4.20E-16 | 2.31E-14 | 441.450387 | 671.770589 | 612.514265 | 664.820505 | 0 |
| 4.28E-16 | 2.35E-14 | 444.34409 | 634.178129 | 602.305694 | 691.264865 | 0 |
| 4.31E-16 | 2.36E-14 | 434.698415 | 769.64445 | 740.58543 | 819.311243 | 0 |
| 4.33E-16 | 2.37E-14 | 447.880837 | 2370.99427 | 2270.94304 | 2461.64522 | 0 |
| 4.37E-16 | 2.39E-14 | 531.476686 | 1760.17241 | 1269.57502 | 1605.68302 | 7.12865473 |
| 4.45E-16 | 2.43E-14 | 460.098692 | 1515.26532 | 1044.98646 | 1208.55367 | 0 |
| 4.52E-16 | 2.46E-14 | 348.530386 | 571.005001 | 441.752712 | 554.403701 | 16.6335277 |
| 4.52E-16 | 2.46E-14 | 440.164297 | 1101.97069 | 892.785944 | 1048.4957 | 0 |
| 4.53E-16 | 2.46E-14 | 438.556685 | 1284.59442 | 1079.32438 | 1164.47974 | 0 |
| 4.65E-16 | 2.52E-14 | 470.708934 | 564.999105 | 496.507776 | 613.323592 | 0 |
| 4.64E-16 | 2.52E-14 | 434.376892 | 1211.85635 | 1071.89996 | 1155.66495 | 0 |
| 4.68E-16 | 2.53E-14 | 1027.26438 | 695.794173 | 512.284658 | 615.643273 | 0 |
| 4.70E-16 | 2.54E-14 | 784.836413 | 2554.06288 | 2395.302 | 2780.36935 | 2.37621824 |
| 4.76E-16 | 2.56E-14 | 430.518622 | 1083.06324 | 995.799707 | 1001.63815 | 0 |
| 4.85E-16 | 2.61E-14 | 430.518622 | 697.796138 | 748.009845 | 781.732415 | 0 |
| 5.03E-16 | 2.70E-14 | 434.698415 | 1131.55529 | 915.987242 | 1009.989 | 0 |
| 5.04E-16 | 2.70E-14 | 486.785059 | 509.833838 | 611.586213 | 736.266672 | 0 |
| 5.13E-16 | 2.75E-14 | 578.740493 | 1217.86224 | 1064.47555 | 1222.00782 | 2.37621824 |
| 5.15E-16 | 2.76E-14 | 430.1971 | 1272.58263 | 1143.35996 | 1245.66857 | 0 |
| 5.25E-16 | 2.80E-14 | 427.62492 | 743.841341 | 668.19738 | 726.524013 | 0 |
| 5.34E-16 | 2.85E-14 | 426.33883 | 1208.2973 | 1137.79165 | 1145.92229 | 0 |
| 5.36E-16 | 2.85E-14 | 427.62492 | 1270.13578 | 1160.99295 | 1171.90272 | 0 |
| 5.40E-16 | 2.87E-14 | 431.161667 | 649.748971 | 721.096339 | 776.165181 | 0 |
| 5.44E-16 | 2.89E-14 | 438.878207 | 1605.5762 | 1841.255 | 1460.93494 | 0 |
| 5.51E-16 | 2.92E-14 | 447.559315 | 2351.19706 | 1939.62851 | 2648.14755 | 0 |
| 5.56E-16 | 2.94E-14 | 443.701045 | 706.02644 | 558.687254 | 640.231889 | 0 |
| 5.63E-16 | 2.97E-14 | 435.984505 | 2047.56565 | 1986.0311 | 2303.90693 | 0 |
| 5.66E-16 | 2.98E-14 | 424.088172 | 1104.41754 | 1159.13684 | 1202.05857 | 0 |
| 5.70E-16 | 3.00E-14 | 448.845405 | 540.53064 | 594.881279 | 509.401894 | 0 |
| 5.71E-16 | 3.00E-14 | 391.935923 | 556.546362 | 540.126216 | 611.931784 | 7.12865473 |
| 5.85E-16 | 3.07E-14 | 436.949072 | 869.742716 | 685.830367 | 692.656674 | 0 |
| 5.92E-16 | 3.10E-14 | 431.48319 | 665.764693 | 579.104396 | 682.914015 | 0 |
| 5.99E-16 | 3.13E-14 | 419.90838 | 907.112736 | 874.224906 | 807.712839 | 0 |
| 5.99E-16 | 3.13E-14 | 828.24195 | 1363.11595 | 1104.38178 | 1263.76208 | 0 |

| | | | | | | |
|---|---|---|---|---|---|---|
| 6.04E-16 | 3.15E-14 | 513.471426 | 629.284436 | 453.817387 | 530.742957 | 0 |
| 6.06E-16 | 3.16E-14 | 433.412325 | 1258.12399 | 978.16672 | 1248.45218 | 0 |
| 6.08E-16 | 3.16E-14 | 467.172187 | 720.485079 | 515.068814 | 593.374338 | 0 |
| 6.12E-16 | 3.18E-14 | 225.708792 | 391.273002 | 449.177128 | 409.191685 | 26.1384007 |
| 6.20E-16 | 3.21E-14 | 483.248312 | 762.30391 | 500.219983 | 675.0271 | 0 |
| 6.26E-16 | 3.24E-14 | 426.660352 | 988.081112 | 779.56361 | 886.581985 | 0 |
| 6.31E-16 | 3.25E-14 | 417.657722 | 861.289974 | 772.139195 | 846.683476 | 0 |
| 6.31E-16 | 3.25E-14 | 418.943812 | 1084.62033 | 1058.90724 | 1163.55187 | 0 |
| 6.30E-16 | 3.25E-14 | 425.695785 | 1157.3584 | 923.411657 | 1016.02017 | 0 |
| 6.32E-16 | 3.25E-14 | 458.812602 | 636.847416 | 491.867516 | 559.506998 | 0 |
| 6.31E-16 | 3.25E-14 | 463.313917 | 817.469177 | 607.874006 | 948.749429 | 0 |
| 6.37E-16 | 3.26E-14 | 416.05011 | 825.92192 | 761.930624 | 814.671882 | 0 |
| 6.38E-16 | 3.27E-14 | 425.374262 | 1574.43451 | 1635.22748 | 1760.63769 | 0 |
| 6.43E-16 | 3.29E-14 | 312.841389 | 545.201892 | 446.392972 | 509.86583 | 14.2573095 |
| 6.48E-16 | 3.31E-14 | 813.773438 | 448.662675 | 574.464137 | 712.141992 | 0 |
| 6.54E-16 | 3.33E-14 | 415.085542 | 777.874752 | 733.161014 | 797.506244 | 0 |
| 6.62E-16 | 3.37E-14 | 2646.77319 | 2454.85438 | 1389.29372 | 1871.0545 | 21.3859642 |
| 6.67E-16 | 3.39E-14 | 429.875577 | 927.132389 | 697.895042 | 831.373583 | 0 |
| 6.72E-16 | 3.41E-14 | 428.589487 | 1329.74986 | 1008.79243 | 1108.8074 | 0 |
| 6.74E-16 | 3.42E-14 | 432.76928 | 1295.49401 | 935.476332 | 1096.74506 | 0 |
| 6.81E-16 | 3.45E-14 | 422.48056 | 623.056099 | 564.255566 | 613.323592 | 0 |
| 6.87E-16 | 3.47E-14 | 419.265335 | 1538.62158 | 1439.40852 | 1523.10238 | 0 |
| 6.88E-16 | 3.47E-14 | 425.05274 | 2274.01018 | 1986.0311 | 2111.83736 | 0 |
| 6.94E-16 | 3.50E-14 | 392.257445 | 1453.64927 | 1427.34385 | 1440.05781 | 19.0097459 |
| 6.98E-16 | 3.51E-14 | 429.554055 | 1113.53761 | 825.966206 | 894.932836 | 0 |
| 7.10E-16 | 3.56E-14 | 675.840287 | 829.036088 | 432.472193 | 623.066251 | 0 |
| 7.24E-16 | 3.63E-14 | 415.407065 | 1233.65553 | 1071.89996 | 1103.7041 | 0 |
| 7.52E-16 | 3.76E-14 | 412.513363 | 701.355188 | 687.68647 | 782.660287 | 0 |
| 7.51E-16 | 3.76E-14 | 409.61966 | 1058.37234 | 1018.07295 | 1078.65155 | 0 |
| 7.54E-16 | 3.76E-14 | 438.878207 | 809.238875 | 1175.84178 | 1152.4174 | 0 |
| 7.75E-16 | 3.86E-14 | 468.779799 | 1289.933 | 1067.2597 | 1158.91251 | 2.37621824 |
| 7.88E-16 | 3.92E-14 | 430.1971 | 850.167944 | 735.017118 | 617.035081 | 0 |
| 7.90E-16 | 3.92E-14 | 415.407065 | 1117.98642 | 890.001788 | 1009.52507 | 0 |
| 7.90E-16 | 3.92E-14 | 406.082913 | 900.661959 | 840.815037 | 861.065497 | 0 |
| 8.05E-16 | 3.99E-14 | 5659.11745 | 26818.55 | 23695.0215 | 25594.4296 | 0 |
| 8.10E-16 | 4.01E-14 | 408.655093 | 1152.24227 | 1024.56932 | 1110.19921 | 0 |
| 8.23E-16 | 4.07E-14 | 411.870318 | 771.868856 | 645.924134 | 704.719014 | 0 |
| 8.24E-16 | 4.07E-14 | 438.556685 | 527.851526 | 507.644399 | 633.272847 | 0 |
| 8.38E-16 | 4.13E-14 | 624.075165 | 969.173661 | 740.58543 | 858.28188 | 142.573095 |
| 8.39E-16 | 4.13E-14 | 411.548795 | 1536.17473 | 1336.39476 | 1422.42824 | 0 |
| 8.50E-16 | 4.18E-14 | 412.19184 | 2167.01625 | 2134.51941 | 2151.73587 | 0 |
| 8.53E-16 | 4.18E-14 | 722.139526 | 2923.31427 | 2785.0838 | 3253.12029 | 0 |
| 8.55E-16 | 4.19E-14 | 462.027827 | 991.862602 | 594.881279 | 785.90784 | 0 |

| | | | | | | |
|---|---|---|---|---|---|---|
| 9.04E-16 | 4.42E-14 | 422.802082 | 1373.34822 | 961.461786 | 1181.64538 | 0 |
| 9.18E-16 | 4.48E-14 | 405.76139 | 732.27443 | 654.276601 | 637.912208 | 0 |
| 9.24E-16 | 4.51E-14 | 419.265335 | 540.308199 | 508.57245 | 587.807104 | 0 |
| 9.32E-16 | 4.54E-14 | 474.245682 | 618.162406 | 695.110886 | 457.441045 | 0 |
| 9.44E-16 | 4.59E-14 | 412.19184 | 792.555831 | 690.470626 | 903.283687 | 0 |
| 9.86E-16 | 4.78E-14 | 420.229902 | 674.662317 | 780.491662 | 575.744764 | 0 |
| 9.86E-16 | 4.78E-14 | 400.61703 | 1190.94693 | 1251.94204 | 1295.30973 | 0 |
| 9.88E-16 | 4.78E-14 | 409.298138 | 1892.30212 | 1537.78203 | 1666.92259 | 0 |
| 1.01E-15 | 4.87E-14 | 437.270595 | 758.29998 | 521.565177 | 682.450079 | 0 |
| 1.01E-15 | 4.87E-14 | 415.085542 | 815.912093 | 625.506992 | 657.397526 | 0 |
| 1.03E-15 | 4.98E-14 | 403.510733 | 1913.65642 | 1818.98176 | 2013.01896 | 0 |
| 1.04E-15 | 5.00E-14 | 402.867688 | 1225.42522 | 991.159447 | 1115.76644 | 0 |
| 1.04E-15 | 5.01E-14 | 400.61703 | 699.575663 | 620.866732 | 621.674443 | 0 |
| 1.05E-15 | 5.02E-14 | 0 | 0 | 0 | 0 | 71.2865473 |
| 1.08E-15 | 5.18E-14 | 424.731217 | 736.723242 | 553.118943 | 573.425083 | 0 |
| 1.10E-15 | 5.28E-14 | 397.080283 | 1479.67482 | 1466.32203 | 1510.57611 | 0 |
| 1.10E-15 | 5.28E-14 | 283.261319 | 532.077897 | 487.227256 | 527.031468 | 4.75243649 |
| 1.11E-15 | 5.28E-14 | 395.151148 | 1021.00232 | 931.764125 | 1068.90889 | 0 |
| 1.15E-15 | 5.50E-14 | 232.139242 | 308.525102 | 295.12051 | 288.568286 | 11.8810912 |
| 1.16E-15 | 5.51E-14 | 398.04485 | 1179.38002 | 956.821526 | 1013.23655 | 0 |
| 1.16E-15 | 5.51E-14 | 539.514748 | 877.083256 | 783.275818 | 891.221347 | 2.37621824 |
| 1.17E-15 | 5.56E-14 | 397.401805 | 836.376628 | 678.405951 | 775.701245 | 0 |
| 1.18E-15 | 5.58E-14 | 390.971355 | 1016.33107 | 926.195813 | 920.91326 | 0 |
| 1.18E-15 | 5.58E-14 | 395.151148 | 1043.24638 | 866.80049 | 1002.56602 | 0 |
| 1.19E-15 | 5.64E-14 | 349.494953 | 593.24906 | 492.795568 | 537.238063 | 4.75243649 |
| 1.20E-15 | 5.68E-14 | 459.134124 | 512.948006 | 484.443101 | 419.398281 | 0 |
| 1.24E-15 | 5.85E-14 | 396.75876 | 844.82937 | 1079.32438 | 931.583792 | 0 |
| 1.25E-15 | 5.87E-14 | 402.546165 | 1645.17062 | 1242.66152 | 1517.53515 | 0 |
| 1.26E-15 | 5.90E-14 | 446.91627 | 454.001249 | 440.824661 | 445.378705 | 0 |
| 1.26E-15 | 5.93E-14 | 459.777169 | 542.977486 | 452.889335 | 436.099982 | 0 |
| 1.27E-15 | 5.97E-14 | 390.649833 | 1376.23995 | 1238.94931 | 1291.59825 | 0 |
| 1.28E-15 | 6.01E-14 | 427.946442 | 1258.79131 | 1989.74331 | 2018.58619 | 0 |
| 1.29E-15 | 6.03E-14 | 396.75876 | 1364.45059 | 1062.61944 | 1204.37825 | 0 |
| 1.30E-15 | 6.05E-14 | 2607.54744 | 1234.10041 | 657.060757 | 817.919435 | 0 |
| 1.31E-15 | 6.10E-14 | 386.791563 | 1086.84473 | 998.583863 | 993.2873 | 0 |
| 1.31E-15 | 6.11E-14 | 1170.66341 | 786.105054 | 436.184401 | 717.709226 | 0 |
| 1.32E-15 | 6.12E-14 | 400.295508 | 1098.85653 | 810.189324 | 909.778793 | 0 |
| 1.32E-15 | 6.13E-14 | 454.311287 | 1803.32588 | 927.123865 | 1335.67218 | 0 |
| 1.33E-15 | 6.15E-14 | 409.941183 | 1579.55065 | 1065.4036 | 1377.89037 | 0 |
| 1.35E-15 | 6.25E-14 | 402.224643 | 769.64445 | 585.60076 | 645.799123 | 0 |
| 1.42E-15 | 6.54E-14 | 412.513363 | 479.137036 | 451.033232 | 476.462427 | 0 |
| 1.42E-15 | 6.55E-14 | 394.829625 | 797.894405 | 624.57894 | 771.061883 | 0 |
| 1.42E-15 | 6.56E-14 | 381.32568 | 880.197424 | 858.448023 | 919.057516 | 0 |

| | | | | | | |
|---|---|---|---|---|---|---|
| 1.45E-15 | 6.69E-14 | 515.079039 | 521.84563 | 392.565961 | 507.082213 | 0 |
| 1.46E-15 | 6.70E-14 | 376.502843 | 706.693762 | 513.21271 | 647.190931 | 0 |
| 1.49E-15 | 6.84E-14 | 384.540905 | 1086.84473 | 954.965423 | 1125.5091 | 0 |
| 1.52E-15 | 6.95E-14 | 390.649833 | 1024.56137 | 781.419714 | 938.542834 | 0 |
| 1.53E-15 | 6.98E-14 | 551.411081 | 1509.48186 | 1028.28152 | 1217.36846 | 2.37621824 |
| 1.58E-15 | 7.21E-14 | 440.807342 | 784.103088 | 511.356606 | 554.867637 | 0 |
| 1.59E-15 | 7.26E-14 | 380.682635 | 843.939608 | 838.958933 | 727.915821 | 0 |
| 1.60E-15 | 7.27E-14 | 388.077653 | 775.205465 | 613.442317 | 724.668268 | 0 |
| 1.62E-15 | 7.35E-14 | 384.862428 | 1111.75808 | 878.865165 | 1027.15464 | 0 |
| 1.62E-15 | 7.35E-14 | 429.875577 | 801.675895 | 504.860243 | 700.079652 | 0 |
| 1.62E-15 | 7.35E-14 | 413.799453 | 468.682328 | 617.154525 | 633.736783 | 0 |
| 1.62E-15 | 7.35E-14 | 378.7535 | 774.983024 | 683.046211 | 764.566777 | 0 |
| 1.62E-15 | 7.35E-14 | 722.461049 | 1715.90673 | 1346.60333 | 1616.35355 | 0 |
| 1.63E-15 | 7.39E-14 | 515.722084 | 1172.7068 | 1177.69788 | 1282.78346 | 2.37621824 |
| 1.66E-15 | 7.51E-14 | 387.75613 | 1192.05913 | 903.922567 | 985.864322 | 0 |
| 1.67E-15 | 7.55E-14 | 471.995024 | 719.150435 | 429.688038 | 592.446465 | 0 |
| 1.68E-15 | 7.56E-14 | 388.077653 | 576.566016 | 574.464137 | 692.192738 | 0 |
| 1.69E-15 | 7.62E-14 | 384.219383 | 609.709664 | 530.845697 | 589.662848 | 0 |
| 1.74E-15 | 7.82E-14 | 380.682635 | 974.512235 | 777.707506 | 886.581985 | 0 |
| 1.74E-15 | 7.82E-14 | 397.723328 | 1013.43934 | 725.736599 | 1035.96943 | 0 |
| 1.75E-15 | 7.83E-14 | 375.538275 | 1251.67321 | 1260.2945 | 1195.56346 | 0 |
| 1.78E-15 | 7.97E-14 | 380.361113 | 631.953723 | 569.823877 | 674.563164 | 0 |
| 1.79E-15 | 7.98E-14 | 1577.38937 | 2995.8299 | 2267.23083 | 2691.29361 | 38.0194919 |
| 1.83E-15 | 8.19E-14 | 378.431978 | 1420.50562 | 1195.33087 | 1394.12813 | 0 |
| 1.85E-15 | 8.22E-14 | 459.134124 | 827.479004 | 474.23453 | 608.684231 | 0 |
| 1.86E-15 | 8.27E-14 | 392.90049 | 816.579415 | 623.650888 | 876.37539 | 0 |
| 1.87E-15 | 8.30E-14 | 434.05537 | 797.894405 | 537.34206 | 527.495404 | 0 |
| 1.87E-15 | 8.30E-14 | 385.826995 | 638.4045 | 518.781022 | 574.816891 | 0 |
| 1.88E-15 | 8.31E-14 | 643.688037 | 2398.79935 | 2249.59785 | 2555.82426 | 21.3859642 |
| 1.89E-15 | 8.34E-14 | 378.431978 | 1366.89744 | 1099.74152 | 1192.77985 | 0 |
| 1.89E-15 | 8.34E-14 | 279.403049 | 442.211898 | 328.530379 | 415.222855 | 4.75243649 |
| 1.90E-15 | 8.39E-14 | 384.540905 | 546.981417 | 675.621796 | 683.841887 | 0 |
| 1.91E-15 | 8.44E-14 | 381.32568 | 770.756653 | 603.233746 | 695.904227 | 0 |
| 1.92E-15 | 8.46E-14 | 415.407065 | 745.398425 | 491.867516 | 616.107209 | 0 |
| 1.94E-15 | 8.54E-14 | 370.072393 | 628.839555 | 620.866732 | 624.45806 | 0 |
| 1.95E-15 | 8.57E-14 | 402.867688 | 1876.73128 | 1165.63321 | 1393.20026 | 0 |
| 1.99E-15 | 8.72E-14 | 395.151148 | 586.575842 | 750.794001 | 854.570391 | 0 |
| 2.01E-15 | 8.80E-14 | 379.396545 | 723.599247 | 676.549847 | 574.816891 | 0 |
| 2.02E-15 | 8.83E-14 | 370.715438 | 773.42594 | 670.053484 | 756.215927 | 0 |
| 2.07E-15 | 9.03E-14 | 33.4383396 | 20.2420939 | 17.6329864 | 14.8459568 | 199.602332 |
| 2.07E-15 | 9.05E-14 | 288.727202 | 346.117562 | 386.997649 | 398.057218 | 9.50487297 |
| 2.08E-15 | 9.05E-14 | 372.001528 | 1248.55905 | 1036.63399 | 1132.46815 | 0 |
| 2.10E-15 | 9.15E-14 | 373.60914 | 1375.79506 | 1666.78124 | 1566.24845 | 0 |

| | | | | | | |
|---|---|---|---|---|---|---|
| 2.11E-15 | 9.17E-14 | 1078.70797 | 712.477218 | 464.025958 | 550.228275 | 0 |
| 2.17E-15 | 9.42E-14 | 367.821736 | 1016.33107 | 1098.81347 | 1170.51091 | 0 |
| 2.18E-15 | 9.45E-14 | 431.48319 | 435.316239 | 681.190107 | 580.848061 | 0 |
| 2.21E-15 | 9.57E-14 | 372.323051 | 610.376986 | 530.845697 | 590.126784 | 0 |
| 2.22E-15 | 9.60E-14 | 790.623818 | 1286.81883 | 1120.15866 | 1245.20463 | 19.0097459 |
| 2.25E-15 | 9.72E-14 | 369.750871 | 691.345361 | 585.60076 | 670.851675 | 0 |
| 2.33E-15 | 1.01E-13 | 366.535646 | 613.491154 | 576.32024 | 655.541782 | 0 |
| 2.34E-15 | 1.01E-13 | 399.973985 | 1091.96087 | 682.118159 | 801.681669 | 0 |
| 2.36E-15 | 1.01E-13 | 498.359869 | 1075.05538 | 1065.4036 | 1086.53847 | 14.2573095 |
| 2.38E-15 | 1.02E-13 | 376.18132 | 1372.6809 | 991.159447 | 1171.43878 | 0 |
| 2.41E-15 | 1.03E-13 | 368.143258 | 1176.71073 | 955.893474 | 972.410173 | 0 |
| 2.40E-15 | 1.03E-13 | 368.464781 | 548.538501 | 640.355823 | 580.384125 | 0 |
| 2.41E-15 | 1.03E-13 | 366.535646 | 1606.24352 | 1354.02775 | 1487.84324 | 0 |
| 2.42E-15 | 1.03E-13 | 856.857452 | 1254.56494 | 783.275818 | 1014.16443 | 0 |
| 2.48E-15 | 1.06E-13 | 363.963466 | 666.432015 | 594.881279 | 687.08944 | 0 |
| 2.50E-15 | 1.06E-13 | 371.358483 | 1667.41468 | 1261.22256 | 1392.73633 | 0 |
| 2.59E-15 | 1.10E-13 | 359.140628 | 1127.55136 | 1029.20958 | 1103.24017 | 0 |
| 2.59E-15 | 1.10E-13 | 358.819106 | 1010.10273 | 916.915294 | 1007.66932 | 0 |
| 2.64E-15 | 1.12E-13 | 446.273225 | 682.002856 | 417.623363 | 527.495404 | 0 |
| 2.64E-15 | 1.12E-13 | 356.889971 | 809.906197 | 746.153741 | 797.97018 | 0 |
| 2.67E-15 | 1.13E-13 | 376.18132 | 694.014648 | 521.565177 | 609.612103 | 0 |
| 2.70E-15 | 1.14E-13 | 408.655093 | 1016.99839 | 601.377642 | 717.24529 | 0 |
| 2.72E-15 | 1.15E-13 | 197.414813 | 298.515275 | 343.379209 | 326.611051 | 19.0097459 |
| 2.76E-15 | 1.16E-13 | 356.246926 | 816.579415 | 915.05919 | 899.108261 | 0 |
| 2.76E-15 | 1.16E-13 | 384.862428 | 589.46757 | 450.10518 | 523.319979 | 0 |
| 2.76E-15 | 1.16E-13 | 380.682635 | 888.427726 | 607.874006 | 712.605928 | 0 |
| 2.76E-15 | 1.16E-13 | 2.25065747 | 2.00196533 | 0.92805192 | 2.31968076 | 52.2768014 |
| 2.77E-15 | 1.16E-13 | 360.105196 | 1023.44917 | 842.671141 | 891.221347 | 0 |
| 2.81E-15 | 1.18E-13 | 370.072393 | 559.215649 | 473.306478 | 542.341361 | 0 |
| 2.81E-15 | 1.18E-13 | 361.069763 | 598.587634 | 531.773748 | 573.425083 | 0 |
| 2.81E-15 | 1.18E-13 | 362.998898 | 512.058244 | 534.557904 | 573.425083 | 0 |
| 2.84E-15 | 1.19E-13 | 354.639313 | 735.166158 | 758.218416 | 816.06369 | 0 |
| 2.85E-15 | 1.19E-13 | 357.211493 | 566.556189 | 558.687254 | 567.857849 | 0 |
| 2.89E-15 | 1.21E-13 | 544.659108 | 588.800248 | 364.724403 | 496.411682 | 0 |
| 2.91E-15 | 1.21E-13 | 422.159037 | 799.229049 | 476.090633 | 583.167742 | 0 |
| 2.91E-15 | 1.21E-13 | 526.010804 | 1060.3743 | 860.304127 | 970.090492 | 4.75243649 |
| 2.92E-15 | 1.21E-13 | 355.925403 | 1812.22351 | 1678.84592 | 1818.62971 | 0 |
| 2.94E-15 | 1.22E-13 | 359.462151 | 1029.23262 | 1073.75607 | 864.776986 | 0 |
| 2.97E-15 | 1.23E-13 | 1074.8497 | 3839.76951 | 3474.62638 | 3948.09665 | 19.0097459 |
| 3.02E-15 | 1.25E-13 | 355.282358 | 688.898515 | 612.514265 | 701.007525 | 0 |
| 3.03E-15 | 1.25E-13 | 390.971355 | 570.56012 | 552.190891 | 424.965515 | 0 |
| 3.03E-15 | 1.25E-13 | 768.117243 | 2759.59799 | 1415.27917 | 2679.23127 | 0 |
| 3.04E-15 | 1.25E-13 | 352.710178 | 628.394674 | 619.938681 | 664.820505 | 0 |

| | | | | | | |
|---|---|---|---|---|---|---|
| 3.09E-15 | 1.27E-13 | 355.603881 | 538.306234 | 548.478683 | 572.497211 | 0 |
| 3.11E-15 | 1.28E-13 | 374.89523 | 530.298372 | 749.865949 | 642.55157 | 0 |
| 3.12E-15 | 1.28E-13 | 353.996268 | 834.152222 | 701.607249 | 767.350394 | 0 |
| 3.13E-15 | 1.28E-13 | 426.981875 | 623.500981 | 546.622579 | 399.449026 | 0 |
| 3.13E-15 | 1.28E-13 | 362.355853 | 868.630513 | 660.772965 | 724.204332 | 0 |
| 3.16E-15 | 1.30E-13 | 395.151148 | 969.173661 | 578.176344 | 799.825925 | 0 |
| 3.17E-15 | 1.30E-13 | 366.214123 | 649.081649 | 560.543358 | 750.184757 | 0 |
| 3.18E-15 | 1.30E-13 | 376.824365 | 787.662138 | 541.054268 | 686.161568 | 0 |
| 3.20E-15 | 1.30E-13 | 369.107826 | 604.59353 | 523.421281 | 703.327205 | 0 |
| 3.21E-15 | 1.31E-13 | 437.270595 | 549.205823 | 374.932974 | 506.618277 | 0 |
| 3.21E-15 | 1.31E-13 | 349.494953 | 826.811682 | 754.506208 | 812.816137 | 0 |
| 3.25E-15 | 1.32E-13 | 348.208863 | 2281.57316 | 2330.33836 | 2319.21682 | 0 |
| 3.26E-15 | 1.32E-13 | 355.603881 | 1534.39521 | 1271.43113 | 1529.13355 | 0 |
| 3.26E-15 | 1.32E-13 | 515.400561 | 652.418258 | 613.442317 | 690.336993 | 4.75243649 |
| 3.27E-15 | 1.32E-13 | 348.530386 | 821.695548 | 754.506208 | 798.434116 | 0 |
| 3.31E-15 | 1.34E-13 | 359.462151 | 602.146684 | 504.860243 | 551.620084 | 0 |
| 3.32E-15 | 1.34E-13 | 356.246926 | 927.799711 | 726.664651 | 860.137625 | 0 |
| 3.31E-15 | 1.34E-13 | 498.359869 | 539.863318 | 358.22804 | 486.205087 | 0 |
| 3.32E-15 | 1.34E-13 | 354.317791 | 1079.05931 | 903.922567 | 1111.59102 | 0 |
| 3.39E-15 | 1.37E-13 | 389.363743 | 1619.58995 | 1414.35112 | 1705.42929 | 0 |
| 3.40E-15 | 1.37E-13 | 378.431978 | 476.467749 | 457.529595 | 409.191685 | 0 |
| 3.45E-15 | 1.38E-13 | 359.462151 | 996.088973 | 778.635558 | 1043.85634 | 0 |
| 3.51E-15 | 1.41E-13 | 347.565818 | 873.079325 | 761.002572 | 818.847307 | 0 |
| 3.53E-15 | 1.41E-13 | 544.980631 | 2444.84455 | 2355.39577 | 2536.33894 | 0 |
| 3.55E-15 | 1.42E-13 | 355.282358 | 920.904053 | 708.103613 | 782.196351 | 0 |
| 3.55E-15 | 1.42E-13 | 371.036961 | 1388.0293 | 914.131138 | 1309.22782 | 0 |
| 3.56E-15 | 1.42E-13 | 615.072535 | 631.731283 | 356.371936 | 515.433064 | 0 |
| 3.57E-15 | 1.42E-13 | 354.639313 | 770.534212 | 980.022824 | 790.547202 | 0 |
| 3.57E-15 | 1.42E-13 | 346.922773 | 902.886365 | 794.412441 | 890.757411 | 0 |
| 3.58E-15 | 1.42E-13 | 441.77191 | 637.069857 | 544.766475 | 641.623697 | 21.3859642 |
| 3.63E-15 | 1.44E-13 | 394.829625 | 461.78667 | 730.376859 | 592.910401 | 0 |
| 3.62E-15 | 1.44E-13 | 344.350593 | 776.540108 | 722.024391 | 743.225714 | 0 |
| 3.65E-15 | 1.45E-13 | 445.951702 | 846.831335 | 611.586213 | 760.855288 | 14.2573095 |
| 3.66E-15 | 1.45E-13 | 349.816476 | 925.575305 | 776.779454 | 945.965813 | 0 |
| 3.69E-15 | 1.46E-13 | 353.353223 | 796.782202 | 622.722836 | 721.420715 | 0 |
| 3.71E-15 | 1.47E-13 | 351.424088 | 495.59764 | 512.284658 | 521.464234 | 0 |
| 3.74E-15 | 1.48E-13 | 437.91364 | 892.876538 | 463.097907 | 651.366356 | 0 |
| 3.75E-15 | 1.48E-13 | 347.244296 | 1099.30141 | 919.69945 | 981.22496 | 0 |
| 3.75E-15 | 1.48E-13 | 349.494953 | 551.87511 | 535.485956 | 602.653061 | 0 |
| 3.79E-15 | 1.49E-13 | 378.110455 | 422.192244 | 564.255566 | 538.165936 | 0 |
| 3.83E-15 | 1.50E-13 | 350.137998 | 951.378414 | 746.153741 | 863.385178 | 0 |
| 3.87E-15 | 1.52E-13 | 442.736477 | 1357.55494 | 1714.11189 | 1454.90377 | 7.12865473 |
| 3.88E-15 | 1.52E-13 | 359.140628 | 561.662496 | 461.241803 | 519.608489 | 0 |

| | | | | | | |
|---|---|---|---|---|---|---|
| 3.90E-15 | 1.52E-13 | 379.075023 | 688.231193 | 465.882062 | 581.775934 | 0 |
| 3.90E-15 | 1.53E-13 | 345.958206 | 694.45953 | 596.737383 | 652.294229 | 0 |
| 3.93E-15 | 1.53E-13 | 350.459521 | 1428.95837 | 1186.05035 | 1532.84504 | 0 |
| 3.99E-15 | 1.55E-13 | 349.173431 | 786.327494 | 647.780238 | 645.799123 | 0 |
| 4.05E-15 | 1.57E-13 | 397.080283 | 1764.84366 | 914.131138 | 1225.71931 | 0 |
| 4.05E-15 | 1.57E-13 | 398.04485 | 385.711987 | 396.278169 | 432.388493 | 0 |
| 4.06E-15 | 1.58E-13 | 586.13551 | 453.333927 | 613.442317 | 368.365304 | 0 |
| 4.09E-15 | 1.59E-13 | 374.252185 | 543.867249 | 414.839207 | 494.555937 | 0 |
| 4.12E-15 | 1.60E-13 | 0 | 0 | 0 | 0 | 392.07601 |
| 4.13E-15 | 1.60E-13 | 560.413711 | 1839.36126 | 1613.88228 | 1697.07844 | 45.1481466 |
| 4.23E-15 | 1.64E-13 | 344.350593 | 942.25835 | 1124.79892 | 1146.85017 | 0 |
| 4.24E-15 | 1.64E-13 | 348.530386 | 843.494727 | 773.995299 | 1013.70049 | 0 |
| 4.25E-15 | 1.64E-13 | 337.920143 | 770.089331 | 732.232962 | 740.906034 | 0 |
| 4.30E-15 | 1.66E-13 | 362.677376 | 582.12703 | 446.392972 | 522.856043 | 0 |
| 4.37E-15 | 1.68E-13 | 359.462151 | 692.902445 | 503.004139 | 623.530187 | 0 |
| 4.37E-15 | 1.68E-13 | 340.170801 | 751.181881 | 656.132705 | 677.346781 | 0 |
| 4.38E-15 | 1.68E-13 | 373.287618 | 832.595137 | 787.916077 | 829.517839 | 7.12865473 |
| 4.46E-15 | 1.71E-13 | 335.991008 | 769.86689 | 738.729326 | 744.617523 | 0 |
| 4.52E-15 | 1.73E-13 | 338.884711 | 621.276575 | 614.370369 | 685.697632 | 0 |
| 4.53E-15 | 1.73E-13 | 349.816476 | 571.005001 | 505.788295 | 638.84008 | 0 |
| 4.56E-15 | 1.74E-13 | 712.172329 | 1606.91084 | 985.591136 | 1254.01942 | 0 |
| 4.60E-15 | 1.76E-13 | 349.816476 | 584.796318 | 491.867516 | 498.267427 | 0 |
| 4.61E-15 | 1.76E-13 | 362.034331 | 498.711808 | 416.695311 | 515.897 | 0 |
| 4.65E-15 | 1.77E-13 | 367.500213 | 655.309985 | 651.492446 | 751.112629 | 0 |
| 4.67E-15 | 1.78E-13 | 355.925403 | 459.562264 | 431.544141 | 508.937958 | 0 |
| 4.68E-15 | 1.78E-13 | 436.306027 | 501.825977 | 351.731677 | 509.401894 | 0 |
| 4.72E-15 | 1.79E-13 | 341.135368 | 581.90459 | 541.98232 | 516.824873 | 0 |
| 4.72E-15 | 1.79E-13 | 596.102708 | 2030.21529 | 2096.46928 | 2216.68693 | 0 |
| 4.75E-15 | 1.80E-13 | 335.669486 | 598.365194 | 593.025175 | 619.354762 | 0 |
| 4.81E-15 | 1.82E-13 | 334.704918 | 1284.59442 | 1214.81996 | 1280.46378 | 0 |
| 4.82E-15 | 1.82E-13 | 608.642085 | 1327.97034 | 1182.33814 | 1347.27058 | 73.6627655 |
| 4.82E-15 | 1.82E-13 | 345.958206 | 684.004822 | 542.910371 | 579.456253 | 0 |
| 4.84E-15 | 1.82E-13 | 190.984363 | 298.070394 | 322.962067 | 351.199667 | 4.75243649 |
| 4.87E-15 | 1.83E-13 | 342.742981 | 588.355367 | 707.175561 | 579.920189 | 0 |
| 4.91E-15 | 1.85E-13 | 339.206233 | 1008.32321 | 806.477116 | 939.00677 | 0 |
| 4.92E-15 | 1.85E-13 | 753.005686 | 2466.19885 | 1165.63321 | 1636.76674 | 0 |
| 4.93E-15 | 1.85E-13 | 357.211493 | 443.10166 | 539.198164 | 450.018067 | 0 |
| 4.93E-15 | 1.85E-13 | 369.750871 | 766.975163 | 509.500502 | 574.352955 | 0 |
| 4.94E-15 | 1.85E-13 | 334.704918 | 701.577629 | 658.916861 | 747.865076 | 0 |
| 4.97E-15 | 1.86E-13 | 519.580354 | 1370.01161 | 1009.72049 | 1238.24559 | 4.75243649 |
| 4.98E-15 | 1.86E-13 | 344.029071 | 774.760583 | 592.097123 | 697.296035 | 0 |
| 5.02E-15 | 1.87E-13 | 341.778413 | 495.375199 | 473.306478 | 517.752745 | 0 |
| 5.08E-15 | 1.90E-13 | 332.132738 | 883.311593 | 801.836856 | 848.539221 | 0 |

| | | | | | | |
|---|---|---|---|---|---|---|
| 5.10E-15 | 1.90E-13 | 336.312531 | 909.559582 | 765.642831 | 766.886458 | 0 |
| 5.10E-15 | 1.90E-13 | 404.153778 | 897.992672 | 740.58543 | 836.012945 | 19.0097459 |
| 5.14E-15 | 1.91E-13 | 330.846649 | 851.057707 | 817.613739 | 863.849114 | 0 |
| 5.15E-15 | 1.91E-13 | 333.418828 | 1503.6984 | 1544.27839 | 1714.24408 | 0 |
| 5.16E-15 | 1.92E-13 | 487.749627 | 618.829728 | 345.235313 | 556.259445 | 0 |
| 5.18E-15 | 1.92E-13 | 340.170801 | 476.69019 | 480.730893 | 484.349342 | 0 |
| 5.29E-15 | 1.96E-13 | 331.811216 | 706.248881 | 640.355823 | 651.830293 | 0 |
| 5.43E-15 | 2.01E-13 | 479.390042 | 1602.23959 | 1508.08437 | 1636.30281 | 11.8810912 |
| 5.43E-15 | 2.01E-13 | 343.064503 | 456.003214 | 443.608816 | 474.142747 | 0 |
| 5.45E-15 | 2.01E-13 | 200.630038 | 254.694478 | 250.574018 | 239.85499 | 11.8810912 |
| 5.58E-15 | 2.06E-13 | 609.606653 | 1589.33803 | 1204.61139 | 1382.06579 | 2.37621824 |
| 5.60E-15 | 2.06E-13 | 339.206233 | 1019.44524 | 758.218416 | 867.096667 | 0 |
| 5.62E-15 | 2.07E-13 | 335.026441 | 667.544218 | 554.046994 | 621.210507 | 0 |
| 5.66E-15 | 2.08E-13 | 219.278342 | 312.751473 | 289.552198 | 334.034029 | 21.3859642 |
| 5.68E-15 | 2.08E-13 | 337.277098 | 1097.52188 | 838.958933 | 896.78858 | 0 |
| 5.75E-15 | 2.11E-13 | 328.917514 | 1461.65713 | 1400.43034 | 1588.51738 | 0 |
| 5.76E-15 | 2.11E-13 | 334.383396 | 744.063782 | 599.521538 | 686.161568 | 0 |
| 5.78E-15 | 2.11E-13 | 351.745611 | 2205.94336 | 1390.22177 | 1544.90738 | 0 |
| 5.81E-15 | 2.12E-13 | 207.38201 | 419.522957 | 408.342843 | 429.14094 | 7.12865473 |
| 5.85E-15 | 2.13E-13 | 380.361113 | 448.662675 | 738.729326 | 580.384125 | 0 |
| 5.88E-15 | 2.14E-13 | 350.137998 | 965.614612 | 666.341276 | 703.791142 | 0 |
| 5.98E-15 | 2.18E-13 | 335.347963 | 1006.0988 | 761.930624 | 859.209752 | 0 |
| 6.05E-15 | 2.20E-13 | 205.452875 | 241.570483 | 288.624146 | 268.155095 | 35.6432737 |
| 6.06E-15 | 2.20E-13 | 332.132738 | 1204.96069 | 945.684903 | 1039.21698 | 0 |
| 6.11E-15 | 2.22E-13 | 362.034331 | 1167.36823 | 687.68647 | 909.778793 | 0 |
| 6.19E-15 | 2.24E-13 | 325.380766 | 796.782202 | 891.857892 | 865.704858 | 0 |
| 6.26E-15 | 2.27E-13 | 0 | 0 | 0 | 0 | 85.5438568 |
| 6.33E-15 | 2.29E-13 | 585.813988 | 1011.43737 | 612.514265 | 803.537414 | 0 |
| 6.35E-15 | 2.30E-13 | 325.059244 | 1663.85563 | 1539.63813 | 1801.46408 | 0 |
| 6.41E-15 | 2.32E-13 | 332.132738 | 1267.4665 | 1023.64126 | 976.585599 | 0 |
| 6.44E-15 | 2.32E-13 | 42215.2607 | 10725.1956 | 9362.18774 | 11233.286 | 40.3957101 |
| 6.44E-15 | 2.32E-13 | 327.631424 | 705.359119 | 595.809331 | 657.397526 | 0 |
| 6.44E-15 | 2.32E-13 | 211.883325 | 310.527067 | 222.73246 | 275.578074 | 4.75243649 |
| 6.45E-15 | 2.32E-13 | 431.161667 | 1227.20475 | 1033.84984 | 1188.60442 | 11.8810912 |
| 6.51E-15 | 2.34E-13 | 321.844019 | 771.868856 | 792.556337 | 781.268479 | 0 |
| 6.54E-15 | 2.34E-13 | 326.666856 | 634.40057 | 557.759202 | 588.734976 | 0 |
| 6.57E-15 | 2.36E-13 | 324.416199 | 721.374841 | 635.715563 | 678.738589 | 0 |
| 6.61E-15 | 2.36E-13 | 324.416199 | 1003.87439 | 868.656594 | 982.616768 | 0 |
| 6.63E-15 | 2.37E-13 | 337.277098 | 602.369124 | 503.932191 | 486.669023 | 0 |
| 6.66E-15 | 2.38E-13 | 332.775783 | 1737.26103 | 1290.92022 | 1339.38367 | 0 |
| 6.68E-15 | 2.38E-13 | 332.454261 | 754.296049 | 588.384915 | 630.025293 | 0 |
| 6.70E-15 | 2.39E-13 | 323.130109 | 913.118632 | 800.908804 | 834.1572 | 0 |
| 6.79E-15 | 2.42E-13 | 334.704918 | 494.707878 | 449.177128 | 448.162322 | 0 |

| | | | | | | |
|---|---|---|---|---|---|---|
| 6.84E-15 | 2.43E-13 | 326.666856 | 960.053597 | 771.211143 | 911.634537 | 0 |
| 6.84E-15 | 2.43E-13 | 320.236406 | 815.022331 | 793.484389 | 829.981775 | 0 |
| 6.92E-15 | 2.46E-13 | 218.635297 | 510.946041 | 412.055051 | 474.142747 | 11.8810912 |
| 6.96E-15 | 2.47E-13 | 327.309901 | 869.965157 | 684.902315 | 787.299649 | 0 |
| 6.99E-15 | 2.48E-13 | 388.720698 | 778.542074 | 439.896609 | 579.456253 | 0 |
| 7.04E-15 | 2.49E-13 | 511.542291 | 474.910665 | 349.875573 | 395.737537 | 0 |
| 7.08E-15 | 2.50E-13 | 319.914884 | 1084.84277 | 1028.28152 | 998.390598 | 0 |
| 7.13E-15 | 2.52E-13 | 382.290248 | 460.229586 | 367.508559 | 381.355516 | 0 |
| 7.22E-15 | 2.55E-13 | 332.454261 | 784.992851 | 584.672708 | 735.3388 | 0 |
| 7.27E-15 | 2.56E-13 | 327.631424 | 1529.05663 | 1135.00749 | 1330.56888 | 0 |
| 7.31E-15 | 2.57E-13 | 319.593361 | 793.890474 | 706.247509 | 751.576565 | 0 |
| 7.31E-15 | 2.57E-13 | 338.241666 | 579.902624 | 446.392972 | 525.175723 | 0 |
| 7.41E-15 | 2.60E-13 | 385.505473 | 543.867249 | 361.940248 | 474.142747 | 0 |
| 7.46E-15 | 2.62E-13 | 317.664226 | 1356.22029 | 1282.56775 | 1358.40505 | 0 |
| 7.47E-15 | 2.62E-13 | 332.132738 | 901.774162 | 649.636342 | 751.112629 | 0 |
| 7.59E-15 | 2.66E-13 | 290.334814 | 502.715739 | 446.392972 | 493.628065 | 9.50487297 |
| 7.61E-15 | 2.66E-13 | 320.879451 | 718.705554 | 616.226473 | 642.087633 | 0 |
| 7.62E-15 | 2.66E-13 | 320.557929 | 554.544397 | 526.205437 | 538.629872 | 0 |
| 7.64E-15 | 2.67E-13 | 585.492465 | 1660.51902 | 1376.30099 | 1538.41228 | 2.37621824 |
| 7.68E-15 | 2.68E-13 | 318.628794 | 683.115059 | 643.139978 | 611.931784 | 0 |
| 7.72E-15 | 2.69E-13 | 0 | 0 | 0 | 0 | 49.9005831 |
| 7.72E-15 | 2.69E-13 | 335.991008 | 570.78256 | 456.601543 | 471.35913 | 0 |
| 7.84E-15 | 2.72E-13 | 315.735091 | 687.563871 | 669.125432 | 684.769759 | 0 |
| 7.86E-15 | 2.73E-13 | 605.105338 | 1101.74825 | 641.283875 | 744.153587 | 0 |
| 7.86E-15 | 2.73E-13 | 386.791563 | 381.708057 | 496.507776 | 372.076793 | 0 |
| 7.88E-15 | 2.73E-13 | 585.492465 | 2188.37055 | 1571.1919 | 1910.02513 | 2.37621824 |
| 8.07E-15 | 2.79E-13 | 807.66451 | 607.485258 | 674.693744 | 326.611051 | 0 |
| 8.11E-15 | 2.80E-13 | 341.778413 | 702.24495 | 481.658945 | 585.023487 | 0 |
| 8.11E-15 | 2.80E-13 | 558.163053 | 389.048596 | 799.052701 | 441.667216 | 0 |
| 8.14E-15 | 2.81E-13 | 314.127479 | 999.870463 | 999.511915 | 1037.82517 | 0 |
| 8.23E-15 | 2.84E-13 | 314.770524 | 1152.46471 | 1172.12957 | 1271.18505 | 0 |
| 8.26E-15 | 2.84E-13 | 371.358483 | 830.148291 | 473.306478 | 612.39572 | 0 |
| 8.29E-15 | 2.85E-13 | 320.879451 | 663.317847 | 547.550631 | 623.530187 | 0 |
| 8.36E-15 | 2.87E-13 | 424.409695 | 336.552616 | 440.824661 | 390.634239 | 0 |
| 8.38E-15 | 2.88E-13 | 218.635297 | 493.818115 | 372.148819 | 438.419663 | 2.37621824 |
| 8.45E-15 | 2.90E-13 | 344.993638 | 599.254956 | 422.263622 | 521.000298 | 0 |
| 8.46E-15 | 2.90E-13 | 269.114329 | 688.453634 | 567.967773 | 641.159761 | 7.12865473 |
| 8.49E-15 | 2.91E-13 | 739.823264 | 835.264425 | 489.08336 | 664.356569 | 0 |
| 8.67E-15 | 2.96E-13 | 314.127479 | 969.840983 | 838.958933 | 925.552622 | 0 |
| 8.67E-15 | 2.96E-13 | 317.664226 | 656.867069 | 567.967773 | 670.387739 | 0 |
| 8.72E-15 | 2.98E-13 | 315.092046 | 852.837231 | 722.024391 | 766.422522 | 0 |
| 8.78E-15 | 2.99E-13 | 366.535646 | 874.191528 | 491.867516 | 656.005718 | 0 |
| 8.85E-15 | 3.01E-13 | 324.094676 | 999.870463 | 732.232962 | 802.145606 | 0 |

| | | | | | | |
|---|---|---|---|---|---|---|
| 8.84E-15 | 3.01E-13 | 335.991008 | 534.524744 | 424.119726 | 443.522961 | 0 |
| 8.91E-15 | 3.03E-13 | 314.127479 | 925.797746 | 791.628285 | 810.496456 | 0 |
| 8.96E-15 | 3.04E-13 | 319.593361 | 474.910665 | 461.241803 | 496.411682 | 0 |
| 9.01E-15 | 3.06E-13 | 317.664226 | 874.636409 | 696.038938 | 840.18837 | 0 |
| 9.16E-15 | 3.10E-13 | 399.973985 | 501.381095 | 356.371936 | 608.684231 | 0 |
| 9.17E-15 | 3.10E-13 | 315.413569 | 730.272465 | 605.08985 | 666.676249 | 0 |
| 9.21E-15 | 3.11E-13 | 340.492323 | 1381.57852 | 844.527244 | 968.234748 | 0 |
| 9.22E-15 | 3.11E-13 | 319.914884 | 368.361621 | 296.048562 | 450.945939 | 9.50487297 |
| 9.24E-15 | 3.12E-13 | 327.952946 | 1511.70627 | 1006.93633 | 1156.12889 | 0 |
| 9.39E-15 | 3.16E-13 | 320.557929 | 440.654814 | 443.608816 | 459.29679 | 0 |
| 9.39E-15 | 3.16E-13 | 309.947686 | 1301.94479 | 1275.14333 | 1430.77909 | 0 |
| 9.40E-15 | 3.16E-13 | 313.162911 | 633.288367 | 558.687254 | 587.807104 | 0 |
| 9.45E-15 | 3.17E-13 | 312.198344 | 1004.54172 | 843.599193 | 898.644325 | 0 |
| 9.45E-15 | 3.17E-13 | 348.530386 | 999.425582 | 587.456863 | 749.256884 | 0 |
| 9.61E-15 | 3.22E-13 | 704.777312 | 1896.97337 | 1152.64048 | 1456.29558 | 16.6335277 |
| 9.62E-15 | 3.23E-13 | 401.581598 | 634.40057 | 467.738166 | 963.13145 | 0 |
| 9.65E-15 | 3.23E-13 | 316.056614 | 580.569946 | 496.507776 | 535.382319 | 0 |
| 9.73E-15 | 3.25E-13 | 362.998898 | 636.847416 | 399.990376 | 499.195299 | 0 |
| 9.79E-15 | 3.27E-13 | 313.805956 | 493.373234 | 510.428554 | 542.341361 | 0 |
| 9.81E-15 | 3.27E-13 | 322.808586 | 816.356974 | 615.298421 | 624.45806 | 0 |
| 9.84E-15 | 3.28E-13 | 374.89523 | 406.176522 | 344.307261 | 365.117751 | 0 |
| 9.87E-15 | 3.29E-13 | 307.697029 | 653.530461 | 666.341276 | 663.892633 | 0 |
| 9.91E-15 | 3.30E-13 | 317.664226 | 668.43398 | 528.989593 | 631.881038 | 0 |
| 1.00E-14 | 3.33E-13 | 252.395159 | 507.164551 | 506.716347 | 546.05285 | 7.12865473 |
| 1.00E-14 | 3.34E-13 | 313.484434 | 497.377165 | 490.011412 | 475.070619 | 0 |
| 1.01E-14 | 3.36E-13 | 306.410939 | 801.008573 | 761.002572 | 784.516032 | 0 |
| 1.02E-14 | 3.37E-13 | 308.983119 | 572.784526 | 547.550631 | 574.352955 | 0 |
| 1.02E-14 | 3.38E-13 | 345.636683 | 411.515096 | 389.781805 | 354.911156 | 0 |
| 1.03E-14 | 3.41E-13 | 328.595991 | 418.633195 | 420.407518 | 382.283389 | 0 |
| 1.03E-14 | 3.41E-13 | 659.121117 | 500.046452 | 357.299988 | 413.831047 | 0 |
| 1.04E-14 | 3.45E-13 | 322.487064 | 558.103446 | 439.896609 | 527.495404 | 0 |
| 1.05E-14 | 3.45E-13 | 315.413569 | 677.776485 | 540.126216 | 579.920189 | 0 |
| 1.06E-14 | 3.48E-13 | 321.200974 | 480.026798 | 412.983103 | 452.337748 | 0 |
| 1.07E-14 | 3.53E-13 | 425.374262 | 440.654814 | 331.314534 | 372.076793 | 0 |
| 1.07E-14 | 3.53E-13 | 324.737721 | 506.71967 | 447.321024 | 593.838274 | 0 |
| 1.08E-14 | 3.54E-13 | 312.841389 | 469.572091 | 456.601543 | 463.008279 | 0 |
| 1.09E-14 | 3.56E-13 | 307.375506 | 745.620866 | 735.94517 | 858.28188 | 0 |
| 1.09E-14 | 3.58E-13 | 172.979103 | 294.956226 | 283.983887 | 270.938712 | 19.0097459 |
| 1.09E-14 | 3.58E-13 | 344.672116 | 647.524565 | 438.040505 | 475.070619 | 0 |
| 1.10E-14 | 3.59E-13 | 311.555299 | 1105.52974 | 1445.90489 | 1435.41845 | 0 |
| 1.11E-14 | 3.64E-13 | 313.162911 | 598.365194 | 571.679981 | 489.916576 | 0 |
| 1.11E-14 | 3.64E-13 | 337.277098 | 418.855636 | 460.313751 | 587.343168 | 0 |
| 1.12E-14 | 3.64E-13 | 376.502843 | 384.599784 | 669.125432 | 514.041256 | 0 |

| | | | | | | |
|---|---|---|---|---|---|---|
| 1.12E-14 | 3.64E-13 | 316.056614 | 440.654814 | 500.219983 | 528.887212 | 0 |
| 1.12E-14 | 3.66E-13 | 304.803326 | 812.353044 | 720.168288 | 810.496456 | 0 |
| 1.13E-14 | 3.67E-13 | 306.410939 | 686.229228 | 594.881279 | 662.96476 | 0 |
| 1.13E-14 | 3.68E-13 | 302.552669 | 887.982845 | 851.023608 | 836.476881 | 0 |
| 1.13E-14 | 3.68E-13 | 307.697029 | 556.991243 | 503.932191 | 545.124978 | 0 |
| 1.14E-14 | 3.70E-13 | 319.593361 | 879.752543 | 1267.71892 | 1261.90633 | 0 |
| 1.14E-14 | 3.70E-13 | 306.732461 | 966.059493 | 868.656594 | 786.835713 | 0 |
| 1.16E-14 | 3.76E-13 | 317.342704 | 497.599605 | 451.961284 | 420.790089 | 0 |
| 1.18E-14 | 3.81E-13 | 304.160281 | 644.632837 | 713.671924 | 747.865076 | 0 |
| 1.19E-14 | 3.85E-13 | 315.735091 | 519.843665 | 692.32673 | 668.068058 | 0 |
| 1.20E-14 | 3.87E-13 | 304.481804 | 617.495085 | 722.952443 | 702.863269 | 0 |
| 1.20E-14 | 3.87E-13 | 578.740493 | 341.66875 | 522.493229 | 390.170303 | 0 |
| 1.20E-14 | 3.88E-13 | 374.573708 | 536.081828 | 341.523105 | 469.967321 | 0 |
| 1.21E-14 | 3.89E-13 | 327.952946 | 1232.54332 | 753.578157 | 939.470706 | 0 |
| 1.21E-14 | 3.90E-13 | 302.874191 | 656.199748 | 588.384915 | 644.407314 | 0 |
| 1.21E-14 | 3.90E-13 | 517.329696 | 521.400749 | 541.98232 | 561.362743 | 16.6335277 |
| 1.22E-14 | 3.92E-13 | 302.874191 | 1079.5042 | 902.066463 | 1002.10209 | 0 |
| 1.23E-14 | 3.93E-13 | 322.808586 | 445.770947 | 386.069597 | 402.696579 | 0 |
| 1.23E-14 | 3.95E-13 | 309.947686 | 926.465067 | 706.247509 | 913.490282 | 0 |
| 1.23E-14 | 3.95E-13 | 290.656336 | 484.920492 | 451.033232 | 481.565725 | 11.8810912 |
| 1.24E-14 | 3.96E-13 | 286.798067 | 396.611576 | 289.552198 | 346.560305 | 11.8810912 |
| 1.24E-14 | 3.96E-13 | 305.446371 | 513.392887 | 478.874789 | 512.185511 | 0 |
| 1.25E-14 | 3.98E-13 | 359.140628 | 780.766479 | 450.10518 | 539.093808 | 0 |
| 1.25E-14 | 4.00E-13 | 300.302011 | 764.973197 | 691.398678 | 749.720821 | 0 |
| 1.26E-14 | 4.01E-13 | 305.124849 | 702.689832 | 577.248292 | 635.128591 | 0 |
| 1.26E-14 | 4.01E-13 | 314.770524 | 696.906376 | 512.284658 | 631.881038 | 0 |
| 1.26E-14 | 4.01E-13 | 415.728587 | 521.84563 | 344.307261 | 373.468602 | 0 |
| 1.28E-14 | 4.06E-13 | 0 | 0 | 0 | 0 | 57.0292378 |
| 1.29E-14 | 4.08E-13 | 322.808586 | 969.618542 | 624.57894 | 758.535607 | 0 |
| 1.33E-14 | 4.21E-13 | 301.909624 | 567.445951 | 510.428554 | 551.620084 | 0 |
| 1.33E-14 | 4.21E-13 | 361.391286 | 524.959798 | 343.379209 | 477.854236 | 0 |
| 1.34E-14 | 4.24E-13 | 1287.37607 | 469.572091 | 537.34206 | 576.672636 | 0 |
| 1.36E-14 | 4.29E-13 | 308.340074 | 497.599605 | 498.363879 | 431.460621 | 0 |
| 1.36E-14 | 4.29E-13 | 299.337444 | 541.197961 | 565.183617 | 585.951359 | 0 |
| 1.36E-14 | 4.29E-13 | 823.740635 | 437.540645 | 372.148819 | 552.547956 | 0 |
| 1.37E-14 | 4.31E-13 | 299.658966 | 760.969267 | 652.420498 | 751.576565 | 0 |
| 1.37E-14 | 4.32E-13 | 519.580354 | 882.866711 | 337.810898 | 821.166988 | 0 |
| 1.37E-14 | 4.32E-13 | 307.697029 | 462.231551 | 465.882062 | 541.877425 | 0 |
| 1.39E-14 | 4.36E-13 | 503.825751 | 2473.31695 | 2416.64719 | 2649.07542 | 2.37621824 |
| 1.39E-14 | 4.37E-13 | 332.454261 | 442.211898 | 673.765692 | 507.54615 | 0 |
| 1.39E-14 | 4.37E-13 | 0 | 0 | 0 | 0 | 68.9103291 |
| 1.42E-14 | 4.47E-13 | 296.122219 | 623.27854 | 647.780238 | 679.202526 | 0 |
| 1.44E-14 | 4.51E-13 | 296.122219 | 681.780416 | 622.722836 | 634.664655 | 0 |

| | | | | | | |
|---|---|---|---|---|---|---|
| 1.44E-14 | 4.51E-13 | 312.519866 | 831.038053 | 907.634775 | 621.674443 | 0 |
| 1.44E-14 | 4.51E-13 | 296.443741 | 619.941931 | 591.169071 | 651.366356 | 0 |
| 1.45E-14 | 4.54E-13 | 350.459521 | 520.955867 | 352.659728 | 424.501578 | 0 |
| 1.46E-14 | 4.54E-13 | 298.694399 | 1042.13418 | 841.743089 | 894.004964 | 0 |
| 1.47E-14 | 4.59E-13 | 298.372876 | 632.398604 | 547.550631 | 574.816891 | 0 |
| 1.48E-14 | 4.61E-13 | 321.844019 | 894.211182 | 577.248292 | 650.438484 | 0 |
| 1.50E-14 | 4.66E-13 | 399.652463 | 408.178487 | 315.537652 | 366.973496 | 0 |
| 1.51E-14 | 4.69E-13 | 659.764162 | 643.965515 | 688.614522 | 670.387739 | 2.37621824 |
| 1.51E-14 | 4.69E-13 | 297.086786 | 504.272823 | 503.004139 | 504.298596 | 0 |
| 1.53E-14 | 4.76E-13 | 353.353223 | 378.149007 | 350.803625 | 329.394667 | 0 |
| 1.55E-14 | 4.80E-13 | 308.661596 | 427.308378 | 532.7018 | 536.310191 | 0 |
| 1.55E-14 | 4.81E-13 | 330.846649 | 635.957654 | 411.126999 | 580.848061 | 0 |
| 1.55E-14 | 4.81E-13 | 295.157651 | 616.382882 | 575.392189 | 656.469654 | 0 |
| 1.58E-14 | 4.89E-13 | 337.920143 | 1037.68536 | 563.327514 | 797.506244 | 0 |
| 1.59E-14 | 4.90E-13 | 295.800696 | 727.380737 | 610.658161 | 711.21412 | 0 |
| 1.59E-14 | 4.91E-13 | 301.266579 | 473.13114 | 430.616089 | 483.42147 | 0 |
| 1.59E-14 | 4.91E-13 | 299.337444 | 1001.64999 | 759.146468 | 939.00677 | 0 |
| 1.60E-14 | 4.92E-13 | 312.519866 | 617.272644 | 444.536868 | 532.134766 | 0 |
| 1.60E-14 | 4.94E-13 | 359.140628 | 430.422546 | 319.249859 | 447.698386 | 0 |
| 1.63E-14 | 5.00E-13 | 290.334814 | 851.280147 | 790.700233 | 827.662094 | 0 |
| 1.63E-14 | 5.02E-13 | 293.871561 | 725.378772 | 615.298421 | 670.387739 | 0 |
| 1.64E-14 | 5.02E-13 | 296.122219 | 690.233158 | 567.039721 | 628.633485 | 0 |
| 1.65E-14 | 5.07E-13 | 301.266579 | 415.519027 | 427.831934 | 435.17211 | 0 |
| 1.67E-14 | 5.12E-13 | 298.694399 | 464.455957 | 555.903098 | 512.185511 | 0 |
| 1.67E-14 | 5.12E-13 | 295.157651 | 601.701803 | 519.709073 | 597.085827 | 0 |
| 1.69E-14 | 5.16E-13 | 305.446371 | 682.670178 | 499.291931 | 571.105402 | 0 |
| 1.70E-14 | 5.18E-13 | 366.214123 | 351.233695 | 322.034015 | 354.44722 | 0 |
| 1.70E-14 | 5.19E-13 | 292.263949 | 1038.79757 | 865.872439 | 883.798368 | 0 |
| 1.71E-14 | 5.22E-13 | 327.952946 | 614.158476 | 652.420498 | 417.542536 | 0 |
| 1.71E-14 | 5.23E-13 | 292.906994 | 763.860994 | 751.722053 | 905.603367 | 0 |
| 1.72E-14 | 5.25E-13 | 289.691769 | 1181.38199 | 1002.29607 | 1084.21879 | 0 |
| 1.73E-14 | 5.25E-13 | 292.585471 | 707.138643 | 598.593486 | 632.344974 | 0 |
| 1.73E-14 | 5.25E-13 | 562.342846 | 361.910844 | 389.781805 | 377.180091 | 7.12865473 |
| 1.74E-14 | 5.27E-13 | 332.132738 | 484.25317 | 354.515832 | 396.201473 | 0 |
| 1.74E-14 | 5.27E-13 | 289.370247 | 593.693941 | 619.938681 | 602.653061 | 0 |
| 1.74E-14 | 5.29E-13 | 297.729831 | 594.583704 | 481.658945 | 547.444659 | 0 |
| 1.75E-14 | 5.30E-13 | 322.165541 | 1019.44524 | 594.881279 | 822.558796 | 0 |
| 1.75E-14 | 5.30E-13 | 297.408309 | 668.43398 | 525.277385 | 621.674443 | 0 |
| 1.75E-14 | 5.30E-13 | 291.942426 | 503.16062 | 499.291931 | 509.86583 | 0 |
| 1.76E-14 | 5.31E-13 | 439.842775 | 999.870463 | 510.428554 | 708.894439 | 11.8810912 |
| 1.78E-14 | 5.38E-13 | 294.836129 | 701.800069 | 676.549847 | 857.354008 | 0 |
| 1.80E-14 | 5.43E-13 | 300.302011 | 938.031978 | 662.629069 | 816.06369 | 0 |
| 1.82E-14 | 5.47E-13 | 308.340074 | 1002.98463 | 678.405951 | 712.141992 | 0 |

| | | | | | | |
|---|---|---|---|---|---|---|
| 1.82E-14 | 5.48E-13 | 453.025197 | 384.599784 | 324.818171 | 349.807858 | 0 |
| 1.83E-14 | 5.50E-13 | 289.691769 | 1166.47847 | 951.253215 | 1002.10209 | 0 |
| 1.83E-14 | 5.51E-13 | 289.048724 | 724.48901 | 635.715563 | 724.204332 | 0 |
| 1.84E-14 | 5.51E-13 | 331.811216 | 708.695728 | 443.608816 | 493.628065 | 0 |
| 1.85E-14 | 5.54E-13 | 285.833499 | 989.193315 | 909.490879 | 971.946237 | 0 |
| 1.85E-14 | 5.55E-13 | 463.313917 | 896.880469 | 627.363096 | 768.278267 | 4.75243649 |
| 1.86E-14 | 5.56E-13 | 320.557929 | 343.670715 | 382.35739 | 353.983283 | 0 |
| 1.86E-14 | 5.57E-13 | 289.691769 | 791.666068 | 669.125432 | 787.763585 | 0 |
| 1.87E-14 | 5.58E-13 | 285.511977 | 928.911914 | 896.498152 | 857.354008 | 0 |
| 1.87E-14 | 5.58E-13 | 341.456891 | 758.077539 | 425.97583 | 570.17753 | 0 |
| 1.88E-14 | 5.60E-13 | 371.680006 | 616.605322 | 563.327514 | 490.844448 | 0 |
| 1.92E-14 | 5.73E-13 | 299.658966 | 517.619259 | 433.400245 | 552.08402 | 0 |
| 1.93E-14 | 5.73E-13 | 305.124849 | 542.755046 | 410.198947 | 481.565725 | 0 |
| 1.93E-14 | 5.74E-13 | 441.128865 | 318.534928 | 345.235313 | 366.045623 | 14.2573095 |
| 1.95E-14 | 5.80E-13 | 289.370247 | 482.028764 | 486.299204 | 480.637853 | 0 |
| 1.97E-14 | 5.84E-13 | 255.931907 | 673.105233 | 553.118943 | 626.313804 | 11.8810912 |
| 1.99E-14 | 5.92E-13 | 281.975229 | 1158.02572 | 1134.07944 | 1207.6258 | 0 |
| 2.02E-14 | 5.98E-13 | 286.798067 | 562.774699 | 605.08985 | 647.654867 | 0 |
| 2.02E-14 | 5.98E-13 | 283.582842 | 652.640698 | 640.355823 | 646.263059 | 0 |
| 2.03E-14 | 6.00E-13 | 308.983119 | 856.17384 | 674.693744 | 820.239116 | 2.37621824 |
| 2.03E-14 | 6.01E-13 | 288.084157 | 792.33339 | 641.283875 | 718.637098 | 0 |
| 2.04E-14 | 6.01E-13 | 292.263949 | 445.103625 | 442.680764 | 492.236257 | 0 |
| 2.06E-14 | 6.09E-13 | 286.476544 | 744.063782 | 634.787511 | 751.576565 | 0 |
| 2.07E-14 | 6.11E-13 | 344.029071 | 513.615328 | 404.630636 | 339.601263 | 0 |
| 2.08E-14 | 6.11E-13 | 287.762634 | 742.951579 | 650.564394 | 813.744009 | 0 |
| 2.08E-14 | 6.11E-13 | 283.261319 | 851.725028 | 968.886201 | 964.059322 | 0 |
| 2.08E-14 | 6.11E-13 | 283.261319 | 784.54797 | 704.391405 | 738.586353 | 0 |
| 2.08E-14 | 6.12E-13 | 285.190454 | 525.849561 | 519.709073 | 536.774127 | 0 |
| 2.09E-14 | 6.14E-13 | 293.550039 | 929.356795 | 668.19738 | 808.176776 | 0 |
| 2.09E-14 | 6.14E-13 | 325.380766 | 337.664819 | 333.170638 | 355.839028 | 0 |
| 2.10E-14 | 6.16E-13 | 490.000284 | 1477.00553 | 1367.02047 | 1477.63664 | 0 |
| 2.11E-14 | 6.18E-13 | 302.874191 | 1332.41915 | 834.318673 | 998.390598 | 0 |
| 2.13E-14 | 6.22E-13 | 292.906994 | 584.796318 | 474.23453 | 594.766146 | 0 |
| 2.13E-14 | 6.22E-13 | 302.552669 | 1517.71216 | 997.655811 | 1003.4939 | 0 |
| 2.13E-14 | 6.22E-13 | 292.585471 | 719.817757 | 543.838423 | 616.107209 | 0 |
| 2.13E-14 | 6.23E-13 | 290.977859 | 1117.54154 | 805.549064 | 971.946237 | 0 |
| 2.14E-14 | 6.24E-13 | 441.128865 | 831.705375 | 644.996082 | 787.763585 | 4.75243649 |
| 2.14E-14 | 6.24E-13 | 288.084157 | 451.999284 | 469.59427 | 490.380512 | 0 |
| 2.15E-14 | 6.25E-13 | 440.164297 | 498.711808 | 309.041288 | 370.221049 | 0 |
| 2.15E-14 | 6.26E-13 | 353.674746 | 352.345898 | 308.113236 | 363.725943 | 0 |
| 2.17E-14 | 6.31E-13 | 294.514606 | 759.857064 | 1078.39633 | 819.311243 | 0 |
| 2.18E-14 | 6.34E-13 | 344.029071 | 1273.47239 | 783.275818 | 823.022732 | 14.2573095 |
| 2.19E-14 | 6.35E-13 | 309.626164 | 359.019116 | 471.450374 | 422.645834 | 0 |

| | | | | | | |
|---|---|---|---|---|---|---|
| 2.19E-14 | 6.35E-13 | 283.261319 | 799.006608 | 687.68647 | 796.114436 | 0 |
| 2.20E-14 | 6.36E-13 | 299.337444 | 399.503304 | 386.997649 | 457.441045 | 0 |
| 2.20E-14 | 6.36E-13 | 346.279728 | 468.904769 | 323.890119 | 379.963708 | 0 |
| 2.21E-14 | 6.40E-13 | 293.871561 | 506.94211 | 458.457647 | 591.054657 | 0 |
| 2.22E-14 | 6.40E-13 | 282.939797 | 735.83348 | 633.859459 | 709.822312 | 0 |
| 2.23E-14 | 6.45E-13 | 286.476544 | 529.40861 | 622.722836 | 640.231889 | 0 |
| 2.24E-14 | 6.45E-13 | 285.833499 | 668.21154 | 551.262839 | 642.087633 | 0 |
| 2.25E-14 | 6.49E-13 | 284.547409 | 492.261031 | 490.939464 | 474.606683 | 0 |
| 2.27E-14 | 6.54E-13 | 476.817862 | 1023.89405 | 725.736599 | 939.00677 | 0 |
| 2.29E-14 | 6.58E-13 | 301.266579 | 583.239233 | 419.479466 | 510.793703 | 0 |
| 2.31E-14 | 6.65E-13 | 305.446371 | 1097.74432 | 660.772965 | 911.634537 | 0 |
| 2.32E-14 | 6.67E-13 | 277.795437 | 1183.16151 | 1210.1797 | 1364.43622 | 0 |
| 2.38E-14 | 6.82E-13 | 12.8608998 | 11.3444702 | 12.0646749 | 12.5262761 | 152.077968 |
| 2.38E-14 | 6.83E-13 | 282.618274 | 600.367159 | 541.98232 | 639.304017 | 0 |
| 2.41E-14 | 6.91E-13 | 306.410939 | 714.701623 | 571.679981 | 628.169549 | 9.50487297 |
| 2.42E-14 | 6.92E-13 | 287.762634 | 465.345719 | 417.623363 | 432.388493 | 0 |
| 2.42E-14 | 6.92E-13 | 287.119589 | 645.077718 | 504.860243 | 572.961147 | 0 |
| 2.42E-14 | 6.92E-13 | 277.473914 | 713.811861 | 771.211143 | 739.050289 | 0 |
| 2.46E-14 | 7.01E-13 | 276.830869 | 859.95533 | 834.318673 | 920.91326 | 0 |
| 2.46E-14 | 7.01E-13 | 284.225887 | 490.703947 | 474.23453 | 549.300403 | 0 |
| 2.48E-14 | 7.07E-13 | 280.367617 | 555.211719 | 520.637125 | 583.631678 | 0 |
| 2.48E-14 | 7.07E-13 | 241.463395 | 587.020723 | 490.011412 | 546.05285 | 9.50487297 |
| 2.49E-14 | 7.09E-13 | 283.582842 | 740.059851 | 603.233746 | 762.247097 | 0 |
| 2.50E-14 | 7.11E-13 | 276.830869 | 904.665889 | 870.512698 | 994.679108 | 0 |
| 2.52E-14 | 7.15E-13 | 51.4435994 | 29.3621582 | 27.8415575 | 39.898509 | 244.750479 |
| 2.53E-14 | 7.18E-13 | 624.396687 | 1404.4899 | 984.663084 | 1201.59463 | 26.1384007 |
| 2.56E-14 | 7.24E-13 | 317.021181 | 421.302482 | 339.667002 | 346.560305 | 0 |
| 2.59E-14 | 7.33E-13 | 275.223257 | 1059.92942 | 928.051917 | 986.328258 | 0 |
| 2.59E-14 | 7.33E-13 | 289.370247 | 426.196175 | 382.35739 | 423.10977 | 0 |
| 2.59E-14 | 7.33E-13 | 302.231146 | 690.455599 | 454.745439 | 554.867637 | 0 |
| 2.60E-14 | 7.34E-13 | 290.977859 | 377.259245 | 438.040505 | 426.357323 | 0 |
| 2.60E-14 | 7.34E-13 | 289.048724 | 404.841878 | 400.918428 | 455.585301 | 0 |
| 2.60E-14 | 7.34E-13 | 277.795437 | 591.691976 | 580.032448 | 650.438484 | 0 |
| 2.62E-14 | 7.38E-13 | 293.228516 | 644.187956 | 459.385699 | 562.290615 | 0 |
| 2.68E-14 | 7.55E-13 | 281.332184 | 477.579952 | 453.817387 | 515.897 | 0 |
| 2.69E-14 | 7.57E-13 | 274.580212 | 702.467391 | 651.492446 | 681.05827 | 0 |
| 2.69E-14 | 7.57E-13 | 354.639313 | 436.428442 | 306.257133 | 356.302964 | 0 |
| 2.71E-14 | 7.60E-13 | 306.732461 | 1550.63337 | 833.390621 | 1218.76027 | 0 |
| 2.74E-14 | 7.68E-13 | 317.342704 | 1036.57316 | 554.046994 | 810.960392 | 0 |
| 2.74E-14 | 7.68E-13 | 276.187824 | 658.201713 | 592.097123 | 587.807104 | 0 |
| 2.77E-14 | 7.76E-13 | 333.740351 | 870.187598 | 451.033232 | 614.251464 | 0 |
| 2.77E-14 | 7.76E-13 | 604.14077 | 1676.3123 | 1567.47969 | 1567.64026 | 16.6335277 |
| 2.78E-14 | 7.77E-13 | 407.369003 | 320.092013 | 334.09869 | 330.786476 | 0 |

| | | | | | | |
|---|---|---|---|---|---|---|
| 2.79E-14 | 7.79E-13 | 276.187824 | 829.258529 | 708.103613 | 701.935397 | 0 |
| 2.79E-14 | 7.79E-13 | 271.364987 | 1021.89208 | 1017.1449 | 1093.96144 | 0 |
| 2.79E-14 | 7.79E-13 | 274.901734 | 783.880648 | 937.332436 | 894.004964 | 0 |
| 2.80E-14 | 7.79E-13 | 284.225887 | 755.185811 | 555.903098 | 672.707419 | 0 |
| 2.80E-14 | 7.81E-13 | 949.455931 | 1115.76201 | 1360.52411 | 1389.48877 | 21.3859642 |
| 2.83E-14 | 7.86E-13 | 310.912254 | 505.829907 | 349.875573 | 430.532748 | 0 |
| 2.83E-14 | 7.87E-13 | 281.975229 | 828.146326 | 614.370369 | 719.564971 | 0 |
| 2.83E-14 | 7.87E-13 | 272.329554 | 1025.67357 | 900.210359 | 951.06911 | 0 |
| 2.84E-14 | 7.88E-13 | 280.046094 | 650.416292 | 582.816604 | 750.648693 | 0 |
| 2.84E-14 | 7.89E-13 | 509.934679 | 424.41665 | 306.257133 | 359.086581 | 0 |
| 2.87E-14 | 7.94E-13 | 275.223257 | 952.713057 | 778.635558 | 826.270285 | 0 |
| 2.91E-14 | 8.06E-13 | 277.152392 | 590.357332 | 506.716347 | 587.807104 | 0 |
| 2.94E-14 | 8.13E-13 | 338.241666 | 344.782918 | 305.329081 | 326.611051 | 0 |
| 2.95E-14 | 8.16E-13 | 328.917514 | 301.851884 | 380.501286 | 355.375092 | 0 |
| 2.96E-14 | 8.18E-13 | 276.509347 | 887.315523 | 692.32673 | 796.578372 | 0 |
| 2.97E-14 | 8.20E-13 | 268.792807 | 1027.23066 | 1019.001 | 1056.38262 | 0 |
| 2.98E-14 | 8.20E-13 | 265.899104 | 1391.58835 | 1369.80463 | 1413.14952 | 0 |
| 2.97E-14 | 8.20E-13 | 272.972599 | 617.272644 | 564.255566 | 583.631678 | 0 |
| 2.98E-14 | 8.21E-13 | 272.651077 | 604.37109 | 592.097123 | 652.758165 | 0 |
| 2.99E-14 | 8.23E-13 | 248.53689 | 476.69019 | 386.997649 | 450.018067 | 14.2573095 |
| 3.00E-14 | 8.23E-13 | 268.149762 | 1115.98445 | 1155.42464 | 1213.19304 | 0 |
| 3.00E-14 | 8.24E-13 | 287.762634 | 509.166516 | 735.017118 | 644.407314 | 0 |
| 3.00E-14 | 8.24E-13 | 284.547409 | 417.743433 | 377.71713 | 416.614664 | 0 |
| 3.01E-14 | 8.26E-13 | 271.686509 | 1389.80882 | 1154.49658 | 1119.94187 | 0 |
| 3.02E-14 | 8.28E-13 | 282.939797 | 492.261031 | 426.903882 | 417.542536 | 0 |
| 3.04E-14 | 8.33E-13 | 315.735091 | 2013.3098 | 947.541007 | 1350.0542 | 0 |
| 3.04E-14 | 8.33E-13 | 17.3622148 | 17.7952474 | 17.6329864 | 18.0935099 | 330.294336 |
| 3.05E-14 | 8.34E-13 | 289.370247 | 502.048417 | 386.069597 | 467.183704 | 0 |
| 3.06E-14 | 8.37E-13 | 276.830869 | 499.601571 | 494.651672 | 445.378705 | 0 |
| 3.07E-14 | 8.38E-13 | 289.048724 | 841.04788 | 558.687254 | 748.792948 | 0 |
| 3.11E-14 | 8.47E-13 | 275.544779 | 634.40057 | 527.133489 | 558.579126 | 0 |
| 3.12E-14 | 8.50E-13 | 412.513363 | 462.453992 | 280.271679 | 407.335941 | 0 |
| 3.13E-14 | 8.52E-13 | 295.800696 | 737.168123 | 473.306478 | 596.157954 | 0 |
| 3.15E-14 | 8.58E-13 | 302.874191 | 618.829728 | 409.270895 | 465.32796 | 0 |
| 3.17E-14 | 8.63E-13 | 280.689139 | 483.585848 | 433.400245 | 410.583494 | 0 |
| 3.18E-14 | 8.63E-13 | 273.937167 | 564.776664 | 500.219983 | 506.154341 | 0 |
| 3.19E-14 | 8.66E-13 | 275.866302 | 546.758976 | 532.7018 | 648.118803 | 0 |
| 3.20E-14 | 8.67E-13 | 305.446371 | 1612.91674 | 855.663867 | 1051.74326 | 0 |
| 3.21E-14 | 8.70E-13 | 272.008032 | 500.936214 | 505.788295 | 491.308384 | 0 |
| 3.25E-14 | 8.78E-13 | 272.972599 | 679.333569 | 563.327514 | 650.438484 | 0 |
| 3.27E-14 | 8.82E-13 | 271.043464 | 1064.15579 | 1282.56775 | 1424.28398 | 0 |
| 3.26E-14 | 8.82E-13 | 277.473914 | 519.621224 | 434.328297 | 496.411682 | 0 |
| 3.27E-14 | 8.82E-13 | 268.471284 | 650.416292 | 620.866732 | 629.561357 | 0 |

| | | | | | | |
|---|---|---|---|---|---|---|
| 3.27E-14 | 8.82E-13 | 362.998898 | 312.751473 | 337.810898 | 308.981477 | 0 |
| 3.30E-14 | 8.90E-13 | 294.514606 | 461.78667 | 370.292715 | 430.068812 | 7.12865473 |
| 3.32E-14 | 8.94E-13 | 326.988379 | 356.79471 | 300.688821 | 328.930731 | 0 |
| 3.32E-14 | 8.95E-13 | 299.658966 | 413.517061 | 404.630636 | 326.611051 | 0 |
| 3.33E-14 | 8.96E-13 | 275.223257 | 629.951758 | 508.57245 | 615.643273 | 0 |
| 3.36E-14 | 9.03E-13 | 272.008032 | 624.613184 | 554.046994 | 670.851675 | 0 |
| 3.44E-14 | 9.23E-13 | 240.498827 | 314.086117 | 282.127783 | 327.538923 | 11.8810912 |
| 3.45E-14 | 9.25E-13 | 264.934537 | 864.626583 | 841.743089 | 867.560603 | 0 |
| 3.45E-14 | 9.26E-13 | 317.985749 | 780.099158 | 509.500502 | 1003.95783 | 0 |
| 3.48E-14 | 9.32E-13 | 272.972599 | 555.6566 | 468.666218 | 514.505192 | 0 |
| 3.50E-14 | 9.35E-13 | 268.471284 | 521.623189 | 529.917645 | 554.867637 | 0 |
| 3.50E-14 | 9.35E-13 | 284.225887 | 785.437732 | 529.917645 | 640.695825 | 0 |
| 3.52E-14 | 9.41E-13 | 273.615644 | 668.43398 | 525.277385 | 623.066251 | 0 |
| 3.53E-14 | 9.42E-13 | 278.760004 | 685.561906 | 600.44959 | 500.123171 | 0 |
| 3.53E-14 | 9.42E-13 | 268.792807 | 694.014648 | 589.312967 | 649.510612 | 0 |
| 3.54E-14 | 9.44E-13 | 268.471284 | 564.109342 | 628.291148 | 649.974548 | 0 |
| 3.55E-14 | 9.47E-13 | 271.686509 | 484.698051 | 446.392972 | 498.731363 | 0 |
| 3.56E-14 | 9.47E-13 | 265.256059 | 663.762728 | 658.916861 | 676.418909 | 0 |
| 3.57E-14 | 9.48E-13 | 385.18395 | 325.208146 | 293.264406 | 364.189879 | 0 |
| 3.58E-14 | 9.51E-13 | 271.364987 | 715.591386 | 573.536085 | 616.571145 | 0 |
| 3.63E-14 | 9.63E-13 | 280.046094 | 435.983561 | 383.285442 | 383.675197 | 0 |
| 3.65E-14 | 9.67E-13 | 295.800696 | 409.068249 | 464.95401 | 340.993071 | 0 |
| 3.66E-14 | 9.71E-13 | 267.828239 | 684.004822 | 587.456863 | 682.914015 | 0 |
| 3.67E-14 | 9.73E-13 | 360.748241 | 494.485437 | 290.48025 | 428.677004 | 0 |
| 3.70E-14 | 9.78E-13 | 385.826995 | 316.977844 | 300.688821 | 342.848816 | 0 |
| 3.73E-14 | 9.87E-13 | 271.364987 | 812.353044 | 617.154525 | 707.038695 | 0 |
| 3.74E-14 | 9.88E-13 | 714.744509 | 334.995532 | 409.270895 | 508.010086 | 9.50487297 |
| 3.75E-14 | 9.90E-13 | 268.471284 | 646.189921 | 679.334003 | 798.898053 | 0 |
| 3.78E-14 | 9.97E-13 | 265.256059 | 544.757011 | 542.910371 | 552.547956 | 0 |
| 3.78E-14 | 9.97E-13 | 669.731359 | 2630.58245 | 2320.12979 | 2605.46543 | 16.6335277 |
| 3.79E-14 | 1.00E-12 | 279.724572 | 424.861532 | 428.759986 | 535.382319 | 0 |
| 3.85E-14 | 1.01E-12 | 267.185194 | 711.365015 | 589.312967 | 689.409121 | 0 |
| 3.87E-14 | 1.02E-12 | 318.628794 | 436.650883 | 444.536868 | 496.411682 | 28.5146189 |
| 3.89E-14 | 1.02E-12 | 281.332184 | 505.162585 | 402.774532 | 405.944132 | 0 |
| 3.89E-14 | 1.02E-12 | 269.114329 | 512.503125 | 449.177128 | 486.669023 | 0 |
| 3.92E-14 | 1.03E-12 | 493.215509 | 1076.39003 | 968.886201 | 1046.17602 | 2.37621824 |
| 3.97E-14 | 1.04E-12 | 264.291492 | 695.794173 | 601.377642 | 659.717207 | 0 |
| 4.02E-14 | 1.05E-12 | 261.719312 | 1220.30909 | 991.159447 | 1128.75666 | 0 |
| 4.04E-14 | 1.06E-12 | 259.790177 | 890.207251 | 880.721269 | 931.119856 | 0 |
| 4.05E-14 | 1.06E-12 | 275.223257 | 401.282829 | 414.839207 | 368.365304 | 0 |
| 4.05E-14 | 1.06E-12 | 271.686509 | 495.152759 | 633.859459 | 522.856043 | 0 |
| 4.05E-14 | 1.06E-12 | 320.236406 | 702.689832 | 621.794784 | 693.584546 | 11.8810912 |
| 4.04E-14 | 1.06E-12 | 266.863672 | 475.577987 | 451.033232 | 493.164129 | 0 |

| | | | | | | |
|---|---|---|---|---|---|---|
| 4.08E-14 | 1.06E-12 | 271.686509 | 387.713953 | 413.911155 | 420.326153 | 0 |
| 4.15E-14 | 1.08E-12 | 260.433222 | 793.668034 | 726.664651 | 750.184757 | 0 |
| 4.24E-14 | 1.10E-12 | 271.043464 | 472.463818 | 499.291931 | 602.653061 | 0 |
| 4.26E-14 | 1.11E-12 | 260.433222 | 811.240841 | 714.599976 | 792.866883 | 0 |
| 4.26E-14 | 1.11E-12 | 256.896474 | 1044.35858 | 1010.64854 | 1046.17602 | 0 |
| 4.28E-14 | 1.11E-12 | 259.790177 | 653.752901 | 638.499719 | 663.428696 | 0 |
| 4.32E-14 | 1.12E-12 | 270.078897 | 503.16062 | 417.623363 | 499.195299 | 0 |
| 4.37E-14 | 1.13E-12 | 271.686509 | 691.122921 | 528.061541 | 725.596141 | 0 |
| 4.42E-14 | 1.15E-12 | 323.773154 | 353.680542 | 283.983887 | 338.67339 | 0 |
| 4.45E-14 | 1.15E-12 | 292.906994 | 317.422725 | 417.623363 | 374.396474 | 0 |
| 4.47E-14 | 1.16E-12 | 254.324294 | 1217.86224 | 1409.71086 | 1409.43803 | 0 |
| 4.51E-14 | 1.17E-12 | 264.291492 | 537.638912 | 462.169855 | 496.875618 | 0 |
| 4.54E-14 | 1.17E-12 | 400.61703 | 425.306413 | 270.99116 | 367.901368 | 0 |
| 4.55E-14 | 1.18E-12 | 291.942426 | 622.166337 | 390.709857 | 536.774127 | 0 |
| 4.56E-14 | 1.18E-12 | 258.825609 | 1483.23387 | 1107.16594 | 1299.48516 | 0 |
| 4.62E-14 | 1.19E-12 | 258.825609 | 611.711629 | 687.68647 | 678.274653 | 0 |
| 4.63E-14 | 1.19E-12 | 1041.73289 | 3684.95085 | 3320.56976 | 3548.18369 | 21.3859642 |
| 4.64E-14 | 1.19E-12 | 268.471284 | 538.973555 | 427.831934 | 501.051043 | 0 |
| 4.67E-14 | 1.20E-12 | 257.539519 | 612.601392 | 599.521538 | 606.828486 | 0 |
| 4.69E-14 | 1.21E-12 | 263.969969 | 543.422367 | 490.011412 | 604.044869 | 0 |
| 4.70E-14 | 1.21E-12 | 259.147132 | 798.339286 | 798.124649 | 677.346781 | 0 |
| 4.70E-14 | 1.21E-12 | 2.57217997 | 2.66928711 | 0 | 0 | 239.998043 |
| 4.76E-14 | 1.22E-12 | 275.866302 | 453.111487 | 360.084144 | 450.482003 | 0 |
| 4.81E-14 | 1.23E-12 | 274.580212 | 398.613542 | 412.983103 | 346.096369 | 0 |
| 4.82E-14 | 1.23E-12 | 970.997938 | 2217.95515 | 1642.65189 | 1747.64748 | 21.3859642 |
| 4.84E-14 | 1.24E-12 | 257.217997 | 774.315702 | 681.190107 | 689.873057 | 0 |
| 4.92E-14 | 1.26E-12 | 267.185194 | 544.31213 | 426.903882 | 501.051043 | 0 |
| 5.01E-14 | 1.28E-12 | 567.808728 | 996.756295 | 741.513482 | 620.282634 | 26.1384007 |
| 5.03E-14 | 1.29E-12 | 256.253429 | 633.288367 | 579.104396 | 633.272847 | 0 |
| 5.05E-14 | 1.29E-12 | 308.018551 | 363.245487 | 438.040505 | 301.558498 | 0 |
| 5.07E-14 | 1.29E-12 | 864.25247 | 602.814006 | 292.336354 | 491.77232 | 0 |
| 5.10E-14 | 1.30E-12 | 253.038204 | 1008.76809 | 897.426204 | 1008.13326 | 0 |
| 5.11E-14 | 1.30E-12 | 301.588101 | 441.767017 | 360.084144 | 594.30221 | 0 |
| 5.14E-14 | 1.31E-12 | 270.078897 | 856.396281 | 574.464137 | 773.8455 | 0 |
| 5.15E-14 | 1.31E-12 | 289.048724 | 570.115238 | 382.35739 | 499.659235 | 9.50487297 |
| 5.16E-14 | 1.31E-12 | 155.295366 | 204.645345 | 178.185968 | 201.34829 | 803.161766 |
| 5.21E-14 | 1.32E-12 | 259.147132 | 625.725387 | 715.528028 | 804.00135 | 0 |
| 5.22E-14 | 1.32E-12 | 261.397789 | 607.485258 | 547.550631 | 485.277214 | 0 |
| 5.26E-14 | 1.33E-12 | 260.111699 | 569.670357 | 472.378426 | 536.310191 | 0 |
| 5.26E-14 | 1.33E-12 | 351.424088 | 351.011255 | 286.768042 | 301.094562 | 0 |
| 5.31E-14 | 1.34E-12 | 255.931907 | 809.461316 | 664.485173 | 738.586353 | 0 |
| 5.33E-14 | 1.35E-12 | 272.651077 | 371.030908 | 332.242586 | 365.581687 | 0 |
| 5.35E-14 | 1.35E-12 | 274.901734 | 344.115596 | 326.674275 | 359.550517 | 0 |

| | | | | | | |
|---|---|---|---|---|---|---|
| 5.36E-14 | 1.35E-12 | 270.078897 | 583.684115 | 426.903882 | 475.998491 | 0 |
| 5.40E-14 | 1.36E-12 | 299.658966 | 315.643201 | 294.192458 | 356.302964 | 0 |
| 5.43E-14 | 1.37E-12 | 256.253429 | 477.579952 | 504.860243 | 519.608489 | 0 |
| 5.53E-14 | 1.39E-12 | 255.288862 | 603.481327 | 540.126216 | 627.241677 | 0 |
| 5.57E-14 | 1.40E-12 | 257.539519 | 803.010539 | 619.938681 | 725.596141 | 0 |
| 5.57E-14 | 1.40E-12 | 274.580212 | 415.963908 | 333.170638 | 379.499772 | 0 |
| 5.58E-14 | 1.40E-12 | 290.977859 | 503.827942 | 331.314534 | 451.873811 | 0 |
| 5.63E-14 | 1.41E-12 | 281.653707 | 361.465963 | 467.738166 | 540.02168 | 0 |
| 5.64E-14 | 1.41E-12 | 257.217997 | 666.209574 | 535.485956 | 595.230082 | 0 |
| 5.68E-14 | 1.42E-12 | 323.130109 | 628.172233 | 336.882846 | 614.7154 | 0 |
| 5.68E-14 | 1.42E-12 | 401.90312 | 503.16062 | 325.746223 | 433.780301 | 0 |
| 5.71E-14 | 1.43E-12 | 248.53689 | 967.839018 | 910.418931 | 939.00677 | 0 |
| 5.73E-14 | 1.43E-12 | 254.324294 | 935.807572 | 733.161014 | 837.404753 | 0 |
| 5.75E-14 | 1.44E-12 | 155.938411 | 165.05092 | 170.761553 | 166.553078 | 9.50487297 |
| 5.85E-14 | 1.46E-12 | 280.046094 | 695.794173 | 442.680764 | 690.336993 | 0 |
| 5.86E-14 | 1.46E-12 | 251.109069 | 622.388778 | 679.334003 | 690.800929 | 0 |
| 5.87E-14 | 1.46E-12 | 263.005402 | 559.660531 | 464.95401 | 633.736783 | 0 |
| 5.89E-14 | 1.47E-12 | 306.410939 | 481.583883 | 936.404384 | 946.893685 | 0 |
| 6.06E-14 | 1.51E-12 | 244.357097 | 1229.87404 | 1441.26463 | 1275.82442 | 0 |
| 6.10E-14 | 1.52E-12 | 288.084157 | 375.034839 | 293.264406 | 358.158709 | 0 |
| 6.11E-14 | 1.52E-12 | 428.589487 | 2451.07289 | 2056.56305 | 2191.17044 | 0 |
| 6.13E-14 | 1.52E-12 | 251.430592 | 595.473466 | 540.126216 | 545.588914 | 0 |
| 6.17E-14 | 1.53E-12 | 253.359727 | 493.373234 | 453.817387 | 485.277214 | 0 |
| 6.20E-14 | 1.54E-12 | 366.535646 | 653.975342 | 732.232962 | 733.019119 | 0 |
| 6.19E-14 | 1.54E-12 | 279.081527 | 514.50509 | 351.731677 | 423.573706 | 0 |
| 6.21E-14 | 1.54E-12 | 372.323051 | 394.16473 | 270.99116 | 322.435625 | 0 |
| 6.23E-14 | 1.54E-12 | 353.674746 | 339.889225 | 794.412441 | 600.33338 | 0 |
| 6.27E-14 | 1.55E-12 | 459.455647 | 1110.64588 | 1036.63399 | 1105.55985 | 2.37621824 |
| 6.29E-14 | 1.55E-12 | 307.697029 | 650.416292 | 514.140762 | 349.807858 | 0 |
| 6.35E-14 | 1.57E-12 | 248.215367 | 665.319812 | 621.794784 | 659.717207 | 0 |
| 6.36E-14 | 1.57E-12 | 268.149762 | 373.700195 | 325.746223 | 356.7669 | 0 |
| 6.45E-14 | 1.59E-12 | 249.822979 | 563.21958 | 518.781022 | 537.238063 | 0 |
| 6.47E-14 | 1.60E-12 | 244.67862 | 1302.61211 | 1081.18048 | 1139.42719 | 0 |
| 6.48E-14 | 1.60E-12 | 251.752114 | 593.026619 | 504.860243 | 573.889019 | 0 |
| 6.49E-14 | 1.60E-12 | 261.076267 | 1146.01393 | 745.225689 | 872.199964 | 0 |
| 6.51E-14 | 1.60E-12 | 246.607755 | 682.447738 | 678.405951 | 673.635292 | 0 |
| 6.52E-14 | 1.60E-12 | 355.925403 | 284.946399 | 368.436611 | 511.257639 | 0 |
| 6.52E-14 | 1.60E-12 | 256.574952 | 555.211719 | 440.824661 | 504.762533 | 0 |
| 6.52E-14 | 1.60E-12 | 247.572322 | 129.015544 | 156.840774 | 115.520102 | 1043.15981 |
| 6.63E-14 | 1.63E-12 | 247.572322 | 779.654276 | 679.334003 | 701.007525 | 0 |
| 6.65E-14 | 1.63E-12 | 263.326924 | 400.837948 | 366.580507 | 344.240624 | 0 |
| 6.66E-14 | 1.63E-12 | 259.468654 | 389.493477 | 355.443884 | 392.95392 | 0 |
| 6.69E-14 | 1.64E-12 | 276.830869 | 514.28265 | 356.371936 | 401.304771 | 0 |

| | | | | | | |
|---|---|---|---|---|---|---|
| 6.73E-14 | 1.65E-12 | 530.512119 | 330.101839 | 325.746223 | 334.497965 | 0 |
| 6.73E-14 | 1.65E-12 | 306.732461 | 1033.68143 | 485.371153 | 923.232941 | 0 |
| 6.75E-14 | 1.65E-12 | 834.029355 | 3014.07003 | 2454.69732 | 2649.07542 | 0 |
| 6.76E-14 | 1.65E-12 | 245.643187 | 814.79989 | 731.304911 | 759.927416 | 0 |
| 6.77E-14 | 1.65E-12 | 272.972599 | 404.619438 | 454.745439 | 331.714348 | 0 |
| 6.78E-14 | 1.65E-12 | 72.3425616 | 77.6317668 | 115.078438 | 142.892335 | 10008.6312 |
| 6.82E-14 | 1.66E-12 | 286.155022 | 326.097909 | 289.552198 | 307.589668 | 0 |
| 6.83E-14 | 1.66E-12 | 251.109069 | 475.133105 | 436.184401 | 471.823066 | 0 |
| 6.89E-14 | 1.68E-12 | 243.39253 | 874.85885 | 895.5701 | 836.476881 | 0 |
| 6.90E-14 | 1.68E-12 | 308.661596 | 545.201892 | 452.889335 | 533.062638 | 2.37621824 |
| 6.91E-14 | 1.68E-12 | 247.572322 | 596.140788 | 543.838423 | 550.692212 | 0 |
| 6.91E-14 | 1.68E-12 | 255.610384 | 465.345719 | 392.565961 | 433.780301 | 0 |
| 6.97E-14 | 1.69E-12 | 251.430592 | 534.747184 | 464.025958 | 553.939765 | 0 |
| 6.98E-14 | 1.69E-12 | 247.2508 | 520.733427 | 559.615306 | 561.826679 | 0 |
| 6.99E-14 | 1.69E-12 | 346.279728 | 671.99303 | 327.602327 | 472.750938 | 0 |
| 7.01E-14 | 1.70E-12 | 273.937167 | 389.271037 | 309.041288 | 349.807858 | 0 |
| 7.05E-14 | 1.71E-12 | 244.357097 | 798.561727 | 879.793217 | 772.453692 | 0 |
| 7.09E-14 | 1.71E-12 | 248.858412 | 837.26639 | 660.772965 | 715.389545 | 0 |
| 7.09E-14 | 1.71E-12 | 301.909624 | 861.067533 | 422.263622 | 675.0271 | 0 |
| 7.26E-14 | 1.75E-12 | 250.144502 | 524.514917 | 454.745439 | 468.111577 | 0 |
| 7.27E-14 | 1.75E-12 | 469.101322 | 1257.23423 | 747.081793 | 888.43773 | 0 |
| 7.31E-14 | 1.76E-12 | 247.893845 | 524.070036 | 473.306478 | 527.95934 | 0 |
| 7.32E-14 | 1.76E-12 | 238.891215 | 1082.8408 | 1105.30983 | 1081.43517 | 0 |
| 7.54E-14 | 1.81E-12 | 343.707548 | 329.656958 | 262.638692 | 320.579881 | 0 |
| 7.55E-14 | 1.81E-12 | 248.858412 | 559.882971 | 481.658945 | 482.029661 | 0 |
| 7.56E-14 | 1.82E-12 | 278.116959 | 1276.36412 | 658.916861 | 895.860708 | 0 |
| 7.62E-14 | 1.83E-12 | 8012.98365 | 14204.3889 | 7082.89223 | 10312.3728 | 42.7719284 |
| 7.63E-14 | 1.83E-12 | 242.10644 | 855.061637 | 758.218416 | 772.453692 | 0 |
| 7.65E-14 | 1.83E-12 | 254.645817 | 559.660531 | 425.97583 | 491.308384 | 0 |
| 7.64E-14 | 1.83E-12 | 489.035717 | 745.843306 | 659.844913 | 675.954972 | 0 |
| 7.66E-14 | 1.83E-12 | 241.141872 | 768.754687 | 843.599193 | 803.073478 | 0 |
| 7.75E-14 | 1.85E-12 | 246.929277 | 514.060209 | 515.996866 | 598.941571 | 0 |
| 7.74E-14 | 1.85E-12 | 243.714052 | 766.530282 | 648.70829 | 700.543588 | 0 |
| 7.77E-14 | 1.85E-12 | 249.179934 | 408.178487 | 446.392972 | 457.441045 | 0 |
| 7.79E-14 | 1.86E-12 | 243.39253 | 616.605322 | 619.010629 | 692.192738 | 0 |
| 7.88E-14 | 1.88E-12 | 248.858412 | 611.711629 | 490.011412 | 526.103596 | 0 |
| 7.88E-14 | 1.88E-12 | 183.267823 | 397.278898 | 417.623363 | 388.778495 | 11.8810912 |
| 7.90E-14 | 1.88E-12 | 242.749485 | 738.947648 | 639.427771 | 678.274653 | 0 |
| 8.03E-14 | 1.91E-12 | 242.749485 | 646.634802 | 574.464137 | 636.056463 | 0 |
| 8.04E-14 | 1.91E-12 | 256.253429 | 531.855457 | 398.134272 | 502.906788 | 0 |
| 8.04E-14 | 1.91E-12 | 251.109069 | 384.154903 | 438.040505 | 450.945939 | 0 |
| 8.05E-14 | 1.91E-12 | 249.179934 | 534.524744 | 436.184401 | 509.86583 | 0 |
| 8.09E-14 | 1.92E-12 | 276.509347 | 422.414685 | 598.593486 | 383.211261 | 0 |

| | | | | | | |
|---|---|---|---|---|---|---|
| 8.10E-14 | 1.92E-12 | 261.076267 | 496.932284 | 378.645182 | 390.170303 | 0 |
| 8.13E-14 | 1.93E-12 | 245.643187 | 631.953723 | 525.277385 | 630.48923 | 0 |
| 8.16E-14 | 1.93E-12 | 410.584228 | 1061.26407 | 677.477899 | 876.839326 | 4.75243649 |
| 8.15E-14 | 1.93E-12 | 246.929277 | 469.794531 | 443.608816 | 421.717962 | 0 |
| 8.21E-14 | 1.94E-12 | 246.286232 | 478.692155 | 433.400245 | 490.380512 | 0 |
| 8.31E-14 | 1.96E-12 | 241.784917 | 537.194031 | 557.759202 | 581.775934 | 0 |
| 8.31E-14 | 1.96E-12 | 249.501457 | 443.991423 | 386.069597 | 437.955727 | 0 |
| 8.36E-14 | 1.97E-12 | 241.463395 | 733.386633 | 631.075304 | 660.181143 | 0 |
| 8.37E-14 | 1.97E-12 | 238.24817 | 1067.4924 | 905.778671 | 949.213366 | 0 |
| 8.40E-14 | 1.98E-12 | 247.893845 | 825.032157 | 603.233746 | 691.264865 | 0 |
| 8.42E-14 | 1.98E-12 | 400.61703 | 711.142574 | 280.271679 | 532.598702 | 0 |
| 8.45E-14 | 1.99E-12 | 332.454261 | 502.715739 | 273.775316 | 477.854236 | 0 |
| 8.46E-14 | 1.99E-12 | 432.126235 | 1496.58031 | 1341.03502 | 1802.39195 | 0 |
| 8.46E-14 | 1.99E-12 | 267.506717 | 1438.74575 | 771.211143 | 977.977407 | 0 |
| 8.48E-14 | 1.99E-12 | 235.997512 | 1117.76398 | 979.094772 | 1072.62038 | 0 |
| 8.49E-14 | 1.99E-12 | 246.286232 | 476.69019 | 419.479466 | 474.142747 | 0 |
| 8.56E-14 | 2.01E-12 | 281.332184 | 653.975342 | 378.645182 | 482.493597 | 0 |
| 8.60E-14 | 2.01E-12 | 475.853294 | 465.345719 | 264.494796 | 340.529135 | 0 |
| 8.60E-14 | 2.01E-12 | 242.10644 | 511.168481 | 532.7018 | 575.744764 | 0 |
| 8.63E-14 | 2.02E-12 | 245.321665 | 873.301766 | 646.852186 | 794.722627 | 0 |
| 8.64E-14 | 2.02E-12 | 244.357097 | 570.115238 | 477.946737 | 549.300403 | 0 |
| 8.66E-14 | 2.02E-12 | 246.929277 | 912.673751 | 688.614522 | 687.08944 | 0 |
| 8.66E-14 | 2.02E-12 | 239.212737 | 635.067891 | 611.586213 | 603.580933 | 0 |
| 8.69E-14 | 2.03E-12 | 931.450671 | 396.834017 | 430.616089 | 382.747325 | 0 |
| 8.71E-14 | 2.03E-12 | 244.357097 | 464.678398 | 478.874789 | 429.14094 | 0 |
| 8.76E-14 | 2.04E-12 | 274.580212 | 536.74915 | 339.667002 | 472.287002 | 0 |
| 8.95E-14 | 2.08E-12 | 480.033087 | 676.441842 | 647.780238 | 676.882845 | 0 |
| 9.00E-14 | 2.10E-12 | 238.569692 | 760.524386 | 667.269328 | 770.597947 | 0 |
| 9.07E-14 | 2.11E-12 | 243.071007 | 580.125065 | 484.443101 | 570.641466 | 0 |
| 9.11E-14 | 2.12E-12 | 334.383396 | 451.776843 | 322.962067 | 385.994878 | 0 |
| 9.21E-14 | 2.14E-12 | 367.500213 | 593.24906 | 585.60076 | 575.744764 | 4.75243649 |
| 9.27E-14 | 2.15E-12 | 241.784917 | 566.333748 | 481.658945 | 553.011892 | 0 |
| 9.30E-14 | 2.16E-12 | 243.714052 | 1032.56923 | 734.089066 | 841.116242 | 0 |
| 9.38E-14 | 2.17E-12 | 253.038204 | 508.499194 | 388.853753 | 411.975302 | 0 |
| 9.39E-14 | 2.17E-12 | 245.321665 | 473.798462 | 432.472193 | 401.768707 | 0 |
| 9.42E-14 | 2.18E-12 | 542.086928 | 324.763265 | 285.83999 | 404.088388 | 0 |
| 9.43E-14 | 2.18E-12 | 23.4711422 | 9.12006429 | 11.136623 | 11.5984038 | 270.88888 |
| 9.48E-14 | 2.19E-12 | 254.645817 | 355.460067 | 319.249859 | 366.50956 | 0 |
| 9.48E-14 | 2.19E-12 | 240.177305 | 790.331425 | 624.57894 | 738.122417 | 0 |
| 9.47E-14 | 2.19E-12 | 248.215367 | 1197.17527 | 761.930624 | 972.874109 | 0 |
| 9.47E-14 | 2.19E-12 | 244.035575 | 451.109521 | 483.515049 | 554.403701 | 0 |
| 9.51E-14 | 2.19E-12 | 260.111699 | 341.891191 | 318.321808 | 397.593282 | 0 |
| 9.55E-14 | 2.20E-12 | 251.430592 | 349.23173 | 363.796351 | 414.294983 | 0 |

| | | | | | | |
|---|---|---|---|---|---|---|
| 9.66E-14 | 2.22E-12 | 448.845405 | 874.413969 | 838.958933 | 947.357621 | 0 |
| 9.70E-14 | 2.23E-12 | 646.58174 | 3422.47096 | 3179.50587 | 2956.66509 | 16.6335277 |
| 9.71E-14 | 2.23E-12 | 241.784917 | 674.217436 | 528.061541 | 619.354762 | 0 |
| 9.73E-14 | 2.23E-12 | 241.141872 | 647.079683 | 517.85297 | 585.023487 | 0 |
| 9.78E-14 | 2.24E-12 | 245.96471 | 508.721635 | 410.198947 | 438.883599 | 0 |
| 9.85E-14 | 2.26E-12 | 242.10644 | 430.867428 | 400.918428 | 415.686792 | 0 |
| 9.89E-14 | 2.27E-12 | 245.321665 | 516.062174 | 412.055051 | 456.513173 | 0 |
| 9.90E-14 | 2.27E-12 | 254.645817 | 392.385205 | 335.954794 | 333.106157 | 0 |
| 9.95E-14 | 2.28E-12 | 415.085542 | 488.479541 | 377.71713 | 475.070619 | 2.37621824 |
| 9.96E-14 | 2.28E-12 | 422.48056 | 1285.92906 | 1218.53217 | 1323.60984 | 0 |
| 9.99E-14 | 2.28E-12 | 281.010662 | 388.603715 | 337.810898 | 397.129346 | 14.2573095 |
| 1.01E-13 | 2.30E-12 | 263.648447 | 341.891191 | 418.551415 | 508.010086 | 0 |
| 1.01E-13 | 2.30E-12 | 253.038204 | 345.895121 | 324.818171 | 321.507753 | 0 |
| 1.01E-13 | 2.31E-12 | 234.3899 | 637.959619 | 626.435044 | 613.787528 | 0 |
| 1.02E-13 | 2.34E-12 | 240.82035 | 574.786491 | 480.730893 | 483.885406 | 0 |
| 1.03E-13 | 2.34E-12 | 232.782287 | 730.494906 | 706.247509 | 727.915821 | 0 |
| 1.03E-13 | 2.34E-12 | 236.319035 | 658.201713 | 747.081793 | 816.06369 | 0 |
| 1.03E-13 | 2.35E-12 | 240.177305 | 427.308378 | 440.824661 | 484.813278 | 0 |
| 1.04E-13 | 2.36E-12 | 249.822979 | 443.324101 | 350.803625 | 394.345729 | 0 |
| 1.04E-13 | 2.36E-12 | 473.602637 | 433.536715 | 261.710641 | 327.074987 | 0 |
| 1.04E-13 | 2.37E-12 | 233.10381 | 1014.10666 | 852.879712 | 856.426135 | 0 |
| 1.05E-13 | 2.37E-12 | 243.714052 | 485.365373 | 417.623363 | 458.368917 | 2.37621824 |
| 1.05E-13 | 2.38E-12 | 235.032945 | 663.317847 | 584.672708 | 658.789335 | 0 |
| 1.05E-13 | 2.38E-12 | 245.643187 | 436.206002 | 364.724403 | 392.95392 | 0 |
| 1.05E-13 | 2.38E-12 | 254.324294 | 510.056278 | 669.125432 | 442.595088 | 0 |
| 1.06E-13 | 2.40E-12 | 236.319035 | 476.91263 | 477.946737 | 501.978916 | 0 |
| 1.06E-13 | 2.40E-12 | 240.498827 | 712.699658 | 537.34206 | 646.263059 | 0 |
| 1.06E-13 | 2.41E-12 | 251.109069 | 399.280863 | 331.314534 | 351.663603 | 0 |
| 1.07E-13 | 2.42E-12 | 375.538275 | 396.834017 | 240.365446 | 397.593282 | 0 |
| 1.07E-13 | 2.42E-12 | 263.326924 | 389.938359 | 298.832717 | 336.35371 | 0 |
| 1.08E-13 | 2.43E-12 | 242.749485 | 642.630872 | 522.493229 | 486.669023 | 0 |
| 1.08E-13 | 2.43E-12 | 435.019937 | 973.844914 | 917.843346 | 990.967619 | 0 |
| 1.08E-13 | 2.43E-12 | 235.67599 | 488.701982 | 473.306478 | 495.019873 | 0 |
| 1.10E-13 | 2.47E-12 | 370.393916 | 718.705554 | 591.169071 | 707.502631 | 4.75243649 |
| 1.10E-13 | 2.47E-12 | 246.286232 | 419.522957 | 355.443884 | 365.117751 | 0 |
| 1.11E-13 | 2.49E-12 | 245.000142 | 545.646773 | 719.240236 | 516.360936 | 0 |
| 1.11E-13 | 2.49E-12 | 248.858412 | 452.666606 | 349.875573 | 401.768707 | 0 |
| 1.11E-13 | 2.49E-12 | 251.430592 | 375.257279 | 438.040505 | 527.495404 | 0 |
| 1.12E-13 | 2.51E-12 | 241.784917 | 371.030908 | 375.861026 | 360.942326 | 0 |
| 1.13E-13 | 2.52E-12 | 228.280972 | 1027.67554 | 961.461786 | 905.139431 | 0 |
| 1.13E-13 | 2.53E-12 | 413.47793 | 1687.65677 | 1431.98411 | 1627.48802 | 0 |
| 1.14E-13 | 2.55E-12 | 234.068377 | 1017.22083 | 762.858676 | 900.036134 | 0 |
| 1.14E-13 | 2.55E-12 | 236.319035 | 503.383061 | 440.824661 | 484.349342 | 0 |

| | | | | | | |
|---|---|---|---|---|---|---|
| 1.14E-13 | 2.55E-12 | 234.3899 | 564.109342 | 503.932191 | 509.401894 | 0 |
| 1.14E-13 | 2.55E-12 | 287.762634 | 416.85367 | 436.184401 | 276.505946 | 0 |
| 1.14E-13 | 2.56E-12 | 337.598621 | 740.059851 | 606.017902 | 653.222101 | 30.8908372 |
| 1.15E-13 | 2.56E-12 | 223.136612 | 1233.65553 | 1286.27996 | 1428.45941 | 0 |
| 1.16E-13 | 2.58E-12 | 241.463395 | 564.999105 | 429.688038 | 508.474022 | 0 |
| 1.17E-13 | 2.60E-12 | 240.177305 | 416.85367 | 391.637909 | 468.575513 | 0 |
| 1.18E-13 | 2.62E-12 | 244.67862 | 954.937463 | 610.658161 | 794.722627 | 0 |
| 1.20E-13 | 2.67E-12 | 248.858412 | 606.150614 | 410.198947 | 510.329766 | 0 |
| 1.21E-13 | 2.70E-12 | 254.324294 | 425.528853 | 315.537652 | 372.54073 | 0 |
| 1.21E-13 | 2.70E-12 | 237.605125 | 580.125065 | 467.738166 | 479.246044 | 0 |
| 1.21E-13 | 2.70E-12 | 271.043464 | 275.826335 | 330.386482 | 375.788283 | 0 |
| 1.22E-13 | 2.71E-12 | 340.170801 | 871.522241 | 725.736599 | 782.196351 | 2.37621824 |
| 1.23E-13 | 2.73E-12 | 226.994882 | 755.630693 | 768.426987 | 762.711033 | 0 |
| 1.23E-13 | 2.74E-12 | 234.068377 | 669.546183 | 535.485956 | 579.920189 | 0 |
| 1.24E-13 | 2.74E-12 | 246.929277 | 432.424512 | 335.026742 | 405.01626 | 0 |
| 1.24E-13 | 2.74E-12 | 255.610384 | 326.320349 | 392.565961 | 465.32796 | 0 |
| 1.24E-13 | 2.74E-12 | 281.975229 | 441.099695 | 283.055835 | 450.018067 | 0 |
| 1.24E-13 | 2.75E-12 | 224.422702 | 1027.00822 | 1044.98646 | 942.718259 | 0 |
| 1.24E-13 | 2.75E-12 | 235.67599 | 474.465784 | 429.688038 | 414.294983 | 0 |
| 1.25E-13 | 2.76E-12 | 236.640557 | 468.459888 | 399.062324 | 456.049237 | 0 |
| 1.26E-13 | 2.79E-12 | 239.53426 | 549.428263 | 492.795568 | 679.666462 | 0 |
| 1.26E-13 | 2.79E-12 | 256.574952 | 367.249418 | 289.552198 | 337.281582 | 0 |
| 1.27E-13 | 2.79E-12 | 299.980489 | 255.584241 | 387.925701 | 334.034029 | 0 |
| 1.27E-13 | 2.80E-12 | 238.24817 | 393.942289 | 368.436611 | 361.870198 | 0 |
| 1.28E-13 | 2.81E-12 | 236.640557 | 626.615149 | 478.874789 | 532.134766 | 0 |
| 1.28E-13 | 2.81E-12 | 237.283602 | 393.497408 | 398.134272 | 456.513173 | 0 |
| 1.28E-13 | 2.82E-12 | 236.96208 | 930.468998 | 651.492446 | 725.132204 | 0 |
| 1.29E-13 | 2.82E-12 | 267.506717 | 314.308557 | 412.983103 | 502.906788 | 0 |
| 1.29E-13 | 2.82E-12 | 272.972599 | 974.512235 | 490.011412 | 641.623697 | 0 |
| 1.30E-13 | 2.85E-12 | 437.91364 | 830.148291 | 764.71478 | 786.835713 | 2.37621824 |
| 1.30E-13 | 2.85E-12 | 229.567062 | 517.619259 | 544.766475 | 572.033275 | 0 |
| 1.31E-13 | 2.86E-12 | 246.286232 | 582.12703 | 399.062324 | 482.493597 | 0 |
| 1.31E-13 | 2.87E-12 | 242.427962 | 468.459888 | 360.084144 | 416.150728 | 0 |
| 1.31E-13 | 2.87E-12 | 0 | 0 | 0 | 0 | 45.1481466 |
| 1.31E-13 | 2.87E-12 | 228.280972 | 567.890832 | 567.039721 | 608.684231 | 0 |
| 1.32E-13 | 2.88E-12 | 245.000142 | 520.955867 | 374.932974 | 437.955727 | 0 |
| 1.32E-13 | 2.88E-12 | 440.48582 | 978.516166 | 794.412441 | 881.478687 | 0 |
| 1.33E-13 | 2.89E-12 | 231.81772 | 1194.50598 | 838.030881 | 930.65592 | 0 |
| 1.33E-13 | 2.89E-12 | 236.96208 | 367.471859 | 389.781805 | 359.550517 | 0 |
| 1.33E-13 | 2.91E-12 | 260.433222 | 476.245308 | 313.681548 | 432.852429 | 0 |
| 1.34E-13 | 2.91E-12 | 229.24554 | 551.87511 | 617.154525 | 650.90242 | 0 |
| 1.34E-13 | 2.91E-12 | 225.708792 | 686.006787 | 684.902315 | 664.356569 | 0 |
| 1.35E-13 | 2.94E-12 | 249.501457 | 309.414864 | 393.494013 | 400.840835 | 0 |

| | | | | | | |
|---|---|---|---|---|---|---|
| 1.35E-13 | 2.94E-12 | 237.283602 | 495.152759 | 639.427771 | 486.205087 | 0 |
| 1.35E-13 | 2.94E-12 | 340.170801 | 392.607646 | 245.933758 | 317.332327 | 0 |
| 1.35E-13 | 2.94E-12 | 232.782287 | 429.087903 | 399.990376 | 438.419663 | 0 |
| 1.36E-13 | 2.94E-12 | 237.926647 | 535.414506 | 407.414792 | 475.070619 | 0 |
| 1.36E-13 | 2.95E-12 | 229.24554 | 502.938179 | 563.327514 | 573.425083 | 0 |
| 1.38E-13 | 2.99E-12 | 486.142014 | 442.211898 | 254.286225 | 325.683178 | 0 |
| 1.39E-13 | 3.00E-12 | 246.286232 | 324.095943 | 302.544925 | 309.445413 | 0 |
| 1.39E-13 | 3.00E-12 | 311.555299 | 300.072359 | 248.717914 | 293.671584 | 0 |
| 1.39E-13 | 3.01E-12 | 237.926647 | 342.558512 | 346.163365 | 343.776688 | 0 |
| 1.39E-13 | 3.01E-12 | 244.035575 | 321.204215 | 341.523105 | 389.242431 | 0 |
| 1.39E-13 | 3.01E-12 | 272.008032 | 421.524923 | 322.962067 | 288.568286 | 0 |
| 1.40E-13 | 3.02E-12 | 295.479174 | 838.156152 | 658.916861 | 786.371776 | 7.12865473 |
| 1.40E-13 | 3.02E-12 | 382.290248 | 309.414864 | 377.71713 | 245.422224 | 0 |
| 1.41E-13 | 3.03E-12 | 227.637927 | 592.359298 | 530.845697 | 539.093808 | 0 |
| 1.41E-13 | 3.03E-12 | 233.425332 | 482.473645 | 406.48674 | 482.493597 | 0 |
| 1.42E-13 | 3.05E-12 | 334.704918 | 847.943538 | 945.684903 | 982.616768 | 11.8810912 |
| 1.42E-13 | 3.05E-12 | 419.265335 | 1109.97856 | 933.620228 | 1060.09411 | 2.37621824 |
| 1.42E-13 | 3.05E-12 | 234.711422 | 517.174377 | 412.983103 | 448.162322 | 0 |
| 1.42E-13 | 3.05E-12 | 252.716682 | 324.540824 | 282.127783 | 342.38488 | 0 |
| 1.42E-13 | 3.05E-12 | 224.744225 | 758.52242 | 827.82231 | 922.305069 | 0 |
| 1.43E-13 | 3.07E-12 | 541.443883 | 1161.36233 | 592.097123 | 863.849114 | 0 |
| 1.43E-13 | 3.07E-12 | 244.035575 | 429.310343 | 329.458431 | 375.788283 | 0 |
| 1.43E-13 | 3.07E-12 | 226.030315 | 543.644808 | 536.414008 | 559.506998 | 0 |
| 1.44E-13 | 3.09E-12 | 235.354467 | 1040.13221 | 676.549847 | 853.642518 | 0 |
| 1.44E-13 | 3.09E-12 | 282.939797 | 442.434338 | 270.063108 | 390.170303 | 0 |
| 1.45E-13 | 3.10E-12 | 231.496197 | 445.326066 | 399.990376 | 456.977109 | 0 |
| 1.46E-13 | 3.12E-12 | 227.316405 | 655.977307 | 547.550631 | 611.467847 | 0 |
| 1.46E-13 | 3.12E-12 | 227.316405 | 621.054134 | 545.694527 | 538.629872 | 0 |
| 1.46E-13 | 3.12E-12 | 226.030315 | 556.991243 | 524.349333 | 574.816891 | 0 |
| 1.46E-13 | 3.12E-12 | 246.607755 | 387.269071 | 312.753496 | 414.294983 | 0 |
| 1.47E-13 | 3.13E-12 | 222.172045 | 803.232979 | 904.850619 | 810.496456 | 0 |
| 1.47E-13 | 3.14E-12 | 364.606511 | 383.487581 | 232.941031 | 417.542536 | 0 |
| 1.47E-13 | 3.14E-12 | 239.855782 | 411.292655 | 338.73895 | 354.911156 | 0 |
| 1.48E-13 | 3.14E-12 | 227.95945 | 527.406645 | 481.658945 | 562.754552 | 0 |
| 1.49E-13 | 3.17E-12 | 229.888585 | 473.353581 | 437.112453 | 417.0786 | 0 |
| 1.49E-13 | 3.17E-12 | 235.67599 | 385.711987 | 486.299204 | 488.988703 | 0 |
| 1.49E-13 | 3.17E-12 | 225.065747 | 797.004643 | 656.132705 | 767.350394 | 0 |
| 1.51E-13 | 3.20E-12 | 225.065747 | 621.054134 | 553.118943 | 617.962954 | 0 |
| 1.52E-13 | 3.22E-12 | 230.53163 | 430.422546 | 388.853753 | 405.944132 | 0 |
| 1.52E-13 | 3.22E-12 | 244.357097 | 535.636947 | 471.450374 | 365.581687 | 0 |
| 1.52E-13 | 3.22E-12 | 266.220627 | 365.025012 | 380.501286 | 268.619032 | 0 |
| 1.53E-13 | 3.23E-12 | 238.24817 | 434.204036 | 347.091417 | 439.347535 | 0 |
| 1.53E-13 | 3.25E-12 | 415.085542 | 968.951221 | 926.195813 | 1038.28911 | 0 |

| | | | | | | |
|---|---|---|---|---|---|---|
| 1.55E-13 | 3.27E-12 | 243.071007 | 308.525102 | 390.709857 | 361.870198 | 0 |
| 1.55E-13 | 3.27E-12 | 233.425332 | 382.82026 | 365.652455 | 429.14094 | 0 |
| 1.55E-13 | 3.27E-12 | 262.683879 | 857.063603 | 451.033232 | 592.910401 | 0 |
| 1.55E-13 | 3.27E-12 | 227.316405 | 557.881006 | 470.522322 | 518.680617 | 0 |
| 1.56E-13 | 3.29E-12 | 241.784917 | 426.418616 | 331.314534 | 357.694773 | 0 |
| 1.56E-13 | 3.29E-12 | 231.496197 | 623.723421 | 467.738166 | 561.362743 | 0 |
| 1.56E-13 | 3.29E-12 | 222.172045 | 708.918168 | 655.204653 | 648.118803 | 0 |
| 1.57E-13 | 3.30E-12 | 355.603881 | 1343.76362 | 1235.2371 | 1395.05601 | 4.75243649 |
| 1.58E-13 | 3.32E-12 | 224.10118 | 562.107377 | 515.996866 | 555.331573 | 0 |
| 1.58E-13 | 3.33E-12 | 230.210107 | 411.959977 | 375.861026 | 389.242431 | 0 |
| 1.59E-13 | 3.34E-12 | 332.775783 | 447.328031 | 312.753496 | 257.484564 | 0 |
| 1.59E-13 | 3.34E-12 | 252.395159 | 1681.206 | 812.045427 | 1085.14666 | 0 |
| 1.59E-13 | 3.34E-12 | 235.997512 | 591.691976 | 543.838423 | 557.651254 | 4.75243649 |
| 1.60E-13 | 3.36E-12 | 222.81509 | 603.258887 | 563.327514 | 623.530187 | 0 |
| 1.60E-13 | 3.36E-12 | 226.030315 | 426.418616 | 332.242586 | 401.768707 | 19.0097459 |
| 1.61E-13 | 3.36E-12 | 233.10381 | 1375.35018 | 864.016335 | 940.398579 | 0 |
| 1.61E-13 | 3.37E-12 | 235.032945 | 410.847774 | 340.595054 | 376.716155 | 0 |
| 1.61E-13 | 3.37E-12 | 276.509347 | 669.768624 | 577.248292 | 605.436677 | 7.12865473 |
| 1.62E-13 | 3.38E-12 | 224.744225 | 1122.65767 | 825.038154 | 868.952411 | 0 |
| 1.63E-13 | 3.40E-12 | 179.409553 | 336.997498 | 289.552198 | 334.961901 | 14.2573095 |
| 1.63E-13 | 3.40E-12 | 230.210107 | 485.142932 | 453.817387 | 583.167742 | 0 |
| 1.63E-13 | 3.40E-12 | 282.939797 | 398.391101 | 395.350117 | 404.088388 | 0 |
| 1.64E-13 | 3.42E-12 | 229.24554 | 852.837231 | 1361.45216 | 1261.90633 | 0 |
| 1.65E-13 | 3.44E-12 | 227.637927 | 497.154724 | 419.479466 | 461.152534 | 0 |
| 1.66E-13 | 3.46E-12 | 219.921387 | 650.416292 | 671.909588 | 678.738589 | 0 |
| 1.66E-13 | 3.46E-12 | 175.551283 | 1301.49991 | 1303.91294 | 1131.54027 | 4.75243649 |
| 1.67E-13 | 3.47E-12 | 236.96208 | 415.519027 | 329.458431 | 397.129346 | 0 |
| 1.67E-13 | 3.47E-12 | 234.3899 | 398.16866 | 387.925701 | 495.019873 | 0 |
| 1.69E-13 | 3.52E-12 | 220.24291 | 605.038411 | 598.593486 | 611.931784 | 0 |
| 1.69E-13 | 3.52E-12 | 226.994882 | 592.359298 | 478.874789 | 496.875618 | 0 |
| 1.69E-13 | 3.52E-12 | 272.972599 | 276.938538 | 291.408302 | 390.170303 | 0 |
| 1.71E-13 | 3.55E-12 | 239.212737 | 377.926567 | 315.537652 | 324.29137 | 0 |
| 1.72E-13 | 3.57E-12 | 219.278342 | 813.020365 | 761.930624 | 696.832099 | 0 |
| 1.72E-13 | 3.57E-12 | 223.458135 | 667.099337 | 557.759202 | 564.610296 | 0 |
| 1.74E-13 | 3.59E-12 | 232.139242 | 369.028943 | 474.23453 | 447.698386 | 0 |
| 1.75E-13 | 3.63E-12 | 233.10381 | 439.097729 | 630.147252 | 560.898807 | 0 |
| 1.76E-13 | 3.63E-12 | 234.711422 | 644.187956 | 521.565177 | 807.248903 | 0 |
| 1.77E-13 | 3.66E-12 | 217.027685 | 757.410217 | 812.973479 | 856.890071 | 0 |
| 1.78E-13 | 3.67E-12 | 221.850522 | 611.489189 | 545.694527 | 527.95934 | 0 |
| 1.78E-13 | 3.67E-12 | 244.035575 | 654.420223 | 420.407518 | 669.459866 | 0 |
| 1.78E-13 | 3.67E-12 | 215.741595 | 968.72878 | 854.735816 | 874.055709 | 0 |
| 1.79E-13 | 3.69E-12 | 218.635297 | 803.900301 | 830.606466 | 696.368163 | 0 |
| 1.80E-13 | 3.71E-12 | 231.496197 | 341.891191 | 341.523105 | 326.147114 | 0 |

| | | | | | | |
|---|---|---|---|---|---|---|
| 1.81E-13 | 3.72E-12 | 232.139242 | 387.491512 | 328.530379 | 369.293176 | 0 |
| 1.81E-13 | 3.72E-12 | 222.172045 | 591.247095 | 509.500502 | 606.36455 | 0 |
| 1.82E-13 | 3.74E-12 | 228.924017 | 545.646773 | 429.688038 | 439.347535 | 0 |
| 1.83E-13 | 3.77E-12 | 246.607755 | 795.447559 | 463.097907 | 762.711033 | 0 |
| 1.86E-13 | 3.81E-12 | 230.853152 | 471.574056 | 364.724403 | 440.739344 | 0 |
| 1.86E-13 | 3.82E-12 | 236.96208 | 389.048596 | 335.026742 | 314.084774 | 0 |
| 1.86E-13 | 3.83E-12 | 218.95682 | 612.601392 | 550.334787 | 585.023487 | 0 |
| 1.87E-13 | 3.84E-12 | 220.564432 | 519.398783 | 473.306478 | 510.329766 | 0 |
| 1.87E-13 | 3.84E-12 | 246.607755 | 369.918705 | 281.199731 | 366.50956 | 0 |
| 1.90E-13 | 3.89E-12 | 218.635297 | 647.302124 | 567.039721 | 640.695825 | 0 |
| 1.91E-13 | 3.90E-12 | 240.177305 | 391.273002 | 537.34206 | 375.788283 | 0 |
| 1.91E-13 | 3.91E-12 | 220.564432 | 268.263354 | 245.005706 | 308.981477 | 9.50487297 |
| 1.92E-13 | 3.93E-12 | 231.81772 | 461.78667 | 365.652455 | 373.004666 | 0 |
| 1.93E-13 | 3.93E-12 | 226.67336 | 490.926388 | 390.709857 | 447.23445 | 0 |
| 1.94E-13 | 3.95E-12 | 264.934537 | 449.552437 | 720.168288 | 390.170303 | 0 |
| 1.94E-13 | 3.96E-12 | 214.133982 | 643.743075 | 534.557904 | 608.220294 | 4.75243649 |
| 1.95E-13 | 3.97E-12 | 233.746855 | 401.282829 | 322.962067 | 411.04743 | 0 |
| 1.95E-13 | 3.97E-12 | 223.779657 | 391.940324 | 379.573234 | 411.975302 | 0 |
| 1.95E-13 | 3.98E-12 | 257.539519 | 352.123458 | 258.926485 | 319.188072 | 0 |
| 1.96E-13 | 3.99E-12 | 224.744225 | 493.595675 | 400.918428 | 462.544343 | 0 |
| 1.96E-13 | 3.99E-12 | 329.882081 | 510.056278 | 244.077654 | 446.306578 | 0 |
| 1.97E-13 | 4.00E-12 | 251.109069 | 262.92478 | 297.904665 | 326.611051 | 0 |
| 1.97E-13 | 4.00E-12 | 245.321665 | 360.5762 | 276.559471 | 331.250412 | 0 |
| 1.97E-13 | 4.01E-12 | 222.172045 | 461.119348 | 408.342843 | 457.904981 | 0 |
| 1.97E-13 | 4.01E-12 | 163.333428 | 222.885474 | 245.005706 | 219.4418 | 4.75243649 |
| 1.98E-13 | 4.01E-12 | 553.983261 | 863.069499 | 860.304127 | 895.396772 | 19.0097459 |
| 1.99E-13 | 4.02E-12 | 220.564432 | 632.843485 | 537.34206 | 521.000298 | 0 |
| 1.99E-13 | 4.04E-12 | 247.572322 | 402.172591 | 373.076871 | 388.314559 | 7.12865473 |
| 2.00E-13 | 4.05E-12 | 229.24554 | 325.653027 | 348.947521 | 377.180091 | 0 |
| 2.00E-13 | 4.06E-12 | 230.210107 | 488.2571 | 505.788295 | 690.800929 | 0 |
| 2.02E-13 | 4.08E-12 | 228.602495 | 329.656958 | 325.746223 | 334.034029 | 0 |
| 2.02E-13 | 4.09E-12 | 242.749485 | 386.156868 | 286.768042 | 347.024241 | 0 |
| 2.02E-13 | 4.09E-12 | 256.896474 | 259.365731 | 270.063108 | 273.722329 | 0 |
| 2.03E-13 | 4.09E-12 | 341.778413 | 238.678756 | 283.055835 | 338.209454 | 0 |
| 2.03E-13 | 4.10E-12 | 226.994882 | 441.989457 | 425.97583 | 357.694773 | 0 |
| 2.04E-13 | 4.12E-12 | 228.280972 | 421.080041 | 364.724403 | 347.952113 | 0 |
| 2.04E-13 | 4.12E-12 | 379.396545 | 1341.53921 | 1315.04957 | 1286.49495 | 0 |
| 2.05E-13 | 4.13E-12 | 224.422702 | 542.532605 | 430.616089 | 556.259445 | 0 |
| 2.08E-13 | 4.18E-12 | 220.564432 | 483.808289 | 420.407518 | 479.246044 | 0 |
| 2.09E-13 | 4.20E-12 | 214.455505 | 684.894584 | 648.70829 | 621.210507 | 0 |
| 2.09E-13 | 4.20E-12 | 246.286232 | 320.536894 | 263.566744 | 300.630626 | 0 |
| 2.09E-13 | 4.21E-12 | 226.351837 | 568.335714 | 416.695311 | 499.195299 | 0 |
| 2.10E-13 | 4.21E-12 | 231.81772 | 583.016793 | 412.983103 | 438.883599 | 0 |

| | | | | | | |
|---|---|---|---|---|---|---|
| 2.10E-13 | 4.22E-12 | 224.10118 | 411.515096 | 357.299988 | 409.191685 | 0 |
| 2.11E-13 | 4.22E-12 | 218.95682 | 447.328031 | 434.328297 | 478.318172 | 0 |
| 2.11E-13 | 4.24E-12 | 932.415239 | 4391.19974 | 2262.59057 | 2884.29105 | 28.5146189 |
| 2.14E-13 | 4.29E-12 | 214.777027 | 788.5519 | 661.701017 | 683.377951 | 0 |
| 2.14E-13 | 4.29E-12 | 212.847892 | 768.309806 | 685.830367 | 717.24529 | 0 |
| 2.15E-13 | 4.29E-12 | 238.24817 | 324.318384 | 277.487523 | 336.35371 | 0 |
| 2.15E-13 | 4.30E-12 | 392.578968 | 978.961047 | 1023.64126 | 1054.06294 | 2.37621824 |
| 2.16E-13 | 4.31E-12 | 220.564432 | 602.591565 | 476.090633 | 594.766146 | 0 |
| 2.16E-13 | 4.31E-12 | 222.493567 | 543.867249 | 425.047778 | 519.608489 | 0 |
| 2.16E-13 | 4.32E-12 | 226.67336 | 546.536536 | 399.990376 | 509.86583 | 0 |
| 2.17E-13 | 4.32E-12 | 226.994882 | 409.29069 | 335.954794 | 369.757113 | 0 |
| 2.18E-13 | 4.34E-12 | 217.027685 | 458.672502 | 473.306478 | 448.626258 | 0 |
| 2.18E-13 | 4.34E-12 | 221.529 | 434.204036 | 449.177128 | 381.819453 | 0 |
| 2.18E-13 | 4.35E-12 | 272.972599 | 343.225834 | 257.998433 | 423.573706 | 0 |
| 2.19E-13 | 4.36E-12 | 11252.0013 | 13364.0084 | 6849.02315 | 9249.95898 | 28.5146189 |
| 2.19E-13 | 4.36E-12 | 222.493567 | 374.812398 | 359.156092 | 403.160515 | 0 |
| 2.19E-13 | 4.36E-12 | 215.420072 | 730.939787 | 969.814253 | 804.00135 | 0 |
| 2.23E-13 | 4.42E-12 | 321.200974 | 353.680542 | 227.37272 | 313.156902 | 0 |
| 2.24E-13 | 4.45E-12 | 222.81509 | 658.201713 | 490.011412 | 513.113383 | 0 |
| 2.26E-13 | 4.48E-12 | 216.706162 | 498.489368 | 473.306478 | 444.914769 | 0 |
| 2.26E-13 | 4.48E-12 | 218.95682 | 540.085758 | 559.615306 | 450.945939 | 0 |
| 2.27E-13 | 4.51E-12 | 226.67336 | 429.087903 | 374.932974 | 507.54615 | 0 |
| 2.28E-13 | 4.51E-12 | 217.67073 | 448.662675 | 411.126999 | 427.285195 | 0 |
| 2.28E-13 | 4.52E-12 | 222.81509 | 342.336072 | 342.451157 | 360.014453 | 0 |
| 2.29E-13 | 4.54E-12 | 299.337444 | 289.17277 | 296.976613 | 498.267427 | 0 |
| 2.31E-13 | 4.56E-12 | 220.885955 | 394.832052 | 362.8683 | 418.006472 | 0 |
| 2.32E-13 | 4.58E-12 | 233.425332 | 449.552437 | 321.105963 | 392.026048 | 0 |
| 2.34E-13 | 4.62E-12 | 227.316405 | 317.645166 | 303.472977 | 318.724136 | 0 |
| 2.34E-13 | 4.63E-12 | 293.871561 | 560.772734 | 276.559471 | 389.706367 | 0 |
| 2.35E-13 | 4.64E-12 | 256.574952 | 881.976949 | 433.400245 | 599.405507 | 0 |
| 2.37E-13 | 4.66E-12 | 225.065747 | 321.871537 | 265.422848 | 320.115944 | 11.8810912 |
| 2.38E-13 | 4.69E-12 | 302.231146 | 324.095943 | 260.782589 | 313.620838 | 0 |
| 2.39E-13 | 4.71E-12 | 819.882365 | 535.192065 | 509.500502 | 555.331573 | 0 |
| 2.39E-13 | 4.71E-12 | 232.139242 | 372.810433 | 304.401029 | 311.301158 | 0 |
| 2.39E-13 | 4.71E-12 | 346.922773 | 410.402893 | 503.932191 | 542.805297 | 2.37621824 |
| 2.40E-13 | 4.72E-12 | 332.454261 | 652.863139 | 532.7018 | 567.857849 | 64.1578926 |
| 2.42E-13 | 4.75E-12 | 600.925545 | 1207.1851 | 938.260488 | 1128.75666 | 19.0097459 |
| 2.42E-13 | 4.75E-12 | 255.931907 | 801.453455 | 399.062324 | 577.136572 | 0 |
| 2.42E-13 | 4.76E-12 | 214.777027 | 822.807751 | 1057.05113 | 1254.01942 | 0 |
| 2.43E-13 | 4.76E-12 | 219.599865 | 356.57227 | 356.371936 | 375.788283 | 0 |
| 2.44E-13 | 4.79E-12 | 286.155022 | 330.101839 | 324.818171 | 229.648395 | 0 |
| 2.44E-13 | 4.79E-12 | 260.111699 | 406.176522 | 491.867516 | 282.537116 | 0 |
| 2.45E-13 | 4.80E-12 | 214.133982 | 645.74504 | 797.196597 | 624.45806 | 0 |

| | | | | | | |
|---|---|---|---|---|---|---|
| 2.45E-13 | 4.80E-12 | 266.542149 | 530.075932 | 382.35739 | 461.616471 | 7.12865473 |
| 2.48E-13 | 4.84E-12 | 223.136612 | 431.312309 | 372.148819 | 347.952113 | 0 |
| 2.49E-13 | 4.86E-12 | 213.81246 | 458.22762 | 467.738166 | 438.883599 | 0 |
| 2.49E-13 | 4.87E-12 | 228.924017 | 434.426477 | 661.701017 | 491.308384 | 0 |
| 2.51E-13 | 4.89E-12 | 212.847892 | 730.939787 | 580.032448 | 652.758165 | 0 |
| 2.51E-13 | 4.90E-12 | 215.09855 | 506.71967 | 444.536868 | 444.450833 | 0 |
| 2.51E-13 | 4.90E-12 | 260.433222 | 267.818473 | 242.22155 | 279.753499 | 0 |
| 2.51E-13 | 4.90E-12 | 246.286232 | 1158.69305 | 564.255566 | 763.174969 | 0 |
| 2.52E-13 | 4.90E-12 | 322.165541 | 275.159013 | 232.941031 | 338.67339 | 0 |
| 2.52E-13 | 4.91E-12 | 263.648447 | 312.751473 | 308.113236 | 237.071373 | 0 |
| 2.53E-13 | 4.92E-12 | 212.847892 | 513.615328 | 472.378426 | 535.846255 | 0 |
| 2.54E-13 | 4.93E-12 | 210.597235 | 597.697872 | 543.838423 | 581.311998 | 0 |
| 2.55E-13 | 4.95E-12 | 211.883325 | 534.524744 | 510.428554 | 581.775934 | 0 |
| 2.56E-13 | 4.96E-12 | 199.343948 | 1171.5946 | 1367.94853 | 1439.59388 | 0 |
| 2.56E-13 | 4.96E-12 | 348.530386 | 618.162406 | 561.47141 | 669.459866 | 21.3859642 |
| 2.56E-13 | 4.96E-12 | 235.354467 | 491.371269 | 428.759986 | 488.060831 | 2.37621824 |
| 2.56E-13 | 4.96E-12 | 208.6681 | 704.469356 | 696.96699 | 636.056463 | 0 |
| 2.61E-13 | 5.05E-12 | 182.624778 | 276.493656 | 288.624146 | 284.856797 | 11.8810912 |
| 2.61E-13 | 5.05E-12 | 209.95419 | 562.997139 | 644.06803 | 628.169549 | 0 |
| 2.61E-13 | 5.06E-12 | 407.690525 | 1138.22851 | 784.20387 | 954.316663 | 2.37621824 |
| 2.63E-13 | 5.09E-12 | 384.862428 | 1037.24048 | 723.880495 | 833.693264 | 14.2573095 |
| 2.63E-13 | 5.09E-12 | 229.888585 | 355.682507 | 290.48025 | 320.579881 | 4.75243649 |
| 2.64E-13 | 5.09E-12 | 210.918757 | 828.146326 | 643.139978 | 712.141992 | 0 |
| 2.64E-13 | 5.10E-12 | 222.81509 | 406.398962 | 327.602327 | 361.870198 | 0 |
| 2.64E-13 | 5.10E-12 | 259.790177 | 288.727889 | 237.581291 | 277.433818 | 0 |
| 2.67E-13 | 5.14E-12 | 208.025055 | 640.851347 | 609.730109 | 603.116997 | 0 |
| 2.68E-13 | 5.16E-12 | 251.109069 | 556.991243 | 530.845697 | 585.023487 | 9.50487297 |
| 2.68E-13 | 5.16E-12 | 606.71295 | 1397.81668 | 1291.84827 | 1301.80484 | 19.0097459 |
| 2.70E-13 | 5.19E-12 | 238.24817 | 286.725924 | 322.034015 | 400.376899 | 0 |
| 2.71E-13 | 5.21E-12 | 209.632667 | 527.851526 | 590.241019 | 598.477635 | 0 |
| 2.73E-13 | 5.25E-12 | 204.488308 | 818.35894 | 981.878928 | 947.821557 | 0 |
| 2.73E-13 | 5.25E-12 | 746.896758 | 238.901196 | 514.140762 | 425.429451 | 0 |
| 2.73E-13 | 5.25E-12 | 447.559315 | 343.225834 | 244.077654 | 290.887967 | 0 |
| 2.74E-13 | 5.25E-12 | 226.030315 | 372.810433 | 443.608816 | 546.05285 | 0 |
| 2.75E-13 | 5.27E-12 | 216.706162 | 385.934428 | 348.947521 | 364.189879 | 0 |
| 2.75E-13 | 5.27E-12 | 227.316405 | 387.936393 | 298.832717 | 354.911156 | 0 |
| 2.76E-13 | 5.28E-12 | 235.032945 | 587.910486 | 361.940248 | 450.018067 | 0 |
| 2.76E-13 | 5.28E-12 | 219.599865 | 381.263175 | 327.602327 | 383.675197 | 0 |
| 2.76E-13 | 5.28E-12 | 231.81772 | 445.548507 | 315.537652 | 353.983283 | 0 |
| 2.75E-13 | 5.28E-12 | 309.304641 | 344.782918 | 253.358173 | 245.422224 | 0 |
| 2.77E-13 | 5.30E-12 | 253.359727 | 371.030908 | 356.371936 | 250.989458 | 0 |
| 2.78E-13 | 5.31E-12 | 216.063117 | 763.638554 | 534.557904 | 636.984336 | 0 |
| 2.81E-13 | 5.37E-12 | 210.275712 | 546.981417 | 474.23453 | 522.856043 | 0 |

| | | | | | | |
|---|---|---|---|---|---|---|
| 2.82E-13 | 5.38E-12 | 210.597235 | 502.048417 | 449.177128 | 500.123171 | 0 |
| 2.82E-13 | 5.38E-12 | 220.885955 | 548.760942 | 397.20622 | 440.739344 | 0 |
| 2.83E-13 | 5.39E-12 | 241.463395 | 253.137394 | 270.063108 | 269.082968 | 0 |
| 2.83E-13 | 5.39E-12 | 359.783673 | 794.335356 | 769.355039 | 839.724434 | 0 |
| 2.84E-13 | 5.41E-12 | 221.850522 | 343.003394 | 308.113236 | 370.221049 | 0 |
| 2.86E-13 | 5.44E-12 | 202.880695 | 953.157939 | 834.318673 | 847.147412 | 0 |
| 2.86E-13 | 5.45E-12 | 217.027685 | 428.643022 | 352.659728 | 415.686792 | 0 |
| 2.86E-13 | 5.45E-12 | 202.559173 | 805.679826 | 837.102829 | 881.478687 | 0 |
| 2.88E-13 | 5.47E-12 | 217.67073 | 405.064319 | 341.523105 | 410.583494 | 0 |
| 2.89E-13 | 5.49E-12 | 205.131353 | 633.733248 | 648.70829 | 649.046676 | 0 |
| 2.89E-13 | 5.49E-12 | 505.433364 | 372.365552 | 226.444668 | 361.870198 | 0 |
| 2.89E-13 | 5.49E-12 | 541.443883 | 2079.81954 | 1779.07552 | 1930.43833 | 16.6335277 |
| 2.90E-13 | 5.50E-12 | 213.169415 | 512.948006 | 416.695311 | 488.988703 | 0 |
| 2.90E-13 | 5.50E-12 | 210.918757 | 574.786491 | 472.378426 | 510.793703 | 0 |
| 2.91E-13 | 5.50E-12 | 208.989622 | 485.810254 | 461.241803 | 498.267427 | 0 |
| 2.90E-13 | 5.50E-12 | 300.623534 | 406.843844 | 227.37272 | 397.129346 | 0 |
| 2.92E-13 | 5.51E-12 | 215.420072 | 536.304268 | 411.126999 | 507.54615 | 0 |
| 2.92E-13 | 5.51E-12 | 218.313775 | 375.034839 | 323.890119 | 346.560305 | 0 |
| 2.99E-13 | 5.64E-12 | 715.709076 | 1159.58281 | 628.291148 | 909.314857 | 0 |
| 2.99E-13 | 5.64E-12 | 211.24028 | 526.072001 | 439.896609 | 458.368917 | 0 |
| 3.01E-13 | 5.67E-12 | 387.75613 | 1146.23637 | 968.886201 | 1065.1974 | 14.2573095 |
| 3.01E-13 | 5.68E-12 | 230.210107 | 462.453992 | 343.379209 | 329.394667 | 0 |
| 3.01E-13 | 5.68E-12 | 214.455505 | 392.385205 | 368.436611 | 442.131152 | 0 |
| 3.02E-13 | 5.69E-12 | 248.858412 | 347.229765 | 245.933758 | 308.981477 | 0 |
| 3.04E-13 | 5.72E-12 | 207.38201 | 531.188135 | 492.795568 | 557.651254 | 0 |
| 3.04E-13 | 5.73E-12 | 198.37938 | 950.933533 | 1011.57659 | 954.316663 | 0 |
| 3.05E-13 | 5.74E-12 | 219.278342 | 326.987671 | 301.616873 | 317.332327 | 0 |
| 3.06E-13 | 5.74E-12 | 213.169415 | 482.696086 | 533.629852 | 414.294983 | 0 |
| 3.06E-13 | 5.75E-12 | 213.169415 | 349.899052 | 371.220767 | 378.107963 | 0 |
| 3.06E-13 | 5.75E-12 | 244.67862 | 554.099516 | 347.091417 | 353.055411 | 0 |
| 3.09E-13 | 5.80E-12 | 205.452875 | 627.06003 | 774.923351 | 717.709226 | 0 |
| 3.09E-13 | 5.80E-12 | 204.488308 | 643.743075 | 587.456863 | 610.539975 | 0 |
| 3.10E-13 | 5.81E-12 | 268.792807 | 310.971948 | 228.300772 | 282.07318 | 0 |
| 3.10E-13 | 5.81E-12 | 210.597235 | 528.741288 | 440.824661 | 453.26562 | 0 |
| 3.10E-13 | 5.81E-12 | 205.452875 | 527.629085 | 558.687254 | 566.002105 | 0 |
| 3.11E-13 | 5.82E-12 | 197.09329 | 1134.00214 | 987.44724 | 1058.23836 | 0 |
| 3.11E-13 | 5.82E-12 | 208.989622 | 427.9757 | 489.08336 | 480.173917 | 0 |
| 3.12E-13 | 5.83E-12 | 216.38464 | 362.578166 | 318.321808 | 352.591475 | 0 |
| 3.13E-13 | 5.84E-12 | 415.085542 | 432.202071 | 227.37272 | 304.342115 | 0 |
| 3.14E-13 | 5.85E-12 | 229.888585 | 1289.71055 | 656.132705 | 866.632731 | 0 |
| 3.15E-13 | 5.89E-12 | 215.420072 | 562.774699 | 409.270895 | 504.762533 | 0 |
| 3.16E-13 | 5.89E-12 | 221.850522 | 298.515275 | 292.336354 | 288.568286 | 0 |
| 3.17E-13 | 5.90E-12 | 214.455505 | 340.334106 | 361.012196 | 396.665409 | 0 |

| | | | | | | |
|---|---|---|---|---|---|---|
| 3.16E-13 | 5.90E-12 | 360.105196 | 370.141146 | 231.084927 | 272.794457 | 0 |
| 3.18E-13 | 5.92E-12 | 359.783673 | 1413.60996 | 1353.09969 | 1542.12377 | 0 |
| 3.18E-13 | 5.92E-12 | 219.599865 | 693.347327 | 461.241803 | 517.288809 | 0 |
| 3.22E-13 | 5.99E-12 | 206.09592 | 592.804179 | 506.716347 | 572.961147 | 0 |
| 3.22E-13 | 5.99E-12 | 204.166785 | 583.461674 | 561.47141 | 625.385932 | 0 |
| 3.23E-13 | 6.01E-12 | 381.647203 | 907.335177 | 814.829583 | 914.418154 | 2.37621824 |
| 3.26E-13 | 6.06E-12 | 218.313775 | 727.603178 | 469.59427 | 584.559551 | 0 |
| 3.26E-13 | 6.06E-12 | 335.669486 | 251.802751 | 249.645966 | 265.835415 | 0 |
| 3.27E-13 | 6.07E-12 | 248.858412 | 454.44613 | 276.559471 | 457.904981 | 0 |
| 3.29E-13 | 6.09E-12 | 221.207477 | 397.056457 | 305.329081 | 348.41605 | 0 |
| 3.29E-13 | 6.09E-12 | 218.95682 | 345.005359 | 298.832717 | 361.406262 | 0 |
| 3.31E-13 | 6.14E-12 | 230.210107 | 391.495443 | 283.983887 | 320.579881 | 0 |
| 3.32E-13 | 6.15E-12 | 249.179934 | 269.153117 | 309.96934 | 239.391054 | 0 |
| 3.33E-13 | 6.16E-12 | 259.147132 | 317.867607 | 232.012979 | 278.361691 | 0 |
| 3.40E-13 | 6.28E-12 | 217.027685 | 631.953723 | 424.119726 | 518.680617 | 0 |
| 3.40E-13 | 6.29E-12 | 324.737721 | 616.605322 | 443.608816 | 502.906788 | 0 |
| 3.42E-13 | 6.32E-12 | 204.488308 | 470.239412 | 483.515049 | 504.762533 | 0 |
| 3.43E-13 | 6.34E-12 | 205.452875 | 481.583883 | 440.824661 | 460.224662 | 0 |
| 3.44E-13 | 6.34E-12 | 229.567062 | 263.592102 | 275.631419 | 316.404455 | 0 |
| 3.48E-13 | 6.41E-12 | 319.914884 | 1194.0611 | 635.715563 | 903.283687 | 7.12865473 |
| 3.49E-13 | 6.43E-12 | 211.883325 | 330.32428 | 375.861026 | 353.983283 | 0 |
| 3.52E-13 | 6.49E-12 | 205.452875 | 478.914596 | 428.759986 | 456.977109 | 0 |
| 3.53E-13 | 6.50E-12 | 214.133982 | 399.503304 | 349.875573 | 328.466795 | 0 |
| 3.53E-13 | 6.50E-12 | 325.702289 | 290.729854 | 219.020252 | 286.248605 | 0 |
| 3.54E-13 | 6.52E-12 | 216.063117 | 703.802034 | 456.601543 | 574.816891 | 0 |
| 3.57E-13 | 6.55E-12 | 263.005402 | 373.477755 | 352.659728 | 405.01626 | 9.50487297 |
| 3.58E-13 | 6.57E-12 | 206.417443 | 556.323922 | 752.650105 | 621.210507 | 0 |
| 3.58E-13 | 6.58E-12 | 217.992252 | 416.408789 | 317.393756 | 415.222855 | 0 |
| 3.62E-13 | 6.65E-12 | 200.630038 | 580.792387 | 572.608033 | 614.7154 | 0 |
| 3.65E-13 | 6.68E-12 | 206.738965 | 389.493477 | 391.637909 | 373.468602 | 0 |
| 3.67E-13 | 6.73E-12 | 635.328452 | 1519.49169 | 1798.56462 | 1782.90663 | 19.0097459 |
| 3.67E-13 | 6.73E-12 | 202.880695 | 552.764872 | 493.72362 | 575.280828 | 0 |
| 3.68E-13 | 6.73E-12 | 208.989622 | 348.119527 | 343.379209 | 352.127539 | 0 |
| 3.69E-13 | 6.74E-12 | 172.336058 | 202.198499 | 170.761553 | 195.781056 | 33.2670554 |
| 3.69E-13 | 6.74E-12 | 197.09329 | 749.402356 | 722.024391 | 761.78316 | 0 |
| 3.73E-13 | 6.82E-12 | 282.939797 | 378.593888 | 390.709857 | 470.895194 | 33.2670554 |
| 3.73E-13 | 6.82E-12 | 369.429348 | 922.461137 | 890.001788 | 976.585599 | 2.37621824 |
| 3.74E-13 | 6.83E-12 | 220.24291 | 344.560478 | 279.343627 | 308.981477 | 0 |
| 3.77E-13 | 6.88E-12 | 245.000142 | 530.520813 | 565.183617 | 308.053604 | 0 |
| 3.78E-13 | 6.90E-12 | 207.38201 | 411.070215 | 359.156092 | 398.98509 | 0 |
| 3.81E-13 | 6.94E-12 | 209.311145 | 387.491512 | 335.026742 | 378.107963 | 0 |
| 3.82E-13 | 6.96E-12 | 352.067133 | 291.619617 | 490.011412 | 241.710735 | 0 |
| 3.82E-13 | 6.96E-12 | 217.027685 | 322.316418 | 284.911939 | 345.168497 | 0 |

| | | | | | | |
|---|---|---|---|---|---|---|
| 3.84E-13 | 6.97E-12 | 219.278342 | 313.418795 | 399.062324 | 311.765094 | 0 |
| 3.83E-13 | 6.97E-12 | 211.883325 | 346.562443 | 399.062324 | 328.002859 | 0 |
| 3.83E-13 | 6.97E-12 | 211.883325 | 311.194389 | 311.825444 | 333.570093 | 0 |
| 3.89E-13 | 7.07E-12 | 715.387554 | 586.353402 | 413.911155 | 223.153289 | 0 |
| 3.94E-13 | 7.16E-12 | 202.23765 | 448.217794 | 426.903882 | 458.368917 | 0 |
| 3.95E-13 | 7.17E-12 | 262.040834 | 320.536894 | 357.299988 | 226.864778 | 0 |
| 3.96E-13 | 7.18E-12 | 205.774398 | 412.404858 | 366.580507 | 381.819453 | 0 |
| 3.97E-13 | 7.20E-12 | 213.169415 | 422.637126 | 322.962067 | 392.489984 | 0 |
| 3.97E-13 | 7.20E-12 | 271.364987 | 643.298193 | 296.976613 | 418.934345 | 0 |
| 4.00E-13 | 7.25E-12 | 196.450245 | 833.70734 | 845.455296 | 702.863269 | 0 |
| 4.04E-13 | 7.32E-12 | 197.09329 | 623.056099 | 596.737383 | 610.076039 | 0 |
| 4.04E-13 | 7.32E-12 | 203.845263 | 486.255135 | 424.119726 | 521.464234 | 0 |
| 4.05E-13 | 7.32E-12 | 211.24028 | 472.018937 | 351.731677 | 443.986897 | 0 |
| 4.07E-13 | 7.34E-12 | 432.447757 | 657.534391 | 513.21271 | 630.025293 | 2.37621824 |
| 4.06E-13 | 7.34E-12 | 226.994882 | 288.060567 | 245.933758 | 284.856797 | 0 |
| 4.09E-13 | 7.38E-12 | 189.055228 | 1046.80543 | 994.871655 | 1037.82517 | 0 |
| 4.09E-13 | 7.38E-12 | 201.594605 | 551.207788 | 461.241803 | 507.54615 | 0 |
| 4.11E-13 | 7.41E-12 | 216.706162 | 285.16884 | 317.393756 | 277.897755 | 0 |
| 4.12E-13 | 7.42E-12 | 214.777027 | 1358.88958 | 714.599976 | 964.987195 | 0 |
| 4.12E-13 | 7.42E-12 | 259.790177 | 348.564408 | 227.37272 | 293.671584 | 0 |
| 4.12E-13 | 7.42E-12 | 388.077653 | 959.386275 | 721.096339 | 809.568584 | 0 |
| 4.13E-13 | 7.44E-12 | 226.030315 | 256.474003 | 254.286225 | 280.217435 | 0 |
| 4.15E-13 | 7.46E-12 | 291.942426 | 356.79471 | 212.523889 | 334.034029 | 0 |
| 4.16E-13 | 7.47E-12 | 199.343948 | 486.922457 | 473.306478 | 478.318172 | 0 |
| 4.20E-13 | 7.55E-12 | 272.329554 | 257.808647 | 226.444668 | 324.755306 | 0 |
| 4.20E-13 | 7.55E-12 | 210.597235 | 315.643201 | 323.890119 | 373.468602 | 0 |
| 4.23E-13 | 7.59E-12 | 210.597235 | 344.338037 | 299.760769 | 349.807858 | 0 |
| 4.26E-13 | 7.64E-12 | 226.67336 | 395.499373 | 270.063108 | 356.7669 | 0 |
| 4.27E-13 | 7.65E-12 | 201.916128 | 655.532426 | 499.291931 | 572.961147 | 0 |
| 4.27E-13 | 7.65E-12 | 215.420072 | 330.54672 | 284.911939 | 287.640414 | 0 |
| 4.28E-13 | 7.66E-12 | 215.741595 | 326.54279 | 277.487523 | 344.240624 | 0 |
| 4.30E-13 | 7.70E-12 | 328.595991 | 439.542611 | 216.236097 | 329.858604 | 0 |
| 4.32E-13 | 7.73E-12 | 370.393916 | 834.597103 | 985.591136 | 921.841133 | 0 |
| 4.33E-13 | 7.73E-12 | 210.275712 | 329.656958 | 309.96934 | 295.991265 | 0 |
| 4.33E-13 | 7.74E-12 | 219.278342 | 265.149186 | 267.278952 | 268.155095 | 0 |
| 4.39E-13 | 7.83E-12 | 210.918757 | 292.509379 | 297.904665 | 294.599456 | 0 |
| 4.39E-13 | 7.84E-12 | 314.127479 | 591.247095 | 619.010629 | 630.025293 | 0 |
| 4.40E-13 | 7.85E-12 | 212.52637 | 383.265141 | 299.760769 | 340.065199 | 0 |
| 4.41E-13 | 7.86E-12 | 201.594605 | 764.528316 | 596.737383 | 566.929977 | 0 |
| 4.41E-13 | 7.86E-12 | 203.52374 | 746.955509 | 589.312967 | 727.451885 | 9.50487297 |
| 4.44E-13 | 7.92E-12 | 202.559173 | 449.774878 | 384.213494 | 402.696579 | 0 |
| 4.48E-13 | 7.97E-12 | 395.151148 | 788.996781 | 610.658161 | 672.243483 | 0 |
| 4.48E-13 | 7.97E-12 | 199.343948 | 438.207967 | 421.33557 | 421.254025 | 0 |

| | | | | | | |
|---|---|---|---|---|---|---|
| 4.49E-13 | 7.97E-12 | 250.466024 | 507.164551 | 274.703367 | 502.442852 | 0 |
| 4.48E-13 | 7.97E-12 | 265.577582 | 225.554761 | 385.141546 | 338.209454 | 0 |
| 4.49E-13 | 7.97E-12 | 249.822979 | 597.03055 | 466.810114 | 295.991265 | 0 |
| 4.49E-13 | 7.97E-12 | 204.80983 | 588.132926 | 429.688038 | 551.156148 | 0 |
| 4.52E-13 | 8.02E-12 | 230.853152 | 457.782739 | 309.96934 | 307.125732 | 0 |
| 4.53E-13 | 8.03E-12 | 205.774398 | 369.918705 | 329.458431 | 332.178284 | 0 |
| 4.53E-13 | 8.03E-12 | 208.6681 | 499.601571 | 364.724403 | 414.294983 | 0 |
| 4.54E-13 | 8.05E-12 | 204.166785 | 385.044666 | 338.73895 | 388.778495 | 0 |
| 4.55E-13 | 8.06E-12 | 231.174675 | 331.214042 | 250.574018 | 269.082968 | 0 |
| 4.58E-13 | 8.10E-12 | 211.561802 | 280.942468 | 287.696094 | 303.414243 | 0 |
| 4.60E-13 | 8.13E-12 | 197.414813 | 647.969446 | 533.629852 | 558.579126 | 0 |
| 4.61E-13 | 8.15E-12 | 201.273083 | 387.046631 | 453.817387 | 408.727749 | 0 |
| 4.62E-13 | 8.16E-12 | 291.299381 | 297.847953 | 226.444668 | 239.391054 | 0 |
| 4.65E-13 | 8.21E-12 | 211.561802 | 366.359656 | 306.257133 | 301.558498 | 0 |
| 4.67E-13 | 8.23E-12 | 399.973985 | 552.319991 | 742.441534 | 704.719014 | 2.37621824 |
| 4.67E-13 | 8.23E-12 | 235.354467 | 376.814364 | 249.645966 | 376.716155 | 0 |
| 4.70E-13 | 8.29E-12 | 195.164155 | 664.43005 | 567.039721 | 589.662848 | 0 |
| 4.70E-13 | 8.29E-12 | 201.916128 | 537.638912 | 428.759986 | 439.811471 | 0 |
| 4.73E-13 | 8.33E-12 | 178.123463 | 946.26228 | 776.779454 | 881.942624 | 2.37621824 |
| 4.75E-13 | 8.36E-12 | 263.648447 | 792.555831 | 622.722836 | 694.976355 | 9.50487297 |
| 4.76E-13 | 8.38E-12 | 209.95419 | 540.75308 | 367.508559 | 470.895194 | 0 |
| 4.80E-13 | 8.45E-12 | 203.845263 | 366.137215 | 426.903882 | 351.663603 | 0 |
| 4.82E-13 | 8.47E-12 | 196.771768 | 748.512594 | 567.039721 | 689.873057 | 0 |
| 4.83E-13 | 8.48E-12 | 218.313775 | 618.162406 | 373.076871 | 463.472215 | 0 |
| 4.83E-13 | 8.48E-12 | 208.989622 | 521.84563 | 423.191674 | 365.581687 | 0 |
| 4.86E-13 | 8.52E-12 | 191.94893 | 657.08951 | 619.010629 | 664.356569 | 0 |
| 4.89E-13 | 8.58E-12 | 211.883325 | 274.04681 | 297.904665 | 326.611051 | 0 |
| 4.93E-13 | 8.64E-12 | 206.09592 | 1383.1356 | 754.506208 | 962.203578 | 0 |
| 4.93E-13 | 8.65E-12 | 236.319035 | 244.01733 | 231.084927 | 256.092756 | 0 |
| 4.94E-13 | 8.65E-12 | 244.035575 | 524.737358 | 505.788295 | 530.742957 | 7.12865473 |
| 4.96E-13 | 8.68E-12 | 198.37938 | 590.134892 | 462.169855 | 517.288809 | 0 |
| 5.00E-13 | 8.75E-12 | 216.706162 | 630.174198 | 627.363096 | 386.92275 | 0 |
| 5.01E-13 | 8.75E-12 | 269.435852 | 275.159013 | 246.86181 | 218.513927 | 0 |
| 5.01E-13 | 8.76E-12 | 196.128723 | 601.256921 | 596.737383 | 492.700193 | 0 |
| 5.02E-13 | 8.78E-12 | 195.485678 | 626.170268 | 510.428554 | 559.970935 | 0 |
| 5.03E-13 | 8.79E-12 | 226.994882 | 297.180631 | 237.581291 | 315.476583 | 0 |
| 5.04E-13 | 8.80E-12 | 234.068377 | 228.446488 | 303.472977 | 296.919137 | 0 |
| 5.08E-13 | 8.87E-12 | 1265.83407 | 3440.71108 | 3888.53753 | 3714.27283 | 26.1384007 |
| 5.12E-13 | 8.93E-12 | 201.594605 | 332.103805 | 326.674275 | 342.848816 | 0 |
| 5.13E-13 | 8.94E-12 | 233.425332 | 307.635339 | 296.048562 | 228.720523 | 0 |
| 5.14E-13 | 8.95E-12 | 232.782287 | 404.174556 | 253.358173 | 342.38488 | 0 |
| 5.17E-13 | 9.00E-12 | 202.559173 | 466.235482 | 374.004923 | 497.339554 | 0 |
| 5.19E-13 | 9.03E-12 | 200.308515 | 423.971769 | 408.342843 | 353.983283 | 0 |

| | | | | | | |
|---|---|---|---|---|---|---|
| 5.20E-13 | 9.04E-12 | 203.52374 | 361.243522 | 309.96934 | 340.065199 | 0 |
| 5.21E-13 | 9.05E-12 | 205.131353 | 359.463997 | 300.688821 | 336.817646 | 0 |
| 5.21E-13 | 9.05E-12 | 189.055228 | 746.288188 | 691.398678 | 794.258691 | 0 |
| 5.22E-13 | 9.05E-12 | 202.880695 | 401.505269 | 333.170638 | 413.367111 | 0 |
| 5.25E-13 | 9.11E-12 | 232.460765 | 397.501339 | 250.574018 | 339.137327 | 0 |
| 5.27E-13 | 9.15E-12 | 217.349207 | 488.701982 | 325.746223 | 352.591475 | 0 |
| 5.30E-13 | 9.18E-12 | 194.199588 | 691.567802 | 541.98232 | 662.500824 | 0 |
| 5.30E-13 | 9.18E-12 | 198.057858 | 377.481685 | 365.652455 | 360.014453 | 0 |
| 5.30E-13 | 9.19E-12 | 262.362357 | 229.11381 | 239.437395 | 314.548711 | 0 |
| 5.32E-13 | 9.21E-12 | 191.305885 | 492.261031 | 451.961284 | 448.162322 | 7.12865473 |
| 5.35E-13 | 9.26E-12 | 198.700903 | 496.264962 | 477.018685 | 631.417102 | 0 |
| 5.40E-13 | 9.33E-12 | 409.61966 | 414.406824 | 412.055051 | 194.853184 | 0 |
| 5.41E-13 | 9.35E-12 | 198.700903 | 1628.71002 | 895.5701 | 1087.0024 | 0 |
| 5.42E-13 | 9.36E-12 | 227.95945 | 289.395211 | 348.019469 | 243.566479 | 0 |
| 5.46E-13 | 9.42E-12 | 194.199588 | 490.703947 | 439.896609 | 505.226469 | 0 |
| 5.46E-13 | 9.42E-12 | 188.733705 | 693.347327 | 860.304127 | 798.434116 | 0 |
| 5.47E-13 | 9.44E-12 | 193.878065 | 592.804179 | 484.443101 | 562.754552 | 0 |
| 5.50E-13 | 9.48E-12 | 202.23765 | 392.385205 | 322.034015 | 348.879986 | 0 |
| 5.51E-13 | 9.49E-12 | 218.635297 | 372.810433 | 315.537652 | 373.468602 | 2.37621824 |
| 5.52E-13 | 9.50E-12 | 382.61177 | 773.64838 | 602.305694 | 691.264865 | 0 |
| 5.53E-13 | 9.52E-12 | 271.043464 | 280.720028 | 207.883629 | 270.938712 | 0 |
| 5.53E-13 | 9.52E-12 | 354.960836 | 955.159904 | 866.80049 | 953.388791 | 0 |
| 5.54E-13 | 9.52E-12 | 194.842633 | 445.103625 | 401.84648 | 433.780301 | 0 |
| 5.56E-13 | 9.56E-12 | 193.878065 | 418.188314 | 425.047778 | 432.388493 | 0 |
| 5.56E-13 | 9.56E-12 | 187.447615 | 761.191707 | 694.182834 | 693.12061 | 0 |
| 5.59E-13 | 9.60E-12 | 210.597235 | 305.188493 | 259.854537 | 287.176478 | 0 |
| 5.62E-13 | 9.65E-12 | 197.09329 | 702.02251 | 511.356606 | 551.620084 | 0 |
| 5.64E-13 | 9.67E-12 | 238.24817 | 449.997318 | 370.292715 | 255.628819 | 0 |
| 5.69E-13 | 9.75E-12 | 195.8072 | 465.123279 | 390.709857 | 449.554131 | 0 |
| 5.72E-13 | 9.79E-12 | 197.736335 | 353.902983 | 341.523105 | 379.499772 | 0 |
| 5.72E-13 | 9.79E-12 | 330.525126 | 1367.7872 | 1156.35269 | 1299.02122 | 0 |
| 5.75E-13 | 9.84E-12 | 214.455505 | 719.150435 | 413.911155 | 509.401894 | 0 |
| 5.76E-13 | 9.86E-12 | 185.51848 | 772.313737 | 771.211143 | 716.317418 | 0 |
| 5.77E-13 | 9.87E-12 | 508.005544 | 683.559941 | 376.789078 | 425.893387 | 16.6335277 |
| 5.81E-13 | 9.92E-12 | 437.270595 | 552.542432 | 193.962851 | 519.144553 | 0 |
| 5.82E-13 | 9.94E-12 | 198.37938 | 533.857422 | 398.134272 | 469.503385 | 0 |
| 5.83E-13 | 9.95E-12 | 191.94893 | 523.180273 | 604.161798 | 501.978916 | 0 |
| 5.84E-13 | 9.96E-12 | 2686.64198 | 8520.80933 | 7614.66598 | 8484.92827 | 9.50487297 |
| 5.84E-13 | 9.96E-12 | 200.630038 | 383.710022 | 322.034015 | 398.521154 | 0 |
| 5.87E-13 | 1.00E-11 | 223.136612 | 497.599605 | 325.746223 | 321.507753 | 0 |
| 5.89E-13 | 1.00E-11 | 209.311145 | 259.14329 | 289.552198 | 308.981477 | 0 |
| 5.91E-13 | 1.01E-11 | 197.414813 | 334.773092 | 356.371936 | 379.963708 | 0 |
| 5.97E-13 | 1.02E-11 | 251.430592 | 279.830265 | 209.739733 | 290.887967 | 0 |

| | | | | | | |
|---|---|---|---|---|---|---|
| 5.98E-13 | 1.02E-11 | 190.984363 | 751.404321 | 569.823877 | 669.459866 | 0 |
| 5.99E-13 | 1.02E-11 | 196.771768 | 403.729675 | 535.485956 | 489.45264 | 0 |
| 6.00E-13 | 1.02E-11 | 208.346577 | 374.589958 | 389.781805 | 286.248605 | 0 |
| 6.01E-13 | 1.02E-11 | 206.09592 | 300.739681 | 268.207004 | 316.404455 | 0 |
| 6.03E-13 | 1.02E-11 | 187.769138 | 684.004822 | 606.017902 | 604.044869 | 0 |
| 6.03E-13 | 1.02E-11 | 206.417443 | 318.312488 | 267.278952 | 312.22903 | 0 |
| 6.05E-13 | 1.02E-11 | 198.700903 | 670.435946 | 464.95401 | 520.536362 | 0 |
| 6.05E-13 | 1.02E-11 | 190.66284 | 475.133105 | 472.378426 | 448.162322 | 0 |
| 6.04E-13 | 1.02E-11 | 212.204847 | 245.574414 | 281.199731 | 284.392861 | 0 |
| 6.05E-13 | 1.02E-11 | 201.916128 | 327.654993 | 439.896609 | 392.95392 | 0 |
| 6.04E-13 | 1.02E-11 | 197.09329 | 472.018937 | 370.292715 | 435.17211 | 0 |
| 6.05E-13 | 1.02E-11 | 240.177305 | 225.33232 | 247.789862 | 231.968076 | 0 |
| 6.05E-13 | 1.02E-11 | 166.870175 | 216.657137 | 191.178695 | 220.369672 | 21.3859642 |
| 6.07E-13 | 1.03E-11 | 195.164155 | 377.926567 | 383.285442 | 349.807858 | 0 |
| 6.13E-13 | 1.04E-11 | 200.308515 | 437.095764 | 333.170638 | 396.665409 | 0 |
| 6.13E-13 | 1.04E-11 | 364.284988 | 298.737716 | 207.883629 | 281.145308 | 0 |
| 6.16E-13 | 1.04E-11 | 317.342704 | 211.096122 | 495.579724 | 439.811471 | 0 |
| 6.21E-13 | 1.05E-11 | 195.485678 | 382.597819 | 340.595054 | 375.788283 | 0 |
| 6.23E-13 | 1.05E-11 | 190.341318 | 491.593709 | 464.025958 | 536.310191 | 0 |
| 6.23E-13 | 1.05E-11 | 231.174675 | 308.747542 | 222.73246 | 281.145308 | 0 |
| 6.28E-13 | 1.06E-11 | 602.533158 | 1371.12381 | 1050.55477 | 1159.84038 | 33.2670554 |
| 6.28E-13 | 1.06E-11 | 193.878065 | 363.023047 | 414.839207 | 403.160515 | 0 |
| 6.32E-13 | 1.06E-11 | 195.164155 | 411.515096 | 350.803625 | 405.480196 | 0 |
| 6.37E-13 | 1.07E-11 | 302.874191 | 612.601392 | 615.298421 | 658.789335 | 0 |
| 6.42E-13 | 1.08E-11 | 1494.11504 | 1576.21404 | 1127.58308 | 1595.94036 | 0 |
| 6.42E-13 | 1.08E-11 | 241.141872 | 317.645166 | 215.308045 | 308.053604 | 0 |
| 6.45E-13 | 1.08E-11 | 239.53426 | 523.402714 | 268.207004 | 426.357323 | 0 |
| 6.45E-13 | 1.08E-11 | 220.564432 | 389.938359 | 260.782589 | 402.232643 | 0 |
| 6.49E-13 | 1.09E-11 | 181.017165 | 986.079146 | 907.634775 | 813.744009 | 0 |
| 6.54E-13 | 1.10E-11 | 190.341318 | 512.280684 | 433.400245 | 486.669023 | 0 |
| 6.60E-13 | 1.11E-11 | 192.591975 | 389.048596 | 372.148819 | 359.086581 | 0 |
| 6.65E-13 | 1.11E-11 | 206.417443 | 549.650704 | 348.947521 | 445.378705 | 0 |
| 6.65E-13 | 1.11E-11 | 192.913498 | 423.749329 | 362.8683 | 400.376899 | 0 |
| 6.67E-13 | 1.12E-11 | 179.731075 | 825.032157 | 851.95166 | 824.878477 | 0 |
| 6.68E-13 | 1.12E-11 | 195.8072 | 476.022868 | 368.436611 | 473.214874 | 0 |
| 6.72E-13 | 1.12E-11 | 215.741595 | 284.279077 | 232.941031 | 266.299351 | 0 |
| 6.73E-13 | 1.12E-11 | 192.913498 | 531.188135 | 411.126999 | 463.472215 | 0 |
| 6.75E-13 | 1.13E-11 | 358.497583 | 690.455599 | 750.794001 | 797.042308 | 0 |
| 6.75E-13 | 1.13E-11 | 216.706162 | 247.576379 | 241.293498 | 291.815839 | 0 |
| 6.76E-13 | 1.13E-11 | 193.556543 | 447.328031 | 388.853753 | 501.051043 | 0 |
| 6.76E-13 | 1.13E-11 | 190.341318 | 480.47168 | 412.983103 | 483.885406 | 0 |
| 6.77E-13 | 1.13E-11 | 179.409553 | 976.06932 | 1484.88307 | 1281.39165 | 0 |
| 6.78E-13 | 1.13E-11 | 194.52111 | 399.725745 | 423.191674 | 344.70456 | 0 |

| | | | | | | |
|---|---|---|---|---|---|---|
| 6.84E-13 | 1.14E-11 | 386.791563 | 910.226904 | 247.789862 | 958.492089 | 0 |
| 6.84E-13 | 1.14E-11 | 254.645817 | 414.184383 | 309.96934 | 367.901368 | 0 |
| 6.88E-13 | 1.14E-11 | 263.326924 | 210.4288 | 245.933758 | 248.205841 | 0 |
| 6.88E-13 | 1.14E-11 | 271.686509 | 398.16866 | 219.948304 | 283.928925 | 0 |
| 6.89E-13 | 1.14E-11 | 195.485678 | 329.434517 | 408.342843 | 380.89158 | 0 |
| 6.89E-13 | 1.14E-11 | 199.986993 | 414.851705 | 316.465704 | 344.70456 | 0 |
| 6.91E-13 | 1.15E-11 | 190.341318 | 669.101302 | 495.579724 | 578.992317 | 0 |
| 6.91E-13 | 1.15E-11 | 187.447615 | 523.402714 | 467.738166 | 537.238063 | 0 |
| 6.91E-13 | 1.15E-11 | 200.308515 | 339.889225 | 466.810114 | 475.534555 | 0 |
| 6.93E-13 | 1.15E-11 | 213.169415 | 319.20225 | 241.293498 | 298.310945 | 0 |
| 6.94E-13 | 1.15E-11 | 262.040834 | 207.314632 | 253.358173 | 254.237011 | 0 |
| 6.94E-13 | 1.15E-11 | 183.589345 | 613.713595 | 660.772965 | 604.044869 | 0 |
| 6.97E-13 | 1.15E-11 | 187.126093 | 460.007145 | 490.939464 | 513.113383 | 0 |
| 6.97E-13 | 1.15E-11 | 255.288862 | 380.373413 | 343.379209 | 471.823066 | 16.6335277 |
| 6.98E-13 | 1.15E-11 | 192.270453 | 405.731641 | 351.731677 | 379.035836 | 0 |
| 6.99E-13 | 1.15E-11 | 191.305885 | 393.719849 | 359.156092 | 372.076793 | 0 |
| 7.01E-13 | 1.16E-11 | 189.37675 | 431.979631 | 394.422065 | 438.419663 | 0 |
| 7.06E-13 | 1.17E-11 | 224.10118 | 262.257458 | 221.804408 | 289.960095 | 0 |
| 7.06E-13 | 1.17E-11 | 275.544779 | 368.13918 | 218.0922 | 259.340309 | 0 |
| 7.07E-13 | 1.17E-11 | 198.057858 | 353.680542 | 303.472977 | 303.878179 | 0 |
| 7.11E-13 | 1.17E-11 | 185.196958 | 504.940145 | 517.85297 | 507.082213 | 0 |
| 7.12E-13 | 1.17E-11 | 349.173431 | 847.721098 | 771.211143 | 817.455499 | 0 |
| 7.15E-13 | 1.18E-11 | 186.80457 | 522.512952 | 460.313751 | 521.92817 | 0 |
| 7.15E-13 | 1.18E-11 | 199.986993 | 351.901017 | 299.760769 | 397.129346 | 0 |
| 7.15E-13 | 1.18E-11 | 191.305885 | 363.690369 | 405.558688 | 432.388493 | 0 |
| 7.17E-13 | 1.18E-11 | 185.51848 | 544.757011 | 501.148035 | 499.195299 | 0 |
| 7.20E-13 | 1.18E-11 | 192.913498 | 482.473645 | 386.069597 | 395.273601 | 0 |
| 7.21E-13 | 1.18E-11 | 201.273083 | 571.227441 | 371.220767 | 456.513173 | 0 |
| 7.21E-13 | 1.18E-11 | 184.553913 | 522.735392 | 541.054268 | 514.969128 | 0 |
| 7.23E-13 | 1.19E-11 | 189.698273 | 466.457922 | 392.565961 | 430.068812 | 0 |
| 7.26E-13 | 1.19E-11 | 189.055228 | 493.150793 | 439.896609 | 554.867637 | 0 |
| 7.28E-13 | 1.19E-11 | 203.52374 | 272.934607 | 253.358173 | 278.825627 | 0 |
| 7.32E-13 | 1.20E-11 | 310.269209 | 1491.46417 | 1315.97762 | 1731.87365 | 2.37621824 |
| 7.32E-13 | 1.20E-11 | 194.52111 | 515.617293 | 387.925701 | 408.263813 | 0 |
| 7.35E-13 | 1.20E-11 | 190.66284 | 387.269071 | 354.515832 | 407.799877 | 0 |
| 7.35E-13 | 1.20E-11 | 192.270453 | 359.019116 | 325.746223 | 363.725943 | 0 |
| 7.35E-13 | 1.20E-11 | 185.840003 | 496.932284 | 554.046994 | 602.189124 | 0 |
| 7.36E-13 | 1.20E-11 | 185.51848 | 546.314095 | 482.586997 | 555.331573 | 0 |
| 7.38E-13 | 1.21E-11 | 183.589345 | 602.814006 | 552.190891 | 640.231889 | 0 |
| 7.41E-13 | 1.21E-11 | 187.769138 | 505.162585 | 425.97583 | 459.760726 | 0 |
| 7.42E-13 | 1.21E-11 | 198.37938 | 450.4422 | 371.220767 | 328.466795 | 0 |
| 7.43E-13 | 1.21E-11 | 190.66284 | 343.225834 | 347.091417 | 357.694773 | 0 |
| 7.44E-13 | 1.21E-11 | 207.38201 | 306.078255 | 472.378426 | 368.365304 | 0 |

| | | | | | | |
|---|---|---|---|---|---|---|
| 7.49E-13 | 1.22E-11 | 190.341318 | 545.869214 | 425.97583 | 558.579126 | 0 |
| 7.55E-13 | 1.23E-11 | 199.022425 | 324.763265 | 272.847264 | 324.755306 | 0 |
| 7.56E-13 | 1.23E-11 | 209.311145 | 361.688403 | 274.703367 | 416.150728 | 0 |
| 7.59E-13 | 1.24E-11 | 187.126093 | 433.091833 | 510.428554 | 521.000298 | 0 |
| 7.63E-13 | 1.24E-11 | 237.283602 | 315.42076 | 229.228823 | 235.679565 | 0 |
| 7.64E-13 | 1.24E-11 | 194.199588 | 300.739681 | 302.544925 | 302.022435 | 0 |
| 7.65E-13 | 1.24E-11 | 188.412183 | 377.036804 | 368.436611 | 371.612857 | 0 |
| 7.67E-13 | 1.25E-11 | 241.463395 | 345.005359 | 213.451941 | 321.507753 | 0 |
| 7.74E-13 | 1.26E-11 | 206.738965 | 250.912988 | 328.530379 | 328.466795 | 0 |
| 7.76E-13 | 1.26E-11 | 348.851908 | 768.754687 | 721.096339 | 810.960392 | 0 |
| 7.77E-13 | 1.26E-11 | 223.136612 | 326.097909 | 621.794784 | 416.614664 | 0 |
| 7.81E-13 | 1.27E-11 | 188.412183 | 527.851526 | 436.184401 | 478.318172 | 9.50487297 |
| 7.86E-13 | 1.27E-11 | 194.199588 | 403.507235 | 316.465704 | 367.437432 | 0 |
| 7.89E-13 | 1.28E-11 | 187.447615 | 503.16062 | 410.198947 | 455.121364 | 0 |
| 7.93E-13 | 1.28E-11 | 184.553913 | 826.589241 | 592.097123 | 742.761778 | 0 |
| 7.95E-13 | 1.29E-11 | 190.341318 | 471.351615 | 369.364663 | 416.614664 | 0 |
| 7.99E-13 | 1.29E-11 | 184.23239 | 582.349471 | 803.69296 | 720.492843 | 0 |
| 8.02E-13 | 1.30E-11 | 190.66284 | 317.645166 | 332.242586 | 324.755306 | 0 |
| 8.03E-13 | 1.30E-11 | 213.490937 | 289.617651 | 226.444668 | 286.712542 | 0 |
| 8.04E-13 | 1.30E-11 | 183.910868 | 729.827584 | 552.190891 | 671.315611 | 0 |
| 8.06E-13 | 1.30E-11 | 304.803326 | 321.426656 | 228.300772 | 254.700947 | 7.12865473 |
| 8.16E-13 | 1.32E-11 | 214.455505 | 436.428442 | 269.135056 | 344.70456 | 0 |
| 8.19E-13 | 1.32E-11 | 299.015921 | 750.069678 | 699.751145 | 792.402946 | 21.3859642 |
| 8.23E-13 | 1.33E-11 | 181.338688 | 769.86689 | 600.44959 | 679.202526 | 0 |
| 8.35E-13 | 1.35E-11 | 388.720698 | 254.249597 | 255.214277 | 232.432012 | 0 |
| 8.36E-13 | 1.35E-11 | 223.136612 | 482.473645 | 457.529595 | 269.546904 | 0 |
| 8.36E-13 | 1.35E-11 | 228.924017 | 308.525102 | 234.797135 | 393.881792 | 0 |
| 8.37E-13 | 1.35E-11 | 335.991008 | 1416.27925 | 1233.381 | 1451.65622 | 26.1384007 |
| 8.41E-13 | 1.35E-11 | 210.275712 | 307.412899 | 232.012979 | 289.960095 | 0 |
| 8.45E-13 | 1.36E-11 | 294.514606 | 592.581738 | 491.867516 | 546.516786 | 14.2573095 |
| 8.58E-13 | 1.38E-11 | 201.594605 | 260.477934 | 245.005706 | 277.897755 | 0 |
| 8.60E-13 | 1.38E-11 | 439.521252 | 251.135429 | 226.444668 | 325.219242 | 0 |
| 8.65E-13 | 1.39E-11 | 185.196958 | 484.47561 | 661.701017 | 607.292422 | 0 |
| 8.64E-13 | 1.39E-11 | 182.303255 | 452.889046 | 477.946737 | 445.842641 | 0 |
| 8.65E-13 | 1.39E-11 | 311.233776 | 663.317847 | 479.802841 | 529.815085 | 4.75243649 |
| 8.68E-13 | 1.39E-11 | 246.286232 | 416.408789 | 220.876356 | 347.024241 | 0 |
| 8.68E-13 | 1.39E-11 | 244.357097 | 331.214042 | 207.883629 | 274.650202 | 0 |
| 8.77E-13 | 1.40E-11 | 180.695643 | 622.833659 | 618.082577 | 783.58816 | 0 |
| 8.78E-13 | 1.40E-11 | 171.37149 | 1021.4472 | 878.865165 | 1071.69251 | 0 |
| 8.80E-13 | 1.41E-11 | 204.80983 | 321.649097 | 355.443884 | 363.725943 | 9.50487297 |
| 8.83E-13 | 1.41E-11 | 186.161525 | 377.481685 | 388.853753 | 347.488177 | 0 |
| 8.85E-13 | 1.42E-11 | 192.913498 | 342.113631 | 484.443101 | 434.708174 | 0 |
| 8.88E-13 | 1.42E-11 | 231.174675 | 298.737716 | 218.0922 | 237.071373 | 0 |

| | | | | | | |
|---|---|---|---|---|---|---|
| 8.92E-13 | 1.42E-11 | 348.851908 | 933.583166 | 588.384915 | 775.701245 | 0 |
| 8.97E-13 | 1.43E-11 | 269.114329 | 258.253528 | 198.60311 | 246.814032 | 0 |
| 8.99E-13 | 1.43E-11 | 0 | 0 | 0 | 0 | 71.2865473 |
| 9.01E-13 | 1.44E-11 | 211.883325 | 770.979093 | 486.299204 | 405.944132 | 0 |
| 9.01E-13 | 1.44E-11 | 182.624778 | 440.209932 | 468.666218 | 414.758919 | 0 |
| 9.04E-13 | 1.44E-11 | 272.008032 | 212.208325 | 220.876356 | 251.453394 | 0 |
| 9.08E-13 | 1.45E-11 | 193.556543 | 294.288904 | 271.919212 | 278.361691 | 0 |
| 9.08E-13 | 1.45E-11 | 213.169415 | 240.903162 | 219.020252 | 254.237011 | 0 |
| 9.10E-13 | 1.45E-11 | 195.485678 | 478.024833 | 816.685687 | 776.165181 | 0 |
| 9.15E-13 | 1.45E-11 | 180.37412 | 532.745219 | 472.378426 | 497.80349 | 0 |
| 9.15E-13 | 1.45E-11 | 184.23239 | 533.857422 | 461.241803 | 423.573706 | 0 |
| 9.18E-13 | 1.46E-11 | 197.09329 | 268.485795 | 251.502069 | 269.082968 | 0 |
| 9.21E-13 | 1.46E-11 | 204.80983 | 924.907983 | 786.988026 | 804.00135 | 4.75243649 |
| 9.26E-13 | 1.47E-11 | 193.878065 | 383.265141 | 295.12051 | 322.899561 | 0 |
| 9.26E-13 | 1.47E-11 | 187.126093 | 525.404679 | 389.781805 | 488.524767 | 0 |
| 9.30E-13 | 1.47E-11 | 187.126093 | 388.381274 | 334.09869 | 405.480196 | 0 |
| 9.30E-13 | 1.47E-11 | 231.496197 | 662.428084 | 616.226473 | 665.748377 | 2.37621824 |
| 9.35E-13 | 1.48E-11 | 182.624778 | 411.515096 | 447.321024 | 396.665409 | 0 |
| 9.36E-13 | 1.48E-11 | 201.273083 | 374.367517 | 263.566744 | 331.250412 | 0 |
| 9.38E-13 | 1.48E-11 | 312.198344 | 393.942289 | 237.581291 | 296.455201 | 35.6432737 |
| 9.39E-13 | 1.49E-11 | 371.358483 | 647.079683 | 492.795568 | 591.518593 | 2.37621824 |
| 9.45E-13 | 1.49E-11 | 182.624778 | 408.623368 | 382.35739 | 422.645834 | 0 |
| 9.49E-13 | 1.50E-11 | 366.214123 | 634.178129 | 514.140762 | 570.641466 | 0 |
| 9.61E-13 | 1.52E-11 | 225.38727 | 214.877612 | 220.876356 | 237.535309 | 0 |
| 9.60E-13 | 1.52E-11 | 192.270453 | 502.048417 | 344.307261 | 424.965515 | 0 |
| 9.60E-13 | 1.52E-11 | 288.405679 | 455.113452 | 400.918428 | 480.173917 | 2.37621824 |
| 9.60E-13 | 1.52E-11 | 237.926647 | 225.554761 | 206.955577 | 240.782863 | 0 |
| 9.64E-13 | 1.52E-11 | 269.757374 | 719.595317 | 676.549847 | 702.863269 | 0 |
| 9.65E-13 | 1.52E-11 | 353.031701 | 643.743075 | 577.248292 | 603.116997 | 0 |
| 9.71E-13 | 1.53E-11 | 169.442355 | 940.256384 | 843.599193 | 893.541027 | 0 |
| 9.72E-13 | 1.53E-11 | 220.885955 | 220.661068 | 221.804408 | 225.47297 | 0 |
| 9.73E-13 | 1.53E-11 | 551.411081 | 565.888867 | 377.71713 | 452.337748 | 2.37621824 |
| 9.76E-13 | 1.54E-11 | 183.267823 | 468.237447 | 396.278169 | 398.521154 | 0 |
| 9.81E-13 | 1.54E-11 | 180.052598 | 489.146863 | 433.400245 | 454.193492 | 0 |
| 9.82E-13 | 1.54E-11 | 188.412183 | 296.290869 | 294.192458 | 316.404455 | 0 |
| 9.84E-13 | 1.55E-11 | 182.303255 | 559.882971 | 661.701017 | 482.957534 | 0 |
| 9.86E-13 | 1.55E-11 | 202.559173 | 448.662675 | 286.768042 | 357.230837 | 0 |
| 9.88E-13 | 1.55E-11 | 194.842633 | 482.696086 | 323.890119 | 454.193492 | 0 |
| 9.90E-13 | 1.55E-11 | 184.875435 | 332.326245 | 387.925701 | 364.653815 | 0 |
| 1.00E-12 | 1.57E-11 | 195.164155 | 254.916919 | 265.422848 | 294.599456 | 0 |
| 1.00E-12 | 1.57E-11 | 231.81772 | 286.503483 | 217.164149 | 223.153289 | 0 |
| 1.00E-12 | 1.57E-11 | 175.22976 | 599.254956 | 595.809331 | 668.99593 | 0 |
| 1.01E-12 | 1.58E-11 | 183.589345 | 348.341968 | 357.299988 | 334.961901 | 0 |

| | | | | | | |
|---|---|---|---|---|---|---|
| 1.01E-12 | 1.58E-11 | 184.553913 | 762.30391 | 501.148035 | 658.325399 | 0 |
| 1.01E-12 | 1.58E-11 | 182.303255 | 454.668571 | 385.141546 | 401.304771 | 0 |
| 1.01E-12 | 1.59E-11 | 188.733705 | 370.586027 | 313.681548 | 307.125732 | 0 |
| 1.02E-12 | 1.59E-11 | 229.888585 | 287.838127 | 452.889335 | 258.876372 | 0 |
| 1.02E-12 | 1.59E-11 | 180.052598 | 629.506877 | 500.219983 | 510.329766 | 0 |
| 1.02E-12 | 1.59E-11 | 184.23239 | 340.778988 | 333.170638 | 373.932538 | 0 |
| 1.02E-12 | 1.59E-11 | 208.989622 | 228.668929 | 250.574018 | 300.16669 | 0 |
| 1.02E-12 | 1.59E-11 | 196.450245 | 277.605859 | 396.278169 | 360.47839 | 0 |
| 1.02E-12 | 1.60E-11 | 576.489836 | 1710.7906 | 1587.89683 | 1691.51121 | 19.0097459 |
| 1.03E-12 | 1.60E-11 | 173.94367 | 627.949792 | 618.082577 | 594.30221 | 0 |
| 1.03E-12 | 1.61E-11 | 287.762634 | 493.595675 | 206.027526 | 412.439239 | 0 |
| 1.04E-12 | 1.62E-11 | 184.23239 | 403.952116 | 336.882846 | 404.088388 | 0 |
| 1.05E-12 | 1.63E-11 | 476.174817 | 2158.78595 | 2148.44019 | 2396.69416 | 16.6335277 |
| 1.05E-12 | 1.63E-11 | 208.025055 | 294.511344 | 229.228823 | 328.930731 | 0 |
| 1.05E-12 | 1.64E-11 | 191.305885 | 299.405037 | 264.494796 | 274.186265 | 0 |
| 1.07E-12 | 1.66E-11 | 220.885955 | 235.119706 | 226.444668 | 208.771268 | 0 |
| 1.07E-12 | 1.67E-11 | 171.693013 | 694.45953 | 749.865949 | 657.861463 | 0 |
| 1.08E-12 | 1.67E-11 | 186.161525 | 355.904948 | 297.904665 | 335.425837 | 0 |
| 1.08E-12 | 1.68E-11 | 282.296752 | 231.338216 | 204.171422 | 237.535309 | 0 |
| 1.08E-12 | 1.68E-11 | 181.981733 | 387.713953 | 343.379209 | 399.449026 | 0 |
| 1.08E-12 | 1.68E-11 | 249.501457 | 405.286759 | 208.811681 | 368.82924 | 0 |
| 1.08E-12 | 1.68E-11 | 182.624778 | 444.436304 | 359.156092 | 440.275408 | 0 |
| 1.09E-12 | 1.68E-11 | 190.984363 | 325.208146 | 271.919212 | 280.681372 | 0 |
| 1.09E-12 | 1.69E-11 | 209.311145 | 441.989457 | 260.782589 | 383.675197 | 0 |
| 1.09E-12 | 1.69E-11 | 223.136612 | 934.472929 | 839.886985 | 839.260498 | 11.8810912 |
| 1.10E-12 | 1.70E-11 | 199.986993 | 243.572449 | 338.73895 | 299.238818 | 0 |
| 1.10E-12 | 1.70E-11 | 187.126093 | 873.079325 | 534.557904 | 604.508805 | 0 |
| 1.11E-12 | 1.71E-11 | 292.263949 | 320.981775 | 188.394539 | 379.963708 | 0 |
| 1.11E-12 | 1.72E-11 | 203.845263 | 234.452384 | 271.919212 | 323.363497 | 0 |
| 1.11E-12 | 1.72E-11 | 20.5774398 | 8.8976237 | 14.8488307 | 6.49510612 | 166.335277 |
| 1.12E-12 | 1.73E-11 | 217.027685 | 544.757011 | 281.199731 | 404.552324 | 0 |
| 1.13E-12 | 1.75E-11 | 183.267823 | 382.375378 | 350.803625 | 318.2602 | 0 |
| 1.15E-12 | 1.77E-11 | 497.395302 | 388.826156 | 205.099474 | 285.320733 | 0 |
| 1.15E-12 | 1.77E-11 | 195.164155 | 276.271216 | 235.725187 | 265.835415 | 0 |
| 1.15E-12 | 1.77E-11 | 501.896616 | 1407.82651 | 765.642831 | 947.357621 | 16.6335277 |
| 1.15E-12 | 1.77E-11 | 320.879451 | 317.867607 | 184.682331 | 391.562112 | 0 |
| 1.15E-12 | 1.78E-11 | 196.128723 | 352.345898 | 253.358173 | 316.868391 | 0 |
| 1.16E-12 | 1.78E-11 | 259.468654 | 203.978023 | 407.414792 | 316.404455 | 0 |
| 1.16E-12 | 1.78E-11 | 170.406923 | 739.83741 | 652.420498 | 659.717207 | 0 |
| 1.16E-12 | 1.79E-11 | 176.194328 | 431.979631 | 424.119726 | 417.542536 | 0 |
| 1.17E-12 | 1.80E-11 | 178.123463 | 398.391101 | 481.658945 | 473.67881 | 0 |
| 1.17E-12 | 1.80E-11 | 171.693013 | 602.146684 | 558.687254 | 602.653061 | 0 |
| 1.17E-12 | 1.80E-11 | 175.551283 | 554.321956 | 461.241803 | 508.474022 | 0 |

| | | | | | | |
|---|---|---|---|---|---|---|
| 1.18E-12 | 1.81E-11 | 175.22976 | 487.812219 | 446.392972 | 508.937958 | 0 |
| 1.19E-12 | 1.82E-11 | 175.551283 | 427.085937 | 427.831934 | 448.626258 | 0 |
| 1.19E-12 | 1.82E-11 | 355.925403 | 685.339465 | 202.315318 | 565.074232 | 0 |
| 1.19E-12 | 1.82E-11 | 197.414813 | 233.117741 | 291.408302 | 257.020628 | 0 |
| 1.19E-12 | 1.82E-11 | 185.196958 | 304.966052 | 275.631419 | 306.19786 | 0 |
| 1.19E-12 | 1.82E-11 | 226.994882 | 267.373592 | 199.531162 | 295.991265 | 0 |
| 1.19E-12 | 1.83E-11 | 182.624778 | 420.857601 | 329.458431 | 390.634239 | 0 |
| 1.19E-12 | 1.83E-11 | 326.988379 | 888.872607 | 735.017118 | 746.009331 | 2.37621824 |
| 1.20E-12 | 1.83E-11 | 184.875435 | 294.511344 | 299.760769 | 276.04201 | 0 |
| 1.20E-12 | 1.83E-11 | 180.052598 | 453.556368 | 380.501286 | 372.54073 | 0 |
| 1.20E-12 | 1.84E-11 | 184.875435 | 491.593709 | 349.875573 | 405.480196 | 0 |
| 1.21E-12 | 1.85E-11 | 274.258689 | 464.900838 | 229.228823 | 266.763287 | 0 |
| 1.21E-12 | 1.85E-11 | 185.196958 | 502.048417 | 368.436611 | 376.252219 | 0 |
| 1.22E-12 | 1.87E-11 | 163.333428 | 1025.00625 | 921.555554 | 868.952411 | 0 |
| 1.22E-12 | 1.87E-11 | 226.351837 | 203.978023 | 256.142329 | 220.369672 | 0 |
| 1.23E-12 | 1.87E-11 | 188.09066 | 558.325887 | 362.8683 | 434.244238 | 0 |
| 1.23E-12 | 1.87E-11 | 182.303255 | 578.790422 | 402.774532 | 555.795509 | 0 |
| 1.24E-12 | 1.89E-11 | 175.551283 | 470.239412 | 408.342843 | 441.667216 | 0 |
| 1.24E-12 | 1.89E-11 | 176.837373 | 434.426477 | 406.48674 | 496.411682 | 0 |
| 1.24E-12 | 1.89E-11 | 179.731075 | 325.653027 | 322.962067 | 324.755306 | 0 |
| 1.24E-12 | 1.89E-11 | 257.539519 | 684.004822 | 827.82231 | 847.147412 | 2.37621824 |
| 1.25E-12 | 1.90E-11 | 173.300625 | 626.615149 | 559.615306 | 503.370724 | 0 |
| 1.25E-12 | 1.90E-11 | 261.076267 | 456.892977 | 401.84648 | 408.263813 | 14.2573095 |
| 1.26E-12 | 1.92E-11 | 198.700903 | 439.987492 | 278.415575 | 334.497965 | 0 |
| 1.26E-12 | 1.92E-11 | 203.845263 | 281.832231 | 215.308045 | 258.876372 | 0 |
| 1.26E-12 | 1.92E-11 | 677.447899 | 1239.88386 | 1235.2371 | 1339.8476 | 21.3859642 |
| 1.26E-12 | 1.92E-11 | 182.9463 | 1050.36448 | 595.809331 | 737.194544 | 0 |
| 1.27E-12 | 1.92E-11 | 195.8072 | 322.98374 | 238.509343 | 305.269988 | 0 |
| 1.27E-12 | 1.93E-11 | 171.37149 | 511.613363 | 504.860243 | 530.279021 | 0 |
| 1.28E-12 | 1.93E-11 | 179.731075 | 532.300338 | 393.494013 | 439.811471 | 0 |
| 1.28E-12 | 1.93E-11 | 331.168171 | 468.682328 | 329.458431 | 413.831047 | 0 |
| 1.28E-12 | 1.94E-11 | 281.975229 | 304.743612 | 191.178695 | 241.246799 | 0 |
| 1.28E-12 | 1.94E-11 | 233.10381 | 350.566374 | 204.171422 | 286.248605 | 0 |
| 1.29E-12 | 1.95E-11 | 187.126093 | 273.824369 | 256.142329 | 259.804245 | 0 |
| 1.29E-12 | 1.95E-11 | 222.172045 | 518.28658 | 458.457647 | 521.92817 | 9.50487297 |
| 1.29E-12 | 1.95E-11 | 205.452875 | 309.637305 | 225.516616 | 339.601263 | 0 |
| 1.29E-12 | 1.95E-11 | 242.10644 | 330.991602 | 194.890903 | 329.858604 | 0 |
| 1.30E-12 | 1.96E-11 | 421.515992 | 808.349113 | 503.004139 | 591.054657 | 16.6335277 |
| 1.30E-12 | 1.96E-11 | 242.10644 | 225.999642 | 235.725187 | 195.781056 | 0 |
| 1.31E-12 | 1.97E-11 | 208.025055 | 415.296586 | 632.931407 | 327.538923 | 0 |
| 1.31E-12 | 1.97E-11 | 196.450245 | 308.080221 | 237.581291 | 255.628819 | 0 |
| 1.32E-12 | 1.99E-11 | 177.480418 | 412.84974 | 366.580507 | 352.127539 | 0 |
| 1.32E-12 | 1.99E-11 | 174.586715 | 389.048596 | 385.141546 | 394.809665 | 0 |

| | | | | | | |
|---|---|---|---|---|---|---|
| 1.33E-12 | 2.00E-11 | 178.766508 | 388.158834 | 324.818171 | 370.221049 | 0 |
| 1.33E-12 | 2.01E-11 | 176.51585 | 341.446309 | 374.004923 | 375.788283 | 0 |
| 1.34E-12 | 2.01E-11 | 171.693013 | 455.113452 | 464.95401 | 469.039449 | 0 |
| 1.34E-12 | 2.01E-11 | 180.052598 | 363.467928 | 307.185185 | 364.653815 | 0 |
| 1.34E-12 | 2.02E-11 | 261.397789 | 720.485079 | 594.881279 | 631.417102 | 2.37621824 |
| 1.35E-12 | 2.02E-11 | 175.22976 | 480.249239 | 392.565961 | 437.491791 | 0 |
| 1.35E-12 | 2.03E-11 | 192.913498 | 277.8283 | 238.509343 | 313.156902 | 0 |
| 1.35E-12 | 2.03E-11 | 171.693013 | 711.809896 | 542.910371 | 596.157954 | 0 |
| 1.36E-12 | 2.04E-11 | 179.409553 | 310.082186 | 297.904665 | 319.652008 | 0 |
| 1.36E-12 | 2.04E-11 | 337.277098 | 640.406466 | 360.084144 | 502.442852 | 35.6432737 |
| 1.36E-12 | 2.04E-11 | 192.913498 | 237.344112 | 295.12051 | 314.548711 | 0 |
| 1.36E-12 | 2.04E-11 | 170.406923 | 637.737179 | 528.061541 | 556.723382 | 0 |
| 1.36E-12 | 2.04E-11 | 184.23239 | 260.700374 | 270.063108 | 280.217435 | 0 |
| 1.37E-12 | 2.05E-11 | 183.589345 | 491.148828 | 342.451157 | 483.42147 | 0 |
| 1.37E-12 | 2.05E-11 | 172.014535 | 442.87922 | 515.068814 | 476.926364 | 0 |
| 1.37E-12 | 2.05E-11 | 194.52111 | 327.877433 | 240.365446 | 331.250412 | 0 |
| 1.37E-12 | 2.05E-11 | 181.981733 | 331.214042 | 279.343627 | 309.909349 | 0 |
| 1.38E-12 | 2.06E-11 | 63.3399317 | 38.0373413 | 49.1867516 | 30.619786 | 313.660808 |
| 1.38E-12 | 2.07E-11 | 172.336058 | 482.473645 | 609.730109 | 617.499017 | 0 |
| 1.39E-12 | 2.07E-11 | 251.109069 | 554.766838 | 831.534518 | 701.471461 | 7.12865473 |
| 1.39E-12 | 2.07E-11 | 176.51585 | 609.487223 | 446.392972 | 477.854236 | 0 |
| 1.39E-12 | 2.08E-11 | 292.263949 | 223.107914 | 221.804408 | 210.627013 | 0 |
| 1.40E-12 | 2.08E-11 | 172.65758 | 425.751294 | 424.119726 | 398.057218 | 0 |
| 1.40E-12 | 2.09E-11 | 216.38464 | 376.147042 | 253.358173 | 238.927118 | 0 |
| 1.40E-12 | 2.09E-11 | 162.047338 | 299.405037 | 302.544925 | 327.538923 | 9.50487297 |
| 1.41E-12 | 2.09E-11 | 170.406923 | 634.845451 | 507.644399 | 574.352955 | 0 |
| 1.41E-12 | 2.10E-11 | 175.551283 | 515.617293 | 397.20622 | 443.986897 | 0 |
| 1.41E-12 | 2.10E-11 | 170.406923 | 455.113452 | 469.59427 | 482.493597 | 0 |
| 1.41E-12 | 2.10E-11 | 181.017165 | 444.881185 | 337.810898 | 352.127539 | 0 |
| 1.42E-12 | 2.11E-11 | 174.908238 | 416.85367 | 357.299988 | 411.04743 | 0 |
| 1.42E-12 | 2.11E-11 | 302.552669 | 911.783988 | 762.858676 | 886.581985 | 2.37621824 |
| 1.42E-12 | 2.11E-11 | 178.444985 | 351.456136 | 354.515832 | 303.414243 | 0 |
| 1.44E-12 | 2.14E-11 | 172.014535 | 550.095585 | 450.10518 | 472.287002 | 0 |
| 1.44E-12 | 2.14E-11 | 179.409553 | 373.032874 | 304.401029 | 329.394667 | 0 |
| 1.45E-12 | 2.14E-11 | 175.551283 | 360.798641 | 350.803625 | 327.538923 | 0 |
| 1.45E-12 | 2.15E-11 | 163.654951 | 755.185811 | 704.391405 | 687.553376 | 0 |
| 1.45E-12 | 2.15E-11 | 219.921387 | 407.956047 | 223.660512 | 380.89158 | 0 |
| 1.47E-12 | 2.18E-11 | 172.65758 | 474.020902 | 396.278169 | 431.460621 | 0 |
| 1.49E-12 | 2.20E-11 | 205.774398 | 377.481685 | 253.358173 | 255.164883 | 0 |
| 1.49E-12 | 2.21E-11 | 165.905608 | 603.258887 | 750.794001 | 681.05827 | 0 |
| 1.50E-12 | 2.22E-11 | 299.337444 | 677.554045 | 626.435044 | 650.90242 | 14.2573095 |
| 1.50E-12 | 2.22E-11 | 185.840003 | 279.607825 | 242.22155 | 291.351903 | 0 |
| 1.51E-12 | 2.22E-11 | 210.597235 | 271.599963 | 199.531162 | 248.205841 | 0 |

| | | | | | | |
|---|---|---|---|---|---|---|
| 1.51E-12 | 2.23E-11 | 189.055228 | 301.184562 | 238.509343 | 264.443606 | 0 |
| 1.51E-12 | 2.23E-11 | 185.51848 | 280.052706 | 246.86181 | 252.845202 | 0 |
| 1.51E-12 | 2.23E-11 | 182.9463 | 473.576021 | 322.962067 | 386.458814 | 0 |
| 1.52E-12 | 2.24E-11 | 172.979103 | 525.404679 | 414.839207 | 440.739344 | 0 |
| 1.52E-12 | 2.24E-11 | 192.270453 | 362.355725 | 329.458431 | 247.277969 | 0 |
| 1.53E-12 | 2.25E-11 | 171.37149 | 381.930497 | 399.062324 | 413.831047 | 0 |
| 1.53E-12 | 2.25E-11 | 193.878065 | 232.67286 | 281.199731 | 230.576267 | 0 |
| 1.53E-12 | 2.25E-11 | 172.979103 | 354.125423 | 358.22804 | 384.603069 | 0 |
| 1.53E-12 | 2.26E-11 | 165.905608 | 573.229407 | 557.759202 | 546.05285 | 0 |
| 1.54E-12 | 2.26E-11 | 349.494953 | 428.865462 | 296.976613 | 358.622645 | 4.75243649 |
| 1.54E-12 | 2.27E-11 | 168.79931 | 612.378951 | 518.781022 | 502.906788 | 0 |
| 1.54E-12 | 2.27E-11 | 208.6681 | 296.958191 | 204.171422 | 266.763287 | 0 |
| 1.54E-12 | 2.27E-11 | 166.22713 | 583.461674 | 528.989593 | 565.074232 | 0 |
| 1.55E-12 | 2.27E-11 | 173.622148 | 332.993567 | 365.652455 | 356.7669 | 0 |
| 1.55E-12 | 2.27E-11 | 183.910868 | 246.241736 | 284.911939 | 292.279775 | 0 |
| 1.55E-12 | 2.27E-11 | 170.728445 | 530.743254 | 429.688038 | 477.854236 | 0 |
| 1.55E-12 | 2.27E-11 | 178.444985 | 349.676611 | 288.624146 | 324.29137 | 0 |
| 1.55E-12 | 2.27E-11 | 169.763878 | 465.790601 | 420.407518 | 436.099982 | 0 |
| 1.56E-12 | 2.28E-11 | 165.584086 | 779.654276 | 699.751145 | 600.797316 | 0 |
| 1.56E-12 | 2.28E-11 | 164.297996 | 625.725387 | 594.881279 | 600.797316 | 0 |
| 1.57E-12 | 2.30E-11 | 165.262563 | 594.583704 | 561.47141 | 643.479442 | 0 |
| 1.57E-12 | 2.30E-11 | 168.79931 | 514.50509 | 444.536868 | 493.628065 | 0 |
| 1.58E-12 | 2.30E-11 | 163.976473 | 667.989099 | 607.874006 | 611.003911 | 0 |
| 1.58E-12 | 2.30E-11 | 209.632667 | 222.440592 | 202.315318 | 214.802438 | 0 |
| 1.59E-12 | 2.32E-11 | 158.510591 | 962.945325 | 801.836856 | 882.870496 | 0 |
| 1.59E-12 | 2.32E-11 | 169.442355 | 447.995353 | 408.342843 | 448.162322 | 0 |
| 1.59E-12 | 2.32E-11 | 181.981733 | 343.003394 | 266.3509 | 320.115944 | 0 |
| 1.59E-12 | 2.32E-11 | 172.979103 | 418.633195 | 350.803625 | 385.994878 | 0 |
| 1.59E-12 | 2.32E-11 | 174.265193 | 320.759334 | 316.465704 | 334.497965 | 0 |
| 1.60E-12 | 2.33E-11 | 0 | 0 | 0 | 0 | 40.3957101 |
| 1.60E-12 | 2.33E-11 | 245.321665 | 484.47561 | 281.199731 | 237.999246 | 0 |
| 1.60E-12 | 2.33E-11 | 170.728445 | 374.145076 | 387.925701 | 380.89158 | 0 |
| 1.60E-12 | 2.33E-11 | 222.493567 | 477.579952 | 374.932974 | 436.563918 | 7.12865473 |
| 1.61E-12 | 2.34E-11 | 316.699659 | 848.38842 | 705.319457 | 770.134011 | 0 |
| 1.61E-12 | 2.34E-11 | 157.867546 | 894.656063 | 807.405168 | 849.467093 | 0 |
| 1.61E-12 | 2.34E-11 | 202.23765 | 465.790601 | 263.566744 | 342.848816 | 0 |
| 1.61E-12 | 2.35E-11 | 186.483048 | 241.792924 | 236.653239 | 237.071373 | 0 |
| 1.62E-12 | 2.36E-11 | 169.442355 | 447.995353 | 505.788295 | 568.785722 | 0 |
| 1.62E-12 | 2.36E-11 | 176.51585 | 284.279077 | 299.760769 | 312.692966 | 0 |
| 1.63E-12 | 2.36E-11 | 169.763878 | 443.10166 | 395.350117 | 455.585301 | 0 |
| 1.64E-12 | 2.38E-11 | 221.850522 | 220.661068 | 402.774532 | 288.10435 | 0 |
| 1.64E-12 | 2.38E-11 | 174.908238 | 342.558512 | 306.257133 | 360.942326 | 0 |
| 1.64E-12 | 2.38E-11 | 850.427002 | 1207.40754 | 713.671924 | 941.326451 | 21.3859642 |

| | | | | | | |
|---|---|---|---|---|---|---|
| 1.64E-12 | 2.38E-11 | 189.055228 | 276.493656 | 224.588564 | 278.361691 | 0 |
| 1.65E-12 | 2.39E-11 | 184.553913 | 244.01733 | 238.509343 | 262.587862 | 0 |
| 1.66E-12 | 2.40E-11 | 168.477788 | 444.881185 | 407.414792 | 428.213068 | 0 |
| 1.67E-12 | 2.41E-11 | 153.366231 | 1057.0377 | 934.54828 | 1079.11549 | 0 |
| 1.67E-12 | 2.41E-11 | 175.551283 | 365.025012 | 325.746223 | 422.181898 | 0 |
| 1.67E-12 | 2.42E-11 | 216.38464 | 244.462211 | 190.250643 | 257.484564 | 0 |
| 1.68E-12 | 2.43E-11 | 173.300625 | 331.881364 | 311.825444 | 346.560305 | 0 |
| 1.68E-12 | 2.43E-11 | 192.913498 | 339.889225 | 244.077654 | 259.340309 | 0 |
| 1.69E-12 | 2.44E-11 | 187.126093 | 479.804358 | 306.257133 | 484.813278 | 0 |
| 1.70E-12 | 2.45E-11 | 171.693013 | 419.967839 | 357.299988 | 442.131152 | 0 |
| 1.70E-12 | 2.46E-11 | 312.841389 | 927.132389 | 714.599976 | 764.102841 | 2.37621824 |
| 1.70E-12 | 2.46E-11 | 172.014535 | 453.556368 | 363.796351 | 390.634239 | 0 |
| 1.70E-12 | 2.46E-11 | 466.529142 | 225.10988 | 322.034015 | 231.968076 | 0 |
| 1.71E-12 | 2.46E-11 | 171.049968 | 350.121492 | 346.163365 | 379.499772 | 0 |
| 1.71E-12 | 2.46E-11 | 168.477788 | 410.402893 | 485.371153 | 482.029661 | 0 |
| 1.71E-12 | 2.47E-11 | 163.654951 | 624.613184 | 554.975046 | 561.826679 | 0 |
| 1.72E-12 | 2.47E-11 | 308.018551 | 390.60568 | 178.185968 | 302.950307 | 0 |
| 1.72E-12 | 2.47E-11 | 211.561802 | 206.202429 | 252.430121 | 316.404455 | 0 |
| 1.72E-12 | 2.47E-11 | 184.875435 | 229.336251 | 248.717914 | 246.814032 | 0 |
| 1.72E-12 | 2.48E-11 | 171.049968 | 859.510449 | 541.98232 | 679.666462 | 0 |
| 1.73E-12 | 2.48E-11 | 198.057858 | 679.333569 | 335.026742 | 556.259445 | 0 |
| 1.73E-12 | 2.49E-11 | 584.527898 | 366.804537 | 361.940248 | 310.837221 | 19.0097459 |
| 1.74E-12 | 2.50E-11 | 175.551283 | 397.501339 | 374.004923 | 520.072426 | 0 |
| 1.75E-12 | 2.50E-11 | 183.589345 | 284.723958 | 235.725187 | 285.320733 | 0 |
| 1.75E-12 | 2.51E-11 | 167.191698 | 421.302482 | 452.889335 | 416.150728 | 0 |
| 1.75E-12 | 2.51E-11 | 238.24817 | 197.304805 | 229.228823 | 314.084774 | 0 |
| 1.76E-12 | 2.52E-11 | 164.297996 | 526.072001 | 525.277385 | 489.916576 | 0 |
| 1.76E-12 | 2.52E-11 | 211.883325 | 244.684652 | 300.688821 | 200.884354 | 0 |
| 1.77E-12 | 2.53E-11 | 239.53426 | 199.08433 | 201.387266 | 227.328714 | 0 |
| 1.77E-12 | 2.54E-11 | 167.834743 | 862.402177 | 589.312967 | 634.200719 | 0 |
| 1.78E-12 | 2.55E-11 | 173.94367 | 558.548328 | 388.853753 | 450.018067 | 0 |
| 1.78E-12 | 2.55E-11 | 183.267823 | 258.698409 | 361.940248 | 337.281582 | 0 |
| 1.79E-12 | 2.56E-11 | 170.728445 | 377.926567 | 331.314534 | 372.076793 | 0 |
| 1.79E-12 | 2.56E-11 | 197.736335 | 368.361621 | 256.142329 | 249.597649 | 0 |
| 1.79E-12 | 2.56E-11 | 165.905608 | 454.891012 | 426.903882 | 457.441045 | 0 |
| 1.80E-12 | 2.58E-11 | 185.840003 | 339.666785 | 243.149602 | 303.414243 | 0 |
| 1.81E-12 | 2.58E-11 | 179.409553 | 403.062353 | 289.552198 | 336.817646 | 0 |
| 1.81E-12 | 2.59E-11 | 184.553913 | 308.080221 | 265.422848 | 382.747325 | 0 |
| 1.82E-12 | 2.59E-11 | 175.22976 | 273.379488 | 313.681548 | 295.991265 | 0 |
| 1.82E-12 | 2.59E-11 | 230.53163 | 297.625513 | 184.682331 | 289.032222 | 0 |
| 1.82E-12 | 2.60E-11 | 173.622148 | 312.084151 | 284.911939 | 303.414243 | 0 |
| 1.82E-12 | 2.60E-11 | 168.156265 | 426.863497 | 372.148819 | 415.686792 | 0 |
| 1.83E-12 | 2.61E-11 | 172.014535 | 504.272823 | 370.292715 | 429.604876 | 0 |

| | | | | | | |
|---|---|---|---|---|---|---|
| 1.84E-12 | 2.61E-11 | 364.606511 | 256.474003 | 204.171422 | 231.50414 | 0 |
| 1.84E-12 | 2.62E-11 | 170.406923 | 340.334106 | 353.58778 | 321.043817 | 0 |
| 1.84E-12 | 2.62E-11 | 175.872805 | 388.158834 | 294.192458 | 358.622645 | 0 |
| 1.84E-12 | 2.62E-11 | 230.53163 | 386.156868 | 199.531162 | 346.096369 | 0 |
| 1.84E-12 | 2.62E-11 | 486.142014 | 330.32428 | 212.523889 | 249.133713 | 0 |
| 1.85E-12 | 2.63E-11 | 328.595991 | 670.213505 | 643.139978 | 662.500824 | 23.7621824 |
| 1.86E-12 | 2.64E-11 | 179.08803 | 349.676611 | 412.055051 | 289.496158 | 0 |
| 1.86E-12 | 2.64E-11 | 179.08803 | 291.842057 | 246.86181 | 273.722329 | 0 |
| 1.87E-12 | 2.65E-11 | 159.153636 | 694.45953 | 621.794784 | 695.440291 | 0 |
| 1.88E-12 | 2.66E-11 | 168.79931 | 371.69823 | 342.451157 | 353.519347 | 0 |
| 1.89E-12 | 2.67E-11 | 173.622148 | 354.347864 | 447.321024 | 508.010086 | 0 |
| 1.89E-12 | 2.68E-11 | 171.049968 | 387.046631 | 322.962067 | 345.168497 | 0 |
| 1.89E-12 | 2.68E-11 | 173.622148 | 278.940503 | 290.48025 | 309.445413 | 0 |
| 1.90E-12 | 2.69E-11 | 180.052598 | 462.231551 | 335.026742 | 322.435625 | 0 |
| 1.90E-12 | 2.69E-11 | 161.082771 | 624.835624 | 549.406735 | 585.023487 | 0 |
| 1.90E-12 | 2.69E-11 | 175.551283 | 259.810612 | 279.343627 | 282.537116 | 0 |
| 1.90E-12 | 2.69E-11 | 200.308515 | 320.092013 | 217.164149 | 247.741905 | 0 |
| 1.90E-12 | 2.69E-11 | 166.22713 | 546.091654 | 432.472193 | 488.060831 | 0 |
| 1.91E-12 | 2.70E-11 | 193.23502 | 288.283008 | 218.0922 | 317.796264 | 0 |
| 1.94E-12 | 2.74E-11 | 185.51848 | 235.342147 | 245.005706 | 217.586055 | 0 |
| 1.95E-12 | 2.75E-11 | 218.95682 | 261.145256 | 445.46492 | 250.061586 | 0 |
| 1.95E-12 | 2.75E-11 | 163.011906 | 558.770768 | 493.72362 | 610.539975 | 0 |
| 1.97E-12 | 2.77E-11 | 182.9463 | 326.097909 | 271.919212 | 244.494352 | 0 |
| 1.96E-12 | 2.77E-11 | 164.941041 | 441.322135 | 400.918428 | 433.316365 | 0 |
| 1.96E-12 | 2.77E-11 | 170.406923 | 369.251383 | 311.825444 | 344.70456 | 0 |
| 1.97E-12 | 2.77E-11 | 169.763878 | 394.832052 | 326.674275 | 358.622645 | 0 |
| 1.98E-12 | 2.79E-11 | 186.80457 | 442.87922 | 273.775316 | 349.343922 | 0 |
| 1.98E-12 | 2.79E-11 | 168.156265 | 399.280863 | 387.925701 | 340.529135 | 0 |
| 1.98E-12 | 2.79E-11 | 223.458135 | 275.826335 | 207.883629 | 199.028609 | 0 |
| 1.98E-12 | 2.79E-11 | 161.725816 | 537.638912 | 507.644399 | 492.236257 | 0 |
| 2.00E-12 | 2.80E-11 | 208.346577 | 320.314453 | 199.531162 | 283.464988 | 0 |
| 2.00E-12 | 2.81E-11 | 194.199588 | 316.088082 | 331.314534 | 219.4418 | 0 |
| 2.00E-12 | 2.81E-11 | 180.052598 | 299.405037 | 238.509343 | 291.351903 | 0 |
| 2.01E-12 | 2.82E-11 | 172.336058 | 301.184562 | 285.83999 | 335.425837 | 0 |
| 2.01E-12 | 2.82E-11 | 165.262563 | 394.16473 | 463.097907 | 456.049237 | 0 |
| 2.01E-12 | 2.82E-11 | 170.0854 | 355.460067 | 462.169855 | 378.107963 | 0 |
| 2.02E-12 | 2.84E-11 | 171.693013 | 473.798462 | 346.163365 | 466.719768 | 0 |
| 2.03E-12 | 2.84E-11 | 180.695643 | 234.674825 | 236.653239 | 263.515734 | 0 |
| 2.03E-12 | 2.85E-11 | 167.51322 | 356.79471 | 330.386482 | 354.911156 | 0 |
| 2.04E-12 | 2.85E-11 | 164.941041 | 398.391101 | 380.501286 | 411.511366 | 0 |
| 2.04E-12 | 2.86E-11 | 179.08803 | 333.883329 | 252.430121 | 289.496158 | 0 |
| 2.05E-12 | 2.87E-11 | 192.591975 | 229.781132 | 309.96934 | 363.262006 | 0 |
| 2.06E-12 | 2.87E-11 | 168.477788 | 442.656779 | 348.947521 | 392.026048 | 0 |

| | | | | | | |
|---|---|---|---|---|---|---|
| 2.06E-12 | 2.88E-11 | 167.51322 | 373.922636 | 331.314534 | 352.127539 | 0 |
| 2.06E-12 | 2.88E-11 | 179.731075 | 288.283008 | 234.797135 | 263.97967 | 0 |
| 2.07E-12 | 2.89E-11 | 185.51848 | 281.832231 | 223.660512 | 239.85499 | 0 |
| 2.08E-12 | 2.90E-11 | 179.409553 | 230.003573 | 266.3509 | 252.381266 | 0 |
| 2.08E-12 | 2.90E-11 | 172.014535 | 347.674646 | 292.336354 | 298.310945 | 0 |
| 2.09E-12 | 2.91E-11 | 175.22976 | 326.54279 | 263.566744 | 285.784669 | 0 |
| 2.09E-12 | 2.91E-11 | 188.412183 | 273.379488 | 253.358173 | 270.01084 | 4.75243649 |
| 2.09E-12 | 2.92E-11 | 160.761248 | 573.006966 | 491.867516 | 541.413489 | 0 |
| 2.09E-12 | 2.92E-11 | 192.270453 | 324.318384 | 556.83115 | 339.601263 | 0 |
| 2.09E-12 | 2.92E-11 | 161.082771 | 468.237447 | 506.716347 | 511.257639 | 0 |
| 2.10E-12 | 2.92E-11 | 185.51848 | 331.436483 | 233.869083 | 328.930731 | 0 |
| 2.10E-12 | 2.92E-11 | 180.695643 | 348.119527 | 248.717914 | 309.909349 | 0 |
| 2.10E-12 | 2.92E-11 | 284.547409 | 912.45131 | 968.886201 | 1005.81358 | 2.37621824 |
| 2.11E-12 | 2.93E-11 | 153.687753 | 753.628727 | 725.736599 | 794.258691 | 0 |
| 2.12E-12 | 2.95E-11 | 1225.96528 | 4133.16865 | 2307.13707 | 3251.72848 | 26.1384007 |
| 2.13E-12 | 2.95E-11 | 171.049968 | 345.895121 | 287.696094 | 312.22903 | 0 |
| 2.13E-12 | 2.96E-11 | 172.336058 | 506.497229 | 382.35739 | 359.550517 | 0 |
| 2.13E-12 | 2.96E-11 | 160.439726 | 552.09755 | 746.153741 | 701.935397 | 0 |
| 2.14E-12 | 2.96E-11 | 154.973843 | 777.429871 | 667.269328 | 736.730608 | 0 |
| 2.14E-12 | 2.97E-11 | 163.333428 | 615.048238 | 410.198947 | 507.54615 | 4.75243649 |
| 2.15E-12 | 2.97E-11 | 477.782429 | 308.302661 | 203.24337 | 259.804245 | 0 |
| 2.15E-12 | 2.97E-11 | 193.556543 | 290.507414 | 243.149602 | 209.235204 | 0 |
| 2.15E-12 | 2.97E-11 | 172.65758 | 430.422546 | 332.242586 | 401.768707 | 4.75243649 |
| 2.15E-12 | 2.98E-11 | 899.619944 | 5467.58976 | 3062.57133 | 3897.99154 | 33.2670554 |
| 2.15E-12 | 2.98E-11 | 20.2559173 | 24.023584 | 19.4890903 | 21.341063 | 111.682257 |
| 2.16E-12 | 2.98E-11 | 249.501457 | 246.686617 | 175.401812 | 241.246799 | 0 |
| 2.19E-12 | 3.03E-11 | 184.23239 | 277.160978 | 257.070381 | 216.194247 | 0 |
| 2.19E-12 | 3.03E-11 | 172.014535 | 283.389315 | 317.393756 | 360.942326 | 0 |
| 2.19E-12 | 3.03E-11 | 160.118203 | 648.191886 | 510.428554 | 608.220294 | 0 |
| 2.20E-12 | 3.03E-11 | 159.796681 | 525.182239 | 487.227256 | 488.524767 | 0 |
| 2.20E-12 | 3.04E-11 | 167.191698 | 353.235661 | 312.753496 | 354.911156 | 0 |
| 2.21E-12 | 3.04E-11 | 161.404293 | 430.644987 | 489.08336 | 469.967321 | 0 |
| 2.21E-12 | 3.05E-11 | 246.929277 | 350.343933 | 180.970124 | 285.320733 | 0 |
| 2.22E-12 | 3.06E-11 | 215.741595 | 449.329997 | 319.249859 | 435.636046 | 9.50487297 |
| 2.22E-12 | 3.06E-11 | 166.870175 | 475.133105 | 748.009845 | 658.325399 | 0 |
| 2.23E-12 | 3.07E-11 | 156.581456 | 573.451847 | 602.305694 | 607.292422 | 0 |
| 2.23E-12 | 3.07E-11 | 294.836129 | 211.096122 | 197.675058 | 225.47297 | 0 |
| 2.24E-12 | 3.08E-11 | 164.941041 | 464.678398 | 374.004923 | 409.191685 | 0 |
| 2.24E-12 | 3.08E-11 | 158.189068 | 524.737358 | 519.709073 | 549.300403 | 0 |
| 2.25E-12 | 3.09E-11 | 161.082771 | 482.696086 | 557.759202 | 461.616471 | 0 |
| 2.25E-12 | 3.09E-11 | 196.771768 | 214.877612 | 193.962851 | 222.689353 | 0 |
| 2.26E-12 | 3.11E-11 | 160.761248 | 483.140967 | 455.673491 | 433.316365 | 0 |
| 2.27E-12 | 3.12E-11 | 201.594605 | 231.560657 | 232.941031 | 187.894141 | 0 |

| | | | | | | |
|---|---|---|---|---|---|---|
| 2.28E-12 | 3.13E-11 | 175.551283 | 324.095943 | 271.919212 | 256.092756 | 0 |
| 2.28E-12 | 3.13E-11 | 165.584086 | 352.345898 | 334.09869 | 323.363497 | 0 |
| 2.30E-12 | 3.15E-11 | 170.0854 | 373.922636 | 294.192458 | 324.29137 | 0 |
| 2.30E-12 | 3.16E-11 | 165.905608 | 345.45024 | 323.890119 | 317.332327 | 0 |
| 2.30E-12 | 3.16E-11 | 162.690383 | 377.926567 | 380.501286 | 410.119558 | 0 |
| 2.31E-12 | 3.16E-11 | 172.014535 | 316.977844 | 265.422848 | 332.64222 | 0 |
| 2.31E-12 | 3.16E-11 | 3295.92711 | 1000.76023 | 752.650105 | 1009.989 | 14.2573095 |
| 2.31E-12 | 3.16E-11 | 255.288862 | 448.217794 | 449.177128 | 427.285195 | 2.37621824 |
| 2.31E-12 | 3.17E-11 | 303.517236 | 672.21547 | 632.003355 | 635.592527 | 0 |
| 2.32E-12 | 3.17E-11 | 164.941041 | 378.371448 | 341.523105 | 337.745518 | 0 |
| 2.32E-12 | 3.17E-11 | 162.690383 | 574.119169 | 433.400245 | 502.442852 | 0 |
| 2.33E-12 | 3.18E-11 | 153.366231 | 647.079683 | 703.463353 | 706.574758 | 0 |
| 2.33E-12 | 3.18E-11 | 182.303255 | 260.477934 | 215.308045 | 274.650202 | 0 |
| 2.36E-12 | 3.21E-11 | 157.224501 | 657.979272 | 558.687254 | 726.987949 | 0 |
| 2.36E-12 | 3.21E-11 | 219.921387 | 290.507414 | 182.826228 | 247.277969 | 0 |
| 2.35E-12 | 3.21E-11 | 166.870175 | 345.672681 | 305.329081 | 366.50956 | 0 |
| 2.38E-12 | 3.24E-11 | 174.586715 | 267.596033 | 330.386482 | 373.932538 | 0 |
| 2.38E-12 | 3.24E-11 | 250.466024 | 189.074504 | 226.444668 | 201.34829 | 0 |
| 2.38E-12 | 3.24E-11 | 154.973843 | 301.629443 | 267.278952 | 288.10435 | 9.50487297 |
| 2.38E-12 | 3.25E-11 | 738.215651 | 2375.44309 | 1482.09891 | 1802.85588 | 21.3859642 |
| 2.41E-12 | 3.28E-11 | 299.980489 | 705.136678 | 632.003355 | 705.646886 | 2.37621824 |
| 2.42E-12 | 3.30E-11 | 167.834743 | 350.121492 | 501.148035 | 441.20328 | 0 |
| 2.44E-12 | 3.32E-11 | 226.994882 | 308.525102 | 219.020252 | 299.702754 | 7.12865473 |
| 2.44E-12 | 3.32E-11 | 161.404293 | 372.587992 | 411.126999 | 418.470408 | 0 |
| 2.45E-12 | 3.32E-11 | 271.043464 | 372.587992 | 402.774532 | 360.014453 | 4.75243649 |
| 2.45E-12 | 3.33E-11 | 288.405679 | 875.971053 | 832.46257 | 887.973794 | 0 |
| 2.46E-12 | 3.34E-11 | 168.477788 | 294.733785 | 268.207004 | 302.486371 | 0 |
| 2.47E-12 | 3.35E-11 | 180.695643 | 468.015006 | 295.12051 | 329.394667 | 0 |
| 2.47E-12 | 3.35E-11 | 0 | 0 | 0 | 0 | 49.9005831 |
| 2.47E-12 | 3.35E-11 | 261.076267 | 253.359835 | 276.559471 | 170.264568 | 0 |
| 2.47E-12 | 3.35E-11 | 168.156265 | 381.040735 | 297.904665 | 330.786476 | 0 |
| 2.47E-12 | 3.35E-11 | 187.769138 | 213.320528 | 200.459214 | 224.081161 | 0 |
| 2.47E-12 | 3.35E-11 | 177.158895 | 425.083972 | 284.911939 | 319.652008 | 0 |
| 2.48E-12 | 3.36E-11 | 329.560559 | 232.227979 | 183.75428 | 253.773075 | 0 |
| 2.49E-12 | 3.37E-11 | 163.333428 | 424.861532 | 355.443884 | 443.522961 | 0 |
| 2.49E-12 | 3.37E-11 | 192.913498 | 225.10988 | 194.890903 | 209.235204 | 0 |
| 2.50E-12 | 3.38E-11 | 148.221871 | 883.534033 | 788.844129 | 804.929223 | 0 |
| 2.51E-12 | 3.39E-11 | 166.22713 | 345.672681 | 396.278169 | 471.35913 | 0 |
| 2.52E-12 | 3.40E-11 | 174.265193 | 267.596033 | 230.156875 | 258.412436 | 0 |
| 2.53E-12 | 3.42E-11 | 394.18658 | 173.948543 | 358.22804 | 272.794457 | 0 |
| 2.54E-12 | 3.43E-11 | 267.506717 | 368.806502 | 179.11402 | 430.532748 | 0 |
| 2.54E-12 | 3.44E-11 | 362.998898 | 233.785063 | 206.027526 | 225.47297 | 0 |
| 2.55E-12 | 3.45E-11 | 208.6681 | 751.404321 | 329.458431 | 413.831047 | 0 |

| | | | | | | |
|---|---|---|---|---|---|---|
| 2.57E-12 | 3.46E-11 | 183.910868 | 205.535107 | 215.308045 | 217.122119 | 0 |
| 2.56E-12 | 3.46E-11 | 159.475158 | 520.510986 | 422.263622 | 470.431257 | 0 |
| 2.57E-12 | 3.47E-11 | 234.711422 | 196.415043 | 204.171422 | 302.022435 | 0 |
| 2.58E-12 | 3.48E-11 | 164.619518 | 437.763086 | 341.523105 | 367.437432 | 0 |
| 2.59E-12 | 3.49E-11 | 156.902978 | 282.499552 | 261.710641 | 264.443606 | 11.8810912 |
| 2.60E-12 | 3.49E-11 | 163.654951 | 305.855815 | 333.170638 | 326.611051 | 0 |
| 2.60E-12 | 3.50E-11 | 173.622148 | 665.097371 | 402.774532 | 440.275408 | 0 |
| 2.60E-12 | 3.50E-11 | 160.761248 | 470.461853 | 401.84648 | 392.95392 | 0 |
| 2.60E-12 | 3.50E-11 | 163.976473 | 312.973914 | 336.882846 | 369.293176 | 0 |
| 2.61E-12 | 3.51E-11 | 177.480418 | 280.720028 | 223.660512 | 246.350096 | 0 |
| 2.63E-12 | 3.53E-11 | 187.447615 | 317.867607 | 342.451157 | 219.4418 | 0 |
| 2.63E-12 | 3.53E-11 | 164.297996 | 300.2948 | 297.904665 | 310.373285 | 0 |
| 2.63E-12 | 3.53E-11 | 168.477788 | 299.182597 | 257.998433 | 283.928925 | 0 |
| 2.66E-12 | 3.57E-11 | 160.118203 | 375.924601 | 361.012196 | 364.189879 | 0 |
| 2.67E-12 | 3.58E-11 | 166.870175 | 266.48383 | 274.703367 | 277.433818 | 0 |
| 2.68E-12 | 3.59E-11 | 158.189068 | 402.617472 | 426.903882 | 450.482003 | 0 |
| 2.69E-12 | 3.61E-11 | 167.834743 | 371.920671 | 283.055835 | 334.034029 | 0 |
| 2.71E-12 | 3.63E-11 | 171.693013 | 262.035018 | 239.437395 | 290.424031 | 0 |
| 2.72E-12 | 3.64E-11 | 159.153636 | 467.125244 | 527.133489 | 412.903175 | 0 |
| 2.72E-12 | 3.64E-11 | 321.200974 | 926.687508 | 856.591919 | 910.706665 | 0 |
| 2.74E-12 | 3.66E-11 | 174.586715 | 387.713953 | 263.566744 | 317.332327 | 0 |
| 2.74E-12 | 3.67E-11 | 173.300625 | 649.748971 | 658.916861 | 706.574758 | 7.12865473 |
| 2.75E-12 | 3.68E-11 | 171.049968 | 244.23977 | 236.653239 | 262.123925 | 0 |
| 2.76E-12 | 3.68E-11 | 161.082771 | 330.769161 | 334.09869 | 332.64222 | 0 |
| 2.76E-12 | 3.69E-11 | 165.905608 | 307.635339 | 279.343627 | 340.529135 | 0 |
| 2.77E-12 | 3.69E-11 | 182.303255 | 289.17277 | 209.739733 | 260.732117 | 0 |
| 2.77E-12 | 3.69E-11 | 435.662982 | 718.927995 | 403.702584 | 516.360936 | 16.6335277 |
| 2.77E-12 | 3.69E-11 | 175.22976 | 342.558512 | 422.263622 | 272.330521 | 0 |
| 2.78E-12 | 3.70E-11 | 193.23502 | 314.308557 | 201.387266 | 268.155095 | 0 |
| 2.78E-12 | 3.70E-11 | 138.254673 | 1020.11256 | 1227.81269 | 1186.74868 | 0 |
| 2.78E-12 | 3.70E-11 | 202.559173 | 190.409147 | 201.387266 | 244.030416 | 0 |
| 2.78E-12 | 3.70E-11 | 259.468654 | 530.075932 | 492.795568 | 517.752745 | 0 |
| 2.79E-12 | 3.71E-11 | 0 | 0 | 0 | 0 | 116.434694 |
| 2.79E-12 | 3.71E-11 | 158.510591 | 377.259245 | 420.407518 | 390.634239 | 0 |
| 2.80E-12 | 3.73E-11 | 194.842633 | 356.349829 | 219.020252 | 256.556692 | 0 |
| 2.81E-12 | 3.73E-11 | 160.118203 | 469.572091 | 374.004923 | 410.119558 | 0 |
| 2.81E-12 | 3.74E-11 | 83.9173715 | 48.7144897 | 47.3306478 | 51.9608489 | 313.660808 |
| 2.81E-12 | 3.74E-11 | 168.79931 | 254.916919 | 302.544925 | 322.899561 | 0 |
| 2.82E-12 | 3.75E-11 | 165.584086 | 465.790601 | 424.119726 | 328.002859 | 0 |
| 2.83E-12 | 3.76E-11 | 183.267823 | 323.428621 | 218.0922 | 315.012647 | 0 |
| 2.85E-12 | 3.78E-11 | 168.79931 | 356.79471 | 268.207004 | 308.517541 | 0 |
| 2.85E-12 | 3.78E-11 | 182.624778 | 264.036983 | 296.048562 | 209.235204 | 0 |
| 2.87E-12 | 3.80E-11 | 152.723186 | 535.192065 | 592.097123 | 531.670829 | 0 |

| | | | | | | |
|---|---|---|---|---|---|---|
| 2.87E-12 | 3.81E-11 | 174.908238 | 235.564587 | 219.020252 | 222.225416 | 0 |
| 2.88E-12 | 3.81E-11 | 168.156265 | 242.682686 | 258.926485 | 264.907542 | 0 |
| 2.90E-12 | 3.85E-11 | 199.343948 | 546.758976 | 257.070381 | 404.552324 | 0 |
| 2.91E-12 | 3.85E-11 | 154.330798 | 506.052348 | 490.011412 | 456.977109 | 0 |
| 2.92E-12 | 3.86E-11 | 165.584086 | 338.999463 | 272.847264 | 309.445413 | 0 |
| 2.92E-12 | 3.87E-11 | 161.082771 | 375.034839 | 316.465704 | 361.870198 | 0 |
| 2.94E-12 | 3.89E-11 | 158.510591 | 453.556368 | 511.356606 | 636.984336 | 0 |
| 2.94E-12 | 3.89E-11 | 243.714052 | 319.20225 | 534.557904 | 216.194247 | 0 |
| 2.95E-12 | 3.90E-11 | 159.475158 | 328.767196 | 353.58778 | 340.065199 | 0 |
| 2.95E-12 | 3.90E-11 | 151.115573 | 575.231372 | 547.550631 | 577.600508 | 0 |
| 2.97E-12 | 3.92E-11 | 167.191698 | 296.51331 | 251.502069 | 268.155095 | 0 |
| 2.98E-12 | 3.93E-11 | 175.551283 | 270.26532 | 268.207004 | 214.338502 | 0 |
| 2.98E-12 | 3.93E-11 | 197.09329 | 197.527246 | 324.818171 | 308.517541 | 0 |
| 2.98E-12 | 3.93E-11 | 161.404293 | 302.519206 | 300.688821 | 303.878179 | 0 |
| 3.00E-12 | 3.96E-11 | 163.654951 | 291.174735 | 286.768042 | 335.425837 | 0 |
| 3.00E-12 | 3.96E-11 | 217.027685 | 550.095585 | 414.839207 | 439.347535 | 14.2573095 |
| 3.00E-12 | 3.96E-11 | 157.546023 | 481.139001 | 388.853753 | 429.604876 | 0 |
| 3.01E-12 | 3.96E-11 | 178.123463 | 213.765409 | 213.451941 | 249.133713 | 0 |
| 3.02E-12 | 3.98E-11 | 189.055228 | 189.741825 | 246.86181 | 256.556692 | 0 |
| 3.02E-12 | 3.98E-11 | 187.447615 | 633.288367 | 304.401029 | 526.103596 | 0 |
| 3.05E-12 | 4.01E-11 | 188.733705 | 245.796855 | 199.531162 | 201.812226 | 0 |
| 3.05E-12 | 4.02E-11 | 177.158895 | 300.51724 | 217.164149 | 270.474776 | 0 |
| 3.06E-12 | 4.03E-11 | 167.191698 | 341.223869 | 264.494796 | 289.032222 | 0 |
| 3.07E-12 | 4.03E-11 | 185.840003 | 279.162944 | 208.811681 | 217.122119 | 0 |
| 3.08E-12 | 4.05E-11 | 570.702431 | 535.636947 | 274.703367 | 399.912962 | 4.75243649 |
| 3.10E-12 | 4.07E-11 | 163.654951 | 279.385384 | 262.638692 | 280.217435 | 0 |
| 3.11E-12 | 4.08E-11 | 186.161525 | 211.541003 | 193.034799 | 239.85499 | 0 |
| 3.13E-12 | 4.11E-11 | 153.044708 | 504.050382 | 452.889335 | 474.142747 | 0 |
| 3.14E-12 | 4.12E-11 | 164.941041 | 328.767196 | 263.566744 | 318.2602 | 0 |
| 3.17E-12 | 4.16E-11 | 287.119589 | 829.480969 | 655.204653 | 746.937204 | 2.37621824 |
| 3.18E-12 | 4.17E-11 | 208.346577 | 430.200106 | 351.731677 | 431.924557 | 7.12865473 |
| 3.19E-12 | 4.18E-11 | 203.202218 | 176.840271 | 248.717914 | 249.597649 | 0 |
| 3.20E-12 | 4.19E-11 | 151.437096 | 496.932284 | 596.737383 | 591.518593 | 0 |
| 3.20E-12 | 4.19E-11 | 176.837373 | 421.524923 | 258.926485 | 327.538923 | 0 |
| 3.21E-12 | 4.20E-11 | 163.654951 | 416.85367 | 306.257133 | 342.38488 | 0 |
| 3.24E-12 | 4.23E-11 | 187.447615 | 276.271216 | 192.106747 | 254.237011 | 0 |
| 3.24E-12 | 4.24E-11 | 160.118203 | 687.11899 | 443.608816 | 518.216681 | 0 |
| 3.25E-12 | 4.25E-11 | 160.761248 | 297.180631 | 287.696094 | 325.219242 | 0 |
| 3.25E-12 | 4.25E-11 | 160.761248 | 314.308557 | 294.192458 | 284.392861 | 0 |
| 3.25E-12 | 4.25E-11 | 156.902978 | 388.158834 | 340.595054 | 375.788283 | 0 |
| 3.26E-12 | 4.26E-11 | 184.23239 | 336.775057 | 213.451941 | 322.435625 | 0 |
| 3.30E-12 | 4.30E-11 | 321.844019 | 204.867786 | 224.588564 | 199.028609 | 0 |
| 3.31E-12 | 4.32E-11 | 160.439726 | 295.623547 | 287.696094 | 282.07318 | 0 |

| | | | | | | |
|---|---|---|---|---|---|---|
| 3.32E-12 | 4.33E-11 | 233.425332 | 1465.21618 | 1551.70281 | 1524.03026 | 0 |
| 3.33E-12 | 4.34E-11 | 328.595991 | 395.944255 | 317.393756 | 414.758919 | 16.6335277 |
| 3.33E-12 | 4.34E-11 | 201.273083 | 424.861532 | 216.236097 | 405.01626 | 0 |
| 3.36E-12 | 4.37E-11 | 141.791421 | 875.081291 | 757.290364 | 913.026346 | 0 |
| 3.36E-12 | 4.38E-11 | 168.477788 | 355.460067 | 278.415575 | 423.10977 | 0 |
| 3.37E-12 | 4.38E-11 | 187.447615 | 199.974093 | 198.60311 | 191.60563 | 0 |
| 3.37E-12 | 4.38E-11 | 173.94367 | 603.926208 | 336.882846 | 416.150728 | 0 |
| 3.37E-12 | 4.38E-11 | 162.047338 | 417.298551 | 311.825444 | 338.209454 | 0 |
| 3.38E-12 | 4.39E-11 | 199.986993 | 260.033053 | 228.300772 | 280.217435 | 7.12865473 |
| 3.38E-12 | 4.39E-11 | 197.414813 | 254.472038 | 179.11402 | 228.256586 | 0 |
| 3.38E-12 | 4.40E-11 | 228.924017 | 257.141325 | 224.588564 | 169.336695 | 0 |
| 3.39E-12 | 4.40E-11 | 153.044708 | 442.656779 | 405.558688 | 458.368917 | 0 |
| 3.40E-12 | 4.41E-11 | 137.611628 | 992.529923 | 1294.63242 | 968.234748 | 0 |
| 3.40E-12 | 4.41E-11 | 147.257303 | 686.451668 | 589.312967 | 612.39572 | 0 |
| 3.41E-12 | 4.42E-11 | 148.543393 | 530.743254 | 602.305694 | 610.539975 | 0 |
| 3.41E-12 | 4.42E-11 | 155.938411 | 539.640877 | 537.34206 | 403.160515 | 0 |
| 3.42E-12 | 4.43E-11 | 154.652321 | 437.095764 | 379.573234 | 388.314559 | 0 |
| 3.42E-12 | 4.43E-11 | 250.787547 | 233.562622 | 167.977397 | 220.833608 | 0 |
| 3.42E-12 | 4.43E-11 | 235.997512 | 806.347148 | 389.781805 | 585.487423 | 11.8810912 |
| 3.43E-12 | 4.44E-11 | 165.905608 | 252.914954 | 233.869083 | 270.01084 | 0 |
| 3.44E-12 | 4.45E-11 | 207.38201 | 290.284973 | 297.904665 | 587.343168 | 0 |
| 3.45E-12 | 4.46E-11 | 176.837373 | 515.394853 | 285.83999 | 392.026048 | 0 |
| 3.46E-12 | 4.48E-11 | 178.444985 | 403.729675 | 662.629069 | 353.519347 | 0 |
| 3.47E-12 | 4.48E-11 | 188.09066 | 462.00911 | 242.22155 | 351.663603 | 0 |
| 3.48E-12 | 4.49E-11 | 596.42423 | 585.90852 | 358.22804 | 166.553078 | 0 |
| 3.49E-12 | 4.51E-11 | 268.792807 | 766.307841 | 222.73246 | 689.409121 | 0 |
| 3.54E-12 | 4.57E-11 | 183.910868 | 247.131498 | 199.531162 | 200.420417 | 0 |
| 3.54E-12 | 4.57E-11 | 213.169415 | 537.416471 | 407.414792 | 475.534555 | 7.12865473 |
| 3.56E-12 | 4.59E-11 | 156.259933 | 420.63516 | 353.58778 | 351.199667 | 0 |
| 3.57E-12 | 4.60E-11 | 153.366231 | 564.999105 | 425.97583 | 542.805297 | 0 |
| 3.61E-12 | 4.65E-11 | 150.151006 | 447.105591 | 491.867516 | 464.864024 | 0 |
| 3.63E-12 | 4.67E-11 | 169.763878 | 333.216007 | 235.725187 | 320.579881 | 0 |
| 3.63E-12 | 4.67E-11 | 161.082771 | 268.485795 | 257.070381 | 286.712542 | 0 |
| 3.65E-12 | 4.69E-11 | 166.22713 | 305.633374 | 235.725187 | 277.433818 | 0 |
| 3.68E-12 | 4.73E-11 | 245.321665 | 422.192244 | 383.285442 | 417.0786 | 2.37621824 |
| 3.68E-12 | 4.74E-11 | 165.262563 | 262.257458 | 228.300772 | 271.402649 | 0 |
| 3.68E-12 | 4.74E-11 | 167.191698 | 312.751473 | 234.797135 | 275.114138 | 0 |
| 3.71E-12 | 4.77E-11 | 211.561802 | 226.222083 | 171.689605 | 204.595843 | 0 |
| 3.72E-12 | 4.78E-11 | 162.368861 | 279.162944 | 382.35739 | 319.188072 | 0 |
| 3.73E-12 | 4.79E-11 | 158.189068 | 464.900838 | 338.73895 | 381.819453 | 0 |
| 3.73E-12 | 4.79E-11 | 318.628794 | 624.835624 | 336.882846 | 437.491791 | 4.75243649 |
| 3.76E-12 | 4.83E-11 | 177.80194 | 405.286759 | 388.853753 | 410.583494 | 4.75243649 |
| 3.76E-12 | 4.83E-11 | 146.935781 | 707.361084 | 553.118943 | 610.076039 | 0 |

| | | | | | | |
|---|---|---|---|---|---|---|
| 3.79E-12 | 4.87E-11 | 207.060488 | 431.75719 | 352.659728 | 418.934345 | 30.8908372 |
| 3.81E-12 | 4.88E-11 | 174.586715 | 216.434696 | 280.271679 | 218.977863 | 0 |
| 3.82E-12 | 4.89E-11 | 153.687753 | 474.465784 | 370.292715 | 453.26562 | 0 |
| 3.83E-12 | 4.90E-11 | 170.728445 | 273.379488 | 346.163365 | 450.018067 | 0 |
| 3.84E-12 | 4.92E-11 | 159.796681 | 260.033053 | 263.566744 | 287.640414 | 0 |
| 3.85E-12 | 4.93E-11 | 153.366231 | 348.786849 | 343.379209 | 338.67339 | 0 |
| 3.85E-12 | 4.93E-11 | 153.366231 | 402.839913 | 401.84648 | 345.168497 | 0 |
| 3.85E-12 | 4.93E-11 | 152.401663 | 500.713774 | 399.062324 | 424.965515 | 0 |
| 3.85E-12 | 4.93E-11 | 154.652321 | 348.341968 | 345.235313 | 411.04743 | 0 |
| 3.86E-12 | 4.93E-11 | 148.543393 | 555.211719 | 463.097907 | 524.711787 | 0 |
| 3.86E-12 | 4.94E-11 | 152.723186 | 368.13918 | 357.299988 | 348.41605 | 0 |
| 3.87E-12 | 4.94E-11 | 241.463395 | 423.971769 | 420.407518 | 420.326153 | 2.37621824 |
| 3.87E-12 | 4.94E-11 | 303.838759 | 507.386991 | 455.673491 | 516.824873 | 0 |
| 3.89E-12 | 4.96E-11 | 179.409553 | 282.499552 | 203.24337 | 224.545097 | 0 |
| 3.89E-12 | 4.96E-11 | 157.546023 | 272.934607 | 289.552198 | 300.630626 | 0 |
| 3.91E-12 | 4.98E-11 | 150.794051 | 453.111487 | 391.637909 | 428.677004 | 0 |
| 3.96E-12 | 5.05E-11 | 166.22713 | 262.035018 | 219.948304 | 276.505946 | 0 |
| 4.00E-12 | 5.10E-11 | 305.767894 | 665.097371 | 469.59427 | 557.187318 | 2.37621824 |
| 4.01E-12 | 5.11E-11 | 173.622148 | 251.58031 | 215.308045 | 309.909349 | 0 |
| 4.05E-12 | 5.15E-11 | 150.472528 | 417.965873 | 389.781805 | 460.224662 | 0 |
| 4.05E-12 | 5.15E-11 | 157.546023 | 286.725924 | 267.278952 | 271.866585 | 0 |
| 4.09E-12 | 5.20E-11 | 252.395159 | 159.267464 | 278.415575 | 257.9485 | 0 |
| 4.10E-12 | 5.21E-11 | 164.297996 | 413.961943 | 713.671924 | 451.873811 | 0 |
| 4.11E-12 | 5.22E-11 | 164.297996 | 218.659102 | 254.286225 | 255.164883 | 0 |
| 4.11E-12 | 5.23E-11 | 168.477788 | 309.859745 | 223.660512 | 262.123925 | 0 |
| 4.11E-12 | 5.23E-11 | 305.767894 | 903.553686 | 751.722053 | 850.394965 | 14.2573095 |
| 4.12E-12 | 5.23E-11 | 149.829483 | 379.928532 | 413.911155 | 409.191685 | 0 |
| 4.12E-12 | 5.23E-11 | 150.472528 | 417.298551 | 556.83115 | 508.010086 | 0 |
| 4.13E-12 | 5.24E-11 | 161.404293 | 263.147221 | 232.012979 | 263.051798 | 0 |
| 4.17E-12 | 5.29E-11 | 157.867546 | 370.808468 | 283.055835 | 327.538923 | 0 |
| 4.20E-12 | 5.33E-11 | 363.320421 | 333.216007 | 392.565961 | 379.499772 | 0 |
| 4.22E-12 | 5.34E-11 | 160.761248 | 309.414864 | 245.933758 | 275.578074 | 0 |
| 4.22E-12 | 5.35E-11 | 1517.9077 | 2220.84687 | 2135.44746 | 2355.86778 | 23.7621824 |
| 4.22E-12 | 5.35E-11 | 153.044708 | 334.773092 | 309.96934 | 321.507753 | 0 |
| 4.22E-12 | 5.35E-11 | 156.581456 | 278.495622 | 325.746223 | 284.392861 | 0 |
| 4.23E-12 | 5.35E-11 | 147.900348 | 416.408789 | 434.328297 | 451.873811 | 0 |
| 4.23E-12 | 5.36E-11 | 152.401663 | 318.090047 | 340.595054 | 329.858604 | 0 |
| 4.26E-12 | 5.39E-11 | 287.441112 | 554.544397 | 567.039721 | 572.033275 | 0 |
| 4.28E-12 | 5.41E-11 | 153.366231 | 314.975879 | 299.760769 | 322.899561 | 0 |
| 4.28E-12 | 5.41E-11 | 158.189068 | 339.666785 | 268.207004 | 341.457007 | 0 |
| 4.29E-12 | 5.42E-11 | 151.437096 | 416.85367 | 345.235313 | 401.768707 | 0 |
| 4.33E-12 | 5.47E-11 | 174.586715 | 326.320349 | 213.451941 | 264.443606 | 0 |
| 4.34E-12 | 5.48E-11 | 280.689139 | 611.93407 | 578.176344 | 616.571145 | 0 |

| | | | | | | |
|---|---|---|---|---|---|---|
| 4.35E-12 | 5.49E-11 | 529.547551 | 217.76934 | 253.358173 | 260.268181 | 0 |
| 4.36E-12 | 5.49E-11 | 173.300625 | 347.897087 | 245.933758 | 425.429451 | 0 |
| 4.36E-12 | 5.49E-11 | 153.687753 | 298.070394 | 306.257133 | 290.887967 | 0 |
| 4.36E-12 | 5.49E-11 | 161.404293 | 325.875468 | 286.768042 | 322.899561 | 0 |
| 4.36E-12 | 5.50E-11 | 295.479174 | 665.764693 | 486.299204 | 640.231889 | 2.37621824 |
| 4.38E-12 | 5.51E-11 | 148.543393 | 399.948185 | 441.752712 | 484.349342 | 0 |
| 4.38E-12 | 5.52E-11 | 168.156265 | 221.105949 | 206.955577 | 253.309139 | 0 |
| 4.40E-12 | 5.54E-11 | 157.867546 | 256.251562 | 250.574018 | 279.289563 | 0 |
| 4.40E-12 | 5.54E-11 | 175.551283 | 315.198319 | 210.667785 | 328.466795 | 0 |
| 4.41E-12 | 5.54E-11 | 263.969969 | 185.293013 | 181.898176 | 269.546904 | 0 |
| 4.41E-12 | 5.54E-11 | 169.442355 | 709.363049 | 351.731677 | 567.393913 | 0 |
| 4.43E-12 | 5.56E-11 | 166.548653 | 338.109701 | 315.537652 | 336.817646 | 11.8810912 |
| 4.43E-12 | 5.56E-11 | 147.900348 | 449.997318 | 538.270112 | 436.099982 | 0 |
| 4.43E-12 | 5.56E-11 | 148.864916 | 416.63123 | 398.134272 | 371.148921 | 0 |
| 4.43E-12 | 5.57E-11 | 163.333428 | 251.58031 | 221.804408 | 278.361691 | 0 |
| 4.44E-12 | 5.57E-11 | 154.973843 | 285.836161 | 348.947521 | 353.983283 | 0 |
| 4.47E-12 | 5.61E-11 | 145.328168 | 506.71967 | 458.457647 | 475.070619 | 0 |
| 4.48E-12 | 5.62E-11 | 172.979103 | 259.810612 | 201.387266 | 213.41063 | 0 |
| 4.49E-12 | 5.62E-11 | 152.080141 | 478.247274 | 368.436611 | 381.355516 | 0 |
| 4.49E-12 | 5.63E-11 | 152.723186 | 366.804537 | 305.329081 | 350.73573 | 0 |
| 4.49E-12 | 5.63E-11 | 174.908238 | 283.834196 | 228.300772 | 202.276162 | 0 |
| 4.51E-12 | 5.64E-11 | 151.115573 | 348.786849 | 332.242586 | 389.706367 | 0 |
| 4.51E-12 | 5.64E-11 | 158.832113 | 542.532605 | 340.595054 | 514.505192 | 0 |
| 4.52E-12 | 5.65E-11 | 151.115573 | 383.932463 | 325.746223 | 360.014453 | 0 |
| 4.52E-12 | 5.65E-11 | 151.437096 | 345.672681 | 311.825444 | 340.529135 | 0 |
| 4.56E-12 | 5.70E-11 | 135.682493 | 731.162227 | 820.397895 | 843.435923 | 0 |
| 4.57E-12 | 5.72E-11 | 195.485678 | 449.997318 | 412.055051 | 424.501578 | 9.50487297 |
| 4.62E-12 | 5.77E-11 | 158.832113 | 243.127568 | 235.725187 | 237.999246 | 0 |
| 4.63E-12 | 5.78E-11 | 161.082771 | 397.278898 | 345.235313 | 267.227223 | 0 |
| 4.63E-12 | 5.78E-11 | 160.439726 | 344.560478 | 250.574018 | 297.847009 | 0 |
| 4.65E-12 | 5.80E-11 | 152.723186 | 476.91263 | 417.623363 | 342.848816 | 0 |
| 4.65E-12 | 5.80E-11 | 138.897718 | 776.317668 | 634.787511 | 694.048482 | 0 |
| 4.66E-12 | 5.82E-11 | 189.698273 | 269.820439 | 183.75428 | 309.445413 | 0 |
| 4.67E-12 | 5.82E-11 | 140.183808 | 627.282471 | 579.104396 | 617.962954 | 0 |
| 4.67E-12 | 5.82E-11 | 147.900348 | 452.444165 | 382.35739 | 407.799877 | 0 |
| 4.67E-12 | 5.82E-11 | 155.295366 | 251.135429 | 290.48025 | 263.97967 | 21.3859642 |
| 4.68E-12 | 5.83E-11 | 146.935781 | 385.934428 | 419.479466 | 425.893387 | 0 |
| 4.69E-12 | 5.83E-11 | 0 | 0 | 0 | 0 | 85.5438568 |
| 4.69E-12 | 5.83E-11 | 146.614258 | 551.430229 | 428.759986 | 504.762533 | 0 |
| 4.70E-12 | 5.84E-11 | 154.973843 | 330.54672 | 270.99116 | 296.455201 | 0 |
| 4.73E-12 | 5.88E-11 | 235.354467 | 546.536536 | 448.249076 | 514.041256 | 0 |
| 4.74E-12 | 5.90E-11 | 170.728445 | 348.786849 | 220.876356 | 301.094562 | 0 |
| 4.75E-12 | 5.90E-11 | 156.259933 | 248.021261 | 270.063108 | 289.496158 | 0 |

| | | | | | | |
|---|---|---|---|---|---|---|
| 4.75E-12 | 5.90E-11 | 154.009276 | 471.129175 | 328.530379 | 426.821259 | 0 |
| 4.77E-12 | 5.92E-11 | 160.439726 | 502.938179 | 313.681548 | 379.963708 | 0 |
| 4.77E-12 | 5.92E-11 | 306.410939 | 529.631051 | 404.630636 | 508.010086 | 2.37621824 |
| 4.81E-12 | 5.96E-11 | 144.042078 | 446.438269 | 497.435827 | 491.77232 | 0 |
| 4.82E-12 | 5.98E-11 | 243.714052 | 573.006966 | 535.485956 | 581.775934 | 4.75243649 |
| 4.83E-12 | 5.99E-11 | 205.452875 | 290.062533 | 180.042072 | 206.915523 | 0 |
| 4.84E-12 | 6.00E-11 | 158.510591 | 310.527067 | 308.113236 | 242.174671 | 0 |
| 4.85E-12 | 6.01E-11 | 222.493567 | 306.078255 | 167.049345 | 235.215629 | 0 |
| 4.88E-12 | 6.04E-11 | 148.864916 | 373.032874 | 330.386482 | 359.550517 | 0 |
| 4.90E-12 | 6.07E-11 | 170.0854 | 274.26925 | 452.889335 | 340.065199 | 7.12865473 |
| 4.91E-12 | 6.07E-11 | 183.589345 | 637.514738 | 300.688821 | 375.788283 | 0 |
| 4.91E-12 | 6.08E-11 | 0 | 0 | 0 | 0 | 30.8908372 |
| 4.93E-12 | 6.09E-11 | 187.447615 | 328.989636 | 190.250643 | 263.97967 | 0 |
| 4.95E-12 | 6.11E-11 | 137.933151 | 679.77845 | 870.512698 | 682.914015 | 0 |
| 4.95E-12 | 6.11E-11 | 183.589345 | 189.741825 | 194.890903 | 255.628819 | 0 |
| 4.96E-12 | 6.13E-11 | 284.547409 | 647.524565 | 533.629852 | 564.14636 | 0 |
| 4.96E-12 | 6.13E-11 | 150.794051 | 351.901017 | 320.177911 | 301.558498 | 0 |
| 4.98E-12 | 6.14E-11 | 153.366231 | 141.472217 | 133.639476 | 151.707121 | 2.37621824 |
| 4.99E-12 | 6.15E-11 | 145.971213 | 437.763086 | 508.57245 | 411.511366 | 0 |
| 4.99E-12 | 6.15E-11 | 147.578826 | 459.562264 | 369.364663 | 413.831047 | 0 |
| 5.00E-12 | 6.16E-11 | 222.172045 | 337.88726 | 170.761553 | 361.870198 | 0 |
| 5.00E-12 | 6.16E-11 | 167.191698 | 272.044845 | 223.660512 | 208.307332 | 0 |
| 5.02E-12 | 6.18E-11 | 162.690383 | 207.314632 | 246.86181 | 227.328714 | 0 |
| 5.04E-12 | 6.20E-11 | 155.295366 | 249.355904 | 245.005706 | 260.268181 | 0 |
| 5.05E-12 | 6.21E-11 | 161.404293 | 213.765409 | 218.0922 | 237.999246 | 0 |
| 5.06E-12 | 6.22E-11 | 153.687753 | 256.474003 | 262.638692 | 270.938712 | 0 |
| 5.06E-12 | 6.22E-11 | 183.589345 | 328.544755 | 192.106747 | 278.825627 | 0 |
| 5.07E-12 | 6.23E-11 | 143.720556 | 427.753259 | 445.46492 | 451.873811 | 0 |
| 5.12E-12 | 6.30E-11 | 158.510591 | 521.623189 | 326.674275 | 378.107963 | 0 |
| 5.13E-12 | 6.30E-11 | 143.720556 | 450.66464 | 420.407518 | 441.667216 | 0 |
| 5.13E-12 | 6.30E-11 | 160.439726 | 230.893335 | 214.379993 | 251.453394 | 0 |
| 5.13E-12 | 6.30E-11 | 201.916128 | 351.901017 | 206.955577 | 211.090949 | 0 |
| 5.17E-12 | 6.34E-11 | 145.971213 | 364.802572 | 394.422065 | 411.511366 | 0 |
| 5.17E-12 | 6.34E-11 | 157.867546 | 404.396997 | 272.847264 | 346.096369 | 0 |
| 5.17E-12 | 6.35E-11 | 154.652321 | 283.166874 | 396.278169 | 350.271794 | 0 |
| 5.18E-12 | 6.35E-11 | 149.186438 | 454.891012 | 398.134272 | 344.240624 | 0 |
| 5.18E-12 | 6.35E-11 | 159.153636 | 268.708236 | 219.948304 | 252.381266 | 0 |
| 5.20E-12 | 6.37E-11 | 288.405679 | 166.163123 | 229.228823 | 214.802438 | 0 |
| 5.20E-12 | 6.37E-11 | 290.334814 | 520.288546 | 450.10518 | 486.205087 | 2.37621824 |
| 5.21E-12 | 6.37E-11 | 145.006646 | 493.595675 | 593.025175 | 442.131152 | 0 |
| 5.21E-12 | 6.38E-11 | 193.23502 | 169.499731 | 193.962851 | 214.802438 | 0 |
| 5.24E-12 | 6.41E-11 | 153.044708 | 487.144897 | 327.602327 | 427.285195 | 0 |
| 5.26E-12 | 6.43E-11 | 156.902978 | 250.912988 | 246.86181 | 302.950307 | 0 |

| | | | | | | |
|---|---|---|---|---|---|---|
| 5.26E-12 | 6.43E-11 | 179.731075 | 259.365731 | 181.898176 | 215.73031 | 0 |
| 5.26E-12 | 6.43E-11 | 165.905608 | 288.060567 | 236.653239 | 210.627013 | 0 |
| 5.27E-12 | 6.44E-11 | 193.23502 | 245.796855 | 179.11402 | 188.358077 | 0 |
| 5.27E-12 | 6.44E-11 | 141.469898 | 661.983203 | 531.773748 | 528.423276 | 0 |
| 5.29E-12 | 6.45E-11 | 141.791421 | 456.448096 | 482.586997 | 476.462427 | 0 |
| 5.29E-12 | 6.46E-11 | 137.611628 | 663.540287 | 721.096339 | 585.023487 | 0 |
| 5.31E-12 | 6.48E-11 | 151.758618 | 342.113631 | 276.559471 | 322.435625 | 0 |
| 5.31E-12 | 6.48E-11 | 168.477788 | 276.493656 | 198.60311 | 275.578074 | 0 |
| 5.32E-12 | 6.48E-11 | 145.649691 | 382.375378 | 360.084144 | 367.437432 | 0 |
| 5.32E-12 | 6.48E-11 | 154.652321 | 252.025191 | 322.034015 | 304.806051 | 0 |
| 5.37E-12 | 6.54E-11 | 141.148376 | 530.965694 | 465.882062 | 512.185511 | 0 |
| 5.39E-12 | 6.57E-11 | 299.015921 | 440.654814 | 401.84648 | 436.563918 | 0 |
| 5.43E-12 | 6.61E-11 | 273.937167 | 275.603894 | 288.624146 | 273.258393 | 2.37621824 |
| 5.45E-12 | 6.63E-11 | 146.614258 | 480.026798 | 424.119726 | 367.437432 | 0 |
| 5.46E-12 | 6.64E-11 | 147.900348 | 446.215828 | 368.436611 | 351.199667 | 0 |
| 5.47E-12 | 6.65E-11 | 134.396403 | 712.032336 | 893.713996 | 1071.22857 | 0 |
| 5.47E-12 | 6.65E-11 | 238.891215 | 513.170447 | 203.24337 | 277.897755 | 0 |
| 5.47E-12 | 6.65E-11 | 268.149762 | 263.592102 | 150.344411 | 276.969882 | 0 |
| 5.51E-12 | 6.69E-11 | 225.38727 | 225.777201 | 158.696878 | 207.37946 | 0 |
| 5.52E-12 | 6.70E-11 | 158.832113 | 278.050741 | 219.020252 | 274.650202 | 0 |
| 5.52E-12 | 6.70E-11 | 141.791421 | 434.648918 | 446.392972 | 451.409875 | 0 |
| 5.52E-12 | 6.70E-11 | 193.878065 | 367.694299 | 196.747006 | 250.061586 | 0 |
| 5.53E-12 | 6.71E-11 | 143.720556 | 593.471501 | 436.184401 | 529.351149 | 0 |
| 5.54E-12 | 6.71E-11 | 220.885955 | 435.53868 | 180.970124 | 372.54073 | 0 |
| 5.54E-12 | 6.71E-11 | 156.581456 | 440.654814 | 316.465704 | 303.414243 | 0 |
| 5.55E-12 | 6.72E-11 | 151.437096 | 268.708236 | 330.386482 | 300.16669 | 0 |
| 5.55E-12 | 6.72E-11 | 137.933151 | 169.944613 | 196.747006 | 188.358077 | 11.8810912 |
| 5.56E-12 | 6.73E-11 | 157.867546 | 243.572449 | 243.149602 | 307.125732 | 0 |
| 5.56E-12 | 6.73E-11 | 193.556543 | 266.038949 | 173.545708 | 207.37946 | 0 |
| 5.57E-12 | 6.74E-11 | 184.553913 | 400.393066 | 210.667785 | 286.712542 | 0 |
| 5.58E-12 | 6.75E-11 | 146.292736 | 373.255314 | 329.458431 | 378.571899 | 0 |
| 5.59E-12 | 6.75E-11 | 275.866302 | 580.569946 | 521.565177 | 592.446465 | 2.37621824 |
| 5.59E-12 | 6.75E-11 | 2.57217997 | 2.22440592 | 0.92805192 | 2.31968076 | 38.0194919 |
| 5.60E-12 | 6.77E-11 | 169.442355 | 260.477934 | 189.322591 | 234.287756 | 0 |
| 5.61E-12 | 6.77E-11 | 143.077511 | 454.001249 | 400.918428 | 414.758919 | 0 |
| 5.61E-12 | 6.78E-11 | 160.761248 | 245.129533 | 207.883629 | 218.049991 | 0 |
| 5.63E-12 | 6.79E-11 | 149.186438 | 328.989636 | 387.925701 | 308.053604 | 0 |
| 5.63E-12 | 6.79E-11 | 150.472528 | 732.496871 | 444.536868 | 510.329766 | 0 |
| 5.63E-12 | 6.79E-11 | 163.976473 | 226.222083 | 193.962851 | 238.463182 | 0 |
| 5.63E-12 | 6.79E-11 | 160.761248 | 286.725924 | 215.308045 | 285.784669 | 0 |
| 5.64E-12 | 6.80E-11 | 151.437096 | 708.473287 | 421.33557 | 511.721575 | 0 |
| 5.68E-12 | 6.84E-11 | 151.758618 | 292.064498 | 252.430121 | 291.815839 | 0 |
| 5.69E-12 | 6.85E-11 | 151.115573 | 677.998926 | 407.414792 | 506.618277 | 0 |

| | | | | | | |
|---|---|---|---|---|---|---|
| 5.72E-12 | 6.89E-11 | 145.971213 | 510.7236 | 384.213494 | 529.351149 | 0 |
| 5.73E-12 | 6.89E-11 | 152.401663 | 315.42076 | 254.286225 | 287.640414 | 0 |
| 5.73E-12 | 6.90E-11 | 147.900348 | 314.975879 | 292.336354 | 302.950307 | 0 |
| 5.74E-12 | 6.91E-11 | 152.401663 | 249.133464 | 245.005706 | 256.092756 | 0 |
| 5.75E-12 | 6.91E-11 | 151.115573 | 305.410933 | 413.911155 | 329.394667 | 0 |
| 5.76E-12 | 6.92E-11 | 154.009276 | 312.529032 | 251.502069 | 333.570093 | 0 |
| 5.79E-12 | 6.95E-11 | 142.434466 | 399.725745 | 451.961284 | 475.070619 | 0 |
| 5.79E-12 | 6.96E-11 | 146.614258 | 392.607646 | 324.818171 | 349.343922 | 0 |
| 5.80E-12 | 6.97E-11 | 144.363601 | 449.552437 | 386.997649 | 374.86041 | 0 |
| 5.81E-12 | 6.97E-11 | 160.761248 | 392.162764 | 264.494796 | 278.361691 | 0 |
| 5.82E-12 | 6.98E-11 | 142.434466 | 393.497408 | 427.831934 | 401.768707 | 0 |
| 5.84E-12 | 7.00E-11 | 165.262563 | 470.906734 | 262.638692 | 403.624452 | 0 |
| 5.86E-12 | 7.02E-11 | 175.22976 | 302.519206 | 197.675058 | 330.786476 | 0 |
| 5.88E-12 | 7.04E-11 | 149.186438 | 435.53868 | 316.465704 | 375.324346 | 0 |
| 5.88E-12 | 7.04E-11 | 158.189068 | 450.887081 | 277.487523 | 378.107963 | 0 |
| 5.89E-12 | 7.05E-11 | 161.082771 | 197.304805 | 214.379993 | 221.76148 | 0 |
| 5.89E-12 | 7.05E-11 | 142.755988 | 403.062353 | 376.789078 | 418.934345 | 0 |
| 5.91E-12 | 7.07E-11 | 138.897718 | 552.542432 | 490.939464 | 492.236257 | 0 |
| 5.92E-12 | 7.08E-11 | 217.349207 | 388.381274 | 225.516616 | 338.209454 | 9.50487297 |
| 5.93E-12 | 7.09E-11 | 149.186438 | 266.70627 | 293.264406 | 279.289563 | 0 |
| 5.93E-12 | 7.09E-11 | 140.183808 | 520.955867 | 450.10518 | 467.183704 | 0 |
| 5.94E-12 | 7.09E-11 | 183.910868 | 171.056816 | 250.574018 | 256.556692 | 0 |
| 5.97E-12 | 7.13E-11 | 140.183808 | 454.891012 | 440.824661 | 439.811471 | 0 |
| 5.97E-12 | 7.13E-11 | 172.65758 | 545.201892 | 442.680764 | 506.154341 | 4.75243649 |
| 6.01E-12 | 7.17E-11 | 142.112943 | 391.050562 | 423.191674 | 389.242431 | 0 |
| 6.02E-12 | 7.18E-11 | 281.010662 | 543.644808 | 468.666218 | 502.906788 | 0 |
| 6.03E-12 | 7.20E-11 | 260.433222 | 429.087903 | 312.753496 | 393.417856 | 4.75243649 |
| 6.06E-12 | 7.22E-11 | 138.576196 | 540.085758 | 481.658945 | 590.590721 | 0 |
| 6.09E-12 | 7.26E-11 | 138.576196 | 519.843665 | 467.738166 | 491.308384 | 0 |
| 6.10E-12 | 7.26E-11 | 157.224501 | 329.212077 | 233.869083 | 281.609244 | 0 |
| 6.09E-12 | 7.26E-11 | 150.794051 | 243.794889 | 263.566744 | 262.123925 | 0 |
| 6.11E-12 | 7.28E-11 | 158.832113 | 281.832231 | 223.660512 | 225.936906 | 0 |
| 6.11E-12 | 7.28E-11 | 138.897718 | 512.280684 | 453.817387 | 497.80349 | 0 |
| 6.14E-12 | 7.31E-11 | 148.864916 | 324.095943 | 274.703367 | 340.529135 | 0 |
| 6.15E-12 | 7.31E-11 | 163.333428 | 242.460246 | 191.178695 | 226.864778 | 0 |
| 6.15E-12 | 7.31E-11 | 206.417443 | 174.615865 | 262.638692 | 350.73573 | 0 |
| 6.17E-12 | 7.33E-11 | 0 | 0 | 0 | 0 | 123.563349 |
| 6.17E-12 | 7.33E-11 | 136.968583 | 512.503125 | 517.85297 | 519.144553 | 0 |
| 6.18E-12 | 7.33E-11 | 148.543393 | 280.052706 | 350.803625 | 315.012647 | 0 |
| 6.18E-12 | 7.34E-11 | 194.842633 | 179.064677 | 278.415575 | 196.244992 | 0 |
| 6.20E-12 | 7.35E-11 | 193.23502 | 460.452026 | 450.10518 | 427.749132 | 9.50487297 |
| 6.20E-12 | 7.35E-11 | 372.644573 | 1063.71091 | 900.210359 | 1027.61858 | 16.6335277 |
| 6.22E-12 | 7.38E-11 | 144.363601 | 356.57227 | 323.890119 | 335.425837 | 0 |

| | | | | | | |
|---|---|---|---|---|---|---|
| 6.25E-12 | 7.40E-11 | 143.399033 | 336.330176 | 355.443884 | 352.591475 | 0 |
| 6.25E-12 | 7.41E-11 | 153.687753 | 305.855815 | 238.509343 | 311.765094 | 0 |
| 6.26E-12 | 7.42E-11 | 170.406923 | 224.220117 | 199.531162 | 179.543291 | 0 |
| 6.28E-12 | 7.44E-11 | 152.723186 | 326.54279 | 246.86181 | 289.496158 | 0 |
| 6.29E-12 | 7.45E-11 | 157.224501 | 268.708236 | 234.797135 | 213.874566 | 0 |
| 6.29E-12 | 7.45E-11 | 193.23502 | 320.092013 | 176.329864 | 322.899561 | 0 |
| 6.30E-12 | 7.45E-11 | 148.543393 | 356.79471 | 279.343627 | 337.745518 | 0 |
| 6.30E-12 | 7.46E-11 | 130.859656 | 812.130603 | 669.125432 | 752.040501 | 0 |
| 6.34E-12 | 7.49E-11 | 134.717926 | 922.461137 | 622.722836 | 752.040501 | 0 |
| 6.34E-12 | 7.49E-11 | 181.338688 | 273.379488 | 176.329864 | 221.76148 | 0 |
| 6.34E-12 | 7.49E-11 | 156.259933 | 273.824369 | 220.876356 | 232.432012 | 0 |
| 6.36E-12 | 7.51E-11 | 150.794051 | 279.830265 | 271.919212 | 346.560305 | 0 |
| 6.36E-12 | 7.51E-11 | 140.826853 | 393.274967 | 402.774532 | 439.811471 | 0 |
| 6.36E-12 | 7.51E-11 | 156.902978 | 259.588171 | 212.523889 | 227.79265 | 0 |
| 6.37E-12 | 7.52E-11 | 215.420072 | 170.834375 | 203.24337 | 175.367865 | 0 |
| 6.38E-12 | 7.52E-11 | 140.505331 | 508.721635 | 433.400245 | 422.645834 | 0 |
| 6.38E-12 | 7.52E-11 | 150.794051 | 288.505448 | 245.005706 | 258.876372 | 0 |
| 6.40E-12 | 7.55E-11 | 221.529 | 281.832231 | 157.768826 | 223.617225 | 0 |
| 6.44E-12 | 7.59E-11 | 151.115573 | 261.812577 | 247.789862 | 302.950307 | 0 |
| 6.45E-12 | 7.59E-11 | 141.469898 | 376.814364 | 372.148819 | 410.119558 | 0 |
| 6.46E-12 | 7.61E-11 | 154.009276 | 557.881006 | 331.314534 | 412.439239 | 0 |
| 6.50E-12 | 7.65E-11 | 149.507961 | 488.924422 | 322.962067 | 400.376899 | 0 |
| 6.50E-12 | 7.65E-11 | 154.973843 | 256.474003 | 214.379993 | 234.287756 | 0 |
| 6.51E-12 | 7.66E-11 | 270.078897 | 560.105412 | 548.478683 | 523.783915 | 0 |
| 6.51E-12 | 7.66E-11 | 164.941041 | 189.296944 | 219.020252 | 191.141694 | 0 |
| 6.52E-12 | 7.66E-11 | 154.652321 | 215.767375 | 261.710641 | 237.535309 | 0 |
| 6.53E-12 | 7.67E-11 | 230.853152 | 165.27336 | 209.739733 | 184.182652 | 0 |
| 6.57E-12 | 7.71E-11 | 171.049968 | 311.416829 | 193.034799 | 274.650202 | 0 |
| 6.57E-12 | 7.72E-11 | 172.979103 | 210.4288 | 172.617657 | 194.389247 | 0 |
| 6.60E-12 | 7.74E-11 | 144.363601 | 325.653027 | 302.544925 | 312.692966 | 0 |
| 6.60E-12 | 7.75E-11 | 142.755988 | 372.365552 | 351.731677 | 330.786476 | 0 |
| 6.61E-12 | 7.75E-11 | 147.900348 | 333.216007 | 269.135056 | 326.147114 | 0 |
| 6.61E-12 | 7.75E-11 | 139.219241 | 472.9087 | 414.839207 | 499.659235 | 0 |
| 6.65E-12 | 7.80E-11 | 144.685123 | 314.530998 | 296.976613 | 300.16669 | 0 |
| 6.67E-12 | 7.81E-11 | 516.365129 | 1392.03323 | 1124.79892 | 1268.40144 | 28.5146189 |
| 6.68E-12 | 7.82E-11 | 148.864916 | 243.794889 | 271.919212 | 265.835415 | 0 |
| 6.71E-12 | 7.85E-11 | 281.332184 | 443.10166 | 467.738166 | 452.801684 | 0 |
| 6.73E-12 | 7.88E-11 | 142.112943 | 347.452205 | 348.019469 | 333.106157 | 0 |
| 6.73E-12 | 7.88E-11 | 145.971213 | 302.296765 | 283.983887 | 276.04201 | 0 |
| 6.74E-12 | 7.88E-11 | 187.769138 | 274.491691 | 167.049345 | 238.463182 | 0 |
| 6.75E-12 | 7.89E-11 | 146.292736 | 806.347148 | 453.817387 | 605.436677 | 0 |
| 6.75E-12 | 7.89E-11 | 146.614258 | 282.499552 | 270.99116 | 268.155095 | 0 |
| 6.79E-12 | 7.93E-11 | 148.221871 | 464.678398 | 321.105963 | 470.431257 | 0 |

| | | | | | | |
|---|---|---|---|---|---|---|
| 6.81E-12 | 7.95E-11 | 169.120833 | 204.422904 | 179.11402 | 185.110524 | 0 |
| 6.81E-12 | 7.95E-11 | 152.723186 | 275.826335 | 224.588564 | 244.958288 | 0 |
| 6.82E-12 | 7.96E-11 | 154.330798 | 381.708057 | 260.782589 | 292.743711 | 0 |
| 6.85E-12 | 7.99E-11 | 0 | 0 | 0 | 0 | 30.8908372 |
| 6.87E-12 | 8.01E-11 | 148.543393 | 327.432552 | 268.207004 | 271.866585 | 0 |
| 6.88E-12 | 8.02E-11 | 227.316405 | 481.139001 | 402.774532 | 442.131152 | 2.37621824 |
| 6.89E-12 | 8.03E-11 | 170.406923 | 521.400749 | 257.070381 | 380.427644 | 0 |
| 6.92E-12 | 8.07E-11 | 150.794051 | 232.67286 | 290.48025 | 281.609244 | 0 |
| 6.94E-12 | 8.08E-11 | 149.507961 | 265.149186 | 233.869083 | 253.773075 | 0 |
| 6.95E-12 | 8.10E-11 | 148.543393 | 309.414864 | 255.214277 | 268.619032 | 0 |
| 6.96E-12 | 8.10E-11 | 135.360971 | 494.485437 | 519.709073 | 586.415295 | 0 |
| 6.98E-12 | 8.12E-11 | 187.447615 | 182.401286 | 173.545708 | 239.85499 | 0 |
| 7.00E-12 | 8.14E-11 | 135.360971 | 482.696086 | 507.644399 | 488.060831 | 0 |
| 7.00E-12 | 8.14E-11 | 0 | 0 | 0 | 0 | 47.5243649 |
| 7.02E-12 | 8.16E-11 | 141.148376 | 335.662854 | 375.861026 | 378.107963 | 0 |
| 7.05E-12 | 8.19E-11 | 145.328168 | 293.844023 | 278.415575 | 325.683178 | 0 |
| 7.05E-12 | 8.19E-11 | 134.396403 | 532.300338 | 514.140762 | 504.298596 | 0 |
| 7.06E-12 | 8.19E-11 | 134.074881 | 577.233337 | 504.860243 | 572.497211 | 0 |
| 7.06E-12 | 8.19E-11 | 156.581456 | 242.237805 | 198.60311 | 227.79265 | 0 |
| 7.08E-12 | 8.21E-11 | 141.791421 | 348.341968 | 322.034015 | 333.570093 | 0 |
| 7.10E-12 | 8.24E-11 | 146.292736 | 268.263354 | 280.271679 | 316.404455 | 0 |
| 7.14E-12 | 8.28E-11 | 151.437096 | 322.538859 | 238.509343 | 287.640414 | 0 |
| 7.15E-12 | 8.28E-11 | 151.115573 | 296.068429 | 309.96934 | 428.677004 | 0 |
| 7.23E-12 | 8.38E-11 | 142.755988 | 413.07218 | 331.314534 | 346.560305 | 0 |
| 7.25E-12 | 8.40E-11 | 834.350877 | 242.460246 | 234.797135 | 438.419663 | 0 |
| 7.25E-12 | 8.40E-11 | 287.119589 | 444.658744 | 397.20622 | 450.482003 | 0 |
| 7.26E-12 | 8.40E-11 | 137.290106 | 479.137036 | 424.119726 | 424.965515 | 0 |
| 7.27E-12 | 8.40E-11 | 171.049968 | 373.255314 | 210.667785 | 368.365304 | 0 |
| 7.27E-12 | 8.40E-11 | 149.507961 | 255.3618 | 339.667002 | 280.217435 | 0 |
| 7.27E-12 | 8.41E-11 | 253.359727 | 270.042879 | 332.242586 | 339.601263 | 0 |
| 7.28E-12 | 8.42E-11 | 149.829483 | 439.32017 | 314.6096 | 308.053604 | 0 |
| 7.28E-12 | 8.42E-11 | 139.862286 | 387.046631 | 348.019469 | 370.221049 | 0 |
| 7.39E-12 | 8.53E-11 | 194.842633 | 576.121134 | 224.588564 | 414.294983 | 0 |
| 7.42E-12 | 8.56E-11 | 150.151006 | 258.031087 | 221.804408 | 263.97967 | 0 |
| 7.44E-12 | 8.58E-11 | 163.333428 | 368.361621 | 331.314534 | 218.513927 | 0 |
| 7.45E-12 | 8.60E-11 | 0 | 0 | 0 | 0 | 111.682257 |
| 7.46E-12 | 8.60E-11 | 164.297996 | 336.552616 | 222.73246 | 232.432012 | 0 |
| 7.47E-12 | 8.61E-11 | 143.720556 | 319.869572 | 277.487523 | 308.053604 | 0 |
| 7.48E-12 | 8.62E-11 | 141.791421 | 305.855815 | 356.371936 | 335.889774 | 0 |
| 7.52E-12 | 8.66E-11 | 142.434466 | 316.088082 | 306.257133 | 360.014453 | 0 |
| 7.52E-12 | 8.66E-11 | 306.089416 | 151.037162 | 296.048562 | 230.576267 | 0 |
| 7.53E-12 | 8.67E-11 | 151.437096 | 415.741467 | 278.415575 | 309.909349 | 0 |
| 7.54E-12 | 8.68E-11 | 142.755988 | 359.019116 | 301.616873 | 376.252219 | 0 |

| | | | | | | |
|---|---|---|---|---|---|---|
| 7.55E-12 | 8.69E-11 | 204.166785 | 220.438627 | 152.200514 | 213.41063 | 0 |
| 7.57E-12 | 8.70E-11 | 176.837373 | 229.336251 | 164.265189 | 211.554885 | 0 |
| 7.59E-12 | 8.72E-11 | 142.434466 | 322.7613 | 289.552198 | 329.858604 | 0 |
| 7.59E-12 | 8.72E-11 | 142.755988 | 318.97981 | 319.249859 | 388.314559 | 0 |
| 7.62E-12 | 8.75E-11 | 145.971213 | 284.723958 | 250.574018 | 298.310945 | 0 |
| 7.64E-12 | 8.77E-11 | 172.65758 | 256.696444 | 172.617657 | 223.153289 | 0 |
| 7.67E-12 | 8.81E-11 | 147.257303 | 260.255493 | 243.149602 | 292.279775 | 0 |
| 7.68E-12 | 8.82E-11 | 151.437096 | 311.416829 | 228.300772 | 279.289563 | 0 |
| 7.69E-12 | 8.82E-11 | 151.115573 | 382.82026 | 440.824661 | 276.505946 | 0 |
| 7.69E-12 | 8.82E-11 | 416.371632 | 702.467391 | 619.010629 | 621.674443 | 19.0097459 |
| 7.72E-12 | 8.85E-11 | 134.396403 | 539.195996 | 454.745439 | 543.733169 | 0 |
| 7.75E-12 | 8.88E-11 | 149.507961 | 230.670894 | 236.653239 | 276.505946 | 0 |
| 7.76E-12 | 8.89E-11 | 150.794051 | 220.661068 | 262.638692 | 226.400842 | 0 |
| 7.79E-12 | 8.92E-11 | 145.328168 | 314.753438 | 258.926485 | 282.537116 | 0 |
| 7.80E-12 | 8.94E-11 | 165.905608 | 213.542969 | 171.689605 | 216.194247 | 0 |
| 7.82E-12 | 8.95E-11 | 143.399033 | 268.930676 | 275.631419 | 280.681372 | 0 |
| 7.84E-12 | 8.97E-11 | 132.467268 | 529.631051 | 497.435827 | 573.425083 | 0 |
| 7.87E-12 | 9.00E-11 | 148.221871 | 331.881364 | 246.86181 | 293.207648 | 0 |
| 7.87E-12 | 9.00E-11 | 143.720556 | 331.436483 | 280.271679 | 285.784669 | 0 |
| 7.89E-12 | 9.02E-11 | 142.112943 | 282.499552 | 308.113236 | 321.043817 | 0 |
| 7.90E-12 | 9.03E-11 | 164.941041 | 226.666964 | 176.329864 | 199.956481 | 0 |
| 7.91E-12 | 9.04E-11 | 182.303255 | 254.694478 | 165.193241 | 209.69914 | 0 |
| 7.92E-12 | 9.05E-11 | 146.292736 | 259.365731 | 245.933758 | 292.743711 | 0 |
| 7.95E-12 | 9.07E-11 | 143.077511 | 316.310522 | 285.83999 | 275.578074 | 0 |
| 7.95E-12 | 9.07E-11 | 142.112943 | 389.048596 | 357.299988 | 302.486371 | 0 |
| 7.98E-12 | 9.10E-11 | 299.015921 | 383.0427 | 348.947521 | 418.470408 | 2.37621824 |
| 7.98E-12 | 9.10E-11 | 136.968583 | 643.520634 | 448.249076 | 647.190931 | 0 |
| 8.01E-12 | 9.13E-11 | 133.110313 | 599.699837 | 480.730893 | 612.39572 | 0 |
| 8.02E-12 | 9.14E-11 | 177.158895 | 162.381632 | 205.099474 | 191.60563 | 0 |
| 8.04E-12 | 9.17E-11 | 149.507961 | 266.48383 | 218.0922 | 265.371479 | 0 |
| 8.06E-12 | 9.18E-11 | 354.960836 | 281.832231 | 161.481034 | 213.41063 | 0 |
| 8.07E-12 | 9.18E-11 | 137.933151 | 471.574056 | 370.292715 | 425.893387 | 0 |
| 8.07E-12 | 9.18E-11 | 139.219241 | 338.109701 | 322.034015 | 330.32254 | 0 |
| 8.12E-12 | 9.24E-11 | 439.521252 | 268.485795 | 228.300772 | 259.340309 | 9.50487297 |
| 8.12E-12 | 9.24E-11 | 134.074881 | 512.503125 | 449.177128 | 452.801684 | 0 |
| 8.16E-12 | 9.28E-11 | 180.695643 | 189.074504 | 371.220767 | 288.568286 | 0 |
| 8.17E-12 | 9.28E-11 | 139.219241 | 341.446309 | 365.652455 | 319.652008 | 0 |
| 8.17E-12 | 9.29E-11 | 209.95419 | 484.698051 | 219.948304 | 241.246799 | 0 |
| 8.28E-12 | 9.40E-11 | 231.81772 | 302.074325 | 278.415575 | 374.86041 | 14.2573095 |
| 8.29E-12 | 9.41E-11 | 135.682493 | 451.109521 | 397.20622 | 489.45264 | 0 |
| 8.29E-12 | 9.41E-11 | 228.602495 | 370.363586 | 158.696878 | 270.474776 | 0 |
| 8.39E-12 | 9.52E-11 | 129.252043 | 597.252991 | 561.47141 | 553.011892 | 0 |
| 8.40E-12 | 9.53E-11 | 162.690383 | 172.83634 | 214.379993 | 212.946693 | 0 |

| | | | | | | |
|---|---|---|---|---|---|---|
| 8.44E-12 | 9.57E-11 | 153.687753 | 589.690011 | 310.897392 | 463.936151 | 0 |
| 8.45E-12 | 9.58E-11 | 181.017165 | 315.865641 | 178.185968 | 233.82382 | 0 |
| 8.46E-12 | 9.59E-11 | 131.502701 | 598.365194 | 607.874006 | 487.132959 | 0 |
| 8.47E-12 | 9.60E-11 | 148.864916 | 435.093799 | 277.487523 | 409.655622 | 0 |
| 8.48E-12 | 9.60E-11 | 145.328168 | 343.670715 | 261.710641 | 286.712542 | 0 |
| 8.49E-12 | 9.61E-11 | 138.897718 | 355.904948 | 399.990376 | 325.683178 | 0 |
| 8.52E-12 | 9.64E-11 | 175.872805 | 160.602108 | 203.24337 | 194.389247 | 0 |
| 8.53E-12 | 9.65E-11 | 196.771768 | 337.88726 | 166.121293 | 309.445413 | 0 |
| 8.54E-12 | 9.66E-11 | 150.472528 | 269.597998 | 214.379993 | 230.576267 | 0 |
| 8.55E-12 | 9.66E-11 | 141.148376 | 413.517061 | 320.177911 | 332.178284 | 0 |
| 8.56E-12 | 9.68E-11 | 139.862286 | 385.044666 | 361.940248 | 495.019873 | 0 |
| 8.61E-12 | 9.73E-11 | 191.94893 | 387.936393 | 412.055051 | 355.839028 | 35.6432737 |
| 8.66E-12 | 9.78E-11 | 151.758618 | 216.657137 | 202.315318 | 246.350096 | 0 |
| 8.70E-12 | 9.82E-11 | 141.791421 | 274.491691 | 276.559471 | 260.732117 | 0 |
| 8.72E-12 | 9.84E-11 | 135.682493 | 477.357511 | 392.565961 | 402.696579 | 0 |
| 8.72E-12 | 9.84E-11 | 218.95682 | 551.207788 | 438.968557 | 487.132959 | 2.37621824 |
| 8.76E-12 | 9.88E-11 | 142.434466 | 321.871537 | 264.494796 | 325.683178 | 0 |
| 8.79E-12 | 9.91E-11 | 140.826853 | 454.891012 | 330.386482 | 362.79807 | 0 |
| 8.80E-12 | 9.92E-11 | 264.934537 | 1418.50366 | 1109.02204 | 1173.29453 | 0 |
| 8.88E-12 | 1.00E-10 | 313.484434 | 205.312667 | 183.75428 | 184.646588 | 0 |
| 8.89E-12 | 1.00E-10 | 204.488308 | 742.506698 | 914.131138 | 896.324644 | 7.12865473 |
| 8.89E-12 | 1.00E-10 | 290.013292 | 179.287118 | 180.970124 | 200.420417 | 0 |
| 8.90E-12 | 1.00E-10 | 138.254673 | 336.997498 | 303.472977 | 330.32254 | 0 |
| 8.92E-12 | 1.00E-10 | 144.363601 | 266.70627 | 235.725187 | 281.609244 | 0 |
| 8.94E-12 | 1.01E-10 | 167.834743 | 229.558691 | 172.617657 | 188.822014 | 0 |
| 9.00E-12 | 1.01E-10 | 265.577582 | 537.194031 | 462.169855 | 485.74115 | 2.37621824 |
| 9.01E-12 | 1.01E-10 | 125.072251 | 621.943896 | 664.485173 | 681.05827 | 0 |
| 9.02E-12 | 1.01E-10 | 268.149762 | 685.561906 | 484.443101 | 603.116997 | 0 |
| 9.03E-12 | 1.01E-10 | 179.731075 | 340.334106 | 178.185968 | 295.991265 | 0 |
| 9.06E-12 | 1.02E-10 | 146.614258 | 549.428263 | 328.530379 | 399.912962 | 0 |
| 9.09E-12 | 1.02E-10 | 154.330798 | 259.810612 | 197.675058 | 277.897755 | 0 |
| 9.17E-12 | 1.03E-10 | 133.110313 | 396.166695 | 425.97583 | 441.667216 | 0 |
| 9.18E-12 | 1.03E-10 | 146.292736 | 224.664998 | 236.653239 | 215.73031 | 0 |
| 9.19E-12 | 1.03E-10 | 138.576196 | 294.288904 | 348.947521 | 328.002859 | 0 |
| 9.21E-12 | 1.03E-10 | 145.328168 | 278.495622 | 239.437395 | 233.359884 | 0 |
| 9.24E-12 | 1.04E-10 | 140.505331 | 332.548686 | 271.919212 | 314.548711 | 0 |
| 9.31E-12 | 1.04E-10 | 143.720556 | 326.54279 | 248.717914 | 306.19786 | 0 |
| 9.33E-12 | 1.04E-10 | 289.370247 | 373.922636 | 367.508559 | 360.942326 | 0 |
| 9.33E-12 | 1.04E-10 | 152.401663 | 282.721993 | 202.315318 | 262.123925 | 0 |
| 9.35E-12 | 1.05E-10 | 139.219241 | 292.509379 | 322.962067 | 282.07318 | 0 |
| 9.40E-12 | 1.05E-10 | 181.338688 | 179.509558 | 157.768826 | 190.677758 | 0 |
| 9.41E-12 | 1.05E-10 | 201.916128 | 275.826335 | 250.574018 | 272.330521 | 0 |
| 9.43E-12 | 1.05E-10 | 131.824223 | 439.765051 | 564.255566 | 492.236257 | 0 |

| | | | | | | |
|---|---|---|---|---|---|---|
| 9.49E-12 | 1.06E-10 | 136.004016 | 380.373413 | 327.602327 | 386.458814 | 0 |
| 9.49E-12 | 1.06E-10 | 431.161667 | 203.755583 | 175.401812 | 341.920944 | 0 |
| 9.50E-12 | 1.06E-10 | 177.158895 | 213.98785 | 157.768826 | 190.677758 | 0 |
| 9.51E-12 | 1.06E-10 | 146.935781 | 252.914954 | 210.667785 | 232.895948 | 0 |
| 9.52E-12 | 1.06E-10 | 154.009276 | 185.960335 | 233.869083 | 240.782863 | 0 |
| 9.59E-12 | 1.07E-10 | 147.257303 | 242.682686 | 218.0922 | 281.609244 | 0 |
| 9.62E-12 | 1.07E-10 | 139.862286 | 492.928353 | 1009.72049 | 752.504437 | 0 |
| 9.63E-12 | 1.07E-10 | 152.401663 | 206.42487 | 192.106747 | 190.213822 | 0 |
| 9.64E-12 | 1.07E-10 | 147.900348 | 313.863676 | 223.660512 | 282.537116 | 0 |
| 9.68E-12 | 1.08E-10 | 127.322908 | 578.34554 | 541.054268 | 524.711787 | 0 |
| 9.69E-12 | 1.08E-10 | 144.685123 | 262.70234 | 223.660512 | 235.679565 | 0 |
| 9.69E-12 | 1.08E-10 | 531.798209 | 253.137394 | 192.106747 | 250.989458 | 0 |
| 9.71E-12 | 1.08E-10 | 148.543393 | 208.204395 | 225.516616 | 199.956481 | 0 |
| 9.72E-12 | 1.08E-10 | 127.322908 | 547.426298 | 517.85297 | 552.547956 | 0 |
| 9.79E-12 | 1.09E-10 | 150.151006 | 361.021082 | 232.941031 | 298.310945 | 0 |
| 9.81E-12 | 1.09E-10 | 145.006646 | 283.834196 | 353.58778 | 251.91733 | 0 |
| 9.83E-12 | 1.09E-10 | 147.578826 | 208.871716 | 206.955577 | 233.82382 | 0 |
| 9.85E-12 | 1.10E-10 | 130.538133 | 487.589779 | 445.46492 | 433.780301 | 0 |
| 9.90E-12 | 1.10E-10 | 138.254673 | 295.401107 | 274.703367 | 284.392861 | 0 |
| 9.91E-12 | 1.10E-10 | 177.80194 | 173.058781 | 160.552982 | 198.564673 | 0 |
| 9.92E-12 | 1.10E-10 | 139.540763 | 324.985706 | 295.12051 | 267.227223 | 0 |
| 9.93E-12 | 1.10E-10 | 134.074881 | 370.141146 | 343.379209 | 356.302964 | 0 |
| 1.00E-11 | 1.11E-10 | 166.22713 | 172.391459 | 265.422848 | 222.225416 | 0 |
| 1.00E-11 | 1.11E-10 | 239.855782 | 422.192244 | 154.056618 | 328.466795 | 0 |
| 1.01E-11 | 1.11E-10 | 148.543393 | 218.881543 | 296.048562 | 313.620838 | 0 |
| 1.01E-11 | 1.12E-10 | 137.611628 | 282.721993 | 287.696094 | 282.537116 | 0 |
| 1.01E-11 | 1.12E-10 | 251.109069 | 605.705733 | 567.039721 | 585.023487 | 0 |
| 1.01E-11 | 1.12E-10 | 19.6128723 | 22.2440592 | 21.3451941 | 18.5574461 | 218.612078 |
| 1.02E-11 | 1.12E-10 | 136.647061 | 302.296765 | 321.105963 | 290.887967 | 0 |
| 1.02E-11 | 1.13E-10 | 133.753358 | 489.814185 | 370.292715 | 438.419663 | 0 |
| 1.02E-11 | 1.13E-10 | 55.3018693 | 38.4822225 | 56.6111669 | 32.4755306 | 235.245606 |
| 1.03E-11 | 1.14E-10 | 172.979103 | 279.162944 | 570.751929 | 294.13552 | 0 |
| 1.03E-11 | 1.14E-10 | 182.9463 | 215.989815 | 152.200514 | 234.751693 | 0 |
| 1.03E-11 | 1.14E-10 | 132.467268 | 368.584062 | 377.71713 | 413.367111 | 0 |
| 1.03E-11 | 1.14E-10 | 139.540763 | 322.316418 | 280.271679 | 264.443606 | 0 |
| 1.03E-11 | 1.14E-10 | 132.145746 | 433.314274 | 399.062324 | 373.468602 | 0 |
| 1.04E-11 | 1.14E-10 | 139.540763 | 490.036625 | 321.105963 | 429.604876 | 0 |
| 1.04E-11 | 1.15E-10 | 144.042078 | 243.127568 | 211.595837 | 240.318926 | 0 |
| 1.05E-11 | 1.16E-10 | 137.933151 | 345.227799 | 295.12051 | 283.001052 | 0 |
| 1.05E-11 | 1.16E-10 | 286.155022 | 349.23173 | 357.299988 | 374.86041 | 2.37621824 |
| 1.06E-11 | 1.17E-10 | 132.788791 | 375.702161 | 348.019469 | 352.127539 | 0 |
| 1.06E-11 | 1.17E-10 | 145.971213 | 201.753617 | 208.811681 | 217.122119 | 0 |
| 1.06E-11 | 1.17E-10 | 133.431836 | 369.473824 | 329.458431 | 357.230837 | 0 |

| | | | | | | |
|---|---|---|---|---|---|---|
| 1.06E-11 | 1.17E-10 | 275.223257 | 437.318205 | 375.861026 | 423.10977 | 0 |
| 1.06E-11 | 1.17E-10 | 134.074881 | 371.69823 | 324.818171 | 337.281582 | 0 |
| 1.07E-11 | 1.18E-10 | 217.349207 | 292.95426 | 240.365446 | 146.603824 | 0 |
| 1.07E-11 | 1.18E-10 | 127.644431 | 532.300338 | 676.549847 | 512.649447 | 0 |
| 1.07E-11 | 1.18E-10 | 146.292736 | 252.692513 | 384.213494 | 392.026048 | 0 |
| 1.09E-11 | 1.20E-10 | 164.619518 | 164.161157 | 209.739733 | 182.326907 | 0 |
| 1.09E-11 | 1.20E-10 | 2141.33982 | 7106.53205 | 3842.13494 | 4992.88086 | 35.6432737 |
| 1.09E-11 | 1.20E-10 | 191.94893 | 268.485795 | 151.272462 | 228.256586 | 0 |
| 1.09E-11 | 1.20E-10 | 143.720556 | 231.338216 | 208.811681 | 219.905736 | 0 |
| 1.10E-11 | 1.21E-10 | 145.971213 | 238.678756 | 332.242586 | 248.669777 | 0 |
| 1.10E-11 | 1.21E-10 | 128.608998 | 561.217615 | 438.040505 | 504.298596 | 0 |
| 1.10E-11 | 1.21E-10 | 182.624778 | 177.730033 | 358.22804 | 250.989458 | 0 |
| 1.11E-11 | 1.22E-10 | 153.366231 | 212.208325 | 225.516616 | 175.831801 | 0 |
| 1.11E-11 | 1.22E-10 | 127.001386 | 517.841699 | 463.097907 | 474.142747 | 0 |
| 1.12E-11 | 1.22E-10 | 265.899104 | 440.654814 | 403.702584 | 472.287002 | 2.37621824 |
| 1.12E-11 | 1.23E-10 | 134.074881 | 317.867607 | 313.681548 | 361.406262 | 0 |
| 1.13E-11 | 1.24E-10 | 184.23239 | 316.532963 | 221.804408 | 225.009033 | 9.50487297 |
| 1.14E-11 | 1.25E-10 | 132.788791 | 315.643201 | 346.163365 | 337.281582 | 0 |
| 1.14E-11 | 1.25E-10 | 136.004016 | 410.847774 | 315.537652 | 326.147114 | 0 |
| 1.14E-11 | 1.25E-10 | 133.753358 | 365.247453 | 307.185185 | 339.601263 | 0 |
| 1.15E-11 | 1.26E-10 | 128.287476 | 454.668571 | 408.342843 | 473.214874 | 0 |
| 1.15E-11 | 1.26E-10 | 143.720556 | 219.103984 | 219.020252 | 201.34829 | 0 |
| 1.15E-11 | 1.26E-10 | 131.502701 | 372.587992 | 479.802841 | 399.449026 | 0 |
| 1.16E-11 | 1.26E-10 | 128.608998 | 579.235303 | 555.903098 | 429.14094 | 0 |
| 1.16E-11 | 1.27E-10 | 181.017165 | 409.735571 | 185.610383 | 304.342115 | 0 |
| 1.16E-11 | 1.27E-10 | 200.308515 | 285.613721 | 149.416359 | 230.112331 | 0 |
| 1.16E-11 | 1.27E-10 | 139.862286 | 249.800785 | 225.516616 | 238.463182 | 0 |
| 1.17E-11 | 1.27E-10 | 243.714052 | 215.100053 | 143.848047 | 202.276162 | 0 |
| 1.17E-11 | 1.27E-10 | 144.363601 | 268.263354 | 210.667785 | 281.145308 | 0 |
| 1.17E-11 | 1.28E-10 | 171.049968 | 212.208325 | 153.128566 | 203.66797 | 0 |
| 1.17E-11 | 1.28E-10 | 133.431836 | 336.775057 | 312.753496 | 300.16669 | 0 |
| 1.17E-11 | 1.28E-10 | 129.573566 | 425.306413 | 378.645182 | 386.92275 | 0 |
| 1.18E-11 | 1.28E-10 | 148.864916 | 228.668929 | 237.581291 | 184.646588 | 0 |
| 1.18E-11 | 1.28E-10 | 117.677234 | 725.156331 | 692.32673 | 711.678056 | 0 |
| 1.18E-11 | 1.29E-10 | 136.004016 | 462.00911 | 623.650888 | 381.355516 | 0 |
| 1.18E-11 | 1.29E-10 | 125.393773 | 511.835803 | 468.666218 | 491.77232 | 0 |
| 1.19E-11 | 1.29E-10 | 138.576196 | 231.560657 | 254.286225 | 264.443606 | 0 |
| 1.19E-11 | 1.30E-10 | 175.22976 | 240.23584 | 165.193241 | 177.687546 | 0 |
| 1.19E-11 | 1.30E-10 | 131.502701 | 347.007324 | 449.177128 | 428.213068 | 0 |
| 1.20E-11 | 1.30E-10 | 133.110313 | 361.021082 | 314.6096 | 311.765094 | 0 |
| 1.21E-11 | 1.32E-10 | 247.572322 | 740.727173 | 546.622579 | 580.384125 | 2.37621824 |
| 1.22E-11 | 1.33E-10 | 136.004016 | 151.704484 | 147.560255 | 146.603824 | 11.8810912 |
| 1.23E-11 | 1.33E-10 | 207.703532 | 180.17688 | 150.344411 | 181.862971 | 0 |

| | | | | | | |
|---|---|---|---|---|---|---|
| 1.23E-11 | 1.34E-10 | 132.788791 | 404.396997 | 313.681548 | 365.117751 | 0 |
| 1.23E-11 | 1.34E-10 | 136.004016 | 297.847953 | 381.429338 | 295.063392 | 0 |
| 1.23E-11 | 1.34E-10 | 127.322908 | 413.517061 | 446.392972 | 405.480196 | 0 |
| 1.24E-11 | 1.34E-10 | 130.859656 | 353.902983 | 322.034015 | 353.055411 | 0 |
| 1.24E-11 | 1.35E-10 | 128.287476 | 418.855636 | 409.270895 | 373.932538 | 0 |
| 1.25E-11 | 1.35E-10 | 136.325538 | 268.708236 | 311.825444 | 357.230837 | 0 |
| 1.25E-11 | 1.36E-10 | 113.818964 | 782.990885 | 765.642831 | 779.87667 | 0 |
| 1.25E-11 | 1.36E-10 | 156.581456 | 693.124886 | 306.257133 | 445.378705 | 0 |
| 1.25E-11 | 1.36E-10 | 161.404293 | 215.989815 | 181.898176 | 164.233398 | 0 |
| 1.26E-11 | 1.36E-10 | 133.431836 | 621.499015 | 460.313751 | 392.026048 | 0 |
| 1.26E-11 | 1.37E-10 | 139.540763 | 316.532963 | 259.854537 | 241.246799 | 0 |
| 1.27E-11 | 1.37E-10 | 131.181178 | 314.975879 | 315.537652 | 318.724136 | 0 |
| 1.28E-11 | 1.38E-10 | 140.826853 | 269.820439 | 213.451941 | 263.97967 | 0 |
| 1.28E-11 | 1.38E-10 | 129.895088 | 376.591923 | 332.242586 | 387.386686 | 0 |
| 1.28E-11 | 1.39E-10 | 130.538133 | 385.044666 | 322.962067 | 369.293176 | 0 |
| 1.28E-11 | 1.39E-10 | 175.872805 | 239.123637 | 161.481034 | 177.687546 | 0 |
| 1.29E-11 | 1.40E-10 | 232.460765 | 767.420044 | 657.988809 | 778.484862 | 0 |
| 1.30E-11 | 1.40E-10 | 124.429206 | 471.574056 | 499.291931 | 590.590721 | 0 |
| 1.30E-11 | 1.40E-10 | 128.608998 | 433.981596 | 362.8683 | 387.386686 | 0 |
| 1.30E-11 | 1.40E-10 | 183.589345 | 240.23584 | 154.056618 | 183.718716 | 0 |
| 1.30E-11 | 1.40E-10 | 139.540763 | 244.01733 | 211.595837 | 248.205841 | 0 |
| 1.31E-11 | 1.41E-10 | 146.614258 | 210.873682 | 179.11402 | 191.141694 | 14.2573095 |
| 1.31E-11 | 1.42E-10 | 541.122361 | 949.154008 | 868.656594 | 962.203578 | 21.3859642 |
| 1.33E-11 | 1.44E-10 | 134.717926 | 294.733785 | 251.502069 | 299.702754 | 0 |
| 1.33E-11 | 1.44E-10 | 167.191698 | 199.751652 | 158.696878 | 168.408823 | 0 |
| 1.35E-11 | 1.45E-10 | 125.715296 | 401.060388 | 480.730893 | 474.606683 | 0 |
| 1.35E-11 | 1.46E-10 | 132.788791 | 293.621582 | 268.207004 | 277.897755 | 0 |
| 1.36E-11 | 1.47E-10 | 153.044708 | 339.221903 | 198.60311 | 302.486371 | 0 |
| 1.37E-11 | 1.48E-10 | 136.647061 | 267.596033 | 262.638692 | 341.457007 | 0 |
| 1.38E-11 | 1.48E-10 | 128.930521 | 383.0427 | 325.746223 | 374.396474 | 0 |
| 1.38E-11 | 1.48E-10 | 136.968583 | 237.788993 | 221.804408 | 228.720523 | 0 |
| 1.38E-11 | 1.49E-10 | 130.859656 | 325.875468 | 301.616873 | 292.279775 | 0 |
| 1.38E-11 | 1.49E-10 | 138.897718 | 288.95033 | 221.804408 | 251.91733 | 0 |
| 1.39E-11 | 1.50E-10 | 131.502701 | 286.503483 | 351.731677 | 319.652008 | 0 |
| 1.40E-11 | 1.50E-10 | 276.830869 | 679.111129 | 390.709857 | 527.031468 | 0 |
| 1.40E-11 | 1.50E-10 | 227.316405 | 731.829549 | 641.283875 | 723.740396 | 2.37621824 |
| 1.40E-11 | 1.50E-10 | 127.322908 | 379.483651 | 371.220767 | 347.488177 | 0 |
| 1.40E-11 | 1.50E-10 | 128.608998 | 327.210111 | 384.213494 | 353.983283 | 0 |
| 1.41E-11 | 1.51E-10 | 130.216611 | 291.842057 | 310.897392 | 315.476583 | 0 |
| 1.41E-11 | 1.51E-10 | 134.074881 | 391.495443 | 280.271679 | 339.137327 | 0 |
| 1.41E-11 | 1.51E-10 | 115.105054 | 688.453634 | 655.204653 | 750.184757 | 0 |
| 1.41E-11 | 1.51E-10 | 193.878065 | 159.045024 | 365.652455 | 301.094562 | 0 |
| 1.41E-11 | 1.51E-10 | 146.935781 | 174.170984 | 192.106747 | 190.677758 | 0 |

| | | | | | | |
|---|---|---|---|---|---|---|
| 1.41E-11 | 1.51E-10 | 133.753358 | 267.818473 | 244.077654 | 266.299351 | 0 |
| 1.42E-11 | 1.52E-10 | 151.758618 | 413.517061 | 233.869083 | 276.505946 | 0 |
| 1.42E-11 | 1.52E-10 | 131.502701 | 276.271216 | 275.631419 | 288.568286 | 0 |
| 1.42E-11 | 1.52E-10 | 167.834743 | 155.708415 | 194.890903 | 234.751693 | 0 |
| 1.42E-11 | 1.52E-10 | 155.616888 | 290.284973 | 178.185968 | 247.277969 | 0 |
| 1.42E-11 | 1.52E-10 | 135.360971 | 294.288904 | 245.005706 | 320.579881 | 0 |
| 1.42E-11 | 1.52E-10 | 255.610384 | 133.686796 | 319.249859 | 308.053604 | 0 |
| 1.43E-11 | 1.53E-10 | 127.322908 | 349.23173 | 364.724403 | 342.848816 | 0 |
| 1.43E-11 | 1.53E-10 | 146.935781 | 181.066642 | 227.37272 | 247.741905 | 0 |
| 1.43E-11 | 1.53E-10 | 130.538133 | 308.747542 | 282.127783 | 300.16669 | 0 |
| 1.44E-11 | 1.53E-10 | 131.502701 | 533.857422 | 367.508559 | 384.139133 | 0 |
| 1.44E-11 | 1.53E-10 | 128.608998 | 313.196354 | 327.602327 | 332.64222 | 0 |
| 1.45E-11 | 1.54E-10 | 149.507961 | 343.003394 | 230.156875 | 425.893387 | 0 |
| 1.45E-11 | 1.54E-10 | 136.647061 | 288.95033 | 234.797135 | 315.476583 | 0 |
| 1.45E-11 | 1.55E-10 | 127.322908 | 359.463997 | 334.09869 | 369.293176 | 0 |
| 1.45E-11 | 1.55E-10 | 142.434466 | 282.277112 | 202.315318 | 248.205841 | 0 |
| 1.45E-11 | 1.55E-10 | 21.5420072 | 18.017688 | 18.5610383 | 19.0213822 | 76.0389838 |
| 1.46E-11 | 1.56E-10 | 159.475158 | 161.936751 | 174.47376 | 211.090949 | 0 |
| 1.47E-11 | 1.56E-10 | 128.930521 | 316.310522 | 301.616873 | 315.476583 | 0 |
| 1.47E-11 | 1.57E-10 | 138.576196 | 298.292834 | 225.516616 | 243.566479 | 0 |
| 1.47E-11 | 1.57E-10 | 190.019795 | 173.058781 | 337.810898 | 200.884354 | 0 |
| 1.47E-11 | 1.57E-10 | 331.489694 | 263.147221 | 144.776099 | 211.090949 | 0 |
| 1.48E-11 | 1.57E-10 | 128.930521 | 406.176522 | 524.349333 | 664.356569 | 0 |
| 1.48E-11 | 1.57E-10 | 126.036818 | 365.025012 | 368.436611 | 411.975302 | 0 |
| 1.48E-11 | 1.57E-10 | 140.826853 | 258.253528 | 212.523889 | 206.451587 | 0 |
| 1.49E-11 | 1.58E-10 | 127.644431 | 446.66071 | 342.451157 | 402.232643 | 0 |
| 1.49E-11 | 1.58E-10 | 128.608998 | 378.149007 | 322.034015 | 324.29137 | 0 |
| 1.49E-11 | 1.58E-10 | 142.434466 | 260.922815 | 239.437395 | 193.461375 | 0 |
| 1.50E-11 | 1.59E-10 | 132.788791 | 291.397176 | 251.502069 | 309.445413 | 0 |
| 1.50E-11 | 1.59E-10 | 132.145746 | 261.812577 | 274.703367 | 251.453394 | 0 |
| 1.50E-11 | 1.59E-10 | 131.824223 | 253.137394 | 277.487523 | 271.866585 | 0 |
| 1.50E-11 | 1.59E-10 | 146.614258 | 271.155082 | 432.472193 | 262.123925 | 0 |
| 1.50E-11 | 1.59E-10 | 123.464639 | 552.987313 | 861.232179 | 572.497211 | 0 |
| 1.50E-11 | 1.59E-10 | 162.047338 | 314.086117 | 773.067247 | 414.294983 | 0 |
| 1.51E-11 | 1.60E-10 | 161.082771 | 191.076469 | 164.265189 | 159.594036 | 0 |
| 1.51E-11 | 1.60E-10 | 181.981733 | 196.859924 | 145.704151 | 173.976057 | 0 |
| 1.51E-11 | 1.60E-10 | 122.178549 | 461.341789 | 432.472193 | 488.060831 | 0 |
| 1.51E-11 | 1.60E-10 | 124.750728 | 429.087903 | 382.35739 | 387.386686 | 0 |
| 1.51E-11 | 1.60E-10 | 130.216611 | 302.964087 | 272.847264 | 293.207648 | 0 |
| 1.53E-11 | 1.62E-10 | 128.930521 | 306.300696 | 294.192458 | 335.425837 | 0 |
| 1.53E-11 | 1.62E-10 | 207.060488 | 157.71038 | 180.042072 | 158.202228 | 0 |
| 1.53E-11 | 1.62E-10 | 126.679863 | 326.097909 | 352.659728 | 344.70456 | 0 |
| 1.54E-11 | 1.62E-10 | 131.824223 | 252.470072 | 297.904665 | 300.630626 | 0 |

| | | | | | | |
|---|---|---|---|---|---|---|
| 1.54E-11 | 1.63E-10 | 127.965953 | 302.296765 | 324.818171 | 317.332327 | 0 |
| 1.54E-11 | 1.63E-10 | 136.647061 | 318.757369 | 565.183617 | 526.103596 | 0 |
| 1.55E-11 | 1.63E-10 | 146.292736 | 233.562622 | 175.401812 | 229.184459 | 0 |
| 1.55E-11 | 1.63E-10 | 135.682493 | 522.735392 | 316.465704 | 534.454446 | 0 |
| 1.55E-11 | 1.64E-10 | 298.372876 | 312.973914 | 338.73895 | 299.238818 | 0 |
| 1.55E-11 | 1.64E-10 | 353.996268 | 199.974093 | 160.552982 | 224.545097 | 0 |
| 1.55E-11 | 1.64E-10 | 132.145746 | 260.477934 | 249.645966 | 245.422224 | 0 |
| 1.56E-11 | 1.64E-10 | 139.219241 | 316.977844 | 245.005706 | 227.79265 | 0 |
| 1.56E-11 | 1.64E-10 | 140.826853 | 268.263354 | 215.308045 | 205.987651 | 0 |
| 1.57E-11 | 1.65E-10 | 270.400419 | 407.288725 | 332.242586 | 380.427644 | 0 |
| 1.57E-11 | 1.65E-10 | 124.750728 | 383.932463 | 356.371936 | 378.107963 | 0 |
| 1.57E-11 | 1.66E-10 | 136.325538 | 234.229944 | 275.631419 | 219.905736 | 0 |
| 1.57E-11 | 1.66E-10 | 140.505331 | 300.2948 | 209.739733 | 291.815839 | 0 |
| 1.59E-11 | 1.67E-10 | 114.140486 | 634.178129 | 747.081793 | 639.304017 | 0 |
| 1.59E-11 | 1.67E-10 | 165.584086 | 253.137394 | 176.329864 | 169.800631 | 0 |
| 1.60E-11 | 1.69E-10 | 124.750728 | 379.03877 | 348.019469 | 373.932538 | 0 |
| 1.62E-11 | 1.70E-10 | 143.399033 | 287.393245 | 194.890903 | 271.402649 | 0 |
| 1.62E-11 | 1.70E-10 | 256.896474 | 480.47168 | 391.637909 | 401.768707 | 2.37621824 |
| 1.62E-11 | 1.70E-10 | 159.796681 | 146.58835 | 182.826228 | 196.708928 | 14.2573095 |
| 1.62E-11 | 1.71E-10 | 115.748099 | 693.792208 | 577.248292 | 591.982529 | 0 |
| 1.63E-11 | 1.71E-10 | 124.107684 | 402.839913 | 355.443884 | 400.376899 | 0 |
| 1.63E-11 | 1.71E-10 | 143.077511 | 226.222083 | 205.099474 | 178.615418 | 0 |
| 1.64E-11 | 1.72E-10 | 123.464639 | 381.930497 | 378.645182 | 425.893387 | 0 |
| 1.64E-11 | 1.72E-10 | 129.573566 | 312.084151 | 270.063108 | 333.570093 | 0 |
| 1.64E-11 | 1.72E-10 | 131.824223 | 270.26532 | 237.581291 | 279.289563 | 0 |
| 1.64E-11 | 1.72E-10 | 139.862286 | 454.001249 | 303.472977 | 278.361691 | 0 |
| 1.65E-11 | 1.73E-10 | 122.500071 | 409.29069 | 386.069597 | 434.244238 | 0 |
| 1.67E-11 | 1.75E-10 | 134.717926 | 280.720028 | 219.948304 | 252.381266 | 0 |
| 1.67E-11 | 1.75E-10 | 222.81509 | 705.359119 | 710.887768 | 688.481249 | 0 |
| 1.67E-11 | 1.75E-10 | 127.965953 | 293.176701 | 279.343627 | 289.496158 | 0 |
| 1.70E-11 | 1.78E-10 | 140.505331 | 276.938538 | 207.883629 | 314.548711 | 0 |
| 1.70E-11 | 1.78E-10 | 121.213981 | 445.326066 | 435.256349 | 399.449026 | 0 |
| 1.70E-11 | 1.78E-10 | 131.502701 | 353.235661 | 258.926485 | 334.497965 | 0 |
| 1.70E-11 | 1.78E-10 | 156.581456 | 222.885474 | 480.730893 | 344.240624 | 0 |
| 1.71E-11 | 1.79E-10 | 121.213981 | 398.835982 | 459.385699 | 441.20328 | 0 |
| 1.72E-11 | 1.80E-10 | 138.897718 | 314.086117 | 215.308045 | 252.845202 | 0 |
| 1.73E-11 | 1.81E-10 | 690.308799 | 1850.03841 | 1982.31889 | 1997.70907 | 23.7621824 |
| 1.73E-11 | 1.81E-10 | 136.968583 | 284.501518 | 229.228823 | 342.848816 | 0 |
| 1.73E-11 | 1.81E-10 | 141.791421 | 272.934607 | 200.459214 | 209.235204 | 0 |
| 1.74E-11 | 1.82E-10 | 132.145746 | 262.257458 | 243.149602 | 225.009033 | 0 |
| 1.74E-11 | 1.82E-10 | 58.5170943 | 60.2814006 | 70.5319457 | 95.106911 | 693.855727 |
| 1.74E-11 | 1.82E-10 | 310.912254 | 180.39932 | 167.977397 | 192.997439 | 0 |
| 1.75E-11 | 1.83E-10 | 133.753358 | 250.245666 | 213.451941 | 262.587862 | 0 |

| | | | | | | |
|---|---|---|---|---|---|---|
| 1.76E-11 | 1.83E-10 | 132.788791 | 263.369661 | 219.948304 | 242.638607 | 0 |
| 1.76E-11 | 1.84E-10 | 243.071007 | 459.562264 | 431.544141 | 452.801684 | 2.37621824 |
| 1.76E-11 | 1.84E-10 | 114.783531 | 655.977307 | 698.823093 | 540.485616 | 0 |
| 1.76E-11 | 1.84E-10 | 139.862286 | 232.450419 | 182.826228 | 216.194247 | 0 |
| 1.76E-11 | 1.84E-10 | 141.469898 | 203.310701 | 178.185968 | 183.718716 | 0 |
| 1.76E-11 | 1.84E-10 | 191.627408 | 190.631588 | 323.890119 | 166.089142 | 0 |
| 1.78E-11 | 1.85E-10 | 132.145746 | 545.424333 | 320.177911 | 422.181898 | 0 |
| 1.80E-11 | 1.87E-10 | 129.573566 | 386.60175 | 275.631419 | 343.776688 | 0 |
| 1.81E-11 | 1.89E-10 | 117.998756 | 521.84563 | 512.284658 | 445.842641 | 0 |
| 1.81E-11 | 1.89E-10 | 114.783531 | 533.1901 | 555.903098 | 594.30221 | 0 |
| 1.82E-11 | 1.89E-10 | 122.500071 | 361.465963 | 465.882062 | 440.275408 | 0 |
| 1.82E-11 | 1.89E-10 | 221.207477 | 324.763265 | 303.472977 | 289.496158 | 4.75243649 |
| 1.83E-11 | 1.90E-10 | 239.212737 | 715.591386 | 504.860243 | 573.889019 | 0 |
| 1.84E-11 | 1.91E-10 | 114.140486 | 728.048059 | 630.147252 | 549.300403 | 0 |
| 1.84E-11 | 1.92E-10 | 122.821594 | 382.375378 | 336.882846 | 363.262006 | 0 |
| 1.85E-11 | 1.92E-10 | 271.686509 | 135.24388 | 209.739733 | 229.648395 | 0 |
| 1.85E-11 | 1.93E-10 | 1242.36293 | 5365.71197 | 2884.38536 | 3652.56932 | 4.75243649 |
| 1.86E-11 | 1.93E-10 | 135.360971 | 244.684652 | 196.747006 | 233.359884 | 0 |
| 1.86E-11 | 1.93E-10 | 127.322908 | 265.371627 | 277.487523 | 263.97967 | 0 |
| 1.86E-11 | 1.93E-10 | 197.414813 | 183.95837 | 140.135839 | 173.512121 | 0 |
| 1.86E-11 | 1.93E-10 | 134.396403 | 221.55083 | 204.171422 | 250.061586 | 0 |
| 1.86E-11 | 1.93E-10 | 160.761248 | 191.966231 | 145.704151 | 181.399035 | 0 |
| 1.87E-11 | 1.94E-10 | 124.429206 | 376.814364 | 312.753496 | 389.706367 | 0 |
| 1.87E-11 | 1.94E-10 | 131.502701 | 279.162944 | 223.660512 | 271.866585 | 0 |
| 1.88E-11 | 1.95E-10 | 441.77191 | 196.859924 | 193.962851 | 214.802438 | 0 |
| 1.89E-11 | 1.95E-10 | 295.800696 | 199.974093 | 272.847264 | 138.716909 | 0 |
| 1.89E-11 | 1.96E-10 | 135.039448 | 395.944255 | 552.190891 | 304.806051 | 0 |
| 1.89E-11 | 1.96E-10 | 127.644431 | 309.859745 | 257.070381 | 301.558498 | 0 |
| 1.90E-11 | 1.96E-10 | 143.399033 | 270.710201 | 191.178695 | 304.806051 | 0 |
| 1.90E-11 | 1.96E-10 | 122.178549 | 399.503304 | 343.379209 | 368.365304 | 0 |
| 1.90E-11 | 1.96E-10 | 129.573566 | 238.901196 | 234.797135 | 260.268181 | 0 |
| 1.90E-11 | 1.96E-10 | 127.965953 | 286.058602 | 326.674275 | 395.273601 | 0 |
| 1.90E-11 | 1.96E-10 | 207.703532 | 567.00107 | 578.176344 | 545.588914 | 0 |
| 1.90E-11 | 1.97E-10 | 150.472528 | 156.598177 | 191.178695 | 170.728504 | 0 |
| 1.92E-11 | 1.98E-10 | 125.715296 | 377.036804 | 293.264406 | 335.425837 | 0 |
| 1.92E-11 | 1.98E-10 | 211.883325 | 376.591923 | 374.004923 | 395.737537 | 4.75243649 |
| 1.92E-11 | 1.98E-10 | 0 | 0 | 0 | 0 | 28.5146189 |
| 1.92E-11 | 1.98E-10 | 139.540763 | 222.218152 | 176.329864 | 220.833608 | 0 |
| 1.93E-11 | 1.99E-10 | 119.284846 | 513.615328 | 402.774532 | 496.411682 | 0 |
| 1.93E-11 | 1.99E-10 | 135.039448 | 217.991781 | 190.250643 | 223.153289 | 0 |
| 1.93E-11 | 1.99E-10 | 181.017165 | 217.546899 | 167.049345 | 270.474776 | 19.0097459 |
| 1.94E-11 | 2.00E-10 | 137.933151 | 189.296944 | 226.444668 | 189.28595 | 0 |
| 1.94E-11 | 2.00E-10 | 241.463395 | 257.141325 | 152.200514 | 225.009033 | 19.0097459 |

| | | | | | | |
|---|---|---|---|---|---|---|
| 1.94E-11 | 2.00E-10 | 199.343948 | 152.594246 | 151.272462 | 186.038397 | 0 |
| 1.95E-11 | 2.01E-10 | 222.81509 | 686.896549 | 602.305694 | 644.87125 | 0 |
| 1.96E-11 | 2.02E-10 | 146.292736 | 169.944613 | 214.379993 | 172.120312 | 0 |
| 1.96E-11 | 2.02E-10 | 132.467268 | 203.978023 | 232.941031 | 216.658183 | 0 |
| 1.96E-11 | 2.02E-10 | 143.720556 | 703.579594 | 304.401029 | 587.807104 | 0 |
| 1.97E-11 | 2.03E-10 | 129.895088 | 224.664998 | 242.22155 | 225.936906 | 0 |
| 1.97E-11 | 2.03E-10 | 124.429206 | 293.844023 | 321.105963 | 354.44722 | 0 |
| 1.98E-11 | 2.03E-10 | 232.460765 | 723.154366 | 657.060757 | 760.391352 | 0 |
| 1.98E-11 | 2.03E-10 | 123.464639 | 301.407003 | 325.746223 | 312.692966 | 0 |
| 1.98E-11 | 2.04E-10 | 180.37412 | 433.759155 | 180.970124 | 467.647641 | 0 |
| 1.99E-11 | 2.04E-10 | 145.006646 | 457.115417 | 232.941031 | 346.096369 | 0 |
| 2.00E-11 | 2.05E-10 | 136.004016 | 224.664998 | 196.747006 | 189.749886 | 0 |
| 2.00E-11 | 2.05E-10 | 234.711422 | 484.920492 | 505.788295 | 480.173917 | 0 |
| 2.01E-11 | 2.06E-10 | 151.437096 | 243.127568 | 230.156875 | 159.1301 | 0 |
| 2.01E-11 | 2.06E-10 | 134.396403 | 193.968197 | 245.933758 | 236.607437 | 0 |
| 2.02E-11 | 2.07E-10 | 114.783531 | 499.37913 | 486.299204 | 520.072426 | 0 |
| 2.03E-11 | 2.08E-10 | 119.927891 | 386.156868 | 515.068814 | 526.567532 | 0 |
| 2.03E-11 | 2.08E-10 | 124.429206 | 360.35376 | 348.947521 | 483.42147 | 0 |
| 2.03E-11 | 2.08E-10 | 110.282216 | 659.981238 | 643.139978 | 585.487423 | 0 |
| 2.04E-11 | 2.09E-10 | 122.500071 | 310.304626 | 331.314534 | 316.868391 | 0 |
| 2.04E-11 | 2.09E-10 | 140.826853 | 203.533142 | 175.401812 | 174.439993 | 0 |
| 2.04E-11 | 2.09E-10 | 124.429206 | 275.603894 | 294.192458 | 299.238818 | 0 |
| 2.07E-11 | 2.12E-10 | 120.249414 | 360.798641 | 361.940248 | 413.367111 | 0 |
| 2.07E-11 | 2.12E-10 | 140.183808 | 236.67679 | 175.401812 | 205.987651 | 0 |
| 2.08E-11 | 2.13E-10 | 129.252043 | 326.320349 | 244.077654 | 271.866585 | 0 |
| 2.09E-11 | 2.13E-10 | 117.998756 | 416.85367 | 389.781805 | 423.10977 | 0 |
| 2.09E-11 | 2.14E-10 | 236.319035 | 577.678219 | 434.328297 | 571.105402 | 0 |
| 2.11E-11 | 2.15E-10 | 118.963324 | 462.00911 | 380.501286 | 402.696579 | 0 |
| 2.12E-11 | 2.16E-10 | 128.287476 | 256.474003 | 239.437395 | 225.009033 | 0 |
| 2.12E-11 | 2.17E-10 | 131.824223 | 256.029122 | 211.595837 | 214.802438 | 0 |
| 2.13E-11 | 2.17E-10 | 183.910868 | 272.044845 | 143.848047 | 199.956481 | 0 |
| 2.15E-11 | 2.19E-10 | 125.072251 | 269.375557 | 254.286225 | 272.330521 | 0 |
| 2.15E-11 | 2.20E-10 | 626.647345 | 1953.25084 | 1585.11267 | 1818.16578 | 23.7621824 |
| 2.16E-11 | 2.20E-10 | 125.072251 | 327.210111 | 263.566744 | 302.486371 | 0 |
| 2.16E-11 | 2.20E-10 | 129.895088 | 207.537073 | 220.876356 | 217.586055 | 0 |
| 2.16E-11 | 2.20E-10 | 127.644431 | 286.948364 | 234.797135 | 255.628819 | 0 |
| 2.16E-11 | 2.20E-10 | 131.502701 | 232.8953 | 208.811681 | 268.619032 | 0 |
| 2.16E-11 | 2.20E-10 | 114.140486 | 489.814185 | 697.895042 | 591.518593 | 0 |
| 2.18E-11 | 2.21E-10 | 125.072251 | 367.249418 | 274.703367 | 338.209454 | 0 |
| 2.18E-11 | 2.21E-10 | 193.878065 | 1029.89994 | 1011.57659 | 970.554429 | 0 |
| 2.18E-11 | 2.22E-10 | 132.145746 | 274.26925 | 205.099474 | 257.020628 | 0 |
| 2.19E-11 | 2.23E-10 | 142.434466 | 185.515454 | 158.696878 | 188.822014 | 0 |
| 2.19E-11 | 2.23E-10 | 144.363601 | 253.359835 | 168.905449 | 214.338502 | 0 |

| | | | | | | |
|---|---|---|---|---|---|---|
| 2.19E-11 | 2.23E-10 | 207.060488 | 218.881543 | 143.848047 | 159.594036 | 0 |
| 2.20E-11 | 2.23E-10 | 136.004016 | 463.566195 | 264.494796 | 318.724136 | 0 |
| 2.21E-11 | 2.24E-10 | 145.971213 | 233.340181 | 232.941031 | 161.449781 | 0 |
| 2.21E-11 | 2.24E-10 | 117.677234 | 506.497229 | 387.925701 | 471.823066 | 0 |
| 2.21E-11 | 2.24E-10 | 136.325538 | 257.808647 | 187.466487 | 258.876372 | 0 |
| 2.22E-11 | 2.25E-10 | 141.791421 | 290.507414 | 184.682331 | 232.895948 | 0 |
| 2.22E-11 | 2.25E-10 | 120.249414 | 432.424512 | 335.954794 | 415.222855 | 0 |
| 2.22E-11 | 2.26E-10 | 202.23765 | 748.735034 | 812.973479 | 815.599754 | 2.37621824 |
| 2.23E-11 | 2.26E-10 | 120.892459 | 307.635339 | 328.530379 | 324.755306 | 0 |
| 2.23E-11 | 2.26E-10 | 118.641801 | 383.932463 | 373.076871 | 455.121364 | 0 |
| 2.23E-11 | 2.26E-10 | 123.143116 | 305.410933 | 273.775316 | 286.248605 | 0 |
| 2.24E-11 | 2.27E-10 | 119.927891 | 472.9087 | 349.875573 | 424.965515 | 0 |
| 2.24E-11 | 2.27E-10 | 159.796681 | 157.932821 | 282.127783 | 215.266374 | 0 |
| 2.24E-11 | 2.27E-10 | 119.606369 | 379.03877 | 347.091417 | 335.889774 | 0 |
| 2.25E-11 | 2.27E-10 | 116.391144 | 396.611576 | 445.46492 | 473.67881 | 0 |
| 2.25E-11 | 2.27E-10 | 120.570936 | 318.090047 | 381.429338 | 346.096369 | 0 |
| 2.25E-11 | 2.27E-10 | 123.143116 | 323.428621 | 273.775316 | 301.558498 | 0 |
| 2.25E-11 | 2.28E-10 | 132.467268 | 286.281042 | 215.308045 | 224.081161 | 0 |
| 2.26E-11 | 2.28E-10 | 117.998756 | 469.572091 | 370.292715 | 430.996685 | 0 |
| 2.26E-11 | 2.28E-10 | 129.252043 | 220.883508 | 211.595837 | 247.741905 | 0 |
| 2.26E-11 | 2.29E-10 | 124.107684 | 281.164909 | 352.659728 | 289.960095 | 0 |
| 2.27E-11 | 2.29E-10 | 131.181178 | 274.714132 | 207.883629 | 275.578074 | 0 |
| 2.27E-11 | 2.29E-10 | 219.599865 | 298.737716 | 265.422848 | 302.486371 | 4.75243649 |
| 2.27E-11 | 2.29E-10 | 135.360971 | 239.123637 | 354.515832 | 238.927118 | 0 |
| 2.27E-11 | 2.29E-10 | 269.114329 | 312.084151 | 309.041288 | 353.983283 | 0 |
| 2.28E-11 | 2.30E-10 | 127.644431 | 234.007503 | 219.020252 | 251.91733 | 0 |
| 2.28E-11 | 2.30E-10 | 124.750728 | 493.595675 | 340.595054 | 345.168497 | 0 |
| 2.28E-11 | 2.30E-10 | 127.965953 | 324.095943 | 241.293498 | 267.691159 | 0 |
| 2.28E-11 | 2.30E-10 | 119.606369 | 336.107735 | 339.667002 | 384.139133 | 0 |
| 2.28E-11 | 2.30E-10 | 126.036818 | 277.605859 | 233.869083 | 268.619032 | 0 |
| 2.30E-11 | 2.31E-10 | 115.748099 | 409.068249 | 442.680764 | 495.019873 | 0 |
| 2.30E-11 | 2.31E-10 | 145.006646 | 180.844202 | 153.128566 | 191.141694 | 0 |
| 2.29E-11 | 2.31E-10 | 148.543393 | 251.357869 | 165.193241 | 192.997439 | 0 |
| 2.30E-11 | 2.31E-10 | 127.322908 | 222.218152 | 224.588564 | 242.638607 | 0 |
| 2.30E-11 | 2.31E-10 | 252.073637 | 466.457922 | 838.030881 | 810.496456 | 11.8810912 |
| 2.31E-11 | 2.32E-10 | 116.391144 | 393.052527 | 397.20622 | 426.357323 | 0 |
| 2.31E-11 | 2.32E-10 | 112.211351 | 558.103446 | 512.284658 | 484.813278 | 0 |
| 2.32E-11 | 2.33E-10 | 120.570936 | 357.906913 | 309.041288 | 323.363497 | 0 |
| 2.32E-11 | 2.33E-10 | 142.434466 | 170.611934 | 260.782589 | 230.112331 | 0 |
| 2.32E-11 | 2.34E-10 | 110.603739 | 738.280326 | 555.903098 | 579.456253 | 0 |
| 2.33E-11 | 2.34E-10 | 163.654951 | 145.476147 | 167.049345 | 208.771268 | 0 |
| 2.33E-11 | 2.34E-10 | 125.072251 | 282.499552 | 239.437395 | 270.938712 | 0 |
| 2.34E-11 | 2.35E-10 | 122.821594 | 265.594067 | 288.624146 | 274.650202 | 0 |

| | | | | | | |
|---|---|---|---|---|---|---|
| 2.34E-11 | 2.35E-10 | 246.286232 | 364.802572 | 362.8683 | 375.324346 | 0 |
| 2.34E-11 | 2.35E-10 | 172.336058 | 229.11381 | 136.423632 | 218.049991 | 0 |
| 2.36E-11 | 2.37E-10 | 115.105054 | 446.88315 | 406.48674 | 483.42147 | 0 |
| 2.37E-11 | 2.37E-10 | 294.193084 | 240.45828 | 140.135839 | 183.718716 | 0 |
| 2.37E-11 | 2.37E-10 | 147.900348 | 218.214221 | 155.912722 | 180.007227 | 0 |
| 2.37E-11 | 2.38E-10 | 152.080141 | 270.042879 | 165.193241 | 295.527328 | 0 |
| 2.39E-11 | 2.39E-10 | 141.469898 | 72.7380737 | 80.7405168 | 74.6937204 | 470.491212 |
| 2.40E-11 | 2.40E-10 | 232.460765 | 479.137036 | 417.623363 | 463.936151 | 2.37621824 |
| 2.40E-11 | 2.41E-10 | 250.466024 | 129.905306 | 346.163365 | 247.741905 | 0 |
| 2.41E-11 | 2.41E-10 | 128.608998 | 541.197961 | 321.105963 | 360.014453 | 0 |
| 2.41E-11 | 2.41E-10 | 307.053984 | 292.95426 | 317.393756 | 461.616471 | 2.37621824 |
| 2.41E-11 | 2.41E-10 | 131.181178 | 346.784884 | 226.444668 | 325.683178 | 0 |
| 2.41E-11 | 2.41E-10 | 131.824223 | 226.889404 | 187.466487 | 235.679565 | 0 |
| 2.42E-11 | 2.42E-10 | 216.38464 | 1214.08075 | 639.427771 | 878.695071 | 2.37621824 |
| 2.42E-11 | 2.42E-10 | 440.807342 | 320.981775 | 133.639476 | 250.525522 | 0 |
| 2.42E-11 | 2.42E-10 | 150.151006 | 155.708415 | 152.200514 | 164.233398 | 0 |
| 2.43E-11 | 2.42E-10 | 126.036818 | 240.23584 | 226.444668 | 224.081161 | 0 |
| 2.43E-11 | 2.42E-10 | 825.66977 | 345.45024 | 274.703367 | 401.768707 | 0 |
| 2.43E-11 | 2.42E-10 | 144.042078 | 154.373771 | 208.811681 | 202.276162 | 0 |
| 2.43E-11 | 2.43E-10 | 119.927891 | 335.440413 | 304.401029 | 365.581687 | 0 |
| 2.45E-11 | 2.45E-10 | 156.259933 | 263.147221 | 153.128566 | 212.018821 | 0 |
| 2.46E-11 | 2.45E-10 | 162.368861 | 243.350008 | 178.185968 | 220.833608 | 14.2573095 |
| 2.47E-11 | 2.46E-10 | 185.51848 | 530.965694 | 467.738166 | 490.844448 | 2.37621824 |
| 2.47E-11 | 2.46E-10 | 174.265193 | 155.485974 | 315.537652 | 209.235204 | 0 |
| 2.48E-11 | 2.47E-10 | 228.280972 | 530.298372 | 436.184401 | 521.000298 | 2.37621824 |
| 2.49E-11 | 2.48E-10 | 123.786161 | 345.005359 | 274.703367 | 272.794457 | 0 |
| 2.50E-11 | 2.48E-10 | 120.892459 | 344.782918 | 303.472977 | 287.176478 | 0 |
| 2.50E-11 | 2.49E-10 | 178.123463 | 506.052348 | 396.278169 | 395.737537 | 7.12865473 |
| 2.50E-11 | 2.49E-10 | 128.930521 | 262.257458 | 232.941031 | 204.595843 | 0 |
| 2.50E-11 | 2.49E-10 | 119.606369 | 336.552616 | 296.976613 | 316.404455 | 0 |
| 2.50E-11 | 2.49E-10 | 207.060488 | 264.926746 | 129.927268 | 211.090949 | 0 |
| 2.51E-11 | 2.49E-10 | 136.004016 | 203.978023 | 172.617657 | 177.22361 | 0 |
| 2.51E-11 | 2.49E-10 | 118.963324 | 310.749508 | 341.523105 | 315.476583 | 0 |
| 2.51E-11 | 2.49E-10 | 117.355711 | 337.219938 | 389.781805 | 362.334134 | 0 |
| 2.52E-11 | 2.50E-10 | 142.755988 | 160.602108 | 236.653239 | 212.946693 | 0 |
| 2.53E-11 | 2.51E-10 | 123.786161 | 253.359835 | 234.797135 | 254.237011 | 0 |
| 2.53E-11 | 2.51E-10 | 134.074881 | 202.64338 | 176.329864 | 226.864778 | 0 |
| 2.53E-11 | 2.51E-10 | 123.786161 | 274.936572 | 236.653239 | 263.051798 | 0 |
| 2.54E-11 | 2.52E-10 | 120.570936 | 336.775057 | 283.983887 | 302.950307 | 0 |
| 2.54E-11 | 2.52E-10 | 114.140486 | 425.751294 | 434.328297 | 395.273601 | 0 |
| 2.55E-11 | 2.53E-10 | 157.546023 | 164.161157 | 141.991943 | 161.913717 | 0 |
| 2.56E-11 | 2.54E-10 | 149.829483 | 420.857601 | 286.768042 | 203.204034 | 0 |
| 2.57E-11 | 2.54E-10 | 132.467268 | 329.434517 | 210.667785 | 285.784669 | 0 |

| | | | | | | |
|---|---|---|---|---|---|---|
| 2.57E-11 | 2.55E-10 | 133.110313 | 187.072538 | 191.178695 | 233.82382 | 0 |
| 2.58E-11 | 2.55E-10 | 154.973843 | 241.125602 | 154.98467 | 177.687546 | 0 |
| 2.58E-11 | 2.56E-10 | 115.105054 | 373.477755 | 451.033232 | 463.008279 | 0 |
| 2.58E-11 | 2.56E-10 | 113.497441 | 482.251204 | 405.558688 | 449.554131 | 0 |
| 2.59E-11 | 2.56E-10 | 137.933151 | 193.968197 | 210.667785 | 291.351903 | 0 |
| 2.59E-11 | 2.57E-10 | 139.862286 | 443.546541 | 225.516616 | 342.848816 | 0 |
| 2.60E-11 | 2.57E-10 | 127.644431 | 211.318563 | 244.077654 | 277.897755 | 0 |
| 2.60E-11 | 2.57E-10 | 312.841389 | 338.554582 | 267.278952 | 289.960095 | 2.37621824 |
| 2.61E-11 | 2.58E-10 | 126.358341 | 342.558512 | 274.703367 | 439.347535 | 0 |
| 2.61E-11 | 2.58E-10 | 117.355711 | 347.229765 | 418.551415 | 347.488177 | 0 |
| 2.63E-11 | 2.59E-10 | 121.535504 | 269.375557 | 269.135056 | 253.773075 | 0 |
| 2.63E-11 | 2.60E-10 | 122.178549 | 292.73182 | 251.502069 | 308.981477 | 0 |
| 2.64E-11 | 2.61E-10 | 129.573566 | 319.424691 | 215.308045 | 288.568286 | 0 |
| 2.65E-11 | 2.61E-10 | 134.396403 | 171.946578 | 173.545708 | 192.533503 | 0 |
| 2.65E-11 | 2.62E-10 | 125.715296 | 238.011434 | 216.236097 | 214.802438 | 0 |
| 2.66E-11 | 2.63E-10 | 141.791421 | 153.706449 | 176.329864 | 163.769461 | 0 |
| 2.66E-11 | 2.63E-10 | 119.284846 | 288.95033 | 286.768042 | 300.630626 | 0 |
| 2.67E-11 | 2.63E-10 | 215.09855 | 292.064498 | 302.544925 | 310.373285 | 0 |
| 2.67E-11 | 2.63E-10 | 129.573566 | 408.178487 | 250.574018 | 303.414243 | 0 |
| 2.67E-11 | 2.63E-10 | 125.393773 | 253.359835 | 379.573234 | 320.579881 | 0 |
| 2.68E-11 | 2.64E-10 | 108.996126 | 620.831693 | 493.72362 | 564.14636 | 0 |
| 2.68E-11 | 2.64E-10 | 134.074881 | 585.018758 | 492.795568 | 528.423276 | 16.6335277 |
| 2.70E-11 | 2.65E-10 | 123.464639 | 244.907092 | 232.941031 | 229.648395 | 0 |
| 2.71E-11 | 2.67E-10 | 123.786161 | 334.550651 | 256.142329 | 265.371479 | 0 |
| 2.72E-11 | 2.68E-10 | 110.282216 | 673.994995 | 489.08336 | 540.485616 | 0 |
| 2.73E-11 | 2.68E-10 | 112.532874 | 431.75719 | 495.579724 | 417.0786 | 0 |
| 2.73E-11 | 2.68E-10 | 126.358341 | 231.115776 | 202.315318 | 241.246799 | 0 |
| 2.74E-11 | 2.69E-10 | 317.664226 | 536.526709 | 495.579724 | 489.45264 | 45.1481466 |
| 2.74E-11 | 2.70E-10 | 127.965953 | 270.710201 | 341.523105 | 229.184459 | 0 |
| 2.76E-11 | 2.71E-10 | 459.777169 | 745.175985 | 1188.83451 | 1013.23655 | 38.0194919 |
| 2.76E-11 | 2.71E-10 | 130.216611 | 214.432731 | 214.379993 | 179.079354 | 0 |
| 2.77E-11 | 2.72E-10 | 116.712666 | 341.223869 | 314.6096 | 333.570093 | 0 |
| 2.77E-11 | 2.72E-10 | 210.275712 | 128.348222 | 190.250643 | 228.256586 | 0 |
| 2.78E-11 | 2.73E-10 | 128.287476 | 298.960156 | 215.308045 | 238.463182 | 0 |
| 2.78E-11 | 2.73E-10 | 145.006646 | 154.373771 | 221.804408 | 180.007227 | 0 |
| 2.78E-11 | 2.73E-10 | 171.37149 | 311.63927 | 231.084927 | 267.227223 | 7.12865473 |
| 2.79E-11 | 2.73E-10 | 180.37412 | 238.678756 | 202.315318 | 132.221803 | 0 |
| 2.79E-11 | 2.73E-10 | 113.818964 | 397.94622 | 370.292715 | 388.778495 | 0 |
| 2.79E-11 | 2.73E-10 | 119.927891 | 370.808468 | 280.271679 | 338.209454 | 0 |
| 2.79E-11 | 2.73E-10 | 114.783531 | 369.028943 | 365.652455 | 351.199667 | 0 |
| 2.79E-11 | 2.74E-10 | 115.105054 | 412.404858 | 386.997649 | 344.240624 | 0 |
| 2.80E-11 | 2.74E-10 | 190.66284 | 711.142574 | 463.097907 | 563.682424 | 2.37621824 |
| 2.82E-11 | 2.76E-10 | 130.859656 | 202.420939 | 173.545708 | 201.34829 | 0 |

| | | | | | | |
|---|---|---|---|---|---|---|
| 2.83E-11 | 2.76E-10 | 159.796681 | 334.10577 | 172.617657 | 392.489984 | 0 |
| 2.83E-11 | 2.76E-10 | 652.690667 | 601.924243 | 438.040505 | 552.08402 | 54.6530196 |
| 2.83E-11 | 2.76E-10 | 121.535504 | 260.700374 | 243.149602 | 286.712542 | 0 |
| 2.84E-11 | 2.77E-10 | 125.393773 | 290.062533 | 218.0922 | 255.628819 | 0 |
| 2.85E-11 | 2.78E-10 | 148.221871 | 344.115596 | 177.257916 | 262.587862 | 0 |
| 2.85E-11 | 2.78E-10 | 137.933151 | 287.170805 | 214.379993 | 184.182652 | 0 |
| 2.86E-11 | 2.79E-10 | 116.391144 | 327.432552 | 311.825444 | 342.848816 | 0 |
| 2.86E-11 | 2.79E-10 | 136.647061 | 312.306592 | 202.315318 | 214.802438 | 0 |
| 2.87E-11 | 2.80E-10 | 126.358341 | 300.962122 | 215.308045 | 277.433818 | 0 |
| 2.87E-11 | 2.80E-10 | 124.750728 | 211.096122 | 223.660512 | 250.989458 | 0 |
| 2.88E-11 | 2.80E-10 | 121.857026 | 241.125602 | 232.941031 | 241.710735 | 0 |
| 2.88E-11 | 2.81E-10 | 251.109069 | 181.289083 | 143.848047 | 167.480951 | 0 |
| 2.91E-11 | 2.84E-10 | 638.543677 | 321.204215 | 189.322591 | 203.204034 | 0 |
| 2.92E-11 | 2.84E-10 | 152.080141 | 256.029122 | 270.99116 | 153.09893 | 0 |
| 2.92E-11 | 2.84E-10 | 115.105054 | 326.76523 | 369.364663 | 381.355516 | 0 |
| 2.93E-11 | 2.85E-10 | 174.265193 | 251.357869 | 389.781805 | 162.377653 | 0 |
| 2.94E-11 | 2.86E-10 | 233.10381 | 186.850098 | 127.143113 | 212.482757 | 0 |
| 2.95E-11 | 2.87E-10 | 145.971213 | 245.574414 | 154.98467 | 198.100737 | 0 |
| 2.96E-11 | 2.88E-10 | 132.467268 | 233.562622 | 182.826228 | 185.110524 | 0 |
| 2.97E-11 | 2.89E-10 | 135.682493 | 302.741646 | 275.631419 | 291.351903 | 9.50487297 |
| 2.97E-11 | 2.89E-10 | 115.748099 | 361.910844 | 311.825444 | 340.529135 | 0 |
| 2.97E-11 | 2.89E-10 | 114.140486 | 421.747363 | 345.235313 | 386.458814 | 0 |
| 2.98E-11 | 2.89E-10 | 112.211351 | 795.89244 | 475.162581 | 573.889019 | 0 |
| 2.98E-11 | 2.90E-10 | 145.006646 | 158.822583 | 209.739733 | 259.804245 | 0 |
| 2.99E-11 | 2.90E-10 | 167.191698 | 131.46239 | 190.250643 | 192.997439 | 0 |
| 2.99E-11 | 2.90E-10 | 156.902978 | 195.0804 | 161.481034 | 140.572654 | 0 |
| 3.00E-11 | 2.91E-10 | 203.52374 | 797.671964 | 650.564394 | 714.461673 | 0 |
| 3.00E-11 | 2.91E-10 | 117.034189 | 286.725924 | 324.818171 | 340.065199 | 0 |
| 3.00E-11 | 2.91E-10 | 117.034189 | 371.475789 | 310.897392 | 307.589668 | 0 |
| 3.00E-11 | 2.91E-10 | 126.679863 | 211.096122 | 190.250643 | 195.781056 | 0 |
| 3.01E-11 | 2.92E-10 | 105.137856 | 542.977486 | 576.32024 | 570.17753 | 0 |
| 3.01E-11 | 2.92E-10 | 229.24554 | 337.219938 | 244.077654 | 205.987651 | 45.1481466 |
| 3.02E-11 | 2.92E-10 | 146.292736 | 175.950509 | 142.919995 | 193.925311 | 0 |
| 3.03E-11 | 2.94E-10 | 132.145746 | 182.846167 | 164.265189 | 181.399035 | 0 |
| 3.06E-11 | 2.96E-10 | 130.538133 | 201.308736 | 168.905449 | 196.244992 | 0 |
| 3.06E-11 | 2.97E-10 | 165.584086 | 250.468107 | 177.257916 | 145.212015 | 0 |
| 3.06E-11 | 2.97E-10 | 130.216611 | 186.627657 | 173.545708 | 208.771268 | 0 |
| 3.08E-11 | 2.98E-10 | 125.393773 | 205.090226 | 211.595837 | 251.453394 | 0 |
| 3.08E-11 | 2.98E-10 | 120.249414 | 250.245666 | 237.581291 | 267.227223 | 0 |
| 3.08E-11 | 2.98E-10 | 122.500071 | 230.670894 | 214.379993 | 221.76148 | 0 |
| 3.09E-11 | 2.99E-10 | 189.055228 | 497.599605 | 443.608816 | 403.160515 | 0 |
| 3.10E-11 | 3.00E-10 | 195.164155 | 140.582454 | 168.905449 | 151.243185 | 0 |
| 3.12E-11 | 3.01E-10 | 118.641801 | 319.647131 | 262.638692 | 283.001052 | 0 |

| | | | | | | |
|---|---|---|---|---|---|---|
| 3.12E-11 | 3.01E-10 | 123.786161 | 343.893156 | 236.653239 | 283.464988 | 0 |
| 3.13E-11 | 3.02E-10 | 173.622148 | 136.578524 | 272.847264 | 264.907542 | 0 |
| 3.13E-11 | 3.02E-10 | 112.532874 | 383.710022 | 353.58778 | 400.376899 | 0 |
| 3.14E-11 | 3.04E-10 | 118.641801 | 289.617651 | 256.142329 | 263.051798 | 0 |
| 3.16E-11 | 3.05E-10 | 497.395302 | 1879.84545 | 1521.07709 | 1685.0161 | 4.75243649 |
| 3.20E-11 | 3.09E-10 | 108.031559 | 445.326066 | 490.011412 | 547.908595 | 0 |
| 3.22E-11 | 3.10E-10 | 117.034189 | 300.2948 | 266.3509 | 302.486371 | 0 |
| 3.22E-11 | 3.10E-10 | 117.677234 | 269.153117 | 258.926485 | 276.505946 | 0 |
| 3.23E-11 | 3.11E-10 | 125.393773 | 318.757369 | 216.236097 | 276.04201 | 0 |
| 3.23E-11 | 3.11E-10 | 137.933151 | 153.261568 | 196.747006 | 169.336695 | 0 |
| 3.23E-11 | 3.11E-10 | 121.213981 | 228.668929 | 219.020252 | 224.081161 | 0 |
| 3.24E-11 | 3.12E-10 | 116.712666 | 311.861711 | 270.99116 | 296.455201 | 0 |
| 3.27E-11 | 3.15E-10 | 129.573566 | 168.387528 | 186.538435 | 201.34829 | 0 |
| 3.28E-11 | 3.16E-10 | 229.24554 | 377.481685 | 438.040505 | 496.875618 | 0 |
| 3.30E-11 | 3.17E-10 | 106.102424 | 494.930318 | 761.002572 | 720.956779 | 0 |
| 3.31E-11 | 3.18E-10 | 127.965953 | 203.088261 | 175.401812 | 184.646588 | 0 |
| 3.31E-11 | 3.18E-10 | 136.647061 | 182.623726 | 157.768826 | 157.274355 | 0 |
| 3.32E-11 | 3.19E-10 | 117.998756 | 294.511344 | 248.717914 | 290.424031 | 0 |
| 3.32E-11 | 3.19E-10 | 202.559173 | 368.806502 | 318.321808 | 340.993071 | 0 |
| 3.35E-11 | 3.22E-10 | 119.284846 | 295.401107 | 237.581291 | 270.01084 | 0 |
| 3.36E-11 | 3.22E-10 | 122.178549 | 349.899052 | 239.437395 | 289.496158 | 0 |
| 3.36E-11 | 3.22E-10 | 190.984363 | 470.906734 | 389.781805 | 431.924557 | 11.8810912 |
| 3.36E-11 | 3.23E-10 | 126.679863 | 408.400928 | 237.581291 | 356.302964 | 0 |
| 3.37E-11 | 3.23E-10 | 118.963324 | 254.694478 | 232.012979 | 247.741905 | 0 |
| 3.38E-11 | 3.24E-10 | 405.439868 | 654.642664 | 460.313751 | 544.197105 | 0 |
| 3.38E-11 | 3.24E-10 | 133.431836 | 175.728068 | 155.912722 | 190.213822 | 0 |
| 3.38E-11 | 3.24E-10 | 114.783531 | 365.469893 | 329.458431 | 296.919137 | 0 |
| 3.38E-11 | 3.24E-10 | 305.767894 | 406.843844 | 114.150386 | 367.901368 | 0 |
| 3.39E-11 | 3.25E-10 | 114.462009 | 330.101839 | 298.832717 | 303.878179 | 0 |
| 3.40E-11 | 3.26E-10 | 245.643187 | 224.887439 | 118.790645 | 230.576267 | 0 |
| 3.42E-11 | 3.28E-10 | 134.074881 | 315.865641 | 192.106747 | 330.32254 | 0 |
| 3.42E-11 | 3.28E-10 | 154.009276 | 187.294979 | 132.711424 | 186.502333 | 0 |
| 3.44E-11 | 3.29E-10 | 117.677234 | 250.468107 | 264.494796 | 296.919137 | 0 |
| 3.47E-11 | 3.32E-10 | 185.51848 | 153.92889 | 141.063891 | 149.851377 | 0 |
| 3.48E-11 | 3.33E-10 | 115.748099 | 272.044845 | 292.336354 | 270.474776 | 0 |
| 3.49E-11 | 3.34E-10 | 118.641801 | 246.686617 | 279.343627 | 236.143501 | 0 |
| 3.50E-11 | 3.34E-10 | 109.317649 | 379.928532 | 455.673491 | 426.357323 | 0 |
| 3.51E-11 | 3.35E-10 | 162.368861 | 762.748791 | 739.657378 | 758.071671 | 2.37621824 |
| 3.50E-11 | 3.35E-10 | 124.429206 | 302.964087 | 211.595837 | 245.88616 | 0 |
| 3.51E-11 | 3.36E-10 | 116.391144 | 274.714132 | 257.070381 | 263.515734 | 0 |
| 3.52E-11 | 3.36E-10 | 141.791421 | 215.322493 | 145.704151 | 210.163077 | 0 |
| 3.53E-11 | 3.37E-10 | 158.510591 | 201.976058 | 434.328297 | 225.47297 | 0 |
| 3.54E-11 | 3.38E-10 | 120.570936 | 212.430766 | 245.005706 | 220.369672 | 0 |

| | | | | | | |
|---|---|---|---|---|---|---|
| 3.54E-11 | 3.38E-10 | 130.216611 | 203.755583 | 161.481034 | 196.708928 | 0 |
| 3.55E-11 | 3.38E-10 | 219.599865 | 416.186348 | 446.392972 | 478.782108 | 0 |
| 3.56E-11 | 3.39E-10 | 104.173289 | 578.567981 | 496.507776 | 514.041256 | 0 |
| 3.56E-11 | 3.40E-10 | 153.687753 | 424.639091 | 183.75428 | 281.145308 | 0 |
| 3.57E-11 | 3.40E-10 | 117.677234 | 308.747542 | 245.005706 | 280.217435 | 0 |
| 3.57E-11 | 3.40E-10 | 140.183808 | 148.590316 | 202.315318 | 167.017014 | 0 |
| 3.57E-11 | 3.40E-10 | 106.102424 | 483.140967 | 488.155308 | 439.347535 | 0 |
| 3.57E-11 | 3.40E-10 | 127.322908 | 189.964266 | 167.977397 | 188.822014 | 0 |
| 3.57E-11 | 3.40E-10 | 154.973843 | 140.137573 | 145.704151 | 172.584248 | 0 |
| 3.58E-11 | 3.41E-10 | 120.249414 | 216.212256 | 219.948304 | 211.090949 | 0 |
| 3.59E-11 | 3.41E-10 | 130.216611 | 200.196533 | 161.481034 | 207.37946 | 0 |
| 3.60E-11 | 3.42E-10 | 117.998756 | 331.658923 | 251.502069 | 281.145308 | 0 |
| 3.60E-11 | 3.43E-10 | 111.568306 | 337.88726 | 329.458431 | 345.168497 | 0 |
| 3.61E-11 | 3.43E-10 | 118.320279 | 311.194389 | 247.789862 | 257.020628 | 0 |
| 3.61E-11 | 3.43E-10 | 110.925261 | 349.899052 | 429.688038 | 371.148921 | 0 |
| 3.61E-11 | 3.43E-10 | 117.677234 | 268.263354 | 248.717914 | 235.215629 | 0 |
| 3.62E-11 | 3.44E-10 | 133.753358 | 191.52135 | 155.912722 | 214.802438 | 0 |
| 3.63E-11 | 3.45E-10 | 301.588101 | 537.638912 | 444.536868 | 499.195299 | 2.37621824 |
| 3.64E-11 | 3.46E-10 | 187.126093 | 811.0184 | 801.836856 | 951.996982 | 0 |
| 3.65E-11 | 3.46E-10 | 110.282216 | 359.019116 | 351.731677 | 365.581687 | 0 |
| 3.66E-11 | 3.47E-10 | 116.069621 | 276.493656 | 262.638692 | 320.115944 | 0 |
| 3.67E-11 | 3.48E-10 | 299.015921 | 396.166695 | 251.502069 | 319.652008 | 0 |
| 3.67E-11 | 3.48E-10 | 217.349207 | 242.237805 | 254.286225 | 271.866585 | 0 |
| 3.68E-11 | 3.49E-10 | 116.712666 | 244.23977 | 257.070381 | 281.609244 | 0 |
| 3.68E-11 | 3.49E-10 | 117.034189 | 266.70627 | 235.725187 | 266.299351 | 0 |
| 3.70E-11 | 3.51E-10 | 291.942426 | 180.844202 | 144.776099 | 178.151482 | 0 |
| 3.71E-11 | 3.51E-10 | 121.213981 | 398.835982 | 298.832717 | 257.020628 | 0 |
| 3.71E-11 | 3.51E-10 | 112.854396 | 361.243522 | 303.472977 | 386.92275 | 0 |
| 3.73E-11 | 3.53E-10 | 136.325538 | 149.924959 | 163.337137 | 189.28595 | 0 |
| 3.73E-11 | 3.53E-10 | 131.181178 | 205.979989 | 215.308045 | 159.1301 | 0 |
| 3.73E-11 | 3.53E-10 | 120.892459 | 231.115776 | 203.24337 | 251.91733 | 0 |
| 3.74E-11 | 3.53E-10 | 191.94893 | 169.277291 | 128.999216 | 162.377653 | 0 |
| 3.75E-11 | 3.54E-10 | 147.257303 | 151.037162 | 142.919995 | 187.894141 | 0 |
| 3.75E-11 | 3.54E-10 | 113.497441 | 343.003394 | 287.696094 | 308.517541 | 0 |
| 3.75E-11 | 3.54E-10 | 129.252043 | 163.048954 | 169.833501 | 190.213822 | 0 |
| 3.75E-11 | 3.54E-10 | 141.148376 | 189.741825 | 154.056618 | 148.459568 | 0 |
| 3.76E-11 | 3.55E-10 | 113.175919 | 323.873503 | 287.696094 | 301.094562 | 0 |
| 3.77E-11 | 3.56E-10 | 100.315019 | 606.595496 | 566.111669 | 581.775934 | 0 |
| 3.79E-11 | 3.58E-10 | 118.641801 | 373.477755 | 255.214277 | 303.878179 | 0 |
| 3.79E-11 | 3.58E-10 | 192.270453 | 278.273181 | 265.422848 | 283.464988 | 7.12865473 |
| 3.81E-11 | 3.59E-10 | 115.105054 | 275.381453 | 273.775316 | 252.381266 | 0 |
| 3.81E-11 | 3.59E-10 | 115.426576 | 159.489905 | 123.430905 | 166.089142 | 9.50487297 |
| 3.83E-11 | 3.61E-10 | 115.748099 | 275.159013 | 268.207004 | 243.102543 | 0 |

| | | | | | | |
|---|---|---|---|---|---|---|
| 3.83E-11 | 3.61E-10 | 118.641801 | 211.541003 | 228.300772 | 231.968076 | 0 |
| 3.85E-11 | 3.63E-10 | 178.766508 | 629.061995 | 484.443101 | 495.48381 | 2.37621824 |
| 3.87E-11 | 3.64E-10 | 122.500071 | 189.519385 | 204.171422 | 186.502333 | 0 |
| 3.88E-11 | 3.65E-10 | 136.004016 | 158.155261 | 182.826228 | 148.459568 | 0 |
| 3.89E-11 | 3.66E-10 | 276.187824 | 165.940682 | 156.840774 | 162.377653 | 0 |
| 3.90E-11 | 3.67E-10 | 121.857026 | 286.503483 | 205.099474 | 268.619032 | 0 |
| 3.91E-11 | 3.68E-10 | 121.857026 | 243.350008 | 206.027526 | 286.248605 | 0 |
| 3.92E-11 | 3.68E-10 | 121.535504 | 194.413078 | 219.020252 | 193.925311 | 0 |
| 3.92E-11 | 3.69E-10 | 154.652321 | 155.263534 | 139.207788 | 142.892335 | 0 |
| 3.92E-11 | 3.69E-10 | 116.391144 | 253.582275 | 233.869083 | 239.85499 | 0 |
| 3.92E-11 | 3.69E-10 | 123.786161 | 285.836161 | 199.531162 | 286.248605 | 0 |
| 3.94E-11 | 3.70E-10 | 135.360971 | 168.609969 | 146.632203 | 157.274355 | 0 |
| 3.96E-11 | 3.72E-10 | 132.145746 | 246.686617 | 234.797135 | 163.305525 | 0 |
| 3.97E-11 | 3.73E-10 | 170.728445 | 134.798999 | 150.344411 | 200.420417 | 0 |
| 3.98E-11 | 3.74E-10 | 0 | 0.66732178 | 0 | 1.39180845 | 216.23586 |
| 4.01E-11 | 3.76E-10 | 149.507961 | 174.170984 | 133.639476 | 155.882547 | 0 |
| 4.01E-11 | 3.77E-10 | 118.641801 | 230.226013 | 205.099474 | 235.215629 | 0 |
| 4.04E-11 | 3.79E-10 | 153.366231 | 208.649276 | 131.783372 | 212.482757 | 0 |
| 4.07E-11 | 3.82E-10 | 113.175919 | 301.851884 | 377.71713 | 295.527328 | 0 |
| 4.08E-11 | 3.83E-10 | 117.034189 | 222.218152 | 220.876356 | 231.968076 | 0 |
| 4.09E-11 | 3.83E-10 | 197.09329 | 505.385026 | 396.278169 | 415.222855 | 4.75243649 |
| 4.11E-11 | 3.85E-10 | 136.004016 | 143.474182 | 161.481034 | 176.295737 | 0 |
| 4.11E-11 | 3.85E-10 | 113.497441 | 282.277112 | 259.854537 | 267.691159 | 0 |
| 4.11E-11 | 3.85E-10 | 126.036818 | 163.716276 | 193.962851 | 202.740098 | 0 |
| 4.13E-11 | 3.86E-10 | 111.246784 | 297.403072 | 376.789078 | 356.7669 | 0 |
| 4.13E-11 | 3.87E-10 | 154.973843 | 301.629443 | 736.873222 | 296.455201 | 0 |
| 4.13E-11 | 3.87E-10 | 165.584086 | 312.084151 | 154.98467 | 185.574461 | 0 |
| 4.15E-11 | 3.89E-10 | 118.641801 | 218.881543 | 207.883629 | 249.133713 | 0 |
| 4.15E-11 | 3.89E-10 | 187.126093 | 399.058423 | 354.515832 | 388.778495 | 30.8908372 |
| 4.17E-11 | 3.90E-10 | 108.674604 | 367.694299 | 428.759986 | 346.560305 | 0 |
| 4.18E-11 | 3.90E-10 | 117.355711 | 225.554761 | 235.725187 | 210.163077 | 0 |
| 4.19E-11 | 3.91E-10 | 230.53163 | 376.591923 | 322.962067 | 354.44722 | 2.37621824 |
| 4.20E-11 | 3.93E-10 | 126.679863 | 175.950509 | 232.012979 | 260.732117 | 0 |
| 4.21E-11 | 3.93E-10 | 169.763878 | 481.806323 | 449.177128 | 485.277214 | 9.50487297 |
| 4.21E-11 | 3.93E-10 | 125.715296 | 166.608004 | 213.451941 | 189.28595 | 0 |
| 4.21E-11 | 3.93E-10 | 131.181178 | 174.838306 | 156.840774 | 153.09893 | 0 |
| 4.22E-11 | 3.94E-10 | 110.282216 | 314.530998 | 302.544925 | 322.899561 | 0 |
| 4.24E-11 | 3.96E-10 | 108.031559 | 430.644987 | 343.379209 | 410.119558 | 0 |
| 4.27E-11 | 3.98E-10 | 131.502701 | 234.674825 | 197.675058 | 159.1301 | 0 |
| 4.26E-11 | 3.98E-10 | 112.854396 | 263.592102 | 296.976613 | 268.155095 | 0 |
| 4.26E-11 | 3.98E-10 | 176.837373 | 255.584241 | 140.135839 | 160.985845 | 0 |
| 4.27E-11 | 3.98E-10 | 109.960694 | 702.467391 | 414.839207 | 478.318172 | 0 |
| 4.27E-11 | 3.98E-10 | 155.295366 | 189.074504 | 149.416359 | 135.469356 | 0 |

| | | | | | | |
|---|---|---|---|---|---|---|
| 4.28E-11 | 3.98E-10 | 142.112943 | 256.918884 | 152.200514 | 270.938712 | 0 |
| 4.28E-11 | 3.99E-10 | 113.497441 | 263.592102 | 294.192458 | 253.773075 | 0 |
| 4.29E-11 | 3.99E-10 | 114.783531 | 586.575842 | 331.314534 | 438.883599 | 0 |
| 4.29E-11 | 3.99E-10 | 113.175919 | 319.20225 | 257.998433 | 296.919137 | 0 |
| 4.29E-11 | 3.99E-10 | 117.034189 | 245.129533 | 227.37272 | 211.090949 | 0 |
| 4.29E-11 | 3.99E-10 | 122.178549 | 288.505448 | 542.910371 | 578.992317 | 0 |
| 4.30E-11 | 4.00E-10 | 168.477788 | 305.410933 | 171.689605 | 212.946693 | 23.7621824 |
| 4.30E-11 | 4.00E-10 | 116.391144 | 250.245666 | 215.308045 | 249.597649 | 0 |
| 4.30E-11 | 4.00E-10 | 133.753358 | 262.70234 | 163.337137 | 207.37946 | 0 |
| 4.31E-11 | 4.00E-10 | 108.031559 | 384.377344 | 335.026742 | 363.725943 | 0 |
| 4.31E-11 | 4.01E-10 | 176.837373 | 154.373771 | 132.711424 | 148.923505 | 0 |
| 4.32E-11 | 4.01E-10 | 110.282216 | 294.511344 | 319.249859 | 308.517541 | 0 |
| 4.36E-11 | 4.05E-10 | 110.282216 | 324.095943 | 298.832717 | 363.262006 | 0 |
| 4.37E-11 | 4.05E-10 | 109.639171 | 350.788814 | 503.932191 | 388.314559 | 0 |
| 4.37E-11 | 4.06E-10 | 113.818964 | 246.909058 | 251.502069 | 242.638607 | 0 |
| 4.37E-11 | 4.06E-10 | 190.984363 | 246.019295 | 191.178695 | 231.040203 | 0 |
| 4.38E-11 | 4.06E-10 | 117.034189 | 236.231909 | 208.811681 | 252.381266 | 0 |
| 4.38E-11 | 4.06E-10 | 132.788791 | 154.818652 | 157.768826 | 147.06776 | 0 |
| 4.39E-11 | 4.07E-10 | 130.859656 | 190.409147 | 148.488307 | 175.831801 | 0 |
| 4.39E-11 | 4.07E-10 | 145.006646 | 167.275326 | 138.279736 | 144.284143 | 0 |
| 4.40E-11 | 4.08E-10 | 128.287476 | 180.39932 | 181.898176 | 151.707121 | 0 |
| 4.41E-11 | 4.08E-10 | 114.783531 | 258.475968 | 226.444668 | 257.020628 | 0 |
| 4.41E-11 | 4.08E-10 | 178.444985 | 352.345898 | 141.991943 | 230.112331 | 0 |
| 4.43E-11 | 4.10E-10 | 126.036818 | 296.958191 | 188.394539 | 236.143501 | 0 |
| 4.43E-11 | 4.10E-10 | 112.854396 | 357.239591 | 484.443101 | 317.796264 | 0 |
| 4.43E-11 | 4.10E-10 | 105.459379 | 373.032874 | 434.328297 | 423.573706 | 0 |
| 4.43E-11 | 4.10E-10 | 115.105054 | 220.883508 | 246.86181 | 243.566479 | 0 |
| 4.44E-11 | 4.11E-10 | 157.867546 | 356.349829 | 257.070381 | 268.619032 | 0 |
| 4.47E-11 | 4.13E-10 | 119.284846 | 195.747721 | 189.322591 | 212.018821 | 0 |
| 4.48E-11 | 4.14E-10 | 117.034189 | 222.885474 | 202.315318 | 230.112331 | 0 |
| 4.49E-11 | 4.14E-10 | 149.186438 | 433.981596 | 184.682331 | 277.433818 | 0 |
| 4.51E-11 | 4.17E-10 | 129.895088 | 202.198499 | 180.042072 | 151.707121 | 0 |
| 4.52E-11 | 4.17E-10 | 444.665612 | 194.413078 | 178.185968 | 192.069567 | 0 |
| 4.53E-11 | 4.18E-10 | 188.733705 | 590.357332 | 481.658945 | 570.17753 | 4.75243649 |
| 4.53E-11 | 4.18E-10 | 117.998756 | 194.857959 | 209.739733 | 200.884354 | 0 |
| 4.53E-11 | 4.18E-10 | 129.573566 | 178.174915 | 147.560255 | 179.543291 | 0 |
| 4.54E-11 | 4.19E-10 | 111.246784 | 345.227799 | 273.775316 | 318.2602 | 0 |
| 4.54E-11 | 4.19E-10 | 139.540763 | 135.02144 | 164.265189 | 169.336695 | 0 |
| 4.55E-11 | 4.20E-10 | 117.998756 | 206.202429 | 195.818954 | 199.028609 | 0 |
| 4.56E-11 | 4.20E-10 | 136.647061 | 258.475968 | 155.912722 | 261.659989 | 0 |
| 4.56E-11 | 4.20E-10 | 110.925261 | 291.174735 | 289.552198 | 348.879986 | 0 |
| 4.56E-11 | 4.20E-10 | 110.282216 | 300.2948 | 278.415575 | 303.878179 | 0 |
| 4.60E-11 | 4.23E-10 | 107.710036 | 319.20225 | 365.652455 | 351.663603 | 0 |

| | | | | | | |
|---|---|---|---|---|---|---|
| 4.60E-11 | 4.23E-10 | 109.960694 | 411.737537 | 314.6096 | 329.394667 | 0 |
| 4.62E-11 | 4.25E-10 | 106.423946 | 386.82419 | 399.990376 | 521.464234 | 0 |
| 4.62E-11 | 4.25E-10 | 159.475158 | 147.922994 | 133.639476 | 195.781056 | 0 |
| 4.63E-11 | 4.26E-10 | 133.753358 | 141.249776 | 185.610383 | 184.182652 | 0 |
| 4.63E-11 | 4.26E-10 | 139.862286 | 259.588171 | 218.0922 | 232.895948 | 26.1384007 |
| 4.63E-11 | 4.26E-10 | 95.8137038 | 696.906376 | 716.45608 | 563.218488 | 0 |
| 4.63E-11 | 4.26E-10 | 115.748099 | 213.320528 | 212.523889 | 225.47297 | 0 |
| 4.65E-11 | 4.27E-10 | 144.363601 | 161.936751 | 134.567528 | 146.603824 | 0 |
| 4.65E-11 | 4.27E-10 | 158.510591 | 136.356083 | 137.351684 | 164.233398 | 0 |
| 4.65E-11 | 4.27E-10 | 118.320279 | 249.800785 | 203.24337 | 207.37946 | 0 |
| 4.65E-11 | 4.27E-10 | 117.355711 | 651.750936 | 331.314534 | 436.099982 | 0 |
| 4.68E-11 | 4.30E-10 | 161.404293 | 396.166695 | 345.235313 | 381.355516 | 7.12865473 |
| 4.69E-11 | 4.30E-10 | 299.980489 | 293.621582 | 244.077654 | 280.681372 | 0 |
| 4.69E-11 | 4.30E-10 | 110.925261 | 261.367696 | 293.264406 | 283.464988 | 0 |
| 4.70E-11 | 4.31E-10 | 135.360971 | 230.003573 | 147.560255 | 196.244992 | 0 |
| 4.71E-11 | 4.32E-10 | 121.213981 | 186.850098 | 200.459214 | 245.88616 | 0 |
| 4.71E-11 | 4.32E-10 | 135.039448 | 264.259424 | 155.912722 | 238.927118 | 0 |
| 4.73E-11 | 4.33E-10 | 211.561802 | 450.219759 | 392.565961 | 447.698386 | 2.37621824 |
| 4.73E-11 | 4.33E-10 | 98.3858838 | 596.140788 | 498.363879 | 643.015506 | 0 |
| 4.73E-11 | 4.33E-10 | 121.535504 | 205.535107 | 172.617657 | 185.110524 | 0 |
| 4.73E-11 | 4.33E-10 | 151.115573 | 292.064498 | 257.070381 | 274.650202 | 9.50487297 |
| 4.75E-11 | 4.35E-10 | 110.603739 | 364.35769 | 277.487523 | 331.250412 | 0 |
| 4.76E-11 | 4.36E-10 | 130.859656 | 176.39539 | 171.689605 | 249.597649 | 0 |
| 4.78E-11 | 4.37E-10 | 106.102424 | 412.627299 | 359.156092 | 350.271794 | 0 |
| 4.79E-11 | 4.38E-10 | 142.434466 | 185.293013 | 153.128566 | 136.861165 | 0 |
| 4.79E-11 | 4.38E-10 | 133.431836 | 322.98374 | 173.545708 | 276.04201 | 0 |
| 4.81E-11 | 4.40E-10 | 113.175919 | 285.613721 | 232.941031 | 283.928925 | 0 |
| 4.82E-11 | 4.41E-10 | 116.391144 | 221.328389 | 239.437395 | 295.991265 | 0 |
| 4.82E-11 | 4.41E-10 | 106.745469 | 343.893156 | 339.667002 | 326.611051 | 0 |
| 4.83E-11 | 4.41E-10 | 118.641801 | 201.976058 | 235.725187 | 281.609244 | 0 |
| 4.84E-11 | 4.41E-10 | 143.720556 | 132.574593 | 193.962851 | 169.800631 | 0 |
| 4.84E-11 | 4.42E-10 | 114.783531 | 292.73182 | 222.73246 | 280.217435 | 0 |
| 4.85E-11 | 4.43E-10 | 213.490937 | 544.089689 | 717.384132 | 583.631678 | 0 |
| 4.86E-11 | 4.43E-10 | 138.576196 | 388.603715 | 271.919212 | 184.646588 | 0 |
| 4.86E-11 | 4.43E-10 | 109.317649 | 341.66875 | 284.911939 | 305.269988 | 0 |
| 4.89E-11 | 4.45E-10 | 106.745469 | 324.095943 | 335.026742 | 334.961901 | 0 |
| 4.89E-11 | 4.45E-10 | 227.95945 | 360.798641 | 317.393756 | 326.147114 | 2.37621824 |
| 4.89E-11 | 4.46E-10 | 138.576196 | 139.915133 | 143.848047 | 150.315313 | 0 |
| 4.90E-11 | 4.47E-10 | 214.777027 | 257.363765 | 238.509343 | 222.689353 | 4.75243649 |
| 4.91E-11 | 4.47E-10 | 104.494811 | 406.398962 | 359.156092 | 395.737537 | 0 |
| 4.91E-11 | 4.47E-10 | 179.409553 | 229.558691 | 118.790645 | 220.833608 | 0 |
| 4.91E-11 | 4.47E-10 | 137.933151 | 205.090226 | 236.653239 | 236.143501 | 2.37621824 |
| 4.92E-11 | 4.48E-10 | 128.608998 | 186.850098 | 146.632203 | 185.110524 | 0 |

| | | | | | | |
|---|---|---|---|---|---|---|
| 4.93E-11 | 4.49E-10 | 114.783531 | 231.338216 | 206.955577 | 225.936906 | 0 |
| 4.94E-11 | 4.49E-10 | 109.639171 | 319.647131 | 270.063108 | 302.950307 | 0 |
| 4.94E-11 | 4.50E-10 | 110.925261 | 287.838127 | 255.214277 | 312.692966 | 0 |
| 4.95E-11 | 4.50E-10 | 119.927891 | 235.787028 | 188.394539 | 265.835415 | 0 |
| 4.96E-11 | 4.50E-10 | 235.032945 | 191.52135 | 134.567528 | 148.459568 | 0 |
| 4.97E-11 | 4.51E-10 | 106.423946 | 333.883329 | 325.746223 | 361.406262 | 0 |
| 4.98E-11 | 4.52E-10 | 132.145746 | 191.298909 | 141.991943 | 194.389247 | 0 |
| 4.99E-11 | 4.53E-10 | 149.507961 | 347.674646 | 267.278952 | 303.414243 | 2.37621824 |
| 5.00E-11 | 4.54E-10 | 107.388514 | 300.739681 | 323.890119 | 324.755306 | 0 |
| 5.01E-11 | 4.55E-10 | 113.497441 | 250.690548 | 221.804408 | 271.866585 | 0 |
| 5.02E-11 | 4.55E-10 | 101.922631 | 405.064319 | 507.644399 | 466.255832 | 0 |
| 5.04E-11 | 4.57E-10 | 105.137856 | 379.706091 | 361.940248 | 340.529135 | 0 |
| 5.06E-11 | 4.59E-10 | 238.569692 | 304.743612 | 135.49558 | 163.305525 | 0 |
| 5.07E-11 | 4.59E-10 | 125.715296 | 157.487939 | 219.020252 | 195.31712 | 0 |
| 5.07E-11 | 4.60E-10 | 123.786161 | 186.627657 | 193.962851 | 156.346483 | 0 |
| 5.08E-11 | 4.60E-10 | 128.930521 | 156.375736 | 156.840774 | 197.172864 | 0 |
| 5.09E-11 | 4.61E-10 | 117.034189 | 239.568518 | 296.048562 | 383.675197 | 0 |
| 5.10E-11 | 4.62E-10 | 109.639171 | 301.184562 | 261.710641 | 309.445413 | 0 |
| 5.11E-11 | 4.62E-10 | 112.854396 | 306.745577 | 234.797135 | 269.082968 | 0 |
| 5.12E-11 | 4.64E-10 | 109.317649 | 269.153117 | 329.458431 | 334.497965 | 0 |
| 5.14E-11 | 4.65E-10 | 115.748099 | 213.765409 | 288.624146 | 297.847009 | 0 |
| 5.14E-11 | 4.65E-10 | 153.366231 | 305.855815 | 142.919995 | 282.07318 | 0 |
| 5.14E-11 | 4.65E-10 | 110.925261 | 350.121492 | 259.854537 | 311.301158 | 0 |
| 5.15E-11 | 4.66E-10 | 167.834743 | 209.983919 | 122.502853 | 172.120312 | 0 |
| 5.20E-11 | 4.70E-10 | 127.001386 | 332.103805 | 418.551415 | 393.417856 | 7.12865473 |
| 5.21E-11 | 4.71E-10 | 110.282216 | 340.556547 | 261.710641 | 342.848816 | 0 |
| 5.23E-11 | 4.73E-10 | 110.603739 | 344.560478 | 259.854537 | 298.310945 | 0 |
| 5.23E-11 | 4.73E-10 | 115.748099 | 249.355904 | 415.767259 | 374.86041 | 0 |
| 5.24E-11 | 4.73E-10 | 113.818964 | 227.334285 | 208.811681 | 244.494352 | 0 |
| 5.24E-11 | 4.73E-10 | 144.685123 | 236.45435 | 137.351684 | 242.174671 | 0 |
| 5.24E-11 | 4.73E-10 | 115.105054 | 236.899231 | 197.675058 | 235.215629 | 0 |
| 5.27E-11 | 4.76E-10 | 107.066991 | 318.090047 | 308.113236 | 375.324346 | 0 |
| 5.28E-11 | 4.76E-10 | 136.004016 | 699.575663 | 257.070381 | 445.842641 | 0 |
| 5.30E-11 | 4.78E-10 | 191.305885 | 653.975342 | 567.967773 | 611.003911 | 0 |
| 5.31E-11 | 4.79E-10 | 122.178549 | 454.891012 | 238.509343 | 326.611051 | 0 |
| 5.33E-11 | 4.80E-10 | 129.895088 | 200.418974 | 198.60311 | 144.284143 | 0 |
| 5.33E-11 | 4.80E-10 | 116.391144 | 509.166516 | 289.552198 | 335.889774 | 0 |
| 5.36E-11 | 4.83E-10 | 120.570936 | 202.86582 | 164.265189 | 199.028609 | 0 |
| 5.37E-11 | 4.84E-10 | 115.748099 | 250.245666 | 342.451157 | 236.143501 | 0 |
| 5.38E-11 | 4.84E-10 | 114.462009 | 230.003573 | 201.387266 | 209.235204 | 0 |
| 5.39E-11 | 4.85E-10 | 111.889829 | 234.452384 | 223.660512 | 252.381266 | 0 |
| 5.39E-11 | 4.85E-10 | 113.175919 | 387.269071 | 252.430121 | 370.684985 | 0 |
| 5.40E-11 | 4.86E-10 | 117.677234 | 181.956405 | 195.818954 | 178.151482 | 0 |

| | | | | | | |
|---|---|---|---|---|---|---|
| 5.42E-11 | 4.88E-10 | 172.65758 | 467.347685 | 438.968557 | 413.367111 | 2.37621824 |
| 5.43E-11 | 4.89E-10 | 110.282216 | 298.515275 | 245.005706 | 267.691159 | 0 |
| 5.44E-11 | 4.89E-10 | 143.399033 | 130.127747 | 201.387266 | 179.079354 | 0 |
| 5.44E-11 | 4.89E-10 | 101.601109 | 437.095764 | 408.342843 | 386.458814 | 0 |
| 5.45E-11 | 4.90E-10 | 104.816334 | 447.550472 | 335.954794 | 390.170303 | 0 |
| 5.47E-11 | 4.91E-10 | 181.338688 | 320.092013 | 331.314534 | 326.611051 | 2.37621824 |
| 5.47E-11 | 4.91E-10 | 115.426576 | 206.869751 | 190.250643 | 224.545097 | 0 |
| 5.47E-11 | 4.91E-10 | 114.783531 | 219.103984 | 196.747006 | 200.420417 | 0 |
| 5.47E-11 | 4.91E-10 | 581.955718 | 316.532963 | 264.494796 | 322.435625 | 0 |
| 5.48E-11 | 4.92E-10 | 107.710036 | 318.757369 | 275.631419 | 339.137327 | 0 |
| 5.49E-11 | 4.93E-10 | 83.9173715 | 749.847237 | 1132.22334 | 1086.07453 | 0 |
| 5.49E-11 | 4.93E-10 | 106.423946 | 318.312488 | 304.401029 | 294.13552 | 0 |
| 5.54E-11 | 4.96E-10 | 109.317649 | 294.956226 | 247.789862 | 287.176478 | 0 |
| 5.54E-11 | 4.97E-10 | 208.025055 | 819.471143 | 963.31789 | 865.240922 | 11.8810912 |
| 5.55E-11 | 4.97E-10 | 142.755988 | 266.48383 | 142.919995 | 257.020628 | 0 |
| 5.56E-11 | 4.98E-10 | 175.22976 | 598.587634 | 461.241803 | 548.372531 | 14.2573095 |
| 5.56E-11 | 4.98E-10 | 122.178549 | 178.174915 | 179.11402 | 154.026802 | 0 |
| 5.57E-11 | 4.99E-10 | 111.246784 | 387.491512 | 268.207004 | 297.383073 | 0 |
| 5.59E-11 | 5.00E-10 | 118.320279 | 175.060746 | 200.459214 | 225.009033 | 0 |
| 5.59E-11 | 5.01E-10 | 108.031559 | 292.73182 | 267.278952 | 270.474776 | 0 |
| 5.60E-11 | 5.01E-10 | 122.821594 | 164.383598 | 160.552982 | 155.418611 | 0 |
| 5.61E-11 | 5.01E-10 | 127.322908 | 255.3618 | 167.977397 | 186.966269 | 0 |
| 5.61E-11 | 5.01E-10 | 117.998756 | 453.111487 | 247.789862 | 340.993071 | 0 |
| 5.62E-11 | 5.02E-10 | 224.744225 | 426.418616 | 313.681548 | 389.706367 | 0 |
| 5.62E-11 | 5.02E-10 | 99.3504513 | 423.971769 | 449.177128 | 449.090194 | 0 |
| 5.63E-11 | 5.03E-10 | 163.976473 | 263.369661 | 145.704151 | 375.788283 | 0 |
| 5.64E-11 | 5.03E-10 | 123.786161 | 207.759513 | 219.948304 | 156.810419 | 0 |
| 5.65E-11 | 5.04E-10 | 105.459379 | 375.034839 | 305.329081 | 387.386686 | 0 |
| 5.68E-11 | 5.07E-10 | 113.818964 | 210.651241 | 197.675058 | 201.34829 | 0 |
| 5.69E-11 | 5.08E-10 | 206.417443 | 702.689832 | 577.248292 | 628.169549 | 0 |
| 5.72E-11 | 5.11E-10 | 120.570936 | 246.464176 | 171.689605 | 230.112331 | 0 |
| 5.73E-11 | 5.11E-10 | 102.244154 | 354.347864 | 432.472193 | 401.304771 | 0 |
| 5.73E-11 | 5.11E-10 | 107.710036 | 282.721993 | 265.422848 | 268.155095 | 0 |
| 5.74E-11 | 5.12E-10 | 96.4567488 | 645.300159 | 541.054268 | 482.957534 | 0 |
| 5.74E-11 | 5.12E-10 | 124.429206 | 260.922815 | 168.905449 | 206.915523 | 0 |
| 5.75E-11 | 5.12E-10 | 119.927891 | 249.355904 | 178.185968 | 201.812226 | 0 |
| 5.78E-11 | 5.14E-10 | 107.388514 | 302.741646 | 263.566744 | 290.887967 | 0 |
| 5.78E-11 | 5.14E-10 | 145.971213 | 684.004822 | 221.804408 | 575.280828 | 0 |
| 5.80E-11 | 5.16E-10 | 189.698273 | 221.55083 | 281.199731 | 118.767655 | 0 |
| 5.81E-11 | 5.17E-10 | 122.178549 | 242.015365 | 173.545708 | 270.474776 | 0 |
| 5.84E-11 | 5.19E-10 | 103.208721 | 390.160799 | 346.163365 | 340.065199 | 0 |
| 5.84E-11 | 5.19E-10 | 169.120833 | 165.05092 | 157.768826 | 120.623399 | 0 |
| 5.84E-11 | 5.19E-10 | 114.462009 | 220.438627 | 212.523889 | 186.966269 | 0 |

| | | | | | | |
|---|---|---|---|---|---|---|
| 5.84E-11 | 5.19E-10 | 109.639171 | 271.822404 | 232.941031 | 278.825627 | 0 |
| 5.85E-11 | 5.20E-10 | 127.001386 | 220.883508 | 149.416359 | 204.595843 | 0 |
| 5.89E-11 | 5.23E-10 | 109.317649 | 297.625513 | 243.149602 | 260.732117 | 0 |
| 5.90E-11 | 5.24E-10 | 120.249414 | 173.726103 | 156.840774 | 178.615418 | 0 |
| 5.91E-11 | 5.25E-10 | 115.426576 | 219.326424 | 181.898176 | 200.884354 | 0 |
| 5.92E-11 | 5.26E-10 | 116.391144 | 246.909058 | 186.538435 | 208.771268 | 0 |
| 5.94E-11 | 5.28E-10 | 120.892459 | 178.397355 | 163.337137 | 212.482757 | 0 |
| 5.95E-11 | 5.28E-10 | 135.682493 | 192.188672 | 134.567528 | 161.449781 | 0 |
| 5.95E-11 | 5.28E-10 | 0 | 0 | 0 | 0 | 54.6530196 |
| 5.95E-11 | 5.28E-10 | 114.140486 | 225.10988 | 348.947521 | 288.568286 | 0 |
| 5.95E-11 | 5.28E-10 | 108.674604 | 284.056637 | 240.365446 | 282.537116 | 0 |
| 5.98E-11 | 5.30E-10 | 131.824223 | 251.135429 | 150.344411 | 200.420417 | 0 |
| 6.00E-11 | 5.32E-10 | 121.535504 | 182.178845 | 153.128566 | 168.872759 | 0 |
| 6.03E-11 | 5.34E-10 | 118.320279 | 204.200464 | 165.193241 | 187.894141 | 0 |
| 6.03E-11 | 5.34E-10 | 175.22976 | 403.729675 | 372.148819 | 370.221049 | 0 |
| 6.05E-11 | 5.36E-10 | 136.004016 | 320.314453 | 159.62493 | 262.123925 | 0 |
| 6.06E-11 | 5.36E-10 | 106.102424 | 276.493656 | 284.911939 | 279.753499 | 0 |
| 6.06E-11 | 5.36E-10 | 115.105054 | 227.556726 | 181.898176 | 216.194247 | 0 |
| 6.06E-11 | 5.36E-10 | 122.178549 | 193.968197 | 153.128566 | 203.204034 | 0 |
| 6.07E-11 | 5.37E-10 | 111.889829 | 237.121672 | 202.315318 | 226.400842 | 0 |
| 6.08E-11 | 5.38E-10 | 108.031559 | 305.633374 | 252.430121 | 267.691159 | 0 |
| 6.09E-11 | 5.38E-10 | 133.431836 | 131.684831 | 175.401812 | 171.19244 | 0 |
| 6.09E-11 | 5.39E-10 | 80.380624 | 857.063603 | 1047.77061 | 904.675495 | 0 |
| 6.10E-11 | 5.39E-10 | 121.213981 | 210.20636 | 157.768826 | 186.038397 | 0 |
| 6.11E-11 | 5.40E-10 | 114.462009 | 244.684652 | 232.012979 | 188.358077 | 0 |
| 6.11E-11 | 5.40E-10 | 153.687753 | 263.147221 | 216.236097 | 238.463182 | 7.12865473 |
| 6.12E-11 | 5.40E-10 | 119.927891 | 160.824548 | 160.552982 | 176.295737 | 0 |
| 6.12E-11 | 5.40E-10 | 131.502701 | 281.387349 | 276.559471 | 194.389247 | 7.12865473 |
| 6.14E-11 | 5.42E-10 | 527.618416 | 930.024117 | 607.874006 | 863.849114 | 23.7621824 |
| 6.14E-11 | 5.42E-10 | 108.996126 | 234.007503 | 284.911939 | 256.556692 | 0 |
| 6.16E-11 | 5.43E-10 | 111.889829 | 332.993567 | 230.156875 | 330.32254 | 0 |
| 6.19E-11 | 5.46E-10 | 7562.20911 | 23312.219 | 19579.1113 | 20473.9663 | 0 |
| 6.20E-11 | 5.46E-10 | 98.7074063 | 405.5092 | 416.695311 | 448.626258 | 0 |
| 6.21E-11 | 5.47E-10 | 148.864916 | 155.263534 | 125.287009 | 143.820207 | 0 |
| 6.21E-11 | 5.47E-10 | 102.887199 | 315.865641 | 361.940248 | 352.591475 | 0 |
| 6.22E-11 | 5.48E-10 | 144.363601 | 425.083972 | 180.042072 | 257.484564 | 0 |
| 6.23E-11 | 5.49E-10 | 112.532874 | 290.729854 | 213.451941 | 241.710735 | 0 |
| 6.23E-11 | 5.49E-10 | 106.102424 | 314.530998 | 265.422848 | 298.310945 | 0 |
| 6.23E-11 | 5.49E-10 | 109.639171 | 235.342147 | 219.020252 | 237.999246 | 0 |
| 6.25E-11 | 5.50E-10 | 98.7074063 | 590.134892 | 406.48674 | 552.547956 | 0 |
| 6.25E-11 | 5.50E-10 | 99.6719738 | 380.150972 | 397.20622 | 418.006472 | 0 |
| 6.26E-11 | 5.51E-10 | 295.157651 | 711.809896 | 805.549064 | 830.445711 | 16.6335277 |
| 6.27E-11 | 5.51E-10 | 222.81509 | 338.777022 | 393.494013 | 306.661796 | 0 |

| | | | | | | |
|---|---|---|---|---|---|---|
| 6.27E-11 | 5.52E-10 | 150.794051 | 122.564766 | 203.24337 | 187.894141 | 0 |
| 6.32E-11 | 5.55E-10 | 147.257303 | 241.792924 | 135.49558 | 173.048184 | 0 |
| 6.33E-11 | 5.56E-10 | 112.211351 | 277.160978 | 206.027526 | 260.268181 | 0 |
| 6.37E-11 | 5.60E-10 | 106.102424 | 291.174735 | 262.638692 | 270.938712 | 0 |
| 6.39E-11 | 5.61E-10 | 166.22713 | 174.393424 | 136.423632 | 127.582442 | 0 |
| 6.40E-11 | 5.62E-10 | 214.777027 | 329.434517 | 379.573234 | 350.271794 | 0 |
| 6.41E-11 | 5.62E-10 | 118.320279 | 190.854028 | 158.696878 | 193.461375 | 0 |
| 6.41E-11 | 5.62E-10 | 111.246784 | 230.003573 | 202.315318 | 211.090949 | 0 |
| 6.43E-11 | 5.64E-10 | 105.780901 | 285.613721 | 260.782589 | 289.496158 | 0 |
| 6.44E-11 | 5.64E-10 | 117.034189 | 221.328389 | 168.905449 | 217.122119 | 0 |
| 6.44E-11 | 5.64E-10 | 121.857026 | 226.444523 | 156.840774 | 209.69914 | 0 |
| 6.44E-11 | 5.64E-10 | 118.641801 | 169.722172 | 156.840774 | 173.976057 | 0 |
| 6.44E-11 | 5.64E-10 | 139.540763 | 432.869393 | 185.610383 | 271.402649 | 0 |
| 6.44E-11 | 5.65E-10 | 113.175919 | 212.875647 | 271.919212 | 318.2602 | 0 |
| 6.46E-11 | 5.66E-10 | 107.388514 | 246.241736 | 244.077654 | 248.205841 | 0 |
| 6.50E-11 | 5.69E-10 | 150.794051 | 222.663033 | 279.343627 | 133.613612 | 0 |
| 6.50E-11 | 5.69E-10 | 107.710036 | 250.912988 | 236.653239 | 273.258393 | 0 |
| 6.58E-11 | 5.76E-10 | 226.030315 | 328.099874 | 287.696094 | 301.094562 | 0 |
| 6.62E-11 | 5.80E-10 | 107.388514 | 365.247453 | 302.544925 | 261.196053 | 0 |
| 6.64E-11 | 5.81E-10 | 117.034189 | 211.763444 | 164.265189 | 197.6368 | 0 |
| 6.65E-11 | 5.82E-10 | 325.380766 | 271.377523 | 229.228823 | 309.909349 | 0 |
| 6.70E-11 | 5.86E-10 | 132.467268 | 261.367696 | 150.344411 | 194.389247 | 0 |
| 6.71E-11 | 5.87E-10 | 108.353081 | 290.507414 | 227.37272 | 271.402649 | 0 |
| 6.72E-11 | 5.87E-10 | 159.153636 | 182.846167 | 134.567528 | 280.217435 | 0 |
| 6.74E-11 | 5.89E-10 | 129.895088 | 159.489905 | 137.351684 | 192.069567 | 0 |
| 6.75E-11 | 5.90E-10 | 124.750728 | 328.767196 | 179.11402 | 250.525522 | 0 |
| 6.79E-11 | 5.93E-10 | 186.80457 | 199.306771 | 150.344411 | 176.759674 | 7.12865473 |
| 6.81E-11 | 5.94E-10 | 212.52637 | 326.097909 | 404.630636 | 402.232643 | 0 |
| 6.81E-11 | 5.95E-10 | 107.066991 | 256.251562 | 231.084927 | 254.237011 | 0 |
| 6.83E-11 | 5.96E-10 | 141.791421 | 125.234054 | 168.905449 | 147.995632 | 0 |
| 6.83E-11 | 5.96E-10 | 137.611628 | 277.160978 | 147.560255 | 199.028609 | 0 |
| 6.87E-11 | 5.99E-10 | 234.711422 | 668.878861 | 715.528028 | 722.348588 | 14.2573095 |
| 6.88E-11 | 6.00E-10 | 114.462009 | 169.499731 | 180.042072 | 187.430205 | 0 |
| 6.88E-11 | 6.00E-10 | 225.708792 | 127.013578 | 163.337137 | 155.882547 | 0 |
| 6.89E-11 | 6.00E-10 | 136.004016 | 215.100053 | 198.60311 | 197.172864 | 26.1384007 |
| 6.94E-11 | 6.05E-10 | 111.889829 | 278.718062 | 200.459214 | 245.88616 | 0 |
| 6.95E-11 | 6.05E-10 | 106.745469 | 236.231909 | 268.207004 | 243.102543 | 0 |
| 6.95E-11 | 6.05E-10 | 121.535504 | 152.149365 | 148.488307 | 172.120312 | 0 |
| 6.97E-11 | 6.07E-10 | 108.353081 | 329.879399 | 235.725187 | 321.043817 | 0 |
| 7.01E-11 | 6.10E-10 | 166.870175 | 217.102018 | 195.818954 | 203.204034 | 11.8810912 |
| 7.02E-11 | 6.11E-10 | 118.963324 | 169.722172 | 149.416359 | 173.976057 | 0 |
| 7.03E-11 | 6.11E-10 | 158.510591 | 211.985885 | 153.128566 | 181.399035 | 4.75243649 |
| 7.04E-11 | 6.13E-10 | 97.4213163 | 381.485616 | 451.961284 | 419.862217 | 0 |

| | | | | | | |
|---|---|---|---|---|---|---|
| 7.05E-11 | 6.13E-10 | 108.353081 | 213.098088 | 219.948304 | 230.112331 | 0 |
| 7.09E-11 | 6.16E-10 | 210.597235 | 363.912809 | 324.818171 | 347.024241 | 2.37621824 |
| 7.10E-11 | 6.17E-10 | 105.459379 | 275.159013 | 246.86181 | 256.092756 | 0 |
| 7.11E-11 | 6.18E-10 | 103.851766 | 337.219938 | 270.063108 | 328.466795 | 0 |
| 7.13E-11 | 6.19E-10 | 117.355711 | 185.737895 | 240.365446 | 173.976057 | 0 |
| 7.14E-11 | 6.20E-10 | 1442.67144 | 1270.58066 | 686.758419 | 973.338045 | 0 |
| 7.16E-11 | 6.21E-10 | 170.728445 | 161.046989 | 135.49558 | 125.262761 | 0 |
| 7.17E-11 | 6.22E-10 | 118.963324 | 194.635518 | 152.200514 | 173.976057 | 0 |
| 7.26E-11 | 6.30E-10 | 110.925261 | 189.519385 | 219.948304 | 195.781056 | 0 |
| 7.26E-11 | 6.30E-10 | 91.6339114 | 489.369303 | 610.658161 | 550.228275 | 0 |
| 7.27E-11 | 6.31E-10 | 115.426576 | 182.401286 | 234.797135 | 275.578074 | 0 |
| 7.28E-11 | 6.31E-10 | 102.887199 | 326.320349 | 278.415575 | 347.024241 | 0 |
| 7.31E-11 | 6.33E-10 | 96.1352263 | 447.328031 | 394.422065 | 434.244238 | 0 |
| 7.33E-11 | 6.35E-10 | 108.031559 | 230.226013 | 208.811681 | 221.297544 | 0 |
| 7.33E-11 | 6.35E-10 | 105.780901 | 239.790959 | 250.574018 | 236.607437 | 0 |
| 7.34E-11 | 6.36E-10 | 118.963324 | 201.308736 | 155.912722 | 167.944887 | 0 |
| 7.35E-11 | 6.36E-10 | 110.282216 | 199.751652 | 191.178695 | 218.513927 | 0 |
| 7.35E-11 | 6.36E-10 | 110.925261 | 336.552616 | 228.300772 | 250.061586 | 0 |
| 7.35E-11 | 6.36E-10 | 102.887199 | 277.383419 | 292.336354 | 281.145308 | 0 |
| 7.37E-11 | 6.37E-10 | 116.069621 | 166.385563 | 163.337137 | 159.594036 | 0 |
| 7.37E-11 | 6.38E-10 | 116.069621 | 227.111845 | 166.121293 | 225.936906 | 0 |
| 7.42E-11 | 6.41E-10 | 357.854538 | 591.247095 | 669.125432 | 594.30221 | 116.434694 |
| 7.42E-11 | 6.41E-10 | 1066.1686 | 323.651062 | 173.545708 | 310.373285 | 0 |
| 7.45E-11 | 6.43E-10 | 109.317649 | 212.430766 | 207.883629 | 253.773075 | 0 |
| 7.46E-11 | 6.44E-10 | 117.355711 | 280.275146 | 176.329864 | 246.350096 | 0 |
| 7.46E-11 | 6.44E-10 | 102.887199 | 304.07629 | 268.207004 | 295.991265 | 0 |
| 7.52E-11 | 6.49E-10 | 108.996126 | 264.481864 | 204.171422 | 239.391054 | 0 |
| 7.54E-11 | 6.51E-10 | 119.606369 | 576.343575 | 294.192458 | 298.774881 | 0 |
| 7.54E-11 | 6.51E-10 | 111.568306 | 329.212077 | 221.804408 | 241.246799 | 0 |
| 7.55E-11 | 6.51E-10 | 111.246784 | 210.651241 | 187.466487 | 242.174671 | 0 |
| 7.55E-11 | 6.51E-10 | 109.960694 | 232.005538 | 207.883629 | 283.001052 | 0 |
| 7.57E-11 | 6.53E-10 | 99.6719738 | 362.578166 | 317.393756 | 347.488177 | 0 |
| 7.59E-11 | 6.54E-10 | 102.565676 | 292.064498 | 278.415575 | 274.650202 | 0 |
| 7.62E-11 | 6.57E-10 | 108.674604 | 218.659102 | 208.811681 | 196.708928 | 0 |
| 7.64E-11 | 6.58E-10 | 105.137856 | 238.678756 | 241.293498 | 240.782863 | 0 |
| 7.65E-11 | 6.60E-10 | 103.851766 | 253.137394 | 257.070381 | 265.835415 | 0 |
| 7.66E-11 | 6.60E-10 | 118.641801 | 193.745756 | 165.193241 | 245.88616 | 0 |
| 7.66E-11 | 6.60E-10 | 163.011906 | 250.912988 | 181.898176 | 236.143501 | 11.8810912 |
| 7.73E-11 | 6.66E-10 | 113.497441 | 220.438627 | 169.833501 | 196.244992 | 0 |
| 7.75E-11 | 6.67E-10 | 154.330798 | 213.542969 | 146.632203 | 128.510314 | 0 |
| 7.75E-11 | 6.67E-10 | 122.178549 | 141.917098 | 143.848047 | 152.634994 | 0 |
| 7.76E-11 | 6.68E-10 | 105.137856 | 239.123637 | 233.869083 | 259.804245 | 0 |
| 7.80E-11 | 6.70E-10 | 149.829483 | 246.909058 | 209.739733 | 246.814032 | 7.12865473 |

| | | | | | | |
|---|---|---|---|---|---|---|
| 7.81E-11 | 6.71E-10 | 1033.3733 | 1386.02733 | 1296.48853 | 1498.9777 | 28.5146189 |
| 7.81E-11 | 6.71E-10 | 103.530244 | 320.092013 | 255.214277 | 321.507753 | 0 |
| 7.82E-11 | 6.72E-10 | 191.305885 | 397.723779 | 380.501286 | 387.850622 | 0 |
| 7.84E-11 | 6.74E-10 | 117.355711 | 226.444523 | 170.761553 | 170.264568 | 0 |
| 7.85E-11 | 6.74E-10 | 101.279586 | 298.960156 | 294.192458 | 285.320733 | 0 |
| 7.85E-11 | 6.74E-10 | 109.960694 | 197.082365 | 181.898176 | 200.884354 | 0 |
| 7.86E-11 | 6.75E-10 | 99.3504513 | 397.723779 | 318.321808 | 356.7669 | 0 |
| 7.87E-11 | 6.75E-10 | 122.500071 | 139.247811 | 166.121293 | 147.531696 | 0 |
| 7.92E-11 | 6.80E-10 | 107.066991 | 206.869751 | 222.73246 | 240.318926 | 0 |
| 7.97E-11 | 6.83E-10 | 102.887199 | 274.26925 | 256.142329 | 264.907542 | 0 |
| 8.00E-11 | 6.86E-10 | 115.748099 | 164.828479 | 109.510126 | 159.1301 | 7.12865473 |
| 8.01E-11 | 6.87E-10 | 154.009276 | 157.043058 | 124.358957 | 131.757867 | 0 |
| 8.01E-11 | 6.87E-10 | 109.317649 | 246.686617 | 204.171422 | 199.492545 | 0 |
| 8.02E-11 | 6.87E-10 | 127.322908 | 183.95837 | 131.783372 | 161.449781 | 0 |
| 8.05E-11 | 6.90E-10 | 106.102424 | 373.255314 | 258.926485 | 278.825627 | 0 |
| 8.05E-11 | 6.90E-10 | 106.423946 | 226.666964 | 209.739733 | 240.318926 | 0 |
| 8.07E-11 | 6.91E-10 | 138.897718 | 135.02144 | 125.287009 | 148.923505 | 0 |
| 8.07E-11 | 6.91E-10 | 233.10381 | 132.129712 | 384.213494 | 181.399035 | 0 |
| 8.07E-11 | 6.91E-10 | 116.712666 | 156.820618 | 154.98467 | 182.790844 | 0 |
| 8.09E-11 | 6.92E-10 | 100.315019 | 363.690369 | 294.192458 | 379.963708 | 0 |
| 8.10E-11 | 6.93E-10 | 133.431836 | 175.728068 | 127.143113 | 147.531696 | 0 |
| 8.17E-11 | 6.99E-10 | 97.0997938 | 564.331783 | 385.141546 | 424.965515 | 0 |
| 8.18E-11 | 6.99E-10 | 105.137856 | 246.909058 | 267.278952 | 222.225416 | 0 |
| 8.18E-11 | 6.99E-10 | 105.137856 | 253.359835 | 219.948304 | 243.102543 | 0 |
| 8.21E-11 | 7.02E-10 | 100.958064 | 344.338037 | 280.271679 | 310.373285 | 0 |
| 8.22E-11 | 7.02E-10 | 133.753358 | 150.1474 | 141.063891 | 216.658183 | 0 |
| 8.26E-11 | 7.05E-10 | 105.780901 | 469.572091 | 561.47141 | 296.455201 | 0 |
| 8.26E-11 | 7.05E-10 | 107.710036 | 201.976058 | 201.387266 | 195.781056 | 0 |
| 8.27E-11 | 7.06E-10 | 127.322908 | 170.834375 | 130.85532 | 190.213822 | 0 |
| 8.27E-11 | 7.06E-10 | 158.832113 | 736.723242 | 820.397895 | 811.888265 | 2.37621824 |
| 8.31E-11 | 7.09E-10 | 174.265193 | 421.080041 | 347.091417 | 369.757113 | 11.8810912 |
| 8.32E-11 | 7.10E-10 | 111.246784 | 212.430766 | 171.689605 | 192.997439 | 0 |
| 8.36E-11 | 7.13E-10 | 119.606369 | 150.592281 | 184.682331 | 147.995632 | 0 |
| 8.36E-11 | 7.13E-10 | 215.741595 | 548.760942 | 479.802841 | 502.442852 | 40.3957101 |
| 8.37E-11 | 7.14E-10 | 100.315019 | 282.721993 | 322.962067 | 298.310945 | 0 |
| 8.38E-11 | 7.14E-10 | 18.0052598 | 15.1259603 | 13.9207788 | 14.8459568 | 64.1578926 |
| 8.39E-11 | 7.15E-10 | 99.6719738 | 385.044666 | 298.832717 | 398.98509 | 0 |
| 8.40E-11 | 7.15E-10 | 173.622148 | 172.6139 | 116.934542 | 142.892335 | 0 |
| 8.40E-11 | 7.15E-10 | 91.9554339 | 511.168481 | 438.968557 | 464.864024 | 0 |
| 8.41E-11 | 7.16E-10 | 107.388514 | 232.005538 | 194.890903 | 218.049991 | 0 |
| 8.41E-11 | 7.16E-10 | 150.794051 | 204.645345 | 171.689605 | 184.182652 | 7.12865473 |
| 8.45E-11 | 7.19E-10 | 99.6719738 | 372.143111 | 339.667002 | 293.207648 | 0 |
| 8.47E-11 | 7.20E-10 | 108.031559 | 196.637484 | 224.588564 | 193.925311 | 0 |

| | | | | | | |
|---|---|---|---|---|---|---|
| 8.49E-11 | 7.22E-10 | 119.284846 | 143.029301 | 154.056618 | 177.22361 | 0 |
| 8.49E-11 | 7.22E-10 | 104.494811 | 273.157048 | 231.084927 | 235.679565 | 0 |
| 8.53E-11 | 7.25E-10 | 167.191698 | 953.82526 | 672.83764 | 777.55699 | 0 |
| 8.58E-11 | 7.29E-10 | 128.608998 | 176.61783 | 139.207788 | 135.469356 | 0 |
| 8.60E-11 | 7.31E-10 | 180.37412 | 216.212256 | 108.582074 | 191.141694 | 0 |
| 8.63E-11 | 7.33E-10 | 106.102424 | 470.461853 | 280.271679 | 328.466795 | 0 |
| 8.64E-11 | 7.34E-10 | 103.208721 | 353.01322 | 251.502069 | 329.394667 | 0 |
| 8.66E-11 | 7.35E-10 | 95.8137038 | 361.021082 | 373.076871 | 360.47839 | 0 |
| 8.67E-11 | 7.36E-10 | 218.313775 | 108.99589 | 212.523889 | 179.543291 | 0 |
| 8.71E-11 | 7.39E-10 | 129.895088 | 222.440592 | 132.711424 | 196.708928 | 0 |
| 8.72E-11 | 7.40E-10 | 175.551283 | 395.054492 | 414.839207 | 391.562112 | 4.75243649 |
| 8.74E-11 | 7.41E-10 | 172.979103 | 327.877433 | 328.530379 | 292.279775 | 2.37621824 |
| 8.77E-11 | 7.43E-10 | 96.1352263 | 336.330176 | 386.997649 | 398.521154 | 0 |
| 8.77E-11 | 7.43E-10 | 101.601109 | 367.026978 | 279.343627 | 290.424031 | 0 |
| 8.79E-11 | 7.45E-10 | 107.066991 | 295.401107 | 210.667785 | 293.671584 | 0 |
| 8.80E-11 | 7.46E-10 | 107.388514 | 243.127568 | 343.379209 | 239.391054 | 0 |
| 8.83E-11 | 7.48E-10 | 99.3504513 | 283.389315 | 308.113236 | 299.238818 | 0 |
| 8.84E-11 | 7.49E-10 | 97.7428388 | 342.558512 | 309.041288 | 332.178284 | 0 |
| 8.86E-11 | 7.50E-10 | 188.733705 | 559.215649 | 412.983103 | 486.669023 | 2.37621824 |
| 8.86E-11 | 7.50E-10 | 231.496197 | 177.285152 | 125.287009 | 142.892335 | 0 |
| 8.89E-11 | 7.52E-10 | 99.6719738 | 284.723958 | 302.544925 | 280.681372 | 0 |
| 8.91E-11 | 7.53E-10 | 112.532874 | 181.956405 | 156.840774 | 171.19244 | 0 |
| 8.91E-11 | 7.54E-10 | 107.710036 | 205.535107 | 187.466487 | 228.720523 | 0 |
| 8.97E-11 | 7.58E-10 | 103.208721 | 263.147221 | 225.516616 | 250.525522 | 0 |
| 8.98E-11 | 7.58E-10 | 107.388514 | 274.936572 | 202.315318 | 282.537116 | 0 |
| 8.98E-11 | 7.58E-10 | 101.279586 | 288.727889 | 256.142329 | 260.732117 | 0 |
| 8.97E-11 | 7.58E-10 | 5.14435994 | 5.56101481 | 4.64025958 | 3.24755306 | 33.2670554 |
| 9.01E-11 | 7.61E-10 | 104.173289 | 217.76934 | 219.948304 | 237.535309 | 0 |
| 9.02E-11 | 7.61E-10 | 105.459379 | 213.320528 | 222.73246 | 200.420417 | 0 |
| 9.06E-11 | 7.65E-10 | 107.388514 | 196.637484 | 236.653239 | 268.155095 | 0 |
| 9.08E-11 | 7.66E-10 | 109.317649 | 169.722172 | 193.034799 | 198.100737 | 0 |
| 9.12E-11 | 7.69E-10 | 133.753358 | 161.714311 | 151.272462 | 120.623399 | 0 |
| 9.12E-11 | 7.70E-10 | 112.854396 | 208.204395 | 157.768826 | 194.389247 | 0 |
| 9.13E-11 | 7.70E-10 | 99.0289288 | 307.635339 | 278.415575 | 304.342115 | 0 |
| 9.14E-11 | 7.71E-10 | 94.5276139 | 363.245487 | 420.407518 | 377.180091 | 0 |
| 9.14E-11 | 7.71E-10 | 108.674604 | 213.765409 | 176.329864 | 196.244992 | 0 |
| 9.15E-11 | 7.71E-10 | 103.851766 | 223.552795 | 294.192458 | 265.371479 | 0 |
| 9.16E-11 | 7.72E-10 | 189.698273 | 374.145076 | 117.862593 | 316.868391 | 0 |
| 9.17E-11 | 7.73E-10 | 93.5630464 | 381.263175 | 416.695311 | 391.098176 | 0 |
| 9.19E-11 | 7.74E-10 | 110.603739 | 228.001607 | 170.761553 | 202.276162 | 0 |
| 9.21E-11 | 7.76E-10 | 101.922631 | 249.355904 | 257.998433 | 236.143501 | 0 |
| 9.22E-11 | 7.77E-10 | 103.851766 | 337.88726 | 235.725187 | 320.579881 | 0 |
| 9.23E-11 | 7.77E-10 | 98.0643613 | 349.676611 | 299.760769 | 309.909349 | 0 |

| | | | | | | |
|---|---|---|---|---|---|---|
| 9.23E-11 | 7.77E-10 | 153.366231 | 151.037162 | 116.00649 | 141.964462 | 0 |
| 9.26E-11 | 7.78E-10 | 99.0289288 | 317.422725 | 311.825444 | 274.186265 | 0 |
| 9.26E-11 | 7.78E-10 | 102.244154 | 244.462211 | 232.012979 | 258.412436 | 0 |
| 9.26E-11 | 7.79E-10 | 118.320279 | 346.784884 | 247.789862 | 186.502333 | 0 |
| 9.28E-11 | 7.80E-10 | 105.137856 | 282.721993 | 210.667785 | 258.876372 | 0 |
| 9.30E-11 | 7.81E-10 | 104.816334 | 223.330355 | 201.387266 | 229.184459 | 0 |
| 9.30E-11 | 7.82E-10 | 101.922631 | 246.241736 | 235.725187 | 262.123925 | 0 |
| 9.32E-11 | 7.82E-10 | 122.178549 | 169.722172 | 131.783372 | 181.399035 | 0 |
| 9.35E-11 | 7.85E-10 | 110.282216 | 366.137215 | 211.595837 | 304.806051 | 0 |
| 9.36E-11 | 7.86E-10 | 102.565676 | 235.342147 | 255.214277 | 227.79265 | 0 |
| 9.38E-11 | 7.87E-10 | 100.636541 | 343.893156 | 567.967773 | 372.54073 | 0 |
| 9.38E-11 | 7.87E-10 | 115.426576 | 243.572449 | 174.47376 | 173.048184 | 0 |
| 9.38E-11 | 7.87E-10 | 109.960694 | 196.192603 | 172.617657 | 169.800631 | 0 |
| 9.37E-11 | 7.87E-10 | 93.2415239 | 422.859566 | 384.213494 | 387.386686 | 0 |
| 9.38E-11 | 7.87E-10 | 106.423946 | 190.631588 | 240.365446 | 237.071373 | 0 |
| 9.40E-11 | 7.88E-10 | 103.851766 | 231.560657 | 208.811681 | 233.359884 | 0 |
| 9.43E-11 | 7.90E-10 | 100.958064 | 291.619617 | 254.286225 | 331.250412 | 0 |
| 9.45E-11 | 7.91E-10 | 114.140486 | 221.995711 | 155.912722 | 207.37946 | 0 |
| 9.45E-11 | 7.91E-10 | 106.745469 | 270.042879 | 359.156092 | 229.648395 | 0 |
| 9.48E-11 | 7.94E-10 | 125.072251 | 158.822583 | 223.660512 | 145.212015 | 0 |
| 9.49E-11 | 7.94E-10 | 103.530244 | 242.015365 | 214.379993 | 223.617225 | 0 |
| 9.51E-11 | 7.96E-10 | 117.677234 | 140.137573 | 149.416359 | 168.408823 | 0 |
| 9.51E-11 | 7.96E-10 | 105.780901 | 255.806681 | 198.60311 | 258.412436 | 0 |
| 9.55E-11 | 7.98E-10 | 191.305885 | 376.814364 | 405.558688 | 401.768707 | 0 |
| 9.55E-11 | 7.98E-10 | 229.567062 | 201.531177 | 108.582074 | 173.512121 | 0 |
| 9.54E-11 | 7.98E-10 | 113.497441 | 147.478113 | 166.121293 | 173.512121 | 0 |
| 9.58E-11 | 8.00E-10 | 111.246784 | 196.192603 | 218.0922 | 162.377653 | 0 |
| 9.58E-11 | 8.01E-10 | 99.3504513 | 339.666785 | 456.601543 | 614.251464 | 0 |
| 9.60E-11 | 8.02E-10 | 110.282216 | 300.072359 | 261.710641 | 189.749886 | 0 |
| 9.66E-11 | 8.07E-10 | 101.922631 | 344.560478 | 270.99116 | 256.092756 | 0 |
| 9.66E-11 | 8.07E-10 | 123.143116 | 363.245487 | 179.11402 | 369.293176 | 0 |
| 9.67E-11 | 8.07E-10 | 104.173289 | 257.808647 | 206.027526 | 245.88616 | 0 |
| 9.67E-11 | 8.07E-10 | 108.353081 | 231.338216 | 176.329864 | 216.658183 | 0 |
| 9.73E-11 | 8.12E-10 | 96.7782713 | 338.109701 | 300.688821 | 321.507753 | 0 |
| 9.73E-11 | 8.12E-10 | 105.780901 | 218.214221 | 195.818954 | 250.525522 | 0 |
| 9.74E-11 | 8.13E-10 | 106.423946 | 224.220117 | 183.75428 | 221.297544 | 0 |
| 9.74E-11 | 8.13E-10 | 202.23765 | 106.326603 | 242.22155 | 284.856797 | 0 |
| 9.79E-11 | 8.16E-10 | 110.925261 | 201.308736 | 237.581291 | 326.611051 | 0 |
| 9.79E-11 | 8.16E-10 | 95.1706589 | 382.152938 | 326.674275 | 354.911156 | 0 |
| 9.79E-11 | 8.16E-10 | 140.826853 | 130.795068 | 128.071165 | 128.97425 | 0 |
| 9.81E-11 | 8.17E-10 | 116.391144 | 146.36591 | 158.696878 | 192.069567 | 0 |
| 9.82E-11 | 8.18E-10 | 129.252043 | 133.686796 | 125.287009 | 143.820207 | 0 |
| 9.86E-11 | 8.21E-10 | 109.317649 | 176.172949 | 162.409085 | 179.543291 | 0 |

| | | | | | | |
|---|---|---|---|---|---|---|
| 9.87E-11 | 8.21E-10 | 152.401663 | 145.031266 | 115.078438 | 143.356271 | 0 |
| 9.87E-11 | 8.21E-10 | 101.922631 | 270.932642 | 229.228823 | 239.85499 | 0 |
| 9.92E-11 | 8.25E-10 | 101.601109 | 244.23977 | 227.37272 | 236.143501 | 0 |
| 9.92E-11 | 8.25E-10 | 96.1352263 | 324.318384 | 307.185185 | 338.67339 | 0 |
| 9.96E-11 | 8.29E-10 | 100.315019 | 598.810075 | 319.249859 | 475.070619 | 0 |
| 9.99E-11 | 8.30E-10 | 89.0617314 | 526.516882 | 474.23453 | 720.028907 | 0 |
| 1.00E-10 | 8.32E-10 | 91.3123889 | 504.272823 | 407.414792 | 418.470408 | 0 |
| 1.00E-10 | 8.32E-10 | 165.262563 | 202.198499 | 117.862593 | 142.428398 | 0 |
| 1.00E-10 | 8.32E-10 | 246.607755 | 399.948185 | 167.977397 | 349.343922 | 0 |
| 1.00E-10 | 8.33E-10 | 105.459379 | 188.407182 | 204.171422 | 191.141694 | 0 |
| 1.01E-10 | 8.35E-10 | 100.315019 | 289.395211 | 241.293498 | 298.774881 | 0 |
| 1.01E-10 | 8.35E-10 | 113.497441 | 182.401286 | 150.344411 | 204.595843 | 0 |
| 1.01E-10 | 8.35E-10 | 101.922631 | 248.021261 | 226.444668 | 224.545097 | 0 |
| 1.01E-10 | 8.39E-10 | 97.7428388 | 279.607825 | 283.983887 | 302.022435 | 0 |
| 1.01E-10 | 8.39E-10 | 118.963324 | 288.060567 | 203.24337 | 430.532748 | 0 |
| 1.01E-10 | 8.39E-10 | 102.565676 | 228.001607 | 210.667785 | 233.359884 | 0 |
| 1.01E-10 | 8.39E-10 | 108.996126 | 191.076469 | 175.401812 | 163.769461 | 0 |
| 1.02E-10 | 8.42E-10 | 105.459379 | 183.513489 | 210.667785 | 216.658183 | 0 |
| 1.02E-10 | 8.43E-10 | 183.267823 | 545.201892 | 526.205437 | 574.816891 | 11.8810912 |
| 1.02E-10 | 8.44E-10 | 108.353081 | 256.918884 | 180.042072 | 228.720523 | 0 |
| 1.02E-10 | 8.45E-10 | 231.81772 | 192.411112 | 126.215061 | 134.541484 | 0 |
| 1.02E-10 | 8.47E-10 | 111.246784 | 193.523315 | 156.840774 | 167.480951 | 0 |
| 1.02E-10 | 8.48E-10 | 102.887199 | 244.23977 | 206.955577 | 249.597649 | 0 |
| 1.02E-10 | 8.48E-10 | 114.783531 | 143.696623 | 194.890903 | 179.079354 | 0 |
| 1.02E-10 | 8.48E-10 | 100.958064 | 270.710201 | 305.329081 | 235.679565 | 0 |
| 1.03E-10 | 8.48E-10 | 115.426576 | 148.367875 | 143.848047 | 143.820207 | 0 |
| 1.03E-10 | 8.48E-10 | 99.6719738 | 249.578345 | 284.911939 | 254.237011 | 0 |
| 1.03E-10 | 8.48E-10 | 105.459379 | 524.514917 | 276.559471 | 364.189879 | 0 |
| 1.03E-10 | 8.49E-10 | 191.305885 | 713.589421 | 412.983103 | 517.752745 | 2.37621824 |
| 1.03E-10 | 8.51E-10 | 97.0997938 | 415.296586 | 296.048562 | 413.367111 | 0 |
| 1.03E-10 | 8.52E-10 | 127.965953 | 211.318563 | 131.783372 | 162.377653 | 0 |
| 1.04E-10 | 8.57E-10 | 201.273083 | 581.459709 | 359.156092 | 437.027855 | 0 |
| 1.05E-10 | 8.63E-10 | 85.8465065 | 593.471501 | 584.672708 | 475.534555 | 0 |
| 1.05E-10 | 8.65E-10 | 95.4921813 | 374.589958 | 322.962067 | 312.692966 | 0 |
| 1.05E-10 | 8.66E-10 | 106.745469 | 171.724137 | 200.459214 | 186.038397 | 0 |
| 1.05E-10 | 8.67E-10 | 106.423946 | 268.708236 | 193.034799 | 219.905736 | 0 |
| 1.06E-10 | 8.71E-10 | 102.887199 | 368.584062 | 240.365446 | 291.815839 | 0 |
| 1.06E-10 | 8.71E-10 | 124.429206 | 152.816687 | 126.215061 | 177.22361 | 0 |
| 1.06E-10 | 8.72E-10 | 135.360971 | 237.788993 | 141.063891 | 149.851377 | 0 |
| 1.06E-10 | 8.74E-10 | 110.282216 | 239.346077 | 171.689605 | 188.358077 | 0 |
| 1.06E-10 | 8.77E-10 | 135.039448 | 236.899231 | 128.999216 | 252.381266 | 0 |
| 1.07E-10 | 8.80E-10 | 118.320279 | 137.023405 | 149.416359 | 179.543291 | 0 |
| 1.07E-10 | 8.83E-10 | 127.644431 | 260.033053 | 140.135839 | 260.732117 | 0 |

| | | | | | | |
|---|---|---|---|---|---|---|
| 1.08E-10 | 8.86E-10 | 105.780901 | 234.897266 | 180.970124 | 217.122119 | 0 |
| 1.08E-10 | 8.86E-10 | 130.859656 | 140.137573 | 119.718697 | 159.1301 | 0 |
| 1.08E-10 | 8.87E-10 | 101.922631 | 264.259424 | 211.595837 | 244.030416 | 0 |
| 1.08E-10 | 8.91E-10 | 111.568306 | 296.958191 | 182.826228 | 220.833608 | 0 |
| 1.08E-10 | 8.91E-10 | 139.862286 | 222.218152 | 153.128566 | 127.582442 | 0 |
| 1.08E-10 | 8.91E-10 | 116.391144 | 221.773271 | 164.265189 | 154.026802 | 0 |
| 1.08E-10 | 8.92E-10 | 107.066991 | 207.981954 | 181.898176 | 171.656376 | 0 |
| 1.09E-10 | 8.96E-10 | 120.249414 | 135.688761 | 130.85532 | 145.675952 | 0 |
| 1.09E-10 | 8.96E-10 | 99.3504513 | 247.79882 | 261.710641 | 236.143501 | 0 |
| 1.09E-10 | 8.97E-10 | 56.5879593 | 30.9192423 | 38.9781805 | 37.5788283 | 1195.23778 |
| 1.09E-10 | 8.98E-10 | 155.938411 | 822.585311 | 707.175561 | 672.243483 | 2.37621824 |
| 1.10E-10 | 9.01E-10 | 99.3504513 | 261.367696 | 236.653239 | 244.958288 | 0 |
| 1.10E-10 | 9.03E-10 | 97.7428388 | 254.472038 | 293.264406 | 304.806051 | 0 |
| 1.10E-10 | 9.04E-10 | 10.6102424 | 9.12006429 | 10.2085711 | 6.03116997 | 78.415202 |
| 1.10E-10 | 9.04E-10 | 111.568306 | 474.688224 | 238.509343 | 299.702754 | 0 |
| 1.10E-10 | 9.04E-10 | 104.816334 | 181.733964 | 191.178695 | 183.718716 | 0 |
| 1.10E-10 | 9.05E-10 | 101.922631 | 207.759513 | 213.451941 | 235.679565 | 0 |
| 1.10E-10 | 9.05E-10 | 103.851766 | 201.753617 | 208.811681 | 254.700947 | 0 |
| 1.11E-10 | 9.06E-10 | 115.105054 | 357.684473 | 186.538435 | 263.515734 | 0 |
| 1.11E-10 | 9.10E-10 | 363.320421 | 1527.27711 | 1385.58151 | 1534.70079 | 19.0097459 |
| 1.11E-10 | 9.11E-10 | 105.780901 | 210.651241 | 172.617657 | 199.956481 | 0 |
| 1.11E-10 | 9.14E-10 | 106.745469 | 223.775236 | 180.042072 | 181.862971 | 0 |
| 1.12E-10 | 9.14E-10 | 100.315019 | 269.153117 | 224.588564 | 287.640414 | 0 |
| 1.12E-10 | 9.15E-10 | 101.922631 | 246.241736 | 204.171422 | 227.328714 | 0 |
| 1.12E-10 | 9.17E-10 | 114.140486 | 443.10166 | 218.0922 | 285.320733 | 0 |
| 1.12E-10 | 9.19E-10 | 121.535504 | 180.844202 | 130.85532 | 147.995632 | 0 |
| 1.12E-10 | 9.20E-10 | 98.3858838 | 255.3618 | 249.645966 | 290.887967 | 0 |
| 1.13E-10 | 9.21E-10 | 115.748099 | 186.182776 | 177.257916 | 284.856797 | 0 |
| 1.13E-10 | 9.21E-10 | 112.854396 | 173.058781 | 157.768826 | 224.545097 | 0 |
| 1.13E-10 | 9.21E-10 | 116.069621 | 133.464355 | 145.704151 | 154.026802 | 0 |
| 1.13E-10 | 9.22E-10 | 97.4213163 | 359.241557 | 283.983887 | 281.145308 | 0 |
| 1.13E-10 | 9.23E-10 | 184.23239 | 151.037162 | 117.862593 | 135.469356 | 0 |
| 1.13E-10 | 9.23E-10 | 147.900348 | 250.245666 | 121.574801 | 177.22361 | 0 |
| 1.15E-10 | 9.37E-10 | 98.7074063 | 251.357869 | 247.789862 | 233.82382 | 0 |
| 1.15E-10 | 9.38E-10 | 108.996126 | 166.608004 | 151.272462 | 169.800631 | 0 |
| 1.15E-10 | 9.41E-10 | 106.423946 | 171.501697 | 168.905449 | 193.461375 | 0 |
| 1.16E-10 | 9.50E-10 | 111.246784 | 157.71038 | 149.416359 | 186.502333 | 0 |
| 1.17E-10 | 9.53E-10 | 100.315019 | 235.564587 | 213.451941 | 221.297544 | 0 |
| 1.17E-10 | 9.54E-10 | 124.107684 | 127.903341 | 132.711424 | 130.366059 | 0 |
| 1.17E-10 | 9.55E-10 | 143.077511 | 133.686796 | 193.962851 | 123.870952 | 0 |
| 1.17E-10 | 9.55E-10 | 124.107684 | 151.037162 | 128.071165 | 128.510314 | 0 |
| 1.17E-10 | 9.56E-10 | 101.922631 | 274.26925 | 214.379993 | 220.833608 | 0 |
| 1.17E-10 | 9.57E-10 | 109.960694 | 150.592281 | 167.049345 | 151.243185 | 0 |

| | | | | | | |
|---|---|---|---|---|---|---|
| 1.18E-10 | 9.58E-10 | 200.630038 | 431.979631 | 357.299988 | 443.522961 | 0 |
| 1.18E-10 | 9.63E-10 | 111.246784 | 204.867786 | 149.416359 | 176.759674 | 0 |
| 1.18E-10 | 9.63E-10 | 103.530244 | 186.182776 | 187.466487 | 183.718716 | 0 |
| 1.19E-10 | 9.65E-10 | 229.24554 | 125.901375 | 131.783372 | 230.576267 | 0 |
| 1.19E-10 | 9.68E-10 | 104.816334 | 218.214221 | 228.300772 | 175.367865 | 0 |
| 1.19E-10 | 9.69E-10 | 133.753358 | 178.842236 | 156.840774 | 115.984038 | 0 |
| 1.19E-10 | 9.70E-10 | 101.601109 | 228.001607 | 194.890903 | 223.617225 | 0 |
| 1.19E-10 | 9.72E-10 | 190.019795 | 377.704126 | 335.954794 | 372.54073 | 2.37621824 |
| 1.20E-10 | 9.74E-10 | 108.031559 | 181.956405 | 295.12051 | 251.91733 | 0 |
| 1.20E-10 | 9.74E-10 | 175.551283 | 959.608716 | 523.421281 | 682.914015 | 0 |
| 1.20E-10 | 9.76E-10 | 612.178833 | 266.038949 | 142.919995 | 215.266374 | 0 |
| 1.20E-10 | 9.76E-10 | 129.895088 | 194.635518 | 365.652455 | 177.687546 | 0 |
| 1.20E-10 | 9.79E-10 | 95.1706589 | 436.650883 | 295.12051 | 373.468602 | 0 |
| 1.21E-10 | 9.80E-10 | 129.895088 | 190.409147 | 121.574801 | 210.163077 | 0 |
| 1.21E-10 | 9.80E-10 | 106.745469 | 248.688582 | 179.11402 | 195.31712 | 0 |
| 1.21E-10 | 9.81E-10 | 108.353081 | 301.629443 | 207.883629 | 197.172864 | 0 |
| 1.21E-10 | 9.82E-10 | 200.308515 | 215.100053 | 101.157659 | 190.677758 | 0 |
| 1.21E-10 | 9.84E-10 | 142.434466 | 179.509558 | 194.890903 | 111.808612 | 0 |
| 1.21E-10 | 9.84E-10 | 99.0289288 | 617.939966 | 332.242586 | 389.706367 | 0 |
| 1.21E-10 | 9.84E-10 | 350.137998 | 207.759513 | 116.00649 | 194.853184 | 0 |
| 1.21E-10 | 9.84E-10 | 171.693013 | 203.533142 | 126.215061 | 124.334889 | 0 |
| 1.23E-10 | 9.94E-10 | 123.464639 | 226.666964 | 131.783372 | 187.430205 | 0 |
| 1.23E-10 | 9.94E-10 | 97.0997938 | 248.688582 | 248.717914 | 250.525522 | 0 |
| 1.23E-10 | 9.98E-10 | 102.887199 | 192.855994 | 226.444668 | 188.358077 | 0 |
| 1.24E-10 | 1.00E-09 | 107.066991 | 153.92889 | 167.049345 | 168.408823 | 0 |
| 1.24E-10 | 1.01E-09 | 105.459379 | 178.619796 | 189.322591 | 161.913717 | 0 |
| 1.25E-10 | 1.01E-09 | 242.427962 | 198.194568 | 228.300772 | 196.708928 | 4.75243649 |
| 1.25E-10 | 1.01E-09 | 96.4567488 | 377.481685 | 264.494796 | 329.394667 | 0 |
| 1.25E-10 | 1.01E-09 | 146.935781 | 208.204395 | 110.438178 | 183.718716 | 0 |
| 1.25E-10 | 1.01E-09 | 95.4921813 | 285.836161 | 276.559471 | 259.804245 | 0 |
| 1.26E-10 | 1.02E-09 | 105.459379 | 199.306771 | 167.049345 | 219.905736 | 0 |
| 1.26E-10 | 1.02E-09 | 183.910868 | 612.601392 | 495.579724 | 590.590721 | 11.8810912 |
| 1.27E-10 | 1.02E-09 | 106.423946 | 158.155261 | 161.481034 | 166.553078 | 0 |
| 1.27E-10 | 1.03E-09 | 136.325538 | 209.094157 | 466.810114 | 202.276162 | 0 |
| 1.28E-10 | 1.03E-09 | 162.368861 | 249.133464 | 114.150386 | 169.336695 | 0 |
| 1.28E-10 | 1.03E-09 | 93.2415239 | 294.066463 | 301.616873 | 327.538923 | 0 |
| 1.28E-10 | 1.04E-09 | 192.591975 | 259.14329 | 137.351684 | 125.262761 | 0 |
| 1.28E-10 | 1.04E-09 | 192.270453 | 366.582096 | 324.818171 | 339.137327 | 0 |
| 1.29E-10 | 1.04E-09 | 196.450245 | 252.247632 | 103.013763 | 302.022435 | 0 |
| 1.29E-10 | 1.04E-09 | 111.889829 | 154.818652 | 159.62493 | 213.874566 | 0 |
| 1.30E-10 | 1.05E-09 | 93.2415239 | 465.123279 | 351.731677 | 322.435625 | 0 |
| 1.30E-10 | 1.05E-09 | 95.1706589 | 260.033053 | 300.688821 | 332.178284 | 0 |
| 1.30E-10 | 1.05E-09 | 113.497441 | 176.61783 | 140.135839 | 144.284143 | 0 |

| | | | | | | |
|---|---|---|---|---|---|---|
| 1.30E-10 | 1.05E-09 | 94.5276139 | 273.824369 | 273.775316 | 276.04201 | 0 |
| 1.30E-10 | 1.05E-09 | 101.279586 | 242.682686 | 190.250643 | 224.081161 | 0 |
| 1.30E-10 | 1.05E-09 | 112.854396 | 152.594246 | 228.300772 | 169.800631 | 0 |
| 1.31E-10 | 1.05E-09 | 102.244154 | 347.452205 | 248.717914 | 227.328714 | 0 |
| 1.31E-10 | 1.05E-09 | 119.927891 | 235.119706 | 137.351684 | 186.038397 | 0 |
| 1.31E-10 | 1.06E-09 | 126.358341 | 125.901375 | 125.287009 | 128.510314 | 0 |
| 1.32E-10 | 1.06E-09 | 105.137856 | 175.060746 | 157.768826 | 179.543291 | 0 |
| 1.32E-10 | 1.06E-09 | 97.4213163 | 270.48776 | 224.588564 | 250.525522 | 0 |
| 1.32E-10 | 1.07E-09 | 210.918757 | 224.887439 | 208.811681 | 161.449781 | 0 |
| 1.32E-10 | 1.07E-09 | 102.565676 | 199.306771 | 219.020252 | 176.295737 | 0 |
| 1.32E-10 | 1.07E-09 | 126.036818 | 177.952474 | 118.790645 | 152.171058 | 0 |
| 1.33E-10 | 1.07E-09 | 100.636541 | 199.751652 | 231.084927 | 266.299351 | 0 |
| 1.33E-10 | 1.07E-09 | 93.8845689 | 292.064498 | 270.063108 | 294.13552 | 0 |
| 1.33E-10 | 1.07E-09 | 99.0289288 | 229.558691 | 224.588564 | 286.712542 | 0 |
| 1.33E-10 | 1.07E-09 | 174.908238 | 175.283187 | 102.085711 | 194.389247 | 0 |
| 1.33E-10 | 1.07E-09 | 91.9554339 | 335.885295 | 313.681548 | 303.414243 | 0 |
| 1.34E-10 | 1.07E-09 | 109.960694 | 174.838306 | 139.207788 | 172.120312 | 0 |
| 1.34E-10 | 1.08E-09 | 219.599865 | 474.910665 | 266.3509 | 365.117751 | 0 |
| 1.34E-10 | 1.08E-09 | 103.208721 | 193.745756 | 243.149602 | 293.671584 | 0 |
| 1.34E-10 | 1.08E-09 | 102.887199 | 218.659102 | 181.898176 | 182.326907 | 0 |
| 1.34E-10 | 1.08E-09 | 95.1706589 | 280.275146 | 296.048562 | 247.741905 | 0 |
| 1.35E-10 | 1.08E-09 | 351.745611 | 306.078255 | 92.8051917 | 280.681372 | 0 |
| 1.35E-10 | 1.08E-09 | 232.139242 | 385.267106 | 246.86181 | 271.402649 | 0 |
| 1.35E-10 | 1.08E-09 | 222.172045 | 749.847237 | 704.391405 | 714.461673 | 14.2573095 |
| 1.35E-10 | 1.08E-09 | 103.530244 | 200.196533 | 167.977397 | 212.482757 | 0 |
| 1.35E-10 | 1.08E-09 | 98.3858838 | 228.001607 | 206.955577 | 225.009033 | 0 |
| 1.35E-10 | 1.08E-09 | 87.4541189 | 418.633195 | 374.932974 | 408.727749 | 0 |
| 1.37E-10 | 1.10E-09 | 130.538133 | 252.025191 | 128.999216 | 183.718716 | 0 |
| 1.37E-10 | 1.10E-09 | 99.0289288 | 202.86582 | 210.667785 | 234.751693 | 0 |
| 1.37E-10 | 1.10E-09 | 97.4213163 | 255.584241 | 222.73246 | 225.936906 | 0 |
| 1.37E-10 | 1.10E-09 | 292.585471 | 244.462211 | 431.544141 | 205.059779 | 4.75243649 |
| 1.38E-10 | 1.10E-09 | 98.3858838 | 238.233874 | 205.099474 | 225.936906 | 0 |
| 1.38E-10 | 1.10E-09 | 138.897718 | 147.700553 | 109.510126 | 143.356271 | 0 |
| 1.38E-10 | 1.10E-09 | 105.459379 | 185.293013 | 170.761553 | 154.490738 | 0 |
| 1.38E-10 | 1.10E-09 | 181.981733 | 479.581917 | 425.97583 | 448.162322 | 16.6335277 |
| 1.38E-10 | 1.10E-09 | 90.3478214 | 317.200285 | 345.235313 | 367.437432 | 0 |
| 1.38E-10 | 1.10E-09 | 241.141872 | 104.102197 | 172.617657 | 211.554885 | 0 |
| 1.38E-10 | 1.11E-09 | 97.0997938 | 233.562622 | 218.0922 | 239.391054 | 0 |
| 1.39E-10 | 1.11E-09 | 119.606369 | 122.119885 | 175.401812 | 162.841589 | 0 |
| 1.39E-10 | 1.11E-09 | 596.102708 | 2703.32052 | 1910.8589 | 2294.6282 | 0 |
| 1.39E-10 | 1.11E-09 | 595.781185 | 280.497587 | 134.567528 | 210.163077 | 0 |
| 1.39E-10 | 1.11E-09 | 215.420072 | 134.576558 | 123.430905 | 149.387441 | 0 |
| 1.39E-10 | 1.11E-09 | 109.960694 | 164.161157 | 141.063891 | 144.284143 | 0 |

| | | | | | | |
|---|---|---|---|---|---|---|
| 1.40E-10 | 1.12E-09 | 107.388514 | 167.497766 | 196.747006 | 150.315313 | 0 |
| 1.40E-10 | 1.12E-09 | 110.603739 | 134.354118 | 168.905449 | 162.841589 | 0 |
| 1.41E-10 | 1.12E-09 | 96.7782713 | 220.661068 | 270.063108 | 274.650202 | 0 |
| 1.41E-10 | 1.12E-09 | 190.019795 | 347.897087 | 310.897392 | 387.386686 | 2.37621824 |
| 1.41E-10 | 1.13E-09 | 100.958064 | 442.656779 | 280.271679 | 265.835415 | 0 |
| 1.42E-10 | 1.13E-09 | 98.7074063 | 233.562622 | 197.675058 | 220.833608 | 0 |
| 1.42E-10 | 1.13E-09 | 97.7428388 | 252.692513 | 216.236097 | 286.712542 | 0 |
| 1.42E-10 | 1.14E-09 | 97.0997938 | 228.224048 | 213.451941 | 240.318926 | 0 |
| 1.43E-10 | 1.14E-09 | 96.7782713 | 364.13525 | 267.278952 | 259.340309 | 0 |
| 1.43E-10 | 1.14E-09 | 101.279586 | 208.204395 | 174.47376 | 201.812226 | 0 |
| 1.43E-10 | 1.14E-09 | 112.532874 | 164.383598 | 133.639476 | 141.964462 | 0 |
| 1.43E-10 | 1.14E-09 | 108.353081 | 187.294979 | 162.409085 | 245.422224 | 0 |
| 1.44E-10 | 1.14E-09 | 134.717926 | 297.180631 | 131.783372 | 212.482757 | 0 |
| 1.44E-10 | 1.15E-09 | 99.3504513 | 201.308736 | 193.962851 | 192.069567 | 0 |
| 1.44E-10 | 1.15E-09 | 96.7782713 | 231.115776 | 216.236097 | 225.009033 | 0 |
| 1.45E-10 | 1.15E-09 | 169.763878 | 285.836161 | 116.934542 | 371.612857 | 0 |
| 1.45E-10 | 1.15E-09 | 89.7047764 | 319.424691 | 335.026742 | 368.82924 | 0 |
| 1.46E-10 | 1.16E-09 | 119.606369 | 149.257638 | 197.675058 | 129.902122 | 0 |
| 1.46E-10 | 1.16E-09 | 107.066991 | 176.39539 | 142.919995 | 174.439993 | 0 |
| 1.46E-10 | 1.16E-09 | 288.405679 | 325.208146 | 314.6096 | 372.54073 | 0 |
| 1.47E-10 | 1.16E-09 | 115.426576 | 332.771126 | 174.47376 | 212.482757 | 0 |
| 1.47E-10 | 1.17E-09 | 101.279586 | 216.212256 | 185.610383 | 256.092756 | 0 |
| 1.48E-10 | 1.17E-09 | 194.842633 | 289.617651 | 257.998433 | 281.609244 | 19.0097459 |
| 1.48E-10 | 1.18E-09 | 107.388514 | 199.529211 | 146.632203 | 199.028609 | 0 |
| 1.48E-10 | 1.18E-09 | 103.851766 | 246.019295 | 169.833501 | 235.215629 | 0 |
| 1.49E-10 | 1.18E-09 | 119.606369 | 121.007682 | 152.200514 | 132.685739 | 0 |
| 1.50E-10 | 1.19E-09 | 94.5276139 | 280.497587 | 236.653239 | 259.340309 | 0 |
| 1.50E-10 | 1.19E-09 | 86.4895515 | 451.776843 | 374.932974 | 383.675197 | 0 |
| 1.51E-10 | 1.19E-09 | 98.0643613 | 246.686617 | 199.531162 | 219.4418 | 0 |
| 1.51E-10 | 1.20E-09 | 120.249414 | 129.905306 | 121.574801 | 153.09893 | 0 |
| 1.51E-10 | 1.20E-09 | 106.745469 | 251.58031 | 174.47376 | 178.151482 | 0 |
| 1.51E-10 | 1.20E-09 | 111.889829 | 190.854028 | 133.639476 | 167.480951 | 0 |
| 1.52E-10 | 1.20E-09 | 96.1352263 | 259.365731 | 242.22155 | 214.338502 | 0 |
| 1.52E-10 | 1.20E-09 | 161.082771 | 548.760942 | 546.622579 | 513.57732 | 0 |
| 1.52E-10 | 1.21E-09 | 107.388514 | 222.663033 | 151.272462 | 216.658183 | 0 |
| 1.53E-10 | 1.21E-09 | 416.05011 | 284.056637 | 131.783372 | 154.954675 | 0 |
| 1.53E-10 | 1.21E-09 | 596.745753 | 988.303552 | 970.742305 | 974.729854 | 163.959059 |
| 1.53E-10 | 1.21E-09 | 90.9908664 | 314.975879 | 283.983887 | 350.73573 | 0 |
| 1.53E-10 | 1.21E-09 | 123.786161 | 144.586385 | 259.854537 | 161.913717 | 0 |
| 1.54E-10 | 1.22E-09 | 101.279586 | 190.854028 | 165.193241 | 188.822014 | 0 |
| 1.54E-10 | 1.22E-09 | 98.7074063 | 193.968197 | 203.24337 | 238.463182 | 0 |
| 1.56E-10 | 1.23E-09 | 166.22713 | 273.157048 | 105.797919 | 237.071373 | 0 |
| 1.56E-10 | 1.23E-09 | 90.9908664 | 328.544755 | 605.08985 | 492.700193 | 0 |

| | | | | | | |
|---|---|---|---|---|---|---|
| 1.57E-10 | 1.24E-09 | 119.284846 | 197.082365 | 314.6096 | 159.594036 | 0 |
| 1.58E-10 | 1.25E-09 | 118.320279 | 121.675004 | 134.567528 | 158.666164 | 0 |
| 1.58E-10 | 1.25E-09 | 91.6339114 | 276.048775 | 292.336354 | 348.879986 | 0 |
| 1.58E-10 | 1.25E-09 | 93.5630464 | 249.355904 | 238.509343 | 264.443606 | 0 |
| 1.59E-10 | 1.25E-09 | 94.2060914 | 258.253528 | 405.558688 | 383.675197 | 0 |
| 1.59E-10 | 1.25E-09 | 101.922631 | 229.781132 | 173.545708 | 186.966269 | 0 |
| 1.59E-10 | 1.25E-09 | 95.8137038 | 368.13918 | 237.581291 | 329.858604 | 0 |
| 1.59E-10 | 1.26E-09 | 126.679863 | 166.830444 | 113.222334 | 182.790844 | 0 |
| 1.59E-10 | 1.26E-09 | 94.5276139 | 224.442558 | 251.502069 | 236.607437 | 0 |
| 1.59E-10 | 1.26E-09 | 102.565676 | 165.27336 | 161.481034 | 191.141694 | 0 |
| 1.60E-10 | 1.26E-09 | 176.837373 | 100.988029 | 247.789862 | 237.999246 | 0 |
| 1.60E-10 | 1.26E-09 | 85.8465065 | 351.456136 | 431.544141 | 421.254025 | 0 |
| 1.60E-10 | 1.26E-09 | 115.105054 | 197.527246 | 128.071165 | 170.728504 | 0 |
| 1.60E-10 | 1.26E-09 | 105.459379 | 168.609969 | 162.409085 | 141.964462 | 0 |
| 1.60E-10 | 1.26E-09 | 195.485678 | 118.560836 | 128.999216 | 146.603824 | 0 |
| 1.60E-10 | 1.26E-09 | 117.034189 | 156.153296 | 127.143113 | 127.118505 | 0 |
| 1.60E-10 | 1.26E-09 | 187.126093 | 242.015365 | 145.704151 | 188.358077 | 7.12865473 |
| 1.61E-10 | 1.26E-09 | 100.315019 | 175.505627 | 171.689605 | 194.853184 | 0 |
| 1.61E-10 | 1.27E-09 | 125.072251 | 133.241915 | 128.071165 | 115.520102 | 0 |
| 1.61E-10 | 1.27E-09 | 94.5276139 | 266.261389 | 221.804408 | 254.237011 | 0 |
| 1.61E-10 | 1.27E-09 | 93.8845689 | 367.249418 | 250.574018 | 313.156902 | 0 |
| 1.61E-10 | 1.27E-09 | 117.677234 | 243.572449 | 143.848047 | 164.697334 | 0 |
| 1.62E-10 | 1.27E-09 | 96.1352263 | 220.216187 | 212.523889 | 204.131907 | 0 |
| 1.62E-10 | 1.27E-09 | 109.639171 | 137.690727 | 132.711424 | 152.171058 | 0 |
| 1.63E-10 | 1.28E-09 | 91.9554339 | 294.066463 | 279.343627 | 252.845202 | 0 |
| 1.63E-10 | 1.28E-09 | 229.567062 | 300.072359 | 218.0922 | 253.309139 | 2.37621824 |
| 1.64E-10 | 1.29E-09 | 94.2060914 | 260.033053 | 221.804408 | 266.299351 | 0 |
| 1.64E-10 | 1.29E-09 | 92.9200014 | 263.592102 | 236.653239 | 262.587862 | 0 |
| 1.65E-10 | 1.30E-09 | 92.9200014 | 242.015365 | 245.005706 | 265.371479 | 0 |
| 1.66E-10 | 1.30E-09 | 96.4567488 | 201.531177 | 199.531162 | 212.946693 | 0 |
| 1.66E-10 | 1.30E-09 | 154.009276 | 166.830444 | 103.013763 | 147.995632 | 0 |
| 1.67E-10 | 1.31E-09 | 244.035575 | 222.663033 | 233.869083 | 227.79265 | 2.37621824 |
| 1.67E-10 | 1.31E-09 | 241.463395 | 713.589421 | 679.334003 | 627.705613 | 35.6432737 |
| 1.67E-10 | 1.31E-09 | 104.816334 | 201.976058 | 150.344411 | 169.800631 | 0 |
| 1.68E-10 | 1.32E-09 | 112.532874 | 184.180811 | 152.200514 | 129.902122 | 0 |
| 1.68E-10 | 1.32E-09 | 106.745469 | 159.934786 | 135.49558 | 168.872759 | 0 |
| 1.68E-10 | 1.32E-09 | 183.267823 | 322.093978 | 286.768042 | 257.020628 | 14.2573095 |
| 1.68E-10 | 1.32E-09 | 99.9934963 | 326.76523 | 257.070381 | 202.276162 | 0 |
| 1.69E-10 | 1.32E-09 | 95.8137038 | 281.60979 | 208.811681 | 256.556692 | 0 |
| 1.69E-10 | 1.32E-09 | 136.647061 | 223.775236 | 113.222334 | 229.184459 | 0 |
| 1.69E-10 | 1.32E-09 | 93.8845689 | 291.842057 | 232.012979 | 246.350096 | 0 |
| 1.69E-10 | 1.32E-09 | 118.320279 | 116.113989 | 137.351684 | 139.644782 | 0 |
| 1.69E-10 | 1.32E-09 | 96.1352263 | 223.552795 | 206.955577 | 197.6368 | 0 |

| | | | | | | |
|---|---|---|---|---|---|---|
| 1.69E-10 | 1.33E-09 | 85.2034615 | 407.511165 | 366.580507 | 364.189879 | 0 |
| 1.69E-10 | 1.33E-09 | 45.6561944 | 17.3503662 | 30.6257133 | 22.2689353 | 180.592587 |
| 1.70E-10 | 1.33E-09 | 95.4921813 | 216.212256 | 201.387266 | 218.513927 | 0 |
| 1.70E-10 | 1.33E-09 | 200.95156 | 113.444702 | 162.409085 | 131.293931 | 0 |
| 1.70E-10 | 1.33E-09 | 97.7428388 | 362.355725 | 228.300772 | 263.97967 | 0 |
| 1.71E-10 | 1.34E-09 | 110.282216 | 158.822583 | 127.143113 | 144.748079 | 0 |
| 1.71E-10 | 1.34E-09 | 137.290106 | 163.048954 | 181.898176 | 209.69914 | 14.2573095 |
| 1.72E-10 | 1.34E-09 | 151.115573 | 383.710022 | 367.508559 | 369.757113 | 2.37621824 |
| 1.72E-10 | 1.34E-09 | 165.584086 | 175.950509 | 118.790645 | 116.91191 | 0 |
| 1.73E-10 | 1.35E-09 | 93.2415239 | 223.997677 | 243.149602 | 239.85499 | 0 |
| 1.74E-10 | 1.35E-09 | 90.0262989 | 284.279077 | 283.983887 | 270.938712 | 0 |
| 1.74E-10 | 1.36E-09 | 110.282216 | 136.800964 | 142.919995 | 183.25478 | 0 |
| 1.74E-10 | 1.36E-09 | 104.494811 | 145.476147 | 143.848047 | 151.707121 | 0 |
| 1.74E-10 | 1.36E-09 | 100.315019 | 172.391459 | 174.47376 | 159.1301 | 0 |
| 1.74E-10 | 1.36E-09 | 267.185194 | 161.936751 | 108.582074 | 183.718716 | 0 |
| 1.74E-10 | 1.36E-09 | 97.7428388 | 241.125602 | 224.588564 | 183.25478 | 0 |
| 1.75E-10 | 1.36E-09 | 68.1627692 | 682.892619 | 897.426204 | 826.270285 | 0 |
| 1.75E-10 | 1.36E-09 | 191.305885 | 304.743612 | 373.076871 | 293.207648 | 0 |
| 1.75E-10 | 1.36E-09 | 114.140486 | 123.676969 | 130.85532 | 152.634994 | 0 |
| 1.76E-10 | 1.37E-09 | 94.8491364 | 209.094157 | 214.379993 | 207.843396 | 0 |
| 1.76E-10 | 1.37E-09 | 115.426576 | 164.606038 | 124.358957 | 133.149675 | 0 |
| 1.76E-10 | 1.37E-09 | 92.5984789 | 264.704305 | 237.581291 | 235.679565 | 0 |
| 1.76E-10 | 1.37E-09 | 91.9554339 | 432.202071 | 268.207004 | 385.067006 | 0 |
| 1.76E-10 | 1.37E-09 | 137.290106 | 123.454529 | 260.782589 | 301.558498 | 0 |
| 1.76E-10 | 1.37E-09 | 107.066991 | 169.499731 | 180.042072 | 134.077548 | 0 |
| 1.78E-10 | 1.39E-09 | 93.2415239 | 250.245666 | 217.164149 | 248.669777 | 0 |
| 1.79E-10 | 1.39E-09 | 101.922631 | 150.1474 | 190.250643 | 184.182652 | 0 |
| 1.79E-10 | 1.39E-09 | 118.963324 | 121.897445 | 124.358957 | 122.94308 | 0 |
| 1.80E-10 | 1.40E-09 | 196.128723 | 291.619617 | 256.142329 | 278.825627 | 2.37621824 |
| 1.80E-10 | 1.40E-09 | 111.889829 | 510.278719 | 201.387266 | 406.408069 | 0 |
| 1.80E-10 | 1.40E-09 | 96.7782713 | 189.964266 | 185.610383 | 187.430205 | 0 |
| 1.80E-10 | 1.40E-09 | 92.2769564 | 244.907092 | 232.012979 | 239.85499 | 0 |
| 1.81E-10 | 1.40E-09 | 98.3858838 | 254.472038 | 336.882846 | 208.771268 | 0 |
| 1.81E-10 | 1.40E-09 | 104.494811 | 162.381632 | 172.617657 | 138.252973 | 0 |
| 1.81E-10 | 1.41E-09 | 114.462009 | 213.098088 | 163.337137 | 131.757867 | 0 |
| 1.81E-10 | 1.41E-09 | 93.5630464 | 369.696265 | 446.392972 | 257.9485 | 0 |
| 1.82E-10 | 1.42E-09 | 96.4567488 | 299.627478 | 202.315318 | 274.186265 | 0 |
| 1.83E-10 | 1.42E-09 | 164.297996 | 241.792924 | 249.645966 | 260.268181 | 2.37621824 |
| 1.83E-10 | 1.42E-09 | 99.9934963 | 162.381632 | 159.62493 | 174.439993 | 0 |
| 1.83E-10 | 1.42E-09 | 113.497441 | 139.915133 | 127.143113 | 178.151482 | 0 |
| 1.84E-10 | 1.43E-09 | 104.173289 | 145.921029 | 141.991943 | 144.284143 | 0 |
| 1.85E-10 | 1.43E-09 | 406.404435 | 716.481148 | 650.564394 | 640.231889 | 9409.82424 |
| 1.85E-10 | 1.43E-09 | 119.927891 | 252.914954 | 138.279736 | 166.553078 | 0 |

| | | | | | | |
|---|---|---|---|---|---|---|
| 1.85E-10 | 1.43E-09 | 106.745469 | 149.257638 | 137.351684 | 180.471163 | 0 |
| 1.86E-10 | 1.44E-09 | 90.9908664 | 264.036983 | 243.149602 | 250.061586 | 0 |
| 1.86E-10 | 1.44E-09 | 96.1352263 | 183.068608 | 193.034799 | 201.34829 | 0 |
| 1.86E-10 | 1.44E-09 | 102.565676 | 162.381632 | 170.761553 | 142.892335 | 0 |
| 1.87E-10 | 1.45E-09 | 85.524984 | 447.772913 | 389.781805 | 326.147114 | 0 |
| 1.88E-10 | 1.45E-09 | 96.1352263 | 252.247632 | 190.250643 | 254.237011 | 0 |
| 1.89E-10 | 1.46E-09 | 91.9554339 | 323.206181 | 239.437395 | 345.632433 | 0 |
| 1.89E-10 | 1.46E-09 | 99.0289288 | 214.877612 | 164.265189 | 204.595843 | 0 |
| 1.89E-10 | 1.46E-09 | 91.3123889 | 254.027157 | 389.781805 | 357.230837 | 0 |
| 1.90E-10 | 1.47E-09 | 104.494811 | 153.261568 | 221.804408 | 168.872759 | 0 |
| 1.91E-10 | 1.47E-09 | 86.168029 | 309.414864 | 329.458431 | 331.250412 | 0 |
| 1.91E-10 | 1.48E-09 | 99.0289288 | 292.73182 | 189.322591 | 226.400842 | 0 |
| 1.91E-10 | 1.48E-09 | 108.353081 | 166.608004 | 127.143113 | 149.387441 | 0 |
| 1.91E-10 | 1.48E-09 | 88.4186864 | 293.176701 | 275.631419 | 281.145308 | 0 |
| 1.91E-10 | 1.48E-09 | 93.2415239 | 234.229944 | 220.876356 | 207.37946 | 0 |
| 1.92E-10 | 1.48E-09 | 89.7047764 | 312.973914 | 257.998433 | 271.402649 | 0 |
| 1.93E-10 | 1.49E-09 | 77.8084441 | 737.613005 | 504.860243 | 506.154341 | 0 |
| 1.93E-10 | 1.49E-09 | 108.674604 | 171.279256 | 126.215061 | 168.872759 | 0 |
| 1.93E-10 | 1.49E-09 | 88.7402089 | 310.749508 | 308.113236 | 261.196053 | 0 |
| 1.93E-10 | 1.49E-09 | 98.7074063 | 303.408968 | 295.12051 | 190.213822 | 0 |
| 1.94E-10 | 1.49E-09 | 193.556543 | 161.936751 | 100.229607 | 154.954675 | 0 |
| 1.94E-10 | 1.49E-09 | 114.783531 | 137.023405 | 115.078438 | 141.03659 | 0 |
| 1.94E-10 | 1.50E-09 | 154.652321 | 116.33643 | 127.143113 | 207.843396 | 0 |
| 1.94E-10 | 1.50E-09 | 141.791421 | 107.661247 | 172.617657 | 132.685739 | 0 |
| 1.94E-10 | 1.50E-09 | 98.7074063 | 187.73986 | 160.552982 | 200.420417 | 0 |
| 1.94E-10 | 1.50E-09 | 90.9908664 | 250.023226 | 231.084927 | 248.669777 | 0 |
| 1.95E-10 | 1.50E-09 | 96.7782713 | 188.407182 | 175.401812 | 211.554885 | 0 |
| 1.95E-10 | 1.50E-09 | 260.754744 | 726.713415 | 675.621796 | 689.873057 | 16.6335277 |
| 1.95E-10 | 1.50E-09 | 96.4567488 | 264.481864 | 194.890903 | 293.671584 | 0 |
| 1.95E-10 | 1.50E-09 | 17.3622148 | 13.7913167 | 13.9207788 | 10.6705315 | 111.682257 |
| 1.96E-10 | 1.50E-09 | 117.677234 | 125.456494 | 121.574801 | 161.913717 | 0 |
| 1.96E-10 | 1.50E-09 | 155.938411 | 448.217794 | 435.256349 | 465.32796 | 4.75243649 |
| 1.97E-10 | 1.51E-09 | 102.887199 | 310.749508 | 185.610383 | 212.018821 | 0 |
| 1.97E-10 | 1.51E-09 | 95.1706589 | 193.968197 | 221.804408 | 189.28595 | 0 |
| 1.97E-10 | 1.51E-09 | 94.5276139 | 195.525281 | 193.034799 | 211.554885 | 0 |
| 1.97E-10 | 1.52E-09 | 93.5630464 | 225.10988 | 206.955577 | 262.123925 | 0 |
| 1.97E-10 | 1.52E-09 | 109.317649 | 139.915133 | 124.358957 | 159.1301 | 0 |
| 1.98E-10 | 1.52E-09 | 95.1706589 | 279.162944 | 209.739733 | 325.683178 | 0 |
| 1.98E-10 | 1.52E-09 | 99.6719738 | 188.852063 | 164.265189 | 227.328714 | 0 |
| 1.98E-10 | 1.52E-09 | 139.540763 | 167.052885 | 151.272462 | 168.872759 | 7.12865473 |
| 1.99E-10 | 1.52E-09 | 97.0997938 | 252.247632 | 180.042072 | 219.4418 | 0 |
| 1.99E-10 | 1.52E-09 | 100.958064 | 144.808826 | 162.409085 | 157.274355 | 0 |
| 1.99E-10 | 1.53E-09 | 106.745469 | 178.842236 | 129.927268 | 166.089142 | 0 |

| | | | | | | |
|---|---|---|---|---|---|---|
| 1.99E-10 | 1.53E-09 | 103.530244 | 319.20225 | 177.257916 | 312.22903 | 0 |
| 2.00E-10 | 1.53E-09 | 99.0289288 | 174.170984 | 154.056618 | 181.862971 | 0 |
| 2.01E-10 | 1.54E-09 | 156.902978 | 188.629622 | 140.135839 | 194.853184 | 11.8810912 |
| 2.01E-10 | 1.54E-09 | 100.315019 | 155.263534 | 216.236097 | 204.131907 | 0 |
| 2.01E-10 | 1.54E-09 | 456.561944 | 806.124707 | 625.506992 | 696.832099 | 149.701749 |
| 2.01E-10 | 1.54E-09 | 84.238894 | 389.048596 | 334.09869 | 342.848816 | 0 |
| 2.01E-10 | 1.54E-09 | 177.158895 | 151.037162 | 167.977397 | 99.2823364 | 0 |
| 2.02E-10 | 1.54E-09 | 90.0262989 | 269.153117 | 235.725187 | 259.804245 | 0 |
| 2.02E-10 | 1.55E-09 | 92.2769564 | 469.794531 | 270.99116 | 333.106157 | 0 |
| 2.03E-10 | 1.55E-09 | 96.7782713 | 196.859924 | 174.47376 | 172.584248 | 0 |
| 2.03E-10 | 1.55E-09 | 268.792807 | 157.043058 | 128.999216 | 135.00542 | 0 |
| 2.03E-10 | 1.55E-09 | 90.3478214 | 286.058602 | 232.941031 | 262.587862 | 0 |
| 2.03E-10 | 1.55E-09 | 101.279586 | 170.389494 | 141.991943 | 167.944887 | 0 |
| 2.04E-10 | 1.56E-09 | 95.8137038 | 202.198499 | 177.257916 | 223.153289 | 0 |
| 2.04E-10 | 1.56E-09 | 100.315019 | 158.155261 | 227.37272 | 222.225416 | 0 |
| 2.04E-10 | 1.56E-09 | 87.1325964 | 623.500981 | 351.731677 | 406.872005 | 0 |
| 2.04E-10 | 1.56E-09 | 98.0643613 | 170.389494 | 160.552982 | 189.28595 | 0 |
| 2.05E-10 | 1.56E-09 | 105.459379 | 146.36591 | 206.955577 | 238.463182 | 0 |
| 2.05E-10 | 1.57E-09 | 110.925261 | 193.523315 | 135.49558 | 140.108718 | 0 |
| 2.06E-10 | 1.57E-09 | 88.0971639 | 280.497587 | 438.968557 | 379.035836 | 0 |
| 2.06E-10 | 1.57E-09 | 92.2769564 | 208.204395 | 238.509343 | 217.586055 | 0 |
| 2.06E-10 | 1.57E-09 | 96.7782713 | 265.594067 | 183.75428 | 221.297544 | 0 |
| 2.06E-10 | 1.57E-09 | 102.244154 | 250.023226 | 164.265189 | 186.038397 | 0 |
| 2.06E-10 | 1.57E-09 | 100.958064 | 204.645345 | 316.465704 | 198.100737 | 0 |
| 2.06E-10 | 1.57E-09 | 142.755988 | 148.145435 | 168.905449 | 101.602017 | 0 |
| 2.06E-10 | 1.57E-09 | 100.636541 | 158.377702 | 143.848047 | 156.346483 | 0 |
| 2.07E-10 | 1.58E-09 | 84.238894 | 343.225834 | 593.025175 | 494.555937 | 0 |
| 2.07E-10 | 1.58E-09 | 93.5630464 | 269.153117 | 199.531162 | 249.133713 | 0 |
| 2.07E-10 | 1.58E-09 | 134.074881 | 106.104163 | 141.991943 | 173.048184 | 0 |
| 2.07E-10 | 1.58E-09 | 137.290106 | 161.714311 | 102.085711 | 145.675952 | 0 |
| 2.08E-10 | 1.58E-09 | 93.8845689 | 247.79882 | 192.106747 | 230.576267 | 0 |
| 2.08E-10 | 1.58E-09 | 174.908238 | 326.097909 | 269.135056 | 300.630626 | 0 |
| 2.08E-10 | 1.59E-09 | 140.826853 | 329.656958 | 353.58778 | 345.168497 | 9.50487297 |
| 2.09E-10 | 1.59E-09 | 92.2769564 | 250.245666 | 205.099474 | 257.9485 | 0 |
| 2.09E-10 | 1.59E-09 | 110.925261 | 123.676969 | 146.632203 | 124.334889 | 0 |
| 2.10E-10 | 1.60E-09 | 189.698273 | 403.284794 | 348.947521 | 375.324346 | 16.6335277 |
| 2.11E-10 | 1.60E-09 | 96.1352263 | 168.387528 | 197.675058 | 181.399035 | 0 |
| 2.11E-10 | 1.60E-09 | 79.737579 | 502.048417 | 400.918428 | 425.429451 | 0 |
| 2.11E-10 | 1.60E-09 | 2.57217997 | 3.78149007 | 2.78415575 | 6.03116997 | 40.3957101 |
| 2.11E-10 | 1.61E-09 | 85.524984 | 312.084151 | 294.192458 | 331.250412 | 0 |
| 2.12E-10 | 1.61E-09 | 102.244154 | 155.930855 | 146.632203 | 136.861165 | 0 |
| 2.13E-10 | 1.62E-09 | 236.319035 | 202.86582 | 152.200514 | 102.065953 | 0 |
| 2.13E-10 | 1.62E-09 | 77.4869216 | 436.428442 | 761.002572 | 617.499017 | 0 |

| | | | | | | |
|---|---|---|---|---|---|---|
| 2.14E-10 | 1.62E-09 | 96.1352263 | 242.682686 | 176.329864 | 238.463182 | 0 |
| 2.14E-10 | 1.62E-09 | 92.5984789 | 407.956047 | 322.962067 | 240.318926 | 0 |
| 2.14E-10 | 1.62E-09 | 144.685123 | 640.628906 | 684.902315 | 663.892633 | 4.75243649 |
| 2.14E-10 | 1.62E-09 | 108.353081 | 120.562801 | 152.200514 | 153.562866 | 0 |
| 2.14E-10 | 1.62E-09 | 108.031559 | 153.484009 | 133.639476 | 198.100737 | 0 |
| 2.14E-10 | 1.63E-09 | 94.2060914 | 197.972127 | 187.466487 | 229.184459 | 0 |
| 2.15E-10 | 1.63E-09 | 89.3832539 | 233.785063 | 255.214277 | 276.04201 | 0 |
| 2.15E-10 | 1.63E-09 | 87.1325964 | 307.635339 | 261.710641 | 309.909349 | 0 |
| 2.15E-10 | 1.63E-09 | 84.238894 | 343.225834 | 308.113236 | 344.240624 | 0 |
| 2.16E-10 | 1.64E-09 | 98.7074063 | 266.48383 | 177.257916 | 201.812226 | 0 |
| 2.17E-10 | 1.64E-09 | 161.082771 | 166.830444 | 101.157659 | 135.469356 | 0 |
| 2.17E-10 | 1.64E-09 | 92.2769564 | 296.73575 | 227.37272 | 224.545097 | 0 |
| 2.17E-10 | 1.65E-09 | 93.8845689 | 250.690548 | 195.818954 | 205.523715 | 0 |
| 2.18E-10 | 1.65E-09 | 168.79931 | 383.932463 | 377.71713 | 435.636046 | 0 |
| 2.18E-10 | 1.65E-09 | 87.1325964 | 271.599963 | 318.321808 | 272.794457 | 0 |
| 2.18E-10 | 1.65E-09 | 200.95156 | 310.527067 | 294.192458 | 335.425837 | 28.5146189 |
| 2.18E-10 | 1.65E-09 | 134.074881 | 213.542969 | 107.654022 | 216.658183 | 0 |
| 2.18E-10 | 1.65E-09 | 110.925261 | 118.115955 | 167.977397 | 157.274355 | 0 |
| 2.19E-10 | 1.65E-09 | 90.6693439 | 227.779167 | 260.782589 | 219.905736 | 0 |
| 2.19E-10 | 1.65E-09 | 94.8491364 | 176.39539 | 206.027526 | 225.47297 | 0 |
| 2.19E-10 | 1.65E-09 | 90.0262989 | 228.001607 | 228.300772 | 254.237011 | 0 |
| 2.19E-10 | 1.65E-09 | 96.4567488 | 254.249597 | 180.970124 | 202.276162 | 0 |
| 2.19E-10 | 1.66E-09 | 96.4567488 | 224.664998 | 177.257916 | 179.543291 | 0 |
| 2.19E-10 | 1.66E-09 | 144.042078 | 251.802751 | 230.156875 | 179.543291 | 7.12865473 |
| 2.20E-10 | 1.66E-09 | 134.717926 | 170.389494 | 112.294282 | 121.551272 | 0 |
| 2.20E-10 | 1.66E-09 | 128.287476 | 106.104163 | 132.711424 | 152.171058 | 0 |
| 2.20E-10 | 1.66E-09 | 83.595849 | 318.090047 | 366.580507 | 424.037642 | 0 |
| 2.20E-10 | 1.66E-09 | 90.9908664 | 232.227979 | 207.883629 | 229.648395 | 0 |
| 2.20E-10 | 1.66E-09 | 98.7074063 | 148.367875 | 156.840774 | 161.913717 | 0 |
| 2.20E-10 | 1.66E-09 | 103.851766 | 185.737895 | 136.423632 | 154.954675 | 0 |
| 2.20E-10 | 1.66E-09 | 165.262563 | 371.920671 | 285.83999 | 308.517541 | 4.75243649 |
| 2.21E-10 | 1.66E-09 | 117.998756 | 249.355904 | 152.200514 | 141.500526 | 0 |
| 2.21E-10 | 1.67E-09 | 261.076267 | 324.318384 | 230.156875 | 302.022435 | 16.6335277 |
| 2.22E-10 | 1.67E-09 | 0 | 0 | 0 | 0 | 38.0194919 |
| 2.22E-10 | 1.67E-09 | 82.6312815 | 399.948185 | 324.818171 | 430.532748 | 0 |
| 2.22E-10 | 1.68E-09 | 83.2743265 | 379.03877 | 332.242586 | 326.147114 | 0 |
| 2.22E-10 | 1.68E-09 | 253.038204 | 153.261568 | 285.83999 | 106.241379 | 0 |
| 2.23E-10 | 1.68E-09 | 137.290106 | 102.767554 | 174.47376 | 192.069567 | 0 |
| 2.23E-10 | 1.68E-09 | 95.4921813 | 177.730033 | 167.977397 | 194.853184 | 0 |
| 2.23E-10 | 1.68E-09 | 104.494811 | 135.466321 | 133.639476 | 161.913717 | 0 |
| 2.25E-10 | 1.69E-09 | 83.595849 | 324.985706 | 530.845697 | 461.616471 | 0 |
| 2.26E-10 | 1.70E-09 | 87.4541189 | 250.468107 | 280.271679 | 263.97967 | 0 |
| 2.26E-10 | 1.70E-09 | 112.211351 | 195.525281 | 120.646749 | 175.367865 | 0 |

| | | | | | | |
|---|---|---|---|---|---|---|
| 2.28E-10 | 1.71E-09 | 94.5276139 | 204.422904 | 211.595837 | 171.19244 | 0 |
| 2.28E-10 | 1.72E-09 | 105.137856 | 138.358048 | 127.143113 | 131.293931 | 0 |
| 2.29E-10 | 1.72E-09 | 95.4921813 | 196.637484 | 255.214277 | 185.110524 | 0 |
| 2.29E-10 | 1.72E-09 | 95.4921813 | 205.312667 | 358.22804 | 291.815839 | 0 |
| 2.29E-10 | 1.72E-09 | 89.0617314 | 236.231909 | 232.941031 | 271.402649 | 0 |
| 2.30E-10 | 1.73E-09 | 89.7047764 | 237.566553 | 215.308045 | 237.535309 | 0 |
| 2.30E-10 | 1.73E-09 | 89.3832539 | 234.229944 | 240.365446 | 220.369672 | 0 |
| 2.32E-10 | 1.74E-09 | 91.3123889 | 221.995711 | 300.688821 | 231.50414 | 0 |
| 2.32E-10 | 1.74E-09 | 104.173289 | 244.462211 | 145.704151 | 207.37946 | 0 |
| 2.32E-10 | 1.74E-09 | 298.051354 | 199.529211 | 120.646749 | 135.469356 | 0 |
| 2.32E-10 | 1.74E-09 | 88.0971639 | 269.375557 | 247.789862 | 237.999246 | 0 |
| 2.33E-10 | 1.75E-09 | 91.3123889 | 254.694478 | 210.667785 | 210.627013 | 0 |
| 2.33E-10 | 1.75E-09 | 92.5984789 | 267.373592 | 193.962851 | 244.494352 | 0 |
| 2.34E-10 | 1.75E-09 | 96.4567488 | 193.968197 | 160.552982 | 168.408823 | 0 |
| 2.34E-10 | 1.75E-09 | 100.958064 | 194.857959 | 141.991943 | 166.089142 | 0 |
| 2.34E-10 | 1.76E-09 | 121.535504 | 170.167053 | 151.272462 | 107.633187 | 0 |
| 2.37E-10 | 1.78E-09 | 192.913498 | 434.426477 | 258.926485 | 350.271794 | 0 |
| 2.38E-10 | 1.79E-09 | 188.412183 | 148.590316 | 175.401812 | 97.4265918 | 0 |
| 2.38E-10 | 1.79E-09 | 86.811074 | 254.916919 | 265.422848 | 254.700947 | 0 |
| 2.39E-10 | 1.79E-09 | 81.023669 | 482.918526 | 338.73895 | 511.257639 | 0 |
| 2.39E-10 | 1.79E-09 | 95.4921813 | 188.852063 | 158.696878 | 191.60563 | 0 |
| 2.39E-10 | 1.79E-09 | 229.567062 | 240.680721 | 88.1649321 | 221.297544 | 0 |
| 2.39E-10 | 1.79E-09 | 96.4567488 | 237.566553 | 168.905449 | 192.533503 | 0 |
| 2.39E-10 | 1.79E-09 | 86.811074 | 257.808647 | 314.6096 | 263.515734 | 0 |
| 2.41E-10 | 1.80E-09 | 95.8137038 | 209.983919 | 162.409085 | 182.790844 | 0 |
| 2.41E-10 | 1.80E-09 | 90.3478214 | 240.680721 | 215.308045 | 289.496158 | 0 |
| 2.41E-10 | 1.80E-09 | 93.8845689 | 176.172949 | 187.466487 | 170.728504 | 0 |
| 2.42E-10 | 1.81E-09 | 95.4921813 | 164.383598 | 165.193241 | 191.141694 | 0 |
| 2.42E-10 | 1.81E-09 | 69.4488592 | 546.758976 | 675.621796 | 648.118803 | 0 |
| 2.42E-10 | 1.81E-09 | 88.0971639 | 302.074325 | 232.941031 | 266.299351 | 0 |
| 2.43E-10 | 1.81E-09 | 92.2769564 | 254.916919 | 191.178695 | 219.905736 | 0 |
| 2.43E-10 | 1.81E-09 | 0 | 0 | 0 | 0 | 76.0389838 |
| 2.43E-10 | 1.81E-09 | 137.290106 | 100.098267 | 179.11402 | 172.120312 | 0 |
| 2.43E-10 | 1.81E-09 | 91.9554339 | 185.070573 | 230.156875 | 217.122119 | 0 |
| 2.43E-10 | 1.81E-09 | 98.7074063 | 183.291048 | 143.848047 | 163.305525 | 0 |
| 2.44E-10 | 1.82E-09 | 86.4895515 | 264.704305 | 248.717914 | 282.537116 | 0 |
| 2.46E-10 | 1.83E-09 | 86.811074 | 286.503483 | 253.358173 | 248.205841 | 0 |
| 2.46E-10 | 1.84E-09 | 100.636541 | 191.743791 | 146.632203 | 149.851377 | 0 |
| 2.46E-10 | 1.84E-09 | 95.4921813 | 173.726103 | 155.912722 | 167.017014 | 0 |
| 2.47E-10 | 1.84E-09 | 88.7402089 | 232.67286 | 217.164149 | 251.453394 | 0 |
| 2.48E-10 | 1.85E-09 | 132.467268 | 269.153117 | 225.516616 | 206.451587 | 14.2573095 |
| 2.49E-10 | 1.85E-09 | 127.001386 | 208.871716 | 112.294282 | 238.927118 | 0 |
| 2.49E-10 | 1.86E-09 | 217.992252 | 318.97981 | 193.962851 | 233.359884 | 14.2573095 |

| | | | | | | |
|---|---|---|---|---|---|---|
| 2.50E-10 | 1.86E-09 | 102.244154 | 339.221903 | 176.329864 | 238.927118 | 0 |
| 2.51E-10 | 1.87E-09 | 93.5630464 | 169.722172 | 173.545708 | 188.358077 | 0 |
| 2.51E-10 | 1.87E-09 | 129.573566 | 194.635518 | 105.797919 | 152.634994 | 0 |
| 2.52E-10 | 1.88E-09 | 87.1325964 | 255.13936 | 245.933758 | 233.82382 | 0 |
| 2.52E-10 | 1.88E-09 | 117.034189 | 252.247632 | 243.149602 | 280.681372 | 16.6335277 |
| 2.53E-10 | 1.88E-09 | 154.973843 | 96.7616577 | 197.675058 | 164.697334 | 0 |
| 2.54E-10 | 1.89E-09 | 150.151006 | 209.761479 | 102.085711 | 147.995632 | 0 |
| 2.55E-10 | 1.90E-09 | 154.973843 | 470.461853 | 455.673491 | 511.721575 | 0 |
| 2.56E-10 | 1.90E-09 | 84.5604165 | 269.153117 | 288.624146 | 277.897755 | 0 |
| 2.56E-10 | 1.91E-09 | 411.227273 | 757.187777 | 660.772965 | 790.083266 | 21.3859642 |
| 2.56E-10 | 1.91E-09 | 156.902978 | 443.546541 | 414.839207 | 432.388493 | 2.37621824 |
| 2.58E-10 | 1.92E-09 | 177.480418 | 294.066463 | 291.408302 | 305.733924 | 0 |
| 2.59E-10 | 1.92E-09 | 121.213981 | 145.476147 | 104.869867 | 165.16127 | 0 |
| 2.61E-10 | 1.94E-09 | 88.7402089 | 229.781132 | 206.955577 | 221.76148 | 0 |
| 2.62E-10 | 1.94E-09 | 91.6339114 | 228.446488 | 180.970124 | 207.37946 | 0 |
| 2.62E-10 | 1.95E-09 | 401.260075 | 704.691797 | 576.32024 | 675.491036 | 30.8908372 |
| 2.64E-10 | 1.96E-09 | 90.9908664 | 189.296944 | 184.682331 | 198.564673 | 0 |
| 2.64E-10 | 1.96E-09 | 107.388514 | 137.468286 | 171.689605 | 121.551272 | 0 |
| 2.64E-10 | 1.96E-09 | 100.958064 | 163.271395 | 133.639476 | 138.252973 | 0 |
| 2.65E-10 | 1.97E-09 | 130.216611 | 126.791138 | 103.941815 | 122.479144 | 0 |
| 2.65E-10 | 1.97E-09 | 88.7402089 | 240.680721 | 203.24337 | 232.432012 | 0 |
| 2.65E-10 | 1.97E-09 | 0 | 233.562622 | 444.536868 | 0 | 0 |
| 2.65E-10 | 1.97E-09 | 97.7428388 | 141.917098 | 146.632203 | 143.820207 | 0 |
| 2.65E-10 | 1.97E-09 | 90.6693439 | 279.607825 | 203.24337 | 223.617225 | 0 |
| 2.66E-10 | 1.97E-09 | 87.7756414 | 243.350008 | 226.444668 | 217.122119 | 0 |
| 2.66E-10 | 1.97E-09 | 89.0617314 | 260.700374 | 213.451941 | 215.266374 | 0 |
| 2.67E-10 | 1.98E-09 | 99.0289288 | 188.184741 | 162.409085 | 139.180845 | 0 |
| 2.67E-10 | 1.98E-09 | 86.811074 | 242.682686 | 226.444668 | 259.340309 | 0 |
| 2.68E-10 | 1.98E-09 | 91.6339114 | 250.023226 | 184.682331 | 215.73031 | 0 |
| 2.68E-10 | 1.98E-09 | 106.102424 | 146.36591 | 154.056618 | 116.447974 | 0 |
| 2.68E-10 | 1.98E-09 | 86.4895515 | 241.125602 | 232.941031 | 244.494352 | 0 |
| 2.68E-10 | 1.98E-09 | 85.524984 | 484.030729 | 291.408302 | 343.312752 | 0 |
| 2.70E-10 | 2.00E-09 | 91.6339114 | 224.887439 | 176.329864 | 220.833608 | 0 |
| 2.71E-10 | 2.00E-09 | 77.1653991 | 366.804537 | 498.363879 | 447.23445 | 0 |
| 2.71E-10 | 2.00E-09 | 80.380624 | 365.692334 | 341.523105 | 319.188072 | 0 |
| 2.71E-10 | 2.00E-09 | 83.2743265 | 286.281042 | 276.559471 | 295.527328 | 0 |
| 2.71E-10 | 2.00E-09 | 155.295366 | 314.975879 | 352.659728 | 362.79807 | 4.75243649 |
| 2.71E-10 | 2.01E-09 | 87.7756414 | 250.245666 | 210.667785 | 238.927118 | 0 |
| 2.71E-10 | 2.01E-09 | 82.952804 | 361.243522 | 280.271679 | 315.940519 | 0 |
| 2.73E-10 | 2.02E-09 | 85.8465065 | 285.836161 | 235.725187 | 261.196053 | 0 |
| 2.74E-10 | 2.03E-09 | 85.524984 | 270.932642 | 235.725187 | 276.04201 | 0 |
| 2.75E-10 | 2.03E-09 | 197.414813 | 520.510986 | 476.090633 | 520.072426 | 14.2573095 |
| 2.75E-10 | 2.03E-09 | 94.8491364 | 211.541003 | 165.193241 | 167.480951 | 0 |

| | | | | | | |
|---|---|---|---|---|---|---|
| 2.75E-10 | 2.03E-09 | 82.6312815 | 283.834196 | 415.767259 | 395.737537 | 0 |
| 2.75E-10 | 2.03E-09 | 417.657722 | 829.70341 | 632.003355 | 748.329012 | 21.3859642 |
| 2.76E-10 | 2.03E-09 | 105.459379 | 151.482043 | 140.135839 | 116.91191 | 0 |
| 2.76E-10 | 2.04E-09 | 118.641801 | 154.818652 | 131.783372 | 106.241379 | 0 |
| 2.77E-10 | 2.04E-09 | 83.2743265 | 379.928532 | 358.22804 | 270.938712 | 0 |
| 2.77E-10 | 2.04E-09 | 86.4895515 | 248.688582 | 226.444668 | 277.897755 | 0 |
| 2.77E-10 | 2.04E-09 | 93.5630464 | 299.405037 | 244.077654 | 185.574461 | 0 |
| 2.77E-10 | 2.04E-09 | 90.9908664 | 180.17688 | 195.818954 | 222.689353 | 0 |
| 2.77E-10 | 2.04E-09 | 121.535504 | 167.052885 | 105.797919 | 135.00542 | 0 |
| 2.78E-10 | 2.05E-09 | 84.238894 | 308.525102 | 265.422848 | 262.123925 | 0 |
| 2.79E-10 | 2.05E-09 | 81.3451915 | 459.117383 | 313.681548 | 360.942326 | 0 |
| 2.79E-10 | 2.05E-09 | 757.828523 | 414.629264 | 287.696094 | 315.012647 | 0 |
| 2.80E-10 | 2.06E-09 | 92.2769564 | 220.661068 | 168.905449 | 203.204034 | 0 |
| 2.80E-10 | 2.06E-09 | 91.3123889 | 221.105949 | 179.11402 | 242.638607 | 0 |
| 2.80E-10 | 2.06E-09 | 96.7782713 | 217.546899 | 183.75428 | 150.779249 | 0 |
| 2.80E-10 | 2.06E-09 | 97.4213163 | 175.060746 | 137.351684 | 166.553078 | 0 |
| 2.80E-10 | 2.06E-09 | 92.9200014 | 191.298909 | 161.481034 | 203.204034 | 0 |
| 2.80E-10 | 2.06E-09 | 121.857026 | 186.627657 | 112.294282 | 229.184459 | 0 |
| 2.81E-10 | 2.07E-09 | 511.542291 | 1049.69716 | 999.511915 | 1043.3924 | 23.7621824 |
| 2.82E-10 | 2.07E-09 | 101.601109 | 235.564587 | 142.919995 | 233.359884 | 0 |
| 2.82E-10 | 2.07E-09 | 109.639171 | 126.123816 | 126.215061 | 141.500526 | 2.37621824 |
| 2.83E-10 | 2.08E-09 | 162.047338 | 238.456315 | 222.73246 | 208.307332 | 0 |
| 2.85E-10 | 2.09E-09 | 163.654951 | 154.596212 | 99.3015551 | 127.118505 | 0 |
| 2.85E-10 | 2.09E-09 | 97.4213163 | 387.713953 | 201.387266 | 252.845202 | 0 |
| 2.85E-10 | 2.09E-09 | 87.1325964 | 262.479899 | 212.523889 | 239.391054 | 0 |
| 2.86E-10 | 2.10E-09 | 91.6339114 | 255.806681 | 201.387266 | 186.502333 | 0 |
| 2.87E-10 | 2.10E-09 | 168.156265 | 348.786849 | 420.407518 | 390.170303 | 0 |
| 2.87E-10 | 2.11E-09 | 80.0591015 | 356.57227 | 310.897392 | 333.570093 | 0 |
| 2.88E-10 | 2.11E-09 | 522.152534 | 181.066642 | 137.351684 | 206.915523 | 0 |
| 2.89E-10 | 2.12E-09 | 102.244154 | 218.214221 | 136.423632 | 172.584248 | 0 |
| 2.89E-10 | 2.12E-09 | 136.647061 | 397.278898 | 418.551415 | 449.554131 | 2.37621824 |
| 2.91E-10 | 2.13E-09 | 87.4541189 | 248.688582 | 241.293498 | 343.776688 | 0 |
| 2.91E-10 | 2.13E-09 | 123.464639 | 830.815613 | 792.556337 | 870.34422 | 0 |
| 2.91E-10 | 2.13E-09 | 90.6693439 | 171.724137 | 189.322591 | 185.110524 | 0 |
| 2.93E-10 | 2.15E-09 | 83.2743265 | 276.493656 | 257.998433 | 273.722329 | 0 |
| 2.94E-10 | 2.15E-09 | 107.066991 | 132.574593 | 201.387266 | 137.789037 | 0 |
| 2.95E-10 | 2.16E-09 | 83.2743265 | 256.474003 | 272.847264 | 275.114138 | 0 |
| 2.96E-10 | 2.17E-09 | 125.393773 | 179.064677 | 231.084927 | 109.488932 | 0 |
| 2.96E-10 | 2.17E-09 | 90.0262989 | 298.070394 | 211.595837 | 215.73031 | 0 |
| 2.96E-10 | 2.17E-09 | 88.4186864 | 267.151152 | 200.459214 | 228.720523 | 0 |
| 2.97E-10 | 2.17E-09 | 91.9554339 | 164.383598 | 166.121293 | 168.408823 | 0 |
| 2.98E-10 | 2.18E-09 | 78.7730116 | 342.336072 | 339.667002 | 334.497965 | 0 |
| 2.98E-10 | 2.18E-09 | 79.094534 | 433.091833 | 332.242586 | 346.096369 | 0 |

| | | | | | | |
|---|---|---|---|---|---|---|
| 2.99E-10 | 2.19E-09 | 252.395159 | 193.968197 | 103.013763 | 140.572654 | 0 |
| 3.00E-10 | 2.19E-09 | 90.0262989 | 177.730033 | 209.739733 | 186.038397 | 0 |
| 3.00E-10 | 2.20E-09 | 103.530244 | 404.396997 | 180.970124 | 261.659989 | 0 |
| 3.01E-10 | 2.20E-09 | 96.7782713 | 139.02537 | 141.991943 | 164.233398 | 0 |
| 3.01E-10 | 2.20E-09 | 79.094534 | 332.326245 | 323.890119 | 390.634239 | 0 |
| 3.02E-10 | 2.21E-09 | 84.881939 | 274.04681 | 230.156875 | 249.133713 | 0 |
| 3.03E-10 | 2.21E-09 | 93.5630464 | 354.347864 | 204.171422 | 239.85499 | 0 |
| 3.03E-10 | 2.21E-09 | 87.7756414 | 213.320528 | 221.804408 | 281.609244 | 0 |
| 3.03E-10 | 2.21E-09 | 89.7047764 | 180.844202 | 232.012979 | 202.276162 | 0 |
| 3.04E-10 | 2.22E-09 | 84.5604165 | 330.101839 | 263.566744 | 248.669777 | 0 |
| 3.06E-10 | 2.23E-09 | 174.908238 | 307.190458 | 271.919212 | 292.743711 | 0 |
| 3.07E-10 | 2.24E-09 | 112.854396 | 125.234054 | 259.854537 | 214.802438 | 0 |
| 3.07E-10 | 2.24E-09 | 82.952804 | 280.497587 | 249.645966 | 282.537116 | 0 |
| 3.07E-10 | 2.24E-09 | 92.9200014 | 187.517419 | 153.128566 | 196.244992 | 0 |
| 3.07E-10 | 2.24E-09 | 80.0591015 | 560.327852 | 322.962067 | 432.852429 | 0 |
| 3.09E-10 | 2.25E-09 | 82.952804 | 321.871537 | 259.854537 | 271.402649 | 0 |
| 3.09E-10 | 2.25E-09 | 183.267823 | 244.907092 | 260.782589 | 253.773075 | 0 |
| 3.10E-10 | 2.26E-09 | 168.79931 | 155.485974 | 127.143113 | 98.8184002 | 0 |
| 3.10E-10 | 2.26E-09 | 81.9882365 | 298.737716 | 260.782589 | 314.548711 | 0 |
| 3.11E-10 | 2.26E-09 | 126.679863 | 174.170984 | 146.632203 | 184.182652 | 4.75243649 |
| 3.12E-10 | 2.27E-09 | 91.6339114 | 166.163123 | 170.761553 | 203.204034 | 0 |
| 3.12E-10 | 2.27E-09 | 87.1325964 | 231.338216 | 195.818954 | 236.143501 | 0 |
| 3.13E-10 | 2.28E-09 | 98.3858838 | 226.222083 | 141.991943 | 208.307332 | 0 |
| 3.14E-10 | 2.28E-09 | 97.0997938 | 316.977844 | 447.321024 | 201.812226 | 0 |
| 3.14E-10 | 2.29E-09 | 484.534402 | 198.417008 | 125.287009 | 190.213822 | 0 |
| 3.15E-10 | 2.29E-09 | 100.315019 | 138.580489 | 122.502853 | 130.366059 | 0 |
| 3.15E-10 | 2.29E-09 | 91.9554339 | 158.600142 | 160.552982 | 161.449781 | 0 |
| 3.15E-10 | 2.29E-09 | 107.710036 | 281.832231 | 139.207788 | 277.897755 | 0 |
| 3.16E-10 | 2.30E-09 | 116.069621 | 234.007503 | 219.020252 | 244.958288 | 4.75243649 |
| 3.17E-10 | 2.30E-09 | 83.2743265 | 246.464176 | 249.645966 | 271.866585 | 0 |
| 3.17E-10 | 2.30E-09 | 92.5984789 | 168.387528 | 160.552982 | 150.315313 | 0 |
| 3.17E-10 | 2.30E-09 | 122.821594 | 143.919063 | 102.085711 | 121.551272 | 0 |
| 3.19E-10 | 2.32E-09 | 146.292736 | 116.33643 | 186.538435 | 327.538923 | 0 |
| 3.19E-10 | 2.32E-09 | 256.896474 | 114.556905 | 256.142329 | 123.870952 | 0 |
| 3.20E-10 | 2.32E-09 | 180.052598 | 342.558512 | 254.286225 | 342.38488 | 0 |
| 3.20E-10 | 2.32E-09 | 85.524984 | 243.350008 | 208.811681 | 244.958288 | 0 |
| 3.20E-10 | 2.32E-09 | 221.529 | 223.330355 | 201.387266 | 289.032222 | 0 |
| 3.20E-10 | 2.32E-09 | 103.851766 | 198.639449 | 131.783372 | 142.428398 | 0 |
| 3.22E-10 | 2.33E-09 | 90.3478214 | 189.519385 | 184.682331 | 162.377653 | 0 |
| 3.22E-10 | 2.33E-09 | 198.700903 | 169.05485 | 98.3735032 | 132.685739 | 0 |
| 3.22E-10 | 2.33E-09 | 97.7428388 | 133.909237 | 135.49558 | 129.438186 | 0 |
| 3.23E-10 | 2.34E-09 | 84.5604165 | 245.129533 | 221.804408 | 235.215629 | 0 |
| 3.23E-10 | 2.34E-09 | 105.137856 | 112.332499 | 126.215061 | 135.933292 | 0 |

| | | | | | | |
|---|---|---|---|---|---|---|
| 3.24E-10 | 2.35E-09 | 95.1706589 | 139.247811 | 182.826228 | 186.502333 | 0 |
| 3.24E-10 | 2.35E-09 | 117.355711 | 104.99196 | 181.898176 | 160.985845 | 0 |
| 3.24E-10 | 2.35E-09 | 98.0643613 | 179.509558 | 141.991943 | 136.861165 | 0 |
| 3.25E-10 | 2.35E-09 | 87.7756414 | 197.082365 | 184.682331 | 208.307332 | 0 |
| 3.25E-10 | 2.35E-09 | 250.466024 | 200.196533 | 183.75428 | 89.5396772 | 0 |
| 3.25E-10 | 2.35E-09 | 122.178549 | 155.041093 | 159.62493 | 99.2823364 | 0 |
| 3.25E-10 | 2.35E-09 | 97.0997938 | 144.586385 | 128.071165 | 141.03659 | 0 |
| 3.26E-10 | 2.35E-09 | 160.118203 | 379.26121 | 108.582074 | 240.782863 | 0 |
| 3.26E-10 | 2.36E-09 | 82.309759 | 283.389315 | 314.6096 | 249.133713 | 0 |
| 3.29E-10 | 2.38E-09 | 91.3123889 | 161.49187 | 219.948304 | 218.049991 | 0 |
| 3.31E-10 | 2.39E-09 | 77.1653991 | 369.473824 | 352.659728 | 323.827434 | 0 |
| 3.33E-10 | 2.41E-09 | 85.2034615 | 230.448454 | 209.739733 | 218.513927 | 0 |
| 3.34E-10 | 2.41E-09 | 206.738965 | 175.728068 | 92.8051917 | 149.387441 | 0 |
| 3.34E-10 | 2.41E-09 | 83.9173715 | 418.633195 | 249.645966 | 377.644027 | 0 |
| 3.35E-10 | 2.42E-09 | 82.309759 | 282.499552 | 249.645966 | 253.773075 | 0 |
| 3.35E-10 | 2.42E-09 | 90.0262989 | 204.422904 | 163.337137 | 189.749886 | 0 |
| 3.36E-10 | 2.43E-09 | 154.652321 | 374.367517 | 388.853753 | 390.634239 | 2.37621824 |
| 3.36E-10 | 2.43E-09 | 80.7021465 | 315.643201 | 275.631419 | 275.114138 | 0 |
| 3.37E-10 | 2.43E-09 | 152.401663 | 251.135429 | 241.293498 | 325.683178 | 11.8810912 |
| 3.37E-10 | 2.43E-09 | 0 | 0 | 0 | 0 | 140.196876 |
| 3.39E-10 | 2.45E-09 | 497.073779 | 978.293726 | 840.815037 | 982.616768 | 2.37621824 |
| 3.39E-10 | 2.45E-09 | 188.733705 | 272.489726 | 302.544925 | 218.977863 | 0 |
| 3.39E-10 | 2.45E-09 | 153.044708 | 249.800785 | 229.228823 | 328.930731 | 4.75243649 |
| 3.40E-10 | 2.45E-09 | 92.2769564 | 166.163123 | 258.926485 | 250.989458 | 0 |
| 3.41E-10 | 2.45E-09 | 85.8465065 | 212.208325 | 201.387266 | 204.131907 | 0 |
| 3.41E-10 | 2.46E-09 | 84.881939 | 227.111845 | 212.523889 | 214.338502 | 0 |
| 3.41E-10 | 2.46E-09 | 101.279586 | 170.167053 | 147.560255 | 120.159463 | 0 |
| 3.41E-10 | 2.46E-09 | 151.115573 | 222.885474 | 238.509343 | 94.6429749 | 0 |
| 3.44E-10 | 2.47E-09 | 95.4921813 | 424.639091 | 200.459214 | 286.248605 | 0 |
| 3.44E-10 | 2.48E-09 | 84.881939 | 248.466142 | 219.948304 | 211.090949 | 0 |
| 3.45E-10 | 2.48E-09 | 80.380624 | 327.654993 | 261.710641 | 325.219242 | 0 |
| 3.46E-10 | 2.49E-09 | 194.199588 | 100.098267 | 126.215061 | 172.584248 | 0 |
| 3.46E-10 | 2.49E-09 | 87.4541189 | 274.04681 | 191.178695 | 240.318926 | 0 |
| 3.47E-10 | 2.49E-09 | 86.811074 | 185.960335 | 195.818954 | 196.708928 | 0 |
| 3.47E-10 | 2.49E-09 | 111.246784 | 119.228158 | 129.927268 | 105.313506 | 0 |
| 3.49E-10 | 2.51E-09 | 118.641801 | 284.946399 | 528.989593 | 171.656376 | 0 |
| 3.49E-10 | 2.51E-09 | 93.8845689 | 147.478113 | 137.351684 | 161.913717 | 0 |
| 3.50E-10 | 2.52E-09 | 84.881939 | 266.928711 | 206.955577 | 274.186265 | 0 |
| 3.50E-10 | 2.52E-09 | 100.958064 | 169.05485 | 124.358957 | 133.613612 | 0 |
| 3.51E-10 | 2.52E-09 | 114.140486 | 103.879757 | 138.279736 | 120.159463 | 0 |
| 3.53E-10 | 2.53E-09 | 90.9908664 | 201.976058 | 157.768826 | 170.728504 | 0 |
| 3.53E-10 | 2.54E-09 | 209.311145 | 131.46239 | 258.926485 | 105.313506 | 0 |
| 3.53E-10 | 2.54E-09 | 87.1325964 | 223.107914 | 183.75428 | 196.244992 | 0 |

| | | | | | | |
|---|---|---|---|---|---|---|
| 3.54E-10 | 2.54E-09 | 129.895088 | 232.227979 | 102.085711 | 200.884354 | 0 |
| 3.54E-10 | 2.54E-09 | 496.430734 | 825.92192 | 737.801274 | 811.888265 | 30.8908372 |
| 3.55E-10 | 2.55E-09 | 102.565676 | 224.887439 | 128.071165 | 203.66797 | 0 |
| 3.55E-10 | 2.55E-09 | 83.9173715 | 231.560657 | 211.595837 | 235.679565 | 0 |
| 3.57E-10 | 2.56E-09 | 96.7782713 | 134.354118 | 146.632203 | 125.262761 | 0 |
| 3.57E-10 | 2.56E-09 | 107.388514 | 179.731999 | 263.566744 | 133.613612 | 0 |
| 3.57E-10 | 2.56E-09 | 98.0643613 | 141.472217 | 136.423632 | 188.822014 | 0 |
| 3.58E-10 | 2.56E-09 | 91.3123889 | 158.600142 | 218.0922 | 177.687546 | 0 |
| 3.58E-10 | 2.57E-09 | 164.619518 | 157.043058 | 94.6612955 | 128.97425 | 0 |
| 3.58E-10 | 2.57E-09 | 89.3832539 | 199.751652 | 160.552982 | 200.884354 | 0 |
| 3.58E-10 | 2.57E-09 | 85.8465065 | 256.696444 | 196.747006 | 221.76148 | 0 |
| 3.59E-10 | 2.57E-09 | 89.7047764 | 254.249597 | 171.689605 | 221.76148 | 0 |
| 3.59E-10 | 2.57E-09 | 81.3451915 | 281.60979 | 242.22155 | 268.155095 | 0 |
| 3.60E-10 | 2.58E-09 | 239.212737 | 253.137394 | 128.071165 | 220.833608 | 7.12865473 |
| 3.60E-10 | 2.58E-09 | 143.720556 | 574.56405 | 419.479466 | 608.684231 | 2.37621824 |
| 3.61E-10 | 2.58E-09 | 81.3451915 | 362.800606 | 253.358173 | 304.806051 | 0 |
| 3.62E-10 | 2.59E-09 | 94.2060914 | 335.885295 | 177.257916 | 252.845202 | 0 |
| 3.63E-10 | 2.59E-09 | 92.2769564 | 193.078434 | 148.488307 | 161.913717 | 0 |
| 3.63E-10 | 2.60E-09 | 119.284846 | 155.041093 | 100.229607 | 176.295737 | 0 |
| 3.63E-10 | 2.60E-09 | 85.2034615 | 201.086296 | 217.164149 | 198.100737 | 0 |
| 3.64E-10 | 2.60E-09 | 71.0564716 | 461.341789 | 493.72362 | 400.376899 | 0 |
| 3.64E-10 | 2.60E-09 | 93.2415239 | 153.039128 | 199.531162 | 151.707121 | 0 |
| 3.65E-10 | 2.61E-09 | 89.0617314 | 177.730033 | 158.696878 | 175.831801 | 0 |
| 3.65E-10 | 2.61E-09 | 110.282216 | 187.517419 | 186.538435 | 109.952868 | 0 |
| 3.67E-10 | 2.62E-09 | 86.4895515 | 243.350008 | 218.0922 | 185.110524 | 0 |
| 3.68E-10 | 2.62E-09 | 126.679863 | 168.165088 | 96.5173994 | 181.399035 | 0 |
| 3.68E-10 | 2.63E-09 | 82.952804 | 335.662854 | 232.941031 | 276.969882 | 0 |
| 3.69E-10 | 2.63E-09 | 82.309759 | 552.319991 | 285.83999 | 386.458814 | 0 |
| 3.69E-10 | 2.64E-09 | 86.811074 | 243.350008 | 181.898176 | 221.297544 | 0 |
| 3.70E-10 | 2.64E-09 | 106.423946 | 117.448633 | 113.222334 | 112.736485 | 0 |
| 3.70E-10 | 2.64E-09 | 118.963324 | 178.174915 | 120.646749 | 164.697334 | 14.2573095 |
| 3.70E-10 | 2.64E-09 | 80.0591015 | 354.125423 | 270.063108 | 284.392861 | 0 |
| 3.71E-10 | 2.64E-09 | 98.0643613 | 145.253707 | 167.977397 | 161.913717 | 9.50487297 |
| 3.71E-10 | 2.64E-09 | 116.712666 | 204.422904 | 109.510126 | 148.459568 | 0 |
| 3.71E-10 | 2.65E-09 | 90.0262989 | 152.371806 | 179.11402 | 192.069567 | 0 |
| 3.72E-10 | 2.65E-09 | 139.862286 | 503.16062 | 450.10518 | 476.926364 | 9.50487297 |
| 3.73E-10 | 2.66E-09 | 155.295366 | 108.99589 | 394.422065 | 257.9485 | 0 |
| 3.74E-10 | 2.66E-09 | 184.553913 | 263.147221 | 217.164149 | 253.309139 | 0 |
| 3.75E-10 | 2.67E-09 | 91.6339114 | 149.480078 | 142.919995 | 161.449781 | 0 |
| 3.75E-10 | 2.67E-09 | 328.274469 | 900.884399 | 1267.71892 | 1218.76027 | 19.0097459 |
| 3.75E-10 | 2.67E-09 | 82.6312815 | 262.035018 | 217.164149 | 254.237011 | 0 |
| 3.76E-10 | 2.68E-09 | 257.217997 | 243.127568 | 306.257133 | 220.833608 | 49.9005831 |
| 3.77E-10 | 2.68E-09 | 87.1325964 | 197.082365 | 197.675058 | 169.800631 | 0 |

| | | | | | | |
|---|---|---|---|---|---|---|
| 3.77E-10 | 2.68E-09 | 108.996126 | 178.842236 | 108.582074 | 149.851377 | 0 |
| 3.80E-10 | 2.70E-09 | 87.7756414 | 164.383598 | 179.11402 | 187.894141 | 0 |
| 3.80E-10 | 2.70E-09 | 70.0919042 | 441.767017 | 457.529595 | 427.749132 | 0 |
| 3.80E-10 | 2.70E-09 | 80.7021465 | 264.704305 | 255.214277 | 244.030416 | 0 |
| 3.81E-10 | 2.71E-09 | 77.1653991 | 435.093799 | 297.904665 | 367.437432 | 0 |
| 3.82E-10 | 2.72E-09 | 191.305885 | 278.940503 | 507.644399 | 337.281582 | 0 |
| 3.83E-10 | 2.72E-09 | 137.290106 | 121.452563 | 97.4454513 | 122.94308 | 0 |
| 3.83E-10 | 2.72E-09 | 85.2034615 | 355.015186 | 226.444668 | 409.655622 | 0 |
| 3.83E-10 | 2.72E-09 | 84.238894 | 252.025191 | 199.531162 | 227.328714 | 0 |
| 3.84E-10 | 2.73E-09 | 105.137856 | 162.381632 | 108.582074 | 153.562866 | 0 |
| 3.84E-10 | 2.73E-09 | 76.5223541 | 347.007324 | 319.249859 | 304.806051 | 0 |
| 3.86E-10 | 2.74E-09 | 99.3504513 | 348.341968 | 161.481034 | 317.796264 | 0 |
| 3.86E-10 | 2.74E-09 | 97.0997938 | 623.27854 | 217.164149 | 471.823066 | 0 |
| 3.87E-10 | 2.74E-09 | 603.497725 | 1201.1792 | 941.044644 | 1035.96943 | 121.18713 |
| 3.87E-10 | 2.74E-09 | 235.997512 | 387.269071 | 477.018685 | 489.916576 | 16.6335277 |
| 3.87E-10 | 2.74E-09 | 103.208721 | 135.466321 | 119.718697 | 111.344676 | 0 |
| 3.87E-10 | 2.74E-09 | 93.8845689 | 171.501697 | 235.725187 | 319.652008 | 0 |
| 3.87E-10 | 2.74E-09 | 88.4186864 | 210.20636 | 296.976613 | 194.853184 | 0 |
| 3.88E-10 | 2.75E-09 | 127.001386 | 164.606038 | 96.5173994 | 189.749886 | 0 |
| 3.88E-10 | 2.75E-09 | 271.686509 | 142.58442 | 125.287009 | 128.510314 | 0 |
| 3.89E-10 | 2.75E-09 | 77.4869216 | 351.901017 | 291.408302 | 301.558498 | 0 |
| 3.89E-10 | 2.75E-09 | 81.3451915 | 266.48383 | 242.22155 | 232.432012 | 0 |
| 3.88E-10 | 2.75E-09 | 84.5604165 | 199.974093 | 197.675058 | 198.564673 | 0 |
| 3.89E-10 | 2.75E-09 | 124.750728 | 119.228158 | 97.4454513 | 138.252973 | 0 |
| 3.90E-10 | 2.76E-09 | 85.524984 | 518.953902 | 253.358173 | 359.086581 | 0 |
| 3.90E-10 | 2.76E-09 | 89.7047764 | 148.367875 | 180.970124 | 181.399035 | 0 |
| 3.90E-10 | 2.76E-09 | 101.601109 | 399.725745 | 227.37272 | 181.862971 | 0 |
| 3.91E-10 | 2.77E-09 | 93.8845689 | 169.277291 | 138.279736 | 200.420417 | 0 |
| 3.92E-10 | 2.77E-09 | 152.080141 | 413.961943 | 372.148819 | 391.562112 | 0 |
| 3.95E-10 | 2.79E-09 | 94.8491364 | 429.310343 | 195.818954 | 283.464988 | 0 |
| 3.96E-10 | 2.80E-09 | 202.559173 | 815.689652 | 745.225689 | 850.394965 | 14.2573095 |
| 3.96E-10 | 2.80E-09 | 91.3123889 | 155.708415 | 145.704151 | 185.574461 | 0 |
| 3.97E-10 | 2.81E-09 | 80.380624 | 255.3618 | 242.22155 | 250.525522 | 0 |
| 3.97E-10 | 2.81E-09 | 132.467268 | 140.582454 | 97.4454513 | 115.984038 | 0 |
| 3.97E-10 | 2.81E-09 | 94.8491364 | 201.531177 | 138.279736 | 158.666164 | 0 |
| 3.98E-10 | 2.81E-09 | 77.4869216 | 288.95033 | 289.552198 | 300.16669 | 0 |
| 3.98E-10 | 2.81E-09 | 86.811074 | 187.294979 | 166.121293 | 187.430205 | 0 |
| 3.99E-10 | 2.82E-09 | 173.300625 | 282.721993 | 239.437395 | 275.578074 | 2.37621824 |
| 4.00E-10 | 2.82E-09 | 84.881939 | 194.857959 | 246.86181 | 203.204034 | 0 |
| 4.00E-10 | 2.82E-09 | 129.573566 | 160.602108 | 166.121293 | 93.2511664 | 0 |
| 4.00E-10 | 2.82E-09 | 117.355711 | 191.298909 | 196.747006 | 229.648395 | 4.75243649 |
| 4.00E-10 | 2.82E-09 | 92.2769564 | 157.71038 | 164.265189 | 133.149675 | 0 |
| 4.02E-10 | 2.83E-09 | 81.9882365 | 223.330355 | 222.73246 | 225.936906 | 0 |

| | | | | | | |
|---|---|---|---|---|---|---|
| 4.02E-10 | 2.84E-09 | 99.0289288 | 128.125781 | 178.185968 | 131.293931 | 0 |
| 4.02E-10 | 2.84E-09 | 84.238894 | 280.720028 | 198.60311 | 254.700947 | 0 |
| 4.05E-10 | 2.85E-09 | 27.3294122 | 16.2381632 | 24.1293498 | 14.3820207 | 102.177384 |
| 4.05E-10 | 2.85E-09 | 165.905608 | 209.316597 | 91.8771398 | 145.675952 | 0 |
| 4.05E-10 | 2.85E-09 | 141.469898 | 266.261389 | 248.717914 | 253.773075 | 16.6335277 |
| 4.05E-10 | 2.85E-09 | 88.4186864 | 161.26943 | 156.840774 | 159.594036 | 0 |
| 4.07E-10 | 2.87E-09 | 82.952804 | 214.210291 | 314.6096 | 270.474776 | 0 |
| 4.07E-10 | 2.87E-09 | 85.524984 | 203.533142 | 174.47376 | 200.420417 | 0 |
| 4.08E-10 | 2.87E-09 | 85.524984 | 212.208325 | 176.329864 | 221.297544 | 0 |
| 4.08E-10 | 2.87E-09 | 86.4895515 | 195.30284 | 180.970124 | 167.944887 | 0 |
| 4.09E-10 | 2.88E-09 | 92.5984789 | 163.938717 | 132.711424 | 173.976057 | 0 |
| 4.09E-10 | 2.88E-09 | 77.1653991 | 321.649097 | 291.408302 | 283.464988 | 0 |
| 4.09E-10 | 2.88E-09 | 78.4514891 | 264.036983 | 283.983887 | 270.474776 | 0 |
| 4.09E-10 | 2.88E-09 | 163.976473 | 310.971948 | 295.12051 | 287.176478 | 0 |
| 4.09E-10 | 2.88E-09 | 90.0262989 | 176.172949 | 222.73246 | 157.738291 | 0 |
| 4.11E-10 | 2.89E-09 | 86.168029 | 238.233874 | 180.970124 | 195.31712 | 0 |
| 4.11E-10 | 2.89E-09 | 85.2034615 | 179.064677 | 197.675058 | 185.574461 | 0 |
| 4.11E-10 | 2.89E-09 | 106.745469 | 151.482043 | 104.869867 | 160.985845 | 0 |
| 4.11E-10 | 2.89E-09 | 91.6339114 | 213.098088 | 161.481034 | 153.562866 | 0 |
| 4.12E-10 | 2.89E-09 | 73.3071291 | 333.660889 | 379.573234 | 391.098176 | 0 |
| 4.12E-10 | 2.89E-09 | 88.0971639 | 213.542969 | 257.070381 | 170.728504 | 0 |
| 4.13E-10 | 2.90E-09 | 76.2008316 | 438.875289 | 322.034015 | 320.115944 | 0 |
| 4.13E-10 | 2.90E-09 | 90.6693439 | 181.733964 | 146.632203 | 151.707121 | 0 |
| 4.14E-10 | 2.90E-09 | 89.7047764 | 184.403251 | 155.912722 | 222.225416 | 0 |
| 4.14E-10 | 2.90E-09 | 80.0591015 | 241.348043 | 267.278952 | 241.710735 | 0 |
| 4.14E-10 | 2.91E-09 | 85.8465065 | 189.964266 | 170.761553 | 202.740098 | 0 |
| 4.15E-10 | 2.91E-09 | 85.8465065 | 233.562622 | 180.042072 | 195.31712 | 0 |
| 4.15E-10 | 2.91E-09 | 92.9200014 | 145.921029 | 129.927268 | 136.861165 | 0 |
| 4.17E-10 | 2.92E-09 | 80.7021465 | 236.67679 | 228.300772 | 244.958288 | 0 |
| 4.17E-10 | 2.92E-09 | 93.2415239 | 163.938717 | 134.567528 | 136.861165 | 0 |
| 4.18E-10 | 2.93E-09 | 82.309759 | 271.599963 | 230.156875 | 212.946693 | 0 |
| 4.18E-10 | 2.93E-09 | 83.2743265 | 232.005538 | 194.890903 | 215.266374 | 0 |
| 4.18E-10 | 2.93E-09 | 83.595849 | 355.682507 | 227.37272 | 258.876372 | 0 |
| 4.19E-10 | 2.93E-09 | 94.2060914 | 169.944613 | 126.215061 | 159.1301 | 0 |
| 4.20E-10 | 2.94E-09 | 74.9147416 | 302.964087 | 351.731677 | 347.952113 | 0 |
| 4.20E-10 | 2.94E-09 | 195.164155 | 635.957654 | 473.306478 | 556.259445 | 0 |
| 4.20E-10 | 2.94E-09 | 99.6719738 | 173.281222 | 120.646749 | 135.00542 | 0 |
| 4.21E-10 | 2.94E-09 | 92.2769564 | 186.627657 | 138.279736 | 155.418611 | 0 |
| 4.21E-10 | 2.94E-09 | 246.286232 | 98.0963013 | 195.818954 | 136.861165 | 0 |
| 4.23E-10 | 2.96E-09 | 128.608998 | 160.157227 | 91.8771398 | 160.985845 | 0 |
| 4.23E-10 | 2.96E-09 | 82.952804 | 232.005538 | 194.890903 | 223.617225 | 0 |
| 4.23E-10 | 2.96E-09 | 79.4160565 | 249.578345 | 244.077654 | 260.732117 | 0 |
| 4.24E-10 | 2.96E-09 | 80.7021465 | 316.310522 | 232.941031 | 261.659989 | 0 |

| | | | | | | |
|---|---|---|---|---|---|---|
| 4.24E-10 | 2.96E-09 | 85.2034615 | 246.464176 | 180.042072 | 225.009033 | 0 |
| 4.24E-10 | 2.96E-09 | 121.535504 | 202.64338 | 178.185968 | 199.028609 | 9.50487297 |
| 4.27E-10 | 2.98E-09 | 81.023669 | 215.767375 | 241.293498 | 235.679565 | 0 |
| 4.28E-10 | 2.99E-09 | 114.140486 | 212.653206 | 106.72597 | 201.34829 | 0 |
| 4.30E-10 | 3.00E-09 | 82.952804 | 290.729854 | 203.24337 | 258.876372 | 0 |
| 4.32E-10 | 3.01E-09 | 85.524984 | 180.621761 | 168.905449 | 189.749886 | 0 |
| 4.32E-10 | 3.02E-09 | 117.677234 | 226.666964 | 105.797919 | 183.718716 | 0 |
| 4.33E-10 | 3.02E-09 | 141.148376 | 510.7236 | 420.407518 | 444.914769 | 2.37621824 |
| 4.34E-10 | 3.03E-09 | 77.8084441 | 278.495622 | 260.782589 | 271.402649 | 0 |
| 4.34E-10 | 3.03E-09 | 81.666714 | 241.125602 | 205.099474 | 231.968076 | 0 |
| 4.36E-10 | 3.04E-09 | 94.8491364 | 133.241915 | 119.718697 | 140.108718 | 0 |
| 4.37E-10 | 3.05E-09 | 91.9554339 | 137.468286 | 197.675058 | 168.408823 | 0 |
| 4.38E-10 | 3.05E-09 | 84.238894 | 280.275146 | 193.034799 | 234.287756 | 0 |
| 4.39E-10 | 3.06E-09 | 82.6312815 | 195.747721 | 214.379993 | 237.071373 | 0 |
| 4.39E-10 | 3.06E-09 | 87.7756414 | 282.277112 | 175.401812 | 227.328714 | 0 |
| 4.39E-10 | 3.06E-09 | 102.244154 | 198.86189 | 116.00649 | 180.935099 | 0 |
| 4.40E-10 | 3.06E-09 | 88.0971639 | 217.76934 | 156.840774 | 219.4418 | 0 |
| 4.40E-10 | 3.06E-09 | 148.221871 | 440.432373 | 365.652455 | 442.131152 | 0 |
| 4.41E-10 | 3.07E-09 | 78.4514891 | 308.302661 | 245.933758 | 276.04201 | 0 |
| 4.43E-10 | 3.08E-09 | 450.131495 | 1216.30516 | 753.578157 | 975.657726 | 78.415202 |
| 4.44E-10 | 3.09E-09 | 84.5604165 | 366.359656 | 219.948304 | 256.092756 | 0 |
| 4.44E-10 | 3.09E-09 | 81.9882365 | 256.251562 | 205.099474 | 221.76148 | 0 |
| 4.45E-10 | 3.09E-09 | 90.6693439 | 152.594246 | 132.711424 | 148.923505 | 0 |
| 4.45E-10 | 3.09E-09 | 159.796681 | 282.721993 | 321.105963 | 309.909349 | 2.37621824 |
| 4.45E-10 | 3.10E-09 | 83.9173715 | 202.86582 | 178.185968 | 198.100737 | 0 |
| 4.46E-10 | 3.10E-09 | 163.654951 | 288.060567 | 276.559471 | 318.724136 | 0 |
| 4.46E-10 | 3.10E-09 | 102.887199 | 152.594246 | 151.272462 | 106.241379 | 0 |
| 4.46E-10 | 3.10E-09 | 80.380624 | 233.562622 | 219.020252 | 234.751693 | 0 |
| 4.46E-10 | 3.10E-09 | 91.6339114 | 129.460425 | 154.98467 | 142.428398 | 0 |
| 4.47E-10 | 3.10E-09 | 92.2769564 | 126.568697 | 147.560255 | 156.810419 | 0 |
| 4.48E-10 | 3.11E-09 | 99.3504513 | 144.141504 | 109.510126 | 135.00542 | 0 |
| 4.48E-10 | 3.11E-09 | 0 | 0 | 0 | 0 | 154.454186 |
| 4.49E-10 | 3.11E-09 | 93.5630464 | 123.89941 | 128.999216 | 137.325101 | 0 |
| 4.49E-10 | 3.11E-09 | 77.4869216 | 396.611576 | 261.710641 | 352.591475 | 0 |
| 4.50E-10 | 3.12E-09 | 87.4541189 | 177.507593 | 160.552982 | 152.171058 | 0 |
| 4.50E-10 | 3.12E-09 | 84.238894 | 210.651241 | 174.47376 | 212.946693 | 0 |
| 4.52E-10 | 3.13E-09 | 90.3478214 | 138.80293 | 152.200514 | 136.397228 | 0 |
| 4.52E-10 | 3.13E-09 | 85.8465065 | 256.696444 | 193.962851 | 184.182652 | 0 |
| 4.52E-10 | 3.13E-09 | 92.9200014 | 143.919063 | 125.287009 | 132.685739 | 0 |
| 4.54E-10 | 3.15E-09 | 93.8845689 | 554.989278 | 234.797135 | 289.032222 | 0 |
| 4.55E-10 | 3.15E-09 | 146.614258 | 429.755225 | 365.652455 | 386.92275 | 2.37621824 |
| 4.57E-10 | 3.17E-09 | 76.2008316 | 314.753438 | 281.199731 | 276.969882 | 0 |
| 4.57E-10 | 3.17E-09 | 97.7428388 | 212.430766 | 133.639476 | 148.923505 | 0 |

| | | | | | | |
|---|---|---|---|---|---|---|
| 4.58E-10 | 3.17E-09 | 128.287476 | 171.501697 | 97.4454513 | 123.870952 | 0 |
| 4.59E-10 | 3.17E-09 | 85.524984 | 292.064498 | 191.178695 | 217.586055 | 0 |
| 4.59E-10 | 3.17E-09 | 86.168029 | 160.824548 | 162.409085 | 164.697334 | 0 |
| 4.59E-10 | 3.17E-09 | 119.284846 | 224.220117 | 112.294282 | 139.180845 | 0 |
| 4.59E-10 | 3.17E-09 | 43.0840145 | 19.3523315 | 16.7049345 | 25.0525522 | 168.711495 |
| 4.59E-10 | 3.18E-09 | 86.4895515 | 232.005538 | 164.265189 | 202.276162 | 0 |
| 4.62E-10 | 3.19E-09 | 2.25065747 | 2.22440592 | 2.78415575 | 1.85574461 | 26.1384007 |
| 4.63E-10 | 3.20E-09 | 104.494811 | 180.621761 | 184.682331 | 109.488932 | 0 |
| 4.64E-10 | 3.21E-09 | 84.881939 | 177.730033 | 181.898176 | 165.16127 | 0 |
| 4.64E-10 | 3.21E-09 | 108.353081 | 209.761479 | 111.36623 | 158.666164 | 0 |
| 4.65E-10 | 3.21E-09 | 147.257303 | 314.308557 | 277.487523 | 299.702754 | 0 |
| 4.65E-10 | 3.21E-09 | 86.168029 | 170.389494 | 157.768826 | 190.213822 | 0 |
| 4.66E-10 | 3.22E-09 | 92.5984789 | 171.501697 | 126.215061 | 157.274355 | 0 |
| 4.68E-10 | 3.23E-09 | 124.429206 | 110.775415 | 156.840774 | 101.602017 | 0 |
| 4.70E-10 | 3.24E-09 | 94.5276139 | 373.032874 | 168.905449 | 319.188072 | 0 |
| 4.70E-10 | 3.24E-09 | 83.9173715 | 196.637484 | 171.689605 | 189.28595 | 0 |
| 4.70E-10 | 3.24E-09 | 80.380624 | 245.129533 | 238.509343 | 206.915523 | 0 |
| 4.71E-10 | 3.25E-09 | 152.401663 | 312.973914 | 250.574018 | 255.164883 | 4.75243649 |
| 4.72E-10 | 3.26E-09 | 96.1352263 | 151.482043 | 301.616873 | 215.73031 | 0 |
| 4.75E-10 | 3.28E-09 | 295.157651 | 142.80686 | 166.121293 | 107.169251 | 0 |
| 4.77E-10 | 3.29E-09 | 93.5630464 | 138.358048 | 136.423632 | 118.303719 | 0 |
| 4.77E-10 | 3.29E-09 | 86.168029 | 167.942647 | 155.912722 | 158.202228 | 0 |
| 4.78E-10 | 3.29E-09 | 85.524984 | 161.046989 | 188.394539 | 166.089142 | 0 |
| 4.79E-10 | 3.30E-09 | 132.145746 | 243.127568 | 183.75428 | 250.061586 | 7.12865473 |
| 4.79E-10 | 3.30E-09 | 86.4895515 | 244.907092 | 169.833501 | 269.082968 | 0 |
| 4.80E-10 | 3.31E-09 | 81.666714 | 451.109521 | 234.797135 | 408.727749 | 0 |
| 4.81E-10 | 3.31E-09 | 124.429206 | 124.566732 | 93.7332436 | 118.767655 | 0 |
| 4.81E-10 | 3.31E-09 | 147.900348 | 597.920312 | 600.44959 | 127.582442 | 0 |
| 4.81E-10 | 3.31E-09 | 84.238894 | 197.304805 | 179.11402 | 168.872759 | 0 |
| 4.82E-10 | 3.32E-09 | 78.7730116 | 302.296765 | 230.156875 | 259.340309 | 0 |
| 4.82E-10 | 3.32E-09 | 69.7703817 | 377.926567 | 440.824661 | 371.148921 | 0 |
| 4.84E-10 | 3.33E-09 | 79.737579 | 260.700374 | 232.012979 | 214.802438 | 0 |
| 4.84E-10 | 3.33E-09 | 84.238894 | 191.966231 | 166.121293 | 203.66797 | 0 |
| 4.85E-10 | 3.33E-09 | 0 | 0 | 644.996082 | 160.521908 | 0 |
| 4.87E-10 | 3.35E-09 | 192.913498 | 225.33232 | 191.178695 | 241.710735 | 2.37621824 |
| 4.87E-10 | 3.35E-09 | 79.4160565 | 325.653027 | 222.73246 | 298.310945 | 0 |
| 4.88E-10 | 3.36E-09 | 239.53426 | 140.582454 | 106.72597 | 131.293931 | 0 |
| 4.89E-10 | 3.36E-09 | 120.570936 | 123.454529 | 116.934542 | 95.106911 | 0 |
| 4.91E-10 | 3.37E-09 | 87.7756414 | 149.257638 | 182.826228 | 211.554885 | 0 |
| 4.93E-10 | 3.38E-09 | 88.4186864 | 141.027336 | 181.898176 | 156.346483 | 0 |
| 4.93E-10 | 3.39E-09 | 79.4160565 | 264.481864 | 217.164149 | 231.968076 | 0 |
| 4.99E-10 | 3.42E-09 | 88.0971639 | 148.812756 | 139.207788 | 144.284143 | 0 |
| 5.00E-10 | 3.43E-09 | 91.9554339 | 139.470251 | 143.848047 | 120.623399 | 0 |

| | | | | | | |
|---|---|---|---|---|---|---|
| 5.00E-10 | 3.43E-09 | 92.2769564 | 141.917098 | 119.718697 | 140.108718 | 0 |
| 5.00E-10 | 3.43E-09 | 98.3858838 | 116.55887 | 111.36623 | 136.861165 | 0 |
| 5.01E-10 | 3.44E-09 | 92.5984789 | 155.485974 | 121.574801 | 138.252973 | 0 |
| 5.01E-10 | 3.44E-09 | 122.500071 | 577.010897 | 637.571667 | 583.167742 | 2.37621824 |
| 5.02E-10 | 3.44E-09 | 105.137856 | 116.33643 | 122.502853 | 180.935099 | 0 |
| 5.03E-10 | 3.45E-09 | 135.360971 | 96.5392171 | 243.149602 | 175.367865 | 0 |
| 5.03E-10 | 3.45E-09 | 80.0591015 | 213.542969 | 204.171422 | 227.79265 | 0 |
| 5.06E-10 | 3.46E-09 | 78.7730116 | 242.460246 | 215.308045 | 240.782863 | 0 |
| 5.10E-10 | 3.50E-09 | 71.0564716 | 320.981775 | 513.21271 | 499.195299 | 0 |
| 5.10E-10 | 3.50E-09 | 825.026725 | 2648.37769 | 1366.09242 | 1818.16578 | 28.5146189 |
| 5.12E-10 | 3.51E-09 | 156.902978 | 121.007682 | 157.768826 | 91.859358 | 0 |
| 5.13E-10 | 3.51E-09 | 78.1299666 | 229.558691 | 228.300772 | 248.669777 | 0 |
| 5.14E-10 | 3.52E-09 | 74.9147416 | 264.259424 | 302.544925 | 335.889774 | 0 |
| 5.15E-10 | 3.52E-09 | 73.6286516 | 357.462032 | 302.544925 | 292.279775 | 0 |
| 5.16E-10 | 3.53E-09 | 0 | 0 | 0 | 0 | 135.44444 |
| 5.17E-10 | 3.54E-09 | 189.37675 | 123.232088 | 101.157659 | 125.726697 | 0 |
| 5.18E-10 | 3.54E-09 | 85.524984 | 175.060746 | 148.488307 | 171.19244 | 0 |
| 5.19E-10 | 3.55E-09 | 87.4541189 | 143.919063 | 194.890903 | 166.089142 | 0 |
| 5.19E-10 | 3.55E-09 | 69.7703817 | 380.150972 | 356.371936 | 360.942326 | 0 |
| 5.19E-10 | 3.55E-09 | 90.3478214 | 152.371806 | 219.020252 | 152.171058 | 0 |
| 5.20E-10 | 3.55E-09 | 87.1325964 | 166.163123 | 149.416359 | 202.740098 | 0 |
| 5.21E-10 | 3.56E-09 | 117.034189 | 193.300875 | 97.4454513 | 159.594036 | 0 |
| 5.22E-10 | 3.57E-09 | 77.8084441 | 530.298372 | 265.422848 | 420.326153 | 0 |
| 5.23E-10 | 3.57E-09 | 78.4514891 | 224.220117 | 223.660512 | 225.009033 | 0 |
| 5.23E-10 | 3.57E-09 | 94.8491364 | 140.804895 | 204.171422 | 134.541484 | 0 |
| 5.23E-10 | 3.57E-09 | 93.2415239 | 162.604073 | 118.790645 | 144.284143 | 0 |
| 5.24E-10 | 3.58E-09 | 236.96208 | 414.851705 | 258.926485 | 314.084774 | 33.2670554 |
| 5.25E-10 | 3.58E-09 | 0.64304499 | 4.00393066 | 2.78415575 | 5.10329766 | 213.859642 |
| 5.26E-10 | 3.59E-09 | 81.3451915 | 374.145076 | 227.37272 | 258.412436 | 0 |
| 5.26E-10 | 3.59E-09 | 90.0262989 | 177.507593 | 131.783372 | 149.387441 | 0 |
| 5.27E-10 | 3.59E-09 | 0 | 0 | 0 | 0 | 21.3859642 |
| 5.27E-10 | 3.60E-09 | 132.788791 | 511.835803 | 452.889335 | 467.183704 | 2.37621824 |
| 5.31E-10 | 3.62E-09 | 72.6640841 | 413.961943 | 314.6096 | 314.548711 | 0 |
| 5.32E-10 | 3.63E-09 | 136.325538 | 183.513489 | 94.6612955 | 124.798825 | 0 |
| 5.32E-10 | 3.63E-09 | 177.480418 | 280.052706 | 211.595837 | 237.999246 | 2.37621824 |
| 5.38E-10 | 3.67E-09 | 110.603739 | 157.043058 | 97.4454513 | 129.438186 | 0 |
| 5.39E-10 | 3.67E-09 | 91.9554339 | 125.456494 | 186.538435 | 181.399035 | 0 |
| 5.40E-10 | 3.68E-09 | 108.674604 | 182.623726 | 104.869867 | 134.077548 | 0 |
| 5.42E-10 | 3.69E-09 | 80.380624 | 204.645345 | 214.379993 | 186.966269 | 0 |
| 5.42E-10 | 3.69E-09 | 119.606369 | 111.442737 | 113.222334 | 96.4987195 | 0 |
| 5.43E-10 | 3.69E-09 | 195.485678 | 80.7459351 | 222.73246 | 189.749886 | 0 |
| 5.45E-10 | 3.71E-09 | 73.3071291 | 342.113631 | 283.055835 | 296.455201 | 0 |
| 5.46E-10 | 3.72E-09 | 97.0997938 | 208.426835 | 129.927268 | 141.03659 | 0 |

| | | | | | | |
|---|---|---|---|---|---|---|
| 5.47E-10 | 3.72E-09 | 60.7677518 | 527.184204 | 535.485956 | 539.093808 | 0 |
| 5.49E-10 | 3.73E-09 | 82.6312815 | 172.6139 | 171.689605 | 169.800631 | 0 |
| 5.50E-10 | 3.74E-09 | 85.2034615 | 172.6139 | 149.416359 | 154.026802 | 0 |
| 5.51E-10 | 3.74E-09 | 80.380624 | 184.625692 | 205.099474 | 203.66797 | 0 |
| 5.51E-10 | 3.75E-09 | 82.952804 | 179.064677 | 167.049345 | 207.37946 | 0 |
| 5.51E-10 | 3.75E-09 | 104.494811 | 107.438806 | 104.869867 | 109.024996 | 0 |
| 5.52E-10 | 3.75E-09 | 85.524984 | 154.818652 | 173.545708 | 145.675952 | 0 |
| 5.53E-10 | 3.76E-09 | 84.238894 | 163.048954 | 237.581291 | 202.276162 | 0 |
| 5.60E-10 | 3.81E-09 | 105.780901 | 128.125781 | 96.5173994 | 121.551272 | 0 |
| 5.61E-10 | 3.81E-09 | 226.351837 | 195.970162 | 127.143113 | 94.6429749 | 0 |
| 5.62E-10 | 3.81E-09 | 97.4213163 | 162.159192 | 109.510126 | 137.325101 | 0 |
| 5.62E-10 | 3.81E-09 | 86.4895515 | 140.804895 | 162.409085 | 142.428398 | 0 |
| 5.62E-10 | 3.81E-09 | 107.388514 | 185.293013 | 193.034799 | 103.457762 | 0 |
| 5.63E-10 | 3.82E-09 | 174.586715 | 346.784884 | 87.2368802 | 308.981477 | 0 |
| 5.63E-10 | 3.82E-09 | 96.4567488 | 133.241915 | 105.797919 | 129.438186 | 0 |
| 5.64E-10 | 3.83E-09 | 78.4514891 | 295.845988 | 210.667785 | 255.164883 | 0 |
| 5.66E-10 | 3.84E-09 | 105.780901 | 162.381632 | 141.063891 | 153.562866 | 0 |
| 5.66E-10 | 3.84E-09 | 124.750728 | 256.696444 | 136.423632 | 192.069567 | 7.12865473 |
| 5.67E-10 | 3.84E-09 | 120.570936 | 68.0668213 | 66.819738 | 63.5592527 | 411.085756 |
| 5.67E-10 | 3.85E-09 | 80.0591015 | 479.137036 | 237.581291 | 356.7669 | 0 |
| 5.70E-10 | 3.86E-09 | 97.7428388 | 268.040914 | 131.783372 | 252.381266 | 0 |
| 5.71E-10 | 3.87E-09 | 105.459379 | 118.115955 | 97.4454513 | 133.149675 | 0 |
| 5.71E-10 | 3.87E-09 | 77.4869216 | 256.029122 | 231.084927 | 214.338502 | 0 |
| 5.72E-10 | 3.87E-09 | 76.8438766 | 235.342147 | 228.300772 | 277.897755 | 0 |
| 5.72E-10 | 3.87E-09 | 83.9173715 | 163.493835 | 179.11402 | 151.707121 | 0 |
| 5.72E-10 | 3.87E-09 | 91.6339114 | 165.495801 | 121.574801 | 178.615418 | 0 |
| 5.73E-10 | 3.88E-09 | 96.4567488 | 113.667143 | 110.438178 | 113.664357 | 0 |
| 5.75E-10 | 3.89E-09 | 155.295366 | 208.426835 | 84.4527244 | 165.625206 | 0 |
| 5.75E-10 | 3.89E-09 | 77.4869216 | 238.011434 | 222.73246 | 214.338502 | 0 |
| 5.75E-10 | 3.89E-09 | 83.595849 | 183.291048 | 152.200514 | 185.110524 | 0 |
| 5.75E-10 | 3.89E-09 | 99.6719738 | 119.895479 | 128.999216 | 189.28595 | 0 |
| 5.77E-10 | 3.90E-09 | 0 | 0 | 0 | 0 | 61.7816743 |
| 5.77E-10 | 3.90E-09 | 2087.96709 | 6123.34463 | 4914.0349 | 5784.81987 | 0 |
| 5.78E-10 | 3.91E-09 | 134.717926 | 122.119885 | 118.790645 | 140.572654 | 342.175427 |
| 5.79E-10 | 3.91E-09 | 85.524984 | 144.363944 | 176.329864 | 150.779249 | 0 |
| 5.79E-10 | 3.91E-09 | 80.7021465 | 220.216187 | 176.329864 | 223.153289 | 0 |
| 5.80E-10 | 3.92E-09 | 82.309759 | 183.95837 | 170.761553 | 163.305525 | 0 |
| 5.81E-10 | 3.93E-09 | 80.7021465 | 181.289083 | 187.466487 | 179.543291 | 0 |
| 5.85E-10 | 3.95E-09 | 86.4895515 | 143.251742 | 138.279736 | 166.089142 | 0 |
| 5.86E-10 | 3.96E-09 | 88.4186864 | 269.153117 | 152.200514 | 249.133713 | 0 |
| 5.87E-10 | 3.96E-09 | 84.881939 | 172.6139 | 157.768826 | 143.356271 | 0 |
| 5.87E-10 | 3.96E-09 | 80.380624 | 178.174915 | 187.466487 | 193.925311 | 0 |
| 5.89E-10 | 3.97E-09 | 88.0971639 | 191.52135 | 249.645966 | 149.851377 | 0 |

| | | | | | | |
|---|---|---|---|---|---|---|
| 5.90E-10 | 3.98E-09 | 83.595849 | 200.196533 | 271.919212 | 177.22361 | 0 |
| 5.91E-10 | 3.99E-09 | 80.380624 | 227.779167 | 192.106747 | 184.182652 | 0 |
| 5.92E-10 | 3.99E-09 | 168.477788 | 119.228158 | 90.9490879 | 170.728504 | 0 |
| 5.92E-10 | 3.99E-09 | 88.7402089 | 179.509558 | 128.071165 | 163.769461 | 0 |
| 5.93E-10 | 4.00E-09 | 81.3451915 | 207.537073 | 170.761553 | 183.25478 | 0 |
| 5.93E-10 | 4.00E-09 | 162.368861 | 412.182418 | 281.199731 | 346.560305 | 0 |
| 5.94E-10 | 4.00E-09 | 89.7047764 | 148.812756 | 119.718697 | 140.572654 | 0 |
| 5.94E-10 | 4.00E-09 | 87.1325964 | 146.58835 | 137.351684 | 130.829995 | 0 |
| 5.96E-10 | 4.02E-09 | 73.9501741 | 353.01322 | 400.918428 | 257.020628 | 0 |
| 5.98E-10 | 4.03E-09 | 83.2743265 | 153.261568 | 160.552982 | 157.738291 | 0 |
| 5.99E-10 | 4.03E-09 | 83.2743265 | 283.611755 | 175.401812 | 225.47297 | 0 |
| 5.99E-10 | 4.03E-09 | 0 | 0 | 0 | 0 | 59.4054561 |
| 6.00E-10 | 4.04E-09 | 123.464639 | 120.785242 | 95.5893474 | 104.84957 | 0 |
| 6.03E-10 | 4.06E-09 | 77.4869216 | 215.100053 | 213.451941 | 210.163077 | 0 |
| 6.04E-10 | 4.06E-09 | 92.2769564 | 111.442737 | 140.135839 | 140.572654 | 0 |
| 6.04E-10 | 4.06E-09 | 190.66284 | 837.488831 | 910.418931 | 841.116242 | 0 |
| 6.04E-10 | 4.06E-09 | 90.6693439 | 127.903341 | 116.00649 | 129.902122 | 0 |
| 6.04E-10 | 4.06E-09 | 136.968583 | 143.474182 | 133.639476 | 86.2921241 | 0 |
| 6.06E-10 | 4.07E-09 | 81.023669 | 219.326424 | 169.833501 | 221.76148 | 0 |
| 6.06E-10 | 4.07E-09 | 83.595849 | 150.592281 | 153.128566 | 167.017014 | 0 |
| 6.06E-10 | 4.07E-09 | 92.2769564 | 139.247811 | 118.790645 | 116.91191 | 0 |
| 6.06E-10 | 4.07E-09 | 93.8845689 | 113.444702 | 112.294282 | 121.087335 | 0 |
| 6.06E-10 | 4.07E-09 | 83.2743265 | 157.043058 | 164.265189 | 150.779249 | 0 |
| 6.07E-10 | 4.08E-09 | 79.094534 | 186.627657 | 199.531162 | 198.564673 | 0 |
| 6.08E-10 | 4.08E-09 | 92.5984789 | 200.863855 | 123.430905 | 166.089142 | 0 |
| 6.09E-10 | 4.09E-09 | 153.687753 | 109.663212 | 100.229607 | 113.664357 | 0 |
| 6.09E-10 | 4.09E-09 | 81.666714 | 256.696444 | 219.020252 | 173.976057 | 0 |
| 6.11E-10 | 4.10E-09 | 83.9173715 | 173.058781 | 145.704151 | 159.1301 | 0 |
| 6.12E-10 | 4.11E-09 | 79.737579 | 186.182776 | 265.422848 | 264.443606 | 0 |
| 6.13E-10 | 4.11E-09 | 80.0591015 | 194.857959 | 175.401812 | 211.554885 | 0 |
| 6.13E-10 | 4.11E-09 | 95.8137038 | 175.283187 | 114.150386 | 136.861165 | 0 |
| 6.14E-10 | 4.12E-09 | 176.194328 | 402.617472 | 217.164149 | 337.745518 | 2.37621824 |
| 6.15E-10 | 4.12E-09 | 181.338688 | 197.082365 | 409.270895 | 99.7462725 | 0 |
| 6.15E-10 | 4.12E-09 | 79.737579 | 205.090226 | 189.322591 | 179.079354 | 0 |
| 6.16E-10 | 4.13E-09 | 186.161525 | 104.547078 | 116.934542 | 117.839782 | 0 |
| 6.16E-10 | 4.13E-09 | 105.137856 | 166.385563 | 103.941815 | 198.564673 | 0 |
| 6.16E-10 | 4.13E-09 | 75.8793091 | 237.566553 | 221.804408 | 237.999246 | 0 |
| 6.17E-10 | 4.13E-09 | 72.9856066 | 299.849919 | 262.638692 | 272.794457 | 0 |
| 6.19E-10 | 4.14E-09 | 76.8438766 | 210.4288 | 233.869083 | 217.586055 | 0 |
| 6.20E-10 | 4.15E-09 | 81.9882365 | 156.820618 | 187.466487 | 196.244992 | 0 |
| 6.20E-10 | 4.15E-09 | 76.5223541 | 228.668929 | 211.595837 | 232.432012 | 0 |
| 6.23E-10 | 4.17E-09 | 87.7756414 | 181.511523 | 128.999216 | 165.625206 | 0 |
| 6.23E-10 | 4.17E-09 | 394.508103 | 273.379488 | 174.47376 | 230.576267 | 0 |

| | | | | | | |
|---|---|---|---|---|---|---|
| 6.23E-10 | 4.17E-09 | 86.4895515 | 136.356083 | 185.610383 | 154.026802 | 0 |
| 6.24E-10 | 4.17E-09 | 64.9475442 | 385.711987 | 421.33557 | 437.491791 | 0 |
| 6.25E-10 | 4.18E-09 | 78.1299666 | 266.261389 | 194.890903 | 255.628819 | 0 |
| 6.26E-10 | 4.18E-09 | 77.1653991 | 216.434696 | 220.876356 | 203.204034 | 0 |
| 6.26E-10 | 4.18E-09 | 91.9554339 | 132.574593 | 153.128566 | 113.664357 | 0 |
| 6.29E-10 | 4.20E-09 | 100.636541 | 163.493835 | 102.085711 | 161.913717 | 0 |
| 6.30E-10 | 4.21E-09 | 61.7323193 | 430.422546 | 527.133489 | 507.54615 | 0 |
| 6.30E-10 | 4.21E-09 | 149.186438 | 359.686438 | 298.832717 | 323.827434 | 0 |
| 6.30E-10 | 4.21E-09 | 70.0919042 | 362.800606 | 290.48025 | 347.952113 | 0 |
| 6.33E-10 | 4.22E-09 | 79.737579 | 217.991781 | 174.47376 | 202.276162 | 0 |
| 6.34E-10 | 4.23E-09 | 82.952804 | 158.377702 | 151.272462 | 179.543291 | 0 |
| 6.35E-10 | 4.24E-09 | 82.952804 | 195.0804 | 155.912722 | 162.377653 | 0 |
| 6.37E-10 | 4.25E-09 | 219.278342 | 102.100232 | 119.718697 | 141.03659 | 0 |
| 6.38E-10 | 4.26E-09 | 97.4213163 | 161.936751 | 111.36623 | 193.461375 | 0 |
| 6.38E-10 | 4.26E-09 | 76.2008316 | 251.357869 | 214.379993 | 224.081161 | 0 |
| 6.38E-10 | 4.26E-09 | 84.238894 | 173.503662 | 149.416359 | 144.284143 | 0 |
| 6.40E-10 | 4.26E-09 | 3006.87838 | 8743.02749 | 7315.83326 | 8177.80254 | 0 |
| 6.40E-10 | 4.27E-09 | 128.287476 | 110.997856 | 107.654022 | 95.5708472 | 0 |
| 6.41E-10 | 4.27E-09 | 85.524984 | 275.826335 | 158.696878 | 226.400842 | 0 |
| 6.42E-10 | 4.28E-09 | 79.094534 | 178.619796 | 202.315318 | 189.749886 | 0 |
| 6.44E-10 | 4.29E-09 | 72.0210391 | 269.820439 | 299.760769 | 276.505946 | 0 |
| 6.45E-10 | 4.30E-09 | 86.811074 | 195.0804 | 136.423632 | 158.666164 | 0 |
| 6.46E-10 | 4.30E-09 | 124.429206 | 332.103805 | 276.559471 | 329.858604 | 7.12865473 |
| 6.46E-10 | 4.30E-09 | 88.4186864 | 121.230123 | 138.279736 | 126.190633 | 0 |
| 6.47E-10 | 4.30E-09 | 72.6640841 | 258.253528 | 399.062324 | 348.879986 | 0 |
| 6.47E-10 | 4.30E-09 | 76.5223541 | 447.550472 | 258.926485 | 290.887967 | 0 |
| 6.47E-10 | 4.30E-09 | 80.0591015 | 191.52135 | 178.185968 | 169.800631 | 0 |
| 6.48E-10 | 4.31E-09 | 83.595849 | 172.169019 | 143.848047 | 154.490738 | 0 |
| 6.48E-10 | 4.31E-09 | 81.3451915 | 202.64338 | 164.265189 | 171.656376 | 0 |
| 6.51E-10 | 4.33E-09 | 83.595849 | 202.198499 | 147.560255 | 177.22361 | 0 |
| 6.51E-10 | 4.33E-09 | 81.023669 | 181.289083 | 158.696878 | 175.367865 | 0 |
| 6.52E-10 | 4.34E-09 | 134.074881 | 312.529032 | 248.717914 | 305.733924 | 2.37621824 |
| 6.53E-10 | 4.34E-09 | 145.971213 | 662.205644 | 123.430905 | 420.790089 | 0 |
| 6.53E-10 | 4.34E-09 | 80.380624 | 200.641414 | 183.75428 | 165.16127 | 0 |
| 6.53E-10 | 4.34E-09 | 82.309759 | 566.556189 | 229.228823 | 401.304771 | 0 |
| 6.54E-10 | 4.34E-09 | 85.2034615 | 139.247811 | 135.49558 | 137.789037 | 0 |
| 6.56E-10 | 4.35E-09 | 180.37412 | 214.655172 | 192.106747 | 218.049991 | 0 |
| 6.58E-10 | 4.37E-09 | 287.119589 | 152.594246 | 154.056618 | 98.3544641 | 0 |
| 6.59E-10 | 4.37E-09 | 74.2716966 | 233.562622 | 347.091417 | 311.765094 | 0 |
| 6.60E-10 | 4.38E-09 | 0 | 0 | 0 | 0 | 26.1384007 |
| 6.61E-10 | 4.38E-09 | 81.666714 | 161.714311 | 154.056618 | 173.048184 | 0 |
| 6.62E-10 | 4.39E-09 | 0 | 0 | 0 | 0 | 19.0097459 |
| 6.64E-10 | 4.40E-09 | 91.9554339 | 162.604073 | 112.294282 | 150.315313 | 0 |

| | | | | | | |
|---|---|---|---|---|---|---|
| 6.64E-10 | 4.40E-09 | 84.5604165 | 162.159192 | 133.639476 | 160.985845 | 0 |
| 6.64E-10 | 4.40E-09 | 77.8084441 | 279.607825 | 193.962851 | 269.546904 | 0 |
| 6.66E-10 | 4.41E-09 | 119.927891 | 114.334465 | 101.157659 | 96.9626556 | 0 |
| 6.66E-10 | 4.42E-09 | 80.7021465 | 172.391459 | 199.531162 | 163.769461 | 0 |
| 6.67E-10 | 4.42E-09 | 140.826853 | 359.463997 | 330.386482 | 381.819453 | 0 |
| 6.71E-10 | 4.44E-09 | 150.472528 | 121.897445 | 100.229607 | 101.138081 | 0 |
| 6.71E-10 | 4.44E-09 | 87.4541189 | 327.432552 | 161.481034 | 275.114138 | 0 |
| 6.73E-10 | 4.46E-09 | 72.6640841 | 256.251562 | 254.286225 | 277.433818 | 0 |
| 6.75E-10 | 4.47E-09 | 375.538275 | 633.288367 | 85.3807764 | 231.968076 | 0 |
| 6.76E-10 | 4.47E-09 | 78.7730116 | 180.39932 | 186.538435 | 216.194247 | 0 |
| 6.76E-10 | 4.48E-09 | 454.63281 | 992.529923 | 881.649321 | 984.008577 | 33.2670554 |
| 6.77E-10 | 4.48E-09 | 80.380624 | 179.064677 | 162.409085 | 167.944887 | 0 |
| 6.79E-10 | 4.49E-09 | 82.309759 | 165.718241 | 144.776099 | 170.264568 | 0 |
| 6.79E-10 | 4.49E-09 | 79.4160565 | 218.659102 | 169.833501 | 201.812226 | 0 |
| 6.80E-10 | 4.50E-09 | 141.469898 | 376.814364 | 315.537652 | 385.530942 | 2.37621824 |
| 6.81E-10 | 4.50E-09 | 74.5932191 | 256.029122 | 225.516616 | 229.648395 | 0 |
| 6.81E-10 | 4.50E-09 | 67.1982017 | 416.63123 | 445.46492 | 315.012647 | 0 |
| 6.81E-10 | 4.50E-09 | 153.044708 | 154.818652 | 235.725187 | 87.6839326 | 0 |
| 6.88E-10 | 4.54E-09 | 202.23765 | 470.906734 | 478.874789 | 465.32796 | 26.1384007 |
| 6.89E-10 | 4.55E-09 | 127.322908 | 417.076111 | 116.934542 | 257.484564 | 0 |
| 6.89E-10 | 4.55E-09 | 148.543393 | 401.060388 | 302.544925 | 317.332327 | 0 |
| 6.90E-10 | 4.56E-09 | 222.81509 | 464.678398 | 357.299988 | 409.655622 | 16.6335277 |
| 6.90E-10 | 4.56E-09 | 74.2716966 | 249.800785 | 221.804408 | 240.782863 | 0 |
| 6.90E-10 | 4.56E-09 | 86.168029 | 191.966231 | 134.567528 | 204.131907 | 0 |
| 6.91E-10 | 4.56E-09 | 80.0591015 | 163.938717 | 167.049345 | 172.120312 | 0 |
| 6.92E-10 | 4.56E-09 | 76.8438766 | 255.806681 | 193.962851 | 228.720523 | 0 |
| 6.94E-10 | 4.58E-09 | 143.399033 | 223.775236 | 218.0922 | 238.463182 | 35.6432737 |
| 6.96E-10 | 4.59E-09 | 142.112943 | 96.3167765 | 109.510126 | 109.952868 | 0 |
| 7.00E-10 | 4.62E-09 | 80.380624 | 159.934786 | 162.409085 | 176.759674 | 0 |
| 7.00E-10 | 4.62E-09 | 82.309759 | 164.383598 | 143.848047 | 153.09893 | 0 |
| 7.02E-10 | 4.62E-09 | 234.3899 | 210.20636 | 189.322591 | 180.471163 | 0 |
| 7.02E-10 | 4.62E-09 | 88.0971639 | 124.566732 | 118.790645 | 140.108718 | 0 |
| 7.02E-10 | 4.63E-09 | 83.2743265 | 182.401286 | 140.135839 | 162.377653 | 0 |
| 7.03E-10 | 4.63E-09 | 79.737579 | 213.320528 | 431.544141 | 443.059025 | 0 |
| 7.08E-10 | 4.66E-09 | 79.737579 | 191.52135 | 161.481034 | 176.759674 | 0 |
| 7.08E-10 | 4.66E-09 | 73.9501741 | 232.450419 | 225.516616 | 248.205841 | 0 |
| 7.09E-10 | 4.67E-09 | 77.4869216 | 204.422904 | 184.682331 | 233.359884 | 0 |
| 7.10E-10 | 4.68E-09 | 180.052598 | 484.030729 | 441.752712 | 480.173917 | 14.2573095 |
| 7.11E-10 | 4.68E-09 | 81.666714 | 274.491691 | 173.545708 | 303.878179 | 0 |
| 7.12E-10 | 4.68E-09 | 131.502701 | 128.348222 | 84.4527244 | 133.613612 | 0 |
| 7.13E-10 | 4.69E-09 | 79.737579 | 185.737895 | 197.675058 | 159.594036 | 0 |
| 7.13E-10 | 4.69E-09 | 126.679863 | 136.800964 | 90.9490879 | 107.169251 | 0 |
| 7.13E-10 | 4.69E-09 | 96.1352263 | 139.692692 | 100.229607 | 140.108718 | 0 |

| | | | | | | |
|---|---|---|---|---|---|---|
| 7.14E-10 | 4.69E-09 | 76.5223541 | 214.210291 | 191.178695 | 203.66797 | 0 |
| 7.14E-10 | 4.70E-09 | 131.181178 | 249.355904 | 96.5173994 | 155.418611 | 0 |
| 7.15E-10 | 4.70E-09 | 79.094534 | 175.283187 | 167.977397 | 173.976057 | 0 |
| 7.16E-10 | 4.71E-09 | 72.3425616 | 287.170805 | 238.509343 | 314.084774 | 0 |
| 7.22E-10 | 4.74E-09 | 520.544921 | 2313.6046 | 2045.42642 | 2152.66374 | 26.1384007 |
| 7.22E-10 | 4.74E-09 | 135.682493 | 410.847774 | 384.213494 | 383.211261 | 0 |
| 7.25E-10 | 4.76E-09 | 323.451631 | 124.566732 | 126.215061 | 147.995632 | 0 |
| 7.25E-10 | 4.76E-09 | 73.6286516 | 223.330355 | 238.509343 | 241.246799 | 0 |
| 7.26E-10 | 4.76E-09 | 86.4895515 | 142.139539 | 248.717914 | 207.843396 | 0 |
| 7.26E-10 | 4.76E-09 | 85.2034615 | 165.495801 | 132.711424 | 136.397228 | 0 |
| 7.26E-10 | 4.77E-09 | 149.186438 | 209.983919 | 177.257916 | 198.564673 | 2.37621824 |
| 7.27E-10 | 4.77E-09 | 77.1653991 | 180.844202 | 197.675058 | 211.554885 | 0 |
| 7.29E-10 | 4.78E-09 | 170.0854 | 87.864034 | 134.567528 | 221.297544 | 0 |
| 7.31E-10 | 4.79E-09 | 0 | 75.852242 | 460.313751 | 0 | 0 |
| 7.31E-10 | 4.80E-09 | 75.5577866 | 201.753617 | 269.135056 | 231.040203 | 0 |
| 7.33E-10 | 4.81E-09 | 95.8137038 | 103.434875 | 107.654022 | 110.416804 | 0 |
| 7.35E-10 | 4.82E-09 | 75.5577866 | 310.527067 | 529.917645 | 274.650202 | 0 |
| 7.38E-10 | 4.83E-09 | 67.8412467 | 321.871537 | 317.393756 | 312.692966 | 0 |
| 7.38E-10 | 4.83E-09 | 148.221871 | 275.159013 | 473.306478 | 449.090194 | 16.6335277 |
| 7.41E-10 | 4.85E-09 | 78.1299666 | 238.011434 | 180.042072 | 194.389247 | 0 |
| 7.41E-10 | 4.85E-09 | 150.151006 | 377.036804 | 266.3509 | 312.22903 | 0 |
| 7.43E-10 | 4.86E-09 | 83.595849 | 145.031266 | 184.682331 | 222.689353 | 0 |
| 7.44E-10 | 4.87E-09 | 158.832113 | 111.887618 | 99.3015551 | 108.097123 | 0 |
| 7.44E-10 | 4.87E-09 | 108.031559 | 190.854028 | 111.36623 | 160.521908 | 2.37621824 |
| 7.45E-10 | 4.87E-09 | 95.4921813 | 114.334465 | 156.840774 | 110.88074 | 0 |
| 7.46E-10 | 4.88E-09 | 95.4921813 | 183.513489 | 118.790645 | 123.870952 | 0 |
| 7.50E-10 | 4.90E-09 | 63.9829767 | 351.456136 | 414.839207 | 478.782108 | 0 |
| 7.50E-10 | 4.91E-09 | 72.6640841 | 238.901196 | 338.73895 | 270.474776 | 0 |
| 7.52E-10 | 4.92E-09 | 143.077511 | 312.084151 | 322.034015 | 348.41605 | 0 |
| 7.56E-10 | 4.94E-09 | 87.1325964 | 120.785242 | 129.927268 | 155.418611 | 0 |
| 7.56E-10 | 4.94E-09 | 187.769138 | 175.283187 | 80.7405168 | 143.356271 | 0 |
| 7.59E-10 | 4.96E-09 | 69.1273367 | 37.3700195 | 40.8342843 | 19.9492545 | 325.541899 |
| 7.60E-10 | 4.96E-09 | 133.110313 | 148.145435 | 245.933758 | 95.5708472 | 0 |
| 7.60E-10 | 4.96E-09 | 92.9200014 | 104.99196 | 111.36623 | 125.262761 | 0 |
| 7.59E-10 | 4.96E-09 | 166.22713 | 110.552974 | 101.157659 | 109.952868 | 0 |
| 7.61E-10 | 4.97E-09 | 162.047338 | 264.259424 | 211.595837 | 252.845202 | 2.37621824 |
| 7.61E-10 | 4.97E-09 | 116.712666 | 215.544934 | 168.905449 | 188.358077 | 0 |
| 7.61E-10 | 4.97E-09 | 158.832113 | 239.790959 | 185.610383 | 207.37946 | 0 |
| 7.65E-10 | 4.99E-09 | 89.7047764 | 126.568697 | 124.358957 | 107.633187 | 0 |
| 7.69E-10 | 5.02E-09 | 90.9908664 | 201.308736 | 126.215061 | 140.572654 | 0 |
| 7.70E-10 | 5.02E-09 | 82.309759 | 133.686796 | 158.696878 | 157.738291 | 0 |
| 7.71E-10 | 5.03E-09 | 88.4186864 | 163.716276 | 138.279736 | 116.447974 | 0 |
| 7.73E-10 | 5.04E-09 | 72.9856066 | 219.103984 | 245.005706 | 239.391054 | 0 |

| | | | | | | |
|---|---|---|---|---|---|---|
| 7.74E-10 | 5.04E-09 | 88.0971639 | 115.446667 | 133.639476 | 116.91191 | 0 |
| 7.76E-10 | 5.06E-09 | 93.5630464 | 174.170984 | 134.567528 | 267.227223 | 0 |
| 7.77E-10 | 5.06E-09 | 198.057858 | 133.686796 | 95.5893474 | 116.91191 | 0 |
| 7.81E-10 | 5.09E-09 | 83.2743265 | 142.139539 | 132.711424 | 132.685739 | 0 |
| 7.84E-10 | 5.10E-09 | 67.1982017 | 387.269071 | 349.875573 | 291.815839 | 0 |
| 7.84E-10 | 5.10E-09 | 78.1299666 | 201.531177 | 172.617657 | 169.800631 | 0 |
| 7.85E-10 | 5.11E-09 | 75.8793091 | 199.529211 | 217.164149 | 184.646588 | 0 |
| 7.85E-10 | 5.11E-09 | 70.0919042 | 288.283008 | 321.105963 | 253.309139 | 0 |
| 7.88E-10 | 5.13E-09 | 120.892459 | 179.064677 | 87.2368802 | 188.358077 | 0 |
| 7.88E-10 | 5.13E-09 | 147.578826 | 224.442558 | 133.639476 | 225.009033 | 9.50487297 |
| 7.90E-10 | 5.14E-09 | 248.53689 | 124.789172 | 100.229607 | 146.603824 | 0 |
| 7.92E-10 | 5.15E-09 | 79.094534 | 180.39932 | 182.826228 | 152.634994 | 0 |
| 7.92E-10 | 5.15E-09 | 73.3071291 | 234.674825 | 219.020252 | 218.049991 | 0 |
| 7.93E-10 | 5.16E-09 | 75.2362641 | 205.757548 | 190.250643 | 223.153289 | 0 |
| 7.94E-10 | 5.16E-09 | 80.0591015 | 194.190637 | 148.488307 | 188.358077 | 0 |
| 7.96E-10 | 5.18E-09 | 107.066991 | 165.495801 | 105.797919 | 106.241379 | 0 |
| 8.00E-10 | 5.20E-09 | 75.5577866 | 209.094157 | 183.75428 | 205.523715 | 0 |
| 8.01E-10 | 5.20E-09 | 127.965953 | 179.287118 | 86.3088283 | 211.554885 | 0 |
| 8.01E-10 | 5.21E-09 | 98.7074063 | 171.501697 | 187.466487 | 200.884354 | 9.50487297 |
| 8.02E-10 | 5.21E-09 | 85.524984 | 155.930855 | 123.430905 | 176.759674 | 0 |
| 8.04E-10 | 5.22E-09 | 80.380624 | 229.558691 | 160.552982 | 177.22361 | 0 |
| 8.04E-10 | 5.22E-09 | 73.9501741 | 218.436662 | 202.315318 | 233.359884 | 0 |
| 8.08E-10 | 5.25E-09 | 154.330798 | 286.281042 | 230.156875 | 269.082968 | 0 |
| 8.09E-10 | 5.25E-09 | 132.788791 | 190.186707 | 83.5246725 | 153.562866 | 0 |
| 8.10E-10 | 5.25E-09 | 81.3451915 | 144.141504 | 151.272462 | 137.325101 | 0 |
| 8.11E-10 | 5.26E-09 | 141.791421 | 295.401107 | 191.178695 | 234.751693 | 7.12865473 |
| 8.13E-10 | 5.28E-09 | 102.244154 | 125.901375 | 133.639476 | 240.782863 | 0 |
| 8.15E-10 | 5.29E-09 | 90.9908664 | 132.129712 | 207.883629 | 132.685739 | 0 |
| 8.15E-10 | 5.29E-09 | 75.8793091 | 253.582275 | 182.826228 | 237.999246 | 0 |
| 8.16E-10 | 5.29E-09 | 138.254673 | 236.899231 | 97.4454513 | 128.510314 | 0 |
| 8.16E-10 | 5.29E-09 | 72.6640841 | 212.875647 | 270.99116 | 250.989458 | 0 |
| 8.16E-10 | 5.29E-09 | 85.8465065 | 149.924959 | 121.574801 | 125.726697 | 0 |
| 8.16E-10 | 5.29E-09 | 99.0289288 | 149.035197 | 95.5893474 | 130.829995 | 0 |
| 8.21E-10 | 5.32E-09 | 87.7756414 | 374.367517 | 160.552982 | 291.351903 | 0 |
| 8.24E-10 | 5.34E-09 | 71.0564716 | 269.375557 | 233.869083 | 253.773075 | 0 |
| 8.26E-10 | 5.35E-09 | 121.213981 | 104.102197 | 94.6612955 | 102.065953 | 0 |
| 8.27E-10 | 5.36E-09 | 95.1706589 | 120.11792 | 97.4454513 | 112.272549 | 0 |
| 8.28E-10 | 5.36E-09 | 84.881939 | 270.26532 | 154.056618 | 305.269988 | 0 |
| 8.31E-10 | 5.38E-09 | 78.4514891 | 227.111845 | 161.481034 | 197.172864 | 0 |
| 8.33E-10 | 5.39E-09 | 72.6640841 | 210.873682 | 232.012979 | 240.782863 | 0 |
| 8.34E-10 | 5.39E-09 | 77.1653991 | 231.115776 | 173.545708 | 190.677758 | 0 |
| 8.36E-10 | 5.41E-09 | 81.023669 | 137.468286 | 153.128566 | 141.964462 | 0 |
| 8.37E-10 | 5.41E-09 | 43.405537 | 713.36698 | 1065.4036 | 990.039747 | 0 |

| | | | | | | |
|---|---|---|---|---|---|---|
| 8.37E-10 | 5.41E-09 | 112.854396 | 605.928174 | 525.277385 | 563.218488 | 2.37621824 |
| 8.40E-10 | 5.43E-09 | 84.5604165 | 155.485974 | 123.430905 | 173.976057 | 0 |
| 8.42E-10 | 5.44E-09 | 81.3451915 | 132.129712 | 151.272462 | 153.562866 | 0 |
| 8.43E-10 | 5.45E-09 | 155.616888 | 264.036983 | 83.5246725 | 305.269988 | 0 |
| 8.48E-10 | 5.48E-09 | 184.553913 | 371.69823 | 187.466487 | 340.993071 | 0 |
| 8.51E-10 | 5.50E-09 | 76.5223541 | 172.391459 | 173.545708 | 179.543291 | 0 |
| 8.51E-10 | 5.50E-09 | 70.0919042 | 236.009469 | 298.832717 | 289.032222 | 0 |
| 8.53E-10 | 5.51E-09 | 83.9173715 | 251.135429 | 154.056618 | 170.264568 | 0 |
| 8.54E-10 | 5.51E-09 | 71.0564716 | 231.783097 | 245.933758 | 243.102543 | 0 |
| 8.55E-10 | 5.52E-09 | 86.811074 | 114.334465 | 130.85532 | 115.520102 | 0 |
| 8.55E-10 | 5.52E-09 | 100.636541 | 146.36591 | 106.72597 | 101.138081 | 0 |
| 8.59E-10 | 5.54E-09 | 108.353081 | 140.804895 | 95.5893474 | 102.993826 | 0 |
| 8.60E-10 | 5.55E-09 | 82.952804 | 141.249776 | 124.358957 | 134.077548 | 0 |
| 8.61E-10 | 5.55E-09 | 87.1325964 | 115.224227 | 112.294282 | 121.551272 | 0 |
| 8.63E-10 | 5.56E-09 | 77.8084441 | 215.767375 | 171.689605 | 265.371479 | 0 |
| 8.66E-10 | 5.58E-09 | 137.611628 | 97.8738607 | 95.5893474 | 115.984038 | 0 |
| 8.66E-10 | 5.59E-09 | 76.2008316 | 200.641414 | 167.977397 | 190.213822 | 0 |
| 8.69E-10 | 5.60E-09 | 126.679863 | 690.233158 | 680.262055 | 726.524013 | 9.50487297 |
| 8.70E-10 | 5.61E-09 | 144.363601 | 290.952295 | 278.415575 | 312.692966 | 0 |
| 8.72E-10 | 5.62E-09 | 80.380624 | 190.409147 | 155.912722 | 145.212015 | 0 |
| 8.75E-10 | 5.63E-09 | 88.7402089 | 111.665177 | 114.150386 | 140.108718 | 0 |
| 8.76E-10 | 5.65E-09 | 91.6339114 | 102.545113 | 126.215061 | 145.212015 | 0 |
| 8.79E-10 | 5.66E-09 | 71.0564716 | 288.283008 | 250.574018 | 225.47297 | 0 |
| 8.80E-10 | 5.67E-09 | 68.4842917 | 263.369661 | 292.336354 | 269.082968 | 0 |
| 8.80E-10 | 5.67E-09 | 117.998756 | 203.533142 | 206.027526 | 232.895948 | 7.12865473 |
| 8.81E-10 | 5.67E-09 | 93.2415239 | 156.820618 | 169.833501 | 103.457762 | 0 |
| 8.84E-10 | 5.69E-09 | 106.423946 | 157.043058 | 145.704151 | 89.5396772 | 0 |
| 8.87E-10 | 5.70E-09 | 78.7730116 | 225.554761 | 165.193241 | 169.336695 | 0 |
| 8.87E-10 | 5.71E-09 | 79.4160565 | 142.58442 | 153.128566 | 147.06776 | 0 |
| 8.89E-10 | 5.71E-09 | 74.2716966 | 383.932463 | 219.020252 | 276.505946 | 0 |
| 8.90E-10 | 5.72E-09 | 102.887199 | 115.891549 | 103.013763 | 92.7872303 | 0 |
| 8.90E-10 | 5.72E-09 | 131.824223 | 397.278898 | 254.286225 | 315.940519 | 14.2573095 |
| 8.94E-10 | 5.75E-09 | 78.1299666 | 235.564587 | 164.265189 | 185.574461 | 0 |
| 8.95E-10 | 5.75E-09 | 79.094534 | 223.330355 | 152.200514 | 192.069567 | 0 |
| 8.96E-10 | 5.76E-09 | 80.380624 | 237.121672 | 155.912722 | 177.22361 | 0 |
| 8.97E-10 | 5.76E-09 | 82.6312815 | 242.237805 | 168.905449 | 342.38488 | 0 |
| 8.97E-10 | 5.76E-09 | 68.1627692 | 356.79471 | 352.659728 | 252.845202 | 0 |
| 8.97E-10 | 5.76E-09 | 141.791421 | 359.463997 | 280.271679 | 342.38488 | 0 |
| 8.98E-10 | 5.77E-09 | 82.309759 | 206.869751 | 135.49558 | 176.759674 | 0 |
| 9.00E-10 | 5.77E-09 | 66.5551567 | 384.154903 | 275.631419 | 342.848816 | 0 |
| 9.00E-10 | 5.77E-09 | 72.9856066 | 270.48776 | 199.531162 | 237.999246 | 0 |
| 9.01E-10 | 5.78E-09 | 84.238894 | 133.909237 | 128.071165 | 115.984038 | 0 |
| 9.03E-10 | 5.79E-09 | 82.6312815 | 122.564766 | 131.783372 | 139.180845 | 0 |

| | | | | | | |
|---|---|---|---|---|---|---|
| 9.06E-10 | 5.81E-09 | 110.603739 | 112.77738 | 86.3088283 | 123.870952 | 0 |
| 9.08E-10 | 5.82E-09 | 87.1325964 | 209.761479 | 122.502853 | 195.31712 | 0 |
| 9.11E-10 | 5.84E-09 | 75.8793091 | 178.174915 | 184.682331 | 165.625206 | 0 |
| 9.11E-10 | 5.84E-09 | 143.077511 | 395.499373 | 350.803625 | 368.365304 | 11.8810912 |
| 9.13E-10 | 5.85E-09 | 477.782429 | 534.969625 | 562.399462 | 622.138379 | 33.2670554 |
| 9.14E-10 | 5.86E-09 | 79.737579 | 142.58442 | 168.905449 | 142.892335 | 0 |
| 9.14E-10 | 5.86E-09 | 84.5604165 | 133.909237 | 119.718697 | 117.839782 | 0 |
| 9.15E-10 | 5.86E-09 | 81.3451915 | 155.263534 | 132.711424 | 134.077548 | 0 |
| 9.15E-10 | 5.86E-09 | 70.0919042 | 270.26532 | 237.581291 | 237.071373 | 0 |
| 9.15E-10 | 5.86E-09 | 74.9147416 | 410.180452 | 245.933758 | 240.318926 | 0 |
| 9.17E-10 | 5.87E-09 | 80.0591015 | 219.993746 | 148.488307 | 175.831801 | 0 |
| 9.18E-10 | 5.87E-09 | 96.1352263 | 163.048954 | 119.718697 | 103.457762 | 0 |
| 9.18E-10 | 5.87E-09 | 130.538133 | 323.651062 | 379.573234 | 370.684985 | 4.75243649 |
| 9.21E-10 | 5.90E-09 | 90.6693439 | 103.879757 | 107.654022 | 108.097123 | 0 |
| 9.22E-10 | 5.90E-09 | 118.641801 | 134.576558 | 90.0210359 | 101.602017 | 0 |
| 9.22E-10 | 5.90E-09 | 176.194328 | 125.678935 | 84.4527244 | 205.987651 | 0 |
| 9.27E-10 | 5.93E-09 | 79.094534 | 165.05092 | 139.207788 | 154.490738 | 0 |
| 9.30E-10 | 5.95E-09 | 106.745469 | 111.220296 | 87.2368802 | 119.231591 | 0 |
| 9.30E-10 | 5.95E-09 | 81.023669 | 166.385563 | 136.423632 | 196.708928 | 0 |
| 9.32E-10 | 5.95E-09 | 83.595849 | 263.814543 | 225.516616 | 142.892335 | 0 |
| 9.33E-10 | 5.96E-09 | 159.153636 | 228.001607 | 207.883629 | 226.864778 | 0 |
| 9.33E-10 | 5.96E-09 | 206.09592 | 100.765588 | 116.934542 | 119.695527 | 0 |
| 9.34E-10 | 5.97E-09 | 88.0971639 | 287.838127 | 138.279736 | 212.482757 | 0 |
| 9.35E-10 | 5.97E-09 | 181.017165 | 145.921029 | 112.294282 | 417.542536 | 0 |
| 9.35E-10 | 5.97E-09 | 84.5604165 | 117.003752 | 116.934542 | 125.726697 | 0 |
| 9.36E-10 | 5.98E-09 | 90.0262989 | 173.948543 | 431.544141 | 236.143501 | 0 |
| 9.37E-10 | 5.98E-09 | 66.8766792 | 303.853849 | 347.091417 | 267.227223 | 0 |
| 9.38E-10 | 5.98E-09 | 80.0591015 | 180.39932 | 177.257916 | 135.00542 | 0 |
| 9.37E-10 | 5.98E-09 | 78.1299666 | 236.899231 | 173.545708 | 167.944887 | 0 |
| 9.38E-10 | 5.98E-09 | 134.717926 | 329.434517 | 365.652455 | 317.332327 | 2.37621824 |
| 9.38E-10 | 5.98E-09 | 163.333428 | 272.489726 | 199.531162 | 236.143501 | 0 |
| 9.39E-10 | 5.99E-09 | 87.7756414 | 148.145435 | 106.72597 | 132.685739 | 0 |
| 9.42E-10 | 6.01E-09 | 80.380624 | 129.015544 | 148.488307 | 145.675952 | 0 |
| 9.43E-10 | 6.01E-09 | 115.426576 | 122.119885 | 179.11402 | 91.3954218 | 0 |
| 9.49E-10 | 6.04E-09 | 77.8084441 | 146.810791 | 166.121293 | 154.026802 | 0 |
| 9.49E-10 | 6.04E-09 | 70.0919042 | 234.229944 | 231.084927 | 238.463182 | 0 |
| 9.49E-10 | 6.04E-09 | 100.315019 | 221.328389 | 104.869867 | 162.841589 | 0 |
| 9.50E-10 | 6.05E-09 | 89.7047764 | 140.360014 | 236.653239 | 139.180845 | 0 |
| 9.52E-10 | 6.06E-09 | 78.7730116 | 154.818652 | 140.135839 | 170.728504 | 0 |
| 9.52E-10 | 6.06E-09 | 114.462009 | 110.997856 | 151.272462 | 89.0757411 | 0 |
| 9.53E-10 | 6.07E-09 | 75.5577866 | 235.342147 | 236.653239 | 406.872005 | 0 |
| 9.54E-10 | 6.07E-09 | 166.22713 | 314.753438 | 180.970124 | 225.47297 | 4.75243649 |
| 9.54E-10 | 6.07E-09 | 691.273367 | 185.737895 | 134.567528 | 197.6368 | 0 |

| | | | | | | |
|---|---|---|---|---|---|---|
| 9.55E-10 | 6.07E-09 | 76.2008316 | 173.948543 | 164.265189 | 206.915523 | 0 |
| 9.58E-10 | 6.09E-09 | 131.502701 | 275.826335 | 100.229607 | 388.314559 | 0 |
| 9.59E-10 | 6.10E-09 | 73.6286516 | 180.621761 | 213.451941 | 198.100737 | 0 |
| 9.63E-10 | 6.12E-09 | 78.4514891 | 158.822583 | 199.531162 | 150.779249 | 0 |
| 9.65E-10 | 6.13E-09 | 71.0564716 | 229.336251 | 217.164149 | 214.802438 | 0 |
| 9.65E-10 | 6.13E-09 | 50.4790319 | 110.330534 | 110.438178 | 165.625206 | 772.270929 |
| 9.65E-10 | 6.13E-09 | 122.821594 | 493.595675 | 377.71713 | 493.164129 | 0 |
| 9.66E-10 | 6.14E-09 | 84.881939 | 151.926925 | 114.150386 | 130.829995 | 0 |
| 9.66E-10 | 6.14E-09 | 76.8438766 | 302.296765 | 280.271679 | 635.592527 | 0 |
| 9.67E-10 | 6.14E-09 | 84.881939 | 155.041093 | 116.934542 | 127.118505 | 0 |
| 9.68E-10 | 6.15E-09 | 65.9121117 | 315.643201 | 291.408302 | 283.001052 | 0 |
| 9.69E-10 | 6.15E-09 | 81.666714 | 145.476147 | 181.898176 | 131.293931 | 0 |
| 9.70E-10 | 6.16E-09 | 608.642085 | 1431.18277 | 1142.43191 | 1334.28037 | 61.7816743 |
| 9.71E-10 | 6.16E-09 | 91.9554339 | 143.029301 | 159.62493 | 100.674145 | 0 |
| 9.73E-10 | 6.17E-09 | 128.930521 | 399.280863 | 351.731677 | 366.973496 | 2.37621824 |
| 9.73E-10 | 6.17E-09 | 79.4160565 | 168.83241 | 141.063891 | 199.492545 | 0 |
| 9.79E-10 | 6.21E-09 | 76.8438766 | 217.546899 | 167.049345 | 167.017014 | 0 |
| 9.82E-10 | 6.23E-09 | 98.0643613 | 199.08433 | 103.013763 | 147.06776 | 0 |
| 9.82E-10 | 6.23E-09 | 142.112943 | 205.979989 | 204.171422 | 192.533503 | 4.75243649 |
| 9.86E-10 | 6.25E-09 | 82.309759 | 207.314632 | 130.85532 | 180.007227 | 0 |
| 9.87E-10 | 6.26E-09 | 174.908238 | 254.027157 | 183.75428 | 205.523715 | 0 |
| 9.88E-10 | 6.26E-09 | 294.514606 | 501.825977 | 447.321024 | 430.996685 | 0 |
| 9.90E-10 | 6.27E-09 | 82.6312815 | 248.466142 | 141.991943 | 188.822014 | 0 |
| 9.91E-10 | 6.28E-09 | 118.641801 | 327.654993 | 270.063108 | 312.692966 | 7.12865473 |
| 9.93E-10 | 6.29E-09 | 159.153636 | 303.408968 | 268.207004 | 276.505946 | 14.2573095 |
| 9.93E-10 | 6.29E-09 | 97.7428388 | 124.566732 | 117.862593 | 90.4675495 | 0 |
| 9.95E-10 | 6.30E-09 | 213.490937 | 439.542611 | 348.019469 | 398.98509 | 16.6335277 |
| 9.98E-10 | 6.32E-09 | 74.9147416 | 201.976058 | 204.171422 | 162.841589 | 0 |
| 9.99E-10 | 6.32E-09 | 111.568306 | 180.17688 | 184.682331 | 173.048184 | 9.50487297 |
| 1.00E-09 | 6.32E-09 | 78.4514891 | 149.035197 | 145.704151 | 182.326907 | 0 |
| 1.01E-09 | 6.36E-09 | 80.0591015 | 146.58835 | 127.143113 | 139.644782 | 0 |
| 1.01E-09 | 6.38E-09 | 83.2743265 | 185.960335 | 125.287009 | 147.06776 | 0 |
| 1.01E-09 | 6.38E-09 | 70.4134267 | 222.218152 | 227.37272 | 213.41063 | 0 |
| 1.01E-09 | 6.39E-09 | 78.1299666 | 150.1474 | 141.063891 | 142.892335 | 0 |
| 1.01E-09 | 6.39E-09 | 72.3425616 | 235.119706 | 189.322591 | 209.235204 | 0 |
| 1.01E-09 | 6.39E-09 | 88.4186864 | 112.77738 | 176.329864 | 131.293931 | 0 |
| 1.01E-09 | 6.40E-09 | 81.666714 | 121.452563 | 143.848047 | 127.582442 | 0 |
| 1.02E-09 | 6.41E-09 | 100.315019 | 109.885653 | 125.287009 | 89.5396772 | 0 |
| 1.02E-09 | 6.41E-09 | 71.6995166 | 261.812577 | 201.387266 | 217.122119 | 0 |
| 1.02E-09 | 6.42E-09 | 67.1982017 | 265.149186 | 330.386482 | 261.659989 | 0 |
| 1.02E-09 | 6.42E-09 | 82.952804 | 145.476147 | 173.545708 | 120.623399 | 0 |
| 1.02E-09 | 6.43E-09 | 89.0617314 | 113.667143 | 100.229607 | 110.416804 | 0 |
| 1.02E-09 | 6.43E-09 | 113.497441 | 98.5411824 | 100.229607 | 93.2511664 | 0 |

| | | | | | | |
|---|---|---|---|---|---|---|
| 1.02E-09 | 6.44E-09 | 79.4160565 | 140.582454 | 150.344411 | 129.438186 | 0 |
| 1.03E-09 | 6.47E-09 | 84.881939 | 133.241915 | 108.582074 | 122.015208 | 0 |
| 1.03E-09 | 6.48E-09 | 92.9200014 | 139.692692 | 95.5893474 | 122.94308 | 0 |
| 1.03E-09 | 6.50E-09 | 108.996126 | 140.137573 | 599.521538 | 489.45264 | 0 |
| 1.03E-09 | 6.51E-09 | 134.396403 | 241.348043 | 287.696094 | 276.04201 | 11.8810912 |
| 1.04E-09 | 6.55E-09 | 158.189068 | 167.052885 | 171.689605 | 215.73031 | 4.75243649 |
| 1.04E-09 | 6.56E-09 | 76.5223541 | 183.291048 | 148.488307 | 163.769461 | 0 |
| 1.04E-09 | 6.57E-09 | 73.6286516 | 236.009469 | 174.47376 | 207.37946 | 0 |
| 1.05E-09 | 6.59E-09 | 148.221871 | 127.013578 | 85.3807764 | 107.633187 | 0 |
| 1.05E-09 | 6.59E-09 | 78.7730116 | 181.733964 | 172.617657 | 278.825627 | 0 |
| 1.05E-09 | 6.59E-09 | 132.467268 | 124.789172 | 98.3735032 | 89.0757411 | 0 |
| 1.05E-09 | 6.59E-09 | 139.862286 | 95.4270142 | 136.423632 | 92.3232941 | 0 |
| 1.05E-09 | 6.59E-09 | 116.069621 | 154.818652 | 280.271679 | 105.777442 | 0 |
| 1.05E-09 | 6.60E-09 | 71.0564716 | 203.755583 | 206.955577 | 207.37946 | 0 |
| 1.05E-09 | 6.60E-09 | 84.5604165 | 192.411112 | 144.776099 | 123.407016 | 0 |
| 1.05E-09 | 6.61E-09 | 78.7730116 | 131.907271 | 165.193241 | 172.120312 | 0 |
| 1.05E-09 | 6.62E-09 | 138.576196 | 325.430587 | 234.797135 | 273.258393 | 4.75243649 |
| 1.05E-09 | 6.63E-09 | 80.380624 | 136.800964 | 168.905449 | 211.090949 | 0 |
| 1.05E-09 | 6.63E-09 | 93.8845689 | 124.789172 | 99.3015551 | 100.210209 | 0 |
| 1.06E-09 | 6.64E-09 | 160.439726 | 384.154903 | 375.861026 | 411.04743 | 2.37621824 |
| 1.06E-09 | 6.64E-09 | 72.0210391 | 182.623726 | 217.164149 | 212.946693 | 0 |
| 1.06E-09 | 6.65E-09 | 80.0591015 | 134.576558 | 128.071165 | 127.118505 | 0 |
| 1.06E-09 | 6.66E-09 | 74.5932191 | 212.653206 | 169.833501 | 173.976057 | 0 |
| 1.06E-09 | 6.66E-09 | 80.7021465 | 178.174915 | 208.811681 | 132.685739 | 0 |
| 1.07E-09 | 6.70E-09 | 96.4567488 | 96.3167765 | 166.121293 | 130.829995 | 0 |
| 1.07E-09 | 6.70E-09 | 217.027685 | 246.241736 | 198.60311 | 241.246799 | 16.6335277 |
| 1.07E-09 | 6.72E-09 | 80.7021465 | 133.019474 | 120.646749 | 129.438186 | 0 |
| 1.07E-09 | 6.73E-09 | 79.094534 | 126.568697 | 150.344411 | 152.634994 | 0 |
| 1.08E-09 | 6.76E-09 | 78.7730116 | 142.80686 | 177.257916 | 137.789037 | 0 |
| 1.08E-09 | 6.79E-09 | 76.8438766 | 188.407182 | 144.776099 | 163.769461 | 0 |
| 1.08E-09 | 6.80E-09 | 84.881939 | 133.241915 | 113.222334 | 165.16127 | 0 |
| 1.08E-09 | 6.80E-09 | 83.9173715 | 108.106128 | 140.135839 | 133.149675 | 0 |
| 1.08E-09 | 6.80E-09 | 87.4541189 | 100.988029 | 115.078438 | 127.118505 | 0 |
| 1.09E-09 | 6.82E-09 | 280.046094 | 152.594246 | 78.8844129 | 229.184459 | 0 |
| 1.09E-09 | 6.82E-09 | 61.0892743 | 358.796676 | 342.451157 | 347.488177 | 0 |
| 1.09E-09 | 6.83E-09 | 151.115573 | 149.702519 | 92.8051917 | 95.5708472 | 0 |
| 1.09E-09 | 6.84E-09 | 73.3071291 | 230.003573 | 172.617657 | 199.028609 | 0 |
| 1.09E-09 | 6.85E-09 | 98.7074063 | 186.627657 | 188.394539 | 96.9626556 | 0 |
| 1.09E-09 | 6.85E-09 | 80.380624 | 147.478113 | 123.430905 | 129.438186 | 0 |
| 1.09E-09 | 6.85E-09 | 59.4816618 | 392.607646 | 387.925701 | 358.158709 | 0 |
| 1.09E-09 | 6.85E-09 | 3.53674746 | 6.45077718 | 12.0646749 | 1.85574461 | 99.8011662 |
| 1.10E-09 | 6.87E-09 | 116.712666 | 662.428084 | 531.773748 | 588.734976 | 0 |
| 1.10E-09 | 6.88E-09 | 100.636541 | 164.161157 | 91.8771398 | 128.510314 | 0 |

| | | | | | | |
|---|---|---|---|---|---|---|
| 1.10E-09 | 6.90E-09 | 101.279586 | 97.8738607 | 108.582074 | 160.057972 | 0 |
| 1.10E-09 | 6.91E-09 | 74.5932191 | 204.422904 | 157.768826 | 212.946693 | 0 |
| 1.11E-09 | 6.95E-09 | 133.110313 | 336.107735 | 319.249859 | 317.332327 | 0 |
| 1.12E-09 | 6.98E-09 | 81.3451915 | 120.11792 | 143.848047 | 121.551272 | 0 |
| 1.12E-09 | 6.99E-09 | 84.5604165 | 119.450598 | 107.654022 | 111.344676 | 0 |
| 1.12E-09 | 7.03E-09 | 77.8084441 | 193.300875 | 138.279736 | 199.492545 | 0 |
| 1.12E-09 | 7.03E-09 | 97.0997938 | 572.339644 | 580.032448 | 554.403701 | 9.50487297 |
| 1.13E-09 | 7.04E-09 | 197.414813 | 237.566553 | 190.250643 | 220.833608 | 28.5146189 |
| 1.13E-09 | 7.04E-09 | 193.23502 | 719.372876 | 412.983103 | 497.339554 | 0 |
| 1.13E-09 | 7.06E-09 | 78.1299666 | 142.361979 | 128.999216 | 141.03659 | 0 |
| 1.13E-09 | 7.08E-09 | 72.0210391 | 230.226013 | 176.329864 | 233.82382 | 0 |
| 1.14E-09 | 7.10E-09 | 78.4514891 | 167.497766 | 129.927268 | 177.22361 | 0 |
| 1.14E-09 | 7.11E-09 | 107.710036 | 85.8620687 | 152.200514 | 167.017014 | 0 |
| 1.14E-09 | 7.12E-09 | 103.530244 | 203.088261 | 189.322591 | 93.7151026 | 0 |
| 1.15E-09 | 7.15E-09 | 67.1982017 | 229.558691 | 282.127783 | 254.237011 | 0 |
| 1.15E-09 | 7.17E-09 | 71.6995166 | 206.64731 | 376.789078 | 283.464988 | 0 |
| 1.15E-09 | 7.18E-09 | 73.3071291 | 164.606038 | 177.257916 | 169.800631 | 0 |
| 1.15E-09 | 7.18E-09 | 74.9147416 | 152.149365 | 162.409085 | 153.562866 | 0 |
| 1.15E-09 | 7.18E-09 | 95.8137038 | 90.9782023 | 122.502853 | 105.313506 | 0 |
| 1.15E-09 | 7.19E-09 | 81.3451915 | 135.911202 | 113.222334 | 128.510314 | 0 |
| 1.15E-09 | 7.19E-09 | 260.111699 | 477.135071 | 356.371936 | 440.739344 | 0 |
| 1.16E-09 | 7.22E-09 | 78.7730116 | 127.458459 | 128.999216 | 143.820207 | 0 |
| 1.16E-09 | 7.22E-09 | 81.3451915 | 112.999821 | 142.919995 | 144.284143 | 0 |
| 1.16E-09 | 7.25E-09 | 115.426576 | 297.180631 | 258.926485 | 279.753499 | 7.12865473 |
| 1.16E-09 | 7.25E-09 | 142.112943 | 267.596033 | 242.22155 | 261.659989 | 2.37621824 |
| 1.17E-09 | 7.26E-09 | 75.5577866 | 183.291048 | 194.890903 | 144.748079 | 0 |
| 1.17E-09 | 7.27E-09 | 151.437096 | 179.731999 | 77.956361 | 250.525522 | 0 |
| 1.17E-09 | 7.27E-09 | 93.5630464 | 369.473824 | 180.970124 | 156.346483 | 0 |
| 1.17E-09 | 7.28E-09 | 66.5551567 | 234.897266 | 261.710641 | 260.268181 | 0 |
| 1.17E-09 | 7.30E-09 | 69.1273367 | 228.89137 | 202.315318 | 224.081161 | 0 |
| 1.17E-09 | 7.30E-09 | 82.952804 | 181.956405 | 147.560255 | 117.839782 | 0 |
| 1.17E-09 | 7.30E-09 | 77.8084441 | 139.470251 | 128.071165 | 141.500526 | 0 |
| 1.18E-09 | 7.31E-09 | 126.358341 | 109.885653 | 221.804408 | 101.602017 | 0 |
| 1.18E-09 | 7.32E-09 | 77.1653991 | 267.596033 | 164.265189 | 184.646588 | 0 |
| 1.18E-09 | 7.33E-09 | 83.2743265 | 170.167053 | 115.078438 | 136.861165 | 0 |
| 1.18E-09 | 7.34E-09 | 107.388514 | 177.730033 | 189.322591 | 88.1478687 | 0 |
| 1.18E-09 | 7.34E-09 | 76.8438766 | 201.308736 | 176.329864 | 140.108718 | 0 |
| 1.18E-09 | 7.35E-09 | 83.2743265 | 122.787207 | 113.222334 | 108.097123 | 0 |
| 1.18E-09 | 7.36E-09 | 146.935781 | 275.603894 | 79.8124649 | 204.131907 | 0 |
| 1.18E-09 | 7.36E-09 | 157.867546 | 130.350187 | 80.7405168 | 117.375846 | 0 |
| 1.19E-09 | 7.37E-09 | 80.0591015 | 121.007682 | 120.646749 | 124.798825 | 0 |
| 1.19E-09 | 7.37E-09 | 152.723186 | 237.344112 | 201.387266 | 225.009033 | 2.37621824 |
| 1.19E-09 | 7.40E-09 | 77.4869216 | 129.905306 | 152.200514 | 167.017014 | 0 |

| | | | | | | |
|---|---|---|---|---|---|---|
| 1.19E-09 | 7.40E-09 | 126.679863 | 128.570662 | 123.430905 | 78.8691457 | 0 |
| 1.20E-09 | 7.42E-09 | 59.8031843 | 367.249418 | 328.530379 | 377.644027 | 0 |
| 1.20E-09 | 7.43E-09 | 121.213981 | 186.182776 | 166.121293 | 188.358077 | 11.8810912 |
| 1.20E-09 | 7.44E-09 | 76.5223541 | 138.580489 | 138.279736 | 141.964462 | 0 |
| 1.20E-09 | 7.45E-09 | 78.1299666 | 160.157227 | 321.105963 | 263.051798 | 0 |
| 1.20E-09 | 7.45E-09 | 71.0564716 | 181.733964 | 193.034799 | 187.894141 | 0 |
| 1.20E-09 | 7.46E-09 | 90.6693439 | 195.747721 | 105.797919 | 196.708928 | 0 |
| 1.20E-09 | 7.47E-09 | 96.4567488 | 122.119885 | 89.092984 | 105.777442 | 0 |
| 1.20E-09 | 7.47E-09 | 99.3504513 | 113.889583 | 281.199731 | 175.831801 | 0 |
| 1.21E-09 | 7.48E-09 | 75.5577866 | 149.480078 | 141.063891 | 149.387441 | 0 |
| 1.21E-09 | 7.50E-09 | 68.4842917 | 265.816508 | 203.24337 | 254.237011 | 0 |
| 1.21E-09 | 7.51E-09 | 59.8031843 | 341.891191 | 357.299988 | 346.560305 | 0 |
| 1.22E-09 | 7.57E-09 | 76.2008316 | 159.934786 | 133.639476 | 154.490738 | 0 |
| 1.23E-09 | 7.59E-09 | 85.2034615 | 109.218331 | 103.941815 | 106.241379 | 0 |
| 1.23E-09 | 7.61E-09 | 70.0919042 | 204.645345 | 193.962851 | 190.213822 | 0 |
| 1.23E-09 | 7.62E-09 | 80.0591015 | 199.974093 | 128.071165 | 205.987651 | 0 |
| 1.23E-09 | 7.64E-09 | 116.391144 | 259.810612 | 218.0922 | 236.143501 | 9.50487297 |
| 1.23E-09 | 7.64E-09 | 75.5577866 | 155.708415 | 144.776099 | 187.894141 | 0 |
| 1.23E-09 | 7.64E-09 | 62.6968867 | 280.052706 | 303.472977 | 321.971689 | 0 |
| 1.24E-09 | 7.65E-09 | 80.380624 | 176.61783 | 121.574801 | 185.110524 | 0 |
| 1.24E-09 | 7.65E-09 | 81.9882365 | 115.001786 | 120.646749 | 109.024996 | 0 |
| 1.24E-09 | 7.67E-09 | 73.3071291 | 326.097909 | 180.970124 | 324.29137 | 0 |
| 1.24E-09 | 7.67E-09 | 94.8491364 | 188.407182 | 106.72597 | 120.623399 | 0 |
| 1.24E-09 | 7.67E-09 | 79.094534 | 139.470251 | 125.287009 | 170.264568 | 0 |
| 1.24E-09 | 7.69E-09 | 69.7703817 | 205.312667 | 228.300772 | 188.358077 | 0 |
| 1.24E-09 | 7.69E-09 | 77.8084441 | 147.033232 | 123.430905 | 152.171058 | 0 |
| 1.25E-09 | 7.70E-09 | 66.2336342 | 254.472038 | 235.725187 | 240.318926 | 0 |
| 1.25E-09 | 7.70E-09 | 74.5932191 | 144.141504 | 163.337137 | 179.079354 | 0 |
| 1.25E-09 | 7.75E-09 | 81.3451915 | 446.438269 | 189.322591 | 243.566479 | 0 |
| 1.26E-09 | 7.75E-09 | 74.5932191 | 250.023226 | 193.962851 | 160.985845 | 0 |
| 1.26E-09 | 7.81E-09 | 75.8793091 | 169.277291 | 134.567528 | 174.903929 | 0 |
| 1.27E-09 | 7.82E-09 | 81.666714 | 158.600142 | 117.862593 | 123.407016 | 0 |
| 1.27E-09 | 7.87E-09 | 151.758618 | 235.342147 | 198.60311 | 235.215629 | 0 |
| 1.27E-09 | 7.87E-09 | 65.5905892 | 242.237805 | 246.86181 | 251.91733 | 0 |
| 1.28E-09 | 7.87E-09 | 87.1325964 | 103.212435 | 103.013763 | 129.902122 | 0 |
| 1.28E-09 | 7.89E-09 | 71.0564716 | 195.747721 | 169.833501 | 207.37946 | 0 |
| 1.28E-09 | 7.90E-09 | 68.4842917 | 252.914954 | 195.818954 | 238.463182 | 0 |
| 1.28E-09 | 7.90E-09 | 136.325538 | 406.176522 | 276.559471 | 317.332327 | 0 |
| 1.28E-09 | 7.91E-09 | 138.576196 | 271.822404 | 260.782589 | 261.659989 | 0 |
| 1.28E-09 | 7.92E-09 | 112.854396 | 128.348222 | 78.8844129 | 130.829995 | 0 |
| 1.29E-09 | 7.92E-09 | 134.074881 | 355.904948 | 329.458431 | 271.866585 | 0 |
| 1.30E-09 | 8.00E-09 | 72.6640841 | 294.733785 | 179.11402 | 219.905736 | 0 |
| 1.30E-09 | 8.01E-09 | 65.9121117 | 237.121672 | 238.509343 | 290.424031 | 0 |

| | | | | | | |
|---|---|---|---|---|---|---|
| 1.30E-09 | 8.01E-09 | 156.902978 | 96.0943359 | 90.9490879 | 125.262761 | 0 |
| 1.30E-09 | 8.01E-09 | 96.4567488 | 165.940682 | 209.739733 | 98.3544641 | 0 |
| 1.30E-09 | 8.02E-09 | 57.8740493 | 429.532784 | 497.435827 | 329.394667 | 0 |
| 1.30E-09 | 8.03E-09 | 64.9475442 | 355.015186 | 245.005706 | 283.464988 | 0 |
| 1.30E-09 | 8.03E-09 | 68.4842917 | 243.794889 | 196.747006 | 216.658183 | 0 |
| 1.31E-09 | 8.03E-09 | 64.6260217 | 280.720028 | 238.509343 | 285.784669 | 0 |
| 1.31E-09 | 8.06E-09 | 86.811074 | 142.80686 | 105.797919 | 107.169251 | 0 |
| 1.31E-09 | 8.08E-09 | 90.6693439 | 176.61783 | 174.47376 | 99.2823364 | 0 |
| 1.31E-09 | 8.09E-09 | 152.723186 | 317.867607 | 214.379993 | 387.850622 | 0 |
| 1.32E-09 | 8.11E-09 | 71.6995166 | 172.169019 | 247.789862 | 198.100737 | 0 |
| 1.32E-09 | 8.12E-09 | 100.636541 | 184.848132 | 177.257916 | 90.4675495 | 0 |
| 1.32E-09 | 8.13E-09 | 286.476544 | 162.604073 | 78.8844129 | 172.120312 | 0 |
| 1.32E-09 | 8.14E-09 | 65.2690667 | 254.472038 | 233.869083 | 254.700947 | 0 |
| 1.32E-09 | 8.14E-09 | 85.8465065 | 170.611934 | 106.72597 | 131.757867 | 0 |
| 1.33E-09 | 8.14E-09 | 80.0591015 | 162.159192 | 117.862593 | 132.221803 | 0 |
| 1.33E-09 | 8.15E-09 | 68.1627692 | 375.47972 | 256.142329 | 229.184459 | 0 |
| 1.33E-09 | 8.16E-09 | 95.4921813 | 154.373771 | 89.092984 | 139.180845 | 0 |
| 1.33E-09 | 8.16E-09 | 79.737579 | 124.121851 | 131.783372 | 111.344676 | 0 |
| 1.33E-09 | 8.17E-09 | 63.9829767 | 269.153117 | 251.502069 | 314.084774 | 0 |
| 1.33E-09 | 8.18E-09 | 70.0919042 | 193.523315 | 177.257916 | 186.966269 | 0 |
| 1.33E-09 | 8.18E-09 | 67.1982017 | 234.897266 | 215.308045 | 213.41063 | 0 |
| 1.33E-09 | 8.18E-09 | 72.0210391 | 177.285152 | 157.768826 | 169.800631 | 0 |
| 1.34E-09 | 8.21E-09 | 154.330798 | 224.220117 | 193.962851 | 201.34829 | 2.37621824 |
| 1.34E-09 | 8.21E-09 | 3.53674746 | 1.77952474 | 0.92805192 | 3.71148921 | 23.7621824 |
| 1.34E-09 | 8.22E-09 | 81.9882365 | 137.245846 | 107.654022 | 154.026802 | 0 |
| 1.35E-09 | 8.26E-09 | 78.7730116 | 159.489905 | 303.472977 | 192.997439 | 0 |
| 1.35E-09 | 8.28E-09 | 88.0971639 | 107.661247 | 95.5893474 | 128.510314 | 0 |
| 1.35E-09 | 8.28E-09 | 72.3425616 | 185.960335 | 154.056618 | 169.800631 | 0 |
| 1.35E-09 | 8.28E-09 | 312.519866 | 163.048954 | 109.510126 | 111.808612 | 0 |
| 1.35E-09 | 8.29E-09 | 79.4160565 | 119.673039 | 128.071165 | 112.736485 | 0 |
| 1.35E-09 | 8.29E-09 | 93.5630464 | 265.594067 | 110.438178 | 222.225416 | 0 |
| 1.35E-09 | 8.30E-09 | 122.821594 | 422.637126 | 351.731677 | 338.209454 | 2.37621824 |
| 1.36E-09 | 8.30E-09 | 73.9501741 | 230.893335 | 153.128566 | 187.894141 | 0 |
| 1.36E-09 | 8.31E-09 | 83.9173715 | 112.332499 | 101.157659 | 128.510314 | 0 |
| 1.36E-09 | 8.32E-09 | 71.6995166 | 185.515454 | 156.840774 | 180.471163 | 0 |
| 1.36E-09 | 8.32E-09 | 128.930521 | 115.669108 | 91.8771398 | 89.0757411 | 0 |
| 1.36E-09 | 8.32E-09 | 81.3451915 | 133.241915 | 108.582074 | 115.520102 | 0 |
| 1.36E-09 | 8.33E-09 | 64.3044992 | 241.792924 | 267.278952 | 312.22903 | 0 |
| 1.36E-09 | 8.33E-09 | 96.1352263 | 101.655351 | 160.552982 | 102.529889 | 0 |
| 1.36E-09 | 8.33E-09 | 59.1601393 | 322.093978 | 491.867516 | 362.79807 | 0 |
| 1.36E-09 | 8.34E-09 | 166.870175 | 140.804895 | 80.7405168 | 111.808612 | 0 |
| 1.37E-09 | 8.35E-09 | 72.3425616 | 190.631588 | 154.98467 | 167.944887 | 0 |
| 1.37E-09 | 8.36E-09 | 78.4514891 | 166.163123 | 326.674275 | 398.521154 | 0 |

| | | | | | | |
|---|---|---|---|---|---|---|
| 1.38E-09 | 8.41E-09 | 99.0289288 | 197.527246 | 102.085711 | 256.556692 | 0 |
| 1.38E-09 | 8.42E-09 | 124.750728 | 109.440771 | 82.5966206 | 102.529889 | 0 |
| 1.38E-09 | 8.42E-09 | 94.8491364 | 90.7557617 | 101.157659 | 96.0347833 | 0 |
| 1.38E-09 | 8.43E-09 | 69.1273367 | 229.11381 | 179.11402 | 223.617225 | 0 |
| 1.38E-09 | 8.43E-09 | 60.7677518 | 311.416829 | 312.753496 | 295.063392 | 0 |
| 1.39E-09 | 8.47E-09 | 70.0919042 | 174.838306 | 178.185968 | 184.646588 | 0 |
| 1.39E-09 | 8.47E-09 | 138.254673 | 305.410933 | 274.703367 | 310.837221 | 0 |
| 1.39E-09 | 8.49E-09 | 197.414813 | 276.716097 | 273.775316 | 322.899561 | 16.6335277 |
| 1.40E-09 | 8.52E-09 | 62.3753642 | 310.304626 | 258.926485 | 295.527328 | 0 |
| 1.40E-09 | 8.52E-09 | 240.177305 | 123.232088 | 116.934542 | 100.674145 | 0 |
| 1.40E-09 | 8.52E-09 | 99.0289288 | 91.2006429 | 120.646749 | 92.7872303 | 0 |
| 1.40E-09 | 8.53E-09 | 76.8438766 | 280.720028 | 148.488307 | 250.061586 | 0 |
| 1.40E-09 | 8.54E-09 | 71.6995166 | 195.970162 | 156.840774 | 212.482757 | 0 |
| 1.40E-09 | 8.54E-09 | 68.8058142 | 342.558512 | 238.509343 | 212.018821 | 0 |
| 1.40E-09 | 8.56E-09 | 78.1299666 | 152.371806 | 180.042072 | 122.015208 | 0 |
| 1.40E-09 | 8.56E-09 | 76.2008316 | 132.797034 | 145.704151 | 124.798825 | 0 |
| 1.41E-09 | 8.56E-09 | 76.8438766 | 137.913167 | 119.718697 | 140.572654 | 0 |
| 1.41E-09 | 8.57E-09 | 106.102424 | 90.3108805 | 113.222334 | 89.5396772 | 0 |
| 1.41E-09 | 8.57E-09 | 89.3832539 | 103.212435 | 90.9490879 | 110.416804 | 0 |
| 1.41E-09 | 8.58E-09 | 65.2690667 | 279.162944 | 232.012979 | 233.359884 | 0 |
| 1.41E-09 | 8.60E-09 | 130.538133 | 551.430229 | 464.95401 | 484.813278 | 0 |
| 1.41E-09 | 8.61E-09 | 210.275712 | 261.145256 | 155.912722 | 188.822014 | 0 |
| 1.41E-09 | 8.61E-09 | 72.9856066 | 156.598177 | 145.704151 | 151.243185 | 0 |
| 1.42E-09 | 8.62E-09 | 71.6995166 | 191.298909 | 172.617657 | 255.628819 | 0 |
| 1.42E-09 | 8.65E-09 | 69.4488592 | 202.86582 | 172.617657 | 196.708928 | 0 |
| 1.42E-09 | 8.67E-09 | 69.4488592 | 203.088261 | 171.689605 | 204.131907 | 0 |
| 1.43E-09 | 8.72E-09 | 102.244154 | 500.713774 | 372.148819 | 438.419663 | 2.37621824 |
| 1.43E-09 | 8.72E-09 | 168.156265 | 148.367875 | 86.3088283 | 99.7462725 | 0 |
| 1.44E-09 | 8.74E-09 | 118.641801 | 368.13918 | 370.292715 | 382.283389 | 2.37621824 |
| 1.44E-09 | 8.74E-09 | 63.3399317 | 369.028943 | 252.430121 | 287.176478 | 0 |
| 1.44E-09 | 8.75E-09 | 56.5879593 | 365.914775 | 361.940248 | 378.571899 | 0 |
| 1.44E-09 | 8.75E-09 | 145.328168 | 271.155082 | 169.833501 | 208.307332 | 0 |
| 1.44E-09 | 8.77E-09 | 74.9147416 | 127.6809 | 146.632203 | 147.995632 | 0 |
| 1.45E-09 | 8.79E-09 | 81.9882365 | 135.466321 | 107.654022 | 110.416804 | 0 |
| 1.45E-09 | 8.82E-09 | 77.1653991 | 123.454529 | 118.790645 | 128.046378 | 0 |
| 1.45E-09 | 8.83E-09 | 68.1627692 | 272.489726 | 226.444668 | 417.0786 | 0 |
| 1.46E-09 | 8.85E-09 | 88.4186864 | 127.6809 | 91.8771398 | 110.88074 | 0 |
| 1.46E-09 | 8.85E-09 | 72.9856066 | 167.275326 | 163.337137 | 139.644782 | 0 |
| 1.46E-09 | 8.88E-09 | 70.4134267 | 302.296765 | 196.747006 | 203.204034 | 0 |
| 1.46E-09 | 8.88E-09 | 71.6995166 | 159.489905 | 227.37272 | 245.88616 | 0 |
| 1.46E-09 | 8.88E-09 | 64.6260217 | 274.26925 | 233.869083 | 351.663603 | 0 |
| 1.46E-09 | 8.88E-09 | 75.2362641 | 337.442379 | 170.761553 | 226.864778 | 0 |
| 1.46E-09 | 8.89E-09 | 83.2743265 | 135.688761 | 100.229607 | 117.839782 | 0 |

| | | | | | | |
|---|---|---|---|---|---|---|
| 1.47E-09 | 8.90E-09 | 70.0919042 | 171.501697 | 168.905449 | 193.925311 | 0 |
| 1.47E-09 | 8.90E-09 | 81.666714 | 112.110059 | 104.869867 | 131.757867 | 0 |
| 1.47E-09 | 8.90E-09 | 114.783531 | 161.936751 | 82.5966206 | 114.128293 | 0 |
| 1.47E-09 | 8.92E-09 | 105.780901 | 142.58442 | 150.344411 | 80.7248903 | 0 |
| 1.48E-09 | 8.96E-09 | 212.52637 | 1100.63605 | 1016.21685 | 1103.24017 | 16.6335277 |
| 1.48E-09 | 8.96E-09 | 76.5223541 | 139.470251 | 119.718697 | 128.97425 | 0 |
| 1.48E-09 | 8.97E-09 | 77.4869216 | 149.257638 | 115.078438 | 146.139888 | 0 |
| 1.48E-09 | 8.97E-09 | 73.3071291 | 169.499731 | 137.351684 | 158.202228 | 0 |
| 1.48E-09 | 8.97E-09 | 76.5223541 | 173.948543 | 126.215061 | 189.28595 | 0 |
| 1.48E-09 | 8.99E-09 | 88.4186864 | 170.611934 | 97.4454513 | 142.428398 | 0 |
| 1.48E-09 | 8.99E-09 | 73.3071291 | 139.470251 | 150.344411 | 142.428398 | 0 |
| 1.49E-09 | 9.01E-09 | 69.1273367 | 200.863855 | 170.761553 | 191.141694 | 0 |
| 1.49E-09 | 9.03E-09 | 69.7703817 | 210.651241 | 284.911939 | 185.110524 | 0 |
| 1.49E-09 | 9.04E-09 | 215.420072 | 907.112736 | 785.131922 | 846.21954 | 16.6335277 |
| 1.50E-09 | 9.05E-09 | 68.8058142 | 198.639449 | 204.171422 | 172.120312 | 0 |
| 1.50E-09 | 9.08E-09 | 75.5577866 | 127.458459 | 150.344411 | 128.510314 | 0 |
| 1.50E-09 | 9.09E-09 | 231.81772 | 110.775415 | 105.797919 | 85.828188 | 719.994128 |
| 1.50E-09 | 9.09E-09 | 70.7349491 | 162.159192 | 190.250643 | 226.864778 | 0 |
| 1.50E-09 | 9.09E-09 | 407.04748 | 627.282471 | 633.859459 | 647.654867 | 30.8908372 |
| 1.51E-09 | 9.12E-09 | 63.9829767 | 222.885474 | 279.343627 | 271.402649 | 0 |
| 1.51E-09 | 9.12E-09 | 74.5932191 | 245.351973 | 169.833501 | 156.346483 | 0 |
| 1.51E-09 | 9.13E-09 | 117.355711 | 394.609611 | 375.861026 | 428.213068 | 0 |
| 1.51E-09 | 9.14E-09 | 107.066991 | 257.586206 | 92.8051917 | 228.256586 | 0 |
| 1.52E-09 | 9.16E-09 | 105.459379 | 523.180273 | 296.048562 | 293.207648 | 7.12865473 |
| 1.52E-09 | 9.16E-09 | 61.0892743 | 270.042879 | 284.911939 | 304.806051 | 0 |
| 1.52E-09 | 9.20E-09 | 64.6260217 | 234.007503 | 220.876356 | 252.381266 | 0 |
| 1.54E-09 | 9.28E-09 | 83.9173715 | 187.73986 | 171.689605 | 106.705315 | 0 |
| 1.54E-09 | 9.28E-09 | 61.4107968 | 297.403072 | 276.559471 | 412.903175 | 0 |
| 1.55E-09 | 9.33E-09 | 69.1273367 | 260.477934 | 173.545708 | 256.092756 | 0 |
| 1.55E-09 | 9.35E-09 | 83.2743265 | 155.708415 | 101.157659 | 134.077548 | 0 |
| 1.55E-09 | 9.36E-09 | 183.589345 | 158.377702 | 90.9490879 | 95.106911 | 0 |
| 1.56E-09 | 9.39E-09 | 80.7021465 | 132.352152 | 104.869867 | 113.200421 | 0 |
| 1.56E-09 | 9.41E-09 | 75.2362641 | 198.86189 | 130.85532 | 173.048184 | 0 |
| 1.56E-09 | 9.41E-09 | 88.7402089 | 93.6474894 | 145.704151 | 114.592229 | 0 |
| 1.56E-09 | 9.42E-09 | 74.5932191 | 185.737895 | 188.394539 | 130.829995 | 0 |
| 1.56E-09 | 9.42E-09 | 72.9856066 | 134.354118 | 147.560255 | 154.954675 | 0 |
| 1.57E-09 | 9.43E-09 | 116.069621 | 176.840271 | 95.5893474 | 299.238818 | 0 |
| 1.57E-09 | 9.44E-09 | 225.065747 | 445.326066 | 463.097907 | 558.579126 | 28.5146189 |
| 1.57E-09 | 9.45E-09 | 81.3451915 | 222.885474 | 126.215061 | 150.315313 | 0 |
| 1.57E-09 | 9.46E-09 | 102.565676 | 109.218331 | 80.7405168 | 104.84957 | 0 |
| 1.57E-09 | 9.46E-09 | 85.8465065 | 153.039128 | 96.5173994 | 128.97425 | 0 |
| 1.57E-09 | 9.48E-09 | 76.5223541 | 169.944613 | 161.481034 | 119.231591 | 0 |
| 1.57E-09 | 9.48E-09 | 180.052598 | 316.755404 | 295.12051 | 321.043817 | 21.3859642 |

| | | | | | | |
|---|---|---|---|---|---|---|
| 1.58E-09 | 9.50E-09 | 62.6968867 | 255.13936 | 240.365446 | 271.866585 | 0 |
| 1.58E-09 | 9.50E-09 | 72.0210391 | 257.586206 | 156.840774 | 255.164883 | 0 |
| 1.58E-09 | 9.50E-09 | 62.6968867 | 322.538859 | 245.933758 | 259.340309 | 0 |
| 1.58E-09 | 9.51E-09 | 55.6233918 | 380.595854 | 342.451157 | 484.813278 | 0 |
| 1.58E-09 | 9.51E-09 | 67.1982017 | 212.430766 | 249.645966 | 186.966269 | 0 |
| 1.58E-09 | 9.53E-09 | 74.5932191 | 269.375557 | 152.200514 | 196.708928 | 0 |
| 1.59E-09 | 9.55E-09 | 125.715296 | 108.99589 | 77.956361 | 112.272549 | 0 |
| 1.59E-09 | 9.57E-09 | 67.5197242 | 182.846167 | 193.962851 | 191.141694 | 0 |
| 1.59E-09 | 9.58E-09 | 1005.07932 | 2576.75182 | 2419.43135 | 2481.59447 | 0 |
| 1.60E-09 | 9.61E-09 | 72.3425616 | 137.468286 | 147.560255 | 155.418611 | 0 |
| 1.60E-09 | 9.61E-09 | 69.7703817 | 160.157227 | 166.121293 | 170.728504 | 0 |
| 1.60E-09 | 9.61E-09 | 111.568306 | 191.076469 | 169.833501 | 163.305525 | 7.12865473 |
| 1.60E-09 | 9.62E-09 | 140.505331 | 234.674825 | 309.96934 | 295.063392 | 0 |
| 1.60E-09 | 9.62E-09 | 75.8793091 | 153.039128 | 117.862593 | 154.026802 | 0 |
| 1.60E-09 | 9.62E-09 | 122.500071 | 310.082186 | 313.681548 | 312.692966 | 0 |
| 1.61E-09 | 9.63E-09 | 77.8084441 | 120.11792 | 115.078438 | 146.603824 | 0 |
| 1.61E-09 | 9.63E-09 | 80.380624 | 229.336251 | 124.358957 | 168.408823 | 0 |
| 1.61E-09 | 9.63E-09 | 72.9856066 | 196.859924 | 142.919995 | 157.274355 | 0 |
| 1.61E-09 | 9.66E-09 | 80.380624 | 262.70234 | 133.639476 | 179.079354 | 0 |
| 1.62E-09 | 9.70E-09 | 346.279728 | 529.186169 | 461.241803 | 522.392106 | 21.3859642 |
| 1.62E-09 | 9.70E-09 | 79.4160565 | 109.440771 | 110.438178 | 133.613612 | 0 |
| 1.62E-09 | 9.73E-09 | 63.9829767 | 259.810612 | 214.379993 | 248.205841 | 0 |
| 1.62E-09 | 9.74E-09 | 108.996126 | 288.060567 | 241.293498 | 265.371479 | 11.8810912 |
| 1.63E-09 | 9.80E-09 | 87.1325964 | 300.962122 | 124.358957 | 202.276162 | 0 |
| 1.64E-09 | 9.82E-09 | 59.1601393 | 294.956226 | 297.904665 | 302.022435 | 0 |
| 1.64E-09 | 9.82E-09 | 72.6640841 | 147.255672 | 134.567528 | 140.572654 | 0 |
| 1.64E-09 | 9.82E-09 | 67.5197242 | 218.881543 | 174.47376 | 226.864778 | 0 |
| 1.64E-09 | 9.83E-09 | 71.6995166 | 141.694657 | 180.042072 | 157.274355 | 0 |
| 1.64E-09 | 9.84E-09 | 74.5932191 | 137.245846 | 124.358957 | 126.654569 | 0 |
| 1.64E-09 | 9.84E-09 | 75.2362641 | 151.037162 | 150.344411 | 118.303719 | 0 |
| 1.65E-09 | 9.89E-09 | 327.631424 | 982.520097 | 994.871655 | 974.265918 | 21.3859642 |
| 1.65E-09 | 9.89E-09 | 73.9501741 | 129.905306 | 128.999216 | 130.829995 | 0 |
| 1.65E-09 | 9.89E-09 | 127.644431 | 299.627478 | 267.278952 | 302.486371 | 0 |
| 1.66E-09 | 9.91E-09 | 63.6614542 | 223.775236 | 232.012979 | 237.999246 | 0 |
| 1.66E-09 | 9.92E-09 | 103.530244 | 86.3069499 | 160.552982 | 199.028609 | 0 |
| 1.66E-09 | 9.93E-09 | 73.3071291 | 170.389494 | 142.919995 | 133.613612 | 0 |
| 1.67E-09 | 9.97E-09 | 63.3399317 | 242.015365 | 226.444668 | 236.143501 | 0 |
| 1.67E-09 | 9.97E-09 | 86.811074 | 160.824548 | 209.739733 | 104.84957 | 0 |
| 1.67E-09 | 9.97E-09 | 65.5905892 | 223.330355 | 192.106747 | 226.400842 | 0 |
| 1.67E-09 | 1.00E-08 | 61.7323193 | 320.314453 | 267.278952 | 244.494352 | 0 |
| 1.69E-09 | 1.01E-08 | 70.0919042 | 173.948543 | 154.056618 | 154.954675 | 0 |
| 1.69E-09 | 1.01E-08 | 128.287476 | 182.846167 | 79.8124649 | 117.375846 | 0 |
| 1.69E-09 | 1.01E-08 | 90.3478214 | 306.968018 | 120.646749 | 333.106157 | 0 |

| | | | | | | |
|---|---|---|---|---|---|---|
| 1.71E-09 | 1.02E-08 | 64.6260217 | 205.757548 | 260.782589 | 221.297544 | 0 |
| 1.71E-09 | 1.02E-08 | 79.737579 | 181.956405 | 111.36623 | 147.06776 | 0 |
| 1.72E-09 | 1.03E-08 | 445.63018 | 136.800964 | 116.00649 | 163.305525 | 0 |
| 1.72E-09 | 1.03E-08 | 102.565676 | 193.523315 | 132.711424 | 167.480951 | 9.50487297 |
| 1.72E-09 | 1.03E-08 | 92.5984789 | 95.8718953 | 154.056618 | 102.529889 | 0 |
| 1.73E-09 | 1.03E-08 | 69.4488592 | 185.515454 | 158.696878 | 159.594036 | 0 |
| 1.73E-09 | 1.03E-08 | 72.9856066 | 148.590316 | 133.639476 | 180.471163 | 0 |
| 1.73E-09 | 1.03E-08 | 69.7703817 | 166.608004 | 149.416359 | 161.913717 | 0 |
| 1.74E-09 | 1.04E-08 | 129.573566 | 462.898873 | 268.207004 | 340.529135 | 2.37621824 |
| 1.74E-09 | 1.04E-08 | 79.4160565 | 187.517419 | 116.934542 | 136.861165 | 0 |
| 1.74E-09 | 1.04E-08 | 742.395444 | 1279.25585 | 1235.2371 | 1372.32314 | 0 |
| 1.75E-09 | 1.04E-08 | 203.52374 | 223.107914 | 193.034799 | 211.554885 | 16.6335277 |
| 1.76E-09 | 1.05E-08 | 143.720556 | 193.300875 | 252.430121 | 269.546904 | 2.37621824 |
| 1.76E-09 | 1.05E-08 | 184.875435 | 153.92889 | 299.760769 | 228.256586 | 2.37621824 |
| 1.76E-09 | 1.05E-08 | 71.6995166 | 188.184741 | 136.423632 | 184.182652 | 0 |
| 1.76E-09 | 1.05E-08 | 77.4869216 | 160.824548 | 118.790645 | 197.6368 | 0 |
| 1.77E-09 | 1.05E-08 | 131.824223 | 168.165088 | 189.322591 | 184.646588 | 16.6335277 |
| 1.77E-09 | 1.05E-08 | 145.649691 | 230.670894 | 190.250643 | 263.515734 | 2.37621824 |
| 1.77E-09 | 1.05E-08 | 68.1627692 | 192.633553 | 160.552982 | 206.451587 | 0 |
| 1.77E-09 | 1.05E-08 | 85.8465065 | 193.300875 | 119.718697 | 280.217435 | 0 |
| 1.77E-09 | 1.06E-08 | 103.208721 | 83.6376628 | 99.3015551 | 94.1790387 | 0 |
| 1.77E-09 | 1.06E-08 | 74.2716966 | 176.840271 | 128.071165 | 140.108718 | 0 |
| 1.78E-09 | 1.06E-08 | 77.8084441 | 148.367875 | 122.502853 | 109.488932 | 0 |
| 1.79E-09 | 1.07E-08 | 72.3425616 | 154.596212 | 127.143113 | 155.882547 | 0 |
| 1.80E-09 | 1.07E-08 | 73.3071291 | 169.722172 | 124.358957 | 159.1301 | 0 |
| 1.81E-09 | 1.07E-08 | 110.603739 | 185.737895 | 78.8844129 | 180.471163 | 0 |
| 1.81E-09 | 1.07E-08 | 70.0919042 | 189.074504 | 147.560255 | 160.521908 | 0 |
| 1.81E-09 | 1.07E-08 | 135.682493 | 494.930318 | 249.645966 | 343.776688 | 2.37621824 |
| 1.82E-09 | 1.08E-08 | 107.710036 | 155.041093 | 80.7405168 | 191.141694 | 0 |
| 1.82E-09 | 1.08E-08 | 111.568306 | 85.4171875 | 94.6612955 | 143.820207 | 0 |
| 1.82E-09 | 1.08E-08 | 72.3425616 | 166.608004 | 145.704151 | 129.438186 | 0 |
| 1.82E-09 | 1.08E-08 | 142.755988 | 230.893335 | 193.962851 | 217.122119 | 0 |
| 1.82E-09 | 1.08E-08 | 69.7703817 | 191.076469 | 144.776099 | 185.574461 | 0 |
| 1.82E-09 | 1.08E-08 | 69.1273367 | 217.102018 | 180.042072 | 156.346483 | 0 |
| 1.83E-09 | 1.08E-08 | 213.490937 | 356.127388 | 247.789862 | 264.907542 | 16.6335277 |
| 1.83E-09 | 1.09E-08 | 98.3858838 | 130.127747 | 78.8844129 | 128.97425 | 0 |
| 1.83E-09 | 1.09E-08 | 72.3425616 | 175.505627 | 137.351684 | 212.018821 | 0 |
| 1.83E-09 | 1.09E-08 | 69.1273367 | 167.275326 | 161.481034 | 148.923505 | 0 |
| 1.84E-09 | 1.09E-08 | 74.5932191 | 189.964266 | 131.783372 | 228.256586 | 0 |
| 1.84E-09 | 1.09E-08 | 104.816334 | 115.224227 | 103.941815 | 78.4052096 | 0 |
| 1.84E-09 | 1.09E-08 | 72.3425616 | 147.478113 | 127.143113 | 135.469356 | 0 |
| 1.84E-09 | 1.09E-08 | 91.6339114 | 98.0963013 | 153.128566 | 212.482757 | 0 |
| 1.85E-09 | 1.10E-08 | 164.619518 | 222.885474 | 156.840774 | 234.751693 | 2.37621824 |

| | | | | | | |
|---|---|---|---|---|---|---|
| 1.85E-09 | 1.10E-08 | 66.5551567 | 173.058781 | 185.610383 | 206.915523 | 0 |
| 1.86E-09 | 1.10E-08 | 92.9200014 | 114.334465 | 87.2368802 | 91.859358 | 0 |
| 1.87E-09 | 1.11E-08 | 179.08803 | 149.035197 | 99.3015551 | 83.9724434 | 0 |
| 1.87E-09 | 1.11E-08 | 63.9829767 | 194.190637 | 257.998433 | 242.174671 | 0 |
| 1.87E-09 | 1.11E-08 | 82.6312815 | 114.779346 | 100.229607 | 151.707121 | 0 |
| 1.87E-09 | 1.11E-08 | 882.579252 | 404.619438 | 320.177911 | 62.6313804 | 0 |
| 1.88E-09 | 1.12E-08 | 68.1627692 | 189.519385 | 161.481034 | 161.449781 | 0 |
| 1.88E-09 | 1.12E-08 | 199.343948 | 248.911023 | 142.919995 | 202.276162 | 2.37621824 |
| 1.89E-09 | 1.12E-08 | 66.2336342 | 174.170984 | 254.286225 | 248.669777 | 0 |
| 1.89E-09 | 1.12E-08 | 76.8438766 | 114.334465 | 133.639476 | 107.169251 | 0 |
| 1.89E-09 | 1.12E-08 | 71.3779941 | 150.814722 | 130.85532 | 166.553078 | 0 |
| 1.90E-09 | 1.12E-08 | 71.6995166 | 127.6809 | 153.128566 | 139.644782 | 0 |
| 1.91E-09 | 1.13E-08 | 62.0538418 | 257.808647 | 213.451941 | 253.773075 | 0 |
| 1.92E-09 | 1.13E-08 | 79.094534 | 106.549044 | 137.351684 | 105.777442 | 0 |
| 1.92E-09 | 1.14E-08 | 54.9803468 | 317.422725 | 346.163365 | 356.302964 | 0 |
| 1.92E-09 | 1.14E-08 | 63.9829767 | 191.076469 | 256.142329 | 235.679565 | 0 |
| 1.93E-09 | 1.14E-08 | 103.208721 | 84.0825439 | 148.488307 | 102.065953 | 0 |
| 1.93E-09 | 1.14E-08 | 72.3425616 | 142.139539 | 121.574801 | 146.603824 | 0 |
| 1.93E-09 | 1.14E-08 | 64.3044992 | 251.135429 | 200.459214 | 201.812226 | 0 |
| 1.93E-09 | 1.14E-08 | 66.5551567 | 207.092192 | 188.394539 | 166.089142 | 0 |
| 1.94E-09 | 1.15E-08 | 65.2690667 | 183.95837 | 185.610383 | 199.492545 | 0 |
| 1.95E-09 | 1.15E-08 | 72.0210391 | 128.570662 | 149.416359 | 128.510314 | 0 |
| 1.95E-09 | 1.15E-08 | 71.3779941 | 132.129712 | 130.85532 | 137.789037 | 0 |
| 1.95E-09 | 1.15E-08 | 513.471426 | 219.103984 | 83.5246725 | 187.430205 | 0 |
| 1.95E-09 | 1.15E-08 | 71.0564716 | 134.131677 | 141.063891 | 166.553078 | 0 |
| 1.95E-09 | 1.15E-08 | 60.1247068 | 237.788993 | 268.207004 | 266.299351 | 0 |
| 1.95E-09 | 1.15E-08 | 99.3504513 | 111.665177 | 88.1649321 | 170.728504 | 0 |
| 1.96E-09 | 1.16E-08 | 65.5905892 | 185.515454 | 180.042072 | 219.4418 | 0 |
| 1.98E-09 | 1.17E-08 | 78.4514891 | 158.155261 | 166.121293 | 104.84957 | 0 |
| 1.98E-09 | 1.17E-08 | 72.0210391 | 160.602108 | 193.034799 | 130.366059 | 0 |
| 1.99E-09 | 1.17E-08 | 73.6286516 | 119.450598 | 161.481034 | 175.831801 | 0 |
| 1.99E-09 | 1.17E-08 | 0 | 325.430587 | 432.472193 | 0 | 0 |
| 1.99E-09 | 1.17E-08 | 83.595849 | 140.804895 | 90.9490879 | 134.541484 | 0 |
| 1.99E-09 | 1.17E-08 | 63.3399317 | 224.887439 | 200.459214 | 207.37946 | 0 |
| 1.99E-09 | 1.17E-08 | 63.0184092 | 210.873682 | 204.171422 | 229.648395 | 0 |
| 1.99E-09 | 1.17E-08 | 71.6995166 | 157.71038 | 134.567528 | 128.510314 | 0 |
| 1.99E-09 | 1.17E-08 | 161.725816 | 86.7518311 | 128.071165 | 92.3232941 | 0 |
| 1.99E-09 | 1.17E-08 | 67.8412467 | 180.844202 | 165.193241 | 152.171058 | 0 |
| 2.01E-09 | 1.19E-08 | 2292.13387 | 555.6566 | 623.650888 | 746.009331 | 28.5146189 |
| 2.02E-09 | 1.19E-08 | 100.315019 | 88.3089152 | 88.1649321 | 90.4675495 | 0 |
| 2.02E-09 | 1.19E-08 | 66.2336342 | 170.834375 | 192.106747 | 168.872759 | 0 |
| 2.03E-09 | 1.19E-08 | 127.965953 | 269.820439 | 246.86181 | 288.10435 | 0 |
| 2.03E-09 | 1.20E-08 | 133.753358 | 117.893514 | 94.6612955 | 80.2609542 | 0 |

| | | | | | | |
|---|---|---|---|---|---|---|
| 2.04E-09 | 1.20E-08 | 63.0184092 | 195.970162 | 284.911939 | 254.237011 | 0 |
| 2.05E-09 | 1.20E-08 | 37.618132 | 85.4171875 | 63.1075304 | 70.518295 | 1306.92003 |
| 2.06E-09 | 1.21E-08 | 75.8793091 | 134.798999 | 111.36623 | 109.488932 | 0 |
| 2.07E-09 | 1.22E-08 | 125.393773 | 324.540824 | 282.127783 | 287.640414 | 30.8908372 |
| 2.07E-09 | 1.22E-08 | 465.886097 | 926.242627 | 638.499719 | 877.767198 | 0 |
| 2.07E-09 | 1.22E-08 | 21.8635297 | 24.4684652 | 22.273246 | 22.7328714 | 95.0487297 |
| 2.08E-09 | 1.22E-08 | 77.4869216 | 101.655351 | 103.941815 | 117.375846 | 0 |
| 2.08E-09 | 1.22E-08 | 123.786161 | 98.0963013 | 90.9490879 | 84.9003157 | 0 |
| 2.09E-09 | 1.23E-08 | 84.5604165 | 97.8738607 | 88.1649321 | 96.9626556 | 0 |
| 2.09E-09 | 1.23E-08 | 63.6614542 | 219.993746 | 186.538435 | 204.595843 | 0 |
| 2.09E-09 | 1.23E-08 | 269.435852 | 411.070215 | 505.788295 | 528.423276 | 0 |
| 2.10E-09 | 1.23E-08 | 57.5525268 | 318.090047 | 286.768042 | 261.659989 | 0 |
| 2.10E-09 | 1.23E-08 | 74.9147416 | 109.218331 | 128.999216 | 111.808612 | 0 |
| 2.10E-09 | 1.23E-08 | 66.8766792 | 153.484009 | 185.610383 | 173.048184 | 0 |
| 2.10E-09 | 1.23E-08 | 87.1325964 | 164.828479 | 128.999216 | 91.3954218 | 0 |
| 2.11E-09 | 1.24E-08 | 63.3399317 | 208.871716 | 189.322591 | 208.307332 | 0 |
| 2.11E-09 | 1.24E-08 | 57.5525268 | 266.70627 | 674.693744 | 627.705613 | 0 |
| 2.11E-09 | 1.24E-08 | 58.8386168 | 262.70234 | 255.214277 | 257.020628 | 0 |
| 2.11E-09 | 1.24E-08 | 62.0538418 | 219.993746 | 204.171422 | 239.85499 | 0 |
| 2.12E-09 | 1.24E-08 | 76.2008316 | 112.110059 | 103.941815 | 106.241379 | 0 |
| 2.12E-09 | 1.24E-08 | 69.1273367 | 139.915133 | 162.409085 | 190.677758 | 0 |
| 2.12E-09 | 1.24E-08 | 68.1627692 | 159.489905 | 141.991943 | 167.944887 | 0 |
| 2.12E-09 | 1.24E-08 | 88.7402089 | 88.7537964 | 169.833501 | 150.779249 | 0 |
| 2.13E-09 | 1.25E-08 | 60.1247068 | 270.932642 | 247.789862 | 224.545097 | 0 |
| 2.14E-09 | 1.25E-08 | 66.8766792 | 266.038949 | 164.265189 | 220.833608 | 0 |
| 2.14E-09 | 1.25E-08 | 75.2362641 | 103.434875 | 128.071165 | 132.685739 | 0 |
| 2.15E-09 | 1.26E-08 | 103.851766 | 183.95837 | 193.034799 | 222.689353 | 11.8810912 |
| 2.15E-09 | 1.26E-08 | 56.2664368 | 373.032874 | 262.638692 | 400.376899 | 0 |
| 2.15E-09 | 1.26E-08 | 88.4186864 | 93.6474894 | 86.3088283 | 92.3232941 | 0 |
| 2.16E-09 | 1.26E-08 | 70.7349491 | 136.133643 | 123.430905 | 138.252973 | 0 |
| 2.17E-09 | 1.27E-08 | 60.4462293 | 225.777201 | 230.156875 | 243.566479 | 0 |
| 2.17E-09 | 1.27E-08 | 74.5932191 | 119.005717 | 106.72597 | 133.613612 | 0 |
| 2.17E-09 | 1.27E-08 | 112.854396 | 131.46239 | 73.3161014 | 111.808612 | 0 |
| 2.17E-09 | 1.27E-08 | 74.5932191 | 112.77738 | 112.294282 | 138.252973 | 0 |
| 2.17E-09 | 1.27E-08 | 384.219383 | 145.253707 | 98.3735032 | 148.459568 | 0 |
| 2.17E-09 | 1.27E-08 | 74.5932191 | 143.251742 | 107.654022 | 150.779249 | 0 |
| 2.18E-09 | 1.27E-08 | 91.6339114 | 131.23995 | 104.869867 | 83.0445711 | 0 |
| 2.18E-09 | 1.28E-08 | 84.5604165 | 213.542969 | 107.654022 | 138.252973 | 0 |
| 2.19E-09 | 1.28E-08 | 70.0919042 | 156.153296 | 129.927268 | 135.933292 | 0 |
| 2.20E-09 | 1.28E-08 | 78.4514891 | 95.6494547 | 103.013763 | 103.457762 | 0 |
| 2.20E-09 | 1.28E-08 | 236.640557 | 470.239412 | 459.385699 | 473.214874 | 0 |
| 2.20E-09 | 1.28E-08 | 20.8989622 | 14.4586385 | 13.9207788 | 17.6295737 | 85.5438568 |
| 2.20E-09 | 1.28E-08 | 0 | 0 | 0 | 0 | 11.8810912 |

| | | | | | | |
|---|---|---|---|---|---|---|
| 2.20E-09 | 1.29E-08 | 94.2060914 | 165.940682 | 125.287009 | 131.293931 | 11.8810912 |
| 2.21E-09 | 1.29E-08 | 66.5551567 | 237.788993 | 157.768826 | 233.359884 | 0 |
| 2.21E-09 | 1.29E-08 | 68.4842917 | 149.702519 | 172.617657 | 139.180845 | 0 |
| 2.21E-09 | 1.29E-08 | 137.290106 | 332.103805 | 213.451941 | 250.989458 | 0 |
| 2.22E-09 | 1.30E-08 | 70.7349491 | 143.251742 | 120.646749 | 144.284143 | 0 |
| 2.23E-09 | 1.30E-08 | 72.0210391 | 242.905127 | 136.423632 | 186.502333 | 0 |
| 2.23E-09 | 1.30E-08 | 117.998756 | 338.332141 | 279.343627 | 317.332327 | 0 |
| 2.23E-09 | 1.30E-08 | 93.2415239 | 88.0864746 | 85.3807764 | 92.7872303 | 0 |
| 2.23E-09 | 1.30E-08 | 91.9554339 | 146.36591 | 103.941815 | 245.88616 | 0 |
| 2.24E-09 | 1.31E-08 | 85.2034615 | 93.4250488 | 89.092984 | 116.91191 | 0 |
| 2.24E-09 | 1.31E-08 | 65.5905892 | 172.6139 | 160.552982 | 192.533503 | 0 |
| 2.25E-09 | 1.31E-08 | 69.1273367 | 151.704484 | 133.639476 | 137.325101 | 0 |
| 2.26E-09 | 1.32E-08 | 64.9475442 | 165.940682 | 180.970124 | 175.367865 | 0 |
| 2.27E-09 | 1.32E-08 | 69.1273367 | 204.645345 | 149.416359 | 260.268181 | 0 |
| 2.27E-09 | 1.32E-08 | 69.1273367 | 153.484009 | 155.912722 | 128.510314 | 0 |
| 2.27E-09 | 1.32E-08 | 59.4816618 | 283.834196 | 292.336354 | 217.122119 | 0 |
| 2.27E-09 | 1.32E-08 | 68.8058142 | 148.812756 | 154.98467 | 130.829995 | 0 |
| 2.28E-09 | 1.33E-08 | 54.9803468 | 283.611755 | 328.530379 | 315.940519 | 0 |
| 2.28E-09 | 1.33E-08 | 97.7428388 | 327.654993 | 199.531162 | 259.340309 | 4.75243649 |
| 2.28E-09 | 1.33E-08 | 560.413711 | 858.175806 | 753.578157 | 746.937204 | 26.1384007 |
| 2.28E-09 | 1.33E-08 | 122.500071 | 247.131498 | 218.0922 | 250.525522 | 4.75243649 |
| 2.28E-09 | 1.33E-08 | 66.2336342 | 170.834375 | 164.265189 | 151.707121 | 0 |
| 2.29E-09 | 1.33E-08 | 81.3451915 | 111.665177 | 87.2368802 | 112.272549 | 0 |
| 2.29E-09 | 1.33E-08 | 56.2664368 | 256.251562 | 335.026742 | 315.940519 | 0 |
| 2.29E-09 | 1.33E-08 | 73.9501741 | 168.165088 | 111.36623 | 146.603824 | 0 |
| 2.30E-09 | 1.33E-08 | 72.0210391 | 114.556905 | 154.98467 | 155.418611 | 0 |
| 2.30E-09 | 1.34E-08 | 65.5905892 | 177.730033 | 154.98467 | 184.182652 | 0 |
| 2.30E-09 | 1.34E-08 | 70.4134267 | 127.013578 | 123.430905 | 140.108718 | 0 |
| 2.31E-09 | 1.34E-08 | 65.2690667 | 296.958191 | 189.322591 | 199.492545 | 0 |
| 2.31E-09 | 1.34E-08 | 67.5197242 | 150.1474 | 141.991943 | 168.408823 | 0 |
| 2.32E-09 | 1.35E-08 | 149.186438 | 168.609969 | 180.970124 | 198.100737 | 2.37621824 |
| 2.32E-09 | 1.35E-08 | 123.786161 | 270.48776 | 255.214277 | 280.217435 | 0 |
| 2.33E-09 | 1.35E-08 | 81.023669 | 98.763623 | 130.85532 | 171.19244 | 0 |
| 2.34E-09 | 1.36E-08 | 75.8793091 | 145.031266 | 103.013763 | 120.159463 | 0 |
| 2.34E-09 | 1.36E-08 | 53.0512119 | 321.204215 | 328.530379 | 344.240624 | 0 |
| 2.34E-09 | 1.36E-08 | 140.183808 | 278.940503 | 301.616873 | 283.464988 | 2.37621824 |
| 2.35E-09 | 1.36E-08 | 78.7730116 | 223.330355 | 172.617657 | 111.808612 | 0 |
| 2.35E-09 | 1.36E-08 | 75.5577866 | 161.49187 | 106.72597 | 171.19244 | 0 |
| 2.35E-09 | 1.36E-08 | 70.4134267 | 195.30284 | 129.927268 | 207.843396 | 0 |
| 2.36E-09 | 1.37E-08 | 71.6995166 | 133.019474 | 122.502853 | 169.800631 | 0 |
| 2.36E-09 | 1.37E-08 | 63.9829767 | 169.277291 | 205.099474 | 184.646588 | 0 |
| 2.38E-09 | 1.38E-08 | 67.5197242 | 138.135608 | 146.632203 | 154.026802 | 0 |
| 2.38E-09 | 1.38E-08 | 142.434466 | 265.149186 | 302.544925 | 281.145308 | 0 |

| | | | | | | |
|---|---|---|---|---|---|---|
| 2.38E-09 | 1.38E-08 | 115.105054 | 350.121492 | 304.401029 | 301.558498 | 2.37621824 |
| 2.38E-09 | 1.38E-08 | 62.6968867 | 199.306771 | 180.042072 | 212.018821 | 0 |
| 2.39E-09 | 1.38E-08 | 72.9856066 | 148.367875 | 117.862593 | 115.056166 | 0 |
| 2.39E-09 | 1.39E-08 | 62.0538418 | 219.326424 | 199.531162 | 189.749886 | 0 |
| 2.39E-09 | 1.39E-08 | 72.6640841 | 157.043058 | 116.00649 | 124.798825 | 0 |
| 2.40E-09 | 1.39E-08 | 83.595849 | 91.6455241 | 98.3735032 | 87.2199964 | 0 |
| 2.41E-09 | 1.39E-08 | 73.6286516 | 158.377702 | 119.718697 | 204.131907 | 0 |
| 2.41E-09 | 1.40E-08 | 68.8058142 | 143.029301 | 139.207788 | 185.110524 | 0 |
| 2.41E-09 | 1.40E-08 | 57.8740493 | 24.9133464 | 22.273246 | 17.1656376 | 318.413245 |
| 2.42E-09 | 1.40E-08 | 57.5525268 | 353.680542 | 244.077654 | 277.433818 | 0 |
| 2.42E-09 | 1.40E-08 | 68.1627692 | 164.383598 | 130.85532 | 154.954675 | 0 |
| 2.43E-09 | 1.41E-08 | 63.9829767 | 177.062712 | 232.941031 | 180.935099 | 0 |
| 2.43E-09 | 1.41E-08 | 69.7703817 | 235.787028 | 150.344411 | 157.738291 | 0 |
| 2.43E-09 | 1.41E-08 | 90.9908664 | 205.979989 | 90.0210359 | 160.521908 | 0 |
| 2.43E-09 | 1.41E-08 | 88.7402089 | 115.001786 | 78.8844129 | 112.736485 | 0 |
| 2.43E-09 | 1.41E-08 | 82.309759 | 110.330534 | 105.797919 | 84.9003157 | 0 |
| 2.44E-09 | 1.41E-08 | 68.4842917 | 142.58442 | 128.999216 | 156.346483 | 0 |
| 2.44E-09 | 1.41E-08 | 605.105338 | 189.296944 | 105.797919 | 166.089142 | 0 |
| 2.44E-09 | 1.41E-08 | 81.3451915 | 101.655351 | 86.3088283 | 109.952868 | 0 |
| 2.45E-09 | 1.42E-08 | 72.6640841 | 120.785242 | 106.72597 | 131.293931 | 0 |
| 2.45E-09 | 1.42E-08 | 80.7021465 | 139.247811 | 90.0210359 | 121.551272 | 0 |
| 2.45E-09 | 1.42E-08 | 85.2034615 | 98.763623 | 86.3088283 | 125.726697 | 0 |
| 2.46E-09 | 1.42E-08 | 189.698273 | 128.348222 | 82.5966206 | 101.138081 | 0 |
| 2.46E-09 | 1.42E-08 | 72.6640841 | 167.497766 | 113.222334 | 173.048184 | 0 |
| 2.47E-09 | 1.43E-08 | 95.1706589 | 150.36984 | 84.4527244 | 102.065953 | 0 |
| 2.47E-09 | 1.43E-08 | 74.2716966 | 180.17688 | 113.222334 | 138.252973 | 0 |
| 2.47E-09 | 1.43E-08 | 76.8438766 | 106.993925 | 93.7332436 | 115.520102 | 0 |
| 2.48E-09 | 1.43E-08 | 73.3071291 | 153.92889 | 223.660512 | 128.046378 | 0 |
| 2.48E-09 | 1.43E-08 | 119.927891 | 359.686438 | 253.358173 | 314.084774 | 4.75243649 |
| 2.48E-09 | 1.43E-08 | 80.0591015 | 258.475968 | 115.078438 | 185.574461 | 0 |
| 2.49E-09 | 1.43E-08 | 118.963324 | 182.178845 | 335.954794 | 90.0036134 | 0 |
| 2.50E-09 | 1.44E-08 | 135.360971 | 218.659102 | 78.8844129 | 117.375846 | 0 |
| 2.50E-09 | 1.44E-08 | 79.094534 | 114.334465 | 122.502853 | 89.0757411 | 0 |
| 2.50E-09 | 1.44E-08 | 75.5577866 | 143.251742 | 99.3015551 | 126.654569 | 0 |
| 2.50E-09 | 1.44E-08 | 49.5144644 | 355.682507 | 474.23453 | 378.571899 | 0 |
| 2.51E-09 | 1.44E-08 | 74.9147416 | 99.2085042 | 128.071165 | 112.272549 | 0 |
| 2.53E-09 | 1.46E-08 | 111.568306 | 219.771305 | 77.956361 | 156.346483 | 0 |
| 2.53E-09 | 1.46E-08 | 67.5197242 | 141.027336 | 141.991943 | 133.613612 | 0 |
| 2.53E-09 | 1.46E-08 | 80.380624 | 122.342326 | 95.5893474 | 93.7151026 | 0 |
| 2.54E-09 | 1.46E-08 | 114.462009 | 216.434696 | 239.437395 | 238.463182 | 2.37621824 |
| 2.54E-09 | 1.46E-08 | 75.2362641 | 422.637126 | 256.142329 | 322.899561 | 2.37621824 |
| 2.54E-09 | 1.46E-08 | 67.8412467 | 160.379667 | 175.401812 | 130.366059 | 0 |
| 2.54E-09 | 1.46E-08 | 72.6640841 | 140.360014 | 139.207788 | 105.313506 | 0 |

| | | | | | | |
|---|---|---|---|---|---|---|
| 2.55E-09 | 1.46E-08 | 62.6968867 | 208.426835 | 184.682331 | 175.831801 | 0 |
| 2.55E-09 | 1.46E-08 | 74.5932191 | 136.578524 | 100.229607 | 141.964462 | 0 |
| 2.55E-09 | 1.47E-08 | 64.9475442 | 170.611934 | 171.689605 | 150.315313 | 0 |
| 2.55E-09 | 1.47E-08 | 54.0157793 | 296.73575 | 288.624146 | 315.940519 | 0 |
| 2.56E-09 | 1.47E-08 | 55.6233918 | 262.035018 | 644.996082 | 509.401894 | 0 |
| 2.56E-09 | 1.47E-08 | 73.9501741 | 105.2144 | 103.013763 | 118.767655 | 0 |
| 2.56E-09 | 1.47E-08 | 63.9829767 | 190.854028 | 159.62493 | 176.759674 | 0 |
| 2.57E-09 | 1.47E-08 | 70.4134267 | 134.354118 | 115.078438 | 122.94308 | 0 |
| 2.57E-09 | 1.48E-08 | 69.1273367 | 184.625692 | 127.143113 | 155.882547 | 0 |
| 2.58E-09 | 1.48E-08 | 59.8031843 | 254.027157 | 204.171422 | 221.76148 | 0 |
| 2.59E-09 | 1.49E-08 | 89.3832539 | 85.6396281 | 90.9490879 | 85.828188 | 0 |
| 2.60E-09 | 1.49E-08 | 79.737579 | 86.9742716 | 120.646749 | 116.447974 | 0 |
| 2.62E-09 | 1.50E-08 | 63.0184092 | 178.619796 | 169.833501 | 179.079354 | 0 |
| 2.62E-09 | 1.51E-08 | 68.4842917 | 122.564766 | 134.567528 | 141.500526 | 0 |
| 2.63E-09 | 1.51E-08 | 76.2008316 | 161.49187 | 103.013763 | 123.870952 | 0 |
| 2.63E-09 | 1.51E-08 | 68.8058142 | 147.033232 | 121.574801 | 134.541484 | 0 |
| 2.63E-09 | 1.51E-08 | 55.3018693 | 273.379488 | 318.321808 | 260.732117 | 0 |
| 2.63E-09 | 1.51E-08 | 74.2716966 | 97.8738607 | 127.143113 | 129.902122 | 0 |
| 2.63E-09 | 1.51E-08 | 74.2716966 | 125.456494 | 97.4454513 | 121.551272 | 0 |
| 2.64E-09 | 1.51E-08 | 66.5551567 | 157.487939 | 140.135839 | 140.572654 | 0 |
| 2.64E-09 | 1.51E-08 | 84.881939 | 182.623726 | 130.85532 | 349.807858 | 0 |
| 2.64E-09 | 1.51E-08 | 115.748099 | 309.192423 | 283.983887 | 348.879986 | 0 |
| 2.66E-09 | 1.52E-08 | 66.5551567 | 196.859924 | 143.848047 | 156.810419 | 0 |
| 2.66E-09 | 1.53E-08 | 72.9856066 | 102.767554 | 107.654022 | 113.200421 | 0 |
| 2.67E-09 | 1.53E-08 | 72.9856066 | 148.367875 | 104.869867 | 128.510314 | 0 |
| 2.67E-09 | 1.53E-08 | 72.3425616 | 112.999821 | 108.582074 | 136.861165 | 0 |
| 2.68E-09 | 1.53E-08 | 60.7677518 | 192.188672 | 214.379993 | 195.31712 | 0 |
| 2.70E-09 | 1.54E-08 | 66.8766792 | 325.208146 | 165.193241 | 225.47297 | 0 |
| 2.70E-09 | 1.54E-08 | 69.4488592 | 130.350187 | 136.423632 | 114.592229 | 0 |
| 2.70E-09 | 1.55E-08 | 82.6312815 | 114.779346 | 82.5966206 | 104.385634 | 0 |
| 2.71E-09 | 1.55E-08 | 101.601109 | 82.970341 | 130.85532 | 199.956481 | 0 |
| 2.71E-09 | 1.55E-08 | 69.1273367 | 173.948543 | 128.999216 | 133.613612 | 0 |
| 2.71E-09 | 1.55E-08 | 138.897718 | 174.615865 | 253.358173 | 250.989458 | 2.37621824 |
| 2.71E-09 | 1.55E-08 | 117.677234 | 100.320707 | 84.4527244 | 82.1166988 | 0 |
| 2.71E-09 | 1.55E-08 | 93.8845689 | 87.4191528 | 83.5246725 | 86.7560603 | 0 |
| 2.71E-09 | 1.55E-08 | 130.538133 | 215.989815 | 263.566744 | 211.090949 | 0 |
| 2.72E-09 | 1.56E-08 | 63.6614542 | 309.859745 | 176.329864 | 229.648395 | 0 |
| 2.72E-09 | 1.56E-08 | 152.080141 | 157.043058 | 240.365446 | 208.771268 | 0 |
| 2.73E-09 | 1.56E-08 | 72.0210391 | 139.02537 | 112.294282 | 110.88074 | 0 |
| 2.74E-09 | 1.56E-08 | 56.2664368 | 271.599963 | 261.710641 | 238.463182 | 0 |
| 2.75E-09 | 1.57E-08 | 59.1601393 | 274.714132 | 196.747006 | 275.578074 | 0 |
| 2.75E-09 | 1.57E-08 | 68.8058142 | 151.704484 | 117.862593 | 142.892335 | 0 |
| 2.75E-09 | 1.57E-08 | 79.737579 | 84.9723063 | 109.510126 | 110.88074 | 0 |

| | | | | | | |
|---|---|---|---|---|---|---|
| 2.75E-09 | 1.57E-08 | 74.9147416 | 125.011613 | 94.6612955 | 127.582442 | 0 |
| 2.76E-09 | 1.57E-08 | 64.3044992 | 185.070573 | 147.560255 | 180.471163 | 0 |
| 2.76E-09 | 1.57E-08 | 59.8031843 | 205.312667 | 253.358173 | 202.740098 | 0 |
| 2.76E-09 | 1.57E-08 | 76.5223541 | 115.224227 | 92.8051917 | 100.674145 | 0 |
| 2.76E-09 | 1.57E-08 | 63.9829767 | 166.163123 | 154.98467 | 192.069567 | 0 |
| 2.76E-09 | 1.58E-08 | 69.1273367 | 144.363944 | 132.711424 | 115.984038 | 0 |
| 2.76E-09 | 1.58E-08 | 63.3399317 | 221.55083 | 163.337137 | 182.326907 | 0 |
| 2.76E-09 | 1.58E-08 | 66.5551567 | 135.466321 | 148.488307 | 135.469356 | 0 |
| 2.77E-09 | 1.58E-08 | 241.141872 | 112.332499 | 95.5893474 | 110.88074 | 0 |
| 2.77E-09 | 1.58E-08 | 63.3399317 | 154.596212 | 189.322591 | 184.646588 | 0 |
| 2.78E-09 | 1.58E-08 | 74.2716966 | 123.89941 | 158.696878 | 103.921698 | 0 |
| 2.78E-09 | 1.59E-08 | 87.7756414 | 80.9683756 | 145.704151 | 125.726697 | 0 |
| 2.80E-09 | 1.60E-08 | 71.0564716 | 149.035197 | 173.545708 | 112.736485 | 0 |
| 2.80E-09 | 1.60E-08 | 63.9829767 | 151.037162 | 166.121293 | 162.377653 | 0 |
| 2.80E-09 | 1.60E-08 | 75.2362641 | 196.859924 | 116.934542 | 245.88616 | 0 |
| 2.82E-09 | 1.61E-08 | 182.303255 | 125.901375 | 180.970124 | 182.790844 | 19.0097459 |
| 2.83E-09 | 1.61E-08 | 67.5197242 | 123.676969 | 157.768826 | 165.16127 | 0 |
| 2.87E-09 | 1.63E-08 | 1.28608998 | 0.88976237 | 0 | 0 | 23.7621824 |
| 2.87E-09 | 1.63E-08 | 100.315019 | 74.0727173 | 155.912722 | 138.252973 | 0 |
| 2.88E-09 | 1.64E-08 | 66.8766792 | 148.812756 | 127.143113 | 136.397228 | 0 |
| 2.89E-09 | 1.64E-08 | 69.4488592 | 194.190637 | 129.927268 | 236.607437 | 0 |
| 2.89E-09 | 1.65E-08 | 60.1247068 | 193.523315 | 207.883629 | 189.28595 | 0 |
| 2.90E-09 | 1.65E-08 | 99.3504513 | 357.684473 | 592.097123 | 469.039449 | 0 |
| 2.91E-09 | 1.65E-08 | 72.3425616 | 114.779346 | 108.582074 | 148.923505 | 0 |
| 2.91E-09 | 1.65E-08 | 60.4462293 | 236.899231 | 341.523105 | 201.812226 | 0 |
| 2.91E-09 | 1.65E-08 | 72.3425616 | 52.2735392 | 45.4745439 | 50.1051043 | 263.760225 |
| 2.91E-09 | 1.66E-08 | 63.0184092 | 156.820618 | 201.387266 | 173.976057 | 0 |
| 2.91E-09 | 1.66E-08 | 72.9856066 | 99.6533854 | 114.150386 | 102.529889 | 0 |
| 2.92E-09 | 1.66E-08 | 340.492323 | 173.058781 | 98.3735032 | 106.705315 | 0 |
| 2.93E-09 | 1.66E-08 | 130.859656 | 107.438806 | 70.5319457 | 154.026802 | 0 |
| 2.93E-09 | 1.67E-08 | 72.0210391 | 122.564766 | 183.75428 | 122.94308 | 0 |
| 2.94E-09 | 1.67E-08 | 66.2336342 | 148.145435 | 199.531162 | 260.732117 | 0 |
| 2.94E-09 | 1.67E-08 | 63.3399317 | 193.523315 | 150.344411 | 179.543291 | 0 |
| 2.95E-09 | 1.67E-08 | 108.353081 | 75.6298014 | 99.3015551 | 91.859358 | 0 |
| 2.95E-09 | 1.68E-08 | 64.6260217 | 233.562622 | 149.416359 | 192.533503 | 0 |
| 2.95E-09 | 1.68E-08 | 72.3425616 | 102.767554 | 102.085711 | 109.488932 | 0 |
| 2.96E-09 | 1.68E-08 | 65.9121117 | 144.586385 | 131.783372 | 139.180845 | 0 |
| 2.96E-09 | 1.68E-08 | 66.8766792 | 151.482043 | 130.85532 | 128.510314 | 0 |
| 2.96E-09 | 1.68E-08 | 65.2690667 | 195.525281 | 150.344411 | 147.531696 | 0 |
| 2.97E-09 | 1.68E-08 | 144.363601 | 76.5195638 | 92.8051917 | 124.334889 | 0 |
| 2.97E-09 | 1.68E-08 | 74.9147416 | 263.147221 | 121.574801 | 185.110524 | 0 |
| 2.98E-09 | 1.69E-08 | 121.535504 | 239.346077 | 193.034799 | 225.936906 | 4.75243649 |
| 2.98E-09 | 1.69E-08 | 60.4462293 | 179.954439 | 189.322591 | 202.276162 | 0 |

| | | | | | | |
|---|---|---|---|---|---|---|
| 2.98E-09 | 1.69E-08 | 67.5197242 | 131.907271 | 124.358957 | 122.479144 | 0 |
| 3.00E-09 | 1.70E-08 | 69.7703817 | 235.564587 | 128.071165 | 184.182652 | 0 |
| 3.00E-09 | 1.70E-08 | 71.3779941 | 112.110059 | 101.157659 | 115.520102 | 0 |
| 3.00E-09 | 1.70E-08 | 54.6588243 | 270.710201 | 253.358173 | 262.123925 | 0 |
| 3.01E-09 | 1.70E-08 | 125.393773 | 213.320528 | 307.185185 | 247.277969 | 2.37621824 |
| 3.01E-09 | 1.70E-08 | 141.469898 | 278.050741 | 265.422848 | 317.332327 | 0 |
| 3.01E-09 | 1.71E-08 | 120.249414 | 133.686796 | 107.654022 | 137.325101 | 7.12865473 |
| 3.01E-09 | 1.71E-08 | 66.8766792 | 148.812756 | 127.143113 | 175.831801 | 0 |
| 3.02E-09 | 1.71E-08 | 65.5905892 | 136.356083 | 138.279736 | 136.397228 | 0 |
| 3.02E-09 | 1.71E-08 | 94.2060914 | 113.222262 | 76.1002572 | 92.3232941 | 0 |
| 3.03E-09 | 1.71E-08 | 83.9173715 | 144.586385 | 173.545708 | 172.584248 | 4.75243649 |
| 3.03E-09 | 1.72E-08 | 653.655235 | 1049.25227 | 912.275034 | 991.895492 | 83.1676385 |
| 3.03E-09 | 1.72E-08 | 73.3071291 | 121.230123 | 94.6612955 | 114.128293 | 0 |
| 3.03E-09 | 1.72E-08 | 67.5197242 | 216.879578 | 134.567528 | 163.769461 | 0 |
| 3.04E-09 | 1.72E-08 | 56.5879593 | 280.497587 | 219.020252 | 243.102543 | 0 |
| 3.04E-09 | 1.72E-08 | 132.467268 | 113.667143 | 69.6038938 | 167.944887 | 0 |
| 3.05E-09 | 1.72E-08 | 63.9829767 | 162.826514 | 141.063891 | 163.769461 | 0 |
| 3.05E-09 | 1.73E-08 | 65.2690667 | 165.940682 | 167.977397 | 131.293931 | 0 |
| 3.06E-09 | 1.73E-08 | 73.3071291 | 105.2144 | 94.6612955 | 108.097123 | 0 |
| 3.06E-09 | 1.73E-08 | 146.935781 | 205.090226 | 154.056618 | 207.843396 | 0 |
| 3.07E-09 | 1.74E-08 | 61.0892743 | 181.066642 | 173.545708 | 175.831801 | 0 |
| 3.07E-09 | 1.74E-08 | 129.573566 | 280.720028 | 219.948304 | 205.523715 | 0 |
| 3.08E-09 | 1.74E-08 | 66.8766792 | 123.009648 | 167.049345 | 171.656376 | 0 |
| 3.08E-09 | 1.74E-08 | 65.2690667 | 219.103984 | 139.207788 | 198.100737 | 0 |
| 3.08E-09 | 1.74E-08 | 63.6614542 | 145.031266 | 156.840774 | 154.954675 | 0 |
| 3.10E-09 | 1.75E-08 | 99.0289288 | 229.781132 | 82.5966206 | 165.16127 | 0 |
| 3.11E-09 | 1.75E-08 | 202.23765 | 104.769519 | 85.3807764 | 111.344676 | 0 |
| 3.12E-09 | 1.76E-08 | 76.8438766 | 136.800964 | 96.5173994 | 100.674145 | 0 |
| 3.12E-09 | 1.76E-08 | 59.1601393 | 194.413078 | 193.034799 | 203.66797 | 0 |
| 3.12E-09 | 1.76E-08 | 85.2034615 | 115.224227 | 77.956361 | 101.602017 | 0 |
| 3.12E-09 | 1.76E-08 | 163.654951 | 231.560657 | 70.5319457 | 411.04743 | 0 |
| 3.13E-09 | 1.77E-08 | 177.158895 | 141.027336 | 100.229607 | 134.541484 | 4.75243649 |
| 3.14E-09 | 1.77E-08 | 68.4842917 | 124.789172 | 110.438178 | 125.726697 | 0 |
| 3.14E-09 | 1.77E-08 | 63.6614542 | 159.489905 | 142.919995 | 154.026802 | 0 |
| 3.15E-09 | 1.78E-08 | 111.889829 | 103.879757 | 105.797919 | 117.839782 | 14.2573095 |
| 3.15E-09 | 1.78E-08 | 55.3018693 | 248.021261 | 304.401029 | 234.751693 | 0 |
| 3.16E-09 | 1.78E-08 | 64.9475442 | 139.470251 | 163.337137 | 136.397228 | 0 |
| 3.17E-09 | 1.78E-08 | 64.9475442 | 144.586385 | 139.207788 | 178.615418 | 0 |
| 3.17E-09 | 1.79E-08 | 66.2336342 | 153.261568 | 123.430905 | 139.644782 | 0 |
| 3.17E-09 | 1.79E-08 | 545.302153 | 1637.16276 | 1134.07944 | 1325.92952 | 4.75243649 |
| 3.18E-09 | 1.79E-08 | 47.5853294 | 308.525102 | 498.363879 | 504.298596 | 0 |
| 3.18E-09 | 1.79E-08 | 326.345334 | 458.894942 | 433.400245 | 516.360936 | 21.3859642 |
| 3.18E-09 | 1.79E-08 | 73.3071291 | 93.2026082 | 111.36623 | 103.457762 | 0 |

| | | | | | | |
|---|---|---|---|---|---|---|
| 3.19E-09 | 1.80E-08 | 117.677234 | 257.141325 | 73.3161014 | 215.73031 | 0 |
| 3.19E-09 | 1.80E-08 | 417.657722 | 309.414864 | 53.8270112 | 245.88616 | 0 |
| 3.20E-09 | 1.80E-08 | 54.6588243 | 242.682686 | 249.645966 | 299.702754 | 0 |
| 3.20E-09 | 1.80E-08 | 165.905608 | 176.39539 | 129.927268 | 196.708928 | 0 |
| 3.21E-09 | 1.80E-08 | 57.5525268 | 222.663033 | 211.595837 | 206.451587 | 0 |
| 3.21E-09 | 1.81E-08 | 308.340074 | 778.542074 | 574.464137 | 615.179337 | 23.7621824 |
| 3.22E-09 | 1.81E-08 | 129.573566 | 189.519385 | 217.164149 | 238.927118 | 0 |
| 3.23E-09 | 1.81E-08 | 71.0564716 | 182.846167 | 110.438178 | 184.182652 | 0 |
| 3.23E-09 | 1.82E-08 | 74.5932191 | 168.387528 | 99.3015551 | 169.800631 | 0 |
| 3.23E-09 | 1.82E-08 | 72.3425616 | 115.446667 | 103.013763 | 96.9626556 | 0 |
| 3.26E-09 | 1.83E-08 | 66.5551567 | 141.249776 | 117.862593 | 139.180845 | 0 |
| 3.27E-09 | 1.84E-08 | 98.3858838 | 253.137394 | 129.927268 | 181.862971 | 4.75243649 |
| 3.28E-09 | 1.84E-08 | 106.423946 | 88.0864746 | 77.0283091 | 90.4675495 | 0 |
| 3.28E-09 | 1.84E-08 | 611.535788 | 1745.71377 | 1517.36488 | 1760.63769 | 0 |
| 3.29E-09 | 1.85E-08 | 109.317649 | 358.796676 | 315.537652 | 442.131152 | 4.75243649 |
| 3.29E-09 | 1.85E-08 | 69.1273367 | 132.352152 | 104.869867 | 131.757867 | 0 |
| 3.29E-09 | 1.85E-08 | 58.8386168 | 202.198499 | 187.466487 | 193.461375 | 0 |
| 3.30E-09 | 1.85E-08 | 66.8766792 | 126.568697 | 146.632203 | 187.430205 | 0 |
| 3.30E-09 | 1.85E-08 | 73.9501741 | 112.55494 | 119.718697 | 90.0036134 | 0 |
| 3.31E-09 | 1.86E-08 | 246.286232 | 97.8738607 | 99.3015551 | 119.695527 | 0 |
| 3.32E-09 | 1.86E-08 | 58.5170943 | 53.3857422 | 57.5392189 | 51.0329766 | 149.701749 |
| 3.32E-09 | 1.86E-08 | 63.6614542 | 136.356083 | 152.200514 | 154.954675 | 0 |
| 3.32E-09 | 1.86E-08 | 70.4134267 | 119.005717 | 111.36623 | 158.666164 | 0 |
| 3.33E-09 | 1.87E-08 | 183.910868 | 143.029301 | 94.6612955 | 78.4052096 | 0 |
| 3.33E-09 | 1.87E-08 | 65.5905892 | 321.871537 | 153.128566 | 244.958288 | 0 |
| 3.34E-09 | 1.87E-08 | 69.7703817 | 135.911202 | 103.941815 | 119.695527 | 0 |
| 3.34E-09 | 1.87E-08 | 72.0210391 | 107.216366 | 96.5173994 | 123.407016 | 0 |
| 3.34E-09 | 1.87E-08 | 64.6260217 | 339.444344 | 187.466487 | 188.822014 | 0 |
| 3.34E-09 | 1.87E-08 | 87.7756414 | 110.108093 | 77.956361 | 89.5396772 | 0 |
| 3.34E-09 | 1.87E-08 | 67.5197242 | 127.013578 | 114.150386 | 117.839782 | 0 |
| 3.35E-09 | 1.87E-08 | 61.4107968 | 176.172949 | 154.056618 | 185.110524 | 0 |
| 3.35E-09 | 1.87E-08 | 63.9829767 | 171.946578 | 154.98467 | 134.541484 | 0 |
| 3.35E-09 | 1.87E-08 | 58.5170943 | 210.651241 | 195.818954 | 187.430205 | 0 |
| 3.35E-09 | 1.87E-08 | 61.7323193 | 179.287118 | 151.272462 | 172.120312 | 0 |
| 3.35E-09 | 1.87E-08 | 62.3753642 | 174.170984 | 154.98467 | 150.315313 | 0 |
| 3.35E-09 | 1.88E-08 | 60.7677518 | 292.73182 | 204.171422 | 182.790844 | 0 |
| 3.36E-09 | 1.88E-08 | 63.6614542 | 137.690727 | 146.632203 | 162.841589 | 0 |
| 3.36E-09 | 1.88E-08 | 91.6339114 | 172.83634 | 79.8124649 | 183.718716 | 0 |
| 3.38E-09 | 1.89E-08 | 61.4107968 | 161.49187 | 163.337137 | 161.913717 | 0 |
| 3.38E-09 | 1.89E-08 | 62.0538418 | 193.968197 | 154.056618 | 231.50414 | 0 |
| 3.38E-09 | 1.89E-08 | 77.1653991 | 608.15258 | 585.60076 | 643.015506 | 0 |
| 3.39E-09 | 1.90E-08 | 110.282216 | 370.363586 | 296.048562 | 328.930731 | 9.50487297 |
| 3.40E-09 | 1.90E-08 | 139.219241 | 240.23584 | 232.012979 | 219.905736 | 14.2573095 |

| | | | | | | |
|---|---|---|---|---|---|---|
| 3.44E-09 | 1.92E-08 | 67.8412467 | 108.99589 | 122.502853 | 116.91191 | 0 |
| 3.44E-09 | 1.92E-08 | 68.4842917 | 123.232088 | 132.711424 | 105.313506 | 0 |
| 3.45E-09 | 1.92E-08 | 58.1955718 | 194.413078 | 191.178695 | 199.956481 | 0 |
| 3.45E-09 | 1.93E-08 | 92.5984789 | 154.596212 | 76.1002572 | 119.231591 | 0 |
| 3.45E-09 | 1.93E-08 | 45.6561944 | 358.351794 | 401.84648 | 405.944132 | 0 |
| 3.48E-09 | 1.94E-08 | 87.4541189 | 187.072538 | 120.646749 | 88.6118049 | 0 |
| 3.48E-09 | 1.94E-08 | 292.263949 | 893.321419 | 853.807764 | 896.78858 | 40.3957101 |
| 3.49E-09 | 1.95E-08 | 64.9475442 | 146.143469 | 126.215061 | 167.480951 | 0 |
| 3.49E-09 | 1.95E-08 | 88.0971639 | 81.858138 | 178.185968 | 138.252973 | 0 |
| 3.49E-09 | 1.95E-08 | 115.748099 | 340.556547 | 234.797135 | 287.640414 | 2.37621824 |
| 3.50E-09 | 1.95E-08 | 116.391144 | 364.802572 | 232.941031 | 315.012647 | 9.50487297 |
| 3.50E-09 | 1.95E-08 | 68.8058142 | 103.657316 | 124.358957 | 113.664357 | 0 |
| 3.51E-09 | 1.96E-08 | 84.238894 | 166.830444 | 86.3088283 | 116.91191 | 0 |
| 3.53E-09 | 1.97E-08 | 105.780901 | 122.564766 | 67.7477899 | 138.716909 | 0 |
| 3.53E-09 | 1.97E-08 | 76.2008316 | 87.6415934 | 90.0210359 | 93.7151026 | 0 |
| 3.53E-09 | 1.97E-08 | 71.3779941 | 118.115955 | 93.7332436 | 120.159463 | 0 |
| 3.54E-09 | 1.97E-08 | 80.380624 | 117.003752 | 83.5246725 | 92.3232941 | 0 |
| 3.55E-09 | 1.98E-08 | 62.3753642 | 176.61783 | 140.135839 | 168.408823 | 0 |
| 3.55E-09 | 1.98E-08 | 67.8412467 | 110.775415 | 130.85532 | 112.272549 | 0 |
| 3.55E-09 | 1.98E-08 | 369.750871 | 415.741467 | 458.457647 | 513.113383 | 28.5146189 |
| 3.55E-09 | 1.98E-08 | 87.1325964 | 155.708415 | 217.164149 | 88.6118049 | 0 |
| 3.56E-09 | 1.98E-08 | 115.748099 | 165.495801 | 75.1722053 | 98.8184002 | 0 |
| 3.56E-09 | 1.98E-08 | 64.9475442 | 135.911202 | 122.502853 | 134.541484 | 0 |
| 3.58E-09 | 1.99E-08 | 64.9475442 | 130.127747 | 147.560255 | 124.334889 | 0 |
| 3.58E-09 | 1.99E-08 | 184.553913 | 197.082365 | 134.567528 | 165.625206 | 0 |
| 3.58E-09 | 1.99E-08 | 68.4842917 | 193.078434 | 120.646749 | 140.108718 | 0 |
| 3.59E-09 | 2.00E-08 | 82.6312815 | 135.02144 | 82.5966206 | 101.138081 | 0 |
| 3.59E-09 | 2.00E-08 | 105.137856 | 145.476147 | 67.7477899 | 140.108718 | 0 |
| 3.60E-09 | 2.00E-08 | 68.4842917 | 131.017509 | 114.150386 | 105.777442 | 0 |
| 3.62E-09 | 2.01E-08 | 66.2336342 | 112.77738 | 142.919995 | 136.861165 | 0 |
| 3.63E-09 | 2.02E-08 | 76.8438766 | 111.887618 | 83.5246725 | 123.870952 | 0 |
| 3.63E-09 | 2.02E-08 | 67.1982017 | 140.582454 | 125.287009 | 193.925311 | 0 |
| 3.63E-09 | 2.02E-08 | 80.7021465 | 85.1947469 | 152.200514 | 115.984038 | 0 |
| 3.63E-09 | 2.02E-08 | 62.3753642 | 160.602108 | 146.632203 | 142.428398 | 0 |
| 3.63E-09 | 2.02E-08 | 71.6995166 | 102.322673 | 93.7332436 | 115.520102 | 0 |
| 3.63E-09 | 2.02E-08 | 63.0184092 | 137.913167 | 154.98467 | 139.644782 | 0 |
| 3.64E-09 | 2.02E-08 | 147.900348 | 102.545113 | 167.049345 | 70.518295 | 0 |
| 3.64E-09 | 2.02E-08 | 58.8386168 | 356.79471 | 196.747006 | 238.927118 | 0 |
| 3.65E-09 | 2.03E-08 | 70.0919042 | 191.743791 | 161.481034 | 109.488932 | 0 |
| 3.65E-09 | 2.03E-08 | 91.3123889 | 133.909237 | 83.5246725 | 86.2921241 | 0 |
| 3.65E-09 | 2.03E-08 | 67.1982017 | 108.328569 | 118.790645 | 118.303719 | 0 |
| 3.66E-09 | 2.03E-08 | 24.4357097 | 20.4645345 | 17.6329864 | 21.8049991 | 76.0389838 |
| 3.67E-09 | 2.04E-08 | 65.9121117 | 119.228158 | 122.502853 | 118.303719 | 0 |

| | | | | | | |
|---|---|---|---|---|---|---|
| 3.68E-09 | 2.04E-08 | 108.674604 | 377.926567 | 330.386482 | 288.10435 | 4.75243649 |
| 3.69E-09 | 2.05E-08 | 62.6968867 | 139.247811 | 188.394539 | 209.69914 | 0 |
| 3.69E-09 | 2.05E-08 | 74.2716966 | 92.9801676 | 88.1649321 | 96.9626556 | 0 |
| 3.69E-09 | 2.05E-08 | 78.7730116 | 93.6474894 | 107.654022 | 160.057972 | 0 |
| 3.70E-09 | 2.05E-08 | 63.6614542 | 137.690727 | 128.999216 | 144.284143 | 0 |
| 3.70E-09 | 2.05E-08 | 61.7323193 | 189.074504 | 154.056618 | 150.315313 | 0 |
| 3.72E-09 | 2.06E-08 | 102.244154 | 323.651062 | 423.191674 | 361.870198 | 0 |
| 3.72E-09 | 2.06E-08 | 75.8793091 | 83.8601033 | 94.6612955 | 101.602017 | 0 |
| 3.73E-09 | 2.07E-08 | 130.216611 | 185.960335 | 179.11402 | 211.090949 | 2.37621824 |
| 3.74E-09 | 2.08E-08 | 138.576196 | 187.294979 | 156.840774 | 190.677758 | 0 |
| 3.75E-09 | 2.08E-08 | 151.758618 | 182.623726 | 167.049345 | 286.712542 | 0 |
| 3.76E-09 | 2.08E-08 | 83.9173715 | 161.49187 | 136.423632 | 154.954675 | 4.75243649 |
| 3.77E-09 | 2.09E-08 | 127.001386 | 76.7420044 | 87.2368802 | 98.8184002 | 0 |
| 3.78E-09 | 2.09E-08 | 94.8491364 | 212.208325 | 83.5246725 | 132.685739 | 0 |
| 3.78E-09 | 2.10E-08 | 88.4186864 | 76.2971232 | 135.49558 | 153.09893 | 0 |
| 3.79E-09 | 2.10E-08 | 72.0210391 | 103.657316 | 90.9490879 | 98.3544641 | 0 |
| 3.79E-09 | 2.10E-08 | 74.2716966 | 86.3069499 | 119.718697 | 123.870952 | 0 |
| 3.79E-09 | 2.10E-08 | 61.4107968 | 241.125602 | 165.193241 | 168.408823 | 0 |
| 3.79E-09 | 2.10E-08 | 264.934537 | 86.5293905 | 110.438178 | 130.829995 | 0 |
| 3.79E-09 | 2.10E-08 | 97.7428388 | 326.76523 | 303.472977 | 353.983283 | 0 |
| 3.79E-09 | 2.10E-08 | 36.0105196 | 35.8129354 | 43.6184401 | 31.0837221 | 133.068222 |
| 3.80E-09 | 2.10E-08 | 66.8766792 | 119.450598 | 119.718697 | 158.202228 | 0 |
| 3.82E-09 | 2.11E-08 | 93.5630464 | 701.800069 | 355.443884 | 506.618277 | 2.37621824 |
| 3.82E-09 | 2.11E-08 | 56.5879593 | 217.546899 | 187.466487 | 215.73031 | 0 |
| 3.83E-09 | 2.12E-08 | 55.3018693 | 286.058602 | 210.667785 | 224.545097 | 0 |
| 3.84E-09 | 2.12E-08 | 57.2310043 | 197.749687 | 259.854537 | 193.461375 | 0 |
| 3.85E-09 | 2.13E-08 | 92.2769564 | 99.875826 | 229.228823 | 111.808612 | 0 |
| 3.85E-09 | 2.13E-08 | 63.0184092 | 172.391459 | 282.127783 | 168.408823 | 0 |
| 3.85E-09 | 2.13E-08 | 115.105054 | 239.790959 | 242.22155 | 242.174671 | 0 |
| 3.87E-09 | 2.14E-08 | 63.6614542 | 148.367875 | 123.430905 | 149.851377 | 0 |
| 3.87E-09 | 2.14E-08 | 67.1982017 | 135.24388 | 104.869867 | 129.902122 | 0 |
| 3.88E-09 | 2.14E-08 | 63.0184092 | 127.458459 | 161.481034 | 161.913717 | 0 |
| 3.90E-09 | 2.15E-08 | 93.8845689 | 130.572628 | 72.3880495 | 101.138081 | 0 |
| 3.90E-09 | 2.15E-08 | 61.4107968 | 180.844202 | 139.207788 | 175.367865 | 0 |
| 3.90E-09 | 2.15E-08 | 143.399033 | 208.204395 | 68.6758419 | 118.767655 | 0 |
| 3.91E-09 | 2.16E-08 | 81.023669 | 88.0864746 | 84.4527244 | 81.1888265 | 0 |
| 3.91E-09 | 2.16E-08 | 60.7677518 | 157.487939 | 146.632203 | 165.625206 | 0 |
| 3.91E-09 | 2.16E-08 | 75.5577866 | 124.566732 | 100.229607 | 86.7560603 | 0 |
| 3.92E-09 | 2.16E-08 | 67.5197242 | 141.472217 | 103.941815 | 135.00542 | 0 |
| 3.92E-09 | 2.16E-08 | 71.0564716 | 151.926925 | 101.157659 | 114.592229 | 0 |
| 3.92E-09 | 2.16E-08 | 61.4107968 | 177.730033 | 162.409085 | 139.644782 | 0 |
| 3.92E-09 | 2.16E-08 | 84.5604165 | 78.7439697 | 82.5966206 | 94.1790387 | 0 |
| 3.93E-09 | 2.16E-08 | 64.9475442 | 164.606038 | 117.862593 | 148.459568 | 0 |

| | | | | | | |
|---|---|---|---|---|---|---|
| 3.93E-09 | 2.17E-08 | 92.2769564 | 103.212435 | 80.7405168 | 77.4773373 | 0 |
| 3.93E-09 | 2.17E-08 | 61.4107968 | 156.153296 | 150.344411 | 141.03659 | 0 |
| 3.94E-09 | 2.17E-08 | 64.3044992 | 174.615865 | 123.430905 | 184.182652 | 0 |
| 3.96E-09 | 2.18E-08 | 59.8031843 | 189.074504 | 151.272462 | 179.543291 | 0 |
| 3.96E-09 | 2.18E-08 | 73.3071291 | 95.8718953 | 99.3015551 | 86.2921241 | 0 |
| 3.96E-09 | 2.18E-08 | 74.5932191 | 95.2045736 | 92.8051917 | 84.9003157 | 0 |
| 3.96E-09 | 2.18E-08 | 215.420072 | 152.149365 | 163.337137 | 149.851377 | 2.37621824 |
| 3.96E-09 | 2.18E-08 | 59.4816618 | 163.716276 | 183.75428 | 159.594036 | 0 |
| 3.98E-09 | 2.19E-08 | 65.2690667 | 126.123816 | 193.034799 | 221.297544 | 0 |
| 3.99E-09 | 2.20E-08 | 127.322908 | 177.507593 | 217.164149 | 230.576267 | 0 |
| 4.00E-09 | 2.20E-08 | 63.3399317 | 170.389494 | 154.98467 | 125.262761 | 0 |
| 4.01E-09 | 2.20E-08 | 64.3044992 | 148.590316 | 120.646749 | 129.438186 | 0 |
| 4.04E-09 | 2.22E-08 | 112.532874 | 250.690548 | 219.020252 | 242.638607 | 4.75243649 |
| 4.04E-09 | 2.22E-08 | 56.2664368 | 189.074504 | 199.531162 | 210.627013 | 0 |
| 4.04E-09 | 2.22E-08 | 64.9475442 | 137.245846 | 189.322591 | 128.97425 | 0 |
| 4.05E-09 | 2.23E-08 | 59.1601393 | 177.285152 | 157.768826 | 203.66797 | 0 |
| 4.06E-09 | 2.23E-08 | 65.9121117 | 126.791138 | 107.654022 | 123.407016 | 0 |
| 4.06E-09 | 2.23E-08 | 81.9882365 | 94.7596924 | 94.6612955 | 159.1301 | 0 |
| 4.06E-09 | 2.23E-08 | 115.426576 | 114.334465 | 75.1722053 | 81.6527626 | 0 |
| 4.08E-09 | 2.24E-08 | 106.423946 | 185.293013 | 68.6758419 | 173.976057 | 0 |
| 4.08E-09 | 2.24E-08 | 75.5577866 | 179.509558 | 93.7332436 | 180.935099 | 0 |
| 4.09E-09 | 2.25E-08 | 56.5879593 | 192.633553 | 185.610383 | 201.34829 | 0 |
| 4.10E-09 | 2.25E-08 | 60.7677518 | 143.919063 | 157.768826 | 148.923505 | 0 |
| 4.13E-09 | 2.27E-08 | 95.4921813 | 159.045024 | 71.4599976 | 160.521908 | 0 |
| 4.13E-09 | 2.27E-08 | 57.5525268 | 197.082365 | 168.905449 | 191.60563 | 0 |
| 4.15E-09 | 2.27E-08 | 60.4462293 | 144.141504 | 154.056618 | 168.408823 | 0 |
| 4.15E-09 | 2.28E-08 | 66.8766792 | 225.33232 | 128.071165 | 152.171058 | 0 |
| 4.16E-09 | 2.28E-08 | 551.089558 | 1374.9053 | 1269.57502 | 1266.54569 | 28.5146189 |
| 4.17E-09 | 2.29E-08 | 58.8386168 | 157.487939 | 180.042072 | 168.408823 | 0 |
| 4.18E-09 | 2.29E-08 | 92.2769564 | 80.3010539 | 169.833501 | 106.241379 | 0 |
| 4.18E-09 | 2.29E-08 | 66.2336342 | 115.001786 | 114.150386 | 147.06776 | 0 |
| 4.18E-09 | 2.29E-08 | 73.6286516 | 112.999821 | 84.4527244 | 100.674145 | 0 |
| 4.18E-09 | 2.29E-08 | 81.9882365 | 141.917098 | 78.8844129 | 149.387441 | 0 |
| 4.19E-09 | 2.29E-08 | 68.1627692 | 116.55887 | 96.5173994 | 114.592229 | 0 |
| 4.19E-09 | 2.29E-08 | 65.2690667 | 112.332499 | 128.071165 | 114.592229 | 0 |
| 4.19E-09 | 2.29E-08 | 63.6614542 | 127.236019 | 120.646749 | 128.046378 | 0 |
| 4.20E-09 | 2.30E-08 | 65.5905892 | 149.257638 | 113.222334 | 170.728504 | 0 |
| 4.20E-09 | 2.30E-08 | 99.0289288 | 282.944434 | 289.552198 | 277.897755 | 7.12865473 |
| 4.20E-09 | 2.30E-08 | 59.8031843 | 221.55083 | 187.466487 | 150.779249 | 0 |
| 4.21E-09 | 2.30E-08 | 64.9475442 | 117.893514 | 116.00649 | 139.644782 | 0 |
| 4.21E-09 | 2.30E-08 | 73.3071291 | 92.0904053 | 96.5173994 | 85.3642518 | 0 |
| 4.22E-09 | 2.31E-08 | 157.546023 | 248.243701 | 141.991943 | 186.502333 | 0 |
| 4.23E-09 | 2.31E-08 | 64.9475442 | 239.346077 | 128.071165 | 204.595843 | 0 |

| | | | | | | |
|---|---|---|---|---|---|---|
| 4.23E-09 | 2.32E-08 | 63.3399317 | 122.342326 | 126.215061 | 132.221803 | 0 |
| 4.24E-09 | 2.32E-08 | 60.7677518 | 142.58442 | 143.848047 | 153.562866 | 0 |
| 4.25E-09 | 2.33E-08 | 66.2336342 | 110.552974 | 119.718697 | 148.923505 | 0 |
| 4.26E-09 | 2.33E-08 | 408.976615 | 768.087366 | 615.298421 | 695.904227 | 0 |
| 4.27E-09 | 2.33E-08 | 66.5551567 | 111.665177 | 103.013763 | 114.592229 | 0 |
| 4.27E-09 | 2.34E-08 | 0 | 0 | 0 | 0 | 59.4054561 |
| 4.30E-09 | 2.35E-08 | 77.1653991 | 133.686796 | 143.848047 | 82.5806349 | 0 |
| 4.30E-09 | 2.35E-08 | 517.972741 | 1290.60032 | 1144.28801 | 1218.29633 | 4.75243649 |
| 4.34E-09 | 2.37E-08 | 66.2336342 | 111.442737 | 112.294282 | 103.921698 | 0 |
| 4.35E-09 | 2.38E-08 | 62.6968867 | 294.288904 | 177.257916 | 163.305525 | 0 |
| 4.35E-09 | 2.38E-08 | 57.5525268 | 191.076469 | 186.538435 | 162.841589 | 0 |
| 4.37E-09 | 2.38E-08 | 64.3044992 | 131.017509 | 111.36623 | 128.510314 | 0 |
| 4.37E-09 | 2.39E-08 | 62.0538418 | 171.946578 | 158.696878 | 126.654569 | 0 |
| 4.37E-09 | 2.39E-08 | 62.6968867 | 199.529211 | 131.783372 | 222.689353 | 0 |
| 4.38E-09 | 2.39E-08 | 67.5197242 | 135.24388 | 109.510126 | 103.457762 | 0 |
| 4.38E-09 | 2.39E-08 | 62.6968867 | 181.511523 | 128.071165 | 147.995632 | 0 |
| 4.42E-09 | 2.41E-08 | 64.3044992 | 137.913167 | 111.36623 | 145.212015 | 0 |
| 4.42E-09 | 2.41E-08 | 58.5170943 | 209.094157 | 190.250643 | 152.634994 | 0 |
| 4.44E-09 | 2.42E-08 | 68.8058142 | 130.127747 | 94.6612955 | 114.128293 | 0 |
| 4.46E-09 | 2.43E-08 | 51.4435994 | 285.16884 | 245.005706 | 235.215629 | 0 |
| 4.46E-09 | 2.43E-08 | 86.168029 | 119.005717 | 72.3880495 | 98.8184002 | 0 |
| 4.47E-09 | 2.44E-08 | 78.1299666 | 94.0923706 | 81.6685687 | 123.870952 | 0 |
| 4.48E-09 | 2.44E-08 | 131.181178 | 198.194568 | 178.185968 | 164.233398 | 0 |
| 4.49E-09 | 2.45E-08 | 78.1299666 | 188.407182 | 91.8771398 | 128.510314 | 0 |
| 4.49E-09 | 2.45E-08 | 75.2362641 | 89.6435588 | 82.5966206 | 86.7560603 | 0 |
| 4.49E-09 | 2.45E-08 | 63.9829767 | 113.444702 | 149.416359 | 130.829995 | 0 |
| 4.50E-09 | 2.45E-08 | 168.477788 | 237.788993 | 146.632203 | 190.677758 | 14.2573095 |
| 4.50E-09 | 2.45E-08 | 68.1627692 | 102.322673 | 140.135839 | 107.169251 | 0 |
| 4.51E-09 | 2.45E-08 | 78.1299666 | 127.903341 | 85.3807764 | 90.9314857 | 0 |
| 4.51E-09 | 2.46E-08 | 56.9094818 | 199.529211 | 164.265189 | 195.781056 | 0 |
| 4.52E-09 | 2.46E-08 | 75.8793091 | 139.692692 | 151.272462 | 84.4363795 | 0 |
| 4.52E-09 | 2.46E-08 | 116.391144 | 205.979989 | 232.941031 | 224.081161 | 2.37621824 |
| 4.53E-09 | 2.46E-08 | 60.7677518 | 131.684831 | 164.265189 | 152.171058 | 0 |
| 4.53E-09 | 2.46E-08 | 60.7677518 | 153.261568 | 308.113236 | 240.782863 | 0 |
| 4.53E-09 | 2.46E-08 | 69.7703817 | 115.446667 | 103.941815 | 90.0036134 | 0 |
| 4.54E-09 | 2.47E-08 | 63.6614542 | 193.745756 | 123.430905 | 154.490738 | 0 |
| 4.54E-09 | 2.47E-08 | 73.3071291 | 96.3167765 | 83.5246725 | 111.808612 | 0 |
| 4.55E-09 | 2.47E-08 | 64.9475442 | 158.822583 | 111.36623 | 133.149675 | 0 |
| 4.56E-09 | 2.48E-08 | 55.6233918 | 218.659102 | 179.11402 | 197.6368 | 0 |
| 4.57E-09 | 2.48E-08 | 66.5551567 | 278.495622 | 126.215061 | 235.679565 | 0 |
| 4.57E-09 | 2.48E-08 | 114.462009 | 555.87904 | 283.055835 | 372.076793 | 11.8810912 |
| 4.57E-09 | 2.49E-08 | 62.3753642 | 145.031266 | 131.783372 | 121.551272 | 0 |
| 4.58E-09 | 2.49E-08 | 58.1955718 | 152.816687 | 180.042072 | 167.944887 | 0 |

| | | | | | | |
|---|---|---|---|---|---|---|
| 4.60E-09 | 2.50E-08 | 71.0564716 | 85.1947469 | 102.085711 | 104.84957 | 0 |
| 4.60E-09 | 2.50E-08 | 56.2664368 | 197.527246 | 170.761553 | 191.141694 | 0 |
| 4.62E-09 | 2.51E-08 | 63.0184092 | 146.143469 | 123.430905 | 121.087335 | 0 |
| 4.63E-09 | 2.51E-08 | 186.80457 | 137.468286 | 192.106747 | 207.843396 | 16.6335277 |
| 4.63E-09 | 2.51E-08 | 50.4790319 | 285.613721 | 237.581291 | 260.268181 | 0 |
| 4.63E-09 | 2.51E-08 | 60.7677518 | 139.247811 | 133.639476 | 141.500526 | 0 |
| 4.64E-09 | 2.52E-08 | 59.8031843 | 163.716276 | 136.423632 | 168.408823 | 0 |
| 4.65E-09 | 2.52E-08 | 115.105054 | 80.0786133 | 74.2441534 | 105.313506 | 0 |
| 4.69E-09 | 2.54E-08 | 68.8058142 | 107.216366 | 90.9490879 | 100.674145 | 0 |
| 4.69E-09 | 2.54E-08 | 96.1352263 | 156.153296 | 105.797919 | 137.325101 | 9.50487297 |
| 4.70E-09 | 2.55E-08 | 59.8031843 | 179.954439 | 142.919995 | 218.049991 | 0 |
| 4.70E-09 | 2.55E-08 | 64.9475442 | 133.909237 | 122.502853 | 104.84957 | 0 |
| 4.72E-09 | 2.56E-08 | 64.6260217 | 106.549044 | 123.430905 | 113.664357 | 0 |
| 4.73E-09 | 2.57E-08 | 69.1273367 | 140.582454 | 100.229607 | 103.921698 | 0 |
| 4.74E-09 | 2.57E-08 | 79.737579 | 98.763623 | 77.0283091 | 124.334889 | 0 |
| 4.74E-09 | 2.57E-08 | 54.0157793 | 229.781132 | 191.178695 | 275.114138 | 0 |
| 4.75E-09 | 2.58E-08 | 103.851766 | 401.282829 | 267.278952 | 308.981477 | 2.37621824 |
| 4.75E-09 | 2.58E-08 | 78.4514891 | 93.2026082 | 127.143113 | 81.1888265 | 0 |
| 4.76E-09 | 2.58E-08 | 2348.72183 | 6120.23046 | 4308.94505 | 5705.48679 | 0 |
| 4.78E-09 | 2.59E-08 | 64.6260217 | 154.818652 | 109.510126 | 164.697334 | 0 |
| 4.78E-09 | 2.59E-08 | 135.039448 | 949.376449 | 1013.43269 | 1060.09411 | 0 |
| 4.79E-09 | 2.59E-08 | 64.9475442 | 117.003752 | 103.013763 | 121.087335 | 0 |
| 4.79E-09 | 2.60E-08 | 132.467268 | 68.7341431 | 107.654022 | 95.106911 | 0 |
| 4.80E-09 | 2.60E-08 | 62.6968867 | 196.415043 | 150.344411 | 127.118505 | 0 |
| 4.80E-09 | 2.60E-08 | 60.7677518 | 127.236019 | 152.200514 | 141.964462 | 0 |
| 4.81E-09 | 2.60E-08 | 112.211351 | 200.641414 | 217.164149 | 242.174671 | 4.75243649 |
| 4.82E-09 | 2.61E-08 | 84.881939 | 107.216366 | 120.646749 | 109.024996 | 9.50487297 |
| 4.82E-09 | 2.61E-08 | 65.9121117 | 136.356083 | 121.574801 | 100.674145 | 0 |
| 4.82E-09 | 2.61E-08 | 91.9554339 | 379.706091 | 389.781805 | 415.222855 | 0 |
| 4.82E-09 | 2.61E-08 | 72.0210391 | 147.922994 | 88.1649321 | 141.03659 | 0 |
| 4.85E-09 | 2.62E-08 | 57.2310043 | 162.381632 | 164.265189 | 169.336695 | 0 |
| 4.87E-09 | 2.63E-08 | 61.4107968 | 158.822583 | 128.999216 | 130.829995 | 0 |
| 4.88E-09 | 2.64E-08 | 63.6614542 | 108.106128 | 141.063891 | 135.00542 | 0 |
| 4.88E-09 | 2.64E-08 | 60.4462293 | 177.730033 | 128.999216 | 172.584248 | 0 |
| 4.89E-09 | 2.64E-08 | 52.4081669 | 209.316597 | 223.660512 | 235.215629 | 0 |
| 4.89E-09 | 2.64E-08 | 70.0919042 | 126.123816 | 148.488307 | 91.3954218 | 0 |
| 4.89E-09 | 2.64E-08 | 68.1627692 | 152.149365 | 97.4454513 | 157.274355 | 0 |
| 4.89E-09 | 2.64E-08 | 73.6286516 | 97.6514201 | 81.6685687 | 87.6839326 | 0 |
| 4.89E-09 | 2.64E-08 | 227.316405 | 576.343575 | 50.1148035 | 509.401894 | 0 |
| 4.91E-09 | 2.65E-08 | 108.996126 | 90.3108805 | 68.6758419 | 94.1790387 | 0 |
| 4.92E-09 | 2.65E-08 | 65.9121117 | 124.566732 | 98.3735032 | 114.592229 | 0 |
| 4.92E-09 | 2.66E-08 | 103.530244 | 271.155082 | 276.559471 | 295.991265 | 0 |
| 4.92E-09 | 2.66E-08 | 109.317649 | 112.999821 | 63.1075304 | 115.056166 | 0 |

| | | | | | | |
|---|---|---|---|---|---|---|
| 4.92E-09 | 2.66E-08 | 76.8438766 | 125.678935 | 77.956361 | 107.169251 | 0 |
| 4.93E-09 | 2.66E-08 | 57.5525268 | 153.039128 | 199.531162 | 171.19244 | 0 |
| 4.93E-09 | 2.66E-08 | 55.3018693 | 178.397355 | 187.466487 | 186.966269 | 0 |
| 4.94E-09 | 2.66E-08 | 61.4107968 | 156.375736 | 120.646749 | 149.851377 | 0 |
| 4.94E-09 | 2.66E-08 | 57.5525268 | 163.938717 | 160.552982 | 207.37946 | 0 |
| 4.94E-09 | 2.67E-08 | 59.4816618 | 153.261568 | 185.610383 | 139.180845 | 0 |
| 4.96E-09 | 2.67E-08 | 111.889829 | 209.316597 | 190.250643 | 205.523715 | 11.8810912 |
| 4.96E-09 | 2.67E-08 | 71.3779941 | 88.0864746 | 87.2368802 | 90.0036134 | 0 |
| 4.96E-09 | 2.67E-08 | 64.3044992 | 109.218331 | 122.502853 | 148.459568 | 0 |
| 4.97E-09 | 2.68E-08 | 107.066991 | 261.812577 | 252.430121 | 252.845202 | 0 |
| 4.98E-09 | 2.69E-08 | 52.0866444 | 239.790959 | 220.876356 | 212.946693 | 0 |
| 4.98E-09 | 2.69E-08 | 63.6614542 | 232.67286 | 126.215061 | 176.759674 | 0 |
| 5.00E-09 | 2.69E-08 | 55.3018693 | 323.873503 | 180.042072 | 252.381266 | 0 |
| 5.00E-09 | 2.69E-08 | 59.4816618 | 172.391459 | 134.567528 | 154.490738 | 0 |
| 5.02E-09 | 2.70E-08 | 67.8412467 | 112.999821 | 92.8051917 | 99.2823364 | 0 |
| 5.03E-09 | 2.71E-08 | 61.0892743 | 157.932821 | 121.574801 | 149.387441 | 0 |
| 5.04E-09 | 2.71E-08 | 62.0538418 | 168.165088 | 119.718697 | 142.892335 | 0 |
| 5.05E-09 | 2.72E-08 | 64.6260217 | 103.879757 | 143.848047 | 126.654569 | 0 |
| 5.05E-09 | 2.72E-08 | 117.355711 | 80.3010539 | 77.956361 | 87.6839326 | 0 |
| 5.05E-09 | 2.72E-08 | 607.99904 | 2104.51045 | 1712.25579 | 2008.84354 | 2.37621824 |
| 5.06E-09 | 2.72E-08 | 53.6942569 | 191.743791 | 207.883629 | 202.276162 | 0 |
| 5.07E-09 | 2.73E-08 | 84.5604165 | 115.446667 | 127.143113 | 70.9822312 | 0 |
| 5.07E-09 | 2.73E-08 | 49.8359869 | 232.450419 | 277.487523 | 257.020628 | 0 |
| 5.07E-09 | 2.73E-08 | 66.2336342 | 279.830265 | 133.639476 | 171.19244 | 0 |
| 5.08E-09 | 2.73E-08 | 11.8963324 | 7.78542074 | 12.0646749 | 7.88691457 | 85.5438568 |
| 5.08E-09 | 2.73E-08 | 61.0892743 | 125.011613 | 147.560255 | 127.582442 | 0 |
| 5.08E-09 | 2.73E-08 | 94.2060914 | 227.334285 | 77.0283091 | 162.377653 | 0 |
| 5.10E-09 | 2.74E-08 | 66.2336342 | 113.889583 | 98.3735032 | 100.674145 | 0 |
| 5.11E-09 | 2.74E-08 | 62.0538418 | 131.23995 | 116.934542 | 121.551272 | 0 |
| 5.11E-09 | 2.74E-08 | 57.5525268 | 167.052885 | 150.344411 | 159.1301 | 0 |
| 5.11E-09 | 2.74E-08 | 66.2336342 | 158.600142 | 103.013763 | 127.118505 | 0 |
| 5.11E-09 | 2.74E-08 | 56.9094818 | 157.932821 | 173.545708 | 161.913717 | 0 |
| 5.11E-09 | 2.74E-08 | 56.2664368 | 156.375736 | 197.675058 | 197.172864 | 0 |
| 5.13E-09 | 2.75E-08 | 62.0538418 | 167.720207 | 117.862593 | 145.675952 | 0 |
| 5.14E-09 | 2.76E-08 | 63.9829767 | 129.015544 | 104.869867 | 140.572654 | 0 |
| 5.14E-09 | 2.76E-08 | 55.9449143 | 168.609969 | 206.955577 | 172.120312 | 0 |
| 5.15E-09 | 2.76E-08 | 2743.22994 | 385.044666 | 115.078438 | 237.071373 | 0 |
| 5.16E-09 | 2.77E-08 | 62.6968867 | 127.013578 | 118.790645 | 161.913717 | 0 |
| 5.16E-09 | 2.77E-08 | 443.058 | 114.334465 | 221.804408 | 86.7560603 | 0 |
| 5.16E-09 | 2.77E-08 | 56.2664368 | 176.840271 | 158.696878 | 184.646588 | 0 |
| 5.16E-09 | 2.77E-08 | 54.0157793 | 203.310701 | 180.970124 | 212.482757 | 0 |
| 5.17E-09 | 2.77E-08 | 684.521394 | 268.708236 | 144.776099 | 227.79265 | 2.37621824 |
| 5.18E-09 | 2.77E-08 | 81.9882365 | 78.0766479 | 77.0283091 | 94.6429749 | 0 |

| | | | | | | |
|---|---|---|---|---|---|---|
| 5.18E-09 | 2.78E-08 | 71.0564716 | 241.570483 | 104.869867 | 210.627013 | 0 |
| 5.19E-09 | 2.78E-08 | 0 | 0 | 0 | 0 | 49.9005831 |
| 5.19E-09 | 2.78E-08 | 75.2362641 | 182.623726 | 89.092984 | 136.861165 | 0 |
| 5.21E-09 | 2.79E-08 | 51.7651219 | 199.751652 | 315.537652 | 274.186265 | 0 |
| 5.22E-09 | 2.80E-08 | 58.1955718 | 143.696623 | 163.337137 | 147.995632 | 0 |
| 5.23E-09 | 2.80E-08 | 126.679863 | 233.562622 | 159.62493 | 219.905736 | 2.37621824 |
| 5.23E-09 | 2.80E-08 | 70.0919042 | 102.989994 | 85.3807764 | 92.3232941 | 0 |
| 5.25E-09 | 2.81E-08 | 57.5525268 | 205.312667 | 146.632203 | 179.543291 | 0 |
| 5.27E-09 | 2.82E-08 | 65.9121117 | 131.684831 | 101.157659 | 157.274355 | 0 |
| 5.28E-09 | 2.83E-08 | 61.7323193 | 156.153296 | 115.078438 | 145.675952 | 0 |
| 5.30E-09 | 2.83E-08 | 51.4435994 | 275.381453 | 206.955577 | 234.287756 | 0 |
| 5.30E-09 | 2.84E-08 | 72.6640841 | 146.58835 | 84.4527244 | 131.293931 | 0 |
| 5.31E-09 | 2.84E-08 | 63.3399317 | 131.46239 | 104.869867 | 134.077548 | 0 |
| 5.34E-09 | 2.85E-08 | 81.9882365 | 127.013578 | 225.516616 | 94.1790387 | 0 |
| 5.35E-09 | 2.86E-08 | 100.636541 | 193.078434 | 160.552982 | 152.171058 | 7.12865473 |
| 5.35E-09 | 2.86E-08 | 57.5525268 | 244.23977 | 150.344411 | 201.34829 | 0 |
| 5.37E-09 | 2.87E-08 | 69.1273367 | 191.52135 | 99.3015551 | 181.862971 | 0 |
| 5.42E-09 | 2.90E-08 | 63.0184092 | 126.346257 | 104.869867 | 124.334889 | 0 |
| 5.44E-09 | 2.91E-08 | 62.6968867 | 131.46239 | 106.72597 | 124.798825 | 0 |
| 5.44E-09 | 2.91E-08 | 125.393773 | 70.958549 | 83.5246725 | 109.024996 | 0 |
| 5.44E-09 | 2.91E-08 | 591.922915 | 850.167944 | 688.614522 | 800.289861 | 42.7719284 |
| 5.45E-09 | 2.91E-08 | 55.6233918 | 173.281222 | 165.193241 | 169.800631 | 0 |
| 5.45E-09 | 2.91E-08 | 97.4213163 | 288.060567 | 349.875573 | 311.765094 | 2.37621824 |
| 5.45E-09 | 2.91E-08 | 61.7323193 | 144.586385 | 173.545708 | 117.375846 | 0 |
| 5.48E-09 | 2.93E-08 | 124.429206 | 156.153296 | 148.488307 | 149.387441 | 0 |
| 5.49E-09 | 2.93E-08 | 66.8766792 | 101.877791 | 101.157659 | 90.0036134 | 0 |
| 5.49E-09 | 2.93E-08 | 85.8465065 | 165.495801 | 77.956361 | 108.561059 | 0 |
| 5.51E-09 | 2.94E-08 | 212.847892 | 121.452563 | 72.3880495 | 108.097123 | 0 |
| 5.52E-09 | 2.95E-08 | 59.8031843 | 122.119885 | 154.98467 | 153.562866 | 0 |
| 5.53E-09 | 2.95E-08 | 63.9829767 | 135.688761 | 107.654022 | 108.561059 | 0 |
| 5.54E-09 | 2.95E-08 | 79.094534 | 95.2045736 | 75.1722053 | 81.1888265 | 0 |
| 5.54E-09 | 2.96E-08 | 61.0892743 | 154.818652 | 116.00649 | 139.644782 | 0 |
| 5.55E-09 | 2.96E-08 | 120.570936 | 130.795068 | 62.1794784 | 104.84957 | 0 |
| 5.55E-09 | 2.96E-08 | 605.42686 | 124.566732 | 108.582074 | 188.358077 | 0 |
| 5.59E-09 | 2.98E-08 | 129.573566 | 260.922815 | 269.135056 | 252.381266 | 0 |
| 5.59E-09 | 2.98E-08 | 61.0892743 | 118.338395 | 136.423632 | 119.231591 | 0 |
| 5.59E-09 | 2.98E-08 | 51.4435994 | 226.889404 | 204.171422 | 218.513927 | 0 |
| 5.62E-09 | 3.00E-08 | 57.2310043 | 205.535107 | 219.948304 | 143.356271 | 0 |
| 5.62E-09 | 3.00E-08 | 53.0512119 | 186.182776 | 193.962851 | 216.194247 | 0 |
| 5.62E-09 | 3.00E-08 | 54.0157793 | 185.960335 | 180.970124 | 181.862971 | 0 |
| 5.63E-09 | 3.00E-08 | 54.3373018 | 203.755583 | 247.789862 | 170.264568 | 0 |
| 5.64E-09 | 3.00E-08 | 77.1653991 | 125.901375 | 199.531162 | 91.3954218 | 0 |
| 5.64E-09 | 3.00E-08 | 64.3044992 | 106.993925 | 98.3735032 | 114.592229 | 0 |

| | | | | | | |
|---|---|---|---|---|---|---|
| 5.64E-09 | 3.00E-08 | 62.6968867 | 104.769519 | 133.639476 | 138.252973 | 0 |
| 5.65E-09 | 3.00E-08 | 118.641801 | 93.86993 | 71.4599976 | 83.0445711 | 0 |
| 5.65E-09 | 3.00E-08 | 67.1982017 | 125.456494 | 90.0210359 | 131.293931 | 0 |
| 5.67E-09 | 3.02E-08 | 54.9803468 | 179.064677 | 162.409085 | 181.862971 | 0 |
| 5.68E-09 | 3.02E-08 | 65.5905892 | 111.442737 | 96.5173994 | 97.4265918 | 0 |
| 5.69E-09 | 3.02E-08 | 61.0892743 | 120.562801 | 115.078438 | 129.902122 | 0 |
| 5.70E-09 | 3.03E-08 | 139.540763 | 78.0766479 | 88.1649321 | 86.7560603 | 0 |
| 5.70E-09 | 3.03E-08 | 67.5197242 | 137.690727 | 90.9490879 | 117.375846 | 0 |
| 5.71E-09 | 3.03E-08 | 0 | 0 | 0 | 0 | 19.0097459 |
| 5.71E-09 | 3.04E-08 | 69.1273367 | 149.702519 | 283.983887 | 129.902122 | 0 |
| 5.71E-09 | 3.04E-08 | 70.0919042 | 128.348222 | 83.5246725 | 120.159463 | 0 |
| 5.72E-09 | 3.04E-08 | 60.4462293 | 176.840271 | 126.215061 | 137.325101 | 0 |
| 5.74E-09 | 3.05E-08 | 106.745469 | 234.897266 | 257.070381 | 254.237011 | 0 |
| 5.75E-09 | 3.05E-08 | 54.3373018 | 167.052885 | 182.826228 | 194.389247 | 0 |
| 5.76E-09 | 3.06E-08 | 62.6968867 | 159.267464 | 113.222334 | 122.94308 | 0 |
| 5.77E-09 | 3.06E-08 | 64.6260217 | 116.781311 | 96.5173994 | 105.777442 | 0 |
| 5.77E-09 | 3.06E-08 | 57.8740493 | 153.261568 | 136.423632 | 144.284143 | 0 |
| 5.78E-09 | 3.07E-08 | 48.8714194 | 366.804537 | 257.070381 | 239.391054 | 0 |
| 5.78E-09 | 3.07E-08 | 49.5144644 | 228.224048 | 229.228823 | 265.835415 | 0 |
| 5.78E-09 | 3.07E-08 | 72.3425616 | 171.056816 | 121.574801 | 90.9314857 | 0 |
| 5.79E-09 | 3.07E-08 | 88.0971639 | 131.23995 | 67.7477899 | 110.416804 | 0 |
| 5.80E-09 | 3.08E-08 | 72.6640841 | 201.753617 | 92.8051917 | 155.882547 | 0 |
| 5.81E-09 | 3.08E-08 | 75.5577866 | 76.2971232 | 83.5246725 | 86.2921241 | 0 |
| 5.81E-09 | 3.08E-08 | 41.476402 | 302.296765 | 503.932191 | 441.667216 | 0 |
| 5.83E-09 | 3.09E-08 | 215.09855 | 448.885116 | 389.781805 | 429.14094 | 0 |
| 5.84E-09 | 3.09E-08 | 61.0892743 | 115.669108 | 127.143113 | 115.056166 | 0 |
| 5.84E-09 | 3.09E-08 | 96.1352263 | 84.3049845 | 70.5319457 | 123.407016 | 0 |
| 5.84E-09 | 3.10E-08 | 127.644431 | 87.6415934 | 70.5319457 | 93.7151026 | 0 |
| 5.87E-09 | 3.11E-08 | 131.824223 | 270.48776 | 219.948304 | 205.987651 | 0 |
| 5.88E-09 | 3.11E-08 | 63.9829767 | 126.791138 | 97.4454513 | 130.829995 | 0 |
| 5.89E-09 | 3.12E-08 | 99.3504513 | 389.271037 | 298.832717 | 345.168497 | 4.75243649 |
| 5.90E-09 | 3.12E-08 | 71.3779941 | 109.885653 | 108.582074 | 78.4052096 | 0 |
| 5.91E-09 | 3.13E-08 | 130.216611 | 147.700553 | 176.329864 | 178.151482 | 0 |
| 5.91E-09 | 3.13E-08 | 74.5932191 | 208.871716 | 90.0210359 | 169.800631 | 0 |
| 5.91E-09 | 3.13E-08 | 147.578826 | 184.625692 | 113.222334 | 175.831801 | 19.0097459 |
| 5.92E-09 | 3.13E-08 | 64.6260217 | 94.3148112 | 104.869867 | 111.344676 | 0 |
| 5.93E-09 | 3.14E-08 | 56.9094818 | 169.944613 | 142.919995 | 150.315313 | 0 |
| 5.93E-09 | 3.14E-08 | 83.2743265 | 171.724137 | 75.1722053 | 136.397228 | 0 |
| 5.93E-09 | 3.14E-08 | 55.6233918 | 183.95837 | 148.488307 | 189.28595 | 0 |
| 5.98E-09 | 3.17E-08 | 59.1601393 | 149.480078 | 122.502853 | 136.861165 | 0 |
| 5.99E-09 | 3.17E-08 | 43.405537 | 36.4802572 | 39.9062324 | 33.4034029 | 163.959059 |
| 6.00E-09 | 3.17E-08 | 127.001386 | 211.318563 | 154.056618 | 186.966269 | 0 |
| 6.00E-09 | 3.17E-08 | 51.1220769 | 346.340002 | 200.459214 | 259.340309 | 0 |

| | | | | | | |
|---|---|---|---|---|---|---|
| 6.00E-09 | 3.17E-08 | 60.7677518 | 143.029301 | 111.36623 | 145.675952 | 0 |
| 6.02E-09 | 3.18E-08 | 86.168029 | 74.2951579 | 75.1722053 | 90.0036134 | 0 |
| 6.02E-09 | 3.18E-08 | 65.2690667 | 139.915133 | 153.128566 | 96.4987195 | 0 |
| 6.02E-09 | 3.18E-08 | 73.3071291 | 305.188493 | 230.156875 | 105.777442 | 0 |
| 6.03E-09 | 3.18E-08 | 66.5551567 | 105.881722 | 87.2368802 | 112.736485 | 0 |
| 6.03E-09 | 3.19E-08 | 75.5577866 | 85.4171875 | 103.941815 | 75.6215927 | 0 |
| 6.04E-09 | 3.19E-08 | 62.3753642 | 103.879757 | 124.358957 | 111.344676 | 0 |
| 6.04E-09 | 3.19E-08 | 56.5879593 | 140.804895 | 174.47376 | 157.738291 | 0 |
| 6.05E-09 | 3.19E-08 | 66.5551567 | 88.7537964 | 94.6612955 | 96.4987195 | 0 |
| 6.05E-09 | 3.19E-08 | 79.094534 | 122.787207 | 74.2441534 | 144.748079 | 0 |
| 6.07E-09 | 3.20E-08 | 201.916128 | 411.070215 | 325.746223 | 376.716155 | 0 |
| 6.09E-09 | 3.21E-08 | 64.6260217 | 105.659281 | 96.5173994 | 96.9626556 | 0 |
| 6.11E-09 | 3.23E-08 | 186.483048 | 214.210291 | 55.683115 | 144.284143 | 0 |
| 6.13E-09 | 3.23E-08 | 55.6233918 | 159.489905 | 150.344411 | 177.687546 | 0 |
| 6.14E-09 | 3.24E-08 | 61.7323193 | 125.011613 | 145.704151 | 107.169251 | 0 |
| 6.16E-09 | 3.25E-08 | 488.071149 | 155.930855 | 89.092984 | 143.820207 | 0 |
| 6.20E-09 | 3.27E-08 | 90.0262989 | 74.5175985 | 168.905449 | 106.241379 | 0 |
| 6.20E-09 | 3.27E-08 | 50.4790319 | 192.188672 | 246.86181 | 261.196053 | 0 |
| 6.22E-09 | 3.28E-08 | 65.2690667 | 125.234054 | 273.775316 | 168.872759 | 0 |
| 6.24E-09 | 3.29E-08 | 60.7677518 | 121.452563 | 109.510126 | 131.293931 | 0 |
| 6.24E-09 | 3.29E-08 | 58.1955718 | 171.279256 | 154.056618 | 126.190633 | 0 |
| 6.24E-09 | 3.29E-08 | 39.8687895 | 304.29873 | 614.370369 | 521.000298 | 0 |
| 6.26E-09 | 3.30E-08 | 109.317649 | 88.3089152 | 69.6038938 | 83.0445711 | 0 |
| 6.27E-09 | 3.30E-08 | 1477.39587 | 3220.05002 | 2155.8646 | 2699.18053 | 42.7719284 |
| 6.27E-09 | 3.30E-08 | 53.3727344 | 237.121672 | 248.717914 | 163.769461 | 0 |
| 6.27E-09 | 3.30E-08 | 62.0538418 | 117.448633 | 102.085711 | 114.128293 | 0 |
| 6.27E-09 | 3.30E-08 | 57.5525268 | 129.015544 | 167.049345 | 175.831801 | 0 |
| 6.30E-09 | 3.32E-08 | 57.5525268 | 143.029301 | 129.927268 | 148.459568 | 0 |
| 6.32E-09 | 3.33E-08 | 104.816334 | 107.216366 | 68.6758419 | 79.797018 | 0 |
| 6.33E-09 | 3.33E-08 | 59.1601393 | 145.698588 | 118.790645 | 132.221803 | 0 |
| 6.34E-09 | 3.34E-08 | 123.786161 | 96.3167765 | 73.3161014 | 78.4052096 | 0 |
| 6.34E-09 | 3.34E-08 | 70.7349491 | 136.133643 | 82.5966206 | 141.964462 | 0 |
| 6.37E-09 | 3.35E-08 | 113.175919 | 194.635518 | 193.034799 | 210.627013 | 2.37621824 |
| 6.38E-09 | 3.35E-08 | 60.7677518 | 107.438806 | 120.646749 | 120.623399 | 0 |
| 6.38E-09 | 3.36E-08 | 27.9724572 | 764.528316 | 541.054268 | 797.506244 | 0 |
| 6.40E-09 | 3.37E-08 | 64.9475442 | 90.0884399 | 115.078438 | 103.921698 | 0 |
| 6.41E-09 | 3.37E-08 | 72.9856066 | 231.338216 | 238.509343 | 96.0347833 | 0 |
| 6.42E-09 | 3.37E-08 | 58.8386168 | 132.352152 | 136.423632 | 118.767655 | 0 |
| 6.42E-09 | 3.37E-08 | 111.246784 | 84.7498657 | 128.071165 | 67.7346781 | 0 |
| 6.42E-09 | 3.37E-08 | 138.897718 | 224.887439 | 141.063891 | 170.264568 | 0 |
| 6.42E-09 | 3.37E-08 | 138.897718 | 66.064856 | 153.128566 | 92.7872303 | 0 |
| 6.43E-09 | 3.38E-08 | 48.8714194 | 226.889404 | 221.804408 | 243.566479 | 0 |
| 6.44E-09 | 3.38E-08 | 63.9829767 | 127.458459 | 184.682331 | 110.416804 | 0 |

| | | | | | | |
|---|---|---|---|---|---|---|
| 6.44E-09 | 3.38E-08 | 60.1247068 | 125.678935 | 120.646749 | 111.344676 | 0 |
| 6.47E-09 | 3.39E-08 | 117.677234 | 148.145435 | 147.560255 | 163.305525 | 4.75243649 |
| 6.49E-09 | 3.41E-08 | 57.5525268 | 129.237984 | 138.279736 | 155.882547 | 0 |
| 6.50E-09 | 3.41E-08 | 58.1955718 | 189.964266 | 131.783372 | 226.400842 | 0 |
| 6.50E-09 | 3.41E-08 | 67.1982017 | 94.3148112 | 111.36623 | 85.3642518 | 0 |
| 6.50E-09 | 3.41E-08 | 108.031559 | 208.204395 | 241.293498 | 220.369672 | 0 |
| 6.51E-09 | 3.42E-08 | 54.0157793 | 179.287118 | 179.11402 | 155.882547 | 0 |
| 6.53E-09 | 3.43E-08 | 106.102424 | 66.5097371 | 234.797135 | 199.492545 | 0 |
| 6.54E-09 | 3.43E-08 | 66.2336342 | 86.7518311 | 96.5173994 | 109.488932 | 0 |
| 6.54E-09 | 3.43E-08 | 81.666714 | 83.1927816 | 167.977397 | 98.3544641 | 0 |
| 6.55E-09 | 3.43E-08 | 54.3373018 | 258.031087 | 163.337137 | 192.997439 | 0 |
| 6.55E-09 | 3.44E-08 | 69.4488592 | 111.220296 | 78.8844129 | 104.84957 | 0 |
| 6.56E-09 | 3.44E-08 | 89.7047764 | 194.635518 | 128.999216 | 191.141694 | 7.12865473 |
| 6.56E-09 | 3.44E-08 | 90.3478214 | 151.926925 | 96.5173994 | 72.8379758 | 0 |
| 6.57E-09 | 3.44E-08 | 57.5525268 | 140.137573 | 127.143113 | 139.644782 | 0 |
| 6.57E-09 | 3.44E-08 | 55.9449143 | 145.698588 | 182.826228 | 148.459568 | 0 |
| 6.57E-09 | 3.44E-08 | 71.0564716 | 79.1888509 | 83.5246725 | 85.3642518 | 0 |
| 6.58E-09 | 3.45E-08 | 533.405821 | 528.963729 | 458.457647 | 431.460621 | 0 |
| 6.61E-09 | 3.46E-08 | 37.618132 | 376.369482 | 526.205437 | 401.304771 | 0 |
| 6.62E-09 | 3.46E-08 | 165.262563 | 296.290869 | 248.717914 | 266.763287 | 40.3957101 |
| 6.62E-09 | 3.47E-08 | 93.2415239 | 79.4112915 | 70.5319457 | 82.1166988 | 0 |
| 6.63E-09 | 3.47E-08 | 75.8793091 | 90.7557617 | 194.890903 | 117.375846 | 0 |
| 6.65E-09 | 3.48E-08 | 54.9803468 | 176.840271 | 181.898176 | 143.356271 | 0 |
| 6.66E-09 | 3.49E-08 | 81.9882365 | 92.9801676 | 87.2368802 | 68.1986142 | 0 |
| 6.66E-09 | 3.49E-08 | 59.4816618 | 175.283187 | 117.862593 | 186.966269 | 0 |
| 6.67E-09 | 3.49E-08 | 48.8714194 | 234.007503 | 207.883629 | 254.237011 | 0 |
| 6.68E-09 | 3.49E-08 | 125.072251 | 228.668929 | 164.265189 | 171.656376 | 0 |
| 6.72E-09 | 3.52E-08 | 72.9856066 | 161.936751 | 81.6685687 | 135.00542 | 0 |
| 6.74E-09 | 3.52E-08 | 85.524984 | 206.202429 | 75.1722053 | 169.336695 | 0 |
| 6.75E-09 | 3.53E-08 | 55.3018693 | 160.602108 | 148.488307 | 147.06776 | 0 |
| 6.78E-09 | 3.54E-08 | 256.896474 | 148.367875 | 167.049345 | 76.549465 | 4.75243649 |
| 6.78E-09 | 3.54E-08 | 136.004016 | 76.0746826 | 77.0283091 | 99.2823364 | 0 |
| 6.80E-09 | 3.55E-08 | 73.3071291 | 205.979989 | 137.351684 | 91.859358 | 0 |
| 6.80E-09 | 3.55E-08 | 225.38727 | 603.036446 | 528.061541 | 533.526574 | 19.0097459 |
| 6.81E-09 | 3.56E-08 | 71.6995166 | 217.546899 | 104.869867 | 122.015208 | 0 |
| 6.81E-09 | 3.56E-08 | 62.3753642 | 165.27336 | 104.869867 | 132.221803 | 0 |
| 6.88E-09 | 3.59E-08 | 102.887199 | 220.661068 | 272.847264 | 256.556692 | 0 |
| 6.89E-09 | 3.60E-08 | 54.3373018 | 147.478113 | 174.47376 | 167.017014 | 0 |
| 6.91E-09 | 3.61E-08 | 57.2310043 | 145.698588 | 203.24337 | 139.644782 | 0 |
| 6.92E-09 | 3.61E-08 | 87.4541189 | 137.913167 | 137.351684 | 66.3428696 | 0 |
| 6.92E-09 | 3.61E-08 | 61.4107968 | 144.363944 | 197.675058 | 115.056166 | 0 |
| 6.92E-09 | 3.61E-08 | 164.619518 | 793.223153 | 384.213494 | 569.249658 | 0 |
| 6.93E-09 | 3.61E-08 | 91.9554339 | 178.619796 | 67.7477899 | 141.03659 | 0 |

| | | | | | | |
|---|---|---|---|---|---|---|
| 6.94E-09 | 3.62E-08 | 73.6286516 | 115.001786 | 103.941815 | 73.3019119 | 0 |
| 6.95E-09 | 3.62E-08 | 82.309759 | 71.8483114 | 83.5246725 | 77.0134011 | 0 |
| 6.95E-09 | 3.63E-08 | 53.3727344 | 192.633553 | 165.193241 | 162.377653 | 0 |
| 6.98E-09 | 3.64E-08 | 63.3399317 | 119.450598 | 115.078438 | 91.3954218 | 0 |
| 6.99E-09 | 3.65E-08 | 61.4107968 | 107.438806 | 112.294282 | 144.284143 | 0 |
| 7.00E-09 | 3.65E-08 | 50.8005544 | 189.964266 | 189.322591 | 222.225416 | 0 |
| 7.01E-09 | 3.65E-08 | 77.1653991 | 98.9860636 | 86.3088283 | 70.518295 | 0 |
| 7.02E-09 | 3.66E-08 | 123.143116 | 191.52135 | 150.344411 | 192.533503 | 0 |
| 7.05E-09 | 3.67E-08 | 65.2690667 | 109.218331 | 136.423632 | 209.235204 | 0 |
| 7.06E-09 | 3.68E-08 | 89.3832539 | 116.55887 | 74.2441534 | 75.6215927 | 0 |
| 7.07E-09 | 3.68E-08 | 60.7677518 | 117.893514 | 101.157659 | 121.087335 | 0 |
| 7.08E-09 | 3.69E-08 | 63.0184092 | 108.328569 | 91.8771398 | 105.313506 | 0 |
| 7.12E-09 | 3.70E-08 | 82.952804 | 119.895479 | 77.0283091 | 183.718716 | 0 |
| 7.14E-09 | 3.72E-08 | 94.5276139 | 185.293013 | 167.977397 | 167.480951 | 7.12865473 |
| 7.15E-09 | 3.72E-08 | 93.8845689 | 99.6533854 | 64.0355823 | 87.2199964 | 0 |
| 7.17E-09 | 3.73E-08 | 69.7703817 | 76.7420044 | 89.092984 | 86.7560603 | 0 |
| 7.20E-09 | 3.75E-08 | 60.4462293 | 161.714311 | 108.582074 | 132.221803 | 0 |
| 7.23E-09 | 3.76E-08 | 312.519866 | 496.264962 | 476.090633 | 488.524767 | 28.5146189 |
| 7.23E-09 | 3.76E-08 | 57.8740493 | 141.917098 | 124.358957 | 120.623399 | 0 |
| 7.24E-09 | 3.77E-08 | 61.0892743 | 192.411112 | 119.718697 | 247.277969 | 0 |
| 7.24E-09 | 3.77E-08 | 69.7703817 | 81.1908162 | 83.5246725 | 80.2609542 | 0 |
| 7.25E-09 | 3.77E-08 | 38.5826995 | 367.694299 | 395.350117 | 358.158709 | 0 |
| 7.26E-09 | 3.77E-08 | 90.3478214 | 157.265499 | 69.6038938 | 103.921698 | 0 |
| 7.35E-09 | 3.82E-08 | 72.0210391 | 157.265499 | 84.4527244 | 108.561059 | 0 |
| 7.35E-09 | 3.82E-08 | 144.685123 | 141.027336 | 139.207788 | 191.60563 | 0 |
| 7.36E-09 | 3.82E-08 | 60.7677518 | 106.993925 | 101.157659 | 116.447974 | 0 |
| 7.38E-09 | 3.83E-08 | 98.0643613 | 72.2931925 | 87.2368802 | 73.3019119 | 0 |
| 7.40E-09 | 3.85E-08 | 57.2310043 | 118.560836 | 154.98467 | 157.738291 | 0 |
| 7.42E-09 | 3.85E-08 | 58.1955718 | 128.570662 | 114.150386 | 122.94308 | 0 |
| 7.42E-09 | 3.85E-08 | 123.143116 | 256.474003 | 157.768826 | 192.997439 | 2.37621824 |
| 7.42E-09 | 3.85E-08 | 61.0892743 | 118.338395 | 104.869867 | 100.210209 | 0 |
| 7.43E-09 | 3.86E-08 | 54.6588243 | 184.625692 | 142.919995 | 223.153289 | 0 |
| 7.43E-09 | 3.86E-08 | 77.4869216 | 73.1829549 | 82.5966206 | 76.0855288 | 0 |
| 7.43E-09 | 3.86E-08 | 69.7703817 | 90.5333211 | 75.1722053 | 98.3544641 | 0 |
| 7.43E-09 | 3.86E-08 | 66.5551567 | 99.2085042 | 101.157659 | 155.882547 | 0 |
| 7.47E-09 | 3.88E-08 | 61.7323193 | 181.289083 | 106.72597 | 139.644782 | 0 |
| 7.47E-09 | 3.88E-08 | 58.1955718 | 115.001786 | 122.502853 | 120.159463 | 0 |
| 7.48E-09 | 3.88E-08 | 54.9803468 | 142.80686 | 149.416359 | 140.572654 | 0 |
| 7.49E-09 | 3.88E-08 | 61.7323193 | 94.3148112 | 105.797919 | 109.952868 | 0 |
| 7.49E-09 | 3.88E-08 | 50.8005544 | 220.883508 | 193.962851 | 174.903929 | 0 |
| 7.51E-09 | 3.90E-08 | 70.4134267 | 85.4171875 | 75.1722053 | 96.4987195 | 0 |
| 7.52E-09 | 3.90E-08 | 34.7244296 | 46.2676432 | 53.8270112 | 64.9510612 | 453.857685 |
| 7.52E-09 | 3.90E-08 | 123.786161 | 119.005717 | 73.3161014 | 72.8379758 | 0 |

| | | | | | | |
|---|---|---|---|---|---|---|
| 7.53E-09 | 3.90E-08 | 50.4790319 | 182.401286 | 187.466487 | 212.482757 | 0 |
| 7.58E-09 | 3.93E-08 | 54.9803468 | 135.911202 | 195.818954 | 212.018821 | 0 |
| 7.59E-09 | 3.93E-08 | 100.958064 | 236.009469 | 246.86181 | 269.546904 | 0 |
| 7.59E-09 | 3.93E-08 | 60.4462293 | 98.5411824 | 117.862593 | 121.551272 | 0 |
| 7.60E-09 | 3.93E-08 | 70.0919042 | 99.6533854 | 76.1002572 | 116.91191 | 0 |
| 7.62E-09 | 3.95E-08 | 239.53426 | 193.968197 | 67.7477899 | 98.8184002 | 0 |
| 7.63E-09 | 3.95E-08 | 55.6233918 | 136.133643 | 139.207788 | 134.077548 | 0 |
| 7.68E-09 | 3.97E-08 | 76.8438766 | 92.757727 | 71.4599976 | 121.087335 | 0 |
| 7.68E-09 | 3.98E-08 | 99.6719738 | 60.5038411 | 130.85532 | 137.325101 | 0 |
| 7.71E-09 | 3.99E-08 | 423.123605 | 138.135608 | 68.6758419 | 197.172864 | 0 |
| 7.72E-09 | 3.99E-08 | 76.5223541 | 73.8502767 | 75.1722053 | 81.1888265 | 0 |
| 7.72E-09 | 4.00E-08 | 61.4107968 | 114.112024 | 93.7332436 | 112.272549 | 0 |
| 7.73E-09 | 4.00E-08 | 121.213981 | 156.375736 | 55.683115 | 157.274355 | 0 |
| 7.74E-09 | 4.00E-08 | 62.6968867 | 92.5352865 | 93.7332436 | 101.602017 | 0 |
| 7.75E-09 | 4.01E-08 | 52.7296894 | 162.604073 | 161.481034 | 160.521908 | 0 |
| 7.77E-09 | 4.02E-08 | 52.0866444 | 161.046989 | 186.538435 | 168.408823 | 0 |
| 7.78E-09 | 4.02E-08 | 57.5525268 | 125.011613 | 176.329864 | 131.293931 | 0 |
| 7.80E-09 | 4.03E-08 | 233.10381 | 97.6514201 | 85.3807764 | 102.065953 | 0 |
| 7.81E-09 | 4.03E-08 | 47.9068519 | 256.029122 | 202.315318 | 221.76148 | 0 |
| 7.81E-09 | 4.04E-08 | 70.4134267 | 85.8620687 | 110.438178 | 77.4773373 | 0 |
| 7.85E-09 | 4.05E-08 | 66.5551567 | 114.779346 | 79.8124649 | 106.705315 | 0 |
| 7.85E-09 | 4.06E-08 | 55.6233918 | 153.706449 | 125.287009 | 149.851377 | 0 |
| 7.85E-09 | 4.06E-08 | 60.1247068 | 111.220296 | 108.582074 | 99.7462725 | 0 |
| 7.86E-09 | 4.06E-08 | 69.7703817 | 155.263534 | 111.36623 | 84.9003157 | 0 |
| 7.87E-09 | 4.06E-08 | 52.4081669 | 172.6139 | 157.768826 | 212.018821 | 0 |
| 7.88E-09 | 4.07E-08 | 337.598621 | 1253.2303 | 1747.52176 | 1618.67323 | 23.7621824 |
| 7.88E-09 | 4.07E-08 | 67.5197242 | 94.7596924 | 77.0283091 | 90.9314857 | 0 |
| 7.90E-09 | 4.08E-08 | 77.8084441 | 86.3069499 | 81.6685687 | 68.6625504 | 0 |
| 7.90E-09 | 4.08E-08 | 84.238894 | 64.0628906 | 102.085711 | 99.2823364 | 0 |
| 7.91E-09 | 4.08E-08 | 57.8740493 | 154.596212 | 113.222334 | 133.613612 | 0 |
| 7.94E-09 | 4.09E-08 | 60.1247068 | 108.328569 | 116.934542 | 154.490738 | 0 |
| 7.98E-09 | 4.12E-08 | 56.9094818 | 118.338395 | 136.423632 | 123.870952 | 0 |
| 7.99E-09 | 4.12E-08 | 58.1955718 | 160.379667 | 111.36623 | 139.180845 | 0 |
| 7.99E-09 | 4.12E-08 | 155.938411 | 53.8306234 | 237.581291 | 166.553078 | 0 |
| 7.99E-09 | 4.12E-08 | 48.5498969 | 215.544934 | 205.099474 | 198.100737 | 0 |
| 8.04E-09 | 4.14E-08 | 63.3399317 | 89.4211182 | 90.0210359 | 100.674145 | 0 |
| 8.05E-09 | 4.15E-08 | 50.8005544 | 199.306771 | 248.717914 | 168.872759 | 0 |
| 8.05E-09 | 4.15E-08 | 59.1601393 | 108.77345 | 104.869867 | 119.695527 | 0 |
| 8.07E-09 | 4.16E-08 | 64.3044992 | 141.249776 | 92.8051917 | 104.84957 | 0 |
| 8.07E-09 | 4.16E-08 | 100.958064 | 203.533142 | 66.819738 | 126.654569 | 0 |
| 8.13E-09 | 4.18E-08 | 44.691627 | 326.54279 | 425.97583 | 223.617225 | 0 |
| 8.13E-09 | 4.18E-08 | 54.3373018 | 169.722172 | 140.135839 | 141.964462 | 0 |
| 8.13E-09 | 4.18E-08 | 62.6968867 | 97.4289795 | 89.092984 | 94.6429749 | 0 |

| | | | | | | |
|---|---|---|---|---|---|---|
| 8.13E-09 | 4.18E-08 | 64.3044992 | 90.3108805 | 103.941815 | 84.9003157 | 0 |
| 8.13E-09 | 4.19E-08 | 92.9200014 | 73.1829549 | 75.1722053 | 75.6215927 | 0 |
| 8.14E-09 | 4.19E-08 | 138.576196 | 268.708236 | 310.897392 | 327.538923 | 14.2573095 |
| 8.14E-09 | 4.19E-08 | 54.3373018 | 143.919063 | 149.416359 | 135.933292 | 0 |
| 8.14E-09 | 4.19E-08 | 66.8766792 | 84.0825439 | 79.8124649 | 95.5708472 | 0 |
| 8.16E-09 | 4.20E-08 | 372.323051 | 552.542432 | 567.967773 | 666.212313 | 23.7621824 |
| 8.18E-09 | 4.20E-08 | 282.618274 | 563.442021 | 490.939464 | 533.526574 | 21.3859642 |
| 8.18E-09 | 4.21E-08 | 137.933151 | 340.334106 | 301.616873 | 325.219242 | 14.2573095 |
| 8.19E-09 | 4.21E-08 | 58.1955718 | 119.228158 | 106.72597 | 124.798825 | 0 |
| 8.22E-09 | 4.23E-08 | 45.6561944 | 221.55083 | 267.278952 | 245.88616 | 0 |
| 8.23E-09 | 4.23E-08 | 56.5879593 | 127.6809 | 117.862593 | 127.582442 | 0 |
| 8.26E-09 | 4.24E-08 | 52.4081669 | 158.155261 | 173.545708 | 152.634994 | 0 |
| 8.26E-09 | 4.24E-08 | 43.405537 | 285.613721 | 245.005706 | 313.156902 | 0 |
| 8.27E-09 | 4.25E-08 | 53.3727344 | 142.139539 | 154.98467 | 154.490738 | 0 |
| 8.28E-09 | 4.25E-08 | 48.2283744 | 200.418974 | 269.135056 | 204.595843 | 0 |
| 8.30E-09 | 4.26E-08 | 108.996126 | 138.80293 | 97.4454513 | 81.6527626 | 2.37621824 |
| 8.32E-09 | 4.27E-08 | 61.4107968 | 103.879757 | 193.034799 | 155.882547 | 0 |
| 8.32E-09 | 4.27E-08 | 507.040976 | 142.361979 | 99.3015551 | 128.510314 | 0 |
| 8.33E-09 | 4.27E-08 | 163.976473 | 76.5195638 | 90.0210359 | 88.6118049 | 0 |
| 8.33E-09 | 4.27E-08 | 60.4462293 | 95.8718953 | 103.941815 | 115.056166 | 0 |
| 8.34E-09 | 4.28E-08 | 235.032945 | 780.321598 | 580.9605 | 657.861463 | 35.6432737 |
| 8.35E-09 | 4.28E-08 | 53.6942569 | 153.261568 | 157.768826 | 224.081161 | 0 |
| 8.36E-09 | 4.29E-08 | 105.459379 | 93.86993 | 60.3233746 | 121.551272 | 0 |
| 8.36E-09 | 4.29E-08 | 77.4869216 | 72.9605143 | 85.3807764 | 72.3740396 | 0 |
| 8.37E-09 | 4.29E-08 | 67.5197242 | 86.3069499 | 92.8051917 | 76.0855288 | 0 |
| 8.41E-09 | 4.31E-08 | 74.2716966 | 112.55494 | 68.6758419 | 108.097123 | 0 |
| 8.42E-09 | 4.32E-08 | 249.179934 | 526.072001 | 43.6184401 | 326.147114 | 0 |
| 8.42E-09 | 4.32E-08 | 77.8084441 | 78.7439697 | 71.4599976 | 75.1576565 | 0 |
| 8.42E-09 | 4.32E-08 | 54.6588243 | 134.131677 | 136.423632 | 137.325101 | 0 |
| 8.46E-09 | 4.33E-08 | 50.1575094 | 205.312667 | 169.833501 | 184.182652 | 0 |
| 8.49E-09 | 4.35E-08 | 61.0892743 | 172.169019 | 103.941815 | 127.582442 | 0 |
| 8.50E-09 | 4.36E-08 | 50.4790319 | 270.710201 | 171.689605 | 202.276162 | 0 |
| 8.51E-09 | 4.36E-08 | 59.1601393 | 101.43291 | 111.36623 | 105.313506 | 0 |
| 8.52E-09 | 4.37E-08 | 82.6312815 | 82.0805786 | 64.9636342 | 90.9314857 | 0 |
| 8.53E-09 | 4.37E-08 | 95.8137038 | 159.045024 | 67.7477899 | 96.0347833 | 0 |
| 8.53E-09 | 4.37E-08 | 93.8845689 | 89.4211182 | 65.8916861 | 77.0134011 | 0 |
| 8.54E-09 | 4.37E-08 | 59.4816618 | 183.735929 | 110.438178 | 137.789037 | 0 |
| 8.54E-09 | 4.37E-08 | 99.6719738 | 186.627657 | 154.056618 | 151.243185 | 11.8810912 |
| 8.54E-09 | 4.37E-08 | 76.5223541 | 177.507593 | 185.610383 | 77.0134011 | 0 |
| 8.55E-09 | 4.38E-08 | 77.8084441 | 88.3089152 | 83.5246725 | 66.8068058 | 0 |
| 8.61E-09 | 4.40E-08 | 58.5170943 | 107.883687 | 104.869867 | 112.272549 | 0 |
| 8.62E-09 | 4.41E-08 | 54.6588243 | 138.580489 | 140.135839 | 128.046378 | 0 |
| 8.64E-09 | 4.42E-08 | 97.7428388 | 466.235482 | 243.149602 | 302.486371 | 2.37621824 |

| | | | | | | |
|---|---|---|---|---|---|---|
| 8.67E-09 | 4.43E-08 | 69.4488592 | 84.5274251 | 183.75428 | 162.377653 | 0 |
| 8.67E-09 | 4.43E-08 | 56.5879593 | 131.017509 | 112.294282 | 139.180845 | 0 |
| 8.68E-09 | 4.44E-08 | 114.140486 | 453.333927 | 232.012979 | 345.632433 | 9.50487297 |
| 8.70E-09 | 4.45E-08 | 77.1653991 | 67.8443807 | 126.215061 | 113.664357 | 0 |
| 8.71E-09 | 4.45E-08 | 49.8359869 | 202.420939 | 174.47376 | 177.687546 | 0 |
| 8.72E-09 | 4.45E-08 | 62.0538418 | 123.676969 | 90.0210359 | 103.921698 | 0 |
| 8.72E-09 | 4.45E-08 | 108.996126 | 201.976058 | 232.941031 | 216.194247 | 11.8810912 |
| 8.72E-09 | 4.45E-08 | 58.1955718 | 106.771484 | 112.294282 | 107.633187 | 0 |
| 8.72E-09 | 4.45E-08 | 51.7651219 | 149.924959 | 166.121293 | 175.831801 | 0 |
| 8.72E-09 | 4.46E-08 | 146.292736 | 163.716276 | 154.98467 | 122.015208 | 0 |
| 8.75E-09 | 4.47E-08 | 51.4435994 | 365.025012 | 269.135056 | 166.553078 | 0 |
| 8.77E-09 | 4.48E-08 | 115.105054 | 91.8679647 | 77.0283091 | 68.1986142 | 0 |
| 8.79E-09 | 4.49E-08 | 117.677234 | 183.735929 | 159.62493 | 158.202228 | 0 |
| 8.81E-09 | 4.50E-08 | 45.6561944 | 228.446488 | 246.86181 | 219.905736 | 0 |
| 8.82E-09 | 4.50E-08 | 123.464639 | 271.822404 | 187.466487 | 59.3838274 | 0 |
| 8.82E-09 | 4.50E-08 | 42.4409695 | 255.806681 | 279.343627 | 305.269988 | 0 |
| 8.83E-09 | 4.50E-08 | 72.9856066 | 152.149365 | 76.1002572 | 114.128293 | 0 |
| 8.84E-09 | 4.51E-08 | 51.4435994 | 167.275326 | 154.056618 | 167.480951 | 0 |
| 8.84E-09 | 4.51E-08 | 87.4541189 | 190.186707 | 67.7477899 | 175.831801 | 0 |
| 8.86E-09 | 4.52E-08 | 50.4790319 | 209.539038 | 155.912722 | 208.307332 | 0 |
| 8.88E-09 | 4.53E-08 | 54.6588243 | 144.141504 | 123.430905 | 145.675952 | 0 |
| 8.88E-09 | 4.53E-08 | 57.2310043 | 147.255672 | 114.150386 | 118.303719 | 0 |
| 8.90E-09 | 4.53E-08 | 53.0512119 | 154.596212 | 140.135839 | 146.139888 | 0 |
| 8.91E-09 | 4.54E-08 | 79.094534 | 115.446667 | 65.8916861 | 128.046378 | 0 |
| 8.95E-09 | 4.56E-08 | 55.3018693 | 131.46239 | 122.502853 | 126.654569 | 0 |
| 8.95E-09 | 4.56E-08 | 60.7677518 | 89.4211182 | 105.797919 | 108.097123 | 0 |
| 8.96E-09 | 4.56E-08 | 59.1601393 | 96.3167765 | 114.150386 | 109.024996 | 0 |
| 9.01E-09 | 4.59E-08 | 90.6693439 | 98.3187419 | 89.092984 | 61.7035081 | 0 |
| 9.01E-09 | 4.59E-08 | 133.110313 | 268.040914 | 135.49558 | 199.956481 | 2.37621824 |
| 9.02E-09 | 4.59E-08 | 52.4081669 | 151.926925 | 147.560255 | 150.315313 | 0 |
| 9.03E-09 | 4.60E-08 | 160.439726 | 183.068608 | 94.6612955 | 141.03659 | 7.12865473 |
| 9.04E-09 | 4.60E-08 | 59.4816618 | 95.8718953 | 105.797919 | 102.529889 | 0 |
| 9.05E-09 | 4.61E-08 | 54.3373018 | 139.915133 | 125.287009 | 145.212015 | 0 |
| 9.07E-09 | 4.62E-08 | 50.1575094 | 162.604073 | 197.675058 | 175.831801 | 0 |
| 9.09E-09 | 4.62E-08 | 55.6233918 | 151.037162 | 117.862593 | 131.293931 | 0 |
| 9.09E-09 | 4.62E-08 | 58.5170943 | 104.547078 | 106.72597 | 102.529889 | 0 |
| 9.09E-09 | 4.62E-08 | 59.8031843 | 158.600142 | 102.085711 | 122.94308 | 0 |
| 9.14E-09 | 4.65E-08 | 64.9475442 | 147.478113 | 134.567528 | 84.4363795 | 0 |
| 9.14E-09 | 4.65E-08 | 68.8058142 | 88.976237 | 171.689605 | 108.561059 | 0 |
| 9.18E-09 | 4.67E-08 | 70.0919042 | 78.0766479 | 73.3161014 | 81.1888265 | 0 |
| 9.19E-09 | 4.67E-08 | 70.4134267 | 109.885653 | 76.1002572 | 82.1166988 | 0 |
| 9.20E-09 | 4.68E-08 | 61.0892743 | 117.671073 | 90.0210359 | 101.602017 | 0 |
| 9.22E-09 | 4.69E-08 | 85.8465065 | 191.52135 | 168.905449 | 183.718716 | 2.37621824 |

| | | | | | | |
|---|---|---|---|---|---|---|
| 9.26E-09 | 4.71E-08 | 0 | 0 | 0 | 0 | 42.7719284 |
| 9.27E-09 | 4.71E-08 | 51.4435994 | 169.05485 | 147.560255 | 164.233398 | 0 |
| 9.27E-09 | 4.71E-08 | 54.6588243 | 122.342326 | 132.711424 | 133.613612 | 0 |
| 9.28E-09 | 4.71E-08 | 54.0157793 | 154.151331 | 128.071165 | 136.861165 | 0 |
| 9.29E-09 | 4.72E-08 | 62.6968867 | 154.596212 | 90.0210359 | 147.995632 | 0 |
| 9.30E-09 | 4.72E-08 | 54.9803468 | 154.373771 | 123.430905 | 128.97425 | 0 |
| 9.32E-09 | 4.73E-08 | 229.567062 | 393.942289 | 292.336354 | 360.014453 | 19.0097459 |
| 9.33E-09 | 4.74E-08 | 54.6588243 | 151.704484 | 128.999216 | 125.262761 | 0 |
| 9.34E-09 | 4.74E-08 | 119.284846 | 142.80686 | 190.250643 | 185.110524 | 0 |
| 9.36E-09 | 4.75E-08 | 79.4160565 | 106.104163 | 202.315318 | 85.3642518 | 0 |
| 9.36E-09 | 4.75E-08 | 68.4842917 | 88.0864746 | 71.4599976 | 89.0757411 | 0 |
| 9.39E-09 | 4.76E-08 | 40.5118345 | 323.206181 | 296.048562 | 277.433818 | 0 |
| 9.40E-09 | 4.77E-08 | 71.6995166 | 123.232088 | 70.5319457 | 118.303719 | 0 |
| 9.41E-09 | 4.77E-08 | 58.1955718 | 98.0963013 | 116.934542 | 112.272549 | 0 |
| 9.42E-09 | 4.77E-08 | 54.0157793 | 138.135608 | 127.143113 | 134.541484 | 0 |
| 9.42E-09 | 4.77E-08 | 48.2283744 | 251.357869 | 173.545708 | 217.122119 | 0 |
| 9.46E-09 | 4.79E-08 | 53.3727344 | 156.153296 | 130.85532 | 141.03659 | 0 |
| 9.47E-09 | 4.80E-08 | 44.691627 | 261.812577 | 210.667785 | 256.092756 | 0 |
| 9.48E-09 | 4.80E-08 | 78.7730116 | 78.9664103 | 66.819738 | 99.7462725 | 0 |
| 9.49E-09 | 4.81E-08 | 104.816334 | 301.629443 | 279.343627 | 315.476583 | 0 |
| 9.50E-09 | 4.81E-08 | 76.8438766 | 73.8502767 | 128.071165 | 83.5085072 | 0 |
| 9.54E-09 | 4.83E-08 | 73.6286516 | 73.8502767 | 78.8844129 | 71.9101035 | 0 |
| 9.56E-09 | 4.84E-08 | 128.608998 | 145.031266 | 88.1649321 | 62.1674443 | 0 |
| 9.57E-09 | 4.84E-08 | 51.1220769 | 190.854028 | 147.560255 | 167.480951 | 0 |
| 9.58E-09 | 4.85E-08 | 62.3753642 | 98.5411824 | 90.0210359 | 129.902122 | 0 |
| 9.61E-09 | 4.86E-08 | 41.476402 | 279.385384 | 287.696094 | 265.371479 | 0 |
| 9.63E-09 | 4.87E-08 | 90.9908664 | 111.887618 | 87.2368802 | 62.1674443 | 0 |
| 9.65E-09 | 4.88E-08 | 713.458419 | 1697.88904 | 1482.09891 | 1607.53876 | 64.1578926 |
| 9.65E-09 | 4.88E-08 | 56.9094818 | 115.891549 | 122.502853 | 103.921698 | 0 |
| 9.67E-09 | 4.89E-08 | 56.2664368 | 118.115955 | 126.215061 | 168.408823 | 0 |
| 9.67E-09 | 4.89E-08 | 51.7651219 | 163.271395 | 142.919995 | 151.707121 | 0 |
| 9.72E-09 | 4.91E-08 | 57.2310043 | 124.566732 | 106.72597 | 107.169251 | 0 |
| 9.73E-09 | 4.92E-08 | 214.777027 | 731.607109 | 1101.59763 | 944.110068 | 0 |
| 9.73E-09 | 4.92E-08 | 102.565676 | 140.137573 | 59.3953227 | 184.182652 | 0 |
| 9.73E-09 | 4.92E-08 | 66.2336342 | 120.11792 | 76.1002572 | 108.561059 | 0 |
| 9.75E-09 | 4.93E-08 | 228.924017 | 226.444523 | 147.560255 | 195.31712 | 45.1481466 |
| 9.77E-09 | 4.94E-08 | 160.761248 | 92.9801676 | 71.4599976 | 87.2199964 | 0 |
| 9.78E-09 | 4.94E-08 | 54.6588243 | 198.194568 | 130.85532 | 142.892335 | 0 |
| 9.79E-09 | 4.94E-08 | 135.039448 | 238.456315 | 134.567528 | 164.233398 | 2.37621824 |
| 9.80E-09 | 4.95E-08 | 50.1575094 | 250.912988 | 159.62493 | 191.60563 | 0 |
| 9.81E-09 | 4.96E-08 | 61.4107968 | 255.3618 | 107.654022 | 212.018821 | 0 |
| 9.82E-09 | 4.96E-08 | 51.7651219 | 144.586385 | 145.704151 | 162.841589 | 0 |
| 9.83E-09 | 4.96E-08 | 170.0854 | 93.4250488 | 217.164149 | 65.4149973 | 0 |

| | | | | | | |
|---|---|---|---|---|---|---|
| 9.83E-09 | 4.96E-08 | 60.4462293 | 121.897445 | 93.7332436 | 96.9626556 | 0 |
| 9.84E-09 | 4.97E-08 | 58.5170943 | 103.212435 | 179.11402 | 157.274355 | 0 |
| 9.86E-09 | 4.97E-08 | 59.8031843 | 92.0904053 | 101.157659 | 94.1790387 | 0 |
| 9.87E-09 | 4.98E-08 | 59.1601393 | 158.600142 | 99.3015551 | 128.97425 | 0 |
| 9.87E-09 | 4.98E-08 | 70.4134267 | 82.7479004 | 70.5319457 | 78.4052096 | 0 |
| 9.89E-09 | 4.99E-08 | 68.8058142 | 110.775415 | 74.2441534 | 86.7560603 | 0 |
| 9.96E-09 | 5.02E-08 | 76.2008316 | 87.864034 | 66.819738 | 112.272549 | 0 |
| 9.98E-09 | 5.03E-08 | 58.1955718 | 179.064677 | 103.941815 | 164.697334 | 0 |
| 9.98E-09 | 5.03E-08 | 61.7323193 | 101.655351 | 86.3088283 | 88.1478687 | 0 |
| 9.99E-09 | 5.04E-08 | 52.4081669 | 194.413078 | 135.49558 | 224.545097 | 0 |
| 1.00E-08 | 5.06E-08 | 60.4462293 | 115.891549 | 273.775316 | 307.589668 | 0 |
| 1.00E-08 | 5.06E-08 | 45.6561944 | 211.985885 | 219.948304 | 207.843396 | 0 |
| 1.01E-08 | 5.09E-08 | 54.6588243 | 130.795068 | 119.718697 | 119.695527 | 0 |
| 1.01E-08 | 5.10E-08 | 48.2283744 | 167.497766 | 324.818171 | 288.568286 | 0 |
| 1.01E-08 | 5.10E-08 | 58.5170943 | 103.879757 | 94.6612955 | 102.529889 | 0 |
| 1.01E-08 | 5.10E-08 | 45.0131495 | 211.763444 | 220.876356 | 230.112331 | 0 |
| 1.01E-08 | 5.11E-08 | 83.595849 | 152.816687 | 64.0355823 | 135.933292 | 0 |
| 1.02E-08 | 5.11E-08 | 43.405537 | 306.300696 | 209.739733 | 294.599456 | 0 |
| 1.02E-08 | 5.13E-08 | 103.530244 | 74.9624797 | 65.8916861 | 86.7560603 | 0 |
| 1.02E-08 | 5.14E-08 | 89.0617314 | 70.2912272 | 68.6758419 | 82.1166988 | 0 |
| 1.02E-08 | 5.14E-08 | 68.1627692 | 87.1967122 | 70.5319457 | 96.4987195 | 0 |
| 1.02E-08 | 5.14E-08 | 61.7323193 | 120.340361 | 93.7332436 | 88.6118049 | 0 |
| 1.02E-08 | 5.15E-08 | 59.8031843 | 98.0963013 | 93.7332436 | 122.94308 | 0 |
| 1.02E-08 | 5.15E-08 | 57.2310043 | 248.688582 | 128.071165 | 155.418611 | 0 |
| 1.02E-08 | 5.15E-08 | 100.958064 | 58.2794352 | 96.5173994 | 107.169251 | 0 |
| 1.03E-08 | 5.17E-08 | 65.5905892 | 154.596212 | 81.6685687 | 141.03659 | 0 |
| 1.03E-08 | 5.17E-08 | 70.0919042 | 127.236019 | 74.2441534 | 94.1790387 | 0 |
| 1.03E-08 | 5.18E-08 | 99.0289288 | 16.2381632 | 23.2012979 | 22.7328714 | 520.391795 |
| 1.03E-08 | 5.18E-08 | 58.5170943 | 144.586385 | 103.013763 | 108.097123 | 0 |
| 1.03E-08 | 5.19E-08 | 105.459379 | 177.952474 | 186.538435 | 199.492545 | 2.37621824 |
| 1.03E-08 | 5.19E-08 | 51.4435994 | 144.363944 | 142.919995 | 160.985845 | 0 |
| 1.03E-08 | 5.19E-08 | 52.7296894 | 179.287118 | 128.071165 | 155.882547 | 0 |
| 1.03E-08 | 5.20E-08 | 372.001528 | 771.646415 | 784.20387 | 789.61933 | 11.8810912 |
| 1.03E-08 | 5.20E-08 | 54.9803468 | 117.671073 | 114.150386 | 126.654569 | 0 |
| 1.04E-08 | 5.20E-08 | 54.9803468 | 125.901375 | 143.848047 | 112.272549 | 0 |
| 1.04E-08 | 5.20E-08 | 69.1273367 | 105.436841 | 167.977397 | 84.9003157 | 0 |
| 1.04E-08 | 5.22E-08 | 62.6968867 | 82.7479004 | 129.927268 | 109.488932 | 0 |
| 1.04E-08 | 5.22E-08 | 63.6614542 | 91.8679647 | 78.8844129 | 83.5085072 | 0 |
| 1.04E-08 | 5.23E-08 | 53.3727344 | 147.255672 | 121.574801 | 138.252973 | 0 |
| 1.04E-08 | 5.24E-08 | 54.6588243 | 181.289083 | 122.502853 | 136.861165 | 0 |
| 1.05E-08 | 5.25E-08 | 114.140486 | 376.369482 | 297.904665 | 349.807858 | 11.8810912 |
| 1.05E-08 | 5.26E-08 | 81.023669 | 75.4073608 | 82.5966206 | 65.4149973 | 0 |
| 1.05E-08 | 5.26E-08 | 51.7651219 | 139.692692 | 148.488307 | 186.038397 | 0 |

| | | | | | | |
|---|---|---|---|---|---|---|
| 1.05E-08 | 5.26E-08 | 52.0866444 | 149.702519 | 283.055835 | 190.677758 | 0 |
| 1.05E-08 | 5.26E-08 | 78.7730116 | 65.6199748 | 140.135839 | 128.97425 | 0 |
| 1.05E-08 | 5.26E-08 | 69.4488592 | 72.9605143 | 103.013763 | 79.3330819 | 0 |
| 1.05E-08 | 5.27E-08 | 58.8386168 | 109.218331 | 92.8051917 | 97.4265918 | 0 |
| 1.05E-08 | 5.27E-08 | 71.0564716 | 139.247811 | 72.3880495 | 142.892335 | 0 |
| 1.05E-08 | 5.28E-08 | 83.595849 | 320.314453 | 299.760769 | 321.507753 | 0 |
| 1.05E-08 | 5.28E-08 | 57.8740493 | 117.893514 | 103.941815 | 96.4987195 | 0 |
| 1.05E-08 | 5.28E-08 | 1.60761248 | 1.11220296 | 0 | 0.46393615 | 26.1384007 |
| 1.06E-08 | 5.29E-08 | 63.3399317 | 88.0864746 | 78.8844129 | 83.9724434 | 0 |
| 1.06E-08 | 5.29E-08 | 60.1247068 | 92.757727 | 136.423632 | 157.738291 | 0 |
| 1.06E-08 | 5.31E-08 | 53.6942569 | 119.673039 | 128.071165 | 139.644782 | 0 |
| 1.06E-08 | 5.31E-08 | 52.4081669 | 131.46239 | 158.696878 | 135.933292 | 0 |
| 1.06E-08 | 5.33E-08 | 56.2664368 | 106.326603 | 108.582074 | 107.633187 | 0 |
| 1.06E-08 | 5.33E-08 | 54.3373018 | 119.228158 | 121.574801 | 119.231591 | 0 |
| 1.07E-08 | 5.33E-08 | 60.1247068 | 102.767554 | 154.98467 | 102.993826 | 0 |
| 1.07E-08 | 5.33E-08 | 55.6233918 | 133.909237 | 113.222334 | 109.952868 | 0 |
| 1.07E-08 | 5.34E-08 | 177.480418 | 724.044128 | 525.277385 | 528.887212 | 16.6335277 |
| 1.07E-08 | 5.34E-08 | 138.897718 | 72.9605143 | 95.5893474 | 73.7658481 | 0 |
| 1.07E-08 | 5.34E-08 | 443.058 | 1060.59674 | 786.059974 | 894.004964 | 35.6432737 |
| 1.07E-08 | 5.35E-08 | 108.674604 | 182.846167 | 160.552982 | 184.646588 | 2.37621824 |
| 1.07E-08 | 5.36E-08 | 51.4435994 | 152.149365 | 140.135839 | 141.03659 | 0 |
| 1.08E-08 | 5.38E-08 | 56.2664368 | 191.52135 | 109.510126 | 173.048184 | 0 |
| 1.08E-08 | 5.38E-08 | 109.317649 | 93.4250488 | 101.157659 | 109.488932 | 14.2573095 |
| 1.08E-08 | 5.38E-08 | 54.6588243 | 125.234054 | 112.294282 | 119.695527 | 0 |
| 1.08E-08 | 5.38E-08 | 68.4842917 | 130.350187 | 449.177128 | 195.781056 | 0 |
| 1.08E-08 | 5.40E-08 | 82.6312815 | 242.015365 | 179.11402 | 73.7658481 | 0 |
| 1.08E-08 | 5.41E-08 | 52.7296894 | 141.249776 | 125.287009 | 134.541484 | 0 |
| 1.08E-08 | 5.41E-08 | 60.4462293 | 98.5411824 | 101.157659 | 83.0445711 | 0 |
| 1.08E-08 | 5.42E-08 | 54.6588243 | 128.793103 | 109.510126 | 134.541484 | 0 |
| 1.09E-08 | 5.42E-08 | 50.1575094 | 159.045024 | 165.193241 | 145.675952 | 0 |
| 1.09E-08 | 5.42E-08 | 75.2362641 | 175.728068 | 72.3880495 | 136.861165 | 0 |
| 1.09E-08 | 5.42E-08 | 51.4435994 | 182.178845 | 131.783372 | 189.749886 | 0 |
| 1.09E-08 | 5.42E-08 | 63.0184092 | 78.7439697 | 83.5246725 | 89.0757411 | 0 |
| 1.09E-08 | 5.43E-08 | 27.6509347 | 443.324101 | 810.189324 | 553.475829 | 0 |
| 1.09E-08 | 5.43E-08 | 64.3044992 | 98.5411824 | 75.1722053 | 109.024996 | 0 |
| 1.09E-08 | 5.45E-08 | 51.7651219 | 156.598177 | 130.85532 | 145.212015 | 0 |
| 1.10E-08 | 5.47E-08 | 56.9094818 | 153.261568 | 103.013763 | 123.407016 | 0 |
| 1.10E-08 | 5.47E-08 | 54.3373018 | 115.001786 | 117.862593 | 122.94308 | 0 |
| 1.10E-08 | 5.47E-08 | 72.0210391 | 108.328569 | 65.8916861 | 100.674145 | 0 |
| 1.10E-08 | 5.47E-08 | 45.6561944 | 197.972127 | 207.883629 | 201.34829 | 0 |
| 1.10E-08 | 5.48E-08 | 103.208721 | 202.64338 | 187.466487 | 192.533503 | 0 |
| 1.10E-08 | 5.49E-08 | 59.1601393 | 158.600142 | 98.3735032 | 117.375846 | 0 |
| 1.10E-08 | 5.49E-08 | 57.8740493 | 177.507593 | 103.941815 | 134.541484 | 0 |

| | | | | | | |
|---|---|---|---|---|---|---|
| 1.10E-08 | 5.49E-08 | 55.9449143 | 132.574593 | 142.919995 | 102.065953 | 0 |
| 1.10E-08 | 5.50E-08 | 57.5525268 | 119.228158 | 95.5893474 | 135.469356 | 0 |
| 1.11E-08 | 5.52E-08 | 52.0866444 | 191.743791 | 196.747006 | 127.118505 | 0 |
| 1.11E-08 | 5.54E-08 | 171.693013 | 266.038949 | 209.739733 | 209.235204 | 16.6335277 |
| 1.11E-08 | 5.54E-08 | 37.2966096 | 326.76523 | 378.645182 | 291.815839 | 0 |
| 1.11E-08 | 5.54E-08 | 50.4790319 | 156.598177 | 159.62493 | 139.644782 | 0 |
| 1.12E-08 | 5.55E-08 | 52.0866444 | 140.804895 | 153.128566 | 126.190633 | 0 |
| 1.12E-08 | 5.55E-08 | 66.5551567 | 128.793103 | 74.2441534 | 111.808612 | 0 |
| 1.12E-08 | 5.56E-08 | 89.3832539 | 237.121672 | 240.365446 | 233.359884 | 0 |
| 1.12E-08 | 5.56E-08 | 162.368861 | 119.228158 | 68.6758419 | 77.9412734 | 0 |
| 1.12E-08 | 5.56E-08 | 66.5551567 | 99.6533854 | 110.438178 | 71.4461673 | 0 |
| 1.12E-08 | 5.57E-08 | 100.315019 | 219.993746 | 283.055835 | 192.069567 | 0 |
| 1.12E-08 | 5.59E-08 | 50.4790319 | 155.263534 | 139.207788 | 158.666164 | 0 |
| 1.12E-08 | 5.59E-08 | 269.757374 | 541.642843 | 271.919212 | 444.914769 | 2991.65877 |
| 1.12E-08 | 5.59E-08 | 52.0866444 | 126.346257 | 141.063891 | 139.180845 | 0 |
| 1.13E-08 | 5.60E-08 | 48.2283744 | 202.64338 | 156.840774 | 185.110524 | 0 |
| 1.13E-08 | 5.60E-08 | 78.7730116 | 73.4053955 | 71.4599976 | 113.664357 | 0 |
| 1.13E-08 | 5.61E-08 | 77.1653991 | 88.0864746 | 64.9636342 | 75.1576565 | 0 |
| 1.13E-08 | 5.61E-08 | 336.634053 | 1037.01804 | 723.880495 | 875.447518 | 54.6530196 |
| 1.13E-08 | 5.61E-08 | 70.0919042 | 123.676969 | 74.2441534 | 156.346483 | 0 |
| 1.13E-08 | 5.62E-08 | 47.9068519 | 163.716276 | 176.329864 | 190.677758 | 0 |
| 1.13E-08 | 5.64E-08 | 101.601109 | 123.009648 | 56.6111669 | 158.202228 | 0 |
| 1.13E-08 | 5.64E-08 | 44.3701045 | 257.586206 | 233.869083 | 193.461375 | 0 |
| 1.14E-08 | 5.64E-08 | 128.287476 | 148.145435 | 176.329864 | 95.106911 | 7.12865473 |
| 1.14E-08 | 5.66E-08 | 104.494811 | 254.472038 | 170.761553 | 211.554885 | 2.37621824 |
| 1.14E-08 | 5.68E-08 | 58.8386168 | 121.452563 | 88.1649321 | 108.561059 | 0 |
| 1.14E-08 | 5.68E-08 | 768.438766 | 2841.67857 | 4029.60142 | 3627.9807 | 285.146189 |
| 1.15E-08 | 5.70E-08 | 51.4435994 | 154.596212 | 128.071165 | 169.800631 | 0 |
| 1.15E-08 | 5.70E-08 | 67.5197242 | 190.854028 | 84.4527244 | 132.685739 | 0 |
| 1.15E-08 | 5.70E-08 | 50.4790319 | 213.765409 | 140.135839 | 172.584248 | 0 |
| 1.15E-08 | 5.71E-08 | 89.3832539 | 77.8542074 | 86.3088283 | 181.862971 | 0 |
| 1.15E-08 | 5.71E-08 | 363.320421 | 684.004822 | 559.615306 | 579.456253 | 0 |
| 1.15E-08 | 5.72E-08 | 171.37149 | 295.401107 | 234.797135 | 314.548711 | 16.6335277 |
| 1.16E-08 | 5.74E-08 | 85.524984 | 64.0628906 | 130.85532 | 88.6118049 | 0 |
| 1.16E-08 | 5.77E-08 | 63.9829767 | 98.763623 | 252.430121 | 171.19244 | 0 |
| 1.16E-08 | 5.77E-08 | 47.9068519 | 178.397355 | 160.552982 | 179.079354 | 0 |
| 1.16E-08 | 5.77E-08 | 93.2415239 | 315.865641 | 215.308045 | 317.796264 | 0 |
| 1.17E-08 | 5.79E-08 | 127.965953 | 62.2833659 | 79.8124649 | 80.7248903 | 556.035069 |
| 1.17E-08 | 5.79E-08 | 64.3044992 | 118.115955 | 75.1722053 | 102.065953 | 0 |
| 1.17E-08 | 5.79E-08 | 60.7677518 | 102.322673 | 80.7405168 | 112.272549 | 0 |
| 1.17E-08 | 5.79E-08 | 46.9422844 | 176.172949 | 186.538435 | 178.615418 | 0 |
| 1.17E-08 | 5.80E-08 | 109.960694 | 163.938717 | 53.8270112 | 139.644782 | 0 |
| 1.17E-08 | 5.80E-08 | 47.9068519 | 171.501697 | 170.761553 | 166.089142 | 0 |

| | | | | | | |
|---|---|---|---|---|---|---|
| 1.17E-08 | 5.80E-08 | 67.8412467 | 108.106128 | 71.4599976 | 85.3642518 | 0 |
| 1.17E-08 | 5.80E-08 | 52.7296894 | 119.005717 | 167.977397 | 143.820207 | 0 |
| 1.17E-08 | 5.81E-08 | 51.4435994 | 130.350187 | 206.027526 | 169.336695 | 0 |
| 1.18E-08 | 5.85E-08 | 127.644431 | 246.019295 | 221.804408 | 235.679565 | 14.2573095 |
| 1.18E-08 | 5.85E-08 | 114.140486 | 69.4014648 | 68.6758419 | 94.1790387 | 0 |
| 1.18E-08 | 5.85E-08 | 48.8714194 | 152.816687 | 157.768826 | 172.120312 | 0 |
| 1.18E-08 | 5.85E-08 | 61.7323193 | 91.2006429 | 77.0283091 | 95.106911 | 0 |
| 1.19E-08 | 5.87E-08 | 54.3373018 | 195.970162 | 116.00649 | 152.634994 | 0 |
| 1.19E-08 | 5.88E-08 | 54.9803468 | 119.673039 | 138.279736 | 103.921698 | 0 |
| 1.19E-08 | 5.88E-08 | 108.353081 | 199.974093 | 144.776099 | 173.048184 | 0 |
| 1.19E-08 | 5.90E-08 | 44.3701045 | 193.968197 | 214.379993 | 230.112331 | 0 |
| 1.20E-08 | 5.92E-08 | 80.7021465 | 292.286938 | 332.242586 | 340.065199 | 2.37621824 |
| 1.20E-08 | 5.92E-08 | 51.4435994 | 139.470251 | 128.999216 | 135.469356 | 0 |
| 1.21E-08 | 5.96E-08 | 102.887199 | 211.985885 | 160.552982 | 157.738291 | 0 |
| 1.21E-08 | 5.96E-08 | 51.7651219 | 129.015544 | 138.279736 | 127.118505 | 0 |
| 1.21E-08 | 5.96E-08 | 51.7651219 | 129.237984 | 128.071165 | 134.541484 | 0 |
| 1.21E-08 | 5.98E-08 | 54.0157793 | 114.334465 | 138.279736 | 112.272549 | 0 |
| 1.21E-08 | 6.00E-08 | 60.4462293 | 186.405216 | 110.438178 | 107.169251 | 0 |
| 1.21E-08 | 6.00E-08 | 66.8766792 | 104.102197 | 76.1002572 | 141.03659 | 0 |
| 1.22E-08 | 6.01E-08 | 48.2283744 | 187.73986 | 151.272462 | 171.656376 | 0 |
| 1.22E-08 | 6.01E-08 | 61.7323193 | 95.8718953 | 147.560255 | 92.7872303 | 0 |
| 1.22E-08 | 6.02E-08 | 59.1601393 | 130.127747 | 95.5893474 | 93.7151026 | 0 |
| 1.22E-08 | 6.02E-08 | 40.5118345 | 227.779167 | 556.83115 | 375.324346 | 0 |
| 1.22E-08 | 6.02E-08 | 72.9856066 | 86.3069499 | 70.5319457 | 68.6625504 | 0 |
| 1.22E-08 | 6.03E-08 | 261.397789 | 861.512415 | 684.902315 | 777.093053 | 21.3859642 |
| 1.22E-08 | 6.03E-08 | 164.297996 | 286.503483 | 238.509343 | 278.825627 | 16.6335277 |
| 1.22E-08 | 6.04E-08 | 142.755988 | 266.038949 | 50.1148035 | 193.461375 | 0 |
| 1.22E-08 | 6.04E-08 | 73.6286516 | 62.9506877 | 94.6612955 | 90.9314857 | 0 |
| 1.23E-08 | 6.06E-08 | 46.2992394 | 194.857959 | 169.833501 | 192.997439 | 0 |
| 1.23E-08 | 6.06E-08 | 89.7047764 | 295.401107 | 244.077654 | 239.391054 | 2.37621824 |
| 1.23E-08 | 6.06E-08 | 72.3425616 | 70.2912272 | 112.294282 | 147.995632 | 0 |
| 1.23E-08 | 6.06E-08 | 48.8714194 | 161.936751 | 147.560255 | 193.925311 | 0 |
| 1.24E-08 | 6.10E-08 | 38.904222 | 262.70234 | 282.127783 | 298.310945 | 0 |
| 1.24E-08 | 6.11E-08 | 54.3373018 | 103.212435 | 119.718697 | 118.303719 | 0 |
| 1.24E-08 | 6.11E-08 | 52.4081669 | 362.800606 | 150.344411 | 217.122119 | 0 |
| 1.24E-08 | 6.12E-08 | 71.0564716 | 86.7518311 | 75.1722053 | 66.8068058 | 0 |
| 1.24E-08 | 6.12E-08 | 53.0512119 | 127.458459 | 117.862593 | 115.056166 | 0 |
| 1.24E-08 | 6.12E-08 | 61.4107968 | 108.106128 | 82.5966206 | 84.9003157 | 0 |
| 1.24E-08 | 6.13E-08 | 104.816334 | 200.863855 | 158.696878 | 204.131907 | 2.37621824 |
| 1.25E-08 | 6.13E-08 | 52.7296894 | 162.381632 | 117.862593 | 132.685739 | 0 |
| 1.25E-08 | 6.13E-08 | 138.254673 | 90.3108805 | 66.819738 | 79.797018 | 0 |
| 1.25E-08 | 6.17E-08 | 54.9803468 | 103.434875 | 103.941815 | 115.520102 | 0 |
| 1.26E-08 | 6.18E-08 | 79.4160565 | 87.6415934 | 65.8916861 | 69.5904227 | 0 |

| | | | | | | |
|---|---|---|---|---|---|---|
| 1.26E-08 | 6.18E-08 | 92.2769564 | 71.4034302 | 186.538435 | 92.7872303 | 0 |
| 1.26E-08 | 6.18E-08 | 49.8359869 | 137.245846 | 168.905449 | 146.139888 | 0 |
| 1.26E-08 | 6.19E-08 | 52.0866444 | 221.55083 | 124.358957 | 184.646588 | 0 |
| 1.26E-08 | 6.19E-08 | 79.737579 | 105.881722 | 87.2368802 | 61.239572 | 0 |
| 1.26E-08 | 6.19E-08 | 137.611628 | 808.126672 | 509.500502 | 542.341361 | 0 |
| 1.26E-08 | 6.20E-08 | 59.8031843 | 341.66875 | 112.294282 | 285.320733 | 0 |
| 1.26E-08 | 6.21E-08 | 47.9068519 | 169.722172 | 158.696878 | 159.594036 | 0 |
| 1.26E-08 | 6.21E-08 | 53.6942569 | 121.452563 | 127.143113 | 105.777442 | 0 |
| 1.27E-08 | 6.22E-08 | 54.6588243 | 170.834375 | 143.848047 | 106.241379 | 0 |
| 1.27E-08 | 6.25E-08 | 69.1273367 | 152.816687 | 87.2368802 | 85.3642518 | 0 |
| 1.27E-08 | 6.25E-08 | 57.5525268 | 98.3187419 | 98.3735032 | 134.541484 | 0 |
| 1.27E-08 | 6.25E-08 | 54.9803468 | 141.917098 | 156.840774 | 101.602017 | 0 |
| 1.27E-08 | 6.25E-08 | 54.3373018 | 128.125781 | 219.948304 | 136.397228 | 0 |
| 1.27E-08 | 6.25E-08 | 49.1929419 | 144.808826 | 151.272462 | 149.387441 | 0 |
| 1.27E-08 | 6.25E-08 | 109.317649 | 54.2755046 | 149.416359 | 117.375846 | 0 |
| 1.27E-08 | 6.26E-08 | 55.6233918 | 95.8718953 | 128.071165 | 114.128293 | 0 |
| 1.28E-08 | 6.27E-08 | 109.639171 | 110.330534 | 54.7550631 | 95.5708472 | 0 |
| 1.28E-08 | 6.30E-08 | 4.82283744 | 6.45077718 | 4.64025958 | 6.03116997 | 71.2865473 |
| 1.29E-08 | 6.32E-08 | 49.8359869 | 136.133643 | 150.344411 | 141.964462 | 0 |
| 1.29E-08 | 6.34E-08 | 48.8714194 | 150.814722 | 146.632203 | 153.562866 | 0 |
| 1.29E-08 | 6.35E-08 | 57.5525268 | 108.77345 | 93.7332436 | 141.03659 | 0 |
| 1.30E-08 | 6.36E-08 | 108.996126 | 188.852063 | 151.272462 | 161.913717 | 2.37621824 |
| 1.30E-08 | 6.36E-08 | 48.2283744 | 191.743791 | 149.416359 | 160.521908 | 0 |
| 1.30E-08 | 6.36E-08 | 94.2060914 | 159.045024 | 117.862593 | 141.03659 | 9.50487297 |
| 1.30E-08 | 6.36E-08 | 87.7756414 | 146.36591 | 58.4672708 | 122.015208 | 0 |
| 1.30E-08 | 6.37E-08 | 60.7677518 | 158.600142 | 85.3807764 | 142.892335 | 0 |
| 1.30E-08 | 6.37E-08 | 103.208721 | 198.194568 | 169.833501 | 181.399035 | 0 |
| 1.30E-08 | 6.37E-08 | 85.2034615 | 161.26943 | 191.178695 | 63.5592527 | 0 |
| 1.30E-08 | 6.38E-08 | 53.0512119 | 130.795068 | 108.582074 | 122.94308 | 0 |
| 1.30E-08 | 6.39E-08 | 58.5170943 | 91.2006429 | 109.510126 | 85.828188 | 0 |
| 1.31E-08 | 6.40E-08 | 43.405537 | 216.657137 | 193.962851 | 236.143501 | 0 |
| 1.31E-08 | 6.44E-08 | 112.532874 | 81.858138 | 83.5246725 | 47.3214874 | 389.699792 |
| 1.32E-08 | 6.45E-08 | 64.3044992 | 137.690727 | 77.0283091 | 105.777442 | 0 |
| 1.32E-08 | 6.45E-08 | 44.691627 | 43.8207967 | 38.9781805 | 48.2493597 | 144.949313 |
| 1.32E-08 | 6.45E-08 | 79.737579 | 95.4270142 | 59.3953227 | 85.3642518 | 0 |
| 1.32E-08 | 6.46E-08 | 51.1220769 | 149.035197 | 126.215061 | 127.582442 | 0 |
| 1.32E-08 | 6.48E-08 | 0 | 0 | 0 | 0 | 14.2573095 |
| 1.32E-08 | 6.48E-08 | 301.266579 | 53.6081828 | 148.488307 | 160.985845 | 0 |
| 1.33E-08 | 6.52E-08 | 54.9803468 | 187.73986 | 106.72597 | 199.956481 | 0 |
| 1.33E-08 | 6.53E-08 | 47.2638069 | 171.946578 | 152.200514 | 194.389247 | 0 |
| 1.33E-08 | 6.53E-08 | 56.5879593 | 131.46239 | 98.3735032 | 99.2823364 | 0 |
| 1.33E-08 | 6.53E-08 | 59.8031843 | 311.416829 | 138.279736 | 140.572654 | 0 |
| 1.34E-08 | 6.54E-08 | 46.2992394 | 294.956226 | 170.761553 | 205.523715 | 0 |

| | | | | | | |
|---|---|---|---|---|---|---|
| 1.34E-08 | 6.56E-08 | 62.3753642 | 80.0786133 | 78.8844129 | 75.6215927 | 0 |
| 1.35E-08 | 6.58E-08 | 57.8740493 | 102.322673 | 85.3807764 | 91.859358 | 0 |
| 1.35E-08 | 6.59E-08 | 62.0538418 | 90.7557617 | 74.2441534 | 82.1166988 | 0 |
| 1.35E-08 | 6.60E-08 | 64.6260217 | 111.665177 | 71.4599976 | 120.159463 | 0 |
| 1.35E-08 | 6.60E-08 | 113.497441 | 207.092192 | 137.351684 | 183.718716 | 0 |
| 1.35E-08 | 6.61E-08 | 53.0512119 | 122.342326 | 109.510126 | 147.06776 | 0 |
| 1.35E-08 | 6.61E-08 | 85.524984 | 86.3069499 | 59.3953227 | 78.4052096 | 0 |
| 1.36E-08 | 6.63E-08 | 61.7323193 | 98.3187419 | 75.1722053 | 84.9003157 | 0 |
| 1.36E-08 | 6.63E-08 | 111.568306 | 122.342326 | 52.8989593 | 161.449781 | 0 |
| 1.36E-08 | 6.63E-08 | 53.6942569 | 108.328569 | 136.423632 | 109.488932 | 0 |
| 1.36E-08 | 6.66E-08 | 93.5630464 | 255.806681 | 221.804408 | 270.938712 | 0 |
| 1.37E-08 | 6.69E-08 | 75.2362641 | 340.111666 | 84.4527244 | 330.32254 | 0 |
| 1.37E-08 | 6.69E-08 | 96.4567488 | 97.4289795 | 61.2514265 | 72.8379758 | 0 |
| 1.37E-08 | 6.70E-08 | 99.3504513 | 258.698409 | 210.667785 | 180.935099 | 0 |
| 1.37E-08 | 6.70E-08 | 52.0866444 | 118.783276 | 168.905449 | 128.97425 | 0 |
| 1.38E-08 | 6.74E-08 | 50.1575094 | 171.279256 | 123.430905 | 179.079354 | 0 |
| 1.38E-08 | 6.74E-08 | 51.4435994 | 167.497766 | 167.049345 | 115.056166 | 0 |
| 1.38E-08 | 6.74E-08 | 48.2283744 | 153.706449 | 281.199731 | 378.571899 | 0 |
| 1.38E-08 | 6.75E-08 | 61.4107968 | 82.970341 | 74.2441534 | 86.7560603 | 0 |
| 1.39E-08 | 6.77E-08 | 126.358341 | 72.2931925 | 65.8916861 | 95.5708472 | 0 |
| 1.39E-08 | 6.78E-08 | 75.8793091 | 97.4289795 | 72.3880495 | 65.4149973 | 0 |
| 1.39E-08 | 6.79E-08 | 48.5498969 | 148.812756 | 143.848047 | 146.603824 | 0 |
| 1.40E-08 | 6.81E-08 | 58.5170943 | 103.657316 | 84.4527244 | 86.7560603 | 0 |
| 1.40E-08 | 6.81E-08 | 84.5604165 | 133.241915 | 152.200514 | 145.675952 | 19.0097459 |
| 1.40E-08 | 6.83E-08 | 43.0840145 | 209.316597 | 210.667785 | 197.172864 | 0 |
| 1.40E-08 | 6.84E-08 | 49.1929419 | 140.137573 | 134.567528 | 150.779249 | 0 |
| 1.41E-08 | 6.85E-08 | 52.4081669 | 137.023405 | 108.582074 | 154.026802 | 0 |
| 1.41E-08 | 6.85E-08 | 50.8005544 | 117.893514 | 159.62493 | 171.656376 | 0 |
| 1.41E-08 | 6.86E-08 | 61.4107968 | 93.6474894 | 90.0210359 | 72.8379758 | 0 |
| 1.41E-08 | 6.86E-08 | 64.6260217 | 84.0825439 | 72.3880495 | 72.8379758 | 0 |
| 1.41E-08 | 6.86E-08 | 132.145746 | 343.225834 | 323.890119 | 373.932538 | 14.2573095 |
| 1.41E-08 | 6.87E-08 | 96.7782713 | 201.976058 | 218.0922 | 189.28595 | 0 |
| 1.41E-08 | 6.87E-08 | 84.5604165 | 271.377523 | 289.552198 | 247.277969 | 0 |
| 1.41E-08 | 6.87E-08 | 52.7296894 | 127.6809 | 105.797919 | 141.964462 | 0 |
| 1.42E-08 | 6.89E-08 | 53.6942569 | 154.151331 | 116.934542 | 108.561059 | 0 |
| 1.42E-08 | 6.90E-08 | 27.6509347 | 353.235661 | 697.895042 | 624.921996 | 0 |
| 1.42E-08 | 6.92E-08 | 54.6588243 | 98.5411824 | 107.654022 | 131.293931 | 0 |
| 1.42E-08 | 6.92E-08 | 54.3373018 | 124.789172 | 96.5173994 | 113.200421 | 0 |
| 1.43E-08 | 6.93E-08 | 6.43044992 | 6.67321777 | 6.49636342 | 3.71148921 | 64.1578926 |
| 1.43E-08 | 6.96E-08 | 85.8465065 | 170.611934 | 374.004923 | 85.828188 | 0 |
| 1.43E-08 | 6.97E-08 | 48.8714194 | 147.478113 | 133.639476 | 147.995632 | 0 |
| 1.43E-08 | 6.97E-08 | 59.4816618 | 111.220296 | 82.5966206 | 87.2199964 | 0 |
| 1.43E-08 | 6.97E-08 | 69.4488592 | 171.946578 | 72.3880495 | 154.954675 | 0 |

| | | | | | | |
|---|---|---|---|---|---|---|
| 1.44E-08 | 6.99E-08 | 48.8714194 | 143.919063 | 141.063891 | 137.325101 | 0 |
| 1.45E-08 | 7.04E-08 | 56.9094818 | 92.3128459 | 87.2368802 | 109.024996 | 0 |
| 1.46E-08 | 7.08E-08 | 56.9094818 | 95.2045736 | 87.2368802 | 88.1478687 | 0 |
| 1.46E-08 | 7.09E-08 | 53.3727344 | 112.77738 | 102.085711 | 106.705315 | 0 |
| 1.46E-08 | 7.09E-08 | 59.4816618 | 91.6455241 | 79.8124649 | 80.7248903 | 0 |
| 1.46E-08 | 7.09E-08 | 69.7703817 | 72.9605143 | 106.72597 | 70.9822312 | 0 |
| 1.46E-08 | 7.09E-08 | 63.6614542 | 74.7400391 | 73.3161014 | 92.7872303 | 0 |
| 1.46E-08 | 7.11E-08 | 45.0131495 | 158.600142 | 216.236097 | 215.73031 | 0 |
| 1.47E-08 | 7.12E-08 | 61.7323193 | 75.4073608 | 108.582074 | 85.3642518 | 0 |
| 1.47E-08 | 7.13E-08 | 61.7323193 | 85.4171875 | 141.991943 | 92.7872303 | 0 |
| 1.47E-08 | 7.13E-08 | 47.5853294 | 148.367875 | 341.523105 | 277.897755 | 0 |
| 1.47E-08 | 7.14E-08 | 129.573566 | 637.069857 | 1180.48204 | 936.223153 | 19.0097459 |
| 1.47E-08 | 7.15E-08 | 65.5905892 | 111.442737 | 70.5319457 | 85.3642518 | 0 |
| 1.47E-08 | 7.15E-08 | 57.8740493 | 80.9683756 | 115.078438 | 106.705315 | 0 |
| 1.48E-08 | 7.16E-08 | 46.9422844 | 183.068608 | 158.696878 | 149.387441 | 0 |
| 1.48E-08 | 7.16E-08 | 123.143116 | 180.844202 | 124.358957 | 232.432012 | 2.37621824 |
| 1.48E-08 | 7.17E-08 | 54.3373018 | 97.2065389 | 144.776099 | 121.551272 | 0 |
| 1.48E-08 | 7.17E-08 | 78.1299666 | 93.6474894 | 60.3233746 | 77.0134011 | 0 |
| 1.48E-08 | 7.17E-08 | 110.603739 | 280.275146 | 227.37272 | 307.589668 | 4.75243649 |
| 1.48E-08 | 7.19E-08 | 161.404293 | 228.224048 | 77.956361 | 70.9822312 | 0 |
| 1.49E-08 | 7.21E-08 | 60.1247068 | 99.875826 | 75.1722053 | 88.6118049 | 0 |
| 1.49E-08 | 7.22E-08 | 88.0971639 | 156.820618 | 58.4672708 | 173.976057 | 0 |
| 1.50E-08 | 7.25E-08 | 52.0866444 | 113.889583 | 108.582074 | 129.438186 | 0 |
| 1.50E-08 | 7.26E-08 | 46.2992394 | 288.95033 | 149.416359 | 276.04201 | 0 |
| 1.51E-08 | 7.29E-08 | 49.1929419 | 213.542969 | 134.567528 | 157.738291 | 0 |
| 1.51E-08 | 7.30E-08 | 41.476402 | 221.105949 | 229.228823 | 205.059779 | 0 |
| 1.51E-08 | 7.30E-08 | 51.4435994 | 116.781311 | 122.502853 | 112.272549 | 0 |
| 1.51E-08 | 7.30E-08 | 57.8740493 | 92.0904053 | 85.3807764 | 81.6527626 | 0 |
| 1.51E-08 | 7.30E-08 | 111.889829 | 193.078434 | 167.977397 | 179.079354 | 0 |
| 1.51E-08 | 7.30E-08 | 49.5144644 | 124.121851 | 148.488307 | 134.541484 | 0 |
| 1.51E-08 | 7.30E-08 | 60.1247068 | 81.1908162 | 76.1002572 | 93.7151026 | 0 |
| 1.51E-08 | 7.30E-08 | 44.048582 | 195.0804 | 180.970124 | 182.790844 | 0 |
| 1.52E-08 | 7.36E-08 | 48.2283744 | 311.861711 | 146.632203 | 218.513927 | 0 |
| 1.52E-08 | 7.37E-08 | 49.5144644 | 144.363944 | 348.947521 | 231.040203 | 0 |
| 1.53E-08 | 7.38E-08 | 66.5551567 | 95.8718953 | 73.3161014 | 70.0543588 | 0 |
| 1.53E-08 | 7.38E-08 | 67.5197242 | 189.296944 | 90.9490879 | 97.8905279 | 0 |
| 1.53E-08 | 7.39E-08 | 93.2415239 | 114.779346 | 55.683115 | 88.6118049 | 0 |
| 1.53E-08 | 7.39E-08 | 97.0997938 | 177.062712 | 204.171422 | 229.184459 | 0 |
| 1.54E-08 | 7.42E-08 | 52.0866444 | 142.139539 | 103.941815 | 142.428398 | 0 |
| 1.54E-08 | 7.43E-08 | 77.8084441 | 108.77345 | 113.222334 | 58.4559551 | 0 |
| 1.54E-08 | 7.43E-08 | 60.7677518 | 71.8483114 | 93.7332436 | 96.9626556 | 0 |
| 1.54E-08 | 7.43E-08 | 66.8766792 | 79.8561727 | 65.8916861 | 96.9626556 | 0 |
| 1.54E-08 | 7.43E-08 | 70.7349491 | 68.5117025 | 157.768826 | 123.407016 | 0 |

| | | | | | | |
|---|---|---|---|---|---|---|
| 1.54E-08 | 7.44E-08 | 50.4790319 | 158.600142 | 143.848047 | 114.128293 | 0 |
| 1.54E-08 | 7.45E-08 | 53.6942569 | 224.887439 | 111.36623 | 245.88616 | 0 |
| 1.54E-08 | 7.45E-08 | 50.8005544 | 149.480078 | 117.862593 | 122.015208 | 0 |
| 1.55E-08 | 7.49E-08 | 64.9475442 | 95.2045736 | 128.999216 | 236.607437 | 0 |
| 1.55E-08 | 7.50E-08 | 53.3727344 | 121.007682 | 95.5893474 | 119.695527 | 0 |
| 1.55E-08 | 7.50E-08 | 105.137856 | 183.735929 | 146.632203 | 167.017014 | 2.37621824 |
| 1.56E-08 | 7.51E-08 | 64.6260217 | 102.100232 | 75.1722053 | 73.3019119 | 0 |
| 1.56E-08 | 7.51E-08 | 53.0512119 | 121.452563 | 115.078438 | 97.4265918 | 0 |
| 1.56E-08 | 7.51E-08 | 72.9856066 | 78.7439697 | 60.3233746 | 83.9724434 | 0 |
| 1.56E-08 | 7.53E-08 | 78.4514891 | 111.665177 | 57.5392189 | 109.952868 | 0 |
| 1.56E-08 | 7.53E-08 | 128.287476 | 64.0628906 | 78.8844129 | 173.048184 | 0 |
| 1.56E-08 | 7.53E-08 | 45.6561944 | 161.936751 | 259.854537 | 183.718716 | 0 |
| 1.56E-08 | 7.53E-08 | 117.355711 | 230.893335 | 171.689605 | 203.204034 | 30.8908372 |
| 1.56E-08 | 7.54E-08 | 49.1929419 | 176.39539 | 121.574801 | 164.233398 | 0 |
| 1.57E-08 | 7.55E-08 | 136.004016 | 213.320528 | 209.739733 | 218.049991 | 19.0097459 |
| 1.57E-08 | 7.55E-08 | 50.8005544 | 122.342326 | 130.85532 | 111.808612 | 0 |
| 1.57E-08 | 7.55E-08 | 79.4160565 | 144.363944 | 143.848047 | 169.336695 | 4.75243649 |
| 1.57E-08 | 7.55E-08 | 42.4409695 | 212.208325 | 198.60311 | 193.461375 | 0 |
| 1.57E-08 | 7.57E-08 | 80.380624 | 104.769519 | 60.3233746 | 77.4773373 | 0 |
| 1.57E-08 | 7.57E-08 | 43.0840145 | 261.812577 | 366.580507 | 179.079354 | 0 |
| 1.57E-08 | 7.57E-08 | 0 | 0 | 0 | 0 | 144.949313 |
| 1.57E-08 | 7.57E-08 | 77.1653991 | 72.0707519 | 61.2514265 | 89.0757411 | 0 |
| 1.57E-08 | 7.57E-08 | 52.4081669 | 103.657316 | 108.582074 | 109.952868 | 0 |
| 1.57E-08 | 7.58E-08 | 57.5525268 | 153.261568 | 88.1649321 | 118.303719 | 0 |
| 1.58E-08 | 7.59E-08 | 64.3044992 | 168.387528 | 78.8844129 | 118.303719 | 0 |
| 1.58E-08 | 7.60E-08 | 68.4842917 | 73.1829549 | 64.9636342 | 88.1478687 | 0 |
| 1.59E-08 | 7.63E-08 | 57.2310043 | 98.0963013 | 160.552982 | 102.993826 | 0 |
| 1.59E-08 | 7.65E-08 | 86.811074 | 153.039128 | 67.7477899 | 81.1888265 | 0 |
| 1.59E-08 | 7.65E-08 | 45.9777169 | 187.072538 | 144.776099 | 183.25478 | 0 |
| 1.59E-08 | 7.66E-08 | 48.2283744 | 241.348043 | 136.423632 | 181.862971 | 0 |
| 1.60E-08 | 7.68E-08 | 118.963324 | 151.926925 | 134.567528 | 223.153289 | 0 |
| 1.60E-08 | 7.68E-08 | 96.1352263 | 96.9840983 | 58.4672708 | 74.2297842 | 0 |
| 1.60E-08 | 7.70E-08 | 58.5170943 | 75.6298014 | 102.085711 | 96.9626556 | 0 |
| 1.61E-08 | 7.74E-08 | 37.2966096 | 252.914954 | 292.336354 | 262.123925 | 0 |
| 1.62E-08 | 7.77E-08 | 53.3727344 | 102.989994 | 170.761553 | 129.902122 | 0 |
| 1.62E-08 | 7.77E-08 | 48.5498969 | 59.1691976 | 50.1148035 | 44.5378705 | 140.196876 |
| 1.62E-08 | 7.78E-08 | 609.28513 | 644.187956 | 730.376859 | 751.576565 | 28.5146189 |
| 1.62E-08 | 7.78E-08 | 49.5144644 | 179.731999 | 117.862593 | 167.944887 | 0 |
| 1.62E-08 | 7.79E-08 | 45.0131495 | 170.834375 | 159.62493 | 175.831801 | 0 |
| 1.62E-08 | 7.79E-08 | 57.8740493 | 81.6356974 | 87.2368802 | 79.797018 | 0 |
| 1.62E-08 | 7.79E-08 | 75.5577866 | 58.5018758 | 88.1649321 | 90.0036134 | 0 |
| 1.62E-08 | 7.80E-08 | 64.6260217 | 81.4132568 | 65.8916861 | 92.3232941 | 0 |
| 1.63E-08 | 7.81E-08 | 52.4081669 | 117.003752 | 98.3735032 | 116.91191 | 0 |

| | | | | | | |
|---|---|---|---|---|---|---|
| 1.63E-08 | 7.82E-08 | 65.9121117 | 74.9624797 | 67.7477899 | 70.9822312 | 0 |
| 1.63E-08 | 7.84E-08 | 98.7074063 | 211.985885 | 167.977397 | 184.646588 | 0 |
| 1.63E-08 | 7.84E-08 | 265.577582 | 303.853849 | 90.0210359 | 189.28595 | 0 |
| 1.63E-08 | 7.84E-08 | 48.8714194 | 127.903341 | 157.768826 | 129.438186 | 0 |
| 1.63E-08 | 7.84E-08 | 75.2362641 | 73.4053955 | 60.3233746 | 84.4363795 | 0 |
| 1.64E-08 | 7.84E-08 | 63.0184092 | 77.4093262 | 78.8844129 | 68.1986142 | 0 |
| 1.64E-08 | 7.85E-08 | 54.6588243 | 91.2006429 | 99.3015551 | 91.3954218 | 0 |
| 1.65E-08 | 7.90E-08 | 68.4842917 | 89.8659993 | 61.2514265 | 91.3954218 | 0 |
| 1.65E-08 | 7.90E-08 | 61.7323193 | 139.02537 | 320.177911 | 123.407016 | 0 |
| 1.65E-08 | 7.90E-08 | 125.715296 | 220.883508 | 215.308045 | 198.564673 | 28.5146189 |
| 1.65E-08 | 7.90E-08 | 51.4435994 | 133.686796 | 103.941815 | 122.015208 | 0 |
| 1.65E-08 | 7.91E-08 | 76.8438766 | 303.408968 | 301.616873 | 294.599456 | 0 |
| 1.65E-08 | 7.91E-08 | 95.4921813 | 219.548865 | 174.47376 | 216.194247 | 0 |
| 1.65E-08 | 7.91E-08 | 54.0157793 | 179.287118 | 103.013763 | 129.902122 | 0 |
| 1.65E-08 | 7.92E-08 | 64.9475442 | 170.611934 | 75.1722053 | 135.933292 | 0 |
| 1.66E-08 | 7.96E-08 | 37.2966096 | 225.33232 | 404.630636 | 310.837221 | 0 |
| 1.66E-08 | 7.96E-08 | 57.2310043 | 84.3049845 | 81.6685687 | 100.210209 | 0 |
| 1.67E-08 | 7.97E-08 | 99.9934963 | 184.403251 | 148.488307 | 166.089142 | 4.75243649 |
| 1.67E-08 | 7.98E-08 | 52.0866444 | 100.098267 | 134.567528 | 118.303719 | 0 |
| 1.67E-08 | 7.99E-08 | 51.4435994 | 120.11792 | 103.941815 | 114.128293 | 0 |
| 1.67E-08 | 8.00E-08 | 86.168029 | 110.330534 | 61.2514265 | 72.8379758 | 0 |
| 1.68E-08 | 8.04E-08 | 47.9068519 | 148.145435 | 130.85532 | 136.861165 | 0 |
| 1.68E-08 | 8.05E-08 | 54.0157793 | 118.338395 | 89.092984 | 117.839782 | 0 |
| 1.68E-08 | 8.05E-08 | 56.2664368 | 146.58835 | 92.8051917 | 183.25478 | 0 |
| 1.68E-08 | 8.05E-08 | 48.8714194 | 141.694657 | 118.790645 | 141.500526 | 0 |
| 1.69E-08 | 8.07E-08 | 50.1575094 | 141.694657 | 118.790645 | 115.520102 | 0 |
| 1.69E-08 | 8.09E-08 | 102.244154 | 161.49187 | 189.322591 | 156.346483 | 0 |
| 1.69E-08 | 8.09E-08 | 61.0892743 | 85.1947469 | 186.538435 | 128.510314 | 0 |
| 1.69E-08 | 8.09E-08 | 51.1220769 | 154.596212 | 106.72597 | 130.829995 | 0 |
| 1.69E-08 | 8.09E-08 | 60.1247068 | 78.2990885 | 72.3880495 | 81.6527626 | 0 |
| 1.70E-08 | 8.11E-08 | 58.1955718 | 124.566732 | 77.956361 | 114.592229 | 0 |
| 1.70E-08 | 8.11E-08 | 40.833357 | 285.16884 | 283.983887 | 180.935099 | 0 |
| 1.70E-08 | 8.12E-08 | 131.824223 | 124.789172 | 130.85532 | 121.087335 | 2.37621824 |
| 1.70E-08 | 8.13E-08 | 134.396403 | 170.834375 | 257.998433 | 117.839782 | 0 |
| 1.70E-08 | 8.13E-08 | 5.46588243 | 1.55708415 | 2.78415575 | 4.17542536 | 45.1481466 |
| 1.70E-08 | 8.13E-08 | 110.925261 | 194.413078 | 128.999216 | 173.512121 | 0 |
| 1.70E-08 | 8.14E-08 | 93.5630464 | 76.5195638 | 61.2514265 | 71.4461673 | 0 |
| 1.71E-08 | 8.15E-08 | 50.1575094 | 132.129712 | 109.510126 | 126.190633 | 0 |
| 1.71E-08 | 8.16E-08 | 514.114471 | 781.211361 | 731.304911 | 725.596141 | 9.50487297 |
| 1.71E-08 | 8.16E-08 | 54.3373018 | 86.9742716 | 107.654022 | 99.7462725 | 0 |
| 1.71E-08 | 8.16E-08 | 63.6614542 | 65.8424154 | 90.0210359 | 96.9626556 | 0 |
| 1.72E-08 | 8.20E-08 | 55.3018693 | 102.545113 | 83.5246725 | 112.272549 | 0 |
| 1.72E-08 | 8.21E-08 | 116.391144 | 165.27336 | 119.718697 | 160.057972 | 2.37621824 |

| | | | | | | |
|---|---|---|---|---|---|---|
| 1.72E-08 | 8.21E-08 | 102.565676 | 74.7400391 | 120.646749 | 59.3838274 | 0 |
| 1.72E-08 | 8.22E-08 | 99.6719738 | 170.611934 | 158.696878 | 175.367865 | 2.37621824 |
| 1.73E-08 | 8.24E-08 | 63.6614542 | 68.2892619 | 72.3880495 | 84.4363795 | 0 |
| 1.73E-08 | 8.24E-08 | 54.9803468 | 123.009648 | 87.2368802 | 135.933292 | 0 |
| 1.73E-08 | 8.27E-08 | 45.334672 | 179.731999 | 147.560255 | 163.769461 | 0 |
| 1.74E-08 | 8.28E-08 | 43.7270595 | 175.950509 | 166.121293 | 214.802438 | 0 |
| 1.74E-08 | 8.29E-08 | 49.5144644 | 132.129712 | 121.574801 | 115.520102 | 0 |
| 1.74E-08 | 8.29E-08 | 57.5525268 | 94.0923706 | 128.999216 | 83.9724434 | 0 |
| 1.74E-08 | 8.29E-08 | 579.062016 | 151.926925 | 90.0210359 | 119.231591 | 0 |
| 1.74E-08 | 8.30E-08 | 31.1876821 | 16.2381632 | 25.9854537 | 15.7738291 | 92.6725115 |
| 1.74E-08 | 8.30E-08 | 51.4435994 | 117.226192 | 101.157659 | 130.829995 | 0 |
| 1.74E-08 | 8.30E-08 | 63.0184092 | 77.4093262 | 173.545708 | 134.541484 | 0 |
| 1.74E-08 | 8.30E-08 | 46.9422844 | 131.017509 | 153.128566 | 160.521908 | 0 |
| 1.74E-08 | 8.31E-08 | 94.8491364 | 188.852063 | 193.034799 | 183.25478 | 0 |
| 1.75E-08 | 8.32E-08 | 67.8412467 | 92.757727 | 62.1794784 | 79.797018 | 0 |
| 1.75E-08 | 8.32E-08 | 64.3044992 | 84.3049845 | 116.00649 | 70.0543588 | 0 |
| 1.75E-08 | 8.34E-08 | 61.0892743 | 101.43291 | 189.322591 | 97.4265918 | 0 |
| 1.75E-08 | 8.35E-08 | 54.9803468 | 162.159192 | 119.718697 | 94.6429749 | 0 |
| 1.76E-08 | 8.36E-08 | 48.8714194 | 131.907271 | 123.430905 | 121.551272 | 0 |
| 1.76E-08 | 8.37E-08 | 52.0866444 | 201.753617 | 118.790645 | 128.046378 | 0 |
| 1.76E-08 | 8.37E-08 | 90.9908664 | 55.3877075 | 166.121293 | 162.377653 | 0 |
| 1.76E-08 | 8.37E-08 | 51.1220769 | 110.997856 | 104.869867 | 128.97425 | 0 |
| 1.76E-08 | 8.38E-08 | 49.5144644 | 129.015544 | 115.078438 | 120.159463 | 0 |
| 1.77E-08 | 8.41E-08 | 52.7296894 | 123.232088 | 94.6612955 | 107.169251 | 0 |
| 1.78E-08 | 8.45E-08 | 53.0512119 | 99.2085042 | 110.438178 | 92.3232941 | 0 |
| 1.78E-08 | 8.45E-08 | 56.5879593 | 80.9683756 | 81.6685687 | 90.0036134 | 0 |
| 1.78E-08 | 8.46E-08 | 62.0538418 | 125.678935 | 71.4599976 | 101.138081 | 0 |
| 1.78E-08 | 8.48E-08 | 48.5498969 | 129.682865 | 120.646749 | 131.293931 | 0 |
| 1.78E-08 | 8.48E-08 | 48.8714194 | 146.143469 | 116.934542 | 129.902122 | 0 |
| 1.79E-08 | 8.49E-08 | 47.5853294 | 125.678935 | 140.135839 | 146.139888 | 0 |
| 1.79E-08 | 8.50E-08 | 122.178549 | 76.2971232 | 64.0355823 | 78.4052096 | 0 |
| 1.79E-08 | 8.52E-08 | 52.7296894 | 126.791138 | 103.941815 | 96.4987195 | 0 |
| 1.80E-08 | 8.56E-08 | 104.173289 | 73.4053955 | 79.8124649 | 60.7756358 | 0 |
| 1.81E-08 | 8.57E-08 | 40.833357 | 185.070573 | 216.236097 | 241.246799 | 0 |
| 1.81E-08 | 8.58E-08 | 40.5118345 | 238.456315 | 191.178695 | 215.266374 | 0 |
| 1.81E-08 | 8.61E-08 | 60.1247068 | 72.2931925 | 75.1722053 | 77.0134011 | 0 |
| 1.81E-08 | 8.61E-08 | 42.762492 | 170.389494 | 181.898176 | 205.523715 | 0 |
| 1.81E-08 | 8.61E-08 | 55.3018693 | 85.6396281 | 137.351684 | 113.664357 | 0 |
| 1.82E-08 | 8.62E-08 | 78.7730116 | 60.9487223 | 162.409085 | 165.16127 | 0 |
| 1.82E-08 | 8.62E-08 | 111.889829 | 198.86189 | 151.272462 | 204.131907 | 30.8908372 |
| 1.83E-08 | 8.67E-08 | 49.5144644 | 127.236019 | 109.510126 | 126.654569 | 0 |
| 1.83E-08 | 8.69E-08 | 57.8740493 | 96.7616577 | 75.1722053 | 111.344676 | 0 |
| 1.84E-08 | 8.70E-08 | 82.309759 | 279.162944 | 241.293498 | 263.051798 | 0 |

| | | | | | | |
|---|---|---|---|---|---|---|
| 1.84E-08 | 8.70E-08 | 38.5826995 | 241.348043 | 220.876356 | 232.895948 | 0 |
| 1.84E-08 | 8.71E-08 | 122.178549 | 91.6455241 | 65.8916861 | 66.8068058 | 0 |
| 1.84E-08 | 8.71E-08 | 1998.90536 | 3434.70519 | 3177.64976 | 3426.63241 | 4.75243649 |
| 1.84E-08 | 8.73E-08 | 172.65758 | 165.718241 | 50.1148035 | 105.313506 | 0 |
| 1.85E-08 | 8.76E-08 | 41.7979245 | 168.609969 | 266.3509 | 226.400842 | 0 |
| 1.86E-08 | 8.79E-08 | 48.8714194 | 132.129712 | 112.294282 | 137.789037 | 0 |
| 1.86E-08 | 8.80E-08 | 40.5118345 | 191.52135 | 207.883629 | 225.47297 | 0 |
| 1.86E-08 | 8.81E-08 | 39.547267 | 224.664998 | 209.739733 | 216.658183 | 0 |
| 1.86E-08 | 8.81E-08 | 112.532874 | 171.946578 | 143.848047 | 49.177232 | 0 |
| 1.86E-08 | 8.81E-08 | 52.0866444 | 100.543148 | 96.5173994 | 101.602017 | 0 |
| 1.86E-08 | 8.83E-08 | 100.958064 | 282.277112 | 220.876356 | 263.97967 | 0 |
| 1.87E-08 | 8.85E-08 | 50.4790319 | 167.052885 | 104.869867 | 149.387441 | 0 |
| 1.87E-08 | 8.86E-08 | 48.2283744 | 128.125781 | 119.718697 | 130.829995 | 0 |
| 1.87E-08 | 8.87E-08 | 55.9449143 | 111.887618 | 78.8844129 | 114.128293 | 0 |
| 1.88E-08 | 8.91E-08 | 53.0512119 | 90.3108805 | 94.6612955 | 99.7462725 | 0 |
| 1.89E-08 | 8.92E-08 | 50.4790319 | 104.102197 | 113.222334 | 132.221803 | 0 |
| 1.89E-08 | 8.93E-08 | 44.691627 | 164.161157 | 167.049345 | 146.139888 | 0 |
| 1.89E-08 | 8.93E-08 | 91.3123889 | 175.950509 | 223.660512 | 214.802438 | 2.37621824 |
| 1.89E-08 | 8.96E-08 | 51.4435994 | 142.80686 | 107.654022 | 105.313506 | 0 |
| 1.89E-08 | 8.96E-08 | 80.380624 | 273.601929 | 250.574018 | 287.176478 | 0 |
| 1.90E-08 | 8.97E-08 | 54.3373018 | 164.606038 | 113.222334 | 97.4265918 | 0 |
| 1.90E-08 | 8.97E-08 | 56.9094818 | 86.7518311 | 77.0283091 | 80.2609542 | 0 |
| 1.90E-08 | 8.98E-08 | 48.8714194 | 122.342326 | 113.222334 | 126.654569 | 0 |
| 1.90E-08 | 8.99E-08 | 72.3425616 | 90.0884399 | 57.5392189 | 82.5806349 | 0 |
| 1.90E-08 | 8.99E-08 | 55.3018693 | 97.8738607 | 80.7405168 | 112.272549 | 0 |
| 1.90E-08 | 9.00E-08 | 104.816334 | 83.6376628 | 66.819738 | 63.0953166 | 0 |
| 1.91E-08 | 9.01E-08 | 43.7270595 | 242.015365 | 153.128566 | 192.997439 | 0 |
| 1.91E-08 | 9.03E-08 | 84.5604165 | 54.4979451 | 98.3735032 | 87.6839326 | 0 |
| 1.91E-08 | 9.04E-08 | 54.6588243 | 99.2085042 | 80.7405168 | 101.138081 | 0 |
| 1.91E-08 | 9.04E-08 | 65.2690667 | 83.4152222 | 122.502853 | 69.5904227 | 0 |
| 1.92E-08 | 9.06E-08 | 48.5498969 | 118.560836 | 147.560255 | 120.623399 | 0 |
| 1.92E-08 | 9.06E-08 | 58.1955718 | 83.4152222 | 71.4599976 | 84.9003157 | 0 |
| 1.92E-08 | 9.07E-08 | 96.7782713 | 170.167053 | 168.905449 | 175.367865 | 0 |
| 1.92E-08 | 9.07E-08 | 59.1601393 | 109.663212 | 72.3880495 | 90.4675495 | 0 |
| 1.92E-08 | 9.08E-08 | 52.4081669 | 97.8738607 | 91.8771398 | 102.529889 | 0 |
| 1.93E-08 | 9.10E-08 | 74.2716966 | 70.7361084 | 105.797919 | 63.5592527 | 0 |
| 1.93E-08 | 9.10E-08 | 43.7270595 | 285.613721 | 216.236097 | 154.026802 | 0 |
| 1.93E-08 | 9.12E-08 | 75.5577866 | 70.0687866 | 81.6685687 | 148.923505 | 0 |
| 1.94E-08 | 9.14E-08 | 54.3373018 | 99.6533854 | 81.6685687 | 105.313506 | 0 |
| 1.94E-08 | 9.14E-08 | 48.2283744 | 126.568697 | 118.790645 | 123.407016 | 0 |
| 1.94E-08 | 9.14E-08 | 98.7074063 | 138.358048 | 50.1148035 | 140.572654 | 0 |
| 1.94E-08 | 9.15E-08 | 60.4462293 | 84.3049845 | 77.0283091 | 68.1986142 | 0 |
| 1.94E-08 | 9.16E-08 | 71.6995166 | 400.837948 | 468.666218 | 430.996685 | 4.75243649 |

| | | | | | | |
|---|---|---|---|---|---|---|
| 1.94E-08 | 9.16E-08 | 40.190312 | 240.45828 | 220.876356 | 182.790844 | 0 |
| 1.95E-08 | 9.18E-08 | 77.8084441 | 60.7262817 | 107.654022 | 73.7658481 | 0 |
| 1.96E-08 | 9.22E-08 | 57.8740493 | 96.3167765 | 71.4599976 | 97.8905279 | 0 |
| 1.96E-08 | 9.22E-08 | 58.1955718 | 76.5195638 | 73.3161014 | 85.3642518 | 0 |
| 1.96E-08 | 9.23E-08 | 74.9147416 | 66.064856 | 64.9636342 | 67.7346781 | 0 |
| 1.96E-08 | 9.25E-08 | 92.2769564 | 205.757548 | 174.47376 | 190.213822 | 0 |
| 1.98E-08 | 9.32E-08 | 47.9068519 | 116.781311 | 132.711424 | 154.490738 | 0 |
| 1.99E-08 | 9.35E-08 | 105.459379 | 72.5156331 | 154.98467 | 64.487125 | 0 |
| 1.99E-08 | 9.35E-08 | 57.5525268 | 105.2144 | 85.3807764 | 154.954675 | 0 |
| 1.99E-08 | 9.35E-08 | 47.2638069 | 141.249776 | 169.833501 | 121.551272 | 0 |
| 1.99E-08 | 9.35E-08 | 53.6942569 | 106.326603 | 93.7332436 | 84.9003157 | 0 |
| 1.99E-08 | 9.35E-08 | 60.7677518 | 145.698588 | 73.3161014 | 125.262761 | 0 |
| 1.99E-08 | 9.35E-08 | 54.6588243 | 114.556905 | 105.797919 | 193.461375 | 0 |
| 1.99E-08 | 9.36E-08 | 465.243052 | 346.340002 | 355.443884 | 421.254025 | 78.415202 |
| 1.99E-08 | 9.36E-08 | 76.5223541 | 124.789172 | 67.7477899 | 71.4461673 | 0 |
| 1.99E-08 | 9.38E-08 | 88.7402089 | 141.249776 | 116.00649 | 149.851377 | 7.12865473 |
| 2.00E-08 | 9.42E-08 | 51.7651219 | 119.228158 | 93.7332436 | 138.252973 | 0 |
| 2.00E-08 | 9.42E-08 | 87.7756414 | 203.533142 | 209.739733 | 248.669777 | 0 |
| 2.00E-08 | 9.42E-08 | 79.737579 | 110.997856 | 277.487523 | 86.2921241 | 0 |
| 2.01E-08 | 9.43E-08 | 64.9475442 | 137.023405 | 83.5246725 | 230.576267 | 0 |
| 2.01E-08 | 9.43E-08 | 53.6942569 | 84.0825439 | 106.72597 | 93.2511664 | 0 |
| 2.01E-08 | 9.44E-08 | 47.5853294 | 141.027336 | 130.85532 | 117.375846 | 0 |
| 2.02E-08 | 9.47E-08 | 658.15655 | 2900.40288 | 2556.78303 | 2792.43169 | 9.50487297 |
| 2.02E-08 | 9.49E-08 | 53.6942569 | 102.545113 | 82.5966206 | 96.9626556 | 0 |
| 2.02E-08 | 9.49E-08 | 93.8845689 | 200.641414 | 160.552982 | 203.204034 | 0 |
| 2.03E-08 | 9.51E-08 | 44.3701045 | 183.068608 | 146.632203 | 154.026802 | 0 |
| 2.03E-08 | 9.52E-08 | 45.9777169 | 158.600142 | 127.143113 | 151.243185 | 0 |
| 2.03E-08 | 9.54E-08 | 88.4186864 | 64.0628906 | 63.1075304 | 102.529889 | 0 |
| 2.03E-08 | 9.55E-08 | 113.175919 | 80.0786133 | 133.639476 | 55.6723382 | 0 |
| 2.03E-08 | 9.55E-08 | 53.0512119 | 105.881722 | 84.4527244 | 102.529889 | 0 |
| 2.04E-08 | 9.55E-08 | 51.4435994 | 100.988029 | 103.941815 | 93.2511664 | 0 |
| 2.04E-08 | 9.59E-08 | 52.4081669 | 116.113989 | 87.2368802 | 115.984038 | 0 |
| 2.05E-08 | 9.60E-08 | 41.1548795 | 177.507593 | 194.890903 | 190.213822 | 0 |
| 2.05E-08 | 9.60E-08 | 79.4160565 | 286.058602 | 238.509343 | 253.309139 | 0 |
| 2.05E-08 | 9.60E-08 | 55.6233918 | 101.655351 | 79.8124649 | 123.870952 | 0 |
| 2.05E-08 | 9.60E-08 | 261.076267 | 625.502946 | 599.521538 | 554.867637 | 59.4054561 |
| 2.05E-08 | 9.60E-08 | 48.5498969 | 116.113989 | 113.222334 | 135.933292 | 0 |
| 2.06E-08 | 9.67E-08 | 152.080141 | 226.666964 | 239.437395 | 235.679565 | 16.6335277 |
| 2.06E-08 | 9.67E-08 | 50.4790319 | 112.55494 | 113.222334 | 96.0347833 | 0 |
| 2.06E-08 | 9.68E-08 | 24.7572322 | 434.426477 | 465.882062 | 506.154341 | 0 |
| 2.07E-08 | 9.69E-08 | 49.5144644 | 161.49187 | 103.941815 | 162.841589 | 0 |
| 2.07E-08 | 9.71E-08 | 45.6561944 | 130.795068 | 143.848047 | 162.841589 | 0 |
| 2.07E-08 | 9.72E-08 | 40.833357 | 174.838306 | 200.459214 | 200.884354 | 0 |

| | | | | | | |
|---|---|---|---|---|---|---|
| 2.07E-08 | 9.72E-08 | 96.7782713 | 151.037162 | 210.667785 | 205.059779 | 4.75243649 |
| 2.08E-08 | 9.74E-08 | 383.254815 | 1013.88422 | 1205.53944 | 1208.55367 | 28.5146189 |
| 2.08E-08 | 9.74E-08 | 44.691627 | 161.936751 | 140.135839 | 149.851377 | 0 |
| 2.08E-08 | 9.74E-08 | 50.8005544 | 173.726103 | 108.582074 | 120.159463 | 0 |
| 2.08E-08 | 9.74E-08 | 52.4081669 | 137.468286 | 89.092984 | 128.046378 | 0 |
| 2.08E-08 | 9.75E-08 | 45.6561944 | 142.58442 | 144.776099 | 131.757867 | 0 |
| 2.08E-08 | 9.76E-08 | 36.0105196 | 209.316597 | 396.278169 | 321.971689 | 0 |
| 2.09E-08 | 9.77E-08 | 38.904222 | 185.960335 | 243.149602 | 247.277969 | 0 |
| 2.09E-08 | 9.78E-08 | 45.9777169 | 140.137573 | 131.783372 | 135.469356 | 0 |
| 2.09E-08 | 9.78E-08 | 51.4435994 | 105.436841 | 98.3735032 | 138.252973 | 0 |
| 2.09E-08 | 9.79E-08 | 67.5197242 | 112.77738 | 81.6685687 | 212.482757 | 0 |
| 2.09E-08 | 9.79E-08 | 52.7296894 | 180.844202 | 103.941815 | 117.375846 | 0 |
| 2.09E-08 | 9.79E-08 | 48.5498969 | 253.804716 | 151.272462 | 134.077548 | 0 |
| 2.09E-08 | 9.79E-08 | 45.9777169 | 127.013578 | 203.24337 | 167.017014 | 0 |
| 2.10E-08 | 9.80E-08 | 48.8714194 | 143.919063 | 119.718697 | 108.561059 | 0 |
| 2.09E-08 | 9.80E-08 | 366.214123 | 495.820081 | 270.99116 | 486.669023 | 21.3859642 |
| 2.10E-08 | 9.83E-08 | 59.1601393 | 74.5175985 | 77.956361 | 69.1264865 | 0 |
| 2.11E-08 | 9.86E-08 | 45.334672 | 141.694657 | 151.272462 | 213.41063 | 0 |
| 2.11E-08 | 9.86E-08 | 85.8465065 | 238.233874 | 241.293498 | 231.040203 | 4.75243649 |
| 2.11E-08 | 9.87E-08 | 47.5853294 | 143.919063 | 117.862593 | 122.015208 | 0 |
| 2.11E-08 | 9.87E-08 | 142.434466 | 127.903341 | 45.4745439 | 134.077548 | 0 |
| 2.11E-08 | 9.87E-08 | 41.476402 | 192.188672 | 175.401812 | 175.367865 | 0 |
| 2.12E-08 | 9.88E-08 | 46.9422844 | 134.354118 | 134.567528 | 118.303719 | 0 |
| 2.12E-08 | 9.91E-08 | 371.036961 | 540.308199 | 409.270895 | 487.132959 | 0 |
| 2.13E-08 | 9.93E-08 | 138.254673 | 352.345898 | 352.659728 | 315.940519 | 4.75243649 |
| 2.13E-08 | 9.96E-08 | 94.2060914 | 177.285152 | 160.552982 | 204.595843 | 0 |
| 2.13E-08 | 9.97E-08 | 51.1220769 | 124.121851 | 103.013763 | 95.106911 | 0 |
| 2.14E-08 | 9.98E-08 | 79.094534 | 69.6239054 | 57.5392189 | 87.6839326 | 0 |
| 2.14E-08 | 9.98E-08 | 70.4134267 | 266.70627 | 85.3807764 | 128.97425 | 0 |
| 2.14E-08 | 9.99E-08 | 49.5144644 | 124.121851 | 100.229607 | 115.056166 | 0 |
| 2.14E-08 | 1.00E-07 | 50.4790319 | 95.4270142 | 125.287009 | 139.180845 | 0 |
| 2.14E-08 | 1.00E-07 | 46.2992394 | 153.484009 | 156.840774 | 119.231591 | 0 |
| 2.15E-08 | 1.00E-07 | 47.9068519 | 134.131677 | 115.078438 | 116.447974 | 0 |
| 2.15E-08 | 1.00E-07 | 66.5551567 | 365.247453 | 346.163365 | 332.64222 | 2.37621824 |
| 2.15E-08 | 1.00E-07 | 50.4790319 | 102.989994 | 99.3015551 | 99.2823364 | 0 |
| 2.15E-08 | 1.00E-07 | 66.8766792 | 83.8601033 | 58.4672708 | 84.4363795 | 0 |
| 2.16E-08 | 1.00E-07 | 68.8058142 | 193.523315 | 85.3807764 | 91.3954218 | 0 |
| 2.16E-08 | 1.00E-07 | 59.1601393 | 79.8561727 | 67.7477899 | 76.0855288 | 0 |
| 2.16E-08 | 1.01E-07 | 79.737579 | 80.9683756 | 58.4672708 | 67.2707419 | 0 |
| 2.16E-08 | 1.01E-07 | 31.1876821 | 305.855815 | 355.443884 | 314.548711 | 0 |
| 2.16E-08 | 1.01E-07 | 30.2231146 | 277.605859 | 500.219983 | 392.489984 | 0 |
| 2.16E-08 | 1.01E-07 | 82.6312815 | 72.5156331 | 64.9636342 | 61.7035081 | 0 |
| 2.16E-08 | 1.01E-07 | 51.4435994 | 124.789172 | 93.7332436 | 101.602017 | 0 |

| | | | | | | |
|---|---|---|---|---|---|---|
| 2.16E-08 | 1.01E-07 | 47.5853294 | 254.249597 | 128.071165 | 170.728504 | 0 |
| 2.16E-08 | 1.01E-07 | 52.4081669 | 144.586385 | 93.7332436 | 107.633187 | 0 |
| 2.16E-08 | 1.01E-07 | 46.2992394 | 135.688761 | 125.287009 | 130.829995 | 0 |
| 2.17E-08 | 1.01E-07 | 630.184092 | 679.333569 | 525.277385 | 641.159761 | 23.7621824 |
| 2.17E-08 | 1.01E-07 | 156.259933 | 173.726103 | 41.7623363 | 197.172864 | 0 |
| 2.17E-08 | 1.01E-07 | 86.168029 | 115.001786 | 163.337137 | 55.6723382 | 0 |
| 2.18E-08 | 1.01E-07 | 40.5118345 | 224.664998 | 180.970124 | 182.790844 | 0 |
| 2.18E-08 | 1.01E-07 | 72.6640841 | 61.3936035 | 59.3953227 | 55.208402 | 182.968805 |
| 2.18E-08 | 1.02E-07 | 41.1548795 | 170.611934 | 179.11402 | 214.802438 | 0 |
| 2.19E-08 | 1.02E-07 | 87.7756414 | 110.108093 | 111.36623 | 128.97425 | 273.265098 |
| 2.19E-08 | 1.02E-07 | 39.8687895 | 281.164909 | 186.538435 | 199.956481 | 0 |
| 2.19E-08 | 1.02E-07 | 142.112943 | 179.287118 | 44.546492 | 135.469356 | 0 |
| 2.20E-08 | 1.02E-07 | 54.6588243 | 97.4289795 | 79.8124649 | 82.5806349 | 0 |
| 2.20E-08 | 1.02E-07 | 62.0538418 | 86.5293905 | 63.1075304 | 96.9626556 | 0 |
| 2.20E-08 | 1.02E-07 | 58.8386168 | 77.6317668 | 73.3161014 | 69.1264865 | 0 |
| 2.21E-08 | 1.03E-07 | 48.2283744 | 200.863855 | 128.071165 | 127.582442 | 0 |
| 2.21E-08 | 1.03E-07 | 368.786303 | 669.101302 | 424.119726 | 710.750184 | 23.7621824 |
| 2.21E-08 | 1.03E-07 | 59.4816618 | 254.694478 | 131.783372 | 100.210209 | 0 |
| 2.22E-08 | 1.03E-07 | 55.9449143 | 74.7400391 | 100.229607 | 114.592229 | 0 |
| 2.22E-08 | 1.03E-07 | 39.8687895 | 286.281042 | 165.193241 | 294.599456 | 0 |
| 2.22E-08 | 1.03E-07 | 45.9777169 | 134.576558 | 123.430905 | 150.315313 | 0 |
| 2.22E-08 | 1.03E-07 | 45.334672 | 135.688761 | 197.675058 | 144.284143 | 0 |
| 2.22E-08 | 1.03E-07 | 35.6889971 | 228.446488 | 297.904665 | 245.88616 | 0 |
| 2.23E-08 | 1.03E-07 | 59.8031843 | 69.4014648 | 76.1002572 | 69.1264865 | 0 |
| 2.23E-08 | 1.04E-07 | 47.9068519 | 174.170984 | 124.358957 | 119.695527 | 0 |
| 2.23E-08 | 1.04E-07 | 56.9094818 | 70.0687866 | 90.9490879 | 96.0347833 | 0 |
| 2.25E-08 | 1.04E-07 | 69.7703817 | 136.800964 | 61.2514265 | 104.385634 | 0 |
| 2.26E-08 | 1.05E-07 | 157.224501 | 48.4920492 | 322.034015 | 150.779249 | 0 |
| 2.28E-08 | 1.06E-07 | 48.5498969 | 129.682865 | 101.157659 | 128.510314 | 0 |
| 2.28E-08 | 1.06E-07 | 63.3399317 | 61.1711629 | 94.6612955 | 93.7151026 | 0 |
| 2.28E-08 | 1.06E-07 | 43.405537 | 156.598177 | 149.416359 | 211.090949 | 0 |
| 2.29E-08 | 1.06E-07 | 49.5144644 | 100.765588 | 101.157659 | 108.097123 | 0 |
| 2.29E-08 | 1.06E-07 | 128.608998 | 93.2026082 | 58.4672708 | 73.7658481 | 0 |
| 2.29E-08 | 1.06E-07 | 44.048582 | 155.485974 | 134.567528 | 162.377653 | 0 |
| 2.29E-08 | 1.06E-07 | 44.048582 | 193.968197 | 134.567528 | 164.233398 | 0 |
| 2.30E-08 | 1.06E-07 | 42.762492 | 178.174915 | 153.128566 | 154.490738 | 0 |
| 2.30E-08 | 1.06E-07 | 103.851766 | 183.291048 | 130.85532 | 157.274355 | 0 |
| 2.30E-08 | 1.07E-07 | 104.494811 | 234.452384 | 139.207788 | 163.769461 | 0 |
| 2.30E-08 | 1.07E-07 | 46.2992394 | 176.39539 | 120.646749 | 140.572654 | 0 |
| 2.30E-08 | 1.07E-07 | 94.8491364 | 146.810791 | 127.143113 | 50.1051043 | 0 |
| 2.30E-08 | 1.07E-07 | 49.5144644 | 132.797034 | 132.711424 | 95.5708472 | 0 |
| 2.31E-08 | 1.07E-07 | 64.9475442 | 88.7537964 | 60.3233746 | 103.457762 | 0 |
| 2.31E-08 | 1.07E-07 | 49.5144644 | 96.0943359 | 110.438178 | 107.169251 | 0 |

| | | | | | | |
|---|---|---|---|---|---|---|
| 2.32E-08 | 1.07E-07 | 92.5984789 | 81.1908162 | 94.6612955 | 52.4247851 | 0 |
| 2.32E-08 | 1.07E-07 | 52.0866444 | 96.5392171 | 88.1649321 | 86.2921241 | 0 |
| 2.32E-08 | 1.08E-07 | 52.0866444 | 112.332499 | 112.294282 | 83.0445711 | 0 |
| 2.32E-08 | 1.08E-07 | 36.3320421 | 219.771305 | 249.645966 | 233.82382 | 0 |
| 2.32E-08 | 1.08E-07 | 49.5144644 | 123.89941 | 105.797919 | 98.3544641 | 0 |
| 2.33E-08 | 1.08E-07 | 30.8661596 | 275.603894 | 433.400245 | 329.858604 | 0 |
| 2.33E-08 | 1.08E-07 | 55.6233918 | 78.2990885 | 75.1722053 | 77.4773373 | 0 |
| 2.34E-08 | 1.08E-07 | 52.0866444 | 114.112024 | 90.0210359 | 89.5396772 | 0 |
| 2.35E-08 | 1.09E-07 | 48.8714194 | 104.769519 | 102.085711 | 111.344676 | 0 |
| 2.35E-08 | 1.09E-07 | 43.0840145 | 139.247811 | 175.401812 | 165.625206 | 0 |
| 2.35E-08 | 1.09E-07 | 91.6339114 | 175.505627 | 164.265189 | 189.749886 | 4.75243649 |
| 2.36E-08 | 1.09E-07 | 58.5170943 | 80.9683756 | 70.5319457 | 69.5904227 | 0 |
| 2.37E-08 | 1.09E-07 | 73.3071291 | 82.3030192 | 57.5392189 | 68.6625504 | 0 |
| 2.37E-08 | 1.09E-07 | 49.8359869 | 93.4250488 | 103.013763 | 110.416804 | 0 |
| 2.37E-08 | 1.09E-07 | 46.2992394 | 127.6809 | 116.00649 | 136.397228 | 0 |
| 2.37E-08 | 1.10E-07 | 45.9777169 | 126.791138 | 136.423632 | 119.695527 | 0 |
| 2.37E-08 | 1.10E-07 | 58.8386168 | 69.4014648 | 74.2441534 | 69.5904227 | 0 |
| 2.37E-08 | 1.10E-07 | 46.2992394 | 123.454529 | 123.430905 | 121.551272 | 0 |
| 2.37E-08 | 1.10E-07 | 93.5630464 | 152.816687 | 191.178695 | 184.646588 | 0 |
| 2.37E-08 | 1.10E-07 | 206.417443 | 556.323922 | 444.536868 | 529.815085 | 7.12865473 |
| 2.37E-08 | 1.10E-07 | 57.2310043 | 77.6317668 | 78.8844129 | 68.6625504 | 0 |
| 2.37E-08 | 1.10E-07 | 155.938411 | 94.3148112 | 52.8989593 | 210.163077 | 0 |
| 2.37E-08 | 1.10E-07 | 53.6942569 | 90.5333211 | 79.8124649 | 81.1888265 | 0 |
| 2.38E-08 | 1.10E-07 | 47.5853294 | 106.104163 | 125.287009 | 118.303719 | 0 |
| 2.38E-08 | 1.10E-07 | 62.0538418 | 108.99589 | 70.5319457 | 73.7658481 | 0 |
| 2.40E-08 | 1.11E-07 | 69.4488592 | 67.3994995 | 62.1794784 | 65.4149973 | 0 |
| 2.40E-08 | 1.11E-07 | 53.3727344 | 91.6455241 | 77.0283091 | 95.5708472 | 0 |
| 2.40E-08 | 1.11E-07 | 53.3727344 | 112.332499 | 189.322591 | 101.138081 | 0 |
| 2.41E-08 | 1.11E-07 | 253.359727 | 61.8384847 | 101.157659 | 114.592229 | 0 |
| 2.41E-08 | 1.11E-07 | 87.7756414 | 208.426835 | 177.257916 | 223.617225 | 0 |
| 2.41E-08 | 1.11E-07 | 48.5498969 | 146.810791 | 99.3015551 | 143.356271 | 0 |
| 2.41E-08 | 1.11E-07 | 43.405537 | 141.249776 | 162.409085 | 145.675952 | 0 |
| 2.41E-08 | 1.11E-07 | 48.8714194 | 102.322673 | 152.200514 | 180.007227 | 0 |
| 2.42E-08 | 1.12E-07 | 38.904222 | 238.901196 | 177.257916 | 205.987651 | 0 |
| 2.43E-08 | 1.12E-07 | 47.2638069 | 128.793103 | 105.797919 | 127.118505 | 0 |
| 2.43E-08 | 1.12E-07 | 49.5144644 | 101.877791 | 94.6612955 | 105.777442 | 0 |
| 2.44E-08 | 1.12E-07 | 38.5826995 | 212.430766 | 190.250643 | 196.708928 | 0 |
| 2.44E-08 | 1.12E-07 | 43.0840145 | 143.696623 | 147.560255 | 173.048184 | 0 |
| 2.45E-08 | 1.13E-07 | 51.4435994 | 117.003752 | 101.157659 | 85.3642518 | 0 |
| 2.45E-08 | 1.13E-07 | 85.524984 | 255.3618 | 185.610383 | 206.915523 | 0 |
| 2.46E-08 | 1.13E-07 | 59.8031843 | 85.6396281 | 100.229607 | 64.487125 | 0 |
| 2.47E-08 | 1.14E-07 | 39.2257445 | 217.991781 | 219.948304 | 167.480951 | 0 |
| 2.48E-08 | 1.14E-07 | 58.1955718 | 84.9723063 | 84.4527244 | 64.9510612 | 0 |

| | | | | | | |
|---|---|---|---|---|---|---|
| 2.48E-08 | 1.14E-07 | 293.228516 | 126.568697 | 65.8916861 | 96.0347833 | 0 |
| 2.48E-08 | 1.14E-07 | 153.366231 | 83.6376628 | 59.3953227 | 87.2199964 | 0 |
| 2.48E-08 | 1.14E-07 | 46.9422844 | 135.466321 | 121.574801 | 108.097123 | 0 |
| 2.49E-08 | 1.15E-07 | 51.1220769 | 114.556905 | 86.3088283 | 98.3544641 | 0 |
| 2.49E-08 | 1.15E-07 | 50.1575094 | 120.11792 | 89.092984 | 108.561059 | 0 |
| 2.50E-08 | 1.15E-07 | 47.5853294 | 107.216366 | 108.582074 | 115.984038 | 0 |
| 2.50E-08 | 1.15E-07 | 46.2992394 | 114.556905 | 149.416359 | 123.407016 | 0 |
| 2.50E-08 | 1.15E-07 | 110.282216 | 226.444523 | 202.315318 | 218.049991 | 0 |
| 2.50E-08 | 1.15E-07 | 49.8359869 | 142.58442 | 94.6612955 | 114.592229 | 0 |
| 2.51E-08 | 1.15E-07 | 72.9856066 | 66.2872965 | 72.3880495 | 128.046378 | 0 |
| 2.51E-08 | 1.16E-07 | 51.7651219 | 102.322673 | 80.7405168 | 101.602017 | 0 |
| 2.52E-08 | 1.16E-07 | 51.1220769 | 85.1947469 | 117.862593 | 101.138081 | 0 |
| 2.52E-08 | 1.16E-07 | 4.50131495 | 5.33857422 | 3.71220767 | 4.17542536 | 28.5146189 |
| 2.52E-08 | 1.16E-07 | 50.8005544 | 134.354118 | 87.2368802 | 128.046378 | 0 |
| 2.53E-08 | 1.16E-07 | 68.4842917 | 66.7321777 | 190.250643 | 193.925311 | 0 |
| 2.53E-08 | 1.16E-07 | 47.9068519 | 104.547078 | 189.322591 | 167.944887 | 0 |
| 2.54E-08 | 1.16E-07 | 65.2690667 | 94.5372518 | 58.4672708 | 80.7248903 | 0 |
| 2.54E-08 | 1.17E-07 | 53.6942569 | 84.3049845 | 109.510126 | 79.797018 | 0 |
| 2.54E-08 | 1.17E-07 | 47.9068519 | 112.55494 | 108.582074 | 102.065953 | 0 |
| 2.55E-08 | 1.17E-07 | 74.2716966 | 55.6101481 | 76.1002572 | 73.7658481 | 0 |
| 2.56E-08 | 1.17E-07 | 46.2992394 | 123.232088 | 125.287009 | 110.88074 | 0 |
| 2.56E-08 | 1.17E-07 | 66.8766792 | 94.5372518 | 56.6111669 | 85.828188 | 0 |
| 2.56E-08 | 1.17E-07 | 63.0184092 | 108.99589 | 115.078438 | 61.7035081 | 0 |
| 2.56E-08 | 1.17E-07 | 54.9803468 | 100.543148 | 73.3161014 | 115.984038 | 0 |
| 2.56E-08 | 1.18E-07 | 44.3701045 | 148.145435 | 122.502853 | 149.387441 | 0 |
| 2.57E-08 | 1.18E-07 | 53.6942569 | 88.7537964 | 76.1002572 | 80.2609542 | 0 |
| 2.58E-08 | 1.18E-07 | 98.0643613 | 120.785242 | 93.7332436 | 96.9626556 | 7.12865473 |
| 2.58E-08 | 1.19E-07 | 88.0971639 | 115.891549 | 63.1075304 | 63.0953166 | 0 |
| 2.59E-08 | 1.19E-07 | 75.2362641 | 100.765588 | 52.8989593 | 88.6118049 | 0 |
| 2.60E-08 | 1.19E-07 | 65.5905892 | 123.89941 | 74.2441534 | 206.915523 | 0 |
| 2.61E-08 | 1.19E-07 | 39.8687895 | 171.279256 | 171.689605 | 215.73031 | 0 |
| 2.61E-08 | 1.19E-07 | 76.2008316 | 126.791138 | 129.927268 | 151.243185 | 4.75243649 |
| 2.62E-08 | 1.20E-07 | 80.7021465 | 60.7262817 | 74.2441534 | 129.902122 | 0 |
| 2.62E-08 | 1.20E-07 | 50.8005544 | 91.4230835 | 97.4454513 | 130.366059 | 0 |
| 2.65E-08 | 1.21E-07 | 64.3044992 | 58.946757 | 73.3161014 | 72.8379758 | 0 |
| 2.65E-08 | 1.21E-07 | 38.261177 | 182.846167 | 193.962851 | 209.69914 | 0 |
| 2.66E-08 | 1.22E-07 | 58.1955718 | 74.9624797 | 77.956361 | 117.375846 | 0 |
| 2.66E-08 | 1.22E-07 | 68.4842917 | 70.7361084 | 56.6111669 | 74.2297842 | 0 |
| 2.67E-08 | 1.22E-07 | 44.048582 | 134.354118 | 128.999216 | 136.397228 | 0 |
| 2.67E-08 | 1.22E-07 | 120.249414 | 77.1868856 | 52.8989593 | 98.8184002 | 0 |
| 2.67E-08 | 1.22E-07 | 60.1247068 | 68.9565837 | 65.8916861 | 67.2707419 | 0 |
| 2.68E-08 | 1.23E-07 | 46.6207619 | 135.466321 | 107.654022 | 115.984038 | 0 |
| 2.68E-08 | 1.23E-07 | 45.0131495 | 130.127747 | 116.934542 | 134.077548 | 0 |

| | | | | | | |
|---|---|---|---|---|---|---|
| 2.68E-08 | 1.23E-07 | 59.4816618 | 141.472217 | 124.358957 | 68.6625504 | 0 |
| 2.69E-08 | 1.23E-07 | 86.168029 | 132.797034 | 51.9709073 | 167.480951 | 0 |
| 2.69E-08 | 1.23E-07 | 58.8386168 | 142.139539 | 159.62493 | 71.9101035 | 0 |
| 2.70E-08 | 1.23E-07 | 53.6942569 | 81.1908162 | 75.1722053 | 96.9626556 | 0 |
| 2.71E-08 | 1.24E-07 | 45.0131495 | 127.458459 | 116.934542 | 134.541484 | 0 |
| 2.71E-08 | 1.24E-07 | 66.8766792 | 179.064677 | 161.481034 | 64.9510612 | 0 |
| 2.72E-08 | 1.24E-07 | 59.4816618 | 111.887618 | 65.8916861 | 86.7560603 | 0 |
| 2.72E-08 | 1.24E-07 | 44.3701045 | 156.375736 | 140.135839 | 119.231591 | 0 |
| 2.72E-08 | 1.24E-07 | 101.279586 | 123.454529 | 57.5392189 | 256.092756 | 0 |
| 2.73E-08 | 1.25E-07 | 50.1575094 | 113.222262 | 88.1649321 | 94.1790387 | 0 |
| 2.73E-08 | 1.25E-07 | 49.8359869 | 110.775415 | 91.8771398 | 142.428398 | 0 |
| 2.73E-08 | 1.25E-07 | 36.0105196 | 193.078434 | 235.725187 | 265.371479 | 0 |
| 2.73E-08 | 1.25E-07 | 101.279586 | 291.842057 | 239.437395 | 274.186265 | 11.8810912 |
| 2.73E-08 | 1.25E-07 | 32.7952946 | 270.48776 | 237.581291 | 302.950307 | 0 |
| 2.74E-08 | 1.25E-07 | 46.6207619 | 121.897445 | 103.013763 | 121.551272 | 0 |
| 2.75E-08 | 1.26E-07 | 52.4081669 | 94.7596924 | 240.365446 | 241.710735 | 0 |
| 2.75E-08 | 1.26E-07 | 218.313775 | 145.031266 | 105.797919 | 132.221803 | 4.75243649 |
| 2.75E-08 | 1.26E-07 | 45.334672 | 186.850098 | 116.934542 | 143.820207 | 0 |
| 2.76E-08 | 1.26E-07 | 50.4790319 | 107.438806 | 94.6612955 | 83.9724434 | 0 |
| 2.76E-08 | 1.26E-07 | 36.9750871 | 218.881543 | 212.523889 | 404.552324 | 0 |
| 2.77E-08 | 1.27E-07 | 246.607755 | 920.236731 | 864.944387 | 974.729854 | 9.50487297 |
| 2.78E-08 | 1.27E-07 | 89.3832539 | 172.169019 | 168.905449 | 175.367865 | 0 |
| 2.78E-08 | 1.27E-07 | 48.8714194 | 117.448633 | 89.092984 | 118.303719 | 0 |
| 2.78E-08 | 1.27E-07 | 16.7191698 | 12.9015544 | 10.2085711 | 6.49510612 | 64.1578926 |
| 2.79E-08 | 1.27E-07 | 43.0840145 | 147.478113 | 133.639476 | 136.861165 | 0 |
| 2.79E-08 | 1.27E-07 | 50.1575094 | 102.100232 | 109.510126 | 171.656376 | 0 |
| 2.79E-08 | 1.27E-07 | 163.654951 | 123.676969 | 110.438178 | 121.551272 | 0 |
| 2.79E-08 | 1.27E-07 | 804.127763 | 244.907092 | 557.759202 | 641.623697 | 21.3859642 |
| 2.80E-08 | 1.27E-07 | 45.6561944 | 133.241915 | 124.358957 | 108.561059 | 0 |
| 2.80E-08 | 1.27E-07 | 57.2310043 | 141.917098 | 96.5173994 | 267.227223 | 0 |
| 2.80E-08 | 1.28E-07 | 79.4160565 | 215.322493 | 209.739733 | 278.825627 | 2.37621824 |
| 2.81E-08 | 1.28E-07 | 56.5879593 | 85.1947469 | 66.819738 | 74.2297842 | 0 |
| 2.83E-08 | 1.29E-07 | 106.423946 | 72.2931925 | 56.6111669 | 75.6215927 | 0 |
| 2.83E-08 | 1.29E-07 | 53.0512119 | 113.444702 | 75.1722053 | 97.4265918 | 0 |
| 2.83E-08 | 1.29E-07 | 202.23765 | 368.806502 | 369.364663 | 361.870198 | 42.7719284 |
| 2.83E-08 | 1.29E-07 | 36.6535646 | 176.172949 | 266.3509 | 248.669777 | 0 |
| 2.83E-08 | 1.29E-07 | 71.3779941 | 98.9860636 | 54.7550631 | 78.4052096 | 0 |
| 2.84E-08 | 1.29E-07 | 48.5498969 | 101.43291 | 91.8771398 | 99.7462725 | 0 |
| 2.84E-08 | 1.29E-07 | 47.9068519 | 92.757727 | 112.294282 | 113.664357 | 0 |
| 2.84E-08 | 1.29E-07 | 73.3071291 | 106.104163 | 54.7550631 | 79.797018 | 0 |
| 2.85E-08 | 1.30E-07 | 43.7270595 | 129.905306 | 128.999216 | 132.685739 | 0 |
| 2.85E-08 | 1.30E-07 | 46.9422844 | 128.793103 | 125.287009 | 97.8905279 | 0 |
| 2.85E-08 | 1.30E-07 | 49.5144644 | 85.4171875 | 114.150386 | 120.159463 | 0 |

| | | | | | | |
|---|---|---|---|---|---|---|
| 2.86E-08 | 1.30E-07 | 45.6561944 | 139.470251 | 143.848047 | 105.313506 | 0 |
| 2.86E-08 | 1.30E-07 | 80.380624 | 199.974093 | 215.308045 | 220.833608 | 14.2573095 |
| 2.86E-08 | 1.30E-07 | 157.867546 | 94.5372518 | 60.3233746 | 75.1576565 | 0 |
| 2.86E-08 | 1.30E-07 | 36.3320421 | 217.546899 | 328.530379 | 198.100737 | 0 |
| 2.86E-08 | 1.30E-07 | 127.001386 | 83.1927816 | 170.761553 | 53.3526574 | 0 |
| 2.88E-08 | 1.31E-07 | 86.4895515 | 222.885474 | 181.898176 | 197.6368 | 4.75243649 |
| 2.88E-08 | 1.31E-07 | 48.5498969 | 116.33643 | 90.0210359 | 105.313506 | 0 |
| 2.89E-08 | 1.31E-07 | 50.8005544 | 100.098267 | 83.5246725 | 84.9003157 | 0 |
| 2.90E-08 | 1.32E-07 | 38.261177 | 185.737895 | 186.538435 | 179.543291 | 0 |
| 2.92E-08 | 1.33E-07 | 85.524984 | 264.481864 | 259.854537 | 187.430205 | 4.75243649 |
| 2.92E-08 | 1.33E-07 | 86.811074 | 210.651241 | 163.337137 | 185.574461 | 2.37621824 |
| 2.93E-08 | 1.33E-07 | 47.5853294 | 102.322673 | 102.085711 | 96.9626556 | 0 |
| 2.93E-08 | 1.33E-07 | 47.9068519 | 123.454529 | 95.5893474 | 102.993826 | 0 |
| 2.94E-08 | 1.33E-07 | 45.0131495 | 115.669108 | 126.215061 | 115.056166 | 0 |
| 2.94E-08 | 1.33E-07 | 50.1575094 | 86.7518311 | 138.279736 | 153.562866 | 0 |
| 2.94E-08 | 1.34E-07 | 49.5144644 | 90.0884399 | 164.265189 | 144.284143 | 0 |
| 2.96E-08 | 1.34E-07 | 93.5630464 | 65.8424154 | 59.3953227 | 69.1264865 | 0 |
| 2.96E-08 | 1.34E-07 | 43.7270595 | 205.757548 | 177.257916 | 118.767655 | 0 |
| 2.97E-08 | 1.35E-07 | 45.334672 | 280.052706 | 274.703367 | 116.447974 | 0 |
| 2.98E-08 | 1.35E-07 | 36.9750871 | 225.999642 | 179.11402 | 205.523715 | 0 |
| 2.98E-08 | 1.35E-07 | 43.7270595 | 119.228158 | 132.711424 | 150.315313 | 0 |
| 2.99E-08 | 1.36E-07 | 64.9475442 | 79.4112915 | 60.3233746 | 111.808612 | 0 |
| 2.99E-08 | 1.36E-07 | 26.3648447 | 10.6771484 | 23.2012979 | 14.8459568 | 204.354769 |
| 3.00E-08 | 1.36E-07 | 77.8084441 | 76.2971232 | 58.4672708 | 60.7756358 | 0 |
| 3.00E-08 | 1.36E-07 | 71.3779941 | 90.5333211 | 54.7550631 | 71.4461673 | 0 |
| 3.00E-08 | 1.36E-07 | 45.6561944 | 146.143469 | 261.710641 | 133.149675 | 0 |
| 3.01E-08 | 1.36E-07 | 77.1653991 | 210.4288 | 229.228823 | 237.535309 | 0 |
| 3.01E-08 | 1.37E-07 | 48.2283744 | 95.6494547 | 97.4454513 | 92.3232941 | 0 |
| 3.02E-08 | 1.37E-07 | 92.9200014 | 223.775236 | 146.632203 | 197.172864 | 4.75243649 |
| 3.02E-08 | 1.37E-07 | 44.3701045 | 112.77738 | 128.071165 | 131.757867 | 0 |
| 3.02E-08 | 1.37E-07 | 28.2939797 | 405.064319 | 343.379209 | 276.969882 | 0 |
| 3.03E-08 | 1.37E-07 | 230.53163 | 153.92889 | 120.646749 | 109.952868 | 2.37621824 |
| 3.04E-08 | 1.37E-07 | 55.9449143 | 76.964445 | 71.4599976 | 66.3428696 | 0 |
| 3.04E-08 | 1.38E-07 | 59.4816618 | 60.05896 | 85.3807764 | 90.4675495 | 0 |
| 3.04E-08 | 1.38E-07 | 45.0131495 | 128.570662 | 106.72597 | 137.789037 | 0 |
| 3.05E-08 | 1.38E-07 | 45.9777169 | 181.289083 | 125.287009 | 113.664357 | 0 |
| 3.06E-08 | 1.39E-07 | 39.8687895 | 143.029301 | 252.430121 | 212.946693 | 0 |
| 3.07E-08 | 1.39E-07 | 48.8714194 | 151.037162 | 95.5893474 | 192.533503 | 0 |
| 3.07E-08 | 1.39E-07 | 46.6207619 | 127.236019 | 115.078438 | 96.9626556 | 0 |
| 3.07E-08 | 1.39E-07 | 62.0538418 | 80.3010539 | 58.4672708 | 96.4987195 | 0 |
| 3.08E-08 | 1.39E-07 | 47.5853294 | 178.842236 | 99.3015551 | 137.789037 | 0 |
| 3.09E-08 | 1.40E-07 | 48.2283744 | 106.104163 | 88.1649321 | 100.210209 | 0 |
| 3.09E-08 | 1.40E-07 | 64.6260217 | 157.487939 | 84.4527244 | 73.3019119 | 0 |

| | | | | | | |
|---|---|---|---|---|---|---|
| 3.09E-08 | 1.40E-07 | 83.2743265 | 107.883687 | 99.3015551 | 99.7462725 | 19.0097459 |
| 3.10E-08 | 1.40E-07 | 54.3373018 | 214.210291 | 88.1649321 | 136.397228 | 0 |
| 3.10E-08 | 1.40E-07 | 43.7270595 | 161.046989 | 113.222334 | 153.09893 | 0 |
| 3.10E-08 | 1.40E-07 | 52.7296894 | 112.332499 | 73.3161014 | 96.0347833 | 0 |
| 3.11E-08 | 1.41E-07 | 45.6561944 | 108.551009 | 108.582074 | 134.077548 | 0 |
| 3.11E-08 | 1.41E-07 | 52.7296894 | 93.2026082 | 75.1722053 | 76.549465 | 0 |
| 3.11E-08 | 1.41E-07 | 65.2690667 | 66.064856 | 62.1794784 | 61.239572 | 0 |
| 3.11E-08 | 1.41E-07 | 38.904222 | 176.61783 | 179.11402 | 159.594036 | 0 |
| 3.12E-08 | 1.41E-07 | 53.0512119 | 97.6514201 | 73.3161014 | 79.797018 | 0 |
| 3.12E-08 | 1.41E-07 | 45.0131495 | 132.797034 | 106.72597 | 121.087335 | 0 |
| 3.13E-08 | 1.41E-07 | 47.9068519 | 110.775415 | 88.1649321 | 109.952868 | 0 |
| 3.13E-08 | 1.41E-07 | 44.691627 | 107.661247 | 128.071165 | 125.262761 | 0 |
| 3.13E-08 | 1.41E-07 | 49.5144644 | 106.326603 | 81.6685687 | 114.128293 | 0 |
| 3.13E-08 | 1.41E-07 | 77.8084441 | 84.0825439 | 93.7332436 | 50.5690405 | 0 |
| 3.14E-08 | 1.42E-07 | 50.1575094 | 166.608004 | 94.6612955 | 109.488932 | 0 |
| 3.14E-08 | 1.42E-07 | 45.9777169 | 131.23995 | 108.582074 | 181.862971 | 0 |
| 3.15E-08 | 1.42E-07 | 53.6942569 | 77.4093262 | 68.6758419 | 80.2609542 | 0 |
| 3.15E-08 | 1.42E-07 | 50.1575094 | 80.5234945 | 126.215061 | 116.91191 | 0 |
| 3.16E-08 | 1.43E-07 | 45.0131495 | 168.165088 | 116.934542 | 119.695527 | 0 |
| 3.16E-08 | 1.43E-07 | 46.2992394 | 97.8738607 | 144.776099 | 154.954675 | 0 |
| 3.17E-08 | 1.43E-07 | 40.190312 | 159.045024 | 144.776099 | 175.367865 | 0 |
| 3.17E-08 | 1.43E-07 | 54.6588243 | 96.7616577 | 72.3880495 | 127.582442 | 0 |
| 3.17E-08 | 1.43E-07 | 45.9777169 | 114.334465 | 107.654022 | 101.138081 | 0 |
| 3.18E-08 | 1.43E-07 | 48.5498969 | 91.6455241 | 93.7332436 | 120.623399 | 0 |
| 3.18E-08 | 1.43E-07 | 47.9068519 | 147.255672 | 91.8771398 | 122.479144 | 0 |
| 3.19E-08 | 1.44E-07 | 48.5498969 | 177.730033 | 94.6612955 | 197.6368 | 0 |
| 3.19E-08 | 1.44E-07 | 45.334672 | 149.035197 | 105.797919 | 121.551272 | 0 |
| 3.19E-08 | 1.44E-07 | 65.5905892 | 148.145435 | 70.5319457 | 80.2609542 | 0 |
| 3.19E-08 | 1.44E-07 | 47.9068519 | 118.783276 | 232.012979 | 127.118505 | 0 |
| 3.19E-08 | 1.44E-07 | 42.119447 | 222.885474 | 125.287009 | 186.502333 | 0 |
| 3.20E-08 | 1.44E-07 | 47.9068519 | 96.0943359 | 95.5893474 | 90.4675495 | 0 |
| 3.20E-08 | 1.44E-07 | 54.0157793 | 108.99589 | 374.932974 | 186.038397 | 0 |
| 3.20E-08 | 1.44E-07 | 90.9908664 | 254.249597 | 153.128566 | 186.502333 | 0 |
| 3.22E-08 | 1.45E-07 | 42.4409695 | 158.377702 | 120.646749 | 150.315313 | 0 |
| 3.22E-08 | 1.45E-07 | 66.8766792 | 298.737716 | 267.278952 | 301.558498 | 2.37621824 |
| 3.22E-08 | 1.45E-07 | 48.8714194 | 132.129712 | 153.128566 | 87.6839326 | 0 |
| 3.22E-08 | 1.45E-07 | 45.0131495 | 146.810791 | 154.056618 | 102.993826 | 0 |
| 3.23E-08 | 1.45E-07 | 261.076267 | 940.033944 | 685.830367 | 779.87667 | 21.3859642 |
| 3.25E-08 | 1.46E-07 | 44.048582 | 163.938717 | 108.582074 | 156.810419 | 0 |
| 3.25E-08 | 1.46E-07 | 52.7296894 | 80.7459351 | 69.6038938 | 85.3642518 | 0 |
| 3.26E-08 | 1.47E-07 | 40.5118345 | 208.426835 | 144.776099 | 154.954675 | 0 |
| 3.26E-08 | 1.47E-07 | 50.1575094 | 92.3128459 | 77.0283091 | 96.4987195 | 0 |
| 3.26E-08 | 1.47E-07 | 51.7651219 | 120.340361 | 75.1722053 | 103.457762 | 0 |

| | | | | | | |
|---|---|---|---|---|---|---|
| 3.28E-08 | 1.47E-07 | 89.3832539 | 168.165088 | 149.416359 | 170.264568 | 0 |
| 3.29E-08 | 1.48E-07 | 52.0866444 | 84.9723063 | 72.3880495 | 77.9412734 | 0 |
| 3.29E-08 | 1.48E-07 | 64.9475442 | 100.988029 | 96.5173994 | 55.208402 | 0 |
| 3.29E-08 | 1.48E-07 | 40.190312 | 205.757548 | 139.207788 | 267.227223 | 0 |
| 3.30E-08 | 1.48E-07 | 36.6535646 | 184.848132 | 206.955577 | 183.718716 | 0 |
| 3.31E-08 | 1.49E-07 | 47.2638069 | 129.237984 | 90.0210359 | 134.077548 | 0 |
| 3.32E-08 | 1.49E-07 | 87.7756414 | 162.826514 | 172.617657 | 154.490738 | 2.37621824 |
| 3.32E-08 | 1.49E-07 | 45.0131495 | 110.330534 | 120.646749 | 104.84957 | 0 |
| 3.33E-08 | 1.49E-07 | 47.9068519 | 127.236019 | 95.5893474 | 94.6429749 | 0 |
| 3.34E-08 | 1.50E-07 | 95.8137038 | 121.230123 | 45.4745439 | 137.325101 | 0 |
| 3.34E-08 | 1.50E-07 | 49.1929419 | 98.763623 | 79.8124649 | 95.106911 | 0 |
| 3.34E-08 | 1.50E-07 | 47.2638069 | 159.934786 | 92.8051917 | 155.882547 | 0 |
| 3.35E-08 | 1.51E-07 | 40.190312 | 137.023405 | 162.409085 | 185.574461 | 0 |
| 3.36E-08 | 1.51E-07 | 45.334672 | 127.458459 | 99.3015551 | 118.303719 | 0 |
| 3.37E-08 | 1.51E-07 | 55.6233918 | 66.064856 | 77.0283091 | 67.2707419 | 0 |
| 3.37E-08 | 1.51E-07 | 45.6561944 | 163.938717 | 272.847264 | 120.159463 | 0 |
| 3.37E-08 | 1.51E-07 | 49.1929419 | 89.6435588 | 81.6685687 | 103.921698 | 0 |
| 3.38E-08 | 1.52E-07 | 32.7952946 | 299.849919 | 208.811681 | 250.061586 | 0 |
| 3.39E-08 | 1.52E-07 | 36.0105196 | 183.95837 | 192.106747 | 223.617225 | 0 |
| 3.39E-08 | 1.52E-07 | 87.1325964 | 157.265499 | 161.481034 | 172.120312 | 2.37621824 |
| 3.40E-08 | 1.52E-07 | 48.5498969 | 95.8718953 | 100.229607 | 81.6527626 | 0 |
| 3.40E-08 | 1.52E-07 | 56.2664368 | 73.8502767 | 82.5966206 | 62.1674443 | 0 |
| 3.40E-08 | 1.52E-07 | 46.6207619 | 118.115955 | 121.574801 | 90.4675495 | 0 |
| 3.41E-08 | 1.53E-07 | 94.5276139 | 169.944613 | 129.927268 | 139.180845 | 4.75243649 |
| 3.41E-08 | 1.53E-07 | 46.6207619 | 140.582454 | 112.294282 | 95.5708472 | 0 |
| 3.42E-08 | 1.53E-07 | 97.7428388 | 213.320528 | 128.071165 | 167.017014 | 0 |
| 3.42E-08 | 1.53E-07 | 48.5498969 | 80.9683756 | 102.085711 | 102.529889 | 0 |
| 3.42E-08 | 1.53E-07 | 51.1220769 | 84.5274251 | 84.4527244 | 72.3740396 | 0 |
| 3.44E-08 | 1.54E-07 | 59.4816618 | 100.765588 | 59.3953227 | 84.9003157 | 0 |
| 3.45E-08 | 1.54E-07 | 57.8740493 | 60.7262817 | 79.8124649 | 93.2511664 | 0 |
| 3.46E-08 | 1.55E-07 | 47.9068519 | 125.901375 | 89.092984 | 99.7462725 | 0 |
| 3.46E-08 | 1.55E-07 | 50.8005544 | 81.4132568 | 94.6612955 | 74.6937204 | 0 |
| 3.46E-08 | 1.55E-07 | 43.405537 | 139.247811 | 112.294282 | 123.870952 | 0 |
| 3.46E-08 | 1.55E-07 | 81.9882365 | 74.5175985 | 167.977397 | 65.4149973 | 0 |
| 3.47E-08 | 1.55E-07 | 42.762492 | 138.580489 | 119.718697 | 180.935099 | 0 |
| 3.48E-08 | 1.56E-07 | 91.9554339 | 219.103984 | 132.711424 | 228.720523 | 0 |
| 3.49E-08 | 1.56E-07 | 54.3373018 | 80.9683756 | 128.071165 | 77.9412734 | 0 |
| 3.49E-08 | 1.56E-07 | 54.9803468 | 89.6435588 | 67.7477899 | 70.518295 | 0 |
| 3.49E-08 | 1.56E-07 | 51.7651219 | 166.830444 | 98.3735032 | 91.859358 | 0 |
| 3.50E-08 | 1.57E-07 | 39.8687895 | 163.938717 | 141.063891 | 152.171058 | 0 |
| 3.50E-08 | 1.57E-07 | 142.434466 | 55.3877075 | 81.6685687 | 83.0445711 | 0 |
| 3.51E-08 | 1.57E-07 | 89.7047764 | 366.804537 | 307.185185 | 281.609244 | 0 |
| 3.52E-08 | 1.57E-07 | 53.0512119 | 74.9624797 | 125.287009 | 88.6118049 | 0 |

| | | | | | | |
|---|---|---|---|---|---|---|
| 3.52E-08 | 1.57E-07 | 55.3018693 | 122.564766 | 77.956361 | 76.0855288 | 0 |
| 3.52E-08 | 1.57E-07 | 202.880695 | 46.934965 | 148.488307 | 100.210209 | 0 |
| 3.52E-08 | 1.58E-07 | 48.8714194 | 110.775415 | 81.6685687 | 93.2511664 | 0 |
| 3.53E-08 | 1.58E-07 | 58.1955718 | 59.3916382 | 95.5893474 | 87.2199964 | 0 |
| 3.54E-08 | 1.58E-07 | 44.691627 | 103.434875 | 111.36623 | 111.344676 | 0 |
| 3.54E-08 | 1.58E-07 | 78.1299666 | 61.6160441 | 107.654022 | 59.8477635 | 0 |
| 3.55E-08 | 1.59E-07 | 48.5498969 | 99.6533854 | 81.6685687 | 89.5396772 | 0 |
| 3.55E-08 | 1.59E-07 | 46.6207619 | 109.440771 | 102.085711 | 89.5396772 | 0 |
| 3.56E-08 | 1.59E-07 | 44.3701045 | 110.330534 | 106.72597 | 130.366059 | 0 |
| 3.56E-08 | 1.59E-07 | 152.080141 | 91.8679647 | 57.5392189 | 73.7658481 | 0 |
| 3.56E-08 | 1.59E-07 | 53.6942569 | 104.547078 | 66.819738 | 107.169251 | 0 |
| 3.56E-08 | 1.59E-07 | 128.930521 | 134.576558 | 97.4454513 | 125.726697 | 0 |
| 3.59E-08 | 1.60E-07 | 51.1220769 | 82.3030192 | 77.956361 | 110.88074 | 0 |
| 3.60E-08 | 1.60E-07 | 79.4160565 | 371.475789 | 182.826228 | 251.91733 | 2.37621824 |
| 3.62E-08 | 1.62E-07 | 74.9147416 | 50.0491333 | 90.9490879 | 102.065953 | 0 |
| 3.62E-08 | 1.62E-07 | 38.904222 | 138.80293 | 181.898176 | 186.502333 | 0 |
| 3.63E-08 | 1.62E-07 | 40.833357 | 147.255672 | 129.927268 | 141.500526 | 0 |
| 3.63E-08 | 1.62E-07 | 46.9422844 | 103.657316 | 96.5173994 | 144.748079 | 0 |
| 3.63E-08 | 1.62E-07 | 56.2664368 | 147.255672 | 79.8124649 | 83.9724434 | 0 |
| 3.63E-08 | 1.62E-07 | 60.7677518 | 58.946757 | 64.0355823 | 69.1264865 | 0 |
| 3.64E-08 | 1.63E-07 | 68.8058142 | 103.212435 | 67.7477899 | 197.6368 | 0 |
| 3.65E-08 | 1.63E-07 | 37.2966096 | 169.944613 | 169.833501 | 178.615418 | 0 |
| 3.66E-08 | 1.63E-07 | 72.3425616 | 100.988029 | 127.143113 | 122.479144 | 9.50487297 |
| 3.68E-08 | 1.64E-07 | 54.9803468 | 77.4093262 | 65.8916861 | 66.3428696 | 0 |
| 3.69E-08 | 1.64E-07 | 41.7979245 | 299.405037 | 146.632203 | 162.377653 | 0 |
| 3.69E-08 | 1.64E-07 | 37.2966096 | 166.163123 | 192.106747 | 166.089142 | 0 |
| 3.69E-08 | 1.65E-07 | 36.6535646 | 199.529211 | 196.747006 | 160.521908 | 0 |
| 3.72E-08 | 1.66E-07 | 75.8793091 | 220.216187 | 216.236097 | 196.708928 | 2.37621824 |
| 3.73E-08 | 1.66E-07 | 45.334672 | 158.377702 | 108.582074 | 107.633187 | 0 |
| 3.73E-08 | 1.66E-07 | 85.2034615 | 169.05485 | 176.329864 | 153.09893 | 0 |
| 3.73E-08 | 1.66E-07 | 34.0813846 | 225.10988 | 206.027526 | 199.028609 | 0 |
| 3.74E-08 | 1.67E-07 | 44.3701045 | 108.328569 | 102.085711 | 115.056166 | 0 |
| 3.74E-08 | 1.67E-07 | 1742.97345 | 2109.40414 | 1773.50721 | 2079.36183 | 0 |
| 3.75E-08 | 1.67E-07 | 51.1220769 | 71.1809896 | 104.869867 | 110.416804 | 0 |
| 3.75E-08 | 1.67E-07 | 41.1548795 | 147.255672 | 129.927268 | 128.046378 | 0 |
| 3.75E-08 | 1.67E-07 | 53.6942569 | 225.554761 | 115.078438 | 95.5708472 | 0 |
| 3.75E-08 | 1.67E-07 | 194.52111 | 83.1927816 | 57.5392189 | 100.210209 | 0 |
| 3.75E-08 | 1.67E-07 | 44.048582 | 102.767554 | 180.970124 | 164.233398 | 0 |
| 3.76E-08 | 1.67E-07 | 43.7270595 | 126.346257 | 105.797919 | 155.418611 | 0 |
| 3.78E-08 | 1.68E-07 | 43.0840145 | 128.570662 | 106.72597 | 128.510314 | 0 |
| 3.78E-08 | 1.68E-07 | 54.3373018 | 116.33643 | 74.2441534 | 76.549465 | 0 |
| 3.78E-08 | 1.68E-07 | 80.380624 | 146.143469 | 128.999216 | 160.985845 | 7.12865473 |
| 3.79E-08 | 1.68E-07 | 35.3674746 | 159.267464 | 290.48025 | 270.01084 | 0 |

| | | | | | | |
|---|---|---|---|---|---|---|
| 3.79E-08 | 1.68E-07 | 51.7651219 | 76.5195638 | 71.4599976 | 95.5708472 | 0 |
| 3.80E-08 | 1.69E-07 | 78.4514891 | 169.499731 | 175.401812 | 151.243185 | 7.12865473 |
| 3.80E-08 | 1.69E-07 | 46.2992394 | 96.7616577 | 89.092984 | 103.921698 | 0 |
| 3.80E-08 | 1.69E-07 | 48.8714194 | 90.7557617 | 76.1002572 | 92.3232941 | 0 |
| 3.80E-08 | 1.69E-07 | 68.8058142 | 237.121672 | 211.595837 | 236.607437 | 2.37621824 |
| 3.81E-08 | 1.69E-07 | 35.6889971 | 177.285152 | 290.48025 | 195.31712 | 0 |
| 3.81E-08 | 1.69E-07 | 98.3858838 | 115.891549 | 137.351684 | 144.284143 | 0 |
| 3.83E-08 | 1.70E-07 | 50.4790319 | 78.2990885 | 136.423632 | 153.562866 | 0 |
| 3.84E-08 | 1.70E-07 | 36.3320421 | 211.763444 | 163.337137 | 185.574461 | 0 |
| 3.84E-08 | 1.71E-07 | 172.014535 | 212.653206 | 203.24337 | 212.946693 | 0 |
| 3.85E-08 | 1.71E-07 | 20.2559173 | 13.123995 | 12.9927268 | 15.309893 | 61.7816743 |
| 3.86E-08 | 1.71E-07 | 45.9777169 | 90.7557617 | 96.5173994 | 100.210209 | 0 |
| 3.86E-08 | 1.71E-07 | 66.5551567 | 265.594067 | 277.487523 | 297.383073 | 0 |
| 3.86E-08 | 1.72E-07 | 88.0971639 | 55.3877075 | 61.2514265 | 77.0134011 | 0 |
| 3.87E-08 | 1.72E-07 | 91.3123889 | 129.015544 | 91.8771398 | 136.397228 | 7.12865473 |
| 3.88E-08 | 1.72E-07 | 81.3451915 | 227.111845 | 165.193241 | 186.966269 | 2.37621824 |
| 3.88E-08 | 1.72E-07 | 63.0184092 | 70.0687866 | 157.768826 | 83.9724434 | 0 |
| 3.89E-08 | 1.73E-07 | 81.023669 | 189.296944 | 139.207788 | 173.048184 | 7.12865473 |
| 3.90E-08 | 1.73E-07 | 43.0840145 | 117.448633 | 109.510126 | 115.520102 | 0 |
| 3.90E-08 | 1.73E-07 | 59.8031843 | 106.326603 | 57.5392189 | 90.0036134 | 0 |
| 3.91E-08 | 1.73E-07 | 330.846649 | 632.621045 | 612.514265 | 627.241677 | 4.75243649 |
| 3.92E-08 | 1.74E-07 | 45.6561944 | 102.989994 | 93.7332436 | 128.510314 | 0 |
| 3.92E-08 | 1.74E-07 | 66.2336342 | 177.952474 | 158.696878 | 197.6368 | 2.37621824 |
| 3.92E-08 | 1.74E-07 | 32.7952946 | 246.909058 | 226.444668 | 200.420417 | 0 |
| 3.92E-08 | 1.74E-07 | 50.1575094 | 74.7400391 | 78.8844129 | 74.6937204 | 0 |
| 3.92E-08 | 1.74E-07 | 48.5498969 | 98.5411824 | 76.1002572 | 102.529889 | 0 |
| 3.93E-08 | 1.74E-07 | 38.904222 | 163.271395 | 135.49558 | 183.25478 | 0 |
| 3.93E-08 | 1.74E-07 | 72.9856066 | 61.3936035 | 65.8916861 | 122.015208 | 0 |
| 3.93E-08 | 1.74E-07 | 193.556543 | 520.288546 | 489.08336 | 536.310191 | 23.7621824 |
| 3.93E-08 | 1.74E-07 | 43.0840145 | 130.350187 | 164.265189 | 107.633187 | 0 |
| 3.95E-08 | 1.75E-07 | 49.1929419 | 98.9860636 | 74.2441534 | 90.0036134 | 0 |
| 3.95E-08 | 1.75E-07 | 45.0131495 | 94.5372518 | 115.078438 | 101.602017 | 0 |
| 3.95E-08 | 1.75E-07 | 43.405537 | 131.017509 | 117.862593 | 102.993826 | 0 |
| 3.96E-08 | 1.75E-07 | 0 | 0 | 0 | 0 | 99.8011662 |
| 3.96E-08 | 1.75E-07 | 47.5853294 | 80.3010539 | 93.7332436 | 89.0757411 | 0 |
| 3.96E-08 | 1.75E-07 | 47.5853294 | 80.9683756 | 93.7332436 | 87.2199964 | 0 |
| 3.97E-08 | 1.76E-07 | 58.1955718 | 68.7341431 | 160.552982 | 188.358077 | 0 |
| 3.97E-08 | 1.76E-07 | 54.6588243 | 101.655351 | 111.36623 | 63.5592527 | 0 |
| 3.98E-08 | 1.76E-07 | 52.0866444 | 66.7321777 | 93.7332436 | 102.065953 | 0 |
| 3.98E-08 | 1.76E-07 | 44.048582 | 100.765588 | 122.502853 | 151.707121 | 0 |
| 3.99E-08 | 1.76E-07 | 65.5905892 | 54.7203857 | 78.8844129 | 64.487125 | 0 |
| 4.00E-08 | 1.77E-07 | 340.492323 | 1876.06396 | 1010.64854 | 1312.93931 | 47.5243649 |
| 4.00E-08 | 1.77E-07 | 48.8714194 | 95.2045736 | 75.1722053 | 86.2921241 | 0 |

| | | | | | | |
|---|---|---|---|---|---|---|
| 4.01E-08 | 1.77E-07 | 53.3727344 | 80.0786133 | 63.1075304 | 74.6937204 | 0 |
| 4.02E-08 | 1.78E-07 | 45.6561944 | 105.881722 | 90.0210359 | 124.798825 | 0 |
| 4.02E-08 | 1.78E-07 | 52.4081669 | 128.125781 | 69.6038938 | 111.344676 | 0 |
| 4.03E-08 | 1.78E-07 | 53.0512119 | 110.108093 | 143.848047 | 71.4461673 | 0 |
| 4.06E-08 | 1.79E-07 | 46.6207619 | 109.218331 | 82.5966206 | 110.416804 | 0 |
| 4.06E-08 | 1.79E-07 | 50.1575094 | 146.36591 | 131.783372 | 76.549465 | 0 |
| 4.06E-08 | 1.80E-07 | 69.1273367 | 65.3975342 | 52.8989593 | 68.6625504 | 0 |
| 4.07E-08 | 1.80E-07 | 0 | 0 | 0 | 0 | 42.7719284 |
| 4.07E-08 | 1.80E-07 | 43.405537 | 108.328569 | 118.790645 | 103.921698 | 0 |
| 4.07E-08 | 1.80E-07 | 43.7270595 | 113.667143 | 125.287009 | 98.8184002 | 0 |
| 4.08E-08 | 1.80E-07 | 118.963324 | 79.6337321 | 51.9709073 | 75.1576565 | 0 |
| 4.08E-08 | 1.80E-07 | 47.5853294 | 94.7596924 | 91.8771398 | 78.4052096 | 0 |
| 4.10E-08 | 1.81E-07 | 70.4134267 | 112.55494 | 50.1148035 | 107.169251 | 0 |
| 4.10E-08 | 1.81E-07 | 44.048582 | 128.348222 | 97.4454513 | 112.272549 | 0 |
| 4.11E-08 | 1.81E-07 | 38.904222 | 195.525281 | 176.329864 | 129.438186 | 0 |
| 4.11E-08 | 1.81E-07 | 47.5853294 | 113.889583 | 80.7405168 | 95.5708472 | 0 |
| 4.11E-08 | 1.82E-07 | 94.2060914 | 145.476147 | 122.502853 | 138.252973 | 0 |
| 4.12E-08 | 1.82E-07 | 52.0866444 | 66.064856 | 90.0210359 | 101.138081 | 0 |
| 4.12E-08 | 1.82E-07 | 78.7730116 | 145.921029 | 180.970124 | 154.490738 | 7.12865473 |
| 4.13E-08 | 1.82E-07 | 133.753358 | 218.881543 | 122.502853 | 178.615418 | 2.37621824 |
| 4.14E-08 | 1.82E-07 | 59.1601393 | 77.6317668 | 61.2514265 | 60.3116997 | 0 |
| 4.14E-08 | 1.83E-07 | 44.048582 | 102.767554 | 103.013763 | 102.529889 | 0 |
| 4.14E-08 | 1.83E-07 | 51.4435994 | 74.9624797 | 72.3880495 | 100.210209 | 0 |
| 4.15E-08 | 1.83E-07 | 39.8687895 | 128.348222 | 206.955577 | 154.954675 | 0 |
| 4.16E-08 | 1.83E-07 | 36.3320421 | 232.450419 | 495.579724 | 184.182652 | 0 |
| 4.16E-08 | 1.83E-07 | 53.0512119 | 91.6455241 | 145.704151 | 76.549465 | 0 |
| 4.16E-08 | 1.83E-07 | 82.309759 | 171.279256 | 137.351684 | 159.594036 | 9.50487297 |
| 4.17E-08 | 1.84E-07 | 33.1168171 | 242.682686 | 196.747006 | 204.595843 | 0 |
| 4.17E-08 | 1.84E-07 | 66.2336342 | 66.7321777 | 110.438178 | 59.8477635 | 0 |
| 4.18E-08 | 1.84E-07 | 153.366231 | 364.580131 | 348.947521 | 342.38488 | 49.9005831 |
| 4.18E-08 | 1.84E-07 | 49.1929419 | 71.4034302 | 90.0210359 | 86.2921241 | 0 |
| 4.19E-08 | 1.84E-07 | 38.261177 | 152.371806 | 141.063891 | 178.151482 | 0 |
| 4.19E-08 | 1.85E-07 | 56.5879593 | 94.5372518 | 60.3233746 | 111.808612 | 0 |
| 4.20E-08 | 1.85E-07 | 44.048582 | 112.77738 | 94.6612955 | 118.767655 | 0 |
| 4.20E-08 | 1.85E-07 | 44.048582 | 103.657316 | 97.4454513 | 114.128293 | 0 |
| 4.21E-08 | 1.85E-07 | 45.6561944 | 124.789172 | 88.1649321 | 105.313506 | 0 |
| 4.23E-08 | 1.86E-07 | 45.9777169 | 93.86993 | 87.2368802 | 91.3954218 | 0 |
| 4.23E-08 | 1.86E-07 | 44.048582 | 235.119706 | 120.646749 | 133.149675 | 0 |
| 4.24E-08 | 1.86E-07 | 79.094534 | 127.903341 | 119.718697 | 122.479144 | 2.37621824 |
| 4.24E-08 | 1.86E-07 | 49.8359869 | 150.1474 | 77.956361 | 116.447974 | 0 |
| 4.24E-08 | 1.87E-07 | 38.5826995 | 136.356083 | 153.128566 | 155.882547 | 0 |
| 4.24E-08 | 1.87E-07 | 315.735091 | 440.654814 | 432.472193 | 473.214874 | 23.7621824 |
| 4.24E-08 | 1.87E-07 | 47.9068519 | 76.5195638 | 96.5173994 | 108.097123 | 0 |

| | | | | | | |
|---|---|---|---|---|---|---|
| 4.25E-08 | 1.87E-07 | 300.302011 | 730.717346 | 504.860243 | 736.266672 | 23.7621824 |
| 4.27E-08 | 1.87E-07 | 43.405537 | 102.545113 | 193.034799 | 154.954675 | 0 |
| 4.27E-08 | 1.88E-07 | 0 | 0 | 0 | 0 | 30.8908372 |
| 4.28E-08 | 1.88E-07 | 42.119447 | 120.11792 | 109.510126 | 117.839782 | 0 |
| 4.28E-08 | 1.88E-07 | 49.5144644 | 94.3148112 | 158.696878 | 90.0036134 | 0 |
| 4.30E-08 | 1.89E-07 | 41.1548795 | 197.972127 | 118.790645 | 144.748079 | 0 |
| 4.31E-08 | 1.89E-07 | 79.4160565 | 59.1691976 | 61.2514265 | 59.8477635 | 0 |
| 4.31E-08 | 1.89E-07 | 50.8005544 | 68.9565837 | 77.956361 | 70.9822312 | 0 |
| 4.31E-08 | 1.89E-07 | 110.282216 | 133.686796 | 42.6903882 | 101.602017 | 0 |
| 4.33E-08 | 1.90E-07 | 464.921529 | 955.382345 | 838.958933 | 979.833152 | 0 |
| 4.34E-08 | 1.91E-07 | 94.8491364 | 85.1947469 | 57.5392189 | 56.6002105 | 0 |
| 4.35E-08 | 1.91E-07 | 90.3478214 | 78.7439697 | 118.790645 | 123.407016 | 4.75243649 |
| 4.36E-08 | 1.91E-07 | 38.904222 | 133.686796 | 156.840774 | 141.03659 | 0 |
| 4.36E-08 | 1.91E-07 | 83.595849 | 54.4979451 | 65.8916861 | 64.0231889 | 0 |
| 4.37E-08 | 1.92E-07 | 49.8359869 | 78.7439697 | 73.3161014 | 71.4461673 | 0 |
| 4.37E-08 | 1.92E-07 | 152.080141 | 302.964087 | 258.926485 | 366.50956 | 16.6335277 |
| 4.38E-08 | 1.92E-07 | 71.6995166 | 132.574593 | 116.934542 | 50.5690405 | 0 |
| 4.39E-08 | 1.93E-07 | 116.391144 | 192.411112 | 100.229607 | 140.572654 | 0 |
| 4.40E-08 | 1.93E-07 | 48.8714194 | 79.6337321 | 115.078438 | 150.315313 | 0 |
| 4.40E-08 | 1.93E-07 | 52.0866444 | 64.952653 | 84.4527244 | 71.4461673 | 0 |
| 4.40E-08 | 1.93E-07 | 111.568306 | 46.2676432 | 108.582074 | 81.1888265 | 0 |
| 4.40E-08 | 1.93E-07 | 65.2690667 | 308.747542 | 252.430121 | 272.794457 | 0 |
| 4.41E-08 | 1.93E-07 | 54.0157793 | 104.102197 | 65.8916861 | 75.6215927 | 0 |
| 4.42E-08 | 1.94E-07 | 49.5144644 | 76.7420044 | 70.5319457 | 78.8691457 | 0 |
| 4.42E-08 | 1.94E-07 | 51.1220769 | 74.2951579 | 107.654022 | 74.6937204 | 0 |
| 4.42E-08 | 1.94E-07 | 78.4514891 | 78.5215291 | 49.1867516 | 68.6625504 | 0 |
| 4.43E-08 | 1.94E-07 | 48.5498969 | 76.2971232 | 128.999216 | 105.777442 | 0 |
| 4.43E-08 | 1.94E-07 | 59.8031843 | 82.7479004 | 160.552982 | 73.3019119 | 0 |
| 4.44E-08 | 1.94E-07 | 40.190312 | 185.515454 | 115.078438 | 170.728504 | 0 |
| 4.45E-08 | 1.95E-07 | 75.5577866 | 227.334285 | 185.610383 | 190.677758 | 0 |
| 4.45E-08 | 1.95E-07 | 89.7047764 | 57.6121134 | 116.934542 | 59.8477635 | 0 |
| 4.45E-08 | 1.95E-07 | 31.8307271 | 186.405216 | 254.286225 | 263.97967 | 0 |
| 4.45E-08 | 1.95E-07 | 43.0840145 | 117.226192 | 100.229607 | 107.169251 | 0 |
| 4.45E-08 | 1.95E-07 | 59.8031843 | 66.064856 | 58.4672708 | 59.8477635 | 0 |
| 4.46E-08 | 1.95E-07 | 41.476402 | 138.358048 | 110.438178 | 121.087335 | 0 |
| 4.46E-08 | 1.95E-07 | 62.6968867 | 101.43291 | 68.6758419 | 57.9920189 | 0 |
| 4.47E-08 | 1.95E-07 | 0 | 0 | 0 | 0 | 30.8908372 |
| 4.48E-08 | 1.96E-07 | 259.468654 | 48.2696086 | 289.552198 | 92.3232941 | 0 |
| 4.48E-08 | 1.96E-07 | 85.2034615 | 96.9840983 | 46.4025958 | 78.8691457 | 0 |
| 4.49E-08 | 1.96E-07 | 41.7979245 | 164.383598 | 182.826228 | 105.313506 | 0 |
| 4.50E-08 | 1.97E-07 | 40.833357 | 153.92889 | 115.078438 | 127.582442 | 0 |
| 4.50E-08 | 1.97E-07 | 66.5551567 | 97.4289795 | 154.056618 | 57.9920189 | 0 |
| 4.52E-08 | 1.98E-07 | 47.5853294 | 80.7459351 | 86.3088283 | 76.549465 | 0 |

| | | | | | | |
|---|---|---|---|---|---|---|
| 4.52E-08 | 1.98E-07 | 46.2992394 | 120.340361 | 83.5246725 | 141.03659 | 0 |
| 4.55E-08 | 1.99E-07 | 79.094534 | 135.911202 | 130.85532 | 122.479144 | 9.50487297 |
| 4.56E-08 | 1.99E-07 | 14.4685123 | 637.069857 | 718.312184 | 522.856043 | 0 |
| 4.56E-08 | 1.99E-07 | 73.3071291 | 100.320707 | 53.8270112 | 64.487125 | 0 |
| 4.57E-08 | 2.00E-07 | 128.608998 | 103.879757 | 43.6184401 | 93.2511664 | 0 |
| 4.57E-08 | 2.00E-07 | 30.2231146 | 248.021261 | 334.09869 | 209.69914 | 0 |
| 4.58E-08 | 2.00E-07 | 46.2992394 | 79.8561727 | 94.6612955 | 89.0757411 | 0 |
| 4.58E-08 | 2.00E-07 | 83.2743265 | 89.4211182 | 68.6758419 | 49.177232 | 0 |
| 4.58E-08 | 2.00E-07 | 118.320279 | 518.731462 | 374.932974 | 436.099982 | 23.7621824 |
| 4.60E-08 | 2.01E-07 | 47.2638069 | 124.344291 | 82.5966206 | 92.3232941 | 0 |
| 4.61E-08 | 2.01E-07 | 0 | 0 | 0 | 0 | 61.7816743 |
| 4.61E-08 | 2.01E-07 | 46.9422844 | 78.0766479 | 95.5893474 | 83.9724434 | 0 |
| 4.61E-08 | 2.01E-07 | 41.7979245 | 223.330355 | 187.466487 | 107.633187 | 0 |
| 4.62E-08 | 2.02E-07 | 46.9422844 | 135.911202 | 86.3088283 | 96.0347833 | 0 |
| 4.64E-08 | 2.02E-07 | 42.119447 | 130.350187 | 112.294282 | 104.385634 | 0 |
| 4.66E-08 | 2.03E-07 | 43.7270595 | 146.810791 | 92.8051917 | 128.046378 | 0 |
| 4.67E-08 | 2.04E-07 | 47.9068519 | 92.3128459 | 100.229607 | 72.3740396 | 0 |
| 4.68E-08 | 2.04E-07 | 43.0840145 | 104.102197 | 97.4454513 | 114.592229 | 0 |
| 4.69E-08 | 2.05E-07 | 51.4435994 | 261.812577 | 121.574801 | 101.602017 | 0 |
| 4.69E-08 | 2.05E-07 | 40.5118345 | 113.667143 | 130.85532 | 158.666164 | 0 |
| 4.69E-08 | 2.05E-07 | 73.3071291 | 307.635339 | 234.797135 | 260.268181 | 4.75243649 |
| 4.69E-08 | 2.05E-07 | 41.1548795 | 111.887618 | 116.00649 | 137.325101 | 0 |
| 4.70E-08 | 2.05E-07 | 60.7677518 | 76.7420044 | 137.351684 | 65.8789335 | 0 |
| 4.70E-08 | 2.05E-07 | 46.2992394 | 86.3069499 | 82.5966206 | 106.241379 | 0 |
| 4.70E-08 | 2.05E-07 | 53.3727344 | 163.938717 | 76.1002572 | 99.2823364 | 0 |
| 4.71E-08 | 2.05E-07 | 233.425332 | 86.3069499 | 73.3161014 | 76.549465 | 0 |
| 4.71E-08 | 2.05E-07 | 23.1496197 | 327.654993 | 392.565961 | 353.055411 | 0 |
| 4.71E-08 | 2.05E-07 | 38.261177 | 170.167053 | 129.927268 | 148.459568 | 0 |
| 4.71E-08 | 2.05E-07 | 46.6207619 | 84.5274251 | 84.4527244 | 79.3330819 | 0 |
| 4.71E-08 | 2.05E-07 | 41.7979245 | 219.326424 | 127.143113 | 127.118505 | 0 |
| 4.72E-08 | 2.06E-07 | 51.1220769 | 72.5156331 | 67.7477899 | 66.8068058 | 0 |
| 4.73E-08 | 2.06E-07 | 59.1601393 | 64.7302124 | 62.1794784 | 56.6002105 | 0 |
| 4.73E-08 | 2.06E-07 | 67.8412467 | 68.0668213 | 50.1148035 | 79.3330819 | 0 |
| 4.74E-08 | 2.06E-07 | 43.7270595 | 110.108093 | 90.0210359 | 115.520102 | 0 |
| 4.75E-08 | 2.07E-07 | 36.6535646 | 161.49187 | 151.272462 | 155.882547 | 0 |
| 4.76E-08 | 2.07E-07 | 42.762492 | 100.765588 | 175.401812 | 205.987651 | 0 |
| 4.78E-08 | 2.08E-07 | 45.6561944 | 106.104163 | 159.62493 | 93.2511664 | 0 |
| 4.80E-08 | 2.09E-07 | 77.4869216 | 74.0727173 | 91.8771398 | 47.7854236 | 0 |
| 4.81E-08 | 2.09E-07 | 146.292736 | 95.6494547 | 55.683115 | 66.8068058 | 0 |
| 4.81E-08 | 2.09E-07 | 56.2664368 | 88.0864746 | 61.2514265 | 64.487125 | 0 |
| 4.82E-08 | 2.09E-07 | 45.6561944 | 107.883687 | 79.8124649 | 109.024996 | 0 |
| 4.83E-08 | 2.10E-07 | 50.4790319 | 76.2971232 | 65.8916861 | 70.9822312 | 0 |
| 4.83E-08 | 2.10E-07 | 48.2283744 | 72.5156331 | 77.0283091 | 88.6118049 | 0 |

| | | | | | | |
|---|---|---|---|---|---|---|
| 4.83E-08 | 2.10E-07 | 71.6995166 | 205.979989 | 135.49558 | 173.048184 | 0 |
| 4.85E-08 | 2.11E-07 | 43.0840145 | 95.4270142 | 102.085711 | 108.097123 | 0 |
| 4.85E-08 | 2.11E-07 | 41.7979245 | 110.108093 | 110.438178 | 148.459568 | 0 |
| 4.86E-08 | 2.11E-07 | 68.1627692 | 99.875826 | 51.9709073 | 134.541484 | 0 |
| 4.89E-08 | 2.12E-07 | 36.9750871 | 173.726103 | 143.848047 | 147.995632 | 0 |
| 4.90E-08 | 2.13E-07 | 44.3701045 | 95.2045736 | 89.092984 | 92.3232941 | 0 |
| 4.90E-08 | 2.13E-07 | 49.5144644 | 82.5254598 | 66.819738 | 76.0855288 | 0 |
| 4.91E-08 | 2.13E-07 | 292.263949 | 801.008573 | 794.412441 | 778.484862 | 0 |
| 4.91E-08 | 2.13E-07 | 50.8005544 | 87.1967122 | 77.0283091 | 64.0231889 | 0 |
| 4.91E-08 | 2.13E-07 | 35.3674746 | 148.367875 | 189.322591 | 239.391054 | 0 |
| 4.92E-08 | 2.14E-07 | 44.691627 | 171.946578 | 89.092984 | 154.954675 | 0 |
| 4.94E-08 | 2.14E-07 | 49.5144644 | 70.958549 | 100.229607 | 76.0855288 | 0 |
| 4.94E-08 | 2.14E-07 | 75.8793091 | 309.637305 | 176.329864 | 225.009033 | 0 |
| 4.95E-08 | 2.15E-07 | 37.9396545 | 185.737895 | 126.215061 | 160.057972 | 0 |
| 4.95E-08 | 2.15E-07 | 117.998756 | 58.0569946 | 57.5392189 | 57.5280828 | 460.986339 |
| 4.95E-08 | 2.15E-07 | 47.9068519 | 85.1947469 | 189.322591 | 128.97425 | 0 |
| 4.96E-08 | 2.15E-07 | 5.46588243 | 5.33857422 | 3.71220767 | 5.56723382 | 19.0097459 |
| 4.98E-08 | 2.16E-07 | 46.2992394 | 92.757727 | 77.0283091 | 85.828188 | 0 |
| 4.98E-08 | 2.16E-07 | 49.8359869 | 75.4073608 | 65.8916861 | 85.3642518 | 0 |
| 4.98E-08 | 2.16E-07 | 67.1982017 | 64.5077718 | 51.9709073 | 63.5592527 | 0 |
| 4.99E-08 | 2.16E-07 | 56.5879593 | 183.95837 | 76.1002572 | 96.9626556 | 0 |
| 5.02E-08 | 2.18E-07 | 103.851766 | 48.047168 | 70.5319457 | 99.2823364 | 0 |
| 5.02E-08 | 2.18E-07 | 142.112943 | 79.1888509 | 77.956361 | 53.3526574 | 0 |
| 5.03E-08 | 2.18E-07 | 30.8661596 | 159.489905 | 545.694527 | 569.713594 | 0 |
| 5.03E-08 | 2.18E-07 | 76.2008316 | 183.291048 | 170.761553 | 188.358077 | 0 |
| 5.03E-08 | 2.18E-07 | 49.5144644 | 140.804895 | 75.1722053 | 100.210209 | 0 |
| 5.05E-08 | 2.19E-07 | 691.594889 | 134.354118 | 162.409085 | 61.7035081 | 0 |
| 5.06E-08 | 2.19E-07 | 449.48845 | 678.666248 | 761.930624 | 737.658481 | 28.5146189 |
| 5.07E-08 | 2.20E-07 | 38.261177 | 195.970162 | 122.502853 | 161.913717 | 0 |
| 5.08E-08 | 2.20E-07 | 44.691627 | 137.690727 | 84.4527244 | 122.479144 | 0 |
| 5.08E-08 | 2.20E-07 | 35.3674746 | 188.852063 | 251.502069 | 147.06776 | 0 |
| 5.08E-08 | 2.20E-07 | 59.1601393 | 73.4053955 | 75.1722053 | 52.4247851 | 0 |
| 5.09E-08 | 2.20E-07 | 36.3320421 | 143.919063 | 202.315318 | 156.810419 | 0 |
| 5.10E-08 | 2.21E-07 | 40.190312 | 164.161157 | 106.72597 | 170.728504 | 0 |
| 5.10E-08 | 2.21E-07 | 38.5826995 | 142.139539 | 127.143113 | 130.829995 | 0 |
| 5.10E-08 | 2.21E-07 | 44.048582 | 133.464355 | 87.2368802 | 113.664357 | 0 |
| 5.11E-08 | 2.21E-07 | 45.334672 | 90.7557617 | 86.3088283 | 81.1888265 | 0 |
| 5.11E-08 | 2.21E-07 | 49.8359869 | 67.8443807 | 68.6758419 | 71.9101035 | 0 |
| 5.12E-08 | 2.22E-07 | 60.1247068 | 57.3896728 | 64.9636342 | 57.5280828 | 0 |
| 5.12E-08 | 2.22E-07 | 30.5446371 | 232.227979 | 200.459214 | 239.391054 | 0 |
| 5.13E-08 | 2.22E-07 | 158.832113 | 94.982133 | 53.8270112 | 72.3740396 | 0 |
| 5.14E-08 | 2.22E-07 | 35.6889971 | 168.83241 | 175.401812 | 145.675952 | 0 |
| 5.14E-08 | 2.22E-07 | 79.4160565 | 244.907092 | 210.667785 | 252.845202 | 2.37621824 |

| | | | | | | |
|---|---|---|---|---|---|---|
| 5.14E-08 | 2.22E-07 | 81.666714 | 118.560836 | 47.3306478 | 80.2609542 | 0 |
| 5.15E-08 | 2.22E-07 | 28.9370247 | 222.885474 | 259.854537 | 250.989458 | 0 |
| 5.15E-08 | 2.23E-07 | 165.584086 | 280.720028 | 254.286225 | 272.794457 | 0 |
| 5.16E-08 | 2.23E-07 | 94.2060914 | 231.783097 | 44.546492 | 172.584248 | 0 |
| 5.16E-08 | 2.23E-07 | 50.1575094 | 106.326603 | 69.6038938 | 78.4052096 | 0 |
| 5.16E-08 | 2.23E-07 | 50.8005544 | 88.0864746 | 62.1794784 | 81.1888265 | 0 |
| 5.17E-08 | 2.23E-07 | 44.691627 | 84.0825439 | 143.848047 | 134.541484 | 0 |
| 5.17E-08 | 2.23E-07 | 39.547267 | 115.891549 | 128.999216 | 126.654569 | 0 |
| 5.18E-08 | 2.24E-07 | 53.6942569 | 144.363944 | 78.8844129 | 77.9412734 | 0 |
| 5.18E-08 | 2.24E-07 | 90.6693439 | 121.007682 | 107.654022 | 131.293931 | 0 |
| 5.18E-08 | 2.24E-07 | 37.2966096 | 128.793103 | 154.98467 | 167.017014 | 0 |
| 5.19E-08 | 2.24E-07 | 46.9422844 | 123.89941 | 75.1722053 | 107.633187 | 0 |
| 5.20E-08 | 2.25E-07 | 49.5144644 | 90.9782023 | 159.62493 | 85.3642518 | 0 |
| 5.20E-08 | 2.25E-07 | 49.5144644 | 88.976237 | 72.3880495 | 68.1986142 | 0 |
| 5.21E-08 | 2.25E-07 | 74.5932191 | 69.4014648 | 48.2586997 | 70.0543588 | 0 |
| 5.22E-08 | 2.25E-07 | 155.616888 | 48.047168 | 103.013763 | 82.1166988 | 0 |
| 5.23E-08 | 2.25E-07 | 51.1220769 | 68.9565837 | 79.8124649 | 113.664357 | 0 |
| 5.23E-08 | 2.26E-07 | 43.0840145 | 91.6455241 | 107.654022 | 96.9626556 | 0 |
| 5.23E-08 | 2.26E-07 | 48.5498969 | 95.2045736 | 67.7477899 | 84.4363795 | 0 |
| 5.24E-08 | 2.26E-07 | 41.7979245 | 108.328569 | 98.3735032 | 116.447974 | 0 |
| 5.24E-08 | 2.26E-07 | 41.1548795 | 173.503662 | 115.078438 | 115.984038 | 0 |
| 5.24E-08 | 2.26E-07 | 45.6561944 | 111.220296 | 77.956361 | 120.623399 | 0 |
| 5.24E-08 | 2.26E-07 | 37.618132 | 155.708415 | 133.639476 | 135.933292 | 0 |
| 5.26E-08 | 2.27E-07 | 64.6260217 | 58.0569946 | 53.8270112 | 72.3740396 | 0 |
| 5.27E-08 | 2.27E-07 | 52.0866444 | 124.789172 | 64.0355823 | 112.736485 | 0 |
| 5.27E-08 | 2.27E-07 | 80.0591015 | 67.8443807 | 48.2586997 | 71.4461673 | 0 |
| 5.27E-08 | 2.27E-07 | 36.0105196 | 160.602108 | 158.696878 | 145.212015 | 0 |
| 5.30E-08 | 2.28E-07 | 36.9750871 | 177.730033 | 126.215061 | 188.822014 | 0 |
| 5.30E-08 | 2.28E-07 | 0 | 0 | 0 | 0 | 45.1481466 |
| 5.30E-08 | 2.28E-07 | 47.9068519 | 67.8443807 | 85.3807764 | 86.7560603 | 0 |
| 5.30E-08 | 2.28E-07 | 44.3701045 | 120.785242 | 90.0210359 | 90.9314857 | 0 |
| 5.31E-08 | 2.28E-07 | 46.9422844 | 96.0943359 | 71.4599976 | 100.210209 | 0 |
| 5.32E-08 | 2.29E-07 | 85.2034615 | 194.190637 | 272.847264 | 157.738291 | 4.75243649 |
| 5.32E-08 | 2.29E-07 | 53.3727344 | 104.547078 | 62.1794784 | 77.4773373 | 0 |
| 5.33E-08 | 2.29E-07 | 77.4869216 | 161.046989 | 203.24337 | 167.944887 | 2.37621824 |
| 5.34E-08 | 2.30E-07 | 36.0105196 | 133.909237 | 180.042072 | 187.430205 | 0 |
| 5.34E-08 | 2.30E-07 | 226.351837 | 96.3167765 | 55.683115 | 90.4675495 | 0 |
| 5.35E-08 | 2.30E-07 | 54.3373018 | 159.934786 | 64.9636342 | 133.149675 | 0 |
| 5.35E-08 | 2.30E-07 | 34.0813846 | 189.741825 | 178.185968 | 160.057972 | 0 |
| 5.35E-08 | 2.30E-07 | 38.261177 | 131.907271 | 128.071165 | 133.613612 | 0 |
| 5.35E-08 | 2.30E-07 | 245.96471 | 197.082365 | 180.970124 | 182.326907 | 544.153978 |
| 5.36E-08 | 2.31E-07 | 41.7979245 | 103.879757 | 128.071165 | 100.210209 | 0 |
| 5.37E-08 | 2.31E-07 | 38.904222 | 123.454529 | 144.776099 | 211.090949 | 0 |

| | | | | | | |
|---|---|---|---|---|---|---|
| 5.37E-08 | 2.31E-07 | 125.072251 | 78.5215291 | 54.7550631 | 65.8789335 | 0 |
| 5.38E-08 | 2.31E-07 | 74.9147416 | 53.8306234 | 64.9636342 | 58.4559551 | 0 |
| 5.38E-08 | 2.31E-07 | 48.2283744 | 73.8502767 | 76.1002572 | 102.529889 | 0 |
| 5.39E-08 | 2.32E-07 | 80.380624 | 115.446667 | 131.783372 | 141.964462 | 14.2573095 |
| 5.42E-08 | 2.33E-07 | 186.161525 | 331.436483 | 217.164149 | 257.484564 | 0 |
| 5.43E-08 | 2.33E-07 | 43.405537 | 93.6474894 | 88.1649321 | 100.210209 | 0 |
| 5.44E-08 | 2.34E-07 | 45.0131495 | 94.0923706 | 78.8844129 | 88.1478687 | 0 |
| 5.44E-08 | 2.34E-07 | 40.5118345 | 103.879757 | 116.934542 | 127.118505 | 0 |
| 5.44E-08 | 2.34E-07 | 71.3779941 | 195.970162 | 190.250643 | 212.482757 | 0 |
| 5.46E-08 | 2.34E-07 | 40.833357 | 115.224227 | 102.085711 | 116.91191 | 0 |
| 5.46E-08 | 2.34E-07 | 55.6233918 | 58.946757 | 87.2368802 | 64.0231889 | 0 |
| 5.48E-08 | 2.35E-07 | 576.168313 | 1554.41486 | 1521.07709 | 1628.41589 | 0 |
| 5.48E-08 | 2.35E-07 | 43.7270595 | 99.4309448 | 138.279736 | 91.859358 | 0 |
| 5.49E-08 | 2.35E-07 | 43.0840145 | 165.940682 | 184.682331 | 91.859358 | 0 |
| 5.50E-08 | 2.36E-07 | 85.8465065 | 210.4288 | 126.215061 | 212.018821 | 0 |
| 5.51E-08 | 2.37E-07 | 49.8359869 | 72.0707519 | 67.7477899 | 64.9510612 | 0 |
| 5.53E-08 | 2.37E-07 | 51.1220769 | 93.6474894 | 72.3880495 | 64.0231889 | 0 |
| 5.54E-08 | 2.38E-07 | 38.261177 | 129.237984 | 124.358957 | 133.613612 | 0 |
| 5.55E-08 | 2.38E-07 | 45.9777169 | 100.988029 | 77.0283091 | 82.5806349 | 0 |
| 5.55E-08 | 2.38E-07 | 142.434466 | 36.2578166 | 646.852186 | 353.983283 | 0 |
| 5.57E-08 | 2.39E-07 | 102.887199 | 192.633553 | 39.9062324 | 180.007227 | 0 |
| 5.58E-08 | 2.39E-07 | 43.0840145 | 123.009648 | 100.229607 | 89.5396772 | 0 |
| 5.58E-08 | 2.39E-07 | 36.0105196 | 163.493835 | 151.272462 | 140.108718 | 0 |
| 5.58E-08 | 2.39E-07 | 51.1220769 | 176.39539 | 72.3880495 | 127.582442 | 0 |
| 5.60E-08 | 2.40E-07 | 24.4357097 | 314.753438 | 291.408302 | 306.19786 | 0 |
| 5.62E-08 | 2.41E-07 | 46.9422844 | 98.0963013 | 74.2441534 | 77.9412734 | 0 |
| 5.64E-08 | 2.42E-07 | 37.618132 | 124.566732 | 135.49558 | 154.490738 | 0 |
| 5.67E-08 | 2.43E-07 | 47.2638069 | 121.452563 | 71.4599976 | 118.303719 | 0 |
| 5.69E-08 | 2.44E-07 | 59.4816618 | 259.810612 | 270.063108 | 291.351903 | 0 |
| 5.69E-08 | 2.44E-07 | 82.952804 | 141.027336 | 142.919995 | 153.09893 | 0 |
| 5.70E-08 | 2.44E-07 | 55.3018693 | 78.7439697 | 53.8270112 | 83.9724434 | 0 |
| 5.71E-08 | 2.44E-07 | 47.2638069 | 94.982133 | 71.4599976 | 77.4773373 | 0 |
| 5.71E-08 | 2.44E-07 | 422.159037 | 65.8424154 | 121.574801 | 101.602017 | 0 |
| 5.71E-08 | 2.45E-07 | 277.152392 | 303.631409 | 250.574018 | 322.899561 | 102.177384 |
| 5.72E-08 | 2.45E-07 | 41.1548795 | 110.997856 | 105.797919 | 98.8184002 | 0 |
| 5.72E-08 | 2.45E-07 | 43.0840145 | 104.324638 | 195.818954 | 116.91191 | 0 |
| 5.73E-08 | 2.45E-07 | 41.7979245 | 145.921029 | 99.3015551 | 108.097123 | 0 |
| 5.73E-08 | 2.45E-07 | 172.65758 | 360.798641 | 326.674275 | 359.086581 | 4.75243649 |
| 5.74E-08 | 2.46E-07 | 39.2257445 | 119.005717 | 157.768826 | 115.984038 | 0 |
| 5.75E-08 | 2.46E-07 | 0 | 0 | 0 | 0 | 61.7816743 |
| 5.76E-08 | 2.46E-07 | 37.2966096 | 262.257458 | 139.207788 | 156.346483 | 0 |
| 5.77E-08 | 2.47E-07 | 45.0131495 | 79.6337321 | 79.8124649 | 87.2199964 | 0 |
| 5.77E-08 | 2.47E-07 | 39.2257445 | 109.885653 | 219.948304 | 254.237011 | 0 |

| | | | | | | |
|---|---|---|---|---|---|---|
| 5.78E-08 | 2.47E-07 | 76.8438766 | 83.6376628 | 49.1867516 | 61.7035081 | 0 |
| 5.79E-08 | 2.48E-07 | 78.4514891 | 175.728068 | 149.416359 | 174.439993 | 0 |
| 5.80E-08 | 2.48E-07 | 82.309759 | 339.666785 | 270.063108 | 322.435625 | 0 |
| 5.80E-08 | 2.48E-07 | 42.119447 | 114.334465 | 94.6612955 | 96.0347833 | 0 |
| 5.80E-08 | 2.48E-07 | 45.334672 | 92.757727 | 98.3735032 | 73.7658481 | 0 |
| 5.81E-08 | 2.48E-07 | 33.7598621 | 177.952474 | 161.481034 | 170.264568 | 0 |
| 5.81E-08 | 2.48E-07 | 84.881939 | 163.048954 | 47.3306478 | 96.9626556 | 0 |
| 5.81E-08 | 2.48E-07 | 53.3727344 | 68.5117025 | 194.890903 | 155.418611 | 0 |
| 5.82E-08 | 2.48E-07 | 44.048582 | 96.3167765 | 89.092984 | 80.7248903 | 0 |
| 5.86E-08 | 2.50E-07 | 61.0892743 | 137.913167 | 65.8916861 | 70.9822312 | 0 |
| 5.88E-08 | 2.51E-07 | 44.048582 | 96.0943359 | 78.8844129 | 95.106911 | 0 |
| 5.88E-08 | 2.51E-07 | 51.1220769 | 85.4171875 | 59.3953227 | 74.6937204 | 0 |
| 5.90E-08 | 2.52E-07 | 41.7979245 | 105.881722 | 91.8771398 | 102.993826 | 0 |
| 5.90E-08 | 2.52E-07 | 52.4081669 | 58.5018758 | 65.8916861 | 77.4773373 | 0 |
| 5.93E-08 | 2.53E-07 | 79.4160565 | 56.9447917 | 58.4672708 | 57.9920189 | 0 |
| 5.94E-08 | 2.54E-07 | 150.794051 | 281.832231 | 237.581291 | 271.402649 | 21.3859642 |
| 5.96E-08 | 2.54E-07 | 47.5853294 | 125.234054 | 70.5319457 | 103.457762 | 0 |
| 5.96E-08 | 2.54E-07 | 4.17979245 | 2.22440592 | 1.85610383 | 1.39180845 | 40.3957101 |
| 5.98E-08 | 2.55E-07 | 105.137856 | 121.230123 | 96.5173994 | 116.91191 | 0 |
| 5.99E-08 | 2.55E-07 | 42.119447 | 88.976237 | 99.3015551 | 104.84957 | 0 |
| 5.99E-08 | 2.55E-07 | 112.854396 | 56.9447917 | 60.3233746 | 71.9101035 | 0 |
| 5.99E-08 | 2.56E-07 | 53.6942569 | 130.795068 | 62.1794784 | 147.531696 | 0 |
| 6.00E-08 | 2.56E-07 | 67.5197242 | 125.234054 | 53.8270112 | 74.6937204 | 0 |
| 6.00E-08 | 2.56E-07 | 390.32831 | 662.205644 | 559.615306 | 574.352955 | 40.3957101 |
| 6.04E-08 | 2.57E-07 | 39.547267 | 136.578524 | 103.941815 | 165.16127 | 0 |
| 6.05E-08 | 2.58E-07 | 44.048582 | 134.576558 | 83.5246725 | 102.993826 | 0 |
| 6.06E-08 | 2.58E-07 | 111.889829 | 90.5333211 | 221.804408 | 212.946693 | 2.37621824 |
| 6.06E-08 | 2.58E-07 | 43.0840145 | 81.6356974 | 113.222334 | 107.633187 | 0 |
| 6.10E-08 | 2.60E-07 | 0 | 0 | 0 | 0 | 9.50487297 |
| 6.13E-08 | 2.61E-07 | 60.7677518 | 56.4999105 | 57.5392189 | 57.5280828 | 0 |
| 6.15E-08 | 2.62E-07 | 39.8687895 | 98.9860636 | 129.927268 | 122.015208 | 0 |
| 6.15E-08 | 2.62E-07 | 41.1548795 | 98.5411824 | 98.3735032 | 102.065953 | 0 |
| 6.16E-08 | 2.62E-07 | 38.261177 | 114.779346 | 123.430905 | 130.366059 | 0 |
| 6.16E-08 | 2.62E-07 | 70.7349491 | 75.1849202 | 46.4025958 | 95.5708472 | 0 |
| 6.16E-08 | 2.62E-07 | 42.4409695 | 84.9723063 | 116.934542 | 128.046378 | 0 |
| 6.16E-08 | 2.62E-07 | 93.8845689 | 294.288904 | 222.73246 | 263.051798 | 11.8810912 |
| 6.20E-08 | 2.64E-07 | 102.565676 | 142.361979 | 95.5893474 | 116.447974 | 4.75243649 |
| 6.21E-08 | 2.64E-07 | 45.334672 | 74.0727173 | 82.5966206 | 98.8184002 | 0 |
| 6.24E-08 | 2.66E-07 | 52.7296894 | 76.2971232 | 55.683115 | 68.6625504 | 0 |
| 6.26E-08 | 2.66E-07 | 41.476402 | 102.322673 | 100.229607 | 91.3954218 | 0 |
| 6.28E-08 | 2.67E-07 | 35.0459521 | 153.484009 | 140.135839 | 158.666164 | 0 |
| 6.29E-08 | 2.67E-07 | 47.2638069 | 69.6239054 | 123.430905 | 100.674145 | 0 |
| 6.29E-08 | 2.68E-07 | 41.476402 | 117.671073 | 91.8771398 | 100.210209 | 0 |

| | | | | | | |
|---|---|---|---|---|---|---|
| 6.31E-08 | 2.68E-07 | 60.7677518 | 90.9782023 | 52.8989593 | 123.870952 | 0 |
| 6.31E-08 | 2.68E-07 | 45.9777169 | 82.0805786 | 73.3161014 | 72.3740396 | 0 |
| 6.31E-08 | 2.68E-07 | 45.334672 | 87.6415934 | 71.4599976 | 83.0445711 | 0 |
| 6.32E-08 | 2.69E-07 | 69.1273367 | 63.84045 | 54.7550631 | 53.8165936 | 0 |
| 6.32E-08 | 2.69E-07 | 53.0512119 | 58.0569946 | 60.3233746 | 71.9101035 | 0 |
| 6.33E-08 | 2.69E-07 | 87.4541189 | 149.702519 | 156.840774 | 157.274355 | 11.8810912 |
| 6.35E-08 | 2.70E-07 | 48.5498969 | 116.781311 | 85.3807764 | 68.6625504 | 0 |
| 6.36E-08 | 2.70E-07 | 41.476402 | 90.7557617 | 101.157659 | 98.3544641 | 0 |
| 6.38E-08 | 2.71E-07 | 74.2716966 | 135.466321 | 132.711424 | 126.190633 | 7.12865473 |
| 6.39E-08 | 2.71E-07 | 15216.6952 | 3390.88439 | 2918.72328 | 3455.86039 | 4.75243649 |
| 6.40E-08 | 2.72E-07 | 46.9422844 | 68.2892619 | 79.8124649 | 70.0543588 | 0 |
| 6.41E-08 | 2.72E-07 | 60.4462293 | 68.2892619 | 192.106747 | 93.2511664 | 0 |
| 6.45E-08 | 2.74E-07 | 399.009418 | 113.222262 | 73.3161014 | 86.7560603 | 0 |
| 6.45E-08 | 2.74E-07 | 61.7323193 | 79.1888509 | 49.1867516 | 66.3428696 | 0 |
| 6.47E-08 | 2.75E-07 | 51.1220769 | 58.0569946 | 73.3161014 | 64.9510612 | 0 |
| 6.48E-08 | 2.75E-07 | 39.2257445 | 135.02144 | 103.941815 | 115.056166 | 0 |
| 6.48E-08 | 2.75E-07 | 42.119447 | 109.885653 | 83.5246725 | 111.808612 | 0 |
| 6.48E-08 | 2.75E-07 | 69.7703817 | 149.702519 | 125.287009 | 126.190633 | 9.50487297 |
| 6.49E-08 | 2.75E-07 | 93.2415239 | 54.4979451 | 62.1794784 | 61.239572 | 0 |
| 6.49E-08 | 2.76E-07 | 70.4134267 | 112.332499 | 77.956361 | 48.2493597 | 0 |
| 6.52E-08 | 2.77E-07 | 38.5826995 | 121.007682 | 149.416359 | 108.097123 | 0 |
| 6.55E-08 | 2.78E-07 | 46.2992394 | 111.220296 | 69.6038938 | 93.2511664 | 0 |
| 6.55E-08 | 2.78E-07 | 45.0131495 | 125.456494 | 76.1002572 | 97.8905279 | 0 |
| 6.56E-08 | 2.78E-07 | 33.7598621 | 142.361979 | 173.545708 | 177.687546 | 0 |
| 6.57E-08 | 2.79E-07 | 77.8084441 | 193.078434 | 155.912722 | 186.502333 | 4.75243649 |
| 6.60E-08 | 2.80E-07 | 48.8714194 | 118.783276 | 68.6758419 | 82.5806349 | 0 |
| 6.61E-08 | 2.80E-07 | 488.071149 | 1376.68483 | 1420.84748 | 1502.68919 | 33.2670554 |
| 6.62E-08 | 2.81E-07 | 0 | 0 | 0 | 0 | 87.920075 |
| 6.63E-08 | 2.81E-07 | 40.833357 | 173.726103 | 104.869867 | 111.808612 | 0 |
| 6.63E-08 | 2.81E-07 | 32.1522496 | 231.338216 | 161.481034 | 177.687546 | 0 |
| 6.66E-08 | 2.82E-07 | 39.2257445 | 100.098267 | 116.934542 | 129.438186 | 0 |
| 6.69E-08 | 2.83E-07 | 58.8386168 | 286.503483 | 261.710641 | 264.907542 | 0 |
| 6.69E-08 | 2.83E-07 | 85.2034615 | 88.976237 | 88.1649321 | 79.3330819 | 9.50487297 |
| 6.70E-08 | 2.84E-07 | 47.5853294 | 98.3187419 | 64.0355823 | 86.7560603 | 0 |
| 6.71E-08 | 2.84E-07 | 64.6260217 | 200.196533 | 81.6685687 | 71.9101035 | 0 |
| 6.72E-08 | 2.84E-07 | 111.568306 | 60.7262817 | 67.7477899 | 56.6002105 | 0 |
| 6.74E-08 | 2.85E-07 | 46.2992394 | 75.6298014 | 106.72597 | 148.923505 | 0 |
| 6.74E-08 | 2.85E-07 | 46.2992394 | 73.6278361 | 72.3880495 | 68.1986142 | 0 |
| 6.75E-08 | 2.85E-07 | 32.4737721 | 163.716276 | 182.826228 | 167.017014 | 0 |
| 6.75E-08 | 2.85E-07 | 41.1548795 | 123.89941 | 97.4454513 | 93.2511664 | 0 |
| 6.75E-08 | 2.86E-07 | 50.4790319 | 114.112024 | 73.3161014 | 184.646588 | 0 |
| 6.76E-08 | 2.86E-07 | 821.489978 | 121.007682 | 79.8124649 | 133.149675 | 0 |
| 6.78E-08 | 2.87E-07 | 41.1548795 | 108.551009 | 93.7332436 | 91.859358 | 0 |

| | | | | | | |
|---|---|---|---|---|---|---|
| 6.79E-08 | 2.87E-07 | 72.9856066 | 225.33232 | 190.250643 | 212.482757 | 4.75243649 |
| 6.80E-08 | 2.88E-07 | 77.1653991 | 159.712345 | 156.840774 | 143.820207 | 2.37621824 |
| 6.85E-08 | 2.90E-07 | 176.194328 | 320.314453 | 298.832717 | 289.032222 | 19.0097459 |
| 6.86E-08 | 2.90E-07 | 34.0813846 | 205.312667 | 140.135839 | 158.666164 | 0 |
| 6.86E-08 | 2.90E-07 | 46.2992394 | 85.1947469 | 65.8916861 | 77.9412734 | 0 |
| 6.87E-08 | 2.90E-07 | 44.3701045 | 96.7616577 | 73.3161014 | 83.9724434 | 0 |
| 6.89E-08 | 2.91E-07 | 38.261177 | 101.655351 | 158.696878 | 162.841589 | 0 |
| 6.90E-08 | 2.91E-07 | 55.9449143 | 311.63927 | 252.430121 | 281.609244 | 2.37621824 |
| 6.90E-08 | 2.92E-07 | 41.1548795 | 239.346077 | 96.5173994 | 178.151482 | 0 |
| 6.91E-08 | 2.92E-07 | 40.5118345 | 165.27336 | 94.6612955 | 126.654569 | 0 |
| 6.92E-08 | 2.92E-07 | 38.904222 | 114.112024 | 106.72597 | 106.705315 | 0 |
| 6.94E-08 | 2.93E-07 | 87.1325964 | 256.251562 | 43.6184401 | 224.545097 | 0 |
| 6.95E-08 | 2.93E-07 | 30.5446371 | 203.755583 | 183.75428 | 184.646588 | 0 |
| 6.95E-08 | 2.94E-07 | 44.3701045 | 84.3049845 | 81.6685687 | 71.9101035 | 0 |
| 6.96E-08 | 2.94E-07 | 39.8687895 | 126.123816 | 92.8051917 | 116.91191 | 0 |
| 6.96E-08 | 2.94E-07 | 71.3779941 | 189.741825 | 168.905449 | 195.31712 | 0 |
| 6.98E-08 | 2.94E-07 | 63.3399317 | 105.436841 | 47.3306478 | 94.6429749 | 0 |
| 6.98E-08 | 2.95E-07 | 42.119447 | 111.887618 | 85.3807764 | 90.4675495 | 0 |
| 7.02E-08 | 2.96E-07 | 98.3858838 | 61.3936035 | 81.6685687 | 48.7132959 | 0 |
| 7.03E-08 | 2.96E-07 | 42.762492 | 82.7479004 | 81.6685687 | 87.6839326 | 0 |
| 7.03E-08 | 2.97E-07 | 92.2769564 | 78.9664103 | 50.1148035 | 57.5280828 | 0 |
| 7.04E-08 | 2.97E-07 | 48.8714194 | 140.582454 | 67.7477899 | 101.602017 | 0 |
| 7.05E-08 | 2.97E-07 | 43.0840145 | 86.5293905 | 78.8844129 | 82.5806349 | 0 |
| 7.06E-08 | 2.98E-07 | 43.405537 | 102.100232 | 154.056618 | 86.2921241 | 0 |
| 7.07E-08 | 2.98E-07 | 171.693013 | 61.8384847 | 77.956361 | 67.7346781 | 0 |
| 7.07E-08 | 2.98E-07 | 90.6693439 | 262.035018 | 403.702584 | 296.455201 | 9.50487297 |
| 7.08E-08 | 2.98E-07 | 43.405537 | 84.5274251 | 76.1002572 | 83.9724434 | 0 |
| 7.10E-08 | 2.99E-07 | 104.494811 | 172.391459 | 157.768826 | 207.37946 | 0 |
| 7.12E-08 | 3.00E-07 | 245.321665 | 584.573877 | 483.515049 | 547.444659 | 85.5438568 |
| 7.15E-08 | 3.01E-07 | 52.0866444 | 56.2774699 | 61.2514265 | 62.6313804 | 0 |
| 7.15E-08 | 3.01E-07 | 36.0105196 | 131.46239 | 122.502853 | 149.851377 | 0 |
| 7.15E-08 | 3.01E-07 | 34.7244296 | 172.169019 | 127.143113 | 156.810419 | 0 |
| 7.17E-08 | 3.02E-07 | 44.048582 | 80.9683756 | 107.654022 | 152.171058 | 0 |
| 7.18E-08 | 3.02E-07 | 40.190312 | 125.234054 | 124.358957 | 88.6118049 | 0 |
| 7.19E-08 | 3.03E-07 | 49.8359869 | 64.0628906 | 70.5319457 | 57.9920189 | 0 |
| 7.19E-08 | 3.03E-07 | 55.9449143 | 69.6239054 | 234.797135 | 128.97425 | 0 |
| 7.21E-08 | 3.03E-07 | 76.2008316 | 94.0923706 | 100.229607 | 92.3232941 | 9.50487297 |
| 7.23E-08 | 3.04E-07 | 51.7651219 | 84.0825439 | 125.287009 | 62.1674443 | 0 |
| 7.24E-08 | 3.05E-07 | 54.0157793 | 54.2755046 | 59.3953227 | 69.5904227 | 0 |
| 7.24E-08 | 3.05E-07 | 40.833357 | 122.119885 | 97.4454513 | 90.4675495 | 0 |
| 7.25E-08 | 3.05E-07 | 45.334672 | 80.0786133 | 100.229607 | 69.5904227 | 0 |
| 7.25E-08 | 3.05E-07 | 24.1141872 | 268.263354 | 314.6096 | 257.9485 | 0 |
| 7.26E-08 | 3.05E-07 | 52.0866444 | 55.6101481 | 87.2368802 | 69.5904227 | 0 |

| | | | | | | |
|---|---|---|---|---|---|---|
| 7.27E-08 | 3.06E-07 | 112.854396 | 136.133643 | 90.9490879 | 103.921698 | 4.75243649 |
| 7.29E-08 | 3.07E-07 | 44.691627 | 185.737895 | 78.8844129 | 185.110524 | 0 |
| 7.29E-08 | 3.07E-07 | 43.7270595 | 93.6474894 | 74.2441534 | 81.1888265 | 0 |
| 7.30E-08 | 3.07E-07 | 36.3320421 | 142.58442 | 117.862593 | 128.97425 | 0 |
| 7.33E-08 | 3.08E-07 | 42.119447 | 118.560836 | 92.8051917 | 83.5085072 | 0 |
| 7.34E-08 | 3.09E-07 | 34.7244296 | 145.476147 | 132.711424 | 145.212015 | 0 |
| 7.35E-08 | 3.09E-07 | 6.10892743 | 6.89565837 | 8.35246725 | 6.49510612 | 33.2670554 |
| 7.37E-08 | 3.10E-07 | 31.1876821 | 191.743791 | 220.876356 | 154.954675 | 0 |
| 7.37E-08 | 3.10E-07 | 41.1548795 | 96.7616577 | 91.8771398 | 85.828188 | 0 |
| 7.39E-08 | 3.10E-07 | 34.4029071 | 144.141504 | 154.98467 | 134.541484 | 0 |
| 7.39E-08 | 3.10E-07 | 51.4435994 | 113.889583 | 58.4672708 | 89.0757411 | 0 |
| 7.40E-08 | 3.11E-07 | 62.0538418 | 51.6062174 | 73.3161014 | 55.6723382 | 0 |
| 7.40E-08 | 3.11E-07 | 105.137856 | 101.43291 | 237.581291 | 147.06776 | 0 |
| 7.41E-08 | 3.11E-07 | 39.547267 | 116.33643 | 98.3735032 | 97.8905279 | 0 |
| 7.42E-08 | 3.12E-07 | 46.2992394 | 113.889583 | 69.6038938 | 83.9724434 | 0 |
| 7.44E-08 | 3.12E-07 | 290.656336 | 140.582454 | 174.47376 | 80.7248903 | 2.37621824 |
| 7.46E-08 | 3.13E-07 | 29.2585471 | 181.733964 | 203.24337 | 221.76148 | 0 |
| 7.46E-08 | 3.13E-07 | 35.3674746 | 196.415043 | 116.00649 | 224.545097 | 0 |
| 7.56E-08 | 3.17E-07 | 35.3674746 | 157.487939 | 127.143113 | 129.438186 | 0 |
| 7.56E-08 | 3.17E-07 | 94.5276139 | 151.926925 | 38.9781805 | 184.182652 | 0 |
| 7.57E-08 | 3.17E-07 | 23.4711422 | 250.690548 | 287.696094 | 326.611051 | 0 |
| 7.57E-08 | 3.18E-07 | 36.9750871 | 130.350187 | 111.36623 | 120.159463 | 0 |
| 7.60E-08 | 3.19E-07 | 79.737579 | 140.360014 | 128.071165 | 143.820207 | 2.37621824 |
| 7.61E-08 | 3.19E-07 | 29.5800696 | 191.298909 | 219.948304 | 179.079354 | 0 |
| 7.62E-08 | 3.20E-07 | 45.6561944 | 71.6258708 | 65.8916861 | 80.7248903 | 0 |
| 7.64E-08 | 3.20E-07 | 55.9449143 | 102.767554 | 103.941815 | 51.4969128 | 0 |
| 7.68E-08 | 3.22E-07 | 45.9777169 | 77.8542074 | 63.1075304 | 77.0134011 | 0 |
| 7.70E-08 | 3.23E-07 | 293.871561 | 76.964445 | 56.6111669 | 131.293931 | 0 |
| 7.70E-08 | 3.23E-07 | 69.4488592 | 182.846167 | 166.121293 | 191.60563 | 2.37621824 |
| 7.71E-08 | 3.23E-07 | 43.7270595 | 70.2912272 | 97.4454513 | 102.065953 | 0 |
| 7.71E-08 | 3.23E-07 | 38.261177 | 119.228158 | 104.869867 | 103.921698 | 0 |
| 7.72E-08 | 3.23E-07 | 72.6640841 | 156.820618 | 168.905449 | 179.543291 | 0 |
| 7.72E-08 | 3.24E-07 | 82.6312815 | 143.474182 | 40.8342843 | 125.262761 | 0 |
| 7.73E-08 | 3.24E-07 | 68.4842917 | 141.917098 | 131.783372 | 163.769461 | 9.50487297 |
| 7.74E-08 | 3.24E-07 | 90.0262989 | 64.5077718 | 51.9709073 | 57.5280828 | 0 |
| 7.74E-08 | 3.24E-07 | 54.6588243 | 222.663033 | 64.0355823 | 240.782863 | 0 |
| 7.75E-08 | 3.25E-07 | 53.6942569 | 84.9723063 | 52.8989593 | 67.7346781 | 0 |
| 7.75E-08 | 3.25E-07 | 41.476402 | 82.7479004 | 90.0210359 | 86.2921241 | 0 |
| 7.77E-08 | 3.25E-07 | 48.8714194 | 83.4152222 | 61.2514265 | 64.487125 | 0 |
| 7.77E-08 | 3.25E-07 | 47.5853294 | 85.8620687 | 88.1649321 | 59.3838274 | 0 |
| 7.79E-08 | 3.26E-07 | 41.476402 | 220.438627 | 97.4454513 | 132.685739 | 0 |
| 7.79E-08 | 3.26E-07 | 40.833357 | 99.875826 | 85.3807764 | 90.4675495 | 0 |
| 7.79E-08 | 3.26E-07 | 32.7952946 | 247.576379 | 136.423632 | 186.038397 | 0 |

| | | | | | | |
|---|---|---|---|---|---|---|
| 7.80E-08 | 3.26E-07 | 32.7952946 | 142.58442 | 171.689605 | 158.666164 | 0 |
| 7.80E-08 | 3.26E-07 | 41.1548795 | 87.4191528 | 84.4527244 | 92.7872303 | 0 |
| 7.82E-08 | 3.27E-07 | 44.048582 | 91.2006429 | 74.2441534 | 73.3019119 | 0 |
| 7.82E-08 | 3.27E-07 | 36.9750871 | 159.267464 | 103.013763 | 152.171058 | 0 |
| 7.82E-08 | 3.27E-07 | 76.8438766 | 167.275326 | 140.135839 | 146.603824 | 0 |
| 7.83E-08 | 3.27E-07 | 22.1850522 | 253.582275 | 345.235313 | 334.034029 | 0 |
| 7.88E-08 | 3.29E-07 | 32.4737721 | 165.27336 | 144.776099 | 168.408823 | 0 |
| 7.88E-08 | 3.29E-07 | 62.6968867 | 127.013578 | 63.1075304 | 60.7756358 | 0 |
| 7.88E-08 | 3.30E-07 | 44.048582 | 78.7439697 | 77.956361 | 69.1264865 | 0 |
| 7.88E-08 | 3.30E-07 | 93.5630464 | 122.119885 | 98.3735032 | 117.839782 | 0 |
| 7.89E-08 | 3.30E-07 | 37.9396545 | 100.988029 | 116.934542 | 111.808612 | 0 |
| 7.90E-08 | 3.30E-07 | 42.762492 | 96.0943359 | 73.3161014 | 88.6118049 | 0 |
| 7.91E-08 | 3.30E-07 | 45.334672 | 65.8424154 | 77.956361 | 92.3232941 | 0 |
| 7.92E-08 | 3.31E-07 | 50.1575094 | 55.1652669 | 64.9636342 | 69.5904227 | 0 |
| 7.92E-08 | 3.31E-07 | 79.737579 | 123.454529 | 40.8342843 | 126.190633 | 0 |
| 7.93E-08 | 3.31E-07 | 52.4081669 | 123.676969 | 57.5392189 | 134.077548 | 0 |
| 7.94E-08 | 3.32E-07 | 39.547267 | 110.108093 | 97.4454513 | 91.859358 | 0 |
| 7.95E-08 | 3.32E-07 | 529.226029 | 872.412004 | 696.038938 | 836.476881 | 30.8908372 |
| 7.96E-08 | 3.32E-07 | 37.2966096 | 116.781311 | 104.869867 | 123.407016 | 0 |
| 7.97E-08 | 3.33E-07 | 43.0840145 | 80.5234945 | 75.1722053 | 76.549465 | 0 |
| 7.98E-08 | 3.33E-07 | 29.2585471 | 197.304805 | 204.171422 | 179.079354 | 0 |
| 7.99E-08 | 3.33E-07 | 37.618132 | 157.932821 | 99.3015551 | 138.252973 | 0 |
| 8.01E-08 | 3.34E-07 | 34.0813846 | 217.324459 | 153.128566 | 131.293931 | 0 |
| 8.04E-08 | 3.36E-07 | 44.691627 | 98.3187419 | 71.4599976 | 74.6937204 | 0 |
| 8.07E-08 | 3.37E-07 | 239.212737 | 586.798283 | 768.426987 | 772.917628 | 23.7621824 |
| 8.08E-08 | 3.37E-07 | 242.749485 | 170.611934 | 28.7696094 | 200.884354 | 0 |
| 8.08E-08 | 3.37E-07 | 46.9422844 | 65.1750936 | 64.0355823 | 82.1166988 | 0 |
| 8.11E-08 | 3.38E-07 | 38.261177 | 97.6514201 | 145.704151 | 183.718716 | 0 |
| 8.13E-08 | 3.39E-07 | 47.5853294 | 93.4250488 | 83.5246725 | 59.3838274 | 0 |
| 8.13E-08 | 3.39E-07 | 38.261177 | 113.444702 | 100.229607 | 102.993826 | 0 |
| 8.13E-08 | 3.39E-07 | 127.322908 | 73.4053955 | 51.0428554 | 66.8068058 | 0 |
| 8.14E-08 | 3.39E-07 | 42.762492 | 122.564766 | 90.0210359 | 78.8691457 | 0 |
| 8.14E-08 | 3.39E-07 | 322.808586 | 618.162406 | 253.358173 | 479.246044 | 0 |
| 8.14E-08 | 3.39E-07 | 45.6561944 | 66.7321777 | 118.790645 | 123.870952 | 0 |
| 8.18E-08 | 3.41E-07 | 37.618132 | 98.3187419 | 133.639476 | 120.623399 | 0 |
| 8.18E-08 | 3.41E-07 | 40.5118345 | 94.7596924 | 88.1649321 | 86.2921241 | 0 |
| 8.18E-08 | 3.41E-07 | 41.476402 | 85.6396281 | 81.6685687 | 83.5085072 | 0 |
| 8.22E-08 | 3.42E-07 | 84.5604165 | 119.005717 | 129.927268 | 115.520102 | 0 |
| 8.23E-08 | 3.43E-07 | 41.476402 | 169.722172 | 90.0210359 | 110.416804 | 0 |
| 8.23E-08 | 3.43E-07 | 49.8359869 | 61.8384847 | 137.351684 | 102.065953 | 0 |
| 8.24E-08 | 3.43E-07 | 38.904222 | 90.5333211 | 152.200514 | 159.594036 | 0 |
| 8.26E-08 | 3.44E-07 | 35.0459521 | 112.110059 | 170.761553 | 189.28595 | 0 |
| 8.27E-08 | 3.44E-07 | 36.0105196 | 145.698588 | 106.72597 | 157.274355 | 0 |

| | | | | | | |
|---|---|---|---|---|---|---|
| 8.27E-08 | 3.44E-07 | 44.691627 | 71.1809896 | 71.4599976 | 94.6429749 | 0 |
| 8.28E-08 | 3.44E-07 | 41.7979245 | 93.86993 | 77.956361 | 83.0445711 | 0 |
| 8.29E-08 | 3.45E-07 | 692.880979 | 3967.67285 | 3187.85833 | 3505.50156 | 9.50487297 |
| 8.35E-08 | 3.47E-07 | 45.0131495 | 81.6356974 | 64.9636342 | 72.3740396 | 0 |
| 8.35E-08 | 3.48E-07 | 33.1168171 | 136.578524 | 149.416359 | 156.346483 | 0 |
| 8.40E-08 | 3.50E-07 | 32.1522496 | 201.531177 | 133.639476 | 183.25478 | 0 |
| 8.41E-08 | 3.50E-07 | 190.984363 | 217.546899 | 213.451941 | 208.771268 | 19.0097459 |
| 8.41E-08 | 3.50E-07 | 69.7703817 | 188.629622 | 159.62493 | 235.679565 | 0 |
| 8.42E-08 | 3.50E-07 | 44.3701045 | 117.893514 | 69.6038938 | 131.293931 | 0 |
| 8.44E-08 | 3.51E-07 | 140.826853 | 95.2045736 | 124.358957 | 93.7151026 | 0 |
| 8.45E-08 | 3.51E-07 | 45.6561944 | 62.5058065 | 69.6038938 | 71.9101035 | 0 |
| 8.46E-08 | 3.52E-07 | 35.6889971 | 123.676969 | 116.00649 | 126.654569 | 0 |
| 8.47E-08 | 3.52E-07 | 34.0813846 | 135.24388 | 130.85532 | 142.428398 | 0 |
| 8.47E-08 | 3.52E-07 | 40.833357 | 96.3167765 | 88.1649321 | 80.7248903 | 0 |
| 8.49E-08 | 3.53E-07 | 93.2415239 | 89.8659993 | 106.72597 | 89.0757411 | 11.8810912 |
| 8.51E-08 | 3.53E-07 | 36.3320421 | 128.793103 | 106.72597 | 123.407016 | 0 |
| 8.51E-08 | 3.53E-07 | 59.1601393 | 356.57227 | 282.127783 | 336.35371 | 4.75243649 |
| 8.51E-08 | 3.54E-07 | 39.8687895 | 87.864034 | 127.143113 | 98.3544641 | 0 |
| 8.53E-08 | 3.54E-07 | 44.691627 | 88.3089152 | 65.8916861 | 74.6937204 | 0 |
| 8.53E-08 | 3.54E-07 | 41.7979245 | 27.3601929 | 25.0574018 | 29.2279775 | 168.711495 |
| 8.53E-08 | 3.54E-07 | 102.887199 | 111.442737 | 90.0210359 | 110.88074 | 2.37621824 |
| 8.54E-08 | 3.55E-07 | 39.8687895 | 94.982133 | 91.8771398 | 86.7560603 | 0 |
| 8.55E-08 | 3.55E-07 | 141.469898 | 60.7262817 | 50.1148035 | 106.241379 | 0 |
| 8.56E-08 | 3.55E-07 | 65.2690667 | 67.3994995 | 85.3807764 | 45.4657428 | 0 |
| 8.57E-08 | 3.56E-07 | 35.3674746 | 132.352152 | 132.711424 | 113.664357 | 0 |
| 8.57E-08 | 3.56E-07 | 36.9750871 | 17.5728068 | 18.5610383 | 13.4541484 | 144.949313 |
| 8.59E-08 | 3.56E-07 | 37.618132 | 237.788993 | 106.72597 | 154.026802 | 0 |
| 8.60E-08 | 3.57E-07 | 60.7677518 | 164.161157 | 161.481034 | 175.831801 | 7.12865473 |
| 8.62E-08 | 3.57E-07 | 43.405537 | 68.2892619 | 104.869867 | 97.4265918 | 0 |
| 8.64E-08 | 3.58E-07 | 39.8687895 | 97.6514201 | 84.4527244 | 94.6429749 | 0 |
| 8.67E-08 | 3.60E-07 | 32.4737721 | 169.722172 | 131.783372 | 172.584248 | 0 |
| 8.70E-08 | 3.61E-07 | 37.9396545 | 98.3187419 | 112.294282 | 101.138081 | 0 |
| 8.70E-08 | 3.61E-07 | 49.8359869 | 93.4250488 | 55.683115 | 75.1576565 | 0 |
| 8.71E-08 | 3.61E-07 | 36.6535646 | 122.787207 | 103.941815 | 117.375846 | 0 |
| 8.76E-08 | 3.63E-07 | 43.405537 | 80.5234945 | 68.6758419 | 94.6429749 | 0 |
| 8.81E-08 | 3.65E-07 | 38.261177 | 114.334465 | 104.869867 | 92.3232941 | 0 |
| 8.84E-08 | 3.66E-07 | 46.9422844 | 80.7459351 | 95.5893474 | 181.399035 | 0 |
| 8.88E-08 | 3.68E-07 | 38.904222 | 87.864034 | 108.582074 | 128.046378 | 0 |
| 8.93E-08 | 3.70E-07 | 78.7730116 | 49.3818115 | 88.1649321 | 54.2805297 | 0 |
| 8.94E-08 | 3.70E-07 | 69.4488592 | 217.102018 | 181.898176 | 149.851377 | 0 |
| 8.94E-08 | 3.70E-07 | 42.4409695 | 109.885653 | 75.1722053 | 83.9724434 | 0 |
| 8.95E-08 | 3.71E-07 | 39.8687895 | 97.8738607 | 82.5966206 | 93.2511664 | 0 |
| 8.96E-08 | 3.71E-07 | 77.8084441 | 135.02144 | 222.73246 | 155.418611 | 7.12865473 |

| | | | | | | |
|---|---|---|---|---|---|---|
| 8.96E-08 | 3.71E-07 | 49.1929419 | 58.0569946 | 118.790645 | 111.344676 | 0 |
| 8.98E-08 | 3.72E-07 | 69.1273367 | 42.9310343 | 77.956361 | 80.2609542 | 0 |
| 9.01E-08 | 3.73E-07 | 32.7952946 | 144.141504 | 149.416359 | 137.325101 | 0 |
| 9.03E-08 | 3.74E-07 | 22.1850522 | 207.314632 | 409.270895 | 396.665409 | 0 |
| 9.05E-08 | 3.74E-07 | 435.34146 | 342.336072 | 343.379209 | 332.64222 | 772.270929 |
| 9.07E-08 | 3.75E-07 | 61.7323193 | 124.566732 | 52.8989593 | 188.822014 | 0 |
| 9.07E-08 | 3.75E-07 | 65.2690667 | 47.3798462 | 64.0355823 | 93.2511664 | 0 |
| 9.08E-08 | 3.75E-07 | 124.750728 | 69.846346 | 63.1075304 | 66.8068058 | 320.789463 |
| 9.08E-08 | 3.75E-07 | 76.5223541 | 145.476147 | 128.999216 | 147.995632 | 0 |
| 9.09E-08 | 3.76E-07 | 42.762492 | 94.3148112 | 99.3015551 | 68.1986142 | 0 |
| 9.11E-08 | 3.77E-07 | 44.048582 | 65.6199748 | 84.4527244 | 71.9101035 | 0 |
| 9.14E-08 | 3.78E-07 | 46.9422844 | 64.2853312 | 59.3953227 | 63.0953166 | 0 |
| 9.15E-08 | 3.78E-07 | 107.066991 | 84.3049845 | 47.3306478 | 59.3838274 | 0 |
| 9.16E-08 | 3.78E-07 | 39.547267 | 105.881722 | 106.72597 | 81.1888265 | 0 |
| 9.16E-08 | 3.79E-07 | 36.0105196 | 105.436841 | 124.358957 | 118.767655 | 0 |
| 9.19E-08 | 3.80E-07 | 38.261177 | 147.033232 | 133.639476 | 88.6118049 | 0 |
| 9.22E-08 | 3.81E-07 | 69.7703817 | 417.965873 | 321.105963 | 433.316365 | 9.50487297 |
| 9.23E-08 | 3.81E-07 | 215.741595 | 123.454529 | 228.300772 | 32.0115944 | 0 |
| 9.27E-08 | 3.83E-07 | 43.7270595 | 72.2931925 | 70.5319457 | 66.8068058 | 0 |
| 9.30E-08 | 3.84E-07 | 37.618132 | 108.99589 | 96.5173994 | 101.138081 | 0 |
| 9.33E-08 | 3.85E-07 | 41.7979245 | 98.0963013 | 74.2441534 | 121.551272 | 0 |
| 9.33E-08 | 3.85E-07 | 41.7979245 | 95.6494547 | 74.2441534 | 119.231591 | 0 |
| 9.34E-08 | 3.86E-07 | 38.261177 | 94.3148112 | 102.085711 | 93.7151026 | 0 |
| 9.36E-08 | 3.86E-07 | 78.7730116 | 114.334465 | 166.121293 | 155.418611 | 2.37621824 |
| 9.38E-08 | 3.87E-07 | 39.8687895 | 90.9782023 | 94.6612955 | 80.2609542 | 0 |
| 9.39E-08 | 3.88E-07 | 35.3674746 | 120.562801 | 109.510126 | 141.03659 | 0 |
| 9.39E-08 | 3.88E-07 | 42.119447 | 128.793103 | 72.3880495 | 116.91191 | 0 |
| 9.41E-08 | 3.88E-07 | 42.4409695 | 74.2951579 | 70.5319457 | 79.797018 | 0 |
| 9.42E-08 | 3.89E-07 | 39.547267 | 80.3010539 | 116.934542 | 122.015208 | 0 |
| 9.45E-08 | 3.90E-07 | 41.1548795 | 74.0727173 | 100.229607 | 86.7560603 | 0 |
| 9.46E-08 | 3.90E-07 | 40.833357 | 76.0746826 | 83.5246725 | 92.7872303 | 0 |
| 9.49E-08 | 3.91E-07 | 104.494811 | 253.359835 | 251.502069 | 260.732117 | 19.0097459 |
| 9.50E-08 | 3.92E-07 | 40.833357 | 89.1986776 | 76.1002572 | 83.0445711 | 0 |
| 9.54E-08 | 3.93E-07 | 66.8766792 | 182.401286 | 179.11402 | 173.976057 | 0 |
| 9.54E-08 | 3.93E-07 | 44.048582 | 95.2045736 | 67.7477899 | 122.479144 | 0 |
| 9.58E-08 | 3.95E-07 | 38.904222 | 118.115955 | 87.2368802 | 95.5708472 | 0 |
| 9.63E-08 | 3.97E-07 | 72.9856066 | 158.822583 | 136.423632 | 147.995632 | 2.37621824 |
| 9.64E-08 | 3.97E-07 | 27.6509347 | 254.027157 | 185.610383 | 183.25478 | 0 |
| 9.64E-08 | 3.97E-07 | 50.1575094 | 58.0569946 | 65.8916861 | 53.3526574 | 0 |
| 9.64E-08 | 3.97E-07 | 35.6889971 | 117.893514 | 104.869867 | 135.469356 | 0 |
| 9.65E-08 | 3.98E-07 | 42.4409695 | 124.566732 | 90.9490879 | 74.2297842 | 0 |
| 9.65E-08 | 3.98E-07 | 37.2966096 | 130.350187 | 96.5173994 | 104.84957 | 0 |
| 9.66E-08 | 3.98E-07 | 69.1273367 | 75.852242 | 45.4745439 | 57.9920189 | 0 |

| | | | | | | |
|---|---|---|---|---|---|---|
| 9.69E-08 | 3.99E-07 | 35.0459521 | 117.226192 | 114.150386 | 151.707121 | 0 |
| 9.69E-08 | 3.99E-07 | 67.1982017 | 73.1829549 | 69.6038938 | 43.1460621 | 0 |
| 9.70E-08 | 3.99E-07 | 35.0459521 | 106.771484 | 149.416359 | 128.97425 | 0 |
| 9.70E-08 | 3.99E-07 | 32.4737721 | 161.936751 | 141.991943 | 131.293931 | 0 |
| 9.72E-08 | 4.00E-07 | 126.036818 | 172.83634 | 135.49558 | 152.634994 | 23.7621824 |
| 9.73E-08 | 4.00E-07 | 73.6286516 | 114.334465 | 51.9709073 | 57.0641466 | 0 |
| 9.75E-08 | 4.01E-07 | 38.5826995 | 86.0845093 | 95.5893474 | 106.705315 | 0 |
| 9.76E-08 | 4.01E-07 | 45.9777169 | 78.0766479 | 60.3233746 | 64.487125 | 0 |
| 9.78E-08 | 4.02E-07 | 40.190312 | 88.0864746 | 77.956361 | 85.828188 | 0 |
| 9.79E-08 | 4.03E-07 | 370.393916 | 391.495443 | 442.680764 | 425.429451 | 68.9103291 |
| 9.80E-08 | 4.03E-07 | 46.6207619 | 69.846346 | 88.1649321 | 58.4559551 | 0 |
| 9.82E-08 | 4.04E-07 | 80.380624 | 181.066642 | 173.545708 | 165.625206 | 0 |
| 9.82E-08 | 4.04E-07 | 73.6286516 | 139.692692 | 135.49558 | 162.377653 | 2.37621824 |
| 9.91E-08 | 4.08E-07 | 56.2664368 | 61.6160441 | 90.0210359 | 50.5690405 | 0 |
| 9.92E-08 | 4.08E-07 | 63.0184092 | 78.2990885 | 46.4025958 | 58.4559551 | 0 |
| 9.93E-08 | 4.08E-07 | 38.5826995 | 100.543148 | 128.071165 | 86.7560603 | 0 |
| 9.94E-08 | 4.08E-07 | 35.6889971 | 106.993925 | 119.718697 | 109.952868 | 0 |
| 9.95E-08 | 4.09E-07 | 32.7952946 | 174.838306 | 207.883629 | 117.375846 | 0 |
| 9.95E-08 | 4.09E-07 | 227.637927 | 544.31213 | 534.557904 | 545.124978 | 97.424948 |
| 9.99E-08 | 4.10E-07 | 45.6561944 | 235.119706 | 72.3880495 | 183.718716 | 0 |
| 9.99E-08 | 4.10E-07 | 42.4409695 | 69.6239054 | 80.7405168 | 101.602017 | 0 |
| 1.00E-07 | 4.12E-07 | 62.6968867 | 69.1790242 | 47.3306478 | 54.7444659 | 0 |
| 1.00E-07 | 4.12E-07 | 43.7270595 | 95.2045736 | 64.9636342 | 111.808612 | 0 |
| 1.00E-07 | 4.12E-07 | 31.5092046 | 136.800964 | 153.128566 | 159.1301 | 0 |
| 1.01E-07 | 4.13E-07 | 94.5276139 | 66.9546183 | 49.1867516 | 56.6002105 | 0 |
| 1.01E-07 | 4.15E-07 | 69.1273367 | 200.641414 | 148.488307 | 164.233398 | 2.37621824 |
| 1.01E-07 | 4.15E-07 | 154.973843 | 80.5234945 | 55.683115 | 60.7756358 | 0 |
| 1.01E-07 | 4.16E-07 | 95.1706589 | 91.2006429 | 37.1220767 | 109.024996 | 0 |
| 1.01E-07 | 4.16E-07 | 15.7546023 | 10.6771484 | 11.136623 | 15.7738291 | 128.315785 |
| 1.02E-07 | 4.17E-07 | 41.1548795 | 92.0904053 | 71.4599976 | 82.5806349 | 0 |
| 1.02E-07 | 4.17E-07 | 36.6535646 | 99.6533854 | 112.294282 | 100.210209 | 0 |
| 1.02E-07 | 4.18E-07 | 56.2664368 | 85.4171875 | 50.1148035 | 59.8477635 | 0 |
| 1.02E-07 | 4.19E-07 | 113.497441 | 80.0786133 | 136.423632 | 167.017014 | 0 |
| 1.03E-07 | 4.24E-07 | 52.7296894 | 57.834554 | 180.970124 | 183.718716 | 0 |
| 1.03E-07 | 4.24E-07 | 36.6535646 | 92.5352865 | 157.768826 | 133.149675 | 0 |
| 1.04E-07 | 4.25E-07 | 32.7952946 | 151.926925 | 125.287009 | 133.149675 | 0 |
| 1.04E-07 | 4.25E-07 | 32.1522496 | 206.202429 | 137.351684 | 137.789037 | 0 |
| 1.04E-07 | 4.25E-07 | 36.9750871 | 110.108093 | 101.157659 | 93.7151026 | 0 |
| 1.04E-07 | 4.26E-07 | 44.3701045 | 144.808826 | 67.7477899 | 108.561059 | 0 |
| 1.04E-07 | 4.27E-07 | 74.2716966 | 78.0766479 | 41.7623363 | 65.8789335 | 0 |
| 1.04E-07 | 4.27E-07 | 67.1982017 | 159.712345 | 174.47376 | 178.151482 | 0 |
| 1.04E-07 | 4.27E-07 | 19.6128723 | 407.511165 | 242.22155 | 371.148921 | 0 |
| 1.05E-07 | 4.29E-07 | 35.6889971 | 103.657316 | 107.654022 | 116.447974 | 0 |

| | | | | | | |
|---|---|---|---|---|---|---|
| 1.05E-07 | 4.29E-07 | 36.6535646 | 112.77738 | 91.8771398 | 116.91191 | 0 |
| 1.05E-07 | 4.30E-07 | 25.0787547 | 254.472038 | 213.451941 | 206.451587 | 0 |
| 1.05E-07 | 4.30E-07 | 39.8687895 | 76.7420044 | 90.9490879 | 82.5806349 | 0 |
| 1.05E-07 | 4.31E-07 | 56.9094818 | 138.80293 | 50.1148035 | 145.675952 | 0 |
| 1.06E-07 | 4.32E-07 | 34.7244296 | 104.769519 | 129.927268 | 124.334889 | 0 |
| 1.06E-07 | 4.33E-07 | 53.0512119 | 80.0786133 | 60.3233746 | 143.820207 | 0 |
| 1.06E-07 | 4.33E-07 | 41.7979245 | 138.358048 | 89.092984 | 79.3330819 | 0 |
| 1.06E-07 | 4.34E-07 | 43.405537 | 96.3167765 | 64.0355823 | 78.8691457 | 0 |
| 1.06E-07 | 4.34E-07 | 42.762492 | 87.864034 | 64.9636342 | 101.602017 | 0 |
| 1.06E-07 | 4.35E-07 | 40.5118345 | 123.89941 | 73.3161014 | 111.344676 | 0 |
| 1.07E-07 | 4.36E-07 | 45.9777169 | 116.55887 | 62.1794784 | 83.9724434 | 0 |
| 1.07E-07 | 4.36E-07 | 54.6588243 | 48.2696086 | 35.2659728 | 41.2903175 | 125.939567 |
| 1.07E-07 | 4.37E-07 | 52.7296894 | 77.8542074 | 47.3306478 | 73.7658481 | 0 |
| 1.07E-07 | 4.37E-07 | 50.1575094 | 124.121851 | 66.819738 | 213.874566 | 0 |
| 1.07E-07 | 4.37E-07 | 48.8714194 | 82.7479004 | 110.438178 | 55.6723382 | 0 |
| 1.07E-07 | 4.37E-07 | 91.9554339 | 189.519385 | 262.638692 | 38.0427644 | 0 |
| 1.07E-07 | 4.38E-07 | 59.1601393 | 242.905127 | 53.8270112 | 250.525522 | 0 |
| 1.07E-07 | 4.38E-07 | 51.1220769 | 78.0766479 | 52.8989593 | 106.241379 | 0 |
| 1.07E-07 | 4.39E-07 | 32.1522496 | 137.913167 | 142.919995 | 130.366059 | 0 |
| 1.08E-07 | 4.39E-07 | 37.618132 | 95.6494547 | 92.8051917 | 90.9314857 | 0 |
| 1.08E-07 | 4.40E-07 | 46.6207619 | 89.1986776 | 99.3015551 | 56.1362743 | 0 |
| 1.08E-07 | 4.40E-07 | 34.4029071 | 162.159192 | 103.013763 | 144.284143 | 0 |
| 1.08E-07 | 4.40E-07 | 37.9396545 | 95.4270142 | 89.092984 | 89.5396772 | 0 |
| 1.08E-07 | 4.40E-07 | 43.0840145 | 82.0805786 | 64.0355823 | 70.518295 | 0 |
| 1.08E-07 | 4.40E-07 | 32.4737721 | 130.350187 | 136.423632 | 132.685739 | 0 |
| 1.08E-07 | 4.41E-07 | 419.265335 | 335.885295 | 38.9781805 | 460.224662 | 0 |
| 1.08E-07 | 4.42E-07 | 43.0840145 | 114.779346 | 68.6758419 | 144.284143 | 0 |
| 1.09E-07 | 4.43E-07 | 66.5551567 | 268.040914 | 159.62493 | 188.822014 | 0 |
| 1.09E-07 | 4.44E-07 | 310.590731 | 896.213147 | 601.377642 | 754.360182 | 26.1384007 |
| 1.09E-07 | 4.44E-07 | 3.21522496 | 2.44684652 | 2.78415575 | 3.24755306 | 19.0097459 |
| 1.09E-07 | 4.44E-07 | 58.5170943 | 329.212077 | 191.178695 | 221.297544 | 2.37621824 |
| 1.09E-07 | 4.45E-07 | 61.7323193 | 49.1593709 | 52.8989593 | 57.0641466 | 0 |
| 1.09E-07 | 4.46E-07 | 48.2283744 | 78.0766479 | 51.9709073 | 69.5904227 | 0 |
| 1.09E-07 | 4.46E-07 | 33.4383396 | 131.684831 | 119.718697 | 122.015208 | 0 |
| 1.09E-07 | 4.46E-07 | 5.78740493 | 9.34250488 | 7.42441534 | 12.0623399 | 76.0389838 |
| 1.10E-07 | 4.47E-07 | 35.3674746 | 99.4309448 | 119.718697 | 147.531696 | 0 |
| 1.10E-07 | 4.48E-07 | 33.1168171 | 135.24388 | 131.783372 | 117.375846 | 0 |
| 1.10E-07 | 4.48E-07 | 35.0459521 | 109.218331 | 126.215061 | 104.84957 | 0 |
| 1.10E-07 | 4.48E-07 | 43.405537 | 65.3975342 | 77.956361 | 103.457762 | 0 |
| 1.10E-07 | 4.50E-07 | 35.6889971 | 99.875826 | 118.790645 | 104.385634 | 0 |
| 1.11E-07 | 4.51E-07 | 63.0184092 | 70.2912272 | 49.1867516 | 50.5690405 | 0 |
| 1.11E-07 | 4.51E-07 | 38.904222 | 149.035197 | 81.6685687 | 115.520102 | 0 |
| 1.11E-07 | 4.52E-07 | 126.036818 | 224.664998 | 90.0210359 | 336.35371 | 0 |

| | | | | | | |
|---|---|---|---|---|---|---|
| 1.11E-07 | 4.52E-07 | 42.762492 | 69.846346 | 64.9636342 | 84.4363795 | 0 |
| 1.11E-07 | 4.52E-07 | 84.5604165 | 128.125781 | 97.4454513 | 121.551272 | 2.37621824 |
| 1.11E-07 | 4.53E-07 | 40.5118345 | 80.3010539 | 71.4599976 | 79.797018 | 0 |
| 1.11E-07 | 4.54E-07 | 60.4462293 | 100.765588 | 177.257916 | 52.8887212 | 0 |
| 1.11E-07 | 4.54E-07 | 40.833357 | 76.0746826 | 70.5319457 | 84.4363795 | 0 |
| 1.12E-07 | 4.54E-07 | 69.1273367 | 206.64731 | 244.077654 | 169.336695 | 4.75243649 |
| 1.12E-07 | 4.55E-07 | 41.7979245 | 104.769519 | 66.819738 | 112.272549 | 0 |
| 1.12E-07 | 4.55E-07 | 42.762492 | 78.5215291 | 209.739733 | 129.902122 | 0 |
| 1.12E-07 | 4.57E-07 | 37.618132 | 119.228158 | 98.3735032 | 85.828188 | 0 |
| 1.12E-07 | 4.57E-07 | 198.37938 | 479.581917 | 455.673491 | 474.606683 | 54.6530196 |
| 1.12E-07 | 4.57E-07 | 33.7598621 | 108.77345 | 132.711424 | 167.480951 | 0 |
| 1.13E-07 | 4.58E-07 | 42.762492 | 76.0746826 | 68.6758419 | 63.5592527 | 0 |
| 1.13E-07 | 4.58E-07 | 42.4409695 | 118.783276 | 152.200514 | 70.0543588 | 0 |
| 1.13E-07 | 4.61E-07 | 57.2310043 | 58.946757 | 45.4745439 | 68.1986142 | 0 |
| 1.14E-07 | 4.62E-07 | 94.2060914 | 169.277291 | 86.3088283 | 147.531696 | 2.37621824 |
| 1.14E-07 | 4.64E-07 | 39.8687895 | 129.905306 | 79.8124649 | 91.3954218 | 0 |
| 1.14E-07 | 4.64E-07 | 53.6942569 | 85.8620687 | 46.4025958 | 86.7560603 | 0 |
| 1.14E-07 | 4.65E-07 | 513.792949 | 712.477218 | 709.031665 | 743.689651 | 42.7719284 |
| 1.14E-07 | 4.65E-07 | 96.4567488 | 55.1652669 | 52.8989593 | 59.3838274 | 0 |
| 1.14E-07 | 4.65E-07 | 40.5118345 | 119.895479 | 76.1002572 | 86.2921241 | 0 |
| 1.14E-07 | 4.65E-07 | 40.190312 | 82.0805786 | 87.2368802 | 133.613612 | 0 |
| 1.14E-07 | 4.65E-07 | 28.9370247 | 255.13936 | 286.768042 | 130.366059 | 0 |
| 1.15E-07 | 4.66E-07 | 87.7756414 | 136.800964 | 38.9781805 | 86.7560603 | 0 |
| 1.15E-07 | 4.66E-07 | 37.618132 | 110.997856 | 161.481034 | 90.0036134 | 0 |
| 1.15E-07 | 4.66E-07 | 185.840003 | 635.957654 | 700.679197 | 793.794755 | 21.3859642 |
| 1.15E-07 | 4.66E-07 | 37.9396545 | 85.6396281 | 86.3088283 | 101.138081 | 0 |
| 1.15E-07 | 4.66E-07 | 12.2178549 | 347.452205 | 608.802058 | 703.791142 | 0 |
| 1.15E-07 | 4.67E-07 | 46.2992394 | 57.6121134 | 65.8916861 | 56.6002105 | 0 |
| 1.15E-07 | 4.67E-07 | 1022.12002 | 130.127747 | 63.1075304 | 145.675952 | 0 |
| 1.15E-07 | 4.68E-07 | 290.334814 | 62.9506877 | 178.185968 | 53.3526574 | 0 |
| 1.16E-07 | 4.69E-07 | 41.476402 | 69.1790242 | 81.6685687 | 105.777442 | 0 |
| 1.16E-07 | 4.70E-07 | 39.547267 | 80.0786133 | 78.8844129 | 104.84957 | 0 |
| 1.16E-07 | 4.70E-07 | 38.904222 | 83.1927816 | 81.6685687 | 80.7248903 | 0 |
| 1.16E-07 | 4.70E-07 | 53.3727344 | 76.5195638 | 62.1794784 | 47.7854236 | 0 |
| 1.16E-07 | 4.70E-07 | 44.3701045 | 109.218331 | 64.0355823 | 77.0134011 | 0 |
| 1.16E-07 | 4.71E-07 | 39.8687895 | 71.6258708 | 100.229607 | 87.2199964 | 0 |
| 1.16E-07 | 4.71E-07 | 40.190312 | 79.8561727 | 72.3880495 | 76.549465 | 0 |
| 1.16E-07 | 4.71E-07 | 41.1548795 | 78.9664103 | 66.819738 | 89.0757411 | 0 |
| 1.16E-07 | 4.71E-07 | 41.1548795 | 211.096122 | 79.8124649 | 141.03659 | 0 |
| 1.17E-07 | 4.73E-07 | 146.614258 | 215.767375 | 257.998433 | 222.689353 | 9.50487297 |
| 1.17E-07 | 4.74E-07 | 38.5826995 | 77.8542074 | 87.2368802 | 92.7872303 | 0 |
| 1.17E-07 | 4.74E-07 | 123.143116 | 71.4034302 | 63.1075304 | 65.4149973 | 327.918118 |
| 1.17E-07 | 4.75E-07 | 45.334672 | 107.883687 | 104.869867 | 57.9920189 | 0 |

| | | | | | | |
|---|---|---|---|---|---|---|
| 1.17E-07 | 4.75E-07 | 52.4081669 | 196.637484 | 176.329864 | 173.048184 | 7.12865473 |
| 1.17E-07 | 4.75E-07 | 143.399033 | 221.328389 | 207.883629 | 226.864778 | 35.6432737 |
| 1.17E-07 | 4.76E-07 | 41.7979245 | 70.2912272 | 67.7477899 | 68.1986142 | 0 |
| 1.18E-07 | 4.77E-07 | 34.4029071 | 125.234054 | 106.72597 | 109.488932 | 0 |
| 1.18E-07 | 4.77E-07 | 43.405537 | 64.952653 | 75.1722053 | 60.7756358 | 0 |
| 1.18E-07 | 4.78E-07 | 39.2257445 | 83.4152222 | 80.7405168 | 76.0855288 | 0 |
| 1.18E-07 | 4.78E-07 | 33.1168171 | 127.236019 | 116.00649 | 122.94308 | 0 |
| 1.19E-07 | 4.81E-07 | 45.334672 | 71.4034302 | 74.2441534 | 54.2805297 | 0 |
| 1.19E-07 | 4.81E-07 | 100.315019 | 575.008931 | 459.385699 | 417.542536 | 14.2573095 |
| 1.20E-07 | 4.86E-07 | 32.4737721 | 128.570662 | 120.646749 | 129.902122 | 0 |
| 1.20E-07 | 4.86E-07 | 32.1522496 | 159.267464 | 114.150386 | 143.820207 | 0 |
| 1.20E-07 | 4.87E-07 | 48.2283744 | 72.2931925 | 51.9709073 | 59.8477635 | 0 |
| 1.20E-07 | 4.88E-07 | 45.334672 | 63.84045 | 56.6111669 | 59.8477635 | 0 |
| 1.21E-07 | 4.88E-07 | 38.5826995 | 76.964445 | 86.3088283 | 98.8184002 | 0 |
| 1.21E-07 | 4.89E-07 | 38.261177 | 123.676969 | 77.0283091 | 114.592229 | 0 |
| 1.21E-07 | 4.90E-07 | 53.0512119 | 74.2951579 | 69.6038938 | 45.929679 | 0 |
| 1.21E-07 | 4.90E-07 | 30.2231146 | 136.800964 | 271.919212 | 167.017014 | 0 |
| 1.21E-07 | 4.91E-07 | 33.7598621 | 110.997856 | 208.811681 | 135.469356 | 0 |
| 1.21E-07 | 4.91E-07 | 66.2336342 | 167.275326 | 150.344411 | 159.1301 | 2.37621824 |
| 1.22E-07 | 4.92E-07 | 35.0459521 | 168.387528 | 96.5173994 | 129.902122 | 0 |
| 1.22E-07 | 4.92E-07 | 16.7191698 | 10.2322673 | 14.8488307 | 8.35085072 | 85.5438568 |
| 1.22E-07 | 4.93E-07 | 40.833357 | 77.1868856 | 167.049345 | 109.488932 | 0 |
| 1.22E-07 | 4.94E-07 | 36.0105196 | 87.1967122 | 114.150386 | 116.447974 | 0 |
| 1.22E-07 | 4.94E-07 | 33.7598621 | 115.001786 | 116.934542 | 109.952868 | 0 |
| 1.23E-07 | 4.96E-07 | 60.7677518 | 52.940861 | 46.4025958 | 60.3116997 | 0 |
| 1.23E-07 | 4.96E-07 | 20.8989622 | 235.342147 | 368.436611 | 259.340309 | 0 |
| 1.23E-07 | 4.96E-07 | 101.279586 | 82.970341 | 40.8342843 | 65.8789335 | 0 |
| 1.23E-07 | 4.97E-07 | 40.833357 | 79.4112915 | 145.704151 | 89.0757411 | 0 |
| 1.23E-07 | 4.97E-07 | 58.5170943 | 207.092192 | 183.75428 | 208.771268 | 2.37621824 |
| 1.23E-07 | 4.98E-07 | 46.6207619 | 55.8325887 | 70.5319457 | 56.1362743 | 0 |
| 1.23E-07 | 4.98E-07 | 78.4514891 | 130.572628 | 77.0283091 | 100.210209 | 0 |
| 1.24E-07 | 4.99E-07 | 108.996126 | 177.285152 | 79.8124649 | 40.8263813 | 0 |
| 1.24E-07 | 5.00E-07 | 35.6889971 | 111.665177 | 90.9490879 | 105.313506 | 0 |
| 1.24E-07 | 5.00E-07 | 31.5092046 | 121.007682 | 142.919995 | 141.964462 | 0 |
| 1.24E-07 | 5.00E-07 | 23.7926647 | 204.200464 | 209.739733 | 316.404455 | 0 |
| 1.24E-07 | 5.00E-07 | 14.4685123 | 6.67321777 | 3.71220767 | 6.95904227 | 149.701749 |
| 1.25E-07 | 5.06E-07 | 48.2283744 | 128.125781 | 56.6111669 | 90.4675495 | 0 |
| 1.25E-07 | 5.06E-07 | 35.0459521 | 101.21047 | 104.869867 | 101.138081 | 0 |
| 1.25E-07 | 5.06E-07 | 49.8359869 | 115.446667 | 52.8989593 | 84.4363795 | 0 |
| 1.26E-07 | 5.07E-07 | 141.148376 | 115.446667 | 31.5537652 | 134.077548 | 0 |
| 1.26E-07 | 5.07E-07 | 31.8307271 | 154.818652 | 122.502853 | 126.190633 | 0 |
| 1.26E-07 | 5.09E-07 | 40.190312 | 84.5274251 | 66.819738 | 89.5396772 | 0 |
| 1.26E-07 | 5.10E-07 | 243.39253 | 744.508663 | 589.312967 | 617.499017 | 0 |

| | | | | | | |
|---|---|---|---|---|---|---|
| 1.26E-07 | 5.10E-07 | 157.546023 | 229.558691 | 203.24337 | 191.141694 | 23.7621824 |
| 1.27E-07 | 5.12E-07 | 35.0459521 | 114.779346 | 153.128566 | 96.0347833 | 0 |
| 1.27E-07 | 5.13E-07 | 35.3674746 | 107.438806 | 98.3735032 | 96.0347833 | 0 |
| 1.27E-07 | 5.14E-07 | 124.750728 | 53.3857422 | 412.055051 | 90.9314857 | 0 |
| 1.28E-07 | 5.14E-07 | 71.6995166 | 199.529211 | 180.042072 | 184.646588 | 9.50487297 |
| 1.28E-07 | 5.15E-07 | 36.9750871 | 102.100232 | 100.229607 | 80.7248903 | 0 |
| 1.28E-07 | 5.17E-07 | 76.8438766 | 99.4309448 | 130.85532 | 38.5067006 | 0 |
| 1.28E-07 | 5.17E-07 | 44.3701045 | 68.2892619 | 74.2441534 | 123.407016 | 0 |
| 1.28E-07 | 5.17E-07 | 66.2336342 | 54.4979451 | 49.1867516 | 51.0329766 | 0 |
| 1.28E-07 | 5.18E-07 | 40.190312 | 141.249776 | 71.4599976 | 113.200421 | 0 |
| 1.29E-07 | 5.18E-07 | 35.3674746 | 102.767554 | 106.72597 | 92.3232941 | 0 |
| 1.29E-07 | 5.19E-07 | 38.5826995 | 74.2951579 | 94.6612955 | 82.5806349 | 0 |
| 1.29E-07 | 5.20E-07 | 54.0157793 | 54.9428263 | 48.2586997 | 53.8165936 | 0 |
| 1.29E-07 | 5.21E-07 | 46.9422844 | 76.2971232 | 73.3161014 | 50.5690405 | 0 |
| 1.30E-07 | 5.22E-07 | 45.6561944 | 61.8384847 | 53.8270112 | 73.3019119 | 0 |
| 1.30E-07 | 5.23E-07 | 84.5604165 | 133.019474 | 117.862593 | 102.993826 | 11.8810912 |
| 1.30E-07 | 5.23E-07 | 46.9422844 | 60.7262817 | 51.0428554 | 63.0953166 | 0 |
| 1.30E-07 | 5.24E-07 | 71.6995166 | 146.143469 | 118.790645 | 153.09893 | 2.37621824 |
| 1.30E-07 | 5.24E-07 | 40.5118345 | 93.2026082 | 64.9636342 | 86.2921241 | 0 |
| 1.30E-07 | 5.24E-07 | 31.1876821 | 164.383598 | 113.222334 | 167.944887 | 0 |
| 1.30E-07 | 5.24E-07 | 73.6286516 | 246.909058 | 175.401812 | 187.894141 | 11.8810912 |
| 1.31E-07 | 5.25E-07 | 46.6207619 | 58.0569946 | 52.8989593 | 58.9198912 | 0 |
| 1.31E-07 | 5.25E-07 | 42.762492 | 131.684831 | 64.0355823 | 103.457762 | 0 |
| 1.31E-07 | 5.25E-07 | 36.9750871 | 196.192603 | 84.4527244 | 159.1301 | 0 |
| 1.31E-07 | 5.26E-07 | 50.1575094 | 59.6140788 | 50.1148035 | 53.3526574 | 0 |
| 1.31E-07 | 5.26E-07 | 57.2310043 | 78.5215291 | 44.546492 | 60.7756358 | 0 |
| 1.31E-07 | 5.27E-07 | 34.7244296 | 117.226192 | 98.3735032 | 101.138081 | 0 |
| 1.31E-07 | 5.27E-07 | 72.9856066 | 55.3877075 | 88.1649321 | 44.5378705 | 0 |
| 1.31E-07 | 5.28E-07 | 36.3320421 | 163.716276 | 116.00649 | 89.5396772 | 0 |
| 1.31E-07 | 5.28E-07 | 46.6207619 | 110.330534 | 58.4672708 | 145.212015 | 0 |
| 1.32E-07 | 5.29E-07 | 39.547267 | 69.4014648 | 77.0283091 | 78.4052096 | 0 |
| 1.32E-07 | 5.31E-07 | 42.762492 | 74.9624797 | 62.1794784 | 61.7035081 | 0 |
| 1.32E-07 | 5.32E-07 | 82.6312815 | 248.688582 | 277.487523 | 162.377653 | 0 |
| 1.33E-07 | 5.33E-07 | 47.5853294 | 71.1809896 | 51.0428554 | 59.3838274 | 0 |
| 1.33E-07 | 5.34E-07 | 71.6995166 | 131.017509 | 128.071165 | 121.551272 | 2.37621824 |
| 1.33E-07 | 5.34E-07 | 67.8412467 | 188.407182 | 154.98467 | 132.685739 | 2.37621824 |
| 1.33E-07 | 5.34E-07 | 41.7979245 | 94.5372518 | 61.2514265 | 99.7462725 | 0 |
| 1.33E-07 | 5.35E-07 | 30.8661596 | 133.464355 | 153.128566 | 124.334889 | 0 |
| 1.33E-07 | 5.36E-07 | 36.3320421 | 89.6435588 | 114.150386 | 88.6118049 | 0 |
| 1.34E-07 | 5.37E-07 | 37.9396545 | 90.0884399 | 77.956361 | 80.7248903 | 0 |
| 1.34E-07 | 5.38E-07 | 40.833357 | 63.3955688 | 77.0283091 | 71.9101035 | 0 |
| 1.34E-07 | 5.38E-07 | 45.334672 | 54.7203857 | 101.157659 | 96.9626556 | 0 |
| 1.34E-07 | 5.38E-07 | 34.0813846 | 102.322673 | 105.797919 | 136.861165 | 0 |

| | | | | | | |
|---|---|---|---|---|---|---|
| 1.34E-07 | 5.39E-07 | 36.6535646 | 90.7557617 | 83.5246725 | 93.7151026 | 0 |
| 1.34E-07 | 5.40E-07 | 36.0105196 | 91.6455241 | 89.092984 | 110.88074 | 0 |
| 1.35E-07 | 5.42E-07 | 60.7677518 | 60.5038411 | 97.4454513 | 46.3936151 | 0 |
| 1.35E-07 | 5.42E-07 | 31.1876821 | 132.352152 | 141.063891 | 120.623399 | 0 |
| 1.35E-07 | 5.43E-07 | 34.4029071 | 182.178845 | 97.4454513 | 130.366059 | 0 |
| 1.36E-07 | 5.44E-07 | 40.833357 | 86.7518311 | 67.7477899 | 68.1986142 | 0 |
| 1.36E-07 | 5.45E-07 | 55.9449143 | 50.2715739 | 84.4527244 | 52.8887212 | 0 |
| 1.36E-07 | 5.45E-07 | 57.5525268 | 181.733964 | 210.667785 | 250.989458 | 0 |
| 1.36E-07 | 5.45E-07 | 43.0840145 | 136.356083 | 63.1075304 | 146.139888 | 0 |
| 1.36E-07 | 5.46E-07 | 41.7979245 | 79.6337321 | 63.1075304 | 65.4149973 | 0 |
| 1.36E-07 | 5.46E-07 | 42.119447 | 81.4132568 | 58.4672708 | 82.1166988 | 0 |
| 1.36E-07 | 5.47E-07 | 49.5144644 | 90.3108805 | 49.1867516 | 70.9822312 | 0 |
| 1.36E-07 | 5.47E-07 | 252.073637 | 442.211898 | 511.356606 | 521.464234 | 30.8908372 |
| 1.37E-07 | 5.48E-07 | 39.8687895 | 66.2872965 | 83.5246725 | 95.5708472 | 0 |
| 1.37E-07 | 5.48E-07 | 38.904222 | 72.5156331 | 135.49558 | 143.356271 | 0 |
| 1.37E-07 | 5.48E-07 | 34.4029071 | 138.80293 | 105.797919 | 98.3544641 | 0 |
| 1.37E-07 | 5.49E-07 | 40.833357 | 67.6219401 | 64.9636342 | 70.9822312 | 0 |
| 1.37E-07 | 5.49E-07 | 87.1325964 | 48.9369303 | 49.1867516 | 96.9626556 | 0 |
| 1.38E-07 | 5.51E-07 | 35.3674746 | 191.076469 | 106.72597 | 107.633187 | 0 |
| 1.38E-07 | 5.52E-07 | 35.6889971 | 102.989994 | 86.3088283 | 123.870952 | 0 |
| 1.38E-07 | 5.53E-07 | 45.9777169 | 190.631588 | 62.1794784 | 142.892335 | 0 |
| 1.38E-07 | 5.53E-07 | 144.685123 | 195.525281 | 144.776099 | 160.985845 | 16.6335277 |
| 1.38E-07 | 5.53E-07 | 27.3294122 | 160.157227 | 165.193241 | 175.367865 | 0 |
| 1.38E-07 | 5.53E-07 | 127.965953 | 179.509558 | 197.675058 | 186.038397 | 16.6335277 |
| 1.39E-07 | 5.54E-07 | 44.691627 | 68.5117025 | 54.7550631 | 59.3838274 | 0 |
| 1.39E-07 | 5.54E-07 | 58.8386168 | 64.7302124 | 44.546492 | 53.8165936 | 0 |
| 1.39E-07 | 5.55E-07 | 44.048582 | 72.2931925 | 53.8270112 | 68.1986142 | 0 |
| 1.39E-07 | 5.55E-07 | 51.4435994 | 119.895479 | 50.1148035 | 87.2199964 | 0 |
| 1.39E-07 | 5.55E-07 | 67.1982017 | 198.86189 | 129.927268 | 164.697334 | 0 |
| 1.39E-07 | 5.56E-07 | 80.380624 | 84.0825439 | 61.2514265 | 40.8263813 | 0 |
| 1.39E-07 | 5.56E-07 | 112.854396 | 65.1750936 | 58.4672708 | 49.177232 | 0 |
| 1.39E-07 | 5.56E-07 | 77.1653991 | 140.582454 | 118.790645 | 107.169251 | 0 |
| 1.39E-07 | 5.56E-07 | 27.6509347 | 189.519385 | 160.552982 | 151.243185 | 0 |
| 1.39E-07 | 5.56E-07 | 45.9777169 | 78.2990885 | 71.4599976 | 51.0329766 | 0 |
| 1.39E-07 | 5.56E-07 | 35.6889971 | 89.4211182 | 98.3735032 | 91.3954218 | 0 |
| 1.39E-07 | 5.57E-07 | 74.9147416 | 66.2872965 | 42.6903882 | 117.839782 | 0 |
| 1.39E-07 | 5.57E-07 | 73.9501741 | 84.5274251 | 40.8342843 | 138.716909 | 0 |
| 1.40E-07 | 5.60E-07 | 35.0459521 | 110.108093 | 110.438178 | 87.2199964 | 0 |
| 1.40E-07 | 5.61E-07 | 91.9554339 | 99.2085042 | 44.546492 | 54.2805297 | 0 |
| 1.41E-07 | 5.61E-07 | 42.119447 | 78.7439697 | 70.5319457 | 58.4559551 | 0 |
| 1.41E-07 | 5.62E-07 | 33.4383396 | 107.438806 | 106.72597 | 109.488932 | 0 |
| 1.41E-07 | 5.62E-07 | 30.2231146 | 19.5747721 | 24.1293498 | 20.4131907 | 87.920075 |
| 1.41E-07 | 5.62E-07 | 36.3320421 | 113.889583 | 84.4527244 | 91.3954218 | 0 |

| | | | | | | |
|---|---|---|---|---|---|---|
| 1.41E-07 | 5.62E-07 | 35.6889971 | 103.879757 | 83.5246725 | 109.952868 | 0 |
| 1.41E-07 | 5.63E-07 | 38.5826995 | 88.0864746 | 70.5319457 | 80.7248903 | 0 |
| 1.41E-07 | 5.64E-07 | 41.1548795 | 66.7321777 | 62.1794784 | 71.4461673 | 0 |
| 1.41E-07 | 5.64E-07 | 86.4895515 | 43.3759155 | 78.8844129 | 55.6723382 | 0 |
| 1.41E-07 | 5.64E-07 | 48.2283744 | 52.2735392 | 61.2514265 | 51.4969128 | 0 |
| 1.42E-07 | 5.65E-07 | 49.1929419 | 48.9369303 | 56.6111669 | 56.1362743 | 0 |
| 1.42E-07 | 5.65E-07 | 37.9396545 | 84.0825439 | 106.72597 | 75.6215927 | 0 |
| 1.42E-07 | 5.65E-07 | 531.476686 | 111.442737 | 56.6111669 | 109.024996 | 0 |
| 1.42E-07 | 5.66E-07 | 118.963324 | 169.277291 | 29.6976613 | 229.184459 | 0 |
| 1.42E-07 | 5.67E-07 | 106.745469 | 34.7007324 | 93.7332436 | 100.210209 | 0 |
| 1.42E-07 | 5.67E-07 | 37.2966096 | 96.3167765 | 77.0283091 | 85.3642518 | 0 |
| 1.42E-07 | 5.67E-07 | 33.7598621 | 143.251742 | 123.430905 | 94.1790387 | 0 |
| 1.42E-07 | 5.67E-07 | 89.3832539 | 93.2026082 | 99.3015551 | 35.2591475 | 0 |
| 1.42E-07 | 5.68E-07 | 35.6889971 | 93.2026082 | 89.092984 | 91.859358 | 0 |
| 1.42E-07 | 5.68E-07 | 37.2966096 | 91.6455241 | 75.1722053 | 91.859358 | 0 |
| 1.43E-07 | 5.71E-07 | 79.737579 | 48.9369303 | 49.1867516 | 61.7035081 | 0 |
| 1.44E-07 | 5.73E-07 | 44.691627 | 66.7321777 | 61.2514265 | 52.8887212 | 0 |
| 1.44E-07 | 5.73E-07 | 99.9934963 | 117.893514 | 32.4818171 | 127.582442 | 0 |
| 1.44E-07 | 5.73E-07 | 32.4737721 | 169.277291 | 107.654022 | 123.407016 | 0 |
| 1.44E-07 | 5.74E-07 | 39.8687895 | 68.5117025 | 77.0283091 | 66.8068058 | 0 |
| 1.44E-07 | 5.75E-07 | 38.261177 | 95.2045736 | 111.36623 | 70.9822312 | 0 |
| 1.45E-07 | 5.76E-07 | 30.2231146 | 141.027336 | 123.430905 | 142.892335 | 0 |
| 1.45E-07 | 5.76E-07 | 63.9829767 | 159.934786 | 144.776099 | 170.728504 | 2.37621824 |
| 1.45E-07 | 5.76E-07 | 81.666714 | 82.5254598 | 77.0283091 | 81.1888265 | 182.968805 |
| 1.45E-07 | 5.77E-07 | 70.0919042 | 133.464355 | 39.9062324 | 93.2511664 | 0 |
| 1.45E-07 | 5.77E-07 | 40.833357 | 78.9664103 | 66.819738 | 63.5592527 | 0 |
| 1.46E-07 | 5.79E-07 | 73.9501741 | 121.452563 | 117.862593 | 115.520102 | 2.37621824 |
| 1.46E-07 | 5.79E-07 | 31.8307271 | 162.604073 | 190.250643 | 102.529889 | 0 |
| 1.46E-07 | 5.80E-07 | 42.762492 | 84.7498657 | 143.848047 | 69.5904227 | 0 |
| 1.46E-07 | 5.80E-07 | 255.931907 | 92.757727 | 47.3306478 | 88.1478687 | 0 |
| 1.46E-07 | 5.80E-07 | 455.9189 | 57.6121134 | 103.013763 | 101.138081 | 0 |
| 1.46E-07 | 5.81E-07 | 42.762492 | 68.2892619 | 59.3953227 | 58.9198912 | 0 |
| 1.47E-07 | 5.84E-07 | 30.8661596 | 134.798999 | 131.783372 | 118.303719 | 0 |
| 1.47E-07 | 5.84E-07 | 326.988379 | 796.559762 | 1086.74879 | 1016.48411 | 28.5146189 |
| 1.47E-07 | 5.86E-07 | 604.462293 | 855.061637 | 837.102829 | 895.396772 | 54.6530196 |
| 1.47E-07 | 5.86E-07 | 33.4383396 | 124.566732 | 97.4454513 | 109.952868 | 0 |
| 1.48E-07 | 5.88E-07 | 138.576196 | 98.9860636 | 35.2659728 | 87.6839326 | 0 |
| 1.48E-07 | 5.89E-07 | 39.2257445 | 92.0904053 | 66.819738 | 78.8691457 | 0 |
| 1.48E-07 | 5.89E-07 | 61.7323193 | 80.3010539 | 45.4745439 | 52.8887212 | 0 |
| 1.48E-07 | 5.89E-07 | 166.22713 | 238.901196 | 273.775316 | 240.782863 | 19.0097459 |
| 1.49E-07 | 5.90E-07 | 40.833357 | 63.6180094 | 67.7477899 | 64.0231889 | 0 |
| 1.49E-07 | 5.90E-07 | 28.6155022 | 202.64338 | 163.337137 | 128.510314 | 0 |
| 1.49E-07 | 5.91E-07 | 82.309759 | 96.5392171 | 106.72597 | 102.065953 | 2.37621824 |

| | | | | | | |
|---|---|---|---|---|---|---|
| 1.49E-07 | 5.91E-07 | 34.0813846 | 164.828479 | 93.7332436 | 122.479144 | 0 |
| 1.49E-07 | 5.92E-07 | 113.175919 | 185.293013 | 174.47376 | 157.738291 | 14.2573095 |
| 1.49E-07 | 5.93E-07 | 55.9449143 | 53.8306234 | 52.8989593 | 98.3544641 | 0 |
| 1.50E-07 | 5.96E-07 | 206.417443 | 342.113631 | 331.314534 | 348.41605 | 21.3859642 |
| 1.50E-07 | 5.97E-07 | 281.010662 | 97.2065389 | 51.9709073 | 79.797018 | 0 |
| 1.51E-07 | 5.99E-07 | 64.3044992 | 46.2676432 | 51.0428554 | 81.1888265 | 0 |
| 1.51E-07 | 6.00E-07 | 38.5826995 | 71.6258708 | 72.3880495 | 74.6937204 | 0 |
| 1.51E-07 | 6.00E-07 | 36.0105196 | 78.5215291 | 101.157659 | 97.4265918 | 0 |
| 1.52E-07 | 6.01E-07 | 34.7244296 | 88.7537964 | 112.294282 | 96.4987195 | 0 |
| 1.52E-07 | 6.01E-07 | 33.7598621 | 115.224227 | 94.6612955 | 104.385634 | 0 |
| 1.52E-07 | 6.02E-07 | 181.338688 | 473.353581 | 420.407518 | 489.916576 | 28.5146189 |
| 1.52E-07 | 6.03E-07 | 28.9370247 | 161.714311 | 123.430905 | 176.295737 | 0 |
| 1.52E-07 | 6.05E-07 | 49.8359869 | 57.834554 | 54.7550631 | 47.3214874 | 0 |
| 1.52E-07 | 6.05E-07 | 86.168029 | 176.39539 | 169.833501 | 181.399035 | 16.6335277 |
| 1.53E-07 | 6.05E-07 | 57.2310043 | 52.940861 | 44.546492 | 57.5280828 | 0 |
| 1.53E-07 | 6.05E-07 | 32.7952946 | 115.669108 | 104.869867 | 106.705315 | 0 |
| 1.53E-07 | 6.06E-07 | 37.9396545 | 77.8542074 | 71.4599976 | 80.7248903 | 0 |
| 1.53E-07 | 6.07E-07 | 32.4737721 | 111.442737 | 103.941815 | 130.366059 | 0 |
| 1.53E-07 | 6.07E-07 | 42.119447 | 87.864034 | 95.5893474 | 57.0641466 | 0 |
| 1.53E-07 | 6.07E-07 | 49.5144644 | 56.4999105 | 49.1867516 | 77.4773373 | 0 |
| 1.53E-07 | 6.08E-07 | 83.2743265 | 78.2990885 | 114.150386 | 83.5085072 | 11.8810912 |
| 1.54E-07 | 6.09E-07 | 35.6889971 | 105.436841 | 92.8051917 | 171.19244 | 0 |
| 1.54E-07 | 6.10E-07 | 45.334672 | 60.05896 | 51.9709073 | 56.1362743 | 0 |
| 1.54E-07 | 6.11E-07 | 37.2966096 | 104.324638 | 71.4599976 | 102.993826 | 0 |
| 1.54E-07 | 6.11E-07 | 70.0919042 | 60.7262817 | 64.0355823 | 40.8263813 | 0 |
| 1.55E-07 | 6.12E-07 | 50.4790319 | 45.6003214 | 59.3953227 | 66.3428696 | 0 |
| 1.55E-07 | 6.13E-07 | 67.8412467 | 57.834554 | 99.3015551 | 44.0739344 | 0 |
| 1.55E-07 | 6.14E-07 | 38.261177 | 67.3994995 | 84.4527244 | 82.1166988 | 0 |
| 1.55E-07 | 6.15E-07 | 37.9396545 | 75.6298014 | 106.72597 | 148.923505 | 0 |
| 1.56E-07 | 6.16E-07 | 42.4409695 | 57.834554 | 66.819738 | 58.4559551 | 0 |
| 1.56E-07 | 6.17E-07 | 36.0105196 | 88.7537964 | 82.5966206 | 84.9003157 | 0 |
| 1.56E-07 | 6.17E-07 | 46.9422844 | 62.0609253 | 52.8989593 | 51.0329766 | 0 |
| 1.56E-07 | 6.18E-07 | 34.4029071 | 97.4289795 | 90.0210359 | 102.529889 | 0 |
| 1.56E-07 | 6.18E-07 | 34.4029071 | 88.976237 | 107.654022 | 97.4265918 | 0 |
| 1.57E-07 | 6.19E-07 | 57.2310043 | 52.2735392 | 46.4025958 | 51.9608489 | 0 |
| 1.57E-07 | 6.20E-07 | 76.5223541 | 49.6042521 | 60.3233746 | 147.531696 | 0 |
| 1.57E-07 | 6.20E-07 | 42.762492 | 58.7243164 | 85.3807764 | 62.6313804 | 0 |
| 1.57E-07 | 6.21E-07 | 32.1522496 | 114.556905 | 105.797919 | 117.375846 | 0 |
| 1.57E-07 | 6.21E-07 | 167.834743 | 177.285152 | 360.084144 | 25.9804245 | 0 |
| 1.57E-07 | 6.22E-07 | 32.7952946 | 163.493835 | 93.7332436 | 142.428398 | 0 |
| 1.57E-07 | 6.22E-07 | 110.925261 | 123.89941 | 30.6257133 | 153.562866 | 0 |
| 1.58E-07 | 6.24E-07 | 36.6535646 | 83.8601033 | 81.6685687 | 77.9412734 | 0 |
| 1.58E-07 | 6.24E-07 | 68.8058142 | 131.907271 | 140.135839 | 122.94308 | 2.37621824 |

| | | | | | | |
|---|---|---|---|---|---|---|
| 1.58E-07 | 6.24E-07 | 37.9396545 | 70.7361084 | 75.1722053 | 84.4363795 | 0 |
| 1.58E-07 | 6.25E-07 | 2.89370247 | 2.8917277 | 0.92805192 | 2.78361691 | 19.0097459 |
| 1.58E-07 | 6.25E-07 | 35.6889971 | 89.8659993 | 85.3807764 | 84.4363795 | 0 |
| 1.59E-07 | 6.27E-07 | 180.052598 | 206.202429 | 287.696094 | 227.328714 | 19.0097459 |
| 1.59E-07 | 6.28E-07 | 33.1168171 | 106.104163 | 96.5173994 | 119.231591 | 0 |
| 1.60E-07 | 6.31E-07 | 41.7979245 | 67.6219401 | 73.3161014 | 55.6723382 | 0 |
| 1.60E-07 | 6.32E-07 | 75.8793091 | 79.1888509 | 36.1940248 | 85.828188 | 0 |
| 1.60E-07 | 6.32E-07 | 93.8845689 | 119.895479 | 78.8844129 | 111.808612 | 0 |
| 1.60E-07 | 6.33E-07 | 38.5826995 | 84.0825439 | 64.9636342 | 83.9724434 | 0 |
| 1.61E-07 | 6.35E-07 | 75.2362641 | 125.456494 | 102.085711 | 117.839782 | 2.37621824 |
| 1.61E-07 | 6.36E-07 | 141.148376 | 191.743791 | 25.9854537 | 192.069567 | 0 |
| 1.61E-07 | 6.36E-07 | 426.33883 | 386.379309 | 279.343627 | 339.601263 | 23.7621824 |
| 1.62E-07 | 6.37E-07 | 157.546023 | 757.632658 | 607.874006 | 712.605928 | 9.50487297 |
| 1.62E-07 | 6.38E-07 | 42.4409695 | 62.5058065 | 54.7550631 | 64.9510612 | 0 |
| 1.62E-07 | 6.39E-07 | 444.665612 | 63.3955688 | 92.8051917 | 93.7151026 | 0 |
| 1.62E-07 | 6.40E-07 | 36.3320421 | 76.2971232 | 90.0210359 | 83.9724434 | 0 |
| 1.62E-07 | 6.41E-07 | 36.9750871 | 80.5234945 | 164.265189 | 108.561059 | 0 |
| 1.63E-07 | 6.42E-07 | 35.0459521 | 83.6376628 | 90.9490879 | 102.065953 | 0 |
| 1.63E-07 | 6.42E-07 | 36.9750871 | 71.1809896 | 98.3735032 | 107.169251 | 0 |
| 1.63E-07 | 6.42E-07 | 33.7598621 | 127.013578 | 99.3015551 | 93.7151026 | 0 |
| 1.63E-07 | 6.42E-07 | 39.2257445 | 67.1770589 | 73.3161014 | 65.4149973 | 0 |
| 1.63E-07 | 6.42E-07 | 38.904222 | 102.767554 | 77.0283091 | 167.944887 | 0 |
| 1.63E-07 | 6.42E-07 | 39.2257445 | 64.952653 | 73.3161014 | 67.7346781 | 0 |
| 1.64E-07 | 6.45E-07 | 34.0813846 | 185.737895 | 95.5893474 | 118.767655 | 0 |
| 1.64E-07 | 6.46E-07 | 41.476402 | 119.005717 | 64.0355823 | 80.7248903 | 0 |
| 1.64E-07 | 6.46E-07 | 71.6995166 | 168.609969 | 130.85532 | 133.149675 | 4.75243649 |
| 1.64E-07 | 6.46E-07 | 37.618132 | 74.7400391 | 152.200514 | 109.488932 | 0 |
| 1.65E-07 | 6.49E-07 | 61.7323193 | 62.0609253 | 40.8342843 | 58.4559551 | 0 |
| 1.65E-07 | 6.49E-07 | 35.6889971 | 99.4309448 | 80.7405168 | 85.3642518 | 0 |
| 1.65E-07 | 6.49E-07 | 30.2231146 | 153.039128 | 119.718697 | 121.551272 | 0 |
| 1.65E-07 | 6.50E-07 | 30.8661596 | 190.631588 | 108.582074 | 138.252973 | 0 |
| 1.65E-07 | 6.51E-07 | 74.9147416 | 127.458459 | 103.941815 | 112.272549 | 2.37621824 |
| 1.66E-07 | 6.53E-07 | 88.0971639 | 88.3089152 | 99.3015551 | 95.106911 | 2.37621824 |
| 1.66E-07 | 6.54E-07 | 36.3320421 | 72.9605143 | 105.797919 | 97.8905279 | 0 |
| 1.67E-07 | 6.56E-07 | 31.1876821 | 141.472217 | 103.941815 | 126.654569 | 0 |
| 1.67E-07 | 6.58E-07 | 66.5551567 | 168.165088 | 128.071165 | 143.820207 | 0 |
| 1.67E-07 | 6.58E-07 | 70.4134267 | 164.161157 | 183.75428 | 108.097123 | 0 |
| 1.67E-07 | 6.59E-07 | 35.6889971 | 92.0904053 | 82.5966206 | 81.1888265 | 0 |
| 1.68E-07 | 6.59E-07 | 30.5446371 | 165.718241 | 110.438178 | 129.438186 | 0 |
| 1.68E-07 | 6.59E-07 | 42.4409695 | 56.9447917 | 56.6111669 | 64.9510612 | 0 |
| 1.68E-07 | 6.62E-07 | 54.3373018 | 143.029301 | 80.7405168 | 51.0329766 | 0 |
| 1.68E-07 | 6.62E-07 | 89.7047764 | 52.7184204 | 121.574801 | 46.3936151 | 0 |
| 1.69E-07 | 6.65E-07 | 30.5446371 | 215.322493 | 117.862593 | 132.685739 | 0 |

| | | | | | | |
|---|---|---|---|---|---|---|
| 1.69E-07 | 6.66E-07 | 259.790177 | 80.5234945 | 41.7623363 | 122.015208 | 0 |
| 1.70E-07 | 6.67E-07 | 34.7244296 | 110.108093 | 79.8124649 | 111.808612 | 0 |
| 1.70E-07 | 6.67E-07 | 33.4383396 | 97.2065389 | 93.7332436 | 104.84957 | 0 |
| 1.70E-07 | 6.68E-07 | 232.782287 | 716.258708 | 744.297637 | 826.734222 | 0 |
| 1.71E-07 | 6.72E-07 | 219.599865 | 266.038949 | 293.264406 | 309.445413 | 21.3859642 |
| 1.71E-07 | 6.73E-07 | 48.2283744 | 53.1633016 | 48.2586997 | 53.3526574 | 0 |
| 1.71E-07 | 6.73E-07 | 32.1522496 | 125.901375 | 120.646749 | 95.106911 | 0 |
| 1.71E-07 | 6.74E-07 | 76.8438766 | 65.6199748 | 39.9062324 | 58.9198912 | 0 |
| 1.72E-07 | 6.75E-07 | 42.119447 | 58.7243164 | 129.927268 | 109.024996 | 0 |
| 1.72E-07 | 6.76E-07 | 83.2743265 | 141.472217 | 120.646749 | 149.851377 | 16.6335277 |
| 1.72E-07 | 6.77E-07 | 42.762492 | 55.1652669 | 64.9636342 | 83.9724434 | 0 |
| 1.72E-07 | 6.77E-07 | 51.4435994 | 113.222262 | 45.4745439 | 104.84957 | 0 |
| 1.73E-07 | 6.77E-07 | 44.691627 | 68.9565837 | 70.5319457 | 48.7132959 | 0 |
| 1.73E-07 | 6.78E-07 | 148.864916 | 445.326066 | 460.313751 | 447.698386 | 19.0097459 |
| 1.73E-07 | 6.78E-07 | 35.6889971 | 105.2144 | 77.956361 | 87.2199964 | 0 |
| 1.73E-07 | 6.78E-07 | 74.5932191 | 165.05092 | 107.654022 | 121.087335 | 0 |
| 1.73E-07 | 6.81E-07 | 33.1168171 | 108.328569 | 95.5893474 | 96.9626556 | 0 |
| 1.74E-07 | 6.84E-07 | 42.762492 | 121.897445 | 58.4672708 | 87.6839326 | 0 |
| 1.75E-07 | 6.85E-07 | 36.0105196 | 88.5313558 | 79.8124649 | 77.0134011 | 0 |
| 1.75E-07 | 6.86E-07 | 34.0813846 | 91.8679647 | 88.1649321 | 113.200421 | 0 |
| 1.75E-07 | 6.86E-07 | 32.1522496 | 97.2065389 | 109.510126 | 121.551272 | 0 |
| 1.75E-07 | 6.86E-07 | 40.190312 | 57.6121134 | 71.4599976 | 68.6625504 | 0 |
| 1.75E-07 | 6.87E-07 | 33.1168171 | 130.572628 | 89.092984 | 111.344676 | 0 |
| 1.75E-07 | 6.88E-07 | 171.37149 | 387.713953 | 253.358173 | 295.063392 | 19.0097459 |
| 1.75E-07 | 6.88E-07 | 176.51585 | 389.938359 | 361.940248 | 398.521154 | 26.1384007 |
| 1.76E-07 | 6.88E-07 | 39.2257445 | 99.4309448 | 61.2514265 | 93.2511664 | 0 |
| 1.76E-07 | 6.88E-07 | 34.0813846 | 88.3089152 | 90.0210359 | 103.921698 | 0 |
| 1.76E-07 | 6.89E-07 | 37.618132 | 153.484009 | 87.2368802 | 83.9724434 | 0 |
| 1.76E-07 | 6.91E-07 | 37.618132 | 98.3187419 | 127.143113 | 69.5904227 | 0 |
| 1.77E-07 | 6.94E-07 | 40.190312 | 68.2892619 | 57.5392189 | 71.9101035 | 0 |
| 1.77E-07 | 6.94E-07 | 76.2008316 | 100.988029 | 36.1940248 | 143.356271 | 0 |
| 1.78E-07 | 6.98E-07 | 39.2257445 | 70.2912272 | 61.2514265 | 68.1986142 | 0 |
| 1.79E-07 | 7.00E-07 | 36.6535646 | 80.7459351 | 70.5319457 | 83.9724434 | 0 |
| 1.79E-07 | 7.03E-07 | 35.3674746 | 83.1927816 | 80.7405168 | 83.5085072 | 0 |
| 1.80E-07 | 7.06E-07 | 19.6128723 | 418.410754 | 886.289581 | 887.509857 | 2.37621824 |
| 1.81E-07 | 7.07E-07 | 37.2966096 | 157.71038 | 71.4599976 | 126.654569 | 0 |
| 1.81E-07 | 7.10E-07 | 0 | 0 | 0 | 0 | 28.5146189 |
| 1.82E-07 | 7.12E-07 | 66.8766792 | 114.334465 | 47.3306478 | 55.208402 | 0 |
| 1.82E-07 | 7.12E-07 | 33.4383396 | 91.4230835 | 104.869867 | 91.3954218 | 0 |
| 1.82E-07 | 7.12E-07 | 36.6535646 | 92.757727 | 73.3161014 | 75.6215927 | 0 |
| 1.82E-07 | 7.12E-07 | 66.5551567 | 67.3994995 | 38.9781805 | 59.3838274 | 0 |
| 1.83E-07 | 7.14E-07 | 27.3294122 | 170.167053 | 157.768826 | 128.046378 | 0 |
| 1.83E-07 | 7.15E-07 | 36.9750871 | 103.212435 | 67.7477899 | 92.3232941 | 0 |

| | | | | | | |
|---|---|---|---|---|---|---|
| 1.83E-07 | 7.15E-07 | 67.1982017 | 143.474182 | 146.632203 | 166.553078 | 4.75243649 |
| 1.83E-07 | 7.18E-07 | 41.1548795 | 70.958549 | 57.5392189 | 58.9198912 | 0 |
| 1.84E-07 | 7.19E-07 | 51.4435994 | 43.1534749 | 72.3880495 | 60.7756358 | 0 |
| 1.84E-07 | 7.20E-07 | 418.62229 | 764.528316 | 836.174777 | 844.363795 | 30.8908372 |
| 1.84E-07 | 7.20E-07 | 31.1876821 | 135.02144 | 121.574801 | 98.3544641 | 0 |
| 1.84E-07 | 7.21E-07 | 44.691627 | 78.0766479 | 48.2586997 | 70.518295 | 0 |
| 1.84E-07 | 7.21E-07 | 37.618132 | 81.4132568 | 65.8916861 | 73.7658481 | 0 |
| 1.85E-07 | 7.22E-07 | 12.5393773 | 23.3562622 | 22.273246 | 27.3722329 | 140.196876 |
| 1.85E-07 | 7.23E-07 | 59.8031843 | 70.5136678 | 42.6903882 | 51.4969128 | 0 |
| 1.85E-07 | 7.24E-07 | 83.9173715 | 227.779167 | 150.344411 | 191.141694 | 14.2573095 |
| 1.85E-07 | 7.25E-07 | 41.7979245 | 107.883687 | 57.5392189 | 79.797018 | 0 |
| 1.86E-07 | 7.25E-07 | 50.1575094 | 53.1633016 | 224.588564 | 164.233398 | 0 |
| 1.87E-07 | 7.29E-07 | 40.5118345 | 60.2814006 | 57.5392189 | 67.7346781 | 0 |
| 1.87E-07 | 7.30E-07 | 144.685123 | 323.428621 | 272.847264 | 304.806051 | 0 |
| 1.87E-07 | 7.31E-07 | 33.4383396 | 118.560836 | 93.7332436 | 89.0757411 | 0 |
| 1.88E-07 | 7.32E-07 | 34.4029071 | 82.3030192 | 87.2368802 | 91.859358 | 0 |
| 1.88E-07 | 7.34E-07 | 34.4029071 | 88.0864746 | 81.6685687 | 92.7872303 | 0 |
| 1.88E-07 | 7.36E-07 | 65.5905892 | 40.4841878 | 55.683115 | 67.2707419 | 0 |
| 1.89E-07 | 7.36E-07 | 27.3294122 | 19.7972127 | 23.2012979 | 24.1246799 | 99.8011662 |
| 1.89E-07 | 7.38E-07 | 55.9449143 | 144.586385 | 412.055051 | 66.3428696 | 0 |
| 1.90E-07 | 7.40E-07 | 50.4790319 | 82.5254598 | 42.6903882 | 75.1576565 | 0 |
| 1.90E-07 | 7.40E-07 | 31.1876821 | 163.938717 | 95.5893474 | 135.00542 | 0 |
| 1.90E-07 | 7.40E-07 | 38.5826995 | 94.7596924 | 160.552982 | 74.2297842 | 0 |
| 1.90E-07 | 7.40E-07 | 40.833357 | 82.5254598 | 54.7550631 | 72.8379758 | 0 |
| 1.90E-07 | 7.41E-07 | 41.1548795 | 64.2853312 | 58.4672708 | 55.6723382 | 0 |
| 1.90E-07 | 7.43E-07 | 48.5498969 | 273.824369 | 224.588564 | 216.658183 | 0 |
| 1.91E-07 | 7.45E-07 | 56.2664368 | 69.4014648 | 55.683115 | 41.7542536 | 0 |
| 1.91E-07 | 7.46E-07 | 46.6207619 | 60.9487223 | 45.4745439 | 69.1264865 | 0 |
| 1.91E-07 | 7.46E-07 | 36.9750871 | 92.9801676 | 66.819738 | 107.633187 | 0 |
| 1.92E-07 | 7.48E-07 | 62.0538418 | 42.2637126 | 58.4672708 | 91.859358 | 0 |
| 1.92E-07 | 7.48E-07 | 36.0105196 | 70.7361084 | 90.0210359 | 106.241379 | 0 |
| 1.92E-07 | 7.49E-07 | 65.9121117 | 121.452563 | 115.078438 | 108.561059 | 7.12865473 |
| 1.92E-07 | 7.50E-07 | 41.7979245 | 56.0550293 | 116.00649 | 93.7151026 | 0 |
| 1.93E-07 | 7.51E-07 | 73.3071291 | 105.881722 | 86.3088283 | 80.2609542 | 11.8810912 |
| 1.93E-07 | 7.51E-07 | 28.2939797 | 177.952474 | 182.826228 | 109.488932 | 0 |
| 1.93E-07 | 7.51E-07 | 162.047338 | 63.3955688 | 51.0428554 | 68.6625504 | 0 |
| 1.93E-07 | 7.52E-07 | 131.181178 | 270.26532 | 302.544925 | 289.496158 | 28.5146189 |
| 1.94E-07 | 7.55E-07 | 166.548653 | 147.700553 | 25.0574018 | 173.512121 | 0 |
| 1.94E-07 | 7.55E-07 | 31.5092046 | 137.468286 | 90.9490879 | 136.397228 | 0 |
| 1.94E-07 | 7.57E-07 | 38.5826995 | 89.1986776 | 61.2514265 | 74.6937204 | 0 |
| 1.95E-07 | 7.59E-07 | 94.8491364 | 84.7498657 | 47.3306478 | 45.929679 | 0 |
| 1.97E-07 | 7.65E-07 | 77.4869216 | 53.3857422 | 83.5246725 | 40.8263813 | 0 |
| 1.97E-07 | 7.65E-07 | 47.9068519 | 120.562801 | 51.9709073 | 74.2297842 | 0 |

| | | | | | | |
|---|---|---|---|---|---|---|
| 1.97E-07 | 7.67E-07 | 42.4409695 | 56.9447917 | 60.3233746 | 51.4969128 | 0 |
| 1.97E-07 | 7.67E-07 | 64.9475442 | 164.606038 | 110.438178 | 145.212015 | 0 |
| 1.97E-07 | 7.67E-07 | 31.5092046 | 101.877791 | 111.36623 | 100.210209 | 0 |
| 1.98E-07 | 7.69E-07 | 41.476402 | 74.2951579 | 51.9709073 | 68.1986142 | 0 |
| 1.98E-07 | 7.69E-07 | 36.9750871 | 66.064856 | 87.2368802 | 103.457762 | 0 |
| 1.98E-07 | 7.69E-07 | 106.745469 | 86.3069499 | 79.8124649 | 34.3312752 | 0 |
| 1.98E-07 | 7.72E-07 | 81.9882365 | 191.966231 | 120.646749 | 142.428398 | 0 |
| 1.99E-07 | 7.74E-07 | 103.208721 | 114.779346 | 37.1220767 | 64.0231889 | 0 |
| 1.99E-07 | 7.74E-07 | 37.2966096 | 91.6455241 | 63.1075304 | 89.5396772 | 0 |
| 1.99E-07 | 7.74E-07 | 172.014535 | 201.976058 | 282.127783 | 244.030416 | 19.0097459 |
| 1.99E-07 | 7.75E-07 | 27.3294122 | 127.903341 | 143.848047 | 208.307332 | 0 |
| 2.00E-07 | 7.78E-07 | 24.4357097 | 217.546899 | 140.135839 | 199.956481 | 0 |
| 2.00E-07 | 7.78E-07 | 47.2638069 | 44.9329997 | 56.6111669 | 58.9198912 | 0 |
| 2.01E-07 | 7.80E-07 | 30.5446371 | 142.361979 | 96.5173994 | 129.902122 | 0 |
| 2.01E-07 | 7.83E-07 | 60.4462293 | 161.936751 | 144.776099 | 155.418611 | 0 |
| 2.02E-07 | 7.84E-07 | 44.048582 | 228.89137 | 274.703367 | 304.342115 | 0 |
| 2.02E-07 | 7.85E-07 | 39.547267 | 95.8718953 | 57.5392189 | 81.6527626 | 0 |
| 2.02E-07 | 7.85E-07 | 0 | 0 | 0 | 0.9278723 | 14.2573095 |
| 2.02E-07 | 7.86E-07 | 37.2966096 | 70.5136678 | 65.8916861 | 69.5904227 | 0 |
| 2.03E-07 | 7.88E-07 | 27.6509347 | 133.464355 | 129.927268 | 142.892335 | 0 |
| 2.04E-07 | 7.91E-07 | 38.904222 | 59.1691976 | 99.3015551 | 82.5806349 | 0 |
| 2.04E-07 | 7.91E-07 | 3.85826995 | 557.213684 | 1391.14982 | 1177.93389 | 0 |
| 2.04E-07 | 7.91E-07 | 86.811074 | 68.2892619 | 36.1940248 | 113.200421 | 0 |
| 2.04E-07 | 7.91E-07 | 31.5092046 | 104.769519 | 103.013763 | 98.3544641 | 0 |
| 2.04E-07 | 7.93E-07 | 224.744225 | 646.412362 | 377.71713 | 469.503385 | 30.8908372 |
| 2.05E-07 | 7.95E-07 | 36.6535646 | 70.0687866 | 72.3880495 | 94.1790387 | 0 |
| 2.06E-07 | 7.98E-07 | 26.0433222 | 311.194389 | 707.175561 | 521.000298 | 2.37621824 |
| 2.07E-07 | 8.04E-07 | 399.009418 | 1170.70484 | 596.737383 | 882.40656 | 0 |
| 2.08E-07 | 8.05E-07 | 83.9173715 | 58.7243164 | 49.1867516 | 45.929679 | 0 |
| 2.08E-07 | 8.07E-07 | 40.5118345 | 122.787207 | 57.5392189 | 100.674145 | 0 |
| 2.08E-07 | 8.07E-07 | 27.6509347 | 142.58442 | 133.639476 | 126.190633 | 0 |
| 2.08E-07 | 8.09E-07 | 148.543393 | 296.290869 | 235.725187 | 251.91733 | 42.7719284 |
| 2.10E-07 | 8.13E-07 | 35.3674746 | 80.5234945 | 108.582074 | 74.6937204 | 0 |
| 2.10E-07 | 8.13E-07 | 56.9094818 | 57.834554 | 325.746223 | 131.293931 | 0 |
| 2.10E-07 | 8.13E-07 | 31.1876821 | 102.322673 | 97.4454513 | 116.447974 | 0 |
| 2.10E-07 | 8.14E-07 | 50.1575094 | 103.434875 | 43.6184401 | 105.313506 | 0 |
| 2.11E-07 | 8.18E-07 | 33.1168171 | 110.997856 | 143.848047 | 82.1166988 | 0 |
| 2.11E-07 | 8.18E-07 | 37.2966096 | 104.102197 | 63.1075304 | 88.6118049 | 0 |
| 2.11E-07 | 8.18E-07 | 73.6286516 | 153.706449 | 95.5893474 | 122.479144 | 2.37621824 |
| 2.11E-07 | 8.18E-07 | 33.7598621 | 87.6415934 | 81.6685687 | 86.2921241 | 0 |
| 2.12E-07 | 8.20E-07 | 36.0105196 | 125.011613 | 67.7477899 | 115.520102 | 0 |
| 2.12E-07 | 8.20E-07 | 1.60761248 | 2.22440592 | 2.78415575 | 5.10329766 | 26.1384007 |
| 2.12E-07 | 8.21E-07 | 331.489694 | 346.784884 | 271.919212 | 388.778495 | 0 |

| | | | | | | |
|---|---|---|---|---|---|---|
| 2.13E-07 | 8.24E-07 | 58.1955718 | 53.8306234 | 40.8342843 | 74.2297842 | 0 |
| 2.13E-07 | 8.24E-07 | 29.9015921 | 136.800964 | 105.797919 | 111.344676 | 0 |
| 2.13E-07 | 8.25E-07 | 0 | 0 | 0 | 0 | 16.6335277 |
| 2.13E-07 | 8.25E-07 | 29.9015921 | 120.785242 | 135.49558 | 100.210209 | 0 |
| 2.13E-07 | 8.25E-07 | 130.216611 | 630.396639 | 683.046211 | 790.083266 | 19.0097459 |
| 2.14E-07 | 8.27E-07 | 43.0840145 | 161.936751 | 55.683115 | 134.541484 | 0 |
| 2.14E-07 | 8.28E-07 | 35.6889971 | 68.0668213 | 90.0210359 | 79.797018 | 0 |
| 2.14E-07 | 8.29E-07 | 51.1220769 | 56.4999105 | 65.8916861 | 41.7542536 | 0 |
| 2.14E-07 | 8.29E-07 | 33.1168171 | 81.4132568 | 97.4454513 | 93.2511664 | 0 |
| 2.15E-07 | 8.30E-07 | 200.95156 | 406.398962 | 354.515832 | 373.932538 | 64.1578926 |
| 2.15E-07 | 8.31E-07 | 72.3425616 | 39.816866 | 53.8270112 | 68.6625504 | 0 |
| 2.15E-07 | 8.33E-07 | 30.2231146 | 109.218331 | 103.013763 | 118.303719 | 0 |
| 2.15E-07 | 8.34E-07 | 74.5932191 | 56.7223511 | 260.782589 | 74.6937204 | 0 |
| 2.16E-07 | 8.34E-07 | 49.5144644 | 70.5136678 | 44.546492 | 52.8887212 | 0 |
| 2.16E-07 | 8.35E-07 | 54.3373018 | 56.4999105 | 41.7623363 | 51.4969128 | 0 |
| 2.16E-07 | 8.35E-07 | 34.7244296 | 119.228158 | 71.4599976 | 107.169251 | 0 |
| 2.16E-07 | 8.36E-07 | 1095.10562 | 494.485437 | 412.983103 | 478.318172 | 7.12865473 |
| 2.16E-07 | 8.36E-07 | 31.5092046 | 119.228158 | 160.552982 | 90.4675495 | 0 |
| 2.17E-07 | 8.37E-07 | 68.1627692 | 78.0766479 | 60.3233746 | 82.1166988 | 2.37621824 |
| 2.17E-07 | 8.39E-07 | 25.4002772 | 136.356083 | 189.322591 | 156.346483 | 0 |
| 2.18E-07 | 8.41E-07 | 54.0157793 | 65.6199748 | 45.4745439 | 46.3936151 | 0 |
| 2.18E-07 | 8.41E-07 | 34.4029071 | 71.8483114 | 115.078438 | 107.169251 | 0 |
| 2.18E-07 | 8.43E-07 | 75.2362641 | 82.5254598 | 90.0210359 | 120.159463 | 7.12865473 |
| 2.18E-07 | 8.43E-07 | 35.3674746 | 84.5274251 | 205.099474 | 112.272549 | 0 |
| 2.19E-07 | 8.44E-07 | 80.7021465 | 159.045024 | 32.4818171 | 129.438186 | 0 |
| 2.19E-07 | 8.44E-07 | 36.3320421 | 110.997856 | 66.819738 | 131.757867 | 0 |
| 2.20E-07 | 8.48E-07 | 43.0840145 | 67.8443807 | 48.2586997 | 58.4559551 | 0 |
| 2.20E-07 | 8.51E-07 | 0 | 0 | 0 | 0 | 11.8810912 |
| 2.21E-07 | 8.52E-07 | 40.190312 | 53.6081828 | 59.3953227 | 61.7035081 | 0 |
| 2.21E-07 | 8.52E-07 | 222.81509 | 300.2948 | 275.631419 | 317.796264 | 33.2670554 |
| 2.21E-07 | 8.54E-07 | 33.4383396 | 79.8561727 | 86.3088283 | 100.210209 | 0 |
| 2.22E-07 | 8.55E-07 | 33.7598621 | 79.4112915 | 84.4527244 | 84.9003157 | 0 |
| 2.22E-07 | 8.56E-07 | 26.3648447 | 138.580489 | 133.639476 | 158.202228 | 0 |
| 2.22E-07 | 8.57E-07 | 36.6535646 | 61.8384847 | 87.2368802 | 87.2199964 | 0 |
| 2.23E-07 | 8.59E-07 | 58.5170943 | 82.0805786 | 77.0283091 | 37.5788283 | 0 |
| 2.23E-07 | 8.61E-07 | 41.1548795 | 106.549044 | 105.797919 | 53.3526574 | 0 |
| 2.24E-07 | 8.64E-07 | 29.9015921 | 106.104163 | 116.00649 | 106.705315 | 0 |
| 2.24E-07 | 8.65E-07 | 137.611628 | 121.897445 | 182.826228 | 26.9082968 | 0 |
| 2.25E-07 | 8.66E-07 | 67.8412467 | 85.1947469 | 69.6038938 | 90.4675495 | 9.50487297 |
| 2.25E-07 | 8.68E-07 | 35.0459521 | 73.1829549 | 73.3161014 | 85.3642518 | 0 |
| 2.26E-07 | 8.71E-07 | 27.0078897 | 146.143469 | 120.646749 | 144.748079 | 0 |
| 2.26E-07 | 8.72E-07 | 62.3753642 | 175.283187 | 154.056618 | 170.264568 | 4.75243649 |
| 2.27E-07 | 8.74E-07 | 41.476402 | 89.6435588 | 109.510126 | 52.8887212 | 0 |

| | | | | | | |
|---|---|---|---|---|---|---|
| 2.27E-07 | 8.76E-07 | 36.0105196 | 69.1790242 | 68.6758419 | 84.4363795 | 0 |
| 2.28E-07 | 8.79E-07 | 54.9803468 | 17.7952474 | 32.4818171 | 28.7640414 | 178.216368 |
| 2.28E-07 | 8.80E-07 | 69.7703817 | 99.4309448 | 35.2659728 | 134.541484 | 0 |
| 2.29E-07 | 8.81E-07 | 37.2966096 | 71.1809896 | 62.1794784 | 64.487125 | 0 |
| 2.30E-07 | 8.85E-07 | 31.5092046 | 104.324638 | 88.1649321 | 131.293931 | 0 |
| 2.30E-07 | 8.87E-07 | 38.904222 | 62.0609253 | 63.1075304 | 55.6723382 | 0 |
| 2.31E-07 | 8.91E-07 | 756.542433 | 2112.29587 | 1509.01242 | 1775.94759 | 7.12865473 |
| 2.32E-07 | 8.92E-07 | 34.0813846 | 96.5392171 | 98.3735032 | 70.9822312 | 0 |
| 2.32E-07 | 8.92E-07 | 68.8058142 | 158.600142 | 166.121293 | 167.480951 | 0 |
| 2.32E-07 | 8.95E-07 | 502.861184 | 437.985527 | 473.306478 | 405.01626 | 38.0194919 |
| 2.33E-07 | 8.95E-07 | 58.1955718 | 73.4053955 | 41.7623363 | 50.1051043 | 0 |
| 2.33E-07 | 8.96E-07 | 31.8307271 | 110.108093 | 108.582074 | 82.5806349 | 0 |
| 2.33E-07 | 8.96E-07 | 32.7952946 | 90.0884399 | 90.9490879 | 81.1888265 | 0 |
| 2.34E-07 | 8.99E-07 | 38.261177 | 112.332499 | 58.4672708 | 99.7462725 | 0 |
| 2.34E-07 | 9.02E-07 | 54.6588243 | 166.163123 | 155.912722 | 186.966269 | 0 |
| 2.35E-07 | 9.03E-07 | 44.3701045 | 66.2872965 | 100.229607 | 50.5690405 | 0 |
| 2.35E-07 | 9.06E-07 | 34.0813846 | 110.552974 | 84.4527244 | 75.6215927 | 0 |
| 2.36E-07 | 9.07E-07 | 36.9750871 | 64.0628906 | 64.9636342 | 64.9510612 | 0 |
| 2.36E-07 | 9.07E-07 | 32.4737721 | 86.7518311 | 95.5893474 | 84.4363795 | 0 |
| 2.36E-07 | 9.09E-07 | 38.904222 | 74.9624797 | 59.3953227 | 57.9920189 | 0 |
| 2.38E-07 | 9.13E-07 | 34.0813846 | 78.2990885 | 74.2441534 | 87.6839326 | 0 |
| 2.38E-07 | 9.15E-07 | 33.7598621 | 95.2045736 | 72.3880495 | 107.633187 | 0 |
| 2.39E-07 | 9.18E-07 | 27.9724572 | 174.615865 | 104.869867 | 142.892335 | 0 |
| 2.39E-07 | 9.19E-07 | 45.0131495 | 69.4014648 | 131.783372 | 57.5280828 | 0 |
| 2.39E-07 | 9.19E-07 | 32.1522496 | 123.232088 | 78.8844129 | 116.91191 | 0 |
| 2.39E-07 | 9.19E-07 | 43.0840145 | 68.7341431 | 48.2586997 | 55.208402 | 0 |
| 2.39E-07 | 9.20E-07 | 31.8307271 | 98.763623 | 86.3088283 | 93.2511664 | 0 |
| 2.40E-07 | 9.21E-07 | 60.1247068 | 50.2715739 | 62.1794784 | 146.139888 | 0 |
| 2.40E-07 | 9.22E-07 | 23.7926647 | 175.283187 | 140.135839 | 202.276162 | 0 |
| 2.41E-07 | 9.24E-07 | 56.2664368 | 89.8659993 | 38.0501286 | 101.138081 | 0 |
| 2.41E-07 | 9.26E-07 | 35.3674746 | 88.5313558 | 69.6038938 | 70.9822312 | 0 |
| 2.41E-07 | 9.26E-07 | 40.833357 | 56.0550293 | 66.819738 | 51.0329766 | 0 |
| 2.41E-07 | 9.26E-07 | 34.7244296 | 69.846346 | 77.0283091 | 77.0134011 | 0 |
| 2.42E-07 | 9.27E-07 | 58.5170943 | 153.706449 | 143.848047 | 151.243185 | 0 |
| 2.42E-07 | 9.28E-07 | 40.5118345 | 67.1770589 | 50.1148035 | 61.7035081 | 0 |
| 2.43E-07 | 9.31E-07 | 34.7244296 | 124.121851 | 77.956361 | 80.7248903 | 0 |
| 2.43E-07 | 9.34E-07 | 56.2664368 | 157.932821 | 163.337137 | 147.531696 | 2.37621824 |
| 2.44E-07 | 9.36E-07 | 57.2310043 | 154.151331 | 45.4745439 | 80.7248903 | 0 |
| 2.44E-07 | 9.37E-07 | 38.261177 | 58.5018758 | 59.3953227 | 72.8379758 | 0 |
| 2.44E-07 | 9.37E-07 | 37.2966096 | 60.7262817 | 62.1794784 | 66.3428696 | 0 |
| 2.45E-07 | 9.38E-07 | 154.652321 | 274.714132 | 241.293498 | 254.237011 | 19.0097459 |
| 2.46E-07 | 9.43E-07 | 61.4107968 | 158.377702 | 152.200514 | 186.966269 | 7.12865473 |
| 2.46E-07 | 9.43E-07 | 39.547267 | 98.0963013 | 53.8270112 | 102.993826 | 0 |

| | | | | | | |
|---|---|---|---|---|---|---|
| 2.46E-07 | 9.43E-07 | 54.0157793 | 259.14329 | 151.272462 | 178.151482 | 2.37621824 |
| 2.46E-07 | 9.44E-07 | 46.9422844 | 152.149365 | 51.9709073 | 89.5396772 | 0 |
| 2.46E-07 | 9.45E-07 | 14.1469898 | 6.22833659 | 7.42441534 | 11.1344676 | 66.5341108 |
| 2.47E-07 | 9.46E-07 | 34.0813846 | 76.0746826 | 73.3161014 | 86.7560603 | 0 |
| 2.47E-07 | 9.47E-07 | 35.3674746 | 146.810791 | 68.6758419 | 111.808612 | 0 |
| 2.47E-07 | 9.48E-07 | 39.2257445 | 65.3975342 | 70.5319457 | 51.9608489 | 0 |
| 2.47E-07 | 9.48E-07 | 95.4921813 | 203.088261 | 211.595837 | 232.895948 | 14.2573095 |
| 2.47E-07 | 9.48E-07 | 32.7952946 | 103.879757 | 84.4527244 | 80.2609542 | 0 |
| 2.48E-07 | 9.52E-07 | 35.6889971 | 67.1770589 | 76.1002572 | 102.065953 | 0 |
| 2.49E-07 | 9.53E-07 | 379.718068 | 685.561906 | 597.665435 | 701.471461 | 0 |
| 2.49E-07 | 9.53E-07 | 83.595849 | 66.064856 | 38.9781805 | 53.8165936 | 0 |
| 2.49E-07 | 9.53E-07 | 41.7979245 | 56.9447917 | 48.2586997 | 55.6723382 | 0 |
| 2.49E-07 | 9.53E-07 | 23.1496197 | 199.08433 | 159.62493 | 157.738291 | 0 |
| 2.49E-07 | 9.54E-07 | 44.3701045 | 77.8542074 | 44.546492 | 68.6625504 | 0 |
| 2.49E-07 | 9.55E-07 | 40.5118345 | 104.99196 | 78.8844129 | 53.8165936 | 0 |
| 2.50E-07 | 9.57E-07 | 29.5800696 | 125.234054 | 92.8051917 | 123.407016 | 0 |
| 2.50E-07 | 9.57E-07 | 43.0840145 | 52.4959798 | 66.819738 | 102.993826 | 0 |
| 2.50E-07 | 9.57E-07 | 60.4462293 | 69.6239054 | 64.9636342 | 36.650956 | 0 |
| 2.50E-07 | 9.58E-07 | 33.7598621 | 75.4073608 | 98.3735032 | 77.4773373 | 0 |
| 2.51E-07 | 9.62E-07 | 58.1955718 | 249.800785 | 242.22155 | 185.110524 | 7.12865473 |
| 2.51E-07 | 9.62E-07 | 29.5800696 | 107.661247 | 112.294282 | 98.8184002 | 0 |
| 2.52E-07 | 9.63E-07 | 37.2966096 | 62.0609253 | 64.9636342 | 59.3838274 | 0 |
| 2.52E-07 | 9.63E-07 | 26.3648447 | 116.113989 | 163.337137 | 143.820207 | 0 |
| 2.52E-07 | 9.65E-07 | 31.1876821 | 101.655351 | 84.4527244 | 114.128293 | 0 |
| 2.53E-07 | 9.66E-07 | 44.691627 | 62.9506877 | 44.546492 | 77.0134011 | 0 |
| 2.53E-07 | 9.67E-07 | 34.0813846 | 176.61783 | 74.2441534 | 127.118505 | 0 |
| 2.53E-07 | 9.67E-07 | 33.7598621 | 88.5313558 | 88.1649321 | 70.0543588 | 0 |
| 2.54E-07 | 9.70E-07 | 65.2690667 | 114.779346 | 123.430905 | 97.8905279 | 7.12865473 |
| 2.54E-07 | 9.71E-07 | 35.6889971 | 72.0707519 | 63.1075304 | 81.1888265 | 0 |
| 2.54E-07 | 9.73E-07 | 108.031559 | 239.123637 | 155.912722 | 174.439993 | 21.3859642 |
| 2.55E-07 | 9.74E-07 | 54.3373018 | 72.9605143 | 38.0501286 | 62.1674443 | 0 |
| 2.55E-07 | 9.74E-07 | 48.8714194 | 65.3975342 | 96.5173994 | 44.0739344 | 0 |
| 2.55E-07 | 9.76E-07 | 32.1522496 | 93.86993 | 121.574801 | 80.7248903 | 0 |
| 2.56E-07 | 9.77E-07 | 43.405537 | 62.9506877 | 49.1867516 | 49.177232 | 0 |
| 2.56E-07 | 9.78E-07 | 48.2283744 | 62.2833659 | 44.546492 | 47.7854236 | 0 |
| 2.57E-07 | 9.82E-07 | 29.5800696 | 188.184741 | 95.5893474 | 128.97425 | 0 |
| 2.57E-07 | 9.82E-07 | 36.0105196 | 79.8561727 | 60.3233746 | 86.2921241 | 0 |
| 2.58E-07 | 9.84E-07 | 32.7952946 | 95.6494547 | 78.8844129 | 81.1888265 | 0 |
| 2.58E-07 | 9.85E-07 | 43.0840145 | 218.881543 | 320.177911 | 57.9920189 | 0 |
| 2.58E-07 | 9.85E-07 | 450.77454 | 432.424512 | 524.349333 | 501.978916 | 35.6432737 |
| 2.59E-07 | 9.87E-07 | 35.0459521 | 70.0687866 | 68.6758419 | 70.518295 | 0 |
| 2.60E-07 | 9.93E-07 | 38.904222 | 71.4034302 | 290.48025 | 149.387441 | 0 |
| 2.60E-07 | 9.94E-07 | 95.4921813 | 72.9605143 | 51.9709073 | 40.3624452 | 0 |

| | | | | | | |
|---|---|---|---|---|---|---|
| 2.60E-07 | 9.94E-07 | 33.1168171 | 82.5254598 | 79.8124649 | 77.0134011 | 0 |
| 2.61E-07 | 9.98E-07 | 36.3320421 | 64.7302124 | 61.2514265 | 69.5904227 | 0 |
| 2.62E-07 | 1.00E-06 | 35.3674746 | 98.0963013 | 62.1794784 | 94.1790387 | 0 |
| 2.62E-07 | 1.00E-06 | 43.7270595 | 238.233874 | 53.8270112 | 210.163077 | 0 |
| 2.63E-07 | 1.00E-06 | 62.6968867 | 51.3837769 | 42.6903882 | 47.3214874 | 0 |
| 2.63E-07 | 1.00E-06 | 29.2585471 | 105.2144 | 111.36623 | 100.674145 | 0 |
| 2.63E-07 | 1.00E-06 | 35.6889971 | 66.5097371 | 64.9636342 | 82.1166988 | 0 |
| 2.63E-07 | 1.00E-06 | 90.6693439 | 107.216366 | 35.2659728 | 62.1674443 | 0 |
| 2.65E-07 | 1.01E-06 | 57.2310043 | 70.2912272 | 37.1220767 | 59.3838274 | 0 |
| 2.65E-07 | 1.01E-06 | 261.397789 | 457.115417 | 322.034015 | 392.026048 | 0 |
| 2.65E-07 | 1.01E-06 | 42.4409695 | 115.224227 | 49.1867516 | 102.993826 | 0 |
| 2.66E-07 | 1.01E-06 | 37.2966096 | 55.8325887 | 65.8916861 | 71.9101035 | 0 |
| 2.66E-07 | 1.02E-06 | 55.9449143 | 46.7125244 | 63.1075304 | 41.7542536 | 0 |
| 2.67E-07 | 1.02E-06 | 50.4790319 | 50.7164551 | 40.8342843 | 61.239572 | 0 |
| 2.67E-07 | 1.02E-06 | 0 | 0 | 0 | 0 | 49.9005831 |
| 2.67E-07 | 1.02E-06 | 497.716824 | 72.9605143 | 52.8989593 | 132.685739 | 0 |
| 2.67E-07 | 1.02E-06 | 33.4383396 | 185.293013 | 90.9490879 | 94.6429749 | 0 |
| 2.68E-07 | 1.02E-06 | 32.7952946 | 79.6337321 | 77.0283091 | 89.0757411 | 0 |
| 2.69E-07 | 1.02E-06 | 37.2966096 | 59.8365194 | 75.1722053 | 58.9198912 | 0 |
| 2.69E-07 | 1.02E-06 | 25.4002772 | 159.045024 | 121.574801 | 148.923505 | 0 |
| 2.69E-07 | 1.03E-06 | 412.19184 | 695.571733 | 514.140762 | 609.612103 | 0 |
| 2.70E-07 | 1.03E-06 | 32.4737721 | 114.334465 | 76.1002572 | 91.859358 | 0 |
| 2.70E-07 | 1.03E-06 | 199.66547 | 376.814364 | 417.623363 | 389.706367 | 28.5146189 |
| 2.71E-07 | 1.03E-06 | 58.1955718 | 109.885653 | 101.157659 | 121.551272 | 9.50487297 |
| 2.71E-07 | 1.03E-06 | 25.7217997 | 137.690727 | 128.071165 | 137.325101 | 0 |
| 2.71E-07 | 1.03E-06 | 70.4134267 | 120.562801 | 90.9490879 | 121.551272 | 0 |
| 2.72E-07 | 1.03E-06 | 45.0131495 | 67.6219401 | 46.4025958 | 96.9626556 | 0 |
| 2.73E-07 | 1.04E-06 | 28.2939797 | 94.0923706 | 145.704151 | 141.03659 | 0 |
| 2.73E-07 | 1.04E-06 | 123.786161 | 266.48383 | 374.932974 | 262.587862 | 16.6335277 |
| 2.74E-07 | 1.04E-06 | 36.6535646 | 61.6160441 | 59.3953227 | 65.8789335 | 0 |
| 2.74E-07 | 1.04E-06 | 32.4737721 | 92.9801676 | 74.2441534 | 88.6118049 | 0 |
| 2.74E-07 | 1.04E-06 | 35.3674746 | 67.1770589 | 64.9636342 | 67.2707419 | 0 |
| 2.74E-07 | 1.04E-06 | 29.2585471 | 94.5372518 | 128.071165 | 106.241379 | 0 |
| 2.74E-07 | 1.04E-06 | 9.96719738 | 10.4547078 | 8.35246725 | 12.5262761 | 111.682257 |
| 2.75E-07 | 1.05E-06 | 28.9370247 | 115.001786 | 110.438178 | 96.9626556 | 0 |
| 2.76E-07 | 1.05E-06 | 36.3320421 | 62.9506877 | 64.9636342 | 60.3116997 | 0 |
| 2.76E-07 | 1.05E-06 | 35.0459521 | 91.6455241 | 76.1002572 | 63.5592527 | 0 |
| 2.77E-07 | 1.05E-06 | 61.7323193 | 130.350187 | 104.869867 | 107.169251 | 0 |
| 2.77E-07 | 1.05E-06 | 32.1522496 | 84.7498657 | 80.7405168 | 82.5806349 | 0 |
| 2.77E-07 | 1.05E-06 | 71.6995166 | 76.7420044 | 37.1220767 | 53.3526574 | 0 |
| 2.78E-07 | 1.06E-06 | 40.5118345 | 82.970341 | 80.7405168 | 48.2493597 | 0 |
| 2.78E-07 | 1.06E-06 | 134.074881 | 84.9723063 | 33.409869 | 76.549465 | 0 |
| 2.78E-07 | 1.06E-06 | 32.4737721 | 104.324638 | 99.3015551 | 71.9101035 | 0 |

| | | | | | | |
|---|---|---|---|---|---|---|
| 2.78E-07 | 1.06E-06 | 380.361113 | 162.159192 | 161.481034 | 63.5592527 | 0 |
| 2.78E-07 | 1.06E-06 | 46.6207619 | 83.4152222 | 48.2586997 | 51.4969128 | 0 |
| 2.78E-07 | 1.06E-06 | 69.4488592 | 96.5392171 | 79.8124649 | 75.1576565 | 11.8810912 |
| 2.78E-07 | 1.06E-06 | 26.0433222 | 129.460425 | 141.991943 | 121.087335 | 0 |
| 2.79E-07 | 1.06E-06 | 27.6509347 | 118.115955 | 105.797919 | 129.438186 | 0 |
| 2.79E-07 | 1.06E-06 | 33.1168171 | 80.7459351 | 74.2441534 | 76.549465 | 0 |
| 2.79E-07 | 1.06E-06 | 38.5826995 | 62.0609253 | 195.818954 | 130.366059 | 0 |
| 2.79E-07 | 1.06E-06 | 27.9724572 | 219.771305 | 108.582074 | 128.510314 | 0 |
| 2.80E-07 | 1.06E-06 | 35.0459521 | 62.0609253 | 90.0210359 | 77.0134011 | 0 |
| 2.80E-07 | 1.06E-06 | 35.3674746 | 77.8542074 | 60.3233746 | 74.2297842 | 0 |
| 2.81E-07 | 1.07E-06 | 77.1653991 | 82.7479004 | 173.545708 | 129.438186 | 2.37621824 |
| 2.82E-07 | 1.07E-06 | 88.4186864 | 82.970341 | 30.6257133 | 97.8905279 | 0 |
| 2.82E-07 | 1.07E-06 | 32.1522496 | 103.212435 | 73.3161014 | 97.8905279 | 0 |
| 2.83E-07 | 1.07E-06 | 51.7651219 | 169.722172 | 60.3233746 | 481.565725 | 0 |
| 2.83E-07 | 1.07E-06 | 48.8714194 | 51.1613363 | 40.8342843 | 53.3526574 | 0 |
| 2.83E-07 | 1.07E-06 | 308.983119 | 603.703768 | 412.983103 | 499.195299 | 26.1384007 |
| 2.84E-07 | 1.08E-06 | 279.724572 | 564.776664 | 496.507776 | 554.403701 | 26.1384007 |
| 2.84E-07 | 1.08E-06 | 45.9777169 | 59.6140788 | 45.4745439 | 46.3936151 | 0 |
| 2.86E-07 | 1.09E-06 | 27.3294122 | 168.609969 | 115.078438 | 111.808612 | 0 |
| 2.87E-07 | 1.09E-06 | 34.4029071 | 67.8443807 | 103.013763 | 75.6215927 | 0 |
| 2.87E-07 | 1.09E-06 | 48.8714194 | 218.881543 | 159.62493 | 200.420417 | 2.37621824 |
| 2.87E-07 | 1.09E-06 | 120.892459 | 608.15258 | 618.082577 | 631.417102 | 0 |
| 2.87E-07 | 1.09E-06 | 30.5446371 | 83.1927816 | 98.3735032 | 115.520102 | 0 |
| 2.88E-07 | 1.09E-06 | 62.3753642 | 38.9271037 | 144.776099 | 83.5085072 | 0 |
| 2.89E-07 | 1.09E-06 | 41.1548795 | 86.0845093 | 138.279736 | 55.208402 | 0 |
| 2.89E-07 | 1.10E-06 | 26.3648447 | 15.1259603 | 25.0574018 | 17.6295737 | 114.058476 |
| 2.90E-07 | 1.10E-06 | 32.7952946 | 74.9624797 | 77.956361 | 77.9412734 | 0 |
| 2.90E-07 | 1.10E-06 | 127.322908 | 62.5058065 | 40.8342843 | 65.4149973 | 0 |
| 2.90E-07 | 1.10E-06 | 79.4160565 | 163.716276 | 121.574801 | 138.252973 | 11.8810912 |
| 2.91E-07 | 1.10E-06 | 29.9015921 | 104.99196 | 156.840774 | 91.3954218 | 0 |
| 2.91E-07 | 1.10E-06 | 42.4409695 | 98.9860636 | 48.2586997 | 71.4461673 | 0 |
| 2.91E-07 | 1.10E-06 | 17.3622148 | 349.899052 | 238.509343 | 191.60563 | 0 |
| 2.92E-07 | 1.10E-06 | 42.762492 | 77.8542074 | 45.4745439 | 61.239572 | 0 |
| 2.92E-07 | 1.10E-06 | 29.2585471 | 145.476147 | 103.013763 | 95.5708472 | 0 |
| 2.93E-07 | 1.11E-06 | 1.92913498 | 739.170089 | 1684.41423 | 1437.73813 | 0 |
| 2.93E-07 | 1.11E-06 | 33.1168171 | 73.8502767 | 87.2368802 | 71.9101035 | 0 |
| 2.94E-07 | 1.11E-06 | 57.2310043 | 69.4014648 | 36.1940248 | 59.3838274 | 0 |
| 2.94E-07 | 1.11E-06 | 31.5092046 | 100.765588 | 157.768826 | 83.5085072 | 0 |
| 2.94E-07 | 1.11E-06 | 2.25065747 | 0.66732178 | 0.92805192 | 0.46393615 | 73.6627655 |
| 2.94E-07 | 1.11E-06 | 46.9422844 | 71.6258708 | 43.6184401 | 51.0329766 | 0 |
| 2.95E-07 | 1.11E-06 | 37.2966096 | 98.3187419 | 55.683115 | 78.4052096 | 0 |
| 2.95E-07 | 1.11E-06 | 56.5879593 | 55.3877075 | 39.9062324 | 89.0757411 | 0 |
| 2.96E-07 | 1.12E-06 | 21.8635297 | 133.019474 | 275.631419 | 207.843396 | 0 |

| | | | | | | |
|---|---|---|---|---|---|---|
| 2.96E-07 | 1.12E-06 | 461.063259 | 591.914416 | 494.651672 | 491.77232 | 0 |
| 2.97E-07 | 1.12E-06 | 48.2283744 | 214.210291 | 191.178695 | 179.079354 | 0 |
| 2.97E-07 | 1.12E-06 | 84.881939 | 161.49187 | 30.6257133 | 104.84957 | 0 |
| 2.97E-07 | 1.12E-06 | 26.6863672 | 126.346257 | 117.862593 | 115.520102 | 0 |
| 2.97E-07 | 1.12E-06 | 66.8766792 | 121.897445 | 47.3306478 | 49.177232 | 0 |
| 2.98E-07 | 1.12E-06 | 83.595849 | 67.3994995 | 37.1220767 | 53.8165936 | 0 |
| 3.00E-07 | 1.13E-06 | 40.190312 | 60.5038411 | 79.8124649 | 49.177232 | 0 |
| 3.01E-07 | 1.14E-06 | 35.0459521 | 66.5097371 | 61.2514265 | 71.9101035 | 0 |
| 3.01E-07 | 1.14E-06 | 63.0184092 | 162.381632 | 104.869867 | 130.829995 | 2.37621824 |
| 3.01E-07 | 1.14E-06 | 34.7244296 | 91.8679647 | 61.2514265 | 79.3330819 | 0 |
| 3.01E-07 | 1.14E-06 | 58.1955718 | 130.795068 | 130.85532 | 189.749886 | 2.37621824 |
| 3.02E-07 | 1.14E-06 | 14.1469898 | 6.89565837 | 16.7049345 | 7.42297842 | 54.6530196 |
| 3.02E-07 | 1.14E-06 | 28.2939797 | 91.6455241 | 125.287009 | 150.779249 | 0 |
| 3.02E-07 | 1.14E-06 | 28.6155022 | 110.552974 | 93.7332436 | 112.736485 | 0 |
| 3.02E-07 | 1.14E-06 | 30.2231146 | 87.4191528 | 95.5893474 | 91.3954218 | 0 |
| 3.02E-07 | 1.14E-06 | 40.190312 | 60.05896 | 46.4025958 | 60.7756358 | 0 |
| 3.03E-07 | 1.14E-06 | 32.7952946 | 69.4014648 | 139.207788 | 115.056166 | 0 |
| 3.03E-07 | 1.14E-06 | 18.9698273 | 220.438627 | 208.811681 | 187.430205 | 0 |
| 3.03E-07 | 1.14E-06 | 117.355711 | 217.76934 | 211.595837 | 242.174671 | 16.6335277 |
| 3.04E-07 | 1.15E-06 | 38.5826995 | 51.3837769 | 54.7550631 | 58.4559551 | 0 |
| 3.04E-07 | 1.15E-06 | 31.1876821 | 128.348222 | 122.502853 | 74.2297842 | 0 |
| 3.05E-07 | 1.15E-06 | 32.1522496 | 119.895479 | 125.287009 | 70.0543588 | 0 |
| 3.05E-07 | 1.15E-06 | 42.4409695 | 84.5274251 | 44.546492 | 85.828188 | 0 |
| 3.05E-07 | 1.15E-06 | 43.405537 | 81.1908162 | 48.2586997 | 119.231591 | 0 |
| 3.07E-07 | 1.16E-06 | 37.2966096 | 74.0727173 | 51.9709073 | 77.4773373 | 0 |
| 3.07E-07 | 1.16E-06 | 45.9777169 | 45.3778809 | 52.8989593 | 44.5378705 | 0 |
| 3.07E-07 | 1.16E-06 | 36.6535646 | 60.9487223 | 55.683115 | 61.239572 | 0 |
| 3.08E-07 | 1.16E-06 | 36.9750871 | 138.135608 | 127.143113 | 57.5280828 | 0 |
| 3.08E-07 | 1.16E-06 | 40.190312 | 59.1691976 | 88.1649321 | 51.9608489 | 0 |
| 3.08E-07 | 1.16E-06 | 28.9370247 | 117.003752 | 98.3735032 | 94.1790387 | 0 |
| 3.08E-07 | 1.16E-06 | 37.9396545 | 62.7282471 | 51.9709073 | 57.0641466 | 0 |
| 3.08E-07 | 1.16E-06 | 67.5197242 | 153.484009 | 115.078438 | 135.00542 | 9.50487297 |
| 3.09E-07 | 1.16E-06 | 32.7952946 | 84.7498657 | 67.7477899 | 85.828188 | 0 |
| 3.09E-07 | 1.16E-06 | 41.1548795 | 60.2814006 | 45.4745439 | 71.4461673 | 0 |
| 3.11E-07 | 1.17E-06 | 85.8465065 | 62.2833659 | 36.1940248 | 132.685739 | 0 |
| 3.11E-07 | 1.17E-06 | 42.762492 | 86.5293905 | 139.207788 | 51.4969128 | 0 |
| 3.11E-07 | 1.17E-06 | 50.4790319 | 70.2912272 | 38.0501286 | 84.4363795 | 0 |
| 3.11E-07 | 1.17E-06 | 35.6889971 | 83.1927816 | 103.941815 | 58.4559551 | 0 |
| 3.12E-07 | 1.17E-06 | 74.2716966 | 109.440771 | 111.36623 | 116.91191 | 11.8810912 |
| 3.12E-07 | 1.17E-06 | 29.5800696 | 101.21047 | 284.911939 | 134.077548 | 0 |
| 3.12E-07 | 1.17E-06 | 64.6260217 | 34.0334106 | 71.4599976 | 84.4363795 | 0 |
| 3.12E-07 | 1.18E-06 | 31.5092046 | 72.2931925 | 99.3015551 | 97.8905279 | 0 |
| 3.13E-07 | 1.18E-06 | 72.3425616 | 119.673039 | 90.9490879 | 130.366059 | 0 |

| | | | | | | |
|---|---|---|---|---|---|---|
| 3.13E-07 | 1.18E-06 | 28.9370247 | 102.989994 | 90.0210359 | 117.839782 | 0 |
| 3.13E-07 | 1.18E-06 | 37.618132 | 50.7164551 | 64.9636342 | 68.6625504 | 0 |
| 3.13E-07 | 1.18E-06 | 64.3044992 | 129.237984 | 98.3735032 | 122.94308 | 2.37621824 |
| 3.13E-07 | 1.18E-06 | 34.7244296 | 62.0609253 | 64.9636342 | 71.4461673 | 0 |
| 3.14E-07 | 1.18E-06 | 149.507961 | 447.328031 | 317.393756 | 375.788283 | 26.1384007 |
| 3.14E-07 | 1.18E-06 | 50.1575094 | 39.1495443 | 55.683115 | 52.8887212 | 0 |
| 3.14E-07 | 1.18E-06 | 45.6561944 | 57.834554 | 72.3880495 | 41.2903175 | 0 |
| 3.14E-07 | 1.18E-06 | 56.5879593 | 218.214221 | 38.9781805 | 228.256586 | 0 |
| 3.15E-07 | 1.18E-06 | 29.9015921 | 135.466321 | 148.488307 | 78.8691457 | 0 |
| 3.16E-07 | 1.19E-06 | 209.95419 | 578.34554 | 551.262839 | 582.703806 | 33.2670554 |
| 3.16E-07 | 1.19E-06 | 127.322908 | 59.8365194 | 44.546492 | 57.9920189 | 0 |
| 3.16E-07 | 1.19E-06 | 39.2257445 | 47.6022868 | 62.1794784 | 68.6625504 | 0 |
| 3.16E-07 | 1.19E-06 | 39.8687895 | 136.578524 | 83.5246725 | 56.6002105 | 0 |
| 3.16E-07 | 1.19E-06 | 274.258689 | 527.406645 | 488.155308 | 519.144553 | 30.8908372 |
| 3.17E-07 | 1.19E-06 | 45.334672 | 40.7066284 | 58.4672708 | 56.6002105 | 0 |
| 3.17E-07 | 1.19E-06 | 42.4409695 | 52.7184204 | 44.546492 | 50.5690405 | 0 |
| 3.17E-07 | 1.19E-06 | 31.1876821 | 96.3167765 | 98.3735032 | 73.7658481 | 0 |
| 3.17E-07 | 1.19E-06 | 89.0617314 | 44.0432373 | 42.6903882 | 77.9412734 | 0 |
| 3.18E-07 | 1.19E-06 | 50.8005544 | 80.0786133 | 123.430905 | 41.7542536 | 0 |
| 3.18E-07 | 1.19E-06 | 33.1168171 | 86.5293905 | 71.4599976 | 69.5904227 | 0 |
| 3.18E-07 | 1.19E-06 | 50.4790319 | 144.141504 | 40.8342843 | 126.654569 | 0 |
| 3.19E-07 | 1.20E-06 | 145.006646 | 58.946757 | 59.3953227 | 48.2493597 | 0 |
| 3.20E-07 | 1.20E-06 | 46.9422844 | 64.952653 | 51.9709073 | 131.293931 | 0 |
| 3.20E-07 | 1.20E-06 | 31.8307271 | 104.547078 | 72.3880495 | 87.2199964 | 0 |
| 3.20E-07 | 1.20E-06 | 33.1168171 | 81.6356974 | 81.6685687 | 64.9510612 | 0 |
| 3.20E-07 | 1.20E-06 | 0 | 0 | 0.92805192 | 0 | 83.1676385 |
| 3.20E-07 | 1.20E-06 | 38.904222 | 111.887618 | 114.150386 | 51.4969128 | 0 |
| 3.20E-07 | 1.20E-06 | 45.0131495 | 68.7341431 | 80.7405168 | 41.2903175 | 0 |
| 3.21E-07 | 1.20E-06 | 27.0078897 | 97.2065389 | 132.711424 | 131.757867 | 0 |
| 3.21E-07 | 1.20E-06 | 0 | 1.33464355 | 4.64025958 | 0 | 80.7914203 |
| 3.21E-07 | 1.20E-06 | 29.2585471 | 100.098267 | 96.5173994 | 90.0036134 | 0 |
| 3.22E-07 | 1.21E-06 | 25.7217997 | 108.106128 | 133.639476 | 168.872759 | 0 |
| 3.22E-07 | 1.21E-06 | 39.2257445 | 47.8247274 | 63.1075304 | 55.6723382 | 0 |
| 3.24E-07 | 1.21E-06 | 239.53426 | 117.003752 | 132.711424 | 139.180845 | 0 |
| 3.25E-07 | 1.22E-06 | 2.89370247 | 2.8917277 | 0.92805192 | 1.85574461 | 40.3957101 |
| 3.25E-07 | 1.22E-06 | 24.4357097 | 117.671073 | 176.329864 | 142.428398 | 0 |
| 3.26E-07 | 1.22E-06 | 91.9554339 | 202.420939 | 190.250643 | 200.884354 | 14.2573095 |
| 3.26E-07 | 1.22E-06 | 45.334672 | 41.8188314 | 71.4599976 | 87.6839326 | 0 |
| 3.26E-07 | 1.22E-06 | 109.317649 | 185.960335 | 253.358173 | 250.061586 | 4.75243649 |
| 3.26E-07 | 1.22E-06 | 28.2939797 | 97.6514201 | 137.351684 | 99.2823364 | 0 |
| 3.27E-07 | 1.22E-06 | 23.7926647 | 148.590316 | 125.287009 | 226.864778 | 0 |
| 3.27E-07 | 1.22E-06 | 34.7244296 | 92.757727 | 59.3953227 | 106.705315 | 0 |
| 3.28E-07 | 1.23E-06 | 30.2231146 | 100.988029 | 77.956361 | 102.065953 | 0 |

| | | | | | | |
|---|---|---|---|---|---|---|
| 3.28E-07 | 1.23E-06 | 60.7677518 | 35.3680542 | 147.560255 | 157.738291 | 0 |
| 3.28E-07 | 1.23E-06 | 138.897718 | 155.485974 | 50.1148035 | 41.2903175 | 0 |
| 3.29E-07 | 1.23E-06 | 32.1522496 | 69.4014648 | 88.1649321 | 80.2609542 | 0 |
| 3.30E-07 | 1.24E-06 | 30.2231146 | 98.3187419 | 79.8124649 | 92.3232941 | 0 |
| 3.31E-07 | 1.24E-06 | 35.6889971 | 104.324638 | 58.4672708 | 78.8691457 | 0 |
| 3.31E-07 | 1.24E-06 | 41.1548795 | 65.3975342 | 48.2586997 | 49.177232 | 0 |
| 3.33E-07 | 1.24E-06 | 149.186438 | 102.989994 | 37.1220767 | 57.9920189 | 0 |
| 3.33E-07 | 1.24E-06 | 34.0813846 | 133.464355 | 130.85532 | 62.1674443 | 0 |
| 3.33E-07 | 1.25E-06 | 43.7270595 | 84.7498657 | 42.6903882 | 67.7346781 | 0 |
| 3.33E-07 | 1.25E-06 | 87.1325964 | 61.3936035 | 90.9490879 | 33.4034029 | 0 |
| 3.33E-07 | 1.25E-06 | 30.2231146 | 119.673039 | 76.1002572 | 109.952868 | 0 |
| 3.33E-07 | 1.25E-06 | 67.1982017 | 189.519385 | 156.840774 | 173.976057 | 0 |
| 3.34E-07 | 1.25E-06 | 19.9343948 | 226.444523 | 193.962851 | 153.09893 | 0 |
| 3.34E-07 | 1.25E-06 | 36.9750871 | 59.8365194 | 51.9709073 | 60.3116997 | 0 |
| 3.35E-07 | 1.25E-06 | 43.405537 | 49.6042521 | 42.6903882 | 56.1362743 | 0 |
| 3.35E-07 | 1.25E-06 | 25.4002772 | 180.844202 | 115.078438 | 124.798825 | 0 |
| 3.35E-07 | 1.25E-06 | 33.4383396 | 84.9723063 | 64.0355823 | 72.8379758 | 0 |
| 3.35E-07 | 1.25E-06 | 36.9750871 | 51.3837769 | 60.3233746 | 64.0231889 | 0 |
| 3.36E-07 | 1.26E-06 | 32.7952946 | 76.2971232 | 67.7477899 | 72.8379758 | 0 |
| 3.36E-07 | 1.26E-06 | 33.4383396 | 85.1947469 | 81.6685687 | 61.7035081 | 0 |
| 3.37E-07 | 1.26E-06 | 33.4383396 | 103.434875 | 63.1075304 | 86.7560603 | 0 |
| 3.38E-07 | 1.26E-06 | 32.4737721 | 71.1809896 | 73.3161014 | 73.3019119 | 0 |
| 3.38E-07 | 1.26E-06 | 29.5800696 | 86.3069499 | 94.6612955 | 90.4675495 | 0 |
| 3.38E-07 | 1.26E-06 | 46.9422844 | 54.053064 | 38.9781805 | 55.208402 | 0 |
| 3.39E-07 | 1.27E-06 | 77.8084441 | 171.501697 | 129.927268 | 141.03659 | 11.8810912 |
| 3.39E-07 | 1.27E-06 | 62.3753642 | 66.9546183 | 36.1940248 | 51.4969128 | 0 |
| 3.40E-07 | 1.27E-06 | 65.2690667 | 175.728068 | 41.7623363 | 75.1576565 | 0 |
| 3.40E-07 | 1.27E-06 | 541.443883 | 1636.49544 | 1225.95658 | 1412.22164 | 47.5243649 |
| 3.40E-07 | 1.27E-06 | 45.334672 | 48.2696086 | 42.6903882 | 47.7854236 | 0 |
| 3.41E-07 | 1.27E-06 | 33.4383396 | 186.850098 | 103.941815 | 75.6215927 | 0 |
| 3.41E-07 | 1.27E-06 | 34.0813846 | 62.5058065 | 77.0283091 | 64.487125 | 0 |
| 3.41E-07 | 1.27E-06 | 23.7926647 | 15.3484009 | 4.64025958 | 12.5262761 | 97.424948 |
| 3.41E-07 | 1.27E-06 | 80.0591015 | 152.149365 | 71.4599976 | 154.490738 | 0 |
| 3.42E-07 | 1.27E-06 | 61.4107968 | 57.3896728 | 61.2514265 | 35.2591475 | 0 |
| 3.42E-07 | 1.28E-06 | 22.1850522 | 129.905306 | 190.250643 | 173.048184 | 0 |
| 3.42E-07 | 1.28E-06 | 22.5065747 | 134.798999 | 168.905449 | 158.666164 | 0 |
| 3.42E-07 | 1.28E-06 | 118.320279 | 124.121851 | 46.4025958 | 42.2181898 | 0 |
| 3.43E-07 | 1.28E-06 | 28.2939797 | 149.480078 | 89.092984 | 112.736485 | 0 |
| 3.43E-07 | 1.28E-06 | 25.4002772 | 172.6139 | 114.150386 | 122.015208 | 0 |
| 3.43E-07 | 1.28E-06 | 25.7217997 | 97.8738607 | 163.337137 | 160.521908 | 0 |
| 3.44E-07 | 1.28E-06 | 63.9829767 | 194.413078 | 121.574801 | 147.995632 | 4.75243649 |
| 3.45E-07 | 1.29E-06 | 32.7952946 | 63.3955688 | 98.3735032 | 90.0036134 | 0 |
| 3.46E-07 | 1.29E-06 | 33.1168171 | 77.6317668 | 63.1075304 | 75.6215927 | 0 |

| | | | | | | |
|---|---|---|---|---|---|---|
| 3.46E-07 | 1.29E-06 | 63.0184092 | 108.551009 | 74.2441534 | 77.4773373 | 7.12865473 |
| 3.47E-07 | 1.29E-06 | 31.8307271 | 87.864034 | 68.6758419 | 103.921698 | 0 |
| 3.47E-07 | 1.29E-06 | 0 | 0 | 0 | 0 | 23.7621824 |
| 3.49E-07 | 1.30E-06 | 37.618132 | 51.1613363 | 54.7550631 | 54.7444659 | 0 |
| 3.50E-07 | 1.30E-06 | 36.9750871 | 62.0609253 | 50.1148035 | 62.6313804 | 0 |
| 3.51E-07 | 1.31E-06 | 44.3701045 | 43.5983561 | 45.4745439 | 52.4247851 | 0 |
| 3.51E-07 | 1.31E-06 | 37.618132 | 94.0923706 | 133.639476 | 55.208402 | 0 |
| 3.52E-07 | 1.31E-06 | 44.048582 | 118.338395 | 43.6184401 | 104.84957 | 0 |
| 3.52E-07 | 1.31E-06 | 63.0184092 | 289.840092 | 155.912722 | 217.122119 | 11.8810912 |
| 3.53E-07 | 1.32E-06 | 30.8661596 | 90.0884399 | 77.0283091 | 78.8691457 | 0 |
| 3.53E-07 | 1.32E-06 | 26.3648447 | 17.7952474 | 18.5610383 | 18.0935099 | 87.920075 |
| 3.54E-07 | 1.32E-06 | 41.7979245 | 197.972127 | 51.9709073 | 132.221803 | 0 |
| 3.55E-07 | 1.32E-06 | 584.527898 | 1160.47257 | 752.650105 | 934.831345 | 35.6432737 |
| 3.55E-07 | 1.32E-06 | 240.498827 | 360.798641 | 325.746223 | 361.406262 | 45.1481466 |
| 3.55E-07 | 1.32E-06 | 323.773154 | 581.682149 | 473.306478 | 586.879231 | 0 |
| 3.56E-07 | 1.32E-06 | 27.9724572 | 102.989994 | 95.5893474 | 99.7462725 | 0 |
| 3.56E-07 | 1.32E-06 | 36.6535646 | 64.7302124 | 57.5392189 | 51.9608489 | 0 |
| 3.57E-07 | 1.33E-06 | 1121.47047 | 1664.74539 | 1356.8119 | 1270.72112 | 0 |
| 3.57E-07 | 1.33E-06 | 28.6155022 | 125.901375 | 90.9490879 | 93.7151026 | 0 |
| 3.57E-07 | 1.33E-06 | 33.1168171 | 87.6415934 | 77.0283091 | 62.1674443 | 0 |
| 3.58E-07 | 1.33E-06 | 27.9724572 | 124.121851 | 85.3807764 | 121.087335 | 0 |
| 3.59E-07 | 1.33E-06 | 25.4002772 | 169.722172 | 128.999216 | 105.777442 | 0 |
| 3.60E-07 | 1.34E-06 | 49.5144644 | 52.2735392 | 103.013763 | 46.8575513 | 0 |
| 3.63E-07 | 1.35E-06 | 34.7244296 | 64.7302124 | 105.797919 | 67.2707419 | 0 |
| 3.64E-07 | 1.35E-06 | 74.2716966 | 50.9388957 | 49.1867516 | 40.3624452 | 0 |
| 3.65E-07 | 1.36E-06 | 67.5197242 | 73.8502767 | 33.409869 | 58.4559551 | 0 |
| 3.66E-07 | 1.36E-06 | 255.931907 | 494.930318 | 580.032448 | 543.733169 | 35.6432737 |
| 3.66E-07 | 1.36E-06 | 33.1168171 | 64.0628906 | 84.4527244 | 112.272549 | 0 |
| 3.67E-07 | 1.36E-06 | 40.5118345 | 55.1652669 | 45.4745439 | 48.7132959 | 0 |
| 3.67E-07 | 1.36E-06 | 2.25065747 | 1.33464355 | 1.85610383 | 1.85574461 | 19.0097459 |
| 3.68E-07 | 1.37E-06 | 29.2585471 | 100.098267 | 87.2368802 | 84.4363795 | 0 |
| 3.69E-07 | 1.37E-06 | 143.399033 | 183.291048 | 146.632203 | 201.812226 | 28.5146189 |
| 3.70E-07 | 1.37E-06 | 20.8989622 | 179.064677 | 163.337137 | 151.707121 | 0 |
| 3.70E-07 | 1.37E-06 | 22.1850522 | 122.787207 | 189.322591 | 173.512121 | 0 |
| 3.70E-07 | 1.37E-06 | 51.4435994 | 176.61783 | 131.783372 | 167.480951 | 2.37621824 |
| 3.70E-07 | 1.37E-06 | 35.0459521 | 74.9624797 | 53.8270112 | 80.7248903 | 0 |
| 3.71E-07 | 1.38E-06 | 27.6509347 | 86.9742716 | 115.078438 | 135.469356 | 0 |
| 3.71E-07 | 1.38E-06 | 30.5446371 | 90.7557617 | 90.9490879 | 71.9101035 | 0 |
| 3.73E-07 | 1.38E-06 | 42.762492 | 103.657316 | 58.4672708 | 50.5690405 | 0 |
| 3.73E-07 | 1.39E-06 | 209.95419 | 117.893514 | 21.3451941 | 241.246799 | 0 |
| 3.74E-07 | 1.39E-06 | 27.3294122 | 84.7498657 | 198.60311 | 161.449781 | 0 |
| 3.75E-07 | 1.39E-06 | 27.3294122 | 108.106128 | 107.654022 | 93.2511664 | 0 |
| 3.77E-07 | 1.40E-06 | 30.2231146 | 81.1908162 | 83.5246725 | 78.8691457 | 0 |

| | | | | | | |
|---|---|---|---|---|---|---|
| 3.77E-07 | 1.40E-06 | 22.8280972 | 214.210291 | 118.790645 | 158.202228 | 0 |
| 3.79E-07 | 1.40E-06 | 68.4842917 | 58.7243164 | 36.1940248 | 50.1051043 | 0 |
| 3.80E-07 | 1.41E-06 | 33.7598621 | 92.0904053 | 97.4454513 | 57.5280828 | 0 |
| 3.81E-07 | 1.41E-06 | 31.5092046 | 116.33643 | 74.2441534 | 78.8691457 | 0 |
| 3.81E-07 | 1.41E-06 | 36.6535646 | 54.4979451 | 66.819738 | 96.0347833 | 0 |
| 3.81E-07 | 1.41E-06 | 29.5800696 | 118.783276 | 78.8844129 | 91.859358 | 0 |
| 3.81E-07 | 1.41E-06 | 36.9750871 | 50.2715739 | 53.8270112 | 58.9198912 | 0 |
| 3.81E-07 | 1.41E-06 | 39.547267 | 65.6199748 | 50.1148035 | 100.210209 | 0 |
| 3.81E-07 | 1.41E-06 | 36.9750871 | 64.0628906 | 48.2586997 | 61.7035081 | 0 |
| 3.82E-07 | 1.41E-06 | 26.0433222 | 105.659281 | 106.72597 | 139.180845 | 0 |
| 3.82E-07 | 1.42E-06 | 0 | 0 | 0 | 0 | 2.37621824 |
| 3.83E-07 | 1.42E-06 | 32.4737721 | 68.0668213 | 243.149602 | 169.800631 | 0 |
| 3.83E-07 | 1.42E-06 | 33.4383396 | 119.228158 | 60.3233746 | 99.2823364 | 0 |
| 3.83E-07 | 1.42E-06 | 26.3648447 | 119.673039 | 119.718697 | 96.0347833 | 0 |
| 3.83E-07 | 1.42E-06 | 28.2939797 | 84.9723063 | 97.4454513 | 109.488932 | 0 |
| 3.84E-07 | 1.42E-06 | 66.2336342 | 96.7616577 | 97.4454513 | 95.5708472 | 2.37621824 |
| 3.85E-07 | 1.42E-06 | 30.2231146 | 77.8542074 | 90.0210359 | 77.9412734 | 0 |
| 3.85E-07 | 1.43E-06 | 145.971213 | 270.48776 | 347.091417 | 277.433818 | 19.0097459 |
| 3.86E-07 | 1.43E-06 | 176.837373 | 442.87922 | 347.091417 | 388.314559 | 21.3859642 |
| 3.86E-07 | 1.43E-06 | 38.5826995 | 63.3955688 | 77.956361 | 46.3936151 | 0 |
| 3.87E-07 | 1.43E-06 | 197.09329 | 293.399141 | 264.494796 | 255.628819 | 21.3859642 |
| 3.88E-07 | 1.44E-06 | 30.2231146 | 81.1908162 | 76.1002572 | 84.4363795 | 0 |
| 3.88E-07 | 1.44E-06 | 41.7979245 | 48.047168 | 171.689605 | 135.469356 | 0 |
| 3.88E-07 | 1.44E-06 | 136.325538 | 168.609969 | 79.8124649 | 70.518295 | 0 |
| 3.89E-07 | 1.44E-06 | 50.4790319 | 166.830444 | 119.718697 | 40.3624452 | 0 |
| 3.90E-07 | 1.44E-06 | 72.3425616 | 201.308736 | 204.171422 | 96.9626556 | 0 |
| 3.90E-07 | 1.44E-06 | 101.279586 | 166.163123 | 125.287009 | 115.056166 | 14.2573095 |
| 3.91E-07 | 1.45E-06 | 44.048582 | 69.846346 | 44.546492 | 46.8575513 | 0 |
| 3.92E-07 | 1.45E-06 | 326.345334 | 86.9742716 | 54.7550631 | 63.5592527 | 0 |
| 3.93E-07 | 1.45E-06 | 84.5604165 | 265.816508 | 327.602327 | 273.722329 | 0 |
| 3.93E-07 | 1.45E-06 | 47.9068519 | 32.0314453 | 36.1940248 | 30.619786 | 111.682257 |
| 3.93E-07 | 1.45E-06 | 30.8661596 | 101.21047 | 70.5319457 | 82.1166988 | 0 |
| 3.93E-07 | 1.45E-06 | 0 | 0 | 0 | 0 | 71.2865473 |
| 3.94E-07 | 1.46E-06 | 51.1220769 | 167.275326 | 149.416359 | 146.139888 | 0 |
| 3.94E-07 | 1.46E-06 | 28.6155022 | 104.324638 | 94.6612955 | 81.6527626 | 0 |
| 3.96E-07 | 1.46E-06 | 25.4002772 | 144.363944 | 108.582074 | 109.952868 | 0 |
| 3.97E-07 | 1.47E-06 | 48.2283744 | 93.86993 | 38.9781805 | 65.8789335 | 0 |
| 3.97E-07 | 1.47E-06 | 180.37412 | 815.689652 | 733.161014 | 688.017312 | 23.7621824 |
| 4.00E-07 | 1.48E-06 | 23.4711422 | 107.438806 | 161.481034 | 168.408823 | 0 |
| 4.00E-07 | 1.48E-06 | 43.0840145 | 61.6160441 | 40.8342843 | 78.4052096 | 0 |
| 4.00E-07 | 1.48E-06 | 32.1522496 | 84.0825439 | 67.7477899 | 122.94308 | 0 |
| 4.01E-07 | 1.48E-06 | 24.1141872 | 171.946578 | 108.582074 | 134.541484 | 0 |
| 4.02E-07 | 1.48E-06 | 103.851766 | 33.1436483 | 72.3880495 | 64.487125 | 0 |

| | | | | | | |
|---|---|---|---|---|---|---|
| 4.02E-07 | 1.48E-06 | 59.4816618 | 58.7243164 | 38.0501286 | 109.952868 | 0 |
| 4.02E-07 | 1.48E-06 | 38.261177 | 46.7125244 | 67.7477899 | 54.2805297 | 0 |
| 4.03E-07 | 1.49E-06 | 35.6889971 | 54.7203857 | 52.8989593 | 61.7035081 | 0 |
| 4.03E-07 | 1.49E-06 | 57.5525268 | 52.940861 | 41.7623363 | 40.8263813 | 0 |
| 4.04E-07 | 1.49E-06 | 95.8137038 | 117.003752 | 86.3088283 | 70.0543588 | 9.50487297 |
| 4.04E-07 | 1.49E-06 | 27.9724572 | 85.8620687 | 107.654022 | 94.6429749 | 0 |
| 4.05E-07 | 1.49E-06 | 261.397789 | 579.012862 | 490.939464 | 549.300403 | 26.1384007 |
| 4.05E-07 | 1.49E-06 | 40.833357 | 61.3936035 | 45.4745439 | 46.8575513 | 0 |
| 4.06E-07 | 1.50E-06 | 27.3294122 | 127.013578 | 90.0210359 | 100.210209 | 0 |
| 4.07E-07 | 1.50E-06 | 27.6509347 | 169.277291 | 78.8844129 | 166.553078 | 0 |
| 4.07E-07 | 1.50E-06 | 48.2283744 | 63.1731283 | 65.8916861 | 36.1870198 | 0 |
| 4.08E-07 | 1.50E-06 | 503.182706 | 85.4171875 | 46.4025958 | 104.84957 | 0 |
| 4.08E-07 | 1.50E-06 | 39.8687895 | 50.7164551 | 49.1867516 | 44.5378705 | 0 |
| 4.08E-07 | 1.51E-06 | 87.4541189 | 192.633553 | 93.7332436 | 230.112331 | 9.50487297 |
| 4.10E-07 | 1.51E-06 | 74.2716966 | 110.775415 | 74.2441534 | 96.9626556 | 2.37621824 |
| 4.11E-07 | 1.51E-06 | 39.8687895 | 53.1633016 | 45.4745439 | 46.8575513 | 0 |
| 4.11E-07 | 1.52E-06 | 40.833357 | 46.934965 | 43.6184401 | 53.3526574 | 0 |
| 4.11E-07 | 1.52E-06 | 59.8031843 | 117.226192 | 117.862593 | 104.84957 | 0 |
| 4.13E-07 | 1.52E-06 | 31.8307271 | 75.852242 | 64.0355823 | 72.3740396 | 0 |
| 4.14E-07 | 1.52E-06 | 118.641801 | 147.478113 | 204.171422 | 211.090949 | 0 |
| 4.14E-07 | 1.53E-06 | 37.9396545 | 49.3818115 | 50.1148035 | 66.3428696 | 0 |
| 4.15E-07 | 1.53E-06 | 28.6155022 | 93.6474894 | 86.3088283 | 83.9724434 | 0 |
| 4.15E-07 | 1.53E-06 | 133.110313 | 102.767554 | 64.0355823 | 96.0347833 | 2.37621824 |
| 4.15E-07 | 1.53E-06 | 29.9015921 | 97.4289795 | 70.5319457 | 92.3232941 | 0 |
| 4.16E-07 | 1.53E-06 | 34.4029071 | 58.5018758 | 163.337137 | 128.046378 | 0 |
| 4.17E-07 | 1.53E-06 | 32.7952946 | 77.6317668 | 64.0355823 | 62.1674443 | 0 |
| 4.17E-07 | 1.54E-06 | 53.6942569 | 76.7420044 | 65.8916861 | 34.3312752 | 0 |
| 4.18E-07 | 1.54E-06 | 81.023669 | 61.1711629 | 38.0501286 | 46.8575513 | 0 |
| 4.19E-07 | 1.54E-06 | 116.391144 | 28.0275146 | 78.8844129 | 116.447974 | 0 |
| 4.21E-07 | 1.55E-06 | 73.9501741 | 157.932821 | 29.6976613 | 107.169251 | 0 |
| 4.22E-07 | 1.55E-06 | 136.325538 | 183.291048 | 179.11402 | 186.966269 | 28.5146189 |
| 4.22E-07 | 1.55E-06 | 69.4488592 | 30.02948 | 106.72597 | 94.1790387 | 0 |
| 4.24E-07 | 1.56E-06 | 26.3648447 | 134.131677 | 98.3735032 | 99.7462725 | 0 |
| 4.24E-07 | 1.56E-06 | 22.5065747 | 11.7893514 | 18.5610383 | 12.9902122 | 85.5438568 |
| 4.25E-07 | 1.56E-06 | 33.1168171 | 69.846346 | 63.1075304 | 58.9198912 | 0 |
| 4.28E-07 | 1.57E-06 | 49.8359869 | 77.4093262 | 35.2659728 | 86.7560603 | 0 |
| 4.28E-07 | 1.57E-06 | 95.1706589 | 42.7085937 | 33.409869 | 48.7132959 | 204.354769 |
| 4.28E-07 | 1.57E-06 | 31.1876821 | 74.7400391 | 68.6758419 | 70.0543588 | 0 |
| 4.28E-07 | 1.57E-06 | 51.7651219 | 138.135608 | 136.423632 | 143.356271 | 0 |
| 4.29E-07 | 1.58E-06 | 29.9015921 | 87.4191528 | 74.2441534 | 77.4773373 | 0 |
| 4.30E-07 | 1.58E-06 | 0.96456749 | 0.22244059 | 0 | 0.46393615 | 28.5146189 |
| 4.31E-07 | 1.58E-06 | 33.4383396 | 93.86993 | 64.9636342 | 62.1674443 | 0 |
| 4.33E-07 | 1.59E-06 | 32.7952946 | 59.1691976 | 67.7477899 | 64.487125 | 0 |

| | | | | | | |
|---|---|---|---|---|---|---|
| 4.33E-07 | 1.59E-06 | 50.8005544 | 53.6081828 | 45.4745439 | 38.0427644 | 0 |
| 4.34E-07 | 1.59E-06 | 48.2283744 | 58.7243164 | 36.1940248 | 52.4247851 | 0 |
| 4.34E-07 | 1.60E-06 | 58.5170943 | 58.2794352 | 70.5319457 | 33.4034029 | 0 |
| 4.35E-07 | 1.60E-06 | 41.1548795 | 70.0687866 | 105.797919 | 45.4657428 | 0 |
| 4.35E-07 | 1.60E-06 | 270.078897 | 250.690548 | 249.645966 | 302.486371 | 38.0194919 |
| 4.37E-07 | 1.60E-06 | 440.48582 | 1098.1892 | 979.094772 | 1003.02996 | 0 |
| 4.37E-07 | 1.60E-06 | 45.9777169 | 47.3798462 | 55.683115 | 38.5067006 | 0 |
| 4.38E-07 | 1.61E-06 | 44.691627 | 91.2006429 | 40.8342843 | 119.231591 | 0 |
| 4.38E-07 | 1.61E-06 | 55.3018693 | 157.71038 | 113.222334 | 136.397228 | 0 |
| 4.38E-07 | 1.61E-06 | 27.0078897 | 92.5352865 | 92.8051917 | 109.952868 | 0 |
| 4.38E-07 | 1.61E-06 | 70.0919042 | 61.8384847 | 35.2659728 | 48.7132959 | 0 |
| 4.39E-07 | 1.61E-06 | 27.3294122 | 98.763623 | 87.2368802 | 98.3544641 | 0 |
| 4.41E-07 | 1.62E-06 | 38.261177 | 58.0569946 | 43.6184401 | 55.6723382 | 0 |
| 4.41E-07 | 1.62E-06 | 16.3976473 | 283.611755 | 168.905449 | 224.081161 | 0 |
| 4.41E-07 | 1.62E-06 | 50.4790319 | 58.7243164 | 97.4454513 | 38.9706367 | 0 |
| 4.43E-07 | 1.62E-06 | 257.217997 | 75.6298014 | 50.1148035 | 63.0953166 | 0 |
| 4.43E-07 | 1.63E-06 | 29.9015921 | 75.4073608 | 75.1722053 | 78.8691457 | 0 |
| 4.45E-07 | 1.63E-06 | 66.5551567 | 95.4270142 | 87.2368802 | 90.4675495 | 2.37621824 |
| 4.46E-07 | 1.63E-06 | 33.4383396 | 56.0550293 | 86.3088283 | 104.385634 | 0 |
| 4.46E-07 | 1.64E-06 | 47.2638069 | 44.9329997 | 40.8342843 | 68.1986142 | 0 |
| 4.47E-07 | 1.64E-06 | 27.0078897 | 137.913167 | 83.5246725 | 111.344676 | 0 |
| 4.48E-07 | 1.64E-06 | 25.0787547 | 157.932821 | 107.654022 | 105.313506 | 0 |
| 4.48E-07 | 1.64E-06 | 28.2939797 | 94.982133 | 77.0283091 | 102.529889 | 0 |
| 4.48E-07 | 1.64E-06 | 23.4711422 | 98.9860636 | 167.977397 | 174.439993 | 0 |
| 4.48E-07 | 1.64E-06 | 27.0078897 | 102.100232 | 107.654022 | 85.3642518 | 0 |
| 4.48E-07 | 1.64E-06 | 93.2415239 | 215.544934 | 206.955577 | 209.69914 | 19.0097459 |
| 4.50E-07 | 1.65E-06 | 33.4383396 | 62.0609253 | 55.683115 | 71.4461673 | 0 |
| 4.50E-07 | 1.65E-06 | 33.4383396 | 79.8561727 | 110.438178 | 57.9920189 | 0 |
| 4.52E-07 | 1.66E-06 | 63.3399317 | 119.228158 | 90.0210359 | 97.4265918 | 11.8810912 |
| 4.52E-07 | 1.66E-06 | 30.8661596 | 375.034839 | 909.490879 | 697.759972 | 7.12865473 |
| 4.52E-07 | 1.66E-06 | 46.9422844 | 78.9664103 | 42.6903882 | 46.8575513 | 0 |
| 4.53E-07 | 1.66E-06 | 52.4081669 | 139.470251 | 128.071165 | 142.428398 | 9.50487297 |
| 4.55E-07 | 1.66E-06 | 36.9750871 | 45.6003214 | 58.4672708 | 54.2805297 | 0 |
| 4.55E-07 | 1.67E-06 | 53.3727344 | 44.7105591 | 38.0501286 | 48.7132959 | 0 |
| 4.56E-07 | 1.67E-06 | 27.9724572 | 91.6455241 | 79.8124649 | 101.138081 | 0 |
| 4.57E-07 | 1.67E-06 | 38.904222 | 14.4586385 | 23.2012979 | 25.0525522 | 116.434694 |
| 4.58E-07 | 1.68E-06 | 46.2992394 | 76.7420044 | 42.6903882 | 46.3936151 | 0 |
| 4.59E-07 | 1.68E-06 | 111.246784 | 44.9329997 | 56.6111669 | 47.7854236 | 0 |
| 4.60E-07 | 1.68E-06 | 31.5092046 | 78.9664103 | 98.3735032 | 61.7035081 | 0 |
| 4.60E-07 | 1.68E-06 | 40.833357 | 63.1731283 | 42.6903882 | 48.2493597 | 0 |
| 4.61E-07 | 1.68E-06 | 409.61966 | 1221.19885 | 1115.5184 | 1208.55367 | 173.463932 |
| 4.62E-07 | 1.69E-06 | 27.3294122 | 103.434875 | 102.085711 | 81.1888265 | 0 |
| 4.63E-07 | 1.69E-06 | 258.182564 | 207.537073 | 25.0574018 | 83.5085072 | 0 |

| | | | | | | |
|---|---|---|---|---|---|---|
| 4.63E-07 | 1.69E-06 | 78.1299666 | 92.9801676 | 62.1794784 | 83.0445711 | 11.8810912 |
| 4.64E-07 | 1.70E-06 | 285.833499 | 76.2971232 | 40.8342843 | 85.828188 | 0 |
| 4.64E-07 | 1.70E-06 | 30.8661596 | 62.0609253 | 84.4527244 | 84.4363795 | 0 |
| 4.64E-07 | 1.70E-06 | 47.2638069 | 44.2656779 | 58.4672708 | 38.9706367 | 0 |
| 4.65E-07 | 1.70E-06 | 91.6339114 | 64.952653 | 29.6976613 | 86.2921241 | 0 |
| 4.65E-07 | 1.70E-06 | 27.0078897 | 95.6494547 | 104.869867 | 85.828188 | 0 |
| 4.66E-07 | 1.70E-06 | 100.315019 | 121.007682 | 75.1722053 | 73.7658481 | 9.50487297 |
| 4.67E-07 | 1.70E-06 | 32.7952946 | 55.6101481 | 69.6038938 | 65.8789335 | 0 |
| 4.68E-07 | 1.71E-06 | 59.4816618 | 136.133643 | 97.4454513 | 116.91191 | 0 |
| 4.69E-07 | 1.71E-06 | 52.0866444 | 56.4999105 | 57.5392189 | 34.3312752 | 0 |
| 4.70E-07 | 1.72E-06 | 32.7952946 | 64.0628906 | 62.1794784 | 93.2511664 | 0 |
| 4.70E-07 | 1.72E-06 | 59.4816618 | 135.466321 | 120.646749 | 125.262761 | 4.75243649 |
| 4.70E-07 | 1.72E-06 | 37.618132 | 84.5274251 | 120.646749 | 49.177232 | 0 |
| 4.71E-07 | 1.72E-06 | 40.833357 | 40.929069 | 51.9709073 | 48.2493597 | 0 |
| 4.72E-07 | 1.72E-06 | 39.8687895 | 175.728068 | 263.566744 | 226.864778 | 0 |
| 4.74E-07 | 1.73E-06 | 47.9068519 | 175.950509 | 128.071165 | 206.451587 | 2.37621824 |
| 4.74E-07 | 1.73E-06 | 25.0787547 | 135.911202 | 131.783372 | 90.0036134 | 0 |
| 4.74E-07 | 1.73E-06 | 27.6509347 | 99.2085042 | 87.2368802 | 83.0445711 | 0 |
| 4.75E-07 | 1.73E-06 | 80.7021465 | 80.3010539 | 46.4025958 | 36.1870198 | 0 |
| 4.75E-07 | 1.73E-06 | 52.7296894 | 40.2617472 | 44.546492 | 43.6099982 | 0 |
| 4.75E-07 | 1.73E-06 | 36.0105196 | 102.100232 | 77.0283091 | 50.1051043 | 0 |
| 4.75E-07 | 1.73E-06 | 180.37412 | 149.035197 | 24.1293498 | 91.3954218 | 0 |
| 4.75E-07 | 1.73E-06 | 31.5092046 | 73.1829549 | 93.7332436 | 61.7035081 | 0 |
| 4.76E-07 | 1.74E-06 | 27.6509347 | 77.8542074 | 99.3015551 | 99.7462725 | 0 |
| 4.77E-07 | 1.74E-06 | 31.8307271 | 157.932821 | 67.7477899 | 94.6429749 | 0 |
| 4.77E-07 | 1.74E-06 | 38.261177 | 83.4152222 | 77.0283091 | 43.6099982 | 0 |
| 4.77E-07 | 1.74E-06 | 55.9449143 | 147.033232 | 122.502853 | 126.654569 | 0 |
| 4.78E-07 | 1.74E-06 | 26.3648447 | 112.110059 | 88.1649321 | 100.210209 | 0 |
| 4.78E-07 | 1.74E-06 | 56.9094818 | 78.5215291 | 32.4818171 | 62.1674443 | 0 |
| 4.78E-07 | 1.74E-06 | 34.0813846 | 86.5293905 | 51.9709073 | 72.3740396 | 0 |
| 4.78E-07 | 1.74E-06 | 45.0131495 | 44.7105591 | 38.9781805 | 50.1051043 | 0 |
| 4.79E-07 | 1.74E-06 | 44.691627 | 192.188672 | 156.840774 | 166.553078 | 2.37621824 |
| 4.80E-07 | 1.75E-06 | 501.896616 | 74.9624797 | 52.8989593 | 94.6429749 | 0 |
| 4.80E-07 | 1.75E-06 | 32.1522496 | 100.543148 | 62.1794784 | 70.9822312 | 0 |
| 4.80E-07 | 1.75E-06 | 34.4029071 | 106.104163 | 92.8051917 | 51.9608489 | 0 |
| 4.81E-07 | 1.75E-06 | 31.8307271 | 83.4152222 | 71.4599976 | 58.9198912 | 0 |
| 4.81E-07 | 1.75E-06 | 118.320279 | 58.5018758 | 38.9781805 | 56.6002105 | 0 |
| 4.81E-07 | 1.75E-06 | 29.2585471 | 70.5136678 | 81.6685687 | 86.2921241 | 0 |
| 4.82E-07 | 1.75E-06 | 59.1601393 | 32.0314453 | 87.2368802 | 109.488932 | 0 |
| 4.82E-07 | 1.76E-06 | 37.9396545 | 43.3759155 | 51.9709073 | 53.8165936 | 0 |
| 4.83E-07 | 1.76E-06 | 810.558213 | 846.386454 | 787.916077 | 872.663901 | 38.0194919 |
| 4.84E-07 | 1.76E-06 | 21.8635297 | 99.6533854 | 226.444668 | 220.833608 | 0 |
| 4.84E-07 | 1.76E-06 | 114.783531 | 54.9428263 | 48.2586997 | 45.4657428 | 0 |

| | | | | | | |
|---|---|---|---|---|---|---|
| 4.85E-07 | 1.77E-06 | 26.6863672 | 94.0923706 | 88.1649321 | 101.138081 | 0 |
| 4.86E-07 | 1.77E-06 | 48.8714194 | 164.606038 | 126.215061 | 169.336695 | 2.37621824 |
| 4.87E-07 | 1.77E-06 | 335.026441 | 96.7616577 | 62.1794784 | 48.2493597 | 0 |
| 4.89E-07 | 1.78E-06 | 29.9015921 | 94.0923706 | 90.0210359 | 64.487125 | 0 |
| 4.89E-07 | 1.78E-06 | 32.1522496 | 100.098267 | 58.4672708 | 123.870952 | 0 |
| 4.89E-07 | 1.78E-06 | 38.5826995 | 46.934965 | 57.5392189 | 44.0739344 | 0 |
| 4.91E-07 | 1.78E-06 | 29.5800696 | 102.989994 | 118.790645 | 65.8789335 | 0 |
| 4.93E-07 | 1.79E-06 | 59.1601393 | 278.940503 | 197.675058 | 287.640414 | 9.50487297 |
| 4.94E-07 | 1.80E-06 | 32.7952946 | 60.9487223 | 58.4672708 | 81.1888265 | 0 |
| 4.94E-07 | 1.80E-06 | 86.811074 | 231.783097 | 192.106747 | 319.188072 | 0 |
| 4.94E-07 | 1.80E-06 | 34.0813846 | 73.6278361 | 51.0428554 | 62.6313804 | 0 |
| 4.94E-07 | 1.80E-06 | 38.904222 | 114.334465 | 44.546492 | 106.705315 | 0 |
| 4.94E-07 | 1.80E-06 | 51.1220769 | 51.828658 | 34.3379209 | 55.6723382 | 0 |
| 4.94E-07 | 1.80E-06 | 47.9068519 | 47.6022868 | 39.9062324 | 41.7542536 | 0 |
| 4.95E-07 | 1.80E-06 | 38.5826995 | 84.5274251 | 43.6184401 | 96.4987195 | 0 |
| 4.96E-07 | 1.80E-06 | 33.7598621 | 87.1967122 | 55.683115 | 62.6313804 | 0 |
| 4.96E-07 | 1.80E-06 | 35.0459521 | 60.5038411 | 185.610383 | 101.602017 | 0 |
| 4.97E-07 | 1.81E-06 | 18.0052598 | 176.61783 | 166.121293 | 186.502333 | 0 |
| 4.97E-07 | 1.81E-06 | 325.380766 | 502.048417 | 411.126999 | 470.895194 | 52.2768014 |
| 4.98E-07 | 1.81E-06 | 111.246784 | 113.667143 | 30.6257133 | 59.8477635 | 0 |
| 4.99E-07 | 1.81E-06 | 90.3478214 | 173.948543 | 137.351684 | 159.1301 | 14.2573095 |
| 4.99E-07 | 1.81E-06 | 45.9777169 | 38.7046631 | 44.546492 | 48.7132959 | 0 |
| 4.99E-07 | 1.81E-06 | 49.8359869 | 160.602108 | 108.582074 | 161.449781 | 7.12865473 |
| 5.01E-07 | 1.82E-06 | 41.1548795 | 60.5038411 | 86.3088283 | 41.7542536 | 0 |
| 5.01E-07 | 1.82E-06 | 345.636683 | 874.85885 | 744.297637 | 929.728047 | 4.75243649 |
| 5.02E-07 | 1.82E-06 | 76.8438766 | 71.6258708 | 62.1794784 | 30.619786 | 0 |
| 5.02E-07 | 1.82E-06 | 34.0813846 | 49.3818115 | 64.0355823 | 65.4149973 | 0 |
| 5.02E-07 | 1.82E-06 | 43.7270595 | 54.2755046 | 52.8989593 | 37.1148921 | 0 |
| 5.02E-07 | 1.82E-06 | 220.24291 | 304.07629 | 288.624146 | 308.517541 | 49.9005831 |
| 5.03E-07 | 1.82E-06 | 54.0157793 | 66.5097371 | 33.409869 | 53.8165936 | 0 |
| 5.03E-07 | 1.82E-06 | 293.871561 | 922.238696 | 714.599976 | 753.43231 | 0 |
| 5.04E-07 | 1.83E-06 | 31.8307271 | 17.1279256 | 20.4171422 | 17.6295737 | 95.0487297 |
| 5.05E-07 | 1.83E-06 | 68.4842917 | 103.657316 | 86.3088283 | 82.1166988 | 0 |
| 5.05E-07 | 1.83E-06 | 53.6942569 | 128.570662 | 64.0355823 | 39.898509 | 0 |
| 5.06E-07 | 1.84E-06 | 60.1247068 | 199.751652 | 222.73246 | 192.069567 | 0 |
| 5.07E-07 | 1.84E-06 | 66.8766792 | 64.5077718 | 35.2659728 | 45.4657428 | 0 |
| 5.08E-07 | 1.84E-06 | 28.6155022 | 145.476147 | 73.3161014 | 101.602017 | 0 |
| 5.08E-07 | 1.84E-06 | 38.5826995 | 89.8659993 | 51.0428554 | 52.4247851 | 0 |
| 5.09E-07 | 1.85E-06 | 33.1168171 | 111.442737 | 77.0283091 | 57.5280828 | 0 |
| 5.10E-07 | 1.85E-06 | 25.4002772 | 131.017509 | 95.5893474 | 98.3544641 | 0 |
| 5.10E-07 | 1.85E-06 | 42.119447 | 70.2912272 | 39.9062324 | 51.9608489 | 0 |
| 5.11E-07 | 1.85E-06 | 41.7979245 | 43.3759155 | 99.3015551 | 62.1674443 | 0 |
| 5.12E-07 | 1.85E-06 | 26.3648447 | 93.4250488 | 87.2368802 | 106.241379 | 0 |

| | | | | | | |
|---|---|---|---|---|---|---|
| 5.12E-07 | 1.85E-06 | 38.904222 | 46.0452026 | 84.4527244 | 54.7444659 | 0 |
| 5.12E-07 | 1.86E-06 | 39.8687895 | 64.5077718 | 45.4745439 | 101.602017 | 0 |
| 5.13E-07 | 1.86E-06 | 30.2231146 | 62.5058065 | 94.6612955 | 81.1888265 | 0 |
| 5.13E-07 | 1.86E-06 | 43.0840145 | 37.1475789 | 69.6038938 | 65.8789335 | 0 |
| 5.14E-07 | 1.86E-06 | 43.0840145 | 67.3994995 | 207.883629 | 66.8068058 | 0 |
| 5.14E-07 | 1.86E-06 | 67.5197242 | 102.989994 | 77.0283091 | 93.2511664 | 2.37621824 |
| 5.14E-07 | 1.86E-06 | 35.0459521 | 140.804895 | 166.121293 | 52.4247851 | 0 |
| 5.15E-07 | 1.86E-06 | 17.3622148 | 166.163123 | 434.328297 | 178.151482 | 0 |
| 5.16E-07 | 1.87E-06 | 47.9068519 | 178.174915 | 124.358957 | 155.418611 | 4.75243649 |
| 5.16E-07 | 1.87E-06 | 30.8661596 | 88.7537964 | 59.3953227 | 80.7248903 | 0 |
| 5.16E-07 | 1.87E-06 | 28.2939797 | 93.2026082 | 72.3880495 | 86.7560603 | 0 |
| 5.17E-07 | 1.87E-06 | 26.3648447 | 129.460425 | 101.157659 | 83.5085072 | 0 |
| 5.17E-07 | 1.87E-06 | 30.5446371 | 67.8443807 | 64.0355823 | 73.7658481 | 0 |
| 5.18E-07 | 1.87E-06 | 31.1876821 | 58.5018758 | 71.4599976 | 72.8379758 | 0 |
| 5.20E-07 | 1.88E-06 | 29.5800696 | 77.4093262 | 70.5319457 | 70.518295 | 0 |
| 5.21E-07 | 1.89E-06 | 187.126093 | 259.14329 | 250.574018 | 288.568286 | 28.5146189 |
| 5.22E-07 | 1.89E-06 | 27.9724572 | 77.8542074 | 86.3088283 | 81.6527626 | 0 |
| 5.22E-07 | 1.89E-06 | 161.082771 | 182.623726 | 168.905449 | 190.677758 | 19.0097459 |
| 5.23E-07 | 1.89E-06 | 134.717926 | 305.855815 | 328.530379 | 295.527328 | 19.0097459 |
| 5.23E-07 | 1.89E-06 | 29.9015921 | 108.328569 | 65.8916861 | 82.1166988 | 0 |
| 5.24E-07 | 1.89E-06 | 29.9015921 | 80.0786133 | 64.9636342 | 73.7658481 | 0 |
| 5.24E-07 | 1.89E-06 | 41.7979245 | 170.389494 | 189.322591 | 178.615418 | 2.37621824 |
| 5.25E-07 | 1.90E-06 | 125.393773 | 182.846167 | 130.85532 | 154.490738 | 16.6335277 |
| 5.26E-07 | 1.90E-06 | 0 | 0 | 0 | 0 | 49.9005831 |
| 5.26E-07 | 1.90E-06 | 37.2966096 | 114.556905 | 103.013763 | 45.929679 | 0 |
| 5.26E-07 | 1.90E-06 | 36.9750871 | 114.112024 | 130.85532 | 47.3214874 | 0 |
| 5.28E-07 | 1.91E-06 | 29.2585471 | 69.4014648 | 86.3088283 | 73.3019119 | 0 |
| 5.28E-07 | 1.91E-06 | 35.3674746 | 59.8365194 | 46.4025958 | 70.518295 | 0 |
| 5.30E-07 | 1.91E-06 | 74.5932191 | 320.981775 | 25.0574018 | 278.361691 | 0 |
| 5.30E-07 | 1.91E-06 | 50.4790319 | 83.6376628 | 38.0501286 | 49.6411682 | 0 |
| 5.31E-07 | 1.92E-06 | 25.7217997 | 117.893514 | 98.3735032 | 88.1478687 | 0 |
| 5.36E-07 | 1.94E-06 | 38.5826995 | 111.887618 | 46.4025958 | 149.851377 | 0 |
| 5.36E-07 | 1.94E-06 | 31.8307271 | 62.0609253 | 57.5392189 | 63.5592527 | 0 |
| 5.37E-07 | 1.94E-06 | 47.2638069 | 73.1829549 | 37.1220767 | 50.1051043 | 0 |
| 5.37E-07 | 1.94E-06 | 31.8307271 | 59.3916382 | 59.3953227 | 63.0953166 | 0 |
| 5.37E-07 | 1.94E-06 | 32.4737721 | 73.4053955 | 52.8989593 | 82.1166988 | 0 |
| 5.37E-07 | 1.94E-06 | 90.9908664 | 184.625692 | 165.193241 | 23.1968076 | 0 |
| 5.37E-07 | 1.94E-06 | 26.0433222 | 121.007682 | 80.7405168 | 109.488932 | 0 |
| 5.38E-07 | 1.94E-06 | 67.5197242 | 91.8679647 | 77.0283091 | 92.7872303 | 0 |
| 5.38E-07 | 1.94E-06 | 28.2939797 | 81.1908162 | 73.3161014 | 83.5085072 | 0 |
| 5.38E-07 | 1.94E-06 | 74.2716966 | 60.05896 | 36.1940248 | 44.5378705 | 0 |
| 5.38E-07 | 1.94E-06 | 27.6509347 | 95.8718953 | 109.510126 | 72.3740396 | 0 |
| 5.39E-07 | 1.94E-06 | 40.833357 | 64.0628906 | 40.8342843 | 47.7854236 | 0 |

| | | | | | | |
|---|---|---|---|---|---|---|
| 5.39E-07 | 1.94E-06 | 32.4737721 | 116.55887 | 55.683115 | 90.4675495 | 0 |
| 5.39E-07 | 1.94E-06 | 32.1522496 | 85.4171875 | 61.2514265 | 59.8477635 | 0 |
| 5.40E-07 | 1.95E-06 | 448.845405 | 921.793815 | 981.878928 | 983.544641 | 57.0292378 |
| 5.41E-07 | 1.95E-06 | 38.5826995 | 40.929069 | 62.1794784 | 70.9822312 | 0 |
| 5.42E-07 | 1.95E-06 | 0 | 0 | 0 | 0 | 42.7719284 |
| 5.43E-07 | 1.96E-06 | 37.618132 | 42.2637126 | 74.2441534 | 63.0953166 | 0 |
| 5.44E-07 | 1.96E-06 | 1484.46936 | 4592.28603 | 3552.58274 | 4153.15643 | 68.9103291 |
| 5.44E-07 | 1.96E-06 | 28.9370247 | 78.9664103 | 171.689605 | 92.3232941 | 0 |
| 5.47E-07 | 1.97E-06 | 27.0078897 | 22.0216187 | 24.1293498 | 23.6607437 | 166.335277 |
| 5.47E-07 | 1.97E-06 | 44.3701045 | 86.3069499 | 48.2586997 | 43.6099982 | 0 |
| 5.50E-07 | 1.98E-06 | 26.3648447 | 110.997856 | 89.092984 | 84.9003157 | 0 |
| 5.51E-07 | 1.98E-06 | 75.2362641 | 79.8561727 | 29.6976613 | 147.531696 | 0 |
| 5.51E-07 | 1.98E-06 | 26.0433222 | 76.964445 | 128.071165 | 130.829995 | 0 |
| 5.55E-07 | 2.00E-06 | 42.762492 | 43.1534749 | 43.6184401 | 70.0543588 | 0 |
| 5.55E-07 | 2.00E-06 | 28.6155022 | 76.5195638 | 141.063891 | 84.9003157 | 0 |
| 5.56E-07 | 2.00E-06 | 52.7296894 | 130.572628 | 113.222334 | 131.757867 | 0 |
| 5.57E-07 | 2.00E-06 | 87.1325964 | 178.174915 | 166.121293 | 188.358077 | 14.2573095 |
| 5.58E-07 | 2.01E-06 | 85.8465065 | 128.793103 | 77.0283091 | 27.8361691 | 0 |
| 5.58E-07 | 2.01E-06 | 39.547267 | 73.4053955 | 39.9062324 | 59.8477635 | 0 |
| 5.59E-07 | 2.01E-06 | 27.9724572 | 81.6356974 | 74.2441534 | 82.1166988 | 0 |
| 5.60E-07 | 2.02E-06 | 108.353081 | 56.0550293 | 104.869867 | 111.344676 | 0 |
| 5.61E-07 | 2.02E-06 | 4.50131495 | 3.11416829 | 9.28051917 | 6.49510612 | 35.6432737 |
| 5.61E-07 | 2.02E-06 | 37.2966096 | 54.9428263 | 41.7623363 | 51.9608489 | 0 |
| 5.62E-07 | 2.02E-06 | 33.1168171 | 57.834554 | 51.0428554 | 59.8477635 | 0 |
| 5.62E-07 | 2.02E-06 | 140.505331 | 222.885474 | 242.22155 | 237.999246 | 23.7621824 |
| 5.62E-07 | 2.02E-06 | 32.7952946 | 50.9388957 | 87.2368802 | 80.2609542 | 0 |
| 5.63E-07 | 2.03E-06 | 49.8359869 | 312.084151 | 157.768826 | 230.576267 | 4.75243649 |
| 5.64E-07 | 2.03E-06 | 40.833357 | 40.2617472 | 46.4025958 | 45.0018067 | 0 |
| 5.64E-07 | 2.03E-06 | 31.1876821 | 70.5136678 | 96.5173994 | 59.3838274 | 0 |
| 5.65E-07 | 2.03E-06 | 22.1850522 | 98.763623 | 193.962851 | 372.54073 | 0 |
| 5.66E-07 | 2.04E-06 | 41.1548795 | 44.2656779 | 68.6758419 | 43.1460621 | 0 |
| 5.67E-07 | 2.04E-06 | 68.4842917 | 118.338395 | 161.481034 | 28.3001052 | 0 |
| 5.67E-07 | 2.04E-06 | 0 | 0 | 0 | 0 | 54.6530196 |
| 5.67E-07 | 2.04E-06 | 26.6863672 | 119.673039 | 78.8844129 | 92.7872303 | 0 |
| 5.69E-07 | 2.04E-06 | 54.6588243 | 164.161157 | 134.567528 | 156.346483 | 7.12865473 |
| 5.70E-07 | 2.05E-06 | 36.0105196 | 49.8266927 | 49.1867516 | 45.4657428 | 0 |
| 5.70E-07 | 2.05E-06 | 29.2585471 | 86.7518311 | 63.1075304 | 99.7462725 | 0 |
| 5.71E-07 | 2.05E-06 | 22.8280972 | 103.434875 | 142.919995 | 118.767655 | 0 |
| 5.73E-07 | 2.06E-06 | 22.5065747 | 92.9801676 | 165.193241 | 185.574461 | 0 |
| 5.75E-07 | 2.06E-06 | 144.363601 | 127.236019 | 22.273246 | 101.138081 | 0 |
| 5.75E-07 | 2.06E-06 | 0 | 0 | 0 | 0 | 42.7719284 |
| 5.77E-07 | 2.07E-06 | 32.7952946 | 75.852242 | 50.1148035 | 80.7248903 | 0 |
| 5.78E-07 | 2.07E-06 | 31.8307271 | 69.4014648 | 52.8989593 | 73.7658481 | 0 |

| | | | | | | |
|---|---|---|---|---|---|---|
| 5.78E-07 | 2.07E-06 | 46.6207619 | 51.3837769 | 35.2659728 | 47.3214874 | 0 |
| 5.79E-07 | 2.08E-06 | 141.148376 | 60.9487223 | 34.3379209 | 71.4461673 | 0 |
| 5.80E-07 | 2.08E-06 | 27.9724572 | 74.7400391 | 77.956361 | 78.8691457 | 0 |
| 5.80E-07 | 2.08E-06 | 27.9724572 | 78.9664103 | 75.1722053 | 77.4773373 | 0 |
| 5.80E-07 | 2.08E-06 | 35.6889971 | 93.2026082 | 45.4745439 | 78.8691457 | 0 |
| 5.82E-07 | 2.09E-06 | 31.8307271 | 74.9624797 | 52.8989593 | 69.1264865 | 0 |
| 5.82E-07 | 2.09E-06 | 72.3425616 | 36.9251383 | 43.6184401 | 55.6723382 | 0 |
| 5.82E-07 | 2.09E-06 | 54.9803468 | 124.566732 | 31.5537652 | 103.457762 | 0 |
| 5.83E-07 | 2.09E-06 | 49.5144644 | 83.8601033 | 33.409869 | 64.9510612 | 0 |
| 5.83E-07 | 2.09E-06 | 28.9370247 | 63.84045 | 87.2368802 | 81.1888265 | 0 |
| 5.85E-07 | 2.10E-06 | 29.2585471 | 66.5097371 | 80.7405168 | 113.200421 | 0 |
| 5.85E-07 | 2.10E-06 | 132.788791 | 71.8483114 | 29.6976613 | 75.1576565 | 0 |
| 5.87E-07 | 2.10E-06 | 31.8307271 | 66.5097371 | 60.3233746 | 54.2805297 | 0 |
| 5.87E-07 | 2.10E-06 | 60.4462293 | 111.442737 | 90.9490879 | 109.024996 | 0 |
| 5.87E-07 | 2.10E-06 | 45.0131495 | 53.3857422 | 75.1722053 | 36.650956 | 0 |
| 5.87E-07 | 2.10E-06 | 22.8280972 | 116.781311 | 105.797919 | 124.334889 | 0 |
| 5.88E-07 | 2.11E-06 | 25.0787547 | 105.436841 | 226.444668 | 102.529889 | 0 |
| 5.88E-07 | 2.11E-06 | 31.1876821 | 149.480078 | 64.0355823 | 89.0757411 | 0 |
| 5.89E-07 | 2.11E-06 | 36.0105196 | 81.1908162 | 47.3306478 | 112.736485 | 0 |
| 5.90E-07 | 2.11E-06 | 241.784917 | 137.023405 | 85.3807764 | 137.789037 | 11.8810912 |
| 5.90E-07 | 2.11E-06 | 47.5853294 | 163.716276 | 254.286225 | 128.510314 | 0 |
| 5.91E-07 | 2.12E-06 | 44.048582 | 74.0727173 | 62.1794784 | 35.7230837 | 0 |
| 5.91E-07 | 2.12E-06 | 36.0105196 | 49.6042521 | 48.2586997 | 45.0018067 | 0 |
| 5.91E-07 | 2.12E-06 | 55.6233918 | 111.887618 | 104.869867 | 103.921698 | 0 |
| 5.91E-07 | 2.12E-06 | 27.9724572 | 25.8031087 | 64.9636342 | 6.49510612 | 294.651062 |
| 5.93E-07 | 2.12E-06 | 28.9370247 | 87.1967122 | 89.092984 | 62.6313804 | 0 |
| 5.93E-07 | 2.12E-06 | 28.6155022 | 70.7361084 | 72.3880495 | 74.6937204 | 0 |
| 5.94E-07 | 2.13E-06 | 26.0433222 | 173.948543 | 75.1722053 | 135.469356 | 0 |
| 5.95E-07 | 2.13E-06 | 27.6509347 | 85.4171875 | 73.3161014 | 79.797018 | 0 |
| 5.96E-07 | 2.13E-06 | 54.6588243 | 40.0393066 | 47.3306478 | 38.9706367 | 0 |
| 5.96E-07 | 2.13E-06 | 155.938411 | 84.9723063 | 67.7477899 | 92.3232941 | 0 |
| 5.97E-07 | 2.13E-06 | 40.833357 | 159.934786 | 45.4745439 | 93.7151026 | 0 |
| 5.97E-07 | 2.13E-06 | 35.0459521 | 47.6022868 | 56.6111669 | 47.3214874 | 0 |
| 5.97E-07 | 2.14E-06 | 12.5393773 | 7.56298014 | 4.64025958 | 6.03116997 | 118.810912 |
| 5.98E-07 | 2.14E-06 | 29.2585471 | 62.5058065 | 80.7405168 | 96.9626556 | 0 |
| 5.98E-07 | 2.14E-06 | 54.0157793 | 52.2735392 | 179.11402 | 55.208402 | 0 |
| 6.00E-07 | 2.15E-06 | 61.0892743 | 110.108093 | 123.430905 | 105.313506 | 9.50487297 |
| 6.00E-07 | 2.15E-06 | 59.1601393 | 83.1927816 | 90.0210359 | 90.4675495 | 7.12865473 |
| 6.00E-07 | 2.15E-06 | 49.8359869 | 23.5787028 | 11.136623 | 26.4443606 | 173.463932 |
| 6.01E-07 | 2.15E-06 | 35.0459521 | 46.0452026 | 61.2514265 | 50.1051043 | 0 |
| 6.02E-07 | 2.15E-06 | 38.5826995 | 54.4979451 | 53.8270112 | 38.9706367 | 0 |
| 6.03E-07 | 2.16E-06 | 54.0157793 | 104.547078 | 126.215061 | 117.375846 | 0 |
| 6.05E-07 | 2.16E-06 | 23.1496197 | 128.570662 | 113.222334 | 99.2823364 | 0 |

| | | | | | | |
|---|---|---|---|---|---|---|
| 6.05E-07 | 2.16E-06 | 32.7952946 | 58.2794352 | 50.1148035 | 66.8068058 | 0 |
| 6.05E-07 | 2.16E-06 | 31.8307271 | 78.9664103 | 52.8989593 | 66.8068058 | 0 |
| 6.05E-07 | 2.16E-06 | 31.8307271 | 65.6199748 | 111.36623 | 62.6313804 | 0 |
| 6.06E-07 | 2.16E-06 | 46.9422844 | 38.0373413 | 143.848047 | 91.859358 | 0 |
| 6.06E-07 | 2.16E-06 | 553.018693 | 1069.71681 | 1122.94282 | 1165.40761 | 83.1676385 |
| 6.07E-07 | 2.17E-06 | 46.6207619 | 146.36591 | 167.049345 | 128.046378 | 2.37621824 |
| 6.08E-07 | 2.17E-06 | 25.0787547 | 119.005717 | 83.5246725 | 102.993826 | 0 |
| 6.09E-07 | 2.17E-06 | 52.0866444 | 121.007682 | 66.819738 | 36.650956 | 0 |
| 6.12E-07 | 2.19E-06 | 29.9015921 | 80.5234945 | 100.229607 | 59.3838274 | 0 |
| 6.13E-07 | 2.19E-06 | 64.3044992 | 94.3148112 | 93.7332436 | 103.457762 | 2.37621824 |
| 6.14E-07 | 2.19E-06 | 30.5446371 | 255.806681 | 253.358173 | 235.679565 | 0 |
| 6.16E-07 | 2.20E-06 | 60.4462293 | 173.058781 | 166.121293 | 155.882547 | 0 |
| 6.17E-07 | 2.20E-06 | 27.6509347 | 96.7616577 | 106.72597 | 66.8068058 | 0 |
| 6.17E-07 | 2.20E-06 | 28.6155022 | 66.5097371 | 76.1002572 | 73.7658481 | 0 |
| 6.18E-07 | 2.20E-06 | 109.960694 | 681.780416 | 511.356606 | 775.237309 | 19.0097459 |
| 6.18E-07 | 2.20E-06 | 100.636541 | 100.765588 | 98.3735032 | 24.1246799 | 0 |
| 6.20E-07 | 2.21E-06 | 0 | 0 | 0 | 0 | 23.7621824 |
| 6.20E-07 | 2.21E-06 | 25.7217997 | 84.7498657 | 97.4454513 | 87.6839326 | 0 |
| 6.20E-07 | 2.21E-06 | 24.7572322 | 84.3049845 | 114.150386 | 105.313506 | 0 |
| 6.21E-07 | 2.21E-06 | 23.1496197 | 126.568697 | 94.6612955 | 120.623399 | 0 |
| 6.21E-07 | 2.22E-06 | 25.0787547 | 83.4152222 | 100.229607 | 109.488932 | 0 |
| 6.24E-07 | 2.22E-06 | 42.119447 | 76.7420044 | 36.1940248 | 81.6527626 | 0 |
| 6.24E-07 | 2.22E-06 | 54.0157793 | 63.6180094 | 30.6257133 | 64.0231889 | 0 |
| 6.25E-07 | 2.23E-06 | 36.0105196 | 60.9487223 | 122.502853 | 56.6002105 | 0 |
| 6.25E-07 | 2.23E-06 | 36.9750871 | 65.3975342 | 63.1075304 | 40.3624452 | 0 |
| 6.26E-07 | 2.23E-06 | 34.0813846 | 63.1731283 | 47.3306478 | 52.4247851 | 0 |
| 6.27E-07 | 2.24E-06 | 38.904222 | 111.665177 | 43.6184401 | 73.3019119 | 0 |
| 6.28E-07 | 2.24E-06 | 114.783531 | 57.834554 | 50.1148035 | 50.5690405 | 299.403499 |
| 6.28E-07 | 2.24E-06 | 91.3123889 | 181.733964 | 138.279736 | 196.708928 | 14.2573095 |
| 6.29E-07 | 2.24E-06 | 27.3294122 | 87.1967122 | 76.1002572 | 75.1576565 | 0 |
| 6.29E-07 | 2.24E-06 | 34.4029071 | 44.2656779 | 62.1794784 | 59.3838274 | 0 |
| 6.30E-07 | 2.24E-06 | 28.2939797 | 63.6180094 | 87.2368802 | 91.3954218 | 0 |
| 6.31E-07 | 2.25E-06 | 29.2585471 | 61.3936035 | 71.4599976 | 77.9412734 | 0 |
| 6.32E-07 | 2.25E-06 | 28.6155022 | 74.2951579 | 64.9636342 | 87.6839326 | 0 |
| 6.32E-07 | 2.25E-06 | 34.4029071 | 47.1574056 | 69.6038938 | 90.4675495 | 0 |
| 6.33E-07 | 2.25E-06 | 29.5800696 | 60.9487223 | 71.4599976 | 67.2707419 | 0 |
| 6.33E-07 | 2.25E-06 | 158.832113 | 326.097909 | 173.545708 | 208.771268 | 0 |
| 6.34E-07 | 2.26E-06 | 50.4790319 | 181.956405 | 153.128566 | 165.16127 | 4.75243649 |
| 6.34E-07 | 2.26E-06 | 26.0433222 | 78.5215291 | 100.229607 | 88.1478687 | 0 |
| 6.35E-07 | 2.26E-06 | 50.4790319 | 312.084151 | 296.976613 | 302.486371 | 9.50487297 |
| 6.35E-07 | 2.26E-06 | 35.6889971 | 136.578524 | 90.0210359 | 49.177232 | 0 |
| 6.38E-07 | 2.27E-06 | 35.0459521 | 52.7184204 | 44.546492 | 49.6411682 | 0 |
| 6.38E-07 | 2.27E-06 | 31.8307271 | 85.1947469 | 51.0428554 | 75.6215927 | 0 |

| | | | | | | |
|---|---|---|---|---|---|---|
| 6.41E-07 | 2.28E-06 | 48.2283744 | 136.133643 | 133.639476 | 124.798825 | 2.37621824 |
| 6.41E-07 | 2.28E-06 | 27.6509347 | 84.5274251 | 68.6758419 | 80.7248903 | 0 |
| 6.42E-07 | 2.28E-06 | 26.6863672 | 139.915133 | 69.6038938 | 117.375846 | 0 |
| 6.43E-07 | 2.29E-06 | 30.5446371 | 65.3975342 | 55.683115 | 67.7346781 | 0 |
| 6.43E-07 | 2.29E-06 | 57.8740493 | 42.9310343 | 113.222334 | 44.0739344 | 0 |
| 6.44E-07 | 2.29E-06 | 23.1496197 | 99.4309448 | 106.72597 | 118.767655 | 0 |
| 6.44E-07 | 2.29E-06 | 33.4383396 | 54.4979451 | 49.1867516 | 50.5690405 | 0 |
| 6.46E-07 | 2.30E-06 | 24.1141872 | 91.8679647 | 108.582074 | 176.295737 | 0 |
| 6.46E-07 | 2.30E-06 | 31.1876821 | 74.0727173 | 51.9709073 | 75.1576565 | 0 |
| 6.48E-07 | 2.30E-06 | 28.6155022 | 71.4034302 | 104.869867 | 69.1264865 | 0 |
| 6.48E-07 | 2.30E-06 | 38.904222 | 66.9546183 | 51.0428554 | 39.898509 | 0 |
| 6.49E-07 | 2.31E-06 | 44.3701045 | 34.7007324 | 70.5319457 | 58.4559551 | 0 |
| 6.51E-07 | 2.31E-06 | 32.1522496 | 57.834554 | 51.9709073 | 54.7444659 | 0 |
| 6.52E-07 | 2.31E-06 | 65.2690667 | 37.5924601 | 38.9781805 | 63.0953166 | 0 |
| 6.52E-07 | 2.32E-06 | 426.017307 | 931.803642 | 636.643615 | 841.580178 | 33.2670554 |
| 6.52E-07 | 2.32E-06 | 35.3674746 | 75.852242 | 42.6903882 | 68.6625504 | 0 |
| 6.56E-07 | 2.33E-06 | 21.2204847 | 125.234054 | 116.00649 | 127.118505 | 0 |
| 6.57E-07 | 2.33E-06 | 37.9396545 | 59.6140788 | 77.956361 | 40.3624452 | 0 |
| 6.58E-07 | 2.34E-06 | 27.9724572 | 66.5097371 | 89.092984 | 75.6215927 | 0 |
| 6.58E-07 | 2.34E-06 | 27.3294122 | 83.4152222 | 91.8771398 | 68.1986142 | 0 |
| 6.59E-07 | 2.34E-06 | 162.690383 | 181.289083 | 161.481034 | 174.439993 | 19.0097459 |
| 6.60E-07 | 2.34E-06 | 621.502985 | 714.256742 | 612.514265 | 707.038695 | 14.2573095 |
| 6.60E-07 | 2.34E-06 | 31.8307271 | 82.970341 | 52.8989593 | 63.0953166 | 0 |
| 6.61E-07 | 2.34E-06 | 28.9370247 | 70.7361084 | 63.1075304 | 70.0543588 | 0 |
| 6.61E-07 | 2.34E-06 | 35.0459521 | 96.5392171 | 44.546492 | 95.5708472 | 0 |
| 6.61E-07 | 2.35E-06 | 26.3648447 | 85.1947469 | 86.3088283 | 76.549465 | 0 |
| 6.61E-07 | 2.35E-06 | 16.7191698 | 11.7893514 | 12.9927268 | 14.3820207 | 123.563349 |
| 6.62E-07 | 2.35E-06 | 51.4435994 | 153.039128 | 103.941815 | 125.262761 | 2.37621824 |
| 6.62E-07 | 2.35E-06 | 30.2231146 | 79.8561727 | 55.683115 | 95.106911 | 0 |
| 6.63E-07 | 2.35E-06 | 40.190312 | 55.6101481 | 51.9709073 | 36.650956 | 0 |
| 6.63E-07 | 2.35E-06 | 28.2939797 | 73.4053955 | 65.8916861 | 74.6937204 | 0 |
| 6.63E-07 | 2.35E-06 | 26.0433222 | 88.976237 | 112.294282 | 76.0855288 | 0 |
| 6.63E-07 | 2.35E-06 | 62.6968867 | 83.4152222 | 84.4527244 | 89.5396772 | 0 |
| 6.66E-07 | 2.36E-06 | 32.4737721 | 109.885653 | 58.4672708 | 64.487125 | 0 |
| 6.66E-07 | 2.36E-06 | 34.0813846 | 44.9329997 | 53.8270112 | 52.4247851 | 0 |
| 6.68E-07 | 2.37E-06 | 25.7217997 | 94.5372518 | 77.956361 | 91.859358 | 0 |
| 6.71E-07 | 2.38E-06 | 39.2257445 | 90.9782023 | 51.9709073 | 45.4657428 | 0 |
| 6.72E-07 | 2.38E-06 | 56.2664368 | 112.332499 | 103.013763 | 111.344676 | 0 |
| 6.73E-07 | 2.38E-06 | 72.0210391 | 26.4704305 | 92.8051917 | 90.9314857 | 0 |
| 6.75E-07 | 2.39E-06 | 470.065889 | 296.958191 | 296.976613 | 257.484564 | 23.7621824 |
| 6.74E-07 | 2.39E-06 | 47.9068519 | 39.5944255 | 38.9781805 | 64.487125 | 0 |
| 6.74E-07 | 2.39E-06 | 274.258689 | 70.5136678 | 37.1220767 | 85.828188 | 0 |
| 6.76E-07 | 2.39E-06 | 113.175919 | 50.2715739 | 38.9781805 | 55.208402 | 0 |

| | | | | | | |
|---|---|---|---|---|---|---|
| 6.76E-07 | 2.39E-06 | 28.2939797 | 64.0628906 | 75.1722053 | 91.3954218 | 0 |
| 6.77E-07 | 2.40E-06 | 39.547267 | 46.7125244 | 38.0501286 | 57.5280828 | 0 |
| 6.79E-07 | 2.40E-06 | 46.9422844 | 86.9742716 | 70.5319457 | 33.4034029 | 0 |
| 6.79E-07 | 2.40E-06 | 33.4383396 | 121.675004 | 54.7550631 | 71.4461673 | 0 |
| 6.79E-07 | 2.40E-06 | 90.6693439 | 157.71038 | 212.523889 | 21.8049991 | 0 |
| 6.80E-07 | 2.41E-06 | 41.476402 | 73.6278361 | 163.337137 | 48.7132959 | 0 |
| 6.81E-07 | 2.41E-06 | 40.5118345 | 240.45828 | 44.546492 | 140.572654 | 0 |
| 6.82E-07 | 2.41E-06 | 18.3267823 | 8.8976237 | 9.28051917 | 12.9902122 | 68.9103291 |
| 6.82E-07 | 2.41E-06 | 25.0787547 | 87.4191528 | 219.948304 | 113.200421 | 0 |
| 6.82E-07 | 2.41E-06 | 47.5853294 | 142.139539 | 34.3379209 | 155.418611 | 0 |
| 6.82E-07 | 2.41E-06 | 27.6509347 | 88.976237 | 65.8916861 | 79.797018 | 0 |
| 6.83E-07 | 2.41E-06 | 28.6155022 | 64.0628906 | 72.3880495 | 68.1986142 | 0 |
| 6.85E-07 | 2.42E-06 | 14.1469898 | 5.11613363 | 5.5683115 | 4.63936151 | 42.7719284 |
| 6.86E-07 | 2.43E-06 | 53.0512119 | 53.6081828 | 44.546492 | 33.867339 | 0 |
| 6.88E-07 | 2.43E-06 | 25.4002772 | 86.3069499 | 86.3088283 | 86.7560603 | 0 |
| 6.89E-07 | 2.44E-06 | 28.9370247 | 100.320707 | 59.3953227 | 83.5085072 | 0 |
| 6.90E-07 | 2.44E-06 | 64.6260217 | 130.572628 | 108.582074 | 114.128293 | 14.2573095 |
| 6.91E-07 | 2.44E-06 | 30.2231146 | 79.4112915 | 80.7405168 | 53.3526574 | 0 |
| 6.91E-07 | 2.44E-06 | 35.3674746 | 116.781311 | 45.4745439 | 88.6118049 | 0 |
| 6.92E-07 | 2.45E-06 | 126.036818 | 49.8266927 | 90.9490879 | 33.4034029 | 0 |
| 6.93E-07 | 2.45E-06 | 135.039448 | 51.6062174 | 39.9062324 | 59.8477635 | 0 |
| 6.93E-07 | 2.45E-06 | 18.3267823 | 129.682865 | 163.337137 | 162.841589 | 0 |
| 6.94E-07 | 2.45E-06 | 28.9370247 | 109.663212 | 60.3233746 | 84.9003157 | 0 |
| 6.95E-07 | 2.46E-06 | 38.5826995 | 41.1515096 | 49.1867516 | 71.4461673 | 0 |
| 6.96E-07 | 2.46E-06 | 25.4002772 | 81.6356974 | 113.222334 | 84.9003157 | 0 |
| 6.96E-07 | 2.46E-06 | 85.2034615 | 171.946578 | 162.409085 | 150.315313 | 14.2573095 |
| 6.98E-07 | 2.46E-06 | 163.976473 | 147.700553 | 76.1002572 | 24.1246799 | 0 |
| 7.00E-07 | 2.47E-06 | 30.5446371 | 80.3010539 | 103.013763 | 54.2805297 | 0 |
| 7.00E-07 | 2.47E-06 | 107.388514 | 547.871179 | 632.003355 | 635.128591 | 9.50487297 |
| 7.01E-07 | 2.47E-06 | 30.5446371 | 56.9447917 | 65.8916861 | 92.3232941 | 0 |
| 7.02E-07 | 2.48E-06 | 0 | 2.00196533 | 0 | 1.85574461 | 26.1384007 |
| 7.02E-07 | 2.48E-06 | 30.8661596 | 52.2735392 | 66.819738 | 60.3116997 | 0 |
| 7.02E-07 | 2.48E-06 | 29.9015921 | 74.7400391 | 54.7550631 | 84.4363795 | 0 |
| 7.03E-07 | 2.48E-06 | 36.6535646 | 63.6180094 | 40.8342843 | 50.5690405 | 0 |
| 7.03E-07 | 2.48E-06 | 224.422702 | 42.7085937 | 69.6038938 | 61.239572 | 0 |
| 7.04E-07 | 2.48E-06 | 394.829625 | 717.370911 | 715.528028 | 680.594334 | 61.7816743 |
| 7.05E-07 | 2.49E-06 | 38.261177 | 63.3955688 | 40.8342843 | 45.929679 | 0 |
| 7.06E-07 | 2.49E-06 | 24.4357097 | 209.761479 | 79.8124649 | 134.541484 | 0 |
| 7.07E-07 | 2.49E-06 | 29.2585471 | 69.1790242 | 59.3953227 | 65.4149973 | 0 |
| 7.09E-07 | 2.50E-06 | 19.9343948 | 131.684831 | 129.927268 | 128.510314 | 0 |
| 7.09E-07 | 2.50E-06 | 35.6889971 | 84.3049845 | 116.934542 | 45.0018067 | 0 |
| 7.09E-07 | 2.50E-06 | 72.3425616 | 265.594067 | 167.977397 | 217.122119 | 11.8810912 |
| 7.12E-07 | 2.51E-06 | 88.7402089 | 57.6121134 | 61.2514265 | 46.8575513 | 190.097459 |

| | | | | | | |
|---|---|---|---|---|---|---|
| 7.12E-07 | 2.51E-06 | 28.9370247 | 82.970341 | 57.5392189 | 91.3954218 | 0 |
| 7.12E-07 | 2.51E-06 | 37.2966096 | 61.1711629 | 38.0501286 | 61.7035081 | 0 |
| 7.13E-07 | 2.51E-06 | 39.547267 | 49.6042521 | 38.0501286 | 43.6099982 | 0 |
| 7.15E-07 | 2.52E-06 | 55.6233918 | 92.3128459 | 106.72597 | 105.777442 | 0 |
| 7.15E-07 | 2.52E-06 | 36.0105196 | 71.4034302 | 41.7623363 | 88.1478687 | 0 |
| 7.16E-07 | 2.52E-06 | 25.4002772 | 84.0825439 | 88.1649321 | 82.5806349 | 0 |
| 7.16E-07 | 2.52E-06 | 38.261177 | 60.2814006 | 45.4745439 | 106.705315 | 0 |
| 7.17E-07 | 2.53E-06 | 28.9370247 | 74.7400391 | 59.3953227 | 68.1986142 | 0 |
| 7.18E-07 | 2.53E-06 | 61.7323193 | 60.5038411 | 126.215061 | 33.4034029 | 0 |
| 7.19E-07 | 2.53E-06 | 31.5092046 | 78.5215291 | 58.4672708 | 52.8887212 | 0 |
| 7.19E-07 | 2.53E-06 | 267.506717 | 808.571554 | 626.435044 | 651.830293 | 28.5146189 |
| 7.21E-07 | 2.54E-06 | 32.1522496 | 57.6121134 | 52.8989593 | 49.177232 | 0 |
| 7.21E-07 | 2.54E-06 | 25.7217997 | 90.5333211 | 82.5966206 | 77.9412734 | 0 |
| 7.23E-07 | 2.54E-06 | 46.2992394 | 53.8306234 | 32.4818171 | 51.9608489 | 0 |
| 7.24E-07 | 2.55E-06 | 44.691627 | 34.923173 | 57.5392189 | 45.4657428 | 0 |
| 7.25E-07 | 2.55E-06 | 40.5118345 | 56.2774699 | 42.6903882 | 38.5067006 | 0 |
| 7.27E-07 | 2.56E-06 | 37.2966096 | 41.3739502 | 43.6184401 | 44.5378705 | 0 |
| 7.27E-07 | 2.56E-06 | 55.3018693 | 59.1691976 | 36.1940248 | 38.9706367 | 0 |
| 7.28E-07 | 2.56E-06 | 39.8687895 | 36.9251383 | 47.3306478 | 56.1362743 | 0 |
| 7.29E-07 | 2.57E-06 | 30.2231146 | 114.334465 | 57.5392189 | 77.9412734 | 0 |
| 7.32E-07 | 2.57E-06 | 30.5446371 | 76.5195638 | 51.0428554 | 77.0134011 | 0 |
| 7.32E-07 | 2.57E-06 | 21.8635297 | 86.3069499 | 177.257916 | 158.202228 | 0 |
| 7.33E-07 | 2.58E-06 | 46.9422844 | 61.3936035 | 32.4818171 | 51.0329766 | 0 |
| 7.36E-07 | 2.59E-06 | 26.3648447 | 73.1829549 | 186.538435 | 111.344676 | 0 |
| 7.36E-07 | 2.59E-06 | 27.6509347 | 72.0707519 | 65.8916861 | 75.1576565 | 0 |
| 7.37E-07 | 2.59E-06 | 35.6889971 | 42.2637126 | 45.4745439 | 47.3214874 | 0 |
| 7.39E-07 | 2.60E-06 | 37.618132 | 67.6219401 | 42.6903882 | 45.4657428 | 0 |
| 7.40E-07 | 2.60E-06 | 0 | 0 | 0 | 0 | 19.0097459 |
| 7.40E-07 | 2.60E-06 | 24.4357097 | 110.552974 | 88.1649321 | 85.3642518 | 0 |
| 7.41E-07 | 2.60E-06 | 36.9750871 | 74.2951579 | 40.8342843 | 54.2805297 | 0 |
| 7.45E-07 | 2.62E-06 | 50.8005544 | 38.2597819 | 37.1220767 | 48.7132959 | 0 |
| 7.46E-07 | 2.62E-06 | 34.4029071 | 43.3759155 | 48.2586997 | 56.6002105 | 0 |
| 7.47E-07 | 2.62E-06 | 27.3294122 | 72.9605143 | 66.819738 | 78.8691457 | 0 |
| 7.48E-07 | 2.63E-06 | 32.1522496 | 46.2676432 | 59.3953227 | 60.7756358 | 0 |
| 7.49E-07 | 2.63E-06 | 60.1247068 | 54.2755046 | 41.7623363 | 34.3312752 | 0 |
| 7.51E-07 | 2.64E-06 | 22.8280972 | 98.5411824 | 107.654022 | 99.2823364 | 0 |
| 7.52E-07 | 2.64E-06 | 32.1522496 | 52.4959798 | 117.862593 | 76.0855288 | 0 |
| 7.52E-07 | 2.64E-06 | 31.1876821 | 116.781311 | 52.8989593 | 154.026802 | 0 |
| 7.53E-07 | 2.64E-06 | 34.7244296 | 64.0628906 | 45.4745439 | 46.8575513 | 0 |
| 7.55E-07 | 2.65E-06 | 54.9803468 | 98.5411824 | 94.6612955 | 117.839782 | 0 |
| 7.55E-07 | 2.65E-06 | 28.2939797 | 94.7596924 | 65.8916861 | 66.8068058 | 0 |
| 7.55E-07 | 2.65E-06 | 55.9449143 | 69.1790242 | 28.7696094 | 82.5806349 | 0 |
| 7.56E-07 | 2.65E-06 | 90.9908664 | 37.8149007 | 40.8342843 | 62.6313804 | 0 |

| | | | | | | |
|---|---|---|---|---|---|---|
| 7.60E-07 | 2.67E-06 | 304.803326 | 352.123458 | 361.012196 | 391.098176 | 85.5438568 |
| 7.62E-07 | 2.67E-06 | 52.0866444 | 108.328569 | 109.510126 | 107.633187 | 0 |
| 7.63E-07 | 2.68E-06 | 0 | 0 | 0 | 0 | 11.8810912 |
| 7.64E-07 | 2.68E-06 | 20.5774398 | 116.781311 | 127.143113 | 116.447974 | 0 |
| 7.64E-07 | 2.68E-06 | 30.8661596 | 62.7282471 | 51.0428554 | 57.0641466 | 0 |
| 7.65E-07 | 2.68E-06 | 35.3674746 | 42.7085937 | 51.9709073 | 70.518295 | 0 |
| 7.65E-07 | 2.68E-06 | 52.0866444 | 186.850098 | 100.229607 | 128.97425 | 0 |
| 7.67E-07 | 2.69E-06 | 24.4357097 | 89.6435588 | 81.6685687 | 112.736485 | 0 |
| 7.68E-07 | 2.69E-06 | 33.4383396 | 81.6356974 | 46.4025958 | 59.3838274 | 0 |
| 7.68E-07 | 2.69E-06 | 36.9750871 | 80.5234945 | 62.1794784 | 39.898509 | 0 |
| 7.68E-07 | 2.69E-06 | 47.9068519 | 143.474182 | 148.488307 | 136.397228 | 0 |
| 7.70E-07 | 2.70E-06 | 22.1850522 | 88.976237 | 128.071165 | 124.334889 | 0 |
| 7.73E-07 | 2.71E-06 | 25.4002772 | 126.568697 | 68.6758419 | 115.984038 | 0 |
| 7.74E-07 | 2.71E-06 | 23.7926647 | 92.5352865 | 99.3015551 | 88.1478687 | 0 |
| 7.74E-07 | 2.71E-06 | 88.4186864 | 68.7341431 | 72.3880495 | 75.6215927 | 2.37621824 |
| 7.75E-07 | 2.71E-06 | 108.031559 | 341.891191 | 358.22804 | 360.942326 | 40.3957101 |
| 7.75E-07 | 2.71E-06 | 30.5446371 | 64.5077718 | 50.1148035 | 67.7346781 | 0 |
| 7.75E-07 | 2.72E-06 | 32.1522496 | 54.7203857 | 48.2586997 | 70.518295 | 0 |
| 7.77E-07 | 2.72E-06 | 27.6509347 | 60.2814006 | 90.9490879 | 81.6527626 | 0 |
| 7.79E-07 | 2.73E-06 | 29.9015921 | 57.1672323 | 57.5392189 | 57.9920189 | 0 |
| 7.79E-07 | 2.73E-06 | 29.2585471 | 68.0668213 | 81.6685687 | 55.208402 | 0 |
| 7.79E-07 | 2.73E-06 | 130.538133 | 503.827942 | 503.932191 | 535.846255 | 2.37621824 |
| 7.80E-07 | 2.73E-06 | 40.190312 | 124.789172 | 41.7623363 | 201.34829 | 0 |
| 7.81E-07 | 2.73E-06 | 24.7572322 | 127.013578 | 77.0283091 | 94.6429749 | 0 |
| 7.83E-07 | 2.74E-06 | 51.7651219 | 121.007682 | 98.3735032 | 118.303719 | 0 |
| 7.85E-07 | 2.75E-06 | 67.5197242 | 78.7439697 | 25.9854537 | 91.859358 | 0 |
| 7.86E-07 | 2.75E-06 | 28.2939797 | 58.5018758 | 70.5319457 | 75.6215927 | 0 |
| 7.88E-07 | 2.76E-06 | 28.9370247 | 104.99196 | 167.049345 | 60.7756358 | 0 |
| 7.89E-07 | 2.76E-06 | 104.173289 | 161.714311 | 178.185968 | 157.274355 | 4.75243649 |
| 7.91E-07 | 2.77E-06 | 28.2939797 | 92.0904053 | 56.6111669 | 92.3232941 | 0 |
| 7.95E-07 | 2.78E-06 | 16.7191698 | 141.249776 | 159.62493 | 182.326907 | 0 |
| 7.98E-07 | 2.79E-06 | 25.0787547 | 111.887618 | 77.0283091 | 84.4363795 | 0 |
| 7.98E-07 | 2.79E-06 | 12.8608998 | 10.899589 | 25.9854537 | 19.4853184 | 99.8011662 |
| 8.00E-07 | 2.80E-06 | 31.1876821 | 51.3837769 | 74.2441534 | 54.7444659 | 0 |
| 8.00E-07 | 2.80E-06 | 30.2231146 | 108.328569 | 53.8270112 | 78.4052096 | 0 |
| 8.02E-07 | 2.80E-06 | 97.4213163 | 99.2085042 | 22.273246 | 100.210209 | 0 |
| 8.02E-07 | 2.80E-06 | 56.2664368 | 57.6121134 | 179.11402 | 43.6099982 | 0 |
| 8.04E-07 | 2.81E-06 | 65.9121117 | 72.9605143 | 26.9135056 | 103.457762 | 0 |
| 8.10E-07 | 2.83E-06 | 28.6155022 | 68.7341431 | 56.6111669 | 69.5904227 | 0 |
| 8.10E-07 | 2.83E-06 | 53.6942569 | 30.02948 | 27.8415575 | 24.1246799 | 144.949313 |
| 8.10E-07 | 2.83E-06 | 18.0052598 | 12.4566732 | 3.71220767 | 17.6295737 | 83.1676385 |
| 8.11E-07 | 2.83E-06 | 23.7926647 | 86.7518311 | 87.2368802 | 109.024996 | 0 |
| 8.16E-07 | 2.85E-06 | 31.5092046 | 58.0569946 | 53.8270112 | 91.3954218 | 0 |

| | | | | | | |
|---|---|---|---|---|---|---|
| 8.16E-07 | 2.85E-06 | 34.7244296 | 51.828658 | 39.9062324 | 52.8887212 | 0 |
| 8.18E-07 | 2.86E-06 | 24.1141872 | 103.212435 | 168.905449 | 81.1888265 | 0 |
| 8.19E-07 | 2.86E-06 | 23.7926647 | 153.484009 | 79.8124649 | 106.241379 | 0 |
| 8.20E-07 | 2.86E-06 | 438.556685 | 667.099337 | 602.305694 | 718.637098 | 42.7719284 |
| 8.22E-07 | 2.87E-06 | 21.8635297 | 115.224227 | 94.6612955 | 111.344676 | 0 |
| 8.23E-07 | 2.87E-06 | 32.7952946 | 53.3857422 | 49.1867516 | 80.7248903 | 0 |
| 8.23E-07 | 2.87E-06 | 49.5144644 | 38.9271037 | 35.2659728 | 53.8165936 | 0 |
| 8.23E-07 | 2.87E-06 | 27.0078897 | 63.1731283 | 77.956361 | 75.6215927 | 0 |
| 8.25E-07 | 2.88E-06 | 26.0433222 | 91.4230835 | 72.3880495 | 73.3019119 | 0 |
| 8.26E-07 | 2.88E-06 | 40.5118345 | 58.946757 | 39.9062324 | 39.4345729 | 0 |
| 8.28E-07 | 2.89E-06 | 44.691627 | 56.2774699 | 31.5537652 | 57.5280828 | 0 |
| 8.30E-07 | 2.89E-06 | 20.5774398 | 112.55494 | 106.72597 | 139.180845 | 0 |
| 8.30E-07 | 2.89E-06 | 31.1876821 | 52.4959798 | 61.2514265 | 48.7132959 | 0 |
| 8.30E-07 | 2.89E-06 | 69.1273367 | 78.9664103 | 70.5319457 | 72.3740396 | 2.37621824 |
| 8.30E-07 | 2.90E-06 | 23.1496197 | 111.220296 | 90.9490879 | 90.0036134 | 0 |
| 8.34E-07 | 2.91E-06 | 9.00262989 | 2.66928711 | 4.64025958 | 4.63936151 | 45.1481466 |
| 8.34E-07 | 2.91E-06 | 32.4737721 | 48.7144897 | 46.4025958 | 50.1051043 | 0 |
| 8.36E-07 | 2.91E-06 | 31.8307271 | 57.834554 | 49.1867516 | 48.2493597 | 0 |
| 8.36E-07 | 2.92E-06 | 26.3648447 | 108.551009 | 62.1794784 | 108.097123 | 0 |
| 8.37E-07 | 2.92E-06 | 0 | 0 | 0 | 0 | 54.6530196 |
| 8.39E-07 | 2.92E-06 | 33.1168171 | 81.858138 | 107.654022 | 45.4657428 | 0 |
| 8.39E-07 | 2.93E-06 | 32.4737721 | 46.4900838 | 47.3306478 | 54.2805297 | 0 |
| 8.41E-07 | 2.93E-06 | 35.6889971 | 48.7144897 | 47.3306478 | 38.9706367 | 0 |
| 8.42E-07 | 2.94E-06 | 268.149762 | 288.060567 | 245.933758 | 293.671584 | 23.7621824 |
| 8.43E-07 | 2.94E-06 | 130.859656 | 57.6121134 | 35.2659728 | 56.6002105 | 0 |
| 8.44E-07 | 2.94E-06 | 26.3648447 | 77.4093262 | 154.056618 | 81.1888265 | 0 |
| 8.44E-07 | 2.94E-06 | 217.349207 | 215.544934 | 192.106747 | 209.69914 | 28.5146189 |
| 8.44E-07 | 2.94E-06 | 29.9015921 | 61.1711629 | 54.7550631 | 53.8165936 | 0 |
| 8.46E-07 | 2.94E-06 | 70.4134267 | 31.1416829 | 161.481034 | 67.2707419 | 0 |
| 8.46E-07 | 2.94E-06 | 108.996126 | 62.9506877 | 83.5246725 | 87.2199964 | 4.75243649 |
| 8.48E-07 | 2.95E-06 | 179.731075 | 312.529032 | 245.005706 | 262.587862 | 35.6432737 |
| 8.48E-07 | 2.95E-06 | 31.1876821 | 52.4959798 | 48.2586997 | 60.3116997 | 0 |
| 8.50E-07 | 2.96E-06 | 34.7244296 | 48.2696086 | 39.9062324 | 53.3526574 | 0 |
| 8.54E-07 | 2.97E-06 | 25.7217997 | 103.212435 | 69.6038938 | 80.7248903 | 0 |
| 8.54E-07 | 2.97E-06 | 30.8661596 | 76.7420044 | 51.9709073 | 55.208402 | 0 |
| 8.54E-07 | 2.97E-06 | 27.6509347 | 70.5136678 | 76.1002572 | 58.9198912 | 0 |
| 8.56E-07 | 2.98E-06 | 30.8661596 | 99.875826 | 53.8270112 | 64.0231889 | 0 |
| 8.58E-07 | 2.98E-06 | 25.4002772 | 74.7400391 | 86.3088283 | 75.1576565 | 0 |
| 8.58E-07 | 2.99E-06 | 64.3044992 | 25.8031087 | 115.078438 | 136.861165 | 0 |
| 8.59E-07 | 2.99E-06 | 140.505331 | 62.2833659 | 36.1940248 | 53.3526574 | 0 |
| 8.59E-07 | 2.99E-06 | 25.0787547 | 89.4211182 | 70.5319457 | 109.952868 | 0 |
| 8.59E-07 | 2.99E-06 | 47.9068519 | 53.1633016 | 31.5537652 | 46.8575513 | 0 |
| 8.61E-07 | 3.00E-06 | 17.3622148 | 10.4547078 | 21.3451941 | 18.0935099 | 49.9005831 |

| | | | | | | |
|---|---|---|---|---|---|---|
| 8.63E-07 | 3.00E-06 | 27.6509347 | 74.2951579 | 58.4672708 | 87.6839326 | 0 |
| 8.64E-07 | 3.00E-06 | 44.3701045 | 33.3660889 | 92.8051917 | 111.344676 | 0 |
| 8.65E-07 | 3.01E-06 | 76.8438766 | 50.2715739 | 38.0501286 | 39.4345729 | 0 |
| 8.66E-07 | 3.01E-06 | 37.2966096 | 37.1475789 | 58.4672708 | 70.9822312 | 0 |
| 8.69E-07 | 3.02E-06 | 18.9698273 | 108.106128 | 149.416359 | 144.284143 | 0 |
| 8.69E-07 | 3.02E-06 | 33.4383396 | 79.1888509 | 44.546492 | 56.6002105 | 0 |
| 8.69E-07 | 3.02E-06 | 79.4160565 | 42.2637126 | 36.1940248 | 50.5690405 | 0 |
| 8.70E-07 | 3.02E-06 | 32.1522496 | 121.230123 | 59.3953227 | 60.3116997 | 0 |
| 8.70E-07 | 3.02E-06 | 26.0433222 | 86.9742716 | 67.7477899 | 75.1576565 | 0 |
| 8.71E-07 | 3.02E-06 | 11.2532874 | 158.155261 | 309.041288 | 346.560305 | 0 |
| 8.72E-07 | 3.03E-06 | 27.6509347 | 62.2833659 | 64.0355823 | 82.1166988 | 0 |
| 8.73E-07 | 3.03E-06 | 6673.19941 | 40679.0458 | 71894.3259 | 38290.0424 | 582.17347 |
| 8.73E-07 | 3.03E-06 | 35.3674746 | 38.4822225 | 67.7477899 | 70.0543588 | 0 |
| 8.74E-07 | 3.04E-06 | 37.9396545 | 58.7243164 | 35.2659728 | 53.8165936 | 0 |
| 8.78E-07 | 3.05E-06 | 29.2585471 | 63.84045 | 53.8270112 | 57.9920189 | 0 |
| 8.80E-07 | 3.06E-06 | 40.833357 | 33.5885295 | 67.7477899 | 60.7756358 | 0 |
| 8.82E-07 | 3.06E-06 | 27.6509347 | 91.2006429 | 64.9636342 | 63.5592527 | 0 |
| 8.85E-07 | 3.07E-06 | 30.2231146 | 64.2853312 | 51.0428554 | 53.8165936 | 0 |
| 8.86E-07 | 3.08E-06 | 730.499111 | 2145.8844 | 2122.45473 | 2241.27555 | 0 |
| 8.86E-07 | 3.08E-06 | 85.524984 | 228.001607 | 107.654022 | 205.987651 | 11.8810912 |
| 8.87E-07 | 3.08E-06 | 173.300625 | 54.4979451 | 51.0428554 | 47.3214874 | 0 |
| 8.87E-07 | 3.08E-06 | 85.8465065 | 92.5352865 | 46.4025958 | 31.0837221 | 0 |
| 8.87E-07 | 3.08E-06 | 27.6509347 | 65.3975342 | 61.2514265 | 65.4149973 | 0 |
| 8.87E-07 | 3.08E-06 | 81.3451915 | 94.7596924 | 25.0574018 | 67.7346781 | 0 |
| 8.88E-07 | 3.08E-06 | 78.7730116 | 85.6396281 | 25.9854537 | 62.1674443 | 0 |
| 8.88E-07 | 3.08E-06 | 42.119447 | 94.5372518 | 33.409869 | 88.1478687 | 0 |
| 8.89E-07 | 3.08E-06 | 20.8989622 | 113.889583 | 113.222334 | 101.602017 | 0 |
| 8.90E-07 | 3.09E-06 | 56.5879593 | 165.05092 | 143.848047 | 169.336695 | 9.50487297 |
| 8.90E-07 | 3.09E-06 | 26.0433222 | 113.444702 | 87.2368802 | 64.9510612 | 0 |
| 8.90E-07 | 3.09E-06 | 25.0787547 | 86.0845093 | 99.3015551 | 69.1264865 | 0 |
| 8.94E-07 | 3.10E-06 | 28.6155022 | 70.7361084 | 55.683115 | 60.7756358 | 0 |
| 8.95E-07 | 3.10E-06 | 71.0564716 | 122.342326 | 132.711424 | 23.6607437 | 0 |
| 8.96E-07 | 3.10E-06 | 63.6614542 | 71.4034302 | 36.1940248 | 37.1148921 | 0 |
| 8.96E-07 | 3.11E-06 | 29.2585471 | 61.1711629 | 61.2514265 | 102.065953 | 0 |
| 8.97E-07 | 3.11E-06 | 77.1653991 | 28.0275146 | 95.5893474 | 56.1362743 | 0 |
| 8.98E-07 | 3.11E-06 | 71.3779941 | 63.3955688 | 51.0428554 | 29.2279775 | 0 |
| 9.02E-07 | 3.12E-06 | 29.9015921 | 51.1613363 | 73.3161014 | 57.0641466 | 0 |
| 9.02E-07 | 3.12E-06 | 601.56859 | 1714.57209 | 1457.96956 | 1582.48621 | 0 |
| 9.02E-07 | 3.13E-06 | 146.614258 | 439.765051 | 416.695311 | 403.160515 | 21.3859642 |
| 9.03E-07 | 3.13E-06 | 28.9370247 | 51.1613363 | 77.0283091 | 72.8379758 | 0 |
| 9.05E-07 | 3.13E-06 | 81.3451915 | 240.45828 | 158.696878 | 197.6368 | 21.3859642 |
| 9.08E-07 | 3.14E-06 | 92.9200014 | 88.3089152 | 67.7477899 | 68.6625504 | 4.75243649 |
| 9.09E-07 | 3.15E-06 | 260.433222 | 981.185453 | 893.713996 | 971.018365 | 30.8908372 |

| | | | | | | |
|---|---|---|---|---|---|---|
| 9.10E-07 | 3.15E-06 | 37.2966096 | 40.0393066 | 38.9781805 | 49.177232 | 0 |
| 9.12E-07 | 3.16E-06 | 28.6155022 | 74.2951579 | 57.5392189 | 57.9920189 | 0 |
| 9.13E-07 | 3.16E-06 | 90.3478214 | 131.017509 | 198.60311 | 194.853184 | 14.2573095 |
| 9.14E-07 | 3.16E-06 | 29.5800696 | 83.6376628 | 49.1867516 | 78.8691457 | 0 |
| 9.14E-07 | 3.16E-06 | 229.24554 | 45.6003214 | 55.683115 | 64.0231889 | 0 |
| 9.21E-07 | 3.19E-06 | 517.008174 | 115.001786 | 76.1002572 | 97.4265918 | 2.37621824 |
| 9.28E-07 | 3.21E-06 | 55.3018693 | 53.3857422 | 42.6903882 | 32.0115944 | 0 |
| 9.28E-07 | 3.21E-06 | 393.222013 | 659.091475 | 592.097123 | 703.327205 | 45.1481466 |
| 9.29E-07 | 3.21E-06 | 28.9370247 | 64.2853312 | 56.6111669 | 54.2805297 | 0 |
| 9.29E-07 | 3.21E-06 | 28.2939797 | 55.3877075 | 61.2514265 | 66.3428696 | 0 |
| 9.29E-07 | 3.21E-06 | 33.4383396 | 63.84045 | 42.6903882 | 48.7132959 | 0 |
| 9.30E-07 | 3.21E-06 | 39.547267 | 48.4920492 | 41.7623363 | 35.7230837 | 0 |
| 9.30E-07 | 3.22E-06 | 30.5446371 | 49.1593709 | 54.7550631 | 50.5690405 | 0 |
| 9.30E-07 | 3.22E-06 | 23.1496197 | 155.263534 | 82.5966206 | 97.8905279 | 0 |
| 9.30E-07 | 3.22E-06 | 34.7244296 | 49.1593709 | 42.6903882 | 40.8263813 | 0 |
| 9.32E-07 | 3.22E-06 | 28.2939797 | 71.6258708 | 56.6111669 | 59.8477635 | 0 |
| 9.32E-07 | 3.22E-06 | 49.8359869 | 63.1731283 | 62.1794784 | 29.2279775 | 0 |
| 9.33E-07 | 3.22E-06 | 28.6155022 | 95.2045736 | 51.9709073 | 85.3642518 | 0 |
| 9.33E-07 | 3.23E-06 | 93.8845689 | 145.476147 | 107.654022 | 128.97425 | 19.0097459 |
| 9.36E-07 | 3.23E-06 | 27.0078897 | 68.7341431 | 64.9636342 | 63.0953166 | 0 |
| 9.37E-07 | 3.24E-06 | 36.3320421 | 50.0491333 | 52.8989593 | 36.1870198 | 0 |
| 9.38E-07 | 3.24E-06 | 50.4790319 | 52.940861 | 44.546492 | 31.5476583 | 0 |
| 9.44E-07 | 3.26E-06 | 28.9370247 | 61.6160441 | 60.3233746 | 51.9608489 | 0 |
| 9.47E-07 | 3.27E-06 | 23.4711422 | 145.698588 | 74.2441534 | 107.169251 | 0 |
| 9.48E-07 | 3.27E-06 | 22.1850522 | 104.102197 | 87.2368802 | 100.210209 | 0 |
| 9.48E-07 | 3.27E-06 | 30.5446371 | 98.763623 | 95.5893474 | 46.8575513 | 0 |
| 9.48E-07 | 3.27E-06 | 44.048582 | 43.5983561 | 33.409869 | 43.1460621 | 0 |
| 9.49E-07 | 3.28E-06 | 22.8280972 | 129.682865 | 78.8844129 | 101.602017 | 0 |
| 9.49E-07 | 3.28E-06 | 29.9015921 | 51.1613363 | 53.8270112 | 52.8887212 | 0 |
| 9.49E-07 | 3.28E-06 | 28.6155022 | 57.3896728 | 54.7550631 | 65.8789335 | 0 |
| 9.49E-07 | 3.28E-06 | 30.2231146 | 48.2696086 | 60.3233746 | 52.8887212 | 0 |
| 9.51E-07 | 3.28E-06 | 27.9724572 | 57.1672323 | 66.819738 | 59.8477635 | 0 |
| 9.52E-07 | 3.29E-06 | 36.3320421 | 69.6239054 | 38.9781805 | 49.177232 | 0 |
| 9.54E-07 | 3.29E-06 | 85.2034615 | 169.944613 | 198.60311 | 237.535309 | 16.6335277 |
| 9.54E-07 | 3.29E-06 | 25.0787547 | 68.9565837 | 94.6612955 | 77.9412734 | 0 |
| 9.55E-07 | 3.29E-06 | 26.3648447 | 74.7400391 | 61.2514265 | 80.2609542 | 0 |
| 9.56E-07 | 3.30E-06 | 48.2283744 | 46.934965 | 30.6257133 | 52.4247851 | 0 |
| 9.57E-07 | 3.30E-06 | 40.833357 | 64.5077718 | 32.4818171 | 57.0641466 | 0 |
| 9.57E-07 | 3.30E-06 | 25.4002772 | 84.0825439 | 96.5173994 | 64.487125 | 0 |
| 9.60E-07 | 3.31E-06 | 38.904222 | 97.4289795 | 35.2659728 | 92.3232941 | 0 |
| 9.60E-07 | 3.31E-06 | 40.190312 | 146.810791 | 140.135839 | 151.243185 | 2.37621824 |
| 9.66E-07 | 3.33E-06 | 44.691627 | 270.48776 | 168.905449 | 209.69914 | 9.50487297 |
| 9.66E-07 | 3.33E-06 | 31.5092046 | 84.5274251 | 46.4025958 | 61.239572 | 0 |

| | | | | | | |
|---|---|---|---|---|---|---|
| 9.67E-07 | 3.33E-06 | 38.261177 | 177.952474 | 138.279736 | 164.697334 | 2.37621824 |
| 9.68E-07 | 3.34E-06 | 32.7952946 | 50.7164551 | 41.7623363 | 62.1674443 | 0 |
| 9.69E-07 | 3.34E-06 | 22.8280972 | 103.879757 | 87.2368802 | 85.3642518 | 0 |
| 9.70E-07 | 3.34E-06 | 34.4029071 | 59.3916382 | 64.9636342 | 38.0427644 | 0 |
| 9.70E-07 | 3.34E-06 | 287.762634 | 661.760762 | 477.018685 | 538.165936 | 28.5146189 |
| 9.72E-07 | 3.35E-06 | 123.464639 | 83.6376628 | 59.3953227 | 81.1888265 | 9.50487297 |
| 9.73E-07 | 3.35E-06 | 26.0433222 | 140.582454 | 70.5319457 | 80.7248903 | 0 |
| 9.73E-07 | 3.35E-06 | 23.4711422 | 74.2951579 | 95.5893474 | 100.674145 | 0 |
| 9.74E-07 | 3.35E-06 | 100.315019 | 217.102018 | 146.632203 | 175.831801 | 16.6335277 |
| 9.74E-07 | 3.35E-06 | 39.547267 | 33.8109701 | 50.1148035 | 47.3214874 | 0 |
| 9.75E-07 | 3.36E-06 | 24.7572322 | 78.2990885 | 115.078438 | 73.3019119 | 0 |
| 9.76E-07 | 3.36E-06 | 27.6509347 | 74.0727173 | 58.4672708 | 60.7756358 | 0 |
| 9.77E-07 | 3.36E-06 | 31.1876821 | 93.6474894 | 73.3161014 | 45.929679 | 0 |
| 9.77E-07 | 3.36E-06 | 32.7952946 | 41.3739502 | 77.956361 | 65.4149973 | 0 |
| 9.77E-07 | 3.36E-06 | 61.4107968 | 33.8109701 | 36.1940248 | 39.898509 | 168.711495 |
| 9.80E-07 | 3.37E-06 | 40.833357 | 152.371806 | 141.991943 | 144.284143 | 0 |
| 9.82E-07 | 3.38E-06 | 28.6155022 | 90.9782023 | 71.4599976 | 52.4247851 | 0 |
| 9.84E-07 | 3.39E-06 | 28.9370247 | 68.5117025 | 77.956361 | 49.6411682 | 0 |
| 9.86E-07 | 3.39E-06 | 47.9068519 | 48.9369303 | 34.3379209 | 90.0036134 | 0 |
| 9.86E-07 | 3.39E-06 | 32.1522496 | 63.84045 | 44.546492 | 48.7132959 | 0 |
| 9.89E-07 | 3.40E-06 | 186.80457 | 1313.5117 | 644.06803 | 880.550815 | 26.1384007 |
| 9.89E-07 | 3.40E-06 | 21.2204847 | 102.322673 | 115.078438 | 92.3232941 | 0 |
| 9.90E-07 | 3.40E-06 | 24.1141872 | 74.2951579 | 121.574801 | 84.4363795 | 0 |
| 9.90E-07 | 3.41E-06 | 27.0078897 | 84.7498657 | 61.2514265 | 64.487125 | 0 |
| 9.92E-07 | 3.41E-06 | 36.3320421 | 60.05896 | 35.2659728 | 57.9920189 | 0 |
| 9.92E-07 | 3.41E-06 | 23.1496197 | 98.0963013 | 76.1002572 | 97.8905279 | 0 |
| 9.94E-07 | 3.42E-06 | 29.9015921 | 86.0845093 | 53.8270112 | 56.1362743 | 0 |
| 1.00E-06 | 3.44E-06 | 27.6509347 | 69.6239054 | 56.6111669 | 61.239572 | 0 |
| 1.00E-06 | 3.45E-06 | 32.1522496 | 41.3739502 | 53.8270112 | 51.0329766 | 0 |
| 1.00E-06 | 3.45E-06 | 23.1496197 | 85.8620687 | 133.639476 | 82.5806349 | 0 |
| 1.01E-06 | 3.46E-06 | 70.0919042 | 132.797034 | 150.344411 | 160.521908 | 7.12865473 |
| 1.01E-06 | 3.47E-06 | 27.9724572 | 60.7262817 | 94.6612955 | 60.3116997 | 0 |
| 1.01E-06 | 3.47E-06 | 1426.91684 | 3653.80917 | 2883.45731 | 3305.54508 | 0 |
| 1.02E-06 | 3.49E-06 | 31.5092046 | 106.326603 | 47.3306478 | 70.9822312 | 0 |
| 1.02E-06 | 3.49E-06 | 20.8989622 | 85.8620687 | 154.056618 | 116.91191 | 0 |
| 1.02E-06 | 3.49E-06 | 0.64304499 | 0.22244059 | 1.85610383 | 0.46393615 | 40.3957101 |
| 1.02E-06 | 3.50E-06 | 25.0787547 | 62.7282471 | 102.085711 | 91.3954218 | 0 |
| 1.02E-06 | 3.51E-06 | 41.1548795 | 113.889583 | 68.6758419 | 36.650956 | 0 |
| 1.02E-06 | 3.51E-06 | 30.8661596 | 108.77345 | 111.36623 | 45.4657428 | 0 |
| 1.03E-06 | 3.52E-06 | 35.0459521 | 53.8306234 | 36.1940248 | 53.8165936 | 0 |
| 1.03E-06 | 3.52E-06 | 23.1496197 | 96.0943359 | 77.956361 | 87.6839326 | 0 |
| 1.03E-06 | 3.53E-06 | 21.5420072 | 86.7518311 | 100.229607 | 112.736485 | 0 |
| 1.03E-06 | 3.53E-06 | 58.8386168 | 79.8561727 | 25.9854537 | 102.993826 | 0 |

| | | | | | | |
|---|---|---|---|---|---|---|
| 1.03E-06 | 3.54E-06 | 17.6837373 | 117.448633 | 139.207788 | 141.03659 | 0 |
| 1.03E-06 | 3.54E-06 | 35.3674746 | 135.24388 | 39.9062324 | 107.633187 | 0 |
| 1.03E-06 | 3.55E-06 | 30.5446371 | 62.0609253 | 46.4025958 | 51.0329766 | 0 |
| 1.04E-06 | 3.55E-06 | 28.9370247 | 91.4230835 | 49.1867516 | 77.4773373 | 0 |
| 1.04E-06 | 3.56E-06 | 44.3701045 | 117.448633 | 31.5537652 | 94.1790387 | 0 |
| 1.04E-06 | 3.56E-06 | 31.1876821 | 90.5333211 | 43.6184401 | 84.9003157 | 0 |
| 1.04E-06 | 3.56E-06 | 29.9015921 | 56.4999105 | 139.207788 | 74.2297842 | 0 |
| 1.04E-06 | 3.56E-06 | 35.0459521 | 57.834554 | 41.7623363 | 92.7872303 | 0 |
| 1.04E-06 | 3.57E-06 | 16.7191698 | 135.911202 | 147.560255 | 137.789037 | 0 |
| 1.04E-06 | 3.58E-06 | 19.2913498 | 131.684831 | 94.6612955 | 149.387441 | 0 |
| 1.05E-06 | 3.59E-06 | 29.5800696 | 49.8266927 | 53.8270112 | 51.0329766 | 0 |
| 1.05E-06 | 3.60E-06 | 30.8661596 | 61.6160441 | 69.6038938 | 43.1460621 | 0 |
| 1.05E-06 | 3.60E-06 | 20.2559173 | 140.582454 | 127.143113 | 86.7560603 | 0 |
| 1.05E-06 | 3.61E-06 | 33.1168171 | 43.8207967 | 44.546492 | 41.7542536 | 0 |
| 1.05E-06 | 3.61E-06 | 484.534402 | 2009.97319 | 1368.87658 | 1620.52898 | 42.7719284 |
| 1.06E-06 | 3.62E-06 | 29.5800696 | 57.834554 | 47.3306478 | 54.7444659 | 0 |
| 1.06E-06 | 3.64E-06 | 23.1496197 | 152.594246 | 66.819738 | 139.180845 | 0 |
| 1.06E-06 | 3.64E-06 | 21.2204847 | 109.440771 | 92.8051917 | 95.106911 | 0 |
| 1.07E-06 | 3.67E-06 | 28.2939797 | 65.8424154 | 51.9709073 | 57.5280828 | 0 |
| 1.07E-06 | 3.67E-06 | 19.2913498 | 150.36984 | 100.229607 | 115.056166 | 0 |
| 1.07E-06 | 3.67E-06 | 32.4737721 | 73.6278361 | 39.9062324 | 71.9101035 | 0 |
| 1.07E-06 | 3.67E-06 | 26.6863672 | 61.1711629 | 63.1075304 | 63.0953166 | 0 |
| 1.07E-06 | 3.68E-06 | 27.6509347 | 60.9487223 | 53.8270112 | 64.9510612 | 0 |
| 1.07E-06 | 3.68E-06 | 19.2913498 | 143.474182 | 101.157659 | 360.942326 | 0 |
| 1.08E-06 | 3.68E-06 | 27.3294122 | 56.0550293 | 108.582074 | 76.0855288 | 0 |
| 1.08E-06 | 3.69E-06 | 48.5498969 | 112.55494 | 116.00649 | 122.015208 | 2.37621824 |
| 1.08E-06 | 3.69E-06 | 17.6837373 | 108.328569 | 158.696878 | 138.252973 | 0 |
| 1.08E-06 | 3.69E-06 | 48.8714194 | 215.544934 | 33.409869 | 107.633187 | 0 |
| 1.08E-06 | 3.70E-06 | 131.502701 | 199.08433 | 191.178695 | 188.358077 | 23.7621824 |
| 1.08E-06 | 3.70E-06 | 262.683879 | 72.7380737 | 26.9135056 | 101.138081 | 0 |
| 1.08E-06 | 3.70E-06 | 27.0078897 | 80.5234945 | 79.8124649 | 53.3526574 | 0 |
| 1.08E-06 | 3.70E-06 | 15.7546023 | 166.385563 | 127.143113 | 166.553078 | 0 |
| 1.08E-06 | 3.71E-06 | 40.5118345 | 85.8620687 | 81.6685687 | 32.4755306 | 0 |
| 1.09E-06 | 3.71E-06 | 213.490937 | 457.782739 | 421.33557 | 447.23445 | 104.553603 |
| 1.09E-06 | 3.73E-06 | 62.6968867 | 65.1750936 | 53.8270112 | 70.0543588 | 7.12865473 |
| 1.09E-06 | 3.73E-06 | 36.0105196 | 35.3680542 | 53.8270112 | 46.8575513 | 0 |
| 1.09E-06 | 3.74E-06 | 27.3294122 | 55.6101481 | 103.013763 | 72.8379758 | 0 |
| 1.10E-06 | 3.75E-06 | 37.2966096 | 153.706449 | 161.481034 | 161.449781 | 0 |
| 1.10E-06 | 3.75E-06 | 375.216753 | 460.229586 | 596.737383 | 604.044869 | 30.8908372 |
| 1.10E-06 | 3.76E-06 | 85.524984 | 36.7026978 | 36.1940248 | 102.065953 | 0 |
| 1.10E-06 | 3.76E-06 | 74.2716966 | 220.438627 | 203.24337 | 184.646588 | 0 |
| 1.10E-06 | 3.77E-06 | 28.9370247 | 60.5038411 | 47.3306478 | 62.1674443 | 0 |
| 1.10E-06 | 3.77E-06 | 181.66021 | 90.3108805 | 180.042072 | 18.0935099 | 0 |

| | | | | | | |
|---|---|---|---|---|---|---|
| 1.10E-06 | 3.77E-06 | 19.2913498 | 178.174915 | 85.3807764 | 153.562866 | 0 |
| 1.11E-06 | 3.78E-06 | 31.8307271 | 77.6317668 | 41.7623363 | 61.7035081 | 0 |
| 1.11E-06 | 3.79E-06 | 0 | 0 | 0 | 0 | 83.1676385 |
| 1.11E-06 | 3.79E-06 | 23.4711422 | 73.6278361 | 83.5246725 | 84.9003157 | 0 |
| 1.11E-06 | 3.80E-06 | 221.529 | 425.751294 | 450.10518 | 462.080407 | 35.6432737 |
| 1.12E-06 | 3.82E-06 | 78.1299666 | 115.224227 | 53.8270112 | 90.9314857 | 2.37621824 |
| 1.12E-06 | 3.82E-06 | 25.7217997 | 69.6239054 | 81.6685687 | 60.3116997 | 0 |
| 1.12E-06 | 3.82E-06 | 21.2204847 | 22.4664998 | 26.9135056 | 42.2181898 | 194.849896 |
| 1.12E-06 | 3.83E-06 | 36.6535646 | 55.1652669 | 37.1220767 | 40.8263813 | 0 |
| 1.13E-06 | 3.84E-06 | 186.80457 | 251.58031 | 258.926485 | 273.258393 | 30.8908372 |
| 1.13E-06 | 3.84E-06 | 67.1982017 | 33.1436483 | 75.1722053 | 38.5067006 | 0 |
| 1.13E-06 | 3.85E-06 | 42.4409695 | 43.8207967 | 38.9781805 | 88.6118049 | 0 |
| 1.13E-06 | 3.85E-06 | 34.0813846 | 47.3798462 | 37.1220767 | 45.929679 | 0 |
| 1.13E-06 | 3.85E-06 | 24.4357097 | 72.7380737 | 68.6758419 | 86.7560603 | 0 |
| 1.13E-06 | 3.86E-06 | 36.9750871 | 44.9329997 | 65.8916861 | 36.650956 | 0 |
| 1.13E-06 | 3.86E-06 | 26.3648447 | 72.5156331 | 60.3233746 | 61.239572 | 0 |
| 1.13E-06 | 3.86E-06 | 23.4711422 | 88.3089152 | 70.5319457 | 87.2199964 | 0 |
| 1.14E-06 | 3.87E-06 | 16.0761248 | 144.808826 | 128.071165 | 160.057972 | 0 |
| 1.14E-06 | 3.88E-06 | 124.750728 | 67.8443807 | 27.8415575 | 59.8477635 | 0 |
| 1.14E-06 | 3.88E-06 | 48.5498969 | 33.3660889 | 51.9709073 | 37.5788283 | 0 |
| 1.14E-06 | 3.88E-06 | 76.2008316 | 134.798999 | 25.0574018 | 69.1264865 | 0 |
| 1.14E-06 | 3.89E-06 | 32.7952946 | 39.3719849 | 84.4527244 | 80.7248903 | 0 |
| 1.15E-06 | 3.91E-06 | 28.6155022 | 52.4959798 | 78.8844129 | 54.7444659 | 0 |
| 1.15E-06 | 3.91E-06 | 52.4081669 | 84.0825439 | 88.1649321 | 101.138081 | 2.37621824 |
| 1.15E-06 | 3.91E-06 | 29.5800696 | 19.5747721 | 25.9854537 | 19.9492545 | 90.2962933 |
| 1.15E-06 | 3.92E-06 | 26.6863672 | 73.6278361 | 62.1794784 | 56.6002105 | 0 |
| 1.15E-06 | 3.92E-06 | 70.4134267 | 229.11381 | 204.171422 | 208.771268 | 0 |
| 1.15E-06 | 3.93E-06 | 128.930521 | 62.5058065 | 28.7696094 | 61.7035081 | 0 |
| 1.16E-06 | 3.94E-06 | 92.9200014 | 69.6239054 | 88.1649321 | 23.1968076 | 0 |
| 1.16E-06 | 3.95E-06 | 25.4002772 | 64.7302124 | 69.6038938 | 65.8789335 | 0 |
| 1.16E-06 | 3.95E-06 | 27.9724572 | 88.5313558 | 49.1867516 | 77.0134011 | 0 |
| 1.16E-06 | 3.96E-06 | 33.4383396 | 48.9369303 | 43.6184401 | 38.5067006 | 0 |
| 1.16E-06 | 3.96E-06 | 513.471426 | 119.673039 | 18.5610383 | 132.685739 | 0 |
| 1.17E-06 | 3.98E-06 | 49.5144644 | 104.769519 | 101.157659 | 106.705315 | 2.37621824 |
| 1.17E-06 | 3.99E-06 | 11.5748099 | 227.334285 | 210.667785 | 175.831801 | 0 |
| 1.17E-06 | 3.99E-06 | 79.4160565 | 126.123816 | 20.4171422 | 117.839782 | 0 |
| 1.17E-06 | 3.99E-06 | 64.6260217 | 127.458459 | 86.3088283 | 24.588616 | 0 |
| 1.18E-06 | 4.00E-06 | 43.0840145 | 84.5274251 | 48.2586997 | 34.7952113 | 0 |
| 1.18E-06 | 4.00E-06 | 26.3648447 | 58.2794352 | 64.9636342 | 61.7035081 | 0 |
| 1.18E-06 | 4.01E-06 | 28.6155022 | 63.6180094 | 66.819738 | 46.3936151 | 0 |
| 1.18E-06 | 4.01E-06 | 25.4002772 | 64.952653 | 64.9636342 | 89.5396772 | 0 |
| 1.18E-06 | 4.02E-06 | 274.901734 | 160.379667 | 279.343627 | 262.123925 | 21.3859642 |
| 1.19E-06 | 4.03E-06 | 21.5420072 | 104.324638 | 77.0283091 | 138.252973 | 0 |

| | | | | | | |
|---|---|---|---|---|---|---|
| 1.19E-06 | 4.03E-06 | 27.3294122 | 87.1967122 | 57.5392189 | 58.9198912 | 0 |
| 1.19E-06 | 4.04E-06 | 28.2939797 | 62.2833659 | 61.2514265 | 120.159463 | 0 |
| 1.19E-06 | 4.04E-06 | 51.7651219 | 40.929069 | 49.1867516 | 44.0739344 | 180.592587 |
| 1.19E-06 | 4.04E-06 | 21.8635297 | 88.976237 | 81.6685687 | 96.9626556 | 0 |
| 1.19E-06 | 4.05E-06 | 6.10892743 | 244.01733 | 400.918428 | 439.811471 | 0 |
| 1.19E-06 | 4.05E-06 | 31.8307271 | 61.1711629 | 38.9781805 | 57.0641466 | 0 |
| 1.19E-06 | 4.06E-06 | 78.4514891 | 67.6219401 | 79.8124649 | 70.9822312 | 0 |
| 1.19E-06 | 4.06E-06 | 23.4711422 | 114.112024 | 76.1002572 | 75.1576565 | 0 |
| 1.20E-06 | 4.07E-06 | 40.833357 | 62.2833659 | 40.8342843 | 34.7952113 | 0 |
| 1.20E-06 | 4.07E-06 | 24.1141872 | 77.4093262 | 89.092984 | 65.4149973 | 0 |
| 1.20E-06 | 4.07E-06 | 24.7572322 | 70.5136678 | 91.8771398 | 64.487125 | 0 |
| 1.20E-06 | 4.07E-06 | 34.7244296 | 35.3680542 | 54.7550631 | 61.239572 | 0 |
| 1.20E-06 | 4.07E-06 | 27.0078897 | 127.458459 | 51.9709073 | 158.202228 | 0 |
| 1.20E-06 | 4.08E-06 | 41.1548795 | 52.4959798 | 38.0501286 | 104.84957 | 0 |
| 1.20E-06 | 4.08E-06 | 54.0157793 | 71.4034302 | 25.9854537 | 66.3428696 | 0 |
| 1.21E-06 | 4.11E-06 | 24.1141872 | 64.7302124 | 78.8844129 | 94.6429749 | 0 |
| 1.21E-06 | 4.11E-06 | 137.290106 | 51.1613363 | 39.9062324 | 48.2493597 | 0 |
| 1.21E-06 | 4.11E-06 | 34.7244296 | 65.1750936 | 36.1940248 | 51.0329766 | 0 |
| 1.21E-06 | 4.11E-06 | 24.7572322 | 66.5097371 | 69.6038938 | 70.0543588 | 0 |
| 1.21E-06 | 4.12E-06 | 22.1850522 | 123.89941 | 73.3161014 | 96.4987195 | 0 |
| 1.22E-06 | 4.13E-06 | 15.4330798 | 161.26943 | 187.466487 | 118.767655 | 0 |
| 1.22E-06 | 4.13E-06 | 38.5826995 | 36.2578166 | 64.0355823 | 40.8263813 | 0 |
| 1.22E-06 | 4.13E-06 | 270.721942 | 132.352152 | 97.4454513 | 110.88074 | 2.37621824 |
| 1.22E-06 | 4.13E-06 | 138.897718 | 43.1534749 | 46.4025958 | 49.177232 | 0 |
| 1.22E-06 | 4.13E-06 | 47.9068519 | 102.322673 | 93.7332436 | 105.777442 | 2.37621824 |
| 1.22E-06 | 4.14E-06 | 81.666714 | 44.7105591 | 38.9781805 | 38.5067006 | 0 |
| 1.22E-06 | 4.14E-06 | 42.762492 | 26.6928711 | 24.1293498 | 30.1558498 | 166.335277 |
| 1.22E-06 | 4.14E-06 | 20.8989622 | 81.1908162 | 110.438178 | 103.921698 | 0 |
| 1.22E-06 | 4.15E-06 | 157.546023 | 37.5924601 | 60.3233746 | 49.6411682 | 0 |
| 1.23E-06 | 4.16E-06 | 37.9396545 | 40.4841878 | 34.3379209 | 41.7542536 | 0 |
| 1.23E-06 | 4.17E-06 | 12.5393773 | 28.917277 | 43.6184401 | 42.6821259 | 142.573095 |
| 1.23E-06 | 4.17E-06 | 14.7900348 | 147.478113 | 130.85532 | 278.825627 | 0 |
| 1.23E-06 | 4.18E-06 | 32.7952946 | 81.1908162 | 38.9781805 | 63.0953166 | 0 |
| 1.24E-06 | 4.19E-06 | 26.3648447 | 98.5411824 | 56.6111669 | 70.0543588 | 0 |
| 1.24E-06 | 4.20E-06 | 30.2231146 | 41.8188314 | 52.8989593 | 57.0641466 | 0 |
| 1.24E-06 | 4.20E-06 | 17.0406923 | 113.667143 | 171.689605 | 122.479144 | 0 |
| 1.24E-06 | 4.21E-06 | 27.0078897 | 57.1672323 | 53.8270112 | 63.0953166 | 0 |
| 1.24E-06 | 4.21E-06 | 38.5826995 | 48.7144897 | 31.5537652 | 53.3526574 | 0 |
| 1.24E-06 | 4.21E-06 | 24.4357097 | 72.0707519 | 64.0355823 | 81.1888265 | 0 |
| 1.24E-06 | 4.21E-06 | 38.5826995 | 63.3955688 | 38.0501286 | 38.5067006 | 0 |
| 1.24E-06 | 4.21E-06 | 48.2283744 | 94.7596924 | 131.783372 | 95.106911 | 2.37621824 |
| 1.25E-06 | 4.22E-06 | 40.5118345 | 76.5195638 | 31.5537652 | 94.6429749 | 0 |
| 1.25E-06 | 4.22E-06 | 23.1496197 | 94.5372518 | 65.8916861 | 111.808612 | 0 |

| | | | | | | |
|---|---|---|---|---|---|---|
| 1.25E-06 | 4.23E-06 | 42.4409695 | 126.123816 | 112.294282 | 123.407016 | 2.37621824 |
| 1.25E-06 | 4.23E-06 | 42.4409695 | 49.3818115 | 35.2659728 | 35.2591475 | 0 |
| 1.25E-06 | 4.23E-06 | 30.8661596 | 19.1298909 | 21.3451941 | 17.6295737 | 92.6725115 |
| 1.25E-06 | 4.24E-06 | 11.5748099 | 9.78738607 | 10.2085711 | 10.6705315 | 104.553603 |
| 1.25E-06 | 4.25E-06 | 36.0105196 | 61.3936035 | 34.3379209 | 48.7132959 | 0 |
| 1.26E-06 | 4.26E-06 | 27.0078897 | 78.2990885 | 51.0428554 | 66.8068058 | 0 |
| 1.26E-06 | 4.26E-06 | 21.5420072 | 124.344291 | 76.1002572 | 97.4265918 | 0 |
| 1.26E-06 | 4.26E-06 | 60.4462293 | 67.3994995 | 81.6685687 | 74.2297842 | 0 |
| 1.26E-06 | 4.26E-06 | 26.6863672 | 59.6140788 | 53.8270112 | 66.3428696 | 0 |
| 1.26E-06 | 4.26E-06 | 0.96456749 | 0 | 0 | 0 | 57.0292378 |
| 1.26E-06 | 4.27E-06 | 20.8989622 | 94.5372518 | 121.574801 | 82.5806349 | 0 |
| 1.27E-06 | 4.28E-06 | 186.161525 | 150.592281 | 116.934542 | 137.325101 | 35.6432737 |
| 1.27E-06 | 4.29E-06 | 34.4029071 | 68.5117025 | 56.6111669 | 35.7230837 | 0 |
| 1.27E-06 | 4.31E-06 | 32.7952946 | 37.8149007 | 61.2514265 | 48.7132959 | 0 |
| 1.28E-06 | 4.33E-06 | 36.6535646 | 62.0609253 | 35.2659728 | 44.0739344 | 0 |
| 1.28E-06 | 4.33E-06 | 39.547267 | 43.1534749 | 45.4745439 | 32.0115944 | 0 |
| 1.28E-06 | 4.34E-06 | 24.4357097 | 114.556905 | 72.3880495 | 67.7346781 | 0 |
| 1.29E-06 | 4.35E-06 | 99.9934963 | 127.6809 | 64.0355823 | 23.1968076 | 0 |
| 1.29E-06 | 4.36E-06 | 21.2204847 | 88.3089152 | 103.013763 | 82.5806349 | 0 |
| 1.29E-06 | 4.37E-06 | 23.7926647 | 76.0746826 | 69.6038938 | 118.767655 | 0 |
| 1.29E-06 | 4.37E-06 | 7.71653991 | 5.56101481 | 5.5683115 | 5.56723382 | 52.2768014 |
| 1.29E-06 | 4.37E-06 | 28.9370247 | 56.9447917 | 48.2586997 | 47.3214874 | 0 |
| 1.29E-06 | 4.37E-06 | 123.464639 | 165.27336 | 169.833501 | 113.200421 | 363.561391 |
| 1.29E-06 | 4.37E-06 | 25.4002772 | 94.0923706 | 55.683115 | 79.3330819 | 0 |
| 1.30E-06 | 4.38E-06 | 27.3294122 | 70.7361084 | 51.0428554 | 57.5280828 | 0 |
| 1.30E-06 | 4.38E-06 | 53.6942569 | 94.3148112 | 221.804408 | 196.708928 | 4.75243649 |
| 1.30E-06 | 4.39E-06 | 31.8307271 | 62.5058065 | 88.1649321 | 40.8263813 | 0 |
| 1.31E-06 | 4.42E-06 | 120.570936 | 46.4900838 | 30.6257133 | 71.4461673 | 0 |
| 1.31E-06 | 4.43E-06 | 23.4711422 | 70.958549 | 111.36623 | 73.7658481 | 0 |
| 1.31E-06 | 4.44E-06 | 31.8307271 | 74.5175985 | 40.8342843 | 52.8887212 | 0 |
| 1.31E-06 | 4.44E-06 | 29.9015921 | 57.6121134 | 54.7550631 | 41.2903175 | 0 |
| 1.32E-06 | 4.44E-06 | 35.6889971 | 100.543148 | 38.0501286 | 62.1674443 | 0 |
| 1.32E-06 | 4.45E-06 | 27.6509347 | 56.9447917 | 50.1148035 | 53.3526574 | 0 |
| 1.32E-06 | 4.45E-06 | 17.0406923 | 122.119885 | 138.279736 | 114.128293 | 0 |
| 1.32E-06 | 4.45E-06 | 49.1929419 | 34.4782918 | 45.4745439 | 34.7952113 | 0 |
| 1.32E-06 | 4.45E-06 | 37.618132 | 78.5215291 | 51.0428554 | 35.2591475 | 0 |
| 1.32E-06 | 4.47E-06 | 22.8280972 | 176.61783 | 64.0355823 | 124.798825 | 0 |
| 1.32E-06 | 4.47E-06 | 28.9370247 | 45.1554403 | 50.1148035 | 52.8887212 | 0 |
| 1.33E-06 | 4.47E-06 | 20.2559173 | 95.2045736 | 92.8051917 | 96.9626556 | 0 |
| 1.33E-06 | 4.48E-06 | 23.7926647 | 59.8365194 | 141.063891 | 110.88074 | 0 |
| 1.33E-06 | 4.48E-06 | 36.3320421 | 72.0707519 | 51.9709073 | 34.7952113 | 0 |
| 1.33E-06 | 4.48E-06 | 42.119447 | 186.627657 | 171.689605 | 30.619786 | 0 |
| 1.33E-06 | 4.48E-06 | 35.6889971 | 82.5254598 | 34.3379209 | 90.4675495 | 0 |

| | | | | | | |
|---|---|---|---|---|---|---|
| 1.33E-06 | 4.49E-06 | 55.6233918 | 103.434875 | 77.0283091 | 76.549465 | 0 |
| 1.33E-06 | 4.49E-06 | 119.927891 | 140.137573 | 101.157659 | 113.200421 | 4.75243649 |
| 1.33E-06 | 4.50E-06 | 45.334672 | 40.929069 | 37.1220767 | 89.0757411 | 0 |
| 1.33E-06 | 4.50E-06 | 135.039448 | 39.3719849 | 65.8916861 | 38.9706367 | 0 |
| 1.34E-06 | 4.51E-06 | 71.0564716 | 26.0255493 | 52.8989593 | 70.0543588 | 0 |
| 1.34E-06 | 4.52E-06 | 139.219241 | 210.20636 | 184.682331 | 164.697334 | 0 |
| 1.34E-06 | 4.53E-06 | 30.2231146 | 40.929069 | 49.1867516 | 50.1051043 | 0 |
| 1.34E-06 | 4.53E-06 | 42.119447 | 69.846346 | 34.3379209 | 40.8263813 | 0 |
| 1.35E-06 | 4.54E-06 | 23.7926647 | 86.7518311 | 83.5246725 | 61.239572 | 0 |
| 1.35E-06 | 4.55E-06 | 27.0078897 | 71.6258708 | 56.6111669 | 51.9608489 | 0 |
| 1.35E-06 | 4.55E-06 | 32.7952946 | 38.9271037 | 39.9062324 | 46.3936151 | 0 |
| 1.36E-06 | 4.58E-06 | 22.5065747 | 91.2006429 | 71.4599976 | 80.2609542 | 0 |
| 1.36E-06 | 4.58E-06 | 25.7217997 | 107.661247 | 51.9709073 | 100.674145 | 0 |
| 1.36E-06 | 4.58E-06 | 27.6509347 | 49.3818115 | 77.956361 | 57.5280828 | 0 |
| 1.36E-06 | 4.59E-06 | 15.1115573 | 157.932821 | 154.98467 | 122.479144 | 0 |
| 1.37E-06 | 4.60E-06 | 67.1982017 | 21.5767375 | 125.287009 | 131.757867 | 0 |
| 1.37E-06 | 4.60E-06 | 1308.27504 | 3201.80989 | 2701.55913 | 3093.06232 | 0 |
| 1.37E-06 | 4.63E-06 | 30.5446371 | 43.3759155 | 82.5966206 | 55.208402 | 0 |
| 1.38E-06 | 4.63E-06 | 30.2231146 | 101.21047 | 42.6903882 | 75.6215927 | 0 |
| 1.38E-06 | 4.63E-06 | 23.7926647 | 66.2872965 | 75.1722053 | 69.5904227 | 0 |
| 1.38E-06 | 4.65E-06 | 41.476402 | 48.4920492 | 62.1794784 | 30.1558498 | 0 |
| 1.38E-06 | 4.65E-06 | 25.7217997 | 78.9664103 | 88.1649321 | 51.9608489 | 0 |
| 1.39E-06 | 4.66E-06 | 39.547267 | 50.7164551 | 118.790645 | 41.7542536 | 0 |
| 1.39E-06 | 4.67E-06 | 51.4435994 | 65.8424154 | 29.6976613 | 41.7542536 | 0 |
| 1.39E-06 | 4.67E-06 | 116.391144 | 110.997856 | 154.98467 | 129.438186 | 35.6432737 |
| 1.39E-06 | 4.67E-06 | 20.8989622 | 76.964445 | 102.085711 | 97.4265918 | 0 |
| 1.39E-06 | 4.68E-06 | 23.1496197 | 65.8424154 | 85.3807764 | 77.4773373 | 0 |
| 1.39E-06 | 4.68E-06 | 26.3648447 | 66.7321777 | 72.3880495 | 50.1051043 | 0 |
| 1.39E-06 | 4.69E-06 | 27.3294122 | 76.2971232 | 50.1148035 | 57.5280828 | 0 |
| 1.40E-06 | 4.70E-06 | 25.0787547 | 76.7420044 | 55.683115 | 70.0543588 | 0 |
| 1.40E-06 | 4.70E-06 | 22.1850522 | 83.8601033 | 88.1649321 | 71.4461673 | 0 |
| 1.40E-06 | 4.71E-06 | 47.2638069 | 65.1750936 | 26.9135056 | 57.9920189 | 0 |
| 1.40E-06 | 4.71E-06 | 52.4081669 | 50.0491333 | 26.9135056 | 52.8887212 | 0 |
| 1.40E-06 | 4.71E-06 | 79.737579 | 111.220296 | 19.4890903 | 121.551272 | 0 |
| 1.40E-06 | 4.71E-06 | 20.2559173 | 121.007682 | 76.1002572 | 114.128293 | 0 |
| 1.40E-06 | 4.72E-06 | 41.476402 | 58.7243164 | 28.7696094 | 57.0641466 | 0 |
| 1.40E-06 | 4.72E-06 | 26.6863672 | 57.3896728 | 51.0428554 | 59.8477635 | 0 |
| 1.41E-06 | 4.73E-06 | 3.53674746 | 4.67125244 | 2.78415575 | 7.88691457 | 16.6335277 |
| 1.41E-06 | 4.73E-06 | 31.5092046 | 37.8149007 | 46.4025958 | 50.1051043 | 0 |
| 1.41E-06 | 4.75E-06 | 30.8661596 | 38.2597819 | 54.7550631 | 50.5690405 | 0 |
| 1.41E-06 | 4.75E-06 | 59.4816618 | 59.1691976 | 33.409869 | 34.3312752 | 0 |
| 1.42E-06 | 4.75E-06 | 125.715296 | 54.4979451 | 29.6976613 | 59.3838274 | 0 |
| 1.42E-06 | 4.78E-06 | 49.5144644 | 81.6356974 | 90.9490879 | 104.385634 | 2.37621824 |

| | | | | | | |
|---|---|---|---|---|---|---|
| 1.43E-06 | 4.79E-06 | 73.9501741 | 93.86993 | 29.6976613 | 41.7542536 | 0 |
| 1.43E-06 | 4.80E-06 | 24.1141872 | 56.4999105 | 96.5173994 | 92.3232941 | 0 |
| 1.43E-06 | 4.80E-06 | 42.119447 | 45.3778809 | 38.9781805 | 31.5476583 | 0 |
| 1.43E-06 | 4.81E-06 | 32.1522496 | 37.5924601 | 62.1794784 | 47.7854236 | 0 |
| 1.44E-06 | 4.82E-06 | 25.4002772 | 75.1849202 | 52.8989593 | 71.9101035 | 0 |
| 1.44E-06 | 4.82E-06 | 26.0433222 | 56.0550293 | 54.7550631 | 66.3428696 | 0 |
| 1.44E-06 | 4.83E-06 | 17.3622148 | 137.913167 | 123.430905 | 99.7462725 | 0 |
| 1.44E-06 | 4.83E-06 | 26.0433222 | 229.781132 | 223.660512 | 184.646588 | 2.37621824 |
| 1.44E-06 | 4.83E-06 | 33.4383396 | 39.3719849 | 76.1002572 | 47.3214874 | 0 |
| 1.45E-06 | 4.85E-06 | 27.6509347 | 61.3936035 | 51.0428554 | 48.2493597 | 0 |
| 1.45E-06 | 4.87E-06 | 87.7756414 | 113.444702 | 20.4171422 | 78.8691457 | 0 |
| 1.46E-06 | 4.88E-06 | 30.2231146 | 64.5077718 | 51.9709073 | 40.3624452 | 0 |
| 1.46E-06 | 4.88E-06 | 28.6155022 | 66.9546183 | 51.9709073 | 45.0018067 | 0 |
| 1.46E-06 | 4.88E-06 | 22.5065747 | 99.2085042 | 69.6038938 | 77.0134011 | 0 |
| 1.46E-06 | 4.89E-06 | 54.3373018 | 39.3719849 | 32.4818171 | 39.4345729 | 0 |
| 1.46E-06 | 4.89E-06 | 22.5065747 | 87.4191528 | 65.8916861 | 87.6839326 | 0 |
| 1.46E-06 | 4.89E-06 | 25.0787547 | 80.5234945 | 53.8270112 | 72.8379758 | 0 |
| 1.46E-06 | 4.90E-06 | 22.1850522 | 81.4132568 | 70.5319457 | 85.828188 | 0 |
| 1.47E-06 | 4.92E-06 | 71.0564716 | 26.6928711 | 62.1794784 | 48.7132959 | 0 |
| 1.47E-06 | 4.92E-06 | 60.1247068 | 30.9192423 | 52.8989593 | 38.0427644 | 0 |
| 1.47E-06 | 4.93E-06 | 63.6614542 | 23.8011434 | 17.6329864 | 24.588616 | 456.233903 |
| 1.47E-06 | 4.93E-06 | 29.5800696 | 54.9428263 | 41.7623363 | 70.9822312 | 0 |
| 1.47E-06 | 4.94E-06 | 11.2532874 | 126.568697 | 257.070381 | 285.784669 | 0 |
| 1.48E-06 | 4.96E-06 | 75.5577866 | 426.641056 | 562.399462 | 487.596895 | 21.3859642 |
| 1.49E-06 | 4.99E-06 | 50.8005544 | 98.9860636 | 117.862593 | 25.5164883 | 0 |
| 1.49E-06 | 4.99E-06 | 21.2204847 | 75.4073608 | 106.72597 | 83.9724434 | 0 |
| 1.49E-06 | 4.99E-06 | 28.6155022 | 41.8188314 | 59.3953227 | 62.1674443 | 0 |
| 1.49E-06 | 5.00E-06 | 39.547267 | 201.531177 | 107.654022 | 146.139888 | 0 |
| 1.50E-06 | 5.01E-06 | 25.0787547 | 84.3049845 | 52.8989593 | 75.1576565 | 0 |
| 1.50E-06 | 5.01E-06 | 42.4409695 | 30.02948 | 43.6184401 | 45.0018067 | 0 |
| 1.50E-06 | 5.02E-06 | 31.8307271 | 44.0432373 | 37.1220767 | 44.0739344 | 0 |
| 1.50E-06 | 5.02E-06 | 73.3071291 | 48.047168 | 28.7696094 | 46.8575513 | 0 |
| 1.50E-06 | 5.02E-06 | 29.9015921 | 72.7380737 | 39.9062324 | 61.239572 | 0 |
| 1.50E-06 | 5.02E-06 | 28.6155022 | 90.7557617 | 43.6184401 | 71.4461673 | 0 |
| 1.50E-06 | 5.02E-06 | 34.4029071 | 57.834554 | 111.36623 | 41.2903175 | 0 |
| 1.50E-06 | 5.03E-06 | 34.7244296 | 48.9369303 | 32.4818171 | 49.6411682 | 0 |
| 1.51E-06 | 5.04E-06 | 21.2204847 | 77.8542074 | 85.3807764 | 85.828188 | 0 |
| 1.51E-06 | 5.05E-06 | 27.9724572 | 44.2656779 | 52.8989593 | 55.208402 | 0 |
| 1.51E-06 | 5.06E-06 | 21.2204847 | 93.2026082 | 73.3161014 | 141.500526 | 0 |
| 1.51E-06 | 5.06E-06 | 66.5551567 | 36.7026978 | 36.1940248 | 40.8263813 | 0 |
| 1.52E-06 | 5.07E-06 | 72.3425616 | 98.763623 | 20.4171422 | 104.385634 | 0 |
| 1.52E-06 | 5.08E-06 | 15.7546023 | 162.604073 | 103.941815 | 147.06776 | 0 |
| 1.52E-06 | 5.08E-06 | 38.5826995 | 70.958549 | 82.5966206 | 30.619786 | 0 |

| | | | | | | |
|---|---|---|---|---|---|---|
| 1.52E-06 | 5.09E-06 | 60.7677518 | 151.482043 | 115.078438 | 114.592229 | 9.50487297 |
| 1.52E-06 | 5.09E-06 | 146.935781 | 134.576558 | 84.4527244 | 101.138081 | 21.3859642 |
| 1.52E-06 | 5.09E-06 | 33.4383396 | 98.0963013 | 39.9062324 | 164.697334 | 0 |
| 1.52E-06 | 5.09E-06 | 39.547267 | 34.923173 | 34.3379209 | 49.177232 | 0 |
| 1.52E-06 | 5.09E-06 | 26.6863672 | 57.834554 | 61.2514265 | 47.7854236 | 0 |
| 1.52E-06 | 5.10E-06 | 31.5092046 | 43.8207967 | 91.8771398 | 51.9608489 | 0 |
| 1.53E-06 | 5.11E-06 | 270.078897 | 562.552258 | 528.061541 | 542.805297 | 0 |
| 1.53E-06 | 5.12E-06 | 18.0052598 | 87.864034 | 170.761553 | 119.695527 | 0 |
| 1.53E-06 | 5.12E-06 | 58.1955718 | 109.663212 | 95.5893474 | 131.293931 | 0 |
| 1.54E-06 | 5.14E-06 | 68.8058142 | 34.923173 | 32.4818171 | 64.9510612 | 0 |
| 1.54E-06 | 5.15E-06 | 72.9856066 | 171.056816 | 178.185968 | 185.110524 | 2.37621824 |
| 1.54E-06 | 5.15E-06 | 28.2939797 | 58.5018758 | 46.4025958 | 46.3936151 | 0 |
| 1.54E-06 | 5.16E-06 | 33.1168171 | 42.4861532 | 81.6685687 | 44.5378705 | 0 |
| 1.55E-06 | 5.16E-06 | 27.6509347 | 49.8266927 | 51.0428554 | 46.8575513 | 0 |
| 1.55E-06 | 5.16E-06 | 39.8687895 | 54.7203857 | 41.7623363 | 31.0837221 | 0 |
| 1.55E-06 | 5.17E-06 | 58.1955718 | 77.6317668 | 81.6685687 | 73.7658481 | 4.75243649 |
| 1.55E-06 | 5.18E-06 | 24.7572322 | 87.1967122 | 75.1722053 | 53.3526574 | 0 |
| 1.56E-06 | 5.19E-06 | 22.8280972 | 71.1809896 | 78.8844129 | 66.8068058 | 0 |
| 1.56E-06 | 5.20E-06 | 57.5525268 | 381.930497 | 228.300772 | 307.125732 | 11.8810912 |
| 1.56E-06 | 5.21E-06 | 30.2231146 | 43.8207967 | 58.4672708 | 40.8263813 | 0 |
| 1.56E-06 | 5.22E-06 | 39.547267 | 174.170984 | 181.898176 | 163.769461 | 4.75243649 |
| 1.57E-06 | 5.22E-06 | 52.7296894 | 102.545113 | 24.1293498 | 116.447974 | 0 |
| 1.57E-06 | 5.23E-06 | 47.2638069 | 37.3700195 | 38.9781805 | 32.4755306 | 0 |
| 1.57E-06 | 5.24E-06 | 42.4409695 | 100.765588 | 107.654022 | 124.798825 | 0 |
| 1.58E-06 | 5.28E-06 | 0 | 0 | 0 | 0 | 49.9005831 |
| 1.58E-06 | 5.28E-06 | 42.119447 | 46.2676432 | 38.9781805 | 30.619786 | 0 |
| 1.58E-06 | 5.28E-06 | 37.618132 | 33.5885295 | 56.6111669 | 88.6118049 | 0 |
| 1.58E-06 | 5.29E-06 | 29.9015921 | 73.6278361 | 51.9709073 | 41.7542536 | 0 |
| 1.59E-06 | 5.29E-06 | 30.2231146 | 37.8149007 | 60.3233746 | 53.3526574 | 0 |
| 1.59E-06 | 5.29E-06 | 23.4711422 | 74.7400391 | 62.1794784 | 68.1986142 | 0 |
| 1.59E-06 | 5.29E-06 | 22.8280972 | 66.064856 | 76.1002572 | 71.4461673 | 0 |
| 1.59E-06 | 5.32E-06 | 42.119447 | 31.8090047 | 37.1220767 | 42.2181898 | 0 |
| 1.59E-06 | 5.32E-06 | 48.5498969 | 75.4073608 | 94.6612955 | 25.9804245 | 0 |
| 1.60E-06 | 5.32E-06 | 48.5498969 | 40.4841878 | 28.7696094 | 47.3214874 | 0 |
| 1.60E-06 | 5.32E-06 | 18.6483048 | 251.135429 | 83.5246725 | 127.118505 | 0 |
| 1.60E-06 | 5.32E-06 | 31.1876821 | 65.6199748 | 36.1940248 | 67.7346781 | 0 |
| 1.60E-06 | 5.32E-06 | 39.8687895 | 112.999821 | 141.063891 | 30.619786 | 0 |
| 1.60E-06 | 5.32E-06 | 24.1141872 | 78.2990885 | 65.8916861 | 57.9920189 | 0 |
| 1.60E-06 | 5.33E-06 | 41.1548795 | 36.9251383 | 33.409869 | 37.1148921 | 0 |
| 1.60E-06 | 5.33E-06 | 46.6207619 | 211.096122 | 106.72597 | 98.8184002 | 0 |
| 1.60E-06 | 5.33E-06 | 729.213021 | 1572.87743 | 1314.12151 | 1537.02047 | 64.1578926 |
| 1.60E-06 | 5.33E-06 | 36.0105196 | 52.2735392 | 57.5392189 | 31.0837221 | 0 |
| 1.60E-06 | 5.34E-06 | 25.4002772 | 70.2912272 | 53.8270112 | 57.5280828 | 0 |

| | | | | | | |
|---|---|---|---|---|---|---|
| 1.60E-06 | 5.34E-06 | 157.224501 | 348.119527 | 251.502069 | 292.279775 | 16.6335277 |
| 1.60E-06 | 5.34E-06 | 339.849278 | 169.05485 | 125.287009 | 176.295737 | 16.6335277 |
| 1.61E-06 | 5.35E-06 | 27.3294122 | 77.1868856 | 44.546492 | 65.4149973 | 0 |
| 1.61E-06 | 5.35E-06 | 30.5446371 | 42.4861532 | 77.956361 | 48.7132959 | 0 |
| 1.61E-06 | 5.35E-06 | 99.3504513 | 88.7537964 | 49.1867516 | 75.1576565 | 0 |
| 1.61E-06 | 5.35E-06 | 11.2532874 | 139.692692 | 305.329081 | 189.28595 | 0 |
| 1.61E-06 | 5.36E-06 | 48.2283744 | 53.1633016 | 62.1794784 | 26.4443606 | 0 |
| 1.61E-06 | 5.36E-06 | 25.7217997 | 59.6140788 | 56.6111669 | 51.9608489 | 0 |
| 1.61E-06 | 5.37E-06 | 38.904222 | 57.6121134 | 56.6111669 | 29.2279775 | 0 |
| 1.62E-06 | 5.38E-06 | 30.8661596 | 36.4802572 | 61.2514265 | 64.487125 | 0 |
| 1.62E-06 | 5.40E-06 | 29.9015921 | 64.952653 | 38.0501286 | 58.9198912 | 0 |
| 1.62E-06 | 5.40E-06 | 16.3976473 | 147.922994 | 90.0210359 | 228.256586 | 0 |
| 1.62E-06 | 5.40E-06 | 24.1141872 | 87.4191528 | 54.7550631 | 76.0855288 | 0 |
| 1.63E-06 | 5.41E-06 | 51.1220769 | 95.6494547 | 78.8844129 | 76.549465 | 2.37621824 |
| 1.63E-06 | 5.41E-06 | 31.8307271 | 46.7125244 | 38.0501286 | 38.9706367 | 0 |
| 1.63E-06 | 5.41E-06 | 27.9724572 | 42.7085937 | 90.9490879 | 79.3330819 | 0 |
| 1.63E-06 | 5.42E-06 | 143.077511 | 360.798641 | 287.696094 | 292.743711 | 2.37621824 |
| 1.63E-06 | 5.43E-06 | 20.8989622 | 102.989994 | 104.869867 | 67.7346781 | 0 |
| 1.63E-06 | 5.43E-06 | 32.4737721 | 60.2814006 | 36.1940248 | 45.0018067 | 0 |
| 1.63E-06 | 5.43E-06 | 37.2966096 | 30.6968018 | 46.4025958 | 45.4657428 | 0 |
| 1.64E-06 | 5.44E-06 | 0 | 0 | 0 | 0 | 52.2768014 |
| 1.64E-06 | 5.44E-06 | 58.8386168 | 48.2696086 | 58.4672708 | 34.3312752 | 268.512662 |
| 1.64E-06 | 5.45E-06 | 24.7572322 | 67.8443807 | 54.7550631 | 61.239572 | 0 |
| 1.64E-06 | 5.46E-06 | 45.0131495 | 58.2794352 | 26.9135056 | 50.5690405 | 0 |
| 1.64E-06 | 5.46E-06 | 41.1548795 | 87.6415934 | 29.6976613 | 60.3116997 | 0 |
| 1.64E-06 | 5.46E-06 | 28.6155022 | 57.3896728 | 41.7623363 | 48.7132959 | 0 |
| 1.65E-06 | 5.46E-06 | 57.2310043 | 39.5944255 | 32.4818171 | 38.0427644 | 0 |
| 1.65E-06 | 5.47E-06 | 71.0564716 | 45.3778809 | 29.6976613 | 43.6099982 | 0 |
| 1.65E-06 | 5.48E-06 | 22.5065747 | 140.804895 | 58.4672708 | 112.736485 | 0 |
| 1.65E-06 | 5.49E-06 | 48.8714194 | 109.885653 | 76.1002572 | 93.2511664 | 0 |
| 1.66E-06 | 5.50E-06 | 31.1876821 | 58.7243164 | 37.1220767 | 76.0855288 | 0 |
| 1.66E-06 | 5.51E-06 | 25.4002772 | 65.6199748 | 51.9709073 | 89.5396772 | 0 |
| 1.66E-06 | 5.51E-06 | 70.4134267 | 55.1652669 | 24.1293498 | 98.8184002 | 0 |
| 1.66E-06 | 5.51E-06 | 20.5774398 | 89.1986776 | 132.711424 | 74.2297842 | 0 |
| 1.66E-06 | 5.52E-06 | 47.2638069 | 49.8266927 | 34.3379209 | 32.0115944 | 0 |
| 1.67E-06 | 5.52E-06 | 200.630038 | 291.174735 | 229.228823 | 276.04201 | 97.424948 |
| 1.67E-06 | 5.53E-06 | 51.4435994 | 132.352152 | 87.2368802 | 137.789037 | 4.75243649 |
| 1.67E-06 | 5.54E-06 | 21.5420072 | 78.9664103 | 75.1722053 | 75.6215927 | 0 |
| 1.67E-06 | 5.54E-06 | 80.7021465 | 48.2696086 | 64.0355823 | 60.7756358 | 154.454186 |
| 1.67E-06 | 5.55E-06 | 24.4357097 | 58.2794352 | 134.567528 | 79.3330819 | 0 |
| 1.68E-06 | 5.57E-06 | 24.7572322 | 59.1691976 | 61.2514265 | 55.6723382 | 0 |
| 1.68E-06 | 5.58E-06 | 21.8635297 | 71.6258708 | 75.1722053 | 76.0855288 | 0 |
| 1.68E-06 | 5.59E-06 | 449.166927 | 77.6317668 | 35.2659728 | 69.5904227 | 0 |

| | | | | | | |
|---|---|---|---|---|---|---|
| 1.68E-06 | 5.59E-06 | 65.9121117 | 50.9388957 | 45.4745439 | 47.7854236 | 137.820658 |
| 1.68E-06 | 5.59E-06 | 26.3648447 | 56.4999105 | 48.2586997 | 53.3526574 | 0 |
| 1.69E-06 | 5.59E-06 | 41.1548795 | 29.5845988 | 56.6111669 | 43.1460621 | 0 |
| 1.69E-06 | 5.59E-06 | 45.9777169 | 107.438806 | 83.5246725 | 135.00542 | 0 |
| 1.69E-06 | 5.60E-06 | 103.851766 | 27.5826335 | 102.085711 | 43.1460621 | 0 |
| 1.69E-06 | 5.60E-06 | 39.8687895 | 42.2637126 | 45.4745439 | 29.6919137 | 0 |
| 1.69E-06 | 5.61E-06 | 42.762492 | 29.8070394 | 70.5319457 | 45.929679 | 0 |
| 1.70E-06 | 5.63E-06 | 27.3294122 | 105.2144 | 43.6184401 | 114.128293 | 0 |
| 1.70E-06 | 5.63E-06 | 31.1876821 | 57.834554 | 35.2659728 | 52.4247851 | 0 |
| 1.70E-06 | 5.63E-06 | 27.6509347 | 48.7144897 | 64.0355823 | 45.0018067 | 0 |
| 1.70E-06 | 5.63E-06 | 25.7217997 | 62.7282471 | 56.6111669 | 49.6411682 | 0 |
| 1.70E-06 | 5.64E-06 | 18.3267823 | 100.988029 | 90.9490879 | 109.488932 | 0 |
| 1.71E-06 | 5.66E-06 | 30.8661596 | 38.0373413 | 41.7623363 | 52.8887212 | 0 |
| 1.71E-06 | 5.66E-06 | 29.9015921 | 37.5924601 | 71.4599976 | 64.0231889 | 0 |
| 1.71E-06 | 5.66E-06 | 23.1496197 | 85.4171875 | 58.4672708 | 74.2297842 | 0 |
| 1.71E-06 | 5.68E-06 | 23.4711422 | 79.8561727 | 61.2514265 | 63.5592527 | 0 |
| 1.72E-06 | 5.68E-06 | 26.0433222 | 46.2676432 | 64.0355823 | 64.9510612 | 0 |
| 1.72E-06 | 5.71E-06 | 48.8714194 | 53.1633016 | 46.4025958 | 26.9082968 | 0 |
| 1.73E-06 | 5.71E-06 | 68.8058142 | 36.9251383 | 55.683115 | 30.619786 | 0 |
| 1.73E-06 | 5.72E-06 | 31.1876821 | 55.3877075 | 35.2659728 | 63.5592527 | 0 |
| 1.73E-06 | 5.72E-06 | 26.6863672 | 57.3896728 | 62.1794784 | 44.5378705 | 0 |
| 1.74E-06 | 5.75E-06 | 27.3294122 | 42.9310343 | 80.7405168 | 68.1986142 | 0 |
| 1.74E-06 | 5.76E-06 | 23.1496197 | 79.6337321 | 56.6111669 | 83.5085072 | 0 |
| 1.74E-06 | 5.76E-06 | 33.7598621 | 156.153296 | 133.639476 | 145.675952 | 0 |
| 1.74E-06 | 5.76E-06 | 221.207477 | 723.599247 | 682.118159 | 741.36997 | 0 |
| 1.74E-06 | 5.77E-06 | 43.7270595 | 44.0432373 | 30.6257133 | 36.1870198 | 0 |
| 1.74E-06 | 5.77E-06 | 22.8280972 | 69.846346 | 70.5319457 | 63.0953166 | 0 |
| 1.75E-06 | 5.77E-06 | 23.7926647 | 59.3916382 | 73.3161014 | 60.3116997 | 0 |
| 1.75E-06 | 5.80E-06 | 36.0105196 | 34.923173 | 36.1940248 | 37.1148921 | 0 |
| 1.75E-06 | 5.80E-06 | 30.5446371 | 80.7459351 | 38.9781805 | 56.1362743 | 0 |
| 1.76E-06 | 5.81E-06 | 20.8989622 | 68.5117025 | 119.718697 | 88.1478687 | 0 |
| 1.77E-06 | 5.84E-06 | 35.6889971 | 35.5904948 | 42.6903882 | 33.867339 | 0 |
| 1.77E-06 | 5.85E-06 | 66.2336342 | 23.3562622 | 37.1220767 | 14.8459568 | 218.612078 |
| 1.77E-06 | 5.85E-06 | 6.10892743 | 7.56298014 | 8.35246725 | 6.49510612 | 194.849896 |
| 1.77E-06 | 5.85E-06 | 33.4383396 | 125.234054 | 36.1940248 | 189.28595 | 0 |
| 1.77E-06 | 5.86E-06 | 24.7572322 | 109.885653 | 49.1867516 | 96.9626556 | 0 |
| 1.78E-06 | 5.87E-06 | 19.9343948 | 78.7439697 | 97.4454513 | 84.4363795 | 0 |
| 1.78E-06 | 5.89E-06 | 25.0787547 | 56.0550293 | 53.8270112 | 56.1362743 | 0 |
| 1.79E-06 | 5.90E-06 | 33.4383396 | 160.379667 | 219.948304 | 161.449781 | 4.75243649 |
| 1.79E-06 | 5.90E-06 | 38.261177 | 33.3660889 | 34.3379209 | 46.3936151 | 0 |
| 1.79E-06 | 5.91E-06 | 26.6863672 | 61.6160441 | 43.6184401 | 60.7756358 | 0 |
| 1.79E-06 | 5.91E-06 | 53.3727344 | 84.0825439 | 134.567528 | 72.8379758 | 0 |
| 1.79E-06 | 5.92E-06 | 46.2992394 | 85.4171875 | 96.5173994 | 87.6839326 | 2.37621824 |

| | | | | | | |
|---|---|---|---|---|---|---|
| 1.79E-06 | 5.92E-06 | 29.5800696 | 157.265499 | 38.9781805 | 143.356271 | 0 |
| 1.79E-06 | 5.93E-06 | 26.3648447 | 62.0609253 | 44.546492 | 61.239572 | 0 |
| 1.80E-06 | 5.94E-06 | 21.2204847 | 133.686796 | 90.9490879 | 66.3428696 | 0 |
| 1.80E-06 | 5.95E-06 | 61.0892743 | 72.7380737 | 58.4672708 | 95.5708472 | 2.37621824 |
| 1.81E-06 | 5.97E-06 | 29.2585471 | 80.0786133 | 88.1649321 | 38.5067006 | 0 |
| 1.81E-06 | 5.98E-06 | 20.2559173 | 84.7498657 | 75.1722053 | 90.4675495 | 0 |
| 1.81E-06 | 5.99E-06 | 16.7191698 | 92.3128459 | 150.344411 | 116.447974 | 0 |
| 1.82E-06 | 5.99E-06 | 245.96471 | 49.3818115 | 42.6903882 | 59.8477635 | 0 |
| 1.83E-06 | 6.03E-06 | 52.4081669 | 35.3680542 | 38.0501286 | 33.4034029 | 0 |
| 1.83E-06 | 6.04E-06 | 28.6155022 | 103.212435 | 39.9062324 | 108.097123 | 0 |
| 1.83E-06 | 6.05E-06 | 26.6863672 | 65.6199748 | 50.1148035 | 46.8575513 | 0 |
| 1.84E-06 | 6.05E-06 | 73.9501741 | 54.2755046 | 150.344411 | 82.1166988 | 2.37621824 |
| 1.84E-06 | 6.06E-06 | 53.6942569 | 82.0805786 | 113.222334 | 66.3428696 | 2.37621824 |
| 1.84E-06 | 6.06E-06 | 34.0813846 | 37.5924601 | 33.409869 | 45.0018067 | 0 |
| 1.84E-06 | 6.06E-06 | 21.5420072 | 72.2931925 | 69.6038938 | 104.385634 | 0 |
| 1.84E-06 | 6.06E-06 | 30.5446371 | 47.1574056 | 35.2659728 | 48.2493597 | 0 |
| 1.84E-06 | 6.07E-06 | 29.2585471 | 42.2637126 | 40.8342843 | 56.1362743 | 0 |
| 1.85E-06 | 6.09E-06 | 181.338688 | 187.962301 | 257.998433 | 323.363497 | 21.3859642 |
| 1.85E-06 | 6.10E-06 | 25.4002772 | 57.3896728 | 49.1867516 | 56.1362743 | 0 |
| 1.85E-06 | 6.10E-06 | 27.3294122 | 45.1554403 | 45.4745439 | 56.6002105 | 0 |
| 1.85E-06 | 6.10E-06 | 20.5774398 | 78.0766479 | 80.7405168 | 77.9412734 | 0 |
| 1.85E-06 | 6.10E-06 | 49.1929419 | 173.058781 | 180.042072 | 152.634994 | 0 |
| 1.86E-06 | 6.11E-06 | 26.3648447 | 60.7262817 | 74.2441534 | 44.5378705 | 0 |
| 1.86E-06 | 6.12E-06 | 46.2992394 | 70.7361084 | 109.510126 | 26.9082968 | 0 |
| 1.86E-06 | 6.13E-06 | 290.656336 | 285.39128 | 317.393756 | 319.652008 | 26.1384007 |
| 1.86E-06 | 6.14E-06 | 240.82035 | 506.71967 | 485.371153 | 573.889019 | 21.3859642 |
| 1.86E-06 | 6.14E-06 | 20.8989622 | 82.3030192 | 68.6758419 | 86.2921241 | 0 |
| 1.87E-06 | 6.16E-06 | 25.4002772 | 52.0510986 | 55.683115 | 51.0329766 | 0 |
| 1.87E-06 | 6.17E-06 | 32.4737721 | 65.1750936 | 127.143113 | 40.8263813 | 0 |
| 1.87E-06 | 6.17E-06 | 83.2743265 | 73.4053955 | 79.8124649 | 66.3428696 | 14.2573095 |
| 1.88E-06 | 6.17E-06 | 47.2638069 | 28.4723958 | 136.423632 | 71.9101035 | 0 |
| 1.88E-06 | 6.17E-06 | 374.89523 | 1395.59228 | 1094.17321 | 1142.67474 | 66.5341108 |
| 1.88E-06 | 6.19E-06 | 71.3779941 | 32.0314453 | 37.1220767 | 46.3936151 | 0 |
| 1.88E-06 | 6.20E-06 | 59.4816618 | 47.8247274 | 145.704151 | 32.0115944 | 0 |
| 1.89E-06 | 6.21E-06 | 103.530244 | 143.919063 | 130.85532 | 124.334889 | 7.12865473 |
| 1.89E-06 | 6.21E-06 | 27.0078897 | 77.4093262 | 80.7405168 | 41.7542536 | 0 |
| 1.89E-06 | 6.21E-06 | 59.1601393 | 209.094157 | 210.667785 | 103.921698 | 0 |
| 1.89E-06 | 6.22E-06 | 37.2966096 | 52.2735392 | 29.6976613 | 42.6821259 | 0 |
| 1.89E-06 | 6.22E-06 | 46.2992394 | 51.3837769 | 25.9854537 | 48.2493597 | 0 |
| 1.90E-06 | 6.23E-06 | 22.5065747 | 139.02537 | 143.848047 | 52.8887212 | 0 |
| 1.90E-06 | 6.25E-06 | 27.9724572 | 48.2696086 | 60.3233746 | 40.8263813 | 0 |
| 1.90E-06 | 6.25E-06 | 19.6128723 | 86.0845093 | 93.7332436 | 77.9412734 | 0 |
| 1.90E-06 | 6.25E-06 | 29.9015921 | 49.1593709 | 49.1867516 | 35.7230837 | 0 |

| | | | | | | |
|---|---|---|---|---|---|---|
| 1.90E-06 | 6.25E-06 | 25.4002772 | 56.4999105 | 54.7550631 | 48.7132959 | 0 |
| 1.90E-06 | 6.25E-06 | 26.0433222 | 59.8365194 | 58.4672708 | 44.5378705 | 0 |
| 1.91E-06 | 6.26E-06 | 22.8280972 | 65.3975342 | 76.1002572 | 59.3838274 | 0 |
| 1.91E-06 | 6.28E-06 | 22.5065747 | 78.7439697 | 57.5392189 | 78.4052096 | 0 |
| 1.91E-06 | 6.29E-06 | 28.2939797 | 82.3030192 | 39.9062324 | 65.8789335 | 0 |
| 1.91E-06 | 6.29E-06 | 35.6889971 | 40.2617472 | 30.6257133 | 45.0018067 | 0 |
| 1.92E-06 | 6.30E-06 | 15.4330798 | 113.667143 | 157.768826 | 107.633187 | 0 |
| 1.92E-06 | 6.31E-06 | 26.3648447 | 52.7184204 | 56.6111669 | 44.0739344 | 0 |
| 1.93E-06 | 6.34E-06 | 21.2204847 | 64.952653 | 78.8844129 | 87.2199964 | 0 |
| 1.93E-06 | 6.34E-06 | 21.2204847 | 68.9565837 | 84.4527244 | 71.9101035 | 0 |
| 1.93E-06 | 6.35E-06 | 24.4357097 | 66.9546183 | 70.5319457 | 49.177232 | 0 |
| 1.94E-06 | 6.36E-06 | 25.7217997 | 62.2833659 | 46.4025958 | 54.7444659 | 0 |
| 1.94E-06 | 6.36E-06 | 24.7572322 | 60.2814006 | 53.8270112 | 52.4247851 | 0 |
| 1.94E-06 | 6.36E-06 | 60.4462293 | 75.4073608 | 82.5966206 | 102.993826 | 9.50487297 |
| 1.94E-06 | 6.36E-06 | 27.9724572 | 53.1633016 | 53.8270112 | 38.9706367 | 0 |
| 1.94E-06 | 6.38E-06 | 35.0459521 | 31.5865641 | 58.4672708 | 43.6099982 | 0 |
| 1.95E-06 | 6.40E-06 | 22.5065747 | 58.5018758 | 107.654022 | 76.549465 | 0 |
| 1.95E-06 | 6.41E-06 | 37.2966096 | 62.2833659 | 128.071165 | 35.7230837 | 0 |
| 1.96E-06 | 6.42E-06 | 43.0840145 | 123.232088 | 31.5537652 | 314.548711 | 0 |
| 1.96E-06 | 6.44E-06 | 37.618132 | 110.108093 | 70.5319457 | 31.5476583 | 0 |
| 1.97E-06 | 6.45E-06 | 29.5800696 | 44.9329997 | 38.0501286 | 40.8263813 | 0 |
| 1.97E-06 | 6.45E-06 | 30.2231146 | 37.8149007 | 81.6685687 | 56.1362743 | 0 |
| 1.97E-06 | 6.46E-06 | 40.190312 | 67.8443807 | 29.6976613 | 44.0739344 | 0 |
| 1.97E-06 | 6.46E-06 | 274.258689 | 173.503662 | 117.862593 | 161.913717 | 21.3859642 |
| 1.97E-06 | 6.46E-06 | 67.1982017 | 57.1672323 | 23.2012979 | 56.6002105 | 0 |
| 1.97E-06 | 6.48E-06 | 51.1220769 | 84.9723063 | 140.135839 | 135.933292 | 0 |
| 1.98E-06 | 6.49E-06 | 37.618132 | 29.3621582 | 42.6903882 | 48.2493597 | 0 |
| 1.98E-06 | 6.50E-06 | 23.7926647 | 50.7164551 | 71.4599976 | 72.8379758 | 0 |
| 1.98E-06 | 6.51E-06 | 47.5853294 | 41.5963908 | 30.6257133 | 34.7952113 | 0 |
| 1.99E-06 | 6.52E-06 | 21.8635297 | 75.1849202 | 105.797919 | 61.7035081 | 0 |
| 1.99E-06 | 6.53E-06 | 16.3976473 | 70.0687866 | 209.739733 | 226.864778 | 0 |
| 2.01E-06 | 6.58E-06 | 22.1850522 | 77.6317668 | 64.9636342 | 63.0953166 | 0 |
| 2.01E-06 | 6.58E-06 | 52.4081669 | 38.9271037 | 32.4818171 | 34.3312752 | 0 |
| 2.01E-06 | 6.59E-06 | 57.2310043 | 75.6298014 | 58.4672708 | 79.3330819 | 0 |
| 2.01E-06 | 6.59E-06 | 472.959592 | 659.758797 | 531.773748 | 599.405507 | 0 |
| 2.01E-06 | 6.59E-06 | 36.9750871 | 52.2735392 | 28.7696094 | 45.4657428 | 0 |
| 2.02E-06 | 6.60E-06 | 25.4002772 | 54.9428263 | 47.3306478 | 53.8165936 | 0 |
| 2.02E-06 | 6.63E-06 | 22.1850522 | 83.4152222 | 56.6111669 | 81.1888265 | 0 |
| 2.03E-06 | 6.64E-06 | 43.0840145 | 70.2912272 | 25.9854537 | 57.0641466 | 0 |
| 2.03E-06 | 6.65E-06 | 21.5420072 | 89.1986776 | 59.3953227 | 83.9724434 | 0 |
| 2.03E-06 | 6.65E-06 | 29.5800696 | 47.1574056 | 37.1220767 | 41.2903175 | 0 |
| 2.03E-06 | 6.65E-06 | 32.1522496 | 12.2342326 | 22.273246 | 13.9180845 | 90.2962933 |
| 2.04E-06 | 6.67E-06 | 34.0813846 | 51.3837769 | 30.6257133 | 44.5378705 | 0 |

| | | | | | | |
|---|---|---|---|---|---|---|
| 2.04E-06 | 6.69E-06 | 85.524984 | 163.493835 | 154.98467 | 160.057972 | 19.0097459 |
| 2.05E-06 | 6.71E-06 | 102.887199 | 65.8424154 | 98.3735032 | 106.241379 | 0 |
| 2.05E-06 | 6.71E-06 | 18.6483048 | 84.3049845 | 91.8771398 | 90.0036134 | 0 |
| 2.05E-06 | 6.71E-06 | 127.322908 | 192.188672 | 186.538435 | 196.244992 | 33.2670554 |
| 2.05E-06 | 6.71E-06 | 25.7217997 | 55.8325887 | 51.0428554 | 45.929679 | 0 |
| 2.05E-06 | 6.71E-06 | 45.9777169 | 111.665177 | 65.8916861 | 79.3330819 | 297.02728 |
| 2.05E-06 | 6.72E-06 | 463.313917 | 858.398246 | 769.355039 | 849.003157 | 38.0194919 |
| 2.06E-06 | 6.73E-06 | 103.851766 | 362.133284 | 337.810898 | 341.457007 | 19.0097459 |
| 2.06E-06 | 6.73E-06 | 21.2204847 | 84.3049845 | 65.8916861 | 71.4461673 | 0 |
| 2.07E-06 | 6.76E-06 | 13.1824223 | 129.682865 | 243.149602 | 121.551272 | 0 |
| 2.07E-06 | 6.76E-06 | 102.565676 | 53.8306234 | 28.7696094 | 45.4657428 | 0 |
| 2.07E-06 | 6.76E-06 | 67.8412467 | 28.0275146 | 69.6038938 | 38.0427644 | 0 |
| 2.08E-06 | 6.80E-06 | 15.4330798 | 97.8738607 | 126.215061 | 133.613612 | 0 |
| 2.08E-06 | 6.81E-06 | 38.5826995 | 72.5156331 | 31.5537652 | 42.6821259 | 0 |
| 2.09E-06 | 6.82E-06 | 158.189068 | 530.298372 | 380.501286 | 462.080407 | 0 |
| 2.09E-06 | 6.82E-06 | 22.8280972 | 106.104163 | 60.3233746 | 66.3428696 | 0 |
| 2.09E-06 | 6.82E-06 | 17.6837373 | 106.326603 | 116.934542 | 80.2609542 | 0 |
| 2.09E-06 | 6.83E-06 | 30.5446371 | 74.5175985 | 37.1220767 | 109.488932 | 0 |
| 2.10E-06 | 6.85E-06 | 30.8661596 | 74.0727173 | 68.6758419 | 33.867339 | 0 |
| 2.10E-06 | 6.86E-06 | 29.2585471 | 38.7046631 | 38.9781805 | 45.0018067 | 0 |
| 2.10E-06 | 6.86E-06 | 33.4383396 | 22.6889404 | 25.9854537 | 25.9804245 | 190.097459 |
| 2.10E-06 | 6.86E-06 | 25.0787547 | 50.0491333 | 51.0428554 | 52.4247851 | 0 |
| 2.10E-06 | 6.86E-06 | 56.9094818 | 123.454529 | 105.797919 | 102.065953 | 0 |
| 2.11E-06 | 6.88E-06 | 46.9422844 | 34.7007324 | 36.1940248 | 32.4755306 | 0 |
| 2.11E-06 | 6.89E-06 | 17.6837373 | 93.86993 | 101.157659 | 90.4675495 | 0 |
| 2.11E-06 | 6.89E-06 | 32.7952946 | 41.5963908 | 63.1075304 | 138.716909 | 0 |
| 2.11E-06 | 6.90E-06 | 43.405537 | 60.7262817 | 85.3807764 | 26.4443606 | 0 |
| 2.12E-06 | 6.93E-06 | 42.762492 | 104.102197 | 90.9490879 | 90.4675495 | 0 |
| 2.13E-06 | 6.95E-06 | 25.4002772 | 50.2715739 | 48.2586997 | 50.5690405 | 0 |
| 2.13E-06 | 6.96E-06 | 33.7598621 | 39.1495443 | 31.5537652 | 39.898509 | 0 |
| 2.13E-06 | 6.97E-06 | 33.4383396 | 42.9310343 | 30.6257133 | 46.3936151 | 0 |
| 2.14E-06 | 6.97E-06 | 50.4790319 | 66.064856 | 82.5966206 | 81.1888265 | 0 |
| 2.14E-06 | 6.97E-06 | 20.8989622 | 142.58442 | 193.034799 | 55.208402 | 0 |
| 2.14E-06 | 6.98E-06 | 19.9343948 | 92.5352865 | 69.6038938 | 81.6527626 | 0 |
| 2.14E-06 | 6.98E-06 | 18.0052598 | 80.9683756 | 105.797919 | 96.4987195 | 0 |
| 2.14E-06 | 6.99E-06 | 25.4002772 | 50.7164551 | 51.0428554 | 47.3214874 | 0 |
| 2.14E-06 | 6.99E-06 | 25.7217997 | 44.4881185 | 58.4672708 | 50.5690405 | 0 |
| 2.14E-06 | 7.00E-06 | 28.6155022 | 42.9310343 | 51.0428554 | 37.5788283 | 0 |
| 2.14E-06 | 7.00E-06 | 18.6483048 | 87.1967122 | 79.8124649 | 141.964462 | 0 |
| 2.15E-06 | 7.00E-06 | 29.5800696 | 35.3680542 | 49.1867516 | 43.1460621 | 0 |
| 2.15E-06 | 7.00E-06 | 24.1141872 | 53.1633016 | 56.6111669 | 81.1888265 | 0 |
| 2.15E-06 | 7.01E-06 | 27.9724572 | 38.2597819 | 46.4025958 | 46.3936151 | 0 |
| 2.16E-06 | 7.04E-06 | 24.4357097 | 141.472217 | 52.8989593 | 79.3330819 | 0 |

| | | | | | | |
|---|---|---|---|---|---|---|
| 2.16E-06 | 7.04E-06 | 23.7926647 | 84.7498657 | 168.905449 | 56.6002105 | 0 |
| 2.16E-06 | 7.04E-06 | 20.5774398 | 115.224227 | 64.9636342 | 81.6527626 | 0 |
| 2.16E-06 | 7.04E-06 | 27.0078897 | 74.0727173 | 50.1148035 | 44.0739344 | 0 |
| 2.16E-06 | 7.05E-06 | 23.1496197 | 52.2735392 | 66.819738 | 67.7346781 | 0 |
| 2.17E-06 | 7.06E-06 | 73.9501741 | 42.4861532 | 28.7696094 | 44.0739344 | 0 |
| 2.18E-06 | 7.10E-06 | 33.4383396 | 48.9369303 | 30.6257133 | 59.8477635 | 0 |
| 2.18E-06 | 7.11E-06 | 96.1352263 | 36.035376 | 55.683115 | 32.9394667 | 0 |
| 2.19E-06 | 7.13E-06 | 27.6509347 | 57.834554 | 41.7623363 | 43.1460621 | 0 |
| 2.19E-06 | 7.13E-06 | 15.4330798 | 102.767554 | 108.582074 | 137.789037 | 0 |
| 2.19E-06 | 7.14E-06 | 759.114613 | 1643.61354 | 1305.76905 | 1421.03643 | 16.6335277 |
| 2.20E-06 | 7.15E-06 | 22.8280972 | 58.5018758 | 58.4672708 | 64.9510612 | 0 |
| 2.21E-06 | 7.18E-06 | 29.5800696 | 35.5904948 | 49.1867516 | 41.2903175 | 0 |
| 2.21E-06 | 7.18E-06 | 53.0512119 | 76.5195638 | 51.0428554 | 24.588616 | 0 |
| 2.21E-06 | 7.18E-06 | 21.8635297 | 68.9565837 | 65.8916861 | 61.7035081 | 0 |
| 2.21E-06 | 7.19E-06 | 20.8989622 | 150.1474 | 63.1075304 | 90.0036134 | 0 |
| 2.22E-06 | 7.22E-06 | 22.1850522 | 94.3148112 | 83.5246725 | 53.3526574 | 0 |
| 2.22E-06 | 7.23E-06 | 26.6863672 | 64.0628906 | 64.9636342 | 39.4345729 | 0 |
| 2.23E-06 | 7.25E-06 | 25.4002772 | 120.340361 | 42.6903882 | 122.015208 | 0 |
| 2.24E-06 | 7.28E-06 | 11.8963324 | 139.02537 | 142.919995 | 177.22361 | 0 |
| 2.24E-06 | 7.28E-06 | 36.3320421 | 70.0687866 | 68.6758419 | 28.3001052 | 0 |
| 2.24E-06 | 7.28E-06 | 33.4383396 | 31.3641235 | 38.9781805 | 44.5378705 | 0 |
| 2.24E-06 | 7.29E-06 | 0 | 0 | 0 | 0 | 11.8810912 |
| 2.25E-06 | 7.31E-06 | 20.5774398 | 107.216366 | 65.8916861 | 75.1576565 | 0 |
| 2.25E-06 | 7.31E-06 | 24.1141872 | 95.8718953 | 51.9709073 | 62.1674443 | 0 |
| 2.25E-06 | 7.31E-06 | 23.4711422 | 49.6042521 | 64.9636342 | 70.518295 | 0 |
| 2.25E-06 | 7.32E-06 | 19.9343948 | 72.5156331 | 77.956361 | 77.9412734 | 0 |
| 2.26E-06 | 7.36E-06 | 20.5774398 | 80.5234945 | 77.956361 | 64.0231889 | 0 |
| 2.27E-06 | 7.38E-06 | 20.2559173 | 70.7361084 | 167.977397 | 84.4363795 | 0 |
| 2.27E-06 | 7.38E-06 | 45.6561944 | 79.8561727 | 79.8124649 | 90.9314857 | 0 |
| 2.27E-06 | 7.39E-06 | 21.5420072 | 73.4053955 | 61.2514265 | 67.2707419 | 0 |
| 2.27E-06 | 7.39E-06 | 61.4107968 | 90.3108805 | 20.4171422 | 76.0855288 | 0 |
| 2.27E-06 | 7.39E-06 | 24.7572322 | 71.4034302 | 48.2586997 | 53.3526574 | 0 |
| 2.28E-06 | 7.40E-06 | 36.0105196 | 89.4211182 | 28.7696094 | 95.106911 | 0 |
| 2.28E-06 | 7.41E-06 | 25.7217997 | 41.5963908 | 79.8124649 | 71.4461673 | 0 |
| 2.28E-06 | 7.41E-06 | 0 | 2.8917277 | 0 | 0 | 87.920075 |
| 2.29E-06 | 7.44E-06 | 21.8635297 | 105.2144 | 139.207788 | 52.8887212 | 0 |
| 2.29E-06 | 7.44E-06 | 516.686651 | 996.088973 | 835.246725 | 960.811769 | 14.2573095 |
| 2.30E-06 | 7.48E-06 | 31.5092046 | 35.1456136 | 96.5173994 | 62.1674443 | 0 |
| 2.31E-06 | 7.50E-06 | 18.9698273 | 93.86993 | 81.6685687 | 75.1576565 | 0 |
| 2.31E-06 | 7.50E-06 | 24.4357097 | 54.4979451 | 47.3306478 | 64.9510612 | 0 |
| 2.31E-06 | 7.51E-06 | 51.7651219 | 111.665177 | 116.934542 | 106.241379 | 0 |
| 2.32E-06 | 7.53E-06 | 360.426718 | 547.203857 | 575.392189 | 623.066251 | 30.8908372 |
| 2.32E-06 | 7.53E-06 | 6.43044992 | 6.67321777 | 6.49636342 | 4.63936151 | 80.7914203 |

| | | | | | | |
|---|---|---|---|---|---|---|
| 2.32E-06 | 7.55E-06 | 24.1141872 | 51.3837769 | 51.0428554 | 57.0641466 | 0 |
| 2.32E-06 | 7.55E-06 | 46.6207619 | 24.023584 | 147.560255 | 107.633187 | 0 |
| 2.33E-06 | 7.55E-06 | 47.5853294 | 34.7007324 | 35.2659728 | 32.0115944 | 0 |
| 2.33E-06 | 7.56E-06 | 22.8280972 | 70.958549 | 54.7550631 | 58.9198912 | 0 |
| 2.33E-06 | 7.56E-06 | 20.5774398 | 86.3069499 | 69.6038938 | 66.3428696 | 0 |
| 2.34E-06 | 7.58E-06 | 24.7572322 | 56.9447917 | 47.3306478 | 50.1051043 | 0 |
| 2.34E-06 | 7.60E-06 | 9.64567488 | 4.44881185 | 13.9207788 | 6.49510612 | 45.1481466 |
| 2.35E-06 | 7.62E-06 | 22.1850522 | 109.218331 | 56.6111669 | 72.3740396 | 0 |
| 2.35E-06 | 7.63E-06 | 37.2966096 | 42.9310343 | 77.956361 | 32.0115944 | 0 |
| 2.36E-06 | 7.66E-06 | 28.2939797 | 54.2755046 | 38.9781805 | 40.8263813 | 0 |
| 2.37E-06 | 7.69E-06 | 23.1496197 | 82.0805786 | 54.7550631 | 57.5280828 | 0 |
| 2.38E-06 | 7.70E-06 | 17.6837373 | 148.367875 | 72.3880495 | 108.097123 | 0 |
| 2.38E-06 | 7.70E-06 | 21.8635297 | 92.0904053 | 54.7550631 | 74.2297842 | 0 |
| 2.38E-06 | 7.71E-06 | 29.2585471 | 82.7479004 | 46.4025958 | 41.7542536 | 0 |
| 2.38E-06 | 7.71E-06 | 382.290248 | 699.130782 | 623.650888 | 720.028907 | 0 |
| 2.38E-06 | 7.71E-06 | 24.7572322 | 55.1652669 | 45.4745439 | 52.8887212 | 0 |
| 2.38E-06 | 7.71E-06 | 20.5774398 | 71.8483114 | 68.6758419 | 69.5904227 | 0 |
| 2.38E-06 | 7.71E-06 | 21.2204847 | 110.775415 | 55.683115 | 88.1478687 | 0 |
| 2.38E-06 | 7.72E-06 | 120.249414 | 345.227799 | 384.213494 | 412.903175 | 23.7621824 |
| 2.38E-06 | 7.72E-06 | 242.427962 | 922.016256 | 780.491662 | 843.435923 | 4.75243649 |
| 2.38E-06 | 7.73E-06 | 29.9015921 | 34.7007324 | 38.9781805 | 45.4657428 | 0 |
| 2.39E-06 | 7.74E-06 | 19.9343948 | 74.5175985 | 70.5319457 | 77.4773373 | 0 |
| 2.39E-06 | 7.74E-06 | 41.1548795 | 73.6278361 | 47.3306478 | 27.8361691 | 0 |
| 2.39E-06 | 7.75E-06 | 35.0459521 | 64.5077718 | 28.7696094 | 51.9608489 | 0 |
| 2.39E-06 | 7.75E-06 | 97.7428388 | 22.0216187 | 51.0428554 | 102.993826 | 0 |
| 2.39E-06 | 7.75E-06 | 25.0787547 | 49.1593709 | 59.3953227 | 45.0018067 | 0 |
| 2.39E-06 | 7.75E-06 | 41.1548795 | 39.3719849 | 74.2441534 | 30.619786 | 0 |
| 2.40E-06 | 7.77E-06 | 27.6509347 | 36.4802572 | 55.683115 | 61.239572 | 0 |
| 2.40E-06 | 7.77E-06 | 43.405537 | 32.0314453 | 51.9709073 | 31.0837221 | 0 |
| 2.41E-06 | 7.79E-06 | 38.904222 | 39.5944255 | 29.6976613 | 34.7952113 | 0 |
| 2.41E-06 | 7.80E-06 | 143.399033 | 193.745756 | 136.423632 | 168.408823 | 19.0097459 |
| 2.41E-06 | 7.81E-06 | 27.6509347 | 40.929069 | 105.797919 | 65.8789335 | 0 |
| 2.43E-06 | 7.85E-06 | 0 | 0 | 0 | 0 | 97.424948 |
| 2.43E-06 | 7.87E-06 | 24.7572322 | 59.8365194 | 45.4745439 | 81.6527626 | 0 |
| 2.43E-06 | 7.88E-06 | 30.5446371 | 115.224227 | 34.3379209 | 154.490738 | 0 |
| 2.43E-06 | 7.88E-06 | 21.2204847 | 91.8679647 | 70.5319457 | 58.9198912 | 0 |
| 2.44E-06 | 7.88E-06 | 22.8280972 | 62.7282471 | 58.4672708 | 53.3526574 | 0 |
| 2.44E-06 | 7.88E-06 | 23.1496197 | 64.0628906 | 49.1867516 | 67.2707419 | 0 |
| 2.44E-06 | 7.88E-06 | 23.1496197 | 54.053064 | 55.683115 | 57.0641466 | 0 |
| 2.44E-06 | 7.89E-06 | 158.832113 | 799.89637 | 575.392189 | 700.079652 | 73.6627655 |
| 2.44E-06 | 7.89E-06 | 53.6942569 | 40.929069 | 25.9854537 | 44.5378705 | 0 |
| 2.44E-06 | 7.89E-06 | 23.4711422 | 65.6199748 | 117.862593 | 54.7444659 | 0 |
| 2.44E-06 | 7.90E-06 | 30.2231146 | 34.2558512 | 54.7550631 | 40.8263813 | 0 |

| | | | | | | |
|---|---|---|---|---|---|---|
| 2.45E-06 | 7.92E-06 | 16.7191698 | 123.454529 | 99.3015551 | 83.5085072 | 0 |
| 2.45E-06 | 7.92E-06 | 23.1496197 | 88.5313558 | 57.5392189 | 55.208402 | 0 |
| 2.46E-06 | 7.94E-06 | 18.0052598 | 78.7439697 | 92.8051917 | 89.5396772 | 0 |
| 2.46E-06 | 7.94E-06 | 68.8058142 | 32.0314453 | 12.9927268 | 39.4345729 | 280.393753 |
| 2.46E-06 | 7.95E-06 | 17.6837373 | 123.232088 | 67.7477899 | 131.757867 | 0 |
| 2.46E-06 | 7.96E-06 | 179.731075 | 304.743612 | 287.696094 | 335.889774 | 23.7621824 |
| 2.47E-06 | 7.97E-06 | 23.7926647 | 42.041272 | 170.761553 | 200.420417 | 0 |
| 2.47E-06 | 7.98E-06 | 21.8635297 | 58.2794352 | 63.1075304 | 64.487125 | 0 |
| 2.47E-06 | 7.98E-06 | 141.469898 | 276.048775 | 264.494796 | 263.97967 | 21.3859642 |
| 2.47E-06 | 7.98E-06 | 55.3018693 | 22.0216187 | 64.9636342 | 80.2609542 | 0 |
| 2.47E-06 | 7.98E-06 | 89.0617314 | 63.6180094 | 19.4890903 | 139.180845 | 0 |
| 2.48E-06 | 8.00E-06 | 20.8989622 | 77.1868856 | 67.7477899 | 61.7035081 | 0 |
| 2.48E-06 | 8.02E-06 | 72.6640841 | 94.7596924 | 17.6329864 | 134.077548 | 0 |
| 2.49E-06 | 8.04E-06 | 22.5065747 | 53.8306234 | 61.2514265 | 81.6527626 | 0 |
| 2.49E-06 | 8.04E-06 | 19.9343948 | 77.4093262 | 94.6612955 | 63.5592527 | 0 |
| 2.50E-06 | 8.06E-06 | 20.5774398 | 60.7262817 | 82.5966206 | 72.8379758 | 0 |
| 2.50E-06 | 8.07E-06 | 22.5065747 | 64.0628906 | 52.8989593 | 62.1674443 | 0 |
| 2.50E-06 | 8.08E-06 | 24.7572322 | 81.858138 | 46.4025958 | 56.6002105 | 0 |
| 2.50E-06 | 8.08E-06 | 51.4435994 | 34.7007324 | 28.7696094 | 42.2181898 | 0 |
| 2.50E-06 | 8.08E-06 | 17.6837373 | 139.692692 | 90.0210359 | 77.9412734 | 0 |
| 2.51E-06 | 8.09E-06 | 73.9501741 | 77.4093262 | 23.2012979 | 45.4657428 | 0 |
| 2.51E-06 | 8.11E-06 | 28.9370247 | 52.7184204 | 33.409869 | 54.7444659 | 0 |
| 2.51E-06 | 8.11E-06 | 34.4029071 | 108.77345 | 32.4818171 | 63.5592527 | 0 |
| 2.52E-06 | 8.12E-06 | 22.5065747 | 57.6121134 | 54.7550631 | 64.0231889 | 0 |
| 2.52E-06 | 8.12E-06 | 33.4383396 | 46.934965 | 28.7696094 | 50.5690405 | 0 |
| 2.52E-06 | 8.14E-06 | 27.0078897 | 39.5944255 | 42.6903882 | 43.1460621 | 0 |
| 2.53E-06 | 8.16E-06 | 8.6811074 | 250.690548 | 128.999216 | 271.402649 | 0 |
| 2.53E-06 | 8.17E-06 | 189.698273 | 343.670715 | 237.581291 | 314.548711 | 0 |
| 2.54E-06 | 8.19E-06 | 31.1876821 | 46.4900838 | 32.4818171 | 38.0427644 | 0 |
| 2.54E-06 | 8.20E-06 | 25.0787547 | 60.5038411 | 51.9709073 | 42.6821259 | 0 |
| 2.54E-06 | 8.20E-06 | 154.652321 | 463.566195 | 364.724403 | 418.470408 | 0 |
| 2.55E-06 | 8.21E-06 | 23.4711422 | 61.3936035 | 53.8270112 | 49.6411682 | 0 |
| 2.55E-06 | 8.21E-06 | 20.5774398 | 74.7400391 | 69.6038938 | 62.6313804 | 0 |
| 2.55E-06 | 8.22E-06 | 15.1115573 | 87.864034 | 117.862593 | 196.244992 | 0 |
| 2.55E-06 | 8.22E-06 | 26.3648447 | 60.9487223 | 76.1002572 | 38.9706367 | 0 |
| 2.55E-06 | 8.22E-06 | 9.96719738 | 11.3444702 | 9.28051917 | 17.1656376 | 57.0292378 |
| 2.55E-06 | 8.22E-06 | 60.4462293 | 32.2538859 | 40.8342843 | 32.9394667 | 0 |
| 2.56E-06 | 8.26E-06 | 31.1876821 | 52.4959798 | 30.6257133 | 47.7854236 | 0 |
| 2.57E-06 | 8.27E-06 | 45.6561944 | 97.8738607 | 104.869867 | 68.6625504 | 0 |
| 2.57E-06 | 8.27E-06 | 17.3622148 | 108.99589 | 125.287009 | 70.9822312 | 0 |
| 2.57E-06 | 8.27E-06 | 31.1876821 | 175.728068 | 36.1940248 | 95.5708472 | 0 |
| 2.57E-06 | 8.28E-06 | 25.7217997 | 84.5274251 | 203.24337 | 51.0329766 | 0 |
| 2.57E-06 | 8.29E-06 | 67.1982017 | 45.3778809 | 23.2012979 | 63.5592527 | 0 |

| | | | | | | |
|---|---|---|---|---|---|---|
| 2.59E-06 | 8.32E-06 | 37.2966096 | 45.822762 | 31.5537652 | 32.0115944 | 0 |
| 2.59E-06 | 8.34E-06 | 18.6483048 | 101.21047 | 72.3880495 | 79.3330819 | 0 |
| 2.60E-06 | 8.36E-06 | 316.378136 | 494.040556 | 194.890903 | 425.429451 | 52.2768014 |
| 2.60E-06 | 8.38E-06 | 24.7572322 | 46.4900838 | 54.7550631 | 45.4657428 | 0 |
| 2.61E-06 | 8.38E-06 | 265.577582 | 667.766658 | 549.406735 | 605.436677 | 30.8908372 |
| 2.61E-06 | 8.39E-06 | 150.472528 | 400.393066 | 438.040505 | 485.277214 | 23.7621824 |
| 2.62E-06 | 8.43E-06 | 20.5774398 | 58.2794352 | 74.2441534 | 88.6118049 | 0 |
| 2.62E-06 | 8.43E-06 | 385.826995 | 672.660352 | 614.370369 | 615.643273 | 0 |
| 2.62E-06 | 8.43E-06 | 153.044708 | 64.0628906 | 59.3953227 | 73.7658481 | 0 |
| 2.62E-06 | 8.43E-06 | 68.1627692 | 40.0393066 | 25.0574018 | 59.3838274 | 0 |
| 2.63E-06 | 8.46E-06 | 33.1168171 | 95.4270142 | 30.6257133 | 70.0543588 | 0 |
| 2.63E-06 | 8.47E-06 | 33.4383396 | 30.4743612 | 39.9062324 | 36.650956 | 0 |
| 2.63E-06 | 8.47E-06 | 22.1850522 | 68.7341431 | 51.0428554 | 85.3642518 | 0 |
| 2.64E-06 | 8.50E-06 | 36.9750871 | 93.2026082 | 137.351684 | 110.88074 | 0 |
| 2.65E-06 | 8.52E-06 | 97.0997938 | 411.070215 | 263.566744 | 322.435625 | 11.8810912 |
| 2.65E-06 | 8.52E-06 | 40.833357 | 142.361979 | 120.646749 | 124.798825 | 4.75243649 |
| 2.66E-06 | 8.54E-06 | 28.6155022 | 62.2833659 | 33.409869 | 60.3116997 | 0 |
| 2.66E-06 | 8.55E-06 | 24.4357097 | 54.9428263 | 52.8989593 | 43.6099982 | 0 |
| 2.66E-06 | 8.56E-06 | 0 | 0 | 0 | 0 | 38.0194919 |
| 2.66E-06 | 8.56E-06 | 35.6889971 | 42.7085937 | 36.1940248 | 29.2279775 | 0 |
| 2.66E-06 | 8.56E-06 | 0 | 1.77952474 | 0 | 0 | 45.1481466 |
| 2.67E-06 | 8.57E-06 | 42.4409695 | 26.4704305 | 39.9062324 | 48.7132959 | 0 |
| 2.67E-06 | 8.58E-06 | 180.695643 | 377.036804 | 332.242586 | 360.942326 | 42.7719284 |
| 2.67E-06 | 8.59E-06 | 27.6509347 | 48.047168 | 37.1220767 | 40.3624452 | 0 |
| 2.68E-06 | 8.60E-06 | 237.283602 | 343.670715 | 309.96934 | 318.2602 | 26.1384007 |
| 2.68E-06 | 8.60E-06 | 22.5065747 | 58.0569946 | 58.4672708 | 52.8887212 | 0 |
| 2.68E-06 | 8.60E-06 | 19.6128723 | 79.6337321 | 62.1794784 | 97.4265918 | 0 |
| 2.68E-06 | 8.60E-06 | 34.7244296 | 30.4743612 | 77.0283091 | 48.7132959 | 0 |
| 2.68E-06 | 8.60E-06 | 12.5393773 | 93.2026082 | 182.826228 | 183.718716 | 0 |
| 2.68E-06 | 8.61E-06 | 21.2204847 | 96.7616577 | 98.3735032 | 51.0329766 | 0 |
| 2.69E-06 | 8.62E-06 | 18.9698273 | 120.11792 | 65.8916861 | 84.4363795 | 0 |
| 2.69E-06 | 8.63E-06 | 28.9370247 | 42.9310343 | 33.409869 | 42.6821259 | 0 |
| 2.69E-06 | 8.64E-06 | 139.862286 | 289.395211 | 246.86181 | 292.279775 | 21.3859642 |
| 2.69E-06 | 8.65E-06 | 19.2913498 | 60.5038411 | 96.5173994 | 134.541484 | 0 |
| 2.70E-06 | 8.66E-06 | 48.8714194 | 60.05896 | 23.2012979 | 91.3954218 | 0 |
| 2.71E-06 | 8.70E-06 | 27.9724572 | 44.9329997 | 37.1220767 | 64.0231889 | 0 |
| 2.71E-06 | 8.71E-06 | 38.261177 | 26.2479899 | 47.3306478 | 51.0329766 | 0 |
| 2.72E-06 | 8.72E-06 | 23.4711422 | 54.053064 | 49.1867516 | 51.9608489 | 0 |
| 2.73E-06 | 8.75E-06 | 0 | 0 | 0 | 0 | 45.1481466 |
| 2.73E-06 | 8.75E-06 | 101.279586 | 33.5885295 | 35.2659728 | 50.5690405 | 0 |
| 2.76E-06 | 8.85E-06 | 353.674746 | 431.75719 | 293.264406 | 364.189879 | 28.5146189 |
| 2.76E-06 | 8.86E-06 | 57.8740493 | 46.0452026 | 24.1293498 | 45.0018067 | 0 |
| 2.76E-06 | 8.86E-06 | 24.4357097 | 42.4861532 | 53.8270112 | 53.3526574 | 0 |

| | | | | | | |
|---|---|---|---|---|---|---|
| 2.76E-06 | 8.86E-06 | 18.3267823 | 139.470251 | 64.9636342 | 97.8905279 | 0 |
| 2.76E-06 | 8.86E-06 | 57.8740493 | 54.9428263 | 21.3451941 | 70.9822312 | 0 |
| 2.77E-06 | 8.89E-06 | 29.2585471 | 33.5885295 | 39.9062324 | 49.177232 | 0 |
| 2.78E-06 | 8.91E-06 | 29.2585471 | 36.4802572 | 51.0428554 | 35.7230837 | 0 |
| 2.78E-06 | 8.93E-06 | 26.6863672 | 48.7144897 | 51.9709073 | 36.1870198 | 0 |
| 2.78E-06 | 8.93E-06 | 20.2559173 | 78.9664103 | 70.5319457 | 59.8477635 | 0 |
| 2.79E-06 | 8.94E-06 | 21.2204847 | 127.013578 | 48.2586997 | 146.603824 | 0 |
| 2.79E-06 | 8.94E-06 | 17.3622148 | 57.3896728 | 300.688821 | 174.903929 | 0 |
| 2.79E-06 | 8.95E-06 | 88.0971639 | 118.560836 | 125.287009 | 14.8459568 | 0 |
| 2.79E-06 | 8.95E-06 | 207.060488 | 674.884757 | 597.665435 | 672.243483 | 0 |
| 2.79E-06 | 8.95E-06 | 22.8280972 | 110.997856 | 55.683115 | 60.7756358 | 0 |
| 2.80E-06 | 8.96E-06 | 21.8635297 | 55.8325887 | 56.6111669 | 64.0231889 | 0 |
| 2.81E-06 | 8.99E-06 | 25.4002772 | 64.5077718 | 40.8342843 | 50.1051043 | 0 |
| 2.81E-06 | 9.00E-06 | 16.7191698 | 86.7518311 | 96.5173994 | 88.6118049 | 0 |
| 2.81E-06 | 9.00E-06 | 19.6128723 | 69.6239054 | 71.4599976 | 69.5904227 | 0 |
| 2.81E-06 | 9.01E-06 | 54.0157793 | 47.1574056 | 23.2012979 | 49.177232 | 0 |
| 2.82E-06 | 9.02E-06 | 49.5144644 | 135.02144 | 122.502853 | 114.592229 | 9.50487297 |
| 2.82E-06 | 9.03E-06 | 24.7572322 | 78.9664103 | 40.8342843 | 65.8789335 | 0 |
| 2.83E-06 | 9.05E-06 | 21.2204847 | 50.0491333 | 97.4454513 | 90.4675495 | 0 |
| 2.83E-06 | 9.06E-06 | 23.4711422 | 53.1633016 | 96.5173994 | 54.2805297 | 0 |
| 2.83E-06 | 9.07E-06 | 25.7217997 | 58.946757 | 42.6903882 | 95.106911 | 0 |
| 2.83E-06 | 9.07E-06 | 30.8661596 | 37.1475789 | 36.1940248 | 62.6313804 | 0 |
| 2.84E-06 | 9.08E-06 | 10.2887199 | 131.684831 | 279.343627 | 144.284143 | 0 |
| 2.84E-06 | 9.08E-06 | 52.7296894 | 21.7991781 | 189.322591 | 100.210209 | 0 |
| 2.86E-06 | 9.14E-06 | 23.4711422 | 69.4014648 | 47.3306478 | 52.8887212 | 0 |
| 2.86E-06 | 9.15E-06 | 142.112943 | 84.3049845 | 82.5966206 | 88.1478687 | 0 |
| 2.86E-06 | 9.16E-06 | 20.2559173 | 75.852242 | 81.6685687 | 56.6002105 | 0 |
| 2.87E-06 | 9.18E-06 | 20.5774398 | 52.940861 | 91.8771398 | 84.4363795 | 0 |
| 2.88E-06 | 9.20E-06 | 29.9015921 | 50.9388957 | 36.1940248 | 34.7952113 | 0 |
| 2.88E-06 | 9.20E-06 | 72.0210391 | 54.4979451 | 47.3306478 | 23.1968076 | 0 |
| 2.88E-06 | 9.21E-06 | 20.8989622 | 66.5097371 | 64.0355823 | 57.9920189 | 0 |
| 2.89E-06 | 9.23E-06 | 44.3701045 | 82.3030192 | 24.1293498 | 55.6723382 | 0 |
| 2.89E-06 | 9.23E-06 | 4.17979245 | 6.22833659 | 7.42441534 | 9.74265918 | 26.1384007 |
| 2.89E-06 | 9.24E-06 | 130.216611 | 167.052885 | 185.610383 | 185.110524 | 59.4054561 |
| 2.89E-06 | 9.24E-06 | 40.5118345 | 30.02948 | 68.6758419 | 36.1870198 | 0 |
| 2.89E-06 | 9.25E-06 | 47.5853294 | 64.952653 | 75.1722053 | 76.0855288 | 2.37621824 |
| 2.90E-06 | 9.27E-06 | 22.8280972 | 68.9565837 | 45.4745439 | 78.8691457 | 0 |
| 2.91E-06 | 9.31E-06 | 37.9396545 | 64.7302124 | 31.5537652 | 34.7952113 | 0 |
| 2.92E-06 | 9.31E-06 | 299.337444 | 575.231372 | 553.118943 | 577.136572 | 66.5341108 |
| 2.92E-06 | 9.32E-06 | 28.9370247 | 42.9310343 | 32.4818171 | 41.2903175 | 0 |
| 2.92E-06 | 9.32E-06 | 61.7323193 | 42.041272 | 49.1867516 | 24.588616 | 0 |
| 2.92E-06 | 9.32E-06 | 24.1141872 | 57.6121134 | 41.7623363 | 58.4559551 | 0 |
| 2.92E-06 | 9.32E-06 | 28.9370247 | 57.1672323 | 37.1220767 | 37.5788283 | 0 |

| | | | | | | |
|---|---|---|---|---|---|---|
| 2.92E-06 | 9.33E-06 | 64.9475442 | 36.035376 | 25.9854537 | 63.5592527 | 0 |
| 2.93E-06 | 9.35E-06 | 28.6155022 | 56.9447917 | 32.4818171 | 64.487125 | 0 |
| 2.93E-06 | 9.36E-06 | 38.261177 | 44.9329997 | 26.9135056 | 37.1148921 | 0 |
| 2.94E-06 | 9.38E-06 | 39.2257445 | 89.8659993 | 111.36623 | 87.6839326 | 0 |
| 2.94E-06 | 9.39E-06 | 24.1141872 | 42.7085937 | 51.0428554 | 54.2805297 | 0 |
| 2.95E-06 | 9.41E-06 | 13.1824223 | 143.919063 | 165.193241 | 90.9314857 | 0 |
| 2.96E-06 | 9.44E-06 | 14.1469898 | 84.3049845 | 157.768826 | 131.293931 | 0 |
| 2.97E-06 | 9.47E-06 | 23.1496197 | 52.4959798 | 47.3306478 | 54.2805297 | 0 |
| 2.98E-06 | 9.50E-06 | 45.6561944 | 22.911381 | 89.092984 | 68.6625504 | 0 |
| 2.99E-06 | 9.53E-06 | 20.8989622 | 57.3896728 | 64.9636342 | 62.1674443 | 0 |
| 2.99E-06 | 9.54E-06 | 30.8661596 | 52.0510986 | 88.1649321 | 33.867339 | 0 |
| 3.00E-06 | 9.56E-06 | 28.2939797 | 41.8188314 | 60.3233746 | 35.2591475 | 0 |
| 3.00E-06 | 9.57E-06 | 17.0406923 | 86.3069499 | 97.4454513 | 77.4773373 | 0 |
| 3.00E-06 | 9.57E-06 | 25.7217997 | 54.053064 | 44.546492 | 38.9706367 | 0 |
| 3.01E-06 | 9.59E-06 | 24.7572322 | 43.8207967 | 44.546492 | 45.0018067 | 0 |
| 3.01E-06 | 9.59E-06 | 299.658966 | 520.955867 | 449.177128 | 516.360936 | 30.8908372 |
| 3.01E-06 | 9.61E-06 | 22.1850522 | 57.1672323 | 52.8989593 | 53.8165936 | 0 |
| 3.02E-06 | 9.62E-06 | 21.2204847 | 65.3975342 | 55.683115 | 59.3838274 | 0 |
| 3.02E-06 | 9.62E-06 | 20.8989622 | 57.3896728 | 63.1075304 | 62.6313804 | 0 |
| 3.02E-06 | 9.63E-06 | 31.5092046 | 31.8090047 | 38.9781805 | 33.867339 | 0 |
| 3.02E-06 | 9.64E-06 | 29.9015921 | 34.7007324 | 65.8916861 | 40.8263813 | 0 |
| 3.03E-06 | 9.67E-06 | 27.9724572 | 35.3680542 | 37.1220767 | 42.6821259 | 0 |
| 3.05E-06 | 9.73E-06 | 32.4737721 | 36.4802572 | 29.6976613 | 37.5788283 | 0 |
| 3.05E-06 | 9.73E-06 | 22.1850522 | 50.4940145 | 54.7550631 | 60.7756358 | 0 |
| 3.06E-06 | 9.74E-06 | 178.123463 | 634.62301 | 600.44959 | 547.444659 | 47.5243649 |
| 3.06E-06 | 9.74E-06 | 25.7217997 | 79.1888509 | 37.1220767 | 68.6625504 | 0 |
| 3.07E-06 | 9.78E-06 | 19.6128723 | 70.5136678 | 64.0355823 | 68.6625504 | 0 |
| 3.08E-06 | 9.81E-06 | 17.3622148 | 88.3089152 | 95.5893474 | 71.4461673 | 0 |
| 3.08E-06 | 9.82E-06 | 15.1115573 | 81.858138 | 107.654022 | 178.151482 | 0 |
| 3.08E-06 | 9.82E-06 | 42.4409695 | 78.0766479 | 78.8844129 | 87.6839326 | 2.37621824 |
| 3.08E-06 | 9.82E-06 | 58.1955718 | 57.3896728 | 20.4171422 | 71.4461673 | 0 |
| 3.09E-06 | 9.85E-06 | 14.4685123 | 81.4132568 | 126.215061 | 143.820207 | 0 |
| 3.10E-06 | 9.86E-06 | 28.2939797 | 62.5058065 | 32.4818171 | 70.9822312 | 0 |
| 3.10E-06 | 9.86E-06 | 28.2939797 | 85.1947469 | 33.409869 | 67.7346781 | 0 |
| 3.10E-06 | 9.87E-06 | 20.8989622 | 63.3955688 | 54.7550631 | 66.3428696 | 0 |
| 3.10E-06 | 9.88E-06 | 14.1469898 | 91.2006429 | 119.718697 | 123.870952 | 0 |
| 3.11E-06 | 9.90E-06 | 100.636541 | 42.4861532 | 51.0428554 | 27.3722329 | 0 |
| 3.11E-06 | 9.90E-06 | 52.7296894 | 124.344291 | 88.1649321 | 129.902122 | 9.50487297 |
| 3.12E-06 | 9.92E-06 | 29.5800696 | 46.934965 | 30.6257133 | 55.208402 | 0 |
| 3.12E-06 | 9.93E-06 | 919.875861 | 954.047701 | 764.71478 | 916.737835 | 45.1481466 |
| 3.12E-06 | 9.93E-06 | 18.0052598 | 80.0786133 | 97.4454513 | 67.7346781 | 0 |
| 3.13E-06 | 9.95E-06 | 150.794051 | 300.962122 | 266.3509 | 295.991265 | 28.5146189 |
| 3.14E-06 | 9.99E-06 | 21.2204847 | 59.8365194 | 57.5392189 | 56.6002105 | 0 |

| | | | | | | |
|---|---|---|---|---|---|---|
| 3.14E-06 | 1.00E-05 | 50.4790319 | 145.253707 | 99.3015551 | 128.97425 | 0 |
| 3.14E-06 | 1.00E-05 | 15.4330798 | 85.1947469 | 117.862593 | 96.0347833 | 0 |
| 3.15E-06 | 1.00E-05 | 20.8989622 | 58.0569946 | 56.6111669 | 70.9822312 | 0 |
| 3.15E-06 | 1.00E-05 | 0 | 0 | 0 | 0 | 16.6335277 |
| 3.18E-06 | 1.01E-05 | 19.9343948 | 83.6376628 | 56.6111669 | 70.518295 | 0 |
| 3.18E-06 | 1.01E-05 | 21.2204847 | 57.1672323 | 61.2514265 | 55.208402 | 0 |
| 3.18E-06 | 1.01E-05 | 14.1469898 | 81.6356974 | 147.560255 | 129.902122 | 0 |
| 3.19E-06 | 1.01E-05 | 23.4711422 | 46.934965 | 46.4025958 | 61.239572 | 0 |
| 3.19E-06 | 1.01E-05 | 127.644431 | 208.871716 | 212.523889 | 187.894141 | 59.4054561 |
| 3.21E-06 | 1.02E-05 | 20.5774398 | 88.3089152 | 53.8270112 | 67.7346781 | 0 |
| 3.22E-06 | 1.02E-05 | 24.7572322 | 83.8601033 | 60.3233746 | 40.3624452 | 0 |
| 3.23E-06 | 1.03E-05 | 29.2585471 | 34.4782918 | 34.3379209 | 47.3214874 | 0 |
| 3.24E-06 | 1.03E-05 | 45.334672 | 41.5963908 | 77.0283091 | 25.9804245 | 0 |
| 3.25E-06 | 1.03E-05 | 24.7572322 | 41.5963908 | 68.6758419 | 47.3214874 | 0 |
| 3.25E-06 | 1.03E-05 | 27.0078897 | 62.5058065 | 35.2659728 | 46.8575513 | 0 |
| 3.25E-06 | 1.03E-05 | 24.7572322 | 100.765588 | 115.078438 | 38.0427644 | 0 |
| 3.25E-06 | 1.03E-05 | 15.1115573 | 87.1967122 | 128.071165 | 94.1790387 | 0 |
| 3.25E-06 | 1.03E-05 | 27.6509347 | 36.4802572 | 35.2659728 | 44.0739344 | 0 |
| 3.25E-06 | 1.03E-05 | 34.0813846 | 50.2715739 | 28.7696094 | 36.1870198 | 0 |
| 3.27E-06 | 1.04E-05 | 7.39501741 | 9.56494547 | 11.136623 | 10.2065953 | 102.177384 |
| 3.27E-06 | 1.04E-05 | 25.0787547 | 52.7184204 | 43.6184401 | 39.4345729 | 0 |
| 3.28E-06 | 1.04E-05 | 15.1115573 | 107.216366 | 93.7332436 | 92.3232941 | 0 |
| 3.28E-06 | 1.04E-05 | 18.6483048 | 5.33857422 | 7.42441534 | 9.27872303 | 83.1676385 |
| 3.28E-06 | 1.04E-05 | 26.0433222 | 35.1456136 | 48.2586997 | 51.4969128 | 0 |
| 3.28E-06 | 1.04E-05 | 35.6889971 | 34.4782918 | 34.3379209 | 29.2279775 | 0 |
| 3.29E-06 | 1.04E-05 | 30.5446371 | 33.3660889 | 42.6903882 | 32.0115944 | 0 |
| 3.29E-06 | 1.04E-05 | 16.0761248 | 78.9664103 | 102.085711 | 92.3232941 | 0 |
| 3.29E-06 | 1.04E-05 | 46.9422844 | 140.804895 | 77.956361 | 84.4363795 | 2.37621824 |
| 3.30E-06 | 1.05E-05 | 19.9343948 | 82.7479004 | 57.5392189 | 65.4149973 | 0 |
| 3.30E-06 | 1.05E-05 | 21.2204847 | 79.6337321 | 47.3306478 | 90.0036134 | 0 |
| 3.30E-06 | 1.05E-05 | 33.1168171 | 65.3975342 | 26.9135056 | 67.7346781 | 0 |
| 3.31E-06 | 1.05E-05 | 17.3622148 | 62.2833659 | 106.72597 | 108.561059 | 0 |
| 3.31E-06 | 1.05E-05 | 30.8661596 | 58.5018758 | 28.7696094 | 66.3428696 | 0 |
| 3.31E-06 | 1.05E-05 | 52.7296894 | 125.011613 | 51.0428554 | 115.984038 | 0 |
| 3.32E-06 | 1.05E-05 | 21.8635297 | 49.1593709 | 59.3953227 | 55.208402 | 0 |
| 3.32E-06 | 1.05E-05 | 26.0433222 | 60.05896 | 46.4025958 | 36.650956 | 0 |
| 3.32E-06 | 1.05E-05 | 24.7572322 | 46.2676432 | 136.423632 | 66.3428696 | 0 |
| 3.33E-06 | 1.06E-05 | 16.0761248 | 89.4211182 | 148.488307 | 77.9412734 | 0 |
| 3.33E-06 | 1.06E-05 | 25.0787547 | 71.4034302 | 40.8342843 | 48.2493597 | 0 |
| 3.34E-06 | 1.06E-05 | 0 | 0 | 0 | 0 | 239.998043 |
| 3.34E-06 | 1.06E-05 | 27.3294122 | 86.9742716 | 36.1940248 | 56.1362743 | 0 |
| 3.35E-06 | 1.06E-05 | 27.3294122 | 37.3700195 | 40.8342843 | 63.5592527 | 0 |
| 3.35E-06 | 1.06E-05 | 24.1141872 | 52.7184204 | 52.8989593 | 39.4345729 | 0 |

| | | | | | | |
|---|---|---|---|---|---|---|
| 3.36E-06 | 1.06E-05 | 31.8307271 | 37.3700195 | 28.7696094 | 38.5067006 | 0 |
| 3.36E-06 | 1.07E-05 | 21.8635297 | 46.0452026 | 71.4599976 | 91.3954218 | 0 |
| 3.37E-06 | 1.07E-05 | 21.2204847 | 65.3975342 | 51.0428554 | 58.9198912 | 0 |
| 3.37E-06 | 1.07E-05 | 30.2231146 | 43.1534749 | 33.409869 | 32.4755306 | 0 |
| 3.37E-06 | 1.07E-05 | 28.6155022 | 32.4763265 | 44.546492 | 36.650956 | 0 |
| 3.37E-06 | 1.07E-05 | 25.4002772 | 103.879757 | 44.546492 | 54.2805297 | 0 |
| 3.38E-06 | 1.07E-05 | 26.6863672 | 37.1475789 | 55.683115 | 85.828188 | 0 |
| 3.38E-06 | 1.07E-05 | 27.3294122 | 35.5904948 | 64.9636342 | 43.6099982 | 0 |
| 3.38E-06 | 1.07E-05 | 56.2664368 | 49.8266927 | 23.2012979 | 41.2903175 | 0 |
| 3.38E-06 | 1.07E-05 | 24.7572322 | 44.0432373 | 88.1649321 | 50.5690405 | 0 |
| 3.39E-06 | 1.07E-05 | 92.9200014 | 57.6121134 | 63.1075304 | 53.8165936 | 0 |
| 3.39E-06 | 1.07E-05 | 24.4357097 | 45.822762 | 40.8342843 | 44.5378705 | 0 |
| 3.40E-06 | 1.07E-05 | 253.038204 | 341.001428 | 248.717914 | 322.899561 | 38.0194919 |
| 3.40E-06 | 1.08E-05 | 18.9698273 | 74.0727173 | 73.3161014 | 61.239572 | 0 |
| 3.41E-06 | 1.08E-05 | 26.0433222 | 35.3680542 | 44.546492 | 42.6821259 | 0 |
| 3.41E-06 | 1.08E-05 | 35.0459521 | 51.1613363 | 33.409869 | 29.6919137 | 0 |
| 3.41E-06 | 1.08E-05 | 18.3267823 | 90.3108805 | 77.0283091 | 62.6313804 | 0 |
| 3.42E-06 | 1.08E-05 | 21.8635297 | 56.9447917 | 69.6038938 | 47.3214874 | 0 |
| 3.42E-06 | 1.08E-05 | 22.5065747 | 14.6810791 | 9.28051917 | 15.7738291 | 85.5438568 |
| 3.42E-06 | 1.08E-05 | 22.5065747 | 73.1829549 | 47.3306478 | 52.8887212 | 0 |
| 3.42E-06 | 1.08E-05 | 308.018551 | 36.035376 | 71.4599976 | 42.2181898 | 0 |
| 3.42E-06 | 1.08E-05 | 26.0433222 | 82.0805786 | 83.5246725 | 34.7952113 | 0 |
| 3.43E-06 | 1.08E-05 | 21.5420072 | 59.1691976 | 50.1148035 | 83.9724434 | 0 |
| 3.43E-06 | 1.08E-05 | 24.4357097 | 41.8188314 | 74.2441534 | 49.177232 | 0 |
| 3.43E-06 | 1.08E-05 | 18.0052598 | 65.8424154 | 89.092984 | 75.6215927 | 0 |
| 3.43E-06 | 1.08E-05 | 22.5065747 | 59.1691976 | 50.1148035 | 46.8575513 | 0 |
| 3.43E-06 | 1.09E-05 | 21.5420072 | 59.3916382 | 48.2586997 | 63.0953166 | 0 |
| 3.44E-06 | 1.09E-05 | 27.0078897 | 35.5904948 | 37.1220767 | 39.898509 | 0 |
| 3.44E-06 | 1.09E-05 | 24.4357097 | 75.852242 | 38.0501286 | 64.9510612 | 0 |
| 3.44E-06 | 1.09E-05 | 82.952804 | 41.3739502 | 27.8415575 | 41.2903175 | 0 |
| 3.44E-06 | 1.09E-05 | 45.334672 | 66.9546183 | 289.552198 | 34.3312752 | 0 |
| 3.44E-06 | 1.09E-05 | 111.889829 | 218.659102 | 270.99116 | 197.6368 | 19.0097459 |
| 3.44E-06 | 1.09E-05 | 29.2585471 | 35.1456136 | 33.409869 | 34.7952113 | 0 |
| 3.44E-06 | 1.09E-05 | 23.1496197 | 42.9310343 | 51.9709073 | 51.0329766 | 0 |
| 3.45E-06 | 1.09E-05 | 62.3753642 | 24.4684652 | 52.8989593 | 38.5067006 | 0 |
| 3.45E-06 | 1.09E-05 | 83.9173715 | 84.0825439 | 44.546492 | 61.7035081 | 2.37621824 |
| 3.46E-06 | 1.09E-05 | 0 | 0 | 0 | 0 | 45.1481466 |
| 3.47E-06 | 1.09E-05 | 48.8714194 | 105.436841 | 100.229607 | 103.921698 | 0 |
| 3.47E-06 | 1.10E-05 | 20.5774398 | 53.8306234 | 59.3953227 | 67.7346781 | 0 |
| 3.48E-06 | 1.10E-05 | 150.794051 | 215.322493 | 228.300772 | 268.619032 | 33.2670554 |
| 3.48E-06 | 1.10E-05 | 32.4737721 | 43.3759155 | 29.6976613 | 33.4034029 | 0 |
| 3.48E-06 | 1.10E-05 | 98.3858838 | 346.784884 | 167.977397 | 233.359884 | 16.6335277 |
| 3.49E-06 | 1.10E-05 | 13.1824223 | 92.9801676 | 122.502853 | 133.149675 | 0 |

| | | | | | | |
|---|---|---|---|---|---|---|
| 3.49E-06 | 1.10E-05 | 31.8307271 | 38.9271037 | 38.9781805 | 28.3001052 | 0 |
| 3.49E-06 | 1.10E-05 | 27.0078897 | 55.8325887 | 52.8989593 | 32.4755306 | 0 |
| 3.49E-06 | 1.10E-05 | 19.6128723 | 85.1947469 | 82.5966206 | 52.4247851 | 0 |
| 3.50E-06 | 1.10E-05 | 30.2231146 | 42.9310343 | 31.5537652 | 65.8789335 | 0 |
| 3.50E-06 | 1.11E-05 | 21.5420072 | 57.1672323 | 48.2586997 | 66.8068058 | 0 |
| 3.51E-06 | 1.11E-05 | 33.1168171 | 27.1377523 | 62.1794784 | 73.7658481 | 0 |
| 3.51E-06 | 1.11E-05 | 19.2913498 | 60.9487223 | 65.8916861 | 70.0543588 | 0 |
| 3.52E-06 | 1.11E-05 | 304.481804 | 627.949792 | 644.996082 | 639.304017 | 149.701749 |
| 3.52E-06 | 1.11E-05 | 63.6614542 | 34.923173 | 35.2659728 | 30.619786 | 125.939567 |
| 3.54E-06 | 1.12E-05 | 24.4357097 | 49.6042521 | 38.0501286 | 48.7132959 | 0 |
| 3.55E-06 | 1.12E-05 | 24.7572322 | 41.1515096 | 42.6903882 | 61.239572 | 0 |
| 3.55E-06 | 1.12E-05 | 83.2743265 | 63.1731283 | 119.718697 | 17.6295737 | 0 |
| 3.56E-06 | 1.12E-05 | 25.0787547 | 39.816866 | 80.7405168 | 51.4969128 | 0 |
| 3.56E-06 | 1.12E-05 | 31.8307271 | 82.5254598 | 43.6184401 | 33.4034029 | 0 |
| 3.56E-06 | 1.12E-05 | 149.507961 | 237.121672 | 121.574801 | 145.212015 | 0 |
| 3.56E-06 | 1.12E-05 | 17.0406923 | 77.1868856 | 74.2441534 | 88.6118049 | 0 |
| 3.57E-06 | 1.12E-05 | 29.9015921 | 46.4900838 | 28.7696094 | 43.1460621 | 0 |
| 3.57E-06 | 1.13E-05 | 25.4002772 | 47.8247274 | 35.2659728 | 50.5690405 | 0 |
| 3.58E-06 | 1.13E-05 | 40.5118345 | 66.5097371 | 28.7696094 | 163.769461 | 0 |
| 3.58E-06 | 1.13E-05 | 33.7598621 | 89.8659993 | 44.546492 | 32.4755306 | 0 |
| 3.59E-06 | 1.13E-05 | 329.560559 | 316.310522 | 320.177911 | 389.706367 | 16.6335277 |
| 3.59E-06 | 1.13E-05 | 19.9343948 | 78.9664103 | 54.7550631 | 64.0231889 | 0 |
| 3.60E-06 | 1.13E-05 | 25.7217997 | 40.4841878 | 36.1940248 | 47.7854236 | 0 |
| 3.61E-06 | 1.14E-05 | 299.015921 | 353.680542 | 230.156875 | 291.351903 | 26.1384007 |
| 3.61E-06 | 1.14E-05 | 26.0433222 | 46.0452026 | 51.0428554 | 33.867339 | 0 |
| 3.61E-06 | 1.14E-05 | 18.6483048 | 184.848132 | 97.4454513 | 57.9920189 | 0 |
| 3.61E-06 | 1.14E-05 | 37.9396545 | 35.3680542 | 25.9854537 | 45.4657428 | 0 |
| 3.63E-06 | 1.14E-05 | 17.6837373 | 65.1750936 | 80.7405168 | 82.1166988 | 0 |
| 3.63E-06 | 1.14E-05 | 21.2204847 | 48.9369303 | 61.2514265 | 84.4363795 | 0 |
| 3.64E-06 | 1.15E-05 | 39.547267 | 88.3089152 | 78.8844129 | 85.3642518 | 2.37621824 |
| 3.64E-06 | 1.15E-05 | 20.5774398 | 65.1750936 | 50.1148035 | 82.1166988 | 0 |
| 3.64E-06 | 1.15E-05 | 22.8280972 | 93.86993 | 40.8342843 | 78.4052096 | 0 |
| 3.64E-06 | 1.15E-05 | 10.6102424 | 6.005896 | 4.64025958 | 2.78361691 | 59.4054561 |
| 3.65E-06 | 1.15E-05 | 40.190312 | 23.3562622 | 124.358957 | 90.9314857 | 0 |
| 3.65E-06 | 1.15E-05 | 37.2966096 | 32.4763265 | 33.409869 | 29.2279775 | 0 |
| 3.66E-06 | 1.15E-05 | 26.0433222 | 57.1672323 | 33.409869 | 53.8165936 | 0 |
| 3.66E-06 | 1.15E-05 | 18.0052598 | 71.6258708 | 112.294282 | 65.8789335 | 0 |
| 3.67E-06 | 1.15E-05 | 22.5065747 | 44.4881185 | 51.0428554 | 51.9608489 | 0 |
| 3.68E-06 | 1.16E-05 | 41.476402 | 66.7321777 | 22.273246 | 58.9198912 | 0 |
| 3.69E-06 | 1.16E-05 | 21.8635297 | 77.4093262 | 49.1867516 | 52.8887212 | 0 |
| 3.69E-06 | 1.16E-05 | 18.6483048 | 63.1731283 | 65.8916861 | 77.9412734 | 0 |
| 3.69E-06 | 1.16E-05 | 36.3320421 | 25.3582275 | 90.9490879 | 67.2707419 | 0 |
| 3.70E-06 | 1.16E-05 | 27.6509347 | 35.8129354 | 33.409869 | 44.5378705 | 0 |

| | | | | | | |
|---|---|---|---|---|---|---|
| 3.72E-06 | 1.17E-05 | 9.00262989 | 153.261568 | 166.121293 | 157.738291 | 0 |
| 3.72E-06 | 1.17E-05 | 164.941041 | 315.865641 | 283.055835 | 321.971689 | 23.7621824 |
| 3.73E-06 | 1.17E-05 | 34.4029071 | 26.0255493 | 72.3880495 | 56.6002105 | 0 |
| 3.73E-06 | 1.17E-05 | 18.6483048 | 56.7223511 | 85.3807764 | 76.549465 | 0 |
| 3.74E-06 | 1.18E-05 | 352.067133 | 544.979451 | 399.990376 | 484.349342 | 71.2865473 |
| 3.75E-06 | 1.18E-05 | 16.7191698 | 70.7361084 | 83.5246725 | 88.6118049 | 0 |
| 3.75E-06 | 1.18E-05 | 38.904222 | 25.3582275 | 60.3233746 | 40.8263813 | 0 |
| 3.76E-06 | 1.18E-05 | 37.9396545 | 87.1967122 | 100.229607 | 81.6527626 | 2.37621824 |
| 3.76E-06 | 1.18E-05 | 40.190312 | 81.4132568 | 76.1002572 | 85.828188 | 0 |
| 3.76E-06 | 1.18E-05 | 22.1850522 | 60.7262817 | 42.6903882 | 68.6625504 | 0 |
| 3.76E-06 | 1.18E-05 | 33.4383396 | 38.2597819 | 94.6612955 | 37.1148921 | 0 |
| 3.76E-06 | 1.18E-05 | 15.4330798 | 104.102197 | 81.6685687 | 86.7560603 | 0 |
| 3.77E-06 | 1.18E-05 | 24.1141872 | 81.6356974 | 39.9062324 | 56.1362743 | 0 |
| 3.77E-06 | 1.18E-05 | 22.5065747 | 48.4920492 | 44.546492 | 58.9198912 | 0 |
| 3.78E-06 | 1.19E-05 | 27.6509347 | 111.220296 | 166.121293 | 143.820207 | 2.37621824 |
| 3.78E-06 | 1.19E-05 | 37.9396545 | 113.889583 | 23.2012979 | 105.777442 | 0 |
| 3.78E-06 | 1.19E-05 | 36.0105196 | 24.9133464 | 102.085711 | 83.5085072 | 0 |
| 3.78E-06 | 1.19E-05 | 23.7926647 | 57.6121134 | 59.3953227 | 37.5788283 | 0 |
| 3.78E-06 | 1.19E-05 | 33.7598621 | 34.2558512 | 37.1220767 | 27.8361691 | 0 |
| 3.79E-06 | 1.19E-05 | 60.4462293 | 36.7026978 | 24.1293498 | 50.5690405 | 0 |
| 3.79E-06 | 1.19E-05 | 47.9068519 | 228.224048 | 156.840774 | 18.0935099 | 0 |
| 3.80E-06 | 1.19E-05 | 35.6889971 | 79.6337321 | 130.85532 | 125.262761 | 0 |
| 3.81E-06 | 1.19E-05 | 45.334672 | 156.820618 | 124.358957 | 136.397228 | 0 |
| 3.81E-06 | 1.19E-05 | 347.887341 | 469.572091 | 361.012196 | 591.518593 | 21.3859642 |
| 3.81E-06 | 1.19E-05 | 27.9724572 | 33.3660889 | 41.7623363 | 62.1674443 | 0 |
| 3.81E-06 | 1.19E-05 | 24.4357097 | 41.1515096 | 38.9781805 | 47.7854236 | 0 |
| 3.81E-06 | 1.19E-05 | 51.1220769 | 123.676969 | 54.7550631 | 23.6607437 | 0 |
| 3.81E-06 | 1.19E-05 | 536.621046 | 992.529923 | 866.80049 | 893.541027 | 45.1481466 |
| 3.82E-06 | 1.20E-05 | 13.5039448 | 110.997856 | 90.0210359 | 121.087335 | 0 |
| 3.82E-06 | 1.20E-05 | 39.2257445 | 98.763623 | 142.919995 | 169.800631 | 4.75243649 |
| 3.83E-06 | 1.20E-05 | 20.2559173 | 76.2971232 | 57.5392189 | 53.8165936 | 0 |
| 3.83E-06 | 1.20E-05 | 37.618132 | 28.4723958 | 28.7696094 | 27.8361691 | 80.7914203 |
| 3.83E-06 | 1.20E-05 | 44.691627 | 81.6356974 | 87.2368802 | 91.3954218 | 11.8810912 |
| 3.84E-06 | 1.20E-05 | 28.2939797 | 30.02948 | 42.6903882 | 46.8575513 | 0 |
| 3.84E-06 | 1.20E-05 | 26.0433222 | 34.0334106 | 73.3161014 | 56.6002105 | 0 |
| 3.84E-06 | 1.20E-05 | 42.762492 | 22.911381 | 116.00649 | 72.8379758 | 0 |
| 3.84E-06 | 1.20E-05 | 86.168029 | 64.5077718 | 51.0428554 | 55.208402 | 0 |
| 3.87E-06 | 1.21E-05 | 19.9343948 | 70.7361084 | 50.1148035 | 73.3019119 | 0 |
| 3.89E-06 | 1.22E-05 | 112.854396 | 39.1495443 | 33.409869 | 40.8263813 | 0 |
| 3.89E-06 | 1.22E-05 | 34.7244296 | 38.2597819 | 59.3953227 | 27.3722329 | 0 |
| 3.89E-06 | 1.22E-05 | 23.1496197 | 165.05092 | 46.4025958 | 77.4773373 | 0 |
| 3.90E-06 | 1.22E-05 | 10.6102424 | 139.692692 | 167.977397 | 111.344676 | 0 |
| 3.90E-06 | 1.22E-05 | 55.9449143 | 112.55494 | 58.4672708 | 20.8771268 | 0 |

| | | | | | | |
|---|---|---|---|---|---|---|
| 3.90E-06 | 1.22E-05 | 2.89370247 | 1.33464355 | 0.92805192 | 2.78361691 | 4.75243649 |
| 3.91E-06 | 1.22E-05 | 30.2231146 | 72.5156331 | 30.6257133 | 46.3936151 | 0 |
| 3.91E-06 | 1.22E-05 | 19.9343948 | 69.6239054 | 50.1148035 | 71.9101035 | 0 |
| 3.91E-06 | 1.22E-05 | 161.725816 | 308.969983 | 292.336354 | 332.64222 | 23.7621824 |
| 3.92E-06 | 1.23E-05 | 19.9343948 | 58.0569946 | 55.683115 | 62.6313804 | 0 |
| 3.92E-06 | 1.23E-05 | 30.5446371 | 49.1593709 | 29.6976613 | 74.6937204 | 0 |
| 3.92E-06 | 1.23E-05 | 29.9015921 | 30.02948 | 62.1794784 | 42.6821259 | 0 |
| 3.92E-06 | 1.23E-05 | 22.5065747 | 43.8207967 | 47.3306478 | 52.4247851 | 0 |
| 3.93E-06 | 1.23E-05 | 23.4711422 | 45.3778809 | 40.8342843 | 58.4559551 | 0 |
| 3.93E-06 | 1.23E-05 | 23.1496197 | 42.2637126 | 47.3306478 | 44.5378705 | 0 |
| 3.94E-06 | 1.23E-05 | 21.2204847 | 52.940861 | 63.1075304 | 47.7854236 | 0 |
| 3.94E-06 | 1.23E-05 | 168.79931 | 48.047168 | 41.7623363 | 34.7952113 | 0 |
| 3.94E-06 | 1.23E-05 | 19.9343948 | 57.3896728 | 59.3953227 | 57.5280828 | 0 |
| 3.95E-06 | 1.23E-05 | 27.6509347 | 37.1475789 | 31.5537652 | 39.4345729 | 0 |
| 3.95E-06 | 1.23E-05 | 80.7021465 | 52.0510986 | 24.1293498 | 39.4345729 | 0 |
| 3.95E-06 | 1.23E-05 | 30.5446371 | 74.9624797 | 30.6257133 | 46.3936151 | 0 |
| 3.97E-06 | 1.24E-05 | 23.7926647 | 41.5963908 | 41.7623363 | 56.6002105 | 0 |
| 3.97E-06 | 1.24E-05 | 0 | 0 | 0 | 0 | 45.1481466 |
| 3.97E-06 | 1.24E-05 | 38.261177 | 51.6062174 | 42.6903882 | 24.1246799 | 0 |
| 3.97E-06 | 1.24E-05 | 44.048582 | 96.0943359 | 84.4527244 | 89.5396772 | 4.75243649 |
| 3.98E-06 | 1.24E-05 | 47.5853294 | 58.0569946 | 112.294282 | 75.6215927 | 2.37621824 |
| 3.98E-06 | 1.24E-05 | 36.9750871 | 131.46239 | 120.646749 | 127.582442 | 4.75243649 |
| 3.99E-06 | 1.25E-05 | 29.9015921 | 54.7203857 | 34.3379209 | 109.952868 | 0 |
| 3.99E-06 | 1.25E-05 | 33.4383396 | 44.0432373 | 48.2586997 | 25.5164883 | 0 |
| 4.00E-06 | 1.25E-05 | 38.5826995 | 38.0373413 | 26.9135056 | 32.0115944 | 0 |
| 4.00E-06 | 1.25E-05 | 18.0052598 | 70.0687866 | 68.6758419 | 146.139888 | 0 |
| 4.00E-06 | 1.25E-05 | 133.431836 | 345.672681 | 345.235313 | 344.70456 | 19.0097459 |
| 4.01E-06 | 1.25E-05 | 24.1141872 | 58.7243164 | 41.7623363 | 40.8263813 | 0 |
| 4.01E-06 | 1.25E-05 | 19.6128723 | 114.334465 | 65.8916861 | 56.6002105 | 0 |
| 4.01E-06 | 1.25E-05 | 219.278342 | 494.707878 | 425.97583 | 689.409121 | 4.75243649 |
| 4.01E-06 | 1.25E-05 | 37.618132 | 84.0825439 | 87.2368802 | 95.106911 | 0 |
| 4.02E-06 | 1.26E-05 | 19.6128723 | 71.1809896 | 62.1794784 | 53.3526574 | 0 |
| 4.03E-06 | 1.26E-05 | 20.2559173 | 58.7243164 | 56.6111669 | 53.3526574 | 0 |
| 4.03E-06 | 1.26E-05 | 35.0459521 | 113.444702 | 92.8051917 | 89.5396772 | 0 |
| 4.04E-06 | 1.26E-05 | 36.9750871 | 26.9153117 | 49.1867516 | 33.4034029 | 0 |
| 4.04E-06 | 1.26E-05 | 29.5800696 | 29.1397176 | 37.1220767 | 44.0739344 | 0 |
| 4.05E-06 | 1.26E-05 | 14.4685123 | 123.232088 | 87.2368802 | 86.7560603 | 0 |
| 4.07E-06 | 1.27E-05 | 14.7900348 | 79.8561727 | 143.848047 | 88.6118049 | 0 |
| 4.08E-06 | 1.27E-05 | 171.693013 | 436.873324 | 376.789078 | 407.799877 | 59.4054561 |
| 4.08E-06 | 1.27E-05 | 39.2257445 | 48.047168 | 30.6257133 | 27.8361691 | 0 |
| 4.08E-06 | 1.27E-05 | 14.7900348 | 78.2990885 | 108.582074 | 94.6429749 | 0 |
| 4.09E-06 | 1.27E-05 | 45.334672 | 52.7184204 | 27.8415575 | 29.6919137 | 0 |
| 4.09E-06 | 1.27E-05 | 55.3018693 | 87.1967122 | 17.6329864 | 89.0757411 | 0 |

| | | | | | | |
|---|---|---|---|---|---|---|
| 4.09E-06 | 1.28E-05 | 19.6128723 | 51.828658 | 64.9636342 | 67.7346781 | 0 |
| 4.10E-06 | 1.28E-05 | 20.2559173 | 88.976237 | 57.5392189 | 53.3526574 | 0 |
| 4.12E-06 | 1.28E-05 | 28.9370247 | 46.2676432 | 30.6257133 | 35.2591475 | 0 |
| 4.12E-06 | 1.28E-05 | 26.6863672 | 84.5274251 | 32.4818171 | 64.0231889 | 0 |
| 4.13E-06 | 1.29E-05 | 31.8307271 | 136.133643 | 83.5246725 | 185.574461 | 2.37621824 |
| 4.13E-06 | 1.29E-05 | 22.8280972 | 43.3759155 | 69.6038938 | 46.8575513 | 0 |
| 4.13E-06 | 1.29E-05 | 129.252043 | 534.969625 | 500.219983 | 483.42147 | 45.1481466 |
| 4.14E-06 | 1.29E-05 | 36.6535646 | 199.306771 | 255.214277 | 212.018821 | 0 |
| 4.14E-06 | 1.29E-05 | 28.6155022 | 56.4999105 | 73.3161014 | 29.6919137 | 0 |
| 4.14E-06 | 1.29E-05 | 43.405537 | 145.698588 | 141.063891 | 139.644782 | 0 |
| 4.15E-06 | 1.29E-05 | 589.99378 | 50.4940145 | 26.9135056 | 90.9314857 | 0 |
| 4.14E-06 | 1.29E-05 | 25.4002772 | 40.929069 | 97.4454513 | 50.1051043 | 0 |
| 4.15E-06 | 1.29E-05 | 28.2939797 | 46.2676432 | 28.7696094 | 45.929679 | 0 |
| 4.15E-06 | 1.29E-05 | 48.2283744 | 84.7498657 | 57.5392189 | 63.5592527 | 2.37621824 |
| 4.16E-06 | 1.29E-05 | 26.6863672 | 49.1593709 | 34.3379209 | 36.1870198 | 0 |
| 4.17E-06 | 1.30E-05 | 23.1496197 | 38.7046631 | 73.3161014 | 57.0641466 | 0 |
| 4.17E-06 | 1.30E-05 | 38.904222 | 94.3148112 | 73.3161014 | 90.0036134 | 0 |
| 4.17E-06 | 1.30E-05 | 13.5039448 | 119.673039 | 79.8124649 | 124.798825 | 0 |
| 4.18E-06 | 1.30E-05 | 33.7598621 | 129.682865 | 93.7332436 | 99.7462725 | 0 |
| 4.18E-06 | 1.30E-05 | 18.3267823 | 68.9565837 | 63.1075304 | 64.487125 | 0 |
| 4.19E-06 | 1.30E-05 | 22.5065747 | 71.8483114 | 43.6184401 | 49.6411682 | 0 |
| 4.19E-06 | 1.30E-05 | 26.3648447 | 130.795068 | 33.409869 | 89.0757411 | 0 |
| 4.20E-06 | 1.31E-05 | 27.0078897 | 50.9388957 | 45.4745439 | 30.619786 | 0 |
| 4.21E-06 | 1.31E-05 | 37.618132 | 34.923173 | 25.0574018 | 44.0739344 | 0 |
| 4.22E-06 | 1.31E-05 | 13.5039448 | 123.454529 | 82.5966206 | 110.416804 | 0 |
| 4.22E-06 | 1.31E-05 | 33.4383396 | 26.0255493 | 39.9062324 | 38.5067006 | 0 |
| 4.23E-06 | 1.31E-05 | 26.0433222 | 36.4802572 | 36.1940248 | 35.7230837 | 0 |
| 4.24E-06 | 1.32E-05 | 20.8989622 | 45.822762 | 56.6111669 | 62.1674443 | 0 |
| 4.26E-06 | 1.32E-05 | 31.8307271 | 123.676969 | 25.9854537 | 164.233398 | 0 |
| 4.27E-06 | 1.33E-05 | 28.9370247 | 29.8070394 | 39.9062324 | 34.3312752 | 0 |
| 4.27E-06 | 1.33E-05 | 68.1627692 | 39.3719849 | 39.9062324 | 25.5164883 | 0 |
| 4.28E-06 | 1.33E-05 | 27.9724572 | 45.3778809 | 33.409869 | 76.549465 | 0 |
| 4.28E-06 | 1.33E-05 | 112.211351 | 192.633553 | 108.582074 | 223.153289 | 16.6335277 |
| 4.29E-06 | 1.33E-05 | 17.3622148 | 8.45274251 | 15.7768826 | 12.9902122 | 66.5341108 |
| 4.30E-06 | 1.34E-05 | 27.3294122 | 31.3641235 | 51.9709073 | 38.9706367 | 0 |
| 4.31E-06 | 1.34E-05 | 11.8963324 | 70.0687866 | 204.171422 | 196.244992 | 0 |
| 4.32E-06 | 1.34E-05 | 26.0433222 | 32.9212077 | 64.0355823 | 47.3214874 | 0 |
| 4.32E-06 | 1.34E-05 | 58.8386168 | 81.4132568 | 64.0355823 | 46.3936151 | 2.37621824 |
| 4.34E-06 | 1.35E-05 | 16.7191698 | 78.5215291 | 64.0355823 | 108.561059 | 0 |
| 4.34E-06 | 1.35E-05 | 10.9317649 | 132.129712 | 99.3015551 | 158.202228 | 0 |
| 4.35E-06 | 1.35E-05 | 21.5420072 | 78.7439697 | 103.013763 | 41.2903175 | 0 |
| 4.37E-06 | 1.36E-05 | 27.6509347 | 30.02948 | 51.0428554 | 40.8263813 | 0 |
| 4.38E-06 | 1.36E-05 | 33.7598621 | 32.9212077 | 26.9135056 | 44.0739344 | 0 |

| | | | | | | |
|---|---|---|---|---|---|---|
| 4.39E-06 | 1.36E-05 | 35.3674746 | 188.407182 | 21.3451941 | 190.677758 | 0 |
| 4.39E-06 | 1.36E-05 | 29.2585471 | 68.0668213 | 62.1794784 | 27.8361691 | 0 |
| 4.41E-06 | 1.37E-05 | 23.1496197 | 63.3955688 | 60.3233746 | 36.650956 | 0 |
| 4.42E-06 | 1.37E-05 | 24.7572322 | 91.8679647 | 33.409869 | 92.7872303 | 0 |
| 4.42E-06 | 1.37E-05 | 31.5092046 | 47.3798462 | 39.9062324 | 26.4443606 | 0 |
| 4.43E-06 | 1.37E-05 | 18.9698273 | 52.0510986 | 71.4599976 | 67.2707419 | 0 |
| 4.43E-06 | 1.38E-05 | 15.4330798 | 63.84045 | 165.193241 | 103.457762 | 0 |
| 4.43E-06 | 1.38E-05 | 23.4711422 | 60.9487223 | 48.2586997 | 37.5788283 | 0 |
| 4.43E-06 | 1.38E-05 | 67.8412467 | 40.929069 | 25.0574018 | 38.0427644 | 0 |
| 4.45E-06 | 1.38E-05 | 51.4435994 | 53.8306234 | 51.0428554 | 20.4131907 | 0 |
| 4.45E-06 | 1.38E-05 | 16.7191698 | 86.5293905 | 68.6758419 | 70.9822312 | 0 |
| 4.46E-06 | 1.38E-05 | 29.9015921 | 46.934965 | 32.4818171 | 30.619786 | 0 |
| 4.47E-06 | 1.39E-05 | 21.8635297 | 48.9369303 | 43.6184401 | 66.3428696 | 0 |
| 4.47E-06 | 1.39E-05 | 60.4462293 | 41.5963908 | 25.0574018 | 35.2591475 | 0 |
| 4.48E-06 | 1.39E-05 | 18.6483048 | 56.4999105 | 63.1075304 | 67.7346781 | 0 |
| 4.49E-06 | 1.39E-05 | 5.14435994 | 1.77952474 | 1.85610383 | 2.31968076 | 66.5341108 |
| 4.50E-06 | 1.39E-05 | 29.9015921 | 64.5077718 | 52.8989593 | 27.3722329 | 0 |
| 4.50E-06 | 1.40E-05 | 28.6155022 | 34.4782918 | 107.654022 | 53.3526574 | 0 |
| 4.51E-06 | 1.40E-05 | 21.5420072 | 51.828658 | 48.2586997 | 44.5378705 | 0 |
| 4.52E-06 | 1.40E-05 | 157.546023 | 57.1672323 | 54.7550631 | 72.3740396 | 0 |
| 4.52E-06 | 1.40E-05 | 220.564432 | 797.004643 | 451.033232 | 613.787528 | 28.5146189 |
| 4.52E-06 | 1.40E-05 | 24.1141872 | 46.7125244 | 35.2659728 | 43.6099982 | 0 |
| 4.53E-06 | 1.40E-05 | 5.46588243 | 219.326424 | 223.660512 | 204.595843 | 0 |
| 4.54E-06 | 1.41E-05 | 26.0433222 | 31.3641235 | 89.092984 | 88.1478687 | 0 |
| 4.54E-06 | 1.41E-05 | 29.5800696 | 45.3778809 | 49.1867516 | 26.9082968 | 0 |
| 4.54E-06 | 1.41E-05 | 20.5774398 | 92.5352865 | 41.7623363 | 95.106911 | 0 |
| 4.55E-06 | 1.41E-05 | 65.5905892 | 102.545113 | 66.819738 | 92.3232941 | 11.8810912 |
| 4.56E-06 | 1.41E-05 | 41.1548795 | 56.4999105 | 21.3451941 | 71.9101035 | 0 |
| 4.56E-06 | 1.41E-05 | 54.9803468 | 22.911381 | 37.1220767 | 58.9198912 | 0 |
| 4.56E-06 | 1.41E-05 | 21.2204847 | 46.934965 | 51.0428554 | 47.7854236 | 0 |
| 4.56E-06 | 1.41E-05 | 87.1325964 | 773.203499 | 958.67763 | 945.03794 | 21.3859642 |
| 4.57E-06 | 1.41E-05 | 35.3674746 | 31.1416829 | 28.7696094 | 31.5476583 | 0 |
| 4.58E-06 | 1.42E-05 | 19.9343948 | 47.8247274 | 62.1794784 | 60.3116997 | 0 |
| 4.58E-06 | 1.42E-05 | 102.565676 | 467.347685 | 215.308045 | 285.784669 | 0 |
| 4.59E-06 | 1.42E-05 | 58.8386168 | 32.4763265 | 28.7696094 | 35.2591475 | 0 |
| 4.61E-06 | 1.43E-05 | 72.9856066 | 33.5885295 | 26.9135056 | 44.0739344 | 0 |
| 4.63E-06 | 1.43E-05 | 36.9750871 | 83.6376628 | 81.6685687 | 90.9314857 | 0 |
| 4.63E-06 | 1.43E-05 | 18.0052598 | 68.0668213 | 57.5392189 | 71.9101035 | 0 |
| 4.63E-06 | 1.43E-05 | 26.0433222 | 99.2085042 | 76.1002572 | 32.0115944 | 0 |
| 4.64E-06 | 1.44E-05 | 18.0052598 | 55.6101481 | 83.5246725 | 68.6625504 | 0 |
| 4.65E-06 | 1.44E-05 | 197.09329 | 516.062174 | 480.730893 | 528.423276 | 49.9005831 |
| 4.66E-06 | 1.44E-05 | 17.0406923 | 78.9664103 | 58.4672708 | 94.1790387 | 0 |
| 4.66E-06 | 1.44E-05 | 94.2060914 | 197.749687 | 267.278952 | 231.968076 | 0 |

| | | | | | | |
|---|---|---|---|---|---|---|
| 4.66E-06 | 1.44E-05 | 19.9343948 | 54.4979451 | 51.9709073 | 54.7444659 | 0 |
| 4.66E-06 | 1.44E-05 | 0 | 0 | 0 | 0 | 11.8810912 |
| 4.66E-06 | 1.44E-05 | 17.3622148 | 74.5175985 | 61.2514265 | 71.4461673 | 0 |
| 4.67E-06 | 1.44E-05 | 17.0406923 | 74.9624797 | 72.3880495 | 64.0231889 | 0 |
| 4.67E-06 | 1.45E-05 | 18.9698273 | 57.1672323 | 70.5319457 | 54.7444659 | 0 |
| 4.69E-06 | 1.45E-05 | 16.0761248 | 74.5175985 | 86.3088283 | 70.518295 | 0 |
| 4.69E-06 | 1.45E-05 | 70.7349491 | 18.4625692 | 72.3880495 | 52.8887212 | 0 |
| 4.71E-06 | 1.46E-05 | 63.0184092 | 150.814722 | 147.560255 | 246.350096 | 16.6335277 |
| 4.71E-06 | 1.46E-05 | 52.4081669 | 38.2597819 | 25.9854537 | 32.4755306 | 0 |
| 4.71E-06 | 1.46E-05 | 23.1496197 | 49.3818115 | 36.1940248 | 52.4247851 | 0 |
| 4.71E-06 | 1.46E-05 | 23.7926647 | 37.1475789 | 45.4745439 | 38.9706367 | 0 |
| 4.72E-06 | 1.46E-05 | 38.261177 | 23.8011434 | 25.9854537 | 27.8361691 | 133.068222 |
| 4.72E-06 | 1.46E-05 | 9.32415239 | 6.89565837 | 5.5683115 | 5.56723382 | 30.8908372 |
| 4.72E-06 | 1.46E-05 | 84.238894 | 231.338216 | 168.905449 | 235.215629 | 16.6335277 |
| 4.74E-06 | 1.46E-05 | 22.8280972 | 39.816866 | 47.3306478 | 67.7346781 | 0 |
| 4.74E-06 | 1.46E-05 | 168.79931 | 434.426477 | 432.472193 | 420.326153 | 26.1384007 |
| 4.75E-06 | 1.47E-05 | 21.8635297 | 42.2637126 | 45.4745439 | 51.0329766 | 0 |
| 4.76E-06 | 1.47E-05 | 55.6233918 | 31.3641235 | 35.2659728 | 28.7640414 | 0 |
| 4.76E-06 | 1.47E-05 | 68.1627692 | 14.4586385 | 105.797919 | 197.172864 | 0 |
| 4.77E-06 | 1.47E-05 | 26.0433222 | 86.5293905 | 30.6257133 | 85.3642518 | 0 |
| 4.77E-06 | 1.47E-05 | 33.1168171 | 22.911381 | 13.9207788 | 14.8459568 | 68.9103291 |
| 4.77E-06 | 1.47E-05 | 33.7598621 | 41.8188314 | 25.0574018 | 35.7230837 | 0 |
| 4.77E-06 | 1.47E-05 | 31.1876821 | 33.3660889 | 29.6976613 | 55.208402 | 0 |
| 4.78E-06 | 1.47E-05 | 31.5092046 | 44.4881185 | 26.9135056 | 66.8068058 | 0 |
| 4.79E-06 | 1.48E-05 | 34.7244296 | 24.2460246 | 49.1867516 | 64.0231889 | 0 |
| 4.79E-06 | 1.48E-05 | 21.2204847 | 42.4861532 | 55.683115 | 51.0329766 | 0 |
| 4.80E-06 | 1.48E-05 | 21.2204847 | 45.6003214 | 48.2586997 | 48.2493597 | 0 |
| 4.80E-06 | 1.48E-05 | 373.287618 | 586.798283 | 580.9605 | 641.159761 | 35.6432737 |
| 4.81E-06 | 1.48E-05 | 20.2559173 | 51.6062174 | 56.6111669 | 48.2493597 | 0 |
| 4.81E-06 | 1.48E-05 | 20.5774398 | 56.4999105 | 52.8989593 | 45.4657428 | 0 |
| 4.81E-06 | 1.48E-05 | 110.603739 | 96.5392171 | 141.991943 | 11.5984038 | 0 |
| 4.82E-06 | 1.49E-05 | 22.5065747 | 38.2597819 | 51.9709073 | 65.8789335 | 0 |
| 4.83E-06 | 1.49E-05 | 50.8005544 | 57.6121134 | 19.4890903 | 51.0329766 | 0 |
| 4.84E-06 | 1.49E-05 | 29.9015921 | 46.0452026 | 29.6976613 | 32.0115944 | 0 |
| 4.87E-06 | 1.50E-05 | 200.630038 | 94.5372518 | 110.438178 | 104.84957 | 496.629613 |
| 4.87E-06 | 1.50E-05 | 0 | 0 | 0 | 0 | 21.3859642 |
| 4.87E-06 | 1.50E-05 | 60.1247068 | 53.1633016 | 83.5246725 | 61.7035081 | 4.75243649 |
| 4.87E-06 | 1.50E-05 | 43.405537 | 28.2499552 | 28.7696094 | 35.7230837 | 0 |
| 4.88E-06 | 1.50E-05 | 95.1706589 | 28.4723958 | 45.4745439 | 35.2591475 | 0 |
| 4.88E-06 | 1.50E-05 | 64.3044992 | 22.6889404 | 37.1220767 | 53.8165936 | 0 |
| 4.88E-06 | 1.50E-05 | 17.0406923 | 61.6160441 | 72.3880495 | 74.2297842 | 0 |
| 4.88E-06 | 1.50E-05 | 15.4330798 | 74.7400391 | 90.0210359 | 75.6215927 | 0 |
| 4.89E-06 | 1.51E-05 | 20.8989622 | 50.4940145 | 47.3306478 | 47.3214874 | 0 |

| | | | | | | |
|---|---|---|---|---|---|---|
| 4.89E-06 | 1.51E-05 | 29.5800696 | 148.812756 | 116.00649 | 93.7151026 | 2.37621824 |
| 4.90E-06 | 1.51E-05 | 48.2283744 | 108.328569 | 82.5966206 | 99.7462725 | 0 |
| 4.91E-06 | 1.51E-05 | 20.5774398 | 65.1750936 | 44.546492 | 53.8165936 | 0 |
| 4.91E-06 | 1.51E-05 | 24.7572322 | 43.3759155 | 38.0501286 | 33.867339 | 0 |
| 4.91E-06 | 1.51E-05 | 140.505331 | 63.3955688 | 46.4025958 | 101.138081 | 0 |
| 4.92E-06 | 1.51E-05 | 157.867546 | 404.619438 | 435.256349 | 465.32796 | 30.8908372 |
| 4.92E-06 | 1.52E-05 | 39.2257445 | 27.5826335 | 30.6257133 | 32.9394667 | 0 |
| 4.92E-06 | 1.52E-05 | 115.426576 | 141.694657 | 174.47376 | 150.315313 | 40.3957101 |
| 4.93E-06 | 1.52E-05 | 5.46588243 | 126.346257 | 264.494796 | 347.488177 | 0 |
| 4.93E-06 | 1.52E-05 | 28.9370247 | 27.3601929 | 138.279736 | 108.561059 | 0 |
| 4.95E-06 | 1.52E-05 | 11.2532874 | 95.2045736 | 110.438178 | 197.172864 | 0 |
| 4.96E-06 | 1.53E-05 | 19.2913498 | 88.0864746 | 45.4745439 | 76.0855288 | 0 |
| 4.96E-06 | 1.53E-05 | 88.4186864 | 491.81615 | 597.665435 | 547.444659 | 21.3859642 |
| 4.96E-06 | 1.53E-05 | 205.774398 | 485.587813 | 259.854537 | 375.788283 | 33.2670554 |
| 4.96E-06 | 1.53E-05 | 20.8989622 | 42.2637126 | 65.8916861 | 54.2805297 | 0 |
| 4.97E-06 | 1.53E-05 | 78.1299666 | 240.45828 | 211.595837 | 232.895948 | 16.6335277 |
| 5.00E-06 | 1.54E-05 | 22.5065747 | 71.4034302 | 37.1220767 | 95.5708472 | 0 |
| 5.00E-06 | 1.54E-05 | 41.476402 | 60.5038411 | 80.7405168 | 85.3642518 | 0 |
| 5.01E-06 | 1.54E-05 | 21.8635297 | 59.6140788 | 50.1148035 | 39.4345729 | 0 |
| 5.01E-06 | 1.54E-05 | 42.762492 | 140.137573 | 56.6111669 | 96.4987195 | 2.37621824 |
| 5.02E-06 | 1.54E-05 | 6.43044992 | 7.56298014 | 4.64025958 | 8.81478687 | 61.7816743 |
| 5.02E-06 | 1.54E-05 | 26.6863672 | 44.4881185 | 28.7696094 | 45.929679 | 0 |
| 5.02E-06 | 1.54E-05 | 19.2913498 | 60.5038411 | 51.0428554 | 56.1362743 | 0 |
| 5.02E-06 | 1.54E-05 | 9.32415239 | 108.77345 | 149.416359 | 171.19244 | 0 |
| 5.05E-06 | 1.55E-05 | 44.691627 | 100.320707 | 51.9709073 | 129.902122 | 2.37621824 |
| 5.06E-06 | 1.55E-05 | 20.8989622 | 92.5352865 | 51.9709073 | 48.7132959 | 0 |
| 5.07E-06 | 1.56E-05 | 19.9343948 | 47.8247274 | 54.7550631 | 53.8165936 | 0 |
| 5.09E-06 | 1.56E-05 | 40.5118345 | 96.0943359 | 61.2514265 | 85.828188 | 0 |
| 5.09E-06 | 1.56E-05 | 21.2204847 | 55.8325887 | 47.3306478 | 42.6821259 | 0 |
| 5.09E-06 | 1.56E-05 | 1497.65179 | 3566.16758 | 3046.79444 | 3364.92891 | 19.0097459 |
| 5.10E-06 | 1.57E-05 | 18.6483048 | 52.2735392 | 59.3953227 | 75.6215927 | 0 |
| 5.10E-06 | 1.57E-05 | 27.6509347 | 52.2735392 | 37.1220767 | 30.1558498 | 0 |
| 5.12E-06 | 1.57E-05 | 17.6837373 | 14.6810791 | 19.4890903 | 20.4131907 | 47.5243649 |
| 5.13E-06 | 1.58E-05 | 42.119447 | 102.100232 | 90.9490879 | 88.6118049 | 7.12865473 |
| 5.14E-06 | 1.58E-05 | 23.4711422 | 42.9310343 | 36.1940248 | 39.898509 | 0 |
| 5.15E-06 | 1.58E-05 | 105.459379 | 270.26532 | 188.394539 | 226.400842 | 19.0097459 |
| 5.15E-06 | 1.58E-05 | 50.1575094 | 26.6928711 | 56.6111669 | 28.7640414 | 0 |
| 5.15E-06 | 1.58E-05 | 33.1168171 | 29.8070394 | 30.6257133 | 29.6919137 | 0 |
| 5.17E-06 | 1.59E-05 | 518.294264 | 1103.30534 | 914.131138 | 983.080705 | 142.573095 |
| 5.19E-06 | 1.59E-05 | 35.0459521 | 137.245846 | 21.3451941 | 151.707121 | 0 |
| 5.19E-06 | 1.59E-05 | 18.3267823 | 47.1574056 | 84.4527244 | 88.1478687 | 0 |
| 5.19E-06 | 1.59E-05 | 41.7979245 | 155.708415 | 51.0428554 | 158.202228 | 2.37621824 |
| 5.21E-06 | 1.60E-05 | 33.4383396 | 30.2519206 | 27.8415575 | 32.4755306 | 0 |

| | | | | | | |
|---|---|---|---|---|---|---|
| 5.21E-06 | 1.60E-05 | 16.3976473 | 68.5117025 | 72.3880495 | 69.5904227 | 0 |
| 5.21E-06 | 1.60E-05 | 52.7296894 | 19.1298909 | 54.7550631 | 58.9198912 | 0 |
| 5.22E-06 | 1.60E-05 | 32.4737721 | 86.9742716 | 24.1293498 | 108.561059 | 0 |
| 5.23E-06 | 1.60E-05 | 44.3701045 | 38.2597819 | 28.7696094 | 26.9082968 | 0 |
| 5.23E-06 | 1.61E-05 | 24.7572322 | 49.3818115 | 52.8989593 | 31.0837221 | 0 |
| 5.24E-06 | 1.61E-05 | 38.5826995 | 35.5904948 | 23.2012979 | 41.2903175 | 0 |
| 5.25E-06 | 1.61E-05 | 31.1876821 | 85.4171875 | 36.1940248 | 34.7952113 | 0 |
| 5.27E-06 | 1.61E-05 | 261.719312 | 503.383061 | 602.305694 | 613.787528 | 2.37621824 |
| 5.27E-06 | 1.62E-05 | 17.3622148 | 74.2951579 | 96.5173994 | 53.3526574 | 0 |
| 5.29E-06 | 1.62E-05 | 48.8714194 | 51.3837769 | 19.4890903 | 77.9412734 | 0 |
| 5.30E-06 | 1.63E-05 | 31.8307271 | 157.487939 | 91.8771398 | 128.510314 | 2.37621824 |
| 5.30E-06 | 1.63E-05 | 21.2204847 | 51.3837769 | 41.7623363 | 46.8575513 | 0 |
| 5.31E-06 | 1.63E-05 | 256.253429 | 415.519027 | 316.465704 | 350.73573 | 99.8011662 |
| 5.33E-06 | 1.63E-05 | 135.682493 | 503.827942 | 360.084144 | 382.747325 | 23.7621824 |
| 5.33E-06 | 1.63E-05 | 32.7952946 | 50.0491333 | 48.2586997 | 23.6607437 | 0 |
| 5.34E-06 | 1.64E-05 | 80.380624 | 37.5924601 | 46.4025958 | 24.1246799 | 0 |
| 5.34E-06 | 1.64E-05 | 20.8989622 | 45.3778809 | 44.546492 | 50.5690405 | 0 |
| 5.35E-06 | 1.64E-05 | 22.1850522 | 64.0628906 | 38.9781805 | 46.8575513 | 0 |
| 5.36E-06 | 1.64E-05 | 36.0105196 | 36.4802572 | 24.1293498 | 34.7952113 | 0 |
| 5.36E-06 | 1.64E-05 | 54.9803468 | 56.9447917 | 51.0428554 | 19.0213822 | 0 |
| 5.36E-06 | 1.64E-05 | 23.7926647 | 32.6987671 | 84.4527244 | 75.1576565 | 0 |
| 5.38E-06 | 1.65E-05 | 20.8989622 | 39.816866 | 59.3953227 | 56.1362743 | 0 |
| 5.38E-06 | 1.65E-05 | 44.691627 | 28.6948364 | 30.6257133 | 30.619786 | 0 |
| 5.39E-06 | 1.65E-05 | 32.1522496 | 51.1613363 | 25.0574018 | 38.0427644 | 0 |
| 5.39E-06 | 1.65E-05 | 38.5826995 | 23.1338216 | 54.7550631 | 38.5067006 | 0 |
| 5.40E-06 | 1.65E-05 | 168.477788 | 158.822583 | 119.718697 | 138.252973 | 23.7621824 |
| 5.41E-06 | 1.66E-05 | 19.2913498 | 56.7223511 | 47.3306478 | 70.0543588 | 0 |
| 5.42E-06 | 1.66E-05 | 18.6483048 | 78.9664103 | 58.4672708 | 51.0329766 | 0 |
| 5.42E-06 | 1.66E-05 | 229.24554 | 50.7164551 | 36.1940248 | 38.9706367 | 0 |
| 5.42E-06 | 1.66E-05 | 25.7217997 | 52.2735392 | 54.7550631 | 29.2279775 | 0 |
| 5.43E-06 | 1.66E-05 | 29.9015921 | 44.9329997 | 25.0574018 | 49.177232 | 0 |
| 5.43E-06 | 1.66E-05 | 23.1496197 | 40.7066284 | 40.8342843 | 68.6625504 | 0 |
| 5.44E-06 | 1.67E-05 | 19.9343948 | 50.9388957 | 46.4025958 | 55.208402 | 0 |
| 5.46E-06 | 1.67E-05 | 25.4002772 | 50.0491333 | 32.4818171 | 36.650956 | 0 |
| 5.46E-06 | 1.67E-05 | 14.7900348 | 58.946757 | 140.135839 | 106.241379 | 0 |
| 5.47E-06 | 1.67E-05 | 20.5774398 | 37.5924601 | 128.071165 | 241.710735 | 0 |
| 5.47E-06 | 1.67E-05 | 20.2559173 | 13.5688761 | 12.0646749 | 11.5984038 | 76.0389838 |
| 5.47E-06 | 1.67E-05 | 22.1850522 | 51.1613363 | 50.1148035 | 36.1870198 | 0 |
| 5.51E-06 | 1.68E-05 | 17.6837373 | 157.932821 | 45.4745439 | 110.416804 | 0 |
| 5.51E-06 | 1.68E-05 | 13.5039448 | 74.7400391 | 109.510126 | 96.0347833 | 0 |
| 5.51E-06 | 1.68E-05 | 19.6128723 | 42.2637126 | 78.8844129 | 70.518295 | 0 |
| 5.51E-06 | 1.68E-05 | 22.5065747 | 36.035376 | 44.546492 | 46.8575513 | 0 |
| 5.51E-06 | 1.68E-05 | 26.0433222 | 37.3700195 | 36.1940248 | 69.5904227 | 0 |

| | | | | | | |
|---|---|---|---|---|---|---|
| 5.52E-06 | 1.69E-05 | 20.2559173 | 52.4959798 | 49.1867516 | 45.0018067 | 0 |
| 5.53E-06 | 1.69E-05 | 15.1115573 | 147.922994 | 62.1794784 | 90.0036134 | 0 |
| 5.53E-06 | 1.69E-05 | 43.7270595 | 149.924959 | 131.783372 | 123.407016 | 9.50487297 |
| 5.53E-06 | 1.69E-05 | 9.32415239 | 115.891549 | 138.279736 | 143.820207 | 0 |
| 5.54E-06 | 1.69E-05 | 15.4330798 | 72.9605143 | 116.00649 | 67.2707419 | 0 |
| 5.55E-06 | 1.70E-05 | 128.287476 | 88.0864746 | 14.8488307 | 61.7035081 | 0 |
| 5.55E-06 | 1.70E-05 | 53.6942569 | 96.7616577 | 77.0283091 | 43.6099982 | 2.37621824 |
| 5.55E-06 | 1.70E-05 | 21.8635297 | 44.7105591 | 39.9062324 | 42.6821259 | 0 |
| 5.56E-06 | 1.70E-05 | 36.6535646 | 71.8483114 | 89.092984 | 83.5085072 | 0 |
| 5.56E-06 | 1.70E-05 | 36.9750871 | 35.1456136 | 25.0574018 | 61.7035081 | 0 |
| 5.57E-06 | 1.70E-05 | 0 | 0 | 0 | 0 | 11.8810912 |
| 5.59E-06 | 1.71E-05 | 133.753358 | 182.401286 | 181.898176 | 196.244992 | 11.8810912 |
| 5.59E-06 | 1.71E-05 | 18.0052598 | 12.2342326 | 11.136623 | 8.81478687 | 83.1676385 |
| 5.59E-06 | 1.71E-05 | 19.2913498 | 72.2931925 | 43.6184401 | 68.1986142 | 0 |
| 5.59E-06 | 1.71E-05 | 21.8635297 | 9.12006429 | 7.42441534 | 11.1344676 | 59.4054561 |
| 5.60E-06 | 1.71E-05 | 59.8031843 | 188.629622 | 110.438178 | 130.366059 | 14.2573095 |
| 5.60E-06 | 1.71E-05 | 17.0406923 | 59.6140788 | 62.1794784 | 79.797018 | 0 |
| 5.61E-06 | 1.71E-05 | 21.5420072 | 56.7223511 | 39.9062324 | 45.0018067 | 0 |
| 5.62E-06 | 1.71E-05 | 15.4330798 | 87.4191528 | 64.0355823 | 80.2609542 | 0 |
| 5.62E-06 | 1.72E-05 | 43.405537 | 117.448633 | 17.6329864 | 148.459568 | 0 |
| 5.63E-06 | 1.72E-05 | 49.5144644 | 23.5787028 | 33.409869 | 42.6821259 | 0 |
| 5.63E-06 | 1.72E-05 | 476.817862 | 1497.24763 | 1190.69061 | 1242.42101 | 47.5243649 |
| 5.63E-06 | 1.72E-05 | 25.4002772 | 33.1436483 | 34.3379209 | 48.7132959 | 0 |
| 5.64E-06 | 1.72E-05 | 20.2559173 | 42.2637126 | 56.6111669 | 52.4247851 | 0 |
| 5.64E-06 | 1.72E-05 | 16.0761248 | 71.6258708 | 62.1794784 | 83.0445711 | 0 |
| 5.65E-06 | 1.72E-05 | 25.4002772 | 46.7125244 | 29.6976613 | 41.2903175 | 0 |
| 5.65E-06 | 1.72E-05 | 23.1496197 | 44.9329997 | 57.5392189 | 34.3312752 | 0 |
| 5.66E-06 | 1.73E-05 | 24.1141872 | 34.7007324 | 35.2659728 | 44.5378705 | 0 |
| 5.67E-06 | 1.73E-05 | 65.5905892 | 42.7085937 | 41.7623363 | 21.8049991 | 0 |
| 5.67E-06 | 1.73E-05 | 36.0105196 | 72.9605143 | 21.3451941 | 67.2707419 | 0 |
| 5.67E-06 | 1.73E-05 | 16.7191698 | 67.8443807 | 99.3015551 | 57.9920189 | 0 |
| 5.68E-06 | 1.73E-05 | 22.5065747 | 43.5983561 | 36.1940248 | 53.8165936 | 0 |
| 5.68E-06 | 1.73E-05 | 22.8280972 | 109.885653 | 60.3233746 | 38.9706367 | 0 |
| 5.70E-06 | 1.74E-05 | 72.3425616 | 32.4763265 | 29.6976613 | 35.2591475 | 0 |
| 5.70E-06 | 1.74E-05 | 17.6837373 | 81.4132568 | 54.7550631 | 60.3116997 | 0 |
| 5.72E-06 | 1.74E-05 | 45.334672 | 67.3994995 | 64.9636342 | 65.8789335 | 7.12865473 |
| 5.72E-06 | 1.74E-05 | 34.4029071 | 28.0275146 | 28.7696094 | 32.0115944 | 0 |
| 5.73E-06 | 1.74E-05 | 35.6889971 | 54.7203857 | 23.2012979 | 87.6839326 | 0 |
| 5.74E-06 | 1.75E-05 | 23.1496197 | 65.3975342 | 132.711424 | 38.9706367 | 0 |
| 5.75E-06 | 1.75E-05 | 58.1955718 | 177.285152 | 132.711424 | 156.346483 | 0 |
| 5.75E-06 | 1.75E-05 | 159.153636 | 363.690369 | 255.214277 | 287.640414 | 23.7621824 |
| 5.77E-06 | 1.76E-05 | 17.3622148 | 61.6160441 | 55.683115 | 78.8691457 | 0 |
| 5.79E-06 | 1.76E-05 | 228.924017 | 37.5924601 | 31.5537652 | 59.8477635 | 0 |

| | | | | | | |
|---|---|---|---|---|---|---|
| 5.79E-06 | 1.76E-05 | 15.7546023 | 106.771484 | 61.2514265 | 74.6937204 | 0 |
| 5.80E-06 | 1.77E-05 | 37.2966096 | 28.917277 | 39.9062324 | 25.9804245 | 0 |
| 5.80E-06 | 1.77E-05 | 37.9396545 | 49.8266927 | 130.85532 | 27.3722329 | 0 |
| 5.81E-06 | 1.77E-05 | 264.934537 | 48.4920492 | 21.3451941 | 79.3330819 | 0 |
| 5.82E-06 | 1.77E-05 | 10.2887199 | 133.464355 | 96.5173994 | 133.613612 | 0 |
| 5.83E-06 | 1.78E-05 | 14.1469898 | 71.8483114 | 93.7332436 | 86.2921241 | 0 |
| 5.84E-06 | 1.78E-05 | 753.970253 | 1594.23173 | 1335.46671 | 1359.33292 | 54.6530196 |
| 5.84E-06 | 1.78E-05 | 22.1850522 | 45.822762 | 122.502853 | 52.8887212 | 0 |
| 5.85E-06 | 1.78E-05 | 57.5525268 | 18.6850098 | 78.8844129 | 48.7132959 | 0 |
| 5.85E-06 | 1.78E-05 | 57.5525268 | 49.6042521 | 66.819738 | 28.7640414 | 142.573095 |
| 5.87E-06 | 1.79E-05 | 79.737579 | 42.4861532 | 23.2012979 | 39.898509 | 0 |
| 5.89E-06 | 1.79E-05 | 126.358341 | 271.155082 | 237.581291 | 225.009033 | 21.3859642 |
| 5.89E-06 | 1.79E-05 | 21.8635297 | 68.0668213 | 37.1220767 | 51.0329766 | 0 |
| 5.89E-06 | 1.79E-05 | 20.2559173 | 48.2696086 | 53.8270112 | 42.6821259 | 0 |
| 5.91E-06 | 1.80E-05 | 149.186438 | 123.454529 | 9.28051917 | 111.808612 | 0 |
| 5.91E-06 | 1.80E-05 | 259.147132 | 75.852242 | 18.5610383 | 58.4559551 | 0 |
| 5.91E-06 | 1.80E-05 | 281.010662 | 356.127388 | 291.408302 | 335.889774 | 28.5146189 |
| 5.91E-06 | 1.80E-05 | 69.4488592 | 119.228158 | 144.776099 | 175.831801 | 14.2573095 |
| 5.91E-06 | 1.80E-05 | 14.1469898 | 70.7361084 | 97.4454513 | 84.9003157 | 0 |
| 5.91E-06 | 1.80E-05 | 21.8635297 | 13.3464355 | 6.49636342 | 12.0623399 | 57.0292378 |
| 5.92E-06 | 1.80E-05 | 38.904222 | 116.781311 | 103.013763 | 111.344676 | 7.12865473 |
| 5.94E-06 | 1.80E-05 | 22.5065747 | 47.8247274 | 37.1220767 | 38.9706367 | 0 |
| 5.94E-06 | 1.81E-05 | 23.1496197 | 45.6003214 | 87.2368802 | 39.898509 | 0 |
| 5.95E-06 | 1.81E-05 | 21.8635297 | 36.9251383 | 60.3233746 | 46.3936151 | 0 |
| 5.96E-06 | 1.81E-05 | 19.2913498 | 43.8207967 | 65.8916861 | 57.5280828 | 0 |
| 5.97E-06 | 1.81E-05 | 434.698415 | 897.770231 | 815.757635 | 855.962199 | 7.12865473 |
| 5.99E-06 | 1.82E-05 | 312.519866 | 632.176164 | 637.571667 | 738.586353 | 35.6432737 |
| 5.99E-06 | 1.82E-05 | 50.8005544 | 30.6968018 | 24.1293498 | 42.6821259 | 0 |
| 6.00E-06 | 1.82E-05 | 45.0131495 | 29.3621582 | 27.8415575 | 31.5476583 | 0 |
| 6.00E-06 | 1.82E-05 | 204.166785 | 385.934428 | 262.638692 | 333.106157 | 26.1384007 |
| 6.01E-06 | 1.82E-05 | 92.5984789 | 166.608004 | 189.322591 | 189.749886 | 0 |
| 6.03E-06 | 1.83E-05 | 17.3622148 | 91.6455241 | 46.4025958 | 116.447974 | 0 |
| 6.04E-06 | 1.83E-05 | 16.0761248 | 64.952653 | 64.0355823 | 78.4052096 | 0 |
| 6.04E-06 | 1.83E-05 | 155.938411 | 289.840092 | 237.581291 | 279.753499 | 0 |
| 6.04E-06 | 1.83E-05 | 70.7349491 | 63.1731283 | 204.171422 | 17.1656376 | 0 |
| 6.07E-06 | 1.84E-05 | 18.6483048 | 57.3896728 | 57.5392189 | 48.2493597 | 0 |
| 6.07E-06 | 1.84E-05 | 945.597661 | 3016.96176 | 2462.12174 | 2911.19935 | 0 |
| 6.08E-06 | 1.84E-05 | 16.0761248 | 51.828658 | 94.6612955 | 97.8905279 | 0 |
| 6.09E-06 | 1.85E-05 | 15.7546023 | 143.029301 | 252.430121 | 206.915523 | 2.37621824 |
| 6.10E-06 | 1.85E-05 | 20.5774398 | 44.9329997 | 303.472977 | 93.7151026 | 0 |
| 6.11E-06 | 1.85E-05 | 27.3294122 | 166.163123 | 25.9854537 | 120.159463 | 0 |
| 6.13E-06 | 1.86E-05 | 16.0761248 | 78.5215291 | 64.0355823 | 64.0231889 | 0 |
| 6.14E-06 | 1.86E-05 | 9.00262989 | 113.667143 | 211.595837 | 113.664357 | 0 |

| | | | | | | |
|---|---|---|---|---|---|---|
| 6.18E-06 | 1.87E-05 | 18.6483048 | 52.0510986 | 50.1148035 | 58.4559551 | 0 |
| 6.19E-06 | 1.88E-05 | 20.2559173 | 49.3818115 | 43.6184401 | 45.4657428 | 0 |
| 6.19E-06 | 1.88E-05 | 15.7546023 | 49.8266927 | 132.711424 | 108.097123 | 0 |
| 6.19E-06 | 1.88E-05 | 323.451631 | 655.532426 | 612.514265 | 618.42689 | 0 |
| 6.18E-06 | 1.88E-05 | 19.6128723 | 49.3818115 | 45.4745439 | 51.4969128 | 0 |
| 6.19E-06 | 1.88E-05 | 12.8608998 | 94.982133 | 82.5966206 | 93.7151026 | 0 |
| 6.20E-06 | 1.88E-05 | 20.2559173 | 44.4881185 | 45.4745439 | 46.8575513 | 0 |
| 6.20E-06 | 1.88E-05 | 47.5853294 | 44.0432373 | 22.273246 | 33.4034029 | 0 |
| 6.21E-06 | 1.88E-05 | 22.5065747 | 42.2637126 | 34.3379209 | 46.8575513 | 0 |
| 6.22E-06 | 1.88E-05 | 31.8307271 | 47.8247274 | 24.1293498 | 36.650956 | 0 |
| 6.24E-06 | 1.89E-05 | 18.0052598 | 45.6003214 | 77.956361 | 71.9101035 | 0 |
| 6.24E-06 | 1.89E-05 | 16.7191698 | 61.3936035 | 62.1794784 | 64.487125 | 0 |
| 6.25E-06 | 1.89E-05 | 20.2559173 | 48.9369303 | 41.7623363 | 48.7132959 | 0 |
| 6.25E-06 | 1.89E-05 | 28.9370247 | 38.0373413 | 26.9135056 | 57.0641466 | 0 |
| 6.25E-06 | 1.89E-05 | 27.9724572 | 29.3621582 | 29.6976613 | 40.3624452 | 0 |
| 6.26E-06 | 1.90E-05 | 45.0131495 | 112.55494 | 64.0355823 | 94.6429749 | 4.75243649 |
| 6.27E-06 | 1.90E-05 | 13.1824223 | 95.6494547 | 74.2441534 | 98.3544641 | 0 |
| 6.30E-06 | 1.91E-05 | 48.5498969 | 73.1829549 | 23.2012979 | 33.867339 | 0 |
| 6.30E-06 | 1.91E-05 | 188.412183 | 192.633553 | 207.883629 | 202.276162 | 57.0292378 |
| 6.30E-06 | 1.91E-05 | 22.5065747 | 48.047168 | 77.0283091 | 36.650956 | 0 |
| 6.32E-06 | 1.91E-05 | 15.4330798 | 59.6140788 | 77.0283091 | 86.7560603 | 0 |
| 6.35E-06 | 1.92E-05 | 24.7572322 | 38.9271037 | 34.3379209 | 66.8068058 | 0 |
| 6.36E-06 | 1.92E-05 | 38.5826995 | 59.6140788 | 21.3451941 | 41.7542536 | 0 |
| 6.37E-06 | 1.93E-05 | 41.1548795 | 71.4034302 | 87.2368802 | 55.208402 | 0 |
| 6.37E-06 | 1.93E-05 | 63.0184092 | 32.4763265 | 24.1293498 | 41.7542536 | 0 |
| 6.38E-06 | 1.93E-05 | 13.1824223 | 158.600142 | 64.0355823 | 105.777442 | 0 |
| 6.40E-06 | 1.94E-05 | 19.2913498 | 104.102197 | 38.9781805 | 90.0036134 | 0 |
| 6.41E-06 | 1.94E-05 | 14.4685123 | 125.678935 | 64.9636342 | 79.797018 | 0 |
| 6.42E-06 | 1.94E-05 | 16.3976473 | 95.2045736 | 70.5319457 | 54.2805297 | 0 |
| 6.42E-06 | 1.94E-05 | 35.6889971 | 48.2696086 | 24.1293498 | 32.0115944 | 0 |
| 6.43E-06 | 1.95E-05 | 12.5393773 | 71.6258708 | 121.574801 | 104.385634 | 0 |
| 6.44E-06 | 1.95E-05 | 409.61966 | 903.998568 | 835.246725 | 866.168794 | 83.1676385 |
| 6.44E-06 | 1.95E-05 | 23.4711422 | 31.5865641 | 78.8844129 | 61.239572 | 0 |
| 6.45E-06 | 1.95E-05 | 15.7546023 | 77.4093262 | 63.1075304 | 66.3428696 | 0 |
| 6.45E-06 | 1.95E-05 | 34.0813846 | 27.3601929 | 34.3379209 | 27.3722329 | 0 |
| 6.46E-06 | 1.95E-05 | 20.2559173 | 71.4034302 | 40.8342843 | 51.4969128 | 0 |
| 6.48E-06 | 1.96E-05 | 26.0433222 | 72.9605143 | 41.7623363 | 32.0115944 | 0 |
| 6.48E-06 | 1.96E-05 | 22.8280972 | 37.3700195 | 38.9781805 | 34.7952113 | 0 |
| 6.49E-06 | 1.96E-05 | 24.1141872 | 30.6968018 | 62.1794784 | 47.3214874 | 0 |
| 6.51E-06 | 1.97E-05 | 20.8989622 | 79.1888509 | 37.1220767 | 59.8477635 | 0 |
| 6.51E-06 | 1.97E-05 | 60.4462293 | 92.3128459 | 13.9207788 | 125.262761 | 0 |
| 6.53E-06 | 1.97E-05 | 14.1469898 | 69.1790242 | 84.4527244 | 84.4363795 | 0 |
| 6.58E-06 | 1.99E-05 | 80.0591015 | 321.649097 | 283.983887 | 301.558498 | 16.6335277 |

| | | | | | | |
|---|---|---|---|---|---|---|
| 6.58E-06 | 1.99E-05 | 21.5420072 | 52.940861 | 35.2659728 | 47.3214874 | 0 |
| 6.58E-06 | 1.99E-05 | 0 | 0 | 0 | 0 | 19.0097459 |
| 6.59E-06 | 1.99E-05 | 19.9343948 | 55.1652669 | 47.3306478 | 41.7542536 | 0 |
| 6.60E-06 | 1.99E-05 | 33.4383396 | 24.4684652 | 32.4818171 | 45.0018067 | 0 |
| 6.60E-06 | 1.99E-05 | 62.6968867 | 181.289083 | 279.343627 | 332.178284 | 21.3859642 |
| 6.61E-06 | 2.00E-05 | 26.0433222 | 113.889583 | 116.00649 | 109.952868 | 2.37621824 |
| 6.61E-06 | 2.00E-05 | 18.6483048 | 57.3896728 | 51.9709073 | 47.3214874 | 0 |
| 6.62E-06 | 2.00E-05 | 35.3674746 | 89.1986776 | 66.819738 | 21.341063 | 0 |
| 6.62E-06 | 2.00E-05 | 18.6483048 | 54.9428263 | 46.4025958 | 55.6723382 | 0 |
| 6.62E-06 | 2.00E-05 | 13.8254673 | 70.7361084 | 85.3807764 | 87.6839326 | 0 |
| 6.62E-06 | 2.00E-05 | 31.5092046 | 22.2440592 | 18.5610383 | 13.4541484 | 154.454186 |
| 6.64E-06 | 2.00E-05 | 23.4711422 | 119.450598 | 160.552982 | 115.520102 | 0 |
| 6.64E-06 | 2.00E-05 | 13.1824223 | 94.5372518 | 121.574801 | 67.2707419 | 0 |
| 6.64E-06 | 2.01E-05 | 13.1824223 | 150.1474 | 64.0355823 | 101.138081 | 0 |
| 6.65E-06 | 2.01E-05 | 25.0787547 | 17.3503662 | 26.9135056 | 18.5574461 | 66.5341108 |
| 6.65E-06 | 2.01E-05 | 21.8635297 | 41.1515096 | 40.8342843 | 36.1870198 | 0 |
| 6.67E-06 | 2.01E-05 | 49.8359869 | 66.064856 | 50.1148035 | 51.4969128 | 0 |
| 6.70E-06 | 2.02E-05 | 24.7572322 | 80.3010539 | 31.5537652 | 50.1051043 | 0 |
| 6.70E-06 | 2.02E-05 | 28.9370247 | 94.982133 | 24.1293498 | 91.3954218 | 0 |
| 6.72E-06 | 2.03E-05 | 22.8280972 | 39.3719849 | 34.3379209 | 36.650956 | 0 |
| 6.76E-06 | 2.04E-05 | 23.4711422 | 34.2558512 | 33.409869 | 39.4345729 | 0 |
| 6.76E-06 | 2.04E-05 | 23.1496197 | 54.053064 | 54.7550631 | 30.619786 | 0 |
| 6.76E-06 | 2.04E-05 | 16.7191698 | 56.0550293 | 67.7477899 | 60.3116997 | 0 |
| 6.77E-06 | 2.04E-05 | 84.5604165 | 152.816687 | 129.927268 | 168.872759 | 16.6335277 |
| 6.77E-06 | 2.04E-05 | 20.8989622 | 46.2676432 | 47.3306478 | 37.1148921 | 0 |
| 6.78E-06 | 2.04E-05 | 90.9908664 | 206.202429 | 156.840774 | 196.244992 | 28.5146189 |
| 6.78E-06 | 2.04E-05 | 25.4002772 | 30.02948 | 62.1794784 | 40.8263813 | 0 |
| 6.78E-06 | 2.04E-05 | 19.6128723 | 57.3896728 | 39.9062324 | 56.6002105 | 0 |
| 6.78E-06 | 2.04E-05 | 21.8635297 | 146.58835 | 33.409869 | 94.6429749 | 0 |
| 6.79E-06 | 2.05E-05 | 14.4685123 | 65.8424154 | 84.4527244 | 77.9412734 | 0 |
| 6.79E-06 | 2.05E-05 | 14.4685123 | 86.7518311 | 65.8916861 | 77.0134011 | 0 |
| 6.80E-06 | 2.05E-05 | 42.762492 | 31.5865641 | 104.869867 | 30.1558498 | 0 |
| 6.80E-06 | 2.05E-05 | 32.1522496 | 33.5885295 | 24.1293498 | 33.4034029 | 0 |
| 6.81E-06 | 2.05E-05 | 23.7926647 | 83.8601033 | 33.409869 | 50.5690405 | 0 |
| 6.86E-06 | 2.07E-05 | 13.8254673 | 80.7459351 | 70.5319457 | 89.0757411 | 0 |
| 6.86E-06 | 2.07E-05 | 13.5039448 | 78.5215291 | 76.1002572 | 91.3954218 | 0 |
| 6.86E-06 | 2.07E-05 | 22.5065747 | 44.4881185 | 33.409869 | 39.4345729 | 0 |
| 6.89E-06 | 2.07E-05 | 141.791421 | 34.4782918 | 36.1940248 | 38.5067006 | 0 |
| 6.89E-06 | 2.08E-05 | 20.2559173 | 60.9487223 | 248.717914 | 57.9920189 | 0 |
| 6.91E-06 | 2.08E-05 | 20.5774398 | 52.7184204 | 40.8342843 | 41.2903175 | 0 |
| 6.93E-06 | 2.09E-05 | 39.547267 | 26.0255493 | 34.3379209 | 27.8361691 | 0 |
| 6.95E-06 | 2.09E-05 | 44.3701045 | 63.6180094 | 18.5610383 | 49.6411682 | 0 |
| 6.96E-06 | 2.09E-05 | 20.5774398 | 17.7952474 | 6.49636342 | 12.9902122 | 47.5243649 |

| | | | | | | |
|---|---|---|---|---|---|---|
| 6.96E-06 | 2.09E-05 | 207.703532 | 558.548328 | 353.58778 | 416.150728 | 21.3859642 |
| 6.98E-06 | 2.10E-05 | 77.4869216 | 63.3955688 | 49.1867516 | 52.8887212 | 7.12865473 |
| 6.99E-06 | 2.10E-05 | 66.8766792 | 52.7184204 | 16.7049345 | 57.0641466 | 0 |
| 7.00E-06 | 2.11E-05 | 17.3622148 | 50.4940145 | 61.2514265 | 59.8477635 | 0 |
| 7.02E-06 | 2.11E-05 | 32.4737721 | 59.1691976 | 61.2514265 | 21.8049991 | 0 |
| 7.03E-06 | 2.12E-05 | 36.0105196 | 66.064856 | 24.1293498 | 35.7230837 | 0 |
| 7.04E-06 | 2.12E-05 | 37.9396545 | 36.9251383 | 24.1293498 | 29.2279775 | 0 |
| 7.04E-06 | 2.12E-05 | 32.4737721 | 26.2479899 | 27.8415575 | 38.0427644 | 0 |
| 7.05E-06 | 2.12E-05 | 426.660352 | 892.654097 | 601.377642 | 623.994124 | 0 |
| 7.06E-06 | 2.12E-05 | 0 | 0 | 0 | 0 | 52.2768014 |
| 7.07E-06 | 2.13E-05 | 15.4330798 | 67.3994995 | 60.3233746 | 78.8691457 | 0 |
| 7.07E-06 | 2.13E-05 | 26.3648447 | 38.0373413 | 71.4599976 | 31.5476583 | 0 |
| 7.07E-06 | 2.13E-05 | 75.2362641 | 29.1397176 | 36.1940248 | 30.619786 | 0 |
| 7.09E-06 | 2.13E-05 | 18.9698273 | 43.1534749 | 77.0283091 | 53.8165936 | 0 |
| 7.10E-06 | 2.13E-05 | 112.854396 | 38.0373413 | 29.6976613 | 36.650956 | 0 |
| 7.12E-06 | 2.14E-05 | 25.7217997 | 36.4802572 | 27.8415575 | 47.3214874 | 0 |
| 7.13E-06 | 2.14E-05 | 9.64567488 | 5.11613363 | 7.42441534 | 6.03116997 | 66.5341108 |
| 7.15E-06 | 2.15E-05 | 13.8254673 | 89.6435588 | 81.6685687 | 67.2707419 | 0 |
| 7.15E-06 | 2.15E-05 | 55.9449143 | 69.1790242 | 42.6903882 | 55.208402 | 0 |
| 7.15E-06 | 2.15E-05 | 264.613014 | 690.233158 | 609.730109 | 682.914015 | 0 |
| 7.16E-06 | 2.15E-05 | 27.0078897 | 47.6022868 | 27.8415575 | 76.549465 | 0 |
| 7.19E-06 | 2.16E-05 | 21.8635297 | 50.9388957 | 33.409869 | 44.0739344 | 0 |
| 7.20E-06 | 2.16E-05 | 81.3451915 | 43.1534749 | 47.3306478 | 64.0231889 | 187.721241 |
| 7.20E-06 | 2.16E-05 | 9.00262989 | 3.78149007 | 2.78415575 | 2.31968076 | 64.1578926 |
| 7.20E-06 | 2.16E-05 | 22.1850522 | 53.3857422 | 32.4818171 | 45.4657428 | 0 |
| 7.20E-06 | 2.16E-05 | 21.8635297 | 49.8266927 | 43.6184401 | 33.4034029 | 0 |
| 7.20E-06 | 2.16E-05 | 12.8608998 | 108.328569 | 88.1649321 | 70.9822312 | 0 |
| 7.21E-06 | 2.17E-05 | 25.7217997 | 27.5826335 | 41.7623363 | 54.2805297 | 0 |
| 7.22E-06 | 2.17E-05 | 9.00262989 | 92.9801676 | 194.890903 | 125.262761 | 0 |
| 7.22E-06 | 2.17E-05 | 70.4134267 | 121.897445 | 37.1220767 | 20.8771268 | 0 |
| 7.22E-06 | 2.17E-05 | 31.8307271 | 42.4861532 | 32.4818171 | 24.1246799 | 0 |
| 7.22E-06 | 2.17E-05 | 24.1141872 | 38.0373413 | 30.6257133 | 33.867339 | 0 |
| 7.23E-06 | 2.17E-05 | 189.055228 | 46.4900838 | 28.7696094 | 42.2181898 | 0 |
| 7.23E-06 | 2.17E-05 | 27.6509347 | 113.889583 | 93.7332436 | 101.602017 | 0 |
| 7.23E-06 | 2.17E-05 | 46.2992394 | 62.2833659 | 49.1867516 | 56.1362743 | 0 |
| 7.23E-06 | 2.17E-05 | 22.8280972 | 165.05092 | 82.5966206 | 33.867339 | 0 |
| 7.23E-06 | 2.17E-05 | 19.2913498 | 38.9271037 | 99.3015551 | 70.9822312 | 0 |
| 7.24E-06 | 2.17E-05 | 40.190312 | 85.1947469 | 72.3880495 | 70.518295 | 7.12865473 |
| 7.24E-06 | 2.17E-05 | 49.5144644 | 43.3759155 | 38.0501286 | 20.4131907 | 0 |
| 7.24E-06 | 2.17E-05 | 19.2913498 | 40.929069 | 51.9709073 | 66.3428696 | 0 |
| 7.25E-06 | 2.17E-05 | 22.8280972 | 80.7459351 | 31.5537652 | 59.8477635 | 0 |
| 7.27E-06 | 2.18E-05 | 101.922631 | 71.8483114 | 70.5319457 | 58.9198912 | 171.087714 |
| 7.27E-06 | 2.18E-05 | 25.4002772 | 30.4743612 | 30.6257133 | 33.4034029 | 0 |

| | | | | | | |
|---|---|---|---|---|---|---|
| 7.27E-06 | 2.18E-05 | 20.5774398 | 40.7066284 | 42.6903882 | 40.3624452 | 0 |
| 7.28E-06 | 2.18E-05 | 22.1850522 | 68.2892619 | 62.1794784 | 31.5476583 | 0 |
| 7.28E-06 | 2.18E-05 | 26.3648447 | 117.448633 | 25.0574018 | 114.592229 | 0 |
| 7.31E-06 | 2.19E-05 | 40.833357 | 49.3818115 | 106.72597 | 95.106911 | 2.37621824 |
| 7.31E-06 | 2.19E-05 | 372.966096 | 175.283187 | 238.509343 | 203.204034 | 983.754353 |
| 7.31E-06 | 2.19E-05 | 28.9370247 | 53.6081828 | 24.1293498 | 42.2181898 | 0 |
| 7.31E-06 | 2.19E-05 | 25.7217997 | 63.84045 | 62.1794784 | 26.4443606 | 0 |
| 7.34E-06 | 2.20E-05 | 247.2508 | 433.091833 | 353.58778 | 414.294983 | 104.553603 |
| 7.35E-06 | 2.20E-05 | 18.9698273 | 78.0766479 | 87.2368802 | 38.0427644 | 0 |
| 7.35E-06 | 2.20E-05 | 23.4711422 | 30.6968018 | 51.0428554 | 38.9706367 | 0 |
| 7.35E-06 | 2.20E-05 | 69.1273367 | 49.3818115 | 29.6976613 | 24.1246799 | 0 |
| 7.36E-06 | 2.20E-05 | 23.7926647 | 28.6948364 | 47.3306478 | 47.3214874 | 0 |
| 7.36E-06 | 2.20E-05 | 20.8989622 | 48.4920492 | 35.2659728 | 46.8575513 | 0 |
| 7.37E-06 | 2.21E-05 | 22.5065747 | 48.047168 | 33.409869 | 37.5788283 | 0 |
| 7.37E-06 | 2.21E-05 | 56.9094818 | 39.1495443 | 26.9135056 | 26.9082968 | 0 |
| 7.38E-06 | 2.21E-05 | 16.3976473 | 78.5215291 | 48.2586997 | 78.4052096 | 0 |
| 7.39E-06 | 2.21E-05 | 25.0787547 | 57.1672323 | 28.7696094 | 84.9003157 | 0 |
| 7.40E-06 | 2.21E-05 | 15.4330798 | 66.5097371 | 74.2441534 | 59.3838274 | 0 |
| 7.40E-06 | 2.22E-05 | 15.4330798 | 54.7203857 | 75.1722053 | 79.3330819 | 0 |
| 7.43E-06 | 2.22E-05 | 36.6535646 | 58.5018758 | 19.4890903 | 62.1674443 | 0 |
| 7.43E-06 | 2.22E-05 | 38.5826995 | 33.8109701 | 25.9854537 | 26.9082968 | 0 |
| 7.44E-06 | 2.23E-05 | 249.501457 | 216.879578 | 296.048562 | 270.474776 | 26.1384007 |
| 7.45E-06 | 2.23E-05 | 15.1115573 | 69.6239054 | 69.6038938 | 63.0953166 | 0 |
| 7.46E-06 | 2.23E-05 | 204.166785 | 450.66464 | 414.839207 | 437.491791 | 0 |
| 7.47E-06 | 2.23E-05 | 14.7900348 | 80.7459351 | 67.7477899 | 64.0231889 | 0 |
| 7.50E-06 | 2.24E-05 | 17.3622148 | 53.8306234 | 50.1148035 | 68.1986142 | 0 |
| 7.51E-06 | 2.25E-05 | 87.1325964 | 44.0432373 | 23.2012979 | 36.1870198 | 0 |
| 7.53E-06 | 2.25E-05 | 171.049968 | 189.074504 | 140.135839 | 155.418611 | 40.3957101 |
| 7.55E-06 | 2.26E-05 | 57.5525268 | 23.3562622 | 45.4745439 | 30.1558498 | 0 |
| 7.55E-06 | 2.26E-05 | 147.578826 | 228.668929 | 206.027526 | 225.936906 | 71.2865473 |
| 7.56E-06 | 2.26E-05 | 52.7296894 | 102.767554 | 89.092984 | 92.3232941 | 16.6335277 |
| 7.56E-06 | 2.26E-05 | 21.2204847 | 39.816866 | 36.1940248 | 51.4969128 | 0 |
| 7.57E-06 | 2.26E-05 | 31.5092046 | 33.5885295 | 25.9854537 | 27.8361691 | 0 |
| 7.62E-06 | 2.28E-05 | 25.0787547 | 72.0707519 | 26.9135056 | 62.1674443 | 0 |
| 7.63E-06 | 2.28E-05 | 131.181178 | 367.249418 | 333.170638 | 340.993071 | 23.7621824 |
| 7.63E-06 | 2.28E-05 | 246.286232 | 390.38324 | 473.306478 | 570.17753 | 30.8908372 |
| 7.64E-06 | 2.28E-05 | 74.2716966 | 90.3108805 | 144.776099 | 147.06776 | 14.2573095 |
| 7.65E-06 | 2.29E-05 | 185.840003 | 183.291048 | 144.776099 | 167.017014 | 21.3859642 |
| 7.67E-06 | 2.29E-05 | 14.1469898 | 79.8561727 | 60.3233746 | 94.6429749 | 0 |
| 7.67E-06 | 2.29E-05 | 206.09592 | 560.550293 | 826.894258 | 714.925609 | 11.8810912 |
| 7.69E-06 | 2.30E-05 | 42.762492 | 27.5826335 | 24.1293498 | 46.8575513 | 0 |
| 7.69E-06 | 2.30E-05 | 18.3267823 | 64.2853312 | 50.1148035 | 45.0018067 | 0 |
| 7.70E-06 | 2.30E-05 | 136.004016 | 363.912809 | 279.343627 | 341.920944 | 21.3859642 |

| | | | | | | |
|---|---|---|---|---|---|---|
| 7.71E-06 | 2.30E-05 | 20.2559173 | 35.1456136 | 57.5392189 | 54.7444659 | 0 |
| 7.73E-06 | 2.31E-05 | 18.0052598 | 72.7380737 | 48.2586997 | 49.6411682 | 0 |
| 7.73E-06 | 2.31E-05 | 27.9724572 | 27.3601929 | 28.7696094 | 32.9394667 | 0 |
| 7.73E-06 | 2.31E-05 | 80.0591015 | 193.745756 | 39.9062324 | 117.375846 | 0 |
| 7.74E-06 | 2.31E-05 | 45.0131495 | 94.5372518 | 44.546492 | 139.644782 | 0 |
| 7.75E-06 | 2.31E-05 | 19.6128723 | 44.9329997 | 39.9062324 | 51.0329766 | 0 |
| 7.75E-06 | 2.31E-05 | 28.9370247 | 78.0766479 | 23.2012979 | 67.2707419 | 0 |
| 7.76E-06 | 2.32E-05 | 73.9501741 | 29.8070394 | 88.1649321 | 22.7328714 | 0 |
| 7.78E-06 | 2.32E-05 | 13.1824223 | 101.43291 | 71.4599976 | 76.0855288 | 0 |
| 7.78E-06 | 2.32E-05 | 24.1141872 | 32.0314453 | 31.5537652 | 32.9394667 | 0 |
| 7.79E-06 | 2.33E-05 | 70.4134267 | 36.7026978 | 21.3451941 | 42.6821259 | 0 |
| 7.81E-06 | 2.33E-05 | 28.9370247 | 23.5787028 | 52.8989593 | 63.5592527 | 0 |
| 7.83E-06 | 2.33E-05 | 26.6863672 | 34.7007324 | 52.8989593 | 27.3722329 | 0 |
| 7.83E-06 | 2.34E-05 | 28.9370247 | 35.1456136 | 26.9135056 | 27.8361691 | 0 |
| 7.83E-06 | 2.34E-05 | 399.009418 | 1177.15562 | 1014.36075 | 1228.96686 | 0 |
| 7.86E-06 | 2.34E-05 | 17.3622148 | 59.6140788 | 46.4025958 | 61.239572 | 0 |
| 7.91E-06 | 2.36E-05 | 38.904222 | 34.2558512 | 22.273246 | 32.4755306 | 0 |
| 7.91E-06 | 2.36E-05 | 23.4711422 | 60.5038411 | 28.7696094 | 50.5690405 | 0 |
| 7.91E-06 | 2.36E-05 | 23.4711422 | 38.4822225 | 29.6976613 | 49.177232 | 0 |
| 7.91E-06 | 2.36E-05 | 53.3727344 | 93.86993 | 96.5173994 | 85.3642518 | 11.8810912 |
| 7.93E-06 | 2.36E-05 | 23.1496197 | 32.4763265 | 32.4818171 | 37.5788283 | 0 |
| 7.97E-06 | 2.38E-05 | 48.8714194 | 47.8247274 | 22.273246 | 29.6919137 | 0 |
| 7.99E-06 | 2.38E-05 | 20.5774398 | 48.047168 | 36.1940248 | 41.7542536 | 0 |
| 8.01E-06 | 2.39E-05 | 131.502701 | 197.972127 | 232.941031 | 226.400842 | 21.3859642 |
| 8.02E-06 | 2.39E-05 | 21.5420072 | 32.9212077 | 42.6903882 | 40.3624452 | 0 |
| 8.02E-06 | 2.39E-05 | 11.8963324 | 99.6533854 | 72.3880495 | 104.385634 | 0 |
| 8.03E-06 | 2.39E-05 | 28.9370247 | 63.1731283 | 25.0574018 | 40.3624452 | 0 |
| 8.03E-06 | 2.39E-05 | 9.32415239 | 105.2144 | 150.344411 | 97.4265918 | 0 |
| 8.05E-06 | 2.40E-05 | 25.4002772 | 44.0432373 | 30.6257133 | 29.6919137 | 0 |
| 8.05E-06 | 2.40E-05 | 62.3753642 | 31.3641235 | 25.9854537 | 33.4034029 | 0 |
| 8.06E-06 | 2.40E-05 | 33.7598621 | 38.0373413 | 28.7696094 | 24.1246799 | 0 |
| 8.06E-06 | 2.40E-05 | 505.433364 | 1020.55744 | 804.621012 | 969.626556 | 45.1481466 |
| 8.07E-06 | 2.40E-05 | 21.5420072 | 42.9310343 | 51.9709073 | 32.4755306 | 0 |
| 8.08E-06 | 2.41E-05 | 23.7926647 | 29.3621582 | 37.1220767 | 33.867339 | 0 |
| 8.09E-06 | 2.41E-05 | 18.3267823 | 53.8306234 | 42.6903882 | 52.8887212 | 0 |
| 8.09E-06 | 2.41E-05 | 25.0787547 | 57.3896728 | 25.9854537 | 58.4559551 | 0 |
| 8.11E-06 | 2.41E-05 | 249.822979 | 182.401286 | 154.98467 | 155.882547 | 21.3859642 |
| 8.11E-06 | 2.42E-05 | 42.762492 | 93.2026082 | 46.4025958 | 83.5085072 | 0 |
| 8.13E-06 | 2.42E-05 | 21.8635297 | 58.946757 | 30.6257133 | 63.5592527 | 0 |
| 8.14E-06 | 2.42E-05 | 44.3701045 | 35.5904948 | 20.4171422 | 38.9706367 | 0 |
| 8.17E-06 | 2.43E-05 | 16.3976473 | 73.1829549 | 61.2514265 | 49.6411682 | 0 |
| 8.18E-06 | 2.43E-05 | 19.2913498 | 37.8149007 | 69.6038938 | 53.8165936 | 0 |
| 8.19E-06 | 2.44E-05 | 29.5800696 | 44.2656779 | 33.409869 | 24.1246799 | 0 |

| | | | | | | |
|---|---|---|---|---|---|---|
| 8.20E-06 | 2.44E-05 | 18.6483048 | 86.5293905 | 49.1867516 | 45.929679 | 0 |
| 8.21E-06 | 2.44E-05 | 19.6128723 | 65.6199748 | 35.2659728 | 65.8789335 | 0 |
| 8.22E-06 | 2.44E-05 | 16.7191698 | 54.4979451 | 61.2514265 | 51.9608489 | 0 |
| 8.22E-06 | 2.45E-05 | 21.8635297 | 40.0393066 | 33.409869 | 36.1870198 | 0 |
| 8.23E-06 | 2.45E-05 | 24.1141872 | 33.5885295 | 28.7696094 | 40.8263813 | 0 |
| 8.25E-06 | 2.45E-05 | 24.4357097 | 38.2597819 | 47.3306478 | 27.3722329 | 0 |
| 8.28E-06 | 2.46E-05 | 32.7952946 | 33.5885295 | 39.9062324 | 22.2689353 | 0 |
| 8.28E-06 | 2.46E-05 | 21.2204847 | 68.2892619 | 78.8844129 | 31.5476583 | 0 |
| 8.30E-06 | 2.47E-05 | 0 | 3.11416829 | 0 | 0 | 78.415202 |
| 8.30E-06 | 2.47E-05 | 0 | 0 | 0 | 0 | 21.3859642 |
| 8.30E-06 | 2.47E-05 | 291.942426 | 659.758797 | 480.730893 | 604.508805 | 0 |
| 8.32E-06 | 2.47E-05 | 50.8005544 | 37.3700195 | 21.3451941 | 33.867339 | 0 |
| 8.32E-06 | 2.47E-05 | 22.5065747 | 29.5845988 | 43.6184401 | 44.0739344 | 0 |
| 8.32E-06 | 2.47E-05 | 43.0840145 | 24.6909058 | 28.7696094 | 32.4755306 | 0 |
| 8.34E-06 | 2.48E-05 | 18.9698273 | 130.795068 | 32.4818171 | 137.325101 | 0 |
| 8.35E-06 | 2.48E-05 | 23.1496197 | 40.929069 | 29.6976613 | 35.7230837 | 0 |
| 8.35E-06 | 2.48E-05 | 12.5393773 | 116.113989 | 63.1075304 | 95.106911 | 0 |
| 8.37E-06 | 2.49E-05 | 0 | 0 | 0 | 0 | 45.1481466 |
| 8.37E-06 | 2.49E-05 | 20.8989622 | 37.5924601 | 38.0501286 | 37.5788283 | 0 |
| 8.39E-06 | 2.49E-05 | 23.4711422 | 27.1377523 | 101.157659 | 115.520102 | 0 |
| 8.39E-06 | 2.49E-05 | 0 | 0 | 0 | 0 | 4.75243649 |
| 8.42E-06 | 2.50E-05 | 91.3123889 | 231.115776 | 85.3807764 | 168.408823 | 14.2573095 |
| 8.42E-06 | 2.50E-05 | 18.6483048 | 46.7125244 | 45.4745439 | 44.5378705 | 0 |
| 8.43E-06 | 2.50E-05 | 54.0157793 | 116.55887 | 12.9927268 | 110.416804 | 0 |
| 8.46E-06 | 2.51E-05 | 13.1824223 | 84.3049845 | 62.1794784 | 101.138081 | 0 |
| 8.45E-06 | 2.51E-05 | 32.1522496 | 33.5885295 | 25.0574018 | 27.3722329 | 0 |
| 8.46E-06 | 2.51E-05 | 16.0761248 | 124.789172 | 47.3306478 | 76.0855288 | 0 |
| 8.48E-06 | 2.52E-05 | 90.0262989 | 248.688582 | 335.954794 | 230.112331 | 19.0097459 |
| 8.50E-06 | 2.52E-05 | 48.2283744 | 27.1377523 | 24.1293498 | 39.898509 | 0 |
| 8.51E-06 | 2.52E-05 | 107.710036 | 96.0943359 | 11.136623 | 75.6215927 | 0 |
| 8.53E-06 | 2.53E-05 | 15.4330798 | 90.0884399 | 60.3233746 | 56.1362743 | 0 |
| 8.53E-06 | 2.53E-05 | 18.3267823 | 61.6160441 | 41.7623363 | 50.1051043 | 0 |
| 8.54E-06 | 2.53E-05 | 38.904222 | 24.2460246 | 26.9135056 | 38.9706367 | 0 |
| 8.55E-06 | 2.54E-05 | 34.0813846 | 20.4645345 | 47.3306478 | 55.6723382 | 0 |
| 8.58E-06 | 2.54E-05 | 20.8989622 | 37.8149007 | 36.1940248 | 38.5067006 | 0 |
| 8.58E-06 | 2.55E-05 | 31.8307271 | 26.4704305 | 28.7696094 | 27.8361691 | 0 |
| 8.59E-06 | 2.55E-05 | 34.7244296 | 80.5234945 | 68.6758419 | 94.6429749 | 4.75243649 |
| 8.60E-06 | 2.55E-05 | 28.2939797 | 24.4684652 | 32.4818171 | 40.3624452 | 0 |
| 8.60E-06 | 2.55E-05 | 16.0761248 | 50.2715739 | 64.9636342 | 63.0953166 | 0 |
| 8.61E-06 | 2.55E-05 | 20.8989622 | 98.3187419 | 187.466487 | 32.0115944 | 0 |
| 8.62E-06 | 2.56E-05 | 23.7926647 | 49.1593709 | 78.8844129 | 29.6919137 | 0 |
| 8.62E-06 | 2.56E-05 | 11.2532874 | 64.7302124 | 115.078438 | 141.03659 | 0 |
| 8.65E-06 | 2.56E-05 | 28.6155022 | 58.946757 | 29.6976613 | 29.6919137 | 0 |

| | | | | | | |
|---|---|---|---|---|---|---|
| 8.67E-06 | 2.57E-05 | 6.10892743 | 8.23030192 | 0 | 4.17542536 | 38.0194919 |
| 8.67E-06 | 2.57E-05 | 6.75197242 | 300.51724 | 155.912722 | 93.2511664 | 0 |
| 8.69E-06 | 2.57E-05 | 23.4711422 | 32.9212077 | 33.409869 | 30.1558498 | 0 |
| 8.69E-06 | 2.57E-05 | 2.57217997 | 1.77952474 | 2.78415575 | 6.03116997 | 14.2573095 |
| 8.71E-06 | 2.58E-05 | 36.3320421 | 68.9565837 | 72.3880495 | 60.7756358 | 0 |
| 8.71E-06 | 2.58E-05 | 25.7217997 | 36.7026978 | 26.9135056 | 30.619786 | 0 |
| 8.74E-06 | 2.59E-05 | 7.07349491 | 285.16884 | 91.8771398 | 147.531696 | 0 |
| 8.74E-06 | 2.59E-05 | 161.404293 | 47.8247274 | 67.7477899 | 61.7035081 | 2.37621824 |
| 8.75E-06 | 2.59E-05 | 22.8280972 | 29.3621582 | 38.9781805 | 36.1870198 | 0 |
| 8.75E-06 | 2.59E-05 | 34.7244296 | 29.3621582 | 27.8415575 | 25.5164883 | 0 |
| 8.78E-06 | 2.60E-05 | 15.1115573 | 79.8561727 | 53.8270112 | 66.3428696 | 0 |
| 8.79E-06 | 2.60E-05 | 8.3595849 | 126.568697 | 179.11402 | 90.9314857 | 0 |
| 8.79E-06 | 2.60E-05 | 34.4029071 | 19.1298909 | 24.1293498 | 16.7017014 | 68.9103291 |
| 8.85E-06 | 2.62E-05 | 45.9777169 | 32.6987671 | 33.409869 | 35.2591475 | 171.087714 |
| 8.86E-06 | 2.62E-05 | 406.725958 | 740.282292 | 628.291148 | 731.163374 | 0 |
| 8.87E-06 | 2.63E-05 | 22.5065747 | 30.4743612 | 64.0355823 | 44.5378705 | 0 |
| 8.88E-06 | 2.63E-05 | 38.261177 | 25.1357869 | 42.6903882 | 25.5164883 | 0 |
| 8.88E-06 | 2.63E-05 | 36.6535646 | 25.8031087 | 21.3451941 | 20.4131907 | 87.920075 |
| 8.91E-06 | 2.64E-05 | 30.8661596 | 63.1731283 | 22.273246 | 43.1460621 | 0 |
| 8.91E-06 | 2.64E-05 | 58.1955718 | 87.864034 | 12.9927268 | 171.19244 | 0 |
| 8.92E-06 | 2.64E-05 | 21.8635297 | 29.5845988 | 75.1722053 | 65.4149973 | 0 |
| 8.96E-06 | 2.65E-05 | 18.9698273 | 44.7105591 | 59.3953227 | 39.4345729 | 0 |
| 8.96E-06 | 2.65E-05 | 21.8635297 | 30.02948 | 93.7332436 | 70.0543588 | 0 |
| 8.98E-06 | 2.66E-05 | 308.661596 | 552.987313 | 437.112453 | 477.3903 | 19.0097459 |
| 8.98E-06 | 2.66E-05 | 18.3267823 | 43.5983561 | 47.3306478 | 45.929679 | 0 |
| 9.00E-06 | 2.66E-05 | 58.5170943 | 169.722172 | 102.085711 | 56.6002105 | 7.12865473 |
| 9.01E-06 | 2.66E-05 | 23.7926647 | 39.816866 | 29.6976613 | 31.0837221 | 0 |
| 9.02E-06 | 2.67E-05 | 26.0433222 | 44.0432373 | 30.6257133 | 27.3722329 | 0 |
| 9.02E-06 | 2.67E-05 | 34.0813846 | 30.2519206 | 24.1293498 | 28.7640414 | 0 |
| 9.03E-06 | 2.67E-05 | 45.334672 | 53.8306234 | 47.3306478 | 54.2805297 | 2.37621824 |
| 9.03E-06 | 2.67E-05 | 31.1876821 | 45.3778809 | 21.3451941 | 60.7756358 | 0 |
| 9.03E-06 | 2.67E-05 | 75.2362641 | 94.3148112 | 60.3233746 | 78.4052096 | 7.12865473 |
| 9.04E-06 | 2.67E-05 | 15.7546023 | 113.667143 | 64.9636342 | 51.0329766 | 0 |
| 9.05E-06 | 2.67E-05 | 44.3701045 | 90.0884399 | 16.7049345 | 60.3116997 | 0 |
| 9.06E-06 | 2.68E-05 | 22.1850522 | 51.6062174 | 28.7696094 | 52.4247851 | 0 |
| 9.08E-06 | 2.68E-05 | 63.9829767 | 33.3660889 | 163.337137 | 93.2511664 | 0 |
| 9.08E-06 | 2.68E-05 | 602.211635 | 1196.73039 | 1062.61944 | 1131.54027 | 0 |
| 9.11E-06 | 2.69E-05 | 30.8661596 | 104.547078 | 126.215061 | 89.0757411 | 2.37621824 |
| 9.12E-06 | 2.69E-05 | 19.6128723 | 51.828658 | 64.0355823 | 34.7952113 | 0 |
| 9.12E-06 | 2.69E-05 | 12.8608998 | 61.8384847 | 96.5173994 | 85.3642518 | 0 |
| 9.12E-06 | 2.69E-05 | 39.8687895 | 34.2558512 | 21.3451941 | 32.4755306 | 0 |
| 9.12E-06 | 2.69E-05 | 37.618132 | 37.5924601 | 34.3379209 | 20.8771268 | 0 |
| 9.13E-06 | 2.70E-05 | 32.7952946 | 66.064856 | 79.8124649 | 19.9492545 | 0 |

| | | | | | | |
|---|---|---|---|---|---|---|
| 9.13E-06 | 2.70E-05 | 5.14435994 | 2.66928711 | 3.71220767 | 2.31968076 | 16.6335277 |
| 9.13E-06 | 2.70E-05 | 19.6128723 | 47.3798462 | 42.6903882 | 36.650956 | 0 |
| 9.15E-06 | 2.70E-05 | 11.2532874 | 97.8738607 | 94.6612955 | 77.4773373 | 0 |
| 9.16E-06 | 2.70E-05 | 18.9698273 | 38.2597819 | 47.3306478 | 46.3936151 | 0 |
| 9.15E-06 | 2.70E-05 | 22.8280972 | 59.6140788 | 29.6976613 | 42.2181898 | 0 |
| 9.16E-06 | 2.70E-05 | 7.39501741 | 78.0766479 | 222.73246 | 183.718716 | 0 |
| 9.19E-06 | 2.71E-05 | 136.647061 | 103.212435 | 99.3015551 | 83.9724434 | 4.75243649 |
| 9.21E-06 | 2.72E-05 | 18.6483048 | 34.7007324 | 90.0210359 | 83.0445711 | 0 |
| 9.23E-06 | 2.72E-05 | 31.1876821 | 86.9742716 | 74.2441534 | 77.0134011 | 0 |
| 9.24E-06 | 2.73E-05 | 29.9015921 | 62.5058065 | 21.3451941 | 52.8887212 | 0 |
| 9.24E-06 | 2.73E-05 | 38.5826995 | 37.5924601 | 19.4890903 | 40.8263813 | 0 |
| 9.26E-06 | 2.73E-05 | 11.5748099 | 78.7439697 | 112.294282 | 78.4052096 | 0 |
| 9.26E-06 | 2.73E-05 | 19.6128723 | 44.7105591 | 36.1940248 | 44.0739344 | 0 |
| 9.28E-06 | 2.74E-05 | 77.1653991 | 334.10577 | 314.6096 | 317.332327 | 19.0097459 |
| 9.28E-06 | 2.74E-05 | 94.2060914 | 27.8050741 | 28.7696094 | 42.6821259 | 0 |
| 9.29E-06 | 2.74E-05 | 18.0052598 | 51.6062174 | 47.3306478 | 42.6821259 | 0 |
| 9.29E-06 | 2.74E-05 | 24.7572322 | 57.1672323 | 25.0574018 | 58.4559551 | 0 |
| 9.31E-06 | 2.74E-05 | 19.6128723 | 36.2578166 | 54.7550631 | 42.6821259 | 0 |
| 9.31E-06 | 2.75E-05 | 3.21522496 | 6.89565837 | 3.71220767 | 4.63936151 | 66.5341108 |
| 9.35E-06 | 2.75E-05 | 42.119447 | 59.1691976 | 64.0355823 | 75.6215927 | 0 |
| 9.40E-06 | 2.77E-05 | 22.8280972 | 30.6968018 | 31.5537652 | 35.7230837 | 0 |
| 9.41E-06 | 2.77E-05 | 69.7703817 | 62.2833659 | 33.409869 | 19.4853184 | 0 |
| 9.43E-06 | 2.78E-05 | 96.4567488 | 239.123637 | 172.617657 | 211.090949 | 19.0097459 |
| 9.43E-06 | 2.78E-05 | 19.9343948 | 34.923173 | 44.546492 | 41.2903175 | 0 |
| 9.47E-06 | 2.79E-05 | 41.476402 | 34.923173 | 19.4890903 | 43.1460621 | 0 |
| 9.48E-06 | 2.79E-05 | 22.5065747 | 55.1652669 | 30.6257133 | 38.0427644 | 0 |
| 9.50E-06 | 2.80E-05 | 27.6509347 | 122.119885 | 80.7405168 | 96.4987195 | 0 |
| 9.50E-06 | 2.80E-05 | 31.8307271 | 30.6968018 | 22.273246 | 35.7230837 | 0 |
| 9.50E-06 | 2.80E-05 | 26.3648447 | 41.3739502 | 31.5537652 | 25.5164883 | 0 |
| 9.52E-06 | 2.80E-05 | 18.9698273 | 41.3739502 | 41.7623363 | 42.2181898 | 0 |
| 9.52E-06 | 2.80E-05 | 211.24028 | 1745.49133 | 1642.65189 | 1572.74355 | 0 |
| 9.54E-06 | 2.81E-05 | 820.846933 | 2014.42201 | 2015.72876 | 2187.92289 | 0 |
| 9.55E-06 | 2.81E-05 | 58.5170943 | 165.27336 | 163.337137 | 179.079354 | 14.2573095 |
| 9.59E-06 | 2.82E-05 | 49.8359869 | 100.098267 | 119.718697 | 13.4541484 | 0 |
| 9.59E-06 | 2.82E-05 | 20.8989622 | 34.4782918 | 36.1940248 | 50.1051043 | 0 |
| 9.60E-06 | 2.82E-05 | 35.6889971 | 35.1456136 | 21.3451941 | 31.5476583 | 0 |
| 9.60E-06 | 2.83E-05 | 25.0787547 | 36.035376 | 40.8342843 | 25.0525522 | 0 |
| 9.61E-06 | 2.83E-05 | 14.4685123 | 110.997856 | 72.3880495 | 52.4247851 | 0 |
| 9.63E-06 | 2.83E-05 | 19.9343948 | 50.9388957 | 38.0501286 | 37.1148921 | 0 |
| 9.64E-06 | 2.84E-05 | 21.5420072 | 38.7046631 | 30.6257133 | 38.5067006 | 0 |
| 9.64E-06 | 2.84E-05 | 63.9829767 | 101.655351 | 110.438178 | 115.056166 | 14.2573095 |
| 9.66E-06 | 2.84E-05 | 6.75197242 | 66.064856 | 327.602327 | 219.905736 | 0 |
| 9.66E-06 | 2.84E-05 | 31.1876821 | 30.9192423 | 23.2012979 | 30.1558498 | 0 |

| | | | | | | |
|---|---|---|---|---|---|---|
| 9.67E-06 | 2.85E-05 | 197.414813 | 311.63927 | 296.976613 | 290.424031 | 4.75243649 |
| 9.67E-06 | 2.85E-05 | 112.211351 | 292.286938 | 348.019469 | 291.815839 | 42.7719284 |
| 9.71E-06 | 2.86E-05 | 41.1548795 | 27.1377523 | 25.9854537 | 72.3740396 | 0 |
| 9.71E-06 | 2.86E-05 | 9.96719738 | 128.125781 | 75.1722053 | 113.200421 | 0 |
| 9.72E-06 | 2.86E-05 | 70.7349491 | 131.23995 | 122.502853 | 126.654569 | 7.12865473 |
| 9.75E-06 | 2.87E-05 | 25.4002772 | 37.8149007 | 61.2514265 | 26.9082968 | 0 |
| 9.77E-06 | 2.87E-05 | 32.1522496 | 28.4723958 | 25.9854537 | 57.5280828 | 0 |
| 9.79E-06 | 2.88E-05 | 21.8635297 | 34.923173 | 59.3953227 | 34.3312752 | 0 |
| 9.81E-06 | 2.88E-05 | 19.6128723 | 33.1436483 | 57.5392189 | 52.4247851 | 0 |
| 9.82E-06 | 2.89E-05 | 59.1601393 | 84.7498657 | 44.546492 | 62.1674443 | 0 |
| 9.82E-06 | 2.89E-05 | 41.7979245 | 158.600142 | 122.502853 | 72.8379758 | 7.12865473 |
| 9.82E-06 | 2.89E-05 | 41.7979245 | 24.2460246 | 25.9854537 | 35.7230837 | 0 |
| 9.83E-06 | 2.89E-05 | 29.5800696 | 89.1986776 | 69.6038938 | 106.241379 | 2.37621824 |
| 9.84E-06 | 2.89E-05 | 31.5092046 | 42.041272 | 21.3451941 | 33.867339 | 0 |
| 9.86E-06 | 2.90E-05 | 35.6889971 | 30.2519206 | 22.273246 | 31.5476583 | 0 |
| 9.87E-06 | 2.90E-05 | 0 | 0 | 0 | 0 | 66.5341108 |
| 9.88E-06 | 2.90E-05 | 20.2559173 | 34.7007324 | 37.1220767 | 43.1460621 | 0 |
| 9.89E-06 | 2.90E-05 | 16.0761248 | 47.8247274 | 65.8916861 | 55.6723382 | 0 |
| 9.89E-06 | 2.91E-05 | 39.547267 | 30.6968018 | 22.273246 | 31.0837221 | 0 |
| 9.91E-06 | 2.91E-05 | 14.4685123 | 53.3857422 | 66.819738 | 86.2921241 | 0 |
| 9.91E-06 | 2.91E-05 | 30.8661596 | 84.0825439 | 77.0283091 | 76.549465 | 0 |
| 9.92E-06 | 2.91E-05 | 17.3622148 | 68.0668213 | 38.9781805 | 94.6429749 | 0 |
| 9.93E-06 | 2.91E-05 | 21.5420072 | 29.8070394 | 43.6184401 | 38.5067006 | 0 |
| 9.94E-06 | 2.92E-05 | 112.211351 | 91.4230835 | 109.510126 | 106.241379 | 358.808955 |
| 9.94E-06 | 2.92E-05 | 74.5932191 | 28.4723958 | 26.9135056 | 36.650956 | 0 |
| 9.94E-06 | 2.92E-05 | 17.6837373 | 101.655351 | 112.294282 | 34.3312752 | 0 |
| 9.95E-06 | 2.92E-05 | 18.0052598 | 48.7144897 | 43.6184401 | 86.2921241 | 0 |
| 9.96E-06 | 2.92E-05 | 15.4330798 | 92.9801676 | 59.3953227 | 51.0329766 | 0 |
| 9.96E-06 | 2.92E-05 | 23.7926647 | 31.5865641 | 27.8415575 | 40.8263813 | 0 |
| 9.97E-06 | 2.92E-05 | 15.1115573 | 78.5215291 | 48.2586997 | 69.5904227 | 0 |
| 1.00E-05 | 2.93E-05 | 21.5420072 | 47.1574056 | 30.6257133 | 37.5788283 | 0 |
| 1.00E-05 | 2.93E-05 | 16.0761248 | 70.958549 | 60.3233746 | 45.4657428 | 0 |
| 1.00E-05 | 2.94E-05 | 11.2532874 | 86.3069499 | 77.0283091 | 95.5708472 | 0 |
| 1.00E-05 | 2.94E-05 | 26.3648447 | 197.972127 | 27.8415575 | 73.7658481 | 0 |
| 1.00E-05 | 2.94E-05 | 17.0406923 | 42.041272 | 66.819738 | 54.2805297 | 0 |
| 1.00E-05 | 2.94E-05 | 14.4685123 | 61.1711629 | 56.6111669 | 81.6527626 | 0 |
| 1.00E-05 | 2.94E-05 | 28.6155022 | 25.3582275 | 28.7696094 | 28.7640414 | 0 |
| 1.01E-05 | 2.95E-05 | 43.405537 | 55.8325887 | 18.5610383 | 37.1148921 | 0 |
| 1.01E-05 | 2.95E-05 | 20.2559173 | 40.2617472 | 36.1940248 | 35.2591475 | 0 |
| 1.01E-05 | 2.97E-05 | 20.5774398 | 30.9192423 | 123.430905 | 78.4052096 | 0 |
| 1.01E-05 | 2.97E-05 | 44.691627 | 20.0196533 | 55.683115 | 32.9394667 | 0 |
| 1.01E-05 | 2.97E-05 | 28.2939797 | 21.5767375 | 77.956361 | 73.3019119 | 0 |
| 1.02E-05 | 2.98E-05 | 55.6233918 | 16.0157227 | 96.5173994 | 49.6411682 | 0 |

| | | | | | | |
|---|---|---|---|---|---|---|
| 1.02E-05 | 2.98E-05 | 58.1955718 | 75.852242 | 14.8488307 | 51.4969128 | 0 |
| 1.02E-05 | 2.99E-05 | 32.4737721 | 87.4191528 | 98.3735032 | 58.4559551 | 2.37621824 |
| 1.02E-05 | 2.99E-05 | 9.32415239 | 93.6474894 | 110.438178 | 104.385634 | 0 |
| 1.02E-05 | 2.99E-05 | 24.1141872 | 57.1672323 | 28.7696094 | 35.2591475 | 0 |
| 1.02E-05 | 3.00E-05 | 33.7598621 | 36.4802572 | 23.2012979 | 76.0855288 | 0 |
| 1.02E-05 | 3.00E-05 | 28.2939797 | 60.05896 | 31.5537652 | 26.9082968 | 0 |
| 1.03E-05 | 3.00E-05 | 16.7191698 | 40.0393066 | 79.8124649 | 66.8068058 | 0 |
| 1.03E-05 | 3.01E-05 | 14.7900348 | 59.3916382 | 64.0355823 | 57.0641466 | 0 |
| 1.03E-05 | 3.01E-05 | 10.9317649 | 90.7557617 | 100.229607 | 75.6215927 | 0 |
| 1.03E-05 | 3.01E-05 | 11.5748099 | 80.0786133 | 75.1722053 | 93.2511664 | 0 |
| 1.03E-05 | 3.03E-05 | 268.149762 | 425.751294 | 382.35739 | 443.986897 | 21.3859642 |
| 1.03E-05 | 3.03E-05 | 12.8608998 | 82.3030192 | 67.7477899 | 69.1264865 | 0 |
| 1.04E-05 | 3.04E-05 | 21.2204847 | 48.2696086 | 30.6257133 | 38.5067006 | 0 |
| 1.04E-05 | 3.04E-05 | 15.1115573 | 82.3030192 | 47.3306478 | 67.7346781 | 0 |
| 1.04E-05 | 3.04E-05 | 63.9829767 | 37.3700195 | 25.9854537 | 26.4443606 | 0 |
| 1.04E-05 | 3.04E-05 | 84.238894 | 272.489726 | 243.149602 | 245.88616 | 19.0097459 |
| 1.04E-05 | 3.04E-05 | 18.3267823 | 41.8188314 | 47.3306478 | 40.3624452 | 0 |
| 1.04E-05 | 3.05E-05 | 21.2204847 | 29.3621582 | 51.0428554 | 43.6099982 | 0 |
| 1.04E-05 | 3.05E-05 | 17.0406923 | 59.6140788 | 39.9062324 | 64.487125 | 0 |
| 1.05E-05 | 3.06E-05 | 20.8989622 | 51.3837769 | 30.6257133 | 41.7542536 | 0 |
| 1.05E-05 | 3.06E-05 | 4.82283744 | 3.33660889 | 2.78415575 | 6.49510612 | 19.0097459 |
| 1.05E-05 | 3.06E-05 | 32.1522496 | 77.8542074 | 99.3015551 | 63.5592527 | 0 |
| 1.05E-05 | 3.07E-05 | 18.3267823 | 38.2597819 | 52.8989593 | 44.0739344 | 0 |
| 1.05E-05 | 3.08E-05 | 20.8989622 | 35.1456136 | 33.409869 | 34.7952113 | 0 |
| 1.05E-05 | 3.08E-05 | 34.4029071 | 86.7518311 | 80.7405168 | 82.1166988 | 0 |
| 1.06E-05 | 3.09E-05 | 36.6535646 | 66.9546183 | 54.7550631 | 64.487125 | 0 |
| 1.06E-05 | 3.09E-05 | 24.7572322 | 24.6909058 | 27.8415575 | 24.588616 | 57.0292378 |
| 1.06E-05 | 3.10E-05 | 19.2913498 | 11.3444702 | 0 | 0 | 594.054561 |
| 1.06E-05 | 3.10E-05 | 32.4737721 | 71.1809896 | 67.7477899 | 76.0855288 | 0 |
| 1.06E-05 | 3.11E-05 | 13.8254673 | 62.0609253 | 66.819738 | 65.4149973 | 0 |
| 1.06E-05 | 3.11E-05 | 48.5498969 | 20.2420939 | 33.409869 | 38.0427644 | 0 |
| 1.06E-05 | 3.11E-05 | 24.4357097 | 94.982133 | 116.00649 | 23.6607437 | 0 |
| 1.07E-05 | 3.12E-05 | 77.4869216 | 155.485974 | 129.927268 | 147.995632 | 1751.27285 |
| 1.07E-05 | 3.12E-05 | 123.464639 | 181.956405 | 172.617657 | 175.831801 | 35.6432737 |
| 1.07E-05 | 3.12E-05 | 16.7191698 | 79.1888509 | 39.9062324 | 64.0231889 | 0 |
| 1.07E-05 | 3.13E-05 | 204.80983 | 223.997677 | 161.481034 | 245.88616 | 23.7621824 |
| 1.07E-05 | 3.14E-05 | 121.213981 | 393.052527 | 271.919212 | 297.847009 | 23.7621824 |
| 1.08E-05 | 3.14E-05 | 16.7191698 | 53.8306234 | 42.6903882 | 57.0641466 | 0 |
| 1.08E-05 | 3.14E-05 | 226.030315 | 143.029301 | 105.797919 | 122.479144 | 0 |
| 1.08E-05 | 3.14E-05 | 19.9343948 | 40.929069 | 33.409869 | 38.9706367 | 0 |
| 1.08E-05 | 3.15E-05 | 18.0052598 | 45.1554403 | 38.9781805 | 52.4247851 | 0 |
| 1.08E-05 | 3.16E-05 | 0 | 1860.938 | 2120.59863 | 2765.98733 | 0 |
| 1.08E-05 | 3.16E-05 | 459.455647 | 423.749329 | 429.688038 | 463.008279 | 35.6432737 |

| | | | | | | |
|---|---|---|---|---|---|---|
| 1.09E-05 | 3.17E-05 | 94.8491364 | 38.0373413 | 23.2012979 | 36.1870198 | 0 |
| 1.09E-05 | 3.17E-05 | 22.5065747 | 60.05896 | 27.8415575 | 43.6099982 | 0 |
| 1.09E-05 | 3.17E-05 | 24.1141872 | 51.1613363 | 65.8916861 | 25.0525522 | 0 |
| 1.09E-05 | 3.19E-05 | 15.4330798 | 49.6042521 | 65.8916861 | 54.7444659 | 0 |
| 1.09E-05 | 3.19E-05 | 23.7926647 | 27.8050741 | 29.6976613 | 39.898509 | 0 |
| 1.09E-05 | 3.19E-05 | 126.679863 | 226.666964 | 215.308045 | 225.009033 | 35.6432737 |
| 1.09E-05 | 3.19E-05 | 18.3267823 | 71.4034302 | 40.8342843 | 43.6099982 | 0 |
| 1.10E-05 | 3.20E-05 | 19.9343948 | 38.7046631 | 34.3379209 | 55.208402 | 0 |
| 1.10E-05 | 3.20E-05 | 23.1496197 | 42.7085937 | 25.9854537 | 37.5788283 | 0 |
| 1.10E-05 | 3.20E-05 | 16.7191698 | 75.4073608 | 43.6184401 | 51.4969128 | 0 |
| 1.10E-05 | 3.22E-05 | 20.8989622 | 34.2558512 | 31.5537652 | 38.5067006 | 0 |
| 1.10E-05 | 3.22E-05 | 18.3267823 | 39.5944255 | 48.2586997 | 39.898509 | 0 |
| 1.11E-05 | 3.22E-05 | 27.0078897 | 22.6889404 | 49.1867516 | 38.5067006 | 0 |
| 1.11E-05 | 3.23E-05 | 23.7926647 | 48.4920492 | 38.9781805 | 25.5164883 | 0 |
| 1.11E-05 | 3.23E-05 | 24.7572322 | 177.952474 | 90.0210359 | 24.588616 | 0 |
| 1.11E-05 | 3.23E-05 | 14.1469898 | 60.2814006 | 69.6038938 | 57.5280828 | 0 |
| 1.11E-05 | 3.23E-05 | 36.9750871 | 32.4763265 | 22.273246 | 27.8361691 | 0 |
| 1.11E-05 | 3.24E-05 | 14.4685123 | 100.320707 | 84.4527244 | 44.5378705 | 0 |
| 1.11E-05 | 3.24E-05 | 10.9317649 | 86.7518311 | 77.956361 | 88.6118049 | 0 |
| 1.11E-05 | 3.24E-05 | 28.6155022 | 26.0255493 | 25.0574018 | 31.5476583 | 0 |
| 1.11E-05 | 3.25E-05 | 23.7926647 | 49.1593709 | 25.9854537 | 36.650956 | 0 |
| 1.12E-05 | 3.25E-05 | 10.9317649 | 14.9035197 | 19.4890903 | 12.9902122 | 45.1481466 |
| 1.12E-05 | 3.27E-05 | 12.5393773 | 79.6337321 | 87.2368802 | 59.3838274 | 0 |
| 1.13E-05 | 3.28E-05 | 38.261177 | 26.6928711 | 57.5392189 | 23.6607437 | 0 |
| 1.13E-05 | 3.28E-05 | 17.6837373 | 42.041272 | 52.8989593 | 41.2903175 | 0 |
| 1.13E-05 | 3.28E-05 | 16.0761248 | 55.1652669 | 47.3306478 | 51.4969128 | 0 |
| 1.13E-05 | 3.28E-05 | 32.1522496 | 181.511523 | 197.675058 | 189.28595 | 9.50487297 |
| 1.13E-05 | 3.30E-05 | 32.4737721 | 169.05485 | 153.128566 | 162.841589 | 0 |
| 1.13E-05 | 3.30E-05 | 100.958064 | 46.0452026 | 22.273246 | 32.9394667 | 0 |
| 1.14E-05 | 3.31E-05 | 20.8989622 | 32.2538859 | 32.4818171 | 37.5788283 | 0 |
| 1.14E-05 | 3.31E-05 | 21.8635297 | 44.0432373 | 26.9135056 | 45.4657428 | 0 |
| 1.14E-05 | 3.32E-05 | 44.3701045 | 24.023584 | 25.0574018 | 35.7230837 | 0 |
| 1.14E-05 | 3.32E-05 | 13.5039448 | 65.1750936 | 58.4672708 | 75.6215927 | 0 |
| 1.14E-05 | 3.32E-05 | 57.8740493 | 46.934965 | 108.582074 | 44.0739344 | 2.37621824 |
| 1.14E-05 | 3.32E-05 | 30.8661596 | 40.2617472 | 22.273246 | 28.7640414 | 0 |
| 1.14E-05 | 3.32E-05 | 62.0538418 | 38.2597819 | 35.2659728 | 19.9492545 | 0 |
| 1.14E-05 | 3.32E-05 | 16.7191698 | 58.7243164 | 41.7623363 | 51.0329766 | 0 |
| 1.14E-05 | 3.33E-05 | 20.8989622 | 55.6101481 | 30.6257133 | 39.898509 | 0 |
| 1.15E-05 | 3.34E-05 | 624.71821 | 421.080041 | 374.004923 | 369.757113 | 99.8011662 |
| 1.15E-05 | 3.34E-05 | 33.1168171 | 81.858138 | 63.1075304 | 63.0953166 | 2.37621824 |
| 1.15E-05 | 3.34E-05 | 396.75876 | 739.392529 | 687.68647 | 745.081459 | 49.9005831 |
| 1.15E-05 | 3.35E-05 | 23.1496197 | 34.7007324 | 25.9854537 | 42.2181898 | 0 |
| 1.15E-05 | 3.35E-05 | 216.063117 | 628.394674 | 712.743872 | 702.863269 | 0 |

| | | | | | | |
|---|---|---|---|---|---|---|
| 1.15E-05 | 3.35E-05 | 209.632667 | 273.824369 | 169.833501 | 219.4418 | 0 |
| 1.15E-05 | 3.36E-05 | 33.7598621 | 99.875826 | 53.8270112 | 77.9412734 | 0 |
| 1.16E-05 | 3.36E-05 | 88.4186864 | 625.280505 | 467.738166 | 577.600508 | 23.7621824 |
| 1.16E-05 | 3.36E-05 | 16.0761248 | 111.220296 | 38.9781805 | 77.0134011 | 0 |
| 1.16E-05 | 3.37E-05 | 19.6128723 | 31.5865641 | 47.3306478 | 42.6821259 | 0 |
| 1.16E-05 | 3.37E-05 | 31.8307271 | 75.852242 | 63.1075304 | 72.3740396 | 2.37621824 |
| 1.16E-05 | 3.37E-05 | 139.219241 | 37.5924601 | 33.409869 | 29.6919137 | 0 |
| 1.16E-05 | 3.38E-05 | 15.1115573 | 103.879757 | 40.8342843 | 83.9724434 | 0 |
| 1.16E-05 | 3.38E-05 | 25.7217997 | 54.9428263 | 24.1293498 | 37.5788283 | 0 |
| 1.16E-05 | 3.38E-05 | 20.8989622 | 37.5924601 | 29.6976613 | 46.8575513 | 0 |
| 1.16E-05 | 3.38E-05 | 19.6128723 | 33.8109701 | 67.7477899 | 43.6099982 | 0 |
| 1.16E-05 | 3.38E-05 | 22.8280972 | 48.4920492 | 29.6976613 | 93.2511664 | 0 |
| 1.17E-05 | 3.39E-05 | 8.0380624 | 102.100232 | 116.934542 | 112.736485 | 0 |
| 1.17E-05 | 3.39E-05 | 37.9396545 | 113.667143 | 18.5610383 | 55.208402 | 0 |
| 1.17E-05 | 3.40E-05 | 61.4107968 | 24.9133464 | 28.7696094 | 32.4755306 | 0 |
| 1.17E-05 | 3.40E-05 | 28.6155022 | 85.8620687 | 75.1722053 | 84.9003157 | 0 |
| 1.17E-05 | 3.40E-05 | 16.7191698 | 93.86993 | 59.3953227 | 40.3624452 | 0 |
| 1.17E-05 | 3.41E-05 | 36.9750871 | 59.3916382 | 71.4599976 | 51.0329766 | 2.37621824 |
| 1.17E-05 | 3.41E-05 | 36.6535646 | 38.2597819 | 20.4171422 | 29.6919137 | 0 |
| 1.18E-05 | 3.41E-05 | 160.439726 | 222.885474 | 187.466487 | 208.771268 | 0 |
| 1.18E-05 | 3.42E-05 | 76.2008316 | 166.608004 | 134.567528 | 156.346483 | 16.6335277 |
| 1.18E-05 | 3.42E-05 | 22.5065747 | 39.3719849 | 27.8415575 | 58.9198912 | 0 |
| 1.18E-05 | 3.42E-05 | 1.92913498 | 2.22440592 | 0 | 0 | 76.0389838 |
| 1.18E-05 | 3.43E-05 | 4.17979245 | 2.00196533 | 3.71220767 | 3.71148921 | 42.7719284 |
| 1.18E-05 | 3.44E-05 | 12.2178549 | 87.1967122 | 72.3880495 | 64.0231889 | 0 |
| 1.19E-05 | 3.45E-05 | 14.1469898 | 49.6042521 | 132.711424 | 70.0543588 | 0 |
| 1.19E-05 | 3.45E-05 | 16.0761248 | 49.3818115 | 55.683115 | 45.929679 | 0 |
| 1.19E-05 | 3.46E-05 | 6.43044992 | 126.568697 | 142.919995 | 124.798825 | 0 |
| 1.19E-05 | 3.46E-05 | 31.8307271 | 28.0275146 | 25.0574018 | 25.5164883 | 0 |
| 1.19E-05 | 3.46E-05 | 52.4081669 | 295.845988 | 195.818954 | 212.018821 | 14.2573095 |
| 1.19E-05 | 3.46E-05 | 314.127479 | 484.47561 | 536.414008 | 566.466041 | 35.6432737 |
| 1.19E-05 | 3.46E-05 | 26.3648447 | 29.1397176 | 25.0574018 | 27.8361691 | 0 |
| 1.19E-05 | 3.46E-05 | 105.780901 | 25.5806681 | 25.0574018 | 57.0641466 | 0 |
| 1.20E-05 | 3.47E-05 | 51.4435994 | 100.765588 | 104.869867 | 57.5280828 | 9.50487297 |
| 1.20E-05 | 3.47E-05 | 176.194328 | 285.836161 | 248.717914 | 262.123925 | 38.0194919 |
| 1.20E-05 | 3.47E-05 | 52.4081669 | 23.3562622 | 31.5537652 | 29.2279775 | 0 |
| 1.20E-05 | 3.47E-05 | 35.6889971 | 23.1338216 | 57.5392189 | 27.8361691 | 0 |
| 1.20E-05 | 3.48E-05 | 263.326924 | 366.137215 | 313.681548 | 348.41605 | 35.6432737 |
| 1.20E-05 | 3.48E-05 | 272.651077 | 327.432552 | 266.3509 | 297.383073 | 0 |
| 1.20E-05 | 3.48E-05 | 68.8058142 | 398.391101 | 192.106747 | 241.246799 | 0 |
| 1.20E-05 | 3.49E-05 | 30.5446371 | 72.7380737 | 28.7696094 | 28.3001052 | 0 |
| 1.20E-05 | 3.49E-05 | 26.0433222 | 29.5845988 | 35.2659728 | 23.6607437 | 0 |
| 1.21E-05 | 3.50E-05 | 31.5092046 | 16.4606038 | 9.28051917 | 11.5984038 | 71.2865473 |

| | | | | | | |
|---|---|---|---|---|---|---|
| 1.21E-05 | 3.50E-05 | 46.6207619 | 124.566732 | 83.5246725 | 91.3954218 | 0 |
| 1.21E-05 | 3.50E-05 | 94.2060914 | 57.6121134 | 19.4890903 | 32.0115944 | 0 |
| 1.21E-05 | 3.51E-05 | 51.7651219 | 26.4704305 | 31.5537652 | 25.0525522 | 0 |
| 1.21E-05 | 3.51E-05 | 63.0184092 | 25.5806681 | 41.7623363 | 24.1246799 | 0 |
| 1.21E-05 | 3.51E-05 | 26.0433222 | 75.4073608 | 25.0574018 | 42.2181898 | 0 |
| 1.21E-05 | 3.52E-05 | 89.3832539 | 93.4250488 | 9.28051917 | 96.4987195 | 0 |
| 1.22E-05 | 3.52E-05 | 11.5748099 | 71.6258708 | 78.8844129 | 77.9412734 | 0 |
| 1.22E-05 | 3.52E-05 | 15.4330798 | 55.3877075 | 50.1148035 | 51.4969128 | 0 |
| 1.22E-05 | 3.53E-05 | 22.8280972 | 56.0550293 | 27.8415575 | 36.1870198 | 0 |
| 1.22E-05 | 3.53E-05 | 419.265335 | 1393.59031 | 1231.52489 | 1338.4558 | 23.7621824 |
| 1.22E-05 | 3.54E-05 | 79.094534 | 176.840271 | 154.056618 | 182.326907 | 14.2573095 |
| 1.22E-05 | 3.54E-05 | 19.6128723 | 37.1475789 | 32.4818171 | 41.7542536 | 0 |
| 1.22E-05 | 3.54E-05 | 97.4213163 | 310.749508 | 303.472977 | 284.856797 | 21.3859642 |
| 1.22E-05 | 3.54E-05 | 17.6837373 | 53.1633016 | 38.9781805 | 42.2181898 | 0 |
| 1.23E-05 | 3.55E-05 | 31.5092046 | 37.3700195 | 19.4890903 | 46.8575513 | 0 |
| 1.23E-05 | 3.55E-05 | 129.252043 | 180.17688 | 170.761553 | 177.687546 | 21.3859642 |
| 1.23E-05 | 3.55E-05 | 18.6483048 | 46.2676432 | 37.1220767 | 37.1148921 | 0 |
| 1.23E-05 | 3.55E-05 | 33.1168171 | 26.4704305 | 27.8415575 | 24.1246799 | 0 |
| 1.23E-05 | 3.56E-05 | 38.904222 | 67.8443807 | 85.3807764 | 45.4657428 | 0 |
| 1.23E-05 | 3.56E-05 | 18.0052598 | 44.2656779 | 38.9781805 | 39.898509 | 0 |
| 1.23E-05 | 3.57E-05 | 17.0406923 | 68.2892619 | 55.683115 | 37.1148921 | 0 |
| 1.23E-05 | 3.57E-05 | 72.6640841 | 214.655172 | 206.955577 | 194.853184 | 2.37621824 |
| 1.24E-05 | 3.58E-05 | 20.2559173 | 56.9447917 | 36.1940248 | 33.4034029 | 0 |
| 1.24E-05 | 3.60E-05 | 136.325538 | 37.5924601 | 21.3451941 | 47.3214874 | 0 |
| 1.25E-05 | 3.60E-05 | 97.4213163 | 47.8247274 | 116.00649 | 12.5262761 | 0 |
| 1.25E-05 | 3.61E-05 | 15.7546023 | 48.9369303 | 65.8916861 | 45.929679 | 0 |
| 1.26E-05 | 3.63E-05 | 34.4029071 | 47.8247274 | 20.4171422 | 30.619786 | 0 |
| 1.26E-05 | 3.63E-05 | 48.8714194 | 31.5865641 | 23.2012979 | 27.3722329 | 0 |
| 1.26E-05 | 3.64E-05 | 51.4435994 | 98.5411824 | 12.0646749 | 92.3232941 | 0 |
| 1.26E-05 | 3.64E-05 | 24.7572322 | 56.9447917 | 65.8916861 | 22.7328714 | 0 |
| 1.26E-05 | 3.64E-05 | 16.7191698 | 41.5963908 | 47.3306478 | 48.7132959 | 0 |
| 1.26E-05 | 3.64E-05 | 18.3267823 | 36.9251383 | 38.9781805 | 52.4247851 | 0 |
| 1.26E-05 | 3.64E-05 | 72.3425616 | 264.481864 | 245.005706 | 370.684985 | 0 |
| 1.26E-05 | 3.65E-05 | 81.023669 | 47.3798462 | 19.4890903 | 32.9394667 | 0 |
| 1.26E-05 | 3.66E-05 | 34.7244296 | 93.86993 | 49.1867516 | 77.4773373 | 2.37621824 |
| 1.27E-05 | 3.66E-05 | 19.6128723 | 37.8149007 | 31.5537652 | 45.0018067 | 0 |
| 1.27E-05 | 3.66E-05 | 14.7900348 | 49.8266927 | 56.6111669 | 58.9198912 | 0 |
| 1.27E-05 | 3.66E-05 | 45.6561944 | 38.4822225 | 17.6329864 | 38.0427644 | 0 |
| 1.27E-05 | 3.66E-05 | 16.7191698 | 77.4093262 | 47.3306478 | 42.6821259 | 0 |
| 1.27E-05 | 3.66E-05 | 13.8254673 | 56.2774699 | 70.5319457 | 57.0641466 | 0 |
| 1.27E-05 | 3.67E-05 | 23.7926647 | 36.035376 | 24.1293498 | 33.867339 | 0 |
| 1.27E-05 | 3.67E-05 | 29.5800696 | 44.7105591 | 19.4890903 | 51.9608489 | 0 |
| 1.27E-05 | 3.67E-05 | 20.2559173 | 37.5924601 | 36.1940248 | 30.1558498 | 0 |

| | | | | | | |
|---|---|---|---|---|---|---|
| 1.27E-05 | 3.67E-05 | 23.1496197 | 55.8325887 | 38.9781805 | 25.9804245 | 0 |
| 1.27E-05 | 3.68E-05 | 38.904222 | 109.440771 | 103.013763 | 14.3820207 | 0 |
| 1.28E-05 | 3.69E-05 | 14.1469898 | 116.33643 | 44.546492 | 76.0855288 | 0 |
| 1.28E-05 | 3.69E-05 | 13.8254673 | 16.4606038 | 12.9927268 | 16.2377653 | 64.1578926 |
| 1.28E-05 | 3.69E-05 | 18.6483048 | 53.8306234 | 37.1220767 | 37.1148921 | 0 |
| 1.28E-05 | 3.70E-05 | 13.1824223 | 113.222262 | 54.7550631 | 65.4149973 | 0 |
| 1.28E-05 | 3.70E-05 | 26.6863672 | 20.9094157 | 135.49558 | 85.828188 | 0 |
| 1.28E-05 | 3.70E-05 | 54.3373018 | 176.172949 | 208.811681 | 179.543291 | 2.37621824 |
| 1.28E-05 | 3.70E-05 | 22.8280972 | 33.1436483 | 27.8415575 | 52.4247851 | 0 |
| 1.28E-05 | 3.70E-05 | 18.0052598 | 38.0373413 | 54.7550631 | 38.9706367 | 0 |
| 1.28E-05 | 3.70E-05 | 46.9422844 | 62.7282471 | 83.5246725 | 49.177232 | 0 |
| 1.28E-05 | 3.70E-05 | 8.6811074 | 96.7616577 | 111.36623 | 92.3232941 | 0 |
| 1.29E-05 | 3.71E-05 | 15.4330798 | 59.3916382 | 43.6184401 | 57.9920189 | 0 |
| 1.29E-05 | 3.71E-05 | 47.5853294 | 36.4802572 | 44.546492 | 33.867339 | 97.424948 |
| 1.29E-05 | 3.71E-05 | 36.6535646 | 45.1554403 | 89.092984 | 80.2609542 | 2.37621824 |
| 1.29E-05 | 3.72E-05 | 18.6483048 | 54.2755046 | 31.5537652 | 51.9608489 | 0 |
| 1.29E-05 | 3.73E-05 | 28.6155022 | 21.3542969 | 32.4818171 | 34.3312752 | 0 |
| 1.30E-05 | 3.74E-05 | 151.115573 | 259.365731 | 200.459214 | 244.958288 | 23.7621824 |
| 1.30E-05 | 3.74E-05 | 41.7979245 | 281.60979 | 341.523105 | 290.887967 | 14.2573095 |
| 1.30E-05 | 3.76E-05 | 17.6837373 | 34.2558512 | 64.0355823 | 51.9608489 | 0 |
| 1.31E-05 | 3.76E-05 | 81.023669 | 190.854028 | 172.617657 | 170.728504 | 0 |
| 1.31E-05 | 3.78E-05 | 17.3622148 | 44.2656779 | 38.0501286 | 46.3936151 | 0 |
| 1.31E-05 | 3.78E-05 | 18.6483048 | 62.7282471 | 30.6257133 | 62.6313804 | 0 |
| 1.31E-05 | 3.78E-05 | 22.1850522 | 32.6987671 | 38.0501286 | 82.5806349 | 0 |
| 1.31E-05 | 3.78E-05 | 30.2231146 | 52.4959798 | 22.273246 | 30.619786 | 0 |
| 1.31E-05 | 3.78E-05 | 22.8280972 | 47.8247274 | 69.6038938 | 25.5164883 | 0 |
| 1.31E-05 | 3.78E-05 | 0.64304499 | 0.22244059 | 0.92805192 | 1.39180845 | 7.12865473 |
| 1.31E-05 | 3.79E-05 | 30.5446371 | 25.3582275 | 23.2012979 | 32.0115944 | 0 |
| 1.32E-05 | 3.81E-05 | 23.4711422 | 36.4802572 | 45.4745439 | 24.1246799 | 0 |
| 1.32E-05 | 3.81E-05 | 12.8608998 | 63.6180094 | 69.6038938 | 61.239572 | 0 |
| 1.33E-05 | 3.83E-05 | 81.3451915 | 40.929069 | 14.8488307 | 65.4149973 | 0 |
| 1.33E-05 | 3.84E-05 | 74.2716966 | 27.5826335 | 31.5537652 | 27.8361691 | 0 |
| 1.34E-05 | 3.85E-05 | 21.5420072 | 90.5333211 | 57.5392189 | 27.3722329 | 0 |
| 1.34E-05 | 3.86E-05 | 39.8687895 | 16.905485 | 106.72597 | 55.208402 | 0 |
| 1.35E-05 | 3.87E-05 | 17.6837373 | 41.5963908 | 38.9781805 | 39.898509 | 0 |
| 1.35E-05 | 3.88E-05 | 16.3976473 | 51.6062174 | 61.2514265 | 38.5067006 | 0 |
| 1.35E-05 | 3.88E-05 | 18.0052598 | 37.5924601 | 63.1075304 | 39.898509 | 0 |
| 1.35E-05 | 3.89E-05 | 47.5853294 | 32.9212077 | 24.1293498 | 24.588616 | 0 |
| 1.35E-05 | 3.89E-05 | 30.2231146 | 127.458459 | 128.999216 | 128.046378 | 7.12865473 |
| 1.35E-05 | 3.89E-05 | 3.21522496 | 161.714311 | 205.099474 | 227.328714 | 0 |
| 1.35E-05 | 3.90E-05 | 71.3779941 | 145.921029 | 74.2441534 | 28.3001052 | 2.37621824 |
| 1.36E-05 | 3.90E-05 | 17.6837373 | 34.7007324 | 51.0428554 | 45.0018067 | 0 |
| 1.36E-05 | 3.91E-05 | 23.4711422 | 53.6081828 | 24.1293498 | 40.3624452 | 0 |

| | | | | | | |
|---|---|---|---|---|---|---|
| 1.36E-05 | 3.92E-05 | 25.7217997 | 46.934965 | 25.0574018 | 95.106911 | 0 |
| 1.36E-05 | 3.92E-05 | 38.5826995 | 58.946757 | 20.4171422 | 29.6919137 | 0 |
| 1.36E-05 | 3.92E-05 | 33.4383396 | 20.2420939 | 12.9927268 | 16.2377653 | 61.7816743 |
| 1.37E-05 | 3.93E-05 | 169.763878 | 85.1947469 | 53.8270112 | 88.6118049 | 9.50487297 |
| 1.37E-05 | 3.93E-05 | 29.9015921 | 126.346257 | 108.582074 | 90.4675495 | 4.75243649 |
| 1.37E-05 | 3.94E-05 | 17.6837373 | 47.3798462 | 37.1220767 | 39.898509 | 0 |
| 1.37E-05 | 3.94E-05 | 20.2559173 | 42.2637126 | 29.6976613 | 34.3312752 | 0 |
| 1.37E-05 | 3.95E-05 | 31.8307271 | 104.102197 | 56.6111669 | 75.1576565 | 0 |
| 1.37E-05 | 3.95E-05 | 38.261177 | 24.9133464 | 64.9636342 | 24.588616 | 0 |
| 1.38E-05 | 3.95E-05 | 12.5393773 | 60.9487223 | 60.3233746 | 95.5708472 | 0 |
| 1.38E-05 | 3.96E-05 | 19.2913498 | 51.6062174 | 32.4818171 | 37.1148921 | 0 |
| 1.38E-05 | 3.96E-05 | 20.5774398 | 26.6928711 | 59.3953227 | 65.8789335 | 0 |
| 1.38E-05 | 3.96E-05 | 18.0052598 | 86.7518311 | 99.3015551 | 29.6919137 | 0 |
| 1.38E-05 | 3.96E-05 | 18.9698273 | 43.8207967 | 30.6257133 | 45.4657428 | 0 |
| 1.38E-05 | 3.96E-05 | 111.568306 | 150.814722 | 154.98467 | 70.518295 | 7.12865473 |
| 1.38E-05 | 3.96E-05 | 62.3753642 | 36.2578166 | 35.2659728 | 37.5788283 | 192.473678 |
| 1.38E-05 | 3.96E-05 | 17.0406923 | 45.3778809 | 51.0428554 | 37.1148921 | 0 |
| 1.38E-05 | 3.97E-05 | 18.9698273 | 59.1691976 | 30.6257133 | 47.3214874 | 0 |
| 1.38E-05 | 3.97E-05 | 80.380624 | 34.2558512 | 22.273246 | 33.867339 | 0 |
| 1.38E-05 | 3.97E-05 | 55.3018693 | 13.7913167 | 54.7550631 | 69.5904227 | 0 |
| 1.38E-05 | 3.97E-05 | 18.0052598 | 66.7321777 | 31.5537652 | 59.3838274 | 0 |
| 1.39E-05 | 3.98E-05 | 18.3267823 | 35.5904948 | 71.4599976 | 42.6821259 | 0 |
| 1.39E-05 | 3.98E-05 | 18.6483048 | 37.5924601 | 33.409869 | 41.7542536 | 0 |
| 1.39E-05 | 3.98E-05 | 23.4711422 | 42.9310343 | 23.2012979 | 38.9706367 | 0 |
| 1.39E-05 | 3.98E-05 | 17.0406923 | 33.1436483 | 75.1722053 | 68.6625504 | 0 |
| 1.39E-05 | 3.98E-05 | 15.1115573 | 70.5136678 | 41.7623363 | 59.8477635 | 0 |
| 1.39E-05 | 3.99E-05 | 20.5774398 | 34.0334106 | 34.3379209 | 28.7640414 | 0 |
| 1.39E-05 | 4.00E-05 | 16.0761248 | 10.4547078 | 10.2085711 | 11.5984038 | 59.4054561 |
| 1.39E-05 | 4.00E-05 | 35.0459521 | 36.9251383 | 23.2012979 | 106.241379 | 0 |
| 1.39E-05 | 4.00E-05 | 22.8280972 | 28.917277 | 57.5392189 | 31.5476583 | 0 |
| 1.40E-05 | 4.01E-05 | 12.8608998 | 55.3877075 | 68.6758419 | 69.1264865 | 0 |
| 1.40E-05 | 4.01E-05 | 36.6535646 | 27.8050741 | 20.4171422 | 46.8575513 | 0 |
| 1.40E-05 | 4.03E-05 | 29.9015921 | 105.436841 | 84.4527244 | 84.9003157 | 2.37621824 |
| 1.41E-05 | 4.03E-05 | 46.2992394 | 119.228158 | 130.85532 | 129.438186 | 11.8810912 |
| 1.41E-05 | 4.03E-05 | 57.2310043 | 182.178845 | 140.135839 | 147.995632 | 14.2573095 |
| 1.41E-05 | 4.05E-05 | 36.9750871 | 60.2814006 | 52.8989593 | 49.6411682 | 0 |
| 1.41E-05 | 4.06E-05 | 85.524984 | 40.7066284 | 28.7696094 | 23.1968076 | 0 |
| 1.42E-05 | 4.06E-05 | 14.1469898 | 51.1613363 | 54.7550631 | 62.1674443 | 0 |
| 1.42E-05 | 4.07E-05 | 26.6863672 | 28.4723958 | 22.273246 | 36.1870198 | 0 |
| 1.42E-05 | 4.08E-05 | 15.7546023 | 47.6022868 | 42.6903882 | 57.0641466 | 0 |
| 1.42E-05 | 4.08E-05 | 22.1850522 | 48.047168 | 25.0574018 | 65.4149973 | 0 |
| 1.43E-05 | 4.10E-05 | 15.1115573 | 49.6042521 | 45.4745439 | 68.1986142 | 0 |
| 1.43E-05 | 4.10E-05 | 51.4435994 | 68.2892619 | 118.790645 | 41.2903175 | 2.37621824 |

| | | | | | | |
|---|---|---|---|---|---|---|
| 1.43E-05 | 4.11E-05 | 18.6483048 | 30.9192423 | 53.8270112 | 44.0739344 | 0 |
| 1.43E-05 | 4.11E-05 | 19.6128723 | 46.934965 | 35.2659728 | 106.705315 | 0 |
| 1.44E-05 | 4.11E-05 | 15.7546023 | 60.2814006 | 38.0501286 | 64.487125 | 0 |
| 1.44E-05 | 4.12E-05 | 93.8845689 | 218.214221 | 154.98467 | 179.079354 | 19.0097459 |
| 1.44E-05 | 4.12E-05 | 612.821878 | 687.786312 | 489.08336 | 209.69914 | 33.2670554 |
| 1.44E-05 | 4.12E-05 | 17.0406923 | 77.4093262 | 38.9781805 | 45.929679 | 0 |
| 1.44E-05 | 4.12E-05 | 225.38727 | 103.657316 | 26.9135056 | 59.8477635 | 2.37621824 |
| 1.44E-05 | 4.13E-05 | 23.4711422 | 46.934965 | 23.2012979 | 60.7756358 | 0 |
| 1.45E-05 | 4.14E-05 | 18.9698273 | 31.5865641 | 48.2586997 | 37.1148921 | 0 |
| 1.45E-05 | 4.16E-05 | 20.2559173 | 36.7026978 | 29.6976613 | 54.7444659 | 0 |
| 1.46E-05 | 4.17E-05 | 529.226029 | 801.231014 | 885.361529 | 938.078898 | 239.998043 |
| 1.46E-05 | 4.18E-05 | 443.701045 | 430.644987 | 263.566744 | 340.065199 | 35.6432737 |
| 1.47E-05 | 4.20E-05 | 14.1469898 | 56.2774699 | 52.8989593 | 53.8165936 | 0 |
| 1.47E-05 | 4.20E-05 | 48.8714194 | 34.4782918 | 22.273246 | 25.5164883 | 0 |
| 1.47E-05 | 4.21E-05 | 16.0761248 | 47.8247274 | 45.4745439 | 42.2181898 | 0 |
| 1.47E-05 | 4.21E-05 | 22.8280972 | 26.4704305 | 28.7696094 | 28.7640414 | 0 |
| 1.47E-05 | 4.22E-05 | 17.0406923 | 47.8247274 | 55.683115 | 34.7952113 | 0 |
| 1.47E-05 | 4.22E-05 | 43.7270595 | 39.3719849 | 81.6685687 | 57.0641466 | 0 |
| 1.48E-05 | 4.23E-05 | 21.2204847 | 27.8050741 | 35.2659728 | 29.6919137 | 0 |
| 1.48E-05 | 4.24E-05 | 21.8635297 | 31.3641235 | 61.2514265 | 31.0837221 | 0 |
| 1.48E-05 | 4.24E-05 | 29.5800696 | 14.4586385 | 24.1293498 | 21.341063 | 57.0292378 |
| 1.48E-05 | 4.25E-05 | 24.7572322 | 106.104163 | 20.4171422 | 87.2199964 | 0 |
| 1.49E-05 | 4.25E-05 | 0 | 0 | 0 | 0 | 289.898626 |
| 1.49E-05 | 4.26E-05 | 20.2559173 | 27.3601929 | 40.8342843 | 50.1051043 | 0 |
| 1.49E-05 | 4.26E-05 | 232.460765 | 31.3641235 | 25.9854537 | 51.9608489 | 0 |
| 1.49E-05 | 4.26E-05 | 46.9422844 | 303.186527 | 408.342843 | 457.904981 | 16.6335277 |
| 1.49E-05 | 4.26E-05 | 50.1575094 | 203.310701 | 193.034799 | 213.874566 | 14.2573095 |
| 1.49E-05 | 4.27E-05 | 4.17979245 | 3.78149007 | 1.85610383 | 3.71148921 | 30.8908372 |
| 1.49E-05 | 4.27E-05 | 24.1141872 | 182.846167 | 154.056618 | 197.6368 | 7.12865473 |
| 1.50E-05 | 4.28E-05 | 51.7651219 | 66.5097371 | 70.5319457 | 40.8263813 | 4.75243649 |
| 1.50E-05 | 4.29E-05 | 295.800696 | 1047.91763 | 961.461786 | 1100.92049 | 61.7816743 |
| 1.51E-05 | 4.31E-05 | 21.5420072 | 25.5806681 | 34.3379209 | 40.8263813 | 0 |
| 1.51E-05 | 4.31E-05 | 84.5604165 | 82.970341 | 10.2085711 | 67.7346781 | 0 |
| 1.51E-05 | 4.32E-05 | 10.6102424 | 78.0766479 | 73.3161014 | 77.4773373 | 0 |
| 1.51E-05 | 4.32E-05 | 39.8687895 | 15.7932821 | 52.8989593 | 51.0329766 | 0 |
| 1.52E-05 | 4.33E-05 | 14.4685123 | 91.4230835 | 40.8342843 | 66.8068058 | 0 |
| 1.52E-05 | 4.34E-05 | 374.573708 | 696.016614 | 605.08985 | 652.758165 | 40.3957101 |
| 1.52E-05 | 4.35E-05 | 14.4685123 | 49.6042521 | 48.2586997 | 65.8789335 | 0 |
| 1.53E-05 | 4.36E-05 | 18.3267823 | 51.828658 | 32.4818171 | 39.4345729 | 0 |
| 1.53E-05 | 4.36E-05 | 15.1115573 | 35.8129354 | 103.013763 | 79.3330819 | 0 |
| 1.53E-05 | 4.37E-05 | 19.2913498 | 43.5983561 | 38.0501286 | 29.2279775 | 0 |
| 1.53E-05 | 4.37E-05 | 30.2231146 | 32.0314453 | 19.4890903 | 33.4034029 | 0 |
| 1.53E-05 | 4.38E-05 | 23.1496197 | 28.0275146 | 25.0574018 | 30.619786 | 0 |

| | | | | | | |
|---|---|---|---|---|---|---|
| 1.53E-05 | 4.38E-05 | 25.7217997 | 40.929069 | 21.3451941 | 32.9394667 | 0 |
| 1.54E-05 | 4.39E-05 | 19.2913498 | 36.035376 | 29.6976613 | 43.1460621 | 0 |
| 1.54E-05 | 4.40E-05 | 73.3071291 | 32.9212077 | 22.273246 | 31.5476583 | 0 |
| 1.55E-05 | 4.41E-05 | 19.6128723 | 61.8384847 | 36.1940248 | 32.4755306 | 0 |
| 1.55E-05 | 4.42E-05 | 33.1168171 | 68.7341431 | 59.3953227 | 16.7017014 | 0 |
| 1.55E-05 | 4.43E-05 | 32.4737721 | 36.035376 | 19.4890903 | 29.2279775 | 0 |
| 1.55E-05 | 4.44E-05 | 26.0433222 | 24.9133464 | 58.4672708 | 30.1558498 | 0 |
| 1.55E-05 | 4.44E-05 | 20.2559173 | 41.3739502 | 37.1220767 | 26.9082968 | 0 |
| 1.56E-05 | 4.44E-05 | 21.5420072 | 28.917277 | 26.9135056 | 33.867339 | 0 |
| 1.56E-05 | 4.44E-05 | 80.0591015 | 26.0255493 | 30.6257133 | 29.6919137 | 0 |
| 1.56E-05 | 4.45E-05 | 16.0761248 | 108.77345 | 153.128566 | 135.00542 | 2.37621824 |
| 1.56E-05 | 4.45E-05 | 252.073637 | 310.082186 | 230.156875 | 274.650202 | 78.415202 |
| 1.56E-05 | 4.45E-05 | 9.00262989 | 78.9664103 | 84.4527244 | 111.808612 | 0 |
| 1.56E-05 | 4.46E-05 | 45.6561944 | 46.934965 | 55.683115 | 69.5904227 | 7.12865473 |
| 1.57E-05 | 4.48E-05 | 49.1929419 | 20.9094157 | 157.768826 | 35.2591475 | 0 |
| 1.57E-05 | 4.48E-05 | 23.4711422 | 32.6987671 | 29.6976613 | 23.6607437 | 0 |
| 1.57E-05 | 4.49E-05 | 104.816334 | 43.3759155 | 23.2012979 | 28.7640414 | 0 |
| 1.57E-05 | 4.49E-05 | 15.7546023 | 37.8149007 | 59.3953227 | 51.4969128 | 0 |
| 1.58E-05 | 4.50E-05 | 10.2887199 | 55.1652669 | 100.229607 | 112.272549 | 0 |
| 1.58E-05 | 4.50E-05 | 12.5393773 | 57.834554 | 59.3953227 | 70.9822312 | 0 |
| 1.59E-05 | 4.52E-05 | 35.3674746 | 33.5885295 | 38.9781805 | 18.0935099 | 0 |
| 1.59E-05 | 4.52E-05 | 15.1115573 | 55.1652669 | 47.3306478 | 44.0739344 | 0 |
| 1.59E-05 | 4.53E-05 | 31.1876821 | 22.6889404 | 39.9062324 | 24.1246799 | 0 |
| 1.59E-05 | 4.54E-05 | 25.4002772 | 61.6160441 | 100.229607 | 132.685739 | 2.37621824 |
| 1.59E-05 | 4.54E-05 | 23.1496197 | 77.6317668 | 33.409869 | 30.619786 | 0 |
| 1.59E-05 | 4.54E-05 | 23.1496197 | 50.9388957 | 22.273246 | 45.0018067 | 0 |
| 1.60E-05 | 4.55E-05 | 24.1141872 | 21.5767375 | 56.6111669 | 48.7132959 | 0 |
| 1.60E-05 | 4.55E-05 | 33.7598621 | 29.1397176 | 50.1148035 | 19.9492545 | 0 |
| 1.60E-05 | 4.56E-05 | 18.3267823 | 47.3798462 | 29.6976613 | 46.8575513 | 0 |
| 1.60E-05 | 4.56E-05 | 21.5420072 | 43.5983561 | 26.9135056 | 30.619786 | 0 |
| 1.60E-05 | 4.57E-05 | 11.8963324 | 72.7380737 | 69.6038938 | 57.5280828 | 0 |
| 1.61E-05 | 4.58E-05 | 17.6837373 | 35.1456136 | 36.1940248 | 44.5378705 | 0 |
| 1.61E-05 | 4.59E-05 | 424.731217 | 481.583883 | 465.882062 | 528.887212 | 38.0194919 |
| 1.61E-05 | 4.59E-05 | 24.7572322 | 28.917277 | 71.4599976 | 30.1558498 | 0 |
| 1.62E-05 | 4.60E-05 | 26.0433222 | 27.8050741 | 29.6976613 | 22.2689353 | 0 |
| 1.62E-05 | 4.61E-05 | 22.1850522 | 41.5963908 | 23.2012979 | 39.4345729 | 0 |
| 1.62E-05 | 4.61E-05 | 42.762492 | 65.6199748 | 14.8488307 | 46.3936151 | 0 |
| 1.63E-05 | 4.63E-05 | 15.1115573 | 51.3837769 | 51.9709073 | 41.7542536 | 0 |
| 1.63E-05 | 4.65E-05 | 7.71653991 | 199.08433 | 114.150386 | 66.3428696 | 0 |
| 1.63E-05 | 4.65E-05 | 30.5446371 | 20.2420939 | 36.1940248 | 28.3001052 | 0 |
| 1.64E-05 | 4.66E-05 | 142.755988 | 35.5904948 | 20.4171422 | 46.3936151 | 0 |
| 1.64E-05 | 4.66E-05 | 46.2992394 | 79.4112915 | 12.0646749 | 92.3232941 | 0 |
| 1.64E-05 | 4.66E-05 | 176.51585 | 195.30284 | 135.49558 | 177.22361 | 61.7816743 |

| | | | | | | |
|---|---|---|---|---|---|---|
| 1.64E-05 | 4.66E-05 | 10.6102424 | 50.7164551 | 141.991943 | 94.6429749 | 0 |
| 1.64E-05 | 4.67E-05 | 14.7900348 | 41.8188314 | 60.3233746 | 51.9608489 | 0 |
| 1.65E-05 | 4.70E-05 | 12.5393773 | 50.0491333 | 76.1002572 | 66.8068058 | 0 |
| 1.66E-05 | 4.71E-05 | 46.6207619 | 191.966231 | 148.488307 | 157.274355 | 0 |
| 1.66E-05 | 4.71E-05 | 14.4685123 | 40.0393066 | 62.1794784 | 66.3428696 | 0 |
| 1.66E-05 | 4.72E-05 | 18.9698273 | 29.3621582 | 54.7550631 | 39.4345729 | 0 |
| 1.66E-05 | 4.72E-05 | 288.405679 | 385.934428 | 374.004923 | 430.532748 | 30.8908372 |
| 1.67E-05 | 4.74E-05 | 42.119447 | 110.330534 | 56.6111669 | 14.8459568 | 0 |
| 1.67E-05 | 4.74E-05 | 7.71653991 | 100.765588 | 93.7332436 | 102.993826 | 0 |
| 1.67E-05 | 4.74E-05 | 30.2231146 | 72.2931925 | 17.6329864 | 53.3526574 | 0 |
| 1.67E-05 | 4.76E-05 | 413.47793 | 348.564408 | 336.882846 | 355.839028 | 33.2670554 |
| 1.68E-05 | 4.77E-05 | 27.9724572 | 24.9133464 | 186.538435 | 53.3526574 | 0 |
| 1.68E-05 | 4.78E-05 | 163.654951 | 29.8070394 | 22.273246 | 53.8165936 | 0 |
| 1.69E-05 | 4.80E-05 | 30.2231146 | 58.2794352 | 66.819738 | 66.3428696 | 2.37621824 |
| 1.69E-05 | 4.80E-05 | 24.1141872 | 52.7184204 | 25.0574018 | 30.1558498 | 0 |
| 1.69E-05 | 4.80E-05 | 10.6102424 | 54.7203857 | 99.3015551 | 88.6118049 | 0 |
| 1.69E-05 | 4.80E-05 | 39.2257445 | 65.8424154 | 103.941815 | 59.3838274 | 4.75243649 |
| 1.69E-05 | 4.80E-05 | 91.6339114 | 54.053064 | 76.1002572 | 56.1362743 | 201.978551 |
| 1.69E-05 | 4.81E-05 | 18.9698273 | 40.929069 | 39.9062324 | 118.303719 | 0 |
| 1.70E-05 | 4.83E-05 | 20.5774398 | 27.3601929 | 38.9781805 | 29.6919137 | 0 |
| 1.71E-05 | 4.85E-05 | 21.2204847 | 29.5845988 | 53.8270112 | 29.6919137 | 0 |
| 1.71E-05 | 4.86E-05 | 24.1141872 | 119.228158 | 84.4527244 | 71.4461673 | 2.37621824 |
| 1.71E-05 | 4.86E-05 | 18.3267823 | 28.6948364 | 49.1867516 | 57.9920189 | 0 |
| 1.71E-05 | 4.86E-05 | 33.7598621 | 47.3798462 | 16.7049345 | 40.3624452 | 0 |
| 1.71E-05 | 4.86E-05 | 27.0078897 | 45.3778809 | 22.273246 | 27.8361691 | 0 |
| 1.72E-05 | 4.87E-05 | 16.3976473 | 41.3739502 | 62.1794784 | 37.5788283 | 0 |
| 1.72E-05 | 4.88E-05 | 17.3622148 | 34.2558512 | 66.819738 | 41.7542536 | 0 |
| 1.72E-05 | 4.89E-05 | 14.1469898 | 54.2755046 | 64.9636342 | 43.1460621 | 0 |
| 1.72E-05 | 4.89E-05 | 161.082771 | 252.914954 | 177.257916 | 203.204034 | 23.7621824 |
| 1.72E-05 | 4.89E-05 | 15.4330798 | 44.2656779 | 44.546492 | 44.0739344 | 0 |
| 1.73E-05 | 4.90E-05 | 47.2638069 | 121.230123 | 74.2441534 | 86.2921241 | 2.37621824 |
| 1.73E-05 | 4.91E-05 | 66.2336342 | 41.8188314 | 57.5392189 | 49.6411682 | 7.12865473 |
| 1.73E-05 | 4.91E-05 | 17.6837373 | 35.3680542 | 36.1940248 | 35.7230837 | 0 |
| 1.73E-05 | 4.91E-05 | 98.3858838 | 140.360014 | 132.711424 | 170.264568 | 19.0097459 |
| 1.73E-05 | 4.91E-05 | 49.5144644 | 133.909237 | 119.718697 | 109.488932 | 0 |
| 1.73E-05 | 4.91E-05 | 28.9370247 | 92.0904053 | 52.8989593 | 101.602017 | 0 |
| 1.74E-05 | 4.93E-05 | 10.6102424 | 147.255672 | 40.8342843 | 141.964462 | 0 |
| 1.74E-05 | 4.93E-05 | 25.7217997 | 27.1377523 | 28.7696094 | 22.2689353 | 0 |
| 1.74E-05 | 4.94E-05 | 17.0406923 | 38.4822225 | 35.2659728 | 41.7542536 | 0 |
| 1.74E-05 | 4.95E-05 | 34.4029071 | 30.02948 | 25.0574018 | 21.341063 | 0 |
| 1.75E-05 | 4.96E-05 | 21.2204847 | 58.7243164 | 24.1293498 | 45.4657428 | 0 |
| 1.75E-05 | 4.96E-05 | 20.8989622 | 48.047168 | 32.4818171 | 26.9082968 | 0 |
| 1.75E-05 | 4.97E-05 | 17.6837373 | 40.2617472 | 31.5537652 | 40.8263813 | 0 |

| | | | | | | |
|---|---|---|---|---|---|---|
| 1.75E-05 | 4.97E-05 | 15.4330798 | 56.7223511 | 46.4025958 | 38.9706367 | 0 |
| 1.75E-05 | 4.97E-05 | 35.0459521 | 63.1731283 | 18.5610383 | 33.4034029 | 0 |
| 1.76E-05 | 4.99E-05 | 14.7900348 | 82.0805786 | 42.6903882 | 49.177232 | 0 |
| 1.76E-05 | 4.99E-05 | 10.6102424 | 9.34250488 | 11.136623 | 7.88691457 | 52.2768014 |
| 1.76E-05 | 4.99E-05 | 18.3267823 | 37.3700195 | 30.6257133 | 36.1870198 | 0 |
| 1.77E-05 | 5.01E-05 | 37.618132 | 23.1338216 | 43.6184401 | 21.8049991 | 0 |
| 1.77E-05 | 5.02E-05 | 55.3018693 | 11.3444702 | 79.8124649 | 73.3019119 | 0 |
| 1.77E-05 | 5.02E-05 | 43.405537 | 102.545113 | 90.0210359 | 82.1166988 | 0 |
| 1.77E-05 | 5.02E-05 | 14.4685123 | 45.822762 | 47.3306478 | 52.8887212 | 0 |
| 1.77E-05 | 5.02E-05 | 4.82283744 | 3.78149007 | 2.78415575 | 2.78361691 | 66.5341108 |
| 1.78E-05 | 5.03E-05 | 8.0380624 | 75.6298014 | 90.0210359 | 142.892335 | 0 |
| 1.78E-05 | 5.04E-05 | 17.6837373 | 44.7105591 | 33.409869 | 35.2591475 | 0 |
| 1.78E-05 | 5.05E-05 | 32.4737721 | 40.4841878 | 25.9854537 | 20.4131907 | 0 |
| 1.78E-05 | 5.05E-05 | 212.52637 | 179.731999 | 177.257916 | 65.4149973 | 16.6335277 |
| 1.79E-05 | 5.06E-05 | 39.2257445 | 10.899589 | 13.9207788 | 12.9902122 | 102.177384 |
| 1.79E-05 | 5.07E-05 | 18.3267823 | 88.0864746 | 29.6976613 | 53.3526574 | 0 |
| 1.79E-05 | 5.07E-05 | 16.7191698 | 57.3896728 | 79.8124649 | 31.5476583 | 0 |
| 1.80E-05 | 5.09E-05 | 45.0131495 | 28.6948364 | 20.4171422 | 28.7640414 | 0 |
| 1.80E-05 | 5.09E-05 | 466.207619 | 1103.75022 | 1010.64854 | 1071.22857 | 0 |
| 1.80E-05 | 5.09E-05 | 22.5065747 | 27.5826335 | 35.2659728 | 24.1246799 | 0 |
| 1.80E-05 | 5.09E-05 | 15.1115573 | 69.1790242 | 34.3379209 | 75.6215927 | 0 |
| 1.80E-05 | 5.10E-05 | 60.4462293 | 76.7420044 | 131.783372 | 43.1460621 | 7.12865473 |
| 1.80E-05 | 5.10E-05 | 20.5774398 | 30.9192423 | 26.9135056 | 29.6919137 | 0 |
| 1.81E-05 | 5.11E-05 | 11.8963324 | 69.6239054 | 56.6111669 | 62.6313804 | 0 |
| 1.81E-05 | 5.11E-05 | 17.0406923 | 53.8306234 | 30.6257133 | 53.3526574 | 0 |
| 1.81E-05 | 5.11E-05 | 16.3976473 | 44.2656779 | 34.3379209 | 50.1051043 | 0 |
| 1.81E-05 | 5.12E-05 | 15.7546023 | 41.8188314 | 38.9781805 | 57.0641466 | 0 |
| 1.81E-05 | 5.12E-05 | 78.7730116 | 219.548865 | 231.084927 | 214.802438 | 19.0097459 |
| 1.81E-05 | 5.13E-05 | 16.7191698 | 80.3010539 | 55.683115 | 32.0115944 | 0 |
| 1.82E-05 | 5.15E-05 | 13.8254673 | 50.0491333 | 47.3306478 | 57.9920189 | 0 |
| 1.83E-05 | 5.17E-05 | 88.7402089 | 37.8149007 | 18.5610383 | 36.1870198 | 0 |
| 1.83E-05 | 5.18E-05 | 21.8635297 | 39.5944255 | 25.0574018 | 28.7640414 | 0 |
| 1.83E-05 | 5.18E-05 | 376.824365 | 621.499015 | 490.939464 | 615.643273 | 35.6432737 |
| 1.83E-05 | 5.19E-05 | 22.8280972 | 28.2499552 | 23.2012979 | 31.5476583 | 0 |
| 1.84E-05 | 5.19E-05 | 33.1168171 | 35.3680542 | 19.4890903 | 26.4443606 | 0 |
| 1.84E-05 | 5.19E-05 | 24.1141872 | 20.2420939 | 60.3233746 | 60.3116997 | 0 |
| 1.84E-05 | 5.20E-05 | 15.4330798 | 42.7085937 | 40.8342843 | 46.8575513 | 0 |
| 1.84E-05 | 5.21E-05 | 19.6128723 | 57.3896728 | 37.1220767 | 28.3001052 | 0 |
| 1.84E-05 | 5.21E-05 | 13.5039448 | 6.22833659 | 9.28051917 | 7.88691457 | 64.1578926 |
| 1.85E-05 | 5.22E-05 | 56.9094818 | 46.4900838 | 20.4171422 | 24.1246799 | 0 |
| 1.85E-05 | 5.23E-05 | 0 | 7.11809896 | 0 | 0 | 178.216368 |
| 1.85E-05 | 5.23E-05 | 17.3622148 | 66.064856 | 33.409869 | 41.2903175 | 0 |
| 1.85E-05 | 5.23E-05 | 16.3976473 | 35.8129354 | 45.4745439 | 39.4345729 | 0 |

| | | | | | | |
|---|---|---|---|---|---|---|
| 1.85E-05 | 5.23E-05 | 24.7572322 | 32.6987671 | 20.4171422 | 37.5788283 | 0 |
| 1.86E-05 | 5.24E-05 | 37.9396545 | 46.0452026 | 65.8916861 | 15.309893 | 0 |
| 1.86E-05 | 5.25E-05 | 110.282216 | 593.24906 | 409.270895 | 464.400087 | 26.1384007 |
| 1.86E-05 | 5.25E-05 | 15.7546023 | 43.8207967 | 40.8342843 | 40.3624452 | 0 |
| 1.86E-05 | 5.26E-05 | 27.9724572 | 28.2499552 | 64.9636342 | 24.1246799 | 0 |
| 1.86E-05 | 5.26E-05 | 19.6128723 | 51.3837769 | 25.9854537 | 40.8263813 | 0 |
| 1.87E-05 | 5.28E-05 | 0 | 0 | 0 | 0 | 7.12865473 |
| 1.87E-05 | 5.29E-05 | 35.0459521 | 83.1927816 | 35.2659728 | 19.4853184 | 0 |
| 1.87E-05 | 5.29E-05 | 13.5039448 | 66.5097371 | 41.7623363 | 64.0231889 | 0 |
| 1.88E-05 | 5.30E-05 | 183.589345 | 144.808826 | 64.9636342 | 94.1790387 | 14.2573095 |
| 1.88E-05 | 5.30E-05 | 117.034189 | 248.688582 | 273.775316 | 279.753499 | 0 |
| 1.88E-05 | 5.32E-05 | 33.7598621 | 70.2912272 | 14.8488307 | 106.705315 | 0 |
| 1.89E-05 | 5.34E-05 | 193.556543 | 611.489189 | 599.521538 | 501.978916 | 33.2670554 |
| 1.89E-05 | 5.34E-05 | 18.0052598 | 33.5885295 | 52.8989593 | 32.9394667 | 0 |
| 1.90E-05 | 5.35E-05 | 16.3976473 | 32.4763265 | 48.2586997 | 49.177232 | 0 |
| 1.90E-05 | 5.35E-05 | 16.7191698 | 41.8188314 | 43.6184401 | 32.9394667 | 0 |
| 1.90E-05 | 5.35E-05 | 422.802082 | 891.097013 | 771.211143 | 860.601561 | 23.7621824 |
| 1.90E-05 | 5.36E-05 | 23.1496197 | 24.023584 | 25.9854537 | 35.2591475 | 0 |
| 1.90E-05 | 5.37E-05 | 39.547267 | 103.879757 | 71.4599976 | 128.510314 | 0 |
| 1.91E-05 | 5.39E-05 | 18.9698273 | 46.4900838 | 25.9854537 | 50.1051043 | 0 |
| 1.91E-05 | 5.39E-05 | 16.3976473 | 33.5885295 | 51.0428554 | 42.2181898 | 0 |
| 1.91E-05 | 5.39E-05 | 2490.83478 | 13273.0302 | 23110.3488 | 11609.5383 | 413.461974 |
| 1.92E-05 | 5.40E-05 | 23.4711422 | 24.023584 | 25.9854537 | 27.3722329 | 0 |
| 1.92E-05 | 5.40E-05 | 25.4002772 | 22.6889404 | 30.6257133 | 53.8165936 | 0 |
| 1.92E-05 | 5.40E-05 | 16.7191698 | 45.6003214 | 32.4818171 | 42.6821259 | 0 |
| 1.92E-05 | 5.42E-05 | 41.7979245 | 56.2774699 | 15.7768826 | 35.2591475 | 0 |
| 1.92E-05 | 5.42E-05 | 90.0262989 | 10.899589 | 4.64025958 | 10.6705315 | 527.52045 |
| 1.93E-05 | 5.43E-05 | 16.7191698 | 39.816866 | 33.409869 | 43.1460621 | 0 |
| 1.93E-05 | 5.43E-05 | 124.750728 | 338.777022 | 302.544925 | 301.558498 | 2.37621824 |
| 1.93E-05 | 5.44E-05 | 19.2913498 | 39.5944255 | 38.0501286 | 25.9804245 | 0 |
| 1.93E-05 | 5.44E-05 | 10.6102424 | 70.0687866 | 67.7477899 | 68.1986142 | 0 |
| 1.93E-05 | 5.44E-05 | 15.7546023 | 43.5983561 | 36.1940248 | 48.7132959 | 0 |
| 1.93E-05 | 5.44E-05 | 11.5748099 | 57.1672323 | 66.819738 | 64.0231889 | 0 |
| 1.93E-05 | 5.44E-05 | 31.1876821 | 76.7420044 | 87.2368802 | 83.9724434 | 4.75243649 |
| 1.94E-05 | 5.45E-05 | 18.9698273 | 36.4802572 | 26.9135056 | 42.6821259 | 0 |
| 1.94E-05 | 5.46E-05 | 14.4685123 | 48.4920492 | 41.7623363 | 51.9608489 | 0 |
| 1.94E-05 | 5.46E-05 | 16.3976473 | 44.4881185 | 33.409869 | 44.0739344 | 0 |
| 1.95E-05 | 5.49E-05 | 86.168029 | 127.458459 | 93.7332436 | 96.4987195 | 16.6335277 |
| 1.95E-05 | 5.50E-05 | 20.5774398 | 37.8149007 | 40.8342843 | 23.6607437 | 0 |
| 1.95E-05 | 5.50E-05 | 20.2559173 | 26.6928711 | 32.4818171 | 28.3001052 | 0 |
| 1.95E-05 | 5.50E-05 | 9.64567488 | 172.6139 | 58.4672708 | 77.9412734 | 0 |
| 1.95E-05 | 5.50E-05 | 16.0761248 | 57.834554 | 53.8270112 | 32.0115944 | 0 |
| 1.96E-05 | 5.51E-05 | 15.4330798 | 38.4822225 | 47.3306478 | 42.2181898 | 0 |

| | | | | | | |
|---|---|---|---|---|---|---|
| 1.96E-05 | 5.51E-05 | 16.7191698 | 44.9329997 | 31.5537652 | 46.3936151 | 0 |
| 1.96E-05 | 5.52E-05 | 15.7546023 | 35.5904948 | 47.3306478 | 72.8379758 | 0 |
| 1.97E-05 | 5.53E-05 | 48.8714194 | 60.2814006 | 66.819738 | 32.0115944 | 0 |
| 1.97E-05 | 5.55E-05 | 22.8280972 | 24.6909058 | 25.0574018 | 28.7640414 | 0 |
| 1.97E-05 | 5.55E-05 | 36.6535646 | 93.86993 | 59.3953227 | 66.3428696 | 4.75243649 |
| 1.98E-05 | 5.56E-05 | 20.5774398 | 33.3660889 | 24.1293498 | 34.3312752 | 0 |
| 1.98E-05 | 5.56E-05 | 24.4357097 | 67.8443807 | 88.1649321 | 81.1888265 | 2.37621824 |
| 1.98E-05 | 5.58E-05 | 20.2559173 | 29.3621582 | 26.9135056 | 42.2181898 | 0 |
| 1.99E-05 | 5.59E-05 | 11.8963324 | 53.1633016 | 77.956361 | 57.5280828 | 0 |
| 1.99E-05 | 5.59E-05 | 13.5039448 | 85.1947469 | 36.1940248 | 82.5806349 | 0 |
| 1.99E-05 | 5.59E-05 | 43.405537 | 32.4763265 | 25.0574018 | 20.4131907 | 0 |
| 1.99E-05 | 5.61E-05 | 205.452875 | 334.773092 | 283.983887 | 307.125732 | 0 |
| 2.00E-05 | 5.61E-05 | 56.2664368 | 132.352152 | 106.72597 | 149.851377 | 16.6335277 |
| 2.00E-05 | 5.61E-05 | 0 | 0 | 0 | 0 | 114.058476 |
| 2.00E-05 | 5.61E-05 | 24.4357097 | 42.041272 | 19.4890903 | 43.6099982 | 0 |
| 2.00E-05 | 5.61E-05 | 56.2664368 | 32.6987671 | 24.1293498 | 22.7328714 | 0 |
| 2.00E-05 | 5.62E-05 | 54.3373018 | 23.3562622 | 28.7696094 | 25.5164883 | 0 |
| 2.00E-05 | 5.62E-05 | 17.0406923 | 33.8109701 | 35.2659728 | 48.7132959 | 0 |
| 2.00E-05 | 5.62E-05 | 35.3674746 | 68.5117025 | 13.9207788 | 87.6839326 | 0 |
| 2.00E-05 | 5.62E-05 | 24.4357097 | 24.2460246 | 23.2012979 | 30.1558498 | 0 |
| 2.00E-05 | 5.63E-05 | 16.3976473 | 42.4861532 | 41.7623363 | 111.344676 | 0 |
| 2.00E-05 | 5.63E-05 | 16.3976473 | 40.0393066 | 38.9781805 | 82.1166988 | 0 |
| 2.01E-05 | 5.64E-05 | 21.2204847 | 48.4920492 | 23.2012979 | 70.0543588 | 0 |
| 2.01E-05 | 5.64E-05 | 17.0406923 | 40.4841878 | 34.3379209 | 34.3312752 | 0 |
| 2.01E-05 | 5.64E-05 | 32.7952946 | 102.100232 | 63.1075304 | 92.7872303 | 4.75243649 |
| 2.01E-05 | 5.64E-05 | 20.8989622 | 60.9487223 | 31.5537652 | 28.7640414 | 0 |
| 2.02E-05 | 5.66E-05 | 1.92913498 | 4.22637126 | 2.78415575 | 8.35085072 | 14.2573095 |
| 2.02E-05 | 5.66E-05 | 13.1824223 | 37.1475789 | 93.7332436 | 78.4052096 | 0 |
| 2.02E-05 | 5.66E-05 | 30.8661596 | 23.3562622 | 21.3451941 | 30.619786 | 0 |
| 2.02E-05 | 5.68E-05 | 39.8687895 | 19.1298909 | 14.8488307 | 27.8361691 | 182.968805 |
| 2.03E-05 | 5.69E-05 | 27.0078897 | 58.7243164 | 17.6329864 | 76.0855288 | 0 |
| 2.03E-05 | 5.69E-05 | 15.7546023 | 99.2085042 | 32.4818171 | 61.239572 | 0 |
| 2.03E-05 | 5.69E-05 | 12.2178549 | 73.1829549 | 46.4025958 | 65.4149973 | 0 |
| 2.03E-05 | 5.71E-05 | 29.5800696 | 27.8050741 | 19.4890903 | 28.7640414 | 0 |
| 2.04E-05 | 5.72E-05 | 17.6837373 | 20.2420939 | 18.5610383 | 25.5164883 | 47.5243649 |
| 2.04E-05 | 5.74E-05 | 10.2887199 | 66.7321777 | 68.6758419 | 72.8379758 | 0 |
| 2.04E-05 | 5.74E-05 | 29.2585471 | 68.2892619 | 58.4672708 | 58.4559551 | 2.37621824 |
| 2.05E-05 | 5.74E-05 | 0.3215225 | 0 | 0 | 0.46393615 | 7.12865473 |
| 2.05E-05 | 5.74E-05 | 30.8661596 | 28.917277 | 18.5610383 | 31.0837221 | 0 |
| 2.05E-05 | 5.74E-05 | 14.1469898 | 51.1613363 | 54.7550631 | 40.3624452 | 0 |
| 2.05E-05 | 5.76E-05 | 148.864916 | 125.901375 | 105.797919 | 114.128293 | 0 |
| 2.05E-05 | 5.76E-05 | 10.6102424 | 55.3877075 | 77.0283091 | 76.549465 | 0 |
| 2.06E-05 | 5.77E-05 | 26.3648447 | 26.4704305 | 28.7696094 | 20.8771268 | 0 |

| | | | | | | |
|---|---|---|---|---|---|---|
| 2.06E-05 | 5.77E-05 | 19.2913498 | 28.4723958 | 29.6976613 | 30.619786 | 0 |
| 2.06E-05 | 5.78E-05 | 36.9750871 | 117.448633 | 37.1220767 | 78.8691457 | 0 |
| 2.06E-05 | 5.78E-05 | 44.691627 | 41.1515096 | 43.6184401 | 29.2279775 | 249.502916 |
| 2.06E-05 | 5.79E-05 | 46.2992394 | 45.822762 | 52.8989593 | 49.6411682 | 7.12865473 |
| 2.07E-05 | 5.80E-05 | 15.1115573 | 37.3700195 | 69.6038938 | 45.4657428 | 0 |
| 2.08E-05 | 5.83E-05 | 36.9750871 | 32.6987671 | 17.6329864 | 29.6919137 | 0 |
| 2.08E-05 | 5.83E-05 | 31.1876821 | 46.0452026 | 79.8124649 | 17.1656376 | 0 |
| 2.08E-05 | 5.84E-05 | 19.9343948 | 82.0805786 | 22.273246 | 80.2609542 | 0 |
| 2.09E-05 | 5.85E-05 | 89.3832539 | 22.0216187 | 27.8415575 | 37.5788283 | 0 |
| 2.09E-05 | 5.86E-05 | 20.8989622 | 22.911381 | 35.2659728 | 37.1148921 | 0 |
| 2.09E-05 | 5.86E-05 | 33.7598621 | 25.1357869 | 20.4171422 | 27.8361691 | 0 |
| 2.09E-05 | 5.86E-05 | 11.5748099 | 90.9782023 | 55.683115 | 55.208402 | 0 |
| 2.09E-05 | 5.86E-05 | 74.5932191 | 262.035018 | 227.37272 | 222.689353 | 19.0097459 |
| 2.09E-05 | 5.87E-05 | 103.851766 | 41.8188314 | 22.273246 | 26.9082968 | 0 |
| 2.10E-05 | 5.88E-05 | 15.4330798 | 36.4802572 | 41.7623363 | 54.2805297 | 0 |
| 2.11E-05 | 5.92E-05 | 21.2204847 | 23.3562622 | 36.1940248 | 29.6919137 | 0 |
| 2.12E-05 | 5.92E-05 | 21.5420072 | 22.2440592 | 34.3379209 | 38.9706367 | 0 |
| 2.12E-05 | 5.93E-05 | 21.5420072 | 23.8011434 | 35.2659728 | 54.2805297 | 0 |
| 2.12E-05 | 5.93E-05 | 21.5420072 | 23.8011434 | 35.2659728 | 27.3722329 | 0 |
| 2.12E-05 | 5.94E-05 | 35.3674746 | 14.0137573 | 66.819738 | 83.5085072 | 0 |
| 2.12E-05 | 5.94E-05 | 330.846649 | 722.487044 | 691.398678 | 701.935397 | 26.1384007 |
| 2.13E-05 | 5.95E-05 | 243.39253 | 418.410754 | 431.544141 | 443.059025 | 102.177384 |
| 2.13E-05 | 5.97E-05 | 26.3648447 | 55.1652669 | 24.1293498 | 25.9804245 | 0 |
| 2.13E-05 | 5.97E-05 | 21.5420072 | 22.6889404 | 33.409869 | 30.1558498 | 0 |
| 2.13E-05 | 5.97E-05 | 151.758618 | 166.163123 | 124.358957 | 159.1301 | 21.3859642 |
| 2.14E-05 | 5.99E-05 | 35.3674746 | 14.9035197 | 59.3953227 | 50.1051043 | 0 |
| 2.14E-05 | 5.99E-05 | 191.94893 | 397.94622 | 265.422848 | 321.507753 | 28.5146189 |
| 2.14E-05 | 6.00E-05 | 136.004016 | 81.6356974 | 87.2368802 | 87.6839326 | 301.779717 |
| 2.15E-05 | 6.00E-05 | 20.8989622 | 36.2578166 | 25.0574018 | 61.239572 | 0 |
| 2.15E-05 | 6.00E-05 | 16.0761248 | 36.035376 | 51.0428554 | 36.1870198 | 0 |
| 2.15E-05 | 6.00E-05 | 15.7546023 | 45.6003214 | 35.2659728 | 78.4052096 | 0 |
| 2.15E-05 | 6.01E-05 | 22.1850522 | 31.5865641 | 179.11402 | 44.5378705 | 0 |
| 2.16E-05 | 6.03E-05 | 20.5774398 | 54.9428263 | 22.273246 | 65.4149973 | 0 |
| 2.16E-05 | 6.04E-05 | 11.2532874 | 44.7105591 | 91.8771398 | 79.797018 | 0 |
| 2.17E-05 | 6.06E-05 | 30.8661596 | 22.2440592 | 25.9854537 | 61.239572 | 0 |
| 2.17E-05 | 6.06E-05 | 178.444985 | 544.757011 | 348.019469 | 366.50956 | 28.5146189 |
| 2.17E-05 | 6.06E-05 | 57.8740493 | 117.893514 | 125.287009 | 123.870952 | 21.3859642 |
| 2.17E-05 | 6.07E-05 | 19.6128723 | 35.1456136 | 29.6976613 | 25.9804245 | 0 |
| 2.17E-05 | 6.08E-05 | 40.190312 | 206.869751 | 267.278952 | 209.235204 | 0 |
| 2.17E-05 | 6.08E-05 | 24.4357097 | 40.2617472 | 36.1940248 | 19.4853184 | 0 |
| 2.18E-05 | 6.08E-05 | 17.6837373 | 41.8188314 | 27.8415575 | 39.898509 | 0 |
| 2.18E-05 | 6.09E-05 | 34.7244296 | 24.9133464 | 26.9135056 | 20.8771268 | 0 |
| 2.18E-05 | 6.09E-05 | 55.6233918 | 123.454529 | 111.36623 | 127.118505 | 14.2573095 |

| | | | | | | |
|---|---|---|---|---|---|---|
| 2.18E-05 | 6.09E-05 | 18.0052598 | 34.923173 | 39.9062324 | 27.8361691 | 0 |
| 2.18E-05 | 6.10E-05 | 21.2204847 | 50.2715739 | 23.2012979 | 35.2591475 | 0 |
| 2.19E-05 | 6.12E-05 | 436.949072 | 1922.99892 | 1087.67685 | 1363.04441 | 16.6335277 |
| 2.19E-05 | 6.13E-05 | 33.1168171 | 34.7007324 | 17.6329864 | 28.7640414 | 0 |
| 2.20E-05 | 6.14E-05 | 18.0052598 | 32.6987671 | 28.7696094 | 38.9706367 | 0 |
| 2.20E-05 | 6.14E-05 | 10.9317649 | 84.3049845 | 55.683115 | 60.7756358 | 0 |
| 2.20E-05 | 6.15E-05 | 18.3267823 | 42.4861532 | 27.8415575 | 33.4034029 | 0 |
| 2.20E-05 | 6.16E-05 | 18.3267823 | 38.0373413 | 26.9135056 | 50.1051043 | 0 |
| 2.21E-05 | 6.16E-05 | 24.7572322 | 36.035376 | 20.4171422 | 57.9920189 | 0 |
| 2.21E-05 | 6.16E-05 | 10.9317649 | 63.1731283 | 90.0210359 | 52.8887212 | 0 |
| 2.21E-05 | 6.17E-05 | 16.3976473 | 36.035376 | 54.7550631 | 34.3312752 | 0 |
| 2.22E-05 | 6.19E-05 | 32.7952946 | 20.2420939 | 25.0574018 | 28.3001052 | 0 |
| 2.22E-05 | 6.19E-05 | 608.642085 | 1112.4254 | 1100.66957 | 1153.34527 | 0 |
| 2.22E-05 | 6.20E-05 | 16.7191698 | 34.923173 | 33.409869 | 37.1148921 | 0 |
| 2.23E-05 | 6.22E-05 | 14.1469898 | 37.8149007 | 50.1148035 | 63.5592527 | 0 |
| 2.24E-05 | 6.25E-05 | 28.9370247 | 26.4704305 | 19.4890903 | 28.3001052 | 0 |
| 2.24E-05 | 6.25E-05 | 11.8963324 | 62.9506877 | 56.6111669 | 51.9608489 | 0 |
| 2.24E-05 | 6.25E-05 | 12.8608998 | 32.4763265 | 116.00649 | 102.529889 | 0 |
| 2.24E-05 | 6.25E-05 | 14.7900348 | 46.2676432 | 36.1940248 | 52.4247851 | 0 |
| 2.24E-05 | 6.26E-05 | 11.8963324 | 64.7302124 | 50.1148035 | 58.4559551 | 0 |
| 2.25E-05 | 6.27E-05 | 48.5498969 | 40.0393066 | 52.8989593 | 52.8887212 | 7.12865473 |
| 2.25E-05 | 6.27E-05 | 15.7546023 | 35.1456136 | 38.9781805 | 41.7542536 | 0 |
| 2.25E-05 | 6.28E-05 | 18.6483048 | 31.3641235 | 45.4745439 | 27.8361691 | 0 |
| 2.25E-05 | 6.28E-05 | 39.547267 | 29.8070394 | 190.250643 | 124.334889 | 0 |
| 2.25E-05 | 6.28E-05 | 31.8307271 | 19.1298909 | 16.7049345 | 19.4853184 | 54.6530196 |
| 2.26E-05 | 6.30E-05 | 133.110313 | 466.235482 | 434.328297 | 488.988703 | 28.5146189 |
| 2.27E-05 | 6.32E-05 | 42.4409695 | 94.3148112 | 45.4745439 | 65.8789335 | 4.75243649 |
| 2.28E-05 | 6.35E-05 | 20.5774398 | 30.4743612 | 24.1293498 | 28.3001052 | 0 |
| 2.28E-05 | 6.35E-05 | 25.7217997 | 22.2440592 | 23.2012979 | 28.3001052 | 0 |
| 2.28E-05 | 6.35E-05 | 12.8608998 | 56.2774699 | 41.7623363 | 79.797018 | 0 |
| 2.28E-05 | 6.36E-05 | 34.0813846 | 85.4171875 | 13.9207788 | 62.6313804 | 0 |
| 2.28E-05 | 6.37E-05 | 42.762492 | 39.1495443 | 14.8488307 | 37.5788283 | 0 |
| 2.29E-05 | 6.37E-05 | 16.7191698 | 33.1436483 | 33.409869 | 44.5378705 | 0 |
| 2.29E-05 | 6.37E-05 | 0 | 0 | 0 | 0 | 121.18713 |
| 2.29E-05 | 6.37E-05 | 45.0131495 | 69.846346 | 12.0646749 | 56.1362743 | 0 |
| 2.29E-05 | 6.37E-05 | 20.2559173 | 36.9251383 | 62.1794784 | 25.0525522 | 0 |
| 2.29E-05 | 6.37E-05 | 12.2178549 | 83.6376628 | 42.6903882 | 63.0953166 | 0 |
| 2.29E-05 | 6.37E-05 | 15.7546023 | 53.6081828 | 31.5537652 | 46.8575513 | 0 |
| 2.29E-05 | 6.37E-05 | 47.2638069 | 34.4782918 | 62.1794784 | 14.8459568 | 0 |
| 2.29E-05 | 6.38E-05 | 138.897718 | 300.2948 | 289.552198 | 353.519347 | 26.1384007 |
| 2.30E-05 | 6.40E-05 | 15.4330798 | 47.6022868 | 32.4818171 | 51.9608489 | 0 |
| 2.30E-05 | 6.41E-05 | 14.1469898 | 44.0432373 | 45.4745439 | 43.1460621 | 0 |
| 2.30E-05 | 6.41E-05 | 10.9317649 | 53.8306234 | 69.6038938 | 65.4149973 | 0 |

| | | | | | | |
|---|---|---|---|---|---|---|
| 2.30E-05 | 6.41E-05 | 11.5748099 | 74.0727173 | 50.1148035 | 58.4559551 | 0 |
| 2.30E-05 | 6.41E-05 | 55.3018693 | 37.1475789 | 17.6329864 | 28.3001052 | 0 |
| 2.31E-05 | 6.42E-05 | 41.476402 | 25.3582275 | 22.273246 | 24.588616 | 0 |
| 2.31E-05 | 6.42E-05 | 9.32415239 | 56.0550293 | 90.0210359 | 88.6118049 | 0 |
| 2.31E-05 | 6.43E-05 | 5.78740493 | 4.22637126 | 1.85610383 | 2.78361691 | 30.8908372 |
| 2.32E-05 | 6.45E-05 | 70.0919042 | 23.5787028 | 48.2586997 | 19.9492545 | 0 |
| 2.33E-05 | 6.47E-05 | 16.7191698 | 52.4959798 | 33.409869 | 34.3312752 | 0 |
| 2.33E-05 | 6.49E-05 | 22.8280972 | 28.4723958 | 21.3451941 | 28.3001052 | 0 |
| 2.33E-05 | 6.50E-05 | 19.9343948 | 70.0687866 | 172.617657 | 23.6607437 | 0 |
| 2.33E-05 | 6.50E-05 | 29.5800696 | 16.0157227 | 54.7550631 | 53.3526574 | 0 |
| 2.34E-05 | 6.50E-05 | 86.4895515 | 52.940861 | 36.1940248 | 65.8789335 | 4.75243649 |
| 2.34E-05 | 6.50E-05 | 40.190312 | 23.8011434 | 24.1293498 | 23.6607437 | 0 |
| 2.34E-05 | 6.51E-05 | 36.6535646 | 50.9388957 | 13.9207788 | 50.1051043 | 0 |
| 2.34E-05 | 6.52E-05 | 60.1247068 | 339.889225 | 334.09869 | 346.560305 | 26.1384007 |
| 2.35E-05 | 6.52E-05 | 13.5039448 | 44.7105591 | 47.3306478 | 48.7132959 | 0 |
| 2.35E-05 | 6.52E-05 | 55.3018693 | 38.7046631 | 16.7049345 | 29.6919137 | 0 |
| 2.35E-05 | 6.54E-05 | 16.7191698 | 83.4152222 | 26.9135056 | 63.5592527 | 0 |
| 2.35E-05 | 6.54E-05 | 17.6837373 | 32.6987671 | 28.7696094 | 35.7230837 | 0 |
| 2.35E-05 | 6.54E-05 | 56.5879593 | 30.02948 | 19.4890903 | 28.7640414 | 0 |
| 2.35E-05 | 6.54E-05 | 95.8137038 | 101.21047 | 131.783372 | 102.065953 | 28.5146189 |
| 2.36E-05 | 6.55E-05 | 7.07349491 | 59.1691976 | 178.185968 | 109.952868 | 0 |
| 2.36E-05 | 6.56E-05 | 24.1141872 | 23.3562622 | 23.2012979 | 26.4443606 | 0 |
| 2.36E-05 | 6.57E-05 | 13.5039448 | 65.6199748 | 50.1148035 | 39.898509 | 0 |
| 2.36E-05 | 6.57E-05 | 17.3622148 | 43.5983561 | 38.0501286 | 27.8361691 | 0 |
| 2.37E-05 | 6.58E-05 | 65.9121117 | 42.9310343 | 34.3379209 | 19.9492545 | 106.929821 |
| 2.37E-05 | 6.58E-05 | 10.2887199 | 70.958549 | 79.8124649 | 53.8165936 | 0 |
| 2.37E-05 | 6.59E-05 | 7.07349491 | 110.775415 | 113.222334 | 69.5904227 | 0 |
| 2.37E-05 | 6.59E-05 | 102.887199 | 225.999642 | 187.466487 | 186.502333 | 2.37621824 |
| 2.37E-05 | 6.60E-05 | 7.71653991 | 72.7380737 | 107.654022 | 85.828188 | 0 |
| 2.38E-05 | 6.60E-05 | 28.2939797 | 42.9310343 | 21.3451941 | 23.6607437 | 0 |
| 2.38E-05 | 6.60E-05 | 16.7191698 | 30.02948 | 70.5319457 | 44.0739344 | 0 |
| 2.38E-05 | 6.61E-05 | 12.8608998 | 39.3719849 | 67.7477899 | 60.7756358 | 0 |
| 2.38E-05 | 6.61E-05 | 55.3018693 | 32.0314453 | 16.7049345 | 34.3312752 | 0 |
| 2.38E-05 | 6.61E-05 | 12.8608998 | 67.1770589 | 61.2514265 | 39.4345729 | 0 |
| 2.38E-05 | 6.61E-05 | 27.3294122 | 71.1809896 | 29.6976613 | 22.7328714 | 0 |
| 2.38E-05 | 6.62E-05 | 197.414813 | 565.221545 | 451.961284 | 522.392106 | 33.2670554 |
| 2.38E-05 | 6.62E-05 | 10.2887199 | 67.1770589 | 55.683115 | 79.3330819 | 0 |
| 2.39E-05 | 6.62E-05 | 21.2204847 | 85.6396281 | 78.8844129 | 96.9626556 | 0 |
| 2.39E-05 | 6.63E-05 | 17.3622148 | 45.822762 | 26.9135056 | 42.6821259 | 0 |
| 2.39E-05 | 6.64E-05 | 47.5853294 | 60.05896 | 36.1940248 | 54.7444659 | 2.37621824 |
| 2.40E-05 | 6.66E-05 | 18.3267823 | 25.5806681 | 57.5392189 | 42.2181898 | 0 |
| 2.40E-05 | 6.67E-05 | 15.7546023 | 35.5904948 | 35.2659728 | 44.0739344 | 0 |
| 2.40E-05 | 6.67E-05 | 22.5065747 | 35.3680542 | 45.4745439 | 20.4131907 | 0 |

| | | | | | | |
|---|---|---|---|---|---|---|
| 2.41E-05 | 6.69E-05 | 10.6102424 | 59.1691976 | 72.3880495 | 58.9198912 | 0 |
| 2.41E-05 | 6.70E-05 | 15.1115573 | 31.8090047 | 68.6758419 | 51.9608489 | 0 |
| 2.42E-05 | 6.70E-05 | 16.3976473 | 33.8109701 | 35.2659728 | 34.7952113 | 0 |
| 2.42E-05 | 6.71E-05 | 26.3648447 | 27.3601929 | 58.4672708 | 22.2689353 | 0 |
| 2.42E-05 | 6.71E-05 | 72.3425616 | 236.45435 | 231.084927 | 239.391054 | 21.3859642 |
| 2.42E-05 | 6.72E-05 | 20.5774398 | 32.6987671 | 33.409869 | 22.2689353 | 0 |
| 2.43E-05 | 6.73E-05 | 24.7572322 | 18.6850098 | 50.1148035 | 39.4345729 | 0 |
| 2.43E-05 | 6.73E-05 | 25.4002772 | 35.3680542 | 47.3306478 | 18.5574461 | 0 |
| 2.43E-05 | 6.74E-05 | 11.2532874 | 65.8424154 | 49.1867516 | 66.3428696 | 0 |
| 2.43E-05 | 6.75E-05 | 268.149762 | 681.113094 | 646.852186 | 731.627311 | 0 |
| 2.44E-05 | 6.75E-05 | 27.3294122 | 21.5767375 | 22.273246 | 30.1558498 | 0 |
| 2.44E-05 | 6.76E-05 | 27.3294122 | 30.4743612 | 34.3379209 | 18.0935099 | 0 |
| 2.44E-05 | 6.76E-05 | 34.0813846 | 46.7125244 | 15.7768826 | 33.867339 | 0 |
| 2.44E-05 | 6.77E-05 | 17.6837373 | 42.2637126 | 61.2514265 | 26.9082968 | 0 |
| 2.45E-05 | 6.79E-05 | 11.5748099 | 53.6081828 | 53.8270112 | 62.6313804 | 0 |
| 2.45E-05 | 6.79E-05 | 263.005402 | 359.908879 | 269.135056 | 315.012647 | 30.8908372 |
| 2.47E-05 | 6.84E-05 | 19.9343948 | 22.6889404 | 36.1940248 | 34.3312752 | 0 |
| 2.47E-05 | 6.85E-05 | 11.8963324 | 56.0550293 | 49.1867516 | 58.9198912 | 0 |
| 2.47E-05 | 6.85E-05 | 18.0052598 | 32.9212077 | 120.646749 | 43.1460621 | 0 |
| 2.47E-05 | 6.85E-05 | 98.0643613 | 24.2460246 | 25.9854537 | 34.3312752 | 0 |
| 2.47E-05 | 6.85E-05 | 12.5393773 | 52.2735392 | 43.6184401 | 67.2707419 | 0 |
| 2.48E-05 | 6.86E-05 | 22.1850522 | 77.8542074 | 103.941815 | 68.1986142 | 2.37621824 |
| 2.48E-05 | 6.87E-05 | 18.6483048 | 30.6968018 | 25.9854537 | 32.4755306 | 0 |
| 2.48E-05 | 6.87E-05 | 68.8058142 | 41.1515096 | 107.654022 | 11.5984038 | 0 |
| 2.48E-05 | 6.88E-05 | 27.9724572 | 55.1652669 | 17.6329864 | 35.7230837 | 0 |
| 2.49E-05 | 6.88E-05 | 153.044708 | 171.501697 | 82.5966206 | 113.664357 | 0 |
| 2.49E-05 | 6.90E-05 | 0 | 0 | 0 | 0 | 38.0194919 |
| 2.50E-05 | 6.93E-05 | 110.282216 | 28.0275146 | 26.9135056 | 29.6919137 | 0 |
| 2.50E-05 | 6.93E-05 | 11.8963324 | 44.0432373 | 77.956361 | 57.9920189 | 0 |
| 2.51E-05 | 6.95E-05 | 15.4330798 | 36.9251383 | 35.2659728 | 41.7542536 | 0 |
| 2.51E-05 | 6.95E-05 | 22.8280972 | 37.1475789 | 32.4818171 | 19.9492545 | 0 |
| 2.51E-05 | 6.95E-05 | 53.6942569 | 29.3621582 | 15.7768826 | 44.0739344 | 0 |
| 2.51E-05 | 6.96E-05 | 28.6155022 | 32.0314453 | 19.4890903 | 24.1246799 | 0 |
| 2.53E-05 | 6.99E-05 | 23.4711422 | 31.1416829 | 25.0574018 | 21.341063 | 0 |
| 2.53E-05 | 7.00E-05 | 95.4921813 | 264.926746 | 199.531162 | 241.710735 | 21.3859642 |
| 2.53E-05 | 7.00E-05 | 13.8254673 | 60.9487223 | 36.1940248 | 51.0329766 | 0 |
| 2.54E-05 | 7.01E-05 | 21.2204847 | 78.9664103 | 77.0283091 | 90.4675495 | 2.37621824 |
| 2.54E-05 | 7.02E-05 | 15.7546023 | 28.4723958 | 56.6111669 | 63.0953166 | 0 |
| 2.54E-05 | 7.03E-05 | 22.1850522 | 20.6869751 | 38.0501286 | 30.619786 | 0 |
| 2.55E-05 | 7.06E-05 | 20.8989622 | 108.551009 | 139.207788 | 121.551272 | 7.12865473 |
| 2.55E-05 | 7.06E-05 | 78.7730116 | 38.9271037 | 41.7623363 | 44.0739344 | 0 |
| 2.56E-05 | 7.07E-05 | 13.5039448 | 35.3680542 | 74.2441534 | 57.0641466 | 0 |
| 2.56E-05 | 7.07E-05 | 49.1929419 | 24.4684652 | 22.273246 | 26.4443606 | 0 |

| | | | | | | |
|---|---|---|---|---|---|---|
| 2.56E-05 | 7.07E-05 | 25.0787547 | 69.4014648 | 19.4890903 | 39.4345729 | 0 |
| 2.56E-05 | 7.08E-05 | 16.7191698 | 32.2538859 | 34.3379209 | 32.0115944 | 0 |
| 2.57E-05 | 7.09E-05 | 141.469898 | 5.7834554 | 231.084927 | 55.208402 | 0 |
| 2.57E-05 | 7.10E-05 | 21.8635297 | 55.6101481 | 23.2012979 | 31.5476583 | 0 |
| 2.58E-05 | 7.12E-05 | 14.1469898 | 45.1554403 | 40.8342843 | 40.8263813 | 0 |
| 2.58E-05 | 7.13E-05 | 89.0617314 | 26.0255493 | 25.9854537 | 29.2279775 | 0 |
| 2.58E-05 | 7.14E-05 | 15.4330798 | 75.6298014 | 93.7332436 | 26.9082968 | 0 |
| 2.58E-05 | 7.14E-05 | 17.6837373 | 44.4881185 | 25.0574018 | 47.7854236 | 0 |
| 2.59E-05 | 7.14E-05 | 14.1469898 | 54.7203857 | 33.409869 | 73.3019119 | 0 |
| 2.60E-05 | 7.17E-05 | 12.8608998 | 60.05896 | 42.6903882 | 48.2493597 | 0 |
| 2.60E-05 | 7.17E-05 | 35.3674746 | 49.1593709 | 54.7550631 | 13.9180845 | 0 |
| 2.60E-05 | 7.17E-05 | 17.6837373 | 29.8070394 | 28.7696094 | 39.898509 | 0 |
| 2.60E-05 | 7.17E-05 | 195.8072 | 577.233337 | 446.392972 | 489.916576 | 33.2670554 |
| 2.60E-05 | 7.18E-05 | 23.7926647 | 23.5787028 | 27.8415575 | 21.8049991 | 0 |
| 2.60E-05 | 7.19E-05 | 45.9777169 | 39.1495443 | 42.6903882 | 38.9706367 | 2.37621824 |
| 2.60E-05 | 7.19E-05 | 210.275712 | 454.001249 | 458.457647 | 457.441045 | 33.2670554 |
| 2.61E-05 | 7.20E-05 | 152.080141 | 368.584062 | 446.392972 | 361.406262 | 0 |
| 2.61E-05 | 7.20E-05 | 14.7900348 | 35.3680542 | 40.8342843 | 44.0739344 | 0 |
| 2.61E-05 | 7.20E-05 | 25.0787547 | 22.911381 | 21.3451941 | 28.7640414 | 0 |
| 2.61E-05 | 7.21E-05 | 252.716682 | 329.212077 | 277.487523 | 325.683178 | 30.8908372 |
| 2.62E-05 | 7.22E-05 | 16.3976473 | 70.958549 | 318.321808 | 27.8361691 | 0 |
| 2.62E-05 | 7.24E-05 | 44.048582 | 51.1613363 | 69.6038938 | 54.7444659 | 9.50487297 |
| 2.63E-05 | 7.25E-05 | 8.0380624 | 92.5352865 | 83.5246725 | 67.7346781 | 0 |
| 2.63E-05 | 7.25E-05 | 208.989622 | 279.830265 | 282.127783 | 248.205841 | 28.5146189 |
| 2.64E-05 | 7.28E-05 | 3.85826995 | 99.4309448 | 170.761553 | 152.171058 | 0 |
| 2.64E-05 | 7.28E-05 | 25.0787547 | 30.2519206 | 73.3161014 | 23.1968076 | 0 |
| 2.64E-05 | 7.29E-05 | 20.8989622 | 49.8266927 | 21.3451941 | 37.1148921 | 0 |
| 2.64E-05 | 7.30E-05 | 24.4357097 | 77.8542074 | 59.3953227 | 73.7658481 | 2.37621824 |
| 2.65E-05 | 7.30E-05 | 47.5853294 | 163.716276 | 167.977397 | 178.615418 | 14.2573095 |
| 2.66E-05 | 7.33E-05 | 13.1824223 | 43.3759155 | 59.3953227 | 42.6821259 | 0 |
| 2.66E-05 | 7.33E-05 | 15.4330798 | 55.6101481 | 28.7696094 | 58.9198912 | 0 |
| 2.67E-05 | 7.36E-05 | 16.0761248 | 32.4763265 | 38.0501286 | 33.4034029 | 0 |
| 2.67E-05 | 7.37E-05 | 156.902978 | 714.701623 | 550.334787 | 619.818698 | 33.2670554 |
| 2.67E-05 | 7.37E-05 | 52.0866444 | 28.4723958 | 22.273246 | 23.1968076 | 0 |
| 2.68E-05 | 7.38E-05 | 49.1929419 | 76.7420044 | 100.229607 | 91.3954218 | 0 |
| 2.68E-05 | 7.38E-05 | 18.0052598 | 26.9153117 | 32.4818171 | 29.6919137 | 0 |
| 2.68E-05 | 7.38E-05 | 141.469898 | 739.170089 | 837.102829 | 864.31305 | 2.37621824 |
| 2.68E-05 | 7.38E-05 | 15.7546023 | 47.6022868 | 82.5966206 | 30.619786 | 0 |
| 2.68E-05 | 7.39E-05 | 162.690383 | 409.29069 | 289.552198 | 367.901368 | 28.5146189 |
| 2.68E-05 | 7.40E-05 | 15.1115573 | 35.8129354 | 36.1940248 | 42.2181898 | 0 |
| 2.69E-05 | 7.42E-05 | 21.8635297 | 21.5767375 | 35.2659728 | 26.4443606 | 0 |
| 2.69E-05 | 7.42E-05 | 27.0078897 | 24.023584 | 27.8415575 | 19.9492545 | 0 |
| 2.70E-05 | 7.43E-05 | 96.7782713 | 178.842236 | 182.826228 | 180.471163 | 2.37621824 |

| | | | | | | |
|---|---|---|---|---|---|---|
| 2.70E-05 | 7.44E-05 | 22.8280972 | 23.3562622 | 24.1293498 | 24.1246799 | 0 |
| 2.70E-05 | 7.44E-05 | 15.4330798 | 44.4881185 | 39.9062324 | 31.5476583 | 0 |
| 2.70E-05 | 7.45E-05 | 20.2559173 | 51.6062174 | 82.5966206 | 21.8049991 | 0 |
| 2.71E-05 | 7.46E-05 | 2174.77816 | 691.790242 | 528.989593 | 787.299649 | 0 |
| 2.72E-05 | 7.48E-05 | 26.0433222 | 40.7066284 | 17.6329864 | 32.0115944 | 0 |
| 2.72E-05 | 7.48E-05 | 36.6535646 | 53.8306234 | 38.0501286 | 46.8575513 | 0 |
| 2.72E-05 | 7.50E-05 | 112.211351 | 290.952295 | 241.293498 | 269.546904 | 23.7621824 |
| 2.73E-05 | 7.51E-05 | 10.9317649 | 8.00786133 | 13.9207788 | 5.56723382 | 45.1481466 |
| 2.73E-05 | 7.52E-05 | 49.1929419 | 28.0275146 | 17.6329864 | 31.0837221 | 0 |
| 2.74E-05 | 7.53E-05 | 16.3976473 | 35.3680542 | 30.6257133 | 33.867339 | 0 |
| 2.75E-05 | 7.58E-05 | 109.960694 | 211.763444 | 243.149602 | 251.91733 | 19.0097459 |
| 2.76E-05 | 7.59E-05 | 21.5420072 | 42.2637126 | 19.4890903 | 44.0739344 | 0 |
| 2.76E-05 | 7.59E-05 | 18.9698273 | 23.3562622 | 35.2659728 | 32.0115944 | 0 |
| 2.76E-05 | 7.61E-05 | 9.96719738 | 77.4093262 | 108.582074 | 45.0018067 | 0 |
| 2.77E-05 | 7.61E-05 | 13.1824223 | 40.929069 | 61.2514265 | 43.6099982 | 0 |
| 2.77E-05 | 7.62E-05 | 18.6483048 | 41.8188314 | 35.2659728 | 23.6607437 | 0 |
| 2.77E-05 | 7.63E-05 | 16.3976473 | 29.5845988 | 34.3379209 | 44.5378705 | 0 |
| 2.77E-05 | 7.63E-05 | 33.1168171 | 23.1338216 | 19.4890903 | 28.3001052 | 0 |
| 2.78E-05 | 7.64E-05 | 15.7546023 | 31.5865641 | 35.2659728 | 42.6821259 | 0 |
| 2.78E-05 | 7.65E-05 | 73.6286516 | 20.4645345 | 19.4890903 | 70.518295 | 0 |
| 2.78E-05 | 7.65E-05 | 12.8608998 | 40.7066284 | 76.1002572 | 47.3214874 | 0 |
| 2.78E-05 | 7.65E-05 | 11.5748099 | 72.9605143 | 46.4025958 | 52.8887212 | 0 |
| 2.78E-05 | 7.66E-05 | 31.5092046 | 74.5175985 | 64.9636342 | 76.549465 | 0 |
| 2.80E-05 | 7.71E-05 | 18.6483048 | 35.5904948 | 23.2012979 | 38.9706367 | 0 |
| 2.81E-05 | 7.72E-05 | 56.9094818 | 268.040914 | 264.494796 | 283.464988 | 0 |
| 2.81E-05 | 7.73E-05 | 8.3595849 | 126.346257 | 76.1002572 | 57.5280828 | 0 |
| 2.82E-05 | 7.74E-05 | 76.2008316 | 139.470251 | 129.927268 | 132.221803 | 0 |
| 2.82E-05 | 7.74E-05 | 27.9724572 | 24.6909058 | 43.6184401 | 19.4853184 | 0 |
| 2.82E-05 | 7.74E-05 | 32.7952946 | 27.3601929 | 18.5610383 | 25.0525522 | 0 |
| 2.82E-05 | 7.75E-05 | 12.5393773 | 49.3818115 | 42.6903882 | 54.2805297 | 0 |
| 2.82E-05 | 7.76E-05 | 11.8963324 | 44.4881185 | 90.9490879 | 51.0329766 | 0 |
| 2.83E-05 | 7.78E-05 | 12.8608998 | 37.1475789 | 75.1722053 | 53.3526574 | 0 |
| 2.83E-05 | 7.78E-05 | 16.3976473 | 40.2617472 | 32.4818171 | 30.1558498 | 0 |
| 2.83E-05 | 7.79E-05 | 25.4002772 | 71.4034302 | 65.8916861 | 56.6002105 | 2.37621824 |
| 2.83E-05 | 7.79E-05 | 17.6837373 | 40.2617472 | 32.4818171 | 25.9804245 | 0 |
| 2.85E-05 | 7.84E-05 | 19.9343948 | 62.9506877 | 25.0574018 | 33.4034029 | 0 |
| 2.85E-05 | 7.84E-05 | 16.0761248 | 31.8090047 | 54.7550631 | 33.867339 | 0 |
| 2.86E-05 | 7.85E-05 | 28.6155022 | 61.6160441 | 47.3306478 | 61.239572 | 0 |
| 2.86E-05 | 7.85E-05 | 39.8687895 | 36.2578166 | 18.5610383 | 36.1870198 | 83.1676385 |
| 2.87E-05 | 7.88E-05 | 10.6102424 | 73.1829549 | 66.819738 | 46.3936151 | 0 |
| 2.87E-05 | 7.88E-05 | 11.5748099 | 50.7164551 | 53.8270112 | 52.4247851 | 0 |
| 2.87E-05 | 7.89E-05 | 15.7546023 | 51.3837769 | 27.8415575 | 47.3214874 | 0 |
| 2.88E-05 | 7.90E-05 | 36.3320421 | 67.6219401 | 33.409869 | 65.8789335 | 2.37621824 |

| | | | | | | |
|---|---|---|---|---|---|---|
| 2.88E-05 | 7.91E-05 | 15.1115573 | 67.3994995 | 32.4818171 | 41.7542536 | 0 |
| 2.88E-05 | 7.92E-05 | 12.2178549 | 38.9271037 | 64.0355823 | 58.4559551 | 0 |
| 2.89E-05 | 7.94E-05 | 244.035575 | 448.662675 | 469.59427 | 521.464234 | 66.5341108 |
| 2.90E-05 | 7.95E-05 | 19.9343948 | 32.2538859 | 28.7696094 | 22.2689353 | 0 |
| 2.90E-05 | 7.95E-05 | 12.8608998 | 39.1495443 | 49.1867516 | 54.7444659 | 0 |
| 2.90E-05 | 7.95E-05 | 0 | 1975.4949 | 1069.11581 | 1095.35325 | 0 |
| 2.90E-05 | 7.97E-05 | 94.8491364 | 49.8266927 | 54.7550631 | 59.8477635 | 278.017534 |
| 2.91E-05 | 7.97E-05 | 9.96719738 | 107.438806 | 38.0501286 | 108.097123 | 0 |
| 2.91E-05 | 7.98E-05 | 11.2532874 | 62.5058065 | 45.4745439 | 60.3116997 | 0 |
| 2.91E-05 | 7.98E-05 | 12.5393773 | 43.5983561 | 52.8989593 | 46.3936151 | 0 |
| 2.91E-05 | 7.98E-05 | 56.5879593 | 98.9860636 | 110.438178 | 144.748079 | 14.2573095 |
| 2.91E-05 | 7.99E-05 | 13.5039448 | 41.3739502 | 48.2586997 | 39.898509 | 0 |
| 2.91E-05 | 7.99E-05 | 82.6312815 | 38.0373413 | 35.2659728 | 54.2805297 | 0 |
| 2.92E-05 | 8.01E-05 | 141.148376 | 408.178487 | 365.652455 | 390.170303 | 28.5146189 |
| 2.92E-05 | 8.02E-05 | 9.32415239 | 68.2892619 | 66.819738 | 63.0953166 | 0 |
| 2.93E-05 | 8.03E-05 | 15.1115573 | 39.1495443 | 31.5537652 | 44.0739344 | 0 |
| 2.93E-05 | 8.03E-05 | 20.8989622 | 20.6869751 | 31.5537652 | 30.619786 | 0 |
| 2.93E-05 | 8.05E-05 | 14.7900348 | 43.1534749 | 32.4818171 | 41.2903175 | 0 |
| 2.94E-05 | 8.06E-05 | 194.199588 | 413.961943 | 357.299988 | 403.624452 | 111.682257 |
| 2.94E-05 | 8.06E-05 | 25.0787547 | 51.3837769 | 21.3451941 | 25.9804245 | 0 |
| 2.94E-05 | 8.07E-05 | 16.3976473 | 73.8502767 | 71.4599976 | 24.1246799 | 0 |
| 2.95E-05 | 8.07E-05 | 39.547267 | 20.2420939 | 21.3451941 | 31.5476583 | 0 |
| 2.95E-05 | 8.07E-05 | 24.4357097 | 38.7046631 | 29.6976613 | 18.5574461 | 0 |
| 2.95E-05 | 8.08E-05 | 31.1876821 | 17.1279256 | 27.8415575 | 42.2181898 | 0 |
| 2.96E-05 | 8.10E-05 | 37.2966096 | 16.2381632 | 44.546492 | 27.8361691 | 0 |
| 2.96E-05 | 8.10E-05 | 230.53163 | 86.0845093 | 89.092984 | 74.6937204 | 356.432737 |
| 2.96E-05 | 8.12E-05 | 13.5039448 | 43.8207967 | 37.1220767 | 63.5592527 | 0 |
| 2.97E-05 | 8.12E-05 | 500.932049 | 733.164193 | 529.917645 | 653.686037 | 0 |
| 2.97E-05 | 8.13E-05 | 6.43044992 | 66.9546183 | 116.934542 | 107.633187 | 0 |
| 2.97E-05 | 8.13E-05 | 63.6614542 | 270.26532 | 154.056618 | 175.367865 | 16.6335277 |
| 2.97E-05 | 8.13E-05 | 266.220627 | 886.425761 | 1195.33087 | 1156.59283 | 0 |
| 2.97E-05 | 8.14E-05 | 28.2939797 | 112.77738 | 91.8771398 | 102.529889 | 7.12865473 |
| 2.97E-05 | 8.14E-05 | 28.6155022 | 37.5924601 | 19.4890903 | 105.313506 | 0 |
| 2.97E-05 | 8.14E-05 | 64.6260217 | 48.2696086 | 31.5537652 | 42.6821259 | 0 |
| 2.98E-05 | 8.16E-05 | 10.9317649 | 100.988029 | 35.2659728 | 94.6429749 | 0 |
| 2.98E-05 | 8.17E-05 | 19.6128723 | 26.2479899 | 24.1293498 | 26.9082968 | 0 |
| 2.99E-05 | 8.18E-05 | 18.0052598 | 52.2735392 | 50.1148035 | 22.7328714 | 0 |
| 3.00E-05 | 8.21E-05 | 11.2532874 | 52.7184204 | 60.3233746 | 48.7132959 | 0 |
| 3.00E-05 | 8.22E-05 | 18.6483048 | 90.9782023 | 94.6612955 | 77.4773373 | 2.37621824 |
| 3.00E-05 | 8.22E-05 | 54.6588243 | 16.905485 | 27.8415575 | 139.180845 | 0 |
| 3.02E-05 | 8.25E-05 | 327.631424 | 59.1691976 | 9.28051917 | 54.2805297 | 0 |
| 3.02E-05 | 8.26E-05 | 8.0380624 | 66.064856 | 77.0283091 | 85.3642518 | 0 |
| 3.02E-05 | 8.26E-05 | 14.1469898 | 45.6003214 | 33.409869 | 45.929679 | 0 |

| | | | | | | |
|---|---|---|---|---|---|---|
| 3.02E-05 | 8.26E-05 | 16.3976473 | 37.3700195 | 27.8415575 | 34.7952113 | 0 |
| 3.03E-05 | 8.28E-05 | 18.6483048 | 40.2617472 | 25.0574018 | 28.3001052 | 0 |
| 3.03E-05 | 8.28E-05 | 15.7546023 | 33.8109701 | 31.5537652 | 35.2591475 | 0 |
| 3.03E-05 | 8.29E-05 | 16.0761248 | 60.7262817 | 34.3379209 | 32.0115944 | 0 |
| 3.04E-05 | 8.30E-05 | 107.710036 | 22.2440592 | 26.9135056 | 34.3312752 | 0 |
| 3.04E-05 | 8.31E-05 | 17.6837373 | 31.3641235 | 25.0574018 | 34.7952113 | 0 |
| 3.04E-05 | 8.31E-05 | 180.37412 | 22.911381 | 25.9854537 | 42.6821259 | 0 |
| 3.04E-05 | 8.32E-05 | 19.6128723 | 38.0373413 | 28.7696094 | 22.7328714 | 0 |
| 3.04E-05 | 8.32E-05 | 19.6128723 | 32.0314453 | 25.0574018 | 24.1246799 | 0 |
| 3.05E-05 | 8.33E-05 | 222.493567 | 423.971769 | 350.803625 | 394.809665 | 85.5438568 |
| 3.05E-05 | 8.33E-05 | 13.8254673 | 42.041272 | 51.0428554 | 35.2591475 | 0 |
| 3.05E-05 | 8.34E-05 | 121.213981 | 265.816508 | 234.797135 | 263.051798 | 30.8908372 |
| 3.05E-05 | 8.35E-05 | 13.5039448 | 54.053064 | 36.1940248 | 44.0739344 | 0 |
| 3.06E-05 | 8.36E-05 | 27.6509347 | 52.0510986 | 54.7550631 | 71.9101035 | 0 |
| 3.06E-05 | 8.37E-05 | 13.8254673 | 44.7105591 | 44.546492 | 35.2591475 | 0 |
| 3.07E-05 | 8.38E-05 | 20.8989622 | 36.4802572 | 22.273246 | 25.5164883 | 0 |
| 3.07E-05 | 8.40E-05 | 10.9317649 | 41.5963908 | 63.1075304 | 106.705315 | 0 |
| 3.08E-05 | 8.40E-05 | 60.1247068 | 20.0196533 | 232.012979 | 23.1968076 | 0 |
| 3.08E-05 | 8.42E-05 | 24.1141872 | 32.2538859 | 17.6329864 | 38.9706367 | 0 |
| 3.09E-05 | 8.45E-05 | 14.4685123 | 46.0452026 | 34.3379209 | 37.5788283 | 0 |
| 3.09E-05 | 8.45E-05 | 25.4002772 | 29.1397176 | 35.2659728 | 17.6295737 | 0 |
| 3.10E-05 | 8.46E-05 | 101.601109 | 54.9428263 | 13.9207788 | 31.5476583 | 0 |
| 3.10E-05 | 8.47E-05 | 51.4435994 | 116.113989 | 137.351684 | 146.139888 | 14.2573095 |
| 3.10E-05 | 8.47E-05 | 27.6509347 | 17.3503662 | 41.7623363 | 28.7640414 | 0 |
| 3.11E-05 | 8.49E-05 | 118.320279 | 337.442379 | 235.725187 | 256.092756 | 7.12865473 |
| 3.12E-05 | 8.51E-05 | 114.783531 | 274.26925 | 279.343627 | 349.343922 | 2.37621824 |
| 3.12E-05 | 8.51E-05 | 55.6233918 | 153.484009 | 17.6329864 | 153.562866 | 0 |
| 3.13E-05 | 8.54E-05 | 30.5446371 | 35.5904948 | 15.7768826 | 62.6313804 | 0 |
| 3.13E-05 | 8.54E-05 | 20.5774398 | 60.7262817 | 109.510126 | 86.2921241 | 2.37621824 |
| 3.13E-05 | 8.54E-05 | 16.7191698 | 41.8188314 | 25.9854537 | 61.239572 | 0 |
| 3.13E-05 | 8.55E-05 | 27.6509347 | 14.9035197 | 19.4890903 | 11.1344676 | 33.2670554 |
| 3.14E-05 | 8.56E-05 | 190.66284 | 183.513489 | 147.560255 | 189.749886 | 54.6530196 |
| 3.14E-05 | 8.57E-05 | 15.4330798 | 30.02948 | 36.1940248 | 39.4345729 | 0 |
| 3.14E-05 | 8.58E-05 | 20.8989622 | 40.929069 | 19.4890903 | 52.4247851 | 0 |
| 3.15E-05 | 8.59E-05 | 19.6128723 | 21.7991781 | 31.5537652 | 28.7640414 | 0 |
| 3.16E-05 | 8.61E-05 | 19.9343948 | 25.1357869 | 61.2514265 | 30.1558498 | 0 |
| 3.16E-05 | 8.61E-05 | 27.9724572 | 44.9329997 | 66.819738 | 73.3019119 | 0 |
| 3.16E-05 | 8.62E-05 | 17.3622148 | 31.5865641 | 28.7696094 | 26.9082968 | 0 |
| 3.17E-05 | 8.65E-05 | 13.8254673 | 30.2519206 | 65.8916861 | 57.0641466 | 0 |
| 3.18E-05 | 8.67E-05 | 17.0406923 | 33.3660889 | 27.8415575 | 28.7640414 | 0 |
| 3.18E-05 | 8.67E-05 | 16.7191698 | 30.4743612 | 29.6976613 | 29.6919137 | 0 |
| 3.18E-05 | 8.67E-05 | 16.7191698 | 34.2558512 | 28.7696094 | 29.2279775 | 0 |
| 3.19E-05 | 8.69E-05 | 172.014535 | 242.460246 | 210.667785 | 191.60563 | 35.6432737 |

| | | | | | | |
|---|---|---|---|---|---|---|
| 3.21E-05 | 8.75E-05 | 28.9370247 | 51.3837769 | 49.1867516 | 79.797018 | 0 |
| 3.22E-05 | 8.77E-05 | 17.0406923 | 28.2499552 | 27.8415575 | 36.1870198 | 0 |
| 3.22E-05 | 8.77E-05 | 11.8963324 | 64.2853312 | 42.6903882 | 47.7854236 | 0 |
| 3.23E-05 | 8.79E-05 | 5.46588243 | 79.4112915 | 94.6612955 | 144.748079 | 0 |
| 3.23E-05 | 8.80E-05 | 15.7546023 | 50.7164551 | 28.7696094 | 36.650956 | 0 |
| 3.23E-05 | 8.81E-05 | 39.547267 | 22.2440592 | 240.365446 | 29.2279775 | 0 |
| 3.24E-05 | 8.82E-05 | 17.6837373 | 30.02948 | 25.0574018 | 30.619786 | 0 |
| 3.24E-05 | 8.82E-05 | 32.4737721 | 63.1731283 | 82.5966206 | 52.8887212 | 0 |
| 3.24E-05 | 8.82E-05 | 51.4435994 | 139.692692 | 117.862593 | 128.97425 | 23.7621824 |
| 3.24E-05 | 8.84E-05 | 95.8137038 | 12.4566732 | 49.1867516 | 39.4345729 | 0 |
| 3.26E-05 | 8.89E-05 | 31.1876821 | 50.7164551 | 40.8342843 | 57.5280828 | 2.37621824 |
| 3.28E-05 | 8.93E-05 | 17.6837373 | 25.3582275 | 38.0501286 | 28.3001052 | 0 |
| 3.28E-05 | 8.94E-05 | 12.2178549 | 40.4841878 | 46.4025958 | 71.9101035 | 0 |
| 3.28E-05 | 8.94E-05 | 17.3622148 | 42.041272 | 44.546492 | 23.1968076 | 0 |
| 3.29E-05 | 8.95E-05 | 55.6233918 | 129.015544 | 138.279736 | 136.397228 | 9.50487297 |
| 3.29E-05 | 8.95E-05 | 12.5393773 | 60.05896 | 46.4025958 | 38.5067006 | 0 |
| 3.29E-05 | 8.95E-05 | 9.64567488 | 50.7164551 | 125.287009 | 58.4559551 | 0 |
| 3.30E-05 | 8.97E-05 | 35.0459521 | 49.8266927 | 73.3161014 | 47.7854236 | 4.75243649 |
| 3.30E-05 | 8.97E-05 | 16.0761248 | 72.9605143 | 23.2012979 | 135.469356 | 0 |
| 3.30E-05 | 8.99E-05 | 38.904222 | 24.9133464 | 19.4890903 | 24.1246799 | 0 |
| 3.31E-05 | 9.00E-05 | 12.8608998 | 32.0314453 | 90.9490879 | 60.7756358 | 0 |
| 3.31E-05 | 9.01E-05 | 11.2532874 | 58.946757 | 73.3161014 | 39.898509 | 0 |
| 3.31E-05 | 9.01E-05 | 7.07349491 | 64.7302124 | 126.215061 | 77.0134011 | 0 |
| 3.31E-05 | 9.02E-05 | 14.7900348 | 27.8050741 | 52.8989593 | 71.9101035 | 0 |
| 3.32E-05 | 9.04E-05 | 35.3674746 | 12.4566732 | 59.3953227 | 132.685739 | 0 |
| 3.33E-05 | 9.06E-05 | 357.211493 | 47.3798462 | 18.5610383 | 29.2279775 | 0 |
| 3.33E-05 | 9.06E-05 | 28.2939797 | 59.1691976 | 46.4025958 | 54.2805297 | 0 |
| 3.33E-05 | 9.06E-05 | 5.46588243 | 3.78149007 | 1.85610383 | 2.31968076 | 40.3957101 |
| 3.33E-05 | 9.07E-05 | 21.5420072 | 28.6948364 | 20.4171422 | 44.5378705 | 0 |
| 3.34E-05 | 9.07E-05 | 155.295366 | 337.219938 | 327.602327 | 287.176478 | 19.0097459 |
| 3.34E-05 | 9.09E-05 | 23.4711422 | 45.822762 | 16.7049345 | 50.1051043 | 0 |
| 3.34E-05 | 9.10E-05 | 33.7598621 | 70.0687866 | 69.6038938 | 70.0543588 | 7.12865473 |
| 3.35E-05 | 9.10E-05 | 31.1876821 | 54.4979451 | 135.49558 | 14.3820207 | 0 |
| 3.35E-05 | 9.10E-05 | 109.317649 | 46.934965 | 31.5537652 | 48.2493597 | 2.37621824 |
| 3.35E-05 | 9.12E-05 | 15.7546023 | 31.5865641 | 30.6257133 | 33.867339 | 0 |
| 3.35E-05 | 9.12E-05 | 23.7926647 | 98.9860636 | 47.3306478 | 86.2921241 | 2.37621824 |
| 3.36E-05 | 9.12E-05 | 11.5748099 | 10.899589 | 3.71220767 | 6.49510612 | 21.3859642 |
| 3.36E-05 | 9.13E-05 | 11.2532874 | 4.22637126 | 4.64025958 | 7.88691457 | 57.0292378 |
| 3.36E-05 | 9.14E-05 | 27.6509347 | 57.834554 | 57.5392189 | 47.3214874 | 0 |
| 3.36E-05 | 9.14E-05 | 12.5393773 | 45.3778809 | 43.6184401 | 43.1460621 | 0 |
| 3.37E-05 | 9.15E-05 | 19.2913498 | 25.8031087 | 28.7696094 | 22.7328714 | 0 |
| 3.38E-05 | 9.19E-05 | 200.95156 | 64.952653 | 85.3807764 | 86.7560603 | 0 |
| 3.39E-05 | 9.21E-05 | 208.346577 | 741.616935 | 686.758419 | 754.824118 | 28.5146189 |

| | | | | | | |
|---|---|---|---|---|---|---|
| 3.39E-05 | 9.21E-05 | 265.899104 | 449.552437 | 410.198947 | 439.811471 | 35.6432737 |
| 3.39E-05 | 9.21E-05 | 24.7572322 | 51.1613363 | 15.7768826 | 64.0231889 | 0 |
| 3.40E-05 | 9.22E-05 | 55.3018693 | 18.9074504 | 26.9135056 | 27.8361691 | 0 |
| 3.40E-05 | 9.22E-05 | 204.166785 | 732.05199 | 624.57894 | 629.561357 | 2.37621824 |
| 3.40E-05 | 9.22E-05 | 93.5630464 | 27.1377523 | 25.0574018 | 25.9804245 | 0 |
| 3.40E-05 | 9.22E-05 | 23.1496197 | 20.6869751 | 25.0574018 | 23.1968076 | 0 |
| 3.40E-05 | 9.24E-05 | 18.6483048 | 23.5787028 | 32.4818171 | 25.9804245 | 0 |
| 3.41E-05 | 9.25E-05 | 7.39501741 | 94.0923706 | 71.4599976 | 70.518295 | 0 |
| 3.42E-05 | 9.29E-05 | 18.0052598 | 33.3660889 | 26.9135056 | 68.1986142 | 0 |
| 3.42E-05 | 9.29E-05 | 4.17979245 | 185.515454 | 94.6612955 | 100.210209 | 0 |
| 3.42E-05 | 9.29E-05 | 15.4330798 | 54.9428263 | 33.409869 | 31.5476583 | 0 |
| 3.42E-05 | 9.29E-05 | 20.8989622 | 19.1298909 | 50.1148035 | 38.9706367 | 0 |
| 3.43E-05 | 9.31E-05 | 21.8635297 | 94.3148112 | 18.5610383 | 52.4247851 | 0 |
| 3.43E-05 | 9.31E-05 | 18.3267823 | 30.9192423 | 25.0574018 | 53.8165936 | 0 |
| 3.43E-05 | 9.31E-05 | 7.39501741 | 54.9428263 | 113.222334 | 88.1478687 | 0 |
| 3.43E-05 | 9.32E-05 | 45.334672 | 109.218331 | 89.092984 | 77.4773373 | 11.8810912 |
| 3.44E-05 | 9.34E-05 | 28.2939797 | 27.5826335 | 23.2012979 | 18.5574461 | 0 |
| 3.44E-05 | 9.34E-05 | 117.677234 | 60.2814006 | 57.5392189 | 90.0036134 | 16.6335277 |
| 3.45E-05 | 9.35E-05 | 16.3976473 | 28.917277 | 28.7696094 | 40.3624452 | 0 |
| 3.47E-05 | 9.41E-05 | 14.7900348 | 51.1613363 | 27.8415575 | 79.797018 | 0 |
| 3.47E-05 | 9.41E-05 | 22.8280972 | 33.8109701 | 27.8415575 | 18.5574461 | 0 |
| 3.47E-05 | 9.42E-05 | 7.07349491 | 51.828658 | 128.071165 | 99.2823364 | 0 |
| 3.48E-05 | 9.43E-05 | 11.2532874 | 90.5333211 | 34.3379209 | 73.7658481 | 0 |
| 3.48E-05 | 9.44E-05 | 11.2532874 | 36.2578166 | 69.6038938 | 72.8379758 | 0 |
| 3.48E-05 | 9.44E-05 | 81.9882365 | 149.035197 | 143.848047 | 185.574461 | 19.0097459 |
| 3.49E-05 | 9.45E-05 | 18.3267823 | 51.6062174 | 34.3379209 | 23.1968076 | 0 |
| 3.49E-05 | 9.45E-05 | 32.4737721 | 20.4645345 | 25.0574018 | 21.341063 | 0 |
| 3.49E-05 | 9.47E-05 | 19.6128723 | 22.0216187 | 25.9854537 | 27.3722329 | 0 |
| 3.49E-05 | 9.47E-05 | 47.5853294 | 21.5767375 | 36.1940248 | 20.8771268 | 92.6725115 |
| 3.50E-05 | 9.48E-05 | 179.731075 | 656.644629 | 438.968557 | 573.425083 | 45.1481466 |
| 3.50E-05 | 9.48E-05 | 18.3267823 | 35.3680542 | 25.9854537 | 24.588616 | 0 |
| 3.51E-05 | 9.50E-05 | 15.1115573 | 34.4782918 | 30.6257133 | 51.0329766 | 0 |
| 3.51E-05 | 9.51E-05 | 19.9343948 | 37.3700195 | 19.4890903 | 43.1460621 | 0 |
| 3.51E-05 | 9.51E-05 | 11.5748099 | 77.4093262 | 42.6903882 | 46.8575513 | 0 |
| 3.51E-05 | 9.52E-05 | 19.6128723 | 34.2558512 | 20.4171422 | 30.619786 | 0 |
| 3.52E-05 | 9.52E-05 | 21.5420072 | 41.1515096 | 18.5610383 | 32.9394667 | 0 |
| 3.52E-05 | 9.53E-05 | 19.6128723 | 27.1377523 | 21.3451941 | 28.3001052 | 0 |
| 3.53E-05 | 9.55E-05 | 99.6719738 | 262.479899 | 180.042072 | 220.369672 | 0 |
| 3.53E-05 | 9.55E-05 | 24.4357097 | 26.0255493 | 80.7405168 | 25.0525522 | 0 |
| 3.53E-05 | 9.55E-05 | 29.9015921 | 49.1593709 | 42.6903882 | 55.6723382 | 2.37621824 |
| 3.53E-05 | 9.56E-05 | 128.287476 | 32.9212077 | 18.5610383 | 33.867339 | 0 |
| 3.53E-05 | 9.57E-05 | 27.0078897 | 33.8109701 | 15.7768826 | 47.7854236 | 0 |
| 3.54E-05 | 9.59E-05 | 18.9698273 | 22.0216187 | 28.7696094 | 27.8361691 | 0 |

| | | | | | | |
|---|---|---|---|---|---|---|
| 3.54E-05 | 9.59E-05 | 12.8608998 | 53.3857422 | 32.4818171 | 64.487125 | 0 |
| 3.55E-05 | 9.61E-05 | 25.7217997 | 37.1475789 | 15.7768826 | 47.3214874 | 0 |
| 3.55E-05 | 9.61E-05 | 13.1824223 | 67.3994995 | 28.7696094 | 117.839782 | 0 |
| 3.55E-05 | 9.62E-05 | 19.6128723 | 45.822762 | 19.4890903 | 57.0641466 | 0 |
| 3.57E-05 | 9.67E-05 | 89.0617314 | 227.111845 | 211.595837 | 214.338502 | 21.3859642 |
| 3.58E-05 | 9.68E-05 | 16.3976473 | 27.1377523 | 33.409869 | 52.4247851 | 0 |
| 3.58E-05 | 9.68E-05 | 23.1496197 | 33.5885295 | 17.6329864 | 29.2279775 | 0 |
| 3.59E-05 | 9.73E-05 | 171.049968 | 37.5924601 | 21.3451941 | 27.8361691 | 0 |
| 3.59E-05 | 9.73E-05 | 21.8635297 | 31.1416829 | 18.5610383 | 28.3001052 | 0 |
| 3.60E-05 | 9.74E-05 | 8.6811074 | 61.8384847 | 64.9636342 | 68.1986142 | 0 |
| 3.60E-05 | 9.74E-05 | 34.0813846 | 67.8443807 | 35.2659728 | 50.1051043 | 2.37621824 |
| 3.61E-05 | 9.75E-05 | 9.96719738 | 56.2774699 | 49.1867516 | 70.518295 | 0 |
| 3.61E-05 | 9.76E-05 | 32.1522496 | 50.2715739 | 37.1220767 | 59.8477635 | 2.37621824 |
| 3.61E-05 | 9.77E-05 | 51.1220769 | 44.2656779 | 29.6976613 | 97.4265918 | 0 |
| 3.62E-05 | 9.78E-05 | 17.3622148 | 34.2558512 | 29.6976613 | 24.1246799 | 0 |
| 3.62E-05 | 9.80E-05 | 21.5420072 | 67.8443807 | 17.6329864 | 50.1051043 | 0 |
| 3.63E-05 | 9.82E-05 | 10.6102424 | 71.8483114 | 38.0501286 | 74.6937204 | 0 |
| 3.64E-05 | 9.85E-05 | 10.6102424 | 63.3955688 | 44.546492 | 55.6723382 | 0 |
| 3.64E-05 | 9.85E-05 | 13.8254673 | 32.0314453 | 84.4527244 | 45.4657428 | 0 |
| 3.66E-05 | 9.90E-05 | 909.265619 | 2477.76576 | 1929.41994 | 2158.23098 | 0 |
| 3.67E-05 | 9.92E-05 | 14.1469898 | 40.929069 | 30.6257133 | 43.6099982 | 0 |
| 3.68E-05 | 9.93E-05 | 105.137856 | 260.255493 | 266.3509 | 244.030416 | 23.7621824 |
| 3.68E-05 | 9.94E-05 | 9.96719738 | 49.1593709 | 74.2441534 | 53.8165936 | 0 |
| 3.68E-05 | 9.95E-05 | 270.078897 | 405.954081 | 400.918428 | 448.626258 | 35.6432737 |
| 3.70E-05 | 0.00010001 | 104.494811 | 123.009648 | 112.294282 | 145.212015 | 21.3859642 |
| 3.70E-05 | 0.00010002 | 53.3727344 | 11.3444702 | 46.4025958 | 46.3936151 | 0 |
| 3.71E-05 | 0.00010014 | 23.7926647 | 19.5747721 | 51.9709073 | 27.3722329 | 0 |
| 3.71E-05 | 0.00010014 | 61.7323193 | 45.3778809 | 76.1002572 | 9.74265918 | 0 |
| 3.72E-05 | 0.00010051 | 19.6128723 | 31.1416829 | 20.4171422 | 42.6821259 | 0 |
| 3.72E-05 | 0.00010056 | 13.5039448 | 47.6022868 | 34.3379209 | 38.5067006 | 0 |
| 3.72E-05 | 0.00010059 | 142.112943 | 338.554582 | 158.696878 | 276.505946 | 11.8810912 |
| 3.73E-05 | 0.00010076 | 21.2204847 | 34.0334106 | 18.5610383 | 29.6919137 | 0 |
| 3.74E-05 | 0.00010112 | 91.9554339 | 45.6003214 | 31.5537652 | 45.0018067 | 0 |
| 3.75E-05 | 0.00010114 | 15.4330798 | 31.5865641 | 30.6257133 | 31.5476583 | 0 |
| 3.75E-05 | 0.00010121 | 11.5748099 | 44.9329997 | 63.1075304 | 41.7542536 | 0 |
| 3.75E-05 | 0.00010125 | 145.328168 | 229.558691 | 247.789862 | 257.9485 | 26.1384007 |
| 3.76E-05 | 0.00010154 | 22.8280972 | 59.1691976 | 73.3161014 | 61.7035081 | 2.37621824 |
| 3.77E-05 | 0.00010179 | 27.9724572 | 54.2755046 | 43.6184401 | 58.4559551 | 2.37621824 |
| 3.77E-05 | 0.00010179 | 17.3622148 | 81.6356974 | 33.409869 | 29.2279775 | 0 |
| 3.77E-05 | 0.00010186 | 20.2559173 | 26.4704305 | 22.273246 | 22.7328714 | 0 |
| 3.78E-05 | 0.0001019 | 16.7191698 | 36.9251383 | 48.2586997 | 23.6607437 | 0 |
| 3.78E-05 | 0.00010194 | 38.904222 | 64.7302124 | 10.2085711 | 77.9412734 | 0 |
| 3.79E-05 | 0.0001022 | 11.5748099 | 46.934965 | 46.4025958 | 45.4657428 | 0 |

| | | | | | | |
|---|---|---|---|---|---|---|
| 3.79E-05 | 0.00010226 | 12.8608998 | 39.3719849 | 39.9062324 | 41.7542536 | 0 |
| 3.79E-05 | 0.00010234 | 69.1273367 | 25.8031087 | 19.4890903 | 29.2279775 | 0 |
| 3.79E-05 | 0.00010238 | 26.6863672 | 34.7007324 | 16.7049345 | 25.9804245 | 0 |
| 3.80E-05 | 0.00010239 | 171.37149 | 371.920671 | 285.83999 | 266.299351 | 0 |
| 3.80E-05 | 0.00010239 | 14.7900348 | 34.2558512 | 35.2659728 | 30.619786 | 0 |
| 3.81E-05 | 0.00010264 | 33.1168171 | 51.3837769 | 20.4171422 | 19.9492545 | 0 |
| 3.81E-05 | 0.00010265 | 11.2532874 | 58.946757 | 45.4745439 | 44.5378705 | 0 |
| 3.81E-05 | 0.00010287 | 11.5748099 | 61.6160441 | 38.9781805 | 49.177232 | 0 |
| 3.86E-05 | 0.00010395 | 38.261177 | 88.5313558 | 14.8488307 | 32.0115944 | 0 |
| 3.86E-05 | 0.00010402 | 197.09329 | 35.3680542 | 13.9207788 | 47.3214874 | 0 |
| 3.86E-05 | 0.00010404 | 9.64567488 | 61.8384847 | 48.2586997 | 64.9510612 | 0 |
| 3.86E-05 | 0.00010407 | 16.3976473 | 61.3936035 | 126.215061 | 22.7328714 | 0 |
| 3.87E-05 | 0.00010433 | 40.5118345 | 47.1574056 | 12.0646749 | 39.4345729 | 0 |
| 3.87E-05 | 0.00010435 | 15.1115573 | 28.0275146 | 37.1220767 | 34.3312752 | 0 |
| 3.89E-05 | 0.00010472 | 27.0078897 | 68.2892619 | 41.7623363 | 64.0231889 | 2.37621824 |
| 3.89E-05 | 0.00010479 | 23.7926647 | 18.4625692 | 18.5610383 | 18.0935099 | 38.0194919 |
| 3.89E-05 | 0.0001048 | 10.9317649 | 75.1849202 | 47.3306478 | 43.6099982 | 0 |
| 3.89E-05 | 0.00010489 | 6.75197242 | 84.7498657 | 81.6685687 | 70.518295 | 0 |
| 3.90E-05 | 0.00010502 | 12.2178549 | 44.2656779 | 47.3306478 | 38.5067006 | 0 |
| 3.91E-05 | 0.00010541 | 114.783531 | 414.629264 | 301.616873 | 307.125732 | 26.1384007 |
| 3.91E-05 | 0.00010543 | 43.0840145 | 134.131677 | 142.919995 | 123.407016 | 7.12865473 |
| 3.93E-05 | 0.00010588 | 14.7900348 | 30.02948 | 36.1940248 | 32.9394667 | 0 |
| 3.93E-05 | 0.00010592 | 12.5393773 | 50.9388957 | 32.4818171 | 59.8477635 | 0 |
| 3.95E-05 | 0.00010634 | 13.5039448 | 77.6317668 | 40.8342843 | 33.867339 | 0 |
| 3.95E-05 | 0.00010636 | 23.7926647 | 62.0609253 | 32.4818171 | 18.5574461 | 0 |
| 3.97E-05 | 0.00010692 | 16.0761248 | 33.1436483 | 28.7696094 | 65.4149973 | 0 |
| 3.98E-05 | 0.00010722 | 32.7952946 | 42.7085937 | 13.9207788 | 32.0115944 | 0 |
| 3.98E-05 | 0.00010722 | 14.1469898 | 29.3621582 | 38.9781805 | 46.3936151 | 0 |
| 3.99E-05 | 0.00010728 | 27.0078897 | 117.671073 | 14.8488307 | 52.4247851 | 0 |
| 3.99E-05 | 0.00010732 | 27.6509347 | 28.2499552 | 17.6329864 | 22.7328714 | 0 |
| 3.99E-05 | 0.00010733 | 49.1929419 | 32.0314453 | 25.9854537 | 16.2377653 | 0 |
| 3.99E-05 | 0.00010736 | 147.578826 | 329.212077 | 333.170638 | 341.920944 | 0 |
| 3.99E-05 | 0.0001075 | 38.904222 | 13.123995 | 54.7550631 | 34.3312752 | 0 |
| 4.00E-05 | 0.00010774 | 4.82283744 | 91.2006429 | 110.438178 | 97.4265918 | 0 |
| 4.01E-05 | 0.00010777 | 9.00262989 | 53.6081828 | 67.7477899 | 60.7756358 | 0 |
| 4.01E-05 | 0.00010784 | 13.8254673 | 32.0314453 | 46.4025958 | 35.7230837 | 0 |
| 4.02E-05 | 0.00010813 | 13.8254673 | 29.1397176 | 43.6184401 | 56.1362743 | 0 |
| 4.02E-05 | 0.00010815 | 89.0617314 | 33.8109701 | 36.1940248 | 49.6411682 | 2.37621824 |
| 4.02E-05 | 0.0001082 | 21.2204847 | 33.3660889 | 17.6329864 | 34.3312752 | 0 |
| 4.03E-05 | 0.00010828 | 17.3622148 | 22.0216187 | 52.8989593 | 38.5067006 | 0 |
| 4.04E-05 | 0.00010862 | 14.7900348 | 33.8109701 | 29.6976613 | 34.3312752 | 0 |
| 4.04E-05 | 0.00010866 | 69.7703817 | 95.8718953 | 104.869867 | 115.056166 | 16.6335277 |
| 4.04E-05 | 0.00010866 | 192.270453 | 576.343575 | 195.818954 | 401.304771 | 0 |

| | | | | | | |
|---|---|---|---|---|---|---|
| 4.04E-05 | 0.00010873 | 63.3399317 | 93.6474894 | 77.0283091 | 89.5396772 | 2.37621824 |
| 4.05E-05 | 0.00010876 | 20.8989622 | 18.2401286 | 34.3379209 | 42.6821259 | 0 |
| 4.05E-05 | 0.00010878 | 28.9370247 | 41.5963908 | 23.2012979 | 17.6295737 | 0 |
| 4.05E-05 | 0.00010888 | 22.8280972 | 63.1731283 | 59.3953227 | 62.1674443 | 0 |
| 4.05E-05 | 0.00010894 | 12.8608998 | 52.2735392 | 36.1940248 | 38.0427644 | 0 |
| 4.05E-05 | 0.00010894 | 24.1141872 | 104.102197 | 81.6685687 | 83.5085072 | 4.75243649 |
| 4.06E-05 | 0.00010899 | 85.524984 | 33.8109701 | 33.409869 | 15.309893 | 0 |
| 4.06E-05 | 0.00010903 | 14.1469898 | 35.1456136 | 31.5537652 | 54.2805297 | 0 |
| 4.07E-05 | 0.00010926 | 64.9475442 | 29.8070394 | 135.49558 | 153.09893 | 7.12865473 |
| 4.07E-05 | 0.00010932 | 1786.70051 | 8434.50238 | 7887.51324 | 8284.97179 | 0 |
| 4.07E-05 | 0.00010944 | 9.64567488 | 50.7164551 | 61.2514265 | 55.6723382 | 0 |
| 4.08E-05 | 0.00010948 | 472.638069 | 1540.84598 | 1322.47398 | 1259.12271 | 0 |
| 4.08E-05 | 0.00010949 | 12.8608998 | 34.0334106 | 42.6903882 | 44.5378705 | 0 |
| 4.10E-05 | 0.00011018 | 25.7217997 | 84.7498657 | 71.4599976 | 62.6313804 | 2.37621824 |
| 4.11E-05 | 0.0001103 | 67.5197242 | 221.995711 | 191.178695 | 211.090949 | 19.0097459 |
| 4.11E-05 | 0.00011048 | 28.9370247 | 29.5845988 | 14.8488307 | 38.0427644 | 0 |
| 4.12E-05 | 0.00011055 | 12.8608998 | 51.6062174 | 47.3306478 | 31.5476583 | 0 |
| 4.12E-05 | 0.00011067 | 24.4357097 | 83.1927816 | 40.8342843 | 17.1656376 | 0 |
| 4.13E-05 | 0.00011101 | 22.5065747 | 29.8070394 | 33.409869 | 17.1656376 | 0 |
| 4.14E-05 | 0.00011115 | 185.51848 | 352.568339 | 326.674275 | 343.776688 | 73.6627655 |
| 4.14E-05 | 0.00011115 | 27.0078897 | 35.8129354 | 15.7768826 | 27.3722329 | 0 |
| 4.14E-05 | 0.00011121 | 30.8661596 | 100.320707 | 15.7768826 | 36.650956 | 0 |
| 4.15E-05 | 0.0001115 | 12.5393773 | 61.1711629 | 30.6257133 | 54.7444659 | 0 |
| 4.15E-05 | 0.0001115 | 66.5551567 | 107.216366 | 141.063891 | 118.303719 | 16.6335277 |
| 4.16E-05 | 0.00011167 | 11.2532874 | 51.828658 | 41.7623363 | 46.8575513 | 0 |
| 4.16E-05 | 0.00011172 | 15.1115573 | 4.44881185 | 4.64025958 | 9.27872303 | 78.415202 |
| 4.17E-05 | 0.0001118 | 30.2231146 | 44.4881185 | 12.9927268 | 46.8575513 | 0 |
| 4.17E-05 | 0.00011187 | 9.96719738 | 46.0452026 | 64.0355823 | 53.8165936 | 0 |
| 4.17E-05 | 0.0001119 | 16.7191698 | 32.6987671 | 28.7696094 | 24.1246799 | 0 |
| 4.17E-05 | 0.00011196 | 158.832113 | 493.818115 | 407.414792 | 492.700193 | 2.37621824 |
| 4.18E-05 | 0.00011211 | 16.7191698 | 46.0452026 | 22.273246 | 38.5067006 | 0 |
| 4.18E-05 | 0.00011212 | 13.8254673 | 38.2597819 | 35.2659728 | 32.4755306 | 0 |
| 4.18E-05 | 0.00011223 | 16.0761248 | 44.4881185 | 23.2012979 | 39.898509 | 0 |
| 4.19E-05 | 0.00011224 | 18.6483048 | 34.7007324 | 44.546492 | 19.9492545 | 0 |
| 4.20E-05 | 0.00011255 | 18.9698273 | 27.1377523 | 25.9854537 | 62.6313804 | 0 |
| 4.21E-05 | 0.00011299 | 236.96208 | 708.250846 | 609.730109 | 682.914015 | 40.3957101 |
| 4.21E-05 | 0.000113 | 0 | 0 | 0 | 0 | 21.3859642 |
| 4.21E-05 | 0.000113 | 17.3622148 | 46.2676432 | 38.0501286 | 21.341063 | 0 |
| 4.23E-05 | 0.00011327 | 10.9317649 | 41.5963908 | 54.7550631 | 49.177232 | 0 |
| 4.23E-05 | 0.00011332 | 20.5774398 | 21.5767375 | 21.3451941 | 29.2279775 | 0 |
| 4.23E-05 | 0.00011332 | 78.1299666 | 27.5826335 | 22.273246 | 23.6607437 | 0 |
| 4.23E-05 | 0.00011349 | 12.2178549 | 40.4841878 | 39.9062324 | 43.1460621 | 0 |
| 4.25E-05 | 0.00011385 | 27.9724572 | 74.7400391 | 100.229607 | 102.529889 | 7.12865473 |

| | | | | | | |
|---|---|---|---|---|---|---|
| 4.25E-05 | 0.00011388 | 98.3858838 | 193.300875 | 149.416359 | 173.512121 | 26.1384007 |
| 4.25E-05 | 0.00011389 | 31.8307271 | 26.9153117 | 14.8488307 | 35.7230837 | 0 |
| 4.26E-05 | 0.00011422 | 13.1824223 | 44.4881185 | 32.4818171 | 39.4345729 | 0 |
| 4.26E-05 | 0.00011422 | 33.4383396 | 82.5254598 | 97.4454513 | 91.859358 | 9.50487297 |
| 4.28E-05 | 0.00011452 | 23.4711422 | 30.02948 | 16.7049345 | 27.8361691 | 0 |
| 4.28E-05 | 0.00011452 | 17.3622148 | 46.0452026 | 20.4171422 | 52.8887212 | 0 |
| 4.28E-05 | 0.00011457 | 40.5118345 | 8.23030192 | 109.510126 | 89.0757411 | 0 |
| 4.29E-05 | 0.00011485 | 15.4330798 | 34.2558512 | 31.5537652 | 26.4443606 | 0 |
| 4.29E-05 | 0.00011497 | 314.770524 | 295.623547 | 200.459214 | 278.361691 | 40.3957101 |
| 4.30E-05 | 0.00011502 | 17.3622148 | 37.5924601 | 31.5537652 | 21.8049991 | 0 |
| 4.30E-05 | 0.0001151 | 90.6693439 | 20.2420939 | 68.6758419 | 16.2377653 | 0 |
| 4.30E-05 | 0.00011516 | 15.1115573 | 58.2794352 | 24.1293498 | 47.3214874 | 0 |
| 4.30E-05 | 0.00011516 | 9.96719738 | 52.0510986 | 45.4745439 | 66.8068058 | 0 |
| 4.30E-05 | 0.00011523 | 9.00262989 | 58.5018758 | 62.1794784 | 54.2805297 | 0 |
| 4.31E-05 | 0.00011544 | 17.6837373 | 54.2755046 | 19.4890903 | 55.6723382 | 0 |
| 4.32E-05 | 0.00011556 | 16.7191698 | 24.6909058 | 26.9135056 | 29.6919137 | 0 |
| 4.32E-05 | 0.00011567 | 34.7244296 | 75.6298014 | 12.0646749 | 43.6099982 | 0 |
| 4.33E-05 | 0.00011585 | 84.881939 | 29.1397176 | 15.7768826 | 35.2591475 | 0 |
| 4.33E-05 | 0.00011587 | 13.5039448 | 38.7046631 | 30.6257133 | 51.9608489 | 0 |
| 4.34E-05 | 0.00011601 | 9.00262989 | 72.5156331 | 41.7623363 | 84.4363795 | 0 |
| 4.34E-05 | 0.00011622 | 16.3976473 | 28.6948364 | 27.8415575 | 25.5164883 | 0 |
| 4.35E-05 | 0.00011629 | 13.8254673 | 38.0373413 | 34.3379209 | 32.0115944 | 0 |
| 4.35E-05 | 0.00011635 | 21.8635297 | 53.1633016 | 102.085711 | 64.487125 | 2.37621824 |
| 4.36E-05 | 0.0001165 | 15.1115573 | 30.4743612 | 31.5537652 | 28.7640414 | 0 |
| 4.36E-05 | 0.00011656 | 28.6155022 | 66.7321777 | 93.7332436 | 64.487125 | 0 |
| 4.37E-05 | 0.00011692 | 18.9698273 | 22.0216187 | 24.1293498 | 24.588616 | 0 |
| 4.38E-05 | 0.00011705 | 1.60761248 | 0.88976237 | 0 | 4.63936151 | 0 |
| 4.38E-05 | 0.00011717 | 111.246784 | 460.229586 | 398.134272 | 428.213068 | 23.7621824 |
| 4.39E-05 | 0.00011725 | 49.1929419 | 38.4822225 | 34.3379209 | 34.7952113 | 85.5438568 |
| 4.39E-05 | 0.00011732 | 15.4330798 | 27.5826335 | 28.7696094 | 38.5067006 | 0 |
| 4.40E-05 | 0.00011772 | 42.4409695 | 55.3877075 | 35.2659728 | 12.0623399 | 0 |
| 4.41E-05 | 0.0001178 | 16.7191698 | 27.3601929 | 38.0501286 | 24.1246799 | 0 |
| 4.41E-05 | 0.00011784 | 28.9370247 | 32.4763265 | 37.1220767 | 14.3820207 | 0 |
| 4.43E-05 | 0.0001184 | 16.3976473 | 31.8090047 | 23.2012979 | 39.4345729 | 0 |
| 4.44E-05 | 0.00011853 | 15.7546023 | 28.0275146 | 30.6257133 | 26.9082968 | 0 |
| 4.44E-05 | 0.00011856 | 13.1824223 | 38.2597819 | 47.3306478 | 31.5476583 | 0 |
| 4.44E-05 | 0.00011862 | 27.6509347 | 129.905306 | 65.8916861 | 96.4987195 | 2.37621824 |
| 4.45E-05 | 0.00011887 | 54.3373018 | 21.5767375 | 30.6257133 | 19.0213822 | 0 |
| 4.45E-05 | 0.00011899 | 19.6128723 | 30.9192423 | 19.4890903 | 25.9804245 | 0 |
| 4.46E-05 | 0.00011925 | 37.9396545 | 64.5077718 | 24.1293498 | 16.2377653 | 0 |
| 4.48E-05 | 0.00011955 | 105.137856 | 438.652848 | 454.745439 | 492.236257 | 28.5146189 |
| 4.49E-05 | 0.00011984 | 12.5393773 | 38.4822225 | 37.1220767 | 40.8263813 | 0 |
| 4.49E-05 | 0.00011994 | 12.5393773 | 36.9251383 | 40.8342843 | 38.5067006 | 0 |

| | | | | | | |
|---|---|---|---|---|---|---|
| 4.51E-05 | 0.00012056 | 7.71653991 | 58.2794352 | 82.5966206 | 63.0953166 | 0 |
| 4.52E-05 | 0.00012064 | 13.5039448 | 50.4940145 | 28.7696094 | 41.7542536 | 0 |
| 4.52E-05 | 0.00012076 | 90.0262989 | 64.2853312 | 66.819738 | 69.1264865 | 23.7621824 |
| 4.52E-05 | 0.00012077 | 7.07349491 | 10.899589 | 4.64025958 | 9.74265918 | 33.2670554 |
| 4.53E-05 | 0.0001208 | 9.00262989 | 38.7046631 | 77.956361 | 88.1478687 | 0 |
| 4.53E-05 | 0.00012081 | 15.7546023 | 42.9310343 | 31.5537652 | 25.0525522 | 0 |
| 4.53E-05 | 0.0001209 | 49.5144644 | 45.3778809 | 27.8415575 | 43.6099982 | 0 |
| 4.54E-05 | 0.00012112 | 45.0131495 | 37.1475789 | 34.3379209 | 37.1148921 | 0 |
| 4.54E-05 | 0.00012116 | 14.7900348 | 26.6928711 | 33.409869 | 38.9706367 | 0 |
| 4.55E-05 | 0.00012146 | 19.9343948 | 57.3896728 | 17.6329864 | 43.1460621 | 0 |
| 4.56E-05 | 0.00012161 | 13.8254673 | 32.2538859 | 32.4818171 | 37.1148921 | 0 |
| 4.56E-05 | 0.00012178 | 272.972599 | 605.483293 | 514.140762 | 621.674443 | 40.3957101 |
| 4.57E-05 | 0.00012186 | 23.4711422 | 66.9546183 | 48.2586997 | 63.0953166 | 0 |
| 4.57E-05 | 0.00012201 | 25.0787547 | 49.8266927 | 13.9207788 | 62.6313804 | 0 |
| 4.58E-05 | 0.00012206 | 39.2257445 | 176.172949 | 162.409085 | 117.375846 | 0 |
| 4.58E-05 | 0.00012214 | 247.2508 | 600.367159 | 545.694527 | 608.684231 | 54.6530196 |
| 4.58E-05 | 0.00012223 | 14.1469898 | 38.9271037 | 44.546492 | 26.9082968 | 0 |
| 4.61E-05 | 0.00012283 | 115.748099 | 56.0550293 | 64.9636342 | 70.9822312 | 403.957101 |
| 4.61E-05 | 0.00012284 | 18.9698273 | 18.9074504 | 34.3379209 | 32.0115944 | 0 |
| 4.62E-05 | 0.00012322 | 18.9698273 | 26.4704305 | 20.4171422 | 38.5067006 | 0 |
| 4.62E-05 | 0.00012322 | 21.2204847 | 19.3523315 | 44.546492 | 119.695527 | 0 |
| 4.63E-05 | 0.00012353 | 28.2939797 | 59.3916382 | 12.0646749 | 87.2199964 | 0 |
| 4.64E-05 | 0.0001238 | 707.349491 | 1673.64302 | 1607.38592 | 1692.43908 | 0 |
| 4.67E-05 | 0.00012442 | 21.5420072 | 25.1357869 | 21.3451941 | 19.9492545 | 0 |
| 4.67E-05 | 0.00012452 | 8.6811074 | 38.2597819 | 93.7332436 | 82.1166988 | 0 |
| 4.67E-05 | 0.00012454 | 25.4002772 | 53.8306234 | 64.0355823 | 45.929679 | 0 |
| 4.72E-05 | 0.00012585 | 14.1469898 | 30.4743612 | 30.6257133 | 39.4345729 | 0 |
| 4.74E-05 | 0.00012625 | 27.6509347 | 45.822762 | 15.7768826 | 26.4443606 | 0 |
| 4.75E-05 | 0.00012668 | 23.7926647 | 22.4664998 | 18.5610383 | 39.898509 | 0 |
| 4.76E-05 | 0.00012674 | 25.7217997 | 19.1298909 | 25.9854537 | 65.4149973 | 0 |
| 4.76E-05 | 0.00012681 | 113.175919 | 105.881722 | 240.365446 | 43.1460621 | 11.8810912 |
| 4.79E-05 | 0.00012759 | 260.433222 | 407.511165 | 407.414792 | 412.903175 | 0 |
| 4.79E-05 | 0.0001277 | 20.8989622 | 16.4606038 | 44.546492 | 50.1051043 | 0 |
| 4.80E-05 | 0.00012795 | 21.2204847 | 19.7972127 | 41.7623363 | 23.6607437 | 0 |
| 4.81E-05 | 0.000128 | 33.7598621 | 73.1829549 | 62.1794784 | 60.3116997 | 0 |
| 4.83E-05 | 0.00012861 | 22.8280972 | 29.5845988 | 38.0501286 | 16.2377653 | 0 |
| 4.83E-05 | 0.0001287 | 18.6483048 | 22.0216187 | 25.9854537 | 22.2689353 | 0 |
| 4.84E-05 | 0.00012891 | 14.1469898 | 26.9153117 | 40.8342843 | 35.7230837 | 0 |
| 4.85E-05 | 0.00012906 | 14.1469898 | 28.0275146 | 38.9781805 | 32.9394667 | 0 |
| 4.85E-05 | 0.00012915 | 64.3044992 | 44.4881185 | 12.0646749 | 31.0837221 | 0 |
| 4.85E-05 | 0.0001292 | 14.7900348 | 56.0550293 | 25.0574018 | 38.9706367 | 0 |
| 4.86E-05 | 0.00012928 | 10.9317649 | 34.0334106 | 73.3161014 | 53.3526574 | 0 |
| 4.86E-05 | 0.00012935 | 20.8989622 | 33.5885295 | 16.7049345 | 33.4034029 | 0 |

| | | | | | | |
|---|---|---|---|---|---|---|
| 4.86E-05 | 0.00012943 | 14.4685123 | 28.0275146 | 53.8270112 | 32.9394667 | 0 |
| 4.87E-05 | 0.00012963 | 574.560701 | 908.002498 | 772.139195 | 845.755604 | 128.315785 |
| 4.87E-05 | 0.00012964 | 137.290106 | 163.938717 | 110.438178 | 138.252973 | 21.3859642 |
| 4.88E-05 | 0.00012974 | 51.4435994 | 29.5845988 | 29.6976613 | 30.1558498 | 185.345023 |
| 4.88E-05 | 0.00012978 | 12.8608998 | 51.3837769 | 29.6976613 | 42.2181898 | 0 |
| 4.89E-05 | 0.00012996 | 23.4711422 | 16.0157227 | 30.6257133 | 30.619786 | 0 |
| 4.90E-05 | 0.00013021 | 9.96719738 | 38.2597819 | 58.4672708 | 66.3428696 | 0 |
| 4.90E-05 | 0.00013023 | 20.8989622 | 25.8031087 | 18.5610383 | 23.6607437 | 0 |
| 4.90E-05 | 0.00013042 | 127.644431 | 245.351973 | 292.336354 | 247.741905 | 26.1384007 |
| 4.92E-05 | 0.00013072 | 10.9317649 | 48.4920492 | 38.0501286 | 51.0329766 | 0 |
| 4.93E-05 | 0.00013099 | 11.8963324 | 49.1593709 | 31.5537652 | 54.2805297 | 0 |
| 4.93E-05 | 0.0001312 | 21.5420072 | 58.7243164 | 53.8270112 | 91.859358 | 0 |
| 4.94E-05 | 0.00013125 | 5.14435994 | 58.946757 | 122.502853 | 106.705315 | 0 |
| 4.94E-05 | 0.00013129 | 29.9015921 | 24.9133464 | 17.6329864 | 69.5904227 | 0 |
| 4.94E-05 | 0.00013131 | 23.4711422 | 29.5845988 | 36.1940248 | 15.7738291 | 0 |
| 4.94E-05 | 0.00013131 | 13.8254673 | 29.5845988 | 32.4818171 | 43.1460621 | 0 |
| 4.94E-05 | 0.00013136 | 11.5748099 | 11.3444702 | 9.28051917 | 7.42297842 | 30.8908372 |
| 4.95E-05 | 0.00013145 | 13.1824223 | 28.0275146 | 42.6903882 | 52.4247851 | 0 |
| 4.95E-05 | 0.00013147 | 7.39501741 | 48.9369303 | 76.1002572 | 90.0036134 | 0 |
| 4.95E-05 | 0.00013165 | 20.2559173 | 38.7046631 | 16.7049345 | 39.4345729 | 0 |
| 4.96E-05 | 0.00013189 | 16.0761248 | 41.3739502 | 51.0428554 | 21.341063 | 0 |
| 4.96E-05 | 0.00013191 | 19.2913498 | 39.3719849 | 57.5392189 | 18.0935099 | 0 |
| 4.97E-05 | 0.00013197 | 20.2559173 | 31.1416829 | 17.6329864 | 27.8361691 | 0 |
| 4.99E-05 | 0.00013261 | 3.21522496 | 92.757727 | 123.430905 | 141.03659 | 0 |
| 4.99E-05 | 0.00013265 | 13.8254673 | 24.4684652 | 58.4672708 | 51.9608489 | 0 |
| 5.00E-05 | 0.00013285 | 35.6889971 | 29.1397176 | 16.7049345 | 21.341063 | 0 |
| 5.01E-05 | 0.00013297 | 15.7546023 | 55.6101481 | 25.0574018 | 32.4755306 | 0 |
| 5.01E-05 | 0.00013298 | 21.8635297 | 25.5806681 | 33.409869 | 17.1656376 | 0 |
| 5.01E-05 | 0.00013298 | 12.5393773 | 49.3818115 | 30.6257133 | 42.2181898 | 0 |
| 5.01E-05 | 0.00013298 | 51.1220769 | 29.3621582 | 14.8488307 | 27.8361691 | 0 |
| 5.01E-05 | 0.00013314 | 29.2585471 | 82.5254598 | 51.0428554 | 70.9822312 | 4.75243649 |
| 5.02E-05 | 0.00013327 | 14.1469898 | 51.3837769 | 24.1293498 | 49.6411682 | 0 |
| 5.02E-05 | 0.00013333 | 19.2913498 | 38.2597819 | 17.6329864 | 34.7952113 | 0 |
| 5.03E-05 | 0.00013341 | 243.071007 | 350.121492 | 308.113236 | 372.54073 | 33.2670554 |
| 5.03E-05 | 0.00013351 | 322.165541 | 1173.15168 | 800.908804 | 943.646132 | 0 |
| 5.03E-05 | 0.00013354 | 15.7546023 | 37.8149007 | 50.1148035 | 22.2689353 | 0 |
| 5.05E-05 | 0.00013393 | 14.4685123 | 51.828658 | 27.8415575 | 32.4755306 | 0 |
| 5.05E-05 | 0.00013393 | 14.4685123 | 34.923173 | 29.6976613 | 28.3001052 | 0 |
| 5.07E-05 | 0.0001344 | 15.4330798 | 25.8031087 | 29.6976613 | 27.8361691 | 0 |
| 5.08E-05 | 0.00013484 | 9.64567488 | 56.0550293 | 86.3088283 | 38.5067006 | 0 |
| 5.09E-05 | 0.00013508 | 223.136612 | 269.375557 | 203.24337 | 227.328714 | 0 |
| 5.10E-05 | 0.00013526 | 26.6863672 | 28.917277 | 56.6111669 | 15.7738291 | 0 |
| 5.10E-05 | 0.00013529 | 27.3294122 | 140.360014 | 106.72597 | 90.9314857 | 0 |

| | | | | | | |
|---|---|---|---|---|---|---|
| 5.10E-05 | 0.00013529 | 201.273083 | 564.554224 | 425.97583 | 471.823066 | 35.6432737 |
| 5.11E-05 | 0.00013565 | 27.9724572 | 55.3877075 | 13.9207788 | 33.867339 | 0 |
| 5.13E-05 | 0.00013594 | 11.8963324 | 62.0609253 | 40.8342843 | 33.867339 | 0 |
| 5.13E-05 | 0.00013612 | 14.7900348 | 27.1377523 | 44.546492 | 29.2279775 | 0 |
| 5.13E-05 | 0.00013614 | 8.6811074 | 42.9310343 | 0 | 0 | 796.033112 |
| 5.15E-05 | 0.00013657 | 17.0406923 | 24.4684652 | 24.1293498 | 39.898509 | 0 |
| 5.15E-05 | 0.00013657 | 21.8635297 | 15.3484009 | 62.1794784 | 46.8575513 | 0 |
| 5.16E-05 | 0.00013669 | 18.0052598 | 37.5924601 | 103.013763 | 23.1968076 | 0 |
| 5.16E-05 | 0.00013679 | 129.252043 | 157.043058 | 145.704151 | 132.221803 | 47.5243649 |
| 5.16E-05 | 0.00013683 | 17.0406923 | 32.6987671 | 24.1293498 | 23.1968076 | 0 |
| 5.16E-05 | 0.00013683 | 13.1824223 | 74.5175985 | 29.6976613 | 40.3624452 | 0 |
| 5.17E-05 | 0.00013697 | 166.548653 | 260.033053 | 267.278952 | 261.196053 | 28.5146189 |
| 5.17E-05 | 0.00013699 | 13.5039448 | 32.2538859 | 37.1220767 | 30.619786 | 0 |
| 5.18E-05 | 0.00013718 | 18.9698273 | 20.6869751 | 40.8342843 | 24.588616 | 0 |
| 5.18E-05 | 0.00013735 | 10.2887199 | 120.11792 | 33.409869 | 59.3838274 | 0 |
| 5.19E-05 | 0.00013736 | 85.8465065 | 207.759513 | 207.883629 | 183.25478 | 21.3859642 |
| 5.20E-05 | 0.00013785 | 15.1115573 | 29.1397176 | 34.3379209 | 25.0525522 | 0 |
| 5.22E-05 | 0.00013837 | 18.0052598 | 20.9094157 | 24.1293498 | 26.9082968 | 0 |
| 5.23E-05 | 0.00013849 | 16.0761248 | 25.8031087 | 24.1293498 | 30.1558498 | 0 |
| 5.23E-05 | 0.00013857 | 9.64567488 | 45.3778809 | 61.2514265 | 47.3214874 | 0 |
| 5.24E-05 | 0.0001388 | 20.8989622 | 54.2755046 | 73.3161014 | 61.239572 | 2.37621824 |
| 5.26E-05 | 0.00013918 | 4.17979245 | 2.00196533 | 4.64025958 | 3.71148921 | 19.0097459 |
| 5.26E-05 | 0.00013918 | 160.761248 | 313.641235 | 397.20622 | 390.170303 | 30.8908372 |
| 5.26E-05 | 0.00013919 | 10.6102424 | 50.9388957 | 209.739733 | 36.1870198 | 0 |
| 5.26E-05 | 0.00013925 | 63.3399317 | 28.2499552 | 11.136623 | 84.4363795 | 0 |
| 5.26E-05 | 0.00013928 | 1961.93027 | 2706.87957 | 1817.12565 | 2168.90151 | 4.75243649 |
| 5.27E-05 | 0.00013951 | 101.601109 | 247.353939 | 208.811681 | 261.659989 | 23.7621824 |
| 5.27E-05 | 0.00013958 | 42.4409695 | 270.710201 | 378.645182 | 305.269988 | 0 |
| 5.28E-05 | 0.00013964 | 34.4029071 | 35.8129354 | 91.8771398 | 12.9902122 | 0 |
| 5.29E-05 | 0.00013987 | 13.8254673 | 30.4743612 | 39.9062324 | 29.2279775 | 0 |
| 5.29E-05 | 0.00013988 | 12.2178549 | 117.448633 | 108.582074 | 134.541484 | 2.37621824 |
| 5.33E-05 | 0.00014092 | 15.7546023 | 37.8149007 | 22.273246 | 57.0641466 | 0 |
| 5.33E-05 | 0.00014107 | 18.6483048 | 18.4625692 | 33.409869 | 29.2279775 | 0 |
| 5.34E-05 | 0.00014117 | 28.2939797 | 39.1495443 | 43.6184401 | 59.8477635 | 2.37621824 |
| 5.34E-05 | 0.00014125 | 46.2992394 | 119.673039 | 130.85532 | 125.726697 | 14.2573095 |
| 5.34E-05 | 0.00014128 | 17.3622148 | 24.2460246 | 25.9854537 | 51.0329766 | 0 |
| 5.34E-05 | 0.00014132 | 9.64567488 | 39.5944255 | 64.0355823 | 54.7444659 | 0 |
| 5.35E-05 | 0.00014145 | 42.4409695 | 129.460425 | 76.1002572 | 109.952868 | 11.8810912 |
| 5.35E-05 | 0.00014156 | 53.3727344 | 36.2578166 | 38.0501286 | 31.0837221 | 147.325531 |
| 5.35E-05 | 0.00014156 | 10.6102424 | 48.047168 | 42.6903882 | 42.6821259 | 0 |
| 5.36E-05 | 0.00014166 | 12.5393773 | 58.946757 | 29.6976613 | 41.2903175 | 0 |
| 5.37E-05 | 0.00014196 | 30.8661596 | 41.5963908 | 35.2659728 | 45.929679 | 2.37621824 |
| 5.37E-05 | 0.00014204 | 21.5420072 | 47.6022868 | 43.6184401 | 15.309893 | 0 |

| | | | | | | |
|---|---|---|---|---|---|---|
| 5.38E-05 | 0.00014227 | 13.1824223 | 48.7144897 | 30.6257133 | 33.4034029 | 0 |
| 5.39E-05 | 0.00014249 | 8.6811074 | 87.1967122 | 50.1148035 | 48.2493597 | 0 |
| 5.40E-05 | 0.00014281 | 8.3595849 | 42.4861532 | 60.3233746 | 109.952868 | 0 |
| 5.41E-05 | 0.00014287 | 18.0052598 | 39.5944255 | 29.6976613 | 19.4853184 | 0 |
| 5.41E-05 | 0.00014287 | 31.5092046 | 42.7085937 | 17.6329864 | 19.4853184 | 0 |
| 5.41E-05 | 0.00014303 | 10.2887199 | 110.552974 | 23.2012979 | 125.262761 | 0 |
| 5.42E-05 | 0.00014321 | 19.2913498 | 45.1554403 | 16.7049345 | 58.9198912 | 0 |
| 5.42E-05 | 0.0001433 | 26.0433222 | 62.0609253 | 169.833501 | 12.0623399 | 0 |
| 5.43E-05 | 0.00014336 | 0 | 0 | 0 | 0 | 4.75243649 |
| 5.43E-05 | 0.00014336 | 17.0406923 | 45.1554403 | 38.0501286 | 19.4853184 | 0 |
| 5.44E-05 | 0.00014373 | 28.9370247 | 20.4645345 | 16.7049345 | 31.0837221 | 0 |
| 5.44E-05 | 0.0001438 | 125.393773 | 285.836161 | 265.422848 | 283.464988 | 52.2768014 |
| 5.45E-05 | 0.00014384 | 74.9147416 | 35.1456136 | 15.7768826 | 24.588616 | 0 |
| 5.46E-05 | 0.00014417 | 33.4383396 | 85.8620687 | 89.092984 | 8.81478687 | 0 |
| 5.46E-05 | 0.00014426 | 186.483048 | 439.097729 | 361.940248 | 440.739344 | 4.75243649 |
| 5.48E-05 | 0.00014465 | 68.8058142 | 14.0137573 | 32.4818171 | 30.619786 | 0 |
| 5.48E-05 | 0.00014465 | 13.1824223 | 39.3719849 | 27.8415575 | 39.898509 | 0 |
| 5.49E-05 | 0.0001448 | 17.6837373 | 38.7046631 | 18.5610383 | 35.7230837 | 0 |
| 5.49E-05 | 0.00014485 | 232.139242 | 463.343754 | 477.018685 | 499.195299 | 57.0292378 |
| 5.50E-05 | 0.00014509 | 170.0854 | 179.954439 | 124.358957 | 143.356271 | 28.5146189 |
| 5.51E-05 | 0.00014553 | 11.5748099 | 40.0393066 | 38.9781805 | 38.5067006 | 0 |
| 5.52E-05 | 0.00014556 | 10.9317649 | 66.064856 | 39.9062324 | 38.5067006 | 0 |
| 5.52E-05 | 0.00014561 | 32.4737721 | 25.8031087 | 13.9207788 | 33.867339 | 0 |
| 5.52E-05 | 0.00014563 | 9.96719738 | 45.822762 | 53.8270112 | 43.1460621 | 0 |
| 5.52E-05 | 0.00014569 | 67.8412467 | 102.989994 | 89.092984 | 108.561059 | 16.6335277 |
| 5.53E-05 | 0.00014582 | 16.3976473 | 21.3542969 | 28.7696094 | 30.619786 | 0 |
| 5.53E-05 | 0.00014585 | 33.1168171 | 43.8207967 | 56.6111669 | 49.177232 | 2.37621824 |
| 5.53E-05 | 0.00014588 | 20.5774398 | 52.940861 | 15.7768826 | 41.7542536 | 0 |
| 5.53E-05 | 0.00014588 | 19.6128723 | 19.5747721 | 33.409869 | 22.2689353 | 0 |
| 5.56E-05 | 0.00014657 | 41.476402 | 76.2971232 | 57.5392189 | 8.81478687 | 0 |
| 5.57E-05 | 0.00014678 | 21.5420072 | 55.3877075 | 97.4454513 | 54.2805297 | 0 |
| 5.58E-05 | 0.00014702 | 11.8963324 | 42.041272 | 36.1940248 | 36.1870198 | 0 |
| 5.60E-05 | 0.00014767 | 19.9343948 | 20.4645345 | 27.8415575 | 19.9492545 | 0 |
| 5.60E-05 | 0.00014773 | 14.4685123 | 34.923173 | 37.1220767 | 24.1246799 | 0 |
| 5.64E-05 | 0.00014858 | 13.1824223 | 35.5904948 | 28.7696094 | 37.1148921 | 0 |
| 5.64E-05 | 0.00014859 | 19.2913498 | 31.3641235 | 20.4171422 | 21.341063 | 0 |
| 5.64E-05 | 0.00014859 | 75.2362641 | 279.607825 | 211.595837 | 242.638607 | 21.3859642 |
| 5.64E-05 | 0.00014877 | 27.9724572 | 15.1259603 | 27.8415575 | 25.9804245 | 0 |
| 5.65E-05 | 0.00014888 | 35.0459521 | 26.9153117 | 25.9854537 | 14.8459568 | 0 |
| 5.65E-05 | 0.00014898 | 10.6102424 | 64.5077718 | 32.4818171 | 54.7444659 | 0 |
| 5.66E-05 | 0.00014912 | 219.599865 | 656.199748 | 554.046994 | 568.321785 | 0 |
| 5.67E-05 | 0.00014933 | 20.2559173 | 28.0275146 | 16.7049345 | 32.0115944 | 0 |
| 5.68E-05 | 0.00014973 | 11.5748099 | 62.7282471 | 34.3379209 | 38.9706367 | 0 |

| | | | | | | |
|---|---|---|---|---|---|---|
| 5.69E-05 | 0.00014991 | 106.745469 | 399.948185 | 244.077654 | 372.54073 | 26.1384007 |
| 5.69E-05 | 0.00014994 | 12.8608998 | 40.2617472 | 28.7696094 | 38.0427644 | 0 |
| 5.69E-05 | 0.00015 | 117.998756 | 270.932642 | 252.430121 | 284.856797 | 26.1384007 |
| 5.70E-05 | 0.00015002 | 19.2913498 | 30.4743612 | 18.5610383 | 51.4969128 | 0 |
| 5.70E-05 | 0.00015018 | 33.7598621 | 43.5983561 | 36.1940248 | 34.3312752 | 0 |
| 5.70E-05 | 0.00015024 | 16.7191698 | 43.3759155 | 21.3451941 | 99.7462725 | 0 |
| 5.71E-05 | 0.00015025 | 20.8989622 | 24.4684652 | 17.6329864 | 23.6607437 | 0 |
| 5.72E-05 | 0.00015048 | 13.1824223 | 98.5411824 | 28.7696094 | 42.2181898 | 0 |
| 5.72E-05 | 0.00015064 | 25.0787547 | 43.5983561 | 21.3451941 | 18.0935099 | 0 |
| 5.73E-05 | 0.00015074 | 16.3976473 | 24.023584 | 23.2012979 | 27.8361691 | 0 |
| 5.74E-05 | 0.00015104 | 23.1496197 | 68.2892619 | 77.0283091 | 72.8379758 | 0 |
| 5.74E-05 | 0.0001511 | 147.257303 | 267.596033 | 213.451941 | 126.654569 | 16.6335277 |
| 5.75E-05 | 0.00015131 | 25.4002772 | 42.4861532 | 12.9927268 | 49.177232 | 0 |
| 5.75E-05 | 0.00015144 | 16.0761248 | 31.5865641 | 33.409869 | 20.8771268 | 0 |
| 5.76E-05 | 0.00015171 | 12.5393773 | 31.5865641 | 36.1940248 | 37.1148921 | 0 |
| 5.77E-05 | 0.00015175 | 10.9317649 | 43.3759155 | 98.3735032 | 35.7230837 | 0 |
| 5.77E-05 | 0.00015179 | 13.8254673 | 31.3641235 | 31.5537652 | 28.3001052 | 0 |
| 5.78E-05 | 0.00015201 | 185.840003 | 479.804358 | 346.163365 | 384.603069 | 33.2670554 |
| 5.78E-05 | 0.00015212 | 122.178549 | 251.135429 | 192.106747 | 226.864778 | 30.8908372 |
| 5.79E-05 | 0.00015222 | 14.1469898 | 23.5787028 | 40.8342843 | 41.2903175 | 0 |
| 5.79E-05 | 0.00015228 | 27.9724572 | 118.560836 | 110.438178 | 110.88074 | 9.50487297 |
| 5.80E-05 | 0.00015253 | 10.9317649 | 75.852242 | 33.409869 | 45.929679 | 0 |
| 5.85E-05 | 0.00015383 | 6.75197242 | 71.1809896 | 124.358957 | 46.3936151 | 0 |
| 5.88E-05 | 0.00015472 | 15.1115573 | 31.5865641 | 28.7696094 | 23.6607437 | 0 |
| 5.88E-05 | 0.00015473 | 17.3622148 | 42.2637126 | 23.2012979 | 23.6607437 | 0 |
| 5.89E-05 | 0.00015478 | 136.647061 | 264.704305 | 340.595054 | 351.663603 | 28.5146189 |
| 5.89E-05 | 0.00015483 | 15.1115573 | 27.5826335 | 24.1293498 | 31.5476583 | 0 |
| 5.90E-05 | 0.00015514 | 13.5039448 | 22.911381 | 53.8270112 | 74.2297842 | 0 |
| 5.91E-05 | 0.0001553 | 16.0761248 | 21.7991781 | 26.9135056 | 32.0115944 | 0 |
| 5.93E-05 | 0.00015591 | 9.96719738 | 96.3167765 | 28.7696094 | 72.3740396 | 0 |
| 5.94E-05 | 0.00015618 | 62.3753642 | 20.0196533 | 20.4171422 | 27.3722329 | 0 |
| 5.97E-05 | 0.0001569 | 105.137856 | 37.8149007 | 8.35246725 | 61.7035081 | 0 |
| 5.98E-05 | 0.00015705 | 0.3215225 | 0 | 0 | 0 | 47.5243649 |
| 6.00E-05 | 0.00015762 | 157.546023 | 425.306413 | 352.659728 | 457.441045 | 7.12865473 |
| 6.00E-05 | 0.00015768 | 93.8845689 | 27.3601929 | 15.7768826 | 32.0115944 | 0 |
| 6.01E-05 | 0.0001579 | 10.6102424 | 57.1672323 | 40.8342843 | 37.5788283 | 0 |
| 6.01E-05 | 0.0001579 | 49.1929419 | 262.257458 | 199.531162 | 212.482757 | 21.3859642 |
| 6.02E-05 | 0.00015809 | 40.5118345 | 63.84045 | 8.35246725 | 65.8789335 | 0 |
| 6.02E-05 | 0.00015815 | 24.7572322 | 10.2322673 | 11.136623 | 7.88691457 | 61.7816743 |
| 6.04E-05 | 0.00015857 | 28.6155022 | 59.8365194 | 52.8989593 | 58.9198912 | 4.75243649 |
| 6.05E-05 | 0.0001589 | 11.8963324 | 30.4743612 | 43.6184401 | 41.2903175 | 0 |
| 6.06E-05 | 0.00015913 | 78.4514891 | 271.377523 | 193.034799 | 194.389247 | 26.1384007 |
| 6.06E-05 | 0.00015918 | 69.1273367 | 13.3464355 | 20.4171422 | 119.695527 | 0 |

| | | | | | | |
|---|---|---|---|---|---|---|
| 6.07E-05 | 0.00015929 | 25.0787547 | 17.5728068 | 57.5392189 | 24.1246799 | 0 |
| 6.07E-05 | 0.00015933 | 18.3267823 | 28.6948364 | 136.423632 | 28.7640414 | 0 |
| 6.07E-05 | 0.00015933 | 16.0761248 | 25.1357869 | 22.273246 | 31.5476583 | 0 |
| 6.08E-05 | 0.00015949 | 14.4685123 | 23.1338216 | 36.1940248 | 34.7952113 | 0 |
| 6.08E-05 | 0.0001595 | 38.5826995 | 93.6474894 | 92.8051917 | 93.2511664 | 11.8810912 |
| 6.08E-05 | 0.00015963 | 0 | 0 | 0 | 0 | 4.75243649 |
| 6.08E-05 | 0.00015965 | 11.5748099 | 52.4959798 | 32.4818171 | 38.9706367 | 0 |
| 6.09E-05 | 0.00015985 | 6.75197242 | 5.7834554 | 11.136623 | 10.6705315 | 54.6530196 |
| 6.11E-05 | 0.0001603 | 7.07349491 | 70.0687866 | 52.8989593 | 69.5904227 | 0 |
| 6.12E-05 | 0.00016056 | 32.4737721 | 45.1554403 | 32.4818171 | 38.9706367 | 0 |
| 6.14E-05 | 0.0001611 | 62.3753642 | 141.917098 | 166.121293 | 147.531696 | 0 |
| 6.14E-05 | 0.0001612 | 15.1115573 | 45.3778809 | 55.683115 | 20.4131907 | 0 |
| 6.15E-05 | 0.00016143 | 32.1522496 | 53.6081828 | 11.136623 | 39.898509 | 0 |
| 6.16E-05 | 0.00016158 | 16.0761248 | 28.4723958 | 21.3451941 | 27.3722329 | 0 |
| 6.18E-05 | 0.00016205 | 24.7572322 | 21.7991781 | 20.4171422 | 65.8789335 | 0 |
| 6.19E-05 | 0.0001623 | 412.834885 | 18.4625692 | 12.0646749 | 65.4149973 | 0 |
| 6.19E-05 | 0.00016235 | 24.7572322 | 14.9035197 | 13.9207788 | 18.5574461 | 90.2962933 |
| 6.19E-05 | 0.00016243 | 68.4842917 | 85.8620687 | 9.28051917 | 31.0837221 | 0 |
| 6.20E-05 | 0.00016244 | 13.5039448 | 22.911381 | 54.7550631 | 49.177232 | 0 |
| 6.20E-05 | 0.00016261 | 19.9343948 | 20.0196533 | 22.273246 | 20.4131907 | 0 |
| 6.20E-05 | 0.00016265 | 0 | 0.22244059 | 0 | 0 | 0 |
| 6.21E-05 | 0.00016279 | 15.7546023 | 22.4664998 | 25.9854537 | 33.867339 | 0 |
| 6.22E-05 | 0.00016309 | 0 | 0 | 0 | 0 | 47.5243649 |
| 6.23E-05 | 0.00016317 | 12.5393773 | 43.3759155 | 99.3015551 | 28.3001052 | 0 |
| 6.23E-05 | 0.00016329 | 14.1469898 | 51.1613363 | 56.6111669 | 21.341063 | 0 |
| 6.23E-05 | 0.00016332 | 13.5039448 | 28.0275146 | 30.6257133 | 32.9394667 | 0 |
| 6.24E-05 | 0.00016346 | 11.8963324 | 50.9388957 | 29.6976613 | 40.3624452 | 0 |
| 6.24E-05 | 0.00016358 | 13.8254673 | 30.9192423 | 44.546492 | 25.9804245 | 0 |
| 6.25E-05 | 0.00016373 | 32.1522496 | 176.39539 | 78.8844129 | 110.88074 | 9.50487297 |
| 6.26E-05 | 0.00016391 | 9.32415239 | 49.6042521 | 64.9636342 | 39.4345729 | 0 |
| 6.26E-05 | 0.00016394 | 43.0840145 | 50.2715739 | 31.5537652 | 60.3116997 | 4.75243649 |
| 6.26E-05 | 0.00016395 | 19.9343948 | 94.5372518 | 13.9207788 | 63.0953166 | 0 |
| 6.27E-05 | 0.00016419 | 12.5393773 | 29.3621582 | 50.1148035 | 34.3312752 | 0 |
| 6.28E-05 | 0.00016449 | 13.1824223 | 38.2597819 | 53.8270112 | 25.5164883 | 0 |
| 6.29E-05 | 0.00016474 | 21.5420072 | 53.8306234 | 58.4672708 | 51.9608489 | 0 |
| 6.29E-05 | 0.00016482 | 12.8608998 | 35.8129354 | 32.4818171 | 29.6919137 | 0 |
| 6.30E-05 | 0.00016503 | 28.6155022 | 17.5728068 | 85.3807764 | 25.0525522 | 0 |
| 6.30E-05 | 0.00016504 | 4.82283744 | 55.3877075 | 111.36623 | 105.313506 | 0 |
| 6.31E-05 | 0.00016511 | 14.4685123 | 52.940861 | 20.4171422 | 60.3116997 | 0 |
| 6.31E-05 | 0.0001652 | 26.0433222 | 105.881722 | 54.7550631 | 37.5788283 | 2.37621824 |
| 6.31E-05 | 0.00016521 | 57.5525268 | 122.119885 | 108.582074 | 121.087335 | 0 |
| 6.31E-05 | 0.00016524 | 15.4330798 | 27.5826335 | 22.273246 | 30.1558498 | 0 |
| 6.32E-05 | 0.00016539 | 25.0787547 | 21.3542969 | 34.3379209 | 16.2377653 | 0 |

| | | | | | | |
|---|---|---|---|---|---|---|
| 6.32E-05 | 0.0001654 | 26.0433222 | 68.0668213 | 56.6111669 | 87.6839326 | 4.75243649 |
| 6.32E-05 | 0.0001654 | 22.8280972 | 23.5787028 | 17.6329864 | 20.4131907 | 0 |
| 6.33E-05 | 0.00016563 | 13.5039448 | 33.1436483 | 25.9854537 | 45.0018067 | 0 |
| 6.33E-05 | 0.0001657 | 18.0052598 | 27.5826335 | 24.1293498 | 19.0213822 | 0 |
| 6.34E-05 | 0.00016586 | 10.9317649 | 32.6987671 | 71.4599976 | 42.6821259 | 0 |
| 6.34E-05 | 0.00016596 | 25.0787547 | 26.2479899 | 14.8488307 | 25.5164883 | 0 |
| 6.36E-05 | 0.00016627 | 13.8254673 | 23.5787028 | 38.9781805 | 38.5067006 | 0 |
| 6.37E-05 | 0.00016653 | 16.0761248 | 36.4802572 | 36.1940248 | 19.4853184 | 0 |
| 6.37E-05 | 0.00016665 | 11.2532874 | 45.3778809 | 30.6257133 | 50.1051043 | 0 |
| 6.37E-05 | 0.00016671 | 27.6509347 | 36.2578166 | 47.3306478 | 12.5262761 | 0 |
| 6.39E-05 | 0.00016704 | 26.3648447 | 69.6239054 | 11.136623 | 58.4559551 | 0 |
| 6.39E-05 | 0.00016709 | 42.762492 | 40.7066284 | 34.3379209 | 50.1051043 | 4.75243649 |
| 6.39E-05 | 0.00016719 | 22.8280972 | 25.5806681 | 45.4745439 | 16.2377653 | 0 |
| 6.40E-05 | 0.00016728 | 11.2532874 | 76.5195638 | 101.157659 | 156.346483 | 0 |
| 6.41E-05 | 0.00016769 | 39.2257445 | 42.7085937 | 41.7623363 | 27.3722329 | 0 |
| 6.43E-05 | 0.00016802 | 33.7598621 | 46.934965 | 47.3306478 | 49.177232 | 7.12865473 |
| 6.44E-05 | 0.00016832 | 32.1522496 | 20.6869751 | 21.3451941 | 18.0935099 | 0 |
| 6.44E-05 | 0.00016839 | 14.4685123 | 35.8129354 | 22.273246 | 37.5788283 | 0 |
| 6.45E-05 | 0.00016856 | 12.5393773 | 26.6928711 | 39.9062324 | 46.3936151 | 0 |
| 6.46E-05 | 0.00016886 | 10.9317649 | 62.0609253 | 28.7696094 | 54.2805297 | 0 |
| 6.46E-05 | 0.00016887 | 37.2966096 | 16.0157227 | 19.4890903 | 48.2493597 | 0 |
| 6.47E-05 | 0.00016891 | 30.5446371 | 27.1377523 | 14.8488307 | 22.7328714 | 0 |
| 6.47E-05 | 0.00016912 | 25.4002772 | 52.0510986 | 12.0646749 | 49.177232 | 0 |
| 6.48E-05 | 0.00016916 | 183.589345 | 418.188314 | 397.20622 | 410.119558 | 0 |
| 6.48E-05 | 0.00016925 | 7.39501741 | 50.2715739 | 113.222334 | 51.4969128 | 0 |
| 6.49E-05 | 0.00016954 | 13.1824223 | 30.02948 | 29.6976613 | 32.4755306 | 0 |
| 6.49E-05 | 0.00016958 | 90.9908664 | 15.1259603 | 79.8124649 | 16.7017014 | 0 |
| 6.49E-05 | 0.00016959 | 18.0052598 | 29.3621582 | 20.4171422 | 21.341063 | 0 |
| 6.50E-05 | 0.00016981 | 14.1469898 | 39.816866 | 22.273246 | 41.7542536 | 0 |
| 6.52E-05 | 0.00017023 | 6.43044992 | 56.9447917 | 109.510126 | 57.0641466 | 0 |
| 6.52E-05 | 0.00017029 | 12.8608998 | 60.7262817 | 64.9636342 | 22.2689353 | 0 |
| 6.53E-05 | 0.0001705 | 55.9449143 | 22.4664998 | 24.1293498 | 18.5574461 | 0 |
| 6.53E-05 | 0.0001705 | 43.405537 | 58.5018758 | 7.42441534 | 102.065953 | 0 |
| 6.53E-05 | 0.00017052 | 167.834743 | 334.995532 | 205.099474 | 260.268181 | 9.50487297 |
| 6.53E-05 | 0.00017054 | 88.7402089 | 93.6474894 | 58.4672708 | 97.8905279 | 19.0097459 |
| 6.55E-05 | 0.00017088 | 145.971213 | 59.3916382 | 9.28051917 | 31.5476583 | 0 |
| 6.56E-05 | 0.00017118 | 18.0052598 | 30.02948 | 17.6329864 | 28.3001052 | 0 |
| 6.57E-05 | 0.00017142 | 10.9317649 | 41.8188314 | 32.4818171 | 68.1986142 | 0 |
| 6.59E-05 | 0.00017205 | 28.2939797 | 22.911381 | 15.7768826 | 22.7328714 | 0 |
| 6.62E-05 | 0.00017259 | 18.3267823 | 17.1279256 | 38.9781805 | 31.5476583 | 0 |
| 6.62E-05 | 0.00017276 | 17.3622148 | 28.6948364 | 18.5610383 | 26.9082968 | 0 |
| 6.65E-05 | 0.00017353 | 21.8635297 | 15.7932821 | 33.409869 | 25.0525522 | 0 |
| 6.66E-05 | 0.00017381 | 15.1115573 | 46.7125244 | 22.273246 | 31.0837221 | 0 |

| | | | | | | |
|---|---|---|---|---|---|---|
| 6.66E-05 | 0.00017381 | 30.8661596 | 47.8247274 | 13.9207788 | 24.588616 | 0 |
| 6.67E-05 | 0.00017393 | 12.5393773 | 24.9133464 | 45.4745439 | 60.3116997 | 0 |
| 6.68E-05 | 0.00017407 | 18.0052598 | 28.2499552 | 20.4171422 | 20.8771268 | 0 |
| 6.68E-05 | 0.00017418 | 81.023669 | 417.076111 | 289.552198 | 322.435625 | 2.37621824 |
| 6.69E-05 | 0.00017436 | 15.7546023 | 24.2460246 | 22.273246 | 30.1558498 | 0 |
| 6.69E-05 | 0.00017451 | 19.2913498 | 58.0569946 | 29.6976613 | 18.5574461 | 0 |
| 6.71E-05 | 0.00017484 | 17.0406923 | 28.2499552 | 18.5610383 | 33.4034029 | 0 |
| 6.73E-05 | 0.00017552 | 9.00262989 | 39.1495443 | 64.9636342 | 50.5690405 | 0 |
| 6.74E-05 | 0.00017562 | 10.6102424 | 37.8149007 | 66.819738 | 36.1870198 | 0 |
| 6.74E-05 | 0.00017572 | 7.39501741 | 62.9506877 | 61.2514265 | 51.4969128 | 0 |
| 6.74E-05 | 0.00017572 | 28.6155022 | 98.0963013 | 100.229607 | 103.457762 | 9.50487297 |
| 6.76E-05 | 0.00017605 | 31.1876821 | 36.9251383 | 13.9207788 | 23.1968076 | 0 |
| 6.76E-05 | 0.00017607 | 0 | 0 | 0 | 0 | 16.6335277 |
| 6.78E-05 | 0.00017664 | 45.334672 | 60.2814006 | 7.42441534 | 70.0543588 | 0 |
| 6.79E-05 | 0.00017686 | 9.32415239 | 40.4841878 | 56.6111669 | 45.929679 | 0 |
| 6.79E-05 | 0.00017696 | 0 | 0 | 0 | 0 | 0 |
| 6.80E-05 | 0.00017701 | 5.46588243 | 51.1613363 | 99.3015551 | 90.0036134 | 0 |
| 6.81E-05 | 0.00017743 | 10.6102424 | 32.0314453 | 45.4745439 | 52.8887212 | 0 |
| 6.82E-05 | 0.00017756 | 10.6102424 | 46.2676432 | 38.9781805 | 36.650956 | 0 |
| 6.82E-05 | 0.00017757 | 6.43044992 | 43.8207967 | 117.862593 | 78.8691457 | 0 |
| 6.83E-05 | 0.00017783 | 49.5144644 | 21.1318563 | 22.273246 | 19.4853184 | 0 |
| 6.84E-05 | 0.000178 | 28.2939797 | 15.3484009 | 11.136623 | 19.0213822 | 78.415202 |
| 6.86E-05 | 0.00017862 | 14.7900348 | 48.047168 | 21.3451941 | 35.2591475 | 0 |
| 6.86E-05 | 0.00017864 | 11.8963324 | 30.02948 | 43.6184401 | 35.2591475 | 0 |
| 6.86E-05 | 0.00017864 | 19.9343948 | 48.4920492 | 14.8488307 | 40.8263813 | 0 |
| 6.88E-05 | 0.00017897 | 19.9343948 | 15.5708415 | 33.409869 | 38.9706367 | 0 |
| 6.90E-05 | 0.00017962 | 14.7900348 | 27.8050741 | 24.1293498 | 25.9804245 | 0 |
| 6.91E-05 | 0.00017979 | 17.3622148 | 123.454529 | 113.222334 | 83.0445711 | 4.75243649 |
| 6.91E-05 | 0.00017982 | 87.1325964 | 119.005717 | 132.711424 | 97.4265918 | 19.0097459 |
| 6.91E-05 | 0.00017982 | 15.1115573 | 30.2519206 | 21.3451941 | 30.1558498 | 0 |
| 6.93E-05 | 0.00018032 | 25.7217997 | 34.2558512 | 14.8488307 | 22.7328714 | 0 |
| 6.93E-05 | 0.00018032 | 22.1850522 | 17.3503662 | 90.9490879 | 32.4755306 | 0 |
| 6.95E-05 | 0.00018079 | 18.3267823 | 56.2774699 | 16.7049345 | 37.5788283 | 0 |
| 6.95E-05 | 0.00018079 | 13.1824223 | 52.940861 | 57.5392189 | 21.8049991 | 0 |
| 6.96E-05 | 0.0001811 | 15.4330798 | 30.4743612 | 21.3451941 | 47.3214874 | 0 |
| 6.96E-05 | 0.00018112 | 85.524984 | 32.9212077 | 12.9927268 | 30.1558498 | 0 |
| 6.97E-05 | 0.00018117 | 26.0433222 | 104.102197 | 25.0574018 | 19.4853184 | 0 |
| 6.97E-05 | 0.00018128 | 235.032945 | 382.152938 | 333.170638 | 350.271794 | 0 |
| 6.98E-05 | 0.00018164 | 8.3595849 | 39.816866 | 114.150386 | 51.0329766 | 0 |
| 7.00E-05 | 0.00018191 | 32.1522496 | 47.6022868 | 19.4890903 | 16.2377653 | 0 |
| 7.01E-05 | 0.00018235 | 11.2532874 | 27.8050741 | 82.5966206 | 47.3214874 | 0 |
| 7.01E-05 | 0.00018235 | 17.0406923 | 23.5787028 | 22.273246 | 20.8771268 | 0 |
| 7.02E-05 | 0.00018262 | 24.7572322 | 29.8070394 | 19.4890903 | 16.7017014 | 0 |

| | | | | | | |
|---|---|---|---|---|---|---|
| 7.03E-05 | 0.00018287 | 13.8254673 | 24.4684652 | 34.3379209 | 29.2279775 | 0 |
| 7.04E-05 | 0.00018304 | 153.044708 | 288.95033 | 234.797135 | 240.318926 | 57.0292378 |
| 7.06E-05 | 0.00018339 | 10.9317649 | 57.6121134 | 25.9854537 | 78.8691457 | 0 |
| 7.08E-05 | 0.0001839 | 15.1115573 | 61.6160441 | 19.4890903 | 43.6099982 | 0 |
| 7.08E-05 | 0.00018391 | 18.3267823 | 103.879757 | 51.9709073 | 65.4149973 | 0 |
| 7.08E-05 | 0.0001841 | 22.1850522 | 14.0137573 | 41.7623363 | 35.7230837 | 0 |
| 7.09E-05 | 0.00018434 | 14.1469898 | 34.923173 | 22.273246 | 34.3312752 | 0 |
| 7.11E-05 | 0.00018485 | 22.5065747 | 30.02948 | 14.8488307 | 25.0525522 | 0 |
| 7.12E-05 | 0.00018495 | 218.313775 | 544.53457 | 438.968557 | 488.988703 | 38.0194919 |
| 7.12E-05 | 0.00018505 | 12.8608998 | 34.4782918 | 36.1940248 | 25.9804245 | 0 |
| 7.13E-05 | 0.00018531 | 8.6811074 | 50.4940145 | 77.956361 | 38.5067006 | 0 |
| 7.14E-05 | 0.00018543 | 33.7598621 | 54.2755046 | 13.9207788 | 23.1968076 | 0 |
| 7.15E-05 | 0.00018563 | 18.3267823 | 34.923173 | 36.1940248 | 16.2377653 | 0 |
| 7.16E-05 | 0.00018598 | 23.4711422 | 49.6042521 | 53.8270112 | 42.6821259 | 0 |
| 7.16E-05 | 0.00018598 | 10.9317649 | 45.1554403 | 36.1940248 | 34.7952113 | 0 |
| 7.16E-05 | 0.00018599 | 18.6483048 | 21.1318563 | 18.5610383 | 25.0525522 | 0 |
| 7.18E-05 | 0.00018645 | 12.8608998 | 5.33857422 | 7.42441534 | 8.81478687 | 21.3859642 |
| 7.18E-05 | 0.00018649 | 9.00262989 | 46.7125244 | 47.3306478 | 45.929679 | 0 |
| 7.20E-05 | 0.00018684 | 13.8254673 | 22.6889404 | 34.3379209 | 37.5788283 | 0 |
| 7.20E-05 | 0.0001869 | 211.883325 | 92.757727 | 96.5173994 | 104.385634 | 11.8810912 |
| 7.20E-05 | 0.00018692 | 45.9777169 | 24.2460246 | 16.7049345 | 21.8049991 | 0 |
| 7.21E-05 | 0.0001872 | 16.7191698 | 27.3601929 | 25.9854537 | 19.0213822 | 0 |
| 7.22E-05 | 0.00018731 | 18.3267823 | 27.3601929 | 20.4171422 | 19.4853184 | 0 |
| 7.22E-05 | 0.00018745 | 15.7546023 | 65.8424154 | 77.0283091 | 77.9412734 | 0 |
| 7.23E-05 | 0.00018758 | 22.5065747 | 52.4959798 | 13.9207788 | 34.7952113 | 0 |
| 7.23E-05 | 0.00018764 | 16.0761248 | 34.2558512 | 18.5610383 | 42.2181898 | 0 |
| 7.24E-05 | 0.0001879 | 12.8608998 | 39.1495443 | 24.1293498 | 48.7132959 | 0 |
| 7.24E-05 | 0.00018794 | 94.5276139 | 77.8542074 | 188.394539 | 109.488932 | 30.8908372 |
| 7.25E-05 | 0.00018804 | 16.7191698 | 17.5728068 | 38.9781805 | 48.2493597 | 0 |
| 7.25E-05 | 0.00018804 | 35.0459521 | 35.1456136 | 25.0574018 | 12.9902122 | 0 |
| 7.25E-05 | 0.00018804 | 36.3320421 | 44.2656779 | 9.28051917 | 66.8068058 | 0 |
| 7.25E-05 | 0.00018808 | 14.1469898 | 25.5806681 | 29.6976613 | 25.5164883 | 0 |
| 7.26E-05 | 0.00018813 | 12.5393773 | 33.3660889 | 52.8989593 | 27.3722329 | 0 |
| 7.27E-05 | 0.00018859 | 14.4685123 | 38.7046631 | 24.1293498 | 26.4443606 | 0 |
| 7.28E-05 | 0.00018874 | 41.7979245 | 37.1475789 | 26.9135056 | 40.8263813 | 0 |
| 7.28E-05 | 0.00018877 | 22.5065747 | 24.2460246 | 65.8916861 | 18.5574461 | 0 |
| 7.29E-05 | 0.00018905 | 22.8280972 | 12.011792 | 76.1002572 | 57.5280828 | 0 |
| 7.31E-05 | 0.00018938 | 243.714052 | 410.180452 | 417.623363 | 421.717962 | 30.8908372 |
| 7.31E-05 | 0.00018953 | 0 | 982.742537 | 638.499719 | 845.291668 | 0 |
| 7.32E-05 | 0.00018975 | 27.3294122 | 54.053064 | 53.8270112 | 62.6313804 | 4.75243649 |
| 7.35E-05 | 0.00019055 | 10.2887199 | 43.3759155 | 38.9781805 | 38.0427644 | 0 |
| 7.36E-05 | 0.00019072 | 14.1469898 | 29.8070394 | 23.2012979 | 42.6821259 | 0 |
| 7.36E-05 | 0.00019072 | 7.71653991 | 33.3660889 | 78.8844129 | 91.859358 | 0 |

| | | | | | | |
|---|---|---|---|---|---|---|
| 7.36E-05 | 0.00019076 | 121.535504 | 512.280684 | 470.522322 | 496.875618 | 76.0389838 |
| 7.40E-05 | 0.00019174 | 34.7244296 | 74.0727173 | 25.9854537 | 50.1051043 | 0 |
| 7.41E-05 | 0.00019187 | 15.1115573 | 77.8542074 | 23.2012979 | 33.867339 | 0 |
| 7.41E-05 | 0.00019205 | 386.47004 | 742.729138 | 581.888552 | 591.518593 | 47.5243649 |
| 7.42E-05 | 0.00019211 | 31.5092046 | 26.9153117 | 12.9927268 | 60.7756358 | 0 |
| 7.42E-05 | 0.00019229 | 0 | 0 | 0 | 0 | 54.6530196 |
| 7.45E-05 | 0.00019304 | 321.522496 | 419.078076 | 268.207004 | 336.817646 | 4.75243649 |
| 7.47E-05 | 0.00019345 | 48.5498969 | 21.3542969 | 16.7049345 | 25.0525522 | 0 |
| 7.47E-05 | 0.0001935 | 10.9317649 | 55.3877075 | 40.8342843 | 30.1558498 | 0 |
| 7.48E-05 | 0.00019353 | 7.71653991 | 43.3759155 | 55.683115 | 70.9822312 | 0 |
| 7.48E-05 | 0.00019354 | 110.925261 | 182.623726 | 180.042072 | 185.110524 | 23.7621824 |
| 7.50E-05 | 0.00019411 | 202.23765 | 381.708057 | 318.321808 | 374.86041 | 80.7914203 |
| 7.51E-05 | 0.00019437 | 32.4737721 | 18.9074504 | 35.2659728 | 28.7640414 | 90.2962933 |
| 7.52E-05 | 0.00019468 | 0 | 0 | 0 | 0 | 42.7719284 |
| 7.52E-05 | 0.00019468 | 58.5170943 | 193.968197 | 137.351684 | 252.381266 | 0 |
| 7.54E-05 | 0.00019502 | 28.2939797 | 48.7144897 | 55.683115 | 54.2805297 | 4.75243649 |
| 7.55E-05 | 0.00019531 | 10.9317649 | 51.1613363 | 26.9135056 | 57.9920189 | 0 |
| 7.57E-05 | 0.00019585 | 58.8386168 | 432.869393 | 132.711424 | 458.368917 | 19.0097459 |
| 7.58E-05 | 0.00019607 | 16.7191698 | 22.6889404 | 22.273246 | 20.8771268 | 0 |
| 7.58E-05 | 0.00019616 | 17.3622148 | 26.6928711 | 22.273246 | 19.0213822 | 0 |
| 7.59E-05 | 0.00019627 | 77.1653991 | 19.3523315 | 15.7768826 | 39.898509 | 0 |
| 7.59E-05 | 0.00019634 | 4.17979245 | 95.4270142 | 68.6758419 | 94.1790387 | 0 |
| 7.60E-05 | 0.00019647 | 13.1824223 | 22.4664998 | 44.546492 | 40.3624452 | 0 |
| 7.61E-05 | 0.0001967 | 43.7270595 | 31.3641235 | 12.0646749 | 27.8361691 | 0 |
| 7.61E-05 | 0.00019682 | 11.5748099 | 29.3621582 | 38.9781805 | 37.1148921 | 0 |
| 7.62E-05 | 0.00019713 | 14.1469898 | 21.7991781 | 32.4818171 | 32.4755306 | 0 |
| 7.64E-05 | 0.00019742 | 22.5065747 | 24.2460246 | 28.7696094 | 14.8459568 | 0 |
| 7.67E-05 | 0.00019832 | 15.4330798 | 24.023584 | 21.3451941 | 30.1558498 | 0 |
| 7.68E-05 | 0.00019853 | 218.313775 | 269.153117 | 167.977397 | 233.359884 | 0 |
| 7.69E-05 | 0.00019886 | 15.1115573 | 32.0314453 | 20.4171422 | 28.3001052 | 0 |
| 7.69E-05 | 0.0001989 | 21.5420072 | 20.0196533 | 20.4171422 | 18.0935099 | 0 |
| 7.70E-05 | 0.0001989 | 16.7191698 | 9.34250488 | 10.2085711 | 12.0623399 | 33.2670554 |
| 7.71E-05 | 0.00019926 | 9.64567488 | 43.5983561 | 40.8342843 | 41.7542536 | 0 |
| 7.71E-05 | 0.00019926 | 16.7191698 | 19.1298909 | 27.8415575 | 23.1968076 | 0 |
| 7.73E-05 | 0.00019963 | 22.5065747 | 72.0707519 | 61.2514265 | 63.0953166 | 2.37621824 |
| 7.73E-05 | 0.00019964 | 4.82283744 | 99.4309448 | 61.2514265 | 77.4773373 | 0 |
| 7.75E-05 | 0.0002002 | 97.7428388 | 130.127747 | 115.078438 | 139.644782 | 33.2670554 |
| 7.78E-05 | 0.00020091 | 36.0105196 | 62.5058065 | 7.42441534 | 108.097123 | 0 |
| 7.80E-05 | 0.00020147 | 20.5774398 | 40.7066284 | 13.9207788 | 37.1148921 | 0 |
| 7.82E-05 | 0.00020206 | 11.8963324 | 53.8306234 | 27.8415575 | 35.2591475 | 0 |
| 7.84E-05 | 0.00020237 | 14.4685123 | 21.7991781 | 29.6976613 | 27.8361691 | 0 |
| 7.85E-05 | 0.00020267 | 15.1115573 | 19.5747721 | 34.3379209 | 45.929679 | 0 |
| 7.85E-05 | 0.00020279 | 19.2913498 | 27.1377523 | 15.7768826 | 37.1148921 | 0 |

| | | | | | | |
|---|---|---|---|---|---|---|
| 7.85E-05 | 0.00020283 | 32.7952946 | 17.1279256 | 19.4890903 | 22.2689353 | 0 |
| 7.88E-05 | 0.00020342 | 14.1469898 | 30.4743612 | 46.4025958 | 22.2689353 | 0 |
| 7.88E-05 | 0.00020352 | 39.2257445 | 20.9094157 | 18.5610383 | 29.6919137 | 197.226114 |
| 7.89E-05 | 0.00020359 | 10.9317649 | 33.1436483 | 35.2659728 | 41.7542536 | 0 |
| 7.89E-05 | 0.00020359 | 38.5826995 | 49.3818115 | 12.0646749 | 24.588616 | 0 |
| 7.91E-05 | 0.0002042 | 47.2638069 | 41.5963908 | 116.934542 | 7.88691457 | 0 |
| 7.93E-05 | 0.00020461 | 87.7756414 | 28.6948364 | 15.7768826 | 25.0525522 | 0 |
| 7.93E-05 | 0.00020474 | 11.8963324 | 39.3719849 | 29.6976613 | 31.5476583 | 0 |
| 7.94E-05 | 0.00020488 | 19.2913498 | 36.4802572 | 15.7768826 | 68.6625504 | 0 |
| 7.95E-05 | 0.00020518 | 28.6155022 | 6.67321777 | 5.5683115 | 8.35085072 | 111.682257 |
| 7.97E-05 | 0.00020578 | 7.07349491 | 12.2342326 | 0 | 12.0623399 | 87.920075 |
| 7.99E-05 | 0.00020622 | 9.64567488 | 39.5944255 | 40.8342843 | 44.0739344 | 0 |
| 8.00E-05 | 0.00020639 | 10.6102424 | 46.934965 | 28.7696094 | 49.6411682 | 0 |
| 8.01E-05 | 0.00020672 | 11.5748099 | 34.0334106 | 32.4818171 | 32.9394667 | 0 |
| 8.01E-05 | 0.00020672 | 13.8254673 | 49.6042521 | 20.4171422 | 41.2903175 | 0 |
| 8.02E-05 | 0.00020681 | 14.4685123 | 29.8070394 | 21.3451941 | 42.2181898 | 0 |
| 8.02E-05 | 0.00020681 | 6.43044992 | 9.78738607 | 16.7049345 | 16.2377653 | 99.8011662 |
| 8.02E-05 | 0.0002069 | 208.346577 | 667.321777 | 412.055051 | 473.214874 | 38.0194919 |
| 8.04E-05 | 0.00020735 | 17.0406923 | 67.6219401 | 65.8916861 | 57.0641466 | 0 |
| 8.04E-05 | 0.00020738 | 13.5039448 | 42.041272 | 27.8415575 | 25.0525522 | 0 |
| 8.05E-05 | 0.0002076 | 15.7546023 | 29.5845988 | 18.5610383 | 32.4755306 | 0 |
| 8.07E-05 | 0.00020799 | 22.5065747 | 17.7952474 | 20.4171422 | 19.4853184 | 0 |
| 8.08E-05 | 0.00020829 | 28.2939797 | 29.8070394 | 14.8488307 | 19.9492545 | 0 |
| 8.08E-05 | 0.0002083 | 19.9343948 | 58.7243164 | 52.8989593 | 50.1051043 | 0 |
| 8.08E-05 | 0.00020836 | 318.628794 | 681.557975 | 642.211927 | 718.173162 | 73.6627655 |
| 8.08E-05 | 0.00020837 | 27.6509347 | 19.5747721 | 35.2659728 | 15.309893 | 0 |
| 8.10E-05 | 0.00020862 | 259.790177 | 409.29069 | 389.781805 | 429.604876 | 38.0194919 |
| 8.10E-05 | 0.00020884 | 34.7244296 | 40.929069 | 25.9854537 | 47.7854236 | 2.37621824 |
| 8.11E-05 | 0.00020887 | 221.850522 | 409.513131 | 391.637909 | 451.409875 | 52.2768014 |
| 8.12E-05 | 0.00020911 | 127.965953 | 41.3739502 | 27.8415575 | 38.9706367 | 2.37621824 |
| 8.13E-05 | 0.00020936 | 21.8635297 | 58.0569946 | 13.9207788 | 34.3312752 | 0 |
| 8.14E-05 | 0.0002097 | 54.9803468 | 16.4606038 | 22.273246 | 25.0525522 | 0 |
| 8.14E-05 | 0.00020973 | 10.6102424 | 52.0510986 | 35.2659728 | 33.4034029 | 0 |
| 8.16E-05 | 0.00021006 | 1144.94161 | 2799.41486 | 2400.87031 | 2588.76372 | 0 |
| 8.16E-05 | 0.00021029 | 13.8254673 | 34.2558512 | 21.3451941 | 35.2591475 | 0 |
| 8.17E-05 | 0.0002103 | 19.2913498 | 45.6003214 | 13.9207788 | 49.177232 | 0 |
| 8.17E-05 | 0.00021032 | 17.0406923 | 62.9506877 | 57.5392189 | 77.9412734 | 0 |
| 8.19E-05 | 0.00021077 | 34.7244296 | 34.7007324 | 10.2085711 | 44.5378705 | 0 |
| 8.20E-05 | 0.00021107 | 29.9015921 | 42.041272 | 31.5537652 | 11.5984038 | 0 |
| 8.20E-05 | 0.00021115 | 9.64567488 | 44.7105591 | 35.2659728 | 46.3936151 | 0 |
| 8.20E-05 | 0.00021116 | 7.71653991 | 72.2931925 | 41.7623363 | 52.4247851 | 0 |
| 8.22E-05 | 0.00021151 | 36.3320421 | 30.4743612 | 49.1867516 | 30.1558498 | 73.6627655 |
| 8.24E-05 | 0.00021208 | 107.710036 | 27.5826335 | 14.8488307 | 29.2279775 | 0 |

| | | | | | | |
|---|---|---|---|---|---|---|
| 8.24E-05 | 0.0002121 | 12.2178549 | 33.8109701 | 26.9135056 | 33.867339 | 0 |
| 8.24E-05 | 0.00021217 | 9.64567488 | 33.5885295 | 64.0355823 | 42.2181898 | 0 |
| 8.25E-05 | 0.0002122 | 7.07349491 | 28.6948364 | 105.797919 | 103.921698 | 0 |
| 8.26E-05 | 0.00021256 | 15.1115573 | 31.8090047 | 21.3451941 | 24.1246799 | 0 |
| 8.26E-05 | 0.00021256 | 27.0078897 | 27.1377523 | 70.5319457 | 93.7151026 | 2.37621824 |
| 8.29E-05 | 0.00021328 | 25.0787547 | 19.3523315 | 16.7049345 | 21.341063 | 0 |
| 8.31E-05 | 0.00021373 | 12.2178549 | 25.1357869 | 36.1940248 | 38.5067006 | 0 |
| 8.32E-05 | 0.00021399 | 16.0761248 | 31.8090047 | 17.6329864 | 32.4755306 | 0 |
| 8.33E-05 | 0.00021429 | 19.9343948 | 37.5924601 | 13.9207788 | 46.8575513 | 0 |
| 8.35E-05 | 0.00021475 | 10.9317649 | 50.7164551 | 25.0574018 | 66.3428696 | 0 |
| 8.35E-05 | 0.00021479 | 20.5774398 | 23.8011434 | 19.4890903 | 17.1656376 | 0 |
| 8.35E-05 | 0.00021484 | 28.9370247 | 43.3759155 | 43.6184401 | 10.6705315 | 0 |
| 8.36E-05 | 0.00021488 | 461.384782 | 441.322135 | 449.177128 | 499.195299 | 97.424948 |
| 8.36E-05 | 0.00021499 | 13.5039448 | 34.4782918 | 22.273246 | 53.3526574 | 0 |
| 8.36E-05 | 0.00021499 | 36.9750871 | 96.3167765 | 128.071165 | 141.500526 | 0 |
| 8.36E-05 | 0.00021499 | 147.257303 | 142.80686 | 138.279736 | 162.841589 | 23.7621824 |
| 8.39E-05 | 0.00021572 | 79.094534 | 336.997498 | 237.581291 | 277.433818 | 23.7621824 |
| 8.39E-05 | 0.00021573 | 12.2178549 | 56.2774699 | 25.0574018 | 36.650956 | 0 |
| 8.41E-05 | 0.0002162 | 121.857026 | 452.666606 | 382.35739 | 432.388493 | 52.2768014 |
| 8.41E-05 | 0.00021627 | 22.8280972 | 35.3680542 | 104.869867 | 14.8459568 | 0 |
| 8.43E-05 | 0.0002166 | 18.0052598 | 46.0452026 | 26.9135056 | 17.6295737 | 0 |
| 8.43E-05 | 0.00021663 | 22.1850522 | 77.4093262 | 12.0646749 | 48.2493597 | 0 |
| 8.43E-05 | 0.00021669 | 36.3320421 | 81.1908162 | 23.2012979 | 53.3526574 | 0 |
| 8.46E-05 | 0.00021724 | 44.048582 | 13.3464355 | 21.3451941 | 38.0427644 | 0 |
| 8.47E-05 | 0.0002175 | 16.7191698 | 37.8149007 | 16.7049345 | 32.4755306 | 0 |
| 8.47E-05 | 0.00021766 | 46.6207619 | 25.1357869 | 37.1220767 | 43.1460621 | 0 |
| 8.49E-05 | 0.00021802 | 28.2939797 | 33.5885295 | 24.1293498 | 77.0134011 | 163.959059 |
| 8.49E-05 | 0.00021803 | 14.7900348 | 52.4959798 | 19.4890903 | 35.2591475 | 0 |
| 8.49E-05 | 0.00021817 | 11.5748099 | 37.3700195 | 26.9135056 | 40.3624452 | 0 |
| 8.51E-05 | 0.00021862 | 47.9068519 | 24.023584 | 13.9207788 | 26.4443606 | 0 |
| 8.53E-05 | 0.00021904 | 16.7191698 | 38.7046631 | 17.6329864 | 80.7248903 | 0 |
| 8.53E-05 | 0.00021906 | 1.28608998 | 108.551009 | 189.322591 | 154.954675 | 0 |
| 8.54E-05 | 0.00021933 | 13.1824223 | 34.0334106 | 29.6976613 | 23.6607437 | 0 |
| 8.54E-05 | 0.00021933 | 41.7979245 | 153.92889 | 77.956361 | 119.231591 | 0 |
| 8.56E-05 | 0.00021965 | 18.6483048 | 25.3582275 | 15.7768826 | 31.5476583 | 0 |
| 8.56E-05 | 0.00021965 | 43.405537 | 14.9035197 | 34.3379209 | 19.0213822 | 0 |
| 8.57E-05 | 0.00022001 | 84.5604165 | 79.4112915 | 85.3807764 | 58.4559551 | 133.068222 |
| 8.57E-05 | 0.00022003 | 12.8608998 | 28.917277 | 25.0574018 | 38.5067006 | 0 |
| 8.58E-05 | 0.00022014 | 18.0052598 | 7.56298014 | 0 | 10.6705315 | 71.2865473 |
| 8.60E-05 | 0.00022071 | 103.530244 | 225.554761 | 195.818954 | 224.081161 | 9.50487297 |
| 8.61E-05 | 0.00022101 | 102.565676 | 26.4704305 | 28.7696094 | 23.6607437 | 171.087714 |
| 8.62E-05 | 0.00022109 | 194.199588 | 906.667855 | 1267.71892 | 1105.09591 | 0 |
| 8.65E-05 | 0.00022178 | 13.5039448 | 23.1338216 | 27.8415575 | 32.9394667 | 0 |

| | | | | | | |
|---|---|---|---|---|---|---|
| 8.64E-05 | 0.00022178 | 41.476402 | 42.9310343 | 51.0428554 | 54.7444659 | 0 |
| 8.65E-05 | 0.00022193 | 12.8608998 | 51.1613363 | 20.4171422 | 51.4969128 | 0 |
| 8.65E-05 | 0.00022196 | 0 | 4.22637126 | 0 | 0 | 123.563349 |
| 8.69E-05 | 0.00022284 | 25.4002772 | 15.5708415 | 26.9135056 | 83.0445711 | 0 |
| 8.72E-05 | 0.00022374 | 12.5393773 | 28.0275146 | 30.6257133 | 27.8361691 | 0 |
| 8.75E-05 | 0.00022431 | 6.75197242 | 50.0491333 | 49.1867516 | 85.828188 | 0 |
| 8.76E-05 | 0.00022468 | 59.4816618 | 110.997856 | 32.4818171 | 112.736485 | 9.50487297 |
| 8.76E-05 | 0.00022471 | 19.2913498 | 20.9094157 | 17.6329864 | 20.4131907 | 0 |
| 8.77E-05 | 0.00022489 | 3.85826995 | 100.320707 | 77.956361 | 76.0855288 | 0 |
| 8.78E-05 | 0.00022497 | 13.1824223 | 27.1377523 | 26.9135056 | 26.4443606 | 0 |
| 8.78E-05 | 0.00022502 | 197.414813 | 348.341968 | 358.22804 | 279.753499 | 33.2670554 |
| 8.79E-05 | 0.00022519 | 23.4711422 | 45.822762 | 40.8342843 | 45.0018067 | 0 |
| 8.80E-05 | 0.0002256 | 7.07349491 | 54.4979451 | 51.0428554 | 56.6002105 | 0 |
| 8.82E-05 | 0.00022595 | 243.39253 | 555.87904 | 395.350117 | 451.409875 | 19.0097459 |
| 8.82E-05 | 0.00022596 | 12.8608998 | 26.9153117 | 25.9854537 | 33.4034029 | 0 |
| 8.82E-05 | 0.00022606 | 11.8963324 | 28.0275146 | 35.2659728 | 30.619786 | 0 |
| 8.83E-05 | 0.00022632 | 116.712666 | 539.640877 | 322.962067 | 389.242431 | 49.9005831 |
| 8.84E-05 | 0.00022661 | 13.8254673 | 23.5787028 | 59.3953227 | 29.6919137 | 0 |
| 8.85E-05 | 0.00022661 | 19.2913498 | 16.2381632 | 25.9854537 | 21.8049991 | 0 |
| 8.85E-05 | 0.00022661 | 35.3674746 | 22.0216187 | 24.1293498 | 17.1656376 | 64.1578926 |
| 8.88E-05 | 0.00022739 | 2.89370247 | 59.8365194 | 161.481034 | 110.88074 | 0 |
| 8.89E-05 | 0.00022769 | 23.1496197 | 20.6869751 | 14.8488307 | 26.9082968 | 0 |
| 8.89E-05 | 0.00022769 | 21.2204847 | 22.911381 | 18.5610383 | 17.1656376 | 0 |
| 8.90E-05 | 0.00022784 | 15.1115573 | 34.923173 | 36.1940248 | 18.0935099 | 0 |
| 8.91E-05 | 0.00022827 | 37.2966096 | 20.4645345 | 13.9207788 | 29.6919137 | 0 |
| 8.92E-05 | 0.00022833 | 6.43044992 | 68.0668213 | 76.1002572 | 43.1460621 | 0 |
| 8.92E-05 | 0.00022842 | 22.1850522 | 22.4664998 | 14.8488307 | 36.650956 | 0 |
| 8.94E-05 | 0.0002289 | 15.7546023 | 28.0275146 | 24.1293498 | 19.0213822 | 0 |
| 8.94E-05 | 0.0002289 | 10.9317649 | 45.6003214 | 33.409869 | 30.619786 | 0 |
| 8.94E-05 | 0.00022894 | 12.5393773 | 32.2538859 | 24.1293498 | 36.1870198 | 0 |
| 8.95E-05 | 0.00022919 | 14.1469898 | 29.5845988 | 38.0501286 | 20.4131907 | 0 |
| 8.97E-05 | 0.00022956 | 55.9449143 | 21.7991781 | 14.8488307 | 27.8361691 | 0 |
| 9.00E-05 | 0.00023032 | 15.1115573 | 26.2479899 | 26.9135056 | 19.4853184 | 0 |
| 9.00E-05 | 0.00023032 | 13.5039448 | 24.6909058 | 29.6976613 | 24.588616 | 0 |
| 9.02E-05 | 0.0002307 | 16.7191698 | 26.4704305 | 24.1293498 | 17.6295737 | 0 |
| 9.02E-05 | 0.00023079 | 30.5446371 | 29.3621582 | 12.9927268 | 22.7328714 | 0 |
| 9.02E-05 | 0.00023079 | 18.6483048 | 20.2420939 | 17.6329864 | 21.8049991 | 0 |
| 9.03E-05 | 0.00023089 | 29.5800696 | 136.800964 | 98.3735032 | 98.8184002 | 11.8810912 |
| 9.03E-05 | 0.00023089 | 28.6155022 | 20.4645345 | 15.7768826 | 20.8771268 | 142.573095 |
| 9.04E-05 | 0.00023117 | 8.6811074 | 47.8247274 | 33.409869 | 87.2199964 | 0 |
| 9.05E-05 | 0.00023151 | 11.8963324 | 56.0550293 | 22.273246 | 46.8575513 | 0 |
| 9.08E-05 | 0.00023227 | 53.6942569 | 22.0216187 | 12.9927268 | 34.3312752 | 0 |
| 9.09E-05 | 0.00023237 | 13.5039448 | 26.2479899 | 25.0574018 | 47.7854236 | 0 |

| | | | | | | |
|---|---|---|---|---|---|---|
| 9.09E-05 | 0.00023245 | 9.64567488 | 51.828658 | 30.6257133 | 46.8575513 | 0 |
| 9.09E-05 | 0.00023246 | 5.78740493 | 40.0393066 | 80.7405168 | 97.4265918 | 0 |
| 9.10E-05 | 0.00023261 | 0.3215225 | 0.44488118 | 0.92805192 | 0.9278723 | 11.8810912 |
| 9.10E-05 | 0.00023267 | 39.8687895 | 87.4191528 | 76.1002572 | 77.9412734 | 0 |
| 9.11E-05 | 0.00023284 | 19.6128723 | 25.3582275 | 18.5610383 | 17.6295737 | 0 |
| 9.11E-05 | 0.00023296 | 24.7572322 | 12.2342326 | 33.409869 | 48.7132959 | 0 |
| 9.11E-05 | 0.00023296 | 20.2559173 | 30.6968018 | 34.3379209 | 13.9180845 | 0 |
| 9.13E-05 | 0.00023341 | 19.2913498 | 14.2361979 | 35.2659728 | 42.6821259 | 0 |
| 9.15E-05 | 0.0002337 | 5.14435994 | 53.6081828 | 86.3088283 | 72.8379758 | 0 |
| 9.16E-05 | 0.00023403 | 13.8254673 | 26.0255493 | 37.1220767 | 22.2689353 | 0 |
| 9.19E-05 | 0.00023475 | 10.9317649 | 39.5944255 | 45.4745439 | 26.9082968 | 0 |
| 9.22E-05 | 0.00023557 | 253.359727 | 224.664998 | 132.711424 | 171.656376 | 54.6530196 |
| 9.23E-05 | 0.00023572 | 13.1824223 | 27.1377523 | 28.7696094 | 24.1246799 | 0 |
| 9.23E-05 | 0.00023582 | 11.8963324 | 33.3660889 | 29.6976613 | 28.3001052 | 0 |
| 9.24E-05 | 0.00023601 | 9.96719738 | 59.6140788 | 28.7696094 | 44.5378705 | 0 |
| 9.24E-05 | 0.00023603 | 22.8280972 | 13.123995 | 29.6976613 | 38.5067006 | 0 |
| 9.26E-05 | 0.00023644 | 20.8989622 | 26.0255493 | 13.9207788 | 29.6919137 | 0 |
| 9.27E-05 | 0.00023674 | 203.845263 | 718.038232 | 0 | 836.012945 | 0 |
| 9.28E-05 | 0.00023699 | 7.07349491 | 7.56298014 | 5.5683115 | 6.49510612 | 57.0292378 |
| 9.30E-05 | 0.00023745 | 35.3674746 | 92.9801676 | 91.8771398 | 97.8905279 | 11.8810912 |
| 9.37E-05 | 0.00023915 | 17.6837373 | 27.5826335 | 55.683115 | 17.6295737 | 0 |
| 9.39E-05 | 0.00023977 | 12.5393773 | 27.8050741 | 30.6257133 | 25.9804245 | 0 |
| 9.40E-05 | 0.00023994 | 15.4330798 | 26.2479899 | 19.4890903 | 23.1968076 | 0 |
| 9.41E-05 | 0.0002401 | 15.1115573 | 24.023584 | 20.4171422 | 23.6607437 | 0 |
| 9.41E-05 | 0.00024013 | 22.5065747 | 61.8384847 | 63.1075304 | 33.4034029 | 0 |
| 9.42E-05 | 0.00024048 | 29.2585471 | 36.4802572 | 53.8270112 | 62.1674443 | 2.37621824 |
| 9.43E-05 | 0.00024057 | 79.4160565 | 120.562801 | 99.3015551 | 127.118505 | 0 |
| 9.45E-05 | 0.00024122 | 35.3674746 | 25.1357869 | 12.9927268 | 24.1246799 | 0 |
| 9.47E-05 | 0.0002416 | 14.7900348 | 59.6140788 | 15.7768826 | 62.1674443 | 0 |
| 9.48E-05 | 0.00024176 | 10.6102424 | 50.0491333 | 29.6976613 | 35.2591475 | 0 |
| 9.48E-05 | 0.00024193 | 220.24291 | 666.876896 | 0 | 800.753797 | 0 |
| 9.49E-05 | 0.00024216 | 16.0761248 | 26.2479899 | 25.9854537 | 17.6295737 | 0 |
| 9.50E-05 | 0.00024221 | 23.1496197 | 18.4625692 | 19.4890903 | 57.0641466 | 0 |
| 9.50E-05 | 0.00024221 | 11.8963324 | 30.2519206 | 27.8415575 | 31.5476583 | 0 |
| 9.50E-05 | 0.00024233 | 48.2283744 | 29.5845988 | 17.6329864 | 16.2377653 | 0 |
| 9.51E-05 | 0.00024241 | 27.0078897 | 17.7952474 | 15.7768826 | 25.5164883 | 0 |
| 9.52E-05 | 0.00024268 | 10.6102424 | 57.1672323 | 29.6976613 | 35.2591475 | 0 |
| 9.52E-05 | 0.00024275 | 9.96719738 | 54.4979451 | 28.7696094 | 43.6099982 | 0 |
| 9.53E-05 | 0.00024296 | 20.2559173 | 45.822762 | 49.1867516 | 58.4559551 | 0 |
| 9.54E-05 | 0.00024323 | 27.9724572 | 36.9251383 | 26.9135056 | 12.0623399 | 0 |
| 9.54E-05 | 0.00024329 | 24.1141872 | 18.017688 | 19.4890903 | 19.9492545 | 61.7816743 |
| 9.55E-05 | 0.00024337 | 13.5039448 | 44.2656779 | 22.273246 | 28.7640414 | 0 |
| 9.58E-05 | 0.0002442 | 249.822979 | 76.2971232 | 313.681548 | 475.534555 | 49.9005831 |

| | | | | | | |
|---|---|---|---|---|---|---|
| 9.59E-05 | 0.00024451 | 10.6102424 | 25.8031087 | 54.7550631 | 43.6099982 | 0 |
| 9.59E-05 | 0.00024451 | 12.2178549 | 23.8011434 | 34.3379209 | 34.3312752 | 0 |
| 9.62E-05 | 0.00024507 | 9.00262989 | 57.834554 | 33.409869 | 44.0739344 | 0 |
| 9.66E-05 | 0.00024613 | 7.39501741 | 47.3798462 | 69.6038938 | 42.6821259 | 0 |
| 9.66E-05 | 0.00024622 | 31.1876821 | 50.4940145 | 46.4025958 | 25.5164883 | 2.37621824 |
| 9.67E-05 | 0.00024637 | 9.96719738 | 25.5806681 | 78.8844129 | 50.5690405 | 0 |
| 9.68E-05 | 0.00024672 | 8.3595849 | 31.5865641 | 54.7550631 | 70.518295 | 0 |
| 9.70E-05 | 0.00024701 | 14.1469898 | 45.6003214 | 28.7696094 | 20.8771268 | 0 |
| 9.70E-05 | 0.00024704 | 27.6509347 | 365.692334 | 247.789862 | 316.868391 | 16.6335277 |
| 9.71E-05 | 0.00024721 | 9.32415239 | 4.22637126 | 0 | 4.17542536 | 109.306039 |
| 9.72E-05 | 0.00024743 | 10.6102424 | 40.2617472 | 28.7696094 | 38.0427644 | 0 |
| 9.72E-05 | 0.0002476 | 27.6509347 | 42.7085937 | 10.2085711 | 41.2903175 | 0 |
| 9.73E-05 | 0.00024767 | 11.2532874 | 32.0314453 | 28.7696094 | 35.7230837 | 0 |
| 9.73E-05 | 0.0002477 | 25.7217997 | 28.4723958 | 17.6329864 | 15.7738291 | 0 |
| 9.74E-05 | 0.00024787 | 114.140486 | 199.974093 | 154.056618 | 158.666164 | 23.7621824 |
| 9.76E-05 | 0.00024837 | 3.85826995 | 114.112024 | 48.2586997 | 109.952868 | 0 |
| 9.76E-05 | 0.00024846 | 24.7572322 | 14.9035197 | 46.4025958 | 21.341063 | 0 |
| 9.77E-05 | 0.00024868 | 45.334672 | 166.608004 | 99.3015551 | 125.262761 | 21.3859642 |
| 9.78E-05 | 0.00024897 | 11.8963324 | 28.6948364 | 26.9135056 | 38.0427644 | 0 |
| 9.78E-05 | 0.00024897 | 60.7677518 | 105.2144 | 168.905449 | 141.500526 | 2.37621824 |
| 9.82E-05 | 0.0002499 | 10.6102424 | 31.3641235 | 35.2659728 | 35.2591475 | 0 |
| 9.83E-05 | 0.00025015 | 325.380766 | 568.113273 | 529.917645 | 577.600508 | 26.1384007 |
| 9.83E-05 | 0.00025016 | 13.1824223 | 24.023584 | 25.0574018 | 30.1558498 | 0 |
| 9.87E-05 | 0.00025122 | 460.420214 | 968.506339 | 853.807764 | 957.10028 | 7.12865473 |
| 9.90E-05 | 0.00025176 | 11.8963324 | 31.3641235 | 25.0574018 | 41.7542536 | 0 |
| 9.91E-05 | 0.00025201 | 8.0380624 | 78.9664103 | 29.6976613 | 63.5592527 | 0 |
| 9.94E-05 | 0.00025275 | 25.7217997 | 14.2361979 | 21.3451941 | 20.8771268 | 80.7914203 |
| 9.94E-05 | 0.00025284 | 38.904222 | 38.2597819 | 10.2085711 | 28.7640414 | 0 |
| 9.95E-05 | 0.00025308 | 124.750728 | 399.948185 | 322.034015 | 388.778495 | 30.8908372 |
| 9.96E-05 | 0.00025322 | 15.1115573 | 30.9192423 | 17.6329864 | 31.0837221 | 0 |
| 0.00010005 | 0.0002544 | 23.1496197 | 31.1416829 | 12.0646749 | 38.5067006 | 0 |
| 0.00010011 | 0.00025455 | 22.8280972 | 20.2420939 | 18.5610383 | 16.7017014 | 0 |
| 0.00010036 | 0.00025518 | 13.8254673 | 16.905485 | 81.6685687 | 52.4247851 | 0 |
| 0.00010044 | 0.00025535 | 137.290106 | 487.144897 | 418.551415 | 440.275408 | 0 |
| 0.00010055 | 0.00025561 | 13.1824223 | 44.0432373 | 23.2012979 | 27.3722329 | 0 |
| 0.00010064 | 0.00025581 | 70.4134267 | 94.0923706 | 106.72597 | 85.828188 | 2.37621824 |
| 0.00010064 | 0.00025581 | 29.2585471 | 65.6199748 | 53.8270112 | 61.7035081 | 0 |
| 0.00010077 | 0.00025612 | 24.7572322 | 24.2460246 | 13.9207788 | 21.341063 | 0 |
| 0.00010083 | 0.00025623 | 9.00262989 | 61.1711629 | 29.6976613 | 50.1051043 | 0 |
| 0.00010084 | 0.00025624 | 14.7900348 | 24.2460246 | 23.2012979 | 20.4131907 | 0 |
| 0.00010094 | 0.00025648 | 6.43044992 | 49.1593709 | 56.6111669 | 61.7035081 | 0 |
| 0.00010131 | 0.0002574 | 72.3425616 | 22.911381 | 18.5610383 | 20.8771268 | 0 |
| 0.00010152 | 0.00025791 | 22.5065747 | 41.8188314 | 14.8488307 | 21.8049991 | 0 |

| | | | | | | |
|---|---|---|---|---|---|---|
| 0.00010157 | 0.00025804 | 14.7900348 | 20.0196533 | 23.2012979 | 31.5476583 | 0 |
| 0.00010183 | 0.00025867 | 18.6483048 | 86.9742716 | 45.4745439 | 50.5690405 | 0 |
| 0.0001019 | 0.00025882 | 20.2559173 | 43.5983561 | 46.4025958 | 57.5280828 | 0 |
| 0.00010195 | 0.00025894 | 468.458277 | 1502.80864 | 1412.49502 | 1515.67941 | 239.998043 |
| 0.00010198 | 0.00025899 | 74.9147416 | 24.4684652 | 20.4171422 | 18.0935099 | 0 |
| 0.00010199 | 0.000259 | 1.60761248 | 109.218331 | 125.287009 | 137.789037 | 0 |
| 0.00010242 | 0.00026006 | 15.4330798 | 19.3523315 | 28.7696094 | 21.341063 | 0 |
| 0.00010242 | 0.00026006 | 24.4357097 | 24.2460246 | 19.4890903 | 14.8459568 | 0 |
| 0.00010254 | 0.00026035 | 12.2178549 | 36.7026978 | 68.6758419 | 22.7328714 | 0 |
| 0.00010262 | 0.00026053 | 21.2204847 | 20.6869751 | 14.8488307 | 32.0115944 | 0 |
| 0.00010268 | 0.00026065 | 21.8635297 | 40.7066284 | 41.7623363 | 54.2805297 | 0 |
| 0.00010271 | 0.00026071 | 145.971213 | 79.8561727 | 85.3807764 | 51.0329766 | 508.510704 |
| 0.00010287 | 0.0002611 | 9.96719738 | 57.3896728 | 30.6257133 | 36.1870198 | 0 |
| 0.000103 | 0.00026141 | 9.96719738 | 71.1809896 | 175.401812 | 72.3740396 | 0 |
| 0.00010313 | 0.00026171 | 32.7952946 | 16.4606038 | 17.6329864 | 22.7328714 | 0 |
| 0.00010328 | 0.00026207 | 22.1850522 | 19.1298909 | 64.0355823 | 19.4853184 | 0 |
| 0.00010357 | 0.00026278 | 19.9343948 | 42.9310343 | 52.8989593 | 55.208402 | 0 |
| 0.00010378 | 0.00026331 | 24.7572322 | 23.3562622 | 13.9207788 | 21.341063 | 0 |
| 0.00010408 | 0.00026405 | 32.1522496 | 14.0137573 | 63.1075304 | 20.4131907 | 0 |
| 0.00010414 | 0.00026417 | 54.3373018 | 42.7085937 | 12.0646749 | 20.4131907 | 0 |
| 0.00010416 | 0.00026421 | 13.1824223 | 26.9153117 | 30.6257133 | 21.8049991 | 0 |
| 0.00010432 | 0.00026459 | 10.9317649 | 48.047168 | 30.6257133 | 29.2279775 | 0 |
| 0.00010433 | 0.00026461 | 19.6128723 | 12.011792 | 76.1002572 | 50.5690405 | 0 |
| 0.00010442 | 0.00026481 | 33.7598621 | 65.3975342 | 57.5392189 | 37.1148921 | 7.12865473 |
| 0.00010447 | 0.00026492 | 14.7900348 | 25.5806681 | 19.4890903 | 23.6607437 | 0 |
| 0.00010452 | 0.00026502 | 15.7546023 | 20.0196533 | 25.0574018 | 19.4853184 | 0 |
| 0.00010454 | 0.00026505 | 24.4357097 | 31.3641235 | 12.0646749 | 27.3722329 | 0 |
| 0.00010455 | 0.00026505 | 6.43044992 | 57.1672323 | 63.1075304 | 46.3936151 | 0 |
| 0.00010457 | 0.00026508 | 13.5039448 | 19.5747721 | 33.409869 | 33.4034029 | 0 |
| 0.00010465 | 0.00026527 | 24.7572322 | 20.0196533 | 13.9207788 | 27.8361691 | 0 |
| 0.00010519 | 0.00026661 | 4.82283744 | 76.2971232 | 60.3233746 | 67.2707419 | 0 |
| 0.00010525 | 0.00026672 | 17.3622148 | 15.7932821 | 20.4171422 | 13.9180845 | 83.1676385 |
| 0.00010524 | 0.00026672 | 18.3267823 | 44.4881185 | 59.3953227 | 57.9920189 | 0 |
| 0.00010551 | 0.00026738 | 15.7546023 | 56.2774699 | 18.5610383 | 28.7640414 | 0 |
| 0.00010554 | 0.00026742 | 22.8280972 | 65.3975342 | 62.1794784 | 65.4149973 | 0 |
| 0.00010558 | 0.00026751 | 36.3320421 | 34.4782918 | 48.2586997 | 37.1148921 | 0 |
| 0.00010562 | 0.00026756 | 9.32415239 | 35.1456136 | 44.546492 | 36.650956 | 0 |
| 0.00010562 | 0.00026756 | 11.8963324 | 27.8050741 | 25.9854537 | 38.0427644 | 0 |
| 0.00010562 | 0.00026756 | 110.925261 | 107.216366 | 112.294282 | 109.024996 | 185.345023 |
| 0.00010577 | 0.0002679 | 12.2178549 | 41.8188314 | 90.0210359 | 21.341063 | 0 |
| 0.00010582 | 0.00026802 | 13.8254673 | 50.7164551 | 17.6329864 | 42.2181898 | 0 |
| 0.00010608 | 0.00026864 | 18.3267823 | 32.2538859 | 16.7049345 | 20.4131907 | 0 |
| 0.00010637 | 0.00026937 | 11.5748099 | 37.3700195 | 24.1293498 | 34.7952113 | 0 |

| | | | | | | |
|---|---|---|---|---|---|---|
| 0.00010646 | 0.00026959 | 10.2887199 | 32.9212077 | 33.409869 | 35.2591475 | 0 |
| 0.00010661 | 0.00026993 | 20.2559173 | 68.0668213 | 12.0646749 | 43.6099982 | 0 |
| 0.00010662 | 0.00026994 | 10.6102424 | 33.8109701 | 31.5537652 | 32.4755306 | 0 |
| 0.00010671 | 0.00027014 | 25.0787547 | 18.2401286 | 14.8488307 | 25.9804245 | 0 |
| 0.00010693 | 0.00027069 | 22.1850522 | 13.7913167 | 23.2012979 | 33.4034029 | 0 |
| 0.00010695 | 0.00027071 | 16.7191698 | 38.7046631 | 17.6329864 | 23.1968076 | 0 |
| 0.00010699 | 0.00027078 | 24.7572322 | 50.2715739 | 10.2085711 | 44.0739344 | 0 |
| 0.000107 | 0.00027079 | 11.8963324 | 56.7223511 | 28.7696094 | 26.9082968 | 0 |
| 0.00010708 | 0.00027098 | 8.3595849 | 36.2578166 | 57.5392189 | 41.7542536 | 0 |
| 0.0001072 | 0.00027127 | 11.2532874 | 36.035376 | 25.0574018 | 36.650956 | 0 |
| 0.0001076 | 0.00027225 | 22.8280972 | 46.7125244 | 12.0646749 | 31.0837221 | 0 |
| 0.0001079 | 0.000273 | 17.0406923 | 25.3582275 | 15.7768826 | 25.0525522 | 0 |
| 0.00010807 | 0.0002734 | 50.8005544 | 56.9447917 | 57.5392189 | 50.5690405 | 173.463932 |
| 0.00010813 | 0.00027354 | 21.5420072 | 19.1298909 | 12.9927268 | 17.6295737 | 83.1676385 |
| 0.00010842 | 0.00027426 | 9.00262989 | 45.822762 | 29.6976613 | 58.9198912 | 0 |
| 0.00010849 | 0.0002744 | 15.7546023 | 48.4920492 | 34.3379209 | 16.2377653 | 0 |
| 0.00010872 | 0.00027496 | 25.0787547 | 17.3503662 | 18.5610383 | 18.0935099 | 0 |
| 0.00010888 | 0.00027534 | 9.32415239 | 34.2558512 | 38.9781805 | 40.3624452 | 0 |
| 0.00010892 | 0.00027544 | 11.8963324 | 46.7125244 | 23.2012979 | 32.9394667 | 0 |
| 0.0001091 | 0.00027586 | 56.9094818 | 32.9212077 | 44.546492 | 31.5476583 | 159.206622 |
| 0.0001093 | 0.00027634 | 16.3976473 | 16.905485 | 34.3379209 | 23.6607437 | 0 |
| 0.0001093 | 0.00027634 | 73.3071291 | 108.99589 | 125.287009 | 110.416804 | 33.2670554 |
| 0.00010936 | 0.00027646 | 10.9317649 | 47.3798462 | 22.273246 | 54.2805297 | 0 |
| 0.00010942 | 0.00027659 | 12.2178549 | 34.923173 | 32.4818171 | 22.2689353 | 0 |
| 0.00010992 | 0.00027784 | 12.2178549 | 20.0196533 | 46.4025958 | 43.6099982 | 0 |
| 0.00011007 | 0.0002782 | 22.5065747 | 35.1456136 | 49.1867516 | 45.4657428 | 0 |
| 0.00011012 | 0.00027829 | 12.8608998 | 26.9153117 | 103.941815 | 29.2279775 | 0 |
| 0.00011031 | 0.00027875 | 15.4330798 | 38.0373413 | 21.3451941 | 20.4131907 | 0 |
| 0.00011042 | 0.00027902 | 12.8608998 | 33.1436483 | 30.6257133 | 20.8771268 | 0 |
| 0.00011066 | 0.0002796 | 12.5393773 | 8.8976237 | 6.49636342 | 13.4541484 | 28.5146189 |
| 0.00011083 | 0.00028 | 26.0433222 | 32.4763265 | 11.136623 | 30.1558498 | 0 |
| 0.00011086 | 0.00028006 | 36.6535646 | 27.5826335 | 40.8342843 | 29.6919137 | 0 |
| 0.00011114 | 0.00028075 | 12.5393773 | 38.9271037 | 33.409869 | 20.8771268 | 0 |
| 0.00011126 | 0.00028102 | 137.611628 | 300.962122 | 348.947521 | 309.445413 | 30.8908372 |
| 0.00011131 | 0.00028115 | 0 | 0 | 0 | 0 | 33.2670554 |
| 0.00011149 | 0.00028158 | 242.10644 | 281.164909 | 267.278952 | 307.589668 | 0 |
| 0.00011167 | 0.000282 | 36.0105196 | 21.1318563 | 27.8415575 | 19.4853184 | 64.1578926 |
| 0.00011179 | 0.00028227 | 8.3595849 | 36.7026978 | 51.9709073 | 40.8263813 | 0 |
| 0.00011209 | 0.00028301 | 30.2231146 | 13.5688761 | 21.3451941 | 24.1246799 | 0 |
| 0.00011228 | 0.00028349 | 30.2231146 | 28.2499552 | 11.136623 | 27.8361691 | 0 |
| 0.00011232 | 0.00028355 | 13.8254673 | 24.023584 | 22.273246 | 44.0739344 | 0 |
| 0.00011249 | 0.00028396 | 17.6837373 | 16.905485 | 20.4171422 | 20.8771268 | 0 |
| 0.00011261 | 0.00028424 | 25.4002772 | 17.3503662 | 15.7768826 | 22.2689353 | 0 |

| | | | | | | |
|---|---|---|---|---|---|---|
| 0.00011287 | 0.0002849 | 101.279586 | 23.3562622 | 22.273246 | 18.0935099 | 0 |
| 0.00011305 | 0.0002853 | 13.5039448 | 25.1357869 | 21.3451941 | 25.5164883 | 0 |
| 0.00011305 | 0.0002853 | 13.5039448 | 35.5904948 | 20.4171422 | 26.4443606 | 0 |
| 0.00011307 | 0.00028532 | 13.1824223 | 25.5806681 | 21.3451941 | 31.0837221 | 0 |
| 0.0001131 | 0.00028539 | 18.0052598 | 21.3542969 | 31.5537652 | 15.7738291 | 0 |
| 0.00011358 | 0.00028658 | 14.7900348 | 25.5806681 | 18.5610383 | 23.6607437 | 0 |
| 0.00011365 | 0.00028673 | 25.4002772 | 72.0707519 | 94.6612955 | 72.3740396 | 0 |
| 0.00011378 | 0.00028703 | 28.9370247 | 9.78738607 | 38.0501286 | 46.8575513 | 0 |
| 0.00011412 | 0.00028787 | 49.5144644 | 271.155082 | 195.818954 | 147.06776 | 0 |
| 0.00011429 | 0.00028826 | 17.6837373 | 47.6022868 | 13.9207788 | 34.7952113 | 0 |
| 0.00011436 | 0.00028844 | 9.64567488 | 45.6003214 | 31.5537652 | 34.7952113 | 0 |
| 0.00011451 | 0.00028879 | 14.7900348 | 28.0275146 | 18.5610383 | 23.1968076 | 0 |
| 0.00011481 | 0.00028952 | 29.2585471 | 19.1298909 | 13.9207788 | 48.7132959 | 0 |
| 0.00011488 | 0.00028968 | 14.1469898 | 23.8011434 | 19.4890903 | 29.6919137 | 0 |
| 0.00011493 | 0.00028978 | 8.3595849 | 30.9192423 | 66.819738 | 46.3936151 | 0 |
| 0.00011495 | 0.00028981 | 19.6128723 | 42.7085937 | 12.0646749 | 70.0543588 | 0 |
| 0.00011507 | 0.00029008 | 24.4357097 | 22.911381 | 12.9927268 | 23.6607437 | 0 |
| 0.00011511 | 0.00029016 | 90.3478214 | 225.554761 | 278.415575 | 243.566479 | 40.3957101 |
| 0.00011522 | 0.00029041 | 15.7546023 | 17.3503662 | 26.9135056 | 44.5378705 | 0 |
| 0.00011558 | 0.00029132 | 6.43044992 | 42.7085937 | 63.1075304 | 56.1362743 | 0 |
| 0.00011561 | 0.00029135 | 7.07349491 | 38.9271037 | 46.4025958 | 78.8691457 | 0 |
| 0.00011561 | 0.00029135 | 11.2532874 | 36.7026978 | 23.2012979 | 40.8263813 | 0 |
| 0.00011564 | 0.00029138 | 13.8254673 | 27.1377523 | 20.4171422 | 23.6607437 | 0 |
| 0.00011583 | 0.00029186 | 8.3595849 | 39.3719849 | 43.6184401 | 40.3624452 | 0 |
| 0.00011588 | 0.00029195 | 122.178549 | 465.345719 | 569.823877 | 507.082213 | 49.9005831 |
| 0.00011589 | 0.00029196 | 16.7191698 | 28.2499552 | 15.7768826 | 23.6607437 | 0 |
| 0.00011605 | 0.00029235 | 17.0406923 | 35.5904948 | 18.5610383 | 19.4853184 | 0 |
| 0.0001161 | 0.00029244 | 296.765264 | 456.670536 | 0 | 617.499017 | 0 |
| 0.00011625 | 0.00029281 | 46.2992394 | 10.6771484 | 30.6257133 | 28.3001052 | 0 |
| 0.00011646 | 0.00029331 | 11.2532874 | 58.0569946 | 21.3451941 | 43.1460621 | 0 |
| 0.0001165 | 0.00029338 | 13.1824223 | 29.8070394 | 25.9854537 | 20.8771268 | 0 |
| 0.00011655 | 0.00029349 | 450.131495 | 334.10577 | 0 | 550.692212 | 0 |
| 0.00011724 | 0.0002952 | 7.71653991 | 46.934965 | 40.8342843 | 45.4657428 | 0 |
| 0.00011764 | 0.00029618 | 325.059244 | 515.172412 | 411.126999 | 468.111577 | 7.12865473 |
| 0.00011844 | 0.00029818 | 10.9317649 | 25.5806681 | 32.4818171 | 37.1148921 | 0 |
| 0.00011875 | 0.00029894 | 307.053984 | 821.028227 | 788.844129 | 868.952411 | 0 |
| 0.00011904 | 0.00029964 | 138.576196 | 602.814006 | 633.859459 | 655.077846 | 0 |
| 0.00011918 | 0.00029998 | 14.1469898 | 26.4704305 | 19.4890903 | 23.1968076 | 0 |
| 0.00011955 | 0.00030089 | 30.2231146 | 12.2342326 | 90.0210359 | 25.5164883 | 0 |
| 0.00011982 | 0.00030154 | 28.9370247 | 75.6298014 | 38.9781805 | 9.27872303 | 0 |
| 0.00011984 | 0.00030156 | 13.5039448 | 34.2558512 | 18.5610383 | 30.619786 | 0 |
| 0.00012001 | 0.00030198 | 202.880695 | 27.8050741 | 12.9927268 | 26.9082968 | 0 |
| 0.00012013 | 0.00030226 | 36.6535646 | 37.1475789 | 25.0574018 | 32.9394667 | 0 |

| | | | | | | |
|---|---|---|---|---|---|---|
| 0.00012019 | 0.00030239 | 38.261177 | 81.1908162 | 39.9062324 | 117.839782 | 7.12865473 |
| 0.00012036 | 0.00030279 | 7.71653991 | 46.2676432 | 50.1148035 | 37.5788283 | 0 |
| 0.00012056 | 0.00030327 | 37.618132 | 44.9329997 | 11.136623 | 21.8049991 | 0 |
| 0.00012079 | 0.00030382 | 20.2559173 | 19.7972127 | 15.7768826 | 19.0213822 | 0 |
| 0.00012091 | 0.00030411 | 5.46588243 | 86.9742716 | 151.272462 | 127.582442 | 2.37621824 |
| 0.00012125 | 0.00030495 | 57.8740493 | 39.3719849 | 38.0501286 | 40.3624452 | 130.692003 |
| 0.0001215 | 0.00030553 | 75.5577866 | 175.728068 | 171.689605 | 161.913717 | 21.3859642 |
| 0.00012155 | 0.00030565 | 13.5039448 | 20.2420939 | 24.1293498 | 30.619786 | 0 |
| 0.00012164 | 0.00030583 | 10.2887199 | 31.5865641 | 29.6976613 | 35.7230837 | 0 |
| 0.00012164 | 0.00030583 | 13.1824223 | 37.8149007 | 23.2012979 | 22.7328714 | 0 |
| 0.00012171 | 0.00030597 | 11.8963324 | 18.017688 | 64.9636342 | 52.4247851 | 0 |
| 0.00012178 | 0.00030613 | 9.64567488 | 25.5806681 | 42.6903882 | 64.0231889 | 0 |
| 0.00012187 | 0.00030634 | 9.64567488 | 62.7282471 | 30.6257133 | 33.4034029 | 0 |
| 0.000122 | 0.00030665 | 11.5748099 | 28.917277 | 28.7696094 | 25.5164883 | 0 |
| 0.00012213 | 0.00030694 | 32.4737721 | 12.9015544 | 22.273246 | 24.1246799 | 0 |
| 0.00012219 | 0.00030707 | 14.1469898 | 8.45274251 | 11.136623 | 3.24755306 | 42.7719284 |
| 0.00012225 | 0.0003072 | 22.8280972 | 46.4900838 | 10.2085711 | 46.3936151 | 0 |
| 0.00012239 | 0.00030753 | 10.6102424 | 24.2460246 | 39.9062324 | 38.5067006 | 0 |
| 0.00012263 | 0.0003081 | 9.32415239 | 49.1593709 | 26.9135056 | 45.4657428 | 0 |
| 0.00012268 | 0.00030821 | 13.5039448 | 23.5787028 | 20.4171422 | 27.8361691 | 0 |
| 0.00012279 | 0.00030847 | 8.0380624 | 40.0393066 | 41.7623363 | 43.1460621 | 0 |
| 0.00012289 | 0.00030868 | 490.000284 | 450.66464 | 1025.49737 | 342.38488 | 28.5146189 |
| 0.0001231 | 0.0003092 | 11.2532874 | 33.1436483 | 24.1293498 | 32.9394667 | 0 |
| 0.00012358 | 0.00031038 | 95.4921813 | 26.2479899 | 18.5610383 | 18.0935099 | 0 |
| 0.00012373 | 0.00031073 | 33.7598621 | 82.5254598 | 112.294282 | 97.4265918 | 11.8810912 |
| 0.00012399 | 0.00031136 | 25.4002772 | 100.765588 | 63.1075304 | 74.6937204 | 9.50487297 |
| 0.00012413 | 0.0003117 | 12.8608998 | 59.8365194 | 64.0355823 | 15.7738291 | 0 |
| 0.00012417 | 0.00031177 | 147.257303 | 896.435588 | 830.606466 | 884.726241 | 45.1481466 |
| 0.0001242 | 0.00031183 | 16.3976473 | 29.1397176 | 15.7768826 | 23.1968076 | 0 |
| 0.00012444 | 0.0003124 | 11.2532874 | 28.4723958 | 41.7623363 | 25.0525522 | 0 |
| 0.00012447 | 0.00031245 | 0 | 0 | 0 | 0 | 2.37621824 |
| 0.00012531 | 0.00031454 | 109.639171 | 288.95033 | 270.99116 | 332.64222 | 0 |
| 0.00012543 | 0.0003148 | 6.75197242 | 29.8070394 | 111.36623 | 59.8477635 | 0 |
| 0.00012546 | 0.00031486 | 104.173289 | 98.0963013 | 125.287009 | 101.602017 | 30.8908372 |
| 0.00012566 | 0.00031535 | 11.2532874 | 31.3641235 | 27.8415575 | 26.4443606 | 0 |
| 0.00012571 | 0.00031545 | 9.64567488 | 40.4841878 | 27.8415575 | 39.898509 | 0 |
| 0.00012597 | 0.00031608 | 44.048582 | 21.5767375 | 12.0646749 | 28.7640414 | 0 |
| 0.00012667 | 0.0003178 | 5.46588243 | 49.6042521 | 110.438178 | 45.4657428 | 0 |
| 0.0001267 | 0.00031785 | 12.2178549 | 24.2460246 | 28.7696094 | 24.1246799 | 0 |
| 0.0001267 | 0.00031785 | 9.00262989 | 32.6987671 | 51.0428554 | 34.3312752 | 0 |
| 0.00012687 | 0.00031824 | 8.6811074 | 46.0452026 | 49.1867516 | 29.2279775 | 0 |
| 0.0001271 | 0.0003188 | 12.5393773 | 26.9153117 | 33.409869 | 20.8771268 | 0 |
| 0.00012739 | 0.00031947 | 18.3267823 | 26.4704305 | 17.6329864 | 16.7017014 | 0 |

| | | | | | | |
|---|---|---|---|---|---|---|
| 0.00012739 | 0.00031947 | 30.5446371 | 24.023584 | 12.0646749 | 23.1968076 | 0 |
| 0.00012783 | 0.00032056 | 11.8963324 | 29.1397176 | 45.4745439 | 22.2689353 | 0 |
| 0.00012787 | 0.00032062 | 54.3373018 | 16.4606038 | 12.9927268 | 47.3214874 | 0 |
| 0.00012824 | 0.00032154 | 14.4685123 | 37.3700195 | 24.1293498 | 18.5574461 | 0 |
| 0.00012842 | 0.00032196 | 6.43044992 | 61.3936035 | 58.4672708 | 38.5067006 | 0 |
| 0.00012852 | 0.00032219 | 5.46588243 | 57.3896728 | 46.4025958 | 73.7658481 | 0 |
| 0.00012857 | 0.00032229 | 21.2204847 | 49.3818115 | 42.6903882 | 36.1870198 | 0 |
| 0.00012886 | 0.00032297 | 0 | 0 | 0 | 0 | 7.12865473 |
| 0.00012886 | 0.00032297 | 55.9449143 | 101.655351 | 101.157659 | 104.84957 | 14.2573095 |
| 0.00012896 | 0.0003232 | 15.7546023 | 72.7380737 | 44.546492 | 13.9180845 | 0 |
| 0.00012914 | 0.00032362 | 25.4002772 | 17.3503662 | 21.3451941 | 15.309893 | 0 |
| 0.00012918 | 0.00032369 | 11.8963324 | 38.4822225 | 26.9135056 | 23.1968076 | 0 |
| 0.00012921 | 0.00032375 | 16.3976473 | 35.5904948 | 13.9207788 | 39.4345729 | 0 |
| 0.00012931 | 0.00032398 | 239.212737 | 426.418616 | 396.278169 | 410.119558 | 4.75243649 |
| 0.00012934 | 0.00032403 | 11.5748099 | 22.911381 | 43.6184401 | 29.2279775 | 0 |
| 0.00012937 | 0.00032408 | 10.2887199 | 94.982133 | 23.2012979 | 41.7542536 | 0 |
| 0.00012946 | 0.00032428 | 21.8635297 | 14.9035197 | 9.28051917 | 13.4541484 | 45.1481466 |
| 0.00012951 | 0.00032437 | 10.6102424 | 34.923173 | 60.3233746 | 24.1246799 | 0 |
| 0.00012952 | 0.00032437 | 179.08803 | 545.424333 | 541.054268 | 557.651254 | 28.5146189 |
| 0.0001296 | 0.00032456 | 17.3622148 | 62.9506877 | 13.9207788 | 34.7952113 | 0 |
| 0.00013013 | 0.00032586 | 10.9317649 | 35.8129354 | 25.9854537 | 28.7640414 | 0 |
| 0.00013029 | 0.00032623 | 14.1469898 | 20.4645345 | 21.3451941 | 21.8049991 | 0 |
| 0.00013063 | 0.00032706 | 15.7546023 | 15.7932821 | 27.8415575 | 40.8263813 | 0 |
| 0.00013104 | 0.00032805 | 15.7546023 | 54.7203857 | 13.9207788 | 41.2903175 | 0 |
| 0.00013137 | 0.00032887 | 123.143116 | 559.660531 | 392.565961 | 418.006472 | 0 |
| 0.00013171 | 0.00032969 | 790.94534 | 1318.62783 | 594.881279 | 963.595386 | 0 |
| 0.00013214 | 0.00033074 | 10.6102424 | 55.6101481 | 43.6184401 | 21.341063 | 0 |
| 0.00013269 | 0.0003321 | 78.4514891 | 43.8207967 | 31.5537652 | 46.3936151 | 144.949313 |
| 0.00013278 | 0.0003323 | 96.4567488 | 341.001428 | 318.321808 | 319.652008 | 28.5146189 |
| 0.00013293 | 0.00033265 | 15.1115573 | 18.6850098 | 19.4890903 | 22.7328714 | 0 |
| 0.00013303 | 0.00033288 | 46.6207619 | 24.4684652 | 16.7049345 | 16.2377653 | 0 |
| 0.00013309 | 0.00033299 | 92.5984789 | 189.074504 | 210.667785 | 167.017014 | 23.7621824 |
| 0.0001337 | 0.00033449 | 28.9370247 | 22.911381 | 51.0428554 | 11.5984038 | 0 |
| 0.00013395 | 0.00033509 | 0 | 679.77845 | 380.501286 | 779.412734 | 0 |
| 0.0001342 | 0.0003357 | 58.8386168 | 76.2971232 | 105.797919 | 92.7872303 | 401.580883 |
| 0.00013428 | 0.00033586 | 0 | 0 | 0 | 0 | 42.7719284 |
| 0.00013441 | 0.00033617 | 24.1141872 | 34.923173 | 33.409869 | 42.2181898 | 0 |
| 0.00013486 | 0.00033727 | 16.3976473 | 19.7972127 | 12.0646749 | 14.3820207 | 35.6432737 |
| 0.00013558 | 0.00033905 | 26.0433222 | 41.5963908 | 48.2586997 | 58.9198912 | 4.75243649 |
| 0.0001357 | 0.00033932 | 44.691627 | 22.4664998 | 18.5610383 | 15.309893 | 0 |
| 0.00013571 | 0.00033934 | 16.7191698 | 28.6948364 | 18.5610383 | 17.1656376 | 0 |
| 0.00013582 | 0.00033957 | 14.1469898 | 5.56101481 | 9.28051917 | 5.56723382 | 64.1578926 |
| 0.00013597 | 0.00033993 | 9.96719738 | 40.4841878 | 34.3379209 | 26.4443606 | 0 |

| | | | | | | |
|---|---|---|---|---|---|---|
| 0.00013605 | 0.0003401 | 13.8254673 | 22.0216187 | 20.4171422 | 21.8049991 | 0 |
| 0.00013613 | 0.00034028 | 15.1115573 | 69.846346 | 23.2012979 | 21.8049991 | 0 |
| 0.00013663 | 0.00034151 | 9.00262989 | 32.2538859 | 41.7623363 | 34.3312752 | 0 |
| 0.00013674 | 0.00034174 | 17.0406923 | 23.3562622 | 16.7049345 | 18.0935099 | 0 |
| 0.000137 | 0.00034238 | 13.8254673 | 8.8976237 | 11.136623 | 14.3820207 | 49.9005831 |
| 0.00013734 | 0.0003432 | 9.96719738 | 26.2479899 | 34.3379209 | 39.898509 | 0 |
| 0.00013826 | 0.00034547 | 18.6483048 | 70.958549 | 55.683115 | 36.1870198 | 0 |
| 0.00013827 | 0.00034548 | 20.5774398 | 24.2460246 | 48.2586997 | 13.4541484 | 0 |
| 0.00013832 | 0.00034556 | 241.784917 | 336.330176 | 374.932974 | 377.644027 | 38.0194919 |
| 0.00013837 | 0.00034567 | 10.9317649 | 32.9212077 | 26.9135056 | 26.4443606 | 0 |
| 0.00013868 | 0.00034639 | 10.9317649 | 22.4664998 | 37.1220767 | 33.867339 | 0 |
| 0.00013867 | 0.00034639 | 8.6811074 | 34.2558512 | 37.1220767 | 38.5067006 | 0 |
| 0.00013903 | 0.00034724 | 0 | 2.8917277 | 3.71220767 | 1.85574461 | 26.1384007 |
| 0.00013941 | 0.00034817 | 2.25065747 | 64.952653 | 130.85532 | 99.7462725 | 0 |
| 0.00013951 | 0.0003484 | 209.95419 | 249.578345 | 242.22155 | 233.359884 | 0 |
| 0.0001396 | 0.0003486 | 18.6483048 | 26.2479899 | 12.9927268 | 26.9082968 | 0 |
| 0.00013997 | 0.00034947 | 7.71653991 | 44.9329997 | 38.0501286 | 40.8263813 | 0 |
| 0.00014007 | 0.0003497 | 15.7546023 | 18.9074504 | 19.4890903 | 18.5574461 | 0 |
| 0.00014012 | 0.00034981 | 37.9396545 | 54.9428263 | 7.42441534 | 37.1148921 | 0 |
| 0.00014014 | 0.00034982 | 36.9750871 | 15.7932821 | 16.7049345 | 22.2689353 | 0 |
| 0.00014017 | 0.00034988 | 0 | 0 | 0.92805192 | 0 | 0 |
| 0.00014058 | 0.00035088 | 77.4869216 | 79.6337321 | 57.5392189 | 91.859358 | 19.0097459 |
| 0.00014065 | 0.00035103 | 10.9317649 | 54.4979451 | 26.9135056 | 26.9082968 | 0 |
| 0.00014069 | 0.00035109 | 9.32415239 | 30.02948 | 36.1940248 | 35.7230837 | 0 |
| 0.00014074 | 0.00035119 | 9.64567488 | 24.2460246 | 50.1148035 | 40.3624452 | 0 |
| 0.00014077 | 0.00035126 | 10.2887199 | 28.2499552 | 28.7696094 | 34.3312752 | 0 |
| 0.00014106 | 0.00035196 | 129.252043 | 639.739144 | 1588.82488 | 1241.0292 | 47.5243649 |
| 0.00014108 | 0.00035196 | 1.28608998 | 1.55708415 | 0.92805192 | 0.46393615 | 0 |
| 0.00014116 | 0.00035213 | 17.3622148 | 54.053064 | 11.136623 | 64.487125 | 0 |
| 0.00014126 | 0.00035235 | 5.78740493 | 28.4723958 | 81.6685687 | 94.1790387 | 0 |
| 0.00014136 | 0.00035259 | 10.2887199 | 24.9133464 | 38.9781805 | 32.4755306 | 0 |
| 0.00014156 | 0.00035307 | 30.5446371 | 7.11809896 | 63.1075304 | 57.5280828 | 0 |
| 0.00014161 | 0.00035316 | 8.3595849 | 55.8325887 | 28.7696094 | 43.1460621 | 0 |
| 0.00014162 | 0.00035316 | 64.9475442 | 63.6180094 | 50.1148035 | 63.0953166 | 14.2573095 |
| 0.00014165 | 0.0003532 | 16.3976473 | 12.6791138 | 58.4672708 | 45.0018067 | 0 |
| 0.00014182 | 0.0003536 | 10.2887199 | 29.1397176 | 26.9135056 | 39.4345729 | 0 |
| 0.00014189 | 0.00035375 | 11.5748099 | 24.9133464 | 25.0574018 | 27.8361691 | 0 |
| 0.00014204 | 0.0003541 | 19.9343948 | 27.3601929 | 12.0646749 | 33.4034029 | 0 |
| 0.00014238 | 0.00035492 | 32.1522496 | 32.4763265 | 22.273246 | 11.1344676 | 0 |
| 0.00014245 | 0.00035507 | 26.3648447 | 20.9094157 | 12.9927268 | 20.8771268 | 0 |
| 0.00014247 | 0.00035509 | 11.8963324 | 15.7932821 | 83.5246725 | 54.7444659 | 0 |
| 0.00014274 | 0.00035573 | 12.8608998 | 20.4645345 | 28.7696094 | 22.7328714 | 0 |
| 0.00014306 | 0.00035651 | 14.4685123 | 20.0196533 | 19.4890903 | 20.8771268 | 0 |

| | | | | | | |
|---|---|---|---|---|---|---|
| 0.00014314 | 0.00035669 | 52.4081669 | 27.8050741 | 12.9927268 | 19.4853184 | 0 |
| 0.0001433 | 0.00035705 | 89.3832539 | 129.460425 | 78.8844129 | 95.106911 | 7.12865473 |
| 0.0001436 | 0.00035777 | 312.519866 | 292.509379 | 279.343627 | 285.784669 | 494.253395 |
| 0.00014397 | 0.00035867 | 15.1115573 | 19.3523315 | 18.5610383 | 20.4131907 | 0 |
| 0.00014412 | 0.00035901 | 5.46588243 | 25.5806681 | 130.85532 | 83.9724434 | 0 |
| 0.00014415 | 0.00035907 | 5.14435994 | 46.0452026 | 94.6612955 | 50.1051043 | 0 |
| 0.00014421 | 0.00035919 | 25.4002772 | 12.9015544 | 21.3451941 | 22.2689353 | 0 |
| 0.00014431 | 0.0003594 | 10.9317649 | 58.7243164 | 18.5610383 | 49.177232 | 0 |
| 0.00014449 | 0.00035982 | 26.0433222 | 14.2361979 | 35.2659728 | 16.7017014 | 0 |
| 0.0001446 | 0.00036009 | 13.8254673 | 25.3582275 | 17.6329864 | 25.0525522 | 0 |
| 0.00014485 | 0.00036067 | 13.8254673 | 21.1318563 | 24.1293498 | 19.0213822 | 0 |
| 0.00014495 | 0.00036091 | 110.603739 | 215.767375 | 193.962851 | 208.307332 | 26.1384007 |
| 0.00014497 | 0.00036092 | 10.6102424 | 23.5787028 | 36.1940248 | 32.0115944 | 0 |
| 0.00014525 | 0.0003616 | 69.7703817 | 54.4979451 | 12.9927268 | 14.3820207 | 0 |
| 0.00014566 | 0.00036258 | 9.00262989 | 42.041272 | 26.9135056 | 43.6099982 | 0 |
| 0.00014567 | 0.00036258 | 35.3674746 | 21.3542969 | 12.0646749 | 24.1246799 | 0 |
| 0.00014616 | 0.00036377 | 9.96719738 | 35.1456136 | 34.3379209 | 25.9804245 | 0 |
| 0.0001463 | 0.00036407 | 17.3622148 | 12.6791138 | 38.9781805 | 37.1148921 | 0 |
| 0.0001463 | 0.00036407 | 15.7546023 | 29.8070394 | 25.9854537 | 14.8459568 | 0 |
| 0.00014665 | 0.00036492 | 18.9698273 | 95.2045736 | 8.35246725 | 72.8379758 | 0 |
| 0.00014704 | 0.00036587 | 6.75197242 | 36.7026978 | 55.683115 | 49.177232 | 0 |
| 0.00014715 | 0.00036611 | 50.8005544 | 208.871716 | 180.970124 | 363.262006 | 26.1384007 |
| 0.00014753 | 0.00036703 | 73.3071291 | 119.228158 | 93.7332436 | 136.861165 | 19.0097459 |
| 0.00014763 | 0.00036724 | 14.1469898 | 36.7026978 | 26.9135056 | 16.7017014 | 0 |
| 0.00014764 | 0.00036725 | 9.96719738 | 24.9133464 | 33.409869 | 51.0329766 | 0 |
| 0.00014772 | 0.00036741 | 52.0866444 | 10.6771484 | 26.9135056 | 26.9082968 | 0 |
| 0.00014775 | 0.00036747 | 9.00262989 | 44.4881185 | 32.4818171 | 31.0837221 | 0 |
| 0.00014799 | 0.00036802 | 9.32415239 | 38.9271037 | 51.0428554 | 25.0525522 | 0 |
| 0.00014924 | 0.00037113 | 20.2559173 | 28.0275146 | 16.7049345 | 15.309893 | 0 |
| 0.00014939 | 0.00037146 | 16.0761248 | 20.6869751 | 16.7049345 | 19.0213822 | 0 |
| 0.00014944 | 0.00037155 | 63.9829767 | 48.7144897 | 61.2514265 | 51.4969128 | 116.434694 |
| 0.00015004 | 0.00037302 | 16.3976473 | 21.1318563 | 15.7768826 | 19.9492545 | 0 |
| 0.00015049 | 0.00037412 | 11.2532874 | 22.6889404 | 29.6976613 | 28.3001052 | 0 |
| 0.00015065 | 0.00037449 | 12.5393773 | 26.4704305 | 20.4171422 | 24.1246799 | 0 |
| 0.00015071 | 0.00037458 | 12.5393773 | 28.2499552 | 33.409869 | 18.5574461 | 0 |
| 0.00015071 | 0.00037458 | 9.32415239 | 41.8188314 | 25.9854537 | 38.5067006 | 0 |
| 0.0001513 | 0.00037602 | 18.6483048 | 27.3601929 | 35.2659728 | 12.5262761 | 0 |
| 0.00015179 | 0.0003772 | 19.2913498 | 41.1515096 | 46.4025958 | 40.3624452 | 2.37621824 |
| 0.00015186 | 0.00037734 | 1.92913498 | 102.545113 | 87.2368802 | 101.138081 | 0 |
| 0.00015187 | 0.00037734 | 20.2559173 | 16.2381632 | 35.2659728 | 16.2377653 | 0 |
| 0.000152 | 0.00037761 | 28.6155022 | 49.1593709 | 41.7623363 | 23.1968076 | 0 |
| 0.00015199 | 0.00037761 | 13.8254673 | 24.6909058 | 20.4171422 | 19.0213822 | 0 |
| 0.0001521 | 0.00037783 | 5.46588243 | 87.1967122 | 44.546492 | 46.3936151 | 0 |

| | | | | | | |
|---|---|---|---|---|---|---|
| 0.00015221 | 0.00037807 | 8.6811074 | 44.4881185 | 31.5537652 | 33.867339 | 0 |
| 0.00015295 | 0.00037989 | 119.284846 | 174.615865 | 164.265189 | 161.913717 | 2.37621824 |
| 0.00015335 | 0.00038086 | 15.7546023 | 22.4664998 | 32.4818171 | 15.309893 | 0 |
| 0.00015344 | 0.00038106 | 9.64567488 | 24.9133464 | 53.8270112 | 34.3312752 | 0 |
| 0.00015346 | 0.00038108 | 11.2532874 | 32.4763265 | 26.9135056 | 22.7328714 | 0 |
| 0.00015371 | 0.00038166 | 7.07349491 | 39.3719849 | 42.6903882 | 46.8575513 | 0 |
| 0.00015381 | 0.0003819 | 488.714194 | 258.253528 | 323.890119 | 345.632433 | 163.959059 |
| 0.000154 | 0.00038233 | 70.0919042 | 475.800427 | 345.235313 | 446.306578 | 0 |
| 0.00015428 | 0.00038299 | 13.5039448 | 34.923173 | 15.7768826 | 44.0739344 | 0 |
| 0.00015436 | 0.00038316 | 21.2204847 | 48.4920492 | 9.28051917 | 59.8477635 | 0 |
| 0.00015473 | 0.00038405 | 11.2532874 | 22.911381 | 43.6184401 | 26.4443606 | 0 |
| 0.00015514 | 0.00038503 | 161.082771 | 359.686438 | 283.055835 | 307.589668 | 33.2670554 |
| 0.00015514 | 0.00038503 | 25.0787547 | 19.1298909 | 12.9927268 | 21.8049991 | 0 |
| 0.00015522 | 0.0003852 | 18.9698273 | 42.4861532 | 12.9927268 | 25.0525522 | 0 |
| 0.0001556 | 0.00038611 | 57.5525268 | 75.4073608 | 81.6685687 | 77.9412734 | 19.0097459 |
| 0.00015563 | 0.00038615 | 24.1141872 | 19.3523315 | 21.3451941 | 19.0213822 | 52.2768014 |
| 0.00015588 | 0.00038673 | 51.1220769 | 23.5787028 | 12.0646749 | 23.1968076 | 0 |
| 0.00015619 | 0.00038747 | 75.2362641 | 311.194389 | 224.588564 | 245.422224 | 30.8908372 |
| 0.00015628 | 0.00038769 | 200.95156 | 533.857422 | 322.034015 | 403.624452 | 38.0194919 |
| 0.00015633 | 0.00038778 | 9.96719738 | 37.3700195 | 29.6976613 | 26.4443606 | 0 |
| 0.00015635 | 0.00038779 | 24.7572322 | 141.249776 | 92.8051917 | 48.7132959 | 7.12865473 |
| 0.00015672 | 0.00038869 | 64.6260217 | 41.5963908 | 6.49636342 | 37.5788283 | 0 |
| 0.0001568 | 0.00038885 | 0 | 520.733427 | 473.306478 | 638.84008 | 0 |
| 0.00015695 | 0.00038918 | 198.37938 | 332.548686 | 374.004923 | 375.788283 | 0 |
| 0.000157 | 0.0003893 | 10.6102424 | 30.02948 | 34.3379209 | 23.1968076 | 0 |
| 0.00015748 | 0.00039045 | 60.1247068 | 34.2558512 | 19.4890903 | 42.2181898 | 2.37621824 |
| 0.00015757 | 0.00039063 | 13.1824223 | 41.8188314 | 34.3379209 | 16.2377653 | 0 |
| 0.00015761 | 0.00039072 | 19.6128723 | 12.9015544 | 25.9854537 | 22.2689353 | 0 |
| 0.00015766 | 0.00039081 | 9.00262989 | 28.6948364 | 37.1220767 | 36.1870198 | 0 |
| 0.00015774 | 0.00039097 | 6.75197242 | 35.3680542 | 76.1002572 | 41.7542536 | 0 |
| 0.00015782 | 0.00039115 | 24.7572322 | 24.4684652 | 12.9927268 | 18.0935099 | 0 |
| 0.0001579 | 0.00039131 | 24.4357097 | 17.7952474 | 18.5610383 | 14.8459568 | 0 |
| 0.00015811 | 0.00039181 | 9.32415239 | 38.9271037 | 25.0574018 | 41.2903175 | 0 |
| 0.00015816 | 0.00039191 | 9.32415239 | 48.2696086 | 21.3451941 | 60.7756358 | 0 |
| 0.00015824 | 0.00039208 | 136.325538 | 544.31213 | 581.888552 | 577.600508 | 0 |
| 0.0001583 | 0.00039218 | 5.46588243 | 48.2696086 | 52.8989593 | 55.6723382 | 0 |
| 0.00015833 | 0.00039223 | 11.2532874 | 42.2637126 | 20.4171422 | 31.5476583 | 0 |
| 0.00015837 | 0.00039231 | 1.60761248 | 1.55708415 | 2.78415575 | 1.85574461 | 111.682257 |
| 0.00015877 | 0.00039328 | 189.698273 | 357.017151 | 227.37272 | 292.279775 | 33.2670554 |
| 0.00015896 | 0.00039372 | 77.4869216 | 37.1475789 | 41.7623363 | 36.650956 | 7.12865473 |
| 0.00015898 | 0.00039374 | 5.78740493 | 36.2578166 | 62.1794784 | 62.1674443 | 0 |
| 0.000159 | 0.00039376 | 179.731075 | 281.164909 | 257.070381 | 267.227223 | 57.0292378 |
| 0.0001593 | 0.00039447 | 22.8280972 | 33.8109701 | 33.409869 | 42.6821259 | 0 |

| | | | | | | |
|---|---|---|---|---|---|---|
| 0.00015935 | 0.00039456 | 12.8608998 | 24.2460246 | 18.5610383 | 31.0837221 | 0 |
| 0.00016019 | 0.00039661 | 11.8963324 | 23.1338216 | 24.1293498 | 23.6607437 | 0 |
| 0.00016075 | 0.00039798 | 16.3976473 | 28.6948364 | 38.9781805 | 13.4541484 | 0 |
| 0.00016085 | 0.00039819 | 2.25065747 | 61.1711629 | 144.776099 | 83.0445711 | 0 |
| 0.00016119 | 0.00039901 | 10.9317649 | 29.3621582 | 24.1293498 | 26.4443606 | 0 |
| 0.00016151 | 0.00039976 | 106.745469 | 252.247632 | 204.171422 | 233.359884 | 35.6432737 |
| 0.00016164 | 0.00040005 | 1.92913498 | 67.8443807 | 121.574801 | 104.84957 | 0 |
| 0.00016192 | 0.00040072 | 16.3976473 | 30.9192423 | 20.4171422 | 15.309893 | 0 |
| 0.00016209 | 0.00040112 | 30.2231146 | 15.7932821 | 20.4171422 | 15.7738291 | 0 |
| 0.00016258 | 0.0004023 | 27.6509347 | 73.6278361 | 6.49636342 | 60.3116997 | 0 |
| 0.00016346 | 0.00040444 | 8.3595849 | 54.7203857 | 27.8415575 | 38.5067006 | 0 |
| 0.00016361 | 0.00040478 | 7.71653991 | 53.6081828 | 40.8342843 | 30.619786 | 0 |
| 0.0001638 | 0.00040521 | 43.7270595 | 216.212256 | 184.682331 | 259.340309 | 19.0097459 |
| 0.00016418 | 0.00040613 | 22.8280972 | 16.2381632 | 16.7049345 | 17.1656376 | 0 |
| 0.00016422 | 0.00040621 | 12.8608998 | 20.6869751 | 20.4171422 | 30.619786 | 0 |
| 0.00016442 | 0.00040668 | 71.6995166 | 205.312667 | 270.99116 | 219.4418 | 0 |
| 0.00016511 | 0.00040834 | 48.8714194 | 12.011792 | 45.4745439 | 16.2377653 | 0 |
| 0.0001652 | 0.00040853 | 865.53856 | 1644.28086 | 1329.8984 | 1507.79249 | 0 |
| 0.00016538 | 0.00040896 | 12.2178549 | 25.5806681 | 31.5537652 | 19.0213822 | 0 |
| 0.00016549 | 0.0004092 | 26.0433222 | 9.56494547 | 33.409869 | 35.7230837 | 0 |
| 0.00016553 | 0.00040923 | 9.32415239 | 29.8070394 | 38.0501286 | 28.7640414 | 0 |
| 0.00016553 | 0.00040923 | 98.3858838 | 174.170984 | 140.135839 | 185.574461 | 23.7621824 |
| 0.00016557 | 0.0004093 | 20.8989622 | 12.9015544 | 20.4171422 | 32.9394667 | 0 |
| 0.00016603 | 0.00041041 | 53.0512119 | 480.026798 | 960.533734 | 866.632731 | 33.2670554 |
| 0.00016634 | 0.00041115 | 27.6509347 | 40.929069 | 25.9854537 | 32.0115944 | 0 |
| 0.00016637 | 0.00041119 | 9.32415239 | 65.1750936 | 19.4890903 | 54.2805297 | 0 |
| 0.00016673 | 0.00041206 | 13.1824223 | 23.5787028 | 18.5610383 | 22.2689353 | 0 |
| 0.00016708 | 0.00041288 | 17.3622148 | 30.2519206 | 23.2012979 | 13.4541484 | 0 |
| 0.0001671 | 0.0004129 | 56.2664368 | 17.3503662 | 17.6329864 | 19.9492545 | 0 |
| 0.00016729 | 0.00041334 | 11.5748099 | 40.0393066 | 19.4890903 | 29.6919137 | 0 |
| 0.00016743 | 0.00041365 | 12.2178549 | 21.5767375 | 21.3451941 | 30.1558498 | 0 |
| 0.00016784 | 0.00041463 | 71.3779941 | 35.1456136 | 25.9854537 | 32.0115944 | 268.512662 |
| 0.00016786 | 0.00041466 | 16.7191698 | 61.1711629 | 12.0646749 | 38.0427644 | 0 |
| 0.00016794 | 0.0004148 | 411.227273 | 1170.4824 | 933.620228 | 1099.06474 | 14.2573095 |
| 0.00016793 | 0.0004148 | 8.3595849 | 40.2617472 | 32.4818171 | 33.867339 | 0 |
| 0.00016797 | 0.00041484 | 16.3976473 | 31.3641235 | 12.9927268 | 29.2279775 | 0 |
| 0.00016813 | 0.0004152 | 11.2532874 | 29.3621582 | 20.4171422 | 43.6099982 | 0 |
| 0.00016834 | 0.0004157 | 9.32415239 | 40.0393066 | 28.7696094 | 29.2279775 | 0 |
| 0.00016855 | 0.00041619 | 25.7217997 | 16.4606038 | 27.8415575 | 13.4541484 | 0 |
| 0.00016877 | 0.00041669 | 14.7900348 | 32.9212077 | 19.4890903 | 17.6295737 | 0 |
| 0.00016901 | 0.00041726 | 23.4711422 | 85.4171875 | 60.3233746 | 59.3838274 | 7.12865473 |
| 0.0001691 | 0.00041744 | 7.39501741 | 56.7223511 | 30.6257133 | 42.6821259 | 0 |
| 0.00016912 | 0.00041747 | 44.691627 | 18.4625692 | 14.8488307 | 20.4131907 | 0 |

| | | | | | | |
|---|---|---|---|---|---|---|
| 0.00016913 | 0.00041747 | 20.2559173 | 46.934965 | 9.28051917 | 61.7035081 | 0 |
| 0.00016923 | 0.00041769 | 9.96719738 | 31.1416829 | 33.409869 | 24.588616 | 0 |
| 0.00017049 | 0.00042075 | 14.7900348 | 58.2794352 | 12.0646749 | 51.4969128 | 0 |
| 0.00017076 | 0.0004214 | 22.1850522 | 42.7085937 | 51.0428554 | 53.3526574 | 2.37621824 |
| 0.00017127 | 0.00042258 | 13.1824223 | 21.1318563 | 21.3451941 | 19.4853184 | 0 |
| 0.00017126 | 0.00042258 | 157.546023 | 266.261389 | 189.322591 | 270.01084 | 38.0194919 |
| 0.00017151 | 0.00042314 | 8.6811074 | 27.1377523 | 49.1867516 | 34.7952113 | 0 |
| 0.00017171 | 0.00042362 | 119.927891 | 25.1357869 | 17.6329864 | 16.7017014 | 0 |
| 0.00017233 | 0.00042511 | 19.2913498 | 36.2578166 | 12.9927268 | 21.341063 | 0 |
| 0.0001729 | 0.00042649 | 9.64567488 | 28.917277 | 26.9135056 | 35.2591475 | 0 |
| 0.00017305 | 0.00042681 | 9.96719738 | 26.2479899 | 26.9135056 | 37.1148921 | 0 |
| 0.00017306 | 0.00042681 | 16.7191698 | 34.7007324 | 12.0646749 | 43.6099982 | 0 |
| 0.00017346 | 0.00042777 | 117.677234 | 164.828479 | 264.494796 | 204.131907 | 26.1384007 |
| 0.00017381 | 0.00042859 | 10.6102424 | 25.8031087 | 33.409869 | 23.6607437 | 0 |
| 0.00017388 | 0.00042875 | 18.9698273 | 42.9310343 | 39.9062324 | 39.4345729 | 0 |
| 0.00017441 | 0.00043001 | 9.32415239 | 24.6909058 | 35.2659728 | 38.5067006 | 0 |
| 0.00017483 | 0.00043103 | 219.599865 | 643.298193 | 684.902315 | 670.851675 | 47.5243649 |
| 0.00017508 | 0.00043162 | 247.2508 | 407.511165 | 0 | 439.347535 | 0 |
| 0.00017541 | 0.00043239 | 14.4685123 | 16.4606038 | 23.2012979 | 19.9492545 | 0 |
| 0.00017572 | 0.00043312 | 115.105054 | 209.094157 | 167.049345 | 191.60563 | 26.1384007 |
| 0.00017631 | 0.00043455 | 13.5039448 | 18.4625692 | 21.3451941 | 20.4131907 | 0 |
| 0.00017649 | 0.00043495 | 155.616888 | 225.10988 | 158.696878 | 234.287756 | 28.5146189 |
| 0.0001765 | 0.00043496 | 11.8963324 | 25.1357869 | 21.3451941 | 22.7328714 | 0 |
| 0.00017655 | 0.00043505 | 41.476402 | 50.2715739 | 37.1220767 | 26.4443606 | 4.75243649 |
| 0.00017661 | 0.00043516 | 47.2638069 | 50.0491333 | 6.49636342 | 33.867339 | 0 |
| 0.00017719 | 0.00043656 | 7.71653991 | 35.5904948 | 33.409869 | 45.4657428 | 0 |
| 0.0001776 | 0.00043752 | 30.2231146 | 212.208325 | 202.315318 | 243.566479 | 16.6335277 |
| 0.00017765 | 0.00043763 | 8.3595849 | 10.4547078 | 10.2085711 | 8.35085072 | 123.563349 |
| 0.00017785 | 0.00043808 | 7.71653991 | 31.1416829 | 45.4745439 | 38.5067006 | 0 |
| 0.00017797 | 0.00043835 | 16.3976473 | 21.1318563 | 13.9207788 | 24.1246799 | 0 |
| 0.00017832 | 0.00043916 | 18.0052598 | 22.4664998 | 12.9927268 | 21.8049991 | 0 |
| 0.00017864 | 0.00043993 | 17.3622148 | 21.7991781 | 24.1293498 | 28.3001052 | 57.0292378 |
| 0.0001788 | 0.00044029 | 8.0380624 | 36.035376 | 50.1148035 | 29.2279775 | 0 |
| 0.00017895 | 0.00044062 | 135.039448 | 257.808647 | 262.638692 | 304.806051 | 30.8908372 |
| 0.00017932 | 0.00044151 | 9.64567488 | 26.4704305 | 36.1940248 | 27.8361691 | 0 |
| 0.00017939 | 0.00044165 | 6.10892743 | 40.2617472 | 41.7623363 | 60.3116997 | 0 |
| 0.00017957 | 0.00044205 | 7.07349491 | 32.0314453 | 44.546492 | 49.177232 | 0 |
| 0.00017995 | 0.00044296 | 10.2887199 | 33.5885295 | 21.3451941 | 35.2591475 | 0 |
| 0.00018007 | 0.0004432 | 6.75197242 | 44.0432373 | 38.9781805 | 42.2181898 | 0 |
| 0.00018006 | 0.0004432 | 11.8963324 | 28.0275146 | 22.273246 | 20.8771268 | 0 |
| 0.00018009 | 0.0004432 | 12.5393773 | 25.8031087 | 17.6329864 | 27.3722329 | 0 |
| 0.00018017 | 0.00044336 | 20.2559173 | 93.86993 | 115.078438 | 136.861165 | 9.50487297 |
| 0.0001802 | 0.0004434 | 157.546023 | 219.993746 | 174.47376 | 191.141694 | 0 |

| | | | | | | |
|---|---|---|---|---|---|---|
| 0.00018027 | 0.00044355 | 9.32415239 | 33.3660889 | 24.1293498 | 47.3214874 | 0 |
| 0.00018037 | 0.00044378 | 13.1824223 | 22.2440592 | 18.5610383 | 20.8771268 | 0 |
| 0.00018042 | 0.00044385 | 21.8635297 | 24.4684652 | 11.136623 | 24.588616 | 0 |
| 0.00018062 | 0.00044432 | 11.8963324 | 23.1338216 | 20.4171422 | 25.9804245 | 0 |
| 0.00018077 | 0.00044466 | 27.3294122 | 51.1613363 | 6.49636342 | 82.1166988 | 0 |
| 0.00018081 | 0.00044472 | 11.5748099 | 27.1377523 | 24.1293498 | 20.8771268 | 0 |
| 0.00018082 | 0.00044472 | 15.7546023 | 26.6928711 | 44.546492 | 13.9180845 | 0 |
| 0.00018085 | 0.00044475 | 4.50131495 | 53.3857422 | 74.2441534 | 48.2493597 | 0 |
| 0.00018117 | 0.00044552 | 17.3622148 | 45.6003214 | 39.9062324 | 47.3214874 | 2.37621824 |
| 0.00018146 | 0.0004462 | 8.3595849 | 29.1397176 | 39.9062324 | 35.2591475 | 0 |
| 0.00018207 | 0.00044767 | 23.4711422 | 89.4211182 | 67.7477899 | 63.5592527 | 0 |
| 0.00018234 | 0.00044829 | 9.96719738 | 41.8188314 | 22.273246 | 32.0115944 | 0 |
| 0.00018238 | 0.00044835 | 10.2887199 | 29.3621582 | 28.7696094 | 23.6607437 | 0 |
| 0.00018239 | 0.00044835 | 28.2939797 | 14.9035197 | 13.9207788 | 28.3001052 | 0 |
| 0.00018301 | 0.00044984 | 7.71653991 | 49.8266927 | 19.4890903 | 177.22361 | 0 |
| 0.0001831 | 0.00045001 | 6.75197242 | 39.1495443 | 38.0501286 | 49.177232 | 0 |
| 0.00018352 | 0.00045101 | 30.8661596 | 27.8050741 | 25.0574018 | 47.7854236 | 0 |
| 0.00018402 | 0.00045223 | 83.9173715 | 21.5767375 | 11.136623 | 29.2279775 | 0 |
| 0.00018433 | 0.00045296 | 21.5420072 | 12.2342326 | 23.2012979 | 52.8887212 | 0 |
| 0.00018437 | 0.00045301 | 9.00262989 | 38.9271037 | 33.409869 | 25.9804245 | 0 |
| 0.00018469 | 0.00045376 | 13.5039448 | 7.11809896 | 5.5683115 | 8.35085072 | 85.5438568 |
| 0.00018472 | 0.00045382 | 207.38201 | 788.774341 | 570.751929 | 634.200719 | 21.3859642 |
| 0.00018505 | 0.00045457 | 0 | 0 | 0 | 0 | 2.37621824 |
| 0.00018519 | 0.00045489 | 45.9777169 | 71.6258708 | 64.0355823 | 4.17542536 | 0 |
| 0.00018523 | 0.00045494 | 13.8254673 | 17.1279256 | 20.4171422 | 30.1558498 | 0 |
| 0.00018523 | 0.00045494 | 4.50131495 | 56.7223511 | 66.819738 | 47.7854236 | 0 |
| 0.0001855 | 0.00045556 | 24.1141872 | 19.5747721 | 43.6184401 | 12.0623399 | 0 |
| 0.00018557 | 0.00045566 | 15.7546023 | 19.7972127 | 19.4890903 | 15.309893 | 0 |
| 0.00018557 | 0.00045566 | 690.630322 | 2372.55136 | 2348.8994 | 1901.67428 | 2.37621824 |
| 0.00018573 | 0.00045602 | 137.611628 | 14.0137573 | 25.9854537 | 19.4853184 | 0 |
| 0.00018613 | 0.00045698 | 9.96719738 | 24.9133464 | 27.8415575 | 31.0837221 | 0 |
| 0.00018627 | 0.00045729 | 58.1955718 | 42.4861532 | 34.3379209 | 6.49510612 | 0 |
| 0.00018639 | 0.00045754 | 10.6102424 | 27.3601929 | 22.273246 | 29.6919137 | 0 |
| 0.0001865 | 0.00045778 | 4.50131495 | 30.9192423 | 107.654022 | 72.3740396 | 0 |
| 0.00018699 | 0.00045896 | 14.4685123 | 29.8070394 | 13.9207788 | 31.5476583 | 0 |
| 0.00018717 | 0.00045935 | 14.7900348 | 14.6810791 | 23.2012979 | 26.9082968 | 0 |
| 0.00018718 | 0.00045935 | 9.64567488 | 27.1377523 | 25.9854537 | 46.3936151 | 0 |
| 0.00018778 | 0.00046076 | 95.8137038 | 10.899589 | 21.3451941 | 31.0837221 | 0 |
| 0.00018778 | 0.00046076 | 182.9463 | 32.9212077 | 22.273246 | 35.2591475 | 2.37621824 |
| 0.00018805 | 0.00046137 | 7.39501741 | 32.0314453 | 51.0428554 | 36.650956 | 0 |
| 0.00018859 | 0.00046267 | 9.32415239 | 52.4959798 | 22.273246 | 36.1870198 | 0 |
| 0.00018875 | 0.00046304 | 84.5604165 | 338.554582 | 370.292715 | 320.115944 | 38.0194919 |
| 0.00018901 | 0.00046365 | 25.7217997 | 13.7913167 | 16.7049345 | 20.4131907 | 0 |

| | | | | | | |
|---|---|---|---|---|---|---|
| 0.00018925 | 0.0004642 | 7.71653991 | 112.55494 | 12.0646749 | 123.870952 | 0 |
| 0.00018941 | 0.00046456 | 18.6483048 | 24.023584 | 12.0646749 | 23.1968076 | 0 |
| 0.00018945 | 0.00046462 | 0 | 478.914596 | 544.766475 | 446.770514 | 0 |
| 0.00018966 | 0.0004651 | 25.4002772 | 18.2401286 | 12.0646749 | 23.6607437 | 0 |
| 0.00018978 | 0.00046536 | 4.82283744 | 38.7046631 | 74.2441534 | 57.9920189 | 0 |
| 0.00018993 | 0.0004657 | 0 | 0 | 1.85610383 | 3.71148921 | 4.75243649 |
| 0.00019024 | 0.00046641 | 170.406923 | 300.962122 | 348.947521 | 396.201473 | 35.6432737 |
| 0.00019063 | 0.00046733 | 9.00262989 | 37.1475789 | 40.8342843 | 23.6607437 | 0 |
| 0.00019083 | 0.00046778 | 14.4685123 | 20.9094157 | 25.0574018 | 15.309893 | 0 |
| 0.00019103 | 0.00046824 | 45.0131495 | 22.911381 | 27.8415575 | 18.0935099 | 228.116951 |
| 0.00019148 | 0.00046931 | 27.0078897 | 35.1456136 | 39.9062324 | 48.2493597 | 4.75243649 |
| 0.00019185 | 0.00047018 | 12.8608998 | 24.6909058 | 16.7049345 | 33.4034029 | 0 |
| 0.00019204 | 0.00047063 | 15.4330798 | 18.4625692 | 18.5610383 | 16.2377653 | 0 |
| 0.00019219 | 0.00047096 | 7.39501741 | 56.2774699 | 28.7696094 | 40.3624452 | 0 |
| 0.00019273 | 0.00047225 | 10.9317649 | 22.911381 | 27.8415575 | 23.1968076 | 0 |
| 0.00019282 | 0.00047242 | 8.0380624 | 30.02948 | 48.2586997 | 32.4755306 | 0 |
| 0.00019283 | 0.00047243 | 102.244154 | 81.1908162 | 98.3735032 | 80.2609542 | 19.0097459 |
| 0.0001931 | 0.00047304 | 15.7546023 | 24.023584 | 25.9854537 | 13.4541484 | 0 |
| 0.00019337 | 0.00047368 | 7.39501741 | 61.8384847 | 32.4818171 | 33.867339 | 0 |
| 0.00019342 | 0.00047377 | 73.3071291 | 159.045024 | 323.890119 | 274.650202 | 23.7621824 |
| 0.00019356 | 0.00047407 | 7.07349491 | 38.7046631 | 51.0428554 | 32.4755306 | 0 |
| 0.00019388 | 0.00047481 | 12.2178549 | 23.5787028 | 26.9135056 | 18.0935099 | 0 |
| 0.00019411 | 0.00047534 | 73.6286516 | 44.7105591 | 29.6976613 | 31.5476583 | 4.75243649 |
| 0.00019419 | 0.0004755 | 173.622148 | 391.940324 | 421.33557 | 412.903175 | 106.929821 |
| 0.00019423 | 0.00047558 | 90.3478214 | 191.298909 | 142.919995 | 186.502333 | 23.7621824 |
| 0.00019431 | 0.00047575 | 25.7217997 | 56.4999105 | 51.9709073 | 35.2591475 | 0 |
| 0.00019442 | 0.00047598 | 21.2204847 | 38.7046631 | 12.0646749 | 20.4131907 | 0 |
| 0.00019448 | 0.00047609 | 9.96719738 | 21.1318563 | 32.4818171 | 41.2903175 | 0 |
| 0.00019472 | 0.00047664 | 82.6312815 | 143.251742 | 93.7332436 | 94.1790387 | 4.75243649 |
| 0.00019509 | 0.00047751 | 103.530244 | 20.6869751 | 12.9927268 | 25.0525522 | 0 |
| 0.0001952 | 0.00047773 | 4.50131495 | 39.3719849 | 49.1867516 | 116.447974 | 0 |
| 0.00019539 | 0.00047816 | 27.9724572 | 12.9015544 | 52.8989593 | 16.7017014 | 0 |
| 0.00019593 | 0.00047946 | 10.6102424 | 23.8011434 | 23.2012979 | 33.4034029 | 0 |
| 0.00019619 | 0.00048007 | 10.2887199 | 34.4782918 | 20.4171422 | 32.9394667 | 0 |
| 0.00019635 | 0.00048041 | 26.3648447 | 46.2676432 | 64.9636342 | 60.3116997 | 7.12865473 |
| 0.00019637 | 0.00048043 | 62.0538418 | 296.290869 | 343.379209 | 360.47839 | 26.1384007 |
| 0.00019681 | 0.00048148 | 18.0052598 | 18.9074504 | 13.9207788 | 18.5574461 | 0 |
| 0.00019714 | 0.00048225 | 17.6837373 | 52.0510986 | 38.0501286 | 41.7542536 | 0 |
| 0.00019739 | 0.00048281 | 10.9317649 | 26.2479899 | 25.9854537 | 21.341063 | 0 |
| 0.00019749 | 0.00048303 | 15.4330798 | 75.852242 | 9.28051917 | 65.8789335 | 0 |
| 0.00019789 | 0.00048396 | 15.4330798 | 24.2460246 | 13.9207788 | 21.8049991 | 0 |
| 0.00019893 | 0.00048647 | 13.5039448 | 26.0255493 | 16.7049345 | 20.4131907 | 0 |
| 0.00019944 | 0.00048769 | 99.3504513 | 168.83241 | 89.092984 | 140.108718 | 21.3859642 |

| | | | | | | |
|---|---|---|---|---|---|---|
| 0.00019987 | 0.00048871 | 0.3215225 | 0.66732178 | 3.71220767 | 0.9278723 | 2.37621824 |
| 0.00019995 | 0.00048886 | 371.358483 | 467.792566 | 465.882062 | 451.873811 | 23.7621824 |
| 0.0002005 | 0.00049018 | 20.8989622 | 36.7026978 | 32.4818171 | 35.7230837 | 2.37621824 |
| 0.00020072 | 0.00049065 | 104.173289 | 187.962301 | 118.790645 | 157.738291 | 35.6432737 |
| 0.00020072 | 0.00049065 | 9.96719738 | 22.4664998 | 30.6257133 | 30.1558498 | 0 |
| 0.0002011 | 0.00049155 | 12.8608998 | 26.0255493 | 18.5610383 | 19.4853184 | 0 |
| 0.0002013 | 0.00049199 | 23.4711422 | 11.5669108 | 24.1293498 | 20.4131907 | 0 |
| 0.00020152 | 0.00049249 | 10.6102424 | 4.00393066 | 8.35246725 | 6.03116997 | 26.1384007 |
| 0.00020182 | 0.00049318 | 17.0406923 | 18.9074504 | 43.6184401 | 15.309893 | 0 |
| 0.00020212 | 0.00049388 | 20.2559173 | 18.4625692 | 14.8488307 | 15.7738291 | 0 |
| 0.00020227 | 0.00049422 | 24.4357097 | 52.940861 | 51.0428554 | 36.1870198 | 4.75243649 |
| 0.00020256 | 0.00049488 | 20.8989622 | 121.007682 | 55.683115 | 76.0855288 | 7.12865473 |
| 0.00020298 | 0.00049588 | 12.8608998 | 20.4645345 | 17.6329864 | 26.4443606 | 0 |
| 0.00020304 | 0.00049599 | 13.8254673 | 18.9074504 | 16.7049345 | 25.5164883 | 0 |
| 0.00020317 | 0.00049626 | 299.337444 | 356.127388 | 158.696878 | 356.302964 | 45.1481466 |
| 0.00020365 | 0.00049741 | 33.4383396 | 204.200464 | 109.510126 | 138.716909 | 21.3859642 |
| 0.00020495 | 0.00050056 | 11.8963324 | 24.6909058 | 18.5610383 | 24.1246799 | 0 |
| 0.00020571 | 0.00050236 | 7.39501741 | 44.4881185 | 27.8415575 | 44.5378705 | 0 |
| 0.00020576 | 0.00050246 | 13.8254673 | 12.6791138 | 55.683115 | 41.2903175 | 0 |
| 0.00020609 | 0.00050321 | 17.6837373 | 21.5767375 | 12.9927268 | 19.4853184 | 0 |
| 0.00020625 | 0.00050358 | 21.2204847 | 16.6830444 | 28.7696094 | 12.9902122 | 0 |
| 0.00020628 | 0.00050362 | 29.9015921 | 12.9015544 | 25.9854537 | 15.7738291 | 0 |
| 0.00020642 | 0.00050392 | 28.2939797 | 50.4940145 | 26.9135056 | 44.5378705 | 2.37621824 |
| 0.00020709 | 0.00050551 | 9.00262989 | 14.6810791 | 15.7768826 | 13.9180845 | 118.810912 |
| 0.00020736 | 0.00050614 | 90.9908664 | 242.460246 | 247.789862 | 242.174671 | 33.2670554 |
| 0.00020747 | 0.00050638 | 15.1115573 | 35.8129354 | 12.0646749 | 42.6821259 | 0 |
| 0.00020751 | 0.00050643 | 11.2532874 | 24.6909058 | 21.3451941 | 22.7328714 | 0 |
| 0.00020843 | 0.00050864 | 29.9015921 | 38.4822225 | 49.1867516 | 80.2609542 | 7.12865473 |
| 0.00020913 | 0.00051031 | 6.75197242 | 45.3778809 | 52.8989593 | 29.2279775 | 0 |
| 0.00020968 | 0.00051161 | 11.8963324 | 16.905485 | 20.4171422 | 16.2377653 | 102.177384 |
| 0.00020985 | 0.000512 | 18.9698273 | 16.6830444 | 55.683115 | 16.7017014 | 0 |
| 0.00021027 | 0.00051299 | 6.10892743 | 58.946757 | 49.1867516 | 31.0837221 | 0 |
| 0.00021064 | 0.00051386 | 14.7900348 | 19.3523315 | 14.8488307 | 22.7328714 | 0 |
| 0.00021104 | 0.00051479 | 6.43044992 | 41.8188314 | 33.409869 | 52.4247851 | 0 |
| 0.00021124 | 0.00051524 | 10.2887199 | 27.5826335 | 24.1293498 | 24.1246799 | 0 |
| 0.00021126 | 0.00051524 | 87.4541189 | 319.424691 | 264.494796 | 270.938712 | 23.7621824 |
| 0.00021148 | 0.00051574 | 11.8963324 | 22.911381 | 18.5610383 | 24.588616 | 0 |
| 0.00021173 | 0.00051632 | 13.8254673 | 16.6830444 | 28.7696094 | 18.5574461 | 0 |
| 0.00021201 | 0.00051695 | 24.1141872 | 52.0510986 | 51.0428554 | 55.6723382 | 7.12865473 |
| 0.00021212 | 0.00051721 | 55.9449143 | 33.3660889 | 47.3306478 | 33.4034029 | 73.6627655 |
| 0.00021245 | 0.00051795 | 8.0380624 | 29.5845988 | 47.3306478 | 30.1558498 | 0 |
| 0.00021279 | 0.00051876 | 28.6155022 | 47.8247274 | 36.1940248 | 56.6002105 | 4.75243649 |
| 0.00021319 | 0.00051968 | 103.208721 | 17.7952474 | 27.8415575 | 12.9902122 | 0 |

| | | | | | | |
|---|---|---|---|---|---|---|
| 0.0002132 | 0.00051968 | 14.1469898 | 22.6889404 | 16.7049345 | 17.6295737 | 0 |
| 0.00021342 | 0.00052017 | 46.2992394 | 24.4684652 | 9.28051917 | 26.4443606 | 0 |
| 0.00021345 | 0.0005202 | 8.6811074 | 32.6987671 | 25.0574018 | 38.9706367 | 0 |
| 0.00021389 | 0.00052125 | 18.0052598 | 93.4250488 | 49.1867516 | 66.8068058 | 4.75243649 |
| 0.00021446 | 0.00052259 | 31.5092046 | 30.9192423 | 22.273246 | 31.5476583 | 2.37621824 |
| 0.00021496 | 0.00052377 | 43.0840145 | 82.5254598 | 67.7477899 | 83.5085072 | 14.2573095 |
| 0.00021513 | 0.00052416 | 8.3595849 | 79.6337321 | 13.9207788 | 93.7151026 | 0 |
| 0.00021526 | 0.00052444 | 16.0761248 | 27.5826335 | 15.7768826 | 16.2377653 | 0 |
| 0.00021553 | 0.00052504 | 3.85826995 | 4.22637126 | 0 | 2.31968076 | 45.1481466 |
| 0.00021586 | 0.00052581 | 12.8608998 | 25.1357869 | 21.3451941 | 16.7017014 | 0 |
| 0.00021639 | 0.00052707 | 20.2559173 | 49.1593709 | 10.2085711 | 28.3001052 | 0 |
| 0.00021662 | 0.00052758 | 29.9015921 | 123.232088 | 151.272462 | 159.594036 | 14.2573095 |
| 0.00021667 | 0.00052768 | 19.9343948 | 15.3484009 | 13.9207788 | 26.4443606 | 0 |
| 0.00021683 | 0.00052803 | 20.8989622 | 25.1357869 | 10.2085711 | 31.0837221 | 0 |
| 0.00021694 | 0.00052827 | 9.32415239 | 33.5885295 | 26.9135056 | 25.5164883 | 0 |
| 0.00021726 | 0.000529 | 248.215367 | 435.093799 | 497.435827 | 500.587107 | 0 |
| 0.00021842 | 0.0005318 | 311.233776 | 316.977844 | 0 | 318.2602 | 0 |
| 0.00021893 | 0.00053299 | 47.2638069 | 167.497766 | 193.962851 | 161.913717 | 0 |
| 0.00021924 | 0.00053368 | 9.00262989 | 27.5826335 | 36.1940248 | 25.9804245 | 0 |
| 0.00021923 | 0.00053368 | 18.6483048 | 36.7026978 | 110.438178 | 9.74265918 | 0 |
| 0.0002198 | 0.00053499 | 117.677234 | 92.9801676 | 103.941815 | 114.592229 | 21.3859642 |
| 0.00021988 | 0.00053512 | 16.7191698 | 28.0275146 | 12.0646749 | 23.6607437 | 0 |
| 0.00021988 | 0.00053512 | 11.2532874 | 30.6968018 | 17.6329864 | 28.7640414 | 0 |
| 0.00021993 | 0.0005352 | 268.149762 | 304.29873 | 318.321808 | 353.055411 | 68.9103291 |
| 0.00022023 | 0.00053587 | 16.7191698 | 64.7302124 | 11.136623 | 33.867339 | 0 |
| 0.00022048 | 0.00053646 | 82.952804 | 208.426835 | 146.632203 | 186.502333 | 30.8908372 |
| 0.00022065 | 0.00053682 | 24.7572322 | 87.4191528 | 69.6038938 | 75.1576565 | 9.50487297 |
| 0.00022104 | 0.00053773 | 14.7900348 | 30.9192423 | 12.9927268 | 26.4443606 | 0 |
| 0.00022135 | 0.00053844 | 16.0761248 | 15.5708415 | 16.7049345 | 18.0935099 | 0 |
| 0.00022207 | 0.00054016 | 12.5393773 | 16.4606038 | 31.5537652 | 22.7328714 | 0 |
| 0.00022218 | 0.0005404 | 9.00262989 | 34.4782918 | 59.3953227 | 21.341063 | 0 |
| 0.0002222 | 0.0005404 | 13.5039448 | 23.1338216 | 16.7049345 | 18.5574461 | 0 |
| 0.00022269 | 0.00054156 | 25.0787547 | 16.0157227 | 14.8488307 | 16.7017014 | 0 |
| 0.00022304 | 0.00054237 | 261.076267 | 280.275146 | 0 | 416.150728 | 0 |
| 0.00022307 | 0.00054239 | 10.6102424 | 36.4802572 | 20.4171422 | 25.0525522 | 0 |
| 0.0002239 | 0.00054439 | 5.78740493 | 32.2538859 | 51.9709073 | 53.8165936 | 0 |
| 0.0002241 | 0.00054482 | 0 | 0.22244059 | 0.92805192 | 0.9278723 | 11.8810912 |
| 0.00022432 | 0.00054531 | 61.7323193 | 52.4959798 | 47.3306478 | 52.8887212 | 190.097459 |
| 0.00022452 | 0.00054577 | 140.505331 | 59.3916382 | 85.3807764 | 63.0953166 | 23.7621824 |
| 0.00022466 | 0.00054608 | 52.0866444 | 48.4920492 | 95.5893474 | 25.5164883 | 7.12865473 |
| 0.00022471 | 0.00054616 | 38.261177 | 18.4625692 | 18.5610383 | 13.4541484 | 0 |
| 0.00022507 | 0.00054698 | 9.00262989 | 20.6869751 | 88.1649321 | 35.7230837 | 0 |
| 0.00022542 | 0.0005478 | 4.17979245 | 52.4959798 | 51.9709073 | 58.9198912 | 0 |

| | | | | | | |
|---|---|---|---|---|---|---|
| 0.00022573 | 0.00054852 | 9.00262989 | 50.4940145 | 29.6976613 | 23.6607437 | 0 |
| 0.00022596 | 0.00054902 | 157.224501 | 344.782918 | 330.386482 | 349.343922 | 35.6432737 |
| 0.00022626 | 0.00054972 | 10.9317649 | 58.0569946 | 12.9927268 | 67.2707419 | 0 |
| 0.00022649 | 0.00055024 | 10.6102424 | 28.2499552 | 20.4171422 | 25.0525522 | 0 |
| 0.00022663 | 0.00055055 | 7.71653991 | 30.6968018 | 32.4818171 | 38.0427644 | 0 |
| 0.00022666 | 0.00055057 | 27.9724572 | 85.1947469 | 53.8270112 | 66.3428696 | 9.50487297 |
| 0.00022691 | 0.00055115 | 610.249698 | 228.668929 | 110.438178 | 237.535309 | 3628.48526 |
| 0.000227 | 0.00055133 | 11.8963324 | 80.0786133 | 14.8488307 | 36.650956 | 0 |
| 0.00022704 | 0.00055137 | 0 | 0 | 0 | 0 | 90.2962933 |
| 0.00022728 | 0.00055192 | 11.2532874 | 55.8325887 | 32.4818171 | 16.7017014 | 0 |
| 0.0002273 | 0.00055192 | 10.2887199 | 35.1456136 | 18.5610383 | 32.9394667 | 0 |
| 0.00022736 | 0.000552 | 31.5092046 | 48.2696086 | 7.42441534 | 29.2279775 | 0 |
| 0.00022736 | 0.000552 | 6.43044992 | 116.33643 | 19.4890903 | 59.8477635 | 0 |
| 0.00022839 | 0.00055445 | 10.9317649 | 29.1397176 | 18.5610383 | 61.7035081 | 0 |
| 0.0002284 | 0.00055445 | 9.00262989 | 32.0314453 | 34.3379209 | 23.1968076 | 0 |
| 0.00022856 | 0.00055479 | 25.4002772 | 12.6791138 | 16.7049345 | 20.8771268 | 0 |
| 0.00022959 | 0.00055726 | 0 | 0 | 0 | 0 | 33.2670554 |
| 0.00022993 | 0.00055803 | 0.3215225 | 0 | 0 | 0 | 4.75243649 |
| 0.00023078 | 0.00056005 | 11.5748099 | 18.9074504 | 22.273246 | 23.1968076 | 0 |
| 0.00023088 | 0.00056025 | 23.1496197 | 14.9035197 | 20.4171422 | 13.9180845 | 0 |
| 0.00023107 | 0.00056067 | 57.5525268 | 34.7007324 | 33.409869 | 38.5067006 | 159.206622 |
| 0.0002311 | 0.00056071 | 21.5420072 | 39.3719849 | 26.9135056 | 36.1870198 | 2.37621824 |
| 0.00023131 | 0.00056118 | 18.3267823 | 50.4940145 | 66.819738 | 61.239572 | 4.75243649 |
| 0.00023154 | 0.00056168 | 326.345334 | 448.885116 | 390.709857 | 417.0786 | 45.1481466 |
| 0.00023156 | 0.00056171 | 40.5118345 | 27.1377523 | 33.409869 | 22.2689353 | 0 |
| 0.00023162 | 0.0005618 | 14.4685123 | 19.7972127 | 17.6329864 | 15.7738291 | 0 |
| 0.00023188 | 0.00056239 | 41.7979245 | 67.6219401 | 25.9854537 | 69.5904227 | 0 |
| 0.00023189 | 0.00056239 | 18.9698273 | 13.7913167 | 15.7768826 | 20.8771268 | 0 |
| 0.00023256 | 0.00056396 | 61.4107968 | 38.7046631 | 45.4745439 | 34.3312752 | 9.50487297 |
| 0.00023262 | 0.00056408 | 106.745469 | 20.4645345 | 12.9927268 | 22.2689353 | 0 |
| 0.00023267 | 0.00056415 | 159.153636 | 317.422725 | 306.257133 | 290.887967 | 42.7719284 |
| 0.00023377 | 0.00056677 | 85.524984 | 97.2065389 | 141.991943 | 154.026802 | 21.3859642 |
| 0.00023402 | 0.00056735 | 252.073637 | 395.054492 | 553.118943 | 522.392106 | 45.1481466 |
| 0.00023413 | 0.00056756 | 21.8635297 | 30.02948 | 13.9207788 | 14.3820207 | 0 |
| 0.00023423 | 0.00056777 | 22.8280972 | 55.8325887 | 72.3880495 | 65.4149973 | 0 |
| 0.00023461 | 0.00056866 | 8.6811074 | 26.6928711 | 32.4818171 | 28.7640414 | 0 |
| 0.0002347 | 0.00056883 | 6.75197242 | 34.0334106 | 32.4818171 | 55.208402 | 0 |
| 0.00023536 | 0.00057038 | 12.5393773 | 14.0137573 | 43.6184401 | 33.867339 | 0 |
| 0.00023542 | 0.00057048 | 8.3595849 | 36.2578166 | 31.5537652 | 25.9804245 | 0 |
| 0.00023556 | 0.00057079 | 18.3267823 | 51.3837769 | 68.6758419 | 27.3722329 | 0 |
| 0.00023591 | 0.00057159 | 10.2887199 | 22.0216187 | 25.9854537 | 24.1246799 | 0 |
| 0.00023632 | 0.00057255 | 0 | 498.934249 | 461.241803 | 355.839028 | 0 |
| 0.00023634 | 0.00057255 | 13.1824223 | 29.1397176 | 14.8488307 | 23.1968076 | 0 |

| | | | | | | |
|---|---|---|---|---|---|---|
| 0.00023643 | 0.00057274 | 18.6483048 | 25.3582275 | 13.9207788 | 15.309893 | 0 |
| 0.00023676 | 0.00057349 | 16.7191698 | 7.11809896 | 15.7768826 | 9.74265918 | 19.0097459 |
| 0.00023692 | 0.00057384 | 3.85826995 | 37.5924601 | 76.1002572 | 68.6625504 | 0 |
| 0.00023701 | 0.00057399 | 4.17979245 | 87.864034 | 25.0574018 | 98.8184002 | 0 |
| 0.00023702 | 0.00057399 | 40.833357 | 26.4704305 | 19.4890903 | 19.4853184 | 64.1578926 |
| 0.00023762 | 0.0005754 | 3.21522496 | 61.3936035 | 63.1075304 | 62.1674443 | 0 |
| 0.00023775 | 0.00057569 | 85.524984 | 13.123995 | 13.9207788 | 33.4034029 | 0 |
| 0.00023928 | 0.00057935 | 9.00262989 | 44.0432373 | 20.4171422 | 35.7230837 | 0 |
| 0.00023943 | 0.00057967 | 97.4213163 | 113.222262 | 116.934542 | 111.344676 | 14.2573095 |
| 0.00023951 | 0.00057982 | 15.4330798 | 15.7932821 | 38.9781805 | 17.6295737 | 0 |
| 0.00024021 | 0.00058147 | 5.14435994 | 44.9329997 | 79.8124649 | 33.867339 | 0 |
| 0.00024053 | 0.0005822 | 86.4895515 | 66.9546183 | 141.063891 | 123.407016 | 19.0097459 |
| 0.00024081 | 0.00058284 | 10.6102424 | 23.1338216 | 21.3451941 | 46.3936151 | 0 |
| 0.00024091 | 0.00058305 | 3.53674746 | 5.56101481 | 4.64025958 | 2.78361691 | 61.7816743 |
| 0.0002416 | 0.00058466 | 13.1824223 | 14.2361979 | 42.6903882 | 27.3722329 | 0 |
| 0.00024161 | 0.00058466 | 15.1115573 | 29.8070394 | 12.0646749 | 28.3001052 | 0 |
| 0.00024178 | 0.00058501 | 30.8661596 | 26.9153117 | 32.4818171 | 23.1968076 | 2.37621824 |
| 0.0002418 | 0.00058502 | 1006.68694 | 82.0805786 | 41.7623363 | 206.915523 | 16.6335277 |
| 0.00024238 | 0.00058638 | 7.71653991 | 32.6987671 | 36.1940248 | 29.2279775 | 0 |
| 0.00024257 | 0.00058681 | 10.9317649 | 20.6869751 | 34.3379209 | 20.8771268 | 0 |
| 0.00024303 | 0.00058789 | 19.6128723 | 24.023584 | 19.4890903 | 11.5984038 | 0 |
| 0.00024312 | 0.00058805 | 1.92913498 | 51.828658 | 102.085711 | 102.065953 | 0 |
| 0.00024342 | 0.00058872 | 0.3215225 | 0.44488118 | 0 | 0 | 57.0292378 |
| 0.00024388 | 0.0005898 | 21.2204847 | 17.1279256 | 12.9927268 | 17.6295737 | 0 |
| 0.00024396 | 0.00058995 | 10.6102424 | 27.5826335 | 32.4818171 | 18.0935099 | 0 |
| 0.00024436 | 0.00059087 | 9.96719738 | 34.0334106 | 18.5610383 | 33.867339 | 0 |
| 0.00024468 | 0.0005916 | 10.6102424 | 32.2538859 | 18.5610383 | 26.4443606 | 0 |
| 0.00024484 | 0.00059196 | 43.405537 | 35.5904948 | 5.5683115 | 47.7854236 | 0 |
| 0.00024496 | 0.0005922 | 10.2887199 | 35.5904948 | 61.2514265 | 16.7017014 | 0 |
| 0.00024501 | 0.00059228 | 14.4685123 | 68.7341431 | 79.8124649 | 8.35085072 | 0 |
| 0.00024523 | 0.00059277 | 14.4685123 | 19.7972127 | 13.9207788 | 25.0525522 | 0 |
| 0.00024535 | 0.00059302 | 10.6102424 | 18.2401286 | 27.8415575 | 29.6919137 | 0 |
| 0.00024555 | 0.00059347 | 23.4711422 | 76.5195638 | 113.222334 | 72.8379758 | 9.50487297 |
| 0.00024575 | 0.00059389 | 33.1168171 | 31.8090047 | 12.9927268 | 13.4541484 | 0 |
| 0.0002458 | 0.00059399 | 130.859656 | 66.9546183 | 38.0501286 | 62.1674443 | 11.8810912 |
| 0.00024607 | 0.00059459 | 12.5393773 | 21.1318563 | 16.7049345 | 20.8771268 | 0 |
| 0.00024634 | 0.00059521 | 9.32415239 | 42.7085937 | 19.4890903 | 34.3312752 | 0 |
| 0.00024657 | 0.00059567 | 51.1220769 | 148.145435 | 134.567528 | 161.913717 | 19.0097459 |
| 0.00024656 | 0.00059567 | 72.3425616 | 18.9074504 | 12.9927268 | 21.8049991 | 0 |
| 0.00024691 | 0.00059645 | 45.6561944 | 17.7952474 | 14.8488307 | 17.1656376 | 0 |
| 0.00024703 | 0.00059669 | 14.1469898 | 14.9035197 | 19.4890903 | 19.4853184 | 0 |
| 0.00024726 | 0.00059719 | 5.78740493 | 46.7125244 | 45.4745439 | 34.7952113 | 0 |
| 0.00024732 | 0.0005973 | 26.3648447 | 13.3464355 | 15.7768826 | 11.5984038 | 161.582841 |

| | | | | | | |
|---|---|---|---|---|---|---|
| 0.00024757 | 0.00059786 | 9.64567488 | 58.5018758 | 13.9207788 | 67.7346781 | 0 |
| 0.00024793 | 0.00059869 | 77.8084441 | 468.904769 | 475.162581 | 483.885406 | 2.37621824 |
| 0.00024862 | 0.00060032 | 22.8280972 | 232.67286 | 232.012979 | 233.359884 | 0 |
| 0.00024954 | 0.00060249 | 7.07349491 | 23.8011434 | 47.3306478 | 50.1051043 | 0 |
| 0.00025003 | 0.00060363 | 126.036818 | 505.829907 | 0 | 412.439239 | 0 |
| 0.0002504 | 0.00060448 | 21.5420072 | 21.3542969 | 11.136623 | 57.5280828 | 0 |
| 0.00025048 | 0.00060462 | 11.8963324 | 28.4723958 | 25.9854537 | 15.7738291 | 0 |
| 0.00025056 | 0.00060479 | 19.2913498 | 35.1456136 | 31.5537652 | 37.5788283 | 2.37621824 |
| 0.00025084 | 0.00060541 | 61.7323193 | 29.8070394 | 7.42441534 | 27.8361691 | 0 |
| 0.00025147 | 0.00060688 | 16.0761248 | 28.917277 | 11.136623 | 31.0837221 | 0 |
| 0.00025189 | 0.00060786 | 9.00262989 | 22.4664998 | 34.3379209 | 29.2279775 | 0 |
| 0.00025315 | 0.00061086 | 80.0591015 | 228.89137 | 232.941031 | 237.071373 | 26.1384007 |
| 0.00025351 | 0.00061168 | 7.39501741 | 57.1672323 | 21.3451941 | 47.3214874 | 0 |
| 0.00025388 | 0.00061252 | 16.3976473 | 27.8050741 | 13.9207788 | 16.7017014 | 0 |
| 0.0002539 | 0.00061253 | 4.82283744 | 75.4073608 | 19.4890903 | 128.046378 | 0 |
| 0.00025409 | 0.00061295 | 256.574952 | 456.225655 | 237.581291 | 467.647641 | 40.3957101 |
| 0.00025424 | 0.00061326 | 42.119447 | 49.3818115 | 6.49636342 | 26.9082968 | 0 |
| 0.00025439 | 0.0006136 | 11.5748099 | 18.017688 | 21.3451941 | 32.9394667 | 0 |
| 0.00025523 | 0.00061556 | 10.9317649 | 26.4704305 | 37.1220767 | 17.1656376 | 0 |
| 0.0002557 | 0.00061665 | 41.7979245 | 25.3582275 | 8.35246725 | 26.4443606 | 0 |
| 0.00025623 | 0.0006179 | 11.2532874 | 28.917277 | 23.2012979 | 17.6295737 | 0 |
| 0.00025728 | 0.00062034 | 16.0761248 | 28.0275146 | 11.136623 | 29.6919137 | 0 |
| 0.00025726 | 0.00062034 | 18.6483048 | 21.1318563 | 32.4818171 | 11.1344676 | 0 |
| 0.00025865 | 0.00062359 | 7.07349491 | 32.2538859 | 38.0501286 | 32.9394667 | 0 |
| 0.00025872 | 0.00062371 | 414.76402 | 719.595317 | 396.278169 | 558.11519 | 52.2768014 |
| 0.00025894 | 0.0006242 | 25.4002772 | 15.5708415 | 12.9927268 | 19.0213822 | 0 |
| 0.00025901 | 0.0006243 | 35.3674746 | 58.946757 | 22.273246 | 49.6411682 | 4.75243649 |
| 0.00025901 | 0.0006243 | 8.6811074 | 44.4881185 | 23.2012979 | 28.7640414 | 0 |
| 0.00025919 | 0.00062467 | 21.8635297 | 15.7932821 | 21.3451941 | 12.5262761 | 0 |
| 0.00025971 | 0.00062589 | 94.8491364 | 21.1318563 | 12.0646749 | 21.341063 | 0 |
| 0.00025978 | 0.00062595 | 858.786587 | 1190.05717 | 1334.53866 | 1440.05781 | 73.6627655 |
| 0.00025976 | 0.00062595 | 122.500071 | 239.123637 | 172.617657 | 222.689353 | 28.5146189 |
| 0.00025982 | 0.00062601 | 6.75197242 | 46.0452026 | 38.0501286 | 28.7640414 | 0 |
| 0.00026035 | 0.00062725 | 18.9698273 | 11.3444702 | 7.42441534 | 10.6705315 | 33.2670554 |
| 0.00026096 | 0.00062867 | 12.8608998 | 17.1279256 | 19.4890903 | 18.0935099 | 0 |
| 0.00026115 | 0.00062909 | 105.780901 | 69.846346 | 39.9062324 | 117.375846 | 14.2573095 |
| 0.00026162 | 0.00063017 | 22.5065747 | 20.6869751 | 10.2085711 | 42.2181898 | 0 |
| 0.00026207 | 0.00063121 | 15.4330798 | 20.9094157 | 25.0574018 | 12.5262761 | 0 |
| 0.00026218 | 0.00063144 | 2.57217997 | 44.2656779 | 69.6038938 | 110.416804 | 0 |
| 0.00026235 | 0.0006318 | 257.861042 | 306.745577 | 0 | 299.702754 | 0 |
| 0.00026267 | 0.00063253 | 36.9750871 | 39.5944255 | 6.49636342 | 31.0837221 | 0 |
| 0.00026289 | 0.00063301 | 11.5748099 | 24.023584 | 16.7049345 | 24.1246799 | 0 |
| 0.00026309 | 0.00063345 | 9.96719738 | 21.5767375 | 24.1293498 | 25.9804245 | 0 |

| | | | | | | |
|---|---|---|---|---|---|---|
| 0.00026352 | 0.00063442 | 9.00262989 | 34.2558512 | 24.1293498 | 25.5164883 | 0 |
| 0.00026409 | 0.00063574 | 6.43044992 | 36.4802572 | 45.4745439 | 32.0115944 | 0 |
| 0.00026418 | 0.00063592 | 206.738965 | 379.26121 | 0 | 300.630626 | 0 |
| 0.00026604 | 0.00064037 | 13.5039448 | 27.1377523 | 19.4890903 | 14.8459568 | 0 |
| 0.0002661 | 0.00064046 | 14.1469898 | 73.1829549 | 60.3233746 | 60.3116997 | 2.37621824 |
| 0.00026668 | 0.00064176 | 57.5525268 | 123.676969 | 113.222334 | 122.015208 | 0 |
| 0.00026667 | 0.00064176 | 27.9724572 | 44.0432373 | 31.5537652 | 49.6411682 | 2.37621824 |
| 0.00026684 | 0.00064205 | 553.983261 | 188.184741 | 0 | 226.864778 | 0 |
| 0.00026683 | 0.00064205 | 12.5393773 | 18.4625692 | 18.5610383 | 18.0935099 | 0 |
| 0.00026736 | 0.00064326 | 0.3215225 | 0 | 0 | 0 | 26.1384007 |
| 0.00026752 | 0.00064361 | 239.855782 | 361.243522 | 296.048562 | 317.796264 | 0 |
| 0.00026769 | 0.00064396 | 80.0591015 | 249.800785 | 348.019469 | 314.548711 | 28.5146189 |
| 0.00026777 | 0.0006441 | 235.354467 | 387.713953 | 0 | 254.237011 | 0 |
| 0.00026783 | 0.00064421 | 25.0787547 | 32.6987671 | 41.7623363 | 41.2903175 | 4.75243649 |
| 0.00026801 | 0.0006446 | 56.2664368 | 14.9035197 | 12.0646749 | 28.7640414 | 0 |
| 0.0002688 | 0.00064646 | 369.429348 | 460.674467 | 384.213494 | 424.037642 | 192.473678 |
| 0.00026904 | 0.00064698 | 14.7900348 | 54.9428263 | 75.1722053 | 84.9003157 | 4.75243649 |
| 0.00026933 | 0.00064763 | 21.8635297 | 39.1495443 | 28.7696094 | 8.81478687 | 0 |
| 0.00026937 | 0.00064767 | 10.9317649 | 36.035376 | 26.9135056 | 16.7017014 | 0 |
| 0.00026954 | 0.00064803 | 20.5774398 | 46.4900838 | 55.683115 | 43.1460621 | 4.75243649 |
| 0.00026984 | 0.00064872 | 25.4002772 | 30.9192423 | 25.0574018 | 8.81478687 | 0 |
| 0.00026994 | 0.0006489 | 8.0380624 | 22.2440592 | 34.3379209 | 48.7132959 | 0 |
| 0.0002702 | 0.00064948 | 9.32415239 | 28.917277 | 22.273246 | 26.4443606 | 0 |
| 0.00027033 | 0.00064975 | 68.8058142 | 54.053064 | 14.8488307 | 8.81478687 | 0 |
| 0.00027061 | 0.00065038 | 8.3595849 | 46.4900838 | 30.6257133 | 22.7328714 | 0 |
| 0.00027143 | 0.0006523 | 78.1299666 | 97.6514201 | 106.72597 | 127.582442 | 30.8908372 |
| 0.00027166 | 0.00065282 | 22.1850522 | 34.4782918 | 27.8415575 | 28.3001052 | 0 |
| 0.00027209 | 0.0006538 | 3.21522496 | 72.5156331 | 73.3161014 | 41.2903175 | 0 |
| 0.00027252 | 0.00065478 | 59.8031843 | 26.4704305 | 28.7696094 | 36.1870198 | 2.37621824 |
| 0.00027255 | 0.00065481 | 66.5551567 | 82.3030192 | 32.4818171 | 45.929679 | 0 |
| 0.00027283 | 0.00065543 | 18.3267823 | 12.9015544 | 16.7049345 | 32.4755306 | 0 |
| 0.000273 | 0.00065579 | 30.8661596 | 35.5904948 | 35.2659728 | 32.0115944 | 7.12865473 |
| 0.00027324 | 0.00065631 | 52.7296894 | 44.4881185 | 43.6184401 | 30.619786 | 118.810912 |
| 0.00027335 | 0.00065654 | 23.4711422 | 33.5885295 | 25.0574018 | 29.2279775 | 2.37621824 |
| 0.00027348 | 0.00065681 | 13.1824223 | 55.8325887 | 12.9927268 | 30.619786 | 0 |
| 0.00027361 | 0.00065708 | 11.2532874 | 23.5787028 | 20.4171422 | 18.5574461 | 0 |
| 0.0002737 | 0.00065724 | 6.10892743 | 54.7203857 | 28.7696094 | 41.2903175 | 0 |
| 0.00027378 | 0.00065739 | 219.599865 | 373.477755 | 550.334787 | 0 | 0 |
| 0.00027422 | 0.00065839 | 21.2204847 | 38.9271037 | 7.42441534 | 62.6313804 | 0 |
| 0.00027446 | 0.00065892 | 12.5393773 | 16.0157227 | 20.4171422 | 19.9492545 | 0 |
| 0.00027464 | 0.0006593 | 7.39501741 | 54.2755046 | 26.9135056 | 30.619786 | 0 |
| 0.00027471 | 0.00065943 | 26.6863672 | 62.5058065 | 14.8488307 | 99.7462725 | 0 |
| 0.00027486 | 0.00065976 | 31.1876821 | 17.1279256 | 25.9854537 | 10.6705315 | 0 |

| | | | | | | |
|---|---|---|---|---|---|---|
| 0.00027593 | 0.00066227 | 94.8491364 | 180.844202 | 189.322591 | 201.812226 | 26.1384007 |
| 0.00027637 | 0.00066327 | 0 | 0.88976237 | 1.85610383 | 0.46393615 | 49.9005831 |
| 0.0002764 | 0.0006633 | 8.0380624 | 28.4723958 | 35.2659728 | 26.4443606 | 0 |
| 0.00027674 | 0.00066407 | 25.7217997 | 37.1475789 | 6.49636342 | 64.487125 | 0 |
| 0.00027707 | 0.00066482 | 20.2559173 | 29.8070394 | 32.4818171 | 32.4755306 | 0 |
| 0.00027733 | 0.00066539 | 15.7546023 | 16.6830444 | 21.3451941 | 13.4541484 | 0 |
| 0.00027745 | 0.00066564 | 17.6837373 | 32.4763265 | 34.3379209 | 9.74265918 | 0 |
| 0.00027786 | 0.00066657 | 11.8963324 | 27.1377523 | 14.8488307 | 26.9082968 | 0 |
| 0.00027807 | 0.00066702 | 25.4002772 | 46.7125244 | 11.136623 | 16.7017014 | 0 |
| 0.00027897 | 0.00066915 | 4.82283744 | 54.2755046 | 46.4025958 | 38.0427644 | 0 |
| 0.00027901 | 0.00066919 | 24.4357097 | 18.2401286 | 12.9927268 | 15.309893 | 0 |
| 0.00027922 | 0.00066963 | 163.333428 | 46.0452026 | 33.409869 | 42.2181898 | 7.12865473 |
| 0.00027947 | 0.00067019 | 19.6128723 | 14.0137573 | 16.7049345 | 15.309893 | 0 |
| 0.00027971 | 0.00067073 | 40.5118345 | 24.2460246 | 9.28051917 | 21.341063 | 0 |
| 0.00027983 | 0.00067096 | 48.8714194 | 98.763623 | 89.092984 | 92.3232941 | 16.6335277 |
| 0.00028006 | 0.00067145 | 9.32415239 | 36.2578166 | 21.3451941 | 25.5164883 | 0 |
| 0.00028032 | 0.00067203 | 12.5393773 | 18.017688 | 20.4171422 | 16.7017014 | 0 |
| 0.00028061 | 0.00067269 | 8.0380624 | 27.8050741 | 26.9135056 | 34.7952113 | 0 |
| 0.0002811 | 0.00067382 | 9.00262989 | 10.2322673 | 7.42441534 | 9.27872303 | 49.9005831 |
| 0.00028155 | 0.00067484 | 32.4737721 | 12.011792 | 20.4171422 | 16.7017014 | 0 |
| 0.00028213 | 0.00067619 | 20.8989622 | 9.78738607 | 25.9854537 | 25.0525522 | 0 |
| 0.00028259 | 0.00067723 | 7.71653991 | 30.4743612 | 31.5537652 | 28.7640414 | 0 |
| 0.00028299 | 0.00067814 | 58.5170943 | 23.8011434 | 45.4745439 | 25.9804245 | 61.7816743 |
| 0.00028307 | 0.00067829 | 323.130109 | 274.26925 | 232.012979 | 160.985845 | 47.5243649 |
| 0.00028345 | 0.00067916 | 14.1469898 | 19.3523315 | 19.4890903 | 13.9180845 | 0 |
| 0.00028353 | 0.00067929 | 18.0052598 | 110.997856 | 68.6758419 | 84.4363795 | 0 |
| 0.00028377 | 0.00067983 | 11.8963324 | 60.7262817 | 12.0646749 | 41.7542536 | 0 |
| 0.00028458 | 0.00068172 | 21.5420072 | 50.7164551 | 58.4672708 | 63.0953166 | 7.12865473 |
| 0.00028554 | 0.00068397 | 21.2204847 | 32.6987671 | 26.9135056 | 32.9394667 | 0 |
| 0.00028559 | 0.00068404 | 10.2887199 | 52.0510986 | 17.6329864 | 27.8361691 | 0 |
| 0.00028756 | 0.0006887 | 23.7926647 | 34.2558512 | 8.35246725 | 25.9804245 | 0 |
| 0.00028784 | 0.00068932 | 0 | 617.272644 | 243.149602 | 392.489984 | 0 |
| 0.000288 | 0.00068965 | 0 | 3.11416829 | 0 | 0.9278723 | 26.1384007 |
| 0.00028821 | 0.00069011 | 63.6614542 | 358.129354 | 349.875573 | 324.755306 | 0 |
| 0.00028836 | 0.00069042 | 13.1824223 | 16.4606038 | 17.6329864 | 17.6295737 | 0 |
| 0.00028941 | 0.00069284 | 15.1115573 | 82.7479004 | 21.3451941 | 15.309893 | 0 |
| 0.00028941 | 0.00069284 | 275.544779 | 436.206002 | 445.46492 | 496.411682 | 0 |
| 0.0002899 | 0.00069396 | 16.0761248 | 17.1279256 | 54.7550631 | 15.7738291 | 0 |
| 0.00029004 | 0.00069425 | 19.6128723 | 30.2519206 | 10.2085711 | 20.8771268 | 0 |
| 0.00029018 | 0.00069454 | 130.859656 | 252.914954 | 229.228823 | 228.720523 | 61.7816743 |
| 0.00029059 | 0.00069546 | 15.7546023 | 8.23030192 | 7.42441534 | 7.88691457 | 35.6432737 |
| 0.00029071 | 0.00069571 | 87.7756414 | 159.934786 | 135.49558 | 155.882547 | 23.7621824 |
| 0.00029122 | 0.00069688 | 8.3595849 | 30.9192423 | 23.2012979 | 31.5476583 | 0 |

| | | | | | | |
|---|---|---|---|---|---|---|
| 0.00029176 | 0.00069811 | 9.64567488 | 38.7046631 | 15.7768826 | 47.3214874 | 0 |
| 0.00029182 | 0.00069821 | 26.6863672 | 93.2026082 | 57.5392189 | 59.3838274 | 9.50487297 |
| 0.00029241 | 0.00069958 | 184.23239 | 117.226192 | 86.3088283 | 104.84957 | 0 |
| 0.00029262 | 0.00070003 | 201.594605 | 297.403072 | 225.516616 | 299.238818 | 35.6432737 |
| 0.00029277 | 0.00070033 | 64.9475442 | 121.675004 | 127.143113 | 135.00542 | 30.8908372 |
| 0.00029343 | 0.00070187 | 25.0787547 | 14.9035197 | 17.6329864 | 13.4541484 | 0 |
| 0.00029352 | 0.00070202 | 10.6102424 | 25.8031087 | 16.7049345 | 32.9394667 | 0 |
| 0.00029357 | 0.00070211 | 9.32415239 | 26.6928711 | 24.1293498 | 22.7328714 | 0 |
| 0.00029408 | 0.00070328 | 205.131353 | 262.479899 | 0 | 373.932538 | 0 |
| 0.00029475 | 0.00070482 | 27.9724572 | 163.271395 | 138.279736 | 116.91191 | 14.2573095 |
| 0.00029492 | 0.00070513 | 5.14435994 | 60.05896 | 42.6903882 | 32.9394667 | 0 |
| 0.00029491 | 0.00070513 | 365.892601 | 718.038232 | 787.916077 | 752.504437 | 0 |
| 0.00029498 | 0.00070521 | 21.8635297 | 17.7952474 | 12.0646749 | 16.7017014 | 0 |
| 0.00029588 | 0.00070732 | 10.9317649 | 17.7952474 | 31.5537652 | 20.8771268 | 0 |
| 0.00029608 | 0.00070774 | 10.2887199 | 39.816866 | 14.8488307 | 39.4345729 | 0 |
| 0.00029617 | 0.00070791 | 13.1824223 | 19.7972127 | 36.1940248 | 14.8459568 | 0 |
| 0.0002964 | 0.00070841 | 42.4409695 | 131.23995 | 38.0501286 | 95.106911 | 7.12865473 |
| 0.00029741 | 0.00071079 | 20.2559173 | 28.917277 | 9.28051917 | 24.588616 | 0 |
| 0.00029803 | 0.00071221 | 11.2532874 | 24.023584 | 15.7768826 | 28.3001052 | 0 |
| 0.00029807 | 0.00071226 | 16.3976473 | 21.3542969 | 11.136623 | 27.8361691 | 0 |
| 0.00029818 | 0.00071246 | 10.6102424 | 22.911381 | 17.6329864 | 27.8361691 | 0 |
| 0.00029854 | 0.00071328 | 16.0761248 | 29.5845988 | 21.3451941 | 11.5984038 | 0 |
| 0.00029859 | 0.00071335 | 12.5393773 | 17.3503662 | 21.3451941 | 16.2377653 | 0 |
| 0.00029862 | 0.00071337 | 5.78740493 | 55.8325887 | 38.0501286 | 30.619786 | 0 |
| 0.00029865 | 0.00071339 | 5.78740493 | 36.4802572 | 64.0355823 | 30.1558498 | 0 |
| 0.0002988 | 0.00071371 | 25.4002772 | 7.78542074 | 28.7696094 | 40.8263813 | 0 |
| 0.00029911 | 0.00071438 | 7.39501741 | 27.5826335 | 31.5537652 | 34.3312752 | 0 |
| 0.00030002 | 0.00071651 | 8.6811074 | 27.5826335 | 24.1293498 | 26.9082968 | 0 |
| 0.00030032 | 0.00071718 | 17.3622148 | 36.035376 | 31.5537652 | 41.2903175 | 2.37621824 |
| 0.00030048 | 0.0007175 | 22.8280972 | 27.5826335 | 20.4171422 | 9.74265918 | 0 |
| 0.00030068 | 0.00071793 | 14.4685123 | 43.1534749 | 10.2085711 | 38.9706367 | 0 |
| 0.0003008 | 0.00071816 | 11.5748099 | 27.8050741 | 40.8342843 | 14.3820207 | 0 |
| 0.0003024 | 0.00072192 | 50.8005544 | 72.9605143 | 57.5392189 | 102.065953 | 7.12865473 |
| 0.00030249 | 0.00072209 | 6.10892743 | 67.3994995 | 23.2012979 | 45.929679 | 0 |
| 0.00030284 | 0.00072289 | 9.96719738 | 22.4664998 | 22.273246 | 21.8049991 | 0 |
| 0.00030304 | 0.0007233 | 5.46588243 | 34.0334106 | 51.0428554 | 38.5067006 | 0 |
| 0.00030307 | 0.00072333 | 9.64567488 | 33.1436483 | 18.5610383 | 26.9082968 | 0 |
| 0.00030334 | 0.00072392 | 6.75197242 | 19.3523315 | 65.8916861 | 51.0329766 | 0 |
| 0.00030364 | 0.00072458 | 13.1824223 | 58.7243164 | 11.136623 | 36.650956 | 0 |
| 0.00030398 | 0.00072535 | 5.14435994 | 24.9133464 | 71.4599976 | 55.208402 | 0 |
| 0.0003048 | 0.00072725 | 65.2690667 | 56.2774699 | 34.3379209 | 48.2493597 | 14.2573095 |
| 0.00030562 | 0.00072916 | 9.00262989 | 30.6968018 | 19.4890903 | 32.4755306 | 0 |
| 0.00030574 | 0.00072938 | 75.2362641 | 28.0275146 | 18.5610383 | 34.7952113 | 2.37621824 |

| | | | | | | |
|---|---|---|---|---|---|---|
| 0.00030639 | 0.00073088 | 0.64304499 | 0.66732178 | 0 | 0 | 30.8908372 |
| 0.0003072 | 0.00073278 | 12.2178549 | 83.6376628 | 83.5246725 | 65.4149973 | 4.75243649 |
| 0.00030751 | 0.00073345 | 270.721942 | 712.477218 | 679.334003 | 739.514225 | 80.7914203 |
| 0.0003084 | 0.00073553 | 59.8031843 | 32.2538859 | 29.6976613 | 31.5476583 | 85.5438568 |
| 0.0003086 | 0.00073595 | 8.6811074 | 23.8011434 | 25.0574018 | 29.6919137 | 0 |
| 0.00030918 | 0.0007373 | 42.762492 | 35.5904948 | 33.409869 | 28.3001052 | 144.949313 |
| 0.00030957 | 0.00073816 | 16.7191698 | 33.8109701 | 34.3379209 | 44.0739344 | 2.37621824 |
| 0.00030959 | 0.00073817 | 34.4029071 | 14.0137573 | 11.136623 | 32.0115944 | 0 |
| 0.00030983 | 0.00073868 | 6.75197242 | 33.3660889 | 38.0501286 | 29.2279775 | 0 |
| 0.00031037 | 0.00073991 | 9.96719738 | 28.917277 | 18.5610383 | 24.1246799 | 0 |
| 0.00031065 | 0.00074053 | 33.7598621 | 27.8050741 | 7.42441534 | 26.9082968 | 0 |
| 0.00031092 | 0.00074112 | 9.00262989 | 38.2597819 | 20.4171422 | 26.4443606 | 0 |
| 0.00031109 | 0.00074147 | 0.3215225 | 78.9664103 | 193.034799 | 183.25478 | 0 |
| 0.00031145 | 0.00074228 | 11.8963324 | 11.1220296 | 12.0646749 | 11.5984038 | 45.1481466 |
| 0.00031204 | 0.00074363 | 9.00262989 | 18.9074504 | 35.2659728 | 30.1558498 | 0 |
| 0.00031292 | 0.00074568 | 50.8005544 | 186.405216 | 154.98467 | 156.346483 | 28.5146189 |
| 0.00031347 | 0.00074694 | 8.6811074 | 19.3523315 | 31.5537652 | 38.0427644 | 0 |
| 0.00031371 | 0.00074744 | 5.14435994 | 47.8247274 | 45.4745439 | 33.4034029 | 0 |
| 0.00031392 | 0.00074788 | 0 | 0 | 0 | 0 | 30.8908372 |
| 0.00031393 | 0.00074788 | 54.3373018 | 129.905306 | 119.718697 | 116.447974 | 0 |
| 0.000314 | 0.00074798 | 12.5393773 | 60.2814006 | 36.1940248 | 54.2805297 | 2.37621824 |
| 0.00031411 | 0.0007482 | 0 | 0 | 0 | 4.63936151 | 66.5341108 |
| 0.00031476 | 0.00074969 | 69.1273367 | 169.944613 | 157.768826 | 174.439993 | 0 |
| 0.00031493 | 0.00075004 | 86.811074 | 93.2026082 | 66.819738 | 101.138081 | 19.0097459 |
| 0.00031499 | 0.00075012 | 11.5748099 | 24.4684652 | 16.7049345 | 19.0213822 | 0 |
| 0.00031552 | 0.00075134 | 6.43044992 | 29.5845988 | 38.0501286 | 36.650956 | 0 |
| 0.00031565 | 0.0007516 | 8.6811074 | 43.5983561 | 43.6184401 | 16.7017014 | 0 |
| 0.00031579 | 0.00075181 | 26.6863672 | 92.5352865 | 25.9854537 | 7.42297842 | 0 |
| 0.00031577 | 0.00075181 | 46.9422844 | 93.6474894 | 80.7405168 | 101.602017 | 4.75243649 |
| 0.00031615 | 0.00075263 | 6.43044992 | 133.686796 | 19.4890903 | 39.898509 | 0 |
| 0.00031641 | 0.00075315 | 160.118203 | 456.003214 | 450.10518 | 467.183704 | 121.18713 |
| 0.00031642 | 0.00075315 | 0 | 251.135429 | 503.004139 | 404.088388 | 0 |
| 0.00031682 | 0.00075405 | 19.6128723 | 67.3994995 | 38.0501286 | 53.8165936 | 4.75243649 |
| 0.00031783 | 0.00075641 | 9.96719738 | 24.023584 | 19.4890903 | 23.1968076 | 0 |
| 0.00031795 | 0.00075664 | 14.1469898 | 42.4861532 | 10.2085711 | 38.0427644 | 0 |
| 0.00031874 | 0.00075847 | 27.9724572 | 18.9074504 | 28.7696094 | 9.27872303 | 0 |
| 0.00031967 | 0.00076062 | 62.0538418 | 16.6830444 | 13.9207788 | 18.5574461 | 0 |
| 0.00032038 | 0.00076226 | 8.0380624 | 38.2597819 | 43.6184401 | 19.0213822 | 0 |
| 0.00032047 | 0.00076243 | 8.6811074 | 23.8011434 | 40.8342843 | 22.7328714 | 0 |
| 0.00032098 | 0.00076359 | 14.1469898 | 22.2440592 | 12.9927268 | 18.5574461 | 0 |
| 0.00032154 | 0.00076487 | 375.859798 | 93.4250488 | 97.4454513 | 92.7872303 | 700.984382 |
| 0.00032162 | 0.000765 | 8.6811074 | 22.911381 | 24.1293498 | 33.4034029 | 0 |
| 0.00032213 | 0.00076616 | 45.6561944 | 18.017688 | 10.2085711 | 23.1968076 | 0 |

| | | | | | | |
|---|---|---|---|---|---|---|
| 0.00032246 | 0.0007669 | 7.07349491 | 26.6928711 | 38.9781805 | 31.0837221 | 0 |
| 0.00032261 | 0.00076719 | 330.525126 | 318.757369 | 262.638692 | 295.063392 | 0 |
| 0.00032344 | 0.00076906 | 122.821594 | 138.135608 | 118.790645 | 125.262761 | 40.3957101 |
| 0.00032344 | 0.00076906 | 29.5800696 | 15.3484009 | 13.9207788 | 15.7738291 | 0 |
| 0.00032469 | 0.00077196 | 4.82283744 | 33.1436483 | 62.1794784 | 41.7542536 | 0 |
| 0.00032491 | 0.00077245 | 35.6889971 | 11.7893514 | 12.0646749 | 73.3019119 | 0 |
| 0.00032521 | 0.00077309 | 3.53674746 | 68.7341431 | 36.1940248 | 61.239572 | 0 |
| 0.00032567 | 0.00077413 | 3.85826995 | 37.3700195 | 145.704151 | 33.867339 | 0 |
| 0.0003261 | 0.0007751 | 45.6561944 | 166.385563 | 94.6612955 | 118.303719 | 26.1384007 |
| 0.00032613 | 0.00077512 | 9.32415239 | 27.3601929 | 30.6257133 | 18.5574461 | 0 |
| 0.00032679 | 0.00077663 | 15.4330798 | 35.3680542 | 10.2085711 | 28.7640414 | 0 |
| 0.00032705 | 0.0007772 | 9.64567488 | 25.5806681 | 18.5610383 | 27.3722329 | 0 |
| 0.00032712 | 0.00077732 | 9.96719738 | 27.1377523 | 17.6329864 | 25.5164883 | 0 |
| 0.00032746 | 0.00077807 | 6.10892743 | 28.2499552 | 36.1940248 | 48.2493597 | 0 |
| 0.00032815 | 0.00077966 | 5.46588243 | 49.3818115 | 56.6111669 | 25.5164883 | 0 |
| 0.00032892 | 0.00078143 | 5.78740493 | 49.8266927 | 55.683115 | 23.6607437 | 0 |
| 0.00032935 | 0.00078239 | 49.1929419 | 125.234054 | 40.8342843 | 77.9412734 | 0 |
| 0.00032941 | 0.00078249 | 56.9094818 | 124.121851 | 104.869867 | 109.952868 | 19.0097459 |
| 0.00032971 | 0.00078313 | 0 | 0 | 0 | 0 | 99.8011662 |
| 0.00033006 | 0.00078392 | 72.9856066 | 16.2381632 | 14.8488307 | 18.0935099 | 0 |
| 0.000331 | 0.00078608 | 13.5039448 | 14.9035197 | 18.5610383 | 16.2377653 | 0 |
| 0.00033166 | 0.00078759 | 0 | 0 | 0 | 0 | 0 |
| 0.0003318 | 0.00078787 | 27.6509347 | 54.2755046 | 8.35246725 | 20.4131907 | 0 |
| 0.00033187 | 0.00078792 | 4.50131495 | 64.7302124 | 47.3306478 | 32.0115944 | 0 |
| 0.00033186 | 0.00078792 | 10.9317649 | 16.905485 | 38.9781805 | 21.341063 | 0 |
| 0.00033202 | 0.00078822 | 8.6811074 | 56.7223511 | 87.2368802 | 49.177232 | 2.37621824 |
| 0.00033275 | 0.0007899 | 6.75197242 | 32.2538859 | 35.2659728 | 29.2279775 | 0 |
| 0.00033295 | 0.00079034 | 11.2532874 | 74.0727173 | 59.3953227 | 149.387441 | 4.75243649 |
| 0.00033311 | 0.00079066 | 35.3674746 | 31.3641235 | 26.9135056 | 37.1148921 | 90.2962933 |
| 0.00033319 | 0.00079073 | 23.1496197 | 19.5747721 | 16.7049345 | 11.1344676 | 0 |
| 0.00033319 | 0.00079073 | 16.0761248 | 19.1298909 | 16.7049345 | 12.5262761 | 0 |
| 0.00033325 | 0.00079081 | 16.3976473 | 17.5728068 | 12.9927268 | 15.7738291 | 0 |
| 0.0003338 | 0.00079206 | 177.480418 | 326.54279 | 320.177911 | 338.67339 | 38.0194919 |
| 0.00033421 | 0.00079298 | 112.211351 | 331.658923 | 320.177911 | 346.096369 | 9.50487297 |
| 0.00033435 | 0.00079325 | 20.5774398 | 20.6869751 | 11.136623 | 16.2377653 | 0 |
| 0.00033476 | 0.00079418 | 11.8963324 | 106.326603 | 8.35246725 | 53.8165936 | 0 |
| 0.0003349 | 0.00079439 | 5.78740493 | 26.4704305 | 48.2586997 | 44.0739344 | 0 |
| 0.00033489 | 0.00079439 | 13.1824223 | 19.5747721 | 44.546492 | 14.8459568 | 0 |
| 0.00033497 | 0.00079451 | 143.720556 | 340.778988 | 0 | 337.745518 | 0 |
| 0.00033501 | 0.00079455 | 27.6509347 | 26.0255493 | 11.136623 | 14.3820207 | 0 |
| 0.00033556 | 0.0007958 | 13.8254673 | 10.899589 | 43.6184401 | 31.5476583 | 0 |
| 0.00033564 | 0.00079592 | 54.9803468 | 26.2479899 | 5.5683115 | 41.2903175 | 0 |
| 0.00033579 | 0.00079623 | 13.1824223 | 16.6830444 | 17.6329864 | 15.309893 | 0 |

| | | | | | | |
|---|---|---|---|---|---|---|
| 0.00033597 | 0.00079659 | 21.2204847 | 54.4979451 | 6.49636342 | 42.2181898 | 0 |
| 0.00033628 | 0.00079727 | 8.6811074 | 39.1495443 | 38.0501286 | 17.1656376 | 0 |
| 0.00033716 | 0.0007993 | 4.82283744 | 66.5097371 | 29.6976613 | 44.5378705 | 0 |
| 0.00033746 | 0.00079997 | 15.1115573 | 13.123995 | 17.6329864 | 17.1656376 | 0 |
| 0.00033832 | 0.00080194 | 11.8963324 | 53.6081828 | 46.4025958 | 10.6705315 | 0 |
| 0.00033863 | 0.00080262 | 246.286232 | 296.068429 | 0 | 220.833608 | 0 |
| 0.00033867 | 0.00080267 | 8.0380624 | 23.8011434 | 30.6257133 | 27.3722329 | 0 |
| 0.00033905 | 0.0008035 | 5.14435994 | 26.2479899 | 55.683115 | 52.4247851 | 0 |
| 0.00033929 | 0.00080402 | 34.0813846 | 14.2361979 | 22.273246 | 12.0623399 | 0 |
| 0.00034001 | 0.00080567 | 13.1824223 | 46.7125244 | 11.136623 | 32.0115944 | 0 |
| 0.00034045 | 0.00080665 | 117.034189 | 323.206181 | 209.739733 | 243.102543 | 30.8908372 |
| 0.00034052 | 0.00080676 | 1.28608998 | 50.2715739 | 119.718697 | 98.8184002 | 0 |
| 0.00034146 | 0.00080892 | 36.9750871 | 21.5767375 | 11.136623 | 15.7738291 | 0 |
| 0.00034224 | 0.00081073 | 98.3858838 | 193.300875 | 204.171422 | 198.100737 | 45.1481466 |
| 0.00034253 | 0.00081136 | 13.5039448 | 48.4920492 | 64.9636342 | 8.81478687 | 0 |
| 0.00034258 | 0.0008114 | 31.8307271 | 23.5787028 | 16.7049345 | 10.2065953 | 0 |
| 0.00034271 | 0.00081165 | 13.1824223 | 18.4625692 | 17.6329864 | 14.3820207 | 0 |
| 0.00034285 | 0.00081194 | 9.96719738 | 27.1377523 | 19.4890903 | 20.4131907 | 0 |
| 0.00034462 | 0.00081608 | 10.6102424 | 28.917277 | 24.1293498 | 15.7738291 | 0 |
| 0.00034507 | 0.00081707 | 12.8608998 | 17.3503662 | 15.7768826 | 16.7017014 | 0 |
| 0.00034539 | 0.00081779 | 8.6811074 | 49.8266927 | 56.6111669 | 79.797018 | 2.37621824 |
| 0.00034564 | 0.00081832 | 5.14435994 | 32.6987671 | 40.8342843 | 48.2493597 | 0 |
| 0.0003458 | 0.00081863 | 21.2204847 | 17.3503662 | 10.2085711 | 23.1968076 | 0 |
| 0.00034606 | 0.0008192 | 8.0380624 | 21.7991781 | 31.5537652 | 30.1558498 | 0 |
| 0.00034642 | 0.00082 | 74.9147416 | 13.5688761 | 12.9927268 | 25.0525522 | 0 |
| 0.0003465 | 0.00082005 | 5.78740493 | 104.102197 | 16.7049345 | 56.1362743 | 0 |
| 0.00034649 | 0.00082005 | 4.50131495 | 59.8365194 | 58.4672708 | 27.8361691 | 0 |
| 0.00034658 | 0.00082019 | 10.9317649 | 18.6850098 | 17.6329864 | 30.1558498 | 0 |
| 0.00034779 | 0.000823 | 32.7952946 | 34.4782918 | 10.2085711 | 14.3820207 | 0 |
| 0.00034783 | 0.00082303 | 9.64567488 | 18.6850098 | 30.6257133 | 22.2689353 | 0 |
| 0.00034794 | 0.00082323 | 6.10892743 | 43.1534749 | 31.5537652 | 31.5476583 | 0 |
| 0.00034816 | 0.0008237 | 10.2887199 | 26.2479899 | 15.7768826 | 29.2279775 | 0 |
| 0.00034859 | 0.00082465 | 14.1469898 | 14.9035197 | 14.8488307 | 18.0935099 | 0 |
| 0.0003487 | 0.00082485 | 19.6128723 | 16.0157227 | 13.9207788 | 13.9180845 | 0 |
| 0.00034983 | 0.00082749 | 6.43044992 | 36.7026978 | 21.3451941 | 90.4675495 | 0 |
| 0.00035053 | 0.00082907 | 21.2204847 | 14.4586385 | 25.9854537 | 11.5984038 | 0 |
| 0.00035087 | 0.00082982 | 8.6811074 | 26.4704305 | 20.4171422 | 30.1558498 | 0 |
| 0.00035095 | 0.00082996 | 4.17979245 | 29.8070394 | 74.2441534 | 50.5690405 | 0 |
| 0.00035122 | 0.00083053 | 8.6811074 | 10.899589 | 88.1649321 | 61.239572 | 0 |
| 0.0003515 | 0.00083114 | 7.39501741 | 54.9428263 | 25.9854537 | 25.5164883 | 0 |
| 0.0003521 | 0.0008325 | 16.0761248 | 20.0196533 | 11.136623 | 19.9492545 | 0 |
| 0.00035219 | 0.00083264 | 26.3648447 | 17.7952474 | 12.9927268 | 13.9180845 | 0 |
| 0.00035283 | 0.00083411 | 3.21522496 | 2.22440592 | 1.85610383 | 3.24755306 | 16.6335277 |

| | | | | | | |
|---|---|---|---|---|---|---|
| 0.00035286 | 0.00083412 | 24.1141872 | 71.4034302 | 21.3451941 | 8.81478687 | 0 |
| 0.00035411 | 0.000837 | 14.1469898 | 43.3759155 | 8.35246725 | 78.8691457 | 0 |
| 0.00035472 | 0.0008384 | 13.1824223 | 25.5806681 | 12.0646749 | 24.1246799 | 0 |
| 0.00035475 | 0.00083841 | 10.6102424 | 16.6830444 | 22.273246 | 21.8049991 | 0 |
| 0.00035485 | 0.00083859 | 9.00262989 | 19.5747721 | 28.7696094 | 25.0525522 | 0 |
| 0.00035488 | 0.00083859 | 16.7191698 | 19.5747721 | 13.9207788 | 13.4541484 | 0 |
| 0.000355 | 0.00083883 | 9.96719738 | 18.9074504 | 28.7696094 | 19.9492545 | 0 |
| 0.00035634 | 0.00084192 | 14.7900348 | 27.1377523 | 11.136623 | 21.8049991 | 0 |
| 0.0003564 | 0.000842 | 3.85826995 | 6.005896 | 8.35246725 | 4.63936151 | 106.929821 |
| 0.00035648 | 0.00084214 | 6.75197242 | 30.2519206 | 28.7696094 | 35.2591475 | 0 |
| 0.00035693 | 0.00084313 | 9.32415239 | 35.1456136 | 19.4890903 | 22.7328714 | 0 |
| 0.00035735 | 0.00084408 | 353.996268 | 729.160262 | 644.06803 | 658.325399 | 21.3859642 |
| 0.00035752 | 0.00084441 | 16.3976473 | 16.4606038 | 13.9207788 | 14.3820207 | 0 |
| 0.00035782 | 0.00084506 | 27.0078897 | 19.7972127 | 15.7768826 | 11.1344676 | 0 |
| 0.00035799 | 0.00084542 | 17.6837373 | 15.5708415 | 13.9207788 | 14.3820207 | 0 |
| 0.00035852 | 0.00084661 | 3.85826995 | 60.5038411 | 38.9781805 | 46.8575513 | 0 |
| 0.00035897 | 0.0008476 | 9.96719738 | 21.3542969 | 21.3451941 | 19.4853184 | 0 |
| 0.00035908 | 0.00084782 | 81.023669 | 148.367875 | 141.991943 | 160.057972 | 23.7621824 |
| 0.00035933 | 0.00084828 | 12.8608998 | 21.1318563 | 14.8488307 | 16.2377653 | 0 |
| 0.00035932 | 0.00084828 | 112.532874 | 141.694657 | 109.510126 | 109.952868 | 2.37621824 |
| 0.00035938 | 0.00084835 | 11.5748099 | 28.0275146 | 36.1940248 | 12.9902122 | 0 |
| 0.00035955 | 0.00084867 | 17.0406923 | 17.1279256 | 11.136623 | 22.2689353 | 0 |
| 0.00035976 | 0.00084912 | 10.2887199 | 17.5728068 | 20.4171422 | 26.9082968 | 0 |
| 0.00035981 | 0.00084917 | 12.2178549 | 28.6948364 | 12.9927268 | 23.6607437 | 0 |
| 0.0003603 | 0.00085027 | 71.3779941 | 30.4743612 | 17.6329864 | 31.0837221 | 0 |
| 0.00036081 | 0.00085141 | 110.603739 | 20.4645345 | 8.35246725 | 25.5164883 | 0 |
| 0.00036135 | 0.00085263 | 10.6102424 | 21.1318563 | 22.273246 | 16.7017014 | 0 |
| 0.0003615 | 0.00085293 | 0 | 385.934428 | 242.22155 | 442.595088 | 0 |
| 0.00036197 | 0.00085397 | 11.2532874 | 54.7203857 | 11.136623 | 41.7542536 | 0 |
| 0.00036221 | 0.00085448 | 24.4357097 | 16.6830444 | 0 | 25.0525522 | 316.037026 |
| 0.00036258 | 0.0008553 | 264.613014 | 343.225834 | 280.271679 | 339.601263 | 4.75243649 |
| 0.00036321 | 0.00085672 | 20.8989622 | 34.0334106 | 47.3306478 | 38.5067006 | 2.37621824 |
| 0.00036328 | 0.00085681 | 17.3622148 | 16.905485 | 11.136623 | 20.8771268 | 0 |
| 0.00036382 | 0.00085804 | 33.1168171 | 22.4664998 | 26.9135056 | 16.2377653 | 109.306039 |
| 0.00036421 | 0.00085889 | 23.7926647 | 21.5767375 | 12.0646749 | 13.4541484 | 0 |
| 0.00036567 | 0.00086227 | 10.2887199 | 21.5767375 | 18.5610383 | 19.9492545 | 0 |
| 0.00036615 | 0.00086334 | 12.5393773 | 11.7893514 | 31.5537652 | 30.619786 | 0 |
| 0.00036619 | 0.00086339 | 9.00262989 | 31.3641235 | 19.4890903 | 24.1246799 | 0 |
| 0.00036632 | 0.00086363 | 19.2913498 | 11.5669108 | 16.7049345 | 17.6295737 | 0 |
| 0.00036691 | 0.00086495 | 0 | 248.911023 | 847.3114 | 176.295737 | 0 |
| 0.00036707 | 0.00086528 | 61.0892743 | 98.9860636 | 35.2659728 | 52.8887212 | 11.8810912 |
| 0.0003681 | 0.00086764 | 77.4869216 | 259.810612 | 280.271679 | 264.443606 | 28.5146189 |
| 0.00036813 | 0.00086765 | 220.24291 | 270.48776 | 259.854537 | 326.611051 | 38.0194919 |

| | | | | | | |
|---|---|---|---|---|---|---|
| 0.00036851 | 0.00086849 | 7.71653991 | 31.1416829 | 29.6976613 | 22.7328714 | 0 |
| 0.00036935 | 0.0008704 | 0 | 0 | 0 | 0 | 42.7719284 |
| 0.00036938 | 0.00087042 | 5.46588243 | 3.55904948 | 0 | 0.9278723 | 11.8810912 |
| 0.00037017 | 0.00087223 | 26.0433222 | 54.9428263 | 35.2659728 | 69.5904227 | 0 |
| 0.00037031 | 0.00087248 | 118.320279 | 215.544934 | 127.143113 | 298.774881 | 28.5146189 |
| 0.00037093 | 0.00087388 | 955.564858 | 291.842057 | 349.875573 | 340.529135 | 68.9103291 |
| 0.00037104 | 0.00087408 | 259.468654 | 351.456136 | 268.207004 | 296.919137 | 45.1481466 |
| 0.0003721 | 0.00087646 | 9.00262989 | 42.2637126 | 17.6329864 | 27.3722329 | 0 |
| 0.0003721 | 0.00087646 | 6.75197242 | 35.3680542 | 26.9135056 | 31.0837221 | 0 |
| 0.00037233 | 0.00087694 | 54.6588243 | 196.859924 | 150.344411 | 158.666164 | 21.3859642 |
| 0.00037263 | 0.00087759 | 16.7191698 | 39.816866 | 9.28051917 | 26.4443606 | 0 |
| 0.00037297 | 0.00087831 | 10.9317649 | 17.5728068 | 18.5610383 | 19.4853184 | 0 |
| 0.00037303 | 0.00087841 | 30.8661596 | 72.7380737 | 14.8488307 | 9.74265918 | 0 |
| 0.00037311 | 0.00087853 | 14.4685123 | 13.3464355 | 33.409869 | 16.7017014 | 0 |
| 0.00037361 | 0.00087965 | 5.78740493 | 29.5845988 | 34.3379209 | 44.5378705 | 0 |
| 0.00037416 | 0.00088088 | 50.4790319 | 13.5688761 | 13.9207788 | 19.9492545 | 0 |
| 0.00037428 | 0.0008811 | 2.89370247 | 0 | 0 | 0 | 204.354769 |
| 0.00037526 | 0.00088334 | 4.17979245 | 2.00196533 | 1.85610383 | 5.10329766 | 49.9005831 |
| 0.00037537 | 0.00088354 | 5.78740493 | 51.6062174 | 40.8342843 | 24.588616 | 0 |
| 0.00037584 | 0.00088458 | 5.46588243 | 36.035376 | 36.1940248 | 36.650956 | 0 |
| 0.00037635 | 0.00088572 | 58.5170943 | 83.8601033 | 72.3880495 | 77.0134011 | 0 |
| 0.00037719 | 0.00088765 | 10.2887199 | 28.2499552 | 18.5610383 | 18.5574461 | 0 |
| 0.0003776 | 0.00088854 | 7.07349491 | 26.9153117 | 28.7696094 | 32.0115944 | 0 |
| 0.00037763 | 0.00088854 | 7.39501741 | 21.3542969 | 40.8342843 | 31.0837221 | 0 |
| 0.00037782 | 0.00088894 | 14.1469898 | 18.017688 | 12.9927268 | 35.2591475 | 0 |
| 0.00037798 | 0.00088924 | 59.8031843 | 30.6968018 | 6.49636342 | 23.6607437 | 0 |
| 0.00037805 | 0.00088935 | 805.413853 | 1085.73253 | 1152.64048 | 1334.74431 | 0 |
| 0.00037837 | 0.00089004 | 39.8687895 | 64.952653 | 66.819738 | 77.4773373 | 14.2573095 |
| 0.00037883 | 0.00089105 | 10.6102424 | 23.1338216 | 15.7768826 | 22.2689353 | 0 |
| 0.00037911 | 0.00089165 | 49.5144644 | 12.9015544 | 17.6329864 | 16.2377653 | 0 |
| 0.00037925 | 0.00089192 | 116.712666 | 274.936572 | 311.825444 | 323.363497 | 0 |
| 0.00037942 | 0.00089225 | 53.0512119 | 14.4586385 | 15.7768826 | 16.2377653 | 0 |
| 0.00038046 | 0.00089466 | 784.514891 | 232.005538 | 139.207788 | 0 | 0 |
| 0.00038105 | 0.00089597 | 242.427962 | 251.58031 | 0 | 220.369672 | 0 |
| 0.0003813 | 0.0008965 | 23.1496197 | 65.8424154 | 15.7768826 | 49.6411682 | 0 |
| 0.00038156 | 0.00089704 | 12.2178549 | 19.1298909 | 13.9207788 | 20.4131907 | 0 |
| 0.00038185 | 0.00089767 | 65.2690667 | 20.2420939 | 13.9207788 | 13.9180845 | 0 |
| 0.00038198 | 0.00089791 | 3.85826995 | 41.3739502 | 45.4745439 | 52.4247851 | 0 |
| 0.00038202 | 0.00089793 | 15.4330798 | 58.0569946 | 58.4672708 | 6.95904227 | 0 |
| 0.00038231 | 0.00089856 | 12.2178549 | 22.0216187 | 15.7768826 | 15.7738291 | 0 |
| 0.00038343 | 0.00090111 | 8.3595849 | 29.3621582 | 21.3451941 | 25.0525522 | 0 |
| 0.00038353 | 0.00090129 | 79.094534 | 36.2578166 | 9.28051917 | 62.6313804 | 2.37621824 |
| 0.00038437 | 0.00090321 | 14.4685123 | 32.0314453 | 39.9062324 | 55.6723382 | 0 |

| | | | | | | |
|---|---|---|---|---|---|---|
| 0.0003846 | 0.00090368 | 6.75197242 | 61.1711629 | 13.9207788 | 76.0855288 | 0 |
| 0.00038494 | 0.00090443 | 248.858412 | 186.405216 | 276.559471 | 214.802438 | 19.0097459 |
| 0.00038503 | 0.00090458 | 61.0892743 | 53.3857422 | 45.4745439 | 57.9920189 | 11.8810912 |
| 0.00038518 | 0.00090485 | 19.6128723 | 13.7913167 | 12.9927268 | 16.7017014 | 0 |
| 0.00038558 | 0.00090574 | 8.3595849 | 30.02948 | 33.409869 | 18.5574461 | 0 |
| 0.00038616 | 0.00090704 | 26.0433222 | 399.725745 | 605.08985 | 541.877425 | 40.3957101 |
| 0.00038638 | 0.00090749 | 8.6811074 | 31.1416829 | 19.4890903 | 25.0525522 | 0 |
| 0.00038708 | 0.00090908 | 0 | 0 | 0 | 0 | 66.5341108 |
| 0.00038718 | 0.00090924 | 13.1824223 | 31.5865641 | 22.273246 | 12.0623399 | 0 |
| 0.00038788 | 0.00091081 | 5.14435994 | 26.0255493 | 83.5246725 | 36.650956 | 0 |
| 0.00038838 | 0.00091193 | 25.4002772 | 45.1554403 | 35.2659728 | 65.8789335 | 7.12865473 |
| 0.00038891 | 0.00091312 | 13.8254673 | 30.4743612 | 10.2085711 | 33.4034029 | 0 |
| 0.00038959 | 0.00091465 | 4.17979245 | 49.6042521 | 41.7623363 | 39.898509 | 0 |
| 0.00038981 | 0.00091509 | 7.07349491 | 24.6909058 | 39.9062324 | 27.3722329 | 0 |
| 0.00039028 | 0.00091615 | 28.9370247 | 45.822762 | 21.3451941 | 50.5690405 | 0 |
| 0.00039068 | 0.00091701 | 231.81772 | 274.491691 | 0 | 203.66797 | 0 |
| 0.00039079 | 0.00091721 | 16.3976473 | 12.2342326 | 15.7768826 | 31.0837221 | 0 |
| 0.00039095 | 0.00091745 | 23.1496197 | 74.2951579 | 103.941815 | 77.0134011 | 0 |
| 0.00039094 | 0.00091745 | 18.0052598 | 9.12006429 | 34.3379209 | 24.1246799 | 0 |
| 0.00039106 | 0.00091764 | 28.9370247 | 91.2006429 | 23.2012979 | 49.177232 | 4.75243649 |
| 0.00039133 | 0.00091821 | 182.624778 | 26.2479899 | 13.9207788 | 9.27872303 | 0 |
| 0.00039136 | 0.00091823 | 0 | 0 | 0 | 0 | 0 |
| 0.00039158 | 0.00091868 | 6.10892743 | 35.1456136 | 90.0210359 | 21.341063 | 0 |
| 0.00039226 | 0.0009202 | 8.0380624 | 26.2479899 | 25.0574018 | 24.1246799 | 0 |
| 0.00039275 | 0.00092123 | 8.6811074 | 69.1790242 | 59.3953227 | 11.1344676 | 0 |
| 0.00039275 | 0.00092123 | 37.618132 | 12.4566732 | 18.5610383 | 17.6295737 | 90.2962933 |
| 0.00039287 | 0.00092144 | 22.1850522 | 8.8976237 | 24.1293498 | 21.8049991 | 0 |
| 0.00039295 | 0.00092156 | 3.21522496 | 39.816866 | 60.3233746 | 56.1362743 | 0 |
| 0.00039398 | 0.00092393 | 8.3595849 | 41.8188314 | 13.9207788 | 67.7346781 | 0 |
| 0.00039511 | 0.0009265 | 127.001386 | 226.889404 | 206.027526 | 206.451587 | 40.3957101 |
| 0.00039525 | 0.00092676 | 0 | 0 | 0 | 0 | 2.37621824 |
| 0.00039578 | 0.00092794 | 8.3595849 | 26.2479899 | 24.1293498 | 22.2689353 | 0 |
| 0.00039591 | 0.00092818 | 226.994882 | 343.670715 | 327.602327 | 0 | 0 |
| 0.00039654 | 0.0009296 | 12.2178549 | 38.9271037 | 69.6038938 | 9.74265918 | 0 |
| 0.00039659 | 0.00092965 | 33.7598621 | 7.34053955 | 25.9854537 | 25.5164883 | 0 |
| 0.00039801 | 0.00093292 | 29.2585471 | 24.6909058 | 34.3379209 | 33.4034029 | 0 |
| 0.00039876 | 0.00093461 | 19.2913498 | 20.0196533 | 9.28051917 | 23.6607437 | 0 |
| 0.00039907 | 0.00093521 | 98.7074063 | 171.056816 | 257.998433 | 276.505946 | 2.37621824 |
| 0.00039905 | 0.00093521 | 224.744225 | 978.961047 | 521.565177 | 701.471461 | 0 |
| 0.00039936 | 0.00093582 | 6.43044992 | 49.1593709 | 22.273246 | 35.7230837 | 0 |
| 0.00040032 | 0.00093801 | 46.6207619 | 19.1298909 | 11.136623 | 16.7017014 | 0 |
| 0.00040037 | 0.00093805 | 7.39501741 | 29.5845988 | 24.1293498 | 28.7640414 | 0 |
| 0.0004006 | 0.00093852 | 8.3595849 | 29.1397176 | 18.5610383 | 32.0115944 | 0 |

| | | | | | | |
|---|---|---|---|---|---|---|
| 0.00040075 | 0.00093881 | 94.2060914 | 180.17688 | 145.704151 | 150.779249 | 33.2670554 |
| 0.00040094 | 0.00093919 | 178.123463 | 278.273181 | 0 | 251.91733 | 0 |
| 0.00040213 | 0.00094192 | 8.3595849 | 32.2538859 | 17.6329864 | 33.4034029 | 0 |
| 0.00040226 | 0.00094215 | 38.904222 | 17.5728068 | 11.136623 | 17.1656376 | 0 |
| 0.00040244 | 0.00094252 | 158.189068 | 277.383419 | 248.717914 | 294.13552 | 45.1481466 |
| 0.00040272 | 0.0009431 | 11.5748099 | 24.4684652 | 15.7768826 | 16.7017014 | 0 |
| 0.00040303 | 0.00094375 | 349.816476 | 541.865283 | 430.616089 | 475.534555 | 206.730987 |
| 0.00040372 | 0.0009453 | 26.6863672 | 33.1436483 | 60.3233746 | 5.56723382 | 0 |
| 0.00040399 | 0.00094589 | 7.71653991 | 24.6909058 | 26.9135056 | 25.9804245 | 0 |
| 0.00040421 | 0.00094632 | 22.1850522 | 15.1259603 | 12.0646749 | 15.7738291 | 0 |
| 0.00040442 | 0.00094675 | 174.908238 | 315.42076 | 240.365446 | 253.773075 | 68.9103291 |
| 0.00040448 | 0.00094682 | 56.2664368 | 150.1474 | 157.768826 | 151.243185 | 21.3859642 |
| 0.00040517 | 0.00094837 | 49.1929419 | 40.2617472 | 19.4890903 | 37.5788283 | 4.75243649 |
| 0.00040609 | 0.00095045 | 17.6837373 | 31.1416829 | 64.9636342 | 26.9082968 | 2.37621824 |
| 0.00040654 | 0.00095144 | 7.39501741 | 44.9329997 | 24.1293498 | 24.588616 | 0 |
| 0.0004066 | 0.00095152 | 6.10892743 | 22.911381 | 42.6903882 | 40.8263813 | 0 |
| 0.00040671 | 0.00095171 | 30.2231146 | 40.7066284 | 44.546492 | 43.6099982 | 7.12865473 |
| 0.00040702 | 0.00095237 | 8.3595849 | 31.8090047 | 19.4890903 | 25.9804245 | 0 |
| 0.00040815 | 0.00095495 | 461.706304 | 635.067891 | 1129.43918 | 789.155393 | 7.12865473 |
| 0.00040846 | 0.00095561 | 5.14435994 | 26.2479899 | 38.9781805 | 60.7756358 | 0 |
| 0.00040856 | 0.0009557 | 10.2887199 | 30.9192423 | 13.9207788 | 28.7640414 | 0 |
| 0.00040854 | 0.0009557 | 16.0761248 | 28.6948364 | 17.6329864 | 11.1344676 | 0 |
| 0.00040877 | 0.00095613 | 9.00262989 | 452.666606 | 600.44959 | 576.2087 | 0 |
| 0.00040894 | 0.00095648 | 246.607755 | 434.871358 | 319.249859 | 396.665409 | 59.4054561 |
| 0.00040962 | 0.000958 | 39.2257445 | 172.391459 | 145.704151 | 199.028609 | 26.1384007 |
| 0.0004097 | 0.00095811 | 8.6811074 | 23.8011434 | 23.2012979 | 21.341063 | 0 |
| 0.00040982 | 0.00095832 | 11.5748099 | 42.7085937 | 26.9135056 | 12.5262761 | 0 |
| 0.00041 | 0.00095868 | 26.3648447 | 12.4566732 | 19.4890903 | 12.9902122 | 0 |
| 0.00041035 | 0.00095944 | 194.199588 | 710.252812 | 533.629852 | 524.711787 | 0 |
| 0.00041109 | 0.0009611 | 45.334672 | 93.6474894 | 81.6685687 | 105.313506 | 16.6335277 |
| 0.0004112 | 0.00096128 | 56.5879593 | 13.7913167 | 11.136623 | 25.0525522 | 0 |
| 0.00041286 | 0.00096511 | 78.7730116 | 240.013399 | 158.696878 | 227.328714 | 26.1384007 |
| 0.00041294 | 0.00096522 | 18.0052598 | 20.4645345 | 9.28051917 | 30.1558498 | 0 |
| 0.00041447 | 0.00096872 | 19.9343948 | 17.5728068 | 12.0646749 | 13.9180845 | 0 |
| 0.00041463 | 0.00096904 | 30.5446371 | 40.0393066 | 27.8415575 | 41.7542536 | 0 |
| 0.00041506 | 0.00096997 | 9.96719738 | 21.1318563 | 27.8415575 | 16.2377653 | 0 |
| 0.00041639 | 0.00097302 | 9.96719738 | 18.4625692 | 18.5610383 | 33.867339 | 0 |
| 0.00041657 | 0.00097337 | 8.3595849 | 24.6909058 | 19.4890903 | 34.3312752 | 0 |
| 0.00041672 | 0.00097364 | 7.39501741 | 24.023584 | 30.6257133 | 26.4443606 | 0 |
| 0.00041706 | 0.00097436 | 9.64567488 | 25.3582275 | 15.7768826 | 27.8361691 | 0 |
| 0.00041708 | 0.00097436 | 22.8280972 | 17.1279256 | 14.8488307 | 11.5984038 | 0 |
| 0.00041721 | 0.00097458 | 41.7979245 | 64.952653 | 77.0283091 | 79.3330819 | 19.0097459 |
| 0.00041734 | 0.00097482 | 0.3215225 | 0.22244059 | 0.92805192 | 0.46393615 | 9.50487297 |

| | | | | | | |
|---|---|---|---|---|---|---|
| 0.00041745 | 0.00097501 | 48.8714194 | 4.89369303 | 195.818954 | 12.9902122 | 0 |
| 0.00041889 | 0.00097832 | 9.00262989 | 39.5944255 | 74.2441534 | 12.5262761 | 0 |
| 0.00041965 | 0.00098001 | 13.5039448 | 13.3464355 | 23.2012979 | 15.309893 | 0 |
| 0.00042076 | 0.00098253 | 48.5498969 | 24.9133464 | 19.4890903 | 29.2279775 | 142.573095 |
| 0.00042093 | 0.00098288 | 184.23239 | 343.448275 | 288.624146 | 315.940519 | 64.1578926 |
| 0.00042116 | 0.00098333 | 20.8989622 | 34.923173 | 41.7623363 | 39.4345729 | 4.75243649 |
| 0.00042152 | 0.0009841 | 0 | 0 | 0 | 0 | 21.3859642 |
| 0.00042167 | 0.0009844 | 6.10892743 | 13.7913167 | 70.5319457 | 70.9822312 | 0 |
| 0.00042172 | 0.00098444 | 5.46588243 | 38.7046631 | 30.6257133 | 35.7230837 | 0 |
| 0.00042186 | 0.0009847 | 4.82283744 | 37.8149007 | 44.546492 | 33.4034029 | 0 |
| 0.00042231 | 0.00098568 | 37.9396545 | 157.487939 | 66.819738 | 103.457762 | 16.6335277 |
| 0.00042269 | 0.00098651 | 9.64567488 | 20.2420939 | 18.5610383 | 21.8049991 | 0 |
| 0.00042278 | 0.00098665 | 66.8766792 | 272.712166 | 320.177911 | 275.578074 | 28.5146189 |
| 0.00042329 | 0.00098776 | 14.1469898 | 47.3798462 | 27.8415575 | 54.7444659 | 0 |
| 0.00042334 | 0.00098781 | 11.2532874 | 38.7046631 | 12.0646749 | 29.2279775 | 0 |
| 0.000424 | 0.00098927 | 17.3622148 | 46.934965 | 13.9207788 | 13.9180845 | 0 |
| 0.00042558 | 0.00099291 | 10.6102424 | 48.4920492 | 10.2085711 | 51.4969128 | 0 |
| 0.00042569 | 0.00099309 | 23.1496197 | 18.6850098 | 25.9854537 | 8.81478687 | 0 |
| 0.00042603 | 0.00099383 | 10.2887199 | 25.8031087 | 14.8488307 | 23.1968076 | 0 |
| 0.00042641 | 0.00099463 | 25.0787547 | 53.1633016 | 28.7696094 | 19.4853184 | 0 |
| 0.00042692 | 0.00099575 | 1175.80777 | 1340.87189 | 1147.07217 | 1235.46197 | 2.37621824 |
| 0.00042711 | 0.00099607 | 14.1469898 | 23.1338216 | 24.1293498 | 10.6705315 | 0 |
| 0.00042712 | 0.00099607 | 0 | 21.5767375 | 18.5610383 | 18.5574461 | 171.087714 |
| 0.00042763 | 0.00099719 | 216.38464 | 148.590316 | 110.438178 | 136.397228 | 313.660808 |
| 0.00042767 | 0.00099722 | 10.2887199 | 24.6909058 | 26.9135056 | 14.3820207 | 0 |
| 0.00042782 | 0.00099752 | 299.980489 | 470.239412 | 444.536868 | 440.275408 | 0 |
| 0.0004279 | 0.00099762 | 149.507961 | 388.381274 | 373.076871 | 404.088388 | 2.37621824 |
| 0.00042819 | 0.00099822 | 11.5748099 | 26.9153117 | 12.0646749 | 30.1558498 | 0 |
| 0.00042924 | 0.00100061 | 11.2532874 | 14.4586385 | 26.9135056 | 18.5574461 | 0 |
| 0.00043056 | 0.00100361 | 9.96719738 | 38.0373413 | 12.9927268 | 33.867339 | 0 |
| 0.00043137 | 0.00100544 | 11.2532874 | 20.9094157 | 16.7049345 | 15.7738291 | 0 |
| 0.00043169 | 0.00100611 | 30.2231146 | 15.3484009 | 9.28051917 | 26.4443606 | 0 |
| 0.0004328 | 0.00100858 | 11.8963324 | 25.3582275 | 66.819738 | 12.5262761 | 0 |
| 0.00043281 | 0.00100858 | 85.2034615 | 14.4586385 | 12.0646749 | 21.341063 | 0 |
| 0.00043314 | 0.00100928 | 15.4330798 | 8.8976237 | 29.6976613 | 57.5280828 | 0 |
| 0.00043346 | 0.00100995 | 9.32415239 | 20.4645345 | 18.5610383 | 24.588616 | 0 |
| 0.00043363 | 0.00101028 | 6.75197242 | 26.9153117 | 30.6257133 | 28.3001052 | 0 |
| 0.0004342 | 0.00101155 | 55.6233918 | 39.5944255 | 22.273246 | 62.1674443 | 0 |
| 0.00043426 | 0.00101161 | 9.00262989 | 31.3641235 | 14.8488307 | 40.3624452 | 0 |
| 0.00043454 | 0.00101219 | 32.1522496 | 33.5885295 | 35.2659728 | 26.9082968 | 109.306039 |
| 0.00043472 | 0.00101253 | 56.5879593 | 22.0216187 | 12.9927268 | 12.5262761 | 0 |
| 0.000435 | 0.00101313 | 4.82283744 | 33.1436483 | 45.4745439 | 36.1870198 | 0 |
| 0.00043544 | 0.00101407 | 4.50131495 | 30.9192423 | 58.4672708 | 38.0427644 | 0 |

| | | | | | | |
|---|---|---|---|---|---|---|
| 0.0004355 | 0.00101414 | 80.380624 | 32.4763265 | 30.6257133 | 27.3722329 | 4.75243649 |
| 0.00043593 | 0.00101508 | 464.600007 | 1051.0318 | 1309.48125 | 1244.74069 | 0 |
| 0.00043664 | 0.00101666 | 7.71653991 | 29.3621582 | 19.4890903 | 33.4034029 | 0 |
| 0.00043702 | 0.00101747 | 81.666714 | 133.909237 | 119.718697 | 146.139888 | 28.5146189 |
| 0.00043708 | 0.00101755 | 25.7217997 | 29.1397176 | 51.0428554 | 6.03116997 | 0 |
| 0.0004373 | 0.00101798 | 23.4711422 | 3.33660889 | 76.1002572 | 77.4773373 | 0 |
| 0.0004379 | 0.00101932 | 13.1824223 | 20.0196533 | 24.1293498 | 11.5984038 | 0 |
| 0.00043848 | 0.0010206 | 8.0380624 | 26.4704305 | 23.2012979 | 22.7328714 | 0 |
| 0.00043957 | 0.00102306 | 114.783531 | 215.544934 | 227.37272 | 237.071373 | 30.8908372 |
| 0.00044044 | 0.001025 | 610.57122 | 435.53868 | 414.839207 | 457.904981 | 54.6530196 |
| 0.00044124 | 0.00102681 | 0.3215225 | 0.88976237 | 0 | 0 | 16.6335277 |
| 0.00044193 | 0.00102834 | 59.1601393 | 19.7972127 | 10.2085711 | 17.6295737 | 0 |
| 0.00044235 | 0.00102925 | 8.6811074 | 22.4664998 | 19.4890903 | 25.9804245 | 0 |
| 0.00044284 | 0.00103031 | 0 | 0 | 0 | 0 | 2.37621824 |
| 0.00044308 | 0.00103079 | 42.119447 | 24.2460246 | 18.5610383 | 29.6919137 | 0 |
| 0.00044425 | 0.00103345 | 17.3622148 | 20.2420939 | 9.28051917 | 24.588616 | 0 |
| 0.00044434 | 0.00103358 | 38.904222 | 82.970341 | 40.8342843 | 72.8379758 | 0 |
| 0.00044473 | 0.00103443 | 12.2178549 | 14.2361979 | 15.7768826 | 20.8771268 | 0 |
| 0.00044488 | 0.00103471 | 6.75197242 | 25.5806681 | 28.7696094 | 31.0837221 | 0 |
| 0.00044496 | 0.00103482 | 7.07349491 | 26.2479899 | 26.9135056 | 27.8361691 | 0 |
| 0.00044537 | 0.00103571 | 104.173289 | 161.046989 | 128.071165 | 181.399035 | 26.1384007 |
| 0.0004457 | 0.00103639 | 39.8687895 | 12.011792 | 15.7768826 | 16.7017014 | 0 |
| 0.00044595 | 0.00103691 | 9.64567488 | 22.911381 | 15.7768826 | 36.650956 | 0 |
| 0.00044605 | 0.00103707 | 1.60761248 | 0.66732178 | 0.92805192 | 0.46393615 | 21.3859642 |
| 0.00044635 | 0.00103768 | 0 | 342.113631 | 415.767259 | 398.521154 | 0 |
| 0.00044704 | 0.00103923 | 36.0105196 | 162.604073 | 170.761553 | 186.502333 | 19.0097459 |
| 0.00044786 | 0.00104105 | 2.89370247 | 0 | 0 | 0 | 49.9005831 |
| 0.00044789 | 0.00104105 | 26.0433222 | 27.5826335 | 7.42441534 | 21.8049991 | 0 |
| 0.00044886 | 0.00104324 | 5.14435994 | 81.4132568 | 14.8488307 | 68.1986142 | 0 |
| 0.00044908 | 0.00104367 | 18.0052598 | 17.3503662 | 11.136623 | 15.309893 | 0 |
| 0.00044913 | 0.00104372 | 18.9698273 | 25.3582275 | 36.1940248 | 27.3722329 | 0 |
| 0.00044936 | 0.00104411 | 10.6102424 | 19.7972127 | 22.273246 | 14.8459568 | 0 |
| 0.00044934 | 0.00104411 | 73.3071291 | 15.5708415 | 9.28051917 | 25.9804245 | 0 |
| 0.00045004 | 0.00104562 | 0 | 0 | 0 | 0 | 9.50487297 |
| 0.00045015 | 0.00104579 | 3.21522496 | 36.4802572 | 67.7477899 | 47.7854236 | 0 |
| 0.00045078 | 0.00104719 | 203.52374 | 131.46239 | 0 | 392.95392 | 0 |
| 0.00045184 | 0.00104958 | 10.6102424 | 44.9329997 | 24.1293498 | 13.9180845 | 0 |
| 0.0004519 | 0.00104966 | 10.9317649 | 18.017688 | 19.4890903 | 15.309893 | 0 |
| 0.00045194 | 0.00104966 | 16.3976473 | 17.7952474 | 10.2085711 | 29.2279775 | 0 |
| 0.00045312 | 0.00105234 | 5.14435994 | 62.9506877 | 74.2441534 | 16.2377653 | 0 |
| 0.00045369 | 0.00105358 | 7.39501741 | 28.2499552 | 23.2012979 | 26.4443606 | 0 |
| 0.00045419 | 0.00105467 | 18.3267823 | 22.911381 | 8.35246725 | 31.5476583 | 0 |
| 0.00045476 | 0.00105593 | 10.6102424 | 20.2420939 | 25.0574018 | 14.3820207 | 0 |

| | | | | | | |
|---|---|---|---|---|---|---|
| 0.00045505 | 0.00105654 | 35.6889971 | 205.090226 | 158.696878 | 165.625206 | 19.0097459 |
| 0.00045511 | 0.0010566 | 24.1141872 | 11.1220296 | 21.3451941 | 13.4541484 | 0 |
| 0.00045557 | 0.0010576 | 9.00262989 | 27.8050741 | 15.7768826 | 29.6919137 | 0 |
| 0.0004557 | 0.00105782 | 0 | 370.586027 | 273.775316 | 282.537116 | 0 |
| 0.000456 | 0.00105843 | 18.6483048 | 20.6869751 | 34.3379209 | 8.81478687 | 0 |
| 0.00045617 | 0.00105877 | 1.28608998 | 0 | 0 | 0 | 9.50487297 |
| 0.00045646 | 0.00105936 | 36.6535646 | 11.3444702 | 33.409869 | 11.1344676 | 0 |
| 0.00045672 | 0.00105989 | 0.96456749 | 3.33660889 | 2.78415575 | 2.78361691 | 83.1676385 |
| 0.00045741 | 0.00106143 | 19.6128723 | 52.4959798 | 5.5683115 | 46.3936151 | 0 |
| 0.00045752 | 0.0010616 | 9.32415239 | 24.023584 | 41.7623363 | 15.7738291 | 0 |
| 0.00045791 | 0.00106243 | 190.019795 | 657.979272 | 0 | 138.252973 | 2.37621824 |
| 0.00045814 | 0.0010629 | 10.9317649 | 18.4625692 | 18.5610383 | 15.309893 | 0 |
| 0.00045827 | 0.00106312 | 3.21522496 | 34.2558512 | 65.8916861 | 51.4969128 | 0 |
| 0.00045831 | 0.00106315 | 317.664226 | 345.005359 | 299.760769 | 350.73573 | 35.6432737 |
| 0.00045879 | 0.00106418 | 30.8661596 | 60.05896 | 55.683115 | 57.5280828 | 11.8810912 |
| 0.00045888 | 0.00106431 | 299.980489 | 400.837948 | 419.479466 | 371.612857 | 175.84015 |
| 0.00045903 | 0.0010646 | 9.32415239 | 20.0196533 | 34.3379209 | 17.6295737 | 0 |
| 0.00045917 | 0.00106486 | 7.07349491 | 21.7991781 | 34.3379209 | 28.3001052 | 0 |
| 0.00045954 | 0.00106563 | 115.748099 | 18.017688 | 7.42441534 | 26.4443606 | 0 |
| 0.000461 | 0.00106893 | 7.71653991 | 30.4743612 | 17.6329864 | 40.8263813 | 0 |
| 0.00046124 | 0.00106943 | 10.9317649 | 16.4606038 | 15.7768826 | 23.6607437 | 0 |
| 0.00046148 | 0.00106991 | 247.572322 | 526.739323 | 676.549847 | 678.274653 | 0 |
| 0.00046239 | 0.00107194 | 25.0787547 | 30.2519206 | 10.2085711 | 13.4541484 | 0 |
| 0.00046317 | 0.00107367 | 14.1469898 | 11.1220296 | 8.35246725 | 12.9902122 | 42.7719284 |
| 0.00046328 | 0.00107386 | 173.300625 | 441.767017 | 417.623363 | 439.347535 | 2.37621824 |
| 0.00046353 | 0.00107436 | 33.7598621 | 15.3484009 | 12.9927268 | 14.3820207 | 0 |
| 0.0004636 | 0.00107445 | 13.5039448 | 26.2479899 | 10.2085711 | 25.9804245 | 0 |
| 0.00046381 | 0.00107486 | 277.473914 | 374.145076 | 403.702584 | 345.168497 | 178.216368 |
| 0.00046386 | 0.00107491 | 9.96719738 | 50.4940145 | 40.8342843 | 54.7444659 | 2.37621824 |
| 0.00046422 | 0.00107567 | 24.1141872 | 38.2597819 | 29.6976613 | 31.0837221 | 4.75243649 |
| 0.00046446 | 0.00107616 | 0.3215225 | 0.22244059 | 0.92805192 | 0.46393615 | 14.2573095 |
| 0.00046462 | 0.00107645 | 129.895088 | 34.7007324 | 21.3451941 | 35.7230837 | 0 |
| 0.00046556 | 0.00107847 | 11.5748099 | 24.023584 | 12.0646749 | 25.9804245 | 0 |
| 0.00046554 | 0.00107847 | 27.3294122 | 39.5944255 | 22.273246 | 64.487125 | 0 |
| 0.00046645 | 0.00108046 | 119.927891 | 103.657316 | 88.1649321 | 101.138081 | 35.6432737 |
| 0.00046725 | 0.00108224 | 6.75197242 | 37.1475789 | 25.0574018 | 25.5164883 | 0 |
| 0.00046759 | 0.00108294 | 20.2559173 | 43.1534749 | 36.1940248 | 36.650956 | 4.75243649 |
| 0.00046807 | 0.00108398 | 21.8635297 | 48.7144897 | 40.8342843 | 5.56723382 | 0 |
| 0.00046853 | 0.00108499 | 10.6102424 | 36.9251383 | 16.7049345 | 17.6295737 | 0 |
| 0.00046874 | 0.00108531 | 5.14435994 | 31.3641235 | 33.409869 | 40.8263813 | 0 |
| 0.00046871 | 0.00108531 | 163.011906 | 277.8283 | 224.588564 | 243.566479 | 0 |
| 0.00046958 | 0.00108718 | 13.8254673 | 43.3759155 | 9.28051917 | 29.2279775 | 0 |
| 0.00047001 | 0.0010881 | 113.175919 | 216.212256 | 149.416359 | 180.935099 | 0 |

| | | | | | | |
|---|---|---|---|---|---|---|
| 0.00047135 | 0.00109114 | 6.43044992 | 40.4841878 | 14.8488307 | 101.138081 | 0 |
| 0.00047168 | 0.00109183 | 10.6102424 | 19.5747721 | 23.2012979 | 14.3820207 | 0 |
| 0.00047173 | 0.00109187 | 81.9882365 | 405.731641 | 283.983887 | 345.168497 | 0 |
| 0.00047255 | 0.00109369 | 5.14435994 | 2.66928711 | 1.85610383 | 0.9278723 | 9.50487297 |
| 0.00047269 | 0.00109393 | 15.1115573 | 5.7834554 | 86.3088283 | 49.6411682 | 0 |
| 0.00047295 | 0.00109447 | 11.5748099 | 65.3975342 | 25.9854537 | 64.0231889 | 2.37621824 |
| 0.00047347 | 0.0010956 | 27.3294122 | 27.5826335 | 23.2012979 | 20.4131907 | 2.37621824 |
| 0.00047465 | 0.00109824 | 0 | 0 | 0 | 0 | 16.6335277 |
| 0.00047592 | 0.00110111 | 7.39501741 | 26.6928711 | 21.3451941 | 30.1558498 | 0 |
| 0.0004764 | 0.00110215 | 108.353081 | 414.184383 | 0 | 215.266374 | 0 |
| 0.00047651 | 0.00110232 | 9.32415239 | 36.2578166 | 13.9207788 | 29.6919137 | 0 |
| 0.00047718 | 0.00110379 | 10.6102424 | 19.1298909 | 16.7049345 | 16.7017014 | 0 |
| 0.00047743 | 0.00110429 | 52.4081669 | 199.306771 | 84.4527244 | 151.243185 | 19.0097459 |
| 0.00047799 | 0.00110552 | 7.07349491 | 19.7972127 | 40.8342843 | 29.2279775 | 0 |
| 0.00048 | 0.0011101 | 17.0406923 | 32.4763265 | 28.7696094 | 28.3001052 | 0 |
| 0.00048062 | 0.00111145 | 1.28608998 | 1.77952474 | 1.85610383 | 2.31968076 | 0 |
| 0.00048129 | 0.00111291 | 7.71653991 | 20.6869751 | 25.0574018 | 28.3001052 | 0 |
| 0.00048138 | 0.00111305 | 9.32415239 | 65.8424154 | 31.5537652 | 12.9902122 | 0 |
| 0.00048291 | 0.00111651 | 14.7900348 | 16.6830444 | 16.7049345 | 11.5984038 | 0 |
| 0.00048296 | 0.00111654 | 19.2913498 | 28.917277 | 11.136623 | 13.4541484 | 0 |
| 0.0004831 | 0.0011168 | 140.505331 | 227.111845 | 0 | 295.991265 | 0 |
| 0.00048373 | 0.00111818 | 7.71653991 | 23.1338216 | 23.2012979 | 25.5164883 | 0 |
| 0.00048465 | 0.00112022 | 9.00262989 | 15.5708415 | 46.4025958 | 25.0525522 | 0 |
| 0.00048471 | 0.00112029 | 8.0380624 | 22.6889404 | 25.0574018 | 21.341063 | 0 |
| 0.00048488 | 0.00112061 | 7.07349491 | 28.917277 | 28.7696094 | 22.2689353 | 0 |
| 0.00048554 | 0.00112206 | 10.6102424 | 21.7991781 | 13.9207788 | 21.341063 | 0 |
| 0.00048595 | 0.00112292 | 7.71653991 | 20.6869751 | 23.2012979 | 35.2591475 | 0 |
| 0.00048604 | 0.00112306 | 7.07349491 | 24.2460246 | 25.0574018 | 29.6919137 | 0 |
| 0.00048678 | 0.0011247 | 261.397789 | 162.159192 | 0 | 219.4418 | 0 |
| 0.00048686 | 0.00112479 | 15.7546023 | 13.7913167 | 12.0646749 | 18.0935099 | 0 |
| 0.0004874 | 0.00112596 | 13.1824223 | 15.5708415 | 12.9927268 | 16.7017014 | 0 |
| 0.00048761 | 0.00112638 | 189.37675 | 108.551009 | 104.869867 | 103.921698 | 23.7621824 |
| 0.00048808 | 0.00112738 | 9.32415239 | 20.4645345 | 16.7049345 | 32.4755306 | 0 |
| 0.00048869 | 0.00112871 | 18.9698273 | 21.1318563 | 9.28051917 | 17.6295737 | 0 |
| 0.00048957 | 0.00113067 | 14.7900348 | 17.3503662 | 11.136623 | 34.3312752 | 0 |
| 0.00049042 | 0.00113256 | 35.6889971 | 21.1318563 | 11.136623 | 12.9902122 | 0 |
| 0.00049121 | 0.00113431 | 9.96719738 | 41.5963908 | 17.6329864 | 18.0935099 | 0 |
| 0.00049272 | 0.00113771 | 23.4711422 | 8.45274251 | 17.6329864 | 34.7952113 | 0 |
| 0.00049332 | 0.00113901 | 9.32415239 | 31.5865641 | 12.9927268 | 44.0739344 | 0 |
| 0.00049353 | 0.00113943 | 7.07349491 | 21.5767375 | 40.8342843 | 25.0525522 | 0 |
| 0.00049371 | 0.00113977 | 129.252043 | 373.700195 | 296.048562 | 330.786476 | 9.50487297 |
| 0.00049388 | 0.00114008 | 11.5748099 | 25.3582275 | 12.9927268 | 18.5574461 | 0 |
| 0.00049422 | 0.00114078 | 13.1824223 | 23.1338216 | 12.9927268 | 14.8459568 | 0 |

| | | | | | | |
|---|---|---|---|---|---|---|
| 0.00049428 | 0.00114084 | 2.89370247 | 52.4959798 | 31.5537652 | 80.2609542 | 0 |
| 0.00049456 | 0.00114142 | 102.244154 | 109.663212 | 201.387266 | 206.451587 | 26.1384007 |
| 0.00049496 | 0.00114225 | 11.8963324 | 19.3523315 | 16.7049345 | 13.4541484 | 0 |
| 0.00049517 | 0.00114265 | 203.202218 | 387.269071 | 394.422065 | 401.768707 | 0 |
| 0.00049539 | 0.0011431 | 10.2887199 | 6.005896 | 4.64025958 | 1.85574461 | 7.12865473 |
| 0.00049563 | 0.00114349 | 12.2178549 | 14.2361979 | 21.3451941 | 14.3820207 | 0 |
| 0.00049561 | 0.00114349 | 4.50131495 | 2.66928711 | 2.78415575 | 3.71148921 | 68.9103291 |
| 0.00049645 | 0.00114531 | 188.09066 | 263.147221 | 308.113236 | 311.765094 | 0 |
| 0.00049674 | 0.00114589 | 18.6483048 | 37.8149007 | 21.3451941 | 34.3312752 | 0 |
| 0.00049681 | 0.00114597 | 25.0787547 | 28.4723958 | 14.8488307 | 9.27872303 | 0 |
| 0.00049819 | 0.00114907 | 41.7979245 | 42.7085937 | 4.64025958 | 26.9082968 | 0 |
| 0.00049878 | 0.00115035 | 7.71653991 | 20.0196533 | 25.9854537 | 26.9082968 | 0 |
| 0.00049951 | 0.00115196 | 9.96719738 | 14.9035197 | 37.1220767 | 20.8771268 | 0 |
| 0.00050051 | 0.00115419 | 9.96719738 | 25.5806681 | 13.9207788 | 24.1246799 | 0 |
| 0.0005007 | 0.00115454 | 8.3595849 | 22.2440592 | 23.2012979 | 19.9492545 | 0 |
| 0.00050079 | 0.00115467 | 42.762492 | 13.7913167 | 13.9207788 | 15.309893 | 0 |
| 0.00050148 | 0.00115619 | 5.78740493 | 2.8917277 | 3.71220767 | 3.71148921 | 9.50487297 |
| 0.00050172 | 0.00115666 | 11.2532874 | 22.0216187 | 13.9207788 | 17.1656376 | 0 |
| 0.00050195 | 0.00115711 | 12.2178549 | 22.4664998 | 11.136623 | 27.8361691 | 0 |
| 0.00050208 | 0.00115733 | 15.7546023 | 29.5845988 | 33.409869 | 31.0837221 | 2.37621824 |
| 0.00050326 | 0.00115998 | 18.9698273 | 29.5845988 | 22.273246 | 46.3936151 | 0 |
| 0.00050441 | 0.00116255 | 135.039448 | 320.981775 | 350.803625 | 349.807858 | 38.0194919 |
| 0.00050655 | 0.00116741 | 18.9698273 | 36.9251383 | 38.0501286 | 50.5690405 | 4.75243649 |
| 0.00050718 | 0.00116878 | 165.584086 | 243.572449 | 0 | 216.194247 | 0 |
| 0.00050763 | 0.00116972 | 6.75197242 | 44.4881185 | 12.9927268 | 86.2921241 | 0 |
| 0.00050868 | 0.00117207 | 0.3215225 | 0.22244059 | 0 | 0 | 19.0097459 |
| 0.00051041 | 0.00117597 | 5.78740493 | 0.22244059 | 0.92805192 | 1.85574461 | 11.8810912 |
| 0.00051151 | 0.00117842 | 22.5065747 | 20.2420939 | 16.7049345 | 9.27872303 | 0 |
| 0.00051154 | 0.00117843 | 17.0406923 | 12.9015544 | 12.9927268 | 15.309893 | 0 |
| 0.00051251 | 0.00118057 | 17.3622148 | 49.3818115 | 45.4745439 | 42.6821259 | 0 |
| 0.00051297 | 0.00118156 | 15.7546023 | 20.2420939 | 9.28051917 | 24.1246799 | 0 |
| 0.00051387 | 0.00118354 | 4.82283744 | 56.2774699 | 23.2012979 | 41.2903175 | 0 |
| 0.00051515 | 0.00118642 | 12.5393773 | 15.3484009 | 12.9927268 | 24.1246799 | 0 |
| 0.00051541 | 0.00118693 | 9.00262989 | 20.0196533 | 53.8270112 | 18.0935099 | 0 |
| 0.00051704 | 0.0011906 | 36.6535646 | 36.2578166 | 7.42441534 | 15.7738291 | 0 |
| 0.0005178 | 0.00119218 | 81.023669 | 4.22637126 | 19.4890903 | 52.4247851 | 0 |
| 0.00051778 | 0.00119218 | 10.9317649 | 14.4586385 | 16.7049345 | 23.6607437 | 0 |
| 0.00051818 | 0.00119297 | 5.46588243 | 26.2479899 | 38.9781805 | 32.9394667 | 0 |
| 0.00051864 | 0.00119396 | 150.794051 | 142.361979 | 139.207788 | 173.048184 | 47.5243649 |
| 0.00051929 | 0.00119529 | 6.75197242 | 16.0157227 | 68.6758419 | 35.2591475 | 0 |
| 0.00051928 | 0.00119529 | 9.00262989 | 20.9094157 | 17.6329864 | 21.8049991 | 0 |
| 0.00051955 | 0.0011958 | 9.32415239 | 0 | 10.2085711 | 13.4541484 | 197.226114 |
| 0.0005196 | 0.00119583 | 116.712666 | 291.397176 | 256.142329 | 283.001052 | 42.7719284 |

| | | | | | | |
|---|---|---|---|---|---|---|
| 0.00052003 | 0.00119674 | 8.0380624 | 61.3936035 | 20.4171422 | 20.8771268 | 0 |
| 0.00052108 | 0.00119907 | 6.43044992 | 26.0255493 | 31.5537652 | 25.9804245 | 0 |
| 0.00052168 | 0.00120037 | 8.6811074 | 27.3601929 | 20.4171422 | 18.0935099 | 0 |
| 0.00052172 | 0.00120039 | 21.8635297 | 20.0196533 | 16.7049345 | 9.27872303 | 0 |
| 0.00052247 | 0.00120202 | 6.10892743 | 33.8109701 | 28.7696094 | 25.5164883 | 0 |
| 0.00052289 | 0.00120292 | 50.4790319 | 311.861711 | 363.796351 | 366.045623 | 52.2768014 |
| 0.00052395 | 0.00120527 | 30.2231146 | 17.3503662 | 10.2085711 | 21.341063 | 64.1578926 |
| 0.00052473 | 0.00120699 | 29.2585471 | 1.77952474 | 7.42441534 | 11.5984038 | 80.7914203 |
| 0.00052486 | 0.00120719 | 0 | 0 | 0 | 0 | 9.50487297 |
| 0.00052558 | 0.00120878 | 70.7349491 | 383.0427 | 0 | 303.878179 | 0 |
| 0.00052629 | 0.00121033 | 8.6811074 | 22.2440592 | 43.6184401 | 16.7017014 | 0 |
| 0.00052709 | 0.00121209 | 27.9724572 | 18.4625692 | 15.7768826 | 9.74265918 | 0 |
| 0.00052713 | 0.00121209 | 25.0787547 | 24.2460246 | 8.35246725 | 16.7017014 | 0 |
| 0.00052726 | 0.00121229 | 34.7244296 | 51.1613363 | 53.8270112 | 52.8887212 | 14.2573095 |
| 0.00052729 | 0.00121229 | 3.53674746 | 36.2578166 | 44.546492 | 48.2493597 | 0 |
| 0.00052738 | 0.00121241 | 3.53674746 | 2.66928711 | 1.85610383 | 3.71148921 | 38.0194919 |
| 0.00052761 | 0.00121286 | 14.4685123 | 11.7893514 | 73.3161014 | 18.5574461 | 0 |
| 0.00052896 | 0.00121588 | 24.4357097 | 23.8011434 | 6.49636342 | 32.9394667 | 0 |
| 0.00052908 | 0.00121608 | 10.9317649 | 15.7932821 | 14.8488307 | 21.8049991 | 0 |
| 0.00052921 | 0.0012162 | 183.267823 | 110.108093 | 71.4599976 | 64.9510612 | 19.0097459 |
| 0.0005292 | 0.0012162 | 11.5748099 | 50.9388957 | 38.9781805 | 35.2591475 | 0 |
| 0.00052958 | 0.00121697 | 48.5498969 | 11.5669108 | 12.9927268 | 20.4131907 | 0 |
| 0.0005303 | 0.00121855 | 1.92913498 | 52.2735392 | 53.8270112 | 70.518295 | 0 |
| 0.00053062 | 0.00121919 | 9.64567488 | 19.5747721 | 24.1293498 | 15.309893 | 0 |
| 0.00053095 | 0.00121987 | 68.4842917 | 75.852242 | 48.2586997 | 82.1166988 | 16.6335277 |
| 0.00053167 | 0.00122145 | 118.963324 | 232.227979 | 0 | 295.991265 | 0 |
| 0.0005327 | 0.00122374 | 120.892459 | 257.586206 | 0 | 261.659989 | 0 |
| 0.00053332 | 0.00122507 | 56.2664368 | 374.812398 | 248.717914 | 307.125732 | 28.5146189 |
| 0.00053338 | 0.00122511 | 74.5932191 | 114.779346 | 96.5173994 | 101.138081 | 21.3859642 |
| 0.00053537 | 0.00122961 | 14.1469898 | 14.6810791 | 43.6184401 | 13.4541484 | 0 |
| 0.00053574 | 0.00123038 | 88.7402089 | 225.554761 | 198.60311 | 197.6368 | 28.5146189 |
| 0.00053592 | 0.0012307 | 0 | 186.405216 | 296.048562 | 395.737537 | 0 |
| 0.00053774 | 0.00123481 | 8.3595849 | 40.2617472 | 16.7049345 | 23.6607437 | 0 |
| 0.00053866 | 0.00123682 | 56.2664368 | 27.3601929 | 49.1867516 | 3.71148921 | 0 |
| 0.00053922 | 0.001238 | 21.5420072 | 44.7105591 | 4.64025958 | 54.2805297 | 0 |
| 0.00053924 | 0.001238 | 66.5551567 | 103.879757 | 90.9490879 | 100.674145 | 2.37621824 |
| 0.00053938 | 0.00123823 | 11.5748099 | 171.279256 | 70.5319457 | 180.007227 | 9.50487297 |
| 0.00054013 | 0.00123986 | 11.2532874 | 29.8070394 | 11.136623 | 25.9804245 | 0 |
| 0.0005416 | 0.00124316 | 14.7900348 | 35.8129354 | 33.409869 | 29.6919137 | 0 |
| 0.00054176 | 0.00124344 | 36.0105196 | 32.0314453 | 5.5683115 | 24.1246799 | 0 |
| 0.00054187 | 0.0012436 | 18.0052598 | 21.7991781 | 8.35246725 | 21.8049991 | 0 |
| 0.000542 | 0.00124373 | 12.8608998 | 19.7972127 | 19.4890903 | 11.1344676 | 0 |
| 0.00054198 | 0.00124373 | 9.32415239 | 15.5708415 | 20.4171422 | 25.9804245 | 0 |

| | | | | | | |
|---|---|---|---|---|---|---|
| 0.00054245 | 0.00124467 | 36.9750871 | 17.3503662 | 49.1867516 | 50.5690405 | 2.37621824 |
| 0.00054251 | 0.00124473 | 9.64567488 | 16.4606038 | 20.4171422 | 18.0935099 | 0 |
| 0.00054269 | 0.00124507 | 24.4357097 | 17.1279256 | 9.28051917 | 17.1656376 | 0 |
| 0.00054273 | 0.00124508 | 24.1141872 | 14.4586385 | 11.136623 | 15.309893 | 0 |
| 0.00054321 | 0.00124608 | 48.8714194 | 46.7125244 | 9.28051917 | 10.2065953 | 0 |
| 0.00054339 | 0.00124642 | 27.3294122 | 11.7893514 | 27.8415575 | 5.56723382 | 68.9103291 |
| 0.0005441 | 0.00124797 | 28.9370247 | 46.0452026 | 38.0501286 | 42.2181898 | 316.037026 |
| 0.00054481 | 0.00124949 | 44.048582 | 21.3542969 | 28.7696094 | 30.1558498 | 2.37621824 |
| 0.0005453 | 0.00125053 | 307.697029 | 527.629085 | 544.766475 | 555.331573 | 59.4054561 |
| 0.00054602 | 0.00125211 | 2.25065747 | 45.1554403 | 64.0355823 | 53.3526574 | 0 |
| 0.00054611 | 0.00125222 | 138.897718 | 198.639449 | 0 | 283.928925 | 0 |
| 0.00054637 | 0.00125265 | 5.78740493 | 38.9271037 | 34.3379209 | 22.2689353 | 0 |
| 0.00054634 | 0.00125265 | 11.2532874 | 53.8306234 | 60.3233746 | 7.42297842 | 0 |
| 0.00054691 | 0.0012538 | 11.2532874 | 24.9133464 | 11.136623 | 33.867339 | 0 |
| 0.00054811 | 0.00125647 | 11.8963324 | 13.3464355 | 28.7696094 | 15.7738291 | 0 |
| 0.00054837 | 0.00125698 | 0 | 109.663212 | 646.852186 | 270.474776 | 0 |
| 0.00054842 | 0.00125702 | 4.50131495 | 27.3601929 | 32.4818171 | 65.4149973 | 0 |
| 0.00054854 | 0.00125718 | 4.82283744 | 39.5944255 | 25.9854537 | 41.7542536 | 0 |
| 0.00054872 | 0.00125743 | 10.2887199 | 21.7991781 | 106.72597 | 13.4541484 | 0 |
| 0.0005487 | 0.00125743 | 45.334672 | 62.2833659 | 4.64025958 | 19.9492545 | 0 |
| 0.0005493 | 0.00125868 | 27.6509347 | 19.3523315 | 13.9207788 | 10.2065953 | 0 |
| 0.00055051 | 0.00126136 | 124.750728 | 274.936572 | 214.379993 | 272.794457 | 0 |
| 0.00055219 | 0.00126512 | 17.6837373 | 11.7893514 | 15.7768826 | 13.4541484 | 0 |
| 0.00055346 | 0.00126795 | 12.8608998 | 9.78738607 | 24.1293498 | 107.169251 | 0 |
| 0.00055421 | 0.00126959 | 13.1824223 | 36.035376 | 9.28051917 | 26.4443606 | 0 |
| 0.00055611 | 0.00127385 | 2.57217997 | 0 | 0 | 0 | 30.8908372 |
| 0.00055623 | 0.00127404 | 34.7244296 | 16.905485 | 24.1293498 | 21.341063 | 47.5243649 |
| 0.00055758 | 0.00127705 | 144.685123 | 119.895479 | 78.8844129 | 102.529889 | 197.226114 |
| 0.00055786 | 0.0012776 | 10.9317649 | 13.7913167 | 16.7049345 | 22.7328714 | 0 |
| 0.00055923 | 0.00128065 | 3.21522496 | 1.55708415 | 2.78415575 | 1.39180845 | 99.8011662 |
| 0.00055957 | 0.00128134 | 28.6155022 | 16.4606038 | 10.2085711 | 15.309893 | 0 |
| 0.00056154 | 0.00128577 | 0 | 0 | 0 | 0 | 0 |
| 0.0005619 | 0.0012865 | 41.7979245 | 98.3187419 | 80.7405168 | 85.828188 | 0 |
| 0.00056222 | 0.00128715 | 15.7546023 | 10.0098267 | 17.6329864 | 34.7952113 | 0 |
| 0.00056513 | 0.00129373 | 60.1247068 | 116.781311 | 201.387266 | 212.482757 | 40.3957101 |
| 0.00056553 | 0.00129455 | 13.8254673 | 2.66928711 | 6.49636342 | 2.78361691 | 4.75243649 |
| 0.00056568 | 0.0012948 | 223.136612 | 141.027336 | 0 | 235.215629 | 0 |
| 0.00056626 | 0.00129604 | 53.6942569 | 13.3464355 | 16.7049345 | 12.9902122 | 0 |
| 0.00056657 | 0.00129666 | 10.2887199 | 18.9074504 | 13.9207788 | 31.0837221 | 0 |
| 0.00056835 | 0.00130065 | 13.5039448 | 11.1220296 | 16.7049345 | 19.4853184 | 0 |
| 0.00056839 | 0.00130066 | 0 | 422.859566 | 153.128566 | 302.022435 | 0 |
| 0.0005689 | 0.00130173 | 22.5065747 | 123.89941 | 32.4818171 | 51.4969128 | 2.37621824 |
| 0.00056944 | 0.00130288 | 8.3595849 | 54.4979451 | 16.7049345 | 22.7328714 | 0 |

| | | | | | | |
|---|---|---|---|---|---|---|
| 0.00056951 | 0.00130294 | 11.2532874 | 14.9035197 | 14.8488307 | 17.1656376 | 0 |
| 0.0005697 | 0.00130329 | 94.5276139 | 12.9015544 | 6.49636342 | 39.4345729 | 0 |
| 0.00057123 | 0.00130671 | 6.43044992 | 42.2637126 | 18.5610383 | 32.4755306 | 0 |
| 0.00057169 | 0.00130767 | 12.8608998 | 12.9015544 | 13.9207788 | 22.2689353 | 0 |
| 0.00057173 | 0.00130767 | 176.837373 | 676.886723 | 398.134272 | 472.750938 | 0 |
| 0.00057202 | 0.00130815 | 9.32415239 | 16.905485 | 24.1293498 | 17.1656376 | 0 |
| 0.00057201 | 0.00130815 | 40.5118345 | 179.954439 | 113.222334 | 152.171058 | 0 |
| 0.0005722 | 0.00130849 | 6.43044992 | 22.4664998 | 38.9781805 | 25.0525522 | 0 |
| 0.00057252 | 0.00130912 | 17.3622148 | 20.0196533 | 8.35246725 | 37.1148921 | 0 |
| 0.00057256 | 0.00130913 | 32.4737721 | 45.6003214 | 51.9709073 | 66.3428696 | 11.8810912 |
| 0.0005751 | 0.00131485 | 14.4685123 | 14.9035197 | 13.9207788 | 12.5262761 | 0 |
| 0.00057772 | 0.00132075 | 11.2532874 | 16.6830444 | 12.9927268 | 20.4131907 | 0 |
| 0.00058027 | 0.00132649 | 289.048724 | 137.468286 | 99.3015551 | 112.272549 | 19.0097459 |
| 0.00058194 | 0.00133022 | 14.4685123 | 39.1495443 | 51.0428554 | 23.1968076 | 2.37621824 |
| 0.00058414 | 0.00133515 | 19.2913498 | 10.899589 | 30.6257133 | 12.5262761 | 0 |
| 0.00058426 | 0.00133533 | 52.4081669 | 30.2519206 | 34.3379209 | 12.9902122 | 0 |
| 0.00058435 | 0.00133544 | 11.2532874 | 27.1377523 | 11.136623 | 23.1968076 | 0 |
| 0.00058441 | 0.00133549 | 15.4330798 | 126.346257 | 71.4599976 | 96.9626556 | 9.50487297 |
| 0.00058485 | 0.00133641 | 9.64567488 | 16.4606038 | 20.4171422 | 16.7017014 | 0 |
| 0.00058544 | 0.00133766 | 16.0761248 | 38.9271037 | 23.2012979 | 34.3312752 | 0 |
| 0.00058576 | 0.00133831 | 80.0591015 | 86.9742716 | 145.704151 | 90.4675495 | 16.6335277 |
| 0.00058583 | 0.00133837 | 8.0380624 | 46.4900838 | 21.3451941 | 18.0935099 | 0 |
| 0.00058621 | 0.00133915 | 10.9317649 | 45.3778809 | 9.28051917 | 36.1870198 | 0 |
| 0.00058633 | 0.00133933 | 8.3595849 | 26.9153117 | 15.7768826 | 25.0525522 | 0 |
| 0.00058653 | 0.00133969 | 22.1850522 | 15.3484009 | 12.9927268 | 9.74265918 | 49.9005831 |
| 0.00058685 | 0.00134035 | 0 | 0 | 0 | 0 | 87.920075 |
| 0.00058952 | 0.00134634 | 3.21522496 | 42.9310343 | 53.8270112 | 35.2591475 | 0 |
| 0.00058968 | 0.00134663 | 17.3622148 | 16.2381632 | 9.28051917 | 20.4131907 | 0 |
| 0.00059057 | 0.00134856 | 6.43044992 | 34.2558512 | 19.4890903 | 32.0115944 | 0 |
| 0.00059078 | 0.00134895 | 9.32415239 | 6.67321777 | 13.9207788 | 7.88691457 | 175.84015 |
| 0.00059168 | 0.00135091 | 9.32415239 | 18.2401286 | 19.4890903 | 16.7017014 | 0 |
| 0.00059279 | 0.00135337 | 235.997512 | 128.125781 | 0 | 228.256586 | 0 |
| 0.00059342 | 0.00135469 | 10.2887199 | 20.2420939 | 12.9927268 | 22.7328714 | 0 |
| 0.00059424 | 0.00135648 | 12.2178549 | 12.6791138 | 15.7768826 | 16.2377653 | 0 |
| 0.00059465 | 0.00135733 | 28.2939797 | 28.4723958 | 7.42441534 | 16.7017014 | 0 |
| 0.00059495 | 0.00135791 | 15.1115573 | 17.5728068 | 42.6903882 | 10.2065953 | 0 |
| 0.00059513 | 0.00135819 | 11.2532874 | 12.2342326 | 33.409869 | 19.0213822 | 0 |
| 0.00059515 | 0.00135819 | 5.46588243 | 47.1574056 | 22.273246 | 31.5476583 | 0 |
| 0.0005952 | 0.0013582 | 20.8989622 | 30.2519206 | 11.136623 | 11.5984038 | 0 |
| 0.0005956 | 0.00135903 | 17.3622148 | 9.78738607 | 20.4171422 | 11.1344676 | 97.424948 |
| 0.00059581 | 0.00135941 | 5.14435994 | 40.929069 | 90.9490879 | 15.7738291 | 0 |
| 0.00059709 | 0.00136224 | 36.3320421 | 83.4152222 | 55.683115 | 54.2805297 | 4.75243649 |
| 0.00059754 | 0.00136318 | 7.71653991 | 18.6850098 | 22.273246 | 27.8361691 | 0 |

| | | | | | | |
|---|---|---|---|---|---|---|
| 0.00059822 | 0.00136465 | 14.1469898 | 24.4684652 | 9.28051917 | 20.8771268 | 0 |
| 0.0005984 | 0.00136497 | 13.5039448 | 17.7952474 | 11.136623 | 15.7738291 | 0 |
| 0.000599 | 0.00136623 | 367.178691 | 903.108805 | 962.389838 | 983.080705 | 95.0487297 |
| 0.00059978 | 0.00136791 | 10.6102424 | 21.3542969 | 12.9927268 | 18.5574461 | 0 |
| 0.00060182 | 0.00137248 | 5.78740493 | 32.2538859 | 25.9854537 | 27.8361691 | 0 |
| 0.00060337 | 0.00137591 | 7.39501741 | 33.5885295 | 18.5610383 | 23.6607437 | 0 |
| 0.00060388 | 0.00137699 | 4.82283744 | 22.0216187 | 63.1075304 | 34.3312752 | 0 |
| 0.00060442 | 0.00137814 | 11.8963324 | 16.0157227 | 12.9927268 | 15.7738291 | 0 |
| 0.00060514 | 0.00137969 | 18.6483048 | 24.6909058 | 23.2012979 | 29.2279775 | 2.37621824 |
| 0.0006055 | 0.00138042 | 51.1220769 | 25.5806681 | 12.9927268 | 37.5788283 | 2.37621824 |
| 0.00060601 | 0.00138129 | 94.5276139 | 48.4920492 | 71.4599976 | 69.1264865 | 225.740733 |
| 0.00060599 | 0.00138129 | 183.267823 | 84.5274251 | 66.819738 | 82.5806349 | 14.2573095 |
| 0.00060599 | 0.00138129 | 32.7952946 | 24.2460246 | 14.8488307 | 17.6295737 | 64.1578926 |
| 0.00060612 | 0.00138144 | 22.8280972 | 12.9015544 | 11.136623 | 16.2377653 | 0 |
| 0.00060692 | 0.00138317 | 30.2231146 | 16.6830444 | 14.8488307 | 19.0213822 | 11.8810912 |
| 0.00060773 | 0.00138493 | 9.00262989 | 56.0550293 | 38.9781805 | 50.1051043 | 0 |
| 0.0006079 | 0.00138522 | 0 | 362.800606 | 194.890903 | 421.254025 | 0 |
| 0.00060882 | 0.00138723 | 17.0406923 | 22.911381 | 21.3451941 | 8.35085072 | 0 |
| 0.00060915 | 0.00138787 | 32.1522496 | 218.881543 | 178.185968 | 192.997439 | 26.1384007 |
| 0.00060943 | 0.00138842 | 17.3622148 | 16.4606038 | 10.2085711 | 14.8459568 | 0 |
| 0.00060956 | 0.00138863 | 9.00262989 | 13.7913167 | 25.9854537 | 25.9804245 | 0 |
| 0.00061007 | 0.00138965 | 8.0380624 | 52.2735392 | 41.7623363 | 63.0953166 | 0 |
| 0.00061009 | 0.00138965 | 5.46588243 | 154.151331 | 45.4745439 | 9.74265918 | 0 |
| 0.00061181 | 0.00139347 | 215.09855 | 243.127568 | 247.789862 | 0 | 0 |
| 0.00061215 | 0.00139416 | 13.1824223 | 9.12006429 | 30.6257133 | 25.9804245 | 0 |
| 0.00061276 | 0.00139545 | 23.4711422 | 24.2460246 | 10.2085711 | 12.0623399 | 0 |
| 0.00061497 | 0.00140037 | 31.1876821 | 15.7932821 | 17.6329864 | 13.4541484 | 40.3957101 |
| 0.00061514 | 0.00140067 | 148.221871 | 284.723958 | 349.875573 | 368.365304 | 11.8810912 |
| 0.00061573 | 0.00140193 | 13.8254673 | 10.6771484 | 35.2659728 | 17.1656376 | 0 |
| 0.00061595 | 0.00140232 | 8.3595849 | 23.3562622 | 18.5610383 | 19.0213822 | 0 |
| 0.00061653 | 0.00140356 | 18.6483048 | 20.0196533 | 59.3953227 | 7.88691457 | 0 |
| 0.00061824 | 0.00140736 | 10.9317649 | 11.7893514 | 21.3451941 | 21.8049991 | 0 |
| 0.00061851 | 0.00140788 | 15.7546023 | 25.8031087 | 8.35246725 | 21.341063 | 0 |
| 0.00061862 | 0.00140804 | 1.28608998 | 40.2617472 | 88.1649321 | 79.3330819 | 0 |
| 0.00061892 | 0.00140861 | 113.497441 | 233.117741 | 0 | 244.958288 | 0 |
| 0.00061923 | 0.00140922 | 157.867546 | 328.322314 | 273.775316 | 263.97967 | 38.0194919 |
| 0.00061938 | 0.00140948 | 22.8280972 | 37.5924601 | 14.8488307 | 41.2903175 | 0 |
| 0.0006199 | 0.00141057 | 311.555299 | 144.363944 | 0 | 143.820207 | 0 |
| 0.00062129 | 0.00141362 | 4.17979245 | 28.6948364 | 37.1220767 | 45.0018067 | 0 |
| 0.0006218 | 0.00141469 | 20.8989622 | 14.2361979 | 50.1148035 | 48.2493597 | 0 |
| 0.00062274 | 0.00141673 | 10.2887199 | 21.3542969 | 19.4890903 | 12.9902122 | 0 |
| 0.00062309 | 0.00141744 | 18.3267823 | 12.2342326 | 13.9207788 | 12.9902122 | 0 |
| 0.00062445 | 0.00142043 | 6.43044992 | 19.7972127 | 35.2659728 | 27.8361691 | 0 |

| | | | | | | |
|---|---|---|---|---|---|---|
| 0.0006247 | 0.00142091 | 7.39501741 | 35.1456136 | 14.8488307 | 34.3312752 | 0 |
| 0.00062482 | 0.00142108 | 6.10892743 | 42.7085937 | 23.2012979 | 24.1246799 | 0 |
| 0.00062522 | 0.00142189 | 0.64304499 | 0.44488118 | 0 | 0.9278723 | 0 |
| 0.00062567 | 0.00142281 | 62.0538418 | 136.578524 | 111.36623 | 134.541484 | 33.2670554 |
| 0.00062589 | 0.00142323 | 45.6561944 | 63.84045 | 49.1867516 | 47.3214874 | 14.2573095 |
| 0.00062603 | 0.00142345 | 2.25065747 | 4.89369303 | 1.85610383 | 5.10329766 | 0 |
| 0.00062629 | 0.00142394 | 7.07349491 | 41.8188314 | 19.4890903 | 22.7328714 | 0 |
| 0.00062643 | 0.00142416 | 165.584086 | 181.511523 | 113.222334 | 147.06776 | 28.5146189 |
| 0.00062703 | 0.00142543 | 17.3622148 | 14.2361979 | 15.7768826 | 10.6705315 | 0 |
| 0.00062796 | 0.00142746 | 8.3595849 | 33.8109701 | 12.9927268 | 33.4034029 | 0 |
| 0.0006281 | 0.00142767 | 30.8661596 | 30.9192423 | 29.6976613 | 44.5378705 | 4.75243649 |
| 0.00062886 | 0.0014293 | 11.8963324 | 28.6948364 | 41.7623363 | 45.929679 | 2.37621824 |
| 0.00062905 | 0.00142964 | 0.64304499 | 0.44488118 | 0.92805192 | 0.46393615 | 2.37621824 |
| 0.00063016 | 0.00143207 | 169.120833 | 178.397355 | 0 | 207.843396 | 0 |
| 0.00063051 | 0.00143277 | 10.9317649 | 10.0098267 | 36.1940248 | 29.6919137 | 0 |
| 0.00063074 | 0.00143319 | 10.6102424 | 11.1220296 | 27.8415575 | 25.5164883 | 0 |
| 0.00063162 | 0.00143509 | 50.8005544 | 12.4566732 | 25.9854537 | 9.27872303 | 0 |
| 0.00063241 | 0.0014368 | 31.5092046 | 24.4684652 | 30.6257133 | 5.56723382 | 0 |
| 0.00063271 | 0.00143737 | 12.5393773 | 22.4664998 | 28.7696094 | 9.74265918 | 0 |
| 0.00063415 | 0.00144054 | 525.367759 | 848.833301 | 702.535301 | 790.547202 | 0 |
| 0.00063471 | 0.00144173 | 11.8963324 | 4.00393066 | 8.35246725 | 4.63936151 | 47.5243649 |
| 0.00063538 | 0.00144314 | 9.32415239 | 19.1298909 | 19.4890903 | 15.309893 | 0 |
| 0.00063672 | 0.00144609 | 14.7900348 | 13.7913167 | 12.9927268 | 12.9902122 | 0 |
| 0.00063679 | 0.00144615 | 5.46588243 | 14.9035197 | 140.135839 | 29.6919137 | 0 |
| 0.00063717 | 0.00144692 | 18.3267823 | 14.2361979 | 11.136623 | 13.9180845 | 0 |
| 0.00063745 | 0.00144746 | 9.64567488 | 16.4606038 | 16.7049345 | 17.6295737 | 0 |
| 0.00063912 | 0.00145116 | 26.6863672 | 15.5708415 | 29.6976613 | 30.1558498 | 2.37621824 |
| 0.00064005 | 0.00145316 | 9.32415239 | 30.2519206 | 35.2659728 | 11.5984038 | 0 |
| 0.00064105 | 0.00145534 | 25.0787547 | 22.4664998 | 22.273246 | 18.5574461 | 54.6530196 |
| 0.00064136 | 0.00145595 | 38.5826995 | 17.3503662 | 11.136623 | 12.9902122 | 0 |
| 0.00064228 | 0.00145793 | 48.5498969 | 19.3523315 | 25.0574018 | 6.49510612 | 0 |
| 0.00064257 | 0.0014584 | 13.8254673 | 22.911381 | 33.409869 | 8.81478687 | 0 |
| 0.00064256 | 0.0014584 | 7.39501741 | 33.3660889 | 20.4171422 | 19.9492545 | 0 |
| 0.00064393 | 0.00146138 | 10.2887199 | 22.2440592 | 12.0646749 | 45.929679 | 0 |
| 0.00064436 | 0.00146227 | 6.43044992 | 19.7972127 | 27.8415575 | 32.9394667 | 0 |
| 0.00064548 | 0.00146471 | 443.701045 | 68.0668213 | 0 | 195.781056 | 0 |
| 0.00064705 | 0.00146818 | 7.39501741 | 32.2538859 | 16.7049345 | 25.9804245 | 0 |
| 0.00064754 | 0.00146918 | 220.24291 | 323.428621 | 370.292715 | 329.858604 | 57.0292378 |
| 0.00064817 | 0.0014705 | 225.065747 | 216.212256 | 0 | 123.870952 | 0 |
| 0.00064847 | 0.0014711 | 18.6483048 | 20.0196533 | 27.8415575 | 48.7132959 | 0 |
| 0.00064922 | 0.00147269 | 124.107684 | 423.082007 | 464.95401 | 437.491791 | 42.7719284 |
| 0.00065009 | 0.00147457 | 16.7191698 | 15.7932821 | 21.3451941 | 9.27872303 | 0 |
| 0.00065155 | 0.00147777 | 21.5420072 | 31.8090047 | 36.1940248 | 5.56723382 | 0 |

| | | | | | | |
|---|---|---|---|---|---|---|
| 0.00065259 | 0.00148004 | 0 | 222.440592 | 315.537652 | 231.968076 | 0 |
| 0.00065436 | 0.00148394 | 21.2204847 | 9.56494547 | 28.7696094 | 12.9902122 | 0 |
| 0.00065583 | 0.00148718 | 66.2336342 | 29.1397176 | 38.9781805 | 38.0427644 | 111.682257 |
| 0.00065667 | 0.00148898 | 250.144502 | 264.481864 | 262.638692 | 256.092756 | 403.957101 |
| 0.00065811 | 0.00149216 | 14.7900348 | 16.2381632 | 20.4171422 | 9.74265918 | 0 |
| 0.0006586 | 0.00149317 | 189.37675 | 442.434338 | 425.047778 | 381.819453 | 30.8908372 |
| 0.00065949 | 0.00149509 | 12.5393773 | 33.5885295 | 8.35246725 | 32.0115944 | 0 |
| 0.00066 | 0.00149613 | 55.9449143 | 50.2715739 | 42.6903882 | 51.4969128 | 21.3859642 |
| 0.00066035 | 0.00149682 | 129.895088 | 317.200285 | 274.703367 | 0 | 0 |
| 0.00066062 | 0.00149734 | 53.3727344 | 15.3484009 | 7.42441534 | 25.5164883 | 0 |
| 0.00066148 | 0.00149918 | 38.261177 | 22.4664998 | 14.8488307 | 8.35085072 | 0 |
| 0.00066152 | 0.00149919 | 23.7926647 | 16.4606038 | 10.2085711 | 13.4541484 | 0 |
| 0.00066209 | 0.00150038 | 5.46588243 | 32.0314453 | 25.0574018 | 29.6919137 | 0 |
| 0.00066257 | 0.00150136 | 36.3320421 | 28.6948364 | 20.4171422 | 33.4034029 | 4.75243649 |
| 0.00066363 | 0.00150361 | 24.7572322 | 5.7834554 | 27.8415575 | 30.1558498 | 0 |
| 0.00066365 | 0.00150361 | 11.8963324 | 13.123995 | 13.9207788 | 16.7017014 | 0 |
| 0.00066453 | 0.00150549 | 5.14435994 | 26.9153117 | 36.1940248 | 28.3001052 | 0 |
| 0.00066487 | 0.00150616 | 12.2178549 | 21.3542969 | 10.2085711 | 20.4131907 | 0 |
| 0.0006668 | 0.00151042 | 63.9829767 | 37.8149007 | 39.9062324 | 42.2181898 | 142.573095 |
| 0.00066691 | 0.00151057 | 19.2913498 | 13.3464355 | 12.0646749 | 12.9902122 | 0 |
| 0.00066707 | 0.00151084 | 7.71653991 | 20.2420939 | 19.4890903 | 23.1968076 | 0 |
| 0.00066734 | 0.00151135 | 14.4685123 | 17.1279256 | 14.8488307 | 16.2377653 | 54.6530196 |
| 0.0006683 | 0.00151343 | 10.6102424 | 11.1220296 | 23.2012979 | 36.650956 | 0 |
| 0.00067026 | 0.00151776 | 25.4002772 | 12.9015544 | 13.9207788 | 12.0623399 | 0 |
| 0.00067042 | 0.00151801 | 12.5393773 | 11.1220296 | 19.4890903 | 15.7738291 | 0 |
| 0.000671 | 0.00151922 | 0 | 569.225476 | 163.337137 | 152.634994 | 0 |
| 0.0006726 | 0.00152275 | 0.96456749 | 2.22440592 | 1.85610383 | 1.39180845 | 40.3957101 |
| 0.00067287 | 0.00152326 | 15.7546023 | 16.0157227 | 10.2085711 | 14.8459568 | 0 |
| 0.00067296 | 0.00152336 | 9.00262989 | 18.017688 | 31.5537652 | 15.309893 | 0 |
| 0.00067527 | 0.0015285 | 11.5748099 | 18.9074504 | 45.4745439 | 11.5984038 | 0 |
| 0.00067559 | 0.00152912 | 7.71653991 | 19.3523315 | 23.2012979 | 19.9492545 | 0 |
| 0.0006762 | 0.00153039 | 0.96456749 | 45.3778809 | 90.9490879 | 83.0445711 | 0 |
| 0.00067727 | 0.0015327 | 0.96456749 | 0.66732178 | 0.92805192 | 0.46393615 | 0 |
| 0.00067733 | 0.00153274 | 4.50131495 | 39.3719849 | 56.6111669 | 20.4131907 | 0 |
| 0.00067743 | 0.00153286 | 104.816334 | 232.005538 | 0 | 231.968076 | 0 |
| 0.00067878 | 0.00153571 | 11.8963324 | 18.9074504 | 12.0646749 | 14.8459568 | 0 |
| 0.00067875 | 0.00153571 | 147.578826 | 551.652669 | 234.797135 | 376.252219 | 40.3957101 |
| 0.00067987 | 0.00153808 | 275.866302 | 276.716097 | 388.853753 | 431.460621 | 19.0097459 |
| 0.0006802 | 0.00153871 | 87.1325964 | 657.534391 | 161.481034 | 632.344974 | 2.37621824 |
| 0.00068213 | 0.00154297 | 111.568306 | 231.783097 | 174.47376 | 192.069567 | 9.50487297 |
| 0.00068283 | 0.00154446 | 9.32415239 | 24.2460246 | 15.7768826 | 16.2377653 | 0 |
| 0.0006831 | 0.00154496 | 295.479174 | 31.8090047 | 16.7049345 | 1.85574461 | 0 |
| 0.00068425 | 0.00154747 | 4.82283744 | 32.2538859 | 33.409869 | 27.8361691 | 0 |

| | | | | | | |
|---|---|---|---|---|---|---|
| 0.00068472 | 0.00154843 | 16.7191698 | 15.3484009 | 16.7049345 | 9.74265918 | 0 |
| 0.00068526 | 0.00154955 | 11.5748099 | 11.3444702 | 18.5610383 | 18.0935099 | 0 |
| 0.00068599 | 0.00155108 | 240.177305 | 152.149365 | 0 | 151.243185 | 0 |
| 0.00068677 | 0.00155275 | 10.2887199 | 21.1318563 | 12.9927268 | 17.1656376 | 0 |
| 0.00068689 | 0.0015529 | 130.216611 | 141.472217 | 0 | 300.16669 | 0 |
| 0.0006873 | 0.00155374 | 62.3753642 | 122.787207 | 129.927268 | 129.902122 | 9.50487297 |
| 0.00068745 | 0.00155396 | 4.50131495 | 14.2361979 | 77.0283091 | 52.8887212 | 0 |
| 0.00068871 | 0.00155671 | 9.64567488 | 26.6928711 | 11.136623 | 36.650956 | 0 |
| 0.0006905 | 0.00156065 | 18.3267823 | 11.5669108 | 12.0646749 | 19.0213822 | 21.3859642 |
| 0.00069129 | 0.00156232 | 585.813988 | 435.983561 | 242.22155 | 263.515734 | 2.37621824 |
| 0.00069171 | 0.00156319 | 11.8963324 | 21.3542969 | 10.2085711 | 20.8771268 | 0 |
| 0.00069244 | 0.00156472 | 48.2283744 | 11.5669108 | 11.136623 | 19.9492545 | 0 |
| 0.00069318 | 0.00156628 | 11.2532874 | 22.4664998 | 10.2085711 | 31.0837221 | 0 |
| 0.00069448 | 0.00156912 | 51.7651219 | 49.3818115 | 21.3451941 | 32.0115944 | 7.12865473 |
| 0.00069475 | 0.00156961 | 4.50131495 | 92.3128459 | 54.7550631 | 60.7756358 | 0 |
| 0.00069506 | 0.00157012 | 82.6312815 | 140.137573 | 159.62493 | 148.459568 | 33.2670554 |
| 0.00069503 | 0.00157012 | 9.64567488 | 9.56494547 | 38.9781805 | 40.3624452 | 0 |
| 0.00069605 | 0.00157225 | 5.14435994 | 62.2833659 | 17.6329864 | 35.7230837 | 0 |
| 0.00069622 | 0.00157253 | 19.9343948 | 5.33857422 | 44.546492 | 32.9394667 | 0 |
| 0.00069648 | 0.001573 | 42.4409695 | 66.9546183 | 88.1649321 | 91.3954218 | 19.0097459 |
| 0.00069779 | 0.00157587 | 12.2178549 | 25.8031087 | 13.9207788 | 12.5262761 | 0 |
| 0.00069869 | 0.00157779 | 125.393773 | 193.745756 | 0 | 221.297544 | 0 |
| 0.0006999 | 0.00158041 | 7.71653991 | 10.0098267 | 50.1148035 | 50.5690405 | 0 |
| 0.00070163 | 0.00158411 | 5.46588243 | 29.8070394 | 25.0574018 | 29.2279775 | 0 |
| 0.0007016 | 0.00158411 | 130.859656 | 197.527246 | 178.185968 | 191.141694 | 33.2670554 |
| 0.00070407 | 0.0015895 | 7.71653991 | 6.005896 | 2.78415575 | 6.03116997 | 4.75243649 |
| 0.00070437 | 0.00159008 | 7.71653991 | 19.7972127 | 27.8415575 | 17.6295737 | 0 |
| 0.00070449 | 0.00159023 | 433.090802 | 1316.84831 | 1256.5823 | 1306.4442 | 0 |
| 0.00070498 | 0.00159123 | 47.2638069 | 283.389315 | 245.005706 | 249.597649 | 4.75243649 |
| 0.0007058 | 0.00159298 | 9.32415239 | 6.45077718 | 7.42441534 | 4.17542536 | 38.0194919 |
| 0.00070889 | 0.00159985 | 9.96719738 | 40.0393066 | 77.0283091 | 7.42297842 | 0 |
| 0.00070942 | 0.00160095 | 16.3976473 | 21.7991781 | 7.42441534 | 31.5476583 | 0 |
| 0.00070964 | 0.00160134 | 6.10892743 | 22.4664998 | 24.1293498 | 31.5476583 | 0 |
| 0.00071013 | 0.00160232 | 4.50131495 | 1.77952474 | 0 | 0 | 159.206622 |
| 0.0007138 | 0.0016105 | 6.75197242 | 28.4723958 | 16.7049345 | 33.4034029 | 0 |
| 0.00071393 | 0.00161068 | 0 | 184.848132 | 361.012196 | 208.771268 | 0 |
| 0.0007143 | 0.00161142 | 10.9317649 | 48.2696086 | 87.2368802 | 56.1362743 | 4.75243649 |
| 0.00071499 | 0.00161286 | 127.001386 | 199.974093 | 204.171422 | 210.627013 | 35.6432737 |
| 0.00071713 | 0.00161759 | 6.75197242 | 18.6850098 | 37.1220767 | 22.7328714 | 0 |
| 0.00071725 | 0.00161775 | 8.6811074 | 33.3660889 | 14.8488307 | 19.4853184 | 0 |
| 0.00071754 | 0.00161829 | 13.5039448 | 15.1259603 | 14.8488307 | 11.1344676 | 0 |
| 0.00071883 | 0.00162106 | 8.6811074 | 14.6810791 | 19.4890903 | 27.3722329 | 0 |
| 0.0007189 | 0.00162106 | 12.5393773 | 12.6791138 | 5.5683115 | 3.71148921 | 64.1578926 |

| | | | | | | |
|---|---|---|---|---|---|---|
| 0.00071891 | 0.00162106 | 90.6693439 | 62.7282471 | 49.1867516 | 57.9920189 | 21.3859642 |
| 0.00071901 | 0.00162118 | 13.5039448 | 30.9192423 | 10.2085711 | 16.2377653 | 0 |
| 0.00072022 | 0.00162379 | 10.2887199 | 23.8011434 | 12.0646749 | 18.5574461 | 0 |
| 0.00072069 | 0.00162475 | 15.4330798 | 7.11809896 | 35.2659728 | 27.8361691 | 0 |
| 0.00072128 | 0.00162596 | 13.5039448 | 10.2322673 | 16.7049345 | 17.1656376 | 0 |
| 0.00072135 | 0.00162602 | 8.3595849 | 21.1318563 | 15.7768826 | 20.8771268 | 0 |
| 0.00072331 | 0.00163025 | 13.5039448 | 31.5865641 | 8.35246725 | 23.1968076 | 0 |
| 0.00072333 | 0.00163025 | 0 | 281.832231 | 287.696094 | 169.336695 | 0 |
| 0.0007234 | 0.00163031 | 277.473914 | 216.434696 | 165.193241 | 0 | 0 |
| 0.00072418 | 0.00163195 | 8.6811074 | 26.9153117 | 13.9207788 | 21.341063 | 0 |
| 0.00072429 | 0.00163209 | 8.3595849 | 23.5787028 | 16.7049345 | 18.0935099 | 0 |
| 0.00072594 | 0.00163569 | 20.5774398 | 18.017688 | 8.35246725 | 16.2377653 | 0 |
| 0.00072605 | 0.00163585 | 0 | 0 | 0 | 0 | 76.0389838 |
| 0.00072645 | 0.00163662 | 93.5630464 | 231.115776 | 0 | 234.751693 | 0 |
| 0.0007275 | 0.00163889 | 9.32415239 | 16.4606038 | 17.6329864 | 15.7738291 | 0 |
| 0.00073039 | 0.00164527 | 4.50131495 | 27.5826335 | 25.0574018 | 59.8477635 | 0 |
| 0.00073072 | 0.0016458 | 9.00262989 | 14.0137573 | 19.4890903 | 22.2689353 | 0 |
| 0.0007307 | 0.0016458 | 121.535504 | 199.751652 | 157.768826 | 174.439993 | 33.2670554 |
| 0.00073095 | 0.00164621 | 6.75197242 | 24.023584 | 29.6976613 | 18.5574461 | 0 |
| 0.00073109 | 0.00164642 | 25.4002772 | 11.7893514 | 11.136623 | 16.2377653 | 0 |
| 0.00073225 | 0.00164893 | 9.64567488 | 22.911381 | 12.9927268 | 18.5574461 | 0 |
| 0.00073439 | 0.00165363 | 0 | 1.55708415 | 0 | 2.31968076 | 97.424948 |
| 0.00073449 | 0.00165374 | 0 | 0.44488118 | 0.92805192 | 0.46393615 | 0 |
| 0.0007351 | 0.001655 | 10.2887199 | 22.4664998 | 11.136623 | 23.1968076 | 0 |
| 0.00073515 | 0.001655 | 7.71653991 | 25.8031087 | 18.5610383 | 18.5574461 | 0 |
| 0.00073523 | 0.00165508 | 5.14435994 | 18.2401286 | 43.6184401 | 38.5067006 | 0 |
| 0.00073661 | 0.00165807 | 32.4737721 | 17.1279256 | 5.5683115 | 46.3936151 | 0 |
| 0.00073785 | 0.00166074 | 8.6811074 | 20.2420939 | 20.4171422 | 14.8459568 | 0 |
| 0.00073865 | 0.00166244 | 205.452875 | 259.588171 | 224.588564 | 276.04201 | 54.6530196 |
| 0.00073914 | 0.00166333 | 30.5446371 | 8.45274251 | 12.0646749 | 32.0115944 | 0 |
| 0.00073914 | 0.00166333 | 8.0380624 | 15.5708415 | 20.4171422 | 31.0837221 | 0 |
| 0.00074098 | 0.00166735 | 71.3779941 | 11.7893514 | 14.8488307 | 13.9180845 | 0 |
| 0.00074198 | 0.00166948 | 9.00262989 | 28.2499552 | 13.9207788 | 19.0213822 | 0 |
| 0.00074237 | 0.00167025 | 13.1824223 | 11.7893514 | 20.4171422 | 12.5262761 | 0 |
| 0.00074272 | 0.00167073 | 8.3595849 | 14.9035197 | 20.4171422 | 24.588616 | 0 |
| 0.00074273 | 0.00167073 | 6.43044992 | 29.5845988 | 15.7768826 | 45.0018067 | 0 |
| 0.00074273 | 0.00167073 | 8.0380624 | 44.4881185 | 16.7049345 | 19.0213822 | 0 |
| 0.00074466 | 0.00167494 | 7.39501741 | 19.5747721 | 18.5610383 | 27.8361691 | 0 |
| 0.0007447 | 0.00167494 | 23.4711422 | 28.2499552 | 14.8488307 | 43.1460621 | 0 |
| 0.00074508 | 0.00167569 | 23.4711422 | 12.4566732 | 10.2085711 | 16.7017014 | 0 |
| 0.00074542 | 0.00167633 | 12.5393773 | 13.3464355 | 12.9927268 | 13.9180845 | 0 |
| 0.00074557 | 0.00167656 | 0 | 375.47972 | 162.409085 | 211.554885 | 0 |
| 0.00074734 | 0.00168031 | 107.066991 | 31.5865641 | 16.7049345 | 29.2279775 | 2.37621824 |

| | | | | | | |
|---|---|---|---|---|---|---|
| 0.0007473 | 0.00168031 | 24.4357097 | 33.1436483 | 5.5683115 | 22.7328714 | 0 |
| 0.00075016 | 0.00168654 | 7.39501741 | 26.0255493 | 16.7049345 | 23.1968076 | 0 |
| 0.00075146 | 0.00168935 | 20.8989622 | 11.7893514 | 15.7768826 | 11.1344676 | 0 |
| 0.00075158 | 0.0016895 | 31.1876821 | 9.12006429 | 12.0646749 | 23.6607437 | 0 |
| 0.00075164 | 0.00168953 | 13.5039448 | 47.1574056 | 31.5537652 | 24.588616 | 2.37621824 |
| 0.0007519 | 0.00169 | 4.50131495 | 0 | 0 | 0 | 102.177384 |
| 0.00075231 | 0.0016907 | 12.8608998 | 29.5845988 | 32.4818171 | 32.9394667 | 2.37621824 |
| 0.00075226 | 0.0016907 | 12.2178549 | 41.5963908 | 10.2085711 | 19.0213822 | 0 |
| 0.00075321 | 0.0016926 | 0 | 205.090226 | 263.566744 | 239.391054 | 0 |
| 0.0007543 | 0.00169494 | 1.60761248 | 65.3975342 | 48.2586997 | 51.4969128 | 0 |
| 0.00075452 | 0.00169534 | 6.75197242 | 19.3523315 | 22.273246 | 28.7640414 | 0 |
| 0.00075478 | 0.00169567 | 0.64304499 | 0.88976237 | 0.92805192 | 0.46393615 | 0 |
| 0.00075473 | 0.00169567 | 7.71653991 | 17.3503662 | 19.4890903 | 25.0525522 | 0 |
| 0.00075596 | 0.00169822 | 15.7546023 | 4.22637126 | 11.136623 | 7.42297842 | 135.44444 |
| 0.00075657 | 0.00169949 | 179.409553 | 423.526888 | 313.681548 | 352.127539 | 80.7914203 |
| 0.00075692 | 0.00170015 | 10.2887199 | 16.0157227 | 14.8488307 | 14.3820207 | 0 |
| 0.00075767 | 0.00170172 | 16.3976473 | 9.56494547 | 28.7696094 | 13.9180845 | 0 |
| 0.00075988 | 0.00170658 | 9.96719738 | 32.9212077 | 44.546492 | 41.2903175 | 2.37621824 |
| 0.00076088 | 0.00170871 | 10.6102424 | 11.7893514 | 19.4890903 | 71.9101035 | 0 |
| 0.00076135 | 0.00170964 | 6.10892743 | 21.1318563 | 28.7696094 | 25.0525522 | 0 |
| 0.00076155 | 0.00170997 | 6.43044992 | 38.2597819 | 14.8488307 | 34.7952113 | 0 |
| 0.00076213 | 0.00171116 | 430.1971 | 568.780595 | 592.097123 | 599.869444 | 7.12865473 |
| 0.00076301 | 0.00171294 | 14.1469898 | 20.6869751 | 9.28051917 | 16.2377653 | 0 |
| 0.00076307 | 0.00171294 | 64.9475442 | 201.308736 | 264.494796 | 221.76148 | 0 |
| 0.00076305 | 0.00171294 | 28.2939797 | 13.3464355 | 69.6038938 | 6.95904227 | 0 |
| 0.00076456 | 0.00171616 | 37.618132 | 70.2912272 | 55.683115 | 55.208402 | 14.2573095 |
| 0.00076557 | 0.00171831 | 0 | 208.426835 | 285.83999 | 211.090949 | 0 |
| 0.00076581 | 0.00171873 | 6.75197242 | 44.0432373 | 17.6329864 | 22.7328714 | 0 |
| 0.00076622 | 0.00171954 | 7.39501741 | 21.3542969 | 19.4890903 | 20.4131907 | 0 |
| 0.00076648 | 0.00172001 | 26.0433222 | 14.9035197 | 10.2085711 | 13.4541484 | 0 |
| 0.00076825 | 0.00172387 | 4.82283744 | 24.9133464 | 25.0574018 | 52.4247851 | 0 |
| 0.00076988 | 0.00172741 | 0 | 0 | 0 | 0 | 23.7621824 |
| 0.00077076 | 0.00172927 | 123.143116 | 155.708415 | 0 | 241.710735 | 0 |
| 0.00077175 | 0.00173138 | 54.0157793 | 28.917277 | 15.7768826 | 60.7756358 | 4.75243649 |
| 0.00077215 | 0.00173216 | 6.10892743 | 36.4802572 | 21.3451941 | 22.2689353 | 0 |
| 0.00077233 | 0.00173245 | 32.4737721 | 54.2755046 | 27.8415575 | 12.0623399 | 0 |
| 0.00077286 | 0.00173353 | 5.78740493 | 33.5885295 | 23.2012979 | 23.1968076 | 0 |
| 0.00077317 | 0.0017341 | 114.462009 | 181.956405 | 410.198947 | 0 | 0 |
| 0.00077553 | 0.00173929 | 9.64567488 | 23.1338216 | 23.2012979 | 11.5984038 | 0 |
| 0.00077567 | 0.00173948 | 192.913498 | 320.981775 | 282.127783 | 286.712542 | 0 |
| 0.00077596 | 0.00174002 | 15.4330798 | 22.2440592 | 27.8415575 | 38.5067006 | 0 |
| 0.00077671 | 0.00174158 | 9.64567488 | 14.9035197 | 14.8488307 | 18.5574461 | 0 |
| 0.00077687 | 0.00174182 | 16.3976473 | 23.5787028 | 23.2012979 | 7.42297842 | 0 |

| | | | | | | |
|---|---|---|---|---|---|---|
| 0.00077697 | 0.00174194 | 7.71653991 | 21.1318563 | 19.4890903 | 18.0935099 | 0 |
| 0.0007771 | 0.00174212 | 118.641801 | 133.019474 | 134.567528 | 135.469356 | 68.9103291 |
| 0.00077789 | 0.00174376 | 16.3976473 | 25.3582275 | 6.49636342 | 38.0427644 | 0 |
| 0.0007794 | 0.00174703 | 25.7217997 | 81.6356974 | 55.683115 | 70.0543588 | 11.8810912 |
| 0.00078038 | 0.00174911 | 22.1850522 | 11.1220296 | 15.7768826 | 11.5984038 | 0 |
| 0.00078195 | 0.00175251 | 12.2178549 | 20.4645345 | 27.8415575 | 9.27872303 | 0 |
| 0.0007822 | 0.00175295 | 0 | 192.633553 | 344.307261 | 181.399035 | 0 |
| 0.00078419 | 0.0017573 | 6.10892743 | 58.2794352 | 88.1649321 | 7.88691457 | 0 |
| 0.00078494 | 0.00175886 | 8.6811074 | 20.4645345 | 72.3880495 | 13.4541484 | 0 |
| 0.00078596 | 0.00176102 | 170.406923 | 378.149007 | 335.026742 | 384.139133 | 2.37621824 |
| 0.00078666 | 0.00176248 | 98.7074063 | 126.568697 | 117.862593 | 94.6429749 | 0 |
| 0.00078767 | 0.00176464 | 8.3595849 | 27.1377523 | 17.6329864 | 15.7738291 | 0 |
| 0.00078801 | 0.00176516 | 182.9463 | 523.625155 | 475.162581 | 507.082213 | 7.12865473 |
| 0.00078801 | 0.00176516 | 23.1496197 | 30.02948 | 7.42441534 | 14.8459568 | 0 |
| 0.00078889 | 0.001767 | 52.4081669 | 17.7952474 | 31.5537652 | 26.4443606 | 140.196876 |
| 0.00078939 | 0.00176801 | 8.6811074 | 51.3837769 | 35.2659728 | 46.8575513 | 0 |
| 0.0007896 | 0.00176828 | 49.1929419 | 33.5885295 | 4.64025958 | 19.4853184 | 0 |
| 0.00078962 | 0.00176828 | 8.3595849 | 37.5924601 | 52.8989593 | 42.2181898 | 2.37621824 |
| 0.00079025 | 0.00176958 | 656.227415 | 1234.10041 | 1115.5184 | 1293.45399 | 64.1578926 |
| 0.00079062 | 0.00177029 | 0.3215225 | 0 | 0 | 0 | 0 |
| 0.0007912 | 0.00177147 | 84.5604165 | 225.10988 | 0 | 233.82382 | 0 |
| 0.00079167 | 0.00177241 | 31.1876821 | 46.4900838 | 6.49636342 | 13.9180845 | 0 |
| 0.00079243 | 0.001774 | 0 | 0 | 0 | 0 | 144.949313 |
| 0.0007929 | 0.00177493 | 13.1824223 | 26.4704305 | 7.42441534 | 47.3214874 | 0 |
| 0.00079329 | 0.00177567 | 5.46588243 | 36.035376 | 26.9135056 | 21.341063 | 0 |
| 0.00079618 | 0.00178202 | 15.1115573 | 15.5708415 | 13.9207788 | 10.2065953 | 0 |
| 0.00079655 | 0.00178274 | 361.712808 | 981.630334 | 880.721269 | 955.244536 | 21.3859642 |
| 0.00079668 | 0.00178292 | 10.9317649 | 13.5688761 | 12.9927268 | 18.0935099 | 0 |
| 0.00079772 | 0.00178513 | 49.8359869 | 114.779346 | 82.5966206 | 99.7462725 | 19.0097459 |
| 0.00079927 | 0.00178846 | 0 | 0 | 0 | 0 | 52.2768014 |
| 0.00080279 | 0.00179623 | 14.7900348 | 13.123995 | 12.0646749 | 12.5262761 | 0 |
| 0.00080519 | 0.00180147 | 7.07349491 | 22.2440592 | 16.7049345 | 30.619786 | 0 |
| 0.00080614 | 0.00180347 | 21.5420072 | 11.7893514 | 14.8488307 | 11.1344676 | 0 |
| 0.00080685 | 0.00180496 | 5.78740493 | 30.6968018 | 22.273246 | 24.1246799 | 0 |
| 0.00080883 | 0.00180926 | 54.6588243 | 31.1416829 | 60.3233746 | 59.8477635 | 87.920075 |
| 0.00080912 | 0.0018098 | 0 | 243.127568 | 160.552982 | 293.671584 | 0 |
| 0.00080958 | 0.0018107 | 8.0380624 | 13.7913167 | 29.6976613 | 22.2689353 | 0 |
| 0.0008109 | 0.00181353 | 3.21522496 | 59.8365194 | 80.7405168 | 15.309893 | 0 |
| 0.00081142 | 0.00181457 | 8.0380624 | 15.5708415 | 23.2012979 | 19.0213822 | 0 |
| 0.00081161 | 0.00181487 | 1.92913498 | 55.6101481 | 35.2659728 | 60.7756358 | 0 |
| 0.00081217 | 0.00181602 | 22.5065747 | 143.029301 | 106.72597 | 99.2823364 | 14.2573095 |
| 0.00081301 | 0.00181776 | 18.0052598 | 41.5963908 | 6.49636342 | 21.341063 | 0 |
| 0.00081492 | 0.00182191 | 80.0591015 | 248.688582 | 0 | 213.41063 | 0 |

| | | | | | | |
|---|---|---|---|---|---|---|
| 0.00081506 | 0.0018221 | 42.762492 | 13.7913167 | 14.8488307 | 10.6705315 | 0 |
| 0.00081574 | 0.0018235 | 262.683879 | 93.6474894 | 0 | 172.584248 | 0 |
| 0.00081762 | 0.00182758 | 11.2532874 | 16.0157227 | 11.136623 | 17.6295737 | 0 |
| 0.00081773 | 0.00182772 | 496.109212 | 447.105591 | 239.437395 | 300.630626 | 0 |
| 0.00081809 | 0.0018284 | 16.3976473 | 36.7026978 | 18.5610383 | 33.4034029 | 0 |
| 0.00081875 | 0.00182974 | 91.3123889 | 83.1927816 | 25.9854537 | 154.954675 | 14.2573095 |
| 0.00081958 | 0.00183149 | 11.2532874 | 37.1475789 | 27.8415575 | 46.3936151 | 0 |
| 0.00082035 | 0.00183308 | 4.50131495 | 27.8050741 | 24.1293498 | 48.2493597 | 0 |
| 0.0008205 | 0.00183329 | 95.8137038 | 218.436662 | 0 | 201.34829 | 0 |
| 0.00082179 | 0.00183605 | 5.78740493 | 22.4664998 | 30.6257133 | 23.1968076 | 0 |
| 0.00082207 | 0.00183656 | 0 | 0 | 0 | 0.46393615 | 26.1384007 |
| 0.00082259 | 0.00183747 | 28.9370247 | 11.7893514 | 13.9207788 | 12.0623399 | 0 |
| 0.00082257 | 0.00183747 | 4.82283744 | 44.4881185 | 21.3451941 | 29.6919137 | 0 |
| 0.0008228 | 0.00183783 | 9.64567488 | 18.4625692 | 12.0646749 | 28.3001052 | 0 |
| 0.0008241 | 0.00184061 | 14.4685123 | 20.4645345 | 36.1940248 | 7.88691457 | 0 |
| 0.00082433 | 0.00184093 | 153.366231 | 192.188672 | 152.200514 | 141.964462 | 30.8908372 |
| 0.00082436 | 0.00184093 | 12.5393773 | 14.0137573 | 11.136623 | 15.309893 | 0 |
| 0.00082462 | 0.00184139 | 12.5393773 | 10.0098267 | 23.2012979 | 15.7738291 | 0 |
| 0.00082472 | 0.00184151 | 7.71653991 | 16.2381632 | 25.0574018 | 19.0213822 | 0 |
| 0.00082554 | 0.00184322 | 80.7021465 | 214.877612 | 0 | 240.318926 | 0 |
| 0.00082586 | 0.00184381 | 18.3267823 | 34.0334106 | 30.6257133 | 38.0427644 | 4.75243649 |
| 0.00082802 | 0.00184851 | 22.5065747 | 19.3523315 | 7.42441534 | 16.7017014 | 0 |
| 0.00082931 | 0.00185127 | 9.96719738 | 27.5826335 | 11.136623 | 19.9492545 | 0 |
| 0.0008334 | 0.00186027 | 8.3595849 | 15.7932821 | 20.4171422 | 17.1656376 | 0 |
| 0.00083474 | 0.00186301 | 3.53674746 | 35.5904948 | 40.8342843 | 30.1558498 | 0 |
| 0.00083472 | 0.00186301 | 26.0433222 | 17.3503662 | 13.9207788 | 8.81478687 | 0 |
| 0.00083495 | 0.00186336 | 9.64567488 | 15.7932821 | 13.9207788 | 16.7017014 | 0 |
| 0.00083518 | 0.00186374 | 8.6811074 | 26.9153117 | 12.0646749 | 24.1246799 | 0 |
| 0.00083606 | 0.00186557 | 8.0380624 | 37.1475789 | 12.0646749 | 26.9082968 | 0 |
| 0.0008381 | 0.00187001 | 7.07349491 | 22.0216187 | 23.2012979 | 17.6295737 | 0 |
| 0.00084056 | 0.00187538 | 42.762492 | 19.3523315 | 8.35246725 | 13.9180845 | 0 |
| 0.0008434 | 0.00188158 | 4.17979245 | 28.2499552 | 30.6257133 | 36.1870198 | 0 |
| 0.00084483 | 0.00188465 | 8.6811074 | 21.7991781 | 16.7049345 | 14.8459568 | 0 |
| 0.00084747 | 0.00189043 | 1.60761248 | 0.88976237 | 6.49636342 | 1.85574461 | 16.6335277 |
| 0.00084771 | 0.00189083 | 207.38201 | 257.586206 | 309.96934 | 312.22903 | 0 |
| 0.00084858 | 0.00189264 | 7.07349491 | 39.3719849 | 12.0646749 | 35.2591475 | 0 |
| 0.00085063 | 0.00189695 | 33.4383396 | 32.6987671 | 18.5610383 | 29.6919137 | 4.75243649 |
| 0.00085059 | 0.00189695 | 10.2887199 | 24.023584 | 12.0646749 | 15.7738291 | 0 |
| 0.0008526 | 0.00190122 | 19.2913498 | 53.6081828 | 21.3451941 | 79.3330819 | 4.75243649 |
| 0.00085488 | 0.0019062 | 4.17979245 | 21.1318563 | 49.1867516 | 34.7952113 | 0 |
| 0.00085563 | 0.00190773 | 9.32415239 | 13.3464355 | 18.5610383 | 16.7017014 | 0 |
| 0.00085673 | 0.00191006 | 5.14435994 | 27.3601929 | 26.9135056 | 25.9804245 | 0 |
| 0.00085792 | 0.00191258 | 2.57217997 | 34.0334106 | 100.229607 | 25.9804245 | 0 |

| | | | | | | |
|---|---|---|---|---|---|---|
| 0.00085807 | 0.00191279 | 9.32415239 | 30.9192423 | 11.136623 | 22.2689353 | 0 |
| 0.00085825 | 0.00191307 | 15.1115573 | 22.4664998 | 7.42441534 | 22.2689353 | 0 |
| 0.00085858 | 0.00191367 | 19.6128723 | 4.44881185 | 0 | 6.95904227 | 104.553603 |
| 0.00085966 | 0.00191595 | 22.5065747 | 20.0196533 | 16.7049345 | 7.42297842 | 0 |
| 0.00086034 | 0.00191721 | 4.82283744 | 24.2460246 | 38.0501286 | 25.9804245 | 0 |
| 0.00086034 | 0.00191721 | 8.3595849 | 42.7085937 | 46.4025958 | 37.1148921 | 0 |
| 0.00086188 | 0.00192039 | 180.052598 | 44.7105591 | 13.9207788 | 30.1558498 | 4.75243649 |
| 0.00086187 | 0.00192039 | 9.96719738 | 13.123995 | 28.7696094 | 14.8459568 | 0 |
| 0.00086456 | 0.00192624 | 175.872805 | 837.933712 | 865.872439 | 842.044115 | 0 |
| 0.00086509 | 0.00192729 | 158.832113 | 210.651241 | 200.459214 | 203.204034 | 30.8908372 |
| 0.00086623 | 0.0019297 | 151.115573 | 321.204215 | 249.645966 | 263.97967 | 0 |
| 0.00086758 | 0.00193258 | 24.7572322 | 21.1318563 | 38.0501286 | 31.0837221 | 2.37621824 |
| 0.00086877 | 0.0019351 | 108.674604 | 191.298909 | 180.970124 | 176.295737 | 40.3957101 |
| 0.00086904 | 0.00193558 | 0 | 191.966231 | 196.747006 | 272.330521 | 0 |
| 0.00086972 | 0.00193697 | 8.6811074 | 11.1220296 | 26.9135056 | 31.0837221 | 0 |
| 0.00087187 | 0.00194163 | 130.216611 | 346.562443 | 342.451157 | 314.548711 | 2.37621824 |
| 0.0008723 | 0.00194245 | 8.0380624 | 14.2361979 | 24.1293498 | 19.9492545 | 0 |
| 0.00087244 | 0.00194264 | 16.7191698 | 16.4606038 | 13.9207788 | 9.27872303 | 0 |
| 0.00087267 | 0.00194302 | 7.71653991 | 11.3444702 | 52.8989593 | 28.3001052 | 0 |
| 0.00087332 | 0.00194433 | 0.3215225 | 0.66732178 | 0 | 1.39180845 | 26.1384007 |
| 0.00087349 | 0.00194458 | 224.422702 | 181.289083 | 0 | 94.1790387 | 0 |
| 0.00087361 | 0.00194474 | 19.9343948 | 15.3484009 | 32.4818171 | 7.42297842 | 0 |
| 0.00087616 | 0.00195028 | 12.8608998 | 80.7459351 | 38.0501286 | 46.3936151 | 4.75243649 |
| 0.00087794 | 0.00195412 | 97.4213163 | 214.655172 | 0 | 181.862971 | 0 |
| 0.00087865 | 0.00195556 | 631.791705 | 2177.47096 | 1975.82253 | 2012.09109 | 99.8011662 |
| 0.00087918 | 0.0019566 | 25.4002772 | 72.2931925 | 68.6758419 | 59.8477635 | 0 |
| 0.0008799 | 0.00195808 | 5.46588243 | 23.5787028 | 27.8415575 | 24.588616 | 0 |
| 0.00088073 | 0.0019598 | 5.78740493 | 29.8070394 | 29.6976613 | 18.0935099 | 0 |
| 0.00088086 | 0.00195996 | 17.3622148 | 3.55904948 | 67.7477899 | 45.929679 | 0 |
| 0.00088102 | 0.00196005 | 209.95419 | 446.438269 | 309.96934 | 366.045623 | 30.8908372 |
| 0.00088096 | 0.00196005 | 8.6811074 | 18.9074504 | 18.5610383 | 13.9180845 | 0 |
| 0.00088332 | 0.00196505 | 8.0380624 | 39.3719849 | 39.9062324 | 49.177232 | 0 |
| 0.00088369 | 0.00196573 | 32.1522496 | 18.017688 | 22.273246 | 19.9492545 | 2.37621824 |
| 0.00088409 | 0.00196649 | 143.720556 | 153.484009 | 155.912722 | 164.233398 | 66.5341108 |
| 0.0008853 | 0.00196905 | 3.21522496 | 35.3680542 | 49.1867516 | 28.7640414 | 0 |
| 0.00088694 | 0.00197257 | 81.666714 | 316.977844 | 390.709857 | 425.429451 | 23.7621824 |
| 0.00088755 | 0.0019738 | 237.926647 | 371.69823 | 253.358173 | 289.960095 | 0 |
| 0.00088804 | 0.00197476 | 10.9317649 | 9.78738607 | 25.9854537 | 19.9492545 | 0 |
| 0.00088901 | 0.00197679 | 41.476402 | 36.9251383 | 3.71220767 | 23.6607437 | 0 |
| 0.00088926 | 0.0019772 | 6.75197242 | 27.1377523 | 15.7768826 | 25.9804245 | 0 |
| 0.00089064 | 0.00198016 | 0 | 17.3503662 | 0 | 30.1558498 | 824.54773 |
| 0.00089072 | 0.00198021 | 4.17979245 | 53.8306234 | 18.5610383 | 37.5788283 | 0 |
| 0.00089242 | 0.00198384 | 31.1876821 | 16.0157227 | 7.42441534 | 18.0935099 | 0 |

| | | | | | | |
|---|---|---|---|---|---|---|
| 0.00089385 | 0.00198689 | 25.4002772 | 15.1259603 | 10.2085711 | 12.0623399 | 0 |
| 0.00089421 | 0.00198757 | 203.845263 | 42.9310343 | 0 | 393.417856 | 0 |
| 0.00089468 | 0.00198848 | 10.9317649 | 30.4743612 | 41.7623363 | 31.5476583 | 0 |
| 0.00089547 | 0.0019901 | 12.2178549 | 8.23030192 | 28.7696094 | 24.1246799 | 0 |
| 0.0008956 | 0.00199024 | 14.1469898 | 18.6850098 | 10.2085711 | 12.5262761 | 0 |
| 0.0008958 | 0.00199057 | 0 | 217.546899 | 162.409085 | 276.04201 | 0 |
| 0.00089649 | 0.00199198 | 14.4685123 | 21.1318563 | 8.35246725 | 16.7017014 | 0 |
| 0.00089671 | 0.00199232 | 51.7651219 | 92.3128459 | 86.3088283 | 74.2297842 | 11.8810912 |
| 0.00090312 | 0.00200644 | 58.1955718 | 117.003752 | 240.365446 | 155.418611 | 23.7621824 |
| 0.00090374 | 0.00200769 | 322.487064 | 443.991423 | 409.270895 | 424.965515 | 54.6530196 |
| 0.00090478 | 0.00200973 | 12.5393773 | 46.2676432 | 20.4171422 | 39.898509 | 0 |
| 0.00090476 | 0.00200973 | 190.66284 | 205.090226 | 135.49558 | 186.966269 | 78.415202 |
| 0.00090571 | 0.00201167 | 3.21522496 | 53.3857422 | 21.3451941 | 47.3214874 | 0 |
| 0.00090656 | 0.00201342 | 4.50131495 | 38.9271037 | 17.6329864 | 45.4657428 | 0 |
| 0.0009068 | 0.00201369 | 18.6483048 | 3.55904948 | 71.4599976 | 38.9706367 | 0 |
| 0.00090676 | 0.00201369 | 7.07349491 | 22.2440592 | 22.273246 | 16.7017014 | 0 |
| 0.00090718 | 0.0020144 | 76.2008316 | 247.131498 | 144.776099 | 205.059779 | 0 |
| 0.00090946 | 0.00201933 | 11.8963324 | 17.5728068 | 10.2085711 | 15.309893 | 0 |
| 0.00091 | 0.00202039 | 18.0052598 | 11.7893514 | 15.7768826 | 10.2065953 | 0 |
| 0.00091371 | 0.00202849 | 9.00262989 | 8.8976237 | 47.3306478 | 32.0115944 | 0 |
| 0.00091397 | 0.00202876 | 25.7217997 | 20.9094157 | 5.5683115 | 24.588616 | 0 |
| 0.00091401 | 0.00202876 | 0 | 0 | 0 | 0 | 40.3957101 |
| 0.00091391 | 0.00202876 | 25.0787547 | 18.2401286 | 15.7768826 | 12.0623399 | 35.6432737 |
| 0.00091462 | 0.00202998 | 12.5393773 | 15.3484009 | 12.9927268 | 11.1344676 | 0 |
| 0.0009157 | 0.00203225 | 10.6102424 | 9.34250488 | 25.9854537 | 24.1246799 | 0 |
| 0.00091606 | 0.00203291 | 161.404293 | 137.468286 | 143.848047 | 110.88074 | 28.5146189 |
| 0.0009166 | 0.00203397 | 4.50131495 | 26.6928711 | 24.1293498 | 40.8263813 | 0 |
| 0.00091675 | 0.00203416 | 134.396403 | 466.013041 | 454.745439 | 494.555937 | 16.6335277 |
| 0.00091706 | 0.00203472 | 38.261177 | 29.8070394 | 24.1293498 | 19.9492545 | 2.37621824 |
| 0.00091806 | 0.00203681 | 26.6863672 | 12.9015544 | 29.6976613 | 7.42297842 | 0 |
| 0.00091817 | 0.00203691 | 21.2204847 | 28.2499552 | 4.64025958 | 37.5788283 | 0 |
| 0.00091899 | 0.00203847 | 8.3595849 | 15.1259603 | 27.8415575 | 15.7738291 | 0 |
| 0.00091895 | 0.00203847 | 6.75197242 | 28.4723958 | 16.7049345 | 21.8049991 | 0 |
| 0.00091908 | 0.00203853 | 27.9724572 | 37.8149007 | 29.6976613 | 39.898509 | 9.50487297 |
| 0.00092089 | 0.00204241 | 6.75197242 | 79.1888509 | 7.42441534 | 48.7132959 | 0 |
| 0.00092542 | 0.00205232 | 5.46588243 | 21.1318563 | 40.8342843 | 21.8049991 | 0 |
| 0.00092896 | 0.00206004 | 742.073921 | 511.168481 | 461.241803 | 552.547956 | 42.7719284 |
| 0.00093022 | 0.0020627 | 9.64567488 | 39.3719849 | 11.136623 | 19.4853184 | 0 |
| 0.00093054 | 0.00206327 | 81.023669 | 118.338395 | 120.646749 | 108.561059 | 38.0194919 |
| 0.00093165 | 0.00206557 | 5.14435994 | 35.1456136 | 27.8415575 | 19.9492545 | 0 |
| 0.0009317 | 0.00206557 | 21.8635297 | 14.9035197 | 22.273246 | 7.42297842 | 0 |
| 0.00093653 | 0.00207614 | 605.748383 | 1303.94675 | 941.972696 | 1200.66676 | 0 |
| 0.00094066 | 0.00208516 | 20.2559173 | 18.9074504 | 41.7623363 | 6.03116997 | 0 |

| | | | | | | |
|---|---|---|---|---|---|---|
| 0.0009429 | 0.00208999 | 14.1469898 | 9.12006429 | 20.4171422 | 14.3820207 | 0 |
| 0.00094361 | 0.00209141 | 14.4685123 | 25.3582275 | 21.3451941 | 7.42297842 | 0 |
| 0.00094425 | 0.0020927 | 32.1522496 | 45.6003214 | 58.4672708 | 38.0427644 | 11.8810912 |
| 0.00094581 | 0.00209602 | 38.5826995 | 78.7439697 | 83.5246725 | 79.797018 | 16.6335277 |
| 0.00094641 | 0.00209722 | 0 | 0.22244059 | 0 | 0 | 0 |
| 0.00094817 | 0.00210098 | 9.32415239 | 18.6850098 | 16.7049345 | 12.5262761 | 0 |
| 0.000949 | 0.00210266 | 22.1850522 | 24.9133464 | 17.6329864 | 6.49510612 | 0 |
| 0.00095013 | 0.00210503 | 25.4002772 | 16.905485 | 7.42441534 | 16.2377653 | 0 |
| 0.0009503 | 0.00210527 | 14.4685123 | 18.017688 | 64.9636342 | 7.88691457 | 0 |
| 0.00095113 | 0.00210698 | 33.7598621 | 103.657316 | 99.3015551 | 87.6839326 | 16.6335277 |
| 0.00095251 | 0.00210989 | 22.8280972 | 13.7913167 | 9.28051917 | 13.9180845 | 0 |
| 0.00095385 | 0.00211272 | 119.606369 | 167.720207 | 0 | 167.017014 | 0 |
| 0.00095484 | 0.00211477 | 39.2257445 | 201.086296 | 78.8844129 | 167.017014 | 19.0097459 |
| 0.00095572 | 0.00211658 | 80.380624 | 14.4586385 | 8.35246725 | 16.2377653 | 0 |
| 0.00095636 | 0.00211759 | 9.64567488 | 5.7834554 | 9.28051917 | 5.56723382 | 30.8908372 |
| 0.00095631 | 0.00211759 | 10.9317649 | 10.4547078 | 18.5610383 | 16.2377653 | 0 |
| 0.00095624 | 0.00211759 | 10.2887199 | 16.2381632 | 11.136623 | 18.0935099 | 0 |
| 0.00095757 | 0.00212012 | 4.17979245 | 29.1397176 | 26.9135056 | 33.867339 | 0 |
| 0.0009581 | 0.00212117 | 9.00262989 | 38.0373413 | 12.9927268 | 17.1656376 | 0 |
| 0.00095893 | 0.00212285 | 0 | 0 | 0 | 0 | 21.3859642 |
| 0.0009613 | 0.00212796 | 15.1115573 | 11.7893514 | 11.136623 | 13.4541484 | 0 |
| 0.00096197 | 0.00212931 | 2.25065747 | 68.0668213 | 51.0428554 | 24.1246799 | 0 |
| 0.00096493 | 0.00213572 | 14.4685123 | 10.2322673 | 16.7049345 | 12.0623399 | 0 |
| 0.00096507 | 0.00213588 | 7.39501741 | 39.1495443 | 12.0646749 | 26.4443606 | 0 |
| 0.00096602 | 0.00213786 | 4.50131495 | 15.7932821 | 38.9781805 | 51.0329766 | 0 |
| 0.00096683 | 0.00213951 | 255.931907 | 148.145435 | 0 | 86.2921241 | 0 |
| 0.00096943 | 0.00214511 | 3.53674746 | 2.44684652 | 3.71220767 | 2.78361691 | 23.7621824 |
| 0.00096968 | 0.00214554 | 5.46588243 | 26.0255493 | 22.273246 | 24.588616 | 0 |
| 0.00097078 | 0.00214781 | 24.1141872 | 17.7952474 | 8.35246725 | 12.9902122 | 0 |
| 0.00097149 | 0.00214925 | 94.5276139 | 261.145256 | 216.236097 | 254.700947 | 0 |
| 0.00097178 | 0.00214974 | 8.6811074 | 20.2420939 | 14.8488307 | 14.8459568 | 0 |
| 0.00097196 | 0.00215002 | 0 | 235.787028 | 293.264406 | 121.551272 | 0 |
| 0.00097209 | 0.00215016 | 2.57217997 | 1.55708415 | 0 | 2.31968076 | 59.4054561 |
| 0.00097276 | 0.0021515 | 29.5800696 | 12.9015544 | 7.42441534 | 23.1968076 | 0 |
| 0.00097457 | 0.00215535 | 11.5748099 | 11.3444702 | 14.8488307 | 13.4541484 | 0 |
| 0.0009759 | 0.00215816 | 19.2913498 | 26.0255493 | 30.6257133 | 29.6919137 | 2.37621824 |
| 0.00097646 | 0.00215925 | 11.2532874 | 26.9153117 | 9.28051917 | 18.5574461 | 0 |
| 0.00097799 | 0.00216249 | 13.5039448 | 26.6928711 | 28.7696094 | 6.95904227 | 0 |
| 0.0009789 | 0.00216436 | 7.39501741 | 26.0255493 | 30.6257133 | 12.5262761 | 0 |
| 0.00097904 | 0.00216452 | 9.96719738 | 3.55904948 | 3.71220767 | 5.56723382 | 42.7719284 |
| 0.00097964 | 0.00216572 | 0 | 160.379667 | 257.070381 | 205.523715 | 0 |
| 0.00098019 | 0.00216678 | 7.07349491 | 29.5845988 | 11.136623 | 51.0329766 | 0 |
| 0.0009803 | 0.00216688 | 65.5905892 | 12.9015544 | 10.2085711 | 14.8459568 | 0 |

| | | | | | | |
|---|---|---|---|---|---|---|
| 0.00098386 | 0.00217461 | 77.4869216 | 12.2342326 | 10.2085711 | 15.309893 | 0 |
| 0.0009864 | 0.00218008 | 9.32415239 | 61.8384847 | 5.5683115 | 51.0329766 | 0 |
| 0.00098884 | 0.00218528 | 77.1653991 | 114.779346 | 78.8844129 | 109.952868 | 28.5146189 |
| 0.00098888 | 0.00218528 | 8.6811074 | 21.5767375 | 12.9927268 | 17.1656376 | 0 |
| 0.00098931 | 0.00218608 | 15.4330798 | 34.923173 | 72.3880495 | 4.17542536 | 0 |
| 0.00099258 | 0.00219317 | 0 | 0 | 0 | 0 | 23.7621824 |
| 0.00099467 | 0.00219764 | 103.208721 | 167.942647 | 0 | 181.399035 | 0 |
| 0.00099517 | 0.0021986 | 11.5748099 | 15.5708415 | 12.0646749 | 12.0623399 | 0 |
| 0.00099671 | 0.00220186 | 7.07349491 | 12.9015544 | 28.7696094 | 28.3001052 | 0 |
| 0.00099715 | 0.00220268 | 24.7572322 | 20.0196533 | 28.7696094 | 33.867339 | 0 |
| 0.00099778 | 0.00220393 | 0 | 0 | 0 | 0 | 30.8908372 |
| 0.00099811 | 0.00220452 | 605.748383 | 37.1475789 | 0 | 109.024996 | 0 |
| 0.00099848 | 0.0022052 | 16.3976473 | 8.00786133 | 32.4818171 | 14.8459568 | 0 |
| 0.00099867 | 0.00220546 | 27.3294122 | 74.9624797 | 87.2368802 | 86.7560603 | 14.2573095 |
| 0.00099883 | 0.00220567 | 0 | 193.300875 | 178.185968 | 240.318926 | 0 |
| 0.0009999 | 0.00220788 | 0 | 0 | 0 | 0 | 7.12865473 |
| 0.00100129 | 0.0022108 | 4.82283744 | 37.5924601 | 25.0574018 | 21.8049991 | 0 |
| 0.00100135 | 0.0022108 | 8.3595849 | 21.7991781 | 12.0646749 | 22.7328714 | 0 |
| 0.00100156 | 0.00221113 | 308.018551 | 782.990885 | 648.70829 | 746.473267 | 0 |
| 0.0010018 | 0.00221152 | 17.6837373 | 13.7913167 | 12.0646749 | 10.2065953 | 0 |
| 0.0010019 | 0.00221159 | 39.8687895 | 80.9683756 | 105.797919 | 83.0445711 | 0 |
| 0.00100244 | 0.00221262 | 16.3976473 | 16.6830444 | 17.6329864 | 7.88691457 | 0 |
| 0.00100552 | 0.00221927 | 9.32415239 | 10.899589 | 19.4890903 | 27.8361691 | 0 |
| 0.00100676 | 0.00222187 | 10.9317649 | 19.3523315 | 10.2085711 | 15.7738291 | 0 |
| 0.00100778 | 0.00222397 | 14.1469898 | 25.8031087 | 22.273246 | 32.9394667 | 0 |
| 0.00100805 | 0.00222443 | 6.75197242 | 27.8050741 | 18.5610383 | 17.6295737 | 0 |
| 0.00100873 | 0.00222579 | 0 | 171.056816 | 379.573234 | 118.303719 | 0 |
| 0.00100895 | 0.00222611 | 27.9724572 | 18.4625692 | 6.49636342 | 17.6295737 | 0 |
| 0.00100908 | 0.00222627 | 108.996126 | 18.9074504 | 20.4171422 | 19.0213822 | 0 |
| 0.00101017 | 0.00222852 | 2.57217997 | 72.2931925 | 32.4818171 | 28.3001052 | 0 |
| 0.0010108 | 0.00222976 | 69.4488592 | 204.422904 | 200.459214 | 248.205841 | 28.5146189 |
| 0.00101106 | 0.00223019 | 9.96719738 | 26.9153117 | 41.7623363 | 8.35085072 | 0 |
| 0.0010118 | 0.00223167 | 9.00262989 | 72.7380737 | 8.35246725 | 26.9082968 | 0 |
| 0.00101186 | 0.00223167 | 4.17979245 | 5.56101481 | 0 | 4.63936151 | 190.097459 |
| 0.00101259 | 0.00223312 | 7.71653991 | 18.6850098 | 14.8488307 | 20.4131907 | 0 |
| 0.0010133 | 0.00223455 | 7.07349491 | 12.9015544 | 27.8415575 | 28.3001052 | 0 |
| 0.00101738 | 0.00224339 | 17.3622148 | 10.4547078 | 32.4818171 | 10.2065953 | 0 |
| 0.00102007 | 0.00224919 | 6.10892743 | 16.2381632 | 25.0574018 | 29.2279775 | 0 |
| 0.00102028 | 0.0022495 | 50.1575094 | 38.7046631 | 25.0574018 | 23.6607437 | 7.12865473 |
| 0.00102227 | 0.00225374 | 9.00262989 | 33.3660889 | 9.28051917 | 28.3001052 | 0 |
| 0.00102239 | 0.00225385 | 0.3215225 | 0.44488118 | 0 | 0 | 7.12865473 |
| 0.00102371 | 0.00225661 | 11.2532874 | 21.5767375 | 8.35246725 | 31.0837221 | 0 |
| 0.00102476 | 0.00225877 | 19.9343948 | 9.34250488 | 12.9927268 | 14.3820207 | 0 |

| | | | | | | |
|---|---|---|---|---|---|---|
| 0.00102616 | 0.00226171 | 22.8280972 | 19.1298909 | 15.7768826 | 16.2377653 | 78.415202 |
| 0.00102762 | 0.0022648 | 10.2887199 | 17.5728068 | 12.0646749 | 13.4541484 | 0 |
| 0.00102791 | 0.00226527 | 8.0380624 | 12.2342326 | 27.8415575 | 20.8771268 | 0 |
| 0.00102815 | 0.00226566 | 0 | 119.228158 | 290.48025 | 220.833608 | 0 |
| 0.00102842 | 0.00226612 | 8.6811074 | 20.0196533 | 16.7049345 | 12.9902122 | 0 |
| 0.00103079 | 0.00227119 | 14.1469898 | 14.9035197 | 16.7049345 | 8.81478687 | 0 |
| 0.00103144 | 0.00227247 | 2.57217997 | 26.9153117 | 44.546492 | 49.6411682 | 0 |
| 0.00103186 | 0.00227325 | 70.0919042 | 45.6003214 | 35.2659728 | 38.5067006 | 0 |
| 0.00103553 | 0.00228118 | 30.2231146 | 57.1672323 | 58.4672708 | 57.0641466 | 0 |
| 0.00103654 | 0.00228324 | 18.9698273 | 30.4743612 | 6.49636342 | 16.7017014 | 0 |
| 0.00103771 | 0.00228568 | 56.5879593 | 18.2401286 | 3.71220767 | 35.2591475 | 0 |
| 0.00103866 | 0.00228761 | 6.43044992 | 29.8070394 | 17.6329864 | 19.4853184 | 0 |
| 0.00103938 | 0.00228906 | 0.3215225 | 0.44488118 | 2.78415575 | 1.39180845 | 4.75243649 |
| 0.00104067 | 0.00229176 | 19.6128723 | 24.6909058 | 12.0646749 | 8.35085072 | 0 |
| 0.00104091 | 0.00229212 | 6.75197242 | 22.4664998 | 16.7049345 | 20.8771268 | 0 |
| 0.00104111 | 0.00229241 | 9.96719738 | 15.5708415 | 11.136623 | 25.5164883 | 0 |
| 0.00104125 | 0.00229257 | 201.273083 | 259.365731 | 232.012979 | 257.020628 | 66.5341108 |
| 0.00104313 | 0.00229657 | 117.034189 | 181.956405 | 0 | 304.342115 | 0 |
| 0.00104478 | 0.00230005 | 0 | 0 | 0 | 0 | 33.2670554 |
| 0.00104822 | 0.00230748 | 34.0813846 | 105.436841 | 48.2586997 | 27.8361691 | 9.50487297 |
| 0.00104927 | 0.00230963 | 38.261177 | 27.8050741 | 20.4171422 | 19.4853184 | 57.0292378 |
| 0.00105178 | 0.002315 | 4.50131495 | 31.3641235 | 18.5610383 | 41.2903175 | 0 |
| 0.00105185 | 0.00231501 | 8.6811074 | 20.0196533 | 13.9207788 | 14.8459568 | 0 |
| 0.00105495 | 0.00232168 | 35.3674746 | 103.434875 | 81.6685687 | 83.0445711 | 16.6335277 |
| 0.00105545 | 0.00232247 | 62.3753642 | 12.9015544 | 6.49636342 | 23.6607437 | 0 |
| 0.00105542 | 0.00232247 | 0 | 1.77952474 | 0 | 0 | 30.8908372 |
| 0.00105625 | 0.00232407 | 18.0052598 | 61.3936035 | 42.6903882 | 23.1968076 | 4.75243649 |
| 0.00105808 | 0.00232795 | 36.3320421 | 11.3444702 | 11.136623 | 9.74265918 | 35.6432737 |
| 0.00105886 | 0.00232951 | 16.3976473 | 26.2479899 | 19.4890903 | 23.6607437 | 2.37621824 |
| 0.00106333 | 0.00233921 | 2.25065747 | 34.0334106 | 48.2586997 | 40.8263813 | 0 |
| 0.00106354 | 0.00233951 | 4.50131495 | 14.6810791 | 44.546492 | 42.6821259 | 0 |
| 0.00106489 | 0.00234234 | 24.4357097 | 29.8070394 | 5.5683115 | 17.1656376 | 0 |
| 0.00106569 | 0.00234393 | 91.3123889 | 163.271395 | 195.818954 | 133.149675 | 28.5146189 |
| 0.00106682 | 0.00234626 | 25.4002772 | 16.4606038 | 22.273246 | 18.0935099 | 23.7621824 |
| 0.00106972 | 0.00235248 | 41.1548795 | 136.800964 | 0 | 441.20328 | 0 |
| 0.00107295 | 0.00235944 | 9.96719738 | 15.5708415 | 11.136623 | 17.1656376 | 0 |
| 0.0010772 | 0.00236862 | 5.78740493 | 21.7991781 | 22.273246 | 21.8049991 | 0 |
| 0.00107744 | 0.00236901 | 77.8084441 | 132.797034 | 0 | 266.763287 | 0 |
| 0.00107788 | 0.00236982 | 39.2257445 | 25.1357869 | 3.71220767 | 27.8361691 | 0 |
| 0.00108247 | 0.00237975 | 7.07349491 | 20.6869751 | 13.9207788 | 31.5476583 | 0 |
| 0.00108465 | 0.0023844 | 17.6837373 | 17.7952474 | 6.49636342 | 23.6607437 | 0 |
| 0.00108613 | 0.00238749 | 216.706162 | 104.324638 | 0 | 121.551272 | 0 |
| 0.00108698 | 0.0023892 | 29.5800696 | 17.7952474 | 20.4171422 | 13.9180845 | 19.0097459 |

| | | | | | | |
|---|---|---|---|---|---|---|
| 0.00109057 | 0.00239692 | 10.9317649 | 13.5688761 | 12.9927268 | 12.0623399 | 0 |
| 0.00109173 | 0.00239933 | 7.39501741 | 3.78149007 | 2.78415575 | 3.24755306 | 11.8810912 |
| 0.00109311 | 0.0024022 | 11.8963324 | 24.6909058 | 7.42441534 | 26.4443606 | 0 |
| 0.00109697 | 0.00241053 | 4.50131495 | 50.2715739 | 12.9927268 | 45.0018067 | 0 |
| 0.00109858 | 0.00241391 | 26.0433222 | 12.6791138 | 14.8488307 | 11.5984038 | 21.3859642 |
| 0.00109916 | 0.00241503 | 11.5748099 | 31.5865641 | 38.0501286 | 23.6607437 | 2.37621824 |
| 0.00110046 | 0.00241773 | 12.8608998 | 41.8188314 | 48.2586997 | 16.7017014 | 2.37621824 |
| 0.00110086 | 0.00241843 | 13.8254673 | 18.6850098 | 7.42441534 | 21.341063 | 0 |
| 0.00110133 | 0.00241932 | 4.50131495 | 21.1318563 | 38.0501286 | 25.9804245 | 0 |
| 0.0011016 | 0.00241975 | 10.6102424 | 12.9015544 | 15.7768826 | 7.88691457 | 35.6432737 |
| 0.00110428 | 0.00242549 | 9.32415239 | 14.6810791 | 34.3379209 | 12.5262761 | 0 |
| 0.00110634 | 0.00242984 | 9.64567488 | 14.2361979 | 12.0646749 | 16.7017014 | 0 |
| 0.00110673 | 0.00243054 | 115.105054 | 0 | 280.271679 | 278.825627 | 0 |
| 0.00110767 | 0.00243246 | 2.89370247 | 11.3444702 | 68.6758419 | 76.549465 | 0 |
| 0.00111002 | 0.00243745 | 94.8491364 | 294.733785 | 304.401029 | 356.7669 | 38.0194919 |
| 0.00111086 | 0.00243914 | 231.496197 | 228.224048 | 141.063891 | 209.69914 | 40.3957101 |
| 0.00111513 | 0.00244836 | 10.6102424 | 43.3759155 | 25.0574018 | 32.4755306 | 2.37621824 |
| 0.00111593 | 0.00244995 | 6.75197242 | 29.5845988 | 14.8488307 | 20.4131907 | 0 |
| 0.00111743 | 0.00245308 | 23.7926647 | 8.00786133 | 13.9207788 | 16.2377653 | 0 |
| 0.00111938 | 0.0024572 | 80.7021465 | 91.6455241 | 84.4527244 | 108.097123 | 28.5146189 |
| 0.00112092 | 0.00246043 | 20.8989622 | 24.9133464 | 35.2659728 | 4.63936151 | 0 |
| 0.00112153 | 0.0024616 | 3.21522496 | 32.2538859 | 36.1940248 | 30.1558498 | 0 |
| 0.00112254 | 0.00246366 | 75.2362641 | 341.66875 | 222.73246 | 258.876372 | 33.2670554 |
| 0.00112418 | 0.00246709 | 12.8608998 | 15.7932821 | 12.9927268 | 9.74265918 | 0 |
| 0.00112569 | 0.00247025 | 0 | 176.172949 | 200.459214 | 193.925311 | 0 |
| 0.00112647 | 0.0024718 | 53.3727344 | 125.234054 | 90.9490879 | 109.952868 | 30.8908372 |
| 0.00112887 | 0.00247691 | 3.21522496 | 23.5787028 | 39.9062324 | 39.4345729 | 0 |
| 0.0011293 | 0.00247762 | 4.50131495 | 42.4861532 | 18.5610383 | 28.3001052 | 0 |
| 0.00112934 | 0.00247762 | 12.5393773 | 10.6771484 | 12.9927268 | 12.9902122 | 0 |
| 0.00112978 | 0.00247842 | 51.1220769 | 56.7223511 | 62.1794784 | 57.0641466 | 2.37621824 |
| 0.00113012 | 0.002479 | 9.00262989 | 13.123995 | 15.7768826 | 14.8459568 | 0 |
| 0.00113123 | 0.00248126 | 0 | 162.381632 | 256.142329 | 161.913717 | 0 |
| 0.00113136 | 0.0024814 | 74.9147416 | 133.909237 | 147.560255 | 142.428398 | 33.2670554 |
| 0.00113145 | 0.00248143 | 24.4357097 | 16.6830444 | 16.7049345 | 6.95904227 | 0 |
| 0.00113414 | 0.00248716 | 15.7546023 | 23.8011434 | 15.7768826 | 7.42297842 | 0 |
| 0.00113431 | 0.00248738 | 7.39501741 | 21.3542969 | 12.9927268 | 24.1246799 | 0 |
| 0.00113462 | 0.00248789 | 8.6811074 | 14.4586385 | 15.7768826 | 14.3820207 | 0 |
| 0.00113739 | 0.00249382 | 15.7546023 | 18.017688 | 7.42441534 | 16.2377653 | 0 |
| 0.00113849 | 0.00249607 | 6.10892743 | 15.3484009 | 23.2012979 | 30.619786 | 0 |
| 0.00113891 | 0.00249681 | 6.43044992 | 24.2460246 | 14.8488307 | 25.0525522 | 0 |
| 0.00113954 | 0.00249796 | 9.96719738 | 12.011792 | 16.7049345 | 12.9902122 | 0 |
| 0.00113958 | 0.00249796 | 9.32415239 | 21.1318563 | 12.0646749 | 14.3820207 | 0 |
| 0.00114112 | 0.00250118 | 0 | 0 | 0 | 0 | 42.7719284 |

| | | | | | | |
|---|---|---|---|---|---|---|
| 0.00114343 | 0.00250608 | 91.9554339 | 71.4034302 | 76.1002572 | 86.7560603 | 116.434694 |
| 0.00114404 | 0.00250725 | 12.5393773 | 35.3680542 | 23.2012979 | 6.95904227 | 0 |
| 0.00114587 | 0.00251109 | 5.78740493 | 14.6810791 | 31.5537652 | 27.8361691 | 0 |
| 0.00114825 | 0.00251615 | 5.46588243 | 23.8011434 | 25.0574018 | 19.4853184 | 0 |
| 0.00115137 | 0.00252281 | 11.8963324 | 14.6810791 | 9.28051917 | 28.7640414 | 0 |
| 0.00115177 | 0.00252348 | 15.4330798 | 63.6180094 | 8.35246725 | 13.4541484 | 0 |
| 0.00115182 | 0.00252348 | 86.168029 | 199.529211 | 0 | 146.139888 | 0 |
| 0.00115505 | 0.00253038 | 10.9317649 | 10.6771484 | 13.9207788 | 14.8459568 | 0 |
| 0.00115634 | 0.00253305 | 8.3595849 | 26.2479899 | 12.9927268 | 15.7738291 | 0 |
| 0.00115863 | 0.0025379 | 2.25065747 | 40.7066284 | 42.6903882 | 34.3312752 | 0 |
| 0.00116 | 0.00254065 | 7.71653991 | 24.6909058 | 13.9207788 | 16.7017014 | 0 |
| 0.00116004 | 0.00254065 | 7.07349491 | 20.2420939 | 14.8488307 | 20.4131907 | 0 |
| 0.00116018 | 0.0025408 | 70.7349491 | 255.584241 | 0 | 135.933292 | 0 |
| 0.00116045 | 0.00254122 | 33.7598621 | 36.2578166 | 24.1293498 | 23.1968076 | 2.37621824 |
| 0.00116138 | 0.00254309 | 10.6102424 | 22.0216187 | 9.28051917 | 17.1656376 | 0 |
| 0.00116256 | 0.00254551 | 169.763878 | 105.881722 | 0 | 138.252973 | 0 |
| 0.00116318 | 0.00254671 | 0 | 184.848132 | 306.257133 | 109.488932 | 0 |
| 0.00116343 | 0.00254709 | 13.5039448 | 16.0157227 | 12.0646749 | 9.74265918 | 0 |
| 0.00116354 | 0.00254717 | 119.927891 | 280.052706 | 401.84648 | 218.977863 | 0 |
| 0.00116413 | 0.00254829 | 12.8608998 | 18.6850098 | 7.42441534 | 29.6919137 | 0 |
| 0.00116607 | 0.00255238 | 4.82283744 | 41.5963908 | 12.9927268 | 42.6821259 | 0 |
| 0.00116642 | 0.00255297 | 6.43044992 | 24.023584 | 13.9207788 | 29.2279775 | 0 |
| 0.00116695 | 0.00255395 | 42.762492 | 38.0373413 | 30.6257133 | 38.5067006 | 40.3957101 |
| 0.00116997 | 0.00256034 | 7.71653991 | 13.123995 | 14.8488307 | 13.4541484 | 35.6432737 |
| 0.00117004 | 0.00256034 | 2.25065747 | 62.7282471 | 58.4672708 | 18.5574461 | 0 |
| 0.00117009 | 0.00256034 | 4.17979245 | 38.4822225 | 21.3451941 | 27.3722329 | 0 |
| 0.00117091 | 0.00256197 | 99.9934963 | 0 | 99.3015551 | 633.736783 | 0 |
| 0.00117167 | 0.00256345 | 11.8963324 | 8.23030192 | 20.4171422 | 19.4853184 | 0 |
| 0.00117333 | 0.00256692 | 25.7217997 | 76.2971232 | 50.1148035 | 58.4559551 | 11.8810912 |
| 0.00117342 | 0.00256694 | 77.8084441 | 144.586385 | 0 | 216.658183 | 0 |
| 0.00117509 | 0.00257043 | 11.5748099 | 64.2853312 | 56.6111669 | 108.097123 | 0 |
| 0.0011756 | 0.00257138 | 11.8963324 | 8.45274251 | 19.4890903 | 18.0935099 | 0 |
| 0.00117949 | 0.00257973 | 29.2585471 | 22.4664998 | 5.5683115 | 16.7017014 | 0 |
| 0.00118007 | 0.00258083 | 6.10892743 | 23.3562622 | 15.7768826 | 25.5164883 | 0 |
| 0.00118056 | 0.00258172 | 7.39501741 | 28.4723958 | 30.6257133 | 10.6705315 | 0 |
| 0.00118093 | 0.00258237 | 9.00262989 | 13.7913167 | 20.4171422 | 12.5262761 | 0 |
| 0.00118234 | 0.0025853 | 18.0052598 | 18.9074504 | 7.42441534 | 13.9180845 | 0 |
| 0.00118402 | 0.00258862 | 20.2559173 | 38.9271037 | 32.4818171 | 31.0837221 | 7.12865473 |
| 0.00118395 | 0.00258862 | 1.60761248 | 1.33464355 | 1.85610383 | 0.9278723 | 0 |
| 0.00118527 | 0.00259119 | 22.1850522 | 14.0137573 | 10.2085711 | 10.6705315 | 0 |
| 0.00118566 | 0.00259187 | 53.3727344 | 123.89941 | 94.6612955 | 62.1674443 | 19.0097459 |
| 0.00118812 | 0.00259708 | 0 | 174.615865 | 293.264406 | 118.303719 | 0 |
| 0.00118887 | 0.00259856 | 11.5748099 | 10.6771484 | 15.7768826 | 12.0623399 | 0 |

| | | | | | | |
|---|---|---|---|---|---|---|
| 0.0011899 | 0.00260064 | 163.333428 | 232.8953 | 271.919212 | 289.496158 | 2.37621824 |
| 0.00119223 | 0.0026054 | 7.39501741 | 38.4822225 | 22.273246 | 11.5984038 | 0 |
| 0.00119223 | 0.0026054 | 35.6889971 | 13.5688761 | 63.1075304 | 4.17542536 | 0 |
| 0.00119343 | 0.00260784 | 4.17979245 | 45.6003214 | 39.9062324 | 14.8459568 | 0 |
| 0.00119466 | 0.00261036 | 11.2532874 | 15.3484009 | 10.2085711 | 13.4541484 | 0 |
| 0.00119582 | 0.00261273 | 89.7047764 | 181.956405 | 0 | 145.675952 | 0 |
| 0.00120125 | 0.00262442 | 11.2532874 | 14.9035197 | 12.0646749 | 11.1344676 | 0 |
| 0.00120261 | 0.00262722 | 19.2913498 | 8.23030192 | 36.1940248 | 11.1344676 | 0 |
| 0.00120328 | 0.0026285 | 7.39501741 | 14.9035197 | 21.3451941 | 16.2377653 | 0 |
| 0.0012037 | 0.00262925 | 0 | 237.121672 | 316.465704 | 74.2297842 | 0 |
| 0.00120402 | 0.00262979 | 54.6588243 | 20.0196533 | 19.4890903 | 16.7017014 | 2.37621824 |
| 0.00120636 | 0.00263472 | 5.46588243 | 33.3660889 | 12.0646749 | 45.4657428 | 0 |
| 0.00120701 | 0.00263598 | 12.5393773 | 6.67321777 | 25.9854537 | 29.2279775 | 0 |
| 0.00120829 | 0.0026386 | 35.6889971 | 292.73182 | 337.810898 | 323.363497 | 35.6432737 |
| 0.00120895 | 0.00263987 | 82.309759 | 115.001786 | 0 | 244.958288 | 0 |
| 0.00120905 | 0.00263991 | 7.07349491 | 19.7972127 | 17.6329864 | 15.7738291 | 0 |
| 0.00120939 | 0.00264049 | 10.2887199 | 14.9035197 | 10.2085711 | 27.3722329 | 0 |
| 0.00121164 | 0.00264524 | 14.1469898 | 11.1220296 | 12.0646749 | 9.74265918 | 14.2573095 |
| 0.00121258 | 0.00264697 | 9.96719738 | 14.0137573 | 12.0646749 | 13.4541484 | 0 |
| 0.0012126 | 0.00264697 | 21.5420072 | 251.58031 | 277.487523 | 160.985845 | 21.3859642 |
| 0.00121394 | 0.00264973 | 6.43044992 | 22.6889404 | 17.6329864 | 18.0935099 | 0 |
| 0.00121574 | 0.00265349 | 90.9908664 | 227.111845 | 207.883629 | 0 | 0 |
| 0.00121702 | 0.0026561 | 9.64567488 | 45.822762 | 25.0574018 | 7.88691457 | 0 |
| 0.0012173 | 0.00265656 | 12.8608998 | 57.6121134 | 49.1867516 | 33.4034029 | 0 |
| 0.00121861 | 0.00265924 | 150.472528 | 409.29069 | 348.947521 | 413.367111 | 0 |
| 0.00122075 | 0.00266374 | 18.0052598 | 11.7893514 | 12.9927268 | 9.74265918 | 0 |
| 0.00122098 | 0.00266405 | 175.22976 | 164.383598 | 119.718697 | 151.707121 | 9.50487297 |
| 0.00122176 | 0.00266558 | 73.9501741 | 188.629622 | 0 | 164.233398 | 0 |
| 0.00122283 | 0.00266776 | 5.14435994 | 20.2420939 | 25.0574018 | 24.1246799 | 0 |
| 0.00122299 | 0.00266793 | 12.2178549 | 18.2401286 | 11.136623 | 10.6705315 | 0 |
| 0.00122463 | 0.00267133 | 0 | 200.196533 | 141.063891 | 210.627013 | 0 |
| 0.00122532 | 0.00267267 | 110.603739 | 120.562801 | 0 | 172.120312 | 0 |
| 0.00122728 | 0.00267677 | 14.1469898 | 8.67518311 | 12.9927268 | 18.5574461 | 0 |
| 0.00122749 | 0.00267705 | 210.918757 | 101.655351 | 199.531162 | 0 | 0 |
| 0.00122837 | 0.0026788 | 44.691627 | 23.3562622 | 31.5537652 | 21.341063 | 97.424948 |
| 0.00122898 | 0.00267996 | 14.1469898 | 26.4704305 | 7.42441534 | 15.7738291 | 0 |
| 0.00123357 | 0.00268979 | 78.4514891 | 95.6494547 | 47.3306478 | 83.9724434 | 19.0097459 |
| 0.0012343 | 0.0026912 | 7.07349491 | 24.4684652 | 12.0646749 | 25.9804245 | 0 |
| 0.00123448 | 0.00269143 | 2.57217997 | 5.56101481 | 7.42441534 | 1.85574461 | 49.9005831 |
| 0.00123461 | 0.00269153 | 21.8635297 | 11.7893514 | 10.2085711 | 12.0623399 | 0 |
| 0.00123667 | 0.00269584 | 6.75197242 | 15.3484009 | 22.273246 | 18.5574461 | 0 |
| 0.00123686 | 0.00269609 | 11.2532874 | 9.34250488 | 17.6329864 | 14.8459568 | 0 |
| 0.00123886 | 0.00270029 | 21.2204847 | 43.8207967 | 47.3306478 | 41.2903175 | 9.50487297 |

| | | | | | | |
|---|---|---|---|---|---|---|
| 0.00123932 | 0.00270111 | 37.2966096 | 12.4566732 | 7.42441534 | 18.5574461 | 0 |
| 0.00123942 | 0.00270116 | 10.2887199 | 24.023584 | 8.35246725 | 21.8049991 | 0 |
| 0.00124065 | 0.00270364 | 57.8740493 | 105.881722 | 99.3015551 | 89.5396772 | 21.3859642 |
| 0.00124087 | 0.00270395 | 95.4921813 | 238.901196 | 0 | 97.8905279 | 0 |
| 0.00124178 | 0.00270574 | 6.43044992 | 17.5728068 | 21.3451941 | 18.0935099 | 0 |
| 0.00124185 | 0.00270574 | 32.4737721 | 19.1298909 | 8.35246725 | 10.6705315 | 0 |
| 0.00124205 | 0.002706 | 23.4711422 | 20.4645345 | 5.5683115 | 18.5574461 | 0 |
| 0.00124217 | 0.00270608 | 9.00262989 | 26.2479899 | 27.8415575 | 8.81478687 | 0 |
| 0.00124279 | 0.00270709 | 93.5630464 | 167.275326 | 158.696878 | 174.439993 | 57.0292378 |
| 0.00124277 | 0.00270709 | 22.8280972 | 23.1338216 | 14.8488307 | 20.8771268 | 2.37621824 |
| 0.00124344 | 0.00270833 | 0.64304499 | 0.44488118 | 0 | 1.39180845 | 78.415202 |
| 0.00124409 | 0.00270956 | 77.1653991 | 159.934786 | 322.034015 | 0 | 0 |
| 0.00124732 | 0.00271642 | 186.483048 | 217.102018 | 219.948304 | 231.50414 | 64.1578926 |
| 0.00124743 | 0.00271645 | 212.847892 | 69.846346 | 0 | 148.923505 | 0 |
| 0.00124749 | 0.00271645 | 11.8963324 | 16.0157227 | 12.0646749 | 10.2065953 | 0 |
| 0.00124855 | 0.00271857 | 4.82283744 | 16.2381632 | 35.2659728 | 29.2279775 | 0 |
| 0.00124947 | 0.00272041 | 17.3622148 | 67.8443807 | 47.3306478 | 59.3838274 | 7.12865473 |
| 0.00125099 | 0.00272353 | 94.5276139 | 104.102197 | 41.7623363 | 70.9822312 | 28.5146189 |
| 0.00125165 | 0.00272479 | 55.9449143 | 40.929069 | 58.4672708 | 38.0427644 | 2.37621824 |
| 0.00125209 | 0.00272558 | 202.880695 | 113.667143 | 0 | 96.0347833 | 0 |
| 0.00125258 | 0.00272647 | 190.66284 | 89.4211182 | 0 | 129.902122 | 0 |
| 0.00125282 | 0.00272681 | 0 | 0 | 0 | 0 | 0 |
| 0.00125335 | 0.0027278 | 7.39501741 | 27.1377523 | 11.136623 | 14.3820207 | 40.3957101 |
| 0.00125386 | 0.00272874 | 76.2008316 | 80.5234945 | 87.2368802 | 69.5904227 | 168.711495 |
| 0.00125509 | 0.00273122 | 10.6102424 | 19.3523315 | 8.35246725 | 24.588616 | 0 |
| 0.00125536 | 0.00273164 | 18.9698273 | 6.89565837 | 15.7768826 | 19.0213822 | 0 |
| 0.00125668 | 0.00273434 | 0 | 0 | 0 | 0 | 33.2670554 |
| 0.0012587 | 0.00273854 | 419.586857 | 971.398067 | 852.879712 | 906.995176 | 0 |
| 0.00125966 | 0.00274047 | 178.123463 | 371.69823 | 382.35739 | 407.335941 | 26.1384007 |
| 0.00126014 | 0.00274133 | 8.3595849 | 16.6830444 | 12.0646749 | 18.5574461 | 0 |
| 0.00126026 | 0.00274142 | 40.190312 | 50.7164551 | 40.8342843 | 65.4149973 | 9.50487297 |
| 0.00126132 | 0.00274354 | 5.78740493 | 47.1574056 | 29.6976613 | 70.518295 | 2.37621824 |
| 0.00126164 | 0.00274405 | 5.46588243 | 31.1416829 | 21.3451941 | 17.6295737 | 0 |
| 0.00126239 | 0.00274552 | 143.399033 | 217.76934 | 0 | 69.5904227 | 0 |
| 0.0012625 | 0.00274558 | 6.75197242 | 40.4841878 | 36.1940248 | 9.27872303 | 0 |
| 0.00126718 | 0.00275558 | 56.9094818 | 135.24388 | 129.927268 | 115.056166 | 4.75243649 |
| 0.00126807 | 0.00275734 | 113.497441 | 88.3089152 | 96.5173994 | 73.7658481 | 299.403499 |
| 0.00127042 | 0.00276226 | 3.85826995 | 37.5924601 | 18.5610383 | 35.2591475 | 0 |
| 0.00127137 | 0.00276416 | 85.2034615 | 125.011613 | 153.128566 | 150.779249 | 0 |
| 0.00127256 | 0.00276655 | 263.648447 | 539.863318 | 454.745439 | 510.329766 | 0 |
| 0.00127386 | 0.0027692 | 8.6811074 | 32.9212077 | 12.0646749 | 15.309893 | 0 |
| 0.0012767 | 0.0027752 | 9.32415239 | 15.1259603 | 14.8488307 | 11.5984038 | 0 |
| 0.00127726 | 0.00277623 | 5.46588243 | 44.7105591 | 11.136623 | 35.7230837 | 0 |

| | | | | | | |
|---|---|---|---|---|---|---|
| 0.00127953 | 0.00278099 | 9.96719738 | 18.2401286 | 11.136623 | 12.9902122 | 0 |
| 0.00128071 | 0.00278337 | 107.710036 | 124.566732 | 0 | 160.057972 | 0 |
| 0.00128107 | 0.00278398 | 7.71653991 | 16.905485 | 12.9927268 | 20.8771268 | 0 |
| 0.0012814 | 0.00278451 | 3.85826995 | 30.9192423 | 24.1293498 | 28.7640414 | 0 |
| 0.00128148 | 0.00278451 | 12.8608998 | 3.78149007 | 17.6329864 | 0 | 128.315785 |
| 0.00128156 | 0.00278451 | 0 | 0 | 0 | 0 | 23.7621824 |
| 0.00128373 | 0.00278903 | 363.641943 | 1362.67107 | 1166.56126 | 1250.77186 | 16.6335277 |
| 0.00128463 | 0.00279082 | 8.0380624 | 26.9153117 | 9.28051917 | 31.5476583 | 0 |
| 0.00128622 | 0.00279409 | 66.2336342 | 11.5669108 | 7.42441534 | 18.5574461 | 0 |
| 0.00128688 | 0.00279522 | 7.71653991 | 25.3582275 | 35.2659728 | 9.74265918 | 0 |
| 0.00128691 | 0.00279522 | 0 | 0.22244059 | 0 | 0 | 21.3859642 |
| 0.00129042 | 0.00280268 | 12.8608998 | 23.5787028 | 25.9854537 | 26.9082968 | 0 |
| 0.00129216 | 0.00280627 | 0 | 0 | 0 | 0 | 0 |
| 0.00129306 | 0.00280805 | 9.00262989 | 16.905485 | 18.5610383 | 10.6705315 | 0 |
| 0.00129349 | 0.00280878 | 97.7428388 | 185.515454 | 154.98467 | 184.646588 | 30.8908372 |
| 0.0012964 | 0.00281493 | 15.4330798 | 4.67125244 | 2.78415575 | 5.56723382 | 47.5243649 |
| 0.001299 | 0.0028204 | 0 | 248.466142 | 158.696878 | 136.397228 | 0 |
| 0.00130089 | 0.00282431 | 0 | 232.005538 | 376.789078 | 51.4969128 | 0 |
| 0.00130194 | 0.0028264 | 0 | 217.76934 | 90.9490879 | 260.268181 | 0 |
| 0.0013037 | 0.00283006 | 0 | 171.946578 | 166.121293 | 189.749886 | 0 |
| 0.00130399 | 0.00283051 | 0.3215225 | 0.44488118 | 0.92805192 | 0 | 0 |
| 0.00130488 | 0.00283225 | 139.219241 | 101.655351 | 270.063108 | 0 | 0 |
| 0.00130519 | 0.00283273 | 11.8963324 | 48.7144897 | 6.49636342 | 22.2689353 | 0 |
| 0.00130621 | 0.00283477 | 6.10892743 | 30.02948 | 14.8488307 | 20.8771268 | 0 |
| 0.0013071 | 0.00283652 | 0 | 316.310522 | 91.8771398 | 173.512121 | 0 |
| 0.00130829 | 0.00283893 | 29.5800696 | 18.017688 | 16.7049345 | 19.9492545 | 0 |
| 0.00131348 | 0.00285 | 4.17979245 | 26.0255493 | 19.4890903 | 39.898509 | 0 |
| 0.00131642 | 0.00285619 | 2.25065747 | 92.0904053 | 46.4025958 | 68.1986142 | 0 |
| 0.00131662 | 0.00285644 | 334.383396 | 524.959798 | 588.384915 | 604.508805 | 0 |
| 0.00131943 | 0.00286235 | 11.2532874 | 22.911381 | 24.1293498 | 7.42297842 | 0 |
| 0.00132149 | 0.00286664 | 0 | 192.188672 | 167.977397 | 164.233398 | 0 |
| 0.00132169 | 0.00286688 | 177.480418 | 114.334465 | 0 | 228.720523 | 0 |
| 0.00132291 | 0.00286935 | 306.089416 | 797.894405 | 633.859459 | 772.453692 | 28.5146189 |
| 0.00132314 | 0.00286967 | 20.5774398 | 13.5688761 | 10.2085711 | 10.2065953 | 0 |
| 0.00132337 | 0.00286997 | 44.048582 | 18.6850098 | 26.9135056 | 25.5164883 | 4.75243649 |
| 0.00132416 | 0.00287149 | 23.4711422 | 14.9035197 | 5.5683115 | 28.3001052 | 0 |
| 0.00132542 | 0.00287404 | 18.6483048 | 23.8011434 | 6.49636342 | 14.3820207 | 0 |
| 0.0013277 | 0.00287881 | 1.60761248 | 0.88976237 | 1.85610383 | 0.9278723 | 2.37621824 |
| 0.0013283 | 0.00287992 | 33.7598621 | 45.822762 | 34.3379209 | 30.619786 | 2.37621824 |
| 0.00132962 | 0.0028826 | 6.75197242 | 26.4704305 | 12.0646749 | 24.588616 | 0 |
| 0.00133055 | 0.00288442 | 14.4685123 | 7.78542074 | 18.5610383 | 14.3820207 | 0 |
| 0.0013308 | 0.00288478 | 24.4357097 | 16.2381632 | 7.42441534 | 12.9902122 | 0 |
| 0.0013332 | 0.00288981 | 47.5853294 | 34.923173 | 5.5683115 | 10.2065953 | 0 |

| | | | | | | |
|---|---|---|---|---|---|---|
| 0.00133685 | 0.00289754 | 4.50131495 | 33.8109701 | 16.7049345 | 30.619786 | 0 |
| 0.00134176 | 0.00290798 | 6.43044992 | 18.4625692 | 15.7768826 | 22.2689353 | 0 |
| 0.00134565 | 0.00291622 | 10.9317649 | 20.9094157 | 15.7768826 | 8.81478687 | 0 |
| 0.00134734 | 0.00291971 | 6.10892743 | 18.017688 | 21.3451941 | 18.0935099 | 0 |
| 0.00134813 | 0.00292122 | 11.5748099 | 10.2322673 | 15.7768826 | 11.5984038 | 0 |
| 0.00134826 | 0.00292131 | 29.5800696 | 15.5708415 | 8.35246725 | 11.5984038 | 0 |
| 0.00134934 | 0.00292346 | 4.82283744 | 24.023584 | 21.3451941 | 24.1246799 | 0 |
| 0.00135235 | 0.00292981 | 6.75197242 | 16.4606038 | 21.3451941 | 15.7738291 | 0 |
| 0.00135297 | 0.00293097 | 18.0052598 | 26.9153117 | 4.64025958 | 26.9082968 | 0 |
| 0.00135525 | 0.0029357 | 0 | 2.00196533 | 0 | 0 | 16.6335277 |
| 0.00135578 | 0.00293667 | 7.07349491 | 12.011792 | 22.273246 | 23.6607437 | 0 |
| 0.00135624 | 0.00293747 | 11.8963324 | 13.5688761 | 12.0646749 | 10.2065953 | 0 |
| 0.00135728 | 0.00293946 | 218.313775 | 103.434875 | 0 | 86.2921241 | 0 |
| 0.00135733 | 0.00293946 | 77.1653991 | 157.265499 | 0 | 161.449781 | 0 |
| 0.00135769 | 0.00294004 | 60.1247068 | 164.161157 | 0 | 196.244992 | 0 |
| 0.00135845 | 0.00294145 | 148.543393 | 271.155082 | 312.753496 | 316.868391 | 42.7719284 |
| 0.00135851 | 0.00294145 | 43.405537 | 8.8976237 | 15.7768826 | 11.1344676 | 0 |
| 0.00135887 | 0.00294202 | 232.460765 | 390.828121 | 409.270895 | 398.521154 | 19.0097459 |
| 0.0013591 | 0.00294234 | 237.926647 | 715.813826 | 632.003355 | 712.605928 | 64.1578926 |
| 0.00136184 | 0.00294808 | 76.8438766 | 62.2833659 | 67.7477899 | 68.6625504 | 97.424948 |
| 0.00136509 | 0.00295493 | 12.2178549 | 17.3503662 | 7.42441534 | 23.1968076 | 0 |
| 0.00136556 | 0.00295577 | 0 | 332.993567 | 557.759202 | 185.110524 | 4.75243649 |
| 0.00136659 | 0.00295778 | 0 | 108.551009 | 193.034799 | 234.287756 | 0 |
| 0.00136933 | 0.00296354 | 53.3727344 | 153.92889 | 0 | 229.648395 | 0 |
| 0.00137189 | 0.00296887 | 20.8989622 | 13.7913167 | 25.9854537 | 23.6607437 | 2.37621824 |
| 0.0013729 | 0.00297088 | 7.71653991 | 15.7932821 | 12.9927268 | 19.9492545 | 0 |
| 0.00137407 | 0.00297322 | 11.8963324 | 10.4547078 | 12.0646749 | 47.7854236 | 0 |
| 0.00137521 | 0.00297548 | 7.71653991 | 24.023584 | 12.0646749 | 17.1656376 | 0 |
| 0.00137624 | 0.00297735 | 0 | 157.932821 | 181.898176 | 173.048184 | 0 |
| 0.0013762 | 0.00297735 | 4.50131495 | 22.911381 | 23.2012979 | 26.4443606 | 0 |
| 0.00137642 | 0.00297754 | 27.6509347 | 77.6317668 | 75.1722053 | 63.0953166 | 14.2573095 |
| 0.00138008 | 0.00298527 | 8.3595849 | 7.78542074 | 15.7768826 | 7.88691457 | 42.7719284 |
| 0.00138055 | 0.0029861 | 269.757374 | 42.4861532 | 14.8488307 | 32.0115944 | 7.12865473 |
| 0.00138294 | 0.00299108 | 14.1469898 | 10.6771484 | 9.28051917 | 23.1968076 | 0 |
| 0.00138322 | 0.00299148 | 3.53674746 | 20.2420939 | 63.1075304 | 23.6607437 | 0 |
| 0.00138791 | 0.00300143 | 4.82283744 | 16.4606038 | 28.7696094 | 28.7640414 | 0 |
| 0.00138931 | 0.00300426 | 8.6811074 | 12.011792 | 13.9207788 | 17.6295737 | 0 |
| 0.00138945 | 0.00300438 | 30.5446371 | 80.7459351 | 55.683115 | 62.6313804 | 14.2573095 |
| 0.00138963 | 0.00300445 | 8.0380624 | 14.4586385 | 13.9207788 | 15.7738291 | 0 |
| 0.00138967 | 0.00300445 | 6.43044992 | 28.4723958 | 15.7768826 | 16.7017014 | 0 |
| 0.00139339 | 0.00301231 | 23.4711422 | 28.917277 | 25.0574018 | 27.8361691 | 7.12865473 |
| 0.00139725 | 0.00302045 | 6.43044992 | 20.0196533 | 13.9207788 | 25.0525522 | 0 |
| 0.00139874 | 0.0030235 | 9.96719738 | 62.5058065 | 62.1794784 | 73.3019119 | 7.12865473 |

| | | | | | | |
|---|---|---|---|---|---|---|
| 0.00139885 | 0.00302354 | 36.6535646 | 32.9212077 | 21.3451941 | 25.0525522 | 7.12865473 |
| 0.00140095 | 0.00302787 | 9.32415239 | 30.2519206 | 31.5537652 | 32.0115944 | 2.37621824 |
| 0.0014039 | 0.00303406 | 9.64567488 | 12.6791138 | 11.136623 | 16.7017014 | 0 |
| 0.00140464 | 0.00303547 | 3.53674746 | 35.5904948 | 21.3451941 | 31.0837221 | 0 |
| 0.00140494 | 0.00303592 | 1.92913498 | 2.66928711 | 0.92805192 | 0.9278723 | 11.8810912 |
| 0.00140555 | 0.00303704 | 7.07349491 | 31.1416829 | 19.4890903 | 12.0623399 | 0 |
| 0.00140833 | 0.00304285 | 96.1352263 | 159.267464 | 0 | 236.607437 | 0 |
| 0.00141075 | 0.00304787 | 144.363601 | 100.765588 | 0 | 127.582442 | 0 |
| 0.00141372 | 0.0030541 | 90.9908664 | 139.02537 | 0 | 146.139888 | 0 |
| 0.00141482 | 0.00305629 | 9.32415239 | 15.7932821 | 12.9927268 | 11.5984038 | 0 |
| 0.00141496 | 0.00305638 | 22.5065747 | 14.9035197 | 14.8488307 | 20.8771268 | 49.9005831 |
| 0.00141567 | 0.0030577 | 63.0184092 | 84.7498657 | 102.085711 | 93.7151026 | 38.0194919 |
| 0.00141575 | 0.0030577 | 32.1522496 | 12.4566732 | 8.35246725 | 13.9180845 | 0 |
| 0.00141612 | 0.00305831 | 8.6811074 | 35.5904948 | 14.8488307 | 11.5984038 | 0 |
| 0.0014181 | 0.00306238 | 46.9422844 | 24.4684652 | 25.0574018 | 25.9804245 | 49.9005831 |
| 0.00141888 | 0.00306388 | 20.8989622 | 8.45274251 | 12.9927268 | 12.9902122 | 0 |
| 0.00142024 | 0.00306661 | 108.674604 | 173.948543 | 144.776099 | 150.779249 | 38.0194919 |
| 0.00142051 | 0.00306699 | 4.17979245 | 26.4704305 | 24.1293498 | 24.588616 | 0 |
| 0.00142345 | 0.00307316 | 37.618132 | 19.5747721 | 17.6329864 | 5.10329766 | 0 |
| 0.00142555 | 0.00307749 | 7.07349491 | 22.0216187 | 11.136623 | 28.7640414 | 0 |
| 0.00142708 | 0.00308058 | 209.311145 | 248.688582 | 61.2514265 | 0 | 0 |
| 0.00142749 | 0.00308129 | 6.10892743 | 23.3562622 | 13.9207788 | 23.6607437 | 0 |
| 0.00142785 | 0.00308187 | 0 | 418.188314 | 32.4818171 | 249.597649 | 0 |
| 0.00143 | 0.00308629 | 39.547267 | 58.0569946 | 21.3451941 | 38.9706367 | 9.50487297 |
| 0.00143073 | 0.00308768 | 52.0866444 | 129.460425 | 90.0210359 | 122.015208 | 26.1384007 |
| 0.00143109 | 0.00308826 | 0 | 180.17688 | 318.321808 | 73.7658481 | 0 |
| 0.00143212 | 0.00309029 | 7.71653991 | 16.905485 | 43.6184401 | 11.5984038 | 0 |
| 0.00143302 | 0.00309203 | 6.75197242 | 41.3739502 | 32.4818171 | 8.81478687 | 0 |
| 0.00143542 | 0.00309701 | 13.5039448 | 27.3601929 | 20.4171422 | 23.1968076 | 0 |
| 0.00143562 | 0.00309723 | 3.53674746 | 0.88976237 | 1.85610383 | 2.78361691 | 26.1384007 |
| 0.00143858 | 0.00310342 | 10.2887199 | 8.8976237 | 21.3451941 | 15.7738291 | 0 |
| 0.00143889 | 0.00310391 | 15.1115573 | 6.89565837 | 30.6257133 | 14.8459568 | 0 |
| 0.00143943 | 0.00310486 | 207.703532 | 69.846346 | 0 | 122.479144 | 0 |
| 0.00143997 | 0.00310584 | 215.420072 | 71.8483114 | 0 | 114.592229 | 0 |
| 0.0014404 | 0.00310656 | 24.1141872 | 14.4586385 | 13.9207788 | 7.42297842 | 0 |
| 0.00144114 | 0.00310796 | 49.1929419 | 140.804895 | 57.5392189 | 113.664357 | 19.0097459 |
| 0.00144201 | 0.00310963 | 81.023669 | 139.692692 | 0 | 158.202228 | 0 |
| 0.00144705 | 0.0031203 | 63.3399317 | 114.779346 | 119.718697 | 104.84957 | 28.5146189 |
| 0.00144739 | 0.00312083 | 84.238894 | 241.348043 | 0 | 194.853184 | 0 |
| 0.00144896 | 0.00312401 | 7.39501741 | 15.5708415 | 13.9207788 | 18.0935099 | 0 |
| 0.00145041 | 0.00312695 | 35.0459521 | 12.6791138 | 12.0646749 | 17.6295737 | 156.830404 |
| 0.00145183 | 0.00312981 | 5.14435994 | 19.7972127 | 22.273246 | 21.8049991 | 0 |
| 0.00145263 | 0.00313133 | 10.2887199 | 21.1318563 | 9.28051917 | 14.3820207 | 0 |

| | | | | | | |
|---|---|---|---|---|---|---|
| 0.00145323 | 0.00313241 | 3.85826995 | 25.3582275 | 21.3451941 | 35.7230837 | 0 |
| 0.00145339 | 0.00313257 | 81.023669 | 139.692692 | 0 | 156.346483 | 0 |
| 0.00145402 | 0.00313372 | 2.89370247 | 48.7144897 | 42.6903882 | 17.1656376 | 0 |
| 0.00145626 | 0.00313834 | 7.07349491 | 20.2420939 | 25.9854537 | 11.5984038 | 0 |
| 0.00145672 | 0.00313914 | 47.9068519 | 3.33660889 | 14.8488307 | 42.6821259 | 0 |
| 0.00145738 | 0.00314036 | 374.89523 | 543.867249 | 492.795568 | 540.02168 | 73.6627655 |
| 0.00145786 | 0.00314119 | 7.71653991 | 14.4586385 | 14.8488307 | 15.309893 | 0 |
| 0.00145947 | 0.00314446 | 7.07349491 | 22.2440592 | 17.6329864 | 12.9902122 | 0 |
| 0.00146044 | 0.00314635 | 102.565676 | 187.73986 | 138.279736 | 186.502333 | 0 |
| 0.00146079 | 0.0031469 | 9.96719738 | 5.33857422 | 1.85610383 | 5.56723382 | 11.8810912 |
| 0.0014642 | 0.00315404 | 14.7900348 | 23.8011434 | 17.6329864 | 27.3722329 | 0 |
| 0.00146473 | 0.00315498 | 4.17979245 | 42.4861532 | 11.136623 | 50.5690405 | 0 |
| 0.00146487 | 0.00315507 | 20.8989622 | 18.2401286 | 22.273246 | 16.2377653 | 0 |
| 0.00146755 | 0.00316065 | 241.141872 | 369.473824 | 270.063108 | 292.279775 | 0 |
| 0.00146868 | 0.00316289 | 0 | 0 | 0 | 0 | 16.6335277 |
| 0.00146966 | 0.00316478 | 14.1469898 | 20.9094157 | 8.35246725 | 11.5984038 | 0 |
| 0.00147085 | 0.00316716 | 271.686509 | 468.237447 | 361.012196 | 449.554131 | 30.8908372 |
| 0.00147326 | 0.00317214 | 8.3595849 | 22.0216187 | 34.3379209 | 8.81478687 | 0 |
| 0.00147548 | 0.00317671 | 59.1601393 | 174.170984 | 0 | 165.625206 | 0 |
| 0.00147652 | 0.00317874 | 187.769138 | 471.351615 | 486.299204 | 394.345729 | 0 |
| 0.00147793 | 0.00318159 | 17.3622148 | 34.2558512 | 23.2012979 | 32.9394667 | 4.75243649 |
| 0.00147868 | 0.00318299 | 72.0210391 | 81.6356974 | 116.934542 | 115.056166 | 45.1481466 |
| 0.00147966 | 0.00318491 | 6.43044992 | 21.7991781 | 23.2012979 | 12.9902122 | 0 |
| 0.00148078 | 0.0031871 | 43.7270595 | 36.035376 | 33.409869 | 32.9394667 | 0 |
| 0.00148124 | 0.0031879 | 35.6889971 | 85.6396281 | 83.5246725 | 81.1888265 | 0 |
| 0.00148321 | 0.00319193 | 3.53674746 | 94.5372518 | 44.546492 | 138.252973 | 4.75243649 |
| 0.0014839 | 0.0031932 | 8.6811074 | 19.1298909 | 11.136623 | 14.3820207 | 0 |
| 0.00148616 | 0.00319766 | 34.4029071 | 15.7932821 | 32.4818171 | 4.17542536 | 0 |
| 0.00148616 | 0.00319766 | 110.925261 | 72.2931925 | 86.3088283 | 74.2297842 | 190.097459 |
| 0.00148692 | 0.0031991 | 5.78740493 | 2.8917277 | 6.49636342 | 4.63936151 | 9.50487297 |
| 0.00148882 | 0.00320277 | 5.14435994 | 45.822762 | 19.4890903 | 16.2377653 | 0 |
| 0.00148874 | 0.00320277 | 39.8687895 | 15.5708415 | 11.136623 | 8.35085072 | 0 |
| 0.00148953 | 0.00320409 | 235.032945 | 85.4171875 | 0 | 83.5085072 | 0 |
| 0.00148989 | 0.00320467 | 143.399033 | 51.3837769 | 36.1940248 | 37.5788283 | 14.2573095 |
| 0.0014904 | 0.00320556 | 57.5525268 | 140.360014 | 126.215061 | 137.325101 | 16.6335277 |
| 0.00149144 | 0.00320759 | 22.5065747 | 10.899589 | 9.28051917 | 12.9902122 | 0 |
| 0.00149329 | 0.00321137 | 9.00262989 | 26.9153117 | 5.5683115 | 80.2609542 | 0 |
| 0.00149406 | 0.00321281 | 11.5748099 | 7.78542074 | 16.7049345 | 21.8049991 | 0 |
| 0.00149475 | 0.00321408 | 6.43044992 | 12.9015544 | 40.8342843 | 18.5574461 | 0 |
| 0.00149689 | 0.00321848 | 17.0406923 | 42.9310343 | 6.49636342 | 13.4541484 | 0 |
| 0.00149744 | 0.00321946 | 65.9121117 | 82.3030192 | 456.601543 | 0 | 0 |
| 0.00149777 | 0.00321996 | 5.78740493 | 21.3542969 | 14.8488307 | 25.0525522 | 0 |
| 0.00149971 | 0.00322394 | 4.82283744 | 37.1475789 | 46.4025958 | 10.6705315 | 0 |

| | | | | | | |
|---|---|---|---|---|---|---|
| 0.00150003 | 0.00322441 | 2.89370247 | 46.4900838 | 50.1148035 | 15.309893 | 0 |
| 0.00150264 | 0.00322961 | 0 | 161.26943 | 162.409085 | 165.16127 | 0 |
| 0.00150259 | 0.00322961 | 3.85826995 | 24.9133464 | 26.9135056 | 25.5164883 | 0 |
| 0.0015042 | 0.00323276 | 18.0052598 | 44.9329997 | 56.6111669 | 50.1051043 | 9.50487297 |
| 0.00150505 | 0.00323438 | 33.7598621 | 61.8384847 | 46.4025958 | 31.0837221 | 11.8810912 |
| 0.00150576 | 0.00323549 | 9.00262989 | 13.123995 | 34.3379209 | 11.5984038 | 0 |
| 0.00150568 | 0.00323549 | 20.5774398 | 8.00786133 | 58.4672708 | 8.81478687 | 0 |
| 0.00150713 | 0.00323822 | 14.7900348 | 13.3464355 | 7.42441534 | 17.1656376 | 0 |
| 0.00150751 | 0.00323884 | 27.0078897 | 25.8031087 | 17.6329864 | 4.63936151 | 0 |
| 0.00150963 | 0.00324319 | 324.737721 | 730.050024 | 496.507776 | 631.881038 | 0 |
| 0.00151323 | 0.00325072 | 5.78740493 | 17.3503662 | 16.7049345 | 27.8361691 | 0 |
| 0.00151348 | 0.00325103 | 120.892459 | 181.511523 | 142.919995 | 0 | 0 |
| 0.00151475 | 0.00325356 | 5.14435994 | 25.8031087 | 17.6329864 | 21.341063 | 0 |
| 0.00152066 | 0.00326605 | 6.10892743 | 17.1279256 | 16.7049345 | 21.341063 | 0 |
| 0.00152202 | 0.00326876 | 0 | 2.66928711 | 0 | 0 | 49.9005831 |
| 0.00152244 | 0.00326944 | 3.85826995 | 25.5806681 | 33.409869 | 20.8771268 | 0 |
| 0.00152253 | 0.00326944 | 22.1850522 | 20.4645345 | 3.71220767 | 45.0018067 | 0 |
| 0.00152405 | 0.00327249 | 7.39501741 | 25.8031087 | 16.7049345 | 12.0623399 | 0 |
| 0.00152523 | 0.00327482 | 3.53674746 | 38.0373413 | 25.9854537 | 21.8049991 | 0 |
| 0.00152599 | 0.00327624 | 16.0761248 | 11.1220296 | 9.28051917 | 12.5262761 | 0 |
| 0.00152631 | 0.00327672 | 26.6863672 | 16.905485 | 21.3451941 | 18.5574461 | 33.2670554 |
| 0.0015271 | 0.0032782 | 13.8254673 | 22.6889404 | 45.4745439 | 5.10329766 | 0 |
| 0.00152808 | 0.00328009 | 118.641801 | 164.606038 | 163.337137 | 163.305525 | 54.6530196 |
| 0.00152918 | 0.00328226 | 5.46588243 | 19.3523315 | 20.4171422 | 19.4853184 | 0 |
| 0.00152948 | 0.00328269 | 6.43044992 | 20.9094157 | 19.4890903 | 13.9180845 | 0 |
| 0.00153143 | 0.00328665 | 4.82283744 | 18.2401286 | 25.0574018 | 23.6607437 | 0 |
| 0.00153241 | 0.00328855 | 12.5393773 | 8.23030192 | 0 | 0 | 99.8011662 |
| 0.00153378 | 0.00329129 | 5.46588243 | 12.011792 | 25.9854537 | 41.2903175 | 0 |
| 0.00153504 | 0.00329378 | 2.89370247 | 35.8129354 | 18.5610383 | 48.7132959 | 0 |
| 0.00153602 | 0.00329567 | 18.0052598 | 15.3484009 | 11.136623 | 13.4541484 | 97.424948 |
| 0.00153673 | 0.00329698 | 6.10892743 | 17.1279256 | 16.7049345 | 20.8771268 | 0 |
| 0.00153812 | 0.00329976 | 314.449001 | 568.335714 | 486.299204 | 591.518593 | 90.2962933 |
| 0.00153953 | 0.00330257 | 0 | 0 | 0 | 0 | 19.0097459 |
| 0.0015414 | 0.00330636 | 33.4383396 | 20.4645345 | 6.49636342 | 11.5984038 | 0 |
| 0.00154273 | 0.00330901 | 104.494811 | 90.5333211 | 44.546492 | 70.518295 | 19.0097459 |
| 0.00154582 | 0.00331543 | 138.576196 | 77.1868856 | 41.7623363 | 55.6723382 | 21.3859642 |
| 0.00154791 | 0.00331969 | 29.2585471 | 210.20636 | 198.60311 | 200.420417 | 23.7621824 |
| 0.00155022 | 0.00332443 | 130.538133 | 34.923173 | 24.1293498 | 30.1558498 | 0 |
| 0.00155158 | 0.00332714 | 506.076409 | 1172.26192 | 979.094772 | 1054.99081 | 4.75243649 |
| 0.00155284 | 0.00332963 | 16.3976473 | 22.0216187 | 20.4171422 | 5.56723382 | 0 |
| 0.00155552 | 0.00333517 | 0 | 0 | 0 | 0 | 73.6627655 |
| 0.00155664 | 0.00333715 | 6.75197242 | 18.4625692 | 13.9207788 | 18.0935099 | 0 |
| 0.00155655 | 0.00333715 | 78.4514891 | 143.696623 | 0 | 141.500526 | 0 |

| | | | | | | |
|---|---|---|---|---|---|---|
| 0.00155782 | 0.00333926 | 4.17979245 | 23.1338216 | 23.2012979 | 25.9804245 | 0 |
| 0.00155783 | 0.00333926 | 49.5144644 | 64.5077718 | 52.8989593 | 61.7035081 | 0 |
| 0.0015596 | 0.00334285 | 8.0380624 | 11.5669108 | 14.8488307 | 20.8771268 | 0 |
| 0.0015601 | 0.00334372 | 4.17979245 | 30.02948 | 23.2012979 | 20.8771268 | 0 |
| 0.00156117 | 0.0033458 | 137.290106 | 288.727889 | 164.265189 | 263.97967 | 38.0194919 |
| 0.00156146 | 0.00334619 | 18.6483048 | 62.0609253 | 10.2085711 | 29.6919137 | 2.37621824 |
| 0.00156492 | 0.00335319 | 138.254673 | 175.283187 | 259.854537 | 295.527328 | 0 |
| 0.00156489 | 0.00335319 | 14.7900348 | 9.12006429 | 10.2085711 | 16.2377653 | 0 |
| 0.00156587 | 0.00335502 | 7.07349491 | 19.1298909 | 12.9927268 | 17.1656376 | 0 |
| 0.00156692 | 0.00335684 | 182.9463 | 361.688403 | 237.581291 | 263.515734 | 90.2962933 |
| 0.0015669 | 0.00335684 | 13.1824223 | 23.8011434 | 43.6184401 | 5.10329766 | 0 |
| 0.00156886 | 0.00336076 | 33.4383396 | 49.1593709 | 55.683115 | 30.1558498 | 0 |
| 0.00156946 | 0.00336184 | 0 | 0 | 0 | 0 | 23.7621824 |
| 0.00156956 | 0.00336184 | 6.43044992 | 31.1416829 | 15.7768826 | 14.8459568 | 0 |
| 0.00156966 | 0.00336185 | 6.43044992 | 32.9212077 | 10.2085711 | 26.9082968 | 0 |
| 0.00157009 | 0.00336254 | 22.5065747 | 60.5038411 | 46.4025958 | 36.650956 | 2.37621824 |
| 0.00157058 | 0.00336338 | 16.0761248 | 67.6219401 | 91.8771398 | 113.200421 | 14.2573095 |
| 0.00157161 | 0.00336523 | 222.81509 | 349.676611 | 195.818954 | 248.669777 | 0 |
| 0.00157164 | 0.00336523 | 0 | 35.5904948 | 438.040505 | 185.574461 | 0 |
| 0.00157205 | 0.00336589 | 3.85826995 | 10.4547078 | 36.1940248 | 70.9822312 | 0 |
| 0.00157553 | 0.00337314 | 20.2559173 | 8.8976237 | 14.8488307 | 10.2065953 | 0 |
| 0.0015761 | 0.00337413 | 6.43044992 | 18.9074504 | 16.7049345 | 15.7738291 | 0 |
| 0.00157656 | 0.0033749 | 125.072251 | 318.090047 | 264.494796 | 294.599456 | 23.7621824 |
| 0.00157709 | 0.00337583 | 22.1850522 | 20.0196533 | 59.3953227 | 26.4443606 | 4.75243649 |
| 0.00157748 | 0.00337618 | 0 | 103.434875 | 236.653239 | 158.202228 | 0 |
| 0.00157751 | 0.00337618 | 55.9449143 | 12.6791138 | 6.49636342 | 16.7017014 | 0 |
| 0.00157756 | 0.00337618 | 108.996126 | 29.1397176 | 13.9207788 | 30.619786 | 4.75243649 |
| 0.00158222 | 0.00338593 | 255.288862 | 121.897445 | 56.6111669 | 95.5708472 | 28.5146189 |
| 0.00158242 | 0.00338616 | 14.4685123 | 17.1279256 | 19.4890903 | 6.49510612 | 0 |
| 0.00158397 | 0.00338925 | 4.50131495 | 28.2499552 | 19.4890903 | 22.2689353 | 0 |
| 0.0015859 | 0.00339296 | 10.6102424 | 17.3503662 | 12.9927268 | 9.27872303 | 0 |
| 0.0015859 | 0.00339296 | 45.334672 | 16.6830444 | 6.49636342 | 12.9902122 | 0 |
| 0.00158667 | 0.00339438 | 8.3595849 | 18.2401286 | 17.6329864 | 10.2065953 | 0 |
| 0.00158779 | 0.00339656 | 6.43044992 | 46.2676432 | 69.6038938 | 57.9920189 | 0 |
| 0.00158951 | 0.00340004 | 12.5393773 | 16.4606038 | 24.1293498 | 6.95904227 | 0 |
| 0.00159029 | 0.00340148 | 0 | 182.401286 | 122.502853 | 175.367865 | 0 |
| 0.00159061 | 0.00340173 | 100.315019 | 208.871716 | 136.423632 | 0 | 0 |
| 0.00159053 | 0.00340173 | 24.1141872 | 69.6239054 | 49.1867516 | 53.8165936 | 0 |
| 0.00159079 | 0.0034019 | 3.85826995 | 63.3955688 | 9.28051917 | 45.0018067 | 0 |
| 0.00159148 | 0.00340316 | 6.75197242 | 24.6909058 | 15.7768826 | 13.9180845 | 0 |
| 0.00159634 | 0.00341333 | 18.3267823 | 13.123995 | 16.7049345 | 12.0623399 | 61.7816743 |
| 0.00159694 | 0.0034144 | 75.8793091 | 118.115955 | 0 | 170.728504 | 0 |
| 0.00159944 | 0.00341953 | 69.1273367 | 9.78738607 | 11.136623 | 11.5984038 | 0 |

| | | | | | | |
|---|---|---|---|---|---|---|
| 0.00160136 | 0.0034232 | 27.3294122 | 40.929069 | 2.78415575 | 24.588616 | 0 |
| 0.00160127 | 0.0034232 | 1.60761248 | 36.035376 | 40.8342843 | 42.2181898 | 0 |
| 0.00160704 | 0.00343512 | 7.07349491 | 40.4841878 | 16.7049345 | 12.0623399 | 0 |
| 0.00160857 | 0.00343817 | 80.0591015 | 113.667143 | 0 | 166.553078 | 0 |
| 0.00161246 | 0.00344627 | 0.3215225 | 0 | 0 | 0 | 30.8908372 |
| 0.0016148 | 0.00345083 | 54.6588243 | 30.4743612 | 24.1293498 | 30.619786 | 185.345023 |
| 0.00161472 | 0.00345083 | 55.9449143 | 166.608004 | 155.912722 | 146.139888 | 26.1384007 |
| 0.00161559 | 0.00345231 | 28.9370247 | 21.5767375 | 26.9135056 | 3.71148921 | 0 |
| 0.00161834 | 0.00345797 | 1.60761248 | 38.0373413 | 51.0428554 | 32.0115944 | 0 |
| 0.00161887 | 0.00345888 | 0 | 191.076469 | 103.941815 | 189.28595 | 0 |
| 0.00162153 | 0.00346434 | 6.10892743 | 14.6810791 | 18.5610383 | 21.341063 | 0 |
| 0.00162189 | 0.00346489 | 3.85826995 | 24.9133464 | 26.9135056 | 23.1968076 | 0 |
| 0.00162236 | 0.00346567 | 9.32415239 | 9.78738607 | 24.1293498 | 13.9180845 | 0 |
| 0.0016239 | 0.00346873 | 314.449001 | 95.2045736 | 149.416359 | 132.685739 | 648.70758 |
| 0.00162841 | 0.00347816 | 18.0052598 | 40.0393066 | 36.1940248 | 29.6919137 | 7.12865473 |
| 0.00162881 | 0.00347878 | 14.7900348 | 35.3680542 | 3.71220767 | 37.1148921 | 0 |
| 0.00162992 | 0.00348093 | 75.8793091 | 137.690727 | 118.790645 | 143.356271 | 4.75243649 |
| 0.00163004 | 0.00348097 | 18.3267823 | 12.6791138 | 17.6329864 | 12.5262761 | 71.2865473 |
| 0.00163178 | 0.00348446 | 5.78740493 | 20.4645345 | 13.9207788 | 26.4443606 | 0 |
| 0.00163309 | 0.00348681 | 0 | 0 | 0 | 0 | 26.1384007 |
| 0.00163302 | 0.00348681 | 9.96719738 | 9.12006429 | 19.4890903 | 13.9180845 | 0 |
| 0.0016372 | 0.00349537 | 21.2204847 | 8.67518311 | 14.8488307 | 5.56723382 | 28.5146189 |
| 0.00163829 | 0.00349748 | 4.50131495 | 17.3503662 | 29.6976613 | 23.6607437 | 0 |
| 0.0016393 | 0.00349942 | 361.069763 | 864.626583 | 585.60076 | 732.091247 | 130.692003 |
| 0.00164039 | 0.00350151 | 259.147132 | 234.674825 | 0 | 19.9492545 | 0 |
| 0.00164072 | 0.00350199 | 16.3976473 | 18.2401286 | 4.64025958 | 54.7444659 | 0 |
| 0.00164118 | 0.00350275 | 0 | 0 | 0 | 0 | 0 |
| 0.00164183 | 0.00350393 | 89.3832539 | 135.911202 | 0 | 121.551272 | 0 |
| 0.00164454 | 0.00350949 | 17.6837373 | 8.45274251 | 29.6976613 | 9.27872303 | 0 |
| 0.00164622 | 0.00351284 | 5.78740493 | 24.6909058 | 12.0646749 | 29.2279775 | 0 |
| 0.00164662 | 0.00351347 | 9.64567488 | 24.2460246 | 15.7768826 | 8.81478687 | 0 |
| 0.0016486 | 0.00351747 | 7.71653991 | 21.7991781 | 12.0646749 | 14.3820207 | 0 |
| 0.00164975 | 0.00351949 | 0 | 125.678935 | 71.4599976 | 355.375092 | 0 |
| 0.00164969 | 0.00351949 | 8.6811074 | 18.017688 | 9.28051917 | 19.4853184 | 0 |
| 0.00165091 | 0.00352174 | 11.2532874 | 25.8031087 | 20.4171422 | 35.7230837 | 2.37621824 |
| 0.00165299 | 0.00352596 | 93.2415239 | 160.379667 | 160.552982 | 173.048184 | 40.3957101 |
| 0.00165331 | 0.00352641 | 4.50131495 | 13.123995 | 46.4025958 | 26.9082968 | 0 |
| 0.00165367 | 0.00352696 | 7.07349491 | 19.7972127 | 12.9927268 | 15.7738291 | 0 |
| 0.00165407 | 0.00352759 | 116.391144 | 365.914775 | 346.163365 | 366.973496 | 45.1481466 |
| 0.00165505 | 0.00352945 | 60.4462293 | 203.310701 | 159.62493 | 186.502333 | 28.5146189 |
| 0.00165604 | 0.00353134 | 18.3267823 | 13.7913167 | 11.136623 | 8.35085072 | 0 |
| 0.0016566 | 0.00353231 | 15.4330798 | 17.7952474 | 5.5683115 | 24.588616 | 0 |
| 0.00165733 | 0.00353365 | 55.9449143 | 134.798999 | 0 | 188.822014 | 0 |

| | | | | | | |
|---|---|---|---|---|---|---|
| 0.00165758 | 0.00353395 | 103.530244 | 260.255493 | 139.207788 | 203.66797 | 33.2670554 |
| 0.00166266 | 0.00354455 | 1.60761248 | 30.6968018 | 33.409869 | 59.8477635 | 0 |
| 0.00166649 | 0.0035525 | 63.6614542 | 68.2892619 | 40.8342843 | 55.208402 | 16.6335277 |
| 0.00166663 | 0.00355257 | 0 | 154.151331 | 151.272462 | 157.274355 | 0 |
| 0.0016688 | 0.00355696 | 95.8137038 | 124.789172 | 130.85532 | 115.056166 | 21.3859642 |
| 0.00166911 | 0.00355741 | 19.2913498 | 10.899589 | 17.6329864 | 11.1344676 | 76.0389838 |
| 0.00167024 | 0.00355958 | 8.0380624 | 12.4566732 | 16.7049345 | 12.9902122 | 0 |
| 0.0016706 | 0.00356012 | 14.1469898 | 11.5669108 | 30.6257133 | 7.88691457 | 0 |
| 0.00167256 | 0.00356408 | 16.3976473 | 15.7932821 | 6.49636342 | 15.309893 | 0 |
| 0.00167595 | 0.00357109 | 4.50131495 | 7.34053955 | 5.5683115 | 6.03116997 | 19.0097459 |
| 0.0016776 | 0.00357436 | 18.3267823 | 18.6850098 | 6.49636342 | 12.9902122 | 0 |
| 0.00168136 | 0.00358192 | 11.8963324 | 14.0137573 | 8.35246725 | 13.4541484 | 0 |
| 0.00168126 | 0.00358192 | 9.32415239 | 10.899589 | 13.9207788 | 12.5262761 | 0 |
| 0.00168167 | 0.00358235 | 3.21522496 | 47.3798462 | 12.0646749 | 49.6411682 | 0 |
| 0.00168311 | 0.00358496 | 7.07349491 | 23.1338216 | 13.9207788 | 13.9180845 | 0 |
| 0.00168303 | 0.00358496 | 157.867546 | 658.646594 | 640.355823 | 636.984336 | 83.1676385 |
| 0.0016833 | 0.00358514 | 6.10892743 | 4.67125244 | 0 | 6.49510612 | 16.6335277 |
| 0.0016896 | 0.00359824 | 0 | 113.222262 | 165.193241 | 189.28595 | 0 |
| 0.00168966 | 0.00359824 | 7.39501741 | 62.0609253 | 19.4890903 | 45.929679 | 0 |
| 0.00169005 | 0.00359885 | 48.2283744 | 100.988029 | 90.9490879 | 97.4265918 | 28.5146189 |
| 0.00169122 | 0.00360111 | 16.7191698 | 6.89565837 | 12.0646749 | 30.619786 | 0 |
| 0.00169424 | 0.00360731 | 0 | 391.273002 | 18.5610383 | 298.774881 | 0 |
| 0.00169478 | 0.00360823 | 0 | 0 | 0 | 0 | 106.929821 |
| 0.00169555 | 0.00360964 | 38.904222 | 12.2342326 | 8.35246725 | 12.5262761 | 0 |
| 0.00169821 | 0.00361507 | 48.8714194 | 144.141504 | 0 | 192.997439 | 0 |
| 0.00170486 | 0.003629 | 19.9343948 | 25.3582275 | 25.0574018 | 27.3722329 | 4.75243649 |
| 0.00170828 | 0.00363606 | 6.10892743 | 22.0216187 | 18.5610383 | 13.9180845 | 0 |
| 0.00170869 | 0.0036367 | 10.2887199 | 18.6850098 | 8.35246725 | 14.8459568 | 0 |
| 0.00170981 | 0.00363884 | 27.9724572 | 25.1357869 | 21.3451941 | 11.5984038 | 0 |
| 0.00171098 | 0.00364111 | 7.39501741 | 19.5747721 | 12.0646749 | 15.309893 | 0 |
| 0.00171115 | 0.00364124 | 3.53674746 | 27.8050741 | 60.3233746 | 14.3820207 | 0 |
| 0.00171257 | 0.00364402 | 117.034189 | 455.780774 | 835.246725 | 694.976355 | 0 |
| 0.00171704 | 0.0036533 | 19.9343948 | 24.9133464 | 22.273246 | 24.588616 | 4.75243649 |
| 0.00171837 | 0.00365592 | 67.8412467 | 59.6140788 | 54.7550631 | 65.4149973 | 11.8810912 |
| 0.00172024 | 0.00365966 | 8.6811074 | 14.0137573 | 12.9927268 | 11.5984038 | 0 |
| 0.00172436 | 0.00366818 | 98.3858838 | 112.55494 | 0 | 123.870952 | 0 |
| 0.00172628 | 0.00367205 | 5.14435994 | 34.2558512 | 12.9927268 | 24.1246799 | 0 |
| 0.00172687 | 0.00367306 | 14.7900348 | 24.023584 | 6.49636342 | 13.9180845 | 0 |
| 0.00172702 | 0.00367315 | 90.0262989 | 41.8188314 | 30.6257133 | 27.8361691 | 118.810912 |
| 0.00172786 | 0.0036747 | 73.6286516 | 15.5708415 | 5.5683115 | 13.4541484 | 0 |
| 0.00172936 | 0.00367766 | 4.82283744 | 1.77952474 | 2.78415575 | 3.71148921 | 26.1384007 |
| 0.0017344 | 0.00368816 | 4.50131495 | 33.3660889 | 19.4890903 | 18.5574461 | 0 |
| 0.00173489 | 0.00368897 | 4.17979245 | 40.929069 | 25.0574018 | 15.309893 | 0 |

| | | | | | | |
|---|---|---|---|---|---|---|
| 0.00173723 | 0.00369369 | 12.2178549 | 13.123995 | 9.28051917 | 11.1344676 | 0 |
| 0.00173858 | 0.00369634 | 1.28608998 | 2.00196533 | 4.64025958 | 1.39180845 | 0 |
| 0.00174081 | 0.00370084 | 122.500071 | 146.143469 | 189.322591 | 216.194247 | 42.7719284 |
| 0.00174194 | 0.00370302 | 0 | 168.165088 | 190.250643 | 104.84957 | 0 |
| 0.00174239 | 0.00370373 | 18.6483048 | 71.8483114 | 69.6038938 | 64.487125 | 11.8810912 |
| 0.00174456 | 0.003708 | 55.6233918 | 195.747721 | 176.329864 | 148.923505 | 38.0194919 |
| 0.00174462 | 0.003708 | 0 | 53.8306234 | 180.970124 | 301.094562 | 0 |
| 0.00174492 | 0.00370841 | 126.679863 | 99.6533854 | 94.6612955 | 100.210209 | 26.1384007 |
| 0.00174776 | 0.00371422 | 16.7191698 | 17.1279256 | 25.0574018 | 17.6295737 | 0 |
| 0.00174994 | 0.0037186 | 4.50131495 | 29.1397176 | 17.6329864 | 21.8049991 | 0 |
| 0.00175216 | 0.00372308 | 16.7191698 | 15.1259603 | 8.35246725 | 10.2065953 | 0 |
| 0.00175602 | 0.00373083 | 42.4409695 | 9.34250488 | 29.6976613 | 21.8049991 | 2.37621824 |
| 0.00175603 | 0.00373083 | 10.9317649 | 11.7893514 | 11.136623 | 10.6705315 | 0 |
| 0.0017564 | 0.00373138 | 147.257303 | 367.249418 | 355.443884 | 379.499772 | 4.75243649 |
| 0.00175657 | 0.00373152 | 8.6811074 | 6.22833659 | 39.9062324 | 29.6919137 | 0 |
| 0.00175683 | 0.00373184 | 87.4541189 | 99.6533854 | 0 | 152.634994 | 0 |
| 0.00175716 | 0.00373231 | 8.6811074 | 22.0216187 | 13.9207788 | 10.2065953 | 0 |
| 0.00175835 | 0.0037346 | 0 | 0 | 0 | 0 | 54.6530196 |
| 0.00176046 | 0.00373883 | 9.64567488 | 23.3562622 | 21.3451941 | 7.42297842 | 0 |
| 0.00176214 | 0.00374217 | 29.5800696 | 252.025191 | 391.637909 | 305.269988 | 0 |
| 0.00176264 | 0.00374299 | 31.5092046 | 9.78738607 | 11.136623 | 11.1344676 | 0 |
| 0.00176872 | 0.00375566 | 0 | 78.5215291 | 298.832717 | 128.046378 | 0 |
| 0.00176884 | 0.00375568 | 5.46588243 | 31.8090047 | 25.9854537 | 11.5984038 | 0 |
| 0.00176989 | 0.00375768 | 61.4107968 | 20.9094157 | 18.5610383 | 23.6607437 | 111.682257 |
| 0.00177023 | 0.00375816 | 5.46588243 | 17.5728068 | 24.1293498 | 16.2377653 | 0 |
| 0.00177674 | 0.00377175 | 42.762492 | 7.78542074 | 6.49636342 | 30.619786 | 0 |
| 0.00177746 | 0.00377304 | 97.7428388 | 88.976237 | 230.156875 | 97.8905279 | 26.1384007 |
| 0.00177787 | 0.00377367 | 54.3373018 | 185.070573 | 0 | 127.118505 | 0 |
| 0.00177823 | 0.0037742 | 22.5065747 | 38.9271037 | 31.5537652 | 36.1870198 | 9.50487297 |
| 0.00178264 | 0.00378331 | 7.07349491 | 20.2420939 | 13.9207788 | 13.4541484 | 0 |
| 0.00178305 | 0.00378394 | 3.21522496 | 2.22440592 | 3.71220767 | 3.71148921 | 14.2573095 |
| 0.00178687 | 0.00379182 | 9.96719738 | 17.5728068 | 37.1220767 | 7.42297842 | 0 |
| 0.00178782 | 0.00379359 | 5.46588243 | 21.5767375 | 16.7049345 | 18.0935099 | 0 |
| 0.00179401 | 0.00380648 | 76.5223541 | 60.9487223 | 76.1002572 | 60.3116997 | 116.434694 |
| 0.00179483 | 0.00380798 | 7.07349491 | 22.4664998 | 17.6329864 | 11.1344676 | 0 |
| 0.00179607 | 0.00381037 | 7.07349491 | 13.123995 | 13.9207788 | 21.8049991 | 0 |
| 0.00179763 | 0.00381346 | 5.14435994 | 26.9153117 | 15.7768826 | 19.4853184 | 0 |
| 0.00179844 | 0.00381492 | 3.53674746 | 92.0904053 | 15.7768826 | 18.0935099 | 0 |
| 0.00179894 | 0.00381575 | 39.8687895 | 68.0668213 | 55.683115 | 61.7035081 | 19.0097459 |
| 0.00180125 | 0.00382041 | 32.4737721 | 154.596212 | 127.143113 | 170.264568 | 0 |
| 0.00180231 | 0.00382241 | 60.4462293 | 187.517419 | 158.696878 | 171.19244 | 42.7719284 |
| 0.00180568 | 0.00382932 | 7.07349491 | 19.5747721 | 18.5610383 | 11.1344676 | 0 |
| 0.00180832 | 0.00383468 | 45.0131495 | 6.89565837 | 17.6329864 | 10.6705315 | 0 |

| | | | | | | |
|---|---|---|---|---|---|---|
| 0.00181028 | 0.00383858 | 41.1548795 | 8.23030192 | 8.35246725 | 19.0213822 | 0 |
| 0.00181376 | 0.00384548 | 12.5393773 | 10.4547078 | 16.7049345 | 8.81478687 | 0 |
| 0.00181368 | 0.00384548 | 19.9343948 | 27.5826335 | 17.6329864 | 4.63936151 | 0 |
| 0.00181538 | 0.00384868 | 4.50131495 | 21.1318563 | 17.6329864 | 28.3001052 | 0 |
| 0.0018163 | 0.00385014 | 5.46588243 | 35.5904948 | 12.0646749 | 22.2689353 | 0 |
| 0.00181626 | 0.00385014 | 106.745469 | 52.4959798 | 106.72597 | 51.0329766 | 133.068222 |
| 0.00181911 | 0.00385561 | 6.43044992 | 17.5728068 | 15.7768826 | 14.8459568 | 0 |
| 0.00181903 | 0.00385561 | 0 | 0 | 0.92805192 | 0 | 0 |
| 0.00182167 | 0.0038608 | 123.786161 | 200.418974 | 92.8051917 | 0 | 0 |
| 0.0018234 | 0.00386421 | 3.21522496 | 20.2420939 | 25.0574018 | 41.2903175 | 0 |
| 0.00182443 | 0.00386615 | 5.46588243 | 43.8207967 | 9.28051917 | 29.6919137 | 0 |
| 0.00182939 | 0.00387642 | 3.21522496 | 40.4841878 | 36.1940248 | 15.309893 | 0 |
| 0.00183297 | 0.00388377 | 0 | 0 | 0 | 0 | 9.50487297 |
| 0.00183851 | 0.00389525 | 0 | 235.787028 | 47.3306478 | 240.318926 | 0 |
| 0.00183923 | 0.00389654 | 12.2178549 | 15.5708415 | 14.8488307 | 7.42297842 | 0 |
| 0.00183993 | 0.00389778 | 11.5748099 | 11.7893514 | 10.2085711 | 10.6705315 | 0 |
| 0.00184364 | 0.00390538 | 44.3701045 | 7.34053955 | 7.42441534 | 25.5164883 | 0 |
| 0.00184531 | 0.00390867 | 19.9343948 | 31.8090047 | 3.71220767 | 21.8049991 | 0 |
| 0.00184625 | 0.00391042 | 106.745469 | 142.361979 | 153.128566 | 154.954675 | 40.3957101 |
| 0.00184928 | 0.00391659 | 49.5144644 | 184.625692 | 0 | 130.829995 | 0 |
| 0.00185167 | 0.00392142 | 11.8963324 | 8.67518311 | 12.0646749 | 13.4541484 | 0 |
| 0.00185635 | 0.00393107 | 0 | 160.602108 | 106.72597 | 177.687546 | 0 |
| 0.00185646 | 0.00393107 | 0 | 3.55904948 | 0 | 10.6705315 | 311.28459 |
| 0.00185976 | 0.00393779 | 2.25065747 | 1.33464355 | 0.92805192 | 1.85574461 | 4.75243649 |
| 0.00186203 | 0.00394235 | 92.9200014 | 81.1908162 | 0 | 160.985845 | 0 |
| 0.00186335 | 0.0039449 | 50.1575094 | 107.883687 | 63.1075304 | 54.7444659 | 11.8810912 |
| 0.00186378 | 0.00394555 | 72.9856066 | 111.665177 | 0 | 148.923505 | 0 |
| 0.00186389 | 0.00394555 | 3.21522496 | 0.22244059 | 0 | 0 | 0 |
| 0.0018642 | 0.00394596 | 33.4383396 | 262.035018 | 247.789862 | 246.814032 | 28.5146189 |
| 0.00186537 | 0.00394819 | 148.864916 | 193.523315 | 182.826228 | 201.812226 | 68.9103291 |
| 0.00186569 | 0.00394861 | 6.10892743 | 19.1298909 | 20.4171422 | 12.9902122 | 0 |
| 0.00186715 | 0.00395146 | 4.82283744 | 20.9094157 | 21.3451941 | 18.0935099 | 0 |
| 0.00186921 | 0.00395557 | 5.78740493 | 22.0216187 | 12.9927268 | 21.341063 | 0 |
| 0.00187044 | 0.00395793 | 6.43044992 | 14.4586385 | 14.8488307 | 19.4853184 | 0 |
| 0.00187243 | 0.00396189 | 0 | 7.11809896 | 0 | 0 | 26.1384007 |
| 0.00187403 | 0.00396503 | 6.75197242 | 15.5708415 | 25.0574018 | 12.0623399 | 0 |
| 0.00187417 | 0.00396506 | 18.9698273 | 18.9074504 | 4.64025958 | 21.8049991 | 0 |
| 0.00187976 | 0.00397664 | 10.9317649 | 7.78542074 | 23.2012979 | 13.9180845 | 0 |
| 0.0018851 | 0.00398769 | 10.2887199 | 34.923173 | 5.5683115 | 25.5164883 | 0 |
| 0.00188526 | 0.00398778 | 0 | 0 | 0 | 6.49510612 | 28.5146189 |
| 0.00188623 | 0.00398957 | 5.78740493 | 15.7932821 | 14.8488307 | 27.8361691 | 0 |
| 0.00188777 | 0.00399259 | 33.1168171 | 135.02144 | 173.545708 | 132.221803 | 26.1384007 |
| 0.00188857 | 0.00399402 | 0 | 173.726103 | 138.279736 | 124.334889 | 0 |

| | | | | | | |
|---|---|---|---|---|---|---|
| 0.00188943 | 0.00399559 | 0 | 87.864034 | 199.531162 | 165.16127 | 0 |
| 0.00189187 | 0.0040005 | 13.1824223 | 13.123995 | 12.9927268 | 7.88691457 | 0 |
| 0.00189327 | 0.00400321 | 26.0433222 | 3.11416829 | 25.9854537 | 28.3001052 | 0 |
| 0.00189799 | 0.00401294 | 7.39501741 | 11.5669108 | 14.8488307 | 17.6295737 | 0 |
| 0.00189834 | 0.00401342 | 22.5065747 | 7.34053955 | 10.2085711 | 17.6295737 | 0 |
| 0.00190032 | 0.00401736 | 18.0052598 | 20.0196533 | 12.0646749 | 6.49510612 | 0 |
| 0.00190124 | 0.00401905 | 128.930521 | 173.726103 | 111.36623 | 264.443606 | 0 |
| 0.00190168 | 0.00401973 | 11.8963324 | 35.3680542 | 51.9709073 | 41.2903175 | 4.75243649 |
| 0.00190431 | 0.00402504 | 11.5748099 | 12.2342326 | 8.35246725 | 13.9180845 | 0 |
| 0.00190512 | 0.0040265 | 77.4869216 | 165.940682 | 166.121293 | 0 | 0 |
| 0.00190579 | 0.00402765 | 10.9317649 | 18.2401286 | 10.2085711 | 9.74265918 | 0 |
| 0.00190709 | 0.00403016 | 14.1469898 | 11.1220296 | 7.42441534 | 23.1968076 | 0 |
| 0.00190839 | 0.0040325 | 36.0105196 | 136.578524 | 122.502853 | 107.633187 | 2.37621824 |
| 0.00190844 | 0.0040325 | 7.71653991 | 9.12006429 | 30.6257133 | 18.0935099 | 0 |
| 0.00191216 | 0.00404011 | 11.8963324 | 9.34250488 | 10.2085711 | 14.3820207 | 0 |
| 0.00191329 | 0.00404224 | 21.8635297 | 21.5767375 | 25.0574018 | 32.4755306 | 0 |
| 0.00191465 | 0.00404486 | 10.9317649 | 19.3523315 | 25.9854537 | 32.0115944 | 0 |
| 0.00191548 | 0.00404635 | 190.984363 | 336.552616 | 397.20622 | 355.839028 | 2.37621824 |
| 0.00191665 | 0.00404856 | 0 | 170.611934 | 129.927268 | 131.757867 | 0 |
| 0.00191875 | 0.00405275 | 21.5420072 | 15.1259603 | 4.64025958 | 26.4443606 | 0 |
| 0.0019201 | 0.00405536 | 0 | 0 | 0 | 0 | 14.2573095 |
| 0.00192186 | 0.00405882 | 9.00262989 | 7.56298014 | 19.4890903 | 29.6919137 | 0 |
| 0.00192279 | 0.00406053 | 22.1850522 | 53.6081828 | 51.0428554 | 54.7444659 | 11.8810912 |
| 0.00192334 | 0.00406143 | 3.21522496 | 18.4625692 | 25.0574018 | 44.0739344 | 0 |
| 0.00192731 | 0.00406955 | 8.0380624 | 13.7913167 | 16.7049345 | 10.6705315 | 0 |
| 0.0019285 | 0.00407183 | 8.6811074 | 19.3523315 | 14.8488307 | 9.27872303 | 0 |
| 0.00192893 | 0.00407248 | 12.5393773 | 25.5806681 | 5.5683115 | 19.9492545 | 0 |
| 0.00193201 | 0.00407872 | 39.8687895 | 80.0786133 | 14.8488307 | 75.6215927 | 9.50487297 |
| 0.00193289 | 0.00408033 | 67.8412467 | 33.8109701 | 25.9854537 | 38.9706367 | 11.8810912 |
| 0.00193496 | 0.00408442 | 3.85826995 | 44.0432373 | 13.9207788 | 26.9082968 | 0 |
| 0.00193799 | 0.00409057 | 17.6837373 | 9.78738607 | 7.42441534 | 20.8771268 | 0 |
| 0.00193956 | 0.00409362 | 190.019795 | 186.627657 | 136.423632 | 171.656376 | 254.255352 |
| 0.0019417 | 0.00409789 | 28.9370247 | 16.6830444 | 16.7049345 | 14.3820207 | 71.2865473 |
| 0.00194186 | 0.00409797 | 11.8963324 | 7.78542074 | 14.8488307 | 13.4541484 | 0 |
| 0.00194603 | 0.00410651 | 112.211351 | 72.0707519 | 85.3807764 | 93.2511664 | 23.7621824 |
| 0.00194653 | 0.0041073 | 172.65758 | 100.543148 | 94.6612955 | 102.993826 | 178.216368 |
| 0.00194777 | 0.00410967 | 68.1627692 | 144.586385 | 0 | 115.056166 | 0 |
| 0.00194799 | 0.00410986 | 10.2887199 | 12.4566732 | 14.8488307 | 8.81478687 | 0 |
| 0.00194994 | 0.00411373 | 0 | 0 | 0 | 0 | 33.2670554 |
| 0.00195049 | 0.00411464 | 0 | 243.127568 | 185.610383 | 56.1362743 | 0 |
| 0.00195094 | 0.00411533 | 0 | 130.127747 | 167.977397 | 129.902122 | 0 |
| 0.00195577 | 0.00412526 | 4.17979245 | 28.2499552 | 22.273246 | 17.6295737 | 0 |
| 0.00195679 | 0.00412714 | 72.3425616 | 123.009648 | 113.222334 | 122.94308 | 26.1384007 |

| | | | | | | |
|---|---|---|---|---|---|---|
| 0.00195966 | 0.00413293 | 5.14435994 | 29.3621582 | 26.9135056 | 11.5984038 | 0 |
| 0.00196029 | 0.00413401 | 124.107684 | 22.4664998 | 31.5537652 | 32.4755306 | 268.512662 |
| 0.00196224 | 0.00413786 | 127.965953 | 213.542969 | 225.516616 | 223.153289 | 0 |
| 0.00196756 | 0.00414882 | 3.21522496 | 21.5767375 | 25.9854537 | 31.0837221 | 0 |
| 0.00196987 | 0.00415342 | 30.2231146 | 32.2538859 | 8.35246725 | 23.6607437 | 2.37621824 |
| 0.00197258 | 0.00415889 | 330.525126 | 425.751294 | 374.932974 | 353.519347 | 59.4054561 |
| 0.00197332 | 0.00416018 | 113.818964 | 70.7361084 | 38.9781805 | 51.9608489 | 16.6335277 |
| 0.00197514 | 0.00416351 | 43.7270595 | 147.478113 | 0 | 166.553078 | 0 |
| 0.00197506 | 0.00416351 | 6.43044992 | 10.2322673 | 17.6329864 | 53.8165936 | 0 |
| 0.0019757 | 0.00416441 | 43.405537 | 107.438806 | 0 | 225.009033 | 0 |
| 0.00197763 | 0.00416823 | 22.8280972 | 177.062712 | 324.818171 | 252.845202 | 26.1384007 |
| 0.00198047 | 0.00417395 | 139.540763 | 239.346077 | 162.409085 | 230.112331 | 38.0194919 |
| 0.00198071 | 0.00417419 | 0 | 106.104163 | 183.75428 | 140.572654 | 0 |
| 0.00198742 | 0.00418807 | 11.2532874 | 14.4586385 | 7.42441534 | 15.7738291 | 0 |
| 0.00198913 | 0.0041914 | 9.32415239 | 16.0157227 | 12.9927268 | 9.27872303 | 0 |
| 0.00199036 | 0.00419375 | 15.1115573 | 10.4547078 | 11.136623 | 9.27872303 | 0 |
| 0.00199057 | 0.00419392 | 6.43044992 | 26.6928711 | 14.8488307 | 12.9902122 | 0 |
| 0.00199192 | 0.0041965 | 7.07349491 | 12.6791138 | 15.7768826 | 14.3820207 | 0 |
| 0.00199456 | 0.00420179 | 3.53674746 | 28.917277 | 22.273246 | 22.2689353 | 0 |
| 0.00199531 | 0.00420312 | 19.9343948 | 30.02948 | 37.1220767 | 47.7854236 | 4.75243649 |
| 0.00200165 | 0.00421621 | 7.71653991 | 16.2381632 | 10.2085711 | 16.7017014 | 0 |
| 0.00200238 | 0.00421748 | 7.07349491 | 16.2381632 | 12.0646749 | 15.309893 | 0 |
| 0.00200338 | 0.00421933 | 3.21522496 | 5.33857422 | 7.42441534 | 9.74265918 | 23.7621824 |
| 0.00201049 | 0.00423403 | 2.89370247 | 32.2538859 | 33.409869 | 19.4853184 | 0 |
| 0.00201084 | 0.00423451 | 9.64567488 | 7.11809896 | 35.2659728 | 17.6295737 | 0 |
| 0.00201244 | 0.0042376 | 21.2204847 | 36.9251383 | 47.3306478 | 57.0641466 | 11.8810912 |
| 0.00201656 | 0.00424602 | 109.639171 | 92.5352865 | 74.2441534 | 81.6527626 | 23.7621824 |
| 0.00201698 | 0.00424665 | 261.397789 | 399.948185 | 372.148819 | 480.637853 | 85.5438568 |
| 0.00202268 | 0.00425838 | 0 | 171.279256 | 431.544141 | 341.920944 | 0 |
| 0.00202283 | 0.00425841 | 25.7217997 | 13.5688761 | 5.5683115 | 16.7017014 | 0 |
| 0.00202347 | 0.0042595 | 13.8254673 | 14.4586385 | 6.49636342 | 15.7738291 | 0 |
| 0.00202863 | 0.00427011 | 57.5525268 | 186.405216 | 0 | 97.4265918 | 0 |
| 0.00202897 | 0.00427054 | 27.6509347 | 15.3484009 | 15.7768826 | 13.9180845 | 35.6432737 |
| 0.0020303 | 0.00427307 | 56.2664368 | 106.104163 | 66.819738 | 105.313506 | 21.3859642 |
| 0.00203348 | 0.00427951 | 4.17979245 | 17.3503662 | 20.4171422 | 31.0837221 | 0 |
| 0.00203407 | 0.00428047 | 224.422702 | 181.066642 | 167.049345 | 180.471163 | 0 |
| 0.00203766 | 0.00428776 | 54.9803468 | 199.529211 | 161.481034 | 139.644782 | 0 |
| 0.00204035 | 0.00429317 | 53.6942569 | 117.448633 | 83.5246725 | 98.3544641 | 33.2670554 |
| 0.00204727 | 0.00430745 | 0 | 254.472038 | 50.1148035 | 179.543291 | 0 |
| 0.00205053 | 0.00431377 | 18.3267823 | 14.9035197 | 6.49636342 | 12.5262761 | 0 |
| 0.00205044 | 0.00431377 | 7.39501741 | 14.0137573 | 13.9207788 | 12.5262761 | 0 |
| 0.00205068 | 0.0043138 | 0 | 221.995711 | 73.3161014 | 151.707121 | 0 |
| 0.00205366 | 0.0043198 | 6.43044992 | 12.011792 | 16.7049345 | 20.4131907 | 0 |

| | | | | | | |
|---|---|---|---|---|---|---|
| 0.0020566 | 0.00432572 | 7.71653991 | 23.1338216 | 25.9854537 | 7.88691457 | 0 |
| 0.00205997 | 0.00433254 | 11.2532874 | 14.4586385 | 10.2085711 | 9.27872303 | 0 |
| 0.0020616 | 0.0043357 | 4.17979245 | 19.1298909 | 25.9854537 | 19.9492545 | 0 |
| 0.00206203 | 0.00433633 | 6.75197242 | 24.9133464 | 13.9207788 | 12.5262761 | 0 |
| 0.00206508 | 0.00434247 | 125.715296 | 96.0943359 | 0 | 86.2921241 | 0 |
| 0.00206544 | 0.00434296 | 36.0105196 | 32.6987671 | 8.35246725 | 5.56723382 | 0 |
| 0.00206802 | 0.00434784 | 7.39501741 | 24.023584 | 7.42441534 | 34.7952113 | 0 |
| 0.00206799 | 0.00434784 | 0 | 0 | 0 | 0 | 16.6335277 |
| 0.00207171 | 0.00435533 | 11.5748099 | 18.9074504 | 5.5683115 | 34.3312752 | 0 |
| 0.002072 | 0.00435567 | 75.8793091 | 114.112024 | 123.430905 | 115.056166 | 47.5243649 |
| 0.00207726 | 0.00436645 | 16.7191698 | 25.5806681 | 36.1940248 | 22.2689353 | 4.75243649 |
| 0.0020799 | 0.00437173 | 47.5853294 | 37.1475789 | 47.3306478 | 41.2903175 | 130.692003 |
| 0.0020821 | 0.00437608 | 49.1929419 | 12.011792 | 5.5683115 | 16.7017014 | 0 |
| 0.00208328 | 0.00437828 | 0.96456749 | 0.44488118 | 1.85610383 | 0.9278723 | 11.8810912 |
| 0.00208763 | 0.0043869 | 24.1141872 | 47.3798462 | 48.2586997 | 42.2181898 | 11.8810912 |
| 0.00208764 | 0.0043869 | 119.284846 | 404.174556 | 679.334003 | 414.758919 | 0 |
| 0.00208791 | 0.0043872 | 185.51848 | 72.2931925 | 50.1148035 | 61.7035081 | 28.5146189 |
| 0.00208884 | 0.00438888 | 0 | 99.2085042 | 149.416359 | 169.800631 | 0 |
| 0.00209006 | 0.00439116 | 6.75197242 | 16.0157227 | 21.3451941 | 11.1344676 | 0 |
| 0.00209348 | 0.00439807 | 6.43044992 | 18.017688 | 36.1940248 | 10.2065953 | 0 |
| 0.00209567 | 0.0044024 | 14.1469898 | 10.2322673 | 34.3379209 | 60.3116997 | 0 |
| 0.00209814 | 0.00440732 | 9.00262989 | 5.33857422 | 28.7696094 | 39.898509 | 0 |
| 0.0021038 | 0.00441893 | 9.00262989 | 16.2381632 | 12.9927268 | 9.27872303 | 0 |
| 0.00210452 | 0.00441989 | 9.00262989 | 11.5669108 | 11.136623 | 12.0623399 | 0 |
| 0.00210446 | 0.00441989 | 18.0052598 | 4.89369303 | 15.7768826 | 26.9082968 | 0 |
| 0.0021077 | 0.00442628 | 6.43044992 | 18.2401286 | 18.5610383 | 11.5984038 | 0 |
| 0.00210927 | 0.00442932 | 32.1522496 | 73.6278361 | 125.287009 | 15.7738291 | 9.50487297 |
| 0.00210966 | 0.00442985 | 82.6312815 | 159.712345 | 139.207788 | 0 | 0 |
| 0.00211173 | 0.00443391 | 12.5393773 | 38.0373413 | 9.28051917 | 9.27872303 | 0 |
| 0.00211263 | 0.00443553 | 114.783531 | 129.015544 | 126.215061 | 0 | 0 |
| 0.00211583 | 0.00444198 | 9.64567488 | 7.34053955 | 19.4890903 | 18.0935099 | 0 |
| 0.00211598 | 0.00444201 | 0 | 65.6199748 | 199.531162 | 177.687546 | 0 |
| 0.00211762 | 0.00444518 | 70.7349491 | 110.997856 | 222.73246 | 0 | 0 |
| 0.00212089 | 0.00445176 | 51.7651219 | 46.4900838 | 37.1220767 | 38.9706367 | 0 |
| 0.00212282 | 0.00445555 | 8.3595849 | 11.7893514 | 12.9927268 | 11.5984038 | 0 |
| 0.00212409 | 0.00445765 | 81.9882365 | 102.989994 | 0 | 118.303719 | 0 |
| 0.00212403 | 0.00445765 | 6.10892743 | 24.023584 | 10.2085711 | 23.1968076 | 0 |
| 0.00212594 | 0.00446125 | 51.4435994 | 195.525281 | 0 | 95.5708472 | 0 |
| 0.00212885 | 0.00446707 | 10.6102424 | 16.6830444 | 7.42441534 | 14.3820207 | 0 |
| 0.00213414 | 0.00447763 | 0 | 186.405216 | 137.351684 | 94.1790387 | 0 |
| 0.00213411 | 0.00447763 | 30.8661596 | 21.5767375 | 27.8415575 | 25.9804245 | 7.12865473 |
| 0.00213976 | 0.00448913 | 6.10892743 | 16.2381632 | 21.3451941 | 12.5262761 | 0 |
| 0.00214492 | 0.00449967 | 22.5065747 | 22.0216187 | 58.4672708 | 8.81478687 | 2.37621824 |

| | | | | | | |
|---|---|---|---|---|---|---|
| 0.00214522 | 0.00450002 | 300.623534 | 992.752364 | 864.016335 | 935.295281 | 0 |
| 0.00214568 | 0.00450071 | 0 | 21.1318563 | 268.207004 | 284.856797 | 0 |
| 0.00214582 | 0.00450073 | 8.0380624 | 39.1495443 | 28.7696094 | 24.588616 | 0 |
| 0.00214692 | 0.00450275 | 0 | 103.212435 | 145.704151 | 160.985845 | 0 |
| 0.00214741 | 0.0045035 | 0 | 151.037162 | 119.718697 | 135.00542 | 0 |
| 0.00214944 | 0.00450748 | 8.0380624 | 18.4625692 | 31.5537652 | 7.88691457 | 0 |
| 0.00215135 | 0.00451121 | 6.75197242 | 8.23030192 | 33.409869 | 23.1968076 | 0 |
| 0.00215307 | 0.00451453 | 0 | 0 | 0 | 0 | 0 |
| 0.00215446 | 0.00451716 | 0 | 0 | 0 | 0 | 38.0194919 |
| 0.00215679 | 0.00452159 | 64.9475442 | 92.0904053 | 84.4527244 | 83.9724434 | 0 |
| 0.00215685 | 0.00452159 | 0 | 0 | 0 | 0 | 9.50487297 |
| 0.0021599 | 0.00452772 | 13.8254673 | 10.2322673 | 7.42441534 | 11.1344676 | 35.6432737 |
| 0.00216303 | 0.00453399 | 0 | 130.795068 | 104.869867 | 173.976057 | 0 |
| 0.00216464 | 0.00453709 | 15.1115573 | 79.6337321 | 38.9781805 | 66.8068058 | 9.50487297 |
| 0.00216591 | 0.00453946 | 1.60761248 | 0.44488118 | 2.78415575 | 1.39180845 | 16.6335277 |
| 0.00217214 | 0.00455195 | 0 | 164.161157 | 136.423632 | 106.241379 | 0 |
| 0.00217208 | 0.00455195 | 0.3215225 | 0 | 0.92805192 | 0 | 0 |
| 0.00217291 | 0.00455328 | 2.57217997 | 32.2538859 | 19.4890903 | 36.650956 | 0 |
| 0.00217457 | 0.00455647 | 92.5984789 | 74.5175985 | 0 | 139.180845 | 0 |
| 0.0021844 | 0.0045768 | 12.2178549 | 10.6771484 | 25.9854537 | 7.88691457 | 0 |
| 0.00218486 | 0.00457747 | 63.0184092 | 85.8620687 | 0 | 173.512121 | 0 |
| 0.0021884 | 0.00458459 | 0 | 0 | 0 | 0 | 23.7621824 |
| 0.00218968 | 0.00458699 | 6.43044992 | 18.2401286 | 12.0646749 | 16.2377653 | 0 |
| 0.00219375 | 0.00459524 | 460.098692 | 27.3601929 | 0 | 46.8575513 | 0 |
| 0.00219445 | 0.00459642 | 7.39501741 | 10.6771484 | 13.9207788 | 19.0213822 | 0 |
| 0.00219467 | 0.00459659 | 0 | 66.7321777 | 225.516616 | 144.748079 | 0 |
| 0.00219716 | 0.00460152 | 36.6535646 | 80.5234945 | 56.6111669 | 88.1478687 | 0 |
| 0.00219912 | 0.00460534 | 0 | 0.66732178 | 0.92805192 | 0.46393615 | 26.1384007 |
| 0.00220033 | 0.0046076 | 126.358341 | 0 | 312.753496 | 66.3428696 | 0 |
| 0.00220468 | 0.00461642 | 10.9317649 | 23.1338216 | 12.0646749 | 7.88691457 | 0 |
| 0.00220614 | 0.00461918 | 57.2310043 | 26.6928711 | 30.6257133 | 29.6919137 | 26.1384007 |
| 0.00220666 | 0.00461999 | 0 | 0 | 9.28051917 | 0 | 21.3859642 |
| 0.00220769 | 0.00462186 | 127.001386 | 68.0668213 | 0 | 108.561059 | 0 |
| 0.00221216 | 0.00463092 | 64.3044992 | 113.444702 | 0 | 128.046378 | 0 |
| 0.00221299 | 0.00463237 | 687.415097 | 1539.73378 | 1046.84256 | 1234.5341 | 0 |
| 0.00221346 | 0.00463305 | 7.39501741 | 10.0098267 | 14.8488307 | 23.6607437 | 0 |
| 0.00221518 | 0.00463638 | 68.4842917 | 75.6298014 | 0 | 177.22361 | 0 |
| 0.00221838 | 0.00464279 | 11.5748099 | 11.7893514 | 11.136623 | 8.81478687 | 0 |
| 0.00221939 | 0.00464462 | 3.53674746 | 19.3523315 | 21.3451941 | 31.5476583 | 0 |
| 0.00222052 | 0.00464669 | 59.8031843 | 82.0805786 | 91.8771398 | 88.1478687 | 30.8908372 |
| 0.00222339 | 0.00465239 | 70.7349491 | 70.7361084 | 0 | 181.862971 | 0 |
| 0.0022245 | 0.00465443 | 164.941041 | 127.013578 | 0 | 89.5396772 | 0 |
| 0.00222496 | 0.00465511 | 46.2992394 | 178.619796 | 187.466487 | 0 | 0 |

| | | | | | | |
|---|---|---|---|---|---|---|
| 0.00223146 | 0.00466842 | 0 | 118.560836 | 134.567528 | 144.284143 | 0 |
| 0.00223402 | 0.00467347 | 11.2532874 | 7.56298014 | 24.1293498 | 12.0623399 | 0 |
| 0.00223728 | 0.00468001 | 33.7598621 | 23.1338216 | 18.5610383 | 19.9492545 | 4.75243649 |
| 0.00224087 | 0.00468722 | 21.8635297 | 27.3601929 | 4.64025958 | 12.9902122 | 0 |
| 0.00224243 | 0.0046902 | 6.75197242 | 36.9251383 | 41.7623363 | 22.7328714 | 2.37621824 |
| 0.00224494 | 0.00469516 | 9.64567488 | 21.3542969 | 8.35246725 | 12.5262761 | 0 |
| 0.00224642 | 0.00469797 | 57.5525268 | 169.944613 | 0 | 91.859358 | 0 |
| 0.00225249 | 0.00471037 | 18.0052598 | 11.1220296 | 6.49636342 | 16.2377653 | 0 |
| 0.00225699 | 0.00471949 | 9.00262989 | 16.6830444 | 12.0646749 | 9.27872303 | 0 |
| 0.00225716 | 0.00471954 | 17.3622148 | 26.0255493 | 2.78415575 | 42.6821259 | 0 |
| 0.00226034 | 0.0047259 | 262.683879 | 430.867428 | 356.371936 | 413.831047 | 0 |
| 0.00226474 | 0.00473474 | 90.6693439 | 0 | 180.970124 | 174.903929 | 0 |
| 0.00226485 | 0.00473474 | 7.07349491 | 61.3936035 | 38.0501286 | 41.2903175 | 2.37621824 |
| 0.00226583 | 0.0047365 | 5.14435994 | 8.23030192 | 12.9927268 | 6.03116997 | 14.2573095 |
| 0.00226951 | 0.00474351 | 8.3595849 | 8.00786133 | 17.6329864 | 20.8771268 | 0 |
| 0.0022695 | 0.00474351 | 27.6509347 | 234.897266 | 0 | 122.015208 | 0 |
| 0.00226961 | 0.00474351 | 5.46588243 | 18.9074504 | 50.1148035 | 10.2065953 | 0 |
| 0.00227105 | 0.00474622 | 15.1115573 | 9.78738607 | 8.35246725 | 12.5262761 | 0 |
| 0.0022733 | 0.00475062 | 10.9317649 | 14.6810791 | 6.49636342 | 38.9706367 | 0 |
| 0.00227376 | 0.00475129 | 133.431836 | 321.871537 | 236.653239 | 273.722329 | 11.8810912 |
| 0.00227655 | 0.00475684 | 53.3727344 | 148.145435 | 193.962851 | 0 | 0 |
| 0.00227963 | 0.00476298 | 4.50131495 | 16.4606038 | 23.2012979 | 19.4853184 | 0 |
| 0.00228533 | 0.00477459 | 100.636541 | 539.640877 | 266.3509 | 360.942326 | 4.75243649 |
| 0.00228601 | 0.0047757 | 3.85826995 | 16.4606038 | 27.8415575 | 23.1968076 | 0 |
| 0.00228637 | 0.00477616 | 23.4711422 | 17.7952474 | 3.71220767 | 23.6607437 | 0 |
| 0.00228668 | 0.00477653 | 6.43044992 | 23.1338216 | 9.28051917 | 22.7328714 | 0 |
| 0.00228786 | 0.00477869 | 80.380624 | 102.767554 | 0 | 108.097123 | 0 |
| 0.00228882 | 0.0047804 | 36.0105196 | 68.9565837 | 80.7405168 | 62.1674443 | 0 |
| 0.00228901 | 0.00478049 | 22.8280972 | 63.6180094 | 73.3161014 | 80.7248903 | 14.2573095 |
| 0.00228962 | 0.00478141 | 59.1601393 | 126.346257 | 0 | 118.303719 | 0 |
| 0.00228973 | 0.00478141 | 0 | 1.11220296 | 0 | 0 | 35.6432737 |
| 0.00229032 | 0.00478234 | 19.9343948 | 16.6830444 | 4.64025958 | 18.0935099 | 0 |
| 0.00229135 | 0.00478419 | 6.75197242 | 27.3601929 | 9.28051917 | 18.5574461 | 0 |
| 0.00229414 | 0.00478973 | 8.6811074 | 9.12006429 | 12.9927268 | 16.2377653 | 0 |
| 0.00229446 | 0.0047901 | 7.71653991 | 10.4547078 | 29.6976613 | 12.5262761 | 0 |
| 0.00230472 | 0.00481121 | 0 | 347.897087 | 180.970124 | 22.7328714 | 0 |
| 0.00230882 | 0.00481947 | 15.1115573 | 6.45077718 | 26.9135056 | 11.1344676 | 0 |
| 0.00230946 | 0.00482052 | 6.43044992 | 23.1338216 | 11.136623 | 15.7738291 | 0 |
| 0.00231431 | 0.00483034 | 4.50131495 | 17.3503662 | 20.4171422 | 19.9492545 | 0 |
| 0.00231462 | 0.00483068 | 9.32415239 | 9.56494547 | 11.136623 | 13.9180845 | 0 |
| 0.00231632 | 0.00483392 | 8.6811074 | 20.0196533 | 7.42441534 | 18.5574461 | 0 |
| 0.00231762 | 0.00483634 | 7.71653991 | 24.6909058 | 52.8989593 | 5.56723382 | 0 |
| 0.00232446 | 0.00485032 | 254.324294 | 537.194031 | 432.472193 | 485.74115 | 9.50487297 |

| | | | | | | |
|---|---|---|---|---|---|---|
| 0.00232548 | 0.00485214 | 117.355711 | 74.2951579 | 0 | 99.7462725 | 0 |
| 0.00233221 | 0.00486589 | 3.85826995 | 21.3542969 | 16.7049345 | 29.6919137 | 0 |
| 0.00233352 | 0.00486826 | 43.7270595 | 114.334465 | 97.4454513 | 90.9314857 | 21.3859642 |
| 0.00233364 | 0.00486826 | 56.2664368 | 11.7893514 | 2.78415575 | 32.9394667 | 0 |
| 0.00233434 | 0.00486943 | 34.7244296 | 25.5806681 | 24.1293498 | 22.7328714 | 21.3859642 |
| 0.0023363 | 0.00487321 | 32.7952946 | 17.3503662 | 8.35246725 | 7.42297842 | 0 |
| 0.0023381 | 0.00487665 | 94.8491364 | 0 | 191.178695 | 151.243185 | 0 |
| 0.00234165 | 0.00488376 | 24.4357097 | 80.7459351 | 56.6111669 | 68.6625504 | 14.2573095 |
| 0.00234239 | 0.004885 | 7.39501741 | 34.4782918 | 25.0574018 | 30.1558498 | 2.37621824 |
| 0.00234267 | 0.00488528 | 11.2532874 | 5.11613363 | 23.2012979 | 27.3722329 | 0 |
| 0.00234345 | 0.00488661 | 0 | 7.56298014 | 0 | 4.63936151 | 71.2865473 |
| 0.00234414 | 0.00488774 | 0 | 135.911202 | 121.574801 | 128.97425 | 0 |
| 0.00234567 | 0.00489062 | 10.9317649 | 6.45077718 | 12.9927268 | 2.78361691 | 7.12865473 |
| 0.00235636 | 0.00491261 | 32.1522496 | 14.9035197 | 6.49636342 | 10.6705315 | 0 |
| 0.00236021 | 0.00492033 | 16.7191698 | 125.234054 | 77.0283091 | 91.859358 | 14.2573095 |
| 0.0023617 | 0.00492314 | 6.10892743 | 14.2361979 | 12.9927268 | 20.4131907 | 0 |
| 0.00236194 | 0.00492333 | 4.50131495 | 28.6948364 | 12.9927268 | 22.7328714 | 0 |
| 0.00236337 | 0.00492601 | 103.851766 | 101.43291 | 0 | 80.7248903 | 0 |
| 0.00236436 | 0.00492777 | 3.53674746 | 4.89369303 | 3.71220767 | 4.63936151 | 49.9005831 |
| 0.00236537 | 0.00492955 | 10.9317649 | 48.9369303 | 65.8916861 | 46.8575513 | 0 |
| 0.00236855 | 0.00493588 | 6.10892743 | 13.123995 | 23.2012979 | 13.4541484 | 0 |
| 0.00236997 | 0.00493854 | 78.7730116 | 0 | 307.185185 | 96.4987195 | 0 |
| 0.00237094 | 0.00494025 | 48.8714194 | 130.127747 | 125.287009 | 102.529889 | 23.7621824 |
| 0.00237155 | 0.00494122 | 3.53674746 | 2.00196533 | 1.85610383 | 2.78361691 | 16.6335277 |
| 0.00237243 | 0.00494264 | 9.96719738 | 20.9094157 | 25.9854537 | 25.9804245 | 0 |
| 0.00237252 | 0.00494264 | 51.1220769 | 97.4289795 | 93.7332436 | 109.488932 | 0 |
| 0.00237694 | 0.00495121 | 248.53689 | 323.428621 | 239.437395 | 291.351903 | 49.9005831 |
| 0.00237687 | 0.00495121 | 161.082771 | 70.7361084 | 0 | 72.8379758 | 0 |
| 0.00237867 | 0.00495452 | 33.4383396 | 14.0137573 | 6.49636342 | 11.1344676 | 0 |
| 0.00238183 | 0.00496079 | 8.3595849 | 12.9015544 | 11.136623 | 11.1344676 | 0 |
| 0.00238271 | 0.00496231 | 86.4895515 | 98.0963013 | 177.257916 | 0 | 0 |
| 0.00238504 | 0.00496687 | 244.035575 | 418.188314 | 676.549847 | 472.287002 | 116.434694 |
| 0.00238674 | 0.0049701 | 28.2939797 | 18.2401286 | 3.71220767 | 19.0213822 | 0 |
| 0.00240107 | 0.00499963 | 1.60761248 | 37.3700195 | 44.546492 | 22.7328714 | 0 |
| 0.00240204 | 0.00500134 | 0 | 62.0609253 | 202.315318 | 150.315313 | 0 |
| 0.00240412 | 0.00500537 | 37.618132 | 10.899589 | 7.42441534 | 12.0623399 | 0 |
| 0.00240442 | 0.00500568 | 10.2887199 | 16.4606038 | 11.136623 | 8.35085072 | 0 |
| 0.00240683 | 0.00501038 | 8.6811074 | 4.22637126 | 3.71220767 | 7.88691457 | 42.7719284 |
| 0.00240857 | 0.0050137 | 6.75197242 | 18.6850098 | 12.9927268 | 12.0623399 | 0 |
| 0.00240993 | 0.00501622 | 46.6207619 | 35.3680542 | 40.8342843 | 26.9082968 | 68.9103291 |
| 0.0024105 | 0.0050171 | 126.679863 | 165.495801 | 190.250643 | 178.615418 | 54.6530196 |
| 0.00241123 | 0.00501831 | 99.3504513 | 94.7596924 | 0 | 87.6839326 | 0 |
| 0.00241223 | 0.00502008 | 151.758618 | 176.172949 | 162.409085 | 167.480951 | 47.5243649 |

| | | | | | | |
|---|---|---|---|---|---|---|
| 0.00241258 | 0.00502048 | 9.00262989 | 10.2322673 | 12.0646749 | 11.1344676 | 0 |
| 0.00241284 | 0.00502072 | 7.71653991 | 14.2361979 | 11.136623 | 12.0623399 | 0 |
| 0.0024147 | 0.00502427 | 23.7926647 | 15.5708415 | 5.5683115 | 12.5262761 | 0 |
| 0.00241631 | 0.00502732 | 3.85826995 | 24.6909058 | 13.9207788 | 32.0115944 | 0 |
| 0.00241776 | 0.00503002 | 5.78740493 | 27.5826335 | 10.2085711 | 19.9492545 | 0 |
| 0.00241804 | 0.0050303 | 3.85826995 | 16.0157227 | 33.409869 | 19.9492545 | 0 |
| 0.00242055 | 0.00503521 | 11.2532874 | 19.5747721 | 14.8488307 | 6.49510612 | 0 |
| 0.0024216 | 0.00503707 | 96.1352263 | 356.127388 | 418.551415 | 382.747325 | 47.5243649 |
| 0.0024233 | 0.0050401 | 8.0380624 | 5.56101481 | 5.5683115 | 9.27872303 | 85.5438568 |
| 0.00242335 | 0.0050401 | 6.43044992 | 19.1298909 | 11.136623 | 15.7738291 | 0 |
| 0.00242401 | 0.00504116 | 27.0078897 | 18.017688 | 6.49636342 | 9.27872303 | 0 |
| 0.00242433 | 0.00504152 | 10.6102424 | 219.771305 | 172.617657 | 209.235204 | 14.2573095 |
| 0.00242461 | 0.00504179 | 4.82283744 | 18.017688 | 13.9207788 | 26.9082968 | 0 |
| 0.00242703 | 0.00504651 | 53.6942569 | 111.442737 | 0 | 134.541484 | 0 |
| 0.00242916 | 0.00505061 | 8.0380624 | 15.7932821 | 15.7768826 | 8.81478687 | 0 |
| 0.00243243 | 0.0050571 | 76.8438766 | 40.2617472 | 98.3735032 | 69.1264865 | 19.0097459 |
| 0.0024347 | 0.00506151 | 3.85826995 | 27.1377523 | 22.273246 | 16.2377653 | 0 |
| 0.00243486 | 0.00506153 | 9.96719738 | 6.67321777 | 4.64025958 | 4.17542536 | 35.6432737 |
| 0.00243544 | 0.00506243 | 0 | 0 | 0 | 0 | 0 |
| 0.00244676 | 0.00508563 | 78.7730116 | 82.7479004 | 0 | 123.407016 | 0 |
| 0.00244758 | 0.00508671 | 10.9317649 | 8.67518311 | 24.1293498 | 9.74265918 | 0 |
| 0.00244755 | 0.00508671 | 82.309759 | 0 | 257.070381 | 112.272549 | 0 |
| 0.00244893 | 0.0050892 | 0.96456749 | 0 | 1.85610383 | 0.46393615 | 7.12865473 |
| 0.00245099 | 0.00509296 | 7.07349491 | 23.3562622 | 7.42441534 | 27.3722329 | 0 |
| 0.00245104 | 0.00509296 | 0 | 131.907271 | 118.790645 | 126.654569 | 0 |
| 0.00245233 | 0.00509532 | 39.2257445 | 11.5669108 | 7.42441534 | 11.1344676 | 0 |
| 0.00245637 | 0.00510341 | 0 | 0 | 0 | 0 | 2.37621824 |
| 0.00246132 | 0.00511338 | 137.290106 | 67.6219401 | 153.128566 | 0 | 0 |
| 0.00246271 | 0.00511594 | 166.22713 | 177.062712 | 125.287009 | 155.418611 | 73.6627655 |
| 0.00246456 | 0.00511948 | 9.32415239 | 16.905485 | 7.42441534 | 15.309893 | 0 |
| 0.0024656 | 0.00512132 | 127.965953 | 52.4959798 | 200.459214 | 0 | 0 |
| 0.00246779 | 0.00512555 | 25.0787547 | 10.4547078 | 8.35246725 | 10.6705315 | 0 |
| 0.00247234 | 0.00513468 | 152.401663 | 83.1927816 | 0 | 61.7035081 | 0 |
| 0.00247267 | 0.00513505 | 11.8963324 | 9.12006429 | 9.28051917 | 12.5262761 | 0 |
| 0.00247352 | 0.00513618 | 4.17979245 | 42.2637126 | 12.0646749 | 21.8049991 | 0 |
| 0.00247346 | 0.00513618 | 5.46588243 | 22.4664998 | 19.4890903 | 11.5984038 | 0 |
| 0.00247672 | 0.00514252 | 12.8608998 | 18.4625692 | 6.49636342 | 12.0623399 | 0 |
| 0.00247908 | 0.00514709 | 12.5393773 | 33.8109701 | 28.7696094 | 34.7952113 | 0 |
| 0.00248066 | 0.00515006 | 0 | 79.1888509 | 182.826228 | 128.97425 | 0 |
| 0.00248241 | 0.00515336 | 8.6811074 | 8.45274251 | 13.9207788 | 15.7738291 | 0 |
| 0.00248891 | 0.00516655 | 27.3294122 | 17.5728068 | 10.2085711 | 6.03116997 | 0 |
| 0.00249178 | 0.00517218 | 7.71653991 | 35.8129354 | 28.7696094 | 5.56723382 | 0 |
| 0.00249532 | 0.00517921 | 6.10892743 | 8.45274251 | 5.5683115 | 5.56723382 | 40.3957101 |

| | | | | | | |
|---|---|---|---|---|---|---|
| 0.00249751 | 0.00518344 | 153.687753 | 147.033232 | 146.632203 | 133.149675 | 61.7816743 |
| 0.00250371 | 0.00519579 | 8.0380624 | 17.3503662 | 9.28051917 | 12.9902122 | 0 |
| 0.00250377 | 0.00519579 | 132.788791 | 69.846346 | 70.5319457 | 74.6937204 | 0 |
| 0.00250447 | 0.00519691 | 6.75197242 | 27.5826335 | 29.6976613 | 6.95904227 | 0 |
| 0.00250604 | 0.00519985 | 25.0787547 | 44.4881185 | 8.35246725 | 5.10329766 | 0 |
| 0.00250757 | 0.00520271 | 80.7021465 | 183.513489 | 175.401812 | 194.853184 | 33.2670554 |
| 0.00251173 | 0.00521102 | 0 | 0 | 0 | 0 | 23.7621824 |
| 0.00251369 | 0.00521476 | 8.0380624 | 10.4547078 | 12.9927268 | 12.0623399 | 0 |
| 0.00251765 | 0.00522267 | 0 | 158.377702 | 112.294282 | 105.777442 | 0 |
| 0.00251978 | 0.00522676 | 9.64567488 | 21.3542969 | 6.49636342 | 17.1656376 | 0 |
| 0.00252058 | 0.0052281 | 7.71653991 | 18.4625692 | 10.2085711 | 12.0623399 | 0 |
| 0.00252089 | 0.00522841 | 60.1247068 | 127.458459 | 0 | 99.7462725 | 0 |
| 0.00252122 | 0.00522878 | 0 | 123.009648 | 130.85532 | 117.839782 | 0 |
| 0.00252317 | 0.0052325 | 0.64304499 | 22.2440592 | 74.2441534 | 55.208402 | 0 |
| 0.00252414 | 0.00523419 | 5.46588243 | 15.1259603 | 14.8488307 | 18.0935099 | 0 |
| 0.00252473 | 0.0052351 | 20.5774398 | 20.4645345 | 7.42441534 | 7.88691457 | 0 |
| 0.00252731 | 0.00524011 | 27.0078897 | 30.9192423 | 10.2085711 | 4.63936151 | 0 |
| 0.00252787 | 0.00524095 | 29.2585471 | 20.4645345 | 24.1293498 | 14.8459568 | 114.058476 |
| 0.00253286 | 0.00525098 | 55.3018693 | 229.558691 | 189.322591 | 176.295737 | 30.8908372 |
| 0.0025334 | 0.00525176 | 84.238894 | 84.7498657 | 0 | 107.169251 | 0 |
| 0.00253433 | 0.00525337 | 5.46588243 | 8.00786133 | 62.1794784 | 19.9492545 | 0 |
| 0.00253455 | 0.00525352 | 9.64567488 | 7.78542074 | 12.9927268 | 16.7017014 | 0 |
| 0.00253741 | 0.0052591 | 73.9501741 | 190.854028 | 174.47376 | 208.771268 | 2.37621824 |
| 0.00253867 | 0.00526141 | 104.173289 | 123.232088 | 0 | 58.9198912 | 0 |
| 0.00254364 | 0.00527137 | 0 | 43.8207967 | 289.552198 | 118.303719 | 0 |
| 0.00254443 | 0.0052727 | 47.9068519 | 211.763444 | 0 | 69.5904227 | 0 |
| 0.00254481 | 0.00527314 | 364.928033 | 1399.15133 | 1086.74879 | 1213.19304 | 0 |
| 0.002546 | 0.00527497 | 22.5065747 | 40.7066284 | 0 | 8.35085072 | 87.920075 |
| 0.00254594 | 0.00527497 | 3.85826995 | 22.6889404 | 26.9135056 | 15.309893 | 0 |
| 0.00254834 | 0.00527949 | 7.71653991 | 14.2361979 | 15.7768826 | 9.27872303 | 0 |
| 0.002549 | 0.00528053 | 88.7402089 | 149.035197 | 103.941815 | 0 | 0 |
| 0.00254968 | 0.00528161 | 32.4737721 | 4.22637126 | 16.7049345 | 16.2377653 | 0 |
| 0.00255053 | 0.00528305 | 3.53674746 | 24.6909058 | 19.4890903 | 21.341063 | 0 |
| 0.00255093 | 0.00528355 | 5.46588243 | 17.3503662 | 13.9207788 | 16.7017014 | 0 |
| 0.00255118 | 0.00528374 | 69.1273367 | 232.67286 | 167.977397 | 216.194247 | 33.2670554 |
| 0.00255255 | 0.00528626 | 0 | 101.21047 | 129.927268 | 140.572654 | 0 |
| 0.00256165 | 0.00530413 | 3.21522496 | 28.6948364 | 39.9062324 | 12.9902122 | 0 |
| 0.00256151 | 0.00530413 | 18.9698273 | 10.4547078 | 8.35246725 | 10.2065953 | 0 |
| 0.00256146 | 0.00530413 | 13.5039448 | 15.1259603 | 20.4171422 | 22.2689353 | 2.37621824 |
| 0.00256233 | 0.0053052 | 3.53674746 | 23.3562622 | 19.4890903 | 22.2689353 | 0 |
| 0.00256255 | 0.00530533 | 70.7349491 | 56.4999105 | 0 | 180.935099 | 0 |
| 0.00256416 | 0.00530835 | 0 | 203.088261 | 74.2441534 | 115.056166 | 0 |
| 0.00256561 | 0.00531102 | 8.3595849 | 10.0098267 | 12.9927268 | 11.5984038 | 0 |

| | | | | | | |
|---|---|---|---|---|---|---|
| 0.00256624 | 0.005312 | 18.0052598 | 12.011792 | 12.0646749 | 17.1656376 | 68.9103291 |
| 0.00256751 | 0.00531431 | 2.89370247 | 27.5826335 | 38.9781805 | 15.309893 | 0 |
| 0.00256854 | 0.0053161 | 31.1876821 | 97.8738607 | 187.466487 | 128.046378 | 21.3859642 |
| 0.0025694 | 0.00531755 | 116.712666 | 71.8483114 | 0 | 89.0757411 | 0 |
| 0.00257232 | 0.00532326 | 6.75197242 | 6.67321777 | 3.71220767 | 8.35085072 | 2.37621824 |
| 0.00257342 | 0.00532522 | 67.8412467 | 225.777201 | 222.73246 | 266.299351 | 52.2768014 |
| 0.00258138 | 0.00534137 | 17.0406923 | 3.78149007 | 7.42441534 | 6.49510612 | 28.5146189 |
| 0.00258283 | 0.00534403 | 30.2231146 | 78.2990885 | 63.1075304 | 79.797018 | 16.6335277 |
| 0.0025837 | 0.00534537 | 0 | 0 | 0 | 4.63936151 | 64.1578926 |
| 0.0025838 | 0.00534537 | 51.4435994 | 103.657316 | 0 | 136.861165 | 0 |
| 0.00258457 | 0.00534664 | 19.9343948 | 10.2322673 | 7.42441534 | 12.0623399 | 0 |
| 0.00258679 | 0.00535058 | 9.64567488 | 16.905485 | 8.35246725 | 11.1344676 | 0 |
| 0.00258676 | 0.00535058 | 2.25065747 | 61.3936035 | 10.2085711 | 41.2903175 | 0 |
| 0.00258933 | 0.0053555 | 76.5223541 | 12.6791138 | 4.64025958 | 12.5262761 | 0 |
| 0.00259082 | 0.00535825 | 29.2585471 | 9.12006429 | 8.35246725 | 12.0623399 | 0 |
| 0.00259184 | 0.00536004 | 9.64567488 | 4.89369303 | 6.49636342 | 5.10329766 | 57.0292378 |
| 0.00259329 | 0.00536271 | 54.6588243 | 137.023405 | 125.287009 | 133.149675 | 26.1384007 |
| 0.00259539 | 0.00536673 | 57.2310043 | 129.015544 | 0 | 98.8184002 | 0 |
| 0.00259756 | 0.00537087 | 6.43044992 | 16.2381632 | 11.136623 | 15.7738291 | 0 |
| 0.00260289 | 0.00538156 | 7.39501741 | 38.4822225 | 4.64025958 | 36.1870198 | 0 |
| 0.00260502 | 0.00538564 | 13.1824223 | 12.6791138 | 8.35246725 | 9.27872303 | 0 |
| 0.00260718 | 0.00538978 | 0 | 73.1829549 | 149.416359 | 157.738291 | 0 |
| 0.00260858 | 0.00539233 | 21.2204847 | 189.296944 | 0 | 154.954675 | 0 |
| 0.00261427 | 0.00540377 | 63.6614542 | 92.9801676 | 0 | 122.479144 | 0 |
| 0.00261545 | 0.00540588 | 3.53674746 | 35.3680542 | 27.8415575 | 12.5262761 | 0 |
| 0.00261782 | 0.00541044 | 14.7900348 | 32.4763265 | 11.136623 | 24.1246799 | 0 |
| 0.00261851 | 0.00541152 | 123.786161 | 35.5904948 | 0 | 155.882547 | 0 |
| 0.00261933 | 0.00541289 | 10.2887199 | 4.44881185 | 24.1293498 | 38.5067006 | 0 |
| 0.00262135 | 0.00541674 | 8.3595849 | 12.2342326 | 17.6329864 | 8.81478687 | 0 |
| 0.00262575 | 0.00542551 | 9.64567488 | 15.3484009 | 8.35246725 | 11.1344676 | 0 |
| 0.00263271 | 0.00543954 | 8.0380624 | 24.6909058 | 23.2012979 | 6.49510612 | 0 |
| 0.0026343 | 0.00544249 | 3.85826995 | 5.56101481 | 9.28051917 | 6.03116997 | 128.315785 |
| 0.002636 | 0.00544566 | 16.0761248 | 22.0216187 | 5.5683115 | 11.5984038 | 0 |
| 0.0026384 | 0.00545029 | 5.46588243 | 12.4566732 | 18.5610383 | 17.6295737 | 0 |
| 0.00264057 | 0.00545444 | 17.3622148 | 28.0275146 | 45.4745439 | 27.3722329 | 7.12865473 |
| 0.00264107 | 0.00545514 | 145.328168 | 44.7105591 | 57.5392189 | 33.867339 | 470.491212 |
| 0.00264693 | 0.00546692 | 9.00262989 | 12.9015544 | 9.28051917 | 11.1344676 | 0 |
| 0.00264803 | 0.00546884 | 3.85826995 | 23.5787028 | 19.4890903 | 18.0935099 | 0 |
| 0.00264932 | 0.00547117 | 11.5748099 | 13.123995 | 39.9062324 | 6.03116997 | 0 |
| 0.0026502 | 0.00547265 | 0 | 0 | 0 | 0 | 52.2768014 |
| 0.00265263 | 0.00547734 | 2.57217997 | 22.6889404 | 31.5537652 | 23.6607437 | 0 |
| 0.00265557 | 0.00548306 | 83.2743265 | 326.54279 | 345.235313 | 276.04201 | 42.7719284 |
| 0.00265711 | 0.00548592 | 208.025055 | 104.324638 | 38.0501286 | 205.059779 | 30.8908372 |

| | | | | | | |
|---|---|---|---|---|---|---|
| 0.00266249 | 0.00549668 | 18.3267823 | 6.89565837 | 11.136623 | 12.5262761 | 0 |
| 0.00266376 | 0.00549897 | 34.0813846 | 211.096122 | 0 | 89.0757411 | 0 |
| 0.00266523 | 0.00550166 | 13.1824223 | 18.6850098 | 37.1220767 | 4.17542536 | 0 |
| 0.00266573 | 0.00550236 | 36.9750871 | 26.4704305 | 20.4171422 | 26.4443606 | 2.37621824 |
| 0.0026716 | 0.00551415 | 6.10892743 | 15.3484009 | 10.2085711 | 33.867339 | 0 |
| 0.00267313 | 0.00551695 | 4.50131495 | 21.1318563 | 14.8488307 | 19.4853184 | 0 |
| 0.00267331 | 0.00551698 | 56.2664368 | 108.328569 | 0 | 114.592229 | 0 |
| 0.0026738 | 0.00551766 | 0 | 82.970341 | 155.912722 | 130.366059 | 0 |
| 0.00267758 | 0.00552512 | 51.7651219 | 50.4940145 | 35.2659728 | 38.9706367 | 16.6335277 |
| 0.00268058 | 0.00553097 | 3.85826995 | 25.1357869 | 51.9709073 | 9.74265918 | 0 |
| 0.00268122 | 0.00553196 | 10.2887199 | 16.905485 | 6.49636342 | 15.309893 | 0 |
| 0.00268365 | 0.00553663 | 12.5393773 | 16.2381632 | 19.4890903 | 24.588616 | 0 |
| 0.00268571 | 0.00554054 | 6.43044992 | 13.3464355 | 11.136623 | 23.1968076 | 0 |
| 0.00269095 | 0.00555101 | 19.9343948 | 10.2322673 | 45.4745439 | 4.63936151 | 0 |
| 0.00269116 | 0.00555111 | 34.4029071 | 13.3464355 | 5.5683115 | 12.5262761 | 0 |
| 0.00269266 | 0.00555386 | 8.0380624 | 35.1456136 | 4.64025958 | 32.4755306 | 0 |
| 0.00269466 | 0.00555764 | 6.43044992 | 10.4547078 | 19.4890903 | 14.8459568 | 0 |
| 0.00269766 | 0.00556349 | 4.17979245 | 4.89369303 | 3.71220767 | 6.95904227 | 19.0097459 |
| 0.00269827 | 0.00556441 | 6.75197242 | 11.5669108 | 17.6329864 | 12.0623399 | 0 |
| 0.002699 | 0.00556558 | 1.60761248 | 1.11220296 | 1.85610383 | 2.31968076 | 7.12865473 |
| 0.00269983 | 0.00556693 | 4.82283744 | 13.3464355 | 18.5610383 | 21.8049991 | 0 |
| 0.00270192 | 0.00557092 | 56.5879593 | 33.5885295 | 36.1940248 | 30.619786 | 35.6432737 |
| 0.00270816 | 0.00558343 | 35.0459521 | 35.1456136 | 30.6257133 | 20.4131907 | 9.50487297 |
| 0.00271216 | 0.00559134 | 0 | 80.5234945 | 149.416359 | 136.861165 | 0 |
| 0.00271273 | 0.00559216 | 10.6102424 | 3.55904948 | 4.64025958 | 0.9278723 | 30.8908372 |
| 0.00271416 | 0.00559477 | 6.10892743 | 24.2460246 | 7.42441534 | 32.4755306 | 0 |
| 0.0027208 | 0.00560811 | 0 | 0 | 0 | 0 | 0 |
| 0.00272278 | 0.00561186 | 52.4081669 | 24.6909058 | 21.3451941 | 1.85574461 | 0 |
| 0.00272446 | 0.00561497 | 24.7572322 | 17.1279256 | 17.6329864 | 15.7738291 | 19.0097459 |
| 0.00273019 | 0.00562643 | 34.4029071 | 69.6239054 | 33.409869 | 63.5592527 | 14.2573095 |
| 0.00273372 | 0.00563336 | 7.71653991 | 11.3444702 | 10.2085711 | 15.7738291 | 0 |
| 0.00273626 | 0.00563825 | 0 | 96.5392171 | 183.75428 | 90.0036134 | 0 |
| 0.00273678 | 0.00563887 | 0 | 143.251742 | 81.6685687 | 139.180845 | 0 |
| 0.00273689 | 0.00563887 | 3.85826995 | 22.6889404 | 20.4171422 | 17.1656376 | 0 |
| 0.00274428 | 0.00565375 | 8.0380624 | 30.6968018 | 22.273246 | 6.03116997 | 0 |
| 0.00274661 | 0.0056582 | 36.0105196 | 200.418974 | 422.263622 | 448.626258 | 2.37621824 |
| 0.00274694 | 0.00565853 | 81.666714 | 300.2948 | 149.416359 | 227.79265 | 35.6432737 |
| 0.0027499 | 0.00566428 | 1.60761248 | 23.3562622 | 47.3306478 | 28.7640414 | 0 |
| 0.00275255 | 0.00566939 | 28.6155022 | 135.911202 | 0 | 158.202228 | 0 |
| 0.00275333 | 0.00567065 | 93.8845689 | 3.11416829 | 16.7049345 | 11.1344676 | 0 |
| 0.00275586 | 0.00567552 | 9.64567488 | 20.0196533 | 25.9854537 | 5.56723382 | 0 |
| 0.00275609 | 0.00567564 | 6.10892743 | 14.0137573 | 12.0646749 | 17.1656376 | 0 |
| 0.00276098 | 0.00568537 | 7.39501741 | 13.123995 | 9.28051917 | 21.341063 | 0 |

| | | | | | | |
|---|---|---|---|---|---|---|
| 0.00276533 | 0.00569397 | 10.2887199 | 9.78738607 | 8.35246725 | 16.7017014 | 0 |
| 0.00277371 | 0.00571087 | 5.46588243 | 18.2401286 | 17.6329864 | 12.0623399 | 0 |
| 0.00277542 | 0.00571404 | 22.8280972 | 128.125781 | 0 | 195.781056 | 0 |
| 0.00277704 | 0.00571682 | 0 | 2.00196533 | 13.9207788 | 0 | 59.4054561 |
| 0.00277711 | 0.00571682 | 4.82283744 | 12.6791138 | 28.7696094 | 16.7017014 | 0 |
| 0.00277953 | 0.00572146 | 77.4869216 | 154.151331 | 22.273246 | 174.439993 | 28.5146189 |
| 0.00278083 | 0.00572378 | 18.3267823 | 40.4841878 | 46.4025958 | 14.3820207 | 0 |
| 0.00278194 | 0.00572573 | 3.85826995 | 16.4606038 | 17.6329864 | 29.6919137 | 0 |
| 0.00278666 | 0.005735 | 24.1141872 | 117.003752 | 0 | 202.740098 | 0 |
| 0.00278679 | 0.005735 | 122.178549 | 80.0786133 | 0 | 67.2707419 | 0 |
| 0.0027873 | 0.00573534 | 5.46588243 | 14.9035197 | 13.9207788 | 17.1656376 | 0 |
| 0.00278713 | 0.00573534 | 2.89370247 | 19.1298909 | 14.8488307 | 60.7756358 | 0 |
| 0.00278929 | 0.00573908 | 25.4002772 | 24.023584 | 18.5610383 | 36.650956 | 0 |
| 0.00278972 | 0.00573964 | 18.6483048 | 9.12006429 | 8.35246725 | 11.1344676 | 0 |
| 0.00279315 | 0.00574633 | 4.50131495 | 13.123995 | 21.3451941 | 21.8049991 | 0 |
| 0.0028037 | 0.00576768 | 10.6102424 | 17.7952474 | 6.49636342 | 13.4541484 | 0 |
| 0.00280644 | 0.00577297 | 8.6811074 | 18.017688 | 12.0646749 | 8.35085072 | 0 |
| 0.00280683 | 0.0057731 | 1.92913498 | 28.0275146 | 35.2659728 | 24.1246799 | 0 |
| 0.00280685 | 0.0057731 | 0 | 0 | 0 | 0 | 0 |
| 0.00280738 | 0.00577384 | 35.0459521 | 137.468286 | 111.36623 | 133.149675 | 0 |
| 0.00281119 | 0.00578133 | 19.6128723 | 9.78738607 | 8.35246725 | 10.2065953 | 0 |
| 0.00281888 | 0.00579678 | 8.3595849 | 10.899589 | 12.9927268 | 9.74265918 | 0 |
| 0.00282113 | 0.00580105 | 0 | 142.58442 | 70.5319457 | 151.243185 | 0 |
| 0.00282479 | 0.00580823 | 4.17979245 | 17.3503662 | 33.409869 | 13.9180845 | 0 |
| 0.0028272 | 0.00581283 | 64.3044992 | 110.330534 | 0 | 90.9314857 | 0 |
| 0.00282828 | 0.0058147 | 57.8740493 | 287.393245 | 290.48025 | 288.568286 | 49.9005831 |
| 0.00283053 | 0.00581897 | 13.5039448 | 11.3444702 | 8.35246725 | 9.27872303 | 0 |
| 0.00283071 | 0.00581897 | 0 | 158.822583 | 244.077654 | 31.5476583 | 0 |
| 0.00283407 | 0.00582553 | 15.7546023 | 12.9015544 | 5.5683115 | 14.8459568 | 0 |
| 0.00283436 | 0.00582576 | 28.9370247 | 4.67125244 | 13.9207788 | 15.7738291 | 0 |
| 0.00283673 | 0.00583028 | 7.39501741 | 14.2361979 | 9.28051917 | 14.8459568 | 0 |
| 0.0028381 | 0.00583274 | 10.9317649 | 9.12006429 | 8.35246725 | 17.1656376 | 0 |
| 0.00283837 | 0.00583294 | 33.1168171 | 9.12006429 | 22.273246 | 5.10329766 | 0 |
| 0.00284109 | 0.00583817 | 27.0078897 | 64.5077718 | 42.6903882 | 53.3526574 | 9.50487297 |
| 0.00284777 | 0.00585155 | 102.565676 | 164.383598 | 0 | 35.7230837 | 0 |
| 0.00284839 | 0.0058521 | 8.0380624 | 29.1397176 | 5.5683115 | 24.588616 | 0 |
| 0.00284825 | 0.0058521 | 7.39501741 | 29.8070394 | 28.7696094 | 23.1968076 | 2.37621824 |
| 0.00284884 | 0.00585268 | 4.17979245 | 31.5865641 | 57.5392189 | 6.49510612 | 0 |
| 0.00285098 | 0.00585672 | 8.0380624 | 9.56494547 | 14.8488307 | 11.1344676 | 0 |
| 0.00285541 | 0.00586546 | 12.8608998 | 7.34053955 | 9.28051917 | 23.1968076 | 0 |
| 0.0028581 | 0.00587061 | 20.5774398 | 9.12006429 | 9.28051917 | 9.74265918 | 0 |
| 0.00286017 | 0.00587452 | 119.927891 | 225.33232 | 223.660512 | 224.545097 | 104.553603 |
| 0.00286393 | 0.00588187 | 29.9015921 | 10.899589 | 9.28051917 | 8.35085072 | 0 |

| | | | | | | |
|---|---|---|---|---|---|---|
| 0.00286618 | 0.00588614 | 20.2559173 | 14.9035197 | 5.5683115 | 11.5984038 | 0 |
| 0.00286945 | 0.00589249 | 5.14435994 | 21.5767375 | 18.5610383 | 11.5984038 | 0 |
| 0.00287877 | 0.00591128 | 21.5420072 | 20.0196533 | 3.71220767 | 16.7017014 | 0 |
| 0.00288172 | 0.00591697 | 47.2638069 | 176.840271 | 0 | 71.4461673 | 0 |
| 0.00288291 | 0.00591905 | 5.78740493 | 12.9015544 | 12.9927268 | 19.9492545 | 0 |
| 0.00289468 | 0.00594286 | 3.21522496 | 34.7007324 | 25.9854537 | 13.4541484 | 0 |
| 0.00289542 | 0.00594401 | 45.334672 | 115.224227 | 0 | 116.91191 | 0 |
| 0.00289835 | 0.00594966 | 18.0052598 | 16.4606038 | 31.5537652 | 23.6607437 | 2.37621824 |
| 0.00289888 | 0.0059504 | 8.3595849 | 5.56101481 | 35.2659728 | 20.8771268 | 0 |
| 0.00290199 | 0.00595641 | 83.595849 | 165.27336 | 112.294282 | 132.685739 | 16.6335277 |
| 0.00290229 | 0.00595665 | 10.9317649 | 29.8070394 | 20.4171422 | 4.63936151 | 0 |
| 0.00290549 | 0.00596286 | 4.50131495 | 17.1279256 | 15.7768826 | 19.4853184 | 0 |
| 0.00290656 | 0.0059647 | 83.9173715 | 189.741825 | 162.409085 | 183.25478 | 9.50487297 |
| 0.00290965 | 0.00597067 | 8.0380624 | 3.55904948 | 5.5683115 | 5.10329766 | 23.7621824 |
| 0.00291049 | 0.00597204 | 8.3595849 | 4.22637126 | 6.49636342 | 2.31968076 | 35.6432737 |
| 0.00291149 | 0.00597373 | 3.85826995 | 13.123995 | 28.7696094 | 22.2689353 | 0 |
| 0.00291476 | 0.00598007 | 6.75197242 | 10.0098267 | 13.9207788 | 16.7017014 | 0 |
| 0.00293138 | 0.0060138 | 49.5144644 | 138.358048 | 0 | 157.274355 | 1123.95123 |
| 0.00294427 | 0.00603987 | 4.50131495 | 25.1357869 | 12.0646749 | 20.4131907 | 0 |
| 0.00294603 | 0.00604312 | 21.2204847 | 10.4547078 | 8.35246725 | 9.27872303 | 0 |
| 0.00294813 | 0.00604706 | 9.96719738 | 14.6810791 | 7.42441534 | 11.5984038 | 0 |
| 0.00294996 | 0.00605043 | 76.5223541 | 94.5372518 | 0 | 83.9724434 | 0 |
| 0.0029587 | 0.00606799 | 86.4895515 | 158.377702 | 0 | 42.2181898 | 0 |
| 0.00295928 | 0.00606881 | 4.17979245 | 23.5787028 | 13.9207788 | 19.9492545 | 0 |
| 0.00296164 | 0.00607328 | 48.8714194 | 9.34250488 | 18.5610383 | 20.4131907 | 0 |
| 0.00296241 | 0.00607449 | 197.736335 | 693.347327 | 561.47141 | 585.487423 | 106.929821 |
| 0.00296483 | 0.00607909 | 78.7730116 | 82.0805786 | 0 | 93.2511664 | 0 |
| 0.00296694 | 0.00608305 | 14.7900348 | 5.33857422 | 8.35246725 | 2.78361691 | 66.5341108 |
| 0.00297019 | 0.00608933 | 14.1469898 | 27.5826335 | 21.3451941 | 3.71148921 | 0 |
| 0.00297658 | 0.00610206 | 21.5420072 | 10.4547078 | 6.49636342 | 12.5262761 | 0 |
| 0.00298504 | 0.00611902 | 4.17979245 | 12.9015544 | 22.273246 | 23.1968076 | 0 |
| 0.00298683 | 0.00612233 | 83.9173715 | 61.6160441 | 53.8270112 | 64.0231889 | 11.8810912 |
| 0.00298768 | 0.00612371 | 17.6837373 | 6.005896 | 13.9207788 | 11.5984038 | 0 |
| 0.00298843 | 0.00612486 | 137.290106 | 337.664819 | 286.768042 | 315.476583 | 109.306039 |
| 0.00298985 | 0.00612739 | 3.85826995 | 14.6810791 | 18.5610383 | 29.2279775 | 0 |
| 0.0029901 | 0.00612753 | 54.6588243 | 64.952653 | 51.0428554 | 63.0953166 | 19.0097459 |
| 0.00299103 | 0.00612908 | 91.6339114 | 114.556905 | 0 | 56.1362743 | 0 |
| 0.00299162 | 0.00612991 | 69.7703817 | 129.905306 | 105.797919 | 124.334889 | 0 |
| 0.00299307 | 0.00613249 | 13.5039448 | 51.3837769 | 29.6976613 | 38.9706367 | 7.12865473 |
| 0.00299438 | 0.00613482 | 5.78740493 | 13.7913167 | 11.136623 | 27.8361691 | 0 |
| 0.00299735 | 0.00614053 | 3.53674746 | 7.34053955 | 47.3306478 | 37.1148921 | 0 |
| 0.00299831 | 0.00614212 | 24.1141872 | 144.363944 | 0 | 150.779249 | 0 |
| 0.00299856 | 0.00614225 | 2.89370247 | 18.4625692 | 24.1293498 | 26.9082968 | 0 |

| | | | | | | |
|---|---|---|---|---|---|---|
| 0.00300122 | 0.00614733 | 2.89370247 | 24.023584 | 32.4818171 | 16.2377653 | 0 |
| 0.00300168 | 0.0061479 | 0 | 0 | 0 | 0 | 0 |
| 0.00300213 | 0.00614846 | 46.9422844 | 10.0098267 | 9.28051917 | 8.35085072 | 0 |
| 0.00300272 | 0.00614928 | 104.173289 | 76.7420044 | 54.7550631 | 64.9510612 | 21.3859642 |
| 0.00300441 | 0.00615237 | 3.53674746 | 12.9015544 | 22.273246 | 35.2591475 | 0 |
| 0.00300681 | 0.0061569 | 4.82283744 | 17.7952474 | 29.6976613 | 11.1344676 | 0 |
| 0.00300761 | 0.00615817 | 82.309759 | 77.6317668 | 0 | 92.3232941 | 0 |
| 0.00300939 | 0.00616145 | 116.712666 | 118.115955 | 64.9636342 | 99.2823364 | 26.1384007 |
| 0.00301516 | 0.00617288 | 38.904222 | 108.328569 | 0 | 134.077548 | 0 |
| 0.00301755 | 0.00617739 | 4.50131495 | 19.3523315 | 31.5537652 | 11.1344676 | 0 |
| 0.00301802 | 0.00617798 | 232.139242 | 148.145435 | 188.394539 | 134.541484 | 249.502916 |
| 0.00301866 | 0.00617891 | 0 | 68.7341431 | 139.207788 | 142.892335 | 0 |
| 0.00302025 | 0.0061818 | 0 | 101.43291 | 151.272462 | 90.9314857 | 0 |
| 0.00303685 | 0.00621539 | 6.43044992 | 17.1279256 | 8.35246725 | 25.9804245 | 0 |
| 0.00303761 | 0.00621658 | 64.6260217 | 61.3936035 | 70.5319457 | 61.7035081 | 33.2670554 |
| 0.00304052 | 0.00622214 | 13.5039448 | 17.1279256 | 5.5683115 | 12.5262761 | 0 |
| 0.00304313 | 0.00622711 | 9.00262989 | 15.7932821 | 33.409869 | 6.03116997 | 0 |
| 0.00304533 | 0.00623123 | 0 | 191.52135 | 76.1002572 | 90.0036134 | 0 |
| 0.00304585 | 0.00623192 | 0 | 99.2085042 | 186.538435 | 70.9822312 | 0 |
| 0.00304949 | 0.00623899 | 2.57217997 | 33.3660889 | 18.5610383 | 23.6607437 | 0 |
| 0.0030545 | 0.00624886 | 0 | 0 | 0 | 0 | 47.5243649 |
| 0.00305873 | 0.00625713 | 67.8412467 | 79.8561727 | 0 | 105.777442 | 0 |
| 0.00305956 | 0.00625844 | 13.5039448 | 10.4547078 | 15.7768826 | 11.1344676 | 16.6335277 |
| 0.0030598 | 0.00625854 | 44.048582 | 147.478113 | 0 | 85.3642518 | 0 |
| 0.00306519 | 0.0062692 | 95.1706589 | 90.9782023 | 119.718697 | 0 | 0 |
| 0.00307056 | 0.0062798 | 15.7546023 | 22.4664998 | 14.8488307 | 14.3820207 | 0 |
| 0.00307111 | 0.00628053 | 17.6837373 | 10.6771484 | 5.5683115 | 16.7017014 | 0 |
| 0.00307561 | 0.00628937 | 7.71653991 | 12.011792 | 16.7049345 | 8.81478687 | 0 |
| 0.00308578 | 0.00630979 | 4.50131495 | 10.0098267 | 25.9854537 | 25.5164883 | 0 |
| 0.00308952 | 0.00631704 | 3.21522496 | 19.3523315 | 26.9135056 | 18.5574461 | 0 |
| 0.00309093 | 0.00631953 | 65.2690667 | 96.0943359 | 0 | 90.0036134 | 0 |
| 0.00309351 | 0.00632443 | 78.7730116 | 81.858138 | 0 | 87.6839326 | 0 |
| 0.003094 | 0.00632505 | 21.2204847 | 13.5688761 | 16.7049345 | 13.9180845 | 42.7719284 |
| 0.00309643 | 0.00632964 | 0 | 108.99589 | 138.279736 | 89.5396772 | 0 |
| 0.00309859 | 0.00633366 | 2.89370247 | 1.33464355 | 0 | 1.39180845 | 16.6335277 |
| 0.00310004 | 0.00633624 | 4.82283744 | 3.11416829 | 11.136623 | 4.17542536 | 28.5146189 |
| 0.00311042 | 0.00635706 | 17.6837373 | 22.4664998 | 25.0574018 | 9.74265918 | 0 |
| 0.00311376 | 0.00636351 | 1.92913498 | 27.5826335 | 68.6758419 | 12.5262761 | 0 |
| 0.00311554 | 0.00636677 | 1.60761248 | 37.1475789 | 41.7623363 | 17.6295737 | 0 |
| 0.00311578 | 0.00636687 | 5.14435994 | 29.1397176 | 10.2085711 | 17.6295737 | 0 |
| 0.00313022 | 0.00639599 | 12.2178549 | 13.3464355 | 5.5683115 | 17.1656376 | 0 |
| 0.00313288 | 0.00640103 | 10.2887199 | 24.023584 | 18.5610383 | 20.4131907 | 0 |
| 0.00313366 | 0.00640223 | 7.71653991 | 15.5708415 | 10.2085711 | 10.2065953 | 0 |

| | | | | | | |
|---|---|---|---|---|---|---|
| 0.00313516 | 0.00640491 | 104.816334 | 88.0864746 | 108.582074 | 0 | 0 |
| 0.0031395 | 0.00641261 | 58.8386168 | 50.9388957 | 56.6111669 | 63.5592527 | 19.0097459 |
| 0.00313949 | 0.00641261 | 7.39501741 | 8.23030192 | 16.7049345 | 14.8459568 | 0 |
| 0.00313935 | 0.00641261 | 5.46588243 | 13.5688761 | 12.9927268 | 18.0935099 | 0 |
| 0.00314197 | 0.00641648 | 235.032945 | 45.822762 | 0 | 43.6099982 | 0 |
| 0.00314191 | 0.00641648 | 82.952804 | 94.5372518 | 0 | 143.356271 | 2.37621824 |
| 0.00314196 | 0.00641648 | 141.791421 | 37.8149007 | 0 | 98.3544641 | 0 |
| 0.00314252 | 0.00641721 | 26.3648447 | 14.6810791 | 12.0646749 | 19.0213822 | 0 |
| 0.00314369 | 0.00641921 | 6.43044992 | 12.6791138 | 14.8488307 | 11.1344676 | 0 |
| 0.00314484 | 0.00642117 | 262.040834 | 324.540824 | 311.825444 | 333.106157 | 26.1384007 |
| 0.00314698 | 0.00642515 | 35.6889971 | 90.9782023 | 0 | 159.1301 | 0 |
| 0.00314719 | 0.0064252 | 21.5420072 | 0 | 4.64025958 | 4.63936151 | 61.7816743 |
| 0.00314889 | 0.00642828 | 14.1469898 | 78.0766479 | 69.6038938 | 61.7035081 | 11.8810912 |
| 0.00315057 | 0.00643131 | 0 | 2.22440592 | 0 | 0 | 0 |
| 0.00315111 | 0.00643204 | 12.2178549 | 90.5333211 | 40.8342843 | 59.3838274 | 9.50487297 |
| 0.00315217 | 0.00643381 | 4.82283744 | 12.6791138 | 26.9135056 | 14.8459568 | 0 |
| 0.00315316 | 0.00643543 | 123.143116 | 215.100053 | 183.75428 | 186.966269 | 52.2768014 |
| 0.00315798 | 0.00644488 | 18.6483048 | 17.1279256 | 21.3451941 | 3.71148921 | 0 |
| 0.00315839 | 0.00644532 | 21.5420072 | 8.45274251 | 12.0646749 | 7.88691457 | 0 |
| 0.00315944 | 0.00644708 | 94.8491364 | 277.605859 | 300.688821 | 271.402649 | 0 |
| 0.00316024 | 0.00644831 | 0 | 105.436841 | 115.078438 | 200.884354 | 0 |
| 0.00316312 | 0.0064538 | 5.46588243 | 38.2597819 | 31.5537652 | 25.0525522 | 0 |
| 0.00316471 | 0.00645665 | 9.32415239 | 18.2401286 | 14.8488307 | 6.49510612 | 0 |
| 0.00317586 | 0.00647901 | 2.57217997 | 1.77952474 | 0.92805192 | 1.39180845 | 16.6335277 |
| 0.00317658 | 0.00648007 | 0 | 52.7184204 | 137.351684 | 165.625206 | 0 |
| 0.00317677 | 0.00648007 | 65.9121117 | 70.7361084 | 0 | 115.520102 | 0 |
| 0.0031808 | 0.0064879 | 6.75197242 | 10.899589 | 11.136623 | 18.0935099 | 0 |
| 0.00318197 | 0.00648951 | 57.8740493 | 89.4211182 | 0 | 103.921698 | 0 |
| 0.00318189 | 0.00648951 | 133.753358 | 272.489726 | 250.574018 | 248.205841 | 78.415202 |
| 0.00318304 | 0.00649129 | 11.8963324 | 6.67321777 | 10.2085711 | 25.0525522 | 0 |
| 0.00320134 | 0.00652822 | 5.78740493 | 16.0157227 | 9.28051917 | 28.7640414 | 0 |
| 0.00320178 | 0.00652872 | 3.53674746 | 19.3523315 | 12.0646749 | 45.0018067 | 0 |
| 0.00320357 | 0.00653198 | 3.21522496 | 20.9094157 | 32.4818171 | 14.8459568 | 0 |
| 0.00320517 | 0.00653483 | 57.5525268 | 59.3916382 | 53.8270112 | 24.588616 | 14.2573095 |
| 0.00320615 | 0.00653605 | 5.46588243 | 17.1279256 | 11.136623 | 17.1656376 | 0 |
| 0.00320604 | 0.00653605 | 17.0406923 | 11.5669108 | 11.136623 | 6.49510612 | 0 |
| 0.00321307 | 0.00654976 | 7.71653991 | 13.5688761 | 9.28051917 | 11.5984038 | 0 |
| 0.00321453 | 0.00655234 | 5.14435994 | 10.2322673 | 20.4171422 | 20.4131907 | 0 |
| 0.00321485 | 0.00655259 | 13.5039448 | 16.905485 | 23.2012979 | 14.3820207 | 0 |
| 0.00321534 | 0.0065532 | 3.21522496 | 28.917277 | 25.9854537 | 13.4541484 | 0 |
| 0.00321574 | 0.00655361 | 76.2008316 | 81.1908162 | 151.272462 | 0 | 0 |
| 0.00322412 | 0.00657029 | 42.762492 | 72.9605143 | 0 | 161.449781 | 0 |
| 0.0032257 | 0.00657312 | 6.43044992 | 14.6810791 | 10.2085711 | 14.3820207 | 0 |

| | | | | | | |
|---|---|---|---|---|---|---|
| 0.00322593 | 0.00657318 | 331.168171 | 456.448096 | 412.983103 | 432.388493 | 130.692003 |
| 0.00322648 | 0.0065739 | 61.7323193 | 111.220296 | 135.49558 | 0 | 0 |
| 0.00322688 | 0.00657393 | 0 | 0 | 0 | 0 | 2.37621824 |
| 0.0032268 | 0.00657393 | 0 | 112.332499 | 188.394539 | 54.7444659 | 0 |
| 0.0032307 | 0.0065813 | 8.3595849 | 8.23030192 | 17.6329864 | 11.1344676 | 0 |
| 0.00323451 | 0.00658868 | 4.82283744 | 24.9133464 | 20.4171422 | 10.2065953 | 0 |
| 0.00323514 | 0.00658955 | 6.10892743 | 10.4547078 | 21.3451941 | 12.9902122 | 0 |
| 0.00323664 | 0.00659222 | 17.6837373 | 22.4664998 | 26.9135056 | 30.619786 | 0 |
| 0.00323786 | 0.00659429 | 52.0866444 | 116.33643 | 0 | 85.828188 | 0 |
| 0.00323826 | 0.00659472 | 8.0380624 | 17.3503662 | 12.9927268 | 7.88691457 | 0 |
| 0.00323892 | 0.00659536 | 104.173289 | 393.942289 | 280.271679 | 481.101789 | 9.50487297 |
| 0.00323897 | 0.00659536 | 63.9829767 | 66.064856 | 75.1722053 | 75.1576565 | 33.2670554 |
| 0.00324704 | 0.0066114 | 22.5065747 | 20.9094157 | 26.9135056 | 32.0115944 | 38.0194919 |
| 0.00325023 | 0.00661748 | 199.986993 | 314.753438 | 331.314534 | 340.993071 | 59.4054561 |
| 0.00325347 | 0.00662369 | 0.3215225 | 0.44488118 | 1.85610383 | 0 | 38.0194919 |
| 0.00325939 | 0.00663534 | 11.2532874 | 7.78542074 | 9.28051917 | 13.4541484 | 0 |
| 0.00325994 | 0.00663605 | 4.82283744 | 1.55708415 | 4.64025958 | 1.39180845 | 7.12865473 |
| 0.00326066 | 0.00663712 | 13.5039448 | 6.67321777 | 9.28051917 | 19.0213822 | 0 |
| 0.00326115 | 0.00663771 | 13.5039448 | 23.5787028 | 10.2085711 | 6.03116997 | 0 |
| 0.00326264 | 0.00664034 | 6.43044992 | 12.2342326 | 12.0646749 | 12.9902122 | 0 |
| 0.0032638 | 0.0066423 | 15.7546023 | 20.2420939 | 7.42441534 | 7.42297842 | 0 |
| 0.00326785 | 0.00665014 | 5.14435994 | 39.816866 | 7.42441534 | 22.2689353 | 0 |
| 0.00326902 | 0.00665213 | 13.5039448 | 17.7952474 | 23.2012979 | 4.17542536 | 0 |
| 0.00327411 | 0.00666208 | 0 | 89.4211182 | 67.7477899 | 190.213822 | 0 |
| 0.0032748 | 0.00666308 | 80.7021465 | 72.0707519 | 0 | 89.0757411 | 0 |
| 0.0032756 | 0.00666429 | 9.96719738 | 13.5688761 | 8.35246725 | 9.27872303 | 0 |
| 0.00327806 | 0.0066689 | 0 | 0 | 0 | 0 | 2.37621824 |
| 0.00327876 | 0.00666993 | 22.8280972 | 3.33660889 | 50.1148035 | 10.2065953 | 0 |
| 0.00328274 | 0.00667761 | 4.50131495 | 23.3562622 | 19.4890903 | 11.5984038 | 0 |
| 0.00328466 | 0.00668111 | 6.75197242 | 12.6791138 | 13.9207788 | 10.2065953 | 0 |
| 0.00328554 | 0.00668251 | 7.39501741 | 20.4645345 | 6.49636342 | 21.8049991 | 0 |
| 0.00328601 | 0.00668306 | 115.426576 | 84.0825439 | 0 | 52.4247851 | 0 |
| 0.00328704 | 0.00668475 | 218.95682 | 292.064498 | 330.386482 | 328.002859 | 14.2573095 |
| 0.00328903 | 0.00668808 | 5.46588243 | 23.8011434 | 12.0646749 | 12.9902122 | 0 |
| 0.00328908 | 0.00668808 | 0 | 3.55904948 | 3.71220767 | 0 | 54.6530196 |
| 0.00329176 | 0.00669313 | 0 | 265.149186 | 80.7405168 | 45.4657428 | 0 |
| 0.0032936 | 0.00669647 | 3.53674746 | 3.55904948 | 2.78415575 | 1.85574461 | 33.2670554 |
| 0.00329497 | 0.00669843 | 54.6588243 | 109.885653 | 123.430905 | 128.510314 | 26.1384007 |
| 0.00329485 | 0.00669843 | 4.50131495 | 13.123995 | 15.7768826 | 25.0525522 | 0 |
| 0.00329644 | 0.00670104 | 0 | 141.917098 | 90.9490879 | 94.1790387 | 0 |
| 0.00329716 | 0.00670208 | 7.39501741 | 11.7893514 | 11.136623 | 10.6705315 | 0 |
| 0.00329812 | 0.00670364 | 59.1601393 | 120.11792 | 126.215061 | 0 | 0 |
| 0.00330233 | 0.00671179 | 13.1824223 | 17.7952474 | 12.9927268 | 12.0623399 | 35.6432737 |

| | | | | | | |
|---|---|---|---|---|---|---|
| 0.00330444 | 0.00671568 | 60.7677518 | 116.113989 | 127.143113 | 0 | 0 |
| 0.00330758 | 0.00672165 | 61.4107968 | 6.45077718 | 9.28051917 | 10.6705315 | 0 |
| 0.00331015 | 0.00672647 | 23.1496197 | 24.9133464 | 22.273246 | 13.9180845 | 2.37621824 |
| 0.00331137 | 0.00672853 | 3.85826995 | 17.7952474 | 15.7768826 | 22.2689353 | 0 |
| 0.00331238 | 0.00673019 | 52.0866444 | 0 | 296.048562 | 151.243185 | 2.37621824 |
| 0.00331359 | 0.00673223 | 0 | 114.779346 | 100.229607 | 106.241379 | 0 |
| 0.00331606 | 0.00673684 | 11.2532874 | 16.905485 | 4.64025958 | 25.0525522 | 0 |
| 0.00331797 | 0.00674032 | 0.3215225 | 0 | 0 | 0 | 59.4054561 |
| 0.00331917 | 0.00674234 | 316.699659 | 766.0854 | 661.701017 | 701.471461 | 4.75243649 |
| 0.00332007 | 0.00674377 | 4.82283744 | 18.6850098 | 12.9927268 | 16.2377653 | 0 |
| 0.00332179 | 0.00674684 | 8.3595849 | 34.4782918 | 3.71220767 | 32.4755306 | 0 |
| 0.00332406 | 0.00675105 | 3.85826995 | 26.9153117 | 12.9927268 | 19.9492545 | 0 |
| 0.00332578 | 0.00675415 | 2.25065747 | 24.9133464 | 22.273246 | 27.3722329 | 0 |
| 0.00332957 | 0.00676144 | 17.0406923 | 8.23030192 | 12.9927268 | 9.74265918 | 14.2573095 |
| 0.00333215 | 0.00676627 | 11.8963324 | 28.6948364 | 18.5610383 | 4.17542536 | 0 |
| 0.00333348 | 0.00676856 | 3.85826995 | 26.0255493 | 18.5610383 | 13.9180845 | 0 |
| 0.0033367 | 0.00677469 | 10.2887199 | 9.56494547 | 9.28051917 | 9.74265918 | 0 |
| 0.00333765 | 0.00677619 | 25.0787547 | 6.89565837 | 8.35246725 | 13.9180845 | 0 |
| 0.00333809 | 0.00677668 | 4.82283744 | 10.2322673 | 22.273246 | 20.8771268 | 0 |
| 0.00333986 | 0.00677986 | 4.50131495 | 22.911381 | 8.35246725 | 38.0427644 | 0 |
| 0.00334174 | 0.00678328 | 63.3399317 | 125.456494 | 109.510126 | 151.707121 | 0 |
| 0.00334322 | 0.00678587 | 148.221871 | 144.586385 | 360.084144 | 275.578074 | 21.3859642 |
| 0.00334618 | 0.00679147 | 54.3373018 | 155.041093 | 0 | 57.0641466 | 0 |
| 0.00335049 | 0.0067998 | 0.96456749 | 0 | 1.85610383 | 0 | 47.5243649 |
| 0.00335118 | 0.00680079 | 7.71653991 | 10.899589 | 9.28051917 | 13.9180845 | 0 |
| 0.00335402 | 0.00680616 | 0.3215225 | 0 | 0.92805192 | 0 | 0 |
| 0.00335568 | 0.00680911 | 2.89370247 | 0.66732178 | 0.92805192 | 3.71148921 | 35.6432737 |
| 0.00335927 | 0.00681599 | 4.17979245 | 26.9153117 | 14.8488307 | 14.8459568 | 0 |
| 0.00336156 | 0.00682021 | 8.6811074 | 22.0216187 | 4.64025958 | 32.0115944 | 0 |
| 0.00336218 | 0.00682106 | 56.5879593 | 85.1947469 | 0 | 102.529889 | 0 |
| 0.00336384 | 0.00682402 | 1.28608998 | 0.44488118 | 0 | 0 | 0 |
| 0.00336451 | 0.00682498 | 5.46588243 | 14.0137573 | 12.0646749 | 17.1656376 | 0 |
| 0.00336515 | 0.00682503 | 16.0761248 | 40.4841878 | 15.7768826 | 3.24755306 | 0 |
| 0.003365 | 0.00682503 | 52.7296894 | 0 | 199.531162 | 130.829995 | 0 |
| 0.00336476 | 0.00682503 | 388.399175 | 783.880648 | 755.43426 | 814.207946 | 168.711495 |
| 0.00336566 | 0.00682565 | 5.46588243 | 59.8365194 | 4.64025958 | 26.4443606 | 0 |
| 0.00336969 | 0.00683341 | 0 | 162.381632 | 69.6038938 | 100.674145 | 0 |
| 0.00337466 | 0.00684308 | 2.89370247 | 18.017688 | 20.4171422 | 28.3001052 | 0 |
| 0.00337591 | 0.00684521 | 21.5420072 | 10.0098267 | 12.0646749 | 6.49510612 | 0 |
| 0.00337625 | 0.00684547 | 84.5604165 | 11.7893514 | 3.71220767 | 11.1344676 | 0 |
| 0.00338012 | 0.0068525 | 56.9094818 | 100.988029 | 0 | 85.3642518 | 0 |
| 0.00338006 | 0.0068525 | 42.762492 | 30.9192423 | 376.789078 | 0 | 0 |
| 0.00338095 | 0.00685378 | 218.635297 | 258.475968 | 287.696094 | 302.950307 | 59.4054561 |

| | | | | | | |
|---|---|---|---|---|---|---|
| 0.00338126 | 0.006854 | 0 | 102.767554 | 134.567528 | 84.4363795 | 0 |
| 0.00338772 | 0.00686667 | 40.5118345 | 36.9251383 | 16.7049345 | 37.5788283 | 9.50487297 |
| 0.00338844 | 0.00686772 | 21.5420072 | 9.34250488 | 0 | 0 | 147.325531 |
| 0.00338874 | 0.00686791 | 0 | 0 | 0 | 0 | 0 |
| 0.00339007 | 0.00687015 | 16.0761248 | 22.6889404 | 36.1940248 | 38.9706367 | 7.12865473 |
| 0.00339025 | 0.00687015 | 11.2532874 | 29.3621582 | 4.64025958 | 15.7738291 | 0 |
| 0.00339068 | 0.00687019 | 58.5170943 | 68.9565837 | 0 | 120.159463 | 0 |
| 0.00339059 | 0.00687019 | 49.8359869 | 14.0137573 | 14.8488307 | 17.1656376 | 42.7719284 |
| 0.00339295 | 0.00687437 | 8.0380624 | 12.9015544 | 20.4171422 | 7.42297842 | 0 |
| 0.00339683 | 0.00688183 | 53.3727344 | 37.8149007 | 52.8989593 | 62.6313804 | 116.434694 |
| 0.00339793 | 0.00688363 | 56.2664368 | 77.4093262 | 0 | 111.344676 | 0 |
| 0.00339961 | 0.00688663 | 0 | 115.446667 | 114.150386 | 88.6118049 | 0 |
| 0.00340555 | 0.00689823 | 63.6614542 | 99.2085042 | 88.1649321 | 89.5396772 | 45.1481466 |
| 0.00341119 | 0.00690925 | 9.96719738 | 28.917277 | 12.9927268 | 6.03116997 | 0 |
| 0.00341424 | 0.006915 | 26.6863672 | 61.1711629 | 48.2586997 | 47.7854236 | 14.2573095 |
| 0.00341706 | 0.00691961 | 19.9343948 | 76.0746826 | 56.6111669 | 45.4657428 | 0 |
| 0.00341713 | 0.00691961 | 4.17979245 | 15.3484009 | 14.8488307 | 23.6607437 | 0 |
| 0.00341693 | 0.00691961 | 11.8963324 | 16.0157227 | 50.1148035 | 3.71148921 | 0 |
| 0.00341886 | 0.0069227 | 2.25065747 | 67.6219401 | 28.7696094 | 9.27872303 | 0 |
| 0.00342285 | 0.00693035 | 4.17979245 | 14.2361979 | 18.5610383 | 19.0213822 | 0 |
| 0.00342405 | 0.00693237 | 18.3267823 | 5.33857422 | 18.5610383 | 10.2065953 | 0 |
| 0.00342804 | 0.00694004 | 27.9724572 | 42.2637126 | 33.409869 | 42.2181898 | 97.424948 |
| 0.00342966 | 0.00694289 | 174.586715 | 355.682507 | 304.401029 | 324.755306 | 0 |
| 0.0034383 | 0.00695997 | 27.0078897 | 111.220296 | 0 | 145.675952 | 0 |
| 0.0034397 | 0.00696238 | 6.10892743 | 30.2519206 | 6.49636342 | 22.7328714 | 0 |
| 0.00344378 | 0.00697022 | 4.82283744 | 1.77952474 | 1.85610383 | 2.78361691 | 21.3859642 |
| 0.00344433 | 0.00697091 | 0 | 132.574593 | 80.7405168 | 105.777442 | 0 |
| 0.00345025 | 0.00698248 | 0 | 92.3128459 | 116.00649 | 106.705315 | 0 |
| 0.00345436 | 0.00699036 | 70.0919042 | 101.21047 | 54.7550631 | 85.3642518 | 0 |
| 0.00345476 | 0.00699076 | 19.2913498 | 12.9015544 | 5.5683115 | 5.56723382 | 30.8908372 |
| 0.00345647 | 0.0069938 | 47.5853294 | 144.586385 | 0 | 66.8068058 | 0 |
| 0.00345814 | 0.00699659 | 189.698273 | 396.166695 | 393.494013 | 396.201473 | 106.929821 |
| 0.00345826 | 0.00699659 | 29.2585471 | 98.9860636 | 82.5966206 | 91.859358 | 19.0097459 |
| 0.00345994 | 0.00699957 | 50.1575094 | 110.775415 | 96.5173994 | 93.7151026 | 23.7621824 |
| 0.00346085 | 0.00700098 | 8.0380624 | 36.9251383 | 5.5683115 | 17.1656376 | 0 |
| 0.00346168 | 0.00700224 | 12.8608998 | 24.023584 | 28.7696094 | 3.24755306 | 0 |
| 0.00346327 | 0.00700503 | 53.6942569 | 10.4547078 | 5.5683115 | 11.1344676 | 0 |
| 0.00346557 | 0.00700927 | 28.6155022 | 84.5274251 | 21.3451941 | 82.1166988 | 11.8810912 |
| 0.00346746 | 0.00701266 | 89.7047764 | 7.34053955 | 34.3379209 | 9.27872303 | 0 |
| 0.00346836 | 0.00701406 | 69.1273367 | 194.190637 | 154.98467 | 158.666164 | 28.5146189 |
| 0.0034726 | 0.00702222 | 47.2638069 | 104.769519 | 0 | 94.1790387 | 0 |
| 0.00347612 | 0.00702892 | 13.1824223 | 8.00786133 | 9.28051917 | 10.2065953 | 0 |
| 0.00347816 | 0.00703262 | 25.7217997 | 22.911381 | 21.3451941 | 29.6919137 | 0 |

| | | | | | | |
|---|---|---|---|---|---|---|
| 0.00348091 | 0.00703776 | 0 | 0 | 0 | 0 | 33.2670554 |
| 0.00348134 | 0.00703821 | 156.902978 | 283.389315 | 248.717914 | 295.063392 | 0 |
| 0.00348199 | 0.00703909 | 352.710178 | 718.260673 | 525.277385 | 614.251464 | 133.068222 |
| 0.00348567 | 0.0070461 | 1.92913498 | 27.1377523 | 28.7696094 | 22.7328714 | 0 |
| 0.00348838 | 0.00705116 | 245.321665 | 23.8011434 | 0 | 62.6313804 | 0 |
| 0.00349181 | 0.00705767 | 3.85826995 | 24.9133464 | 14.8488307 | 16.7017014 | 0 |
| 0.00349885 | 0.00707148 | 67.5197242 | 92.757727 | 0 | 74.6937204 | 0 |
| 0.00350143 | 0.00707627 | 0 | 102.767554 | 103.013763 | 105.777442 | 0 |
| 0.00350545 | 0.00708395 | 4.82283744 | 27.5826335 | 29.6976613 | 7.42297842 | 0 |
| 0.00350983 | 0.00709239 | 47.9068519 | 197.749687 | 101.157659 | 126.654569 | 26.1384007 |
| 0.00351831 | 0.0071091 | 23.7926647 | 15.5708415 | 24.1293498 | 13.4541484 | 114.058476 |
| 0.00351867 | 0.00710939 | 228.602495 | 43.3759155 | 56.6111669 | 47.3214874 | 23.7621824 |
| 0.00352633 | 0.00712445 | 30.8661596 | 41.3739502 | 40.8342843 | 17.6295737 | 9.50487297 |
| 0.00353342 | 0.00713834 | 31.8307271 | 100.098267 | 0 | 135.933292 | 0 |
| 0.00353454 | 0.00714017 | 5.14435994 | 14.2361979 | 12.9927268 | 16.2377653 | 0 |
| 0.0035354 | 0.00714148 | 7.71653991 | 14.2361979 | 6.49636342 | 38.9706367 | 0 |
| 0.00354122 | 0.0071528 | 122.178549 | 466.680363 | 451.961284 | 444.914769 | 0 |
| 0.00354229 | 0.00715454 | 8.0380624 | 8.8976237 | 10.2085711 | 14.3820207 | 0 |
| 0.00354736 | 0.00716435 | 1.60761248 | 8.23030192 | 68.6758419 | 41.7542536 | 0 |
| 0.00355102 | 0.00717132 | 9.00262989 | 11.7893514 | 8.35246725 | 10.2065953 | 0 |
| 0.00355757 | 0.00718412 | 81.023669 | 172.169019 | 153.128566 | 158.202228 | 33.2670554 |
| 0.0035588 | 0.00718617 | 0 | 93.6474894 | 141.063891 | 80.7248903 | 0 |
| 0.00356515 | 0.00719855 | 14.4685123 | 26.0255493 | 38.0501286 | 21.8049991 | 2.37621824 |
| 0.0035668 | 0.00720144 | 15.4330798 | 24.9133464 | 20.4171422 | 21.8049991 | 4.75243649 |
| 0.00356742 | 0.00720227 | 33.1168171 | 25.8031087 | 23.2012979 | 24.1246799 | 64.1578926 |
| 0.00356814 | 0.00720329 | 11.2532874 | 11.3444702 | 5.5683115 | 35.7230837 | 0 |
| 0.00356868 | 0.00720396 | 14.1469898 | 8.00786133 | 18.5610383 | 6.95904227 | 0 |
| 0.00357694 | 0.0072202 | 6.75197242 | 47.3798462 | 16.7049345 | 29.2279775 | 0 |
| 0.00357733 | 0.00722055 | 126.036818 | 61.3936035 | 51.0428554 | 67.2707419 | 232.869388 |
| 0.00357848 | 0.00722244 | 10.6102424 | 15.5708415 | 9.28051917 | 7.42297842 | 0 |
| 0.00358003 | 0.00722513 | 161.725816 | 13.7913167 | 9.28051917 | 33.4034029 | 4.75243649 |
| 0.00358123 | 0.00722712 | 62.3753642 | 84.7498657 | 0 | 85.3642518 | 0 |
| 0.00358411 | 0.0072325 | 18.3267823 | 17.5728068 | 32.4818171 | 2.78361691 | 0 |
| 0.00358917 | 0.00724226 | 0 | 164.161157 | 140.135839 | 40.8263813 | 0 |
| 0.00359373 | 0.00725104 | 1.60761248 | 26.0255493 | 36.1940248 | 23.1968076 | 0 |
| 0.00359789 | 0.00725899 | 7.07349491 | 5.56101481 | 5.5683115 | 4.17542536 | 47.5243649 |
| 0.00359947 | 0.00726174 | 106.102424 | 53.6081828 | 80.7405168 | 46.3936151 | 28.5146189 |
| 0.00360029 | 0.00726297 | 59.1601393 | 76.7420044 | 166.121293 | 0 | 0 |
| 0.00360349 | 0.00726854 | 98.3858838 | 78.0766479 | 0 | 57.9920189 | 0 |
| 0.00360347 | 0.00726854 | 70.4134267 | 104.324638 | 108.582074 | 0 | 0 |
| 0.00360947 | 0.00728018 | 246.607755 | 976.06932 | 505.788295 | 665.284441 | 0 |
| 0.00361247 | 0.0072858 | 7.39501741 | 12.4566732 | 9.28051917 | 11.5984038 | 0 |
| 0.00361602 | 0.00729252 | 4.50131495 | 16.0157227 | 25.9854537 | 11.5984038 | 0 |

| | | | | | | |
|---|---|---|---|---|---|---|
| 0.00361996 | 0.00730003 | 2.89370247 | 26.0255493 | 44.546492 | 10.2065953 | 0 |
| 0.00362048 | 0.00730063 | 85.524984 | 255.584241 | 223.660512 | 235.679565 | 52.2768014 |
| 0.00362187 | 0.00730299 | 16.0761248 | 27.5826335 | 7.42441534 | 6.49510612 | 0 |
| 0.00362382 | 0.00730649 | 213.490937 | 614.158476 | 841.743089 | 803.537414 | 16.6335277 |
| 0.00362477 | 0.00730797 | 229.24554 | 278.050741 | 213.451941 | 288.568286 | 149.701749 |
| 0.00362722 | 0.00731247 | 70.7349491 | 59.6140788 | 0 | 104.385634 | 0 |
| 0.00362749 | 0.00731259 | 0 | 61.6160441 | 110.438178 | 147.995632 | 0 |
| 0.00363482 | 0.00732691 | 5.46588243 | 17.3503662 | 9.28051917 | 20.4131907 | 0 |
| 0.00363845 | 0.00733379 | 89.0617314 | 0 | 91.8771398 | 161.913717 | 0 |
| 0.00363912 | 0.0073347 | 157.224501 | 407.288725 | 219.948304 | 273.258393 | 49.9005831 |
| 0.00364192 | 0.00733991 | 3.85826995 | 18.9074504 | 13.9207788 | 21.341063 | 0 |
| 0.00364445 | 0.00734456 | 60.1247068 | 115.224227 | 66.819738 | 117.839782 | 4.75243649 |
| 0.00365046 | 0.00735624 | 0 | 101.21047 | 107.654022 | 96.0347833 | 0 |
| 0.0036512 | 0.0073573 | 0 | 73.8502767 | 128.999216 | 107.633187 | 0 |
| 0.00366269 | 0.00737999 | 212.847892 | 44.2656779 | 70.5319457 | 0 | 0 |
| 0.00366409 | 0.00738237 | 95.4921813 | 54.053064 | 0 | 83.9724434 | 0 |
| 0.00366578 | 0.00738535 | 16.0761248 | 2.8917277 | 20.4171422 | 29.6919137 | 0 |
| 0.00366954 | 0.00739248 | 4.50131495 | 26.0255493 | 10.2085711 | 19.0213822 | 0 |
| 0.0036735 | 0.00740001 | 8.0380624 | 14.6810791 | 15.7768826 | 6.95904227 | 0 |
| 0.00368135 | 0.00741538 | 5.78740493 | 20.6869751 | 7.42441534 | 24.1246799 | 0 |
| 0.00368697 | 0.00742625 | 37.618132 | 59.1691976 | 43.6184401 | 56.6002105 | 7.12865473 |
| 0.00368828 | 0.00742845 | 10.2887199 | 6.89565837 | 10.2085711 | 16.7017014 | 0 |
| 0.00368987 | 0.00743121 | 22.5065747 | 15.5708415 | 13.9207788 | 13.4541484 | 0 |
| 0.00369112 | 0.00743282 | 51.1220769 | 143.251742 | 0 | 56.6002105 | 0 |
| 0.003691 | 0.00743282 | 7.39501741 | 7.34053955 | 20.4171422 | 13.9180845 | 0 |
| 0.00369167 | 0.00743349 | 5.78740493 | 5.11613363 | 3.71220767 | 4.17542536 | 2.37621824 |
| 0.00369214 | 0.007434 | 43.7270595 | 94.3148112 | 0 | 102.529889 | 0 |
| 0.00369237 | 0.00743401 | 2.57217997 | 7.78542074 | 39.9062324 | 45.0018067 | 0 |
| 0.00369368 | 0.0074362 | 64.6260217 | 73.6278361 | 0 | 90.4675495 | 0 |
| 0.00369593 | 0.00744029 | 60.1247068 | 81.1908162 | 57.5392189 | 76.549465 | 2.37621824 |
| 0.00369681 | 0.00744162 | 0 | 129.905306 | 97.4454513 | 79.797018 | 0 |
| 0.00369761 | 0.00744273 | 4.82283744 | 14.9035197 | 12.0646749 | 19.9492545 | 0 |
| 0.00369781 | 0.00744273 | 5.78740493 | 9.34250488 | 18.5610383 | 15.309893 | 0 |
| 0.00370515 | 0.00745707 | 0 | 123.232088 | 158.696878 | 47.3214874 | 0 |
| 0.0037093 | 0.00746498 | 52.0866444 | 79.4112915 | 0 | 102.529889 | 0 |
| 0.00371087 | 0.00746769 | 3.21522496 | 32.9212077 | 8.35246725 | 37.1148921 | 0 |
| 0.00371479 | 0.00747513 | 7.07349491 | 27.5826335 | 9.28051917 | 10.6705315 | 0 |
| 0.00371851 | 0.00748217 | 0 | 91.2006429 | 204.171422 | 46.8575513 | 0 |
| 0.0037203 | 0.00748533 | 49.8359869 | 98.0963013 | 0 | 86.2921241 | 0 |
| 0.00372264 | 0.00748958 | 36.6535646 | 114.779346 | 0 | 97.4265918 | 0 |
| 0.00372883 | 0.00750157 | 6.43044992 | 17.3503662 | 13.9207788 | 8.81478687 | 0 |
| 0.00372913 | 0.00750173 | 8.3595849 | 12.4566732 | 9.28051917 | 9.27872303 | 0 |
| 0.00373314 | 0.00750936 | 18.3267823 | 6.89565837 | 19.4890903 | 6.95904227 | 0 |

| | | | | | | |
|---|---|---|---|---|---|---|
| 0.00373423 | 0.00751072 | 135.039448 | 169.277291 | 93.7332436 | 138.716909 | 33.2670554 |
| 0.00373426 | 0.00751072 | 2.89370247 | 51.1613363 | 9.28051917 | 25.0525522 | 0 |
| 0.00373659 | 0.00751478 | 25.0787547 | 132.129712 | 0 | 115.520102 | 0 |
| 0.00373673 | 0.00751478 | 0 | 84.5274251 | 99.3015551 | 119.231591 | 0 |
| 0.0037403 | 0.00752152 | 5.14435994 | 17.1279256 | 11.136623 | 15.7738291 | 0 |
| 0.00374353 | 0.00752756 | 35.6889971 | 118.115955 | 0 | 96.0347833 | 0 |
| 0.00374501 | 0.00753008 | 153.366231 | 141.917098 | 127.143113 | 126.190633 | 320.789463 |
| 0.00374776 | 0.0075347 | 44.048582 | 148.590316 | 156.840774 | 158.202228 | 35.6432737 |
| 0.00374763 | 0.0075347 | 4.50131495 | 43.5983561 | 24.1293498 | 31.5476583 | 2.37621824 |
| 0.00374798 | 0.00753471 | 24.4357097 | 10.6771484 | 16.7049345 | 4.63936151 | 0 |
| 0.00374965 | 0.0075376 | 216.38464 | 442.656779 | 375.861026 | 436.099982 | 0 |
| 0.00375009 | 0.00753805 | 25.4002772 | 131.684831 | 0 | 114.128293 | 0 |
| 0.00375666 | 0.00755079 | 2.57217997 | 26.9153117 | 19.4890903 | 20.4131907 | 0 |
| 0.00375702 | 0.00755108 | 26.3648447 | 120.785242 | 0 | 120.623399 | 0 |
| 0.00375975 | 0.0075561 | 0.96456749 | 0 | 0 | 0.46393615 | 23.7621824 |
| 0.00376469 | 0.00756558 | 10.6102424 | 7.34053955 | 17.6329864 | 8.81478687 | 0 |
| 0.00377098 | 0.00757777 | 3.21522496 | 1.55708415 | 1.85610383 | 1.39180845 | 7.12865473 |
| 0.00377279 | 0.00758096 | 105.459379 | 186.850098 | 112.294282 | 156.346483 | 21.3859642 |
| 0.00377468 | 0.0075843 | 103.851766 | 142.361979 | 141.063891 | 135.00542 | 30.8908372 |
| 0.00378059 | 0.00759571 | 63.6614542 | 144.363944 | 77.956361 | 0 | 0 |
| 0.0037813 | 0.00759669 | 6.75197242 | 24.4684652 | 43.6184401 | 18.0935099 | 2.37621824 |
| 0.00378256 | 0.00759878 | 63.6614542 | 98.0963013 | 67.7477899 | 83.0445711 | 19.0097459 |
| 0.00378361 | 0.00760043 | 14.4685123 | 18.2401286 | 7.42441534 | 12.5262761 | 7.12865473 |
| 0.00378847 | 0.00760974 | 67.5197242 | 146.36591 | 111.36623 | 145.675952 | 11.8810912 |
| 0.00379923 | 0.0076309 | 0 | 69.4014648 | 191.178695 | 66.3428696 | 0 |
| 0.0038022 | 0.00763641 | 4.50131495 | 31.8090047 | 35.2659728 | 27.8361691 | 0 |
| 0.00380474 | 0.00764105 | 5.46588243 | 15.3484009 | 11.136623 | 14.3820207 | 0 |
| 0.00380517 | 0.00764146 | 42.4409695 | 118.338395 | 0 | 79.797018 | 0 |
| 0.00380752 | 0.00764572 | 18.3267823 | 20.9094157 | 12.9927268 | 12.5262761 | 0 |
| 0.0038087 | 0.00764726 | 7.07349491 | 9.56494547 | 12.0646749 | 11.5984038 | 0 |
| 0.00380874 | 0.00764726 | 11.2532874 | 18.2401286 | 34.3379209 | 3.71148921 | 0 |
| 0.00380987 | 0.00764908 | 77.4869216 | 153.92889 | 58.4672708 | 0 | 0 |
| 0.00381549 | 0.0076599 | 307.375506 | 583.016793 | 569.823877 | 618.42689 | 16.6335277 |
| 0.003821 | 0.0076705 | 8.0380624 | 8.23030192 | 10.2085711 | 25.9804245 | 0 |
| 0.00382338 | 0.00767483 | 50.4790319 | 107.661247 | 109.510126 | 98.8184002 | 7.12865473 |
| 0.00382628 | 0.00768017 | 10.2887199 | 13.5688761 | 10.2085711 | 6.95904227 | 0 |
| 0.00383529 | 0.0076978 | 13.8254673 | 36.4802572 | 4.64025958 | 10.6705315 | 0 |
| 0.00383633 | 0.00769944 | 9.32415239 | 38.7046631 | 10.2085711 | 34.3312752 | 0 |
| 0.00383882 | 0.00770398 | 0 | 96.0943359 | 154.056618 | 61.7035081 | 0 |
| 0.0038436 | 0.0077131 | 92.5984789 | 71.6258708 | 54.7550631 | 59.8477635 | 21.3859642 |
| 0.00384584 | 0.00771715 | 8.6811074 | 6.45077718 | 24.1293498 | 12.5262761 | 0 |
| 0.00384721 | 0.00771943 | 23.7926647 | 18.017688 | 12.0646749 | 17.1656376 | 87.920075 |
| 0.00384972 | 0.00772401 | 2.57217997 | 0.66732178 | 0 | 0.46393615 | 4.75243649 |

| | | | | | | |
|---|---|---|---|---|---|---|
| 0.00385273 | 0.00772958 | 111.246784 | 23.5787028 | 0 | 137.325101 | 0 |
| 0.00385668 | 0.00773705 | 54.9803468 | 139.470251 | 90.0210359 | 0 | 0 |
| 0.00386115 | 0.00774555 | 11.2532874 | 32.0314453 | 51.9709073 | 44.5378705 | 0 |
| 0.00387124 | 0.00776533 | 7.39501741 | 22.4664998 | 6.49636342 | 15.7738291 | 0 |
| 0.00387519 | 0.00777279 | 41.476402 | 88.0864746 | 174.47376 | 0 | 0 |
| 0.00388293 | 0.00778785 | 0 | 0 | 0 | 0 | 7.12865473 |
| 0.00388478 | 0.00779109 | 3.85826995 | 13.7913167 | 11.136623 | 53.3526574 | 0 |
| 0.00388974 | 0.00780058 | 188.412183 | 466.902804 | 375.861026 | 404.088388 | 57.0292378 |
| 0.00390645 | 0.00783362 | 5.46588243 | 15.5708415 | 12.9927268 | 11.5984038 | 0 |
| 0.00391013 | 0.0078405 | 13.1824223 | 14.6810791 | 6.49636342 | 8.81478687 | 0 |
| 0.00391034 | 0.0078405 | 6.43044992 | 15.5708415 | 9.28051917 | 12.5262761 | 0 |
| 0.0039168 | 0.00785297 | 78.4514891 | 199.751652 | 122.502853 | 165.625206 | 0 |
| 0.00391843 | 0.00785577 | 7.39501741 | 13.123995 | 7.42441534 | 17.1656376 | 0 |
| 0.00391989 | 0.0078578 | 0 | 107.216366 | 72.3880495 | 118.303719 | 0 |
| 0.00391991 | 0.0078578 | 36.3320421 | 18.4625692 | 23.2012979 | 26.4443606 | 104.553603 |
| 0.003922 | 0.00786105 | 52.0866444 | 10.0098267 | 6.49636342 | 8.81478687 | 0 |
| 0.00392199 | 0.00786105 | 12.2178549 | 4.67125244 | 20.4171422 | 14.3820207 | 0 |
| 0.00392357 | 0.00786373 | 146.614258 | 0 | 109.510126 | 72.8379758 | 0 |
| 0.003931 | 0.00787817 | 8.3595849 | 12.011792 | 10.2085711 | 8.35085072 | 0 |
| 0.00393408 | 0.00788386 | 9.32415239 | 10.899589 | 10.2085711 | 7.88691457 | 0 |
| 0.00393638 | 0.00788799 | 11.2532874 | 25.8031087 | 4.64025958 | 13.9180845 | 0 |
| 0.00394311 | 0.00790101 | 10.6102424 | 7.11809896 | 11.136623 | 9.74265918 | 49.9005831 |
| 0.00395109 | 0.00791653 | 24.4357097 | 8.23030192 | 5.5683115 | 16.2377653 | 0 |
| 0.0039524 | 0.00791869 | 131.502701 | 345.227799 | 336.882846 | 362.79807 | 0 |
| 0.00395387 | 0.00792116 | 42.119447 | 50.2715739 | 158.696878 | 75.1576565 | 14.2573095 |
| 0.00395757 | 0.0079281 | 15.4330798 | 132.797034 | 0 | 151.243185 | 0 |
| 0.00396388 | 0.00794026 | 7.07349491 | 10.2322673 | 9.28051917 | 16.7017014 | 0 |
| 0.00397089 | 0.00795384 | 5.78740493 | 23.1338216 | 42.6903882 | 5.10329766 | 0 |
| 0.0039767 | 0.007965 | 0 | 101.43291 | 144.776099 | 58.9198912 | 0 |
| 0.00397815 | 0.00796742 | 35.6889971 | 140.360014 | 0 | 72.3740396 | 0 |
| 0.00397942 | 0.00796949 | 42.762492 | 105.436841 | 0 | 83.5085072 | 0 |
| 0.00398456 | 0.00797931 | 399.33094 | 457.115417 | 0 | 199.492545 | 7.12865473 |
| 0.00398668 | 0.00798309 | 47.2638069 | 40.7066284 | 47.3306478 | 43.6099982 | 14.2573095 |
| 0.00399077 | 0.0079908 | 0 | 121.675004 | 58.4672708 | 122.015208 | 0 |
| 0.00399189 | 0.00799258 | 0 | 110.108093 | 103.013763 | 79.3330819 | 0 |
| 0.0039993 | 0.00800693 | 27.9724572 | 0 | 0 | 0 | 460.986339 |
| 0.00400457 | 0.00801699 | 11.5748099 | 15.1259603 | 5.5683115 | 11.5984038 | 0 |
| 0.00400827 | 0.00802393 | 31.5092046 | 30.6968018 | 27.8415575 | 42.6821259 | 11.8810912 |
| 0.00400859 | 0.0080241 | 18.0052598 | 8.23030192 | 67.7477899 | 39.898509 | 2.37621824 |
| 0.00401211 | 0.00803066 | 0 | 113.222262 | 167.049345 | 41.7542536 | 0 |
| 0.00401547 | 0.00803692 | 89.7047764 | 62.7282471 | 0 | 67.2707419 | 0 |
| 0.00401685 | 0.0080392 | 9.32415239 | 6.22833659 | 6.49636342 | 6.49510612 | 35.6432737 |
| 0.00401938 | 0.00804379 | 57.8740493 | 84.9723063 | 0 | 153.09893 | 0 |

| | | | | | | |
|---|---|---|---|---|---|---|
| 0.00402414 | 0.00805283 | 61.7323193 | 77.6317668 | 0 | 78.8691457 | 0 |
| 0.00402866 | 0.00806139 | 131.502701 | 83.1927816 | 41.7623363 | 86.7560603 | 152.077968 |
| 0.0040324 | 0.0080684 | 9.32415239 | 6.005896 | 17.6329864 | 12.9902122 | 0 |
| 0.00403309 | 0.0080693 | 83.2743265 | 250.023226 | 261.710641 | 306.661796 | 0 |
| 0.00403417 | 0.00807097 | 58.8386168 | 21.3542969 | 11.136623 | 22.2689353 | 2.37621824 |
| 0.00403601 | 0.00807419 | 1.60761248 | 20.4645345 | 24.1293498 | 38.5067006 | 0 |
| 0.00403826 | 0.0080782 | 0 | 119.228158 | 130.85532 | 53.8165936 | 0 |
| 0.00403926 | 0.00807972 | 0 | 0.22244059 | 1.85610383 | 1.39180845 | 40.3957101 |
| 0.00404739 | 0.0080955 | 11.2532874 | 41.1515096 | 3.71220767 | 15.309893 | 0 |
| 0.00405711 | 0.00811446 | 35.6889971 | 46.934965 | 25.9854537 | 41.2903175 | 85.5438568 |
| 0.00406293 | 0.00812543 | 12.8608998 | 14.0137573 | 6.49636342 | 8.81478687 | 0 |
| 0.00406308 | 0.00812543 | 43.405537 | 56.2774699 | 62.1794784 | 64.9510612 | 19.0097459 |
| 0.00406414 | 0.00812708 | 7.71653991 | 14.2361979 | 7.42441534 | 12.5262761 | 0 |
| 0.00406528 | 0.00812887 | 8.0380624 | 11.1220296 | 7.42441534 | 15.7738291 | 0 |
| 0.00406779 | 0.0081334 | 0 | 72.2931925 | 101.157659 | 118.303719 | 0 |
| 0.00407069 | 0.00813872 | 0 | 1.77952474 | 0 | 1.39180845 | 28.5146189 |
| 0.00407113 | 0.00813912 | 121.535504 | 199.974093 | 239.437395 | 217.122119 | 66.5341108 |
| 0.00407183 | 0.00814003 | 63.3399317 | 54.4979451 | 0 | 106.241379 | 0 |
| 0.00407645 | 0.00814878 | 2.89370247 | 37.1475789 | 23.2012979 | 11.1344676 | 0 |
| 0.00407842 | 0.00815224 | 31.8307271 | 78.5215291 | 0 | 138.252973 | 0 |
| 0.00407916 | 0.00815323 | 6.43044992 | 15.3484009 | 10.2085711 | 10.6705315 | 0 |
| 0.00408705 | 0.00816852 | 116.391144 | 185.960335 | 0 | 12.5262761 | 0 |
| 0.00409661 | 0.00818714 | 0 | 97.8738607 | 88.1649321 | 100.674145 | 0 |
| 0.00409933 | 0.00819209 | 4.82283744 | 38.7046631 | 9.28051917 | 14.3820207 | 0 |
| 0.00410057 | 0.00819407 | 8.6811074 | 14.4586385 | 7.42441534 | 10.2065953 | 0 |
| 0.00410086 | 0.00819417 | 0 | 0.44488118 | 0 | 0 | 0 |
| 0.00410177 | 0.00819551 | 61.0892743 | 50.4940145 | 0 | 116.447974 | 0 |
| 0.00410222 | 0.00819591 | 7.39501741 | 8.45274251 | 12.9927268 | 11.1344676 | 0 |
| 0.00410425 | 0.00819948 | 5.14435994 | 8.23030192 | 23.2012979 | 19.0213822 | 0 |
| 0.00410834 | 0.0082071 | 20.5774398 | 4.89369303 | 36.1940248 | 6.95904227 | 0 |
| 0.00410855 | 0.0082071 | 0 | 0 | 0 | 0 | 16.6335277 |
| 0.00411507 | 0.00821964 | 36.9750871 | 150.1474 | 0 | 61.239572 | 0 |
| 0.00411713 | 0.00822326 | 38.261177 | 66.064856 | 0 | 136.861165 | 0 |
| 0.00413114 | 0.00825076 | 320.557929 | 287.838127 | 325.746223 | 309.909349 | 406.33332 |
| 0.00414152 | 0.008271 | 0 | 68.2892619 | 128.071165 | 95.106911 | 0 |
| 0.0041419 | 0.00827126 | 19.2913498 | 31.8090047 | 12.0646749 | 3.24755306 | 0 |
| 0.00414277 | 0.0082725 | 33.4383396 | 31.8090047 | 30.6257133 | 32.4755306 | 49.9005831 |
| 0.00414385 | 0.00827417 | 2.89370247 | 0 | 0 | 0 | 9.50487297 |
| 0.0041502 | 0.00828637 | 70.0919042 | 107.883687 | 0 | 46.8575513 | 0 |
| 0.00415106 | 0.0082876 | 11.2532874 | 19.3523315 | 4.64025958 | 14.3820207 | 0 |
| 0.00415266 | 0.00829029 | 20.2559173 | 8.67518311 | 6.49636342 | 11.5984038 | 0 |
| 0.00415561 | 0.00829569 | 141.791421 | 55.8325887 | 77.0283091 | 0 | 0 |
| 0.00415794 | 0.00829984 | 2.89370247 | 10.0098267 | 21.3451941 | 47.3214874 | 0 |

| | | | | | | |
|---|---|---|---|---|---|---|
| 0.00416263 | 0.00830872 | 11.2532874 | 39.1495443 | 36.1940248 | 37.1148921 | 7.12865473 |
| 0.00416523 | 0.00831341 | 18.3267823 | 22.911381 | 2.78415575 | 17.6295737 | 0 |
| 0.00416711 | 0.00831666 | 113.818964 | 321.426656 | 281.199731 | 280.681372 | 0 |
| 0.00416831 | 0.00831858 | 63.9829767 | 121.897445 | 0 | 44.5378705 | 0 |
| 0.0041712 | 0.00832385 | 39.547267 | 74.5175985 | 0 | 117.375846 | 0 |
| 0.00417171 | 0.00832436 | 55.9449143 | 40.4841878 | 56.6111669 | 45.0018067 | 102.177384 |
| 0.00417286 | 0.00832616 | 0 | 101.655351 | 50.1148035 | 152.634994 | 0 |
| 0.0041757 | 0.00833133 | 6.43044992 | 22.6889404 | 14.8488307 | 7.42297842 | 0 |
| 0.00417654 | 0.00833252 | 9.32415239 | 21.1318563 | 3.71220767 | 31.5476583 | 0 |
| 0.00418263 | 0.00834418 | 6.75197242 | 10.0098267 | 24.1293498 | 9.27872303 | 0 |
| 0.00418464 | 0.00834769 | 93.5630464 | 61.3936035 | 0 | 61.7035081 | 0 |
| 0.00418541 | 0.00834823 | 9.96719738 | 58.7243164 | 32.4818171 | 42.6821259 | 7.12865473 |
| 0.00418532 | 0.00834823 | 5.46588243 | 13.5688761 | 12.0646749 | 12.5262761 | 0 |
| 0.00418874 | 0.00835439 | 5.46588243 | 23.5787028 | 22.273246 | 6.95904227 | 0 |
| 0.0041895 | 0.00835491 | 8.0380624 | 9.78738607 | 11.136623 | 8.81478687 | 0 |
| 0.00418931 | 0.00835491 | 20.8989622 | 15.5708415 | 14.8488307 | 3.71148921 | 0 |
| 0.004194 | 0.00836339 | 0 | 92.5352865 | 103.013763 | 87.6839326 | 0 |
| 0.00419987 | 0.00837459 | 41.476402 | 12.011792 | 4.64025958 | 10.6705315 | 0 |
| 0.00420014 | 0.00837464 | 8.3595849 | 20.6869751 | 19.4890903 | 22.2689353 | 2.37621824 |
| 0.00420286 | 0.00837956 | 25.0787547 | 8.8976237 | 11.136623 | 6.49510612 | 0 |
| 0.00420415 | 0.00838164 | 10.6102424 | 25.3582275 | 9.28051917 | 6.49510612 | 0 |
| 0.0042044 | 0.00838164 | 0 | 10.899589 | 0 | 0 | 49.9005831 |
| 0.00420877 | 0.00838987 | 4.17979245 | 14.9035197 | 24.1293498 | 12.0623399 | 0 |
| 0.00421552 | 0.00840282 | 34.4029071 | 101.877791 | 0 | 96.4987195 | 0 |
| 0.00421718 | 0.00840563 | 157.867546 | 463.566195 | 508.57245 | 186.038397 | 2.37621824 |
| 0.00422348 | 0.0084177 | 15.1115573 | 16.6830444 | 23.2012979 | 24.588616 | 2.37621824 |
| 0.00423354 | 0.00843724 | 11.2532874 | 8.67518311 | 7.42441534 | 11.1344676 | 0 |
| 0.00423668 | 0.008443 | 62.6968867 | 60.7262817 | 0 | 91.3954218 | 0 |
| 0.00423835 | 0.00844583 | 4.82283744 | 15.5708415 | 10.2085711 | 20.4131907 | 0 |
| 0.00423937 | 0.00844735 | 56.9094818 | 138.580489 | 154.056618 | 134.077548 | 14.2573095 |
| 0.004244 | 0.00845609 | 24.4357097 | 21.3542969 | 19.4890903 | 16.7017014 | 4.75243649 |
| 0.00424687 | 0.0084613 | 31.1876821 | 150.36984 | 114.150386 | 0 | 0 |
| 0.0042493 | 0.00846563 | 2.89370247 | 24.2460246 | 17.6329864 | 17.6295737 | 0 |
| 0.00425988 | 0.00848594 | 12.2178549 | 12.4566732 | 4.64025958 | 19.9492545 | 0 |
| 0.00426 | 0.00848594 | 106.745469 | 117.226192 | 103.941815 | 94.1790387 | 118.810912 |
| 0.00426186 | 0.00848915 | 0 | 52.940861 | 162.409085 | 86.7560603 | 0 |
| 0.00426393 | 0.00849278 | 0 | 147.478113 | 123.430905 | 39.4345729 | 0 |
| 0.00428333 | 0.00853091 | 9.64567488 | 27.5826335 | 12.0646749 | 5.56723382 | 0 |
| 0.00428649 | 0.0085367 | 9.32415239 | 8.23030192 | 17.6329864 | 7.88691457 | 0 |
| 0.00428857 | 0.00854033 | 0 | 52.940861 | 259.854537 | 41.7542536 | 0 |
| 0.00428946 | 0.0085416 | 49.1929419 | 55.6101481 | 0 | 121.551272 | 0 |
| 0.0042906 | 0.00854336 | 4.82283744 | 13.3464355 | 12.9927268 | 15.309893 | 0 |
| 0.00429358 | 0.0085488 | 0 | 77.8542074 | 133.639476 | 75.1576565 | 0 |

| | | | | | | |
|---|---|---|---|---|---|---|
| 0.00429873 | 0.00855854 | 16.3976473 | 39.5944255 | 20.4171422 | 30.619786 | 0 |
| 0.00430129 | 0.00856312 | 4.50131495 | 13.7913167 | 14.8488307 | 14.8459568 | 0 |
| 0.00430357 | 0.00856716 | 11.8963324 | 12.6791138 | 18.5610383 | 22.7328714 | 0 |
| 0.00430416 | 0.00856779 | 63.6614542 | 40.7066284 | 0 | 126.190633 | 0 |
| 0.00430439 | 0.00856779 | 6.43044992 | 34.0334106 | 16.7049345 | 31.0837221 | 0 |
| 0.00430559 | 0.00856966 | 40.190312 | 7.78542074 | 8.35246725 | 8.81478687 | 0 |
| 0.00430624 | 0.00857045 | 2.25065747 | 22.4664998 | 22.273246 | 21.341063 | 0 |
| 0.00430773 | 0.00857291 | 6.43044992 | 20.0196533 | 14.8488307 | 7.42297842 | 0 |
| 0.00430987 | 0.00857666 | 0 | 0 | 0 | 0 | 59.4054561 |
| 0.0043103 | 0.00857701 | 5.14435994 | 8.23030192 | 17.6329864 | 23.6607437 | 0 |
| 0.00431125 | 0.0085784 | 149.507961 | 299.849919 | 176.329864 | 214.338502 | 57.0292378 |
| 0.0043128 | 0.00858096 | 57.2310043 | 102.545113 | 0 | 132.685739 | 0 |
| 0.0043265 | 0.00860771 | 89.0617314 | 112.999821 | 86.3088283 | 109.952868 | 28.5146189 |
| 0.00432974 | 0.00861365 | 14.7900348 | 6.67321777 | 9.28051917 | 10.6705315 | 0 |
| 0.00434684 | 0.00864716 | 48.8714194 | 117.893514 | 180.042072 | 120.623399 | 0 |
| 0.00434829 | 0.00864953 | 298.051354 | 567.00107 | 518.781022 | 598.477635 | 270.88888 |
| 0.0043588 | 0.00866993 | 15.1115573 | 29.8070394 | 2.78415575 | 17.1656376 | 0 |
| 0.00436156 | 0.00867491 | 155.938411 | 240.680721 | 206.955577 | 227.79265 | 0 |
| 0.00436324 | 0.00867773 | 9.00262989 | 19.1298909 | 23.2012979 | 17.6295737 | 0 |
| 0.00436449 | 0.0086797 | 44.3701045 | 113.667143 | 0 | 64.487125 | 0 |
| 0.00436907 | 0.0086883 | 14.1469898 | 86.5293905 | 180.042072 | 160.057972 | 19.0097459 |
| 0.00437043 | 0.00869037 | 0 | 0 | 0 | 0 | 23.7621824 |
| 0.00437063 | 0.00869037 | 29.9015921 | 15.1259603 | 31.5537652 | 1.85574461 | 0 |
| 0.00437215 | 0.00869245 | 26.0433222 | 55.8325887 | 26.9135056 | 40.8263813 | 11.8810912 |
| 0.00437219 | 0.00869245 | 1.28608998 | 3.33660889 | 2.78415575 | 5.10329766 | 0 |
| 0.00437479 | 0.00869711 | 10.2887199 | 22.6889404 | 10.2085711 | 6.03116997 | 0 |
| 0.00437749 | 0.00870145 | 9.00262989 | 14.6810791 | 10.2085711 | 6.95904227 | 0 |
| 0.00437739 | 0.00870145 | 100.636541 | 146.58835 | 251.502069 | 169.336695 | 38.0194919 |
| 0.00438359 | 0.00871306 | 8.6811074 | 26.6928711 | 2.78415575 | 40.8263813 | 0 |
| 0.00438508 | 0.00871551 | 83.2743265 | 70.7361084 | 0 | 56.1362743 | 0 |
| 0.00439257 | 0.00872988 | 8.0380624 | 13.7913167 | 9.28051917 | 8.35085072 | 0 |
| 0.00439767 | 0.00873949 | 4.17979245 | 12.4566732 | 18.5610383 | 15.7738291 | 0 |
| 0.00440263 | 0.00874884 | 8.6811074 | 8.23030192 | 8.35246725 | 18.5574461 | 0 |
| 0.00440602 | 0.00875504 | 11.2532874 | 16.0157227 | 4.64025958 | 14.8459568 | 0 |
| 0.00440703 | 0.00875653 | 12.2178549 | 11.7893514 | 39.9062324 | 16.2377653 | 0 |
| 0.00441447 | 0.00877081 | 17.3622148 | 12.2342326 | 16.7049345 | 4.17542536 | 0 |
| 0.00441501 | 0.00877137 | 5.46588243 | 16.0157227 | 13.9207788 | 9.74265918 | 0 |
| 0.00442026 | 0.00878127 | 0 | 18.6850098 | 142.919995 | 196.244992 | 0 |
| 0.00442251 | 0.00878523 | 6.43044992 | 8.23030192 | 25.0574018 | 11.5984038 | 0 |
| 0.00442475 | 0.00878916 | 0.96456749 | 23.3562622 | 57.5392189 | 21.8049991 | 0 |
| 0.00443199 | 0.00880302 | 33.4383396 | 10.899589 | 13.9207788 | 4.17542536 | 0 |
| 0.0044336 | 0.0088057 | 8.0380624 | 9.34250488 | 10.2085711 | 9.27872303 | 0 |
| 0.00443721 | 0.00881234 | 53.0512119 | 7.78542074 | 4.64025958 | 13.9180845 | 0 |

| | | | | | | |
|---|---|---|---|---|---|---|
| 0.00443812 | 0.00881312 | 5.78740493 | 11.1220296 | 14.8488307 | 10.6705315 | 0 |
| 0.00443796 | 0.00881312 | 68.1627692 | 65.6199748 | 0 | 72.8379758 | 0 |
| 0.004439 | 0.00881434 | 11.2532874 | 20.2420939 | 3.71220767 | 19.9492545 | 0 |
| 0.00445357 | 0.00884276 | 113.175919 | 106.993925 | 111.36623 | 117.839782 | 61.7816743 |
| 0.00445417 | 0.00884341 | 309.947686 | 545.424333 | 458.457647 | 539.093808 | 0 |
| 0.00445465 | 0.00884385 | 52.7296894 | 286.281042 | 23.2012979 | 0 | 0 |
| 0.00445883 | 0.00885163 | 67.8412467 | 83.8601033 | 97.4454513 | 104.84957 | 7.12865473 |
| 0.00446225 | 0.00885789 | 42.119447 | 106.104163 | 0 | 70.518295 | 0 |
| 0.00447103 | 0.00887479 | 4.82283744 | 34.4782918 | 12.9927268 | 9.74265918 | 0 |
| 0.00447197 | 0.00887613 | 79.094534 | 29.3621582 | 53.8270112 | 27.8361691 | 244.750479 |
| 0.00447257 | 0.00887681 | 0 | 140.804895 | 95.5893474 | 52.4247851 | 0 |
| 0.00447647 | 0.00888402 | 0 | 0 | 0 | 0 | 0 |
| 0.00447874 | 0.008888 | 4.50131495 | 13.123995 | 17.6329864 | 12.9902122 | 0 |
| 0.00448689 | 0.00890366 | 7.07349491 | 15.7932821 | 6.49636342 | 17.1656376 | 0 |
| 0.00448778 | 0.00890489 | 2.89370247 | 19.7972127 | 24.1293498 | 14.8459568 | 0 |
| 0.00448819 | 0.00890517 | 61.4107968 | 99.4309448 | 102.085711 | 96.0347833 | 26.1384007 |
| 0.00448894 | 0.00890576 | 0 | 1.55708415 | 0 | 0 | 21.3859642 |
| 0.00448901 | 0.00890576 | 8.3595849 | 10.2322673 | 9.28051917 | 8.81478687 | 0 |
| 0.0044938 | 0.00891474 | 78.7730116 | 44.4881185 | 148.488307 | 0 | 0 |
| 0.00449591 | 0.0089184 | 19.2913498 | 14.9035197 | 13.9207788 | 12.5262761 | 0 |
| 0.00449806 | 0.00892213 | 65.5905892 | 103.657316 | 87.2368802 | 97.4265918 | 26.1384007 |
| 0.00450285 | 0.0089311 | 3.85826995 | 12.4566732 | 22.273246 | 15.7738291 | 0 |
| 0.0045047 | 0.00893425 | 14.4685123 | 24.9133464 | 4.64025958 | 9.74265918 | 0 |
| 0.00450529 | 0.0089349 | 0 | 100.098267 | 82.5966206 | 90.0036134 | 0 |
| 0.00451322 | 0.00895009 | 1.60761248 | 1.55708415 | 0.92805192 | 0.46393615 | 40.3957101 |
| 0.00451716 | 0.00895737 | 84.881939 | 205.979989 | 303.472977 | 274.186265 | 0 |
| 0.00451894 | 0.00896038 | 0 | 0 | 0 | 0 | 47.5243649 |
| 0.00451926 | 0.00896048 | 33.7598621 | 12.011792 | 6.49636342 | 7.42297842 | 0 |
| 0.00452048 | 0.00896238 | 6.43044992 | 13.7913167 | 9.28051917 | 11.1344676 | 0 |
| 0.00452516 | 0.00897112 | 4.50131495 | 22.2440592 | 9.28051917 | 18.0935099 | 0 |
| 0.00452702 | 0.00897428 | 76.2008316 | 70.2912272 | 0 | 58.9198912 | 0 |
| 0.00452768 | 0.00897507 | 10.6102424 | 18.2401286 | 6.49636342 | 8.81478687 | 0 |
| 0.0045292 | 0.00897756 | 8.3595849 | 34.7007324 | 30.6257133 | 38.9706367 | 0 |
| 0.00453479 | 0.0089881 | 20.5774398 | 124.344291 | 0 | 107.633187 | 0 |
| 0.00453575 | 0.00898947 | 2.89370247 | 13.5688761 | 29.6976613 | 18.5574461 | 0 |
| 0.00454583 | 0.00900876 | 0 | 110.997856 | 31.5537652 | 172.120312 | 0 |
| 0.00454602 | 0.00900876 | 11.8963324 | 15.5708415 | 4.64025958 | 13.4541484 | 0 |
| 0.00454728 | 0.00901074 | 6.43044992 | 8.23030192 | 16.7049345 | 12.0623399 | 0 |
| 0.00454772 | 0.00901108 | 11.8963324 | 177.507593 | 0 | 107.633187 | 0 |
| 0.00455177 | 0.00901856 | 12.2178549 | 10.6771484 | 18.5610383 | 5.10329766 | 0 |
| 0.0045551 | 0.00902464 | 4.50131495 | 10.4547078 | 14.8488307 | 22.7328714 | 0 |
| 0.00455787 | 0.00902959 | 69.7703817 | 135.24388 | 129.927268 | 115.984038 | 45.1481466 |
| 0.00456136 | 0.00903598 | 15.1115573 | 109.885653 | 0 | 149.851377 | 0 |

| | | | | | | |
|---|---|---|---|---|---|---|
| 0.00456238 | 0.00903746 | 4.82283744 | 10.6771484 | 21.3451941 | 12.9902122 | 0 |
| 0.00456307 | 0.0090383 | 48.2283744 | 10.899589 | 5.5683115 | 8.35085072 | 0 |
| 0.00456873 | 0.00904898 | 12.2178549 | 7.56298014 | 3.71220767 | 5.10329766 | 99.8011662 |
| 0.0045695 | 0.00904997 | 58.5170943 | 72.0707519 | 0 | 73.7658481 | 0 |
| 0.0045735 | 0.00905735 | 54.3373018 | 79.4112915 | 78.8844129 | 104.385634 | 23.7621824 |
| 0.00457852 | 0.00906677 | 0 | 0 | 0 | 0 | 2.37621824 |
| 0.00458128 | 0.00907169 | 114.462009 | 48.9369303 | 0 | 53.8165936 | 0 |
| 0.00459516 | 0.00909865 | 63.0184092 | 100.543148 | 86.3088283 | 91.859358 | 49.9005831 |
| 0.00460428 | 0.00911617 | 91.6339114 | 118.115955 | 48.2586997 | 0 | 0 |
| 0.00461187 | 0.00913065 | 7.07349491 | 12.9015544 | 13.9207788 | 7.42297842 | 0 |
| 0.00461452 | 0.00913537 | 7.07349491 | 10.4547078 | 8.35246725 | 13.9180845 | 0 |
| 0.00461509 | 0.00913596 | 13.5039448 | 22.4664998 | 24.1293498 | 2.78361691 | 0 |
| 0.00461714 | 0.00913947 | 100.958064 | 107.216366 | 62.1794784 | 84.4363795 | 35.6432737 |
| 0.00461837 | 0.00914137 | 79.094534 | 156.153296 | 116.00649 | 112.736485 | 30.8908372 |
| 0.00461914 | 0.00914236 | 0 | 77.6317668 | 103.013763 | 89.0757411 | 0 |
| 0.00462858 | 0.00916051 | 62.0538418 | 49.8266927 | 0 | 97.4265918 | 0 |
| 0.00463153 | 0.00916581 | 434.05537 | 328.544755 | 346.163365 | 379.035836 | 116.434694 |
| 0.00463475 | 0.00917164 | 40.5118345 | 120.562801 | 0 | 59.8477635 | 0 |
| 0.00463931 | 0.00918013 | 15.7546023 | 13.7913167 | 12.9927268 | 10.6705315 | 40.3957101 |
| 0.00464579 | 0.0091924 | 0 | 0 | 0 | 0 | 14.2573095 |
| 0.00464705 | 0.00919436 | 42.762492 | 100.988029 | 135.49558 | 120.159463 | 2.37621824 |
| 0.00464804 | 0.00919578 | 99.9934963 | 56.0550293 | 0 | 53.3526574 | 0 |
| 0.00465905 | 0.00921702 | 0 | 52.940861 | 91.8771398 | 136.397228 | 0 |
| 0.00466352 | 0.0092253 | 125.072251 | 251.357869 | 245.933758 | 261.659989 | 14.2573095 |
| 0.00466483 | 0.00922737 | 11.5748099 | 22.911381 | 6.49636342 | 7.88691457 | 0 |
| 0.00466842 | 0.00923392 | 28.9370247 | 78.0766479 | 0 | 123.870952 | 0 |
| 0.0046774 | 0.00925115 | 0 | 0.22244059 | 0 | 0 | 0 |
| 0.0046826 | 0.00926089 | 9.32415239 | 38.9271037 | 6.49636342 | 8.81478687 | 0 |
| 0.00468487 | 0.00926483 | 140.505331 | 318.757369 | 262.638692 | 315.476583 | 52.2768014 |
| 0.00468638 | 0.00926726 | 8.3595849 | 3.78149007 | 9.28051917 | 2.78361691 | 38.0194919 |
| 0.00469348 | 0.00928075 | 3.53674746 | 15.7932821 | 12.0646749 | 28.3001052 | 0 |
| 0.00469468 | 0.00928258 | 5.46588243 | 19.5747721 | 8.35246725 | 14.8459568 | 0 |
| 0.00469951 | 0.00929105 | 0 | 0 | 0 | 0 | 0 |
| 0.00469951 | 0.00929105 | 0 | 0 | 0 | 0 | 0 |
| 0.00469982 | 0.00929111 | 11.5748099 | 10.2322673 | 7.42441534 | 8.35085072 | 0 |
| 0.00470139 | 0.00929367 | 17.3622148 | 9.34250488 | 4.64025958 | 17.6295737 | 0 |
| 0.0047142 | 0.00931845 | 10.6102424 | 20.9094157 | 12.0646749 | 22.7328714 | 2.37621824 |
| 0.00471538 | 0.00932022 | 0 | 68.9565837 | 87.2368802 | 113.200421 | 0 |
| 0.00472048 | 0.00932977 | 3.53674746 | 12.011792 | 15.7768826 | 28.3001052 | 0 |
| 0.00472386 | 0.00933589 | 48.5498969 | 44.0432373 | 0 | 130.829995 | 0 |
| 0.00472775 | 0.00934302 | 24.7572322 | 39.816866 | 51.0428554 | 36.650956 | 14.2573095 |
| 0.00473516 | 0.00935713 | 43.0840145 | 100.320707 | 0 | 66.8068058 | 0 |
| 0.00474284 | 0.00937175 | 11.5748099 | 12.011792 | 5.5683115 | 11.1344676 | 0 |

| | | | | | | |
|---|---|---|---|---|---|---|
| 0.00475595 | 0.00939711 | 0 | 161.714311 | 79.8124649 | 47.3214874 | 0 |
| 0.00475816 | 0.00940092 | 3.21522496 | 11.1220296 | 64.9636342 | 11.1344676 | 0 |
| 0.00476195 | 0.00940787 | 6.43044992 | 14.2361979 | 11.136623 | 8.81478687 | 0 |
| 0.00476652 | 0.00941632 | 203.202218 | 613.491154 | 630.147252 | 640.695825 | 2.37621824 |
| 0.00477341 | 0.0094291 | 6.10892743 | 12.011792 | 10.2085711 | 11.1344676 | 0 |
| 0.00477354 | 0.0094291 | 5.14435994 | 25.8031087 | 7.42441534 | 17.1656376 | 0 |
| 0.00477409 | 0.00942963 | 3.21522496 | 13.5688761 | 28.7696094 | 15.309893 | 0 |
| 0.00477592 | 0.00943268 | 1.28608998 | 24.4684652 | 56.6111669 | 14.3820207 | 0 |
| 0.00478763 | 0.00945526 | 27.9724572 | 7.34053955 | 26.9135056 | 19.0213822 | 47.5243649 |
| 0.00479082 | 0.00946101 | 45.6561944 | 84.5274251 | 0 | 74.2297842 | 0 |
| 0.00479166 | 0.00946211 | 101.601109 | 190.854028 | 179.11402 | 166.553078 | 7.12865473 |
| 0.00480033 | 0.00947867 | 10.2887199 | 13.3464355 | 4.64025958 | 19.9492545 | 0 |
| 0.00480429 | 0.00948594 | 0 | 58.7243164 | 91.8771398 | 120.159463 | 0 |
| 0.00480555 | 0.00948786 | 5.14435994 | 12.9015544 | 11.136623 | 13.9180845 | 0 |
| 0.00480989 | 0.00949588 | 10.9317649 | 19.5747721 | 20.4171422 | 12.5262761 | 2.37621824 |
| 0.00481263 | 0.00950017 | 6.10892743 | 29.5845988 | 11.136623 | 8.35085072 | 0 |
| 0.00481243 | 0.00950017 | 137.933151 | 286.725924 | 257.998433 | 262.587862 | 49.9005831 |
| 0.00481894 | 0.00951208 | 0 | 50.7164551 | 150.344411 | 80.2609542 | 0 |
| 0.00482063 | 0.00951485 | 226.351837 | 103.212435 | 45.4745439 | 51.0329766 | 0 |
| 0.00482158 | 0.00951617 | 145.328168 | 793.890474 | 566.111669 | 634.664655 | 2.37621824 |
| 0.00482615 | 0.00952458 | 0 | 67.8443807 | 132.711424 | 70.9822312 | 0 |
| 0.00482641 | 0.00952458 | 3.53674746 | 13.5688761 | 19.4890903 | 16.7017014 | 0 |
| 0.00482675 | 0.00952469 | 0 | 8.8976237 | 0 | 4.63936151 | 21.3859642 |
| 0.00482971 | 0.00952997 | 0 | 56.0550293 | 132.711424 | 84.9003157 | 0 |
| 0.00483096 | 0.00953187 | 32.7952946 | 10.0098267 | 8.35246725 | 6.49510612 | 0 |
| 0.0048388 | 0.00954679 | 2.25065747 | 0.88976237 | 0.92805192 | 1.39180845 | 7.12865473 |
| 0.00484239 | 0.00955332 | 5.78740493 | 22.911381 | 6.49636342 | 18.0935099 | 0 |
| 0.00484791 | 0.00956364 | 37.9396545 | 42.4861532 | 0 | 157.274355 | 0 |
| 0.00484846 | 0.00956416 | 0 | 22.2440592 | 318.321808 | 47.7854236 | 0 |
| 0.00485087 | 0.00956836 | 8.0380624 | 9.78738607 | 11.136623 | 7.88691457 | 0 |
| 0.0048546 | 0.00957516 | 25.7217997 | 68.9565837 | 78.8844129 | 64.487125 | 0 |
| 0.00485905 | 0.00958336 | 5.78740493 | 15.5708415 | 7.42441534 | 19.0213822 | 0 |
| 0.00485988 | 0.00958446 | 0 | 151.926925 | 69.6038938 | 130.366059 | 0 |
| 0.0048756 | 0.00961488 | 1.28608998 | 23.3562622 | 38.0501286 | 21.341063 | 0 |
| 0.00487604 | 0.00961519 | 202.23765 | 412.84974 | 438.968557 | 410.583494 | 61.7816743 |
| 0.00487996 | 0.00962235 | 50.4790319 | 102.989994 | 67.7477899 | 80.7248903 | 28.5146189 |
| 0.00488157 | 0.00962496 | 11.8963324 | 2.8917277 | 29.6976613 | 19.9492545 | 0 |
| 0.00488694 | 0.00963498 | 38.261177 | 19.3523315 | 29.6976613 | 18.0935099 | 102.177384 |
| 0.00488961 | 0.0096397 | 5.14435994 | 15.1259603 | 15.7768826 | 9.27872303 | 0 |
| 0.0048908 | 0.00964147 | 0 | 145.253707 | 97.4454513 | 41.2903175 | 0 |
| 0.00489144 | 0.00964217 | 9.32415239 | 28.917277 | 12.0646749 | 22.2689353 | 2.37621824 |
| 0.00489674 | 0.00965205 | 14.4685123 | 21.1318563 | 8.35246725 | 24.588616 | 2.37621824 |
| 0.00491111 | 0.0096798 | 20.8989622 | 38.0373413 | 7.42441534 | 3.71148921 | 0 |

| | | | | | | |
|---|---|---|---|---|---|---|
| 0.00491146 | 0.00967993 | 41.1548795 | 101.655351 | 110.438178 | 0 | 0 |
| 0.00491216 | 0.00968074 | 76.2008316 | 42.9310343 | 0 | 83.9724434 | 0 |
| 0.00491293 | 0.00968169 | 2.89370247 | 28.917277 | 42.6903882 | 32.4755306 | 2.37621824 |
| 0.00491493 | 0.00968507 | 4.82283744 | 14.2361979 | 18.5610383 | 9.74265918 | 0 |
| 0.00492369 | 0.00970176 | 0 | 78.0766479 | 105.797919 | 77.4773373 | 0 |
| 0.00493891 | 0.00973119 | 3.85826995 | 0 | 0 | 0 | 199.602332 |
| 0.00493964 | 0.00973205 | 9.00262989 | 174.838306 | 0 | 115.984038 | 0 |
| 0.00494138 | 0.00973491 | 24.7572322 | 20.9094157 | 22.273246 | 14.3820207 | 64.1578926 |
| 0.00494234 | 0.00973623 | 5.78740493 | 17.5728068 | 8.35246725 | 12.9902122 | 0 |
| 0.00494279 | 0.00973654 | 2.89370247 | 39.816866 | 10.2085711 | 19.0213822 | 0 |
| 0.00494429 | 0.00973892 | 11.5748099 | 11.3444702 | 6.49636342 | 8.81478687 | 0 |
| 0.00495435 | 0.00975818 | 0 | 99.6533854 | 32.4818171 | 164.233398 | 0 |
| 0.00495853 | 0.00976583 | 12.8608998 | 10.6771484 | 4.64025958 | 17.6295737 | 0 |
| 0.0049594 | 0.00976697 | 49.8359869 | 78.7439697 | 0 | 69.5904227 | 0 |
| 0.00496127 | 0.00977008 | 11.5748099 | 15.5708415 | 6.49636342 | 7.88691457 | 0 |
| 0.00496254 | 0.00977201 | 68.1627692 | 64.2853312 | 0 | 62.6313804 | 0 |
| 0.00496964 | 0.00978542 | 0 | 70.7361084 | 56.6111669 | 147.06776 | 0 |
| 0.0049795 | 0.00980426 | 0.64304499 | 0.44488118 | 0 | 0 | 0 |
| 0.00498002 | 0.00980471 | 120.892459 | 209.316597 | 125.287009 | 163.305525 | 0 |
| 0.00498448 | 0.00981292 | 0 | 82.7479004 | 90.9490879 | 83.9724434 | 0 |
| 0.00499265 | 0.00982842 | 17.0406923 | 27.5826335 | 18.5610383 | 26.9082968 | 0 |
| 0.00499316 | 0.00982886 | 9.32415239 | 10.0098267 | 6.49636342 | 12.0623399 | 0 |
| 0.00499628 | 0.00983443 | 3.85826995 | 5.11613363 | 39.9062324 | 29.6919137 | 0 |
| 0.00500027 | 0.00984171 | 1.60761248 | 0.88976237 | 0.92805192 | 0.46393615 | 54.6530196 |
| 0.00500667 | 0.00985372 | 0.3215225 | 0.22244059 | 0 | 0 | 7.12865473 |
| 0.00500777 | 0.00985532 | 3.53674746 | 9.56494547 | 23.2012979 | 22.7328714 | 0 |
| 0.00500975 | 0.00985863 | 13.5039448 | 140.360014 | 166.121293 | 0 | 0 |
| 0.00501539 | 0.00986915 | 65.5905892 | 57.1672323 | 0 | 71.9101035 | 0 |
| 0.00502345 | 0.00988437 | 18.6483048 | 131.907271 | 0 | 93.2511664 | 0 |
| 0.00502371 | 0.00988437 | 9.00262989 | 9.34250488 | 8.35246725 | 8.81478687 | 0 |
| 0.0050247 | 0.00988574 | 181.017165 | 583.239233 | 555.903098 | 619.354762 | 2.37621824 |
| 0.00502559 | 0.0098864 | 5.14435994 | 28.917277 | 7.42441534 | 15.309893 | 0 |
| 0.00502563 | 0.0098864 | 0 | 82.5254598 | 74.2441534 | 101.138081 | 0 |
| 0.00502668 | 0.00988791 | 73.9501741 | 104.769519 | 60.3233746 | 0 | 0 |
| 0.00503437 | 0.00990246 | 64.3044992 | 104.99196 | 99.3015551 | 90.0036134 | 11.8810912 |
| 0.00503763 | 0.00990828 | 187.126093 | 333.883329 | 334.09869 | 332.64222 | 59.4054561 |
| 0.00503885 | 0.0099101 | 4.17979245 | 8.00786133 | 21.3451941 | 24.1246799 | 0 |
| 0.00504351 | 0.00991869 | 9.96719738 | 12.4566732 | 7.42441534 | 7.88691457 | 0 |
| 0.0050466 | 0.00992419 | 108.031559 | 100.543148 | 57.5392189 | 57.9920189 | 23.7621824 |
| 0.0050483 | 0.00992695 | 0 | 80.7459351 | 104.869867 | 72.3740396 | 0 |
| 0.00504997 | 0.00992965 | 129.895088 | 52.4959798 | 103.941815 | 12.0623399 | 14.2573095 |
| 0.00505041 | 0.00992994 | 5.14435994 | 8.00786133 | 12.9927268 | 34.7952113 | 0 |
| 0.0050516 | 0.0099317 | 18.3267823 | 157.043058 | 0 | 76.0855288 | 0 |

| | | | | | | |
|---|---|---|---|---|---|---|
| 0.00505447 | 0.00993676 | 0 | 92.0904053 | 82.5966206 | 81.1888265 | 0 |
| 0.00505492 | 0.00993706 | 40.190312 | 59.6140788 | 0 | 107.633187 | 0 |
| 0.005058 | 0.00994245 | 19.2913498 | 12.4566732 | 18.5610383 | 12.5262761 | 30.8908372 |
| 0.00505825 | 0.00994245 | 0 | 40.4841878 | 116.934542 | 118.303719 | 0 |
| 0.00506095 | 0.00994716 | 39.547267 | 29.3621582 | 23.2012979 | 7.88691457 | 99.8011662 |
| 0.00506298 | 0.00995059 | 24.4357097 | 117.448633 | 134.567528 | 0 | 0 |
| 0.00506379 | 0.00995159 | 0 | 0 | 0 | 0 | 28.5146189 |
| 0.0050659 | 0.00995516 | 8.3595849 | 6.22833659 | 20.4171422 | 10.6705315 | 0 |
| 0.00506631 | 0.00995538 | 18.0052598 | 15.5708415 | 12.9927268 | 12.0623399 | 0 |
| 0.00507481 | 0.0099715 | 3.85826995 | 11.3444702 | 16.7049345 | 19.4853184 | 0 |
| 0.00508369 | 0.00998836 | 5.14435994 | 10.4547078 | 11.136623 | 19.4853184 | 0 |
| 0.00510243 | 0.0100246 | 5.14435994 | 9.34250488 | 17.6329864 | 12.9902122 | 0 |
| 0.00511495 | 0.01004862 | 11.5748099 | 6.22833659 | 7.42441534 | 29.2279775 | 0 |
| 0.00512065 | 0.01005923 | 4.17979245 | 37.8149007 | 11.136623 | 11.1344676 | 0 |
| 0.00512095 | 0.01005923 | 57.5525268 | 79.4112915 | 0 | 204.131907 | 2.37621824 |
| 0.00512627 | 0.01006908 | 33.4383396 | 11.5669108 | 7.42441534 | 6.03116997 | 0 |
| 0.00512778 | 0.01007146 | 9.96719738 | 20.0196533 | 4.64025958 | 12.9902122 | 0 |
| 0.0051281 | 0.0100715 | 24.4357097 | 16.905485 | 7.42441534 | 20.8771268 | 0 |
| 0.00513245 | 0.01007946 | 10.6102424 | 6.45077718 | 10.2085711 | 10.2065953 | 0 |
| 0.00513923 | 0.01009218 | 14.4685123 | 11.3444702 | 6.49636342 | 7.88691457 | 0 |
| 0.00514078 | 0.01009464 | 0 | 81.858138 | 112.294282 | 64.0231889 | 0 |
| 0.00514206 | 0.01009656 | 185.196958 | 228.446488 | 225.516616 | 240.782863 | 49.9005831 |
| 0.00514809 | 0.01010723 | 54.9803468 | 63.3955688 | 0 | 74.2297842 | 0 |
| 0.00514801 | 0.01010723 | 15.7546023 | 11.3444702 | 12.9927268 | 12.9902122 | 14.2573095 |
| 0.00514901 | 0.01010843 | 15.7546023 | 10.6771484 | 16.7049345 | 8.81478687 | 61.7816743 |
| 0.00515141 | 0.01011256 | 22.5065747 | 38.9271037 | 35.2659728 | 31.0837221 | 0 |
| 0.0051533 | 0.01011568 | 0.3215225 | 0.66732178 | 0 | 0 | 33.2670554 |
| 0.00515462 | 0.01011769 | 123.786161 | 117.448633 | 54.7550631 | 107.169251 | 0 |
| 0.00515886 | 0.01012542 | 126.358341 | 72.5156331 | 68.6758419 | 74.2297842 | 0 |
| 0.00516068 | 0.0101284 | 3.85826995 | 16.0157227 | 13.9207788 | 14.8459568 | 0 |
| 0.00516806 | 0.0101423 | 47.9068519 | 69.4014648 | 0 | 77.0134011 | 0 |
| 0.0051762 | 0.01015767 | 4.50131495 | 7.56298014 | 29.6976613 | 17.1656376 | 0 |
| 0.00517841 | 0.0101613 | 5.14435994 | 14.4586385 | 16.7049345 | 8.81478687 | 0 |
| 0.00517865 | 0.0101613 | 11.2532874 | 29.5845988 | 12.9927268 | 14.3820207 | 0 |
| 0.00518026 | 0.01016386 | 51.4435994 | 88.976237 | 96.5173994 | 0 | 0 |
| 0.00518063 | 0.01016399 | 45.9777169 | 78.5215291 | 0 | 70.518295 | 0 |
| 0.00518142 | 0.01016495 | 5.14435994 | 10.6771484 | 18.5610383 | 10.6705315 | 0 |
| 0.00518489 | 0.01017117 | 30.2231146 | 92.0904053 | 0 | 87.6839326 | 0 |
| 0.00518713 | 0.01017496 | 5.78740493 | 12.011792 | 24.1293498 | 7.88691457 | 0 |
| 0.00518848 | 0.01017703 | 49.5144644 | 0 | 115.078438 | 124.334889 | 0 |
| 0.00518894 | 0.01017734 | 38.261177 | 93.2026082 | 0 | 70.0543588 | 0 |
| 0.00519022 | 0.01017925 | 0 | 0 | 0.92805192 | 0 | 57.0292378 |
| 0.00519152 | 0.0101812 | 74.2716966 | 107.216366 | 124.358957 | 146.603824 | 30.8908372 |

| | | | | | | |
|---|---|---|---|---|---|---|
| 0.00519208 | 0.01018171 | 0.3215225 | 2.00196533 | 2.78415575 | 2.78361691 | 7.12865473 |
| 0.00519495 | 0.01018663 | 3.21522496 | 12.4566732 | 26.9135056 | 15.7738291 | 0 |
| 0.00519519 | 0.01018663 | 0 | 104.324638 | 62.1794784 | 89.5396772 | 0 |
| 0.00520447 | 0.01020423 | 59.8031843 | 72.2931925 | 0 | 58.9198912 | 0 |
| 0.00520654 | 0.01020709 | 383.254815 | 517.396818 | 209.739733 | 647.190931 | 0 |
| 0.00520626 | 0.01020709 | 3.53674746 | 16.0157227 | 14.8488307 | 16.2377653 | 0 |
| 0.00521515 | 0.01022276 | 283.582842 | 286.725924 | 276.559471 | 283.001052 | 59.4054561 |
| 0.00521505 | 0.01022276 | 8.3595849 | 7.56298014 | 8.35246725 | 17.1656376 | 0 |
| 0.00521545 | 0.01022276 | 0 | 0 | 0 | 0 | 85.5438568 |
| 0.00521691 | 0.01022504 | 17.3622148 | 32.0314453 | 95.5893474 | 49.6411682 | 11.8810912 |
| 0.0052192 | 0.01022893 | 9.96719738 | 25.3582275 | 16.7049345 | 13.9180845 | 0 |
| 0.00522204 | 0.01023389 | 61.7323193 | 3.78149007 | 17.6329864 | 5.56723382 | 0 |
| 0.00522409 | 0.01023733 | 310.590731 | 1481.45435 | 2502.02797 | 2018.12226 | 144.949313 |
| 0.00522545 | 0.01023939 | 135.039448 | 352.79078 | 331.314534 | 302.486371 | 2.37621824 |
| 0.00522923 | 0.0102462 | 21.5420072 | 139.02537 | 0 | 73.3019119 | 0 |
| 0.00524079 | 0.01026825 | 82.6312815 | 0 | 156.840774 | 52.4247851 | 0 |
| 0.00524495 | 0.01027581 | 3.85826995 | 20.0196533 | 12.9927268 | 13.4541484 | 0 |
| 0.0052522 | 0.01028881 | 50.8005544 | 79.1888509 | 0 | 62.1674443 | 0 |
| 0.00525191 | 0.01028881 | 6.10892743 | 22.2440592 | 22.273246 | 24.588616 | 2.37621824 |
| 0.0052556 | 0.01029486 | 6.75197242 | 1.77952474 | 7.42441534 | 3.24755306 | 47.5243649 |
| 0.005258 | 0.01029897 | 0 | 50.2715739 | 113.222334 | 96.9626556 | 0 |
| 0.00526202 | 0.01030625 | 162.047338 | 210.873682 | 172.617657 | 209.69914 | 45.1481466 |
| 0.00526279 | 0.01030714 | 6.43044992 | 10.4547078 | 8.35246725 | 14.8459568 | 0 |
| 0.00526402 | 0.01030836 | 109.639171 | 10.6771484 | 1.85610383 | 8.35085072 | 0 |
| 0.00526375 | 0.01030836 | 64.6260217 | 72.5156331 | 0 | 53.3526574 | 0 |
| 0.00526512 | 0.01030991 | 31.1876821 | 94.3148112 | 131.783372 | 0 | 0 |
| 0.00526825 | 0.01031544 | 6.43044992 | 10.6771484 | 8.35246725 | 13.9180845 | 0 |
| 0.0052764 | 0.01033079 | 29.5800696 | 116.33643 | 0 | 67.7346781 | 0 |
| 0.00528378 | 0.01034407 | 8.6811074 | 8.45274251 | 8.35246725 | 9.74265918 | 0 |
| 0.00528379 | 0.01034407 | 0.3215225 | 26.0255493 | 56.6111669 | 40.8263813 | 0 |
| 0.00528611 | 0.01034801 | 48.2283744 | 81.1908162 | 0 | 63.0953166 | 0 |
| 0.00528868 | 0.01035243 | 4.82283744 | 12.4566732 | 17.6329864 | 10.2065953 | 0 |
| 0.0052908 | 0.01035597 | 1.60761248 | 30.2519206 | 18.5610383 | 23.1968076 | 0 |
| 0.00530042 | 0.01037402 | 3.85826995 | 18.6850098 | 16.7049345 | 11.1344676 | 0 |
| 0.00530063 | 0.01037402 | 67.1982017 | 74.5175985 | 52.8989593 | 32.4755306 | 23.7621824 |
| 0.00531087 | 0.01039345 | 9.64567488 | 9.12006429 | 7.42441534 | 9.27872303 | 0 |
| 0.00531283 | 0.01039669 | 0.64304499 | 2.66928711 | 0 | 0.9278723 | 0 |
| 0.00531834 | 0.01040686 | 8.0380624 | 25.5806681 | 3.71220767 | 21.8049991 | 0 |
| 0.00532179 | 0.01041301 | 44.691627 | 78.9664103 | 0 | 69.1264865 | 0 |
| 0.00532492 | 0.01041853 | 47.5853294 | 20.0196533 | 34.3379209 | 27.3722329 | 59.4054561 |
| 0.0053345 | 0.01043666 | 90.0262989 | 186.850098 | 103.013763 | 130.366059 | 33.2670554 |
| 0.00533885 | 0.01044456 | 2.25065747 | 0.44488118 | 3.71220767 | 1.39180845 | 52.2768014 |
| 0.00534838 | 0.01046255 | 86.811074 | 38.0373413 | 0 | 72.3740396 | 0 |

| | | | | | | |
|---|---|---|---|---|---|---|
| 0.00534866 | 0.01046255 | 13.5039448 | 9.56494547 | 6.49636342 | 8.81478687 | 0 |
| 0.00535149 | 0.01046746 | 79.094534 | 75.852242 | 50.1148035 | 68.6625504 | 0 |
| 0.00535226 | 0.01046836 | 42.762492 | 84.3049845 | 0 | 66.8068058 | 0 |
| 0.00535561 | 0.0104743 | 4.17979245 | 12.6791138 | 20.4171422 | 11.5984038 | 0 |
| 0.00536391 | 0.01048993 | 33.4383396 | 90.0884399 | 127.143113 | 0 | 0 |
| 0.00536503 | 0.01049151 | 8.3595849 | 9.78738607 | 9.28051917 | 7.88691457 | 0 |
| 0.00536575 | 0.01049231 | 4.82283744 | 2.66928711 | 8.35246725 | 4.17542536 | 16.6335277 |
| 0.00536634 | 0.01049286 | 4.50131495 | 16.4606038 | 14.8488307 | 10.2065953 | 0 |
| 0.00536745 | 0.01049441 | 9.96719738 | 6.005896 | 10.2085711 | 4.63936151 | 38.0194919 |
| 0.0053704 | 0.01049957 | 22.8280972 | 123.454529 | 0 | 77.0134011 | 0 |
| 0.00537356 | 0.01050515 | 6.43044992 | 11.1220296 | 28.7696094 | 47.7854236 | 0 |
| 0.00537462 | 0.0105066 | 5.78740493 | 15.5708415 | 7.42441534 | 15.309893 | 0 |
| 0.00537713 | 0.01051089 | 43.0840145 | 108.328569 | 0 | 50.1051043 | 0 |
| 0.00538658 | 0.01052876 | 45.334672 | 49.1593709 | 46.4025958 | 47.3214874 | 73.6627655 |
| 0.0053918 | 0.01053797 | 20.2559173 | 78.9664103 | 17.6329864 | 126.190633 | 9.50487297 |
| 0.00539192 | 0.01053797 | 14.7900348 | 30.4743612 | 24.1293498 | 24.588616 | 7.12865473 |
| 0.0053923 | 0.01053811 | 103.530244 | 220.216187 | 217.164149 | 217.586055 | 38.0194919 |
| 0.00539392 | 0.01054003 | 3.21522496 | 21.7991781 | 14.8488307 | 14.3820207 | 0 |
| 0.00539378 | 0.01054003 | 12.2178549 | 6.89565837 | 10.2085711 | 9.74265918 | 11.8810912 |
| 0.00539894 | 0.01054924 | 5.14435994 | 7.78542074 | 16.7049345 | 17.6295737 | 0 |
| 0.00540616 | 0.01056274 | 29.2585471 | 43.1534749 | 36.1940248 | 26.4443606 | 9.50487297 |
| 0.00541074 | 0.0105707 | 17.6837373 | 114.112024 | 101.157659 | 105.777442 | 0 |
| 0.00541087 | 0.0105707 | 7.07349491 | 6.67321777 | 6.49636342 | 9.27872303 | 147.325531 |
| 0.0054113 | 0.01057093 | 18.9698273 | 14.0137573 | 16.7049345 | 13.9180845 | 111.682257 |
| 0.00541539 | 0.01057831 | 4.17979245 | 23.3562622 | 10.2085711 | 13.9180845 | 0 |
| 0.00542341 | 0.01059335 | 0 | 93.2026082 | 146.632203 | 34.7952113 | 0 |
| 0.00542644 | 0.01059866 | 21.2204847 | 104.769519 | 0 | 96.0347833 | 0 |
| 0.00542749 | 0.0105997 | 64.6260217 | 6.005896 | 12.0646749 | 4.63936151 | 0 |
| 0.0054276 | 0.0105997 | 0.64304499 | 0 | 0 | 0 | 4.75243649 |
| 0.00543656 | 0.01061659 | 114.462009 | 51.6062174 | 45.4745439 | 48.7132959 | 156.830404 |
| 0.00543712 | 0.01061705 | 124.107684 | 409.29069 | 363.796351 | 349.807858 | 0 |
| 0.00544827 | 0.01063821 | 99.9934963 | 45.3778809 | 0 | 51.0329766 | 0 |
| 0.00545061 | 0.01064217 | 9.64567488 | 12.9015544 | 23.2012979 | 18.5574461 | 2.37621824 |
| 0.0054517 | 0.01064367 | 12.2178549 | 21.3542969 | 10.2085711 | 18.0935099 | 2.37621824 |
| 0.0054536 | 0.01064677 | 13.8254673 | 16.4606038 | 18.5610383 | 3.24755306 | 0 |
| 0.00545743 | 0.01065362 | 7.07349491 | 9.34250488 | 8.35246725 | 12.0623399 | 0 |
| 0.00546258 | 0.01066284 | 7.07349491 | 28.0275146 | 39.9062324 | 34.7952113 | 4.75243649 |
| 0.00546278 | 0.01066284 | 0 | 4.00393066 | 0 | 0 | 78.415202 |
| 0.00547426 | 0.01068461 | 22.1850522 | 15.3484009 | 10.2085711 | 10.2065953 | 23.7621824 |
| 0.00548393 | 0.01070288 | 0 | 76.7420044 | 125.287009 | 53.3526574 | 0 |
| 0.0054869 | 0.01070805 | 113.818964 | 339.444344 | 282.127783 | 345.632433 | 0 |
| 0.00548934 | 0.01071218 | 65.5905892 | 38.2597819 | 51.9709073 | 34.7952113 | 42.7719284 |
| 0.00549142 | 0.01071562 | 3.85826995 | 22.6889404 | 10.2085711 | 15.7738291 | 0 |

| | | | | | | |
|---|---|---|---|---|---|---|
| 0.00549426 | 0.01072054 | 64.3044992 | 73.1829549 | 0 | 49.6411682 | 0 |
| 0.00549584 | 0.010723 | 6.75197242 | 17.7952474 | 4.64025958 | 28.3001052 | 0 |
| 0.00549648 | 0.01072362 | 27.0078897 | 19.5747721 | 18.5610383 | 21.8049991 | 92.6725115 |
| 0.00549687 | 0.01072377 | 58.5170943 | 27.1377523 | 0 | 132.685739 | 0 |
| 0.00549868 | 0.01072669 | 15.1115573 | 8.8976237 | 10.2085711 | 6.03116997 | 0 |
| 0.00550071 | 0.01073002 | 0 | 92.757727 | 77.0283091 | 75.1576565 | 0 |
| 0.00550468 | 0.01073714 | 3.53674746 | 17.7952474 | 11.136623 | 19.9492545 | 0 |
| 0.00550578 | 0.01073867 | 29.5800696 | 20.0196533 | 5.5683115 | 5.10329766 | 0 |
| 0.0055085 | 0.01074334 | 3.85826995 | 27.8050741 | 6.49636342 | 25.9804245 | 0 |
| 0.00550937 | 0.01074443 | 0 | 0.22244059 | 0 | 0 | 4.75243649 |
| 0.00551269 | 0.01075027 | 81.3451915 | 41.1515096 | 0 | 68.6625504 | 0 |
| 0.00553082 | 0.01078501 | 34.4029071 | 148.145435 | 69.6038938 | 0 | 0 |
| 0.00553688 | 0.01079619 | 17.3622148 | 15.5708415 | 18.5610383 | 16.2377653 | 42.7719284 |
| 0.00554097 | 0.01080354 | 163.654951 | 437.763086 | 423.191674 | 420.326153 | 66.5341108 |
| 0.00556229 | 0.01084449 | 18.9698273 | 15.1259603 | 16.7049345 | 10.6705315 | 26.1384007 |
| 0.00556271 | 0.01084468 | 6.10892743 | 9.12006429 | 13.9207788 | 9.74265918 | 0 |
| 0.0055635 | 0.01084558 | 39.547267 | 74.2951579 | 0 | 77.0134011 | 0 |
| 0.00556508 | 0.01084803 | 229.24554 | 539.863318 | 523.421281 | 546.980722 | 30.8908372 |
| 0.00557036 | 0.01085771 | 7.39501741 | 11.7893514 | 13.9207788 | 6.49510612 | 0 |
| 0.00557734 | 0.01087067 | 5.78740493 | 12.2342326 | 13.9207788 | 8.35085072 | 0 |
| 0.0055831 | 0.01088127 | 312.519866 | 456.670536 | 626.435044 | 600.33338 | 23.7621824 |
| 0.00558396 | 0.01088232 | 39.8687895 | 78.5215291 | 0 | 71.9101035 | 0 |
| 0.00558592 | 0.01088551 | 5.14435994 | 44.2656779 | 12.0646749 | 6.95904227 | 0 |
| 0.00559083 | 0.01089444 | 9.96719738 | 126.791138 | 183.75428 | 0 | 0 |
| 0.00561696 | 0.01094472 | 9.00262989 | 10.4547078 | 6.49636342 | 10.2065953 | 0 |
| 0.00562275 | 0.01095538 | 31.5092046 | 91.6455241 | 0 | 75.1576565 | 0 |
| 0.00562898 | 0.01096689 | 0.64304499 | 0 | 0 | 0 | 11.8810912 |
| 0.00562975 | 0.0109671 | 2.25065747 | 19.7972127 | 21.3451941 | 17.6295737 | 0 |
| 0.00562969 | 0.0109671 | 0.3215225 | 0 | 0 | 0 | 0 |
| 0.0056308 | 0.01096852 | 48.5498969 | 57.834554 | 0 | 79.797018 | 0 |
| 0.00563954 | 0.01098492 | 44.3701045 | 98.763623 | 59.3953227 | 43.1460621 | 19.0097459 |
| 0.00564687 | 0.01099855 | 23.4711422 | 98.9860636 | 109.510126 | 126.654569 | 21.3859642 |
| 0.00564875 | 0.01100158 | 84.881939 | 162.381632 | 122.502853 | 115.520102 | 2.37621824 |
| 0.00564969 | 0.01100276 | 10.2887199 | 141.027336 | 0 | 110.416804 | 0 |
| 0.00565077 | 0.01100424 | 2.57217997 | 16.0157227 | 18.5610383 | 20.4131907 | 0 |
| 0.00565616 | 0.01101408 | 71.3779941 | 54.4979451 | 0 | 57.5280828 | 0 |
| 0.00567169 | 0.0110437 | 4.82283744 | 12.2342326 | 10.2085711 | 15.7738291 | 0 |
| 0.00567606 | 0.01105157 | 0 | 110.108093 | 59.3953227 | 76.0855288 | 0 |
| 0.00567715 | 0.01105305 | 10.2887199 | 15.5708415 | 5.5683115 | 9.27872303 | 0 |
| 0.00568009 | 0.01105775 | 2.25065747 | 21.5767375 | 14.8488307 | 23.6607437 | 0 |
| 0.00568022 | 0.01105775 | 22.5065747 | 95.8718953 | 0 | 93.7151026 | 0 |
| 0.0056815 | 0.01105961 | 45.334672 | 45.3778809 | 161.481034 | 0 | 0 |
| 0.0056821 | 0.01106012 | 6.75197242 | 9.56494547 | 12.0646749 | 8.35085072 | 0 |

| | | | | | | |
|---|---|---|---|---|---|---|
| 0.00568696 | 0.01106894 | 0 | 80.0786133 | 59.3953227 | 104.84957 | 0 |
| 0.00569285 | 0.01107976 | 34.0813846 | 84.0825439 | 0 | 75.1576565 | 0 |
| 0.00569536 | 0.01108402 | 24.7572322 | 27.8050741 | 34.3379209 | 33.4034029 | 19.0097459 |
| 0.00569694 | 0.01108645 | 37.9396545 | 71.4034302 | 0 | 80.2609542 | 0 |
| 0.00570015 | 0.01109206 | 8.3595849 | 8.23030192 | 8.35246725 | 9.74265918 | 0 |
| 0.00571525 | 0.01112078 | 13.8254673 | 18.6850098 | 12.0646749 | 12.5262761 | 2.37621824 |
| 0.00571662 | 0.01112282 | 2.25065747 | 23.5787028 | 27.8415575 | 12.0623399 | 0 |
| 0.00572111 | 0.01113091 | 7.07349491 | 17.5728068 | 5.5683115 | 14.8459568 | 0 |
| 0.00572875 | 0.01114513 | 30.5446371 | 33.1436483 | 27.8415575 | 39.4345729 | 106.929821 |
| 0.00573848 | 0.01116341 | 8.3595849 | 15.1259603 | 19.4890903 | 4.63936151 | 0 |
| 0.00574018 | 0.01116608 | 6.10892743 | 12.6791138 | 7.42441534 | 14.3820207 | 0 |
| 0.00574142 | 0.01116745 | 203.202218 | 138.80293 | 126.215061 | 122.94308 | 0 |
| 0.00574155 | 0.01116745 | 0 | 12.9015544 | 0 | 0 | 118.810912 |
| 0.0057423 | 0.01116826 | 51.1220769 | 94.982133 | 0 | 44.0739344 | 0 |
| 0.005743 | 0.01116897 | 10.2887199 | 8.8976237 | 9.28051917 | 6.95904227 | 0 |
| 0.00575366 | 0.01118906 | 9.96719738 | 13.3464355 | 12.0646749 | 5.10329766 | 0 |
| 0.00575928 | 0.01119934 | 83.2743265 | 20.4645345 | 14.8488307 | 20.4131907 | 49.9005831 |
| 0.00576867 | 0.01121696 | 46.9422844 | 75.1849202 | 0 | 61.239572 | 0 |
| 0.00577224 | 0.01122324 | 52.4081669 | 64.0628906 | 58.4672708 | 64.9510612 | 71.2865473 |
| 0.0057838 | 0.01124507 | 39.547267 | 78.5215291 | 0 | 68.6625504 | 0 |
| 0.00578701 | 0.01125065 | 17.6837373 | 65.3975342 | 212.523889 | 0 | 0 |
| 0.00578851 | 0.01125294 | 8.3595849 | 3.55904948 | 0 | 0 | 35.6432737 |
| 0.00578942 | 0.01125404 | 123.786161 | 179.064677 | 216.236097 | 190.677758 | 0 |
| 0.00579723 | 0.01126857 | 13.5039448 | 97.6514201 | 216.236097 | 202.740098 | 0 |
| 0.00580044 | 0.01127417 | 63.3399317 | 39.816866 | 0 | 83.9724434 | 0 |
| 0.0058016 | 0.01127577 | 27.0078897 | 21.1318563 | 19.4890903 | 20.8771268 | 33.2670554 |
| 0.00580324 | 0.0112783 | 73.9501741 | 70.7361084 | 72.3880495 | 0 | 0 |
| 0.00580692 | 0.01128481 | 24.4357097 | 33.8109701 | 58.4672708 | 74.2297842 | 14.2573095 |
| 0.00580812 | 0.01128648 | 0.96456749 | 0.88976237 | 0.92805192 | 0.46393615 | 0 |
| 0.00580948 | 0.01128848 | 43.405537 | 64.5077718 | 0 | 76.0855288 | 0 |
| 0.00581189 | 0.0112925 | 6.10892743 | 7.78542074 | 13.9207788 | 11.5984038 | 0 |
| 0.00581372 | 0.01129541 | 1.92913498 | 2.8917277 | 2.78415575 | 3.71148921 | 21.3859642 |
| 0.005818 | 0.01130307 | 10.6102424 | 38.7046631 | 39.9062324 | 0 | 259.007789 |
| 0.00582401 | 0.01131409 | 3.21522496 | 16.6830444 | 24.1293498 | 11.1344676 | 0 |
| 0.00582582 | 0.01131696 | 0.3215225 | 0 | 0 | 0.46393615 | 0 |
| 0.00582959 | 0.01132363 | 11.8963324 | 10.6771484 | 8.35246725 | 7.88691457 | 85.5438568 |
| 0.00583423 | 0.01133198 | 0 | 55.3877075 | 82.5966206 | 103.921698 | 0 |
| 0.00583511 | 0.01133304 | 1.60761248 | 27.1377523 | 21.3451941 | 19.4853184 | 0 |
| 0.00583655 | 0.01133518 | 36.9750871 | 57.1672323 | 152.200514 | 0 | 0 |
| 0.00583919 | 0.01133965 | 17.0406923 | 6.67321777 | 5.5683115 | 16.7017014 | 0 |
| 0.00584409 | 0.01134852 | 0 | 88.976237 | 81.6685687 | 66.8068058 | 0 |
| 0.0058448 | 0.01134923 | 10.6102424 | 5.11613363 | 8.35246725 | 34.7952113 | 0 |
| 0.00584534 | 0.01134964 | 126.036818 | 190.854028 | 188.394539 | 197.172864 | 0 |

| | | | | | | |
|---|---|---|---|---|---|---|
| 0.00584955 | 0.01135715 | 30.2231146 | 79.4112915 | 0 | 84.9003157 | 0 |
| 0.00585586 | 0.01136874 | 64.3044992 | 51.1613363 | 0 | 64.487125 | 0 |
| 0.00585669 | 0.0113697 | 0 | 121.452563 | 80.7405168 | 46.3936151 | 0 |
| 0.00586089 | 0.01137719 | 40.5118345 | 59.3916382 | 138.279736 | 0 | 0 |
| 0.00586462 | 0.01138378 | 0 | 78.7439697 | 180.970124 | 25.5164883 | 0 |
| 0.0058708 | 0.01139513 | 75.8793091 | 79.6337321 | 61.2514265 | 0 | 0 |
| 0.00587252 | 0.01139781 | 4.82283744 | 18.6850098 | 17.6329864 | 7.42297842 | 0 |
| 0.00587588 | 0.01140366 | 18.6483048 | 29.8070394 | 2.78415575 | 10.2065953 | 0 |
| 0.00587684 | 0.01140488 | 34.0813846 | 59.6140788 | 0 | 99.7462725 | 0 |
| 0.00588032 | 0.01141098 | 0 | 0 | 0 | 0 | 0 |
| 0.00588701 | 0.0114233 | 52.4081669 | 8.67518311 | 5.5683115 | 7.42297842 | 0 |
| 0.00588864 | 0.01142579 | 3.53674746 | 17.1279256 | 11.136623 | 18.5574461 | 0 |
| 0.00589093 | 0.01142958 | 0 | 109.663212 | 51.9709073 | 81.1888265 | 0 |
| 0.00589367 | 0.01143424 | 52.7296894 | 83.1927816 | 102.085711 | 77.0134011 | 0 |
| 0.00589423 | 0.01143466 | 18.0052598 | 11.7893514 | 4.64025958 | 9.74265918 | 0 |
| 0.00589503 | 0.01143555 | 0.96456749 | 26.9153117 | 25.0574018 | 29.2279775 | 0 |
| 0.00589826 | 0.01144116 | 8.3595849 | 9.56494547 | 7.42441534 | 9.27872303 | 0 |
| 0.00589952 | 0.01144294 | 293.871561 | 435.983561 | 450.10518 | 506.618277 | 2.37621824 |
| 0.00589988 | 0.01144299 | 97.0997938 | 63.84045 | 0 | 32.4755306 | 0 |
| 0.00590474 | 0.01145174 | 4.82283744 | 18.2401286 | 8.35246725 | 14.3820207 | 0 |
| 0.00590645 | 0.0114544 | 9.64567488 | 6.89565837 | 8.35246725 | 10.6705315 | 0 |
| 0.00590921 | 0.01145909 | 54.9803468 | 8.00786133 | 3.71220767 | 11.5984038 | 0 |
| 0.00591238 | 0.01146459 | 13.5039448 | 5.11613363 | 36.1940248 | 6.49510612 | 0 |
| 0.00591354 | 0.01146617 | 10.2887199 | 24.6909058 | 25.0574018 | 24.588616 | 4.75243649 |
| 0.005915 | 0.01146834 | 286.476544 | 86.7518311 | 36.1940248 | 108.097123 | 26.1384007 |
| 0.00592575 | 0.01148852 | 0 | 114.779346 | 65.8916861 | 60.7756358 | 0 |
| 0.0059293 | 0.01149474 | 5.46588243 | 10.0098267 | 9.28051917 | 18.5574461 | 0 |
| 0.00593424 | 0.01150367 | 7.71653991 | 9.56494547 | 8.35246725 | 8.81478687 | 0 |
| 0.00593577 | 0.01150596 | 55.6233918 | 66.064856 | 0 | 56.6002105 | 0 |
| 0.00593752 | 0.0115087 | 7.07349491 | 11.3444702 | 5.5683115 | 33.4034029 | 0 |
| 0.00595031 | 0.01153282 | 54.9803468 | 69.4014648 | 40.8342843 | 50.1051043 | 19.0097459 |
| 0.00595595 | 0.01154309 | 4.50131495 | 10.2322673 | 15.7768826 | 12.9902122 | 0 |
| 0.00595979 | 0.01154987 | 0 | 60.5038411 | 103.013763 | 74.2297842 | 0 |
| 0.00596737 | 0.01156388 | 2.57217997 | 36.9251383 | 7.42441534 | 25.9804245 | 0 |
| 0.00597855 | 0.01158488 | 3.53674746 | 10.4547078 | 27.8415575 | 13.9180845 | 0 |
| 0.00598221 | 0.01159131 | 32.4737721 | 82.970341 | 119.718697 | 0 | 0 |
| 0.00598423 | 0.01159456 | 18.3267823 | 10.899589 | 6.49636342 | 6.95904227 | 0 |
| 0.00599715 | 0.01161892 | 5.46588243 | 11.5669108 | 9.28051917 | 12.5262761 | 0 |
| 0.00600024 | 0.01162423 | 4.82283744 | 9.56494547 | 13.9207788 | 13.4541484 | 0 |
| 0.00600717 | 0.01163699 | 55.3018693 | 50.4940145 | 0 | 72.8379758 | 0 |
| 0.00600891 | 0.0116397 | 3.85826995 | 5.33857422 | 0 | 0 | 16.6335277 |
| 0.00601276 | 0.0116458 | 9.32415239 | 8.45274251 | 6.49636342 | 12.0623399 | 0 |
| 0.00601243 | 0.0116458 | 45.6561944 | 28.2499552 | 28.7696094 | 25.9804245 | 121.18713 |

| | | | | | | |
|---|---|---|---|---|---|---|
| 0.00601701 | 0.01165337 | 7.71653991 | 30.4743612 | 2.78415575 | 25.5164883 | 0 |
| 0.00602317 | 0.01166437 | 8.3595849 | 18.2401286 | 3.71220767 | 23.1968076 | 0 |
| 0.00602338 | 0.01166437 | 7.39501741 | 22.6889404 | 18.5610383 | 17.1656376 | 2.37621824 |
| 0.00602761 | 0.01167188 | 48.5498969 | 58.7243164 | 0 | 70.9822312 | 0 |
| 0.00603089 | 0.01167757 | 10.2887199 | 11.7893514 | 5.5683115 | 9.74265918 | 0 |
| 0.00603611 | 0.01168699 | 62.6968867 | 80.0786133 | 70.5319457 | 0 | 0 |
| 0.00604613 | 0.01170572 | 2.89370247 | 29.3621582 | 13.9207788 | 12.9902122 | 0 |
| 0.00605155 | 0.01171554 | 37.618132 | 60.05896 | 138.279736 | 0 | 0 |
| 0.00605338 | 0.0117184 | 8.3595849 | 13.5688761 | 5.5683115 | 11.5984038 | 0 |
| 0.00605794 | 0.01172657 | 5.14435994 | 13.5688761 | 31.5537652 | 25.5164883 | 68.9103291 |
| 0.00606499 | 0.01173953 | 272.008032 | 622.388778 | 267.278952 | 454.193492 | 0 |
| 0.00606777 | 0.01174425 | 5.14435994 | 8.67518311 | 9.28051917 | 38.9706367 | 0 |
| 0.00607089 | 0.01174961 | 11.5748099 | 9.78738607 | 9.28051917 | 6.03116997 | 0 |
| 0.00607188 | 0.01175085 | 135.360971 | 56.9447917 | 73.3161014 | 65.4149973 | 11.8810912 |
| 0.0060757 | 0.01175756 | 0 | 82.7479004 | 66.819738 | 82.5806349 | 0 |
| 0.00607781 | 0.01176096 | 7.07349491 | 10.0098267 | 10.2085711 | 7.88691457 | 0 |
| 0.00608516 | 0.01177452 | 126.358341 | 39.3719849 | 18.5610383 | 37.5788283 | 0 |
| 0.00608822 | 0.01177947 | 10.2887199 | 10.6771484 | 19.4890903 | 4.63936151 | 0 |
| 0.00608842 | 0.01177947 | 50.8005544 | 35.5904948 | 0 | 105.313506 | 0 |
| 0.00609306 | 0.01178776 | 0 | 88.7537964 | 77.0283091 | 66.3428696 | 0 |
| 0.00609532 | 0.01179145 | 8.3595849 | 14.6810791 | 4.64025958 | 16.7017014 | 0 |
| 0.00610031 | 0.01180043 | 6.10892743 | 8.67518311 | 11.136623 | 10.6705315 | 0 |
| 0.00610582 | 0.01181041 | 2.89370247 | 30.4743612 | 8.35246725 | 22.2689353 | 0 |
| 0.00613179 | 0.01185997 | 24.1141872 | 91.8679647 | 0 | 82.5806349 | 0 |
| 0.00613296 | 0.01186155 | 7.07349491 | 9.34250488 | 9.28051917 | 8.81478687 | 0 |
| 0.00613704 | 0.01186875 | 6.43044992 | 26.9153117 | 43.6184401 | 34.3312752 | 4.75243649 |
| 0.00613798 | 0.0118699 | 1.28608998 | 25.5806681 | 17.6329864 | 30.619786 | 0 |
| 0.00614227 | 0.01187668 | 123.786161 | 33.8109701 | 0 | 43.1460621 | 0 |
| 0.00614205 | 0.01187668 | 114.462009 | 46.4900838 | 61.2514265 | 0 | 0 |
| 0.00614255 | 0.01187668 | 0.3215225 | 0 | 0 | 0 | 28.5146189 |
| 0.00614456 | 0.01187988 | 66.8766792 | 117.671073 | 217.164149 | 57.0641466 | 0 |
| 0.00614667 | 0.01188327 | 17.3622148 | 7.34053955 | 12.9927268 | 5.56723382 | 0 |
| 0.00614775 | 0.01188469 | 0 | 88.0864746 | 77.0283091 | 65.8789335 | 0 |
| 0.00615401 | 0.01189611 | 154.330798 | 258.253528 | 227.37272 | 249.133713 | 106.929821 |
| 0.00617237 | 0.01193091 | 21.8635297 | 26.6928711 | 19.4890903 | 5.56723382 | 0 |
| 0.00618421 | 0.01195311 | 5.14435994 | 21.1318563 | 5.5683115 | 21.8049991 | 0 |
| 0.00619197 | 0.01196742 | 36.9750871 | 96.9840983 | 0 | 52.4247851 | 0 |
| 0.00619511 | 0.0119728 | 2.25065747 | 22.0216187 | 35.2659728 | 9.74265918 | 0 |
| 0.00621945 | 0.01201914 | 0 | 41.3739502 | 208.811681 | 35.7230837 | 0 |
| 0.00622373 | 0.01202605 | 128.930521 | 224.887439 | 212.523889 | 230.576267 | 83.1676385 |
| 0.00622343 | 0.01202605 | 80.0591015 | 58.946757 | 0 | 40.3624452 | 0 |
| 0.00623451 | 0.01204618 | 32.4737721 | 20.2420939 | 4.64025958 | 5.10329766 | 0 |
| 0.00623598 | 0.01204832 | 13.8254673 | 15.1259603 | 9.28051917 | 20.8771268 | 2.37621824 |

| | | | | | | |
|---|---|---|---|---|---|---|
| 0.00623656 | 0.01204876 | 1.28608998 | 27.1377523 | 25.9854537 | 19.0213822 | 0 |
| 0.00623909 | 0.01205295 | 0 | 0 | 0 | 0 | 14.2573095 |
| 0.00624995 | 0.01207324 | 90.9908664 | 87.1967122 | 103.013763 | 81.6527626 | 28.5146189 |
| 0.00625252 | 0.01207751 | 0 | 124.344291 | 35.2659728 | 88.6118049 | 0 |
| 0.00625639 | 0.0120843 | 172.336058 | 537.416471 | 460.313751 | 600.797316 | 0 |
| 0.00625734 | 0.01208544 | 5.14435994 | 8.23030192 | 17.6329864 | 12.0623399 | 0 |
| 0.00627076 | 0.01211066 | 13.8254673 | 27.3601929 | 3.71220767 | 9.27872303 | 0 |
| 0.00627118 | 0.01211077 | 5.14435994 | 8.00786133 | 19.4890903 | 12.0623399 | 0 |
| 0.00627283 | 0.01211326 | 386.791563 | 1274.36215 | 962.389838 | 1022.97921 | 0 |
| 0.00627992 | 0.01212621 | 27.0078897 | 33.5885295 | 58.4672708 | 43.1460621 | 14.2573095 |
| 0.00628026 | 0.01212621 | 10.9317649 | 4.67125244 | 15.7768826 | 10.6705315 | 0 |
| 0.00629105 | 0.01214635 | 0.64304499 | 0.88976237 | 0.92805192 | 0.46393615 | 2.37621824 |
| 0.0062943 | 0.01215123 | 225.065747 | 0 | 24.1293498 | 65.4149973 | 0 |
| 0.00629403 | 0.01215123 | 58.1955718 | 44.4881185 | 0 | 72.8379758 | 0 |
| 0.00629622 | 0.01215425 | 4.82283744 | 10.4547078 | 13.9207788 | 11.1344676 | 0 |
| 0.00629712 | 0.01215528 | 8.0380624 | 29.5845988 | 3.71220767 | 15.7738291 | 0 |
| 0.00630101 | 0.01216209 | 141.469898 | 426.863497 | 394.422065 | 356.7669 | 0 |
| 0.00630662 | 0.01217222 | 2.57217997 | 13.3464355 | 27.8415575 | 15.309893 | 0 |
| 0.00631766 | 0.01219283 | 3.21522496 | 10.2322673 | 19.4890903 | 19.9492545 | 0 |
| 0.00633376 | 0.01222321 | 0.96456749 | 1.33464355 | 1.85610383 | 1.85574461 | 26.1384007 |
| 0.0063462 | 0.0122465 | 4.50131495 | 22.2440592 | 8.35246725 | 13.4541484 | 0 |
| 0.00634735 | 0.01224803 | 46.9422844 | 76.2971232 | 89.092984 | 0 | 0 |
| 0.00634961 | 0.01225169 | 7.39501741 | 9.78738607 | 6.49636342 | 14.8459568 | 0 |
| 0.00635492 | 0.01226122 | 0 | 64.7302124 | 73.3161014 | 89.0757411 | 0 |
| 0.00635615 | 0.01226289 | 0 | 92.0904053 | 128.999216 | 30.619786 | 0 |
| 0.0063605 | 0.01227059 | 15.4330798 | 24.023584 | 12.9927268 | 25.0525522 | 4.75243649 |
| 0.00637632 | 0.0123004 | 28.6155022 | 11.5669108 | 8.35246725 | 4.63936151 | 0 |
| 0.00637694 | 0.01230089 | 115.105054 | 223.552795 | 126.215061 | 170.264568 | 0 |
| 0.0064017 | 0.01234794 | 4.17979245 | 27.3601929 | 5.5683115 | 23.1968076 | 0 |
| 0.00640598 | 0.0123555 | 3.21522496 | 13.7913167 | 20.4171422 | 12.9902122 | 0 |
| 0.00641246 | 0.01236727 | 112.211351 | 201.976058 | 202.315318 | 199.028609 | 33.2670554 |
| 0.00641405 | 0.01236964 | 1.60761248 | 17.1279256 | 17.6329864 | 33.867339 | 0 |
| 0.00641708 | 0.01237478 | 8.0380624 | 40.0393066 | 5.5683115 | 48.7132959 | 0 |
| 0.0064307 | 0.01240032 | 275.866302 | 386.379309 | 358.22804 | 406.408069 | 0 |
| 0.00643558 | 0.01240903 | 164.941041 | 132.797034 | 93.7332436 | 125.726697 | 35.6432737 |
| 0.00643759 | 0.01241219 | 0 | 12.011792 | 13.9207788 | 10.2065953 | 102.177384 |
| 0.00644193 | 0.01241985 | 2.89370247 | 12.9015544 | 16.7049345 | 19.9492545 | 0 |
| 0.00644465 | 0.01242437 | 69.7703817 | 38.4822225 | 0 | 67.2707419 | 0 |
| 0.00644778 | 0.01242969 | 209.632667 | 0 | 92.8051917 | 16.2377653 | 0 |
| 0.00645392 | 0.01244083 | 1.60761248 | 25.1357869 | 15.7768826 | 25.0525522 | 0 |
| 0.00645722 | 0.01244647 | 15.1115573 | 163.048954 | 0 | 110.88074 | 2.37621824 |
| 0.00645972 | 0.01245058 | 5.14435994 | 16.4606038 | 12.9927268 | 7.88691457 | 0 |
| 0.00646221 | 0.01245466 | 83.9173715 | 114.112024 | 111.36623 | 119.231591 | 40.3957101 |

| | | | | | | |
|---|---|---|---|---|---|---|
| 0.00647714 | 0.01248271 | 0 | 56.0550293 | 88.1649321 | 82.1166988 | 0 |
| 0.00647759 | 0.01248288 | 27.9724572 | 16.0157227 | 3.71220767 | 7.88691457 | 0 |
| 0.00648412 | 0.01249474 | 0 | 58.946757 | 132.711424 | 47.7854236 | 0 |
| 0.00648755 | 0.01250064 | 0 | 74.2951579 | 75.1722053 | 73.7658481 | 0 |
| 0.00648955 | 0.01250378 | 29.2585471 | 8.23030192 | 9.28051917 | 5.56723382 | 0 |
| 0.00650041 | 0.01252399 | 51.7651219 | 85.6396281 | 0 | 39.898509 | 0 |
| 0.00650554 | 0.01253315 | 84.238894 | 41.5963908 | 86.3088283 | 0 | 0 |
| 0.00651227 | 0.01254539 | 10.6102424 | 15.5708415 | 4.64025958 | 10.2065953 | 0 |
| 0.00651482 | 0.01254959 | 71.0564716 | 111.665177 | 81.6685687 | 106.705315 | 28.5146189 |
| 0.00652335 | 0.0125653 | 22.8280972 | 26.6928711 | 29.6976613 | 35.7230837 | 11.8810912 |
| 0.00653337 | 0.01258387 | 15.4330798 | 88.0864746 | 44.546492 | 58.4559551 | 14.2573095 |
| 0.00653566 | 0.01258758 | 6.43044992 | 13.3464355 | 10.2085711 | 7.42297842 | 0 |
| 0.00654557 | 0.01260522 | 0 | 24.6909058 | 100.229607 | 131.757867 | 0 |
| 0.00654535 | 0.01260522 | 0 | 0 | 0 | 0 | 73.6627655 |
| 0.00655234 | 0.01261752 | 23.1496197 | 115.891549 | 96.5173994 | 0 | 0 |
| 0.00655429 | 0.01262056 | 1.60761248 | 10.899589 | 24.1293498 | 38.9706367 | 0 |
| 0.00656052 | 0.01263184 | 49.1929419 | 85.8620687 | 0 | 41.2903175 | 0 |
| 0.00656238 | 0.01263469 | 301.588101 | 605.705733 | 490.939464 | 550.228275 | 87.920075 |
| 0.00656347 | 0.01263606 | 66.2336342 | 50.7164551 | 0 | 52.8887212 | 0 |
| 0.00657681 | 0.01266104 | 16.0761248 | 57.3896728 | 0 | 149.851377 | 0 |
| 0.00658176 | 0.01266983 | 0 | 0.22244059 | 0 | 0 | 19.0097459 |
| 0.00658238 | 0.01267031 | 190.66284 | 79.1888509 | 62.1794784 | 92.7872303 | 38.0194919 |
| 0.00658309 | 0.01267094 | 14.4685123 | 27.5826335 | 19.4890903 | 29.2279775 | 4.75243649 |
| 0.00658823 | 0.01268012 | 45.9777169 | 57.3896728 | 0 | 66.8068058 | 0 |
| 0.00658974 | 0.01268229 | 28.6155022 | 68.2892619 | 0 | 86.2921241 | 0 |
| 0.00659496 | 0.01269088 | 8.6811074 | 21.1318563 | 9.28051917 | 5.56723382 | 0 |
| 0.0065949 | 0.01269088 | 3.53674746 | 13.123995 | 13.9207788 | 15.309893 | 0 |
| 0.00659771 | 0.01269544 | 0 | 72.9605143 | 93.7332436 | 57.5280828 | 0 |
| 0.00660045 | 0.01269999 | 9.32415239 | 12.6791138 | 3.71220767 | 28.3001052 | 0 |
| 0.00660343 | 0.01270501 | 10.9317649 | 88.7537964 | 0 | 133.149675 | 0 |
| 0.00660966 | 0.01271626 | 96.7782713 | 88.7537964 | 77.0283091 | 96.0347833 | 28.5146189 |
| 0.00661313 | 0.0127222 | 0 | 94.982133 | 115.078438 | 32.0115944 | 0 |
| 0.00662682 | 0.01274782 | 6.75197242 | 12.2342326 | 8.35246725 | 8.35085072 | 0 |
| 0.00662911 | 0.0127515 | 7.07349491 | 16.2381632 | 5.5683115 | 12.5262761 | 0 |
| 0.0066301 | 0.01275267 | 20.5774398 | 30.02948 | 37.1220767 | 45.4657428 | 11.8810912 |
| 0.00663269 | 0.01275692 | 26.0433222 | 77.6317668 | 0 | 81.6527626 | 0 |
| 0.00663557 | 0.01276172 | 1.28608998 | 24.2460246 | 21.3451941 | 23.6607437 | 0 |
| 0.00664339 | 0.01277604 | 17.6837373 | 2.66928711 | 12.9927268 | 20.8771268 | 0 |
| 0.00664987 | 0.01278777 | 72.9856066 | 30.6968018 | 0 | 75.1576565 | 0 |
| 0.00665118 | 0.01278957 | 4.82283744 | 11.3444702 | 12.0646749 | 10.6705315 | 0 |
| 0.00665314 | 0.01279261 | 203.52374 | 377.481685 | 296.976613 | 308.981477 | 140.196876 |
| 0.00666676 | 0.01281806 | 18.0052598 | 20.2420939 | 6.49636342 | 19.0213822 | 2.37621824 |
| 0.00666838 | 0.01282044 | 0 | 101.43291 | 61.2514265 | 61.7035081 | 0 |

| | | | | | | |
|---|---|---|---|---|---|---|
| 0.00666929 | 0.01282145 | 8.0380624 | 43.8207967 | 33.409869 | 43.1460621 | 7.12865473 |
| 0.00667349 | 0.0128288 | 4.50131495 | 13.3464355 | 14.8488307 | 9.27872303 | 0 |
| 0.00668193 | 0.01284428 | 10.6102424 | 12.4566732 | 4.64025958 | 11.1344676 | 0 |
| 0.00668527 | 0.01284997 | 5.46588243 | 10.6771484 | 10.2085711 | 10.2065953 | 0 |
| 0.00668771 | 0.01285393 | 6.75197242 | 10.899589 | 9.28051917 | 7.88691457 | 0 |
| 0.00669212 | 0.01286167 | 36.6535646 | 12.2342326 | 7.42441534 | 19.0213822 | 0 |
| 0.00670063 | 0.01287728 | 5.14435994 | 12.9015544 | 11.136623 | 9.27872303 | 0 |
| 0.00670238 | 0.01287917 | 94.5276139 | 451.331962 | 654.276601 | 565.074232 | 71.2865473 |
| 0.0067021 | 0.01287917 | 2.89370247 | 19.5747721 | 11.136623 | 19.4853184 | 0 |
| 0.00670807 | 0.01288938 | 249.179934 | 258.031087 | 245.933758 | 281.145308 | 0 |
| 0.00673145 | 0.01293356 | 0 | 74.0727173 | 85.3807764 | 60.7756358 | 0 |
| 0.00673202 | 0.01293393 | 118.320279 | 72.9605143 | 30.6257133 | 0 | 0 |
| 0.00673769 | 0.01294407 | 11.5748099 | 14.4586385 | 12.9927268 | 13.9180845 | 0 |
| 0.00674246 | 0.0129525 | 4.50131495 | 10.2322673 | 19.4890903 | 10.2065953 | 0 |
| 0.00674552 | 0.01295763 | 24.7572322 | 8.00786133 | 12.0646749 | 4.63936151 | 0 |
| 0.00676338 | 0.0129912 | 12.2178549 | 21.7991781 | 2.78415575 | 15.309893 | 0 |
| 0.006768 | 0.01299934 | 81.666714 | 58.946757 | 0 | 34.3312752 | 0 |
| 0.00677097 | 0.01300429 | 104.173289 | 103.212435 | 90.0210359 | 101.138081 | 30.8908372 |
| 0.00677909 | 0.01301916 | 4.50131495 | 10.4547078 | 14.8488307 | 11.1344676 | 0 |
| 0.00680015 | 0.01305885 | 5.14435994 | 11.7893514 | 16.7049345 | 7.88691457 | 0 |
| 0.00680225 | 0.01306213 | 3.85826995 | 12.4566732 | 12.0646749 | 15.309893 | 0 |
| 0.0068048 | 0.01306629 | 5.14435994 | 9.12006429 | 14.8488307 | 10.2065953 | 0 |
| 0.00680658 | 0.01306753 | 27.9724572 | 94.7596924 | 0 | 59.8477635 | 0 |
| 0.00680661 | 0.01306753 | 24.7572322 | 13.123995 | 5.5683115 | 6.03116997 | 0 |
| 0.00680602 | 0.01306753 | 2.89370247 | 15.1259603 | 13.9207788 | 18.5574461 | 0 |
| 0.00681888 | 0.01309033 | 55.6233918 | 0 | 82.5966206 | 102.529889 | 0 |
| 0.0068277 | 0.01310651 | 25.4002772 | 8.67518311 | 10.2085711 | 16.7017014 | 114.058476 |
| 0.00683506 | 0.0131199 | 40.5118345 | 73.8502767 | 0 | 55.208402 | 0 |
| 0.00684134 | 0.0131312 | 8.3595849 | 10.2322673 | 9.28051917 | 6.49510612 | 0 |
| 0.00684646 | 0.01314028 | 18.0052598 | 56.7223511 | 25.0574018 | 41.7542536 | 11.8810912 |
| 0.00685872 | 0.01316306 | 281.975229 | 1419.39342 | 1006.00828 | 1159.37644 | 0 |
| 0.00685987 | 0.01316451 | 3.21522496 | 16.2381632 | 18.5610383 | 11.1344676 | 0 |
| 0.00687732 | 0.01319725 | 24.7572322 | 101.21047 | 98.3735032 | 0 | 0 |
| 0.00689106 | 0.01322286 | 6.43044992 | 14.9035197 | 10.2085711 | 6.95904227 | 0 |
| 0.00689728 | 0.01323404 | 0 | 154.373771 | 141.991943 | 28.7640414 | 0 |
| 0.00689812 | 0.01323491 | 45.334672 | 21.1318563 | 0 | 141.500526 | 0 |
| 0.00690314 | 0.01324378 | 0 | 0 | 0 | 0 | 57.0292378 |
| 0.00690494 | 0.01324647 | 4.17979245 | 12.6791138 | 25.0574018 | 8.35085072 | 0 |
| 0.00690559 | 0.01324696 | 0 | 52.4959798 | 83.5246725 | 83.0445711 | 0 |
| 0.00691079 | 0.01325618 | 8.3595849 | 7.78542074 | 6.49636342 | 16.7017014 | 0 |
| 0.00691558 | 0.01326462 | 30.8661596 | 13.123995 | 3.71220767 | 8.35085072 | 0 |
| 0.00691843 | 0.01326932 | 28.9370247 | 68.5117025 | 0 | 79.3330819 | 0 |
| 0.00692047 | 0.01327248 | 0 | 134.798999 | 38.0501286 | 122.94308 | 2.37621824 |

| | | | | | | |
|---|---|---|---|---|---|---|
| 0.00693263 | 0.01329505 | 9.00262989 | 11.1220296 | 17.6329864 | 4.63936151 | 0 |
| 0.00693359 | 0.01329614 | 48.2283744 | 73.8502767 | 0 | 45.4657428 | 0 |
| 0.00693427 | 0.01329669 | 0 | 101.43291 | 32.4818171 | 99.7462725 | 0 |
| 0.00693698 | 0.01330112 | 20.8989622 | 79.6337321 | 52.8989593 | 77.9412734 | 16.6335277 |
| 0.00694385 | 0.01331353 | 34.4029071 | 67.1770589 | 0 | 69.1264865 | 0 |
| 0.00694884 | 0.01332233 | 10.2887199 | 7.78542074 | 8.35246725 | 8.81478687 | 54.6530196 |
| 0.00695406 | 0.01333159 | 109.317649 | 0 | 27.8415575 | 125.726697 | 0 |
| 0.00695581 | 0.01333418 | 3.21522496 | 29.1397176 | 5.5683115 | 27.8361691 | 0 |
| 0.00696375 | 0.01334864 | 46.2992394 | 56.2774699 | 0 | 62.1674443 | 0 |
| 0.00697513 | 0.0133697 | 5.14435994 | 22.911381 | 58.4672708 | 38.5067006 | 4.75243649 |
| 0.00697693 | 0.01337237 | 9.96719738 | 6.89565837 | 22.273246 | 6.03116997 | 0 |
| 0.0069777 | 0.0133731 | 13.5039448 | 3.78149007 | 7.42441534 | 36.650956 | 0 |
| 0.00697894 | 0.01337472 | 41.476402 | 130.127747 | 99.3015551 | 98.3544641 | 26.1384007 |
| 0.00698267 | 0.01338109 | 122.821594 | 25.3582275 | 20.4171422 | 30.619786 | 0 |
| 0.00698564 | 0.01338603 | 8.0380624 | 21.3542969 | 6.49636342 | 7.88691457 | 0 |
| 0.00699129 | 0.01339608 | 54.9803468 | 81.1908162 | 0 | 35.2591475 | 0 |
| 0.00699572 | 0.01340382 | 0 | 39.5944255 | 128.071165 | 64.0231889 | 0 |
| 0.0070043 | 0.01341949 | 19.6128723 | 5.7834554 | 12.0646749 | 6.49510612 | 0 |
| 0.00700682 | 0.01342356 | 78.4514891 | 203.533142 | 171.689605 | 201.34829 | 40.3957101 |
| 0.00701046 | 0.01342977 | 7.39501741 | 7.56298014 | 7.42441534 | 17.1656376 | 0 |
| 0.00701521 | 0.01343809 | 0 | 1.77952474 | 0 | 0 | 0 |
| 0.00702112 | 0.01344865 | 63.6614542 | 29.8070394 | 0 | 81.1888265 | 0 |
| 0.00702616 | 0.01345755 | 7.39501741 | 16.2381632 | 29.6976613 | 14.3820207 | 2.37621824 |
| 0.00702751 | 0.01345937 | 5.78740493 | 26.0255493 | 17.6329864 | 21.341063 | 0 |
| 0.00703445 | 0.01347188 | 6.75197242 | 8.67518311 | 6.49636342 | 11.5984038 | 0 |
| 0.00703626 | 0.01347458 | 9.00262989 | 10.6771484 | 4.64025958 | 17.6295737 | 0 |
| 0.00703676 | 0.01347478 | 49.1929419 | 18.017688 | 15.7768826 | 13.4541484 | 4.75243649 |
| 0.00704474 | 0.01348928 | 48.5498969 | 36.035376 | 0 | 88.1478687 | 0 |
| 0.00704695 | 0.01349275 | 4.50131495 | 7.78542074 | 14.8488307 | 17.1656376 | 0 |
| 0.00706373 | 0.0135241 | 10.2887199 | 8.23030192 | 5.5683115 | 8.81478687 | 14.2573095 |
| 0.0070667 | 0.01352903 | 22.8280972 | 63.1731283 | 0 | 100.210209 | 0 |
| 0.00707267 | 0.01353969 | 8.6811074 | 5.7834554 | 32.4818171 | 7.42297842 | 0 |
| 0.00707383 | 0.01354113 | 37.618132 | 76.964445 | 84.4527244 | 80.2609542 | 0 |
| 0.00707641 | 0.01354531 | 14.4685123 | 16.4606038 | 10.2085711 | 11.1344676 | 16.6335277 |
| 0.00708088 | 0.01355309 | 8.6811074 | 4.44881185 | 1.85610383 | 4.63936151 | 7.12865473 |
| 0.00708242 | 0.01355526 | 2.57217997 | 27.3601929 | 8.35246725 | 23.1968076 | 0 |
| 0.00708749 | 0.01356419 | 3.53674746 | 28.4723958 | 45.4745439 | 3.24755306 | 0 |
| 0.00708793 | 0.01356426 | 111.889829 | 109.440771 | 101.157659 | 106.241379 | 168.711495 |
| 0.00709006 | 0.01356757 | 50.4790319 | 18.9074504 | 2.78415575 | 5.56723382 | 0 |
| 0.00709677 | 0.01357964 | 4.17979245 | 9.34250488 | 23.2012979 | 11.1344676 | 0 |
| 0.00709996 | 0.01358375 | 207.703532 | 619.941931 | 516.924918 | 569.713594 | 0 |
| 0.00709968 | 0.01358375 | 10.6102424 | 8.67518311 | 4.64025958 | 20.4131907 | 0 |
| 0.00710013 | 0.01358375 | 23.4711422 | 72.9605143 | 0 | 85.3642518 | 0 |

| | | | | | | |
|---|---|---|---|---|---|---|
| 0.00710115 | 0.01358494 | 2.57217997 | 22.0216187 | 10.2085711 | 21.8049991 | 0 |
| 0.00710534 | 0.01359218 | 53.0512119 | 42.2637126 | 0 | 69.5904227 | 0 |
| 0.00710723 | 0.01359502 | 26.6863672 | 74.5175985 | 0 | 75.1576565 | 0 |
| 0.00711446 | 0.01360808 | 33.1168171 | 13.123995 | 15.7768826 | 17.1656376 | 2.37621824 |
| 0.0071226 | 0.01362287 | 13.1824223 | 150.592281 | 0 | 57.9920189 | 0 |
| 0.00712326 | 0.01362336 | 19.2913498 | 29.1397176 | 38.0501286 | 31.0837221 | 125.939567 |
| 0.00712551 | 0.01362689 | 14.7900348 | 9.34250488 | 6.49636342 | 10.2065953 | 57.0292378 |
| 0.0071321 | 0.01363871 | 1.92913498 | 12.011792 | 31.5537652 | 19.0213822 | 0 |
| 0.00713353 | 0.01364067 | 10.6102424 | 19.3523315 | 4.64025958 | 8.81478687 | 0 |
| 0.00713856 | 0.01364874 | 46.2992394 | 42.2637126 | 22.273246 | 22.7328714 | 92.6725115 |
| 0.00713835 | 0.01364874 | 15.4330798 | 6.005896 | 13.9207788 | 6.03116997 | 0 |
| 0.00714546 | 0.01366115 | 75.8793091 | 39.816866 | 26.9135056 | 38.5067006 | 199.602332 |
| 0.00714702 | 0.01366336 | 0.3215225 | 0.44488118 | 0.92805192 | 0 | 0 |
| 0.00715216 | 0.01367241 | 6.43044992 | 6.67321777 | 10.2085711 | 15.7738291 | 0 |
| 0.00715542 | 0.01367787 | 11.8963324 | 6.45077718 | 8.35246725 | 8.35085072 | 0 |
| 0.0071573 | 0.01368068 | 40.833357 | 57.3896728 | 0 | 65.8789335 | 0 |
| 0.00716262 | 0.01368929 | 38.904222 | 83.1927816 | 0 | 46.8575513 | 0 |
| 0.00716251 | 0.01368929 | 3.85826995 | 2.44684652 | 3.71220767 | 1.39180845 | 0 |
| 0.00716746 | 0.01369777 | 0 | 94.5372518 | 92.8051917 | 36.650956 | 0 |
| 0.00716818 | 0.01369836 | 48.5498969 | 46.2676432 | 0 | 68.6625504 | 0 |
| 0.00717195 | 0.0137048 | 189.698273 | 270.48776 | 243.149602 | 261.659989 | 2.37621824 |
| 0.00717651 | 0.01371273 | 74.2716966 | 104.769519 | 90.0210359 | 28.7640414 | 21.3859642 |
| 0.00717692 | 0.01371273 | 27.3294122 | 6.67321777 | 8.35246725 | 6.95904227 | 0 |
| 0.00718099 | 0.01371972 | 7.71653991 | 7.34053955 | 6.49636342 | 7.88691457 | 38.0194919 |
| 0.00718404 | 0.01372478 | 5.46588243 | 22.2440592 | 6.49636342 | 12.0623399 | 0 |
| 0.00718548 | 0.01372674 | 8.0380624 | 87.864034 | 0 | 142.428398 | 0 |
| 0.00719442 | 0.01374305 | 22.5065747 | 11.7893514 | 0 | 0 | 54.6530196 |
| 0.00719538 | 0.01374409 | 31.8307271 | 84.0825439 | 91.8771398 | 0 | 0 |
| 0.00721081 | 0.01377279 | 32.7952946 | 74.0727173 | 88.1649321 | 77.0134011 | 26.1384007 |
| 0.00721868 | 0.01378705 | 25.7217997 | 13.7913167 | 8.35246725 | 13.9180845 | 144.949313 |
| 0.00722789 | 0.01380385 | 21.8635297 | 0 | 16.7049345 | 20.4131907 | 78.415202 |
| 0.007229 | 0.01380518 | 53.3727344 | 53.8306234 | 0 | 53.3526574 | 0 |
| 0.00723269 | 0.01381144 | 22.5065747 | 100.765588 | 0 | 61.239572 | 0 |
| 0.00723423 | 0.01381359 | 96.1352263 | 17.3503662 | 8.35246725 | 19.4853184 | 4.75243649 |
| 0.00723659 | 0.01381733 | 39.8687895 | 77.4093262 | 0 | 48.7132959 | 0 |
| 0.00723954 | 0.01382217 | 5.46588243 | 6.005896 | 10.2085711 | 40.3624452 | 0 |
| 0.00724097 | 0.01382412 | 22.5065747 | 15.3484009 | 22.273246 | 14.3820207 | 45.1481466 |
| 0.00724302 | 0.01382725 | 22.5065747 | 7.34053955 | 8.35246725 | 6.49510612 | 0 |
| 0.0072496 | 0.01383902 | 4.82283744 | 6.89565837 | 8.35246725 | 51.9608489 | 0 |
| 0.00725862 | 0.01385545 | 2.89370247 | 23.5787028 | 17.6329864 | 9.74265918 | 0 |
| 0.00726252 | 0.01386211 | 18.0052598 | 24.023584 | 18.5610383 | 19.0213822 | 7.12865473 |
| 0.00726727 | 0.01387039 | 77.1653991 | 77.6317668 | 0 | 23.6607437 | 0 |
| 0.00727724 | 0.01388863 | 1.92913498 | 12.4566732 | 35.2659728 | 16.2377653 | 0 |

| | | | | | | |
|---|---|---|---|---|---|---|
| 0.00728014 | 0.01389337 | 0 | 71.8483114 | 67.7477899 | 70.0543588 | 0 |
| 0.00728379 | 0.01389957 | 69.7703817 | 168.165088 | 193.962851 | 166.553078 | 38.0194919 |
| 0.00729942 | 0.0139286 | 89.7047764 | 110.997856 | 90.9490879 | 80.7248903 | 121.18713 |
| 0.00730493 | 0.01393831 | 22.5065747 | 1.55708415 | 30.6257133 | 12.9902122 | 0 |
| 0.00730889 | 0.01394508 | 3.21522496 | 17.1279256 | 10.2085711 | 18.0935099 | 0 |
| 0.00731865 | 0.01396292 | 67.1982017 | 34.2558512 | 0 | 64.0231889 | 0 |
| 0.00732238 | 0.01396924 | 149.829483 | 488.479541 | 234.797135 | 409.655622 | 0 |
| 0.00732576 | 0.0139749 | 4.50131495 | 16.905485 | 11.136623 | 9.27872303 | 0 |
| 0.00733451 | 0.0139908 | 3.53674746 | 12.4566732 | 20.4171422 | 10.6705315 | 0 |
| 0.00734306 | 0.0140063 | 87.4541189 | 211.763444 | 129.927268 | 184.182652 | 28.5146189 |
| 0.00734689 | 0.01401283 | 0 | 70.0687866 | 61.2514265 | 77.9412734 | 0 |
| 0.00735024 | 0.01401841 | 70.7349491 | 57.3896728 | 0 | 36.1870198 | 0 |
| 0.00736417 | 0.01404418 | 26.0433222 | 98.9860636 | 41.7623363 | 58.9198912 | 16.6335277 |
| 0.00737014 | 0.01405477 | 23.7926647 | 72.5156331 | 52.8989593 | 56.6002105 | 16.6335277 |
| 0.00737094 | 0.0140555 | 4.82283744 | 9.34250488 | 12.0646749 | 11.5984038 | 0 |
| 0.00737229 | 0.01405728 | 28.9370247 | 51.3837769 | 0 | 94.1790387 | 0 |
| 0.00737673 | 0.01406494 | 168.477788 | 12.4566732 | 0 | 47.7854236 | 0 |
| 0.00738471 | 0.01407936 | 40.190312 | 44.9329997 | 0 | 80.2609542 | 0 |
| 0.00739037 | 0.01408936 | 2.89370247 | 22.6889404 | 10.2085711 | 16.7017014 | 0 |
| 0.00739233 | 0.01409229 | 0.96456749 | 2.66928711 | 0 | 1.85574461 | 0 |
| 0.00739425 | 0.01409516 | 63.3399317 | 62.2833659 | 64.9636342 | 0 | 0 |
| 0.00739826 | 0.01410201 | 22.1850522 | 8.8976237 | 4.64025958 | 9.74265918 | 0 |
| 0.00739996 | 0.01410444 | 12.8608998 | 8.8976237 | 16.7049345 | 4.17542536 | 0 |
| 0.0074014 | 0.0141056 | 7.07349491 | 17.1279256 | 6.49636342 | 8.81478687 | 0 |
| 0.00740141 | 0.0141056 | 37.618132 | 24.6909058 | 28.7696094 | 22.7328714 | 90.2962933 |
| 0.00740306 | 0.01410795 | 37.618132 | 53.3857422 | 0 | 72.3740396 | 0 |
| 0.00740872 | 0.01411794 | 10.2887199 | 5.11613363 | 19.4890903 | 7.88691457 | 0 |
| 0.007414 | 0.0141272 | 2.89370247 | 8.23030192 | 26.9135056 | 19.0213822 | 0 |
| 0.00742092 | 0.01413959 | 7.07349491 | 10.2322673 | 5.5683115 | 19.9492545 | 0 |
| 0.00743623 | 0.01416795 | 6.75197242 | 18.6850098 | 14.8488307 | 24.1246799 | 2.37621824 |
| 0.00744742 | 0.01418848 | 5.46588243 | 10.2322673 | 12.0646749 | 8.35085072 | 0 |
| 0.0074495 | 0.01419163 | 7.07349491 | 10.0098267 | 13.9207788 | 6.03116997 | 0 |
| 0.00745028 | 0.01419232 | 21.8635297 | 52.4959798 | 0 | 112.272549 | 0 |
| 0.00745214 | 0.01419506 | 74.2716966 | 181.289083 | 165.193241 | 239.85499 | 40.3957101 |
| 0.00745829 | 0.01420597 | 40.833357 | 45.6003214 | 43.6184401 | 42.6821259 | 35.6432737 |
| 0.00746118 | 0.01421067 | 0 | 0 | 0 | 12.9902122 | 57.0292378 |
| 0.00746924 | 0.01422522 | 6.43044992 | 7.78542074 | 11.136623 | 8.81478687 | 0 |
| 0.00747437 | 0.01423417 | 49.1929419 | 48.9369303 | 0 | 60.3116997 | 0 |
| 0.00747817 | 0.01423981 | 10.6102424 | 6.45077718 | 7.42441534 | 9.74265918 | 0 |
| 0.00747783 | 0.01423981 | 4.50131495 | 14.0137573 | 10.2085711 | 10.6705315 | 0 |
| 0.00747932 | 0.01424119 | 0 | 58.2794352 | 41.7623363 | 121.087335 | 0 |
| 0.00748068 | 0.01424296 | 10.2887199 | 13.5688761 | 6.49636342 | 6.49510612 | 0 |
| 0.00748547 | 0.01425128 | 5.46588243 | 4.67125244 | 2.78415575 | 3.24755306 | 23.7621824 |

| | | | | | | |
|---|---|---|---|---|---|---|
| 0.00748978 | 0.01425807 | 16.0761248 | 10.0098267 | 12.0646749 | 16.2377653 | 0 |
| 0.00748989 | 0.01425807 | 77.1653991 | 52.940861 | 35.2659728 | 42.6821259 | 0 |
| 0.00749455 | 0.01426614 | 41.1548795 | 59.3916382 | 0 | 58.9198912 | 0 |
| 0.00749705 | 0.0142701 | 3.21522496 | 21.1318563 | 13.9207788 | 10.6705315 | 0 |
| 0.0075074 | 0.014289 | 43.0840145 | 49.8266927 | 0 | 66.8068058 | 0 |
| 0.00751197 | 0.01429689 | 13.8254673 | 13.5688761 | 11.136623 | 12.9902122 | 0 |
| 0.0075298 | 0.01433001 | 6.10892743 | 11.5669108 | 9.28051917 | 7.88691457 | 0 |
| 0.0075324 | 0.01433415 | 66.5551567 | 51.3837769 | 32.4818171 | 69.1264865 | 19.0097459 |
| 0.00753312 | 0.01433471 | 4.17979245 | 57.834554 | 10.2085711 | 5.56723382 | 0 |
| 0.00753629 | 0.01433993 | 0 | 31.8090047 | 108.582074 | 82.5806349 | 0 |
| 0.00753961 | 0.01434543 | 3.85826995 | 0.66732178 | 0 | 0 | 7.12865473 |
| 0.00754662 | 0.01435795 | 31.1876821 | 55.6101481 | 0 | 79.797018 | 0 |
| 0.00754945 | 0.01436252 | 16.7191698 | 11.3444702 | 14.8488307 | 14.3820207 | 4.75243649 |
| 0.00756037 | 0.01438248 | 47.5853294 | 84.5274251 | 0 | 34.3312752 | 0 |
| 0.00757203 | 0.01440386 | 18.9698273 | 4.44881185 | 7.42441534 | 13.9180845 | 0 |
| 0.00757506 | 0.01440881 | 7.71653991 | 10.2322673 | 12.9927268 | 5.56723382 | 0 |
| 0.00758125 | 0.01441976 | 32.4737721 | 50.2715739 | 0 | 83.9724434 | 0 |
| 0.00758493 | 0.01442595 | 13.1824223 | 3.33660889 | 9.28051917 | 27.3722329 | 0 |
| 0.00759054 | 0.01443579 | 0 | 86.3069499 | 51.9709073 | 69.5904227 | 0 |
| 0.00759317 | 0.01443998 | 64.9475442 | 30.9192423 | 0 | 68.6625504 | 0 |
| 0.00759499 | 0.01444264 | 47.2638069 | 33.3660889 | 26.9135056 | 30.619786 | 26.1384007 |
| 0.00760033 | 0.01445196 | 3.21522496 | 5.56101481 | 1.85610383 | 2.78361691 | 83.1676385 |
| 0.00760375 | 0.01445765 | 11.5748099 | 214.432731 | 179.11402 | 182.790844 | 26.1384007 |
| 0.00761262 | 0.0144737 | 50.4790319 | 50.7164551 | 0 | 55.208402 | 0 |
| 0.00761696 | 0.01448113 | 23.4711422 | 12.011792 | 12.9927268 | 19.0213822 | 19.0097459 |
| 0.00765611 | 0.01455474 | 17.3622148 | 32.9212077 | 18.5610383 | 23.1968076 | 7.12865473 |
| 0.0076665 | 0.01457367 | 56.2664368 | 35.5904948 | 0 | 68.6625504 | 0 |
| 0.0076678 | 0.01457532 | 0 | 91.8679647 | 47.3306478 | 69.5904227 | 0 |
| 0.00766897 | 0.01457671 | 9.96719738 | 12.2342326 | 18.5610383 | 3.71148921 | 0 |
| 0.0076781 | 0.01459324 | 6.43044992 | 8.8976237 | 10.2085711 | 7.88691457 | 0 |
| 0.0076818 | 0.01459945 | 5.78740493 | 12.4566732 | 6.49636342 | 13.4541484 | 0 |
| 0.00768526 | 0.01460521 | 10.6102424 | 9.12006429 | 4.64025958 | 13.4541484 | 0 |
| 0.00768663 | 0.01460698 | 4.17979245 | 0 | 2.78415575 | 3.71148921 | 35.6432737 |
| 0.00768961 | 0.01461183 | 36.3320421 | 63.1731283 | 0 | 59.8477635 | 0 |
| 0.00769661 | 0.01462405 | 56.2664368 | 45.822762 | 0 | 53.8165936 | 0 |
| 0.00769692 | 0.01462405 | 4.82283744 | 9.12006429 | 13.9207788 | 10.2065953 | 0 |
| 0.00770046 | 0.01462995 | 45.0131495 | 56.7223511 | 0 | 54.2805297 | 0 |
| 0.00771592 | 0.0146585 | 7.71653991 | 7.34053955 | 9.28051917 | 10.2065953 | 38.0194919 |
| 0.00771806 | 0.01466173 | 8.3595849 | 4.67125244 | 11.136623 | 3.71148921 | 23.7621824 |
| 0.00771912 | 0.01466294 | 26.6863672 | 62.5058065 | 0 | 78.8691457 | 0 |
| 0.00772282 | 0.01466912 | 0 | 0 | 9.28051917 | 8.81478687 | 211.483424 |
| 0.00773431 | 0.01469012 | 3.85826995 | 7.56298014 | 17.6329864 | 18.0935099 | 0 |
| 0.00773628 | 0.01469304 | 2.25065747 | 19.1298909 | 12.0646749 | 21.8049991 | 0 |

| | | | | | | |
|---|---|---|---|---|---|---|
| 0.00773943 | 0.0146982 | 1.92913498 | 14.6810791 | 13.9207788 | 31.5476583 | 0 |
| 0.00774294 | 0.01470402 | 77.8084441 | 25.8031087 | 0 | 64.9510612 | 0 |
| 0.00774491 | 0.01470694 | 29.5800696 | 84.0825439 | 92.8051917 | 75.1576565 | 21.3859642 |
| 0.00775518 | 0.0147256 | 0 | 0 | 0 | 0 | 23.7621824 |
| 0.00775857 | 0.01473122 | 0 | 75.4073608 | 135.49558 | 23.6607437 | 0 |
| 0.00776125 | 0.01473547 | 89.3832539 | 144.363944 | 117.862593 | 150.315313 | 42.7719284 |
| 0.00776688 | 0.01474533 | 20.2559173 | 102.989994 | 99.3015551 | 61.7035081 | 21.3859642 |
| 0.00777022 | 0.01475083 | 214.777027 | 272.489726 | 0 | 322.899561 | 16.6335277 |
| 0.0077765 | 0.01476193 | 10.6102424 | 17.3503662 | 19.4890903 | 2.78361691 | 0 |
| 0.00777823 | 0.01476438 | 118.320279 | 32.2538859 | 0 | 31.5476583 | 0 |
| 0.00778594 | 0.01477817 | 4.82283744 | 13.5688761 | 29.6976613 | 29.2279775 | 0 |
| 0.00779854 | 0.01480126 | 2.25065747 | 32.2538859 | 19.4890903 | 8.81478687 | 0 |
| 0.00780308 | 0.01480904 | 14.4685123 | 8.45274251 | 4.64025958 | 11.1344676 | 0 |
| 0.00780394 | 0.01480983 | 8.3595849 | 5.11613363 | 12.0646749 | 11.1344676 | 0 |
| 0.00780867 | 0.01481797 | 257.539519 | 203.755583 | 99.3015551 | 92.7872303 | 14.2573095 |
| 0.0078179 | 0.01483466 | 17.3622148 | 51.3837769 | 48.2586997 | 72.8379758 | 14.2573095 |
| 0.00782081 | 0.01483934 | 0.64304499 | 0.22244059 | 0.92805192 | 0 | 30.8908372 |
| 0.00782143 | 0.01483967 | 16.3976473 | 35.1456136 | 0 | 164.233398 | 0 |
| 0.00782682 | 0.01484907 | 41.1548795 | 39.3719849 | 0 | 81.1888265 | 0 |
| 0.0078281 | 0.01484981 | 6.43044992 | 13.5688761 | 4.64025958 | 22.2689353 | 0 |
| 0.00782774 | 0.01484981 | 11.5748099 | 63.84045 | 143.848047 | 113.200421 | 0 |
| 0.00783111 | 0.01485469 | 77.8084441 | 43.5983561 | 0 | 38.9706367 | 0 |
| 0.0078445 | 0.01487925 | 5.46588243 | 18.6850098 | 7.42441534 | 9.74265918 | 0 |
| 0.00784626 | 0.01488175 | 48.5498969 | 9.56494547 | 3.71220767 | 7.42297842 | 0 |
| 0.00785128 | 0.01489043 | 7.71653991 | 103.879757 | 0 | 111.344676 | 0 |
| 0.00785595 | 0.01489844 | 6.10892743 | 8.00786133 | 10.2085711 | 9.27872303 | 0 |
| 0.00785686 | 0.01489933 | 52.0866444 | 88.976237 | 0 | 27.3722329 | 0 |
| 0.00786337 | 0.01491084 | 14.7900348 | 10.2322673 | 11.136623 | 10.6705315 | 38.0194919 |
| 0.00787303 | 0.01492831 | 0 | 0 | 0 | 0 | 0 |
| 0.0078741 | 0.0149295 | 6.75197242 | 14.9035197 | 8.35246725 | 6.95904227 | 0 |
| 0.00787546 | 0.01493125 | 2.57217997 | 15.3484009 | 14.8488307 | 16.7017014 | 0 |
| 0.00787822 | 0.01493563 | 0 | 72.7380737 | 88.1649321 | 45.0018067 | 0 |
| 0.00787984 | 0.01493786 | 0 | 74.5175985 | 65.8916861 | 60.7756358 | 0 |
| 0.00789545 | 0.01496661 | 0 | 85.4171875 | 70.5319457 | 48.2493597 | 0 |
| 0.00790626 | 0.01498611 | 0 | 52.0510986 | 45.4745439 | 115.056166 | 0 |
| 0.00790662 | 0.01498611 | 26.3648447 | 11.7893514 | 22.273246 | 11.1344676 | 90.2962933 |
| 0.0079124 | 0.0149962 | 6.10892743 | 6.45077718 | 23.2012979 | 9.27872303 | 0 |
| 0.00791679 | 0.01500369 | 6.75197242 | 5.56101481 | 12.9927268 | 12.5262761 | 0 |
| 0.00791813 | 0.01500537 | 6.43044992 | 29.1397176 | 27.8415575 | 33.867339 | 4.75243649 |
| 0.0079186 | 0.01500543 | 5.78740493 | 9.34250488 | 8.35246725 | 10.6705315 | 0 |
| 0.00792261 | 0.01501217 | 0 | 0 | 0 | 0 | 2.37621824 |
| 0.00792581 | 0.01501739 | 0 | 0.44488118 | 0 | 0 | 54.6530196 |
| 0.00792829 | 0.01502126 | 174.908238 | 0 | 25.9854537 | 58.9198912 | 0 |

| | | | | | | |
|---|---|---|---|---|---|---|
| 0.00793177 | 0.01502615 | 0 | 0 | 0 | 0 | 0 |
| 0.00793177 | 0.01502615 | 0 | 0 | 0 | 0 | 0 |
| 0.00793427 | 0.01503004 | 27.3294122 | 75.1849202 | 0 | 61.7035081 | 0 |
| 0.00794692 | 0.01505231 | 39.547267 | 6.22833659 | 6.49636342 | 7.42297842 | 0 |
| 0.00794666 | 0.01505231 | 307.697029 | 849.945504 | 695.110886 | 771.52582 | 0 |
| 0.0079474 | 0.01505236 | 5.78740493 | 11.7893514 | 6.49636342 | 13.4541484 | 0 |
| 0.0079545 | 0.01506496 | 136.647061 | 12.9015544 | 0 | 55.6723382 | 0 |
| 0.00795554 | 0.01506609 | 3.53674746 | 15.3484009 | 16.7049345 | 9.27872303 | 0 |
| 0.00795938 | 0.01507251 | 56.2664368 | 47.1574056 | 0 | 49.6411682 | 0 |
| 0.00796087 | 0.01507448 | 22.8280972 | 45.3778809 | 0 | 111.808612 | 0 |
| 0.00796664 | 0.01508371 | 66.5551567 | 92.9801676 | 34.3379209 | 0 | 0 |
| 0.00796656 | 0.01508371 | 14.4685123 | 3.78149007 | 21.3451941 | 7.88691457 | 0 |
| 0.00797039 | 0.01508996 | 6.75197242 | 5.33857422 | 24.1293498 | 10.2065953 | 0 |
| 0.00797193 | 0.01509204 | 28.6155022 | 45.3778809 | 0 | 94.6429749 | 0 |
| 0.00798511 | 0.01511614 | 44.691627 | 62.9506877 | 0 | 46.3936151 | 0 |
| 0.00798712 | 0.01511908 | 10.2887199 | 17.7952474 | 21.3451941 | 24.588616 | 2.37621824 |
| 0.00799602 | 0.01513508 | 1.92913498 | 1.55708415 | 8.35246725 | 1.85574461 | 28.5146189 |
| 0.00800367 | 0.01514871 | 49.5144644 | 112.999821 | 0 | 52.4247851 | 0 |
| 0.00800833 | 0.01515668 | 2.89370247 | 23.8011434 | 9.28051917 | 16.2377653 | 0 |
| 0.00801155 | 0.01516192 | 8.0380624 | 10.4547078 | 4.64025958 | 18.5574461 | 0 |
| 0.00803089 | 0.01519767 | 97.0997938 | 50.7164551 | 48.2586997 | 39.898509 | 66.5341108 |
| 0.00803197 | 0.01519885 | 9.64567488 | 28.4723958 | 34.3379209 | 19.4853184 | 4.75243649 |
| 0.00803926 | 0.01521179 | 13.5039448 | 8.00786133 | 6.49636342 | 7.42297842 | 0 |
| 0.00805666 | 0.01524386 | 11.2532874 | 93.86993 | 0 | 94.1790387 | 0 |
| 0.00805949 | 0.01524835 | 36.3320421 | 64.7302124 | 0 | 54.2805297 | 0 |
| 0.00806086 | 0.0152501 | 0 | 81.1908162 | 95.5893474 | 33.867339 | 0 |
| 0.00807168 | 0.0152697 | 163.333428 | 373.255314 | 236.653239 | 419.398281 | 0 |
| 0.00807793 | 0.01528067 | 9.96719738 | 7.78542074 | 25.0574018 | 4.63936151 | 0 |
| 0.00808532 | 0.01529379 | 253.038204 | 25.5806681 | 13.9207788 | 0 | 0 |
| 0.00808755 | 0.01529715 | 25.0787547 | 5.7834554 | 3.71220767 | 19.4853184 | 0 |
| 0.00809645 | 0.01531312 | 54.0157793 | 119.673039 | 94.6612955 | 101.602017 | 28.5146189 |
| 0.00809957 | 0.01531816 | 52.4081669 | 94.0923706 | 98.3735032 | 0 | 0 |
| 0.00810697 | 0.0153313 | 23.7926647 | 33.1436483 | 0 | 132.685739 | 0 |
| 0.00811141 | 0.01533883 | 36.0105196 | 52.0510986 | 0 | 67.2707419 | 0 |
| 0.00812104 | 0.01535617 | 106.423946 | 169.944613 | 275.631419 | 231.50414 | 47.5243649 |
| 0.00813347 | 0.01537881 | 182.303255 | 323.428621 | 316.465704 | 302.486371 | 0 |
| 0.00814402 | 0.01539789 | 7.71653991 | 6.45077718 | 10.2085711 | 5.56723382 | 9.50487297 |
| 0.00815183 | 0.01541179 | 10.2887199 | 9.34250488 | 10.2085711 | 5.10329766 | 0 |
| 0.00815635 | 0.01541948 | 70.4134267 | 72.7380737 | 0 | 23.1968076 | 0 |
| 0.00815938 | 0.01542434 | 100.958064 | 22.6889404 | 17.6329864 | 15.7738291 | 0 |
| 0.00817364 | 0.01545043 | 2.57217997 | 1.11220296 | 0.92805192 | 1.85574461 | 26.1384007 |
| 0.00817519 | 0.01545249 | 42.119447 | 33.5885295 | 29.6976613 | 30.1558498 | 377.818701 |
| 0.00817877 | 0.01545838 | 2.89370247 | 13.5688761 | 13.9207788 | 15.7738291 | 0 |

| | | | | | | |
|---|---|---|---|---|---|---|
| 0.00820048 | 0.01549559 | 4.82283744 | 18.4625692 | 26.9135056 | 21.341063 | 0 |
| 0.00820076 | 0.01549559 | 8.3595849 | 90.9782023 | 0 | 113.200421 | 0 |
| 0.00819898 | 0.01549559 | 81.023669 | 174.170984 | 111.36623 | 154.954675 | 71.2865473 |
| 0.00819941 | 0.01549559 | 7.71653991 | 10.0098267 | 5.5683115 | 11.1344676 | 0 |
| 0.00819986 | 0.01549559 | 2.57217997 | 4.89369303 | 0.92805192 | 3.71148921 | 21.3859642 |
| 0.0082066 | 0.01550481 | 14.7900348 | 21.1318563 | 26.9135056 | 18.0935099 | 0 |
| 0.00820702 | 0.01550481 | 84.238894 | 114.556905 | 104.869867 | 128.97425 | 33.2670554 |
| 0.00820663 | 0.01550481 | 124.429206 | 121.897445 | 133.639476 | 134.541484 | 38.0194919 |
| 0.00822355 | 0.01553516 | 3.53674746 | 14.6810791 | 10.2085711 | 14.3820207 | 0 |
| 0.00822762 | 0.01554199 | 26.6863672 | 98.763623 | 0 | 43.6099982 | 0 |
| 0.00824656 | 0.0155769 | 0 | 61.8384847 | 68.6758419 | 65.4149973 | 0 |
| 0.00825107 | 0.01558454 | 4.82283744 | 12.6791138 | 7.42441534 | 13.9180845 | 0 |
| 0.0082655 | 0.01561091 | 50.4790319 | 41.5963908 | 0 | 58.9198912 | 0 |
| 0.00827405 | 0.01562619 | 5.14435994 | 10.899589 | 10.2085711 | 8.81478687 | 0 |
| 0.00827701 | 0.01563089 | 3.53674746 | 11.1220296 | 14.8488307 | 12.5262761 | 0 |
| 0.00828089 | 0.01563734 | 3.21522496 | 11.7893514 | 12.9927268 | 16.2377653 | 0 |
| 0.00831458 | 0.01570007 | 2.89370247 | 50.2715739 | 17.6329864 | 4.63936151 | 0 |
| 0.00833109 | 0.01573037 | 3.21522496 | 35.3680542 | 22.273246 | 4.63936151 | 0 |
| 0.00833375 | 0.01573452 | 23.4711422 | 80.9683756 | 95.5893474 | 0 | 0 |
| 0.00833486 | 0.01573572 | 45.0131495 | 45.822762 | 96.5173994 | 0 | 0 |
| 0.00833646 | 0.01573786 | 177.80194 | 384.377344 | 304.401029 | 346.096369 | 97.424948 |
| 0.00834056 | 0.01574471 | 14.7900348 | 8.67518311 | 15.7768826 | 3.71148921 | 0 |
| 0.00834188 | 0.01574633 | 54.3373018 | 0 | 61.2514265 | 100.210209 | 0 |
| 0.00834888 | 0.01575866 | 56.9094818 | 0 | 166.121293 | 26.4443606 | 0 |
| 0.0083504 | 0.01576064 | 90.0262989 | 86.9742716 | 48.2586997 | 70.9822312 | 140.196876 |
| 0.00835279 | 0.01576426 | 22.5065747 | 72.7380737 | 0 | 69.5904227 | 0 |
| 0.00836237 | 0.01578125 | 8.0380624 | 12.4566732 | 5.5683115 | 8.81478687 | 0 |
| 0.00836273 | 0.01578125 | 43.405537 | 37.8149007 | 0 | 72.8379758 | 0 |
| 0.00836393 | 0.01578264 | 19.9343948 | 15.7932821 | 6.49636342 | 17.1656376 | 0 |
| 0.00836641 | 0.01578643 | 0 | 11.7893514 | 6.49636342 | 0 | 64.1578926 |
| 0.00837208 | 0.01579625 | 0 | 76.2971232 | 58.4672708 | 60.3116997 | 0 |
| 0.00837458 | 0.01580007 | 11.8963324 | 10.6771484 | 9.28051917 | 4.63936151 | 0 |
| 0.0083751 | 0.01580018 | 55.9449143 | 160.824548 | 153.128566 | 138.716909 | 7.12865473 |
| 0.0083851 | 0.01581815 | 169.120833 | 222.663033 | 262.638692 | 280.681372 | 80.7914203 |
| 0.00839505 | 0.01583603 | 53.6942569 | 40.0393066 | 0 | 56.1362743 | 0 |
| 0.00842522 | 0.01589206 | 49.8359869 | 60.9487223 | 0 | 39.4345729 | 0 |
| 0.00843047 | 0.01590106 | 41.476402 | 46.0452026 | 0 | 62.6313804 | 0 |
| 0.00844221 | 0.01592232 | 1.28608998 | 14.4586385 | 28.7696094 | 21.341063 | 0 |
| 0.0084437 | 0.01592423 | 3.53674746 | 1.55708415 | 2.78415575 | 3.24755306 | 7.12865473 |
| 0.00844807 | 0.01593158 | 114.783531 | 28.4723958 | 0 | 32.0115944 | 0 |
| 0.00845087 | 0.01593598 | 2.57217997 | 22.4664998 | 7.42441534 | 25.0525522 | 0 |
| 0.00845257 | 0.01593827 | 0.3215225 | 0.22244059 | 0.92805192 | 0 | 21.3859642 |
| 0.00845417 | 0.0159404 | 41.7979245 | 7.34053955 | 4.64025958 | 7.88691457 | 0 |

| | | | | | | |
|---|---|---|---|---|---|---|
| 0.00845645 | 0.01594203 | 3.85826995 | 33.1436483 | 22.273246 | 19.4853184 | 0 |
| 0.00845572 | 0.01594203 | 281.010662 | 242.015365 | 177.257916 | 204.131907 | 76.0389838 |
| 0.00845628 | 0.01594203 | 0 | 12.9015544 | 0 | 4.63936151 | 180.592587 |
| 0.00846235 | 0.01595226 | 9.96719738 | 8.45274251 | 5.5683115 | 9.27872303 | 0 |
| 0.00846978 | 0.01596536 | 8.3595849 | 4.00393066 | 13.9207788 | 15.309893 | 0 |
| 0.00847368 | 0.01597182 | 7.07349491 | 4.89369303 | 23.2012979 | 10.2065953 | 0 |
| 0.00848591 | 0.01599398 | 0 | 108.328569 | 81.6685687 | 65.4149973 | 0 |
| 0.00849522 | 0.01601062 | 5.78740493 | 12.9015544 | 27.8415575 | 21.8049991 | 0 |
| 0.00849867 | 0.01601622 | 20.2559173 | 20.6869751 | 6.49636342 | 12.9902122 | 0 |
| 0.00850134 | 0.01602036 | 6.43044992 | 15.7932821 | 8.35246725 | 7.42297842 | 35.6432737 |
| 0.00850415 | 0.01602477 | 4.82283744 | 12.011792 | 13.9207788 | 7.42297842 | 0 |
| 0.00850926 | 0.01603349 | 3.21522496 | 12.011792 | 12.0646749 | 16.7017014 | 0 |
| 0.0085137 | 0.01604097 | 49.1929419 | 11.3444702 | 38.9781805 | 17.1656376 | 7.12865473 |
| 0.00852677 | 0.01606469 | 7.71653991 | 19.5747721 | 4.64025958 | 10.2065953 | 0 |
| 0.00852791 | 0.01606594 | 0.96456749 | 17.7952474 | 29.6976613 | 22.2689353 | 0 |
| 0.00852884 | 0.01606679 | 8.0380624 | 8.00786133 | 7.42441534 | 7.88691457 | 0 |
| 0.0085308 | 0.01606958 | 2.57217997 | 22.2440592 | 29.6976613 | 6.49510612 | 0 |
| 0.00853607 | 0.01607861 | 0 | 69.1790242 | 116.934542 | 70.0543588 | 0 |
| 0.00853701 | 0.01607949 | 90.3478214 | 0 | 47.3306478 | 75.1576565 | 0 |
| 0.0085377 | 0.01607988 | 6.10892743 | 7.56298014 | 8.35246725 | 12.0623399 | 0 |
| 0.00854395 | 0.01609075 | 26.3648447 | 152.816687 | 37.1220767 | 0 | 0 |
| 0.00855399 | 0.01610875 | 9.96719738 | 5.56101481 | 25.9854537 | 6.03116997 | 0 |
| 0.00855741 | 0.0161143 | 5.14435994 | 0.88976237 | 0 | 2.31968076 | 4.75243649 |
| 0.00856042 | 0.01611906 | 102.565676 | 17.3503662 | 0 | 56.6002105 | 0 |
| 0.00857132 | 0.01613868 | 73.9501741 | 94.7596924 | 116.934542 | 100.210209 | 30.8908372 |
| 0.00857326 | 0.01614143 | 5.78740493 | 10.2322673 | 8.35246725 | 8.81478687 | 0 |
| 0.00858394 | 0.01616064 | 2.89370247 | 14.4586385 | 22.273246 | 9.74265918 | 0 |
| 0.00858446 | 0.0161607 | 0 | 38.0373413 | 125.287009 | 46.8575513 | 0 |
| 0.0085866 | 0.01616383 | 15.4330798 | 9.78738607 | 12.9927268 | 12.9902122 | 0 |
| 0.00859356 | 0.01617566 | 51.4435994 | 31.3641235 | 0 | 70.518295 | 0 |
| 0.00859385 | 0.01617566 | 5.46588243 | 8.67518311 | 9.28051917 | 10.2065953 | 0 |
| 0.00861326 | 0.0162113 | 45.0131495 | 22.4664998 | 17.6329864 | 17.1656376 | 161.582841 |
| 0.0086143 | 0.01621234 | 108.353081 | 45.6003214 | 0 | 20.4131907 | 0 |
| 0.00861501 | 0.01621277 | 0 | 0 | 0 | 0 | 0 |
| 0.00862215 | 0.0162253 | 3.21522496 | 14.4586385 | 17.6329864 | 9.74265918 | 0 |
| 0.00862294 | 0.01622588 | 46.2992394 | 87.6415934 | 47.3306478 | 0 | 0 |
| 0.00862631 | 0.01623132 | 19.2913498 | 88.7537964 | 0 | 60.7756358 | 0 |
| 0.00863421 | 0.01624528 | 10.6102424 | 16.905485 | 12.9927268 | 11.1344676 | 2.37621824 |
| 0.00863623 | 0.01624816 | 11.5748099 | 15.3484009 | 22.273246 | 31.5476583 | 0 |
| 0.00863808 | 0.01625074 | 8.6811074 | 9.34250488 | 8.35246725 | 6.03116997 | 0 |
| 0.00864609 | 0.01626489 | 27.6509347 | 38.9271037 | 0 | 98.3544641 | 0 |
| 0.00866284 | 0.0162955 | 5.46588243 | 3.11416829 | 2.78415575 | 1.85574461 | 19.0097459 |
| 0.00867605 | 0.01631942 | 12.2178549 | 2.66928711 | 10.2085711 | 31.0837221 | 0 |

| | | | | | | |
|---|---|---|---|---|---|---|
| 0.00867761 | 0.01632074 | 5.46588243 | 6.89565837 | 9.28051917 | 20.8771268 | 0 |
| 0.00867772 | 0.01632074 | 152.401663 | 221.55083 | 163.337137 | 208.771268 | 71.2865473 |
| 0.00868281 | 0.01632941 | 4.82283744 | 7.78542074 | 11.136623 | 13.4541484 | 0 |
| 0.00868462 | 0.01633191 | 2.57217997 | 12.6791138 | 13.9207788 | 19.4853184 | 0 |
| 0.0086999 | 0.01635973 | 49.1929419 | 53.3857422 | 0 | 43.6099982 | 0 |
| 0.00870046 | 0.01635986 | 6.10892743 | 13.7913167 | 4.64025958 | 19.0213822 | 0 |
| 0.00871009 | 0.01637705 | 16.7191698 | 102.989994 | 60.3233746 | 75.1576565 | 0 |
| 0.00871205 | 0.01637983 | 57.5525268 | 60.9487223 | 0 | 32.0115944 | 0 |
| 0.00872284 | 0.0163992 | 20.8989622 | 107.661247 | 36.1940248 | 104.385634 | 0 |
| 0.00873837 | 0.01642747 | 32.4737721 | 16.6830444 | 6.49636342 | 12.5262761 | 0 |
| 0.00874546 | 0.01643987 | 17.6837373 | 10.6771484 | 4.64025958 | 7.42297842 | 0 |
| 0.00874765 | 0.01644307 | 17.0406923 | 81.6356974 | 108.582074 | 0 | 0 |
| 0.00874986 | 0.01644632 | 4.17979245 | 8.8976237 | 12.0646749 | 13.4541484 | 0 |
| 0.00875219 | 0.01644978 | 10.9317649 | 12.6791138 | 7.42441534 | 5.10329766 | 0 |
| 0.00875431 | 0.01645285 | 0 | 48.047168 | 81.6685687 | 62.6313804 | 0 |
| 0.00875586 | 0.01645484 | 3.21522496 | 2.00196533 | 1.85610383 | 4.63936151 | 26.1384007 |
| 0.00875685 | 0.01645577 | 20.8989622 | 78.5215291 | 0 | 63.5592527 | 0 |
| 0.00875795 | 0.01645693 | 7.07349491 | 12.2342326 | 20.4171422 | 20.4131907 | 2.37621824 |
| 0.00876027 | 0.01646037 | 24.7572322 | 87.864034 | 78.8844129 | 0 | 0 |
| 0.00876124 | 0.01646128 | 9.64567488 | 5.11613363 | 6.49636342 | 27.8361691 | 0 |
| 0.00876224 | 0.01646223 | 18.9698273 | 203.310701 | 173.545708 | 94.6429749 | 30.8908372 |
| 0.00876813 | 0.01647238 | 11.2532874 | 6.67321777 | 10.2085711 | 6.03116997 | 0 |
| 0.00877137 | 0.01647753 | 34.4029071 | 8.45274251 | 3.71220767 | 9.27872303 | 0 |
| 0.00877499 | 0.01648247 | 32.1522496 | 50.0491333 | 0 | 68.6625504 | 0 |
| 0.00877546 | 0.01648247 | 0 | 0 | 0 | 0 | 2.37621824 |
| 0.00877478 | 0.01648247 | 3.85826995 | 2.00196533 | 0 | 1.85574461 | 14.2573095 |
| 0.00879361 | 0.01651563 | 3.85826995 | 9.34250488 | 18.5610383 | 10.6705315 | 0 |
| 0.00879656 | 0.01651932 | 0 | 45.6003214 | 109.510126 | 45.4657428 | 0 |
| 0.00879649 | 0.01651932 | 32.7952946 | 48.9369303 | 0 | 68.6625504 | 0 |
| 0.0087974 | 0.01651999 | 23.7926647 | 73.6278361 | 95.5893474 | 0 | 0 |
| 0.0088121 | 0.01654667 | 2.89370247 | 14.6810791 | 12.0646749 | 15.309893 | 0 |
| 0.00881671 | 0.01655439 | 5.78740493 | 10.899589 | 6.49636342 | 12.0623399 | 0 |
| 0.00883663 | 0.01659087 | 9.96719738 | 6.22833659 | 6.49636342 | 11.1344676 | 0 |
| 0.00884162 | 0.01659932 | 44.048582 | 57.3896728 | 0 | 44.0739344 | 0 |
| 0.00884512 | 0.01660495 | 14.7900348 | 61.6160441 | 0 | 101.602017 | 0 |
| 0.00884759 | 0.01660867 | 60.1247068 | 0 | 74.2441534 | 70.0543588 | 0 |
| 0.00884874 | 0.0166099 | 0.64304499 | 0.88976237 | 0 | 0 | 16.6335277 |
| 0.00885143 | 0.01661403 | 37.9396545 | 66.5097371 | 0 | 43.6099982 | 0 |
| 0.00885478 | 0.01661938 | 26.3648447 | 31.1416829 | 21.3451941 | 43.1460621 | 11.8810912 |
| 0.00885906 | 0.01662649 | 23.1496197 | 156.598177 | 37.1220767 | 0 | 0 |
| 0.00886159 | 0.01663031 | 3.53674746 | 18.9074504 | 20.4171422 | 6.49510612 | 0 |
| 0.00886846 | 0.01664227 | 3.53674746 | 16.905485 | 11.136623 | 10.6705315 | 0 |
| 0.00887237 | 0.01664867 | 2.57217997 | 4.00393066 | 32.4818171 | 32.4755306 | 0 |

| | | | | | | |
|---|---|---|---|---|---|---|
| 0.00888378 | 0.01666917 | 29.2585471 | 40.2617472 | 33.409869 | 29.6919137 | 0 |
| 0.00889309 | 0.0166857 | 5.78740493 | 8.45274251 | 8.35246725 | 10.2065953 | 0 |
| 0.00889784 | 0.01669369 | 30.8661596 | 173.948543 | 113.222334 | 133.149675 | 28.5146189 |
| 0.00890509 | 0.01670636 | 14.1469898 | 20.4645345 | 6.49636342 | 4.17542536 | 0 |
| 0.00891039 | 0.01671536 | 11.2532874 | 5.33857422 | 18.5610383 | 6.03116997 | 0 |
| 0.0089114 | 0.01671633 | 17.0406923 | 4.00393066 | 9.28051917 | 10.6705315 | 0 |
| 0.00891309 | 0.01671764 | 9.32415239 | 11.1220296 | 35.2659728 | 2.78361691 | 0 |
| 0.0089126 | 0.01671764 | 77.1653991 | 219.326424 | 186.538435 | 204.595843 | 54.6530196 |
| 0.00891586 | 0.0167204 | 0 | 54.9428263 | 99.3015551 | 41.7542536 | 0 |
| 0.00891606 | 0.0167204 | 72.6640841 | 37.1475789 | 0 | 39.898509 | 0 |
| 0.00891517 | 0.0167204 | 67.5197242 | 53.6081828 | 51.9709073 | 0 | 0 |
| 0.00892037 | 0.01672755 | 46.2992394 | 20.2420939 | 24.1293498 | 20.8771268 | 78.415202 |
| 0.00892439 | 0.01673416 | 4.50131495 | 14.0137573 | 7.42441534 | 12.9902122 | 0 |
| 0.00893572 | 0.01675447 | 4.82283744 | 11.1220296 | 9.28051917 | 9.74265918 | 0 |
| 0.00894114 | 0.01676371 | 1.92913498 | 18.6850098 | 17.6329864 | 14.8459568 | 0 |
| 0.00894379 | 0.01676774 | 0 | 56.0550293 | 91.8771398 | 45.0018067 | 0 |
| 0.00894502 | 0.01676863 | 36.6535646 | 50.4940145 | 94.6612955 | 0 | 0 |
| 0.00894526 | 0.01676863 | 3.21522496 | 16.0157227 | 8.35246725 | 19.0213822 | 0 |
| 0.00895077 | 0.01677719 | 0.96456749 | 10.6771484 | 42.6903882 | 23.6607437 | 0 |
| 0.00895083 | 0.01677719 | 5.78740493 | 7.78542074 | 46.4025958 | 5.10329766 | 0 |
| 0.00895955 | 0.0167926 | 50.4790319 | 45.3778809 | 0 | 47.7854236 | 0 |
| 0.00896854 | 0.01680852 | 72.0210391 | 62.0609253 | 52.8989593 | 64.487125 | 85.5438568 |
| 0.00897034 | 0.01681009 | 52.7296894 | 21.1318563 | 25.0574018 | 25.5164883 | 68.9103291 |
| 0.00897038 | 0.01681009 | 42.762492 | 149.480078 | 0 | 72.8379758 | 2.37621824 |
| 0.00897372 | 0.01681541 | 0 | 57.1672323 | 64.9636342 | 64.9510612 | 0 |
| 0.00897563 | 0.01681806 | 1.92913498 | 13.5688761 | 18.5610383 | 19.4853184 | 0 |
| 0.00898278 | 0.01683052 | 66.8766792 | 49.8266927 | 0 | 32.0115944 | 0 |
| 0.00899066 | 0.01684434 | 20.5774398 | 121.452563 | 0 | 35.7230837 | 0 |
| 0.00899223 | 0.01684634 | 117.355711 | 30.6968018 | 0 | 25.5164883 | 0 |
| 0.00899882 | 0.01685776 | 0.96456749 | 0.66732178 | 0 | 0.46393615 | 33.2670554 |
| 0.00901209 | 0.01688167 | 2.89370247 | 11.7893514 | 10.2085711 | 25.9804245 | 0 |
| 0.00901406 | 0.01688352 | 0 | 0 | 0 | 0 | 7.12865473 |
| 0.00901408 | 0.01688352 | 0 | 33.3660889 | 84.4527244 | 78.4052096 | 0 |
| 0.00902522 | 0.01690345 | 9.00262989 | 8.00786133 | 5.5683115 | 10.2065953 | 0 |
| 0.0090415 | 0.01693299 | 0 | 57.3896728 | 49.1867516 | 82.5806349 | 0 |
| 0.00904258 | 0.01693407 | 31.1876821 | 69.1790242 | 0 | 48.7132959 | 0 |
| 0.00904623 | 0.01693969 | 3.21522496 | 14.0137573 | 13.9207788 | 11.1344676 | 0 |
| 0.00904659 | 0.01693969 | 0 | 74.9624797 | 55.683115 | 56.6002105 | 0 |
| 0.00907817 | 0.01699788 | 0 | 24.4684652 | 128.999216 | 59.3838274 | 0 |
| 0.00908337 | 0.01700668 | 32.7952946 | 45.1554403 | 0 | 70.518295 | 0 |
| 0.00908513 | 0.01700902 | 0 | 66.5097371 | 73.3161014 | 47.7854236 | 0 |
| 0.00908642 | 0.01701049 | 0 | 35.5904948 | 90.0210359 | 68.6625504 | 0 |
| 0.00909137 | 0.01701797 | 3.85826995 | 22.2440592 | 24.1293498 | 4.63936151 | 0 |

| | | | | | | |
|---|---|---|---|---|---|---|
| 0.00909143 | 0.01701797 | 1.92913498 | 1.77952474 | 0 | 2.78361691 | 30.8908372 |
| 0.00909361 | 0.01702111 | 5.78740493 | 9.12006429 | 6.49636342 | 17.1656376 | 0 |
| 0.00910553 | 0.01704247 | 148.864916 | 309.859745 | 239.437395 | 269.546904 | 0 |
| 0.00910745 | 0.01704512 | 6.10892743 | 16.0157227 | 5.5683115 | 10.6705315 | 0 |
| 0.00910854 | 0.01704621 | 106.423946 | 277.383419 | 232.941031 | 280.681372 | 0 |
| 0.00911514 | 0.01705762 | 52.0866444 | 22.4664998 | 0 | 83.9724434 | 0 |
| 0.0091297 | 0.01708391 | 47.5853294 | 56.2774699 | 79.8124649 | 70.9822312 | 9.50487297 |
| 0.00913094 | 0.01708527 | 3.53674746 | 13.7913167 | 10.2085711 | 12.9902122 | 0 |
| 0.00913316 | 0.01708848 | 2.57217997 | 12.2342326 | 19.4890903 | 13.4541484 | 0 |
| 0.00915386 | 0.01712626 | 41.1548795 | 99.4309448 | 120.646749 | 131.293931 | 28.5146189 |
| 0.00916037 | 0.01713749 | 12.5393773 | 8.8976237 | 8.35246725 | 5.10329766 | 0 |
| 0.00916939 | 0.0171534 | 3.85826995 | 12.6791138 | 24.1293498 | 6.95904227 | 0 |
| 0.00916997 | 0.01715355 | 41.476402 | 70.7361084 | 60.3233746 | 0 | 0 |
| 0.00917335 | 0.01715892 | 0 | 55.8325887 | 58.4672708 | 70.9822312 | 0 |
| 0.0091798 | 0.01717002 | 105.137856 | 14.9035197 | 0 | 55.208402 | 0 |
| 0.00918073 | 0.0171708 | 0 | 0 | 0 | 0 | 0 |
| 0.00918215 | 0.0171725 | 0 | 80.9683756 | 50.1148035 | 56.1362743 | 0 |
| 0.00918327 | 0.01717366 | 40.833357 | 70.2912272 | 54.7550631 | 36.1870198 | 16.6335277 |
| 0.00918642 | 0.01717858 | 70.7349491 | 56.0550293 | 33.409869 | 46.8575513 | 19.0097459 |
| 0.00919103 | 0.01718625 | 2.57217997 | 26.0255493 | 8.35246725 | 16.7017014 | 0 |
| 0.00919757 | 0.01719752 | 13.5039448 | 6.005896 | 7.42441534 | 7.88691457 | 0 |
| 0.00920185 | 0.01720457 | 5.14435994 | 5.56101481 | 12.9927268 | 19.0213822 | 0 |
| 0.00920358 | 0.01720684 | 123.143116 | 159.489905 | 153.128566 | 193.925311 | 78.415202 |
| 0.00921932 | 0.01723532 | 0.3215225 | 0.88976237 | 0.92805192 | 1.85574461 | 4.75243649 |
| 0.00922349 | 0.01724216 | 0 | 76.7420044 | 39.9062324 | 72.3740396 | 0 |
| 0.00922659 | 0.01724699 | 9.00262989 | 1.55708415 | 29.6976613 | 24.588616 | 0 |
| 0.00922844 | 0.01724853 | 6.10892743 | 8.23030192 | 25.9854537 | 6.03116997 | 0 |
| 0.00922838 | 0.01724853 | 20.8989622 | 6.005896 | 8.35246725 | 6.49510612 | 0 |
| 0.00924246 | 0.01727281 | 4.82283744 | 6.45077718 | 14.8488307 | 12.9902122 | 0 |
| 0.00924209 | 0.01727281 | 74.2716966 | 69.846346 | 85.3807764 | 110.416804 | 28.5146189 |
| 0.00925501 | 0.01729531 | 17.6837373 | 7.11809896 | 15.7768826 | 3.71148921 | 0 |
| 0.00925667 | 0.01729745 | 3.85826995 | 14.0137573 | 26.9135056 | 6.03116997 | 0 |
| 0.00926052 | 0.01730369 | 0.64304499 | 0 | 0 | 0 | 4.75243649 |
| 0.00927251 | 0.01732514 | 3.21522496 | 15.1259603 | 12.0646749 | 11.5984038 | 0 |
| 0.00928285 | 0.01734349 | 4.82283744 | 30.9192423 | 9.28051917 | 6.49510612 | 0 |
| 0.00928633 | 0.01734902 | 1.60761248 | 12.6791138 | 20.4171422 | 23.1968076 | 0 |
| 0.00928784 | 0.01735089 | 5.46588243 | 8.00786133 | 9.28051917 | 10.2065953 | 0 |
| 0.0092911 | 0.01735573 | 0 | 40.7066284 | 91.8771398 | 57.5280828 | 0 |
| 0.00929147 | 0.01735573 | 0 | 0 | 0 | 0 | 40.3957101 |
| 0.00929368 | 0.0173589 | 5.46588243 | 12.4566732 | 6.49636342 | 11.1344676 | 0 |
| 0.00929652 | 0.01736325 | 0 | 69.1790242 | 66.819738 | 48.7132959 | 0 |
| 0.00929878 | 0.0173665 | 6.10892743 | 7.11809896 | 7.42441534 | 19.4853184 | 0 |
| 0.00930223 | 0.01737198 | 3.21522496 | 2.66928711 | 23.2012979 | 49.177232 | 0 |

| | | | | | | |
|---|---|---|---|---|---|---|
| 0.0093098 | 0.01738515 | 3.53674746 | 17.3503662 | 9.28051917 | 12.0623399 | 0 |
| 0.00931041 | 0.01738532 | 70.0919042 | 143.251742 | 170.761553 | 159.594036 | 0 |
| 0.00931241 | 0.0173881 | 1.92913498 | 7.11809896 | 31.5537652 | 23.1968076 | 0 |
| 0.00931804 | 0.01739763 | 0.3215225 | 1.33464355 | 0.92805192 | 0.46393615 | 0 |
| 0.00933158 | 0.01742195 | 11.2532874 | 19.5747721 | 11.136623 | 10.6705315 | 2.37621824 |
| 0.00933482 | 0.01742704 | 5.78740493 | 10.6771484 | 8.35246725 | 7.88691457 | 0 |
| 0.00934687 | 0.01744857 | 37.9396545 | 44.7105591 | 0 | 59.8477635 | 0 |
| 0.00934931 | 0.01745215 | 8.3595849 | 4.22637126 | 11.136623 | 14.8459568 | 0 |
| 0.00935256 | 0.01745724 | 34.4029071 | 70.2912272 | 0 | 41.2903175 | 0 |
| 0.00935822 | 0.01746684 | 20.5774398 | 62.0609253 | 0 | 73.7658481 | 0 |
| 0.00936828 | 0.01748465 | 7.71653991 | 6.005896 | 2.78415575 | 3.71148921 | 71.2865473 |
| 0.00937473 | 0.01749481 | 0 | 120.785242 | 44.546492 | 76.549465 | 2.37621824 |
| 0.00937477 | 0.01749481 | 27.3294122 | 15.1259603 | 9.28051917 | 9.27872303 | 0 |
| 0.00937716 | 0.01749791 | 8.3595849 | 9.78738607 | 1.85610383 | 0.9278723 | 14.2573095 |
| 0.00937747 | 0.01749791 | 608.642085 | 406.843844 | 388.853753 | 427.749132 | 0 |
| 0.00937839 | 0.01749866 | 63.3399317 | 42.2637126 | 0 | 37.5788283 | 0 |
| 0.00937971 | 0.01750015 | 0.64304499 | 0.66732178 | 0.92805192 | 1.39180845 | 0 |
| 0.00938347 | 0.0175062 | 4.82283744 | 12.4566732 | 14.8488307 | 6.49510612 | 0 |
| 0.00938914 | 0.01751581 | 7.71653991 | 14.4586385 | 3.71220767 | 15.309893 | 0 |
| 0.00939816 | 0.01753166 | 54.0157793 | 31.3641235 | 30.6257133 | 29.6919137 | 299.403499 |
| 0.00940897 | 0.01755085 | 0 | 0 | 0 | 0 | 59.4054561 |
| 0.00941119 | 0.01755402 | 117.355711 | 140.360014 | 103.941815 | 140.572654 | 0 |
| 0.00941542 | 0.01756094 | 0 | 77.8542074 | 23.2012979 | 102.529889 | 0 |
| 0.00941726 | 0.01756341 | 56.5879593 | 100.320707 | 81.6685687 | 83.9724434 | 0 |
| 0.00942141 | 0.01757017 | 7.39501741 | 5.56101481 | 12.0646749 | 8.81478687 | 0 |
| 0.00943621 | 0.01759582 | 0 | 76.5195638 | 89.092984 | 64.9510612 | 0 |
| 0.00943569 | 0.01759582 | 6.43044992 | 9.12006429 | 8.35246725 | 7.42297842 | 0 |
| 0.00944227 | 0.01760614 | 3.85826995 | 10.0098267 | 9.28051917 | 18.5574461 | 0 |
| 0.00944297 | 0.01760647 | 4.17979245 | 12.2342326 | 8.35246725 | 12.9902122 | 0 |
| 0.00945587 | 0.01762954 | 5.78740493 | 11.5669108 | 5.5683115 | 14.3820207 | 0 |
| 0.00946351 | 0.01764282 | 1.28608998 | 0.22244059 | 0 | 0.9278723 | 0 |
| 0.00946961 | 0.01765321 | 43.0840145 | 31.3641235 | 0 | 71.9101035 | 0 |
| 0.00947103 | 0.01765487 | 45.9777169 | 96.9840983 | 35.2659728 | 0 | 0 |
| 0.00947879 | 0.01766836 | 6.43044992 | 3.33660889 | 3.71220767 | 2.31968076 | 16.6335277 |
| 0.00948125 | 0.01767197 | 9.00262989 | 25.5806681 | 23.2012979 | 21.341063 | 0 |
| 0.00948324 | 0.01767469 | 2.89370247 | 19.5747721 | 14.8488307 | 9.27872303 | 0 |
| 0.00949032 | 0.01768691 | 51.1220769 | 187.517419 | 74.2441534 | 146.139888 | 30.8908372 |
| 0.00949364 | 0.01769213 | 6.10892743 | 12.011792 | 8.35246725 | 6.95904227 | 0 |
| 0.00949658 | 0.01769661 | 2.89370247 | 15.1259603 | 13.9207788 | 11.5984038 | 0 |
| 0.00949868 | 0.01769955 | 0.64304499 | 10.4547078 | 33.409869 | 39.898509 | 0 |
| 0.00950572 | 0.01771168 | 9.96719738 | 11.7893514 | 4.64025958 | 8.35085072 | 0 |
| 0.00951886 | 0.01773519 | 5.46588243 | 21.5767375 | 5.5683115 | 10.6705315 | 0 |
| 0.00951948 | 0.01773536 | 9.00262989 | 8.8976237 | 6.49636342 | 6.95904227 | 0 |

| | | | | | | |
|---|---|---|---|---|---|---|
| 0.00952003 | 0.0177354 | 5.78740493 | 14.4586385 | 7.42441534 | 7.88691457 | 0 |
| 0.00952692 | 0.01774726 | 17.6837373 | 35.5904948 | 39.9062324 | 31.5476583 | 11.8810912 |
| 0.00953677 | 0.01776462 | 37.618132 | 29.3621582 | 21.3451941 | 17.6295737 | 140.196876 |
| 0.00955858 | 0.01780427 | 6.75197242 | 13.123995 | 17.6329864 | 4.17542536 | 0 |
| 0.00956491 | 0.01781507 | 7.71653991 | 56.7223511 | 0 | 139.644782 | 0 |
| 0.00956838 | 0.01782055 | 4.50131495 | 10.899589 | 8.35246725 | 12.0623399 | 0 |
| 0.00957036 | 0.01782325 | 5.14435994 | 11.1220296 | 2.78415575 | 0 | 28.5146189 |
| 0.00957787 | 0.01783624 | 351.102566 | 1039.68733 | 871.44075 | 849.467093 | 130.692003 |
| 0.00958009 | 0.01783939 | 5.46588243 | 1.77952474 | 3.71220767 | 1.39180845 | 26.1384007 |
| 0.00958676 | 0.01785082 | 6.43044992 | 2.8917277 | 1.85610383 | 5.56723382 | 2.37621824 |
| 0.00958883 | 0.01785369 | 29.2585471 | 38.9271037 | 0 | 81.6527626 | 0 |
| 0.0095895 | 0.01785395 | 0 | 91.4230835 | 71.4599976 | 29.6919137 | 0 |
| 0.00959368 | 0.01786075 | 5.14435994 | 14.6810791 | 0 | 0 | 92.6725115 |
| 0.00959708 | 0.01786609 | 0 | 138.358048 | 89.092984 | 10.2065953 | 0 |
| 0.00960413 | 0.01787822 | 4.82283744 | 25.1357869 | 6.49636342 | 9.74265918 | 0 |
| 0.00960575 | 0.01788024 | 7.71653991 | 40.929069 | 34.3379209 | 30.1558498 | 7.12865473 |
| 0.00961538 | 0.01789719 | 45.0131495 | 109.218331 | 128.071165 | 147.531696 | 23.7621824 |
| 0.00961896 | 0.01790286 | 11.2532874 | 6.45077718 | 13.9207788 | 5.10329766 | 0 |
| 0.00964393 | 0.01794833 | 97.7428388 | 151.482043 | 73.3161014 | 109.952868 | 33.2670554 |
| 0.00964658 | 0.01795228 | 26.3648447 | 44.2656779 | 0 | 78.8691457 | 0 |
| 0.00965204 | 0.01796084 | 11.5748099 | 7.56298014 | 3.71220767 | 21.8049991 | 0 |
| 0.00965225 | 0.01796084 | 72.3425616 | 23.5787028 | 0 | 53.8165936 | 0 |
| 0.00966078 | 0.01797573 | 2.57217997 | 11.5669108 | 26.9135056 | 10.6705315 | 0 |
| 0.00967381 | 0.01799897 | 2.25065747 | 3.33660889 | 0.92805192 | 0.9278723 | 0 |
| 0.00967527 | 0.01800068 | 38.904222 | 54.053064 | 0 | 45.929679 | 0 |
| 0.009683 | 0.01801407 | 0 | 78.0766479 | 70.5319457 | 36.650956 | 0 |
| 0.00969172 | 0.0180293 | 28.9370247 | 102.989994 | 97.4454513 | 87.2199964 | 23.7621824 |
| 0.00969476 | 0.01803396 | 39.2257445 | 123.676969 | 270.063108 | 119.695527 | 30.8908372 |
| 0.0097002 | 0.01804308 | 9.32415239 | 5.56101481 | 8.35246725 | 8.81478687 | 0 |
| 0.00970698 | 0.01805469 | 1.28608998 | 19.3523315 | 16.7049345 | 22.2689353 | 0 |
| 0.00971543 | 0.01806942 | 3.21522496 | 27.3601929 | 12.9927268 | 7.42297842 | 0 |
| 0.00973787 | 0.01811014 | 262.683879 | 21.1318563 | 0 | 90.9314857 | 4.75243649 |
| 0.0097411 | 0.01811515 | 10.9317649 | 40.0393066 | 187.466487 | 0 | 0 |
| 0.00976942 | 0.01816681 | 0 | 0 | 0 | 0 | 2.37621824 |
| 0.00978109 | 0.01818751 | 2.57217997 | 10.899589 | 19.4890903 | 13.9180845 | 0 |
| 0.00980205 | 0.01822547 | 24.1141872 | 16.4606038 | 21.3451941 | 17.6295737 | 80.7914203 |
| 0.00980453 | 0.01822907 | 12.2178549 | 14.2361979 | 25.0574018 | 18.5574461 | 4.75243649 |
| 0.00980641 | 0.01823156 | 1.60761248 | 5.33857422 | 52.8989593 | 19.4853184 | 0 |
| 0.00980797 | 0.01823346 | 19.6128723 | 11.3444702 | 3.71220767 | 7.88691457 | 0 |
| 0.00981874 | 0.01825248 | 23.7926647 | 66.2872965 | 0 | 57.0641466 | 0 |
| 0.00982789 | 0.01826847 | 221.207477 | 442.87922 | 412.055051 | 408.263813 | 0 |
| 0.00983816 | 0.01828655 | 7.07349491 | 7.56298014 | 8.35246725 | 7.42297842 | 0 |
| 0.00984589 | 0.01829992 | 300.623534 | 430.422546 | 370.292715 | 449.090194 | 0 |

| | | | | | | |
|---|---|---|---|---|---|---|
| 0.00984755 | 0.01830097 | 3.21522496 | 13.3464355 | 9.28051917 | 16.7017014 | 0 |
| 0.00984721 | 0.01830097 | 1.92913498 | 31.1416829 | 19.4890903 | 7.88691457 | 0 |
| 0.00985553 | 0.01831479 | 312.519866 | 617.272644 | 333.170638 | 521.000298 | 2.37621824 |
| 0.0098604 | 0.01832283 | 0 | 19.5747721 | 0 | 0 | 125.939567 |
| 0.00986364 | 0.01832785 | 3.53674746 | 2.66928711 | 48.2586997 | 19.9492545 | 0 |
| 0.00988364 | 0.01836399 | 16.3976473 | 93.4250488 | 0 | 52.4247851 | 0 |
| 0.00988465 | 0.01836484 | 1.60761248 | 22.6889404 | 16.7049345 | 14.3820207 | 0 |
| 0.00989329 | 0.01837989 | 0 | 59.1691976 | 60.3233746 | 57.5280828 | 0 |
| 0.00990084 | 0.01839289 | 0 | 90.7557617 | 85.3807764 | 21.8049991 | 0 |
| 0.00990435 | 0.01839839 | 0 | 57.834554 | 55.683115 | 63.5592527 | 0 |
| 0.00990714 | 0.01840258 | 0.64304499 | 0 | 0 | 0 | 0 |
| 0.00991038 | 0.01840757 | 4.50131495 | 9.12006429 | 10.2085711 | 10.6705315 | 0 |
| 0.00992436 | 0.01843251 | 35.6889971 | 23.1338216 | 54.7550631 | 19.4853184 | 59.4054561 |
| 0.00993993 | 0.01846041 | 3.85826995 | 18.4625692 | 6.49636342 | 14.8459568 | 0 |
| 0.00995893 | 0.01849468 | 4.50131495 | 20.4645345 | 4.64025958 | 17.6295737 | 0 |
| 0.00997133 | 0.01851668 | 10.9317649 | 15.5708415 | 8.35246725 | 16.7017014 | 0 |
| 0.00997931 | 0.01853049 | 11.8963324 | 12.4566732 | 24.1293498 | 26.4443606 | 4.75243649 |
| 0.00999082 | 0.01855083 | 0.96456749 | 20.9094157 | 16.7049345 | 26.9082968 | 0 |
| 0.00999303 | 0.01855392 | 8.6811074 | 12.9015544 | 3.71220767 | 12.5262761 | 0 |
| 0.00999444 | 0.01855449 | 5.78740493 | 9.34250488 | 14.8488307 | 6.03116997 | 0 |
| 0.009994 | 0.01855449 | 0 | 15.3484009 | 0 | 0 | 90.2962933 |
| 0.00999798 | 0.01856004 | 0 | 0 | 0 | 0 | 4.75243649 |
| 0.01001396 | 0.01858867 | 14.1469898 | 40.929069 | 0 | 120.623399 | 0 |
| 0.01003161 | 0.01862041 | 10.6102424 | 8.8976237 | 4.64025958 | 9.27872303 | 0 |
| 0.01003592 | 0.01862739 | 190.984363 | 354.792745 | 291.408302 | 328.466795 | 109.306039 |
| 0.01003979 | 0.01863353 | 6.43044992 | 14.6810791 | 9.28051917 | 5.56723382 | 0 |
| 0.01005803 | 0.01866635 | 7.39501741 | 17.7952474 | 8.35246725 | 5.10329766 | 0 |
| 0.01007301 | 0.0186921 | 0 | 0 | 0 | 0 | 57.0292378 |
| 0.01007301 | 0.0186921 | 0 | 0 | 0 | 0 | 57.0292378 |
| 0.01007519 | 0.01869512 | 31.5092046 | 45.1554403 | 0 | 62.6313804 | 0 |
| 0.01008225 | 0.01870718 | 4.50131495 | 7.56298014 | 23.2012979 | 0 | 38.0194919 |
| 0.01008503 | 0.01871027 | 0.3215225 | 0.22244059 | 0 | 0.46393615 | 45.1481466 |
| 0.0100845 | 0.01871027 | 6.75197242 | 17.5728068 | 6.49636342 | 6.95904227 | 0 |
| 0.0100899 | 0.01871828 | 74.2716966 | 129.015544 | 127.143113 | 134.541484 | 35.6432737 |
| 0.01010679 | 0.01874858 | 89.7047764 | 42.4861532 | 0 | 21.341063 | 0 |
| 0.01011086 | 0.01875509 | 22.1850522 | 89.1986776 | 0 | 41.2903175 | 0 |
| 0.01011536 | 0.0187624 | 6.10892743 | 8.67518311 | 6.49636342 | 11.1344676 | 0 |
| 0.01014091 | 0.01880875 | 28.6155022 | 65.1750936 | 0 | 46.8575513 | 0 |
| 0.01017121 | 0.01886391 | 5.78740493 | 16.0157227 | 6.49636342 | 8.35085072 | 0 |
| 0.01017629 | 0.01887229 | 0 | 60.05896 | 63.1075304 | 51.4969128 | 0 |
| 0.0101801 | 0.01887832 | 22.8280972 | 21.5767375 | 16.7049345 | 17.6295737 | 21.3859642 |
| 0.01018809 | 0.01889211 | 1.92913498 | 4.44881185 | 1.85610383 | 2.78361691 | 11.8810912 |
| 0.01019244 | 0.01889912 | 26.3648447 | 50.7164551 | 33.409869 | 42.6821259 | 4.75243649 |

| | | | | | | |
|---|---|---|---|---|---|---|
| 0.0101934 | 0.01889986 | 59.4816618 | 157.932821 | 119.718697 | 141.964462 | 59.4054561 |
| 0.01020339 | 0.01891734 | 13.8254673 | 2.8917277 | 39.9062324 | 5.56723382 | 0 |
| 0.01021755 | 0.01894255 | 0 | 0 | 0.92805192 | 0.46393615 | 0 |
| 0.01022033 | 0.01894666 | 0 | 0 | 0 | 0 | 9.50487297 |
| 0.01022933 | 0.0189623 | 0 | 6.89565837 | 13.9207788 | 0 | 57.0292378 |
| 0.01023648 | 0.01897359 | 6.75197242 | 10.0098267 | 4.64025958 | 17.6295737 | 0 |
| 0.01023655 | 0.01897359 | 18.6483048 | 12.2342326 | 12.9927268 | 8.81478687 | 21.3859642 |
| 0.01023778 | 0.01897483 | 4.50131495 | 88.7537964 | 0 | 117.839782 | 0 |
| 0.01023858 | 0.01897526 | 25.4002772 | 6.22833659 | 4.64025958 | 9.74265918 | 0 |
| 0.01024021 | 0.01897725 | 1.28608998 | 25.3582275 | 11.136623 | 24.1246799 | 0 |
| 0.0102438 | 0.01898285 | 80.7021465 | 129.460425 | 118.790645 | 111.808612 | 61.7816743 |
| 0.01026236 | 0.0190162 | 211.24028 | 59.8365194 | 46.4025958 | 51.4969128 | 23.7621824 |
| 0.01026796 | 0.01902534 | 7.39501741 | 12.4566732 | 13.9207788 | 22.2689353 | 2.37621824 |
| 0.01026843 | 0.01902534 | 0.3215225 | 0.44488118 | 0 | 0.46393615 | 0 |
| 0.01028035 | 0.01904639 | 17.3622148 | 58.7243164 | 0 | 77.0134011 | 0 |
| 0.01029611 | 0.01907454 | 0 | 71.1809896 | 81.6685687 | 30.1558498 | 0 |
| 0.01029692 | 0.01907499 | 0 | 104.102197 | 32.4818171 | 50.1051043 | 0 |
| 0.01030366 | 0.01908642 | 3.21522496 | 40.7066284 | 12.0646749 | 5.56723382 | 0 |
| 0.01031788 | 0.01911172 | 4.82283744 | 7.56298014 | 12.0646749 | 9.74265918 | 0 |
| 0.01032121 | 0.01911682 | 43.405537 | 84.9723063 | 38.0501286 | 0 | 0 |
| 0.01032749 | 0.0191274 | 189.37675 | 324.318384 | 241.293498 | 329.394667 | 80.7914203 |
| 0.01033357 | 0.0191376 | 7.39501741 | 11.3444702 | 7.42441534 | 6.03116997 | 0 |
| 0.0103739 | 0.01921118 | 20.5774398 | 119.450598 | 0 | 27.8361691 | 0 |
| 0.01037444 | 0.01921118 | 87.4541189 | 102.100232 | 201.387266 | 196.244992 | 40.3957101 |
| 0.01038479 | 0.01922929 | 0.3215225 | 14.2361979 | 59.3953227 | 23.6607437 | 0 |
| 0.01039229 | 0.01924213 | 2.57217997 | 10.899589 | 13.9207788 | 17.6295737 | 0 |
| 0.01039509 | 0.01924625 | 139.862286 | 4.44881185 | 0 | 69.5904227 | 0 |
| 0.01041611 | 0.01928199 | 50.1575094 | 38.0373413 | 74.2441534 | 0 | 0 |
| 0.01041542 | 0.01928199 | 35.3674746 | 56.7223511 | 70.5319457 | 0 | 0 |
| 0.01041564 | 0.01928199 | 27.6509347 | 88.976237 | 53.8270112 | 0 | 0 |
| 0.01041936 | 0.01928694 | 3.21522496 | 21.3542969 | 10.2085711 | 9.74265918 | 0 |
| 0.01042024 | 0.01928751 | 46.6207619 | 114.112024 | 98.3735032 | 167.017014 | 30.8908372 |
| 0.01042957 | 0.01930363 | 30.2231146 | 103.212435 | 0 | 23.6607437 | 0 |
| 0.0104301 | 0.01930363 | 6.10892743 | 6.22833659 | 8.35246725 | 13.4541484 | 0 |
| 0.01043104 | 0.01930431 | 46.9422844 | 42.041272 | 0 | 43.6099982 | 0 |
| 0.01044306 | 0.01932549 | 0 | 21.1318563 | 71.4599976 | 100.674145 | 0 |
| 0.01045086 | 0.01933886 | 23.4711422 | 52.7184204 | 0 | 65.8789335 | 0 |
| 0.01046358 | 0.01936133 | 37.618132 | 47.6022868 | 37.1220767 | 35.2591475 | 7.12865473 |
| 0.01048493 | 0.01939977 | 2.57217997 | 18.9074504 | 15.7768826 | 9.27872303 | 0 |
| 0.01048906 | 0.01940634 | 104.816334 | 30.02948 | 16.7049345 | 34.3312752 | 0 |
| 0.01049554 | 0.01941727 | 357.854538 | 616.382882 | 585.60076 | 633.272847 | 0 |
| 0.01050152 | 0.01942726 | 6.10892743 | 22.4664998 | 2.78415575 | 19.9492545 | 0 |
| 0.01050741 | 0.01943708 | 51.7651219 | 129.905306 | 73.3161014 | 96.0347833 | 14.2573095 |

| | | | | | | |
|---|---|---|---|---|---|---|
| 0.01051615 | 0.01945218 | 68.8058142 | 186.627657 | 287.696094 | 246.814032 | 38.0194919 |
| 0.01051704 | 0.01945277 | 31.8307271 | 56.7223511 | 0 | 46.3936151 | 0 |
| 0.01052105 | 0.01945911 | 38.5826995 | 32.0314453 | 0 | 66.8068058 | 0 |
| 0.0105261 | 0.01946524 | 6.10892743 | 9.34250488 | 13.9207788 | 5.56723382 | 0 |
| 0.01052598 | 0.01946524 | 51.7651219 | 37.8149007 | 30.6257133 | 37.5788283 | 66.5341108 |
| 0.01052519 | 0.01946524 | 7.39501741 | 5.7834554 | 7.42441534 | 12.5262761 | 0 |
| 0.01054128 | 0.01949224 | 13.5039448 | 94.3148112 | 0 | 54.2805297 | 0 |
| 0.01054324 | 0.0194948 | 22.8280972 | 9.78738607 | 3.71220767 | 7.88691457 | 0 |
| 0.01055536 | 0.01951613 | 31.8307271 | 62.2833659 | 0 | 41.7542536 | 0 |
| 0.01056117 | 0.01952493 | 0 | 0 | 0 | 0 | 7.12865473 |
| 0.01056128 | 0.01952493 | 29.2585471 | 8.23030192 | 3.71220767 | 8.35085072 | 0 |
| 0.01056782 | 0.01953595 | 0 | 108.551009 | 139.207788 | 78.8691457 | 2.37621824 |
| 0.01057192 | 0.01954246 | 26.3648447 | 5.7834554 | 3.71220767 | 12.9902122 | 0 |
| 0.0105731 | 0.01954357 | 4.50131495 | 10.899589 | 11.136623 | 7.88691457 | 0 |
| 0.01057964 | 0.01955458 | 26.3648447 | 25.8031087 | 0 | 104.84957 | 0 |
| 0.0105985 | 0.01958837 | 29.5800696 | 39.5944255 | 0 | 69.1264865 | 0 |
| 0.01060089 | 0.01959102 | 0 | 38.2597819 | 46.4025958 | 94.6429749 | 0 |
| 0.0106011 | 0.01959102 | 0 | 0 | 0 | 5.10329766 | 90.2962933 |
| 0.01060204 | 0.01959167 | 1.92913498 | 27.3601929 | 18.5610383 | 8.35085072 | 0 |
| 0.01060658 | 0.01959899 | 14.7900348 | 42.7085937 | 0 | 106.241379 | 0 |
| 0.01062066 | 0.01962393 | 60.7677518 | 0 | 78.8844129 | 46.8575513 | 0 |
| 0.01062209 | 0.01962549 | 131.502701 | 288.283008 | 215.308045 | 250.525522 | 42.7719284 |
| 0.0106251 | 0.01962998 | 164.619518 | 594.361263 | 686.758419 | 782.196351 | 106.929821 |
| 0.01062667 | 0.0196318 | 3.53674746 | 8.8976237 | 15.7768826 | 11.1344676 | 0 |
| 0.01062972 | 0.01963635 | 2.25065747 | 7.56298014 | 23.2012979 | 19.9492545 | 0 |
| 0.01063665 | 0.01964807 | 31.5092046 | 43.8207967 | 0 | 59.3838274 | 0 |
| 0.0106495 | 0.01966965 | 5.46588243 | 8.45274251 | 12.0646749 | 6.95904227 | 0 |
| 0.01064945 | 0.01966965 | 7.39501741 | 8.23030192 | 5.5683115 | 10.2065953 | 0 |
| 0.01065287 | 0.0196748 | 7.39501741 | 7.34053955 | 5.5683115 | 13.9180845 | 0 |
| 0.01067964 | 0.01972316 | 17.6837373 | 21.7991781 | 8.35246725 | 2.31968076 | 0 |
| 0.01068095 | 0.01972449 | 33.1168171 | 34.4782918 | 0 | 70.0543588 | 0 |
| 0.01068581 | 0.01973239 | 32.1522496 | 20.4645345 | 23.2012979 | 55.208402 | 0 |
| 0.01069891 | 0.0197555 | 25.7217997 | 101.877791 | 96.5173994 | 95.5708472 | 23.7621824 |
| 0.01070053 | 0.01975739 | 25.4002772 | 77.4093262 | 0 | 39.4345729 | 0 |
| 0.01070645 | 0.01976724 | 1.60761248 | 15.7932821 | 17.6329864 | 17.1656376 | 0 |
| 0.01071711 | 0.01978584 | 37.9396545 | 72.2931925 | 0 | 28.7640414 | 0 |
| 0.01071956 | 0.01978929 | 38.261177 | 37.3700195 | 0 | 57.0641466 | 0 |
| 0.01072076 | 0.0197904 | 23.1496197 | 18.6850098 | 17.6329864 | 16.7017014 | 7.12865473 |
| 0.01072278 | 0.01979306 | 130.216611 | 145.698588 | 164.265189 | 130.829995 | 76.0389838 |
| 0.01073179 | 0.0198086 | 716.352121 | 1606.91084 | 1399.50229 | 1532.84504 | 0 |
| 0.01073911 | 0.01982101 | 2.57217997 | 13.5688761 | 17.6329864 | 10.6705315 | 0 |
| 0.01074163 | 0.01982458 | 15.7546023 | 7.11809896 | 5.5683115 | 7.42297842 | 0 |
| 0.01074458 | 0.01982894 | 79.4160565 | 292.064498 | 242.22155 | 263.051798 | 11.8810912 |

| | | | | | | |
|---|---|---|---|---|---|---|
| 0.01075016 | 0.01983814 | 240.177305 | 591.469535 | 478.874789 | 527.95934 | 0 |
| 0.01077634 | 0.01988537 | 0 | 69.6239054 | 35.2659728 | 69.1264865 | 0 |
| 0.01077918 | 0.01988952 | 19.6128723 | 71.1809896 | 0 | 53.8165936 | 0 |
| 0.01078256 | 0.01989467 | 4.50131495 | 13.123995 | 29.6976613 | 20.8771268 | 2.37621824 |
| 0.01078862 | 0.01990476 | 0 | 57.3896728 | 49.1867516 | 62.6313804 | 0 |
| 0.01079481 | 0.01991509 | 0 | 69.846346 | 68.6758419 | 35.2591475 | 0 |
| 0.01079614 | 0.01991645 | 0 | 31.3641235 | 98.3735032 | 51.0329766 | 0 |
| 0.01080395 | 0.01992977 | 0.3215225 | 0 | 1.85610383 | 0.46393615 | 0 |
| 0.01081325 | 0.01994583 | 3.53674746 | 13.123995 | 8.35246725 | 13.9180845 | 0 |
| 0.01081682 | 0.01995132 | 13.1824223 | 5.11613363 | 25.0574018 | 4.17542536 | 0 |
| 0.0108232 | 0.01996198 | 7.71653991 | 11.1220296 | 4.64025958 | 9.74265918 | 0 |
| 0.01082604 | 0.01996613 | 47.2638069 | 60.9487223 | 0 | 27.3722329 | 0 |
| 0.01083128 | 0.0199747 | 41.7979245 | 99.4309448 | 90.0210359 | 101.602017 | 7.12865473 |
| 0.01083463 | 0.01997978 | 86.168029 | 120.340361 | 135.49558 | 131.757867 | 45.1481466 |
| 0.01084222 | 0.01999269 | 19.2913498 | 65.6199748 | 0 | 58.9198912 | 0 |
| 0.01086276 | 0.02002946 | 63.6614542 | 30.2519206 | 0 | 40.8263813 | 0 |
| 0.01086806 | 0.02003814 | 32.1522496 | 16.905485 | 28.7696094 | 14.3820207 | 80.7914203 |
| 0.01087711 | 0.02005372 | 7.07349491 | 67.1770589 | 146.632203 | 0 | 0 |
| 0.0108933 | 0.02008246 | 9.00262989 | 178.397355 | 39.9062324 | 0 | 0 |
| 0.01089896 | 0.0200918 | 2.25065747 | 18.017688 | 13.9207788 | 12.0623399 | 0 |
| 0.01090056 | 0.02009366 | 46.6207619 | 15.3484009 | 6.49636342 | 8.81478687 | 0 |
| 0.01091693 | 0.02012272 | 54.6588243 | 41.3739502 | 59.3953227 | 0 | 0 |
| 0.01091761 | 0.02012287 | 41.1548795 | 54.7203857 | 0 | 35.2591475 | 0 |
| 0.01092364 | 0.02013288 | 0 | 0 | 0 | 0 | 0 |
| 0.01093358 | 0.0201501 | 0 | 56.4999105 | 42.6903882 | 70.518295 | 0 |
| 0.01093453 | 0.02015076 | 25.4002772 | 62.7282471 | 0 | 48.2493597 | 0 |
| 0.01093685 | 0.02015392 | 6.43044992 | 9.78738607 | 7.42441534 | 16.7017014 | 64.1578926 |
| 0.01094931 | 0.02017467 | 5.78740493 | 10.0098267 | 12.9927268 | 5.56723382 | 0 |
| 0.01094874 | 0.02017467 | 9.64567488 | 92.757727 | 0 | 66.3428696 | 0 |
| 0.01095232 | 0.02017911 | 2.25065747 | 4.00393066 | 24.1293498 | 36.650956 | 0 |
| 0.01097045 | 0.02021091 | 3.85826995 | 4.22637126 | 3.71220767 | 2.78361691 | 16.6335277 |
| 0.01097078 | 0.02021091 | 27.0078897 | 11.5669108 | 8.35246725 | 12.5262761 | 121.18713 |
| 0.01097202 | 0.0202121 | 7.07349491 | 13.123995 | 9.28051917 | 5.10329766 | 66.5341108 |
| 0.01097382 | 0.02021429 | 3.21522496 | 17.7952474 | 9.28051917 | 11.1344676 | 0 |
| 0.01097965 | 0.02022392 | 18.6483048 | 8.45274251 | 8.35246725 | 4.17542536 | 0 |
| 0.01098037 | 0.02022414 | 8.6811074 | 4.89369303 | 0 | 1.85574461 | 40.3957101 |
| 0.01098475 | 0.0202311 | 2.89370247 | 10.899589 | 9.28051917 | 23.1968076 | 0 |
| 0.01099094 | 0.0202414 | 10.2887199 | 8.23030192 | 4.64025958 | 9.27872303 | 0 |
| 0.01099215 | 0.02024252 | 0.64304499 | 0.66732178 | 0 | 0.46393615 | 0 |
| 0.01101749 | 0.02028807 | 73.3071291 | 52.7184204 | 0 | 18.5574461 | 0 |
| 0.01102011 | 0.02029179 | 2.25065747 | 10.0098267 | 15.7768826 | 19.4853184 | 0 |
| 0.01104277 | 0.02033241 | 0 | 32.9212077 | 56.6111669 | 84.9003157 | 0 |
| 0.01104891 | 0.0203426 | 47.5853294 | 63.3955688 | 43.6184401 | 0 | 0 |

| | | | | | | |
|---|---|---|---|---|---|---|
| 0.01105146 | 0.02034617 | 6.10892743 | 10.2322673 | 5.5683115 | 10.6705315 | 0 |
| 0.01105414 | 0.02035 | 11.2532874 | 4.44881185 | 12.9927268 | 6.95904227 | 0 |
| 0.01106118 | 0.02036185 | 0 | 53.3857422 | 26.9135056 | 101.602017 | 0 |
| 0.01106373 | 0.02036543 | 30.8661596 | 74.2951579 | 0 | 32.4755306 | 0 |
| 0.01107047 | 0.02037672 | 3.21522496 | 14.0137573 | 13.9207788 | 8.81478687 | 0 |
| 0.01107739 | 0.02038833 | 159.153636 | 189.741825 | 166.121293 | 205.523715 | 38.0194919 |
| 0.01108128 | 0.02039437 | 30.5446371 | 60.05896 | 0 | 41.7542536 | 0 |
| 0.01108666 | 0.02040317 | 0 | 109.663212 | 30.6257133 | 42.6821259 | 0 |
| 0.01108974 | 0.02040772 | 10.6102424 | 8.8976237 | 3.71220767 | 12.5262761 | 0 |
| 0.0110922 | 0.02041112 | 3.21522496 | 19.3523315 | 6.49636342 | 16.2377653 | 0 |
| 0.01109777 | 0.02042026 | 4.50131495 | 15.3484009 | 7.42441534 | 9.27872303 | 0 |
| 0.01109953 | 0.02042238 | 9.00262989 | 9.78738607 | 3.71220767 | 13.9180845 | 0 |
| 0.01110148 | 0.02042484 | 0 | 68.7341431 | 40.8342843 | 58.9198912 | 0 |
| 0.01111232 | 0.02044368 | 62.6968867 | 82.7479004 | 117.862593 | 109.488932 | 30.8908372 |
| 0.01111743 | 0.02045133 | 30.8661596 | 210.651241 | 282.127783 | 237.999246 | 0 |
| 0.0111181 | 0.02045133 | 28.6155022 | 51.828658 | 81.6685687 | 0 | 0 |
| 0.0111183 | 0.02045133 | 145.971213 | 451.109521 | 369.364663 | 344.70456 | 16.6335277 |
| 0.01112228 | 0.02045752 | 6.43044992 | 10.6771484 | 5.5683115 | 9.27872303 | 0 |
| 0.01112357 | 0.02045878 | 13.8254673 | 2.22440592 | 14.8488307 | 14.3820207 | 0 |
| 0.01112669 | 0.02046341 | 3.21522496 | 21.5767375 | 23.2012979 | 4.63936151 | 0 |
| 0.0111314 | 0.02047095 | 54.3373018 | 224.664998 | 128.071165 | 162.841589 | 38.0194919 |
| 0.01113494 | 0.02047634 | 153.044708 | 262.035018 | 291.408302 | 297.383073 | 78.415202 |
| 0.01113784 | 0.02048055 | 0 | 95.6494547 | 66.819738 | 21.8049991 | 0 |
| 0.01114171 | 0.02048655 | 0 | 45.3778809 | 77.956361 | 45.929679 | 0 |
| 0.01114471 | 0.02048982 | 3.85826995 | 20.2420939 | 7.42441534 | 10.2065953 | 0 |
| 0.01114419 | 0.02048982 | 46.9422844 | 29.1397176 | 25.0574018 | 34.7952113 | 19.0097459 |
| 0.01115505 | 0.02050772 | 201.916128 | 491.81615 | 385.141546 | 473.67881 | 0 |
| 0.01116624 | 0.02052716 | 67.5197242 | 0 | 49.1867516 | 63.0953166 | 0 |
| 0.01116689 | 0.02052724 | 471.351979 | 774.760583 | 701.607249 | 770.597947 | 128.315785 |
| 0.01117452 | 0.02054014 | 0 | 63.3955688 | 50.1148035 | 52.4247851 | 0 |
| 0.01118432 | 0.02055704 | 3.85826995 | 14.6810791 | 12.0646749 | 7.42297842 | 0 |
| 0.01119123 | 0.02056862 | 1.28608998 | 0.22244059 | 0 | 0.46393615 | 33.2670554 |
| 0.01121145 | 0.02060464 | 10.2887199 | 12.011792 | 4.64025958 | 6.95904227 | 0 |
| 0.01121866 | 0.02061677 | 36.0105196 | 73.1829549 | 47.3306478 | 0 | 0 |
| 0.01122454 | 0.02062644 | 49.8359869 | 50.2715739 | 0 | 30.1558498 | 0 |
| 0.01122649 | 0.0206289 | 2.89370247 | 11.3444702 | 13.9207788 | 12.0623399 | 0 |
| 0.01122966 | 0.0206336 | 21.8635297 | 2.22440592 | 34.3379209 | 4.63936151 | 0 |
| 0.01123067 | 0.02063433 | 38.5826995 | 48.4920492 | 0 | 40.8263813 | 0 |
| 0.01123164 | 0.020635 | 48.8714194 | 16.6830444 | 0 | 82.1166988 | 0 |
| 0.01124231 | 0.02065346 | 0 | 38.9271037 | 85.3807764 | 46.8575513 | 0 |
| 0.0112464 | 0.02065985 | 7.07349491 | 14.0137573 | 4.64025958 | 9.27872303 | 0 |
| 0.01125 | 0.02066534 | 3.21522496 | 8.8976237 | 15.7768826 | 12.0623399 | 0 |
| 0.01125572 | 0.02067471 | 27.0078897 | 44.0432373 | 45.4745439 | 41.2903175 | 114.058476 |

| | | | | | | |
|---|---|---|---|---|---|---|
| 0.011262 | 0.02068511 | 29.2585471 | 68.0668213 | 89.092984 | 64.0231889 | 21.3859642 |
| 0.01126264 | 0.02068517 | 21.5420072 | 30.2519206 | 0 | 99.2823364 | 0 |
| 0.01126683 | 0.02069173 | 2.57217997 | 13.3464355 | 11.136623 | 15.7738291 | 0 |
| 0.01127339 | 0.02070265 | 12.5393773 | 9.56494547 | 6.49636342 | 5.10329766 | 0 |
| 0.01127881 | 0.02071148 | 13.8254673 | 43.5983561 | 0 | 100.674145 | 0 |
| 0.01128072 | 0.02071384 | 129.895088 | 308.747542 | 212.523889 | 246.350096 | 7.12865473 |
| 0.01129448 | 0.02073798 | 16.3976473 | 2.22440592 | 10.2085711 | 18.0935099 | 0 |
| 0.01129755 | 0.02074248 | 0 | 36.4802572 | 64.0355823 | 67.7346781 | 0 |
| 0.01131546 | 0.02077423 | 8.6811074 | 8.67518311 | 5.5683115 | 7.42297842 | 0 |
| 0.01131745 | 0.02077676 | 8.6811074 | 10.899589 | 14.8488307 | 7.88691457 | 16.6335277 |
| 0.01135009 | 0.02083554 | 7.07349491 | 5.33857422 | 11.136623 | 5.56723382 | 49.9005831 |
| 0.01135284 | 0.02083945 | 0 | 47.8247274 | 25.9854537 | 108.561059 | 0 |
| 0.01135441 | 0.02084119 | 34.7244296 | 10.899589 | 30.6257133 | 16.2377653 | 35.6432737 |
| 0.01136201 | 0.02085401 | 20.2559173 | 60.7262817 | 0 | 57.0641466 | 0 |
| 0.0113636 | 0.0208558 | 6.43044992 | 32.4763265 | 7.42441534 | 4.63936151 | 0 |
| 0.0113893 | 0.02090181 | 4.82283744 | 11.3444702 | 9.28051917 | 7.42297842 | 0 |
| 0.01139407 | 0.02090943 | 11.8963324 | 41.8188314 | 0 | 111.344676 | 0 |
| 0.01139795 | 0.0209154 | 271.364987 | 425.306413 | 365.652455 | 379.963708 | 0 |
| 0.01140315 | 0.02092381 | 47.2638069 | 0 | 111.36623 | 32.9394667 | 0 |
| 0.01141291 | 0.02094057 | 7.39501741 | 106.326603 | 0 | 62.6313804 | 0 |
| 0.0114232 | 0.02095832 | 23.1496197 | 62.0609253 | 77.956361 | 0 | 0 |
| 0.01143295 | 0.02097505 | 4.17979245 | 15.5708415 | 9.28051917 | 7.88691457 | 0 |
| 0.01144084 | 0.02098839 | 0 | 69.6239054 | 90.0210359 | 21.341063 | 0 |
| 0.01145137 | 0.02100559 | 0 | 49.8266927 | 47.3306478 | 67.2707419 | 0 |
| 0.01145147 | 0.02100559 | 38.5826995 | 43.1534749 | 0 | 44.5378705 | 0 |
| 0.01145325 | 0.02100772 | 2.25065747 | 1.55708415 | 0 | 0.46393615 | 30.8908372 |
| 0.0114605 | 0.02101987 | 25.7217997 | 3.78149007 | 3.71220767 | 19.9492545 | 0 |
| 0.01146161 | 0.02102076 | 28.9370247 | 14.9035197 | 1.85610383 | 8.81478687 | 0 |
| 0.01146574 | 0.02102718 | 3.53674746 | 12.2342326 | 14.8488307 | 7.88691457 | 0 |
| 0.01147193 | 0.0210374 | 4.82283744 | 7.56298014 | 9.28051917 | 10.6705315 | 0 |
| 0.01148248 | 0.0210556 | 17.3622148 | 39.816866 | 0 | 91.3954218 | 0 |
| 0.01148962 | 0.02106753 | 40.833357 | 63.84045 | 0 | 27.3722329 | 0 |
| 0.01149236 | 0.02107141 | 0 | 27.3601929 | 99.3015551 | 50.5690405 | 0 |
| 0.01150158 | 0.02108718 | 0 | 25.3582275 | 84.4527244 | 65.4149973 | 0 |
| 0.01150986 | 0.0211012 | 21.2204847 | 32.9212077 | 0 | 91.859358 | 0 |
| 0.01151325 | 0.02110627 | 203.52374 | 255.584241 | 223.660512 | 228.720523 | 7.12865473 |
| 0.0115194 | 0.02111638 | 45.6561944 | 4.22637126 | 0 | 5.56723382 | 104.553603 |
| 0.01153808 | 0.02114948 | 772.940081 | 2220.40199 | 1876.52098 | 2160.55066 | 0 |
| 0.01155438 | 0.02117821 | 6.10892743 | 8.67518311 | 10.2085711 | 6.03116997 | 0 |
| 0.01155553 | 0.02117917 | 49.8359869 | 93.4250488 | 116.934542 | 109.024996 | 19.0097459 |
| 0.0115607 | 0.02118749 | 217.992252 | 6.005896 | 28.7696094 | 0 | 0 |
| 0.01156736 | 0.02119853 | 6.10892743 | 11.5669108 | 4.64025958 | 12.9902122 | 0 |
| 0.01158879 | 0.02123664 | 23.1496197 | 41.5963908 | 0 | 70.9822312 | 0 |

| | | | | | | |
|---|---|---|---|---|---|---|
| 0.01159816 | 0.02125265 | 3.21522496 | 1.55708415 | 0 | 1.39180845 | 23.7621824 |
| 0.01165464 | 0.021355 | 2.89370247 | 3.33660889 | 3.71220767 | 2.78361691 | 66.5341108 |
| 0.01167706 | 0.02139491 | 20.8989622 | 7.78542074 | 6.49636342 | 5.10329766 | 0 |
| 0.01168943 | 0.0214164 | 9.96719738 | 8.23030192 | 4.64025958 | 8.81478687 | 0 |
| 0.0117024 | 0.02143798 | 2.57217997 | 28.2499552 | 9.28051917 | 10.2065953 | 0 |
| 0.01170248 | 0.02143798 | 3.21522496 | 10.2322673 | 14.8488307 | 10.2065953 | 0 |
| 0.0117114 | 0.02145315 | 7.39501741 | 8.67518311 | 5.5683115 | 8.35085072 | 0 |
| 0.01172997 | 0.02148599 | 20.2559173 | 60.2814006 | 0 | 54.7444659 | 0 |
| 0.01173578 | 0.02149548 | 144.042078 | 368.13918 | 269.135056 | 267.691159 | 4.75243649 |
| 0.01174379 | 0.02150897 | 25.0787547 | 14.2361979 | 12.9927268 | 12.9902122 | 19.0097459 |
| 0.01174757 | 0.02151472 | 0 | 60.5038411 | 50.1148035 | 50.5690405 | 0 |
| 0.01177826 | 0.02156976 | 33.7598621 | 88.5313558 | 0 | 20.8771268 | 0 |
| 0.01178105 | 0.02157369 | 42.4409695 | 68.0668213 | 0 | 23.1968076 | 0 |
| 0.01178573 | 0.02158108 | 85.524984 | 52.940861 | 47.3306478 | 45.929679 | 190.097459 |
| 0.01179007 | 0.02158785 | 92.2769564 | 18.9074504 | 0 | 34.7952113 | 0 |
| 0.01179129 | 0.02158892 | 217.349207 | 995.644092 | 924.339709 | 999.782406 | 0 |
| 0.01180539 | 0.02161355 | 0 | 56.2774699 | 67.7477899 | 38.9706367 | 0 |
| 0.01180697 | 0.02161527 | 25.7217997 | 49.6042521 | 0 | 53.8165936 | 0 |
| 0.01181675 | 0.02163082 | 52.7296894 | 34.0334106 | 0 | 38.9706367 | 0 |
| 0.01181625 | 0.02163082 | 0 | 95.6494547 | 49.1867516 | 28.3001052 | 0 |
| 0.01182022 | 0.021636 | 27.9724572 | 64.7302124 | 0 | 37.5788283 | 0 |
| 0.0118236 | 0.02164101 | 14.7900348 | 16.0157227 | 21.3451941 | 22.7328714 | 7.12865473 |
| 0.01185097 | 0.02168992 | 3.21522496 | 21.5767375 | 10.2085711 | 8.35085072 | 0 |
| 0.01185966 | 0.02170465 | 15.7546023 | 4.00393066 | 6.49636342 | 12.0623399 | 0 |
| 0.01186235 | 0.02170809 | 3.21522496 | 15.1259603 | 11.136623 | 9.27872303 | 0 |
| 0.01186283 | 0.02170809 | 4.17979245 | 3.78149007 | 0 | 5.56723382 | 7.12865473 |
| 0.01186451 | 0.02170998 | 23.4711422 | 22.0216187 | 144.776099 | 0 | 0 |
| 0.01186665 | 0.02171271 | 20.5774398 | 51.3837769 | 0 | 62.1674443 | 0 |
| 0.01187169 | 0.02172076 | 8.3595849 | 15.3484009 | 18.5610383 | 13.9180845 | 26.1384007 |
| 0.01188273 | 0.02173859 | 33.4383396 | 48.7144897 | 0 | 42.6821259 | 0 |
| 0.0118825 | 0.02173859 | 23.1496197 | 26.0255493 | 23.2012979 | 32.9394667 | 11.8810912 |
| 0.01188697 | 0.02174516 | 0 | 74.0727173 | 38.0501286 | 51.4969128 | 0 |
| 0.01188853 | 0.02174684 | 13.8254673 | 54.9428263 | 0 | 77.9412734 | 0 |
| 0.01188972 | 0.02174782 | 9.96719738 | 4.89369303 | 5.5683115 | 18.5574461 | 0 |
| 0.01189151 | 0.02174992 | 45.0131495 | 16.2381632 | 116.934542 | 0 | 0 |
| 0.01190559 | 0.0217745 | 21.8635297 | 48.9369303 | 0 | 61.7035081 | 0 |
| 0.01193863 | 0.02183373 | 6.10892743 | 11.1220296 | 17.6329864 | 4.17542536 | 0 |
| 0.0119447 | 0.02184365 | 0 | 37.8149007 | 33.409869 | 102.065953 | 0 |
| 0.01196682 | 0.0218829 | 0 | 23.5787028 | 57.5392189 | 93.2511664 | 0 |
| 0.01198269 | 0.02191075 | 209.632667 | 369.918705 | 386.069597 | 346.096369 | 104.553603 |
| 0.01198348 | 0.02191099 | 16.3976473 | 61.1711629 | 0 | 61.7035081 | 0 |
| 0.01199212 | 0.0219256 | 1.92913498 | 21.3542969 | 12.9927268 | 12.0623399 | 0 |
| 0.01199797 | 0.02193437 | 4.50131495 | 7.78542074 | 9.28051917 | 11.1344676 | 0 |

| | | | | | | |
|---|---|---|---|---|---|---|
| 0.01199822 | 0.02193437 | 3.53674746 | 3.33660889 | 13.9207788 | 42.6821259 | 0 |
| 0.01200491 | 0.02194541 | 0 | 50.4940145 | 39.9062324 | 71.4461673 | 0 |
| 0.01201258 | 0.02195824 | 41.476402 | 45.1554403 | 0 | 36.650956 | 0 |
| 0.01201785 | 0.02196667 | 108.353081 | 287.170805 | 247.789862 | 282.537116 | 80.7914203 |
| 0.012028 | 0.02198404 | 28.2939797 | 31.5865641 | 0 | 72.3740396 | 0 |
| 0.01202968 | 0.02198591 | 41.1548795 | 87.4191528 | 64.0355823 | 51.9608489 | 26.1384007 |
| 0.01205318 | 0.02202766 | 4.50131495 | 3.33660889 | 23.2012979 | 19.0213822 | 0 |
| 0.01205445 | 0.02202878 | 5.14435994 | 7.34053955 | 8.35246725 | 10.2065953 | 0 |
| 0.01205543 | 0.02202939 | 0 | 24.6909058 | 112.294282 | 42.2181898 | 0 |
| 0.01205655 | 0.02203023 | 51.7651219 | 21.5767375 | 0 | 57.9920189 | 0 |
| 0.01206159 | 0.02203824 | 14.1469898 | 31.5865641 | 28.7696094 | 23.6607437 | 9.50487297 |
| 0.01206443 | 0.02204223 | 0 | 55.8325887 | 68.6758419 | 37.1148921 | 0 |
| 0.01207026 | 0.02205169 | 511.863814 | 135.911202 | 115.078438 | 159.1301 | 836.428822 |
| 0.01207643 | 0.02206177 | 17.0406923 | 6.67321777 | 3.71220767 | 11.5984038 | 0 |
| 0.01208598 | 0.02207801 | 51.7651219 | 19.1298909 | 27.8415575 | 21.341063 | 35.6432737 |
| 0.01209306 | 0.02208975 | 34.7244296 | 26.4704305 | 0 | 70.0543588 | 0 |
| 0.01209705 | 0.02209584 | 5.14435994 | 10.0098267 | 7.42441534 | 8.35085072 | 0 |
| 0.01210251 | 0.02210423 | 51.7651219 | 31.1416829 | 67.7477899 | 0 | 0 |
| 0.01210296 | 0.02210423 | 6.43044992 | 7.34053955 | 8.35246725 | 6.95904227 | 0 |
| 0.01210718 | 0.02211073 | 258.182564 | 145.253707 | 133.639476 | 121.087335 | 254.255352 |
| 0.01211396 | 0.02212191 | 9.00262989 | 3.78149007 | 10.2085711 | 4.63936151 | 2.37621824 |
| 0.01212137 | 0.02213304 | 7.71653991 | 13.5688761 | 12.9927268 | 13.9180845 | 2.37621824 |
| 0.0121211 | 0.02213304 | 111.889829 | 384.154903 | 309.96934 | 344.70456 | 26.1384007 |
| 0.01212471 | 0.02213794 | 7.07349491 | 6.005896 | 16.7049345 | 6.03116997 | 0 |
| 0.01213509 | 0.0221557 | 54.3373018 | 150.36984 | 151.272462 | 169.800631 | 0 |
| 0.0121455 | 0.0221735 | 4.17979245 | 9.56494547 | 9.28051917 | 9.74265918 | 0 |
| 0.012158 | 0.02219446 | 32.7952946 | 30.2519206 | 0 | 65.4149973 | 0 |
| 0.0121583 | 0.02219446 | 16.7191698 | 44.9329997 | 0 | 78.8691457 | 0 |
| 0.01216679 | 0.02220876 | 6.10892743 | 16.4606038 | 28.7696094 | 2.31968076 | 0 |
| 0.01217118 | 0.02221555 | 0 | 0 | 0 | 0 | 0 |
| 0.01218316 | 0.02223621 | 0 | 0 | 0 | 0 | 52.2768014 |
| 0.01218963 | 0.02224682 | 14.4685123 | 50.4940145 | 111.36623 | 0 | 0 |
| 0.0122001 | 0.02226473 | 59.8031843 | 112.999821 | 98.3735032 | 90.0036134 | 30.8908372 |
| 0.01220927 | 0.02227903 | 76.5223541 | 66.5097371 | 0 | 48.7132959 | 297.02728 |
| 0.01220887 | 0.02227903 | 5.14435994 | 11.5669108 | 5.5683115 | 11.5984038 | 0 |
| 0.01221131 | 0.02228155 | 12.2178549 | 10.4547078 | 13.9207788 | 2.78361691 | 0 |
| 0.01221248 | 0.02228247 | 72.9856066 | 122.119885 | 125.287009 | 111.808612 | 21.3859642 |
| 0.01221621 | 0.02228807 | 24.1141872 | 52.0510986 | 0 | 51.4969128 | 0 |
| 0.01222458 | 0.02230213 | 73.3071291 | 195.0804 | 110.438178 | 125.726697 | 0 |
| 0.01222698 | 0.0223053 | 60.7677518 | 44.0432373 | 30.6257133 | 67.7346781 | 87.920075 |
| 0.01225658 | 0.0223576 | 9.64567488 | 5.56101481 | 6.49636342 | 8.81478687 | 0 |
| 0.01225764 | 0.0223576 | 8.6811074 | 4.67125244 | 8.35246725 | 9.74265918 | 0 |
| 0.01225745 | 0.0223576 | 55.3018693 | 0 | 37.1220767 | 82.5806349 | 0 |

| | | | | | | |
|---|---|---|---|---|---|---|
| 0.01226071 | 0.02236198 | 19.9343948 | 7.34053955 | 5.5683115 | 6.03116997 | 0 |
| 0.01227185 | 0.0223811 | 4.82283744 | 2.00196533 | 3.71220767 | 0.9278723 | 19.0097459 |
| 0.01227432 | 0.02238316 | 0 | 0 | 0 | 0 | 0 |
| 0.01227432 | 0.02238316 | 0 | 0 | 0 | 0 | 0 |
| 0.01228739 | 0.02240579 | 9.32415239 | 8.67518311 | 7.42441534 | 5.10329766 | 0 |
| 0.01228815 | 0.02240597 | 142.755988 | 299.627478 | 253.358173 | 301.094562 | 147.325531 |
| 0.01230626 | 0.02243533 | 11.2532874 | 80.3010539 | 0 | 58.4559551 | 0 |
| 0.01230515 | 0.02243533 | 14.7900348 | 56.0550293 | 0 | 69.5904227 | 0 |
| 0.01230616 | 0.02243533 | 0 | 46.7125244 | 69.6038938 | 42.6821259 | 0 |
| 0.01230844 | 0.0224381 | 0 | 49.6042521 | 41.7623363 | 67.2707419 | 0 |
| 0.0123134 | 0.02244592 | 23.4711422 | 41.1515096 | 0 | 64.9510612 | 0 |
| 0.01234088 | 0.02249479 | 3.85826995 | 19.5747721 | 6.49636342 | 10.6705315 | 0 |
| 0.01235097 | 0.02251196 | 0 | 81.1908162 | 28.7696094 | 55.208402 | 0 |
| 0.01235687 | 0.02252028 | 9.00262989 | 13.123995 | 11.136623 | 12.9902122 | 0 |
| 0.01235684 | 0.02252028 | 18.9698273 | 79.4112915 | 62.1794784 | 0 | 0 |
| 0.01236399 | 0.02253081 | 179.409553 | 5.11613363 | 0 | 27.3722329 | 0 |
| 0.01236334 | 0.02253081 | 3.53674746 | 13.3464355 | 8.35246725 | 11.1344676 | 0 |
| 0.01237117 | 0.02254268 | 82.309759 | 133.686796 | 109.510126 | 107.169251 | 52.2768014 |
| 0.01237343 | 0.02254557 | 0 | 42.4861532 | 22.273246 | 36.650956 | 206.730987 |
| 0.01238369 | 0.02256304 | 0.96456749 | 27.8050741 | 5.5683115 | 40.3624452 | 0 |
| 0.01239108 | 0.02257528 | 0 | 51.3837769 | 41.7623363 | 64.487125 | 0 |
| 0.01239433 | 0.02257999 | 4.50131495 | 8.8976237 | 12.0646749 | 7.42297842 | 0 |
| 0.01240333 | 0.02259517 | 6.43044992 | 34.2558512 | 6.49636342 | 21.8049991 | 0 |
| 0.01240459 | 0.02259624 | 1.92913498 | 19.7972127 | 9.28051917 | 17.6295737 | 0 |
| 0.01240617 | 0.02259789 | 12.2178549 | 7.34053955 | 5.5683115 | 9.74265918 | 28.5146189 |
| 0.01240848 | 0.02260088 | 0 | 34.2558512 | 78.8844129 | 48.7132959 | 0 |
| 0.0124101 | 0.0226026 | 0 | 0.22244059 | 0 | 0.46393615 | 0 |
| 0.0124437 | 0.02266257 | 10.6102424 | 24.023584 | 8.35246725 | 2.78361691 | 0 |
| 0.01244606 | 0.02266564 | 0 | 34.2558512 | 76.1002572 | 50.5690405 | 0 |
| 0.01245309 | 0.02267722 | 0 | 80.9683756 | 24.1293498 | 62.6313804 | 0 |
| 0.01246315 | 0.02269431 | 37.9396545 | 15.7932821 | 11.136623 | 12.0623399 | 2.37621824 |
| 0.01246672 | 0.02269958 | 0 | 35.8129354 | 78.8844129 | 46.3936151 | 0 |
| 0.01246784 | 0.02270039 | 6.43044992 | 17.3503662 | 3.71220767 | 11.5984038 | 0 |
| 0.01247029 | 0.02270363 | 3.21522496 | 9.78738607 | 12.0646749 | 11.5984038 | 0 |
| 0.01247431 | 0.02270971 | 0 | 61.6160441 | 41.7623363 | 53.3526574 | 0 |
| 0.0124764 | 0.02271229 | 5.14435994 | 8.67518311 | 6.49636342 | 11.5984038 | 0 |
| 0.01248689 | 0.02273015 | 9.96719738 | 79.1888509 | 0 | 63.0953166 | 0 |
| 0.01248834 | 0.02273156 | 5.46588243 | 5.11613363 | 11.136623 | 12.5262761 | 0 |
| 0.01248914 | 0.02273178 | 23.4711422 | 8.45274251 | 8.35246725 | 7.88691457 | 123.563349 |
| 0.01250255 | 0.02275472 | 16.7191698 | 42.2637126 | 0 | 79.797018 | 0 |
| 0.01250309 | 0.02275472 | 18.0052598 | 11.3444702 | 2.78415575 | 8.81478687 | 0 |
| 0.01251119 | 0.02276823 | 30.2231146 | 57.834554 | 59.3953227 | 0 | 0 |
| 0.01251786 | 0.02277913 | 139.540763 | 357.239591 | 348.947521 | 370.684985 | 71.2865473 |

| | | | | | | |
|---|---|---|---|---|---|---|
| 0.01251935 | 0.02278062 | 2.89370247 | 14.0137573 | 12.9927268 | 9.27872303 | 0 |
| 0.01252822 | 0.02279551 | 17.6837373 | 9.78738607 | 2.78415575 | 10.2065953 | 0 |
| 0.01254253 | 0.02282033 | 0 | 45.1554403 | 59.3953227 | 51.0329766 | 0 |
| 0.01254458 | 0.02282282 | 20.2559173 | 46.934965 | 0 | 63.0953166 | 0 |
| 0.01255239 | 0.02283579 | 56.9094818 | 22.0216187 | 0 | 48.7132959 | 0 |
| 0.01255312 | 0.02283589 | 5.46588243 | 6.22833659 | 6.49636342 | 25.5164883 | 0 |
| 0.012571 | 0.02286718 | 23.1496197 | 52.0510986 | 0 | 51.0329766 | 0 |
| 0.01257209 | 0.02286746 | 0 | 42.7085937 | 62.1794784 | 51.0329766 | 0 |
| 0.01257252 | 0.02286746 | 28.6155022 | 9.12006429 | 2.78415575 | 8.35085072 | 0 |
| 0.01257484 | 0.02287045 | 23.7926647 | 52.7184204 | 0 | 49.177232 | 0 |
| 0.01257647 | 0.02287219 | 0 | 56.2774699 | 25.0574018 | 85.3642518 | 0 |
| 0.01257882 | 0.02287397 | 52.4081669 | 30.4743612 | 0 | 39.4345729 | 0 |
| 0.01257829 | 0.02287397 | 47.2638069 | 33.8109701 | 0 | 97.8905279 | 415.838193 |
| 0.01258338 | 0.02288104 | 38.5826995 | 25.8031087 | 92.8051917 | 0 | 0 |
| 0.01260189 | 0.02291346 | 6.75197242 | 9.78738607 | 21.3451941 | 3.71148921 | 0 |
| 0.0126044 | 0.02291678 | 7.71653991 | 18.2401286 | 1.85610383 | 23.1968076 | 0 |
| 0.01261352 | 0.02293212 | 57.5525268 | 36.9251383 | 0 | 29.2279775 | 0 |
| 0.01261509 | 0.02293374 | 8.0380624 | 5.56101481 | 8.35246725 | 7.42297842 | 0 |
| 0.01262025 | 0.02294188 | 2.89370247 | 8.23030192 | 15.7768826 | 13.4541484 | 0 |
| 0.01262808 | 0.02295488 | 9.00262989 | 6.005896 | 5.5683115 | 10.2065953 | 0 |
| 0.01264615 | 0.02298582 | 24.1141872 | 13.123995 | 165.193241 | 0 | 0 |
| 0.01264647 | 0.02298582 | 11.8963324 | 88.5313558 | 72.3880495 | 0 | 0 |
| 0.01264881 | 0.02298883 | 5.78740493 | 9.78738607 | 6.49636342 | 7.88691457 | 0 |
| 0.01266311 | 0.02301357 | 5.78740493 | 3.33660889 | 5.5683115 | 3.71148921 | 61.7816743 |
| 0.01266759 | 0.02302047 | 0 | 0 | 0 | 0 | 23.7621824 |
| 0.01266939 | 0.0230225 | 0 | 84.9723063 | 64.9636342 | 20.4131907 | 0 |
| 0.0126737 | 0.0230291 | 0 | 86.0845093 | 26.9135056 | 51.9608489 | 0 |
| 0.01267599 | 0.023032 | 8.3595849 | 8.00786133 | 3.71220767 | 19.0213822 | 0 |
| 0.01268895 | 0.02305431 | 13.8254673 | 14.0137573 | 13.9207788 | 14.3820207 | 11.8810912 |
| 0.01269337 | 0.02306109 | 0 | 0 | 0 | 0 | 19.0097459 |
| 0.0126957 | 0.02306409 | 17.6837373 | 6.005896 | 7.42441534 | 5.56723382 | 0 |
| 0.01269658 | 0.02306444 | 1.60761248 | 21.5767375 | 17.6329864 | 10.2065953 | 0 |
| 0.01269744 | 0.02306475 | 18.3267823 | 34.0334106 | 35.2659728 | 45.0018067 | 9.50487297 |
| 0.01269972 | 0.02306766 | 0 | 0 | 0 | 0 | 28.5146189 |
| 0.0127195 | 0.02310233 | 5.14435994 | 9.56494547 | 13.9207788 | 5.56723382 | 0 |
| 0.01272177 | 0.0231052 | 1.28608998 | 2.22440592 | 0 | 1.85574461 | 14.2573095 |
| 0.01273382 | 0.02312317 | 16.3976473 | 0 | 50.1148035 | 133.613612 | 0 |
| 0.01273358 | 0.02312317 | 2.57217997 | 19.3523315 | 21.3451941 | 29.6919137 | 2.37621824 |
| 0.01273265 | 0.02312317 | 0 | 65.3975342 | 38.9781805 | 51.4969128 | 0 |
| 0.01273441 | 0.02312317 | 1.92913498 | 1.55708415 | 0 | 1.39180845 | 49.9005831 |
| 0.01273792 | 0.02312669 | 2.57217997 | 26.0255493 | 10.2085711 | 8.81478687 | 0 |
| 0.01273841 | 0.02312669 | 41.476402 | 79.1888509 | 73.3161014 | 106.705315 | 26.1384007 |
| 0.01273723 | 0.02312669 | 9.32415239 | 17.5728068 | 17.6329864 | 20.8771268 | 0 |

| | | | | | | |
|---|---|---|---|---|---|---|
| 0.01274702 | 0.02314106 | 103.851766 | 5.11613363 | 94.6612955 | 0 | 0 |
| 0.01275924 | 0.02316201 | 6.75197242 | 5.11613363 | 8.35246725 | 11.1344676 | 0 |
| 0.01277252 | 0.02318487 | 189.055228 | 300.51724 | 258.926485 | 264.443606 | 475.243649 |
| 0.01277493 | 0.02318799 | 6.43044992 | 19.5747721 | 30.6257133 | 7.88691457 | 0 |
| 0.01279208 | 0.02321786 | 0 | 0 | 0 | 0 | 0 |
| 0.01279825 | 0.0232278 | 3.21522496 | 11.5669108 | 10.2085711 | 11.1344676 | 0 |
| 0.01279923 | 0.02322833 | 70.4134267 | 29.8070394 | 0 | 27.8361691 | 0 |
| 0.01280507 | 0.02323769 | 20.5774398 | 9.78738607 | 8.35246725 | 12.0623399 | 2.37621824 |
| 0.0128067 | 0.02323939 | 0.3215225 | 0 | 0 | 0 | 0 |
| 0.01281485 | 0.02325291 | 72.0210391 | 24.6909058 | 0 | 32.4755306 | 0 |
| 0.01281817 | 0.02325675 | 76.8438766 | 149.035197 | 69.6038938 | 77.0134011 | 30.8908372 |
| 0.01281834 | 0.02325675 | 2.57217997 | 10.6771484 | 15.7768826 | 11.5984038 | 0 |
| 0.0128324 | 0.02327974 | 6.43044992 | 7.11809896 | 5.5683115 | 13.9180845 | 0 |
| 0.01283239 | 0.02327974 | 1.60761248 | 23.1338216 | 38.9781805 | 4.17542536 | 0 |
| 0.01287074 | 0.02334711 | 87.4541189 | 8.00786133 | 89.092984 | 0 | 0 |
| 0.01287092 | 0.02334711 | 5.78740493 | 8.8976237 | 9.28051917 | 6.03116997 | 0 |
| 0.0128744 | 0.02335217 | 10.2887199 | 8.67518311 | 5.5683115 | 6.03116997 | 0 |
| 0.01287565 | 0.02335316 | 4.82283744 | 10.899589 | 6.49636342 | 9.74265918 | 0 |
| 0.01288358 | 0.02336629 | 0 | 18.6850098 | 127.143113 | 38.0427644 | 0 |
| 0.01288588 | 0.0233692 | 181.66021 | 64.5077718 | 34.3379209 | 55.6723382 | 23.7621824 |
| 0.01289289 | 0.02338065 | 3.85826995 | 8.67518311 | 11.136623 | 9.74265918 | 0 |
| 0.01289432 | 0.02338198 | 50.4790319 | 120.785242 | 116.934542 | 106.705315 | 45.1481466 |
| 0.01289962 | 0.02339034 | 4.50131495 | 22.911381 | 13.9207788 | 18.5574461 | 2.37621824 |
| 0.01291758 | 0.02342163 | 3.85826995 | 9.78738607 | 9.28051917 | 10.2065953 | 0 |
| 0.01292657 | 0.02343668 | 11.2532874 | 8.23030192 | 5.5683115 | 6.95904227 | 19.0097459 |
| 0.01293211 | 0.02344546 | 9.32415239 | 14.9035197 | 7.42441534 | 19.9492545 | 2.37621824 |
| 0.0129486 | 0.0234741 | 2.89370247 | 6.67321777 | 19.4890903 | 14.3820207 | 0 |
| 0.01295802 | 0.02348934 | 0 | 55.3877075 | 47.3306478 | 49.6411682 | 0 |
| 0.01295841 | 0.02348934 | 165.262563 | 315.198319 | 295.12051 | 310.837221 | 0 |
| 0.01296076 | 0.02349233 | 0 | 49.8266927 | 55.683115 | 46.8575513 | 0 |
| 0.01296554 | 0.02349973 | 17.0406923 | 103.879757 | 131.783372 | 100.210209 | 23.7621824 |
| 0.01298414 | 0.02353217 | 13.1824223 | 4.44881185 | 4.64025958 | 18.0935099 | 0 |
| 0.01300987 | 0.02357754 | 54.9803468 | 34.2558512 | 0 | 31.5476583 | 0 |
| 0.01301219 | 0.02358047 | 74.5932191 | 0 | 32.4818171 | 62.6313804 | 0 |
| 0.01301552 | 0.02358397 | 5.46588243 | 10.899589 | 5.5683115 | 9.74265918 | 0 |
| 0.01301507 | 0.02358397 | 201.273083 | 4.00393066 | 0 | 21.341063 | 0 |
| 0.01301724 | 0.02358454 | 36.3320421 | 36.4802572 | 0 | 45.4657428 | 0 |
| 0.01301676 | 0.02358454 | 43.405537 | 38.9271037 | 0 | 35.7230837 | 0 |
| 0.01301872 | 0.02358596 | 35.6889971 | 46.0452026 | 0 | 36.650956 | 0 |
| 0.01303005 | 0.02360522 | 184.875435 | 329.212077 | 321.105963 | 332.64222 | 42.7719284 |
| 0.01303678 | 0.02361613 | 18.9698273 | 82.3030192 | 0 | 33.867339 | 0 |
| 0.0130414 | 0.02362323 | 1.92913498 | 10.6771484 | 25.9854537 | 11.1344676 | 0 |
| 0.01306269 | 0.02366052 | 52.0866444 | 51.6062174 | 37.1220767 | 0 | 0 |

| | | | | | | |
|---|---|---|---|---|---|---|
| 0.01306706 | 0.02366716 | 23.7926647 | 115.891549 | 113.222334 | 106.241379 | 26.1384007 |
| 0.01307412 | 0.02367868 | 95.1706589 | 9.78738607 | 0 | 47.3214874 | 0 |
| 0.0130935 | 0.02371251 | 0 | 0.88976237 | 0.92805192 | 0.9278723 | 0 |
| 0.01309626 | 0.02371623 | 11.8963324 | 30.02948 | 11.136623 | 18.5574461 | 4.75243649 |
| 0.01310054 | 0.0237227 | 12.8608998 | 9.12006429 | 6.49636342 | 4.63936151 | 0 |
| 0.01310716 | 0.02373341 | 27.9724572 | 19.3523315 | 11.136623 | 21.341063 | 125.939567 |
| 0.01311035 | 0.02373791 | 110.603739 | 236.009469 | 193.034799 | 233.359884 | 9.50487297 |
| 0.01311481 | 0.0237447 | 10.2887199 | 11.7893514 | 5.5683115 | 5.10329766 | 0 |
| 0.01312811 | 0.02376749 | 27.6509347 | 117.226192 | 123.430905 | 124.334889 | 28.5146189 |
| 0.01313052 | 0.02377058 | 365.892601 | 695.126851 | 657.060757 | 617.499017 | 83.1676385 |
| 0.0131562 | 0.02381579 | 27.6509347 | 32.2538859 | 94.6612955 | 0 | 0 |
| 0.0131627 | 0.02382628 | 22.8280972 | 46.0452026 | 0 | 54.2805297 | 0 |
| 0.0131757 | 0.02384853 | 23.4711422 | 47.3798462 | 0 | 51.4969128 | 0 |
| 0.01317775 | 0.02385096 | 2.89370247 | 11.7893514 | 9.28051917 | 14.3820207 | 0 |
| 0.01317856 | 0.02385113 | 0 | 0 | 0 | 0 | 49.9005831 |
| 0.01321883 | 0.02392274 | 9.96719738 | 8.8976237 | 7.42441534 | 12.9902122 | 33.2670554 |
| 0.01322085 | 0.02392511 | 2.89370247 | 19.7972127 | 8.35246725 | 10.6705315 | 0 |
| 0.01322458 | 0.02393057 | 58.8386168 | 16.6830444 | 9.28051917 | 23.1968076 | 0 |
| 0.01322606 | 0.02393195 | 0 | 51.1613363 | 43.6184401 | 56.1362743 | 0 |
| 0.01323147 | 0.02394046 | 74.9147416 | 215.544934 | 174.47376 | 244.030416 | 0 |
| 0.01323469 | 0.023945 | 17.6837373 | 48.2696086 | 0 | 63.0953166 | 0 |
| 0.01324249 | 0.02395721 | 12.8608998 | 54.2755046 | 0 | 70.9822312 | 0 |
| 0.01324286 | 0.02395721 | 11.2532874 | 4.22637126 | 7.42441534 | 9.74265918 | 0 |
| 0.01325986 | 0.02398666 | 1.92913498 | 13.5688761 | 12.9927268 | 15.7738291 | 0 |
| 0.0132736 | 0.02401023 | 89.0617314 | 0 | 35.2659728 | 45.4657428 | 0 |
| 0.01328829 | 0.0240355 | 0 | 6.67321777 | 0 | 1.39180845 | 0 |
| 0.01331415 | 0.02408099 | 5.14435994 | 3.33660889 | 4.64025958 | 4.17542536 | 4.75243649 |
| 0.01331567 | 0.02408244 | 0 | 38.7046631 | 83.5246725 | 35.2591475 | 0 |
| 0.01332414 | 0.02409646 | 163.333428 | 751.404321 | 374.004923 | 587.807104 | 0 |
| 0.01334085 | 0.0241254 | 10.6102424 | 29.8070394 | 19.4890903 | 25.5164883 | 2.37621824 |
| 0.01335188 | 0.02414145 | 789.980773 | 232.67286 | 218.0922 | 209.69914 | 76.0389838 |
| 0.01335069 | 0.02414145 | 37.618132 | 110.997856 | 99.3015551 | 100.674145 | 28.5146189 |
| 0.01335146 | 0.02414145 | 11.8963324 | 20.4645345 | 20.4171422 | 31.5476583 | 0 |
| 0.01335641 | 0.02414705 | 6.75197242 | 25.8031087 | 46.4025958 | 33.4034029 | 0 |
| 0.01335596 | 0.02414705 | 218.635297 | 196.415043 | 141.991943 | 176.295737 | 102.177384 |
| 0.01335907 | 0.02415055 | 0.96456749 | 0 | 0 | 0 | 7.12865473 |
| 0.01336167 | 0.02415395 | 42.762492 | 24.023584 | 0 | 41.7542536 | 135.44444 |
| 0.01336398 | 0.02415683 | 54.0157793 | 54.7203857 | 42.6903882 | 55.6723382 | 201.978551 |
| 0.01336913 | 0.02416484 | 15.7546023 | 10.0098267 | 11.136623 | 9.27872303 | 0 |
| 0.01338748 | 0.0241967 | 66.2336342 | 297.403072 | 170.761553 | 230.576267 | 73.6627655 |
| 0.01340116 | 0.02422014 | 9.00262989 | 8.23030192 | 6.49636342 | 5.56723382 | 0 |
| 0.01340842 | 0.02423196 | 9.00262989 | 38.0373413 | 144.776099 | 0 | 0 |
| 0.01341218 | 0.02423745 | 33.4383396 | 134.576558 | 124.358957 | 168.408823 | 0 |

| | | | | | | |
|---|---|---|---|---|---|---|
| 0.01343533 | 0.02427537 | 67.8412467 | 14.4586385 | 0 | 51.0329766 | 0 |
| 0.01343529 | 0.02427537 | 1.28608998 | 10.2322673 | 24.1293498 | 18.5574461 | 0 |
| 0.01343522 | 0.02427537 | 6.10892743 | 8.00786133 | 6.49636342 | 7.88691457 | 0 |
| 0.01343634 | 0.02427588 | 16.0761248 | 86.3069499 | 0 | 34.7952113 | 0 |
| 0.01344469 | 0.02428967 | 5.46588243 | 12.2342326 | 12.0646749 | 4.63936151 | 0 |
| 0.01344674 | 0.02429207 | 17.3622148 | 46.2676432 | 0 | 64.9510612 | 0 |
| 0.01344829 | 0.02429357 | 34.7244296 | 28.6948364 | 0 | 56.1362743 | 0 |
| 0.01345221 | 0.02429935 | 7.71653991 | 8.45274251 | 3.71220767 | 17.1656376 | 0 |
| 0.01346563 | 0.02432227 | 22.1850522 | 61.3936035 | 89.092984 | 51.0329766 | 19.0097459 |
| 0.01347609 | 0.02433987 | 0.96456749 | 12.011792 | 41.7623363 | 11.5984038 | 0 |
| 0.01348946 | 0.0243627 | 19.2913498 | 0 | 86.3088283 | 72.8379758 | 0 |
| 0.01349282 | 0.02436746 | 0.64304499 | 10.899589 | 26.9135056 | 29.2279775 | 0 |
| 0.01349361 | 0.02436758 | 4.50131495 | 2.66928711 | 12.9927268 | 38.0427644 | 0 |
| 0.01349889 | 0.02437581 | 63.3399317 | 108.328569 | 94.6612955 | 117.375846 | 0 |
| 0.01350957 | 0.02439379 | 5.14435994 | 12.4566732 | 4.64025958 | 12.9902122 | 0 |
| 0.01352054 | 0.02441228 | 5.14435994 | 78.5215291 | 0 | 83.5085072 | 0 |
| 0.01352711 | 0.02442223 | 8.3595849 | 9.78738607 | 0.92805192 | 5.10329766 | 9.50487297 |
| 0.0135275 | 0.02442223 | 0 | 41.5963908 | 32.4818171 | 82.1166988 | 0 |
| 0.01354531 | 0.02445308 | 30.2231146 | 3.33660889 | 5.5683115 | 10.2065953 | 0 |
| 0.01354997 | 0.02446017 | 0 | 40.2617472 | 52.8989593 | 56.1362743 | 0 |
| 0.01355777 | 0.02447188 | 4.17979245 | 10.6771484 | 17.6329864 | 5.56723382 | 0 |
| 0.01355791 | 0.02447188 | 6.43044992 | 9.34250488 | 4.64025958 | 11.1344676 | 0 |
| 0.01356126 | 0.02447662 | 7.07349491 | 2.00196533 | 25.0574018 | 16.2377653 | 0 |
| 0.01359184 | 0.02453 | 0.3215225 | 1.55708415 | 0 | 1.85574461 | 2.37621824 |
| 0.0135923 | 0.02453 | 5.78740493 | 16.0157227 | 2.78415575 | 19.9492545 | 0 |
| 0.01359677 | 0.02453676 | 0 | 0 | 0 | 0 | 47.5243649 |
| 0.01360336 | 0.02454734 | 0 | 0 | 25.0574018 | 0 | 384.947355 |
| 0.01362695 | 0.02458726 | 6.43044992 | 7.78542074 | 6.49636342 | 7.42297842 | 0 |
| 0.01362645 | 0.02458726 | 5.46588243 | 8.45274251 | 16.7049345 | 5.10329766 | 0 |
| 0.01364091 | 0.02461114 | 74.5932191 | 220.216187 | 215.308045 | 211.554885 | 4.75243649 |
| 0.0136425 | 0.02461269 | 3.21522496 | 10.2322673 | 13.9207788 | 8.81478687 | 0 |
| 0.01366931 | 0.02465973 | 22.8280972 | 6.89565837 | 3.71220767 | 8.35085072 | 0 |
| 0.01367992 | 0.02467755 | 1.60761248 | 14.4586385 | 16.7049345 | 13.9180845 | 0 |
| 0.01368097 | 0.02467811 | 24.1141872 | 0 | 50.1148035 | 100.674145 | 0 |
| 0.01368726 | 0.02468813 | 464.600007 | 952.935498 | 875.152958 | 933.439536 | 0 |
| 0.01369213 | 0.0246956 | 3.53674746 | 12.011792 | 9.28051917 | 9.27872303 | 0 |
| 0.01370099 | 0.02471025 | 0 | 81.1908162 | 27.8415575 | 47.3214874 | 0 |
| 0.01370731 | 0.02472033 | 5.46588243 | 13.3464355 | 8.35246725 | 5.56723382 | 0 |
| 0.01371932 | 0.02474065 | 0 | 13.3464355 | 115.078438 | 51.0329766 | 0 |
| 0.01372499 | 0.02474956 | 4.82283744 | 17.3503662 | 4.64025958 | 11.1344676 | 0 |
| 0.0137429 | 0.02478054 | 14.4685123 | 60.5038411 | 82.5966206 | 0 | 0 |
| 0.01374791 | 0.02478823 | 31.1876821 | 0 | 91.8771398 | 45.4657428 | 0 |
| 0.01375046 | 0.02479151 | 0 | 6.89565837 | 6.49636342 | 11.5984038 | 23.7621824 |

| | | | | | | |
|---|---|---|---|---|---|---|
| 0.01376095 | 0.02480908 | 25.4002772 | 3.78149007 | 6.49636342 | 8.35085072 | 0 |
| 0.01376286 | 0.02481121 | 136.647061 | 18.4625692 | 0 | 12.5262761 | 0 |
| 0.01376614 | 0.02481579 | 37.2966096 | 0 | 54.7550631 | 69.5904227 | 0 |
| 0.0137678 | 0.02481744 | 4.82283744 | 10.4547078 | 8.35246725 | 6.95904227 | 0 |
| 0.01377311 | 0.02482568 | 18.0052598 | 18.2401286 | 2.78415575 | 5.56723382 | 0 |
| 0.01377453 | 0.02482692 | 52.7296894 | 37.3700195 | 46.4025958 | 0 | 0 |
| 0.01378864 | 0.02485101 | 3.21522496 | 13.123995 | 6.49636342 | 16.7017014 | 0 |
| 0.01378955 | 0.02485133 | 0 | 38.2597819 | 52.8989593 | 57.0641466 | 0 |
| 0.01379053 | 0.02485176 | 4.50131495 | 22.0216187 | 3.71220767 | 13.9180845 | 0 |
| 0.01383811 | 0.02493616 | 72.3425616 | 39.816866 | 37.1220767 | 17.6295737 | 0 |
| 0.01383897 | 0.02493638 | 145.649691 | 431.75719 | 425.97583 | 418.006472 | 40.3957101 |
| 0.01384361 | 0.02494341 | 52.7296894 | 16.905485 | 0 | 56.1362743 | 0 |
| 0.01384843 | 0.02495076 | 97.7428388 | 23.3562622 | 0 | 19.0213822 | 0 |
| 0.01387109 | 0.02499024 | 9.96719738 | 19.1298909 | 1.85610383 | 13.9180845 | 0 |
| 0.01387239 | 0.02499124 | 164.619518 | 119.450598 | 79.8124649 | 81.1888265 | 118.810912 |
| 0.01388161 | 0.02500652 | 29.2585471 | 90.5333211 | 48.2586997 | 85.828188 | 19.0097459 |
| 0.01388551 | 0.0250122 | 22.5065747 | 50.2715739 | 0 | 46.3936151 | 0 |
| 0.01389793 | 0.02503324 | 6.75197242 | 11.5669108 | 4.64025958 | 8.35085072 | 0 |
| 0.01389914 | 0.02503409 | 24.4357097 | 66.7321777 | 101.157659 | 76.549465 | 0 |
| 0.01390461 | 0.02504142 | 5.46588243 | 10.2322673 | 5.5683115 | 9.27872303 | 0 |
| 0.01390471 | 0.02504142 | 144.685123 | 328.767196 | 327.602327 | 372.076793 | 42.7719284 |
| 0.01391068 | 0.02504896 | 26.0433222 | 50.9388957 | 64.0355823 | 0 | 0 |
| 0.01391113 | 0.02504896 | 47.2638069 | 112.55494 | 129.927268 | 127.582442 | 33.2670554 |
| 0.01390967 | 0.02504896 | 111.246784 | 157.487939 | 199.531162 | 171.656376 | 0 |
| 0.01391664 | 0.02505754 | 57.2310043 | 0 | 52.8989593 | 47.7854236 | 0 |
| 0.01393785 | 0.0250944 | 4.17979245 | 8.8976237 | 8.35246725 | 9.74265918 | 0 |
| 0.01394719 | 0.02510988 | 131.502701 | 298.070394 | 316.465704 | 331.250412 | 118.810912 |
| 0.01394967 | 0.02511299 | 17.3622148 | 96.3167765 | 121.574801 | 88.1478687 | 16.6335277 |
| 0.01395104 | 0.02511411 | 4.50131495 | 7.56298014 | 10.2085711 | 8.35085072 | 0 |
| 0.01395267 | 0.02511436 | 0 | 0 | 0 | 0 | 0 |
| 0.01395267 | 0.02511436 | 0 | 0 | 0 | 0 | 0 |
| 0.01395362 | 0.02511472 | 0 | 94.7596924 | 21.3451941 | 46.3936151 | 0 |
| 0.01395512 | 0.02511608 | 0 | 77.6317668 | 51.9709073 | 25.5164883 | 0 |
| 0.01397657 | 0.02515335 | 39.8687895 | 29.8070394 | 0 | 88.1478687 | 2.37621824 |
| 0.01398259 | 0.02516282 | 9.96719738 | 48.7144897 | 0 | 84.4363795 | 0 |
| 0.01398362 | 0.02516334 | 15.4330798 | 72.9605143 | 0 | 41.7542536 | 0 |
| 0.01398578 | 0.02516588 | 53.0512119 | 22.911381 | 0 | 42.6821259 | 0 |
| 0.01398654 | 0.0251659 | 0 | 16.6830444 | 62.1794784 | 86.2921241 | 0 |
| 0.0140042 | 0.02519634 | 99.9934963 | 596.363228 | 1306.6971 | 560.898807 | 156.830404 |
| 0.01400638 | 0.02519756 | 0.64304499 | 2.44684652 | 0.92805192 | 5.10329766 | 38.0194919 |
| 0.0140063 | 0.02519756 | 2.89370247 | 8.67518311 | 3.71220767 | 9.74265918 | 2.37621824 |
| 0.01400832 | 0.0251997 | 3.21522496 | 9.34250488 | 11.136623 | 11.1344676 | 0 |
| 0.01402821 | 0.02523412 | 0 | 58.5018758 | 37.1220767 | 51.4969128 | 0 |

| | | | | | | |
|---|---|---|---|---|---|---|
| 0.01403983 | 0.02525368 | 0.64304499 | 0.22244059 | 0 | 1.39180845 | 0 |
| 0.01406662 | 0.02530052 | 4.50131495 | 5.33857422 | 5.5683115 | 6.49510612 | 0 |
| 0.01407363 | 0.02531177 | 32.7952946 | 60.05896 | 0 | 25.9804245 | 0 |
| 0.01407654 | 0.02531369 | 4.50131495 | 3.78149007 | 31.5537652 | 10.2065953 | 0 |
| 0.01407644 | 0.02531369 | 3.85826995 | 9.78738607 | 8.35246725 | 10.2065953 | 0 |
| 0.01407695 | 0.02531369 | 4.50131495 | 9.56494547 | 8.35246725 | 7.88691457 | 0 |
| 0.014086 | 0.02532861 | 378.7535 | 625.725387 | 554.046994 | 626.77774 | 14.2573095 |
| 0.01410484 | 0.02536022 | 5.46588243 | 13.7913167 | 16.7049345 | 16.7017014 | 0 |
| 0.01410509 | 0.02536022 | 15.7546023 | 7.11809896 | 4.64025958 | 6.95904227 | 0 |
| 0.01411348 | 0.02537395 | 27.3294122 | 138.358048 | 115.078438 | 115.984038 | 38.0194919 |
| 0.01412915 | 0.02540076 | 85.524984 | 93.86993 | 111.36623 | 86.7560603 | 392.07601 |
| 0.01413021 | 0.02540132 | 3.85826995 | 8.00786133 | 10.2085711 | 10.2065953 | 0 |
| 0.01413257 | 0.0254042 | 45.0131495 | 162.159192 | 91.8771398 | 118.303719 | 40.3957101 |
| 0.01414129 | 0.02541851 | 6.43044992 | 22.0216187 | 3.71220767 | 8.81478687 | 0 |
| 0.01414601 | 0.02542564 | 0 | 60.2814006 | 30.6257133 | 57.9920189 | 0 |
| 0.01415196 | 0.02543498 | 5.78740493 | 5.56101481 | 11.136623 | 7.88691457 | 0 |
| 0.0141549 | 0.0254389 | 3.53674746 | 8.45274251 | 13.9207788 | 8.81478687 | 0 |
| 0.0141581 | 0.02544329 | 7.07349491 | 90.5333211 | 0 | 55.208402 | 0 |
| 0.01415918 | 0.02544387 | 129.895088 | 390.828121 | 48.2586997 | 154.490738 | 42.7719284 |
| 0.01416664 | 0.02545591 | 103.208721 | 258.475968 | 244.077654 | 243.566479 | 57.0292378 |
| 0.01417001 | 0.02546061 | 0 | 42.4861532 | 49.1867516 | 53.3526574 | 0 |
| 0.01417817 | 0.02547392 | 3.21522496 | 12.9015544 | 31.5537652 | 4.17542536 | 0 |
| 0.01418788 | 0.02549 | 15.1115573 | 52.7184204 | 0 | 58.9198912 | 0 |
| 0.01419561 | 0.02550253 | 1.60761248 | 16.4606038 | 17.6329864 | 11.1344676 | 0 |
| 0.01421225 | 0.02552863 | 96.4567488 | 17.7952474 | 0 | 24.1246799 | 0 |
| 0.01421242 | 0.02552863 | 165.905608 | 218.436662 | 189.322591 | 218.049991 | 57.0292378 |
| 0.01421106 | 0.02552863 | 0.64304499 | 9.56494547 | 32.4818171 | 25.0525522 | 0 |
| 0.01421737 | 0.02553617 | 108.674604 | 511.390922 | 444.536868 | 484.813278 | 2.37621824 |
| 0.01422624 | 0.02555073 | 36.6535646 | 38.0373413 | 0 | 37.5788283 | 0 |
| 0.01422714 | 0.02555099 | 6.43044992 | 7.78542074 | 5.5683115 | 8.81478687 | 0 |
| 0.01423447 | 0.02556279 | 0 | 37.8149007 | 53.8270112 | 53.8165936 | 0 |
| 0.01424158 | 0.02557418 | 64.6260217 | 20.2420939 | 0 | 37.1148921 | 0 |
| 0.01425292 | 0.02559318 | 28.2939797 | 6.45077718 | 4.64025958 | 6.03116997 | 0 |
| 0.01425901 | 0.02560219 | 110.603739 | 191.076469 | 186.538435 | 206.451587 | 23.7621824 |
| 0.01425946 | 0.02560219 | 77.4869216 | 99.2085042 | 0 | 55.208402 | 4.75243649 |
| 0.01426767 | 0.02561556 | 0 | 65.3975342 | 38.9781805 | 83.5085072 | 0 |
| 0.01426879 | 0.02561621 | 40.190312 | 64.0628906 | 32.4818171 | 0 | 0 |
| 0.01427551 | 0.02562692 | 90.6693439 | 150.36984 | 147.560255 | 160.985845 | 16.6335277 |
| 0.01427883 | 0.02563149 | 5.78740493 | 8.8976237 | 4.64025958 | 14.3820207 | 0 |
| 0.01431123 | 0.02568692 | 0 | 0 | 0 | 0 | 0 |
| 0.01431123 | 0.02568692 | 0 | 0 | 0 | 0 | 0 |
| 0.01432002 | 0.02570133 | 126.036818 | 44.0432373 | 48.2586997 | 35.7230837 | 178.216368 |
| 0.01433484 | 0.02572655 | 32.4737721 | 38.9271037 | 0 | 40.8263813 | 0 |

| | | | | | | |
|---|---|---|---|---|---|---|
| 0.01435683 | 0.02576464 | 8.0380624 | 4.44881185 | 2.78415575 | 2.31968076 | 14.2573095 |
| 0.01436329 | 0.02577486 | 40.190312 | 39.3719849 | 53.8270112 | 0 | 0 |
| 0.01436814 | 0.02578218 | 55.3018693 | 42.4861532 | 12.0646749 | 31.0837221 | 11.8810912 |
| 0.01439608 | 0.02583096 | 4.82283744 | 5.56101481 | 10.2085711 | 11.5984038 | 0 |
| 0.01439726 | 0.02583169 | 4.17979245 | 12.011792 | 27.8415575 | 3.71148921 | 0 |
| 0.01440242 | 0.02583957 | 0.64304499 | 1.77952474 | 0.92805192 | 0.9278723 | 35.6432737 |
| 0.01440714 | 0.02584665 | 32.4737721 | 17.7952474 | 17.6329864 | 14.8459568 | 190.097459 |
| 0.0144186 | 0.02586584 | 2.57217997 | 10.4547078 | 8.35246725 | 21.341063 | 0 |
| 0.01444142 | 0.02590468 | 40.190312 | 28.2499552 | 0 | 44.5378705 | 0 |
| 0.01444179 | 0.02590468 | 0 | 0 | 0 | 0 | 21.3859642 |
| 0.01445014 | 0.02591827 | 0 | 32.9212077 | 36.1940248 | 82.1166988 | 0 |
| 0.01446865 | 0.02595009 | 10.2887199 | 8.45274251 | 5.5683115 | 5.56723382 | 0 |
| 0.01448284 | 0.02597415 | 3.53674746 | 6.89565837 | 20.4171422 | 8.81478687 | 0 |
| 0.01448528 | 0.02597714 | 11.5748099 | 4.67125244 | 5.5683115 | 10.6705315 | 0 |
| 0.0144929 | 0.02598942 | 27.9724572 | 62.0609253 | 46.4025958 | 0 | 0 |
| 0.0145164 | 0.02603018 | 4.50131495 | 3.55904948 | 5.5683115 | 2.78361691 | 49.9005831 |
| 0.01453505 | 0.02606085 | 7.39501741 | 25.8031087 | 7.42441534 | 15.309893 | 2.37621824 |
| 0.01453468 | 0.02606085 | 21.2204847 | 20.4645345 | 16.7049345 | 24.588616 | 9.50487297 |
| 0.01454984 | 0.02608598 | 22.8280972 | 15.7932821 | 8.35246725 | 24.588616 | 19.0097459 |
| 0.01456602 | 0.0261136 | 10.9317649 | 1.55708415 | 13.9207788 | 21.8049991 | 0 |
| 0.01456894 | 0.02611744 | 103.208721 | 16.2381632 | 0 | 22.2689353 | 0 |
| 0.01458786 | 0.02614995 | 0 | 38.0373413 | 55.683115 | 49.6411682 | 0 |
| 0.0145909 | 0.02615402 | 2.89370247 | 16.905485 | 9.28051917 | 9.27872303 | 0 |
| 0.01459625 | 0.02616221 | 2.57217997 | 1.55708415 | 2.78415575 | 1.85574461 | 16.6335277 |
| 0.01459937 | 0.02616641 | 1.92913498 | 19.5747721 | 8.35246725 | 15.7738291 | 0 |
| 0.01460058 | 0.02616718 | 0.64304499 | 1.11220296 | 0.92805192 | 0.46393615 | 23.7621824 |
| 0.01460257 | 0.02616936 | 20.5774398 | 62.9506877 | 0 | 36.1870198 | 0 |
| 0.01460553 | 0.02617327 | 23.1496197 | 55.6101481 | 37.1220767 | 49.6411682 | 16.6335277 |
| 0.01461399 | 0.02618702 | 6.75197242 | 0 | 116.00649 | 81.1888265 | 0 |
| 0.01461848 | 0.02619368 | 7.71653991 | 35.1456136 | 4.64025958 | 4.17542536 | 0 |
| 0.01463266 | 0.02621769 | 4.17979245 | 12.011792 | 8.35246725 | 7.42297842 | 0 |
| 0.01463679 | 0.02622369 | 7.07349491 | 11.7893514 | 4.64025958 | 7.42297842 | 0 |
| 0.01463963 | 0.02622739 | 0 | 44.9329997 | 43.6184401 | 53.8165936 | 0 |
| 0.01464611 | 0.02623761 | 12.8608998 | 14.2361979 | 7.42441534 | 10.6705315 | 0 |
| 0.01465131 | 0.02624552 | 218.313775 | 237.788993 | 295.12051 | 308.981477 | 61.7816743 |
| 0.01467462 | 0.02628587 | 11.2532874 | 11.1220296 | 11.136623 | 8.81478687 | 26.1384007 |
| 0.0147311 | 0.02638563 | 37.9396545 | 58.2794352 | 20.4171422 | 41.7542536 | 11.8810912 |
| 0.01473885 | 0.02639812 | 33.4383396 | 46.4900838 | 0 | 31.5476583 | 0 |
| 0.01474941 | 0.02641563 | 51.4435994 | 47.1574056 | 33.409869 | 0 | 0 |
| 0.01475055 | 0.02641626 | 2.57217997 | 15.3484009 | 8.35246725 | 12.9902122 | 0 |
| 0.01475262 | 0.02641856 | 5.78740493 | 9.34250488 | 34.3379209 | 2.78361691 | 0 |
| 0.01475795 | 0.0264267 | 6.75197242 | 3.55904948 | 12.9927268 | 11.5984038 | 0 |
| 0.01475918 | 0.02642749 | 0 | 38.2597819 | 43.6184401 | 61.239572 | 0 |

| | | | | | | |
|---|---|---|---|---|---|---|
| 0.01476532 | 0.02643708 | 0 | 50.0491333 | 58.4672708 | 34.7952113 | 0 |
| 0.0147706 | 0.02644512 | 0 | 4.44881185 | 0 | 7.88691457 | 2.37621824 |
| 0.0147876 | 0.02647416 | 40.190312 | 29.8070394 | 0 | 40.8263813 | 0 |
| 0.01480883 | 0.02651076 | 37.618132 | 56.0550293 | 38.0501286 | 42.6821259 | 19.0097459 |
| 0.01481929 | 0.02652807 | 8.3595849 | 8.8976237 | 7.42441534 | 4.63936151 | 0 |
| 0.01482826 | 0.02654272 | 71.0564716 | 203.755583 | 227.37272 | 211.554885 | 49.9005831 |
| 0.01484312 | 0.0265679 | 39.2257445 | 35.8129354 | 0 | 34.7952113 | 0 |
| 0.01485171 | 0.02658187 | 0 | 16.2381632 | 116.00649 | 37.1148921 | 0 |
| 0.01485975 | 0.02659483 | 0 | 50.7164551 | 40.8342843 | 49.6411682 | 0 |
| 0.0148693 | 0.02661051 | 12.2178549 | 22.4664998 | 2.78415575 | 6.03116997 | 0 |
| 0.01490048 | 0.0266649 | 36.0105196 | 95.2045736 | 75.1722053 | 80.2609542 | 14.2573095 |
| 0.01490405 | 0.02666987 | 4.50131495 | 12.6791138 | 5.5683115 | 10.2065953 | 0 |
| 0.01491708 | 0.02669075 | 11.2532874 | 25.8031087 | 16.7049345 | 0 | 80.7914203 |
| 0.01491731 | 0.02669075 | 52.7296894 | 34.4782918 | 0 | 25.9804245 | 0 |
| 0.01492693 | 0.02670654 | 0 | 0 | 0 | 0 | 0 |
| 0.0149296 | 0.0267099 | 17.6837373 | 14.0137573 | 3.71220767 | 4.63936151 | 0 |
| 0.01493078 | 0.02671059 | 0 | 0 | 0 | 0 | 0 |
| 0.01494124 | 0.02672789 | 16.7191698 | 22.6889404 | 31.5537652 | 17.6295737 | 9.50487297 |
| 0.0149483 | 0.02673908 | 1.28608998 | 14.6810791 | 17.6329864 | 14.8459568 | 0 |
| 0.01495619 | 0.02675179 | 0 | 48.2696086 | 61.2514265 | 33.4034029 | 0 |
| 0.01496549 | 0.02676699 | 2.57217997 | 23.8011434 | 17.6329864 | 4.63936151 | 0 |
| 0.01499359 | 0.02681583 | 260.754744 | 541.420402 | 533.629852 | 540.02168 | 4.75243649 |
| 0.01500627 | 0.02683708 | 41.1548795 | 49.6042521 | 38.9781805 | 0 | 0 |
| 0.01503244 | 0.02688245 | 1.92913498 | 20.9094157 | 7.42441534 | 16.2377653 | 0 |
| 0.01503716 | 0.02688947 | 16.3976473 | 16.4606038 | 54.7550631 | 80.7248903 | 0 |
| 0.01503853 | 0.02689048 | 0 | 44.4881185 | 38.9781805 | 57.5280828 | 0 |
| 0.01504252 | 0.02689476 | 5.78740493 | 103.879757 | 0 | 46.8575513 | 0 |
| 0.01504222 | 0.02689476 | 188.733705 | 274.26925 | 167.049345 | 203.66797 | 9.50487297 |
| 0.01504478 | 0.02689622 | 0 | 0 | 0 | 0 | 26.1384007 |
| 0.01504493 | 0.02689622 | 2.89370247 | 7.34053955 | 25.0574018 | 8.81478687 | 0 |
| 0.01505683 | 0.02691605 | 0 | 47.1574056 | 53.8270112 | 39.4345729 | 0 |
| 0.01506084 | 0.02692037 | 0 | 0 | 0 | 0 | 33.2670554 |
| 0.01506084 | 0.02692037 | 0 | 0 | 0 | 0 | 33.2670554 |
| 0.01506317 | 0.02692309 | 550.446513 | 1109.08879 | 918.771398 | 1008.13326 | 0 |
| 0.01507911 | 0.02695015 | 114.140486 | 159.267464 | 151.272462 | 195.31712 | 0 |
| 0.01508644 | 0.02696083 | 53.6942569 | 27.1377523 | 51.9709073 | 0 | 0 |
| 0.01508668 | 0.02696083 | 24.4357097 | 160.602108 | 187.466487 | 135.469356 | 40.3957101 |
| 0.01510075 | 0.02698453 | 4.50131495 | 11.5669108 | 4.64025958 | 15.309893 | 0 |
| 0.01510237 | 0.026986 | 25.7217997 | 12.9015544 | 40.8342843 | 23.1968076 | 9.50487297 |
| 0.0151159 | 0.02700873 | 30.8661596 | 17.3503662 | 12.9927268 | 7.42297842 | 26.1384007 |
| 0.01514306 | 0.02705582 | 7.07349491 | 5.33857422 | 12.0646749 | 6.03116997 | 0 |
| 0.01517056 | 0.02710353 | 4.17979245 | 9.12006429 | 7.42441534 | 9.74265918 | 0 |
| 0.01517424 | 0.02710865 | 1.60761248 | 3.33660889 | 0 | 0 | 9.50487297 |

| | | | | | | |
|---|---|---|---|---|---|---|
| 0.01517777 | 0.02711352 | 22.1850522 | 40.929069 | 0 | 50.1051043 | 0 |
| 0.01518525 | 0.027124 | 0 | 49.6042521 | 45.4745439 | 44.0739344 | 0 |
| 0.01518513 | 0.027124 | 0.64304499 | 0.44488118 | 0 | 0.46393615 | 0 |
| 0.01519926 | 0.02714758 | 9.64567488 | 10.4547078 | 10.2085711 | 8.35085072 | 26.1384007 |
| 0.01520474 | 0.02715594 | 0 | 48.4920492 | 82.5966206 | 20.8771268 | 0 |
| 0.01522223 | 0.02718573 | 8.6811074 | 11.1220296 | 5.5683115 | 5.10329766 | 0 |
| 0.01524612 | 0.02722694 | 9.64567488 | 72.5156331 | 0 | 51.9608489 | 0 |
| 0.0152806 | 0.02728707 | 0 | 61.3936035 | 61.2514265 | 23.6607437 | 0 |
| 0.01528643 | 0.02729604 | 4.82283744 | 4.67125244 | 18.5610383 | 9.27872303 | 0 |
| 0.01531558 | 0.02734518 | 0 | 0 | 0 | 0 | 0 |
| 0.01531558 | 0.02734518 | 0 | 0 | 0 | 0 | 0 |
| 0.01532223 | 0.0273556 | 8.3595849 | 14.9035197 | 7.42441534 | 3.71148921 | 0 |
| 0.01532406 | 0.02735743 | 30.5446371 | 39.816866 | 0 | 38.0427644 | 0 |
| 0.01532525 | 0.0273581 | 32.1522496 | 7.78542074 | 4.64025958 | 4.17542536 | 0 |
| 0.01533873 | 0.0273807 | 6.10892743 | 11.1220296 | 24.1293498 | 2.78361691 | 0 |
| 0.01535722 | 0.02741204 | 0 | 12.2342326 | 0 | 0 | 68.9103291 |
| 0.01535954 | 0.02741204 | 14.4685123 | 5.56101481 | 13.9207788 | 7.88691457 | 2.37621824 |
| 0.01535867 | 0.02741204 | 6.75197242 | 7.34053955 | 5.5683115 | 7.88691457 | 0 |
| 0.01535898 | 0.02741204 | 0 | 5.7834554 | 0 | 0 | 0 |
| 0.01537388 | 0.02743618 | 0 | 38.7046631 | 42.6903882 | 57.9920189 | 0 |
| 0.0153893 | 0.02746224 | 0 | 73.1829549 | 25.9854537 | 46.3936151 | 0 |
| 0.01539884 | 0.02747781 | 19.9343948 | 50.9388957 | 0 | 43.1460621 | 0 |
| 0.01542705 | 0.02752668 | 8.3595849 | 64.952653 | 0 | 63.0953166 | 0 |
| 0.01542986 | 0.02752731 | 0 | 34.4782918 | 72.3880495 | 36.1870198 | 0 |
| 0.01542983 | 0.02752731 | 0 | 0.22244059 | 0 | 0 | 0 |
| 0.01542829 | 0.02752731 | 25.4002772 | 8.45274251 | 10.2085711 | 8.81478687 | 0 |
| 0.01543546 | 0.02753585 | 1.60761248 | 9.34250488 | 31.5537652 | 10.2065953 | 0 |
| 0.01543688 | 0.02753691 | 33.1168171 | 34.2558512 | 0 | 40.3624452 | 0 |
| 0.015446 | 0.02755172 | 30.5446371 | 58.2794352 | 0 | 24.588616 | 0 |
| 0.01544742 | 0.02755281 | 34.0813846 | 34.4782918 | 0 | 38.9706367 | 0 |
| 0.0154524 | 0.02756022 | 34.4029071 | 42.041272 | 0 | 31.5476583 | 0 |
| 0.01545379 | 0.02756124 | 7.71653991 | 6.22833659 | 8.35246725 | 5.56723382 | 0 |
| 0.01545867 | 0.02756847 | 0 | 37.8149007 | 23.2012979 | 90.9314857 | 0 |
| 0.01547427 | 0.02759483 | 14.1469898 | 51.828658 | 0 | 55.208402 | 0 |
| 0.01547792 | 0.02759989 | 2.57217997 | 7.34053955 | 17.6329864 | 12.5262761 | 0 |
| 0.01548057 | 0.02760314 | 0 | 38.0373413 | 48.2586997 | 51.9608489 | 0 |
| 0.01549029 | 0.02761902 | 0 | 61.8384847 | 31.5537652 | 47.3214874 | 0 |
| 0.01550296 | 0.02764014 | 34.4029071 | 29.1397176 | 0 | 45.0018067 | 0 |
| 0.01550432 | 0.0276411 | 6.10892743 | 5.33857422 | 1.85610383 | 1.85574461 | 26.1384007 |
| 0.0155069 | 0.02764423 | 22.8280972 | 46.934965 | 0 | 41.2903175 | 0 |
| 0.01550775 | 0.02764428 | 0 | 0 | 0 | 0 | 7.12865473 |
| 0.01551343 | 0.02765295 | 9.64567488 | 7.34053955 | 2.78415575 | 20.8771268 | 0 |
| 0.015518 | 0.02765963 | 36.0105196 | 58.0569946 | 74.2441534 | 70.0543588 | 204.354769 |

| | | | | | | |
|---|---|---|---|---|---|---|
| 0.01551943 | 0.02766019 | 15.7546023 | 8.00786133 | 7.42441534 | 3.71148921 | 0 |
| 0.01551996 | 0.02766019 | 0.96456749 | 14.4586385 | 20.4171422 | 16.7017014 | 0 |
| 0.01552623 | 0.0276699 | 0 | 73.6278361 | 57.5392189 | 19.0213822 | 0 |
| 0.01553047 | 0.02767598 | 362.034331 | 639.294263 | 566.111669 | 600.33338 | 19.0097459 |
| 0.01553227 | 0.02767773 | 3.53674746 | 11.3444702 | 12.9927268 | 6.49510612 | 0 |
| 0.01553866 | 0.02768683 | 0 | 67.6219401 | 64.0355823 | 18.5574461 | 0 |
| 0.01553902 | 0.02768683 | 6.43044992 | 3.33660889 | 10.2085711 | 18.0935099 | 0 |
| 0.01554772 | 0.02769939 | 15.1115573 | 82.970341 | 0 | 29.6919137 | 0 |
| 0.01554757 | 0.02769939 | 2.25065747 | 8.8976237 | 8.35246725 | 28.3001052 | 0 |
| 0.01555058 | 0.02770302 | 3.21522496 | 11.7893514 | 9.28051917 | 9.27872303 | 0 |
| 0.01555926 | 0.02771702 | 0 | 1.55708415 | 0 | 0 | 0 |
| 0.01556803 | 0.02773118 | 78.7730116 | 172.6139 | 135.49558 | 154.490738 | 0 |
| 0.01558384 | 0.02775786 | 31.8307271 | 66.064856 | 74.2441534 | 55.208402 | 23.7621824 |
| 0.01559067 | 0.02776856 | 142.112943 | 135.911202 | 183.75428 | 227.79265 | 9.50487297 |
| 0.01559355 | 0.02777222 | 0 | 53.8306234 | 27.8415575 | 59.8477635 | 0 |
| 0.01559832 | 0.02777777 | 0 | 24.4684652 | 103.013763 | 28.7640414 | 0 |
| 0.01559796 | 0.02777777 | 9.64567488 | 10.2322673 | 1.85610383 | 24.1246799 | 0 |
| 0.01562604 | 0.02782566 | 0.96456749 | 15.5708415 | 24.1293498 | 12.9902122 | 0 |
| 0.01565893 | 0.02788276 | 56.9094818 | 60.7262817 | 0 | 104.385634 | 4.75243649 |
| 0.01566355 | 0.02788952 | 142.434466 | 145.253707 | 116.00649 | 132.685739 | 66.5341108 |
| 0.01566782 | 0.02789564 | 5.14435994 | 8.8976237 | 37.1220767 | 2.78361691 | 0 |
| 0.01567177 | 0.0279012 | 0 | 37.3700195 | 45.4745439 | 54.7444659 | 0 |
| 0.01567378 | 0.02790329 | 17.0406923 | 5.33857422 | 8.35246725 | 10.6705315 | 61.7816743 |
| 0.01568068 | 0.02791409 | 8.6811074 | 6.89565837 | 7.42441534 | 5.10329766 | 0 |
| 0.01568834 | 0.02792627 | 103.851766 | 26.4704305 | 0 | 11.1344676 | 0 |
| 0.01570479 | 0.02795406 | 15.7546023 | 8.23030192 | 3.71220767 | 6.95904227 | 0 |
| 0.01570674 | 0.02795605 | 106.102424 | 258.475968 | 206.955577 | 197.6368 | 49.9005831 |
| 0.01570828 | 0.02795732 | 0 | 139.02537 | 180.042072 | 38.0427644 | 7.12865473 |
| 0.01570923 | 0.02795752 | 33.7598621 | 45.822762 | 47.3306478 | 0 | 0 |
| 0.01571151 | 0.02796011 | 0 | 2.44684652 | 23.2012979 | 21.8049991 | 49.9005831 |
| 0.01574546 | 0.02801904 | 5.78740493 | 76.0746826 | 0 | 64.487125 | 0 |
| 0.01574851 | 0.02802298 | 91.9554339 | 28.2499552 | 0 | 12.9902122 | 0 |
| 0.01575615 | 0.02803437 | 0 | 23.5787028 | 72.3880495 | 48.7132959 | 0 |
| 0.01575732 | 0.02803437 | 4.82283744 | 8.67518311 | 6.49636342 | 8.81478687 | 0 |
| 0.01575741 | 0.02803437 | 4.17979245 | 0.88976237 | 0 | 0 | 4.75243649 |
| 0.01576034 | 0.02803663 | 117.034189 | 153.706449 | 155.912722 | 169.336695 | 11.8810912 |
| 0.01576031 | 0.02803663 | 0.96456749 | 18.4625692 | 30.6257133 | 8.35085072 | 0 |
| 0.01576175 | 0.02803765 | 198.057858 | 520.066105 | 438.968557 | 475.070619 | 95.0487297 |
| 0.01578284 | 0.02807368 | 30.5446371 | 12.4566732 | 8.35246725 | 18.5574461 | 54.6530196 |
| 0.01578952 | 0.02808408 | 26.3648447 | 8.00786133 | 9.28051917 | 2.31968076 | 0 |
| 0.01579223 | 0.02808742 | 103.530244 | 5.56101481 | 0 | 44.0739344 | 0 |
| 0.01582079 | 0.02813671 | 41.1548795 | 69.4014648 | 23.2012979 | 0 | 0 |
| 0.01582807 | 0.02814818 | 0.96456749 | 10.6771484 | 24.1293498 | 18.5574461 | 0 |

| | | | | | | |
|---|---|---|---|---|---|---|
| 0.01583236 | 0.02815433 | 50.8005544 | 14.6810791 | 0 | 52.8887212 | 0 |
| 0.01583657 | 0.02815923 | 23.7926647 | 63.1731283 | 0 | 57.9920189 | 2.37621824 |
| 0.01583679 | 0.02815923 | 0 | 0 | 0 | 0 | 11.8810912 |
| 0.01584704 | 0.02817596 | 31.8307271 | 36.2578166 | 0 | 38.0427644 | 0 |
| 0.01584919 | 0.02817829 | 137.933151 | 399.280863 | 352.659728 | 392.489984 | 0 |
| 0.0158551 | 0.0281873 | 1.28608998 | 14.4586385 | 12.0646749 | 20.4131907 | 0 |
| 0.01586027 | 0.02819501 | 72.0210391 | 46.934965 | 18.5610383 | 0 | 0 |
| 0.01589402 | 0.02825352 | 0 | 57.6121134 | 27.8415575 | 54.2805297 | 0 |
| 0.01590583 | 0.02827302 | 21.2204847 | 15.1259603 | 12.0646749 | 14.8459568 | 38.0194919 |
| 0.0159164 | 0.02829032 | 6.43044992 | 16.905485 | 11.136623 | 12.9902122 | 2.37621824 |
| 0.01591946 | 0.02829425 | 25.0787547 | 48.2696086 | 0 | 35.2591475 | 0 |
| 0.01593774 | 0.02832525 | 5.78740493 | 4.44881185 | 11.136623 | 9.74265918 | 0 |
| 0.01595922 | 0.02836193 | 2.25065747 | 10.6771484 | 7.42441534 | 25.0525522 | 0 |
| 0.0159752 | 0.02838882 | 160.118203 | 14.9035197 | 12.0646749 | 0 | 0 |
| 0.01598149 | 0.0283985 | 32.4737721 | 10.6771484 | 6.49636342 | 9.27872303 | 0 |
| 0.01598437 | 0.02840213 | 155.295366 | 189.074504 | 183.75428 | 193.461375 | 140.196876 |
| 0.01598526 | 0.02840219 | 0 | 0 | 0 | 0 | 4.75243649 |
| 0.0160038 | 0.02843213 | 6.75197242 | 8.8976237 | 5.5683115 | 6.49510612 | 0 |
| 0.01600348 | 0.02843213 | 10.9317649 | 8.00786133 | 9.28051917 | 5.10329766 | 114.058476 |
| 0.01602197 | 0.02846275 | 38.5826995 | 19.5747721 | 79.8124649 | 0 | 0 |
| 0.01602272 | 0.02846275 | 88.7402089 | 21.3542969 | 0 | 18.0935099 | 0 |
| 0.01602498 | 0.02846525 | 268.471284 | 356.79471 | 366.580507 | 395.273601 | 59.4054561 |
| 0.01603208 | 0.02847636 | 87.1325964 | 174.170984 | 151.272462 | 183.25478 | 66.5341108 |
| 0.01603575 | 0.0284799 | 45.9777169 | 121.230123 | 122.502853 | 143.356271 | 14.2573095 |
| 0.01603576 | 0.0284799 | 27.0078897 | 7.34053955 | 2.78415575 | 7.88691457 | 0 |
| 0.01604758 | 0.02849938 | 6.43044992 | 8.67518311 | 9.28051917 | 4.63936151 | 0 |
| 0.01605383 | 0.02850898 | 73.6286516 | 18.017688 | 0 | 28.3001052 | 0 |
| 0.01607954 | 0.02855313 | 0 | 18.4625692 | 59.3953227 | 69.1264865 | 0 |
| 0.01608405 | 0.02855962 | 4.82283744 | 5.56101481 | 13.9207788 | 7.88691457 | 0 |
| 0.01608967 | 0.02856208 | 0 | 0 | 0 | 0 | 0 |
| 0.01608967 | 0.02856208 | 0 | 0 | 0 | 0 | 0 |
| 0.01608967 | 0.02856208 | 0 | 0 | 0 | 0 | 0 |
| 0.01608967 | 0.02856208 | 0 | 0 | 0 | 0 | 0 |
| 0.01608967 | 0.02856208 | 0 | 0 | 0 | 0 | 0 |
| 0.01609524 | 0.02857045 | 14.4685123 | 29.8070394 | 0 | 81.1888265 | 0 |
| 0.01609978 | 0.028577 | 0.64304499 | 1.33464355 | 3.71220767 | 0.46393615 | 2.37621824 |
| 0.01610253 | 0.02858038 | 4.50131495 | 13.5688761 | 7.42441534 | 6.49510612 | 0 |
| 0.01611045 | 0.02859293 | 106.745469 | 10.2322673 | 0 | 25.5164883 | 0 |
| 0.01611277 | 0.02859554 | 27.3294122 | 37.1475789 | 0 | 41.7542536 | 0 |
| 0.01611659 | 0.0286008 | 32.1522496 | 48.4920492 | 0 | 26.9082968 | 0 |
| 0.01612938 | 0.028622 | 5.78740493 | 73.1829549 | 89.092984 | 0 | 0 |
| 0.01617006 | 0.02869267 | 18.3267823 | 58.0569946 | 0 | 37.1148921 | 0 |
| 0.01617668 | 0.02870291 | 0 | 0 | 0 | 0 | 0 |

| | | | | | | |
|---|---|---|---|---|---|---|
| 0.01618209 | 0.02871099 | 0 | 26.9153117 | 42.6903882 | 70.9822312 | 0 |
| 0.01619655 | 0.02873513 | 37.2966096 | 26.9153117 | 35.2659728 | 23.6607437 | 78.415202 |
| 0.01620079 | 0.02874113 | 6.43044992 | 8.23030192 | 25.0574018 | 15.309893 | 0 |
| 0.01622144 | 0.02877625 | 16.0761248 | 19.7972127 | 17.6329864 | 21.341063 | 28.5146189 |
| 0.01622242 | 0.02877648 | 72.9856066 | 148.590316 | 162.409085 | 153.562866 | 42.7719284 |
| 0.01622483 | 0.02877922 | 8.3595849 | 6.67321777 | 3.71220767 | 15.309893 | 0 |
| 0.01623739 | 0.02879999 | 0 | 68.7341431 | 22.273246 | 51.4969128 | 0 |
| 0.01623895 | 0.02880069 | 0 | 30.2519206 | 44.546492 | 62.6313804 | 0 |
| 0.0162395 | 0.02880069 | 9.32415239 | 64.0628906 | 0 | 55.208402 | 0 |
| 0.01624679 | 0.02881059 | 5.46588243 | 7.11809896 | 6.49636342 | 8.35085072 | 0 |
| 0.01624669 | 0.02881059 | 66.5551567 | 117.671073 | 74.2441534 | 100.210209 | 0 |
| 0.01625207 | 0.02881843 | 169.442355 | 272.267285 | 273.775316 | 149.387441 | 21.3859642 |
| 0.01625419 | 0.02882068 | 66.2336342 | 132.797034 | 88.1649321 | 98.3544641 | 42.7719284 |
| 0.01625965 | 0.02882883 | 36.6535646 | 10.2322673 | 6.49636342 | 8.81478687 | 2.37621824 |
| 0.0162633 | 0.02883378 | 6.75197242 | 5.7834554 | 7.42441534 | 6.95904227 | 0 |
| 0.01626454 | 0.02883446 | 0 | 23.1338216 | 94.6612955 | 32.0115944 | 0 |
| 0.01629358 | 0.02888443 | 1.92913498 | 14.6810791 | 10.2085711 | 13.9180845 | 0 |
| 0.016303 | 0.0288996 | 19.9343948 | 40.929069 | 38.0501286 | 31.0837221 | 14.2573095 |
| 0.01630501 | 0.02890165 | 13.5039448 | 47.1574056 | 46.4025958 | 50.1051043 | 14.2573095 |
| 0.01631377 | 0.02891564 | 13.8254673 | 7.34053955 | 9.28051917 | 6.49510612 | 73.6627655 |
| 0.01633268 | 0.02894764 | 0 | 55.3877075 | 81.6685687 | 14.8459568 | 0 |
| 0.01637353 | 0.02901851 | 36.6535646 | 20.9094157 | 0 | 51.9608489 | 0 |
| 0.01637643 | 0.02902212 | 19.6128723 | 0 | 68.6758419 | 67.2707419 | 0 |
| 0.01638021 | 0.02902729 | 33.4383396 | 0 | 76.1002572 | 39.4345729 | 0 |
| 0.01639403 | 0.02905026 | 27.6509347 | 48.9369303 | 49.1867516 | 0 | 0 |
| 0.01640254 | 0.02906294 | 14.7900348 | 7.34053955 | 2.78415575 | 11.1344676 | 0 |
| 0.01640292 | 0.02906294 | 70.7349491 | 187.072538 | 174.47376 | 159.1301 | 45.1481466 |
| 0.0164083 | 0.02907095 | 30.8661596 | 64.5077718 | 0 | 19.0213822 | 0 |
| 0.01642914 | 0.02910633 | 2.57217997 | 8.23030192 | 18.5610383 | 9.74265918 | 0 |
| 0.01643622 | 0.02911734 | 59.4816618 | 257.808647 | 344.307261 | 240.318926 | 2.37621824 |
| 0.01643945 | 0.02912154 | 5.14435994 | 5.56101481 | 9.28051917 | 8.81478687 | 0 |
| 0.01644457 | 0.02912907 | 14.4685123 | 47.6022868 | 0 | 53.8165936 | 0 |
| 0.01647234 | 0.02917673 | 72.9856066 | 0.66732178 | 9.28051917 | 5.10329766 | 0 |
| 0.01647571 | 0.02918116 | 17.6837373 | 11.7893514 | 9.28051917 | 10.6705315 | 33.2670554 |
| 0.01648194 | 0.0291865 | 22.1850522 | 38.2597819 | 12.0646749 | 31.5476583 | 30.8908372 |
| 0.01648217 | 0.0291865 | 3.53674746 | 10.899589 | 9.28051917 | 7.88691457 | 0 |
| 0.01648219 | 0.0291865 | 20.5774398 | 29.8070394 | 0 | 62.1674443 | 0 |
| 0.01648025 | 0.0291865 | 7.07349491 | 12.2342326 | 4.64025958 | 6.49510612 | 0 |
| 0.01648412 | 0.02918836 | 226.994882 | 318.534928 | 354.515832 | 340.065199 | 0 |
| 0.01649238 | 0.02920147 | 2.25065747 | 3.78149007 | 34.3379209 | 14.8459568 | 0 |
| 0.01649985 | 0.02921316 | 4.82283744 | 13.5688761 | 28.7696094 | 2.31968076 | 0 |
| 0.01650937 | 0.02922846 | 49.5144644 | 23.1338216 | 0 | 34.7952113 | 0 |
| 0.0165248 | 0.02925425 | 0 | 0 | 0 | 0 | 0 |

| | | | | | | |
|---|---|---|---|---|---|---|
| 0.01653024 | 0.02926233 | 0 | 40.929069 | 41.7623363 | 50.1051043 | 0 |
| 0.01653389 | 0.02926726 | 2.25065747 | 12.011792 | 11.136623 | 12.0623399 | 0 |
| 0.0165363 | 0.02926999 | 3.21522496 | 7.56298014 | 10.2085711 | 12.5262761 | 0 |
| 0.01653849 | 0.02927232 | 2.57217997 | 10.899589 | 16.7049345 | 7.88691457 | 0 |
| 0.01654025 | 0.0292739 | 45.0131495 | 24.023584 | 0 | 37.1148921 | 0 |
| 0.01655773 | 0.0293033 | 3.85826995 | 11.1220296 | 12.9927268 | 5.56723382 | 0 |
| 0.01656818 | 0.02932025 | 5.78740493 | 4.89369303 | 8.35246725 | 10.2065953 | 0 |
| 0.01657359 | 0.02932828 | 0 | 27.3601929 | 50.1148035 | 58.9198912 | 0 |
| 0.01657772 | 0.02933404 | 15.7546023 | 46.4900838 | 0 | 51.0329766 | 0 |
| 0.01657897 | 0.0293347 | 6.43044992 | 3.55904948 | 9.28051917 | 15.7738291 | 0 |
| 0.0166055 | 0.0293801 | 8.6811074 | 7.11809896 | 9.28051917 | 4.17542536 | 0 |
| 0.01661002 | 0.02938501 | 0.3215225 | 0.88976237 | 0 | 0.46393615 | 0 |
| 0.0166092 | 0.02938501 | 3.21522496 | 12.9015544 | 10.2085711 | 7.42297842 | 0 |
| 0.01661591 | 0.02939388 | 66.5551567 | 5.7834554 | 0.92805192 | 7.88691457 | 0 |
| 0.016625 | 0.02940842 | 2.57217997 | 13.5688761 | 8.35246725 | 12.0623399 | 0 |
| 0.01663102 | 0.02941751 | 3.53674746 | 3.78149007 | 14.8488307 | 20.8771268 | 0 |
| 0.01664287 | 0.02943693 | 6.75197242 | 6.45077718 | 7.42441534 | 6.03116997 | 0 |
| 0.01664431 | 0.02943794 | 11.8963324 | 3.78149007 | 5.5683115 | 12.5262761 | 0 |
| 0.01665923 | 0.02946278 | 0 | 39.816866 | 57.5392189 | 36.1870198 | 0 |
| 0.01666293 | 0.02946776 | 1.92913498 | 14.4586385 | 12.9927268 | 10.6705315 | 0 |
| 0.01666679 | 0.02947305 | 6.10892743 | 15.1259603 | 4.64025958 | 6.95904227 | 0 |
| 0.01667401 | 0.02948426 | 4.17979245 | 24.6909058 | 11.136623 | 18.0935099 | 0 |
| 0.01667968 | 0.02949274 | 17.0406923 | 96.9840983 | 65.8916861 | 90.9314857 | 21.3859642 |
| 0.01668883 | 0.02950737 | 56.2664368 | 18.017688 | 12.9927268 | 18.0935099 | 68.9103291 |
| 0.01669351 | 0.02951408 | 194.52111 | 383.0427 | 371.220767 | 379.963708 | 54.6530196 |
| 0.01669482 | 0.02951486 | 11.5748099 | 28.6948364 | 5.5683115 | 2.31968076 | 0 |
| 0.01670009 | 0.02952262 | 24.1141872 | 29.3621582 | 17.6329864 | 28.3001052 | 0 |
| 0.01670132 | 0.02952324 | 163.333428 | 130.795068 | 93.7332436 | 117.375846 | 64.1578926 |
| 0.0167075 | 0.02953262 | 4.82283744 | 8.45274251 | 9.28051917 | 6.03116997 | 0 |
| 0.01671086 | 0.029537 | 54.9803468 | 20.9094157 | 52.8989593 | 0 | 0 |
| 0.01671302 | 0.02953926 | 18.6483048 | 3.33660889 | 0 | 3.71148921 | 52.2768014 |
| 0.01671962 | 0.02954939 | 3.53674746 | 88.0864746 | 0 | 64.9510612 | 0 |
| 0.0167412 | 0.02958597 | 0 | 0 | 0 | 5.56723382 | 9.50487297 |
| 0.01676937 | 0.02963419 | 39.2257445 | 35.3680542 | 0 | 28.7640414 | 0 |
| 0.01679209 | 0.02967279 | 78.7730116 | 26.9153117 | 26.9135056 | 0 | 0 |
| 0.01679566 | 0.02967754 | 4.50131495 | 2.44684652 | 0 | 6.03116997 | 71.2865473 |
| 0.01681705 | 0.02971377 | 9.32415239 | 81.4132568 | 0 | 38.9706367 | 0 |
| 0.01682983 | 0.02973479 | 28.6155022 | 64.2853312 | 45.4745439 | 33.4034029 | 19.0097459 |
| 0.0168322 | 0.02973586 | 0.3215225 | 0 | 0 | 0 | 14.2573095 |
| 0.01683205 | 0.02973586 | 0 | 42.7085937 | 57.5392189 | 32.9394667 | 0 |
| 0.01683406 | 0.02973757 | 0 | 69.846346 | 14.8488307 | 63.0953166 | 0 |
| 0.01684337 | 0.02975246 | 19.9343948 | 9.78738607 | 12.0646749 | 11.5984038 | 90.2962933 |
| 0.0168514 | 0.02976508 | 3.85826995 | 12.6791138 | 11.136623 | 5.56723382 | 0 |

| | | | | | | |
|---|---|---|---|---|---|---|
| 0.01685519 | 0.02976865 | 1.28608998 | 6.67321777 | 17.6329864 | 27.8361691 | 0 |
| 0.01685513 | 0.02976865 | 6.10892743 | 13.5688761 | 10.2085711 | 19.4853184 | 0 |
| 0.01687018 | 0.02979356 | 0 | 28.4723958 | 75.1722053 | 34.7952113 | 0 |
| 0.01689347 | 0.02983314 | 55.6233918 | 28.2499552 | 36.1940248 | 29.6919137 | 154.454186 |
| 0.01690028 | 0.0298436 | 4.17979245 | 7.78542074 | 26.9135056 | 4.63936151 | 0 |
| 0.01690444 | 0.02984937 | 27.3294122 | 57.834554 | 0 | 23.6607437 | 0 |
| 0.01691066 | 0.02985879 | 12.5393773 | 28.917277 | 31.5537652 | 24.588616 | 7.12865473 |
| 0.01691494 | 0.02986477 | 11.8963324 | 16.0157227 | 15.7768826 | 18.0935099 | 2.37621824 |
| 0.01692969 | 0.02988925 | 26.0433222 | 29.8070394 | 74.2441534 | 0 | 0 |
| 0.016935 | 0.02989705 | 3.21522496 | 3.78149007 | 18.5610383 | 18.0935099 | 0 |
| 0.01693695 | 0.02989737 | 0 | 0 | 0 | 0 | 0 |
| 0.01693695 | 0.02989737 | 0 | 0 | 0 | 0 | 0 |
| 0.01693793 | 0.02989752 | 5.78740493 | 16.0157227 | 11.136623 | 15.309893 | 0 |
| 0.01696737 | 0.02994791 | 55.6233918 | 57.3896728 | 46.4025958 | 61.239572 | 26.1384007 |
| 0.01698521 | 0.02997783 | 0 | 187.072538 | 177.257916 | 114.128293 | 9.50487297 |
| 0.0170136 | 0.03002637 | 0 | 0 | 0 | 0 | 95.0487297 |
| 0.01701777 | 0.03003211 | 4.82283744 | 18.4625692 | 12.9927268 | 3.24755306 | 0 |
| 0.01701864 | 0.03003211 | 34.4029071 | 71.1809896 | 0 | 13.4541484 | 0 |
| 0.01702142 | 0.03003544 | 58.5170943 | 116.55887 | 103.013763 | 79.797018 | 40.3957101 |
| 0.01703522 | 0.03005663 | 0 | 0 | 0 | 0 | 42.7719284 |
| 0.01703522 | 0.03005663 | 0 | 0 | 0 | 0 | 42.7719284 |
| 0.01704249 | 0.03006788 | 0 | 0 | 0 | 0 | 0 |
| 0.01704353 | 0.03006814 | 13.1824223 | 33.1436483 | 15.7768826 | 17.1656376 | 7.12865473 |
| 0.01705446 | 0.03008586 | 2.89370247 | 10.0098267 | 8.35246725 | 12.9902122 | 0 |
| 0.01708638 | 0.03014059 | 2.25065747 | 14.9035197 | 8.35246725 | 12.9902122 | 0 |
| 0.01709141 | 0.03014661 | 0 | 25.8031087 | 45.4745439 | 64.0231889 | 0 |
| 0.01709159 | 0.03014661 | 0 | 0 | 0 | 0 | 23.7621824 |
| 0.01710809 | 0.03017255 | 13.1824223 | 0 | 131.783372 | 31.5476583 | 0 |
| 0.01710752 | 0.03017255 | 0.96456749 | 26.2479899 | 12.0646749 | 13.4541484 | 0 |
| 0.01711975 | 0.03019153 | 36.3320421 | 28.2499552 | 0 | 37.5788283 | 0 |
| 0.0171243 | 0.03019797 | 28.9370247 | 45.822762 | 0 | 28.7640414 | 0 |
| 0.01713282 | 0.03021141 | 111.889829 | 245.796855 | 364.724403 | 320.115944 | 0 |
| 0.01714571 | 0.03023256 | 1.92913498 | 26.2479899 | 4.64025958 | 17.6295737 | 0 |
| 0.01715571 | 0.0302486 | 26.0433222 | 20.2420939 | 14.8488307 | 26.9082968 | 11.8810912 |
| 0.01716146 | 0.03025716 | 0.96456749 | 0.88976237 | 0 | 1.39180845 | 2.37621824 |
| 0.01716604 | 0.03026365 | 17.0406923 | 1.33464355 | 12.0646749 | 14.8459568 | 0 |
| 0.01717426 | 0.03027656 | 11.8963324 | 4.44881185 | 10.2085711 | 5.10329766 | 0 |
| 0.01719389 | 0.03030957 | 101.279586 | 16.4606038 | 0 | 15.7738291 | 0 |
| 0.01720902 | 0.03033207 | 21.2204847 | 22.4664998 | 20.4171422 | 20.4131907 | 66.5341108 |
| 0.01720913 | 0.03033207 | 15.7546023 | 67.6219401 | 0 | 31.5476583 | 0 |
| 0.01720936 | 0.03033207 | 9.00262989 | 18.9074504 | 1.85610383 | 11.5984038 | 0 |
| 0.01721794 | 0.03034561 | 1.92913498 | 17.7952474 | 8.35246725 | 13.4541484 | 0 |
| 0.01721979 | 0.03034727 | 0 | 90.7557617 | 12.9927268 | 48.2493597 | 0 |

| | | | | | | |
|---|---|---|---|---|---|---|
| 0.01723453 | 0.03037031 | 0 | 63.6180094 | 41.7623363 | 28.3001052 | 0 |
| 0.01723467 | 0.03037031 | 6.43044992 | 6.005896 | 4.64025958 | 17.6295737 | 0 |
| 0.01723665 | 0.03037222 | 2.25065747 | 9.56494547 | 21.3451941 | 8.35085072 | 0 |
| 0.01724249 | 0.03038091 | 4.17979245 | 6.89565837 | 8.35246725 | 9.74265918 | 0 |
| 0.01725415 | 0.03039986 | 0 | 80.0786133 | 21.3451941 | 39.4345729 | 0 |
| 0.01727634 | 0.03043577 | 5.78740493 | 6.89565837 | 5.5683115 | 9.27872303 | 0 |
| 0.01727616 | 0.03043577 | 103.851766 | 69.846346 | 71.4599976 | 69.1264865 | 259.007789 |
| 0.01727749 | 0.03043621 | 5.78740493 | 7.78542074 | 5.5683115 | 7.88691457 | 0 |
| 0.01728977 | 0.03045466 | 1.60761248 | 9.78738607 | 14.8488307 | 16.7017014 | 0 |
| 0.01728963 | 0.03045466 | 0.3215225 | 0.66732178 | 0 | 0 | 0 |
| 0.0172951 | 0.03046244 | 0 | 39.1495443 | 26.9135056 | 69.1264865 | 0 |
| 0.0173034 | 0.03047546 | 0 | 68.0668213 | 28.7696094 | 37.5788283 | 0 |
| 0.01731397 | 0.03049249 | 3.85826995 | 5.11613363 | 10.2085711 | 16.2377653 | 0 |
| 0.01733391 | 0.03052602 | 10.9317649 | 13.3464355 | 14.8488307 | 17.6295737 | 4.75243649 |
| 0.01733648 | 0.03052893 | 7.71653991 | 16.6830444 | 8.35246725 | 12.5262761 | 2.37621824 |
| 0.01734165 | 0.03053644 | 0.96456749 | 14.9035197 | 19.4890903 | 14.3820207 | 0 |
| 0.01734875 | 0.03054735 | 9.32415239 | 44.4881185 | 0 | 70.518295 | 0 |
| 0.01735796 | 0.03056197 | 8.0380624 | 63.6180094 | 0 | 55.208402 | 0 |
| 0.01738999 | 0.03061676 | 20.2559173 | 34.4782918 | 0 | 51.4969128 | 0 |
| 0.01739191 | 0.03061853 | 5.14435994 | 8.00786133 | 6.49636342 | 7.42297842 | 0 |
| 0.0174017 | 0.03063417 | 1.60761248 | 3.33660889 | 3.71220767 | 3.71148921 | 21.3859642 |
| 0.01741236 | 0.03065133 | 5.46588243 | 7.56298014 | 4.64025958 | 14.3820207 | 0 |
| 0.01743738 | 0.03069376 | 6.10892743 | 14.6810791 | 13.9207788 | 11.1344676 | 2.37621824 |
| 0.01745844 | 0.03072923 | 3.85826995 | 12.2342326 | 3.71220767 | 21.341063 | 0 |
| 0.01746015 | 0.03073063 | 42.119447 | 34.2558512 | 42.6903882 | 0 | 0 |
| 0.01746437 | 0.03073644 | 2.25065747 | 1.33464355 | 0 | 1.39180845 | 26.1384007 |
| 0.01746788 | 0.03074102 | 0 | 74.0727173 | 39.9062324 | 23.1968076 | 0 |
| 0.01746884 | 0.03074109 | 8.3595849 | 66.7321777 | 0 | 50.5690405 | 0 |
| 0.01747018 | 0.03074185 | 15.7546023 | 44.4881185 | 0 | 49.177232 | 0 |
| 0.01748183 | 0.03076075 | 26.6863672 | 42.7085937 | 0 | 32.4755306 | 0 |
| 0.01749451 | 0.03078144 | 5.14435994 | 10.6771484 | 5.5683115 | 7.42297842 | 0 |
| 0.01750489 | 0.0307981 | 1.60761248 | 11.3444702 | 27.8415575 | 7.88691457 | 0 |
| 0.01751643 | 0.03081679 | 7.07349491 | 3.11416829 | 9.28051917 | 15.7738291 | 0 |
| 0.01752974 | 0.03083704 | 0 | 40.0393066 | 46.4025958 | 41.7542536 | 0 |
| 0.01752977 | 0.03083704 | 0 | 42.2637126 | 68.6758419 | 24.1246799 | 0 |
| 0.01753517 | 0.03084492 | 18.3267823 | 11.5669108 | 10.2085711 | 11.5984038 | 47.5243649 |
| 0.01753658 | 0.03084579 | 27.6509347 | 41.1515096 | 0 | 32.4755306 | 0 |
| 0.01753947 | 0.03084926 | 50.4790319 | 30.6968018 | 0 | 23.1968076 | 0 |
| 0.01755299 | 0.03087142 | 219.921387 | 383.265141 | 313.681548 | 389.242431 | 57.0292378 |
| 0.01755605 | 0.0308752 | 99.6719738 | 14.4586385 | 0 | 17.6295737 | 0 |
| 0.01755844 | 0.03087778 | 0 | 0 | 19.4890903 | 16.2377653 | 173.463932 |
| 0.01758429 | 0.03092162 | 9.32415239 | 13.5688761 | 14.8488307 | 6.95904227 | 2.37621824 |
| 0.0175876 | 0.03092583 | 6.43044992 | 1.55708415 | 3.71220767 | 3.24755306 | 7.12865473 |

| | | | | | | |
|---|---|---|---|---|---|---|
| 0.01759012 | 0.03092626 | 0 | 0 | 0 | 0 | 0 |
| 0.0175906 | 0.03092626 | 0 | 49.3818115 | 38.9781805 | 39.898509 | 0 |
| 0.01759012 | 0.03092626 | 0 | 0 | 0 | 0 | 0 |
| 0.01761656 | 0.03097028 | 32.4737721 | 4.44881185 | 3.71220767 | 7.42297842 | 0 |
| 0.0176229 | 0.03097981 | 0 | 10.0098267 | 0 | 0 | 42.7719284 |
| 0.0176302 | 0.03099101 | 4.50131495 | 10.2322673 | 5.5683115 | 9.27872303 | 0 |
| 0.01764776 | 0.03102026 | 0 | 0 | 0 | 0 | 40.3957101 |
| 0.01765076 | 0.0310234 | 38.261177 | 40.0393066 | 0 | 23.6607437 | 0 |
| 0.01765139 | 0.0310234 | 13.8254673 | 44.4881185 | 0 | 53.3526574 | 0 |
| 0.01766473 | 0.03104522 | 280.046094 | 800.118811 | 786.988026 | 778.948798 | 199.602332 |
| 0.01767274 | 0.03105768 | 0 | 0 | 0 | 0.46393615 | 0 |
| 0.01767822 | 0.03106569 | 43.0840145 | 60.2814006 | 0 | 12.0623399 | 0 |
| 0.01768306 | 0.03107257 | 3.21522496 | 12.2342326 | 6.49636342 | 11.5984038 | 0 |
| 0.01769313 | 0.03108864 | 2.89370247 | 9.12006429 | 18.5610383 | 6.95904227 | 0 |
| 0.01771154 | 0.03111937 | 6.43044992 | 4.22637126 | 11.136623 | 7.88691457 | 0 |
| 0.01771765 | 0.03112848 | 31.8307271 | 52.4959798 | 0 | 20.8771268 | 0 |
| 0.01772836 | 0.03114567 | 14.7900348 | 14.4586385 | 11.136623 | 12.9902122 | 2.37621824 |
| 0.01774147 | 0.03116707 | 34.7244296 | 10.0098267 | 0 | 79.3330819 | 0 |
| 0.01777586 | 0.03122586 | 6.43044992 | 15.3484009 | 0 | 142.892335 | 0 |
| 0.01778523 | 0.03124069 | 48.5498969 | 20.6869751 | 0 | 34.7952113 | 0 |
| 0.01778701 | 0.03124218 | 146.614258 | 405.731641 | 309.96934 | 347.952113 | 0 |
| 0.01779092 | 0.03124742 | 0 | 66.9546183 | 39.9062324 | 25.9804245 | 0 |
| 0.01780424 | 0.03126917 | 1.28608998 | 13.3464355 | 12.0646749 | 18.5574461 | 0 |
| 0.01782271 | 0.03129998 | 25.0787547 | 22.2440592 | 3.71220767 | 1.85574461 | 0 |
| 0.01782555 | 0.03130333 | 27.3294122 | 18.4625692 | 3.71220767 | 0 | 111.682257 |
| 0.01784803 | 0.03134117 | 54.0157793 | 11.1220296 | 0 | 50.5690405 | 0 |
| 0.0178607 | 0.03136179 | 0 | 36.4802572 | 64.9636342 | 29.6919137 | 0 |
| 0.01786804 | 0.03137305 | 0 | 0 | 0 | 0 | 0 |
| 0.01788678 | 0.03140431 | 6.10892743 | 9.34250488 | 3.71220767 | 12.0623399 | 0 |
| 0.01789845 | 0.03142315 | 4.82283744 | 1.33464355 | 1.85610383 | 3.24755306 | 14.2573095 |
| 0.01791627 | 0.03145199 | 25.7217997 | 41.3739502 | 0 | 33.4034029 | 0 |
| 0.01791674 | 0.03145199 | 0.96456749 | 11.7893514 | 14.8488307 | 22.7328714 | 0 |
| 0.01793037 | 0.03147427 | 0 | 48.047168 | 33.409869 | 45.929679 | 0 |
| 0.01794603 | 0.03149988 | 4.17979245 | 7.56298014 | 13.9207788 | 6.03116997 | 0 |
| 0.01794683 | 0.03149988 | 12.5393773 | 66.5097371 | 0 | 36.1870198 | 0 |
| 0.01796444 | 0.03152913 | 4.50131495 | 18.9074504 | 3.71220767 | 10.6705315 | 0 |
| 0.01798056 | 0.03155578 | 16.7191698 | 7.56298014 | 2.78415575 | 8.81478687 | 0 |
| 0.01799228 | 0.03157306 | 5.78740493 | 13.7913167 | 3.71220767 | 9.27872303 | 0 |
| 0.01799201 | 0.03157306 | 25.4002772 | 75.852242 | 0 | 15.309893 | 0 |
| 0.0180018 | 0.03158811 | 13.1824223 | 28.2499552 | 0 | 77.0134011 | 0 |
| 0.01802052 | 0.03161932 | 120.570936 | 150.814722 | 90.9490879 | 83.0445711 | 251.879134 |
| 0.01802415 | 0.03162404 | 16.3976473 | 47.8247274 | 0 | 42.2181898 | 0 |
| 0.01803499 | 0.0316414 | 0 | 36.035376 | 35.2659728 | 56.6002105 | 0 |

| | | | | | | |
|---|---|---|---|---|---|---|
| 0.0180396 | 0.03164784 | 4.17979245 | 70.958549 | 0 | 67.2707419 | 0 |
| 0.01804311 | 0.03165234 | 4.17979245 | 10.6771484 | 7.42441534 | 6.95904227 | 0 |
| 0.01807855 | 0.03170956 | 59.4816618 | 87.4191528 | 63.1075304 | 58.4559551 | 0 |
| 0.01807764 | 0.03170956 | 4.17979245 | 8.00786133 | 9.28051917 | 6.95904227 | 0 |
| 0.01807681 | 0.03170956 | 68.1627692 | 38.9271037 | 70.5319457 | 39.898509 | 30.8908372 |
| 0.01808266 | 0.03171513 | 43.0840145 | 12.011792 | 2.78415575 | 13.4541484 | 0 |
| 0.01809472 | 0.03173462 | 13.8254673 | 48.7144897 | 69.6038938 | 59.8477635 | 16.6335277 |
| 0.01809875 | 0.03174003 | 5.46588243 | 0 | 102.085711 | 73.3019119 | 0 |
| 0.01812017 | 0.03177594 | 19.6128723 | 157.265499 | 50.1148035 | 74.2297842 | 0 |
| 0.01813282 | 0.03179646 | 3.53674746 | 11.5669108 | 5.5683115 | 12.5262761 | 0 |
| 0.01814107 | 0.03180926 | 36.0105196 | 34.4782918 | 0 | 28.3001052 | 0 |
| 0.01815043 | 0.03182402 | 9.96719738 | 19.5747721 | 9.28051917 | 32.9394667 | 4.75243649 |
| 0.01815353 | 0.0318278 | 0 | 40.4841878 | 57.5392189 | 30.1558498 | 0 |
| 0.01817032 | 0.03185059 | 0 | 0 | 0 | 0 | 0 |
| 0.01817032 | 0.03185059 | 0 | 0 | 0 | 0 | 0 |
| 0.01817032 | 0.03185059 | 0 | 0 | 0 | 0 | 0 |
| 0.01817032 | 0.03185059 | 0 | 0 | 0 | 0 | 0 |
| 0.01817379 | 0.03185502 | 121.535504 | 140.360014 | 137.351684 | 124.798825 | 0 |
| 0.01818173 | 0.03186728 | 0 | 51.3837769 | 29.6976613 | 46.3936151 | 0 |
| 0.01818481 | 0.03187101 | 9.32415239 | 36.2578166 | 0 | 77.4773373 | 0 |
| 0.01818683 | 0.03187289 | 2.57217997 | 12.011792 | 6.49636342 | 16.7017014 | 0 |
| 0.01819026 | 0.03187724 | 4.50131495 | 17.7952474 | 4.64025958 | 8.35085072 | 0 |
| 0.01819432 | 0.03188271 | 17.6837373 | 17.1279256 | 24.1293498 | 20.4131907 | 9.50487297 |
| 0.01820057 | 0.031892 | 0 | 31.1416829 | 53.8270112 | 42.2181898 | 0 |
| 0.01820421 | 0.03189671 | 11.8963324 | 4.67125244 | 6.49636342 | 6.49510612 | 0 |
| 0.01822585 | 0.03193296 | 27.3294122 | 13.5688761 | 12.9927268 | 13.4541484 | 38.0194919 |
| 0.01823579 | 0.03194872 | 9.96719738 | 51.6062174 | 0 | 55.208402 | 0 |
| 0.01824353 | 0.0319606 | 39.8687895 | 0 | 33.409869 | 64.487125 | 0 |
| 0.01825751 | 0.03198301 | 37.618132 | 44.9329997 | 33.409869 | 39.4345729 | 33.2670554 |
| 0.01825821 | 0.03198301 | 3.53674746 | 8.00786133 | 15.7768826 | 6.49510612 | 0 |
| 0.01826369 | 0.03199093 | 9.32415239 | 43.5983561 | 0 | 66.8068058 | 0 |
| 0.01826628 | 0.0319938 | 0 | 28.4723958 | 44.546492 | 54.7444659 | 0 |
| 0.01829546 | 0.03204324 | 0 | 22.4664998 | 59.3953227 | 49.177232 | 0 |
| 0.01830354 | 0.03205573 | 15.4330798 | 0 | 76.1002572 | 58.4559551 | 0 |
| 0.01830995 | 0.03206528 | 7.07349491 | 24.023584 | 67.7477899 | 37.5788283 | 9.50487297 |
| 0.0183155 | 0.03207334 | 12.2178549 | 3.33660889 | 7.42441534 | 8.81478687 | 0 |
| 0.01832032 | 0.0320801 | 0 | 58.5018758 | 13.9207788 | 68.6625504 | 0 |
| 0.01832626 | 0.03208884 | 5.78740493 | 6.89565837 | 5.5683115 | 8.35085072 | 0 |
| 0.01834095 | 0.03211289 | 165.905608 | 512.725566 | 521.565177 | 538.629872 | 28.5146189 |
| 0.01834411 | 0.03211675 | 1.28608998 | 11.3444702 | 10.2085711 | 25.0525522 | 0 |
| 0.01835159 | 0.03212817 | 7.39501741 | 5.33857422 | 5.5683115 | 8.81478687 | 0 |
| 0.01835665 | 0.03213537 | 5.46588243 | 6.89565837 | 9.28051917 | 5.56723382 | 0 |
| 0.01838028 | 0.03217505 | 7.07349491 | 9.56494547 | 9.28051917 | 4.17542536 | 7.12865473 |

| | | | | | | |
|---|---|---|---|---|---|---|
| 0.01839773 | 0.03220239 | 2.89370247 | 8.00786133 | 11.136623 | 10.6705315 | 0 |
| 0.01839781 | 0.03220239 | 10.2887199 | 51.6062174 | 0 | 53.3526574 | 0 |
| 0.01841959 | 0.03223883 | 7.39501741 | 7.34053955 | 4.64025958 | 7.42297842 | 0 |
| 0.01842251 | 0.03224226 | 0.64304499 | 0.22244059 | 0 | 0 | 0 |
| 0.01842514 | 0.03224519 | 1.60761248 | 8.23030192 | 20.4171422 | 13.4541484 | 0 |
| 0.01843677 | 0.03226386 | 53.6942569 | 21.7991781 | 15.7768826 | 17.1656376 | 9.50487297 |
| 0.01846714 | 0.03231534 | 12.2178549 | 10.6771484 | 12.0646749 | 6.03116997 | 23.7621824 |
| 0.01846827 | 0.03231563 | 23.1496197 | 35.1456136 | 63.1075304 | 0 | 0 |
| 0.0184791 | 0.0323329 | 12.2178549 | 65.6199748 | 43.6184401 | 41.7542536 | 14.2573095 |
| 0.01849039 | 0.03235096 | 0 | 6.005896 | 0 | 5.56723382 | 57.0292378 |
| 0.01851233 | 0.03238767 | 12.2178549 | 14.6810791 | 12.0646749 | 16.7017014 | 0 |
| 0.01851714 | 0.0323944 | 26.3648447 | 62.0609253 | 40.8342843 | 51.0329766 | 19.0097459 |
| 0.01851865 | 0.03239536 | 18.3267823 | 31.5865641 | 0 | 54.7444659 | 0 |
| 0.01852317 | 0.03240158 | 0 | 26.9153117 | 57.5392189 | 43.1460621 | 0 |
| 0.01852668 | 0.03240604 | 62.0538418 | 49.3818115 | 100.229607 | 141.500526 | 47.5243649 |
| 0.01854448 | 0.03243381 | 30.8661596 | 31.8090047 | 54.7550631 | 0 | 0 |
| 0.01854366 | 0.03243381 | 9.96719738 | 38.4822225 | 0 | 69.5904227 | 0 |
| 0.01856809 | 0.0324734 | 10.2887199 | 9.34250488 | 9.28051917 | 11.1344676 | 0 |
| 0.0185711 | 0.03247699 | 5.46588243 | 8.00786133 | 4.64025958 | 10.6705315 | 0 |
| 0.01857779 | 0.03248699 | 5.46588243 | 10.4547078 | 3.71220767 | 12.0623399 | 0 |
| 0.01858743 | 0.03250217 | 33.1168171 | 19.1298909 | 0 | 50.5690405 | 0 |
| 0.01860592 | 0.0325328 | 9.32415239 | 7.78542074 | 3.71220767 | 7.88691457 | 0 |
| 0.01860775 | 0.03253431 | 6.75197242 | 8.45274251 | 2.78415575 | 20.4131907 | 0 |
| 0.01861189 | 0.03253986 | 0 | 0 | 0 | 0 | 0 |
| 0.01862673 | 0.03256411 | 25.4002772 | 49.1593709 | 42.6903882 | 0 | 0 |
| 0.01863858 | 0.03258313 | 5.46588243 | 7.78542074 | 4.64025958 | 11.1344676 | 0 |
| 0.01864519 | 0.03259299 | 0 | 29.8070394 | 54.7550631 | 41.2903175 | 0 |
| 0.01864876 | 0.03259585 | 51.4435994 | 21.1318563 | 17.6329864 | 15.7738291 | 9.50487297 |
| 0.01864798 | 0.03259585 | 7.07349491 | 11.7893514 | 16.7049345 | 2.31968076 | 0 |
| 0.01865081 | 0.03259774 | 27.0078897 | 0 | 59.3953227 | 50.5690405 | 0 |
| 0.01865489 | 0.03260317 | 15.1115573 | 13.3464355 | 12.9927268 | 14.3820207 | 116.434694 |
| 0.01865641 | 0.03260414 | 0 | 34.923173 | 43.6184401 | 45.4657428 | 0 |
| 0.01866877 | 0.03262405 | 8.3595849 | 83.8601033 | 0 | 33.4034029 | 0 |
| 0.01870285 | 0.0326802 | 5.46588243 | 6.67321777 | 4.64025958 | 16.2377653 | 0 |
| 0.01870211 | 0.0326802 | 98.0643613 | 422.192244 | 240.365446 | 281.145308 | 0 |
| 0.01872234 | 0.03271257 | 5.78740493 | 15.3484009 | 3.71220767 | 8.35085072 | 0 |
| 0.01874989 | 0.03275899 | 2.25065747 | 9.12006429 | 11.136623 | 13.4541484 | 0 |
| 0.01878633 | 0.03281896 | 1.60761248 | 31.8090047 | 5.5683115 | 12.5262761 | 0 |
| 0.01878602 | 0.03281896 | 0 | 0 | 0 | 0 | 0 |
| 0.01878769 | 0.03281896 | 2.57217997 | 6.22833659 | 12.9927268 | 15.309893 | 0 |
| 0.01878811 | 0.03281896 | 0 | 0.44488118 | 0.92805192 | 0 | 0 |
| 0.01878972 | 0.03282007 | 14.1469898 | 5.11613363 | 10.2085711 | 10.6705315 | 99.8011662 |
| 0.01879628 | 0.03282982 | 19.9343948 | 46.0452026 | 0 | 34.7952113 | 0 |

| | | | | | | |
|---|---|---|---|---|---|---|
| 0.01880523 | 0.03284375 | 31.8307271 | 24.9133464 | 0 | 41.2903175 | 0 |
| 0.0188153 | 0.03285962 | 36.3320421 | 41.3739502 | 36.1940248 | 0 | 0 |
| 0.01881992 | 0.03286599 | 0 | 14.6810791 | 80.7405168 | 44.0739344 | 0 |
| 0.01883783 | 0.03289557 | 4.82283744 | 2.8917277 | 13.9207788 | 17.1656376 | 0 |
| 0.01883954 | 0.03289684 | 13.8254673 | 59.6140788 | 53.8270112 | 0 | 0 |
| 0.01884379 | 0.03290255 | 0 | 0 | 0 | 0 | 0 |
| 0.01885069 | 0.0329129 | 0 | 33.8109701 | 37.1220767 | 53.3526574 | 0 |
| 0.01885249 | 0.03291433 | 9.32415239 | 7.11809896 | 5.5683115 | 5.10329766 | 9.50487297 |
| 0.01889584 | 0.03297975 | 36.0105196 | 67.6219401 | 0 | 26.9082968 | 0 |
| 0.01889581 | 0.03297975 | 0 | 0 | 0 | 0 | 0 |
| 0.01889581 | 0.03297975 | 0 | 0 | 0 | 0 | 0 |
| 0.01889581 | 0.03297975 | 0 | 0 | 0 | 0 | 0 |
| 0.01889581 | 0.03297975 | 0 | 0 | 0 | 0 | 0 |
| 0.01889581 | 0.03297975 | 0 | 0 | 0 | 0 | 0 |
| 0.0188988 | 0.03298319 | 3.21522496 | 1.33464355 | 25.9854537 | 28.7640414 | 0 |
| 0.01890594 | 0.03299396 | 5.46588243 | 4.22637126 | 6.49636342 | 6.49510612 | 2.37621824 |
| 0.01891081 | 0.03300074 | 8.0380624 | 4.22637126 | 7.42441534 | 7.88691457 | 0 |
| 0.01893461 | 0.03304056 | 14.4685123 | 56.7223511 | 0 | 35.7230837 | 0 |
| 0.01894197 | 0.03304878 | 32.7952946 | 46.2676432 | 0 | 20.8771268 | 0 |
| 0.01894227 | 0.03304878 | 89.7047764 | 12.4566732 | 0 | 20.8771268 | 0 |
| 0.01894223 | 0.03304878 | 3.85826995 | 7.11809896 | 11.136623 | 11.5984038 | 0 |
| 0.01894851 | 0.03305795 | 2.25065747 | 12.2342326 | 11.136623 | 9.74265918 | 0 |
| 0.01895126 | 0.03306104 | 34.4029071 | 34.0334106 | 30.6257133 | 38.0427644 | 0 |
| 0.01895922 | 0.03307286 | 8.3595849 | 19.5747721 | 0 | 110.416804 | 0 |
| 0.01896 | 0.03307286 | 2.89370247 | 5.7834554 | 4.64025958 | 2.78361691 | 9.50487297 |
| 0.01896159 | 0.03307392 | 3.53674746 | 13.7913167 | 4.64025958 | 12.9902122 | 0 |
| 0.01896854 | 0.03308432 | 0.3215225 | 2.22440592 | 0 | 2.31968076 | 19.0097459 |
| 0.01897662 | 0.03309498 | 22.1850522 | 8.45274251 | 9.28051917 | 12.9902122 | 54.6530196 |
| 0.0189765 | 0.03309498 | 31.5092046 | 19.3523315 | 16.7049345 | 16.2377653 | 21.3859642 |
| 0.01899426 | 0.03312402 | 69.4488592 | 63.1731283 | 77.0283091 | 60.3116997 | 0 |
| 0.01899909 | 0.03313073 | 13.8254673 | 57.6121134 | 0 | 36.1870198 | 0 |
| 0.01901067 | 0.03314922 | 8.3595849 | 16.2381632 | 12.9927268 | 1.85574461 | 0 |
| 0.01901835 | 0.03316088 | 0 | 40.4841878 | 49.1867516 | 33.4034029 | 0 |
| 0.01903394 | 0.03318635 | 2.89370247 | 7.78542074 | 8.35246725 | 15.309893 | 0 |
| 0.01904741 | 0.03320812 | 13.1824223 | 92.0904053 | 30.6257133 | 0 | 0 |
| 0.01904908 | 0.0332093 | 1.60761248 | 1.55708415 | 1.85610383 | 1.85574461 | 7.12865473 |
| 0.01905178 | 0.03321229 | 3.53674746 | 9.12006429 | 5.5683115 | 15.7738291 | 0 |
| 0.01905567 | 0.03321735 | 51.1220769 | 82.3030192 | 34.3379209 | 135.469356 | 35.6432737 |
| 0.01907016 | 0.03324089 | 23.4711422 | 0 | 42.6903882 | 72.8379758 | 0 |
| 0.01907826 | 0.03325327 | 1.60761248 | 11.7893514 | 13.9207788 | 12.5262761 | 0 |
| 0.01908528 | 0.03326379 | 0 | 48.4920492 | 43.6184401 | 31.0837221 | 0 |
| 0.01909893 | 0.03328586 | 0 | 18.4625692 | 48.2586997 | 64.487125 | 0 |
| 0.01910084 | 0.03328747 | 0 | 47.1574056 | 26.9135056 | 50.5690405 | 0 |

| | | | | | | |
|---|---|---|---|---|---|---|
| 0.01912984 | 0.03333628 | 4.82283744 | 10.2322673 | 12.0646749 | 4.17542536 | 0 |
| 0.01913823 | 0.03334917 | 171.693013 | 529.631051 | 505.788295 | 523.319979 | 0 |
| 0.01915383 | 0.03337462 | 2.89370247 | 12.6791138 | 9.28051917 | 7.88691457 | 0 |
| 0.01917175 | 0.03340395 | 8.0380624 | 8.8976237 | 13.9207788 | 11.1344676 | 0 |
| 0.01917264 | 0.03340395 | 52.0866444 | 28.0275146 | 0 | 20.8771268 | 0 |
| 0.01917563 | 0.03340743 | 23.7926647 | 25.5806681 | 0 | 50.5690405 | 0 |
| 0.01918801 | 0.03342726 | 5.46588243 | 5.56101481 | 7.42441534 | 7.88691457 | 0 |
| 0.01922098 | 0.03348296 | 3.85826995 | 6.67321777 | 12.0646749 | 7.42297842 | 0 |
| 0.01924801 | 0.0335283 | 2.57217997 | 8.23030192 | 10.2085711 | 12.9902122 | 0 |
| 0.01925243 | 0.03353428 | 18.3267823 | 42.7085937 | 0 | 38.9706367 | 0 |
| 0.01925611 | 0.03353895 | 43.405537 | 24.6909058 | 0 | 29.2279775 | 0 |
| 0.01926648 | 0.03355526 | 4.50131495 | 5.56101481 | 13.9207788 | 6.95904227 | 0 |
| 0.01927937 | 0.03357598 | 4.17979245 | 10.2322673 | 9.28051917 | 5.56723382 | 0 |
| 0.01928576 | 0.03358538 | 0.3215225 | 0.22244059 | 0 | 0 | 0 |
| 0.01930138 | 0.03360562 | 14.1469898 | 52.4959798 | 0 | 38.5067006 | 0 |
| 0.01929994 | 0.03360562 | 0 | 23.5787028 | 47.3306478 | 54.7444659 | 0 |
| 0.01930091 | 0.03360562 | 0 | 46.0452026 | 19.4890903 | 64.0231889 | 0 |
| 0.01929889 | 0.03360562 | 5.14435994 | 8.8976237 | 4.64025958 | 9.74265918 | 0 |
| 0.01931656 | 0.03362952 | 95.8137038 | 151.926925 | 138.279736 | 147.531696 | 85.5438568 |
| 0.01931711 | 0.03362952 | 18.3267823 | 42.4861532 | 0 | 38.9706367 | 0 |
| 0.01932108 | 0.0336347 | 0 | 60.7262817 | 17.6329864 | 52.4247851 | 0 |
| 0.01934566 | 0.03367575 | 21.2204847 | 55.6101481 | 0 | 25.0525522 | 0 |
| 0.01935426 | 0.03368898 | 4.50131495 | 16.0157227 | 2.78415575 | 15.7738291 | 0 |
| 0.01936647 | 0.03370847 | 0.96456749 | 0.22244059 | 0 | 0.46393615 | 0 |
| 0.01936811 | 0.03370958 | 0 | 27.1377523 | 56.6111669 | 40.3624452 | 0 |
| 0.01938339 | 0.03373444 | 4.17979245 | 9.78738607 | 4.64025958 | 13.4541484 | 0 |
| 0.0193982 | 0.03375847 | 1.28608998 | 15.3484009 | 18.5610383 | 9.27872303 | 0 |
| 0.01939966 | 0.03375925 | 104.494811 | 27.8050741 | 16.7049345 | 19.4853184 | 0 |
| 0.01940813 | 0.03377225 | 28.2939797 | 24.9133464 | 25.0574018 | 27.3722329 | 11.8810912 |
| 0.0194102 | 0.0337741 | 9.96719738 | 4.89369303 | 3.71220767 | 14.8459568 | 0 |
| 0.01943376 | 0.03381335 | 41.476402 | 44.7105591 | 0 | 15.7738291 | 0 |
| 0.01946733 | 0.03387002 | 8.6811074 | 22.6889404 | 22.273246 | 30.619786 | 0 |
| 0.01949251 | 0.03391208 | 99.3504513 | 9.78738607 | 0 | 19.9492545 | 0 |
| 0.01949364 | 0.03391229 | 15.4330798 | 24.023584 | 16.7049345 | 2.78361691 | 0 |
| 0.01949706 | 0.03391648 | 2.89370247 | 6.67321777 | 11.136623 | 12.5262761 | 0 |
| 0.01949829 | 0.03391687 | 0 | 49.6042521 | 44.546492 | 28.3001052 | 0 |
| 0.01950162 | 0.0339209 | 2.57217997 | 2.66928711 | 2.78415575 | 2.31968076 | 11.8810912 |
| 0.01957776 | 0.03404982 | 17.6837373 | 14.6810791 | 19.4890903 | 13.4541484 | 7.12865473 |
| 0.01957683 | 0.03404982 | 15.4330798 | 8.45274251 | 9.28051917 | 11.1344676 | 16.6335277 |
| 0.01960482 | 0.03409511 | 19.2913498 | 15.7932821 | 0 | 79.3330819 | 0 |
| 0.01961283 | 0.03410729 | 23.1496197 | 24.6909058 | 0 | 51.4969128 | 0 |
| 0.01961508 | 0.03410943 | 54.6588243 | 97.2065389 | 158.696878 | 140.572654 | 38.0194919 |
| 0.01962299 | 0.03412143 | 25.0787547 | 73.8502767 | 0 | 13.4541484 | 0 |

| | | | | | | |
|---|---|---|---|---|---|---|
| 0.01965386 | 0.03417334 | 0 | 5.56101481 | 3.71220767 | 0 | 2.37621824 |
| 0.01965981 | 0.03418184 | 21.5420072 | 29.1397176 | 0 | 47.3214874 | 0 |
| 0.01966078 | 0.03418184 | 1.92913498 | 1.11220296 | 0 | 3.71148921 | 9.50487297 |
| 0.01966196 | 0.03418212 | 2.25065747 | 17.3503662 | 34.3379209 | 2.31968076 | 0 |
| 0.01966442 | 0.03418358 | 49.1929419 | 183.95837 | 157.768826 | 159.594036 | 42.7719284 |
| 0.01966483 | 0.03418358 | 6.10892743 | 5.56101481 | 8.35246725 | 6.03116997 | 0 |
| 0.01967141 | 0.03419325 | 6.43044992 | 4.44881185 | 8.35246725 | 1.85574461 | 23.7621824 |
| 0.01967972 | 0.03420594 | 19.2913498 | 14.0137573 | 21.3451941 | 13.9180845 | 133.068222 |
| 0.01968865 | 0.03421968 | 0 | 13.123995 | 52.8989593 | 69.5904227 | 0 |
| 0.01971723 | 0.03426759 | 25.0787547 | 33.1436483 | 0 | 36.650956 | 0 |
| 0.01972194 | 0.034274 | 23.1496197 | 40.7066284 | 0 | 32.0115944 | 0 |
| 0.01973294 | 0.03429135 | 2.25065747 | 18.017688 | 12.9927268 | 6.03116997 | 0 |
| 0.01973819 | 0.03429869 | 0 | 26.9153117 | 50.1148035 | 45.0018067 | 0 |
| 0.01974064 | 0.03430118 | 1.92913498 | 9.78738607 | 11.136623 | 14.3820207 | 0 |
| 0.01975101 | 0.03431742 | 7.39501741 | 10.4547078 | 10.2085711 | 12.9902122 | 2.37621824 |
| 0.01977218 | 0.03435244 | 6.75197242 | 5.11613363 | 5.5683115 | 9.74265918 | 0 |
| 0.01977964 | 0.03436362 | 3.85826995 | 16.4606038 | 3.71220767 | 12.5262761 | 0 |
| 0.01978259 | 0.03436698 | 0 | 32.4763265 | 22.273246 | 74.2297842 | 0 |
| 0.01981755 | 0.03442593 | 15.4330798 | 8.23030192 | 15.7768826 | 9.74265918 | 47.5243649 |
| 0.01982195 | 0.03443179 | 25.0787547 | 37.5924601 | 0 | 32.0115944 | 0 |
| 0.01983665 | 0.03445556 | 36.9750871 | 30.9192423 | 0 | 26.4443606 | 0 |
| 0.01984572 | 0.03446953 | 0 | 260.033053 | 0 | 354.44722 | 0 |
| 0.01985349 | 0.03448125 | 0 | 27.5826335 | 53.8270112 | 40.3624452 | 0 |
| 0.01988162 | 0.03452831 | 4.82283744 | 4.67125244 | 4.64025958 | 5.10329766 | 23.7621824 |
| 0.0198843 | 0.03453017 | 2.89370247 | 6.67321777 | 10.2085711 | 13.4541484 | 0 |
| 0.01988474 | 0.03453017 | 15.4330798 | 11.3444702 | 1.85610383 | 8.81478687 | 0 |
| 0.01988892 | 0.03453565 | 164.619518 | 184.625692 | 119.718697 | 145.675952 | 7.12865473 |
| 0.01989559 | 0.03454545 | 1.92913498 | 28.4723958 | 12.9927268 | 4.63936151 | 0 |
| 0.01993964 | 0.03462015 | 26.3648447 | 0 | 31.5537652 | 78.8691457 | 0 |
| 0.01994069 | 0.03462018 | 0 | 23.5787028 | 48.2586997 | 51.0329766 | 0 |
| 0.01994443 | 0.03462488 | 0.64304499 | 0.22244059 | 0 | 0.9278723 | 0 |
| 0.01995864 | 0.03464778 | 17.3622148 | 30.4743612 | 26.9135056 | 23.6607437 | 11.8810912 |
| 0.01997446 | 0.03467345 | 0 | 8.67518311 | 131.783372 | 23.1968076 | 0 |
| 0.01997965 | 0.03468067 | 157.867546 | 487.144897 | 365.652455 | 410.583494 | 99.8011662 |
| 0.01998208 | 0.03468309 | 9.64567488 | 9.56494547 | 2.78415575 | 8.81478687 | 0 |
| 0.0199971 | 0.03470507 | 3.85826995 | 4.89369303 | 0 | 6.03116997 | 7.12865473 |
| 0.01999625 | 0.03470507 | 13.5039448 | 4.89369303 | 3.71220767 | 10.2065953 | 0 |
| 0.01999784 | 0.03470507 | 1.28608998 | 0.66732178 | 0 | 0.46393615 | 26.1384007 |
| 0.02003569 | 0.03475105 | 0 | 0 | 0 | 0 | 0 |
| 0.02003569 | 0.03475105 | 0 | 0 | 0 | 0 | 0 |
| 0.02003569 | 0.03475105 | 0 | 0 | 0 | 0 | 0 |
| 0.02003569 | 0.03475105 | 0 | 0 | 0 | 0 | 0 |
| 0.02003569 | 0.03475105 | 0 | 0 | 0 | 0 | 0 |

| | | | | | | |
|---|---|---|---|---|---|---|
| 0.02003569 | 0.03475105 | 0 | 0 | 0 | 0 | 0 |
| 0.02002968 | 0.03475105 | 0 | 45.1554403 | 92.8051917 | 8.81478687 | 0 |
| 0.02003569 | 0.03475105 | 0 | 0 | 0 | 0 | 0 |
| 0.02003569 | 0.03475105 | 0 | 0 | 0 | 0 | 0 |
| 0.02003569 | 0.03475105 | 0 | 0 | 0 | 0 | 0 |
| 0.02002625 | 0.03475105 | 3.53674746 | 8.00786133 | 11.136623 | 6.95904227 | 0 |
| 0.02003795 | 0.03475317 | 27.6509347 | 39.1495443 | 0 | 27.3722329 | 0 |
| 0.02003986 | 0.0347547 | 27.9724572 | 48.2696086 | 0 | 45.929679 | 2.37621824 |
| 0.02004982 | 0.03477017 | 3.53674746 | 11.1220296 | 6.49636342 | 8.81478687 | 0 |
| 0.02006361 | 0.03479229 | 4.82283744 | 12.9015544 | 4.64025958 | 7.88691457 | 0 |
| 0.02007543 | 0.034811 | 0 | 81.6356974 | 17.6329864 | 33.4034029 | 0 |
| 0.02009857 | 0.03484932 | 4.50131495 | 8.23030192 | 5.5683115 | 9.27872303 | 0 |
| 0.02010801 | 0.03486389 | 9.96719738 | 51.3837769 | 0 | 47.7854236 | 0 |
| 0.02012535 | 0.03489216 | 34.0813846 | 33.1436483 | 42.6903882 | 0 | 0 |
| 0.02013254 | 0.03490283 | 54.0157793 | 70.0687866 | 67.7477899 | 83.0445711 | 28.5146189 |
| 0.02013386 | 0.03490332 | 12.8608998 | 35.1456136 | 0 | 57.0641466 | 0 |
| 0.02013752 | 0.03490607 | 0 | 0 | 0 | 0 | 0 |
| 0.02013752 | 0.03490607 | 0 | 0 | 0 | 0 | 0 |
| 0.02013911 | 0.03490703 | 31.8307271 | 47.3798462 | 31.5537652 | 0 | 0 |
| 0.02014539 | 0.03491611 | 27.9724572 | 18.017688 | 26.9135056 | 19.0213822 | 4.75243649 |
| 0.02014673 | 0.03491664 | 9.64567488 | 12.4566732 | 9.28051917 | 9.27872303 | 0 |
| 0.02016937 | 0.03495408 | 2.25065747 | 12.4566732 | 11.136623 | 8.81478687 | 0 |
| 0.02017655 | 0.03496293 | 37.2966096 | 43.1534749 | 29.6976613 | 0 | 0 |
| 0.02017587 | 0.03496293 | 22.5065747 | 51.828658 | 49.1867516 | 57.5280828 | 19.0097459 |
| 0.02018671 | 0.03497873 | 21.5420072 | 45.3778809 | 0 | 29.2279775 | 0 |
| 0.02018925 | 0.03498078 | 13.5039448 | 49.1593709 | 0 | 39.898509 | 0 |
| 0.02018997 | 0.03498078 | 2.89370247 | 4.67125244 | 18.5610383 | 12.0623399 | 0 |
| 0.02019393 | 0.03498582 | 0 | 68.9565837 | 30.6257133 | 25.5164883 | 0 |
| 0.02020236 | 0.03499862 | 0 | 43.3759155 | 27.8415575 | 48.7132959 | 0 |
| 0.02020865 | 0.03500773 | 27.6509347 | 6.45077718 | 6.49636342 | 2.78361691 | 0 |
| 0.02021976 | 0.03502517 | 7.39501741 | 8.45274251 | 17.6329864 | 10.6705315 | 2.37621824 |
| 0.02024981 | 0.03507541 | 0 | 19.5747721 | 66.819738 | 39.898509 | 0 |
| 0.02026008 | 0.0350914 | 42.4409695 | 19.1298909 | 0 | 34.7952113 | 0 |
| 0.02028115 | 0.03512609 | 0 | 0 | 0 | 0 | 80.7914203 |
| 0.02030427 | 0.03516431 | 25.0787547 | 28.2499552 | 27.8415575 | 13.4541484 | 38.0194919 |
| 0.02032793 | 0.03520348 | 16.0761248 | 60.7262817 | 0 | 26.4443606 | 0 |
| 0.020331 | 0.03520698 | 0.3215225 | 0 | 0 | 0 | 0 |
| 0.02035202 | 0.03524156 | 19.6128723 | 12.6791138 | 0 | 84.4363795 | 0 |
| 0.02035607 | 0.03524677 | 54.9803468 | 35.8129354 | 0 | 12.9902122 | 0 |
| 0.02036687 | 0.03526365 | 64.3044992 | 21.7991781 | 0 | 18.0935099 | 0 |
| 0.02038038 | 0.03528523 | 79.4160565 | 47.3798462 | 0 | 0 | 387.323574 |
| 0.02038263 | 0.03528731 | 13.1824223 | 12.6791138 | 0 | 0 | 52.2768014 |
| 0.02039182 | 0.0353014 | 50.8005544 | 26.4704305 | 0 | 20.4131907 | 0 |

| | | | | | | |
|---|---|---|---|---|---|---|
| 0.0204004 | 0.03531444 | 25.7217997 | 25.5806681 | 0 | 43.1460621 | 0 |
| 0.02042188 | 0.0353498 | 14.1469898 | 32.6987671 | 0 | 55.6723382 | 0 |
| 0.02042322 | 0.0353503 | 1.28608998 | 11.7893514 | 15.7768826 | 12.9902122 | 0 |
| 0.02042565 | 0.03535211 | 5.14435994 | 6.22833659 | 6.49636342 | 7.88691457 | 0 |
| 0.02042636 | 0.03535211 | 191.627408 | 255.806681 | 206.955577 | 259.804245 | 68.9103291 |
| 0.02043973 | 0.03537342 | 18.0052598 | 10.4547078 | 7.42441534 | 8.35085072 | 0 |
| 0.02046952 | 0.03542316 | 19.2913498 | 5.33857422 | 6.49636342 | 4.17542536 | 0 |
| 0.02048263 | 0.0354422 | 14.7900348 | 22.6889404 | 0 | 70.518295 | 0 |
| 0.02048165 | 0.0354422 | 52.7296894 | 40.0393066 | 29.6976613 | 33.4034029 | 133.068222 |
| 0.02049039 | 0.0354538 | 23.7926647 | 54.9428263 | 0 | 20.4131907 | 0 |
| 0.02049264 | 0.03545588 | 2.57217997 | 10.4547078 | 11.136623 | 8.35085072 | 0 |
| 0.0204985 | 0.03546419 | 4.82283744 | 11.7893514 | 6.49636342 | 5.56723382 | 0 |
| 0.020528 | 0.0355134 | 29.9015921 | 14.9035197 | 14.8488307 | 15.7738291 | 47.5243649 |
| 0.0205707 | 0.03558543 | 0 | 20.6869751 | 85.3807764 | 25.5164883 | 0 |
| 0.02058238 | 0.03560381 | 20.5774398 | 17.1279256 | 14.8488307 | 12.5262761 | 85.5438568 |
| 0.02058439 | 0.03560547 | 13.5039448 | 20.6869751 | 25.0574018 | 22.7328714 | 9.50487297 |
| 0.02060438 | 0.03563821 | 12.5393773 | 58.0569946 | 76.1002572 | 67.7346781 | 2.37621824 |
| 0.02060708 | 0.03564105 | 33.4383396 | 8.45274251 | 11.136623 | 0.9278723 | 0 |
| 0.02065014 | 0.03571368 | 0 | 0 | 0 | 0 | 38.0194919 |
| 0.02065766 | 0.03572486 | 0 | 35.1456136 | 28.7696094 | 55.208402 | 0 |
| 0.02066772 | 0.03574041 | 15.1115573 | 60.2814006 | 0 | 27.3722329 | 0 |
| 0.02069462 | 0.03578509 | 8.0380624 | 8.23030192 | 3.71220767 | 7.42297842 | 0 |
| 0.02069721 | 0.03578773 | 8.3595849 | 44.7105591 | 0 | 58.4559551 | 0 |
| 0.02070265 | 0.03579529 | 1.28608998 | 11.5669108 | 11.136623 | 18.5574461 | 0 |
| 0.02070392 | 0.03579565 | 2.89370247 | 12.9015544 | 7.42441534 | 8.81478687 | 0 |
| 0.02071454 | 0.03581218 | 5.14435994 | 10.0098267 | 2.78415575 | 19.9492545 | 0 |
| 0.02075642 | 0.03588273 | 0.64304499 | 0.44488118 | 0 | 0.46393615 | 0 |
| 0.02078429 | 0.03592906 | 32.1522496 | 0 | 59.3953227 | 35.7230837 | 0 |
| 0.02079211 | 0.03594074 | 3.21522496 | 7.56298014 | 8.35246725 | 10.6705315 | 0 |
| 0.02079678 | 0.03594697 | 3.85826995 | 5.7834554 | 11.136623 | 8.35085072 | 0 |
| 0.02081245 | 0.0359722 | 13.1824223 | 41.5963908 | 0 | 45.929679 | 0 |
| 0.02082683 | 0.0359952 | 5.46588243 | 74.7400391 | 0 | 87.2199964 | 0 |
| 0.0208351 | 0.03600765 | 0 | 47.3798462 | 60.3233746 | 17.1656376 | 0 |
| 0.02083936 | 0.03601045 | 0 | 28.2499552 | 45.4745439 | 43.6099982 | 0 |
| 0.02083993 | 0.03601045 | 2.89370247 | 4.67125244 | 1.85610383 | 2.78361691 | 2.37621824 |
| 0.02083985 | 0.03601045 | 10.6102424 | 16.0157227 | 2.78415575 | 5.56723382 | 0 |
| 0.02084628 | 0.03601812 | 24.4357097 | 28.4723958 | 7.42441534 | 68.6625504 | 9.50487297 |
| 0.02084651 | 0.03601812 | 3.53674746 | 18.2401286 | 9.28051917 | 4.63936151 | 0 |
| 0.02085882 | 0.03603754 | 7.39501741 | 8.23030192 | 12.0646749 | 0 | 42.7719284 |
| 0.0208624 | 0.03604155 | 0 | 0.44488118 | 0 | 0 | 2.37621824 |
| 0.02086329 | 0.03604155 | 0 | 30.4743612 | 78.8844129 | 19.0213822 | 0 |
| 0.02086954 | 0.0360505 | 7.07349491 | 6.89565837 | 11.136623 | 3.71148921 | 0 |
| 0.02092022 | 0.03613619 | 4.50131495 | 4.89369303 | 8.35246725 | 11.1344676 | 0 |

| | | | | | | |
|---|---|---|---|---|---|---|
| 0.0209267 | 0.03614552 | 26.6863672 | 11.3444702 | 0.92805192 | 10.2065953 | 0 |
| 0.02093099 | 0.03615108 | 42.119447 | 40.7066284 | 57.5392189 | 52.8887212 | 0 |
| 0.02094302 | 0.03617 | 0.3215225 | 0.88976237 | 1.85610383 | 0 | 0 |
| 0.02094528 | 0.03617206 | 0 | 42.9310343 | 56.6111669 | 21.341063 | 0 |
| 0.02094817 | 0.03617519 | 18.3267823 | 46.934965 | 0 | 30.619786 | 0 |
| 0.02095095 | 0.03617813 | 1.28608998 | 2.44684652 | 2.78415575 | 1.39180845 | 2.37621824 |
| 0.02097037 | 0.0362098 | 11.8963324 | 7.78542074 | 3.71220767 | 6.03116997 | 0 |
| 0.02098418 | 0.03623179 | 0 | 0 | 0 | 0 | 2.37621824 |
| 0.02102905 | 0.0363074 | 10.9317649 | 12.9015544 | 1.85610383 | 9.74265918 | 0 |
| 0.02103707 | 0.03631938 | 22.5065747 | 12.9015544 | 97.4454513 | 0 | 0 |
| 0.02104368 | 0.03632893 | 0.64304499 | 0.66732178 | 0 | 0 | 0 |
| 0.02104764 | 0.0363339 | 0.96456749 | 0.22244059 | 0 | 0.46393615 | 0 |
| 0.02106386 | 0.03636004 | 8.3595849 | 53.6081828 | 0 | 47.7854236 | 0 |
| 0.02106821 | 0.03636568 | 1.28608998 | 21.3542969 | 38.0501286 | 14.3820207 | 0 |
| 0.02107579 | 0.0363769 | 71.6995166 | 17.7952474 | 10.2085711 | 13.9180845 | 0 |
| 0.02107811 | 0.03637726 | 8.0380624 | 14.6810791 | 1.85610383 | 12.0623399 | 0 |
| 0.02107816 | 0.03637726 | 1.92913498 | 13.7913167 | 9.28051917 | 11.1344676 | 0 |
| 0.0210972 | 0.03640826 | 0 | 18.4625692 | 57.5392189 | 46.3936151 | 0 |
| 0.02110928 | 0.03642723 | 26.3648447 | 16.4606038 | 8.35246725 | 4.17542536 | 0 |
| 0.0211151 | 0.0364354 | 31.8307271 | 30.9192423 | 0 | 27.8361691 | 0 |
| 0.02112707 | 0.03645419 | 13.8254673 | 6.67321777 | 6.49636342 | 7.88691457 | 0 |
| 0.02113284 | 0.03646227 | 10.6102424 | 12.6791138 | 6.49636342 | 2.78361691 | 0 |
| 0.02114224 | 0.03647662 | 0.64304499 | 16.6830444 | 8.35246725 | 30.1558498 | 0 |
| 0.02117094 | 0.03652425 | 6.43044992 | 44.4881185 | 87.2368802 | 0 | 0 |
| 0.02117201 | 0.03652425 | 20.8989622 | 39.816866 | 0 | 32.0115944 | 0 |
| 0.02120507 | 0.0365794 | 32.4737721 | 158.377702 | 59.3953227 | 110.88074 | 0 |
| 0.021235 | 0.03662915 | 0.96456749 | 2.22440592 | 1.85610383 | 2.31968076 | 2.37621824 |
| 0.02124416 | 0.03664308 | 3.85826995 | 12.011792 | 9.28051917 | 5.10329766 | 0 |
| 0.02125003 | 0.03665132 | 129.895088 | 127.236019 | 34.3379209 | 168.408823 | 52.2768014 |
| 0.02126215 | 0.03667034 | 4.50131495 | 7.56298014 | 11.136623 | 5.10329766 | 0 |
| 0.02127376 | 0.03668848 | 9.00262989 | 13.123995 | 2.78415575 | 6.95904227 | 0 |
| 0.0212759 | 0.0366903 | 0 | 44.7105591 | 28.7696094 | 42.2181898 | 0 |
| 0.02127819 | 0.03669236 | 3.85826995 | 7.11809896 | 6.49636342 | 11.1344676 | 0 |
| 0.02128605 | 0.03670403 | 23.1496197 | 45.822762 | 0 | 24.588616 | 0 |
| 0.02129305 | 0.03671422 | 0 | 24.6909058 | 51.9709073 | 40.8263813 | 0 |
| 0.02130677 | 0.03672847 | 0 | 0 | 0 | 0 | 0 |
| 0.02130677 | 0.03672847 | 0 | 0 | 0 | 0 | 0 |
| 0.02130677 | 0.03672847 | 0 | 0 | 0 | 0 | 0 |
| 0.02130677 | 0.03672847 | 0 | 0 | 0 | 0 | 0 |
| 0.02130677 | 0.03672847 | 0 | 0 | 0 | 0 | 0 |
| 0.02131829 | 0.03674644 | 4.17979245 | 9.56494547 | 6.49636342 | 6.95904227 | 0 |
| 0.02133419 | 0.03677196 | 6.75197242 | 8.23030192 | 2.78415575 | 15.7738291 | 0 |
| 0.02133979 | 0.03677973 | 42.762492 | 26.4704305 | 0 | 23.1968076 | 0 |

| | | | | | | |
|---|---|---|---|---|---|---|
| 0.02135778 | 0.03680885 | 89.7047764 | 175.505627 | 189.322591 | 188.358077 | 52.2768014 |
| 0.02137164 | 0.03683086 | 0 | 0 | 0 | 28.3001052 | 230.49317 |
| 0.02139034 | 0.03686119 | 0.3215225 | 2.44684652 | 1.85610383 | 1.85574461 | 52.2768014 |
| 0.02139453 | 0.03686464 | 109.960694 | 417.743433 | 352.659728 | 403.624452 | 0 |
| 0.02139381 | 0.03686464 | 19.6128723 | 39.3719849 | 51.9709073 | 0 | 0 |
| 0.02140123 | 0.0368743 | 0 | 0 | 0 | 0 | 26.1384007 |
| 0.02140797 | 0.03688401 | 26.0433222 | 22.911381 | 18.5610383 | 17.1656376 | 40.3957101 |
| 0.02141587 | 0.03689574 | 0 | 0 | 0 | 0 | 23.7621824 |
| 0.0214465 | 0.03694661 | 68.8058142 | 32.9212077 | 15.7768826 | 0 | 0 |
| 0.02145679 | 0.03696245 | 47.9068519 | 65.1750936 | 51.0428554 | 46.8575513 | 0 |
| 0.02147675 | 0.03699495 | 27.9724572 | 37.1475789 | 83.5246725 | 0 | 0 |
| 0.02149393 | 0.03702265 | 0 | 46.7125244 | 29.6976613 | 38.5067006 | 0 |
| 0.02149896 | 0.03702942 | 0 | 3.11416829 | 2.78415575 | 3.71148921 | 0 |
| 0.02150845 | 0.03704386 | 13.5039448 | 11.5669108 | 10.2085711 | 11.5984038 | 111.682257 |
| 0.02151329 | 0.03705029 | 65.9121117 | 310.304626 | 236.653239 | 266.763287 | 28.5146189 |
| 0.02158858 | 0.03717616 | 13.5039448 | 34.2558512 | 0 | 51.0329766 | 0 |
| 0.02158843 | 0.03717616 | 6.43044992 | 32.6987671 | 0 | 80.2609542 | 0 |
| 0.02159252 | 0.03718104 | 2.25065747 | 6.89565837 | 13.9207788 | 12.0623399 | 0 |
| 0.02162491 | 0.03723491 | 95.8137038 | 12.011792 | 0 | 13.9180845 | 0 |
| 0.02163475 | 0.03724995 | 0 | 0 | 0 | 0 | 76.0389838 |
| 0.02163938 | 0.03725601 | 23.4711422 | 71.1809896 | 21.3451941 | 0 | 0 |
| 0.02164234 | 0.0372592 | 0 | 41.8188314 | 29.6976613 | 42.6821259 | 0 |
| 0.02165426 | 0.03727781 | 4.82283744 | 5.56101481 | 5.5683115 | 14.8459568 | 0 |
| 0.02166228 | 0.03728971 | 308.983119 | 481.361442 | 392.565961 | 453.26562 | 125.939567 |
| 0.02166451 | 0.03728973 | 28.9370247 | 34.2558512 | 25.0574018 | 27.3722329 | 68.9103291 |
| 0.02166451 | 0.03728973 | 15.7546023 | 11.3444702 | 8.35246725 | 6.95904227 | 0 |
| 0.02168645 | 0.03732553 | 1.92913498 | 0 | 2.78415575 | 0.9278723 | 26.1384007 |
| 0.02168752 | 0.03732553 | 6.43044992 | 13.123995 | 7.42441534 | 18.0935099 | 0 |
| 0.0217112 | 0.03736399 | 11.5748099 | 13.7913167 | 5.5683115 | 2.78361691 | 0 |
| 0.02171209 | 0.03736399 | 1.28608998 | 8.8976237 | 29.6976613 | 8.35085072 | 0 |
| 0.02171557 | 0.03736806 | 0.96456749 | 1.33464355 | 0.92805192 | 0.46393615 | 49.9005831 |
| 0.02172961 | 0.03739031 | 107.710036 | 122.342326 | 177.257916 | 62.1674443 | 206.730987 |
| 0.02173605 | 0.03739947 | 17.0406923 | 44.0432373 | 0 | 64.9510612 | 0 |
| 0.02174383 | 0.03741094 | 2.25065747 | 7.56298014 | 13.9207788 | 10.6705315 | 0 |
| 0.02176156 | 0.03743954 | 8.3595849 | 2.00196533 | 12.9927268 | 12.0623399 | 0 |
| 0.02180978 | 0.03752059 | 0 | 32.0314453 | 42.6903882 | 38.5067006 | 0 |
| 0.02181206 | 0.03752259 | 3.85826995 | 6.005896 | 10.2085711 | 7.88691457 | 0 |
| 0.02183415 | 0.03755866 | 101.922631 | 12.6791138 | 0 | 11.1344676 | 0 |
| 0.02185121 | 0.0375861 | 208.346577 | 327.210111 | 315.537652 | 299.702754 | 646.331362 |
| 0.02185417 | 0.03758927 | 15.4330798 | 25.5806681 | 79.8124649 | 0 | 0 |
| 0.02187883 | 0.03762975 | 17.6837373 | 7.34053955 | 3.71220767 | 5.10329766 | 0 |
| 0.02188126 | 0.03763201 | 0 | 267.596033 | 347.091417 | 0 | 0 |
| 0.02188322 | 0.03763346 | 0 | 45.6003214 | 18.5610383 | 55.208402 | 0 |

| | | | | | | |
|---|---|---|---|---|---|---|
| 0.02188779 | 0.03763939 | 35.6889971 | 16.4606038 | 61.2514265 | 0 | 0 |
| 0.02189281 | 0.03764611 | 20.8989622 | 0 | 56.6111669 | 49.177232 | 0 |
| 0.02189913 | 0.03765505 | 48.5498969 | 8.45274251 | 5.5683115 | 0.9278723 | 0 |
| 0.02190429 | 0.03766199 | 78.4514891 | 200.196533 | 210.667785 | 174.439993 | 0 |
| 0.02191275 | 0.03767461 | 4.82283744 | 4.67125244 | 3.71220767 | 2.31968076 | 47.5243649 |
| 0.02191933 | 0.03768399 | 0 | 10.4547078 | 31.5537652 | 96.0347833 | 0 |
| 0.02193636 | 0.03771136 | 23.4711422 | 143.919063 | 154.98467 | 169.800631 | 47.5243649 |
| 0.02193983 | 0.0377154 | 14.7900348 | 56.0550293 | 50.1148035 | 48.7132959 | 16.6335277 |
| 0.02197694 | 0.03777725 | 402.867688 | 1269.24602 | 1219.46022 | 1225.71931 | 16.6335277 |
| 0.0219832 | 0.03778607 | 0.3215225 | 0.22244059 | 0 | 0 | 40.3957101 |
| 0.02198521 | 0.0377876 | 0 | 34.0334106 | 25.9854537 | 55.6723382 | 0 |
| 0.02199149 | 0.03779647 | 0 | 4.89369303 | 0 | 0 | 76.0389838 |
| 0.02199941 | 0.03780816 | 0.3215225 | 0 | 0 | 0 | 0 |
| 0.02200563 | 0.03781691 | 64.9475442 | 220.216187 | 180.042072 | 224.545097 | 0 |
| 0.02204464 | 0.03788202 | 0 | 6.005896 | 0 | 0 | 59.4054561 |
| 0.02205473 | 0.03789741 | 39.547267 | 28.6948364 | 36.1940248 | 0 | 0 |
| 0.02206455 | 0.03791236 | 0 | 38.4822225 | 30.6257133 | 43.6099982 | 0 |
| 0.02206854 | 0.03791727 | 135.360971 | 321.426656 | 318.321808 | 314.548711 | 73.6627655 |
| 0.02207392 | 0.03792458 | 0 | 1.55708415 | 0 | 0 | 30.8908372 |
| 0.022086 | 0.0379434 | 0 | 23.5787028 | 57.5392189 | 35.2591475 | 0 |
| 0.02210267 | 0.03797009 | 0.64304499 | 1.77952474 | 0.92805192 | 0.46393615 | 7.12865473 |
| 0.02211691 | 0.03799261 | 4.82283744 | 11.3444702 | 3.71220767 | 10.2065953 | 0 |
| 0.02212776 | 0.03800349 | 26.0433222 | 25.1357869 | 13.9207788 | 6.03116997 | 0 |
| 0.02212715 | 0.03800349 | 2.25065747 | 8.45274251 | 12.9927268 | 9.74265918 | 0 |
| 0.02212611 | 0.03800349 | 26.3648447 | 21.5767375 | 0 | 43.1460621 | 0 |
| 0.0221249 | 0.03800349 | 5.78740493 | 5.33857422 | 9.28051917 | 5.56723382 | 0 |
| 0.02214481 | 0.03803084 | 32.7952946 | 35.5904948 | 0 | 21.341063 | 0 |
| 0.02215186 | 0.03804099 | 3.21522496 | 10.0098267 | 16.7049345 | 4.63936151 | 0 |
| 0.02216961 | 0.03806954 | 24.7572322 | 36.9251383 | 0 | 27.3722329 | 0 |
| 0.02218188 | 0.03808866 | 17.3622148 | 19.5747721 | 18.5610383 | 15.7738291 | 64.1578926 |
| 0.02219308 | 0.03810596 | 43.7270595 | 25.5806681 | 0 | 21.8049991 | 0 |
| 0.02221918 | 0.03814882 | 0 | 24.4684652 | 78.8844129 | 20.8771268 | 0 |
| 0.02222177 | 0.03815132 | 3.85826995 | 9.56494547 | 6.49636342 | 7.42297842 | 0 |
| 0.02222731 | 0.03815889 | 6.43044992 | 4.67125244 | 10.2085711 | 5.56723382 | 0 |
| 0.0222308 | 0.03816294 | 8.0380624 | 9.12006429 | 13.9207788 | 2.31968076 | 0 |
| 0.02223486 | 0.03816795 | 5.46588243 | 9.34250488 | 4.64025958 | 6.95904227 | 0 |
| 0.02224487 | 0.03818318 | 60.4462293 | 8.67518311 | 0 | 37.1148921 | 0 |
| 0.02224742 | 0.03818562 | 355.282358 | 505.385026 | 593.953227 | 532.598702 | 0 |
| 0.02225986 | 0.03820503 | 0 | 6.89565837 | 0 | 0 | 26.1384007 |
| 0.02228681 | 0.03824933 | 0 | 37.5924601 | 38.0501286 | 35.7230837 | 0 |
| 0.02231957 | 0.03830359 | 18.3267823 | 28.4723958 | 0 | 45.4657428 | 0 |
| 0.0223306 | 0.03832057 | 7.07349491 | 11.1220296 | 12.9927268 | 2.31968076 | 0 |
| 0.02234115 | 0.03833672 | 1.28608998 | 12.2342326 | 14.8488307 | 11.5984038 | 0 |

| | | | | | | |
|---|---|---|---|---|---|---|
| 0.02234378 | 0.03833928 | 13.5039448 | 4.22637126 | 5.5683115 | 6.49510612 | 0 |
| 0.0223507 | 0.0383492 | 16.3976473 | 24.4684652 | 0 | 55.6723382 | 0 |
| 0.0223519 | 0.03834929 | 4.50131495 | 4.67125244 | 1.85610383 | 2.31968076 | 59.4054561 |
| 0.02236129 | 0.03836345 | 2.25065747 | 2.8917277 | 29.6976613 | 13.9180845 | 0 |
| 0.0223649 | 0.03836768 | 4.17979245 | 16.0157227 | 1.85610383 | 21.8049991 | 0 |
| 0.02238762 | 0.03840471 | 29.5800696 | 0 | 46.4025958 | 44.5378705 | 0 |
| 0.02239097 | 0.03840849 | 4.82283744 | 44.4881185 | 0 | 114.592229 | 2.37621824 |
| 0.02241377 | 0.03844564 | 42.4409695 | 50.7164551 | 16.7049345 | 0 | 0 |
| 0.02242196 | 0.03845635 | 15.4330798 | 35.5904948 | 0 | 42.2181898 | 0 |
| 0.0224223 | 0.03845635 | 0 | 55.8325887 | 27.8415575 | 30.619786 | 0 |
| 0.02243151 | 0.03846663 | 12.8608998 | 45.822762 | 23.2012979 | 36.650956 | 11.8810912 |
| 0.02243172 | 0.03846663 | 0.64304499 | 1.55708415 | 0.92805192 | 0 | 0 |
| 0.02243043 | 0.03846663 | 0.3215225 | 0.22244059 | 0 | 0 | 0 |
| 0.02247821 | 0.03854438 | 0 | 20.4645345 | 12.0646749 | 23.6607437 | 83.1676385 |
| 0.02248781 | 0.03855888 | 54.6588243 | 29.5845988 | 40.8342843 | 31.5476583 | 78.415202 |
| 0.02250514 | 0.03858663 | 11.2532874 | 57.3896728 | 0 | 32.0115944 | 0 |
| 0.02250984 | 0.03859272 | 45.9777169 | 0 | 42.6903882 | 31.0837221 | 0 |
| 0.02251791 | 0.03860459 | 4.82283744 | 7.78542074 | 5.5683115 | 6.95904227 | 11.8810912 |
| 0.02251946 | 0.03860529 | 1.92913498 | 9.12006429 | 17.6329864 | 8.35085072 | 0 |
| 0.02252505 | 0.03861289 | 25.4002772 | 28.4723958 | 0 | 33.867339 | 0 |
| 0.02252856 | 0.03861694 | 0 | 0 | 0 | 0 | 0 |
| 0.0225389 | 0.0386327 | 2.57217997 | 10.899589 | 9.28051917 | 8.35085072 | 0 |
| 0.02255593 | 0.03865993 | 14.4685123 | 5.7834554 | 0 | 117.839782 | 0 |
| 0.02258846 | 0.03871174 | 55.9449143 | 52.4959798 | 0 | 5.56723382 | 0 |
| 0.02258832 | 0.03871174 | 32.4737721 | 47.1574056 | 32.4818171 | 47.3214874 | 19.0097459 |
| 0.02259814 | 0.03872437 | 21.8635297 | 52.940861 | 51.9709073 | 49.6411682 | 16.6335277 |
| 0.02259777 | 0.03872437 | 16.3976473 | 65.3975342 | 0 | 19.0213822 | 0 |
| 0.02260324 | 0.03873114 | 16.3976473 | 10.2322673 | 7.42441534 | 12.0623399 | 125.939567 |
| 0.02262364 | 0.03876413 | 3.85826995 | 4.89369303 | 8.35246725 | 13.4541484 | 0 |
| 0.02262615 | 0.03876645 | 124.429206 | 33.8109701 | 38.0501286 | 44.5378705 | 28.5146189 |
| 0.02263401 | 0.03877795 | 48.5498969 | 0 | 46.4025958 | 25.9804245 | 0 |
| 0.02263638 | 0.03878003 | 50.4790319 | 127.903341 | 54.7550631 | 123.407016 | 0 |
| 0.02264855 | 0.03879891 | 17.6837373 | 4.44881185 | 6.49636342 | 4.63936151 | 0 |
| 0.02265022 | 0.03879978 | 71.0564716 | 125.234054 | 85.3807764 | 43.6099982 | 30.8908372 |
| 0.02265294 | 0.03880247 | 18.6483048 | 37.5924601 | 32.4818171 | 30.1558498 | 14.2573095 |
| 0.02265454 | 0.03880323 | 6.75197242 | 9.56494547 | 10.2085711 | 13.9180845 | 0 |
| 0.02265718 | 0.03880578 | 0 | 32.2538859 | 81.6685687 | 13.4541484 | 0 |
| 0.02266118 | 0.03881066 | 124.429206 | 8.23030192 | 0 | 8.81478687 | 0 |
| 0.02266375 | 0.03881309 | 3.53674746 | 10.2322673 | 6.49636342 | 7.88691457 | 0 |
| 0.02266821 | 0.03881875 | 65.9121117 | 123.009648 | 98.3735032 | 122.479144 | 57.0292378 |
| 0.02267322 | 0.03882534 | 29.9015921 | 88.3089152 | 125.287009 | 140.108718 | 38.0194919 |
| 0.0226798 | 0.03883463 | 3.53674746 | 41.8188314 | 0 | 82.5806349 | 0 |
| 0.02269509 | 0.03885884 | 0 | 59.1691976 | 25.0574018 | 30.619786 | 0 |

| | | | | | | |
|---|---|---|---|---|---|---|
| 0.02270047 | 0.03886608 | 0 | 0.44488118 | 0.92805192 | 0 | 0 |
| 0.022711 | 0.03888213 | 21.8635297 | 13.7913167 | 4.64025958 | 8.35085072 | 2.37621824 |
| 0.02271866 | 0.03889327 | 0 | 25.3582275 | 40.8342843 | 45.929679 | 0 |
| 0.02272348 | 0.03889756 | 170.406923 | 184.180811 | 145.704151 | 182.326907 | 45.1481466 |
| 0.02272334 | 0.03889756 | 2.57217997 | 6.005896 | 12.0646749 | 12.5262761 | 0 |
| 0.02273259 | 0.03890523 | 0 | 0 | 0 | 0 | 0 |
| 0.02273259 | 0.03890523 | 0 | 0 | 0 | 0 | 0 |
| 0.02273259 | 0.03890523 | 0 | 0 | 0 | 0 | 0 |
| 0.02273259 | 0.03890523 | 0 | 0 | 0 | 0 | 0 |
| 0.02274079 | 0.0389173 | 25.4002772 | 5.7834554 | 2.78415575 | 6.49510612 | 0 |
| 0.02274413 | 0.03892102 | 89.3832539 | 99.875826 | 0 | 208.771268 | 4.75243649 |
| 0.02274906 | 0.03892748 | 15.7546023 | 37.1475789 | 0 | 38.9706367 | 0 |
| 0.02275811 | 0.03894099 | 14.1469898 | 7.34053955 | 2.78415575 | 7.42297842 | 0 |
| 0.02276464 | 0.03895018 | 13.8254673 | 4.89369303 | 10.2085711 | 7.88691457 | 21.3859642 |
| 0.02276953 | 0.03895656 | 39.8687895 | 50.4940145 | 17.6329864 | 0 | 0 |
| 0.02277619 | 0.03896597 | 4.82283744 | 0 | 0 | 3.24755306 | 0 |
| 0.02277792 | 0.03896696 | 4.17979245 | 68.0668213 | 0 | 48.7132959 | 0 |
| 0.02277992 | 0.0389684 | 7.07349491 | 31.3641235 | 0 | 73.7658481 | 0 |
| 0.0227887 | 0.03898143 | 2.25065747 | 10.899589 | 11.136623 | 8.35085072 | 0 |
| 0.02279664 | 0.03899303 | 37.2966096 | 74.2951579 | 84.4527244 | 90.0036134 | 28.5146189 |
| 0.02282156 | 0.03903367 | 29.2585471 | 0 | 38.0501286 | 52.4247851 | 0 |
| 0.02285835 | 0.03909461 | 89.7047764 | 23.5787028 | 40.8342843 | 40.3624452 | 19.0097459 |
| 0.02286709 | 0.03910757 | 119.284846 | 374.367517 | 312.753496 | 350.73573 | 2.37621824 |
| 0.02287323 | 0.03911608 | 0 | 33.8109701 | 28.7696094 | 48.7132959 | 0 |
| 0.0228794 | 0.03912465 | 32.7952946 | 49.6042521 | 0 | 30.619786 | 0 |
| 0.02288238 | 0.03912776 | 3.21522496 | 11.3444702 | 10.2085711 | 5.56723382 | 0 |
| 0.02289558 | 0.03914834 | 17.6837373 | 56.9447917 | 42.6903882 | 42.6821259 | 0 |
| 0.02291403 | 0.03917788 | 8.6811074 | 54.053064 | 0 | 40.3624452 | 0 |
| 0.02291893 | 0.03918246 | 38.904222 | 19.5747721 | 47.3306478 | 0 | 0 |
| 0.02291903 | 0.03918246 | 2.25065747 | 19.3523315 | 5.5683115 | 10.6705315 | 0 |
| 0.02292516 | 0.03919095 | 0 | 46.0452026 | 39.9062324 | 25.5164883 | 0 |
| 0.02294164 | 0.03921713 | 69.4488592 | 30.02948 | 0 | 8.35085072 | 0 |
| 0.02294534 | 0.03922062 | 4.50131495 | 10.6771484 | 15.7768826 | 14.8459568 | 0 |
| 0.02294601 | 0.03922062 | 4.17979245 | 6.67321777 | 7.42441534 | 7.42297842 | 0 |
| 0.02294727 | 0.03922077 | 0 | 36.7026978 | 38.9781805 | 33.867339 | 0 |
| 0.02295652 | 0.03923458 | 0 | 97.4289795 | 16.7049345 | 18.5574461 | 0 |
| 0.02295904 | 0.03923691 | 27.6509347 | 36.7026978 | 0 | 23.1968076 | 0 |
| 0.02296078 | 0.03923789 | 11.5748099 | 46.7125244 | 0 | 38.5067006 | 0 |
| 0.02296628 | 0.03924529 | 0 | 44.7105591 | 32.4818171 | 32.9394667 | 0 |
| 0.02297063 | 0.03925073 | 10.6102424 | 6.67321777 | 9.28051917 | 13.4541484 | 116.434694 |
| 0.022975 | 0.03925621 | 1.60761248 | 9.34250488 | 12.0646749 | 13.9180845 | 0 |
| 0.02301698 | 0.03932593 | 14.7900348 | 30.2519206 | 68.6758419 | 0 | 0 |
| 0.02303509 | 0.03935487 | 8.3595849 | 7.11809896 | 7.42441534 | 3.71148921 | 0 |

| | | | | | | |
|---|---|---|---|---|---|---|
| 0.02303853 | 0.03935876 | 34.4029071 | 4.44881185 | 2.78415575 | 6.03116997 | 0 |
| 0.0230467 | 0.03937072 | 18.9698273 | 44.9329997 | 41.7623363 | 0 | 0 |
| 0.02306388 | 0.03939806 | 0 | 22.6889404 | 58.4672708 | 32.9394667 | 0 |
| 0.02310773 | 0.03947096 | 0.96456749 | 0.44488118 | 0.92805192 | 0 | 54.6530196 |
| 0.02311504 | 0.03948144 | 82.952804 | 11.1220296 | 27.8415575 | 0 | 0 |
| 0.0231208 | 0.03948928 | 0 | 26.2479899 | 17.6329864 | 78.8691457 | 0 |
| 0.02312641 | 0.03949685 | 10.9317649 | 46.4900838 | 0 | 39.898509 | 0 |
| 0.02315403 | 0.03954202 | 32.4737721 | 90.3108805 | 121.574801 | 109.488932 | 9.50487297 |
| 0.02315761 | 0.03954613 | 0.64304499 | 0 | 0 | 0.46393615 | 0 |
| 0.02316268 | 0.03954675 | 12.5393773 | 46.7125244 | 52.8989593 | 42.6821259 | 14.2573095 |
| 0.02316232 | 0.03954675 | 0 | 0 | 0 | 0 | 71.2865473 |
| 0.02316232 | 0.03954675 | 0 | 0 | 0 | 0 | 71.2865473 |
| 0.0231598 | 0.03954675 | 13.8254673 | 0 | 59.3953227 | 55.6723382 | 0 |
| 0.0231729 | 0.0395622 | 3.21522496 | 15.1259603 | 6.49636342 | 6.95904227 | 0 |
| 0.02319088 | 0.03959089 | 97.4213163 | 139.470251 | 112.294282 | 124.334889 | 0 |
| 0.02319713 | 0.03959894 | 3.85826995 | 4.00393066 | 14.8488307 | 9.74265918 | 0 |
| 0.02319795 | 0.03959894 | 44.3701045 | 24.2460246 | 0 | 20.8771268 | 0 |
| 0.0232047 | 0.03960844 | 114.140486 | 158.822583 | 161.481034 | 176.295737 | 0 |
| 0.02321444 | 0.03962306 | 18.0052598 | 30.9192423 | 0 | 40.3624452 | 0 |
| 0.02324638 | 0.03967558 | 168.79931 | 64.5077718 | 179.11402 | 88.6118049 | 0 |
| 0.02327777 | 0.03972712 | 5.46588243 | 8.45274251 | 13.9207788 | 3.24755306 | 0 |
| 0.02328469 | 0.03973691 | 18.0052598 | 19.7972127 | 0 | 57.5280828 | 0 |
| 0.02329609 | 0.03975437 | 42.119447 | 59.1691976 | 12.0646749 | 0 | 0 |
| 0.02329833 | 0.03975617 | 18.9698273 | 18.6850098 | 25.0574018 | 7.42297842 | 61.7816743 |
| 0.02331083 | 0.03977548 | 0 | 22.911381 | 54.7550631 | 34.7952113 | 0 |
| 0.02334592 | 0.03983334 | 4.50131495 | 4.00393066 | 23.2012979 | 6.03116997 | 0 |
| 0.0233483 | 0.03983538 | 7.39501741 | 8.23030192 | 2.78415575 | 11.1344676 | 0 |
| 0.02335473 | 0.03984433 | 15.7546023 | 18.2401286 | 0 | 65.8789335 | 0 |
| 0.02335924 | 0.03984999 | 2.57217997 | 6.67321777 | 12.9927268 | 9.74265918 | 0 |
| 0.02336335 | 0.03985498 | 3.21522496 | 2.22440592 | 0.92805192 | 4.63936151 | 21.3859642 |
| 0.0233661 | 0.03985766 | 5.46588243 | 6.22833659 | 4.64025958 | 10.2065953 | 0 |
| 0.02338196 | 0.0398827 | 8.0380624 | 7.11809896 | 5.5683115 | 4.63936151 | 0 |
| 0.02339544 | 0.03990366 | 12.2178549 | 18.6850098 | 0 | 74.6937204 | 0 |
| 0.0234287 | 0.03995837 | 126.679863 | 39.1495443 | 91.8771398 | 45.4657428 | 21.3859642 |
| 0.02343991 | 0.03997546 | 34.0813846 | 19.1298909 | 29.6976613 | 19.9492545 | 154.454186 |
| 0.02345707 | 0.04000269 | 0 | 25.1357869 | 29.6976613 | 57.5280828 | 0 |
| 0.02347188 | 0.04002592 | 3.21522496 | 11.5669108 | 4.64025958 | 12.5262761 | 0 |
| 0.02348558 | 0.04004523 | 13.8254673 | 42.9310343 | 29.6976613 | 57.5280828 | 14.2573095 |
| 0.0234849 | 0.04004523 | 0.96456749 | 12.9015544 | 29.6976613 | 6.49510612 | 0 |
| 0.02349247 | 0.04005494 | 291.942426 | 146.143469 | 170.761553 | 202.276162 | 9.50487297 |
| 0.023505 | 0.04007427 | 5.14435994 | 6.22833659 | 6.49636342 | 6.49510612 | 0 |
| 0.02354965 | 0.04014837 | 24.1141872 | 32.2538859 | 0 | 29.2279775 | 0 |
| 0.02356361 | 0.04016809 | 0 | 27.1377523 | 33.409869 | 49.177232 | 0 |

| | | | | | | |
|---|---|---|---|---|---|---|
| 0.02356278 | 0.04016809 | 2.57217997 | 0 | 5.5683115 | 3.71148921 | 68.9103291 |
| 0.02362184 | 0.04026532 | 64.6260217 | 161.714311 | 0 | 105.313506 | 9.50487297 |
| 0.02362453 | 0.04026787 | 0 | 29.3621582 | 51.9709073 | 28.7640414 | 0 |
| 0.02364449 | 0.04029985 | 16.0761248 | 42.041272 | 48.2586997 | 0 | 0 |
| 0.02365035 | 0.04030779 | 4.17979245 | 6.89565837 | 4.64025958 | 15.7738291 | 0 |
| 0.02365344 | 0.04031101 | 12.8608998 | 13.5688761 | 26.9135056 | 11.1344676 | 2.37621824 |
| 0.02367349 | 0.0403411 | 37.9396545 | 0 | 45.4745439 | 32.4755306 | 0 |
| 0.02367237 | 0.0403411 | 0 | 0 | 0.92805192 | 0.46393615 | 0 |
| 0.02368597 | 0.04036031 | 0 | 54.9428263 | 25.0574018 | 31.0837221 | 0 |
| 0.0237264 | 0.04042716 | 0 | 0 | 0 | 0 | 4.75243649 |
| 0.02373505 | 0.04043986 | 2.25065747 | 10.899589 | 7.42441534 | 12.0623399 | 0 |
| 0.02374106 | 0.04044804 | 2.25065747 | 8.23030192 | 6.49636342 | 20.4131907 | 0 |
| 0.02375635 | 0.04047204 | 10.2887199 | 2.8917277 | 10.2085711 | 6.49510612 | 0 |
| 0.02375778 | 0.04047243 | 1.92913498 | 8.67518311 | 8.35246725 | 17.1656376 | 0 |
| 0.02376497 | 0.04048263 | 2.25065747 | 19.5747721 | 17.6329864 | 3.24755306 | 0 |
| 0.023777 | 0.04050107 | 16.0761248 | 24.9133464 | 0 | 51.0329766 | 0 |
| 0.02379209 | 0.04052472 | 0 | 18.4625692 | 50.1148035 | 44.0739344 | 0 |
| 0.02379723 | 0.04053143 | 0 | 8.00786133 | 0 | 1.85574461 | 42.7719284 |
| 0.02380539 | 0.04054328 | 277.473914 | 96.3167765 | 65.8916861 | 148.923505 | 449.105248 |
| 0.0238197 | 0.04056558 | 0.3215225 | 3.11416829 | 2.78415575 | 4.17542536 | 19.0097459 |
| 0.02384006 | 0.0405982 | 90.9908664 | 78.2990885 | 0 | 69.1264865 | 7.12865473 |
| 0.02384936 | 0.04061199 | 0 | 0 | 0 | 0 | 33.2670554 |
| 0.02385822 | 0.04062502 | 0 | 0.66732178 | 0.92805192 | 0.46393615 | 49.9005831 |
| 0.0238673 | 0.04063843 | 3.85826995 | 14.0137573 | 9.28051917 | 4.17542536 | 0 |
| 0.02388126 | 0.04066013 | 28.2939797 | 15.1259603 | 0 | 47.7854236 | 0 |
| 0.02389253 | 0.04067726 | 0 | 0 | 2.78415575 | 1.85574461 | 33.2670554 |
| 0.02389499 | 0.04067939 | 36.9750871 | 28.917277 | 0 | 20.4131907 | 0 |
| 0.02390381 | 0.04068928 | 0.3215225 | 0 | 0 | 0 | 0 |
| 0.02390442 | 0.04068928 | 4.82283744 | 5.33857422 | 7.42441534 | 7.42297842 | 0 |
| 0.0239024 | 0.04068928 | 3.53674746 | 7.78542074 | 9.28051917 | 6.49510612 | 0 |
| 0.02391587 | 0.04070671 | 9.32415239 | 34.4782918 | 0 | 55.6723382 | 0 |
| 0.02392164 | 0.0407124 | 7.39501741 | 7.56298014 | 7.42441534 | 3.71148921 | 0 |
| 0.02392043 | 0.0407124 | 3.85826995 | 20.2420939 | 2.78415575 | 11.1344676 | 0 |
| 0.02394419 | 0.04074872 | 2.89370247 | 8.23030192 | 7.42441534 | 10.6705315 | 0 |
| 0.02394776 | 0.04075274 | 7.39501741 | 0 | 79.8124649 | 52.8887212 | 0 |
| 0.02396135 | 0.0407738 | 45.6561944 | 141.249776 | 142.919995 | 138.716909 | 40.3957101 |
| 0.02397738 | 0.04079901 | 9.32415239 | 6.89565837 | 4.64025958 | 5.10329766 | 0 |
| 0.02399859 | 0.04083303 | 3.85826995 | 4.44881185 | 4.64025958 | 31.0837221 | 0 |
| 0.02400204 | 0.04083684 | 0 | 0 | 0 | 0 | 68.9103291 |
| 0.02400894 | 0.04084652 | 13.1824223 | 38.7046631 | 57.5392189 | 0 | 0 |
| 0.02401284 | 0.0408511 | 168.156265 | 278.718062 | 356.371936 | 300.16669 | 116.434694 |
| 0.02402126 | 0.04086334 | 89.3832539 | 219.103984 | 197.675058 | 211.554885 | 0 |
| 0.02402893 | 0.04087432 | 0 | 22.2440592 | 18.5610383 | 80.2609542 | 0 |

| | | | | | | |
|---|---|---|---|---|---|---|
| 0.0240339 | 0.04088072 | 0 | 25.1357869 | 43.6184401 | 39.4345729 | 0 |
| 0.02403542 | 0.04088124 | 3.85826995 | 10.2322673 | 26.9135056 | 2.31968076 | 0 |
| 0.02407242 | 0.04094211 | 16.3976473 | 27.5826335 | 64.9636342 | 0 | 0 |
| 0.02408064 | 0.04095401 | 18.9698273 | 25.3582275 | 0 | 43.6099982 | 0 |
| 0.02408211 | 0.04095443 | 31.5092046 | 22.2440592 | 0 | 31.0837221 | 0 |
| 0.02408958 | 0.04096507 | 3.21522496 | 10.0098267 | 12.0646749 | 5.10329766 | 0 |
| 0.02413138 | 0.04103408 | 0 | 45.3778809 | 43.6184401 | 20.8771268 | 0 |
| 0.02413275 | 0.04103433 | 26.6863672 | 0 | 56.6111669 | 33.867339 | 0 |
| 0.02414559 | 0.04105409 | 5.46588243 | 3.55904948 | 44.546492 | 2.31968076 | 0 |
| 0.02416449 | 0.04108415 | 333.740351 | 440.209932 | 500.219983 | 509.401894 | 0 |
| 0.02418327 | 0.04111401 | 18.3267823 | 9.12006429 | 11.136623 | 8.35085072 | 35.6432737 |
| 0.02419721 | 0.04113563 | 556.876963 | 2186.36858 | 1509.01242 | 1718.88344 | 235.245606 |
| 0.02422132 | 0.04117453 | 1.92913498 | 3.55904948 | 4.64025958 | 2.78361691 | 35.6432737 |
| 0.02423178 | 0.04119023 | 32.4737721 | 42.041272 | 51.0428554 | 43.6099982 | 21.3859642 |
| 0.02423515 | 0.04119387 | 222.81509 | 504.940145 | 447.321024 | 567.393913 | 0 |
| 0.02423783 | 0.04119428 | 2.25065747 | 10.6771484 | 38.0501286 | 2.31968076 | 0 |
| 0.02423695 | 0.04119428 | 15.1115573 | 4.89369303 | 4.64025958 | 7.88691457 | 0 |
| 0.0242797 | 0.04126334 | 5.46588243 | 9.56494547 | 2.78415575 | 13.9180845 | 0 |
| 0.02428968 | 0.04127614 | 6.10892743 | 6.45077718 | 7.42441534 | 4.63936151 | 0 |
| 0.02428857 | 0.04127614 | 4.17979245 | 6.22833659 | 6.49636342 | 8.81478687 | 0 |
| 0.02429408 | 0.04128152 | 6.10892743 | 7.34053955 | 3.71220767 | 9.27872303 | 0 |
| 0.02429679 | 0.04128358 | 83.595849 | 55.6101481 | 73.3161014 | 51.9608489 | 28.5146189 |
| 0.02429774 | 0.04128358 | 53.6942569 | 141.694657 | 116.00649 | 123.870952 | 35.6432737 |
| 0.02432092 | 0.04132088 | 0 | 28.4723958 | 55.683115 | 25.5164883 | 0 |
| 0.02432419 | 0.04132436 | 79.4160565 | 69.6239054 | 88.1649321 | 74.2297842 | 87.920075 |
| 0.02434269 | 0.04132657 | 0 | 0 | 0 | 0 | 0 |
| 0.02434269 | 0.04132657 | 0 | 0 | 0 | 0 | 0 |
| 0.02434269 | 0.04132657 | 0 | 0 | 0 | 0 | 0 |
| 0.02434269 | 0.04132657 | 0 | 0 | 0 | 0 | 0 |
| 0.02434269 | 0.04132657 | 0 | 0 | 0 | 0 | 0 |
| 0.02434269 | 0.04132657 | 0 | 0 | 0 | 0 | 0 |
| 0.02434269 | 0.04132657 | 0 | 0 | 0 | 0 | 0 |
| 0.02434269 | 0.04132657 | 0 | 0 | 0 | 0 | 0 |
| 0.02434269 | 0.04132657 | 0 | 0 | 0 | 0 | 0 |
| 0.02434191 | 0.04132657 | 10.9317649 | 44.4881185 | 0 | 38.5067006 | 0 |
| 0.02432726 | 0.04132657 | 35.6889971 | 25.8031087 | 0 | 23.1968076 | 0 |
| 0.02434269 | 0.04132657 | 0 | 0 | 0 | 0 | 0 |
| 0.02434269 | 0.04132657 | 0 | 0 | 0 | 0 | 0 |
| 0.02434269 | 0.04132657 | 0 | 0 | 0 | 0 | 0 |
| 0.0243509 | 0.04133842 | 16.3976473 | 7.11809896 | 10.2085711 | 7.88691457 | 0 |
| 0.02435513 | 0.04134352 | 0 | 25.8031087 | 15.7768826 | 78.8691457 | 0 |
| 0.02435795 | 0.04134621 | 54.3373018 | 27.3601929 | 0 | 12.5262761 | 0 |
| 0.02436382 | 0.04135409 | 25.4002772 | 23.8011434 | 0 | 35.2591475 | 0 |

| | | | | | | |
|---|---|---|---|---|---|---|
| 0.02436818 | 0.0413594 | 3.85826995 | 6.005896 | 12.9927268 | 6.03116997 | 0 |
| 0.02437203 | 0.04136386 | 28.2939797 | 26.4704305 | 0 | 28.7640414 | 0 |
| 0.02438297 | 0.04138033 | 19.6128723 | 30.4743612 | 0 | 35.2591475 | 0 |
| 0.02438839 | 0.04138671 | 23.4711422 | 5.33857422 | 3.71220767 | 5.10329766 | 0 |
| 0.02438919 | 0.04138671 | 51.1220769 | 173.058781 | 0 | 42.2181898 | 7.12865473 |
| 0.02439859 | 0.04140058 | 22.8280972 | 35.8129354 | 0 | 25.9804245 | 0 |
| 0.02442948 | 0.04145089 | 0 | 0 | 0 | 0 | 0 |
| 0.02443865 | 0.04146436 | 2.57217997 | 15.7932821 | 4.64025958 | 12.0623399 | 0 |
| 0.02444518 | 0.04147335 | 27.3294122 | 84.0825439 | 67.7477899 | 87.6839326 | 26.1384007 |
| 0.02447502 | 0.04152189 | 0 | 36.4802572 | 16.7049345 | 60.3116997 | 0 |
| 0.02448222 | 0.04152992 | 43.405537 | 41.1515096 | 76.1002572 | 66.8068058 | 28.5146189 |
| 0.02448178 | 0.04152992 | 5.46588243 | 14.0137573 | 8.35246725 | 14.8459568 | 0 |
| 0.02448639 | 0.0415349 | 3.85826995 | 10.6771484 | 4.64025958 | 9.27872303 | 0 |
| 0.02448918 | 0.04153753 | 20.5774398 | 36.035376 | 0 | 28.3001052 | 0 |
| 0.02451605 | 0.04157891 | 8.6811074 | 7.78542074 | 3.71220767 | 6.03116997 | 0 |
| 0.02451507 | 0.04157891 | 9.96719738 | 43.3759155 | 59.3953227 | 0 | 0 |
| 0.02451962 | 0.04158287 | 6.75197242 | 6.67321777 | 7.42441534 | 4.17542536 | 0 |
| 0.02455249 | 0.04163651 | 163.654951 | 198.417008 | 190.250643 | 192.533503 | 142.573095 |
| 0.02455936 | 0.04164607 | 6.43044992 | 14.4586385 | 125.287009 | 0 | 0 |
| 0.02456455 | 0.04165277 | 10.9317649 | 43.3759155 | 22.273246 | 49.6411682 | 11.8810912 |
| 0.02461362 | 0.04173387 | 0.96456749 | 19.5747721 | 7.42441534 | 16.7017014 | 0 |
| 0.02464865 | 0.04179116 | 4.50131495 | 41.8188314 | 0 | 66.8068058 | 0 |
| 0.02466319 | 0.04181371 | 15.1115573 | 47.6022868 | 0 | 26.9082968 | 0 |
| 0.02468701 | 0.04185199 | 5.14435994 | 63.84045 | 0 | 40.8263813 | 0 |
| 0.02471919 | 0.04190442 | 115.105054 | 0 | 17.6329864 | 12.0623399 | 0 |
| 0.02473108 | 0.04192247 | 4.82283744 | 1.77952474 | 21.3451941 | 12.9902122 | 0 |
| 0.02474456 | 0.04194321 | 18.6483048 | 1.77952474 | 27.8415575 | 2.31968076 | 0 |
| 0.0247469 | 0.04194505 | 2.89370247 | 2.44684652 | 12.0646749 | 27.3722329 | 0 |
| 0.02476032 | 0.0419657 | 0 | 265.149186 | 308.113236 | 247.741905 | 19.0097459 |
| 0.02477549 | 0.04198929 | 16.7191698 | 28.4723958 | 0 | 41.7542536 | 0 |
| 0.02477776 | 0.04199102 | 6.75197242 | 4.89369303 | 6.49636342 | 6.03116997 | 0 |
| 0.02479185 | 0.04201066 | 12.2178549 | 25.8031087 | 76.1002572 | 0 | 0 |
| 0.02479169 | 0.04201066 | 27.9724572 | 0 | 32.4818171 | 54.2805297 | 0 |
| 0.02479507 | 0.04201401 | 2.25065747 | 9.34250488 | 17.6329864 | 6.03116997 | 0 |
| 0.02480072 | 0.04202147 | 0 | 45.6003214 | 22.273246 | 39.4345729 | 0 |
| 0.0248279 | 0.04206541 | 38.5826995 | 44.7105591 | 0 | 25.0525522 | 0 |
| 0.0248454 | 0.04209293 | 0.3215225 | 0.88976237 | 2.78415575 | 1.85574461 | 0 |
| 0.02484734 | 0.0420941 | 27.3294122 | 5.56101481 | 3.71220767 | 4.17542536 | 0 |
| 0.02485419 | 0.04210358 | 24.7572322 | 36.4802572 | 0 | 22.7328714 | 0 |
| 0.02487619 | 0.04213872 | 15.7546023 | 28.6948364 | 0 | 43.1460621 | 0 |
| 0.02487828 | 0.04214014 | 23.7926647 | 34.2558512 | 16.7049345 | 83.5085072 | 2.37621824 |
| 0.0248805 | 0.04214179 | 30.8661596 | 18.9074504 | 0 | 34.7952113 | 0 |
| 0.0248992 | 0.04217134 | 0 | 0 | 0 | 0 | 66.5341108 |

| | | | | | | |
|---|---|---|---|---|---|---|
| 0.02491627 | 0.042196 | 0 | 7.11809896 | 0 | 0 | 85.5438568 |
| 0.02491549 | 0.042196 | 1.60761248 | 0 | 0.92805192 | 1.85574461 | 2.37621824 |
| 0.02493828 | 0.04223115 | 29.5800696 | 15.7932821 | 0 | 41.7542536 | 0 |
| 0.02494628 | 0.04224256 | 1.60761248 | 7.56298014 | 14.8488307 | 12.5262761 | 0 |
| 0.02495134 | 0.04224901 | 119.606369 | 222.663033 | 207.883629 | 218.049991 | 0 |
| 0.02497124 | 0.04228057 | 24.7572322 | 13.3464355 | 13.9207788 | 14.8459568 | 19.0097459 |
| 0.02497344 | 0.04228217 | 65.5905892 | 73.1829549 | 140.135839 | 100.210209 | 30.8908372 |
| 0.02497887 | 0.04228924 | 38.904222 | 26.4704305 | 0 | 19.4853184 | 0 |
| 0.02498177 | 0.04229202 | 19.6128723 | 47.1574056 | 0 | 20.8771268 | 0 |
| 0.02499598 | 0.04231395 | 2.57217997 | 6.67321777 | 12.9927268 | 8.81478687 | 0 |
| 0.02499783 | 0.04231495 | 15.4330798 | 27.5826335 | 0 | 45.0018067 | 0 |
| 0.02500786 | 0.04232769 | 3.85826995 | 14.9035197 | 8.35246725 | 4.17542536 | 0 |
| 0.02500787 | 0.04232769 | 31.5092046 | 12.011792 | 66.819738 | 0 | 0 |
| 0.02504915 | 0.04239542 | 0 | 10.6771484 | 122.502853 | 12.0623399 | 0 |
| 0.02506216 | 0.04241319 | 0.3215225 | 0 | 0 | 0.9278723 | 0 |
| 0.02506214 | 0.04241319 | 32.4737721 | 18.4625692 | 0 | 33.4034029 | 0 |
| 0.02506518 | 0.04241616 | 5.14435994 | 4.22637126 | 1.85610383 | 3.24755306 | 11.8810912 |
| 0.02507534 | 0.04243121 | 71.3779941 | 10.899589 | 2.78415575 | 3.24755306 | 0 |
| 0.02508235 | 0.04244094 | 0 | 37.1475789 | 15.7768826 | 59.3838274 | 0 |
| 0.02509994 | 0.04246858 | 5.78740493 | 12.2342326 | 3.71220767 | 6.49510612 | 0 |
| 0.02511783 | 0.0424967 | 4.17979245 | 5.11613363 | 6.49636342 | 8.35085072 | 64.1578926 |
| 0.02512328 | 0.04250378 | 0.96456749 | 1.11220296 | 0 | 0.46393615 | 0 |
| 0.02513099 | 0.04251469 | 0 | 30.9192423 | 68.6758419 | 15.309893 | 0 |
| 0.0251323 | 0.04251477 | 44.048582 | 0 | 24.1293498 | 45.4657428 | 0 |
| 0.0251371 | 0.04251647 | 7.07349491 | 63.84045 | 0 | 63.0953166 | 2.37621824 |
| 0.02513559 | 0.04251647 | 0 | 208.649276 | 328.530379 | 0 | 0 |
| 0.02513696 | 0.04251647 | 175.872805 | 242.237805 | 251.502069 | 239.391054 | 35.6432737 |
| 0.02514855 | 0.0425337 | 21.8635297 | 36.035376 | 0 | 25.5164883 | 0 |
| 0.02515141 | 0.04253641 | 3.85826995 | 8.00786133 | 5.5683115 | 8.81478687 | 0 |
| 0.02516524 | 0.04255766 | 4.50131495 | 7.78542074 | 5.5683115 | 6.95904227 | 0 |
| 0.02516742 | 0.0425592 | 117.677234 | 25.8031087 | 12.0646749 | 17.1656376 | 11.8810912 |
| 0.02518851 | 0.04259272 | 55.3018693 | 131.684831 | 131.783372 | 91.3954218 | 14.2573095 |
| 0.02519565 | 0.04260266 | 40.190312 | 39.1495443 | 39.9062324 | 37.1148921 | 59.4054561 |
| 0.02521547 | 0.04263402 | 18.3267823 | 44.7105591 | 37.1220767 | 0 | 0 |
| 0.02523105 | 0.04265822 | 11.5748099 | 45.6003214 | 0 | 33.867339 | 0 |
| 0.0252634 | 0.04271077 | 0 | 43.8207967 | 71.4599976 | 8.35085072 | 0 |
| 0.02528128 | 0.04273886 | 9.96719738 | 7.11809896 | 2.78415575 | 6.95904227 | 57.0292378 |
| 0.02528798 | 0.04274803 | 3.53674746 | 5.33857422 | 8.35246725 | 10.6705315 | 0 |
| 0.02530237 | 0.0427702 | 38.5826995 | 12.4566732 | 0 | 38.0427644 | 0 |
| 0.02530559 | 0.0427735 | 4.50131495 | 2.66928711 | 12.0646749 | 14.8459568 | 0 |
| 0.02530704 | 0.0427738 | 0.96456749 | 16.0157227 | 15.7768826 | 9.27872303 | 0 |
| 0.02534837 | 0.04284151 | 7.39501741 | 6.67321777 | 3.71220767 | 5.56723382 | 92.6725115 |
| 0.02535885 | 0.04285706 | 5.78740493 | 6.89565837 | 5.5683115 | 5.56723382 | 0 |

| | | | | | | |
|---|---|---|---|---|---|---|
| 0.0253672 | 0.04286902 | 3.21522496 | 8.23030192 | 12.0646749 | 5.56723382 | 0 |
| 0.02537168 | 0.04287444 | 3.21522496 | 9.34250488 | 8.35246725 | 6.49510612 | 0 |
| 0.0253795 | 0.0428855 | 11.8963324 | 29.3621582 | 0 | 50.1051043 | 0 |
| 0.02538193 | 0.04288745 | 0 | 26.4704305 | 21.3451941 | 62.1674443 | 0 |
| 0.02538981 | 0.04289861 | 12.2178549 | 30.9192423 | 33.409869 | 29.6919137 | 11.8810912 |
| 0.02539593 | 0.0429068 | 82.309759 | 78.2990885 | 110.438178 | 73.7658481 | 175.84015 |
| 0.02541111 | 0.04293029 | 20.2559173 | 39.1495443 | 0 | 24.588616 | 0 |
| 0.0254127 | 0.04293082 | 2.57217997 | 6.005896 | 12.9927268 | 9.74265918 | 0 |
| 0.02541788 | 0.04293742 | 10.6102424 | 51.3837769 | 0 | 31.0837221 | 0 |
| 0.02542425 | 0.04294603 | 4.50131495 | 6.005896 | 7.42441534 | 6.49510612 | 0 |
| 0.02545836 | 0.04300149 | 81.666714 | 97.4289795 | 83.5246725 | 103.457762 | 7.12865473 |
| 0.02547953 | 0.04303322 | 5.14435994 | 6.67321777 | 6.49636342 | 5.56723382 | 0 |
| 0.02547971 | 0.04303322 | 27.6509347 | 23.1338216 | 0 | 31.0837221 | 0 |
| 0.02548928 | 0.04304723 | 8.3595849 | 33.3660889 | 38.0501286 | 51.4969128 | 0 |
| 0.02549378 | 0.0430505 | 0 | 0 | 0 | 0 | 0 |
| 0.02549378 | 0.0430505 | 0 | 0 | 0 | 0 | 0 |
| 0.02549966 | 0.04305827 | 35.0459521 | 19.1298909 | 8.35246725 | 16.7017014 | 4.75243649 |
| 0.02550435 | 0.04306403 | 21.2204847 | 36.9251383 | 26.9135056 | 19.4853184 | 104.553603 |
| 0.02551804 | 0.04308498 | 3.85826995 | 4.67125244 | 11.136623 | 8.35085072 | 0 |
| 0.02552578 | 0.04309589 | 30.2231146 | 6.005896 | 0 | 70.518295 | 0 |
| 0.02553344 | 0.04310666 | 3.21522496 | 12.011792 | 3.71220767 | 14.3820207 | 0 |
| 0.02554 | 0.04311557 | 23.1496197 | 29.3621582 | 80.7405168 | 0 | 0 |
| 0.02555594 | 0.04313914 | 29.5800696 | 23.3562622 | 44.546492 | 0 | 0 |
| 0.02555653 | 0.04313914 | 83.2743265 | 114.779346 | 102.085711 | 113.200421 | 14.2573095 |
| 0.02556876 | 0.04315762 | 0 | 15.3484009 | 77.956361 | 24.1246799 | 0 |
| 0.02557804 | 0.04317113 | 45.334672 | 0 | 77.956361 | 8.35085072 | 0 |
| 0.02559201 | 0.04319253 | 11.2532874 | 22.4664998 | 9.28051917 | 19.4853184 | 2.37621824 |
| 0.02560191 | 0.04320707 | 2.57217997 | 17.7952474 | 17.6329864 | 2.78361691 | 0 |
| 0.02560884 | 0.0432166 | 40.833357 | 58.5018758 | 0 | 5.56723382 | 0 |
| 0.02562348 | 0.04323914 | 12.2178549 | 7.34053955 | 2.78415575 | 6.95904227 | 0 |
| 0.02563003 | 0.04324802 | 17.3622148 | 20.4645345 | 31.5537652 | 24.588616 | 11.8810912 |
| 0.02564383 | 0.04326697 | 3.53674746 | 46.7125244 | 0 | 64.487125 | 0 |
| 0.02564371 | 0.04326697 | 0 | 0 | 0 | 0 | 0 |
| 0.02565009 | 0.04327535 | 2.57217997 | 11.7893514 | 5.5683115 | 11.5984038 | 0 |
| 0.02565561 | 0.04328049 | 1.28608998 | 0.66732178 | 0 | 0 | 0 |
| 0.0256557 | 0.04328049 | 0.96456749 | 0 | 0 | 0 | 21.3859642 |
| 0.02566255 | 0.04328987 | 15.1115573 | 14.4586385 | 3.71220767 | 11.5984038 | 0 |
| 0.02566584 | 0.04329326 | 30.2231146 | 21.5767375 | 21.3451941 | 59.3838274 | 14.2573095 |
| 0.02566967 | 0.04329754 | 2.57217997 | 7.11809896 | 12.0646749 | 8.35085072 | 0 |
| 0.02567708 | 0.04330786 | 0.64304499 | 0 | 0 | 0.46393615 | 2.37621824 |
| 0.02570199 | 0.04334772 | 13.1824223 | 0 | 27.8415575 | 0 | 152.077968 |
| 0.02572782 | 0.0433891 | 43.405537 | 40.929069 | 0 | 22.7328714 | 2.37621824 |
| 0.02576959 | 0.04345736 | 0.3215225 | 0.44488118 | 0 | 0 | 0 |

| | | | | | | |
|---|---|---|---|---|---|---|
| 0.02579223 | 0.04349337 | 6.10892743 | 59.3916382 | 59.3953227 | 28.7640414 | 73.6627655 |
| 0.02579803 | 0.04350097 | 0 | 30.2519206 | 40.8342843 | 31.5476583 | 0 |
| 0.02582987 | 0.04355246 | 11.2532874 | 6.67321777 | 8.35246725 | 2.78361691 | 0 |
| 0.0258376 | 0.04356332 | 7.39501741 | 28.2499552 | 18.5610383 | 23.1968076 | 7.12865473 |
| 0.02584575 | 0.04357488 | 3.85826995 | 6.67321777 | 9.28051917 | 6.03116997 | 0 |
| 0.02585986 | 0.04359211 | 0 | 0 | 0 | 0 | 64.1578926 |
| 0.02585986 | 0.04359211 | 0 | 0 | 0 | 0 | 64.1578926 |
| 0.02585986 | 0.04359211 | 0 | 0 | 0 | 0 | 64.1578926 |
| 0.02586839 | 0.04360431 | 4.17979245 | 5.56101481 | 7.42441534 | 7.88691457 | 0 |
| 0.02588434 | 0.04362901 | 66.2336342 | 174.838306 | 92.8051917 | 136.397228 | 0 |
| 0.0258905 | 0.04363721 | 18.3267823 | 24.2460246 | 0 | 41.7542536 | 0 |
| 0.02589927 | 0.0436498 | 2.57217997 | 8.45274251 | 12.0646749 | 6.95904227 | 0 |
| 0.02591532 | 0.04367465 | 27.0078897 | 23.5787028 | 23.2012979 | 36.1870198 | 14.2573095 |
| 0.0259267 | 0.04368947 | 0 | 0 | 0 | 0 | 4.75243649 |
| 0.0259267 | 0.04368947 | 0 | 0 | 0 | 0 | 4.75243649 |
| 0.02594541 | 0.04371879 | 17.3622148 | 17.5728068 | 0 | 55.208402 | 0 |
| 0.02595005 | 0.04372442 | 2.57217997 | 24.6909058 | 8.35246725 | 4.17542536 | 0 |
| 0.02595845 | 0.04373639 | 24.7572322 | 34.0334106 | 0 | 22.7328714 | 0 |
| 0.02597473 | 0.04376162 | 63.3399317 | 137.468286 | 196.747006 | 148.459568 | 40.3957101 |
| 0.0260259 | 0.04384565 | 24.1141872 | 6.005896 | 4.64025958 | 3.24755306 | 0 |
| 0.02603221 | 0.04385408 | 2.25065747 | 13.123995 | 34.3379209 | 1.85574461 | 0 |
| 0.02604874 | 0.04387972 | 0.96456749 | 2.00196533 | 2.78415575 | 0.46393615 | 21.3859642 |
| 0.02605696 | 0.04389138 | 8.3595849 | 32.4763265 | 0 | 55.208402 | 0 |
| 0.0260647 | 0.04390221 | 506.076409 | 308.525102 | 314.6096 | 305.733924 | 11.8810912 |
| 0.02607498 | 0.04391734 | 3.53674746 | 6.89565837 | 7.42441534 | 7.88691457 | 0 |
| 0.02608106 | 0.04392318 | 0.3215225 | 0 | 0 | 0 | 0 |
| 0.02608106 | 0.04392318 | 0.3215225 | 0 | 0 | 0 | 0 |
| 0.02609071 | 0.04393723 | 4.82283744 | 11.5669108 | 3.71220767 | 7.88691457 | 0 |
| 0.02609979 | 0.04394813 | 3.21522496 | 14.2361979 | 10.2085711 | 4.17542536 | 0 |
| 0.0260988 | 0.04394813 | 4.50131495 | 35.8129354 | 7.42441534 | 9.74265918 | 0 |
| 0.0261059 | 0.04395622 | 0 | 44.4881185 | 9.28051917 | 64.487125 | 0 |
| 0.02612447 | 0.04398528 | 196.128723 | 339.444344 | 275.631419 | 432.388493 | 0 |
| 0.02613106 | 0.04399417 | 5.14435994 | 5.33857422 | 7.42441534 | 6.03116997 | 0 |
| 0.02613562 | 0.04399964 | 3.53674746 | 8.67518311 | 10.2085711 | 5.10329766 | 0 |
| 0.02613955 | 0.04400406 | 13.1824223 | 0 | 77.0283091 | 32.9394667 | 0 |
| 0.0261748 | 0.0440433 | 0 | 0 | 0 | 0 | 0 |
| 0.0261748 | 0.0440433 | 0 | 0 | 0 | 0 | 0 |
| 0.02616622 | 0.0440433 | 5.14435994 | 23.5787028 | 0 | 84.4363795 | 0 |
| 0.0261748 | 0.0440433 | 0 | 0 | 0 | 0 | 0 |
| 0.0261748 | 0.0440433 | 0 | 0 | 0 | 0 | 0 |
| 0.0261748 | 0.0440433 | 0 | 0 | 0 | 0 | 0 |
| 0.0261748 | 0.0440433 | 0 | 0 | 0 | 0 | 0 |
| 0.0261748 | 0.0440433 | 0 | 0 | 0 | 0 | 0 |

| | | | | | | |
|---|---|---|---|---|---|---|
| 0.02616909 | 0.0440433 | 4.17979245 | 47.3798462 | 10.2085711 | 22.2689353 | 4.75243649 |
| 0.02617595 | 0.0440433 | 3.85826995 | 10.0098267 | 5.5683115 | 6.95904227 | 0 |
| 0.02618397 | 0.04405459 | 5.78740493 | 12.9015544 | 3.71220767 | 6.03116997 | 0 |
| 0.02621943 | 0.04411204 | 29.5800696 | 44.4881185 | 22.273246 | 0 | 0 |
| 0.02623784 | 0.04414081 | 0 | 24.9133464 | 45.4745439 | 32.4755306 | 0 |
| 0.0262451 | 0.04415081 | 9.64567488 | 6.89565837 | 129.927268 | 0 | 0 |
| 0.02625282 | 0.0441616 | 117.034189 | 62.5058065 | 40.8342843 | 49.6411682 | 0 |
| 0.02629335 | 0.04422756 | 21.8635297 | 16.905485 | 65.8916861 | 0 | 0 |
| 0.02629941 | 0.04423334 | 0 | 35.5904948 | 31.5537652 | 33.867339 | 0 |
| 0.02629845 | 0.04423334 | 0.3215225 | 0.66732178 | 0 | 0 | 0 |
| 0.02632234 | 0.04426969 | 0 | 59.3916382 | 16.7049345 | 32.4755306 | 0 |
| 0.02632832 | 0.04427753 | 26.6863672 | 41.3739502 | 26.9135056 | 0 | 0 |
| 0.02633754 | 0.04429083 | 2.25065747 | 11.7893514 | 10.2085711 | 6.95904227 | 0 |
| 0.0263417 | 0.04429561 | 27.3294122 | 18.017688 | 0 | 37.1148921 | 0 |
| 0.0263578 | 0.04432046 | 2.89370247 | 2.8917277 | 21.3451941 | 12.0623399 | 0 |
| 0.02636009 | 0.0443221 | 11.5748099 | 22.2440592 | 0 | 59.3838274 | 0 |
| 0.0263632 | 0.04432512 | 0 | 32.0314453 | 41.7623363 | 27.8361691 | 0 |
| 0.02636484 | 0.04432565 | 16.3976473 | 9.56494547 | 12.0646749 | 5.56723382 | 23.7621824 |
| 0.02639045 | 0.04436649 | 49.5144644 | 25.5806681 | 33.409869 | 26.4443606 | 16.6335277 |
| 0.02642446 | 0.04442145 | 140.826853 | 141.027336 | 158.696878 | 164.233398 | 149.701749 |
| 0.02643957 | 0.04444464 | 2.57217997 | 8.45274251 | 9.28051917 | 8.35085072 | 0 |
| 0.0264425 | 0.04444733 | 29.5800696 | 26.2479899 | 0 | 24.1246799 | 0 |
| 0.02644428 | 0.04444811 | 0.3215225 | 0 | 1.85610383 | 3.24755306 | 0 |
| 0.02645975 | 0.04447189 | 4.50131495 | 6.005896 | 12.9927268 | 4.63936151 | 0 |
| 0.02646252 | 0.04447211 | 54.0157793 | 21.5767375 | 0 | 0 | 401.580883 |
| 0.02646186 | 0.04447211 | 0 | 0 | 0 | 0 | 16.6335277 |
| 0.02646794 | 0.04447899 | 1.60761248 | 13.123995 | 8.35246725 | 11.5984038 | 0 |
| 0.02647724 | 0.0444924 | 15.4330798 | 4.89369303 | 3.71220767 | 6.49510612 | 0 |
| 0.02648137 | 0.04449711 | 9.32415239 | 50.0491333 | 0 | 32.9394667 | 0 |
| 0.02649272 | 0.04451395 | 1.60761248 | 4.22637126 | 12.0646749 | 25.5164883 | 0 |
| 0.02650499 | 0.04453235 | 0.64304499 | 17.3503662 | 10.2085711 | 17.6295737 | 0 |
| 0.02651123 | 0.04454028 | 84.5604165 | 88.3089152 | 48.2586997 | 69.1264865 | 30.8908372 |
| 0.02651236 | 0.04454028 | 0.64304499 | 1.55708415 | 0 | 2.78361691 | 0 |
| 0.02652151 | 0.04455343 | 34.7244296 | 19.5747721 | 0 | 26.9082968 | 0 |
| 0.02652407 | 0.04455552 | 53.0512119 | 17.5728068 | 0 | 17.6295737 | 0 |
| 0.02655667 | 0.04460804 | 11.5748099 | 38.7046631 | 0 | 37.1148921 | 0 |
| 0.02656183 | 0.04461449 | 33.7598621 | 19.7972127 | 0 | 27.3722329 | 0 |
| 0.02656425 | 0.04461474 | 3.21522496 | 14.6810791 | 5.5683115 | 6.95904227 | 0 |
| 0.02656463 | 0.04461474 | 12.2178549 | 8.8976237 | 2.78415575 | 5.56723382 | 71.2865473 |
| 0.02658104 | 0.04464006 | 3.85826995 | 8.00786133 | 3.71220767 | 16.7017014 | 0 |
| 0.02658794 | 0.04464943 | 18.3267823 | 59.6140788 | 23.2012979 | 32.4755306 | 14.2573095 |
| 0.02659963 | 0.04466682 | 14.4685123 | 62.7282471 | 41.7623363 | 46.8575513 | 0 |
| 0.02661213 | 0.04468455 | 6.43044992 | 34.4782918 | 0 | 58.9198912 | 0 |

| | | | | | | |
|---|---|---|---|---|---|---|
| 0.02661284 | 0.04468455 | 0 | 39.5944255 | 27.8415575 | 33.4034029 | 0 |
| 0.02662106 | 0.04469612 | 71.0564716 | 7.56298014 | 0 | 23.1968076 | 0 |
| 0.02662874 | 0.04470679 | 2.25065747 | 6.22833659 | 7.42441534 | 19.9492545 | 0 |
| 0.02664641 | 0.04472975 | 0 | 0 | 0 | 0 | 0 |
| 0.02664641 | 0.04472975 | 0 | 0 | 0 | 0 | 0 |
| 0.02664641 | 0.04472975 | 0 | 0 | 0 | 0 | 0 |
| 0.02665803 | 0.04474479 | 0 | 0 | 0 | 0 | 30.8908372 |
| 0.02665803 | 0.04474479 | 0 | 0 | 0 | 0 | 30.8908372 |
| 0.0266999 | 0.04481283 | 5.14435994 | 44.7105591 | 0 | 52.8887212 | 0 |
| 0.02670441 | 0.04481817 | 16.7191698 | 7.56298014 | 10.2085711 | 6.49510612 | 2.37621824 |
| 0.02672376 | 0.04484841 | 0.3215225 | 0 | 0 | 0.9278723 | 0 |
| 0.02675395 | 0.04489482 | 6.43044992 | 64.5077718 | 0 | 30.1558498 | 0 |
| 0.02675408 | 0.04489482 | 8.6811074 | 27.8050741 | 31.5537652 | 16.7017014 | 7.12865473 |
| 0.02676382 | 0.04490892 | 92.2769564 | 188.407182 | 191.178695 | 197.172864 | 4.75243649 |
| 0.02677166 | 0.04491983 | 21.5420072 | 74.5175985 | 42.6903882 | 83.5085072 | 0 |
| 0.026776 | 0.04492488 | 427.946442 | 745.620866 | 689.542574 | 688.945185 | 216.23586 |
| 0.02679808 | 0.04495968 | 54.0157793 | 46.0452026 | 69.6038938 | 27.3722329 | 0 |
| 0.02681773 | 0.0449904 | 1.92913498 | 10.2322673 | 10.2085711 | 9.27872303 | 0 |
| 0.026823 | 0.044997 | 1.92913498 | 14.9035197 | 6.49636342 | 10.6705315 | 0 |
| 0.02686338 | 0.0450625 | 20.8989622 | 12.4566732 | 20.4171422 | 17.6295737 | 49.9005831 |
| 0.02686542 | 0.04506367 | 4.17979245 | 8.00786133 | 5.5683115 | 6.95904227 | 0 |
| 0.02686999 | 0.04506909 | 0 | 215.767375 | 0 | 240.318926 | 0 |
| 0.02688456 | 0.04509128 | 110.925261 | 160.602108 | 160.552982 | 178.151482 | 54.6530196 |
| 0.02689163 | 0.04510089 | 6.75197242 | 7.78542074 | 4.64025958 | 5.10329766 | 0 |
| 0.02689977 | 0.04511229 | 18.0052598 | 28.6948364 | 0 | 34.3312752 | 0 |
| 0.02690631 | 0.04512101 | 33.4383396 | 0 | 36.1940248 | 37.1148921 | 0 |
| 0.02690822 | 0.04512196 | 3.21522496 | 10.6771484 | 5.5683115 | 8.35085072 | 0 |
| 0.0269367 | 0.04516748 | 2.89370247 | 9.12006429 | 6.49636342 | 9.27872303 | 0 |
| 0.02693994 | 0.04517065 | 74.9147416 | 104.102197 | 103.941815 | 92.7872303 | 0 |
| 0.02695407 | 0.04519209 | 5.46588243 | 8.23030192 | 2.78415575 | 13.9180845 | 0 |
| 0.02698427 | 0.04524047 | 2.25065747 | 9.12006429 | 10.2085711 | 8.35085072 | 0 |
| 0.02702733 | 0.04531041 | 25.7217997 | 14.9035197 | 0 | 43.6099982 | 0 |
| 0.02703239 | 0.04531663 | 0 | 36.7026978 | 9.28051917 | 70.9822312 | 0 |
| 0.02703836 | 0.04532438 | 0 | 0 | 0 | 0 | 2.37621824 |
| 0.02704195 | 0.04532814 | 3.21522496 | 6.89565837 | 13.9207788 | 5.56723382 | 0 |
| 0.02705271 | 0.04534392 | 27.6509347 | 17.1279256 | 10.2085711 | 18.0935099 | 123.563349 |
| 0.02706575 | 0.04536351 | 0 | 0.22244059 | 0 | 0 | 0 |
| 0.02706727 | 0.04536381 | 50.4790319 | 74.7400391 | 77.0283091 | 83.9724434 | 38.0194919 |
| 0.02708278 | 0.04538754 | 14.7900348 | 28.2499552 | 0 | 40.3624452 | 0 |
| 0.02709182 | 0.04540042 | 0 | 42.2637126 | 31.5537652 | 26.4443606 | 0 |
| 0.02716747 | 0.04552492 | 5.14435994 | 4.89369303 | 11.136623 | 5.10329766 | 0 |
| 0.0271823 | 0.04554752 | 16.3976473 | 17.5728068 | 36.1940248 | 20.4131907 | 4.75243649 |
| 0.02719274 | 0.04556274 | 5.14435994 | 5.7834554 | 5.5683115 | 6.95904227 | 0 |

| | | | | | | |
|---|---|---|---|---|---|---|
| 0.02721811 | 0.04560298 | 52.7296894 | 127.6809 | 46.4025958 | 86.2921241 | 30.8908372 |
| 0.02722382 | 0.04561026 | 30.5446371 | 31.5865641 | 0 | 18.0935099 | 0 |
| 0.02723145 | 0.04562078 | 319.593361 | 269.375557 | 0 | 289.960095 | 33.2670554 |
| 0.02727538 | 0.04568982 | 27.0078897 | 17.3503662 | 0 | 36.650956 | 0 |
| 0.02727468 | 0.04568982 | 138.254673 | 69.6239054 | 104.869867 | 77.4773373 | 254.255352 |
| 0.02728989 | 0.04571186 | 1.28608998 | 12.9015544 | 9.28051917 | 12.9902122 | 0 |
| 0.02730468 | 0.04573436 | 132.788791 | 240.013399 | 111.36623 | 178.615418 | 57.0292378 |
| 0.02731869 | 0.04575554 | 34.7244296 | 0 | 61.2514265 | 17.1656376 | 0 |
| 0.02732582 | 0.0457652 | 60.7677518 | 102.322673 | 102.085711 | 108.561059 | 11.8810912 |
| 0.02735239 | 0.04580743 | 1.28608998 | 1.55708415 | 0 | 1.85574461 | 0 |
| 0.02736708 | 0.04582885 | 8.0380624 | 28.0275146 | 0 | 58.9198912 | 0 |
| 0.02736791 | 0.04582885 | 32.4737721 | 25.8031087 | 0 | 20.8771268 | 0 |
| 0.02737629 | 0.0458406 | 0 | 7.56298014 | 25.0574018 | 93.7151026 | 0 |
| 0.02738547 | 0.0458537 | 6.43044992 | 52.2735392 | 0 | 38.0427644 | 0 |
| 0.02739839 | 0.04587304 | 3.85826995 | 7.78542074 | 5.5683115 | 7.88691457 | 0 |
| 0.02741397 | 0.04589396 | 54.0157793 | 75.1849202 | 20.4171422 | 33.4034029 | 19.0097459 |
| 0.02741497 | 0.04589396 | 37.9396545 | 23.1338216 | 31.5537652 | 0 | 0 |
| 0.02741352 | 0.04589396 | 31.8307271 | 28.0275146 | 0 | 19.4853184 | 0 |
| 0.02741818 | 0.04589705 | 183.267823 | 287.838127 | 257.998433 | 251.91733 | 118.810912 |
| 0.02743658 | 0.04592556 | 0.64304499 | 0.22244059 | 0 | 0.46393615 | 30.8908372 |
| 0.02743958 | 0.04592829 | 6.43044992 | 65.1750936 | 0 | 28.3001052 | 0 |
| 0.02744684 | 0.04593815 | 4.82283744 | 12.2342326 | 4.64025958 | 5.56723382 | 0 |
| 0.0274624 | 0.04596191 | 5.78740493 | 10.4547078 | 5.5683115 | 4.17542536 | 0 |
| 0.02746521 | 0.04596434 | 16.0761248 | 34.4782918 | 86.3088283 | 0 | 0 |
| 0.02747306 | 0.04597519 | 23.7926647 | 4.44881185 | 3.71220767 | 5.10329766 | 0 |
| 0.02749649 | 0.0460121 | 6.75197242 | 6.67321777 | 3.71220767 | 7.42297842 | 0 |
| 0.02750485 | 0.04602175 | 28.6155022 | 36.2578166 | 26.9135056 | 0 | 0 |
| 0.02750636 | 0.04602175 | 2.57217997 | 8.45274251 | 9.28051917 | 7.88691457 | 0 |
| 0.02750557 | 0.04602175 | 198.057858 | 443.10166 | 411.126999 | 421.254025 | 45.1481466 |
| 0.02750838 | 0.04602284 | 2.25065747 | 11.7893514 | 8.35246725 | 7.88691457 | 0 |
| 0.02752877 | 0.04605467 | 0.64304499 | 2.00196533 | 3.71220767 | 3.24755306 | 2.37621824 |
| 0.02753927 | 0.04606994 | 22.8280972 | 30.4743612 | 0 | 25.0525522 | 0 |
| 0.02755485 | 0.04609371 | 7.71653991 | 22.2440592 | 25.0574018 | 17.1656376 | 0 |
| 0.02755645 | 0.0460941 | 0 | 23.8011434 | 37.1220767 | 38.5067006 | 0 |
| 0.02757101 | 0.04611388 | 3.21522496 | 5.56101481 | 14.8488307 | 6.49510612 | 0 |
| 0.02757101 | 0.04611388 | 2.57217997 | 2.22440592 | 0 | 5.10329766 | 0 |
| 0.02759191 | 0.04614578 | 16.3976473 | 37.3700195 | 41.7623363 | 0 | 0 |
| 0.02759283 | 0.04614578 | 0 | 211.318563 | 218.0922 | 216.658183 | 21.3859642 |
| 0.02760211 | 0.04615901 | 2.89370247 | 6.005896 | 10.2085711 | 8.81478687 | 0 |
| 0.02761184 | 0.04617299 | 3.53674746 | 7.56298014 | 8.35246725 | 6.03116997 | 0 |
| 0.02761741 | 0.0461777 | 28.2939797 | 47.8247274 | 0 | 11.1344676 | 0 |
| 0.0276174 | 0.0461777 | 6.75197242 | 5.7834554 | 7.42441534 | 4.17542536 | 0 |
| 0.02765405 | 0.04623667 | 0 | 37.1475789 | 21.3451941 | 41.7542536 | 0 |

| | | | | | | |
|---|---|---|---|---|---|---|
| 0.02765641 | 0.04623832 | 68.4842917 | 22.4664998 | 6.49636342 | 0 | 216.23586 |
| 0.02767592 | 0.04626864 | 23.1496197 | 27.3601929 | 0 | 27.3722329 | 0 |
| 0.02768975 | 0.04628946 | 1.28608998 | 2.00196533 | 0 | 0.46393615 | 0 |
| 0.02770661 | 0.04631533 | 8.6811074 | 3.33660889 | 6.49636342 | 7.42297842 | 0 |
| 0.02771263 | 0.04632311 | 49.8359869 | 19.5747721 | 25.9854537 | 0 | 0 |
| 0.02774181 | 0.04636957 | 11.2532874 | 37.8149007 | 0 | 36.1870198 | 0 |
| 0.02775815 | 0.04639459 | 23.7926647 | 60.05896 | 0 | 9.27872303 | 0 |
| 0.02780377 | 0.04646853 | 0 | 156.153296 | 153.128566 | 171.656376 | 19.0097459 |
| 0.02780819 | 0.04647361 | 58.1955718 | 97.6514201 | 83.5246725 | 104.84957 | 35.6432737 |
| 0.02781245 | 0.04647841 | 0 | 44.9329997 | 31.5537652 | 23.1968076 | 0 |
| 0.02786375 | 0.04656182 | 0 | 16.6830444 | 52.8989593 | 33.867339 | 0 |
| 0.02789134 | 0.04660562 | 99.9934963 | 113.222262 | 106.72597 | 94.1790387 | 396.828447 |
| 0.02789872 | 0.04661101 | 0 | 0 | 0 | 0 | 0 |
| 0.02789872 | 0.04661101 | 0 | 0 | 0 | 0 | 0 |
| 0.02789872 | 0.04661101 | 0 | 0 | 0 | 0 | 0 |
| 0.02793524 | 0.0466697 | 79.737579 | 183.735929 | 163.337137 | 162.377653 | 52.2768014 |
| 0.02793679 | 0.04666997 | 26.6863672 | 18.4625692 | 0 | 33.867339 | 0 |
| 0.02794594 | 0.04668294 | 37.618132 | 98.5411824 | 103.013763 | 98.3544641 | 33.2670554 |
| 0.02795802 | 0.04670081 | 21.8635297 | 68.0668213 | 0 | 7.88691457 | 0 |
| 0.02798206 | 0.04673864 | 5.46588243 | 9.78738607 | 2.78415575 | 0 | 30.8908372 |
| 0.02800015 | 0.04676653 | 0 | 0 | 0 | 0 | 59.4054561 |
| 0.02804403 | 0.0468375 | 27.0078897 | 17.7952474 | 23.2012979 | 12.5262761 | 68.9103291 |
| 0.02804727 | 0.04684058 | 3.85826995 | 6.005896 | 8.35246725 | 6.49510612 | 0 |
| 0.02810198 | 0.04692962 | 38.5826995 | 17.1279256 | 38.0501286 | 0 | 0 |
| 0.02810598 | 0.04693397 | 136.647061 | 119.005717 | 96.5173994 | 108.097123 | 45.1481466 |
| 0.02810893 | 0.04693657 | 0 | 0 | 0 | 0 | 21.3859642 |
| 0.028167 | 0.0470312 | 2.57217997 | 7.11809896 | 10.2085711 | 8.35085072 | 0 |
| 0.02817642 | 0.04704459 | 3.53674746 | 7.78542074 | 5.5683115 | 8.81478687 | 0 |
| 0.0281829 | 0.04705309 | 105.459379 | 192.633553 | 155.912722 | 184.646588 | 28.5146189 |
| 0.02819961 | 0.04707864 | 32.4737721 | 27.3601929 | 0 | 18.5574461 | 0 |
| 0.02822622 | 0.04712072 | 0 | 12.4566732 | 53.8270112 | 39.898509 | 0 |
| 0.02823018 | 0.04712311 | 24.4357097 | 0 | 47.3306478 | 33.867339 | 0 |
| 0.02823045 | 0.04712311 | 0 | 24.2460246 | 10.2085711 | 82.5806349 | 0 |
| 0.0282397 | 0.04713622 | 39.2257445 | 28.4723958 | 36.1940248 | 30.619786 | 57.0292378 |
| 0.02824641 | 0.04714507 | 4.50131495 | 5.56101481 | 5.5683115 | 8.81478687 | 0 |
| 0.02825974 | 0.04716499 | 10.2887199 | 9.56494547 | 10.2085711 | 5.56723382 | 2.37621824 |
| 0.02827735 | 0.04717097 | 0 | 0 | 0 | 0 | 0 |
| 0.02827735 | 0.04717097 | 0 | 0 | 0 | 0 | 0 |
| 0.02827207 | 0.04717097 | 1.60761248 | 4.22637126 | 14.8488307 | 19.0213822 | 0 |
| 0.02827735 | 0.04717097 | 0 | 0 | 0 | 0 | 0 |
| 0.02827735 | 0.04717097 | 0 | 0 | 0 | 0 | 0 |
| 0.02827735 | 0.04717097 | 0 | 0 | 0 | 0 | 0 |
| 0.02827735 | 0.04717097 | 0 | 0 | 0 | 0 | 0 |

| | | | | | | |
|---|---|---|---|---|---|---|
| 0.02827735 | 0.04717097 | 0 | 0 | 0 | 0 | 0 |
| 0.02827735 | 0.04717097 | 0 | 0 | 0 | 0 | 0 |
| 0.02827735 | 0.04717097 | 0 | 0 | 0 | 0 | 0 |
| 0.02828419 | 0.04718004 | 11.2532874 | 61.8384847 | 0 | 18.5574461 | 0 |
| 0.02828839 | 0.04718121 | 47.2638069 | 16.4606038 | 0 | 19.4853184 | 0 |
| 0.02828789 | 0.04718121 | 2.57217997 | 6.005896 | 13.9207788 | 7.88691457 | 0 |
| 0.02828909 | 0.04718121 | 3.21522496 | 12.2342326 | 4.64025958 | 8.81478687 | 0 |
| 0.02831138 | 0.04721605 | 31.1876821 | 15.5708415 | 0 | 32.9394667 | 0 |
| 0.02832713 | 0.04723996 | 1.28608998 | 10.6771484 | 9.28051917 | 14.8459568 | 0 |
| 0.02834136 | 0.04726135 | 13.1824223 | 31.1416829 | 0 | 37.5788283 | 0 |
| 0.02839571 | 0.04734963 | 3.85826995 | 6.89565837 | 6.49636342 | 6.95904227 | 0 |
| 0.02840287 | 0.04735922 | 8.0380624 | 4.00393066 | 3.71220767 | 3.24755306 | 83.1676385 |
| 0.02841735 | 0.04738103 | 9.32415239 | 13.7913167 | 9.28051917 | 6.03116997 | 0 |
| 0.02842232 | 0.04738697 | 0 | 60.05896 | 37.1220767 | 11.1344676 | 0 |
| 0.02843433 | 0.04740464 | 12.8608998 | 1.77952474 | 22.273246 | 3.71148921 | 0 |
| 0.0285065 | 0.04752261 | 13.8254673 | 9.78738607 | 6.49636342 | 7.88691457 | 0 |
| 0.02851645 | 0.04753684 | 4.82283744 | 4.00393066 | 12.9927268 | 6.03116997 | 0 |
| 0.02853157 | 0.04755968 | 30.2231146 | 8.45274251 | 5.5683115 | 6.95904227 | 0 |
| 0.02853532 | 0.04756358 | 5.14435994 | 1.77952474 | 12.0646749 | 17.1656376 | 0 |
| 0.02854673 | 0.04758024 | 16.7191698 | 0 | 38.0501286 | 54.2805297 | 0 |
| 0.02855 | 0.04758333 | 16.7191698 | 4.00393066 | 5.5683115 | 4.63936151 | 0 |
| 0.02856595 | 0.0476052 | 29.2585471 | 25.1357869 | 36.1940248 | 72.8379758 | 11.8810912 |
| 0.02856485 | 0.0476052 | 5.14435994 | 28.0275146 | 21.3451941 | 22.2689353 | 4.75243649 |
| 0.02856886 | 0.04760758 | 27.0078897 | 29.5845988 | 0 | 20.4131907 | 0 |
| 0.02857021 | 0.04760758 | 0.64304499 | 10.0098267 | 20.4171422 | 13.4541484 | 0 |
| 0.02857965 | 0.04762096 | 0 | 33.5885295 | 34.3379209 | 28.3001052 | 0 |
| 0.02859357 | 0.04764179 | 1.28608998 | 12.6791138 | 10.2085711 | 11.1344676 | 0 |
| 0.02861453 | 0.04767435 | 56.2664368 | 56.7223511 | 52.8989593 | 54.2805297 | 85.5438568 |
| 0.02863228 | 0.04770058 | 19.6128723 | 67.8443807 | 61.2514265 | 61.7035081 | 0 |
| 0.02863311 | 0.04770058 | 81.3451915 | 153.261568 | 171.689605 | 146.139888 | 26.1384007 |
| 0.02864064 | 0.04771076 | 3.85826995 | 16.4606038 | 7.42441534 | 3.71148921 | 0 |
| 0.02866037 | 0.04774127 | 6.43044992 | 8.00786133 | 2.78415575 | 9.74265918 | 0 |
| 0.02866659 | 0.04774927 | 31.8307271 | 27.8050741 | 0 | 18.0935099 | 0 |
| 0.02867199 | 0.0477559 | 0 | 0 | 0 | 0 | 30.8908372 |
| 0.02867493 | 0.04775843 | 48.8714194 | 27.5826335 | 0 | 10.2065953 | 0 |
| 0.0287323 | 0.04785161 | 4.17979245 | 2.8917277 | 8.35246725 | 18.0935099 | 0 |
| 0.02874438 | 0.04786936 | 0 | 46.7125244 | 37.1220767 | 16.7017014 | 0 |
| 0.02876082 | 0.04789437 | 4.50131495 | 3.55904948 | 30.6257133 | 3.71148921 | 0 |
| 0.02876784 | 0.04790368 | 20.5774398 | 36.035376 | 0 | 21.341063 | 0 |
| 0.02877503 | 0.04791328 | 0 | 29.8070394 | 33.409869 | 32.4755306 | 0 |
| 0.02881457 | 0.04797675 | 0 | 34.7007324 | 23.2012979 | 38.9706367 | 0 |
| 0.02881741 | 0.04797911 | 0 | 219.326424 | 0 | 206.451587 | 0 |
| 0.02882774 | 0.04799393 | 4.50131495 | 44.7105591 | 0 | 50.5690405 | 0 |

| | | | | | | |
|---|---|---|---|---|---|---|
| 0.02884565 | 0.04802137 | 153.366231 | 252.247632 | 261.710641 | 347.024241 | 85.5438568 |
| 0.02884971 | 0.04802338 | 21.8635297 | 34.7007324 | 0 | 20.8771268 | 0 |
| 0.02884898 | 0.04802338 | 44.691627 | 92.9801676 | 85.3807764 | 109.488932 | 33.2670554 |
| 0.02886679 | 0.04804943 | 19.9343948 | 40.4841878 | 0 | 19.0213822 | 0 |
| 0.02889416 | 0.0480926 | 8.6811074 | 4.89369303 | 8.35246725 | 3.71148921 | 0 |
| 0.02889661 | 0.04809431 | 1.28608998 | 0 | 1.85610383 | 0.9278723 | 0 |
| 0.0289445 | 0.04817164 | 1.92913498 | 11.7893514 | 19.4890903 | 4.17542536 | 0 |
| 0.02895097 | 0.04818001 | 18.0052598 | 8.00786133 | 10.2085711 | 9.74265918 | 26.1384007 |
| 0.02902502 | 0.04830086 | 15.7546023 | 25.3582275 | 0 | 38.0427644 | 0 |
| 0.02902925 | 0.04830552 | 366.857168 | 380.595854 | 355.443884 | 296.919137 | 313.660808 |
| 0.02903101 | 0.04830605 | 27.9724572 | 146.143469 | 154.056618 | 119.231591 | 38.0194919 |
| 0.02905754 | 0.0483478 | 16.0761248 | 10.4547078 | 13.9207788 | 11.5984038 | 57.0292378 |
| 0.02906211 | 0.04835302 | 22.8280972 | 48.7144897 | 0 | 12.5262761 | 0 |
| 0.02908778 | 0.04839333 | 0 | 23.3562622 | 25.9854537 | 48.7132959 | 0 |
| 0.02909369 | 0.04840076 | 4.82283744 | 23.8011434 | 93.7332436 | 0 | 0 |
| 0.02910172 | 0.04841174 | 6.75197242 | 11.5669108 | 12.9927268 | 6.95904227 | 0 |
| 0.02910732 | 0.04841866 | 4.82283744 | 7.34053955 | 5.5683115 | 5.56723382 | 0 |
| 0.02913282 | 0.04845867 | 11.5748099 | 1.55708415 | 7.42441534 | 13.4541484 | 0 |
| 0.02913821 | 0.04846526 | 6.10892743 | 17.7952474 | 33.409869 | 25.0525522 | 0 |
| 0.02915446 | 0.04848989 | 9.00262989 | 7.11809896 | 8.35246725 | 11.5984038 | 2.37621824 |
| 0.02919695 | 0.04855815 | 0 | 0 | 0 | 0 | 57.0292378 |
| 0.02920795 | 0.04857404 | 2.89370247 | 8.8976237 | 6.49636342 | 8.35085072 | 0 |
| 0.02924492 | 0.04863312 | 102.565676 | 146.143469 | 119.718697 | 125.726697 | 28.5146189 |
| 0.02924982 | 0.04863887 | 0 | 39.1495443 | 32.4818171 | 24.1246799 | 0 |
| 0.02926087 | 0.04865483 | 9.64567488 | 10.0098267 | 4.64025958 | 3.24755306 | 0 |
| 0.02926409 | 0.04865539 | 0 | 0 | 0 | 0 | 0 |
| 0.02926409 | 0.04865539 | 0 | 0 | 0 | 0 | 0 |
| 0.0293084 | 0.04872664 | 8.0380624 | 56.0550293 | 0 | 26.4443606 | 0 |
| 0.02931205 | 0.0487303 | 14.4685123 | 30.02948 | 0 | 34.3312752 | 0 |
| 0.02931481 | 0.04873249 | 12.8608998 | 21.3542969 | 0 | 49.6411682 | 0 |
| 0.02934775 | 0.04878484 | 5.78740493 | 7.34053955 | 2.78415575 | 6.49510612 | 28.5146189 |
| 0.02936301 | 0.04880779 | 191.94893 | 441.099695 | 373.076871 | 353.055411 | 99.8011662 |
| 0.02937181 | 0.04882001 | 11.2532874 | 43.3759155 | 55.683115 | 59.3838274 | 16.6335277 |
| 0.02941985 | 0.04889502 | 4.82283744 | 20.6869751 | 98.3735032 | 0 | 0 |
| 0.02941883 | 0.04889502 | 12.8608998 | 13.7913167 | 13.9207788 | 19.0213822 | 2.37621824 |
| 0.02942546 | 0.04890193 | 20.5774398 | 25.8031087 | 0 | 29.2279775 | 0 |
| 0.02948624 | 0.04900052 | 1.60761248 | 13.123995 | 13.9207788 | 6.03116997 | 0 |
| 0.02949047 | 0.04900513 | 4.50131495 | 5.11613363 | 13.9207788 | 4.63936151 | 0 |
| 0.02951466 | 0.0490429 | 32.1522496 | 16.0157227 | 23.2012979 | 19.9492545 | 140.196876 |
| 0.02955355 | 0.0491051 | 19.2913498 | 10.0098267 | 6.49636342 | 16.7017014 | 57.0292378 |
| 0.02957112 | 0.04913187 | 85.8465065 | 222.663033 | 206.955577 | 224.545097 | 11.8810912 |
| 0.02959071 | 0.04915957 | 9.00262989 | 4.67125244 | 8.35246725 | 3.71148921 | 0 |
| 0.02959021 | 0.04915957 | 2.25065747 | 51.1613363 | 0 | 58.9198912 | 0 |

| | | | | | | |
|---|---|---|---|---|---|---|
| 0.02961535 | 0.04919745 | 0 | 18.9074504 | 50.1148035 | 29.2279775 | 0 |
| 0.02961644 | 0.04919745 | 13.8254673 | 0 | 69.6038938 | 28.7640414 | 0 |
| 0.02962807 | 0.04921434 | 14.7900348 | 3.55904948 | 6.49636342 | 4.63936151 | 0 |
| 0.02962965 | 0.04921454 | 24.1141872 | 23.8011434 | 0 | 26.9082968 | 0 |
| 0.0296494 | 0.04924491 | 0.96456749 | 18.6850098 | 6.49636342 | 14.8459568 | 0 |
| 0.02965842 | 0.04925746 | 193.878065 | 49.8266927 | 37.1220767 | 51.4969128 | 28.5146189 |
| 0.02971342 | 0.04934637 | 1.28608998 | 19.7972127 | 8.35246725 | 8.35085072 | 0 |
| 0.02975648 | 0.04941544 | 225.065747 | 235.119706 | 280.271679 | 206.451587 | 78.415202 |
| 0.02976218 | 0.04942247 | 65.9121117 | 50.4940145 | 128.071165 | 20.8771268 | 125.939567 |
| 0.02978306 | 0.0494547 | 0 | 0 | 331.314534 | 169.336695 | 0 |
| 0.02979745 | 0.04947614 | 67.8412467 | 116.33643 | 131.783372 | 117.375846 | 313.660808 |
| 0.02981252 | 0.04949873 | 17.6837373 | 11.1220296 | 0 | 59.8477635 | 0 |
| 0.02981842 | 0.0495043 | 3.85826995 | 1.77952474 | 8.35246725 | 29.2279775 | 0 |
| 0.02981882 | 0.0495043 | 30.8661596 | 26.2479899 | 0 | 18.5574461 | 0 |
| 0.02983979 | 0.04953668 | 31.1876821 | 68.0668213 | 66.819738 | 61.7035081 | 30.8908372 |
| 0.02984242 | 0.0495386 | 8.3595849 | 7.56298014 | 3.71220767 | 5.10329766 | 0 |
| 0.0298558 | 0.04955836 | 0.96456749 | 12.011792 | 12.9927268 | 11.5984038 | 0 |
| 0.02988367 | 0.04960218 | 8.3595849 | 27.8050741 | 0 | 51.4969128 | 0 |
| 0.02992706 | 0.04967175 | 9.96719738 | 4.67125244 | 6.49636342 | 4.17542536 | 0 |
| 0.02994705 | 0.04970248 | 0 | 20.4645345 | 53.8270112 | 24.1246799 | 0 |
| 0.02996385 | 0.0497279 | 3.53674746 | 7.34053955 | 4.64025958 | 11.5984038 | 0 |
| 0.02997307 | 0.04974074 | 0 | 375.034839 | 97.4454513 | 0 | 0 |
| 0.0299859 | 0.04975959 | 1.92913498 | 8.23030192 | 12.0646749 | 8.35085072 | 0 |
| 0.02999647 | 0.04977466 | 0 | 0 | 0 | 0 | 28.5146189 |
| 0.0300106 | 0.04979565 | 33.1168171 | 136.133643 | 154.056618 | 145.675952 | 0 |
| 0.03001414 | 0.04979908 | 129.573566 | 249.133464 | 215.308045 | 267.227223 | 0 |
| 0.0300311 | 0.04982477 | 3.21522496 | 14.6810791 | 6.49636342 | 5.10329766 | 0 |
| 0.03003702 | 0.04983213 | 5.46588243 | 45.822762 | 0 | 41.7542536 | 0 |
| 0.03005153 | 0.04985375 | 29.5800696 | 0 | 46.4025958 | 25.5164883 | 0 |
| 0.03009889 | 0.04992984 | 26.0433222 | 26.4704305 | 29.6976613 | 32.0115944 | 49.9005831 |
| 0.03012359 | 0.04996836 | 50.8005544 | 69.4014648 | 72.3880495 | 85.3642518 | 7.12865473 |

| GFP (-) 2 | GFP (-) 3 |
|---|---|
| 5900.95837 | 5779.47349 |
| 1194.3062 | 1217.21294 |
| 762.72737 | 558.270598 |
| 187.288927 | 86.9437817 |
| 18256.5989 | 9165.70499 |
| 24.4289905 | 41.1838966 |
| 16057.9898 | 12181.2814 |
| 11880.6324 | 7449.7093 |
| 4212.6437 | 3070.48829 |
| 8615.29065 | 6131.8246 |
| 9201.58642 | 6218.76839 |
| 1397.88112 | 1148.57312 |
| 3639.91958 | 2960.66457 |
| 2.71433228 | 0 |
| 5.42866456 | 0 |
| 2201.32348 | 1139.42114 |
| 439.721829 | 370.655069 |
| 2434.75605 | 1166.87707 |
| 445.150494 | 736.73415 |
| 610.724763 | 851.133863 |
| 0 | 0 |
| 0 | 0 |
| 8.14299683 | 4.57598851 |
| 0 | 0 |
| 0 | 0 |
| 5.42866456 | 0 |
| 0 | 0 |
| 0 | 0 |
| 0 | 0 |
| 16.2859937 | 0 |
| 3181.19743 | 5381.36249 |
| 0 | 18.303954 |
| 0 | 0 |
| 0 | 13.7279655 |
| 1224.16386 | 544.542633 |
| 1001.58861 | 1478.04429 |
| 10.8573291 | 0 |
| 0 | 0 |
| 0 | 0 |
| 0 | 0 |
| 0 | 0 |
| 0 | 0 |

| | |
|---|---|
| 0 | 0 |
| 2.71433228 | 0 |
| 13.5716614 | 0 |
| 0 | 0 |
| 5.42866456 | 0 |
| 0 | 0 |
| 0 | 0 |
| 0 | 0 |
| 306.719547 | 160.159598 |
| 5.42866456 | 22.8799426 |
| 1131.87656 | 398.111 |
| 0 | 0 |
| 1848.46028 | 2562.55357 |
| 0 | 0 |
| 0 | 0 |
| 798.01369 | 297.439253 |
| 5567.0955 | 3509.78319 |
| 469.579484 | 283.711288 |
| 719.298054 | 530.814667 |
| 0 | 0 |
| 0 | 0 |
| 0 | 0 |
| 0 | 0 |
| 301.290883 | 169.311575 |
| 757.298706 | 356.927104 |
| 0 | 0 |
| 0 | 0 |
| 0 | 0 |
| 0 | 0 |
| 29.8576551 | 0 |
| 2079.17852 | 1601.59598 |
| 0 | 0 |
| 0 | 0 |
| 0 | 0 |
| 0 | 0 |
| 0 | 0 |
| 0 | 0 |
| 0 | 0 |
| 0 | 0 |
| 0 | 0 |
| 0 | 0 |
| 0 | 0 |
| 0 | 0 |

| | |
|---|---|
| 1311.02249 | 1157.72509 |
| 0 | 0 |
| 0 | 4.57598851 |
| 0 | 0 |
| 0 | 0 |
| 0 | 0 |
| 0 | 27.4559311 |
| 0 | 0 |
| 0 | 0 |
| 0 | 0 |
| 0 | 0 |
| 0 | 0 |
| 13.5716614 | 0 |
| 0 | 0 |
| 0 | 0 |
| 0 | 0 |
| 0 | 9.15197702 |
| 0 | 0 |
| 0 | 0 |
| 149.288275 | 370.655069 |
| 0 | 4.57598851 |
| 0 | 4.57598851 |
| 0 | 0 |
| 0 | 0 |
| 0 | 0 |
| 0 | 0 |
| 0 | 0 |
| 2.71433228 | 0 |
| 0 | 9.15197702 |
| 415.292839 | 256.255357 |
| 1552.59806 | 960.957587 |
| 0 | 0 |
| 0 | 0 |
| 0 | 0 |
| 0 | 0 |
| 0 | 0 |
| 0 | 9.15197702 |
| 0 | 0 |
| 0 | 0 |
| 0 | 0 |
| 0 | 0 |
| 0 | 0 |
| 0 | 0 |

| | |
|---|---|
| 5.42866456 | 45.7598851 |
| 0 | 0 |
| 0 | 0 |
| 0 | 0 |
| 0 | 0 |
| 0 | 0 |
| 0 | 0 |
| 5.42866456 | 9.15197702 |
| 8.14299683 | 4.57598851 |
| 0 | 0 |
| 0 | 0 |
| 0 | 0 |
| 0 | 0 |
| 0 | 0 |
| 209.003585 | 237.951403 |
| 0 | 0 |
| 2.71433228 | 9.15197702 |
| 0 | 0 |
| 0 | 0 |
| 0 | 0 |
| 0 | 0 |
| 0 | 0 |
| 0 | 0 |
| 0 | 13.7279655 |
| 0 | 0 |
| 0 | 0 |
| 0 | 0 |
| 0 | 0 |
| 0 | 0 |
| 713.869389 | 581.150541 |
| 0 | 4.57598851 |
| 0 | 0 |
| 0 | 0 |
| 0 | 0 |
| 0 | 0 |
| 0 | 0 |
| 0 | 4.57598851 |
| 13.5716614 | 4.57598851 |
| 0 | 9.15197702 |
| 0 | 0 |
| 0 | 9.15197702 |
| 2.71433228 | 0 |
| 0 | 0 |

| | |
|---|---|
| 0 | 0 |
| 0 | 0 |
| 0 | 0 |
| 0 | 0 |
| 0 | 0 |
| 0 | 0 |
| 0 | 0 |
| 0 | 0 |
| 0 | 0 |
| 0 | 0 |
| 0 | 0 |
| 0 | 0 |
| 0 | 0 |
| 0 | 0 |
| 0 | 0 |
| 0 | 0 |
| 0 | 0 |
| 0 | 0 |
| 0 | 0 |
| 249.71857 | 59.4878506 |
| 0 | 0 |
| 5.42866456 | 9.15197702 |
| 0 | 0 |
| 0 | 0 |
| 0 | 0 |
| 0 | 0 |
| 0 | 0 |
| 0 | 0 |
| 0 | 0 |
| 2.71433228 | 9.15197702 |
| 0 | 0 |
| 0 | 0 |
| 407.149842 | 521.66269 |
| 0 | 0 |
| 0 | 0 |
| 0 | 0 |
| 0 | 0 |
| 0 | 0 |
| 0 | 0 |
| 0 | 0 |
| 0 | 0 |
| 0 | 0 |
| 29.8576551 | 27.4559311 |

| | |
|---|---|
| 0 | 0 |
| 0 | 0 |
| 133.002282 | 105.247736 |
| 0 | 0 |
| 0 | 0 |
| 0 | 0 |
| 0 | 0 |
| 0 | 4.57598851 |
| 0 | 0 |
| 0 | 0 |
| 0 | 0 |
| 0 | 0 |
| 0 | 0 |
| 0 | 0 |
| 0 | 0 |
| 0 | 0 |
| 0 | 0 |
| 0 | 0 |
| 0 | 0 |
| 2263.75312 | 590.302518 |
| 1620.45637 | 260.831345 |
| 0 | 0 |
| 0 | 0 |
| 0 | 0 |
| 2.71433228 | 0 |
| 0 | 0 |
| 0 | 0 |
| 0 | 0 |
| 0 | 0 |
| 0 | 0 |
| 0 | 0 |
| 0 | 0 |
| 0 | 0 |
| 0 | 0 |
| 0 | 0 |
| 437.007497 | 256.255357 |
| 0 | 0 |
| 0 | 0 |
| 0 | 0 |
| 0 | 0 |
| 0 | 0 |
| 0 | 0 |
| 0 | 0 |

| | |
|---|---|
| 241.575573 | 137.279655 |
| 0 | 0 |
| 0 | 0 |
| 0 | 0 |
| 0 | 0 |
| 5.42866456 | 22.8799426 |
| 10.8573291 | 0 |
| 0 | 0 |
| 0 | 0 |
| 0 | 0 |
| 0 | 0 |
| 0 | 0 |
| 0 | 13.7279655 |
| 0 | 0 |
| 0 | 0 |
| 0 | 0 |
| 0 | 0 |
| 0 | 0 |
| 236.146908 | 324.895184 |
| 0 | 0 |
| 0 | 0 |
| 0 | 0 |
| 0 | 0 |
| 0 | 0 |
| 103.144627 | 128.127678 |
| 0 | 0 |
| 0 | 0 |
| 0 | 0 |
| 0 | 0 |
| 0 | 0 |
| 179.14593 | 242.527391 |
| 0 | 0 |
| 0 | 0 |
| 2.71433228 | 13.7279655 |
| 0 | 0 |
| 0 | 0 |
| 0 | 0 |
| 0 | 0 |
| 0 | 0 |
| 0 | 0 |
| 0 | 0 |
| 0 | 0 |
| 0 | 0 |

| | |
|---|---|
| 2.71433228 | 4.57598851 |
| 0 | 0 |
| 0 | 0 |
| 0 | 0 |
| 0 | 0 |
| 0 | 0 |
| 0 | 13.7279655 |
| 5.42866456 | 18.303954 |
| 168.288601 | 196.767506 |
| 0 | 0 |
| 0 | 0 |
| 0 | 0 |
| 48.857981 | 100.671747 |
| 0 | 0 |
| 0 | 0 |
| 0 | 0 |
| 0 | 0 |
| 0 | 0 |
| 0 | 0 |
| 0 | 0 |
| 0 | 0 |
| 0 | 0 |
| 0 | 0 |
| 0 | 0 |
| 0 | 0 |
| 0 | 0 |
| 16.2859937 | 9.15197702 |
| 0 | 0 |
| 0 | 0 |
| 0 | 0 |
| 0 | 0 |
| 0 | 0 |
| 0 | 0 |
| 0 | 0 |
| 0 | 0 |
| 0 | 0 |
| 0 | 0 |
| 0 | 0 |
| 0 | 0 |
| 0 | 0 |
| 0 | 0 |
| 0 | 0 |
| 5.42866456 | 0 |

| | |
|---|---|
| 0 | 0 |
| 0 | 0 |
| 0 | 0 |
| 0 | 0 |
| 0 | 0 |
| 0 | 0 |
| 0 | 0 |
| 0 | 0 |
| 0 | 0 |
| 0 | 0 |
| 0 | 0 |
| 0 | 0 |
| 105.858959 | 292.863265 |
| 8.14299683 | 0 |
| 0 | 0 |
| 0 | 0 |
| 0 | 0 |
| 0 | 0 |
| 0 | 0 |
| 5.42866456 | 0 |
| 0 | 0 |
| 0 | 0 |
| 0 | 0 |
| 95.0016297 | 160.159598 |
| 2.71433228 | 4.57598851 |
| 0 | 4.57598851 |
| 0 | 18.303954 |
| 0 | 0 |
| 0 | 0 |
| 0 | 0 |
| 0 | 0 |
| 0 | 0 |
| 0 | 0 |
| 0 | 0 |
| 97.715962 | 224.223437 |
| 0 | 0 |
| 146.573943 | 91.5197702 |
| 0 | 0 |
| 0 | 0 |
| 0 | 0 |
| 0 | 0 |
| 0 | 0 |
| 0 | 0 |

| | |
|---|---|
| 0 | 0 |
| 0 | 0 |
| 0 | 0 |
| 0 | 0 |
| 0 | 0 |
| 0 | 0 |
| 0 | 0 |
| 0 | 0 |
| 0 | 0 |
| 0 | 0 |
| 0 | 0 |
| 0 | 0 |
| 0 | 0 |
| 0 | 0 |
| 0 | 0 |
| 0 | 0 |
| 0 | 0 |
| 0 | 0 |
| 0 | 0 |
| 0 | 0 |
| 0 | 4.57598851 |
| 0 | 0 |
| 0 | 0 |
| 0 | 0 |
| 21.7146582 | 18.303954 |
| 0 | 0 |
| 0 | 0 |
| 0 | 4.57598851 |
| 0 | 0 |
| 0 | 4.57598851 |
| 13.5716614 | 0 |
| 0 | 0 |
| 0 | 0 |
| 0 | 0 |
| 0 | 0 |
| 0 | 0 |
| 0 | 0 |
| 0 | 0 |
| 0 | 0 |
| 5.42866456 | 0 |
| 43.4293164 | 36.6079081 |
| 0 | 0 |
| 0 | 0 |

| | |
|---|---|
| 2.71433228 | 13.7279655 |
| 0 | 0 |
| 0 | 9.15197702 |
| 0 | 0 |
| 5.42866456 | 13.7279655 |
| 168.288601 | 91.5197702 |
| 0 | 0 |
| 0 | 0 |
| 0 | 0 |
| 0 | 0 |
| 76.0013038 | 151.007621 |
| 0 | 0 |
| 0 | 0 |
| 0 | 0 |
| 0 | 0 |
| 0 | 0 |
| 0 | 0 |
| 0 | 0 |
| 0 | 0 |
| 152.002608 | 155.583609 |
| 73.2869715 | 36.6079081 |
| 0 | 0 |
| 0 | 0 |
| 0 | 4.57598851 |
| 0 | 0 |
| 0 | 0 |
| 0 | 0 |
| 0 | 0 |
| 0 | 0 |
| 0 | 0 |
| 0 | 0 |
| 0 | 0 |
| 0 | 0 |
| 0 | 0 |
| 0 | 0 |
| 0 | 0 |
| 0 | 0 |
| 0 | 0 |
| 0 | 0 |
| 0 | 0 |
| 0 | 0 |
| 0 | 0 |
| 0 | 0 |

| | |
|---|---|
| 0 | 0 |
| 0 | 0 |
| 0 | 0 |
| 0 | 0 |
| 0 | 0 |
| 0 | 0 |
| 0 | 0 |
| 0 | 0 |
| 0 | 0 |
| 0 | 0 |
| 0 | 0 |
| 0 | 0 |
| 5.42866456 | 0 |
| 0 | 0 |
| 10.8573291 | 50.3358736 |
| 0 | 0 |
| 0 | 0 |
| 0 | 0 |
| 0 | 0 |
| 0 | 0 |
| 0 | 0 |
| 0 | 0 |
| 0 | 0 |
| 0 | 0 |
| 0 | 0 |
| 0 | 0 |
| 2.71433228 | 13.7279655 |
| 0 | 0 |
| 0 | 0 |
| 0 | 0 |
| 0 | 0 |
| 0 | 0 |
| 0 | 0 |
| 0 | 0 |
| 0 | 0 |
| 0 | 0 |
| 195.431924 | 96.0957587 |
| 0 | 0 |
| 0 | 0 |
| 564.581114 | 192.191517 |
| 0 | 0 |
| 0 | 0 |
| 0 | 0 |

| | |
|---|---|
| 0 | 0 |
| 0 | 0 |
| 0 | 0 |
| 0 | 0 |
| 0 | 4.57598851 |
| 0 | 0 |
| 0 | 0 |
| 0 | 0 |
| 0 | 0 |
| 0 | 0 |
| 0 | 0 |
| 0 | 0 |
| 0 | 0 |
| 0 | 0 |
| 0 | 0 |
| 0 | 0 |
| 0 | 0 |
| 0 | 0 |
| 0 | 0 |
| 0 | 0 |
| 0 | 0 |
| 0 | 0 |
| 16.2859937 | 0 |
| 0 | 0 |
| 0 | 0 |
| 1598.74171 | 878.589794 |
| 0 | 0 |
| 0 | 0 |
| 0 | 0 |
| 0 | 0 |
| 0 | 0 |
| 0 | 0 |
| 24.4289905 | 9.15197702 |
| 355.577528 | 183.03954 |
| 0 | 0 |
| 0 | 0 |
| 0 | 0 |
| 0 | 0 |
| 0 | 0 |
| 0 | 0 |
| 0 | 0 |
| 0 | 0 |
| 0 | 0 |

| | |
|---|---|
| 0 | 0 |
| 2.71433228 | 0 |
| 0 | 0 |
| 0 | 0 |
| 0 | 0 |
| 0 | 0 |
| 0 | 0 |
| 0 | 0 |
| 0 | 0 |
| 0 | 0 |
| 0 | 0 |
| 0 | 4.57598851 |
| 0 | 0 |
| 0 | 0 |
| 0 | 0 |
| 0 | 0 |
| 0 | 0 |
| 0 | 0 |
| 0 | 0 |
| 0 | 0 |
| 0 | 0 |
| 0 | 0 |
| 165.574269 | 242.527391 |
| 0 | 0 |
| 0 | 0 |
| 0 | 0 |
| 0 | 0 |
| 0 | 0 |
| 0 | 0 |
| 0 | 0 |
| 43.4293164 | 100.671747 |
| 0 | 0 |
| 0 | 0 |
| 0 | 0 |
| 0 | 4.57598851 |
| 0 | 0 |
| 0 | 0 |
| 27.1433228 | 13.7279655 |
| 0 | 0 |
| 0 | 0 |
| 2.71433228 | 0 |
| 0 | 0 |
| 0 | 0 |

| | |
|---|---|
| 0 | 0 |
| 0 | 0 |
| 0 | 0 |
| 0 | 0 |
| 0 | 0 |
| 0 | 0 |
| 0 | 0 |
| 0 | 0 |
| 0 | 0 |
| 0 | 0 |
| 0 | 0 |
| 0 | 0 |
| 0 | 0 |
| 0 | 0 |
| 0 | 0 |
| 0 | 13.7279655 |
| 0 | 0 |
| 0 | 0 |
| 0 | 0 |
| 0 | 0 |
| 0 | 0 |
| 0 | 0 |
| 0 | 0 |
| 0 | 0 |
| 0 | 0 |
| 0 | 0 |
| 0 | 4.57598851 |
| 0 | 0 |
| 0 | 0 |
| 0 | 0 |
| 0 | 0 |
| 0 | 0 |
| 0 | 0 |
| 0 | 0 |
| 0 | 0 |
| 0 | 0 |
| 0 | 0 |
| 0 | 0 |
| 0 | 0 |
| 0 | 0 |
| 0 | 0 |
| 0 | 0 |
| 0 | 4.57598851 |

| | |
|---|---|
| 0 | 0 |
| 0 | 0 |
| 0 | 0 |
| 19.0003259 | 22.8799426 |
| 0 | 0 |
| 0 | 0 |
| 0 | 0 |
| 0 | 0 |
| 0 | 0 |
| 0 | 0 |
| 0 | 0 |
| 0 | 0 |
| 0 | 0 |
| 0 | 4.57598851 |
| 0 | 0 |
| 0 | 0 |
| 0 | 0 |
| 0 | 0 |
| 0 | 0 |
| 0 | 0 |
| 0 | 0 |
| 0 | 0 |
| 0 | 0 |
| 2.71433228 | 18.303954 |
| 0 | 0 |
| 0 | 0 |
| 0 | 0 |
| 0 | 0 |
| 0 | 0 |
| 0 | 0 |
| 2.71433228 | 22.8799426 |
| 0 | 0 |
| 0 | 0 |
| 0 | 0 |
| 0 | 32.0319196 |
| 0 | 0 |
| 0 | 0 |
| 0 | 0 |
| 100.430294 | 151.007621 |
| 0 | 0 |
| 0 | 0 |
| 2.71433228 | 0 |
| 0 | 0 |

| | |
|---|---|
| 0 | 0 |
| 0 | 0 |
| 0 | 0 |
| 0 | 0 |
| 0 | 0 |
| 0 | 0 |
| 0 | 0 |
| 0 | 0 |
| 0 | 0 |
| 0 | 0 |
| 0 | 0 |
| 0 | 0 |
| 0 | 0 |
| 122.144953 | 68.6398277 |
| 0 | 0 |
| 0 | 0 |
| 0 | 9.15197702 |
| 0 | 0 |
| 0 | 4.57598851 |
| 0 | 0 |
| 0 | 4.57598851 |
| 0 | 0 |
| 0 | 0 |
| 0 | 0 |
| 0 | 4.57598851 |
| 0 | 0 |
| 0 | 0 |
| 0 | 0 |
| 0 | 0 |
| 0 | 0 |
| 0 | 0 |
| 0 | 0 |
| 0 | 0 |
| 0 | 0 |
| 0 | 0 |
| 0 | 0 |
| 0 | 0 |
| 0 | 0 |
| 0 | 0 |
| 0 | 0 |
| 0 | 0 |
| 0 | 0 |
| 0 | 0 |

| | |
|---|---|
| 0 | 0 |
| 10.8573291 | 0 |
| 0 | 0 |
| 0 | 0 |
| 0 | 4.57598851 |
| 0 | 0 |
| 0 | 0 |
| 0 | 0 |
| 0 | 0 |
| 0 | 0 |
| 0 | 0 |
| 0 | 0 |
| 0 | 4.57598851 |
| 0 | 4.57598851 |
| 0 | 0 |
| 0 | 0 |
| 0 | 0 |
| 0 | 0 |
| 0 | 0 |
| 0 | 0 |
| 0 | 0 |
| 0 | 0 |
| 0 | 22.8799426 |
| 0 | 0 |
| 0 | 0 |
| 0 | 0 |
| 0 | 0 |
| 0 | 0 |
| 0 | 9.15197702 |
| 0 | 0 |
| 0 | 9.15197702 |
| 0 | 0 |
| 0 | 0 |
| 0 | 0 |
| 0 | 0 |
| 0 | 0 |
| 0 | 0 |
| 0 | 0 |
| 0 | 0 |
| 233.432576 | 219.647449 |
| 0 | 0 |
| 0 | 0 |
| 0 | 0 |

| | |
|---|---|
| 0 | 0 |
| 0 | 0 |
| 0 | 0 |
| 0 | 0 |
| 0 | 0 |
| 0 | 0 |
| 0 | 0 |
| 0 | 0 |
| 0 | 0 |
| 0 | 0 |
| 0 | 0 |
| 0 | 0 |
| 0 | 0 |
| 0 | 4.57598851 |
| 0 | 0 |
| 0 | 0 |
| 0 | 0 |
| 0 | 0 |
| 0 | 0 |
| 0 | 0 |
| 0 | 0 |
| 0 | 0 |
| 16.2859937 | 27.4559311 |
| 0 | 0 |
| 0 | 0 |
| 0 | 0 |
| 138.430946 | 324.895184 |
| 0 | 0 |
| 0 | 0 |
| 0 | 0 |
| 0 | 0 |
| 0 | 0 |
| 0 | 0 |
| 0 | 0 |
| 0 | 0 |
| 0 | 0 |
| 0 | 0 |
| 0 | 0 |
| 0 | 0 |
| 0 | 0 |
| 0 | 0 |
| 0 | 4.57598851 |
| 0 | 0 |

| | |
|---|---|
| 0 | 0 |
| 0 | 0 |
| 0 | 0 |
| 0 | 0 |
| 0 | 0 |
| 0 | 0 |
| 0 | 0 |
| 0 | 0 |
| 0 | 0 |
| 0 | 0 |
| 0 | 0 |
| 0 | 0 |
| 0 | 0 |
| 0 | 0 |
| 0 | 0 |
| 0 | 0 |
| 0 | 0 |
| 0 | 0 |
| 0 | 0 |
| 2.71433228 | 9.15197702 |
| 0 | 0 |
| 0 | 0 |
| 0 | 0 |
| 5.42866456 | 0 |
| 0 | 0 |
| 0 | 0 |
| 0 | 0 |
| 0 | 0 |
| 0 | 0 |
| 2.71433228 | 0 |
| 0 | 0 |
| 0 | 0 |
| 0 | 0 |
| 0 | 0 |
| 0 | 0 |
| 0 | 0 |
| 0 | 0 |
| 0 | 0 |
| 0 | 0 |
| 0 | 0 |
| 0 | 0 |
| 0 | 0 |
| 0 | 0 |

| | |
|---|---|
| 0 | 0 |
| 0 | 0 |
| 0 | 0 |
| 0 | 0 |
| 0 | 0 |
| 0 | 0 |
| 0 | 0 |
| 0 | 0 |
| 206.289253 | 64.0638392 |
| 27.1433228 | 77.7918047 |
| 0 | 0 |
| 0 | 0 |
| 0 | 0 |
| 0 | 0 |
| 0 | 0 |
| 0 | 0 |
| 0 | 0 |
| 0 | 0 |
| 0 | 0 |
| 0 | 0 |
| 0 | 4.57598851 |
| 0 | 0 |
| 0 | 0 |
| 16.2859937 | 4.57598851 |
| 0 | 0 |
| 0 | 0 |
| 0 | 0 |
| 0 | 9.15197702 |
| 0 | 0 |
| 0 | 0 |
| 19.0003259 | 36.6079081 |
| 0 | 0 |
| 5.42866456 | 13.7279655 |
| 0 | 0 |
| 0 | 0 |
| 2.71433228 | 0 |
| 0 | 0 |
| 0 | 0 |
| 0 | 0 |
| 0 | 0 |
| 0 | 0 |
| 0 | 0 |
| 0 | 0 |

| | |
|---|---|
| 0 | 0 |
| 0 | 9.15197702 |
| 0 | 0 |
| 0 | 0 |
| 0 | 0 |
| 0 | 0 |
| 0 | 0 |
| 0 | 0 |
| 0 | 0 |
| 0 | 13.7279655 |
| 13.5716614 | 36.6079081 |
| 0 | 0 |
| 0 | 0 |
| 5.42866456 | 22.8799426 |
| 0 | 0 |
| 0 | 0 |
| 0 | 0 |
| 0 | 0 |
| 8.14299683 | 0 |
| 0 | 0 |
| 0 | 0 |
| 0 | 0 |
| 32.5719873 | 22.8799426 |
| 0 | 0 |
| 0 | 0 |
| 0 | 0 |
| 51.5723133 | 114.399713 |
| 0 | 9.15197702 |
| 0 | 0 |
| 0 | 0 |
| 230.718244 | 22.8799426 |
| 0 | 0 |
| 2.71433228 | 13.7279655 |
| 0 | 0 |
| 0 | 0 |
| 0 | 0 |
| 0 | 0 |
| 0 | 0 |
| 0 | 0 |
| 0 | 0 |
| 0 | 0 |
| 0 | 0 |
| 0 | 0 |

| | |
|---|---|
| 0 | 0 |
| 0 | 0 |
| 2.71433228 | 4.57598851 |
| 0 | 0 |
| 0 | 0 |
| 0 | 0 |
| 0 | 0 |
| 0 | 0 |
| 0 | 0 |
| 0 | 0 |
| 0 | 0 |
| 0 | 0 |
| 0 | 0 |
| 0 | 0 |
| 0 | 0 |
| 0 | 0 |
| 0 | 0 |
| 0 | 0 |
| 0 | 0 |
| 0 | 0 |
| 81.4299683 | 82.3677932 |
| 0 | 0 |
| 0 | 0 |
| 0 | 9.15197702 |
| 0 | 0 |
| 0 | 0 |
| 0 | 0 |
| 0 | 0 |
| 0 | 0 |
| 0 | 0 |
| 0 | 0 |
| 0 | 0 |
| 0 | 0 |
| 0 | 0 |
| 8.14299683 | 4.57598851 |
| 0 | 0 |
| 0 | 4.57598851 |
| 0 | 9.15197702 |
| 0 | 0 |
| 0 | 0 |
| 0 | 0 |
| 0 | 0 |
| 0 | 0 |

| | |
|---|---|
| 0 | 0 |
| 0 | 0 |
| 0 | 0 |
| 0 | 0 |
| 0 | 0 |
| 0 | 0 |
| 0 | 0 |
| 0 | 0 |
| 0 | 0 |
| 0 | 0 |
| 0 | 0 |
| 0 | 0 |
| 0 | 0 |
| 0 | 0 |
| 0 | 0 |
| 0 | 0 |
| 0 | 0 |
| 0 | 0 |
| 0 | 0 |
| 0 | 0 |
| 0 | 0 |
| 0 | 0 |
| 0 | 0 |
| 0 | 4.57598851 |
| 0 | 0 |
| 0 | 0 |
| 0 | 0 |
| 0 | 0 |
| 0 | 0 |
| 0 | 0 |
| 0 | 0 |
| 0 | 0 |
| 0 | 0 |
| 0 | 0 |
| 0 | 0 |
| 0 | 0 |
| 0 | 0 |
| 19.0003259 | 18.303954 |
| 0 | 0 |
| 0 | 0 |
| 0 | 0 |
| 0 | 0 |
| 0 | 0 |

| | |
|---|---|
| 0 | 0 |
| 0 | 0 |
| 0 | 0 |
| 0 | 0 |
| 0 | 0 |
| 0 | 0 |
| 0 | 0 |
| 0 | 0 |
| 0 | 0 |
| 0 | 0 |
| 0 | 0 |
| 0 | 0 |
| 0 | 0 |
| 0 | 0 |
| 0 | 0 |
| 57.0009778 | 59.4878506 |
| 0 | 0 |
| 0 | 0 |
| 0 | 0 |
| 0 | 0 |
| 0 | 0 |
| 0 | 0 |
| 0 | 0 |
| 0 | 0 |
| 0 | 0 |
| 0 | 0 |
| 0 | 0 |
| 135.716614 | 73.2158162 |
| 0 | 0 |
| 0 | 0 |
| 0 | 0 |
| 0 | 0 |
| 0 | 0 |
| 0 | 0 |
| 0 | 0 |
| 0 | 0 |
| 0 | 0 |
| 0 | 0 |
| 0 | 0 |
| 0 | 0 |
| 67.8583069 | 50.3358736 |
| 0 | 0 |
| 0 | 0 |

| | |
|---|---|
| 0 | 0 |
| 0 | 0 |
| 0 | 0 |
| 0 | 0 |
| 0 | 0 |
| 0 | 0 |
| 0 | 0 |
| 0 | 0 |
| 0 | 0 |
| 0 | 0 |
| 0 | 0 |
| 0 | 0 |
| 0 | 0 |
| 0 | 0 |
| 0 | 0 |
| 0 | 0 |
| 0 | 0 |
| 0 | 0 |
| 0 | 0 |
| 0 | 0 |
| 0 | 0 |
| 0 | 0 |
| 0 | 0 |
| 0 | 0 |
| 0 | 0 |
| 0 | 0 |
| 0 | 0 |
| 0 | 0 |
| 0 | 0 |
| 0 | 0 |
| 0 | 0 |
| 0 | 0 |
| 0 | 0 |
| 0 | 0 |
| 0 | 0 |
| 0 | 0 |
| 0 | 0 |
| 0 | 0 |
| 0 | 0 |
| 0 | 0 |
| 0 | 0 |
| 0 | 0 |
| 0 | 0 |

| | |
|---|---|
| 0 | 0 |
| 0 | 0 |
| 0 | 0 |
| 0 | 0 |
| 0 | 0 |
| 0 | 0 |
| 0 | 0 |
| 0 | 0 |
| 0 | 0 |
| 0 | 0 |
| 0 | 13.7279655 |
| 0 | 0 |
| 0 | 0 |
| 0 | 0 |
| 0 | 0 |
| 2.71433228 | 0 |
| 0 | 0 |
| 0 | 0 |
| 0 | 0 |
| 0 | 9.15197702 |
| 0 | 0 |
| 0 | 0 |
| 0 | 0 |
| 0 | 0 |
| 0 | 0 |
| 0 | 0 |
| 0 | 0 |
| 0 | 0 |
| 0 | 0 |
| 0 | 0 |
| 0 | 9.15197702 |
| 0 | 0 |
| 0 | 0 |
| 0 | 0 |
| 0 | 0 |
| 0 | 0 |
| 0 | 0 |
| 0 | 0 |
| 0 | 0 |
| 0 | 0 |
| 0 | 0 |
| 2.71433228 | 0 |
| 0 | 0 |

| | |
|---|---|
| 0 | 0 |
| 0 | 0 |
| 0 | 0 |
| 0 | 0 |
| 0 | 0 |
| 0 | 0 |
| 0 | 0 |
| 0 | 0 |
| 0 | 9.15197702 |
| 0 | 0 |
| 0 | 0 |
| 0 | 0 |
| 382.720851 | 118.975701 |
| 0 | 0 |
| 0 | 0 |
| 0 | 0 |
| 0 | 0 |
| 0 | 0 |
| 0 | 0 |
| 0 | 0 |
| 0 | 0 |
| 0 | 4.57598851 |
| 0 | 0 |
| 0 | 0 |
| 548.29512 | 448.446874 |
| 0 | 13.7279655 |
| 0 | 0 |
| 0 | 0 |
| 0 | 0 |
| 0 | 0 |
| 0 | 0 |
| 0 | 0 |
| 0 | 0 |
| 0 | 0 |
| 0 | 0 |
| 0 | 0 |
| 0 | 0 |
| 0 | 0 |
| 0 | 0 |
| 0 | 0 |
| 0 | 0 |
| 0 | 0 |
| 0 | 0 |

| | |
|---|---|
| 0 | 0 |
| 0 | 0 |
| 0 | 0 |
| 0 | 0 |
| 0 | 0 |
| 0 | 0 |
| 0 | 0 |
| 0 | 0 |
| 0 | 0 |
| 0 | 0 |
| 0 | 0 |
| 0 | 0 |
| 0 | 0 |
| 0 | 0 |
| 0 | 0 |
| 0 | 0 |
| 0 | 0 |
| 0 | 0 |
| 0 | 0 |
| 21.7146582 | 0 |
| 0 | 0 |
| 0 | 0 |
| 0 | 0 |
| 0 | 0 |
| 0 | 0 |
| 0 | 0 |
| 141.145278 | 141.855644 |
| 0 | 0 |
| 0 | 0 |
| 0 | 0 |
| 0 | 0 |
| 0 | 0 |
| 0 | 0 |
| 0 | 0 |
| 0 | 0 |
| 0 | 0 |
| 0 | 0 |
| 0 | 0 |
| 0 | 0 |
| 0 | 0 |
| 0 | 0 |
| 0 | 0 |
| 0 | 0 |

| | |
|---|---|
| 0 | 0 |
| 0 | 0 |
| 0 | 0 |
| 0 | 0 |
| 0 | 0 |
| 0 | 0 |
| 0 | 0 |
| 0 | 0 |
| 0 | 0 |
| 0 | 0 |
| 0 | 0 |
| 0 | 0 |
| 0 | 0 |
| 0 | 0 |
| 0 | 0 |
| 0 | 0 |
| 0 | 0 |
| 0 | 0 |
| 0 | 0 |
| 0 | 0 |
| 0 | 0 |
| 0 | 0 |
| 0 | 0 |
| 0 | 0 |
| 0 | 0 |
| 0 | 0 |
| 0 | 0 |
| 0 | 0 |
| 0 | 0 |
| 0 | 0 |
| 0 | 0 |
| 0 | 0 |
| 13.5716614 | 0 |
| 0 | 0 |
| 0 | 0 |
| 0 | 0 |
| 0 | 0 |
| 0 | 0 |
| 0 | 0 |
| 0 | 0 |
| 0 | 0 |
| 0 | 0 |
| 0 | 0 |

| | |
|---|---|
| 0 | 0 |
| 0 | 0 |
| 0 | 0 |
| 0 | 0 |
| 0 | 0 |
| 0 | 0 |
| 0 | 0 |
| 0 | 0 |
| 0 | 0 |
| 0 | 0 |
| 0 | 0 |
| 0 | 0 |
| 0 | 0 |
| 0 | 0 |
| 0 | 0 |
| 0 | 0 |
| 0 | 0 |
| 0 | 0 |
| 0 | 0 |
| 0 | 0 |
| 0 | 0 |
| 228.003911 | 169.311575 |
| 0 | 0 |
| 0 | 0 |
| 0 | 0 |
| 0 | 0 |
| 0 | 0 |
| 81.4299683 | 41.1838966 |
| 0 | 0 |
| 0 | 0 |
| 0 | 0 |
| 0 | 0 |
| 0 | 0 |
| 0 | 0 |
| 0 | 0 |
| 1563.45539 | 983.83753 |
| 0 | 0 |
| 0 | 0 |
| 0 | 0 |
| 0 | 0 |
| 0 | 0 |
| 0 | 0 |
| 0 | 0 |

| | |
|---|---|
| 0 | 0 |
| 0 | 0 |
| 0 | 0 |
| 0 | 0 |
| 0 | 0 |
| 0 | 0 |
| 0 | 0 |
| 0 | 0 |
| 0 | 0 |
| 0 | 0 |
| 0 | 13.7279655 |
| 0 | 0 |
| 0 | 0 |
| 2.71433228 | 13.7279655 |
| 0 | 0 |
| 0 | 0 |
| 0 | 0 |
| 0 | 0 |
| 0 | 0 |
| 0 | 0 |
| 0 | 4.57598851 |
| 0 | 0 |
| 0 | 0 |
| 2.71433228 | 4.57598851 |
| 0 | 0 |
| 0 | 0 |
| 0 | 0 |
| 0 | 0 |
| 0 | 0 |
| 0 | 0 |
| 0 | 0 |
| 0 | 0 |
| 0 | 0 |
| 0 | 0 |
| 0 | 0 |
| 0 | 0 |
| 0 | 0 |
| 0 | 0 |
| 1908.17559 | 1665.65982 |
| 0 | 0 |
| 0 | 0 |
| 0 | 0 |
| 0 | 0 |

| | |
|---|---|
| 0 | 0 |
| 0 | 0 |
| 0 | 18.303954 |
| 0 | 0 |
| 0 | 0 |
| 31195.8209 | 1143.99713 |
| 0 | 0 |
| 0 | 0 |
| 0 | 0 |
| 0 | 18.303954 |
| 0 | 0 |
| 0 | 0 |
| 0 | 0 |
| 0 | 0 |
| 5.42866456 | 4.57598851 |
| 0 | 0 |
| 0 | 0 |
| 0 | 0 |
| 0 | 0 |
| 0 | 0 |
| 0 | 9.15197702 |
| 0 | 0 |
| 0 | 0 |
| 0 | 0 |
| 0 | 0 |
| 0 | 0 |
| 0 | 0 |
| 0 | 0 |
| 0 | 0 |
| 2.71433228 | 0 |
| 0 | 0 |
| 0 | 0 |
| 0 | 0 |
| 0 | 0 |
| 0 | 0 |
| 0 | 0 |
| 2.71433228 | 4.57598851 |
| 0 | 0 |
| 0 | 0 |
| 0 | 0 |
| 0 | 0 |
| 0 | 0 |
| 0 | 0 |

| | |
|---|---|
| 0 | 0 |
| 0 | 0 |
| 0 | 0 |
| 0 | 0 |
| 0 | 0 |
| 0 | 0 |
| 0 | 0 |
| 0 | 0 |
| 0 | 0 |
| 0 | 0 |
| 0 | 0 |
| 0 | 0 |
| 2.71433228 | 0 |
| 0 | 0 |
| 0 | 0 |
| 0 | 0 |
| 0 | 0 |
| 0 | 0 |
| 0 | 0 |
| 0 | 0 |
| 0 | 0 |
| 0 | 0 |
| 0 | 0 |
| 0 | 0 |
| 0 | 0 |
| 0 | 0 |
| 0 | 4.57598851 |
| 0 | 0 |
| 0 | 0 |
| 0 | 13.7279655 |
| 0 | 0 |
| 0 | 0 |
| 0 | 0 |
| 0 | 0 |
| 0 | 0 |
| 0 | 0 |
| 564.581114 | 2498.48973 |
| 0 | 0 |
| 0 | 0 |
| 0 | 0 |
| 0 | 0 |
| 0 | 0 |
| 0 | 0 |

| | |
|---|---|
| 2.71433228 | 0 |
| 0 | 0 |
| 0 | 0 |
| 0 | 0 |
| 0 | 0 |
| 0 | 0 |
| 0 | 0 |
| 0 | 0 |
| 2.71433228 | 0 |
| 2.71433228 | 0 |
| 0 | 0 |
| 0 | 0 |
| 0 | 0 |
| 0 | 0 |
| 0 | 0 |
| 0 | 0 |
| 0 | 0 |
| 0 | 0 |
| 0 | 0 |
| 0 | 0 |
| 0 | 0 |
| 0 | 13.7279655 |
| 0 | 0 |
| 0 | 0 |
| 0 | 0 |
| 0 | 0 |
| 0 | 0 |
| 0 | 0 |
| 0 | 0 |
| 0 | 0 |
| 0 | 0 |
| 2.71433228 | 0 |
| 0 | 0 |
| 0 | 0 |
| 0 | 0 |
| 0 | 0 |
| 0 | 0 |
| 0 | 0 |
| 0 | 0 |
| 0 | 0 |
| 0 | 4.57598851 |
| 0 | 0 |
| 0 | 0 |

| | |
|---|---|
| 0 | 0 |
| 0 | 0 |
| 43.4293164 | 68.6398277 |
| 0 | 0 |
| 0 | 0 |
| 0 | 0 |
| 0 | 0 |
| 0 | 0 |
| 0 | 0 |
| 0 | 0 |
| 0 | 0 |
| 0 | 4.57598851 |
| 0 | 0 |
| 0 | 0 |
| 0 | 0 |
| 0 | 0 |
| 0 | 0 |
| 0 | 0 |
| 0 | 0 |
| 0 | 0 |
| 0 | 0 |
| 0 | 0 |
| 0 | 0 |
| 0 | 0 |
| 0 | 0 |
| 0 | 0 |
| 0 | 0 |
| 0 | 0 |
| 0 | 0 |
| 0 | 0 |
| 0 | 0 |
| 0 | 0 |
| 0 | 0 |
| 116.716288 | 68.6398277 |
| 0 | 0 |
| 0 | 0 |
| 0 | 4.57598851 |
| 0 | 0 |
| 0 | 0 |
| 0 | 0 |
| 0 | 0 |
| 0 | 0 |
| 0 | 0 |

| | |
|---|---|
| 0 | 0 |
| 0 | 0 |
| 0 | 0 |
| 0 | 0 |
| 0 | 0 |
| 0 | 0 |
| 0 | 0 |
| 0 | 0 |
| 0 | 0 |
| 0 | 0 |
| 0 | 0 |
| 0 | 0 |
| 0 | 0 |
| 0 | 0 |
| 0 | 0 |
| 0 | 0 |
| 0 | 0 |
| 0 | 0 |
| 0 | 0 |
| 0 | 0 |
| 0 | 4.57598851 |
| 0 | 0 |
| 0 | 0 |
| 0 | 0 |
| 0 | 0 |
| 0 | 0 |
| 0 | 0 |
| 0 | 0 |
| 0 | 0 |
| 0 | 0 |
| 0 | 0 |
| 0 | 0 |
| 0 | 0 |
| 0 | 0 |
| 0 | 0 |
| 0 | 0 |
| 0 | 0 |
| 0 | 0 |
| 0 | 0 |
| 0 | 0 |
| 0 | 0 |
| 0 | 0 |
| 2.71433228 | 0 |

| | |
|---|---|
| 0 | 0 |
| 0 | 0 |
| 0 | 0 |
| 0 | 0 |
| 0 | 0 |
| 0 | 0 |
| 0 | 0 |
| 0 | 0 |
| 0 | 0 |
| 0 | 0 |
| 0 | 0 |
| 0 | 0 |
| 2.71433228 | 0 |
| 0 | 0 |
| 10.8573291 | 36.6079081 |
| 0 | 0 |
| 0 | 0 |
| 0 | 0 |
| 0 | 0 |
| 2.71433228 | 4.57598851 |
| 0 | 0 |
| 0 | 13.7279655 |
| 0 | 0 |
| 0 | 0 |
| 0 | 0 |
| 8.14299683 | 13.7279655 |
| 0 | 0 |
| 0 | 0 |
| 0 | 0 |
| 0 | 0 |
| 0 | 0 |
| 0 | 0 |
| 0 | 0 |
| 0 | 0 |
| 0 | 0 |
| 0 | 0 |
| 0 | 0 |
| 0 | 0 |
| 0 | 0 |
| 0 | 0 |
| 0 | 0 |
| 0 | 0 |
| 0 | 0 |

| | |
|---|---|
| 0 | 0 |
| 0 | 0 |
| 0 | 0 |
| 0 | 0 |
| 0 | 0 |
| 0 | 0 |
| 0 | 0 |
| 0 | 0 |
| 0 | 0 |
| 0 | 0 |
| 0 | 0 |
| 0 | 0 |
| 0 | 0 |
| 0 | 0 |
| 0 | 0 |
| 0 | 9.15197702 |
| 0 | 0 |
| 0 | 0 |
| 0 | 0 |
| 0 | 0 |
| 0 | 0 |
| 0 | 0 |
| 0 | 0 |
| 0 | 0 |
| 13.5716614 | 0 |
| 0 | 0 |
| 0 | 0 |
| 0 | 0 |
| 0 | 0 |
| 0 | 0 |
| 0 | 0 |
| 0 | 0 |
| 0 | 0 |
| 0 | 0 |
| 0 | 0 |
| 0 | 0 |
| 2.71433228 | 0 |
| 0 | 0 |
| 0 | 0 |
| 0 | 0 |
| 0 | 0 |
| 0 | 0 |
| 0 | 0 |

| | |
|---|---|
| 0 | 0 |
| 0 | 0 |
| 0 | 4.57598851 |
| 0 | 0 |
| 0 | 0 |
| 0 | 0 |
| 0 | 0 |
| 0 | 0 |
| 0 | 0 |
| 0 | 0 |
| 0 | 0 |
| 0 | 0 |
| 0 | 0 |
| 0 | 0 |
| 176.431598 | 4.57598851 |
| 0 | 0 |
| 0 | 0 |
| 0 | 0 |
| 0 | 0 |
| 0 | 0 |
| 0 | 0 |
| 0 | 0 |
| 0 | 0 |
| 0 | 0 |
| 0 | 0 |
| 0 | 0 |
| 0 | 0 |
| 0 | 0 |
| 0 | 0 |
| 0 | 0 |
| 0 | 0 |
| 0 | 13.7279655 |
| 0 | 4.57598851 |
| 0 | 0 |
| 0 | 4.57598851 |
| 29.8576551 | 27.4559311 |
| 0 | 0 |
| 0 | 0 |
| 0 | 0 |
| 0 | 0 |
| 0 | 0 |
| 0 | 0 |
| 0 | 0 |

| | |
|---|---|
| 0 | 0 |
| 0 | 0 |
| 0 | 0 |
| 0 | 0 |
| 0 | 0 |
| 0 | 0 |
| 0 | 0 |
| 0 | 0 |
| 0 | 0 |
| 0 | 0 |
| 0 | 0 |
| 0 | 0 |
| 0 | 0 |
| 0 | 0 |
| 0 | 9.15197702 |
| 0 | 9.15197702 |
| 0 | 0 |
| 0 | 4.57598851 |
| 0 | 0 |
| 0 | 0 |
| 0 | 0 |
| 0 | 4.57598851 |
| 0 | 0 |
| 0 | 0 |
| 0 | 0 |
| 0 | 0 |
| 0 | 0 |
| 0 | 0 |
| 0 | 0 |
| 0 | 0 |
| 0 | 0 |
| 0 | 0 |
| 0 | 0 |
| 0 | 0 |
| 0 | 0 |
| 0 | 0 |
| 0 | 0 |
| 0 | 0 |
| 0 | 0 |
| 0 | 0 |
| 0 | 0 |
| 0 | 0 |
| 0 | 0 |

| | |
|---|---|
| 0 | 0 |
| 0 | 0 |
| 0 | 0 |
| 0 | 13.7279655 |
| 0 | 0 |
| 0 | 0 |
| 0 | 0 |
| 0 | 0 |
| 0 | 0 |
| 0 | 0 |
| 0 | 0 |
| 0 | 0 |
| 0 | 0 |
| 0 | 0 |
| 0 | 0 |
| 0 | 0 |
| 0 | 0 |
| 0 | 0 |
| 0 | 18.303954 |
| 0 | 0 |
| 0 | 0 |
| 0 | 0 |
| 0 | 0 |
| 0 | 0 |
| 0 | 0 |
| 0 | 0 |
| 0 | 0 |
| 0 | 0 |
| 0 | 0 |
| 0 | 0 |
| 0 | 0 |
| 0 | 0 |
| 0 | 0 |
| 0 | 0 |
| 0 | 0 |
| 0 | 0 |
| 0 | 0 |
| 0 | 0 |
| 0 | 0 |
| 0 | 0 |
| 0 | 0 |
| 0 | 0 |
| 0 | 0 |

| | |
|---|---|
| 0 | 0 |
| 0 | 4.57598851 |
| 0 | 0 |
| 0 | 0 |
| 0 | 0 |
| 0 | 0 |
| 0 | 0 |
| 0 | 0 |
| 0 | 0 |
| 0 | 0 |
| 0 | 0 |
| 0 | 0 |
| 0 | 0 |
| 0 | 0 |
| 0 | 0 |
| 0 | 13.7279655 |
| 0 | 0 |
| 0 | 0 |
| 0 | 0 |
| 0 | 0 |
| 0 | 0 |
| 0 | 0 |
| 0 | 0 |
| 0 | 0 |
| 0 | 0 |
| 32.5719873 | 4.57598851 |
| 0 | 0 |
| 0 | 0 |
| 0 | 0 |
| 0 | 0 |
| 0 | 0 |
| 0 | 0 |
| 0 | 0 |
| 21.7146582 | 18.303954 |
| 0 | 0 |
| 10.8573291 | 4.57598851 |
| 0 | 0 |
| 0 | 0 |
| 0 | 0 |
| 0 | 0 |
| 0 | 0 |
| 0 | 0 |
| 0 | 0 |

| | |
|---|---|
| 0 | 0 |
| 0 | 0 |
| 0 | 0 |
| 0 | 0 |
| 0 | 0 |
| 0 | 0 |
| 0 | 0 |
| 0 | 0 |
| 0 | 0 |
| 0 | 0 |
| 0 | 0 |
| 0 | 0 |
| 0 | 0 |
| 0 | 22.8799426 |
| 0 | 0 |
| 0 | 0 |
| 0 | 0 |
| 0 | 0 |
| 0 | 0 |
| 0 | 0 |
| 0 | 4.57598851 |
| 0 | 0 |
| 0 | 0 |
| 0 | 0 |
| 0 | 0 |
| 0 | 0 |
| 0 | 0 |
| 0 | 0 |
| 0 | 0 |
| 0 | 0 |
| 0 | 0 |
| 0 | 0 |
| 0 | 4.57598851 |
| 0 | 0 |
| 0 | 0 |
| 0 | 0 |
| 5.42866456 | 0 |
| 0 | 0 |
| 0 | 0 |
| 2.71433228 | 0 |
| 0 | 0 |
| 0 | 4.57598851 |
| 0 | 0 |

| | |
|---|---|
| 0 | 0 |
| 0 | 0 |
| 0 | 0 |
| 0 | 0 |
| 0 | 0 |
| 0 | 0 |
| 0 | 0 |
| 0 | 0 |
| 0 | 0 |
| 0 | 0 |
| 0 | 0 |
| 0 | 0 |
| 0 | 0 |
| 0 | 0 |
| 0 | 0 |
| 0 | 0 |
| 0 | 0 |
| 0 | 0 |
| 2.71433228 | 9.15197702 |
| 0 | 0 |
| 0 | 0 |
| 0 | 0 |
| 0 | 0 |
| 0 | 0 |
| 0 | 0 |
| 0 | 0 |
| 0 | 0 |
| 0 | 0 |
| 0 | 0 |
| 0 | 0 |
| 0 | 0 |
| 0 | 0 |
| 0 | 0 |
| 0 | 0 |
| 0 | 0 |
| 0 | 0 |
| 0 | 0 |
| 0 | 0 |
| 0 | 0 |
| 0 | 0 |
| 0 | 0 |
| 0 | 0 |
| 0 | 0 |

| | |
|---|---|
| 0 | 0 |
| 0 | 0 |
| 0 | 0 |
| 0 | 0 |
| 0 | 0 |
| 0 | 0 |
| 0 | 0 |
| 0 | 0 |
| 0 | 0 |
| 0 | 4.57598851 |
| 0 | 0 |
| 0 | 0 |
| 0 | 0 |
| 0 | 0 |
| 0 | 0 |
| 0 | 0 |
| 0 | 0 |
| 0 | 0 |
| 0 | 9.15197702 |
| 0 | 4.57598851 |
| 0 | 0 |
| 0 | 4.57598851 |
| 0 | 0 |
| 0 | 0 |
| 0 | 0 |
| 0 | 0 |
| 0 | 0 |
| 0 | 0 |
| 0 | 0 |
| 0 | 0 |
| 2.71433228 | 0 |
| 0 | 0 |
| 0 | 0 |
| 0 | 22.8799426 |
| 0 | 0 |
| 0 | 0 |
| 0 | 0 |
| 76.0013038 | 0 |
| 0 | 0 |
| 0 | 0 |
| 0 | 0 |
| 0 | 0 |
| 0 | 0 |

| | |
|---|---|
| 0 | 0 |
| 0 | 0 |
| 0 | 0 |
| 0 | 0 |
| 0 | 0 |
| 0 | 0 |
| 0 | 0 |
| 0 | 0 |
| 0 | 0 |
| 0 | 0 |
| 0 | 0 |
| 0 | 0 |
| 8.14299683 | 27.4559311 |
| 0 | 0 |
| 0 | 0 |
| 0 | 0 |
| 0 | 0 |
| 0 | 0 |
| 0 | 0 |
| 0 | 0 |
| 10.8573291 | 73.2158162 |
| 0 | 0 |
| 0 | 0 |
| 5.42866456 | 0 |
| 24.4289905 | 0 |
| 0 | 0 |
| 0 | 0 |
| 0 | 0 |
| 0 | 0 |
| 0 | 0 |
| 0 | 0 |
| 0 | 0 |
| 0 | 0 |
| 0 | 0 |
| 0 | 0 |
| 0 | 0 |
| 0 | 0 |
| 0 | 4.57598851 |
| 0 | 0 |
| 0 | 0 |
| 0 | 0 |
| 0 | 0 |
| 0 | 0 |

| | |
|---|---|
| 0 | 0 |
| 13.5716614 | 0 |
| 0 | 0 |
| 0 | 0 |
| 0 | 0 |
| 0 | 0 |
| 0 | 0 |
| 0 | 0 |
| 0 | 0 |
| 0 | 0 |
| 0 | 0 |
| 0 | 0 |
| 0 | 0 |
| 0 | 4.57598851 |
| 0 | 0 |
| 0 | 0 |
| 0 | 0 |
| 0 | 0 |
| 0 | 0 |
| 0 | 0 |
| 0 | 0 |
| 0 | 4.57598851 |
| 0 | 0 |
| 0 | 0 |
| 0 | 0 |
| 0 | 0 |
| 0 | 0 |
| 0 | 0 |
| 0 | 0 |
| 0 | 0 |
| 0 | 0 |
| 0 | 0 |
| 0 | 0 |
| 0 | 0 |
| 0 | 0 |
| 0 | 0 |
| 0 | 0 |
| 0 | 0 |
| 0 | 0 |
| 0 | 0 |
| 0 | 0 |
| 0 | 0 |
| 0 | 0 |

| | |
|---|---|
| 0 | 0 |
| 0 | 0 |
| 0 | 0 |
| 0 | 0 |
| 0 | 0 |
| 0 | 0 |
| 0 | 0 |
| 0 | 0 |
| 0 | 0 |
| 0 | 4.57598851 |
| 0 | 0 |
| 0 | 9.15197702 |
| 0 | 0 |
| 0 | 0 |
| 0 | 0 |
| 0 | 0 |
| 0 | 0 |
| 0 | 0 |
| 0 | 0 |
| 0 | 0 |
| 0 | 0 |
| 0 | 0 |
| 2.71433228 | 27.4559311 |
| 0 | 0 |
| 0 | 0 |
| 0 | 0 |
| 0 | 0 |
| 2.71433228 | 22.8799426 |
| 0 | 0 |
| 0 | 0 |
| 0 | 0 |
| 0 | 0 |
| 0 | 0 |
| 0 | 0 |
| 0 | 0 |
| 0 | 0 |
| 0 | 0 |
| 0 | 0 |
| 0 | 0 |
| 0 | 0 |
| 0 | 0 |
| 0 | 0 |
| 0 | 0 |

| | |
|---|---|
| 0 | 9.15197702 |
| 0 | 0 |
| 420.721503 | 77.7918047 |
| 0 | 0 |
| 0 | 0 |
| 0 | 0 |
| 0 | 0 |
| 0 | 0 |
| 0 | 0 |
| 0 | 0 |
| 0 | 0 |
| 0 | 0 |
| 0 | 4.57598851 |
| 0 | 0 |
| 0 | 0 |
| 0 | 0 |
| 0 | 13.7279655 |
| 0 | 0 |
| 0 | 0 |
| 13.5716614 | 9.15197702 |
| 0 | 0 |
| 0 | 0 |
| 0 | 4.57598851 |
| 0 | 0 |
| 0 | 0 |
| 0 | 4.57598851 |
| 0 | 0 |
| 2.71433228 | 4.57598851 |
| 0 | 4.57598851 |
| 0 | 0 |
| 0 | 0 |
| 0 | 0 |
| 0 | 0 |
| 0 | 0 |
| 0 | 0 |
| 0 | 0 |
| 0 | 0 |
| 0 | 0 |
| 0 | 0 |
| 0 | 0 |
| 0 | 0 |
| 0 | 0 |
| 0 | 0 |

| | |
|---|---|
| 0 | 0 |
| 0 | 0 |
| 0 | 0 |
| 0 | 0 |
| 0 | 0 |
| 0 | 0 |
| 0 | 0 |
| 0 | 0 |
| 0 | 0 |
| 0 | 0 |
| 0 | 0 |
| 0 | 0 |
| 32.5719873 | 73.2158162 |
| 0 | 0 |
| 0 | 0 |
| 0 | 0 |
| 0 | 0 |
| 0 | 0 |
| 0 | 0 |
| 0 | 0 |
| 0 | 0 |
| 0 | 0 |
| 0 | 0 |
| 0 | 0 |
| 0 | 4.57598851 |
| 0 | 0 |
| 0 | 0 |
| 0 | 0 |
| 0 | 0 |
| 184.574595 | 155.583609 |
| 0 | 0 |
| 0 | 0 |
| 0 | 0 |
| 0 | 0 |
| 0 | 0 |
| 0 | 0 |
| 0 | 0 |
| 0 | 0 |
| 0 | 0 |
| 0 | 0 |
| 0 | 0 |
| 0 | 0 |
| 0 | 0 |

| | |
|---|---|
| 0 | 0 |
| 0 | 0 |
| 0 | 0 |
| 0 | 0 |
| 0 | 0 |
| 0 | 0 |
| 0 | 0 |
| 0 | 0 |
| 0 | 0 |
| 0 | 0 |
| 0 | 0 |
| 0 | 0 |
| 0 | 0 |
| 0 | 0 |
| 0 | 0 |
| 0 | 0 |
| 0 | 0 |
| 0 | 0 |
| 0 | 0 |
| 0 | 0 |
| 2.71433228 | 0 |
| 0 | 0 |
| 0 | 0 |
| 0 | 0 |
| 0 | 0 |
| 0 | 0 |
| 0 | 0 |
| 0 | 0 |
| 0 | 0 |
| 0 | 0 |
| 0 | 9.15197702 |
| 0 | 0 |
| 0 | 0 |
| 0 | 0 |
| 0 | 0 |
| 0 | 0 |
| 0 | 0 |
| 0 | 0 |
| 0 | 0 |
| 0 | 0 |
| 0 | 0 |
| 0 | 0 |
| 0 | 0 |

| | |
|---|---|
| 0 | 0 |
| 0 | 0 |
| 0 | 0 |
| 0 | 0 |
| 2.71433228 | 0 |
| 0 | 0 |
| 0 | 0 |
| 0 | 0 |
| 0 | 0 |
| 8.14299683 | 9.15197702 |
| 0 | 0 |
| 0 | 0 |
| 0 | 0 |
| 0 | 0 |
| 0 | 0 |
| 0 | 0 |
| 0 | 0 |
| 268.718896 | 439.294897 |
| 0 | 0 |
| 0 | 0 |
| 0 | 0 |
| 0 | 0 |
| 0 | 0 |
| 0 | 0 |
| 0 | 9.15197702 |
| 0 | 0 |
| 0 | 0 |
| 0 | 0 |
| 0 | 0 |
| 0 | 0 |
| 0 | 18.303954 |
| 0 | 0 |
| 0 | 0 |
| 0 | 0 |
| 0 | 0 |
| 0 | 0 |
| 0 | 0 |
| 0 | 0 |
| 0 | 0 |
| 0 | 0 |
| 0 | 0 |
| 0 | 0 |
| 0 | 0 |

| | |
|---|---|
| 0 | 0 |
| 0 | 0 |
| 0 | 0 |
| 0 | 0 |
| 0 | 0 |
| 0 | 0 |
| 0 | 0 |
| 0 | 0 |
| 0 | 0 |
| 0 | 0 |
| 0 | 0 |
| 0 | 0 |
| 0 | 0 |
| 0 | 0 |
| 0 | 0 |
| 0 | 0 |
| 0 | 0 |
| 0 | 0 |
| 0 | 0 |
| 0 | 0 |
| 0 | 0 |
| 0 | 0 |
| 0 | 0 |
| 0 | 0 |
| 0 | 0 |
| 0 | 0 |
| 0 | 0 |
| 0 | 0 |
| 0 | 0 |
| 388.149516 | 137.279655 |
| 0 | 0 |
| 0 | 0 |
| 0 | 0 |
| 0 | 4.57598851 |
| 0 | 0 |
| 0 | 0 |
| 0 | 0 |
| 0 | 0 |
| 0 | 0 |
| 0 | 0 |
| 0 | 0 |
| 0 | 0 |
| 0 | 0 |

| | |
|---|---|
| 0 | 0 |
| 0 | 0 |
| 0 | 0 |
| 0 | 0 |
| 0 | 0 |
| 0 | 0 |
| 0 | 0 |
| 0 | 0 |
| 0 | 0 |
| 0 | 0 |
| 0 | 0 |
| 0 | 0 |
| 0 | 0 |
| 0 | 0 |
| 0 | 0 |
| 0 | 0 |
| 0 | 0 |
| 0 | 0 |
| 0 | 0 |
| 0 | 0 |
| 0 | 0 |
| 2.71433228 | 0 |
| 0 | 0 |
| 0 | 0 |
| 0 | 0 |
| 0 | 0 |
| 0 | 0 |
| 0 | 0 |
| 0 | 0 |
| 0 | 0 |
| 0 | 0 |
| 0 | 0 |
| 0 | 0 |
| 0 | 0 |
| 0 | 0 |
| 0 | 0 |
| 0 | 0 |
| 0 | 0 |
| 0 | 0 |
| 0 | 0 |
| 0 | 0 |
| 0 | 0 |
| 0 | 0 |

| | |
|---|---|
| 0 | 0 |
| 0 | 0 |
| 0 | 0 |
| 0 | 0 |
| 0 | 0 |
| 43.4293164 | 4.57598851 |
| 0 | 0 |
| 0 | 0 |
| 0 | 0 |
| 0 | 0 |
| 0 | 0 |
| 0 | 0 |
| 0 | 0 |
| 0 | 0 |
| 0 | 0 |
| 0 | 0 |
| 0 | 0 |
| 0 | 0 |
| 0 | 0 |
| 0 | 0 |
| 0 | 0 |
| 0 | 0 |
| 0 | 0 |
| 0 | 0 |
| 0 | 0 |
| 0 | 0 |
| 0 | 0 |
| 0 | 0 |
| 0 | 0 |
| 0 | 0 |
| 0 | 0 |
| 0 | 0 |
| 0 | 0 |
| 0 | 0 |
| 0 | 0 |
| 0 | 0 |
| 0 | 0 |
| 0 | 0 |
| 0 | 0 |
| 0 | 0 |
| 0 | 0 |
| 0 | 0 |
| 0 | 0 |

| | |
|---|---|
| 0 | 0 |
| 0 | 0 |
| 0 | 0 |
| 0 | 0 |
| 0 | 0 |
| 0 | 0 |
| 2.71433228 | 0 |
| 0 | 0 |
| 0 | 0 |
| 0 | 0 |
| 0 | 0 |
| 0 | 0 |
| 0 | 0 |
| 0 | 0 |
| 0 | 0 |
| 0 | 0 |
| 0 | 0 |
| 0 | 0 |
| 0 | 0 |
| 8.14299683 | 0 |
| 0 | 0 |
| 0 | 0 |
| 0 | 9.15197702 |
| 0 | 4.57598851 |
| 179.14593 | 100.671747 |
| 0 | 0 |
| 0 | 0 |
| 0 | 0 |
| 0 | 0 |
| 0 | 0 |
| 0 | 0 |
| 0 | 0 |
| 0 | 0 |
| 0 | 0 |
| 0 | 0 |
| 0 | 0 |
| 0 | 0 |
| 0 | 0 |
| 0 | 0 |
| 0 | 0 |
| 0 | 0 |
| 0 | 0 |
| 0 | 0 |

| | |
|---|---|
| 0 | 0 |
| 0 | 0 |
| 0 | 0 |
| 0 | 0 |
| 0 | 0 |
| 0 | 0 |
| 0 | 0 |
| 0 | 4.57598851 |
| 2.71433228 | 0 |
| 0 | 0 |
| 0 | 0 |
| 0 | 0 |
| 0 | 0 |
| 0 | 0 |
| 0 | 0 |
| 0 | 0 |
| 0 | 0 |
| 0 | 0 |
| 0 | 9.15197702 |
| 0 | 0 |
| 0 | 0 |
| 0 | 0 |
| 0 | 0 |
| 0 | 0 |
| 0 | 0 |
| 0 | 4.57598851 |
| 0 | 0 |
| 0 | 0 |
| 252.432902 | 68.6398277 |
| 0 | 0 |
| 0 | 0 |
| 0 | 0 |
| 0 | 0 |
| 0 | 0 |
| 0 | 0 |
| 0 | 0 |
| 0 | 0 |
| 0 | 0 |
| 0 | 0 |
| 0 | 0 |
| 0 | 0 |
| 0 | 0 |
| 0 | 0 |

| | |
|---|---|
| 0 | 0 |
| 0 | 0 |
| 0 | 0 |
| 0 | 0 |
| 2.71433228 | 0 |
| 0 | 0 |
| 0 | 0 |
| 0 | 0 |
| 0 | 0 |
| 0 | 0 |
| 0 | 0 |
| 0 | 0 |
| 0 | 0 |
| 0 | 0 |
| 0 | 0 |
| 0 | 0 |
| 0 | 0 |
| 0 | 0 |
| 0 | 0 |
| 0 | 13.7279655 |
| 0 | 0 |
| 0 | 4.57598851 |
| 0 | 0 |
| 0 | 0 |
| 0 | 0 |
| 0 | 0 |
| 0 | 0 |
| 0 | 0 |
| 0 | 0 |
| 0 | 0 |
| 0 | 0 |
| 5.42866456 | 0 |
| 76.0013038 | 32.0319196 |
| 0 | 0 |
| 0 | 0 |
| 0 | 0 |
| 304.005215 | 306.59123 |
| 0 | 0 |
| 0 | 0 |
| 0 | 0 |
| 0 | 0 |
| 0 | 0 |
| 0 | 0 |

| | |
|---|---|
| 0 | 0 |
| 0 | 0 |
| 0 | 0 |
| 0 | 0 |
| 0 | 0 |
| 0 | 0 |
| 0 | 0 |
| 0 | 0 |
| 0 | 0 |
| 0 | 0 |
| 0 | 0 |
| 0 | 0 |
| 0 | 0 |
| 0 | 0 |
| 0 | 0 |
| 21.7146582 | 0 |
| 0 | 0 |
| 0 | 0 |
| 0 | 0 |
| 0 | 0 |
| 0 | 0 |
| 0 | 0 |
| 0 | 0 |
| 0 | 0 |
| 0 | 0 |
| 0 | 0 |
| 0 | 0 |
| 0 | 0 |
| 0 | 0 |
| 0 | 0 |
| 0 | 0 |
| 0 | 0 |
| 0 | 0 |
| 0 | 0 |
| 0 | 0 |
| 0 | 0 |
| 0 | 0 |
| 0 | 0 |
| 0 | 0 |
| 0 | 0 |
| 0 | 0 |
| 0 | 0 |
| 0 | 0 |

| | |
|---|---|
| 2.71433228 | 0 |
| 0 | 0 |
| 0 | 0 |
| 0 | 0 |
| 0 | 0 |
| 0 | 0 |
| 0 | 0 |
| 0 | 0 |
| 0 | 0 |
| 0 | 0 |
| 0 | 0 |
| 0 | 0 |
| 0 | 0 |
| 0 | 0 |
| 0 | 0 |
| 0 | 0 |
| 0 | 0 |
| 0 | 0 |
| 0 | 4.57598851 |
| 0 | 0 |
| 0 | 0 |
| 0 | 0 |
| 0 | 0 |
| 0 | 0 |
| 0 | 0 |
| 0 | 0 |
| 0 | 0 |
| 0 | 0 |
| 0 | 0 |
| 0 | 0 |
| 0 | 0 |
| 0 | 0 |
| 0 | 0 |
| 0 | 0 |
| 2.71433228 | 0 |
| 0 | 0 |
| 0 | 0 |
| 0 | 0 |
| 0 | 0 |
| 0 | 0 |
| 0 | 0 |
| 0 | 4.57598851 |
| 0 | 0 |

| | |
|---|---|
| 8.14299683 | 13.7279655 |
| 0 | 0 |
| 0 | 0 |
| 0 | 0 |
| 0 | 0 |
| 0 | 0 |
| 0 | 0 |
| 0 | 0 |
| 0 | 0 |
| 0 | 0 |
| 0 | 0 |
| 0 | 4.57598851 |
| 0 | 4.57598851 |
| 0 | 0 |
| 0 | 0 |
| 0 | 0 |
| 0 | 0 |
| 0 | 0 |
| 0 | 0 |
| 0 | 0 |
| 0 | 0 |
| 0 | 0 |
| 0 | 0 |
| 0 | 0 |
| 0 | 0 |
| 0 | 0 |
| 0 | 0 |
| 0 | 0 |
| 0 | 0 |
| 0 | 0 |
| 0 | 4.57598851 |
| 0 | 0 |
| 0 | 54.9118621 |
| 0 | 0 |
| 0 | 0 |
| 0 | 0 |
| 0 | 0 |
| 0 | 4.57598851 |
| 0 | 0 |
| 0 | 0 |
| 0 | 0 |
| 0 | 0 |
| 2.71433228 | 0 |

| | |
|---|---|
| 0 | 0 |
| 0 | 0 |
| 0 | 0 |
| 8.14299683 | 4.57598851 |
| 0 | 0 |
| 0 | 0 |
| 0 | 0 |
| 0 | 0 |
| 0 | 0 |
| 0 | 0 |
| 0 | 0 |
| 0 | 4.57598851 |
| 0 | 0 |
| 0 | 0 |
| 0 | 0 |
| 0 | 0 |
| 0 | 0 |
| 0 | 0 |
| 0 | 0 |
| 0 | 0 |
| 0 | 0 |
| 0 | 0 |
| 0 | 0 |
| 0 | 0 |
| 0 | 0 |
| 0 | 0 |
| 0 | 0 |
| 0 | 0 |
| 0 | 0 |
| 0 | 0 |
| 0 | 0 |
| 0 | 0 |
| 0 | 0 |
| 0 | 0 |
| 19.0003259 | 4.57598851 |
| 0 | 0 |
| 35.2863196 | 210.495472 |
| 0 | 0 |
| 0 | 0 |
| 2.71433228 | 4.57598851 |
| 0 | 0 |
| 0 | 0 |

| | |
|---|---|
| 0 | 0 |
| 0 | 0 |
| 0 | 0 |
| 0 | 0 |
| 0 | 0 |
| 0 | 0 |
| 0 | 0 |
| 0 | 0 |
| 0 | 0 |
| 0 | 9.15197702 |
| 0 | 0 |
| 65.1439747 | 96.0957587 |
| 0 | 0 |
| 0 | 0 |
| 0 | 0 |
| 0 | 4.57598851 |
| 0 | 0 |
| 13.5716614 | 13.7279655 |
| 0 | 0 |
| 0 | 0 |
| 0 | 0 |
| 0 | 0 |
| 0 | 0 |
| 0 | 0 |
| 0 | 0 |
| 0 | 0 |
| 0 | 0 |
| 0 | 0 |
| 0 | 0 |
| 0 | 0 |
| 0 | 0 |
| 0 | 0 |
| 0 | 0 |
| 0 | 0 |
| 0 | 0 |
| 0 | 0 |
| 0 | 0 |
| 0 | 0 |
| 0 | 0 |
| 0 | 0 |
| 0 | 0 |
| 0 | 0 |
| 0 | 0 |

| | |
|---|---|
| 0 | 0 |
| 0 | 0 |
| 0 | 0 |
| 0 | 0 |
| 0 | 0 |
| 0 | 0 |
| 0 | 0 |
| 0 | 0 |
| 0 | 0 |
| 0 | 0 |
| 0 | 0 |
| 2.71433228 | 0 |
| 0 | 4.57598851 |
| 0 | 0 |
| 0 | 0 |
| 0 | 0 |
| 0 | 0 |
| 0 | 0 |
| 0 | 0 |
| 0 | 0 |
| 0 | 0 |
| 0 | 0 |
| 0 | 0 |
| 0 | 0 |
| 0 | 0 |
| 0 | 0 |
| 10.8573291 | 27.4559311 |
| 0 | 0 |
| 0 | 0 |
| 0 | 0 |
| 0 | 0 |
| 0 | 0 |
| 293.147886 | 91.5197702 |
| 0 | 0 |
| 0 | 0 |
| 0 | 0 |
| 0 | 0 |
| 0 | 0 |
| 0 | 0 |
| 0 | 0 |
| 0 | 0 |
| 0 | 0 |
| 0 | 0 |

| | |
|---|---|
| 0 | 0 |
| 0 | 0 |
| 0 | 0 |
| 0 | 0 |
| 0 | 0 |
| 0 | 0 |
| 0 | 0 |
| 0 | 0 |
| 0 | 0 |
| 0 | 0 |
| 0 | 0 |
| 0 | 0 |
| 0 | 0 |
| 0 | 0 |
| 0 | 0 |
| 0 | 0 |
| 0 | 0 |
| 0 | 0 |
| 0 | 0 |
| 0 | 0 |
| 0 | 0 |
| 0 | 0 |
| 0 | 0 |
| 0 | 4.57598851 |
| 0 | 0 |
| 2.71433228 | 0 |
| 0 | 0 |
| 0 | 0 |
| 0 | 0 |
| 0 | 0 |
| 0 | 0 |
| 0 | 0 |
| 0 | 0 |
| 0 | 0 |
| 0 | 0 |
| 0 | 0 |
| 309.43388 | 32.0319196 |
| 0 | 0 |
| 0 | 0 |
| 0 | 0 |
| 0 | 0 |
| 0 | 0 |
| 0 | 0 |

| | |
|---|---|
| 0 | 0 |
| 0 | 0 |
| 0 | 0 |
| 0 | 0 |
| 0 | 0 |
| 0 | 0 |
| 0 | 0 |
| 0 | 0 |
| 0 | 0 |
| 0 | 0 |
| 0 | 0 |
| 0 | 0 |
| 0 | 0 |
| 0 | 0 |
| 0 | 0 |
| 0 | 0 |
| 0 | 0 |
| 0 | 0 |
| 0 | 0 |
| 0 | 0 |
| 0 | 0 |
| 0 | 0 |
| 0 | 0 |
| 2.71433228 | 0 |
| 0 | 0 |
| 0 | 0 |
| 0 | 0 |
| 0 | 0 |
| 0 | 0 |
| 0 | 0 |
| 0 | 0 |
| 0 | 0 |
| 0 | 0 |
| 0 | 0 |
| 2.71433228 | 0 |
| 0 | 0 |
| 2.71433228 | 0 |
| 0 | 0 |
| 0 | 0 |
| 0 | 0 |
| 0 | 0 |
| 0 | 0 |
| 0 | 0 |

| | |
|---|---|
| 0 | 0 |
| 0 | 0 |
| 0 | 0 |
| 57.0009778 | 86.9437817 |
| 0 | 0 |
| 0 | 4.57598851 |
| 0 | 0 |
| 0 | 0 |
| 0 | 0 |
| 0 | 0 |
| 0 | 0 |
| 0 | 0 |
| 0 | 0 |
| 157.431272 | 45.7598851 |
| 0 | 0 |
| 0 | 0 |
| 0 | 0 |
| 0 | 0 |
| 0 | 0 |
| 0 | 0 |
| 0 | 0 |
| 0 | 0 |
| 0 | 0 |
| 0 | 0 |
| 0 | 0 |
| 0 | 4.57598851 |
| 0 | 0 |
| 0 | 0 |
| 0 | 0 |
| 2.71433228 | 4.57598851 |
| 0 | 0 |
| 0 | 0 |
| 0 | 0 |
| 0 | 0 |
| 0 | 0 |
| 171.002934 | 27.4559311 |
| 0 | 0 |
| 0 | 0 |
| 0 | 0 |
| 0 | 0 |
| 0 | 0 |
| 0 | 0 |
| 0 | 0 |

| | |
|---|---|
| 0 | 0 |
| 0 | 0 |
| 0 | 0 |
| 0 | 0 |
| 0 | 0 |
| 0 | 0 |
| 0 | 0 |
| 0 | 0 |
| 0 | 0 |
| 0 | 0 |
| 0 | 0 |
| 0 | 0 |
| 0 | 0 |
| 0 | 0 |
| 0 | 0 |
| 0 | 0 |
| 0 | 0 |
| 0 | 0 |
| 0 | 0 |
| 0 | 0 |
| 0 | 0 |
| 0 | 0 |
| 0 | 0 |
| 0 | 0 |
| 0 | 0 |
| 0 | 0 |
| 0 | 0 |
| 0 | 0 |
| 0 | 0 |
| 0 | 0 |
| 0 | 0 |
| 0 | 0 |
| 0 | 0 |
| 0 | 18.303954 |
| 0 | 0 |
| 0 | 0 |
| 0 | 0 |
| 0 | 0 |
| 0 | 0 |
| 0 | 0 |
| 0 | 0 |
| 0 | 0 |
| 0 | 0 |

| | |
|---|---|
| 0 | 0 |
| 0 | 0 |
| 0 | 0 |
| 0 | 0 |
| 0 | 0 |
| 0 | 0 |
| 0 | 0 |
| 0 | 0 |
| 0 | 0 |
| 0 | 0 |
| 0 | 0 |
| 13.5716614 | 22.8799426 |
| 0 | 0 |
| 0 | 0 |
| 0 | 0 |
| 0 | 4.57598851 |
| 0 | 0 |
| 0 | 0 |
| 0 | 9.15197702 |
| 0 | 0 |
| 0 | 0 |
| 0 | 0 |
| 0 | 0 |
| 0 | 0 |
| 0 | 0 |
| 0 | 0 |
| 0 | 0 |
| 0 | 4.57598851 |
| 0 | 0 |
| 0 | 0 |
| 0 | 0 |
| 0 | 0 |
| 0 | 0 |
| 0 | 0 |
| 0 | 0 |
| 0 | 0 |
| 0 | 0 |
| 0 | 4.57598851 |
| 0 | 0 |
| 0 | 0 |
| 0 | 0 |
| 0 | 13.7279655 |
| 0 | 0 |

| | |
|---|---|
| 0 | 0 |
| 0 | 0 |
| 0 | 0 |
| 0 | 0 |
| 0 | 0 |
| 0 | 0 |
| 0 | 0 |
| 0 | 0 |
| 0 | 0 |
| 0 | 0 |
| 0 | 0 |
| 0 | 0 |
| 0 | 0 |
| 0 | 0 |
| 0 | 0 |
| 0 | 0 |
| 0 | 0 |
| 0 | 0 |
| 0 | 0 |
| 0 | 0 |
| 0 | 0 |
| 0 | 0 |
| 0 | 0 |
| 0 | 0 |
| 0 | 0 |
| 0 | 0 |
| 0 | 4.57598851 |
| 100.430294 | 329.471173 |
| 0 | 0 |
| 0 | 0 |
| 673.154405 | 530.814667 |
| 0 | 0 |
| 0 | 0 |
| 0 | 0 |
| 0 | 0 |
| 0 | 0 |
| 0 | 0 |
| 0 | 0 |
| 0 | 0 |
| 0 | 0 |
| 0 | 0 |
| 0 | 0 |
| 0 | 0 |

| | |
|---|---|
| 0 | 4.57598851 |
| 0 | 0 |
| 0 | 0 |
| 0 | 0 |
| 0 | 0 |
| 0 | 0 |
| 0 | 0 |
| 0 | 0 |
| 0 | 0 |
| 0 | 0 |
| 0 | 0 |
| 0 | 0 |
| 0 | 0 |
| 0 | 0 |
| 0 | 0 |
| 0 | 0 |
| 0 | 0 |
| 0 | 0 |
| 0 | 0 |
| 0 | 0 |
| 0 | 0 |
| 0 | 0 |
| 0 | 0 |
| 0 | 0 |
| 0 | 0 |
| 0 | 0 |
| 0 | 0 |
| 0 | 0 |
| 0 | 0 |
| 0 | 0 |
| 0 | 0 |
| 0 | 0 |
| 0 | 0 |
| 0 | 0 |
| 0 | 0 |
| 0 | 0 |
| 0 | 0 |
| 0 | 0 |
| 0 | 0 |
| 0 | 0 |
| 0 | 0 |
| 0 | 4.57598851 |
| 0 | 0 |

| | |
|---|---|
| 0 | 0 |
| 0 | 0 |
| 0 | 0 |
| 0 | 0 |
| 0 | 0 |
| 0 | 0 |
| 0 | 0 |
| 0 | 0 |
| 0 | 0 |
| 0 | 0 |
| 0 | 0 |
| 0 | 0 |
| 0 | 0 |
| 0 | 0 |
| 0 | 0 |
| 0 | 0 |
| 0 | 4.57598851 |
| 0 | 0 |
| 0 | 0 |
| 0 | 0 |
| 0 | 0 |
| 2.71433228 | 0 |
| 0 | 0 |
| 0 | 0 |
| 0 | 0 |
| 0 | 0 |
| 0 | 0 |
| 0 | 0 |
| 0 | 0 |
| 0 | 0 |
| 0 | 0 |
| 0 | 0 |
| 0 | 0 |
| 0 | 0 |
| 2.71433228 | 0 |
| 0 | 0 |
| 0 | 0 |
| 0 | 0 |
| 0 | 0 |
| 0 | 0 |
| 0 | 0 |
| 0 | 0 |
| 0 | 0 |

| | |
|---|---|
| 0 | 0 |
| 0 | 0 |
| 0 | 0 |
| 0 | 0 |
| 0 | 0 |
| 0 | 0 |
| 0 | 0 |
| 0 | 0 |
| 0 | 0 |
| 0 | 0 |
| 0 | 0 |
| 0 | 0 |
| 0 | 0 |
| 0 | 0 |
| 0 | 0 |
| 0 | 0 |
| 119.43062 | 114.399713 |
| 0 | 0 |
| 0 | 0 |
| 0 | 0 |
| 0 | 0 |
| 0 | 0 |
| 0 | 0 |
| 0 | 0 |
| 0 | 0 |
| 0 | 0 |
| 0 | 0 |
| 0 | 0 |
| 0 | 0 |
| 0 | 0 |
| 0 | 0 |
| 0 | 0 |
| 0 | 0 |
| 0 | 0 |
| 0 | 0 |
| 0 | 0 |
| 0 | 0 |
| 0 | 0 |
| 0 | 0 |
| 0 | 0 |
| 0 | 0 |
| 0 | 0 |
| 0 | 0 |

| | |
|---|---|
| 0 | 0 |
| 0 | 0 |
| 0 | 0 |
| 0 | 0 |
| 2.71433228 | 0 |
| 0 | 0 |
| 0 | 0 |
| 0 | 0 |
| 0 | 0 |
| 2.71433228 | 0 |
| 0 | 0 |
| 0 | 0 |
| 0 | 0 |
| 0 | 0 |
| 0 | 0 |
| 0 | 0 |
| 0 | 0 |
| 0 | 0 |
| 2.71433228 | 0 |
| 0 | 0 |
| 0 | 0 |
| 0 | 0 |
| 0 | 0 |
| 0 | 0 |
| 0 | 0 |
| 0 | 0 |
| 0 | 0 |
| 0 | 0 |
| 0 | 4.57598851 |
| 0 | 0 |
| 0 | 0 |
| 0 | 0 |
| 0 | 0 |
| 0 | 0 |
| 0 | 0 |
| 0 | 0 |
| 0 | 0 |
| 0 | 0 |
| 0 | 0 |
| 0 | 0 |
| 2326.18276 | 599.454495 |
| 0 | 0 |
| 0 | 0 |

| | |
|---|---|
| 0 | 0 |
| 0 | 0 |
| 0 | 0 |
| 0 | 0 |
| 0 | 0 |
| 0 | 0 |
| 0 | 0 |
| 0 | 0 |
| 0 | 0 |
| 0 | 0 |
| 0 | 0 |
| 0 | 0 |
| 0 | 4.57598851 |
| 0 | 0 |
| 0 | 0 |
| 0 | 0 |
| 2.71433228 | 64.0638392 |
| 0 | 0 |
| 0 | 0 |
| 0 | 0 |
| 0 | 0 |
| 0 | 0 |
| 0 | 0 |
| 0 | 0 |
| 0 | 0 |
| 0 | 0 |
| 0 | 0 |
| 0 | 0 |
| 0 | 0 |
| 0 | 0 |
| 0 | 0 |
| 0 | 0 |
| 5.42866456 | 0 |
| 0 | 0 |
| 0 | 0 |
| 0 | 0 |
| 480.436813 | 146.431632 |
| 0 | 0 |
| 0 | 0 |
| 0 | 0 |
| 10.8573291 | 32.0319196 |
| 0 | 0 |
| 8.14299683 | 0 |

| | |
|---|---|
| 0 | 0 |
| 0 | 4.57598851 |
| 0 | 0 |
| 0 | 0 |
| 0 | 0 |
| 0 | 0 |
| 0 | 0 |
| 0 | 9.15197702 |
| 0 | 0 |
| 0 | 0 |
| 0 | 0 |
| 0 | 0 |
| 0 | 4.57598851 |
| 0 | 0 |
| 0 | 0 |
| 0 | 0 |
| 0 | 0 |
| 0 | 0 |
| 0 | 0 |
| 0 | 0 |
| 0 | 0 |
| 0 | 0 |
| 0 | 0 |
| 0 | 0 |
| 0 | 0 |
| 0 | 0 |
| 2.71433228 | 0 |
| 0 | 0 |
| 0 | 0 |
| 0 | 0 |
| 0 | 0 |
| 0 | 0 |
| 0 | 36.6079081 |
| 0 | 0 |
| 0 | 0 |
| 0 | 0 |
| 0 | 0 |
| 0 | 0 |
| 0 | 0 |
| 0 | 4.57598851 |
| 0 | 0 |
| 0 | 0 |
| 0 | 0 |

| | |
|---|---|
| 0 | 0 |
| 0 | 0 |
| 0 | 0 |
| 0 | 0 |
| 0 | 0 |
| 0 | 0 |
| 0 | 0 |
| 0 | 0 |
| 0 | 0 |
| 0 | 0 |
| 0 | 0 |
| 0 | 0 |
| 0 | 0 |
| 0 | 0 |
| 0 | 0 |
| 0 | 0 |
| 0 | 0 |
| 0 | 0 |
| 0 | 0 |
| 0 | 0 |
| 0 | 0 |
| 0 | 0 |
| 0 | 0 |
| 0 | 0 |
| 0 | 4.57598851 |
| 0 | 0 |
| 0 | 0 |
| 0 | 0 |
| 0 | 0 |
| 0 | 0 |
| 0 | 0 |
| 0 | 0 |
| 0 | 0 |
| 0 | 0 |
| 0 | 0 |
| 0 | 0 |
| 0 | 0 |
| 0 | 0 |
| 0 | 0 |
| 0 | 0 |
| 0 | 0 |
| 0 | 0 |
| 0 | 0 |

| | |
|---|---|
| 2.71433228 | 0 |
| 0 | 0 |
| 0 | 0 |
| 0 | 0 |
| 0 | 0 |
| 0 | 0 |
| 559.152449 | 695.550254 |
| 0 | 0 |
| 0 | 0 |
| 0 | 0 |
| 0 | 0 |
| 0 | 0 |
| 0 | 0 |
| 0 | 0 |
| 0 | 0 |
| 0 | 0 |
| 0 | 0 |
| 2.71433228 | 4.57598851 |
| 0 | 0 |
| 0 | 0 |
| 0 | 0 |
| 0 | 4.57598851 |
| 2.71433228 | 0 |
| 0 | 0 |
| 0 | 0 |
| 0 | 0 |
| 0 | 0 |
| 0 | 0 |
| 0 | 0 |
| 0 | 0 |
| 0 | 0 |
| 0 | 0 |
| 0 | 0 |
| 0 | 0 |
| 0 | 0 |
| 0 | 0 |
| 0 | 0 |
| 0 | 0 |
| 0 | 0 |
| 0 | 0 |
| 0 | 0 |
| 0 | 0 |
| 0 | 0 |

| | |
|---|---|
| 0 | 0 |
| 0 | 0 |
| 0 | 0 |
| 0 | 0 |
| 0 | 0 |
| 0 | 0 |
| 0 | 0 |
| 0 | 0 |
| 0 | 0 |
| 0 | 0 |
| 0 | 0 |
| 0 | 0 |
| 0 | 0 |
| 0 | 0 |
| 0 | 0 |
| 0 | 0 |
| 0 | 0 |
| 0 | 9.15197702 |
| 0 | 0 |
| 0 | 0 |
| 0 | 0 |
| 10.8573291 | 4.57598851 |
| 0 | 0 |
| 0 | 0 |
| 0 | 0 |
| 0 | 0 |
| 0 | 0 |
| 76.0013038 | 91.5197702 |
| 0 | 0 |
| 27.1433228 | 0 |
| 0 | 0 |
| 0 | 0 |
| 0 | 0 |
| 0 | 0 |
| 0 | 0 |
| 0 | 0 |
| 0 | 0 |
| 0 | 0 |
| 0 | 0 |
| 0 | 0 |
| 0 | 0 |
| 0 | 4.57598851 |
| 0 | 0 |

| | |
|---|---|
| 0 | 0 |
| 119.43062 | 50.3358736 |
| 0 | 0 |
| 0 | 0 |
| 0 | 0 |
| 0 | 0 |
| 0 | 0 |
| 0 | 0 |
| 0 | 0 |
| 0 | 0 |
| 0 | 0 |
| 0 | 0 |
| 0 | 0 |
| 0 | 0 |
| 0 | 0 |
| 0 | 0 |
| 0 | 0 |
| 0 | 0 |
| 0 | 0 |
| 0 | 4.57598851 |
| 0 | 0 |
| 0 | 0 |
| 0 | 0 |
| 0 | 0 |
| 0 | 0 |
| 0 | 0 |
| 0 | 4.57598851 |
| 0 | 0 |
| 0 | 0 |
| 0 | 0 |
| 0 | 0 |
| 32.5719873 | 22.8799426 |
| 0 | 0 |
| 0 | 0 |
| 0 | 0 |
| 0 | 0 |
| 0 | 0 |
| 0 | 0 |
| 0 | 0 |
| 0 | 0 |
| 2.71433228 | 4.57598851 |
| 0 | 0 |
| 0 | 0 |

| | |
|---|---|
| 0 | 0 |
| 0 | 0 |
| 0 | 0 |
| 0 | 0 |
| 0 | 32.0319196 |
| 0 | 0 |
| 0 | 0 |
| 0 | 0 |
| 0 | 0 |
| 0 | 0 |
| 0 | 0 |
| 0 | 0 |
| 0 | 0 |
| 2.71433228 | 0 |
| 0 | 0 |
| 0 | 0 |
| 0 | 0 |
| 0 | 0 |
| 5.42866456 | 0 |
| 0 | 0 |
| 0 | 0 |
| 0 | 0 |
| 0 | 0 |
| 0 | 0 |
| 19.0003259 | 0 |
| 0 | 0 |
| 0 | 0 |
| 0 | 0 |
| 0 | 0 |
| 0 | 0 |
| 0 | 0 |
| 0 | 0 |
| 0 | 0 |
| 0 | 0 |
| 0 | 0 |
| 0 | 0 |
| 0 | 0 |
| 0 | 4.57598851 |
| 0 | 0 |
| 0 | 0 |
| 0 | 0 |
| 0 | 0 |
| 0 | 0 |

| | |
|---|---|
| 0 | 0 |
| 0 | 13.7279655 |
| 0 | 0 |
| 0 | 0 |
| 0 | 0 |
| 0 | 0 |
| 0 | 0 |
| 0 | 0 |
| 0 | 0 |
| 0 | 0 |
| 0 | 0 |
| 0 | 0 |
| 0 | 0 |
| 0 | 0 |
| 0 | 0 |
| 0 | 0 |
| 0 | 0 |
| 0 | 22.8799426 |
| 0 | 4.57598851 |
| 0 | 0 |
| 0 | 0 |
| 2.71433228 | 0 |
| 0 | 9.15197702 |
| 0 | 0 |
| 0 | 0 |
| 0 | 0 |
| 0 | 0 |
| 0 | 0 |
| 0 | 0 |
| 0 | 0 |
| 0 | 0 |
| 0 | 0 |
| 0 | 0 |
| 0 | 0 |
| 0 | 0 |
| 0 | 0 |
| 0 | 0 |
| 0 | 0 |
| 0 | 0 |
| 0 | 22.8799426 |
| 0 | 0 |
| 13.5716614 | 13.7279655 |
| 0 | 0 |

| | |
|---|---|
| 0 | 0 |
| 0 | 4.57598851 |
| 0 | 0 |
| 0 | 0 |
| 0 | 0 |
| 0 | 0 |
| 0 | 0 |
| 0 | 0 |
| 0 | 0 |
| 0 | 0 |
| 0 | 0 |
| 0 | 0 |
| 0 | 0 |
| 0 | 0 |
| 32.5719873 | 132.703667 |
| 0 | 0 |
| 0 | 0 |
| 0 | 0 |
| 0 | 0 |
| 0 | 0 |
| 0 | 0 |
| 0 | 0 |
| 0 | 0 |
| 0 | 0 |
| 0 | 0 |
| 0 | 0 |
| 0 | 0 |
| 0 | 0 |
| 8.14299683 | 9.15197702 |
| 0 | 0 |
| 0 | 0 |
| 0 | 0 |
| 0 | 0 |
| 0 | 0 |
| 0 | 0 |
| 0 | 0 |
| 0 | 0 |
| 0 | 0 |
| 0 | 0 |
| 0 | 0 |
| 0 | 0 |
| 0 | 0 |
| 0 | 0 |

| | |
|---|---|
| 0 | 0 |
| 0 | 0 |
| 0 | 0 |
| 0 | 0 |
| 0 | 0 |
| 0 | 0 |
| 10.8573291 | 9.15197702 |
| 0 | 0 |
| 0 | 0 |
| 0 | 0 |
| 0 | 0 |
| 0 | 0 |
| 0 | 0 |
| 0 | 0 |
| 0 | 0 |
| 0 | 13.7279655 |
| 0 | 0 |
| 0 | 0 |
| 0 | 0 |
| 0 | 0 |
| 0 | 0 |
| 0 | 0 |
| 0 | 0 |
| 0 | 0 |
| 0 | 0 |
| 0 | 0 |
| 0 | 0 |
| 2.71433228 | 9.15197702 |
| 0 | 0 |
| 0 | 0 |
| 0 | 0 |
| 0 | 0 |
| 0 | 0 |
| 0 | 0 |
| 0 | 0 |
| 0 | 0 |
| 0 | 0 |
| 0 | 0 |
| 0 | 0 |
| 0 | 0 |
| 0 | 0 |
| 0 | 0 |
| 0 | 0 |

| | |
|---|---|
| 0 | 0 |
| 0 | 0 |
| 0 | 0 |
| 0 | 0 |
| 21.7146582 | 27.4559311 |
| 0 | 0 |
| 0 | 0 |
| 0 | 0 |
| 0 | 0 |
| 0 | 0 |
| 0 | 0 |
| 0 | 0 |
| 0 | 4.57598851 |
| 0 | 0 |
| 0 | 0 |
| 0 | 0 |
| 0 | 0 |
| 0 | 0 |
| 0 | 0 |
| 0 | 0 |
| 0 | 0 |
| 0 | 0 |
| 0 | 0 |
| 0 | 0 |
| 0 | 0 |
| 0 | 0 |
| 0 | 0 |
| 0 | 0 |
| 0 | 0 |
| 0 | 0 |
| 0 | 0 |
| 0 | 0 |
| 0 | 9.15197702 |
| 0 | 0 |
| 0 | 0 |
| 0 | 0 |
| 0 | 0 |
| 0 | 0 |
| 0 | 0 |
| 0 | 0 |
| 0 | 0 |
| 2.71433228 | 4.57598851 |
| 0 | 0 |

| | |
|---|---|
| 0 | 0 |
| 0 | 0 |
| 0 | 0 |
| 0 | 0 |
| 0 | 0 |
| 0 | 0 |
| 0 | 0 |
| 0 | 9.15197702 |
| 0 | 0 |
| 0 | 0 |
| 0 | 0 |
| 0 | 0 |
| 0 | 0 |
| 0 | 0 |
| 0 | 0 |
| 0 | 0 |
| 0 | 0 |
| 0 | 0 |
| 0 | 0 |
| 0 | 0 |
| 0 | 0 |
| 0 | 0 |
| 0 | 0 |
| 0 | 0 |
| 2.71433228 | 0 |
| 0 | 0 |
| 0 | 0 |
| 0 | 0 |
| 0 | 0 |
| 0 | 0 |
| 0 | 0 |
| 0 | 0 |
| 0 | 0 |
| 0 | 4.57598851 |
| 0 | 0 |
| 0 | 0 |
| 0 | 0 |
| 0 | 0 |
| 0 | 0 |
| 0 | 0 |
| 0 | 0 |
| 0 | 0 |
| 0 | 0 |

| | |
|---|---|
| 0 | 4.57598851 |
| 0 | 0 |
| 0 | 0 |
| 0 | 0 |
| 0 | 0 |
| 0 | 4.57598851 |
| 0 | 0 |
| 0 | 0 |
| 0 | 9.15197702 |
| 0 | 0 |
| 0 | 0 |
| 0 | 0 |
| 0 | 0 |
| 0 | 0 |
| 0 | 0 |
| 19.0003259 | 0 |
| 0 | 0 |
| 0 | 0 |
| 0 | 0 |
| 0 | 0 |
| 0 | 0 |
| 0 | 0 |
| 0 | 0 |
| 0 | 4.57598851 |
| 0 | 0 |
| 0 | 0 |
| 0 | 0 |
| 0 | 0 |
| 0 | 0 |
| 0 | 9.15197702 |
| 0 | 0 |
| 0 | 0 |
| 0 | 0 |
| 0 | 0 |
| 0 | 0 |
| 0 | 0 |
| 0 | 0 |
| 0 | 0 |
| 0 | 0 |
| 0 | 0 |
| 0 | 0 |
| 0 | 0 |
| 0 | 0 |

| | |
|---|---|
| 0 | 0 |
| 0 | 0 |
| 0 | 0 |
| 0 | 0 |
| 0 | 0 |
| 0 | 0 |
| 0 | 0 |
| 0 | 0 |
| 29.8576551 | 36.6079081 |
| 0 | 0 |
| 0 | 0 |
| 0 | 0 |
| 0 | 0 |
| 0 | 0 |
| 2.71433228 | 4.57598851 |
| 0 | 0 |
| 0 | 0 |
| 0 | 0 |
| 0 | 0 |
| 0 | 0 |
| 0 | 0 |
| 0 | 0 |
| 0 | 0 |
| 0 | 0 |
| 0 | 0 |
| 0 | 0 |
| 0 | 0 |
| 0 | 9.15197702 |
| 0 | 32.0319196 |
| 0 | 0 |
| 0 | 0 |
| 70.5726392 | 0 |
| 0 | 0 |
| 0 | 0 |
| 0 | 0 |
| 0 | 0 |
| 0 | 0 |
| 0 | 0 |
| 0 | 0 |
| 0 | 0 |
| 0 | 0 |
| 0 | 0 |
| 0 | 4.57598851 |

| | |
|---|---|
| 0 | 0 |
| 0 | 0 |
| 0 | 0 |
| 0 | 0 |
| 0 | 0 |
| 0 | 4.57598851 |
| 0 | 0 |
| 0 | 0 |
| 0 | 0 |
| 0 | 0 |
| 0 | 0 |
| 0 | 0 |
| 0 | 0 |
| 0 | 0 |
| 0 | 0 |
| 0 | 0 |
| 0 | 0 |
| 0 | 4.57598851 |
| 0 | 0 |
| 0 | 0 |
| 0 | 4.57598851 |
| 0 | 0 |
| 0 | 0 |
| 0 | 0 |
| 0 | 0 |
| 0 | 0 |
| 0 | 0 |
| 0 | 4.57598851 |
| 0 | 0 |
| 0 | 0 |
| 0 | 0 |
| 0 | 0 |
| 0 | 0 |
| 0 | 0 |
| 19.0003259 | 32.0319196 |
| 0 | 0 |
| 0 | 0 |
| 0 | 0 |
| 0 | 0 |
| 19.0003259 | 41.1838966 |
| 0 | 0 |
| 0 | 4.57598851 |
| 0 | 0 |

| | |
|---|---|
| 0 | 0 |
| 0 | 0 |
| 0 | 0 |
| 0 | 0 |
| 0 | 0 |
| 0 | 22.8799426 |
| 0 | 0 |
| 0 | 0 |
| 0 | 0 |
| 0 | 0 |
| 0 | 0 |
| 0 | 0 |
| 0 | 0 |
| 0 | 0 |
| 0 | 0 |
| 0 | 0 |
| 0 | 0 |
| 0 | 0 |
| 0 | 0 |
| 0 | 0 |
| 0 | 0 |
| 108.573291 | 91.5197702 |
| 0 | 0 |
| 0 | 0 |
| 0 | 0 |
| 0 | 0 |
| 0 | 0 |
| 0 | 0 |
| 0 | 0 |
| 0 | 0 |
| 0 | 0 |
| 0 | 0 |
| 0 | 0 |
| 0 | 0 |
| 0 | 0 |
| 0 | 0 |
| 0 | 0 |
| 0 | 0 |
| 0 | 0 |
| 0 | 0 |
| 0 | 0 |
| 0 | 0 |
| 0 | 0 |

| | |
|---|---|
| 0 | 0 |
| 0 | 0 |
| 0 | 9.15197702 |
| 0 | 0 |
| 0 | 0 |
| 0 | 0 |
| 0 | 0 |
| 0 | 0 |
| 0 | 0 |
| 0 | 0 |
| 8.14299683 | 4.57598851 |
| 0 | 0 |
| 0 | 0 |
| 0 | 0 |
| 0 | 0 |
| 0 | 0 |
| 0 | 0 |
| 0 | 0 |
| 0 | 0 |
| 0 | 0 |
| 0 | 0 |
| 0 | 0 |
| 0 | 0 |
| 0 | 0 |
| 0 | 0 |
| 0 | 0 |
| 0 | 0 |
| 0 | 0 |
| 0 | 0 |
| 0 | 0 |
| 0 | 0 |
| 0 | 0 |
| 0 | 0 |
| 46.1436487 | 18.303954 |
| 0 | 0 |
| 114.001956 | 105.247736 |
| 0 | 0 |
| 0 | 0 |
| 0 | 0 |
| 0 | 0 |
| 0 | 0 |
| 0 | 0 |
| 0 | 0 |

| | |
|---|---|
| 0 | 0 |
| 0 | 0 |
| 0 | 4.57598851 |
| 0 | 0 |
| 0 | 0 |
| 0 | 0 |
| 0 | 0 |
| 0 | 0 |
| 0 | 0 |
| 0 | 0 |
| 0 | 0 |
| 0 | 4.57598851 |
| 0 | 0 |
| 0 | 0 |
| 0 | 0 |
| 0 | 0 |
| 0 | 0 |
| 0 | 0 |
| 0 | 0 |
| 0 | 0 |
| 0 | 0 |
| 0 | 0 |
| 0 | 0 |
| 0 | 0 |
| 0 | 0 |
| 0 | 0 |
| 152.002608 | 123.55169 |
| 0 | 0 |
| 0 | 0 |
| 0 | 0 |
| 0 | 0 |
| 0 | 0 |
| 0 | 0 |
| 0 | 0 |
| 0 | 0 |
| 0 | 0 |
| 0 | 0 |
| 0 | 0 |
| 0 | 0 |
| 0 | 0 |
| 0 | 0 |
| 0 | 0 |
| 0 | 0 |

| | |
|---|---|
| 0 | 0 |
| 0 | 0 |
| 0 | 0 |
| 0 | 0 |
| 0 | 0 |
| 0 | 0 |
| 0 | 0 |
| 0 | 0 |
| 0 | 0 |
| 0 | 0 |
| 0 | 0 |
| 0 | 0 |
| 0 | 0 |
| 0 | 0 |
| 0 | 0 |
| 0 | 0 |
| 0 | 0 |
| 0 | 0 |
| 0 | 0 |
| 0 | 0 |
| 0 | 0 |
| 0 | 0 |
| 0 | 0 |
| 2.71433228 | 0 |
| 0 | 0 |
| 0 | 0 |
| 0 | 0 |
| 0 | 0 |
| 0 | 0 |
| 0 | 0 |
| 0 | 0 |
| 0 | 0 |
| 0 | 0 |
| 0 | 0 |
| 0 | 0 |
| 0 | 0 |
| 0 | 0 |
| 0 | 0 |
| 0 | 0 |
| 0 | 0 |
| 0 | 0 |
| 0 | 0 |
| 0 | 0 |

| | |
|---|---|
| 0 | 0 |
| 0 | 0 |
| 0 | 0 |
| 0 | 0 |
| 0 | 0 |
| 0 | 0 |
| 0 | 0 |
| 0 | 0 |
| 8.14299683 | 0 |
| 0 | 0 |
| 0 | 0 |
| 0 | 0 |
| 0 | 0 |
| 0 | 0 |
| 0 | 0 |
| 0 | 0 |
| 0 | 0 |
| 0 | 0 |
| 0 | 0 |
| 0 | 0 |
| 0 | 0 |
| 0 | 0 |
| 0 | 0 |
| 0 | 0 |
| 0 | 0 |
| 0 | 0 |
| 0 | 0 |
| 0 | 0 |
| 0 | 0 |
| 0 | 0 |
| 0 | 0 |
| 0 | 4.57598851 |
| 0 | 0 |
| 0 | 0 |
| 0 | 0 |
| 0 | 0 |
| 0 | 0 |
| 0 | 0 |
| 0 | 0 |
| 0 | 0 |
| 0 | 0 |
| 0 | 0 |
| 0 | 0 |

| | |
|---|---|
| 0 | 0 |
| 0 | 0 |
| 0 | 0 |
| 0 | 0 |
| 0 | 0 |
| 0 | 0 |
| 0 | 0 |
| 0 | 0 |
| 0 | 0 |
| 1351.73747 | 201.343494 |
| 0 | 0 |
| 0 | 0 |
| 0 | 0 |
| 282.290557 | 96.0957587 |
| 0 | 0 |
| 0 | 0 |
| 0 | 0 |
| 0 | 0 |
| 0 | 0 |
| 0 | 0 |
| 0 | 0 |
| 0 | 0 |
| 0 | 0 |
| 0 | 0 |
| 0 | 0 |
| 0 | 0 |
| 0 | 0 |
| 0 | 0 |
| 0 | 0 |
| 0 | 0 |
| 0 | 0 |
| 0 | 0 |
| 0 | 0 |
| 0 | 0 |
| 0 | 0 |
| 0 | 0 |
| 0 | 0 |
| 0 | 0 |
| 0 | 0 |
| 0 | 0 |
| 0 | 0 |
| 0 | 0 |
| 0 | 0 |

| | |
|---|---|
| 0 | 13.7279655 |
| 0 | 0 |
| 0 | 0 |
| 0 | 0 |
| 0 | 0 |
| 0 | 0 |
| 0 | 0 |
| 0 | 0 |
| 0 | 0 |
| 0 | 4.57598851 |
| 0 | 0 |
| 0 | 0 |
| 0 | 0 |
| 0 | 0 |
| 0 | 0 |
| 0 | 0 |
| 0 | 0 |
| 0 | 0 |
| 0 | 0 |
| 0 | 0 |
| 0 | 0 |
| 0 | 0 |
| 0 | 0 |
| 0 | 0 |
| 0 | 0 |
| 0 | 0 |
| 0 | 0 |
| 0 | 0 |
| 0 | 0 |
| 0 | 0 |
| 0 | 0 |
| 0 | 0 |
| 0 | 0 |
| 0 | 0 |
| 0 | 0 |
| 0 | 0 |
| 0 | 0 |
| 0 | 4.57598851 |
| 0 | 0 |
| 0 | 0 |
| 0 | 0 |
| 0 | 0 |
| 0 | 0 |

| | |
|---|---|
| 0 | 0 |
| 0 | 0 |
| 0 | 0 |
| 0 | 0 |
| 0 | 0 |
| 0 | 0 |
| 0 | 0 |
| 0 | 0 |
| 0 | 13.7279655 |
| 0 | 0 |
| 0 | 0 |
| 0 | 0 |
| 0 | 0 |
| 0 | 0 |
| 0 | 0 |
| 0 | 0 |
| 0 | 0 |
| 0 | 4.57598851 |
| 0 | 0 |
| 0 | 0 |
| 0 | 0 |
| 0 | 0 |
| 2.71433228 | 0 |
| 0 | 0 |
| 0 | 0 |
| 0 | 0 |
| 0 | 0 |
| 0 | 0 |
| 0 | 0 |
| 0 | 0 |
| 0 | 0 |
| 0 | 0 |
| 0 | 0 |
| 0 | 0 |
| 0 | 4.57598851 |
| 0 | 0 |
| 0 | 0 |
| 0 | 0 |
| 0 | 0 |
| 0 | 22.8799426 |
| 0 | 0 |
| 0 | 0 |
| 0 | 0 |

| | |
|---|---|
| 0 | 0 |
| 0 | 0 |
| 0 | 0 |
| 0 | 0 |
| 0 | 0 |
| 0 | 0 |
| 0 | 0 |
| 0 | 0 |
| 0 | 0 |
| 0 | 0 |
| 0 | 0 |
| 0 | 0 |
| 0 | 0 |
| 0 | 0 |
| 0 | 0 |
| 0 | 0 |
| 0 | 0 |
| 0 | 0 |
| 0 | 0 |
| 2.71433228 | 22.8799426 |
| 0 | 0 |
| 0 | 0 |
| 0 | 4.57598851 |
| 0 | 0 |
| 0 | 0 |
| 0 | 0 |
| 0 | 0 |
| 0 | 0 |
| 0 | 0 |
| 0 | 0 |
| 0 | 0 |
| 0 | 0 |
| 0 | 0 |
| 2.71433228 | 0 |
| 0 | 0 |
| 0 | 0 |
| 86.8586329 | 183.03954 |
| 0 | 0 |
| 0 | 0 |
| 0 | 0 |
| 0 | 0 |
| 0 | 0 |
| 0 | 0 |

| | |
|---|---|
| 0 | 0 |
| 0 | 0 |
| 0 | 0 |
| 0 | 0 |
| 0 | 0 |
| 0 | 0 |
| 0 | 0 |
| 0 | 0 |
| 0 | 0 |
| 0 | 0 |
| 0 | 0 |
| 0 | 0 |
| 0 | 0 |
| 0 | 0 |
| 0 | 0 |
| 0 | 0 |
| 0 | 0 |
| 0 | 0 |
| 0 | 0 |
| 0 | 0 |
| 0 | 0 |
| 0 | 0 |
| 0 | 0 |
| 0 | 0 |
| 0 | 0 |
| 0 | 0 |
| 0 | 0 |
| 0 | 0 |
| 0 | 0 |
| 0 | 0 |
| 0 | 0 |
| 0 | 0 |
| 5.42866456 | 27.4559311 |
| 0 | 0 |
| 0 | 0 |
| 0 | 0 |
| 0 | 0 |
| 0 | 0 |
| 0 | 0 |
| 0 | 0 |
| 0 | 0 |
| 0 | 0 |
| 0 | 0 |

| | |
|---|---|
| 0 | 0 |
| 369.14919 | 311.167219 |
| 0 | 0 |
| 0 | 0 |
| 0 | 0 |
| 0 | 0 |
| 0 | 4.57598851 |
| 0 | 4.57598851 |
| 0 | 0 |
| 0 | 0 |
| 0 | 0 |
| 0 | 0 |
| 0 | 0 |
| 0 | 0 |
| 0 | 0 |
| 0 | 0 |
| 0 | 0 |
| 0 | 4.57598851 |
| 0 | 0 |
| 0 | 0 |
| 0 | 0 |
| 0 | 0 |
| 0 | 0 |
| 0 | 0 |
| 0 | 0 |
| 0 | 0 |
| 0 | 0 |
| 0 | 0 |
| 0 | 0 |
| 0 | 0 |
| 0 | 0 |
| 0 | 0 |
| 0 | 0 |
| 0 | 0 |
| 0 | 0 |
| 0 | 0 |
| 0 | 0 |
| 0 | 4.57598851 |
| 0 | 0 |
| 0 | 0 |
| 0 | 0 |
| 28514.0606 | 3862.1343 |
| 0 | 0 |

| | |
|---|---|
| 0 | 0 |
| 0 | 0 |
| 0 | 0 |
| 0 | 0 |
| 0 | 0 |
| 0 | 0 |
| 0 | 0 |
| 0 | 0 |
| 0 | 0 |
| 0 | 0 |
| 0 | 0 |
| 0 | 0 |
| 0 | 0 |
| 0 | 0 |
| 0 | 0 |
| 0 | 0 |
| 0 | 0 |
| 0 | 0 |
| 0 | 0 |
| 0 | 0 |
| 0 | 0 |
| 0 | 0 |
| 0 | 0 |
| 0 | 0 |
| 0 | 0 |
| 0 | 0 |
| 0 | 0 |
| 0 | 0 |
| 0 | 0 |
| 312.148212 | 100.671747 |
| 0 | 0 |
| 0 | 0 |
| 0 | 0 |
| 0 | 0 |
| 0 | 0 |
| 0 | 0 |
| 0 | 0 |
| 0 | 0 |
| 0 | 0 |
| 0 | 0 |
| 0 | 0 |
| 0 | 0 |
| 0 | 0 |

| | |
|---|---|
| 0 | 0 |
| 0 | 0 |
| 0 | 0 |
| 0 | 0 |
| 86.8586329 | 132.703667 |
| 0 | 0 |
| 0 | 0 |
| 0 | 0 |
| 0 | 0 |
| 0 | 0 |
| 0 | 0 |
| 0 | 0 |
| 0 | 0 |
| 0 | 0 |
| 0 | 0 |
| 0 | 0 |
| 0 | 0 |
| 0 | 0 |
| 0 | 0 |
| 0 | 0 |
| 0 | 0 |
| 0 | 0 |
| 0 | 0 |
| 0 | 0 |
| 0 | 0 |
| 0 | 0 |
| 0 | 0 |
| 0 | 0 |
| 0 | 0 |
| 0 | 0 |
| 0 | 0 |
| 0 | 13.7279655 |
| 0 | 0 |
| 0 | 0 |
| 0 | 0 |
| 0 | 18.303954 |
| 0 | 0 |
| 0 | 0 |
| 57.0009778 | 118.975701 |
| 0 | 0 |
| 0 | 0 |
| 0 | 0 |
| 0 | 0 |

| | |
|---|---|
| 0 | 0 |
| 0 | 0 |
| 0 | 0 |
| 0 | 0 |
| 0 | 0 |
| 0 | 0 |
| 0 | 0 |
| 0 | 0 |
| 0 | 0 |
| 0 | 0 |
| 0 | 0 |
| 0 | 0 |
| 0 | 0 |
| 0 | 4.57598851 |
| 0 | 0 |
| 2.71433228 | 13.7279655 |
| 0 | 0 |
| 0 | 0 |
| 0 | 0 |
| 0 | 0 |
| 0 | 0 |
| 0 | 0 |
| 0 | 0 |
| 0 | 0 |
| 0 | 0 |
| 0 | 0 |
| 0 | 0 |
| 0 | 0 |
| 0 | 0 |
| 0 | 0 |
| 0 | 0 |
| 0 | 0 |
| 0 | 0 |
| 35.2863196 | 27.4559311 |
| 0 | 0 |
| 0 | 0 |
| 0 | 0 |
| 0 | 0 |
| 0 | 0 |
| 0 | 0 |
| 0 | 0 |
| 0 | 0 |
| 0 | 0 |

| | |
|---|---|
| 0 | 0 |
| 0 | 0 |
| 0 | 0 |
| 0 | 0 |
| 0 | 0 |
| 0 | 0 |
| 0 | 0 |
| 0 | 0 |
| 0 | 0 |
| 0 | 0 |
| 0 | 0 |
| 0 | 0 |
| 0 | 0 |
| 0 | 0 |
| 0 | 0 |
| 0 | 0 |
| 2.71433228 | 0 |
| 0 | 0 |
| 0 | 0 |
| 0 | 0 |
| 0 | 0 |
| 0 | 0 |
| 0 | 0 |
| 0 | 0 |
| 0 | 0 |
| 0 | 0 |
| 0 | 0 |
| 0 | 0 |
| 0 | 0 |
| 0 | 0 |
| 0 | 0 |
| 29.8576551 | 32.0319196 |
| 0 | 0 |
| 0 | 0 |
| 0 | 0 |
| 0 | 0 |
| 0 | 0 |
| 0 | 0 |
| 0 | 0 |
| 0 | 0 |
| 2.71433228 | 0 |
| 0 | 0 |
| 0 | 0 |

| | |
|---|---|
| 0 | 0 |
| 0 | 0 |
| 0 | 0 |
| 0 | 0 |
| 10.8573291 | 22.8799426 |
| 0 | 0 |
| 0 | 0 |
| 0 | 4.57598851 |
| 0 | 0 |
| 0 | 0 |
| 0 | 0 |
| 0 | 4.57598851 |
| 0 | 0 |
| 0 | 0 |
| 0 | 0 |
| 0 | 18.303954 |
| 0 | 0 |
| 0 | 0 |
| 0 | 0 |
| 0 | 0 |
| 0 | 0 |
| 0 | 0 |
| 0 | 0 |
| 0 | 0 |
| 0 | 0 |
| 0 | 0 |
| 0 | 0 |
| 0 | 0 |
| 0 | 0 |
| 0 | 0 |
| 0 | 0 |
| 0 | 0 |
| 0 | 0 |
| 0 | 0 |
| 0 | 0 |
| 0 | 0 |
| 0 | 0 |
| 0 | 0 |
| 0 | 0 |
| 0 | 0 |
| 0 | 0 |
| 0 | 0 |
| 0 | 0 |
| 0 | 0 |

| | |
|---|---|
| 0 | 0 |
| 0 | 0 |
| 0 | 0 |
| 0 | 0 |
| 0 | 0 |
| 0 | 0 |
| 0 | 0 |
| 0 | 0 |
| 0 | 0 |
| 0 | 0 |
| 0 | 0 |
| 13.5716614 | 18.303954 |
| 0 | 0 |
| 0 | 0 |
| 0 | 0 |
| 0 | 0 |
| 0 | 0 |
| 0 | 0 |
| 0 | 0 |
| 0 | 0 |
| 2.71433228 | 9.15197702 |
| 2.71433228 | 4.57598851 |
| 0 | 0 |
| 0 | 0 |
| 0 | 0 |
| 0 | 0 |
| 0 | 9.15197702 |
| 0 | 0 |
| 0 | 0 |
| 0 | 0 |
| 0 | 4.57598851 |
| 0 | 0 |
| 0 | 4.57598851 |
| 0 | 0 |
| 0 | 0 |
| 0 | 0 |
| 0 | 0 |
| 0 | 0 |
| 0 | 0 |
| 0 | 0 |
| 0 | 0 |
| 0 | 0 |
| 0 | 0 |

| | |
|---|---|
| 0 | 0 |
| 0 | 0 |
| 0 | 0 |
| 0 | 0 |
| 0 | 0 |
| 0 | 0 |
| 0 | 0 |
| 0 | 0 |
| 0 | 0 |
| 0 | 0 |
| 0 | 4.57598851 |
| 0 | 0 |
| 0 | 0 |
| 0 | 0 |
| 0 | 0 |
| 0 | 0 |
| 0 | 4.57598851 |
| 0 | 0 |
| 0 | 0 |
| 2.71433228 | 0 |
| 0 | 0 |
| 0 | 0 |
| 0 | 0 |
| 0 | 0 |
| 0 | 0 |
| 0 | 0 |
| 0 | 0 |
| 0 | 0 |
| 2.71433228 | 0 |
| 0 | 0 |
| 0 | 0 |
| 0 | 0 |
| 0 | 0 |
| 0 | 0 |
| 0 | 13.7279655 |
| 0 | 0 |
| 0 | 4.57598851 |
| 0 | 0 |
| 0 | 0 |
| 0 | 0 |
| 0 | 0 |
| 0 | 0 |
| 0 | 0 |

| | |
|---|---|
| 0 | 0 |
| 0 | 0 |
| 0 | 0 |
| 0 | 0 |
| 0 | 0 |
| 0 | 0 |
| 0 | 0 |
| 0 | 0 |
| 0 | 0 |
| 0 | 0 |
| 0 | 0 |
| 0 | 0 |
| 0 | 0 |
| 0 | 0 |
| 0 | 0 |
| 0 | 0 |
| 0 | 0 |
| 0 | 0 |
| 0 | 0 |
| 24.4289905 | 50.3358736 |
| 32.5719873 | 0 |
| 2.71433228 | 0 |
| 0 | 18.303954 |
| 0 | 0 |
| 0 | 0 |
| 0 | 0 |
| 0 | 0 |
| 0 | 0 |
| 0 | 0 |
| 0 | 0 |
| 0 | 0 |
| 0 | 0 |
| 0 | 0 |
| 0 | 0 |
| 0 | 0 |
| 0 | 0 |
| 0 | 0 |
| 0 | 0 |
| 0 | 0 |
| 0 | 0 |
| 0 | 0 |
| 0 | 0 |
| 0 | 0 |

| | |
|---|---|
| 0 | 0 |
| 0 | 4.57598851 |
| 0 | 0 |
| 0 | 0 |
| 0 | 0 |
| 0 | 0 |
| 0 | 0 |
| 0 | 0 |
| 0 | 0 |
| 0 | 0 |
| 0 | 0 |
| 0 | 0 |
| 0 | 0 |
| 0 | 0 |
| 0 | 0 |
| 0 | 0 |
| 0 | 0 |
| 0 | 0 |
| 0 | 0 |
| 0 | 0 |
| 0 | 0 |
| 0 | 0 |
| 0 | 0 |
| 0 | 0 |
| 0 | 0 |
| 0 | 0 |
| 0 | 0 |
| 0 | 0 |
| 0 | 0 |
| 0 | 0 |
| 2.71433228 | 4.57598851 |
| 0 | 0 |
| 2.71433228 | 13.7279655 |
| 0 | 0 |
| 0 | 0 |
| 0 | 0 |
| 0 | 0 |
| 2.71433228 | 0 |
| 0 | 0 |
| 0 | 0 |
| 0 | 0 |
| 59.7153101 | 77.7918047 |
| 0 | 0 |

| | |
|---|---|
| 0 | 0 |
| 0 | 0 |
| 0 | 0 |
| 0 | 0 |
| 0 | 0 |
| 0 | 9.15197702 |
| 0 | 0 |
| 0 | 0 |
| 0 | 0 |
| 0 | 0 |
| 0 | 0 |
| 0 | 0 |
| 0 | 0 |
| 27.1433228 | 77.7918047 |
| 0 | 0 |
| 0 | 0 |
| 0 | 0 |
| 0 | 0 |
| 0 | 0 |
| 0 | 0 |
| 0 | 0 |
| 0 | 0 |
| 0 | 0 |
| 0 | 0 |
| 0 | 0 |
| 0 | 0 |
| 0 | 0 |
| 0 | 0 |
| 0 | 4.57598851 |
| 0 | 0 |
| 0 | 0 |
| 0 | 0 |
| 0 | 0 |
| 0 | 0 |
| 0 | 0 |
| 0 | 0 |
| 0 | 0 |
| 0 | 0 |
| 0 | 0 |
| 0 | 0 |
| 0 | 4.57598851 |
| 0 | 0 |
| 0 | 0 |

| | |
|---|---|
| 0 | 0 |
| 0 | 0 |
| 133.002282 | 141.855644 |
| 0 | 0 |
| 2.71433228 | 0 |
| 0 | 0 |
| 0 | 0 |
| 0 | 0 |
| 0 | 0 |
| 0 | 0 |
| 0 | 0 |
| 0 | 0 |
| 0 | 0 |
| 0 | 4.57598851 |
| 0 | 0 |
| 0 | 0 |
| 0 | 0 |
| 0 | 0 |
| 0 | 0 |
| 0 | 0 |
| 0 | 0 |
| 0 | 0 |
| 0 | 0 |
| 0 | 0 |
| 0 | 0 |
| 0 | 0 |
| 0 | 0 |
| 0 | 0 |
| 0 | 0 |
| 0 | 0 |
| 0 | 0 |
| 0 | 0 |
| 0 | 0 |
| 0 | 0 |
| 0 | 0 |
| 0 | 13.7279655 |
| 0 | 0 |
| 0 | 0 |
| 0 | 0 |
| 0 | 0 |
| 0 | 0 |
| 0 | 0 |
| 0 | 0 |

| | |
|---|---|
| 0 | 0 |
| 0 | 0 |
| 0 | 0 |
| 0 | 0 |
| 0 | 0 |
| 0 | 0 |
| 0 | 0 |
| 0 | 0 |
| 0 | 0 |
| 0 | 0 |
| 0 | 0 |
| 0 | 0 |
| 0 | 0 |
| 0 | 0 |
| 0 | 0 |
| 0 | 0 |
| 0 | 0 |
| 0 | 4.57598851 |
| 0 | 0 |
| 119.43062 | 45.7598851 |
| 0 | 0 |
| 0 | 0 |
| 0 | 0 |
| 0 | 0 |
| 0 | 0 |
| 0 | 13.7279655 |
| 0 | 0 |
| 0 | 0 |
| 0 | 0 |
| 0 | 0 |
| 0 | 0 |
| 13.5716614 | 269.983322 |
| 0 | 0 |
| 0 | 0 |
| 0 | 0 |
| 0 | 0 |
| 0 | 0 |
| 0 | 0 |
| 0 | 0 |
| 0 | 0 |
| 0 | 0 |
| 0 | 0 |
| 0 | 0 |

| | |
|---|---|
| 0 | 0 |
| 0 | 0 |
| 0 | 0 |
| 0 | 0 |
| 290.433554 | 215.07146 |
| 0 | 0 |
| 130.287949 | 45.7598851 |
| 0 | 0 |
| 0 | 0 |
| 0 | 0 |
| 5.42866456 | 0 |
| 0 | 0 |
| 0 | 0 |
| 0 | 0 |
| 0 | 0 |
| 0 | 0 |
| 0 | 0 |
| 0 | 0 |
| 0 | 0 |
| 0 | 0 |
| 0 | 0 |
| 0 | 0 |
| 0 | 0 |
| 0 | 0 |
| 0 | 0 |
| 0 | 0 |
| 0 | 0 |
| 0 | 0 |
| 0 | 0 |
| 0 | 0 |
| 0 | 0 |
| 0 | 0 |
| 0 | 0 |
| 0 | 0 |
| 0 | 0 |
| 0 | 0 |
| 0 | 0 |
| 0 | 0 |
| 0 | 0 |
| 0 | 0 |
| 0 | 0 |
| 0 | 0 |
| 0 | 0 |

| | |
|---|---|
| 0 | 0 |
| 0 | 0 |
| 0 | 0 |
| 0 | 0 |
| 0 | 0 |
| 0 | 0 |
| 0 | 0 |
| 0 | 0 |
| 0 | 0 |
| 0 | 0 |
| 0 | 0 |
| 0 | 0 |
| 0 | 0 |
| 0 | 0 |
| 48.857981 | 22.8799426 |
| 0 | 0 |
| 0 | 0 |
| 0 | 0 |
| 0 | 0 |
| 0 | 0 |
| 0 | 0 |
| 0 | 0 |
| 0 | 0 |
| 0 | 0 |
| 0 | 0 |
| 0 | 0 |
| 51.5723133 | 13.7279655 |
| 570.009778 | 36.6079081 |
| 0 | 0 |
| 0 | 0 |
| 84.1443006 | 45.7598851 |
| 0 | 0 |
| 0 | 0 |
| 0 | 0 |
| 0 | 0 |
| 0 | 0 |
| 0 | 0 |
| 0 | 0 |
| 0 | 0 |
| 0 | 0 |
| 0 | 0 |
| 0 | 0 |
| 0 | 0 |

| | |
|---|---|
| 0 | 0 |
| 0 | 0 |
| 0 | 0 |
| 0 | 0 |
| 0 | 0 |
| 0 | 0 |
| 0 | 0 |
| 0 | 0 |
| 0 | 0 |
| 0 | 0 |
| 0 | 0 |
| 0 | 0 |
| 0 | 0 |
| 0 | 0 |
| 0 | 0 |
| 0 | 0 |
| 8.14299683 | 4.57598851 |
| 2.71433228 | 0 |
| 504.865804 | 347.775127 |
| 0 | 0 |
| 0 | 0 |
| 0 | 0 |
| 0 | 0 |
| 0 | 0 |
| 0 | 0 |
| 0 | 0 |
| 0 | 0 |
| 0 | 0 |
| 0 | 0 |
| 0 | 0 |
| 0 | 0 |
| 103.144627 | 18.303954 |
| 0 | 41.1838966 |
| 589.010104 | 439.294897 |
| 0 | 0 |
| 0 | 0 |
| 0 | 0 |
| 0 | 0 |
| 0 | 0 |
| 0 | 0 |
| 0 | 0 |
| 0 | 0 |
| 0 | 0 |

| | |
|---|---|
| 0 | 0 |
| 0 | 0 |
| 0 | 0 |
| 0 | 0 |
| 0 | 0 |
| 0 | 9.15197702 |
| 0 | 0 |
| 0 | 0 |
| 0 | 0 |
| 0 | 0 |
| 0 | 0 |
| 67.8583069 | 18.303954 |
| 0 | 0 |
| 0 | 0 |
| 0 | 0 |
| 0 | 13.7279655 |
| 0 | 0 |
| 0 | 0 |
| 0 | 0 |
| 0 | 0 |
| 0 | 0 |
| 0 | 0 |
| 0 | 0 |
| 0 | 0 |
| 0 | 0 |
| 0 | 0 |
| 0 | 0 |
| 0 | 0 |
| 0 | 0 |
| 0 | 0 |
| 0 | 0 |
| 0 | 0 |
| 0 | 0 |
| 0 | 0 |
| 0 | 0 |
| 0 | 0 |
| 0 | 0 |
| 0 | 0 |
| 0 | 0 |
| 0 | 0 |
| 0 | 0 |
| 0 | 0 |
| 2.71433228 | 0 |

| | |
|---|---|
| 0 | 0 |
| 0 | 0 |
| 0 | 0 |
| 0 | 0 |
| 0 | 0 |
| 0 | 0 |
| 0 | 0 |
| 0 | 4.57598851 |
| 0 | 0 |
| 0 | 0 |
| 0 | 0 |
| 0 | 0 |
| 0 | 0 |
| 0 | 0 |
| 0 | 0 |
| 0 | 0 |
| 92.2872974 | 9.15197702 |
| 0 | 0 |
| 0 | 0 |
| 0 | 0 |
| 0 | 0 |
| 0 | 0 |
| 0 | 0 |
| 0 | 0 |
| 0 | 0 |
| 0 | 0 |
| 0 | 0 |
| 0 | 0 |
| 0 | 0 |
| 0 | 0 |
| 0 | 0 |
| 2.71433228 | 0 |
| 0 | 0 |
| 0 | 0 |
| 0 | 0 |
| 0 | 0 |
| 0 | 4.57598851 |
| 0 | 0 |
| 0 | 0 |
| 54.2866456 | 311.167219 |
| 0 | 0 |
| 146.573943 | 64.0638392 |
| 0 | 0 |

| | |
|---|---|
| 0 | 0 |
| 0 | 0 |
| 0 | 0 |
| 0 | 0 |
| 2.71433228 | 0 |
| 0 | 0 |
| 0 | 0 |
| 0 | 0 |
| 0 | 0 |
| 0 | 0 |
| 2.71433228 | 0 |
| 0 | 0 |
| 0 | 0 |
| 0 | 0 |
| 0 | 0 |
| 0 | 0 |
| 0 | 0 |
| 0 | 0 |
| 38.0006519 | 4.57598851 |
| 0 | 0 |
| 0 | 4.57598851 |
| 0 | 0 |
| 0 | 0 |
| 0 | 0 |
| 0 | 0 |
| 0 | 0 |
| 13.5716614 | 22.8799426 |
| 0 | 0 |
| 0 | 0 |
| 0 | 0 |
| 0 | 4.57598851 |
| 0 | 0 |
| 0 | 0 |
| 0 | 0 |
| 0 | 0 |
| 0 | 0 |
| 0 | 0 |
| 0 | 0 |
| 0 | 0 |
| 0 | 0 |
| 0 | 0 |
| 0 | 0 |
| 0 | 0 |

| | |
|---|---|
| 0 | 0 |
| 0 | 0 |
| 0 | 0 |
| 0 | 0 |
| 0 | 0 |
| 0 | 4.57598851 |
| 0 | 0 |
| 0 | 0 |
| 0 | 0 |
| 0 | 0 |
| 0 | 4.57598851 |
| 0 | 0 |
| 0 | 0 |
| 0 | 0 |
| 0 | 0 |
| 0 | 0 |
| 0 | 0 |
| 0 | 0 |
| 29.8576551 | 18.303954 |
| 0 | 0 |
| 2.71433228 | 0 |
| 0 | 0 |
| 0 | 0 |
| 8.14299683 | 9.15197702 |
| 0 | 0 |
| 0 | 0 |
| 0 | 0 |
| 0 | 0 |
| 0 | 4.57598851 |
| 0 | 0 |
| 0 | 0 |
| 355.577528 | 375.231058 |
| 0 | 0 |
| 0 | 0 |
| 0 | 0 |
| 0 | 0 |
| 2.71433228 | 9.15197702 |
| 0 | 9.15197702 |
| 0 | 0 |
| 0 | 0 |
| 0 | 0 |
| 0 | 0 |
| 0 | 0 |

| | |
|---|---|
| 0 | 0 |
| 0 | 0 |
| 0 | 0 |
| 0 | 0 |
| 0 | 0 |
| 0 | 0 |
| 0 | 0 |
| 0 | 0 |
| 0 | 0 |
| 0 | 0 |
| 0 | 0 |
| 0 | 0 |
| 0 | 0 |
| 0 | 0 |
| 0 | 0 |
| 0 | 0 |
| 0 | 0 |
| 0 | 0 |
| 2.71433228 | 0 |
| 0 | 0 |
| 0 | 0 |
| 0 | 0 |
| 2.71433228 | 0 |
| 0 | 0 |
| 0 | 0 |
| 0 | 18.303954 |
| 0 | 0 |
| 0 | 0 |
| 0 | 0 |
| 0 | 0 |
| 0 | 0 |
| 0 | 0 |
| 0 | 0 |
| 0 | 0 |
| 0 | 0 |
| 0 | 0 |
| 0 | 0 |
| 0 | 0 |
| 0 | 0 |
| 0 | 0 |
| 0 | 0 |
| 0 | 0 |
| 0 | 0 |

| | |
|---|---|
| 0 | 0 |
| 0 | 0 |
| 0 | 0 |
| 0 | 0 |
| 0 | 4.57598851 |
| 0 | 0 |
| 0 | 0 |
| 0 | 0 |
| 0 | 0 |
| 0 | 0 |
| 0 | 0 |
| 0 | 0 |
| 0 | 0 |
| 0 | 0 |
| 0 | 0 |
| 0 | 0 |
| 0 | 0 |
| 0 | 0 |
| 0 | 4.57598851 |
| 0 | 0 |
| 0 | 0 |
| 0 | 0 |
| 0 | 0 |
| 0 | 0 |
| 0 | 0 |
| 0 | 0 |
| 0 | 0 |
| 0 | 0 |
| 0 | 0 |
| 0 | 0 |
| 0 | 0 |
| 2.71433228 | 0 |
| 0 | 0 |
| 0 | 0 |
| 0 | 0 |
| 0 | 0 |
| 0 | 0 |
| 0 | 4.57598851 |
| 0 | 0 |
| 0 | 0 |
| 0 | 0 |
| 0 | 0 |
| 0 | 0 |

| | |
|---|---|
| 0 | 0 |
| 0 | 0 |
| 0 | 0 |
| 0 | 0 |
| 0 | 9.15197702 |
| 0 | 0 |
| 0 | 0 |
| 0 | 0 |
| 0 | 0 |
| 0 | 0 |
| 0 | 0 |
| 0 | 0 |
| 0 | 0 |
| 0 | 0 |
| 0 | 0 |
| 0 | 0 |
| 0 | 0 |
| 0 | 0 |
| 0 | 0 |
| 0 | 0 |
| 0 | 4.57598851 |
| 0 | 0 |
| 0 | 0 |
| 0 | 0 |
| 0 | 0 |
| 0 | 0 |
| 0 | 0 |
| 0 | 0 |
| 0 | 0 |
| 0 | 0 |
| 0 | 4.57598851 |
| 0 | 0 |
| 0 | 0 |
| 0 | 0 |
| 0 | 0 |
| 0 | 0 |
| 0 | 0 |
| 0 | 0 |
| 0 | 0 |
| 0 | 0 |
| 0 | 0 |
| 0 | 0 |
| 0 | 0 |

| | |
|---|---|
| 0 | 0 |
| 0 | 0 |
| 0 | 0 |
| 0 | 0 |
| 0 | 0 |
| 1007.01728 | 741.310139 |
| 2.71433228 | 0 |
| 0 | 0 |
| 0 | 0 |
| 0 | 0 |
| 0 | 0 |
| 0 | 0 |
| 2.71433228 | 32.0319196 |
| 0 | 0 |
| 0 | 0 |
| 0 | 0 |
| 0 | 0 |
| 0 | 0 |
| 0 | 0 |
| 0 | 0 |
| 0 | 4.57598851 |
| 24.4289905 | 4.57598851 |
| 0 | 0 |
| 0 | 0 |
| 0 | 0 |
| 0 | 0 |
| 0 | 0 |
| 0 | 0 |
| 0 | 0 |
| 0 | 0 |
| 0 | 0 |
| 0 | 0 |
| 0 | 0 |
| 0 | 0 |
| 0 | 0 |
| 0 | 0 |
| 0 | 0 |
| 0 | 0 |
| 0 | 0 |
| 0 | 0 |
| 0 | 0 |
| 0 | 0 |
| 0 | 0 |

| | |
|---|---|
| 0 | 0 |
| 0 | 0 |
| 0 | 0 |
| 0 | 0 |
| 0 | 0 |
| 0 | 0 |
| 0 | 0 |
| 0 | 0 |
| 0 | 0 |
| 0 | 0 |
| 0 | 0 |
| 0 | 0 |
| 0 | 0 |
| 0 | 0 |
| 0 | 0 |
| 0 | 0 |
| 0 | 0 |
| 0 | 0 |
| 0 | 0 |
| 19.0003259 | 0 |
| 0 | 0 |
| 0 | 0 |
| 0 | 0 |
| 0 | 0 |
| 0 | 0 |
| 0 | 0 |
| 0 | 0 |
| 0 | 0 |
| 0 | 0 |
| 0 | 0 |
| 0 | 0 |
| 0 | 0 |
| 0 | 0 |
| 0 | 0 |
| 0 | 0 |
| 0 | 0 |
| 0 | 0 |
| 0 | 0 |
| 0 | 0 |
| 0 | 0 |
| 86.8586329 | 169.311575 |
| 5.42866456 | 0 |
| 0 | 0 |

| | |
|---|---|
| 0 | 0 |
| 0 | 0 |
| 0 | 4.57598851 |
| 0 | 0 |
| 0 | 0 |
| 0 | 0 |
| 0 | 4.57598851 |
| 2.71433228 | 13.7279655 |
| 0 | 13.7279655 |
| 0 | 0 |
| 0 | 0 |
| 0 | 0 |
| 0 | 0 |
| 0 | 0 |
| 0 | 0 |
| 0 | 0 |
| 0 | 0 |
| 0 | 0 |
| 0 | 0 |
| 0 | 0 |
| 0 | 18.303954 |
| 0 | 0 |
| 0 | 0 |
| 0 | 0 |
| 0 | 0 |
| 0 | 0 |
| 0 | 0 |
| 0 | 0 |
| 0 | 0 |
| 0 | 0 |
| 0 | 0 |
| 0 | 0 |
| 0 | 0 |
| 0 | 0 |
| 0 | 0 |
| 0 | 0 |
| 0 | 0 |
| 0 | 0 |
| 0 | 0 |
| 0 | 0 |
| 0 | 0 |
| 0 | 0 |
| 0 | 0 |

| | |
|---|---|
| 0 | 0 |
| 0 | 0 |
| 0 | 0 |
| 0 | 0 |
| 0 | 0 |
| 0 | 0 |
| 0 | 0 |
| 0 | 0 |
| 0 | 0 |
| 0 | 0 |
| 0 | 0 |
| 0 | 0 |
| 0 | 0 |
| 0 | 0 |
| 0 | 0 |
| 0 | 0 |
| 0 | 0 |
| 0 | 0 |
| 0 | 0 |
| 0 | 0 |
| 0 | 0 |
| 0 | 0 |
| 0 | 0 |
| 0 | 0 |
| 0 | 0 |
| 0 | 0 |
| 0 | 0 |
| 0 | 0 |
| 0 | 0 |
| 0 | 0 |
| 0 | 0 |
| 0 | 0 |
| 0 | 4.57598851 |
| 0 | 0 |
| 0 | 0 |
| 0 | 0 |
| 0 | 0 |
| 0 | 4.57598851 |
| 0 | 4.57598851 |
| 0 | 0 |
| 0 | 4.57598851 |
| 0 | 0 |
| 0 | 0 |

| | |
|---|---|
| 0 | 0 |
| 0 | 0 |
| 0 | 0 |
| 0 | 0 |
| 0 | 0 |
| 0 | 0 |
| 0 | 0 |
| 0 | 0 |
| 0 | 4.57598851 |
| 0 | 0 |
| 0 | 0 |
| 0 | 0 |
| 0 | 0 |
| 0 | 0 |
| 0 | 0 |
| 0 | 0 |
| 0 | 0 |
| 0 | 0 |
| 0 | 0 |
| 0 | 0 |
| 0 | 0 |
| 0 | 0 |
| 0 | 0 |
| 184.574595 | 109.823724 |
| 0 | 0 |
| 0 | 0 |
| 0 | 0 |
| 0 | 0 |
| 0 | 0 |
| 0 | 0 |
| 0 | 0 |
| 0 | 0 |
| 0 | 0 |
| 0 | 0 |
| 0 | 0 |
| 0 | 0 |
| 0 | 0 |
| 0 | 0 |
| 0 | 0 |
| 0 | 0 |
| 0 | 0 |
| 0 | 0 |
| 0 | 0 |

| | |
|---|---|
| 0 | 0 |
| 0 | 0 |
| 0 | 0 |
| 0 | 0 |
| 0 | 0 |
| 0 | 0 |
| 0 | 9.15197702 |
| 0 | 0 |
| 0 | 0 |
| 0 | 0 |
| 0 | 0 |
| 0 | 0 |
| 0 | 0 |
| 0 | 0 |
| 0 | 0 |
| 0 | 0 |
| 0 | 0 |
| 0 | 0 |
| 0 | 0 |
| 0 | 0 |
| 13.5716614 | 9.15197702 |
| 2.71433228 | 0 |
| 0 | 0 |
| 0 | 0 |
| 0 | 0 |
| 0 | 0 |
| 2.71433228 | 0 |
| 0 | 0 |
| 0 | 0 |
| 0 | 0 |
| 0 | 0 |
| 2.71433228 | 4.57598851 |
| 0 | 0 |
| 0 | 0 |
| 0 | 0 |
| 0 | 0 |
| 0 | 0 |
| 0 | 0 |
| 0 | 0 |
| 0 | 0 |
| 0 | 0 |
| 0 | 0 |
| 0 | 0 |

| | |
|---|---|
| 0 | 0 |
| 0 | 0 |
| 0 | 0 |
| 0 | 0 |
| 0 | 0 |
| 0 | 0 |
| 0 | 0 |
| 0 | 0 |
| 0 | 0 |
| 0 | 0 |
| 0 | 0 |
| 0 | 0 |
| 0 | 0 |
| 0 | 0 |
| 0 | 0 |
| 0 | 0 |
| 2939.62186 | 2141.56262 |
| 0 | 0 |
| 0 | 27.4559311 |
| 0 | 0 |
| 0 | 0 |
| 0 | 4.57598851 |
| 0 | 0 |
| 0 | 4.57598851 |
| 0 | 0 |
| 0 | 0 |
| 0 | 0 |
| 0 | 0 |
| 0 | 0 |
| 0 | 0 |
| 0 | 0 |
| 0 | 0 |
| 0 | 0 |
| 0 | 0 |
| 0 | 0 |
| 0 | 0 |
| 0 | 0 |
| 65.1439747 | 45.7598851 |
| 0 | 0 |
| 0 | 0 |
| 0 | 0 |
| 0 | 0 |
| 0 | 9.15197702 |

| | |
|---|---|
| 0 | 0 |
| 0 | 0 |
| 0 | 0 |
| 0 | 0 |
| 0 | 0 |
| 0 | 0 |
| 0 | 0 |
| 0 | 0 |
| 0 | 32.0319196 |
| 0 | 0 |
| 0 | 0 |
| 0 | 0 |
| 0 | 9.15197702 |
| 0 | 0 |
| 5.42866456 | 0 |
| 0 | 0 |
| 0 | 0 |
| 0 | 0 |
| 0 | 0 |
| 0 | 0 |
| 0 | 0 |
| 0 | 0 |
| 0 | 4.57598851 |
| 0 | 0 |
| 0 | 0 |
| 0 | 0 |
| 0 | 0 |
| 0 | 0 |
| 0 | 0 |
| 0 | 0 |
| 0 | 0 |
| 0 | 0 |
| 2.71433228 | 0 |
| 0 | 0 |
| 0 | 0 |
| 0 | 0 |
| 0 | 0 |
| 0 | 0 |
| 0 | 0 |
| 0 | 0 |
| 0 | 0 |
| 0 | 0 |
| 0 | 0 |

| | |
|---|---|
| 0 | 0 |
| 0 | 0 |
| 0 | 0 |
| 0 | 4.57598851 |
| 0 | 0 |
| 0 | 0 |
| 0 | 0 |
| 0 | 0 |
| 0 | 0 |
| 0 | 0 |
| 0 | 27.4559311 |
| 0 | 0 |
| 0 | 0 |
| 0 | 0 |
| 0 | 0 |
| 0 | 0 |
| 0 | 4.57598851 |
| 0 | 0 |
| 0 | 0 |
| 0 | 0 |
| 0 | 0 |
| 0 | 0 |
| 0 | 0 |
| 0 | 0 |
| 0 | 0 |
| 0 | 0 |
| 0 | 0 |
| 0 | 0 |
| 0 | 0 |
| 0 | 0 |
| 0 | 0 |
| 2.71433228 | 0 |
| 0 | 0 |
| 0 | 0 |
| 0 | 0 |
| 0 | 0 |
| 0 | 0 |
| 0 | 0 |
| 0 | 0 |
| 0 | 0 |
| 0 | 0 |
| 0 | 0 |
| 0 | 0 |

| | |
|---|---|
| 0 | 0 |
| 0 | 0 |
| 0 | 0 |
| 0 | 0 |
| 0 | 0 |
| 0 | 0 |
| 0 | 0 |
| 0 | 0 |
| 0 | 0 |
| 0 | 0 |
| 0 | 0 |
| 0 | 0 |
| 0 | 0 |
| 0 | 0 |
| 0 | 0 |
| 0 | 0 |
| 0 | 0 |
| 0 | 0 |
| 0 | 0 |
| 0 | 0 |
| 0 | 0 |
| 0 | 0 |
| 0 | 0 |
| 0 | 0 |
| 0 | 0 |
| 0 | 0 |
| 0 | 0 |
| 0 | 0 |
| 0 | 0 |
| 0 | 0 |
| 0 | 0 |
| 0 | 0 |
| 0 | 0 |
| 0 | 0 |
| 0 | 0 |
| 0 | 0 |
| 0 | 0 |
| 0 | 0 |
| 2.71433228 | 0 |
| 0 | 0 |
| 0 | 0 |
| 0 | 4.57598851 |
| 0 | 0 |

| | |
|---|---|
| 0 | 0 |
| 4394.50396 | 361.503092 |
| 0 | 0 |
| 19.0003259 | 22.8799426 |
| 0 | 22.8799426 |
| 73.2869715 | 118.975701 |
| 0 | 0 |
| 0 | 0 |
| 0 | 0 |
| 0 | 0 |
| 5.42866456 | 27.4559311 |
| 0 | 0 |
| 0 | 0 |
| 0 | 0 |
| 0 | 0 |
| 0 | 0 |
| 0 | 0 |
| 0 | 0 |
| 0 | 0 |
| 0 | 0 |
| 0 | 0 |
| 0 | 0 |
| 0 | 0 |
| 0 | 0 |
| 0 | 0 |
| 0 | 0 |
| 0 | 4.57598851 |
| 0 | 0 |
| 0 | 0 |
| 0 | 0 |
| 0 | 0 |
| 0 | 0 |
| 0 | 0 |
| 0 | 0 |
| 0 | 0 |
| 0 | 0 |
| 0 | 0 |
| 0 | 0 |
| 0 | 0 |
| 0 | 0 |
| 0 | 18.303954 |
| 203.574921 | 96.0957587 |
| 184.574595 | 105.247736 |

| | |
|---|---|
| 5.42866456 | 0 |
| 0 | 0 |
| 0 | 0 |
| 0 | 4.57598851 |
| 0 | 0 |
| 0 | 0 |
| 2.71433228 | 0 |
| 0 | 0 |
| 0 | 0 |
| 0 | 0 |
| 0 | 0 |
| 0 | 0 |
| 0 | 0 |
| 0 | 0 |
| 0 | 0 |
| 0 | 0 |
| 0 | 0 |
| 0 | 0 |
| 2.71433228 | 0 |
| 0 | 0 |
| 0 | 0 |
| 0 | 0 |
| 0 | 0 |
| 0 | 0 |
| 0 | 0 |
| 0 | 0 |
| 0 | 0 |
| 0 | 0 |
| 0 | 0 |
| 0 | 0 |
| 0 | 0 |
| 0 | 4.57598851 |
| 0 | 0 |
| 0 | 0 |
| 0 | 0 |
| 0 | 18.303954 |
| 0 | 0 |
| 0 | 0 |
| 0 | 0 |
| 0 | 0 |
| 0 | 0 |
| 0 | 0 |
| 0 | 13.7279655 |

| | |
|---|---|
| 0 | 0 |
| 0 | 0 |
| 0 | 0 |
| 0 | 0 |
| 0 | 0 |
| 0 | 0 |
| 0 | 0 |
| 0 | 0 |
| 1433.16744 | 375.231058 |
| 0 | 0 |
| 0 | 0 |
| 0 | 0 |
| 0 | 0 |
| 0 | 0 |
| 0 | 0 |
| 0 | 0 |
| 0 | 0 |
| 0 | 0 |
| 0 | 0 |
| 0 | 0 |
| 0 | 0 |
| 0 | 0 |
| 0 | 0 |
| 0 | 0 |
| 0 | 0 |
| 0 | 0 |
| 0 | 0 |
| 0 | 0 |
| 0 | 0 |
| 0 | 0 |
| 0 | 0 |
| 0 | 0 |
| 0 | 0 |
| 0 | 0 |
| 0 | 0 |
| 0 | 0 |
| 0 | 0 |
| 0 | 0 |
| 0 | 4.57598851 |
| 5.42866456 | 4.57598851 |
| 0 | 0 |
| 0 | 0 |

| | |
|---|---|
| 0 | 0 |
| 0 | 0 |
| 0 | 0 |
| 0 | 0 |
| 0 | 0 |
| 0 | 0 |
| 0 | 0 |
| 0 | 0 |
| 0 | 0 |
| 0 | 0 |
| 0 | 0 |
| 0 | 0 |
| 0 | 0 |
| 0 | 0 |
| 0 | 0 |
| 0 | 0 |
| 0 | 0 |
| 0 | 0 |
| 0 | 0 |
| 0 | 0 |
| 0 | 0 |
| 0 | 4.57598851 |
| 0 | 0 |
| 0 | 0 |
| 0 | 0 |
| 0 | 0 |
| 0 | 0 |
| 0 | 0 |
| 0 | 0 |
| 0 | 0 |
| 0 | 0 |
| 0 | 0 |
| 0 | 0 |
| 0 | 0 |
| 0 | 0 |
| 0 | 4.57598851 |
| 0 | 0 |
| 0 | 4.57598851 |
| 0 | 0 |
| 0 | 0 |
| 0 | 0 |
| 0 | 0 |
| 0 | 0 |

| | |
|---|---|
| 0 | 0 |
| 0 | 0 |
| 0 | 0 |
| 0 | 0 |
| 0 | 0 |
| 0 | 0 |
| 0 | 0 |
| 0 | 0 |
| 0 | 0 |
| 0 | 0 |
| 0 | 0 |
| 0 | 0 |
| 0 | 0 |
| 0 | 0 |
| 0 | 0 |
| 21.7146582 | 41.1838966 |
| 0 | 0 |
| 70.5726392 | 54.9118621 |
| 0 | 0 |
| 0 | 0 |
| 0 | 0 |
| 0 | 0 |
| 2.71433228 | 0 |
| 0 | 0 |
| 0 | 0 |
| 597.153101 | 256.255357 |
| 0 | 0 |
| 0 | 0 |
| 0 | 0 |
| 0 | 0 |
| 0 | 0 |
| 0 | 0 |
| 0 | 0 |
| 0 | 0 |
| 0 | 0 |
| 0 | 0 |
| 0 | 0 |
| 0 | 0 |
| 0 | 0 |
| 0 | 0 |
| 0 | 0 |
| 0 | 0 |
| 0 | 0 |

| | |
|---|---|
| 0 | 0 |
| 0 | 0 |
| 0 | 0 |
| 0 | 0 |
| 0 | 0 |
| 0 | 13.7279655 |
| 0 | 0 |
| 0 | 0 |
| 0 | 0 |
| 0 | 0 |
| 5.42866456 | 0 |
| 8.14299683 | 59.4878506 |
| 0 | 0 |
| 0 | 0 |
| 0 | 0 |
| 0 | 0 |
| 0 | 0 |
| 0 | 0 |
| 0 | 0 |
| 0 | 4.57598851 |
| 0 | 0 |
| 0 | 4.57598851 |
| 0 | 0 |
| 0 | 0 |
| 0 | 0 |
| 0 | 0 |
| 0 | 0 |
| 0 | 0 |
| 0 | 0 |
| 0 | 0 |
| 0 | 0 |
| 0 | 4.57598851 |
| 0 | 0 |
| 0 | 0 |
| 2.71433228 | 0 |
| 0 | 0 |
| 0 | 0 |
| 0 | 0 |
| 0 | 0 |
| 0 | 41.1838966 |
| 0 | 0 |
| 0 | 0 |
| 0 | 0 |

| | |
|---|---|
| 0 | 0 |
| 0 | 0 |
| 0 | 0 |
| 0 | 9.15197702 |
| 0 | 0 |
| 13.5716614 | 73.2158162 |
| 0 | 4.57598851 |
| 0 | 0 |
| 0 | 0 |
| 0 | 0 |
| 0 | 0 |
| 5.42866456 | 0 |
| 0 | 0 |
| 0 | 27.4559311 |
| 0 | 0 |
| 0 | 0 |
| 0 | 0 |
| 0 | 0 |
| 0 | 0 |
| 0 | 0 |
| 160.145604 | 224.223437 |
| 0 | 0 |
| 0 | 0 |
| 0 | 0 |
| 0 | 0 |
| 0 | 0 |
| 0 | 0 |
| 0 | 0 |
| 0 | 0 |
| 0 | 0 |
| 0 | 0 |
| 0 | 0 |
| 0 | 0 |
| 0 | 0 |
| 0 | 0 |
| 0 | 0 |
| 0 | 0 |
| 0 | 0 |
| 0 | 0 |
| 0 | 0 |
| 2.71433228 | 0 |
| 0 | 0 |
| 0 | 0 |

| | |
|---|---|
| 0 | 0 |
| 0 | 0 |
| 0 | 0 |
| 0 | 0 |
| 0 | 0 |
| 0 | 0 |
| 2.71433228 | 32.0319196 |
| 0 | 0 |
| 0 | 0 |
| 0 | 0 |
| 0 | 0 |
| 0 | 0 |
| 0 | 0 |
| 0 | 0 |
| 0 | 0 |
| 0 | 0 |
| 0 | 0 |
| 0 | 0 |
| 0 | 0 |
| 0 | 4.57598851 |
| 0 | 0 |
| 0 | 0 |
| 0 | 0 |
| 0 | 0 |
| 2.71433228 | 0 |
| 0 | 0 |
| 0 | 0 |
| 0 | 0 |
| 0 | 0 |
| 0 | 0 |
| 0 | 0 |
| 0 | 0 |
| 0 | 0 |
| 0 | 0 |
| 0 | 0 |
| 0 | 0 |
| 0 | 0 |
| 0 | 0 |
| 0 | 0 |
| 0 | 0 |
| 0 | 0 |
| 252.432902 | 237.951403 |
| 0 | 0 |

| | |
|---|---|
| 0 | 0 |
| 0 | 0 |
| 0 | 0 |
| 0 | 0 |
| 0 | 0 |
| 0 | 0 |
| 0 | 4.57598851 |
| 0 | 0 |
| 0 | 0 |
| 0 | 4.57598851 |
| 0 | 4.57598851 |
| 10.8573291 | 0 |
| 0 | 0 |
| 0 | 0 |
| 0 | 0 |
| 0 | 0 |
| 0 | 0 |
| 0 | 0 |
| 0 | 0 |
| 2.71433228 | 4.57598851 |
| 111.287623 | 123.55169 |
| 0 | 0 |
| 0 | 0 |
| 0 | 0 |
| 0 | 0 |
| 0 | 0 |
| 0 | 0 |
| 0 | 0 |
| 2.71433228 | 0 |
| 0 | 0 |
| 0 | 0 |
| 0 | 0 |
| 0 | 0 |
| 0 | 0 |
| 0 | 0 |
| 0 | 0 |
| 0 | 0 |
| 0 | 0 |
| 0 | 0 |
| 0 | 0 |
| 0 | 0 |
| 0 | 0 |
| 0 | 0 |

| | |
|---|---|
| 0 | 0 |
| 0 | 0 |
| 0 | 0 |
| 0 | 0 |
| 0 | 0 |
| 0 | 0 |
| 0 | 0 |
| 0 | 0 |
| 0 | 0 |
| 0 | 4.57598851 |
| 0 | 0 |
| 0 | 0 |
| 0 | 0 |
| 0 | 0 |
| 0 | 0 |
| 0 | 0 |
| 0 | 0 |
| 0 | 0 |
| 0 | 0 |
| 0 | 0 |
| 0 | 0 |
| 0 | 0 |
| 0 | 0 |
| 0 | 0 |
| 0 | 0 |
| 0 | 0 |
| 0 | 0 |
| 0 | 0 |
| 0 | 0 |
| 0 | 0 |
| 0 | 0 |
| 0 | 0 |
| 0 | 0 |
| 0 | 0 |
| 0 | 0 |
| 0 | 0 |
| 0 | 0 |
| 0 | 0 |
| 0 | 0 |
| 0 | 0 |
| 0 | 0 |
| 0 | 4.57598851 |
| 0 | 0 |

| | |
|---|---|
| 0 | 0 |
| 0 | 0 |
| 0 | 0 |
| 0 | 22.8799426 |
| 0 | 0 |
| 108.573291 | 13.7279655 |
| 0 | 0 |
| 0 | 41.1838966 |
| 0 | 0 |
| 0 | 0 |
| 0 | 0 |
| 0 | 0 |
| 0 | 0 |
| 0 | 0 |
| 0 | 0 |
| 0 | 0 |
| 0 | 0 |
| 0 | 0 |
| 0 | 0 |
| 0 | 0 |
| 0 | 0 |
| 0 | 0 |
| 0 | 4.57598851 |
| 0 | 0 |
| 0 | 0 |
| 0 | 0 |
| 0 | 0 |
| 0 | 0 |
| 0 | 0 |
| 0 | 0 |
| 0 | 0 |
| 0 | 0 |
| 0 | 0 |
| 0 | 0 |
| 0 | 0 |
| 0 | 0 |
| 0 | 0 |
| 0 | 0 |
| 0 | 0 |
| 0 | 0 |
| 0 | 4.57598851 |
| 0 | 0 |
| 0 | 0 |

| | |
|---|---|
| 0 | 0 |
| 0 | 0 |
| 0 | 0 |
| 0 | 18.303954 |
| 0 | 0 |
| 0 | 0 |
| 0 | 0 |
| 0 | 0 |
| 0 | 0 |
| 0 | 9.15197702 |
| 0 | 0 |
| 0 | 0 |
| 0 | 0 |
| 0 | 0 |
| 0 | 0 |
| 0 | 0 |
| 0 | 0 |
| 0 | 0 |
| 95.0016297 | 13.7279655 |
| 0 | 0 |
| 13.5716614 | 0 |
| 0 | 0 |
| 0 | 0 |
| 0 | 0 |
| 0 | 0 |
| 0 | 0 |
| 2.71433228 | 0 |
| 0 | 0 |
| 0 | 4.57598851 |
| 0 | 0 |
| 0 | 0 |
| 0 | 0 |
| 0 | 0 |
| 0 | 0 |
| 0 | 0 |
| 0 | 0 |
| 0 | 0 |
| 0 | 0 |
| 0 | 0 |
| 0 | 0 |
| 0 | 0 |
| 2.71433228 | 0 |
| 0 | 0 |

| | |
|---|---|
| 0 | 0 |
| 0 | 0 |
| 0 | 0 |
| 0 | 0 |
| 0 | 0 |
| 0 | 0 |
| 0 | 0 |
| 0 | 0 |
| 0 | 0 |
| 2.71433228 | 0 |
| 0 | 0 |
| 0 | 0 |
| 0 | 0 |
| 0 | 0 |
| 0 | 0 |
| 0 | 0 |
| 0 | 0 |
| 0 | 0 |
| 0 | 0 |
| 0 | 41.1838966 |
| 0 | 0 |
| 0 | 0 |
| 0 | 0 |
| 0 | 0 |
| 89.5729652 | 41.1838966 |
| 0 | 0 |
| 0 | 0 |
| 0 | 0 |
| 0 | 0 |
| 0 | 0 |
| 0 | 0 |
| 0 | 0 |
| 0 | 0 |
| 0 | 0 |
| 0 | 0 |
| 0 | 0 |
| 0 | 0 |
| 0 | 0 |
| 0 | 0 |
| 0 | 0 |
| 0 | 0 |
| 0 | 0 |
| 0 | 0 |

| | |
|---|---|
| 0 | 0 |
| 16.2859937 | 224.223437 |
| 0 | 0 |
| 0 | 0 |
| 0 | 0 |
| 0 | 0 |
| 0 | 0 |
| 0 | 0 |
| 0 | 0 |
| 0 | 0 |
| 0 | 0 |
| 0 | 0 |
| 0 | 0 |
| 0 | 0 |
| 0 | 0 |
| 0 | 0 |
| 0 | 0 |
| 0 | 0 |
| 0 | 0 |
| 0 | 0 |
| 54.2866456 | 0 |
| 0 | 0 |
| 0 | 0 |
| 0 | 0 |
| 0 | 9.15197702 |
| 0 | 0 |
| 0 | 0 |
| 0 | 0 |
| 0 | 0 |
| 0 | 0 |
| 0 | 0 |
| 0 | 0 |
| 0 | 0 |
| 0 | 0 |
| 0 | 13.7279655 |
| 0 | 0 |
| 0 | 0 |
| 0 | 0 |
| 0 | 0 |
| 0 | 0 |
| 0 | 0 |
| 0 | 0 |
| 0 | 0 |

| | |
|---|---|
| 0 | 0 |
| 0 | 0 |
| 0 | 0 |
| 0 | 0 |
| 0 | 0 |
| 0 | 0 |
| 0 | 0 |
| 0 | 0 |
| 29.8576551 | 41.1838966 |
| 0 | 0 |
| 0 | 0 |
| 0 | 0 |
| 0 | 4.57598851 |
| 0 | 0 |
| 0 | 0 |
| 0 | 0 |
| 0 | 0 |
| 0 | 0 |
| 0 | 0 |
| 0 | 0 |
| 0 | 0 |
| 0 | 0 |
| 0 | 0 |
| 0 | 0 |
| 0 | 18.303954 |
| 0 | 0 |
| 0 | 0 |
| 0 | 0 |
| 0 | 13.7279655 |
| 0 | 0 |
| 0 | 0 |
| 0 | 0 |
| 0 | 4.57598851 |
| 0 | 0 |
| 2.71433228 | 18.303954 |
| 0 | 0 |
| 0 | 0 |
| 0 | 0 |
| 0 | 0 |
| 0 | 0 |
| 114.001956 | 228.799426 |
| 0 | 4.57598851 |
| 0 | 0 |

| | |
|---|---|
| 0 | 0 |
| 0 | 0 |
| 0 | 0 |
| 0 | 0 |
| 0 | 0 |
| 0 | 0 |
| 0 | 0 |
| 0 | 0 |
| 0 | 0 |
| 0 | 0 |
| 2.71433228 | 22.8799426 |
| 0 | 0 |
| 0 | 0 |
| 0 | 0 |
| 0 | 0 |
| 0 | 0 |
| 0 | 0 |
| 0 | 0 |
| 0 | 0 |
| 0 | 0 |
| 0 | 0 |
| 0 | 0 |
| 0 | 0 |
| 0 | 0 |
| 0 | 0 |
| 0 | 0 |
| 0 | 0 |
| 0 | 0 |
| 0 | 0 |
| 0 | 0 |
| 0 | 0 |
| 0 | 0 |
| 0 | 0 |
| 0 | 0 |
| 0 | 0 |
| 0 | 0 |
| 0 | 0 |
| 0 | 0 |
| 0 | 0 |
| 2.71433228 | 0 |
| 0 | 0 |
| 0 | 0 |
| 0 | 0 |

| | |
|---|---|
| 0 | 0 |
| 0 | 0 |
| 0 | 0 |
| 0 | 0 |
| 0 | 0 |
| 2.71433228 | 0 |
| 0 | 0 |
| 0 | 0 |
| 0 | 0 |
| 0 | 0 |
| 0 | 0 |
| 0 | 0 |
| 0 | 4.57598851 |
| 0 | 0 |
| 0 | 0 |
| 0 | 0 |
| 0 | 0 |
| 0 | 22.8799426 |
| 0 | 0 |
| 13.5716614 | 9.15197702 |
| 0 | 0 |
| 0 | 0 |
| 0 | 0 |
| 0 | 0 |
| 0 | 0 |
| 0 | 0 |
| 0 | 4.57598851 |
| 0 | 0 |
| 0 | 0 |
| 0 | 0 |
| 5.42866456 | 4.57598851 |
| 0 | 0 |
| 0 | 0 |
| 0 | 0 |
| 0 | 0 |
| 0 | 0 |
| 2.71433228 | 0 |
| 0 | 0 |
| 0 | 0 |
| 0 | 0 |
| 0 | 0 |
| 0 | 13.7279655 |
| 0 | 0 |

| | |
|---|---|
| 0 | 0 |
| 0 | 0 |
| 0 | 0 |
| 0 | 0 |
| 0 | 0 |
| 0 | 0 |
| 0 | 0 |
| 0 | 4.57598851 |
| 0 | 0 |
| 0 | 0 |
| 0 | 0 |
| 0 | 0 |
| 0 | 0 |
| 0 | 0 |
| 0 | 0 |
| 0 | 0 |
| 0 | 0 |
| 0 | 4.57598851 |
| 0 | 0 |
| 0 | 0 |
| 0 | 0 |
| 0 | 0 |
| 0 | 0 |
| 0 | 0 |
| 0 | 4.57598851 |
| 0 | 0 |
| 0 | 0 |
| 0 | 0 |
| 0 | 0 |
| 0 | 0 |
| 0 | 0 |
| 0 | 0 |
| 0 | 0 |
| 0 | 0 |
| 0 | 0 |
| 0 | 0 |
| 0 | 0 |
| 0 | 0 |
| 0 | 0 |
| 0 | 0 |
| 0 | 0 |
| 179.14593 | 562.846587 |
| 0 | 0 |

| | |
|---|---|
| 0 | 0 |
| 0 | 0 |
| 0 | 4.57598851 |
| 0 | 0 |
| 0 | 0 |
| 0 | 0 |
| 0 | 0 |
| 0 | 0 |
| 0 | 0 |
| 0 | 0 |
| 0 | 0 |
| 0 | 0 |
| 0 | 0 |
| 0 | 0 |
| 0 | 0 |
| 0 | 0 |
| 0 | 0 |
| 0 | 0 |
| 0 | 0 |
| 0 | 0 |
| 0 | 0 |
| 0 | 0 |
| 0 | 0 |
| 0 | 0 |
| 0 | 0 |
| 46.1436487 | 105.247736 |
| 0 | 0 |
| 0 | 0 |
| 0 | 0 |
| 0 | 0 |
| 0 | 0 |
| 0 | 0 |
| 0 | 0 |
| 0 | 0 |
| 0 | 0 |
| 0 | 0 |
| 0 | 0 |
| 0 | 0 |
| 0 | 0 |
| 0 | 0 |
| 0 | 0 |
| 0 | 0 |
| 0 | 0 |

| | |
|---|---|
| 0 | 0 |
| 0 | 0 |
| 0 | 0 |
| 0 | 0 |
| 0 | 0 |
| 32.5719873 | 100.671747 |
| 0 | 0 |
| 0 | 0 |
| 0 | 0 |
| 0 | 0 |
| 0 | 0 |
| 0 | 0 |
| 0 | 0 |
| 0 | 0 |
| 0 | 0 |
| 5.42866456 | 9.15197702 |
| 0 | 0 |
| 0 | 0 |
| 0 | 0 |
| 0 | 0 |
| 21.7146582 | 0 |
| 0 | 0 |
| 0 | 0 |
| 0 | 0 |
| 0 | 0 |
| 0 | 0 |
| 0 | 0 |
| 0 | 0 |
| 0 | 0 |
| 0 | 0 |
| 0 | 0 |
| 0 | 0 |
| 0 | 0 |
| 0 | 0 |
| 0 | 0 |
| 0 | 0 |
| 0 | 0 |
| 0 | 13.7279655 |
| 0 | 0 |
| 0 | 0 |
| 0 | 0 |
| 0 | 0 |
| 0 | 0 |

| | |
|---|---|
| 0 | 0 |
| 0 | 0 |
| 0 | 0 |
| 0 | 0 |
| 0 | 0 |
| 0 | 0 |
| 0 | 0 |
| 0 | 0 |
| 0 | 0 |
| 0 | 4.57598851 |
| 0 | 0 |
| 0 | 0 |
| 0 | 4.57598851 |
| 0 | 0 |
| 0 | 0 |
| 0 | 0 |
| 0 | 0 |
| 0 | 0 |
| 0 | 0 |
| 0 | 0 |
| 0 | 0 |
| 0 | 0 |
| 0 | 0 |
| 0 | 0 |
| 0 | 0 |
| 0 | 0 |
| 0 | 0 |
| 0 | 0 |
| 0 | 0 |
| 0 | 0 |
| 0 | 0 |
| 0 | 0 |
| 0 | 0 |
| 0 | 0 |
| 0 | 0 |
| 0 | 0 |
| 0 | 0 |
| 0 | 0 |
| 0 | 0 |
| 0 | 0 |
| 0 | 0 |
| 0 | 0 |
| 0 | 0 |
| 0 | 0 |
| 0 | 9.15197702 |

| | |
|---|---|
| 19.0003259 | 27.4559311 |
| 0 | 0 |
| 0 | 0 |
| 0 | 0 |
| 0 | 0 |
| 0 | 0 |
| 0 | 0 |
| 0 | 0 |
| 0 | 4.57598851 |
| 0 | 0 |
| 0 | 0 |
| 0 | 0 |
| 0 | 0 |
| 0 | 0 |
| 0 | 0 |
| 0 | 0 |
| 0 | 0 |
| 0 | 0 |
| 0 | 0 |
| 10.8573291 | 0 |
| 0 | 0 |
| 0 | 0 |
| 0 | 0 |
| 0 | 0 |
| 0 | 0 |
| 0 | 0 |
| 0 | 0 |
| 2.71433228 | 4.57598851 |
| 0 | 0 |
| 0 | 0 |
| 0 | 0 |
| 0 | 0 |
| 0 | 18.303954 |
| 0 | 0 |
| 0 | 0 |
| 40.7149842 | 50.3358736 |
| 0 | 0 |
| 0 | 0 |
| 0 | 0 |
| 0 | 0 |
| 0 | 0 |
| 0 | 0 |
| 0 | 0 |

| | |
|---|---|
| 0 | 0 |
| 0 | 0 |
| 0 | 0 |
| 0 | 0 |
| 0 | 0 |
| 0 | 0 |
| 0 | 0 |
| 0 | 0 |
| 0 | 0 |
| 0 | 0 |
| 0 | 0 |
| 0 | 0 |
| 0 | 0 |
| 0 | 0 |
| 0 | 0 |
| 0 | 0 |
| 0 | 0 |
| 0 | 0 |
| 0 | 0 |
| 0 | 0 |
| 0 | 0 |
| 0 | 0 |
| 0 | 0 |
| 0 | 0 |
| 0 | 0 |
| 0 | 0 |
| 0 | 0 |
| 1606.88471 | 906.045725 |
| 0 | 0 |
| 0 | 0 |
| 0 | 0 |
| 0 | 0 |
| 5.42866456 | 64.0638392 |
| 0 | 0 |
| 0 | 0 |
| 0 | 0 |
| 0 | 0 |
| 0 | 0 |
| 0 | 0 |
| 0 | 0 |
| 0 | 0 |
| 0 | 0 |
| 0 | 0 |

| | |
|---|---|
| 0 | 0 |
| 0 | 0 |
| 0 | 0 |
| 0 | 0 |
| 0 | 0 |
| 2.71433228 | 0 |
| 0 | 0 |
| 181.860263 | 32.0319196 |
| 0 | 0 |
| 0 | 0 |
| 0 | 0 |
| 0 | 0 |
| 0 | 0 |
| 0 | 0 |
| 0 | 0 |
| 0 | 0 |
| 0 | 0 |
| 0 | 0 |
| 0 | 32.0319196 |
| 0 | 0 |
| 0 | 0 |
| 0 | 0 |
| 2.71433228 | 0 |
| 0 | 0 |
| 0 | 0 |
| 0 | 0 |
| 0 | 0 |
| 0 | 0 |
| 0 | 0 |
| 0 | 0 |
| 0 | 0 |
| 0 | 0 |
| 0 | 0 |
| 0 | 0 |
| 0 | 0 |
| 0 | 0 |
| 0 | 0 |
| 0 | 0 |
| 0 | 0 |
| 0 | 0 |
| 0 | 4.57598851 |
| 0 | 0 |
| 0 | 0 |

| | |
|---|---|
| 0 | 0 |
| 0 | 0 |
| 0 | 0 |
| 0 | 0 |
| 0 | 0 |
| 0 | 0 |
| 0 | 0 |
| 0 | 0 |
| 2.71433228 | 4.57598851 |
| 0 | 0 |
| 0 | 0 |
| 2.71433228 | 0 |
| 0 | 0 |
| 8004.56589 | 2018.01093 |
| 0 | 0 |
| 0 | 0 |
| 0 | 0 |
| 0 | 0 |
| 32.5719873 | 109.823724 |
| 0 | 0 |
| 0 | 0 |
| 0 | 0 |
| 0 | 0 |
| 0 | 0 |
| 0 | 0 |
| 0 | 0 |
| 225.289579 | 160.159598 |
| 0 | 0 |
| 0 | 0 |
| 0 | 0 |
| 0 | 0 |
| 0 | 22.8799426 |
| 0 | 0 |
| 0 | 0 |
| 0 | 0 |
| 0 | 0 |
| 0 | 4.57598851 |
| 708.440725 | 315.743207 |
| 0 | 0 |
| 0 | 0 |
| 0 | 0 |
| 0 | 0 |
| 0 | 0 |

| | |
|---|---|
| 0 | 0 |
| 0 | 0 |
| 0 | 0 |
| 0 | 0 |
| 0 | 0 |
| 0 | 0 |
| 0 | 0 |
| 0 | 0 |
| 0 | 0 |
| 5.42866456 | 0 |
| 0 | 0 |
| 0 | 0 |
| 0 | 0 |
| 2.71433228 | 4.57598851 |
| 0 | 0 |
| 0 | 0 |
| 0 | 0 |
| 0 | 0 |
| 0 | 0 |
| 0 | 0 |
| 0 | 0 |
| 0 | 0 |
| 0 | 0 |
| 0 | 0 |
| 0 | 0 |
| 0 | 0 |
| 0 | 0 |
| 0 | 0 |
| 0 | 0 |
| 0 | 0 |
| 0 | 0 |
| 0 | 0 |
| 0 | 0 |
| 0 | 0 |
| 0 | 0 |
| 0 | 0 |
| 0 | 0 |
| 0 | 0 |
| 0 | 0 |
| 0 | 0 |
| 0 | 0 |
| 0 | 0 |
| 0 | 0 |

| | |
|---|---|
| 0 | 0 |
| 0 | 0 |
| 0 | 0 |
| 0 | 0 |
| 0 | 13.7279655 |
| 0 | 0 |
| 0 | 0 |
| 0 | 0 |
| 0 | 0 |
| 0 | 0 |
| 0 | 0 |
| 0 | 0 |
| 0 | 0 |
| 0 | 0 |
| 0 | 0 |
| 0 | 0 |
| 0 | 0 |
| 366.434858 | 41.1838966 |
| 0 | 0 |
| 0 | 0 |
| 0 | 0 |
| 0 | 0 |
| 0 | 0 |
| 0 | 0 |
| 0 | 0 |
| 0 | 0 |
| 0 | 4.57598851 |
| 0 | 0 |
| 0 | 0 |
| 0 | 0 |
| 0 | 0 |
| 1414.16712 | 645.21438 |
| 0 | 0 |
| 426.150168 | 247.10338 |
| 0 | 0 |
| 0 | 0 |
| 103.144627 | 41.1838966 |
| 0 | 0 |
| 0 | 0 |
| 0 | 0 |
| 0 | 0 |
| 0 | 0 |
| 0 | 0 |

| | |
|---|---|
| 0 | 0 |
| 0 | 0 |
| 0 | 0 |
| 0 | 0 |
| 0 | 4.57598851 |
| 0 | 0 |
| 0 | 0 |
| 0 | 0 |
| 0 | 0 |
| 0 | 0 |
| 5.42866456 | 0 |
| 0 | 0 |
| 0 | 0 |
| 0 | 4.57598851 |
| 0 | 0 |
| 0 | 0 |
| 0 | 0 |
| 0 | 0 |
| 0 | 0 |
| 0 | 0 |
| 0 | 0 |
| 0 | 0 |
| 0 | 0 |
| 13.5716614 | 9.15197702 |
| 0 | 0 |
| 0 | 0 |
| 0 | 0 |
| 0 | 0 |
| 0 | 0 |
| 0 | 0 |
| 0 | 0 |
| 0 | 4.57598851 |
| 2.71433228 | 0 |
| 0 | 0 |
| 0 | 0 |
| 0 | 0 |
| 0 | 0 |
| 0 | 0 |
| 135.716614 | 32.0319196 |
| 0 | 0 |
| 0 | 0 |
| 0 | 0 |
| 0 | 0 |

| | |
|---|---|
| 0 | 0 |
| 0 | 0 |
| 0 | 0 |
| 0 | 0 |
| 0 | 0 |
| 0 | 0 |
| 0 | 0 |
| 0 | 0 |
| 0 | 0 |
| 0 | 0 |
| 0 | 0 |
| 0 | 4.57598851 |
| 0 | 0 |
| 0 | 0 |
| 0 | 0 |
| 0 | 0 |
| 0 | 0 |
| 0 | 0 |
| 0 | 18.303954 |
| 0 | 0 |
| 0 | 0 |
| 0 | 0 |
| 0 | 0 |
| 0 | 0 |
| 0 | 0 |
| 0 | 0 |
| 0 | 0 |
| 0 | 0 |
| 0 | 13.7279655 |
| 0 | 0 |
| 0 | 0 |
| 0 | 0 |
| 0 | 0 |
| 0 | 0 |
| 0 | 0 |
| 0 | 0 |
| 0 | 0 |
| 0 | 4.57598851 |
| 0 | 0 |
| 0 | 0 |
| 0 | 0 |
| 0 | 0 |
| 0 | 0 |

| | |
|---|---|
| 0 | 0 |
| 0 | 0 |
| 0 | 0 |
| 0 | 0 |
| 0 | 0 |
| 0 | 0 |
| 0 | 0 |
| 0 | 0 |
| 0 | 0 |
| 0 | 0 |
| 0 | 0 |
| 0 | 0 |
| 16.2859937 | 9.15197702 |
| 0 | 0 |
| 0 | 4.57598851 |
| 0 | 0 |
| 0 | 4.57598851 |
| 0 | 0 |
| 0 | 0 |
| 0 | 0 |
| 10.8573291 | 59.4878506 |
| 0 | 0 |
| 0 | 0 |
| 0 | 0 |
| 0 | 0 |
| 0 | 0 |
| 0 | 0 |
| 0 | 0 |
| 0 | 0 |
| 0 | 0 |
| 2.71433228 | 0 |
| 0 | 0 |
| 0 | 0 |
| 0 | 0 |
| 0 | 0 |
| 219.860915 | 164.735586 |
| 0 | 0 |
| 0 | 0 |
| 0 | 0 |
| 0 | 0 |
| 0 | 0 |
| 0 | 0 |
| 0 | 0 |

| | |
|---|---|
| 0 | 0 |
| 0 | 4.57598851 |
| 2.71433228 | 0 |
| 0 | 0 |
| 0 | 0 |
| 0 | 0 |
| 0 | 0 |
| 0 | 0 |
| 0 | 0 |
| 5.42866456 | 9.15197702 |
| 0 | 0 |
| 2.71433228 | 0 |
| 0 | 4.57598851 |
| 0 | 0 |
| 0 | 0 |
| 0 | 0 |
| 0 | 0 |
| 0 | 0 |
| 0 | 0 |
| 0 | 0 |
| 0 | 0 |
| 0 | 0 |
| 0 | 0 |
| 0 | 0 |
| 0 | 0 |
| 0 | 0 |
| 0 | 4.57598851 |
| 0 | 0 |
| 0 | 0 |
| 0 | 0 |
| 0 | 0 |
| 0 | 0 |
| 0 | 0 |
| 2.71433228 | 0 |
| 304.005215 | 50.3358736 |
| 2.71433228 | 0 |
| 0 | 0 |
| 0 | 0 |
| 0 | 45.7598851 |
| 0 | 0 |
| 0 | 0 |
| 0 | 0 |
| 0 | 0 |

| | |
|---|---|
| 0 | 0 |
| 0 | 0 |
| 0 | 0 |
| 0 | 0 |
| 0 | 0 |
| 0 | 0 |
| 0 | 0 |
| 0 | 0 |
| 0 | 0 |
| 228.003911 | 192.191517 |
| 0 | 0 |
| 0 | 0 |
| 0 | 0 |
| 0 | 4.57598851 |
| 0 | 0 |
| 0 | 0 |
| 0 | 0 |
| 0 | 0 |
| 0 | 0 |
| 0 | 0 |
| 0 | 0 |
| 0 | 0 |
| 0 | 0 |
| 0 | 0 |
| 0 | 0 |
| 0 | 0 |
| 0 | 0 |
| 0 | 0 |
| 0 | 0 |
| 0 | 0 |
| 0 | 0 |
| 0 | 0 |
| 0 | 0 |
| 0 | 0 |
| 0 | 0 |
| 0 | 0 |
| 0 | 0 |
| 0 | 0 |
| 0 | 0 |
| 16.2859937 | 18.303954 |
| 0 | 0 |
| 0 | 0 |
| 0 | 4.57598851 |

| | |
|---|---|
| 0 | 0 |
| 0 | 0 |
| 119.43062 | 4.57598851 |
| 0 | 0 |
| 0 | 0 |
| 0 | 0 |
| 0 | 0 |
| 0 | 0 |
| 0 | 0 |
| 0 | 0 |
| 0 | 9.15197702 |
| 0 | 0 |
| 0 | 0 |
| 0 | 0 |
| 0 | 0 |
| 0 | 0 |
| 0 | 0 |
| 0 | 0 |
| 0 | 0 |
| 0 | 4.57598851 |
| 0 | 0 |
| 0 | 0 |
| 0 | 0 |
| 0 | 0 |
| 0 | 0 |
| 0 | 0 |
| 0 | 0 |
| 0 | 0 |
| 0 | 0 |
| 0 | 0 |
| 0 | 0 |
| 0 | 0 |
| 0 | 4.57598851 |
| 0 | 0 |
| 0 | 0 |
| 0 | 0 |
| 0 | 0 |
| 0 | 0 |
| 0 | 0 |
| 0 | 0 |
| 0 | 0 |
| 0 | 0 |
| 0 | 0 |

| | |
|---|---|
| 0 | 0 |
| 0 | 0 |
| 0 | 0 |
| 0 | 0 |
| 0 | 0 |
| 2.71433228 | 0 |
| 0 | 0 |
| 0 | 0 |
| 0 | 0 |
| 0 | 0 |
| 0 | 0 |
| 0 | 0 |
| 0 | 0 |
| 130.287949 | 128.127678 |
| 0 | 0 |
| 0 | 0 |
| 0 | 0 |
| 0 | 4.57598851 |
| 0 | 0 |
| 0 | 0 |
| 0 | 0 |
| 0 | 0 |
| 0 | 64.0638392 |
| 0 | 0 |
| 2.71433228 | 0 |
| 0 | 0 |
| 0 | 0 |
| 0 | 0 |
| 0 | 0 |
| 0 | 0 |
| 0 | 0 |
| 0 | 0 |
| 0 | 0 |
| 2.71433228 | 0 |
| 0 | 0 |
| 8.14299683 | 22.8799426 |
| 0 | 0 |
| 0 | 0 |
| 0 | 0 |
| 0 | 0 |
| 0 | 0 |
| 0 | 0 |
| 0 | 0 |

| | |
|---|---|
| 0 | 0 |
| 54.2866456 | 0 |
| 0 | 0 |
| 0 | 0 |
| 0 | 0 |
| 0 | 0 |
| 0 | 0 |
| 0 | 0 |
| 0 | 0 |
| 0 | 0 |
| 0 | 0 |
| 0 | 0 |
| 0 | 0 |
| 0 | 0 |
| 0 | 0 |
| 0 | 0 |
| 0 | 0 |
| 0 | 0 |
| 0 | 0 |
| 0 | 0 |
| 0 | 0 |
| 0 | 0 |
| 0 | 0 |
| 0 | 22.8799426 |
| 0 | 22.8799426 |
| 2.71433228 | 0 |
| 0 | 0 |
| 0 | 0 |
| 0 | 0 |
| 0 | 0 |
| 0 | 0 |
| 0 | 0 |
| 0 | 0 |
| 0 | 0 |
| 0 | 0 |
| 0 | 0 |
| 0 | 0 |
| 0 | 0 |
| 0 | 0 |
| 0 | 0 |
| 0 | 0 |
| 0 | 0 |
| 0 | 0 |

| | |
|---|---|
| 0 | 0 |
| 0 | 0 |
| 0 | 0 |
| 0 | 45.7598851 |
| 0 | 0 |
| 0 | 0 |
| 0 | 0 |
| 567.295446 | 420.990943 |
| 0 | 0 |
| 352.863196 | 315.743207 |
| 0 | 0 |
| 0 | 0 |
| 0 | 0 |
| 0 | 0 |
| 0 | 0 |
| 0 | 0 |
| 0 | 0 |
| 0 | 0 |
| 0 | 0 |
| 0 | 0 |
| 0 | 0 |
| 0 | 0 |
| 0 | 0 |
| 0 | 0 |
| 0 | 0 |
| 0 | 0 |
| 0 | 0 |
| 0 | 0 |
| 0 | 0 |
| 0 | 0 |
| 0 | 0 |
| 0 | 0 |
| 0 | 0 |
| 0 | 0 |
| 0 | 0 |
| 0 | 0 |
| 0 | 4.57598851 |
| 2.71433228 | 0 |
| 0 | 0 |
| 0 | 0 |
| 0 | 0 |
| 0 | 0 |
| 0 | 0 |

| | |
|---|---|
| 0 | 0 |
| 0 | 0 |
| 0 | 0 |
| 0 | 0 |
| 0 | 0 |
| 0 | 0 |
| 0 | 0 |
| 0 | 0 |
| 0 | 0 |
| 0 | 0 |
| 0 | 0 |
| 0 | 0 |
| 0 | 0 |
| 0 | 0 |
| 0 | 0 |
| 0 | 0 |
| 0 | 0 |
| 0 | 0 |
| 5.42866456 | 0 |
| 10.8573291 | 50.3358736 |
| 0 | 0 |
| 0 | 0 |
| 0 | 0 |
| 0 | 0 |
| 0 | 0 |
| 0 | 0 |
| 0 | 0 |
| 0 | 0 |
| 0 | 0 |
| 0 | 4.57598851 |
| 0 | 0 |
| 0 | 0 |
| 0 | 0 |
| 0 | 0 |
| 0 | 0 |
| 0 | 0 |
| 0 | 0 |
| 0 | 0 |
| 0 | 0 |
| 2.71433228 | 0 |
| 0 | 0 |
| 0 | 0 |
| 0 | 0 |

| | |
|---|---|
| 0 | 0 |
| 0 | 0 |
| 0 | 0 |
| 0 | 0 |
| 0 | 0 |
| 0 | 0 |
| 0 | 0 |
| 0 | 13.7279655 |
| 0 | 0 |
| 0 | 0 |
| 0 | 0 |
| 0 | 0 |
| 35.2863196 | 54.9118621 |
| 0 | 0 |
| 0 | 0 |
| 0 | 0 |
| 0 | 0 |
| 0 | 0 |
| 0 | 0 |
| 0 | 0 |
| 0 | 0 |
| 0 | 0 |
| 0 | 0 |
| 0 | 0 |
| 0 | 0 |
| 0 | 0 |
| 0 | 0 |
| 0 | 0 |
| 0 | 0 |
| 0 | 0 |
| 0 | 0 |
| 0 | 4.57598851 |
| 0 | 0 |
| 0 | 0 |
| 0 | 0 |
| 0 | 0 |
| 0 | 0 |
| 0 | 0 |
| 0 | 0 |
| 0 | 0 |
| 0 | 0 |
| 0 | 0 |
| 0 | 0 |

| | |
|---|---|
| 0 | 0 |
| 0 | 0 |
| 0 | 0 |
| 0 | 0 |
| 0 | 0 |
| 0 | 0 |
| 0 | 0 |
| 0 | 0 |
| 0 | 0 |
| 0 | 0 |
| 0 | 0 |
| 0 | 0 |
| 0 | 0 |
| 0 | 0 |
| 0 | 0 |
| 0 | 0 |
| 0 | 0 |
| 0 | 0 |
| 0 | 0 |
| 0 | 0 |
| 0 | 36.6079081 |
| 0 | 4.57598851 |
| 0 | 0 |
| 130.287949 | 86.9437817 |
| 0 | 0 |
| 0 | 0 |
| 0 | 4.57598851 |
| 0 | 0 |
| 0 | 0 |
| 0 | 0 |
| 0 | 0 |
| 0 | 0 |
| 0 | 0 |
| 0 | 0 |
| 5.42866456 | 27.4559311 |
| 0 | 0 |
| 0 | 0 |
| 0 | 0 |
| 0 | 0 |
| 0 | 0 |
| 0 | 0 |
| 0 | 0 |
| 0 | 0 |

| | |
|---|---|
| 0 | 0 |
| 19.0003259 | 9.15197702 |
| 0 | 0 |
| 0 | 0 |
| 0 | 0 |
| 0 | 0 |
| 0 | 0 |
| 0 | 0 |
| 0 | 0 |
| 0 | 0 |
| 0 | 0 |
| 0 | 0 |
| 0 | 0 |
| 0 | 0 |
| 0 | 0 |
| 0 | 0 |
| 0 | 0 |
| 0 | 0 |
| 0 | 0 |
| 0 | 0 |
| 0 | 0 |
| 0 | 0 |
| 187.288927 | 82.3677932 |
| 0 | 0 |
| 0 | 0 |
| 0 | 0 |
| 2.71433228 | 0 |
| 0 | 0 |
| 0 | 0 |
| 0 | 0 |
| 0 | 0 |
| 0 | 0 |
| 0 | 0 |
| 0 | 0 |
| 0 | 0 |
| 0 | 0 |
| 0 | 0 |
| 0 | 0 |
| 0 | 0 |
| 0 | 0 |
| 0 | 0 |
| 0 | 0 |
| 0 | 0 |

| | |
|---|---|
| 8.14299683 | 4.57598851 |
| 0 | 0 |
| 0 | 0 |
| 0 | 0 |
| 0 | 0 |
| 0 | 0 |
| 0 | 0 |
| 0 | 0 |
| 0 | 0 |
| 0 | 0 |
| 0 | 0 |
| 0 | 0 |
| 0 | 0 |
| 0 | 0 |
| 0 | 0 |
| 0 | 0 |
| 0 | 0 |
| 0 | 0 |
| 0 | 0 |
| 0 | 0 |
| 0 | 0 |
| 0 | 0 |
| 0 | 0 |
| 0 | 0 |
| 0 | 0 |
| 0 | 0 |
| 0 | 0 |
| 0 | 0 |
| 0 | 0 |
| 0 | 0 |
| 0 | 0 |
| 0 | 0 |
| 0 | 4.57598851 |
| 0 | 0 |
| 0 | 0 |
| 0 | 0 |
| 0 | 0 |
| 59.7153101 | 4.57598851 |
| 0 | 0 |
| 0 | 0 |
| 0 | 0 |
| 0 | 0 |
| 0 | 0 |

| | |
|---|---|
| 0 | 4.57598851 |
| 0 | 0 |
| 0 | 0 |
| 0 | 0 |
| 0 | 0 |
| 0 | 0 |
| 0 | 0 |
| 0 | 0 |
| 0 | 0 |
| 0 | 0 |
| 0 | 0 |
| 0 | 0 |
| 0 | 0 |
| 0 | 0 |
| 0 | 0 |
| 0 | 0 |
| 0 | 0 |
| 0 | 0 |
| 0 | 0 |
| 0 | 0 |
| 0 | 0 |
| 0 | 0 |
| 0 | 0 |
| 0 | 0 |
| 0 | 0 |
| 0 | 4.57598851 |
| 0 | 0 |
| 0 | 0 |
| 0 | 0 |
| 0 | 0 |
| 0 | 0 |
| 0 | 0 |
| 0 | 0 |
| 0 | 0 |
| 0 | 0 |
| 2.71433228 | 0 |
| 0 | 0 |
| 0 | 0 |
| 0 | 0 |
| 0 | 0 |
| 0 | 0 |
| 0 | 9.15197702 |
| 0 | 0 |

| | |
|---|---|
| 0 | 0 |
| 0 | 0 |
| 0 | 0 |
| 0 | 0 |
| 0 | 0 |
| 0 | 0 |
| 0 | 0 |
| 0 | 0 |
| 0 | 0 |
| 0 | 0 |
| 0 | 0 |
| 0 | 4.57598851 |
| 0 | 0 |
| 0 | 0 |
| 0 | 0 |
| 0 | 0 |
| 0 | 0 |
| 0 | 0 |
| 0 | 0 |
| 0 | 0 |
| 0 | 0 |
| 0 | 0 |
| 2.71433228 | 0 |
| 0 | 0 |
| 0 | 0 |
| 0 | 0 |
| 0 | 0 |
| 0 | 0 |
| 0 | 0 |
| 2.71433228 | 0 |
| 0 | 0 |
| 0 | 0 |
| 2.71433228 | 59.4878506 |
| 0 | 0 |
| 0 | 0 |
| 0 | 0 |
| 0 | 0 |
| 0 | 0 |
| 0 | 0 |
| 0 | 0 |
| 0 | 0 |
| 0 | 0 |
| 0 | 0 |

| | |
|---|---|
| 0 | 0 |
| 0 | 0 |
| 0 | 0 |
| 0 | 0 |
| 0 | 9.15197702 |
| 0 | 0 |
| 2.71433228 | 0 |
| 0 | 0 |
| 0 | 0 |
| 0 | 18.303954 |
| 92.2872974 | 73.2158162 |
| 0 | 0 |
| 0 | 4.57598851 |
| 0 | 0 |
| 0 | 0 |
| 0 | 0 |
| 0 | 0 |
| 0 | 0 |
| 0 | 0 |
| 0 | 0 |
| 0 | 36.6079081 |
| 0 | 0 |
| 8.14299683 | 0 |
| 0 | 0 |
| 0 | 0 |
| 0 | 0 |
| 0 | 0 |
| 0 | 0 |
| 0 | 22.8799426 |
| 0 | 0 |
| 0 | 0 |
| 0 | 0 |
| 0 | 0 |
| 5.42866456 | 251.679368 |
| 0 | 0 |
| 0 | 0 |
| 0 | 0 |
| 0 | 0 |
| 0 | 0 |
| 0 | 0 |
| 0 | 0 |
| 114.001956 | 91.5197702 |
| 0 | 0 |

| | |
|---|---|
| 0 | 0 |
| 0 | 0 |
| 0 | 0 |
| 0 | 0 |
| 0 | 0 |
| 0 | 0 |
| 0 | 0 |
| 10.8573291 | 114.399713 |
| 0 | 0 |
| 0 | 0 |
| 0 | 0 |
| 0 | 0 |
| 0 | 0 |
| 0 | 0 |
| 0 | 0 |
| 0 | 0 |
| 0 | 4.57598851 |
| 0 | 0 |
| 0 | 0 |
| 0 | 18.303954 |
| 0 | 0 |
| 0 | 0 |
| 0 | 0 |
| 0 | 0 |
| 0 | 0 |
| 0 | 0 |
| 0 | 0 |
| 0 | 0 |
| 0 | 0 |
| 16.2859937 | 27.4559311 |
| 0 | 0 |
| 0 | 0 |
| 0 | 0 |
| 0 | 0 |
| 0 | 0 |
| 0 | 0 |
| 0 | 0 |
| 0 | 0 |
| 0 | 13.7279655 |
| 0 | 0 |
| 0 | 0 |
| 0 | 0 |
| 0 | 0 |

| | |
|---|---|
| 0 | 0 |
| 0 | 0 |
| 95.0016297 | 13.7279655 |
| 0 | 0 |
| 0 | 0 |
| 0 | 0 |
| 0 | 0 |
| 0 | 0 |
| 0 | 0 |
| 0 | 27.4559311 |
| 0 | 0 |
| 2.71433228 | 0 |
| 0 | 0 |
| 0 | 0 |
| 0 | 0 |
| 0 | 0 |
| 0 | 0 |
| 5.42866456 | 0 |
| 0 | 0 |
| 0 | 0 |
| 0 | 0 |
| 0 | 4.57598851 |
| 0 | 0 |
| 0 | 0 |
| 0 | 0 |
| 0 | 0 |
| 0 | 0 |
| 0 | 0 |
| 0 | 0 |
| 2.71433228 | 0 |
| 0 | 0 |
| 0 | 0 |
| 0 | 0 |
| 0 | 0 |
| 0 | 0 |
| 0 | 0 |
| 13.5716614 | 105.247736 |
| 0 | 0 |
| 0 | 0 |
| 0 | 0 |
| 0 | 0 |
| 0 | 0 |
| 0 | 0 |

| | |
|---|---|
| 0 | 0 |
| 0 | 0 |
| 0 | 0 |
| 0 | 0 |
| 0 | 0 |
| 0 | 0 |
| 0 | 0 |
| 0 | 0 |
| 8.14299683 | 0 |
| 0 | 0 |
| 8.14299683 | 132.703667 |
| 0 | 0 |
| 0 | 0 |
| 0 | 0 |
| 0 | 0 |
| 0 | 0 |
| 0 | 0 |
| 0 | 0 |
| 0 | 0 |
| 0 | 0 |
| 0 | 0 |
| 0 | 0 |
| 0 | 0 |
| 0 | 0 |
| 0 | 0 |
| 0 | 0 |
| 0 | 0 |
| 0 | 0 |
| 0 | 0 |
| 0 | 0 |
| 0 | 0 |
| 0 | 0 |
| 0 | 0 |
| 0 | 0 |
| 0 | 0 |
| 0 | 0 |
| 0 | 0 |
| 0 | 0 |
| 0 | 0 |
| 0 | 0 |
| 0 | 0 |
| 0 | 0 |
| 0 | 0 |

| | |
|---|---|
| 5.42866456 | 4.57598851 |
| 0 | 0 |
| 0 | 0 |
| 0 | 0 |
| 0 | 0 |
| 0 | 0 |
| 0 | 0 |
| 24.4289905 | 0 |
| 0 | 0 |
| 0 | 0 |
| 0 | 0 |
| 0 | 0 |
| 0 | 4.57598851 |
| 0 | 0 |
| 1482.02542 | 425.566932 |
| 0 | 0 |
| 127.573617 | 164.735586 |
| 0 | 0 |
| 0 | 0 |
| 0 | 0 |
| 0 | 0 |
| 0 | 0 |
| 0 | 0 |
| 0 | 0 |
| 2.71433228 | 0 |
| 0 | 0 |
| 0 | 0 |
| 0 | 0 |
| 0 | 0 |
| 0 | 0 |
| 0 | 0 |
| 0 | 0 |
| 0 | 0 |
| 0 | 0 |
| 0 | 0 |
| 0 | 0 |
| 0 | 0 |
| 0 | 0 |
| 0 | 0 |
| 0 | 0 |
| 0 | 0 |
| 0 | 0 |
| 0 | 13.7279655 |

| | |
|---|---|
| 0 | 0 |
| 0 | 0 |
| 0 | 18.303954 |
| 0 | 0 |
| 0 | 0 |
| 0 | 0 |
| 0 | 0 |
| 0 | 0 |
| 0 | 0 |
| 2.71433228 | 4.57598851 |
| 0 | 0 |
| 0 | 0 |
| 0 | 0 |
| 0 | 0 |
| 0 | 0 |
| 0 | 0 |
| 0 | 0 |
| 0 | 0 |
| 0 | 0 |
| 0 | 0 |
| 0 | 0 |
| 0 | 0 |
| 0 | 0 |
| 0 | 0 |
| 0 | 0 |
| 0 | 0 |
| 0 | 0 |
| 0 | 0 |
| 293.147886 | 9.15197702 |
| 0 | 0 |
| 0 | 0 |
| 0 | 0 |
| 0 | 0 |
| 0 | 0 |
| 0 | 0 |
| 0 | 0 |
| 0 | 0 |
| 0 | 0 |
| 0 | 0 |
| 0 | 0 |
| 526.580462 | 443.870886 |
| 0 | 0 |
| 0 | 0 |

| | |
|---|---|
| 0 | 0 |
| 0 | 0 |
| 0 | 0 |
| 13.5716614 | 0 |
| 0 | 18.303954 |
| 0 | 0 |
| 0 | 0 |
| 0 | 0 |
| 2.71433228 | 0 |
| 0 | 0 |
| 0 | 0 |
| 0 | 32.0319196 |
| 0 | 0 |
| 0 | 0 |
| 0 | 4.57598851 |
| 0 | 0 |
| 0 | 0 |
| 0 | 0 |
| 0 | 0 |
| 0 | 0 |
| 0 | 0 |
| 0 | 0 |
| 0 | 0 |
| 0 | 0 |
| 0 | 0 |
| 0 | 0 |
| 0 | 0 |
| 0 | 0 |
| 2.71433228 | 0 |
| 0 | 4.57598851 |
| 0 | 0 |
| 0 | 0 |
| 0 | 0 |
| 67.8583069 | 45.7598851 |
| 0 | 0 |
| 0 | 0 |
| 0 | 0 |
| 0 | 27.4559311 |
| 0 | 0 |
| 59.7153101 | 9.15197702 |
| 0 | 0 |
| 0 | 0 |
| 0 | 0 |

| | |
|---|---|
| 0 | 0 |
| 0 | 4.57598851 |
| 0 | 9.15197702 |
| 0 | 0 |
| 0 | 0 |
| 0 | 0 |
| 0 | 0 |
| 0 | 0 |
| 0 | 0 |
| 0 | 0 |
| 0 | 0 |
| 0 | 0 |
| 0 | 0 |
| 0 | 0 |
| 0 | 0 |
| 0 | 18.303954 |
| 0 | 0 |
| 141.145278 | 22.8799426 |
| 0 | 4.57598851 |
| 0 | 0 |
| 0 | 0 |
| 0 | 0 |
| 0 | 0 |
| 0 | 13.7279655 |
| 0 | 0 |
| 0 | 0 |
| 0 | 0 |
| 0 | 0 |
| 124.859285 | 45.7598851 |
| 0 | 0 |
| 0 | 0 |
| 0 | 0 |
| 0 | 0 |
| 0 | 0 |
| 0 | 0 |
| 0 | 0 |
| 0 | 0 |
| 0 | 0 |
| 0 | 0 |
| 0 | 0 |
| 0 | 0 |
| 0 | 0 |
| 0 | 0 |

| | |
|---|---|
| 0 | 0 |
| 0 | 0 |
| 0 | 0 |
| 0 | 0 |
| 0 | 0 |
| 0 | 0 |
| 0 | 0 |
| 0 | 0 |
| 0 | 0 |
| 0 | 96.0957587 |
| 0 | 0 |
| 0 | 0 |
| 0 | 0 |
| 0 | 0 |
| 0 | 0 |
| 0 | 0 |
| 0 | 0 |
| 0 | 4.57598851 |
| 0 | 0 |
| 0 | 0 |
| 0 | 0 |
| 0 | 0 |
| 0 | 0 |
| 0 | 0 |
| 0 | 0 |
| 0 | 0 |
| 0 | 0 |
| 16.2859937 | 22.8799426 |
| 0 | 0 |
| 0 | 0 |
| 0 | 0 |
| 0 | 4.57598851 |
| 0 | 0 |
| 0 | 0 |
| 0 | 0 |
| 0 | 0 |
| 0 | 0 |
| 0 | 0 |
| 0 | 0 |
| 0 | 0 |
| 0 | 0 |
| 0 | 0 |
| 0 | 0 |

| | |
|---|---|
| 0 | 0 |
| 0 | 0 |
| 0 | 0 |
| 0 | 0 |
| 0 | 0 |
| 0 | 0 |
| 0 | 0 |
| 0 | 0 |
| 0 | 0 |
| 0 | 0 |
| 0 | 0 |
| 0 | 0 |
| 0 | 0 |
| 0 | 0 |
| 0 | 0 |
| 0 | 4.57598851 |
| 0 | 0 |
| 0 | 0 |
| 0 | 0 |
| 0 | 0 |
| 0 | 0 |
| 0 | 0 |
| 0 | 0 |
| 0 | 0 |
| 0 | 0 |
| 24.4289905 | 18.303954 |
| 0 | 0 |
| 16.2859937 | 4.57598851 |
| 29.8576551 | 27.4559311 |
| 0 | 0 |
| 0 | 0 |
| 0 | 0 |
| 0 | 0 |
| 0 | 0 |
| 0 | 0 |
| 0 | 0 |
| 0 | 0 |
| 0 | 0 |
| 0 | 0 |
| 0 | 0 |
| 0 | 0 |
| 0 | 0 |
| 0 | 0 |

| | |
|---|---|
| 0 | 0 |
| 0 | 0 |
| 0 | 0 |
| 0 | 0 |
| 0 | 0 |
| 0 | 0 |
| 181.860263 | 68.6398277 |
| 0 | 0 |
| 0 | 0 |
| 0 | 0 |
| 0 | 0 |
| 0 | 0 |
| 0 | 4.57598851 |
| 0 | 0 |
| 0 | 0 |
| 0 | 0 |
| 0 | 0 |
| 0 | 0 |
| 0 | 0 |
| 0 | 0 |
| 0 | 0 |
| 0 | 0 |
| 0 | 0 |
| 0 | 0 |
| 0 | 0 |
| 0 | 0 |
| 0 | 0 |
| 0 | 0 |
| 0 | 0 |
| 0 | 0 |
| 0 | 4.57598851 |
| 0 | 0 |
| 0 | 4.57598851 |
| 0 | 0 |
| 0 | 0 |
| 0 | 0 |
| 0 | 0 |
| 0 | 0 |
| 0 | 0 |
| 0 | 0 |
| 0 | 0 |
| 0 | 0 |

| | |
|---|---|
| 0 | 0 |
| 0 | 0 |
| 0 | 0 |
| 0 | 0 |
| 0 | 4.57598851 |
| 0 | 0 |
| 0 | 0 |
| 0 | 0 |
| 0 | 0 |
| 0 | 4.57598851 |
| 0 | 0 |
| 0 | 0 |
| 0 | 0 |
| 0 | 0 |
| 0 | 0 |
| 0 | 0 |
| 0 | 0 |
| 0 | 0 |
| 0 | 0 |
| 0 | 0 |
| 0 | 0 |
| 0 | 0 |
| 0 | 0 |
| 0 | 0 |
| 2.71433228 | 50.3358736 |
| 0 | 0 |
| 0 | 0 |
| 0 | 0 |
| 0 | 0 |
| 0 | 0 |
| 0 | 0 |
| 0 | 0 |
| 2.71433228 | 27.4559311 |
| 0 | 0 |
| 0 | 0 |
| 0 | 0 |
| 0 | 0 |
| 0 | 4.57598851 |
| 0 | 0 |
| 0 | 0 |
| 0 | 0 |
| 0 | 0 |
| 0 | 0 |

| | |
|---|---|
| 0 | 0 |
| 0 | 0 |
| 0 | 77.7918047 |
| 0 | 0 |
| 0 | 0 |
| 0 | 0 |
| 0 | 0 |
| 0 | 4.57598851 |
| 0 | 0 |
| 2.71433228 | 0 |
| 0 | 0 |
| 0 | 0 |
| 0 | 0 |
| 0 | 0 |
| 0 | 0 |
| 0 | 0 |
| 0 | 0 |
| 0 | 0 |
| 0 | 0 |
| 580.867107 | 160.159598 |
| 0 | 0 |
| 0 | 0 |
| 0 | 0 |
| 0 | 0 |
| 0 | 0 |
| 181.860263 | 114.399713 |
| 0 | 0 |
| 0 | 4.57598851 |
| 0 | 0 |
| 0 | 0 |
| 0 | 0 |
| 0 | 0 |
| 0 | 0 |
| 0 | 0 |
| 0 | 0 |
| 0 | 0 |
| 0 | 0 |
| 0 | 0 |
| 0 | 0 |
| 2.71433228 | 0 |
| 0 | 0 |
| 0 | 0 |
| 0 | 0 |

| | |
|---|---|
| 0 | 0 |
| 0 | 0 |
| 0 | 0 |
| 0 | 0 |
| 909.301313 | 777.918047 |
| 0 | 0 |
| 0 | 0 |
| 822.44268 | 439.294897 |
| 0 | 4.57598851 |
| 0 | 0 |
| 0 | 0 |
| 0 | 0 |
| 0 | 0 |
| 0 | 0 |
| 0 | 0 |
| 0 | 0 |
| 0 | 0 |
| 0 | 0 |
| 0 | 0 |
| 0 | 0 |
| 0 | 0 |
| 0 | 0 |
| 0 | 0 |
| 0 | 0 |
| 0 | 0 |
| 0 | 0 |
| 0 | 0 |
| 0 | 0 |
| 0 | 0 |
| 0 | 0 |
| 0 | 0 |
| 2.71433228 | 0 |
| 0 | 0 |
| 0 | 4.57598851 |
| 0 | 0 |
| 0 | 0 |
| 0 | 0 |
| 0 | 0 |
| 0 | 0 |
| 0 | 0 |
| 0 | 0 |
| 0 | 0 |
| 0 | 0 |

| | |
|---|---|
| 0 | 0 |
| 0 | 0 |
| 0 | 0 |
| 0 | 0 |
| 2.71433228 | 22.8799426 |
| 0 | 0 |
| 0 | 0 |
| 0 | 0 |
| 0 | 0 |
| 8.14299683 | 32.0319196 |
| 0 | 0 |
| 0 | 9.15197702 |
| 0 | 0 |
| 0 | 0 |
| 0 | 0 |
| 0 | 0 |
| 0 | 0 |
| 0 | 0 |
| 59.7153101 | 50.3358736 |
| 0 | 0 |
| 0 | 0 |
| 0 | 0 |
| 0 | 0 |
| 0 | 0 |
| 0 | 0 |
| 0 | 4.57598851 |
| 0 | 0 |
| 0 | 0 |
| 119.43062 | 41.1838966 |
| 0 | 0 |
| 0 | 0 |
| 0 | 0 |
| 0 | 4.57598851 |
| 0 | 0 |
| 0 | 0 |
| 0 | 0 |
| 0 | 0 |
| 0 | 0 |
| 0 | 0 |
| 0 | 0 |
| 0 | 4.57598851 |
| 0 | 0 |
| 0 | 0 |

| | |
|---|---|
| 0 | 0 |
| 0 | 0 |
| 0 | 0 |
| 0 | 0 |
| 0 | 0 |
| 0 | 0 |
| 0 | 0 |
| 0 | 0 |
| 0 | 0 |
| 0 | 0 |
| 0 | 0 |
| 154.71694 | 169.311575 |
| 0 | 0 |
| 0 | 0 |
| 0 | 0 |
| 0 | 0 |
| 0 | 0 |
| 0 | 0 |
| 0 | 0 |
| 0 | 0 |
| 0 | 0 |
| 0 | 0 |
| 0 | 0 |
| 0 | 0 |
| 0 | 0 |
| 0 | 4.57598851 |
| 0 | 0 |
| 0 | 4.57598851 |
| 0 | 0 |
| 43.4293164 | 215.07146 |
| 0 | 0 |
| 0 | 0 |
| 0 | 0 |
| 0 | 0 |
| 428.8645 | 59.4878506 |
| 0 | 0 |
| 0 | 0 |
| 0 | 0 |
| 0 | 0 |
| 0 | 0 |
| 0 | 0 |
| 0 | 0 |
| 5.42866456 | 0 |

| | |
|---|---|
| 0 | 0 |
| 0 | 0 |
| 0 | 0 |
| 0 | 0 |
| 0 | 0 |
| 0 | 0 |
| 0 | 0 |
| 0 | 0 |
| 0 | 0 |
| 8.14299683 | 22.8799426 |
| 0 | 0 |
| 0 | 0 |
| 0 | 0 |
| 0 | 0 |
| 0 | 0 |
| 0 | 0 |
| 0 | 0 |
| 0 | 36.6079081 |
| 0 | 0 |
| 0 | 0 |
| 0 | 0 |
| 0 | 0 |
| 0 | 0 |
| 0 | 0 |
| 0 | 0 |
| 0 | 0 |
| 0 | 0 |
| 0 | 0 |
| 0 | 0 |
| 0 | 0 |
| 0 | 0 |
| 0 | 0 |
| 0 | 0 |
| 0 | 0 |
| 0 | 0 |
| 0 | 0 |
| 0 | 0 |
| 0 | 0 |
| 0 | 0 |
| 0 | 27.4559311 |
| 0 | 0 |
| 295.862218 | 361.503092 |
| 0 | 0 |

| | |
|---|---|
| 2.71433228 | 9.15197702 |
| 5.42866456 | 22.8799426 |
| 0 | 0 |
| 0 | 0 |
| 0 | 0 |
| 0 | 0 |
| 0 | 0 |
| 0 | 0 |
| 0 | 0 |
| 0 | 0 |
| 0 | 0 |
| 0 | 0 |
| 0 | 0 |
| 0 | 0 |
| 0 | 0 |
| 0 | 0 |
| 0 | 0 |
| 0 | 0 |
| 0 | 0 |
| 0 | 0 |
| 70.5726392 | 54.9118621 |
| 0 | 0 |
| 0 | 0 |
| 0 | 0 |
| 0 | 0 |
| 0 | 0 |
| 0 | 0 |
| 0 | 0 |
| 0 | 0 |
| 0 | 0 |
| 5.42866456 | 9.15197702 |
| 0 | 0 |
| 0 | 0 |
| 0 | 0 |
| 0 | 0 |
| 453.29349 | 41.1838966 |
| 0 | 0 |
| 0 | 0 |
| 0 | 0 |
| 0 | 0 |
| 0 | 0 |
| 0 | 0 |
| 0 | 22.8799426 |

| | |
|---|---|
| 2.71433228 | 0 |
| 0 | 0 |
| 0 | 0 |
| 0 | 0 |
| 0 | 0 |
| 0 | 0 |
| 0 | 0 |
| 0 | 0 |
| 0 | 0 |
| 0 | 0 |
| 0 | 0 |
| 0 | 0 |
| 0 | 0 |
| 0 | 0 |
| 0 | 0 |
| 0 | 0 |
| 0 | 0 |
| 0 | 0 |
| 0 | 0 |
| 0 | 0 |
| 0 | 4.57598851 |
| 0 | 0 |
| 0 | 0 |
| 0 | 0 |
| 0 | 0 |
| 0 | 0 |
| 0 | 0 |
| 0 | 0 |
| 0 | 0 |
| 0 | 0 |
| 0 | 0 |
| 0 | 0 |
| 0 | 9.15197702 |
| 0 | 0 |
| 0 | 4.57598851 |
| 0 | 0 |
| 0 | 0 |
| 0 | 0 |
| 0 | 0 |
| 0 | 0 |
| 0 | 0 |
| 0 | 0 |
| 0 | 0 |

| | |
|---|---|
| 0 | 0 |
| 0 | 0 |
| 0 | 0 |
| 0 | 0 |
| 0 | 0 |
| 0 | 0 |
| 0 | 0 |
| 0 | 4.57598851 |
| 0 | 0 |
| 0 | 0 |
| 0 | 0 |
| 0 | 0 |
| 0 | 4.57598851 |
| 0 | 0 |
| 0 | 0 |
| 0 | 0 |
| 0 | 0 |
| 0 | 0 |
| 0 | 0 |
| 0 | 0 |
| 0 | 0 |
| 0 | 0 |
| 0 | 0 |
| 0 | 0 |
| 0 | 0 |
| 0 | 0 |
| 0 | 0 |
| 0 | 0 |
| 2.71433228 | 0 |
| 0 | 0 |
| 0 | 0 |
| 0 | 4.57598851 |
| 0 | 0 |
| 0 | 0 |
| 0 | 0 |
| 0 | 0 |
| 0 | 0 |
| 0 | 0 |
| 0 | 0 |
| 0 | 0 |
| 0 | 0 |
| 195.431924 | 100.671747 |
| 0 | 0 |

| | |
|---|---|
| 0 | 0 |
| 0 | 0 |
| 0 | 0 |
| 0 | 0 |
| 0 | 0 |
| 0 | 0 |
| 0 | 0 |
| 0 | 0 |
| 0 | 0 |
| 0 | 0 |
| 0 | 0 |
| 0 | 0 |
| 0 | 0 |
| 0 | 0 |
| 0 | 0 |
| 0 | 0 |
| 0 | 0 |
| 0 | 0 |
| 0 | 0 |
| 0 | 0 |
| 0 | 0 |
| 0 | 0 |
| 0 | 0 |
| 173.717266 | 183.03954 |
| 0 | 0 |
| 0 | 0 |
| 0 | 0 |
| 0 | 0 |
| 0 | 0 |
| 0 | 0 |
| 0 | 0 |
| 0 | 0 |
| 0 | 0 |
| 0 | 0 |
| 0 | 22.8799426 |
| 0 | 0 |
| 0 | 0 |
| 0 | 0 |
| 0 | 0 |
| 0 | 0 |
| 0 | 0 |
| 0 | 0 |
| 0 | 0 |

| | |
|---|---|
| 0 | 0 |
| 27.1433228 | 0 |
| 0 | 0 |
| 0 | 0 |
| 0 | 0 |
| 0 | 0 |
| 0 | 0 |
| 0 | 0 |
| 0 | 0 |
| 0 | 0 |
| 2.71433228 | 0 |
| 0 | 0 |
| 0 | 0 |
| 2.71433228 | 0 |
| 0 | 0 |
| 0 | 0 |
| 0 | 0 |
| 0 | 0 |
| 0 | 0 |
| 0 | 0 |
| 10.8573291 | 18.303954 |
| 0 | 0 |
| 0 | 0 |
| 0 | 0 |
| 0 | 0 |
| 0 | 0 |
| 0 | 0 |
| 0 | 0 |
| 0 | 0 |
| 0 | 0 |
| 0 | 0 |
| 0 | 0 |
| 0 | 0 |
| 0 | 0 |
| 0 | 0 |
| 0 | 0 |
| 0 | 0 |
| 0 | 0 |
| 0 | 0 |
| 0 | 0 |
| 0 | 0 |
| 0 | 0 |
| 0 | 0 |

| | |
|---|---|
| 0 | 0 |
| 32.5719873 | 54.9118621 |
| 0 | 0 |
| 0 | 0 |
| 0 | 0 |
| 0 | 0 |
| 0 | 0 |
| 2.71433228 | 0 |
| 0 | 0 |
| 0 | 0 |
| 0 | 0 |
| 0 | 0 |
| 0 | 32.0319196 |
| 0 | 0 |
| 0 | 0 |
| 0 | 0 |
| 0 | 0 |
| 0 | 0 |
| 0 | 0 |
| 0 | 0 |
| 0 | 0 |
| 0 | 0 |
| 0 | 0 |
| 0 | 0 |
| 0 | 0 |
| 0 | 0 |
| 0 | 0 |
| 0 | 0 |
| 0 | 0 |
| 0 | 0 |
| 0 | 0 |
| 0 | 0 |
| 0 | 0 |
| 0 | 0 |
| 0 | 0 |
| 0 | 4.57598851 |
| 2.71433228 | 0 |
| 0 | 0 |
| 0 | 0 |
| 0 | 0 |
| 0 | 0 |
| 0 | 0 |
| 0 | 0 |

| | |
|---|---|
| 0 | 0 |
| 0 | 0 |
| 0 | 0 |
| 2.71433228 | 32.0319196 |
| 0 | 0 |
| 0 | 0 |
| 0 | 0 |
| 0 | 0 |
| 0 | 0 |
| 16.2859937 | 0 |
| 0 | 0 |
| 0 | 0 |
| 0 | 0 |
| 0 | 0 |
| 0 | 0 |
| 0 | 4.57598851 |
| 0 | 0 |
| 0 | 0 |
| 0 | 0 |
| 0 | 0 |
| 0 | 0 |
| 0 | 0 |
| 0 | 0 |
| 0 | 0 |
| 0 | 4.57598851 |
| 0 | 0 |
| 0 | 0 |
| 0 | 0 |
| 0 | 0 |
| 0 | 0 |
| 0 | 0 |
| 0 | 0 |
| 0 | 0 |
| 0 | 0 |
| 0 | 4.57598851 |
| 0 | 0 |
| 10.8573291 | 54.9118621 |
| 0 | 0 |
| 0 | 0 |
| 0 | 0 |
| 0 | 0 |
| 0 | 0 |
| 0 | 0 |

| | |
|---|---|
| 0 | 0 |
| 0 | 0 |
| 0 | 0 |
| 0 | 0 |
| 0 | 0 |
| 0 | 0 |
| 0 | 0 |
| 95.0016297 | 343.199138 |
| 0 | 0 |
| 16.2859937 | 0 |
| 0 | 0 |
| 0 | 0 |
| 0 | 0 |
| 0 | 18.303954 |
| 0 | 0 |
| 0 | 0 |
| 0 | 0 |
| 0 | 0 |
| 350.148864 | 118.975701 |
| 0 | 0 |
| 0 | 0 |
| 0 | 0 |
| 0 | 0 |
| 0 | 0 |
| 0 | 0 |
| 0 | 4.57598851 |
| 0 | 0 |
| 0 | 0 |
| 0 | 0 |
| 0 | 0 |
| 0 | 0 |
| 0 | 0 |
| 0 | 0 |
| 8.14299683 | 18.303954 |
| 0 | 0 |
| 0 | 0 |
| 2.71433228 | 4.57598851 |
| 0 | 0 |
| 0 | 0 |
| 0 | 0 |
| 0 | 0 |
| 0 | 0 |
| 0 | 0 |

| | |
|---|---|
| 0 | 0 |
| 2.71433228 | 9.15197702 |
| 0 | 0 |
| 0 | 0 |
| 0 | 0 |
| 0 | 0 |
| 0 | 9.15197702 |
| 0 | 0 |
| 0 | 0 |
| 0 | 0 |
| 0 | 0 |
| 0 | 0 |
| 0 | 0 |
| 0 | 0 |
| 0 | 4.57598851 |
| 0 | 4.57598851 |
| 0 | 0 |
| 437.007497 | 50.3358736 |
| 0 | 0 |
| 0 | 0 |
| 0 | 0 |
| 0 | 0 |
| 0 | 0 |
| 0 | 0 |
| 0 | 18.303954 |
| 0 | 0 |
| 0 | 0 |
| 0 | 27.4559311 |
| 0 | 0 |
| 0 | 0 |
| 0 | 0 |
| 59.7153101 | 45.7598851 |
| 0 | 0 |
| 0 | 0 |
| 0 | 0 |
| 0 | 0 |
| 0 | 0 |
| 0 | 0 |
| 0 | 0 |
| 0 | 0 |
| 0 | 0 |
| 76.0013038 | 32.0319196 |
| 0 | 22.8799426 |

| | |
|---|---|
| 0 | 0 |
| 0 | 0 |
| 16.2859937 | 77.7918047 |
| 0 | 0 |
| 0 | 0 |
| 0 | 0 |
| 0 | 0 |
| 0 | 0 |
| 0 | 0 |
| 27.1433228 | 73.2158162 |
| 0 | 0 |
| 0 | 0 |
| 0 | 0 |
| 0 | 0 |
| 0 | 0 |
| 0 | 0 |
| 0 | 41.1838966 |
| 0 | 0 |
| 5.42866456 | 4.57598851 |
| 0 | 0 |
| 0 | 0 |
| 0 | 0 |
| 0 | 0 |
| 0 | 0 |
| 0 | 0 |
| 0 | 0 |
| 0 | 0 |
| 133.002282 | 22.8799426 |
| 0 | 0 |
| 5.42866456 | 50.3358736 |
| 0 | 0 |
| 0 | 0 |
| 0 | 0 |
| 0 | 0 |
| 0 | 0 |
| 0 | 0 |
| 0 | 0 |
| 0 | 0 |
| 0 | 4.57598851 |
| 0 | 0 |
| 0 | 0 |
| 0 | 0 |
| 0 | 0 |

| | |
|---|---|
| 0 | 0 |
| 217.146582 | 201.343494 |
| 0 | 0 |
| 0 | 0 |
| 0 | 0 |
| 0 | 0 |
| 0 | 68.6398277 |
| 0 | 0 |
| 2.71433228 | 13.7279655 |
| 2.71433228 | 0 |
| 0 | 0 |
| 0 | 0 |
| 0 | 0 |
| 0 | 0 |
| 2.71433228 | 0 |
| 0 | 0 |
| 0 | 0 |
| 0 | 0 |
| 0 | 0 |
| 0 | 0 |
| 0 | 0 |
| 0 | 0 |
| 0 | 0 |
| 0 | 0 |
| 0 | 0 |
| 0 | 0 |
| 0 | 0 |
| 0 | 0 |
| 0 | 0 |
| 0 | 0 |
| 0 | 0 |
| 0 | 0 |
| 0 | 0 |
| 0 | 4.57598851 |
| 0 | 0 |
| 0 | 0 |
| 0 | 0 |
| 0 | 0 |
| 0 | 0 |
| 0 | 0 |
| 0 | 0 |
| 0 | 0 |
| 0 | 0 |

| | |
|---|---|
| 0 | 0 |
| 0 | 0 |
| 179.14593 | 54.9118621 |
| 0 | 0 |
| 0 | 0 |
| 0 | 0 |
| 0 | 0 |
| 0 | 0 |
| 0 | 0 |
| 0 | 27.4559311 |
| 0 | 0 |
| 0 | 0 |
| 0 | 0 |
| 0 | 0 |
| 0 | 0 |
| 0 | 0 |
| 0 | 0 |
| 0 | 0 |
| 0 | 0 |
| 0 | 0 |
| 0 | 0 |
| 0 | 0 |
| 0 | 0 |
| 0 | 0 |
| 0 | 0 |
| 0 | 0 |
| 0 | 0 |
| 0 | 0 |
| 0 | 0 |
| 19.0003259 | 0 |
| 0 | 0 |
| 0 | 0 |
| 0 | 0 |
| 0 | 0 |
| 0 | 0 |
| 0 | 0 |
| 0 | 0 |
| 0 | 0 |
| 0 | 0 |
| 57.0009778 | 0 |
| 0 | 0 |
| 0 | 0 |
| 0 | 0 |

| | |
|---|---|
| 0 | 0 |
| 0 | 0 |
| 0 | 0 |
| 0 | 0 |
| 0 | 0 |
| 0 | 0 |
| 0 | 0 |
| 0 | 0 |
| 0 | 0 |
| 0 | 22.8799426 |
| 0 | 0 |
| 0 | 0 |
| 0 | 0 |
| 0 | 0 |
| 5.42866456 | 160.159598 |
| 0 | 0 |
| 0 | 0 |
| 0 | 0 |
| 0 | 0 |
| 0 | 0 |
| 0 | 27.4559311 |
| 0 | 0 |
| 0 | 4.57598851 |
| 2.71433228 | 0 |
| 0 | 0 |
| 0 | 4.57598851 |
| 0 | 0 |
| 0 | 0 |
| 0 | 0 |
| 0 | 0 |
| 0 | 0 |
| 0 | 0 |
| 0 | 0 |
| 214.43225 | 32.0319196 |
| 0 | 0 |
| 0 | 0 |
| 0 | 0 |
| 2.71433228 | 9.15197702 |
| 0 | 0 |
| 0 | 0 |
| 0 | 0 |
| 0 | 0 |
| 0 | 0 |

| | |
|---|---|
| 2.71433228 | 4.57598851 |
| 0 | 0 |
| 2.71433228 | 0 |
| 0 | 0 |
| 0 | 0 |
| 0 | 0 |
| 0 | 0 |
| 0 | 0 |
| 0 | 0 |
| 0 | 0 |
| 0 | 0 |
| 0 | 0 |
| 0 | 0 |
| 0 | 0 |
| 0 | 0 |
| 0 | 0 |
| 0 | 0 |
| 0 | 0 |
| 0 | 0 |
| 0 | 0 |
| 0 | 0 |
| 0 | 18.303954 |
| 0 | 0 |
| 0 | 0 |
| 0 | 0 |
| 73.2869715 | 96.0957587 |
| 0 | 0 |
| 0 | 0 |
| 0 | 0 |
| 0 | 0 |
| 0 | 0 |
| 0 | 0 |
| 0 | 0 |
| 0 | 0 |
| 0 | 0 |
| 0 | 0 |
| 0 | 0 |
| 0 | 0 |
| 152.002608 | 4.57598851 |
| 0 | 0 |
| 0 | 0 |
| 0 | 0 |
| 0 | 0 |

| | |
|---|---|
| 0 | 27.4559311 |
| 0 | 4.57598851 |
| 0 | 0 |
| 0 | 0 |
| 0 | 0 |
| 0 | 0 |
| 0 | 0 |
| 0 | 0 |
| 0 | 0 |
| 0 | 0 |
| 0 | 0 |
| 195.431924 | 96.0957587 |
| 0 | 0 |
| 0 | 0 |
| 0 | 0 |
| 0 | 0 |
| 0 | 0 |
| 0 | 0 |
| 0 | 0 |
| 0 | 0 |
| 0 | 0 |
| 0 | 0 |
| 0 | 0 |
| 0 | 0 |
| 0 | 0 |
| 0 | 0 |
| 0 | 0 |
| 0 | 0 |
| 0 | 0 |
| 0 | 0 |
| 0 | 0 |
| 0 | 0 |
| 0 | 0 |
| 0 | 0 |
| 0 | 0 |
| 0 | 0 |
| 0 | 0 |
| 0 | 0 |
| 0 | 0 |
| 0 | 0 |
| 0 | 0 |
| 0 | 0 |
| 5.42866456 | 0 |

| | |
|---|---|
| 0 | 0 |
| 0 | 0 |
| 0 | 0 |
| 0 | 0 |
| 0 | 9.15197702 |
| 0 | 0 |
| 0 | 0 |
| 0 | 0 |
| 0 | 0 |
| 2.71433228 | 0 |
| 0 | 0 |
| 0 | 0 |
| 0 | 0 |
| 67.8583069 | 4.57598851 |
| 0 | 0 |
| 0 | 0 |
| 0 | 0 |
| 0 | 0 |
| 0 | 0 |
| 0 | 0 |
| 0 | 0 |
| 0 | 0 |
| 0 | 0 |
| 0 | 0 |
| 0 | 0 |
| 8.14299683 | 45.7598851 |
| 0 | 0 |
| 0 | 0 |
| 0 | 0 |
| 355.577528 | 128.127678 |
| 0 | 0 |
| 0 | 0 |
| 0 | 0 |
| 2.71433228 | 18.303954 |
| 46.1436487 | 13.7279655 |
| 0 | 0 |
| 0 | 0 |
| 0 | 0 |
| 0 | 22.8799426 |
| 0 | 0 |
| 0 | 0 |
| 0 | 0 |
| 0 | 0 |

| | |
|---|---|
| 0 | 0 |
| 0 | 0 |
| 0 | 0 |
| 0 | 0 |
| 0 | 0 |
| 0 | 0 |
| 0 | 0 |
| 0 | 0 |
| 0 | 0 |
| 0 | 0 |
| 0 | 0 |
| 0 | 9.15197702 |
| 0 | 0 |
| 0 | 0 |
| 0 | 0 |
| 0 | 0 |
| 0 | 0 |
| 0 | 0 |
| 0 | 0 |
| 0 | 0 |
| 0 | 0 |
| 0 | 0 |
| 0 | 0 |
| 0 | 0 |
| 0 | 0 |
| 0 | 0 |
| 0 | 0 |
| 184.574595 | 151.007621 |
| 0 | 0 |
| 0 | 0 |
| 0 | 0 |
| 285.004889 | 123.55169 |
| 2.71433228 | 0 |
| 0 | 0 |
| 0 | 0 |
| 0 | 0 |
| 0 | 0 |
| 0 | 0 |
| 0 | 0 |
| 0 | 0 |
| 0 | 0 |
| 0 | 0 |
| 0 | 0 |

| | |
|---|---|
| 2.71433228 | 0 |
| 0 | 0 |
| 190.003259 | 9.15197702 |
| 0 | 0 |
| 0 | 0 |
| 0 | 0 |
| 0 | 0 |
| 0 | 0 |
| 2.71433228 | 0 |
| 0 | 0 |
| 65.1439747 | 86.9437817 |
| 0 | 0 |
| 0 | 0 |
| 2.71433228 | 13.7279655 |
| 35.2863196 | 4.57598851 |
| 0 | 0 |
| 0 | 0 |
| 0 | 41.1838966 |
| 0 | 0 |
| 0 | 0 |
| 0 | 0 |
| 0 | 0 |
| 0 | 0 |
| 0 | 0 |
| 0 | 0 |
| 0 | 0 |
| 2.71433228 | 0 |
| 0 | 0 |
| 0 | 0 |
| 135.716614 | 18.303954 |
| 0 | 0 |
| 13.5716614 | 50.3358736 |
| 0 | 0 |
| 0 | 0 |
| 0 | 0 |
| 0 | 0 |
| 0 | 0 |
| 0 | 0 |
| 0 | 0 |
| 0 | 0 |
| 0 | 0 |
| 0 | 0 |
| 0 | 0 |

| | |
|---|---|
| 0 | 0 |
| 0 | 0 |
| 0 | 0 |
| 0 | 0 |
| 0 | 0 |
| 0 | 0 |
| 0 | 0 |
| 0 | 0 |
| 0 | 0 |
| 0 | 0 |
| 135.716614 | 151.007621 |
| 0 | 0 |
| 0 | 0 |
| 0 | 0 |
| 0 | 0 |
| 0 | 4.57598851 |
| 0 | 0 |
| 0 | 0 |
| 0 | 0 |
| 0 | 0 |
| 0 | 0 |
| 0 | 0 |
| 0 | 0 |
| 2.71433228 | 0 |
| 0 | 0 |
| 5.42866456 | 0 |
| 0 | 0 |
| 0 | 0 |
| 0 | 0 |
| 2.71433228 | 18.303954 |
| 157.431272 | 118.975701 |
| 0 | 0 |
| 89.5729652 | 4.57598851 |
| 0 | 4.57598851 |
| 0 | 0 |
| 0 | 0 |
| 0 | 0 |
| 0 | 0 |
| 0 | 0 |
| 0 | 0 |
| 0 | 0 |
| 0 | 0 |
| 0 | 0 |

| | |
|---|---|
| 0 | 0 |
| 0 | 0 |
| 0 | 0 |
| 0 | 0 |
| 0 | 13.7279655 |
| 0 | 0 |
| 0 | 0 |
| 0 | 0 |
| 0 | 0 |
| 0 | 0 |
| 0 | 0 |
| 0 | 0 |
| 0 | 0 |
| 0 | 0 |
| 0 | 0 |
| 0 | 0 |
| 0 | 0 |
| 0 | 4.57598851 |
| 0 | 0 |
| 16.2859937 | 0 |
| 0 | 0 |
| 0 | 0 |
| 0 | 4.57598851 |
| 0 | 0 |
| 0 | 0 |
| 0 | 0 |
| 0 | 0 |
| 0 | 0 |
| 0 | 0 |
| 0 | 0 |
| 57.0009778 | 22.8799426 |
| 0 | 0 |
| 0 | 0 |
| 65.1439747 | 86.9437817 |
| 0 | 0 |
| 0 | 0 |
| 542.866456 | 613.18246 |
| 0 | 0 |
| 2.71433228 | 0 |
| 0 | 0 |
| 8.14299683 | 82.3677932 |
| 0 | 0 |
| 0 | 0 |

| | |
|---|---|
| 0 | 0 |
| 0 | 0 |
| 0 | 0 |
| 0 | 0 |
| 8.14299683 | 0 |
| 0 | 32.0319196 |
| 0 | 0 |
| 0 | 0 |
| 0 | 4.57598851 |
| 0 | 0 |
| 0 | 0 |
| 0 | 0 |
| 0 | 0 |
| 0 | 0 |
| 0 | 0 |
| 0 | 0 |
| 0 | 0 |
| 0 | 0 |
| 0 | 0 |
| 0 | 0 |
| 0 | 0 |
| 0 | 0 |
| 0 | 0 |
| 0 | 0 |
| 0 | 0 |
| 10.8573291 | 36.6079081 |
| 0 | 0 |
| 0 | 0 |
| 2.71433228 | 0 |
| 0 | 0 |
| 0 | 0 |
| 2.71433228 | 0 |
| 0 | 0 |
| 0 | 0 |
| 0 | 0 |
| 233.432576 | 132.703667 |
| 0 | 0 |
| 0 | 0 |
| 0 | 0 |
| 0 | 0 |
| 146.573943 | 64.0638392 |
| 0 | 0 |
| 0 | 0 |

| | |
|---|---|
| 2.71433228 | 0 |
| 0 | 0 |
| 0 | 0 |
| 0 | 0 |
| 0 | 0 |
| 0 | 0 |
| 0 | 0 |
| 0 | 0 |
| 0 | 4.57598851 |
| 0 | 0 |
| 0 | 0 |
| 0 | 0 |
| 0 | 0 |
| 0 | 0 |
| 0 | 4.57598851 |
| 0 | 0 |
| 0 | 0 |
| 0 | 0 |
| 0 | 0 |
| 0 | 0 |
| 0 | 0 |
| 0 | 0 |
| 0 | 0 |
| 0 | 0 |
| 0 | 0 |
| 0 | 0 |
| 2.71433228 | 4.57598851 |
| 0 | 0 |
| 0 | 0 |
| 0 | 0 |
| 0 | 0 |
| 0 | 0 |
| 0 | 0 |
| 0 | 0 |
| 0 | 0 |
| 0 | 0 |
| 0 | 0 |
| 0 | 0 |
| 0 | 0 |
| 0 | 0 |
| 0 | 0 |
| 0 | 0 |
| 0 | 0 |

| | |
|---|---|
| 0 | 0 |
| 0 | 0 |
| 0 | 0 |
| 0 | 0 |
| 0 | 0 |
| 0 | 0 |
| 0 | 0 |
| 0 | 0 |
| 0 | 0 |
| 0 | 13.7279655 |
| 0 | 0 |
| 0 | 0 |
| 0 | 0 |
| 0 | 0 |
| 0 | 0 |
| 0 | 0 |
| 0 | 0 |
| 0 | 0 |
| 2.71433228 | 36.6079081 |
| 0 | 0 |
| 0 | 0 |
| 0 | 0 |
| 2.71433228 | 0 |
| 0 | 0 |
| 0 | 45.7598851 |
| 0 | 0 |
| 0 | 0 |
| 0 | 0 |
| 46.1436487 | 4.57598851 |
| 0 | 0 |
| 0 | 27.4559311 |
| 152.002608 | 82.3677932 |
| 2.71433228 | 4.57598851 |
| 0 | 0 |
| 0 | 9.15197702 |
| 0 | 0 |
| 0 | 0 |
| 0 | 0 |
| 0 | 0 |
| 0 | 0 |
| 0 | 0 |
| 0 | 0 |
| 0 | 0 |

| | |
|---|---|
| 0 | 0 |
| 0 | 0 |
| 0 | 0 |
| 0 | 0 |
| 0 | 0 |
| 0 | 0 |
| 0 | 0 |
| 0 | 0 |
| 0 | 4.57598851 |
| 0 | 0 |
| 0 | 0 |
| 0 | 0 |
| 0 | 0 |
| 0 | 0 |
| 0 | 0 |
| 2.71433228 | 0 |
| 0 | 0 |
| 0 | 0 |
| 0 | 0 |
| 0 | 0 |
| 0 | 0 |
| 0 | 0 |
| 0 | 0 |
| 5.42866456 | 82.3677932 |
| 0 | 0 |
| 0 | 0 |
| 0 | 0 |
| 0 | 0 |
| 0 | 0 |
| 0 | 0 |
| 0 | 0 |
| 0 | 0 |
| 0 | 0 |
| 0 | 0 |
| 0 | 0 |
| 0 | 0 |
| 0 | 0 |
| 0 | 0 |
| 0 | 4.57598851 |
| 0 | 0 |
| 0 | 0 |
| 0 | 0 |
| 0 | 0 |

| | |
|---|---|
| 0 | 0 |
| 0 | 0 |
| 2.71433228 | 68.6398277 |
| 0 | 0 |
| 184.574595 | 4.57598851 |
| 0 | 0 |
| 0 | 0 |
| 0 | 0 |
| 62.4296424 | 233.375414 |
| 0 | 0 |
| 0 | 0 |
| 0 | 0 |
| 0 | 0 |
| 0 | 0 |
| 0 | 0 |
| 0 | 4.57598851 |
| 0 | 0 |
| 0 | 0 |
| 0 | 0 |
| 0 | 0 |
| 2.71433228 | 0 |
| 122.144953 | 45.7598851 |
| 0 | 0 |
| 0 | 0 |
| 0 | 4.57598851 |
| 0 | 0 |
| 0 | 0 |
| 0 | 0 |
| 0 | 0 |
| 0 | 0 |
| 0 | 0 |
| 0 | 0 |
| 16.2859937 | 13.7279655 |
| 0 | 0 |
| 0 | 0 |
| 0 | 0 |
| 0 | 0 |
| 0 | 0 |
| 0 | 0 |
| 0 | 0 |
| 173.717266 | 4.57598851 |
| 0 | 0 |
| 0 | 0 |

| | |
|---|---|
| 0 | 0 |
| 0 | 0 |
| 0 | 0 |
| 0 | 0 |
| 0 | 0 |
| 0 | 0 |
| 0 | 0 |
| 0 | 0 |
| 0 | 0 |
| 0 | 0 |
| 0 | 0 |
| 0 | 0 |
| 0 | 0 |
| 2.71433228 | 4.57598851 |
| 0 | 0 |
| 0 | 0 |
| 0 | 0 |
| 0 | 0 |
| 0 | 0 |
| 0 | 0 |
| 0 | 4.57598851 |
| 0 | 0 |
| 0 | 0 |
| 2.71433228 | 0 |
| 1330.02282 | 379.807046 |
| 0 | 0 |
| 0 | 0 |
| 0 | 0 |
| 0 | 0 |
| 0 | 0 |
| 0 | 4.57598851 |
| 0 | 0 |
| 0 | 0 |
| 1115.59057 | 22.8799426 |
| 0 | 0 |
| 0 | 0 |
| 0 | 0 |
| 0 | 0 |
| 469.579484 | 201.343494 |
| 0 | 0 |
| 0 | 0 |
| 0 | 4.57598851 |
| 0 | 0 |

| | |
|---|---|
| 0 | 0 |
| 0 | 0 |
| 0 | 0 |
| 0 | 0 |
| 249.71857 | 109.823724 |
| 0 | 0 |
| 0 | 0 |
| 0 | 0 |
| 0 | 0 |
| 13.5716614 | 13.7279655 |
| 2.71433228 | 0 |
| 0 | 9.15197702 |
| 0 | 0 |
| 0 | 0 |
| 0 | 0 |
| 0 | 0 |
| 8.14299683 | 128.127678 |
| 0 | 0 |
| 0 | 0 |
| 0 | 0 |
| 0 | 0 |
| 0 | 0 |
| 0 | 0 |
| 0 | 0 |
| 0 | 0 |
| 0 | 0 |
| 0 | 0 |
| 705.726392 | 338.62315 |
| 0 | 0 |
| 0 | 0 |
| 0 | 0 |
| 0 | 0 |
| 0 | 0 |
| 0 | 0 |
| 0 | 0 |
| 0 | 0 |
| 0 | 18.303954 |
| 0 | 0 |
| 0 | 0 |
| 0 | 0 |
| 0 | 0 |
| 0 | 0 |
| 0 | 0 |

| | |
|---|---|
| 0 | 0 |
| 0 | 0 |
| 0 | 0 |
| 0 | 0 |
| 0 | 0 |
| 0 | 0 |
| 0 | 0 |
| 0 | 0 |
| 0 | 0 |
| 0 | 0 |
| 0 | 0 |
| 0 | 0 |
| 0 | 0 |
| 0 | 0 |
| 0 | 0 |
| 0 | 0 |
| 0 | 0 |
| 0 | 0 |
| 0 | 0 |
| 0 | 0 |
| 0 | 0 |
| 116.716288 | 73.2158162 |
| 0 | 0 |
| 0 | 0 |
| 0 | 0 |
| 0 | 0 |
| 48.857981 | 64.0638392 |
| 0 | 0 |
| 0 | 0 |
| 0 | 0 |
| 0 | 0 |
| 0 | 0 |
| 0 | 4.57598851 |
| 0 | 0 |
| 0 | 0 |
| 0 | 0 |
| 0 | 0 |
| 2.71433228 | 4.57598851 |
| 0 | 0 |
| 0 | 0 |
| 0 | 0 |
| 0 | 0 |
| 0 | 0 |

| | |
|---|---|
| 0 | 0 |
| 0 | 0 |
| 0 | 0 |
| 0 | 0 |
| 0 | 0 |
| 0 | 0 |
| 0 | 0 |
| 62.4296424 | 187.615529 |
| 0 | 0 |
| 0 | 0 |
| 0 | 0 |
| 0 | 0 |
| 86.8586329 | 96.0957587 |
| 0 | 0 |
| 0 | 0 |
| 0 | 0 |
| 5.42866456 | 18.303954 |
| 0 | 0 |
| 0 | 0 |
| 0 | 0 |
| 0 | 0 |
| 0 | 0 |
| 0 | 0 |
| 0 | 0 |
| 0 | 0 |
| 0 | 0 |
| 0 | 0 |
| 0 | 0 |
| 5.42866456 | 45.7598851 |
| 0 | 0 |
| 290.433554 | 59.4878506 |
| 0 | 0 |
| 0 | 0 |
| 0 | 0 |
| 0 | 0 |
| 13.5716614 | 0 |
| 0 | 0 |
| 0 | 0 |
| 0 | 0 |
| 0 | 0 |
| 0 | 0 |
| 0 | 0 |
| 219.860915 | 265.407334 |

| | |
|---|---|
| 0 | 0 |
| 0 | 0 |
| 0 | 0 |
| 0 | 4.57598851 |
| 0 | 0 |
| 0 | 0 |
| 0 | 0 |
| 0 | 0 |
| 0 | 0 |
| 0 | 0 |
| 0 | 0 |
| 0 | 0 |
| 0 | 0 |
| 0 | 0 |
| 0 | 0 |
| 0 | 0 |
| 0 | 0 |
| 0 | 0 |
| 0 | 0 |
| 0 | 0 |
| 0 | 0 |
| 0 | 0 |
| 0 | 0 |
| 0 | 0 |
| 0 | 0 |
| 0 | 0 |
| 0 | 0 |
| 149.288275 | 9.15197702 |
| 0 | 0 |
| 0 | 0 |
| 0 | 0 |
| 0 | 0 |
| 0 | 0 |
| 0 | 0 |
| 0 | 0 |
| 0 | 0 |
| 0 | 0 |
| 0 | 0 |
| 0 | 0 |
| 0 | 4.57598851 |
| 0 | 0 |
| 0 | 0 |
| 0 | 0 |

| | |
|---|---|
| 152.002608 | 123.55169 |
| 0 | 4.57598851 |
| 168.288601 | 13.7279655 |
| 0 | 0 |
| 0 | 0 |
| 0 | 0 |
| 0 | 4.57598851 |
| 0 | 0 |
| 0 | 0 |
| 0 | 0 |
| 2.71433228 | 4.57598851 |
| 0 | 0 |
| 0 | 0 |
| 0 | 0 |
| 0 | 4.57598851 |
| 8.14299683 | 18.303954 |
| 0 | 0 |
| 0 | 0 |
| 0 | 0 |
| 0 | 0 |
| 0 | 0 |
| 0 | 27.4559311 |
| 0 | 0 |
| 0 | 0 |
| 0 | 4.57598851 |
| 0 | 0 |
| 0 | 0 |
| 0 | 0 |
| 24.4289905 | 0 |
| 0 | 0 |
| 0 | 0 |
| 0 | 0 |
| 70.5726392 | 100.671747 |
| 0 | 0 |
| 0 | 0 |
| 0 | 0 |
| 0 | 0 |
| 0 | 0 |
| 0 | 0 |
| 616.153427 | 210.495472 |
| 119.43062 | 100.671747 |
| 0 | 0 |
| 0 | 0 |

| | |
|---|---|
| 0 | 0 |
| 0 | 0 |
| 0 | 0 |
| 0 | 41.1838966 |
| 0 | 0 |
| 0 | 0 |
| 0 | 0 |
| 0 | 0 |
| 0 | 0 |
| 0 | 0 |
| 0 | 0 |
| 0 | 0 |
| 0 | 0 |
| 0 | 0 |
| 0 | 0 |
| 65.1439747 | 36.6079081 |
| 0 | 0 |
| 0 | 0 |
| 0 | 0 |
| 76.0013038 | 4.57598851 |
| 0 | 0 |
| 0 | 0 |
| 0 | 0 |
| 0 | 0 |
| 0 | 0 |
| 0 | 0 |
| 0 | 18.303954 |
| 0 | 0 |
| 0 | 0 |
| 0 | 0 |
| 8.14299683 | 0 |
| 0 | 0 |
| 0 | 0 |
| 0 | 0 |
| 0 | 0 |
| 0 | 0 |
| 0 | 0 |
| 0 | 0 |
| 0 | 0 |
| 0 | 0 |
| 0 | 0 |
| 0 | 0 |
| 192.717592 | 155.583609 |

| | |
|---|---|
| 0 | 0 |
| 0 | 0 |
| 0 | 0 |
| 0 | 0 |
| 0 | 0 |
| 0 | 0 |
| 0 | 0 |
| 0 | 0 |
| 0 | 0 |
| 0 | 0 |
| 0 | 0 |
| 409.864174 | 0 |
| 0 | 0 |
| 0 | 0 |
| 0 | 0 |
| 0 | 0 |
| 0 | 0 |
| 0 | 0 |
| 0 | 45.7598851 |
| 0 | 0 |
| 0 | 0 |
| 0 | 0 |
| 0 | 0 |
| 0 | 0 |
| 0 | 0 |
| 0 | 0 |
| 0 | 0 |
| 0 | 0 |
| 0 | 0 |
| 0 | 0 |
| 0 | 0 |
| 0 | 0 |
| 0 | 0 |
| 0 | 0 |
| 0 | 0 |
| 0 | 0 |
| 0 | 0 |
| 43.4293164 | 0 |
| 0 | 0 |
| 0 | 0 |
| 2.71433228 | 4.57598851 |
| 0 | 0 |
| 0 | 0 |

| | |
|---|---|
| 0 | 0 |
| 0 | 0 |
| 0 | 0 |
| 0 | 0 |
| 0 | 0 |
| 0 | 0 |
| 0 | 0 |
| 2.71433228 | 0 |
| 0 | 0 |
| 0 | 0 |
| 0 | 0 |
| 0 | 0 |
| 0 | 0 |
| 0 | 0 |
| 0 | 0 |
| 0 | 0 |
| 0 | 0 |
| 2.71433228 | 0 |
| 0 | 0 |
| 0 | 0 |
| 0 | 0 |
| 0 | 0 |
| 0 | 0 |
| 0 | 0 |
| 0 | 0 |
| 0 | 0 |
| 0 | 0 |
| 0 | 0 |
| 0 | 0 |
| 0 | 0 |
| 0 | 0 |
| 0 | 0 |
| 0 | 13.7279655 |
| 0 | 0 |
| 0 | 0 |
| 0 | 0 |
| 0 | 0 |
| 0 | 0 |
| 0 | 0 |
| 0 | 0 |
| 8.14299683 | 0 |
| 0 | 0 |

| | |
|---|---|
| 0 | 0 |
| 0 | 0 |
| 0 | 0 |
| 0 | 0 |
| 0 | 0 |
| 0 | 4.57598851 |
| 0 | 0 |
| 0 | 0 |
| 0 | 18.303954 |
| 0 | 0 |
| 0 | 0 |
| 0 | 0 |
| 0 | 0 |
| 0 | 0 |
| 428.8645 | 169.311575 |
| 0 | 4.57598851 |
| 0 | 0 |
| 0 | 0 |
| 0 | 0 |
| 0 | 0 |
| 0 | 0 |
| 0 | 0 |
| 0 | 0 |
| 0 | 0 |
| 0 | 0 |
| 0 | 0 |
| 0 | 0 |
| 0 | 0 |
| 0 | 0 |
| 0 | 0 |
| 0 | 18.303954 |
| 0 | 0 |
| 59.7153101 | 118.975701 |
| 0 | 0 |
| 0 | 0 |
| 86.8586329 | 4.57598851 |
| 0 | 0 |
| 0 | 0 |
| 0 | 0 |
| 0 | 0 |
| 0 | 0 |
| 0 | 0 |
| 0 | 0 |

| | |
|---|---|
| 0 | 0 |
| 0 | 0 |
| 0 | 0 |
| 0 | 0 |
| 0 | 0 |
| 0 | 0 |
| 0 | 0 |
| 0 | 0 |
| 0 | 0 |
| 0 | 0 |
| 0 | 0 |
| 0 | 0 |
| 0 | 0 |
| 0 | 0 |
| 0 | 0 |
| 0 | 0 |
| 0 | 0 |
| 0 | 0 |
| 0 | 0 |
| 0 | 0 |
| 0 | 0 |
| 0 | 0 |
| 0 | 0 |
| 0 | 0 |
| 0 | 0 |
| 0 | 4.57598851 |
| 0 | 0 |
| 0 | 0 |
| 0 | 18.303954 |
| 0 | 0 |
| 0 | 0 |
| 0 | 0 |
| 0 | 0 |
| 73.2869715 | 50.3358736 |
| 0 | 4.57598851 |
| 0 | 0 |
| 0 | 0 |
| 0 | 4.57598851 |
| 0 | 0 |
| 0 | 0 |
| 13.5716614 | 0 |
| 0 | 0 |
| 0 | 0 |

| | |
|---|---|
| 0 | 0 |
| 0 | 0 |
| 5.42866456 | 36.6079081 |
| 0 | 0 |
| 0 | 22.8799426 |
| 0 | 0 |
| 0 | 0 |
| 409.864174 | 91.5197702 |
| 0 | 0 |
| 62.4296424 | 9.15197702 |
| 0 | 0 |
| 0 | 0 |
| 0 | 0 |
| 0 | 0 |
| 0 | 0 |
| 0 | 0 |
| 0 | 0 |
| 0 | 0 |
| 0 | 0 |
| 0 | 0 |
| 0 | 0 |
| 0 | 0 |
| 0 | 0 |
| 0 | 0 |
| 86.8586329 | 215.07146 |
| 0 | 0 |
| 0 | 13.7279655 |
| 0 | 0 |
| 0 | 0 |
| 0 | 0 |
| 0 | 0 |
| 0 | 0 |
| 0 | 0 |
| 2.71433228 | 0 |
| 0 | 0 |
| 0 | 0 |
| 0 | 0 |
| 0 | 0 |
| 0 | 0 |
| 0 | 0 |
| 0 | 0 |
| 0 | 0 |
| 0 | 0 |

| | |
|---|---|
| 0 | 0 |
| 0 | 0 |
| 111.287623 | 279.135299 |
| 0 | 0 |
| 0 | 0 |
| 0 | 0 |
| 0 | 9.15197702 |
| 0 | 0 |
| 0 | 0 |
| 0 | 0 |
| 0 | 0 |
| 0 | 0 |
| 0 | 0 |
| 0 | 0 |
| 0 | 0 |
| 0 | 0 |
| 0 | 0 |
| 0 | 0 |
| 0 | 0 |
| 0 | 0 |
| 0 | 0 |
| 0 | 0 |
| 13.5716614 | 18.303954 |
| 0 | 0 |
| 0 | 0 |
| 0 | 0 |
| 152.002608 | 82.3677932 |
| 0 | 0 |
| 0 | 0 |
| 0 | 0 |
| 241.575573 | 283.711288 |
| 0 | 0 |
| 0 | 0 |
| 0 | 0 |
| 0 | 0 |
| 0 | 0 |
| 0 | 0 |
| 0 | 0 |
| 0 | 0 |
| 0 | 0 |
| 0 | 0 |
| 0 | 0 |
| 0 | 0 |

| | |
|---|---|
| 0 | 0 |
| 0 | 0 |
| 675.868737 | 146.431632 |
| 852.300335 | 32.0319196 |
| 0 | 0 |
| 0 | 0 |
| 0 | 0 |
| 0 | 4.57598851 |
| 0 | 0 |
| 21.7146582 | 27.4559311 |
| 0 | 0 |
| 27.1433228 | 50.3358736 |
| 0 | 0 |
| 0 | 0 |
| 0 | 0 |
| 0 | 0 |
| 0 | 0 |
| 0 | 0 |
| 0 | 0 |
| 0 | 0 |
| 65.1439747 | 18.303954 |
| 0 | 0 |
| 789.870693 | 494.206759 |
| 0 | 0 |
| 0 | 0 |
| 0 | 0 |
| 0 | 0 |
| 0 | 0 |
| 0 | 0 |
| 0 | 0 |
| 0 | 0 |
| 0 | 0 |
| 0 | 0 |
| 0 | 0 |
| 0 | 0 |
| 0 | 0 |
| 0 | 0 |
| 0 | 0 |
| 0 | 0 |
| 0 | 0 |
| 0 | 0 |
| 0 | 0 |
| 0 | 0 |

| | |
|---|---|
| 0 | 4.57598851 |
| 27.1433228 | 4.57598851 |
| 0 | 0 |
| 0 | 0 |
| 0 | 0 |
| 19.0003259 | 0 |
| 0 | 0 |
| 0 | 0 |
| 0 | 0 |
| 0 | 0 |
| 0 | 0 |
| 0 | 0 |
| 0 | 0 |
| 0 | 0 |
| 0 | 0 |
| 0 | 0 |
| 0 | 59.4878506 |
| 0 | 0 |
| 0 | 0 |
| 0 | 0 |
| 0 | 0 |
| 0 | 0 |
| 0 | 0 |
| 0 | 0 |
| 35.2863196 | 41.1838966 |
| 0 | 0 |
| 0 | 0 |
| 0 | 0 |
| 0 | 0 |
| 0 | 0 |
| 0 | 0 |
| 0 | 0 |
| 0 | 0 |
| 0 | 0 |
| 0 | 0 |
| 0 | 0 |
| 0 | 0 |
| 143.859611 | 86.9437817 |
| 0 | 0 |
| 0 | 0 |
| 0 | 0 |
| 0 | 0 |
| 0 | 0 |

| | |
|---|---|
| 0 | 0 |
| 0 | 0 |
| 0 | 0 |
| 0 | 0 |
| 0 | 0 |
| 0 | 0 |
| 0 | 0 |
| 0 | 0 |
| 0 | 0 |
| 0 | 0 |
| 0 | 0 |
| 0 | 0 |
| 0 | 0 |
| 0 | 0 |
| 0 | 0 |
| 0 | 0 |
| 0 | 0 |
| 0 | 0 |
| 0 | 0 |
| 0 | 0 |
| 119.43062 | 631.486415 |
| 0 | 0 |
| 0 | 0 |
| 0 | 4.57598851 |
| 0 | 0 |
| 0 | 0 |
| 0 | 0 |
| 0 | 4.57598851 |
| 0 | 0 |
| 0 | 0 |
| 0 | 0 |
| 0 | 0 |
| 0 | 0 |
| 0 | 0 |
| 0 | 0 |
| 0 | 0 |
| 0 | 0 |
| 0 | 0 |
| 0 | 0 |
| 0 | 0 |
| 0 | 0 |
| 0 | 0 |
| 0 | 0 |

| | |
|---|---|
| 0 | 13.7279655 |
| 8.14299683 | 0 |
| 0 | 0 |
| 0 | 0 |
| 0 | 0 |
| 0 | 0 |
| 0 | 27.4559311 |
| 0 | 0 |
| 0 | 9.15197702 |
| 0 | 0 |
| 2.71433228 | 22.8799426 |
| 0 | 0 |
| 0 | 0 |
| 0 | 0 |
| 0 | 0 |
| 0 | 0 |
| 0 | 0 |
| 0 | 0 |
| 0 | 0 |
| 0 | 0 |
| 0 | 0 |
| 0 | 0 |
| 0 | 0 |
| 2.71433228 | 0 |
| 2.71433228 | 155.583609 |
| 0 | 0 |
| 0 | 0 |
| 0 | 0 |
| 0 | 0 |
| 0 | 0 |
| 0 | 0 |
| 0 | 0 |
| 0 | 0 |
| 0 | 0 |
| 0 | 0 |
| 0 | 0 |
| 0 | 0 |
| 0 | 0 |
| 0 | 0 |
| 0 | 4.57598851 |
| 293.147886 | 114.399713 |
| 0 | 0 |
| 0 | 0 |

| | |
|---|---|
| 0 | 32.0319196 |
| 0 | 0 |
| 0 | 0 |
| 0 | 0 |
| 2.71433228 | 0 |
| 0 | 0 |
| 0 | 0 |
| 0 | 0 |
| 0 | 0 |
| 0 | 0 |
| 0 | 0 |
| 0 | 0 |
| 0 | 0 |
| 0 | 0 |
| 0 | 0 |
| 0 | 0 |
| 0 | 27.4559311 |
| 0 | 0 |
| 0 | 0 |
| 0 | 0 |
| 2.71433228 | 141.855644 |
| 154.71694 | 636.062403 |
| 0 | 0 |
| 0 | 0 |
| 0 | 0 |
| 0 | 0 |
| 0 | 0 |
| 0 | 0 |
| 0 | 0 |
| 2.71433228 | 0 |
| 0 | 0 |
| 0 | 0 |
| 0 | 0 |
| 0 | 0 |
| 0 | 0 |
| 54.2866456 | 59.4878506 |
| 0 | 0 |
| 0 | 0 |
| 670.440073 | 457.598851 |
| 0 | 0 |
| 0 | 0 |
| 0 | 0 |
| 0 | 0 |

| | |
|---|---|
| 420.721503 | 237.951403 |
| 0 | 0 |
| 0 | 0 |
| 0 | 4.57598851 |
| 0 | 0 |
| 0 | 0 |
| 0 | 0 |
| 0 | 0 |
| 0 | 0 |
| 0 | 0 |
| 0 | 0 |
| 0 | 0 |
| 0 | 0 |
| 0 | 0 |
| 0 | 0 |
| 0 | 0 |
| 0 | 0 |
| 0 | 0 |
| 0 | 0 |
| 0 | 0 |
| 0 | 0 |
| 0 | 0 |
| 0 | 0 |
| 2.71433228 | 0 |
| 0 | 27.4559311 |
| 0 | 0 |
| 0 | 0 |
| 0 | 0 |
| 0 | 0 |
| 0 | 0 |
| 0 | 0 |
| 0 | 0 |
| 293.147886 | 283.711288 |
| 317.576877 | 9.15197702 |
| 0 | 0 |
| 0 | 0 |
| 0 | 0 |
| 0 | 0 |
| 0 | 4.57598851 |
| 0 | 0 |
| 0 | 0 |
| 0 | 4.57598851 |
| 0 | 0 |

| | |
|---|---|
| 0 | 0 |
| 0 | 0 |
| 0 | 0 |
| 0 | 0 |
| 0 | 0 |
| 0 | 0 |
| 0 | 0 |
| 0 | 0 |
| 0 | 0 |
| 0 | 0 |
| 0 | 0 |
| 0 | 0 |
| 0 | 0 |
| 0 | 0 |
| 0 | 0 |
| 0 | 0 |
| 0 | 0 |
| 0 | 0 |
| 0 | 0 |
| 0 | 0 |
| 0 | 0 |
| 0 | 9.15197702 |
| 0 | 0 |
| 0 | 0 |
| 0 | 0 |
| 54.2866456 | 0 |
| 0 | 0 |
| 0 | 0 |
| 0 | 0 |
| 0 | 9.15197702 |
| 0 | 0 |
| 195.431924 | 64.0638392 |
| 0 | 0 |
| 0 | 0 |
| 21.7146582 | 36.6079081 |
| 0 | 0 |
| 0 | 9.15197702 |
| 0 | 0 |
| 0 | 0 |
| 0 | 0 |
| 0 | 0 |
| 0 | 0 |
| 0 | 0 |

| | |
|---|---|
| 0 | 0 |
| 0 | 0 |
| 0 | 0 |
| 0 | 0 |
| 0 | 0 |
| 0 | 0 |
| 0 | 0 |
| 0 | 0 |
| 0 | 0 |
| 0 | 0 |
| 0 | 0 |
| 0 | 0 |
| 0 | 0 |
| 10.8573291 | 18.303954 |
| 0 | 0 |
| 0 | 0 |
| 0 | 0 |
| 0 | 0 |
| 0 | 0 |
| 0 | 0 |
| 0 | 0 |
| 0 | 0 |
| 0 | 0 |
| 0 | 4.57598851 |
| 0 | 0 |
| 5.42866456 | 0 |
| 0 | 0 |
| 0 | 0 |
| 0 | 0 |
| 0 | 0 |
| 0 | 0 |
| 0 | 0 |
| 0 | 0 |
| 0 | 4.57598851 |
| 0 | 32.0319196 |
| 0 | 0 |
| 0 | 0 |
| 0 | 0 |
| 0 | 0 |
| 0 | 0 |
| 0 | 0 |
| 279.576225 | 118.975701 |
| 0 | 0 |

| | |
|---|---|
| 0 | 4.57598851 |
| 0 | 13.7279655 |
| 0 | 0 |
| 21.7146582 | 0 |
| 0 | 0 |
| 260.575899 | 384.383035 |
| 0 | 0 |
| 0 | 0 |
| 0 | 0 |
| 0 | 0 |
| 0 | 0 |
| 0 | 0 |
| 0 | 0 |
| 0 | 0 |
| 0 | 22.8799426 |
| 0 | 0 |
| 0 | 0 |
| 0 | 0 |
| 0 | 0 |
| 0 | 0 |
| 198.146256 | 59.4878506 |
| 0 | 0 |
| 10.8573291 | 0 |
| 0 | 0 |
| 0 | 0 |
| 0 | 0 |
| 0 | 0 |
| 2.71433228 | 0 |
| 0 | 0 |
| 0 | 0 |
| 0 | 0 |
| 2.71433228 | 0 |
| 0 | 0 |
| 0 | 0 |
| 0 | 0 |
| 0 | 0 |
| 0 | 0 |
| 0 | 0 |
| 0 | 0 |
| 0 | 0 |
| 0 | 0 |
| 0 | 0 |
| 0 | 0 |

| | |
|---|---|
| 0 | 0 |
| 0 | 0 |
| 0 | 0 |
| 0 | 0 |
| 0 | 0 |
| 0 | 0 |
| 0 | 0 |
| 0 | 0 |
| 0 | 0 |
| 133.002282 | 4.57598851 |
| 0 | 0 |
| 0 | 0 |
| 0 | 0 |
| 0 | 0 |
| 0 | 0 |
| 0 | 0 |
| 0 | 0 |
| 0 | 0 |
| 0 | 0 |
| 0 | 0 |
| 0 | 0 |
| 10.8573291 | 96.0957587 |
| 0 | 0 |
| 0 | 0 |
| 0 | 0 |
| 0 | 0 |
| 0 | 0 |
| 0 | 0 |
| 2.71433228 | 0 |
| 0 | 0 |
| 0 | 0 |
| 0 | 0 |
| 0 | 0 |
| 0 | 0 |
| 143.859611 | 114.399713 |
| 0 | 0 |
| 0 | 68.6398277 |
| 0 | 0 |
| 0 | 0 |
| 0 | 0 |
| 0 | 9.15197702 |
| 0 | 50.3358736 |
| 146.573943 | 13.7279655 |

| | |
|---|---|
| 0 | 0 |
| 0 | 0 |
| 0 | 0 |
| 0 | 0 |
| 0 | 0 |
| 0 | 0 |
| 314.862544 | 64.0638392 |
| 0 | 0 |
| 0 | 0 |
| 0 | 0 |
| 0 | 0 |
| 0 | 0 |
| 0 | 0 |
| 0 | 0 |
| 19.0003259 | 54.9118621 |
| 0 | 0 |
| 0 | 0 |
| 0 | 0 |
| 0 | 22.8799426 |
| 5.42866456 | 68.6398277 |
| 0 | 0 |
| 0 | 0 |
| 0 | 0 |
| 0 | 0 |
| 0 | 0 |
| 0 | 0 |
| 0 | 0 |
| 0 | 0 |
| 0 | 0 |
| 0 | 0 |
| 0 | 0 |
| 0 | 0 |
| 29.8576551 | 4.57598851 |
| 0 | 0 |
| 0 | 0 |
| 0 | 0 |
| 0 | 0 |
| 0 | 0 |
| 0 | 0 |
| 16.2859937 | 13.7279655 |
| 0 | 0 |
| 0 | 0 |
| 0 | 0 |

| | |
|---|---|
| 0 | 0 |
| 0 | 0 |
| 0 | 0 |
| 331.148538 | 228.799426 |
| 0 | 0 |
| 0 | 0 |
| 0 | 0 |
| 0 | 0 |
| 0 | 0 |
| 0 | 0 |
| 0 | 0 |
| 0 | 0 |
| 0 | 0 |
| 0 | 0 |
| 0 | 0 |
| 0 | 0 |
| 0 | 0 |
| 0 | 0 |
| 0 | 0 |
| 0 | 0 |
| 0 | 0 |
| 0 | 0 |
| 0 | 0 |
| 0 | 0 |
| 0 | 0 |
| 0 | 0 |
| 0 | 0 |
| 0 | 22.8799426 |
| 0 | 0 |
| 0 | 0 |
| 0 | 22.8799426 |
| 0 | 0 |
| 0 | 0 |
| 0 | 0 |
| 0 | 0 |
| 76.0013038 | 41.1838966 |
| 0 | 0 |
| 0 | 0 |
| 0 | 4.57598851 |
| 0 | 0 |
| 0 | 0 |
| 0 | 0 |
| 0 | 0 |

| | |
|---|---|
| 0 | 0 |
| 0 | 0 |
| 86.8586329 | 27.4559311 |
| 0 | 0 |
| 0 | 0 |
| 0 | 0 |
| 0 | 0 |
| 0 | 32.0319196 |
| 0 | 4.57598851 |
| 0 | 0 |
| 0 | 0 |
| 0 | 0 |
| 0 | 0 |
| 2.71433228 | 0 |
| 38.0006519 | 4.57598851 |
| 0 | 0 |
| 0 | 0 |
| 0 | 0 |
| 5.42866456 | 41.1838966 |
| 0 | 0 |
| 89.5729652 | 45.7598851 |
| 0 | 0 |
| 10.8573291 | 0 |
| 0 | 0 |
| 0 | 0 |
| 0 | 0 |
| 0 | 0 |
| 0 | 0 |
| 0 | 0 |
| 0 | 0 |
| 0 | 0 |
| 0 | 0 |
| 0 | 0 |
| 0 | 0 |
| 0 | 0 |
| 0 | 0 |
| 0 | 0 |
| 0 | 0 |
| 46.1436487 | 4.57598851 |
| 0 | 0 |
| 0 | 0 |
| 0 | 0 |
| 0 | 0 |

| | |
|---|---|
| 0 | 0 |
| 0 | 0 |
| 0 | 0 |
| 0 | 0 |
| 0 | 0 |
| 0 | 0 |
| 0 | 0 |
| 0 | 0 |
| 0 | 0 |
| 0 | 27.4559311 |
| 0 | 0 |
| 0 | 0 |
| 0 | 0 |
| 0 | 0 |
| 0 | 0 |
| 0 | 0 |
| 0 | 0 |
| 0 | 0 |
| 0 | 0 |
| 0 | 0 |
| 0 | 0 |
| 0 | 0 |
| 0 | 0 |
| 0 | 0 |
| 0 | 0 |
| 0 | 13.7279655 |
| 0 | 0 |
| 0 | 0 |
| 0 | 0 |
| 0 | 0 |
| 0 | 0 |
| 0 | 0 |
| 62.4296424 | 59.4878506 |
| 32.5719873 | 22.8799426 |
| 0 | 0 |
| 0 | 0 |
| 0 | 0 |
| 0 | 0 |
| 89.5729652 | 164.735586 |
| 0 | 0 |
| 0 | 0 |
| 0 | 0 |
| 0 | 0 |

| | |
|---|---|
| 0 | 0 |
| 0 | 0 |
| 0 | 0 |
| 2.71433228 | 0 |
| 0 | 0 |
| 0 | 0 |
| 0 | 0 |
| 0 | 0 |
| 0 | 0 |
| 0 | 0 |
| 0 | 0 |
| 0 | 0 |
| 0 | 0 |
| 0 | 0 |
| 0 | 0 |
| 0 | 0 |
| 0 | 0 |
| 0 | 0 |
| 0 | 0 |
| 0 | 0 |
| 0 | 0 |
| 0 | 0 |
| 0 | 0 |
| 0 | 0 |
| 27.1433228 | 0 |
| 0 | 0 |
| 0 | 0 |
| 0 | 0 |
| 0 | 0 |
| 0 | 0 |
| 0 | 0 |
| 0 | 0 |
| 0 | 0 |
| 0 | 0 |
| 0 | 0 |
| 0 | 0 |
| 0 | 0 |
| 0 | 0 |
| 0 | 0 |
| 0 | 0 |
| 0 | 0 |
| 0 | 4.57598851 |
| 0 | 0 |

| | |
|---|---|
| 0 | 9.15197702 |
| 0 | 0 |
| 0 | 0 |
| 5.42866456 | 151.007621 |
| 0 | 0 |
| 0 | 0 |
| 0 | 0 |
| 0 | 0 |
| 27.1433228 | 41.1838966 |
| 0 | 0 |
| 0 | 0 |
| 0 | 0 |
| 0 | 0 |
| 0 | 0 |
| 0 | 0 |
| 0 | 0 |
| 0 | 0 |
| 0 | 0 |
| 0 | 0 |
| 442.436161 | 22.8799426 |
| 0 | 0 |
| 0 | 0 |
| 268.718896 | 50.3358736 |
| 0 | 0 |
| 0 | 0 |
| 0 | 0 |
| 0 | 0 |
| 8.14299683 | 0 |
| 0 | 0 |
| 0 | 0 |
| 0 | 0 |
| 0 | 0 |
| 0 | 0 |
| 0 | 4.57598851 |
| 0 | 0 |
| 0 | 0 |
| 0 | 0 |
| 0 | 0 |
| 0 | 0 |
| 751.870041 | 0 |
| 0 | 0 |
| 0 | 0 |
| 0 | 0 |

| | |
|---|---|
| 0 | 0 |
| 0 | 0 |
| 0 | 0 |
| 0 | 0 |
| 0 | 0 |
| 0 | 0 |
| 0 | 0 |
| 0 | 0 |
| 0 | 0 |
| 0 | 0 |
| 0 | 4.57598851 |
| 0 | 0 |
| 2.71433228 | 45.7598851 |
| 0 | 0 |
| 0 | 0 |
| 0 | 0 |
| 0 | 0 |
| 0 | 0 |
| 116.716288 | 50.3358736 |
| 0 | 0 |
| 0 | 0 |
| 0 | 0 |
| 0 | 0 |
| 0 | 0 |
| 0 | 0 |
| 0 | 0 |
| 0 | 0 |
| 0 | 0 |
| 0 | 0 |
| 0 | 0 |
| 0 | 0 |
| 0 | 0 |
| 0 | 0 |
| 0 | 0 |
| 0 | 0 |
| 0 | 0 |
| 40.7149842 | 4.57598851 |
| 0 | 9.15197702 |
| 0 | 0 |
| 73.2869715 | 36.6079081 |
| 0 | 0 |
| 0 | 0 |
| 0 | 0 |

| | |
|---|---|
| 0 | 0 |
| 0 | 0 |
| 0 | 0 |
| 0 | 0 |
| 0 | 0 |
| 0 | 0 |
| 0 | 0 |
| 157.431272 | 151.007621 |
| 198.146256 | 192.191517 |
| 0 | 0 |
| 0 | 0 |
| 0 | 0 |
| 0 | 0 |
| 0 | 0 |
| 0 | 18.303954 |
| 0 | 0 |
| 0 | 0 |
| 0 | 0 |
| 0 | 0 |
| 0 | 0 |
| 92.2872974 | 36.6079081 |
| 0 | 0 |
| 0 | 0 |
| 0 | 0 |
| 0 | 0 |
| 0 | 0 |
| 0 | 0 |
| 0 | 0 |
| 0 | 0 |
| 0 | 0 |
| 0 | 0 |
| 0 | 0 |
| 347.434532 | 27.4559311 |
| 0 | 0 |
| 0 | 0 |
| 0 | 0 |
| 0 | 0 |
| 0 | 0 |
| 0 | 0 |
| 0 | 0 |
| 0 | 0 |
| 0 | 0 |
| 0 | 0 |

| | |
|---|---|
| 0 | 0 |
| 0 | 0 |
| 0 | 0 |
| 0 | 0 |
| 10.8573291 | 82.3677932 |
| 0 | 0 |
| 0 | 0 |
| 0 | 0 |
| 2.71433228 | 0 |
| 0 | 0 |
| 0 | 0 |
| 0 | 0 |
| 0 | 0 |
| 0 | 0 |
| 0 | 0 |
| 0 | 0 |
| 0 | 0 |
| 0 | 0 |
| 0 | 0 |
| 0 | 0 |
| 0 | 0 |
| 0 | 4.57598851 |
| 0 | 9.15197702 |
| 0 | 50.3358736 |
| 0 | 0 |
| 0 | 0 |
| 0 | 0 |
| 0 | 64.0638392 |
| 0 | 0 |
| 0 | 0 |
| 0 | 0 |
| 124.859285 | 86.9437817 |
| 5.42866456 | 4.57598851 |
| 0 | 0 |
| 0 | 0 |
| 0 | 0 |
| 0 | 4.57598851 |
| 0 | 0 |
| 0 | 0 |
| 0 | 0 |
| 0 | 0 |
| 0 | 0 |
| 0 | 0 |

| | |
|---|---|
| 420.721503 | 27.4559311 |
| 0 | 0 |
| 0 | 0 |
| 0 | 0 |
| 0 | 0 |
| 0 | 0 |
| 0 | 0 |
| 0 | 0 |
| 0 | 0 |
| 0 | 0 |
| 0 | 0 |
| 0 | 0 |
| 0 | 0 |
| 0 | 0 |
| 0 | 0 |
| 0 | 0 |
| 0 | 0 |
| 2.71433228 | 91.5197702 |
| 0 | 0 |
| 0 | 0 |
| 0 | 0 |
| 0 | 0 |
| 0 | 0 |
| 0 | 0 |
| 0 | 0 |
| 0 | 0 |
| 62.4296424 | 41.1838966 |
| 0 | 0 |
| 0 | 0 |
| 0 | 41.1838966 |
| 0 | 4.57598851 |
| 0 | 0 |
| 0 | 0 |
| 2.71433228 | 0 |
| 0 | 0 |
| 0 | 0 |
| 0 | 0 |
| 0 | 0 |
| 29.8576551 | 4.57598851 |
| 0 | 0 |
| 0 | 0 |
| 0 | 0 |
| 0 | 0 |

| | |
|---|---|
| 0 | 0 |
| 0 | 0 |
| 0 | 0 |
| 0 | 0 |
| 0 | 0 |
| 0 | 0 |
| 2.71433228 | 9.15197702 |
| 0 | 9.15197702 |
| 0 | 0 |
| 0 | 9.15197702 |
| 0 | 0 |
| 0 | 0 |
| 0 | 0 |
| 0 | 0 |
| 0 | 0 |
| 0 | 0 |
| 0 | 4.57598851 |
| 0 | 0 |
| 0 | 4.57598851 |
| 0 | 0 |
| 0 | 0 |
| 0 | 0 |
| 0 | 0 |
| 0 | 0 |
| 0 | 0 |
| 0 | 0 |
| 0 | 0 |
| 0 | 0 |
| 0 | 0 |
| 0 | 0 |
| 0 | 0 |
| 0 | 0 |
| 119.43062 | 45.7598851 |
| 0 | 0 |
| 0 | 0 |
| 0 | 0 |
| 0 | 0 |
| 0 | 0 |
| 0 | 0 |
| 0 | 0 |
| 0 | 0 |
| 0 | 0 |

| | |
|---|---|
| 0 | 0 |
| 0 | 0 |
| 0 | 0 |
| 0 | 0 |
| 0 | 0 |
| 0 | 0 |
| 0 | 0 |
| 0 | 0 |
| 0 | 0 |
| 0 | 0 |
| 0 | 0 |
| 0 | 0 |
| 0 | 0 |
| 0 | 0 |
| 0 | 0 |
| 67.8583069 | 9.15197702 |
| 0 | 0 |
| 0 | 0 |
| 0 | 0 |
| 0 | 4.57598851 |
| 0 | 41.1838966 |
| 0 | 0 |
| 0 | 0 |
| 0 | 0 |
| 0 | 0 |
| 0 | 0 |
| 16.2859937 | 0 |
| 0 | 0 |
| 0 | 0 |
| 0 | 0 |
| 38.0006519 | 68.6398277 |
| 0 | 0 |
| 0 | 0 |
| 0 | 18.303954 |
| 0 | 0 |
| 0 | 0 |
| 0 | 4.57598851 |
| 0 | 0 |
| 0 | 0 |
| 0 | 0 |
| 84.1443006 | 13.7279655 |
| 0 | 0 |
| 0 | 18.303954 |

| | |
|---|---|
| 0 | 0 |
| 78.7156361 | 9.15197702 |
| 0 | 0 |
| 0 | 0 |
| 0 | 0 |
| 0 | 0 |
| 0 | 0 |
| 13.5716614 | 0 |
| 0 | 0 |
| 0 | 0 |
| 0 | 0 |
| 70.5726392 | 137.279655 |
| 35.2863196 | 96.0957587 |
| 0 | 0 |
| 0 | 0 |
| 0 | 0 |
| 0 | 0 |
| 0 | 0 |
| 0 | 0 |
| 0 | 0 |
| 312.148212 | 155.583609 |
| 0 | 0 |
| 0 | 0 |
| 0 | 0 |
| 0 | 0 |
| 0 | 0 |
| 0 | 0 |
| 0 | 0 |
| 0 | 0 |
| 0 | 0 |
| 0 | 0 |
| 0 | 0 |
| 0 | 0 |
| 0 | 0 |
| 1115.59057 | 434.718909 |
| 2.71433228 | 201.343494 |
| 0 | 0 |
| 0 | 0 |
| 0 | 0 |
| 0 | 0 |
| 0 | 0 |
| 0 | 0 |
| 0 | 0 |

| | |
|---|---|
| 0 | 0 |
| 0 | 9.15197702 |
| 0 | 0 |
| 0 | 0 |
| 2.71433228 | 4.57598851 |
| 0 | 27.4559311 |
| 0 | 0 |
| 54.2866456 | 36.6079081 |
| 0 | 0 |
| 0 | 0 |
| 0 | 0 |
| 0 | 0 |
| 0 | 0 |
| 0 | 4.57598851 |
| 0 | 0 |
| 0 | 0 |
| 0 | 0 |
| 0 | 4.57598851 |
| 0 | 0 |
| 0 | 0 |
| 27.1433228 | 36.6079081 |
| 0 | 0 |
| 0 | 0 |
| 0 | 0 |
| 0 | 0 |
| 0 | 0 |
| 0 | 0 |
| 0 | 4.57598851 |
| 0 | 0 |
| 173.717266 | 0 |
| 0 | 0 |
| 0 | 0 |
| 152.002608 | 86.9437817 |
| 0 | 0 |
| 0 | 0 |
| 0 | 0 |
| 0 | 0 |
| 0 | 0 |
| 203.574921 | 302.015242 |
| 0 | 0 |
| 0 | 0 |
| 0 | 0 |
| 0 | 0 |

| | |
|---|---|
| 0 | 0 |
| 0 | 0 |
| 0 | 0 |
| 0 | 0 |
| 0 | 0 |
| 0 | 0 |
| 0 | 0 |
| 0 | 41.1838966 |
| 0 | 0 |
| 0 | 0 |
| 2.71433228 | 0 |
| 0 | 0 |
| 24.4289905 | 54.9118621 |
| 0 | 0 |
| 0 | 0 |
| 0 | 0 |
| 0 | 0 |
| 0 | 0 |
| 0 | 0 |
| 0 | 0 |
| 0 | 0 |
| 0 | 0 |
| 0 | 0 |
| 0 | 0 |
| 0 | 0 |
| 0 | 18.303954 |
| 0 | 0 |
| 0 | 0 |
| 0 | 0 |
| 0 | 0 |
| 0 | 0 |
| 0 | 0 |
| 0 | 0 |
| 0 | 0 |
| 0 | 0 |
| 0 | 0 |
| 181.860263 | 45.7598851 |
| 0 | 0 |
| 0 | 0 |
| 0 | 0 |
| 0 | 0 |
| 0 | 0 |
| 0 | 0 |

| | |
|---|---|
| 0 | 0 |
| 0 | 0 |
| 0 | 0 |
| 0 | 0 |
| 0 | 0 |
| 0 | 0 |
| 0 | 0 |
| 0 | 0 |
| 0 | 4.57598851 |
| 0 | 0 |
| 10.8573291 | 36.6079081 |
| 0 | 32.0319196 |
| 76.0013038 | 36.6079081 |
| 0 | 0 |
| 195.431924 | 73.2158162 |
| 0 | 4.57598851 |
| 0 | 0 |
| 0 | 0 |
| 0 | 0 |
| 0 | 0 |
| 0 | 0 |
| 0 | 0 |
| 0 | 0 |
| 0 | 0 |
| 0 | 0 |
| 0 | 0 |
| 0 | 0 |
| 0 | 0 |
| 0 | 0 |
| 0 | 0 |
| 0 | 0 |
| 0 | 0 |
| 0 | 0 |
| 0 | 0 |
| 0 | 0 |
| 0 | 0 |
| 0 | 0 |
| 0 | 0 |
| 0 | 0 |
| 0 | 13.7279655 |
| 0 | 0 |
| 0 | 0 |
| 0 | 0 |

| | |
|---|---|
| 0 | 0 |
| 0 | 0 |
| 0 | 0 |
| 0 | 0 |
| 0 | 0 |
| 0 | 0 |
| 0 | 0 |
| 0 | 0 |
| 0 | 0 |
| 276.861892 | 228.799426 |
| 0 | 0 |
| 0 | 27.4559311 |
| 0 | 0 |
| 0 | 0 |
| 0 | 0 |
| 0 | 0 |
| 0 | 0 |
| 0 | 0 |
| 0 | 0 |
| 100.430294 | 82.3677932 |
| 0 | 0 |
| 0 | 0 |
| 0 | 0 |
| 0 | 0 |
| 0 | 0 |
| 70.5726392 | 0 |
| 0 | 0 |
| 0 | 0 |
| 0 | 0 |
| 0 | 0 |
| 0 | 18.303954 |
| 0 | 0 |
| 0 | 0 |
| 0 | 22.8799426 |
| 0 | 0 |
| 0 | 0 |
| 0 | 64.0638392 |
| 0 | 0 |
| 0 | 0 |
| 0 | 0 |
| 0 | 0 |
| 0 | 0 |
| 0 | 0 |

| | |
|---|---|
| 0 | 0 |
| 0 | 0 |
| 0 | 0 |
| 51.5723133 | 22.8799426 |
| 0 | 0 |
| 0 | 0 |
| 0 | 0 |
| 0 | 0 |
| 0 | 0 |
| 0 | 0 |
| 0 | 0 |
| 0 | 0 |
| 0 | 0 |
| 0 | 0 |
| 0 | 0 |
| 0 | 0 |
| 0 | 0 |
| 0 | 0 |
| 21.7146582 | 73.2158162 |
| 0 | 0 |
| 0 | 0 |
| 0 | 0 |
| 0 | 0 |
| 2.71433228 | 0 |
| 0 | 0 |
| 0 | 0 |
| 0 | 0 |
| 0 | 0 |
| 0 | 0 |
| 0 | 0 |
| 0 | 0 |
| 2.71433228 | 54.9118621 |
| 0 | 0 |
| 0 | 0 |
| 0 | 0 |
| 0 | 0 |
| 0 | 0 |
| 0 | 0 |
| 0 | 0 |
| 0 | 0 |
| 0 | 0 |
| 0 | 0 |
| 0 | 22.8799426 |

| | |
|---|---|
| 0 | 0 |
| 97.715962 | 4.57598851 |
| 0 | 0 |
| 0 | 0 |
| 5.42866456 | 0 |
| 0 | 0 |
| 0 | 0 |
| 0 | 0 |
| 0 | 0 |
| 0 | 0 |
| 62.4296424 | 132.703667 |
| 2.71433228 | 0 |
| 0 | 0 |
| 0 | 0 |
| 176.431598 | 82.3677932 |
| 0 | 0 |
| 0 | 4.57598851 |
| 0 | 0 |
| 0 | 0 |
| 0 | 0 |
| 0 | 0 |
| 0 | 0 |
| 0 | 0 |
| 0 | 0 |
| 0 | 0 |
| 13.5716614 | 0 |
| 0 | 0 |
| 0 | 0 |
| 0 | 0 |
| 0 | 0 |
| 0 | 0 |
| 0 | 0 |
| 0 | 0 |
| 0 | 0 |
| 0 | 0 |
| 0 | 0 |
| 0 | 0 |
| 0 | 0 |
| 0 | 0 |
| 0 | 0 |
| 0 | 0 |
| 0 | 0 |
| 198.146256 | 123.55169 |

| | |
|---|---|
| 32.5719873 | 96.0957587 |
| 0 | 0 |
| 0 | 0 |
| 0 | 0 |
| 0 | 0 |
| 0 | 0 |
| 0 | 0 |
| 0 | 0 |
| 38.0006519 | 0 |
| 8.14299683 | 13.7279655 |
| 0 | 0 |
| 0 | 0 |
| 0 | 0 |
| 0 | 0 |
| 0 | 0 |
| 0 | 0 |
| 200.860589 | 18.303954 |
| 0 | 0 |
| 0 | 4.57598851 |
| 32.5719873 | 45.7598851 |
| 0 | 0 |
| 0 | 0 |
| 423.435835 | 196.767506 |
| 16.2859937 | 242.527391 |
| 0 | 0 |
| 0 | 0 |
| 0 | 0 |
| 0 | 0 |
| 0 | 0 |
| 0 | 0 |
| 0 | 0 |
| 0 | 0 |
| 0 | 0 |
| 2.71433228 | 0 |
| 0 | 4.57598851 |
| 0 | 0 |
| 0 | 0 |
| 8.14299683 | 0 |
| 0 | 0 |
| 0 | 0 |
| 0 | 0 |
| 643.29675 | 425.566932 |
| 0 | 0 |

| | |
|---|---|
| 0 | 0 |
| 0 | 0 |
| 0 | 0 |
| 0 | 0 |
| 1623.1707 | 677.2463 |
| 0 | 0 |
| 0 | 0 |
| 27.1433228 | 68.6398277 |
| 0 | 0 |
| 0 | 0 |
| 0 | 0 |
| 0 | 0 |
| 0 | 0 |
| 0 | 0 |
| 0 | 0 |
| 0 | 0 |
| 0 | 0 |
| 0 | 0 |
| 0 | 0 |
| 0 | 0 |
| 0 | 0 |
| 0 | 0 |
| 0 | 0 |
| 0 | 27.4559311 |
| 0 | 0 |
| 0 | 0 |
| 0 | 0 |
| 13.5716614 | 54.9118621 |
| 0 | 0 |
| 46.1436487 | 73.2158162 |
| 2.71433228 | 4.57598851 |
| 0 | 0 |
| 0 | 0 |
| 0 | 0 |
| 0 | 0 |
| 0 | 0 |
| 0 | 0 |
| 0 | 0 |
| 0 | 0 |
| 24.4289905 | 96.0957587 |
| 0 | 0 |
| 0 | 0 |
| 0 | 32.0319196 |

| | |
|---|---|
| 0 | 0 |
| 0 | 0 |
| 0 | 0 |
| 5.42866456 | 0 |
| 2.71433228 | 0 |
| 0 | 0 |
| 0 | 0 |
| 0 | 0 |
| 0 | 0 |
| 0 | 0 |
| 0 | 0 |
| 0 | 0 |
| 0 | 0 |
| 0 | 0 |
| 0 | 41.1838966 |
| 0 | 0 |
| 0 | 0 |
| 0 | 0 |
| 0 | 0 |
| 0 | 0 |
| 0 | 0 |
| 0 | 0 |
| 0 | 0 |
| 48.857981 | 4.57598851 |
| 0 | 0 |
| 0 | 0 |
| 0 | 0 |
| 0 | 0 |
| 0 | 0 |
| 0 | 0 |
| 0 | 0 |
| 0 | 0 |
| 0 | 0 |
| 0 | 0 |
| 0 | 0 |
| 0 | 0 |
| 0 | 0 |
| 0 | 4.57598851 |
| 0 | 0 |
| 0 | 0 |
| 0 | 0 |
| 0 | 0 |
| 0 | 0 |

| | |
|---|---|
| 0 | 0 |
| 0 | 0 |
| 0 | 0 |
| 0 | 0 |
| 0 | 0 |
| 0 | 0 |
| 0 | 0 |
| 0 | 0 |
| 173.717266 | 86.9437817 |
| 2.71433228 | 141.855644 |
| 0 | 32.0319196 |
| 0 | 0 |
| 0 | 0 |
| 0 | 0 |
| 0 | 0 |
| 0 | 0 |
| 0 | 0 |
| 8.14299683 | 13.7279655 |
| 0 | 0 |
| 0 | 0 |
| 29.8576551 | 36.6079081 |
| 0 | 0 |
| 0 | 0 |
| 0 | 0 |
| 0 | 0 |
| 0 | 0 |
| 0 | 0 |
| 0 | 0 |
| 0 | 0 |
| 0 | 0 |
| 0 | 0 |
| 0 | 0 |
| 0 | 0 |
| 0 | 0 |
| 0 | 0 |
| 0 | 0 |
| 5.42866456 | 0 |
| 0 | 0 |
| 0 | 0 |
| 0 | 0 |
| 0 | 0 |
| 0 | 0 |
| 0 | 0 |

| | |
|---|---|
| 187.288927 | 86.9437817 |
| 0 | 0 |
| 0 | 0 |
| 206.289253 | 32.0319196 |
| 0 | 4.57598851 |
| 0 | 0 |
| 0 | 0 |
| 2.71433228 | 0 |
| 0 | 0 |
| 0 | 0 |
| 0 | 0 |
| 0 | 0 |
| 179.14593 | 114.399713 |
| 89.5729652 | 109.823724 |
| 0 | 36.6079081 |
| 0 | 0 |
| 0 | 0 |
| 89.5729652 | 91.5197702 |
| 0 | 0 |
| 0 | 0 |
| 0 | 0 |
| 0 | 0 |
| 0 | 0 |
| 65.1439747 | 0 |
| 0 | 0 |
| 0 | 9.15197702 |
| 0 | 0 |
| 0 | 0 |
| 0 | 0 |
| 0 | 0 |
| 0 | 0 |
| 19.0003259 | 18.303954 |
| 0 | 0 |
| 0 | 0 |
| 0 | 0 |
| 0 | 4.57598851 |
| 0 | 45.7598851 |
| 0 | 4.57598851 |
| 0 | 0 |
| 0 | 0 |
| 0 | 0 |
| 0 | 0 |
| 0 | 0 |

| | |
|---|---|
| 76.0013038 | 27.4559311 |
| 0 | 0 |
| 0 | 0 |
| 0 | 0 |
| 0 | 0 |
| 0 | 0 |
| 0 | 22.8799426 |
| 0 | 0 |
| 2.71433228 | 0 |
| 0 | 0 |
| 0 | 0 |
| 0 | 0 |
| 0 | 0 |
| 0 | 0 |
| 0 | 0 |
| 0 | 0 |
| 0 | 0 |
| 0 | 0 |
| 187.288927 | 9.15197702 |
| 2.71433228 | 4.57598851 |
| 0 | 0 |
| 0 | 0 |
| 0 | 0 |
| 0 | 0 |
| 0 | 0 |
| 0 | 0 |
| 0 | 4.57598851 |
| 0 | 0 |
| 0 | 0 |
| 0 | 0 |
| 59.7153101 | 4.57598851 |
| 0 | 59.4878506 |
| 0 | 0 |
| 0 | 0 |
| 0 | 0 |
| 0 | 0 |
| 0 | 0 |
| 0 | 0 |
| 0 | 0 |
| 0 | 0 |
| 0 | 0 |
| 0 | 0 |
| 0 | 0 |

| | |
|---|---|
| 40.7149842 | 0 |
| 10.8573291 | 45.7598851 |
| 0 | 0 |
| 0 | 0 |
| 0 | 22.8799426 |
| 0 | 0 |
| 0 | 0 |
| 0 | 0 |
| 0 | 0 |
| 5.42866456 | 0 |
| 0 | 0 |
| 0 | 0 |
| 0 | 0 |
| 0 | 0 |
| 0 | 0 |
| 2.71433228 | 36.6079081 |
| 0 | 0 |
| 0 | 0 |
| 0 | 0 |
| 0 | 0 |
| 0 | 4.57598851 |
| 0 | 0 |
| 0 | 0 |
| 209.003585 | 544.542633 |
| 0 | 0 |
| 0 | 0 |
| 0 | 0 |
| 0 | 0 |
| 0 | 0 |
| 0 | 0 |
| 0 | 0 |
| 0 | 0 |
| 0 | 0 |
| 0 | 0 |
| 0 | 0 |
| 0 | 0 |
| 0 | 0 |
| 0 | 0 |
| 0 | 0 |
| 0 | 0 |
| 0 | 0 |
| 0 | 0 |
| 0 | 0 |

| | |
|---|---|
| 0 | 0 |
| 0 | 0 |
| 0 | 0 |
| 0 | 0 |
| 0 | 0 |
| 0 | 0 |
| 0 | 0 |
| 0 | 0 |
| 0 | 0 |
| 0 | 0 |
| 0 | 50.3358736 |
| 0 | 0 |
| 0 | 0 |
| 0 | 0 |
| 0 | 0 |
| 0 | 0 |
| 0 | 0 |
| 0 | 0 |
| 0 | 0 |
| 0 | 0 |
| 32.5719873 | 59.4878506 |
| 0 | 4.57598851 |
| 0 | 0 |
| 0 | 0 |
| 2.71433228 | 4.57598851 |
| 0 | 4.57598851 |
| 537.437791 | 178.463552 |
| 1351.73747 | 663.518334 |
| 0 | 4.57598851 |
| 0 | 0 |
| 0 | 0 |
| 0 | 0 |
| 4701.22351 | 178.463552 |
| 5.42866456 | 36.6079081 |
| 0 | 0 |
| 0 | 0 |
| 0 | 27.4559311 |
| 0 | 0 |
| 2.71433228 | 22.8799426 |
| 0 | 0 |
| 0 | 0 |
| 0 | 0 |
| 0 | 0 |

| | |
|---|---|
| 0 | 0 |
| 0 | 0 |
| 0 | 0 |
| 0 | 0 |
| 0 | 0 |
| 2.71433228 | 4.57598851 |
| 0 | 0 |
| 0 | 0 |
| 0 | 0 |
| 0 | 0 |
| 0 | 0 |
| 0 | 0 |
| 0 | 0 |
| 0 | 0 |
| 0 | 0 |
| 0 | 0 |
| 0 | 0 |
| 0 | 0 |
| 0 | 0 |
| 0 | 0 |
| 0 | 0 |
| 65.1439747 | 54.9118621 |
| 0 | 0 |
| 0 | 0 |
| 0 | 0 |
| 0 | 0 |
| 0 | 0 |
| 0 | 9.15197702 |
| 0 | 0 |
| 0 | 0 |
| 0 | 0 |
| 0 | 0 |
| 0 | 0 |
| 0 | 0 |
| 0 | 0 |
| 0 | 0 |
| 0 | 0 |
| 0 | 0 |
| 176.431598 | 82.3677932 |
| 0 | 0 |
| 76.0013038 | 0 |
| 0 | 0 |
| 0 | 32.0319196 |

| | |
|---|---|
| 29.8576551 | 4.57598851 |
| 2.71433228 | 0 |
| 0 | 0 |
| 0 | 0 |
| 0 | 0 |
| 0 | 0 |
| 0 | 0 |
| 0 | 0 |
| 0 | 0 |
| 0 | 0 |
| 0 | 0 |
| 0 | 0 |
| 0 | 0 |
| 0 | 0 |
| 0 | 0 |
| 0 | 4.57598851 |
| 0 | 0 |
| 0 | 0 |
| 0 | 0 |
| 0 | 22.8799426 |
| 0 | 0 |
| 0 | 0 |
| 16.2859937 | 54.9118621 |
| 48.857981 | 9.15197702 |
| 0 | 0 |
| 0 | 0 |
| 0 | 0 |
| 0 | 0 |
| 0 | 0 |
| 0 | 0 |
| 0 | 0 |
| 0 | 0 |
| 0 | 0 |
| 0 | 0 |
| 2.71433228 | 41.1838966 |
| 0 | 0 |
| 0 | 0 |
| 119.43062 | 64.0638392 |
| 0 | 27.4559311 |
| 16.2859937 | 0 |
| 0 | 0 |
| 0 | 0 |
| 431.578832 | 114.399713 |

| | |
|---|---|
| 2.71433228 | 13.7279655 |
| 0 | 0 |
| 0 | 0 |
| 0 | 0 |
| 0 | 0 |
| 0 | 0 |
| 0 | 0 |
| 0 | 0 |
| 0 | 0 |
| 0 | 86.9437817 |
| 2.71433228 | 32.0319196 |
| 0 | 0 |
| 0 | 0 |
| 0 | 0 |
| 0 | 0 |
| 0 | 0 |
| 0 | 0 |
| 0 | 0 |
| 0 | 4.57598851 |
| 0 | 0 |
| 0 | 0 |
| 0 | 22.8799426 |
| 0 | 0 |
| 0 | 0 |
| 0 | 0 |
| 0 | 0 |
| 0 | 0 |
| 0 | 0 |
| 0 | 0 |
| 0 | 0 |
| 0 | 0 |
| 0 | 0 |
| 0 | 18.303954 |
| 0 | 0 |
| 0 | 0 |
| 0 | 0 |
| 0 | 0 |
| 0 | 0 |
| 0 | 0 |
| 0 | 0 |
| 0 | 0 |
| 0 | 0 |
| 0 | 0 |

| | |
|---|---|
| 0 | 0 |
| 0 | 0 |
| 0 | 0 |
| 385.435183 | 54.9118621 |
| 0 | 0 |
| 0 | 0 |
| 0 | 0 |
| 19.0003259 | 0 |
| 0 | 0 |
| 0 | 0 |
| 5.42866456 | 0 |
| 0 | 0 |
| 0 | 0 |
| 214.43225 | 123.55169 |
| 0 | 0 |
| 0 | 0 |
| 0 | 0 |
| 0 | 0 |
| 0 | 0 |
| 0 | 0 |
| 0 | 18.303954 |
| 0 | 0 |
| 0 | 0 |
| 0 | 0 |
| 0 | 0 |
| 0 | 0 |
| 40.7149842 | 18.303954 |
| 0 | 0 |
| 0 | 0 |
| 0 | 0 |
| 0 | 0 |
| 0 | 0 |
| 0 | 0 |
| 0 | 0 |
| 0 | 0 |
| 0 | 0 |
| 0 | 0 |
| 0 | 0 |
| 0 | 0 |
| 0 | 0 |
| 0 | 0 |
| 0 | 0 |

| | |
|---|---|
| 0 | 0 |
| 0 | 0 |
| 130.287949 | 178.463552 |
| 5.42866456 | 18.303954 |
| 0 | 0 |
| 0 | 0 |
| 0 | 0 |
| 0 | 4.57598851 |
| 0 | 0 |
| 0 | 0 |
| 0 | 0 |
| 0 | 0 |
| 0 | 0 |
| 0 | 0 |
| 0 | 22.8799426 |
| 184.574595 | 100.671747 |
| 0 | 0 |
| 0 | 0 |
| 0 | 0 |
| 0 | 0 |
| 0 | 0 |
| 0 | 0 |
| 0 | 0 |
| 0 | 0 |
| 0 | 0 |
| 0 | 0 |
| 0 | 0 |
| 38.0006519 | 50.3358736 |
| 0 | 0 |
| 0 | 0 |
| 0 | 0 |
| 0 | 0 |
| 0 | 4.57598851 |
| 0 | 0 |
| 0 | 0 |
| 2.71433228 | 0 |
| 0 | 0 |
| 0 | 0 |
| 0 | 0 |
| 0 | 0 |
| 0 | 0 |
| 0 | 0 |
| 2.71433228 | 0 |

| | |
|---|---|
| 0 | 0 |
| 0 | 0 |
| 0 | 0 |
| 0 | 0 |
| 0 | 0 |
| 0 | 0 |
| 0 | 0 |
| 0 | 0 |
| 0 | 0 |
| 0 | 0 |
| 54.2866456 | 86.9437817 |
| 0 | 41.1838966 |
| 0 | 0 |
| 0 | 0 |
| 0 | 0 |
| 0 | 0 |
| 0 | 0 |
| 2.71433228 | 0 |
| 0 | 0 |
| 0 | 0 |
| 225.289579 | 128.127678 |
| 0 | 0 |
| 160.145604 | 0 |
| 0 | 0 |
| 0 | 0 |
| 0 | 0 |
| 0 | 0 |
| 222.575247 | 9.15197702 |
| 0 | 0 |
| 0 | 0 |
| 2.71433228 | 0 |
| 0 | 0 |
| 0 | 0 |
| 0 | 0 |
| 0 | 0 |
| 0 | 0 |
| 0 | 0 |
| 0 | 0 |
| 0 | 0 |
| 0 | 0 |
| 0 | 0 |
| 0 | 0 |
| 0 | 0 |

| | |
|---|---|
| 0 | 0 |
| 0 | 0 |
| 0 | 0 |
| 0 | 0 |
| 0 | 0 |
| 0 | 0 |
| 0 | 0 |
| 0 | 0 |
| 0 | 0 |
| 0 | 0 |
| 0 | 0 |
| 16.2859937 | 59.4878506 |
| 0 | 0 |
| 0 | 0 |
| 0 | 0 |
| 0 | 0 |
| 0 | 0 |
| 0 | 0 |
| 0 | 0 |
| 0 | 0 |
| 0 | 0 |
| 0 | 0 |
| 0 | 0 |
| 0 | 0 |
| 0 | 0 |
| 0 | 0 |
| 0 | 0 |
| 0 | 0 |
| 0 | 0 |
| 0 | 0 |
| 0 | 0 |
| 0 | 0 |
| 0 | 0 |
| 0 | 0 |
| 0 | 0 |
| 0 | 0 |
| 0 | 0 |
| 0 | 0 |
| 0 | 0 |
| 0 | 0 |
| 0 | 0 |
| 0 | 0 |
| 19.0003259 | 22.8799426 |

| | |
|---|---|
| 0 | 0 |
| 0 | 0 |
| 0 | 0 |
| 0 | 13.7279655 |
| 0 | 0 |
| 0 | 0 |
| 103.144627 | 13.7279655 |
| 0 | 0 |
| 0 | 0 |
| 0 | 0 |
| 0 | 0 |
| 0 | 0 |
| 0 | 0 |
| 0 | 0 |
| 0 | 0 |
| 0 | 0 |
| 0 | 0 |
| 187.288927 | 384.383035 |
| 0 | 0 |
| 0 | 0 |
| 0 | 0 |
| 0 | 0 |
| 0 | 0 |
| 0 | 0 |
| 0 | 0 |
| 0 | 0 |
| 0 | 0 |
| 0 | 0 |
| 0 | 0 |
| 0 | 0 |
| 13.5716614 | 0 |
| 0 | 0 |
| 0 | 0 |
| 0 | 0 |
| 0 | 13.7279655 |
| 0 | 4.57598851 |
| 0 | 0 |
| 0 | 0 |
| 0 | 0 |
| 0 | 0 |
| 0 | 0 |
| 0 | 0 |
| 0 | 0 |

| | |
|---|---|
| 0 | 0 |
| 0 | 0 |
| 0 | 0 |
| 24.4289905 | 73.2158162 |
| 0 | 0 |
| 0 | 0 |
| 0 | 0 |
| 10.8573291 | 0 |
| 0 | 0 |
| 105.858959 | 4.57598851 |
| 0 | 0 |
| 0 | 0 |
| 0 | 0 |
| 0 | 0 |
| 179.14593 | 114.399713 |
| 0 | 0 |
| 0 | 0 |
| 0 | 0 |
| 0 | 45.7598851 |
| 0 | 0 |
| 0 | 0 |
| 0 | 0 |
| 0 | 0 |
| 0 | 0 |
| 0 | 0 |
| 0 | 0 |
| 0 | 0 |
| 0 | 0 |
| 0 | 0 |
| 0 | 0 |
| 0 | 0 |
| 0 | 0 |
| 0 | 59.4878506 |
| 0 | 0 |
| 0 | 0 |
| 0 | 0 |
| 0 | 0 |
| 0 | 0 |
| 27.1433228 | 233.375414 |
| 0 | 0 |
| 575.438443 | 457.598851 |
| 0 | 0 |
| 0 | 0 |

| | |
|---|---|
| 0 | 0 |
| 0 | 0 |
| 0 | 0 |
| 0 | 0 |
| 0 | 0 |
| 0 | 0 |
| 13.5716614 | 32.0319196 |
| 0 | 0 |
| 0 | 0 |
| 0 | 0 |
| 0 | 22.8799426 |
| 0 | 0 |
| 0 | 0 |
| 0 | 0 |
| 0 | 0 |
| 0 | 0 |
| 0 | 77.7918047 |
| 0 | 0 |
| 0 | 9.15197702 |
| 0 | 0 |
| 0 | 0 |
| 0 | 169.311575 |
| 0 | 0 |
| 0 | 0 |
| 0 | 0 |
| 0 | 0 |
| 1243.16418 | 370.655069 |
| 0 | 0 |
| 0 | 32.0319196 |
| 0 | 0 |
| 0 | 0 |
| 0 | 0 |
| 0 | 0 |
| 0 | 0 |
| 0 | 0 |
| 0 | 0 |
| 0 | 0 |
| 0 | 0 |
| 0 | 0 |
| 0 | 0 |
| 0 | 0 |
| 0 | 0 |
| 0 | 0 |

| | |
|---|---|
| 0 | 0 |
| 0 | 0 |
| 0 | 4.57598851 |
| 0 | 0 |
| 0 | 0 |
| 0 | 0 |
| 0 | 0 |
| 0 | 0 |
| 0 | 0 |
| 0 | 0 |
| 0 | 0 |
| 0 | 27.4559311 |
| 0 | 4.57598851 |
| 323.005541 | 0 |
| 0 | 0 |
| 0 | 0 |
| 0 | 0 |
| 0 | 0 |
| 0 | 0 |
| 0 | 0 |
| 0 | 0 |
| 0 | 0 |
| 0 | 0 |
| 0 | 0 |
| 0 | 0 |
| 0 | 0 |
| 203.574921 | 50.3358736 |
| 0 | 0 |
| 0 | 0 |
| 95.0016297 | 100.671747 |
| 0 | 0 |
| 0 | 0 |
| 0 | 0 |
| 0 | 0 |
| 103.144627 | 201.343494 |
| 0 | 0 |
| 0 | 0 |
| 103.144627 | 4.57598851 |
| 0 | 0 |
| 0 | 0 |
| 19.0003259 | 0 |
| 0 | 0 |
| 0 | 0 |

| | |
|---|---|
| 0 | 0 |
| 2.71433228 | 0 |
| 100.430294 | 123.55169 |
| 10.8573291 | 0 |
| 0 | 0 |
| 0 | 0 |
| 0 | 0 |
| 0 | 0 |
| 0 | 0 |
| 0 | 0 |
| 0 | 0 |
| 0 | 0 |
| 0 | 0 |
| 0 | 0 |
| 0 | 0 |
| 0 | 0 |
| 0 | 0 |
| 0 | 0 |
| 0 | 0 |
| 0 | 0 |
| 81.4299683 | 0 |
| 48.857981 | 13.7279655 |
| 0 | 0 |
| 0 | 0 |
| 0 | 0 |
| 0 | 0 |
| 0 | 0 |
| 358.291861 | 210.495472 |
| 0 | 0 |
| 0 | 0 |
| 0 | 0 |
| 0 | 0 |
| 0 | 0 |
| 0 | 0 |
| 0 | 0 |
| 43.4293164 | 100.671747 |
| 10.8573291 | 0 |
| 0 | 0 |
| 13.5716614 | 0 |
| 0 | 0 |
| 0 | 0 |
| 0 | 0 |
| 0 | 0 |

| | |
|---|---|
| 0 | 0 |
| 0 | 0 |
| 105.858959 | 0 |
| 0 | 0 |
| 0 | 0 |
| 0 | 0 |
| 0 | 0 |
| 0 | 0 |
| 0 | 0 |
| 0 | 0 |
| 0 | 0 |
| 0 | 0 |
| 0 | 50.3358736 |
| 0 | 0 |
| 0 | 0 |
| 0 | 0 |
| 0 | 0 |
| 0 | 0 |
| 0 | 0 |
| 0 | 0 |
| 0 | 0 |
| 0 | 0 |
| 0 | 0 |
| 0 | 4.57598851 |
| 361.006193 | 132.703667 |
| 0 | 0 |
| 0 | 0 |
| 0 | 0 |
| 0 | 0 |
| 0 | 0 |
| 0 | 0 |
| 0 | 0 |
| 0 | 0 |
| 2.71433228 | 13.7279655 |
| 0 | 0 |
| 0 | 0 |
| 0 | 0 |
| 0 | 0 |
| 0 | 0 |
| 0 | 0 |
| 0 | 0 |
| 0 | 0 |
| 0 | 0 |

| | |
|---|---|
| 0 | 0 |
| 0 | 0 |
| 0 | 0 |
| 0 | 0 |
| 16.2859937 | 68.6398277 |
| 0 | 0 |
| 0 | 0 |
| 0 | 0 |
| 0 | 0 |
| 0 | 0 |
| 0 | 0 |
| 0 | 0 |
| 0 | 0 |
| 2.71433228 | 0 |
| 0 | 0 |
| 0 | 0 |
| 0 | 0 |
| 0 | 0 |
| 0 | 0 |
| 2.71433228 | 9.15197702 |
| 0 | 0 |
| 0 | 0 |
| 0 | 0 |
| 0 | 0 |
| 453.29349 | 320.319196 |
| 0 | 0 |
| 0 | 0 |
| 0 | 27.4559311 |
| 0 | 0 |
| 0 | 0 |
| 0 | 0 |
| 0 | 0 |
| 0 | 0 |
| 0 | 0 |
| 0 | 0 |
| 0 | 0 |
| 0 | 0 |
| 0 | 4.57598851 |
| 0 | 0 |
| 2.71433228 | 0 |
| 0 | 0 |
| 0 | 0 |
| 0 | 0 |

| | |
|---|---|
| 0 | 0 |
| 0 | 0 |
| 0 | 0 |
| 0 | 0 |
| 0 | 18.303954 |
| 0 | 0 |
| 0 | 0 |
| 0 | 0 |
| 0 | 0 |
| 0 | 36.6079081 |
| 0 | 0 |
| 0 | 0 |
| 0 | 0 |
| 0 | 0 |
| 0 | 0 |
| 0 | 0 |
| 0 | 0 |
| 0 | 0 |
| 0 | 0 |
| 0 | 0 |
| 0 | 0 |
| 0 | 0 |
| 0 | 0 |
| 0 | 0 |
| 0 | 0 |
| 0 | 18.303954 |
| 2.71433228 | 36.6079081 |
| 0 | 0 |
| 0 | 0 |
| 0 | 0 |
| 0 | 0 |
| 0 | 0 |
| 0 | 0 |
| 0 | 0 |
| 0 | 0 |
| 0 | 0 |
| 0 | 0 |
| 0 | 0 |
| 0 | 0 |
| 2.71433228 | 4.57598851 |
| 0 | 4.57598851 |
| 0 | 0 |
| 0 | 0 |

| | |
|---|---|
| 0 | 0 |
| 0 | 0 |
| 0 | 0 |
| 0 | 0 |
| 0 | 0 |
| 0 | 0 |
| 0 | 0 |
| 0 | 0 |
| 0 | 0 |
| 0 | 0 |
| 0 | 0 |
| 0 | 0 |
| 0 | 0 |
| 0 | 0 |
| 0 | 0 |
| 0 | 0 |
| 0 | 27.4559311 |
| 0 | 0 |
| 0 | 0 |
| 0 | 0 |
| 0 | 0 |
| 0 | 0 |
| 0 | 0 |
| 0 | 0 |
| 0 | 0 |
| 0 | 0 |
| 0 | 0 |
| 0 | 0 |
| 0 | 0 |
| 0 | 0 |
| 0 | 0 |
| 0 | 0 |
| 0 | 0 |
| 0 | 0 |
| 0 | 13.7279655 |
| 0 | 0 |
| 0 | 45.7598851 |
| 0 | 0 |
| 0 | 0 |
| 0 | 0 |
| 0 | 0 |
| 0 | 0 |
| 27.1433228 | 0 |

| | |
|---|---|
| 0 | 0 |
| 0 | 0 |
| 0 | 0 |
| 0 | 45.7598851 |
| 0 | 0 |
| 0 | 4.57598851 |
| 0 | 0 |
| 43.4293164 | 41.1838966 |
| 0 | 0 |
| 0 | 0 |
| 51.5723133 | 50.3358736 |
| 0 | 0 |
| 0 | 0 |
| 0 | 0 |
| 0 | 0 |
| 0 | 0 |
| 0 | 0 |
| 0 | 0 |
| 0 | 0 |
| 0 | 0 |
| 0 | 0 |
| 0 | 0 |
| 5.42866456 | 0 |
| 0 | 0 |
| 0 | 18.303954 |
| 0 | 0 |
| 0 | 18.303954 |
| 0 | 0 |
| 0 | 0 |
| 0 | 0 |
| 0 | 0 |
| 0 | 0 |
| 0 | 0 |
| 0 | 0 |
| 0 | 0 |
| 0 | 0 |
| 0 | 0 |
| 2.71433228 | 0 |
| 293.147886 | 155.583609 |
| 0 | 0 |
| 0 | 0 |
| 0 | 0 |
| 0 | 0 |

| | |
|---|---|
| 0 | 0 |
| 0 | 0 |
| 2.71433228 | 9.15197702 |
| 0 | 0 |
| 0 | 0 |
| 0 | 0 |
| 285.004889 | 141.855644 |
| 0 | 0 |
| 0 | 0 |
| 0 | 0 |
| 0 | 0 |
| 0 | 0 |
| 0 | 4.57598851 |
| 0 | 0 |
| 0 | 0 |
| 0 | 0 |
| 0 | 0 |
| 0 | 0 |
| 35.2863196 | 50.3358736 |
| 0 | 0 |
| 0 | 0 |
| 0 | 0 |
| 0 | 0 |
| 0 | 0 |
| 0 | 0 |
| 1691.02901 | 787.070024 |
| 0 | 0 |
| 0 | 41.1838966 |
| 0 | 0 |
| 0 | 0 |
| 2.71433228 | 59.4878506 |
| 0 | 0 |
| 0 | 0 |
| 2.71433228 | 0 |
| 0 | 0 |
| 0 | 0 |
| 0 | 0 |
| 0 | 0 |
| 0 | 0 |
| 0 | 0 |
| 0 | 0 |
| 0 | 0 |
| 0 | 0 |

| | |
|---|---|
| 0 | 0 |
| 0 | 0 |
| 0 | 0 |
| 0 | 0 |
| 0 | 0 |
| 0 | 0 |
| 0 | 0 |
| 0 | 0 |
| 0 | 0 |
| 10.8573291 | 45.7598851 |
| 0 | 0 |
| 0 | 4.57598851 |
| 0 | 0 |
| 0 | 4.57598851 |
| 0 | 0 |
| 0 | 0 |
| 0 | 0 |
| 0 | 0 |
| 0 | 0 |
| 0 | 0 |
| 0 | 0 |
| 0 | 0 |
| 0 | 0 |
| 0 | 0 |
| 2.71433228 | 45.7598851 |
| 0 | 32.0319196 |
| 0 | 4.57598851 |
| 0 | 0 |
| 0 | 0 |
| 0 | 0 |
| 447.864826 | 183.03954 |
| 8.14299683 | 22.8799426 |
| 0 | 0 |
| 0 | 0 |
| 0 | 0 |
| 0 | 0 |
| 0 | 4.57598851 |
| 0 | 0 |
| 0 | 0 |
| 0 | 0 |
| 0 | 0 |
| 0 | 0 |
| 0 | 13.7279655 |

| | |
|---|---|
| 0 | 4.57598851 |
| 0 | 0 |
| 0 | 0 |
| 0 | 0 |
| 0 | 0 |
| 0 | 0 |
| 0 | 0 |
| 5.42866456 | 4.57598851 |
| 5.42866456 | 9.15197702 |
| 0 | 0 |
| 0 | 0 |
| 0 | 0 |
| 0 | 0 |
| 0 | 0 |
| 27.1433228 | 4.57598851 |
| 0 | 0 |
| 0 | 0 |
| 0 | 4.57598851 |
| 0 | 0 |
| 0 | 0 |
| 0 | 0 |
| 0 | 0 |
| 0 | 0 |
| 0 | 0 |
| 0 | 0 |
| 0 | 0 |
| 2.71433228 | 0 |
| 73.2869715 | 9.15197702 |
| 0 | 0 |
| 78.7156361 | 18.303954 |
| 0 | 0 |
| 0 | 9.15197702 |
| 0 | 0 |
| 0 | 0 |
| 0 | 0 |
| 0 | 0 |
| 252.432902 | 73.2158162 |
| 217.146582 | 18.303954 |
| 2.71433228 | 0 |
| 0 | 0 |
| 0 | 0 |
| 0 | 13.7279655 |
| 0 | 59.4878506 |

| | |
|---|---|
| 0 | 0 |
| 0 | 0 |
| 0 | 0 |
| 73.2869715 | 13.7279655 |
| 0 | 0 |
| 0 | 0 |
| 0 | 0 |
| 0 | 0 |
| 0 | 0 |
| 0 | 0 |
| 0 | 0 |
| 0 | 0 |
| 0 | 0 |
| 0 | 0 |
| 0 | 0 |
| 0 | 0 |
| 0 | 0 |
| 5.42866456 | 0 |
| 0 | 0 |
| 0 | 0 |
| 0 | 0 |
| 0 | 0 |
| 0 | 0 |
| 0 | 0 |
| 0 | 0 |
| 0 | 0 |
| 0 | 0 |
| 0 | 0 |
| 146.573943 | 160.159598 |
| 0 | 4.57598851 |
| 0 | 0 |
| 0 | 0 |
| 0 | 0 |
| 0 | 0 |
| 0 | 0 |
| 0 | 0 |
| 0 | 0 |
| 21.7146582 | 0 |
| 0 | 0 |
| 0 | 0 |
| 0 | 0 |
| 0 | 0 |
| 0 | 0 |

| | |
|---|---|
| 0 | 0 |
| 0 | 0 |
| 0 | 0 |
| 0 | 0 |
| 0 | 0 |
| 0 | 0 |
| 0 | 0 |
| 0 | 0 |
| 0 | 0 |
| 0 | 0 |
| 0 | 0 |
| 0 | 0 |
| 0 | 0 |
| 2.71433228 | 0 |
| 0 | 0 |
| 0 | 0 |
| 0 | 0 |
| 0 | 0 |
| 0 | 0 |
| 0 | 73.2158162 |
| 0 | 0 |
| 0 | 0 |
| 0 | 0 |
| 0 | 0 |
| 0 | 22.8799426 |
| 0 | 0 |
| 0 | 0 |
| 0 | 0 |
| 0 | 0 |
| 0 | 0 |
| 0 | 36.6079081 |
| 0 | 0 |
| 0 | 4.57598851 |
| 0 | 0 |
| 0 | 0 |
| 0 | 0 |
| 0 | 0 |
| 0 | 0 |
| 0 | 0 |
| 0 | 0 |
| 0 | 0 |
| 0 | 0 |
| 0 | 0 |

| | |
|---|---|
| 0 | 0 |
| 0 | 0 |
| 0 | 0 |
| 0 | 27.4559311 |
| 0 | 0 |
| 0 | 0 |
| 0 | 0 |
| 0 | 0 |
| 0 | 0 |
| 0 | 0 |
| 0 | 0 |
| 0 | 0 |
| 0 | 0 |
| 0 | 0 |
| 0 | 0 |
| 114.001956 | 137.279655 |
| 0 | 0 |
| 0 | 0 |
| 0 | 0 |
| 0 | 0 |
| 0 | 18.303954 |
| 0 | 0 |
| 0 | 0 |
| 0 | 0 |
| 0 | 0 |
| 0 | 0 |
| 0 | 0 |
| 0 | 0 |
| 0 | 0 |
| 0 | 0 |
| 0 | 0 |
| 0 | 27.4559311 |
| 0 | 0 |
| 0 | 0 |
| 0 | 0 |
| 0 | 0 |
| 0 | 0 |
| 0 | 0 |
| 0 | 0 |
| 0 | 0 |
| 0 | 0 |
| 0 | 0 |
| 0 | 27.4559311 |

| | |
|---|---|
| 21.7146582 | 13.7279655 |
| 0 | 0 |
| 0 | 0 |
| 2.71433228 | 0 |
| 0 | 0 |
| 0 | 0 |
| 0 | 0 |
| 0 | 0 |
| 0 | 0 |
| 160.145604 | 0 |
| 0 | 0 |
| 0 | 54.9118621 |
| 0 | 0 |
| 5.42866456 | 0 |
| 0 | 0 |
| 0 | 0 |
| 0 | 0 |
| 0 | 0 |
| 0 | 0 |
| 8.14299683 | 18.303954 |
| 0 | 0 |
| 0 | 0 |
| 0 | 0 |
| 0 | 0 |
| 0 | 0 |
| 214.43225 | 22.8799426 |
| 0 | 0 |
| 0 | 0 |
| 0 | 0 |
| 43.4293164 | 0 |
| 0 | 0 |
| 0 | 0 |
| 0 | 0 |
| 0 | 0 |
| 0 | 0 |
| 0 | 0 |
| 2.71433228 | 50.3358736 |
| 0 | 0 |
| 0 | 0 |
| 0 | 0 |
| 0 | 0 |
| 0 | 0 |
| 0 | 0 |

| | |
|---|---|
| 0 | 4.57598851 |
| 0 | 0 |
| 0 | 0 |
| 0 | 0 |
| 0 | 0 |
| 0 | 0 |
| 0 | 0 |
| 0 | 0 |
| 0 | 27.4559311 |
| 0 | 0 |
| 0 | 0 |
| 0 | 0 |
| 0 | 0 |
| 0 | 0 |
| 0 | 0 |
| 0 | 0 |
| 0 | 0 |
| 0 | 0 |
| 0 | 0 |
| 0 | 0 |
| 0 | 0 |
| 0 | 0 |
| 0 | 0 |
| 0 | 0 |
| 5.42866456 | 0 |
| 0 | 0 |
| 610.724763 | 178.463552 |
| 2.71433228 | 54.9118621 |
| 130.287949 | 233.375414 |
| 0 | 0 |
| 0 | 0 |
| 0 | 0 |
| 0 | 0 |
| 0 | 0 |
| 0 | 0 |
| 0 | 0 |
| 0 | 9.15197702 |
| 0 | 0 |
| 0 | 0 |
| 0 | 0 |
| 0 | 0 |
| 0 | 0 |
| 0 | 0 |

| | |
|---|---|
| 0 | 0 |
| 0 | 0 |
| 13.5716614 | 4.57598851 |
| 276.861892 | 32.0319196 |
| 0 | 0 |
| 0 | 0 |
| 0 | 4.57598851 |
| 0 | 4.57598851 |
| 0 | 0 |
| 0 | 0 |
| 0 | 0 |
| 0 | 0 |
| 2.71433228 | 0 |
| 0 | 0 |
| 0 | 13.7279655 |
| 2.71433228 | 0 |
| 0 | 0 |
| 160.145604 | 503.358736 |
| 0 | 0 |
| 0 | 0 |
| 0 | 0 |
| 0 | 0 |
| 70.5726392 | 0 |
| 0 | 0 |
| 0 | 0 |
| 0 | 4.57598851 |
| 0 | 0 |
| 0 | 0 |
| 0 | 0 |
| 0 | 0 |
| 0 | 0 |
| 35.2863196 | 0 |
| 0 | 0 |
| 0 | 0 |
| 0 | 9.15197702 |
| 0 | 0 |
| 0 | 0 |
| 0 | 0 |
| 0 | 0 |
| 0 | 0 |
| 0 | 0 |
| 0 | 0 |
| 0 | 0 |

| | |
|---|---|
| 0 | 0 |
| 19.0003259 | 173.887563 |
| 0 | 0 |
| 160.145604 | 73.2158162 |
| 0 | 0 |
| 0 | 0 |
| 0 | 0 |
| 0 | 0 |
| 0 | 0 |
| 0 | 0 |
| 0 | 0 |
| 2.71433228 | 0 |
| 0 | 0 |
| 0 | 0 |
| 0 | 0 |
| 0 | 0 |
| 141.145278 | 41.1838966 |
| 0 | 0 |
| 0 | 0 |
| 0 | 0 |
| 0 | 0 |
| 0 | 0 |
| 0 | 0 |
| 0 | 0 |
| 0 | 0 |
| 0 | 0 |
| 0 | 0 |
| 0 | 0 |
| 0 | 0 |
| 0 | 0 |
| 0 | 0 |
| 0 | 0 |
| 0 | 0 |
| 0 | 45.7598851 |
| 0 | 45.7598851 |
| 0 | 0 |
| 0 | 0 |
| 0 | 0 |
| 0 | 0 |
| 0 | 0 |
| 0 | 27.4559311 |
| 0 | 0 |
| 0 | 9.15197702 |

| | |
|---|---|
| 0 | 0 |
| 0 | 0 |
| 0 | 0 |
| 0 | 0 |
| 3922.21014 | 1034.1734 |
| 0 | 0 |
| 0 | 0 |
| 0 | 0 |
| 19.0003259 | 59.4878506 |
| 0 | 0 |
| 0 | 0 |
| 0 | 0 |
| 0 | 0 |
| 0 | 0 |
| 0 | 0 |
| 0 | 0 |
| 0 | 0 |
| 0 | 0 |
| 0 | 0 |
| 0 | 0 |
| 0 | 0 |
| 236.146908 | 9.15197702 |
| 0 | 0 |
| 0 | 0 |
| 0 | 0 |
| 157.431272 | 0 |
| 0 | 0 |
| 0 | 18.303954 |
| 0 | 0 |
| 0 | 0 |
| 2.71433228 | 0 |
| 0 | 0 |
| 0 | 0 |
| 0 | 0 |
| 0 | 0 |
| 0 | 0 |
| 0 | 0 |
| 0 | 0 |
| 105.858959 | 96.0957587 |
| 0 | 0 |
| 0 | 0 |
| 0 | 0 |
| 0 | 0 |

| | |
|---|---|
| 0 | 0 |
| 0 | 0 |
| 0 | 0 |
| 0 | 0 |
| 0 | 0 |
| 0 | 0 |
| 0 | 0 |
| 0 | 0 |
| 35.2863196 | 13.7279655 |
| 0 | 0 |
| 0 | 0 |
| 8.14299683 | 4.57598851 |
| 0 | 0 |
| 0 | 0 |
| 67.8583069 | 9.15197702 |
| 0 | 0 |
| 0 | 0 |
| 0 | 0 |
| 119.43062 | 18.303954 |
| 0 | 4.57598851 |
| 0 | 0 |
| 0 | 0 |
| 0 | 0 |
| 0 | 0 |
| 0 | 0 |
| 0 | 0 |
| 8.14299683 | 4.57598851 |
| 0 | 0 |
| 0 | 0 |
| 0 | 0 |
| 0 | 4.57598851 |
| 0 | 0 |
| 0 | 0 |
| 0 | 0 |
| 0 | 0 |
| 0 | 0 |
| 0 | 0 |
| 0 | 0 |
| 0 | 0 |
| 0 | 0 |
| 0 | 36.6079081 |
| 0 | 0 |
| 0 | 0 |

| | |
|---|---|
| 0 | 0 |
| 0 | 0 |
| 0 | 0 |
| 0 | 0 |
| 0 | 4.57598851 |
| 0 | 0 |
| 0 | 0 |
| 0 | 0 |
| 0 | 0 |
| 0 | 0 |
| 2.71433228 | 4.57598851 |
| 2.71433228 | 45.7598851 |
| 0 | 0 |
| 0 | 0 |
| 0 | 0 |
| 0 | 0 |
| 0 | 0 |
| 0 | 0 |
| 0 | 22.8799426 |
| 0 | 0 |
| 0 | 0 |
| 0 | 0 |
| 0 | 0 |
| 0 | 0 |
| 0 | 0 |
| 0 | 0 |
| 0 | 0 |
| 0 | 0 |
| 0 | 0 |
| 0 | 0 |
| 0 | 0 |
| 0 | 0 |
| 89.5729652 | 0 |
| 16.2859937 | 77.7918047 |
| 0 | 0 |
| 0 | 0 |
| 0 | 4.57598851 |
| 0 | 0 |
| 59.7153101 | 105.247736 |
| 0 | 0 |
| 0 | 0 |
| 0 | 4.57598851 |
| 0 | 0 |

| | |
|---|---|
| 0 | 0 |
| 0 | 0 |
| 0 | 0 |
| 0 | 0 |
| 0 | 0 |
| 13.5716614 | 32.0319196 |
| 0 | 0 |
| 97.715962 | 18.303954 |
| 0 | 0 |
| 0 | 4.57598851 |
| 0 | 18.303954 |
| 0 | 0 |
| 0 | 0 |
| 0 | 0 |
| 0 | 0 |
| 0 | 0 |
| 475.008149 | 41.1838966 |
| 0 | 0 |
| 0 | 0 |
| 0 | 0 |
| 100.430294 | 73.2158162 |
| 0 | 0 |
| 5.42866456 | 9.15197702 |
| 0 | 0 |
| 0 | 0 |
| 0 | 0 |
| 0 | 0 |
| 0 | 0 |
| 0 | 0 |
| 0 | 0 |
| 0 | 0 |
| 0 | 0 |
| 0 | 0 |
| 0 | 0 |
| 2.71433228 | 0 |
| 0 | 0 |
| 0 | 0 |
| 0 | 0 |
| 0 | 0 |
| 0 | 0 |
| 16.2859937 | 0 |
| 0 | 0 |
| 0 | 0 |

| | |
|---|---|
| 0 | 0 |
| 0 | 0 |
| 0 | 0 |
| 0 | 0 |
| 0 | 0 |
| 0 | 0 |
| 0 | 0 |
| 0 | 0 |
| 0 | 0 |
| 0 | 0 |
| 0 | 0 |
| 0 | 0 |
| 0 | 0 |
| 0 | 4.57598851 |
| 2.71433228 | 0 |
| 2.71433228 | 9.15197702 |
| 0 | 0 |
| 0 | 0 |
| 0 | 0 |
| 0 | 0 |
| 0 | 0 |
| 0 | 0 |
| 0 | 0 |
| 0 | 32.0319196 |
| 0 | 0 |
| 0 | 0 |
| 0 | 0 |
| 0 | 0 |
| 0 | 0 |
| 0 | 0 |
| 0 | 0 |
| 0 | 0 |
| 0 | 0 |
| 2.71433228 | 54.9118621 |
| 0 | 9.15197702 |
| 0 | 0 |
| 0 | 0 |
| 0 | 0 |
| 0 | 0 |
| 0 | 0 |
| 0 | 0 |
| 0 | 0 |
| 0 | 0 |

| | |
|---|---|
| 0 | 0 |
| 0 | 0 |
| 0 | 0 |
| 0 | 32.0319196 |
| 0 | 0 |
| 0 | 0 |
| 0 | 0 |
| 0 | 0 |
| 0 | 0 |
| 0 | 0 |
| 0 | 0 |
| 0 | 0 |
| 2.71433228 | 50.3358736 |
| 0 | 0 |
| 0 | 0 |
| 219.860915 | 114.399713 |
| 0 | 0 |
| 0 | 0 |
| 0 | 0 |
| 0 | 0 |
| 0 | 0 |
| 222.575247 | 54.9118621 |
| 0 | 0 |
| 0 | 0 |
| 0 | 0 |
| 0 | 0 |
| 0 | 0 |
| 0 | 0 |
| 0 | 0 |
| 0 | 0 |
| 0 | 0 |
| 0 | 0 |
| 0 | 0 |
| 0 | 0 |
| 0 | 0 |
| 0 | 0 |
| 0 | 0 |
| 8.14299683 | 50.3358736 |
| 0 | 0 |
| 0 | 0 |
| 0 | 0 |
| 0 | 0 |
| 0 | 0 |

| | |
|---|---|
| 0 | 0 |
| 59.7153101 | 9.15197702 |
| 0 | 0 |
| 0 | 0 |
| 0 | 4.57598851 |
| 0 | 0 |
| 0 | 0 |
| 0 | 0 |
| 0 | 0 |
| 0 | 0 |
| 0 | 0 |
| 0 | 0 |
| 0 | 0 |
| 0 | 4.57598851 |
| 0 | 0 |
| 0 | 0 |
| 57.0009778 | 160.159598 |
| 0 | 0 |
| 0 | 0 |
| 2.71433228 | 32.0319196 |
| 0 | 0 |
| 0 | 0 |
| 0 | 0 |
| 0 | 4.57598851 |
| 0 | 0 |
| 0 | 0 |
| 0 | 0 |
| 24.4289905 | 18.303954 |
| 0 | 0 |
| 0 | 0 |
| 0 | 0 |
| 0 | 0 |
| 0 | 0 |
| 0 | 0 |
| 0 | 4.57598851 |
| 0 | 0 |
| 0 | 0 |
| 0 | 54.9118621 |
| 0 | 0 |
| 0 | 0 |
| 0 | 0 |
| 0 | 0 |
| 0 | 0 |

| | |
|---|---|
| 8.14299683 | 18.303954 |
| 0 | 4.57598851 |
| 0 | 0 |
| 0 | 0 |
| 0 | 0 |
| 233.432576 | 0 |
| 0 | 50.3358736 |
| 0 | 0 |
| 0 | 0 |
| 0 | 0 |
| 0 | 0 |
| 8.14299683 | 18.303954 |
| 152.002608 | 68.6398277 |
| 10.8573291 | 4.57598851 |
| 0 | 18.303954 |
| 0 | 0 |
| 0 | 0 |
| 0 | 0 |
| 0 | 0 |
| 0 | 0 |
| 0 | 0 |
| 0 | 0 |
| 0 | 0 |
| 0 | 0 |
| 0 | 0 |
| 0 | 0 |
| 0 | 0 |
| 0 | 0 |
| 0 | 27.4559311 |
| 0 | 0 |
| 0 | 0 |
| 48.857981 | 86.9437817 |
| 0 | 0 |
| 0 | 0 |
| 0 | 4.57598851 |
| 0 | 0 |
| 10.8573291 | 0 |
| 0 | 0 |
| 0 | 0 |
| 0 | 0 |
| 0 | 0 |
| 0 | 0 |
| 0 | 0 |

| | |
|---|---|
| 0 | 0 |
| 0 | 0 |
| 57.0009778 | 155.583609 |
| 0 | 0 |
| 0 | 0 |
| 0 | 0 |
| 0 | 0 |
| 0 | 0 |
| 0 | 0 |
| 54.2866456 | 183.03954 |
| 249.71857 | 68.6398277 |
| 0 | 0 |
| 0 | 0 |
| 0 | 0 |
| 0 | 0 |
| 0 | 0 |
| 46.1436487 | 45.7598851 |
| 0 | 0 |
| 2.71433228 | 0 |
| 0 | 0 |
| 0 | 0 |
| 0 | 0 |
| 0 | 0 |
| 2.71433228 | 0 |
| 0 | 18.303954 |
| 0 | 0 |
| 0 | 0 |
| 0 | 0 |
| 0 | 27.4559311 |
| 0 | 0 |
| 0 | 0 |
| 0 | 0 |
| 0 | 0 |
| 100.430294 | 0 |
| 0 | 0 |
| 0 | 0 |
| 0 | 4.57598851 |
| 0 | 0 |
| 0 | 0 |
| 0 | 0 |
| 0 | 0 |
| 352.863196 | 164.735586 |
| 0 | 0 |

| | |
|---|---|
| 0 | 0 |
| 0 | 0 |
| 0 | 0 |
| 0 | 0 |
| 0 | 0 |
| 0 | 0 |
| 0 | 0 |
| 0 | 0 |
| 0 | 0 |
| 0 | 0 |
| 0 | 0 |
| 0 | 0 |
| 209.003585 | 210.495472 |
| 0 | 0 |
| 0 | 13.7279655 |
| 0 | 0 |
| 0 | 0 |
| 0 | 0 |
| 81.4299683 | 18.303954 |
| 0 | 0 |
| 0 | 0 |
| 0 | 0 |
| 0 | 4.57598851 |
| 0 | 0 |
| 0 | 0 |
| 0 | 4.57598851 |
| 439.721829 | 151.007621 |
| 0 | 0 |
| 0 | 0 |
| 0 | 0 |
| 0 | 0 |
| 0 | 0 |
| 0 | 0 |
| 0 | 13.7279655 |
| 0 | 0 |
| 0 | 0 |
| 306.719547 | 334.047161 |
| 0 | 0 |
| 268.718896 | 151.007621 |
| 0 | 36.6079081 |
| 1167.16288 | 260.831345 |
| 0 | 45.7598851 |
| 0 | 0 |

| | |
|---|---|
| 5.42866456 | 13.7279655 |
| 0 | 0 |
| 78.7156361 | 118.975701 |
| 0 | 0 |
| 0 | 0 |
| 0 | 0 |
| 0 | 0 |
| 0 | 0 |
| 0 | 0 |
| 0 | 0 |
| 0 | 0 |
| 0 | 4.57598851 |
| 0 | 0 |
| 8.14299683 | 100.671747 |
| 0 | 0 |
| 0 | 0 |
| 21.7146582 | 0 |
| 0 | 0 |
| 0 | 0 |
| 173.717266 | 86.9437817 |
| 0 | 0 |
| 0 | 0 |
| 0 | 0 |
| 0 | 0 |
| 0 | 0 |
| 0 | 0 |
| 0 | 0 |
| 0 | 0 |
| 0 | 0 |
| 0 | 0 |
| 0 | 0 |
| 0 | 0 |
| 0 | 0 |
| 0 | 0 |
| 0 | 0 |
| 0 | 0 |
| 0 | 0 |
| 0 | 4.57598851 |
| 127.573617 | 36.6079081 |
| 0 | 0 |
| 0 | 0 |
| 0 | 0 |
| 0 | 0 |

| | |
|---|---|
| 0 | 0 |
| 0 | 0 |
| 97.715962 | 0 |
| 0 | 13.7279655 |
| 0 | 0 |
| 0 | 0 |
| 0 | 0 |
| 0 | 0 |
| 0 | 0 |
| 0 | 0 |
| 0 | 0 |
| 73.2869715 | 27.4559311 |
| 0 | 0 |
| 0 | 0 |
| 0 | 0 |
| 0 | 0 |
| 0 | 0 |
| 0 | 0 |
| 86.8586329 | 9.15197702 |
| 0 | 0 |
| 0 | 0 |
| 0 | 0 |
| 0 | 0 |
| 0 | 0 |
| 2.71433228 | 0 |
| 0 | 4.57598851 |
| 19.0003259 | 0 |
| 0 | 0 |
| 0 | 0 |
| 0 | 0 |
| 0 | 0 |
| 0 | 0 |
| 0 | 0 |
| 0 | 0 |
| 0 | 0 |
| 0 | 0 |
| 0 | 0 |
| 0 | 0 |
| 0 | 4.57598851 |
| 0 | 0 |
| 46.1436487 | 68.6398277 |
| 0 | 0 |
| 38.0006519 | 36.6079081 |

| | |
|---|---|
| 0 | 0 |
| 0 | 0 |
| 0 | 0 |
| 0 | 0 |
| 0 | 0 |
| 0 | 0 |
| 0 | 0 |
| 0 | 0 |
| 0 | 0 |
| 10.8573291 | 315.743207 |
| 0 | 0 |
| 0 | 0 |
| 0 | 0 |
| 0 | 0 |
| 0 | 0 |
| 0 | 0 |
| 0 | 0 |
| 2.71433228 | 4.57598851 |
| 0 | 0 |
| 0 | 22.8799426 |
| 0 | 0 |
| 86.8586329 | 0 |
| 0 | 0 |
| 130.287949 | 105.247736 |
| 0 | 0 |
| 0 | 0 |
| 314.862544 | 50.3358736 |
| 0 | 0 |
| 0 | 0 |
| 0 | 0 |
| 0 | 0 |
| 0 | 0 |
| 0 | 0 |
| 0 | 32.0319196 |
| 0 | 0 |
| 0 | 22.8799426 |
| 0 | 9.15197702 |
| 0 | 0 |
| 0 | 0 |
| 0 | 0 |
| 0 | 0 |
| 0 | 0 |
| 0 | 0 |

| | |
|---|---|
| 0 | 0 |
| 2.71433228 | 0 |
| 2.71433228 | 0 |
| 141.145278 | 370.655069 |
| 0 | 0 |
| 0 | 0 |
| 0 | 0 |
| 0 | 0 |
| 0 | 0 |
| 0 | 0 |
| 2.71433228 | 0 |
| 222.575247 | 398.111 |
| 0 | 0 |
| 2.71433228 | 0 |
| 0 | 0 |
| 0 | 0 |
| 0 | 4.57598851 |
| 0 | 0 |
| 0 | 0 |
| 0 | 0 |
| 0 | 0 |
| 0 | 0 |
| 0 | 0 |
| 0 | 0 |
| 0 | 0 |
| 0 | 0 |
| 0 | 0 |
| 0 | 0 |
| 0 | 0 |
| 0 | 0 |
| 0 | 0 |
| 116.716288 | 27.4559311 |
| 2.71433228 | 0 |
| 0 | 0 |
| 5.42866456 | 0 |
| 5.42866456 | 0 |
| 0 | 0 |
| 0 | 0 |
| 542.866456 | 347.775127 |
| 0 | 0 |
| 0 | 0 |
| 0 | 0 |
| 0 | 0 |

| | |
|---|---|
| 0 | 0 |
| 0 | 0 |
| 0 | 0 |
| 0 | 0 |
| 0 | 0 |
| 0 | 0 |
| 0 | 0 |
| 0 | 0 |
| 0 | 0 |
| 0 | 0 |
| 0 | 0 |
| 0 | 0 |
| 363.720525 | 100.671747 |
| 298.576551 | 36.6079081 |
| 0 | 0 |
| 0 | 0 |
| 0 | 0 |
| 0 | 0 |
| 0 | 0 |
| 317.576877 | 86.9437817 |
| 0 | 0 |
| 5.42866456 | 9.15197702 |
| 0 | 0 |
| 0 | 0 |
| 0 | 0 |
| 2.71433228 | 0 |
| 0 | 0 |
| 0 | 0 |
| 0 | 0 |
| 89.5729652 | 45.7598851 |
| 0 | 0 |
| 2.71433228 | 0 |
| 0 | 0 |
| 0 | 0 |
| 5.42866456 | 9.15197702 |
| 0 | 32.0319196 |
| 247.004237 | 96.0957587 |
| 0 | 0 |
| 0 | 0 |
| 0 | 0 |
| 0 | 0 |
| 0 | 0 |
| 0 | 0 |

| | |
|---|---|
| 0 | 0 |
| 0 | 0 |
| 0 | 0 |
| 0 | 0 |
| 0 | 0 |
| 0 | 0 |
| 0 | 9.15197702 |
| 0 | 0 |
| 0 | 18.303954 |
| 0 | 0 |
| 0 | 0 |
| 0 | 0 |
| 0 | 0 |
| 0 | 0 |
| 0 | 0 |
| 0 | 0 |
| 0 | 0 |
| 2.71433228 | 4.57598851 |
| 0 | 0 |
| 0 | 0 |
| 0 | 0 |
| 0 | 0 |
| 0 | 0 |
| 0 | 0 |
| 0 | 13.7279655 |
| 0 | 0 |
| 0 | 0 |
| 0 | 0 |
| 0 | 0 |
| 0 | 0 |
| 0 | 0 |
| 0 | 0 |
| 0 | 0 |
| 0 | 64.0638392 |
| 0 | 0 |
| 51.5723133 | 0 |
| 0 | 36.6079081 |
| 0 | 0 |
| 0 | 0 |
| 0 | 0 |
| 0 | 0 |
| 0 | 0 |
| 0 | 4.57598851 |

| | |
|---|---|
| 0 | 9.15197702 |
| 0 | 0 |
| 0 | 0 |
| 0 | 0 |
| 0 | 0 |
| 374.577854 | 105.247736 |
| 0 | 0 |
| 0 | 0 |
| 0 | 0 |
| 0 | 0 |
| 0 | 0 |
| 0 | 0 |
| 0 | 0 |
| 0 | 0 |
| 0 | 0 |
| 92.2872974 | 41.1838966 |
| 0 | 0 |
| 0 | 0 |
| 0 | 0 |
| 0 | 0 |
| 0 | 0 |
| 0 | 73.2158162 |
| 0 | 0 |
| 0 | 0 |
| 0 | 0 |
| 5.42866456 | 4.57598851 |
| 0 | 0 |
| 0 | 0 |
| 0 | 0 |
| 0 | 0 |
| 24.4289905 | 27.4559311 |
| 0 | 27.4559311 |
| 0 | 0 |
| 2.71433228 | 4.57598851 |
| 0 | 0 |
| 0 | 0 |
| 0 | 0 |
| 0 | 0 |
| 0 | 0 |
| 0 | 0 |
| 0 | 0 |
| 0 | 0 |
| 0 | 0 |

| | |
|---|---|
| 0 | 0 |
| 0 | 0 |
| 0 | 0 |
| 0 | 0 |
| 0 | 0 |
| 0 | 0 |
| 2.71433228 | 0 |
| 8.14299683 | 22.8799426 |
| 5.42866456 | 32.0319196 |
| 0 | 0 |
| 0 | 0 |
| 0 | 0 |
| 0 | 0 |
| 0 | 54.9118621 |
| 0 | 0 |
| 0 | 0 |
| 244.289905 | 54.9118621 |
| 0 | 0 |
| 24.4289905 | 128.127678 |
| 0 | 0 |
| 0 | 0 |
| 0 | 0 |
| 0 | 0 |
| 0 | 0 |
| 0 | 32.0319196 |
| 0 | 59.4878506 |
| 0 | 0 |
| 236.146908 | 146.431632 |
| 0 | 0 |
| 0 | 0 |
| 0 | 0 |
| 0 | 0 |
| 0 | 0 |
| 0 | 0 |
| 1370.7378 | 146.431632 |
| 8.14299683 | 4.57598851 |
| 0 | 4.57598851 |
| 59.7153101 | 77.7918047 |
| 0 | 0 |
| 0 | 0 |
| 0 | 0 |
| 35.2863196 | 36.6079081 |
| 0 | 0 |

| | |
|---|---|
| 0 | 0 |
| 0 | 0 |
| 0 | 0 |
| 0 | 0 |
| 32.5719873 | 45.7598851 |
| 0 | 0 |
| 0 | 4.57598851 |
| 0 | 0 |
| 0 | 0 |
| 0 | 0 |
| 0 | 0 |
| 0 | 0 |
| 352.863196 | 4.57598851 |
| 0 | 0 |
| 43.4293164 | 13.7279655 |
| 0 | 0 |
| 0 | 0 |
| 0 | 0 |
| 0 | 0 |
| 0 | 0 |
| 301.290883 | 13.7279655 |
| 0 | 4.57598851 |
| 0 | 0 |
| 0 | 0 |
| 0 | 0 |
| 0 | 0 |
| 0 | 0 |
| 97.715962 | 41.1838966 |
| 0 | 0 |
| 0 | 0 |
| 0 | 0 |
| 0 | 0 |
| 0 | 0 |
| 0 | 0 |
| 0 | 0 |
| 0 | 0 |
| 0 | 0 |
| 0 | 0 |
| 0 | 0 |
| 0 | 0 |
| 0 | 0 |
| 0 | 0 |
| 0 | 0 |

| | |
|---|---|
| 0 | 0 |
| 5.42866456 | 36.6079081 |
| 686.726066 | 1116.5412 |
| 0 | 0 |
| 0 | 0 |
| 0 | 0 |
| 0 | 0 |
| 0 | 0 |
| 0 | 0 |
| 0 | 0 |
| 0 | 0 |
| 0 | 0 |
| 0 | 0 |
| 0 | 0 |
| 0 | 0 |
| 0 | 0 |
| 0 | 0 |
| 0 | 0 |
| 0 | 0 |
| 0 | 0 |
| 0 | 0 |
| 2.71433228 | 0 |
| 0 | 0 |
| 0 | 0 |
| 0 | 0 |
| 0 | 0 |
| 0 | 0 |
| 0 | 0 |
| 0 | 0 |
| 0 | 0 |
| 152.002608 | 334.047161 |
| 0 | 0 |
| 0 | 0 |
| 0 | 0 |
| 0 | 0 |
| 0 | 0 |
| 0 | 0 |
| 0 | 0 |
| 0 | 0 |
| 0 | 0 |
| 2.71433228 | 0 |
| 0 | 0 |
| 0 | 0 |

| | |
|---|---|
| 0 | 0 |
| 10.8573291 | 45.7598851 |
| 0 | 0 |
| 0 | 0 |
| 0 | 0 |
| 0 | 0 |
| 111.287623 | 100.671747 |
| 0 | 27.4559311 |
| 0 | 0 |
| 0 | 0 |
| 0 | 0 |
| 0 | 0 |
| 0 | 0 |
| 0 | 0 |
| 0 | 4.57598851 |
| 48.857981 | 96.0957587 |
| 0 | 0 |
| 0 | 4.57598851 |
| 0 | 0 |
| 0 | 0 |
| 0 | 0 |
| 0 | 0 |
| 0 | 0 |
| 5.42866456 | 50.3358736 |
| 0 | 0 |
| 0 | 0 |
| 0 | 0 |
| 0 | 0 |
| 0 | 0 |
| 0 | 0 |
| 0 | 0 |
| 0 | 0 |
| 0 | 0 |
| 0 | 0 |
| 0 | 36.6079081 |
| 0 | 0 |
| 0 | 0 |
| 0 | 54.9118621 |
| 0 | 0 |
| 0 | 0 |
| 0 | 0 |
| 16.2859937 | 0 |
| 0 | 4.57598851 |

| | |
|---|---|
| 0 | 0 |
| 0 | 0 |
| 0 | 0 |
| 0 | 0 |
| 0 | 0 |
| 2.71433228 | 0 |
| 0 | 0 |
| 0 | 0 |
| 0 | 0 |
| 0 | 0 |
| 0 | 0 |
| 0 | 0 |
| 0 | 0 |
| 0 | 0 |
| 0 | 0 |
| 5.42866456 | 32.0319196 |
| 0 | 0 |
| 84.1443006 | 0 |
| 0 | 0 |
| 0 | 0 |
| 0 | 0 |
| 0 | 0 |
| 0 | 0 |
| 0 | 0 |
| 0 | 4.57598851 |
| 0 | 0 |
| 0 | 0 |
| 0 | 0 |
| 0 | 0 |
| 0 | 0 |
| 0 | 0 |
| 133.002282 | 132.703667 |
| 0 | 0 |
| 10.8573291 | 173.887563 |
| 0 | 0 |
| 0 | 0 |
| 0 | 0 |
| 0 | 0 |
| 0 | 0 |
| 0 | 0 |
| 0 | 0 |
| 0 | 0 |
| 0 | 0 |

| | |
|---|---|
| 0 | 0 |
| 0 | 0 |
| 0 | 0 |
| 0 | 4.57598851 |
| 0 | 0 |
| 0 | 27.4559311 |
| 0 | 0 |
| 0 | 0 |
| 0 | 0 |
| 0 | 0 |
| 0 | 0 |
| 0 | 0 |
| 0 | 32.0319196 |
| 0 | 0 |
| 2.71433228 | 0 |
| 0 | 0 |
| 0 | 0 |
| 0 | 0 |
| 0 | 0 |
| 0 | 0 |
| 0 | 0 |
| 0 | 0 |
| 0 | 0 |
| 0 | 0 |
| 0 | 0 |
| 0 | 0 |
| 0 | 0 |
| 255.147234 | 4.57598851 |
| 0 | 0 |
| 0 | 0 |
| 0 | 0 |
| 59.7153101 | 128.127678 |
| 0 | 0 |
| 0 | 0 |
| 0 | 0 |
| 0 | 0 |
| 0 | 0 |
| 0 | 0 |
| 0 | 0 |
| 0 | 0 |
| 0 | 0 |
| 0 | 0 |
| 0 | 27.4559311 |

| | |
|---|---|
| 0 | 0 |
| 0 | 0 |
| 0 | 0 |
| 0 | 0 |
| 0 | 0 |
| 0 | 0 |
| 0 | 0 |
| 0 | 0 |
| 0 | 0 |
| 0 | 0 |
| 0 | 9.15197702 |
| 0 | 0 |
| 0 | 0 |
| 0 | 0 |
| 0 | 0 |
| 0 | 0 |
| 2.71433228 | 0 |
| 0 | 0 |
| 0 | 0 |
| 0 | 0 |
| 78.7156361 | 13.7279655 |
| 141.145278 | 64.0638392 |
| 29.8576551 | 18.303954 |
| 0 | 0 |
| 0 | 0 |
| 0 | 0 |
| 0 | 0 |
| 0 | 0 |
| 138.430946 | 0 |
| 0 | 0 |
| 0 | 0 |
| 0 | 0 |
| 0 | 0 |
| 0 | 0 |
| 0 | 0 |
| 0 | 0 |
| 0 | 0 |
| 0 | 0 |
| 0 | 0 |
| 0 | 0 |
| 0 | 0 |
| 0 | 18.303954 |
| 0 | 0 |

| | |
|---|---|
| 0 | 0 |
| 0 | 0 |
| 0 | 0 |
| 0 | 0 |
| 0 | 0 |
| 594.438769 | 36.6079081 |
| 0 | 0 |
| 0 | 0 |
| 0 | 0 |
| 692.154731 | 32.0319196 |
| 0 | 0 |
| 0 | 0 |
| 0 | 0 |
| 0 | 0 |
| 0 | 0 |
| 0 | 0 |
| 0 | 0 |
| 0 | 0 |
| 0 | 0 |
| 0 | 0 |
| 0 | 0 |
| 0 | 0 |
| 0 | 0 |
| 100.430294 | 36.6079081 |
| 0 | 0 |
| 5.42866456 | 0 |
| 0 | 0 |
| 0 | 0 |
| 0 | 27.4559311 |
| 0 | 0 |
| 0 | 0 |
| 0 | 0 |
| 0 | 0 |
| 0 | 0 |
| 2.71433228 | 9.15197702 |
| 0 | 0 |
| 0 | 0 |
| 0 | 4.57598851 |
| 0 | 0 |
| 0 | 0 |
| 0 | 0 |
| 0 | 0 |
| 0 | 0 |

| | |
|---|---|
| 122.144953 | 18.303954 |
| 2.71433228 | 100.671747 |
| 0 | 0 |
| 27.1433228 | 64.0638392 |
| 0 | 0 |
| 0 | 0 |
| 0 | 0 |
| 92.2872974 | 27.4559311 |
| 0 | 0 |
| 0 | 0 |
| 0 | 0 |
| 0 | 4.57598851 |
| 0 | 0 |
| 0 | 0 |
| 0 | 4.57598851 |
| 13.5716614 | 9.15197702 |
| 0 | 0 |
| 0 | 0 |
| 0 | 0 |
| 0 | 0 |
| 0 | 0 |
| 0 | 0 |
| 2.71433228 | 0 |
| 198.146256 | 13.7279655 |
| 2.71433228 | 41.1838966 |
| 0 | 0 |
| 0 | 0 |
| 0 | 0 |
| 0 | 0 |
| 24.4289905 | 196.767506 |
| 0 | 0 |
| 0 | 0 |
| 0 | 0 |
| 0 | 0 |
| 0 | 0 |
| 2.71433228 | 50.3358736 |
| 0 | 0 |
| 0 | 0 |
| 0 | 0 |
| 358.291861 | 196.767506 |
| 0 | 0 |
| 0 | 9.15197702 |
| 0 | 0 |

| | |
|---|---|
| 0 | 0 |
| 0 | 0 |
| 0 | 0 |
| 0 | 0 |
| 0 | 0 |
| 0 | 0 |
| 0 | 0 |
| 0 | 0 |
| 171.002934 | 4.57598851 |
| 0 | 0 |
| 0 | 0 |
| 0 | 0 |
| 0 | 0 |
| 0 | 0 |
| 0 | 0 |
| 0 | 41.1838966 |
| 0 | 0 |
| 19.0003259 | 0 |
| 0 | 0 |
| 0 | 0 |
| 0 | 0 |
| 0 | 0 |
| 0 | 0 |
| 2.71433228 | 45.7598851 |
| 0 | 0 |
| 2.71433228 | 0 |
| 0 | 0 |
| 0 | 0 |
| 0 | 0 |
| 0 | 0 |
| 0 | 0 |
| 0 | 0 |
| 0 | 0 |
| 0 | 0 |
| 0 | 0 |
| 0 | 0 |
| 135.716614 | 0 |
| 735.584047 | 100.671747 |
| 16.2859937 | 9.15197702 |
| 0 | 0 |
| 0 | 0 |
| 0 | 0 |
| 0 | 0 |

| | |
|---|---|
| 0 | 0 |
| 0 | 0 |
| 0 | 0 |
| 0 | 0 |
| 0 | 0 |
| 0 | 0 |
| 7097.97891 | 352.351115 |
| 0 | 0 |
| 0 | 100.671747 |
| 0 | 0 |
| 0 | 0 |
| 0 | 0 |
| 0 | 0 |
| 0 | 0 |
| 0 | 0 |
| 0 | 0 |
| 160.145604 | 0 |
| 2.71433228 | 54.9118621 |
| 0 | 0 |
| 0 | 0 |
| 100.430294 | 96.0957587 |
| 0 | 0 |
| 0 | 0 |
| 0 | 0 |
| 0 | 4.57598851 |
| 0 | 0 |
| 0 | 9.15197702 |
| 0 | 0 |
| 0 | 0 |
| 0 | 0 |
| 105.858959 | 82.3677932 |
| 0 | 0 |
| 0 | 0 |
| 0 | 0 |
| 0 | 9.15197702 |
| 0 | 0 |
| 0 | 0 |
| 0 | 0 |
| 0 | 0 |
| 0 | 4.57598851 |
| 0 | 0 |
| 0 | 0 |
| 0 | 0 |

| | |
|---|---|
| 0 | 0 |
| 124.859285 | 123.55169 |
| 0 | 0 |
| 0 | 0 |
| 195.431924 | 96.0957587 |
| 0 | 0 |
| 0 | 0 |
| 0 | 0 |
| 0 | 32.0319196 |
| 0 | 0 |
| 0 | 0 |
| 0 | 0 |
| 0 | 0 |
| 257.861566 | 0 |
| 0 | 0 |
| 0 | 0 |
| 0 | 0 |
| 0 | 0 |
| 0 | 0 |
| 0 | 0 |
| 0 | 0 |
| 0 | 0 |
| 0 | 32.0319196 |
| 0 | 0 |
| 0 | 0 |
| 0 | 0 |
| 0 | 0 |
| 0 | 0 |
| 0 | 0 |
| 0 | 0 |
| 0 | 0 |
| 0 | 0 |
| 0 | 0 |
| 0 | 0 |
| 0 | 0 |
| 0 | 0 |
| 0 | 0 |
| 0 | 0 |
| 0 | 0 |
| 0 | 0 |
| 0 | 0 |
| 0 | 0 |
| 19.0003259 | 283.711288 |

| | |
|---|---|
| 0 | 0 |
| 2.71433228 | 54.9118621 |
| 16.2859937 | 0 |
| 0 | 0 |
| 0 | 0 |
| 0 | 0 |
| 0 | 0 |
| 0 | 0 |
| 0 | 0 |
| 0 | 0 |
| 0 | 0 |
| 0 | 0 |
| 0 | 0 |
| 0 | 0 |
| 0 | 0 |
| 0 | 0 |
| 0 | 0 |
| 0 | 0 |
| 0 | 0 |
| 0 | 0 |
| 0 | 0 |
| 0 | 0 |
| 0 | 0 |
| 0 | 0 |
| 0 | 0 |
| 0 | 0 |
| 0 | 0 |
| 0 | 0 |
| 0 | 0 |
| 0 | 0 |
| 0 | 141.855644 |
| 0 | 0 |
| 0 | 0 |
| 198.146256 | 45.7598851 |
| 0 | 0 |
| 0 | 0 |
| 0 | 0 |
| 0 | 0 |
| 0 | 0 |
| 0 | 0 |
| 0 | 0 |
| 0 | 0 |
| 0 | 0 |

| | |
|---|---|
| 0 | 0 |
| 0 | 0 |
| 0 | 0 |
| 0 | 0 |
| 0 | 4.57598851 |
| 19.0003259 | 0 |
| 0 | 9.15197702 |
| 0 | 0 |
| 0 | 0 |
| 0 | 91.5197702 |
| 0 | 36.6079081 |
| 0 | 0 |
| 0 | 0 |
| 0 | 0 |
| 0 | 0 |
| 89.5729652 | 196.767506 |
| 0 | 0 |
| 0 | 0 |
| 0 | 0 |
| 0 | 0 |
| 0 | 0 |
| 0 | 0 |
| 0 | 0 |
| 0 | 0 |
| 0 | 0 |
| 8.14299683 | 0 |
| 2.71433228 | 0 |
| 0 | 0 |
| 2.71433228 | 0 |
| 10.8573291 | 0 |
| 0 | 0 |
| 0 | 0 |
| 114.001956 | 86.9437817 |
| 0 | 0 |
| 0 | 0 |
| 0 | 0 |
| 0 | 0 |
| 0 | 0 |
| 0 | 0 |
| 0 | 0 |
| 0 | 0 |
| 0 | 4.57598851 |
| 0 | 0 |

| | |
|---|---|
| 0 | 0 |
| 2.71433228 | 22.8799426 |
| 0 | 0 |
| 0 | 0 |
| 2.71433228 | 0 |
| 0 | 0 |
| 0 | 0 |
| 0 | 0 |
| 0 | 0 |
| 0 | 0 |
| 0 | 0 |
| 0 | 4.57598851 |
| 0 | 0 |
| 0 | 0 |
| 0 | 0 |
| 0 | 0 |
| 0 | 0 |
| 0 | 0 |
| 16.2859937 | 59.4878506 |
| 0 | 0 |
| 0 | 0 |
| 0 | 0 |
| 342.005867 | 292.863265 |
| 0 | 13.7279655 |
| 0 | 0 |
| 8.14299683 | 0 |
| 0 | 0 |
| 0 | 0 |
| 0 | 4.57598851 |
| 0 | 0 |
| 0 | 0 |
| 0 | 0 |
| 222.575247 | 4.57598851 |
| 0 | 27.4559311 |
| 0 | 0 |
| 0 | 0 |
| 59.7153101 | 4.57598851 |
| 0 | 0 |
| 0 | 0 |
| 5.42866456 | 27.4559311 |
| 59.7153101 | 32.0319196 |
| 0 | 0 |
| 0 | 0 |

| | |
|---|---|
| 0 | 0 |
| 0 | 0 |
| 2.71433228 | 27.4559311 |
| 0 | 0 |
| 16.2859937 | 0 |
| 0 | 0 |
| 0 | 0 |
| 0 | 0 |
| 0 | 0 |
| 0 | 0 |
| 0 | 0 |
| 0 | 54.9118621 |
| 0 | 0 |
| 0 | 0 |
| 0 | 0 |
| 0 | 0 |
| 0 | 13.7279655 |
| 0 | 0 |
| 0 | 0 |
| 0 | 0 |
| 0 | 0 |
| 0 | 0 |
| 0 | 0 |
| 0 | 0 |
| 0 | 0 |
| 0 | 0 |
| 0 | 0 |
| 0 | 0 |
| 0 | 0 |
| 0 | 0 |
| 0 | 0 |
| 0 | 0 |
| 8.14299683 | 27.4559311 |
| 0 | 0 |
| 0 | 0 |
| 0 | 0 |
| 0 | 0 |
| 0 | 0 |
| 0 | 0 |
| 0 | 0 |
| 2.71433228 | 9.15197702 |
| 0 | 0 |
| 0 | 0 |

| | |
|---|---|
| 0 | 91.5197702 |
| 0 | 0 |
| 0 | 9.15197702 |
| 325.719873 | 132.703667 |
| 0 | 0 |
| 333.86287 | 54.9118621 |
| 0 | 0 |
| 0 | 0 |
| 0 | 0 |
| 0 | 0 |
| 0 | 0 |
| 0 | 0 |
| 0 | 0 |
| 21.7146582 | 68.6398277 |
| 0 | 0 |
| 5.42866456 | 0 |
| 0 | 0 |
| 0 | 0 |
| 138.430946 | 0 |
| 19.0003259 | 0 |
| 0 | 0 |
| 358.291861 | 36.6079081 |
| 0 | 22.8799426 |
| 0 | 0 |
| 0 | 0 |
| 0 | 0 |
| 0 | 0 |
| 0 | 0 |
| 0 | 22.8799426 |
| 0 | 0 |
| 40.7149842 | 91.5197702 |
| 0 | 0 |
| 0 | 0 |
| 0 | 0 |
| 0 | 0 |
| 0 | 0 |
| 0 | 0 |
| 0 | 0 |
| 0 | 0 |
| 0 | 0 |
| 808.871019 | 1226.36492 |
| 0 | 0 |
| 0 | 0 |

| | |
|---|---|
| 0 | 0 |
| 46.1436487 | 41.1838966 |
| 13.5716614 | 0 |
| 0 | 0 |
| 0 | 0 |
| 0 | 0 |
| 0 | 0 |
| 0 | 0 |
| 16.2859937 | 4.57598851 |
| 0 | 0 |
| 0 | 0 |
| 0 | 0 |
| 0 | 0 |
| 0 | 0 |
| 0 | 0 |
| 0 | 0 |
| 0 | 13.7279655 |
| 0 | 0 |
| 0 | 68.6398277 |
| 0 | 0 |
| 0 | 0 |
| 160.145604 | 41.1838966 |
| 0 | 0 |
| 0 | 0 |
| 0 | 0 |
| 0 | 0 |
| 0 | 0 |
| 0 | 0 |
| 133.002282 | 54.9118621 |
| 0 | 0 |
| 0 | 0 |
| 0 | 0 |
| 0 | 0 |
| 2.71433228 | 64.0638392 |
| 0 | 0 |
| 0 | 0 |
| 0 | 0 |
| 0 | 0 |
| 0 | 0 |
| 0 | 0 |
| 0 | 0 |
| 0 | 0 |
| 0 | 0 |

| | |
|---|---|
| 0 | 0 |
| 0 | 0 |
| 0 | 0 |
| 0 | 0 |
| 0 | 0 |
| 0 | 0 |
| 0 | 0 |
| 0 | 0 |
| 0 | 0 |
| 0 | 0 |
| 0 | 0 |
| 0 | 0 |
| 0 | 0 |
| 65.1439747 | 18.303954 |
| 0 | 0 |
| 0 | 0 |
| 0 | 0 |
| 0 | 0 |
| 0 | 0 |
| 0 | 0 |
| 0 | 0 |
| 0 | 0 |
| 0 | 0 |
| 0 | 0 |
| 0 | 0 |
| 0 | 0 |
| 0 | 0 |
| 0 | 0 |
| 0 | 0 |
| 0 | 0 |
| 0 | 0 |
| 0 | 0 |
| 0 | 0 |
| 0 | 0 |
| 0 | 0 |
| 0 | 0 |
| 0 | 0 |
| 0 | 0 |
| 0 | 0 |
| 0 | 0 |
| 0 | 0 |
| 0 | 0 |
| 73.2869715 | 4.57598851 |

| | |
|---|---|
| 0 | 0 |
| 0 | 0 |
| 0 | 0 |
| 0 | 0 |
| 0 | 0 |
| 0 | 0 |
| 0 | 0 |
| 0 | 0 |
| 244.289905 | 0 |
| 0 | 0 |
| 0 | 0 |
| 0 | 96.0957587 |
| 0 | 0 |
| 0 | 0 |
| 0 | 0 |
| 0 | 0 |
| 0 | 0 |
| 0 | 0 |
| 0 | 0 |
| 2.71433228 | 36.6079081 |
| 0 | 0 |
| 0 | 0 |
| 0 | 0 |
| 0 | 0 |
| 0 | 4.57598851 |
| 0 | 0 |
| 0 | 0 |
| 0 | 0 |
| 0 | 0 |
| 597.153101 | 73.2158162 |
| 86.8586329 | 22.8799426 |
| 0 | 4.57598851 |
| 0 | 0 |
| 152.002608 | 41.1838966 |
| 0 | 0 |
| 0 | 0 |
| 0 | 0 |
| 0 | 0 |
| 0 | 0 |
| 0 | 0 |
| 0 | 0 |
| 0 | 0 |
| 0 | 0 |

| | |
|---|---|
| 0 | 0 |
| 97.715962 | 0 |
| 146.573943 | 27.4559311 |
| 0 | 9.15197702 |
| 2.71433228 | 45.7598851 |
| 0 | 13.7279655 |
| 135.716614 | 32.0319196 |
| 0 | 0 |
| 0 | 0 |
| 0 | 0 |
| 0 | 0 |
| 0 | 0 |
| 0 | 0 |
| 0 | 0 |
| 0 | 0 |
| 0 | 0 |
| 35.2863196 | 13.7279655 |
| 21.7146582 | 9.15197702 |
| 0 | 0 |
| 0 | 0 |
| 0 | 18.303954 |
| 0 | 0 |
| 0 | 0 |
| 0 | 0 |
| 0 | 0 |
| 0 | 0 |
| 135.716614 | 13.7279655 |
| 0 | 0 |
| 0 | 0 |
| 0 | 0 |
| 0 | 32.0319196 |
| 0 | 0 |
| 0 | 0 |
| 0 | 0 |
| 2.71433228 | 0 |
| 0 | 0 |
| 0 | 0 |
| 0 | 0 |
| 0 | 0 |
| 0 | 0 |
| 0 | 0 |
| 0 | 0 |
| 0 | 4.57598851 |

| | |
|---|---|
| 0 | 0 |
| 5.42866456 | 77.7918047 |
| 0 | 18.303954 |
| 0 | 0 |
| 0 | 0 |
| 16.2859937 | 45.7598851 |
| 0 | 0 |
| 0 | 91.5197702 |
| 0 | 0 |
| 0 | 0 |
| 0 | 0 |
| 0 | 0 |
| 0 | 0 |
| 0 | 0 |
| 2.71433228 | 4.57598851 |
| 0 | 0 |
| 0 | 0 |
| 10.8573291 | 0 |
| 0 | 0 |
| 0 | 0 |
| 0 | 0 |
| 184.574595 | 27.4559311 |
| 0 | 0 |
| 0 | 0 |
| 0 | 0 |
| 390.863848 | 45.7598851 |
| 0 | 0 |
| 0 | 0 |
| 0 | 0 |
| 2.71433228 | 0 |
| 10.8573291 | 32.0319196 |
| 0 | 0 |
| 0 | 0 |
| 0 | 0 |
| 0 | 0 |
| 2.71433228 | 4.57598851 |
| 0 | 0 |
| 0 | 36.6079081 |
| 0 | 50.3358736 |
| 0 | 0 |
| 0 | 0 |
| 0 | 0 |
| 0 | 0 |

| | |
|---|---|
| 0 | 13.7279655 |
| 0 | 0 |
| 0 | 0 |
| 0 | 0 |
| 0 | 4.57598851 |
| 0 | 0 |
| 59.7153101 | 128.127678 |
| 0 | 0 |
| 0 | 0 |
| 0 | 0 |
| 54.2866456 | 128.127678 |
| 0 | 0 |
| 0 | 0 |
| 0 | 0 |
| 0 | 0 |
| 0 | 0 |
| 2.71433228 | 13.7279655 |
| 0 | 0 |
| 0 | 100.671747 |
| 0 | 0 |
| 0 | 0 |
| 0 | 0 |
| 21.7146582 | 0 |
| 27.1433228 | 146.431632 |
| 27.1433228 | 9.15197702 |
| 0 | 0 |
| 0 | 0 |
| 0 | 0 |
| 0 | 45.7598851 |
| 0 | 0 |
| 0 | 0 |
| 0 | 0 |
| 0 | 0 |
| 0 | 0 |
| 8.14299683 | 0 |
| 0 | 0 |
| 0 | 0 |
| 0 | 0 |
| 0 | 0 |
| 0 | 0 |
| 0 | 0 |
| 13.5716614 | 0 |
| 0 | 59.4878506 |

| | |
|---|---|
| 0 | 0 |
| 0 | 0 |
| 0 | 0 |
| 247.004237 | 54.9118621 |
| 2.71433228 | 45.7598851 |
| 0 | 0 |
| 146.573943 | 0 |
| 0 | 0 |
| 0 | 0 |
| 0 | 0 |
| 0 | 4.57598851 |
| 0 | 0 |
| 0 | 0 |
| 0 | 4.57598851 |
| 0 | 0 |
| 0 | 0 |
| 0 | 0 |
| 0 | 0 |
| 0 | 0 |
| 0 | 4.57598851 |
| 8.14299683 | 187.615529 |
| 0 | 0 |
| 352.863196 | 352.351115 |
| 355.577528 | 462.17484 |
| 0 | 0 |
| 0 | 45.7598851 |
| 0 | 50.3358736 |
| 0 | 0 |
| 0 | 0 |
| 0 | 0 |
| 0 | 0 |
| 0 | 0 |
| 0 | 0 |
| 0 | 0 |
| 0 | 0 |
| 0 | 0 |
| 0 | 0 |
| 0 | 9.15197702 |
| 0 | 0 |
| 54.2866456 | 265.407334 |
| 0 | 0 |
| 0 | 0 |
| 0 | 0 |

| | |
|---|---|
| 0 | 0 |
| 108.573291 | 4.57598851 |
| 0 | 0 |
| 0 | 4.57598851 |
| 0 | 0 |
| 0 | 0 |
| 0 | 0 |
| 0 | 0 |
| 0 | 0 |
| 0 | 0 |
| 95.0016297 | 0 |
| 0 | 0 |
| 0 | 0 |
| 2.71433228 | 77.7918047 |
| 0 | 4.57598851 |
| 0 | 0 |
| 0 | 13.7279655 |
| 0 | 0 |
| 0 | 0 |
| 0 | 0 |
| 0 | 0 |
| 0 | 0 |
| 0 | 0 |
| 0 | 68.6398277 |
| 0 | 4.57598851 |
| 0 | 0 |
| 70.5726392 | 27.4559311 |
| 0 | 0 |
| 0 | 0 |
| 0 | 0 |
| 0 | 4.57598851 |
| 0 | 0 |
| 0 | 0 |
| 214.43225 | 0 |
| 0 | 0 |
| 0 | 0 |
| 0 | 0 |
| 0 | 0 |
| 0 | 0 |
| 0 | 0 |
| 0 | 0 |
| 0 | 0 |
| 0 | 0 |

| | |
|---|---|
| 0 | 0 |
| 0 | 0 |
| 0 | 0 |
| 0 | 0 |
| 0 | 0 |
| 127.573617 | 4.57598851 |
| 0 | 0 |
| 48.857981 | 0 |
| 0 | 0 |
| 0 | 0 |
| 0 | 0 |
| 0 | 0 |
| 0 | 0 |
| 0 | 64.0638392 |
| 0 | 0 |
| 133.002282 | 224.223437 |
| 0 | 0 |
| 0 | 0 |
| 0 | 0 |
| 0 | 0 |
| 0 | 0 |
| 0 | 50.3358736 |
| 0 | 0 |
| 24.4289905 | 82.3677932 |
| 0 | 54.9118621 |
| 0 | 0 |
| 0 | 0 |
| 114.001956 | 59.4878506 |
| 0 | 0 |
| 0 | 0 |
| 0 | 41.1838966 |
| 2.71433228 | 4.57598851 |
| 0 | 0 |
| 2.71433228 | 4.57598851 |
| 19.0003259 | 0 |
| 0 | 0 |
| 0 | 0 |
| 0 | 0 |
| 0 | 0 |
| 0 | 0 |
| 43.4293164 | 4.57598851 |
| 0 | 0 |
| 0 | 27.4559311 |

| | |
|---|---|
| 0 | 0 |
| 0 | 0 |
| 2.71433228 | 41.1838966 |
| 95.0016297 | 13.7279655 |
| 0 | 0 |
| 0 | 0 |
| 0 | 0 |
| 157.431272 | 0 |
| 0 | 0 |
| 0 | 0 |
| 0 | 0 |
| 0 | 0 |
| 0 | 0 |
| 0 | 0 |
| 2.71433228 | 0 |
| 57.0009778 | 59.4878506 |
| 0 | 0 |
| 0 | 0 |
| 0 | 0 |
| 0 | 0 |
| 0 | 0 |
| 0 | 0 |
| 0 | 0 |
| 0 | 0 |
| 0 | 0 |
| 0 | 0 |
| 0 | 0 |
| 0 | 0 |
| 0 | 0 |
| 0 | 0 |
| 0 | 0 |
| 0 | 0 |
| 0 | 0 |
| 0 | 0 |
| 0 | 0 |
| 0 | 0 |
| 0 | 0 |
| 0 | 0 |
| 0 | 0 |
| 0 | 0 |
| 0 | 54.9118621 |
| 0 | 0 |
| 0 | 0 |

| | |
|---|---|
| 0 | 0 |
| 0 | 0 |
| 0 | 0 |
| 0 | 41.1838966 |
| 73.2869715 | 128.127678 |
| 0 | 0 |
| 0 | 160.159598 |
| 38.0006519 | 0 |
| 0 | 4.57598851 |
| 0 | 0 |
| 0 | 0 |
| 0 | 0 |
| 0 | 0 |
| 0 | 0 |
| 0 | 0 |
| 0 | 0 |
| 19.0003259 | 86.9437817 |
| 0 | 0 |
| 0 | 0 |
| 0 | 0 |
| 0 | 4.57598851 |
| 0 | 0 |
| 0 | 0 |
| 0 | 0 |
| 0 | 0 |
| 0 | 41.1838966 |
| 54.2866456 | 50.3358736 |
| 0 | 0 |
| 0 | 0 |
| 5.42866456 | 0 |
| 0 | 0 |
| 0 | 0 |
| 0 | 0 |
| 0 | 0 |
| 0 | 0 |
| 0 | 0 |
| 0 | 0 |
| 0 | 0 |
| 2.71433228 | 9.15197702 |
| 0 | 0 |
| 0 | 0 |
| 488.57981 | 32.0319196 |
| 0 | 27.4559311 |

| | |
|---|---|
| 0 | 0 |
| 0 | 0 |
| 0 | 0 |
| 0 | 0 |
| 0 | 0 |
| 46.1436487 | 27.4559311 |
| 458.722155 | 41.1838966 |
| 73.2869715 | 251.679368 |
| 2.71433228 | 27.4559311 |
| 0 | 0 |
| 0 | 0 |
| 0 | 0 |
| 0 | 0 |
| 0 | 4.57598851 |
| 0 | 0 |
| 5.42866456 | 27.4559311 |
| 0 | 0 |
| 0 | 0 |
| 0 | 0 |
| 0 | 0 |
| 2.71433228 | 0 |
| 0 | 0 |
| 0 | 0 |
| 0 | 0 |
| 0 | 0 |
| 0 | 0 |
| 0 | 0 |
| 0 | 0 |
| 0 | 0 |
| 0 | 0 |
| 0 | 0 |
| 0 | 0 |
| 0 | 0 |
| 0 | 0 |
| 0 | 0 |
| 27.1433228 | 22.8799426 |
| 0 | 0 |
| 0 | 0 |
| 0 | 4.57598851 |
| 0 | 0 |
| 0 | 0 |
| 0 | 0 |
| 0 | 0 |

| | |
|---|---|
| 5.42866456 | 0 |
| 0 | 0 |
| 0 | 0 |
| 0 | 0 |
| 0 | 0 |
| 494.008475 | 45.7598851 |
| 76.0013038 | 109.823724 |
| 2.71433228 | 0 |
| 2.71433228 | 105.247736 |
| 0 | 0 |
| 0 | 0 |
| 0 | 0 |
| 0 | 0 |
| 0 | 0 |
| 0 | 0 |
| 0 | 0 |
| 0 | 0 |
| 0 | 0 |
| 0 | 0 |
| 0 | 0 |
| 0 | 0 |
| 0 | 32.0319196 |
| 0 | 0 |
| 0 | 0 |
| 0 | 0 |
| 244.289905 | 4.57598851 |
| 149.288275 | 86.9437817 |
| 298.576551 | 329.471173 |
| 0 | 0 |
| 27.1433228 | 0 |
| 0 | 0 |
| 209.003585 | 13.7279655 |
| 19.0003259 | 4.57598851 |
| 0 | 0 |
| 16.2859937 | 32.0319196 |
| 301.290883 | 64.0638392 |
| 0 | 0 |
| 0 | 0 |
| 0 | 0 |
| 0 | 0 |
| 0 | 0 |
| 2.71433228 | 9.15197702 |
| 0 | 0 |

| | |
|---|---|
| 0 | 0 |
| 0 | 0 |
| 0 | 0 |
| 0 | 0 |
| 65.1439747 | 27.4559311 |
| 0 | 0 |
| 19.0003259 | 0 |
| 0 | 0 |
| 0 | 0 |
| 0 | 0 |
| 0 | 0 |
| 0 | 0 |
| 0 | 32.0319196 |
| 0 | 0 |
| 0 | 0 |
| 0 | 4.57598851 |
| 0 | 0 |
| 0 | 0 |
| 0 | 0 |
| 0 | 4.57598851 |
| 0 | 27.4559311 |
| 0 | 0 |
| 0 | 0 |
| 0 | 0 |
| 78.7156361 | 36.6079081 |
| 0 | 50.3358736 |
| 0 | 0 |
| 0 | 0 |
| 0 | 0 |
| 32.5719873 | 18.303954 |
| 0 | 0 |
| 0 | 0 |
| 0 | 0 |
| 0 | 0 |
| 0 | 0 |
| 0 | 0 |
| 0 | 0 |
| 0 | 0 |
| 0 | 0 |
| 266.004563 | 13.7279655 |
| 0 | 0 |
| 0 | 18.303954 |
| 0 | 0 |

| | |
|---|---|
| 0 | 0 |
| 0 | 0 |
| 0 | 77.7918047 |
| 0 | 0 |
| 0 | 0 |
| 0 | 0 |
| 0 | 0 |
| 0 | 0 |
| 0 | 0 |
| 0 | 0 |
| 366.434858 | 118.975701 |
| 27.1433228 | 0 |
| 119.43062 | 54.9118621 |
| 0 | 0 |
| 382.720851 | 228.799426 |
| 0 | 4.57598851 |
| 2.71433228 | 0 |
| 0 | 0 |
| 2.71433228 | 0 |
| 0 | 0 |
| 0 | 0 |
| 0 | 4.57598851 |
| 0 | 0 |
| 0 | 0 |
| 0 | 0 |
| 0 | 0 |
| 152.002608 | 77.7918047 |
| 0 | 59.4878506 |
| 0 | 0 |
| 0 | 0 |
| 0 | 0 |
| 0 | 0 |
| 0 | 0 |
| 0 | 0 |
| 0 | 0 |
| 0 | 0 |
| 0 | 4.57598851 |
| 0 | 0 |
| 0 | 0 |
| 2.71433228 | 0 |
| 0 | 0 |
| 0 | 0 |
| 0 | 0 |

| | |
|---|---|
| 0 | 0 |
| 0 | 0 |
| 21.7146582 | 45.7598851 |
| 27.1433228 | 41.1838966 |
| 0 | 0 |
| 89.5729652 | 173.887563 |
| 0 | 0 |
| 0 | 0 |
| 0 | 0 |
| 0 | 0 |
| 8.14299683 | 4.57598851 |
| 0 | 0 |
| 0 | 109.823724 |
| 0 | 0 |
| 0 | 0 |
| 0 | 0 |
| 0 | 0 |
| 0 | 0 |
| 0 | 0 |
| 2.71433228 | 91.5197702 |
| 16.2859937 | 68.6398277 |
| 0 | 0 |
| 0 | 0 |
| 0 | 0 |
| 0 | 0 |
| 0 | 0 |
| 0 | 0 |
| 0 | 0 |
| 393.57818 | 45.7598851 |
| 0 | 0 |
| 0 | 0 |
| 0 | 0 |
| 0 | 0 |
| 0 | 0 |
| 0 | 0 |
| 0 | 0 |
| 0 | 0 |
| 0 | 4.57598851 |
| 0 | 0 |
| 2.71433228 | 0 |
| 0 | 0 |
| 0 | 0 |
| 0 | 0 |

| | |
|---|---|
| 0 | 0 |
| 0 | 0 |
| 119.43062 | 132.703667 |
| 855.014668 | 512.510713 |
| 0 | 0 |
| 2.71433228 | 91.5197702 |
| 0 | 0 |
| 8.14299683 | 4.57598851 |
| 0 | 0 |
| 0 | 0 |
| 0 | 0 |
| 0 | 0 |
| 0 | 0 |
| 0 | 0 |
| 0 | 0 |
| 0 | 0 |
| 0 | 0 |
| 0 | 0 |
| 499.437139 | 82.3677932 |
| 0 | 0 |
| 0 | 0 |
| 545.580788 | 13.7279655 |
| 0 | 0 |
| 0 | 0 |
| 0 | 0 |
| 0 | 0 |
| 0 | 54.9118621 |
| 0 | 0 |
| 0 | 0 |
| 0 | 0 |
| 0 | 0 |
| 0 | 0 |
| 29.8576551 | 27.4559311 |
| 0 | 0 |
| 0 | 0 |
| 0 | 0 |
| 0 | 0 |
| 0 | 73.2158162 |
| 0 | 45.7598851 |
| 48.857981 | 0 |
| 0 | 0 |
| 0 | 0 |
| 0 | 0 |

| | |
|---|---|
| 0 | 0 |
| 0 | 0 |
| 0 | 0 |
| 0 | 0 |
| 0 | 0 |
| 8.14299683 | 45.7598851 |
| 0 | 0 |
| 0 | 0 |
| 86.8586329 | 132.703667 |
| 0 | 59.4878506 |
| 0 | 0 |
| 0 | 0 |
| 0 | 0 |
| 0 | 4.57598851 |
| 0 | 4.57598851 |
| 0 | 9.15197702 |
| 0 | 0 |
| 0 | 0 |
| 0 | 0 |
| 0 | 9.15197702 |
| 0 | 0 |
| 0 | 0 |
| 0 | 0 |
| 0 | 0 |
| 5.42866456 | 64.0638392 |
| 244.289905 | 27.4559311 |
| 0 | 0 |
| 0 | 82.3677932 |
| 0 | 36.6079081 |
| 97.715962 | 9.15197702 |
| 0 | 0 |
| 0 | 0 |
| 0 | 0 |
| 57.0009778 | 4.57598851 |
| 0 | 0 |
| 0 | 0 |
| 0 | 0 |
| 89.5729652 | 64.0638392 |
| 0 | 0 |
| 0 | 0 |
| 0 | 0 |
| 0 | 0 |
| 211.717918 | 9.15197702 |

| | |
|---|---|
| 13.5716614 | 0 |
| 0 | 0 |
| 0 | 0 |
| 0 | 0 |
| 0 | 0 |
| 0 | 0 |
| 0 | 0 |
| 0 | 0 |
| 0 | 0 |
| 0 | 0 |
| 0 | 0 |
| 0 | 0 |
| 0 | 50.3358736 |
| 0 | 0 |
| 0 | 0 |
| 0 | 0 |
| 0 | 0 |
| 0 | 32.0319196 |
| 0 | 0 |
| 0 | 0 |
| 0 | 0 |
| 0 | 201.343494 |
| 116.716288 | 4.57598851 |
| 0 | 0 |
| 0 | 0 |
| 0 | 0 |
| 0 | 0 |
| 0 | 0 |
| 0 | 22.8799426 |
| 0 | 0 |
| 0 | 0 |
| 0 | 0 |
| 0 | 0 |
| 0 | 0 |
| 0 | 0 |
| 0 | 0 |
| 0 | 0 |
| 0 | 0 |
| 0 | 4.57598851 |
| 0 | 0 |
| 0 | 0 |
| 0 | 0 |
| 2.71433228 | 0 |

| | |
|---|---|
| 0 | 0 |
| 0 | 0 |
| 0 | 0 |
| 0 | 0 |
| 0 | 0 |
| 19.0003259 | 178.463552 |
| 0 | 0 |
| 0 | 0 |
| 0 | 0 |
| 0 | 0 |
| 0 | 0 |
| 51.5723133 | 13.7279655 |
| 0 | 0 |
| 10.8573291 | 100.671747 |
| 2.71433228 | 13.7279655 |
| 0 | 0 |
| 0 | 0 |
| 0 | 0 |
| 0 | 0 |
| 0 | 0 |
| 0 | 0 |
| 127.573617 | 4.57598851 |
| 0 | 0 |
| 0 | 27.4559311 |
| 0 | 0 |
| 0 | 0 |
| 0 | 0 |
| 0 | 0 |
| 0 | 0 |
| 0 | 0 |
| 0 | 0 |
| 2.71433228 | 9.15197702 |
| 0 | 0 |
| 0 | 0 |
| 0 | 0 |
| 0 | 4.57598851 |
| 0 | 0 |
| 0 | 0 |
| 0 | 0 |
| 40.7149842 | 0 |
| 2.71433228 | 0 |
| 0 | 0 |
| 0 | 0 |

| | |
|---|---|
| 0 | 0 |
| 27.1433228 | 18.303954 |
| 0 | 0 |
| 0 | 0 |
| 0 | 0 |
| 0 | 0 |
| 0 | 0 |
| 0 | 0 |
| 0 | 0 |
| 0 | 54.9118621 |
| 29.8576551 | 13.7279655 |
| 0 | 0 |
| 40.7149842 | 118.975701 |
| 0 | 0 |
| 78.7156361 | 118.975701 |
| 0 | 4.57598851 |
| 0 | 0 |
| 0 | 0 |
| 187.288927 | 0 |
| 138.430946 | 4.57598851 |
| 0 | 0 |
| 0 | 0 |
| 10.8573291 | 0 |
| 0 | 0 |
| 0 | 0 |
| 0 | 0 |
| 141.145278 | 0 |
| 0 | 0 |
| 0 | 0 |
| 0 | 68.6398277 |
| 0 | 0 |
| 0 | 0 |
| 0 | 0 |
| 0 | 0 |
| 390.863848 | 173.887563 |
| 0 | 0 |
| 0 | 0 |
| 0 | 0 |
| 0 | 0 |
| 0 | 0 |
| 0 | 0 |
| 0 | 0 |
| 0 | 0 |

| | |
|---|---|
| 0 | 0 |
| 0 | 0 |
| 0 | 0 |
| 0 | 45.7598851 |
| 0 | 4.57598851 |
| 0 | 0 |
| 0 | 4.57598851 |
| 0 | 0 |
| 0 | 0 |
| 0 | 0 |
| 0 | 45.7598851 |
| 0 | 0 |
| 0 | 0 |
| 0 | 0 |
| 0 | 0 |
| 0 | 0 |
| 0 | 0 |
| 0 | 0 |
| 0 | 0 |
| 0 | 0 |
| 0 | 0 |
| 0 | 0 |
| 0 | 0 |
| 0 | 0 |
| 0 | 0 |
| 0 | 0 |
| 0 | 0 |
| 0 | 0 |
| 0 | 0 |
| 0 | 0 |
| 0 | 0 |
| 0 | 0 |
| 0 | 0 |
| 105.858959 | 4.57598851 |
| 57.0009778 | 9.15197702 |
| 0 | 0 |
| 0 | 0 |
| 0 | 0 |
| 0 | 0 |
| 0 | 0 |
| 0 | 0 |
| 0 | 0 |
| 0 | 0 |

| | |
|---|---|
| 0 | 0 |
| 0 | 0 |
| 371.863522 | 22.8799426 |
| 0 | 0 |
| 0 | 0 |
| 2.71433228 | 0 |
| 0 | 0 |
| 0 | 0 |
| 0 | 59.4878506 |
| 10.8573291 | 86.9437817 |
| 0 | 36.6079081 |
| 2.71433228 | 4.57598851 |
| 0 | 18.303954 |
| 0 | 0 |
| 0 | 0 |
| 65.1439747 | 4.57598851 |
| 0 | 0 |
| 0 | 0 |
| 0 | 0 |
| 0 | 45.7598851 |
| 0 | 0 |
| 0 | 0 |
| 0 | 0 |
| 0 | 0 |
| 236.146908 | 0 |
| 0 | 13.7279655 |
| 0 | 0 |
| 0 | 0 |
| 0 | 0 |
| 0 | 68.6398277 |
| 0 | 0 |
| 2.71433228 | 0 |
| 0 | 0 |
| 59.7153101 | 105.247736 |
| 0 | 0 |
| 76.0013038 | 4.57598851 |
| 0 | 41.1838966 |
| 0 | 0 |
| 0 | 0 |
| 0 | 0 |
| 2733.3326 | 22.8799426 |
| 0 | 0 |
| 0 | 0 |

| | |
|---|---|
| 0 | 0 |
| 0 | 0 |
| 2.71433228 | 0 |
| 0 | 0 |
| 0 | 0 |
| 0 | 0 |
| 0 | 0 |
| 2.71433228 | 32.0319196 |
| 0 | 0 |
| 0 | 0 |
| 2.71433228 | 0 |
| 13.5716614 | 36.6079081 |
| 0 | 0 |
| 0 | 0 |
| 0 | 0 |
| 0 | 0 |
| 0 | 27.4559311 |
| 0 | 0 |
| 0 | 0 |
| 0 | 0 |
| 0 | 0 |
| 0 | 77.7918047 |
| 116.716288 | 59.4878506 |
| 0 | 0 |
| 0 | 0 |
| 0 | 0 |
| 0 | 0 |
| 105.858959 | 77.7918047 |
| 0 | 4.57598851 |
| 0 | 0 |
| 0 | 0 |
| 0 | 0 |
| 0 | 0 |
| 0 | 0 |
| 0 | 0 |
| 0 | 0 |
| 0 | 105.247736 |
| 0 | 0 |
| 13.5716614 | 0 |
| 0 | 0 |
| 0 | 0 |
| 152.002608 | 45.7598851 |
| 0 | 0 |

| | |
|---|---|
| 0 | 0 |
| 0 | 0 |
| 0 | 0 |
| 0 | 0 |
| 86.8586329 | 9.15197702 |
| 0 | 0 |
| 0 | 0 |
| 0 | 0 |
| 0 | 0 |
| 0 | 0 |
| 0 | 0 |
| 0 | 0 |
| 0 | 0 |
| 0 | 0 |
| 290.433554 | 4.57598851 |
| 0 | 0 |
| 0 | 0 |
| 0 | 0 |
| 0 | 0 |
| 5.42866456 | 4.57598851 |
| 0 | 0 |
| 0 | 0 |
| 0 | 0 |
| 0 | 0 |
| 0 | 0 |
| 0 | 0 |
| 2.71433228 | 73.2158162 |
| 0 | 0 |
| 0 | 0 |
| 0 | 32.0319196 |
| 0 | 0 |
| 0 | 0 |
| 13.5716614 | 4.57598851 |
| 0 | 0 |
| 0 | 0 |
| 0 | 4.57598851 |
| 0 | 0 |
| 0 | 0 |
| 0 | 0 |
| 5.42866456 | 141.855644 |
| 0 | 0 |
| 0 | 0 |
| 0 | 0 |

| | |
|---|---|
| 0 | 0 |
| 0 | 0 |
| 2.71433228 | 0 |
| 0 | 0 |
| 0 | 0 |
| 0 | 105.247736 |
| 0 | 0 |
| 0 | 0 |
| 0 | 0 |
| 2.71433228 | 4.57598851 |
| 67.8583069 | 0 |
| 0 | 0 |
| 0 | 0 |
| 0 | 0 |
| 0 | 0 |
| 152.002608 | 0 |
| 0 | 0 |
| 0 | 0 |
| 67.8583069 | 0 |
| 0 | 0 |
| 0 | 18.303954 |
| 0 | 0 |
| 0 | 0 |
| 0 | 0 |
| 2.71433228 | 0 |
| 0 | 0 |
| 0 | 0 |
| 0 | 0 |
| 0 | 4.57598851 |
| 0 | 0 |
| 8.14299683 | 59.4878506 |
| 0 | 0 |
| 0 | 0 |
| 165.574269 | 0 |
| 0 | 0 |
| 0 | 0 |
| 0 | 0 |
| 0 | 0 |
| 0 | 0 |
| 0 | 0 |
| 0 | 73.2158162 |
| 0 | 0 |
| 0 | 0 |

| | |
|---|---|
| 119.43062 | 22.8799426 |
| 0 | 0 |
| 0 | 0 |
| 0 | 0 |
| 0 | 0 |
| 0 | 0 |
| 0 | 0 |
| 13.5716614 | 4.57598851 |
| 13.5716614 | 0 |
| 0 | 0 |
| 0 | 0 |
| 0 | 0 |
| 8.14299683 | 50.3358736 |
| 0 | 0 |
| 0 | 0 |
| 0 | 0 |
| 5.42866456 | 64.0638392 |
| 0 | 4.57598851 |
| 0 | 82.3677932 |
| 0 | 0 |
| 122.144953 | 73.2158162 |
| 0 | 0 |
| 0 | 0 |
| 0 | 0 |
| 0 | 0 |
| 65.1439747 | 4.57598851 |
| 0 | 0 |
| 168.288601 | 114.399713 |
| 0 | 0 |
| 0 | 0 |
| 0 | 0 |
| 0 | 0 |
| 0 | 0 |
| 168.288601 | 114.399713 |
| 0 | 0 |
| 0 | 0 |
| 0 | 0 |
| 0 | 0 |
| 0 | 0 |
| 0 | 0 |
| 0 | 0 |
| 0 | 0 |
| 371.863522 | 109.823724 |

| | |
|---|---|
| 0 | 0 |
| 135.716614 | 9.15197702 |
| 0 | 0 |
| 0 | 0 |
| 73.2869715 | 274.559311 |
| 0 | 0 |
| 0 | 0 |
| 0 | 0 |
| 0 | 0 |
| 127.573617 | 22.8799426 |
| 0 | 0 |
| 0 | 0 |
| 0 | 0 |
| 0 | 0 |
| 0 | 0 |
| 100.430294 | 22.8799426 |
| 0 | 0 |
| 0 | 0 |
| 0 | 0 |
| 0 | 9.15197702 |
| 0 | 0 |
| 0 | 0 |
| 0 | 0 |
| 0 | 0 |
| 0 | 0 |
| 2.71433228 | 4.57598851 |
| 0 | 0 |
| 0 | 0 |
| 0 | 0 |
| 5.42866456 | 27.4559311 |
| 0 | 0 |
| 0 | 0 |
| 0 | 13.7279655 |
| 0 | 0 |
| 0 | 0 |
| 0 | 0 |
| 0 | 0 |
| 0 | 0 |
| 0 | 0 |
| 0 | 0 |
| 0 | 0 |
| 0 | 0 |
| 0 | 64.0638392 |

| | |
|---|---|
| 268.718896 | 215.07146 |
| 0 | 0 |
| 0 | 0 |
| 0 | 0 |
| 0 | 0 |
| 0 | 0 |
| 0 | 0 |
| 0 | 0 |
| 0 | 0 |
| 0 | 0 |
| 0 | 0 |
| 0 | 0 |
| 0 | 0 |
| 2.71433228 | 9.15197702 |
| 0 | 0 |
| 0 | 0 |
| 0 | 0 |
| 0 | 0 |
| 0 | 36.6079081 |
| 0 | 0 |
| 0 | 0 |
| 0 | 0 |
| 350.148864 | 183.03954 |
| 168.288601 | 22.8799426 |
| 0 | 0 |
| 0 | 0 |
| 0 | 0 |
| 0 | 0 |
| 0 | 9.15197702 |
| 0 | 0 |
| 0 | 9.15197702 |
| 0 | 0 |
| 0 | 0 |
| 0 | 0 |
| 0 | 0 |
| 0 | 0 |
| 0 | 0 |
| 0 | 0 |
| 114.001956 | 9.15197702 |
| 0 | 0 |
| 0 | 0 |
| 0 | 0 |
| 0 | 0 |

| | |
|---|---|
| 10.8573291 | 73.2158162 |
| 0 | 0 |
| 0 | 0 |
| 0 | 0 |
| 0 | 0 |
| 0 | 0 |
| 0 | 0 |
| 0 | 0 |
| 0 | 0 |
| 0 | 0 |
| 0 | 0 |
| 0 | 0 |
| 0 | 0 |
| 0 | 18.303954 |
| 0 | 0 |
| 0 | 0 |
| 0 | 0 |
| 176.431598 | 50.3358736 |
| 0 | 0 |
| 0 | 0 |
| 0 | 0 |
| 0 | 0 |
| 0 | 0 |
| 5.42866456 | 0 |
| 46.1436487 | 0 |
| 0 | 0 |
| 0 | 45.7598851 |
| 0 | 0 |
| 0 | 0 |
| 0 | 0 |
| 0 | 0 |
| 0 | 0 |
| 0 | 0 |
| 0 | 0 |
| 65.1439747 | 96.0957587 |
| 0 | 0 |
| 0 | 0 |
| 0 | 0 |
| 51.5723133 | 4.57598851 |
| 0 | 0 |
| 0 | 0 |
| 0 | 0 |
| 0 | 0 |

| | |
|---|---|
| 0 | 0 |
| 0 | 0 |
| 0 | 0 |
| 0 | 0 |
| 0 | 0 |
| 0 | 0 |
| 0 | 0 |
| 0 | 0 |
| 19.0003259 | 59.4878506 |
| 0 | 0 |
| 0 | 9.15197702 |
| 0 | 0 |
| 35.2863196 | 0 |
| 0 | 0 |
| 0 | 0 |
| 0 | 0 |
| 0 | 9.15197702 |
| 8.14299683 | 4.57598851 |
| 19.0003259 | 4.57598851 |
| 0 | 0 |
| 0 | 0 |
| 13.5716614 | 128.127678 |
| 168.288601 | 50.3358736 |
| 901.158316 | 146.431632 |
| 0 | 0 |
| 0 | 0 |
| 0 | 73.2158162 |
| 0 | 77.7918047 |
| 0 | 73.2158162 |
| 0 | 0 |
| 0 | 18.303954 |
| 0 | 0 |
| 0 | 0 |
| 0 | 0 |
| 0 | 0 |
| 0 | 32.0319196 |
| 355.577528 | 123.55169 |
| 0 | 0 |
| 27.1433228 | 0 |
| 0 | 54.9118621 |
| 0 | 0 |
| 0 | 0 |
| 0 | 0 |

| | |
|---|---|
| 0 | 0 |
| 0 | 0 |
| 0 | 0 |
| 0 | 0 |
| 43.4293164 | 292.863265 |
| 70.5726392 | 0 |
| 171.002934 | 0 |
| 0 | 0 |
| 0 | 0 |
| 0 | 0 |
| 2.71433228 | 4.57598851 |
| 0 | 4.57598851 |
| 59.7153101 | 45.7598851 |
| 0 | 0 |
| 0 | 0 |
| 21.7146582 | 68.6398277 |
| 0 | 0 |
| 0 | 0 |
| 0 | 0 |
| 0 | 0 |
| 73.2869715 | 4.57598851 |
| 0 | 0 |
| 0 | 0 |
| 0 | 0 |
| 0 | 0 |
| 2.71433228 | 0 |
| 0 | 0 |
| 2.71433228 | 0 |
| 65.1439747 | 0 |
| 0 | 0 |
| 0 | 0 |
| 0 | 4.57598851 |
| 168.288601 | 4.57598851 |
| 0 | 0 |
| 0 | 0 |
| 0 | 0 |
| 0 | 0 |
| 0 | 164.735586 |
| 13.5716614 | 4.57598851 |
| 0 | 0 |
| 0 | 0 |
| 0 | 0 |
| 0 | 0 |

| | |
|---|---|
| 0 | 0 |
| 0 | 0 |
| 0 | 0 |
| 0 | 0 |
| 0 | 0 |
| 0 | 0 |
| 0 | 0 |
| 0 | 0 |
| 0 | 0 |
| 5.42866456 | 100.671747 |
| 0 | 0 |
| 0 | 0 |
| 0 | 0 |
| 0 | 0 |
| 0 | 0 |
| 0 | 0 |
| 0 | 0 |
| 0 | 82.3677932 |
| 238.86124 | 73.2158162 |
| 437.007497 | 201.343494 |
| 0 | 0 |
| 0 | 0 |
| 0 | 0 |
| 0 | 0 |
| 0 | 0 |
| 0 | 0 |
| 0 | 0 |
| 0 | 0 |
| 0 | 0 |
| 0 | 0 |
| 0 | 0 |
| 16.2859937 | 86.9437817 |
| 0 | 0 |
| 0 | 0 |
| 0 | 0 |
| 0 | 0 |
| 0 | 0 |
| 0 | 0 |
| 0 | 0 |
| 0 | 0 |
| 0 | 0 |
| 0 | 0 |
| 0 | 0 |

| | |
|---|---|
| 0 | 0 |
| 0 | 0 |
| 0 | 0 |
| 0 | 0 |
| 0 | 128.127678 |
| 0 | 0 |
| 2.71433228 | 0 |
| 0 | 0 |
| 0 | 0 |
| 0 | 0 |
| 198.146256 | 32.0319196 |
| 35.2863196 | 22.8799426 |
| 0 | 0 |
| 0 | 0 |
| 252.432902 | 105.247736 |
| 0 | 0 |
| 0 | 4.57598851 |
| 0 | 0 |
| 0 | 0 |
| 0 | 0 |
| 0 | 0 |
| 0 | 0 |
| 0 | 0 |
| 0 | 0 |
| 0 | 4.57598851 |
| 0 | 0 |
| 0 | 0 |
| 0 | 0 |
| 2.71433228 | 0 |
| 0 | 123.55169 |
| 0 | 0 |
| 0 | 0 |
| 0 | 0 |
| 0 | 0 |
| 0 | 0 |
| 0 | 0 |
| 0 | 0 |
| 0 | 4.57598851 |
| 0 | 4.57598851 |
| 0 | 0 |
| 40.7149842 | 77.7918047 |
| 0 | 0 |
| 0 | 0 |

| | |
|---|---|
| 0 | 0 |
| 0 | 0 |
| 0 | 0 |
| 0 | 0 |
| 0 | 0 |
| 0 | 77.7918047 |
| 0 | 0 |
| 0 | 0 |
| 38.0006519 | 4.57598851 |
| 54.2866456 | 45.7598851 |
| 0 | 4.57598851 |
| 0 | 0 |
| 0 | 4.57598851 |
| 65.1439747 | 9.15197702 |
| 81.4299683 | 0 |
| 0 | 0 |
| 127.573617 | 4.57598851 |
| 0 | 0 |
| 0 | 0 |
| 0 | 50.3358736 |
| 0 | 0 |
| 16.2859937 | 9.15197702 |
| 0 | 0 |
| 0 | 13.7279655 |
| 0 | 18.303954 |
| 0 | 0 |
| 0 | 0 |
| 0 | 0 |
| 743.727044 | 160.159598 |
| 78.7156361 | 18.303954 |
| 0 | 0 |
| 0 | 0 |
| 0 | 0 |
| 2.71433228 | 9.15197702 |
| 0 | 32.0319196 |
| 0 | 0 |
| 0 | 0 |
| 0 | 0 |
| 0 | 0 |
| 0 | 0 |
| 0 | 0 |
| 0 | 0 |
| 0 | 0 |

| | |
|---|---|
| 0 | 0 |
| 0 | 0 |
| 0 | 0 |
| 0 | 4.57598851 |
| 0 | 0 |
| 0 | 0 |
| 0 | 0 |
| 0 | 0 |
| 0 | 0 |
| 0 | 64.0638392 |
| 0 | 0 |
| 0 | 0 |
| 0 | 0 |
| 0 | 0 |
| 40.7149842 | 13.7279655 |
| 0 | 0 |
| 0 | 0 |
| 0 | 0 |
| 0 | 0 |
| 0 | 0 |
| 100.430294 | 68.6398277 |
| 0 | 0 |
| 2.71433228 | 18.303954 |
| 0 | 0 |
| 0 | 0 |
| 0 | 0 |
| 73.2869715 | 311.167219 |
| 0 | 0 |
| 0 | 0 |
| 21.7146582 | 54.9118621 |
| 0 | 0 |
| 43.4293164 | 233.375414 |
| 0 | 91.5197702 |
| 0 | 0 |
| 0 | 0 |
| 5.42866456 | 18.303954 |
| 0 | 0 |
| 0 | 9.15197702 |
| 160.145604 | 169.311575 |
| 0 | 0 |
| 0 | 27.4559311 |
| 0 | 0 |
| 0 | 0 |

| | |
|---|---|
| 0 | 0 |
| 0 | 18.303954 |
| 0 | 0 |
| 0 | 0 |
| 0 | 0 |
| 0 | 0 |
| 0 | 36.6079081 |
| 0 | 0 |
| 0 | 0 |
| 46.1436487 | 132.703667 |
| 0 | 0 |
| 0 | 13.7279655 |
| 62.4296424 | 0 |
| 0 | 0 |
| 0 | 0 |
| 0 | 13.7279655 |
| 5.42866456 | 9.15197702 |
| 0 | 41.1838966 |
| 0 | 0 |
| 0 | 0 |
| 0 | 0 |
| 0 | 0 |
| 0 | 59.4878506 |
| 0 | 0 |
| 0 | 0 |
| 0 | 0 |
| 0 | 0 |
| 0 | 4.57598851 |
| 0 | 0 |
| 0 | 0 |
| 0 | 0 |
| 0 | 0 |
| 0 | 0 |
| 2.71433228 | 4.57598851 |
| 0 | 0 |
| 0 | 0 |
| 0 | 0 |
| 0 | 13.7279655 |
| 0 | 0 |
| 0 | 0 |
| 35.2863196 | 32.0319196 |
| 0 | 0 |
| 0 | 0 |

| | |
|---|---|
| 0 | 0 |
| 0 | 0 |
| 0 | 0 |
| 0 | 0 |
| 16.2859937 | 0 |
| 40.7149842 | 297.439253 |
| 0 | 0 |
| 0 | 0 |
| 0 | 0 |
| 0 | 0 |
| 0 | 0 |
| 0 | 0 |
| 0 | 0 |
| 1560.74106 | 297.439253 |
| 0 | 0 |
| 0 | 0 |
| 0 | 36.6079081 |
| 192.717592 | 13.7279655 |
| 0 | 0 |
| 0 | 73.2158162 |
| 0 | 0 |
| 65.1439747 | 86.9437817 |
| 0 | 0 |
| 43.4293164 | 265.407334 |
| 0 | 0 |
| 0 | 0 |
| 67.8583069 | 32.0319196 |
| 0 | 0 |
| 0 | 0 |
| 0 | 0 |
| 0 | 0 |
| 0 | 0 |
| 0 | 0 |
| 0 | 0 |
| 0 | 0 |
| 0 | 9.15197702 |
| 0 | 0 |
| 0 | 0 |
| 0 | 0 |
| 0 | 0 |
| 0 | 0 |
| 0 | 0 |
| 0 | 0 |

| | |
|---|---|
| 0 | 0 |
| 0 | 0 |
| 0 | 0 |
| 0 | 0 |
| 0 | 36.6079081 |
| 16.2859937 | 155.583609 |
| 0 | 0 |
| 0 | 0 |
| 0 | 0 |
| 8.14299683 | 132.703667 |
| 0 | 0 |
| 0 | 0 |
| 0 | 0 |
| 0 | 0 |
| 0 | 0 |
| 0 | 4.57598851 |
| 27.1433228 | 100.671747 |
| 0 | 0 |
| 0 | 4.57598851 |
| 5.42866456 | 0 |
| 0 | 0 |
| 0 | 0 |
| 8.14299683 | 0 |
| 0 | 0 |
| 0 | 0 |
| 0 | 4.57598851 |
| 0 | 0 |
| 0 | 0 |
| 0 | 4.57598851 |
| 0 | 0 |
| 0 | 0 |
| 0 | 54.9118621 |
| 0 | 0 |
| 2.71433228 | 32.0319196 |
| 0 | 0 |
| 0 | 0 |
| 0 | 0 |
| 0 | 0 |
| 551.009452 | 123.55169 |
| 0 | 0 |
| 122.144953 | 0 |
| 0 | 0 |
| 0 | 0 |

| | |
|---|---|
| 0 | 0 |
| 57.0009778 | 59.4878506 |
| 0 | 4.57598851 |
| 0 | 0 |
| 0 | 0 |
| 21.7146582 | 0 |
| 0 | 0 |
| 0 | 0 |
| 0 | 4.57598851 |
| 0 | 0 |
| 0 | 0 |
| 0 | 0 |
| 0 | 0 |
| 2.71433228 | 82.3677932 |
| 0 | 0 |
| 0 | 0 |
| 0 | 0 |
| 0 | 36.6079081 |
| 0 | 0 |
| 29.8576551 | 0 |
| 0 | 0 |
| 0 | 0 |
| 0 | 0 |
| 385.435183 | 59.4878506 |
| 0 | 0 |
| 0 | 0 |
| 0 | 0 |
| 0 | 0 |
| 0 | 0 |
| 0 | 0 |
| 8.14299683 | 32.0319196 |
| 0 | 0 |
| 0 | 0 |
| 244.289905 | 128.127678 |
| 0 | 0 |
| 0 | 0 |
| 0 | 0 |
| 0 | 0 |
| 19.0003259 | 18.303954 |
| 21.7146582 | 0 |
| 2.71433228 | 22.8799426 |
| 0 | 0 |
| 0 | 0 |

| | |
|---|---|
| 0 | 0 |
| 0 | 0 |
| 0 | 0 |
| 0 | 0 |
| 0 | 0 |
| 1175.30588 | 201.343494 |
| 0 | 0 |
| 108.573291 | 4.57598851 |
| 0 | 0 |
| 0 | 0 |
| 0 | 0 |
| 0 | 0 |
| 2.71433228 | 13.7279655 |
| 0 | 0 |
| 0 | 0 |
| 0 | 0 |
| 0 | 0 |
| 0 | 0 |
| 2.71433228 | 0 |
| 0 | 0 |
| 0 | 0 |
| 0 | 0 |
| 19.0003259 | 86.9437817 |
| 0 | 146.431632 |
| 0 | 0 |
| 8.14299683 | 32.0319196 |
| 0 | 0 |
| 314.862544 | 22.8799426 |
| 0 | 0 |
| 5.42866456 | 9.15197702 |
| 0 | 0 |
| 0 | 0 |
| 0 | 0 |
| 0 | 0 |
| 412.578506 | 50.3358736 |
| 0 | 0 |
| 0 | 0 |
| 0 | 0 |
| 0 | 0 |
| 111.287623 | 233.375414 |
| 0 | 0 |
| 0 | 4.57598851 |
| 0 | 0 |

| | |
|---|---|
| 0 | 0 |
| 0 | 0 |
| 0 | 0 |
| 0 | 0 |
| 0 | 0 |
| 0 | 0 |
| 0 | 32.0319196 |
| 0 | 4.57598851 |
| 0 | 0 |
| 0 | 0 |
| 0 | 0 |
| 0 | 0 |
| 27.1433228 | 18.303954 |
| 0 | 0 |
| 0 | 0 |
| 5.42866456 | 4.57598851 |
| 2.71433228 | 0 |
| 0 | 64.0638392 |
| 0 | 0 |
| 0 | 0 |
| 76.0013038 | 0 |
| 0 | 0 |
| 0 | 0 |
| 0 | 0 |
| 0 | 0 |
| 0 | 0 |
| 0 | 0 |
| 0 | 0 |
| 5.42866456 | 0 |
| 0 | 0 |
| 0 | 0 |
| 363.720525 | 585.726529 |
| 344.720199 | 18.303954 |
| 0 | 0 |
| 0 | 0 |
| 711.155057 | 343.199138 |
| 0 | 0 |
| 0 | 0 |
| 0 | 54.9118621 |
| 0 | 0 |
| 0 | 0 |
| 0 | 0 |
| 0 | 0 |

| | |
|---|---|
| 0 | 0 |
| 632.439421 | 114.399713 |
| 0 | 36.6079081 |
| 0 | 0 |
| 0 | 0 |
| 35.2863196 | 196.767506 |
| 0 | 13.7279655 |
| 0 | 0 |
| 0 | 0 |
| 0 | 0 |
| 38.0006519 | 18.303954 |
| 0 | 0 |
| 0 | 0 |
| 0 | 0 |
| 0 | 0 |
| 0 | 0 |
| 0 | 0 |
| 0 | 0 |
| 0 | 0 |
| 8.14299683 | 4.57598851 |
| 0 | 0 |
| 0 | 0 |
| 192.717592 | 4.57598851 |
| 0 | 0 |
| 0 | 0 |
| 2.71433228 | 0 |
| 0 | 0 |
| 130.287949 | 9.15197702 |
| 5.42866456 | 0 |
| 0 | 0 |
| 0 | 0 |
| 0 | 0 |
| 51.5723133 | 132.703667 |
| 0 | 0 |
| 0 | 0 |
| 0 | 36.6079081 |
| 27.1433228 | 0 |
| 2.71433228 | 9.15197702 |
| 0 | 0 |
| 0 | 0 |
| 0 | 0 |
| 0 | 0 |
| 0 | 0 |

| | |
|---|---|
| 0 | 0 |
| 0 | 0 |
| 0 | 0 |
| 0 | 0 |
| 0 | 0 |
| 0 | 0 |
| 0 | 0 |
| 0 | 82.3677932 |
| 0 | 0 |
| 32.5719873 | 54.9118621 |
| 0 | 0 |
| 57.0009778 | 247.10338 |
| 0 | 0 |
| 0 | 50.3358736 |
| 0 | 0 |
| 0 | 50.3358736 |
| 5.42866456 | 4.57598851 |
| 0 | 0 |
| 0 | 0 |
| 0 | 0 |
| 0 | 4.57598851 |
| 0 | 0 |
| 0 | 0 |
| 0 | 0 |
| 0 | 0 |
| 0 | 0 |
| 0 | 0 |
| 0 | 0 |
| 0 | 0 |
| 0 | 0 |
| 0 | 0 |
| 0 | 0 |
| 0 | 0 |
| 8.14299683 | 18.303954 |
| 0 | 0 |
| 0 | 0 |
| 0 | 0 |
| 0 | 0 |
| 0 | 0 |
| 0 | 0 |
| 2.71433228 | 4.57598851 |
| 0 | 0 |
| 0 | 0 |

| | |
|---|---|
| 0 | 82.3677932 |
| 0 | 0 |
| 0 | 4.57598851 |
| 0 | 0 |
| 0 | 0 |
| 0 | 0 |
| 0 | 0 |
| 21.7146582 | 73.2158162 |
| 0 | 45.7598851 |
| 0 | 22.8799426 |
| 86.8586329 | 0 |
| 350.148864 | 4.57598851 |
| 0 | 0 |
| 0 | 0 |
| 0 | 77.7918047 |
| 0 | 0 |
| 76.0013038 | 4.57598851 |
| 0 | 0 |
| 0 | 0 |
| 0 | 0 |
| 0 | 0 |
| 2.71433228 | 4.57598851 |
| 0 | 0 |
| 0 | 0 |
| 0 | 0 |
| 0 | 0 |
| 0 | 18.303954 |
| 0 | 22.8799426 |
| 0 | 0 |
| 0 | 0 |
| 293.147886 | 343.199138 |
| 746.441376 | 91.5197702 |
| 0 | 0 |
| 0 | 0 |
| 119.43062 | 0 |
| 0 | 0 |
| 0 | 0 |
| 0 | 0 |
| 0 | 0 |
| 2.71433228 | 22.8799426 |
| 0 | 0 |
| 2.71433228 | 0 |
| 0 | 0 |

| | |
|---|---|
| 0 | 0 |
| 0 | 0 |
| 0 | 0 |
| 0 | 0 |
| 0 | 0 |
| 0 | 0 |
| 0 | 0 |
| 0 | 0 |
| 0 | 0 |
| 0 | 0 |
| 0 | 54.9118621 |
| 0 | 0 |
| 0 | 4.57598851 |
| 29.8576551 | 91.5197702 |
| 0 | 0 |
| 0 | 0 |
| 0 | 0 |
| 0 | 0 |
| 0 | 0 |
| 0 | 64.0638392 |
| 0 | 0 |
| 0 | 0 |
| 86.8586329 | 32.0319196 |
| 0 | 0 |
| 0 | 0 |
| 0 | 0 |
| 0 | 0 |
| 0 | 18.303954 |
| 0 | 0 |
| 0 | 0 |
| 5.42866456 | 9.15197702 |
| 0 | 0 |
| 0 | 0 |
| 0 | 0 |
| 0 | 0 |
| 0 | 0 |
| 0 | 0 |
| 0 | 0 |
| 0 | 0 |
| 0 | 0 |
| 0 | 0 |
| 0 | 0 |
| 143.859611 | 73.2158162 |

| | |
|---|---|
| 0 | 0 |
| 0 | 0 |
| 0 | 0 |
| 0 | 0 |
| 548.29512 | 86.9437817 |
| 0 | 0 |
| 0 | 0 |
| 0 | 0 |
| 0 | 0 |
| 0 | 0 |
| 108.573291 | 36.6079081 |
| 0 | 0 |
| 225.289579 | 36.6079081 |
| 0 | 0 |
| 0 | 0 |
| 0 | 0 |
| 0 | 0 |
| 0 | 0 |
| 0 | 123.55169 |
| 0 | 9.15197702 |
| 0 | 0 |
| 0 | 0 |
| 0 | 0 |
| 2.71433228 | 50.3358736 |
| 0 | 4.57598851 |
| 0 | 22.8799426 |
| 0 | 59.4878506 |
| 0 | 0 |
| 0 | 0 |
| 0 | 0 |
| 0 | 0 |
| 181.860263 | 32.0319196 |
| 0 | 0 |
| 0 | 0 |
| 0 | 0 |
| 0 | 0 |
| 0 | 0 |
| 13.5716614 | 27.4559311 |
| 0 | 0 |
| 374.577854 | 77.7918047 |
| 8.14299683 | 0 |
| 0 | 0 |
| 2.71433228 | 0 |

| | |
|---|---|
| 0 | 0 |
| 0 | 0 |
| 0 | 0 |
| 0 | 0 |
| 0 | 0 |
| 0 | 0 |
| 0 | 0 |
| 105.858959 | 86.9437817 |
| 2.71433228 | 86.9437817 |
| 0 | 0 |
| 0 | 0 |
| 21.7146582 | 0 |
| 0 | 0 |
| 0 | 0 |
| 0 | 0 |
| 0 | 0 |
| 0 | 0 |
| 5.42866456 | 118.975701 |
| 135.716614 | 4.57598851 |
| 0 | 0 |
| 0 | 0 |
| 0 | 0 |
| 0 | 123.55169 |
| 0 | 0 |
| 0 | 0 |
| 0 | 0 |
| 13.5716614 | 9.15197702 |
| 0 | 0 |
| 92.2872974 | 41.1838966 |
| 0 | 0 |
| 0 | 0 |
| 0 | 0 |
| 0 | 0 |
| 0 | 0 |
| 0 | 0 |
| 0 | 0 |
| 0 | 0 |
| 5.42866456 | 0 |
| 0 | 0 |
| 0 | 0 |
| 0 | 0 |
| 0 | 0 |
| 5.42866456 | 77.7918047 |

| | |
|---|---|
| 65.1439747 | 105.247736 |
| 0 | 0 |
| 0 | 22.8799426 |
| 0 | 0 |
| 0 | 0 |
| 0 | 0 |
| 0 | 59.4878506 |
| 0 | 0 |
| 0 | 0 |
| 0 | 0 |
| 0 | 0 |
| 0 | 0 |
| 0 | 0 |
| 0 | 0 |
| 0 | 0 |
| 0 | 0 |
| 0 | 0 |
| 24.4289905 | 141.855644 |
| 0 | 0 |
| 0 | 0 |
| 0 | 0 |
| 0 | 0 |
| 0 | 41.1838966 |
| 0 | 0 |
| 0 | 0 |
| 0 | 0 |
| 108.573291 | 0 |
| 434.293164 | 224.223437 |
| 0 | 0 |
| 0 | 0 |
| 0 | 0 |
| 0 | 0 |
| 0 | 0 |
| 0 | 0 |
| 0 | 0 |
| 0 | 0 |
| 0 | 0 |
| 0 | 0 |
| 0 | 0 |
| 0 | 0 |
| 0 | 0 |
| 0 | 0 |
| 0 | 0 |

| | |
|---|---|
| 16.2859937 | 114.399713 |
| 0 | 0 |
| 0 | 0 |
| 0 | 0 |
| 149.288275 | 45.7598851 |
| 5.42866456 | 45.7598851 |
| 70.5726392 | 32.0319196 |
| 0 | 0 |
| 184.574595 | 9.15197702 |
| 0 | 0 |
| 0 | 0 |
| 0 | 0 |
| 0 | 0 |
| 0 | 0 |
| 0 | 0 |
| 2.71433228 | 109.823724 |
| 0 | 0 |
| 0 | 0 |
| 0 | 0 |
| 0 | 0 |
| 0 | 0 |
| 0 | 0 |
| 0 | 0 |
| 0 | 0 |
| 0 | 0 |
| 2.71433228 | 0 |
| 0 | 0 |
| 0 | 0 |
| 0 | 0 |
| 0 | 0 |
| 0 | 0 |
| 0 | 118.975701 |
| 0 | 0 |
| 0 | 0 |
| 0 | 0 |
| 192.717592 | 224.223437 |
| 0 | 0 |
| 0 | 0 |
| 0 | 0 |
| 0 | 27.4559311 |
| 0 | 0 |
| 0 | 0 |
| 0 | 9.15197702 |

| | |
|---|---|
| 0 | 0 |
| 0 | 0 |
| 0 | 0 |
| 8.14299683 | 4.57598851 |
| 0 | 0 |
| 0 | 0 |
| 0 | 0 |
| 0 | 0 |
| 0 | 4.57598851 |
| 0 | 0 |
| 0 | 0 |
| 0 | 4.57598851 |
| 0 | 0 |
| 0 | 0 |
| 0 | 0 |
| 0 | 0 |
| 0 | 0 |
| 13.5716614 | 27.4559311 |
| 0 | 0 |
| 0 | 105.247736 |
| 0 | 0 |
| 434.293164 | 173.887563 |
| 0 | 0 |
| 0 | 0 |
| 5.42866456 | 151.007621 |
| 0 | 0 |
| 97.715962 | 9.15197702 |
| 0 | 0 |
| 179.14593 | 82.3677932 |
| 0 | 0 |
| 0 | 0 |
| 0 | 0 |
| 0 | 0 |
| 0 | 0 |
| 0 | 0 |
| 0 | 45.7598851 |
| 0 | 0 |
| 0 | 0 |
| 0 | 0 |
| 0 | 0 |
| 0 | 0 |
| 0 | 0 |
| 0 | 0 |

| | |
|---|---|
| 0 | 0 |
| 0 | 0 |
| 0 | 0 |
| 399.006845 | 41.1838966 |
| 0 | 0 |
| 8.14299683 | 9.15197702 |
| 5.42866456 | 0 |
| 0 | 0 |
| 0 | 0 |
| 0 | 0 |
| 0 | 0 |
| 0 | 0 |
| 0 | 9.15197702 |
| 0 | 0 |
| 0 | 0 |
| 0 | 0 |
| 0 | 0 |
| 0 | 0 |
| 124.859285 | 4.57598851 |
| 0 | 22.8799426 |
| 0 | 0 |
| 0 | 0 |
| 0 | 0 |
| 0 | 0 |
| 0 | 0 |
| 2.71433228 | 0 |
| 0 | 0 |
| 0 | 0 |
| 0 | 0 |
| 0 | 0 |
| 0 | 0 |
| 0 | 0 |
| 0 | 0 |
| 16.2859937 | 9.15197702 |
| 0 | 0 |
| 0 | 0 |
| 0 | 0 |
| 0 | 0 |
| 0 | 0 |
| 0 | 0 |
| 0 | 0 |
| 70.5726392 | 73.2158162 |
| 0 | 0 |

| | |
|---|---|
| 0 | 0 |
| 0 | 0 |
| 0 | 0 |
| 0 | 0 |
| 0 | 0 |
| 0 | 0 |
| 0 | 0 |
| 0 | 4.57598851 |
| 0 | 0 |
| 0 | 41.1838966 |
| 0 | 0 |
| 0 | 0 |
| 2.71433228 | 59.4878506 |
| 40.7149842 | 9.15197702 |
| 35.2863196 | 9.15197702 |
| 0 | 0 |
| 0 | 0 |
| 3067.19547 | 571.998564 |
| 0 | 0 |
| 0 | 0 |
| 0 | 0 |
| 0 | 0 |
| 0 | 0 |
| 0 | 0 |
| 0 | 4.57598851 |
| 0 | 18.303954 |
| 0 | 0 |
| 2.71433228 | 205.919483 |
| 0 | 0 |
| 0 | 0 |
| 0 | 0 |
| 0 | 27.4559311 |
| 0 | 0 |
| 10.8573291 | 27.4559311 |
| 0 | 0 |
| 0 | 0 |
| 0 | 0 |
| 27.1433228 | 0 |
| 0 | 0 |
| 0 | 0 |
| 0 | 0 |
| 0 | 0 |
| 0 | 0 |

| | |
|---|---|
| 0 | 0 |
| 92.2872974 | 9.15197702 |
| 0 | 0 |
| 0 | 0 |
| 0 | 0 |
| 0 | 0 |
| 0 | 0 |
| 0 | 0 |
| 0 | 0 |
| 0 | 0 |
| 464.15082 | 544.542633 |
| 0 | 0 |
| 0 | 0 |
| 0 | 0 |
| 5.42866456 | 9.15197702 |
| 0 | 0 |
| 0 | 0 |
| 0 | 0 |
| 0 | 0 |
| 0 | 0 |
| 0 | 27.4559311 |
| 0 | 0 |
| 48.857981 | 77.7918047 |
| 0 | 0 |
| 0 | 0 |
| 0 | 4.57598851 |
| 0 | 0 |
| 0 | 0 |
| 0 | 0 |
| 0 | 0 |
| 0 | 0 |
| 0 | 0 |
| 27.1433228 | 118.975701 |
| 0 | 0 |
| 81.4299683 | 0 |
| 108.573291 | 4.57598851 |
| 2.71433228 | 0 |
| 0 | 0 |
| 0 | 0 |
| 0 | 0 |
| 0 | 0 |
| 0 | 4.57598851 |
| 0 | 0 |

| | |
|---|---|
| 0 | 0 |
| 0 | 0 |
| 0 | 0 |
| 0 | 0 |
| 0 | 0 |
| 0 | 0 |
| 0 | 0 |
| 0 | 4.57598851 |
| 0 | 0 |
| 152.002608 | 22.8799426 |
| 0 | 0 |
| 374.577854 | 50.3358736 |
| 0 | 0 |
| 304.005215 | 27.4559311 |
| 0 | 0 |
| 0 | 0 |
| 0 | 18.303954 |
| 0 | 0 |
| 0 | 0 |
| 0 | 41.1838966 |
| 2.71433228 | 0 |
| 2.71433228 | 0 |
| 0 | 0 |
| 2.71433228 | 22.8799426 |
| 0 | 0 |
| 0 | 0 |
| 0 | 0 |
| 0 | 0 |
| 13.5716614 | 82.3677932 |
| 0 | 0 |
| 0 | 0 |
| 0 | 0 |
| 0 | 0 |
| 0 | 0 |
| 0 | 0 |
| 0 | 0 |
| 0 | 0 |
| 0 | 0 |
| 0 | 4.57598851 |
| 0 | 0 |
| 0 | 0 |
| 0 | 0 |
| 0 | 0 |

| | |
|---|---|
| 21.7146582 | 164.735586 |
| 0 | 0 |
| 116.716288 | 0 |
| 0 | 0 |
| 0 | 0 |
| 0 | 0 |
| 0 | 0 |
| 0 | 9.15197702 |
| 0 | 0 |
| 0 | 0 |
| 103.144627 | 45.7598851 |
| 19.0003259 | 41.1838966 |
| 76.0013038 | 86.9437817 |
| 5.42866456 | 105.247736 |
| 0 | 18.303954 |
| 0 | 0 |
| 8.14299683 | 82.3677932 |
| 0 | 0 |
| 0 | 0 |
| 0 | 0 |
| 0 | 0 |
| 0 | 0 |
| 0 | 0 |
| 0 | 0 |
| 0 | 0 |
| 0 | 0 |
| 2.71433228 | 27.4559311 |
| 0 | 0 |
| 0 | 0 |
| 0 | 0 |
| 0 | 0 |
| 0 | 0 |
| 0 | 82.3677932 |
| 0 | 0 |
| 76.0013038 | 9.15197702 |
| 0 | 0 |
| 0 | 77.7918047 |
| 0 | 0 |
| 0 | 0 |
| 0 | 0 |
| 0 | 0 |
| 0 | 0 |
| 252.432902 | 13.7279655 |

| | |
|---|---|
| 0 | 0 |
| 0 | 0 |
| 0 | 4.57598851 |
| 0 | 0 |
| 0 | 0 |
| 0 | 0 |
| 0 | 0 |
| 0 | 4.57598851 |
| 187.288927 | 41.1838966 |
| 0 | 0 |
| 0 | 0 |
| 0 | 0 |
| 0 | 0 |
| 241.575573 | 27.4559311 |
| 0 | 0 |
| 0 | 0 |
| 0 | 0 |
| 0 | 0 |
| 0 | 0 |
| 0 | 0 |
| 0 | 27.4559311 |
| 86.8586329 | 4.57598851 |
| 0 | 0 |
| 0 | 73.2158162 |
| 0 | 0 |
| 59.7153101 | 4.57598851 |
| 0 | 68.6398277 |
| 0 | 0 |
| 0 | 0 |
| 0 | 0 |
| 108.573291 | 4.57598851 |
| 0 | 0 |
| 0 | 0 |
| 0 | 0 |
| 380.006519 | 118.975701 |
| 0 | 0 |
| 0 | 0 |
| 0 | 0 |
| 0 | 0 |
| 0 | 0 |
| 0 | 0 |
| 0 | 0 |
| 149.288275 | 27.4559311 |

| | |
|---|---|
| 0 | 0 |
| 0 | 0 |
| 176.431598 | 265.407334 |
| 2.71433228 | 128.127678 |
| 0 | 0 |
| 0 | 0 |
| 0 | 0 |
| 181.860263 | 118.975701 |
| 0 | 0 |
| 92.2872974 | 105.247736 |
| 0 | 0 |
| 0 | 0 |
| 35.2863196 | 22.8799426 |
| 0 | 0 |
| 0 | 0 |
| 8.14299683 | 13.7279655 |
| 0 | 0 |
| 0 | 0 |
| 0 | 0 |
| 0 | 0 |
| 0 | 0 |
| 89.5729652 | 54.9118621 |
| 0 | 50.3358736 |
| 0 | 0 |
| 0 | 0 |
| 8.14299683 | 18.303954 |
| 0 | 0 |
| 0 | 0 |
| 0 | 22.8799426 |
| 241.575573 | 13.7279655 |
| 0 | 0 |
| 57.0009778 | 183.03954 |
| 0 | 0 |
| 0 | 0 |
| 0 | 0 |
| 0 | 0 |
| 0 | 0 |
| 0 | 0 |
| 0 | 4.57598851 |
| 57.0009778 | 4.57598851 |
| 0 | 0 |
| 0 | 0 |
| 5.42866456 | 9.15197702 |

| | |
|---|---|
| 0 | 36.6079081 |
| 0 | 45.7598851 |
| 282.290557 | 54.9118621 |
| 0 | 0 |
| 0 | 0 |
| 0 | 0 |
| 0 | 0 |
| 0 | 0 |
| 0 | 0 |
| 0 | 0 |
| 13.5716614 | 192.191517 |
| 10.8573291 | 4.57598851 |
| 0 | 0 |
| 0 | 0 |
| 0 | 0 |
| 0 | 0 |
| 0 | 54.9118621 |
| 0 | 0 |
| 0 | 0 |
| 0 | 0 |
| 0 | 0 |
| 0 | 0 |
| 5.42866456 | 0 |
| 0 | 0 |
| 40.7149842 | 114.399713 |
| 0 | 0 |
| 0 | 0 |
| 0 | 4.57598851 |
| 133.002282 | 279.135299 |
| 0 | 0 |
| 0 | 0 |
| 0 | 0 |
| 0 | 0 |
| 0 | 0 |
| 0 | 0 |
| 108.573291 | 0 |
| 2.71433228 | 9.15197702 |
| 0 | 0 |
| 0 | 0 |
| 0 | 0 |
| 81.4299683 | 0 |
| 0 | 0 |
| 0 | 0 |

| | |
|---|---|
| 0 | 0 |
| 2.71433228 | 0 |
| 0 | 0 |
| 141.145278 | 0 |
| 40.7149842 | 73.2158162 |
| 0 | 0 |
| 0 | 0 |
| 0 | 0 |
| 0 | 0 |
| 0 | 0 |
| 0 | 0 |
| 38.0006519 | 4.57598851 |
| 0 | 0 |
| 0 | 0 |
| 0 | 0 |
| 0 | 0 |
| 0 | 0 |
| 0 | 0 |
| 0 | 0 |
| 0 | 0 |
| 0 | 22.8799426 |
| 0 | 0 |
| 0 | 4.57598851 |
| 0 | 0 |
| 0 | 0 |
| 67.8583069 | 36.6079081 |
| 0 | 0 |
| 0 | 0 |
| 0 | 0 |
| 0 | 27.4559311 |
| 0 | 0 |
| 0 | 0 |
| 0 | 0 |
| 0 | 0 |
| 0 | 0 |
| 0 | 0 |
| 0 | 0 |
| 0 | 0 |
| 0 | 0 |
| 0 | 0 |
| 0 | 0 |
| 0 | 0 |
| 0 | 0 |

| | |
|---|---|
| 0 | 0 |
| 0 | 0 |
| 0 | 0 |
| 0 | 0 |
| 0 | 0 |
| 0 | 0 |
| 605.296098 | 173.887563 |
| 2.71433228 | 0 |
| 0 | 0 |
| 0 | 0 |
| 0 | 59.4878506 |
| 0 | 0 |
| 0 | 0 |
| 0 | 0 |
| 0 | 36.6079081 |
| 0 | 0 |
| 89.5729652 | 0 |
| 0 | 45.7598851 |
| 78.7156361 | 54.9118621 |
| 0 | 0 |
| 0 | 0 |
| 8.14299683 | 45.7598851 |
| 219.860915 | 73.2158162 |
| 0 | 41.1838966 |
| 0 | 0 |
| 2.71433228 | 0 |
| 0 | 0 |
| 0 | 0 |
| 2.71433228 | 0 |
| 0 | 0 |
| 0 | 0 |
| 0 | 0 |
| 184.574595 | 9.15197702 |
| 0 | 0 |
| 40.7149842 | 4.57598851 |
| 0 | 0 |
| 0 | 0 |
| 2.71433228 | 0 |
| 0 | 0 |
| 0 | 0 |
| 0 | 0 |
| 19.0003259 | 105.247736 |
| 103.144627 | 0 |

| | |
|---|---|
| 0 | 0 |
| 0 | 0 |
| 0 | 9.15197702 |
| 0 | 0 |
| 0 | 0 |
| 0 | 82.3677932 |
| 0 | 0 |
| 168.288601 | 22.8799426 |
| 0 | 0 |
| 0 | 0 |
| 0 | 0 |
| 0 | 32.0319196 |
| 0 | 0 |
| 0 | 0 |
| 347.434532 | 18.303954 |
| 0 | 0 |
| 0 | 0 |
| 0 | 0 |
| 0 | 0 |
| 0 | 0 |
| 0 | 0 |
| 13.5716614 | 36.6079081 |
| 0 | 0 |
| 0 | 50.3358736 |
| 2.71433228 | 27.4559311 |
| 0 | 0 |
| 0 | 0 |
| 0 | 0 |
| 0 | 0 |
| 0 | 0 |
| 0 | 0 |
| 0 | 13.7279655 |
| 10.8573291 | 27.4559311 |
| 0 | 0 |
| 0 | 0 |
| 1384.30946 | 183.03954 |
| 0 | 0 |
| 0 | 9.15197702 |
| 0 | 4.57598851 |
| 0 | 0 |
| 0 | 0 |
| 238.86124 | 0 |
| 2.71433228 | 0 |

| | |
|---|---|
| 0 | 0 |
| 249.71857 | 393.535012 |
| 0 | 0 |
| 0 | 41.1838966 |
| 0 | 4.57598851 |
| 0 | 0 |
| 0 | 0 |
| 0 | 36.6079081 |
| 0 | 0 |
| 412.578506 | 50.3358736 |
| 0 | 0 |
| 0 | 0 |
| 0 | 0 |
| 0 | 0 |
| 0 | 0 |
| 76.0013038 | 0 |
| 2.71433228 | 4.57598851 |
| 0 | 0 |
| 0 | 0 |
| 0 | 0 |
| 0 | 0 |
| 0 | 0 |
| 0 | 0 |
| 0 | 0 |
| 0 | 0 |
| 84.1443006 | 4.57598851 |
| 0 | 0 |
| 0 | 0 |
| 0 | 0 |
| 0 | 0 |
| 2.71433228 | 4.57598851 |
| 0 | 0 |
| 0 | 0 |
| 1370.7378 | 672.670311 |
| 0 | 0 |
| 0 | 0 |
| 331.148538 | 59.4878506 |
| 152.002608 | 4.57598851 |
| 0 | 0 |
| 0 | 0 |
| 0 | 0 |
| 0 | 0 |
| 0 | 0 |

| | |
|---|---|
| 0 | 0 |
| 0 | 0 |
| 0 | 45.7598851 |
| 0 | 0 |
| 0 | 0 |
| 76.0013038 | 224.223437 |
| 0 | 0 |
| 0 | 0 |
| 0 | 0 |
| 0 | 0 |
| 0 | 0 |
| 0 | 0 |
| 0 | 0 |
| 0 | 0 |
| 0 | 0 |
| 0 | 0 |
| 0 | 0 |
| 0 | 0 |
| 0 | 0 |
| 0 | 0 |
| 0 | 0 |
| 179.14593 | 242.527391 |
| 0 | 0 |
| 0 | 0 |
| 81.4299683 | 54.9118621 |
| 0 | 4.57598851 |
| 0 | 0 |
| 0 | 0 |
| 0 | 0 |
| 51.5723133 | 36.6079081 |
| 2.71433228 | 0 |
| 0 | 0 |
| 0 | 0 |
| 0 | 0 |
| 325.719873 | 379.807046 |
| 0 | 0 |
| 0 | 0 |
| 0 | 0 |
| 0 | 0 |
| 0 | 0 |
| 0 | 0 |
| 0 | 0 |
| 0 | 0 |

| | |
|---|---|
| 0 | 9.15197702 |
| 0 | 0 |
| 0 | 4.57598851 |
| 0 | 0 |
| 0 | 4.57598851 |
| 0 | 0 |
| 0 | 0 |
| 0 | 0 |
| 0 | 13.7279655 |
| 0 | 0 |
| 0 | 4.57598851 |
| 0 | 4.57598851 |
| 0 | 0 |
| 0 | 0 |
| 0 | 27.4559311 |
| 73.2869715 | 64.0638392 |
| 0 | 0 |
| 57.0009778 | 4.57598851 |
| 0 | 4.57598851 |
| 0 | 0 |
| 0 | 0 |
| 32.5719873 | 0 |
| 0 | 0 |
| 0 | 0 |
| 192.717592 | 9.15197702 |
| 0 | 0 |
| 0 | 0 |
| 0 | 0 |
| 0 | 0 |
| 0 | 0 |
| 0 | 0 |
| 0 | 0 |
| 0 | 0 |
| 0 | 0 |
| 2.71433228 | 0 |
| 0 | 0 |
| 0 | 0 |
| 0 | 0 |
| 0 | 0 |
| 0 | 0 |
| 0 | 0 |
| 0 | 0 |
| 0 | 0 |

| | |
|---|---|
| 0 | 0 |
| 0 | 0 |
| 0 | 0 |
| 51.5723133 | 77.7918047 |
| 2.71433228 | 96.0957587 |
| 0 | 0 |
| 46.1436487 | 18.303954 |
| 0 | 0 |
| 0 | 0 |
| 81.4299683 | 164.735586 |
| 0 | 0 |
| 95.0016297 | 4.57598851 |
| 0 | 0 |
| 0 | 0 |
| 0 | 0 |
| 0 | 0 |
| 13.5716614 | 9.15197702 |
| 0 | 0 |
| 0 | 0 |
| 0 | 4.57598851 |
| 0 | 0 |
| 0 | 0 |
| 0 | 0 |
| 0 | 0 |
| 8.14299683 | 0 |
| 0 | 41.1838966 |
| 13.5716614 | 0 |
| 0 | 0 |
| 0 | 0 |
| 0 | 0 |
| 0 | 0 |
| 0 | 0 |
| 5.42866456 | 0 |
| 0 | 0 |
| 0 | 0 |
| 0 | 0 |
| 0 | 0 |
| 0 | 0 |
| 0 | 0 |
| 0 | 0 |
| 0 | 0 |
| 16.2859937 | 54.9118621 |
| 0 | 0 |

| | |
|---|---|
| 0 | 0 |
| 0 | 0 |
| 8.14299683 | 32.0319196 |
| 0 | 0 |
| 0 | 0 |
| 2.71433228 | 22.8799426 |
| 0 | 0 |
| 19.0003259 | 18.303954 |
| 0 | 0 |
| 0 | 0 |
| 0 | 0 |
| 0 | 0 |
| 5.42866456 | 0 |
| 0 | 0 |
| 0 | 0 |
| 0 | 0 |
| 130.287949 | 9.15197702 |
| 0 | 0 |
| 124.859285 | 13.7279655 |
| 0 | 54.9118621 |
| 0 | 32.0319196 |
| 0 | 0 |
| 0 | 0 |
| 0 | 68.6398277 |
| 0 | 0 |
| 0 | 0 |
| 48.857981 | 4.57598851 |
| 0 | 0 |
| 0 | 0 |
| 192.717592 | 0 |
| 0 | 0 |
| 0 | 0 |
| 92.2872974 | 4.57598851 |
| 92.2872974 | 4.57598851 |
| 0 | 0 |
| 0 | 0 |
| 0 | 0 |
| 0 | 0 |
| 0 | 0 |
| 0 | 0 |
| 0 | 4.57598851 |
| 0 | 0 |
| 0 | 0 |

| | |
|---|---|
| 0 | 0 |
| 0 | 0 |
| 0 | 0 |
| 0 | 100.671747 |
| 0 | 0 |
| 0 | 0 |
| 0 | 0 |
| 0 | 0 |
| 57.0009778 | 137.279655 |
| 0 | 0 |
| 73.2869715 | 27.4559311 |
| 0 | 0 |
| 0 | 0 |
| 0 | 0 |
| 0 | 0 |
| 0 | 0 |
| 0 | 0 |
| 0 | 0 |
| 0 | 22.8799426 |
| 103.144627 | 0 |
| 0 | 0 |
| 0 | 0 |
| 84.1443006 | 123.55169 |
| 0 | 0 |
| 0 | 0 |
| 48.857981 | 0 |
| 0 | 0 |
| 0 | 0 |
| 0 | 0 |
| 0 | 0 |
| 0 | 18.303954 |
| 0 | 0 |
| 0 | 4.57598851 |
| 0 | 0 |
| 162.859937 | 13.7279655 |
| 2.71433228 | 0 |
| 0 | 0 |
| 95.0016297 | 36.6079081 |
| 0 | 0 |
| 0 | 0 |
| 0 | 0 |
| 0 | 0 |
| 0 | 0 |

| | |
|---|---|
| 0 | 0 |
| 0 | 0 |
| 0 | 0 |
| 0 | 0 |
| 0 | 0 |
| 187.288927 | 0 |
| 0 | 0 |
| 124.859285 | 22.8799426 |
| 0 | 0 |
| 0 | 0 |
| 0 | 0 |
| 0 | 0 |
| 0 | 0 |
| 0 | 0 |
| 0 | 0 |
| 0 | 0 |
| 0 | 0 |
| 51.5723133 | 4.57598851 |
| 0 | 36.6079081 |
| 0 | 0 |
| 0 | 9.15197702 |
| 0 | 0 |
| 0 | 0 |
| 0 | 4.57598851 |
| 0 | 22.8799426 |
| 0 | 0 |
| 0 | 0 |
| 0 | 0 |
| 0 | 0 |
| 0 | 0 |
| 0 | 96.0957587 |
| 0 | 0 |
| 0 | 0 |
| 0 | 0 |
| 2.71433228 | 45.7598851 |
| 0 | 0 |
| 0 | 0 |
| 0 | 0 |
| 0 | 0 |
| 0 | 0 |
| 0 | 0 |
| 0 | 0 |
| 0 | 0 |

| | |
|---|---|
| 0 | 0 |
| 0 | 0 |
| 35.2863196 | 64.0638392 |
| 0 | 0 |
| 81.4299683 | 77.7918047 |
| 0 | 0 |
| 0 | 32.0319196 |
| 0 | 0 |
| 0 | 4.57598851 |
| 0 | 0 |
| 0 | 0 |
| 0 | 0 |
| 0 | 0 |
| 0 | 0 |
| 2.71433228 | 4.57598851 |
| 0 | 0 |
| 0 | 0 |
| 0 | 4.57598851 |
| 0 | 0 |
| 0 | 0 |
| 0 | 0 |
| 0 | 0 |
| 0 | 0 |
| 46.1436487 | 109.823724 |
| 108.573291 | 9.15197702 |
| 0 | 13.7279655 |
| 0 | 9.15197702 |
| 0 | 27.4559311 |
| 27.1433228 | 18.303954 |
| 0 | 0 |
| 0 | 0 |
| 0 | 0 |
| 0 | 0 |
| 2.71433228 | 0 |
| 0 | 0 |
| 0 | 0 |
| 0 | 0 |
| 0 | 0 |
| 0 | 0 |
| 0 | 0 |
| 0 | 0 |
| 0 | 4.57598851 |
| 0 | 0 |

| | |
|---|---|
| 0 | 123.55169 |
| 0 | 0 |
| 0 | 0 |
| 0 | 0 |
| 2.71433228 | 91.5197702 |
| 0 | 0 |
| 0 | 0 |
| 0 | 0 |
| 2.71433228 | 0 |
| 0 | 4.57598851 |
| 0 | 4.57598851 |
| 0 | 0 |
| 532.009126 | 45.7598851 |
| 86.8586329 | 4.57598851 |
| 0 | 0 |
| 0 | 0 |
| 0 | 0 |
| 0 | 0 |
| 0 | 0 |
| 0 | 146.431632 |
| 0 | 0 |
| 0 | 0 |
| 0 | 0 |
| 0 | 0 |
| 0 | 0 |
| 0 | 0 |
| 0 | 0 |
| 0 | 0 |
| 0 | 0 |
| 0 | 0 |
| 0 | 0 |
| 0 | 0 |
| 0 | 0 |
| 0 | 0 |
| 16.2859937 | 4.57598851 |
| 0 | 0 |
| 119.43062 | 4.57598851 |
| 0 | 0 |
| 0 | 0 |
| 103.144627 | 91.5197702 |
| 0 | 0 |
| 0 | 27.4559311 |
| 0 | 0 |

| | |
|---|---|
| 0 | 0 |
| 0 | 22.8799426 |
| 0 | 0 |
| 0 | 0 |
| 0 | 0 |
| 0 | 0 |
| 13.5716614 | 32.0319196 |
| 0 | 0 |
| 27.1433228 | 13.7279655 |
| 0 | 0 |
| 2.71433228 | 0 |
| 0 | 0 |
| 0 | 0 |
| 366.434858 | 82.3677932 |
| 0 | 9.15197702 |
| 0 | 0 |
| 62.4296424 | 0 |
| 0 | 0 |
| 51.5723133 | 45.7598851 |
| 0 | 0 |
| 13.5716614 | 0 |
| 19.0003259 | 0 |
| 19.0003259 | 4.57598851 |
| 35.2863196 | 22.8799426 |
| 0 | 0 |
| 0 | 0 |
| 0 | 0 |
| 0 | 0 |
| 0 | 82.3677932 |
| 116.716288 | 425.566932 |
| 0 | 54.9118621 |
| 0 | 0 |
| 0 | 0 |
| 0 | 0 |
| 0 | 0 |
| 0 | 0 |
| 0 | 0 |
| 5.42866456 | 64.0638392 |
| 198.146256 | 27.4559311 |
| 0 | 0 |
| 0 | 50.3358736 |
| 561.866782 | 141.855644 |
| 0 | 0 |

| | |
|---|---|
| 0 | 0 |
| 0 | 0 |
| 19.0003259 | 146.431632 |
| 0 | 0 |
| 0 | 0 |
| 0 | 0 |
| 0 | 0 |
| 0 | 0 |
| 0 | 0 |
| 2.71433228 | 9.15197702 |
| 0 | 0 |
| 0 | 0 |
| 29.8576551 | 41.1838966 |
| 0 | 0 |
| 40.7149842 | 64.0638392 |
| 0 | 0 |
| 0 | 0 |
| 195.431924 | 13.7279655 |
| 0 | 0 |
| 0 | 0 |
| 0 | 132.703667 |
| 0 | 0 |
| 0 | 0 |
| 0 | 0 |
| 0 | 0 |
| 0 | 0 |
| 0 | 54.9118621 |
| 0 | 0 |
| 2.71433228 | 54.9118621 |
| 0 | 0 |
| 0 | 0 |
| 0 | 0 |
| 48.857981 | 4.57598851 |
| 0 | 0 |
| 0 | 0 |
| 0 | 0 |
| 0 | 0 |
| 0 | 0 |
| 0 | 0 |
| 0 | 0 |
| 0 | 0 |
| 0 | 0 |
| 0 | 0 |

| | |
|---|---|
| 0 | 0 |
| 0 | 0 |
| 173.717266 | 219.647449 |
| 0 | 0 |
| 0 | 0 |
| 0 | 0 |
| 0 | 0 |
| 0 | 0 |
| 67.8583069 | 45.7598851 |
| 0 | 0 |
| 0 | 0 |
| 0 | 36.6079081 |
| 184.574595 | 169.311575 |
| 0 | 0 |
| 0 | 0 |
| 0 | 0 |
| 648.725414 | 96.0957587 |
| 0 | 0 |
| 157.431272 | 196.767506 |
| 0 | 0 |
| 0 | 0 |
| 40.7149842 | 73.2158162 |
| 0 | 41.1838966 |
| 0 | 22.8799426 |
| 0 | 0 |
| 51.5723133 | 91.5197702 |
| 0 | 0 |
| 0 | 0 |
| 89.5729652 | 0 |
| 0 | 0 |
| 0 | 0 |
| 13.5716614 | 4.57598851 |
| 350.148864 | 183.03954 |
| 0 | 0 |
| 0 | 132.703667 |
| 5.42866456 | 64.0638392 |
| 0 | 0 |
| 0 | 0 |
| 0 | 0 |
| 0 | 0 |
| 0 | 0 |
| 0 | 0 |
| 0 | 41.1838966 |

| | |
|---|---|
| 0 | 0 |
| 0 | 0 |
| 0 | 0 |
| 0 | 0 |
| 0 | 0 |
| 0 | 0 |
| 0 | 0 |
| 0 | 0 |
| 0 | 0 |
| 8.14299683 | 64.0638392 |
| 0 | 0 |
| 0 | 0 |
| 0 | 0 |
| 24.4289905 | 0 |
| 0 | 0 |
| 0 | 0 |
| 0 | 0 |
| 0 | 96.0957587 |
| 0 | 0 |
| 0 | 0 |
| 0 | 0 |
| 0 | 0 |
| 0 | 0 |
| 0 | 0 |
| 0 | 0 |
| 0 | 0 |
| 0 | 0 |
| 0 | 0 |
| 0 | 0 |
| 0 | 0 |
| 0 | 0 |
| 0 | 0 |
| 0 | 0 |
| 0 | 0 |
| 0 | 0 |
| 0 | 0 |
| 0 | 0 |
| 0 | 0 |
| 0 | 0 |
| 0 | 0 |
| 149.288275 | 9.15197702 |
| 0 | 0 |
| 35.2863196 | 155.583609 |

| | |
|---|---|
| 0 | 0 |
| 0 | 0 |
| 0 | 0 |
| 0 | 0 |
| 0 | 0 |
| 0 | 0 |
| 0 | 0 |
| 0 | 0 |
| 0 | 0 |
| 200.860589 | 32.0319196 |
| 0 | 68.6398277 |
| 0 | 0 |
| 0 | 0 |
| 0 | 32.0319196 |
| 0 | 0 |
| 0 | 0 |
| 0 | 13.7279655 |
| 0 | 0 |
| 0 | 0 |
| 0 | 0 |
| 0 | 0 |
| 0 | 0 |
| 0 | 0 |
| 0 | 0 |
| 0 | 0 |
| 0 | 0 |
| 0 | 0 |
| 0 | 0 |
| 0 | 0 |
| 0 | 9.15197702 |
| 0 | 27.4559311 |
| 0 | 0 |
| 0 | 0 |
| 130.287949 | 54.9118621 |
| 2.71433228 | 0 |
| 0 | 0 |
| 0 | 0 |
| 0 | 0 |
| 0 | 0 |
| 10.8573291 | 68.6398277 |
| 0 | 0 |
| 0 | 0 |
| 0 | 0 |

| | |
|---|---|
| 0 | 0 |
| 0 | 45.7598851 |
| 0 | 0 |
| 0 | 0 |
| 0 | 73.2158162 |
| 0 | 0 |
| 0 | 0 |
| 0 | 0 |
| 127.573617 | 0 |
| 0 | 0 |
| 0 | 0 |
| 0 | 45.7598851 |
| 0 | 0 |
| 0 | 0 |
| 0 | 0 |
| 0 | 0 |
| 0 | 59.4878506 |
| 0 | 0 |
| 0 | 0 |
| 0 | 18.303954 |
| 0 | 0 |
| 0 | 0 |
| 0 | 0 |
| 0 | 0 |
| 0 | 0 |
| 0 | 0 |
| 0 | 0 |
| 48.857981 | 4.57598851 |
| 0 | 0 |
| 0 | 0 |
| 0 | 54.9118621 |
| 0 | 0 |
| 0 | 4.57598851 |
| 70.5726392 | 0 |
| 0 | 0 |
| 29.8576551 | 173.887563 |
| 0 | 0 |
| 0 | 0 |
| 0 | 0 |
| 0 | 0 |
| 0 | 0 |
| 0 | 0 |
| 0 | 9.15197702 |

| | |
|---|---|
| 0 | 0 |
| 0 | 0 |
| 0 | 0 |
| 0 | 0 |
| 0 | 0 |
| 0 | 0 |
| 0 | 0 |
| 0 | 0 |
| 0 | 0 |
| 0 | 4.57598851 |
| 0 | 4.57598851 |
| 0 | 0 |
| 0 | 0 |
| 0 | 4.57598851 |
| 0 | 0 |
| 0 | 0 |
| 0 | 0 |
| 0 | 0 |
| 0 | 0 |
| 0 | 0 |
| 0 | 13.7279655 |
| 5.42866456 | 0 |
| 5.42866456 | 0 |
| 0 | 0 |
| 0 | 0 |
| 0 | 0 |
| 0 | 0 |
| 0 | 0 |
| 0 | 0 |
| 0 | 0 |
| 0 | 0 |
| 0 | 0 |
| 0 | 0 |
| 0 | 0 |
| 0 | 0 |
| 0 | 0 |
| 0 | 0 |
| 0 | 0 |
| 0 | 0 |
| 0 | 0 |
| 13.5716614 | 64.0638392 |
| 0 | 0 |
| 0 | 0 |

| | |
|---|---|
| 0 | 0 |
| 0 | 0 |
| 0 | 0 |
| 0 | 0 |
| 0 | 0 |
| 0 | 4.57598851 |
| 0 | 0 |
| 0 | 0 |
| 0 | 0 |
| 0 | 54.9118621 |
| 0 | 0 |
| 0 | 0 |
| 2.71433228 | 0 |
| 0 | 0 |
| 0 | 0 |
| 0 | 0 |
| 0 | 0 |
| 0 | 0 |
| 0 | 0 |
| 0 | 0 |
| 0 | 0 |
| 0 | 0 |
| 0 | 0 |
| 0 | 0 |
| 0 | 0 |
| 0 | 0 |
| 59.7153101 | 18.303954 |
| 0 | 0 |
| 0 | 0 |
| 2.71433228 | 0 |
| 130.287949 | 0 |
| 0 | 0 |
| 0 | 0 |
| 0 | 0 |
| 0 | 4.57598851 |
| 0 | 0 |
| 5.42866456 | 0 |
| 0 | 0 |
| 0 | 0 |
| 0 | 0 |
| 0 | 0 |
| 0 | 0 |
| 0 | 0 |

| | |
|---|---|
| 0 | 0 |
| 0 | 0 |
| 0 | 0 |
| 0 | 0 |
| 0 | 0 |
| 27.1433228 | 9.15197702 |
| 0 | 0 |
| 0 | 0 |
| 0 | 0 |
| 0 | 0 |
| 0 | 0 |
| 0 | 0 |
| 0 | 0 |
| 10.8573291 | 0 |
| 0 | 0 |
| 0 | 0 |
| 0 | 0 |
| 0 | 0 |
| 0 | 0 |
| 0 | 0 |
| 84.1443006 | 50.3358736 |
| 0 | 0 |
| 0 | 0 |
| 0 | 4.57598851 |
| 162.859937 | 105.247736 |
| 0 | 0 |
| 0 | 0 |
| 0 | 0 |
| 0 | 0 |
| 32.5719873 | 32.0319196 |
| 0 | 0 |
| 0 | 0 |
| 0 | 22.8799426 |
| 187.288927 | 22.8799426 |
| 5.42866456 | 146.431632 |
| 0 | 0 |
| 0 | 0 |
| 146.573943 | 256.255357 |
| 0 | 0 |
| 0 | 0 |
| 119.43062 | 36.6079081 |
| 0 | 0 |
| 0 | 0 |

| | |
|---|---|
| 0 | 0 |
| 0 | 0 |
| 0 | 0 |
| 0 | 4.57598851 |
| 0 | 0 |
| 651.439747 | 4.57598851 |
| 8.14299683 | 77.7918047 |
| 0 | 0 |
| 0 | 0 |
| 51.5723133 | 160.159598 |
| 0 | 0 |
| 312.148212 | 41.1838966 |
| 78.7156361 | 0 |
| 0 | 0 |
| 0 | 0 |
| 0 | 0 |
| 0 | 0 |
| 103.144627 | 13.7279655 |
| 0 | 0 |
| 0 | 0 |
| 0 | 68.6398277 |
| 0 | 0 |
| 0 | 0 |
| 38.0006519 | 4.57598851 |
| 0 | 0 |
| 0 | 0 |
| 447.864826 | 137.279655 |
| 0 | 0 |
| 16.2859937 | 0 |
| 62.4296424 | 13.7279655 |
| 420.721503 | 96.0957587 |
| 0 | 0 |
| 0 | 0 |
| 0 | 0 |
| 0 | 0 |
| 0 | 0 |
| 133.002282 | 22.8799426 |
| 0 | 0 |
| 0 | 0 |
| 0 | 0 |
| 0 | 0 |
| 0 | 0 |
| 0 | 0 |

| | |
|---|---|
| 0 | 0 |
| 0 | 0 |
| 282.290557 | 187.615529 |
| 0 | 0 |
| 0 | 0 |
| 0 | 0 |
| 0 | 59.4878506 |
| 0 | 0 |
| 0 | 0 |
| 57.0009778 | 0 |
| 0 | 0 |
| 0 | 0 |
| 0 | 0 |
| 0 | 0 |
| 0 | 0 |
| 0 | 0 |
| 0 | 0 |
| 0 | 0 |
| 0 | 0 |
| 95.0016297 | 9.15197702 |
| 0 | 0 |
| 0 | 0 |
| 0 | 0 |
| 0 | 0 |
| 336.577202 | 18.303954 |
| 0 | 0 |
| 0 | 0 |
| 0 | 0 |
| 0 | 0 |
| 0 | 0 |
| 0 | 0 |
| 0 | 0 |
| 0 | 0 |
| 0 | 0 |
| 165.574269 | 247.10338 |
| 431.578832 | 86.9437817 |
| 0 | 0 |
| 0 | 0 |
| 0 | 0 |
| 0 | 0 |
| 0 | 0 |
| 0 | 0 |
| 8.14299683 | 18.303954 |

| | |
|---|---|
| 0 | 4.57598851 |
| 2.71433228 | 22.8799426 |
| 0 | 0 |
| 0 | 0 |
| 0 | 0 |
| 0 | 4.57598851 |
| 0 | 0 |
| 0 | 0 |
| 0 | 0 |
| 0 | 0 |
| 5.42866456 | 77.7918047 |
| 0 | 0 |
| 192.717592 | 77.7918047 |
| 0 | 0 |
| 0 | 0 |
| 0 | 0 |
| 0 | 0 |
| 0 | 0 |
| 0 | 0 |
| 0 | 0 |
| 570.009778 | 68.6398277 |
| 0 | 27.4559311 |
| 0 | 0 |
| 0 | 0 |
| 40.7149842 | 201.343494 |
| 0 | 4.57598851 |
| 0 | 0 |
| 0 | 0 |
| 0 | 0 |
| 0 | 0 |
| 0 | 0 |
| 0 | 0 |
| 5.42866456 | 59.4878506 |
| 0 | 0 |
| 0 | 0 |
| 0 | 0 |
| 0 | 0 |
| 29.8576551 | 9.15197702 |
| 62.4296424 | 9.15197702 |
| 0 | 0 |
| 29.8576551 | 114.399713 |
| 0 | 0 |
| 0 | 0 |

| | |
|---|---|
| 0 | 0 |
| 0 | 0 |
| 0 | 0 |
| 0 | 27.4559311 |
| 0 | 0 |
| 2.71433228 | 45.7598851 |
| 0 | 4.57598851 |
| 5.42866456 | 0 |
| 0 | 0 |
| 0 | 0 |
| 2.71433228 | 0 |
| 0 | 0 |
| 0 | 0 |
| 0 | 0 |
| 59.7153101 | 0 |
| 0 | 0 |
| 0 | 4.57598851 |
| 0 | 0 |
| 0 | 0 |
| 0 | 0 |
| 0 | 18.303954 |
| 0 | 0 |
| 0 | 0 |
| 0 | 0 |
| 0 | 0 |
| 0 | 0 |
| 0 | 0 |
| 388.149516 | 0 |
| 0 | 187.615529 |
| 0 | 0 |
| 0 | 0 |
| 0 | 0 |
| 0 | 0 |
| 0 | 0 |
| 0 | 0 |
| 24.4289905 | 68.6398277 |
| 0 | 0 |
| 0 | 0 |
| 0 | 0 |
| 0 | 0 |
| 0 | 77.7918047 |
| 0 | 4.57598851 |
| 0 | 0 |

| | |
|---|---|
| 35.2863196 | 18.303954 |
| 35.2863196 | 18.303954 |
| 0 | 0 |
| 0 | 0 |
| 108.573291 | 0 |
| 0 | 0 |
| 0 | 0 |
| 0 | 0 |
| 0 | 0 |
| 0 | 0 |
| 0 | 0 |
| 0 | 0 |
| 0 | 0 |
| 0 | 0 |
| 0 | 0 |
| 0 | 4.57598851 |
| 76.0013038 | 0 |
| 0 | 4.57598851 |
| 0 | 0 |
| 0 | 0 |
| 513.008801 | 160.159598 |
| 2.71433228 | 0 |
| 0 | 0 |
| 0 | 0 |
| 0 | 0 |
| 0 | 0 |
| 0 | 59.4878506 |
| 0 | 0 |
| 0 | 0 |
| 0 | 0 |
| 0 | 0 |
| 0 | 4.57598851 |
| 0 | 0 |
| 0 | 0 |
| 0 | 0 |
| 81.4299683 | 36.6079081 |
| 141.145278 | 9.15197702 |
| 0 | 0 |
| 0 | 0 |
| 0 | 9.15197702 |
| 0 | 22.8799426 |
| 732.869715 | 0 |
| 0 | 0 |

| | |
|---|---|
| 0 | 4.57598851 |
| 0 | 0 |
| 13.5716614 | 18.303954 |
| 0 | 0 |
| 0 | 96.0957587 |
| 0 | 9.15197702 |
| 0 | 0 |
| 0 | 0 |
| 0 | 0 |
| 0 | 0 |
| 0 | 0 |
| 0 | 0 |
| 0 | 0 |
| 0 | 0 |
| 0 | 0 |
| 0 | 0 |
| 0 | 0 |
| 0 | 0 |
| 0 | 0 |
| 0 | 0 |
| 54.2866456 | 0 |
| 0 | 0 |
| 0 | 0 |
| 0 | 0 |
| 453.29349 | 109.823724 |
| 0 | 0 |
| 0 | 0 |
| 0 | 0 |
| 0 | 4.57598851 |
| 108.573291 | 100.671747 |
| 0 | 0 |
| 0 | 0 |
| 57.0009778 | 4.57598851 |
| 141.145278 | 32.0319196 |
| 0 | 0 |
| 0 | 0 |
| 0 | 0 |
| 0 | 0 |
| 89.5729652 | 0 |
| 0 | 0 |
| 0 | 0 |
| 0 | 4.57598851 |
| 0 | 0 |

| | |
|---|---|
| 0 | 4.57598851 |
| 0 | 13.7279655 |
| 597.153101 | 0 |
| 0 | 0 |
| 0 | 0 |
| 0 | 0 |
| 0 | 4.57598851 |
| 2.71433228 | 0 |
| 2.71433228 | 0 |
| 5.42866456 | 151.007621 |
| 0 | 0 |
| 0 | 0 |
| 0 | 0 |
| 0 | 0 |
| 0 | 0 |
| 0 | 0 |
| 0 | 0 |
| 0 | 4.57598851 |
| 0 | 0 |
| 0 | 0 |
| 0 | 0 |
| 0 | 0 |
| 92.2872974 | 4.57598851 |
| 0 | 0 |
| 0 | 0 |
| 0 | 0 |
| 0 | 0 |
| 0 | 0 |
| 2.71433228 | 0 |
| 0 | 0 |
| 0 | 0 |
| 287.719221 | 9.15197702 |
| 0 | 0 |
| 0 | 173.887563 |
| 0 | 0 |
| 0 | 0 |
| 0 | 0 |
| 0 | 0 |
| 5.42866456 | 0 |
| 0 | 0 |
| 0 | 0 |
| 65.1439747 | 0 |
| 0 | 0 |

| | |
|---|---|
| 0 | 0 |
| 124.859285 | 41.1838966 |
| 0 | 0 |
| 0 | 0 |
| 0 | 0 |
| 0 | 0 |
| 0 | 18.303954 |
| 0 | 0 |
| 0 | 18.303954 |
| 0 | 4.57598851 |
| 0 | 0 |
| 0 | 0 |
| 0 | 0 |
| 0 | 0 |
| 0 | 0 |
| 32.5719873 | 0 |
| 0 | 0 |
| 0 | 0 |
| 0 | 0 |
| 0 | 0 |
| 0 | 4.57598851 |
| 0 | 0 |
| 0 | 0 |
| 0 | 0 |
| 0 | 73.2158162 |
| 8.14299683 | 18.303954 |
| 0 | 0 |
| 0 | 0 |
| 0 | 0 |
| 0 | 0 |
| 0 | 0 |
| 0 | 0 |
| 0 | 0 |
| 0 | 0 |
| 0 | 0 |
| 0 | 0 |
| 0 | 22.8799426 |
| 0 | 0 |
| 0 | 0 |
| 0 | 0 |
| 0 | 0 |
| 0 | 0 |
| 0 | 0 |

| | |
|---|---|
| 2.71433228 | 18.303954 |
| 0 | 0 |
| 0 | 0 |
| 0 | 0 |
| 0 | 0 |
| 0 | 0 |
| 0 | 0 |
| 0 | 22.8799426 |
| 0 | 0 |
| 0 | 0 |
| 0 | 0 |
| 214.43225 | 32.0319196 |
| 0 | 0 |
| 0 | 0 |
| 0 | 0 |
| 0 | 0 |
| 0 | 0 |
| 0 | 0 |
| 0 | 0 |
| 0 | 0 |
| 0 | 0 |
| 116.716288 | 173.887563 |
| 206.289253 | 50.3358736 |
| 0 | 4.57598851 |
| 0 | 0 |
| 0 | 0 |
| 0 | 0 |
| 0 | 0 |
| 0 | 0 |
| 2.71433228 | 0 |
| 0 | 0 |
| 0 | 50.3358736 |
| 0 | 0 |
| 0 | 0 |
| 0 | 0 |
| 0 | 0 |
| 0 | 0 |
| 0 | 0 |
| 0 | 0 |
| 0 | 0 |
| 0 | 0 |
| 0 | 0 |
| 0 | 0 |

| | |
|---|---|
| 0 | 41.1838966 |
| 0 | 0 |
| 57.0009778 | 0 |
| 0 | 0 |
| 0 | 50.3358736 |
| 0 | 0 |
| 32.5719873 | 0 |
| 0 | 0 |
| 0 | 0 |
| 0 | 0 |
| 0 | 0 |
| 0 | 0 |
| 0 | 0 |
| 0 | 0 |
| 0 | 0 |
| 0 | 164.735586 |
| 0 | 0 |
| 0 | 18.303954 |
| 0 | 0 |
| 0 | 0 |
| 0 | 0 |
| 0 | 0 |
| 100.430294 | 77.7918047 |
| 0 | 45.7598851 |
| 0 | 0 |
| 0 | 0 |
| 0 | 0 |
| 0 | 0 |
| 0 | 0 |
| 0 | 0 |
| 0 | 0 |
| 0 | 0 |
| 0 | 59.4878506 |
| 0 | 0 |
| 0 | 0 |
| 0 | 0 |
| 0 | 0 |
| 0 | 0 |
| 8.14299683 | 0 |
| 0 | 0 |
| 0 | 0 |
| 0 | 0 |
| 0 | 0 |

| | |
|---|---|
| 0 | 0 |
| 0 | 36.6079081 |
| 0 | 0 |
| 76.0013038 | 0 |
| 0 | 0 |
| 0 | 0 |
| 0 | 0 |
| 0 | 0 |
| 0 | 0 |
| 0 | 0 |
| 40.7149842 | 0 |
| 0 | 0 |
| 0 | 4.57598851 |
| 160.145604 | 4.57598851 |
| 130.287949 | 13.7279655 |
| 0 | 0 |
| 76.0013038 | 0 |
| 0 | 0 |
| 0 | 0 |
| 40.7149842 | 86.9437817 |
| 0 | 4.57598851 |
| 0 | 36.6079081 |
| 0 | 0 |
| 27.1433228 | 0 |
| 0 | 0 |
| 2.71433228 | 0 |
| 0 | 0 |
| 0 | 0 |
| 0 | 0 |
| 0 | 0 |
| 105.858959 | 0 |
| 0 | 0 |
| 0 | 0 |
| 2.71433228 | 64.0638392 |
| 5.42866456 | 0 |
| 0 | 0 |
| 0 | 0 |
| 0 | 0 |
| 0 | 0 |
| 0 | 0 |
| 0 | 0 |
| 0 | 0 |
| 0 | 0 |

| | |
|---|---|
| 0 | 0 |
| 0 | 0 |
| 290.433554 | 9.15197702 |
| 0 | 0 |
| 0 | 0 |
| 0 | 0 |
| 67.8583069 | 32.0319196 |
| 0 | 0 |
| 0 | 73.2158162 |
| 27.1433228 | 59.4878506 |
| 0 | 0 |
| 0 | 0 |
| 200.860589 | 18.303954 |
| 0 | 0 |
| 0 | 0 |
| 0 | 0 |
| 40.7149842 | 45.7598851 |
| 0 | 0 |
| 0 | 0 |
| 0 | 0 |
| 0 | 0 |
| 0 | 0 |
| 0 | 0 |
| 95.0016297 | 4.57598851 |
| 0 | 0 |
| 0 | 0 |
| 0 | 0 |
| 2.71433228 | 41.1838966 |
| 0 | 0 |
| 0 | 0 |
| 0 | 0 |
| 0 | 0 |
| 0 | 0 |
| 5.42866456 | 9.15197702 |
| 0 | 0 |
| 21.7146582 | 64.0638392 |
| 0 | 0 |
| 0 | 0 |
| 0 | 0 |
| 0 | 0 |
| 108.573291 | 36.6079081 |
| 0 | 0 |
| 0 | 77.7918047 |

| | |
|---|---|
| 0 | 0 |
| 0 | 0 |
| 116.716288 | 0 |
| 401.721177 | 169.311575 |
| 0 | 0 |
| 0 | 0 |
| 0 | 0 |
| 0 | 0 |
| 0 | 0 |
| 0 | 0 |
| 29.8576551 | 18.303954 |
| 0 | 0 |
| 95.0016297 | 114.399713 |
| 0 | 0 |
| 0 | 0 |
| 2.71433228 | 0 |
| 0 | 0 |
| 0 | 0 |
| 0 | 0 |
| 0 | 0 |
| 510.294468 | 32.0319196 |
| 2.71433228 | 9.15197702 |
| 0 | 0 |
| 0 | 0 |
| 21.7146582 | 0 |
| 0 | 0 |
| 0 | 0 |
| 0 | 4.57598851 |
| 0 | 4.57598851 |
| 0 | 0 |
| 103.144627 | 114.399713 |
| 0 | 0 |
| 0 | 0 |
| 0 | 0 |
| 0 | 0 |
| 0 | 0 |
| 0 | 0 |
| 0 | 0 |
| 0 | 0 |
| 0 | 0 |
| 217.146582 | 36.6079081 |
| 0 | 86.9437817 |
| 8.14299683 | 22.8799426 |

| | |
|---|---|
| 16.2859937 | 4.57598851 |
| 0 | 0 |
| 0 | 128.127678 |
| 2.71433228 | 4.57598851 |
| 344.720199 | 18.303954 |
| 0 | 0 |
| 154.71694 | 18.303954 |
| 0 | 0 |
| 0 | 0 |
| 0 | 0 |
| 8.14299683 | 27.4559311 |
| 0 | 0 |
| 0 | 0 |
| 0 | 59.4878506 |
| 0 | 0 |
| 0 | 0 |
| 0 | 0 |
| 0 | 0 |
| 0 | 0 |
| 0 | 0 |
| 168.288601 | 36.6079081 |
| 0 | 0 |
| 0 | 0 |
| 0 | 0 |
| 0 | 0 |
| 0 | 0 |
| 0 | 0 |
| 0 | 0 |
| 0 | 0 |
| 0 | 0 |
| 0 | 0 |
| 0 | 0 |
| 0 | 0 |
| 0 | 0 |
| 0 | 0 |
| 0 | 0 |
| 0 | 0 |
| 0 | 22.8799426 |
| 0 | 0 |
| 0 | 13.7279655 |
| 105.858959 | 0 |
| 0 | 0 |
| 0 | 36.6079081 |

| | |
|---|---|
| 10.8573291 | 91.5197702 |
| 0 | 0 |
| 0 | 0 |
| 0 | 0 |
| 67.8583069 | 132.703667 |
| 0 | 0 |
| 0 | 0 |
| 0 | 0 |
| 0 | 0 |
| 0 | 45.7598851 |
| 0 | 0 |
| 5.42866456 | 0 |
| 0 | 0 |
| 0 | 0 |
| 0 | 0 |
| 0 | 0 |
| 0 | 0 |
| 10.8573291 | 45.7598851 |
| 0 | 0 |
| 0 | 0 |
| 0 | 0 |
| 81.4299683 | 0 |
| 238.86124 | 187.615529 |
| 0 | 0 |
| 0 | 0 |
| 0 | 0 |
| 0 | 0 |
| 0 | 0 |
| 0 | 0 |
| 0 | 0 |
| 0 | 0 |
| 5.42866456 | 13.7279655 |
| 0 | 0 |
| 0 | 0 |
| 0 | 0 |
| 0 | 0 |
| 0 | 0 |
| 2.71433228 | 0 |
| 40.7149842 | 100.671747 |
| 198.146256 | 0 |
| 0 | 0 |
| 0 | 0 |
| 0 | 73.2158162 |

| | |
|---|---|
| 0 | 86.9437817 |
| 0 | 0 |
| 0 | 0 |
| 0 | 0 |
| 0 | 0 |
| 0 | 0 |
| 0 | 0 |
| 54.2866456 | 4.57598851 |
| 0 | 0 |
| 0 | 0 |
| 0 | 0 |
| 0 | 0 |
| 0 | 36.6079081 |
| 0 | 4.57598851 |
| 0 | 0 |
| 0 | 0 |
| 124.859285 | 22.8799426 |
| 0 | 0 |
| 0 | 0 |
| 0 | 0 |
| 0 | 4.57598851 |
| 0 | 0 |
| 0 | 0 |
| 0 | 141.855644 |
| 0 | 0 |
| 0 | 0 |
| 192.717592 | 0 |
| 0 | 0 |
| 0 | 0 |
| 0 | 0 |
| 0 | 91.5197702 |
| 5.42866456 | 137.279655 |
| 0 | 187.615529 |
| 0 | 0 |
| 0 | 0 |
| 70.5726392 | 4.57598851 |
| 0 | 0 |
| 0 | 0 |
| 0 | 73.2158162 |
| 0 | 0 |
| 0 | 0 |
| 0 | 0 |
| 0 | 0 |

| | |
|---|---|
| 0 | 0 |
| 0 | 0 |
| 0 | 0 |
| 0 | 0 |
| 0 | 0 |
| 95.0016297 | 114.399713 |
| 0 | 0 |
| 0 | 0 |
| 0 | 0 |
| 0 | 0 |
| 0 | 0 |
| 0 | 0 |
| 0 | 0 |
| 0 | 0 |
| 40.7149842 | 0 |
| 0 | 0 |
| 0 | 105.247736 |
| 0 | 0 |
| 0 | 0 |
| 0 | 0 |
| 0 | 0 |
| 0 | 4.57598851 |
| 0 | 0 |
| 0 | 0 |
| 0 | 0 |
| 469.579484 | 86.9437817 |
| 0 | 77.7918047 |
| 0 | 0 |
| 0 | 0 |
| 0 | 0 |
| 0 | 0 |
| 0 | 4.57598851 |
| 0 | 0 |
| 0 | 0 |
| 0 | 0 |
| 0 | 0 |
| 0 | 0 |
| 0 | 0 |
| 0 | 0 |
| 0 | 0 |
| 0 | 0 |
| 0 | 0 |
| 57.0009778 | 82.3677932 |

| | |
|---|---|
| 0 | 0 |
| 0 | 0 |
| 0 | 0 |
| 0 | 0 |
| 0 | 0 |
| 0 | 77.7918047 |
| 0 | 0 |
| 0 | 0 |
| 0 | 0 |
| 65.1439747 | 22.8799426 |
| 0 | 151.007621 |
| 0 | 0 |
| 51.5723133 | 205.919483 |
| 0 | 0 |
| 0 | 0 |
| 0 | 0 |
| 0 | 0 |
| 103.144627 | 4.57598851 |
| 0 | 0 |
| 0 | 0 |
| 0 | 0 |
| 0 | 0 |
| 0 | 0 |
| 0 | 0 |
| 0 | 0 |
| 0 | 22.8799426 |
| 0 | 0 |
| 0 | 0 |
| 0 | 0 |
| 0 | 0 |
| 0 | 0 |
| 0 | 0 |
| 0 | 0 |
| 0 | 0 |
| 0 | 0 |
| 0 | 82.3677932 |
| 483.151145 | 27.4559311 |
| 247.004237 | 0 |
| 0 | 0 |
| 29.8576551 | 54.9118621 |
| 0 | 0 |
| 0 | 0 |
| 0 | 0 |

| | |
|---|---|
| 0 | 45.7598851 |
| 2.71433228 | 4.57598851 |
| 0 | 0 |
| 0 | 0 |
| 0 | 0 |
| 0 | 0 |
| 0 | 0 |
| 0 | 0 |
| 5.42866456 | 109.823724 |
| 29.8576551 | 196.767506 |
| 0 | 0 |
| 0 | 0 |
| 0 | 0 |
| 594.438769 | 32.0319196 |
| 0 | 0 |
| 181.860263 | 9.15197702 |
| 0 | 0 |
| 0 | 0 |
| 0 | 0 |
| 0 | 0 |
| 0 | 0 |
| 0 | 0 |
| 0 | 0 |
| 0 | 0 |
| 0 | 0 |
| 10.8573291 | 77.7918047 |
| 0 | 0 |
| 0 | 0 |
| 173.717266 | 13.7279655 |
| 0 | 0 |
| 0 | 0 |
| 0 | 0 |
| 0 | 0 |
| 0 | 0 |
| 0 | 0 |
| 0 | 0 |
| 0 | 0 |
| 0 | 0 |
| 0 | 0 |
| 0 | 9.15197702 |
| 0 | 0 |
| 0 | 0 |
| 0 | 0 |

| | |
|---|---|
| 0 | 0 |
| 0 | 0 |
| 0 | 0 |
| 5.42866456 | 0 |
| 0 | 0 |
| 0 | 4.57598851 |
| 0 | 0 |
| 0 | 0 |
| 0 | 0 |
| 0 | 0 |
| 0 | 0 |
| 0 | 0 |
| 692.154731 | 626.910426 |
| 0 | 0 |
| 122.144953 | 137.279655 |
| 0 | 0 |
| 0 | 0 |
| 0 | 0 |
| 0 | 0 |
| 806.156687 | 302.015242 |
| 105.858959 | 32.0319196 |
| 0 | 0 |
| 103.144627 | 0 |
| 0 | 0 |
| 0 | 36.6079081 |
| 0 | 0 |
| 0 | 0 |
| 0 | 0 |
| 0 | 0 |
| 8.14299683 | 41.1838966 |
| 2.71433228 | 0 |
| 0 | 0 |
| 0 | 0 |
| 743.727044 | 256.255357 |
| 0 | 0 |
| 0 | 0 |
| 13.5716614 | 73.2158162 |
| 0 | 0 |
| 0 | 36.6079081 |
| 423.435835 | 68.6398277 |
| 0 | 0 |
| 0 | 0 |
| 0 | 0 |

| | |
|---|---|
| 0 | 0 |
| 0 | 73.2158162 |
| 0 | 128.127678 |
| 0 | 128.127678 |
| 0 | 0 |
| 38.0006519 | 22.8799426 |
| 0 | 0 |
| 0 | 0 |
| 0 | 0 |
| 0 | 0 |
| 0 | 0 |
| 0 | 0 |
| 0 | 0 |
| 0 | 4.57598851 |
| 0 | 0 |
| 0 | 0 |
| 0 | 0 |
| 2.71433228 | 4.57598851 |
| 358.291861 | 183.03954 |
| 0 | 0 |
| 0 | 0 |
| 0 | 0 |
| 2.71433228 | 0 |
| 0 | 0 |
| 21.7146582 | 22.8799426 |
| 0 | 0 |
| 62.4296424 | 0 |
| 0 | 0 |
| 0 | 0 |
| 0 | 0 |
| 0 | 4.57598851 |
| 0 | 0 |
| 0 | 0 |
| 0 | 0 |
| 0 | 0 |
| 0 | 0 |
| 0 | 0 |
| 0 | 0 |
| 2.71433228 | 114.399713 |
| 0 | 0 |
| 0 | 0 |
| 0 | 0 |
| 0 | 0 |

| | |
|---|---|
| 0 | 0 |
| 0 | 0 |
| 0 | 0 |
| 0 | 0 |
| 0 | 0 |
| 0 | 0 |
| 0 | 0 |
| 0 | 0 |
| 0 | 0 |
| 0 | 0 |
| 0 | 0 |
| 0 | 0 |
| 827.871345 | 132.703667 |
| 0 | 0 |
| 0 | 0 |
| 0 | 0 |
| 0 | 0 |
| 0 | 0 |
| 0 | 0 |
| 0 | 0 |
| 0 | 0 |
| 0 | 0 |
| 0 | 0 |
| 0 | 41.1838966 |
| 0 | 18.303954 |
| 0 | 0 |
| 0 | 0 |
| 0 | 0 |
| 143.859611 | 32.0319196 |
| 0 | 22.8799426 |
| 0 | 0 |
| 0 | 0 |
| 0 | 13.7279655 |
| 10.8573291 | 0 |
| 0 | 0 |
| 0 | 45.7598851 |
| 0 | 0 |
| 0 | 0 |
| 0 | 0 |
| 0 | 9.15197702 |
| 0 | 0 |
| 0 | 0 |
| 2.71433228 | 4.57598851 |

| | |
|---|---|
| 0 | 0 |
| 0 | 0 |
| 958.159294 | 315.743207 |
| 0 | 4.57598851 |
| 0 | 123.55169 |
| 0 | 0 |
| 0 | 0 |
| 0 | 0 |
| 0 | 118.975701 |
| 0 | 0 |
| 0 | 0 |
| 0 | 0 |
| 0 | 0 |
| 0 | 0 |
| 0 | 0 |
| 0 | 0 |
| 0 | 0 |
| 0 | 0 |
| 0 | 0 |
| 0 | 0 |
| 0 | 0 |
| 10.8573291 | 0 |
| 0 | 0 |
| 0 | 0 |
| 8.14299683 | 96.0957587 |
| 0 | 0 |
| 0 | 0 |
| 0 | 0 |
| 0 | 64.0638392 |
| 0 | 0 |
| 0 | 0 |
| 0 | 0 |
| 0 | 0 |
| 0 | 0 |
| 0 | 0 |
| 0 | 0 |
| 0 | 0 |
| 0 | 0 |
| 0 | 0 |
| 187.288927 | 41.1838966 |
| 0 | 0 |
| 0 | 0 |
| 0 | 0 |

| | |
|---|---|
| 0 | 0 |
| 0 | 0 |
| 0 | 109.823724 |
| 0 | 0 |
| 0 | 0 |
| 21.7146582 | 59.4878506 |
| 78.7156361 | 0 |
| 0 | 0 |
| 0 | 0 |
| 0 | 160.159598 |
| 0 | 0 |
| 0 | 0 |
| 0 | 0 |
| 0 | 0 |
| 2.71433228 | 0 |
| 5.42866456 | 114.399713 |
| 0 | 0 |
| 5.42866456 | 4.57598851 |
| 0 | 0 |
| 0 | 45.7598851 |
| 0 | 0 |
| 0 | 0 |
| 0 | 0 |
| 0 | 0 |
| 0 | 0 |
| 238.86124 | 41.1838966 |
| 24.4289905 | 22.8799426 |
| 0 | 0 |
| 122.144953 | 50.3358736 |
| 0 | 9.15197702 |
| 0 | 0 |
| 0 | 0 |
| 2.71433228 | 9.15197702 |
| 0 | 9.15197702 |
| 13.5716614 | 9.15197702 |
| 0 | 59.4878506 |
| 293.147886 | 178.463552 |
| 51.5723133 | 137.279655 |
| 2.71433228 | 0 |
| 2.71433228 | 4.57598851 |
| 0 | 0 |
| 0 | 0 |
| 0 | 32.0319196 |

| | |
|---|---|
| 0 | 0 |
| 0 | 0 |
| 0 | 0 |
| 0 | 0 |
| 0 | 0 |
| 0 | 0 |
| 0 | 0 |
| 0 | 0 |
| 0 | 0 |
| 0 | 0 |
| 0 | 0 |
| 0 | 0 |
| 0 | 0 |
| 32.5719873 | 0 |
| 0 | 0 |
| 0 | 0 |
| 16.2859937 | 100.671747 |
| 0 | 0 |
| 0 | 0 |
| 0 | 0 |
| 0 | 0 |
| 0 | 0 |
| 0 | 0 |
| 0 | 73.2158162 |
| 0 | 0 |
| 0 | 4.57598851 |
| 390.863848 | 173.887563 |
| 0 | 0 |
| 0 | 0 |
| 0 | 64.0638392 |
| 0 | 0 |
| 0 | 0 |
| 0 | 0 |
| 0 | 0 |
| 149.288275 | 0 |
| 0 | 0 |
| 0 | 0 |
| 0 | 0 |
| 0 | 0 |
| 0 | 0 |
| 0 | 0 |
| 0 | 0 |
| 76.0013038 | 54.9118621 |

| | |
|---|---|
| 0 | 0 |
| 0 | 0 |
| 0 | 0 |
| 0 | 0 |
| 0 | 0 |
| 0 | 0 |
| 0 | 0 |
| 0 | 0 |
| 0 | 0 |
| 2.71433228 | 18.303954 |
| 200.860589 | 73.2158162 |
| 0 | 0 |
| 0 | 0 |
| 0 | 0 |
| 738.29838 | 260.831345 |
| 27.1433228 | 4.57598851 |
| 0 | 0 |
| 0 | 0 |
| 0 | 22.8799426 |
| 0 | 0 |
| 149.288275 | 9.15197702 |
| 0 | 0 |
| 0 | 0 |
| 0 | 45.7598851 |
| 0 | 0 |
| 0 | 0 |
| 5.42866456 | 50.3358736 |
| 0 | 22.8799426 |
| 0 | 0 |
| 0 | 114.399713 |
| 0 | 114.399713 |
| 0 | 0 |
| 0 | 0 |
| 0 | 0 |
| 0 | 0 |
| 0 | 0 |
| 0 | 0 |
| 0 | 0 |
| 59.7153101 | 0 |
| 0 | 36.6079081 |
| 146.573943 | 13.7279655 |
| 0 | 0 |
| 0 | 0 |

| | |
|---|---|
| 0 | 86.9437817 |
| 195.431924 | 9.15197702 |
| 0 | 0 |
| 0 | 0 |
| 0 | 0 |
| 0 | 0 |
| 184.574595 | 0 |
| 0 | 4.57598851 |
| 0 | 0 |
| 5.42866456 | 0 |
| 100.430294 | 205.919483 |
| 0 | 0 |
| 2.71433228 | 0 |
| 0 | 0 |
| 0 | 0 |
| 0 | 0 |
| 0 | 0 |
| 0 | 0 |
| 0 | 0 |
| 0 | 0 |
| 0 | 0 |
| 0 | 0 |
| 0 | 0 |
| 0 | 0 |
| 0 | 0 |
| 0 | 0 |
| 0 | 0 |
| 105.858959 | 0 |
| 0 | 0 |
| 0 | 0 |
| 0 | 0 |
| 0 | 0 |
| 0 | 0 |
| 2.71433228 | 86.9437817 |
| 0 | 0 |
| 2.71433228 | 0 |
| 0 | 0 |
| 2.71433228 | 73.2158162 |
| 0 | 0 |
| 111.287623 | 0 |
| 111.287623 | 0 |
| 328.434206 | 100.671747 |
| 0 | 0 |

| | |
|---|---|
| 13.5716614 | 27.4559311 |
| 0 | 0 |
| 0 | 0 |
| 0 | 0 |
| 0 | 0 |
| 2.71433228 | 0 |
| 40.7149842 | 22.8799426 |
| 0 | 0 |
| 0 | 0 |
| 0 | 18.303954 |
| 0 | 0 |
| 0 | 0 |
| 0 | 0 |
| 0 | 0 |
| 0 | 0 |
| 8.14299683 | 4.57598851 |
| 0 | 0 |
| 0 | 0 |
| 76.0013038 | 91.5197702 |
| 0 | 0 |
| 0 | 0 |
| 0 | 0 |
| 0 | 0 |
| 0 | 32.0319196 |
| 0 | 0 |
| 0 | 4.57598851 |
| 0 | 0 |
| 0 | 0 |
| 0 | 0 |
| 0 | 0 |
| 0 | 0 |
| 0 | 0 |
| 0 | 0 |
| 2.71433228 | 0 |
| 143.859611 | 9.15197702 |
| 13.5716614 | 54.9118621 |
| 16.2859937 | 0 |
| 0 | 0 |
| 0 | 0 |
| 0 | 0 |
| 0 | 0 |
| 0 | 0 |
| 0 | 0 |

| | |
|---|---|
| 0 | 0 |
| 146.573943 | 59.4878506 |
| 0 | 0 |
| 0 | 0 |
| 0 | 0 |
| 10.8573291 | 86.9437817 |
| 0 | 0 |
| 0 | 0 |
| 0 | 0 |
| 0 | 0 |
| 32.5719873 | 32.0319196 |
| 0 | 0 |
| 67.8583069 | 178.463552 |
| 0 | 0 |
| 13.5716614 | 27.4559311 |
| 0 | 0 |
| 46.1436487 | 4.57598851 |
| 2.71433228 | 0 |
| 0 | 0 |
| 0 | 0 |
| 0 | 0 |
| 48.857981 | 205.919483 |
| 0 | 0 |
| 0 | 0 |
| 0 | 13.7279655 |
| 0 | 0 |
| 0 | 0 |
| 0 | 82.3677932 |
| 0 | 13.7279655 |
| 0 | 0 |
| 0 | 0 |
| 0 | 9.15197702 |
| 0 | 9.15197702 |
| 217.146582 | 4.57598851 |
| 0 | 45.7598851 |
| 0 | 0 |
| 0 | 9.15197702 |
| 0 | 0 |
| 0 | 0 |
| 203.574921 | 27.4559311 |
| 0 | 0 |
| 0 | 0 |
| 0 | 132.703667 |

| | |
|---|---|
| 0 | 0 |
| 0 | 0 |
| 0 | 54.9118621 |
| 225.289579 | 0 |
| 0 | 0 |
| 0 | 0 |
| 0 | 0 |
| 0 | 0 |
| 0 | 0 |
| 0 | 105.247736 |
| 0 | 105.247736 |
| 0 | 0 |
| 0 | 0 |
| 0 | 0 |
| 0 | 0 |
| 255.147234 | 27.4559311 |
| 105.858959 | 77.7918047 |
| 0 | 0 |
| 263.290231 | 118.975701 |
| 0 | 0 |
| 0 | 0 |
| 0 | 0 |
| 0 | 0 |
| 0 | 0 |
| 0 | 100.671747 |
| 2.71433228 | 0 |
| 0 | 0 |
| 0 | 0 |
| 0 | 0 |
| 0 | 0 |
| 0 | 0 |
| 0 | 0 |
| 0 | 0 |
| 0 | 0 |
| 0 | 0 |
| 0 | 0 |
| 0 | 0 |
| 0 | 0 |
| 0 | 82.3677932 |
| 0 | 0 |
| 2.71433228 | 4.57598851 |
| 0 | 0 |
| 138.430946 | 73.2158162 |

| | |
|---|---|
| 0 | 0 |
| 0 | 0 |
| 0 | 0 |
| 293.147886 | 64.0638392 |
| 0 | 0 |
| 0 | 0 |
| 0 | 0 |
| 0 | 0 |
| 0 | 0 |
| 0 | 0 |
| 108.573291 | 4.57598851 |
| 0 | 41.1838966 |
| 0 | 13.7279655 |
| 10.8573291 | 100.671747 |
| 0 | 0 |
| 0 | 0 |
| 0 | 0 |
| 0 | 0 |
| 0 | 0 |
| 16.2859937 | 0 |
| 0 | 0 |
| 0 | 0 |
| 10.8573291 | 45.7598851 |
| 0 | 0 |
| 0 | 0 |
| 0 | 0 |
| 70.5726392 | 0 |
| 2.71433228 | 0 |
| 0 | 0 |
| 415.292839 | 32.0319196 |
| 0 | 0 |
| 0 | 0 |
| 0 | 0 |
| 0 | 0 |
| 54.2866456 | 9.15197702 |
| 5.42866456 | 86.9437817 |
| 0 | 0 |
| 198.146256 | 13.7279655 |
| 43.4293164 | 50.3358736 |
| 0 | 0 |
| 0 | 0 |
| 0 | 0 |
| 0 | 0 |

| | |
|---|---|
| 0 | 0 |
| 0 | 0 |
| 0 | 27.4559311 |
| 0 | 0 |
| 0 | 73.2158162 |
| 0 | 0 |
| 0 | 0 |
| 0 | 0 |
| 176.431598 | 9.15197702 |
| 0 | 0 |
| 0 | 0 |
| 0 | 0 |
| 0 | 0 |
| 0 | 0 |
| 2.71433228 | 0 |
| 84.1443006 | 82.3677932 |
| 0 | 41.1838966 |
| 0 | 0 |
| 46.1436487 | 0 |
| 0 | 0 |
| 0 | 0 |
| 209.003585 | 18.303954 |
| 2.71433228 | 36.6079081 |
| 0 | 45.7598851 |
| 0 | 0 |
| 0 | 0 |
| 0 | 0 |
| 0 | 0 |
| 0 | 0 |
| 0 | 100.671747 |
| 0 | 100.671747 |
| 0 | 100.671747 |
| 0 | 100.671747 |
| 0 | 100.671747 |
| 0 | 0 |
| 0 | 73.2158162 |
| 0 | 0 |
| 0 | 0 |
| 0 | 0 |
| 0 | 0 |
| 0 | 0 |
| 0 | 0 |
| 2.71433228 | 50.3358736 |

| | |
|---|---|
| 0 | 0 |
| 27.1433228 | 205.919483 |
| 0 | 4.57598851 |
| 114.001956 | 27.4559311 |
| 0 | 0 |
| 0 | 0 |
| 0 | 0 |
| 0 | 0 |
| 0 | 0 |
| 0 | 0 |
| 0 | 32.0319196 |
| 78.7156361 | 123.55169 |
| 19.0003259 | 59.4878506 |
| 0 | 0 |
| 0 | 0 |
| 0 | 0 |
| 0 | 0 |
| 0 | 0 |
| 0 | 0 |
| 5.42866456 | 41.1838966 |
| 0 | 0 |
| 0 | 0 |
| 0 | 0 |
| 0 | 0 |
| 0 | 0 |
| 0 | 0 |
| 0 | 0 |
| 0 | 0 |
| 0 | 0 |
| 0 | 68.6398277 |
| 0 | 0 |
| 0 | 0 |
| 0 | 0 |
| 13.5716614 | 82.3677932 |
| 152.002608 | 77.7918047 |
| 0 | 0 |
| 0 | 0 |
| 0 | 0 |
| 100.430294 | 4.57598851 |
| 0 | 0 |
| 0 | 0 |
| 0 | 0 |
| 97.715962 | 0 |

| | |
|---|---|
| 0 | 0 |
| 0 | 0 |
| 0 | 0 |
| 0 | 0 |
| 0 | 0 |
| 0 | 0 |
| 0 | 0 |
| 0 | 0 |
| 0 | 0 |
| 0 | 0 |
| 0 | 0 |
| 0 | 59.4878506 |
| 0 | 0 |
| 0 | 0 |
| 0 | 0 |
| 0 | 0 |
| 0 | 0 |
| 0 | 0 |
| 0 | 0 |
| 0 | 0 |
| 0 | 0 |
| 0 | 4.57598851 |
| 0 | 0 |
| 57.0009778 | 146.431632 |
| 206.289253 | 128.127678 |
| 0 | 0 |
| 0 | 13.7279655 |
| 5.42866456 | 41.1838966 |
| 0 | 0 |
| 0 | 0 |
| 10.8573291 | 123.55169 |
| 0 | 0 |
| 10.8573291 | 151.007621 |
| 0 | 0 |
| 0 | 0 |
| 2.71433228 | 18.303954 |
| 0 | 0 |
| 0 | 9.15197702 |
| 0 | 9.15197702 |
| 0 | 0 |
| 0 | 0 |
| 10.8573291 | 45.7598851 |
| 0 | 0 |

| | |
|---|---|
| 0 | 0 |
| 0 | 4.57598851 |
| 0 | 0 |
| 439.721829 | 9.15197702 |
| 0 | 0 |
| 0 | 0 |
| 0 | 9.15197702 |
| 2.71433228 | 4.57598851 |
| 0 | 0 |
| 0 | 0 |
| 0 | 96.0957587 |
| 0 | 96.0957587 |
| 0 | 4.57598851 |
| 8.14299683 | 36.6079081 |
| 0 | 0 |
| 0 | 0 |
| 0 | 0 |
| 0 | 0 |
| 2.71433228 | 0 |
| 2.71433228 | 0 |
| 2.71433228 | 0 |
| 95.0016297 | 0 |
| 0 | 0 |
| 0 | 0 |
| 0 | 0 |
| 0 | 0 |
| 0 | 13.7279655 |
| 0 | 0 |
| 0 | 0 |
| 0 | 0 |
| 0 | 0 |
| 0 | 64.0638392 |
| 0 | 0 |
| 171.002934 | 86.9437817 |
| 0 | 45.7598851 |
| 0 | 0 |
| 0 | 0 |
| 0 | 0 |
| 103.144627 | 13.7279655 |
| 0 | 0 |
| 0 | 0 |
| 0 | 0 |
| 0 | 0 |

| | |
|---|---|
| 0 | 0 |
| 0 | 0 |
| 0 | 0 |
| 0 | 0 |
| 0 | 0 |
| 0 | 0 |
| 48.857981 | 526.238679 |
| 0 | 0 |
| 0 | 0 |
| 70.5726392 | 0 |
| 0 | 0 |
| 0 | 0 |
| 0 | 0 |
| 0 | 0 |
| 0 | 0 |
| 0 | 0 |
| 0 | 0 |
| 0 | 0 |
| 0 | 0 |
| 0 | 0 |
| 10.8573291 | 4.57598851 |
| 0 | 0 |
| 0 | 0 |
| 0 | 0 |
| 0 | 0 |
| 0 | 18.303954 |
| 0 | 0 |
| 0 | 0 |
| 0 | 0 |
| 0 | 0 |
| 0 | 0 |
| 0 | 0 |
| 0 | 0 |
| 0 | 0 |
| 0 | 0 |
| 16.2859937 | 41.1838966 |
| 0 | 0 |
| 0 | 0 |
| 86.8586329 | 233.375414 |
| 0 | 0 |
| 377.292187 | 45.7598851 |
| 0 | 0 |
| 29.8576551 | 13.7279655 |

| | |
|---|---|
| 92.2872974 | 0 |
| 0 | 0 |
| 92.2872974 | 0 |
| 0 | 0 |
| 228.003911 | 13.7279655 |
| 0 | 0 |
| 0 | 4.57598851 |
| 0 | 0 |
| 0 | 0 |
| 407.149842 | 151.007621 |
| 0 | 96.0957587 |
| 0 | 0 |
| 0 | 0 |
| 0 | 0 |
| 0 | 0 |
| 0 | 0 |
| 0 | 4.57598851 |
| 0 | 0 |
| 0 | 0 |
| 0 | 0 |
| 0 | 73.2158162 |
| 0 | 0 |
| 0 | 0 |
| 0 | 0 |
| 65.1439747 | 151.007621 |
| 0 | 0 |
| 0 | 0 |
| 0 | 91.5197702 |
| 0 | 0 |
| 0 | 64.0638392 |
| 0 | 0 |
| 0 | 0 |
| 0 | 0 |
| 0 | 0 |
| 0 | 0 |
| 0 | 0 |
| 0 | 0 |
| 0 | 0 |
| 0 | 0 |
| 0 | 0 |
| 534.723459 | 114.399713 |
| 0 | 0 |
| 0 | 0 |

| | |
|---|---|
| 0 | 0 |
| 0 | 0 |
| 27.1433228 | 0 |
| 0 | 0 |
| 5.42866456 | 13.7279655 |
| 0 | 4.57598851 |
| 0 | 0 |
| 0 | 0 |
| 0 | 22.8799426 |
| 0 | 0 |
| 0 | 0 |
| 0 | 0 |
| 0 | 0 |
| 89.5729652 | 0 |
| 89.5729652 | 0 |
| 89.5729652 | 0 |
| 89.5729652 | 0 |
| 27.1433228 | 54.9118621 |
| 0 | 0 |
| 0 | 0 |
| 0 | 0 |
| 0 | 0 |
| 0 | 0 |
| 0 | 0 |
| 0 | 0 |
| 67.8583069 | 27.4559311 |
| 0 | 0 |
| 0 | 0 |
| 211.717918 | 100.671747 |
| 0 | 0 |
| 0 | 0 |
| 0 | 0 |
| 0 | 0 |
| 0 | 0 |
| 0 | 0 |
| 0 | 0 |
| 0 | 0 |
| 0 | 0 |
| 0 | 146.431632 |
| 0 | 0 |
| 0 | 0 |
| 0 | 0 |
| 116.716288 | 9.15197702 |

| | |
|---|---|
| 0 | 0 |
| 0 | 0 |
| 0 | 0 |
| 0 | 68.6398277 |
| 0 | 0 |
| 0 | 0 |
| 51.5723133 | 13.7279655 |
| 0 | 0 |
| 0 | 0 |
| 5.42866456 | 77.7918047 |
| 5.42866456 | 0 |
| 0 | 0 |
| 0 | 0 |
| 0 | 0 |
| 21.7146582 | 22.8799426 |
| 0 | 0 |
| 0 | 0 |
| 2.71433228 | 0 |
| 0 | 0 |
| 0 | 0 |
| 0 | 0 |
| 0 | 0 |
| 0 | 0 |
| 29.8576551 | 9.15197702 |
| 0 | 0 |
| 0 | 0 |
| 0 | 0 |
| 0 | 0 |
| 0 | 0 |
| 0 | 0 |
| 0 | 196.767506 |
| 0 | 0 |
| 0 | 0 |
| 0 | 0 |
| 0 | 64.0638392 |
| 0 | 0 |
| 0 | 0 |
| 0 | 0 |
| 86.8586329 | 0 |
| 0 | 0 |
| 0 | 77.7918047 |
| 21.7146582 | 9.15197702 |
| 0 | 0 |

| | |
|---|---|
| 0 | 0 |
| 0 | 0 |
| 0 | 0 |
| 0 | 0 |
| 0 | 0 |
| 2.71433228 | 45.7598851 |
| 0 | 0 |
| 187.288927 | 0 |
| 2.71433228 | 0 |
| 0 | 86.9437817 |
| 0 | 86.9437817 |
| 0 | 86.9437817 |
| 0 | 86.9437817 |
| 0 | 86.9437817 |
| 0 | 0 |
| 8.14299683 | 114.399713 |
| 0 | 0 |
| 0 | 0 |
| 0 | 0 |
| 0 | 0 |
| 263.290231 | 18.303954 |
| 0 | 0 |
| 0 | 18.303954 |
| 0 | 0 |
| 86.8586329 | 0 |
| 0 | 0 |
| 0 | 36.6079081 |
| 8.14299683 | 109.823724 |
| 141.145278 | 50.3358736 |
| 32.5719873 | 4.57598851 |
| 0 | 0 |
| 0 | 0 |
| 0 | 0 |
| 0 | 0 |
| 0 | 0 |
| 0 | 36.6079081 |
| 0 | 0 |
| 2.71433228 | 4.57598851 |
| 0 | 0 |
| 0 | 0 |
| 0 | 0 |
| 0 | 0 |
| 0 | 0 |

| | |
|---|---|
| 0 | 0 |
| 127.573617 | 4.57598851 |
| 0 | 0 |
| 0 | 4.57598851 |
| 0 | 0 |
| 0 | 0 |
| 0 | 0 |
| 0 | 0 |
| 0 | 0 |
| 0 | 0 |
| 0 | 0 |
| 0 | 0 |
| 0 | 0 |
| 0 | 50.3358736 |
| 0 | 0 |
| 0 | 0 |
| 0 | 0 |
| 0 | 0 |
| 29.8576551 | 68.6398277 |
| 0 | 0 |
| 0 | 0 |
| 0 | 0 |
| 0 | 0 |
| 0 | 54.9118621 |
| 0 | 0 |
| 0 | 0 |
| 0 | 0 |
| 0 | 13.7279655 |
| 211.717918 | 96.0957587 |
| 0 | 0 |
| 0 | 0 |
| 2.71433228 | 9.15197702 |
| 0 | 0 |
| 0 | 4.57598851 |
| 0 | 0 |
| 0 | 0 |
| 0 | 45.7598851 |
| 0 | 0 |
| 16.2859937 | 100.671747 |
| 0 | 0 |
| 0 | 0 |
| 0 | 0 |
| 0 | 0 |

| | |
|---|---|
| 149.288275 | 13.7279655 |
| 0 | 0 |
| 54.2866456 | 0 |
| 0 | 0 |
| 0 | 0 |
| 0 | 0 |
| 5.42866456 | 41.1838966 |
| 78.7156361 | 4.57598851 |
| 0 | 0 |
| 0 | 0 |
| 0 | 0 |
| 0 | 0 |
| 0 | 0 |
| 0 | 0 |
| 0 | 0 |
| 0 | 0 |
| 0 | 0 |
| 0 | 0 |
| 222.575247 | 0 |
| 0 | 0 |
| 0 | 0 |
| 0 | 0 |
| 0 | 0 |
| 40.7149842 | 0 |
| 0 | 0 |
| 0 | 0 |
| 0 | 68.6398277 |
| 0 | 0 |
| 0 | 0 |
| 0 | 0 |
| 0 | 54.9118621 |
| 0 | 0 |
| 0 | 0 |
| 0 | 73.2158162 |
| 0 | 0 |
| 57.0009778 | 13.7279655 |
| 0 | 0 |
| 2.71433228 | 0 |
| 0 | 82.3677932 |
| 0 | 82.3677932 |
| 0 | 82.3677932 |
| 0 | 82.3677932 |
| 0 | 82.3677932 |

| | |
|---|---|
| 0 | 82.3677932 |
| 0 | 0 |
| 0 | 82.3677932 |
| 0 | 82.3677932 |
| 0 | 82.3677932 |
| 0 | 0 |
| 0 | 0 |
| 0 | 0 |
| 0 | 0 |
| 0 | 0 |
| 0 | 0 |
| 0 | 0 |
| 0 | 0 |
| 0 | 0 |
| 0 | 0 |
| 0 | 0 |
| 81.4299683 | 0 |
| 81.4299683 | 0 |
| 0 | 0 |
| 10.8573291 | 4.57598851 |
| 198.146256 | 50.3358736 |
| 0 | 0 |
| 0 | 0 |
| 0 | 0 |
| 0 | 0 |
| 0 | 0 |
| 0 | 0 |
| 0 | 0 |
| 0 | 0 |
| 0 | 0 |
| 0 | 0 |
| 0 | 0 |
| 0 | 0 |
| 0 | 0 |
| 162.859937 | 32.0319196 |
| 0 | 0 |
| 0 | 86.9437817 |
| 0 | 0 |
| 0 | 0 |
| 0 | 0 |
| 1055.87526 | 457.598851 |
| 111.287623 | 86.9437817 |
| 0 | 0 |

| | |
|---|---|
| 0 | 0 |
| 0 | 0 |
| 0 | 0 |
| 0 | 0 |
| 0 | 0 |
| 0 | 4.57598851 |
| 0 | 0 |
| 0 | 0 |
| 135.716614 | 32.0319196 |
| 0 | 0 |
| 0 | 0 |
| 0 | 0 |
| 46.1436487 | 36.6079081 |
| 0 | 0 |
| 241.575573 | 0 |
| 0 | 0 |
| 0 | 22.8799426 |
| 0 | 0 |
| 2.71433228 | 0 |
| 0 | 0 |
| 0 | 0 |
| 0 | 0 |
| 0 | 0 |
| 0 | 0 |
| 0 | 0 |
| 0 | 0 |
| 0 | 45.7598851 |
| 0 | 0 |
| 0 | 0 |
| 0 | 0 |
| 0 | 0 |
| 0 | 4.57598851 |
| 0 | 0 |
| 0 | 0 |
| 100.430294 | 0 |
| 0 | 0 |
| 0 | 0 |
| 0 | 0 |
| 5.42866456 | 160.159598 |
| 0 | 36.6079081 |
| 0 | 0 |
| 0 | 0 |
| 0 | 0 |

| | |
|---|---|
| 0 | 0 |
| 0 | 22.8799426 |
| 84.1443006 | 0 |
| 0 | 0 |
| 0 | 0 |
| 0 | 68.6398277 |
| 0 | 0 |
| 2.71433228 | 9.15197702 |
| 0 | 0 |
| 0 | 0 |
| 73.2869715 | 0 |
| 46.1436487 | 0 |
| 0 | 0 |
| 0 | 4.57598851 |
| 0 | 9.15197702 |
| 0 | 0 |
| 0 | 0 |
| 0 | 0 |
| 0 | 4.57598851 |
| 0 | 0 |
| 233.432576 | 22.8799426 |
| 0 | 0 |
| 0 | 0 |
| 0 | 0 |
| 0 | 0 |
| 32.5719873 | 13.7279655 |
| 0 | 64.0638392 |
| 0 | 0 |
| 5.42866456 | 27.4559311 |
| 0 | 0 |
| 0 | 0 |
| 0 | 0 |
| 0 | 0 |
| 0 | 0 |
| 0 | 0 |
| 0 | 77.7918047 |
| 0 | 77.7918047 |
| 0 | 77.7918047 |
| 0 | 77.7918047 |
| 0 | 77.7918047 |
| 0 | 0 |
| 0 | 0 |
| 0 | 0 |

| | |
|---|---|
| 0 | 0 |
| 78.7156361 | 581.150541 |
| 0 | 4.57598851 |
| 0 | 77.7918047 |
| 0 | 0 |
| 0 | 9.15197702 |
| 73.2869715 | 27.4559311 |
| 8.14299683 | 0 |
| 0 | 0 |
| 27.1433228 | 9.15197702 |
| 0 | 4.57598851 |
| 0 | 0 |
| 114.001956 | 13.7279655 |
| 2.71433228 | 54.9118621 |
| 86.8586329 | 0 |
| 0 | 0 |
| 0 | 0 |
| 0 | 0 |
| 0 | 0 |
| 0 | 0 |
| 0 | 0 |
| 0 | 0 |
| 0 | 0 |
| 222.575247 | 297.439253 |
| 29.8576551 | 105.247736 |
| 0 | 4.57598851 |
| 0 | 36.6079081 |
| 2.71433228 | 0 |
| 0 | 0 |
| 0 | 0 |
| 0 | 0 |
| 483.151145 | 183.03954 |
| 2.71433228 | 0 |
| 0 | 0 |
| 0 | 0 |
| 0 | 0 |
| 0 | 0 |
| 0 | 0 |
| 46.1436487 | 4836.81986 |
| 0 | 0 |
| 0 | 0 |
| 0 | 0 |
| 0 | 0 |

| | |
|---|---|
| 0 | 0 |
| 0 | 0 |
| 0 | 0 |
| 0 | 50.3358736 |
| 0 | 27.4559311 |
| 0 | 0 |
| 13.5716614 | 0 |
| 0 | 0 |
| 323.005541 | 0 |
| 0 | 0 |
| 0 | 0 |
| 29.8576551 | 4.57598851 |
| 0 | 82.3677932 |
| 0 | 50.3358736 |
| 241.575573 | 0 |
| 0 | 0 |
| 0 | 0 |
| 21.7146582 | 164.735586 |
| 2.71433228 | 4.57598851 |
| 0 | 0 |
| 0 | 22.8799426 |
| 0 | 0 |
| 5.42866456 | 0 |
| 0 | 0 |
| 0 | 0 |
| 0 | 0 |
| 0 | 0 |
| 0 | 0 |
| 0 | 0 |
| 5.42866456 | 178.463552 |
| 0 | 0 |
| 0 | 0 |
| 0 | 0 |
| 0 | 0 |
| 0 | 0 |
| 0 | 0 |
| 0 | 0 |
| 122.144953 | 0 |
| 40.7149842 | 59.4878506 |
| 0 | 0 |
| 0 | 0 |
| 0 | 0 |
| 0 | 0 |

| | |
|---|---|
| 0 | 0 |
| 0 | 0 |
| 10.8573291 | 0 |
| 0 | 0 |
| 0 | 0 |
| 0 | 0 |
| 0 | 0 |
| 0 | 0 |
| 0 | 0 |
| 0 | 0 |
| 0 | 0 |
| 81.4299683 | 0 |
| 0 | 45.7598851 |
| 200.860589 | 54.9118621 |
| 610.724763 | 9.15197702 |
| 0 | 0 |
| 0 | 0 |
| 2.71433228 | 100.671747 |
| 0 | 0 |
| 0 | 0 |
| 73.2869715 | 0 |
| 0 | 0 |
| 0 | 0 |
| 0 | 0 |
| 0 | 0 |
| 8.14299683 | 22.8799426 |
| 0 | 0 |
| 2.71433228 | 50.3358736 |
| 0 | 0 |
| 2.71433228 | 4.57598851 |
| 0 | 0 |
| 0 | 32.0319196 |
| 0 | 0 |
| 0 | 0 |
| 0 | 0 |
| 2.71433228 | 0 |
| 0 | 0 |
| 0 | 0 |
| 0 | 0 |
| 2.71433228 | 9.15197702 |
| 0 | 4.57598851 |
| 0 | 0 |
| 0 | 0 |

| | |
|---|---|
| 0 | 68.6398277 |
| 0 | 0 |
| 0 | 0 |
| 2.71433228 | 86.9437817 |
| 0 | 0 |
| 0 | 73.2158162 |
| 0 | 73.2158162 |
| 0 | 73.2158162 |
| 0 | 73.2158162 |
| 0 | 0 |
| 13.5716614 | 0 |
| 0 | 0 |
| 0 | 0 |
| 16.2859937 | 50.3358736 |
| 0 | 0 |
| 230.718244 | 9.15197702 |
| 0 | 0 |
| 0 | 0 |
| 0 | 0 |
| 0 | 0 |
| 0 | 0 |
| 0 | 0 |
| 100.430294 | 0 |
| 0 | 0 |
| 2.71433228 | 0 |
| 0 | 0 |
| 0 | 18.303954 |
| 0 | 0 |
| 0 | 0 |
| 0 | 0 |
| 0 | 0 |
| 0 | 0 |
| 2.71433228 | 0 |
| 0 | 0 |
| 0 | 0 |
| 0 | 0 |
| 0 | 0 |
| 0 | 0 |
| 0 | 0 |
| 21.7146582 | 4.57598851 |
| 0 | 0 |
| 0 | 0 |
| 0 | 0 |

| | |
|---|---|
| 0 | 0 |
| 0 | 0 |
| 0 | 0 |
| 0 | 0 |
| 0 | 0 |
| 0 | 0 |
| 0 | 0 |
| 0 | 41.1838966 |
| 0 | 73.2158162 |
| 0 | 9.15197702 |
| 0 | 0 |
| 0 | 0 |
| 0 | 0 |
| 0 | 0 |
| 43.4293164 | 0 |
| 0 | 0 |
| 0 | 0 |
| 0 | 50.3358736 |
| 0 | 0 |
| 0 | 41.1838966 |
| 0 | 0 |
| 0 | 0 |
| 0 | 0 |
| 13.5716614 | 118.975701 |
| 0 | 0 |
| 0 | 0 |
| 0 | 0 |
| 0 | 0 |
| 0 | 0 |
| 0 | 41.1838966 |
| 0 | 0 |
| 0 | 0 |
| 0 | 0 |
| 0 | 27.4559311 |
| 10.8573291 | 128.127678 |
| 0 | 0 |
| 0 | 0 |
| 0 | 0 |
| 0 | 0 |
| 2.71433228 | 105.247736 |
| 0 | 0 |
| 0 | 0 |
| 0 | 0 |

| | |
|---|---|
| 0 | 64.0638392 |
| 0 | 0 |
| 0 | 0 |
| 0 | 0 |
| 0 | 0 |
| 2.71433228 | 4.57598851 |
| 0 | 0 |
| 0 | 73.2158162 |
| 0 | 0 |
| 0 | 32.0319196 |
| 0 | 0 |
| 0 | 0 |
| 0 | 0 |
| 0 | 0 |
| 0 | 0 |
| 0 | 0 |
| 0 | 0 |
| 27.1433228 | 22.8799426 |
| 1495.59709 | 77.7918047 |
| 0 | 54.9118621 |
| 0 | 0 |
| 0 | 4.57598851 |
| 0 | 0 |
| 0 | 0 |
| 0 | 0 |
| 0 | 64.0638392 |
| 0 | 0 |
| 0 | 77.7918047 |
| 0 | 0 |
| 0 | 0 |
| 0 | 0 |
| 0 | 0 |
| 0 | 0 |
| 0 | 0 |
| 0 | 0 |
| 0 | 0 |
| 0 | 0 |
| 0 | 0 |
| 0 | 0 |
| 0 | 0 |
| 184.574595 | 132.703667 |
| 0 | 54.9118621 |
| 0 | 0 |

| | |
|---|---|
| 0 | 0 |
| 0 | 0 |
| 0 | 0 |
| 0 | 0 |
| 0 | 0 |
| 0 | 0 |
| 0 | 0 |
| 0 | 0 |
| 0 | 0 |
| 0 | 109.823724 |
| 21.7146582 | 32.0319196 |
| 893.015319 | 118.975701 |
| 0 | 22.8799426 |
| 0 | 13.7279655 |
| 0 | 105.247736 |
| 0 | 0 |
| 138.430946 | 54.9118621 |
| 0 | 0 |
| 0 | 0 |
| 0 | 0 |
| 0 | 0 |
| 0 | 0 |
| 0 | 36.6079081 |
| 0 | 0 |
| 352.863196 | 128.127678 |
| 0 | 68.6398277 |
| 0 | 68.6398277 |
| 0 | 68.6398277 |
| 0 | 68.6398277 |
| 0 | 68.6398277 |
| 0 | 68.6398277 |
| 0 | 68.6398277 |
| 0 | 68.6398277 |
| 0 | 68.6398277 |
| 0 | 0 |
| 0 | 0 |
| 0 | 68.6398277 |
| 0 | 68.6398277 |
| 0 | 68.6398277 |
| 0 | 4.57598851 |
| 0 | 0 |
| 0 | 0 |
| 0 | 0 |

| | |
|---|---|
| 0 | 0 |
| 0 | 0 |
| 0 | 0 |
| 0 | 0 |
| 0 | 0 |
| 0 | 0 |
| 67.8583069 | 0 |
| 0 | 0 |
| 0 | 0 |
| 0 | 0 |
| 0 | 4.57598851 |
| 2.71433228 | 0 |
| 0 | 0 |
| 0 | 0 |
| 0 | 0 |
| 0 | 0 |
| 0 | 0 |
| 62.4296424 | 105.247736 |
| 0 | 0 |
| 0 | 0 |
| 0 | 0 |
| 0 | 0 |
| 0 | 0 |
| 0 | 0 |
| 0 | 0 |
| 0 | 0 |
| 0 | 0 |
| 0 | 0 |
| 0 | 0 |
| 0 | 0 |
| 0 | 0 |
| 0 | 0 |
| 0 | 0 |
| 0 | 0 |
| 0 | 0 |
| 2.71433228 | 0 |
| 43.4293164 | 4.57598851 |
| 0 | 0 |
| 0 | 0 |
| 0 | 0 |
| 0 | 18.303954 |
| 0 | 0 |
| 0 | 0 |

| | |
|---|---|
| 2.71433228 | 86.9437817 |
| 0 | 68.6398277 |
| 0 | 0 |
| 0 | 0 |
| 0 | 59.4878506 |
| 179.14593 | 18.303954 |
| 62.4296424 | 18.303954 |
| 0 | 0 |
| 0 | 0 |
| 0 | 0 |
| 0 | 0 |
| 0 | 0 |
| 0 | 0 |
| 0 | 0 |
| 0 | 77.7918047 |
| 0 | 0 |
| 0 | 77.7918047 |
| 2.71433228 | 0 |
| 0 | 0 |
| 0 | 0 |
| 2.71433228 | 18.303954 |
| 0 | 64.0638392 |
| 0 | 0 |
| 0 | 0 |
| 0 | 0 |
| 0 | 0 |
| 157.431272 | 18.303954 |
| 0 | 0 |
| 0 | 0 |
| 0 | 0 |
| 0 | 0 |
| 0 | 50.3358736 |
| 78.7156361 | 54.9118621 |
| 0 | 0 |
| 0 | 0 |
| 0 | 0 |
| 5.42866456 | 22.8799426 |
| 0 | 0 |
| 0 | 0 |
| 0 | 0 |
| 0 | 0 |
| 2.71433228 | 9.15197702 |
| 0 | 0 |

| | |
|---|---|
| 0 | 0 |
| 0 | 0 |
| 0 | 0 |
| 0 | 0 |
| 0 | 0 |
| 105.858959 | 160.159598 |
| 0 | 0 |
| 0 | 0 |
| 0 | 0 |
| 0 | 0 |
| 62.4296424 | 18.303954 |
| 0 | 0 |
| 0 | 0 |
| 0 | 13.7279655 |
| 65.1439747 | 0 |
| 65.1439747 | 0 |
| 5.42866456 | 0 |
| 27.1433228 | 68.6398277 |
| 0 | 0 |
| 0 | 0 |
| 0 | 0 |
| 2.71433228 | 0 |
| 0 | 0 |
| 0 | 54.9118621 |
| 0 | 0 |
| 0 | 0 |
| 2.71433228 | 4.57598851 |
| 0 | 0 |
| 0 | 0 |
| 0 | 0 |
| 0 | 0 |
| 0 | 0 |
| 32.5719873 | 4.57598851 |
| 0 | 0 |
| 0 | 100.671747 |
| 0 | 18.303954 |
| 0 | 4.57598851 |
| 0 | 0 |
| 0 | 0 |
| 0 | 27.4559311 |
| 130.287949 | 219.647449 |
| 0 | 0 |
| 0 | 50.3358736 |

| | |
|---|---|
| 241.575573 | 123.55169 |
| 0 | 0 |
| 0 | 0 |
| 0 | 0 |
| 0 | 0 |
| 0 | 0 |
| 0 | 0 |
| 0 | 0 |
| 0 | 0 |
| 0 | 41.1838966 |
| 0 | 0 |
| 0 | 0 |
| 0 | 0 |
| 2.71433228 | 4.57598851 |
| 2.71433228 | 4.57598851 |
| 0 | 0 |
| 0 | 0 |
| 0 | 0 |
| 0 | 45.7598851 |
| 0 | 0 |
| 0 | 0 |
| 2.71433228 | 4.57598851 |
| 0 | 0 |
| 152.002608 | 0 |
| 0 | 0 |
| 0 | 73.2158162 |
| 0 | 73.2158162 |
| 0 | 0 |
| 0 | 0 |
| 0 | 4.57598851 |
| 0 | 0 |
| 111.287623 | 4.57598851 |
| 0 | 0 |
| 0 | 0 |
| 0 | 0 |
| 0 | 64.0638392 |
| 0 | 64.0638392 |
| 0 | 0 |
| 0 | 64.0638392 |
| 0 | 64.0638392 |
| 0 | 64.0638392 |
| 0 | 64.0638392 |
| 0 | 64.0638392 |

| | |
|---|---|
| 0 | 0 |
| 0 | 0 |
| 0 | 0 |
| 0 | 0 |
| 0 | 0 |
| 0 | 0 |
| 0 | 27.4559311 |
| 0 | 0 |
| 0 | 0 |
| 0 | 59.4878506 |
| 0 | 0 |
| 0 | 0 |
| 0 | 0 |
| 0 | 0 |
| 0 | 0 |
| 0 | 0 |
| 0 | 0 |
| 233.432576 | 0 |
| 2.71433228 | 13.7279655 |
| 637.868085 | 279.135299 |
| 0 | 0 |
| 0 | 0 |
| 86.8586329 | 4.57598851 |
| 0 | 0 |
| 211.717918 | 260.831345 |
| 0 | 13.7279655 |
| 0 | 0 |
| 0 | 0 |
| 0 | 0 |
| 0 | 0 |
| 0 | 0 |
| 0 | 0 |
| 54.2866456 | 4.57598851 |
| 0 | 0 |
| 0 | 0 |
| 0 | 0 |
| 0 | 0 |
| 0 | 0 |
| 8.14299683 | 18.303954 |
| 0 | 0 |
| 0 | 0 |
| 0 | 18.303954 |
| 0 | 0 |

| | |
|---|---|
| 0 | 0 |
| 0 | 0 |
| 0 | 0 |
| 62.4296424 | 0 |
| 62.4296424 | 0 |
| 62.4296424 | 0 |
| 0 | 4.57598851 |
| 0 | 4.57598851 |
| 0 | 0 |
| 0 | 0 |
| 67.8583069 | 0 |
| 0 | 0 |
| 0 | 4.57598851 |
| 0 | 73.2158162 |
| 0 | 22.8799426 |
| 499.437139 | 224.223437 |
| 0 | 22.8799426 |
| 0 | 0 |
| 0 | 0 |
| 16.2859937 | 77.7918047 |
| 0 | 0 |
| 0 | 0 |
| 0 | 0 |
| 0 | 0 |
| 0 | 0 |
| 0 | 0 |
| 0 | 0 |
| 0 | 0 |
| 38.0006519 | 0 |
| 0 | 0 |
| 0 | 0 |
| 0 | 0 |
| 0 | 0 |
| 0 | 36.6079081 |
| 0 | 0 |
| 16.2859937 | 45.7598851 |
| 0 | 68.6398277 |
| 2.71433228 | 0 |
| 0 | 0 |
| 0 | 0 |
| 0 | 0 |
| 10.8573291 | 0 |
| 0 | 0 |

| | |
|---|---|
| 0 | 0 |
| 0 | 0 |
| 5.42866456 | 22.8799426 |
| 0 | 0 |
| 100.430294 | 311.167219 |
| 0 | 0 |
| 0 | 0 |
| 0 | 0 |
| 0 | 50.3358736 |
| 0 | 105.247736 |
| 0 | 0 |
| 0 | 0 |
| 0 | 0 |
| 0 | 0 |
| 0 | 0 |
| 0 | 0 |
| 0 | 0 |
| 0 | 0 |
| 2.71433228 | 54.9118621 |
| 0 | 0 |
| 0 | 0 |
| 0 | 0 |
| 0 | 0 |
| 0 | 4.57598851 |
| 0 | 0 |
| 0 | 0 |
| 0 | 0 |
| 0 | 0 |
| 230.718244 | 18.303954 |
| 0 | 0 |
| 0 | 82.3677932 |
| 0 | 0 |
| 0 | 9.15197702 |
| 0 | 0 |
| 0 | 0 |
| 119.43062 | 4.57598851 |
| 0 | 0 |
| 10.8573291 | 9.15197702 |
| 0 | 0 |
| 0 | 0 |
| 0 | 0 |
| 0 | 0 |
| 0 | 0 |

| | |
|---|---|
| 81.4299683 | 388.959023 |
| 0 | 0 |
| 84.1443006 | 0 |
| 0 | 0 |
| 0 | 0 |
| 0 | 0 |
| 0 | 0 |
| 2.71433228 | 4.57598851 |
| 0 | 0 |
| 0 | 0 |
| 0 | 0 |
| 396.292513 | 59.4878506 |
| 59.7153101 | 0 |
| 59.7153101 | 0 |
| 59.7153101 | 0 |
| 0 | 0 |
| 0 | 0 |
| 0 | 0 |
| 0 | 0 |
| 162.859937 | 0 |
| 0 | 0 |
| 19.0003259 | 73.2158162 |
| 0 | 0 |
| 0 | 0 |
| 0 | 0 |
| 0 | 9.15197702 |
| 0 | 0 |
| 0 | 0 |
| 0 | 73.2158162 |
| 0 | 0 |
| 0 | 0 |
| 0 | 0 |
| 0 | 0 |
| 336.577202 | 18.303954 |
| 0 | 0 |
| 51.5723133 | 64.0638392 |
| 0 | 59.4878506 |
| 0 | 59.4878506 |
| 0 | 0 |
| 0 | 59.4878506 |
| 0 | 59.4878506 |
| 0 | 59.4878506 |
| 0 | 59.4878506 |

| | |
|---|---|
| 0 | 59.4878506 |
| 0 | 59.4878506 |
| 0 | 59.4878506 |
| 0 | 0 |
| 0 | 0 |
| 0 | 0 |
| 0 | 0 |
| 0 | 0 |
| 0 | 0 |
| 0 | 0 |
| 0 | 0 |
| 0 | 18.303954 |
| 0 | 4.57598851 |
| 0 | 0 |
| 0 | 0 |
| 0 | 4.57598851 |
| 0 | 0 |
| 0 | 4.57598851 |
| 0 | 0 |
| 0 | 0 |
| 0 | 0 |
| 19.0003259 | 0 |
| 0 | 4.57598851 |
| 0 | 0 |
| 0 | 0 |
| 0 | 0 |
| 0 | 0 |
| 67.8583069 | 132.703667 |
| 0 | 22.8799426 |
| 0 | 64.0638392 |
| 0 | 0 |
| 0 | 0 |
| 0 | 0 |
| 2.71433228 | 0 |
| 0 | 0 |
| 0 | 0 |
| 0 | 0 |
| 0 | 0 |
| 0 | 0 |
| 0 | 0 |
| 0 | 0 |
| 0 | 0 |
| 0 | 0 |

| | |
|---|---|
| 21.7146582 | 160.159598 |
| 0 | 0 |
| 48.857981 | 9.15197702 |
| 0 | 0 |
| 0 | 0 |
| 0 | 105.247736 |
| 0 | 0 |
| 35.2863196 | 22.8799426 |
| 0 | 0 |
| 1536.31207 | 690.974265 |
| 0 | 0 |
| 5.42866456 | 77.7918047 |
| 0 | 0 |
| 0 | 0 |
| 0 | 0 |
| 0 | 4.57598851 |
| 0 | 0 |
| 0 | 0 |
| 0 | 9.15197702 |
| 0 | 0 |
| 0 | 0 |
| 0 | 0 |
| 0 | 59.4878506 |
| 0 | 0 |
| 0 | 0 |
| 57.0009778 | 0 |
| 57.0009778 | 0 |
| 0 | 0 |
| 0 | 0 |
| 0 | 0 |
| 0 | 82.3677932 |
| 0 | 0 |
| 0 | 0 |
| 0 | 0 |
| 0 | 9.15197702 |
| 0 | 0 |
| 0 | 0 |
| 0 | 0 |
| 0 | 361.503092 |
| 59.7153101 | 9.15197702 |
| 105.858959 | 54.9118621 |
| 0 | 0 |
| 0 | 0 |

| | |
|---|---|
| 0 | 0 |
| 0 | 0 |
| 0 | 0 |
| 0 | 0 |
| 0 | 0 |
| 0 | 0 |
| 0 | 0 |
| 0 | 0 |
| 222.575247 | 205.919483 |
| 0 | 0 |
| 789.870693 | 50.3358736 |
| 0 | 0 |
| 0 | 0 |
| 0 | 0 |
| 2.71433228 | 0 |
| 0 | 0 |
| 0 | 0 |
| 0 | 0 |
| 0 | 0 |
| 0 | 0 |
| 0 | 0 |
| 0 | 0 |
| 0 | 0 |
| 0 | 4.57598851 |
| 43.4293164 | 22.8799426 |
| 70.5726392 | 0 |
| 0 | 0 |
| 0 | 0 |
| 0 | 0 |
| 141.145278 | 27.4559311 |
| 0 | 41.1838966 |
