## Supplemental Figure 3 for "Downregulation of Neurodevelopmental Gene Expression in iPSC-Derived Cerebral Organoids Upon Infection by Human Cytomegalovirus"

**Supplemental Table S3.** Includes Gene and rank lists from GSEA analyses found in Figure 3. All comparisons were made relative to Mock and each list contained all significant differentially expressed genes as revealed by DESeq2

| NAME | SYMBOL | TITLE | RANK IN GEN | RANK METRI | RUNNING ES | CORE ENRICI |
|---|---|---|---|---|---|---|
| **row_0** | CD47 | CD47 molecu | 10 | 3567.07446 | 0.2492651 | Yes |
| **row_1** | GBP4 | guanylate binc | 150 | 339.55188 | 0.26279798 | Yes |
| **row_2** | LAMP3 | lysosomal ass | 649 | 103.491318 | 0.23327412 | No |
| **row_3** | NCOA7 | nuclear recept | 909 | 74.330658 | 0.2193566 | No |
| **row_4** | EIF2AK2 | eukaryotic trar | 1216 | 50.5981865 | 0.2003048 | No |
| **row_5** | IRF7 | interferon regu | 1620 | -9.9604034 | 0.17124143 | No |
| **row_6** | NMI | N-myc and ST | 1855 | -12.753016 | 0.15485439 | No |
| **row_7** | CMPK2 | cytidine/uridin | 2153 | -15.278153 | 0.1339918 | No |
| **row_8** | SP110 | SP110 nuclea | 2228 | -15.843646 | 0.12963733 | No |
| **row_9** | PSMB8 | proteasome 2 | 2344 | -16.856472 | 0.12232602 | No |
| **row_10** | TXNIP | thioredoxin int | 2376 | -17.118099 | 0.12123642 | No |
| **row_11** | IFI35 | interferon indu | 2772 | -20.530298 | 0.09350465 | No |
| **row_12** | MOV10 | Mov10 RISC c | 2838 | -21.055349 | 0.09018011 | No |
| **row_13** | PARP9 | poly(ADP-ribo | 3230 | -24.617743 | 0.06303021 | No |
| **row_14** | DHX58 | DExH-box hel | 3256 | -24.914341 | 0.06293012 | No |
| **row_15** | PARP12 | poly(ADP-ribo | 3329 | -25.645409 | 0.05941033 | No |
| **row_16** | TRAFD1 | TRAF-type zir | 3357 | -25.921444 | 0.05923313 | No |
| **row_17** | IFIH1 | interferon indu | 3386 | -26.098246 | 0.05899447 | No |
| **row_18** | IFIT2 | interferon indu | 3731 | -29.999317 | 0.03569269 | No |
| **row_19** | USP18 | ubiquitin spec | 4191 | -35.513855 | 0.00428468 | No |
| **row_20** | TMEM140 | transmembrar | 4465 | -38.731377 | -0.0131618 | No |
| **row_21** | IRF2 | interferon regu | 4507 | -39.162704 | -0.0134448 | No |
| **row_22** | IFI30 | IFI30 lysosom | 4530 | -39.44434 | -0.012305 | No |
| **row_23** | RSAD2 | radical S-aden | 4669 | -40.988914 | -0.0196235 | No |
| **row_24** | GMPR | guanosine mc | 4963 | -44.384579 | -0.0381508 | No |
| **row_25** | TRIM21 | tripartite motif | 5174 | -47.212055 | -0.0503503 | No |
| **row_26** | C1S | complement C | 5316 | -49.296707 | -0.0573081 | No |
| **row_27** | TDRD7 | tudor domain | 5393 | -50.331558 | -0.0593931 | No |
| **row_28** | DDX60 | DExD/H-box h | 5466 | -51.355556 | -0.061111 | No |
| **row_29** | IFI44 | interferon indu | 5702 | -55.15657 | -0.0745999 | No |
| **row_30** | ELF1 | E74 like ETS t | 6123 | -63.081059 | -0.1011957 | No |
| **row_31** | TENT5A | terminal nucle | 6177 | -63.952568 | -0.1006275 | No |
| **row_32** | ISG15 | ISG15 ubiquit | 6479 | -70.364784 | -0.1179247 | No |
| **row_33** | NUB1 | negative regul | 6643 | -73.920769 | -0.1247814 | No |
| **row_34** | PLSCR1 | phospholipid | 6686 | -74.892319 | -0.1226341 | No |
| **row_35** | UBA7 | ubiquitin like r | 6764 | -76.634247 | -0.1229496 | No |
| **row_36** | BST2 | bone marrow | 7257 | -89.166603 | -0.1530343 | No |
| **row_37** | TRIM26 | tripartite motif | 7484 | -94.926094 | -0.1630713 | No |
| **row_38** | IFI44L | interferon indu | 8103 | -114.26035 | -0.2007024 | No |
| **row_39** | TAP1 | transporter 1, | 8158 | -116.07171 | -0.1965552 | No |
| **row_40** | UBE2L6 | ubiquitin conju | 8442 | -126.97215 | -0.2085557 | No |
| **row_41** | MVB12A | multivesicular | 8540 | -130.94151 | -0.2065419 | No |
| **row_42** | LY6E | lymphocyte ar | 8551 | -131.21906 | -0.1980837 | No |
| **row_43** | SLC25A28 | solute carrier f | 8838 | -142.59526 | -0.2092107 | No |

| | | | | | | |
|---|---|---|---|---|---|---|
| **row_44** | HLA-C | major histocor | 8881 | -144.32292 | -0.2021973 | No |
| **row_45** | ADAR | adenosine dea | 9061 | -152.67032 | -0.2047163 | No |
| **row_46** | IRF9 | interferon regu | 9127 | -155.57648 | -0.1986128 | No |
| **row_47** | HELZ2 | helicase with z | 9257 | -161.26155 | -0.1968371 | No |
| **row_48** | CSF1 | colony stimula | 9351 | -166.25504 | -0.192053 | No |
| **row_49** | IFITM1 | interferon indu | 9615 | -181.3557 | -0.1987649 | No |
| **row_50** | TRIM25 | tripartite motif | 9719 | -186.39101 | -0.1933079 | No |
| **row_51** | PSME1 | proteasome ac | 10066 | -210.09407 | -0.2041352 | No |
| **row_52** | LAP3 | leucine amino | 10302 | -228.00081 | -0.2055101 | No |
| **row_53** | PSME2 | proteasome ac | 10379 | -233.03122 | -0.1947904 | No |
| **row_54** | IFITM2 | interferon indu | 10833 | -273.94855 | -0.2090443 | No |
| **row_55** | PSMA3 | proteasome 2 | 10903 | -281.48401 | -0.1944117 | No |
| **row_56** | RNF31 | ring finger pro | 10937 | -285.12659 | -0.1768652 | No |
| **row_57** | WARS1 | tryptophanyl-t | 11210 | -315.72379 | -0.1748244 | No |
| **row_58** | CMTR1 | cap methyltrar | 11626 | -367.3056 | -0.1797289 | No |
| **row_59** | CD74 | CD74 molecu | 11806 | -395.2486 | -0.1652464 | No |
| **row_60** | STAT2 | signal transdu | 11860 | -407.09146 | -0.1406289 | No |
| **row_61** | OGFR | opioid growth | 11971 | -428.81546 | -0.1186982 | No |
| **row_62** | CNP | 2',3'-cyclic nu | 12632 | -632.13733 | -0.1231348 | No |
| **row_63** | IFITM3 | interferon indu | 12925 | -789.66852 | -0.0893539 | No |
| **row_64** | B2M | beta-2-microg | 13022 | -870.55383 | -0.0354294 | No |
| **row_65** | LGALS3BP | galectin 3 binc | 13226 | -1119.8126 | 0.02806276 | No |

HMENT

| NAME | SYMBOL | TITLE | RANK IN GEI | RANK METRI | RUNNING ES | CORE ENRICI |
|---|---|---|---|---|---|---|
| **row_0** | IRF1 | interferon regu | 144 | 2.95010591 | -0.0030284 | No |
| **row_1** | GBP4 | guanylate binc | 352 | 1.87184191 | -0.0144334 | No |
| **row_2** | CASP8 | caspase 8 [So | 666 | 1.21339834 | -0.036492 | No |
| **row_3** | CMPK2 | cytidine/uridin | 832 | 0.99999046 | -0.0470476 | No |
| **row_4** | GMPR | guanosine mc | 1177 | -2.0803921 | -0.0690678 | No |
| **row_5** | IFI35 | interferon indu | 1304 | -2.2203798 | -0.0727921 | No |
| **row_6** | PARP12 | poly(ADP-ribo | 1433 | -2.3384011 | -0.0763292 | No |
| **row_7** | IRF7 | interferon regu | 1547 | -2.4566913 | -0.0782839 | No |
| **row_8** | PROCR | protein C rece | 1838 | -2.6856821 | -0.0940736 | No |
| **row_9** | RIPK2 | receptor intera | 2026 | -2.8110082 | -0.1010426 | No |
| **row_10** | IL4R | interleukin 4 re | 2494 | -3.1426797 | -0.1299884 | No |
| **row_11** | TRIM25 | tripartite motif | 2551 | -3.2033882 | -0.1250456 | No |
| **row_12** | BST2 | bone marrow | 3214 | -3.5974441 | -0.1686305 | No |
| **row_13** | TRIM5 | tripartite motif | 3453 | -3.7460012 | -0.1769991 | No |
| **row_14** | IRF9 | interferon regu | 4028 | -4.0084872 | -0.2121432 | No |
| **row_15** | PLSCR1 | phospholipid | 4221 | -4.0928836 | -0.2157066 | No |
| **row_16** | UBA7 | ubiquitin like r | 4272 | -4.1143513 | -0.20756 | No |
| **row_17** | CMTR1 | cap methyltrar | 4466 | -4.1947112 | -0.2109023 | No |
| **row_18** | B2M | beta-2-microg | 4509 | -4.2108688 | -0.2018123 | No |
| **row_19** | C1S | complement C | 5061 | -4.4565287 | -0.2337363 | No |
| **row_20** | MVB12A | multivesicular | 5235 | -4.5333657 | -0.2344303 | No |
| **row_21** | PSME1 | proteasome ac | 5685 | -4.7191563 | -0.2572071 | No |
| **row_22** | WARS1 | tryptophanyl-t | 6339 | -4.9983668 | -0.2958837 | No |
| **row_23** | HELZ2 | helicase with z | 6352 | -5.0030913 | -0.281975 | No |
| **row_24** | USP18 | ubiquitin spec | 6636 | -5.1264262 | -0.2899251 | No |
| **row_25** | STAT2 | signal transdu | 6931 | -5.2488713 | -0.2984129 | Yes |
| **row_26** | TRIM26 | tripartite motif | 7026 | -5.2894111 | -0.2903772 | Yes |
| **row_27** | ELF1 | E74 like ETS t | 7091 | -5.3197832 | -0.2797906 | Yes |
| **row_28** | UBE2L6 | ubiquitin conju | 7102 | -5.3260822 | -0.2647565 | Yes |
| **row_29** | TDRD7 | tudor domain | 7218 | -5.3808684 | -0.2581708 | Yes |
| **row_30** | IFITM3 | interferon indu | 7848 | -5.7201529 | -0.2927305 | Yes |
| **row_31** | TRAFD1 | TRAF-type zir | 8045 | -5.834569 | -0.2914374 | Yes |
| **row_32** | PARP9 | poly(ADP-ribo | 8235 | -5.9630704 | -0.2891877 | Yes |
| **row_33** | PSMA3 | proteasome 2 | 8248 | -5.9710793 | -0.2723977 | Yes |
| **row_34** | TAP1 | transporter 1, | 8329 | -6.0205317 | -0.2610374 | Yes |
| **row_35** | ISG15 | ISG15 ubiquit | 8364 | -6.0509825 | -0.2458138 | Yes |
| **row_36** | HLA-C | major histocor | 8518 | -6.1409802 | -0.240082 | Yes |
| **row_37** | OGFR | opioid growth | 8677 | -6.2536311 | -0.234425 | Yes |
| **row_38** | SLC25A28 | solute carrier f | 8680 | -6.2565346 | -0.2159651 | Yes |
| **row_39** | NCOA7 | nuclear recept | 8874 | -6.4049635 | -0.212728 | Yes |
| **row_40** | PSMB8 | proteasome 2 | 9228 | -6.6906061 | -0.221763 | Yes |
| **row_41** | TMEM140 | transmembrar | 9546 | -6.9973841 | -0.2269323 | Yes |
| **row_42** | CNP | 2',3'-cyclic nu | 9548 | -6.9995503 | -0.2061786 | Yes |
| **row_43** | CD47 | CD47 molecu | 9614 | -7.0569668 | -0.1905029 | Yes |

| | | | | | | |
|---|---|---|---|---|---|---|
| **row_44** | LAP3 | leucine amino | 9776 | -7.2330103 | -0.1821766 | Yes |
| **row_45** | TENT5A | terminal nucle | 9789 | -7.2458577 | -0.1615919 | Yes |
| **row_46** | NUB1 | negative regul | 9855 | -7.3181763 | -0.1451387 | Yes |
| **row_47** | IFITM2 | interferon indu | 9975 | -7.438611 | -0.1327557 | Yes |
| **row_48** | LGALS3BP | galectin 3 binc | 10081 | -7.5652132 | -0.1188477 | Yes |
| **row_49** | PSME2 | proteasome ac | 10336 | -7.9208307 | -0.1161012 | Yes |
| **row_50** | TXNIP | thioredoxin int | 10447 | -8.0962133 | -0.1010227 | Yes |
| **row_51** | EIF2AK2 | eukaryotic trar | 10505 | -8.1854591 | -0.0813317 | Yes |
| **row_52** | CD74 | CD74 molecu | 10656 | -8.4618111 | -0.0684454 | Yes |
| **row_53** | RNF31 | ring finger pro | 10832 | -8.8088646 | -0.0565764 | Yes |
| **row_54** | ADAR | adenosine dea | 10870 | -8.8694983 | -0.0332089 | Yes |
| **row_55** | IFI30 | IFI30 lysosom | 11012 | -9.1798573 | -0.017447 | Yes |
| **row_56** | DHX58 | DExH-box hel | 11358 | -9.9224367 | -0.0162057 | Yes |
| **row_57** | TRIM21 | tripartite motif | 11495 | -9.9948854 | 0.00239236 | Yes |
| **row_58** | IFITM1 | interferon indu | 11601 | -9.9987545 | 0.02354434 | Yes |
| **row_59** | LY6E | lymphocyte ar | 11674 | -9.9991941 | 0.0474041 | Yes |

HMENT

| NAME | SYMBOL | TITLE | RANK IN GEN | RANK METRI | RUNNING ES | CORE ENRICI |
|---|---|---|---|---|---|---|
| row_0 | TSPYL2 | TSPY like 2 [S | 960 | -1.6531411 | -0.0763857 | No |
| row_1 | EIF2S1 | eukaryotic trar | 1104 | -1.9492968 | -0.0850716 | No |
| row_2 | IFIT1 | interferon indu | 1162 | -2.0645001 | -0.0864982 | No |
| row_3 | SLC30A5 | solute carrier f | 1670 | -2.555783 | -0.1241791 | No |
| row_4 | CNOT2 | CCR4-NOT tra | 2100 | -2.8611627 | -0.1549584 | No |
| row_5 | EIF4E | eukaryotic trar | 2519 | -3.171581 | -0.1843418 | No |
| row_6 | EDC4 | enhancer of m | 2582 | -3.2195098 | -0.1843539 | No |
| row_7 | PSAT1 | phosphoserin | 2898 | -3.4305148 | -0.2048519 | No |
| row_8 | DCP2 | decapping mF | 3014 | -3.4895046 | -0.2087985 | No |
| row_9 | ERN 1.00 | endoplasmic r | 3047 | -3.5092735 | -0.2058837 | No |
| row_10 | PAIP1 | poly(A) bindin | 3123 | -3.5508714 | -0.2064417 | No |
| row_11 | ERO1A | endoplasmic r | 3391 | -3.7132041 | -0.2225429 | No |
| row_12 | WIPI1 | WD repeat do | 3627 | -3.8319228 | -0.235823 | No |
| row_13 | KIF5B | kinesin family | 3675 | -3.8584645 | -0.2335905 | No |
| row_14 | ARFGAP1 | ADP ribosylati | 4130 | -4.0611858 | -0.2645299 | No |
| row_15 | IMP3 | IMP U3 small | 4231 | -4.0962133 | -0.266283 | No |
| row_16 | NHP2 | NHP2 ribonuc | 4301 | -4.1229382 | -0.2654428 | No |
| row_17 | CCL2 | C-C motif che | 4315 | -4.1296086 | -0.2599837 | No |
| row_18 | EIF4G1 | eukaryotic trar | 4391 | -4.1632676 | -0.2595734 | No |
| row_19 | EXOSC5 | exosome com | 4484 | -4.2009401 | -0.2605026 | No |
| row_20 | CNOT4 | CCR4-NOT tra | 4610 | -4.2687402 | -0.2640401 | No |
| row_21 | DNAJA4 | DnaJ heat sho | 4790 | -4.3448167 | -0.2719011 | No |
| row_22 | TTC37 | tetratricopeptic | 4795 | -4.3474364 | -0.2653571 | No |
| row_23 | SRPRA | SRP receptor s | 4889 | -4.3859725 | -0.2660759 | No |
| row_24 | GEMIN4 | gem nuclear o | 4914 | -4.3957176 | -0.2611014 | No |
| row_25 | CXXC1 | CXXC finger p | 4918 | -4.3981681 | -0.2543948 | No |
| row_26 | PREB | prolactin regul | 5156 | -4.4918242 | -0.2667962 | No |
| row_27 | ASNS | asparagine syr | 5194 | -4.5087986 | -0.2627127 | No |
| row_28 | ATF4 | activating tran | 5296 | -4.5536489 | -0.2638248 | No |
| row_29 | NFYB | nuclear transc | 5842 | -4.7887979 | -0.3011024 | No |
| row_30 | SEC31A | SEC31 homol | 6004 | -4.8580399 | -0.3066707 | No |
| row_31 | PARN | poly(A)-specifi | 6050 | -4.8793373 | -0.3026596 | No |
| row_32 | HSP90B1 | heat shock pro | 6487 | -5.0604591 | -0.330538 | No |
| row_33 | YWHAZ | tyrosine 3-mo | 6534 | -5.0801768 | -0.3262917 | No |
| row_34 | PDIA5 | protein disulfic | 6683 | -5.1452947 | -0.3303361 | No |
| row_35 | FKBP14 | FKBP prolyl is | 6701 | -5.1493945 | -0.3235939 | No |
| row_36 | EXOSC2 | exosome com | 6950 | -5.2573833 | -0.3356903 | No |
| row_37 | CEBPG | CCAAT enhan | 7065 | -5.3093538 | -0.3366774 | Yes |
| row_38 | DCTN1 | dynactin subu | 7130 | -5.3417311 | -0.3334988 | Yes |
| row_39 | VEGFA | vascular endo | 7262 | -5.4043975 | -0.3357347 | Yes |
| row_40 | EXOC2 | exocyst comp | 7364 | -5.4589329 | -0.3354156 | Yes |
| row_41 | YIF1A | Yip1 interactin | 7397 | -5.4745712 | -0.3293937 | Yes |
| row_42 | POP4 | POP4 homolo | 7467 | -5.5045052 | -0.3263692 | Yes |
| row_43 | SHC1 | SHC adaptor p | 7587 | -5.5706019 | -0.3273548 | Yes |

| | | | | | | |
|---|---|---|---|---|---|---|
| **row_44** | BAG3 | BAG cochape | 7806 | -5.6951056 | -0.3362903 | Yes |
| **row_45** | SRPRB | SRP receptor s | 7898 | -5.750627 | -0.3346871 | Yes |
| **row_46** | KHSRP | KH-type splici | 7949 | -5.781219 | -0.3296617 | Yes |
| **row_47** | FUS | FUS RNA binc | 8042 | -5.8323002 | -0.3280117 | Yes |
| **row_48** | EXOSC9 | exosome com | 8083 | -5.8573675 | -0.3220429 | Yes |
| **row_49** | EDEM1 | ER degradatio | 8099 | -5.8711429 | -0.313995 | Yes |
| **row_50** | H2AX | H2A.X variant | 8200 | -5.9421697 | -0.3128296 | Yes |
| **row_51** | EEF2 | eukaryotic trar | 8210 | -5.9482765 | -0.3041661 | Yes |
| **row_52** | CHAC1 | ChaC glutathi | 8226 | -5.9567823 | -0.2959829 | Yes |
| **row_53** | EIF4A3 | eukaryotic trar | 8254 | -5.9753175 | -0.2887579 | Yes |
| **row_54** | IARS1 | isoleucyl-tRNA | 8447 | -6.098845 | -0.2949156 | Yes |
| **row_55** | DKC1 | dyskerin pseu | 8515 | -6.1385121 | -0.2907241 | Yes |
| **row_56** | CALR | calreticulin [Sc | 8520 | -6.1440992 | -0.2813396 | Yes |
| **row_57** | NOLC1 | nucleolar and | 8540 | -6.1564431 | -0.2731699 | Yes |
| **row_58** | TUBB2A | tubulin beta 2 | 8669 | -6.2463865 | -0.2738276 | Yes |
| **row_59** | DNAJB9 | DnaJ heat shc | 8727 | -6.2892313 | -0.268575 | Yes |
| **row_60** | SDAD1 | SDA1 domain | 8753 | -6.3072352 | -0.2606607 | Yes |
| **row_61** | SPCS1 | signal peptida | 8904 | -6.4264407 | -0.2628442 | Yes |
| **row_62** | PDIA6 | protein disulfi | 8999 | -6.5036678 | -0.2602974 | Yes |
| **row_63** | LSM 1.00 | LSM1 homolc | 9082 | -6.5731597 | -0.2566531 | Yes |
| **row_64** | SERP1 | stress associa | 9169 | -6.6465611 | -0.2532221 | Yes |
| **row_65** | HERPUD1 | homocysteine | 9364 | -6.8067536 | -0.2584251 | Yes |
| **row_66** | EIF4A2 | eukaryotic trar | 9366 | -6.8115916 | -0.2477384 | Yes |
| **row_67** | NPM1 | nucleophosm | 9529 | -6.9757423 | -0.250041 | Yes |
| **row_68** | ATP6V0D1 | ATPase H+ tra | 9537 | -6.9830613 | -0.2395769 | Yes |
| **row_69** | EIF4A1 | eukaryotic trar | 9761 | -7.2195864 | -0.2465137 | Yes |
| **row_70** | LSM 4.00 | LSM4 homolc | 9763 | -7.2206702 | -0.2351802 | Yes |
| **row_71** | HYOU1 | hypoxia up-re | 10097 | -7.584393 | -0.2505923 | Yes |
| **row_72** | WFS1 | wolframin ER | 10144 | -7.6513219 | -0.242281 | Yes |
| **row_73** | KDELR3 | KDEL endopla | 10198 | -7.7319698 | -0.2344183 | Yes |
| **row_74** | EIF2AK3 | eukaryotic trar | 10482 | -8.1443253 | -0.2448306 | Yes |
| **row_75** | SSR1 | signal sequen | 10499 | -8.1674623 | -0.2332346 | Yes |
| **row_76** | EXOSC1 | exosome com | 10506 | -8.1926365 | -0.2207759 | Yes |
| **row_77** | CEBPB | CCAAT enhan | 10590 | -8.3240128 | -0.2144459 | Yes |
| **row_78** | NOP56 | NOP56 ribonu | 10597 | -8.3440247 | -0.2017479 | Yes |
| **row_79** | TARS1 | threonyl-tRNA | 10676 | -8.5048828 | -0.1947205 | Yes |
| **row_80** | DDIT4 | DNA damage | 10745 | -8.6368647 | -0.1866615 | Yes |
| **row_81** | DDX10 | DEAD-box hel | 10874 | -8.8761539 | -0.1831617 | Yes |
| **row_82** | DNAJC3 | DnaJ heat shc | 10973 | -9.0911913 | -0.1768531 | Yes |
| **row_83** | SLC1A4 | solute carrier f | 11060 | -9.3038445 | -0.1692209 | Yes |
| **row_84** | HSPA5 | heat shock pr | 11083 | -9.367403 | -0.1562216 | Yes |
| **row_85** | XBP1 | X-box binding | 11172 | -9.558073 | -0.148352 | Yes |
| **row_86** | EIF4EBP1 | eukaryotic trar | 11234 | -9.7401066 | -0.1379728 | Yes |
| **row_87** | ALDH18A1 | aldehyde deh | 11255 | -9.7755289 | -0.1241636 | Yes |
| **row_88** | SLC7A5 | solute carrier f | 11276 | -9.8239088 | -0.110278 | Yes |

| | | | | | | |
|---|---|---|---|---|---|---|
| **row_89** | MTHFD2 | methylenetetra | 11363 | -9.9324741 | -0.1016519 | Yes |
| **row_90** | EXOSC4 | exosome com | 11591 | -9.9986372 | -0.1045243 | Yes |
| **row_91** | GOSR2 | golgi SNAP re | 11721 | -9.9994326 | -0.0993308 | Yes |
| **row_92** | NFYA | nuclear transc | 11725 | -9.9994459 | -0.0837687 | Yes |
| **row_93** | SPCS3 | signal peptida | 11828 | -9.9997063 | -0.076353 | Yes |
| **row_94** | DCP1A | decapping mF | 11959 | -9.9998598 | -0.0712412 | Yes |
| **row_95** | SEC11A | SEC11 homol | 11998 | -9.9998846 | -0.0585586 | Yes |
| **row_96** | STC2 | stanniocalcin : | 12037 | -9.9999046 | -0.0458759 | Yes |
| **row_97** | BANF1 | BAF nuclear a | 12090 | -9.9999342 | -0.0343453 | Yes |
| **row_98** | XPOT | exportin for tF | 12164 | -9.9999666 | -0.0245427 | Yes |
| **row_99** | RPS14 | ribosomal pro | 12220 | -9.9999895 | -0.0132589 | Yes |
| **row_100** | HSPA9 | heat shock pr | 12239 | -9.9999952 | 0.00106973 | Yes |

HMENT

| NAME | SYMBOL | TITLE | RANK IN GEN | RANK METRI | RUNNING ES | CORE ENRICI |
|---|---|---|---|---|---|---|
| row_0 | PSMB2 | proteasome 2 | 158 | 325.56189 | -0.0090179 | No |
| row_1 | SNRPD3 | small nuclear r | 545 | 123.203529 | -0.0367563 | No |
| row_2 | HDAC2 | histone deace | 929 | 72.678833 | -0.0647 | No |
| row_3 | PSMA4 | proteasome 2 | 1769 | -11.957121 | -0.1271637 | No |
| row_4 | SNRPD2 | small nuclear r | 1866 | -12.868557 | -0.1342133 | No |
| row_5 | HDGF | heparin bindin | 1868 | -12.871877 | -0.1341785 | No |
| row_6 | RNPS1 | RNA binding p | 1981 | -13.831942 | -0.1424131 | No |
| row_7 | ORC2 | origin recogni | 2106 | -14.897097 | -0.1515334 | No |
| row_8 | TXNL4A | thioredoxin lik | 2155 | -15.303273 | -0.1549829 | No |
| row_9 | PSMA6 | proteasome 2 | 2165 | -15.36896 | -0.1555235 | No |
| row_10 | CCT7 | chaperonin cc | 2258 | -16.072845 | -0.1622476 | No |
| row_11 | RPL6 | ribosomal pro | 2398 | -17.264765 | -0.1724664 | No |
| row_12 | PHB | prohibitin [Sou | 2669 | -19.708227 | -0.1924333 | No |
| row_13 | USP1 | ubiquitin spec | 2751 | -20.36705 | -0.1983006 | No |
| row_14 | CCT2 | chaperonin cc | 2806 | -20.802021 | -0.2021509 | No |
| row_15 | UBE2L3 | ubiquitin conju | 2840 | -21.064741 | -0.2044329 | No |
| row_16 | SSBP1 | single strande | 3077 | -23.149006 | -0.2218351 | No |
| row_17 | SLC25A3 | solute carrier f | 3136 | -23.671297 | -0.2259592 | No |
| row_18 | HSPD1 | heat shock pro | 3350 | -25.83353 | -0.2416236 | No |
| row_19 | CTPS1 | CTP synthase | 3377 | -26.044889 | -0.2433413 | No |
| row_20 | PCNA | proliferating ce | 3441 | -26.587105 | -0.2478135 | No |
| row_21 | EXOSC7 | exosome com | 3501 | -27.342031 | -0.2519811 | No |
| row_22 | EIF3J | eukaryotic trar | 3502 | -27.344082 | -0.2517489 | No |
| row_23 | PSMA1 | proteasome 2 | 3514 | -27.445667 | -0.2523361 | No |
| row_24 | PSMB3 | proteasome 2 | 3570 | -28.111855 | -0.2561988 | No |
| row_25 | CLNS1A | chloride nucle | 3706 | -29.756065 | -0.2660133 | No |
| row_26 | MRPL9 | mitochondrial | 3900 | -32.180176 | -0.2801323 | No |
| row_27 | RPL34 | ribosomal pro | 4103 | -34.475685 | -0.294903 | No |
| row_28 | SNRPG | small nuclear r | 4122 | -34.68927 | -0.2959507 | No |
| row_29 | NHP2 | NHP2 ribonuc | 4505 | -39.145458 | -0.3241045 | No |
| row_30 | ACP1 | acid phosphat | 4976 | -44.51976 | -0.358775 | No |
| row_31 | NOP16 | NOP16 nucleo | 5024 | -45.151123 | -0.3618964 | No |
| row_32 | RAN | RAN, member | 5098 | -46.080025 | -0.3669489 | No |
| row_33 | CDC45 | cell division cy | 5123 | -46.428223 | -0.3683444 | No |
| row_34 | PTGES3 | prostaglandin | 5284 | -48.768867 | -0.3798617 | No |
| row_35 | TRA2B | transformer 2 | 5401 | -50.453259 | -0.3880835 | No |
| row_36 | CSTF2 | cleavage stimu | 5676 | -54.802216 | -0.4080507 | No |
| row_37 | GNL3 | G protein nuc | 5727 | -55.468792 | -0.4113083 | No |
| row_38 | AP3S1 | adaptor relatec | 5835 | -57.385777 | -0.4188002 | No |
| row_39 | EIF4E | eukaryotic trar | 5957 | -59.603874 | -0.4273172 | No |
| row_40 | CCT4 | chaperonin cc | 5968 | -59.825325 | -0.4275549 | No |
| row_41 | EIF4G2 | eukaryotic trar | 6291 | -66.239738 | -0.4510044 | No |
| row_42 | SNRPB2 | small nuclear r | 6324 | -66.990746 | -0.4528219 | No |
| row_43 | DUT | deoxyuridine t | 6706 | -75.136436 | -0.4805955 | No |

| | | | | | | |
|---|---|---|---|---|---|---|
| **row_44** | SYNCRIP | synaptotagmi | 6810 | -77.959984 | -0.4876144 | No |
| **row_45** | POLD2 | DNA polymera | 6865 | -79.312126 | -0.4909678 | No |
| **row_46** | APEX1 | apurinic/apyrir | 7056 | -84.207634 | -0.5044214 | No |
| **row_47** | NOLC1 | nucleolar and | 7096 | -85.067085 | -0.5066074 | No |
| **row_48** | AIMP2 | aminoacyl tRN | 7122 | -85.819962 | -0.5077429 | No |
| **row_49** | SET | SET nuclear p | 7285 | -90.025032 | -0.5190591 | No |
| **row_50** | TCP1 | t-complex 1 [S | 7299 | -90.326904 | -0.5192615 | No |
| **row_51** | CCT3 | chaperonin cc | 7346 | -91.233231 | -0.5219172 | No |
| **row_52** | HSP90AB1 | heat shock pr | 7357 | -91.502136 | -0.5218859 | No |
| **row_53** | TFDP1 | transcription fa | 7376 | -92.045364 | -0.5224466 | No |
| **row_54** | PRPS2 | phosphoribos | 7472 | -94.377419 | -0.5287296 | No |
| **row_55** | RPS10 | ribosomal pro | 7501 | -95.295563 | -0.5300084 | No |
| **row_56** | MAD2L1 | mitotic arrest d | 7701 | -100.91022 | -0.5439913 | No |
| **row_57** | SRSF2 | serine and arg | 7917 | -107.8864 | -0.559108 | No |
| **row_58** | FAM120A | family with se | 7961 | -109.31349 | -0.5613864 | No |
| **row_59** | LDHA | lactate dehydr | 7984 | -109.80038 | -0.5620947 | No |
| **row_60** | RFC4 | replication fac | 8028 | -111.42232 | -0.5643551 | No |
| **row_61** | PABPC4 | poly(A) bindin | 8172 | -116.63804 | -0.5740285 | No |
| **row_62** | HPRT1 | hypoxanthine | 8393 | -125.57224 | -0.5893679 | No |
| **row_63** | EEF1B2 | eukaryotic tran | 8415 | -126.29381 | -0.5898616 | No |
| **row_64** | NCBP2 | nuclear cap bi | 8430 | -126.65629 | -0.5898301 | No |
| **row_65** | YWHAE | tyrosine 3-mo | 8450 | -127.25491 | -0.5901665 | No |
| **row_66** | EIF4H | eukaryotic tran | 8456 | -127.44774 | -0.5894572 | No |
| **row_67** | NAP1L1 | nucleosome a | 8702 | -136.94119 | -0.6065643 | No |
| **row_68** | RUVBL2 | RuvB like AAA | 8732 | -138.12292 | -0.6075541 | No |
| **row_69** | SRSF7 | serine and arg | 8785 | -140.40822 | -0.6102396 | No |
| **row_70** | CCNA2 | cyclin A2 [Sou | 8797 | -140.82231 | -0.6098642 | No |
| **row_71** | XRCC6 | X-ray repair cr | 8807 | -141.33958 | -0.6093352 | No |
| **row_72** | BUB3 | BUB3 mitotic | 8887 | -144.62422 | -0.6139984 | No |
| **row_73** | CDK2 | cyclin depend | 8957 | -148.14723 | -0.6178859 | No |
| **row_74** | ODC1 | ornithine deca | 8964 | -148.41666 | -0.6170731 | No |
| **row_75** | POLE3 | DNA polymera | 8977 | -149.05492 | -0.6167023 | No |
| **row_76** | VDAC1 | voltage depen | 9069 | -153.02469 | -0.622189 | No |
| **row_77** | RRP9 | ribosomal RNA | 9134 | -155.76862 | -0.625639 | No |
| **row_78** | LSM 2.00 | LSM2 homolo | 9139 | -156.04254 | -0.6246123 | No |
| **row_79** | U2AF1 | U2 small nucl | 9290 | -162.87425 | -0.634415 | No |
| **row_80** | HDDC2 | HD domain co | 9387 | -168.87025 | -0.64014 | No |
| **row_81** | CDC20 | cell division cy | 9422 | -170.53409 | -0.6412274 | No |
| **row_82** | HNRNPR | heterogeneou | 9466 | -173.03481 | -0.6429647 | No |
| **row_83** | NPM1 | nucleophosm | 9504 | -175.36215 | -0.6442349 | No |
| **row_84** | MRPL23 | mitochondrial | 9512 | -175.8186 | -0.643264 | No |
| **row_85** | NOP56 | NOP56 ribonu | 9542 | -177.35413 | -0.6439207 | No |
| **row_86** | CCT5 | chaperonin cc | 9560 | -178.29921 | -0.6436745 | No |
| **row_87** | HNRNPD | heterogeneou | 9583 | -179.479 | -0.6437911 | No |
| **row_88** | MRPS18B | mitochondrial | 9631 | -182.04585 | -0.6457502 | No |

| | | | | | | |
|---|---|---|---|---|---|---|
| **row_89** | SRSF1 | serine and arg | 9733 | -186.9841 | -0.6516942 | No |
| **row_90** | KARS1 | lysyl-tRNA syr | 9773 | -189.77835 | -0.6529911 | No |
| **row_91** | MCM5 | minichromosc | 9850 | -195.02733 | -0.6570026 | No |
| **row_92** | SERBP1 | SERPINE1 mF | 9867 | -195.87169 | -0.6565325 | No |
| **row_93** | SNRPA1 | small nuclear r | 9989 | -203.7578 | -0.6638256 | No |
| **row_94** | VDAC3 | voltage depen | 10189 | -218.17567 | -0.6768127 | No |
| **row_95** | MCM7 | minichromosc | 10498 | -242.32735 | -0.6977231 | No |
| **row_96** | RANBP1 | RAN binding p | 10646 | -256.05579 | -0.7065109 | No |
| **row_97** | SNRPD1 | small nuclear r | 10902 | -281.40021 | -0.7231372 | No |
| **row_98** | TYMS | thymidylate sy | 10921 | -283.02841 | -0.7220763 | No |
| **row_99** | HNRNPA3 | heterogeneou | 10951 | -286.57162 | -0.7218056 | No |
| **row_100** | SNRPA | small nuclear r | 10998 | -291.08646 | -0.7227643 | No |
| **row_101** | NDUFAB1 | NADH:ubiquir | 11076 | -299.19098 | -0.7259659 | No |
| **row_102** | NCBP1 | nuclear cap bi | 11270 | -322.62869 | -0.7376187 | No |
| **row_103** | COPS5 | COP9 signalo | 11272 | -322.80188 | -0.7349524 | No |
| **row_104** | SRSF3 | serine and arg | 11299 | -326.01709 | -0.7341231 | No |
| **row_105** | G3BP1 | G3BP stress g | 11399 | -336.71732 | -0.7386466 | No |
| **row_106** | CANX | calnexin [Sour | 11513 | -351.39923 | -0.7440894 | Yes |
| **row_107** | IARS1 | isoleucyl-tRN/ | 11527 | -353.64703 | -0.742056 | Yes |
| **row_108** | MCM2 | minichromosc | 11551 | -357.37616 | -0.7407367 | Yes |
| **row_109** | PWP1 | PWP1 homolc | 11573 | -359.78033 | -0.7392478 | Yes |
| **row_110** | LSM 7.00 | LSM7 homolc | 11602 | -363.97513 | -0.7382453 | Yes |
| **row_111** | SRPK1 | SRSF protein | 11629 | -367.62512 | -0.7370627 | Yes |
| **row_112** | PSMD14 | proteasome 2 | 11646 | -370.29834 | -0.7351117 | Yes |
| **row_113** | MCM4 | minichromosc | 11657 | -371.46777 | -0.7327033 | Yes |
| **row_114** | PSMC6 | proteasome 2 | 11712 | -379.70923 | -0.733506 | Yes |
| **row_115** | EIF2S1 | eukaryotic trar | 11769 | -388.26901 | -0.7343853 | Yes |
| **row_116** | IFRD1 | interferon relat | 11777 | -389.44583 | -0.7316005 | Yes |
| **row_117** | PRPF31 | pre-mRNA prc | 11798 | -393.62717 | -0.7297496 | Yes |
| **row_118** | RRM1 | ribonucleotide | 11841 | -403.95349 | -0.7294517 | Yes |
| **row_119** | VBP1 | VHL binding p | 11864 | -408.46436 | -0.727624 | Yes |
| **row_120** | DDX18 | DEAD-box hel | 11874 | -410.20129 | -0.7248122 | Yes |
| **row_121** | UBE2E1 | ubiquitin conju | 11892 | -414.82782 | -0.7225576 | Yes |
| **row_122** | ABCE1 | ATP binding c | 11917 | -419.39563 | -0.7207862 | Yes |
| **row_123** | PSMA2 | proteasome 2 | 12012 | -435.80984 | -0.7240955 | Yes |
| **row_124** | CAD | carbamoyl-ph | 12025 | -439.06625 | -0.7212623 | Yes |
| **row_125** | PRDX4 | peroxiredoxin | 12066 | -447.48541 | -0.7204456 | Yes |
| **row_126** | RPL14 | ribosomal pro | 12067 | -447.95517 | -0.716642 | Yes |
| **row_127** | HSPE1 | heat shock pr | 12097 | -455.03033 | -0.7149409 | Yes |
| **row_128** | SRM | spermidine sy | 12138 | -464.00937 | -0.7139839 | Yes |
| **row_129** | CNBP | CCHC-type zi | 12170 | -471.91888 | -0.7122886 | Yes |
| **row_130** | DDX21 | DExD-box hel | 12192 | -478.05927 | -0.7097954 | Yes |
| **row_131** | ETF1 | eukaryotic trar | 12195 | -478.62149 | -0.7058806 | Yes |
| **row_132** | EIF3B | eukaryotic trar | 12215 | -483.96832 | -0.7031881 | Yes |
| **row_133** | CUL1 | cullin 1 [Sourc | 12227 | -487.68832 | -0.6998674 | Yes |

| | | | | | | |
|---|---|---|---|---|---|---|
| **row_134** | PSMD7 | proteasome 2 | 12229 | -489.30826 | -0.6957873 | Yes |
| **row_135** | CYC1 | cytochrome c | 12331 | -521.38324 | -0.698892 | Yes |
| **row_136** | PRDX3 | peroxiredoxin | 12364 | -530.94818 | -0.69677 | Yes |
| **row_137** | C1QBP | complement ( | 12393 | -538.20697 | -0.6942881 | Yes |
| **row_138** | PSMC4 | proteasome 2 | 12402 | -543.72021 | -0.690268 | Yes |
| **row_139** | FBL | fibrillarin [Sour | 12459 | -563.88635 | -0.6896561 | Yes |
| **row_140** | NME1 | NME/NM23 n | 12467 | -567.15558 | -0.6853624 | Yes |
| **row_141** | COX5A | cytochrome c | 12522 | -583.74994 | -0.6844327 | Yes |
| **row_142** | TOMM70 | translocase of | 12524 | -584.54547 | -0.6795439 | Yes |
| **row_143** | CDK4 | cyclin depend | 12533 | -586.95557 | -0.6751567 | Yes |
| **row_144** | DEK | DEK proto-on | 12543 | -591.48346 | -0.6708056 | Yes |
| **row_145** | GOT2 | glutamic-oxalo | 12655 | -641.40594 | -0.6736369 | Yes |
| **row_146** | EIF2S2 | eukaryotic trar | 12661 | -643.02759 | -0.6685498 | Yes |
| **row_147** | EIF1AX | eukaryotic trar | 12665 | -646.8457 | -0.6632811 | Yes |
| **row_148** | GLO1 | glyoxalase I [S | 12743 | -679.44342 | -0.663254 | Yes |
| **row_149** | DHX15 | DEAH-box hel | 12799 | -703.81989 | -0.6613794 | Yes |
| **row_150** | ERH | ERH mRNA sp | 12807 | -710.5639 | -0.655868 | Yes |
| **row_151** | GSPT1 | G1 to S phase | 12816 | -714.3399 | -0.6503992 | Yes |
| **row_152** | EIF3D | eukaryotic trar | 12835 | -724.224 | -0.6455921 | Yes |
| **row_153** | SF3A1 | splicing factor | 12838 | -725.65424 | -0.6395798 | Yes |
| **row_154** | IMPDH2 | inosine mono | 12846 | -728.1358 | -0.6339192 | Yes |
| **row_155** | RPS3 | ribosomal pro | 12854 | -733.33789 | -0.6282145 | Yes |
| **row_156** | XPOT | exportin for tF | 12872 | -745.11029 | -0.6231555 | Yes |
| **row_157** | SF3B3 | splicing factor | 12873 | -746.48517 | -0.6168172 | Yes |
| **row_158** | SSB | small RNA bin | 12921 | -788.9491 | -0.6136231 | Yes |
| **row_159** | RSL1D1 | ribosomal L1 | 12950 | -808.75079 | -0.608844 | Yes |
| **row_160** | PA2G4 | proliferation-as | 12954 | -810.77576 | -0.6021835 | Yes |
| **row_161** | PSMD8 | proteasome 2 | 12968 | -826.44037 | -0.5961357 | Yes |
| **row_162** | PPM1G | protein phosp | 12975 | -832.98468 | -0.5895103 | Yes |
| **row_163** | STARD7 | StAR related li | 12984 | -835.88245 | -0.5830094 | Yes |
| **row_164** | CBX3 | chromobox 3 | 12987 | -838.76672 | -0.5760366 | Yes |
| **row_165** | PSMD1 | proteasome 2 | 13000 | -848.92377 | -0.5697234 | Yes |
| **row_166** | SMARCC1 | SWI/SNF relat | 13016 | -865.88831 | -0.5634897 | Yes |
| **row_167** | PHB2 | prohibitin 2 [S | 13030 | -879.1936 | -0.556994 | Yes |
| **row_168** | TARDBP | TAR DNA binc | 13034 | -881.14319 | -0.549736 | Yes |
| **row_169** | HNRNPC | heterogeneou | 13056 | -904.54602 | -0.5436215 | Yes |
| **row_170** | KPNA2 | karyopherin su | 13067 | -914.93646 | -0.5365986 | Yes |
| **row_171** | EPRS1 | glutamyl-proly | 13070 | -918.43567 | -0.5289493 | Yes |
| **row_172** | TUFM | Tu translation | 13129 | -978.07751 | -0.5249696 | Yes |
| **row_173** | H2AZ1 | H2A.Z variant | 13153 | -1005.7172 | -0.5181453 | Yes |
| **row_174** | PSMD3 | proteasome 2 | 13161 | -1015.6555 | -0.5100434 | Yes |
| **row_175** | PSMA7 | proteasome 2 | 13165 | -1026.2896 | -0.501553 | Yes |
| **row_176** | RPLP0 | ribosomal pro | 13175 | -1042.8042 | -0.4933698 | Yes |
| **row_177** | ILF2 | interleukin enh | 13215 | -1094.0721 | -0.4869883 | Yes |
| **row_178** | RAD23B | RAD23 homo | 13228 | -1120.9146 | -0.4783656 | Yes |

| | | | | | | |
|---|---|---|---|---|---|---|
| **row_179** | KPNB1 | karyopherin su | 13328 | -1367.2808 | -0.4741387 | Yes |
| **row_180** | XPO1 | exportin 1 [So | 13335 | -1380.3431 | -0.4628657 | Yes |
| **row_181** | HNRNPU | heterogeneou | 13383 | -1549.4451 | -0.4532143 | Yes |
| **row_182** | PCBP1 | poly(rC) bindir | 13397 | -1611.0588 | -0.4405043 | Yes |
| **row_183** | RPL22 | ribosomal pro | 13479 | -2219.9307 | -0.4276953 | Yes |
| **row_184** | PGK 1.00 | phosphoglyce | 13500 | -2467.1528 | -0.4082383 | Yes |
| **row_185** | RPS5 | ribosomal pro | 13502 | -2577.4143 | -0.3864282 | Yes |
| **row_186** | TRIM28 | tripartite motif | 13512 | -2781.1719 | -0.3634846 | Yes |
| **row_187** | RPL18 | ribosomal pro | 13515 | -2819.5486 | -0.3396931 | Yes |
| **row_188** | EIF4A1 | eukaryotic trar | 13517 | -2847.5481 | -0.3155893 | Yes |
| **row_189** | YWHAQ | tyrosine 3-mo | 13543 | -3553.1133 | -0.2872843 | Yes |
| **row_190** | PPIA | peptidylprolyl | 13546 | -3642.3215 | -0.2565068 | Yes |
| **row_191** | HNRNPA2B1 | heterogeneou | 13555 | -3850.2842 | -0.2244108 | Yes |
| **row_192** | RPS6 | ribosomal pro | 13557 | -3874.8772 | -0.1915841 | Yes |
| **row_193** | PABPC1 | poly(A) bindin | 13563 | -4034.8516 | -0.1576973 | Yes |
| **row_194** | RACK1 | receptor for ac | 13568 | -4328.4961 | -0.1212426 | Yes |
| **row_195** | HNRNPA1 | heterogeneou | 13585 | -6266.3975 | -0.0692281 | Yes |
| **row_196** | RPS2 | ribosomal pro | 13593 | -8328.8271 | 9.69E-04 | Yes |

HMENT

| NAME | SYMBOL | TITLE | RANK IN GEN | RANK METRI | RUNNING ES | CORE ENRICI |
|---|---|---|---|---|---|---|
| **row_0** | SRSF10 | serine and arg | 495 | 133.72377 | -0.0347329 | No |
| **row_1** | ARID4A | AT-rich interac | 529 | 126.302185 | -0.0351759 | No |
| **row_2** | SLC38A1 | solute carrier f | 623 | 107.562988 | -0.0403864 | No |
| **row_3** | STMN1 | stathmin 1 [Sc | 647 | 104.083931 | -0.0404391 | No |
| **row_4** | NUP50 | nucleoporin 5 | 752 | 90.4270859 | -0.0467422 | No |
| **row_5** | AURKA | aurora kinase | 759 | 89.5569382 | -0.0457604 | No |
| **row_6** | RBM14 | RNA binding r | 877 | 76.4854202 | -0.0532542 | No |
| **row_7** | PLK4 | polo like kinas | 1145 | 55.819191 | -0.0722491 | No |
| **row_8** | NOTCH2 | notch recepto | 1568 | 22.5436935 | -0.1033187 | No |
| **row_9** | CDK1 | cyclin depend | 1798 | -12.140559 | -0.1201802 | No |
| **row_10** | EXO1 | exonuclease 1 | 1871 | -12.892405 | -0.1253368 | No |
| **row_11** | PRMT5 | protein arginir | 2014 | -14.106836 | -0.1356875 | No |
| **row_12** | DKC1 | dyskerin pseu | 2193 | -15.579227 | -0.1486959 | No |
| **row_13** | H2AZ2 | H2A.Z variant | 2207 | -15.661372 | -0.1494143 | No |
| **row_14** | NUSAP1 | nucleolar and | 2214 | -15.740179 | -0.14961 | No |
| **row_15** | PML | PML nuclear b | 2230 | -15.870028 | -0.150474 | No |
| **row_16** | NEK2 | NIMA related k | 2769 | -20.512587 | -0.1902153 | No |
| **row_17** | FBXO5 | F-box protein | 3163 | -23.94738 | -0.2191027 | No |
| **row_18** | JPT1 | Jupiter microtu | 3170 | -23.992151 | -0.2191669 | No |
| **row_19** | PTTG1 | PTTG1 regula | 3194 | -24.23123 | -0.2204933 | No |
| **row_20** | POLQ | DNA polymera | 3420 | -26.392439 | -0.2368296 | No |
| **row_21** | CENPA | centromere pr | 3439 | -26.564247 | -0.2377464 | No |
| **row_22** | KMT5A | lysine methyltı | 3505 | -27.380651 | -0.2421506 | No |
| **row_23** | BCL3 | BCL3 transcrip | 3599 | -28.47468 | -0.2486228 | No |
| **row_24** | HMGA1 | high mobility g | 3614 | -28.653833 | -0.2492084 | No |
| **row_25** | CENPE | centromere pr | 3725 | -29.917566 | -0.2569236 | No |
| **row_26** | CHEK1 | checkpoint kir | 3734 | -30.025553 | -0.2570404 | No |
| **row_27** | NUP98 | nucleoporin 9 | 3764 | -30.45174 | -0.2587145 | No |
| **row_28** | CDKN3 | cyclin depend | 3845 | -31.46172 | -0.2641708 | No |
| **row_29** | HUS1 | HUS1 checkpo | 4028 | -33.576889 | -0.2771899 | No |
| **row_30** | MTF2 | metal respons | 4037 | -33.728752 | -0.2772477 | No |
| **row_31** | RAD54L | RAD54 like [So | 4194 | -35.597256 | -0.2882983 | No |
| **row_32** | CHMP1A | charged multi | 4431 | -38.242573 | -0.3052647 | No |
| **row_33** | KIF2C | kinesin family | 4508 | -39.165268 | -0.3103002 | No |
| **row_34** | CUL3 | cullin 3 [Sourc | 4719 | -41.466957 | -0.3252789 | No |
| **row_35** | TRAIP | TRAF interacti | 4725 | -41.51012 | -0.3249891 | No |
| **row_36** | DTYMK | deoxythymidy | 4751 | -41.814102 | -0.326184 | No |
| **row_37** | BARD1 | BRCA1 assoc | 4797 | -42.377369 | -0.3288595 | No |
| **row_38** | ODF2 | outer dense fil | 4825 | -42.705204 | -0.3301891 | No |
| **row_39** | TTK | TTK protein ki | 4907 | -43.637138 | -0.3355257 | No |
| **row_40** | CDC45 | cell division cy | 5123 | -46.428223 | -0.3507976 | No |
| **row_41** | CDKN2C | cyclin depend | 5190 | -47.380005 | -0.3549573 | No |
| **row_42** | TLE3 | TLE family me | 5206 | -47.593224 | -0.3553152 | No |
| **row_43** | DMD | dystrophin [So | 5282 | -48.756283 | -0.3601232 | No |

| | | | | | | |
|---|---|---|---|---|---|---|
| **row_44** | AMD 1.00 | adenosylmeth | 5300 | -48.989536 | -0.3606079 | No |
| **row_45** | KIF23 | kinesin family | 5319 | -49.317059 | -0.3611617 | No |
| **row_46** | TRA2B | transformer 2 | 5401 | -50.453259 | -0.3663895 | No |
| **row_47** | BIRC5 | baculoviral IAF | 5531 | -52.503952 | -0.3751595 | No |
| **row_48** | NUMA1 | nuclear mitotic | 5689 | -54.966728 | -0.3859756 | No |
| **row_49** | H2BC12 | H2B clustered | 5741 | -55.701981 | -0.3888854 | No |
| **row_50** | SFPQ | splicing factor | 5777 | -56.340992 | -0.3905933 | No |
| **row_51** | SUV39H1 | suppressor of | 5810 | -56.955692 | -0.392068 | No |
| **row_52** | NDC80 | NDC80 kineto | 5862 | -57.906651 | -0.3949426 | No |
| **row_53** | AURKB | aurora kinase | 5973 | -59.882671 | -0.4021798 | No |
| **row_54** | EZH2 | enhancer of ze | 6091 | -62.582741 | -0.4098953 | No |
| **row_55** | PBK | PDZ binding k | 6184 | -64.066917 | -0.4157252 | No |
| **row_56** | EWSR1 | EWS RNA bin | 6315 | -66.781548 | -0.4243419 | No |
| **row_57** | PRC1 | protein regula | 6429 | -69.195747 | -0.431654 | No |
| **row_58** | MNAT1 | MNAT1 comp | 6452 | -69.743767 | -0.4321799 | No |
| **row_59** | GINS2 | GINS complex | 6561 | -72.005676 | -0.4390748 | No |
| **row_60** | DBF4 | DBF4 zinc fin | 6652 | -74.192108 | -0.4445942 | No |
| **row_61** | SYNCRIP | synaptotagmi | 6810 | -77.959984 | -0.4550435 | No |
| **row_62** | RASAL2 | RAS protein a | 7033 | -83.569229 | -0.4702443 | No |
| **row_63** | KIF20B | kinesin family | 7050 | -83.883072 | -0.4700978 | No |
| **row_64** | NOLC1 | nucleolar and | 7096 | -85.067085 | -0.4720923 | No |
| **row_65** | POLA2 | DNA polymera | 7130 | -85.953621 | -0.4731789 | No |
| **row_66** | TROAP | trophinin asso | 7151 | -86.389275 | -0.4732904 | No |
| **row_67** | UPF1 | UPF1 RNA he | 7228 | -88.318268 | -0.4775418 | No |
| **row_68** | FANCC | FA compleme | 7283 | -89.90239 | -0.4801295 | No |
| **row_69** | CDC7 | cell division cy | 7344 | -91.169502 | -0.4831438 | No |
| **row_70** | ESPL1 | extra spindle p | 7348 | -91.336487 | -0.4819102 | No |
| **row_71** | CDC25A | cell division cy | 7358 | -91.505447 | -0.4811208 | No |
| **row_72** | TFDP1 | transcription fa | 7376 | -92.045364 | -0.4809187 | No |
| **row_73** | WRN | WRN RecQ lik | 7420 | -93.149597 | -0.4826353 | No |
| **row_74** | YTHDC1 | YTH domain c | 7430 | -93.257462 | -0.4818179 | No |
| **row_75** | ORC5 | origin recognit | 7626 | -98.557083 | -0.4947687 | No |
| **row_76** | MAD2L1 | mitotic arrest c | 7701 | -100.91022 | -0.4986703 | No |
| **row_77** | CCNB2 | cyclin B2 [Sou | 7747 | -102.57158 | -0.5003856 | No |
| **row_78** | UBE2C | ubiquitin conju | 7765 | -102.99552 | -0.5000088 | No |
| **row_79** | E2F3 | E2F transcript | 7825 | -104.78731 | -0.5027313 | No |
| **row_80** | CDC25B | cell division cy | 7862 | -106.17209 | -0.5037189 | No |
| **row_81** | CBX1 | chromobox 1 | 7879 | -106.52879 | -0.5032112 | No |
| **row_82** | SRSF2 | serine and arg | 7917 | -107.8864 | -0.5042458 | No |
| **row_83** | PURA | purine rich ele | 7972 | -109.50436 | -0.5065208 | No |
| **row_84** | CCNT1 | cyclin T1 [Sou | 7986 | -109.96906 | -0.5057348 | No |
| **row_85** | SAP30 | Sin3A associa | 8030 | -111.43317 | -0.5071598 | No |
| **row_86** | BUB1 | BUB1 mitotic | 8092 | -113.9671 | -0.5098848 | No |
| **row_87** | KIF4A | kinesin family | 8174 | -116.73381 | -0.5140554 | No |
| **row_88** | MAPK14 | mitogen-activa | 8225 | -119.39951 | -0.5158746 | No |

| | | | | | | |
|---|---|---|---|---|---|---|
| **row_89** | E2F1 | E2F transcript | 8378 | -125.27557 | -0.5251967 | No |
| **row_90** | KIF11 | kinesin family | 8414 | -126.267 | -0.5257892 | No |
| **row_91** | RACGAP1 | Rac GTPase a | 8426 | -126.51835 | -0.5245903 | No |
| **row_92** | CKS1B | CDC28 protei | 8463 | -127.76732 | -0.5252333 | No |
| **row_93** | NASP | nuclear autoa | 8688 | -136.51584 | -0.5397385 | No |
| **row_94** | CCNF | cyclin F [Sour | 8700 | -136.92657 | -0.5383735 | No |
| **row_95** | SMC2 | structural mai | 8705 | -137.05138 | -0.5364852 | No |
| **row_96** | CASP8AP2 | caspase 8 ass | 8716 | -137.65048 | -0.5350342 | No |
| **row_97** | CCNA2 | cyclin A2 [Sou | 8797 | -140.82231 | -0.5387459 | No |
| **row_98** | BUB3 | BUB3 mitotic | 8887 | -144.62422 | -0.5430674 | No |
| **row_99** | CKS2 | CDC28 protei | 8963 | -148.28918 | -0.5462877 | No |
| **row_100** | ODC1 | ornithine deca | 8964 | -148.41666 | -0.5439202 | No |
| **row_101** | LMNB1 | lamin B1 [Sou | 8992 | -149.72861 | -0.5435426 | No |
| **row_102** | HMGB3 | high mobility | 9095 | -154.35699 | -0.548677 | No |
| **row_103** | MAP3K20 | mitogen-activ | 9106 | -154.85654 | -0.5469515 | No |
| **row_104** | CHAF1A | chromatin ass | 9201 | -158.64874 | -0.5514216 | No |
| **row_105** | TGFB1 | transforming | 9212 | -159.09102 | -0.5496286 | No |
| **row_106** | INCENP | inner centrom | 9294 | -163.00812 | -0.553061 | No |
| **row_107** | SMC4 | structural mai | 9411 | -170.12029 | -0.5589865 | No |
| **row_108** | CDC20 | cell division cy | 9422 | -170.53409 | -0.557011 | No |
| **row_109** | HNRNPD | heterogeneou | 9583 | -179.479 | -0.5660643 | No |
| **row_110** | SRSF1 | serine and arg | 9733 | -186.9841 | -0.5741786 | No |
| **row_111** | RPA2 | replication pro | 9848 | -194.95108 | -0.5795592 | No |
| **row_112** | MCM5 | minichromoso | 9850 | -195.02733 | -0.5765226 | No |
| **row_113** | LIG3 | DNA ligase 3 [ | 9861 | -195.5159 | -0.5741485 | No |
| **row_114** | EFNA5 | ephrin A5 [Sou | 9992 | -204.29659 | -0.5805717 | No |
| **row_115** | MYBL2 | MYB proto-or | 10018 | -206.67432 | -0.5791367 | No |
| **row_116** | TACC3 | transforming a | 10151 | -215.93233 | -0.5855232 | No |
| **row_117** | SS18 | SS18 subunit | 10177 | -217.51537 | -0.5839154 | No |
| **row_118** | TENT4A | terminal nucle | 10203 | -219.22391 | -0.5822803 | No |
| **row_119** | SLC12A2 | solute carrier f | 10230 | -221.80647 | -0.5806785 | No |
| **row_120** | TMPO | thymopoietin | 10339 | -230.53242 | -0.5850446 | No |
| **row_121** | MKI67 | marker of prol | 10388 | -233.69356 | -0.5848917 | No |
| **row_122** | STAG1 | stromal antige | 10529 | -244.89174 | -0.591412 | No |
| **row_123** | KIF22 | kinesin family | 10673 | -257.79898 | -0.5979498 | No |
| **row_124** | CUL5 | cullin 5 [Sourc | 10689 | -259.2012 | -0.5949323 | No |
| **row_125** | SMAD3 | SMAD family r | 10796 | -269.55249 | -0.598527 | No |
| **row_126** | UCK2 | uridine-cytidin | 10840 | -274.98001 | -0.597343 | No |
| **row_127** | SNRPD1 | small nuclear r | 10902 | -281.40021 | -0.5973973 | No |
| **row_128** | E2F4 | E2F transcript | 10993 | -290.7146 | -0.5994628 | No |
| **row_129** | MEIS2 | Meis homeob | 11072 | -298.88486 | -0.6005043 | No |
| **row_130** | MCM3 | minichromoso | 11267 | -322.37292 | -0.6098104 | No |
| **row_131** | G3BP1 | G3BP stress g | 11399 | -336.71732 | -0.6141956 | Yes |
| **row_132** | ABL1 | ABL proto-on | 11407 | -338.06058 | -0.6093243 | Yes |
| **row_133** | POLE | DNA polymera | 11436 | -340.27371 | -0.6059817 | Yes |

| | | | | | | |
|---|---|---|---|---|---|---|
| **row_134** | DDX39A | DExD-box hel | 11514 | -351.53784 | -0.6061087 | Yes |
| **row_135** | PDS5B | PDS5 cohesin | 11529 | -353.74277 | -0.6015086 | Yes |
| **row_136** | NSD2 | nuclear recept | 11532 | -354.18997 | -0.5960076 | Yes |
| **row_137** | TPX2 | TPX2 microtul | 11534 | -354.46402 | -0.5904278 | Yes |
| **row_138** | CENPF | centromere pr | 11535 | -354.77417 | -0.5847685 | Yes |
| **row_139** | MCM2 | minichromosc | 11551 | -357.37616 | -0.5801849 | Yes |
| **row_140** | ILF3 | interleukin enh | 11585 | -361.5907 | -0.5768746 | Yes |
| **row_141** | CTCF | CCCTC-bindir | 11643 | -370.00684 | -0.5752175 | Yes |
| **row_142** | CUL4A | cullin 4A [Sou | 11667 | -372.9769 | -0.5709809 | Yes |
| **row_143** | LBR | lamin B recept | 11677 | -374.57886 | -0.5656759 | Yes |
| **row_144** | MT2A | metallothionei | 11755 | -385.98254 | -0.5652536 | Yes |
| **row_145** | DR1 | down-regulato | 11757 | -386.31314 | -0.5591657 | Yes |
| **row_146** | TNPO2 | transportin 2 [ | 11794 | -392.90076 | -0.5555794 | Yes |
| **row_147** | HIRA | histone cell cy | 11944 | -423.70001 | -0.5599177 | Yes |
| **row_148** | FOXN3 | forkhead box | 11948 | -424.52899 | -0.5533691 | Yes |
| **row_149** | CUL1 | cullin 1 [Sourc | 12227 | -487.68832 | -0.5662942 | Yes |
| **row_150** | PRPF4B | pre-mRNA prc | 12247 | -492.95303 | -0.5598458 | Yes |
| **row_151** | SLC7A1 | solute carrier f | 12328 | -521.00195 | -0.557493 | Yes |
| **row_152** | ATF5 | activating tran | 12389 | -537.63489 | -0.5533854 | Yes |
| **row_153** | MEIS1 | Meis homeob | 12427 | -553.81909 | -0.5473067 | Yes |
| **row_154** | H2AX | H2A.X variant | 12475 | -569.98444 | -0.5417149 | Yes |
| **row_155** | CDK4 | cyclin depend | 12533 | -586.95557 | -0.5365971 | Yes |
| **row_156** | CDKN1B | cyclin depend | 12554 | -594.79688 | -0.5285986 | Yes |
| **row_157** | TOP2A | DNA topoison | 12642 | -636.25903 | -0.5249286 | Yes |
| **row_158** | ATRX | ATRX chroma | 12739 | -676.07898 | -0.5212937 | Yes |
| **row_159** | HIF1A | hypoxia induc | 12801 | -705.68658 | -0.5145799 | Yes |
| **row_160** | GSPT1 | G1 to S phase | 12816 | -714.3399 | -0.5042276 | Yes |
| **row_161** | TOP 1.00 | DNA topoison | 12996 | -845.62341 | -0.5040698 | Yes |
| **row_162** | UBE2S | ubiquitin conju | 13006 | -854.74939 | -0.4911053 | Yes |
| **row_163** | SMARCC1 | SWI/SNF relat | 13016 | -865.88831 | -0.4779632 | Yes |
| **row_164** | KPNA2 | karyopherin su | 13067 | -914.93646 | -0.4670922 | Yes |
| **row_165** | CCND1 | cyclin D1 [Sou | 13099 | -947.91846 | -0.45428 | Yes |
| **row_166** | H2AZ1 | H2A.Z variant | 13153 | -1005.7172 | -0.4421843 | Yes |
| **row_167** | RAD23B | RAD23 homo | 13228 | -1120.9146 | -0.4298151 | Yes |
| **row_168** | RAD21 | RAD21 cohes | 13266 | -1197.0808 | -0.4134752 | Yes |
| **row_169** | KIF5B | kinesin family | 13308 | -1318.3112 | -0.3954994 | Yes |
| **row_170** | KPNB1 | karyopherin su | 13328 | -1367.2808 | -0.3751039 | Yes |
| **row_171** | XPO1 | exportin 1 [So | 13335 | -1380.3431 | -0.3535319 | Yes |
| **row_172** | SQLE | squalene epox | 13381 | -1530.3182 | -0.3324721 | Yes |
| **row_173** | HNRNPU | heterogeneou | 13383 | -1549.4451 | -0.3078302 | Yes |
| **row_174** | SLC7A5 | solute carrier f | 13439 | -1887.4026 | -0.2818191 | Yes |
| **row_175** | HMGN2 | high mobility g | 13450 | -2000.1737 | -0.2506576 | Yes |
| **row_176** | PAFAH1B1 | platelet activat | 13454 | -2039.5105 | -0.2183473 | Yes |
| **row_177** | NCL | nucleolin [Sou | 13532 | -3250.1819 | -0.1722359 | Yes |
| **row_178** | MARCKS | myristoylated | 13553 | -3776.4138 | -0.113485 | Yes |
