## Supplemental Table 4 for "Downregulation of Neurodevelopmental Gene Expression in iPSC-Derived Cerebral Organoids Upon Infection by Human Cytomegalovirus"

**Supplemental Table S1. Ingenuity Pathway Analysis (IPA) of GFP (+) versus Mock organoid s**

"Canonical Pathways" are a list of IPA-predicted pathways affected in GFP (+) vs Mock organoid su
"Upstream Regulators" are predicted key transcription factor regulators that distinguish GFP (+) vs
"Diseases and Functions" are predicted diseases or functional outcomes associated with GFI

**ubpopulations. Related to Figure 4.**

ubpopulations.

Mock organoid subpopulations.

ᵓ (+) vs Mock organoid subpopulations

© 2000-2021 QIAGEN. All rights reserved.

| Ingenuity Ca | -log(p-value | Ratio | z-score | Molecules |
|---|---|---|---|---|
| EIF2 Signalin | 47.9 | 0.527 | -7.034 | ACTA2,ACTB,AKT2,AKT3,ATF4,ATF5,CCN |
| Protein Ubiq | 22.9 | 0.356 | No activity p | AMFR,ANAPC5,B2M,BAP1,CDC34,CUL1, |
| Huntington's | 22.1 | 0.349 | -3.024 | AKT2,AKT3,AP2A2,ATF4,ATP5F1A,ATP5I |
| Regulation o | 21.9 | 0.413 | -0.816 | AKT2,AKT3,EIF1,EIF2S2,EIF3B,EIF3C,EIF |
| Inhibition of | 18.7 | 0.404 | -0.973 | AKT2,AKT3,CNOT1,CNOT3,DDX6,EDC4,E |
| BAG2 Signali | 18.1 | 0.524 | 0.243 | ANXA2,ATXN3,CDKN1A,CTSB,HSP90AA1 |
| Estrogen Rec | 17.8 | 0.285 | -6.754 | AKT2,AKT3,ATF4,ATP5F1A,ATP5F1B,ATI |
| mTOR Signal | 17.5 | 0.354 | -0.928 | AKT2,AKT3,CDC42,DDIT4,DGKZ,EIF3B,EI |
| Phagosome I | 16.9 | 0.39 | No activity p | ATP6AP1,ATP6AP2,ATP6V0A1,ATP6V0B |
| Mitochondria | 16.6 | 0.376 | No activity p | ACO2,APP,ATP5F1A,ATP5F1B,ATP5F1C, |
| Synaptogene | 16.4 | 0.301 | -8.102 | ACTR3,AFDN,AKT2,AKT3,AP1B1,AP2A1, |
| Polyamine R | 15.9 | 0.565 | No activity p | APC,AZIN1,CTNNB1,OAZ1,OAZ2,ODC1,F |
| FAT10 Signal | 15.8 | 0.589 | -1.342 | MAP1LC3B,PSMA1,PSMA4,PSMA5,PSM |
| Sirtuin Signa | 15.4 | 0.301 | 0.372 | ACLY,ACSS2,ADAM10,APEX1,APP,ATG4E |
| AMPK Signal | 13.4 | 0.306 | -1.131 | ACACA,ACTB,AK1,AKT2,AKT3,ARID1A,AT |
| Semaphorin | 13 | 0.358 | -0.555 | AKT2,AKT3,CD44,CRMP1,DPYSL2,DPYSL |
| Axonal Guida | 12.9 | 0.241 | No activity p | ABL1,ABLIM1,ACTR3,ADAM10,ADAM23 |
| Integrin Sign | 12.8 | 0.315 | -5.695 | ABL1,ACTA2,ACTB,ACTG1,ACTN1,ACTN |
| Signaling by | 12.8 | 0.291 | -6.018 | ACTA2,ACTB,ACTG1,ACTR3,ARHGEF17, |
| Tight Junctio | 12.5 | 0.331 | No activity p | ACTA2,ACTB,ACTG1,AFDN,AKT2,AKT3,A |
| Remodeling | 11.7 | 0.471 | -2.887 | ACTA2,ACTB,ACTG1,ACTN1,ACTN4,ACTI |
| Actin Cytosk | 11.6 | 0.29 | -5.939 | ABI2,ACTA2,ACTB,ACTG1,ACTN1,ACTN4 |
| Oxidative Ph | 11.5 | 0.385 | -6.481 | ATP5F1A,ATP5F1B,ATP5F1C,ATP5F1E,A |
| RhoGDI Sign | 11.1 | 0.298 | 4.423 | ACTA2,ACTB,ACTG1,ACTR3,ARHGAP5,A |
| Clathrin-mec | 10.8 | 0.306 | No activity p | ACTA2,ACTB,ACTG1,ACTR3,AP1B1,AP1 |
| Coronavirus I | 10.8 | 0.3 | 3.939 | ABL1,ADAM9,ATF4,ATP6AP1,CCND1,CD |
| BEX2 Signali | 10.7 | 0.42 | -0.686 | AKT2,AKT3,BEX1,BEX2,CCND1,CDH2,CD |
| RhoA Signali | 10.6 | 0.355 | -4.938 | ACTA2,ACTB,ACTG1,ACTR3,ARHGAP5,A |
| Ephrin Recep | 10 | 0.294 | -5.603 | ABL1,ACP1,ACTR3,ADAM10,AKT2,AKT3, |
| HIPPO signal | 9.97 | 0.4 | 3.13 | AJUBA,CD44,CSNK1D,CSNK1E,CUL1,DLG |
| Wnt/β-caten | 9.83 | 0.306 | 0 | ACVR1B,AKT2,AKT3,APC,BCL9,CCND1,C |
| Hypoxia Sign | 9.75 | 0.419 | -2.333 | ATF4,CDC34,CREBBP,CSNK1D,EP300,HIF |
| PI3K/AKT Sig | 9.74 | 0.291 | -2.188 | AKT2,AKT3,CCND1,CDC37,CDKN1A,CDKN |
| Epithelial Ad | 9.35 | 0.31 | No activity p | ACTA2,ACTB,ACTG1,ACTN1,ACTN4,ACTI |
| IGF-1 Signali | 9.06 | 0.356 | -5 | AKT2,AKT3,CCN1,CCN2,CSNK2A1,CSNK2 |
| Amyloid Proc | 8.94 | 0.471 | -2.333 | AKT2,AKT3,APP,BACE1,CAPN1,CAPN2,C |
| ILK Signaling | 8.89 | 0.283 | -3.355 | ACTA2,ACTB,ACTG1,ACTN1,ACTN4,AKT |
| ERK/MAPK S | 8.86 | 0.276 | -1.697 | ATF4,BCAR1,CREBBP,DUSP4,FYN,GRB2 |
| Insulin Secre | 8.84 | 0.257 | -7.07 | ASIC1,ATF4,CACNA1H,CACNB1,CACNB4, |
| Ephrin B Sign | 8.69 | 0.403 | -3.8 | ACP1,CAP1,CDC42,CTNNB1,CXCL12,EFN |
| Reelin Signal | 8.62 | 0.325 | -5.754 | ACTR3,AFDN,AKT2,AKT3,APBB1,APOE,A |
| Sertoli Cell-S | 8.2 | 0.272 | No activity p | A2M,ACTA2,ACTB,ACTG1,ACTN1,ACTN4 |
| Autophagy | 8.07 | 0.268 | -4.811 | AKT2,AKT3,ATF4,ATG4B,ATG9A,BNIP3, |

| | | | | |
|---|---|---|---|---|
| CDK5 Signali | 8.05 | 0.33 | -2.959 | ABL1,CAPN1,EGR1,GNAL,GNAS,ITGA2,I |
| RAR Activati | 7.72 | 0.27 | No activity p | ACTB,AKR1C3,AKR1C4,AKT2,AKT3,ARID1 |
| Protein Kinas | 7.64 | 0.221 | -1.053 | ACP1,ADD1,ADD2,AKAP12,AKAP9,ANAP |
| Molecular M | 7.6 | 0.216 | No activity p | ABL1,AKT2,AKT3,APC,ARHGEF17,ARHGI |
| Caveolar-me | 7.55 | 0.373 | No activity p | ABL1,ACTA2,ACTB,ACTG1,ARCN1,B2M, |
| Regulation o | 7.06 | 0.31 | -3.413 | ACTA2,ACTB,ACTR3,ARHGDIA,ARPC1A, |
| Ferroptosis S | 7.05 | 0.302 | -1.622 | ACACA,ACSL4,ARF1,ARF3,ARF4,ARF5,A |
| Virus Entry vi | 6.79 | 0.317 | No activity p | ABL1,ACTA2,ACTB,ACTG1,AP1B1,AP1G1 |
| PTEN Signali | 6.26 | 0.273 | 3.888 | AKT2,AKT3,BCAR1,CCND1,CDC42,CDKN1 |
| Androgen Sig | 6.03 | 0.26 | -3.962 | CACNA1H,CACNB1,CACNB4,CACNG7,CA |
| ATM Signalir | 5.97 | 0.309 | 1 | ABL1,ATF4,BRAT1,CBX1,CBX5,CDKN1A, |
| Rac Signaling | 5.94 | 0.275 | -4.243 | ABI2,ACTR3,ARPC1A,ARPC2,ARPC3,BRK |
| tRNA Chargi | 5.93 | 0.436 | -4.123 | DARS1,FARSA,GARS1,HARS1,IARS1,IAR |
| Endocannabi | 5.89 | 0.285 | -3.922 | AKT2,AKT3,ATF4,CCND1,CDKN1B,CNR1, |
| Germ Cell-Se | 5.88 | 0.257 | No activity p | A2M,ACTA2,ACTB,ACTG1,ACTN1,ACTN4 |
| Renal Cell Ca | 5.71 | 0.325 | -3.207 | AKT2,AKT3,CDC42,CREBBP,ELOB,EP300, |
| Cardiac β-adı | 5.66 | 0.253 | 1.512 | AKAP12,AKAP9,APEX1,ATP2A2,CACNB1, |
| Hepatic Fibro | 5.64 | 0.203 | -5.02 | ACTA2,ACVR1B,AKT2,AKT3,APC,ATF4,BI |
| Superpathwa | 5.63 | 0.483 | -3.742 | ACAT2,CYP51A1,DHCR7,FDFT1,FDPS,HM |
| Spliceosoma | 5.62 | 0.388 | -4.359 | CASC3,DDX39B,DDX46,DHX15,DHX38,EF |
| 14-3-3-medi | 5.54 | 0.276 | -2.746 | AKT2,AKT3,CDKN1B,GRB2,GSK3A,GSK3 |
| Leukocyte Ex | 5.51 | 0.244 | -4.422 | ABL1,ACTA2,ACTB,ACTG1,ACTN1,ACTN |
| Colorectal Ca | 5.49 | 0.223 | -5.864 | AKT2,AKT3,APC,CCND1,CDC42,CTNNB1, |
| Cell Cycle: G2 | 5.46 | 0.38 | -0.535 | ABL1,CDC34,CDKN1A,CUL1,EP300,FBXL5 |
| PFKFB4 Signa | 5.42 | 0.391 | -3.638 | ATF4,CREBBP,GNAS,GPI,HK1,MAP2K2,M |
| HIF1α Signal | 5.28 | 0.236 | -3.938 | AKT2,AKT3,APEX1,CDKN1A,CREBBP,EGL |
| Cyclins and C | 5.27 | 0.31 | 2.357 | ABL1,CCND1,CCND2,CDC34,CDK4,CDKN1 |
| Pancreatic A | 5.18 | 0.27 | -3.674 | ABL1,AKT2,AKT3,CCND1,CDC42,CDK4,CD |
| Unfolded pro | 5.16 | 0.3 | -3.873 | AMFR,ATF4,ATF6,CALR,CANX,CEBPG,DN |
| Cardiac Hype | 5.11 | 0.221 | -5.778 | CACNB1,CACNB4,CACNG7,CALM1 (inclu |
| MSP-RON Sig | 5.08 | 0.261 | -4.564 | ACTA2,AKT2,AKT3,ATF4,CCND1,CREBBP |
| Telomerase S | 5 | 0.28 | -0.816 | ABL1,AKT2,AKT3,CDKN1A,GRB2,HDAC2, |
| HER-2 Signal | 4.95 | 0.235 | -1.474 | AKT2,AKT3,ARF1,ARF3,ARF4,ARF5,CCN |
| Insulin Recep | 4.93 | 0.257 | -2.333 | ACLY,AKT2,AKT3,ASIC1,FYN,GRB2,GSK3 |
| Cell Cycle: G1 | 4.89 | 0.324 | -0.5 | ABL1,CCND1,CCND2,CDC34,CDK4,CDKN1 |
| Sumoylation | 4.89 | 0.282 | 0.655 | ARHGDIA,CBX4,CDC42,CREBBP,CTBP1,C |
| Fcγ Receptor | 4.77 | 0.287 | -4.811 | ACTA2,ACTB,ACTG1,ACTR3,AKT2,AKT3, |
| EGF Signalin | 4.76 | 0.345 | -4.243 | AKT2,AKT3,CSNK2A1,CSNK2B,GRB2,JAK |
| Endocannabi | 4.71 | 0.252 | 1.859 | AKT2,AKT3,ATF4,CAMKK2,CCND1,CCND |
| p70S6K Signa | 4.71 | 0.258 | -1.89 | AKT2,AKT3,EEF2,GNAI1,GRB2,JAK1,MA |
| Glycolysis I | 4.66 | 0.462 | -3.464 | ALDOA,ALDOC,ENO1,ENO2,GPI,PFKL,PF |
| FAK Signaling | 4.61 | 0.265 | No activity p | ACTA2,ACTB,ACTG1,AKT2,AKT3,ARHGE |
| Prostate Can | 4.58 | 0.268 | No activity p | ABL1,AKT2,AKT3,ATF4,CCND1,CDKN1A, |
| 3-phosphoin | 4.54 | 0.227 | -4.564 | ACP1,ATP1A1,CDIPT,DUSP4,DUSP8,G6P |
| G Beta Gam | 4.52 | 0.256 | -5.303 | AKT2,AKT3,CACNA1H,CACNB1,CACNB4, |

| | | | | |
|---|---|---|---|---|
| Semaphorin | 4.48 | 0.323 | No activity p | CDC42,CRMP1,DPYSL2,DPYSL3,DPYSL4,F |
| Paxillin Signa | 4.47 | 0.269 | -3.266 | ACTA2,ACTB,ACTG1,ACTN1,ACTN4,ARF |
| Cholesterol E | 4.41 | 0.615 | -2.828 | CYP51A1,DHCR7,FDFT1,LBR,LSS,MSMO |
| Cholesterol E | 4.41 | 0.615 | -2.828 | CYP51A1,DHCR7,FDFT1,LBR,LSS,MSMO |
| Cholesterol E | 4.41 | 0.615 | -2.828 | CYP51A1,DHCR7,FDFT1,LBR,LSS,MSMO |
| Mouse Embr | 4.36 | 0.269 | -4.315 | AKT2,AKT3,APC,CREBBP,CTNNB1,FZD3, |
| Coronavirus I | 4.28 | 0.356 | -3.5 | ARCN1,COPB1,COPE,COPZ1,GSK3A,GSK |
| Role of NFAT | 4.27 | 0.218 | -4.867 | AKT2,AKT3,CABIN1,CACNA1H,CACNB1,C |
| Opioid Signa | 4.22 | 0.207 | -6.062 | AKT2,AKT3,AP1B1,AP2A1,AP2A2,AP2B1 |
| Superpathwa | 4.22 | 0.215 | -4.964 | ACP1,ATP1A1,CDIPT,DUSP4,DUSP8,G6P |
| RAN Signalir | 4.2 | 0.529 | -3 | CSE1L,IPO5,KPNA2,KPNA3,KPNB1,RANB |
| Cell Cycle Re | 4.15 | 0.378 | 1 | BTG1,CCND1,CDK4,E2F6,PPM1L,PPP2CA |
| PAK Signaling | 4.12 | 0.254 | -3.53 | ARHGEF7,CDC42,DSCAM,GIT1,GRB2,ITG |
| Dopamine-D | 4.12 | 0.227 | -3.452 | ATF4,ATP2A2,CACNB1,CACNB4,CACNG7 |
| Ovarian Canc | 4.09 | 0.234 | -3.771 | ABL1,AKT2,AKT3,APC,CCND1,CD44,CDK4 |
| Aryl Hydroca | 4.03 | 0.233 | -1.347 | ALDH18A1,ALDH2,ALDH3B1,ALDH6A1,A |
| NRF2-media | 4.03 | 0.211 | -4.811 | ACTA2,ACTB,ACTG1,ATF4,CCT7,CDC34,C |
| Hereditary B | 4.02 | 0.239 | No activity p | ACTB,AKT2,AKT3,ARID1A,CCND1,CCND2 |
| Glioblastoma | 4.02 | 0.228 | -3.536 | AKT2,AKT3,APC,CCND1,CCND2,CDC42,C |
| Regulation o | 4 | 0.221 | No activity p | AKT2,AKT3,APC,BCL9,CDH2,CTNNB1,EG |
| GABA Recep | 3.99 | 0.244 | No activity p | ALDH9A1,AP1B1,AP2A1,AP2A2,AP2B1,A |
| Dopamine R | 3.97 | 0.286 | -2.449 | PPM1L,PPP1CA,PPP1CB,PPP1CC,PPP1R1 |
| GP6 Signalin | 3.88 | 0.244 | -4.017 | ADAM10,AKT2,AKT3,CALM1 (includes o |
| Gap Junction | 3.84 | 0.217 | No activity p | ACTA2,ACTB,ACTG1,AKT2,AKT3,CSNK1A |
| Neuregulin S | 3.79 | 0.248 | -2.982 | AKT2,AKT3,CDKN1B,DLG4,ERBB2,GRB2, |
| PPARα/RXRα | 3.79 | 0.218 | 1.521 | ACVR1B,ADIPOR1,ADIPOR2,AP2A2,CAN |
| Chronic Myel | 3.71 | 0.252 | No activity p | ABL1,AKT2,AKT3,CCND1,CCND2,CDK4,C |
| Glioma Signa | 3.7 | 0.242 | -4.123 | ABL1,AKT2,AKT3,CALM1 (includes other |
| D-myo-inosit | 3.68 | 0.22 | -4.158 | ACP1,ATP1A1,DUSP4,DUSP8,G6PC3,ITP |
| D-myo-inosit | 3.68 | 0.22 | -4.158 | ACP1,ATP1A1,DUSP4,DUSP8,G6PC3,ITP |
| Agrin Interac | 3.66 | 0.286 | -3.3 | ACTA2,ACTB,ACTG1,AGRN,ARHGEF7,C |
| CXCR4 Signal | 3.58 | 0.222 | -4.811 | AKT2,AKT3,BCAR1,CDC42,CXCL12,EGR1, |
| Mitotic Roles | 3.55 | 0.288 | -1.342 | ANAPC5,CAPN1,FZR1,HSP90AA1,HSP90 |
| Regulation C | 3.53 | 0.214 | -4.867 | AKT2,AKT3,CDC42,CDH2,EGR1,ERBB2,F |
| IL-22 Signalir | 3.51 | 0.417 | -3 | AKT2,AKT3,JAK1,MAP2K4,MAPK1,MAPK |
| D-myo-inosit | 3.48 | 0.212 | -4.017 | ACP1,ATP1A1,DUSP4,DUSP8,G6PC3,ITP |
| IL-1 Signaling | 3.45 | 0.255 | -2.887 | ELP1,GNA12,GNA13,GNAI1,GNAL,GNA |
| IL-7 Signaling | 3.42 | 0.269 | -2.524 | AKT2,AKT3,CCND1,CDKN1B,FOXG1,FYN, |
| α-Adrenergic | 3.39 | 0.245 | -3.317 | CALM1 (includes others),GNA12,GNA13 |
| 3-phosphoino | 3.38 | 0.212 | -4.271 | ACP1,ATP1A1,DUSP4,DUSP8,G6PC3,INP |
| Oncostatin M | 3.36 | 0.326 | -3.742 | GRB2,IL6ST,JAK1,MAP2K2,MAPK1,MAP |
| Production o | 3.28 | 0.209 | -0.686 | AKT2,AKT3,APOE,CDC42,CLU,CREBBP,E |
| Calcium Sign | 3.27 | 0.204 | -4.041 | ACTA2,ATF4,ATP2A2,ATP2B1,ATP2C1,C |
| Thrombin Sig | 3.26 | 0.205 | -4.796 | AKT2,AKT3,ARHGEF2,ARHGEF9,CDC42,C |
| G Protein Sig | 3.24 | 0.318 | No activity p | ABL1,GNA12,GNA13,GNAI1,GNAL,GNA |

| | | | | |
|---|---|---|---|---|
| PDGF Signali | 3.22 | 0.256 | -3.411 | ABL1,ACP1,CSNK2A1,CSNK2B,EIF2AK2,G |
| Gluconeoger | 3.18 | 0.385 | -3.162 | ALDOA,ALDOC,ENO1,ENO2,GPI,MDH1,M |
| DNA Methyla | 3.18 | 0.343 | No activity pa | CHD3,CHD4,DNMT1,H3-3A/H3-3B,HDAC |
| ErbB2-ErbB3 | 3.18 | 0.277 | -2.5 | CCND1,CDKN1B,ERBB2,GRB2,GSK3A,GS |
| Hepatic Fibro | 3.14 | 0.206 | No activity pa | A2M,ACTA2,CCN2,COL11A1,COL12A1,CO |
| Actin Nuclea | 3.12 | 0.247 | -2.84 | ACTR3,ARPC1A,ARPC2,ARPC3,CDC42,GN |
| Senescence I | 3.11 | 0.189 | -4.061 | ACVR1B,AKT2,AKT3,ANAPC5,ASXL1,CAC |
| Aldosterone | 3.1 | 0.213 | -3.5 | AHCY,ASIC1,DNAJA1,DNAJB1,DNAJB11,I |
| Dilated Cardi | 3.09 | 0.219 | -0.6 | ACTA2,ACTB,ACTG1,AKT2,AKT3,ATP2A2 |
| Regulation o | 3.01 | 0.247 | -2.309 | ACTN1,ACTN4,CAPN1,CAPN2,CAPNS1,C |
| TR/RXR Activ | 2.96 | 0.25 | No activity pa | ACACA,AKR1C3,AKT2,AKT3,COL6A3,ENC |
| CNTF Signali | 2.95 | 0.281 | -4 | CNTFR,GRB2,IL6ST,JAK1,LIFR,MAP2K2,M |
| IL-8 Signaling | 2.95 | 0.199 | -5.488 | AKT2,AKT3,CCND1,CCND2,CDC42,ELP1,G |
| Granzyme A | 2.94 | 0.421 | No activity pa | ANP32A,APEX1,CREBBP,EP300,H1-0,H1- |
| fMLP Signalir | 2.93 | 0.221 | -4.146 | ACTR3,ARPC1A,ARPC2,ARPC3,CALM1 (ir |
| IL-3 Signaling | 2.92 | 0.253 | -4.472 | AKT2,AKT3,GAB2,GRB2,JAK1,JUN,MAP2 |
| FLT3 Signalir | 2.85 | 0.25 | -4.359 | AKT2,AKT3,ATF4,CREBBP,GAB2,GRB2,M |
| NER (Nucleo | 2.84 | 0.233 | -2.4 | CETN2,COPS3,COPS6,COPS8,CUL4A,DDE |
| Iron homeos | 2.82 | 0.216 | No activity pa | ACO2,ATP6AP1,ATP6AP2,ATP6V0A1,ATI |
| p53 Signaling | 2.79 | 0.235 | -0.258 | AKT2,AKT3,CCND1,CCND2,CCNG1,CCNK, |
| Role of Tissu | 2.77 | 0.224 | No activity pa | AKT2,AKT3,CCN1,CCN2,CDC42,EGR1,FGA |
| IL-6 Signaling | 2.77 | 0.219 | -3.922 | A2M,AKT2,AKT3,COL1A1,CRP,CSNK2A1, |
| VEGF Signali | 2.73 | 0.232 | -4.359 | ACTA2,ACTB,ACTG1,ACTN1,ACTN4,AKT2 |
| Inhibition of | 2.73 | 0.324 | -3 | AKT2,AKT3,CD47,FYN,HSPG2,JUN,MAP2 |
| JAK/Stat Sigr | 2.71 | 0.244 | -3.578 | AKT2,AKT3,CDKN1A,GRB2,JAK1,JUN,MA |
| Endometrial | 2.7 | 0.267 | -2.673 | AKT2,AKT3,CCND1,CTNNA1,CTNNA2,CT |
| Non-Small Ce | 2.68 | 0.234 | -3.742 | ABL1,AKT2,AKT3,CCND1,CCND2,CDK4,E2 |
| Role of JAK f | 2.68 | 0.36 | No activity pa | IL6ST,JAK1,MAP2K4,MAPK1,MAPK11,MA |
| Xenobiotic M | 2.68 | 0.198 | -3.244 | ALDH18A1,ALDH2,ALDH3B1,ALDH6A1,AL |
| Netrin Signal | 2.62 | 0.25 | -3.771 | ABLIM1,CACNA1H,CACNB1,CACNB4,CAC |
| Glucocorticoi | 2.61 | 0.164 | No activity pa | A2M,ACTB,AKT2,AKT3,ANXA1,ARID1A,A |
| Gαi Signaling | 2.57 | 0.21 | -1.606 | CNR1,GABBR1,GNA12,GNA13,GNAI1,G |
| HOTAIR Regu | 2.56 | 0.202 | -4.243 | AKT2,AKT3,BRD4,CD44,CDKN1A,COL1A1 |
| Renin-Angiot | 2.55 | 0.217 | -4.379 | GRB2,JUN,MAP2K2,MAP2K4,MAPK1,MA |
| Zymosterol E | 2.55 | 0.667 | -2 | CYP51A1,LBR,MSMO1,TM7SF2 |
| Gαq Signalin | 2.53 | 0.2 | -4.264 | AKT2,AKT3,ARHGEF25,CALM1 (includes |
| Mechanisms | 2.51 | 0.293 | No activity pa | ACTA2,ACTB,ACTG1,CHMP4B,LMNB1,LM |
| Apelin Adipo | 2.45 | 0.233 | -0.728 | GNAI1,GPX1,GPX4,GPX8,GSTP1,HIF1A,M |
| Relaxin Signa | 2.45 | 0.203 | -4.243 | AKT2,AKT3,APEX1,GDE1,GNA12,GNA13 |
| CCR3 Signalir | 2.41 | 0.207 | -1.897 | CALM1 (includes others),CCR3,GNA12,G |
| Adipogenesis | 2.41 | 0.207 | No activity pa | BSCL2,CTBP1,CTBP2,CTNNB1,DGKD,DLK |
| Synaptic Lon | 2.41 | 0.209 | -2.887 | ATF4,CALM1 (includes others),CREBBP,I |
| Pentose Phos | 2.38 | 0.5 | -2.236 | G6PD,H6PD,PGD,TALDO1,TKT |
| Apelin Muscl | 2.35 | 0.271 | No activity pa | AKT2,AKT3,GNA12,GNA13,GNAI1,GNAL |
| Cellular Effec | 2.35 | 0.201 | No activity pa | ACTA2,ACTB,ACTG1,CACNB1,CACNB4,CA |

| | | | | |
|---|---|---|---|---|
| NGF Signalir | 2.34 | 0.212 | -4.69 | AKT2,AKT3,ATF4,CDC42,CREBBP,GRB2,I |
| ErbB Signalir | 2.33 | 0.223 | -2.982 | CDC42,ERBB2,GRB2,GSK3B,JUN,MAP2K |
| GNRH Signal | 2.32 | 0.191 | -4.536 | ATF4,CACNA1H,CACNB1,CACNB4,CACN( |
| Xenobiotic M | 2.28 | 0.19 | -3.333 | ALDH18A1,ALDH2,ALDH3B1,ALDH6A1,AI |
| HGF Signalin | 2.26 | 0.205 | -3.962 | AKT2,AKT3,CCND1,CDC42,CDKN1A,GRB2 |
| Pyridoxal 5'-ị | 2.25 | 0.242 | -3 | AKT2,CDK4,CSNK1A1,CSNK1D,EIF2AK2,F |
| Thioredoxin I | 2.23 | 0.571 | -2 | NXN,SELENOT,TXN,TXNRD1 |
| Inhibition of | 2.21 | 0.282 | 0 | A2M,ADAM10,HSPG2,MMP14,MMP15,M |
| Antigen Pres | 2.21 | 0.282 | No activity pa | B2M,CALR,CANX,CD74,HLA-A,HLA-B,HLA |
| TCA Cycle II ( | 2.2 | 0.333 | -2.828 | ACO2,DLD,DLST,FH,MDH1,MDH2,OGDH, |
| Gαs Signalin | 2.19 | 0.207 | -3.153 | ADD1,ADD2,ATF4,CNR1,CREBBP,GNA12 |
| Acute Phase | 2.19 | 0.189 | -3.272 | A2M,AKT2,AKT3,C4A/C4B,CRABP1,CRAE |
| Sonic Hedgel | 2.19 | 0.31 | 1.667 | GRK2,GSK3B,PRKACA,PRKACB,PRKAR1A |
| Telomere Ex | 2.19 | 0.4 | No activity pa | HNRNPA1,HNRNPA2B1,TERF2IP,TNKS,X |
| Amyotrophic | 2.18 | 0.209 | -2 | AKT3,CACNB1,CACNB4,CACNG7,CAPN1, |
| Pentose Phos | 2.17 | 0.75 | No activity pa | G6PD,H6PD,PGD |
| Role of CHK I | 2.09 | 0.246 | -3.317 | ATMIN,CDKN1A,E2F6,MDC1,PCNA,PPM1 |
| Kinetochore I | 2.07 | 0.21 | 1.528 | ANAPC5,ARPP19,BOD1,CENPT,DSN1,EN |
| Granzyme B | 2.03 | 0.375 | -1.633 | DFFB,LMNB1,LMNB2,NUMA1,PARP1,PF |
| Parkinson's S | 2.03 | 0.375 | No activity pa | MAPK1,MAPK11,PARK7,SEPTIN5,SNCA,U |
| Melanocyte I | 2 | 0.213 | -4.472 | ATF4,CREBBP,EP300,GNAS,GRB2,MAP2 |
| GM-CSF Sigr | 2 | 0.229 | -3.742 | AKT2,AKT3,CCND1,GRB2,MAP2K2,MAPK |
| Gα12/13 Sig | 1.94 | 0.195 | -3.8 | AKT2,AKT3,CDC42,CDH11,CDH2,CDH4,CT |
| Human Embr | 1.92 | 0.187 | No activity pa | AKT2,AKT3,APC,CTNNB1,FGFR1,FGFR3, |
| Endocannabi | 1.9 | 0.19 | -2.6 | CACNA1H,CACNB1,CACNB4,CACNG7,CN |
| PCP pathway | 1.89 | 0.233 | -3.606 | CELSR1,DAAM1,EFNB1,FZD3,FZD7,JUN, |
| Xenobiotic M | 1.89 | 0.17 | No activity pa | ALDH18A1,ALDH2,ALDH3B1,ALDH6A1,AI |
| Melatonin Si | 1.88 | 0.222 | -4 | CALM1 (includes others),GNAI1,GNAO1 |
| P2Y Purigeni | 1.84 | 0.194 | -4.379 | AKT2,AKT3,ATF4,CREBBP,GNAI1,GNB1, |
| Notch Signal | 1.84 | 0.263 | -1.89 | CNTN1,DTX3,DTX4,FURIN,JAG2,NCSTN, |
| Corticotropin | 1.82 | 0.188 | -3.128 | ATF4,CACNA1H,CACNB1,CACNB4,CACN( |
| IL-2 Signaling | 1.82 | 0.23 | -3.317 | AKT2,AKT3,CSNK2A1,CSNK2B,GRB2,JAK |
| White Adipo | 1.82 | 0.191 | -4.2 | ATF4,CACNA1H,CACNB1,CACNB4,CACN( |
| Epoxysqualer | 1.82 | 1 | No activity pa | FDFT1,SQLE |
| Spermine Bio | 1.82 | 1 | No activity pa | AMD1,SMS |
| Formaldehyc | 1.82 | 1 | No activity pa | ADH5,ESD |
| 2-ketoglutara | 1.81 | 0.6 | No activity pa | DLD,DLST,OGDH |
| Prolactin Sigr | 1.78 | 0.209 | -3.5 | CREBBP,EP300,FYN,GRB2,JUN,MAP2K2 |
| Chemokine S | 1.77 | 0.212 | -2 | CALM1 (includes others),CCR3,CXCL12,G |
| Leptin Signal | 1.77 | 0.216 | -2.828 | AKT2,AKT3,GRB2,MAP2K2,MAPK1,MAPI |
| Superpathwa | 1.76 | 0.333 | -2.449 | ACAT2,FDPS,HMGCR,HMGCS1,IDI1,MVD |
| BMP signalir | 1.73 | 0.207 | -4 | CREBBP,GRB2,JUN,MAGED1,MAP2K2,M |
| Xenobiotic M | 1.73 | 0.207 | -0.471 | ALDH18A1,ALDH2,ALDH3B1,ALDH6A1,AI |
| Nitric Oxide S | 1.73 | 0.193 | -3.9 | AKT2,AKT3,ATP2A2,CACNB1,CACNB4,CA |
| Apelin Endot | 1.71 | 0.187 | -4.025 | AKT2,AKT3,CALM1 (includes others),GN |

| | | | | |
|---|---|---|---|---|
| Thrombopoie | 1.71 | 0.222 | -3.742 | GAB2,GRB2,JUN,MAP2K2,MAPK1,MAPK |
| Role of NAN | 1.68 | 0.192 | -3.441 | AKT2,AKT3,APC,CTNNB1,FZD3,FZD7,GR |
| Role of Oct4 | 1.66 | 0.239 | No activity pa | IGF2BP1,KDM5B,NANOG,NR2F1,NR2F2 |
| DNA Double- | 1.66 | 0.357 | No activity pa | DCLRE1C,PARP1,PRKDC,XRCC5,XRCC6 |
| Mevalonate | 1.66 | 0.357 | -2.236 | ACAT2,HMGCR,HMGCS1,IDI1,MVD |
| Cysteine Bios | 1.65 | 0.292 | -2.646 | AHCY,AHCYL1,CBS/CBSL,EHMT1,MAT2A |
| Growth Horn | 1.6 | 0.211 | -3.051 | A2M,GHR,IGF1R,IGF2,IGFBP3,MAPK1,M |
| Ephrin A Sigr | 1.6 | 0.234 | No activity pa | ADAM10,BCAR1,CDC42,EFNA3,EPHA4,F |
| Ceramide Sig | 1.59 | 0.2 | -2.828 | AKT2,AKT3,CTSD,JUN,MAP2K4,MAPK3,P |
| Bladder Canc | 1.59 | 0.19 | -2.646 | ABL1,CCND1,CDK4,CDKN1A,E2F6,ERBB2 |
| Calcium Trar | 1.58 | 0.4 | -2 | ANXA5,ATP2A2,ATP2B1,ATP2C1 |
| PEDF Signali | 1.58 | 0.202 | -1.5 | AKT2,AKT3,MAPK1,MAPK11,MAPK3,PIK3 |
| FGF Signalin | 1.58 | 0.202 | -4 | AKT2,AKT3,ATF4,CREBBP,FGF13,FGFR1 |
| Acute Myeloi | 1.55 | 0.198 | -3.873 | AKT2,AKT3,CCND1,GRB2,IDH1,IDH2,MA |
| Natural Kille | 1.51 | 0.171 | -2.401 | AKT2,AKT3,B2M,CDC42,COL18A1,COL1A |
| Osteoarthriti | 1.51 | 0.167 | -0.756 | ANXA2,ANXA5,ATF4,COL2A1,CREBBP,CT |
| Glioma Invas | 1.5 | 0.205 | -1.508 | CD44,CDC42,ITGAV,MAPK1,MAPK3,MM |
| Role of MAP | 1.49 | 0.188 | -3.9 | ATP6AP1,ATP6AP2,ATP6V0A1,ATP6V0B |
| Superpathwa | 1.49 | 0.243 | -3 | AHCY,AHCYL1,CBS/CBSL,DLD,EHMT1,GC |
| Cardiac Hype | 1.48 | 0.151 | -5.417 | ACVR1B,AKT2,AKT3,APEX1,ATF6,ATP2A |
| Role of JAK1, | 1.47 | 0.269 | No activity pa | JAK1,PTPN2,RELA,STAT1,STAT2,STAT3,T |
| D-myo-inosit | 1.47 | 0.269 | -1.89 | CDIPT,PI4KA,PIP4K2A,PIP4K2B,PIP5K1A,I |
| Apelin Liver S | 1.47 | 0.269 | -2.646 | COL18A1,COL1A1,COL2A1,GSK3A,GSK3B |
| Endoplasmic | 1.43 | 0.286 | -2.236 | ATF4,ATF6,CALR,HSP90B1,MBTPS1,XBP |
| PPAR Signali | 1.43 | 0.187 | 2.828 | CITED2,CREBBP,ELP1,EP300,GRB2,HSP9 |
| Melanoma S | 1.41 | 0.22 | -2.714 | AKT2,AKT3,CCND1,CDK4,CDKN1A,MAP2 |
| Hypusine Bio | 1.38 | 0.667 | No activity pa | DHPS,EIF5A |
| Neurotrophir | 1.36 | 0.197 | -3.873 | ATF4,CDC42,CREBBP,GRB2,JUN,MAP2K |
| Role of PKR i | 1.35 | 0.176 | -1.043 | EIF2AK2,ELP1,HMGB1,HSP90AA1,HSP90 |
| Acetyl-CoA B | 1.35 | 0.429 | No activity pa | DLD,PDHA1,PDHA2 |
| Aspartate De | 1.35 | 0.429 | No activity pa | GOT2,MDH1,MDH2 |
| Methionine D | 1.34 | 0.273 | -2.449 | AHCY,AHCYL1,EHMT1,MAT2A,PRMT1,PR |
| Necroptosis S | 1.34 | 0.172 | -4.041 | CAPN1,CAPN2,CAPNS1,EIF2AK2,FKBP1A |
| Breast Cance | 1.33 | 0.147 | -4.196 | ADGRA3,ADGRB2,ADGRE2,ADGRL1,AN |
| Adrenomedu | 1.33 | 0.166 | -4.131 | AKT2,AKT3,CALM1 (includes others),CSK |
| IL-15 Signali | 1.32 | 0.195 | -3.873 | AKT2,AKT3,JAK1,MAP2K2,MAPK1,MAPK |
| NAD Signalir | 1.31 | 0.172 | -3.71 | AKT2,AKT3,GJA1,GSK3B,H1-0,H1-10,HIF |
| Tumor Micro | 1.3 | 0.168 | -4.382 | AKT2,AKT3,CCND1,CD44,COL1A1,CXCL12 |
| Cleavage and | 1.29 | 0.333 | No activity pa | CPSF1,CPSF2,PABPN1,PAPOLA |
| 4-1BB Signal | 1.29 | 0.235 | -2.646 | ELP1,JUN,MAP2K2,MAP2K4,MAPK1,MA |
| Activation of | 1.28 | 0.2 | -1.941 | ADAR,CREBBP,ELP1,IFNA16,JUN,MAP2K |
| B Cell Recep | 1.27 | 0.165 | -4.271 | ABL1,AKT2,AKT3,ATF4,CALM1 (includes |
| Salvage Path | 1.27 | 0.184 | -3.3 | AK1,AKT2,CDK4,CMPK1,CSNK1A1,CSNK1 |
| ERK5 Signali | 1.26 | 0.194 | -3.742 | ATF4,CREBBP,GNA12,GNA13,IL6ST,RAF |
| Cholecystoki | 1.24 | 0.176 | -4.123 | BCAR1,CDC42,EPHA4,GNA12,GNA13,GR |

| | | | | |
|---|---|---|---|---|
| iNOS Signali | 1.24 | 0.213 | -2.646 | CALM1 (includes others),CREBBP,HMGA |
| nNOS Signali | 1.24 | 0.213 | -2.236 | CALM1 (includes others),CAPN1,CAPN2, |
| Thyroid Canc | 1.24 | 0.19 | -2.84 | AKT2,AKT3,CCND1,CTNNB1,GSK3B,JUN, |
| Estrogen-Dep | 1.24 | 0.19 | -3.464 | AKR1C4,AKT2,AKT3,ATF4,CCND1,CREBB |
| Glutamate R | 1.24 | 0.197 | -1.342 | CALM1 (includes others),DLG4,GLUL,GN |
| Role of p14/p | 1.23 | 0.241 | 2.449 | MDM2,NPM1,PIK3R3,RAC1,SENP3,SF3A |
| Apelin Cardic | 1.23 | 0.182 | -4.243 | AKT2,AKT3,ATP2A2,GNAI1,HIF1A,MAPK |
| Role of Macr | 1.23 | 0.154 | No activity pa | AKT2,AKT3,APC,ATF4,CALM1 (includes o |
| γ-linolenate | 1.21 | 0.278 | -1.342 | ACSL3,ACSL4,ACSL6,CYB5R3,FADS2 |
| Sperm Motili | 1.2 | 0.157 | -3.464 | ABL1,CACNA1H,CALM1 (includes others |
| Role of BRCA | 1.2 | 0.188 | -0.816 | ACTB,ARID1A,CDKN1A,DPF1,E2F6,MDC1 |
| IL-17A Signal | 1.19 | 0.194 | -3.317 | AKT2,AKT3,GSK3B,JAK1,MAP2K2,MAP2I |
| Regulation C | 1.18 | 0.184 | -2.673 | APC,BCL9,CTNNB1,FZD3,FZD7,GSK3B,H |
| Vitamin-C Tr | 1.18 | 0.25 | -1.633 | AKR1C4,NXN,SELENOT,SLC2A1,TXN,TXN |
| Lymphotoxin | 1.15 | 0.2 | -2.828 | AKT2,AKT3,CREBBP,ELP1,EP300,MAPK1, |
| ErbB4 Signal | 1.15 | 0.191 | -2.887 | GRB2,MAP2K2,MAPK1,MAPK3,NCSTN,N |
| Neuroinflam | 1.14 | 0.152 | -4.11 | ACVR1B,AKT2,AKT3,APP,ATF4,B2M,BAC |
| Branched-cha | 1.11 | 0.5 | No activity pa | BCKDHB,DLD |
| Glutathione | 1.11 | 0.5 | No activity pa | GSR,PDIA3 |
| Oxidized GTF | 1.11 | 0.5 | No activity pa | DDX6,RUVBL2 |
| Synaptic Long | 1.1 | 0.16 | -1.671 | CACNA1H,CACNB1,CACNB4,CACNG7,GN |
| TGF-β Signal | 1.1 | 0.177 | -3 | ACVR1B,CDC42,CREBBP,EP300,GRB2,JU |
| Glutathione | 1.1 | 0.24 | -0.816 | GPX1,GPX4,GPX8,GSR,GSTP1,PRDX6 |
| eNOS Signali | 1.09 | 0.164 | -2.837 | AKT2,AKT3,AQP1,CALM1 (includes other |
| FAT10 Cance | 1.09 | 0.2 | -1.897 | ACVR1B,AKT2,AKT3,CTNNB1,GSK3B,PCI |
| Apoptosis Sig | 1.06 | 0.173 | 0.5 | ACIN1,CAPN1,CAPN2,CAPNS1,DFFB,ELP |
| Docosahexae | 1.05 | 0.211 | No activity pa | AKT2,AKT3,APP,GSK3A,GSK3B,PDPK1,PI |
| Sucrose Degr | 1.05 | 0.333 | No activity pa | ALDOA,ALDOC,TPI1 |
| GADD45 Sigr | 1.04 | 0.25 | No activity pa | CCND1,CCND2,CDK4,CDKN1A,PCNA |
| Valine Degra | 1.04 | 0.25 | 0 | ALDH6A1,BCKDHB,DLD,ECHS1,HSD17B4 |
| IL-17A Signal | 1.03 | 0.231 | -2.236 | JUN,MAP2K4,MAPK1,MAPK11,MAPK3,R |
| Estrogen-me | 1.03 | 0.231 | 0.447 | CCND1,CDK4,CDKN1A,CDKN1B,E2F6,TFD |
| NF-κB Activa | 1.02 | 0.179 | -2.673 | AKT2,AKT3,EIF2AK2,ELP1,ITGA2,ITGA3,I |
| Role of Wnt/ | 1.02 | 0.179 | -0.816 | APC,CSNK1A1,CSNK1D,CSNK1E,CSNK1G |
| Erythropoieti | 1 | 0.158 | -4.315 | AKT2,AKT3,CCND1,CCND2,CTNNB1,GRB |
| STAT3 Pathw | 0.979 | 0.163 | -3.441 | CDKN1A,FGFR1,FGFR3,GHR,IGF1R,IL6S |
| Apelin Pancr | 0.971 | 0.196 | 2.333 | MAP2K4,PIK3R3,PRKACA,PRKACB,PRKAI |
| PI3K Signalin | 0.955 | 0.161 | -4.69 | ABL1,AKT2,AKT3,ATF4,ATF5,ATF6,ATF6 |
| Embryonic St | 0.936 | 0.3 | No activity pa | HOXB5,NANOG,SOX2 |
| CD40 Signali | 0.928 | 0.179 | -3.317 | ELP1,JUN,MAP2K2,MAP2K4,MAPK1,MA |
| Role of JAK2 | 0.928 | 0.206 | No activity pa | GHR,JAK1,PTPN1,SHC1,STAT1,STAT3,TY |
| Creatine-pho | 0.928 | 0.4 | No activity pa | CKB,MAP4K4 |
| Lysine Degra | 0.928 | 0.4 | No activity pa | AASDHPPT,ALDH7A1 |
| Lysine Degra | 0.928 | 0.4 | No activity pa | AASDHPPT,ALDH7A1 |
| Trans, trans- | 0.928 | 0.4 | No activity pa | FDPS,IDI1 |

| | | | | |
|---|---|---|---|---|
| dTMP De Nov | 0.928 | 0.4 | No activity pa | DHFR,SHMT2 |
| IL-15 Product | 0.921 | 0.163 | -4.472 | ABL1,CSK,DDR1,EPHA4,EPHB2,ERBB2,F( |
| Acyl Carrier F | 0.91 | 1 | No activity pa | AASDHPPT |
| Acetyl-CoA B | 0.91 | 1 | No activity pa | ACLY |
| Glutamine B | 0.91 | 1 | No activity pa | GLUL |
| Asparagine E | 0.91 | 1 | No activity pa | ASNS |
| Lanosterol Bi | 0.91 | 1 | No activity pa | LSS |
| Systemic Lup | 0.903 | 0.148 | -4.529 | AKT2,AKT3,CALM1 (includes others),CCN |
| Small Cell Lu | 0.886 | 0.167 | -2.121 | ABL1,AKT2,AKT3,CCND1,CCND2,CDK4,CD |
| Death Recep | 0.886 | 0.167 | -2.84 | ACIN1,ACTA2,ACTB,ACTG1,DFFB,ELP1,H |
| GDNF Family | 0.857 | 0.171 | -3.606 | CDC42,GRB2,JUN,MAP2K2,MAP2K4,MA |
| Ethanol Degr | 0.836 | 0.217 | -1.342 | ACSS2,ALDH2,ALDH3B1,ALDH7A1,ALDH9 |
| Interferon Si | 0.83 | 0.194 | -1.89 | IFITM3,JAK1,PTPN2,RELA,STAT1,STAT2, |
| RANK Signali | 0.827 | 0.165 | -3.606 | AKT2,AKT3,CALM1 (includes others),ELP |
| VEGF Family | 0.827 | 0.167 | -3.606 | AKT2,AKT3,GRB2,MAP2K2,MAPK1,MAP |
| Isoleucine De | 0.827 | 0.235 | No activity pa | ACAT2,DLD,ECHS1,HSD17B4 |
| Dermatan Su | 0.827 | 0.235 | -1 | CEMIP2,GM2A,IDS,OGA |
| Histamine De | 0.827 | 0.235 | -1 | ALDH2,ALDH3B1,ALDH7A1,ALDH9A1 |
| Myc Mediate | 0.804 | 0.18 | -1.667 | CREBBP,MCL1,MDM2,PRKACA,PRKACB, |
| LPS-stimulat | 0.796 | 0.165 | -3.464 | CDC42,ELP1,JUN,MAP2K2,MAP2K4,MAP |
| NAD Biosynth | 0.788 | 0.333 | No activity pa | NMNAT2,QPRT |
| Arginine Bios | 0.788 | 0.333 | No activity pa | GLUD1,OAT |
| Pyruvate Fer | 0.788 | 0.333 | No activity pa | LDHA,LDHB |
| Chondroitin a | 0.788 | 0.333 | No activity pa | CHPF,CHPF2 |
| Pentose Phos | 0.788 | 0.333 | No activity pa | TALDO1,TKT |
| Selenocystei | 0.788 | 0.333 | No activity pa | SARS1,SEPHS1 |
| NAD Biosynth | 0.788 | 0.333 | No activity pa | NAMPT,NMNAT2 |
| Cancer Drug | 0.762 | 0.172 | No activity pa | AKT2,AKT3,FOXG1,MAP2K2,MAPK1,MA |
| Neuropathic | 0.752 | 0.158 | -3.5 | GRIA1,GRINA,KCNN3,KCNQ2,MAPK1,M |
| BER pathway | 0.752 | 0.25 | No activity pa | APEX1,PARP1,PCNA |
| Heparan Sulf | 0.738 | 0.163 | -2.496 | B3GAT1,CES3,CHST10,CHST3,EXT1,EXT2 |
| Heparan Sulf | 0.733 | 0.164 | -2.309 | CES3,CHST10,CHST3,EXT1,EXT2,EXTL3,H |
| Role of PI3K/ | 0.73 | 0.167 | -1.897 | AKT2,AKT3,GNAI1,GSK3B,IFNA16,KPNA |
| UVB-Inducec | 0.73 | 0.173 | -2.646 | H3-3A/H3-3B,JUN,MAP2K4,MAPK1,MAF |
| Ethanol Degr | 0.708 | 0.188 | -1.633 | ACSS2,ADH5,ALDH2,ALDH3B1,ALDH7A1, |
| Assembly of | 0.678 | 0.231 | No activity pa | GTF3A,GTF3C1,SF3A1 |
| Superpathwa | 0.678 | 0.286 | No activity pa | PSAT1,SHMT2 |
| NAD Phospho | 0.678 | 0.231 | No activity pa | ACP1,NADK,NNT |
| Circadian Rh | 0.666 | 0.182 | No activity pa | ATF4,CREBBP,CSNK1D,CSNK1E,GRINA,F |
| Phototransdu | 0.664 | 0.167 | No activity pa | GNAT1,GNB1,PDC,PRKACA,PRKACB,PRK |
| Macropinocy | 0.65 | 0.158 | -2.646 | ACTN4,ANKFY1,CDC42,ITGB1,ITGB5,PIK |
| SPINK1 Gene | 0.644 | 0.159 | -2.714 | AKT2,AKT3,JAK1,MAP2K2,MAPK1,MAPK |
| Fatty Acid α- | 0.642 | 0.2 | -1 | ALDH2,ALDH3B1,ALDH7A1,ALDH9A1 |
| UDP-D-xylose | 0.635 | 0.5 | No activity pa | UGDH |
| β-alanine De | 0.635 | 0.5 | No activity pa | ALDH6A1 |

| | | | | |
|---|---|---|---|---|
| Choline Degr | 0.635 | 0.5 | No activity pa | ALDH7A1 |
| Alanine Degr | 0.635 | 0.5 | No activity pa | GPT2 |
| Alanine Biosy | 0.635 | 0.5 | No activity pa | GPT2 |
| Palmitate Bi | 0.635 | 0.5 | No activity pa | FASN |
| Uridine-5'-ph | 0.635 | 0.5 | No activity pa | CAD |
| Spermidine E | 0.635 | 0.5 | No activity pa | AMD1 |
| Sulfate Activ | 0.635 | 0.5 | No activity pa | PAPSS1 |
| Fatty Acid Bi | 0.635 | 0.5 | No activity pa | FASN |
| Cysteine Bios | 0.635 | 0.5 | No activity pa | CBS/CBSL |
| Putrescine Bi | 0.635 | 0.5 | No activity pa | ODC1 |
| Glycine Biosy | 0.635 | 0.5 | No activity pa | SHMT2 |
| Glutamate B | 0.635 | 0.5 | No activity pa | GLUD1 |
| Glutamate D | 0.635 | 0.5 | No activity pa | GLUD1 |
| PD-1, PD-L1 c | 0.635 | 0.151 | 2.673 | AKT2,AKT3,B2M,CDKN1B,CSK,CSNK2A1, |
| Factors Prom | 0.631 | 0.146 | -2.837 | ACVR1B,APC,ATF4,CCND1,CREBBP,CTNI |
| Antiprolifera | 0.625 | 0.156 | -1.667 | CDKN1A,CDKN1B,GNB1,GNB2,GNG2,GI |
| Fatty Acid β- | 0.625 | 0.176 | -1.633 | ACSL3,ACSL4,ACSL6,ECHS1,HSD17B4,IVI |
| IL-4 Signaling | 0.618 | 0.152 | No activity pa | AKT2,AKT3,GRB2,HLA-A,HLA-B,HLA-DPA |
| DNA Double- | 0.611 | 0.214 | No activity pa | ABL1,ATRX,RAD51 |
| Fatty Acid Ac | 0.611 | 0.214 | No activity pa | ACSL3,ACSL4,ACSL6 |
| Chondroitin S | 0.602 | 0.161 | -2.333 | B3GAT1,CHPF,CHPF2,CHST10,CHST3,HS( |
| Putrescine De | 0.592 | 0.19 | -1 | ALDH2,ALDH3B1,ALDH7A1,ALDH9A1 |
| April Mediate | 0.59 | 0.167 | -2.236 | ELP1,JUN,MAP2K4,MAPK1,MAPK11,REL |
| Superoxide R | 0.587 | 0.25 | No activity pa | SOD1,SOD2 |
| Nucleotide E | 0.587 | 0.171 | No activity pa | POLR2A,POLR2B,POLR2C,POLR2E,POLR |
| Wnt/Ca+ pat | 0.582 | 0.156 | -3.162 | ATF4,CREBBP,FZD3,FZD7,GSK3B,PDIA3, |
| GPCR-Media | 0.582 | 0.147 | -3.638 | CACNA1H,CACNB1,CACNB4,CACNG7,GN |
| Fc Epsilon RI | 0.564 | 0.145 | -4 | AKT2,AKT3,FYN,GRB2,MAP2K2,MAP2K4 |
| B Cell Activa | 0.558 | 0.163 | -2.449 | ELP1,JUN,MAP2K4,MAPK1,MAPK11,REL |
| PXR/RXR Act | 0.556 | 0.154 | No activity pa | AKT2,AKT3,CES3,PRKACA,PRKACB,PRKA |
| Sphingosine- | 0.545 | 0.144 | -3.606 | AKT2,AKT3,ASAH1,CDC42,GNA12,GNA1 |
| CDP-diacylgly | 0.545 | 0.182 | -2 | AGPAT1,AGPAT3,AGPAT4,MBOAT7 |
| Pyrimidine D | 0.545 | 0.182 | -2 | AK1,CMPK1,NME2,RRM1 |
| UVC-Induced | 0.538 | 0.157 | -2.646 | JUN,MAP2K2,MAP2K4,MAPK1,MAPK11, |
| MIF Regulati | 0.527 | 0.159 | -2.646 | CD74,JUN,MAP2K4,MAPK1,MAPK3,MIF, |
| Dermatan Su | 0.52 | 0.153 | -2.333 | B3GAT1,CHPF,CHPF2,CHST10,CHST3,HS( |
| Phosphatidyl | 0.513 | 0.222 | No activity pa | CHKA,SELENOI |
| Citrulline Bio | 0.513 | 0.222 | No activity pa | ALDH18A1,OAT |
| Dopamine De | 0.51 | 0.167 | -0.447 | ALDH2,ALDH3B1,ALDH7A1,ALDH9A1,SU |
| Xenobiotic M | 0.509 | 0.14 | -3.578 | ABCA2,AKT2,AKT3,CREBBP,CUL3,EP300, |
| Apelin Cardia | 0.503 | 0.174 | 0 | AKT2,AKT3,CCN2,TGFB2 |
| Mismatch Re | 0.5 | 0.188 | No activity pa | MSH2,MSH6,PCNA |
| Chondroitin S | 0.5 | 0.188 | No activity pa | CEMIP2,GM2A,OGA |
| IL-12 Signalir | 0.498 | 0.14 | No activity pa | AKT2,AKT3,APOE,CLU,ELP1,EP300,IFNA |
| Role of IL-17 | 0.498 | 0.156 | -2.646 | ATF4,CREBBP,MAP2K2,MAPK1,MAPK3,F |

| | | | | |
|---|---|---|---|---|
| UVA-Inducec | 0.492 | 0.143 | -2.828 | JUN,MAP2K4,MAPK1,MAPK11,MAPK3,N |
| IL-17A Signal | 0.487 | 0.158 | No activity pa | GSK3B,JUN,MAPK1,MAPK11,MAPK3,REI |
| L-carnitine B | 0.485 | 0.333 | No activity pa | ALDH9A1 |
| Diphthamide | 0.485 | 0.333 | No activity pa | EEF2 |
| Methylglyoxa | 0.485 | 0.333 | No activity pa | GLO1 |
| Biotin-carbox | 0.485 | 0.333 | No activity pa | ACACA |
| Thyroid Horm | 0.485 | 0.333 | No activity pa | CTSD |
| 1D-myo-inos | 0.485 | 0.333 | No activity pa | ITPK1 |
| Glutamate D | 0.485 | 0.333 | No activity pa | GOT2 |
| S-adenosyl-L | 0.485 | 0.333 | No activity pa | MAT2A |
| Aspartate Bi | 0.485 | 0.333 | No activity pa | GOT2 |
| Stearate Bios | 0.483 | 0.151 | -1.89 | ACOT7,ACSL3,ACSL4,ACSL6,FASN,MBOA |
| IL-23 Signalir | 0.471 | 0.152 | -2.646 | AKT2,AKT3,HIF1A,PIK3R3,RELA,STAT3,T |
| Phosphatidyl | 0.465 | 0.167 | -2 | AGPAT1,AGPAT3,AGPAT4,MBOAT7 |
| Glutaryl-CoA | 0.455 | 0.176 | No activity pa | ACAT2,HSD17B4,PARK7 |
| Toll-like Rece | 0.435 | 0.141 | -1.342 | EIF2AK2,JUN,MAP2K4,MAP4K4,MAPK1, |
| Tryptophan D | 0.428 | 0.16 | -1 | ALDH2,ALDH3B1,ALDH7A1,ALDH9A1 |
| Phagosome I | 0.42 | 0.134 | No activity pa | CDC42,FCGR3A/FCGR3B,FN1,ITGA11,IT |
| Chondroitin S | 0.419 | 0.146 | -1.89 | CHPF,CHST10,CHST3,HS6ST1,NDST1,SU |
| Neuroprotec | 0.416 | 0.136 | 0.905 | APP,HLA-A,HLA-B,HTRA1,LONP1,MAPT, |
| Cell Cycle Co | 0.412 | 0.143 | -2.121 | CDK11B,CDK16,CDK4,CDK9,MCM7,ORC4 |
| 1D-myo-inos | 0.412 | 0.167 | No activity pa | ITPK1,ITPKB,SEC16A |
| Mitochondria | 0.412 | 0.167 | No activity pa | ACSL3,ACSL4,ACSL6 |
| G-Protein Co | 0.411 | 0.13 | No activity pa | AKT2,AKT3,APEX1,ATF4,CNR1,CREBBP,D |
| cAMP-media | 0.406 | 0.131 | -1.89 | AKAP12,AKAP9,APEX1,ATF4,CALM1 (inc |
| IL-10 Signalir | 0.401 | 0.139 | No activity pa | ELP1,JAK1,JUN,MAP2K4,MAP4K4,MAPK |
| Agranulocyte | 0.397 | 0.131 | No activity pa | ACTA2,ACTB,ACTG1,CCR3,CD99,CXCL12, |
| Ketogenesis | 0.396 | 0.182 | No activity pa | ACAT2,HMGCS1 |
| Role of IL-17 | 0.39 | 0.14 | No activity pa | MAP2K2,MAP2K4,MAPK1,MAPK11,MAP |
| Retinoate Bi | 0.387 | 0.25 | No activity pa | RBP1 |
| PRPP Biosynt | 0.387 | 0.25 | No activity pa | PRPS1 |
| Phenylethyla | 0.387 | 0.25 | No activity pa | ALDH2 |
| Proline Biosy | 0.387 | 0.25 | No activity pa | ALDH18A1 |
| Arginine Deg | 0.387 | 0.25 | No activity pa | OAT |
| L-cysteine De | 0.387 | 0.25 | No activity pa | GOT2 |
| Acetate Conv | 0.387 | 0.25 | No activity pa | ACSS2 |
| GPCR-Media | 0.383 | 0.137 | -1.265 | GNAI1,GNAS,PDIA3,PLCG1,PRKACA,PRK |
| Type II Diabe | 0.381 | 0.132 | -2.496 | ACSL3,ACSL4,ACSL6,ADIPOR1,ADIPOR2, |
| Role of Oste | 0.375 | 0.129 | No activity pa | AKT2,AKT3,APC,CALM1 (includes others |
| Assembly of | 0.374 | 0.14 | -2.646 | POLR2A,POLR2B,POLR2C,POLR2E,POLR |
| IL-9 Signaling | 0.359 | 0.143 | -2.236 | JAK1,PIK3R3,RELA,STAT1,STAT3 |
| Noradrenalir | 0.359 | 0.143 | -1.342 | ADH5,ALDH2,ALDH3B1,ALDH7A1,ALDH9 |
| NAD biosynth | 0.35 | 0.167 | No activity pa | NMNAT2,QPRT |
| LXR/RXR Act | 0.345 | 0.13 | -1.667 | ACACA,APOE,C4A/C4B,CLU,CYP51A1,EC |
| The Visual Cy | 0.341 | 0.15 | No activity pa | AKR1C3,RBP1,RDH11 |

| | | | | |
|---|---|---|---|---|
| MIF-mediate | 0.334 | 0.139 | -2.236 | CD74,MAPK1,MAPK3,MIF,RELA |
| Retinoate Bi | 0.334 | 0.139 | 0.447 | AKR1C3,AKR1C4,NT5C1B,RBP1,RDH11 |
| Pyrimidine R | 0.334 | 0.139 | -2.236 | AK1,CAD,CMPK1,DDX3X,NME2 |
| Retinol Biosy | 0.333 | 0.136 | -0.816 | AKR1C3,CES3,DDHD2,ESD,RBP1,RDH11 |
| MSP-RON Si | 0.319 | 0.128 | -1.604 | ATF4,CREBBP,GAB2,GRB2,HLA-DPA1,JU |
| FcγRIIB Signa | 0.317 | 0.129 | -2.646 | CACNA1H,CACNB1,CACNB4,CACNG7,GR |
| Proline Biosy | 0.316 | 0.2 | No activity p | OAT |
| Serine Biosy | 0.316 | 0.2 | No activity p | PSAT1 |
| Eumelanin B | 0.316 | 0.2 | No activity p | MIF |
| Myo-inositol | 0.316 | 0.2 | No activity p | ISYNA1 |
| 2-oxobutano | 0.316 | 0.2 | No activity p | DLD |
| Folate Polygl | 0.316 | 0.2 | No activity p | SHMT2 |
| Role of JAK1 | 0.315 | 0.13 | No activity p | GRB2,JAK1,MAPK1,MAPK3,PIK3R3,RAP1 |
| Acyl-CoA Hyc | 0.31 | 0.154 | No activity p | ACOT7,PPT1 |
| Oleate Biosy | 0.31 | 0.154 | No activity p | ALDH6A1,FADS2 |
| Bile Acid Bio | 0.31 | 0.154 | No activity p | AKR1C3,AKR1C4 |
| UDP-N-acety | 0.31 | 0.154 | No activity p | GPI,HK1 |
| Endothelin-1 | 0.305 | 0.126 | -4.264 | EDNRB,GNA12,GNA13,GNAI1,GNAL,GN |
| VDR/RXR Ac | 0.301 | 0.128 | 0 | CDKN1A,CDKN1B,CSNK2A1,EP300,IGFBI |
| Transcriptior | 0.296 | 0.13 | No activity p | H3-3A/H3-3B,L1CAM,NANOG,SET,SOX2 |
| Role of RIG1 | 0.293 | 0.13 | 0 | CREBBP,EP300,IFNA16,MAVS,RELA,TRA |
| Dermatan Su | 0.293 | 0.13 | -1.633 | CHST10,CHST3,HS6ST1,NDST1,SULT1C4 |
| Antioxidant A | 0.281 | 0.125 | 2.646 | MAP2K4,MAPK1,MAPK11,MAPK3,NXN,F |
| Glycogen De | 0.276 | 0.143 | No activity p | GAA,PYGB |
| Urate Biosyn | 0.276 | 0.143 | No activity p | IMPDH2,NT5C1B |
| Phenylalanin | 0.276 | 0.143 | No activity p | ALDH2,GOT2 |
| Colanic Acid | 0.276 | 0.143 | No activity p | GPI,UGDH |
| Triacylglycer | 0.275 | 0.128 | -2.449 | AGPAT1,AGPAT3,AGPAT4,MBOAT7,PLP |
| Arginine Deg | 0.263 | 0.167 | No activity p | OAT |
| Glycine Clea | 0.263 | 0.167 | No activity p | DLD |
| Rapoport-Lu | 0.263 | 0.167 | No activity p | PGAM1 |
| Lactose Degr | 0.263 | 0.167 | No activity p | PSAP |
| GDP-mannos | 0.263 | 0.167 | No activity p | GPI |
| nNOS Signali | 0.258 | 0.125 | No activity p | CACNA1H,CACNB1,CACNB4,CACNG7,CA |
| Superpathwa | 0.245 | 0.133 | No activity p | ALDH18A1,OAT |
| Tryptophan D | 0.234 | 0.125 | No activity p | ACAT2,HSD17B4,PARK7 |
| Trehalose De | 0.22 | 0.143 | No activity p | HK1 |
| Phosphatidyl | 0.22 | 0.143 | No activity p | CHKA |
| NAD Salvage | 0.22 | 0.143 | No activity p | NMNAT2 |
| Ceramide De | 0.22 | 0.143 | No activity p | ASAH1 |
| Glycoaminog | 0.22 | 0.143 | No activity p | B3GAT1 |
| Inositol Pyro | 0.186 | 0.125 | No activity p | IP6K1 |
| LPS/IL-1 Mec | 0 | 0.103 | 0 | ACSL3,ACSL4,ACSL6,ALDH18A1,ALDH2,A |
| Coagulation | 0 | 0.0857 | No activity p | A2M,FGA,KLKB1 |
| Hepatic Chol | 0 | 0.0688 | No activity p | ELP1,JUN,MAP2K4,PRKACA,PRKACB,PRK |

| | | | | |
|---|---|---|---|---|
| FXR/RXR Act | 0 | 0.111 | No activity pa | AKT2,AKT3,APOE,C4A/C4B,CLU,CREBBP |
| Role of Patte | 0 | 0.0577 | -1.414 | EIF2AK2,MAP2K4,MAPK1,MAPK3,MAVS |
| TREM1 Signa | 0 | 0.117 | -3 | AKT2,AKT3,GRB2,ITGB1,MAPK1,MAPK3 |
| Role of NFAT | 0 | 0.0811 | -3.888 | AKT2,AKT3,CABIN1,CALM1 (includes oth |
| CCR5 Signali | 0 | 0.0461 | -2.828 | CACNA1H,CACNB1,CACNB4,CACNG7,CA |
| Calcium-indu | 0 | 0.0297 | -2.53 | ATP2A2,CABIN1,CALM1 (includes others |
| Cytotoxic T L | 0 | 0.00993 | No activity pa | B2M,DFFB,HLA-A,HLA-B |
| CD27 Signali | 0 | 0.105 | -2.236 | ELP1,JUN,MAP2K2,MAP2K4,MAP3K10,R |
| IL-17 Signali | 0 | 0.102 | -3.441 | AKT2,AKT3,GSK3B,HSP90AA1,HSP90AB1 |
| Airway Patho | 0 | 0.0254 | No activity pa | FGF13,MMP2,TGFB2 |
| CTLA4 Signal | 0 | 0.0614 | No activity pa | AKT2,AKT3,AP1B1,AP1G1,AP1M1,AP1S1 |
| Induction of | 0 | 0.0923 | -1.633 | DFFB,ELP1,MAP2K4,RELA,SLC25A3,SLC2 |
| Airway Inflar | 0 | 0.0312 | No activity pa | TGFB2 |
| T Helper Cell | 0 | 0.0156 | No activity pa | HLA-A,HLA-B,HLA-DPA1,IL6ST,STAT1,ST |
| CD28 Signali | 0 | 0.0505 | -4.146 | ACTR3,AKT2,AKT3,ARPC1A,ARPC2,ARPC |
| Role of Cytok | 0 | 0.0185 | No activity pa | IFNA16 |
| Dendritic Cel | 0 | 0.0423 | -4.082 | AKT2,AKT3,ATF4,B2M,COL18A1,COL1A1 |
| Angiopoietin | 0 | 0.117 | -1.414 | AKT2,AKT3,ELP1,GRB2,PAK3,PIK3R3,RA |
| iCOS-iCOSL S | 0 | 0.0351 | -3.5 | AKT2,AKT3,CALM1 (includes others),CSK |
| Lipid Antiger | 0 | 0.0279 | No activity pa | AP1B1,AP2A1,AP2A2,AP2B1,AP2M1,AP2 |
| HMGB1 Sign | 0 | 0.12 | -4.123 | AKT2,AKT3,CDC42,HMGB1,JUN,KAT2A,M |
| Maturity Ons | 0 | 0.103 | No activity pa | APOE,CACNB1,CACNB4,CACNG7,CREBB |
| CREB Signali | 0 | 0.118 | -6.181 | ADGRA3,ADGRB2,ADGRE2,ADGRL1,AKT |
| Type I Diabet | 0 | 0.0226 | -2.236 | CPE,HLA-A,HLA-B,HLA-DPA1,HSPD1,JAK1 |
| Basal Cell Ca | 0 | 0.111 | -1.89 | APC,CTNNB1,FZD3,FZD7,GSK3B,SMO,TC |
| Primary Imm | 0 | 0.0185 | No activity pa | DCLRE1C |
| Allograft Rej | 0 | 0.00868 | No activity pa | B2M,HLA-A,HLA-B,HLA-DPA1 |
| Autoimmune | 0 | 0.00696 | No activity pa | HLA-A,HLA-B,HLA-DPA1 |
| Graft-versus- | 0 | 0.00708 | No activity pa | HLA-A,HLA-B,HLA-DPA1 |
| Communicat | 0 | 0.00634 | No activity pa | B2M,HLA-A,HLA-B |
| Crosstalk bet | 0 | 0.11 | No activity pa | ACTA2,ACTB,ACTG1,FSCN1,HLA-A,HLA-I |
| Systemic Lup | 0 | 0.0601 | No activity pa | AKT2,AKT3,CD2BP2,EFTUD2,FCGR3A/FC |
| Cdc42 Signal | 0 | 0.0741 | -3.024 | ACTR3,APC,ARPC1A,ARPC2,ARPC3,B2M, |
| Retinoic acid | 0 | 0.1 | -1 | CRABP1,CRABP2,IFNA16,PARP1,TNFRSI |
| Phospholipas | 0 | 0.0844 | -5.337 | ARHGEF17,ARHGEF18,ARHGEF2,ARHGI |
| Altered T Cel | 0 | 0.0106 | No activity pa | HLA-A,HLA-B,HLA-DPA1,RELA,TRAF3 |
| Atherosclero | 0 | 0.0769 | No activity pa | APOE,CCR3,CLU,COL18A1,COL1A1,COL2 |
| B Cell Develo | 0 | 0.0698 | No activity pa | HLA-A,HLA-B,HLA-DPA1 |
| Regulation o | 0 | 0.0412 | -3.441 | CALM1 (includes others),ELP1,FYN,GRB |
| TWEAK Signa | 0 | 0.0541 | No activity pa | RELA,TRAF3 |
| Nur77 Signal | 0 | 0.0327 | -1.155 | B2M,CABIN1,CALM1 (includes others),E |
| PKCθ Signali | 0 | 0.0429 | -4.243 | CACNA1H,CACNB1,CACNB4,CACNG7,FY |
| Role of Lipids | 0 | 0.087 | No activity pa | FDPS,IFNA16 |
| TNFR1 Signa | 0 | 0.0962 | -1.342 | CDC42,JUN,MAP2K4,PAK3,RELA |
| Role of MAP | 0 | 0.111 | No activity pa | AKT2,AKT3,MAP2K2,MAP2K4,MAPK1,MA |

| | | | | |
|---|---|---|---|---|
| TNFR2 Signa | 0 | 0.0938 | No activity p | JUN,MAP2K4,RELA |
| Role of Hype | 0 | 0.0698 | -0.816 | EIF2AK2,IFNA16,JAK1,STAT1,STAT2,TYK |
| Antiprolifera | 0 | 0.0223 | 0.333 | CDC34,CDKN1B,CUL1,DPP7,MAPK1,PABI |
| MSP-RON Si | 0 | 0.103 | No activity p | ACTA2,ACTB,ACTG1,KLKB1,PIK3R3,RPS6 |
| OX40 Signali | 0 | 0.0175 | -2 | B2M,HLA-A,HLA-B,HLA-DPA1,JUN,MAP2 |
| Extrinsic Prot | 0 | 0.0625 | No activity p | FGA |
| Tumoricidal | 0 | 0.0833 | No activity p | DFFB,M6PR |
| Intrinsic Prot | 0 | 0.119 | -0.447 | COL18A1,COL1A1,COL2A1,FGA,KLKB1 |
| Assembly of | 0 | 0.0833 | No activity p | UBTF |
| Bupropion De | 0 | 0.04 | No activity p | CYP51A1 |
| Sphingosine | 0 | 0.111 | No activity p | ASAH1 |
| Glucose and | 0 | 0.0833 | No activity p | HK1 |
| Thyroid Horn | 0 | 0.0769 | No activity p | B4GAT1,EXT2,SULT1C4 |
| Nicotine Deg | 0 | 0.0517 | No activity p | B4GAT1,CYP51A1,EXT2 |
| Prostanoid B | 0 | 0.1 | No activity p | PTGES3 |
| Androgen Bio | 0 | 0.111 | No activity p | AKR1C3,AKR1C4 |
| Ubiquinol-10 | 0 | 0.087 | No activity p | CYP26B1,MICAL1 |
| tRNA Splicin | 0 | 0.0909 | 0 | APEX1,GDE1,PDE11A,PDE7A |
| Melatonin De | 0 | 0.0656 | -1 | B4GAT1,CYP51A1,EXT2,SULT1C4 |
| NAD Salvage | 0 | 0.111 | No activity p | ACP1,NMNAT2,NT5C1B |
| Triacylglycer | 0 | 0.118 | -1.633 | ABHD12,CES3,DDHD2,NDST1,PPME1,PR |
| Phospholipas | 0 | 0.0606 | No activity p | PDIA3,PLCG1,PLD3,PRDX6 |
| Choline Biosy | 0 | 0.0667 | No activity p | PLD3 |
| Estrogen Bio | 0 | 0.114 | -0.447 | AKR1C3,AKR1C4,CYP51A1,HSD17B12,HS |
| Leukotriene | 0 | 0.0714 | No activity p | GGT7 |
| Glutathione- | 0 | 0.121 | No activity p | GSTA4,GSTM3,GSTP1,LANCL1 |
| Glycogen Deg | 0 | 0.0833 | No activity p | PYGB |
| Nicotine Deg | 0 | 0.0455 | No activity p | B4GAT1,CYP51A1,EXT2 |
| Superpathwa | 0 | 0.087 | No activity p | ITPKB,SEC16A |
| Purine Nucle | 0 | 0.105 | No activity p | IMPDH2,NT5C1B |
| D-myo-inosit | 0 | 0.111 | No activity p | ITPKB,SEC16A |
| Serotonin De | 0 | 0.118 | -1.414 | ADH5,ALDH2,ALDH3B1,ALDH7A1,ALDH9 |
| Leucine Degr | 0 | 0.111 | No activity p | IVD |
| D-myo-inosit | 0 | 0.0588 | No activity p | SEC16A |
| γ-glutamyl C | 0 | 0.0909 | No activity p | GGT7 |
| Ketolysis | 0 | 0.1 | No activity p | ACAT2 |
| Pyrimidine R | 0 | 0.118 | -2 | AK1,CMPK1,DDX3X,NME2 |
| Purine Nucle | 0 | 0.0909 | No activity p | IMPDH2 |
| Methylglyoxa | 0 | 0.05 | No activity p | AKR1C3 |
| Guanosine N | 0 | 0.0769 | No activity p | NT5C1B |
| Adenosine N | 0 | 0.0625 | No activity p | NT5C1B |
| Superpathwa | 0 | 0.0606 | -1 | B4GAT1,CYP51A1,EXT2,SULT1C4 |
| Histidine Deg | 0 | 0.1 | No activity p | CYP26B1,MICAL1 |
| Pregnenolon | 0 | 0.111 | No activity p | CYP26B1,MICAL1 |
| Acetone Deg | 0 | 0.0312 | No activity p | CYP51A1 |

| | | | | |
|---|---|---|---|---|
| GDP-glucose | 0 | 0.0909 | No activity pa | HK1 |
| Glycine Beta | 0 | 0.1 | No activity pa | SHMT2 |
| Oxidative Eth | 0 | 0.0909 | -1.342 | ACSS2,ALDH2,ALDH3B1,ALDH7A1,ALDH |
| Folate Trans | 0 | 0.111 | No activity pa | SHMT2 |
| DNA damage | 0 | 0.105 | No activity pa | AKT2,AKT3 |
| Granulocyte | 0 | 0.0794 | No activity pa | CCR3,CD99,CXCL12,EZR,GLG1,GNAI1,IT( |
| Tec Kinase Si | 0 | 0.0719 | -3.838 | ACTA2,ACTB,ACTG1,CDC42,FYN,GNA12, |
| SAPK/JNK Sig | 0 | 0.05 | -3.441 | CDC42,DUSP4,DUSP8,GNA12,GNA13,GN |
| Serotonin Re | 0 | 0.0233 | No activity pa | SLC18A3 |
| p38 MAPK Si | 0 | 0.119 | -3.317 | ATF4,CREBBP,H3-3A/H3-3B,HSPB1,MAP |
| NF-κB Signal | 0 | 0.042 | -3.545 | AKT2,AKT3,CREBBP,CSNK2A1,CSNK2B,E |
| T Cell Recept | 0 | 0.0608 | -3.55 | AKT2,AKT3,B2M,CALM1 (includes others |
| Gustation Pa | 0 | 0.114 | No activity pa | APEX1,ASIC1,CACNA1H,CACNB1,CACNB |
| Inflammasor | 0 | 0.05 | No activity pa | CTSB |
| Th1 and Th2 | 0 | 0.11 | No activity pa | ACVR1B,CCR3,CHD4,GRB2,HLA-A,HLA-B |
| Th1 Pathway | 0 | 0.107 | -3.317 | GRB2,HLA-A,HLA-B,HLA-DPA1,JAK1,KLR |
| Th2 Pathway | 0 | 0.117 | -1.508 | ACVR1B,CCR3,CHD4,GRB2,HLA-A,HLA-B |
| Th17 Activati | 0 | 0.0195 | -1 | HIF1A,HSP90AA1,HSP90AB1,HSP90B1,J |
| SPINK1 Panc | 0 | 0.1 | 1.633 | CPD,CPE,CTSA,CTSB,KLKB1,TGFBR1 |
| T Cell Exhaus | 0 | 0.0607 | -2.6 | ACVR1B,AKT2,AKT3,HLA-A,HLA-B,HLA-D |
| Systemic Lup | 0 | 0.0746 | -3.833 | AKT2,AKT3,ATF4,B2M,CD44,CDC42,CREI |
| Complement | 0 | 0.0811 | No activity pa | C1QBP,C4A/C4B,CD59 |
| Eicosanoid Si | 0 | 0.0299 | No activity pa | AKR1C3,PRDX6 |
| Role of MAP | 0 | 0.105 | -1.89 | EIF2AK2,JUN,MAP2K2,MAP2K4,MAPK1, |

ND1,EIF1,EIF2AK1,EIF2AK2,EIF2S2,EIF3B,EIF3C,EIF3D,EIF3E,EIF3F,EIF3G,EIF3H,EIF3J,EIF3K,EIF
DNAJA1,DNAJB1,DNAJB11,DNAJB2,DNAJC10,DNAJC5,DNAJC8,ELOB,FZR1,HLA-A,HLA-B,HSP90
1B,ATP5F1C,ATP5F1E,CAPN1,CAPN2,CAPNS1,CLTA,CPLX2,CREBBP,CTSD,DCTN1,DLG4,DNAJB
3D,EIF3E,EIF3F,EIF3G,EIF3H,EIF3J,EIF3K,EIF3L,EIF3M,EIF4A1,EIF4A3,EIF4G1,EIF4G2,EIF4G3,F
XOSC10,MAPK1,MAPK11,MAPK3,MAPKAPK2,PABPN1,PAPOLA,PPM1L,PPP2CA,PPP2CB,PPP2R1
,HSPA1A/HSPA1B,HSPA4,MAPK1,MAPK3,MAPKAPK2,MAPT,MDM2,PINK1,PSMA1,PSMA4,PSM
5F1C,ATP5F1E,CACNB1,CACNB4,CACNG7,CARM1,CCND1,CDKN1A,CREBBP,CTBP1,CTBP2,DDX
F3C,EIF3D,EIF3E,EIF3F,EIF3G,EIF3H,EIF3J,EIF3K,EIF3L,EIF3M,EIF4A1,EIF4A3,EIF4B,EIF4G1,EIF
,ATP6V0C,ATP6V0D1,ATP6V0E2,ATP6V1B2,ATP6V1D,ATP6V1E1,ATP6V1F,ATP6V1G1,B2M,CA
ATP5F1E,ATP5MC2,ATP5MC3,ATP5MG,ATP5PD,ATP5PO,BACE1,COX4I1,COX5A,COX5B,COX6A
AP2A2,AP2B1,AP2M1,AP2S1,APOE,ARHGEF7,ARPC1A,ARPC2,ARPC3,ATF4,CACNB1,CACNB4,C
SMA1,PSMA4,PSMA5,PSMA6,PSMA7,PSMB1,PSMB2,PSMB4,PSMB5,PSMB6,PSMC1,PSMC2,P
A6,PSMA7,PSMB1,PSMB2,PSMB4,PSMB5,PSMB6,PSMC1,PSMC2,PSMC3,PSMC4,PSMC5,PSMC
,ATG9A,ATP5F1A,ATP5F1B,ATP5F1C,ATP5F1E,G6PD,GABARAP,GABARAPL1,GABARAPL2,GL
TF4,CAB39,CAMKK2,CCND1,CDKN1A,CHRNA3,CHRNA4,CREBBP,DPF1,EEF2,ELAVL1,EP300,FAS
,DPYSL4,ERBB2,FARP1,FYN,GSK3B,ITGA11,ITGA2,ITGA3,ITGA7,ITGAV,ITGB1,ITGB5,LIMK1,
,ADAM9,ADAMTS1,ADAMTS7,AKT2,AKT3,ARHGEF7,ARPC1A,ARPC2,ARPC3,BCAR1,CDC42,CH
,ACTR3,AKT2,AKT3,ARF1,ARF3,ARF4,ARF5,ARHGAP5,ARHGEF7,ARPC1A,ARPC2,ARPC3,BCAR
ARHGEF18,ARHGEF2,ARHGEF7,ARHGEF9,ARPC1A,ARPC2,ARPC3,CDC42,CDC42EP4,CDH11,CD
RHGEF2,CASK,CDC42,CDK4,CPSF1,CPSF2,CTNNA1,CTNNB1,GOSR2,GPAA1,HSF1,JAM3,JUN,M
R3,APC,ARPC1A,ARPC2,ARPC3,CTNNA1,CTNNA2,CTNNB1,CTNND1,DNM3,HGS,MAPRE1,MAPR
,ACTR3,APC,ARHGAP24,ARHGEF7,ARPC1A,ARPC2,ARPC3,BCAR1,BRK1,CDC42,CSK,CYFIP1,CY
TP5MC2,ATP5MC3,ATP5MG,ATP5PD,ATP5PO,COX4I1,COX5A,COX5B,COX6A1,COX6C,COX7A2,
RHGDIA,ARHGEF17,ARHGEF18,ARHGEF2,ARHGEF7,ARHGEF9,ARPC1A,ARPC2,ARPC3,CD44,C
1,AP1M1,AP1S1,AP2A1,AP2A2,AP2B1,AP2M1,AP2S1,APOE,ARPC1A,ARPC2,ARPC3,CDC42,CLT
K4,E2F6,EEF1A2,EP300,FAU,FURIN,HDAC2,HDAC5,HDAC6,HIF1A,JAK1,JUN,KPNB1,MAP2K4,M
N1A,CTNNB1,ERBB2,GSK3B,HIF1A,JUN,LDB1,LGALS1,MAP2,MAP2K4,MMP2,MTOR,PPM1L,P
RPC1A,ARPC2,ARPC3,CDC42EP4,EZR,GNA12,GNA13,IGF1R,KTN1,LIMK1,LPAR2,MPRIP,MSN,M
ARPC1A,ARPC2,ARPC3,ATF4,BCAR1,CDC42,CREBBP,CXCL12,EFNA3,EFNB1,EFNB2,EFNB3,EPH
1,DLG4,DLG5,FAT4,ITCH,NF2,PPM1L,PPP1CA,PPP1CB,PPP1CC,PPP1R10,PPP1R12A,PPP1R14B,
44,CDH2,CREBBP,CSNK1A1,CSNK1D,CSNK1E,CSNK1G2,CSNK2A1,CSNK2B,CTNNB1,DKK3,EP30
1A,HSP90AA1,HSP90AB1,HSP90B1,JUN,LDHA,MDM2,P4HB,SUMO1,UBE2D2,UBE2D3,UBE2E1
1B,CTNNB1,ELP1,GAB2,GHR,GRB2,GSK3A,GSK3B,HSP90AA1,HSP90AB1,HSP90B1,IL6ST,ITGA
R3,ACVR1B,AFDN,AKT2,AKT3,APC,ARPC1A,ARPC2,ARPC3,CDC42,CDH2,CTNNA1,CTNNA2,CTN
B,GRB2,IGF1R,IGFBP2,IGFBP3,IGFBP4,IGFBP7,JAK1,JUN,MAP2K2,MAPK1,MAPK3,PDPK1,PIK3
APNS1,CSNK1A1,CSNK1D,CSNK1E,CSNK2A1,CSNK2B,GSK3B,MAPK1,MAPK11,MAPK3,MAPT,N
2,AKT3,ATF4,CCND1,CDC42,CREBBP,CTNNB1,FLNA,FLNB,FLNC,FN1,GSK3A,GSK3B,HIF1A,ITGB
,H3-3A/H3-3B,HSPB1,ITGA11,ITGA2,ITGA3,ITGA7,ITGAV,ITGB1,ITGB5,MAP2K2,MAPK1,MAPK
,CACNG7,CLCN3,CPE,CREBBP,DLD,EIF1,EIF2S2,EIF4A1,EIF4A3,EIF4G1,EIF4G2,EIF4G3,FURIN,F
B1,EFNB2,EFNB3,EPHB2,GNA12,GNA13,GNAI1,GNAL,GNAO1,GNAS,GNAT1,GNB1,GNB2,GN
PP,ARHGEF2,ARHGEF9,ARPC1A,ARPC2,ARPC3,CDC42,CDH2,CNR1,DCX,FYN,GRINA,GSK3B,ITG
,AFDN,AKT2,AKT3,BCAR1,CDC42,CTNNA1,CTNNA2,CTNNB1,DLG1,EPN1,GSK3A,GSK3B,ITGA1
ALM1 (includes others),CAMKK2,CDKN1B,CREBBP,DAPK3,GABARAP,GABARAPL1,GABARAPL

TGA3,ITGB1,LAMA5,LAMC1,MAP2K2,MAPK1,MAPK11,MAPK3,MAPT,PPM1L,PPP1CA,PPP1CB,P
1A,CARM1,CITED2,CRABP1,CRABP2,CREBBP,CSK,CSNK2A1,CSNK2B,DPF1,EP300,IGFBP3,JUN,N
C5,APEX1,ATF4,CALM1 (includes others),CREBBP,CTNNB1,DUSP3,DUSP4,DUSP8,FLNA,FLNB,F
EF18,ARHGEF2,ARHGEF7,ARHGEF9,CCND1,CCND2,CDC42,CDK11B,CDK16,CDK4,CDK9,CDKN1A,
COPB1,COPE,COPZ1,FLNA,FLNB,FLNC,FLOT1,FLOT2,FYN,HLA-A,HLA-B,ITGA11,ITGA2,ITGA3,IT
ARPC2,ARPC3,CDC42,ITGA11,ITGA2,ITGA3,ITGA7,ITGAV,ITGB1,ITGB5,LIMK1,MPRIP,MYL12A,N
TF4,BAP1,CBS/CBSL,CDKN1A,CTSB,FDFT1,FTH1,FTL,G3BP1,GPX4,H2AX,H2AZ1,HMGCR,HSPB1
L,AP1M1,AP1S1,AP2A1,AP2A2,AP2B1,AP2M1,AP2S1,B2M,CDC42,CLTA,CXADR,FLNA,FLNB,FLN
A,CDKN1B,CSNK2A1,CSNK2B,ELP1,FGFR1,FGFR3,FOXG1,GHR,GRB2,GSK3A,GSK3B,IGF1R,ITG
LM1 (includes others),CALR,CCND1,CREBBP,DNAJB1,EP300,GNA12,GNA13,GNAI1,GNAL,GNA
CREBBP,H2AX,JUN,MAP2K4,MAPK11,MDC1,MDM2,PPM1L,PPP1CC,PPP2CA,PPP2CB,PPP2R1A,P
1,CD44,CDC42,CYFIP1,CYFIP2,ITGA11,ITGA2,ITGA3,ITGA7,ITGAV,ITGB1,ITGB5,JUN,LIMK1,MA
S2,KARS1,LARS1,MARS1,NARS1,QARS1,RARS1,SARS1,TARS1,VARS1,WARS1,YARS1
CREBBP,CTNNB1,GNAI1,GNAL,GNAO1,GNB1,GNB2,GNG2,GNG4,GSK3B,MAP2K2,MAP2K4,M
4,AFDN,BCAR1,CDC42,CDH2,CTNNA1,CTNNA2,CTNNB1,CTNND1,EPN1,ITGA2,ITGA3,ITGB1,LIM
FH,GRB2,HIF1A,JUN,MAP2K2,MAPK1,MAPK3,PAK3,PIK3R3,RAC1,RAP1B,RAPGEF1,RBX1,RPS2
CACNB4,CACNG7,GDE1,GNA12,GNA13,GNAI1,GNAL,GNAO1,GNAS,GNAT1,GNB1,GNB2,GNG
RD4,CACNB1,CACNB4,CACNG7,CALM1 (includes others),CCDC88A,CCN2,CCND1,CDC42,CDKN1E
MGCR,HMGCS1,IDI1,LBR,LSS,MSMO1,MVD,SQLE,TM7SF2
TUD2,EIF4A3,PLRG1,PRPF19,SF1,SF3A1,SF3A2,SF3A3,SF3B1,SF3B2,SNRNP200,U2AF1/U2AF
B,JUN,MAP2K2,MAP2K4,MAPK1,MAPK3,MAPT,PDIA3,PIK3R3,PLCG1,PRKCI,RAP1B,SNCA,SRPK
4,AFDN,ARHGAP5,BCAR1,CD44,CD99,CDC42,CTNNA1,CTNNA2,CTNNB1,CTNND1,CTTN,CXCL12
FZD3,FZD7,GNA12,GNA13,GNAI1,GNAL,GNAO1,GNAS,GNAT1,GNB1,GNB2,GNG2,GNG4,GRE
5,HIPK2,MDM2,PRKDC,RPRM,SKP1,TOP2B,TRIP12,YWHAB,YWHAE,YWHAG,YWHAH,YWHAQ,Y
MAP2K4,MAPK1,MAPK3,PFKM,PRKACA,PRKACB,PRKAR1A,PRKAR1B,PRKAR2A,PRKAR2B,TGFB
N2,ELOB,EP300,GPI,HIF1A,HK1,HSP90AA1,HSPA1A/HSPA1B,HSPA4,IGF2,JUN,KDM1A,LDHA,LD
A,CDKN1B,CUL1,E2F6,FBXL5,GSK3B,HDAC2,HDAC5,HDAC6,PPM1L,PPP2CA,PPP2CB,PPP2R1A,P
DKN1A,CDKN1B,E2F6,ERBB2,GRB2,HDAC2,HDAC5,HDAC6,JAK1,MAP2K2,MAP2K4,MAPK1,MAPK
NAJA1,DNAJB1,DNAJB11,DNAJB2,DNAJC10,DNAJC5,DNAJC8,HSP90B1,HSPA1A/HSPA1B,HSPA4
des others),CDC42,CREBBP,EP300,GNA12,GNA13,GNAI1,GNAL,GNAO1,GNAS,GNAT1,GNB1,G
,CTNNB1,FLNA,GRB2,GSK3B,HIF1A,ITGB1,JUN,KLKB1,MAP2K2,MAPK1,MAPK3,MTOR,PIK3R3,
HDAC5,HDAC6,HSP90AA1,HSP90AB1,HSP90B1,MAP2K2,MAPK1,MAPK3,PDPK1,PIK3R3,PPM1L,
D1,CDC42,CDK4,CDKN1A,CDKN1B,COPS6,COX4I1,COX5A,COX5B,COX6A1,COX6C,COX7A2,COX7
A,GSK3B,JAK1,MAP2K2,MAPK1,MAPK3,MTOR,PDPK1,PIK3R3,PPP1CA,PPP1CB,PPP1CC,PPP1R1
A,CDKN1B,CUL1,E2F6,FBXL5,GNL3,GSK3B,HDAC2,HDAC5,HDAC6,MDM2,NRG1,RPL11,RPL5,SK
TBP2,EP300,HDAC2,JUN,KDM1A,MAP2K4,MDM2,PCNA,RAC1,RAC3,RANGAP1,RHOA,RHOBTB
ARPC1A,ARPC2,ARPC3,CDC42,EZR,FCGR3A/FCGR3B,FYN,GAB2,MAPK1,MAPK3,NCK2,PIK3R3,P
1,JUN,MAP2K4,MAPK1,MAPK11,MAPK3,MTOR,PIK3R3,PLCG1,SHC1,SOS1,SRF,STAT1,STAT3
2,CDKN1A,CDKN1B,CNR1,CREBBP,CTNNB1,GNA13,GNAI1,GNAL,GNAO1,GSK3B,HIF1A,MAP2K
P2K2,MAPK1,MAPK3,MAPT,MTOR,PDIA3,PDPK1,PIK3R3,PLCG1,PPM1L,PPP2CA,PPP2CB,PPP2R1
KM,PFKP,PGAM1,PGK1,PKM,TPI1
F7,BCAR1,CAPN1,CAPN2,CAPNS1,CSK,FYN,GRB2,ITGA11,ITGA2,ITGA3,ITGA7,ITGAV,ITGB1,ITG
CDKN1B,CREBBP,CTNNB1,E2F6,GRB2,GSK3B,GSTP1,HDAC2,HDAC5,HDAC6,HSP90AA1,HSP90A
C3,ITPK1,MTMR4,NUDT3,NUDT4,PDXP,PI4KA,PIK3R3,PIP4K2A,PIP4K2B,PIP5K1A,PIP5K1C,PPFIA
CACNG7,CDC42,GNA12,GNA13,GNAI1,GNAL,GNAO1,GNAS,GNAT1,GNB1,GNB2,GNG2,GNG4

:YN,ITGB1,LIMK1,MAPK1,MAPK3,PAK3,PLXNA1,PLXNA2,PLXNA3,PLXNB1,RAC1,RAC3,RHOA,RH
1,ARHGEF7,BCAR1,CDC42,CSK,GRB2,ITGA11,ITGA2,ITGA3,ITGA7,ITGAV,ITGB1,ITGB5,MAP2K4
1,SQLE,TM7SF2
1,SQLE,TM7SF2
1,SQLE,TM7SF2
FZD7,GRB2,GSK3B,ID2,ID4,IL6ST,JAK1,LIFR,MAP2K2,MAPK1,MAPK11,MAPK3,NANOG,PIK3R3,
3B,HNRNPA1,IFITM3,TUBA1A,TUBA1B,TUBA1C,TUBB,TUBB2B,TUBB4A,TUBB4B,TUBB6
ACNB4,CACNG7,CALM1 (includes others),CSNK1A1,EP300,GNAI1,GNAS,GNB1,GNB2,GNG2,G
,AP2M1,AP2S1,ATF4,CACNA1H,CACNB1,CACNB4,CACNG7,CALM1 (includes others),CDC42,CDK
C3,IP6K1,ITPK1,ITPKB,MTMR4,NUDT3,NUDT4,PDXP,PI4KA,PIK3R3,PIP4K2A,PIP4K2B,PIP5K1A,PI
P1,RANGAP1,TNPO1,XPO1
,PPP2CB,PPP2R1A,PPP2R2A,PPP2R5B,PPP2R5C,PPP2R5D,PRMT1,PTPA
A11,ITGA2,ITGA3,ITGA7,ITGAV,ITGB1,ITGB5,LIMK1,MAP2K2,MAP2K4,MAPK1,MAPK3,MYL12
7,CALM1 (includes others),CAMKK2,CREBBP,CSNK1A1,CSNK1D,CSNK1E,CSNK1G2,GNAI1,GNAS
,CTNNB1,E2F6,FZD3,FZD7,GJA1,GSK3B,HDAC2,HDAC5,HDAC6,MAP2K2,MAPK1,MAPK3,MMP2
LDH7A1,ALDH9A1,CCND1,CCND2,CDK4,CDKN1A,CDKN1B,CTSD,DHFR,EP300,GSTA4,GSTM3,GS
REBBP,CUL3,DNAJA1,DNAJB1,DNAJB11,DNAJB2,DNAJC10,DNAJC5,DNAJC8,ENC1,EP300,ERP2
,CDK4,CDKN1A,CREBBP,DPF1,EP300,H2AX,HDAC2,HDAC5,HDAC6,MSH2,MSH6,NPM1,PBRM1,P
K4,CDKN1A,CDKN1B,CTNNB1,E2F6,FZD3,FZD7,GRB2,GSK3B,IGF1R,IGF2,MAP2K2,MAPK1,MA
R1,FGF13,FGFR1,FGFR3,FZD3,FZD7,GRB2,GSK3B,HIF1A,ID2,JAG2,JAK1,LOX,MAP2K2,MAP2K4
P2M1,AP2S1,CACNA1H,CACNB1,CACNB4,CACNG7,GABARAP,GABBR1,GABRB3,GNA12,GNA1
0,PPP1R12A,PPP1R14B,PPP2CA,PPP2CB,PPP2R1A,PPP2R2A,PPP2R5B,PPP2R5C,PPP2R5D,PRKA
thers),COL11A1,COL12A1,COL18A1,COL1A1,COL2A1,COL4A2,COL4A5,COL5A1,COL5A2,COL6A1
1,CSNK1D,CSNK1E,CSNK1G2,CTNNB1,DBN1,GJA1,GNAI1,GNAS,GRB2,GRIA1,MAP2K2,MAPK1
,HSP90AA1,HSP90AB1,HSP90B1,ITGA11,ITGA2,ITGA3,ITGA7,ITGAV,ITGB1,ITGB5,MAP2K2,MA
D1,CKAP5,CREBBP,ELP1,EP300,FASN,GHR,GNAS,GOT2,GRB2,HSP90AA1,HSP90AB1,HSP90B1,
KN1A,CDKN1B,CTBP1,CTBP2,E2F6,GAB2,GRB2,HDAC2,HDAC5,HDAC6,MAP2K2,MAPK1,MAPK
s),CCND1,CCND2,CDK4,CDKN1A,E2F6,GRB2,HDAC2,HDAC5,HDAC6,IDH1,IDH2,IGF1R,IGF2,MAP
K1,MTMR4,NUDT3,NUDT4,PDXP,PPFIA2,PPFIA3,PPM1L,PPP1CA,PPP1CB,PPP1CC,PPP1R10,PPP1
K1,MTMR4,NUDT3,NUDT4,PDXP,PPFIA2,PPFIA3,PPM1L,PPP1CA,PPP1CB,PPP1CC,PPP1R10,PPP1
C42,CTTN,DAG1,ERBB2,ITGB1,JUN,LAMC1,MAP2K4,MAPK1,MAPK3,NRG1,PAK3,RAC1,RAC3,R
ELMO2,GNA12,GNA13,GNAI1,GNAL,GNAO1,GNAS,GNAT1,GNB1,GNB2,GNG2,GNG4,JUN,M
AB1,HSP90B1,PLK2,PLK4,PPM1L,PPP2CA,PPP2CB,PPP2R1A,PPP2R2A,PPP2R5B,PPP2R5C,PPP2R
GF13,FGFR1,FGFR3,GRB2,GSK3B,HSF1,ID2,JAK1,JUN,MAP2K2,MAP2K4,MAPK1,MAPK11,MAP
11,MAPK3,STAT1,STAT3,TYK2
K1,MTMR4,NUDT3,NUDT4,PDXP,PIP4K2A,PLCG1,PPFIA2,PPFIA3,PPM1L,PPP1CA,PPP1CB,PPP1C
O1,GNAS,GNAT1,GNB1,GNB2,GNG2,GNG4,JUN,MAP2K4,MAPK1,MAPK11,PRKACA,PRKACB,P
,GRB2,GSK3A,GSK3B,JAK1,JUN,MAPK1,MAPK11,MAPK3,MCL1,PDPK1,PIK3R3,SHC1,SLC2A1,SO
,GNAI1,GNAL,GNAO1,GNAS,GNAT1,GNB1,GNB2,GNG2,GNG4,MAP2K2,MAPK1,MAPK3,PHKB
P4A,ITPK1,MTMR4,NUDT3,NUDT4,PDXP,PPFIA2,PPFIA3,PPM1L,PPP1CA,PPP1CB,PPP1CC,PPP1R
K3,MT2A,RAP1B,SHC1,SOS1,STAT1,STAT3,TIMP3,TYK2
LP1,JAK1,JUN,MAP2K4,MAP3K10,MAPK1,MAPK11,MAPK3,PIK3R3,PLCG1,PPM1L,PPP1CA,PPP1C
BIN1,CACNA1H,CACNB1,CACNB4,CACNG7,CALM1 (includes others),CALR,CAMKK2,CHRNA3,C
GNA12,GNA13,GNAI1,GNAL,GNAO1,GNAS,GNAT1,GNB1,GNB2,GNG2,GNG4,GRB2,MAP2K2,
O1,GNAS,GNAT1,GNB1,GNB2,GNG2,GNG4,PLCG1,TUB

GRB2,JAK1,JUN,MAP2K2,MAP2K4,MAPK1,MAPK3,PDGFRB,PIK3R3,PLCG1,RAP1B,SHC1,SOS1,S
MDH2,ME2,PGAM1,PGK1
2,MBD3,MTA1,MTA2,RBBP4,RBBP7,SAP18,SUDS3
SK3B,JUN,MAP2K2,MAPK1,MAPK3,NRG1,PDPK1,PIK3R3,RAP1B,SHC1,SOS1,STAT3,TYK2
OL18A1,COL1A1,COL2A1,COL4A2,COL4A5,COL5A1,COL5A2,COL6A1,COL6A2,COL6A3,COL7A1,E
NA12,GRB2,ITGA11,ITGA2,ITGA3,ITGA7,ITGAV,ITGB1,ITGB5,NCK2,PPP1R12A,RAC1,RAC3,RAP
NB1,CACNB4,CACNG7,CALM1 (includes others),CAPN1,CAPN2,CAPNS1,CCND1,CCND2,CDK4,C
DNAJB2,DNAJC10,DNAJC5,DNAJC8,HSP90AA1,HSP90AB1,HSP90B1,HSPA12A,HSPA1A/HSPA1B
,CACNA1H,CACNB1,CACNB4,CACNG7,GNAS,GSK3B,KDM1A,KDM5A,LMNA,MAPK1,MAPK11,M
CND1,CCND2,CDK4,CDKN1B,EZR,GRB2,ITGA11,ITGA2,ITGA3,ITGA7,ITGAV,ITGB1,ITGB5,MAPK
01,EP300,FASN,FGA,HIF1A,LDLR,MDM2,MTOR,NCOA4,NCOA6,NCOR2,PFKP,PIK3R3,SLC2A1,TE
MAPK1,MAPK3,MTOR,PIK3R3,RAP1B,RPS6KA2,SOS1,STAT1,STAT3,TYK2
GNA12,GNA13,GNAI1,GNAL,GNAO1,GNAS,GNAT1,GNB1,GNB2,GNG2,GNG4,ITGAV,JUN,LAS
-10,HMGB2,SET
ncludes others),CDC42,GNA12,GNA13,GNAI1,GNAL,GNAO1,GNAS,GNAT1,GNB1,GNB2,GNG2
2K2,MAPK1,MAPK3,PIK3R3,PPP3CA,PPP3CB,PRKCI,RAC1,RAP1B,RAPGEF1,SHC1,SOS1,STAT1,S
MAP2K2,MAPK1,MAPK11,MAPK3,MTOR,PDPK1,PIK3R3,RAP1B,RPS6KA2,SHC1,SOS1,STAT1,STA
31,EP300,GPS1,H3-3A/H3-3B,NEDD8,PCNA,POLD2,POLR2A,POLR2B,POLR2C,POLR2E,POLR2G,
P6V0B,ATP6V0C,ATP6V0D1,ATP6V0E2,ATP6V1B2,ATP6V1D,ATP6V1E1,ATP6V1F,ATP6V1G1,CD
CDK4,CDKN1A,CSNK1D,CTNNB1,EP300,GNL3,GSK3B,HIF1A,HIPK2,JUN,MDM2,PCNA,PIK3R3,P
A,FYN,GNA12,GNA13,ITGA3,ITGAV,ITGB1,LIMK1,MAPK1,MAPK11,MAPK3,MTOR,P4HB,PDIA6,
CSNK2B,ELP1,GRB2,HSPB1,IL6ST,JUN,MAP2K2,MAP2K4,MAP4K4,MAPK1,MAPK11,MAPK3,MA
2,AKT3,EIF1,EIF2S2,ELAVL1,GRB2,HIF1A,MAP2K2,MAPK1,MAPK3,PIK3R3,PLCG1,RAP1B,ROCK2
2K4,MAPK1,MAPK11,TGFBR1,VEGFA
AP2K2,MAPK1,MAPK3,MTOR,PIK3R3,PTPN1,RAP1B,RELA,SHC1,SOS1,STAT1,STAT2,STAT3,TYK
NNB1,ERBB2,GRB2,GSK3B,MAP2K2,MAPK1,MAPK3,PDPK1,PIK3R3,RAP1B,SOS1
2F6,ERBB2,FHIT,GRB2,HDAC2,HDAC5,HDAC6,MAP2K2,MAPK1,MAPK3,PDPK1,PIK3R3,PLCG1,RA
APK3,STAT1,STAT3,TYK2
LDH7A1,ALDH9A1,CES3,CHST10,CHST3,CITED2,CREBBP,EP300,ESD,GSTA4,GSTM3,GSTP1,HS6
CNG7,NCK2,NTN1,PPP3CA,PPP3CB,PRKACA,PRKACB,PRKAR1A,PRKAR1B,PRKAR2A,PRKAR2B,R
ATP5F1A,ATP5F1B,ATP5F1C,ATP5F1E,B2M,CDKN1A,CREBBP,DPF1,ELP1,EP300,FKBP4,GHR,GJA
NAL,GNAO1,GNAS,GNAT1,GNB1,GNB2,GNG2,GNG4,GRB2,MAPK1,MAPK3,PRKACA,PRKACB,
,CREBBP,CTNNB1,ELAVL1,EP300,ERBB2,H3-3A/H3-3B,HDAC2,HSF1,KDM1A,MDM2,MEX3B,M
APK11,MAPK3,PAK3,PIK3R3,PLCG1,PRKACA,PRKACB,PRKAR1A,PRKAR1B,PRKAR2A,PRKAR2B,P

others),CDC42,CSK,GNA12,GNA13,GNAI1,GNAL,GNAO1,GNAS,GNAT1,GNB1,GNB2,GNG2,G
MNB2,PRKCI,SH3GL1,SH3GLB1,VPS28,VPS4A,XPO1
MAPK1,MAPK11,MAPK3,NOX4,PRDX6,PRKACA,PRKACB,PRKAR1A,PRKAR1B,PRKAR2A,PRKAR2
,GNAI1,GNAL,GNAO1,GNAS,GNAT1,GNB1,GNB2,GNG2,GNG4,JUN,MAPK1,MAPK3,PDE11A,P
GNA13,GNAI1,GNAL,GNAO1,GNAS,GNAT1,GNB1,GNB2,GNG2,GNG4,LIMK1,MAP2K2,MAPK1,
1,FGFR1,FGFR3,FZD3,FZD7,HDAC2,HDAC5,HDAC6,HIF1A,KAT2A,KDM1A,NR2F2,RBBP4,RBBP7
EP300,GRIA1,GRINA,MAP2K2,MAPK1,MAPK3,PDIA3,PLCG1,PPP1CA,PPP1CB,PPP1CC,PPP1R10,

,GNAO1,GNAS,GNAT1,GNB1,GNB2,GNG2,GNG4
ACNG7,CALM1 (includes others),GNAS,KCNN3,KCNQ2,MPRIP,MYH10,MYH9,MYL12A,MYL12B,

MAP2K2,MAP2K4,MAP3K10,MAPK1,MAPK3,PDPK1,PIK3R3,PLCG1,RAC1,RAP1B,RELA,RHOA,RC
2,MAP2K4,MAPK1,MAPK11,MAPK3,MTOR,NCK2,NRG1,PAK3,PDPK1,PIK3R3,PLCG1,PRKCI,RAP
G7,CALM1 (includes others),CDC42,CREBBP,EGR1,GNAI1,GNAS,GNB1,GNB2,GNG2,GNG4,GR
LDH7A1,ALDH9A1,CHST10,CHST3,CITED2,CREBBP,EP300,GSTA4,GSTM3,GSTP1,HS6ST1,HSP90
2,ITGA11,ITGA2,ITGA3,ITGA7,ITGAV,ITGB1,ITGB5,JUN,MAP2K2,MAP2K4,MAP3K10,MAPK1,M
AM20B,HIPK1,LIMK1,MAP2K2,MAP2K4,MAPK1,MAPK3,PAK3,PDXK,PKN1,PRPF4B

MMP2,MMP24,THBS2,TIMP1,TIMP2,TIMP3
-DPA1,PDIA3,PSMB5,PSMB6,TAPBP
SDHA
,GNA13,GNAI1,GNAL,GNAO1,GNAS,GNAT1,GNB1,GNB2,GNG2,GNG4,MAP2K2,MAPK1,MAP
P2,CRP,ELP1,FGA,FN1,FTL,GRB2,HNRNPK,IL6ST,JUN,KLKB1,MAP2K2,MAP2K4,MAPK1,MAPK1
,PRKAR1B,PRKAR2A,PRKAR2B,SMO
RCC5,XRCC6
CAPN2,CAPNS1,GLUL,GPX1,GRIA1,GRIK5,GRINA,NEFL,NEFM,PIK3R3,PPP3CA,PRPH,RAB5A,R

1L,PPP2CA,PPP2CB,PPP2R1A,PPP2R2A,PPP2R5B,PPP2R5C,PPP2R5D,PTPA
SA,H2AX,H2AZ1,MACROH2A1,MASTL,NSL1,PPP1CA,PPP1CB,PPP1CC,PPP1R10,PPP1R12A,PPP1
KDC
JCHL1
K2,MAPK1,MAPK3,PIK3R3,PLCG1,PRKACA,PRKACB,PRKAR1A,PRKAR1B,PRKAR2A,PRKAR2B,R
1,MAPK3,PIK3R3,PPP3CA,PPP3CB,RACK1,RAP1B,SHC1,SOS1,STAT1,STAT3
NNB1,ELP1,GNA12,GNA13,JUN,LPAR2,MAP2K2,MAP2K4,MAPK1,MAPK3,MYL12A,MYL12B,M
FZD3,FZD7,GNA12,GNA13,GNAI1,GNAL,GNAO1,GNAS,GNAT1,GNB1,GNB2,GNG2,GNG4,GS
R1,GNAI1,GNAL,GNAO1,GNB1,GNB2,GNG2,GNG4,GRIA1,GRINA,MAPK1,MAPK11,MAPK3,PD
LGR4,MAP2K4,PFN1,PFN2,RAC1,RHOA,ROCK2,SMO
LDH7A1,ALDH9A1,CES3,CHST10,CHST3,CITED2,CREBBP,CUL3,EP300,ESD,FTL,GSTA4,GSTM3,G
,MAP2K2,MAP2K4,MAPK1,MAPK3,PDIA3,PLCG1,PRKACA,PRKACB,PRKAR1A,PRKAR1B,PRKAR2
GNB2,GNG2,GNG4,JUN,MAP2K2,MAPK1,MAPK3,PDIA3,PIK3R3,PLCG1,PRKACA,PRKACB,PRKA
NOTCH1,NOTCH2,NUMBL,RBPJ
G7,CALM1 (includes others),CNR1,CREBBP,GNAI1,GNAO1,GNAS,JUN,MAP2K2,MAPK1,MAPK1
1,JUN,MAP2K2,MAPK1,MAPK3,PIK3R3,RAP1B,SHC1,SOS1
G7,CAMKK2,CREBBP,CTBP1,CTBP2,FGFR1,FGFR3,GNAS,LDHA,LDHB,MAPK1,MAPK11,NDN,PR

,MAPK1,MAPK3,PDPK1,PIK3R3,PLCG1,PRKCI,RAP1B,SHC1,SOS1,STAT1,STAT3,TCF7
NAI1,JUN,LIMK1,MAP2K2,MAPK1,MAPK11,MAPK3,MPRIP,PLCG1,PPP1CB,PPP1R12A,RAP1B,R
K3,PDIA3,PIK3R3,PLCG1,PRKACA,PRKACB,PRKAR1A,PRKAR1B,PRKAR2A,PRKAR2B,STAT3
[illegible]
MAP2K4,MAPK1,MAPK11,MAPK3,PRKACA,PRKACB,PRKAR1A,PRKAR1B,PRKAR2A,PRKAR2B,RA
LDH7A1,ALDH9A1,CREBBP,EP300,GSTA4,GSTM3,GSTP1,HDAC5,HSP90AA1,HSP90AB1,HSP90B
ACNG7,CALM1 (includes others),HSP90AA1,HSP90AB1,HSP90B1,MAP2K2,MAPK1,MAPK3,PIK3
A12,GNA13,GNAI1,GNAL,GNAO1,GNAS,GNAT1,GNB1,GNB2,GNG2,GNG4,HDAC5,HIF1A,JUN

K3,PIK3R3,PLCG1,PRKCI,RAP1B,SHC1,SOS1,STAT1,STAT3
B2,GSK3B,IL6ST,JAK1,LIFR,MAP2K2,MAPK1,MAPK3,NANOG,PIK3R3,RAP1B,SHC1,SMO,SOS1,S
2,NR2F6,PARP1,PHB,SH3GLB1,SOX2,TASOR

,PRMT1,PRMT5
MAPK3,PDPK1,PIK3R3,PLCG1,PRKCI,RPS6KA2,SRF,STAT1,STAT3
YN,LIMK1,PIK3R3,RAC1,RHOA,ROCK2
PIK3R3,PPM1L,PPP2CA,PPP2CB,PPP2R1A,PPP2R2A,PPP2R5B,PPP2R5C,PPP2R5D,PTPA,RAP1B,R
2,FGF13,FGFR3,HDAC2,HDAC5,HDAC6,MAP2K2,MAPK1,MAPK3,MDM2,MMP14,MMP15,MMP2,

3R3,RAC1,RAP1B,RELA,RHOA,ROCK2,SERPINF1,SOD2,SRF,TCF7,WASF2,ZEB1
,FGFR3,GRB2,MAPK1,MAPK11,MAPK3,MAPKAPK2,PIK3R3,PLCG1,RAC1,SOS1,STAT3
P2K2,MAP2K4,MAPK1,MAPK3,MTOR,PIK3R3,RAP1B,RELA,SOS1,STAT3,TCF3,TCF7
A1,COL2A1,FCGR3A/FCGR3B,FYN,GRB2,HLA-A,HLA-B,HSPA1A/HSPA1B,HSPA4,ITGB1,KLRD1,LI
TNNA1,CTNNA2,CTNNB1,DDIT4,FGFR1,FGFR3,FN1,FZD3,FZD7,H19,HIF1A,HMGB1,HTRA1,ITG
P2,PIK3R3,RAC1,RAC3,RAP1B,RHOA,RHOBTB2,TIMP1,TIMP2,TIMP3
,ATP6V0C,ATP6V0D1,ATP6V0E2,ATP6V1B2,ATP6V1D,ATP6V1E1,ATP6V1F,ATP6V1G1,JUN,MA
OT2,MAT2A,PRMT1,PRMT5
2,CACNB1,CACNB4,CACNG7,CALM1 (includes others),CTNNB1,DIAPH3,DLG1,EDNRB,EP300,FG
TYK2
PIP5K1C,PLCG1
3,MAP2K4,PDGFRB
P1
0AA1,HSP90AB1,HSP90B1,JUN,MAP2K2,MAP4K4,MAPK1,MAPK3,NCOR2,NR2F1,PDGFRB,RAP
K2,MAPK1,MAPK3,MDM2,PIK3R3,RAP1B

2,MAP2K4,MAPK1,MAPK3,PDPK1,PIK3R3,PLCG1,RAP1B,SHC1,SOS1
0AB1,HSP90B1,HSPA1A/HSPA1B,HSPA4,JAK1,JUN,MAP2K4,MAPK1,MAPK11,MAPK3,MAVS,ME

RMT5
,GLUD1,GLUL,JAK1,MDM2,PPP3CA,PPP3CB,SLC25A3,SLC25A5,STAT1,STAT2,TIMM13,TIMM23
KHD1/ANKHD1-EIF4EBP3,ARHGEF17,ARHGEF18,ARHGEF2,ARHGEF7,ARHGEF9,ATF4,CCND1,C
K,GNAS,GRB2,GSK3A,GSK3B,HIF1A,KCNN3,KCNQ2,MAP2K2,MAP2K4,MAPK1,MAPK11,MAPK3,
11,MAPK3,MTOR,PIK3R3,RAC3,RAP1B,RELA,SHC1,STAT3,TYK2
1A,HSPD1,IDH2,LDHA,LDHB,NADK,NAMPT,NMNAT2,PARP1,PIK3R3,POLR2A,POLR2B,POLR2C,P
2,FGF13,FN1,FOXG1,HIF1A,HLA-A,HLA-B,IGF2,JUN,MAP2K2,MAPK1,MAPK3,MMP14,MMP15,M

PK11,MAPK3,RELA
K4,MAVS,PIN1,PPIB,RELA,STAT1,STAT2,TRAF3
others),CDC42,CREBBP,CSK,EGR1,ELP1,GAB2,GRB2,GSK3A,GSK3B,JUN,MAP2K2,MAP2K4,MA
D,EIF2AK2,FAM20B,HIPK1,LIMK1,MAP2K2,MAP2K4,MAPK1,MAPK3,NME2,PAK3,PKN1,PRPF4B
P1B,RPS6KA2,WNK1,YWHAB,YWHAE,YWHAG,YWHAH,YWHAQ,YWHAZ
RB2,JUN,MAP2K2,MAP2K4,MAPK1,MAPK3,PRKCI,RAC1,RAC3,RAP1B,RHOA,RHOBTB2,ROCK2,S

\1,JAK1,JUN,MAPK1,MAPK11,RELA,STAT1,TYK2
CAPNS1,DLG4,GRINA,PFKM,PPP3CA,PPP3CB,PRKCI
,MAP2K2,MAPK1,MAPK3,MTOR,PIK3R3,RAP1B,SHC1,TCF3,TCF7
P,HSD17B12,HSD17B4,IGF1R,JUN,MAPK1,MAPK3,PIK3R3,RAP1B,RELA
IB1,GNB2,GNG2,GNG4,GRIA1,GRIK5,GRINA,SLC1A3,SLC1A4,SLC38A1
\1,UBTF
1,MAPK11,MAPK3,MYL12A,MYL12B,MYL6,MYL6B,MYL9,PDIA3,PIK3R3,PLCG1,PRKCI,SLC9A6
others),CCND1,CEBPG,CREBBP,CSNK1A1,CSNK1D,CSNK1E,CSNK1G2,CTNNB1,CXCL12,DAAM1,D

),CSK,DDR1,EPHA4,EPHB2,ERBB2,FGFR1,FGFR3,FYN,GNA12,GNA13,GNAI1,GNAL,GNAO1,GN
L,MSH2,MSH6,PBRM1,RAD51,SMARCA4,SMARCB1,SMARCC1,SMARCD1,STAT1
K4,MAPK1,MAPK11,MAPK3,PIK3R3,RELA,STAT3,TYK2
IF1A,JAG2,LOX,NCSTN,NOTCH1,RBPJ,RELA,SMO,TCF3,TCF7
IRD1
,MAPK3,PDPK1,PIK3R3,RELA,TRAF3
IRG1,PDPK1,PIK3R3,PLCG1,PRKCI,RAP1B,SHC1,SOS1
CE1,CREBBP,CRP,CTNNB1,CXCL12,ELP1,GABBR1,GABRB3,GLUL,GRIA1,GRINA,GSK3B,HLA-A,H

IA12,GNA13,GNAI1,GNAL,GNAO1,GNAS,GNAT1,GRIA1,IGF1R,MAP2K2,MAPK1,MAPK3,PDIA3
IN,MAP2K2,MAP2K4,MAPK1,MAPK11,MAPK3,PMEPA1,RAP1B,SOS1,TGFB2,TGFBR1,TGIF1

rs),CHRNA3,CHRNA4,GNAS,HSP90AA1,HSP90AB1,HSP90B1,HSPA1A/HSPA1B,HSPA4,LPAR2,PD
IA,RELA,STAT3,TGFB2,TGFBR1
1,LMNA,MAP2K2,MAP2K4,MAP4K4,MAPK1,MAPK3,MCL1,PARP1,PLCG1,RAP1B,RELA,SPTAN1
K3R3,SERPINF1

ELA
)P1
TGAV,ITGB1,MAPK1,MAPK3,PIK3R3,PRKCI,RAP1B,RELA
2,CTNNB1,FZD3,FZD7,GSK3B,IFNA16,NCOA4,SMO,TCF3,TCF7
2,GSK3B,HIF1A,JUN,MACROH2A2,MAP2K2,MAPK1,MAPK3,MDM2,MTOR,PDPK1,PIK3R3,PLCG
T,MAP2K2,MAP2K4,MAP3K10,MAPK1,MAPK11,MAPK3,NDUFA13,PDGFRB,PTPN2,RAC1,RAP1E
R1A,PRKAR1B,PRKAR2A,PRKAR2B,RELA
B,CALM1 (includes others),CD81,FYN,JUN,MAP2K2,MAPK1,MAPK3,PDIA3,PDPK1,PLCG1,PPP3C

PK11,MAPK3,MAPKAPK2,PIK3R3,RELA,STAT3,TRAF3
K2

GFR1,FGFR3,FYN,IGF1R,JAK1,LMTK3,MAP2K4,PDGFRB,PRKCI,PTK7,RELA,STAT1,TNK2,TYK2

ND1,CCND2,CSK,CTNNB1,FOXG1,FYN,GRB2,GSK3B,IFNA16,IL6ST,JAK1,JUN,MAP2K2,MAP4K4,
DKN1B,E2F6,FHIT,HDAC2,HDAC5,HDAC6,PIK3R3,RELA,TFDP1,TRAF3
ISPB1,LIMK1,LMNA,MAP2K4,MAP4K4,PARP1,RELA,SPTAN1,TNFRSF10B,TNKS
PK1,MAPK3,PIK3R3,PLCG1,RAC1,RAP1B,SHC1,SOS1
9A1
TYK2
P1,JUN,MAP2K2,MAP2K4,MAP3K10,MAPK1,MAPK11,MAPK3,PIK3R3,PPP3CA,PPP3CB,RELA
K3,NRP2,PIK3R3,PLCG1,PRKCI,RAP1B,SHC1,SOS1,VEGFA

PRKAR1A,PRKAR1B,PRKAR2A,PRKAR2B
K1,MAPK11,MAPK3,PIK3R3,PRKCI,RAC1,RAP1B,RELA,SRF

PK3,MDM2,PIK3R3,RAP1B,YBX1
APK3,PDIA3,PIK3R3,PLCG1,PRKACA,PRKACB,PRKAR1A,PRKAR1B,PRKAR2A,PRKAR2B,PRKCI

2,EXTL3,HS6ST1,NDST1,PPME1,PRDX6,SULT1C4,SULT4A1
IS6ST1,NDST1,PPME1,PRDX6,SULT1C4,SULT4A1
3,MAP2K2,MAPK1,MAPK3,PIK3R3,RELA
K11,MAPK3,MTOR,PIK3R3,PRKCI
,ALDH9A1

ER1
KAR1A,PRKAR1B,PRKAR2A,PRKAR2B
3R3,PLCG1,PRKCI,RAB5A,RAC1,RAP1B,RHOA
3,MT2A,PIK3R3,RAP1B,STAT3,TYK2

,CSNK2B,GSK3B,HLA-A,HLA-B,HLA-DPA1,JAK1,PIK3R3,PLCG1,TGFB2,TYK2
NB1,FZD3,FZD7,GSK3B,LRP6,MAP2K4,NOX4,PDIA3,PLCG1,PRKCI,RHOA,ROCK2,SMO,TCF3,TCF[illegible]
NG4,MAP2K2,MAPK1,MAPK3,PIK3R3,RAP1B,SLC39A7
D
A1,HMGA1,JAK1,MTOR,PIK3R3,RAP1B,SHC1,SOS1,TYK2

6ST1,NDST1,SULT1C4,SULT4A1

A,TRAF3

2G,RAD23B
PLCG1,PPP3CA,RELA,SMO
IAI1,GNAS,GNG2,GNG4,PDIA3,PLCG1,PRKACA,PRKACB,PRKAR1A,PRKAR1B,PRKAR2A,PRKAR2
I,MAPK1,MAPK11,MAPK3,PDPK1,PIK3R3,PLCG1,PRKCI,RAC1,RAC3,RAP1B,SOS1
A,TRAF3
R1A,PRKAR1B,PRKAR2A,PRKAR2B,RELA
3,GNAI1,MAPK1,MAPK3,PDGFRB,PDIA3,PIK3R3,PLCG1,RAC1,RAC3,RHOA,RHOBTB2

MAPK3,PRKCI,RAP1B
RELA
6ST1,NDST1,SULT1C4,SULT4A1

LT1C4
,FTL,GSTA4,GSTM3,GSTP1,MAP2K2,MAP2K4,MAP3K10,MAPK1,MAPK11,MAPK3,PIK3R3,PRKCI

16,JUN,MAP2K2,MAP2K4,MAPK1,MAPK11,MAPK3,PIK3R3,PRKCI,RAB7A,RELA,STAT1,TGFB2
RELA,RPS6KA2

ЛTOR,PARP1,PDIA3,PIK3R3,PLCG1,RAP1B,RPS6KA2,STAT1,TNKS
LA

AT7,PPT1,TECR
YK2

MAPK11,RELA,RPS27A,TOLLIP,UBA52,UBB

GA2,ITGA3,ITGA7,ITGAV,ITGB1,ITGB5,MARCKS,MRC2,PDIA3,PIK3R3,PLCG1,PRKCI,RAC1,RAC3
LT1C4,SULT4A1
PREP,PRKACA,PRKACB,PRKAR1A,PRKAR1B,PRKAR2A,PRKAR2B,PRSS23,TMPRSS4,YWHAE
,PCNA,TOP2B

DUSP4,ELP1,FYN,GABBR1,GDE1,GNAI1,GNAL,GNAO1,GNAS,GRB2,GRK2,MAP2K2,MAPK1,MA
ludes others),CNR1,CREBBP,DUSP4,GABBR1,GDE1,GNAI1,GNAL,GNAO1,GNAS,GRK2,MAP2K
1,MAPK11,RELA,STAT3,TYK2
EZR,FN1,GLG1,GNAI1,ITGA2,ITGA3,ITGB1,JAM3,MMP14,MMP15,MMP2,MMP24,MSN,MYH1

K3,MAPKAPK2,PIK3R3,RELA

ACB,PRKAR1A,PRKAR1B,PRKAR2A,PRKAR2B
AKT2,AKT3,CACNA1H,CACNB1,CACNB4,CACNG7,MAP2K4,MAPK1,MAPK3,MTOR,PDPK1,PIK3R
),COL1A1,CSNK1A1,CSNK1D,CSNK1E,CSNK1G2,CTNNB1,DKK3,ELP1,FZD3,FZD7,GSK3B,JUN,LR
2G,TAF15,TAF7

A1

HS1,FASN,FDFT1,FGA,HMGCR,LDLR,NCOR2,PLTP,RELA,SERPINF1,TTR

JN,KLKB1,MAPK1,MAPK3,PIK3R3,RAP1B,RELA,SOS1,STAT1,STAT3
RB2,MAP2K4,PDPK1,PIK3R3,RAP1B,SHC1,SOS1

1B,SHC1,STAT1,STAT3

NAO1,GNAS,GNAT1,GRB2,JUN,MAPK1,MAPK11,MAPK3,PDIA3,PIK3R3,PLCG1,PLD3,PRDX6,PRK
P3,NCOR2,PRKCI,PSMC5,TGFB2,YY1
,STAT3,TRIM24
AF3
,SULT4A1
PDIA3,PLCG1,PLD3,PRDX6,RELA,SELENOT,SLC2A1,TXN,TXNRD1

PR2,PLPPR3

LM1 (includes others),DAG1

ALDH3B1,ALDH6A1,ALDH7A1,ALDH9A1,APOE,CHST10,CHST3,FABP5,FABP7,GSTA4,GSTM3,GST

KAR1A,PRKAR1B,PRKAR2A,PRKAR2B,PRKCI,RELA,SLC4A2,TGFB2

,FASN,FGA,G6PC3,MAP2K4,PLTP,SERPINF1,SLC4A2,TTR
,PIK3R3,PRKCI,RELA,TGFB2
,PLCG1,RELA,STAT3
hers),CSNK1A1,CSNK1D,CSNK1E,CSNK1G2,ELP1,FCGR3A/FCGR3B,FYN,GNA12,GNA13,GNAI1,(
LM1 (includes others),GNA12,GNA13,GNAI1,GNAL,GNAO1,GNAS,GNAT1,GNB1,GNB2,GNG2
s),CAPN2,EP300,HDAC2,HLA-A,HLA-B,HLA-DPA1,PLCG1,PPP3CA,PPP3CB,PRKCI

ELA
1,HSP90B1,JAK1,JUN,MAP2K4,MAPK1,MAPK11,MAPK3,MMP2,PIK3R3,RAP1B,RELA,SRSF1,TGF

L,AP2A1,AP2A2,AP2B1,AP2M1,AP2S1,B2M,CLTA,FYN,GRB2,HLA-A,HLA-B,PIK3R3,PLCG1,PPM1
25A5

AT3,TGFBR1
C3,CALM1 (includes others),CDC42,CSK,ELP1,FYN,GRB2,HLA-A,HLA-B,HLA-DPA1,JUN,MAP2K2,M

.,COL2A1,CREBBP,ELP1,FCGR3A/FCGR3B,FSCN1,HLA-A,HLA-B,HLA-DPA1,MAP2K4,MAPK1,MAI
P1B,RELA,SOS1
K,GAB2,GRB2,HLA-A,HLA-B,HLA-DPA1,PDPK1,PIK3R3,PLCG1,PPP3CA,PPP3CB,RAC1,RELA,SHC1
2S1,B2M,CALR,CANX,PDIA3,PSAP
MAP2K2,MAP2K4,MAPK1,MAPK11,MAPK3,PIK3R3,RAC1,RAC3,RAP1B,RBBP7,RELA,RHOA,RHOI
P,EP300,KLF11,OGDH
T2,AKT3,ATF4,CACNA1H,CACNB1,CACNB4,CACNG7,CALM1 (includes others),CCR3,CELSR1,CNF
1,MAP2K4,MAPK1,MAPK11,RELA,STAT1
CF3,TCF7

3,KLRD1,NECTIN2,RELA,TLN1
CGR3B,GRB2,HLA-A,HLA-B,HNRNPA2B1,HNRNPC,IFNA16,JUN,LSM14A,LSM14B,LSM4,LSM7,M
,CDC42,CDC42BPA,DIAPH3,EXOC4,EXOC5,EXOC7,FGD1,FNBP1L,GSK3B,HLA-A,HLA-B,HLA-DPA1
F10B,TNKS
EF7,ARHGEF9,ATF4,CALM1 (includes others),CDC42,CREBBP,EP300,FYN,GNA13,GNAS,GNB1,(

A1,CXCL12,GLG1,PRDX6,RELA

2,JUN,MAP2K2,MAP2K4,MAPK1,MAPK3,PABPC4,PLCG1,PPP3CA,PPP3CB,RAC1,RAP1B,RELA,SC

P300,HDAC2,HLA-A,HLA-B,HLA-DPA1,MAP2K2,MAPK1,MAPK3,PCNA,PPP3CA,PPP3CB,PRKCI,SII
N,GRB2,HLA-A,HLA-B,HLA-DPA1,JUN,MAP2K4,MAP3K10,MAPK1,MAPK3,PIK3R3,PLCG1,PPP3C/

APK11,MAPK3,PRDX6,RAP1B

2
C4,SKP1,TGFB2,TGFBR1
KA2
K4,RELA,TRAF3

DX6

D17B4

A1,B4GAT1,EXT2,SULT1C4

9A1

GB1,JAM3,MMP14,MMP15,MMP2,MMP24,MSN,RDX,THY1

,GNA13,GNAI1,GNAL,GNAO1,GNAS,GNAT1,GNB1,GNB2,GNG2,GNG4,GTF2I,ITGA11,ITGA2,IT

NB1,GNB2,GNG2,GNG4,GRB2,HNRNPK,JUN,MAP2K4,MAP3K10,MAP4K4,MAPK8IP1,MAPK8IP

P2K4,MAPK11,MAPKAPK2,MAPT,MKNK2,RPS6KA2,SRF,STAT1,TGFB2,TGFBR1

IF2AK2,EP300,FGFR1,FGFR3,GHR,GSK3B,HDAC2,IGF1R,MAP4K4,PDGFRB,PIK3R3,PRKACA,PR

s),CDKN1B,CSK,CTNNB1,ELP1,FYN,GRB2,GSK3A,GSK3B,HLA-A,HLA-B,HLA-DPA1,ITGA2,ITGA3,

4,CACNG7,GDE1,GNB1,GNB2,GNG2,GNG4,PDE11A,PDE7A,PRKACA,PRKACB,PRKAR1A,PRKAR

,HLA-DPA1,JAG2,JAK1,JUN,KLRD1,NCSTN,NOTCH1,NOTCH2,PIK3R3,STAT1,STAT3,TGFBR1,TY

D1,NCSTN,NOTCH1,NOTCH2,PIK3R3,STAT1,STAT3,TYK2

,HLA-DPA1,JAG2,JAK1,JUN,NCSTN,NOTCH1,NOTCH2,PIK3R3,TGFBR1,TYK2

AK1,MTOR,RELA,STAT3,TYK2

DPA1,IFNA16,JAK1,JUN,MAP2K4,MAPK1,MAPK3,MGAT5,MTOR,PIK3R3,PLCG1,PPM1L,PPP2CA,

3BP,DNMT1,EZR,GNA12,GNA13,GNAI1,GRB2,HLA-A,HLA-B,HLA-DPA1,JUN,MAP2K2,MAP2K4,

MAPK11,MAPK3,PRDX6

3L,EIF3M,EIF4A1,EIF4A3,EIF4G1,EIF4G2,EIF4G3,EIF5,EIF5B,FAU,GRB2,GSK3B,HNRNPA1,IGF1
0AA1,HSP90AB1,HSP90B1,HSPA12A,HSPA1A/HSPA1B,HSPA4,HSPB1,HSPD1,HSPE1,HSPH1,MDN
1,DNAJC5,DNM3,DYNC1I2,EP300,GNB1,GNB2,GNG2,GNG4,GOSR2,GPAA1,GRB2,HDAC2,HDAC
AU,GRB2,ITGA11,ITGA2,ITGA3,ITGA7,ITGAV,ITGB1,ITGB5,MAP2K2,MAPK1,MAPK11,MAPK3,M
A,PPP2R2A,PPP2R5B,PPP2R5C,PPP2R5D,PRKACA,PRKACB,PRKAR1A,PRKAR1B,PRKAR2A,PRKA
IA5,PSMA6,PSMA7,PSMB1,PSMB2,PSMB4,PSMB5,PSMB6,PSMC1,PSMC2,PSMC3,PSMC4,PSMC
5,DLG4,EP300,FOXG1,GNA12,GNA13,GNAI1,GNAL,GNAO1,GNAS,GNAT1,GNB1,GNB2,GNG2,
4G2,EIF4G3,FAU,FKBP1A,HIF1A,MAPK1,MAPK3,MTOR,PDPK1,PIK3R3,PLD3,PPM1L,PPP2CA,PPP
LR,CANX,CTSA,CTSB,CTSC,CTSD,DCTN4,DYNC1I2,DYNC1LI1,DYNC1LI2,DYNLL1,DYNLRB1,DYNLT
L,COX6C,COX7A2,COX7A2L,COX7C,COX8A,CYB5R3,FIS1,FURIN,GPX4,GSR,MAP2K4,NCSTN,NDU
ADM1,CALM1 (includes others),CDC42,CDH11,CDH2,CDH4,CLASP2,CNTNAP1,CPLX2,CREBBP,CT
SMC3,PSMC4,PSMC5,PSMC6,PSMD1,PSMD10,PSMD12,PSMD13,PSMD14,PSMD2,PSMD3,PSM
C6,PSMD1,PSMD10,PSMD12,PSMD13,PSMD14,PSMD2,PSMD3,PSMD4,PSMD5,PSMD7,PSMD8,P
JD1,GOT2,GSK3B,H1-0,H1-10,H3-3A/H3-3B,HIF1A,HSF1,IDH2,JUN,KAT2A,LDHA,LDHB,MAP1LC
N,FOXG1,GNA12,GNA13,GNAI1,GNAL,GNAO1,GNAS,GNAT1,GNB1,GNB2,GNG2,GNG4,HMGC
IAP2K2,MAP2K4,MAPT,MICAL1,MPRIP,MYL12A,MYL12B,MYL6,MYL6B,MYL9,NRP2,PAK3,PIK3R
IP1A,CXCL12,DPYSL2,ECEL1,EFNA3,EFNB1,EFNB2,EFNB3,EPHA4,EPHB2,ERBB2,FYN,FZD3,FZD
1,CAPN1,CAPN2,CAPNS1,CDC42,CTTN,FYN,GIT1,GRB2,GSK3B,ITGA11,ITGA2,ITGA3,ITGA7,ITG
H2,CDH4,CYFIP1,DIAPH3,EZR,GNA12,GNA13,GNAI1,GNAL,GNAO1,GNAS,GNAT1,GNB1,GNB2,G
YH10,MYH9,MYL6,MYL6B,MYL9,MYO18A,NAPA,NAPB,NECTIN1,NECTIN2,NSF,PPM1L,PPP2CA,
E2,MAPRE3,RAB5A,RAB5B,RAB5C,RAB7A,TUBA1A,TUBA1B,TUBA1C,TUBB,TUBB2B,TUBB4A
IP2,DIAPH3,EZR,FGD1,FGF13,FLNA,FN1,GIT1,GNA12,GNA13,GRB2,ITGA11,ITGA2,ITGA3,ITGA
COX7A2L,COX7C,COX8A,NDUFA10,NDUFA11,NDUFA13,NDUFA4,NDUFA5,NDUFB10,NDUFB11
C42,CDH11,CDH2,CDH4,CREBBP,DGKZ,EP300,EZR,GNA12,GNA13,GNAI1,GNAL,GNAO1,GNAS,
A,CLU,CSNK2A1,CSNK2B,CTTN,DNM3,EPHB2,EPN1,FGF13,GAK,GRB2,HGS,HIP1,HIP1R,ITGB1,I
PK1,MAPK11,MAPK3,MAVS,NPM1,NUP98,RAB7A,RELA,RPS10,RPS12,RPS13,RPS14,RPS15,RP
P2CA,PPP2CB,PPP2R1A,PPP2R2A,PPP2R5B,PPP2R5C,PPP2R5D,PTPA,RELA,SMARCB1,STAT3,T
YL12A,MYL12B,MYL6,MYL6B,MYL9,NRP2,PFN1,PFN2,PI4KA,PIP4K2A,PIP4K2B,PIP5K1A,PIP5K1
A4,EPHB2,FYN,GNA12,GNA13,GNAI1,GNAL,GNAO1,GNAS,GNAT1,GNB1,GNB2,GNG2,GNG4,
PPP2CA,PPP2CB,PPP2R1A,PPP2R2A,PPP2R5B,PPP2R5C,PPP2R5D,PTPA,SCRIB,SKP1,YWHAB,YW
0,FZD3,FZD7,GJA1,GNAO1,GSK3A,GSK3B,JUN,LRP6,MDM2,PIN1,PPM1L,PPP2CA,PPP2CB,PPP2
,UBE2G1,UBE2G2,UBE2H,UBE2I,UBE2J1,UBE2K,UBE2L3,UBE2M,UBE2Q1,UBE2R2,UBE2V1,U
11,ITGA2,ITGA3,ITGA7,ITGAV,ITGB1,ITGB5,JAK1,LIMS1,MAP2K2,MAPK1,MAPK3,MAPK8IP1,M
B1,CTNND1,EPN1,FGFR1,MYH10,MYH9,MYL6,MYL6B,MYL9,MYO18A,NECTIN1,NECTIN2,NOT
R3,PRKACA,PRKACB,PRKAR1A,PRKAR1B,PRKAR2A,PRKAR2B,PRKCI,RAP1B,SHC1,SOS1,SRF,ST
STN,PRKACA,PRKACB,PRKAR1A,PRKAR1B,PRKAR2A,PRKAR2B
1,ITGB5,JUN,LIMS1,MAP2K4,MAPK1,MAPK3,MTOR,MYH10,MYH9,MYL6,MYL6B,MYL9,MYO18
3,MKNK2,PAK3,PIK3R3,PLCG1,PPM1L,PPP1CA,PPP1CB,PPP1CC,PPP1R10,PPP1R12A,PPP1R14B,
YN,GHR,GNAS,GOSR2,GPAA1,JAK1,KLF11,MAPK1,MAPK11,MAPK3,MTOR,NAPA,NAPB,NSF,PA
G2,GNG4,HNRNPK,LIMK1,MAPK1,MAPK3,NCK2,RAC1,RAC3,RHOA,ROCK2
A3,ITGB1,LIMK1,MAP1B,MAP2K2,MAP2K4,MAP3K10,MAPK1,MAPK3,MAPK8IP1,MAPK8IP2,MA
1,ITGA2,ITGA3,ITGA7,ITGAV,ITGB1,ITGB5,JAM3,JUN,MAP2K2,MAP2K4,MAP3K10,MAPK1,MA
2,GSK3A,GSK3B,HIF1A,JUN,LAMP1,MAP1LC3A,MAP1LC3B,MAP2K2,MAP2K4,MAPK1,MAPK3,M

'PP1CC,PPP1R10,PPP1R12A,PPP1R14B,PPP2CA,PPP2CB,PPP2R1A,PPP2R2A,PPP2R5B,PPP2R5C,I
/IAP2K4,MAPK1,MAPK11,MAPKAPK2,NCOR2,NR2F1,NR2F2,NR2F6,NSD1,NT5C1B,PARP1,PBRI
LNC,GDE1,GNA13,GNAI1,GNAS,GNB1,GNB2,GNG2,GNG4,GSK3A,GSK3B,H1-0,H1-10,H3-3A/I
CDKN1B,CREBBP,CTNNA1,CTNNA2,CTNNB1,CTNND1,E2F6,EP300,FYN,FZD3,FZD7,GAB2,GNA
GA7,ITGAV,ITGB1,ITGB5,PTPN1,RAB5A,RAB5B,RAB5C
/IYL12B,MYL6,MYL6B,MYL9,PAK3,PFN1,PFN2,PI4KA,PIP4K2A,PIP4K2B,PIP5K1A,PIP5K1C,PPP1CI
,IREB2,MACROH2A1,MAP2K2,MAPK1,MAPK3,NCOA4,NOX4,PEBP1,RAP1B,SLC1A5,SLC38A1,SL
C,FYN,HLA-A,HLA-B,ITGA2,ITGB1,ITGB5,PIK3R3,PLCG1,PRKCI,RAC1,RAC3,RAP1B,TFRC
;A11,ITGA2,ITGA3,ITGA7,ITGAV,ITGB1,ITGB5,MAP2K2,MAPK1,MAPK3,MAST2,MCRS1,PDGFRI
O1,GNAS,GNAT1,GNB1,GNB2,GNG2,GNG4,H3-3A/H3-3B,HSP90AA1,HSPA4,JUN,MAPK1,MAF
'PP2R2A,PPP2R5B,PPP2R5C,PPP2R5D,PTPA,RAD51,SMC3,TP53BP1,TRIM28,TRRAP,USP7,ZEB1
P2K2,MAP2K4,MAPK1,MAPK3,NCKAP1,NOX4,PAK3,PI4KA,PIK3R3,PIP4K2A,PIP4K2B,PIP5K1A,PII

IAPK1,MAPK11,MAPK3,MTOR,PIK3R3,PRKACA,PRKACB,PRKAR1A,PRKAR1B,PRKAR2A,PRKAR2E
IK1,MAP2K2,MAP2K4,MAP3K10,MAPK1,MAPK3,NECTIN2,PAK3,PDPK1,PIK3R3,RAC1,RAC3,RAP
.7A,SLC2A1,SOS1,UBA52,UBB,VEGFA,VHL
;2,GNG4,GRK2,PDE11A,PDE7A,PKIA,PPM1L,PPP1CA,PPP1CB,PPP1CC,PPP1R10,PPP1R12A,PPP1I
3,CNR1,COL18A1,COL1A1,COL2A1,CREBBP,CSNK1A1,CSNK1D,CSNK1E,CSNK1G2,CTNNB1,FGFI

1L5,U2AF2,XAB2
2,TSC2,TUBA1A,TUBA1B,TUBA1C,TUBB,TUBB2B,TUBB4A,TUBB4B,TUBB6,VIM,YWHAB,YWH/
,EDIL3,EZR,GNAI1,ITGA2,ITGA3,ITGB1,JAM3,MAP2K2,MAP2K4,MAPK1,MAPK11,MMP14,MMP
32,GRK2,GSK3B,IL6ST,JAK1,JUN,LRP6,MAP2K2,MAP2K4,MAPK1,MAPK3,MMP14,MMP15,MMP
WHAZ
2,TKT
)HB,MAP2K2,MAP2K4,MAPK1,MAPK3,MDM2,MKNK2,MMP14,MMP15,MMP2,MMP24,MTOR,I
'PP2R2A,PPP2R5B,PPP2R5C,PPP2R5D,PTPA,SKP1,TFDP1,TGFB2
(3,MDM2,NOTCH1,PIK3R3,PLD3,RAC1,RAD51,RALGDS,RELA,STAT1,STAT3,TFDP1,TGFB2,TGFE
I,HSPH1,INSIG1,MBTPS1,OS9,P4HB,PDIA6,PPP1R15A,SCAP,UBXN4,VCP,XBP1
;NB2,GNG2,GNG4,GRB2,GSK3B,HSPB1,IGF1R,JUN,MAP2K2,MAP2K4,MAP3K10,MAPK1,MAPI
.RAP1B,RELA,RPS6KA2,SOS1,STAT3,TCF3,TCF7,TRIM24,VEGFA,VHL,VIM,YWHAB,YWHAE,YWI
PPP2CA,PPP2CB,PPP2R1A,PPP2R2A,PPP2R5B,PPP2R5C,PPP2R5D,PTGES3,PTPA,RAP1B,SHC1,S(
A2L,COX7C,COX8A,ERBB2,FYN,GRB2,GSK3A,GSK3B,ITGB1,ITGB5,JUN,KPNB1,MAP2K2,MAPK1
0,PPP1R12A,PPP1R14B,PRKACA,PRKACB,PRKAR1A,PRKAR1B,PRKAR2A,PRKAR2B,PRKCI,PTPN1
(P1,TFDP1,TGFB2
2,RNF4,SAE1,SENP3,SENP5,SENP6,SERBP1,SNCA,STUB1,SUMO1,UBE2I,ZEB1
'IP5K1A,PLCG1,PLD3,PRKCI,RAB11A,RAB11B,RAC1,RAC3,TLN1

(2,MAP2K4,MAPK1,MAPK3,MMP2,MTOR,PIK3R3,PRKACA,PRKACB,PRKAR1A,PRKAR1B,PRKAR2
LA,PPP2R2A,PPP2R5B,PPP2R5C,PPP2R5D,PRKCI,PTPA,RAP1B,SHC1,SOS1,YWHAB,YWHAE,YWI

GB5,MAP2K2,MAPK1,MAPK3,PAK3,PDPK1,PIK3R3,PLCG1,RAC1,RAP1B,SOS1,TLN1
.B1,HSP90B1,MAP2K2,MAPK1,MAPK3,MDM2,MTOR,PDPK1,PIK3R3,RAP1B,RELA,SOS1,TFDP1
A2,PPFIA3,PPM1L,PPP1CA,PPP1CB,PPP1CC,PPP1R10,PPP1R12A,PPP1R14B,PPP2CA,PPP2CB,PPP2
-,GRB2,MAPK1,MAPK3,PDPK1,PLCG1,PRKACA,PRKACB,PRKAR1A,PRKAR1B,PRKAR2A,PRKAR2B

IOBTB2,ROCK2
,MAPK1,MAPK11,NCK2,PAK3,PIK3R3,PTPN12,RAC1,RAP1B,SOS1,TLN1

RAP1B,SMO,SOS1,SOX2,STAT3,TCF3,TCF7,TYK2

NG4,GRB2,GSK3B,HDAC2,HDAC5,HDAC6,IGF1R,IL6ST,MAP2K2,MAP2K4,MAPK1,MAPK11,MA
N1B,CLTA,CREBBP,CTNNB1,EP300,FYN,GNA12,GNA13,GNAI1,GNAL,GNAO1,GNAS,GNAT1,G
IP5K1C,PLCG1,PPFIA2,PPFIA3,PPM1L,PPP1CA,PPP1CB,PPP1CC,PPP1R10,PPP1R12A,PPP1R14B,P

A,MYL12B,MYL6,MYL6B,MYL9,NCK2,PAK3,PDGFRB,PIK3R3,RAC1,RAP1B,SHC1,SOS1
,GRINA,PDIA3,PLCG1,PPM1L,PPP1CA,PPP1CB,PPP1CC,PPP1R10,PPP1R12A,PPP1R14B,PPP2CA,
,MSH2,MSH6,MTOR,PIK3R3,PRKACA,PRKACB,PRKAR1A,PRKAR1B,PRKAR2A,PRKAR2B,RAD51,
TP1,HSP90AA1,HSP90AB1,HSP90B1,HSPB1,JUN,MAPK1,MAPK3,MCM7,MDM2,NCOR2,NEDD8,
9,FTH1,FTL,GSK3B,GSR,GSTA4,GSTM3,GSTP1,HERPUD1,HSP90AA1,HSP90AB1,HSP90B1,JUN,
IK3R3,POLR2A,POLR2B,POLR2C,POLR2E,POLR2G,RAD51,RAP1B,RPS27A,SMARCA4,SMARCB1
PK3,MDM2,MTOR,NF2,PDGFRB,PDIA3,PIK3R3,PLCG1,RAC1,RAC3,RAP1B,RHOA,RHOBTB2,SH
,MAPK1,MAPK3,MMP2,NCSTN,NOTCH1,NOTCH2,PDGFRB,PIK3R3,RAP1B,RBPJ,RELA,RHOA,S
3,GNAI1,GNAL,GNAO1,GNAS,GNAT1,GNB1,GNB2,GNG2,GNG4,KCNN3,KCNQ2,NSF,RPS27A,
CA,PRKACB,PRKAR1A,PRKAR1B,PRKAR2A,PRKAR2B,PTPA,SLC18A3
,COL6A2,COL6A3,COL7A1,FGA,FYN,GRB2,GSK3A,GSK3B,LAMA5,LAMC1,PDPK1,PIK3R3,PLCG1,
,MAPK3,PDIA3,PIK3R3,PLCG1,PPP3CA,PPP3CB,PRKACA,PRKACB,PRKAR1A,PRKAR1B,PRKAR2A,
PK1,MAPK3,MTOR,NRG1,PDPK1,PIK3R3,PLCG1,PRKCI,RAP1B,RNF41,SHC1,SOS1
ITGB5,JUN,MAP2K2,MAP2K4,MAP4K4,MAPK1,MAPK3,MED12,NCOA6,NCOR2,NR2F1,PDIA3,P
,MDM2,PIK3R3,RAP1B,RELA,SOS1,TFDP1,TGFB2,TGFBR1
2K2,MAPK1,MAPK3,MDM2,MTOR,PDGFRB,PIK3R3,PLCG1,PRKCI,RAP1B,SHC1,SOS1,TFDP1
R12A,PPP1R14B,PPP2CA,PPP2CB,PPP2R1A,PPP2R2A,PPP2R5B,PPP2R5C,PPP2R5D,PPP3CA,PP
R12A,PPP1R14B,PPP2CA,PPP2CB,PPP2R1A,PPP2R2A,PPP2R5B,PPP2R5C,PPP2R5D,PPP3CA,PP
AP1B
AP2K2,MAP2K4,MAPK1,MAPK3,MYL12A,MYL12B,MYL6,MYL6B,MYL9,PAK3,PIK3R3,PRKCI,RAC
5D,PTPA,SLK,SMC3
K3,MEST,MMP2,MTOR,PDGFRB,PIK3R3,RAC1,RAP1B,RELA,RHOA,SHC1,SHC2,SHC3,SOS1,STA

C,PPP1R10,PPP1R12A,PPP1R14B,PPP2CA,PPP2CB,PPP2R1A,PPP2R2A,PPP2R5B,PPP2R5C,PPP2
RKAR1A,PRKAR1B,PRKAR2A,PRKAR2B,RELA,TOLLIP
S1,STAT1
,PLCG1,PRKACA,PRKACB,PRKAR1A,PRKAR1B,PRKAR2A,PRKAR2B,PRKCI,PYGB,RAP1B
10,PPP1R12A,PPP1R14B,PPP2CA,PPP2CB,PPP2R1A,PPP2R2A,PPP2R5B,PPP2R5C,PPP2R5D,PPP

B,PPP1CC,PPP1R10,PPP1R12A,PPP1R14B,PPP2CA,PPP2CB,PPP2R1A,PPP2R2A,PPP2R5B,PPP2R
HRNA4,CREBBP,EP300,GRIA1,GRINA,HDAC2,HDAC5,HDAC6,MAPK1,MAPK3,MICU1,MYH10,M
MAPK1,MAPK11,MAPK3,MPRIP,MYL12A,MYL12B,MYL6,MYL6B,MYL9,PDIA3,PDPK1,PIK3R3,PL

RF,STAT1,STAT3,TYK2

DNRB,FGFR1,FN1,IGF1R,IGF2,IGFBP3,IGFBP4,KLF6,LHX2,MMP2,MYH10,MYH9,MYL6,MYL6B,M
1B,RHOA,RHOBTB2,ROCK2,SOS1
DKN1A,CDKN1B,CREBBP,DLD,E2F6,EP300,EP400,FZR1,HIPK2,JUN,MAP2K2,MAP2K4,MAPK1,MA
,HSPA4,HSPB1,HSPD1,HSPE1,HSPH1,MAP2K2,MAPK1,MAPK3,PDIA3,PDPK1,PI4KA,PIK3R3,PIP4I
APK3,MTOR,MYH10,MYH9,MYL6,MYL6B,MYL9,MYO18A,PRKACA,PRKACB,PRKAR1A,PRKAR1B,
(1,MAPK3,RAP1B,TLN1
3L1XR1,THRA

P1,LIMK1,MAP2K2,MAP2K4,MAP4K4,MAPK1,MAPK3,MMP2,MTOR,MYL12B,MYL9,NOX4,PIK3R3

,GNG4,MAP2K2,MAPK1,MAPK3,NOX4,PIK3R3,PPP3CA,PPP3CB,PRKCI,RAC1,RAC3,RAP1B,RELA
TAT3
T2,STAT3
RAD23B,RBX1,TCEA1,TOP2B,UBE2I,USP7,XAB2
C34,CIAO2B,CUL1,FBXL5,FTH1,FTL,HIF1A,HIF3A,IREB2,JAK1,MAPK1,MAPK3,PDGFRB,SKP1,STA
RKDC,RPRM,SERPINE2,TNFRSF10B
PDXP,PIK3R3,RAC1,RAP1B,RPS6KA2,VEGFA
PKAPK2,MCL1,PIK3R3,RAP1B,RELA,SHC1,SOS1,SRF,STAT3,VEGFA
2,SHC1,SOS1,VEGFA,YWHAE

2

P1B,SOS1,TFDP1

ST1,HSP90AA1,HSP90AB1,HSP90B1,NCOR2,NDST1,PPM1A,PPP1CA,PPP1CB,PPP1CC,PPP1R10,P
AC1,RAC3,UNC5A
1,GRB2,H3-3A/H3-3B,HLA-A,HLA-B,HLA-DPA1,HMGB1,HSP90AA1,HSP90AB1,HSP90B1,HSPA1
PRKAR1A,PRKAR1B,PRKAR2A,PRKAR2B,RALGDS,RAP1B,SHC1,SHC2,SHC3,SOS1,STAT3
MP14,MMP15,MMP2,MMP24,MRTFA,PIK3R3,RBBP4,RBBP7,RELA,ROCK2,SRF,STAT3,TCF3,TC
RKCI,RAC1,RAP1B,RELA,SHC1,SHC2,SHC3,SOS1,STAT1,STAT3

NG4,GRK2,GSK3B,MAP2K2,MAPK1,MAPK3,PIK3R3,PLCG1,PLD3,PPP3CA,PPP3CB,PRKCI,RAC1,RA

3,RAC1,RAC3,SOD1
DE7A,PIK3R3,PRKACA,PRKACB,PRKAR1A,PRKAR1B,PRKAR2A,PRKAR2B,RAP1B,RELA,SLC39A7,
MAPK11,MAPK3,MPRIP,PAK3,PIK3R3,PPP1CB,PPP1R12A,PRKCI,RAC1,RAP1B,RHOA,ROCK2
7,RBP1,SAP18,SIN3B,SMO,SOX9,TBL1XR1,TCF7,TXNIP,XBP1
PPP1R12A,PPP1R14B,PPP3CA,PPP3CB,PRKACA,PRKACB,PRKAR1A,PRKAR1B,PRKAR2A,PRKAR2

MYL6,MYL6B,MYL9,MYO18A,PABPC4,PDIA3,PLCG1,PPP1CB,PPP1R12A,PRKACA,PRKACB,PRKAI

)CK2,RPS6KA2,SHC1,SHC2,SHC3,SOS1,TRIO
1B,SHC1,SOS1
B2,JUN,MAP2K2,MAP2K4,MAP3K10,MAPK1,MAPK11,MAPK3,MMP2,PAK3,PRKACA,PRKACB,PR
AA1,HSP90AB1,HSP90B1,MAP2K2,MAP2K4,MAPK1,MAPK3,NDST1,PPM1L,PPP2CA,PPP2CB,PPI
APK3,PIK3R3,PLCG1,PRKCI,RAC1,RAP1B,RAPGEF1,SOS1,STAT3

K3,PRKACA,PRKACB,PRKAR1A,PRKAR1B,PRKAR2A,PRKAR2B
.1,MAPK3,MTOR,NOLC1,PDPK1,PIK3R3,RAP1B,RBP1,RELA,SERPINF1,SHC1,SOD2,SOS1,STAT3,

AB5B,RAB5C,RAC1,SOD1,SSR4,VEGFA

R14B,PPP2R5B,PPP2R5C,PPP2R5D,SKA2,SMC3

AP1B,RPS6KA2,SHC1,SOS1

YL6,MYL6B,MYL9,PIK3R3,RAP1B,RELA,RHOA,ROCK2
K3A,GSK3B,NANOG,PDGFRB,PDPK1,PIK3R3,SMO,SOX2,TCF3,TCF7,TGFB2,TGFBR1
IA3,PLCG1,PPP3CA,PPP3CB,PRKACA,PRKACB,PRKAR1A,PRKAR1B,PRKAR2A,PRKAR2B,RIMS1

STP1,HDAC5,HS6ST1,HSP90AA1,HSP90AB1,HSP90B1,MAP2K2,MAP2K4,MAP3K10,MAPK1,MAP
A,PRKAR2B,PRKCI
R1A,PRKAR1B,PRKAR2A,PRKAR2B,PRKCI,RAP1B,RELA

.1,MAPK3,PLCG1,PRKACA,PRKACB,PRKAR1A,PRKAR1B,PRKAR2A,PRKAR2B,PRKCI,RAP1B,SLC3

KACA,PRKACB,PRKAR1A,PRKAR1B,PRKAR2A,PRKAR2B,THRA,VEGFA,VGF

HOA,ROCK2

P1B,RELA,SOS1
1,PTGES3,RELA,SUMO1
R3,PRKACA,PRKACB,PRKAR1A,PRKAR1B,PRKAR2A,PRKAR2B,PRKCI,SLC7A1,VEGFA
,MAP2K2,MAP2K4,MAPK1,MAPK3,MTOR,PIK3R3,PRKCI,RAP1B,RELA

OX2,STAT3,TYK2

ELA
,MMP24,RAP1B,TFDP1,VEGFA

MK1,MAP2K2,MAP3K10,MAPK1,MAPK11,MAPK3,MTOR,NECTIN2,PAK3,PIK3R3,PLCG1,RAC1,R
A11,ITGA2,ITGA3,ITGA7,ITGAV,ITGB1,ITGB5,MTOR,MYBBP1A,NAMPT,NOTCH1,RAC1,RBPJ,R

P2K2,MAP2K4,MAPK1,MAPK11,MAPK3,NUP153,PRDX6,RAP1B

F13,FGFR1,FGFR3,FZD3,FZD7,GDE1,GHR,GNA12,GNA13,GNAI1,GNAS,GNB1,GNB2,GNG2,G

1B,RELA,SHC1,SOS1

TAP2,NPM1,PDGFRB,RELA,STAT1,STAT2,STAT3,TRAF3,TYK2

B,TIMM50,TNFRSF10B,TOMM20,TOMM40,TOMM6,TOMM7,TOMM70,VDAC1,VDAC2,VDAC3
CND2,CCR3,CDK4,CDKN1B,CELSR1,CNR1,CREBBP,E2F6,EDNRB,FZD3,FZD7,GABBR1,GNA13,G
MMP2,PDIA3,PIK3R3,PLCG1,PRKACA,PRKACB,PRKAR1A,PRKAR1B,PRKAR2A,PRKAR2B,RAP1B,

POLR2E,POLR2G,SLC7A5,SOD2,TGFB2,TNKS,VEGFA
MMP2,MMP24,PIK3R3,RAC1,RAP1B,RELA,SLC1A4,SLC2A1,STAT3,TGFB2,VEGFA

P3K10,MAPK1,MAPK11,MAPK3,MTOR,PDPK1,PIK3R3,PPP3CA,PPP3CB,RAC1,RAP1B,RELA,SHC1

HC1,SOS1,SRF

)KK3,ELP1,FCGR3A/FCGR3B,FN1,FZD3,FZD7,GNAO1,GSK3B,IL6ST,JUN,LRP6,MAP2K2,MAP2K4

IAS,GNAT1,GNB1,GNB2,GNG2,GNG4,IGF1R,JAK1,LMTK3,MAP2K4,PDGFRB,PDIA3,PLCG1,PRD

LA-B,HLA-DPA1,HMGB1,IFNA16,JAK1,JUN,MAP2K4,MAPK1,MAPK11,MAPK3,MAPT,MFGE8,NC

,PLCG1,PPM1L,PPP2CA,PPP2CB,PPP2R1A,PPP2R2A,PPP2R5B,PPP2R5C,PPP2R5D,PRDX6,PRKCI,I

)PK1,PIK3R3,PLCG1,PRKACA,PRKACB,PRKAR1A,PRKAR1B,PRKAR2A,PRKAR2B,PRKCI,SLC7A1,ST

1,PRKCI,RAC1,RAC3,RAP1B,RELA,SHC1,SHC2,SHC3,SOS1,TGFB2
3,STAT3,TGFB2,TGFBR1,TYK2,VEGFA

:A,PPP3CB,PRKCI,RAC1,RAP1B,RELA

MAPK1,MAPK3,MAVS,MCL1,MTOR,PDPK1,PIK3R3,PPP3CA,PPP3CB,PRKCI,RAC1,RAC3,RAP1B,R

7,TGFB2,TGFBR1

.B,PRKCI

,RAP1B,RBX1

,RHOA,RHOBTB2

PK3,PDE11A,PDE7A,PDPK1,PIK3R3,PRKACA,PRKACB,PRKAR1A,PRKAR1B,PRKAR2A,PRKAR2B,R
2,MAPK1,MAPK3,PDE11A,PDE7A,PKIA,PPP3CA,PPP3CB,PRKACA,PRKACB,PRKAR1A,PRKAR1B,PF

0,MYH9,MYL6,MYL6B,MYL9,MYO18A,PODXL,PODXL2,RDX

3,PKM,PRKCI,RELA
P6,MAP2K4,MAPK1,MAPK3,MMP14,PIK3R3,PPP3CA,PPP3CB,RELA,SFRP1,SMO,TCF3,TCF7

CI,RAP1B,SHC1,SHC2,SHC3,SOS1

P1,HMGCS1,HS6ST1,JUN,MAP2K4,NDST1,PLTP,SULT1C4,SULT4A1,XPO1

GNAL,GNAO1,GNAS,GNAT1,GNB1,GNB2,GNG2,GNG4,GRB2,GSK3A,GSK3B,HLA-A,HLA-B,HLA
,GNG4,JUN,MAP2K4,MAPK1,MAPK11,PLCG1,PRKCI

B2,VEGFA

L,PPP2CA,PPP2CB,PPP2R1A,PPP2R2A,PPP2R5B,PPP2R5C,PPP2R5D,PTPA

MAP2K4,PDPK1,PIK3R3,PLCG1,PPP3CA,PPP3CB,RAC1,RELA

PK11,MAPK3,PDIA3,PIK3R3,PLCG1,RELA,STAT1,STAT2

3TB2,TGFB2

R1,CREBBP,EDNRB,EP300,FGFR1,FGFR3,FZD3,FZD7,GABBR1,GHR,GNA12,GNA13,GNAI1,GNA

MAPK1,MAPK3,MTOR,PIK3R3,PLCG1,PRPF19,PRPF31,PRPF4B,PRPF6,PRPF8,RAP1B,SART1,SNR
,ITGA11,ITGA2,ITGA3,ITGA7,ITGAV,ITGB1,ITGB5,JUN,LIMK1,MAP2K4,MAPK1,MAPK11,MPRIP

GNB2,GNG2,GNG4,GRB2,HDAC2,HDAC5,HDAC6,ITGA11,ITGA2,ITGA3,ITGA7,ITGAV,ITGB1,ITG

OS1,TGFB2,TGFBR1

N3B
A,PPP3CB,RAC1,RAC3,RAP1B,RELA,SOS1

TGA3,ITGA7,ITGAV,ITGB1,ITGB5,JAK1,MAP2K4,PAK3,PIK3R3,PLCG1,PRKCI,RAC1,RAC3,RELA,R
2,MAPK8IP3,PIK3R3,RAC1,RAC3,RAP1B,SHC1,SOS1

ACB,RAP1B,RELA,TGFBR1,TRAF3,UBE2V1
ITGB1,JUN,MAP2K2,MAP2K4,MAPK1,MAPK11,MAPK3,MTOR,PIK3R3,PLCG1,PPP3CA,PPP3CB,R
1B,PRKAR2A,PRKAR2B

2

PP2CB,PPP2R1A,PPP2R2A,PPP2R5B,PPP2R5C,PPP2R5D,PTPA,RAP1B,STAT1,STAT2,STAT3,TCF
MAPK1,MAPK3,MSN,MTOR,PIK3R3,PLCG1,PPM1L,PPP2CA,PPP2CB,PPP2R1A,PPP2R2A,PPP2R5

R,MAP2K2,MAPK1,MAPK3,NOX4,PAIP1,PDPK1,PIK3R3,PPP1CA,PPP1CB,PPP1CC,PPP1R15A,PTB
M2,NEDD4L,PSMA1,PSMA4,PSMA5,PSMA6,PSMA7,PSMB1,PSMB2,PSMB4,PSMB5,PSMB6,PSM
C5,HDAC6,HIP1,HSPA1A/HSPA1B,HSPA4,IGF1R,JUN,MAP2K4,MAP3K10,MAPK1,MAPK3,MTOR,
MTOR,PAIP1,PAIP2,PDPK1,PIK3R3,PPM1L,PPP2CA,PPP2CB,PPP2R1A,PPP2R2A,PPP2R5B,PPP2R5
R2B,PSMA1,PSMA4,PSMA5,PSMA6,PSMA7,PSMB1,PSMB2,PSMB4,PSMB5,PSMB6,PSMC1,PSM
C5,PSMC6,PSMD1,PSMD10,PSMD12,PSMD13,PSMD14,PSMD2,PSMD3,PSMD4,PSMD5,PSMD7,
,GNG4,GRB2,GSK3A,GSK3B,HIF1A,HNRNPD,HSP90AA1,HSP90AB1,HSP90B1,IGF1R,IGF2,JAK1,
P2CB,PPP2R1A,PPP2R2A,PPP2R5B,PPP2R5C,PPP2R5D,PRKCI,PTPA,RAC1,RAC3,RAP1B,RHEB,RH
1,GOSR2,GPAA1,HLA-A,HLA-B,LAMP1,M6PR,NAPA,NAPB,NOX4,NSF,PRDX1,PRDX2,PRDX5,PR
JFA10,NDUFA11,NDUFA13,NDUFA4,NDUFA5,NDUFB10,NDUFB11,NDUFB2,NDUFB4,NDUFB8
NNB1,CTNND1,DLG4,DNAJC5,EFNA3,EFNB1,EFNB2,EFNB3,EPHA4,EPHB2,FARP1,FYN,GOSR2,
D4,PSMD5,PSMD7,PSMD8,PSME3,PSMF1
PSME3,PSMF1,SQSTM1,UBA1,UBA6
3A,MAP1LC3B,MAPK1,MAPK3,MTOR,NAMPT,NDRG1,NDUFA10,NDUFA11,NDUFA13,NDUFA4
CR,MAPK1,MAPK11,MTOR,ORAI2,PBRM1,PFKFB3,PFKL,PFKM,PFKP,PIK3R3,PPM1A,PPM1G,PPM
3,PIP5K1C,PLCG1,PLXNA1,PLXNA2,PLXNA3,PLXNB1,PPP1CB,PPP1R12A,PRKACA,PRKACB,PRKA
7,GIT1,GNA12,GNA13,GNAI1,GNAL,GNAO1,GNAS,GNAT1,GNB1,GNB2,GNG2,GNG4,GRB2,G
AV,ITGB1,ITGB5,LIMS1,MAP2K2,MAP2K4,MAPK1,MAPK3,MPRIP,MYL12A,MYL12B,MYL9,NCK2
GNG2,GNG4,ITGA11,ITGA2,ITGA3,ITGA7,ITGAV,ITGB1,ITGB5,JUN,LIMK1,MAP2K2,MAP2K4,M
PPP2CB,PPP2R1A,PPP2R2A,PPP2R5B,PPP2R5C,PPP2R5D,PRKACA,PRKACB,PRKAR1A,PRKAR1B,
,TUBB4B,TUBB6,ZYX
A7,ITGAV,ITGB1,ITGB5,LIMK1,MAP2K2,MAPK1,MAPK3,MPRIP,MSN,MYH10,MYH9,MYL12A,MY
,NDUFB2,NDUFB4,NDUFB8,NDUFB9,NDUFS1,NDUFS5,NDUFS6,NDUFS7,NDUFS8,NDUFV1,N
,GNAT1,GNB1,GNB2,GNG2,GNG4,ITGA11,ITGA2,ITGA3,ITGA7,ITGAV,ITGB1,ITGB5,LIMK1,MS
TGB5,LDLR,MDM2,PICALM,PIK3R3,PIP5K1C,PPP3CA,PPP3CB,RAB11A,RAB11B,RAB4A,RAB5A,
PS15A,RPS16,RPS17,RPS19,RPS20,RPS21,RPS24,RPS25,RPS26,RPS27,RPS27A,RPS28,RPS29,R
CF3,TCF7,TSC2,VEGFA
C,PKN1,PLXNA1,PPP1CB,PPP1R12A,RDX,RHOA,ROCK2,SEPTIN2,SEPTIN3,SEPTIN5,SEPTIN6,SEP
GRB2,GRINA,ITGA11,ITGA2,ITGA3,ITGA7,ITGAV,ITGB1,ITGB5,LIMK1,MAP2K2,MAP4K4,MAPK
HAE,YWHAG,YWHAH,YWHAQ,YWHAZ
2R1A,PPP2R2A,PPP2R5B,PPP2R5C,PPP2R5D,PTPA,RPS27A,RUVBL2,SFRP1,SMO,SOX12,SOX2,S
JBE2V2,VEGFA,VHL
ICL1,MDM2,MTOR,NANOG,PDPK1,PIK3R3,PPM1L,PPP2CA,PPP2CB,PPP2R1A,PPP2R2A,PPP2R5
CH1,NOTCH2,RAC1,RAP1B,RAPGEF1,RHOA,TCF3,TCF7,TGFB2,TGFBR1,TUBA1A,TUBA1B,TUB
AT3,YWHAB,YWHAE,YWHAG,YWHAH,YWHAQ,YWHAZ

A,NACA,NCK2,PDPK1,PIK3R3,PPM1L,PPP1R12A,PPP1R14B,PPP2CA,PPP2CB,PPP2R1A,PPP2R2A
PPP2CA,PPP2CB,PPP2R1A,PPP2R2A,PPP2R5B,PPP2R5C,PPP2R5D,PRKACA,PRKACB,PRKAR1A,P
AIP1,PDHA1,PDHA2,PDIA3,PIK3R3,PLCG1,PRKACA,PRKACB,PRKAR1A,PRKAR1B,PRKAR2A,PRKA

APK8IP3,MAPT,MTOR,PAFAH1B1,PAFAH1B2,PAFAH1B3,PIK3R3,RAC1,RAP1B,RAPGEF1,RHOA
PK11,MAPK3,NECTIN1,NECTIN2,PRKACA,PRKACB,PRKAR1A,PRKAR1B,PRKAR2A,PRKAR2B,RAC
ITOR,PIK3R3,PPM1L,PPP2CA,PPP2CB,PPP2R1A,PPP2R2A,PPP2R5B,PPP2R5C,PPP2R5D,PPP3CA,

PPP2R5D,PRKACA,PRKACB,PRKAR1A,PRKAR1B,PRKAR2A,PRKAR2B,PTPA,RAP1B
M1,PDPK1,PIK3R3,PRKACA,PRKACB,PRKAR1A,PRKAR1B,PRKAR2A,PRKAR2B,PRKCI,PRMT1,PRM
H3-3B,KDELR1,KDELR2,MAP2K2,MAPK1,MAPK3,MYH10,MYL12A,MYL12B,MYL6,MYL6B,MYL9,M
12,GNA13,GNAI1,GNAL,GNAO1,GNAS,GNAT1,GNB1,GNB2,GNG2,GNG4,GRB2,GSK3A,GSK3E

B,PPP1R12A,RAC1,RAC3,RHOA,RHOBTB2
C3A2,SQSTM1,STAT3,TFRC,TXNRD1,USP7,VDAC2

B,PDPK1,PIK3R3,RAC1,RAC3,RAP1B,RELA,SHC1,SOS1,TGFBR1,YWHAH
PK3,NCOA4,POLR2A,POLR2B,POLR2C,POLR2E,POLR2G,PRKACA,PRKACB,PRKAR1A,PRKAR1B,PI

P5K1C,PRKCI,RAC1,RAC3,RAP1B,RELA,RHOA

B,RAC1,RAC3,RAP1B,RHOA,STAT3,STMN2
1B,RHOA,RHOBTB2,TGFB2,TGFBR1,TUBA1A,TUBA1B,TUBA1C,TUBB,TUBB2B,TUBB4A,TUBB

R14B,PPP2CA,PPP2CB,PPP2R1A,PPP2R2A,PPP2R5B,PPP2R5C,PPP2R5D,PRKACA,PRKACB,PRKAI
R1,FTH1,FTL,FZD3,FZD7,GNAI1,GSK3B,HIF1A,ITGA11,ITGA2,ITGA3,ITGA7,ITGAV,ITGB1,ITGB5

AE,YWHAG,YWHAH,YWHAQ,YWHAZ
15,MMP2,MMP24,MSN,MYL6,PIK3R3,PLCG1,PRKCI,RAC1,RAP1B,RDX,RHOA,ROCK2,THY1,TIMI
2,MMP24,MSH2,MSH6,PIK3R3,PRKACA,PRKACB,PRKAR1A,PRKAR1B,PRKAR2A,PRKAR2B,RAC1

NOX4,PIK3R3,PKM,PLCG1,PPP3CA,PPP3CB,PRKCI,RAC1,RAC3,RACK1,RAP1B,RBX1,SLC2A1,STAT

R1,TYK2,VEGFA

K11,MAPK3,MAPKAPK2,MTOR,MYL12A,MYL12B,MYL6,MYL6B,MYL9,PDIA3,PIK3R3,PLCG1,PPP3
IAG,YWHAH,YWHAQ,YWHAZ
OS1,TERF2IP
,MAPK3,MDM2,MMP2,MTOR,NRG1,PIK3R3,PLCG1,PRKCI,RAP1B,RELA,RHEB,SOS1,STAT3,TSC
,PTPRF,RAP1B,RAPGEF1,SHC1,SOS1,TSC2,VAMP2

2A,PRKAR2B,RHOA,ROCK2,TCF3,TCF7,VEGFA,VIM
IAG,YWHAH,YWHAQ,YWHAZ

2R1A,PPP2R2A,PPP2R5B,PPP2R5C,PPP2R5D,PPP3CA,PPP3CB,PPP4C,PPP5C,PTPA,PTPN1,PTPN1
,PRKCI,RAP1B,SHC1,SOS1

'K3,PDIA3,PIK3R3,PLCG1,PPP3CA,PPP3CB,PRKACA,PRKACB,PRKAR1A,PRKAR1B,PRKAR2A,PRKA
NB1,GNB2,GNG2,GNG4,GRINA,GRK2,GSK3B,MAP2K2,MAP2K4,MAPK1,MAPK3,OGFR,PPP3CA
'PP2CA,PPP2CB,PPP2R1A,PPP2R2A,PPP2R5B,PPP2R5C,PPP2R5D,PPP3CA,PPP3CB,PPP4C,PPP5C,

PPP2CB,PPP2R1A,PPP2R2A,PPP2R5B,PPP2R5C,PPP2R5D,PPP3CA,PPP3CB,PRKACA,PRKACB,PRK
RAP1B,SMO,TCF3,TCF7,TFDP1,VEGFA
NFIA,NFIB,NFIX,NR2F1,PTGES3,RELA,SMARCA4,TFDP1,TGFB2
MAP2K2,MAP2K4,MAPK1,MAPK3,PIK3R3,PPIB,PRDX1,PRKCI,RAP1B,RBX1,SOD1,SOD2,SQSTM
,SMARCC1,SMARCD1,UBA52,UBB
C1,SHC2,SHC3,SMO,SOS1,TCF3,TSC2
MO,SOS1,STAT3,TCF3,TCF7,TGFB2,TGFBR1,TYK2,ZEB1,ZEB2
UBA52,UBB,UBQLN1

,PRKCI,RAC1,RAP1B,TLN1
,PRKAR2B,PRKCI,RAP1B,SGSM3,SLC39A7,SOS1,TUBA1A,TUBA1B,TUBA1C,TUBB,TUBB2B,TUE

LCG1,PRKACA,PRKACB,PRKAR1A,PRKAR1B,PRKAR2A,PRKAR2B,RAP1B,RELA,SHC1,SOS1,TGFB

3CB,PPP4C,PPP5C,PTPA,PTPN1,PTPN12,PTPN2,PTPN20,PTPN23,PTPRF,SET,WBP11
3CB,PPP4C,PPP5C,PTPA,PTPN1,PTPN12,PTPN2,PTPN20,PTPN23,PTPRF,SET,WBP11

,RAC3,RAP1B,RHOA,RHOBTB2,ROCK2

T3,TCF3,TGFB2,TGFBR1,TYK2,VIM,ZEB1,ZEB2

R5D,PPP3CA,PPP3CB,PPP4C,PPP5C,PTPA,PTPN1,PTPN12,PTPN2,PTPN20,PTPN23,PTPRF,SET,W

3CA,PPP3CB,PPP4C,PPP5C,PTPA,PTPN1,PTPN12,PTPN2,PTPN20,PTPN23,PTPRF,SET,WBP11

5C,PPP2R5D,PRKCI,PTPA,RAC1,RAC3,RAP1B,RELA,RHOA,RHOBTB2,STAT1,TYK2
H9,MYL6,MYL6B,MYL9,MYO18A,PPP3CA,PPP3CB,PRKACA,PRKACB,PRKAR1A,PRKAR1B,PRKAR
G1,PPP1CB,PPP1R12A,PRKCI,RAC1,RAC3,RAP1B,RELA,RHOA,RHOBTB2,ROCK2,SHC1,SOS1

MYL9,MYO18A,PDGFRB,RELA,STAT1,TGFB2,TGFBR1,TIMP1,TIMP2,VEGFA

APK3,MAPKAPK2,MDM2,MTOR,PARP1,PDHA1,PDHA2,PHF1,PIK3R3,PPM1L,PPP1CA,PPP2CA,PP
K2A,PIP4K2B,PIP5K1A,PIP5K1C,PLCG1,PRKCI,SMO,SOS1,TRAP1
,PRKAR2A,PRKAR2B,TPM1

3,PLD3,PRKCI,RAC1,RAC3,RAP1B,RELA,RHOA,RHOBTB2,ROCK2,VEGFA

AT3,TFRC,TYK2

PP1R12A,PPP1R14B,PRKACA,PRKACB,PRKAR1A,PRKAR1B,PRKAR2A,PRKAR2B,PRKCI,SULT1C4

A/HSPA1B,HSPA4,IFNA16,IL6ST,JAK1,JUN,KRT36,KRT8,MAP2K2,MAP2K4,MAPK1,MAPK11,MA

F7,VIM,YY1

AC3,RELA,RHOA,RHOBTB2,ROCK2

VEGFA

B,PRKCI,RAP1B

R1A,PRKAR1B,PRKAR2A,PRKAR2B

KAR1A,PRKAR1B,PRKAR2A,PRKAR2B,PRKCI,RAC1,RAP1B,RELA,SOS1
2R1A,PPP2R2A,PPP2R5B,PPP2R5C,PPP2R5D,PRKCI,PTPA,RACK1,SULT1C4,SULT4A1

TCF3,TTR

K11,MAPK3,NCOR2,NDST1,PIK3R3,PPM1L,PPP2CA,PPP2CB,PPP2R1A,PPP2R2A,PPP2R5B,PPP2F

A7,SMO,VEGFA

AC3,RAP1B,RELA,SOS1,TYK2,ULBP1
ELA,SMO,SOX9,TCF3,TCF7,TGFBR1,TIMP3,VEGFA

NG4,GSK3A,GSK3B,HDAC2,HDAC5,HDAC6,HSPB1,IGF1R,IL6ST,ITGA11,ITGA2,ITGA3,ITGA7,ITG

NB1,GNB2,GNG2,GNG4,GPR107,GPRC5B,GRB2,HIF1A,JUN,LGR4,LPAR2,MAP2K2,MAPK1,MAI
RELA,SHC1,SHC2,SHC3,SLC39A7,SOS1

.,SOS1,TCF3

4,MAPK1,MAPK3,MAPKAPK2,MIF,PDIA3,PIK3R3,PLCG1,PPP3CA,PPP3CB,PRKCI,RAC1,RAP1B,REL

X6,PRKACA,PRKACB,PRKAR1A,PRKAR1B,PRKAR2A,PRKAR2B,PRKCI,PTK7,TNK2,TYK2

STN,NOX4,PIK3R3,PLCG1,PPP3CA,PPP3CB,RAC1,RAC3,RELA,SLC1A3,SNCA,SOD2,STAT1,TGFB2

PTPA,RAP1B,SLC39A7

TUB1,VEGFA

ELA,SHC1,SHC2,SHC3,SOS1,STAT1,STAT2,STAT3,TGFB2,TRAF3,TYK2

AP1B,RELA,SHC1,SOS1,STAT3,SYNGAP1
RKAR2A,PRKAR2B,STAT3

-DPA1,IPO5,JUN,KPNA2,KPNA3,KPNB1,MAP2K2,MAPK1,MAPK3,PIK3R3,PLCG1,PPP3CA,PPP3CI

L,GNAO1,GNAS,GNAT1,GNB1,GNB2,GNG2,GNG4,GPR107,GPRC5B,GRB2,GRIA1,GRIK5,IGF1

NP200,SNRNP70,SNRPB,SNRPD2,SNRPD3,SNRPN,SNU13,SOS1
,MYL12A,MYL12B,MYL6,MYL6B,MYL9,PAK3,PPP1CB,PPP1R12A,PRKCI,TNK2

B5,MAP2K2,MAPK1,MAPK3,MARCKS,MPRIP,MYL12A,MYL12B,MYL6,MYL6B,MYL9,PEBP1,PLC

HOA,RHOBTB2,STAT1,STAT2,STAT3,TNFRSF10B,TYK2

AC1,RAP1B,RELA,SHC3,SOS1,TCF3,TCF7

7,TGFBR1,TYK2,VEGFA
B,PPP2R5C,PPP2R5D,PPP3CA,PPP3CB,PTPA,RAB4A,RAC1,RAC3,RAP1B,RDX,RHOA,RHOBTB2,R[illegible]

P1,RAP1B,RPL10,RPL11,RPL12,RPL13,RPL13A,RPL14,RPL15,RPL17,RPL18,RPL19,RPL22,RPL23,
MC1,PSMC2,PSMC3,PSMC4,PSMC5,PSMC6,PSMD1,PSMD10,PSMD12,PSMD13,PSMD14,PSMD2,
NAPA,NAPB,NCOR2,NSF,PDPK1,PIK3R3,POLR2A,POLR2B,POLR2C,POLR2E,POLR2G,PRKCI,PSM
C,PPP2R5D,PTPA,RAP1B,RPS10,RPS12,RPS13,RPS14,RPS15,RPS15A,RPS16,RPS17,RPS19,RPS
MC2,PSMC3,PSMC4,PSMC5,PSMC6,PSMD1,PSMD10,PSMD12,PSMD13,PSMD14,PSMD2,PSMD
PSMD8,PSME3,PSMF1,RELA,STUB1
JUN,LIMK1,MAP2K2,MAPK1,MAPK3,MDK,MED12,MED18,MMP14,MMP15,MMP2,MMP24,MP
OA,RHOBTB2,RPS10,RPS12,RPS13,RPS14,RPS15,RPS15A,RPS16,RPS17,RPS19,RPS20,RPS21,
DX6,RAB5A,RAB5B,RAB5C,RAB7A,RAC1,RAC3,SNAP25,TUBA1A,TUBA1B,TUBA1C,TUBB,TUBE
,NDUFB9,NDUFS1,NDUFS5,NDUFS6,NDUFS7,NDUFS8,NDUFV1,NDUFV2,OGDH,PARK7,PDHA
GPAA1,GRB2,GRIA1,GRINA,GSK3B,LIMK1,MAP1B,MAPK1,MAPK3,MAPT,MARCKS,MTOR,NAP

,NDUFA5,NDUFB10,NDUFB11,NDUFB2,NDUFB4,NDUFB8,NDUFB9,NDUFS1,NDUFS5,NDUFS
M1L,PPP2CA,PPP2CB,PPP2R1A,PPP2R2A,PPP2R5B,PPP2R5C,PPP2R5D,PRKACA,PRKACB,PRKAR
R1A,PRKAR1B,PRKAR2A,PRKAR2B,RAC1,RHOA,ROCK2,SEMA6A,SEMA6C,SEMA6D,SMC3,VCAN
SK3B,ITGA11,ITGA2,ITGA3,ITGA7,ITGAV,ITGB1,ITGB5,KLC1,L1CAM,LIMK1,LINGO1,MAP2K2,M
,PAK3,PFN1,PFN2,PIK3R3,PLCG1,PPP1CB,PPP1R12A,RAC1,RAC3,RAP1B,RAPGEF1,RHOA,RHO
AP3K10,MAPK1,MAPK3,MSN,MYL12A,MYL12B,MYL6,MYL6B,MYL9,NOX4,PAK3,PI4KA,PIK3R3,
PRKAR2A,PRKAR2B,PRKCI,PTPA,RAC1,RELA,RHOA,SAFB,SNAP25,SPTAN1,SYMPK,TGFB2,TGF

L12B,MYL6,MYL6B,MYL9,MYO18A,NCKAP1,PAK3,PFN1,PFN2,PIK3R3,PIP5K1A,PIP5K1C,PPP1CE
DUFV2,SDHA,UQCR11,UQCRC1,UQCRC2,UQCRFS1,UQCRH
N,MYH10,MYH9,MYL12A,MYL12B,MYL6,MYL6B,MYL9,PAK3,PI4KA,PIP4K2A,PIP4K2B,PIP5K1A,
RAB5B,RAB5C,RAB7A,RAC1,RPS27A,SH3BP4,SH3GL1,SH3GLB1,SH3KBP1,SNAP91,TFRC,UBA5
S3,RPS4X,RPS5,RPS7,RPS8,RPS9,STAT1,STAT2,STAT3,TFDP1,TGFBR1,TNPO1,TNPO2,TOMM

TIN7,SEPTIN9
1,MAPK3,NCK2,PAK3,RAC1,RAC3,RAP1B,RAPGEF1,RHOA,ROCK2,SDCBP,SHC1,SOS1,STAT3,VE

OX4,SOX9,TCF3,TCF7,TGFB2,TGFBR1,TLE1,TLE4,UBA52,UBB

,PPP2R5C,PPP2R5D,PTPA,RAP1B,RELA,RHEB,SHC1,SOS1,TSC2,TYK2,YWHAB,YWHAE,YWHAG,
1C,TUBB,TUBB2B,TUBB4A,TUBB4B,TUBB6,ZYX

,PPP2R5B,PPP2R5C,PPP2R5D,PTPA,RAC1,RAC3,RELA,RHOA,RHOBTB2,TMSB10/TMSB4X,VEGF
KAR1B,PRKAR2A,PRKAR2B,PRKCI,PTPA,RAC1,RAC3,RAP1B,RAPGEF1,SHC1,SOS1,SRF,STAT1,
2B,PRKCI,RAP1B,SEC11A,SEC61A1,SEC61B,SLC2A1,SNAP25,SPCS1,SPCS2,SRP14,SRP54,SRP7

1,RAP1B,SPTAN1,SPTBN1,SYMPK,TJAP1,TUBA1A,TUBA1B,TUBA1C,TUBB,TUBB2B,TUBB4A,T
PP3CB,PRKACA,PRKACB,PRKAR1A,PRKAR1B,PRKAR2A,PRKAR2B,PTPA,RAB7A,RB1CC1,SLC1A

MT2,PSMC5,RAC1,RBP1,RDH11,RELA,SMARCA4,SMARCB1,SMARCC1,SMARCD1,TGFB2,TRIM24
NTN1,PDE11A,PDE7A,PDIA3,PHKB,PLCG1,PPP1CA,PPP1CB,PPP1CC,PPP1R10,PPP1R12A,PPP1R14
3,HDAC2,HDAC5,HDAC6,HIF1A,HIPK2,ITGA11,ITGA2,ITGA3,ITGA7,ITGAV,ITGB1,ITGB5,JAK1,JUN

RKAR2A,PRKAR2B,PRKCI,RELA,SHC1,TAF15,TAF7

4B,TUBB6,ZYX

R1A,PRKAR1B,PRKAR2A,PRKAR2B,PTPA
,JAK1,JUN,LRP6,MAP2K2,MAP2K4,MAPK1,MAPK11,MAPK3,MTOR,MYL12A,MYL12B,MYL6,MYL

P1,TIMP2,TIMP3
1,RAC3,RALGDS,RAP1B,RELA,RHOA,RHOBTB2,SMO,SOS1,STAT1,STAT3,TCF3,TCF7,TGFB2,TGF

T3,STUB1,TGFB2,VEGFA,VHL,VIM

3CA,PPP3CB,PRKACA,PRKACB,PRKAR1A,PRKAR1B,PRKAR2A,PRKAR2B,RAC1,RAC3,RAP1B,RHOA

C2

2,PTPN2,PTPN20,PTPN23,PTPRF,SET,WBP11

\AR2B,PRKCI,RAP1B,RCAN1,SHC1,SHC2,SHC3,SOS1,TGFB2,TGFBR1
\,PPP3CB,PRKACA,PRKACB,PRKAR1A,PRKAR1B,PRKAR2A,PRKAR2B,PRKCI,RAC1,RAC3,RAP1B,F
PTPA,PTPN1,PTPN12,PTPN2,PTPN20,PTPN23,PTPRF,SEC16A,SET,WBP11

\AR1A,PRKAR1B,PRKAR2A,PRKAR2B,PRKCI,PTPA

1,STIP1,TXN,TXNRD1,UBB,UBE2K,USP14,VCP

3B4A,TUBB4B,TUBB6

2,TGFBR1

BP11

\2A,PRKAR2B,RAP1B,RCAN1,TPM1,TPM3,TPM4

P2CB,PPP2R1A,PPP2R2A,PPP2R5B,PPP2R5C,PPP2R5D,PPP3CA,PPP3CB,PTPA,RAP1B,SOD2,SQS

,SULT4A1

PK3,MMP2,NCOR2,NDUFA10,NDUFA11,NDUFA13,NDUFA4,NDUFA5,NDUFB10,NDUFB11,ND

R5C,PPP2R5D,PRKCI,PTGES3,PTPA,RAP1B,RBX1,RELA,SULT1C4,SULT4A1,SUMO1 [illegible]AV,ITGB1,ITGB5,JUN,MAP2K2,MAP2K4,MAP3K10,MAPK1,MAPK11,MAPK3,MAPKAPK2,MKNK[illegible]

[illegible]K3,MMP2,MTOR,PIK3R3,PPM1L,PPP1CA,PPP1CB,PPP1CC,PPP1R10,PPP1R12A,PPP1R14B,PPP2

.A,RHOA,ROCK2,SFRP1,SMO,STAT3,TCF3,TCF7,TRAF3,VEGFA

2,TGFBR1,TRAF3,TYK2

3,RAP1B,RCAN1,RELA,SOS1,TNPO1,XPO1

.R,LGR4,LPAR2,MAP2K2,MAPK1,MAPK3,PDGFRB,PDIA3,PIK3R3,PLCG1,POLR2A,POLR2B,POLR2

G1,PLD3,PPP1CB,PPP1R12A,PPP3CA,PPP3CB,PRKCI,RAC1,RAC3,RALGDS,RAP1B,RELA,RHOA,RH

OCK2,SOS1,STAT3

RPL23A,RPL24,RPL26,RPL27,RPL28,RPL29,RPL3,RPL30,RPL31,RPL34,RPL35,RPL35A,RPL36,RPL3
PSMD3,PSMD4,PSMD5,PSMD7,PSMD8,RBX1,RPS27A,SKP1,SMO,STUB1,TRAP1,UBA1,UBA52,
A1,PSMA4,PSMA5,PSMA6,PSMA7,PSMB1,PSMB2,PSMB4,PSMB5,PSMB6,PSMC1,PSMC2,PSM
20,RPS21,RPS24,RPS25,RPS26,RPS27,RPS27A,RPS28,RPS29,RPS3,RPS4X,RPS5,RPS7,RPS8,RPS
3,PSMD4,PSMD5,PSMD7,PSMD8,PSME3,PSMF1,PTPA,YWHAB,YWHAE,YWHAG,YWHAH,YWHA

RIP,MTOR,MYL12A,MYL12B,MYL6,MYL6B,MYL9,NCOR2,NDUFA10,NDUFA11,NDUFA13,NDUF
RPS24,RPS25,RPS26,RPS27,RPS27A,RPS28,RPS29,RPS3,RPS4X,RPS5,RPS6KA2,RPS7,RPS8,RPS
32B,TUBB4A,TUBB4B,TUBB6,VAMP2,VPS16,VPS28,VPS37B,VPS41,VTI1B,YKT6
1,PINK1,PRDX3,PRDX5,SDHA,SNCA,SOD2,TRAK1,UQCR11,UQCRC1,UQCRC2,UQCRFS1,UQCRH,
1L1,NAP1L4,NAPA,NAPB,NECTIN1,NLGN2,NRXN2,NSF,PAFAH1B1,PIK3R3,PLCG1,PRKACA,PRK

6,NDUFS7,NDUFS8,NDUFV1,NDUFV2,PARP1,PDHA1,PFKFB3,PFKM,PGAM1,PGK1,PRKDC,RELA
1A,PRKAR1B,PRKAR2A,PRKAR2B,PTPA,RAB11A,RAB1A,RAB27A,RAB2A,RAB3A,RAB6A,RAB7A
N
IAPK1,MAPK3,MICAL1,MMP14,MMP15,MMP2,MMP24,MYL12A,MYL12B,MYL6,MYL6B,MYL9,N
3TB2,SHC1,SOS1,TLN1,TNK2,TSPAN3,TSPAN5,TSPAN6,TSPAN7,ZYX
PIP4K2A,PIP4K2B,PIP5K1A,PIP5K1C,PKN1,PPP1R12A,PRKCI,RAC1,RAC3,RDX,RELA,RHOA,RHOB
3R1,VAMP2,VTI1B,YBX3,YKT6

3,PPP1R12A,RAC1,RAC3,RAP1B,RDX,RHOA,ROCK2,SHC1,SOS1,TLN1,TMSB10/TMSB4X,TRIO,W

PIP5K1C,PPP1R12A,RAC1,RAC3,RDX,RHOA,RHOBTB2,ROCK2,WASF2
2,UBB,VEGFA
70,TRAF3,TYK2

GFA

YWHAH,YWHAQ,YWHAZ

A,VIM
STAT3,TLN1,YWHAB,YWHAG,YWHAH,YWHAQ,YWHAZ
2,SRP9,SSR1,SSR2,SSR3,SSR4,STAT1,STAT2,STAT3,TCF3,TYK2,VAMP2,VTI1B,YKT6

UBB4B,TUBB6,YBX3
5,SLC3A2,SLC7A5,SQSTM1,TGFB2,TRAF3,TSC2,ULK1,VEGFA,VPS16,VPS41

4,VEGFA
4B,PPP3CA,PPP3CB,PRKACA,PRKACB,PRKAR1A,PRKAR1B,PRKAR2A,PRKAR2B,PRKCI,PTP4A1,PT
N,LRP6,MAP2K2,MAP2K4,MAPK1,MAPK11,MAPK3,MDM2,NCSTN,NOTCH1,PAK3,PIK3R3,PRKAC

L6B,MYL9,NOX4,PDGFRB,PIK3R3,PLCG1,PRKACA,PRKACB,PRKAR1A,PRKAR1B,PRKAR2A,PRKAR

FBR1,TYK2,VEGFA

A,RHOBTB2,ROCK2,SOS1,SRF,TGFB2,TGFBR1

[illegible]PS6KA2,SOS1,SRF,TCF3,TCF7

TM1,TGFB2,TGFBR1,VHL,ZFP36L1

UFB2,NDUFB4,NDUFB8,NDUFB9,NDUFS1,NDUFS5,NDUFS6,NDUFS7,NDUFS8,NDUFV1,NDUI

2,MTOR,PDE11A,PDE7A,PDIA3,PIK3R3,PKN1,PLCG1,PPP3CA,PPP3CB,PRKACA,PRKACB,PRKAR1[illegible]

CA,PPP2CB,PPP2R1A,PPP2R2A,PPP2R5B,PPP2R5C,PPP2R5D,PPP4C,PRKACA,PRKACB,PRKAR1A,

2C,POLR2E,POLR2G,PRKACA,PRKACB,PRKAR1A,PRKAR1B,PRKAR2A,PRKAR2B,PRKCI,RAP1B,SH

HOBTB2,SHC1,SOS1

36A,RPL36AL,RPL37,RPL37A,RPL38,RPL39,RPL4,RPL41,RPL5,RPL6,RPL7,RPL7L1,RPL8,RPL9,RPL
UBB,UBE2D2,UBE2D3,UBE2E1,UBE2G1,UBE2G2,UBE2H,UBE2I,UBE2J1,UBE2K,UBE2L3,UBE2
C3,PSMC4,PSMC5,PSMC6,PSMD1,PSMD10,PSMD12,PSMD13,PSMD14,PSMD2,PSMD3,PSMD4,
9,SHC1,SOS1
Q,YWHAZ,ZFP36L1,ZFP36L2

A4,NDUFA5,NDUFB10,NDUFB11,NDUFB2,NDUFB4,NDUFB8,NDUFB9,NDUFS1,NDUFS5,NDUI
9,TSC2,ULK1,VEGFA

,VDAC1,VDAC2,VDAC3
ACB,PRKAR1A,PRKAR1B,PRKAR2A,PRKAR2B,RAB3A,RAB5A,RAB5B,RAB5C,RAC1,RAP1B,RAPG

,SDHA,SF3A1,SLC25A5,SLC2A1,SOD1,SOD2,STAT3,TIMM13,TIMM23B,TIMM50,TOMM20,TOM
,SLC2A1,SMARCA4,SMARCB1,SMARCC1,SMARCD1,TBC1D1,TSC2,ULK1

ICK2,NRP2,NTN1,PAK3,PDIA3,PFN1,PFN2,PIK3R3,PITRM1,PLCG1,PLXNA1,PLXNA2,PLXNA3,PLXI

TB2,ROCK2,SEPTIN2,SEPTIN3,SEPTIN5,SEPTIN6,SEPTIN7,SEPTIN9,STMN1,VIM,WASF3

'ASF2

TPN1,PTPN12,PTPN2,PTPN23,PTPRA,PTPRF,PTPRS,PTPRU,PTPRZ1,PYGB,RAP1B,RELA,RHOA,R
CA,PRKACB,PRKAR1A,PRKAR1B,PRKAR2A,PRKAR2B,PRKCI,PRKDC,RAC1,RAC3,RALGDS,RAP1B,

R2B,PRKCI,RAC1,RAC3,RAP1B,RELA,RHOA,RHOBTB2,ROCK2,SDHA,SMO,SOD2,SOS1,STAT3,TC

FV2,PBRM1,PIK3R3,POLR2A,POLR2B,POLR2C,POLR2E,POLR2G,PPP3CA,PPP3CB,PRKACA,PRKA[illegible]

A,PRKAR1B,PRKAR2A,PRKAR2B,PRKCI,RAP1B,RCAN1,RELA,RHOA,ROCK2,SMO,SRF,STAT3,TGF

,PRKAR1B,PRKAR2A,PRKAR2B,PRKCI,PTPA,RAC1,RAC3,RAP1B,RB1CC1,RHOA,SHC1,SHC2,SHC3

C1,SHC2,SHC3,SMO,SOS1,TGFB2,TGFBR1,VEGFA

P0,RPLP1,RPLP2,RPS10,RPS12,RPS13,RPS14,RPS15,RPS15A,RPS16,RPS17,RPS19,RPS20,RPS2[illegible]
M,UBE2Q1,UBE2R2,UBE2V1,UBE2V2,UBE3A,UBE4B,UCHL1,USO1,USP10,USP11,USP14,USP1[illegible]
PSMD5,PSMD7,PSMD8,PSME3,PSMF1,RCOR2,RPS27A,SDHA,SHC1,SHC2,SHC3,SNAP25,SNCA,S[illegible]

[illegible]S6,NDUFS7,NDUFS8,NDUFV1,NDUFV2,NOTCH1,PCNA,PDIA3,PELP1,PIK3R3,PLCG1,POLR2B,P[illegible]

[illegible]EF1,RHOA,SGTA,SHC1,SHC2,SHC3,SNAP25,SNCA,SNCG,SOS1,STXBP1,SYNGAP1,SYT1,THBS2,

[illegible]M40,TOMM6,TOMM7,TOMM70,TP53BP1,TRIM28,TUBA1A,TUBA1B,TUBA1C,UQCRC2,UQCR[illegible]

NB1,PLXNB2,PPP3CA,PPP3CB,PRKACA,PRKACB,PRKAR1A,PRKAR1B,PRKAR2A,PRKAR2B,PRKCI,

OCK2,SMO,TCF3,TCF7,TGFB2,TGFBR1,TIMM50,YWHAB,YWHAE,YWHAG,YWHAH,YWHAQ,YW
RAPGEF1,RBPJ,RELA,RHOA,RHOBTB2,SHC1,SHC2,SHC3,SMO,SOS1,SYNGAP1,TCF3,TFDP1,TGF

F3,TCF7,TFRC,TGFB2,TGFBR1,TIMP1,VEGFA

CB,PTGES3,RAC1,RAP1B,RELA,SDHA,SHC1,SMARCA4,SMARCB1,SMARCC1,SMARCD1,SOS1,ST[illegible]

B2,TGFBR1,TSC2

,SMO,SOS1,STAT3,STMN1,TGFB2,TUBA1A,TUBA1B,TUBA1C,TUBB,TUBB2B,TUBB4A,TUBB4

1,RPS24,RPS25,RPS26,RPS27,RPS27A,RPS28,RPS29,RPS3,RPS4X,RPS5,RPS7,RPS8,RPS9,SHC1,
9,USP22,USP24,USP3,USP33,USP45,USP47,USP48,USP5,USP7,VHL
OS1,UBA52,UBB,VAMP2,VTI1B,YKT6,ZDHHC17

PP1CB,PPP1R12A,PRKACA,PRKACB,PRKAR1A,PRKAR1B,PRKAR2A,PRKAR2B,PRKCI,PRKDC,RAP

TLN1,VAMP2,VTI1B,YKT6

FS1,VDAC1,VDAC2,VDAC3,XRCC5,XRCC6

PSMD14,RAC1,RAC3,RAP1B,RHOA,ROCK2,RTN4,SDCBP,SEMA3C,SEMA4C,SEMA6A,SEMA6C,

HAZ
B2,TGFBR1,TYK2

AT1,STAT3,SUMO1,TAF15,TAF7,TGFB2,TGFBR1,TYK2,UBE2I,UQCRC2,UQCRFS1,YWHAH

3,TUBB6,VEGFA

SOS1,UBA52,VEGFA,WARS1

LB,RBFOX2,RELA,RHOA,ROCK2,SDHA,SHC1,SHC2,SHC3,SOD2,SOS1,TBL1XR1,THRAP3,TRRAP,T

EMA6D,SHC1,SLIT1,SLIT2,SMO,SOS1,SRGAP2,SRGAP3,TUBA1A,TUBA1B,TUBA1C,TUBB,TUBB

YK2,UQCRC2,UQCRFS1,VEGFA

2B,TUBB4A,TUBB4B,TUBB6,UNC5A,VEGFA

© 2000-2021 QIAGEN. All rights reserved.

| Upstream Re | Expr Fold Ch | Molecule Ty | Predicted Ac | Activation z- | Flags | p-value of ov |
|---|---|---|---|---|---|---|
| MYCN | -8.56 | transcription | Inhibited | -3.008 | | 2.29E-69 |
| MYC | | transcription | Inhibited | -8.777 | bias | 2.04E-68 |
| torin1 | | chemical rea | Activated | 6.388 | bias | 3.6E-58 |
| ST1926 | | chemical dru | Activated | 9.954 | bias | 2.09E-56 |
| CD 437 | | chemical dru | Activated | 9.779 | bias | 1.16E-55 |
| TP53 | -8.88 | transcription | Inhibited | -3.389 | | 3.16E-49 |
| LARP1 | -11.308 | translation re | Activated | 8.245 | bias | 1.28E-46 |
| MAPT | -10.101 | other | | 0.805 | | 2.03E-46 |
| APP | -12.771 | other | Inhibited | -3.606 | bias | 4.26E-44 |
| beta-estradio | | chemical - er | Inhibited | -9.933 | bias | 5.1E-41 |
| sirolimus | | chemical dru | Activated | 9.528 | bias | 2.18E-38 |
| 5-fluorouraci | | chemical dru | Activated | 6.427 | | 5.99E-36 |
| RICTOR | -7.069 | other | Activated | 9.142 | bias | 1.96E-35 |
| Lh | | complex | Inhibited | -5.305 | bias | 3.58E-34 |
| tazemetosta | | chemical dru | | 1.949 | | 7.63E-31 |
| 1,2-dithiol-3- | | chemical rea | Inhibited | -8.508 | bias | 3.63E-30 |
| SP2509 | | chemical rea | Activated | 2.029 | | 8.41E-30 |
| TGFB1 | -6.676 | growth facto | Inhibited | -9.635 | bias | 1.18E-29 |
| HTT | -4.106 | transcription | Inhibited | -3.252 | | 8.72E-29 |
| PTP4A1 | -10.755 | phosphatase | | -1.166 | | 1.1E-28 |
| YAP1 | -7.629 | transcription | Inhibited | -3.785 | bias | 5.33E-28 |
| metribolone | | chemical rea | Inhibited | -8.916 | bias | 1.34E-27 |
| MKNK1 | -3.873 | kinase | Inhibited | -7.616 | bias | 3.81E-26 |
| NFE2L2 | -6 | transcription | Inhibited | -9.767 | bias | 7.79E-26 |
| CTNNB1 | -10.623 | transcription | Inhibited | -5.759 | bias | 1.11E-25 |
| MLXIPL | | transcription | Inhibited | -7.947 | bias | 1.23E-25 |
| ESR1 | | ligand-deper | Inhibited | -2.656 | | 2.21E-24 |
| decitabine | | chemical dru | Inhibited | -2.992 | bias | 6.67E-23 |
| arsenic trioxi | | chemical dru | | 0.642 | | 6.74E-23 |
| HRAS | -8.678 | enzyme | | -0.302 | bias | 9.07E-23 |
| EGFR | -6.211 | kinase | Inhibited | -6.023 | bias | 9.17E-22 |
| imatinib | | chemical dru | Activated | 3.72 | bias | 9.86E-22 |
| PSEN1 | -8.422 | peptidase | Activated | 2.258 | | 2.07E-21 |
| D-glucose | | chemical - er | Inhibited | -4.527 | bias | 2.49E-21 |
| GnRH analog | | biologic drug | Activated | 2.194 | | 4.4E-21 |
| dihydrotesto: | | chemical - er | Inhibited | -4.268 | bias | 5.79E-21 |
| HNF4A | | transcription | | -0.382 | | 9.38E-21 |
| BDNF | -6.219 | growth facto | Inhibited | -3.388 | bias | 1.15E-20 |
| trichostatin A | | chemical dru | Inhibited | -2.226 | | 1.87E-20 |
| acyline | | biologic drug | | | | 2.68E-20 |
| geldanamyci | | chemical dru | | 0.087 | | 6.21E-20 |
| FN1 | -11.749 | enzyme | Inhibited | -3.216 | bias | 1.11E-19 |
| mibolerone | | chemical dru | Inhibited | -4.258 | bias | 2.65E-18 |

| | | | | | | |
|---|---|---|---|---|---|---|
| ERBB2 | -9.289 | kinase | Inhibited | -3.919 | bias | 3.18E-18 |
| FLT1 | | kinase | Inhibited | -2.897 | bias | 1.11E-17 |
| FSH | | complex | | -1.102 | | 1.6E-17 |
| MMP12 | | peptidase | | -0.739 | bias | 2.93E-17 |
| KRAS | -2.928 | enzyme | | -0.409 | | 7.59E-17 |
| IND S7 | | chemical - ki | | 1.46 | bias | 1.28E-16 |
| XBP1 | -9.262 | transcription | Inhibited | -5.882 | bias | 2.33E-16 |
| MEL S3 | | chemical - ki | | 1.877 | bias | 5.44E-16 |
| CD3 | | complex | Inhibited | -7.578 | bias | 1.37E-15 |
| PAX3-FOXO1 | | fusion gene/ | | 1.786 | | 1.79E-15 |
| miR-1-3p (ar | | mature micr | Activated | 7.637 | bias | 2.3E-15 |
| HSF1 | -9.596 | transcription | Inhibited | -2.718 | | 2.79E-15 |
| uranyl nitrate | | chemical tox | Inhibited | -2.173 | | 3.43E-15 |
| ATF4 | -11.414 | transcription | Inhibited | -6.363 | bias | 3.56E-15 |
| tunicamycin | | chemical - er | Inhibited | -6.066 | bias | 4.64E-15 |
| gentamicin | | chemical dru | Inhibited | -4.246 | | 5.41E-15 |
| CST5 | | other | Activated | 3.205 | bias | 8.59E-15 |
| IL3 | | cytokine | Inhibited | -4.978 | bias | 1.03E-14 |
| ESR2 | | ligand-deper | Inhibited | -5.875 | bias | 1.82E-14 |
| SYVN1 | -8.172 | transporter | Inhibited | -6.981 | bias | 2.76E-14 |
| HSF2 | -5.507 | transcription | Inhibited | -3.598 | bias | 4.01E-14 |
| dexamethas | | chemical dru | Inhibited | -3.033 | | 4.23E-14 |
| tanespimycir | | chemical dru | Activated | 4.437 | bias | 5.25E-14 |
| valproic acid | | chemical dru | | 0.95 | | 6.65E-14 |
| mir-122 | | microRNA | Activated | 6.067 | bias | 7.85E-14 |
| LMNB1 | -9.232 | other | | 1.901 | bias | 7.95E-14 |
| Akt | | group | Inhibited | -3.808 | bias | 9.53E-14 |
| Z-LLL-CHO | | chemical - pr | Inhibited | -5.825 | bias | 1.38E-13 |
| IND S1 | | chemical - ki | | 0.943 | | 3.33E-13 |
| PC-SPES | | chemical dru | | | | 4.39E-13 |
| CAB39L | 10.244 | kinase | Inhibited | -4.887 | bias | 4.98E-13 |
| hydrogen per | | chemical - er | Inhibited | -7.45 | bias | 6.54E-13 |
| MTOR | -9.683 | kinase | | -1.509 | | 8.22E-13 |
| INSR | | kinase | Inhibited | -6.291 | bias | 9.3E-13 |
| OGA | -11.171 | enzyme | | 1.923 | | 9.42E-13 |
| WT1 | | transcription | | -0.913 | | 1.04E-12 |
| E2F1 | -2.807 | transcription | Inhibited | -2.47 | bias | 1.15E-12 |
| ADORA2A | 7.331 | G-protein co | Inhibited | -3.343 | | 2.12E-12 |
| mir-1 | | microRNA | Activated | 6.495 | bias | 2.75E-12 |
| trans-hydrox | | chemical dru | | 1.816 | | 2.93E-12 |
| forskolin | | chemical tox | Inhibited | -5.589 | bias | 3.59E-12 |
| SP1 | 8.964 | transcription | Inhibited | -5.807 | bias | 4.31E-12 |
| SEL1L | -3.478 | other | Inhibited | -2.459 | | 4.9E-12 |
| nitrofurantoi | | chemical dru | Inhibited | -4.006 | | 7.67E-12 |
| tretinoin | | chemical - er | Inhibited | -3.468 | bias | 1.09E-11 |

| | | | | | | |
|---|---|---|---|---|---|---|
| tetraethylam | | chemical dru | | 1.816 | | 2.46E-11 |
| levodopa | | chemical - er | Inhibited | -2.032 | | 2.67E-11 |
| UBQLN2 | -8.699 | other | | | | 2.73E-11 |
| LONP1 | -10.121 | peptidase | Activated | 3.352 | bias | 5.27E-11 |
| mono-(2-eth | | chemical tox | Inhibited | -6.096 | bias | 5.31E-11 |
| CPT1B | -8.016 | enzyme | Activated | 3.461 | bias | 7.56E-11 |
| HIF1A | -10.088 | transcription | Inhibited | -6.169 | bias | 7.71E-11 |
| PDGF BB | | complex | Inhibited | -3.732 | bias | 7.8E-11 |
| MYCL | -6.854 | transcription | Inhibited | -5.857 | bias | 8.03E-11 |
| glutamine | | chemical - er | | 0.836 | bias | 8.47E-11 |
| VHL | -10.528 | transcription | | 1.845 | bias | 8.58E-11 |
| NRG1 | 10.05 | growth facto | Inhibited | -4.808 | bias | 8.94E-11 |
| CDK4/6 | | group | | 0.828 | | 1.13E-10 |
| FOS | -8.603 | transcription | | -1.681 | bias | 1.19E-10 |
| STK11 | -5.591 | kinase | Inhibited | -3.034 | | 1.27E-10 |
| TCR | | complex | Inhibited | -4.808 | bias | 1.4E-10 |
| PTEN | -8.745 | phosphatase | | 0.454 | | 1.92E-10 |
| 8-bromo-cAN | | chemical rea | Inhibited | -7.602 | bias | 2.41E-10 |
| MMP3 | | peptidase | | -1.797 | | 2.52E-10 |
| IGF1R | -9.641 | transmembra | Inhibited | -5.067 | bias | 2.71E-10 |
| PTPRR | | phosphatase | Activated | 5.74 | bias | 3.07E-10 |
| ciprofloxacin | | chemical dru | | 1.025 | | 3.1E-10 |
| Mek | | group | Inhibited | -2.534 | bias | 3.33E-10 |
| diclofenac | | chemical dru | Inhibited | -3.822 | bias | 3.57E-10 |
| TAZ | -6.031 | enzyme | Inhibited | -3.443 | bias | 3.89E-10 |
| PAX3 | | transcription | Inhibited | -2 | bias | 4.2E-10 |
| medroxyprog | | chemical dru | Inhibited | -7.401 | bias | 4.28E-10 |
| let-7a-5p (ar | | mature micr | Activated | 6.403 | bias | 5.19E-10 |
| lactacystin | | chemical - pi | Inhibited | -3.841 | bias | 6.45E-10 |
| OGT | -9.775 | enzyme | | -0.601 | | 6.62E-10 |
| PCGEM1 | | other | Inhibited | -2.936 | bias | 7.67E-10 |
| MEL T1 | | chemical - ki | | 0.277 | | 8.86E-10 |
| EIF2AK3 | -7.695 | kinase | Inhibited | -4.324 | bias | 9.56E-10 |
| IPMK | | kinase | Inhibited | -3.491 | bias | 1.26E-09 |
| KAT2A | -9.387 | enzyme | Inhibited | -4.765 | bias | 1.34E-09 |
| desmopressi | | biologic drug | | -0.523 | bias | 1.42E-09 |
| miR-30c-5p ( | | mature micr | Activated | 6.435 | bias | 1.48E-09 |
| molybdenum | | chemical rea | Activated | 5.814 | bias | 1.78E-09 |
| Insulin | | group | Inhibited | -5.355 | bias | 2.67E-09 |
| mir-210 | | microRNA | Activated | 5.093 | bias | 2.73E-09 |
| ERBB3 | | kinase | | -0.506 | | 3.85E-09 |
| RET | -8.562 | kinase | Inhibited | -3.473 | bias | 3.97E-09 |
| 2-amino-1-m | | chemical tox | Inhibited | -4.67 | bias | 4.18E-09 |
| AGT | -4.916 | growth facto | Inhibited | -7.596 | bias | 4.4E-09 |
| DDX5 | -10.818 | enzyme | Inhibited | -4.745 | bias | 4.46E-09 |

| | | | | | | |
|---|---|---|---|---|---|---|
| IL15 | | cytokine | Inhibited | -4.153 | bias | 4.69E-09 |
| (-)-norephed | | chemical dru | Inhibited | -3.464 | bias | 5.68E-09 |
| CLPP | -8.956 | peptidase | Activated | 5.196 | bias | 6.7E-09 |
| maslinic acic | | chemical - e | Activated | 3.5 | bias | 7.25E-09 |
| salirasib | | chemical dru | Activated | 3.854 | bias | 9.17E-09 |
| mir-133 | | microRNA | Activated | 5.176 | bias | 9.69E-09 |
| pirinixic acid | | chemical tox | Inhibited | -5.992 | bias | 1.09E-08 |
| ICMT | -1.884 | enzyme | | 0 | bias | 1.16E-08 |
| topotecan | | chemical dru | | 0.692 | bias | 1.5E-08 |
| ERN1 | -5.296 | kinase | Inhibited | -4.794 | bias | 1.52E-08 |
| TSC2 | -9.272 | other | Activated | 4.119 | bias | 1.61E-08 |
| HBA1/HBA2 | | transporter | Inhibited | -3.71 | bias | 1.62E-08 |
| SOD1 | -10.315 | enzyme | Activated | 2.132 | bias | 1.73E-08 |
| EGF | | growth facto | Inhibited | -6.367 | bias | 1.89E-08 |
| RTN4 | -11.673 | other | | 1.546 | | 2.09E-08 |
| IGF1 | -6.401 | growth facto | Inhibited | -5.674 | bias | 2.09E-08 |
| PD98059 | | chemical - ki | Activated | 6.992 | bias | 2.16E-08 |
| AKT1 | -8.323 | kinase | Inhibited | -5.039 | bias | 2.34E-08 |
| cisplatin | | chemical dru | Inhibited | -2.119 | bias | 2.4E-08 |
| CSF1 | -8.006 | cytokine | Inhibited | -5.388 | bias | 3.01E-08 |
| Ins1 | | other | Inhibited | -3.924 | bias | 3.04E-08 |
| Firre | | other | Inhibited | -4.626 | bias | 3.33E-08 |
| GAST | | other | Inhibited | -5.276 | bias | 3.33E-08 |
| EPAS1 | -8.458 | transcription | Inhibited | -3.888 | bias | 3.74E-08 |
| SNAI1 | -5.194 | transcription | | -0.44 | | 3.8E-08 |
| PDX1 | 6.115 | transcription | | -0.863 | | 3.92E-08 |
| RB1 | -5.375 | transcription | Inhibited | -3.27 | | 4.01E-08 |
| APOE | -9.363 | transporter | | 1.42 | bias | 4.64E-08 |
| FMR1 | 10.637 | translation r | Activated | 2.794 | bias | 4.7E-08 |
| carbamazepi | | chemical dru | Inhibited | -4.643 | bias | 4.97E-08 |
| benzene | | chemical tox | | -0.91 | | 5.02E-08 |
| bortezomib | | chemical dru | Inhibited | -2.684 | | 6.06E-08 |
| sulindac sulfi | | chemical dru | | 0.974 | | 6.27E-08 |
| elaidic acid | | chemical - e | Inhibited | -4.954 | bias | 6.35E-08 |
| EDN1 | -5.037 | cytokine | Inhibited | -4.023 | bias | 6.49E-08 |
| CCN2 | -11.854 | growth facto | Inhibited | -3.369 | bias | 6.71E-08 |
| phenethyl isc | | chemical dru | | 1.969 | | 6.71E-08 |
| FUS | -10.899 | transcription | | -1 | bias | 7.71E-08 |
| PRL | | cytokine | Inhibited | -5.665 | bias | 7.96E-08 |
| ALKBH5 | -8.763 | enzyme | | | | 8.5E-08 |
| SCAP | -9.705 | other | Inhibited | -4.777 | bias | 8.5E-08 |
| miR-16-5p (a | | mature micr | Activated | 6.242 | bias | 8.76E-08 |
| CD44 | -10.607 | other | Inhibited | -2.085 | bias | 1.04E-07 |
| MAP2K5 | -6.924 | kinase | Inhibited | -4.527 | bias | 1.12E-07 |
| HMGA1 | -10.215 | transcription | Inhibited | -2.638 | bias | 1.13E-07 |

| | | | | | | |
|---|---|---|---|---|---|---|
| POLG | -7.733 | enzyme | | | | 1.16E-07 |
| isoquercitrin | | chemical dru | Inhibited | -3.302 | bias | 1.19E-07 |
| KDM8 | -5.343 | enzyme | Inhibited | -2.782 | bias | 1.23E-07 |
| interferon be | | biologic drug | | | | 1.31E-07 |
| thyroid horm | | chemical - er | | 0.646 | | 1.36E-07 |
| RCE1 | -6.986 | peptidase | | | | 1.42E-07 |
| rifaximin | | chemical dru | | -0.775 | | 1.42E-07 |
| puromycin ar | | chemical rea | | -0.998 | bias | 1.45E-07 |
| NME1 | -8.331 | kinase | | 0.931 | bias | 1.57E-07 |
| MIR17HG | | other | Activated | 3.738 | bias | 1.65E-07 |
| SMYD1 | | transcription | | | | 1.68E-07 |
| NR4A1 | -4.415 | ligand-deper | Activated | 3.313 | | 1.78E-07 |
| ERG | | transcription | Inhibited | -3.502 | bias | 1.85E-07 |
| mir-8 | | microRNA | Activated | 5.059 | bias | 1.85E-07 |
| PRKAG3 | | other | | | | 1.92E-07 |
| methylpredn | | chemical dru | Inhibited | -5.763 | | 1.93E-07 |
| eprenetapop | | chemical dru | | | | 1.99E-07 |
| DDIT3 | -7.673 | transcription | | -1.521 | | 2.02E-07 |
| miR-29b-3p ( | | mature micr | Activated | 4.636 | bias | 2.05E-07 |
| SREBF2 | -6.371 | transcription | Inhibited | -5.14 | bias | 2.05E-07 |
| TRAP1 | -9.898 | enzyme | | -0.171 | | 0.00000023 |
| trichloroethy | | chemical tox | | -0.626 | | 2.41E-07 |
| NR3C1 | | ligand-deper | | -0.039 | | 2.46E-07 |
| NFKBIA | -7.984 | transcription | | -0.397 | | 2.59E-07 |
| thioacetamic | | chemical tox | Inhibited | -3.872 | bias | 2.61E-07 |
| PLA2R1 | | transmembra | Inhibited | -3.825 | bias | 0.00000027 |
| TP73 | 8.94 | transcription | Inhibited | -3.977 | bias | 2.88E-07 |
| Immunoglob | | complex | | -0.276 | | 3.05E-07 |
| isobutylmeth | | chemical tox | Inhibited | -2.363 | bias | 3.33E-07 |
| YY1 | -9.644 | transcription | | 0.424 | | 3.47E-07 |
| clopidogrel | | chemical dru | | -1.569 | | 3.63E-07 |
| MFSD2A | -7.499 | transporter | Activated | 3.592 | bias | 3.72E-07 |
| JUN | -9.376 | transcription | Inhibited | -3.657 | bias | 4.07E-07 |
| miR-155-5p ( | | mature micr | Activated | 6.364 | bias | 4.25E-07 |
| miR-338-3p ( | | mature micr | Activated | 4.065 | bias | 0.00000046 |
| nocodazole | | chemical rea | | -1.146 | | 0.00000047 |
| RRP1B | -9.36 | transcription | | | | 0.00000047 |
| lithium chlor | | chemical dru | Inhibited | -2.17 | bias | 0.0000005 |
| bisphenol A | | chemical - er | Inhibited | -2.04 | bias | 5.18E-07 |
| SPDEF | | transcription | Activated | 2.797 | bias | 5.63E-07 |
| TP63 | 3.791 | transcription | Inhibited | -2.824 | bias | 0.00000057 |
| PRNP | -9.164 | other | | -1.095 | | 0.00000058 |
| TWIST1 | -8.812 | transcription | Inhibited | -2.44 | bias | 6.15E-07 |
| bexarotene | | chemical dru | Inhibited | -2.09 | bias | 6.54E-07 |
| MPZ | -4.459 | other | | | | 8.45E-07 |

| | | | | | | |
|---|---|---|---|---|---|---|
| curcumin | | chemical dru | Activated | 4.471 | bias | 0.00000086 |
| tosedostat | | chemical dru | Inhibited | -3.86 | bias | 8.69E-07 |
| BRAF | -3.935 | kinase | Inhibited | -2.184 | bias | 9.23E-07 |
| KDM5A | -9.116 | transcription | Activated | 5.406 | bias | 9.41E-07 |
| TFRC | -9.461 | transporter | Activated | 2.708 | bias | 9.54E-07 |
| F2 | | peptidase | Inhibited | -6.835 | bias | 0.00000102 |
| PKM | -12.425 | kinase | Inhibited | -2.282 | bias | 0.00000105 |
| OSM | | cytokine | | -1.873 | bias | 0.00000106 |
| bardoxolone | | chemical dru | | -0.648 | | 0.00000106 |
| butyric acid | | chemical - er | | 0.452 | | 0.00000124 |
| KLF3 | -8.925 | transcription | Activated | 4.212 | bias | 0.00000133 |
| VEGFA | -10.85 | growth facto | Inhibited | -6.047 | bias | 0.00000134 |
| L-histidine | | chemical - er | Activated | 2.446 | bias | 0.00000135 |
| FOXO4 | -6.414 | transcription | | -0.194 | | 0.00000146 |
| CREB1 | -8.26 | transcription | | -1.52 | bias | 0.00000147 |
| QKI | -8.218 | other | | -1.406 | bias | 0.00000151 |
| methylnitros | | chemical tox | | -1.172 | bias | 0.00000173 |
| PLN | 6.921 | transporter | | | | 0.00000186 |
| RASSF5 | -6.337 | other | | -0.082 | | 0.00000202 |
| FGF2 | -8.155 | growth facto | Inhibited | -5.107 | bias | 0.00000203 |
| MAPK7 | -7.983 | kinase | Inhibited | -3.112 | bias | 0.00000212 |
| COLQ | -4.744 | other | | -0.607 | | 0.0000022 |
| TEAD | | group | Inhibited | -2.186 | bias | 0.00000246 |
| 6-hydroxydop | | chemical tox | Inhibited | -4.792 | bias | 0.00000262 |
| HEXIM1 | -6.685 | transcription | Activated | 2.085 | bias | 0.00000268 |
| ATP7B | | transporter | Inhibited | -3 | | 0.00000287 |
| HGF | | growth facto | Inhibited | -5.685 | bias | 0.00000288 |
| methapyriler | | chemical dru | Inhibited | -3.545 | | 0.00000306 |
| Pln | | other | | | | 0.00000327 |
| MAX | -7.396 | transcription | | 0.922 | | 0.00000333 |
| KIF3A | -10.07 | enzyme | Activated | 2.601 | bias | 0.00000339 |
| methylmercu | | chemical tox | | -0.626 | | 0.00000343 |
| estrogen rec | | group | Activated | 3.463 | | 0.00000368 |
| raloxifene | | chemical dru | | -0.532 | | 0.00000369 |
| thapsigargin | | chemical tox | Inhibited | -3.551 | bias | 0.00000369 |
| vancomycin | | biologic drug | | -1.074 | | 0.00000375 |
| C4BP | | complex | Inhibited | -2.646 | bias | 0.00000388 |
| TRPC4AP | -9.448 | transporter | Activated | 2.449 | bias | 0.00000388 |
| MTBP | -3.326 | other | | -1.987 | bias | 0.00000388 |
| histidinol | | chemical - er | Inhibited | -2.2 | bias | 0.00000428 |
| IL4 | | cytokine | Inhibited | -6.609 | bias | 0.00000429 |
| 1-methyl-4-p | | chemical tox | | -0.35 | bias | 0.00000472 |
| phytohemagg | | chemical dru | Inhibited | -6.579 | bias | 0.00000488 |
| cigarette sm | | chemical tox | Inhibited | -2.843 | bias | 0.0000049 |
| H1-6 | | other | Inhibited | -2.502 | bias | 0.00000496 |

| | | | | | | |
|---|---|---|---|---|---|---|
| H1f1 | | other | Inhibited | -2.502 | bias | 0.00000496 |
| cyclosporin A | | biologic drug | | 1.935 | bias | 0.00000503 |
| LY294002 | | chemical - ki | Activated | 6.963 | bias | 0.00000514 |
| GLI1 | -6.269 | transcription | Inhibited | -5.007 | bias | 0.00000523 |
| camptotheci | | chemical dru | Inhibited | -2.414 | bias | 0.00000553 |
| MXD1 | -7.56 | transcription | Activated | 4.973 | bias | 0.00000603 |
| Ngf | | group | Inhibited | -4.074 | bias | 0.0000061 |
| ARNT | -8.119 | transcription | Inhibited | -3.933 | bias | 0.00000628 |
| SKP2 | -8.225 | other | | -1.004 | | 0.00000629 |
| TWIST2 | -4.582 | transcription | Inhibited | -2.019 | | 0.00000649 |
| mir-29 | | microRNA | Activated | 4.633 | bias | 0.00000653 |
| CDK19 | -8.595 | kinase | Activated | 3.962 | bias | 0.00000665 |
| estrogen | | chemical dru | Inhibited | -3.895 | bias | 0.00000672 |
| peoniflorin | | chemical dru | | -0.286 | | 0.0000079 |
| SP3 | -8.341 | transcription | | -1.636 | bias | 0.00000797 |
| SNHG20 | 1.392 | other | | -1.335 | bias | 0.00000799 |
| salinomycin | | chemical - er | Activated | 2.16 | bias | 0.00000799 |
| ERBB4 | -5.513 | kinase | | -1.03 | | 0.00000803 |
| NRF1 | -6.678 | transcription | Inhibited | -3.731 | bias | 0.00000806 |
| black raspber | | chemical dru | | | | 0.00000808 |
| ATN1 | -7.153 | transcription | | | | 0.00000845 |
| halofuginone | | chemical dru | | 1.98 | bias | 0.0000085 |
| SREBF1 | -8.94 | transcription | Inhibited | -5.864 | bias | 0.00000876 |
| SRSF1 | -10.699 | other | | 0.008 | | 0.0000092 |
| PPARGC1A | | transcription | Inhibited | -2.26 | | 0.00000921 |
| indomethaci | | chemical dru | | -0.945 | | 0.00000948 |
| FOXO3 | -8.025 | transcription | | -0.221 | | 0.00000954 |
| SNAI2 | -6.614 | transcription | | -1.861 | | 0.00000954 |
| MEG3 | 1.949 | other | | 0.637 | | 0.00000964 |
| HOXA4 | -7.899 | transcription | | 0.389 | | 0.00000964 |
| mir-10 | | microRNA | Activated | 4.035 | bias | 0.00000987 |
| AR | -6.632 | ligand-deper | Inhibited | -3.652 | bias | 0.00001 |
| bromobenzer | | chemical tox | | 1.546 | | 0.0000106 |
| COL4A3 | | other | | | | 0.0000111 |
| IREB2 | -9.719 | translation re | | 1.132 | | 0.0000111 |
| clozapine | | chemical dru | | -1.818 | bias | 0.0000112 |
| CLOCK | -6.292 | transcription | | 0.868 | | 0.0000112 |
| SP110 | -4.609 | transcription | Activated | 3.781 | | 0.0000114 |
| cobalt chloric | | chemical rea | Inhibited | -4.121 | bias | 0.0000125 |
| CYP1A1 | | enzyme | | 0 | | 0.0000128 |
| TXNIP | -9.967 | other | | 1.128 | | 0.000013 |
| miR-335-3p ( | | mature micr | Activated | 3.606 | bias | 0.0000133 |
| n-nitrosomet | | chemical tox | | | | 0.0000144 |
| SPZ1 | | transcription | Inhibited | -2.779 | bias | 0.0000147 |
| CDKN2A | | transcription | | 1.365 | | 0.000015 |

| | | | | | | |
|---|---|---|---|---|---|---|
| tomatidine | | chemical - er | Activated | 2.646 | bias | 0.0000155 |
| TLE3 | -7.938 | other | | | | 0.0000157 |
| doxorubicin | | chemical dru | Inhibited | -3.943 | bias | 0.0000176 |
| silibinin | | chemical dru | Activated | 2.739 | bias | 0.0000178 |
| homocystein | | chemical - er | Inhibited | -2.034 | | 0.0000183 |
| COMMD1 | -6.811 | transporter | Activated | 3.317 | bias | 0.0000188 |
| SMAD7 | -7.094 | transcription | Activated | 2.939 | bias | 0.0000194 |
| ATP5IF1 | -7.714 | other | | | | 0.0000197 |
| M344 | | chemical rea | Inhibited | -3.036 | bias | 0.0000206 |
| fulvestrant | | chemical dru | | 1.574 | | 0.0000208 |
| PROM1 | -7.362 | other | Activated | 3 | bias | 0.0000214 |
| INSIG2 | -7.487 | other | Activated | 2.933 | bias | 0.0000214 |
| KDM3B | -9.952 | enzyme | Activated | 2 | bias | 0.0000229 |
| BCL2 | -8.15 | transporter | Inhibited | -2.489 | bias | 0.000023 |
| CDKN1A | -10.797 | kinase | Activated | 2.377 | bias | 0.0000231 |
| HSP90B1 | -12.689 | other | Inhibited | -3.578 | bias | 0.0000234 |
| EGR2 | | transcription | Inhibited | -4.735 | bias | 0.0000244 |
| Hbb-b1 | | transporter | Inhibited | -4.082 | bias | 0.0000246 |
| NPC1 | -8.328 | transporter | | 1.594 | bias | 0.0000246 |
| CCND1 | -10.515 | transcription | | -0.495 | | 0.0000247 |
| EIF4G2 | -13.559 | translation re | Inhibited | -2.538 | | 0.000026 |
| MTM1 | -6.631 | phosphatase | Activated | 2.75 | bias | 0.000026 |
| ammonium c | | chemical dru | Inhibited | -2.525 | bias | 0.000026 |
| Sos | | group | | | | 0.0000265 |
| MYCBP | -6.158 | transcription | Inhibited | -2.449 | bias | 0.0000267 |
| PTBP3 | -7.812 | other | | -0.835 | bias | 0.0000267 |
| SPP1 | -6.735 | cytokine | Inhibited | -3.854 | bias | 0.0000267 |
| 26s Proteaso | | complex | Activated | 4.807 | bias | 0.0000287 |
| TGFB3 | -8.006 | growth facto | Inhibited | -4.635 | bias | 0.0000307 |
| NRG2 | | growth facto | | | | 0.0000336 |
| Srebp | | group | Inhibited | -2.901 | bias | 0.0000345 |
| RHOJ | 1.405 | enzyme | | | | 0.0000345 |
| EIF4E | -8.033 | translation re | Inhibited | -6.449 | bias | 0.0000355 |
| ATG5 | -7.46 | other | | -1.568 | | 0.0000357 |
| vincristine | | chemical dru | | 0.072 | | 0.0000366 |
| TFAP2A | -4.456 | transcription | | 1.092 | | 0.0000367 |
| ONECUT1 | -6.258 | transcription | | | | 0.0000404 |
| H2AX | -9.78 | transcription | Inhibited | -2.447 | bias | 0.0000413 |
| miR-122-5p ( | | mature micr | Activated | 4.471 | bias | 0.0000435 |
| COPS5 | -8.96 | transcription | | -0.563 | | 0.000044 |
| BRD7 | -8.401 | transcription | Inhibited | -2.416 | bias | 0.000044 |
| EPO | | cytokine | Inhibited | -4.15 | bias | 0.0000445 |
| IFNG | | cytokine | Inhibited | -5.152 | bias | 0.000045 |
| PHF21A | -7.756 | other | Inhibited | -2.623 | bias | 0.000046 |
| STX3 | -6.37 | transporter | | 0.378 | | 0.000046 |

| | | | | | | |
|---|---|---|---|---|---|---|
| IL5 | | cytokine | Inhibited | -7.111 | bias | 0.0000498 |
| TERT | | enzyme | | -1.4 | | 0.0000537 |
| deferoxamin | | chemical dru | Inhibited | -3.169 | bias | 0.0000576 |
| miR-296-5p ( | | mature micr | Activated | 3.105 | bias | 0.0000611 |
| TCF7L2 | -4.744 | transcription | Inhibited | -5.564 | bias | 0.0000632 |
| BRCA1 | 3.148 | transcription | | -1.488 | bias | 0.0000645 |
| INS | | other | | -1.873 | | 0.0000648 |
| CXCL12 | -9.229 | cytokine | | -1.931 | bias | 0.0000725 |
| cadmium chl | | chemical tox | Inhibited | -3.225 | bias | 0.0000728 |
| WNT3A | | cytokine | | -1.586 | bias | 0.0000744 |
| CD28 | | transmembra | Inhibited | -4.887 | bias | 0.000077 |
| IGF2 | -12.39 | growth facto | | -1.77 | bias | 0.0000782 |
| HNRNPH1 | -10.787 | other | | | | 0.0000784 |
| NCAM1 | -11.084 | other | | -1.744 | bias | 0.0000815 |
| glutamyl-Se- | | chemical - er | Activated | 2.333 | bias | 0.0000815 |
| CEBPB | -8.096 | transcription | Inhibited | -6.688 | bias | 0.0000844 |
| ATF6 | -9.286 | transcription | Inhibited | -3.352 | bias | 0.0000871 |
| napabucasin | | chemical dru | Activated | 3.437 | bias | 0.0000871 |
| MITF | | transcription | Inhibited | -4.254 | bias | 0.0000872 |
| HSPA9 | -8.252 | other | Inhibited | -2.197 | bias | 0.0000897 |
| FAS | -6.308 | transmembra | Activated | 2.154 | | 0.0000936 |
| iron | | chemical - er | | -0.374 | | 0.0000996 |
| PGR | 7.293 | ligand-deper | Inhibited | -4.694 | bias | 0.0001 |
| alvocidib | | chemical dru | Activated | 2.957 | bias | 0.0001 |
| SOX4 | -10.339 | transcription | Inhibited | -5.175 | bias | 0.000101 |
| G protein alp | | group | Inhibited | -4.707 | bias | 0.000103 |
| Mir200 | | group | Activated | 2.55 | bias | 0.000103 |
| GF 120918 | | chemical rea | Inhibited | -2.538 | bias | 0.000103 |
| GPER1 | | G-protein co | Inhibited | -2.21 | bias | 0.000106 |
| FOXM1 | -6.453 | transcription | Inhibited | -3.764 | bias | 0.000107 |
| RAF1 | -3.325 | kinase | Inhibited | -5.55 | bias | 0.000107 |
| CD38 | | enzyme | Inhibited | -5.793 | bias | 0.000109 |
| prostaglandi | | chemical - er | Inhibited | -3.07 | bias | 0.000109 |
| ARHGAP31 | -7.443 | other | | 1.677 | | 0.000109 |
| RLIM | -7.827 | enzyme | Inhibited | -2.449 | bias | 0.000112 |
| DYSF | | other | | | | 0.000113 |
| dichlorovinyl | | chemical tox | | 0.487 | | 0.000115 |
| tamoxifen | | chemical dru | | -0.539 | | 0.000115 |
| PI3K (comple | | complex | Inhibited | -4.915 | bias | 0.000124 |
| RASSF1 | -7.683 | other | | -0.128 | | 0.00013 |
| NGF | -4.23 | growth facto | Inhibited | -5.663 | bias | 0.000136 |
| MAP4K4 | -10.307 | kinase | Activated | 4.386 | bias | 0.000138 |
| SPRY2 | -7.975 | other | | 1.663 | bias | 0.00014 |
| NFE2L1 | -11.405 | transcription | Inhibited | -3.592 | bias | 0.00014 |
| TCOF1 | -8.1 | transporter | | | | 0.000142 |

| | | | | | | |
|---|---|---|---|---|---|---|
| EGR1 | -9.716 | transcription | Inhibited | -4.303 | bias | 0.000144 |
| NFASC | -10.049 | other | Inhibited | -2.339 | bias | 0.000145 |
| NOTCH1 | -10.13 | transcription | | -1.484 | | 0.000146 |
| ZBED6 | -6.422 | transcription | | 1.058 | bias | 0.000156 |
| IGF2BP1 | -9.89 | translation re | | -1.212 | | 0.000156 |
| FHIT | 9.371 | enzyme | Activated | 2.992 | bias | 0.000156 |
| mir-182 | | microRNA | Activated | 2.007 | bias | 0.000157 |
| 4-phenylbuty | | chemical - er | | 0.508 | | 0.000157 |
| nicotine | | chemical dru | Inhibited | -3.494 | bias | 0.000167 |
| Laminin (con | | complex | | 1.256 | | 0.000167 |
| SRSF2 | -10.04 | transcription | | 0.083 | bias | 0.000167 |
| Abcb1b | | transporter | Activated | 2.534 | bias | 0.000168 |
| selenomethy | | chemical - er | Activated | 2.53 | bias | 0.000168 |
| PSEN2 | -7.337 | peptidase | | 1.541 | | 0.000169 |
| ramipril | | chemical dru | Activated | 2.887 | bias | 0.000169 |
| MET | -6.398 | kinase | Inhibited | -3.223 | bias | 0.000172 |
| FBXW7 | -7.518 | enzyme | Activated | 3.251 | bias | 0.000176 |
| MUC1 | | other | Inhibited | -3.955 | bias | 0.000176 |
| EIF3M | -9.636 | other | Inhibited | -2.236 | bias | 0.00018 |
| ADRA1D | -4.822 | G-protein co | | | | 0.000188 |
| cetuximab | | biologic drug | | -0.677 | | 0.000194 |
| IGFBP7 | -10.539 | transporter | | 0.174 | | 0.000194 |
| mir-140 | | microRNA | | | | 0.000194 |
| lith-O-Asp | | chemical rea | | 1.083 | | 0.000194 |
| SBDS | -9.164 | other | | | | 0.000198 |
| nitric oxide | | chemical - er | Inhibited | -3.603 | bias | 0.000206 |
| ATM | -1.039 | kinase | Inhibited | -2.672 | bias | 0.000218 |
| interferon al | | biologic drug | Activated | 3.051 | bias | 0.00022 |
| ING1 | -7.632 | transcription | | -0.806 | | 0.00022 |
| ILK | -8.868 | kinase | | -1.219 | bias | 0.000224 |
| tert-butyl-hy | | chemical rea | Inhibited | -3.869 | bias | 0.000227 |
| COL18A1 | -11.962 | other | Activated | 3.608 | bias | 0.000239 |
| AFAP1-AS1 | | other | | 1.157 | | 0.00024 |
| NELFB | -9.894 | other | | 0.378 | | 0.00024 |
| CTNNBIP1 | -7.061 | other | | 1.79 | bias | 0.00024 |
| arsenite | | chemical tox | Inhibited | -3.131 | bias | 0.000241 |
| enalapril | | biologic drug | Activated | 2.582 | | 0.000251 |
| PDGF (family | | group | Inhibited | -2.449 | bias | 0.000255 |
| WNT5B | -8.027 | other | | -1.961 | bias | 0.000255 |
| mir-148 | | microRNA | Activated | 3.267 | bias | 0.00026 |
| VCP | -11.811 | enzyme | | | | 0.00026 |
| alvespimycin | | chemical dru | Activated | 2.428 | bias | 0.000263 |
| AHR | -2.61 | ligand-deper | | 0.851 | | 0.000264 |
| testosterone | | chemical - er | Inhibited | -4.598 | bias | 0.000264 |
| FICD | -5.322 | enzyme | | | | 0.000264 |

| | | | | | | |
|---|---|---|---|---|---|---|
| PDE8A | | enzyme | Activated | 2 | bias | 0.000264 |
| NBR2 | | other | | 1 | bias | 0.000264 |
| RNASEH1 | -3.74 | enzyme | Inhibited | -2 | bias | 0.000264 |
| Collagen type | | complex | Inhibited | -2.737 | bias | 0.000273 |
| methylseleni | | chemical rea | Activated | 2.213 | | 0.000282 |
| RAC1 | -11.205 | enzyme | Inhibited | -4.011 | bias | 0.000282 |
| SIRT1 | -8.124 | transcription | Activated | 2.027 | bias | 0.000284 |
| BMP7 | -4.535 | growth facto | Inhibited | -2.936 | bias | 0.000285 |
| Pdgf (comple | | complex | Inhibited | -4.212 | bias | 0.000302 |
| miR-450a-5p | | mature micr | Activated | 2.496 | bias | 0.000304 |
| SFPQ | -11.117 | other | Inhibited | -2.976 | bias | 0.000304 |
| RPTOR | -8.866 | other | Inhibited | -5.044 | bias | 0.00032 |
| lipopolysacch | | chemical dru | Inhibited | -8.36 | bias | 0.000325 |
| ANGPT2 | | growth facto | Inhibited | -4.669 | bias | 0.000329 |
| SFRP1 | -10.577 | transmembra | | 1.68 | bias | 0.000334 |
| IGFBP2 | -12.715 | other | Inhibited | -2.674 | bias | 0.000334 |
| ITGB3 | -6.529 | transmembra | Inhibited | -2.406 | bias | 0.000337 |
| IDH1 | -10.681 | enzyme | | 0.651 | | 0.000341 |
| mitomycin C | | chemical dru | Inhibited | -3.337 | bias | 0.000342 |
| folic acid | | chemical - er | | -0.098 | | 0.000342 |
| RAPGEF1 | -10.479 | other | Inhibited | -2.8 | bias | 0.000344 |
| spermidine | | chemical - er | | 0.342 | | 0.000344 |
| UCP1 | | transporter | Inhibited | -2.22 | | 0.000346 |
| NORAD | -4.793 | other | Inhibited | -3.687 | bias | 0.000353 |
| cholesterol | | chemical - er | | 0.621 | bias | 0.000357 |
| let-7 | | microRNA | Activated | 5.977 | bias | 0.000361 |
| CCN1 | -10.494 | other | Inhibited | -2.52 | bias | 0.000364 |
| sterol | | chemical - er | Activated | 2.832 | bias | 0.00038 |
| MED1 | -5.155 | transcription | Inhibited | -3.479 | bias | 0.000385 |
| SUCNR1 | -5.53 | G-protein co | Inhibited | -3 | bias | 0.000393 |
| mir-192 | | microRNA | | 0.207 | bias | 0.000393 |
| PPP1R13L | | transcription | | -1.539 | bias | 0.000393 |
| RND3 | -7.984 | enzyme | | -1.369 | | 0.000393 |
| LDL | | complex | Inhibited | -4.225 | bias | 0.000395 |
| SAFB2 | -9.761 | other | | 0.632 | bias | 0.0004 |
| RPS15 | -11.68 | other | | | | 0.000424 |
| IRS1 | -7.799 | enzyme | | -1.998 | bias | 0.000426 |
| FEV | | transcription | Inhibited | -4.796 | bias | 0.000436 |
| LDB1 | -9.976 | transcription | | -1.491 | | 0.000454 |
| ERK1/2 | | group | Inhibited | -3.041 | bias | 0.000459 |
| (±)-2-hydroxy | | chemical dru | | -1.914 | | 0.000463 |
| DANCR | -7.885 | other | | 0.416 | bias | 0.000463 |
| HIF3A | 10.377 | transcription | | 1.166 | bias | 0.000463 |
| ACO1 | -8.683 | enzyme | | 0.769 | | 0.000463 |
| chloroproma | | chemical dru | | -1.491 | bias | 0.000468 |

| | | | | | | |
|---|---|---|---|---|---|---|
| SSRP1 | -10.73 | transcription | | -1.98 | bias | 0.00048 |
| LMO2 | -6.272 | transcription | | -1.491 | | 0.000496 |
| KITLG | -7.642 | growth facto | Inhibited | -3.619 | bias | 0.000506 |
| simvastatin | | chemical dru | | -0.399 | | 0.000507 |
| inosine | | chemical - er | Inhibited | -3.938 | bias | 0.000525 |
| RAC2 | | enzyme | | | | 0.000525 |
| TEAD4 | -5.958 | transcription | Inhibited | -2.409 | bias | 0.000533 |
| SH3TC2 | | other | Inhibited | -2.714 | bias | 0.000542 |
| ROCK1 | -3.274 | kinase | Inhibited | -2.553 | bias | 0.000542 |
| AKT3 | -10.694 | kinase | | -0.594 | | 0.000542 |
| mifepristone | | chemical dru | Activated | 2.638 | | 0.000543 |
| nitrofen | | chemical tox | Activated | 3.592 | bias | 0.000558 |
| olaparib | | chemical dru | | 0.2 | bias | 0.000565 |
| TMPO | -9.205 | other | Inhibited | -2.236 | bias | 0.000565 |
| CAPNS1 | -9.267 | peptidase | Inhibited | -2.195 | bias | 0.000565 |
| HOXA11-AS | | other | | -0.862 | bias | 0.000568 |
| ESRP1 | | other | | 0.199 | | 0.000568 |
| LINC01139 | | other | Inhibited | -2.449 | bias | 0.000568 |
| DAB2IP | -7.736 | other | | 0.606 | bias | 0.000568 |
| ARHGAP5 | -9.392 | enzyme | Inhibited | -2.425 | bias | 0.000568 |
| RANBP9 | -9.475 | other | | 1.741 | bias | 0.000568 |
| PCBP2 | -10.552 | other | | -0.277 | bias | 0.000568 |
| INCB054329 | | chemical dru | | 0.97 | | 0.000568 |
| MXI1 | -8.855 | transcription | Activated | 2.63 | bias | 0.000577 |
| mir-214 | | microRNA | | 0.747 | bias | 0.000591 |
| lysophosphat | | chemical - ot | Inhibited | -3.583 | bias | 0.000613 |
| kanamycin A | | chemical dru | | 1.838 | | 0.000616 |
| MRTFB | -7.899 | transcription | Inhibited | -5.056 | bias | 0.000637 |
| miR-145-5p ( | | mature micr | Activated | 3.977 | bias | 0.000638 |
| ADRA1B | | G-protein co | | | | 0.000647 |
| sodium arser | | chemical dru | Inhibited | -2.446 | bias | 0.000656 |
| CCN5 | | growth facto | | 0.234 | | 0.000664 |
| lovastatin | | chemical dru | Inhibited | -3.849 | bias | 0.000679 |
| fenretinide | | chemical dru | | -0.554 | | 0.000702 |
| EP300 | -9.82 | transcription | Inhibited | -4.389 | bias | 0.000724 |
| GNRH | | group | Inhibited | -2.027 | bias | 0.00074 |
| NSUN6 | -7.059 | enzyme | Inhibited | -3.606 | bias | 0.00074 |
| PRKN | -6.222 | enzyme | | 0.803 | bias | 0.00074 |
| sucrose | | chemical - er | Inhibited | -2.73 | bias | 0.00074 |
| STAT4 | | transcription | Inhibited | -6.64 | bias | 0.000752 |
| ITGB1 | -11.852 | transmembra | | 0.507 | | 0.000754 |
| LLGL2 | | other | Inhibited | -3.317 | bias | 0.000758 |
| HNRNPA2B1 | -9.995 | other | | -0.152 | bias | 0.00077 |
| Growth horm | | group | Inhibited | -3.197 | bias | 0.000784 |
| eflornithine | | chemical dru | | -1.765 | | 0.000795 |

| | | | | | | |
|---|---|---|---|---|---|---|
| ouabain | | chemical dru | | -0.226 | bias | 0.000795 |
| STOX1 | -5.971 | other | | | | 0.000823 |
| manganese ( | | chemical rea | | -0.898 | bias | 0.000823 |
| D-tubocurari | | chemical dru | | | | 0.000823 |
| nilotinib | | chemical dru | | 1.585 | bias | 0.000824 |
| ITGA5 | -8.645 | transmembra | | -1.528 | bias | 0.000824 |
| NOX4 | 10.768 | enzyme | | -0.424 | bias | 0.000824 |
| diethylnitros | | chemical tox | | -1.254 | | 0.000827 |
| SUPT16H | -2.626 | transcription | | | | 0.000851 |
| amino acids | | chemical - er | | 0.824 | | 0.000851 |
| NRAS | -1.458 | enzyme | | -0.47 | bias | 0.000867 |
| progesterone | | chemical - er | Inhibited | -4.358 | | 0.000883 |
| hydrocortisor | | chemical - er | | -0.191 | | 0.000884 |
| S100B | -2.296 | other | | -0.277 | bias | 0.000923 |
| GPI | -11.926 | enzyme | | -1.333 | bias | 0.000923 |
| CREB3 | -8.953 | transcription | | -1.348 | bias | 0.000923 |
| PIK3R1 | -8.011 | kinase | | -1.936 | bias | 0.00093 |
| miR-291a-3p | | mature micr | Activated | 4.543 | bias | 0.000932 |
| ASPSCR1-TFI | | fusion gene/ | Inhibited | -4.082 | bias | 0.000932 |
| lomustine | | chemical dru | Inhibited | -2.644 | bias | 0.000983 |
| VCAN | -11.339 | other | | 0.241 | | 0.00101 |
| phenylbutazo | | chemical dru | Inhibited | -3.674 | bias | 0.00102 |
| RBM5 | -9.66 | other | | 1.056 | bias | 0.00102 |
| miR-483-3p ( | | mature micr | Activated | 3.726 | bias | 0.00102 |
| HBEGF | -7.617 | growth facto | Inhibited | -2.331 | bias | 0.00102 |
| losartan pota | | chemical dru | Activated | 3.079 | bias | 0.00103 |
| bufalin | | chemical rea | | 1.897 | | 0.00104 |
| mir-21 | | microRNA | Activated | 4.494 | bias | 0.00105 |
| colistin | | biologic drug | | 1.633 | | 0.00105 |
| PITX2 | | transcription | | -1.426 | bias | 0.00106 |
| Hif1 | | complex | Inhibited | -3.546 | bias | 0.00107 |
| NUPR1 | -6.638 | transcription | | 1.009 | bias | 0.00109 |
| E2F4 | -8.809 | transcription | | | | 0.00111 |
| MBOAT7 | -9.26 | enzyme | Activated | 2.425 | bias | 0.00111 |
| DGCR5 | -6.769 | other | | 1.709 | | 0.00111 |
| LIMS1 | -9.228 | other | Inhibited | -2.417 | bias | 0.00111 |
| PPBP | | cytokine | Inhibited | -2.425 | bias | 0.00111 |
| clioquinol | | chemical dru | | -1.633 | bias | 0.00111 |
| 2-deoxygluco | | chemical dru | | -1.602 | | 0.00113 |
| CGA | | other | | 1.474 | | 0.00115 |
| crizotinib | | chemical dru | | -1.029 | | 0.00117 |
| FLVCR1 | | transporter | Activated | 2 | bias | 0.00119 |
| DDX21 | -6.956 | enzyme | Inhibited | -2 | bias | 0.00119 |
| GLS | -8.887 | enzyme | | -1.961 | bias | 0.00119 |
| PUF60 | -8.767 | other | | -1.993 | bias | 0.00119 |

| | | | | | | |
|---|---|---|---|---|---|---|
| ARHGEF25 | -9.213 | other | Inhibited | -2 | bias | 0.00119 |
| NUDT21 | -5.233 | other | Activated | 2 | bias | 0.00119 |
| GTF2H4 | -8.258 | transcription | | | | 0.00119 |
| DVL3 | -3.188 | other | | | | 0.00119 |
| ADAM9 | -10.539 | peptidase | | | | 0.00119 |
| ADAM10 | -9.646 | peptidase | | -1.525 | bias | 0.00119 |
| BMI1 | -8.946 | transcription | | 1.819 | | 0.00119 |
| triamterene | | chemical dru | Inhibited | -3.179 | bias | 0.00119 |
| p70 S6k | | group | Inhibited | -2.138 | bias | 0.0012 |
| POR | -8.406 | enzyme | Activated | 4.008 | bias | 0.00125 |
| UCHL1 | -12.491 | peptidase | | 1.579 | | 0.00127 |
| PIAS1 | -7.407 | transcription | | -1.222 | bias | 0.00127 |
| STEAP3 | -4.856 | transporter | Activated | 3.988 | bias | 0.00128 |
| HNF1B | | transcription | Activated | 2.38 | | 0.0013 |
| DGCR8 | -9.229 | enzyme | | 0.399 | | 0.0013 |
| ADRA1A | | G-protein co | | | | 0.0013 |
| CDKN1B | -9.841 | kinase | Activated | 2.757 | bias | 0.00134 |
| Pkg | | group | | -1.349 | bias | 0.00134 |
| FAM3B | | cytokine | | | | 0.00134 |
| pitavastatin | | chemical dru | Inhibited | -2.969 | bias | 0.00134 |
| evofosfamid | | chemical dru | Activated | 2.236 | bias | 0.00135 |
| artemisinin | | chemical dru | | 0.497 | bias | 0.00135 |
| cocaine | | chemical dru | Inhibited | -2.488 | bias | 0.00135 |
| TEAD1 | -8.105 | transcription | | -1.789 | bias | 0.00136 |
| PPARGC1B | | transcription | Inhibited | -3.554 | bias | 0.00136 |
| gemcitabine | | chemical dru | | -1.647 | bias | 0.00136 |
| PRKAA2 | -4.664 | kinase | Activated | 3.886 | bias | 0.00136 |
| CDH2 | -11.268 | other | | 0.603 | bias | 0.00137 |
| PNN | -10.76 | other | | 0.64 | | 0.00137 |
| PDLIM2 | -6.77 | other | | 1.504 | | 0.00138 |
| grape seed e | | chemical dru | | 0.905 | bias | 0.0014 |
| ROR1 | -7.025 | kinase | Inhibited | -2.064 | bias | 0.0014 |
| TRPM2 | | ion channel | | 0.784 | | 0.0014 |
| PRKG1 | -5.841 | kinase | | -0.095 | | 0.0014 |
| mir-193 | | microRNA | Activated | 3.25 | bias | 0.0014 |
| carfilzomib | | biologic drug | Inhibited | -2.06 | bias | 0.00142 |
| ITGA6 | -8.132 | transmembra | | -0.691 | bias | 0.00142 |
| DPP-23 | | chemical rea | Inhibited | -2.828 | bias | 0.00142 |
| GH1 | | growth facto | | -0.698 | | 0.00144 |
| CCL5 | 2.051 | cytokine | Inhibited | -4.28 | bias | 0.00151 |
| NCOA3 | -7.732 | transcription | Inhibited | -3.61 | bias | 0.00151 |
| FLI1 | -3.705 | transcription | Inhibited | -2.961 | bias | 0.00151 |
| RHO | | G-protein co | | 0.5 | | 0.00151 |
| glucosamine | | chemical - er | | 0.197 | | 0.00151 |
| HDL-choleste | | complex | Activated | 2.887 | bias | 0.00154 |

| | | | | | | |
|---|---|---|---|---|---|---|
| ITCH | 9.719 | enzyme | Activated | 2.882 | bias | 0.00154 |
| SOX11 | -4.034 | transcription | | -1.52 | bias | 0.00156 |
| PML | -7.15 | transcription | Activated | 3.32 | | 0.00157 |
| hexachlorobe | | chemical tox | Inhibited | -3.011 | bias | 0.00158 |
| panobinostat | | chemical dru | | 0.929 | | 0.00162 |
| mir-34 | | microRNA | Activated | 3.641 | bias | 0.00162 |
| PIK3CA | -8.351 | kinase | | -1.414 | | 0.00162 |
| romidepsin | | biologic drug | | 0.205 | | 0.00164 |
| WBP2 | -10.26 | transcription | | -1.61 | | 0.00174 |
| MAP2K1/2 | | group | Inhibited | -2.822 | bias | 0.00178 |
| MMP9 | | peptidase | | -1.456 | | 0.00178 |
| seocalcitol | | chemical dru | Inhibited | -3.162 | bias | 0.00183 |
| FOLR1 | -5.755 | transporter | | | | 0.00187 |
| NPPB | -7.706 | other | Activated | 3.283 | bias | 0.00187 |
| NTRK1 | | kinase | Inhibited | -2.901 | bias | 0.0019 |
| CDH1 | -5.744 | other | | 1.291 | | 0.00192 |
| MBNL1 | -6.049 | other | | 1.377 | bias | 0.00199 |
| SERPINA3 | -7.213 | other | | -0.277 | | 0.00199 |
| PAWR | -6.476 | transcription | | 0.6 | bias | 0.00199 |
| MTORC1 | | complex | Inhibited | -2.618 | bias | 0.00199 |
| fenamic acid | | chemical rea | Inhibited | -3.236 | bias | 0.00206 |
| OSMI-1 | | chemical rea | | | | 0.00207 |
| SLC48A1 | -2.963 | transporter | | | | 0.00207 |
| CERS4 | -8.582 | transcription | | | | 0.00207 |
| EPM2A | -3.981 | phosphatase | | | | 0.00207 |
| SERTAD1 | | transcription | | | | 0.00207 |
| TPM1 | -11.528 | other | | | | 0.00207 |
| NPTX1 | -8.852 | other | | | | 0.00207 |
| FTMT | | enzyme | | | | 0.00207 |
| RASSF7 | -7.259 | other | | | | 0.00207 |
| 3K3A-APC pr | | biologic drug | | | | 0.00207 |
| 4-ethylsulfar | | chemical rea | | | | 0.00207 |
| 4-hydroxyphe | | chemical rea | | | | 0.00207 |
| polyhydroxye | | chemical rea | | | | 0.00207 |
| neratinib | | chemical dru | | | | 0.00207 |
| (S)-duloxetin | | chemical dru | | | | 0.00207 |
| 3,4-dihydroxy | | chemical - er | | | | 0.00207 |
| EWSR1-FLI1 | | fusion gene/ | Activated | 3.71 | bias | 0.00209 |
| SAV1 | | other | Activated | 2.813 | bias | 0.0021 |
| IAPP | 4.669 | other | | | | 0.0021 |
| 5-azacytidine | | chemical dru | Inhibited | -3.529 | bias | 0.00215 |
| GFAP | -5.216 | other | | -1.165 | | 0.00218 |
| asciminib | | chemical dru | Activated | 2.643 | bias | 0.00218 |
| SUB1 | -10.252 | transcription | Inhibited | -2.138 | bias | 0.00224 |
| FGFR1 | -9.393 | kinase | | -0.976 | bias | 0.00225 |

| | | | | | | |
|---|---|---|---|---|---|---|
| RELA | -9.122 | transcription | Inhibited | -2.698 | bias | 0.00225 |
| anisomycin | | chemical - er | | -0.867 | | 0.00233 |
| L-glutamic ac | | chemical - er | Inhibited | -4.101 | bias | 0.00238 |
| Alpha catenin | | group | Activated | 4.771 | bias | 0.00242 |
| AKT2 | -9.481 | kinase | Inhibited | -2.063 | bias | 0.00244 |
| 3-nitropropic | | chemical tox | | -0.202 | | 0.00244 |
| IFT88 | -6.77 | other | Activated | 2.779 | | 0.00245 |
| sulforafan | | chemical dru | Inhibited | -2.825 | | 0.00248 |
| SKIL | -8.927 | transcription | | -1.873 | bias | 0.00256 |
| methyl meth | | chemical tox | Inhibited | -2.643 | bias | 0.00256 |
| sunitinib | | chemical dru | | 0.791 | bias | 0.00263 |
| EXOSC3 | -6.425 | enzyme | Activated | 2.975 | bias | 0.00264 |
| SDCBP | -9.513 | enzyme | Inhibited | -2.147 | bias | 0.00264 |
| Y 27632 | | chemical dru | Activated | 3.469 | bias | 0.00267 |
| tetrachlorodi | | chemical tox | | 0.477 | | 0.00269 |
| metronidazo | | chemical dru | | 1.633 | | 0.0027 |
| APC/APC2 | | group | | -0.156 | bias | 0.00271 |
| TACSTD2 | | other | | | | 0.00271 |
| KDM4B | -9.119 | enzyme | | -1.319 | bias | 0.00271 |
| SLC18A2 | | transporter | | -0.555 | bias | 0.00271 |
| genipin | | chemical - er | | 0.447 | bias | 0.00271 |
| PINK1 | -9.931 | kinase | Inhibited | -2.216 | bias | 0.00271 |
| MBD3 | -10.121 | other | | | | 0.00286 |
| U0126 | | chemical - ki | Activated | 5.153 | bias | 0.00289 |
| diethylstilbes | | chemical dru | Inhibited | -2.954 | bias | 0.00293 |
| dithiothreitol | | chemical rea | | -0.426 | bias | 0.00298 |
| fludarabine | | chemical dru | | 0.544 | | 0.00298 |
| Collagen type | | complex | | 0.378 | | 0.00302 |
| EDN3 | -4.961 | other | | -0.113 | bias | 0.00302 |
| PCDH11Y | | other | Inhibited | -2.813 | bias | 0.00302 |
| MYOC | | other | | -1.765 | bias | 0.00303 |
| wortmannin | | chemical - ki | Activated | 5.248 | bias | 0.00313 |
| SETD2 | -4.334 | enzyme | Activated | 2.111 | bias | 0.00316 |
| PAK1 | -4.538 | kinase | | -1.703 | | 0.00316 |
| WDFY2 | -7.368 | other | Inhibited | -2 | bias | 0.0032 |
| CLDN1 | -7.943 | other | | -1.091 | bias | 0.0032 |
| RHEB | -9.829 | enzyme | | 0.218 | bias | 0.0032 |
| miR-3118 (ar | | mature micro | | 0.957 | bias | 0.0032 |
| LY2109761 | | chemical - ki | Activated | 2 | bias | 0.0032 |
| AATF | -8.409 | transcription | | -0.991 | bias | 0.0032 |
| CCG-1423 | | chemical rea | | 1.982 | bias | 0.0032 |
| ECT2 | -5.553 | other | | -0.152 | | 0.0032 |
| ABCB7 | -6.36 | transporter | | -1.131 | bias | 0.0032 |
| Gm35986 | | other | | 1.98 | bias | 0.0032 |
| CDH13 | -8.337 | other | | -1.165 | bias | 0.0032 |

| | | | | | | |
|---|---|---|---|---|---|---|
| SF1 | -9.985 | transcription | | 0.061 | | 0.0032 |
| NET1 | -7.135 | other | | 1.98 | bias | 0.0032 |
| CXXC4 | -7.794 | other | | 1 | bias | 0.0032 |
| risedronic ac | | chemical dru | | -1 | bias | 0.0032 |
| octyl gallate | | chemical rea | | 0.581 | bias | 0.0032 |
| alpha-hydrox | | chemical - er | | -1 | bias | 0.0032 |
| UDP-D-gluco | | chemical - er | Activated | 2 | bias | 0.0032 |
| gefitinib | | chemical dru | | 0.675 | bias | 0.0032 |
| TGFBR2 | -6.617 | kinase | | -1.785 | | 0.00321 |
| poly rI:rC-RN | | biologic drug | Inhibited | -2.986 | bias | 0.00322 |
| bevacizumab | | biologic drug | | 1.134 | bias | 0.00329 |
| PPP1R1B | -6.675 | phosphatase | Inhibited | -2.596 | bias | 0.00329 |
| FANCD2 | | other | | -1.408 | bias | 0.00329 |
| DNAJB6 | -8.567 | transcription | | 1.941 | bias | 0.00329 |
| CREB3L2 | -8.648 | transcription | Inhibited | -2.425 | bias | 0.0033 |
| CHRM1 | | G-protein co | | -1.516 | bias | 0.0033 |
| MBTPS1 | -10.013 | peptidase | Inhibited | -2.449 | bias | 0.0033 |
| FZR1 | -9.745 | kinase | | 1.539 | bias | 0.0033 |
| SERPINB5 | | other | | 0.64 | | 0.0033 |
| PEX5L | 8.125 | ion channel | Activated | 2.425 | bias | 0.0033 |
| vatalanib | | chemical dru | | 0.505 | bias | 0.0033 |
| SOX9 | -9.114 | transcription | | -1.066 | bias | 0.00335 |
| STUB1 | -9.593 | enzyme | Activated | 2.488 | bias | 0.00336 |
| acetaminoph | | chemical dru | Inhibited | -3.837 | bias | 0.00345 |
| KDM5B | -10.461 | transcription | Activated | 2.633 | bias | 0.00349 |
| ruxolitinib | | chemical dru | Activated | 3 | bias | 0.00358 |
| SIRT2 | -8.447 | transcription | Inhibited | -2.132 | bias | 0.00358 |
| mir-144 | | microRNA | | 1.208 | bias | 0.00358 |
| SPRY1 | -6.361 | other | | 1.706 | bias | 0.00358 |
| TBXT | | transcription | Inhibited | -2.138 | bias | 0.00358 |
| cyanocobalar | | chemical - er | | 0.309 | | 0.00358 |
| Ige | | complex | Inhibited | -6.187 | bias | 0.00369 |
| phosphate | | chemical - er | | -1.914 | bias | 0.00373 |
| TIMP3 | -10.662 | other | Activated | 3.051 | bias | 0.00378 |
| TEAD2 | -7.874 | transcription | Inhibited | -3.464 | bias | 0.00378 |
| ACKR3 | -7.399 | G-protein co | | -1.131 | bias | 0.0039 |
| SRC | -4.649 | kinase | Inhibited | -3.433 | bias | 0.00391 |
| MASTL | 10.411 | kinase | Inhibited | -3.578 | bias | 0.00394 |
| miR-200b-3p | | mature micr | Activated | 2.778 | bias | 0.00394 |
| RUVBL1 | -8.959 | transcription | Inhibited | -3.317 | bias | 0.00394 |
| hexamethyle | | chemical rea | | 1.14 | | 0.00394 |
| ZFHX3 | -7.384 | transcription | | | | 0.00402 |
| DSCAML1 | -7.273 | other | Inhibited | -2.837 | bias | 0.00402 |
| miR-221-3p ( | | mature micr | Activated | 2.601 | bias | 0.00403 |
| branched cha | | chemical dru | Activated | 2.795 | bias | 0.00422 |

| | | | | | | |
|---|---|---|---|---|---|---|
| miR-199a-3p | | mature micr | Activated | 2.755 | bias | 0.00422 |
| PDGFC | -8.762 | growth facto | Inhibited | -2.773 | bias | 0.00422 |
| GNA12 | -10.045 | enzyme | Inhibited | -2.528 | bias | 0.00425 |
| ATG7 | -7.323 | enzyme | | -0.187 | | 0.00425 |
| (+)-MK-801 | | chemical dru | | -1.035 | | 0.00425 |
| chloroquine | | chemical dru | Inhibited | -3.115 | | 0.00437 |
| FGF1 | -4.928 | growth facto | | -1.073 | | 0.00439 |
| tributyltin | | chemical rea | | 0 | | 0.00439 |
| SMAD3 | -8.702 | transcription | Inhibited | -5.189 | bias | 0.00442 |
| NEUROD1 | | transcription | Inhibited | -2.308 | bias | 0.00458 |
| TEAD3 | -8.148 | transcription | Inhibited | -3.207 | bias | 0.00458 |
| mir-183 | | microRNA | Activated | 2.158 | | 0.00458 |
| HSPA5 | -6.08 | enzyme | | -0.105 | bias | 0.00458 |
| RHOA | -11.096 | enzyme | Inhibited | -2.736 | bias | 0.00461 |
| ursolic acid | | chemical dru | | 1.678 | | 0.00467 |
| HOXA9 | | transcription | | -0.896 | bias | 0.00468 |
| LIN28A | | other | Inhibited | -2.945 | bias | 0.00477 |
| NTF3 | -5.436 | growth facto | | -1.706 | bias | 0.00477 |
| prednisolone | | chemical dru | Activated | 3.449 | bias | 0.00477 |
| FZD8 | -8.861 | G-protein co | | -1.228 | | 0.0048 |
| p85 (pik3r) | | group | | -1.969 | bias | 0.0048 |
| MDM4 | -7.791 | transcription | | 1.979 | bias | 0.0048 |
| BARX2 | | transcription | Inhibited | -2.2 | bias | 0.0048 |
| acyclic retinc | | chemical dru | | 1.817 | bias | 0.0048 |
| acadesine | | chemical dru | | -0.314 | | 0.0048 |
| SPHK1 | -4.474 | kinase | Inhibited | -2.902 | bias | 0.00482 |
| INSIG1 | -9.475 | other | Activated | 3.864 | bias | 0.00482 |
| TRPM8 | | ion channel | | -1.287 | bias | 0.00486 |
| Sox2ot | | other | | -1.179 | bias | 0.00486 |
| MLH1 | -7.907 | enzyme | | -1.406 | bias | 0.00486 |
| ACTG1 | -15.034 | other | | 1.491 | bias | 0.00486 |
| S100P | | other | | | | 0.00486 |
| CYBA | -6.033 | enzyme | Inhibited | -2.236 | bias | 0.00486 |
| CBR-470-1 | | chemical - ki | | -1.342 | bias | 0.00486 |
| WNT1 | | cytokine | Inhibited | -3.463 | bias | 0.00488 |
| PRKAA1 | -6.249 | kinase | Activated | 3.826 | bias | 0.00491 |
| SB-431542 | | chemical rea | | 0.861 | | 0.00491 |
| KDM3A | -9.613 | transcription | Inhibited | -3.396 | bias | 0.005 |
| IRS2 | -3.023 | enzyme | | -0.572 | | 0.005 |
| ibuprofen | | chemical dru | Activated | 2.797 | bias | 0.005 |
| NCOA4 | -10.172 | transcription | | -0.981 | bias | 0.00504 |
| rosuvastatin | | chemical dru | Inhibited | -2.538 | | 0.00504 |
| betulinic acic | | chemical dru | | 0.765 | bias | 0.00504 |
| amphetamin | | chemical dru | Inhibited | -2.88 | bias | 0.00512 |
| PDGFB | -6.184 | growth facto | | -1.768 | bias | 0.00512 |

| | | | | | | |
|---|---|---|---|---|---|---|
| ELL2 | -4.582 | transcription | Inhibited | -3.742 | bias | 0.00513 |
| sorbitol | | chemical - er | | -0.64 | bias | 0.00515 |
| exisulind | | chemical - er | | -0.243 | | 0.00515 |
| dihydroartem | | chemical dru | | 0.7 | bias | 0.00515 |
| MFN2 | -9.798 | enzyme | Activated | 2.236 | bias | 0.00515 |
| Collagen Alpl | | group | | -1.457 | bias | 0.00515 |
| IDH2 | -9.75 | enzyme | | | | 0.00515 |
| DNAJC3 | -1.761 | other | | | | 0.00515 |
| F10 | | peptidase | Inhibited | -2.412 | bias | 0.00515 |
| RLN2 | | other | | -0.238 | | 0.00515 |
| LAMC1 | -10.482 | other | Activated | 2.431 | bias | 0.00515 |
| delta-aminol | | chemical - er | | -1.732 | bias | 0.00515 |
| SNCA | -9.031 | enzyme | | -0.098 | | 0.00525 |
| fenofibrate | | chemical dru | Inhibited | -2.649 | | 0.00535 |
| triamcinolon | | chemical dru | | 1.427 | | 0.00541 |
| miR-141-3p ( | | mature micr | Activated | 3.113 | bias | 0.00543 |
| paclitaxel | | chemical dru | | -0.068 | bias | 0.00557 |
| gentamicin C | | chemical dru | Inhibited | -3 | bias | 0.00559 |
| HTATIP2 | | transcription | | 1.387 | bias | 0.00575 |
| bleomycin | | biologic drug | Inhibited | -5.18 | bias | 0.00577 |
| aldesleukin | | biologic drug | Inhibited | -2.496 | bias | 0.00585 |
| atorvastatin | | chemical dru | Inhibited | -2.859 | | 0.00588 |
| androgen | | chemical dru | | 1.371 | | 0.00591 |
| DKK1 | -5.565 | growth facto | | 0.965 | bias | 0.00598 |
| ZEB1 | 10.286 | transcription | Inhibited | -2.419 | | 0.00601 |
| PTK2 | -8.869 | kinase | Inhibited | -2.305 | bias | 0.00608 |
| ETV5 | -5.226 | transcription | | 0.144 | | 0.00611 |
| MRTFA | -9.003 | transcription | Inhibited | -4.251 | | 0.00613 |
| everolimus | | chemical dru | | -0.804 | | 0.00615 |
| TRIM37 | -9.12 | enzyme | Activated | 3 | bias | 0.00625 |
| LIN28B | -5.473 | other | | -1.141 | bias | 0.00625 |
| apomorphine | | chemical dru | | -0.541 | | 0.00625 |
| CP-55940 | | chemical rea | | | | 0.00635 |
| monobutyl pl | | chemical tox | | -0.777 | | 0.00635 |
| PCGF2 | -8.371 | transcription | | 1.216 | | 0.00635 |
| C1QA | | other | | -0.64 | bias | 0.00641 |
| PPP3CA | -9.571 | phosphatase | | -0.933 | bias | 0.00653 |
| MMP1 | | peptidase | Inhibited | -2.76 | bias | 0.00658 |
| ADRB | | group | | 1.941 | bias | 0.0066 |
| cinnamaldeh | | chemical tox | | 0 | | 0.00671 |
| PPARA | | ligand-deper | Inhibited | -2.859 | | 0.00671 |
| ST6GALNAC1 | | enzyme | | -1.98 | bias | 0.00671 |
| BCAR4 | | other | | -1.98 | bias | 0.00671 |
| antisense oli | | biologic drug | | | | 0.00671 |
| MED28 | -7.337 | other | | 1.928 | bias | 0.00671 |

| | | | | | | |
|---|---|---|---|---|---|---|
| RAP1GAP | -6.954 | other | Activated | 2 | bias | 0.00671 |
| CBR3-AS1 | | other | | -1.999 | bias | 0.00671 |
| PHIP | -8.521 | other | | -1.963 | bias | 0.00671 |
| CCNA2 | -7.763 | other | | | | 0.00671 |
| LDHB | -12.176 | enzyme | | -1.982 | bias | 0.00671 |
| CCND2 | -11.407 | other | | | | 0.00671 |
| MAD2L2 | -7.904 | enzyme | | | | 0.00671 |
| NKX3-2 | -7.556 | transcription | | -1.982 | bias | 0.00671 |
| SENP2 | -8.493 | peptidase | | 1.951 | bias | 0.00671 |
| SLC6A2 | | transporter | | -1.067 | bias | 0.00671 |
| TNIK | -8.958 | kinase | | 0.152 | bias | 0.00671 |
| jasplakinolid | | chemical - er | | -1.954 | bias | 0.00671 |
| epigallocatec | | chemical dru | | 0.627 | bias | 0.00672 |
| methotrexat | | chemical dru | | -0.591 | | 0.00673 |
| Gli | | group | | | | 0.00678 |
| TFEC | | transcription | | | | 0.00678 |
| MNT | -8.903 | transcription | | | | 0.00678 |
| riboflavin | | chemical - er | | -1.134 | | 0.00678 |
| RUNX3 | 3.468 | transcription | Activated | 4.03 | bias | 0.0068 |
| ID1 | -8.214 | transcription | | -1.546 | bias | 0.00695 |
| JUNB | -8.574 | transcription | Inhibited | -2.206 | bias | 0.00699 |
| 2-bromoethy | | chemical rea | Inhibited | -2.883 | bias | 0.00706 |
| cerivastatin | | chemical dru | | -0.973 | | 0.00725 |
| PTGS2 | | enzyme | Inhibited | -2.626 | bias | 0.00729 |
| Creb | | group | Inhibited | -3.602 | bias | 0.00739 |
| L-alanine | | chemical - er | | | | 0.0075 |
| inecalcitol | | chemical dru | | | | 0.0075 |
| MARCHF5 | -9.101 | enzyme | | | | 0.0075 |
| FDX2 | -7.044 | transporter | | | | 0.0075 |
| DNAJB4 | -6.204 | other | | | | 0.0075 |
| LINC00887 | 2.458 | other | | | | 0.0075 |
| MUCL1 | | other | | | | 0.0075 |
| asparagine | | chemical - er | | | | 0.0075 |
| CTSD | -11.152 | peptidase | | | | 0.0075 |
| PSME3 | -9.627 | peptidase | | | | 0.0075 |
| FBLN2 | -6.842 | other | | | | 0.0075 |
| TIAM1 | | other | | | | 0.0075 |
| BOP1 | -9.421 | other | | | | 0.0075 |
| CHCHD4 | -5.941 | enzyme | | | | 0.0075 |
| AP2A1 | -9.863 | transporter | | | | 0.0075 |
| CHRM4 | -4.196 | G-protein co | | | | 0.0075 |
| CTNNAL1 | -7.691 | other | | | | 0.0075 |
| RNF139 | -8.508 | enzyme | | | | 0.0075 |
| SORL1 | -3.6 | transporter | | | | 0.0075 |
| SNRK | -7.671 | kinase | | | | 0.0075 |

| | | | | | | |
|---|---|---|---|---|---|---|
| FDXR | -7.519 | enzyme | | | | 0.0075 |
| MME | -6.017 | peptidase | | | | 0.0075 |
| PDGFRbeta-r | | chemical rea | | | | 0.0075 |
| benfur | | chemical rea | | | | 0.0075 |
| SRT2183 | | chemical rea | | | | 0.0075 |
| phenylacetic | | chemical - er | | | | 0.0075 |
| phenformin | | chemical dru | | | | 0.0075 |
| NSC719239 | | chemical dru | | | | 0.0075 |
| tipifarnib | | chemical dru | | | | 0.0075 |
| (-)-isoproter | | chemical dru | | | | 0.0075 |
| sertindole | | chemical dru | | | | 0.0075 |
| cyclohexaned | | chemical rea | | | | 0.0075 |
| L-asparagine | | chemical - er | | | | 0.0075 |
| AGN194204 | | chemical dru | Inhibited | -4.031 | bias | 0.00754 |
| THRAP3 | -9.608 | transcription | | 1.131 | bias | 0.00767 |
| TCF/LEF | | group | Inhibited | -2.169 | bias | 0.00767 |
| 48s | | complex | Inhibited | -2.433 | bias | 0.00767 |
| Nkx2-2os | | other | Inhibited | -2.425 | bias | 0.00767 |
| TIMP2 | -10.425 | other | | -1.085 | bias | 0.00767 |
| SPTLC2 | | enzyme | Activated | 2.449 | bias | 0.00767 |
| mir-322 | | microRNA | | 0.393 | bias | 0.00767 |
| ST6GAL1 | -8.781 | enzyme | Inhibited | -2.434 | bias | 0.00767 |
| leupeptin | | chemical - pr | Inhibited | -2.414 | bias | 0.00767 |
| oleoyl-estror | | chemical dru | | 1.937 | bias | 0.00768 |
| vanillin | | chemical - er | | 1.414 | | 0.00768 |
| JAK2 | -6.166 | kinase | Inhibited | -2.279 | bias | 0.00768 |
| MAPK9 | -8.436 | kinase | | -1.02 | bias | 0.00775 |
| JAG1 | -8.815 | growth facto | Inhibited | -2.71 | bias | 0.00781 |
| BRD4 | -9.757 | kinase | Inhibited | -2.915 | bias | 0.00787 |
| IRF4 | | transcription | | 1.714 | | 0.00793 |
| calcitriol | | chemical dru | | 0.768 | | 0.00795 |
| Irp | | group | | 0.927 | bias | 0.00798 |
| L-serine | | chemical - er | | 0.038 | bias | 0.00798 |
| ZFAS1 | -8.574 | other | | 0.328 | | 0.00798 |
| LINC-ROR | | other | | 0.555 | | 0.00798 |
| SART1 | -9.695 | other | | 0.447 | bias | 0.00798 |
| NEUROD2 | | transcription | | -1.131 | bias | 0.00798 |
| MMP2 | -10.767 | peptidase | | -0.629 | | 0.00804 |
| GDNF | | growth facto | | -1.655 | bias | 0.00804 |
| miR-133a-3p | | mature micr | Activated | 2.975 | bias | 0.00805 |
| XAV939 | | chemical rea | | | | 0.00805 |
| NPC2 | -4.48 | transporter | | 0.707 | bias | 0.00805 |
| GRN | -10.338 | growth facto | | 0.296 | | 0.00805 |
| epoxomicin | | chemical - pr | | 0.314 | bias | 0.00805 |
| RAB1B | -9.996 | other | Inhibited | -2.164 | bias | 0.00807 |

| | | | | | | |
|---|---|---|---|---|---|---|
| ANXA2 | -11.828 | other | | 1.188 | | 0.00807 |
| ezetimibe | | chemical dru | Inhibited | -3.133 | | 0.00807 |
| actinomycin | | biologic drug | Activated | 3.424 | bias | 0.00811 |
| 2-(4-amino-1 | | chemical rea | | 0.332 | | 0.00819 |
| CX3CL1 | -7.466 | cytokine | | -0.711 | | 0.00821 |
| bicalutamide | | chemical dru | Activated | 2.043 | bias | 0.00845 |
| FBXO32 | -8.472 | enzyme | | -1.588 | | 0.00853 |
| diphenylenei | | chemical rea | Activated | 2.921 | bias | 0.0089 |
| LMNA | -9.134 | other | | 1.531 | | 0.00891 |
| Tgf beta | | group | Inhibited | -5.398 | bias | 0.00902 |
| dacinostat | | chemical dru | | -0.21 | bias | 0.0093 |
| CDK1 | -7.908 | kinase | | -1.231 | bias | 0.0093 |
| PTP4A3 | -8.19 | phosphatase | | -0.863 | | 0.0093 |
| SMARCD3 | -8.868 | transcription | Inhibited | -2.63 | bias | 0.0093 |
| PKNOX1 | | transcription | | -1.964 | bias | 0.0093 |
| TFAM | -8.519 | transcription | | 0.378 | bias | 0.0093 |
| NAB2 | -9.336 | transcription | | 1.076 | bias | 0.0093 |
| NCD-38 | | chemical rea | | 0.44 | | 0.0093 |
| (1S,2R)-NCL- | | chemical rea | | 0.456 | | 0.0093 |
| THBS1 | -3.454 | other | | -0.224 | | 0.00961 |
| NR1D1 | -6.453 | ligand-deper | | -1.078 | | 0.00961 |
| sorafenib | | chemical dru | | 1.221 | bias | 0.00961 |
| cyclic AMP | | chemical - er | Inhibited | -3.99 | bias | 0.0098 |
| miR-204-5p ( | | mature micr | Activated | 3.357 | bias | 0.00983 |
| F2R | -7.135 | G-protein co | Inhibited | -4.239 | bias | 0.00984 |
| EZR | -9.126 | other | | 0.498 | | 0.01 |
| NGFR | -5.104 | transmembra | | -1.576 | bias | 0.01 |
| valsartan | | chemical dru | | 1.748 | bias | 0.01 |
| ziritaxestat | | chemical dru | | -1.387 | | 0.01 |
| SOX6 | -4.337 | transcription | | 0.039 | | 0.0101 |
| carvedilol | | chemical dru | | 1.498 | bias | 0.0101 |
| BMPR2 | -5.848 | kinase | | | | 0.0102 |
| leucine | | chemical - er | | 0.647 | | 0.0102 |
| PHB | -9.568 | transcription | | 1.872 | bias | 0.0102 |
| trabectedin | | chemical dru | | 0.822 | bias | 0.0102 |
| trovafloxacin | | chemical dru | | 0.5 | bias | 0.0103 |
| CG | | complex | Inhibited | -4.482 | bias | 0.0106 |
| PPARG | | ligand-deper | Inhibited | -2.525 | | 0.0108 |
| temozolomic | | chemical dru | | 0.136 | | 0.0109 |
| PTC-209 | | chemical rea | | 1.772 | | 0.0109 |
| heparin | | chemical - er | | -0.165 | | 0.0109 |
| N-Ac-Leu-Leu | | chemical - pr | | -1.911 | bias | 0.0109 |
| SMARCB1 | -9.597 | transcription | Inhibited | -2.639 | bias | 0.0109 |
| salubrinal | | chemical rea | | 1.741 | bias | 0.011 |
| OMA1 | -5.626 | peptidase | | -1.633 | | 0.011 |

| | | | | | | |
|---|---|---|---|---|---|---|
| DPH5 | -7.454 | enzyme | | | | 0.011 |
| NREP | -10.052 | other | | -1.706 | | 0.011 |
| PMP22 | -8.842 | other | | -0.075 | | 0.011 |
| POU3F2 | -7.991 | transcription | Inhibited | -2 | bias | 0.011 |
| ZNF148 | | transcription | | 0.161 | bias | 0.011 |
| JARID2 | -8.093 | transcription | | 0.9 | | 0.011 |
| mevastatin | | chemical dru | | -1.755 | bias | 0.011 |
| amlodipine | | chemical dru | | 1.633 | bias | 0.011 |
| GDF2 | | growth facto | Inhibited | -4.382 | bias | 0.0111 |
| allopurinol | | chemical dru | Inhibited | -2.535 | bias | 0.0111 |
| CXCR4 | -7.556 | G-protein co | | -1.711 | bias | 0.0114 |
| DACH1 | -8.186 | transcription | | 1.394 | bias | 0.0117 |
| CLDN7 | | other | | 1.53 | | 0.0118 |
| FASN | -12.67 | enzyme | | -0.414 | bias | 0.0119 |
| ANXA7 | -8.275 | ion channel | Inhibited | -2.034 | bias | 0.0119 |
| brefeldin A | | chemical - er | | 1.155 | | 0.0119 |
| L-triiodothyro | | chemical - er | Inhibited | -2.915 | bias | 0.0119 |
| N-Cadherin | | group | | | | 0.0121 |
| Retinoic acid | | complex | Inhibited | -2 | bias | 0.0121 |
| CACNA1C | -7.819 | ion channel | | | | 0.0121 |
| hydrogel | | chemical dru | | | | 0.0121 |
| MIA | | other | | -1.981 | bias | 0.0121 |
| MAP3K12 | -8.382 | kinase | | -1.165 | bias | 0.0121 |
| HMGCR | -11.57 | enzyme | | -1 | | 0.0121 |
| ERCC4 | -3.831 | enzyme | | -1 | bias | 0.0121 |
| miR-192-5p ( | | mature micro | | -0.106 | bias | 0.0121 |
| mir-326 | | microRNA | | 1.982 | bias | 0.0121 |
| EFEMP1 | | enzyme | | 1.067 | | 0.0121 |
| LAMA4 | -6.619 | enzyme | | -0.339 | | 0.0121 |
| SDC2 | -3.857 | other | | -1.4 | bias | 0.0121 |
| PPM1A | -9.334 | phosphatase | Inhibited | -2 | bias | 0.0121 |
| BLACAT1 | -5.541 | other | | -1.969 | bias | 0.0121 |
| protocatechu | | chemical - er | | | | 0.0121 |
| TCF4 | -7.731 | transcription | | -1.16 | | 0.0121 |
| ADCYAP1 | -7.231 | other | Inhibited | -2.16 | | 0.0121 |
| plicamycin | | chemical dru | Activated | 2.703 | bias | 0.0122 |
| corticosteroi | | chemical dru | | | | 0.0122 |
| miR-199a-5p | | mature micro | Activated | 4.068 | bias | 0.0122 |
| SOX2-OT | -5.735 | other | | -1.385 | bias | 0.0123 |
| LAMTOR1 | -8.876 | other | Inhibited | -2.236 | bias | 0.0123 |
| JMJD6 | -6.839 | transmembra | | -0.218 | | 0.0123 |
| CCAT1 | | other | | -0.447 | | 0.0123 |
| CIC | -10.354 | transcription | | 1.387 | bias | 0.0123 |
| miR-451a (ar | | mature micro | Activated | 2.214 | bias | 0.0123 |
| CLCA2 | | ion channel | | 0.692 | bias | 0.0123 |

| | | | | | | |
|---|---|---|---|---|---|---|
| DBI | -8.716 | other | Inhibited | -2.236 | bias | 0.0123 |
| ITGA3 | -9.352 | other | | | | 0.0123 |
| KRIT1 | -6.443 | other | | -0.447 | bias | 0.0123 |
| azetidyl-2-ca | | chemical rea | Inhibited | -2.219 | bias | 0.0123 |
| ferric ammo | | chemical dru | | -0.421 | | 0.0123 |
| putrescine | | chemical - e | | -1.387 | bias | 0.0123 |
| zinostatin | | biologic drug | Inhibited | -2.219 | bias | 0.0123 |
| zalcitabine | | chemical dru | | -1.342 | bias | 0.0123 |
| miR-92a-3p ( | | mature micr | Activated | 2.393 | bias | 0.0123 |
| SUZ12 | -8.461 | enzyme | | | | 0.0124 |
| FUT8 | -8.636 | enzyme | | 1.195 | bias | 0.0124 |
| AChR | | complex | | | | 0.0124 |
| MK2206 | | chemical dru | | 1.937 | bias | 0.0124 |
| SPOP | -8.371 | other | | | | 0.0124 |
| FBN1 | -3.58 | other | Activated | 2.433 | bias | 0.0124 |
| SIN3B | -9.72 | transcription | | | | 0.0124 |
| PPP1R15A | -9.762 | other | | -0.762 | | 0.0124 |
| ST8SIA1 | -8.581 | enzyme | | -1.254 | | 0.0124 |
| MDM2 | -9.316 | transcription | | -1.102 | | 0.0124 |
| REST | -7.21 | transcription | Activated | 4.041 | bias | 0.0124 |
| PLAG1 | | transcription | Inhibited | -2.76 | bias | 0.0126 |
| caspase | | group | Inhibited | -2.621 | bias | 0.0128 |
| NUP98-HOXA | | fusion gene/ | | | | 0.0128 |
| CHI3L1 | -4.523 | enzyme | | | | 0.0129 |
| calyculin A | | chemical tox | | -0.707 | | 0.0129 |
| methyl-beta- | | chemical dru | | -0.816 | | 0.0129 |
| IgG | | complex | Activated | 4.017 | bias | 0.0132 |
| 3-deazanepla | | chemical dru | Inhibited | -3.194 | bias | 0.0132 |
| miR-182-5p ( | | mature micr | Activated | 3.843 | bias | 0.0132 |
| CD247 | | transmembra | Inhibited | -3.873 | bias | 0.0132 |
| ERK | | group | Inhibited | -5.391 | bias | 0.0133 |
| PTH | | other | Inhibited | -2.854 | bias | 0.0133 |
| Histone h3 | | group | | | | 0.0135 |
| miR-21-5p (a | | mature micr | Activated | 3.373 | bias | 0.0136 |
| CASR | | G-protein co | Inhibited | -3.014 | bias | 0.0136 |
| tetracycline | | chemical dru | | -0.336 | | 0.0139 |
| phenacetin | | chemical dru | Inhibited | -2.72 | bias | 0.014 |
| SAHM1 | | chemical rea | Activated | 2.714 | bias | 0.014 |
| haloperidol | | chemical dru | | -1.813 | bias | 0.014 |
| cardiotoxin | | chemical - ot | Inhibited | -3.302 | bias | 0.0142 |
| PRKCA | -2.532 | kinase | Inhibited | -2.501 | bias | 0.0143 |
| ESRRA | -6.983 | transcription | Inhibited | -4.43 | bias | 0.0145 |
| ZAP70 | | kinase | Inhibited | -3.713 | bias | 0.0147 |
| ABCB1 | | transporter | Activated | 2.534 | bias | 0.015 |
| CUL4B | -9.213 | other | Activated | 3 | bias | 0.015 |

| | | | | | | |
|---|---|---|---|---|---|---|
| picropodophy | | chemical dru | | 1.709 | bias | 0.0151 |
| Hmgb1 | | transcription | | -1.342 | bias | 0.0151 |
| KDM4A | -9.792 | transcription | | -1 | bias | 0.0151 |
| mir-137 | | microRNA | | 1.71 | bias | 0.0151 |
| mir-194 | | microRNA | Activated | 2.233 | bias | 0.0151 |
| DDR1 | -10.745 | kinase | | -0.798 | bias | 0.0151 |
| ARG1 | | enzyme | Activated | 2.433 | bias | 0.0151 |
| DDX17 | -11.839 | enzyme | | -1.117 | bias | 0.0151 |
| DDX3X | -10.814 | enzyme | | -1.761 | bias | 0.0151 |
| ECM1 | -6.042 | transporter | | 0.008 | bias | 0.0151 |
| ethidium | | chemical dru | | | | 0.0151 |
| HOXD10 | | transcription | | 1.846 | | 0.0152 |
| NEUROG1 | -4.636 | transcription | | 0.535 | | 0.0152 |
| PRMT1 | -10.47 | enzyme | Inhibited | -2.057 | bias | 0.0152 |
| TOB1 | -8.193 | transcription | | -1.388 | | 0.0153 |
| RAS | | group | | -1.857 | bias | 0.0153 |
| DSCAM | 9.688 | other | Inhibited | -3.144 | bias | 0.0156 |
| TFAP2C | -6.409 | transcription | | -0.039 | | 0.0157 |
| RUNX2 | -4.942 | transcription | | -0.353 | bias | 0.0158 |
| ESRRG | -3.216 | ligand-deper | | -0.682 | | 0.0158 |
| CNTF | | cytokine | | 0.04 | | 0.0158 |
| Stat3-Stat3 | | complex | | -1.614 | bias | 0.0159 |
| ALDH1A2 | -8.015 | enzyme | Activated | 2.333 | bias | 0.0159 |
| mir-23 | | microRNA | | -0.07 | bias | 0.0159 |
| NF1 | -7.821 | other | Activated | 2.156 | bias | 0.0159 |
| selenium | | chemical dru | Inhibited | -2.753 | | 0.0159 |
| N-acetyl-L-cy | | chemical dru | Activated | 4.183 | bias | 0.0159 |
| NFYB | -5.694 | transcription | | | | 0.0162 |
| Npm | | group | | | | 0.0163 |
| 4-aminopyra | | chemical rea | | | | 0.0163 |
| loganin | | chemical - er | | | | 0.0163 |
| Alpha 1 antit | | group | | | | 0.0163 |
| opioid recept | | group | | | | 0.0163 |
| ARHGEF12 | -7.837 | other | | | | 0.0163 |
| chartreusin | | chemical rea | | | | 0.0163 |
| MBNL3 | | other | | | | 0.0163 |
| PLEKHM1 | -7.179 | other | | | | 0.0163 |
| TCHP | | other | | | | 0.0163 |
| USP42 | | peptidase | | | | 0.0163 |
| RBM17 | -9.17 | other | | | | 0.0163 |
| CERS3 | | transcription | | | | 0.0163 |
| RTN3 | -11.904 | other | | | | 0.0163 |
| SLC9A9 | -4.141 | transporter | | | | 0.0163 |
| SETX | -9.151 | enzyme | | | | 0.0163 |
| BATF2 | | transcription | | | | 0.0163 |

| | | | |
|---|---|---|---|
| NAT8 | | transcription | 0.0163 |
| TLN2 | -5.022 | other | 0.0163 |
| MLXIP | -3.422 | transcription | 0.0163 |
| TMEM106B | -9.001 | other | 0.0163 |
| MIR31HG | 7.631 | other | 0.0163 |
| BACE1-AS | | other | 0.0163 |
| PX 478 | | chemical dru | 0.0163 |
| LOXL1 | -8.686 | enzyme | 0.0163 |
| EIF4H | -11.301 | translation re | 0.0163 |
| CTS-1027 | | chemical dru | 0.0163 |
| SH3GLB1 | -9.428 | enzyme | 0.0163 |
| MCF2L | -7.845 | other | 0.0163 |
| ATP1A1 | -11.123 | transporter | 0.0163 |
| CNGA2 | | ion channel | 0.0163 |
| SNX6 | -5.987 | transporter | 0.0163 |
| NEFH | -8.417 | other | 0.0163 |
| CELA1 | | peptidase | 0.0163 |
| DPP10-AS1 | | other | 0.0163 |
| ELOC | -8.484 | transcription | 0.0163 |
| MLIP | | transcription | 0.0163 |
| ACP3 | | phosphatase | 0.0163 |
| TJP1 | -7.231 | other | 0.0163 |
| ELP3 | -8.52 | enzyme | 0.0163 |
| miR-483-5p ( | | mature micr | 0.0163 |
| miR-548h-5p | | mature micr | 0.0163 |
| miR-3189-3p | | mature micr | 0.0163 |
| mir-630 | | microRNA | 0.0163 |
| NUBP1 | -6.89 | other | 0.0163 |
| IDE | -7.313 | peptidase | 0.0163 |
| ITM2B | -11.236 | other | 0.0163 |
| NASP | -10.038 | other | 0.0163 |
| PLPP3 | -8.795 | phosphatase | 0.0163 |
| Dst | | other | 0.0163 |
| NEK4 | 9.84 | kinase | 0.0163 |
| RPS6KA2 | -9.66 | kinase | 0.0163 |
| KHDRBS2 | -5.667 | other | 0.0163 |
| MTHFD2 | -8.938 | enzyme | 0.0163 |
| DPH1 | -8.292 | other | 0.0163 |
| ADAMTS5 | -4.341 | peptidase | 0.0163 |
| FRAT1 | -5.779 | other | 0.0163 |
| RGS20 | | enzyme | 0.0163 |
| MID1 | -8.657 | other | 0.0163 |
| CGGBP1 | -9.827 | transcription | 0.0163 |
| SNRPE | -3.75 | other | 0.0163 |
| NAT8B | | enzyme | 0.0163 |

| | | | | | |
|---|---|---|---|---|---|
| VPS26B | -10.414 | transporter | | | 0.0163 |
| RPS4Y1 | -6.648 | other | | | 0.0163 |
| nexrutine | | chemical dru | | | 0.0163 |
| SEPTIN5 | -10.503 | enzyme | | | 0.0163 |
| MAP4 | -11.264 | other | | | 0.0163 |
| RPS19 | -11.665 | other | | | 0.0163 |
| indotecan | | chemical dru | | | 0.0163 |
| sorafenib de | | chemical rea | | | 0.0163 |
| gamitrinib-T | | chemical rea | | | 0.0163 |
| buformin | | chemical dru | | | 0.0163 |
| telomestatin | | chemical - er | | | 0.0163 |
| inostamycin | | chemical rea | | | 0.0163 |
| clorgyline | | chemical dru | | | 0.0163 |
| 2-hydroxy-im | | chemical rea | | | 0.0163 |
| Z-IL-CHO | | chemical - pr | | | 0.0163 |
| glycoconjuga | | chemical - ot | | | 0.0163 |
| N(6)-carboxy | | chemical - er | | | 0.0163 |
| 2-keto-4-me | | chemical - er | | | 0.0163 |
| DOT1L | -3.766 | phosphatase | | | 0.0163 |
| BPIFB1 | | other | 1.709 | bias | 0.0163 |
| miR-7a-5p (a | | mature micr Activated | 2.605 | bias | 0.0163 |
| miR-218-5p ( | | mature micr | 1.793 | bias | 0.0163 |
| TRPC1 | | ion channel | -0.478 | | 0.0163 |
| HDAC4 | -1.872 | transcription | -0.617 | | 0.0163 |
| CTSB | -10.918 | peptidase | -1.805 | bias | 0.0164 |
| CPE | -11.835 | peptidase | 1.718 | bias | 0.0164 |
| WR 1065 | | chemical dru Inhibited | -2.184 | bias | 0.0164 |
| pituitary ade | | biologic drug | -1.131 | | 0.0167 |
| Cdc42 | | enzyme Inhibited | -3.302 | bias | 0.0167 |
| CDK2AP1 | -9.194 | other | | | 0.0167 |
| NOTCH3 | -2.823 | transcription Inhibited | -2.231 | bias | 0.0169 |
| PAX5 | | transcription | 1.811 | | 0.0169 |
| PTGES | | enzyme | 1.177 | bias | 0.0169 |
| L-aspartic ac | | chemical - er | | | 0.017 |
| MELK | -5.003 | kinase | | | 0.017 |
| PFDN6 | -7.035 | other | | | 0.017 |
| ZMIZ2 | -9.794 | transcription | | | 0.017 |
| RNF5 | -9.549 | enzyme | | | 0.017 |
| PHF6 | -8.522 | transcription | | | 0.017 |
| Cathepsin | | group | | | 0.017 |
| SEMA3F | -8.532 | other | | | 0.017 |
| MCF2 | | other | | | 0.017 |
| EAF2 | | transcription | | | 0.017 |
| miR-382-5p ( | | mature micr | | | 0.017 |
| miR-431-3p ( | | mature micr | | | 0.017 |

| | | | | | | |
|---|---|---|---|---|---|---|
| WSB1 | -11.334 | other | | | | 0.017 |
| FLOT1 | -10.551 | other | | | | 0.017 |
| NRP2 | -9.667 | kinase | | | | 0.017 |
| KCTD11 | -7.726 | other | | | | 0.017 |
| PORCN | -7.459 | enzyme | | | | 0.017 |
| DCX | -11.767 | other | | | | 0.017 |
| H19 | -10.819 | other | | | | 0.017 |
| NME3 | -8.218 | kinase | | | | 0.017 |
| BBC3 | -8.308 | other | | | | 0.017 |
| NUBPL | -6.935 | other | | | | 0.017 |
| PSMB5 | -9.678 | peptidase | | | | 0.017 |
| talazoparib | | chemical dru | | | | 0.017 |
| Cox5b | | other | | | | 0.017 |
| CLDN2 | | other | | | | 0.017 |
| tideglusib | | chemical dru | | | | 0.017 |
| 2,2'-dipyridyl | | chemical tox | | | | 0.017 |
| vinyl carbam | | chemical tox | | | | 0.017 |
| Collagen type | | complex | | | | 0.017 |
| setanaxib | | chemical dru | | | | 0.017 |
| mini-GAGR | | chemical rea | | | | 0.017 |
| angustibalin | | chemical - e | | | | 0.017 |
| Cr6+ | | chemical rea | | | | 0.017 |
| MYOD1 | | transcription | Inhibited | -2.023 | bias | 0.0173 |
| CD40LG | | cytokine | Inhibited | -4.323 | bias | 0.0174 |
| TO-901317 | | chemical rea | Inhibited | -2.776 | bias | 0.0178 |
| PP2/AG1879 | | chemical - ki | Activated | 3.369 | bias | 0.0178 |
| PTTG1 | -7.042 | transcription | | 0.441 | bias | 0.0178 |
| AMPK | | complex | | -0.462 | | 0.0178 |
| digoxin | | chemical dru | | -0.2 | | 0.0179 |
| trypsin | | group | | -1.067 | bias | 0.0179 |
| FOXD2-AS1 | | other | | -1.336 | bias | 0.0179 |
| POLR3G | -5.143 | enzyme | | -0.447 | bias | 0.0179 |
| Laminin1 | | complex | | | | 0.0179 |
| SOX5 | -7.977 | transcription | | -1.387 | bias | 0.0179 |
| L1CAM | -11.828 | other | Inhibited | -2 | bias | 0.0179 |
| MIRLET7 | | group | | 1 | bias | 0.0179 |
| MED16 | -7 | transcription | | 1.987 | bias | 0.0179 |
| BCAR1 | -9.193 | enzyme | | | | 0.0179 |
| DDR2 | -7.364 | kinase | | -1.526 | bias | 0.0179 |
| USP8 | -7.422 | peptidase | | -0.927 | bias | 0.0179 |
| TRRAP | -9.316 | transcription | | | | 0.0179 |
| AK1 | -9.118 | kinase | | | | 0.0179 |
| ATXN3 | 10.499 | peptidase | | | | 0.0179 |
| CYP51A1 | -10.682 | enzyme | Activated | 2.236 | bias | 0.0179 |
| APOC3 | | transporter | Inhibited | -2.236 | bias | 0.0179 |

| | | | | | | |
|---|---|---|---|---|---|---|
| adefovir dipi |  | chemical dru | Activated | 2.2 | bias | 0.0179 |
| etoposide |  | chemical dru | Inhibited | -2.088 | bias | 0.018 |
| SP4 | -7.069 | transcription |  | -1.446 | bias | 0.018 |
| rasagiline |  | chemical dru |  | -1.609 | bias | 0.0181 |
| PAK2 | -3.5 | kinase |  | 1.897 |  | 0.0181 |
| KIN001-043 |  | chemical - ki |  | 1.117 |  | 0.0181 |
| S-adenosylm |  | chemical - er |  | 1.943 | bias | 0.0181 |
| corticosteror |  | chemical - er |  | -1.083 |  | 0.0188 |
| CDK9 | -9.236 | kinase |  |  |  | 0.0189 |
| ELAVL1 | -9.998 | other | Inhibited | -2.977 | bias | 0.0193 |
| BMS-690514 |  | chemical dru | Activated | 2.333 | bias | 0.0194 |
| OVA-8 |  | chemical rea |  |  |  | 0.0194 |
| RUNX1-RUN |  | fusion gene/ |  | -1.98 |  | 0.0194 |
| FOXC2 | -6.789 | transcription | Inhibited | -2.79 | bias | 0.0195 |
| INHBA | -6.052 | growth facto | Inhibited | -2.74 | bias | 0.0195 |
| TGFBR |  | group |  | -1.067 | bias | 0.0195 |
| BMS-387032 |  | chemical dru |  | 1.952 | bias | 0.0195 |
| FA2H |  | enzyme |  | 1.214 |  | 0.0195 |
| LRRK2 |  | kinase |  |  |  | 0.0195 |
| AFF4 | -8.268 | transcription |  |  |  | 0.0195 |
| MEMO1 | -7.676 | other |  | -1.981 | bias | 0.0195 |
| CPEB4 | -8.581 | other |  |  |  | 0.0195 |
| Presenilin |  | group |  |  |  | 0.0195 |
| SPTAN1 | -12.584 | other |  |  |  | 0.0195 |
| AGRN | -11.503 | other |  | -1.97 | bias | 0.0195 |
| ZIC2 | -3.303 | transcription |  | -1 |  | 0.0195 |
| TPM3 | -10.3 | other | Activated | 2 | bias | 0.0195 |
| mir-338 |  | microRNA |  | 1.993 | bias | 0.0195 |
| mir-497 |  | microRNA |  | 1.939 | bias | 0.0195 |
| GUSB | -7.575 | enzyme |  |  |  | 0.0195 |
| UBQLN1 | -10.59 | other |  | 0 |  | 0.0195 |
| MFGE8 | -9.608 | other |  | 0.152 |  | 0.0195 |
| DIAPH1 | -8.971 | other |  | -1.98 |  | 0.0195 |
| TSG101 | -4.297 | transcription |  | 0.218 | bias | 0.0195 |
| PPP2R1A | -12.095 | phosphatase |  | -1.98 |  | 0.0195 |
| YBX3 | -9.104 | transcription |  |  |  | 0.0195 |
| TKT | -11.275 | enzyme |  | -1.98 | bias | 0.0195 |
| GNRH2 |  | other |  | -0.152 | bias | 0.0195 |
| SERPINH1 | -5.621 | other |  | -1.972 | bias | 0.0195 |
| SLC30A3 |  | transporter |  | -1.982 | bias | 0.0195 |
| thiocoraline |  | chemical rea |  | -0.283 | bias | 0.0195 |
| raltitrexed |  | chemical dru |  | -1 | bias | 0.0195 |
| 7,8-dihydro-7 |  | chemical tox |  | -1.969 | bias | 0.0195 |
| MK2 inhibito |  | chemical - ki |  | 1.972 | bias | 0.0195 |
| SP600125 |  | chemical - ki | Activated | 3.924 | bias | 0.0197 |

| | | | | | | |
|---|---|---|---|---|---|---|
| NAMPT | -9.007 | cytokine | | -1.763 | bias | 0.0198 |
| ZNF106 | -8.889 | other | Activated | 2.111 | | 0.0198 |
| TNC | -5.511 | other | | -0.047 | bias | 0.0198 |
| verapamil | | chemical dru | | 1.037 | bias | 0.0198 |
| MM-589 | | chemical rea | Activated | 2.137 | | 0.0198 |
| CXCL8 | | cytokine | | -1.715 | bias | 0.0199 |
| 3,4,5,3',4'-pe | | chemical tox | | -1.766 | bias | 0.0199 |
| mycophenoli | | chemical dru | Inhibited | -2.254 | bias | 0.0199 |
| cycloheximid | | chemical rea | | 1.252 | | 0.02 |
| 15(S)-HETE | | chemical - er | | -0.995 | | 0.0203 |
| glycine | | chemical - er | | -0.928 | | 0.0203 |
| AURKA | 3.34 | kinase | | 0.498 | | 0.0203 |
| DEF6 | | other | | -1.708 | bias | 0.0203 |
| miR-100-5p ( | | mature micr | | 1.725 | bias | 0.0203 |
| miR-205-5p ( | | mature micr | | 1.432 | bias | 0.0203 |
| SLC6A3 | | transporter | | -1.387 | | 0.0203 |
| XPO1 | -11.056 | transporter | | | | 0.0203 |
| latrunculin B | | chemical rea | | 1.664 | bias | 0.0203 |
| Mir122a,b | | group | | 1.537 | | 0.0204 |
| TMPRSS2-ER | | fusion gene/ | | -1.027 | | 0.0208 |
| SEMA7A | -8.156 | transmembra | Inhibited | -2.646 | bias | 0.0209 |
| Pde4 | | group | | 0.956 | bias | 0.0209 |
| ALDH2 | -9.571 | enzyme | Activated | 2.63 | bias | 0.0209 |
| FGF4 | | growth facto | Inhibited | -2.376 | bias | 0.0209 |
| FOXF2 | -6.076 | transcription | | 1.309 | bias | 0.0209 |
| AURK | | group | Activated | 3.45 | bias | 0.021 |
| ANLN | -7.748 | other | Activated | 3.464 | bias | 0.021 |
| LRP1 | -5.518 | transmembra | | 1.077 | | 0.021 |
| cytochalasin | | chemical tox | | 1.718 | bias | 0.021 |
| FOXO1 | -7.493 | transcription | Inhibited | -3.473 | bias | 0.0214 |
| SET | -12.094 | phosphatase | Activated | 2.333 | bias | 0.0216 |
| ethionine | | chemical tox | | -1.897 | bias | 0.0216 |
| BIRC5 | -7.23 | other | | | | 0.0218 |
| CLEC11A | -6.348 | growth facto | | | | 0.0218 |
| TGFB2 | -9.635 | growth facto | Inhibited | -4.13 | bias | 0.0221 |
| EGLN | | group | Activated | 4.623 | bias | 0.0223 |
| SIRT6 | -7.763 | enzyme | Activated | 2.828 | bias | 0.0223 |
| BACH1 | -8.223 | transcription | Activated | 3.208 | bias | 0.0223 |
| PIN1 | -9.522 | enzyme | Inhibited | -2.184 | bias | 0.0225 |
| TBP | -7.448 | transcription | | | | 0.0226 |
| 3,3'-diindolyl | | chemical dru | | 0.475 | | 0.0226 |
| Brd4 | | kinase | Inhibited | -2.883 | bias | 0.0226 |
| benzo(a)pyre | | chemical tox | Inhibited | -2.537 | bias | 0.0232 |
| Collagen(s) | | complex | | -0.396 | bias | 0.0233 |
| A2M | -10.849 | transporter | Inhibited | -3.176 | bias | 0.0233 |

| | | | | | | |
|---|---|---|---|---|---|---|
| GPX1 | -10.115 | enzyme | | 1.555 | bias | 0.0233 |
| U73122 | | chemical rea | Activated | 3.286 | bias | 0.0233 |
| genistein | | chemical dru | Inhibited | -2.173 | | 0.0234 |
| Calcineurin A | | group | | -0.956 | bias | 0.0235 |
| SOX17 | | transcription | | 0.308 | | 0.0235 |
| FGFR4 | -5.02 | kinase | | -0.418 | bias | 0.0235 |
| daunorubicin | | chemical dru | | -0.869 | | 0.0235 |
| RGS2 | -7.598 | enzyme | Activated | 2.236 | bias | 0.0239 |
| EBF1 | -6.408 | transcription | Inhibited | -4.168 | bias | 0.0241 |
| metformin | | chemical dru | Activated | 2.506 | | 0.0242 |
| TFEB | -4.473 | transcription | Inhibited | -3.392 | bias | 0.0244 |
| NFYA | -8.708 | transcription | Inhibited | -3.405 | bias | 0.0249 |
| SMO | -9.497 | G-protein co | Inhibited | -2 | bias | 0.0249 |
| U46619 | | chemical rea | | -1 | bias | 0.0249 |
| COL9A1 | -8.438 | other | | 1.387 | | 0.0249 |
| YWHAG | -12.506 | other | | | | 0.0249 |
| NDUFA13 | -10.189 | enzyme | | | | 0.0249 |
| PTBP1 | -10.342 | enzyme | | 0.49 | | 0.0249 |
| SRSF3 | -10.673 | other | | -1.498 | bias | 0.0249 |
| MGAT3 | -9.898 | enzyme | | -0.562 | bias | 0.0249 |
| SAT1 | -8.693 | enzyme | | 0.106 | bias | 0.0249 |
| COL6A1 | -10.691 | other | | -0.632 | | 0.0249 |
| CAMKK2 | 9.927 | kinase | | -0.651 | bias | 0.0249 |
| ID4 | -10.807 | transcription | | | | 0.0249 |
| aloxistatin | | chemical - pr | | -0.2 | | 0.0249 |
| manganese | | chemical - er | | -1.432 | bias | 0.0249 |
| glucagon | | biologic drug | | -0.653 | bias | 0.025 |
| methylnitron | | chemical tox | Inhibited | -2.183 | bias | 0.0251 |
| letrozole | | chemical dru | | 1.238 | | 0.0251 |
| TNF | | cytokine | Inhibited | -3.278 | bias | 0.0251 |
| E2F6 | 9.75 | transcription | | 1.342 | bias | 0.0254 |
| Raf | | group | | -1.007 | bias | 0.0255 |
| F3 | -7.639 | transmembra | Inhibited | -2.364 | bias | 0.0255 |
| EFNA1 | -5.178 | other | | -0.535 | | 0.0255 |
| BCL6 | -6.678 | transcription | | -0.125 | | 0.0256 |
| PLAUR | 3.716 | transmembra | Inhibited | -2.37 | bias | 0.0257 |
| ABL1 | -9.724 | kinase | | -0.958 | bias | 0.0257 |
| INHA | -4.448 | growth facto | | 0.127 | | 0.0257 |
| ATF3 | -7.475 | transcription | | -1.595 | | 0.0257 |
| Jnk | | group | Inhibited | -3.293 | bias | 0.0259 |
| CAV1 | -6.882 | transmembra | | -0.089 | bias | 0.0261 |
| PDGF-AA | | complex | | -1.98 | bias | 0.0263 |
| UHRF2 | | enzyme | Inhibited | -2.646 | bias | 0.0263 |
| MLX | -7.584 | transcription | Inhibited | -2.357 | bias | 0.0263 |
| LCAT | -6.659 | enzyme | | 1.8 | | 0.0263 |

| | | | | | | |
|---|---|---|---|---|---|---|
| CYP7A1 | | enzyme | | -1.912 | | 0.0263 |
| THZ2 | | chemical - ki | | 1.912 | bias | 0.0263 |
| PARP1 | -10.785 | enzyme | | -0.722 | bias | 0.0265 |
| CREBZF | -8.244 | transcription | Activated | 2.401 | bias | 0.0266 |
| HNF1A-AS1 | 3.418 | other | | -1.039 | bias | 0.0266 |
| CCKBR | -5.141 | G-protein co | | | | 0.0266 |
| EPHA2 | -8.79 | kinase | | 0.11 | | 0.0266 |
| MUC4 | | other | Inhibited | -2.332 | bias | 0.0266 |
| ethosuximide | | chemical dru | Inhibited | -2.449 | bias | 0.0266 |
| thioridazine | | chemical dru | | -0.447 | bias | 0.0266 |
| leuprolide | | biologic drug | | 0.056 | | 0.0266 |
| ACOX1 | -1.917 | enzyme | Activated | 3.68 | | 0.0271 |
| S100A4 | | other | | -0.999 | bias | 0.0272 |
| TERC | | other | | | | 0.0281 |
| UPF1 | -9.926 | enzyme | | 1.687 | bias | 0.0281 |
| mir-17 | | microRNA | Activated | 3.135 | bias | 0.0282 |
| PTHLH | -4.674 | other | | -1.868 | bias | 0.0282 |
| MSTN | | growth facto | | -1.113 | | 0.0288 |
| CHD4 | -10.893 | enzyme | | 1.286 | bias | 0.0288 |
| FGFR2 | -8.866 | kinase | | -0.096 | | 0.0293 |
| HULC | | other | | -1 | bias | 0.0293 |
| PARP | | group | | | | 0.0293 |
| cercosporin | | chemical - er | | 1.982 | bias | 0.0293 |
| DCLK1 | -10.632 | kinase | | -1.117 | bias | 0.0293 |
| PDGFA | 4.116 | growth facto | | -1.964 | bias | 0.0293 |
| mir-185 | | microRNA | | 1.969 | bias | 0.0293 |
| NMNAT1 | -6.027 | enzyme | | -0.898 | bias | 0.0293 |
| LAMP2 | -8.094 | enzyme | | -0.747 | | 0.0293 |
| N4BP1 | -6.436 | other | | 0.232 | | 0.0293 |
| ANGPTL3 | | growth facto | | | | 0.0293 |
| Igkv1-117 | | other | | | | 0.0293 |
| TP53BP2 | -8.44 | other | | -0.347 | bias | 0.0293 |
| COPS8 | -9.325 | other | | 0 | bias | 0.0293 |
| calmidazoliu | | chemical dru | | 1 | bias | 0.0293 |
| selenomethi | | chemical - er | | 0.81 | | 0.0293 |
| E2F3 | -6.916 | transcription | | 0.331 | bias | 0.0296 |
| FGF8 | | growth facto | Inhibited | -2.179 | bias | 0.0299 |
| okadaic acid | | chemical tox | Inhibited | -2.461 | | 0.0299 |
| PP1 | | chemical - ki | | 1.961 | bias | 0.0302 |
| T3-TR-RXR | | complex | | -1.803 | bias | 0.0302 |
| F7 | | peptidase | Inhibited | -3.082 | bias | 0.0302 |
| 4-methylnitr | | chemical tox | | -1.657 | bias | 0.0302 |
| LPIN1 | -8.945 | phosphatase | | -0.662 | bias | 0.0305 |
| BAK1 | -7.462 | other | | -1.82 | bias | 0.0305 |
| GCK | | kinase | | 0.563 | | 0.0305 |

| | | | | | | |
|---|---|---|---|---|---|---|
| PRKCI | -9.107 | kinase | | -1.091 | bias | 0.0305 |
| FAAH | | enzyme | | 1.673 | bias | 0.0305 |
| ferulic acid | | chemical - er | | 1.58 | bias | 0.0305 |
| carbenoxolor | | chemical tox | | 1.257 | bias | 0.0305 |
| miR-17-5p (a | | mature micr | Activated | 3.568 | bias | 0.0307 |
| Rp-8-pCPT-c( | | chemical rea | | | | 0.0307 |
| C9orf116 | -6.061 | other | | | | 0.0307 |
| F Actin | | complex | | | | 0.0307 |
| LINC00842 | | other | | | | 0.0307 |
| NSD3 | -8.428 | enzyme | | | | 0.0307 |
| CPXM1 | -9.591 | peptidase | | | | 0.0307 |
| ZMYND10 | -8.007 | other | | | | 0.0307 |
| lumefantrine | | chemical dru | | | | 0.0307 |
| LDL-choleste | | complex | | | | 0.0307 |
| PFDN5 | -6.983 | transcription | | | | 0.0307 |
| PLSCR1 | -6.858 | enzyme | | | | 0.0307 |
| miR-320b (ar | | mature micr | | | | 0.0307 |
| BIX 01294 | | chemical rea | | | | 0.0307 |
| AMBN | | growth facto | | | | 0.0307 |
| LIMS2 | -5.678 | other | | | | 0.0307 |
| ELP1 | -9.606 | other | | | | 0.0307 |
| erastin | | chemical dru | | | | 0.0307 |
| SRPK1 | -9.147 | kinase | | | | 0.0307 |
| TDG | -7.649 | enzyme | | | | 0.0307 |
| GJA8 | | transporter | | | | 0.0307 |
| EFEMP2 | -8.483 | other | | | | 0.0307 |
| CBLC | | enzyme | | | | 0.0307 |
| RAB7A | -11.043 | enzyme | | | | 0.0307 |
| CTH | -6.101 | enzyme | | | | 0.0307 |
| SRGN | 2.614 | other | | | | 0.0307 |
| dec-RRLL-cm | | chemical rea | | | | 0.0307 |
| PK11007 | | chemical rea | | | | 0.0307 |
| zonisamide | | chemical dru | | | | 0.0307 |
| juglone | | chemical tox | | | | 0.0307 |
| aurintricarbo | | chemical dru | | | | 0.0307 |
| GSI34 | | chemical rea | | | | 0.0307 |
| PDZ1i | | chemical rea | | | | 0.0307 |
| Z-VEID-FMK | | chemical rea | | | | 0.0307 |
| bucladesine | | chemical tox | Inhibited | -4.12 | bias | 0.0308 |
| LRP6 | -9.055 | transmembra | | 0.717 | | 0.0315 |
| dinoprost | | chemical - er | | -1.374 | bias | 0.0319 |
| N-cor | | group | | 1.616 | bias | 0.032 |
| RBL2 | -8.841 | other | Activated | 2.867 | bias | 0.032 |
| TNFSF10 | | cytokine | | -1.002 | | 0.032 |
| AGK | -8.215 | kinase | Inhibited | -3.44 | bias | 0.0322 |

| | | | | | | |
|---|---|---|---|---|---|---|
| BAX | -8.374 | transporter | Inhibited | -2.075 | bias | 0.0322 |
| FOXA1 | -5.349 | transcription | | 1.546 | | 0.0324 |
| GSTO1 | -7.453 | enzyme | | 0.954 | bias | 0.0326 |
| 25-hydroxych | | chemical rea | | 1.833 | | 0.0326 |
| BTRC | -8.194 | enzyme | | 0.765 | bias | 0.0326 |
| P2RY2 | | G-protein co | | -1.937 | bias | 0.0326 |
| berberine | | chemical dru | | 1.279 | bias | 0.0328 |
| ABCB4 | | transporter | Activated | 2.787 | bias | 0.0334 |
| NVP-TAE684 | | chemical - ki | | 1.019 | | 0.0334 |
| ACVRL1 | -4.688 | kinase | | -1.706 | bias | 0.0334 |
| amiodarone | | chemical dru | | -1.414 | bias | 0.0334 |
| mercuric chl | | chemical tox | | | | 0.0334 |
| FUS-DDIT3 | | fusion gene/ | | 1 | bias | 0.0334 |
| phorbol 12,1 | | chemical - er | | -0.479 | bias | 0.0334 |
| Nos | | group | | -1.432 | bias | 0.0335 |
| SIRT7 | -6.1 | enzyme | Inhibited | -2.186 | | 0.0335 |
| CLCN5 | -6.298 | ion channel | | 1.067 | bias | 0.0335 |
| NSD1 | -9.165 | transcription | | | | 0.0335 |
| APBB1 | -10.397 | transcription | | -1.144 | bias | 0.0335 |
| HBP1 | -7.494 | transcription | | 1.331 | bias | 0.0335 |
| Smad2/3/4 | | group | | -1.432 | bias | 0.0335 |
| TOPBP1 | -8.977 | other | Activated | 2.226 | bias | 0.0335 |
| pyridoxine | | chemical - er | | -1.342 | | 0.0335 |
| cobalt | | chemical tox | Inhibited | -2.236 | bias | 0.0335 |
| PKNOX2 | -7.068 | transcription | | -0.816 | bias | 0.034 |
| RFX1 | -7.835 | transcription | | | | 0.034 |
| HNRNPA1 | -10.978 | other | | | | 0.034 |
| HRG | | other | | -1.671 | bias | 0.034 |
| CSNK2A1 | -10.035 | kinase | Inhibited | -2.449 | bias | 0.034 |
| HULC | | other | | -0.933 | | 0.034 |
| EML4-ALK | | fusion gene/ | | -0.946 | | 0.034 |
| SMAD4 | | transcription | Inhibited | -2.314 | bias | 0.0344 |
| staurosporin | | chemical - ki | | -0.387 | | 0.0344 |
| ATF1 | -6.701 | transcription | | -1.231 | bias | 0.0352 |
| PRKCE | -8.642 | kinase | Inhibited | -3.224 | bias | 0.0356 |
| NTRK2 | -8.141 | kinase | Inhibited | -2.837 | bias | 0.0356 |
| l-asparagina | | biologic drug | | 1.027 | bias | 0.0358 |
| Esrra | | transcription | | -1.053 | | 0.0358 |
| TET2 | -2.731 | enzyme | Inhibited | -2.2 | bias | 0.0358 |
| ritonavir | | chemical dru | | | | 0.0358 |
| GnRH-A | | chemical rea | | -0.961 | | 0.0362 |
| LDLR | -10.328 | transporter | | -1.265 | | 0.0363 |
| HNRNPK | -10.554 | other | | -0.342 | | 0.0363 |
| oxaliplatin | | chemical dru | | -1.427 | | 0.0367 |
| Ubiquitin | | group | | 0.896 | bias | 0.0367 |

| | | | | | | |
|---|---|---|---|---|---|---|
| PRKAR1A | -10.65 | kinase | | -0.116 | | 0.0367 |
| THBS4 | -5.519 | other | | -1.409 | bias | 0.0367 |
| ROCK | | group | Inhibited | -2.433 | bias | 0.0368 |
| Am 580 | | chemical rea | | 0.311 | | 0.0368 |
| S-nitroso-N-a | | chemical rea | | -0.851 | | 0.0369 |
| KLF5 | -5.655 | transcription | | -1.418 | bias | 0.0369 |
| carbon tetrac | | chemical tox | Inhibited | -3.632 | bias | 0.0376 |
| SMAD2 | -7.513 | transcription | Inhibited | -3.152 | bias | 0.0385 |
| dopamine | | chemical - er | | -1.596 | bias | 0.0386 |
| HOXA10 | | transcription | Activated | 2.335 | bias | 0.0389 |
| 4-hydroxytan | | chemical dru | Inhibited | -2.807 | bias | 0.0389 |
| tacrolimus | | chemical dru | | 0.671 | bias | 0.0392 |
| CCAR2 | -10.447 | peptidase | Inhibited | -3 | bias | 0.0393 |
| nafenopin | | chemical dru | Inhibited | -2.219 | bias | 0.0393 |
| SH42 | | chemical rea | | -0.333 | bias | 0.0393 |
| captopril | | chemical dru | | -1.22 | | 0.0395 |
| miR-125b-5p | | mature micr | Activated | 3.722 | bias | 0.0398 |
| CDC73 | -5.614 | other | | | | 0.0399 |
| miR-23a-3p ( | | mature micr | | 1.97 | bias | 0.0399 |
| XIAP | -8.03 | enzyme | Inhibited | -2.401 | bias | 0.0399 |
| PSMD10 | -9.283 | transcription | | 0 | bias | 0.0399 |
| BAG1 | -3.344 | other | | -1.969 | bias | 0.0399 |
| diazoxide | | chemical dru | Activated | 2.646 | bias | 0.0399 |
| LCK | | kinase | Inhibited | -3.286 | bias | 0.0404 |
| 2,4,5,2',4',5'- | | chemical tox | | 0.714 | bias | 0.0405 |
| POSTN | -9.729 | other | | -1.505 | bias | 0.0408 |
| fasudil | | chemical dru | Activated | 2.396 | bias | 0.0408 |
| lipid | | chemical - er | | 1.342 | | 0.0408 |
| 17-alpha-eth | | chemical dru | Inhibited | -3.123 | bias | 0.0409 |
| IKBKE | -5.192 | kinase | Inhibited | -3.068 | bias | 0.0412 |
| TCF | | group | | -1.996 | bias | 0.0412 |
| IKZF1 | | transcription | Activated | 3.881 | bias | 0.0414 |
| azathioprine | | chemical dru | Activated | 2 | bias | 0.0415 |
| mGluR | | group | | -1.067 | bias | 0.0415 |
| MSI2 | -9.215 | other | | 0.83 | bias | 0.0415 |
| DLX6-AS1 | | other | | -1 | bias | 0.0415 |
| C9orf72 | -8.203 | other | | 0 | | 0.0415 |
| PPP1R15B | -2.556 | phosphatase | | 1.964 | bias | 0.0415 |
| PBRM1 | -9.471 | other | | | | 0.0415 |
| ZFP57 | | transcription | | | | 0.0415 |
| G2535 | | chemical rea | | 1.086 | bias | 0.0415 |
| USP9X | -7.117 | peptidase | | -0.849 | bias | 0.0415 |
| CCN4 | | other | | -1.969 | bias | 0.0415 |
| FZD7 | -9.007 | G-protein co | | -1.091 | bias | 0.0415 |
| NRBP2 | -5.543 | kinase | | 0.984 | bias | 0.0415 |

| | | | | | | |
|---|---|---|---|---|---|---|
| CHRM3 | -6.678 | G-protein co | | -0.849 | bias | 0.0415 |
| miR-193a-3p | | mature micr | | 1.231 | bias | 0.0415 |
| miR-142-3p | | mature micr | Activated | 2 | bias | 0.0415 |
| miR-378a-3p | | mature micr | | 1.969 | bias | 0.0415 |
| PTCSC3 | 4.69 | other | | 1.987 | bias | 0.0415 |
| LIMA1 | -7.692 | other | | | | 0.0415 |
| AFP | | transporter | | -1.067 | bias | 0.0415 |
| KLB | 1.211 | enzyme | | 0.849 | bias | 0.0415 |
| TNFSF8 | | cytokine | Activated | 2 | bias | 0.0415 |
| ELK3 | | transcription | | | | 0.0415 |
| CNB-001 | | chemical rea | Inhibited | -2 | bias | 0.0415 |
| MYZAP | | other | | -1.982 | | 0.0415 |
| TAS1R3 | -4.438 | G-protein co | | 0 | bias | 0.0415 |
| KMUP-1 | | chemical rea | | 0.038 | bias | 0.0415 |
| sesame oil | | chemical rea | | | | 0.0415 |
| rosmarinic a | | chemical - e | | | | 0.0415 |
| hesperetin | | chemical dru | | 0 | | 0.0415 |
| adaphostin | | chemical dru | | -0.308 | bias | 0.0415 |
| baicalin | | chemical - e | Inhibited | -3.148 | bias | 0.0416 |
| EGR3 | 8.08 | transcription | | -0.588 | | 0.0416 |
| WWTR1 | -8.269 | transcription | | -0.847 | bias | 0.0416 |
| CAT | -7.324 | enzyme | Activated | 2.239 | bias | 0.0418 |
| SAFB | -9.624 | other | | 1.155 | bias | 0.0418 |
| DCN | -6.815 | other | | 0.397 | | 0.0418 |
| EHMT2 | -9.823 | transcription | | 1 | | 0.0418 |
| Ras homolog | | group | | -0.277 | bias | 0.0426 |
| HDAC7 | -8.728 | transcription | | -0.643 | | 0.0426 |
| AXIN1 | -8.271 | other | | 1.481 | bias | 0.0426 |
| PD 169316 | | chemical - ki | | 0 | | 0.0426 |
| epirubicin | | chemical dru | | | | 0.0426 |
| linalool | | chemical - e | | | | 0.0426 |
| AG490 | | chemical - ki | Activated | 3.056 | bias | 0.0436 |
| ITGAV | -9.968 | transmembr | | -1.977 | bias | 0.0437 |
| NF2 | 10.345 | other | Activated | 2.02 | | 0.0437 |
| nelfinavir | | chemical dru | Inhibited | -3.873 | bias | 0.0437 |
| prostaglandi | | chemical - e | | 0.887 | | 0.0438 |
| epothilone B | | chemical dru | | -1.109 | | 0.0438 |
| H1-2 | -6.323 | other | | -0.689 | | 0.0438 |
| miR-128-3p | | mature micr | Activated | 2.2 | bias | 0.0438 |
| ACACB | | enzyme | Activated | 2.216 | bias | 0.0438 |
| 4-nitroquinol | | chemical tox | | | | 0.0438 |
| 2-acetylamir | | chemical tox | | -1.172 | bias | 0.0438 |
| WIN 55,212- | | chemical - ot | | 1.109 | | 0.0438 |
| reserpine | | chemical dru | | -1.22 | bias | 0.0438 |
| mannose | | chemical - e | | 1.342 | | 0.0438 |

| | | | |
|---|---|---|---|
| GY1-22 | | chemical rea | 0.0446 |
| phosphatidyl | | chemical - er | 0.0446 |
| Diap | | group | 0.0446 |
| Ggtase I | | complex | 0.0446 |
| (diaminocycl | | chemical rea | 0.0446 |
| AMOT | -7.971 | other | 0.0446 |
| ANTXR2 | -7.349 | transmembra | 0.0446 |
| aliskiren | | chemical dru | 0.0446 |
| benfotiamine | | chemical dru | 0.0446 |
| Tropomyosin | | group | 0.0446 |
| FKHR | | group | 0.0446 |
| Beta Secreta | | group | 0.0446 |
| TFIIH | | complex | 0.0446 |
| G protein be | | complex | 0.0446 |
| POFUT1 | -2.575 | enzyme | 0.0446 |
| RHOBTB2 | -9.031 | enzyme | 0.0446 |
| RFFL | -7.738 | enzyme | 0.0446 |
| PTPMT1 | -7.587 | phosphatase | 0.0446 |
| NIBAN2 | -9.464 | transcription | 0.0446 |
| OR51E1 | | G-protein co | 0.0446 |
| CHD5 | -5.225 | enzyme | 0.0446 |
| DERL1 | -9.027 | other | 0.0446 |
| BCS1L | -6.379 | other | 0.0446 |
| NDUFAF3 | -8.364 | other | 0.0446 |
| PRMT8 | -6.746 | enzyme | 0.0446 |
| Fascin | | group | 0.0446 |
| SENP8 | | peptidase | 0.0446 |
| NHLRC1 | | enzyme | 0.0446 |
| ITGBL1 | -5.651 | other | 0.0446 |
| IPPK | -7.227 | kinase | 0.0446 |
| ESRP2 | | other | 0.0446 |
| KLHL2 | -7.919 | other | 0.0446 |
| AMOTL1 | -8.852 | other | 0.0446 |
| CLASP2 | -10.065 | other | 0.0446 |
| EXOC3L1 | | other | 0.0446 |
| RADIL | -6.891 | other | 0.0446 |
| PTPN23 | -9.586 | phosphatase | 0.0446 |
| OLA1 | -9.874 | enzyme | 0.0446 |
| VAX2 | -5.331 | transcription | 0.0446 |
| ATP13A2 | -10.037 | transporter | 0.0446 |
| Eotaxin | | group | 0.0446 |
| isoleucine | | chemical - er | 0.0446 |
| ACAT1 | -3.977 | enzyme | 0.0446 |
| TSIX | 1.76 | other | 0.0446 |
| CSNK1E | -11.355 | kinase | 0.0446 |

| | | | |
|---|---|---|---|
| LPXN | | transcription | 0.0446 |
| SRGAP1 | -5.233 | other | 0.0446 |
| GPC3 | -11.109 | other | 0.0446 |
| RRAD | -6.352 | enzyme | 0.0446 |
| PAG1 | -3.172 | other | 0.0446 |
| SEMA3D | -7.58 | other | 0.0446 |
| PSMC5 | -9.595 | transcription | 0.0446 |
| PURB | -2.534 | transcription | 0.0446 |
| U2AF2 | -10.688 | other | 0.0446 |
| CUL4A | -9.169 | other | 0.0446 |
| miR-222-5p ( | | mature micr | 0.0446 |
| miR-515-3p ( | | mature micr | 0.0446 |
| miR-743b-3p | | mature micr | 0.0446 |
| mir-153 | | microRNA | 0.0446 |
| mir-1180 | | microRNA | 0.0446 |
| MAPK15 | -4.568 | kinase | 0.0446 |
| SGI 1776 | | chemical dru | 0.0446 |
| STIP1 | -10.845 | other | 0.0446 |
| RBM4 | -9.9 | other | 0.0446 |
| SEMA3C | -10.474 | other | 0.0446 |
| EIF2B1 | -8.975 | translation re | 0.0446 |
| AHNAK | -8.43 | other | 0.0446 |
| KCNQ2 | -10.993 | ion channel | 0.0446 |
| opaganib | | chemical dru | 0.0446 |
| DLST | -9.085 | enzyme | 0.0446 |
| SRPK2 | -9.19 | kinase | 0.0446 |
| SLC12A6 | -7.48 | transporter | 0.0446 |
| MXD4 | -7.961 | transcription | 0.0446 |
| IGF2BP3 | -8.863 | translation re | 0.0446 |
| Ank2 | | other | 0.0446 |
| KRT5 | | other | 0.0446 |
| MAP2 | -11.698 | other | 0.0446 |
| TRERF1 | -7.16 | transcription | 0.0446 |
| MANF | -8.917 | other | 0.0446 |
| Manf | | other | 0.0446 |
| SYNPO | -5.921 | other | 0.0446 |
| APPL1 | -4.638 | other | 0.0446 |
| VDAC1 | -10.281 | ion channel | 0.0446 |
| GLUD1 | -10.084 | enzyme | 0.0446 |
| RAB3B | | enzyme | 0.0446 |
| RACGAP1 | -7.607 | transporter | 0.0446 |
| Gm33994 | | other | 0.0446 |
| glycosylated | | chemical rea | 0.0446 |
| 8-butyl-10-[( | | chemical rea | 0.0446 |
| MXD3 | -7.009 | transcription | 0.0446 |

| | | | | | | |
|---|---|---|---|---|---|---|
| SRSF5 | -10.308 | other | | | | 0.0446 |
| EXOC7 | -9.633 | transporter | | | | 0.0446 |
| G3BP2 | -10.623 | enzyme | | | | 0.0446 |
| RDH16 | 6.37 | enzyme | | | | 0.0446 |
| GIMAP1-GIM | | other | | | | 0.0446 |
| YY1AP1 | -7.771 | other | | | | 0.0446 |
| WASF1 | -7.044 | other | | | | 0.0446 |
| (+)-R-3-(4-ch | | chemical rea | | | | 0.0446 |
| SB-657510 | | chemical rea | | | | 0.0446 |
| BR-4628 | | chemical rea | | | | 0.0446 |
| FTL | -13.094 | enzyme | | | | 0.0446 |
| (E)-1-(2-nitro | | chemical rea | | | | 0.0446 |
| (E)-9-(2-nitro | | chemical rea | | | | 0.0446 |
| norcantharid | | chemical rea | | | | 0.0446 |
| LY3214996 | | chemical dru | | | | 0.0446 |
| 2,5-di-tert-bu | | chemical tox | | | | 0.0446 |
| GSK-3beta in | | chemical - ki | | | | 0.0446 |
| CVT-313 | | chemical - ki | | | | 0.0446 |
| apomine | | chemical dru | | | | 0.0446 |
| NSC 644221 | | chemical rea | | | | 0.0446 |
| nitromifene | | chemical rea | | | | 0.0446 |
| ethyl-3-triflu | | chemical rea | | | | 0.0446 |
| TTI-101 | | chemical dru | | | | 0.0446 |
| potassium cy | | chemical rea | | | | 0.0446 |
| nickel subsul | | chemical tox | | | | 0.0446 |
| SYUIQ-5 | | chemical rea | | | | 0.0446 |
| ACTG1-MITF | | fusion gene/ | | | | 0.0446 |
| 5-phenylvale | | chemical rea | | | | 0.0446 |
| oleuropein a | | chemical - en | | | | 0.0446 |
| 3,4-methylen | | chemical dru | | | | 0.0446 |
| atosibanum | | biologic drug | | | | 0.0446 |
| NSOP00313 | | chemical rea | | | | 0.0446 |
| R-etodolac | | chemical dru | | | | 0.0446 |
| E64 | | chemical - en | | | | 0.0446 |
| Mn2+ | | chemical - en | | | | 0.0446 |
| mir-15 | | microRNA | Activated | 2.827 | bias | 0.0449 |
| sphingosine- | | chemical - en | Inhibited | -2.272 | bias | 0.0452 |
| PLIN5 | | other | | -0.652 | | 0.0459 |
| mir-221 | | microRNA | | 1.409 | bias | 0.0459 |
| TSC1 | -8.74 | other | | 1.647 | | 0.0459 |
| NDRG1 | -9.851 | kinase | | 0.411 | | 0.047 |
| ETV6-RUNX1 | | fusion gene/ | Activated | 3.794 | bias | 0.0471 |
| PTPN6 | | phosphatase | | 1.719 | bias | 0.0473 |
| BSG | -12.018 | transporter | Inhibited | -2.283 | bias | 0.0474 |
| EOMES | -8.024 | transcription | Activated | 2.728 | | 0.0478 |

| | | | | | | |
|---|---|---|---|---|---|---|
| BCAP31 | -8.798 | transporter | | 1.394 | bias | 0.0482 |
| HEY1 | -8.266 | transcription | | 1.386 | bias | 0.0482 |
| IL2 | | cytokine | Inhibited | -3.187 | bias | 0.0483 |
| L-proline | | chemical - er | | | | 0.0488 |
| menthol | | chemical dru | | | | 0.0488 |
| MRPS18B | -2.99 | other | | | | 0.0488 |
| PDCD10 | -6.549 | other | | | | 0.0488 |
| SCGB3A2 | | other | | | | 0.0488 |
| SH2D5 | -5.947 | other | | | | 0.0488 |
| TRIM29 | | transcription | | | | 0.0488 |
| ATE1 | -7.559 | enzyme | | | | 0.0488 |
| EHT-1864 | | chemical rea | | | | 0.0488 |
| WNT16 | | other | | | | 0.0488 |
| Collagen type | | complex | | | | 0.0488 |
| HNRNPR | -9.511 | other | | | | 0.0488 |
| SERPINC1 | | enzyme | | | | 0.0488 |
| PRKG2 | | kinase | | | | 0.0488 |
| WNT2 | | cytokine | | | | 0.0488 |
| BANP | -5.682 | other | | | | 0.0488 |
| DAB2 | -8.65 | other | | | | 0.0488 |
| SNCB | -8.441 | other | | | | 0.0488 |
| miR-542-3p ( | | mature micr | | | | 0.0488 |
| NUP107 | | other | | | | 0.0488 |
| BANCR | | other | | | | 0.0488 |
| RAD51AP1 | | other | | | | 0.0488 |
| KHDRBS1 | -11.328 | transcription | | | | 0.0488 |
| GABBR2 | -5.653 | G-protein co | | | | 0.0488 |
| PROK1 | | growth facto | | | | 0.0488 |
| sulfinosine | | chemical rea | | | | 0.0488 |
| TP53INP1 | -8.193 | other | | | | 0.0488 |
| SF3B2 | -11.18 | other | | | | 0.0488 |
| SHCBP1 | | other | | | | 0.0488 |
| SUPT5H | -10.169 | transcription | | | | 0.0488 |
| PGK1 | -10.565 | kinase | | | | 0.0488 |
| Gm26313 | | other | | | | 0.0488 |
| temsirolimus | | chemical dru | | | | 0.0488 |
| 2-mercaptoe | | chemical - er | | | | 0.0488 |
| butylideneph | | chemical rea | | | | 0.0488 |
| geranylgerar | | chemical - er | | | | 0.0488 |
| tyrphostin AG | | chemical - ki | | | | 0.0488 |
| 3,5-dihydroxy | | chemical rea | | | | 0.0488 |
| Muscarinic cl | | group | | | | 0.0488 |
| LEP | | growth facto | Inhibited | -2.813 | bias | 0.0489 |
| Vegf | | group | Inhibited | -6.542 | bias | 0.0491 |
| BMP10 | | growth facto | | -1.462 | bias | 0.0495 |

| | | | | | |
|---|---|---|---|---|---|
| POU5F1 | 2.819 | transcription | Inhibited | -2.491 | 0.0505 |
| METTL3 | -8.561 | enzyme | | -1.961 | 0.0562 |
| CREM | -5.581 | transcription | | -1.819 | 0.0564 |
| EZH2 | -7.513 | transcription | Activated | 2.291 | 0.0571 |
| PROX1 | -4.471 | transcription | Inhibited | -2.254 | 0.0695 |
| EGLN1 | -5.987 | enzyme | Activated | 3.075 | 0.0716 |
| CYP19A1 | | enzyme | Inhibited | -2.09 | 0.0719 |
| SLC13A1 | | transporter | Activated | 2.668 | 0.0728 |
| enzalutamid | | chemical dru | | -1.964 | 0.0732 |
| CUL3 | -9.936 | enzyme | | 1.97 | 0.0761 |
| BAP1 | -10.212 | peptidase | Activated | 2.19 | 0.0846 |
| WWOX | -6.773 | enzyme | Activated | 2.2 | 0.102 |
| SOD2 | -9.647 | enzyme | | -1.98 | 0.102 |
| zinc | | chemical dru | | 1.794 | 0.102 |
| SLC2A1 | -10.976 | transporter | | -1.936 | 0.114 |
| BCL2L1 | -7.994 | other | | -1.835 | 0.119 |
| ethanol | | chemical - er | Inhibited | -3.986 | 0.119 |
| docetaxel | | chemical dru | Activated | 2.096 | 0.12 |
| NCOA1 | -8.666 | transcription | Activated | 2.892 | 0.128 |
| RARG | -6.083 | ligand-deper | | -1.905 | 0.139 |
| SOX7 | 2.44 | transcription | Inhibited | -2.714 | 0.14 |
| NGLY1 | -7.647 | enzyme | Inhibited | -2.613 | 0.15 |
| Il3 | | cytokine | | -1.913 | 0.166 |
| 8-(4-chloropl | | chemical rea | | -1.995 | 0.189 |
| NKX2-3 | | transcription | Inhibited | -2.632 | 0.189 |
| DICER1 | -9.664 | enzyme | Activated | 4.27 | 0.19 |
| VDR | | transcription | Inhibited | -2.668 | 0.204 |
| NR1H2 | -8.264 | ligand-deper | Inhibited | -2.988 | 0.236 |
| obeticholic a | | chemical dru | Activated | 2.152 | 0.245 |
| IDO1 | | enzyme | | -1.941 | 0.276 |
| NR6A1 | | ligand-deper | Inhibited | -2 | 0.276 |
| VEGFB | -8.424 | growth facto | Inhibited | -2.397 | 0.281 |
| TCF3 | -9.944 | transcription | Inhibited | -2.89 | 0.293 |
| PKD1 | -11.603 | ion channel | Activated | 2.76 | 0.294 |
| quercetin | | chemical dru | | -1.83 | 0.3 |
| IRAK3 | | kinase | Inhibited | -2.219 | 0.308 |
| ciprofibrate | | chemical dru | Inhibited | -2.849 | 0.312 |
| THRB | | ligand-deper | | -1.891 | 0.327 |
| mir-204 | | microRNA | Inhibited | -2.449 | 0.336 |
| ID2 | -10.042 | transcription | | 1.85 | 0.341 |
| anandamide | | chemical - er | Inhibited | -2.207 | 0.403 |
| THRA | -9.92 | ligand-deper | | -1.754 | 0.423 |
| KCNK9 | | ion channel | Activated | 2.449 | 0.427 |
| PPIF | -8.406 | enzyme | Activated | 3 | 0.438 |
| melatonin | | chemical - er | | -1.824 | 0.465 |

| | | | | | |
|---|---|---|---|---|---|
| CX3CR1 | | G-protein co | Inhibited | -2.584 | 0.484 |
| HDAC2 | -11.226 | transcription | Activated | 2.265 | 0.487 |
| Rxr | | group | | -1.982 | 0.49 |
| NR1H3 | -5.814 | ligand-deper | Inhibited | -3.775 | 0.497 |
| arsenic | | chemical tox | | -1.99 | 0.516 |
| MEX3A | -10.741 | other | Inhibited | -2.236 | 0.543 |
| FFAR3 | | G-protein co | Activated | 2.53 | 1 |
| STAT5a/b | | group | | -1.96 | 1 |
| 3M-002 | | chemical rea | Activated | 2.236 | 1 |
| MALAT1 | -14.16 | other | | -1.96 | 1 |
| NR1H4 | | ligand-deper | | 1.836 | 1 |
| GATA4 | | transcription | Inhibited | -2.484 | 1 |
| AXL | -8.02 | kinase | | -1.98 | 1 |
| ID3 | -8.996 | transcription | | 1.939 | 1 |
| S100A8 | | other | Inhibited | -2.125 | 1 |
| HOXA3 | -8.266 | transcription | Inhibited | -2.236 | 1 |
| CITED2 | -9.366 | transcription | | 1.993 | 1 |
| GHRL | | growth facto | | -1.913 | 1 |
| PPP3R1 | -4.034 | phosphatase | | -1.912 | 1 |
| dehydroisoar | | chemical - er | | -1.756 | 1 |

| **Target Mole** | Mechanistic Network |
|---|---|
| ABCA2,ABCA | 1056 (16) |
| ABCA2,ABCA | 1317 (18) |
| ACSS2,ALDH | 1039 (18) |
| ACTB,AP2M1 | |
| ALDOA,AP2M | |
| A2M,ABCA2, | 1092 (16) |
| EEF1B2,EEF1 | |
| A2M,ACLY,A | 1206 (19) |
| ABCA7,ABCD | 1141 (16) |
| A2M,ABCA3, | 1314 (19) |
| ACACA,ACAD | 1106 (19) |
| ACVR1B,AHS | 1121 (18) |
| ATP5F1A,ATI | 900 (12) |
| ACTA2,ACTB, | 1199 (22) |
| ABCE1,ADAM | |
| ACTG1,AGPA | 194 (3) |
| ABCE1,ADAM | |
| ABCE1,ABCF | 1172 (19) |
| ACAT2,ACTA | 1282 (21) |
| ACLY,ACTN1, | |
| ACACA,ACAT | 1117 (21) |
| ACACA,ACLY, | 1196 (17) |
| ACOT7,AGRN | |
| ACTG1,AGPA | 903 (11) |
| ACACA,ACAD | 1026 (21) |
| ACACA,ACLY, | |
| ABCA3,ABI2, | 1294 (17) |
| ABCE1,ADAM | 1179 (20) |
| ACAT2,ARL6I | 1169 (22) |
| A2M,ACACA, | 1098 (21) |
| ACOT7,ACTA | 1115 (18) |
| ABL1,APP,AR | 1142 (21) |
| ACO2,ACTB,A | 1182 (19) |
| ACACA,ACHE | 1258 (19) |
| ACACA,ACOT | |
| ACACA,ACAD | 1185 (22) |
| ABCE1,ACIN1 | 1186 (15) |
| ACOT7,AGRN | 1167 (22) |
| ABCE1,ACTB, | 1266 (20) |
| EEF1B2,EEF2 | |
| ACACA,ACTA | 1246 (19) |
| ACTA2,ACTN | 921 (17) |
| ACACA,ALCA | 757 (7) |

| | |
|---|---|
| ABL1,ACSL4,. | 1188 (20) |
| ACTN4,ADD1 | 697 (7) |
| ACTA2,ACTB, | 1326 (22) |
| ACTN1,ACTN | 876 (13) |
| ABCA3,ACLY, | 1060 (19) |
| ACTB,ATP5F1 | |
| ACACA,ADAN | 907 (16) |
| ACTB,ATP5F1 | |
| ACTN4,ACTR | 1106 (22) |
| ACTN1,AHCY | 738 (10) |
| ADAR,AGRN, | |
| ACACA,ACLY, | 940 (15) |
| ATP5F1B,CAI | |
| APEX1,APOE, | 859 (16) |
| ACACA,ACTA | 616 (15) |
| ACSL3,AGPA | |
| ABLIM1,ACA | |
| ABCA2,ALDH | 1161 (19) |
| ABCE1,ABLIN | 1298 (17) |
| ACSL3,ADAN | 998 (10) |
| CCT2,CCT3,C | 262 (7) |
| ACACA,ACLY, | 1132 (17) |
| ACACA,AHSA | 1135 (19) |
| ACACA,ADRN | 1378 (23) |
| ACTR1A,ADA | |
| ACTN4,AEBP | 880 (11) |
| A2M,ACTA2,. | 1108 (22) |
| ABL1,ACTB,A | 1097 (21) |
| ACTB,ATP5F1 | |
| ACLY,ANXA1, | |
| ATP5F1E,ATI | 951 (14) |
| ACAT2,ACTA | 1048 (16) |
| ACACA,ACLY, | 1089 (20) |
| ACADVL,ACO | 1101 (22) |
| ABLIM1,ACO | 1299 (23) |
| ACTN1,AHCY | 1029 (15) |
| ACTR1A,ACTI | 1011 (17) |
| AKT2,ALDOC, | 468 (7) |
| ADAR,ADD1,. | |
| ACTN4,AP1G | 1071 (16) |
| ACTA2,ACTB, | 1124 (18) |
| ACTA2,ADAN | 1226 (24) |
| ATP5F1C,ATI | |
| A2M,ACADVI | |
| A2M,ABCE1,. | 1182 (19) |

| | |
|---|---|
| ARHGDIA,AT | |
| ABCA7,ABI2, | |
| ADRM1,FDFT | |
| ANXA2,ASNS | |
| ACADVL,ACA | 929 (17) |
| ACACA,ACAD | |
| ACTA2,ADAM | 782 (14) |
| ACAT2,ACTA: | 1234 (19) |
| ARPP19,ATP5 | |
| ACTA2,ACTB, | 1157 (20) |
| AHCYL1,ARP( | 1048 (22) |
| ACHE,ACTN1 | 1127 (19) |
| AFAP1,ARHG | |
| A2M,ACACA, | 1181 (17) |
| A2M,ADGRE: | 1177 (22) |
| ABCD3,ABLIN | 1100 (20) |
| ACACA,ACO2 | 1162 (19) |
| ABL1,ACSL4,. | 1320 (17) |
| ADAR,AJUBA | |
| ACP1,ACTA2, | 1104 (21) |
| AP1M1,BTBD | |
| ACO2,ACTN1 | 639 (7) |
| ABCE1,APLP2 | 1086 (19) |
| APP,ASNS,AT | 1026 (22) |
| ADAMTS1,AL | 455 (11) |
| ANOS1,ANXA | 844 (7) |
| ABL1,ACTG1, | 1352 (18) |
| ACP1,ACSL6,. | |
| ACTB,APP,AS | 1033 (17) |
| ACLY,ACO2,A | 1185 (20) |
| ACACA,ACLY, | 804 (7) |
| ATP5F1B,C1( | |
| ALDH18A1,AI | 850 (15) |
| ABHD12,ACT | |
| ACTN4,AKAP | 939 (11) |
| ACTB,ATP1A | |
| AP2A1,ATP2A | |
| ABCA3,ACTA | |
| ACACA,ACLY, | 1292 (20) |
| ABL1,ACVR1I | |
| ACTA2,ACTB, | 1147 (19) |
| AKAP9,ANXA | 1016 (20) |
| CCND1,CCT2, | |
| ACAT2,ACTA: | 1169 (18) |
| ACTA2,ATP5I | 1098 (18) |

| | |
|---|---|
| ACTN4,AKT2, | 1107 (17) |
| CYP51A1,DH | 692 (9) |
| ACADVL,ACO | |
| ACTN1,CCND | 692 (7) |
| ALDOC,ATF4, | 1116 (18) |
| ARHGDIA,AT | 735 (10) |
| ABCD3,ACAC | 1014 (19) |
| ACTN4,AEBP | 815 (11) |
| AKAP12,ARH | 963 (14) |
| ACACA,ACTA | 1070 (18) |
| A2M,ACACA, | 1081 (20) |
| ATP5F1C,AT | |
| ACTB,ALDOC | 1063 (16) |
| ACSL4,ACTN | 1416 (19) |
| ACTR1A,AP2 | 989 (14) |
| A2M,ACACA, | 1390 (22) |
| ACAT2,ACOT | 1090 (19) |
| ACAT2,ACLY, | 1247 (19) |
| A2M,ABL1,A | 1219 (16) |
| APOE,APP,C | 1214 (21) |
| ACACA,ACTB | 1063 (17) |
| CAPN1,COTL | |
| ARCN1,ATF4 | 1149 (23) |
| ACACA,ACLY, | 1057 (19) |
| ACTN1,ACTR | 1047 (16) |
| ACTB,ACTG1 | 1012 (18) |
| ACO2,ACTB, | 1070 (18) |
| ABCA2,ACTA | 1150 (23) |
| AGAP1,ALDC | 849 (7) |
| ASNS,ATF4, | 361 (8) |
| ATF5,BNIP3, | |
| ATF4,BNIP3, | 1123 (20) |
| APP,BACE1,C | 1162 (21) |
| ACACA,ACAT | |
| ACTA2,ACTB, | 1172 (22) |
| ABCF1,ADAM | 1146 (19) |
| CAPN2,CCND | 1050 (18) |
| ATF4,ATF5,C | |
| A2M,ACTB,A | 1159 (22) |
| ASAH1,CLU,C | |
| ABCA3,ACAC | 781 (12) |
| ACTR1A,ARH | 408 (7) |
| ABCA2,ACTA | 1284 (22) |
| ACACA,ACLY, | 538 (7) |
| ACTA2,ARL2 | 1082 (20) |

| | |
|---|---|
| COX7A2L,EEF | |
| ACAT2,CYP51 | 536 (5) |
| ACACA,ACLY, | 790 (11) |
| ATP5MC3,AT | |
| APOE,ATF4,A | 1307 (23) |
| ACTN4,AEBP | 692 (9) |
| ANXA5,FUBP | |
| A2M,ACTB,A | |
| ACIN1,BOP1, | 993 (18) |
| ACTA2,APP,C | 847 (7) |
| ACO2,ATP5F | |
| ACACA,ACLY, | 1140 (22) |
| ACTN1,ADD1 | 1060 (17) |
| ACTA2,ACTN | 1056 (19) |
| ACSL6,ACTR1 | |
| A2M,AHCY,A | 1010 (18) |
| AKAP9,CD81, | 819 (7) |
| ASNS,ATF4,A | 1115 (18) |
| ARPC3,BACE | 566 (11) |
| ACACA,ACLY, | 1091 (21) |
| ANXA1,ATP5 | 691 (7) |
| ATF5,BNIP3, | |
| A2M,ABL1,A | 1461 (18) |
| A2M,ABCD3, | 1141 (20) |
| ACTA2,ADAM | 810 (7) |
| ACSL3,ACSL4 | |
| ACTA2,ADAM | 1010 (17) |
| ACACA,ACBD | 1049 (19) |
| ACLY,ACSL3, | 1013 (22) |
| ABCE1,ACTA | 1081 (20) |
| AGRN,ASNS, | 691 (7) |
| ACSL4,ALDO | |
| A2M,ACAT2, | 1239 (20) |
| ANKFY1,CCN | |
| CCND1,COL1 | |
| CCND1,CDK4, | 1197 (22) |
| CCND1,CCNG | |
| ACAT2,ACLY, | 1209 (22) |
| ABL1,APEH,A | 1305 (20) |
| APC,CCN2,CD | 911 (12) |
| ABI2,ACTA2, | 999 (15) |
| APOE,APP,AT | 1070 (17) |
| ACTA2,ADAM | 1000 (17) |
| AKAP8L,AKR | 866 (7) |
| ATF4,CDK2A | |

| | |
|---|---|
| ACTA2,APOE | 1055 (15) |
| ASNS,ATF5, | |
| APLP2,APP,A | 1092 (20) |
| ACO2,ATP1A | |
| ACTA2,ANXA | 263 (5) |
| ACTA2,BSG, | 1098 (16) |
| ACLY,ALDOA, | 1064 (20) |
| A2M,ACSL3, | 1117 (18) |
| ACTA2,CCND | 466 (4) |
| A2M,ACHE,A | 1081 (20) |
| ABCF1,ALDH | |
| ACHE,ACTA2, | 1095 (21) |
| ASNS,ATF4, | 1035 (17) |
| ACACA,ACLY, | 1113 (23) |
| ACHE,AKAP8 | 1290 (22) |
| CHGB,CLTA, | 513 (7) |
| ATP1A1,ATP | 1031 (18) |
| ACTB,ADAM | 843 (11) |
| CD99,CDKN1 | 715 (7) |
| ACTA2,AGAP | 1094 (21) |
| ACACA,ACLY, | 1152 (21) |
| ACHE,AGRN, | |
| CALD1,CCN1, | 58 (6) |
| ATF4,BNIP3L | 1210 (24) |
| ANXA1,CCND | 1150 (22) |
| ACLY,APP,CC | |
| A2M,ACTA2, | 1075 (20) |
| A2M,ACTG1, | |
| ACTB,AK1,AL | |
| APEX1,BCR, | 604 (4) |
| ACAT2,ACLY, | |
| ATF5,BNIP3, | |
| AKT2,AKT3,A | 1301 (16) |
| AP1G1,CBX6, | 1120 (20) |
| ADIPOR1,AD | 763 (19) |
| A2M,ADAMT | |
| CYP51A1,FDF | |
| APP,BZW2,F | |
| CAD,CCND2, | |
| ASNS,ATF4,E | |
| ABCA2,ABLIN | 1111 (18) |
| ATF4,BAG6, | 1000 (16) |
| ADH5,AKT2, | 1162 (18) |
| ACTA2,ATF4, | 1079 (17) |
| ATP6V0E2,B | |

| | |
|---|---|
| ATP6V0E2,B2 | |
| A2M,ABCD3, | 920 (16) |
| ACACA,ACSS | 994 (18) |
| ACADVL,CCN | 1233 (25) |
| ACTA2,AKAP | 1111 (22) |
| ACO2,CCND2 | 813 (7) |
| ACHE,ANXA2 | 1208 (19) |
| AKT2,ALDH7 | 1023 (19) |
| CCND1,CCND | 977 (18) |
| BCL11A,BIN1 | 339 (5) |
| ACTA2,ARPC | 942 (21) |
| ACVR1B,ALD | |
| ABI2,AP3D1, | 1209 (19) |
| ATP5F1A,CLT | |
| ACTA2,ADH5 | 1168 (20) |
| CCND1,CDH2 | |
| ACTA2,CCND | 1051 (15) |
| ACTA2,ADAM | 1137 (19) |
| ATP1A1,CD4 | |
| ACTN4,ANXA | |
| ATP1A1,ATP | |
| AKAP12,BGN | 970 (12) |
| ACACA,ACLY, | 1037 (18) |
| COL1A1,FN1, | |
| ACACA,ACAD | 1220 (20) |
| A2M,ABCA3, | 1062 (18) |
| ACLY,ACTA2, | 1164 (22) |
| BSG,CCN1,C | 830 (21) |
| ACTA2,COL1 | 904 (10) |
| ACTA2,CCN1, | 315 (9) |
| AP1M1,APC, | 1128 (18) |
| ACTR3,ADGR | 1115 (21) |
| ACTN4,CAPN | |
| CCN2,COL1A | 914 (12) |
| ACO2,APP,FT | 251 (5) |
| ACACA,ACAT | 981 (16) |
| ACSL4,AKT3, | |
| ACP1,ANXA5 | |
| ACTA2,ADAM | 953 (14) |
| AP2M1,BANI | 1063 (7) |
| ALDOA,CCND | 1149 (23) |
| COL1A1,COL | |
| ACTN4,ANXA | |
| CCND1,CD44, | 582 (6) |
| A2M,AK1,AM | 1117 (21) |

| | |
|---|---|
| EIF2S2,EIF3C | |
| ATP5F1A,ATI | |
| A2M,ABCA3, | 1022 (17) |
| CCN2,CCND1 | 1059 (18) |
| ABCD3,ACLY, | 1033 (21) |
| ALDOA,BNIP | 598 (7) |
| ACTA2,ACVR | 903 (18) |
| ACACA,ACAD | |
| ABCA7,ADAN | 450 (4) |
| ABL1,AP1G1, | 1197 (23) |
| DHCR7,FDPS, | 325 (7) |
| ACACA,ACLY, | 692 (9) |
| ADAMTS1,AI | |
| ATP2A2,BNII | 1103 (19) |
| ACP1,ACTA2, | 1022 (17) |
| ARF1,ARF3,A | 854 (9) |
| BNIP3L,CDKN | 988 (10) |
| ATP5F1C,ATI | 263 (5) |
| AKR1C4,APLI | 888 (15) |
| AFAP1,ARHG | 1074 (18) |
| ACTB,CCND2 | |
| ATG4B,BNIP | 697 (10) |
| APP,ATP1A1, | 1007 (13) |
| ADD2,ADNP, | |
| CAD,CCND2,C | |
| CDH2,CTNNB | |
| CAPNS1,CD4 | 1009 (17) |
| ADRM1,ATF5 | 1153 (19) |
| ACTA2,CCN1, | 1077 (19) |
| BIN1,BNIP3,C | 1088 (20) |
| ACLY,ACSS2,C | 820 (7) |
| ACTG1,AKAP | |
| B2M,BCLAF1 | 989 (13) |
| APP,BMF,C1 | 797 (9) |
| AKT2,CDKN1 | 1101 (21) |
| ALCAM,ANXA | 1012 (18) |
| AKIRIN2,AKR | |
| ACTB,ATP5F1 | 900 (11) |
| AKT3,ALDOA | |
| CDKN1A,CDK | 1066 (22) |
| CCNG1,CDKN | 1322 (24) |
| ACTB,ATP6V | 1213 (22) |
| A2M,ABCD3, | 1094 (19) |
| CHGB,RAB3A | |
| ANAPC5,CAN | |

| | |
|---|---|
| A2M,ALDOA, | 1108 (21) |
| ASNS,B2M,B | 1035 (20) |
| ALDOA,ANXA | 1023 (16) |
| CCND1,CDKN | |
| ACSL3,ADD1, | 1040 (18) |
| ACTB,AK1,AT | 1233 (20) |
| ACACA,ACAD | 1224 (20) |
| ACTA2,AKR1 | 909 (17) |
| ANXA5,ATF4 | 1246 (19) |
| ACADVL,ACT | 1083 (19) |
| ACTR3,AHCY | 1075 (19) |
| ARHGDIA,AR | 915 (20) |
| ALDH18A1,A | |
| CALR,GLUL,H | |
| ATP1A1,CLU, | |
| ACTA2,AKR1 | 1169 (19) |
| APP,ATP2A2, | 1170 (22) |
| ACTB,ALCAM | |
| ACHE,ACO2,A | 947 (11) |
| CDKN1A,CRA | 966 (12) |
| ACTA2,AKAP | 1039 (20) |
| APP,CCND1,C | 1113 (20) |
| ADAMTS1,AH | 1348 (20) |
| CCND1,CDK9, | 1041 (22) |
| APBB1,BEX3, | 593 (8) |
| APPBP2,CALR | |
| COL1A1,JUN, | 742 (5) |
| ACBD5,APP,A | |
| ACTA2,ASAH | 1313 (21) |
| ANXA1,CCND | 1107 (22) |
| ACAT2,ACOT | 1177 (19) |
| ACTA2,ALDO | 1017 (16) |
| CCND1,CDKN | 824 (7) |
| ARHGEF18,C | 487 (9) |
| CCN2,CDH2,C | 272 (6) |
| ACTG1,ACTR | |
| CCND1,CDK4, | |
| ABCE1,ACAC | 1349 (24) |
| ADIPOR1,ATI | 1140 (18) |
| APOE,CARD8 | 1029 (21) |
| A2M,ACHE,A | 1349 (21) |
| ACACA,ACAD | 1107 (20) |
| CDKN1A,FAB | 1010 (18) |
| GSTP1,PSMA | |
| ANXA2,AZIN | |

| | |
|---|---|
| ACHE,ACTA2, | 1172 (16) |
| DCTN1,DCTN | |
| ACTA2,CAD, | 1197 (22) |
| ANKRD50,AR | |
| ACTB,BCL11 | 668 (7) |
| CDKN1A,IPO | |
| CDH4,FN1,GI | |
| ADIPOR2,ATI | 984 (19) |
| ALDOA,CCN2 | 1114 (20) |
| ACTA2,APP, | 1223 (20) |
| APP,ATF4,CC | 988 (15) |
| ACBD5,APP, | |
| ATP1A1,ATP | 637 (7) |
| ACTB,ANXA5 | 1152 (18) |
| CCN2,COL1A | 660 (7) |
| BMF,CCND1, | 1061 (17) |
| APP,ATP5F1 | 1105 (19) |
| ARID1A,CCN | 1134 (21) |
| EIF3C,EIF3F, | |
| CDKN1B,CSR | 909 (13) |
| BNIP3L,FN1, | 1139 (20) |
| ASNS,CCND1 | 909 (16) |
| BNIP3,HIF1A | |
| CCT6A,HNRN | |
| ADD2,AKR1C | |
| APP,BNIP3, | 1135 (20) |
| ACTB,ADH5, | 1195 (20) |
| ARL4C,CALM | |
| ADNP,CDKN1 | |
| CCND1,CD44, | 1052 (21) |
| CNR1,DDAH1 | 900 (9) |
| ANTXR1,APC | 1089 (21) |
| AFAP1,ARHG | |
| CCN2,CDKN1 | 708 (7) |
| CCND1,CDKN | 1079 (18) |
| APEX1,ARL6I | 1147 (22) |
| A2M,ACTA2, | 954 (12) |
| CALR,EEF1B2 | |
| ACTA2,CCND | 667 (7) |
| ACTA2,ALCAI | |
| HERPUD1,LM | |
| ACACA,CCND | 966 (13) |
| A2M,ACACA, | 1121 (17) |
| A2M,ALDOA, | 1441 (21) |
| DNAJB1,HSP | |

| | |
|---|---|
| CYP51A1,FDF | 540 (7) |
| CDH2,NOTCH | 472 (5) |
| GPAA1,PSME | |
| ACTA2,APOE | 965 (20) |
| ACSL3,ACTN4 | 904 (18) |
| ACTA2,APBB | 1021 (17) |
| ACTB,ADAR,A | 965 (18) |
| ACTA2,AP2A1 | 1039 (22) |
| CCND1,CCND | 997 (19) |
| COL5A2,FZD7 | |
| ACIN1,BEX1,C | |
| ACACA,CCND | 1105 (17) |
| ACLY,ACSS2,A | 1349 (18) |
| ACTA2,ARHG | 1046 (20) |
| ACACA,ACLY, | 1132 (23) |
| CADM1,CCNE | 1202 (20) |
| ACLY,APLP2,C | 873 (17) |
| APOE,ATF4,C | 1031 (20) |
| ALYREF,ATF6 | 1105 (25) |
| APC,APP,CDK | 1018 (19) |
| CDKN1A,CDK | |
| AMD1,CDKN1 | |
| ACACA,AGAF | |
| BASP1,CCN1, | 432 (7) |
| ABCA2,ABCA | 926 (15) |
| ACTA2,APC,B | 940 (17) |
| CCN1,CCND1 | 1055 (20) |
| ACACA,ACLY, | 645 (11) |
| ACACA,ACLY, | 1185 (17) |
| ACTA2,COL1A | |
| ACTA2,ALCAI | 1034 (17) |
| CCND2,CDKN | 946 (13) |
| CCND1,CDKN | 937 (15) |
| AP1G1,APOE | 1173 (19) |
| B2M,CHD3,E | |
| CDKN1A,EIF2 | 883 (9) |
| ACACA,ACTR | 1022 (20) |
| ANXA2,CHGE | |
| ACTB,ACVR1 | |
| AKR1C3,ASAI | 1149 (18) |
| CDKN1A,DHF | |
| CDH2,CTNNB | 98 (3) |
| HIF1A,PDGFI | 625 (8) |
| ACO2,APP,FT | 251 (5) |
| ACAT2,DHCR | |

| | |
|---|---|
| AP1G1,CDKN | |
| ACTB,ACVR1 | |
| ACHE,ACTB,A | 1107 (18) |
| ACTA2,ADAN | 1058 (19) |
| A2M,ALCAM, | |
| ALDOA,ANXA | 626 (7) |
| ACTA2,CALD1 | 318 (7) |
| ACLY,CYP51A | |
| APP,CCN2,CC | 1055 (20) |
| BMF,CDKN1E | 885 (7) |
| AKR1C4,ANX | 1260 (20) |
| AQP1,CLCN7, | |
| CHGA,NCAM | |
| COL11A1,COI | |
| CAPN1,CAPN | 944 (19) |
| CDH2,CDKN1 | 849 (10) |
| CD44,CTNND | |
| ALDOA,EGR1 | 597 (7) |
| CD44,CDKN1, | 1175 (21) |
| ACTA2,CCN2, | 742 (7) |
| DBN1,DLG4,( | 739 (7) |
| CCN1,CCN2,H | 301 (6) |
| CCND1,CCND | 401 (3) |
| APEX1,IARS1 | 179 (3) |
| ALCAM,ATF4 | |
| CD74,CYP51A | 1125 (21) |
| AGPAT3,AHC | |
| ACTA2,ACTB, | 714 (13) |
| ACBD3,ACSS: | 796 (7) |
| CCND1,CDKN | 690 (11) |
| AHCY,CBS/CE | 1076 (19) |
| ACTA2,CCN2, | 961 (14) |
| ACAT2,ADAN | 1124 (19) |
| AFDN,AP2M: | 1104 (15) |
| ACHE,ACTA2, | 1176 (17) |
| CHGA,EGR1,( | 1281 (22) |
| AMOTL2,CCN | |
| ACTA2,CCND | 1213 (22) |
| ACACA,ACLY, | |
| AKAP8L,ALD( | 738 (7) |
| ACTA2,APOE | 1115 (23) |
| AMOTL2,CCN | |
| AKT3,ATRX,C | |
| ACACA,ACTB | 1010 (22) |
| APP,ATF4,CD | 1399 (22) |

| | |
|---|---|
| ACTB,ANXA5 | 1159 (22) |
| ACTG1,AHCY | |
| IGF1R,MTOR | |
| CNTN1,DHFR | |
| ABCD3,CDKN | 943 (13) |
| APOE,CCND1 | 1018 (19) |
| ACTA2,BMF, | 1150 (20) |
| ACTA2,CCND | 1026 (17) |
| AP1G1,CDKN | |
| ACTB,ASNS, | 1060 (20) |
| AGRN,AK1,A | 1079 (23) |
| ACOT7,ACTA | 1243 (21) |
| ACHE,ALDH1 | 1221 (17) |
| APP,CDKN1A | 1035 (17) |
| CCND1,CDKN | 1098 (20) |
| ANXA1,ARF4 | |
| ACTA2,ATF4, | 1221 (20) |
| ACSL6,ADAM | |
| APOE,ASAH1 | |
| A2M,ANXA1, | |
| AEBP1,AGRN | 994 (19) |
| A2M,ADAMT | 888 (10) |
| ADAM9,ANX | |
| ADIPOR2,CTN | |
| ALDOA,CCND | 1123 (21) |
| ADIPOR1,API | 919 (14) |
| CCND1,CDK4, | 1027 (15) |
| ACTA2,ARCN | 1033 (19) |
| AGPAT3,AHC | |
| ATP1A1,CCN | 930 (12) |
| ALDOA,BNIP | 799 (11) |
| AGRN,AKAP1 | 998 (13) |
| ATRX,BNIP3, | 475 (5) |
| ACACA,ACLY, | 309 (6) |
| CDH2,CTNNB | |
| CTNNA1,GJA | 768 (7) |
| COL18A1,CO | |
| APP,CCND1, | 1135 (18) |
| ACACA,APP, | 882 (17) |
| ADAMTS1,BE | 711 (7) |
| COL5A2,COX | 1168 (22) |
| RPL35,RPS14 | |
| GPAA1,PSME | |
| ACTA2,COL1 | 1004 (12) |
| CDKN1B,H2A | |

| | |
|---|---|
| CDC42,JUN,R | 824 (13) |
| COL11A1,COI | |
| CAD,CCND2,C | |
| CTNNB1,CTN | |
| ITGA3,ITGB1 | 786 (11) |
| ACTA2,APP,A | 982 (17) |
| BNIP3,CADM | 1136 (22) |
| A2M,ADAMT | |
| CCND1,CDH2 | 994 (20) |
| ACADVL,ACA | 642 (7) |
| ANXA6,ATP5 | 783 (13) |
| ACACA,ACTA | 1160 (19) |
| ANXA2,CCN2 | 734 (11) |
| ACADVL,ACTI | 846 (17) |
| ACTA2,CCND | 1000 (13) |
| CDKN1B,CSR | 797 (14) |
| B2M,B4GALT | 1061 (19) |
| CTNNB1,EGF | 1008 (17) |
| CDKN1A,FN1 | 491 (11) |
| CYP51A1,FDF | 383 (7) |
| CCND1,CCND | |
| CCND1,CDK4, | 943 (11) |
| ATP5F1B,CDI | 1210 (23) |
| ACTA2,AMO | 602 (11) |
| ACACA,ACAD | 1101 (12) |
| CCND1,CD47, | 1188 (20) |
| ACOT7,ACTA | 1106 (23) |
| ACTA2,AKT3, | 1075 (20) |
| BSG,CDK4,CD | |
| CCND1,CHPF, | |
| APP,CCND1,C | 652 (7) |
| AMOTL2,CCN | 696 (10) |
| BNIP3,COX4I | 1004 (16) |
| ACTA2,CCN2, | 917 (19) |
| ACTN4,CD15 | 932 (11) |
| ATF4,CDK9,E | 30 (3) |
| BFSP1,CCND | 1110 (20) |
| ATF4,ATF6B, | |
| APC,ATF4,CD | 1244 (20) |
| ALCAM,ATF4 | 1133 (19) |
| CCDC80,CCN | 1336 (18) |
| CCN2,CCND1 | |
| AP2A2,APP,A | |
| ACACA,ATP2 | 1093 (21) |
| CYP51A1,DH | |

| | |
|---|---|
| CCN1,CCN2,C | 1034 (20) |
| ADAM10,AD/ | |
| ACACA,ACSL4 | 1103 (20) |
| A2M,ANXA1, | 789 (11) |
| CCND2,CDC42 | 730 (11) |
| ACTR1A,AHC | 918 (15) |
| ALCAM,ATP1 | 1192 (23) |
| ARF4,CCND1 | 1086 (18) |
| ADRM1,BMF | |
| CCN2,CCND1 | 1113 (20) |
| ACTA2,CCND | 1209 (25) |
| ACTR3,CDKN | 973 (17) |
| ABCE1,ACTN | |
| ACAT2,CCN2, | 1220 (15) |
| CDKN1A,DLG | 1114 (22) |
| ACTB,CCND1 | 1108 (19) |
| ABLIM1,NAN | |
| APP,CCN2,CC | 999 (16) |
| APP,CDKN1A | 819 (10) |
| ATF4,CCND1, | 1140 (19) |
| A2M,ADAMT | |
| CCND1,CDKN | |
| IGF1R,ITGB1 | 677 (7) |
| ASAH1,CERS1 | |
| GSK3B,LAMP | |
| CCND1,CDK4, | 845 (11) |
| CALD1,RHOA | |
| DLG4,KCNQ2 | |
| FTH1,FTL,TFI | |
| CCND1,CDKN | 804 (7) |
| APP,BACE1,R | 818 (10) |
| CD44,CTNNB | |
| CD44,CTNNB | |
| CALR,EZR,SR | |
| CCND1,CDKN | 1095 (20) |
| ACTB,HDAC2 | |
| CCND2,HMGI | |
| CALD1,CCN1, | |
| CCN2,CCND1 | |
| CYP51A1,FDF | 339 (7) |
| ANXA2,ATP5 | 1062 (20) |
| ATF6,CD44,IT | 415 (7) |
| DNMT1,HSF1 | |
| ADGRL1,CCN | |
| ATF6,CCND1, | 1057 (18) |

| | |
|---|---|
| A2M,ACTA2,/ | 1108 (18) |
| AQP1,BACE1 | 1103 (24) |
| ACLY,ACTA2, | 1147 (19) |
| ADAMTS1,B( | 949 (13) |
| ACTA2,AKT2, | 1192 (22) |
| APP,ATP6V1I | 961 (14) |
| ACAT2,ACLY, | 566 (7) |
| CCND1,CD44, | 997 (16) |
| ACTA2,CDH1 | |
| APC,BTG1,C( | 1215 (17) |
| CCN2,CCND2 | 973 (18) |
| ANTXR1,CDK | |
| CDC42,CTNN | 1102 (22) |
| ACTA2,ACTB, | 1017 (18) |
| ACHE,ACOT7 | 1207 (20) |
| AGPAT3,AHC | |
| CDKN1A,CDK | 1007 (15) |
| CCND1,CHGA | |
| LOX,LOXL2,N | |
| CLSTN1,DLG4 | |
| CCND1,CDK4, | 723 (10) |
| BNIP3,CCND1 | 597 (7) |
| APC,APOE,C( | 848 (7) |
| ABI2,ACLY,A( | 1064 (18) |
| AKAP8L,ATXI | 1261 (22) |
| ATP2A2,CCNI | 1002 (18) |
| CCND1,CDKN | 1097 (21) |
| ACTA2,ITGB1 | 1135 (21) |
| CCND1,CDH2 | 1170 (25) |
| CCND1,CD44, | 564 (7) |
| ADAMTS1,AI | 1098 (19) |
| ACACA,ACTA | 1030 (18) |
| CCN1,CCN2,( | |
| CCND1,CDKN | 1040 (17) |
| ACO2,AKT2,F | 678 (7) |
| CTNNB1,MN | 563 (7) |
| BACE1,CDKN | |
| GNAO1,LIMK | |
| AKT2,ATF5,C | |
| APP,CDKN1A | 973 (11) |
| ACTA2,CCN2, | |
| CCND1,CDC4 | 942 (11) |
| FTH1,FTL,TFI | 250 (5) |
| FDFT1,HMG( | |
| AKT3,CCND1, | 1066 (18) |

| | |
|---|---|
| ACTA2,COL1/ | |
| CDH2,FN1,IT( | 710 (7) |
| CCND1,CDKN | 692 (10) |
| GJA1,HMGCI | |
| ADAM10,APF | 738 (7) |
| FN1,VIM,ZEE | |
| CALR,HSP90E | |
| ADAMTS1,BI | 1124 (20) |
| ACSL3,ACTA2 | 1078 (17) |
| ABCE1,ACO2 | 1099 (17) |
| CDH2,CXCL12 | 626 (12) |
| CCND1,CD44, | 866 (10) |
| CDKN1A,GPX | |
| CDH2,CDKN1 | 879 (12) |
| ARF4,ATF5,C | |
| ACHE,BACE1, | 1131 (19) |
| ACACA,FASN | 642 (8) |
| CCND1,CDKN | 922 (13) |
| CDC42,CDH2, | 861 (12) |
| DHCR7,FDFT | |
| ACTA2,CCND | 1064 (16) |
| ADAMTS1,AI | 1006 (13) |
| ACTA2,CDKN | 1299 (21) |
| A2M,ACTB,A | 1208 (18) |
| ARL6IP5,CCN | 680 (7) |
| COL1A1,COX | |
| ACLY,CDKN1I | 827 (11) |
| ADAM10,APF | |
| FABP7,FGFR | |
| ALCAM,CD15 | |
| APC,APP,CSN | |
| ACSL4,ACTN: | 1180 (18) |
| A2M,COL5A1 | 634 (7) |
| ADAM10,APF | 1091 (21) |
| CALD1,CCN1, | 312 (7) |
| AKR1C3,CCNI | 1120 (18) |
| CCND1,CCND | 1062 (19) |
| ARHGEF2,AT | |
| BAP1,CDKN1 | |
| CCND2,ERGI( | |
| CDKN1A,CDK | 402 (7) |
| C1QL1,CCND | |
| ARHGAP33,C | |
| BMF,BNIP3L, | |
| ACTA2,CTNN | |

| | |
|---|---|
| CALU,CD44,F | |
| ACTA2,CCND | 1115 (18) |
| ACTA2,ARHG | 1059 (18) |
| ACACA,ACLY, | 1082 (17) |
| ALDOA,APP, | 1137 (20) |
| APP,ATP1A1, | 988 (18) |
| APOE,CCN1, | 1031 (18) |
| ACLY,ATP5M | 684 (7) |
| ABI2,ACTA2, | 1116 (17) |
| ACTN4,ATP2 | 638 (7) |
| ACTA2,AMO | |
| EGR1,FN1,IT | 211 (5) |
| ACADVL,ACL | 611 (15) |
| ACTA2,CCN2, | 898 (17) |
| CCND1,CDKN | 1052 (20) |
| AHCYL1,ARF | |
| CCND1,CDKN | 203 (4) |
| ACHE,CCND2 | 1228 (23) |
| APC,APOE,A | 1164 (16) |
| ACTA2,CCN2, | 880 (12) |
| FASN,HIF1A, | 1184 (21) |
| CDKN1A,HIP | 975 (17) |
| ACTA2,DYNL | |
| CCND1,CDKN | 836 (14) |
| ACACA,CDKN | 1097 (19) |
| ACTA2,APP,C | 1209 (20) |
| ACACA,ACLY, | 781 (11) |
| CIRBP,GPX4, | 328 (7) |
| CD44,CDH2,N | |
| APC,BNIP3,C | 819 (7) |
| ACTA2,ACTB, | 711 (7) |
| ACTB,CCND1 | |
| CDKN1A,FN1 | 974 (17) |
| G6PD,GSR,PI | 189 (3) |
| ACTA2,CCDC | 1075 (21) |
| ACOT7,ACTA | 1220 (25) |
| ACTA2,BGN, | 1165 (23) |
| ACTA2,AJUB | 790 (11) |
| ACACA,ACLY, | 1238 (24) |
| APP,BACE1,C | 1038 (18) |
| ALDH7A1,CC | |
| CYP51A1,DH | |
| CCND1,CXCL | 1005 (16) |
| ATF4,CCNK,C | 1027 (18) |
| ACTA2,ADAM | 1140 (23) |

| | |
|---|---|
| CITED2,CLPTI | |
| AQP1,CCN1,( | 1054 (18) |
| CDKN1A,CDK | 913 (12) |
| CCND1,CDK4, | 1080 (15) |
| ATF4,BNIP3,I | 1059 (19) |
| CDH2,CDKN1 | 970 (19) |
| EGR1,FN1,T( | |
| ATF4,BASP1, | |
| CCN1,CCN2,E | 1067 (21) |
| ACTA2,EDNR | 622 (7) |
| ACTA2,CDH1 | 660 (7) |
| JAK1,RPL35,F | |
| CAMKV,CCD( | 950 (10) |
| ABCD3,ACAD | 1077 (20) |
| ANOS1,AQP1 | 965 (10) |
| BAP1,CDH2,( | 962 (12) |
| ABCA2,ACTA | 1067 (19) |
| A2M,ADAMT | |
| CCND1,CDH2 | 918 (15) |
| ACTA2,CCN2, | 1096 (18) |
| B2M,CALR,C | |
| APOE,APP,C( | 1123 (20) |
| AKR1C3,ALDI | 1123 (21) |
| ACTA2,APP,B | 1234 (17) |
| ABCA3,CCN1 | 983 (15) |
| ACTA2,CCN1, | 1157 (22) |
| AGRN,ALCAN | 1001 (15) |
| ACTA2,ACTB, | 714 (13) |
| CCN2,CCND1 | 1100 (18) |
| CD63,CKB,CU | 547 (5) |
| BCL11A,CD4 | |
| APC,EGR1,GI | 1256 (20) |
| ACTB,ALDOC | |
| BCL9L,COL11 | |
| CCND2,CDH2 | 1119 (19) |
| C4A/C4B,CD4 | 705 (7) |
| APP,ATP2A2, | 1130 (18) |
| CDK4,CDKN1 | 981 (16) |
| ABL1,ACAT2, | 1008 (19) |
| ACACA,ACSL | 201 (3) |
| ABCD3,ACAC | 783 (22) |
| B2M,CDH2,C | |
| CDH2,MTOR, | 805 (7) |
| APP,ATXN3,( | |
| ACTA2,AEBP | |

| | |
|---|---|
| CCND1,CDH2 | 1064 (18) |
| CCND1,CDH2 | |
| CCND1,CCND | 719 (9) |
| CDKN1A,CDK | 1036 (20) |
| ATP5F1A,ATI | |
| CCND1,CCND | 783 (11) |
| CDH2,EGR1,I | 337 (6) |
| COL2A1,FGFI | |
| CTNNB1,LDH | 926 (10) |
| CLSTN1,DLG | 296 (6) |
| CCND1,GRIA | 674 (7) |
| CCN1,CCN2,S | 423 (11) |
| ADAM10,APC | 1008 (15) |
| ABCD3,ACAD | 1092 (19) |
| ACTA2,CCND | |
| BCL11A,COL6 | |
| CAD,CCND2,C | |
| APC,CSNK1A | 317 (5) |
| CCN1,CCN2,C | 816 (17) |
| CCN2,CCND1 | 1164 (18) |
| ACAT2,ACLY, | 981 (13) |
| A2M,ADAMT | |
| CCN2,CDKN1 | 989 (17) |
| ANXA1,ANXA | 1051 (19) |
| ACTA2,ANTX | 1222 (20) |
| CCND1,CDKN | |
| CCND1,CDKN | 1000 (12) |
| COL6A1,HGS | |
| FTH1,IREB2, | |
| CCND1,CDKN | 893 (12) |
| CDH2,VIM,ZE | |
| CCND1,CDKN | 941 (12) |
| ASNS,OAZ1,C | |
| EEF2,ROCK2, | |
| ACACA,FASN | 325 (8) |
| FBLN1,MMP | |
| JUN,RAC1,TI | 461 (7) |
| BOP1,CDKN1 | 863 (12) |
| ATF4,HIF1A,I | |
| APP,MT2A,T | |
| EGR1,ITGAV, | |
| CCND1,FN1,I | 621 (7) |
| FASN,FDFT1, | 540 (7) |
| APP,HYOU1,I | |
| CACYBP,CCN | |

| | |
|---|---|
| FTH1,IREB2, | 249 (5) |
| APP,MAP2,S | 738 (7) |
| ACTA2,COL1 | |
| CDKN1A,CDK | 632 (7) |
| FASN,HMGC | |
| HLA-A,MMP | |
| ATF4,SLC2A1 | 340 (9) |
| ITGA2,ITGA3 | |
| CDKN1A,SQS | 970 (11) |
| CD44,MMP2, | 621 (7) |
| ACTB,APP,M | |
| ENO2,NCAM | |
| CCND1,CDKN | 920 (11) |
| ACAT2,ANAP | |
| APLP2,CCND | |
| CCND1,FN1, | |
| ATF4,ATF5,C | |
| CCND1,CHGB | |
| CDKN1A,CDK | 989 (18) |
| ATF4,CCN2, | 490 (6) |
| CDH2,FGFR1 | |
| CCND2,CDH2 | 950 (17) |
| APP,CD151, | 738 (7) |
| ACACA,ACAD | |
| CCN2,DDIT4, | |
| ACTB,ACTN1 | 1106 (21) |
| ACACA,ACLY, | 1173 (20) |
| ACTA2,CCND | 1105 (18) |
| ABLIM1,ACT | 926 (12) |
| ATF4,B2M,B | 899 (12) |
| ABL1,ABLIM | 1313 (19) |
| ACO2,APP,FT | |
| ATF4,CCND1, | 1102 (19) |
| CDH2,MMP1 | |
| CDKN1A,DCT | 622 (7) |
| DDIT4,HIF1A, | 597 (7) |
| CNTN2,DLG4 | |
| ACO2,ATF6, | 1120 (22) |
| ACTB,CCND1 | 1091 (20) |
| CCN2,FSCN1, | 834 (11) |
| ARGLU1,CRA | |
| AKR1C3,COX | 984 (11) |
| APP,CCND1, | 1035 (19) |
| H2AZ1,NOX4 | 249 (7) |
| ARF4,DERL1, | |

| | |
|---|---|
| CDKN1A,IGFI | 1165 (20) |
| COL1A1,CYP5 | 980 (18) |
| ACTB,APP,AT | 1173 (18) |
| ADAMTS1,AK | 1175 (24) |
| ADAMTS1,CF | 1132 (20) |
| CDH2,CLU,CT | 1279 (19) |
| CADM1,CALR | |
| ACTA2,ATP6/ | 947 (15) |
| ACTA2,AGPA | 1043 (21) |
| ACTA2,BASP | 1196 (19) |
| ABL1,CDKN1 | 1089 (20) |
| CDKN1A,CDK | 998 (20) |
| CSK,FN1,HDA | 872 (10) |
| ALCAM,CD15 | 961 (13) |
| AMD1,COL5A | |
| ECH1,ECHS1, | |
| BCAR1,CDKN | 413 (8) |
| ATF4,DISP3,H | 371 (8) |
| ATF4,DISP3,H | 295 (7) |
| CCN2,COL1A | 1081 (22) |
| ACACA,ACLY, | 790 (11) |
| ACTA2,ATF5, | 1066 (20) |
| ACHE,ACTA2, | 1143 (20) |
| ATF4,ATP2B | |
| AKT3,CCN1,C | 1178 (17) |
| ATF4,BOP1,C | |
| A2M,APP,CCI | |
| APP,AQP1,CA | |
| ABCA3,AKR1 | |
| CCND1,CDKN | |
| ACTA2,APP,A | |
| CTNNB1,CXC | |
| ACACA,ASNS | |
| CCND1,CDK4, | |
| ARL6IP1,CCN | |
| B2M,CDKN1A | |
| ACTA2,ACTB, | |
| ABCA3,ACAC | |
| CDKN1A,H2A | |
| CCDC80,CDKN | |
| ACADVL,ACTA | |
| ABCA7,ACHE | |
| A2M,BNIP3,E | |
| ATF4,ATF6,D | |
| ACADVL,COX | |

CREBBP,EEF
ACTA2,ITGA
APOE,FLOT1,
ALDOC,CDK5
CDKN1A,CTN
CCND1,CCND
ADAMTS1,A
ANXA2,COL1
ATP1A1,CDH
A2M,ADAMT
CCND1,CD44,
ATF6,CDKN1
ABLIM1,APL
ACACA,ACLY,
APC,CDH2,CR
APP,ATP2A2,
A2M,ABCD3,
AKT3,CCND1,
CRABP2,IGF
ANXA2,ENO
CDH2,CTNNB
CDH2,FN1,M
CRP,JUN,MA
CCN1,CCN2,
EP300,FN1,G
DHFR,IGF1R,
CD47,NOTCH
FBLN5,RELA,
HIF1A,ITGB1
ACTA2,CDKN
CDKN1A,ITG
CCND1,CTNN
APP,HIF1A,S
ALCAM,ANX
ACTN1,ARH
ADH5,AKT2,
AQP1,ARL4C,
BGN,CALD1,
CD44,CDH2,
ACACA,FASN
CCND1,GRN,
CDH2,CDKN1
CCND1,DUSP
ADAM10,CC
CDH2,CDKN1

ACSS2,FASN,
FN1,ITGA2,M
EFNB2,JUN,F
ASNS,ATP2A
ELN,FTH1,FT
CDKN1A,IGF
CDKN1A,CRC
CXCL12,EGLN
CCND1,CCND
ATP2A2,CALU
ALDH2,ELN,G
ALDOC,ASNS
ATF4,CCND1,
BRD2,BRD4,I
CCN1,CCN2,C
CAD,CDK4,DH
ATF4,CCND1,
C4A/C4B,CDK
CCND1,CCNG
ARRDC3,B3G
AKAP8L,AQP
APP,CCND1,C
HOXB5,HOXB
ARL4C,CALD1
CCND1,CCND
ARL6IP5,CCN
APOE,APP,AS
CDKN1A,DUS
CDH4,CTTN,F
COL4A5,EDIL
ACTA2,ACTN
ADAMTS1,AT
ADAM10,ADI
ACTA2,ARF4,
ABL1,APC,CC
CCN2,COL1A
A2M,ANXA1,
ABLIM1,BNIP
CTNNB1,DLG
ADAM10,B2M
APP,ARL4C,A
ACADVL,ALD
ANXA1,COL4
ACBD5,APP,A
DARS1,GLO1

CCND2,ELN,
CCND1,CDKN
CDKN1A,FHL
APP,E2F6,KD
ACTA2,CUL4
CDH2,CDKN1
CDKN1A,GSK
CCND1,CDKN
CCND1,CDKN
ADAMTS7,G
CCN1,CCN2,
CD44,EZR,F
BEX1,CCND2,
CCND1,CDH2
AMOTL2,CC
ARHGEF2,B4
ATF5,CNTN1
ALCAM,CCN
ACTA2,C4A/
ACO2,ALDO
A2M,ACHE,A
A2M,CDKN1
BGN,CCN1,C
CELF1,CELF2,
CBS/CBSL,C
CDKN1A,CDK
ACTA2,ADIP
ACLY,ACTA2,
CDKN1A,RPL
APOE,HMGC
CCN2,NOX4
CTSB,MMP2
BACE1,TXN
ACTA2,RHOA
CDKN1A,TNF
ABLIM1,CD4
MAP1LC3B,S
SQSTM1,VE
CDKN1A,MD
ERBB2,FN1
ASAH1,CERS
APP,BACE1
FTH1,TFRC
ACTB,ENSA
CCN1,JUN

APP,BACE1
ITGB1,TLN1
ARRDC4,TXN
SQSTM1,TAF
CDKN1A,HIF1
APP,BACE1
LDHA,SLC2A1
ELN,SNCA
CCND1,EIF4H
ACTA2,COL1A
MCL1,SQSTM
CCND1,RHOA
HMGCR,LDLR
CYP26B1,GSI
BACE1,CDKN
CDK4,NEFL
HLA-A,HSPA1
CTNNB1,TIM
HNRNPA2B1
HK1,SLC2A1
ERBB2,SHC1
MMP14,YBX1
DEK,SOX9
ALCAM,ARHG
ERBB2,MTA1
ARHGEF25,S
CDKN1B,DICE
FTH1,TFRC
APP,SNCA
APP,BACE1
CCND1,CDKN
CTNND1,FN1
MAP2,MAPT
CDKN1A,MDI
EGR1,TRAF3
CDKN1A,STA
NANOG,SOX
CCND1,EEF2
BGN,VCAN
CTNNB1,JUN
CTNNB1,GSK
PDPK1,PPP2C
CDKN1A,FMR
MAP1LC3A,M
APP,BACE1

SORT1,VPS3
CDH2,VIM
RELA,STAT3
SEPTIN2,SEP
HIF1A,VEGF
CD74,MIF
HIF1A,SLC2A
CCND1,MCL1
NES,SOX2
ATF4,XBP1
CDKN1A,CDK
CCND1,TNFR
GNAS,HIF1A
CTBP2,CTNN
APP,TCF3
MGAT3,MGA
CCN2,FN1
CTBP2,CTNN
LMO4,NANO
CCND1,CDH2
IGF1R,MKNK
COL1A1,FCG
CCND1,CCND
CCND1,CDKN
ACTA2,APP,C
CHGA,CHGB,
CDKN1A,GSR
C1QL1,CCND
ACTA2,CAPZ
CCND2,CSNK
ACTA2,ACTN
CCND1,CDKN
ANXA1,ANXA
CCND1,CDKN
CDKN1A,MCL
PFDN1,PFDN
CCND1,CD44,
ACACA,FDFT
H2AX,POLR2
ACTA2,CCND
HIF1A,MMP2
CCND1,ITGA
H2AX,HIF1A,
SEPTIN3,TAG
CD81,DICER1

ALDOC,HIPK2
CCND1,CDKN
CTNNB1,H2A
CCND2,CDKN
CCND1,CTNN
ACTN1,ACTN
COL2A1,IGF2
ITGA2,ITGB1
H2AX,PRDX1,
NDUFA13,NI
GSTP1,PSMB
CHGA,NCAM
POSTN,PPP3
CCND1,ITGA
CTNNB1,GSK
FTH1,LOX,LO
CDKN1A,CDK
ITGA2,ITGAV
ACTA2,CD44,
GPX4,GSR,SC
ACTA2,COL1
HIF1A,MT2A,
ACACA,ACHE
ALCAM,ANX
ACACA,ACLY,
ACSL4,CCN2,
APEX1,ATRX,
ACACA,ACO2
ACACA,FASN
CDKN1A,ELN
CDC42,CDH2,
CDKN1B,EZR
CDH11,CDKN
CDKN1B,COL
ARHGDIA,IDI
ACVR1B,BCL
CCN1,CCN2,N
CCND1,CDKN
COL1A1,ITGB
APP,BACE1,G
CCND2,CDKN
AK1,ENO1,G
APP,CXCL12,I
HMGCR,LSS,
CLPX,HSPD1,

ACTA2,COL1
ACHE,ADAM
ADH5,CCND1
AFDN,ALDO
CKB,H2AX,H
CCND1,COL1
ACACA,ACTA
ACACA,ANXA
CAD,CCND2,
ACHE,ACTA2,
CCND1,CDKN
CSNK2B,FBL,
AGAP1,CAL
ACTA2,ADD1
ACTA2,ACTN
ACTA2,CD44,
CCND1,CDK9,
CCN1,CCN2,
MAP1LC3B,
HNRNPH1,PT
CCND1,CTSD,
AUTS2,CHD2
APP,BACE1,
CDKN1A,ITG
AGRN,MAP1
APOE,CCND1
ITGA3,MMP
CCND1,CDH2
CCND1,CDK4,
APBB1,CCN1
GABRB3,TA
CCND1,CDH2
APC,DIAPH3,
CDKN1A,FA
MCL1,PPP2C
CCND1,ERBB
CCND1,H2AX
CDH11,JUN,
CDH2,COL1A
APP,DCX,DLG
CHGA,HES6,
ANXA2,CDKN
DNMT1,H2A
ACTA2,FN1,
ACAT2,AKR1

ATXN10,CCR
C4A/C4B,CO
CDH11,CDKN
ANXA2,CAPN
CCDC80,COL1
ACTA2,ANXA
CCND2,CD44,
ACTA2,AKAP
ACTB,ALDOA
EGR1,HMGC
ATF4,CCND1,
CDKN1A,CTN
CDH2,FN1,M
CCND1,FGFR
ATP1A1,NCO
CLSTN1,DLG
CCND1,CDKN
ACTA2,ACTB,
ADAM10,CC
ARRDC4,BEX
CCN2,COL1A
CDKN1A,COL
ATF4,ATF5,P
CDKN1A,CTN
ACTA2,CCND
CDKN1A,CY
CDKN1A,CY
APP,CDH2,CT
ACTA2,AMO
A2M,ACACA,
CORO1C,CO
ANXA1,C4A/
BGN,CCN2,C
ALDOA,ANX
ACTA2,CCN2,
A2M,ALDOC,
ACSL3,ACSL4
ATP5F1B,AT
ATF4,ATF6,C
ACTB,ASNS,
CCND1,CDK4,
ACTA2,COL1
ACIN1,AJUB
ACTB,ATF4,C
ACLY,ATF4,A

ACTA2,CCN2,
ATF4,ATP2A2
A2M,AKR1C3
ATP2A2,B3G
AP2M1,CCND
ACACA,ACLY,
ABCA2,ABCA
ATXN10,CHM
AKT2,AKT3,C
ACACA,ACTA
ATP6V0B,AT
ACLY,CALR,C
ALCAM,BGN,
MMP14,NOX
COL12A1,CO
CCT6A,GPI,H
A2M,CCND1,
CD44,CDC42,
CCND1,CD44,
CDH2,DAG1,
CDKN1A,DHF
AK1,ALDOA,
ACACA,CCND
CDKN1A,CDK
APP,CD74,IG
CDKN1A,HSP
CTNNB1,EIF4
APC,BTG1,CC
CCND1,CDKN
A2M,ABR,AC
CBX5,DCTN4,
CCND1,CDKN
CCN1,CCN2,C
ACHE,CCN2,C
A2M,ABCD3,
ACTA2,CCND
ABL1,CCND1,
ACTA2,ADAM
ADIPOR1,AR
A2M,ACTA2,
CCND1,CD27
ACTA2,CDKN
ADIPOR2,CK
ACACA,ARRD
APOE,APP,CC

ACACA,FASN
CCND1,CDKN
ATXN10,CCN
ATF4,FASN,H
CDH2,CTNNB
CDH2,CTNNB
GRN,MAPK1,
CCND1,CDH2
APC,CCND1,C
CDK4,CDKN1
APP,EGR1,M
ABCD3,ACAD
BNIP3,CDKN1
CDKN1A,CDK
ADAM9,CD16
APP,CCN2,CC
ADAMTS1,AD
ACTA2,AKIRI
BCL11A,CDKN
CCND1,CCND
ITGB1,LAMP
APOE,CDKN1
ACTA2,CCN2,
NOTCH1,SNC
EGR1,ELN,N
EPHB2,FDFT1
ATXN10,NEL
FN1,HIF1A,M
HLA-A,HLA-B
ACACA,ACSS
CCND2,CDKN
CCN2,CTNNB
CCND2,CDK4,
EGR1,FN1,N
CTSD,MMP2,
AKR1C3,ARP
CCND1,COL1
ANXA1,APP,B
CCND2,CDKN
ACACA,AKR1
CCN1,CCN2,C
ANXA1,CDKN
ACACA,ACAD
CDKN1B,DNA
ACLY,ATP5F1

CCND1,EZR,F
ATP5PD,ERBI
APP,BACE1,C
ACACA,CCN2
ACTA2,APP,C
CDKN1A,ROC
CDKN1A,EI24
CCN2,FLNA,N
CDH2,FN1,VI
CCND1,CDH2
COL4A2,COL5
MMP14,MMI
CTNNB1,HSP
SHC1,SYP,UL
PFDN1,PFDN
CCND1,CDK4,
AQP1,TAGLN
ROCK2,SOD2
CDKN1A,CDK
CTNNA1,GJA
CTNNB1,RHC
FTH1,GPX4,N
MCL1,RAC1,V
APEX1,CRABI
ATF4,ATF6,C
CCN2,ELN,FB
NOX4,SQSTN
EGR1,MTOR,
ASNS,CDKN1
NANOG,SOX
COL11A1,FN
CDKN1A,MDI
PCNA,SOD1,S
EGR1,FASN,I
APP,IGF1R,PT
CCND1,MTOI
CDC42,RAC1,
CCND1,CCND
ABCA3,ADAN
ACTA2,APP,C
AKR1C4,ATP1
ACACA,ADIPC
CAMKK2,CCN
CSE1L,CTNNI
GOT2,NDUF/

CDKN1A,CDK
ACTA2,ALDH
APLP2,CELSR
ACAT2,ACLY,
CCND1,CDKN
ACHE,CCND1
ACACA,ATP5
ACTA2,CCN2,
CCND1,CCND
ACTB,CDKN1
ASAH1,FADS
CDH2,CDKN1
ASNS,CAMK
APP,CCND2,
ADH5,CD59,
CDKN1A,GPA
B2M,CTSB,P
CTSD,KLF6,P
ACTA2,APP,
CCND1,CDKN
ACTA2,CCN2,
ABL1,CDKN1
APC,CSNK1A
HIF1A,IREB2,
CDKN1A,COL
COL1A1,KIF3
APP,CD44,EI
ARF3,CD74,F
CCND1,FN1,
CDH2,ITGB1,
COL5A2,CTN
ARHGEF2,BG
ANXA5,APP,
ATP1A1,CYP
ACTA2,APP,
APP,CCND1,
ASNS,ATF5,
ALDOA,ALD
AGAP1,ANX
ACAT2,ATP2
AGPAT1,ANX
ACACA,ACLY,
CCN2,CDKN1
ANXA2,CCN
FAM89B,HS

CCND1,CDC4
ATF4,ATF6,C
ACTA2,APP,C
CNR1,CYP26
BNIP3,CDKN
ACTA2,CCND
ACTA2,APOE
ACTA2,BGN,
APEX1,ATP1
ACTB,APOE,
CAPNS1,CCN
ABCD3,ABLI
CCND1,CNN3
ATP5F1B,CC
AKT2,GAB2,
A2M,ANXA1,
ADAMTS1,AJ
APP,CCND1,
ACSL6,ATAT1
CCND1,CDH2
CCND1,CDK4,
CCND1,CDKN
ACACA,EGR1
ANXA1,CSNK
ACSL3,AGPA
ACTA2,ATF4,
ACTA2,CCN2,
ACLY,CLU,DH
ALDH18A1,A
ACLY,ADAR,
CCND1,CCT8,
ACP1,AJUBA,
ACLY,CALR,G
DLG4,ERBB2
CDKN1A,FN1
CDKN1A,NA
EDNRB,GLU
ATF4,CEBPG,
CDKN1A,CRA
DLK1,H19,IG
MTA2,RELA,
CTNNB1,GPS
CCND1,CDH2
CCND1,CD44,
ANXA2,EP30

CCN1,EGR1,I
CCND1,E2F6,
BCLAF1,HMG
CDK4,IGF1R,
CCND1,CTNN
ID2,KRT8,VCA
EGR1,PTP4A
CCND1,EGR1
MSH2,PRKDC
CCT8,EGR1,E
ASNS,ATF4,H
ATP2A2,CDH
DDIT4,SLC1A
HMGCR,LDLR
EGR1,GJA1,F
ACTA2,CCN2,
CHGA,LDLR,N
CCND1,CDK4,
CCNG1,CIRBI
APBB1,EFNB
CCN1,CCN2,C
CDKN1A,CDK
B2M,CD74,CI
BGN,CCN2,C
C8orf44-SGK
CCN2,CCND1
ALCAM,CCND
CCND1,CCND
CCND1,CCND
CCND1,CTNN
DHCR7,ECHS
A2M,ACHE,A
CDKN1A,CDK
ACSL4,CCND
ARHGEF2,AS
CCND1,CDK4,
CDKN1A,CDK
ACTB,CCND1
DCX,LDLR,SN
ACACA,ACLY,
CDKN1A,CTN
CCND1,CDKN
CNR1,DCX,M
CHGA,CNR1,
ATF4,FASN,H

CCND1,CDKN
STAT1,STAT2
ACTA2,SRF
CCN1,CCN2
CDKN1A,CDK
CCN1,CCN2
CTNNB1,VIM
ATP6AP2,FN
NOX4,TKT
TPM1,TPM3
A2M,CCND1
APP,NES
CDKN1A,COL
CCN1,CCN2
NOTCH1,NOT
CDKN1A,MSI
CDKN1A,MDI
CDKN1A,CDK
CCND1,PKM
CDKN1A,CDK
VIM,ZEB1
MMP2,XBP1
ATF4,HIF1A
CCND1,CTSD
EIF4G1,EIF4I
CCN1,CCN2
MDM2,SNCA
LAMP1,MAP
CD44,CDH11
CUL1,CUL4A
ZEB1,ZEB2
CCND1,JUN
CCN1,CCN2
GRIA1,SNAP
CDKN1A,SEP
VIM,ZEB1
CDH2,VIM
CCND1,CDKN
EFNB2,EPHB
ATF4,CTTN
MAPK1,MAPI
CDKN1A,SOD
HMGCR,ITGE
COL1A1,XIST
APP,PER1

ACTA2,TAGL
CCND1,CTNN
CCND1,LDLR
ATP2A2,CCN
FYN,RAD51
CCND1,ERBB
CCND2,CDKN
ACTA2,HTRA
CCND1,CDKN
CDKN1A,CDK
ACTA2,ROCK
BCOR,BTG1
CYFIP1,FMR1
APP,MCL1
CTNNB1,GSK
JUN,PCNA
CCND1,MCL1
GRK2,STAT3
MCL1,ULBP1
PLXNA2,SEM
ATF4,CCND1
CCND1,CCND
DLG4,SYP
APP,MAP2
JUN,PRKDC
CCND1,MAPT
CTNNB1,VIM
ID2,ODC1
CD44,IGF2
L1CAM,SPTB
SERPINF1,VE
PRKAR2A,PR
CDKN1A,CDK
ATF6,XBP1
HSP90B1,PDI
ACTN4,MAPT
SNAP25,VAM
APP,VDAC1
MAPK3,SLC2
SNAP25,SYP
CDC42,CDKN
NANOG,SOX
TOMM40,VD
CDKN1A,TNF
ID2,ODC1

AZIN1,VEGF
SNCA,ZEB2
NANOG,SAR
CCND1,CTNN
HSP90B1,HS
NF2,YY1
APP,EZR
CDKN1A,MD
ATP2A2,MT
CCN2,LOX
FTH1,TFRC
CCND1,SQST
CCND1,SQST
PCNA,SOD2
GPX4,NCOA
ASNS,EIF1
CDKN1A,MD
NANOG,SOX
HMGCR,PITP
CDKN1A,HIF
PPP1R9B,SY
HIF1A,SLC2A
ACTA2,HIF1
TIMP1,TIMP
NDRG1,VEG
CDKN1A,CDK
APEX1,HIF1A
ATF4,XBP1
APP,SQSTM1
EIF4G2,JUN
CDKN1A,GJA
ACTA2,COL1
CCND1,MCL1
APP,MMP2
ATP13A2,GA
APP,BACE1,
ACTA2,ANXA
ACADVL,ACS
BMF,CCN2,C
ATP6V1B2,B
CCND1,CDK4,
ABLIM1,ABR
CCND1,CD44,
BSG,CCND1,
ACTA2,AFAP

ACACA,ATF6,
ACTA2,CDKN
ACVR1B,ANX
CCND1,CDKN
CHRNA4,RHO
ACTA2,SOX2,
CTNNB1,RHO
CTSD,MMP1
CCND1,MCL1
CD44,CDKN1
ACTB,CCND1
ACTA2,APP,
ACADVL,CDK
ITGA2,ITGAV
CANX,GPM6
HSPG2,MAP
GRIA1,NES,S
CTNNB1,GSK
CCND1,CDKN
CCND1,CTNN
MDM2,PDPK
RPL11,RPL22
NUP133,NU
CCND1,CDKN
CD44,NANO
CCND1,CD44,
GABBR1,GL
CCN2,MCL1,
GSTP1,HIF1A
CDKN1A,MD
HOXA5,HOX
CCND1,NAN
CDKN1A,ME
CTNNB1,MN
CANX,GPM6
CCND1,LAM
HERPUD1,KI
CDH2,MMP2,
CD59,HMGC
CDKN1A,ERB
APP,DLG4,F
CCN1,FMR1,
ABCD3,ACAC
A2M,ACTA2,
A2M,AQP1,C

ALDOC,ANKR
BRD4,PSMD7
APOE,ATP2A
ACTB,ADAM1
AKT2,CHGA,
ATP13A2,CD
ACACA,ACAD
ACLY,ACTG1,
CHGA,ENO2,
BRD2,BRD4,
ATF4,CCN1,C
COL1A1,ITG
ATP1A1,CDK
BEX3,BLMH,
ACACA,FN1,
ATP8A2,CD4
ACACA,ACLY,
CCND2,CDH2
ACSL4,AGRN
CCN2,CHST1
COL4A2,CRA
ATG9A,BNIP
ADAM10,ATP
CDKN1A,CDK
ACTA2,AGAP
CCN1,CCND1
ABL1,ACACA,
ABCA2,ACAC
ACTA2,CCN2,
SLC1A4,SLC1
LMNA,NANC
ATP2A2,BMP
AFDN,ALCAM
AMOTL2,ATP
ARRDC4,CAP
ACTA2,CCN2,
ABCD3,ADAM
ABCD3,ACAC
ATF4,GNB1,
ACTA2,CCND
FKBP1A,HSB
ACACA,APP,
A2M,DKK3,E
CITED2,COPS
ACACA,CDKN

ANXA5,APP,A
APEX1,CDKN
ADAM10,APC
ABCA2,ACAC
FTH1,FTL,HIF
CCND1,CD44,
ARHGEF2,B4
CCND1,CCND
ARRDC4,CCN
LTBP3,SRF,V
ACACA,ACTA
ACACA,ATP2
CDH2,MMP2,
CCND2,CCNG
A2M,ADAMT
ANXA2,FAT4
HIF1A,PGK1,
ACACA,CTNN
ABCA3,AQP1
B2M,CACNA

© 2000-2021 QIAGEN. All rights reserved.

| ID | Consistency : | Node Total | Regulator To | Regulators | Target Total | Target Molec |
|---|---|---|---|---|---|---|
| 1 | 3.873 | 17 | 1 | RAC1 | 15 | APBB1,CCN2 |
| 2 | 3.671 | 21 | 1 | F2 | 19 | CCN1,CCN2,C |
| 3 | 3.464 | 14 | 1 | COL18A1 | 12 | CCND1,CTNN |
| 4 | 3.334 | 28 | 1 | IL15 | 26 | ATF4,CCN2,C |
| 5 | 3.317 | 13 | 1 | COL18A1 | 11 | CCND1,CTNN |
| 6 | 3.317 | 13 | 1 | COL18A1 | 11 | CCND1,CTNN |
| 7 | 3.25 | 18 | 1 | RAC1 | 16 | APBB1,CCND |
| 8 | 3.25 | 18 | 1 | RAC1 | 16 | APBB1,CCND |
| 9 | 3.162 | 12 | 1 | SPHK1 | 10 | APP,CCN2,CC |
| 10 | 3.064 | 20 | 1 | EDN1 | 18 | ACTB,CDC42, |
| 11 | 3.064 | 20 | 1 | EDN1 | 18 | CCN1,CCND1 |
| 12 | 3.015 | 13 | 1 | GAST | 11 | ATF4,CCND1, |
| 13 | 2.91 | 19 | 1 | RPTOR | 17 | ACACA,CCND |
| 14 | 2.907 | 22 | 1 | EBF1 | 20 | AKT2,AKT3,C |
| 15 | 2.887 | 14 | 1 | NCOA3 | 12 | CCN1,CCN2,C |
| 16 | 2.858 | 8 | 1 | PKM | 6 | CCND1,CDKN |
| 17 | 2.846 | 12 | 1 | PIN1 | 10 | CCND1,CDKN |
| 18 | 2.846 | 12 | 1 | PRKCE | 10 | APP,CCN1,CC |
| 19 | 2.828 | 10 | 1 | CCN2 | 8 | CCND1,COL1, |
| 20 | 2.828 | 10 | 1 | TGFB2 | 8 | CCN2,CD59,C |
| 21 | 2.828 | 10 | 1 | alvespimycin | 8 | ACACA,CCND |
| 22 | 2.774 | 15 | 1 | EDN1 | 13 | CDC42,CDH2, |
| 23 | 2.774 | 15 | 1 | PCGEM1 | 13 | ACACA,ACLY, |
| 24 | 2.714 | 13 | 1 | miR-16-5p (a | 11 | CCND1,CCND |
| 25 | 2.667 | 11 | 1 | FOXC2 | 9 | CDKN1B,CTN |
| 26 | 2.667 | 11 | 1 | SPZ1 | 9 | CCND1,CD44, |
| 27 | 2.667 | 11 | 1 | miR-133a-3p | 9 | CCN2,FSCN1, |
| 28 | 2.646 | 9 | 1 | F7 | 7 | CCN1,CCN2,C |
| 29 | 2.646 | 9 | 1 | INSIG2 | 7 | ACACA,ACLY, |
| 30 | 2.646 | 9 | 1 | miR-155-5p | 7 | CCN1,CCND1 |
| 31 | 2.598 | 14 | 1 | RAC1 | 12 | APBB1,CDH2 |
| 32 | 2.53 | 12 | 1 | KAT2A | 10 | ACTN4,CCND |
| 33 | 2.53 | 12 | 1 | RHOA | 10 | CCND1,CDKN |
| 34 | 2.53 | 12 | 1 | phytohemag | 10 | APP,CD47,CR |
| 35 | 2.53 | 12 | 1 | sphingosine- | 10 | ACTA2,ANXA |
| 36 | 2.5 | 6 | 1 | A2M | 4 | CCND1,CDH2 |
| 37 | 2.496 | 15 | 1 | SCAP | 13 | ACACA,ACLY, |
| 38 | 2.475 | 10 | 1 | diphenylenei | 8 | EGR1,FN1,HI |
| 39 | 2.475 | 10 | 1 | mir-148 | 8 | ALCAM,COL1 |
| 40 | 2.475 | 10 | 1 | mir-17 | 8 | CCN2,CCND1 |
| 41 | 2.475 | 10 | 1 | mir-193 | 8 | ACTN4,CD15 |
| 42 | 2.449 | 8 | 1 | BSG | 6 | CCND1,CDKN |
| 43 | 2.449 | 8 | 1 | ESRRA | 6 | ASAH1,CHKA |

| | | | | | |
|---|---|---|---|---|---|
| 44 | 2.449 | 8 | 1 | GAST | 6 CTNNB1,MA |
| 45 | 2.449 | 8 | 1 | miR-182-5p | 6 FN1,IGF1R,IT |
| 46 | 2.449 | 8 | 1 | miR-182-5p | 6 FN1,IGF1R,IT |
| 47 | 2.449 | 8 | 1 | napabucasin | 6 DDR1,ERBB2 |
| 48 | 2.449 | 8 | 1 | ramipril | 6 COL1A1,DDR |
| 49 | 2.405 | 16 | 1 | ERK | 14 ADAM10,CCN |
| 50 | 2.333 | 11 | 1 | 26s Proteaso | 9 CDKN1A,CDK |
| 51 | 2.333 | 11 | 1 | IL5 | 9 ANXA2,CCND |
| 52 | 2.333 | 11 | 1 | MKNK1 | 9 APC,ATAT1,F |
| 53 | 2.333 | 11 | 1 | cobalt chloric | 9 ADAMTS1,CD |
| 54 | 2.333 | 11 | 1 | ibuprofen | 9 APP,CCND1,E |
| 55 | 2.309 | 5 | 1 | PINK1 | 3 CCND1,HIF1A |
| 56 | 2.309 | 5 | 1 | PINK1 | 3 CCND1,HIF1A |
| 57 | 2.309 | 14 | 1 | RAC1 | 12 CCND1,CTNN |
| 58 | 2.268 | 9 | 1 | AG490 | 7 CDKN1A,HIF1 |
| 59 | 2.268 | 9 | 1 | BAX | 7 CDKN1A,CDK |
| 60 | 2.268 | 9 | 1 | CD38 | 7 ANXA2,CCND |
| 61 | 2.268 | 9 | 1 | FLI1 | 7 CCN2,CCND1 |
| 62 | 2.268 | 9 | 1 | GPER1 | 7 ASAH1,CCN1 |
| 63 | 2.268 | 9 | 1 | KDM3A | 7 AJUBA,ALCA |
| 64 | 2.268 | 9 | 1 | L-glutamic a | 7 CCND1,CDKN |
| 65 | 2.268 | 9 | 1 | MKNK1 | 7 AGRN,CCN1, |
| 66 | 2.268 | 9 | 1 | MTORC1 | 7 CCND1,FASN |
| 67 | 2.268 | 9 | 1 | PRKAA1 | 7 CDKN1A,CLU |
| 68 | 2.268 | 9 | 1 | ROR1 | 7 AMOTL2,CCN |
| 69 | 2.268 | 9 | 1 | mir-29 | 7 BACE1,CDC4 |
| 70 | 2.236 | 7 | 1 | 2-bromoethy | 5 ADAMTS1,AN |
| 71 | 2.236 | 7 | 1 | EGR2 | 5 EPHA4,MAP2 |
| 72 | 2.236 | 7 | 1 | FBXW7 | 5 CDKN1B,HM |
| 73 | 2.236 | 7 | 1 | GDF2 | 5 CXCL12,FGF |
| 74 | 2.236 | 7 | 1 | HEXIM1 | 5 ANXA1,CCND |
| 75 | 2.236 | 7 | 1 | IKBKE | 5 ACLY,CCND1, |
| 76 | 2.236 | 7 | 1 | KIF3A | 5 ACLY,CDKN1 |
| 77 | 2.236 | 7 | 1 | MBTPS1 | 5 ACACA,FASN |
| 78 | 2.236 | 7 | 1 | MET | 5 CDH2,ITGA3, |
| 79 | 2.236 | 7 | 1 | NF1 | 5 CCND1,CTNN |
| 80 | 2.236 | 7 | 1 | NF1 | 5 CCND1,CTNN |
| 81 | 2.236 | 7 | 1 | PDGFC | 5 CCND1,CTNN |
| 82 | 2.236 | 7 | 1 | PPP1R1B | 5 CCND1,CD44, |
| 83 | 2.236 | 7 | 1 | PPP1R1B | 5 CCND1,CD44, |
| 84 | 2.236 | 7 | 1 | RUNX3 | 5 CDKN1A,IL6S |
| 85 | 2.236 | 7 | 1 | RUNX3 | 5 CDKN1A,IL6S |
| 86 | 2.236 | 7 | 1 | SEMA7A | 5 CCN2,CTSB,E |
| 87 | 2.236 | 7 | 1 | TEAD | 5 CCN1,CCN2,C |
| 88 | 2.236 | 7 | 1 | allopurinol | 5 ADAMTS1,AN |

| | | | | | | |
|---|---|---|---|---|---|---|
| 89 | 2.236 | 7 | 1 | branched cha | 5 | CTNNB1,HIF1 |
| 90 | 2.236 | 7 | 1 | fenamic acid | 5 | ADAMTS1,AN |
| 91 | 2.236 | 7 | 1 | gentamicin C | 5 | ADAMTS1,AN |
| 92 | 2.236 | 7 | 1 | hexachlorobe | 5 | ANXA1,CD44 |
| 93 | 2.236 | 7 | 1 | leupeptin | 5 | APP,IGF1R,IG |
| 94 | 2.236 | 7 | 1 | leupeptin | 5 | APP,IGF1R,IG |
| 95 | 2.236 | 7 | 1 | miR-145-5p | 5 | IGF1R,MDM2 |
| 96 | 2.236 | 7 | 1 | miR-145-5p | 5 | IGF1R,MDM2 |
| 97 | 2.236 | 7 | 1 | miR-199a-3p | 5 | CD44,FN1,MT |
| 98 | 2.236 | 7 | 1 | miR-450a-5p | 5 | GSK3B,HNRN |
| 99 | 2.236 | 7 | 1 | triamterene | 5 | ADAMTS1,AN |
| 100 | 2.219 | 15 | 1 | TCR | 13 | CCND1,CD44, |
| 101 | 2.138 | 16 | 1 | FGF2 | 14 | CCN1,CTNNB |
| 102 | 2.121 | 4 | 1 | ARHGAP5 | 2 | COL1A1,FN1 |
| 103 | 2.121 | 10 | 1 | CCAR2 | 8 | CCND1,KIF1A |
| 104 | 2.121 | 10 | 1 | PRKAA2 | 8 | CDKN1A,CDK |
| 105 | 2.121 | 10 | 1 | SIRT6 | 8 | ACSL4,AKT3, |
| 106 | 2.121 | 10 | 1 | Y 27632 | 8 | ACTB,AMOTI |
| 107 | 2.121 | 10 | 1 | carbamazepi | 8 | CTNNB1,EIF4 |
| 108 | 2.121 | 10 | 1 | let-7 | 8 | ACTA2,CCND |
| 109 | 2.121 | 10 | 1 | miR-141-3p | 8 | CDH2,CTNNB |
| 110 | 2.121 | 10 | 1 | miR-141-3p | 8 | CDH2,CTNNB |
| 111 | 2.121 | 10 | 1 | mir-15 | 8 | APP,CCND1,C |
| 112 | 2.111 | 13 | 1 | NGF | 11 | APP,ELAVL4, |
| 113 | 2.041 | 8 | 1 | ERK1/2 | 6 | ITGA3,MAPK |
| 114 | 2.041 | 8 | 1 | F10 | 6 | CCN1,CCN2,E |
| 115 | 2.041 | 8 | 1 | MRTFB | 6 | CCN1,ITGB1, |
| 116 | 2.041 | 8 | 1 | REST | 6 | CXCL12,DCX, |
| 117 | 2.041 | 8 | 1 | SETD2 | 6 | CCN1,CCN2,C |
| 118 | 2.041 | 8 | 1 | methapyriler | 6 | ANXA2,ATF4 |
| 119 | 2 | 6 | 1 | 4-hydroxytan | 4 | CAPNS1,CCN |
| 120 | 2 | 6 | 1 | ACACB | 4 | ACACA,ACLY, |
| 121 | 2 | 6 | 1 | ARHGEF25 | 4 | CDC42,JUN,R |
| 122 | 2 | 6 | 1 | CSNK2A1 | 4 | CCND1,FN1,F |
| 123 | 2 | 6 | 1 | Cdc42 | 4 | FLNA,GSK3B |
| 124 | 2 | 6 | 1 | IGFBP2 | 4 | CCND1,FN1,I |
| 125 | 2 | 6 | 1 | IPMK | 4 | CAPNS1,CCN |
| 126 | 2 | 6 | 1 | IPMK | 4 | ACTB,CCN1,J |
| 127 | 2 | 6 | 1 | IPMK | 4 | CAPNS1,CCN |
| 128 | 2 | 6 | 1 | ITCH | 4 | CCN1,CCN2,C |
| 129 | 2 | 6 | 1 | MTM1 | 4 | IGF1R,MAP1 |
| 130 | 2 | 6 | 1 | NF2 | 4 | CCND1,CDKN |
| 131 | 2 | 6 | 1 | NF2 | 4 | CCND1,CDKN |
| 132 | 2 | 6 | 1 | PCDH11Y | 4 | CCND1,EP300 |
| 133 | 2 | 6 | 1 | PLAG1 | 4 | EFNB1,IGF2, |

| | | | | | | |
|---|---|---|---|---|---|---|
| 134 | 2 | 6 | 1 | PLAG1 | 4 | EFNB1,IGF2, |
| 135 | 2 | 6 | 1 | RAC1 | 4 | CDH2,FN1,IT( |
| 136 | 2 | 6 | 1 | ST6GAL1 | 4 | CDH2,FN1,HI |
| 137 | 2 | 6 | 1 | ST6GAL1 | 4 | CDH2,FN1,RE |
| 138 | 2 | 6 | 1 | STEAP3 | 4 | ANXA2,CCN2 |
| 139 | 2 | 6 | 1 | STEAP3 | 4 | ANXA2,CCN2 |
| 140 | 2 | 6 | 1 | TAZ | 4 | CDC42,CDH2, |
| 141 | 2 | 6 | 1 | TCF/LEF | 4 | CCND1,FN1,J |
| 142 | 2 | 11 | 1 | TFEB | 9 | ATP6V1B2,C |
| 143 | 2 | 6 | 1 | ethosuximide | 4 | CCND1,CTNN |
| 144 | 2 | 6 | 1 | inosine | 4 | A2M,ALCAM, |
| 145 | 2 | 6 | 1 | inosine | 4 | A2M,ALCAM, |
| 146 | 2 | 6 | 1 | lomustine | 4 | ANXA1,ANXA |
| 147 | 2 | 11 | 1 | maslinic acic | 9 | CCND1,CCND |
| 148 | 2 | 6 | 1 | miR-182-5p | 4 | ITGB1,MTSS |
| 149 | 2 | 6 | 1 | molybdenum | 4 | FN1,L1CAM,I |
| 150 | 2 | 6 | 1 | okadaic acid | 4 | APP,CCND1,J |
| 151 | 2 | 6 | 1 | phenacetin | 4 | ANXA1,ANXA |
| 152 | 2 | 6 | 1 | phenacetin | 4 | ANXA2,CD44 |
| 153 | 2 | 6 | 1 | phenacetin | 4 | ANXA1,CD44 |
| 154 | 2 | 6 | 1 | salinomycin | 4 | CCND1,CTNN |
| 155 | 1.964 | 23 | 1 | IL3 | 21 | CCND1,CCND |
| 156 | 1.897 | 12 | 1 | miR-141-3p | 10 | CDH2,CTNNB |
| 157 | 1.89 | 9 | 1 | CCL5 | 7 | ALCAM,CD44 |
| 158 | 1.89 | 9 | 1 | FGF8 | 7 | CCND1,CTNN |
| 159 | 1.89 | 9 | 1 | TIMP3 | 7 | CREBBP,CTN |
| 160 | 1.89 | 9 | 1 | miR-291a-3p | 7 | CCND1,CCND |
| 161 | 1.89 | 9 | 1 | mir-1 | 7 | CXCL12,FN1, |
| 162 | 1.89 | 9 | 1 | mir-210 | 7 | ABL1,CDH2,C |
| 163 | 1.89 | 9 | 1 | silibinin | 7 | CDH2,CDKN1 |
| 164 | 1.809 | 13 | 1 | phytohemag | 11 | APP,BCR,CCN |
| 165 | 1.809 | 13 | 1 | thioacetamic | 11 | CLU,COL1A1, |
| 166 | 1.807 | 17 | 1 | CD40LG | 15 | CCN2,CD44,C |
| 167 | 1.807 | 17 | 1 | RET | 15 | ANXA1,CCND |
| 168 | 1.789 | 7 | 1 | 6-hydroxydop | 5 | ATF4,JUN,M |
| 169 | 1.789 | 7 | 1 | BMP7 | 5 | CDH2,IGF2,M |
| 170 | 1.789 | 7 | 1 | CYBA | 5 | CDKN1A,FN1 |
| 171 | 1.789 | 7 | 1 | F2R | 5 | CXCL12,FN1, |
| 172 | 1.789 | 7 | 1 | FHIT | 5 | CDKN1A,MM |
| 173 | 1.789 | 7 | 1 | GNRH | 5 | EGR1,JUN,M |
| 174 | 1.789 | 7 | 1 | LAMC1 | 5 | ACTA2,FN1,I |
| 175 | 1.789 | 7 | 1 | LIN28A | 5 | CCND1,CDKN |
| 176 | 1.789 | 7 | 1 | RBL2 | 5 | CCND1,FGFR |
| 177 | 1.789 | 7 | 1 | ROCK1 | 5 | CCN2,CCND1 |
| 178 | 1.789 | 7 | 1 | SDCBP | 5 | CDC42,CTNN |

| | | | | | | |
|---|---|---|---|---|---|---|
| 179 | 1.789 | 7 | 1 | SMAD3 | 5 | CDH11,CDH2, |
| 180 | 1.789 | 7 | 1 | TEAD4 | 5 | CDKN1A,CDK |
| 181 | 1.789 | 7 | 1 | U73122 | 5 | FN1,MMP14, |
| 182 | 1.789 | 7 | 1 | diazoxide | 5 | ACACA,EGR1 |
| 183 | 1.789 | 7 | 1 | glutamyl-Se- | 5 | CLU,EGR1,JU |
| 184 | 1.789 | 7 | 1 | methyl meth | 5 | CDKN1A,CTS |
| 185 | 1.789 | 7 | 1 | miR-17-5p (a | 5 | APP,RHOA,SI |
| 186 | 1.789 | 7 | 1 | nitrofen | 5 | EGR1,FGFR1 |
| 187 | 1.789 | 7 | 1 | nitrofen | 5 | FGFR1,FGFR |
| 188 | 1.789 | 7 | 1 | selenomethy | 5 | CLU,EGR1,JU |
| 189 | 1.768 | 10 | 1 | Collagen type | 8 | APOE,APP,CD |
| 190 | 1.768 | 10 | 1 | F3 | 8 | CCN1,CDC42, |
| 191 | 1.768 | 10 | 1 | F3 | 8 | CDC42,EGR1, |
| 192 | 1.768 | 10 | 1 | FEV | 8 | CNTN1,ITGB |
| 193 | 1.768 | 10 | 1 | FOXM1 | 8 | CCND1,CCND |
| 194 | 1.768 | 10 | 1 | SPDEF | 8 | CCN2,CDH11, |
| 195 | 1.768 | 10 | 1 | U0126 | 8 | CTNNB1,GSK |
| 196 | 1.768 | 10 | 1 | mir-10 | 8 | EPHB2,FGFR |
| 197 | 1.768 | 10 | 1 | sucrose | 8 | ACACA,ACLY, |
| 198 | 1.75 | 18 | 1 | POR | 16 | ACLY,ASNS,A |
| 199 | 1.732 | 5 | 1 | 48s | 3 | CCND1,IGF1F |
| 200 | 1.732 | 5 | 1 | 48s | 3 | CCND1,IGF1F |
| 201 | 1.732 | 5 | 1 | 48s | 3 | ATF4,CCND1, |
| 202 | 1.732 | 5 | 1 | 48s | 3 | CCND1,IGF1F |
| 203 | 1.732 | 5 | 1 | 48s | 3 | CCND1,IGF1F |
| 204 | 1.732 | 5 | 1 | 48s | 3 | IGF1R,PPP1R |
| 205 | 1.732 | 5 | 1 | ALDH1A2 | 3 | CCN1,CCN2,C |
| 206 | 1.732 | 5 | 1 | ARG1 | 3 | GSK3B,MAPT |
| 207 | 1.732 | 5 | 1 | BMS-690514 | 3 | CCND1,CDKN |
| 208 | 1.732 | 5 | 1 | CCN1 | 3 | CTNNB1,ITGA |
| 209 | 1.732 | 5 | 1 | CREBZF | 3 | ATF4,HSPA1A |
| 210 | 1.732 | 5 | 1 | DPP-23 | 3 | ATF4,HSPA1A |
| 211 | 1.732 | 5 | 1 | EIF3M | 3 | EIF3C,EIF3H,I |
| 212 | 1.732 | 5 | 1 | FBN1 | 3 | CCN1,CCN2,C |
| 213 | 1.732 | 14 | 1 | HIF1A | 12 | ADAM10,CDH |
| 214 | 1.732 | 5 | 1 | LIMS1 | 3 | GJA1,HIF1A,I |
| 215 | 1.732 | 5 | 1 | LIMS1 | 3 | GJA1,ITGB1,S |
| 216 | 1.732 | 5 | 1 | LLGL2 | 3 | AMOTL2,CCN |
| 217 | 1.732 | 5 | 1 | MAP2K1/2 | 3 | DUSP4,EGR1 |
| 218 | 1.732 | 5 | 1 | Mir200 | 3 | JUN,SOX2,ZE |
| 219 | 1.732 | 5 | 1 | NPPB | 3 | CCN2,FN1,HN |
| 220 | 1.732 | 5 | 1 | NSUN6 | 3 | FLNA,FSCN1, |
| 221 | 1.732 | 5 | 1 | Nkx2-2os | 3 | CCND1,GNAC |
| 222 | 1.732 | 5 | 1 | Nkx2-2os | 3 | CCND1,GNAC |
| 223 | 1.732 | 5 | 1 | PPBP | 3 | COL6A3,PDG |

| | | | | | | |
|---|---|---|---|---|---|---|
| 224 | 1.732 | 5 | 1 | SUB1 | 3 | CCND1,MDM |
| 225 | 1.732 | 5 | 1 | SUB1 | 3 | CCND1,MDM |
| 226 | 1.732 | 5 | 1 | SUB1 | 3 | CNTFR,MDM |
| 227 | 1.732 | 5 | 1 | TPM3 | 3 | ITGA3,MMP2 |
| 228 | 1.732 | 5 | 1 | TPM3 | 3 | ITGA3,MMP2 |
| 229 | 1.732 | 5 | 1 | ammonium | 3 | CTSB,GRN,V |
| 230 | 1.732 | 5 | 1 | ibuprofen | 3 | APP,BACE1,R |
| 231 | 1.732 | 5 | 1 | miR-142-3p | 3 | BCLAF1,LIFR, |
| 232 | 1.732 | 5 | 1 | miR-296-5p | 3 | CCND1,COL1 |
| 233 | 1.732 | 5 | 1 | mir-133 | 3 | CDC42,FN1,S |
| 234 | 1.701 | 30 | 1 | EGF | 28 | CCND1,CCND |
| 235 | 1.671 | 31 | 1 | genistein | 29 | A2M,ASAH1, |
| 236 | 1.667 | 11 | 1 | ASPSCR1-TFI | 9 | APOE,ASAH1 |
| 237 | 1.667 | 11 | 1 | PP2/AG1879 | 9 | ACSL4,CDKN |
| 238 | 1.667 | 11 | 1 | PRL | 9 | ACTR3,APP,F |
| 239 | 1.667 | 11 | 1 | YAP1 | 9 | CDKN1B,CTN |
| 240 | 1.667 | 11 | 1 | bisphenol A | 9 | ABL1,CDKN1 |
| 241 | 1.667 | 11 | 1 | mir-8 | 9 | CCND1,CCND |
| 242 | 1.664 | 15 | 1 | MITF | 13 | ACHE,ALCAM |
| 243 | 1.633 | 8 | 1 | BRD4 | 6 | CCND1,COL1 |
| 244 | 1.633 | 26 | 1 | EGR1 | 24 | ACHE,APP,AT |
| 245 | 1.633 | 8 | 1 | INHBA | 6 | CCND2,NRG1 |
| 246 | 1.633 | 8 | 1 | Jnk | 6 | APP,ASAH1,C |
| 247 | 1.633 | 8 | 1 | NFYA | 6 | CDKN1A,FAS |
| 248 | 1.633 | 8 | 1 | NOTCH3 | 6 | CDH11,CDKN |
| 249 | 1.633 | 8 | 1 | Ngf | 6 | APP,FGFR1,I |
| 250 | 1.633 | 8 | 1 | SPTLC2 | 6 | ATF4,CCN2,C |
| 251 | 1.633 | 8 | 1 | SRC | 6 | CTNNB1,FN1 |
| 252 | 1.633 | 8 | 1 | TFRC | 6 | ANXA2,CDKN |
| 253 | 1.633 | 8 | 1 | acetaminoph | 6 | A2M,CDKN1 |
| 254 | 1.633 | 8 | 1 | lithium chlor | 6 | CDKN1B,CTN |
| 255 | 1.581 | 12 | 1 | PDGF BB | 10 | CDKN1B,EIF4 |
| 256 | 1.581 | 12 | 1 | PML | 10 | APOE,CCND1 |
| 257 | 1.512 | 9 | 1 | ERG | 7 | CTNNB1,GAI |
| 258 | 1.512 | 9 | 1 | Hif1 | 7 | CDKN1A,FN1 |
| 259 | 1.512 | 30 | 1 | IGF1 | 28 | APP,CCND1,C |
| 260 | 1.512 | 9 | 1 | Ige | 7 | CCND2,COL1 |
| 261 | 1.512 | 9 | 1 | SP110 | 7 | DICER1,HDA |
| 262 | 1.512 | 9 | 1 | diethylstilbe | 7 | ATXN3,EGR1 |
| 263 | 1.512 | 9 | 1 | fasudil | 7 | ACTA2,CCN2, |
| 264 | 1.508 | 13 | 1 | PI3K (comple | 11 | CCND1,CCND |
| 265 | 1.508 | 13 | 1 | SMO | 11 | ALCAM,CCNE |
| 266 | 1.5 | 6 | 1 | ANXA7 | 4 | CDH2,EPHB2, |
| 267 | 1.5 | 6 | 1 | DSCAM | 4 | L1CAM,LHX2, |
| 268 | 1.5 | 6 | 1 | EWSR1-FLI1 | 4 | IGF1R,MAP2 |

| | | | | | | |
|---|---|---|---|---|---|---|
| 269 | 1.5 | 6 | 1 | HSP90B1 | 4 | CDC42,EIF5A |
| 270 | 1.5 | 6 | 1 | JAG1 | 4 | CCND1,CCND |
| 271 | 1.5 | 18 | 1 | LY294002 | 16 | APOE,CCND1 |
| 272 | 1.5 | 6 | 1 | MXD1 | 4 | CCND2,CCNG |
| 273 | 1.5 | 6 | 1 | NRF1 | 4 | CD47,CDH2,F |
| 274 | 1.5 | 6 | 1 | PRMT1 | 4 | CCND1,CDH2 |
| 275 | 1.5 | 6 | 1 | SAV1 | 4 | CCN2,CCND1 |
| 276 | 1.5 | 6 | 1 | SOX4 | 4 | CDH2,DICER1 |
| 277 | 1.5 | 6 | 1 | STUB1 | 4 | CDKN1A,ERB |
| 278 | 1.5 | 6 | 1 | STUB1 | 4 | MAPT,PINK1, |
| 279 | 1.5 | 6 | 1 | WDFY2 | 4 | ACO2,AKT2,F |
| 280 | 1.5 | 6 | 1 | arsenite | 4 | CDH2,EGR1,I |
| 281 | 1.5 | 6 | 1 | caspase | 4 | APP,CCND1,C |
| 282 | 1.5 | 6 | 1 | miR-21-5p (a | 4 | NFIB,SMARC |
| 283 | 1.5 | 6 | 1 | p70 S6k | 4 | CDH2,CDK4,D |
| 284 | 1.46 | 40 | 1 | NFE2L2 | 38 | ARF1,ATF4,C |
| 285 | 1.443 | 14 | 1 | SP1 | 12 | CDH2,CDKN1 |
| 286 | 1.429 | 26 | 1 | Creb | 24 | APBA2,ARHG |
| 287 | 1.414 | 10 | 1 | ARNT | 8 | AKT2,CCND2, |
| 288 | 1.414 | 10 | 1 | GLI1 | 8 | CCND1,CCND |
| 289 | 1.414 | 4 | 1 | L1CAM | 2 | MMP2,STMN |
| 290 | 1.414 | 4 | 1 | L1CAM | 2 | MMP2,STMN |
| 291 | 1.342 | 7 | 1 | DSCAM | 5 | L1CAM,LHX2, |
| 292 | 1.342 | 7 | 1 | ELAVL1 | 5 | CTNNB1,EIF4 |
| 293 | 1.342 | 7 | 1 | EOMES | 5 | ALCAM,GNA |
| 294 | 1.342 | 7 | 1 | FGF4 | 5 | CDKN1A,CTN |
| 295 | 1.342 | 7 | 1 | GNA12 | 5 | EZR,GNA13,I |
| 296 | 1.342 | 7 | 1 | KDM3B | 5 | ADAMTS1,CC |
| 297 | 1.342 | 7 | 1 | N-acetyl-L-cy | 5 | CDKN1B,CXC |
| 298 | 1.342 | 7 | 1 | lovastatin | 5 | CDKN1B,FGF |
| 299 | 1.342 | 7 | 1 | miR-199a-5p | 5 | CDKN1A,FN1 |
| 300 | 1.342 | 7 | 1 | seocalcitol | 5 | ACTR3,CDKN |
| 301 | 1.342 | 22 | 1 | silibinin | 20 | CCND1,CD44, |
| 302 | 1.336 | 16 | 1 | PRKAA2 | 14 | ACTA2,CDKN |
| 303 | 1.333 | 11 | 1 | AKT1 | 9 | CDKN1A,EP3 |
| 304 | 1.333 | 11 | 1 | Growth horm | 9 | APLP1,APOE, |
| 305 | 1.333 | 11 | 1 | LCK | 9 | ANXA1,CSNK |
| 306 | 1.333 | 11 | 1 | PGR | 9 | CDKN1B,DPY |
| 307 | 1.333 | 11 | 1 | PPARA | 9 | APEX1,CCND |
| 308 | 1.333 | 11 | 1 | TCF7L2 | 9 | BACE1,CCN2, |
| 309 | 1.333 | 11 | 1 | plicamycin | 9 | B4GALT5,CA |
| 310 | 1.309 | 23 | 1 | COL18A1 | 21 | ANTXR1,APC |
| 311 | 1.265 | 12 | 1 | methylseleni | 10 | CCND1,CDC4 |
| 312 | 1.25 | 18 | 1 | metformin | 16 | CCND1,CCND |
| 313 | 1.225 | 8 | 1 | Alpha cateni | 6 | CDKN1B,CXC |

| | | | | | | |
|---|---|---|---|---|---|---|
| 314 | 1.225 | 8 | 1 | BCL2 | 6 | CCND1,CDKN |
| 315 | 1.225 | 8 | 1 | CASR | 6 | ABL1,CDKN1I |
| 316 | 1.225 | 8 | 1 | CDK19 | 6 | CCND2,CDKN |
| 317 | 1.225 | 8 | 1 | DSCAM | 6 | CNTNAP1,L1( |
| 318 | 1.225 | 8 | 1 | ETV6-RUNX1 | 6 | CDH2,NOTCH |
| 319 | 1.225 | 8 | 1 | HBEGF | 6 | CCND1,CCND |
| 320 | 1.225 | 8 | 1 | NTRK2 | 6 | CCND1,CCND |
| 321 | 1.225 | 8 | 1 | PTGS2 | 6 | CDKN1B,CXCI |
| 322 | 1.225 | 8 | 1 | ROCK | 6 | CCN2,CCND1 |
| 323 | 1.225 | 8 | 1 | amphetamin | 6 | CD47,CDH2,E |
| 324 | 1.225 | 8 | 1 | lactacystin | 6 | CDKN1B,GJA |
| 325 | 1.225 | 8 | 1 | mir-17 | 6 | APP,CCN2,CC |
| 326 | 1.206 | 13 | 1 | 8-bromo-cAI | 11 | CDKN1B,FGF |
| 327 | 1.206 | 13 | 1 | Z-LLL-CHO | 11 | ABL1,APP,CD |
| 328 | 1.183 | 37 | 1 | CTNNB1 | 35 | APP,CCND1,( |
| 329 | 1.155 | 14 | 1 | 17-alpha-eth | 12 | APOE,ATF4,( |
| 330 | 1.155 | 5 | 1 | ABCB1 | 3 | APP,DDIT4,HI |
| 331 | 1.155 | 5 | 1 | AKT2 | 3 | CTNNB1,ITGI |
| 332 | 1.155 | 5 | 1 | ARG1 | 3 | GSK3B,MAPT |
| 333 | 1.155 | 5 | 1 | Abcb1b | 3 | APP,DDIT4,HI |
| 334 | 1.155 | 5 | 1 | BARX2 | 3 | L1CAM,NCAI |
| 335 | 1.155 | 5 | 1 | BARX2 | 3 | FLNA,L1CAM |
| 336 | 1.155 | 5 | 1 | BRD7 | 3 | DICER1,FN1,' |
| 337 | 1.155 | 5 | 1 | BSG | 3 | CDKN1B,CXCI |
| 338 | 1.155 | 5 | 1 | BSG | 3 | CDKN1B,CXCI |
| 339 | 1.155 | 5 | 1 | CAPNS1 | 3 | CCND1,CCND |
| 340 | 1.155 | 5 | 1 | CAT | 3 | CDKN1B,FN1 |
| 341 | 1.155 | 5 | 1 | COMMD1 | 3 | PFKFB3,PGK: |
| 342 | 1.155 | 14 | 1 | CSF1 | 12 | APOE,APP,CC |
| 343 | 1.155 | 5 | 1 | DSCAML1 | 3 | DCLK1,MAPT |
| 344 | 1.155 | 5 | 1 | GF 120918 | 3 | APP,DDIT4,HI |
| 345 | 1.155 | 5 | 1 | PLAUR | 3 | CDH2,ITGB1, |
| 346 | 1.155 | 5 | 1 | PLAUR | 3 | CDH2,ITGB1, |
| 347 | 1.155 | 5 | 1 | PLAUR | 3 | CCND1,CDH2 |
| 348 | 1.155 | 5 | 1 | POU3F2 | 3 | CDK5R2,FABI |
| 349 | 1.155 | 5 | 1 | RAP1GAP | 3 | CCND1,CDK4, |
| 350 | 1.155 | 5 | 1 | RLIM | 3 | CCN2,CDKN1 |
| 351 | 1.155 | 5 | 1 | SMARCD3 | 3 | ALCAM,CD44 |
| 352 | 1.155 | 5 | 1 | SMARCD3 | 3 | CD44,COL1A: |
| 353 | 1.155 | 5 | 1 | TBXT | 3 | CD44,COL1A: |
| 354 | 1.155 | 5 | 1 | TEAD2 | 3 | CALD1,CCN1, |
| 355 | 1.155 | 5 | 1 | TEAD3 | 3 | CALD1,CCN1, |
| 356 | 1.155 | 5 | 1 | TWIST2 | 3 | CDH2,CTNNB |
| 357 | 1.155 | 5 | 1 | WR 1065 | 3 | CDKN1A,MDI |
| 358 | 1.155 | 5 | 1 | ZEB1 | 3 | FN1,MAP1B, |

| | | | | | | |
|---|---|---|---|---|---|---|
| 359 | 1.155 | 5 | 1 | adefovir dipi | 3 | ACTA2,FN1,T |
| 360 | 1.155 | 5 | 1 | azetidyl-2-ca | 3 | ATP2A2,CLU, |
| 361 | 1.155 | 5 | 1 | cadmium chl | 3 | CDH2,CXCL12 |
| 362 | 1.155 | 14 | 1 | cardiotoxin | 12 | CCND1,CCND |
| 363 | 1.155 | 5 | 1 | chloroquine | 3 | APP,MAPT,VI |
| 364 | 1.155 | 5 | 1 | cobalt | 3 | HIF1A,PFKFB |
| 365 | 1.155 | 14 | 1 | cyclic AMP | 12 | APP,CDC42,C |
| 366 | 1.155 | 5 | 1 | leupeptin | 3 | APP,IGF2,MA |
| 367 | 1.155 | 5 | 1 | methylnitron | 3 | CCND1,CDKN |
| 368 | 1.155 | 5 | 1 | miR-451a (a | 3 | ADAM10,CCN |
| 369 | 1.155 | 5 | 1 | miR-451a (a | 3 | CCND1,CDKN |
| 370 | 1.155 | 5 | 1 | miR-451a (a | 3 | ADAM10,CCN |
| 371 | 1.155 | 5 | 1 | mir-194 | 3 | ACTA2,MDM |
| 372 | 1.155 | 5 | 1 | nafenopin | 3 | CDK4,DLD,HS |
| 373 | 1.155 | 5 | 1 | sterol | 3 | ACLY,HMGCF |
| 374 | 1.155 | 5 | 1 | sterol | 3 | ACACA,IDH1, |
| 375 | 1.155 | 14 | 1 | wortmannin | 12 | CCND1,CCND |
| 376 | 1.155 | 5 | 1 | zinostatin | 3 | CDKN1A,IGF1 |
| 377 | 1.155 | 29 | 1 | bucladesine | 27 | APOE,CD44,C |
| 378 | 1.147 | 21 | 1 | medroxyprog | 19 | AKT3,ALDH18 |
| 379 | 1.134 | 9 | 1 | EPO | 7 | ACTB,CXCL12 |
| 380 | 1.134 | 9 | 1 | HNF1B | 7 | CD44,CTNNB |
| 381 | 1.134 | 9 | 1 | MED1 | 7 | ARID1A,BCL1 |
| 382 | 1.134 | 9 | 1 | MUC1 | 7 | ARID1A,CCNI |
| 383 | 1.134 | 9 | 1 | PRKCA | 7 | APP,BNIP3L,I |
| 384 | 1.118 | 22 | 1 | COL18A1 | 20 | ANTXR1,APC |
| 385 | 1.109 | 15 | 1 | Vegf | 13 | ADAM10,CTN |
| 386 | 1.109 | 15 | 1 | forskolin | 13 | CDH2,CDKN1 |
| 387 | 1.069 | 16 | 1 | MMP1 | 14 | CDK4,CDKN1 |
| 388 | 1.069 | 16 | 1 | estrogen rec | 14 | AKT3,CCN2,C |
| 389 | 1.061 | 10 | 1 | EP300 | 8 | CNN3,COL2A |
| 390 | 1.061 | 34 | 1 | HTT | 32 | AKT2,CCN2,C |
| 391 | 1.061 | 10 | 1 | LDL | 8 | APOE,CBS/CI |
| 392 | 1.061 | 10 | 1 | PLA2R1 | 8 | ALDH2,COX5 |
| 393 | 1.061 | 10 | 1 | RAF1 | 8 | CCND1,CTNN |
| 394 | 1.043 | 25 | 1 | Tgf beta | 23 | ACTA2,CCN1, |
| 395 | 1 | 6 | 1 | ATM | 4 | CDKN1B,FN1 |
| 396 | 1 | 38 | 1 | CEBPB | 36 | APP,ARPP19, |
| 397 | 1 | 6 | 1 | H2AX | 4 | ACTB,HIF1A,I |
| 398 | 1 | 6 | 1 | HOXA10 | 4 | ACTB,APOE,H |
| 399 | 1 | 6 | 1 | M344 | 4 | APBB1,APP,N |
| 400 | 1 | 11 | 1 | M344 | 9 | APBA2,APBB |
| 401 | 1 | 6 | 1 | PHF21A | 4 | RAB3A,SNAP |
| 402 | 1 | 6 | 1 | Pdgf (comple | 4 | CDKN1B,FN1 |
| 403 | 1 | 11 | 1 | RELA | 9 | APP,BNIP3L,C |

| | | | | | |
|---|---|---|---|---|---|
| 404 | 1 | 6 | 1 | alvocidib | 4 CCND1,CDKN |
| 405 | 1 | 6 | 1 | bicalutamide | 4 CDH2,CTNNB |
| 406 | 1 | 18 | 1 | doxorubicin | 16 A2M,ABCA7, |
| 407 | 1 | 6 | 1 | miR-92a-3p | 4 CCND1,CCND |
| 408 | 1 | 11 | 1 | nitric oxide | 9 APP,CCND1,C |
| 409 | 1 | 6 | 1 | salirasib | 4 CDKN1B,MT( |
| 410 | 1 | 6 | 1 | tosedostat | 4 CBS/CBSL,DE |
| 411 | 0.97 | 19 | 1 | EIF2AK3 | 17 ARF4,ATF4,A |
| 412 | 0.97 | 19 | 1 | miR-125b-5p | 17 ADAMTS1,AJ |
| 413 | 0.949 | 12 | 1 | losartan pota | 10 APP,CST3,EG |
| 414 | 0.949 | 12 | 1 | mycophenoli | 10 ACTA2,AKAP |
| 415 | 0.918 | 21 | 1 | ADORA2A | 19 APP,ARHGEF |
| 416 | 0.905 | 13 | 1 | BRAF | 11 CCND1,CDKN |
| 417 | 0.905 | 13 | 1 | BRAF | 11 APP,CCND1,C |
| 418 | 0.905 | 13 | 1 | Ins1 | 11 CCND1,EGR1 |
| 419 | 0.894 | 7 | 1 | PTH | 5 CDKN1B,CTN |
| 420 | 0.894 | 7 | 1 | TET2 | 5 CTNNB1,FOX |
| 421 | 0.894 | 7 | 1 | mir-122 | 5 ADAM10,CSF |
| 422 | 0.894 | 7 | 1 | mir-183 | 5 EGR1,ITGB1, |
| 423 | 0.894 | 7 | 1 | nicotine | 5 CDKN1B,DLG |
| 424 | 0.866 | 14 | 1 | SPP1 | 12 CD44,CDKN1 |
| 425 | 0.853 | 24 | 1 | carbon tetra | 22 ACTA2,APOE |
| 426 | 0.832 | 15 | 1 | testosterone | 13 A2M,BGN,CD |
| 427 | 0.816 | 8 | 1 | CDKN1A | 6 ADAM10,CDK |
| 428 | 0.816 | 8 | 1 | EIF4G2 | 6 ACTB,CTSD,I( |
| 429 | 0.816 | 8 | 1 | JAK2 | 6 CDKN1B,CTN |
| 430 | 0.816 | 8 | 1 | JAK2 | 6 CCND1,CCND |
| 431 | 0.816 | 8 | 1 | STAT4 | 6 CDKN1B,FSCI |
| 432 | 0.816 | 8 | 1 | TWIST1 | 6 CDH2,CXCL12 |
| 433 | 0.816 | 8 | 1 | WNT1 | 6 CDKN1B,CTN |
| 434 | 0.816 | 8 | 1 | miR-204-5p | 6 EFNB1,ID2,PI |
| 435 | 0.816 | 8 | 1 | mir-182 | 6 FN1,LRP6,NC |
| 436 | 0.775 | 17 | 1 | IL2 | 15 ARNT2,CCND |
| 437 | 0.775 | 17 | 1 | SIRT1 | 15 CCND1,CCND |
| 438 | 0.756 | 9 | 1 | homocystein | 7 ATF4,CDKN1 |
| 439 | 0.75 | 18 | 1 | thapsigargin | 16 ATF4,ATP2A |
| 440 | 0.743 | 31 | 1 | NRG1 | 29 ACHE,ARHGE |
| 441 | 0.718 | 33 | 1 | Lh | 31 ACTB,ARL6IP |
| 442 | 0.707 | 20 | 1 | EGFR | 18 APLP2,APP,C |
| 443 | 0.707 | 10 | 1 | ATF6 | 8 APP,ATP2A2, |
| 444 | 0.707 | 10 | 1 | INSIG1 | 8 ACLY,APOE,D |
| 445 | 0.707 | 10 | 1 | IgG | 8 APOE,APP,DH |
| 446 | 0.707 | 10 | 1 | TGFB3 | 8 ACTA2,CCN2, |
| 447 | 0.696 | 35 | 1 | LDL | 33 ATP2A2,CAPI |
| 448 | 0.696 | 35 | 1 | LDL | 33 ATP2A2,CAPI |

| | | | | | | |
|---|---|---|---|---|---|---|
| 449 | 0.696 | 35 | 1 | methylpredn | 33 | A2M,AHCY,A |
| 450 | 0.667 | 11 | 1 | PPARGC1A | 9 | CDKN1A,ERB |
| 451 | 0.667 | 11 | 1 | SMARCB1 | 9 | BNIP3L,CD44 |
| 452 | 0.667 | 11 | 1 | bexarotene | 9 | APOE,APP,CF |
| 453 | 0.603 | 13 | 1 | 5-azacytidine | 11 | ANXA2,CANX |
| 454 | 0.603 | 13 | 1 | ATF4 | 11 | CDKN1A,CDK |
| 455 | 0.603 | 13 | 1 | EGLN | 11 | A2M,CSRP2,I |
| 456 | 0.603 | 13 | 1 | MMP1 | 11 | CDK4,CDKN1 |
| 457 | 0.603 | 13 | 1 | SOD1 | 11 | APEX1,CCND |
| 458 | 0.603 | 13 | 1 | bleomycin | 11 | ACTA2,CCN2, |
| 459 | 0.603 | 13 | 1 | deferoxamin | 11 | ALDOA,CCNE |
| 460 | 0.603 | 13 | 1 | elaidic acid | 11 | ACLY,APOE,A |
| 461 | 0.577 | 5 | 1 | EXOSC3 | 3 | CDKN1A,ITG |
| 462 | 0.577 | 5 | 1 | HSPA9 | 3 | CDKN1A,CTN |
| 463 | 0.577 | 14 | 1 | MIR17HG | 12 | APP,CCND1,C |
| 464 | 0.577 | 5 | 1 | MUC4 | 3 | CDKN1A,CDK |
| 465 | 0.577 | 5 | 1 | MYCBP | 3 | CAD,CCND2,C |
| 466 | 0.577 | 5 | 1 | PDE8A | 3 | FDPS,HMGCF |
| 467 | 0.577 | 5 | 1 | SUCNR1 | 3 | ACTA2,COL1 |
| 468 | 0.577 | 5 | 1 | evofosfamid | 3 | CCND1,CCND |
| 469 | 0.577 | 5 | 1 | miR-128-3p | 3 | LDLR,SNAP2 |
| 470 | 0.577 | 5 | 1 | miR-200b-3p | 3 | CDKN1B,VEG |
| 471 | 0.577 | 5 | 1 | miR-200b-3p | 3 | CDKN1B,RER |
| 472 | 0.577 | 5 | 1 | miR-221-3p | 3 | BNIP3L,CDKN |
| 473 | 0.577 | 5 | 1 | miR-221-3p | 3 | BNIP3L,CDKN |
| 474 | 0.577 | 5 | 1 | sodium arser | 3 | CDKN1B,HDA |
| 475 | 0.557 | 31 | 1 | JUN | 29 | AKAP12,AKT |
| 476 | 0.557 | 31 | 1 | PTGS2 | 29 | ANXA1,ANXA |
| 477 | 0.555 | 15 | 1 | AR | 13 | ATP2A2,CDKI |
| 478 | 0.555 | 15 | 1 | PAX3 | 13 | APP,CDKN1A |
| 479 | 0.555 | 15 | 1 | prednisolone | 13 | APOE,APP,CF |
| 480 | 0.539 | 33 | 1 | AGT | 31 | CCND2,CD44, |
| 481 | 0.535 | 16 | 1 | ANGPT2 | 14 | BCR,CYP51A |
| 482 | 0.535 | 16 | 1 | EPAS1 | 14 | AKAP12,CDKI |
| 483 | 0.535 | 16 | 1 | FOXO1 | 14 | CCND1,CNR1 |
| 484 | 0.535 | 16 | 1 | VEGFA | 14 | ALDH2,CDKN |
| 485 | 0.516 | 17 | 1 | INSR | 15 | AQP1,CCN2,C |
| 486 | 0.5 | 18 | 1 | CG | 16 | CCND2,CDK4, |
| 487 | 0.5 | 18 | 1 | IGF1R | 16 | ALDOA,CDKN |
| 488 | 0.5 | 6 | 1 | NEUROD1 | 4 | DCX,NCAM1, |
| 489 | 0.5 | 6 | 1 | mitomycin C | 4 | CDKN1A,MDI |
| 490 | 0.5 | 6 | 1 | pitavastatin | 4 | FDFT1,FDPS, |
| 491 | 0.459 | 21 | 1 | metribolone | 19 | ACLY,APP,AT |
| 492 | 0.453 | 80 | 1 | APP | 78 | ABCA7,ACHE |
| 493 | 0.447 | 7 | 1 | ABCB4 | 5 | ACTA2,CCN2, |

| | | | | | | |
|---|---|---|---|---|---|---|
| 494 | 0.447 | 7 | 1 | CDKN1B | 5 | B4GALT5,GA |
| 495 | 0.447 | 7 | 1 | JUNB | 5 | CCND1,CDKN |
| 496 | 0.447 | 22 | 1 | PD98059 | 20 | ACTA2,BGN, |
| 497 | 0.447 | 7 | 1 | PEX5L | 5 | DHCR7,FDFT |
| 498 | 0.447 | 7 | 1 | PPARGC1B | 5 | FDFT1,FDPS, |
| 499 | 0.447 | 7 | 1 | SH3TC2 | 5 | ACLY,DHCR7, |
| 500 | 0.447 | 7 | 1 | SIRT2 | 5 | ACLY,DHCR7, |
| 501 | 0.447 | 22 | 1 | SMAD4 | 20 | ARHGEF2,CD |
| 502 | 0.447 | 7 | 1 | ezetimibe | 5 | DHCR7,FDFT |
| 503 | 0.447 | 7 | 1 | isoquercitrin | 5 | DHCR7,FDFT |
| 504 | 0.447 | 7 | 1 | miR-122-5p | 5 | AKT3,FUBP1, |
| 505 | 0.447 | 7 | 1 | rosuvastatin | 5 | DHCR7,FDFT |
| 506 | 0.438 | 49 | 1 | lipopolysacch | 47 | ALCAM,ALDH |
| 507 | 0.436 | 23 | 1 | LEP | 21 | ATP2A2,CDKI |
| 508 | 0.426 | 24 | 1 | Akt | 22 | ATF4,CCND2, |
| 509 | 0.426 | 24 | 1 | ESR2 | 22 | AKAP12,APC, |
| 510 | 0.417 | 25 | 1 | Insulin | 23 | ALDOA,CDKN |
| 511 | 0.417 | 94 | 1 | SP2509 | 92 | ABCE1,ADAM |
| 512 | 0.417 | 25 | 1 | curcumin | 23 | CCND2,CD44, |
| 513 | 0.408 | 8 | 1 | (-)-norephed | 6 | DHCR7,FDFT |
| 514 | 0.408 | 8 | 1 | FMR1 | 6 | APP,DAG1,M |
| 515 | 0.408 | 8 | 1 | IFT88 | 6 | ACLY,FDPS,H |
| 516 | 0.408 | 8 | 1 | PTK2 | 6 | ACTA2,COL1 |
| 517 | 0.408 | 8 | 1 | TO-901317 | 6 | ALDH2,CDK4, |
| 518 | 0.408 | 8 | 1 | aldesleukin | 6 | B2M,CANX,F |
| 519 | 0.408 | 8 | 1 | atorvastatin | 6 | CDKN1B,COL |
| 520 | 0.378 | 9 | 1 | ATP7B | 7 | ACLY,APP,FD |
| 521 | 0.378 | 9 | 1 | SMAD2 | 7 | ACTA2,BGN, |
| 522 | 0.378 | 9 | 1 | SMAD2 | 7 | CDKN1A,FN1 |
| 523 | 0.378 | 9 | 1 | SREBF2 | 7 | ACLY,DHCR7, |
| 524 | 0.378 | 9 | 1 | lysophosphat | 7 | CD74,DHCR7, |
| 525 | 0.371 | 31 | 1 | dihydrotesto | 29 | APOE,APP,BC |
| 526 | 0.359 | 33 | 1 | ERBB2 | 31 | ABL1,AKT2,A |
| 527 | 0.354 | 10 | 1 | AGN194204 | 8 | ARL4C,BNIP3 |
| 528 | 0.354 | 10 | 1 | ERN1 | 8 | ARF4,ATF4,A |
| 529 | 0.354 | 10 | 1 | KDM8 | 8 | ALDOA,GSR, |
| 530 | 0.354 | 10 | 1 | MAP2K5 | 8 | ACLY,DHCR7, |
| 531 | 0.354 | 10 | 1 | TSC2 | 8 | ATF4,CCND2, |
| 532 | 0.354 | 10 | 1 | benzo(a)pyre | 8 | AJUBA,CDKN |
| 533 | 0.333 | 38 | 1 | hydrogen pe | 36 | ACTA2,APP,C |
| 534 | 0.327 | 86 | 1 | camptotheci | 84 | ACTA2,AKAP |
| 535 | 0.316 | 12 | 1 | NR4A1 | 10 | ALDOA,CCND |
| 536 | 0.316 | 12 | 1 | estrogen | 10 | APOE,CCN2, |
| 537 | 0.316 | 12 | 1 | pirinixic acid | 10 | CDK4,CDKN1 |
| 538 | 0.302 | 46 | 1 | D-glucose | 44 | ACHE,ACLY,A |

| | | | | | | |
|---|---|---|---|---|---|---|
| 539 | 0.302 | 13 | 1 | Mek | 11 | B4GAT1,CCN |
| 540 | 0.295 | 48 | 1 | cigarette sm | 46 | ACTA2,ATF4, |
| 541 | 0.277 | 15 | 1 | PPARG | 13 | ATP2A2,CDKI |
| 542 | 0.277 | 15 | 1 | SP600125 | 13 | CD44,CDKN1/ |
| 543 | 0.277 | 15 | 1 | SP600125 | 13 | APOE,AQP1,( |
| 544 | 0.277 | 15 | 1 | actinomycin | 13 | ACTB,APP,AT |
| 545 | 0.277 | 15 | 1 | fenofibrate | 13 | APP,CLU,CRP |
| 546 | 0.277 | 15 | 1 | tanespimycir | 13 | CAPN1,CDK4, |
| 547 | 0.267 | 16 | 1 | CD44 | 14 | CCND2,CDKN |
| 548 | 0.267 | 16 | 1 | SREBF1 | 14 | ACLY,CDK4,C |
| 549 | 0.258 | 17 | 1 | CD 437 | 15 | ATP5F1A,CDI |
| 550 | 0.25 | 18 | 1 | HSF1 | 16 | ACLY,CCN2,C |
| 551 | 0.25 | 18 | 1 | RB1 | 16 | CD44,CDKN1/ |
| 552 | 0.25 | 18 | 1 | mibolerone | 16 | ARF4,C4A/C4 |
| 553 | 0.25 | 18 | 1 | poly rI:rC-RN | 16 | ATF4,CCND2, |
| 554 | 0.243 | 19 | 1 | MRTFA | 17 | ACTA2,ACTB, |
| 555 | 0.243 | 19 | 1 | SMAD7 | 17 | ACTA2,ACVR |
| 556 | 0.239 | 72 | 1 | TGFB1 | 70 | ACLY,ACTA2, |
| 557 | 0.228 | 79 | 1 | dexamethas( | 77 | ACLY,ACTA2, |
| 558 | 0.224 | 22 | 1 | ADCYAP1 | 20 | ARHGEF25,A |
| 559 | 0.224 | 22 | 1 | E2F1 | 20 | CDK4,CTSB,D |
| 560 | 0.224 | 22 | 1 | TP63 | 20 | APC,CCND2,( |
| 561 | 0.224 | 22 | 1 | mifepristone | 20 | APOE,AQP1,( |
| 562 | 0.218 | 23 | 1 | TP73 | 21 | BAP1,CDKN1 |
| 563 | 0.213 | 24 | 1 | HGF | 22 | AKAP12,CD44 |
| 564 | 0.213 | 24 | 1 | IL4 | 22 | ALDH2,BNIP3 |
| 565 | 0.183 | 32 | 1 | PTPRR | 30 | AP1M1,BTBD |
| 566 | 0.183 | 32 | 1 | progesterone | 30 | ACTA2,ACTB, |
| 567 | 0.177 | 34 | 1 | decitabine | 32 | AGRN,ANP32 |
| 568 | 0.177 | 34 | 1 | sirolimus | 32 | ATF4,CCND2, |
| 569 | 0 | 16 | 1 | 1,2-dithiol-3- | 14 | ATP1A1,CTSI |
| 570 | 0 | 5 | 1 | 3-deazanepla | 3 | CDKN1A,IGFI |
| 571 | 0 | 6 | 1 | 3-deazanepla | 4 | HIF1A,IGFBP |
| 572 | 0 | 8 | 1 | ACOX1 | 6 | CD63,CXCL12 |
| 573 | 0 | 4 | 1 | ALDH2 | 2 | ATF4,SHMT2 |
| 574 | 0 | 5 | 1 | ALDH2 | 3 | ATF4,PSAT1, |
| 575 | 0 | 4 | 1 | ALDH2 | 2 | ATF4,SLC1A4 |
| 576 | 0 | 16 | 1 | BDNF | 14 | APC,ATP2A2, |
| 577 | 0 | 16 | 1 | Brd4 | 14 | ACTA2,COL1/ |
| 578 | 0 | 13 | 1 | CD28 | 11 | CCND2,CDK4, |
| 579 | 0 | 10 | 1 | CD28 | 8 | CCND2,CDK4, |
| 580 | 0 | 12 | 1 | CD28 | 10 | BIN1,CDK4,FI |
| 581 | 0 | 14 | 1 | CST5 | 12 | CTSB,HNRNF |
| 582 | 0 | 14 | 1 | DDX5 | 12 | ATP5F1C,ATI |
| 583 | 0 | 10 | 1 | EIF4E | 8 | CDK4,CDKN1/ |

| | | | | | |
|---|---|---|---|---|---|
| 584 | 0 | 9 | 1 | EIF4E | 7 CDK4,CDKN1/ |
| 585 | 0 | 16 | 1 | FAS | 14 ACTA2,BCL11 |
| 586 | 0 | 12 | 1 | FAS | 10 AKT2,CD44,C |
| 587 | 0 | 9 | 1 | FLT1 | 7 APLP2,CANX, |
| 588 | 0 | 9 | 1 | HBA1/HBA2 | 7 ATP5F1C,ATI |
| 589 | 0 | 11 | 1 | HMGA1 | 9 ACTA2,CCND |
| 590 | 0 | 9 | 1 | HMGA1 | 7 CCND2,CD44, |
| 591 | 0 | 7 | 1 | HSF2 | 5 CLU,HIF1A,H |
| 592 | 0 | 13 | 1 | Hbb-b1 | 11 ATP5F1C,ATI |
| 593 | 0 | 12 | 1 | IKZF1 | 10 AJUBA,CCND |
| 594 | 0 | 28 | 1 | IKZF1 | 26 AJUBA,ANKF |
| 595 | 0 | 28 | 1 | IKZF1 | 26 AJUBA,ANKF |
| 596 | 0 | 9 | 1 | IKZF1 | 7 AQP1,CEMIP |
| 597 | 0 | 7 | 1 | ITGB3 | 5 APLP2,FN1,IT |
| 598 | 0 | 9 | 1 | ITGB3 | 7 COL1A1,FN1, |
| 599 | 0 | 9 | 1 | ITGB3 | 7 ACLY,APLP2,I |
| 600 | 0 | 11 | 1 | KDM5A | 9 ATP1A1,ATP! |
| 601 | 0 | 11 | 1 | KITLG | 9 ACHE,ACTB,C |
| 602 | 0 | 5 | 1 | LINC01139 | 3 ALDOA,IGFBI |
| 603 | 0 | 10 | 1 | LONP1 | 8 ATF4,ATP5F1 |
| 604 | 0 | 10 | 1 | MAPK7 | 8 ACLY,DHCR7, |
| 605 | 0 | 7 | 1 | MBOAT7 | 5 ACLY,ACSS2,I |
| 606 | 0 | 5 | 1 | MFN2 | 3 ATF4,HIF1A,) |
| 607 | 0 | 6 | 1 | MFSD2A | 4 DHCR7,FDFT |
| 608 | 0 | 9 | 1 | MM-589 | 7 COL1A1,COL! |
| 609 | 0 | 12 | 1 | MM-589 | 10 CCDC80,COL1 |
| 610 | 0 | 20 | 1 | MYCN | 18 ALDOA,CCND |
| 611 | 0 | 18 | 1 | MYCN | 16 ALDOA,CCND |
| 612 | 0 | 49 | 1 | MYC | 47 ABCA2,ABCA |
| 613 | 0 | 9 | 1 | MYOD1 | 7 ALDOA,CDKN |
| 614 | 0 | 12 | 1 | MYOD1 | 10 ACHE,ACTA2, |
| 615 | 0 | 6 | 1 | NORAD | 4 CCN2,IGFBP3 |
| 616 | 0 | 5 | 1 | NTRK1 | 3 IGF2,LGALS1 |
| 617 | 0 | 6 | 1 | NUDT21 | 4 COL11A1,CO |
| 618 | 0 | 5 | 1 | PDGF (family | 3 HSPD1,PDIA3 |
| 619 | 0 | 5 | 1 | PROM1 | 3 DHCR7,FDPS, |
| 620 | 0 | 19 | 1 | PSEN1 | 17 ALDOA,ARF4 |
| 621 | 0 | 18 | 1 | PSEN1 | 16 ALDOA,ARF4 |
| 622 | 0 | 17 | 1 | PSEN1 | 15 CDH2,CTNNB |
| 623 | 0 | 6 | 1 | RUVBL1 | 4 CCND2,ERGI( |
| 624 | 0 | 5 | 1 | Retinoic acid | 3 IGFBP3,RBP1 |
| 625 | 0 | 5 | 1 | SAHM1 | 3 CD44,FLNA,P |
| 626 | 0 | 15 | 1 | SEL1L | 13 ATP5F1C,ATI |
| 627 | 0 | 7 | 1 | TOPBP1 | 5 ABL1,CDKN1/ |
| 628 | 0 | 7 | 1 | TOPBP1 | 5 ABL1,CDKN1/ |

| | | | | | |
|---|---|---|---|---|---|
| 629 | 0 | 7 | 1 | TOPBP1 | 5 ABL1,CDKN1. |
| 630 | 0 | 7 | 1 | TRIM37 | 5 CKB,DNMT1, |
| 631 | 0 | 6 | 1 | TRPC4AP | 4 APP,FBL,NOI |
| 632 | 0 | 8 | 1 | UCP1 | 6 ARHGAP33,A |
| 633 | 0 | 5 | 1 | XIAP | 3 CDH2,CDK4,S |
| 634 | 0 | 6 | 1 | XIAP | 4 CCND1,CDH2 |
| 635 | 0 | 5 | 1 | XIAP | 3 CDK4,CDKN1. |
| 636 | 0 | 5 | 1 | XIAP | 3 CDK4,RELA,S |
| 637 | 0 | 5 | 1 | ZNF106 | 3 C4A/C4B,LG/ |
| 638 | 0 | 6 | 1 | baicalin | 4 COL1A1,FABI |
| 639 | 0 | 8 | 1 | bortezomib | 6 CXCL12,HSP9 |
| 640 | 0 | 15 | 1 | bortezomib | 13 ATF4,CDK4,C |
| 641 | 0 | 38 | 1 | cisplatin | 36 A2M,ACTA2,. |
| 642 | 0 | 8 | 1 | diclofenac | 6 ATF4,CDKN1. |
| 643 | 0 | 7 | 1 | enalapril | 5 CDKN1B,CLU |
| 644 | 0 | 11 | 1 | enalapril | 9 A2M,ACTA2, |
| 645 | 0 | 8 | 1 | etoposide | 6 CDK4,CDKN1. |
| 646 | 0 | 7 | 1 | etoposide | 5 BIN1,CDKN1I |
| 647 | 0 | 12 | 1 | etoposide | 10 ACHE,ADAM: |
| 648 | 0 | 16 | 1 | gentamicin | 14 ALCAM,CLU, |
| 649 | 0 | 24 | 1 | gentamicin | 22 AQP1,C4A/C |
| 650 | 0 | 13 | 1 | isobutylmeth | 11 CD44,CDKN1. |
| 651 | 0 | 17 | 1 | isobutylmeth | 15 ACLY,CCN2,D |
| 652 | 0 | 12 | 1 | let-7a-5p (ar | 10 ACTA2,CEMII |
| 653 | 0 | 22 | 1 | levodopa | 20 ABCA7,ACTA |
| 654 | 0 | 8 | 1 | miR-30c-5p ( | 6 ATP2A2,ATR: |
| 655 | 0 | 30 | 1 | miR-30c-5p ( | 28 AP2A1,ATP2. |
| 656 | 0 | 14 | 1 | miR-335-3p | 12 COL1A1,COL |
| 657 | 0 | 12 | 1 | miR-335-3p | 10 COL1A1,COL |
| 658 | 0 | 12 | 1 | miR-335-3p | 10 COL1A1,COL |
| 659 | 0 | 17 | 1 | miR-338-3p | 15 CCND1,COL1. |
| 660 | 0 | 14 | 1 | miR-338-3p | 12 COL1A1,COL |
| 661 | 0 | 15 | 1 | mir-21 | 13 ARF4,CDKN1 |
| 662 | 0 | 13 | 1 | mir-21 | 11 ACTA2,CANX |
| 663 | 0 | 6 | 1 | phenylbutazc | 4 A2M,ASNS,C |
| 664 | 0 | 7 | 1 | phenylbutazc | 5 A2M,ASNS,C |
| 665 | 0 | 8 | 1 | phenylbutazc | 6 A2M,C4A/C4 |
| 666 | 0 | 5 | 1 | selenium | 3 CDKN1A,CDK |
| 667 | 0 | 9 | 1 | sulforafan | 7 CHRNA4,HIF: |
| 668 | 0 | 17 | 1 | tunicamycin | 15 ACTA2,ADAN |
| 669 | 0 | 14 | 1 | uranyl nitrate | 12 CANX,CD74, |
| 670 | -1.162 | 76 | 1 | beta-estradic | 74 ABL1,AJUBA, |
| 671 | -1.501 | 77 | 1 | TP53 | 75 A2M,ABCA2, |
| 672 | -1.732 | 14 | 1 | FN1 | 12 ACTA2,CCN2, |
| 673 | -1.82 | 53 | 1 | tretinoin | 51 A2M,ADAM1 |

| | | | | | | |
|---|---|---|---|---|---|---|
| 674 | -1.854 | 59 | 1 | IFNG | 57 | A2M,ACLY,A( |
| 675 | -1.919 | 24 | 1 | ESR1 | 22 | ACVR1B,ARH |
| 676 | -2.183 | 19 | 1 | L-triiodothyr( | 17 | APP,ATP1B2, |
| 677 | -2.2 | 27 | 1 | miR-29b-3p ( | 25 | BACE1,CCND |
| 678 | -2.214 | 12 | 1 | SYVN1 | 10 | AMOTL2,CCN |
| 679 | -2.593 | 20 | 1 | imatinib | 18 | ABL1,APP,AT |
| 680 | -2.694 | 29 | 1 | miR-1-3p (ar | 27 | ANXA2,AP3D |
| 681 | -2.907 | 22 | 1 | XBP1 | 20 | APP,ATF4,AT |
| 682 | -3.13 | 22 | 1 | nitrofurantoi | 20 | A2M,ACTB,A |
| 683 | -3.357 | 17 | 1 | H1-6 | 15 | ATP6V0E2,B( |
| 684 | -3.357 | 17 | 1 | H1f1 | 15 | ATP6V0E2,B( |
| 685 | -3.597 | 67 | 1 | TNF | 65 | A2M,ACTA2,. |
| 686 | -3.638 | 19 | 1 | CD3 | 17 | ATF4,CCND2, |
| 687 | -3.662 | 153 | 1 | trichostatin / | 151 | ABCE1,ACTB, |
| 688 | -3.75 | 18 | 1 | 5-fluorouraci | 16 | ALDOA,ATF4 |
| 689 | -3.753 | 14 | 1 | CPT1B | 12 | AKT3,FLOT2, |
| 690 | -3.78 | 9 | 1 | Srebp | 7 | ACLY,CDKN1/ |
| 691 | -4.009 | 16 | 1 | mono-(2-eth | 14 | COX7A2,GPI, |
| 692 | -4.158 | 9 | 1 | mir-34 | 7 | CCND1,CD47, |
| 693 | -4.743 | 12 | 1 | CAB39L | 10 | ATP5MG,ATF |
| 694 | -4.743 | 12 | 1 | STK11 | 10 | ALCAM,ALDC |
| 695 | -4.743 | 12 | 1 | tert-butyl-hy( | 10 | CNR1,FTH1,F |
| 696 | -4.919 | 7 | 1 | miR-7a-5p (a | 5 | IGF1R,MKNK |
| 697 | -5 | 11 | 1 | cocaine | 9 | CNR1,GABRE |
| 698 | -5.367 | 7 | 1 | RGS2 | 5 | CXCL12,EPN1 |
| 699 | -5.814 | 7 | 1 | ANLN | 5 | CDKN1A,CYF |
| 700 | -5.814 | 7 | 1 | AURK | 5 | CDKN1A,CYF |
| 701 | -6 | 6 | 1 | L-histidine | 4 | EGR1,ELAVL1 |
| 702 | -6 | 6 | 1 | histidinol | 4 | ASNS,ATF4,E |
| 703 | -6 | 6 | 1 | miR-483-3p ( | 4 | CTNNB1,GJA |
| 704 | -6.261 | 7 | 1 | KDM5B | 5 | CDKN1B,ERB |
| 705 | -6.5 | 6 | 1 | PPM1A | 4 | CDKN1A,ITG/ |
| 706 | -6.5 | 6 | 1 | PPM1A | 4 | CDKN1A,ITG/ |
| 707 | -7.506 | 5 | 1 | MLX | 3 | ACACA,IGF2, |
| 708 | -8.083 | 5 | 1 | 2-amino-1-m | 3 | HSP90AA1,R/ |
| 709 | -9.192 | 4 | 1 | NFASC | 2 | KIF5A,NRCAI |
| 710 | -9.827 | 9 | 1 | interferon al | 7 | ARL4C,CTR9, |
| 711 | -9.899 | 4 | 1 | TMPO | 2 | COL1A1,TIMI |
| 712 | -12.06 | 13 | 1 | KLF3 | 11 | ATF4,CAPNS1 |
| 713 | -12.522 | 7 | 1 | G protein alp | 5 | CALR,GPRC5I |
| 714 | -13.522 | 37 | 1 | ST1926 | 35 | ACTB,ARF1,C |
| 715 | -14.5 | 6 | 1 | LY2109761 | 4 | AKT2,ATF5,C |
| 716 | -16.743 | 5 | 1 | prostaglandi | 3 | CCND1,CDKN |
| 717 | -17.321 | 5 | 1 | CUL4B | 3 | IGFBP3,SOD1 |
| 718 | -17.321 | 5 | 1 | CUL4B | 3 | IGFBP3,SOD1 |

| | | | | | | |
|---|---|---|---|---|---|---|
| 719 | -17.321 | 5 | 1 | CUL4B | 3 | IGFBP3,SOD: |
| 720 | -17.321 | 5 | 1 | CUL4B | 3 | IGFBP3,SOD: |
| 721 | -25.981 | 5 | 1 | RAPGEF1 | 3 | CDKN1A,CDK |
| 722 | -28.29 | 15 | 1 | GnRH analog | 13 | ACTN4,FBLN |

| Disease & Fu | Diseases & F | Known Regulator-Disease/Function Relationship |
|---|---|---|
| 1 | Cell prolifera | 100% (1/1) |
| 1 | Cell prolifera | 0% (0/1) |
| 1 | Cell prolifera | 0% (0/1) |
| 1 | Cell prolifera | 100% (1/1) |
| 1 | Colony forma | 0% (0/1) |
| 1 | Colony forma | 0% (0/1) |
| 1 | Outgrowth o | 100% (1/1) |
| 1 | Proliferation | 100% (1/1) |
| 1 | Cell prolifera | 100% (1/1) |
| 1 | Growth of ne | 0% (0/1) |
| 1 | Outgrowth o | 0% (0/1) |
| 1 | Proliferation | 0% (0/1) |
| 1 | Cell prolifera | 100% (1/1) |
| 1 | Cell prolifera | 0% (0/1) |
| 1 | Cell prolifera | 100% (1/1) |
| 1 | Cell prolifera | 100% (1/1) |
| 1 | Cell prolifera | 100% (1/1) |
| 1 | Cell prolifera | 100% (1/1) |
| 1 | Outgrowth o | 0% (0/1) |
| 1 | Cell prolifera | 0% (0/1) |
| 1 | Cell prolifera | 100% (1/1) |
| 1 | Outgrowth o | 0% (0/1) |
| 1 | Cell prolifera | 100% (1/1) |
| 1 | Proliferation | 0% (0/1) |
| 1 | Cell prolifera | 0% (0/1) |
| 1 | Cell prolifera | 100% (1/1) |
| 1 | Cell prolifera | 100% (1/1) |
| 1 | Cell prolifera | 100% (1/1) |
| 1 | Cell prolifera | 0% (0/1) |
| 1 | Cell prolifera | 100% (1/1) |
| 1 | Outgrowth o | 100% (1/1) |
| 1 | Cell prolifera | 0% (0/1) |
| 1 | Colony forma | 100% (1/1) |
| 1 | Growth of ne | 0% (0/1) |
| 1 | Cell prolifera | 100% (1/1) |
| 1 | Proliferation | 100% (1/1) |
| 1 | Cell prolifera | 0% (0/1) |
| 1 | Colony forma | 100% (1/1) |
| 1 | Cell prolifera | 100% (1/1) |
| 1 | Cell prolifera | 100% (1/1) |
| 1 | Cell prolifera | 100% (1/1) |
| 1 | Colony forma | 0% (0/1) |
| 1 | Cell prolifera | 100% (1/1) |

| | | |
|---|---|---|
| 1 | Differentiati | 0% (0/1) |
| 1 | Outgrowth o | 0% (0/1) |
| 1 | Outgrowth o | 0% (0/1) |
| 1 | Outgrowth o | 0% (0/1) |
| 1 | Outgrowth o | 0% (0/1) |
| 1 | Proliferation | 100% (1/1) |
| 1 | Cell prolifera | 0% (0/1) |
| 1 | Cell prolifera | 0% (0/1) |
| 1 | Migration of | 0% (0/1) |
| 1 | Cell prolifera | 0% (0/1) |
| 1 | Cell prolifera | 100% (1/1) |
| 1 | Cell prolifera | 100% (1/1) |
| 1 | Colony forma | 100% (1/1) |
| 1 | Proliferation | 100% (1/1) |
| 1 | Cell prolifera | 100% (1/1) |
| 1 | Proliferation | 100% (1/1) |
| 1 | Cell prolifera | 0% (0/1) |
| 1 | Cell prolifera | 0% (0/1) |
| 1 | Cell prolifera | 100% (1/1) |
| 1 | Cell prolifera | 0% (0/1) |
| 1 | Cell prolifera | 0% (0/1) |
| 1 | Dendritic gro | 0% (0/1) |
| 1 | Cell prolifera | 100% (1/1) |
| 1 | Outgrowth o | 0% (0/1) |
| 1 | Cell prolifera | 100% (1/1) |
| 1 | Branching of | 0% (0/1) |
| 1 | Cell prolifera | 0% (0/1) |
| 1 | Migration of | 100% (1/1) |
| 1 | Colony forma | 0% (0/1) |
| 1 | Colony forma | 0% (0/1) |
| 1 | Colony forma | 0% (0/1) |
| 1 | Colony forma | 0% (0/1) |
| 1 | Colony forma | 100% (1/1) |
| 1 | Cell prolifera | 0% (0/1) |
| 1 | Long-term p | 0% (0/1) |
| 1 | Colony forma | 100% (1/1) |
| 1 | Proliferation | 100% (1/1) |
| 1 | Cell prolifera | 0% (0/1) |
| 1 | Cell prolifera | 100% (1/1) |
| 1 | Proliferation | 0% (0/1) |
| 1 | Quantity of r | 100% (1/1) |
| 1 | Quantity of r | 100% (1/1) |
| 1 | Cell prolifera | 0% (0/1) |
| 1 | Cell prolifera | 0% (0/1) |
| 1 | Cell prolifera | 0% (0/1) |

| | | |
|---|---|---|
| 1 | Cell prolifera | 0% (0/1) |
| 1 | Cell prolifera | 0% (0/1) |
| 1 | Cell prolifera | 0% (0/1) |
| 1 | Colony forma | 0% (0/1) |
| 1 | Outgrowth o | 0% (0/1) |
| 1 | Outgrowth o | 0% (0/1) |
| 1 | Cell prolifera | 0% (0/1) |
| 1 | Colony forma | 0% (0/1) |
| 1 | Cell prolifera | 100% (1/1) |
| 1 | Cell prolifera | 0% (0/1) |
| 1 | Cell prolifera | 0% (0/1) |
| 1 | Cell prolifera | 0% (0/1) |
| 1 | Dendritic gro | 0% (0/1) |
| 1 | Outgrowth o | 100% (1/1) |
| 1 | Cell prolifera | 0% (0/1) |
| 1 | Cell prolifera | 0% (0/1) |
| 1 | Cell prolifera | 100% (1/1) |
| 1 | Cell prolifera | 0% (0/1) |
| 1 | Cell prolifera | 0% (0/1) |
| 1 | Cell prolifera | 0% (0/1) |
| 1 | Morphogene | 0% (0/1) |
| 1 | Neuritogenes | 0% (0/1) |
| 1 | Cell prolifera | 100% (1/1) |
| 1 | Growth of ax | 100% (1/1) |
| 1 | Dendritic gro | 100% (1/1) |
| 1 | Cell prolifera | 0% (0/1) |
| 1 | Outgrowth o | 0% (0/1) |
| 1 | Growth of ne | 100% (1/1) |
| 1 | Proliferation | 0% (0/1) |
| 1 | Cell prolifera | 0% (0/1) |
| 1 | Dendritic gro | 0% (0/1) |
| 1 | Cell prolifera | 0% (0/1) |
| 1 | Cell prolifera | 0% (0/1) |
| 1 | Proliferation | 0% (0/1) |
| 1 | Migration of | 100% (1/1) |
| 1 | Outgrowth o | 0% (0/1) |
| 1 | Branching of | 0% (0/1) |
| 1 | Cell prolifera | 0% (0/1) |
| 1 | Dendritic gro | 0% (0/1) |
| 1 | Proliferation | 0% (0/1) |
| 1 | Cell prolifera | 0% (0/1) |
| 1 | Colony forma | 100% (1/1) |
| 1 | Proliferation | 100% (1/1) |
| 1 | Proliferation | 0% (0/1) |
| 1 | Outgrowth o | 0% (0/1) |

| | | |
|---|---|---|
| 1 | Outgrowth o | 0% (0/1) |
| 1 | Extension of | 100% (1/1) |
| 1 | Growth of ne | 0% (0/1) |
| 1 | Outgrowth o | 0% (0/1) |
| 1 | Cell prolifera | 0% (0/1) |
| 1 | Cell prolifera | 0% (0/1) |
| 1 | Extension of | 0% (0/1) |
| 1 | Colony forma | 0% (0/1) |
| 1 | Motor dysfur | 0% (0/1) |
| 1 | Proliferation | 0% (0/1) |
| 1 | Branching of | 100% (1/1) |
| 1 | Branching of | 100% (1/1) |
| 1 | Cell prolifera | 0% (0/1) |
| 1 | Proliferation | 0% (0/1) |
| 1 | Branching of | 0% (0/1) |
| 1 | Extension of | 0% (0/1) |
| 1 | Outgrowth o | 0% (0/1) |
| 1 | Cell prolifera | 0% (0/1) |
| 1 | Cell prolifera | 0% (0/1) |
| 1 | Colony forma | 0% (0/1) |
| 1 | Cell prolifera | 0% (0/1) |
| 1 | Proliferation | 0% (0/1) |
| 1 | Developmen | 0% (0/1) |
| 1 | Neuritogene | 0% (0/1) |
| 1 | Proliferation | 0% (0/1) |
| 1 | Colony forma | 0% (0/1) |
| 1 | Cell prolifera | 0% (0/1) |
| 1 | Differentiati | 100% (1/1) |
| 1 | Branching of | 0% (0/1) |
| 1 | Migration of | 0% (0/1) |
| 1 | Developmen | 0% (0/1) |
| 1 | Outgrowth o | 0% (0/1) |
| 1 | Cell prolifera | 100% (1/1) |
| 1 | Colony forma | 100% (1/1) |
| 1 | Cell prolifera | 0% (0/1) |
| 1 | Long-term pe | 0% (0/1) |
| 1 | Cell prolifera | 0% (0/1) |
| 1 | Differentiati | 0% (0/1) |
| 1 | Proliferation | 0% (0/1) |
| 1 | Proliferation | 0% (0/1) |
| 1 | Cell prolifera | 0% (0/1) |
| 1 | Cell prolifera | 0% (0/1) |
| 1 | Formation of | 0% (0/1) |
| 1 | Cell prolifera | 0% (0/1) |
| 1 | Differentiati | 0% (0/1) |

| | | |
|---|---|---|
| 1 | Long-term p | 0% (0/1) |
| 1 | Quantity of r | 0% (0/1) |
| 1 | Colony forma | 0% (0/1) |
| 1 | Cell prolifera | 0% (0/1) |
| 1 | Proliferation | 0% (0/1) |
| 1 | Cell prolifera | 0% (0/1) |
| 1 | Differentiati | 0% (0/1) |
| 1 | Developmen | 0% (0/1) |
| 1 | Differentiati | 0% (0/1) |
| 1 | Proliferation | 0% (0/1) |
| 1 | Developmen | 0% (0/1) |
| 1 | Outgrowth o | 0% (0/1) |
| 1 | Proliferation | 0% (0/1) |
| 1 | Developmen | 0% (0/1) |
| 1 | Formation o | 0% (0/1) |
| 1 | Cell prolifera | 100% (1/1) |
| 1 | Dendritic gro | 0% (0/1) |
| 1 | Formation o | 0% (0/1) |
| 1 | Cell prolifera | 0% (0/1) |
| 1 | Cell prolifera | 100% (1/1) |
| 1 | Cell prolifera | 0% (0/1) |
| 1 | Outgrowth o | 0% (0/1) |
| 1 | Proliferation | 0% (0/1) |
| 1 | Proliferation | 0% (0/1) |
| 1 | Proliferation | 0% (0/1) |
| 1 | Quantity of r | 0% (0/1) |
| 1 | Cell prolifera | 0% (0/1) |
| 1 | Growth of a | 0% (0/1) |
| 1 | Cell prolifera | 0% (0/1) |
| 1 | Dendritic gro | 100% (1/1) |
| 1 | Proliferation | 0% (0/1) |
| 1 | Proliferation | 0% (0/1) |
| 1 | Proliferation | 0% (0/1) |
| 1 | Cell prolifera | 0% (0/1) |
| 1 | Migration of | 0% (0/1) |
| 1 | Growth of ne | 0% (0/1) |
| 1 | Outgrowth o | 0% (0/1) |
| 1 | Cell prolifera | 0% (0/1) |
| 1 | Long-term p | 0% (0/1) |
| 1 | Cell prolifera | 0% (0/1) |
| 1 | Cell prolifera | 0% (0/1) |
| 1 | Migration of | 0% (0/1) |
| 1 | Outgrowth o | 0% (0/1) |
| 1 | Proliferation | 0% (0/1) |
| 1 | Cell prolifera | 0% (0/1) |

| | | |
|---|---|---|
| 1 | Cell prolifera | 0% (0/1) |
| 1 | Colony forma | 0% (0/1) |
| 1 | Quantity of r | 100% (1/1) |
| 1 | Growth of ne | 0% (0/1) |
| 1 | Outgrowth o | 0% (0/1) |
| 1 | Cell prolifera | 0% (0/1) |
| 1 | Axonogenesi | 0% (0/1) |
| 1 | Quantity of r | 0% (0/1) |
| 1 | Outgrowth o | 0% (0/1) |
| 1 | Extension of | 0% (0/1) |
| 1 | Formation of | 0% (0/1) |
| 1 | Movement D | 0% (0/1) |
| 1 | Movement D | 0% (0/1) |
| 1 | Differentiatio | 0% (0/1) |
| 1 | Differentiatio | 0% (0/1) |
| 1 | Migration of | 0% (0/1) |
| 1 | Outgrowth o | 0% (0/1) |
| 1 | Formation of | 0% (0/1) |
| 1 | Neuritogenes | 0% (0/1) |
| 1 | Outgrowth o | 0% (0/1) |
| 1 | Motor dysfur | 100% (1/1) |
| 1 | Formation of | 0% (0/1) |
| 1 | Differentiatio | 0% (0/1) |
| 1 | Cell prolifera | 0% (0/1) |
| 1 | Cell prolifera | 100% (1/1) |
| 1 | Differentiatio | 100% (1/1) |
| 1 | Proliferation | 0% (0/1) |
| 1 | Migration of | 0% (0/1) |
| 1 | Cell prolifera | 0% (0/1) |
| 1 | Branching of | 0% (0/1) |
| 1 | Migration of | 0% (0/1) |
| 1 | Differentiatio | 100% (1/1) |
| 1 | Developmen | 0% (0/1) |
| 1 | Proliferation | 100% (1/1) |
| 1 | Cell prolifera | 0% (0/1) |
| 1 | Formation of | 100% (1/1) |
| 1 | Congenital m | 0% (0/1) |
| 1 | Developmen | 0% (0/1) |
| 1 | Long-term po | 0% (0/1) |
| 1 | Cell prolifera | 0% (0/1) |
| 1 | Formation of | 0% (0/1) |
| 1 | Cell prolifera | 100% (1/1) |
| 1 | Outgrowth o | 0% (0/1) |
| 1 | Congenital m | 0% (0/1) |
| 1 | Outgrowth o | 0% (0/1) |

| | | |
|---|---|---|
| 1 | Outgrowth o | 0% (0/1) |
| 1 | Formation of | 0% (0/1) |
| 1 | Formation of | 0% (0/1) |
| 1 | Cell prolifera | 0% (0/1) |
| 1 | Long-term p | 0% (0/1) |
| 1 | Formation of | 100% (1/1) |
| 1 | Cell prolifera | 0% (0/1) |
| 1 | Extension of | 0% (0/1) |
| 1 | Cell prolifera | 0% (0/1) |
| 1 | Long-term p | 0% (0/1) |
| 1 | Cell prolifera | 0% (0/1) |
| 1 | Long-term p | 0% (0/1) |
| 1 | Developmen | 0% (0/1) |
| 1 | Differentiati | 0% (0/1) |
| 1 | Branching of | 100% (1/1) |
| 1 | Motor dysfur | 100% (1/1) |
| 1 | Migration of | 0% (0/1) |
| 1 | Motor dysfur | 100% (1/1) |
| 1 | Cell prolifera | 0% (0/1) |
| 1 | Formation of | 100% (1/1) |
| 1 | Growth of ne | 100% (1/1) |
| 1 | Outgrowth o | 100% (1/1) |
| 1 | Congenital e | 0% (0/1) |
| 1 | Differentiati | 0% (0/1) |
| 1 | Branching of | 0% (0/1) |
| 1 | Cell prolifera | 100% (1/1) |
| 1 | Colony forma | 100% (1/1) |
| 1 | Cell prolifera | 0% (0/1) |
| 1 | Migration of | 0% (0/1) |
| 1 | Differentiati | 0% (0/1) |
| 1 | Cell prolifera | 0% (0/1) |
| 1 | Differentiati | 0% (0/1) |
| 1 | Central nerv | 100% (1/1) |
| 1 | Cerebral disc | 0% (0/1) |
| 1 | Extension of | 0% (0/1) |
| 1 | Formation of | 0% (0/1) |
| 1 | Cell prolifera | 100% (1/1) |
| 1 | Differentiati | 0% (0/1) |
| 1 | Proliferation | 0% (0/1) |
| 1 | Axonogenesi | 0% (0/1) |
| 1 | Motor dysfur | 0% (0/1) |
| 1 | Nervous syst | 100% (1/1) |
| 1 | Proliferation | 0% (0/1) |
| 1 | Developmen | 100% (1/1) |
| 1 | Differentiati | 0% (0/1) |
